# Insights into long non-coding RNA regulation of anthocyanin carrot root pigmentation

**DOI:** 10.1101/2020.10.27.356964

**Authors:** Constanza Chialva, Thomas Blein, Martin Crespi, Diego Lijavetzky

**Author notes:** Constanza Chialva and Thomas Blein contributed equally to this work. Correspondence and requests for materials should be addressed to D.L. or M.C.

## Abstract

Carrot (*Daucus carota* L.) is one of the most cultivated vegetable in the world and of great importance in the human diet. Its storage organs can accumulate large quantities of anthocyanins, metabolites that confer the purple pigmentation to carrot tissues and whose biosynthesis is well characterized. Long non-coding RNAs (lncRNAs) play critical roles in regulating gene expression of various biological processes in plants. In this study, we used a high throughput stranded RNA-seq to identify and analyze the expression profiles of lncRNAs in phloem and xylem root samples using two genotypes with a strong difference in anthocyanin production. We discovered and annotated 8484 new genes, including 2095 new protein-coding and 6373 non-coding transcripts. Moreover, we identified 639 differentially expressed lncRNAs between the phenotypically contrasted genotypes, including certain only detected in a particular tissue. We then established correlations between lncRNAs and anthocyanin biosynthesis genes in order to identify a molecular framework for the differential expression of the pathway between genotypes. A specific natural antisense transcript (NAT) linked to the *DcMYB7* key anthocyanin biosynthetic transcription factor suggested how the regulation of this pathway may have evolved between genotypes.

## INTRODUCTION

Anthocyanins are flavonoids, a class of phenolic compounds synthesized via the phenylpropanoid pathway, a late branch of the shikimic acid pathway^1^. They are secondary metabolites that confer purple, red, and blue pigmentation to several organs and tissues of many plant species^2^. These water-soluble pigments serve in various roles in the plant, including attracting pollinators to flowers and seed dispersers to fruits, protection against UV radiation, amelioration of different abiotic and biotic stresses, such as drought, wounding, cold temperatures, and pathogen attacks^3,4^, as well as participation in physiological processes such as leaf senescence^5,6^. As dietary components, anthocyanins possess various health-promoting effects, mainly due to their antioxidant and anti-inflammatory properties, including protection against cancer, strokes and other chronic human disorders^7^.

Carrot (*Daucus carota* subsp. *carota* L.; 2n = 2x = 18) is a globally important root crop with yellow and purple as the first documented colors for domesticated carrot in Central Asia approximately 1,100 years ago^8^. Orange carrots were not reliably reported until the sixteenth century in Europe^9,10^, where its popularity was fortuitous for modern consumers because the orange pigmentation results from high quantities of α- and β-carotene, making carrots the richest source of provitamin A in the US diet^11^. Additionally, with its great nutrition and economic value, carrot has been well known as a nice model plant for genetic and molecular studies^11^. Carrot is one of the crops that can accumulate large quantities of anthocyanins in its storage roots (up to 17–18 mg/100 g fresh weight)^12^. Purple carrots accumulate almost exclusively derivatives of cyanidin glycosides with five cyanidin pigments reported in most studies^13,14^. The root content of these five anthocyanin pigments vary across carrot genetic backgrounds^12,15^. In addition, anthocyanin pigmentation also varies between root tissues, ranging from fully pigmented roots (i.e., purple color in the root phloem and xylem) to pigmentation only in the outer-most layer of the phloem^16,17^.

Regardless of the plant species, at least two classes of genes are involved in anthocyanin biosynthesis: structural genes encoding the enzymes that directly catalyze the production of anthocyanins, and regulatory genes that control the transcription of structural genes^18,19^. In most cases, the anthocyanin biosynthetic structural genes are regulated by transcription factors (TFs) belonging to the R2R3–<u>M</u>YB, <u>b</u>asic helix-loop-helix (bHLH) and <u>W</u>D-repeat protein families, in the form of the ‘MBW’ complex^19,20^. Recent reports pointed out that gene regulation by TFs may play a key role controlling anthocyanin pigmentation in purple carrots^17,21,22^. Moreover, the broad variation observed among purple carrot root genotypes, regarding both anthocyanin concentration and pigment distribution in the phloem and xylem tissues, suggests independent genetic regulation in these two root tissues^23^. In this sense, Xu et al.^16^ found that the expression pattern of a R2R3–MYB TF, *DcMYB6*, is correlated with anthocyanin production in carrot roots and that the overexpression of this gene in *Arabidopsis thaliana* enhanced anthocyanin accumulation in vegetative and reproductive tissues in this heterologous system. Similarly, Kodama et al.^24^ found that a total of 10 *MYB*, *bHLH* and *WD40* genes were consistently up- or downregulated in a purple color-specific manner, including *DcMYB6*. Iorizzo et al.^25^ identified a cluster of MYB TFs, with *DcMYB7* as a candidate gene for root and petiole pigmentation, and *DcMYB11* as a candidate gene for petiole pigmentation. Bannoud et al.^23^ showed that *DcMYB7* and *DcMYB6* participate in the regulation of phloem pigmentation in purple-rooted samples. Finally, Xu et al.^26^, by means of loss- and gain-of-function mutation experiments, demonstrated that *DcMYB7* is the main determinant that controls purple pigmentation in carrot roots.

Non-coding RNAs with a length higher than 200 nucleotides are defined as long noncoding RNAs (lncRNAs). They were originally considered to be transcriptional byproducts, or transcriptional ‘noise’, and were often dismissed in transcriptome analyses due to their low expression and low sequence conservation compared with protein-coding mRNAs. However, specific lncRNAs were shown to be involved in chromatin modification, epigenetic regulation, genomic imprinting, transcriptional control as well as pre- and post-translational mRNA processing in diverse biological processes in plants^27–30^. Certain lncRNAs can be precursors of small interfering RNA (siRNA) or microRNA (miRNAs), triggering the repression of protein-coding genes at the transcription level (transcriptional gene silencing or TGS) or at post-transcriptional level (PTGS)^27,31^. Additionally, other lncRNAs can act as endogenous target mimics of miRNAs, to fine-tune the miRNA-dependent regulation of target genes^32,33^. It has been suggested that lncRNAs can regulate gene expression in both the *cis*- and *trans*-acting mode^35^. The *cis*-acting lncRNAs can be classified by their relative position to annotated genes^27,34,35^ and notably include long noncoding natural antisense (lncNATs) transcribed in opposite strand of a coding gene, overlapping with at least one of its exons^36,37^. Other so-called intronic lncRNAs are transcribed within introns of a protein-coding gene^38^ whereas long intergenic ncRNAs (lincRNAs) are transcripts located farther than 1 kb from protein-coding genes^27,34,35^. Among these cis-lncRNAs, NATs are of special interest as they have been shown to provide a mechanism for locally regulating the transcription or translation of the target gene on the other strand, providing novel mechanisms involved in the regulation of key biological processes^39^, plant development^40^ and environmentally dependent gene expression^36,37^.

As mentioned above, several differential expression analyses have been performed between purple and non-purple carrot roots allowing the identification of the main structural genes and TFs involved in anthocyanin biosynthesis in whole roots and/or phloem tissues^16,21,23–26^. However, the identification and functional prediction of lncRNA in carrot or putatively involved in carrot anthocyanin biosynthesis regulation has not yet been reported. In the present study, we combined a high throughput stranded RNA-Seq based approach with a dedicated bioinformatic pipeline, to annotate lncRNAs and analyze the expression profiles of lncNATs putatively associated to the carrot root anthocyanin biosynthesis regulation. In addition, we individually analyzed the gene expression patterns in phloem and xylem root of purple and orange *D. carota* genotypes. Our findings point to a role of antisense transcription in the anthocyanin biosynthesis regulation in the carrot root at a tissue-specific level.

## RESULTS

### RNA-seq data mining, identification and annotation of anthocyanin-related lncRNAs

In order to thoroughly identify and annotate lncRNAs related to anthocyanin biosynthesis regulation in carrot roots, we performed a whole transcriptome RNA-seq analysis of specific tissues from the carrot genotypes ‘Nightbird’ (purple phloem and xylem) and ‘Musica’ (orange phloem and xylem) (Supplementary Figure S1). We generated an average of 51.4 million of reads per sample from the 12 carrot root samples (i.e., two phenotypes x two tissues x three biological replicates), ranging from 43.5 million to 60.3 million. The average GC content (%) was 44.8% and the average ratio of bases that have phred^41^ quality score of over 30 (Q30) was 94.1%. The average mapping rate to the carrot genome was 90.9% (Supplementary Table S1). We identified and annotated 8484 new transcripts, including 2095 new protein-coding and 6373 non-coding transcripts (1521 lncNATs, 4852 lincRNAs and 16 structural transcripts) (Supplementary Table S2 and Supplementary File S1). Those were added to the 34263 known carrot transcripts^42^ to complete the final set of 42747 transcripts used for this work. The set contains 34204 coding transcripts and 7288 noncoding transcripts (1521 lncNATs, 5767 lincRNAs) and 1255 structural transcripts (Fig 1A and Supplementary Table S3). As expected, the newly predicted protein-coding genes carry ORFs presenting strong homologies with already annotated ones. In contrary, the great majority of the newly predicted non-coding transcripts present no conservation of their predicted ORFs^43,44^ (Fig 1B). Most non-coding transcripts presented less than 1000 bp long, being 400-800 bp the most frequent length class. Coding transcripts between 500-1000 bp long were the most frequent, while most structural transcripts presented less than 200 bp (Fig 1C). Noncoding transcripts predominantly presented one exon and unexpectedly^45^, only one exon was also the most frequent class for coding transcripts (Fig 1D). Additionally, we found no particular bias for the distribution of the noncoding transcripts along the nine carrot chromosomes (Fig 1E). Finally, the expression level of the coding sequences (measured as normalized counts) was similar within the known, novel and total transcripts. This was also observed for the noncoding transcripts. As expected, the expression level of the coding genes was higher than that of the noncoding ones independently if they were already known or newly predicted (Fig. 1F). Normalized counts for each of the 12 sequenced libraries were included in Supplementary Table S4.

**Figure 1.**
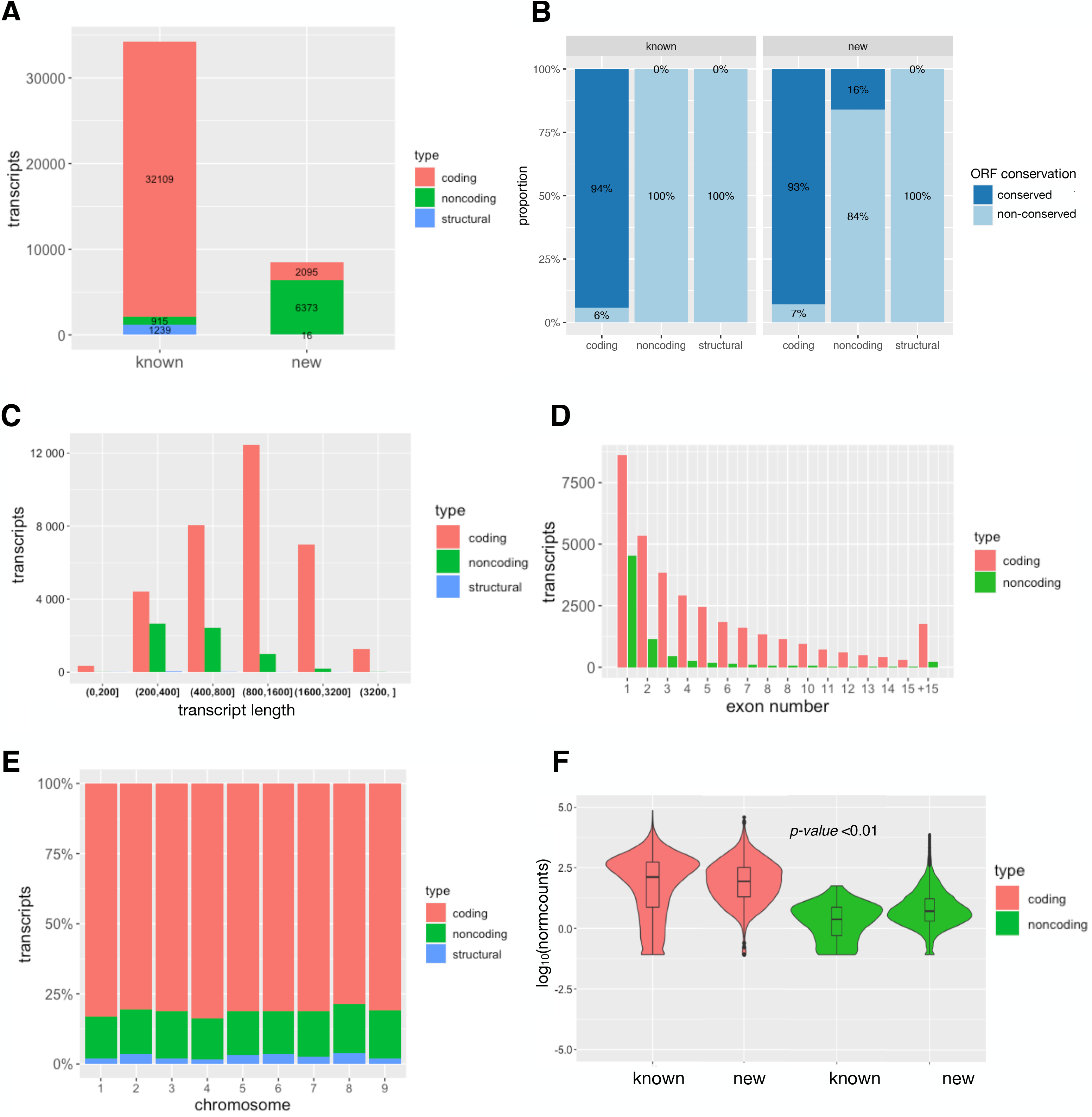
Characteristics of carrot transcripts. (A) Distribution of coding, noncoding and structural sequences between the known and newly annotated transcripts. (B) Conservation of the known and newly predicted protein-coding and non-coding transcripts. (C) Transcript length distributions for the total coding, noncoding and structural RNAs. (D) Number of exons per transcript for the total coding and noncoding RNAs. (E) Proportional distribution of the total coding, noncoding and structural RNAs along each chromosome. (F) Violin plot of the expression levels of carrot total coding and noncoding RNAs. The y-axis represents the average log2 of normalized count values. t-test *p*-value < 0.01 is considered to be significantly different.

### Variation in coding and noncoding expression was mainly explained by the anthocyanin-pigmentation phenotype difference between orange and purple carrots

We sampled phloem and xylem tissues from orange and purple carrot genotypes (Supplementary Figure S1). Considering the global gene variation of the 12 evaluated libraries (i.e., three for each phenotype/tissue combination), the color phenotype was clearly the main source of variation (PC1, 49 %), while the tissue specificity factor was also important albeit less significant (PC2, 18%), (Fig. 2A).

**Figure 2.**
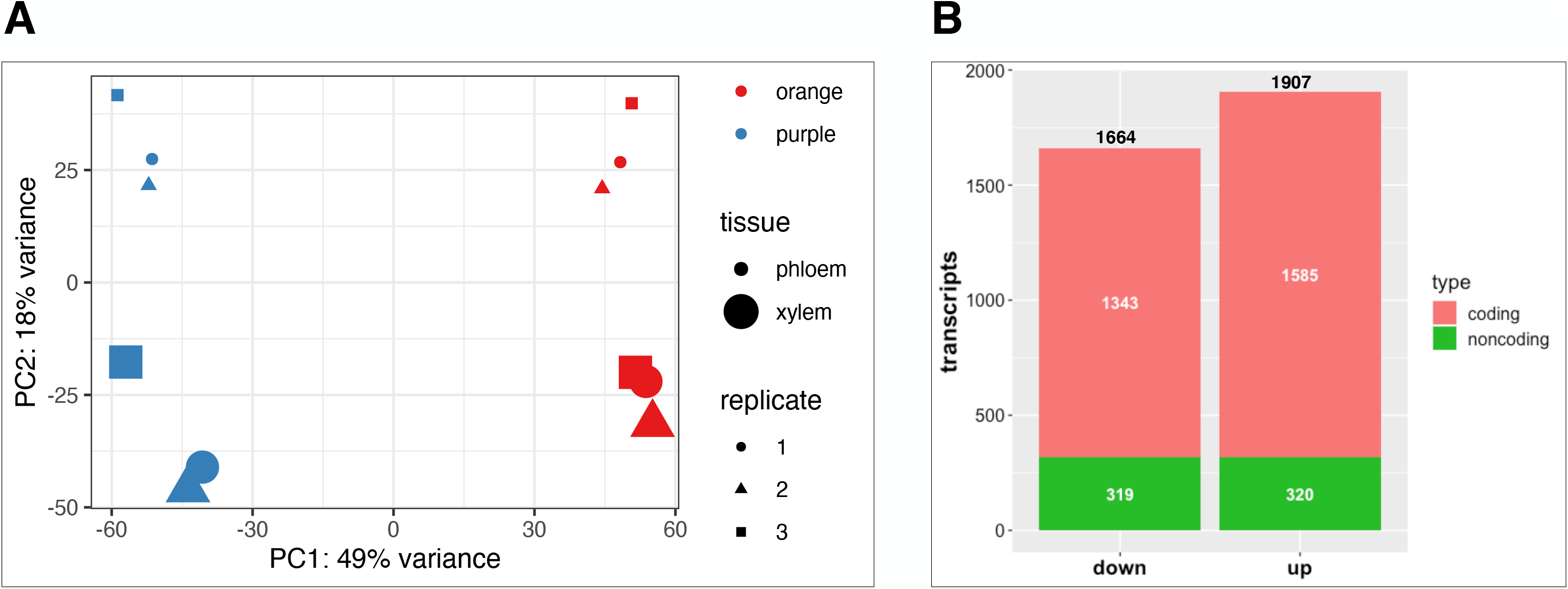
Expression of carrot coding and noncoding RNAs. (A) PCA analysis of the global gene expression of the 12 evaluated libraries (three replicates for each color-phenotype and tissue type combination). (B) Differentially expressed genes (up- and down-regulated) between purple and orange carrots (Bonferroni’s adjusted *p*-value < 0.01) distributed by coding and noncoding transcripts.

We then assessed the variation in mRNA and ncRNA gene expression between purple and orange carrot roots in our RNA-seq analysis. A total of 3567 genes were differentially expressed (DEG) between purple and orange carrots (Bonferroni’s adjusted *p*-value < 0.01), divided in 2928 mRNA and 639 lncRNAs (Fig. 2B) and representing 10% and 15% of the mRNA and lncRNA expressed genes, respectively. Within the 3567 DEGs, we found 1664 downregulated and 1907 upregulated transcripts. In turn, the downregulated transcripts were distributed into 1343 coding and 319 noncoding transcripts, while the upregulated were divided into 1585 and 320 coding and noncoding transcripts, respectively (Fig. 2B). All information concerning the differentially expressed analysis and gene annotation is detailed in Supplementary Table S5.

As expected, we identified several differentially expressed genes (DEG) between the two genotypes known to be involved in carrot root anthocyanin biosynthesis^21,23–26^. Most of the known genes of the pathway and their main regulators were differentially expressed between the two genotypes (Supplementary Table S5). Several genes were induced in purple tissues and they mainly comprised genes representing: i) the early step in the flavonoid/anthocyanin pathway, like chalcone synthase (*DcCHS1*/DCAR_030786); chalcone isomerase (*DcCHI1*/DCAR_027694) and (*DcCHIL*/DCAR_019805); flavanone 3-hydroxylase (*DcF3H1*/DCAR_009483), and flavonoid 3′-hydroxylase (*DcF3’H1*/DCAR_014032); ii) cytochrome P450 (CYP450) proteins, putatively related to the flavonoid and isoflavonoid biosynthesis pathways^23,46^; iii) ATP-binding cassette (ABC) transporters, potentially related to anthocyanin transport^47,48^; and iv) genes from the late steps of the pathway, like dihydro-flavonol 4-reductase (*DcDFR1*/DCAR_021485), leucoanthocyanidin dioxygenase (*DcLDOX1*/ DCAR_006772), and UDP-glycosyltransferase (*DcUFGT*/DCAR_009823) and the recently described *DcUCGXT1*/DCAR_021269 and *DcSAT1*/MSTRG.8365, which were confirmed to be responsible for anthocyanin glycosylation and acylation, respectively^26,49^. Finally, the most significant regulatory genes of the pathway, belonging to the MYB, bHLH and WD40 TF gene families^21,23–26^ were also differentially expressed between purple and orange genotypes (Supplementary Table S5). We further analyzed the tissue differential expression distribution of those 26 ‘MBW’ TFs and found that *DcMYB6* and *DcMYB7,* the two most studied TFs associated with anthocyanin biosynthesis regulation^23–26^, were differentially expressed between purple and orange carrots, both in phloem and xylem tissues (Supplementary Figure S2). Interestingly, three genes recently described to be regulated by *DcMYB7*^26^ (i.e. *DcbHLH3*, *DcUCGXT1* and *DcSAT1*) also displayed no tissue specificity. *DcbHLH3* was described as a co-regulator in anthocyanin biosynthesis, while *DcUCGXT1* and *DcSAT1* participate in anthocyanin glycosylation and acylation, respectively^26,49^. Additionally, seven TFs showed xylem preferential expression-specificity, while only one was preferentially expressed specifically in phloem. Finally, differential expression of 11 TFs was just detected when the 12 libraries were jointly analyzed, presumably because they have significant but low expression differences (Supplementary Figure S2).

### Putative regulation of anthocyanin-related genes by carrot antisense lncRNAs

In order to investigate the putative involvement of carrot lncRNAs in the regulation of the anthocyanin biosynthesis in different carrot root tissues, we predicted the potential targets of lncRNAs in *cis*-regulatory relationship, particularly those classified as natural antisense transcripts (lncNATs). The selection of such lncRNAs was based on three assumptions: i) both, the lncRNA and the putative target were differentially expressed between purple and orange tissues (Supplementary Table S5); ii) the lncRNAs were antisense of the target genes; and iii) the Pearson and Spearman correlation coefficients between the expression levels of these genes were ≥0.70 or ≤−0.70, and *p*□<□0.01.

According to these criteria, we found 19 differentially expressed lncNATs, since the lncRNAs were located in the antisense orientation (in the opposite strand) to a target mRNA, being most of them fully overlapping pairs (Supplementary Table S5 and S6). About 79% of those lncNATs were expressed in concordance with the sense strand transcript, while five out of the 19 presented discordant expression (i.e. when the lncNAT expression increase, the sense strand transcript was repressed) (Supplementary Table S5 and S6). Interestingly, we detected two lncNATs (MSTRG.27767/*asDcMyb6* and MSTRG.9120/*asDcMyb7*) in antisense relationship to the critical regulators *DcMYB6* and *DcMYB7*, respectively, with concordant expression correlation (Fig. 3). *DcMYB6* showed a log_2_ fold-change of 7.6 with an adjusted *p*-value of 4.5 10^−30^, while *DcMYB7* presented a log_2_ fold-change of 11.7 with an adjusted *p*-value of 3.8 10^−37^. Accordingly, the two detected antisense lncRNAs also presented significant differential expression, where *asDcMYB6* displayed a log_2_ fold-change of 6.5 with an adjusted *p*-value of 2.1 10^−13^ and *asDcMYB7* presented a log_2_ fold-change of 6.1 with an adjusted *p*-value of 1.3 10^−04^ (Supplementary Table S5). Finally, the Pearson and Spearman correlation coefficients between the expression levels of each sense/antisense pair were ≥0.79 and *p*-value□< 0.01 (Supplementary Table S6). On the other hand, as also detailed in Supplementary Table S5, two out of the four lncNATs showing discordant expression were found in the antisense relationship with disease resistance related genes (a predicted Catalase, and probable disease resistance protein At5g63020).

**Figure 3.**
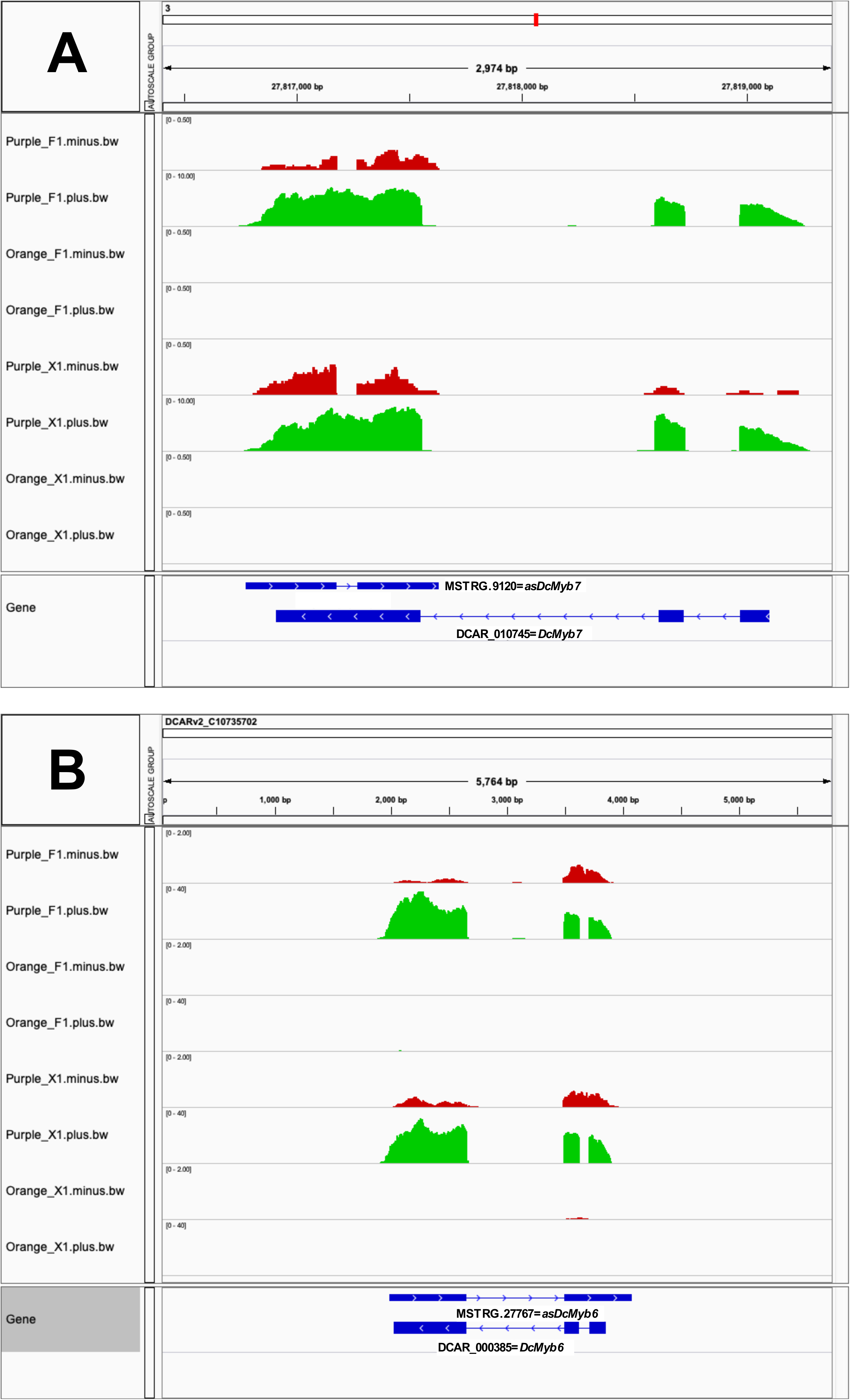
Strand specific expression of R2R3–MYB TFs and their lncNATs. Coverage data for the sense (green) and antisense (red) strands corresponding to *DcMYB7*/*asDcMYB7* (A) and *DcMYB6/as DcMYB6* (B), respectively. Tracks correspond to four carrot libraries: two phloem samples Purple_F1 and Orange F1; and two xylem samples Purple_X1 and Orange_X1. Data Range of each track was set to allow an even visualization of the mRNA and lncRNA transcripts by enlarging the last ones (20x).

### The differential expression of *DcMYB6* and *DcMYB7* and their lncNATs was validated by RT-qPCR

In order to validate the differential expression results obtained by RNA-seq, we performed a RT-qPCR analysis of *DcMYB6* and *DcMYB7* and their corresponding lncNATs (*asDcMYB6*and *asDcMYB7*). As shown in Fig. 4, the expression of the four genes was detected by RNA-seq and RT-qPCR in all purple samples, being mostly undetected in orange tissues. Moreover, both techniques allowed the detection of gene expression in orange tissues only for *DcMYB6*, displaying significantly lower values than in purple tissues. The comparative RT-qPCR expression of the four genes in purple phloem and xylem tissues is presented in Supplementary Figure S3.

**Figure 4.**
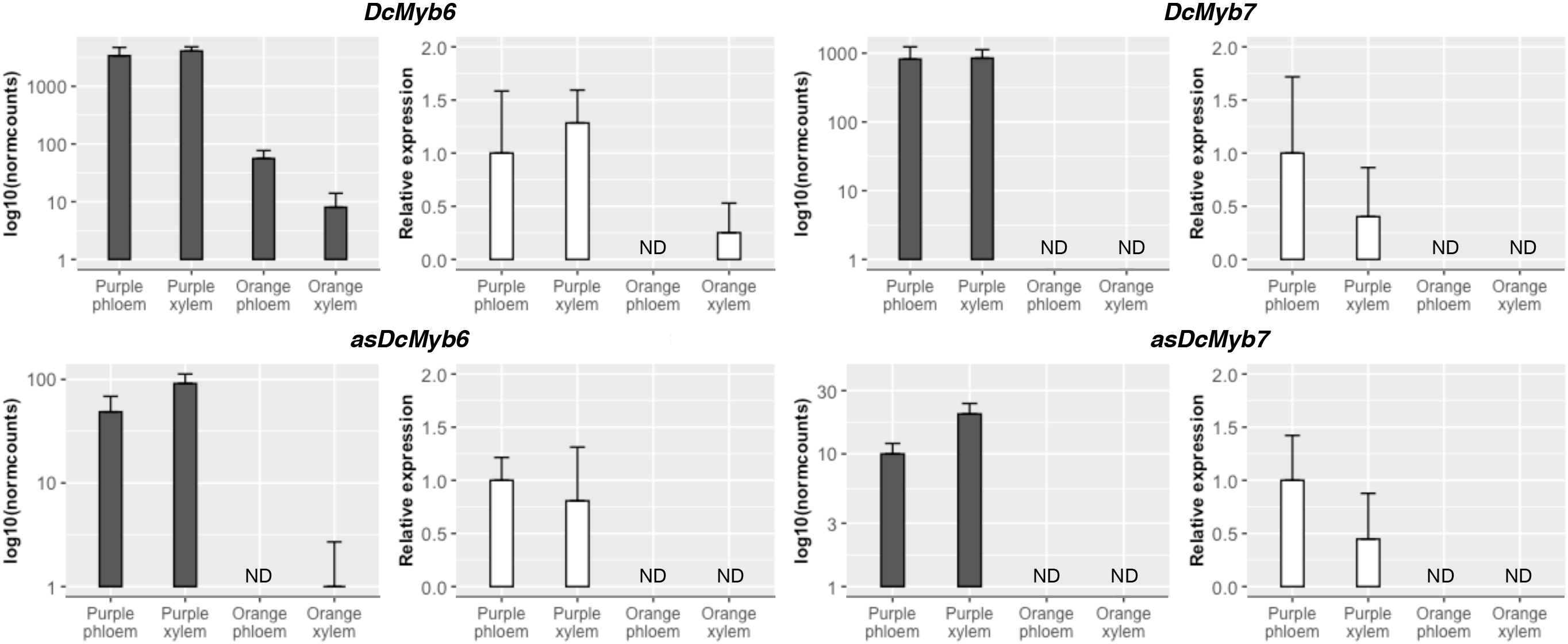
Comparison of expression results from RNA-Seq (log_10_ of normalized counts) and RT-qPCR (Relative expression) methods for *DcMyb6*, *DcMyb7* and their corresponding lncNATs. Data are means ± SD of three biological replicates. For RT-qPCR, carrot *actin-7* was used as reference gene and ‘Purple phloem’ as reference sample. ND= Not detected.

## DISCUSION

The presence of color in flowers, fruits and other organs and tissues, plays several biological functions mostly driven by the adaptive behavior of plants in response to the environment^2,20,50,51^. But in turn, plant organ pigmentation has served as a natural genetic marker since the early works of Mendel^52,53^. Anthocyanins are flavonoid pigments that accumulate in plant cell vacuoles^54^ and are mainly responsible for most tissue and organ coloration^19,20,50^. Genetic analyses using model plant species like Arabidopsis, petunia and maize allowed the identification of most structural genes in the anthocyanin biosynthesis pathway as well as the main regulatory genes controlling pigment synthesis. In carrot, anthocyanin pigmentation is responsible for the purple phenotype^9,55^. Two main genes, *P*_*1*_ and *P*_*3*_, have been identified in chromosome 3 and suggested to be responsible for the two independent mutations underlying the domestication of purple carrots^17^. Despite several carrot structural genes from the anthocyanin biosynthesis pathway have shown expression correlation with the purple phenotype^21,22^, none of them co-localize with *P*_*1*_ and *P*_*3*_. A similar situation occurs in other plants like grapevine, where accumulation of anthocyanins correlated with the expression of several structural genes of the pathway but none of them co-localized with the ‘color locus’ in chromosome 2^56,57^. Finally, this discrepancy was solved by a study describing an insertion mutation in the promotor of a R2R3–MYB TF (i.e. *VviMybA1*)^58^ explaining the lack of color of white grapevine cultivars. In the same direction, several recent works^16,23–25,49^ focused on the role of carrot TFs putatively involved in the regulation of anthocyanin biosynthesis in purple genotypes, particularly those belonging to the ‘MBW’ complex (i.e., R2R3–<u>M</u>YB, basic helix-loop-helix -<u>b</u>HLH- and <u>W</u>D-repeat TFs). Two recent reports showed that three R2R3–MYB TFs are involved in the *P*_*1*_ and *P*_*3*_ loci: *DcMYB113* has been suggested to correspond to *P*_*1*_^49^, while *DcMYB6* and *DcMYB7* were proposed as the two main candidate TFs underlying the carrot root anthocyanin pigmentation in the *P*_*3*_ locus^25^. However, knockdown and overexpression functional analyses demonstrated that *DcMYB7* (but not *DcMYB6*) is the *P*_*3*_ gene controlling purple pigmentation in carrot roots^26^. Likewise described for the grapevine *VviMybA1* gene^58^, non-purple carrot genotypes seems to arise by an insertion mutation in the promoter region of *DcMYB7*^26^, yet the authors imply the existence of an additional genetic factor suppressing the expression of *DcMYB7* in non-purple pigmented peridermal carrot root tissues.

In this work, we performed a thorough transcriptomic analysis by comparing two carrot hybrids with contrasted anthocyanin pigmentation phenotypes (i.e. purple vs. orange), both in phloem and xylem tissues. The study corroborates the involvement of the principal reported structural genes of the anthocyanin biosynthesis pathway^21,22^, but mostly, the key TF genes reported as the main regulators explain the carrot purple phenotype (i.e. *DcMYB6* and *DcMYB7*)^16,25,26^. Interestingly, the performed dissection between phloem and xylem purple samples, allowed us to show that there is no tissue-specific expression of such key genes, contrary to previously suggested for *DcMYB6* and *DcMYB7*^16,23,25^. One possible explanation for such discrepancy is that none of the reported works^16,23,25^ performed phloem and xylem transcriptomic analyses independently.

We showed here a first whole genome identification and annotation of lncRNAs in carrot by combining a high throughput stranded RNA-Seq based approach with a focused bioinformatic pipeline. Through this process, we identified 6373 novel lncRNAs, as compared to the 915 sequences annotated in the original carrot genome assembly^42^. Moreover, 10% of them (641 genes) can be defined as anthocyanin biosynthesis-related lncRNAs since we found them differentially expressed between purple and orange carrots. In order to assess the presumed function of such lncRNAs, we focused on those showing an antisense relationship with differentially expressed protein coding genes, known (or putatively) involved in carrot anthocyanin biosynthesis and depicted in the precedent paragraph. Additionally, the selected lncNATs had to present a statistically significant Pearson and Spearman correlation with their putative targets to further refine our functional predictions. This led us to identify 19 differentially expressed lncNATs between purple and orange carrots. Interestingly, we found two of these lncNATs (*asDcMYB6* and *asDcMYB7*) transcribed in opposite direction to *DcMYB6* and *DcMYB7*, respectively. Moreover, *asDcMYB6* and *asDcMYB7* exhibited concordant expression patterns with their corresponding sense transcripts opening the possibility that non-coding RNA antisense transcription is a new player in the regulation of carrot anthocyanin biosynthesis, through *DcMYB7* (and/or *DcMYB6*). This regulation maybe linked to the previously proposed unknown genetic factors^26^.

Antisense transcripts, particularly lncNATs, present in many genomes of diverse kingdoms, showed either positively or negatively correlated expression with their corresponding sense transcripts. This antisense lncRNAs regulate the expression of their sense transcripts in a negative or positive way, by means of different transcriptional or post-transcriptional mechanisms. In particular cases, upregulation of sense gene expression may be explained by the participation of a lncNAT in the inhibition of other factors at translational level, such as efficient translation initiation or elongation^59–61^.

In plants, both repression and activation roles have been assigned to some lncNATs in response to environmental conditions. While *COOLAIR* and *COLDAIR* negatively regulates *FLC* in vernalization responses^38,62^, and *SVALKA* controls *CBF1* expression to consequently regulate freezing tolerance^37^, the expression of another member of the *FLC* family (*MAF4*) is activated by the lncNAT MAS to fine-tune flowering time^36^. On the other hand, a rice lncNAT (TWISTED LEAF) have shown to maintains leaf blade flattening by regulating its associated sense R2R3-MYB gene^40^.

Anthocyanins are known to participate in abiotic stress responses and adaptation to environmental variations^3,4,63^, so the evolutionary role of the newly identified antisense transcripts *asDcMYB7* and *asDcMYB6* may be linked to the activation of anthocyanin biosynthesis through *DcMYB7* and *DcMYB6*. Hence, our work hints to new antisense regulations potentially involved in the variable expression of anthocyanin genes among carrot ecotypes.

## METHODS

### Sample preparation and plant material

Total RNA was obtained independently from three biological replicates of phloem and xylem root samples of two *Daucus carota* L genotypes: ‘Nightbird’, a purple root hybrid (purple phloem and xylem) and ‘Musica’, a non-anthocyanin pigmentated root hybrid. Plants were germinated from seeds and roots were collected after 12 weeks. Frozen samples were grinded using liquid nitrogen and RNA was extracted using TRI Reagent^®^ (Sigma-Aldrich) and purified using SV Total RNA Isolation System (Promega). RNA samples were quantified, and purity measured using a spectrophotometer (AmpliQuant AQ-07). RNA integrity and potential genomic DNA contaminations were checked through agarose gel electrophoresis.

### Library construction and RNA sequencing

Twelve samples (two genotypes x two tissues x three biological replicates) were sent to the Macrogen sequencing service (Seoul, Korea). Once in destination they were checked for total RNA integrity using a Bioanalyzer RNA Nano 6000 chip. All the samples qualified to proceed with the library construction having an RNA Integrity Number (RIN) ≥7. NGS transcriptomic libraries were constructed using a TruSeq Stranded mRNA LT Sample Prep Kit (Illumina). To verify the size of PCR enriched fragments, the template size distribution was checked on an Agilent Technologies 2100 Bioanalyzer using a DNA 1000 chip. The sequencing of libraries was performed as paired-end 101 bp reads on an Illumina HiSeq 2500 platform. The quality of the raw reads in the FastQ files was checked through FastQC^64^ and were then trimmed for sequencing adaptor and low quality sequences using Trimmomatic^65^ using ‘ILLUMINACLIP:TruSeq3-PE.fa:2:30:10 LEADING:21 TRAILING:21 MINLEN:30’ as parameters. For removing reads corresponding to remaining ribosomal RNA, trimmed reads were mapped to the rRNA reference using SortMeRNA^66^ using ‘--ref silva-bac-16s-id90.fasta --ref silva-bac-23s-id98.fasta --ref silva-euk-18s-id95.fasta --ref silva-euk-28s-id98.fasta --paired_in --fastx --log -e 1e-07 -a 4 -v’ as parameters.

### New transcripts assembly and lncRNA identification

Clean filtered reads were aligned on the *Daucus carota* genome^42^ using the STAR aligner^67^ using ‘--alignIntronMin 20 --alignIntronMax 20000 --outSAMtype BAM SortedByCoordinate --outReadsUnmapped Fastx’ as parameters. Subsequently, the aligned reads were assembled by means of StringTie^68^ and new transcripts were extracted and annotated using the GffCompare^69^ program (GffCompare classes “u”, “x”, to adjust). Only new transcripts whose length was greater than 200 nt were kept. The classification of the newly predicted transcript was performed as follow: i) coding, if their predicted open reading frame (ORF) was greater than 120 aa or if they were predicted as coding by CPC2^70^ calculator; ii) structural, in case of homology with structural RNA (tRNA, rRNA, snRNA or snoRNA) after the analysis against Rfam^71^; and iii) non-coding, if they were predicted as non-coding by CPC2 calculator or in case of homology with known structured non-coding RNA in Rfam (miRNA precursors, lncRNA). For each transcript, the longest ORF on the forward strand with at least 70 amino acid was predicted using TransDecoder (“-S -m 50”, v5.5.0)^72^. Each ORFs was then search against UniRef90 using DIAMOND v2.0.6^73^. Hits with an e-value lower than 1e-10 were considered as positive.

### Differential expression analysis

We performed a strand-specific read counting of coding and non-coding gene using on the carrot official annotation and the newly predicted genes of this study for each of the 12 aligned BAM files by means of the featureCount^74^ software included in the Rsubread package^75^. The resulted normalized counts (median of ratios)^76^ were used for differential expression analysis with DEseq2^77^. Differentially expressed genes were declared as having a Bonferroni’s adjusted *p*-value < 0.01. Reads corresponding to the strand specific expression of mRNAs and their lncNATs were visualized with the Integrative Genomics Viewer (IGV) software^78^. Additional Venn diagrams were performed with Venny v2.1^79^.

### Real-time quantitative PCR (RT-qPCR) expression analysis

One microgram of total RNA from each of the 12 carrot samples described above was used for RT-qPCR. Protocols for cDNA synthesis and RT-qPCR were performed according to Lijavetzky et al. (2008) using a StepOne Plus Real-Time PCR System (Applied Biosystems, Life Technologies). Non-template controls were included for each primer pair, and each RT-qPCR reaction was completed in triplicate. Expression data were normalized against the carrot *actin-7* gene (LOC108202619). Relative quantification was performed by means of the ΔΔCt method using the ‘pcr’ R package^80^. Gene-specific primers were designed using the Primer Blast web tool^81^ and the sequences are described in Supplementary Table S7.

## Supporting information

Supplementary information_Chialva_etal_2020_R1

## ACKNOWLEDGEMENTS

This work was supported by Agencia Nacional de Promoción Científica y Tecnológica (ANPCyT): PICT2015-0822 & PICT-2016-3134; LabEx Sciences des Plantes de Saclay (SPS). Travel suport to Diego Lijavetzky was funded by Programa de Movilidad para Docentes de la UNCuyo, INRA and Universite Paris Sud. We thank to Dr. Pablo Cavagnaro for providing and growing the plant materials.

## AUTHOR CONTRIBUTIONS

C.C. performed the samples collection, laboratory work for library preparation, participated in the draft of the manuscript and figures preparation; T.B. designed the bioinformatic pipeline related to raw data processing, new transcripts identification and differential expression analysis. M.C provided the main insight on the design of the transcriptomic experiment. D.L. wrote the final manuscript, designed and coordinated the experiments. All authors carefully read and helped to improve the final content of the manuscript.

## ADDITIONAL INFORMATION

### Data access

Sequence files generated during this study have been deposited into the NCBI BioProject database accession PRJNA668894.

### Competing Interests

The authors declare no competing interests.

## FIGURE LEGENDS

**Supplementary Figure S1**. Scheme of the performed dissection between xylem and phloem tissues.

**Supplementary Figure S2**. Tissue specific differential expression of the 26 ‘MBW’ TFs identified in the experiment. a) Genes differentially expressed between purple and orange carrots both in xylem and phloem tissues; b) genes differentially expressed between purple and orange carrots just in xylem; c) gene differentially expressed between purple and orange carrots just in phloem; d) genes differentially expressed between purple and orange carrots detected after the join analysis of phloem and xylem samples.

**Supplementary Figure S3**. Comparative RT-qPCR expression of *DcMyb6*, *DcMyb7* and their corresponding lncNATs in purple phloem and xylem. Data are means ± SD of three biological replicates. Carrot *actin-7* was used as reference gene and *asDcMyb7* as reference sample.

**Supplementary Table S1**. Summary of NGS and quality control data regarding the 12 sequenced libraries.

**Supplementary Table S2**. Genome annotation of the newly identified transcripts.

**Supplementary Table S3**. Known and newly annotated carrot genes classified as coding, noncoding and structural transcripts.

**Supplementary Table S4**. Normalized counts for each of the 12 sequenced libraries.

**Supplementary Table S5**. Overall differentially expressed genes (DEGs) list, including statistical tests, *cis*-located sequences, gene lengths and gene products. The 21 identified lncNAT/coding transcript pairs are sorted on top of the list.

**Supplementary Table S6**. Pearson and Spearman correlation coefficients between the expression levels of the 19 identified lncNAT/coding transcript pairs across the 12 analyzed libraries.

**Supplementary Table S7**. Primers used for RT-qPCR.

**Supplementary File S1.** FASTA sequences of the newly annotated transcripts.

## Notes

### Competing Interest Statement

The authors have declared no competing interest.

### Summary of Updates

The Title of the manuscript modified; Section on Results "The differential expression of DcMYB6 and DcMYB7 and their lncNATs was validated by RT-qPCR" added; Section "RNA-seq data mining, identification and annotation of anthocyanin-related lncRNAs" updated with conservation information on new and known lncRNAs and mRNAs

## REFERENCES

1. Herrmann, K. M. & Weaver, L. M. The Shikimate Pathway. Annu. Rev. Plant Physiol. Plant Mol. Biol. 50, 473–503 (1999).

2. Harborne, J. B. & Williams, C. A. Advances in flavonoid research since 1992. Phytochemistry 55, 481–504 (2000).

3. Koes, R. E., Quattrocchio, F. & Mol, J. N. M. M. The flavonoid biosynthetic pathway in plants: Function and evolution. BioEssays 16, 123–132 (1994).

4. Shirley, B. W. Flavonoid biosynthesis: ‘new’ functions for an ‘old’ pathway. Trends Plant Sci. 1, 377–382 (1996).

5. Hatier, J.-H. B. & Gould, K. S. Anthocyanin Function in Vegetative Organs. in Anthocyanins 1–19 (Springer New York, 2008). doi:10.1007/978-0-387-77335-3_1

6. Gould, K. S. & Lister, C. Flavonoid function in plants. Flavonoids Chem. Biochem. Appl. 397–441 (2006).

7. Lila, M. A. Anthocyanins and human health: An in vitro investigative approach. Journal of Biomedicine and Biotechnology 2004, 306–313 (2004).

8. Iorizzo, M. et al. Genetic structure and domestication of carrot (Daucus carota subsp. sativus) (Apiaceae). Am. J. Bot. 100, 930–938 (2013).

9. Simon, P. W. Domestication, Historical Development, and Modern Breeding of Carrot. in Plant Breeding Reviews 157–190 (John Wiley & Sons, Inc., 2010). doi:10.1002/9780470650172.ch5

10. Arscott, S. A. & Tanumihardjo, S. A. Carrots of Many Colors Provide Basic Nutrition and Bioavailable Phytochemicals Acting as a Functional Food. Compr. Rev. Food Sci. Food Saf. 9, 223–239 (2010).

11. Simon, P. W., Pollak, L. M., Clevidence, B. A., Holden, J. M. & Haytowitz, D. B. Plant Breeding for Human Nutritional Quality. in Plant Breeding Reviews 31, 325–392 (John Wiley & Sons, Inc., 2009).

12. Montilla, E. C., Arzaba, M. R., Hillebrand, S. & Winterhalter, P. Anthocyanin composition of black carrot (Daucus carota ssp. sativus var. atrorubens Alef.) Cultivars antonina, beta sweet, deep purple, and purple haze. J. Agric. Food Chem. 59, 3385–3390 (2011).

13. Kammerer, D., Carle, R. & Schieber, A. Quantification of anthocyanins in black carrot extracts (Daucus carota ssp. sativus var. atrorubens Alef.) and evaluation of their color properties. Eur. Food Res. Technol. 219, 479–486 (2004).

14. Kammerer, D., Carle, R. & Schieber, A. Detection of peonidin and pelargonidin glycosides in black carrots (Daucus carota ssp.sativus var.atrorubens Alef.) by high-performance liquid chromatography/electrospray ionization mass spectrometry. Rapid Commun. Mass Spectrom. 17, 2407–2412 (2003).

15. Mazza, G. & Miniati, E. (Enrico). Anthocyanins in fruits, vegetables, and grains. (CRC Press, 1993).

16. Xu, Z. S., Feng, K., Que, F., Wang, F. & Xiong, A. S. A MYB transcription factor, DcMYB6, is involved in regulating anthocyanin biosynthesis in purple carrot taproots. Sci. Rep. 7, 1–9 (2017).

17. Cavagnaro, P. F. et al. A gene-derived SNP-based high resolution linkage map of carrot including the location of QTL conditioning root and leaf anthocyanin pigmentation. BMC Genomics 15, 1118 (2014).

18. Jaakola, L. et al. Expression of genes involved in anthocyanin biosynthesis in relation to anthocyanin, proanthocyanidin, and flavonol levels during bilberry fruit development. Plant Physiol 130, 729–39 (2002).

19. Koes, R., Verweij, W. & Quattrocchio, F. Flavonoids: a colorful model for the regulation and evolution of biochemical pathways. Trends Plant Sci 10, 236–242 (2005).

20. Winkel-Shirley, B. Flavonoid biosynthesis. A colorful model for genetics, biochemistry, cell biology, and biotechnology. Plant Physiol 126, 485–493 (2001).

21. Yildiz, M. et al. Expression and mapping of anthocyanin biosynthesis genes in carrot. Theor. Appl. Genet. 126, 1–14 (2013).

22. Xu, Z. S. et al. Transcript profiling of structural genes involved in cyanidin-based anthocyanin biosynthesis between purple and non-purple carrot (Daucus carota L.) cultivars reveals distinct patterns. BMC Plant Biol. 14, (2014).

23. Bannoud, F. et al. Dissecting the genetic control of root and leaf tissue-specific anthocyanin pigmentation in carrot (Daucus carota L.). Theor. Appl. Genet. 132, 2485–2507 (2019).

24. Kodama, M. et al. Identification of transcription factor genes involved in anthocyanin biosynthesis in carrot (Daucus carota L.) using RNA-Seq. BMC Genomics 19, 811 (2018).

25. Iorizzo, M. et al. A Cluster of MYB Transcription Factors Regulates Anthocyanin Biosynthesis in Carrot (Daucus carota L.) Root and Petiole. Front. Plant Sci. 9, 1927 (2018).

26. Xu, Z.-S., Yang, Q.-Q., Feng, K. & Xiong, A.-S. Changing Carrot Color: Insertions in DcMYB7 Alter the Regulation of Anthocyanin Biosynthesis and Modification. Plant Physiol. pp.00523.2019 (2019). doi:10.1104/pp.19.00523

27. Ariel, F., Romero-Barrios, N., Jégu, T., Benhamed, M. & Crespi, M. Battles and hijacks: noncoding transcription in plants. Trends Plant Sci. 20, 362–371 (2015).

28. Forestan, C. et al. Stress-induced and epigenetic-mediated maize transcriptome regulation study by means of transcriptome reannotation and differential expression analysis. Sci. Rep. 6, 1–20 (2016).

29. Forestan, C. et al. Epigenetic signatures of stress adaptation and flowering regulation in response to extended drought and recovery in Zea mays. Plant Cell Environ. 43, 55–75 (2020).

30. Mercer, T. R. & Mattick, J. S. Structure and function of long noncoding RNAs in epigenetic regulation. Nature Structural and Molecular Biology 20, 300–307 (2013).

31. Wu, H. J., Wang, Z. M., Wang, M. & Wang, X. J. Widespread long noncoding RNAs as endogenous target mimics for microRNAs in plants. Plant Physiol. 161, 1875–1884 (2013).

32. Zhang, G. et al. Transcriptomic and functional analyses unveil the role of long non-coding RNAs in anthocyanin biosynthesis during sea buckthorn fruit ripening. DNA Res. 25, 465–476 (2018).

33. Yang, T. et al. Systematic identification of long noncoding <scp>RNA</scp> s expressed during light-induced anthocyanin accumulation in apple fruit. Plant J. 100, 572–590 (2019).

34. Rinn, J. L. & Chang, H. Y. Genome Regulation by Long Noncoding RNAs. Annu. Rev. Biochem. 81, 145–166 (2012).

35. Bonasio, R. & Shiekhattar, R. Regulation of Transcription by Long Noncoding RNAs. Annu. Rev. Genet. 48, 433–455 (2014).

36. Zhao, X. et al. Global identification of Arabidopsis lncRNAs reveals the regulation of MAF4 by a natural antisense RNA. Nat. Commun. 9, 5056 (2018).

37. Kindgren, P., Ard, R., Ivanov, M. & Marquardt, S. Transcriptional read-through of the long non-coding RNA SVALKA governs plant cold acclimation. Nat. Commun. 9, (2018).

38. Heo, J. B. & Sung, S. Vernalization-mediated epigenetic silencing by a long intronic noncoding RNA. Science (80-.). 331, 76–79 (2011).

39. Thieffry, A. et al. Characterization of Arabidopsis thaliana promoter Bidirectionality and Antisense RNAs by Depletion of Nuclear RNA Decay Pathways. Plant Cell tpc.00815.2019 (2020). doi:10.1105/tpc.19.00815

40. Liu, X. et al. A novel antisense long noncoding RNA, TWISTED LEAF, maintains leaf blade flattening by regulating its associated sense R2R3-MYB gene in rice. New Phytol. 218, 774–788 (2018).

41. Ewing, B., Hillier, L. D., Wendl, M. C. & Green, P. Base-calling of automated sequencer traces using phred. I. Accuracy assessment. Genome Res. 8, 175–185 (1998).

42. Iorizzo, M. et al. A high-quality carrot genome assembly provides new insights into carotenoid accumulation and asterid genome evolution. Nat. Genet. 48, 657–666 (2016).

43. Deng, P., Liu, S., Nie, X., Weining, S. & Wu, L. Conservation analysis of long non-coding RNAs in plants. Sci. China Life Sci. 61, 190–198 (2018).

44. Diederichs, S. The four dimensions of noncoding RNA conservation. Trends in Genetics 30, 121–123 (2014).

45. Ramírez-Sánchez, O., Pérez-Rodríguez, P., Delaye, L. & Tiessen, A. Plant Proteins Are Smaller Because They Are Encoded by Fewer Exons than Animal Proteins. Genomics, Proteomics Bioinforma. 14, 357–370 (2016).

46. Ayabe, S. I. & Akashi, T. Cytochrome P450s in flavonoid metabolism. Phytochemistry Reviews 5, 271–282 (2006).

47. Kovinich, N. et al. Arabidopsis MATE45 antagonizes local abscisic acid signaling to mediate development and abiotic stress responses. Plant Direct 2, e00087 (2018).

48. Francisco, R. M. et al. ABCC1, an ATP Binding Cassette Protein from Grape Berry, Transports Anthocyanidin 3-O-Glucosides. Plant Cell Online (2013). doi:10.1105/tpc.112.102152

49. Xu, Z., Yang, Q., Feng, K., Yu, X. & Xiong, A. DcMYB113, a root-specific R2R3-MYB, conditions anthocyanin biosynthesis and modification in carrot. Plant Biotechnol. J. pbi.13325 (2020). doi:10.1111/pbi.13325

50. Mol, J., Grotewold, E. & Koes, R. How genes paint flowers and seeds. Trends Plant Sci 3, 212–217 (1998).

51. Winkel-Shirley, B. Biosynthesis of flavonoids and effects of stress. Curr Opin Plant Biol 5, 218–223 (2002).

52. Henig, R. M. The monk in the garden: the lost and found genius of Gregor Mendel, the father of genetics. (Houghton Mifflin Harcourt, 2017).

53. Hartl, D. L. & Orel, V. What did Gregor Mendel think he discovered? Genetics 131, 245 (1992).

54. Petrussa, E. et al. Plant Flavonoids—Biosynthesis, Transport and Involvement in Stress Responses. Int. J. Mol. Sci. 14, 14950–14973 (2013).

55. Simon, P. W. Inheritance and expression of purple and yellow storage root color in carrot. J. Hered. 87, 63–66 (1996).

56. Kobayashi, S., Ishimaru, M., Ding, C. K., Yakushiji, H. & Goto, N. Comparison of UDP-glucose:flavonoid 3-O-glucosyltransferase (UFGT) gene sequences between white grapes (Vitis vinifera) and their sports with red skin. Plant Sci 160, 543–550 (2001).

57. Kobayashi, S., Ishimaru, M., Hiraoka, K. & Honda, C. Myb-related genes of the Kyoho grape (Vitis labruscana) regulate anthocyanin biosynthesis. Planta 215, 924–933 (2002).

58. Kobayashi, S., Goto-Yamamoto, N. & Hirochika, H. Retrotransposon-induced mutations in grape skin color. Science (80-.). 304, 982 (2004).

59. Pelechano, V. & Steinmetz, L. M. Gene regulation by antisense transcription. Nature Reviews Genetics 14, 880–893 (2013).

60. Wight, M. & Werner, A. The functions of natural antisense transcripts. Essays Biochem. 54, 91–101 (2013).

61. Faghihi, M. A. & Wahlestedt, C. Regulatory roles of natural antisense transcripts. Nature Reviews Molecular Cell Biology 10, 637–643 (2009).

62. Swiezewski, S., Liu, F., Magusin, A. & Dean, C. Cold-induced silencing by long antisense transcripts of an Arabidopsis Polycomb target. Nature 462, 799–802 (2009).

63. Rienth, M. et al. Day and night heat stress trigger different transcriptomic responses in green and ripening grapevine (vitis vinifera) fruit. BMC Plant Biol. 14, 108 (2014).

64. Andrews, S. FastQC: a quality control tool for high throughput sequence data. (2010). Available at: https://www.bioinformatics.babraham.ac.uk/projects/fastqc/.

65. Bolger, A. M., Lohse, M. & Usadel, B. Trimmomatic: a flexible trimmer for Illumina sequence data. Bioinformatics 30, 2114–2120 (2014).

66. Kopylova, E., Noé, L. & Touzet, H. SortMeRNA: fast and accurate filtering of ribosomal RNAs in metatranscriptomic data. Bioinformatics 28, 3211–3217 (2012).

67. Dobin, A. et al. STAR: ultrafast universal RNA-seq aligner. Bioinformatics 29, 15–21 (2013).

68. Pertea, M. et al. StringTie enables improved reconstruction of a transcriptome from RNA-seq reads. Nat. Biotechnol. 33, 290–295 (2015).

69. GffCompare. A Program for processing GTF/GFF files. Available at: https://ccb.jhu.edu/software/stringtie/gffcompare.shtml.

70. Kang, Y.-J. et al. CPC2: a fast and accurate coding potential calculator based on sequence intrinsic features. Nucleic Acids Res. 45, W12–W16 (2017).

71. Nawrocki, E. P. & Eddy, S. R. Infernal 1.1: 100-fold faster RNA homology searches. Bioinformatics 29, 2933–2935 (2013).

72. Haas, B. J. et al. De novo transcript sequence reconstruction from RNA-seq using the Trinity platform for reference generation and analysis. Nat Protoc (2013).

73. Buchfink, B., Xie, C. & Huson, D. H. Fast and sensitive protein alignment using DIAMOND. Nat. Methods 12, 59–60 (2015).

74. Liao, Y., Smyth, G. K. & Shi, W. featureCounts: an efficient general purpose program for assigning sequence reads to genomic features. Bioinformatics 30, 923–930 (2014).

75. Liao, Y., Smyth, G. K. & Shi, W. The R package Rsubread is easier, faster, cheaper and better for alignment and quantification of RNA sequencing reads. Nucleic Acids Res. 47, e47–e47 (2019).

76. Anders, S. & Huber, W. Differential expression analysis for sequence count data. Genome Biol. 11, R106 (2010).

77. Love, M. I., Huber, W. & Anders, S. Moderated estimation of fold change and dispersion for RNA-seq data with DESeq2. Genome Biol. 15, 550 (2014).

78. Thorvaldsdóttir, H., Robinson, J. T. & Mesirov, J. P. Integrative Genomics Viewer (IGV): high-performance genomics data visualization and exploration. Brief. Bioinform. 14, 178–192 (2012).

79. Oliveros, J. C. Venny. Venny. An interactive tool for comparing lists with Venn’s diagrams. (2015).

80. Ahmed, M. & Kim, D. R. pcr: An R package for quality assessment, analysis and testing of qPCR data. PeerJ 2018, e4473 (2018).

81. Ye, J. et al. Primer-BLAST: a tool to design target-specific primers for polymerase chain reaction. BMC Bioinformatics 13, 134 (2012).

