## Supplemental File 1 for "Insights into long non-coding RNA regulation of anthocyanin carrot root pigmentation"

>MSTRG.3.1 gene=MSTRG.3

CAGTAATCCATTTTTCTTTCTTTACTCAGATCTACTTTCTCTCTCTCCTCGTTTCTCTTACCTCCGTCTT

TCTCTCAACCTGCCCCGATGTAAATCAATATAGTTTGCTACAACTGAATTTTGTTCCCATTGAGATAGAT

ACAAACATATAGATATAATTTAGTGCTGAAGTTGCGGTGATTTTGGTGTTTGATGATACTACGTTTTGGC

AGTGGTGATGTTGGTGATGGGGTGTCAACGGTGGTAGTGCAATGGTAGTGGCAGAGGCAGAGGCGAATTC

ACCCACCATCACGTCTCCACAAGTCCGCTGTTTACGCCTCCTGCTATATACTTTTGTGATTTTCGCTTTG

CAAATTAGTGATTTGTGCTTTATAAATTAGTGATTATCACTTTAGAAATTGGTAAAACAACGGGTACAGC

ACAGAAGCTAGTGACAGTGACACCAGGATGTGGCTGTGTTTTCAGACGACAGAGGTGAGGCTGGAGGTGA

CGGAGGTGAGGCGGG

>MSTRG.8.1 gene=MSTRG.8

GATACTACTACCTCCATTCTGCACCTTCCTGATGATTGTCTCTATTTCATTTTTCAACGCCTTGAATCTG

TTTTTGATCGCAAATCCTTTGGTCTCACTTGCCATCGCTGGCTATTCATAGAAAATACAAGCAGGCGGTC

TCTGCAATTCCCGTGTTCCTTCCGCCACCTACACCGCACTTCATTATCTCGAACCAGCTCAACTGTTGGT

TCTTTCCAGTTGTATACAATGCTTGATCGCTTCCAACACCTAGAGTTATTGTGCCTTTCAGGATGCGTAC

ATCTTCCTGACTCGGGGTTATCTCAATTGCAATATTATGGCTCGAAATTGCAGACGCTGTATCTAGATTG

TTGTTTTGGAATCACTGATGAT

>MSTRG.13.1 gene=MSTRG.13

GTAAATTTATTCTAAATAAAATGTCGATTCTGCCCTCCCCAGTCTACTCCGAGCCTGTTTCCTGACCCCG

GCCTCTGGGATGTAGCGATCATATCTTTCACATTGCATGCTCAGCTCGTTACTTCCTCCACCCACATTCA

TCTTTCACGGCTTATTTAGGCGATCATCATATCTGCGCAGTTTATGTATCAGGACTTAGGAGTGGAGCAG

GTGTCTGTGATCTAAATTCACGGTTCAGGGATGATAATTGGAGGATATGCCCTTCGGTGGTGAGCGCCTT

ATTTGCCTTCTGTGGAGGTTTGACAATGGAGTTTGTGTAATGTAGTGCGTGAAAACTCAACTGGCTTGTT

TCAGGAGATTAAGTGAGTATGATGTTAAGACTATCGGGTGTGTATCCGTGACATGTTGGACAATCTGGTA

GGCTCGAAAAGGAATTGGTATGGAGGGAAAAAAAATCAGTTGCTACAAAAGTGGTAAATAATGCAATTAT

TTTCCTTGATACATGGAATCTTGCTCGAACATATGATTCTAATCATCCGGTTGCCGCTACAAATCATGCT

GAAGTAGCAGAGCGCTGGTCTAAACTTGAAATTGGTAATATCAAGTTTAATTGTGATGCTGCTATTTTTA

GCTCTGAAAGCTGGCCCATGATCATAATTGTGTGTTGTTAGAGACTGCTGCTAATGTCCGTGTAGGGGGA

ATGGAACTGCAACTTGCTGAAGCGATTAGTTTAAAGGTGGTTGTGAATTGGATCAAAGATCTCAACATCA

ATACTTTTGCTGGTGTGCAGACTGATTGTGAGCACACATTCCTTGCTCTCTTCTTCAGGTCTGATTGTTG

CGATTGCAAGGCATTATATTACCAAATGTTTCTATGTCTTTTGTAAAATGGCTCGGAAACAGGACAATCA

ATTAGTCTTCTTGTTCACTACCTGGCCGTAGTCAATGAGGGTTCTGTCCCTATTGAGTTGGAGCCAAACT

TGATAGTTGATTTATCTTAATGAAATTGTTCCATTACCTTTCTAGAAAAATATATGTTTCCTTATATCGG

TATTAATTACACCTACGTTACAGATGTATGAACTACACTTGAC

>MSTRG.13.2 gene=MSTRG.13

CTTTCACATTGCATGCTCAGCTCGTTACTTCCTCCACCCACATTCATCTTTCACGGCTTATTTAGGCGAT

CATCATATCTGCGCAGTTTATGTATCAGGACTTAGGAGTGGAGCAGGTGTCTGTGATCTAAATTCACGGT

TCAGGGATGATAATTGGAGGATATGCCCTTCGGTGGTGAGCGCCTTATTTGCCTTCTGTGGAGGTATAGG

CTCAGCAAACCTGAAACATCAGAAATGCCATAGGGTGTTGCAAAGAACGCGTTCATTACATATACGGCCA

AAGGTTTGACAATGGAGTTTGTGTAATGTAGTGCGTGAAAACTCAACTGGCTTGTTTCAGGAGATTAAGT

GAGTATGATGTTAAGACTATCGGGTGTGTATCCGTGACATGTTGGACAATCTGGTAGGCTCGAAAAGGAA

TTGGTATGGAGGGAAAAAAAATCAGTTGCTACAAAAGTGGTAAATAATGCAATTATTTTCCTTGATACAT

GGAATCTTGCTCGAACATATGATTCTAATCATCCGGTTGCCGCTACAAATCATGCTGAAGTAGCAGAGCG

CTGGTCTAAACTTGAAATTGGTAATATCAAGTTTAATTGTGATGCTGCTATTTTTAGCTCTGAAAGCTGG

CCCATGATCATAATTGTGTGTTGTTAGAGACTGCTGCTAATGTCCGTGTAGGGGGAATGGAACTGCAACT

TGCTGAAGCGATTAGTTTAAAGGTGGTTGTGAATTGGATCAAAGATCTCAACATCAATACTTTTGCTGGT

GTGCAGACTGATTGTGAGCACACATTCCTTGCTCTCTTCTTCAGGTCTGATTGTTGCGATTGCAAGGCAT

TATATTACCAAATGTTTCTATGTCTTTTGTAAAATGGCTCGGAAACAGGACAATCAATTAGTCTTCTTGT

TCACTACCTGGCCGTAGTCAATGAGGGTTCTGTCCCTATTGAGTTGGAGCCAAACTTGATAGTTGATTTA

TCTTAATGAAATTGTTCCATTACCTTTCTAGAAAAATATATGTTTCCTTATATCGGTATTAATTA

>MSTRG.16.1 gene=MSTRG.16

GCTAAATTTTGTGTCACCTCTGGAAGCTCGTCGATAATTTTGTCCACTGGTATGCCGACGATCTGTTCCG

CCTCCTTATTGAAGAGGGTGAAAGTTGTTTGGGCTGTTGAATCTTCCACTCGGACGGTAAGTCTGAACCT

GTATTCAATTGATGAGTATCCTTAGTTTTTTTAATAGAATGCAAATGTGGTTGGCATTCATGCTAAAGGA

TGATTGCAGTTAAATTACCGTGGCTTGAAGTCTCCGGCCTCCTTAGAGCAGAATGTGCAGTAATACTTCC

CCTCAGTAGTCTGTACTTTGCGGGCGCATCGTGGGCAACAATTGTAGTACCATCCCTCGCCTTCCATCAC

CGCCACTATTGTCGCCTCGCATGTCCGTCGTAGAACCTACAATTTAAGTATTTTTTTATAGCAGTTGCTA

GGGGTTAACAGTGAGTTTGTCAGTCCTAAGAATATGCACATTAGGACCTCATGCTTACCAGGTAG

>MSTRG.22.1 gene=MSTRG.22

GGCAAAACAGGCCTCTCCCTCTTCTTCATTGTCTGCATATCCAGCCCGCTATTACTCTCAACCAAGAATT

TGAAATTAATTTCTGTAGTTTTGAAGCACAGATACAAAATTGAGAACAAGCAAGAAGACGAAGCAAAAAA

TACTCTATACTCACTTGAATTAGTTTTGGGTAAATTAACACTTGTTTATATATTTAATTATAGATAGAGA

GGCCTAGCTACAAATCGGTATCGGAAAATTCAAAATCCAGGTAATTCTAAGC

>MSTRG.26.1 gene=MSTRG.26

GTCATCGTGGATCAGTCCTCAATCATCGCATCATTTACCGGAACAGGGAATTAGATGCAATCGATGATTA

TGTTCGAATTGGTGAAAGCACCGCAATTGAAAGTTTGAGGAAATTTGTCAAAGCGATTGTGCAAATATTT

GGGGAACAATATTTAAGAAGGCCACATAGCGGTGATGTTAAAAGATTAATGGAGGTTGCTGAGCAGCGTG

GATTTCCAGGAATGCTAGGCAGTATCGATTGTATGCATTGGAGGTGGAAAAACTGTCCAACTGCGTGGCA

TGGGGCTTTTTCAGGTCGTTACCATGAGCCAACAATCATCCTTGAAGCTGTAGCTTCGTATGATCTTTGG

ATATGGCATGCTTATTTTGGGTTACCTGGATCGTTGAATGATATAAATGTATTGGACCGATCCACTTTAT

TTTCTGAATGGGCAGAGGGTCAAGCACCAGAAGTTAACTATACCATCAATGGGCACAATTACA

>MSTRG.27.1 gene=MSTRG.27

CAGGTGCACTTCAAGATGGATTGTGGTGAAAAAGTGCCATCTTTTTATAATTTGATCAGTGGAGACTGGA

GAGTATTCTGAATATGACGATTTTACACAAAATCAGGAAGAAAGGAATTTAGAAGCTACAATACAAAATT

CAAGTACTCGTTTTACAGAAAGTCAAGAAGAAAGAAATGCAGAATTCATATCAGTTGCATCTCAAGTATT

TTGTTCACAGAAAGTGAAGGAGAAAAGGCATCGAACGAAAAATTTTACAAGGCAAGAGGATGAATTATTG

ATTTCTGCTTGGCAAAATGTGTCACTGGATCCCATCACAGGGGCTGATAAAAAAATGGAACCTATTGGCA

GAGAGTGCAAAGCTACTTTATGAAGCATAAAAACTTTGAATCAGATCATACTTGGGGTTCATTGATGCAT

CGTTGGTCCGTGATTCAACTTGGGGTCAATAAATTTCAAGGATTTTATAATCAAATGGATGGAAGAAGCG

GGTACTCTGAAATTGATAAGATTCAAGGTGCAAAGCATATGTACAAAGAAGTATGCAAACAAAGCTTTTC

TCTCGAGCATTGTTGGAAGTACTTGGAACATATTGAGATATTTGCCTAAGTGGAATGTTGATTTTGCTAC

GAAAAGGACAAAAGTATCTCAAAAGAACAGTCCAACAACTTCTTCACCATCAACTCCAGAGTGTACAATC

GTGGAAAACTCGGACTTGGAAAGGCCTATCGGAAGGAAGGCTGCAAAAGAACTTCAGAAAAAAAAGAAAG

AAATTGGACAATGAGCACGATGATGATGCTGGAGCTGTAATTTTAGAAAAAAATGAGAGTTGATCAGATT

GAATCTAGAAAGCAAAGAAATGAGCACCTTAAAGAGATGTTGCAGTTGGCAAAAGAGAGAGATGAACGTG

AGAAGAGAAAAGAAGCCGCTGAACAAGTTGAAGCTGATGATAAAATTATGGCTATGGATTCAAGTTCTAT

GGGGGTAATTGAGGCTGAGTATTTTAACTTGAGAAAGCAGAAGATTATTGAAAGAAAGCGCAAACTTTTG

GTCAAGTAATTAATGTGTTATACAGCTAGAAATCTTTTGTCATTTAATTT

>MSTRG.29.1 gene=MSTRG.29

GACAGATCTTCAATCCACCAGTCTCCTTCACATTTGGCCTGCCGATTCATCTGAAGTCTTCCAGAACTAG

TATATTCCAAGCGAGAGAAGCTTGAAGCGATCTGTTCAACACACGCTTTGGATACTATAGCGTCAATGCA

TCTGCTAACAATTTTAAGCTCATCAGCAAAAGGAAGTAGGTTTTCACATTGTTGCAGTACTGCAATACAC

ATTTCAAGGTTCTTGCAGACAATAGTCTCCAGATATTCTTCAGCACGGGACCCAAGATTGTTACTTGAGA

ACTCCTCAGTCATTTCAAGGTAGTCGGAGACGCAAAGAAGTTGAGCAACATTAGCCGAT

>MSTRG.31.1 gene=MSTRG.31

CATGGCTCCCCCAGATAACTTCTCCATAAGAGTCGTGGTAGATGCTATCTCTCGTAGTGGAGATATCGTT

GAATCCGTGATCTATTTGGATTAGGAGTCTGAGGTAGTCATACTCAAGACTAATTCTAACAGGAGTAGGA

TTCTCCAAGATGGGCCTTCGGCCTATCTCCGACCTTATCGCCAAGAGGCCTAGTCCTGATCAAACTAGGA

CTCCTGGTTCCATAGAACTACGTACGGCTTGATCCCCTATAAATATAGGGGTACGTAGGCACATTGGGGG

ATCATGAGTTGAGAGCACGTGAGACCAACTCAAAACCCTAATCTCAGCCACCCCCTAAAACAAACAACCA

CCACTCTCCGGCGACCAAAACCAC

>MSTRG.38.1 gene=MSTRG.38

CTCCCTGAACTGCGCAACCAAAAACTCCCCGAAGCAGCAGCTCGGATCCCAACAGTAAACCGCTAACAAA

ATATAAACTGTGTTGCTAACCAGGAGGGGGCTTAGGAGAAGACGGAGGAGGATAGATCTAAGTTTGTGTG

GGCAATTTAGAGAATTTGATTGATGAGAATCATGCTATTGTTTCCAGTTTCTGTTGGTCCCGAGTATTGT

G

>MSTRG.42.1 gene=MSTRG.42

CAAAATTTAAGGTCAAAGAAAGTTTGTAAAAAGTATGAGCGATGATAAATGGGAGTGAAAAGTTTCTATA

ATGAGATTTTTGAAGAAATTGAGTAATGAAAATGATGAATATTTGCCTACAGAATAGTACAAATTTCATT

CCGACCTGAATTACTTTCATCCAACTAAATGGAGCATTGAGGATTCGTGTTTTAAAACTAAAAAATAAAT

CAAGATCATGAAAGTGAAATTAGGGCAAGCAGAAAAATAGGGGCAAATGACCTTTTCATCACATGTTGTA

TGGTCCTTGATTTTACTATTGCCCCTTGACCATCTCTAGAGTTAATATTTGCTATTCATAGTTCAAGCTA

AAATTTAATTCGCTATATATCATCATAAAATTCTATAATTTTTTTCCTGTGGTGTTAGGCATCAAATTAT

GATCACTAAAATCTCTCTTTCCTGTACCAGGGGCTAAACACATTGGTGGTTGTTCAGATACTGTTGAAAT

TTATCAGAAAGGGGAGCTTGAAAATTTGTTATCTGAAGCTGGCGCAAAGAAATAGAGCTAGTCTTTAACC

TCATCAAGGATCTCTATGCTGCTTCACTTTCATCAGAAAGAGTCACGAGTAATCAGTGTTGATTTGTCCA

CGAGTCCATAGGTTTCGGCATTAACACGTTTTGTTTGATATGTTTTTCTTAAGAAGTGTGCAAGTCCAGG

TAAATTTGTAGTGTTGAAAGAGTGAACTTGGTTTTATCCACATATTGCGATGGCATTGTTTTCCCTCAAG

AGGACTAAAATCTTTTTTGATCAAACTCGAGATGAGTTCCTTTTCTTATGGAACTTAGCTGATCTTGATA

AAGATTATATTAAGTACTCCCTCAGTCTCAATGAGTTGTATACATTTGGACGGAGGTGATATGTTATATG

TATACATTTGAGACGGAGGGTTCTGCACGAATTTTAAGGCTCTTATAATAGCATAAATTAGGAGTTGTAC

GGATGCTGTTGGGCCAGCCACATGTTCCCATTTCTGACTACCCAACTGGGATTCAAGTTCGAAACATTTC

CTAGATTTAAAGGCAAAAATGGATATTCTTAT

>MSTRG.43.1 gene=MSTRG.43

GTAGTATTGTGGGGTTTATGATTTTTTGTGGTGTGCAGAGCATTATGGCTCCTACAAGGAAGACAAGCTC

CTCTTTGAATACAACTAAAGTCATTATGGCTCCTACAAGGAAGACAAGATCCTCTTCGAATACAACTAAA

GTCAACCTTATGAAGGCATTAAAAAGAAACCATTCAGCAGGGGAGACAAGTGGGACGTCTAAGAAGCTTA

AGGCATCAACAAAAAACCATGCTGCAGGGGAGACAAGTGGGCTAGCTAAAAAGGCATTAAAAGGAAGCAC

TGTGCAGCCTCCCTCTCAGAGTTCTTCACCTAAATCTGAAAAGAAGAAGGCCATTTCTACAAATCTACAA

CAGCAACTGAGTAAGAAGCGGAAACAAAGGGAAGAAGGGCCTATAGATAACGAACAAGGAGTTAAGCATT

TGAAACGAGGACCTGTTACGATGGTCCGGATTCTTAGGCGTAAGATGTTGGGAGTCAAGTTAGCGGTTTC

TTTCAATGCAAAGGGTGAGCCATATGGTAAAGTTGCCACAGAGATG

>MSTRG.44.1 gene=MSTRG.44

GTACAAGACAGAAAAACTTGTAACCATAGCTGAAGAAATTGAGAAGATGAAGAAGAAAGTTGAAGAGGGG

GAAGTGACAGAGGAAGGCACTAATGATGTCTTGACCAAGGTTCTTGGTAACTCTGAGCACCGAGGACGGG

TGCGTGGTCAGGGGAGTTATGTTAAGCAATCCAACTATTTTAATCTGCCAAGACAGAAGAGAAAAAACCG

ATCAATTGAAGAAAAGATCCAAGAAGGTATTCAAAAGTTTATGGCTGATGAGACTAACAGGATCATCAAA

GAAAGAGATGAATTTTGGGCTGCAGAGATGGCAAAGCTGAAAGAAGCTTTAAGTATAAAGATTGATGGGA

GTCCAAATATTGGATCACAACAAGGAAGTTGTTCAAAAGGAGGACTTGAAAATGTGCTGGAATTGACAGC

TGTTAAAAAGAAATTGGATCTCAATGAATCTCCTAGAGGGGAAGATGCTGAGAATAATTTTCAGGAAGTT

GATGATGAAGAGAAGAACAAGGATAAAGAAAAAGATGTTGCTGAATTAAAGAGCAGTGGTGAAGAGAAAG

GAGTTGAGTATCTTGTAGTTGAGGACAAGGATGTGGAAGTGGCTGAGGAAAATGAAGGTATTCTATGGGA

GTTAGCAATTGGAACCCCAACCAACATTGTTGCTCATGCAACAGTTGACTTTGTCACTGCTGTTCTTCAT

GGGAAGCCACTTGGAGGAGATAACGTACGAGTATCAATTACTCGTGTTATCCAGGGAGCTGCTGAGATCC

CATTCCCAATTGACGATGAAATCATCACAGTAGATCAAGCTGTTGGAACTTTCATTGCATGGCCGAGAAA

CATGTTACGGGAAGTGAAGGCAAACAGTGATCGAGTGAAGTCTCGAGCAGTGAAGGGAGGCAAGAAGAAT

GGCCCAAGAAAAATGAAGAAGGTGAATGACTTGGTCTCGGAACCAGAGCCAGTGGTTAATATGGGCCCTG

ACTTCCCTCCTACTTTGAAAAAATTGTGGCTTTGGGCAAGCGATGCTTTGAAGGATGGTCGTTCACAGGC

GTTCAAGTTATCCGAAGAAGCTTTTGGATCATCGGATAAAAAGTGTTTGTTTAAATCCGACATAAGTGCT

GTGTGCTTTGGAGGTGAAATATCAGGAACTGTCATTTGCATGTTTATCAATATACTCCAGGAAAATTTAA

GAAAGCACAAAATGACAGACATGATATCTTTTGTTGATCCAGCCAAGATTGGGGCTCTTGGTTGTGGTAC

TCCAGCAGCGAGGTCACGCGCCCTTGCTCTCAGATTCAAAAGTGCTAAGCCATCCCAAGTCTTTCTTTTG

CCATATCACCACACGAACCACTGGGCACTTACTGTTGTAAATCCCGATGCACAAATGGTTTATCACTTGG

ACCCGTTAAAGCGGAGAATTGCAAATGAAGAATGGATTGAAGTTGTTAACAATGGCATAAAAATTTACAA

GGAGGATGTGAAGAGGTTTCTGAAGAAGAAAATAAATTGGGAGAATTTAGCGGGTGTTCCTGCCCAGACT

GGGACCACTGATTGTGGTCTTTTTGTGATGCTTTATATGAGAGAAATATGTATAGACAAGGAACTCAAGT

TTGCGTCCAAGTGGGCGCGGAGGAGCAATCTTGTCTTTGATAGCGATGATCTCAATGAGATCAGGAGTGC

CTGGGCAAAGTACTTCATGAGGCAGCATGCTAGTTAGGCTGCATAGAGGATTGATTATTTTCGGATGCAC

ATTTATATATATTGGGACTTATTACATGGTTTATTTTGCCTAAATTTAGACTTAAGTTATATCCATGTAT

GTCTTTTTTGGATATGTGCATCCGAAAATACAATACATTGTCGCGTCAAGATTGGTAGAATCTGTTGGTG

GGTTTTTGGATTGTTAGGTGCGTTTGCCTTGGTTAGTATTGTTGGATGATATGTGAATATTAGTTGGTGG

ATATTAGTTGGTTTTTGAATTGTTTTGGCTTTTATGTTTGGTTGGAAACATAGTTGGTAAATTGTGAAGA

TTGTTGGTATGTGGTGGC

>MSTRG.46.1 gene=MSTRG.46

GTCATACTACAAGAATCACACAATAAACACCACCAAATCAGAGTCCGTCAAAGTCGGTGCACAGCCAAGA

TAACTGCCTCTACTAGCCGTTTAAATCCGGTGAATCTCCGGCATTATATCTCACCGGACTGTCATTTTCC

GGCCAACCCACTCATCACAAGGTGTTAAAATGAATGGAATTCAGAATCACAAAACTCACAATATCGAGAT

ACCTTATCTAGGATGTTAGGGACGTGTAGTCAACGTTTTTGAACCGAGTGCAGGCATGCCAGGAAATAGA

CTGTTGATGGATAAGCCTCATCATAATGG

>MSTRG.49.1 gene=MSTRG.49

TAAGCTTTTCTGTTAAACAAAACAAGAAATGGACATTACATATAGACATGCTACAGTATTACACTATTAC

TGAAGGCAGCTGTAACCCTGATCACAGACAAATAATCAGCCAACACGAAATTAGTAAAGTCCTGGGCTCG

AATAAACCATCCTCTTCTTCCCAATTCAGGTATTTATGTCTATGCTTCTATGCTCGTGTTCATGGTACTG

GAGTTGGATGCCCTCCTTGGAATGTTTTTGTGGCACTCCTACCACTTTAAAGGACTATGATCAGTTCTAT

TTTATGGCATGGAGTGGTGGAACGACGAGTCTAGCTTGATCTGCTTATAGAGGTCAGATCTTCACGGTTG

AGCTTGAACTAATACACATACAGGGTTAGATTATCTGTCAGAAAATTCAGTTGCTTAATTAGTTTCCTCG

GTATCTTTAAGCTTTTTACTATTTCTCTACTCTTCACTGAATTTTGAGTTTTTTTCCGTCCATTTATGTC

ACTGTG

>MSTRG.59.1 gene=MSTRG.59

CGAAGGGTGTAAGGGAGAGATATATGTCTCCTCAACTCTTTACAAACTGCAAGCTGTTTGCATTAGCATA

TCGCTCCATAAATCTCATGAACCTGTCCCAATCTCGTGGAGTACGCATGACATACTTGGCTTCAATCCCT

GCAGGCTTTCCGTTTACAAACCTGGCGTTTACATCAACTGATTGAAGAGTCCCTTCCTCATCAATCATGT

AGAAGCCAGTGATCTCGCCAACCTCACCAGATGAGTCAAAAACTGATGGCTCGTCAAACCTAAATATAGC

CATACCATTTGTTCCATCCCTAGATTTAGTTAGCTTCACATCTGGAACAGTCTGCTCATCAGTCCCTTGG

ATAAATTGAATCGTTGGTTTAACCATCATTACACATAATCTTGACATCTGAATTTTTCTTCGTTGAGGGG

CATATTGTGAAGAGGACAGGTGCAATGTTTGTCCGCTGAATGAGGAGCCTGCACTACGGATG

>MSTRG.63.5 gene=MSTRG.63

TTCCCTTGACCCTAGACTTTTAAGCAGCTCCTCACGCTGTATCAATTACAATCTCAAGTCTCCGTCCATG

AATCCAATCTCGCTCAATTATATTTGATGGTTGGGACTGCACAGGCTGACAAAACGAGGCAAAGTGGTTC

CAATCCAAACATGCAATCTATGCTCAATCATAATCTGTTGATCTTCCCATTTTATACCATGCCAAACTGT

ATGGAGACGATGACAAAAGCTAAAAGCTAACGACTGTTTGCCAGTTTGCCAGCAAGTACTCCTGCTCTTT

CACAAACTAAGGTTAGTCTTGTGTTGACCGTGATGTTC

>MSTRG.65.7 gene=MSTRG.65

GTGATGATTATGATTATTGGAGTAATCTTCCCGCTGATATTCTTGAGCTTATCGTGGGGAAGTTTAGTTA

TCTCCGTGACTATTACAGATTCATTGCTGTCTGCAGATCATGGAGGGATTCACTTAGTCATTTCAAGAAG

CCCTGTCTTCCCCTCACCGAATATCCGTTCCTTCTTCTTGCGGAAGATGTTGCTCCAGGAGCTATGCTTG

AATATGACCCTAGTCAAGATTATAGTGAAGTAGAAGATAATAACGATGAGGAAGATGATAGTGGTGATGA

TGGAGGCGACAATGATGTTAGTGGTGGTGATGAAGAAGAAGAAGATAGCAGTGATGATGATGATGATGGC

AGTGGTGATGGTGATGATAATCACGAAGAAGACAATAATGGTGATGATGAAGACAATGACAAAGTAGATG

ATACCGAGGAAGATGAATATGATCATAATAAAATTTCAGTCGGTAGTCGTCGTGGTTTCTACAGTCTTTT

GACGGGAAAAACGTACTATATTGATTTGCCGGAAGCTGCTGGAAGACTAATAGTGGGGACCAATAAAGGG

TGGCTAGTAACTCTAGG

>MSTRG.73.2 gene=MSTRG.73

CTCAACTAAACCCGTACCAACACTCACTCTAATATTCCTATAAAAACACCCACACACACACTTGCTGTTT

GTTCTTTTCTTGATTCAAAAATAAAACGCAGAAATTAGAAAGCAATGGAGCTTGATCTGGGCGCAATGGC

GTCTATGATCGGCGTATCAGTCCCTGTTCTGAGGTTTCTGTTGTGTTTCATAGCTACTGTGCCGGTGAGT

TTTGTGTGGCGGTTGATTCCCGGCGGGCCGGCGGTGAAGAACTTGTATGCTGCGGTGACTGGGGTGGTTT

TGTCGTATTTGTCGTTTGGGGCGTCGTCGAATCTTCATTTTTTGGTGCCTATGCTGTGTGGGTATGGGGC

GATGGTGGTGTATCGGGAGAGATGTGGGCTTGTGACTTTTGTTTTGGCTATGGGATATCTCATTGGCTGC

CATGTATACTACATGAGTGGGGATGCATGGAAGGAAGGGGGAATTGATGCCACCGGTGCCTTGATGGTTC

TCACACTGAAAGTCATTTCTTGTGCAATTAACTACAATGATGGATTACTAAAGGAGGAAGATTTACGTGA

ATCACAAAAGAAAAATCGTTTACTTGAGTTGCCAACTATAACTGAGTATTTTGGATACTGCCTCTGTTGT

GGTAGTCACTTTGCTGGCCCAGTTTTTGAAATTAAGGATTATCTTAATTGGACAGAAAAGAAGGGGATTT

GGACAGCTTCAGATAAGGGGAAATCCCCATCCCCATATTGGCCAGCTCTCAGAGCAATCCTCCAAGCTGT

TATATGCATGGGCTTGTACCTGTATCTTTCGCCCCACTTCCCACTGACCCGTTTTACGGAACCTATATAT

CAAGAATGGGGTTTTTGGAAACGAATAAGTTACCAGTACATGGCTGGTTTTACGGCACGTTGGAAATATT

ACTTTATATGGTCAATCTCAGAATCTTCTATTATTATTTCTGGTCTGGGTTTTAGTGGCTGGACAAATTC

TTCACCACCCAAAGCACGATGGGACCGTGCAAAAAATGTAGATATCTTGGGTGTTGAGTTGGCAACGAGT

TCAGTTCAGCTGCCACTTGTATGGAACATACAAGTTAGTACCTGGCTTCGTCATTATGTTTATGACAGAC

TCGTTCAGAAGGGAAAGAAACCTGGTTTCTTCCAGTTGCTGGCAACACAGACTGTTACTTACCGCATACA

AGAGCGTATACTTCATAGCAACCATTGTTCCCGTAACATTGATCCTTCTTGGAAAGATTATTAAACCGGC

AAGGCCCGCCAGAACAAAAGCTCGCAAAGAAGAGTGAGATGGAATTGCAATTTCGGATTCTGAAATCCCT

GGATTGAACTTTACAAATCATTAGTATGCTTCTTCCAGCCATATTCCCTTGTCTTGCTATTCCGTAAAGT

GTTTTCTTTTTTATAGTTGAAATTGCTTTGCCTGTCCATGTTTGGACCACTGGCCTCACTTCAGAAATCT

AGGCCAACAGGCAATCGATTAGAAAGTTGTATTCTTACTTAGATCAGAACAAGTGATTGCAAGAATTATA

TTTCGGTTGGAATTTTTCCTCTTATCCGTGACAGTCCAATTAAAATGTGTGAAAATATAACCCTTATCAC

TTTGTGGAGGGAATTTTTTAGTTCTTCGTACCAGTAAATTACAATCCCTGTTGAAA

>MSTRG.73.1 gene=MSTRG.73

CAAAAACACAACAAGCAAACACAACTCAACTAAACCCGTACCAACACTCACTCTAATATTCCTATAAAAA

CACCCACACACACACTTGCTGTTTGTTCTTTTCTTGATTCAAAAATAAAACGCAGAAATTAGAAAGCAAT

GGAGCTTGATCTGGGCGCAATGGCGTCTATGATCGGCGTATCAGTCCCTGTTCTGAGGTTTCTGTTGTGT

TTCATAGCTACTGTGCCGGTGAGTTTTGTGTGGCGGTTGATTCCCGGCGGGCCGGCGGTGAAGAACTTGT

ATGCTGCGGTGACTGGGGTGGTTTTGTCGTATTTGTCGTTTGGGGCGTCGTCGAATCTTCATTTTTTGGT

GCCTATGCTGTGTGGGTATGGGGCGATGGTGGTGTATCGGGAGAGATGTGGGCTTGTGACTTTTGTTTTG

GCTATGGGATATCTCATTGGCTGCCATGTATACTACATGAGTGGGGATGCATGGAAGGAAGGGGGAATTG

ATGCCACCGGTGCCTTGATGGTTCTCACACTGAAAGTCATTTCTTGTGCAATTAACTACAATGATGGATT

ACTAAAGGAGGAAGATTTACGTGAATCACAAAAGAAAAATCGTTTACTTGAGTTGCCAACTATAACTGAG

TATTTTGGATACTGCCTCTGTTGTGGTAGTCACTTTGCTGGCCCAGTTTTTGAAATTAAGGATTATCTTA

ATTGGACAGAAAAGAAGGGGATTTGGACAGCTTCAGATAAGGGGAAATCCCCATCCCCATATTGGCCAGC

TCTCAGAGCAATCCTCCAAGCTGTTATATGCATGGGCTTGTACCTGTATCTTTCGCCCCACTTCCCACTG

ACCCGTTTTACGGAACCTATATATCAAGAATGGGGTTTTTGGAAACGAATAAGTTACCAGTACATGGCTG

GTTTTACGGCACGTTGGAAATATTACTTTATATGGTCAATCTCAGAATCTTCTATTATTATTTCTGGTCT

GGGTTTTAGTGGCTGGACAAATTCTTCACCACCCAAAGCACGATGGGACCGTGCAAAAAATGTAGATATC

TTGGGTGTTGAGTTGGCAACGAGTTCAGTTCAGCTGCCACTTGTATGGAACATACAAGTTAGTACCTGGC

TTCGTCATTATGTTTATGACAGACTCGTTCAGAAGGGAAAGAAACCTGGTTTCTTCCAGTTGCTGGCAAC

ACAGACTGTTAGTGCTGTATGGCATGGTCTTTATCCTGGGTACATGATATTTTTTGTTCAGACAGCTTTG

ATGATTGCTGGATCCAGAGTCCTATACAAATGGCAGCAATCTCTTCCTCCAAAAATGGCCATAGTGAAGA

ATCTATTGGTATTCATTAACTTTTTGTATACACTTATGGTTCTTAACTGCTCTGCCGCTGGTTTCATGGT

ATTAAGCTTCCATGAAACTCTTACCGCATACAAGAGCGTATACTTCATAGCAACCATTGTTCCCGTAACA

TTGATCCTTCTTGGAAAGATTATTAAACCGGCAAGGCCCGCCAGAACAAAAGCTCGCAAAGAAGAGTGAG

ATGGAATTGCAATTTCGGATTCTGAAATCCCTGGATTGAACTTTACAAATCATTAGTATGCTTCTTCCAG

CCATATTCCCTTGTCTTGCTATTCCGTAAAGTGTTTTCTTTTTTATAGTTGAAATTGCTTTGCCTGTCCA

TGTTTGGACCACTGGCCTCACTTCAGAAATCTAGGCCAACAGGCAATCGATTAGAAAGTTGTATTCTTAC

TTAGATCAGAACAAGTGATTGCAAGAATTATATTTCGGTTGGAATTTTTCCTCTTATCCGTGACAGTCCA

ATTAAAATGTGTGAAAATATAACCCTTATCACTTTGTGGAGGGAATTTTTTAGTTCTTCGTACCAGTAAA

TTACAATCCCTGTTGAAA

>MSTRG.84.1 gene=MSTRG.84

TTCTCCTGTACAAACTCACTTCTTCTTTTTCTCCGTAAAAATGAGAGAGATTCTACACATCCAAGGAGGC

CAATGCGGGAACCAGATCGGAGCCAAGTTCTGGGGAGTGGTCTGCGCTGAGCACGGAATTGACTCAACAG

GCCGTTACAGCGGAGACTCTGAGCTCCAACTCGAACGCATCAATGTCTACTACAATGAAGCTAGCTGTGG

CCGCTTTGTTCCACGTGCAGTTCTCATGGACCTGGAACCTGGTACTATGGATAGTGTCCGCTCTGGACCG

TATGGTCAGATTTTTCGACCTGATAACTTTGTTTTCTGACAGTCTGGAGCTGGGAATAACTGGGCTAAAG

GACATTACACGGAAGGAGCTGAGCTTATTGATTCGGTTCTTGATGTTGTTCGTAAAGAAGCTGAAAATTG

TGATTGCCTCCAAGGGTTTCAGGTGTGTCATTCGTTGGGAGGAGGGACGGGGTCAGGAATGGGGACCTTG

CTTATATCAAAGATCAGAGAAGAGTATCCGGATCGAATGATGATGACTTTCTCAGTTTTTCCATCGCCAA

AGGTGTCTGATACGGTGGTGGAGCCTTACAATGCTACATTGTCTGTGCATCAGCTTGTTGAGAATGCCGA

TGAGTGTATGGTCTTGGATAATGAAGCTCTGTATGATATCTGTTTCCGTACTCTCAAGCTCACCACTCCC

AGTTTTGGAGATTTGAATCACCTGATTTCTGCAACCATGAGTGGTGTCACTTGCTGCCTTCGGTTTCCTG

GTCAGCTTAACTCTGATCTGCGCAAGCTTGCTGTTAACCTGATTCCATTTCCTAGACTTCACTTCTTCAT

GGTTGGCTTTGCACCCCTTACGTCCCGTGGCTCACAGCAGTACCGTGCTCTCACTGTTCCTGAACTTACC

CAGCAAATGTGGGACTCGAAGAATATGATGTGTGCTGCAGATCCTAGACATGGTCGGTATCTGACTGCAT

CTGCCATGTTCCGTGGTAAAATGAGCACCAAGGAGGTTGATGAGCAAATGATCAATGTACAGAACAAGAA

CTCATCGTACTTCGTCGAATGGATTCCCAACAATGTGAAGTCAACTGTCTGCGACATCCCCCCTATAGGT

TTGAAGATGTCCTCGACGTTTATTGGGAATTCCACGTCAATTCAAGAGATGTTTAGGAGGGTCAGTGAGC

AATTCACTGCTATGTTCCGTAGGAAAGCTTTCCTGCATTGGTACACCGGAGAGGGGATGGACGAAATGGA

GTTCACGGAAGCTGAGAGCAACATGAATGACCTGGTTTCAGAATACCAGCAATATCAAGATGCAACTGCT

GATGAGGAAGCTGATTACGAGGAAGATGAAGAGAGTTATGAAGACGAGGCTTAAACTTAGACAGATAGAT

ATTTGGTTTTTGTTTTTGGTTCTTTCGCTTGAAGTTGAAACTTGAAACAGTGTGTTTTTCATTTCCTGCC

TCTGGGATATTTGCTTGATCCTGTGTTTATTTCACTTGCTTCAATCTTGTAAAACTAGCA

>MSTRG.88.1 gene=MSTRG.88

ATTTAATTTAAAAAGAAAAAGAAAAAAAACAAGAAGGGAGTTGCCAGTTGCCACTCTCAATTCAAAGTTC

TTGAAATTTTCTTGTTTGTATTTCATCACCATTTTGTCTTCACACACCCCTCCCCCTTCTTCCCCTCTTT

TCAGTTTCAAATCTAGGGTTTCTGTGTACTCTTTACAGCCCATTCTTCCCTTCAAAATTTTTGAGTTTTG

CTCACATTTTTAATCAAACCCTTCTGTTTTCAACCAAGATTTCTCACTCAGCAGCCACTAGAACTATAAA

GATTCAATCTTTATCATTTTTTAGGGCTTTTAGAGTGGTTTTTAATCAAACCCATCTTTTTTCACTCAGC

AGCCACTAGGAATGTAAAGTTTCAATCTTTATCATATTTTAGTGCTTTTAGAGTGGTTTTTTTTTCATAT

TCTCATGTTTTAGAACAACCCCATTAAAGCTTTTGTCAAGATTTTGGTAAATTGTATTGATATAGAATCT

GGGTTGGTGTTATTTTGTCTTTCTTGAGTGAATTGCTGAGTAAATGCATGGATAGTAGAGAAGAAATGAC

ACTTTCAGGGTCATCACCTTACTATTTTAACGGAGGGATAGGTGGGCCCGTCCTTGATTCACCTGAATTT

AAGGACTTCTCAAACCCTAATGATTCAGTTGAGACCAATGTGATGGTTAGTCATCTGGGTCCTACTCAAA

ATGTTGAAAATTTGGGTACTAATTTTGGTGAGGAGATGAACATGGTTATGAATTCTGGTGGTTATGGTGG

TGGTATGGCTTCGGGGGAGGTTGATTTAAGTAAGAAGAAGAGGGGGAGGCCTAGGAAATATGGGCCTGAT

GGGGCCAATGTGGCTTTGGCATTGTCCCCATTGTCGTCGAATCTTTCAGCTGGTTCGGGTACAGAGGTGG

AGAGGAAGAATAGAGGGAGGCCTAAAGGGAGCGGAAGGAAGCAAAGATTAGCATCTCTTGGCGAATGGAT

GAATAGTTCAGCCGGAATGGCCTTTACACCACACGTCATTCACATTGGAGCAGGAGAGGATATTGCATCA

AAGATTTTACTCTTCGCTCAACAGAGACCAAGGGCTTTATGCATTATGTCTGCAAATGGCACAGTTTCTG

CTGTGACTCTAACCCAGCCTGAATCTTCTCATGACTCTGTCACATATGAGGGCCGTTTCCAGATATTGTG

TTTGTCAGGTTCATACTTGCTCTCTGAGGGTGGCGGACCACGTGATCGAACTGGGGGTCTAAGTGTATCT

GTCTGCAGTCCTGATGGCCATGTGATAGGAGGTGCAGTTGGGGGAAGGCTTATTGCAGCGAGCTTGGTTC

AGGTCATTGTTTGCAGCTTTGTATATGGTGATTCAAAGGCAAAGATCAATCCAAATGGGGCAGCGGCCAA

AGCTGACCTTGATTCCGAAGTGCAACCCACAGAAAAACTATCTACTCCAGGTAGTGCAGGGCCCAGTGGA

GATATTACCCCCAACTTGGCACCAGCTGGTGCTGAAGATATTTCCCTTAAACCAGCAATTGCTGATTGTC

CCCCAGATTCTCAGCCGGGTATTACTGATATAAAAACGGAAATTGACCTAGCCCATGGTTGAGGCATTTA

CTGCAACAATGATGTGGAGAAGTTGATCATACCTTGTTACAAGAGCTTAGAAAATGGGTTTAACATAACT

GTTGATGTAACAGAACGCCATTTTGGGGTTCTCAAAGTAGGAATATTAGGAAGTTGGTGTAGTTAAACGA

GTTATCGTATGTTTTGATGACAATGTAGGTTGGTAGTTGTTGGAAGCTGTTTAGTGGAAACTTTTAGCTA

GGCCTAGTAAACTGTTGATGATAATGTTTGAATTTGTTCTAATTTGTGTCGAATTGTGCTTTGTATGGAG

AAACGTTTCCGCATTTTGCAAATTAGGTTGTAAAATACTCCTATTCATCCTG

>MSTRG.88.2 gene=MSTRG.88

GGGAGTTGCCAGTTGCCACTCTCAATTCAAAGTTCTTGAAATTTTCTTGTTTGTATTTCATCACCATTTT

GTCTTCACACACCCCTCCCCCTTCTTCCCCTCTTTTCAGTTTCAAATCTAGGGTTTCTGTGTACTCTTTA

CAGCCCATTCTTCCCTTCAAAATTTTTGAGTTTTGCTCACATTTTTAATCAAACCCTTCTGTTTTCAACC

AAGATTTCTCACTCAGCAGCCACTAGAACTATAAAGATTCAATCTTTATCATTTTTTAGGGCTTTTAGAG

TGTGGTTTTTTTTTCATATTCTCATGTTTTAGAACAACCCCATTAAAGCTTTTGTCAAGATTTTGGTAAA

TTGTATTGATATAGAATCTGGGTTGGTGTTATTTTGTCTTTCTTGAGTGAATTGCTGAGTAAATGCATGG

ATAGTAGAGAAGAAATGACACTTTCAGGGTCATCACCTTACTATTTTAACGGAGGGATAGGTGGGCCCGT

CCTTGATTCACCTGAATTTAAGGACTTCTCAAACCCTAATGATTCAGTTGAGACCAATGTGATGGTTAGT

CATCTGGGTCCTACTCAAAATGTTGAAAATTTGGGTACTAATTTTGGTGAGGAGATGAACATGGTTATGA

ATTCTGGTGGTTATGGTGGTGGTATGGCTTCGGGGGAGGTTGATTTAAGTAAGAAGAAGAGGGGGAGGCC

TAGGAAATATGGGCCTGATGGGGCCAATGTGGCTTTGGCATTGTCCCCATTGTCGTCGAATCTTTCAGCT

GGTTCGGGTACAGAGGTGGAGAGGAAGAATAGAGGGAGGCCTAAAGGGAGCGGAAGGAAGCAAAGATTAG

CATCTCTTGGCGAATGGATGAATAGTTCAGCCGGAATGGCCTTTACACCACACGTCATTCACATTGGAGC

AGGAGAGGATATTGCATCAAAGATTTTACTCTTCGCTCAACAGAGACCAAGGGCTTTATGCATTATGTCT

GCAAATGGCACAGTTTCTGCTGTGACTCTAACCCAGCCTGAATCTTCTCATGACTCTGTCACATATGAGG

GCCGTTTCCAGATATTGTGTTTGTCAGGTTCATACTTGCTCTCTGAGGGTGGCGGACCACGTGATCGAAC

TGGGGGTCTAAGTGTATCTGTCTGCAGTCCTGATGGCCATGTGATAGGAGGTGCAGTTGGGGGAAGGCTT

ATTGCAGCGAGCTTGGTTCAGGTCATTGTTTGCAGCTTTGTATATGGTGATTCAAAGGCAAAGATCAATC

CAAATGGGGCAGCGGCCAAAGCTGACCTTGATTCCGAAGTGCAACCCACAGAAAAACTATCTACTCCAGG

TAGTGCAGGGCCCAGTGGAGATATTACCCCCAACTTGGCACCAGCTGGTGCTGAAGATATTTCCCTTAAA

CCAGCAATTGCTGATTGTCCCCCAGATTCTCAGCCGGGTATTACTGATATAAAAACGGAAATTGACCTAG

CCCATGGTTGAGGCATTTACTGCAACAATGATGTGGAGAAGTTGATCATACCTTGTTACAAGAGCTTAGA

AAATGGGTTTAACATAACTGTTGATGTAACAGAACGCCATTTTGGGGTTCTCAAAGTAGGAATATTAGGA

AGTTGGTGTAGTTAAACGAGTTATCGTATGTTTTGATGACAATGTAGGTTGGTAGTTGTTGGAAGCTGTT

TAGTGGAAACTTTTAGCTAGGCCTAGTAAACTGTTGATGATAATGTTTGAATTTGTTCTAATTTGTGTCG

AATTGTGCTTTGTATGGAGAAACGTTTCCG

>MSTRG.90.1 gene=MSTRG.90

ATATGTGCCCCTTTCAACGATCGCTCATTTTTATACAGTGTTTTGTATAGCTTGCAGAAGCCGAGTCCTC

TGTTCCAAACGTCTACAATTCGAACGTTTACAATCCGTTGGGACAGAGGGAGTATTGCGATGTCAGGGAA

AGATTTCCATTCGAGCACAAGAATCGAACTATCAACAAAGACCGAAAAAAAATGAAAGTATGGTCACTCC

CCTCCTATGCACACATCTGACTCCAAATGACTAGCTCCTTCAGGCTGAAAATCATAGAAGCTTACTACAC

TTAACTCCATTACCATCTACTAGCATCCAAGAGGCGAGACAGAGTTTACGATCATGAAAGGCACAGGTAA

AGTCCCGATTTCTATAATGTTTGGCGTAAAAGGGATGATTCATGGGTTCAACTCTTTATTCAAACATACA

TCAATATGTATGCGATATTCTTGTTTTGCATTTCTACTCTAATATAATAAGTTAC

>MSTRG.96.1 gene=MSTRG.96

CCAAATTTCAAGTTATCCGGCCTGGCTGCTTCATGAAACTGTTATATTGGTAGTTCTAGGTAAAGGTACA

AATTTTGTAGCAAGTAGATAGTTTAGTTCTTATAGTGTCAACTAGAACTGTCGACCGCTTCTCTCTTGGA

GACTAGTGTTCCCGGACTTTCAATGTATATTTGTTCTATACATGTTACTGAAGAACACACGCTATTACAT

CGAAATATGTTCTGAATCATTGTAATCATTTACGAACTGGGCTCTGTACTTAATTTAAGCATTGGTTTGA

GGCTGTTGAACAAGGTTTTCCAGAGCTTTATACTTATCAGAACTTACAGTTGAATTGTTGGATGAAATTT

AATGTTTTTTCAA

>MSTRG.98.1 gene=MSTRG.98

CCCGAAAGCCCATGCAATGCCTTGAATCCCCACAGAAGCACATTTATCAGGTGCATTCGAGACTCCCATA

ACAGTCAAAACAGAAACATACAAGAACAAGAAAGTGGCTATAAATTCTGCAATGCCAGCCCTATAAAAAG

ACCATGAGACTAGTTCACCATGCTCATACAGAGGTGCTGGTGGTGGCTCCACGTAGTCTCGATCGTGCAT

TGCAGAAGCAGATGTTCCAAGGGGAATCCTCTCAGGGTACTTGTTGGTTCCAACTTTGTCATCCTCTTCT

TCAATCCCCATTGATGCTTTATTTTAGCAAACTTTGTTAAGGGATAAGAAGTGCTGTGAGTTGGTGCTTA

AGGTGAGTGTTTTATAGGTGAAAATTTGAGCAAAATGCAGAACACAAATTTATGCAGATTAATGTAAATT

ATTGGTTTTTTGGGAGGGAGTAAGACAATGTCACAATGGTGGTCTGGAACTGTGACTGTTATCAAATATG

TTTGTTTGTATTAATTTGTTCTTATGTGCTTG

>MSTRG.99.1 gene=MSTRG.99

ACAAACAAACATATTTGATAACAGTCACAGTTCCAGACCACCATTGTGACATTGTCTTACTCCCTCCCAA

AAAACCAATAATTTACATTAATCTGCATAAATTTGTGTTCTGCATTTTGCTCAAATTTTCACCTATAAAA

CACTCACCTTAAGCACCAACTCACAGCACTTCTTATCCCTTAACAAAGTTTGCTAAAATAAAGCATCAAT

GGGGATTGAAGAAGAGGATGACAAAGTTGGAACCAACAAGTACCCTGAGAGGATTCCCCTTGGAACATCT

GCTTCTGCAATGCACGATCGAGACTACGTGGAGCCACCACCAGCACCTCTGTATGAGCATGGTGAACTAG

TCTCATGGTCTTTTTATAGGGCTGGCATTGCAGAATTTATAGCCACTTTCTTGTTCTTGTATGTTTCTGT

TTTGACTGTTATGGGAGTCTCGAATGCACCTGATAAATGTGCTTCTGTGGGGATTCAAGGCATTGCATGG

GCTTTCGGGGGTATGATCTTTGCACTTGTCTACTGCACTGCTGGTATTTCAGGAGGACATATCAACCCTG

CTGTGACCTTTGGTTTATTTCTGGGAAGGAAGCTTTCGTTGACCAGGGCATTGTTCTACATAGTGATGCA

GTGCCTTGGTGCTATCTGTGGTGCTGGTGTTATAAAGGGCTTTCAAGGATCCTCAAAATTTGAGCTCAAT

GGTGGGGGAGCTAATGCTGTGAATCATGGCTACACAAAGGGTGATGGTCTTGGTGCTGAGATTGTTGGCA

CCTTTGTCCTTGTTTACACTGTGTTTTCCGCGACTGATGCTAAGGGAAGTGCCAGAGACTCAACTGTCCC

TATTTTGGCCCCTCTTCCCATTGGCTTCGCGGTGTTCTGTGTTCATTTAGCCACCATCCCCATTACAGGG

ACCGGAATCAACCCTGCTAGAAGTCTTGGAGCTGCCATAATTTACAACAAAAGTCACGCTTGGGACGATC

ATTGGATTTTCTGGGTCGGGCCATTCATCGGTGCCTCACTTGCTGCTGCATACCACCTGGTTGTCATCAG

GGCCTTTCCTTTCAAGAGCAAATCTTAAATTTCAGTCTGCATGTACCAGAACTATTTGTCATCCCTTTTC

GATCTCAGTCTTTCATTTTAGTCTATTGTAATTGTCACTGTCATGTGAATTCGTCGGTTTCAGCTATTAA

AGTATCCATGCACATCATTGTTTTATGCTGTATTTAAGCTTGAGTTTGCATCTGAAGAAGATATTCTTTT

TTTATTGTTTGATTCATAATAGCAACATGTTTCACATCTCCACAAAATTATATATACCTTATTCTGGAGA

CAAGATATTAATAATCA

>MSTRG.100.1 gene=MSTRG.100

ACGCCGTCTCCATCAACGTCCGCTGCTCCAACGGCTCCAAGTTTAGTGTCCGAGCGAGTGTGGGATCCAC

TGTGTCGCAGTTCAAACAAGTTGTCGCGGAGAATTGCAATGTTCCAGCTGTGCAGCAGCGGCTGATTTAC

AAGGGACGGATCTTAAAGGATGATCAAACCCTAGATAGTTATGGCTTAATTGGGTTCTGTAGCTGGGTAA

GTTCATGTGAGACCCGACCATGGCTCAGGTGAGGGGTTCCAATAACCCTAAGTGGTTTGTTACATTATTA

ATTCGATTGAATCTAGTATTGATCCCAAGTTTGTTGAATTCGAGTTATTTGATTTAATTTTATTTGTCGT

CGTTTTTTTATATTTTGATATTCATCCATTTTATATATTGTGTTGTTCTGGTATTGCTAATTTTATGGAT

GTAGGAACTTCTTGATATCTATGGTGGTGATGGATAGGTCCAGAAAGGGTGGAGAAGAAGAGCTATTTTG

GCGATAATTGTGCCTTAATTAAGGATACTAATTATTCTTAATTAAGGTTTTAGTACCGTTTATGGCGACC

ATTTATGGCGACCATTATTGGGGCATTATTGGCCCATAAGAGGCATTTAAGTGCCATAATGTTGGCTCGT

TTATGTCTTTTATTTTAATTTTAAATTTGCTCACATTCGTTCATTCTGTTTTGTCTTATTTTCAGGATGT

TTGAAAAAGCATTTAAGAATTTATCTATTAAGGCGTCTTCCGAGAAGACGCGAATTTTGGTGGACAAGAC

AGTTGAGCTACAAACAAAGATAGAAAGGAATAAAGAAGACTTCATTTTTAGTCTATTTTGTAATCGATGT

ATTTTTTTGTAGGGTACTCTTTTTCCCCCTTGAGTTTTTTTCCCACAGGGTTTTGCTCTTGAGGGGTTTT

AACGAGGCCCTTTTTTTGTGGGTTTTCTCTTCGGACTTGAAAGGTTATGTTATCTAAGCGTCCGGAGTAT

TTACATACTTTCGGTTAGATGATATACCTTCTTGAATGAAGGAAACTCTGTCCATCCGGATTGTAATTTT

TTCTTGGTTTAATACAACCGTTTATTGTGACAAAAAAAAAAAAAAAAGCTTTGTTCTTTTTACTCGCATA

ACAACTCTGATTGGGAGTGTACTACCAAACTTCTCTTGATATTGTGATGGACCCCACACGTATTGTATCC

C

>MSTRG.102.1 gene=MSTRG.102

TTGTAGTGTGGATTCTAAACTGATTGTTTCACAGATGGAAGTGAAATAAGTTAGTGCAAGACCAATGCTG

CTTGTCGGTCAGTGATCATCTGTTTACTAATCATTTCTGCAGAAGCATTCCTGCATCTGGAACGGATGAA

ATTCATGTTAGTATCATATTTATATGATTTTACCTTTTGTTTTAGACGGAAATTATGTACATTAACATTT

TTTCCGAGCTATTGGTAGGCATTTGGTATTAGAATATGTTTCTGTTCTTCTGACAGTTTCCTTGTTTAAT

TCCGTTTCATGTTTAATTCCGCTCTTTCGAGAGTTCGGAGAAAAGTACCCGGGCCTTAAAACAGACACTC

AACCGGAAACAGATGAAATTTTCGGCTGGACCTTATGGTCAGTTCAAAACTGTTCTTGTGTAGGCTCTTG

AGACTTGATGAAGTGATGTTCATG

>MSTRG.104.1 gene=MSTRG.104

CAACTACTTTCCTCCACTATCTTCATTCTATAATAATATAAACACTATTACACCCACTACTTTCTTCCAC

TATCTCAAATCTATTATTAAATATAAATGGGTCCCACCACTATACCCATTTTTCATCTAACTTTACTCAT

TTCTGACCCATTTTCTTGGTCTCCGTGTCCCAGCCATTTGTATACAAATGACTGGGACGGAGGGAGTATT

A

>MSTRG.106.1 gene=MSTRG.106

AAATATGTTGAAATACAAGTCCTGTTCATACCCTTACTTGATACTAACAACTATACTAGACTTTTATGGC

AATTGTAGTTGGGAGTTTAGCAGTTGAATAGTACCAGTAACTAGCACACGGTTTATTTATGTAGCATAGA

CTTTCTGTACATTACTTACTCTATATATGTAGCTCTTGGGCATGTAATAGCAATACACACAGAACAGTAA

TACAAGAG

>MSTRG.108.1 gene=MSTRG.108

CTTCCGCGGCCTCTCCTTCATTCTTGGCTTCCGCGGCCTCTCCTTCATCCTCGGCTTCCGCGGCCTCATC

TCTCCTTCATCCTCGGCTTCCGCGGCCTCATCTTCATCAACTCAGTCTTGGCTTCCGCGGCCTCTCCTTC

AGCAACTCAGTCTTGGCTTCCGCGGTCTCTCCTTCAGCCTCGGCTTCCGCGGCCTCATCTTCATCAACTC

AGCTTCGGCTTCCGCGGCCTCATTCTCAGCAACCTCGCCATTGTCTCTATCAAGCAAACACTGCCTTCAC

CCCCGTTAGGCAGAATCATCTTCATCTTCAAACACCTCCTTCAACATCGGCTTCAGAGGAAATTTCTTCA

ACATAAGGCGCGCCAAGTTAAACAACGTCAACATCTTCTGCATCAGCGAAGAACAACCAAGATAGGACAT

CTCAGGAGATGAAGGACATCAAGATCTCGTCGACATCTTCAATGCCACCAAGATCTGGGGGGTAGTTGTT

ATACGCAAAAATCACCATGAAGACCCGGCCCATAAAAGAAGACCATGATGGGCTCAAGAATGGACCAAGC

CCAAGCTAGCCATGTCAGTTAGAAGAGACCACTTCAAGAAGATTGCGCGCCTTATCTTCAGGGCACGTCA

AGTTGAAGCTTACAGAGAAAGATCGT

>MSTRG.109.1 gene=MSTRG.109

GAAAAATCGATCAGAATTAGGAGAAACGTATAAAAATTTTGCCACTATGATTGAAACTCAATTCAAAAAG

AAAATCAAGCTTTTTCGTGCCGATAATGCCAAAGAATATAAAGACTCAGAACTTACCAAATTTTTGGCAT

CTTATGGTACTCTTGCACAAAGCTCTTGTCCTTACACATCACAACAAAATGGTCGTGCTGAACGTAAACA

TAGGCACATTCTTGATACAGTTCGAGCTCTTCTCATTTCTTCTTCTTGTCCAGAATTATTTTGGGGAGAA

GCTGCATTAACTGCTGTTTACACAATAAACATGATTCCTTCTCCCGTTATAGATAATGTGTCACCTTATG

AAAAATTATATGAGGAACCTCCGTCATATGATTCTCTTAAGGTTTTTGGTTCTGCCTGCTTTGTTCTTCT

TCCTCCCCATGAAAGAACAAAACTAGAGCCACGTGCCCGTCTTTGTTGTTTTATTGGTTATGGTAGTGGT

CAAAAGGGATATCGCTGTTGGGATCCTATCTCTCAACGTCTTCGTGTCTCTCGTAATGTGACATTTTGGG

AACACAAAATGTTTTCATCTATGTCAGGATTTAAAATTGACACGTCGACTAGTATTGCCCACTTTACAGA

TCCTTTTGTTGATCTACACCCC

>MSTRG.110.1 gene=MSTRG.110

AGTGTTCAAACTTAACAAGGCTCTTTATGGTCTTAAACAAGCACCTCGAGCTTGGTTTGAAAAATTCAGC

AAAATAGTTCAACAATTGGGATTTTCTTCAAGCCCATATGATCATGCTCTTTTCATACGAAGGTCTGAAC

GAGGAATTGTTTTACTTCTTCTATATGTCGATGACATGATCATTACTGGAGATGACGTCAAAGGCATTTC

TGAACTTAAACAATTTCTCAGCCAGCAATTTGAAATGAAGGATCTTGGTTCACTAAGCTACTTCTTGGGT

CTTGAGGTTTCTTCTGTCCCTACTGGCTACACTCTTTCACAAGTAAAATATGC

>MSTRG.111.1 gene=MSTRG.111

GGTTTTCGTGTTAAAATCAGTTCTCTTCAGTCAGTGATCCCGCGGAAAATGAAGAAGGTCGTCGTGAAGT

TGGATATATTTGATGAAAAAGAAAAGAAGAAGGCCTTAAAAACGGTTTCCAGTCTCTCAGGTACCAATAA

TTATAAATGTAACATCTTGTACGTGGTACAGAAGACTCCCGCGCATAGATTACTGATAAGTGGCACTGTG

TTTGTAGGAATTGCGTCCATATCAGTAGATATGACGGATAGGAAGCTGACAGTGATCGGAGATGTTGATC

CTATTGGCGTGGTGAACAAATTGAGAAAACTATGGCACGCAGAATTAGTATCGGTTGGTCCTGATAAGGA

ACCGGAAAAGAATAAAGATGATGAGAAGAGTAAAGAAGAGAAGAAGAAGATTGAGGACAAGGAGAAGGAA

GAAGCGGAAAAGAAGAAAAAACAAGAAGAACAGATTCAAAATCTTCTCAGTGCCTATGGCTATCAGAATT

ACGCTCCTTATAATGTTATGCCCCAATACGTTGTTCAAAGTGCTGATGATTATCCACATTCTTGTGTTAT

CAGCTGAATAATTGTTTCGAGTTTATGAATGTAAATAAAAAACCACATGTAATTTTTTTTGGTTGTCGGA

AAATTCCATCTGAGTTTAAACAAGTGTTTGATCTCCAATTGATAGGTTTTGGTTGATTTGATCGAATCTT

CATTGGCCATGATGCTATCTGGTCTGCATTGTTGTACGATACAAGTTCCCTATCGGATGATTTTCTTGTA

CAATTCTGTTTCGTCTCTTTTTTGG

>MSTRG.114.1 gene=MSTRG.114

CTATGTCGATATAAGACTGAACTCAATTTCGAGCTCGAACCAGAGGCGATGCAAGAGGAGAATGAGGAGA

CATTATCATATGAGGAGACATCAAATGTGGAGTCACTATTCTTGGTGTCTTCATAATACTCTCATCCCAC

CTTCTGAATGACAAGCTGAGCTGCAGATTCTTATGCCACATGTGCTTCTTAGCTGCCGCTTCACGTTCAT

CAGCCTGAAGCACAAATTTCTTGATCCGTCTCAGCGCGATCCTCATCGTCTCATCATCCATATTTGCATA

GCAGACTCTGAACCACCCTGGCTCTGAGCAATGAAAGGAGGATCCAGGGGACACATTGAGCTTAACTTCA

TTGATAATTACGCGCCATAAGTCCATTTCAGCTTCGACTGTGGACTCTTTGAGTAAACGTCGTAAATCCA

TCCAGAAGAAAAGGCCTGCATTGCTCTGTAGGTTTCCTACCCGGACCTGCATTAGGCCTCTGGTGAAAAC

GCCATGTCGTGCAGCTAGTCTTTTCCTGCTTTCGGATATGTATTTATTTATGAATGAATCGTCTGACAAC

ATGTTTGCGATTAATTGTTGAGTCTGAGTTGAGACTAATCCAAAACTTGACATTTTTCTAGCAGAATTCA

CAACTTTGTCATTGTAGGAGTAAATGATCCCAATTCTGAACCCGGGGAAGCCCATGTCTTTCGACAAGCT

GTACACCAGATGAATCAAATCGCGGTTGCATTTCTTGTCCTCTTCGAGTATCTCAGCAATGCTGACATAC

ATTGGCCTGGTAAAAACCGTGGCAGCATAAATTTCATCGCAGATCAGGTGGATGTTTTTCTCATTGATAA

AATTCACTAGGCCTTGTAACGTGTCACGGTCCAAAATTGTGCCTAATGGATTCGAAGGATTTGTGATTAG

CAAGCCTTTAACGTTGATATTCGACTCCTCTGCTTTCTGATACGCTTCTTCCAGGGCTTCTCTGGTGATC

ATGAAATTATTCGAGCTCTCACAGACAACTGGAAGGAGTTTTACTCCAGTTCGCCACCTCAAATCGCGGT

CAAATCTGCAAAACATCGTAATAAAAATGTAAATTTCTGAGTTTTTTT

>MSTRG.119.1 gene=MSTRG.119

CAGGATGAAGAATAGGCATAGTTATCCTAAAGACGTTGAGGCTTGCATTAAGAAATATGGATCTTCACTA

CCCAAAAGATTCAGATACTCTGCCCTAAAAAAATAACAAATTCATTTAAAGAGGAATTAGGCAAAGGCGG

GTATGGAAACGTATACAAAGGTAAGTTACCAGATGGTTGTGTCGTGGCTGTGAAGGTTTTAATTGAAACC

AAAGGAAACGGGGAGGAATTCTTGAATGAAAATGCAAGTATTGGCAGGACTTCTCATGTCAATGTTGTCT

CTCTGCTCGGGTTTGTCTACGAGGGTAAAAGGCGAGCCCTGATATATGAATACATGCCCAATGGATCTCT

AGAGAAGTTCATCCATGGCACTACTCCGTCGTTGAAAGGACAGAATCTCAGCTGGGAGAAGCTGTACAGC

ATTGCACTTGGAATAGCTCGAGGGCTAGAGTACCTGCATTGTGGCTGCACCGCACAAATCCTACATTTTG

ACATAAAGCCGCATAATATTCTTCTTGACCAAGATTTTAGCCCCAAGATATCAGACTTTGGGCTTGCAAA

GATGTACACCAGAAAAAAAGTGTGTTATTTTCTATAATGCAAGCTAGAGGAACCATTGGCTATATTGCTC

CTGAAGTAATTAGTAGAAAAGAAATTTCGGGCCAGTGTCACATAAGTCTGATGTCTATAGCTACGGAATG

ATGACTCTGGAAATGGTTGGAGGAAGAAAGAATGTGGATGCTAGAGCAGCTCACAGTAGTGAGATATATT

TTCCTAGATGGGCTTACAAGCATATCCATGATAATGAGGATACTACTAGTACTGCAGATAATATAACAGA

AGAAGATAATATAACAGAAGACGAAAACCATGTTGCCAGGAGGATGATGATCGTGGGACTGTGGTGCATT

CAGGCCAATCCGTCACAAAGGCCATCAATAAGCAAGGTGATTGAAATGTTTGAAGTGGATTTAGCAACTT

TGGAAATACCAGCTAAGCCTTACTTGTGTTCTGCTCCCACTTCACCGCGGCATTCTCAGCAGAAGGCTCT

TTCCGGATCTTCTTCACTAAGTGAATCCACTGAAACTAGTTCGAATTAATATCAGATGAGAATTATATAA

CTCCATCGACTCATGTAATGTGTTTCCTTGAATCTTGTCTAGTGCCAAGTTAGCTAAGCTAGGTGATCAA

TTCGAC

>MSTRG.125.1 gene=MSTRG.125

CAGTCACTTAGCCCCCAAGGTAGGGAATTGGTATTGATCTTCGATTATCGGGGCAGGTACATTGTTCCCA

AAGTTTCCAGTGTATCCTCCTCTTGGACCACGGCCGCGTCCACGTCCACGACCAGGGCCATAGTATTTTT

CTCCTTCAGCAGGCTTTAGAAACTCATTAATGCTGAGAGCCTTTCTGGCTTTCTCTTCCTTTTCAGCCTC

CTTACGCTTTTCCTTATCAGACCCCAATTTGACAAAGATCTCCTCTTCATTTTTCTTGCTTGAAAGAAGC

TGCATTGATTCAAAATCTTTGTCCAAGTCAACCTTTCTCTCTTCAGATTTAAGGGCCAGCAGTGCCTTTC

TCTTCTCCTCACGCACCTTCTCATACTCCTCCAGAGTCATTTCCTCAGGCTCCTTCTCTTCAGGCTCAGT

CACAGGATTCTCCTTGCTTGCATCACCAGCATCTTCCTGCTCTGCTTGTTTCTCAACATCAACATTCTTC

TCACCATCATTGACAAGCTCTTCGTTTACC

>MSTRG.129.2 gene=MSTRG.129

ATAACTTTAATGTTGCTTAAAGCCTTCCACTCTATTCCATTCTATTCCACTCATTCTCAACGAAGACAAC

CTGATGATAATTCTCCAAACTACCCTCTTCCAGCCATTATATAAACAAATTGAAATCAGACTCGAAGCAT

TCTTACGACCCAGCTACATATATTCCTCACATCCTCCTCTCATTCCTTCATCTTATCTTCACACTCTTAG

AGTCTGAGAGAGTCATGGCCTTTTCACATGCCATGCTCATGCTAACCTTCTCACTTACGTTCGTTGTCTT

ATCCTCTGGTCAAGCTCCGGCAATGTCACCCTCCACCACCATGCCACCTATGATGATGACACCACCTCCA

ATGGTGACTCCACCTCCTATGATGATGACACCACCTCCAATGATGATGACACCTCCTCCAATGATGATGC

CAGCACCAATAGCTAGTCCACCTGAGATGATGACTCCAGCCATGAGTCCTGATTCGATGAGTTCACCACC

AGCACCAATGGGACCTGCAATGGCTCCGACCCCAACCATGACGGGTATGTCACCCGTGGAGTCTACCCCA

GGGTCAGCAGCTTTCAAGCAGGGAAGTAGCCTAGCCATGCTAGCATTATTGGGAGGAGCTGTACTTCTGG

TTTAAAACTTTTGTGTATTATATTGTGTCTCTTTTGTTTTGGTCCTCGTGGAGTTTTAATCTTTGTGTTT

TATCTTGGAGGTCAGTATGTGGTCTGTTGAAGATTTATTTAATGTTACAACTCTCTTGTAGCATTTTAAA

CTAGGTCAATAAGAATACTGTTATTCAGTTTAATCTCGTAATTGTTACTGAGTCAGGAATTACACAGTTA

TGTAAAGTCTATATATTTTTGCTTTAGAGGTATTGTTTTCAGCTATTTGTTGTGTCTCTCTTACCGTCGT

TCAGTACTGCGAATACAATATCCTTGCTACGTG

>MSTRG.129.3 gene=MSTRG.129

GACAACCTGATGATAATTCTCCAAACTACCCTCTTCCAGCCATTATATAAACAAATTGAAATCAGACTCG

AAGCATTCTTACGACCCAGCTACATATATTCCTCACATCCTCCTCTCATTCCTTCATCTTATCTTCACAC

TCTTAGAGTCTGAGAGAGTCATGGCCTTTTCACATGCCATGCTCATGCTAACCTTCTCACTTACGTTCGT

TGTCTTATCCTCTGGTCAAGCTCCGGCAATGTCACCCTCCACCACCATGCCACCTATGATGATGACACCA

CCTCCAATGGTGACTCCACCTCCTATGATGATGACACCACCTCCAATGATGATGACACCTCCTGAGATGA

TGACACCACCTCCAGTGATGATGACACCTCCTCCGATGATGATGCCAGCACCAATGGCTACTCCACCTGA

GATGATGATGACTCCAGCCATGAGTCCTGATTCGATGAGTTCACCACCAGCACCAATGGGACCTGCAATG

GCTCCGACCCCAACCATGACGGGTATGTCACCCGTGGAGTCTACCCCAGGGTCAGCAGCTTTCAAGCAGG

GAAGTAGCCTAGCCATGCTAGCATTATTGGGAGGAGCTGTACTTCTGGTTTAAAACTTTTGTGTATTATA

TTGTGTCTCTTTTGTTTTGGTCCTCGTGGAGTTTTAATCTTTGTGTTTTATCTTGGAGGTCAGTATGTGG

TCTGTTGAAGATTTATTTAATGTTACAACTCTCTTGTAGCATTTTAAACTAGGTCAATAAGAATACTGTT

ATTCAGTTTAATCTCGTAATTGTTACTGAGTCAGGAATTACACAGTTATGTAAAGTCTATATATTTTTGC

TTTAGAGGTATTGTTTTCAGCTATTTGTTGTGTCTCTCTTACCGTCGTTCAGTACTGC

>MSTRG.132.1 gene=MSTRG.132

CCGCATTTCATACAAACAAACCCCAGGAGTGCCTTCGTTTTTTTGCTTTACCGAGTGCGAGCTTGTAATT

GTTGGGGATGATTTCAGTGGTGAAGAAAGATTGGCCTGATGAGGATGCGATGAAGTTGTCGTTTGAGACT

GGCAGTGCTGTGAAGGTCAAATTTAAGCGCAGGAAAGTACCAGTTGTTAGGGATTTTCCTTTGGGATGTG

GACCAAATGCACTAAAGGTTAGTGATGTTAAGGGTTCTTCGCAGAGTGGATCAAATGCAATGAAGGTCTC

TGCTATTGGGGGATTTCCTGCGGGGAATGGGCAAAATGCAGTAAAGGTCTCTGCTGTTAGAGATTTTCCT

GCGGTGAATGTTTTGAATGCACTAAAGGTCTCTGCTGAGGATTTTTCTTTGAGGAATGGACTTGGTGAAC

AAATGGCATCTGGTGTTAAAGATTCGCCTGCAGGGAGAGGACCGGATGCACCAAATGCCAATGCTGCTGT

AGAATCTCCTACGGAGAGCGGATCAAATGCAATGAAGGGCTCTGCAACTGGCGAATTATCTGTGGGTAAT

GGACCAAATGCACTAAAGGTCCCAGGTGATAGAGATTCTTCTGCAGTGAATGTTTCGAAGGTCTCTTCTT

GGGATATTCCTGCGGGGAATGGATCAATTGAACCATTGGTCTTGCCTTCTAGGGATTCTCCTGCAGGGAG

GAGACTGAATGCACCAAAGGTCAATGTTGTTACAAATTTTCCCACGGTGAGCGGATCAAATGCAATGAAG

GTTTCTGCTATTGGGGAATTCTCTTCAAGAACTGAACCAAGTGCAATGAAGGCTCCTGCTGTTGGAGATG

TTTCTGCAATGAATGCTTCAAATGCTTTAAAGCTCTCTGCTATTGGGGATATTCCAGTGGCGAATGTACC

AAGTGCATTAAAGGCCTCTGCTGTTAAAGATTTCTCTGGAGGGAATGGACCGAAATCATCAAGGGTCTAT

GCTGTTAGACGAGGTTTTCCAGTTGTTCGGGATTTCCCTCCAGGATGTGGACCAGGTGTTAGAGGAACCA

GACTAAAAATACAGTAATGTCAAATGAAAAAACATGTTGTACAGGAGAGAAGTCACAACAGAACTTAGAG

ACCAACTAAGAACTCATGGCATCCGTGGAAAGTAACTGTCTTCTGAAACTTTTAAGATCTTTTGCTAAAT

ATTTCATTACACAGTTCCTCGACAGACAGCTGCTCCAGCGGAATCAAACTTTTATTGGTCAAGCTCAGTA

CAAAAACATATGCAGCTTAATATACCAAAAACATTTCACATCCTGACAATATCCATATCCAGTCAACACA

AGTTATGATCACTACTCTAGACTTCTTATATGATATACATATATGATAGGCTTCTTATATGATTATCCTT

TTAGTTTTGAAATTATGCATATATGATATGGCTGCCGGTC

>MSTRG.132.2 gene=MSTRG.132

CCGCATTTCATACAAACAAACCCCAGGAGTGCCTTCGTTTTTTTGCTTTACCGAGTGCGAGCTTGTAATT

GTTGGGGATGATTTCAGTGGTGAAGAAAGATTGGCCTGATGAGGATGCGATGAAGTTGTCGTTTGAGACT

GGCAGTGCTGTGAAGGTCAAATTTAAGCGCAGGAAAGTACCAGTTGTTAGGGATTTTCCTTTGGGATGTG

GACCAAATGCACTAAAGGTTAGTGATGTTAAGGGTTCTTCGCAGAGTGGATCAAATGCAATGAAGGTCTC

TGCTATTGGGGGATTTCCTGCGGGGAATGGGCAAAATGCAGTAAAGGTCTCTGCTGTTAGAGATTTTCCT

GCGGTGAATGTTTTGAATGCACTAAAGGTCTCTGCTGAGGATTTTTCTTTGAGGAATGGACTTGGTGAAC

AAATGGCATCTGGTGTTAAAGATTCGCCTGCAGGGAGAGGACCGGATGCACCAAATGCCAATGCTGCTGT

AGAATCTCCTACGGAGAGCGGATCAAATGCAATGAAGGGCTCTGCAACTGGCGAATTATCTGTGGGTAAT

GGACCAAATGCACTAAAGGTCCCAGGTGATAGAGATTCTTCTGCAGTGAATGTTTCGAAGGTCTCTTCTT

GGGATATTCCTGCGGGGAATGGATCAATTGAACCATTGGTCTTGCCTTCTAGGGATTCTCCTGCAGGGAG

GAGACTGAATGCACCAAAGGTCAATGTTGTTACAAATTTTCCCACGGTGAGCGGATCAAATGCAATGAAG

GTTTCTGCTATTGGGGAATTCTCTTCAAGAACTGAACCAAGTGCAATGAAGGCTCCTGCTGTTGGAGATG

TTTCTGCAATGAATGCTTCAAATGCTTTAAAGCTCTCTGCTATTGGGGATATTCCAGTGGCGAATGTACC

AAGTGCATTAAAGGCCTCTGCTGTTAAAGATTTCTCTGGAGGGAATGGACCGAAATCATCAAGGGTCTAT

GCTGTTAGACGAGGTTTTCCAGTTGTTCGGGATTTCCCTCCAGGATGTGGACCAGGTGTTAGAGGAACCA

GACTAAAAATACAGTAATGTCAAATGAAAAAACATGTTGTACAGGTAGATTATTGGCGTCACTTTTGTGT

CCTATGATGTTCTACCTCTGAAAATTTTGAACAAAATCCTTACATGCGCCGTTCATATTTCAGGAGAGAA

GTCACAACAGAACTTAGAGACCAACTAAGAACTCATGGCATCCGTGGAAAGTAACTGTCTTCTGAAACTT

TTAAGATCTTTTGCTAAATATTTCATTACACAGTTCCTCGACAGACAGCTGCTCCAGCGGAATCAAACTT

TTATTGGTCAAGCTCAGTACAAAAACATATGCAGCTTAATATACCAAAAACATTTCACATCCTGACAATA

TCCATATCCAGTCAACACAAGTTATGATCACTACTCTAGACTTCTTATATGATATACATATATGATAGGC

TTCTTATATGATTATCCTTTTAGTTTTGAAATTATGCATATATGATATGGCTGCCGGTC

>MSTRG.132.3 gene=MSTRG.132

CTCGAACTCTCGACTTCTTGTTATCGAGACGGAGTGCCTTCGTTTTTTTGCTTTACCGAGTGCGAGCTTG

TAATTGTTGGGGATGATTTCAGTGGTGAAGAAAGATTGGCCTGATGAGGATGCGATGAAGTTGTCGTTTG

AGACTGGCAGTGCTGTGAAGGTCAAATTTAAGCGCAGGAAAGTACCAGTTGTTAGGGATTTTCCTTTGGG

ATGTGGACCAAATGCACTAAAGGTTAGTGATGTTAAGGGTTCTTCGCAGAGTGGATCAAATGCAATGAAG

GTCTCTGCTATTGGGGGATTTCCTGCGGGGAATGGGCAAAATGCAGTAAAGGTCTCTGCTGTTAGAGATT

TTCCTGCGGTGAATGTTTTGAATGCACTAAAGGTCTCTGCTGAGGATTTTTCTTTGAGGAATGGACTTGG

TGAACAAATGGCATCTGGTGTTAAAGATTCGCCTGCAGGGAGAGGACCGGATGCACCAAATGCCAATGCT

GCTGTAGAATCTCCTACGGAGAGCGGATCAAATGCAATGAAGGGCTCTGCAACTGGCGAATTATCTGTGG

GTAATGGACCAAATGCACTAAAGGTCCCAGGTGATAGAGATTCTTCTGCAGTGAATGTTTCGAAGGTCTC

TTCTTGGGATATTCCTGCGGGGAATGGATCAATTGAACCATTGGTCTTGCCTTCTAGGGATTCTCCTGCA

GGGAGGAGACTGAATGCACCAAAGGTCAATGTTGTTACAAATTTTCCCACGGTGAGCGGATCAAATGCAA

TGAAGGTTTCTGCTATTGGGGAATTCTCTTCAAGAACTGAACCAAGTGCAATGAAGGCTCCTGCTGTTGG

AGATGTTTCTGCAATGAATGCTTCAAATGCTTTAAAGCTCTCTGCTATTGGGGATATTCCAGTGGCGAAT

GTACCAAGTGCATTAAAGGCCTCTGCTGTTAAAGATTTCTCTGGAGGGAATGGACCGAAATCATCAAGGG

TCTATGCTGTTAGACGAGGTTTTCCAGTTGTTCGGGATTTCCCTCCAGGATGTGGACCAGGTGTTAGAGG

AACCAGACTAAAAATACAGTAATGTCAAATGAAAAAACATGTTGTACAGGAGAGAAGTCACAACAGAACT

TAGAGACCAACTAAGAACTCATGGCATCCGTGGAAAGTAACTGTCTTCTGAAACTTTTAAGATCTTTTGC

TAAATATTTCATTACACAGTTCCTCGACAGACAGCTGCTCCAGCGGAATCAAACTTTTATTGGTCAAGCT

CAGTACAAAAACATATGCAGCTTAATATACCAAAAACATTTCACATCCTGACAATATCCATATCCAGTCA

ACACAAGTTATGATCACTACTCTAGACTTCTTATATGATATACATATATGATAGGCTTCTTATATGATTA

TCCTTTTAGTTTTGAAATTATGCATATATGATATGGCTGCCGGTCCACAACATACGAGTGATTCTAGCTA

AATCGAAAAACCTCAAAATGAGACAATTTTATGTTGAATGATACTGTAGCATCCTCTATATGTACATAAA

AAATTTATGGTTAATTTGTCCAGCATCTTTTTTT

>MSTRG.147.1 gene=MSTRG.147

CGATCAAGCGGTTAAAACATCAAATGTACCTCCGTCACGCGAGTGGCGTGCCCGAAACCGTGACCGGTAA

CATAATACGCGAAGATCAGCCGATTCGCACCTGCTGAACCATTAACAGCCCGATCTCCTCCTTCAATAAT

CATCCTCTCCGACTCAAACCAGATCAAATTTTCGATACCGATGAAGAAAACGTAGATTTTGTGATCAATG

TGGATGTGAATGCGCAACAATGAGAGCTTATTCGAGTGAATTTATAGAGGGTTTAGGTGAGGGAGACGCT

GACGTGGATTTGGGGCTTATCTGTTATGAAGTTAGTAACTGTTATTTGTGTGTTTAATTGCGTGAATAAA

GAAAATTCTGGTGATGCATCTGCATATTCTCCGGCCCATTCTTCTTGAAAGTGATGCCCGTTGGATTGGA

TCGAGTTTGGATAAGGGGCTGTGGGAATTAATGGGCTTCTTGGTTTTTTGCGGGTCTTGGGTTTGGAGGA

ATAA

>MSTRG.164.1 gene=MSTRG.164

CTAGATTTACAATCTTCCTGACAGATCTCGAGTCCTGAAGTTTGACAGTGGATTTTTTCTCGACACCGAG

AACGATAGTATCGGTGCCTCTGACGCCCACGGCGGCGTTACCTTTGCGGACGGCTTCAAGTGCGTATTCA

ACCTGAAATAGATGACCGTCCGGTGAAAAGACAGTGATTGCTCTATCGTATCTTGCCATCTTGGTTCCCT

TTGTG

>MSTRG.166.1 gene=MSTRG.166

GAAAAAATAAGAGCGGCACTTTAAAAAAATAAGCGGGTTAGGGTTCTCTCTCAAAGGGGGCAACAAATCT

CTCTCAGGTGGCTGTATCTTCAGGTTTTGGGTTTTTGTTAACTTCTTGGTGGATATTCTCGGTTTTCTCT

TCTCTCAGGTTTTGCGTAAGATCAAGTTTAATAGGCTTGGTGTTGAGGATCTTAGGTTTGGAGTAAATTA

ATAGTCCGAAAAGTCGGGTTCTTTTAAATATGAGAATGAACAAATGCTCAAGGAAGGATTATATCAGCGA

TCTGCCCGAGAGCATCGTCGATATCATCCTAACCAAACTGCCAATAAGGGATGCTGTAAGAACCAGCATT

TTGTCTACCAAGTGGAGGTATCAATGGGCGACCATGACACGAGTCGTATTTAAAAACCACGTGCCATATA

AGAGTGACAACAAACTTGCCGAGCAAGAAATTACAAACTTTATCATGCGATTTCTATTTCTTCATCAGGG

CCCTATTTATAAGTTCAAACTATCTACTAACTGCTCCATGAATTCTACTGACATGGATCAATGGCTACTT

TTCCTATCAAGGAAAGATATAAAAGAGTTAGTTCTGATGTTAGACGCCAACTGTCTGAAGATACCCTCAC

ATATATTTTCTTGTCAGAAACTGACCACATTGAAGCTTCAAGCATCGGACGTGAAACCTCCTTTAAGATT

TCGTGGATTGCCATGTTTGAAGTACCTTAACCTATGTAGTGGCTTGTTTACTATTGAGGATATTGAAAAC

CTTATTTCAGGTTGCCCTCTTCTAGAAAAGTTTACATTCACAAATGTTGGGGACCCTATGGGTTTTGCCA

TCCATGGCCCAAATTTGAAGCATCTTATTTTGAATGGGGACTTTTCAAACTTAAATCTTGAGCATTCTCC

GTTTCTGGATGTCTTAAGGGTTGACTTTCGTGCACAGGTTTGGGAAAGTAACGTCCTTATGAAAGTTCCA

GTTACATATGATCGGCTAAAGTTTATTGATCTTCAAGGAATAAATTACGAAGAGGGGAATGCAGTGTTGT

ATGTTCTTCACTTGATTTTGCATTCCCCTAATCTACAAGAACTGGAAATTGGAGCTACACAGTTTGAAAG

TTCTCATGATAAAGCTGTTGACTTGGATTTTTGGGAGAAAGAATGTCCTACCGATTTCACATTTAAGCAT

CTTAAATTAGTTGAGATGTGTGGTTTGCCCAACAAAAGTTGTGTGGAATTTCTCAAATTTGTGCTTGGAT

GTTCTCCGGTGCTTGAAGTAATGAGAGTGTCACTTGATGAGGCTTACGATGGAGAAATGAATATGGCGAA

TGAACTGCTGCATTTTCAATGTGCTTCTCCAAAAGTTGATATAAAATTCTTTGATTAGTCATCTGAAGCA

TACTTATGCGTTTCCTTCCCCTTTTGTTCTTAGCTGCTTTATTTCGTCATGTCTCAATTTTTTATTTTTT

TTGTGAGAGTCAAGTCTCAAATTTAGTTGCAAAAGAAATCTGTACATTAGAAGAATAAGCTTTTGCAAAT

TGGGGACTCTTCAAGACTCTAATGACATGATTTTTTTTTTTTTTATGGATGACGCCTTAACAGGGAGAGC

ATTCCTTAGTCTTTATCAAGACTTCAATCTCTGAACTCTTTCCATGCTTGGTGGATTTTCCGGTAATGAT

GAATTGAATCGACTATTAAAACAGTAAAAACAACTTTGATGATGTATCTCAGAACCACGCTTCCTGACAA

TCACTATTCTTCTGATGTTAGATTGGATATAAACA

>MSTRG.166.2 gene=MSTRG.166

ATATCTACACACACCGTGTAATCCAAGAATATGAGAGCTAAGAATAAAAGAAAAGAAAAACTCCAGGCTG

CGTGGCGAGAAGAAAACTAGAAAAGAGGTTTTGCGTAAGATCAAGTTTAATAGGCTTGGTGTTGAGGATC

TTAGGTTTGGAGTAAATTAATAGTCCGAAAAGTCGGGTTCTTTTAAATATGAGAATGAACAAATGCTCAA

GGAAGGATTATATCAGCGATCTGCCCGAGAGCATCGTCGATATCATCCTAACCAAACTGCCAATAAGGGA

TGCTGTAAGAACCAGCATTTTGTCTACCAAGTGGAGGTATCAATGGGCGACCATGACACGAGTCGTATTT

AAAAACCACGTGCCATATAAGAGTGACAACAAACTTGCCGAGCAAGAAATTACAAACTTTATCATGCGAT

TTCTATTTCTTCATCAGGGCCCTATTTATAAGTTCAAACTATCTACTAACTGCTCCATGAATTCTACTGA

CATGGATCAATGGCTACTTTTCCTATCAAGGAAAGATATAAAAGAGTTAGTTCTGATGTTAGACGCCAAC

TGTCTGAAGATACCCTCACATATATTTTCTTGTCAGAAACTGACCACATTGAAGCTTCAAGCATCGGACG

TGAAACCTCCTTTAAGATTTCGTGGATTGCCATGTTTGAAGTACCTTAACCTATGTAGTGGCTTGTTTAC

TATTGAGGATATTGAAAACCTTATTTCAGGTTGCCCTCTTCTAGAAAAGTTTACATTCACAAATGTTGGG

GACCCTATGGGTTTTGCCATCCATGGCCCAAATTTGAAGCATCTTATTTTGAATGGGGACTTTTCAAACT

TAAATCTTGAGCATTCTCCGTTTCTGGATGTCTTAAGGGTTGACTTTCGTGCACAGGTTTGGGAAAGTAA

CGTCCTTATGAAAGTTCCAGTTACATATGATCGGCTAAAGTTTATTGATCTTCAAGGAATAAATTACGAA

GAGGGGAATGCAGTGTTGTATGTTCTTCACTTGATTTTGCATTCCCCTAATCTACAAGAACTGGAAATTG

GAGCTACACAGTTTGAAAGTTCTCATGATAAAGCTGTTGACTTGGATTTTTGGGAGAAAGAATGTCCTAC

CGATTTCACATTTAAGCATCTTAAATTAGTTGAGATGTGTGGTTTGCCCAACAAAAGTTGTGTGGAATTT

CTCAAATTTGTGCTTGGATGTTCTCCGGTGCTTGAAGTAATGAGAGTGTCACTTGATGAGGCTTACGATG

GAGAAATGAATATGGCGAATGAACTGCTGCATTTTCAATGTGCTTCTCCAAAAGTTGATATAAAATTCTT

TGATTAGTCATCTGAAGCATACTTATGCGTTTCCTTCCCCTTTTGTTCTTAGCTGCTTTATTTCGTCATG

TCTCAATTTTTTATTTTTTTTGTGAGAGTCAAGTCTCAAATTTAGTTGCAAAAGAAATCTGTACATTAGA

AGAATAAGCTTTTGCAAATTGGGGACTCTTCAAGACTCTAATGACATGATTTTTTTTTTTTTTA

>MSTRG.176.1 gene=MSTRG.176

CTTTCCTCTTCATTGCATCATTTCCGCAATTAATAGTTCTTCCTGGTTTCGGAGAAACTGGCCAGTGGCC

ACAAGTCACAGCCATATCACCCAGATGTTGCAGAGCTTCAAGTGTTTGAATGATATGCGACATTTTAGGC

CTATCTTTAGGATTCGGACTCACACAACGCAGTGCCAACTTTGCCATTTCTTTTGCTCCTTTAACCGAAT

ATTGCCCACAAAGCCGGGGGTCCATGATGTAACGTAGCCTCCGGCTACTAGCCAAGTATGGTTTTGCCCA

ACTGACCAGGTTTTGCTCTGTTTTCATTCTTTTTTCAGTAGCTCTTCATCCTGTTAGTAGTTCAAGCAGA

ACAACTCCGAAGCTATAGATCACTCCTGGTTGTCAAGTGTCCTGCAAATCGATGGTCAAAATGAAATATC

TGAATACAGATTGTTACCCATTTCTGGGTCGAGAGTGGATCCCGAGAGCTCTGATACCAGGAAAATGATC

AATTCTCCC

>MSTRG.177.1 gene=MSTRG.177

GAAAACATGAAAAATTTGTCCTACTCTATCGCTTTTCAACGGAAACTAAAATTCATCTCATATAACAGAC

TCAAGAACGAAAAATTTCTGGCGAAATAGCTATTATATGTATCAGGAAATCAAAAAGTGAGTGAGAAAGT

ACAGTTATGTAAGCTTCGAGCCGCAGTACTCAGGCATGGAGTGGCATTAGTATAAGCACTGTTGACTCTA

TACACTGAGTGCTCCTTGAACCAGGTTCCAGCATCTTGTTAAAAGAACACTAAGTTCCGACAGACAGGTG

TTATTACAGCTCTGAAGATCTTCTGTTCTTTTCAAATTGATCGTAAACTGACCGAGTTCTTGCAAACAAA

AGAGCACTTGATGCTAGCCCGAAAACAGATACACATCCCCACCACACGAATGTCCAGAAATAGCATTGTC

TTCCCATGCACACCAATGAGTTGGCTGGCAGCATTTGAATGTCCTCAGTTGCATTGGCATCGTAAACGAG

AGCAGATAGAAGACCATATAAAAGCGATCCAATGGGTATATTAGTTATCAGAATGTTATGGTTGACACCC

ACACTGTTAGGCCCGAAAAGCTCAGATGTTATTGACACCGCAGCTGCAAATATGAACCCAGAGCTTAAGC

CAATCAGAGCTGTGCCTACCTGTAATGCCATCTCACTGCCTGTTCCTGCAAGCAAGAAAAAGGCAACTGG

TGTAGGTAAAAGCGCAAGGGCTAGCCATCCGGTCCTGGCAAAATAAAATTTCCTGCAGAATTAAGAATGT

TGTATCAGTAACGTACAAGCACTCGTGTAAACCAAATTGCTAAAGTCTGTACTTACATCCGCAAGATATC

TGGAGCTGCTGACAGCAAGCGGCCAAAGAAGGAGAAAGCGGAATATAGTGTGATAAGAGTCGAGGTTGTT

GAACTCAGGCCAAGTGACTGTGATATCTGTCCTAAATTGTTGCTATACACTAGTCCTATTGTACCTCCAA

AAAAGTAAGCAAAGTAGTAGAGCCAGAAATCAATCCTACGAATAAGCTTGTTTGCCTTGTGATCTTCTCC

AAGCATTACCAAGCGATCTCCTTCTATGATCCTTACAAAGAACCCTTCTCTAGATTTGGTGACTTCTTCA

ATAATACGATTAGGATATGCATCATCACTACCTGTATACGATACTACATTAATAACTGGACTATTCTCCT

GACTCATGAACTCTTTATGAAGTTGAAGATCATCGTGATCAACAAGAATGAAACTTGTGCTATCAACACG

AATGCTGGAGTAAATTGTACGGTGAAACCAATCACGAGCATACACAACACCAGGGATAGCTAGAGGAAGA

ATCAGAAGTATAATGGCACCACAGAACAGAATACGGGCGGATGATGGATCTCCTGGATGGAGCACGAGAA

GATAGAGGCCAGTGATGACAGCAAGAAAATTTAGCAGCAGAAATATTAGGCGGTCACGTTTAACAGCATC

AGTAGGAAGATGGTTCACCGGGGGTTGCCTTAGAATCGGGATTAGAGCTGCTATGGATGTTAAAAGAGGT

ATAAATGCATTGAGAAGAAGGTATAGGTCACTCGATGATGAATCAAATGATTTGGCAGCAAGGTTGTACA

AGGCTGCGCTTACACCATTGAAGCTTACAGTTAGAGATATTGCTAAAGGTCGATTAGCTGGGAAATTCTT

GATGCAGAGAACAAAACACACAGTGTTGAACCAACAGATGCTACATCCAGCTAACAAACACAGCATGAAA

ACCTGAAACAAGTTACCGGAAGAATAAGCTACCAAATGAGGTGAGTCTGTAGAGTCTCGTATACACTACA

ATGCTATAACACCTGAAAATGTCACAAGTACTTATCCGACAAATGTGGGTGAAGGAGGGATCAGTCCAAT

ATACTTCGTAAATTTGGAAAGATTCAGTTAAGGTGAGCTACCATTAAACTTAATCTTACAGCAAATGCCT

CACATTCTGAATAGTTTTAGTAATTCAGCATCATCACAACTTATACGACTCTATTTCCTGCCCAACCCCG

AAAGAAAGGCTAATTTTCC

>MSTRG.180.1 gene=MSTRG.180

AAGTACTTTCAGGAGAATCAGGTGCACTAGGCTCTTTTTTTGGAGTAGTTGTGATCTTAGTTGTTTTGCT

GTCAGTTTGGTGTAGTAGGGAAGTGGTGTTAGGAATGGCCATGGATATTAATGGGAATTCTCGAAGCGTG

GTTTCAAGTCTGGGAAATCTCAAACACCCTGCGGTTGCTCTTCTTTGTTTTTGCTCAAGTGGAGTTGCAC

TTTCTTTGTTTTAAATGTGATTCAGATTTTGTGTAGTTGCTTTTTTCTGGAGCTTGGTTTTAAAAGAGGT

TTTTGGTTTGTAATTATTTGAGGGGTGCCCACTCTGATATTTATGTATTATATGTTTTCCTTCTTTTTAA

AATTATTC

>MSTRG.187.1 gene=MSTRG.187

AGCATATTTTCTTTACTTGTACTGTTGTACATAATATTACCAATATCAACTTGTCTTCACGCAATATTTT

GAGTCCTTCGTTATACTTGGAGAAGTATACTAGCTGCACAAGATGTAGTAAGAAGAGGGTGCAGGAAGCG

AATTGGCGATGGTAAACAGACTGAAATATGGAAGGTCCCTTGGCTTCCATGTAAAGAGATTGGTCTCGTG

ACTACAGAGATGCCCATGCAAATGGAAGGAGTAAGGTTTGTAGTTTCATGCAGGCAAACCAGAAAAAATG

GGATGAGGATGTGCTTATGGACATTGTAATGAAAGAGATAGAAATTGTATCAGAAAAATTCCACCATCAA

TAAGAGAAGGAAGTGACTCATGGTTGGCTTCTACATGATAAGGGGTGCGTTACGGTTAGGAGTTGTTATA

GAATTTTGCAAGGGGAGGTGGAGGCGCCATATGCGTCGTTTTGGGCGAAACTTTGGGGTTTGAAGTTCAG

GGTAAAGTTACTCAGTATGTATGTCGCGTGTGTTCTTGGTGCTTGCCTACTACAGCAAGGCTTACTACAA

AATGAGTGCAAGTTGACACGATTTGCCAATGGTGTAGACGGTATCACGAAACAGATAATCATGTGTTAGT

CGAATGTGAAATGGCAAATGCAACGTGGATGGAAGTTGGTCCTCAGGCGTGCATTAGTACACTGCCAGGT

GATAATGTGTTTGATGTTATATGTCGATGTTTTAACAGCTGCACCAGATAACAACATACCTTGATAGTTT

TGATGTGTTGGAGTATTTGGACCAGAAGAAACAAGTGGGAGGGAAAAAATTAATATGTCAGAGTTTGGCA

TCAAAGCAGCTGCCCTGAACCTGCTTGCAGACTGGAAGAAGGCACAGGAACAACAGTTAACACCTGGTAC

AAGGTCGAGAACACTGGGTGCAGTATGTAAATGGGAGAAGCCTCAGACTTCTTGGGTGAAAATGAATTTA

GATGCTGCATTTTTTGAAGAGATTGATTGTATTGGTTTGGGCAGTGTGGTTCGAGGTGCAGACGGACAAT

TTATCATGGCTATGCATAGGAGACAACAAGGGCTGATGGCACCAAGAGAGGCTGAGGCATTATGCTTGAA

AGAAGCACTGATCTGGCTTAAGGACAAGAGTTTCAGCAAGTGTAATTTCGAAACTGATTCACAATTTCTC

GCTTGAGCTTGCAAAGGAGCTGGTGGAAGGTCATATTTTGATACAGTTATTGGAGATTGTATTGATTTAT

ATAATAACTTTGATAAAGTGTCATTATGTTTTACACATAGGTCTGCAAATGGAGTGGCTCATGTTTTAGC

AAGGATGGCTTATTCTATGTCAGGCTCTCGGGAATGGCATATTAATGCTCCCGATTTTATTCATCATGTA

ATTTGTTCCGAAGCTTTATAAGAAATGCAAGTACGAT

>MSTRG.188.1 gene=MSTRG.188

TGGAGGCGCCATATGCGTCGTTTTGGGCGAAACTTTGGGGTTTGAAGTTCAGGGTAAAGTTACTCAGTAT

GTATGTCGCGTGTGTTCTTGGTGCTTGCCTACTACAGCAAGGCTTACTACAAAATGAGTGCAAGTTGACA

CGATTTGCCAATGGTGTAGACGGTATCACGAAACAGATAATCATGTGTTAGTCGAATGTGAAATGGCAAA

TGCAACGTGGATGGAAGTTGGTCCTCAGGCGTGCATTAGTACACTGCCAGGTGATAATGTGTTTGATGTT

ATATGTCGATGTTTTAACAGCTGCACCAGATAACAACATACCTTGATAGTTTTGATGTGTTGGAGTATTT

GGACCAGAAGAAACAAGTGGGAGGGAAAAAATTAATATGTCAGAGTTTGGCATCAAAGCAGCTGCCCTGA

ACCTGCTTGCAGACTGGAAGAAGGCACAGGAACAACAGTTAACACCTGGTACAAGGTCGAGAACACTGGG

TGCAGTATGTAAATGGGAGAAGCCTCAGGCATTATGCTTGAAAGAAGCACTGATCTGGCTTAAGGACAAG

AGTTTCAGCAAGTGTAATTTCGAAACTGATTCACAATTTCTCGCTTGAGCTTGCAAAGGAGCTGGTGGAA

GGTCATATTTTGATACAGTTATTGGAGATTGTATTGATTTATATAATAACTTTGATAAAGTGTCATTATG

TTTTACACATAGGTCTGCAAATGGAGTGGCTCATGTTTTAGCAAGGATGGCTTATTCTATGTCAGGCTCT

CGGGAATGGCATATTAATGCTCCCGATTTTATTCATCATGTAATTTGTTCC

>MSTRG.189.1 gene=MSTRG.189

ATTTAACTCAAATCAAAATATTGAAGAACATAACTTGTTCAATTCAGCTAGAATATTCTCACTTAGACAT

TTAATCAAACACCTTGATATCAATTTTCATGTATAAGCAGATAATAAACACGCAGCACCAGCAGATCCAT

AAATTGCATAAATGGTGAGCCAATGCACACAAATATATGCATAAATTTGTGACGAGAGTATAGATATGAG

GAGAGTTTTGTGCAAAGAGTTACCTTGGAAAGAGCCTTCACTATGGAGCAATATGATGCAGAGATGGAGA

GTGGAGAGAGATATTGGGAGAGAGGGAGAGATGGAGAGGAGAGACTGTTAGAGAGAGAGGCGAGATGACG

TGTGAGAGAGAGAGGCGAAATAGACGCACAGGCAGAGAGAATTGAAGAAACGAGGGAGAAGATGT

>MSTRG.190.1 gene=MSTRG.190

CGGGATTGGGCTTCCCATTCCATAGCCTTACGTTTGGATTAAATTTTTAAGACAAGGGAATGGGAACTCT

TGTGCCACTAGCCTGTTAACACAAACCTGACCAATACCCCGGGTTACCAATATTGATTCCCGGTCTCGAC

ATAACATCTTCTCCCTCGTTTCTTCAATTCTCTCTGCCTGTGCGTCTATTTCGCCTCTCTCTCTCACACG

TCATCTCGCCTCTCTCTCTAACAGTCTCTCCTCTCCATCTCTCCCTCTCTCCCAATATCTCTCTCCACTC

TCCATCTCTGCATCATATTGCTCCATAGTGAAGGCTCTTTCCAAGGTAACTCTTTGCACAAAACTCTCCT

CATATCTATACTCTCGT

>MSTRG.194.1 gene=MSTRG.194

TTGATCAAACTTCTTATCTACACATTGTAGTAGATAACCATACCAGTTATATTTAAAGCACTGATTCAAA

TTTCCGGAATACGACAGAATCTTCAAACACACATAAGGTGTTCTGGACCCCTCTATTAGAATATTAGTGA

GGACAATGAGAAAGTTCAATTTAAATCTCAAGTCAACTAATTTCGATGATTAAATAACATCACATAAACT

ATTTGCAGTAATCTTATGAGGACCCTTGTACTCGTTGAACTGTGATCTCCAAACATCTGTATACTCACTG

TTGTAAGATCCTCAAACATTAACTCCCCCTTTCGGCAAGCCAAGTATAATCTGGACATCATTTTCCGAAA

AGTTTATAATATTCTCACCAATTGAAATTCTTGAATCGATTGATTTGTATGACTTCGCAATCAAAAAAGT

CAAAGGTTGAGGGTATTCATTGATGTTGAAAACTAGAACCTTTTCAAAACCAGTCTCTACAACCTATTGC

CTTTGTACATCCGAAAGATTGTCCACGACTCTCTTCAAGAACATAGGTTTTGCTCGAACCAACAAACGGC

CGGTATTCTTATCGAGATTCTTCTCAGCCATCATATCACTGAAAAAGGAATATGTGGTTGCGGAGATCAT

ACAAAAGTCATAACAGATATATTG

>MSTRG.197.5 gene=MSTRG.197

CAAACCAAGTGCCCTTTAACATATAAGCCGTCCCCGATAATTAATTTTATTCTTCCTAAATCTTCGCGTA

CTCAACAAACCAAGTGCATCGCCGCTTTCCAGGTGCCCATATCAGTCATGTAAGAATTGTTGCGCAAAAT

CTCAAAACCCGTGCCATGTACATGGAATTCTCATAGGGTACCCAAACCAGTGAGCAAGAAGAATATGTTA

GAAGTCCAGAAAAGGAGTTCAGGAGTTCAGGAGAGATGAGGCATTGGCTGCGTTTATTGACAAAATTAAC

AAAGCCCAGAATGAGAAAGATCTGAAACTCTGCATGGAGATGAAAGATCAGCTTGTTAATCCGCATACTG

AGAGTACTCAAGCAGAATCCGAAAGTGTTGCAATTCGTATGGCGCTCCAGACTGATGATGTGGTCAAACC

TCTGTTCGCTTAATTTTTTTTATTTTTTATTTTGAAAGGTACCCCAAAACTTGTTTGTTGTTTTCGGAAG

GTACCCCAAAACTTTTGATCCTCCAAATTATAGACAAATTAGTGCATACTCTGATTCCCTGGAGGAGCTG

TAGGAGCTGTAGTCTATGATGAGTTCTTGGTACCAAAAACTTGAATGACTCAGTCGGCGAATTGAATAAG

AATCAAACCATTAATATGCATCCGTCCGTTTAGAACGGTTGCATACTAATGGTTTGCTTGCTAGTCGATT

TGCCAAATGAAACATTCAAGTTTTTTGGTACCAAGAACTCATCATAGTCTACAACTCCTTAATATGCTCC

AGGAAATATAGATCATGGTTTATAAGTAAATAATCATTTATAGTTTTAGTGGAGGATCAACGAACACTTG

ATCACGTCATCTGTCAGAGATTGACCACATCATCAGTTTGGAGGCCCATAAAAATTGCATCACTTTCGGG

TTCTGCATGAGGACTCTCGGTATGCTGATTAAAAAGTTGATCTTTCATCTCCATACAGACTTTCGGATCT

TCCTCATTCTGGGCTTTGTTGATTTTGTCAATACAAGTTATCCACTCAATTAATTATATATCACATATTT

ATATTTTAATTCAGGATTTTTAATATATTTTGGGTTTTTCTTGAAAAATATACTCGCTCTGTTCTAGCTA

>MSTRG.197.6 gene=MSTRG.197

CCAAGTGCCCTTTAACATATAAGCCGTCCCCGATAATTAATTTTATTCTTCCTAAATCTTCGCGTACTCA

ACAAACCAAGTGCATCGCCGCTTTCCAGGTGCCCATATCAGTCATGTAAGAATTGTTGCGCAAAATCTCA

AAACCCGTGCCATGTACATGAACAACATCTCTGCGCTACTTGCTGAGTATATATCTAAATATATCTACGT

CCCACAAGACCCTTCAATCTCACAAGTAACGTTGCCAAGAATCATAACATCACAAACTTAGACATTCAAC

AATTGACACTGTGGAATTCTCATAGGGTACCCAAACCAGTGAGCAAGAAGAATATGTTAGAAGTCCAGAA

AAGGAGTTCAGGAGTTCAGGAGAGATGAGGCATTGGCTGCGTTTATTGACAAAATTAACAAAGCCCAGAA

TGAGAAAGATCTGAAACTCTGCATGGAGATGAAAGATCAGCTTGTTAATCCGCATACTGAGAGTACTCAA

GCAGAATCCGAAAGTGTTGCAATTCGTATGGCGCTCCAGACTGATGATGTGGTCAAACCTCTGTTCGCTT

AATTTTTTTTATTTTTTATTTTGAAAGGTACCCCAAAACTTGTTTGTTGTTTTCGGAAGGTACCCCAAAA

CTTTTGATCCTCCAAATTATAGACAAATTAGTGCATACTCTGATTCCCTGGAGGAGCTGTAGGAGCTGTA

GTCTATGATGAGTTCTTGGTACCAAAAACTTGAATGACTCAGTCGGCGAATTGAATAAGAATCAAACCAT

TAATATGCATCCGTCCGTTTAGAACGGTTGCATACTAATGGTTTGCTTGCTAGTCGATTTGCCAAATGAA

ACATTCAAGTTTTTTGGTACCAAGAACTCATCATAGTCTACAACTCCTTAATATGCTCCAGGAAATATAG

ATCATGGTTTATAAGTAAATAATCATTTATAGTTTTAGTGGAGGATCAACGAACACTTGATCACGTCATC

TGTCAGAGATTGACCACATCATCAGTTTGGAGGCCCATAAAAATTGCATCACTTTCGGGTTCTGCATGAG

GACTCTCGGTATGCTGATTAAAAAGTTGATCTTTCATCTCCATACAGACTTTCGGATCTTCCTCATTCTG

GGCTTTGTTGATTTTGTCAATACAAGTTATCCACTCAATTAATTATATATCACATATTTATATTTTAATT

CAGGATTTTTAATATATTTTGGGTTTTTCTTGAAAAA

>MSTRG.198.1 gene=MSTRG.198

TTCTGATTTCTCTGTTAAACTAAAATAGCCAGATTTGAACCATTTCATCCTCACCTACGATTTGAATCAC

AAAATCGATTAAAAACTCGAGGATTCCTAATTCTTTTTCAGAAATCCCATGGCTGCGTGTTTGTTTTGTG

TGTTTTTTTTCTCAGTCACCTACGATTTGAATCACAAAATCGATTAAAAACTCGAGGATTCCTAATCCTT

TTTCAGAAATCCCATGGCTGCGTGTTTGTTCTGTGTGTTTTTTTTCTCAGGGATATATAAAAACTAAAGA

AAGCTACCGACGAGACTCGGCTAATGACTCCACGATCTCGTACAAGAGAGACATGATAGCGGTGGTAGCC

GAAGAGGATGGTAGTGACGGCGGCAGTTCCGAGATCAGAATGAGATTCGGGTGTCGGGTTCAGAAAAGAT

GCATTAATTGTGCCTTAATTGCGGCGTTAATTACGGCATTAATGTTTTGTTAATTGTGCCTTAATTG

>MSTRG.201.1 gene=MSTRG.201

ATTGCAGTCACAGTCTTGCTACTTGTGCCCGTGGATCGATACATACACGAACACGATACATACATCTCCT

ATTCTTACCGGTACCAACGCTCACTCTCCCCGACCTCCTCTCCTTGTGTCTATACATACATCGTGTATAT

GTATGTATGTATAGATTAAAAGCTTAATTGATAAGCTCTGATCTCAATACAGAAATGAGTAGCAGGAGAG

GAGGTGCGGAGATGTTGAAAGATAGCAGGACCAAACAGATTAATGATGCTCTTGATAAGCATTTGGAGCG

ATCTTCGCCTTCTTCGTCTCGAGGCGGAGGAGGAATAGCTTTTAAGGAGAAGGATCGGGTTTCCGGGTTG

TCCGGGAACGGAAAGGCTTCGTTTCAGAAGAAGCCTATTGAGGAATCTGAAACCGACAGCGAGGAATCAG

ATGTTAGCGGTTCTGATGGGGTAGACTCATCATGGATTTCTTGGTTTTGCAATTTGCGGGGAAATGAATT

CTTTTGTGAAGTTGATGAGGATTACATTCAAGATGATTTTAATTTATGTGGATTAAGCAGCCAAGTTCCA

TACTATGATCATGCTCTTGATCTGATTCTAGATCTCGAGTCCTCTCCTGGTGATGTATTTACAGAGGAAC

AGCATGATTTAGTTGAAACAGCAGCAGAGATGCTATATGGTCTGATACATGTTCGATATGTATTGACAAC

TAAAGGGCAGGCTGCTATGTTGGAGAAGTACAAAAATGCTGAGTTCGGTCGATGCCCTAGAGTTTGCTGC

TCTGGACAACCCTGCTTAGCTGTTGGTCAATCAGATATTCCTCGTCAATCAACTGTGAAAATTTATTGTC

CCAAGTGCGAAGATATATATGCCCCTCGATCCAGGCATCAAGATAACCTTGACGGAGCTTATTTTGGAAC

GACATTTCCTCACCTGTTTTTGATGACCTACGGGCATCTTAAACCACAGAAGAAATTGACGAGTTATATT

CCAAGAGTATTTGGTTTCAAGGTTCACAAACCATGATGCTGTAGTTATGTTACTCAGTTACCTGGGATTA

AATCTTGTGCTGCTGAACGCAAACCCCAATACCATCACTCAGACTCCTCGGATAACAGATGATAGACTAG

AACAGTTATCTACTATGAAAGTCTTTGGTGAGAGCAATTGTATTCAAATGTATGCTAAATTTTACTTATA

AATTTAGGTTCGATAATGAATTTGTTGGGATGTATAATAACTTTGTTGTAGATTAGCTGCAGAGAGAATG

GTCTTCTTCTCAAGTATGTAGTGGATCTTATTGAATGGAATTATGTCCTGAATTTTTTGGTTTAAAA

>MSTRG.204.1 gene=MSTRG.204

CTTAAATTGTTGTTTAACTTGGGATTATTCTGATTGGGTTTGATTCTTTTTTCTGGAATAGCGAAAGTGA

TATTGATGATGAAGCAGATTGCGGAGAGTCCAGTTTGATGCACTTCCCAGTGCATGATCAAGCATCACTG

CACGAAACGATGCAACAATGTATCATCAAGGCAGCAAATTCCCTCCTGCATGACTCTGCCAATGCGGAAC

CTTTACCTTATATCAACCATTGGACTCATAAACACCCGCTGACACTTAGAAACAAGAACGCAACAACTTC

AACCTCTAACTTGAAACATAAATTAGCAGAAACTGAATTGCTAATCTGTGACGGGTGCACTAAACCTATA

TCTTTGGTTGATGATATATCCTACGAATGCAATTTATGCAAATTTTTTCTCCACAAATCATGCGCCCTAT

TTCCAGAAGAGATTGAGCATCATCTAGCAGGCAAGCTGTGGGGAGTACAAGTAGATAACGAACAAATAAT

TTATTGCCAGGGTTGCAGTCATCTTGGTAATGGGATTTTCATGCGCAATGAAACAGCCTGTTTTGATATC

GGGTGTGCTTCGTTACCCAGAATAATCAAACATGAATCTCATCGTCATCCTCTTAAGCAATTACCATATC

CAGATGATTATATTTGCAAAGCATGTCGTTCTGAACTTTTAACAGACAGAGAAATTATGATGTATGGATG

TGAAAGATGCGAATTCTACATACATATATGGTGTGCGTTAAGGCCACGTCGGGTGAATCATCGATGGGAT

CCTCATACTCTGGACCTGATTCTGTCGCTCAACAATGTACCTGATCATCCTCATGAATTCGAGTGTGAAC

TCTGTTCCGAGCAGATAGACCCAAACACTTGGTTCTATCACTGCAATGTTTGTGATCTATCATTTGATAC

TTTCTGTATTGATCCTGATTCCTGGCTCTCCAATATCAAGCTTGGTGCCACCAACATTCATACCGACTCA

CATCCTCACTCACACGGCCTGACATTGGCCGTGAATAAAAAGAAGAGGAAATGCGACAAGTGTGGTATAG

ATGCACCCGGCTGGATAGTCCTCGAATGTTCAGAATGCGAATTTATGGTTCATGTGCGTTGTTGATGAAA

TCAGACTTCAAATCAGATGCTGCACTTGTGTCAAATGTAATGAAGAATGAAAGTGTCTTATTGCAAAGCC

TGATGGAGCATTGCACTAGAATAAAAGGCCTTTGGCTATATTTTTTTCTTTGGAATGCAAAAATTATCAA

CTATGTTCTTGCTCCTCAAAACAATGGTCTTAATTACCATATTTGTGTTGAAAACATATTGTACT

>MSTRG.205.1 gene=MSTRG.205

CTTAAATTGTTGTTTAACTTGGGATTATTCTGATTGGGTTTGATTCTTTTTTCTGGAATAGCGAAAGTGA

TATTGATGATGAAGCAGATTGCGGAGAGTCCAGTTTGATGCACTTCCCAGTGCATGATCAAGCATCACTG

CACGAAACGATGCAACAATGTATCATCAAGGCAGCAAATTCCCTCCTGCATGACTCTGCCAATGCGGAAC

CTTTACCTTATATCAACCATTGGACTCATAAACACCCGCTGACACTTAGAAACAAGAACGCAACAACTTC

AACCTCTAACTTGAAACATAAATTAGCAGAAACTGAATTGCTAATCTGTGACGGGTGCACTAAACCTATA

TCTTTGGTTGATGATATATCCTACGAATGCAATTTATGCAAATTTTTTCTCCACAAATCATGCGCCCTAT

TTCCAGAAGAGATTGAGCATCATCTAGCAGGCAAGCTGTGGGGAGTACAAGTAGATAACGAACAAATAAT

TTATTGCCAGGGTTGCAGTCATCTTGGTAATGGGATTTTCATGCGCAATGAAACAGCCTGTTTTGATATC

GGGTGTGCTTCGTTACCCAGAATAATCAAACATGAATCTCATCGTCATCCTCTTAAGCAATTACCATATC

CAGATGATTATATTTGCAAAGCATGTCGTTCTGAACTTTTAACAGACAGAGAAATTATGATGTATGGATG

TGAAAGATGCGAATTCTACATACATATATGGTGTGCGTTAAGGCCACGTCGGGTGAATCATCGATGGGAT

CCTCATACTCTGGACCTGATTCTGTCGCTCAACAATGTACCTGATCATCCTCATGAATTCGAGTGTGAAC

TCTGTTCCGAGCAGATAGACCCAAACACTTGGTTCTATCACTGCAATGTTTGTGATCTATCATTTGATAC

TTTCTGTATTGATCCTGATTCCTGGCTCTCCAATATCAAGCTTGGTGCCACCAACATTCATACCGACTCA

CATCCTCACTCACACGGCCTGACATTGGCCGTGAATAAAAAGAAGAGGAAATGCGACAAGTGTGGTATAG

ATGCACCCGGCTGGATAGTCCTCGAATGTTCAGAATGCGAATTTATGGTTCATGTGCGTTGTTGATGAAA

TCAGACTTCAAATCAGATGCTGCACTTGTGTCAAATGTAATGAAGAATGAAAGTGTCTTATTGCAAAGCC

TGATGGAGCATTGCACTAGAATAAAAGGCCTTTGGCTATATTTTTTTCTTTGGAATGCAAAAATTATCAA

CTATGTTCTTGCTCCTCAAAACAATGGTCTTAATTACCATATTTGTGTTGAAAACATATTGTACT

>MSTRG.206.1 gene=MSTRG.206

TAATATATATATGAACAGAGTGCAGTGGTCGTGTTGAATCATTCATACTCGCCTTCAGTAAATATACATC

TCCCTTTGTCTCTCTCTCTCTCTCTCTGGTAAACATAGATGGCAGAATTGAAATTTTTTTTTCATGAGCA

TCAGCTGATACTAAATGAAGCTGAGCCTGTTGTCGGTAAGGATGTAGAATGTGTTGGGTGCAAACAGCCA

ATCAACAAACTCATAGACGCATTTTATAGGTGCAACAAGTCCCTTATTGATAGCAGTCCAGCGAGTGATT

GTGTTGGTTTCTATATGCATAAAACTTGTTCTGAGTTGCCCTCAACTTTTACACACCCTATGAACCCGAA

AAAACCTCTAAGCCTCTTTGTGCTTCCATATAAAAAAGAATACTTTTATATTTGTCATGCTTGTGACAGC

CACTCTCAGCACTTTATGTATGGTTCTGATTCTTCTGTCAGTTCCGAATTTCGTGTTTGTTTGAAATGTG

TAATGTCAGAACTTAAATCTCTAGAGGACCGTAACCTTTGTCATCCAGGTCACAATCACCCATTAACTTT

AGTCCAAAGCCCAGCTTTGTTCCTGTGTCATGCTTGTAACACTACAGCTACAGACTTGTCCTATATCTGT

ACAAGATGTTGTTTTTATATACACAAGAATTGTGCCAATGCACCCACCACCTATCAAAGTAAATTCCACA

ATGAACACGCTCTCATCTTGACCTACTCTCTTCCCCAACAATATCGTGAATATCCCTGTTACTGCAGCAT

CTGCGAGAAACATATAAATCCCATCTACTGGGTTTATATTTGTGCGAATTGCAGATTTTTTGCTCACGTG

AAATGTGCTTCATTAACTGAGTTGTTGAGTGAAAGTGATATTGATGATGAAGCAGATTGCGGAGAGTCTA

ATTTGATGCAGTTCCCAGTGCATGATGAAGCATCACTACATGAAACAATGCAAGAATGTATCATCAAGGC

AGCAAATTCTTCCACTGGGGAACCTTCACCTTATATCAATCATTGGGCCCATGGACACCAACTGGCACTC

GGAAACAAGAATGCAAAAACTTTGCCCCTCAACTTGAAGCCGAAAATAGCAGAAACTGAATTGCTAATCT

GTGATGGTTGCATGAATCCTATCTCCTTGGTTGATGTATTCTATGAATGCAGTTTATGCAATTTTTTTCT

CCACAGATCATGCTCCCAATTCCCCGAAAGGATTAAGCATCATCTAGCAGGCGATCTGGAGGGAATTCTA

GCACGAATGTCAGGTGAACTATATACTCTTGAATGCAGAGGTTGCGGCATTCGTGGTAATGGGATTTGCA

TGCACAATGATGAATACGCTTTTGACATCAAGTGCGCTTCATTACCCAGAATAATCAAACATAAAGGTCA

TCGTCACCCTCTTCAGCAATTAAAGACTCCAGATGACTTTCTTTGCAAAGCATGTTGGCGTTACCGTGTA

ACTGAAGAGGTAAGAACCATCGTGTATGGATGTGAAAAATGTGATTTTTTCACACATATAGGGTGTGTGT

TAAGTGCACAAGTGGTGAAACATAGATGGGATCCTCATCCTCTCTACTTGATCCTGTCTCTTAAGAATGT

ACCTGATCATCCCCATGATTTCCATTGTGAATTTTGCCCCGATCAGATAAACACCAACAGTTGGTTTTAT

CACTGCAATGTATGCGATCTATCATTTCATACCAGCTGTATTGATCCAGATGATTGGGTGTCCAATATCA

AGTTTGGTGCTACCGACATTTACAGCGACAAACATCCACATCCACACGGCCTCACATATATTCTCAATAA

AAAGACGAGGAATTGCAATTTATGTGGTAAAGATGCACGCGGTATGCGGGCTCTCCAATGTTCAACATGC

AAATATATAGTGCATGAGAATTGTTTCATTGAAGATTGAAATGAGGCAAAAATCAGATATTGTATTACTG

GATTATTCAGGTTCAGAGGATGTACAGAAAGAAGTAGCCACCGCATTCTTGAAGAATCTCTAGAACTCGT

TGAGCATTAGTAACAGTTCGCTCTATAGCATTAAGAGTTCTAATATTTCAAT

>MSTRG.206.2 gene=MSTRG.206

TAATATATATATGAACAGAGTGCAGTGGTCGTGTTGAATCATTCATACTCGCCTTCAGTAAATATACATC

TCCCTTTGTCTCTCTCTCTCTCTCTCTGGTAAACATAGATGGCAGAATTGAAATTTTTTTTTCATGAGCA

TCAGCTGATACTAAATGAAGCTGAGCCTGTTGTCGGTAAGGATGTAGAATGTGTTGGGTGCAAACAGCCA

ATCAACAAACTCATAGACGCATTTTATAGGTGCAACAAGTCCCTTATTGATAGCAGTCCAGCGAGTGATT

GTGTTGGTTTCTATATGCATAAAACTTGTTCTGAGTTGCCCTCAACTTTTACACACCCTATGAACCCGAA

AAAACCTCTAAGCCTCTTTGTGCTTCCATATAAAAAAGAATACTTTTATATTTGTCATGCTTGTGACAGC

CACTCTCAGCACTTTATGTATGGTTCTGATTCTTCTGTCAGTTCCGAATTTCGTGTTTGTTTGAAATGTG

TAATGTCAGAACTTAAATCTCTAGAGGACCGTAACCTTTGTCATCCAGGTCACAATCACCCATTAACTTT

AGTCCAAAGCCCAGCTTTGTTCCTGTGTCATGCTTGTAACACTACAGCTACAGACTTGTCCTATATCTGT

ACAAGATGTTGTTTTTATATACACAAGAATTGTGCCAATGCACCCACCACCTATCAAAGTAAATTCCACA

ATGAACACGCTCTCATCTTGACCTACTCTCTTCCCCAACAATATCGTCAATATCTTTGTTACTGCAGCAT

CTGCGAGAAACATATAAATCCCATCTACTGGGTTTATATTTGTGCGAATTGCAGATTTTTTGCTCACGTG

AAATGTGCTTCATTAACTGAGATGTTGAGTGAAAGTGATATTGATGATGAAGCAGATTGCGGAGAGTCTA

ATTTGATGCAGTTCCCAGTGCATGATGAAGCATCACTACATGAAACAATGCAAGAATGTATCATCAAGGC

AGCAAATTCTTCCACTGGGGAACCTTCACCTTATATCAATCATTGGGCCCATGGACACCAACTGGCACTC

GGAAACAAGAATGCAAAAACTTTGCCCCTCAACTTGAAGCCGAAAATAGCAGAAACTGAATTGCTAATCT

GTGATGGTTGCATGAATCCTATCTCCTTGGTTGATGTATTCTATGAATGCAGTTTATGCAATTTTTTTCT

CCACAGATCATGCTCCCAATTCCCCGAAAGGATTAAGCATCATCTAGCAGGCGATCTGGAGGGAATTCTA

GCACGAATGTCAGGTGAACTATATACTCTTGAATGCAGAGGTTGCGGCATTCGTGGTAATGGGATTTGCA

TGCACAATGATGAATACGCTTTTGACATCAAGTGCGCTTCATTACCCAGAATAATCAAACATAAAGGTCA

TCGTCACCCTCTTCAGCAATTAAAGACTCCAGATGACTTTCTTTGCAAAGCATGTTGGCGTTACCGTGTA

ACTGAAGAGGTAAGAACCATCGTGTATGGATGTGAAAAATGTGATTTTTTCACACATATAGGGTGTGTGT

TAAGTGCACAAGTGGTGAAACATAGATGGGATCCTCATCCTCTCTACTTGATCCTGTCTCTTAAGAATGT

ACCTGATCATCCCCATGATTTCCATTGTGAATTTTGCCCCGATCAGATAAACACCAACAGTTGGTTTTAT

CACTGCAATGTATGCGATCTATCATTTCATACCAGCTGTATTGATCCAGATGATTGGGTGTCCAATATCA

AGTTTGGTGCTACCGACATTTACAGCGACAAACATCCACATCCACACGGCCTCACATATATTCTCAATAA

AAAGACGAGGAATTGCAATTTATGTGGTAAAGATGCACGCGGTATGCGGGCTCTCCAATGTTCAACATGC

AAATATATAGTGCATGAGAATTGTTTCATTGAAGATTGAAATGAGGCAAAAATCAGATATTGTATTACTG

GATTATTCAGGTTCAGAGGATGTACAGAAAGAAGTAGCCACCGCATTCTTGAAGAATCTCTAGAACTCGT

TGAGCATTAGTAACAGTTCGCTCTATAGCATTAAGAGTTCTAATATTTCAAT

>MSTRG.206.4 gene=MSTRG.206

AGAATTATAATTTTTTTCATAGTCCGAGACAGAGAATTATAAATTTTTAATAGTCCGAAACTTCACACTC

ATACTGGTCTTTATATGTTTCTGGGATGAAATGGTATCACAAAGAATATAAATGTATGAATGATATTGAT

GATGAAGCAGATTGCGGAGAGTCTAATTTGATGCAGTTCCCAGTGCATGATGAAGCATCACTACATGAAA

CAATGCAAGAATGTATCATCAAGGCAGCAAATTCTTCCACTGGGGAACCTTCACCTTATATCAATCATTG

GGCCCATGGACACCAACTGGCACTCGGAAACAAGAATGCAAAAACTTTGCCCCTCAACTTGAAGCCGAAA

ATAGCAGAAACTGAATTGCTAATCTGTGATGGTTGCATGAATCCTATCTCCTTGGTTGATGTATTCTATG

AATGCAGTTTATGCAATTTTTTTCTCCACAGATCATGCTCCCAATTCCCCGAAAGGATTAAGCATCATCT

AGCAGGCGATCTGGAGGGAATTCTAGCACGAATGTCAGGTGAACTATATACTCTTGAATGCAGAGGTTGC

GGCATTCGTGGTAATGGGATTTGCATGCACAATGATGAATACGCTTTTGACATCAAGTGCGCTTCATTAC

CCAGAATAATCAAACATAAAGGTCATCGTCACCCTCTTCAGCAATTAAAGACTCCAGATGACTTTCTTTG

CAAAGCATGTTGGCGTTACCGTGTAACTGAAGAGGTAAGAACCATCGTGTATGGATGTGAAAAATGTGAT

TTTTTCACACATATAGGGTGTGTGTTAAGTGCACAAGTGGTGAAACATAGATGGGATCCTCATCCTCTCT

ACTTGATCCTGTCTCTTAAGAATGTACCTGATCATCCCCATGATTTCCATTGTGAATTTTGCCCCGATCA

GATAAACACCAACAGTTGGTTTTATCACTGCAATGTATGCGATCTATCATTTCATACCAGCTGTATTGAT

CCAGATGATTGGGTGTCCAATATCAAGTTTGGTGCTACCGACATTTACAGCGACAAACATCCACATCCAC

ACGGCCTCACATATATTCTCAATAAAAAGACGAGGAATTGCAATTTATGTGGTAAAGATGCACGCGGTAT

GCGGGCTCTCCAATGTTCAACATGCAAATATATAGTGCATGAGAATTGTTTCATTGAAGATTGAAATGAG

GCAAAAATCAGATATTGTATTACTGGATTATTCAGGTTCAGAGGATGTACAGAAAGAAGTAGCCACCGCA

TTCTTGAAGAATCTCTAGAACTCGTTGAGC

>MSTRG.206.3 gene=MSTRG.206

AAACATAGATGGCAGAATTGAAATTTTTTTTTCATGAGCATCAGCTGATACTAAATGAAGCTGAGCCTGT

TGTCGGTAAGGATGTAGAATGTGTTGGGTGCAAACAGCCAATCAACAAACTCATAGACGCATTTTATAGG

TGCAACAAGTCCCTTATTGATAGCAGTCCAGCGAGTGATTGTGTTGGTTTCTATATGCATAAAACTTGTT

CTGAGTTGCCCTCAACTTTTACACACCCTATGAACCCGAAAAAACCTCTAAGCCTCTTTGTGCTTCCATA

TAAAAAAGAATACTTTTATATTTGTCATGCTTGTGACAGCCACTCTCAGCACTTTATGTATGGTTCTGAT

TCTTCTGTCAGTTCCGAATTTCGTGTTTGTTTGAAATGTGTAATGTCAGAACTTAAATCTCTAGAGGACC

GTAACCTTTGTCATCCAGGTCACAATCACCCATTAACTTTAGTCCAAAGCCCAGCTTTGTTCCTGTGTCA

TGCTTGTAACACTACAGCTACAGACTTGTCCTATATCTGTACAAGATGTTGTTTTTATATACACAAGAAT

TGTGCCAATGCACCCACCACCTATCAAAGTAAATTCCACAATGAACACGCTCTCATCTTGACCTACTCTC

TTCCCCAACAATATCGTCAATATCTTTGTTACTGCAGCATCTGCGAGAAACATATAAATCCCATCTACTG

GGTTTATATTTGTGCGAATTGCAGATTTTTTGCTCACGTGAAATGTGCTTCATTAACTGAGATGTTGAGT

GAAAGTGATATTGATGATGAAGCAGATTGCGGAGAGTCCAGTTTGATGCAGTTCCCAGTGCATGATGAAG

CATCACTACATGAAACAATGCAACAATTGAAAGTGATATTGATGATGAAGCAGATTGCGGAGAGTCTAAT

TTGATGCAGTTCCCAGTGCATGATGAAGCATCACTACATGAAACAATGCAAGAATGTATCATCAAGGCAG

CAAATTCTTCCACTGGGGAACCTTCACCTTATATCAATCATTGGGCCCATGGACACCAACTGGCACTCGG

AAACAAGAATGCAAAAACTTTGCCCCTCAACTTGAAGCCGAAAATAGCAGAAACTGAATTGCTAATCTGT

GATGGTTGCATGAATCCTATCTCCTTGGTTGATGTATTCTATGAATGCAGTTTATGCAATTTTTTTCTCC

ACAGATCATGCTCCCAATTCCCCGAAAGGATTAAGCATCATCTAGCAGGCGATCTGGAGGGAATTCTAGC

ACGAATGTCAGGTGAACTATATACTCTTGAATGCAGAGGTTGCGGCATTCGTGGTAATGGGATTTGCATG

CACAATGATGAATACGCTTTTGACATCAAGTGCGCTTCATTACCCAGAATAATCAAACATAAAGGTCATC

GTCACCCTCTTCAGCAATTAAAGACTCCAGATGACTTTCTTTGCAAAGCATGTTGGCGTTACCGTGTAAC

TGAAGAGGTAAGAACCATCGTGTATGGATGTGAAAAATGTGATTTTTTCACACATATAGGGTGTGTGTTA

AGTGCACAAGTGGTGAAACATAGATGGGATCCTCATCCTCTCTACTTGATCCTGTCTCTTAAGAATGTAC

CTGATCATCCCCATGATTTCCATTGTGAATTTTGCCCCGATCAGATAAACACCAACAGTTGGTTTTATCA

CTGCAATGTATGCGATCTATCATTTCATACCAGCTGTATTGATCCAGATGATTGGGTGTCCAATATCAAG

TTTGGTGCTACCGACATTTACAGCGACAAACATCCACATCCACACGGCCTCACATATATTCTCAATAAAA

AGACGAGGAATTGCAATTTATGTGGTAAAGATGCACGCGGTATGCGGGCTCTCCAATGTTCAACATGCAA

ATATATAGTGCATGAGAATTGTTTCATTGAAGATTGAAATGAGGCAAAAATCAGATATTGTATTACTGGA

TTATTCAGGTTCAGAGGATGTACAGAAAGAAGTAGCCACCGCATTCTTGAAGAATCTCTAGAACTCGTTG

AGC

>MSTRG.206.7 gene=MSTRG.206

CTCTCTCTCTCTCTCTGGTAAACATAGATGGCAGAATTGAAATTTTTTTTTCATGAGCATCAGCTGATAC

TAAATGAAGCTGAGCCTGTTGTCGGTAAGGATGTAGAATGTGTTGGGTGCAAACAGCCAATCAACAAACT

CATAGACGCATTTTATAGGTGCAACAAGTCCCTTATTGATAGCAGTCCAGCGAGTGATTGTGTTGGTTTC

TATATGCATAAAACTTGTTCTGAGTTGCCCTCAACTTTTACACACCCTATGAACCCGAAAAAACCTCTAA

GCCTCTTTGTGCTTCCATATAAAAAAGAATACTTTTATATTTGTCATGCTTGTGACAGCCACTCTCAGCA

CTTTATGTATGGTTCTGATTCTTCTGTCAGTTCCGAATTTCGTGTTTGTTTGAAATGTGTAATGTCAGAA

CTTAAATCTCTAGAGGACCGTAACCTTTGTCATCCAGGTCACAATCACCCATTAACTTTAGTCCAAAGCC

CAGCTTTGTTCCTGTGTCATGCTTGTAACACTACAGCTACAGACTTGTCCTATATCTGTACAAGATGTTG

TTTTTATATACACAAGAATTGTGCCAATGCACCCACCACCTATCAAAGTAAATTCCACAATGAACACGCT

CTCATCTTGACCTACTCTCTTCCCCAACAATATCGTCAATATCTTTGTTACTGCAGCATCTGCGAGAAAC

ATATAAATCCCATCTACTGGGTTTATATTTGTGCGAATTGCAGATTTTTTGCTCACGTGAAATGTGCTTC

ATTAACTGAGATGTTGAGGTTGATGTACTATCTCCTATTATTATTTCTTATATATTTGATATTGAATACT

TATTTAATAAATATTTAGTTTTCTGACATTGTTGTTTTACTTGAAATTATTCTGATTGGATATGAATCTT

TCTTTTGGAATAGTGAAAGTGATATTGATGATGAAGCAGATTGCGGAGAGTCTAATTTGATGCAGTTCCC

AGTGCATGATGAAGCATCACTACATGAAACAATGCAAGAATGTATCATCAAGGCAGCAAATTCTTCCACT

GGGGAACCTTCACCTTATATCAATCATTGGGCCCATGGACACCAACTGGCACTCGGAAACAAGAATGCAA

AAACTTTGCCCCTCAACTTGAAGCCGAAAATAGCAGAAACTGAATTGCTAATCTGTGATGGTTGCATGAA

TCCTATCTCCTTGGTTGATGTATTCTATGAATGCAGTTTATGCAATTTTTTTCTCCACAGATCATGCTCC

CAATTCCCCGAAAGGATTAAGCATCATCTAGCAGGCGATCTGGAGGGAATTCTAGCACGAATGTCAGGTG

AACTATATACTCTTGAATGCAGAGGTTGCGGCATTCGTGGTAATGGGATTTGCATGCACAATGATGAATA

CGCTTTTGACATCAAGTGCGCTTCATTACCCAGAATAATCAAACATAAAGGTCATCGTCACCCTCTTCAG

CAATTAAAGACTCCAGATGACTTTCTTTGCAAAGCATGTTGGCGTTACCGTGTAACTGAAGAGGTAAGAA

CCATCGTGTATGGATGTGAAAAATGTGATTTTTTCACACATATAGGGTGTGTGTTAAGTGCACAAGTGGT

GAAACATAGATGGGATCCTCATCCTCTCTACTTGATCCTGTCTCTTAAGAATGTACCTGATCATCCCCAT

GATTTCCATTGTGAATTTTGCCCCGATCAGATAAACACCAACAGTTGGTTTTATCACTGCAATGTATGCG

ATCTATCATTTCATACCAGCTGTATTGATCCAGATGATTGGGTGTCCAATATCAAGTTTGGTGCTACCGA

CATTTACAGCGACAAACATCCACATCCACACGGCCTCACATATATTCTCAATAAAAAGACGAGGAATTGC

AATTTATGTGGTAAAGATGCACGCGGTATGCGGGCTCTCCAATGTTCAACATGCAAATATATAGTGCATG

AGAATTGTTTCATTGAAGATTGAAATGAGGCAAAAATCAGATATTGTATTACTGGATTATTCAGGTTCAG

AGGATGTACAGAAAGAAGTAGCCACCG

>MSTRG.206.5 gene=MSTRG.206

TAATATATATATGAACAGAGTGCAGTGGTCGTGTTGAATCATTCATACTCGCCTTCAGTAAATATACATC

TCCCTTTGTCTCTCTCTCTCTCTCTCTGGTAAACATAGATGGCAGAATTGAAATTTTTTTTTCATGAGCA

TCAGCTGATACTAAATGAAGCTGAGCCTGTTGTCGGTAAGGATGTAGAATGTGTTGGGTGCAAACAGCCA

ATCAACAAACTCATAGACGCATTTTATAGGTGCAACAAGTCCCTTATTGATAGCAGTCCAGCGAGTGATT

GTGTTGGTTTCTATATGCATAAAACTTGTTCTGAGTTGCCCTCAACTTTTACACACCCTATGAACCCGAA

AAAACCTCTAAGCCTCTTTGTGCTTCCATATAAAAAAGAATACTTTTATATTTGTCATGCTTGTGACAGC

CACTCTCAGCACTTTATGTATGGTTCTGATTCTTCTGTCAGTTCCGAATTTCGTGTTTGTTTGAAATGTG

TAATGTCAGAACTTAAATCTCTAGAGGACCGTAACCTTTGTCATCCAGGTCACAATCACCCATTAACTTT

AGTCCAAAGCCCAGCTTTGTTCCTGTGTCATGCTTGTAACACTACAGCTACAGACTTGTCCTATATCTGT

ACAAGATGTTGTTTTTATATACACAAGAATTGTGCCAATGCACCCACCACCTATCAAAGTAAATTCCACA

ATGAACACGCTCTCATCTTGACCTACTCTCTTCCCCAACAATATCGTGAATATCCCTGTTACTGCAGCAT

CTGCGAGAAACATATAAATCCCATCTACTGGGTTTATATTTGTGCGAATTGCAGATTTTTTGCTCACGTG

AAATGTGCTTCATTAACTGAGTTGTTGAGGTTGATGTACTATCTCCTATTATTATTTCTTATATATTTGA

TATTGAATACTTATTTAATAAATATTTAGTTTTCTGACATTGTTGTTTTACTTGAAATTATTCTGATTGG

ATATGAATCTTTCTTTTGGAATAGTGAAAGTGATATTGATGATGAAGCAGATTGCGGAGAGTCTAATTTG

ATGCAGTTCCCAGTGCATGATGAAGCATCACTACATGAAACAATGCAAGAATGTATCATCAAGGCAGCAA

ATTCTTCCACTGGGGAACCTTCACCTTATATCAATCATTGGGCCCATGGACACCAACTGGCACTCGGAAA

CAAGAATGCAAAAACTTTGCCCCTCAACTTGAAGCCGAAAATAGCAGAAACTGAATTGCTAATCTGTGAT

GGTTGCATGAATCCTATCTCCTTGGTTGATGTATTCTATGAATGCAGTTTATGCAATTTTTTTCTCCACA

GATCATGCTCCCAATTCCCCGAAAGGATTAAGCATCATCTAGCAGGCGATCTGGAGGGAATTCTAGCACG

AATGTCAGGTGAACTATATACTCTTGAATGCAGAGGTTGCGGCATTCGTGGTAATGGGATTTGCATGCAC

AATGATGAATACGCTTTTGACATCAAGTGCGCTTCATTACCCAGAATAATCAAACATAAAGGTCATCGTC

ACCCTCTTCAGCAATTAAAGACTCCAGATGACTTTCTTTGCAAAGCATGTTGGCGTTACCGTGTAACTGA

AGAGGTAAGAACCATCGTGTATGGATGTGAAAAATGTGATTTTTTCACACATATAGGGTGTGTGTTAAGT

GCACAAGTGGTGAAACATAGATGGGATCCTCATCCTCTCTACTTGATCCTGTCTCTTAAGAATGTACCTG

ATCATCCCCATGATTTCCATTGTGAATTTTGCCCCGATCAGATAAACACCAACAGTTGGTTTTATCACTG

CAATGTATGCGATCTATCATTTCATACCAGCTGTATTGATCCAGATGATTGGGTGTCCAATATCAAGTTT

GGTGCTACCGACATTTACAGCGACAAACATCCACATCCACACGGCCTCACATATATTCTCAATAAAAAGA

CGAGGAATTGCAATTTATGTGGTAAAGATGCACGCGGTATGCGGGCTCTCCAATGTTCAACATGCAAATA

TATAGTGCATGAGAATTGTTTCATTGAAGATTGAAATGAGGCAAAAATCAGATATTGTATTACTGGATTA

TTCAGGTTCAGAGGATGTACAGAAAGAAGTAGCCACCG

>MSTRG.206.6 gene=MSTRG.206

TAATATATATATGAACAGAGTGCAGTGGTCGTGTTGAATCATTCATACTCGCCTTCAGTAAATATACATC

TCCCTTTGTCTCTCTCTCTCTCTCTCTGGTAAACATAGATGGCAGAATTGAAATTTTTTTTTCATGAGCA

TCAGCTGATACTAAATGAAGCTGAGCCTGTTGTCGGTAAGGATGTAGAATGTGTTGGGTGCAAACAGCCA

ATCAACAAACTCATAGACGCATTTTATAGGTGCAACAAGTCCCTTATTGATAGCAGTCCAGCGAGTGATT

GTGTTGGTTTCTATATGCATAAAACTTGTTCTGAGTTGCCCTCAACTTTTACACACCCTATGAACCCGAA

AAAACCTCTAAGCCTCTTTGTGCTTCCATATAAAAAAGAATACTTTTATATTTGTCATGCTTGTGACAGC

CACTCTCAGCACTTTATGTATGGTTCTGATTCTTCTGTCAGTTCCGAATTTCGTGTTTGTTTGAAATGTG

TAATGTCAGAACTTAAATCTCTAGAGGACCGTAACCTTTGTCATCCAGGTCACAATCACCCATTAACTTT

AGTCCAAAGCCCAGCTTTGTTCCTGTGTCATGCTTGTAACACTACAGCTACAGACTTGTCCTATATCTGT

ACAAGATGTTGTTTTTATATACACAAGAATTGTGCCAATGCACCCACCACCTATCAAAGTAAATTCCACA

ATGAACACGCTCTCATCTTGACCTACTCTCTTCCCCAACAATATCGTCAATATCTTTGTTACTGCAGCAT

CTGCGAGAAACATATAAATCCCATCTACTGGGTTTATATTTGTGCGAATTGCAGATTTTTTGCTCACGTG

AAATGTGCTTCATTAACTGAGATGTTGAGTGAAAGTGATATTGATGATGAAGCAGATTGCGGAGAGTCCA

GTTTGATGCAGTTCCCAGTGCATGATGAAGCATCACTACATGAAACAATGCAACAATGTATCATCAAGGC

CGTAAATCCTTCCACTGGGGAACCTTCACCTTATATCAATCATTGGGCCCATGGACACCAACTGGCACTC

GGAAACAAGAATGCAAAAACTTTGCCCCTCAACTTGAAGCCGAAAATAGCAGAAACTGAATTGCTAATCT

GTGATGGTTGCATGAATCCTATCTCCTTGGTTGATGTATTCTATGAATGCAGTTTATGCAATTTTTTTCT

CCACAGATCATGCTCCCAATTCCCCAAAAGGATTAAGCATCATCTAGCAGGCGATCTGGAGGCAGTGCTA

GCACGAGAGTCAGGTGAACTATATACTTTTCAATGCAGTGGTTGCGGCATTTATGGTAATGGGATTTGCA

TGCTCAATGATGAATACACTTTTGACATCAAGTGCGCTTCATTACCCAGAATAATCAAACATAAAGGTCA

TCGTCACCCTCTTCAGCAATTAAAGACTCCAGATGACTTTCTTTGCAAAGGTTGTTGGCGTGAGCCTGTA

ACTGAAGAGGTAAGAACCATCGTGTACGGATGTGAAAAATGTGAATTTTACACACATATAGGGTGTGTGT

TAAGTGCACAAGTGGTGAAACATAGGTGGGATCCTCATCCTCTCTACTTGATCCTGTCTCTCAAGAATGT

ACCCGATCATCCCCACGAATTCCCTTGTGAATTTTGCTCCAAAGAGATAAACACCAACAGTTGGTTTTAT

CACTGCAATGTATGCGATCTATCATTTCATATTAGCTGTATTGATCCAGATGATTGGCTGTCCAATATCA

AGTTTGGTGCCACCAACATTTACAGCGACAAACATCCTCATCCACACGGCCTCACATATATTCTCAATAA

AAAGACAAGGAATTGCAATTTATGTGGGTTTGCTTCAGTTGGGATTGCTTTTAGTGGGGATGAATAATTG

TGATTTGGGATTCTTTGACTTGGATAATATGATCTAGGGTTCTTTGGCTTGGAGTCTTGGACAATCTGAG

TTTGAGTGAAAGTGATATTGATGATGAAGCAGATTGCGGAGAGTCTAATTTGATGCAGTTCCCAGTGCAT

GATGAAGCATCACTACATGAAACAATGCAAGAATGTATCATCAAGGCAGCAAATTCTTCCACTGGGGAAC

CTTCACCTTATATCAATCATTGGGCCCATGGACACCAACTGGCACTCGGAAACAAGAATGCAAAAACTTT

GCCCCTCAACTTGAAGCCGAAAATAGCAGAAACTGAATTGCTAATCTGTGATGGTTGCATGAATCCTATC

TCCTTGGTTGATGTATTCTATGAATGCAGTTTATGCAATTTTTTTCTCCACAGATCATGCTCCCAATTCC

CCGAAAGGATTAAGCATCATCTAGCAGGCGATCTGGAGGGAATTCTAGCACGAATGTCAGGTGAACTATA

TACTCTTGAATGCAGAGGTTGCGGCATTCGTGGTAATGGGATTTGCATGCACAATGATGAATACGCTTTT

GACATCAAGTGCGCTTCATTACCCAGAATAATCAAACATAAAGGTCATCGTCACCCTCTTCAGCAATTAA

AGACTCCAGATGACTTTCTTTGCAAAGCATGTTGGCGTTACCGTGTAACTGAAGAGGTAAGAACCATCGT

GTATGGATGTGAAAAATGTGATTTTTTCACACATATAGGGTGTGTGTTAAGTGCACAAGTGGTGAAACAT

AGATGGGATCCTCATCCTCTCTACTTGATCCTGTCTCTTAAGAATGTACCTGATCATCCCCATGATTTCC

ATTGTGAATTTTGCCCCGATCAGATAAACACCAACAGTTGGTTTTATCACTGCAATGTATGCGATCTATC

ATTTCATACCAGCTGTATTGATCCAGATGATTGGGTGTCCAATATCAAGTTTGGTGCTACCGACATTTAC

AGCGACAAACATCCACATCCACACGGCCTCACATATATTCTCAATAAAAAGACGAGGAATTGCAATTTAT

GTGGTAAAGATGCACGCGGTATGCGGGCTCTCCAATGTTCAACATGCAAATATATAGTGCATGAGAATTG

TTTCATTGAAGATTGAAATGAGGCAAAAATCAGATATTGTATTACTGGATTATTCAGGTTCAGAGGATGT

ACAGAAAGAAGTAGCCACCG

>MSTRG.206.8 gene=MSTRG.206

GATTTGTTCAATGCAAACTCAAAACTGGTTTAAAGTCAAAAGCACGAGATTTTCATATCAAGGAGGTGGG

CATTAAGTCTGGGTTTTTCACTATGCGGCCTTCTACTTGAAAGTGATATTGATGATGAAGCAGATTGCGG

AGAGTCCAGTTTGATGCAGTTCCCAGTGCATGATGAAGCATCACTACATGAAACAATGCAACAATGTATC

ATCAAGGCCGTAAATCCTTCCACTGGGGAACCTTCACCTTATATCAATCATTGGGCCCATGGACACCAAC

TGGCACTCGGAAACAAGAATGCAAAAACTTTGCCCCTCAACTTGAAGCCGAAAATAGCAGAAACTGAATT

GCTAATCTGTGATGGTTGCATGAATCCTATCTCCTTGGTTGATGTATTCTATGAATGCAGTTTATGCAAT

TTTTTTCTCCACAGATCATGCTCCCAATTCCCCAAAAGGATTAAGCATCATCTAGCAGGCGATCTGGAGG

CAGTGCTAGCACGAGAGTCAGGTGAACTATATACTTTTCAATGCAGTGGTTGCGGCATTTATGGTAATGG

GATTTGCATGCTCAATGATGAATACACTTTTGACATCAAGTGCGCTTCATTACCCAGAATAATCAAACAT

AAAGGTCATCGTCACCCTCTTCAGCAATTAAAGACTCCAGATGACTTTCTTTGCAAAGGTTGTTGGCGTG

AGCCTGTAACTGAAGAGGTAAGAACCATCGTGTACGGATGTGAAAAATGTGAATTTTACACACATATAGG

GTGTGTGTTAAGTGCACAAGTGGTGAAACATAGGTGGGATCCTCATCCTCTCTACTTGATCCTGTCTCTC

AAGAATGTACCCGATCATCCCCACGAATTCCCTTGTGAATTTTGCTCCAAAGAGATAAACACCAACAGTT

GGTTTTATCACTGCAATGTATGCGATCTATCATTTCATATTAGCTGTATTGATCCAGATGATTGGCTGTC

CAATATCAAGTTTGGTGCCACCAACATTTACAGCGACAAACATCCTCATCCACACGGCCTCACATATATT

CTCAATAAAAAGACAAGGAATTGCAATTTATGTGGGGAAGATGCACGCGATACGCCGGCTCTCCAATGTT

CACCATGCAAATATATAGTGCATCGGAATTGTTTCATTAAAGAATGAAATCAGGGAAAATTATATTGTGT

ATTGCTGGATTGTATAGGTTCAGAAAATTTACAAATCCTTTGTGCTTGTATATAT

>MSTRG.206.9 gene=MSTRG.206

TCTCTCTTAAAAATAGATGGCAGAACTGAAACATTTTTTTCATGAGCATAAGCTGATACTGAATGAAGCT

GAGCCTGTTATCGATAAGGAAGTAGAATGTGTTGGGTGCAGACGGCCAATAAACAAACTCATAGACGCGT

TTTATAAGTGCAACAATTCCCTTATTGATAGCAGTCCATCGAGTGATTGTGTTGGTTTCTATATGCATAA

AACTTGTTCTGAGTTGCCCTCAACTTTTACACGCCCTTTGTTCCCAAAACAACCTCTAAGCCTCTTTGCG

CTTCCATATAAAAAACAAAACTTTTTTTCTTGTGATGCTTGTGACAGCGAATCTCAGTGCTTTATGTATT

GTTCTGAGTCTTCTGCCAATTGGGCCAATTCCGAATTTCTTGTTTGTTTGAAATGTGTAATATCAGAACT

TAAATCTCTAGAGGAACGAAACCGTTATCATCCGGGTCATGATCACCCATTAACCTTAGTCCAAAGCCCA

GCTTTGTTCCTGTGTCATGCTTGTAACACTACAGCAACAGATTTGTCCTATATTTGTACAACTTGTCCTT

TCTGGATACACGAGAGTTGTGCCAATGCACCCATCACCTACCAAAGTAAATTTCACAATGAACACGCTCT

CATCTTGAACTACTCTCTTCCCCAACAATTTCGTCAATTTACCTGTTTCTGCAGCATCTGCGATGAAAAT

ATAAATCCCATCGACTGGGTTTATATTTGTGCAAATTGCAGATTTTTTGCTCACGTGAAATGTGCTTTAT

CAACTTTGATTGAAAGTGATATTGATGATGAAGCAGATTGCGGAGAGTCCAGTTTGATGCAGTTCCCAGT

GCATGATGAAGCATCACTACATGAAACAATGCAACAATGTATCATCAAGGCCGTAAATCCTTCCACTGGG

GAACCTTCACCTTATATCAATCATTGGGCCCATGGACACCAACTGGCACTCGGAAACAAGAATGCAAAAA

CTTTGCCCCTCAACTTGAAGCCGAAAATAGCAGAAACTGAATTGCTAATCTGTGATGGTTGCATGAATCC

TATCTCCTTGGTTGATGTATTCTATGAATGCAGTTTATGCAATTTTTTTCTCCACAGATCATGCTCCCAA

TTCCCCAAAAGGATTAAGCATCATCTAGCAGGCGATCTGGAGGCAGTGCTAGCACGAGAGTCAGGTGAAC

TATATACTTTTCAATGCAGTGGTTGCGGCATTTATGGTAATGGGATTTGCATGCTCAATGATGAATACAC

TTTTGACATCAAGTGCGCTTCATTACCCAGAATAATCAAACATAAAGGTCATCGTCACCCTCTTCAGCAA

TTAAAGACTCCAGATGACTTTCTTTGCAAAGGTTGTTGGCGTGAGCCTGTAACTGAAGAGGTAAGAACCA

TCGTGTACGGATGTGAAAAATGTGAATTTTACACACATATAGGGTGTGTGTTAAGTGCACAAGTGGTGAA

ACATAGGTGGGATCCTCATCCTCTCTACTTGATCCTGTCTCTCAAGAATGTACCCGATCATCCCCACGAA

TTCCCTTGTGAATTTTGCTCCAAAGAGATAAACACCAACAGTTGGTTTTATCACTGCAATGTATGCGATC

TATCATTTCATATTAGCTGTATTGATCCAGATGATTGGCTGTCCAATATCAAGTTTGGTGCCACCAACAT

TTACAGCGACAAACATCCTCATCCACACGGCCTCACATATATTCTCAATAAAAAGACAAGGAATTGCAAT

TTATGTGGGGAAGATGCACGCGATACGCCGGCTCTCCAATGTTCACCATGCAAATATATAGTGCATCGGA

ATTGTTTCATTAAAGAATGAAATCAGGGAAAATTATATTGTGTATTGCTGGATTGTATAGGTTCAGAAAA

TTTACAAATCCTTTGTGCTTGTATATAT

>MSTRG.219.1 gene=MSTRG.219

GAGTGTAAAGTGCTGAATCGTTTCGTGATGCTATCTGTTTCGAAGTCTAAAAACAGTAAACAATGAAGAA

GATGCTCCAGAAAAACAAAACAATGATCTGCCTGATTGGGTTGGCAAGGGAAAACAGTAGTTCTACCAAA

CCCGCCTCAAAGTGGACAACGAGAAATGCCAACATTCTCATTCCGGAATTCCCACTTTGCTTGATATTTG

TCAGACAACAAAAATTATTTGTGGTCATGCAGCTGATATGAGGCAAAAGAACCCAAAGGCTGGCAACCCA

TTTAATGAAATTGCACCTATTAAATGGGAGATAATTTGTGACCACTGCGAGGGATTCCTTGTGACTCGGC

CAACTGTGTGCGAGAGAACATCAGAAGAGCACACAATTGCTGAAGGCGAAAGCATTGGAAGCAGCTATGA

ATGACATTAATAACTCATTTGGAAATGGAGCTGTAACAATATTAGGCAATGCTGGTGGATCTCTTGTTGA

CTCTTTTCCAAGTGGTTGTCTGACATTGGACATTGCATTGGGTGTCGGCCTTCCTAGAGGAAATATTGTT

GAGTAAACAATGAAGATGATGCTCCAAAAAGATGGAAAACAGTAACTAAGGCTGGCTAATCTGCTACTTA

TCTGCTATTAAGAAGCTGCTGCTTGTCTACTTGGTTATTTCGTTGAAGATAAAGCAACGCGACATATCGG

GAACTTTTGAAATTGCAAGAACTAAGTTAAAAGCTGGCAATGTAGATGATTTTTTTCCAAGTAGTTGAAC

CTACCAATGTATTCCTTACCATATTGTCTTATTCTTATTTCTATGTAACATCAATGCTCAAAAGACTTAA

AATCTAAGTTACTCAAGTAGTTTTTATATTTGTCTGTTTATTTATTTACCTATATAATGTATATAATACG

TTTGTGTGATGACAGAGGATCGATTCAGCACTTTGTAAAAAAAATTCATAGTTTAC

>MSTRG.219.2 gene=MSTRG.219

AGGCCTTTTAGTTAACCCAATTTATTTTTAGGTATAAAATCACTCGTCGTGTAGGTTAAGGCGGCAGCTT

CTCAATTACATTAGGGCATCGCCTAGATTAGGGCATCGCCGCAACGATCTATGATAATCGTGTGTGTATG

TATCTAGAGTGTAAAGTGCTGAATCGTTTCGTGATGCTATCTGTTTCGAAGTCTAAAAACAGTAAACAAT

GAAGAAGATGCTCCAGAAAAACAAAACAATGATCTGCCTGATTGGGTTGGCAAGGGAAAACAGTAGTTCT

ACCAAACCCGCCTCAAAGTGGACAACGAGAAATGCCAACATTCTCATTCCGGAATTCCCACTTTGCTTGA

TATTTGTCAGACAACAAAAATTATTTGTTACATTGGTCATGCAGCTGATATGAGGCAAAAGAACCCAAAG

GCTGGCAACCCATTTAATGAAATTGCACCTATTAAATGGGAGATAATTTGTGACCACTGCGAGGGATTCC

TTGTGACTCGGCCAACTGTGTGCGAGAGAACATCAGAAGAGCACACAATTGCTGAAGTGACTCTTTTCCA

AGTGGTTGTCTGACATTGGACATTGCATTGGGTGTCGGCCTTCCTAGAGGAAATATTGTTGAGGTATGTT

TTCGTACACACTTTGAGGTATTGTAAAACAAACTCAGTACAATCTTTTTCACGTGAAGATTATCTTGTTT

AATATCCGGGATACGTATGGCTCAGAGATGGTGAAGTAAACAATGAAGATGATGCTCCAAAAAGATGGAA

AACAGTAACTAAGGCTGGCTAATCTGCTACTTATCTGCTATTAAGAAGCTGCTGCTTGTCTACTTGGTTA

TTTCGTTGAAGATAAAGCAACGCGACATATCGGGAACTTTTGAAATTGCAAGAACTAAGTTAAAAGCTGG

CAATGTAGATGATTTTTTTCCAAGTAGTTGAACCTACCAATGTATTCCTTACCATATTGTCTTATTCTTA

TTTCTATGTAACATCAATGCTCAAAAGACTTAAAATCTAAGTTACTCAAGTAGTTTTTATATTTGTCTGT

TTATTTATTTACCTATATAATGTATATAATACGTTTGTGTGATGACAGAGGATCGATTCAG

>MSTRG.219.3 gene=MSTRG.219

AGGCCTTTTAGTTAACCCAATTTATTTTTAGGTATAAAATCACTCGTCGTGTAGGTTAAGGCGGCAGCTT

CTCAATTACATTAGGGCATCGCCTAGATTAGGGCATCGCCGCAACGATCTATGATAATCGTGTGTGTATG

TATCTAGAGTGTAAAGTGCTGAATCGTTTCGTGATGCTATCTGTTTCGAAGTCTAAAAACAGTAAACAAT

GAAGAAGATGCTCCAGAAAAACAAAACAATGATCTGCCTGATTGGGTTGGCAAGGGAAAACAGTAGTTCT

ACCAAACCCGCCTCAAAGTGGACAACGAGAAATGCCAACATTCTCATTCCGGAATTCCCACTTTGCTTGA

TATTTGTCAGACAACAAAAATTATTTGTGGTCATGCAGCTGATATGAGGCAAAAGAACCCAAAGGCTGGC

AACCCATTTAATGAAATTGCACCTATTAAATGGGAGATAATTTGTGACCACTGCGAGGGATTCCTTGTGA

CTCGGCCAACTGTGTGCGAGAGAACATCAGAAGAGCACACAATTGCTGAAGTGACTCTTTTCCAAGTGGT

TGTCTGACATTGGACATTGCATTGGGTGTCGGCCTTCCTAGAGGAAATATTGTTGAGGTATGTTTTCGTA

CACACTTTGAGGTATTGTAAAACAAACTCAGTACAATCTTTTTCACGTGAAGATTATCTTGTTTAATATC

CGGGATACGTATGGCTCAGAGATGGTGAAGTAAACAATGAAGATGATGCTCCAAAAAGATGGAAAACAGT

AACTAAGGCTGGCTAATCTGCTACTTATCTGCTATTAAGAAGCTGCTGCTTGTCTACTTGGTTATTTCGT

TGAAGATAAAGCAACGCGACATATCGGGAACTTTTGAAATTGCAAGAACTAAGTTAAAAGCTGGCAATGT

AGATGATTTTTTTCCAAGTAGTTGAACCTACCAATGTATTCCTTACCATATTGTCTTATTCTTATTTCTA

TGTAACATCAATGCTCAAAAGACTTAAAATCTAAGTTACTCAAGTAGTTTTTATATTTGTCTGTTTATTT

ATTTACCTATATAATGTATATAATACGTTTGTGTGATGACAGAG

>MSTRG.219.4 gene=MSTRG.219

TCTAAGAGCACATATATTAATATTTTAGTACTCTCATATCTGTCAATTTTTTAATTAATGCATAGAATAT

GTAATTCGTGAAATATTGAAATTCATTATGTCAGGTTGGCAAGGGAAAACAGTAGTTCTACCAAACCCGC

CTCAAAGTGGACAACGAGAAATGCCAACATTCTCATTCCGGAATTCCCACTTTGCTTGATATTTGTCAGA

CAACAAAAATTATTTGTGGTCATGCAGCTGATATGAGGCAAAAGAACCCAAAGGCTGGCAACCCATTTAA

TGAAATTGCACCTATTAAATGGGAGATAATTTGTGACCACTGCGAGGGATTCCTTGTGACTCGGCCAACT

GTGTGCGAGAGAACATCAGAAGAGCACACAATTGCTGAAGGCGAAAGCATTGGAAGCAGCTATGAATGAC

ATTAATAACTCATTTGGAAATGGAGCTGTAACAATATTAGGCAATGCTGGTGGATCTCTTGTGTATGTTG

ACTCTTTTCCAAGTGGTTGTCTGACATTGGACATTGCATTGGGTGTCGGCCTTCCTAGAGGAAATATTGT

TGAGGTATGTTTTCGTACACACTTTGAGGTATTGTAAAACAAACTCAGTACAATCTTTTTCACGTGAAGA

TTATCTTGTTTAATATCCGGGATACGTATGGCTCAGAGATGGTGAAGTAAACAATGAAGATGATGCTCCA

AAAAGATGGAAAACAGTAACTAAGGCTGGCTAATCTGCTACTTATCTGCTATTAAGAAGCTGCTGCTTGT

CTACTTGGTTATTTCGTTGAAGATAAAGCAACGCGACATATCGGGAACTTTTGAAATTGCAAGAACTAAG

TTAAAAGCTGGCAATGTAGATGATTTTTTTCCAAGTAGTTGAACCTACCAATGTATTCCTTACCATATTG

TCTTATTCTTATTTCTATGTAACATC

>MSTRG.219.5 gene=MSTRG.219

TGGGAAATTGGATTTCATTTGAAATTTTGGATTTGATCAAATAATCTGTTTGGGAATGTGGATTTGGATT

TCATTTGAAATCCAGACATTCAAATATTTGTATAAATTTGAGTGTTTGAAATGACAACTTAAATCTTGTC

ATTTGAAATGAAATGCATGTCTCCAAACGGCCTCTTATTTAAATTAAAACATCGAGGGAGGATCCATTTG

AATCAGCGCTATCGTGCAATTGGGATCAATTATATATGTTGGCAAGGGAAAACAGTAGTTCTACCAAACC

CGCCTCAAAGTGGACAACGAGAAATGCCAACATTCTCATTCCGGAATTCCCACTTTGCTTGATATTTTGG

TCATGCAGCTGATATGAGGCAAAAGAACCCAAAGGCTGGCAACCCATTTAATGAAATTGCACCTATTAAA

TGGGAGATAATTTGTGACCACTGCGAGGGATTCCTTGTGACTCGGCCAACTGTGTGCGAGAGAACATCAG

AAGAGCACACAATTGCTGAAGGCGAAAGCATTGGAAGCAGCTATGAATGACATTAATAACTCATTTGGAA

ATGGAGCTGTAACAATATTAGGCAATGCTGGTGGATCTCTTGTGTATGTTGACTCTTTTCCAAGTGGTTG

TCTGACATTGGACATTGCATTGGGTGTCGGCCTTCCTAGAGGAAATATTGTTGAGGTATGTTTTCGTACA

CACTTTGAGGTATTGTAAAACAAACTCAGTACAATCTTTTTCACGTGAAGATTATCTTGTTTAATATCCG

GGATACGTATGGCTCAGAGATGGTGAAGTAAACAATGAAGATGATGCTCCAAAAAGATGGAAAACAGTAA

CTAAGGCTGGCTAATCTGCTACTTATCTGCTATTAAGAAGCTGCTGCTTGTCTACTTGGTTATTTCGTTG

AAGATAAAGCAACGCGACATATCGGGAACTTTTGAAATTGCAAGAACTAAGTTAAAAGCTGGCAATGTAG

ATGATTTTTTTCCAAGTAGTTGAACCTACCAATGTATTCCTTACCATATTGTCTTATTCTTATTTCTATG

TAACATC

>MSTRG.219.6 gene=MSTRG.219

TTTAGGTATAAAATCACTCGTCGTGTAGGTTAAGGCGGCAGCTTCTCAATTACATTAGGGCATCGCCTAG

ATTAGGGCATCGCCGCAACGATCTATGATAATCGTGTGTGTATGTATCTAGAGTGTAAAGTGCTGAATCG

TTTCGTGATGCTATCTGTTTCGAAGTCTAAAAACAGTAAACAATGAAGAAGATGCTCCAGAAAAACAAAA

CAATGATCTGCCTGATTGGGTTGGCAAGGGAAAACAGTAGTTCTACCAAACCCGCCTCAAAGTGGACAAC

GAGAAATGCCAACATTCTCATTCCGGAATTCCCACTTTGCTTGATATTTTGGTCATGCAGCTGATATGAG

GCAAAAGAACCCAAAGGCTGGCAACCCATTTAATGAAATTGCACCTATTAAATGGGAGATAATTTGTGAC

CACTGCGAGGGATTCCTTGTGACTCGGCCAACTGTGTGCGAGAGAACATCAGAAGAGCACACAATTGCTG

AAGTGACTCTTTTCCAAGTGGTTGTCTGACATTGGACATTGCATTGGGTGTCGGCCTTCCTAGAGGAAAT

ATTGTTGAGGTATGTTTTCGTACACACTTTGAGGTATTGTAAAACAAACTCAGTACAATCTTTTTCACGT

GAAGATTATCTTGTTTAATATCCGGGATACGTATGGCTCAGAGATGGTGAAGTAAACAATGAAGATGATG

CTCCAAAAAGATGGAAAACAGTAACTAAGGCTGGCTAATCTGCTACTTATCTGCTATTAAGAAGCTGCTG

CTTGTCTACTTGGTTATTTCGTTGAAGATAAAGCAACGCGACATATCGGGAACTTTTGAAATTGCAAGAA

CTAAGTTAAAAGCTGGCAATGTAGATGATTTTTTTCCAAGTAGTTGAACCTACCAATGTATTCCTTACCA

TATTGTCTTATTCTTATTTCTAT

>MSTRG.219.7 gene=MSTRG.219

AAGTGGACAACGAGAAATGCCAACATTCTCATTGCGGAATTCCCACTTTGCTTGATATTTGTCAGACAAC

AAAAATTATTTGTTACATGTACAAACTTCATACAGTATATGCAAAATCGTGACAGTCAATCTTTATCTCT

CTAACAAGCTTTATATGTCCAATGTACTCTTTGTGTCCAGTGGTCATGCAGCTGATATGAGGCAAAAGAA

CCCAAAGGCTGGCAACCCATTTAATGAAATGCGAAAGCATTGGAAGCAGCTATGAATGACATTAATAACT

CATTTGGAAATGGAGCTGTAACAATATTAGGCAATGCTGGTGGATCTCTTGTTGACTCTTTTCCAAGTGG

TTGTCTGACATTGGACATTGCATTGGGTGTCGGCCTTCCTAGAGGAAATATTGTTGAGGTATGTTTTCGT

ACACACTTTGAGGTATTGTAAAACAAACTCAGTACAATCTTTTTCACGTGAAGATTATCTTGTTTAATAT

CCGGGATACGTATGGCTCAGAGATGGTGAAGTAAACAATGAAGATGATGCTCCAAAAAGATGGAAAACAG

TAACTAAGGCTGGCTAATCTGCTACTTATCTGCTATTAAGAAGCTGCTGCTTGTCTACTTGGTTATTTCG

TTGAAGATAAAGCAACGCGACATATCGGGAACTTTTGAAATTGCAAGAACTAAGTTAAAAGCTGGCAATG

TAGATGATTTTTTTCCAAGTAGTTGAACCTACCAATGTATTCCTTACCATATTGTCTTATTCTTATTTCT

AT

>MSTRG.220.1 gene=MSTRG.220

GCAAGGTCTGCAATTTTTTCTGTCTTCGAGTCTTTGGAATGTTGCTGTTATGGAATGCCCGAGGAAAATA

TAACTTCGGAATCAAGAAGTTTCGAAAGGTAGCTTGGCAAATCAGCCCAACCAAAGCTGCTCAGAAGTAC

AACTTAGATACAAGTTGTTTTTTCAAATATGTGTGAGTCCTTGTATTGATATCTATCAGGCTAACGAGTG

ATGCCACTATGGATATGATCTTCTAGTGAAATTTGTATATACATTCATTTATGCACGTATGTATATTAAT

GTGTTAACTGTACCAAATATTTATCATTTAATATTTTTATTCACGGCAATGTTGGTTTTAA

>MSTRG.220.2 gene=MSTRG.220

GCAAGGTCTGCAATTTTTTCTGTCTTCGAGTCTTTGGAATGTTGCTGTTATGGAATGCCCGAGGAAAATA

TAACTTCGGAATCAAGAAGTTTCGAAAGGTAGCTTGGCAAATCAGCCCAACCAAAGCTGCTCAGAAGTAC

AACTTAGATACAAGTTGTTTTTTCAAATATGTGTGAGTCCTTGTATTGATATCTATCAGGCTAACGAGTG

ATGCCACTATGGATATGATCTTCTAGCACGTATGTATATTAATGTGTTAACTGTACCAAATATTTATCAT

TTAATATTTTTATTCACGGCAATGTTGGTTTTAA

>MSTRG.221.1 gene=MSTRG.221

CTGCCATAAAACAAACTCAATACAATCTTATTCACGTAAAGATTGTTTTGTTTAATAATATCCGGGAGAC

GGATGGCTCATAAATGGTGAAGTAAACAAGGAAGATGATTTGCAGTAAACAATGAAGATGATGCTCCAAA

ATGATGAAAAACAATTATCTAAGGCCCGCTAATTTGCTAGTAACCATATTGTCTTATTCTTATTCCTATG

TAAGATCAGGGCTCAAAAG

>MSTRG.222.1 gene=MSTRG.222

TTTTTCACGTGAAGATTATCTCGTTTAATATCTGGGATACGTATGGCTCAGAGATGGTGAAGGTATGCTT

GTTATATAGTTATAATGTACTTGAAAATCTATTAAGTTTTCTAGTATTCAATTCACTGGTTTTCAAATTT

CTTCTTGCAGTAAACAATGAAGATGATGCTCCAAAAAGATGGAAAACAGTAACTAAGGCTGGCTAATCTG

CTACTTATCTGCTATTAAGAAGCTGCTGCTTGTCTACTTGGTTATTTCGTTGAAGATAAAGCAACGCGAC

ATATCGGGAACTTTTGAAATTGCAAGAACTAAGTTAAAAGCTGGCAATGTAGATGATTTTTTTCCAAGTA

GTTGAACCTACCAATGTATTCCTTACCATATTGTCTTATTCTTATTTCTATGTAACATCAATGCTCAAAA

GACTTAAAATCTAAGTTACTCAAGTAGTTTTTATATTTGTCTGTTTATTTATT

>MSTRG.222.2 gene=MSTRG.222

TTTTTCACGTGAAGATTATCTCGTTTAATATCTGGGATACGTATGGCTCAGAGATGGTGAAGTAAACAAT

GAAGATGATGCTCCAAAAAGATGGAAAACAGTAACTAAGGCTGGCTAATCTGCTACTTATCTGCTATTAA

GAAGCTGCTGCTTGTCTACTTGGTTATTTCGTTGAAGATAAAGCAACGCGACATATCGGGAACTTTTGAA

ATTGCAAGAACTAAGTTAAAAGCTGGCAATGTAGATGATTTTTTTCCAAGTAGTTGAACCTACCAATGTA

TTCCTTACCATATTGTCTTATTCTTATTTCTATGTAACATCAATGCTCAAAAGACTTAAAATCTAAGTTA

CTCAAGTAGTTTTTATATTTGTCTGTTTATTTATT

>MSTRG.223.1 gene=MSTRG.223

CTATCATTGATTAAATATTAAAAAATTTAGAATTTAACTTTAAAGGCGTTTTCTCTTGGCTTTTAATCTG

AAACACTTGCAATATCACTTGTACAAAAGGTAACCGAGAAGGTAATAGGAAAATTAATCCAGCTTCTGCA

AAGAGAGAAGGTATTCTACTTCTTTTGATCAGAGATTATTATATGTTCCTTGGACCCTGCAATAAAACCA

ACTCAATACAATCTTATTCACGTGAAGATTGTCTTGTTTAATAATATCTTTGAGACGGATGGCTCAGAAA

TGGTGAAGTAAACAAGGAAGATGATTTGCAGTAAACAATGAAGATGATGCTCCAAAATGATGAAAAACAA

TTTTCTAAGGC

>MSTRG.224.1 gene=MSTRG.224

CATGGCGTTACTCAGCAACGCTCTCCGGCAAGCCTTCATGCCGAAGCACGAGTACGACAATCTGAGCGAA

GAAGACAAAGCGCTGATCCAATTACAACGCCCCGTCTTGATTTCCCTCTTGTTGTGTATCGTGATTGTGA

TCGTTGTGTCCACCTCGATTAGCGTGAAGATTGTGTTCCCGGCGAAGGACGGGAAACGCGTGTTTTGCCG

GGATTTGAGGATTCAGCCGTTGTCGATAAATGTGAGCTCTGGTGGCGGTGGCGGCGGGGAGGATGTGTTT

CCTGGGGCGTTTTATTTGACGGATCAGCAGACGGTGGATTATTATTGGATGGTGGTGTTTATGCC

>MSTRG.225.1 gene=MSTRG.225

ATTAGATTAGGGCACGCCGCATCGATGGATAATAATCGTGTGTGTATGTATCTATTCGAGTGTATTCATA

ATTCAGTGTTGCTAGATCGATGTTCAAATTCTTTATGTTGTCCGTTTCGAAGCCGAAAACCAGTAAACAA

TGAAGAAGATGCTCCAGCAAAACAAAACTATGTTCTATTGCCACCTCAGTGGGCCGGCAAGGGATAAAAG

TGATTCTACCAAACCTTGCTCTAAGTGGACAAAAAGAAATGCCAACATTTCTCGATATGTCTTGCCGCTG

CTTCTTTTGGTGGAAAAGAACCCATAGGCTGGCACCACTGCTAGGGATTCCTTGTGACTCGGCTCACCGA

AATAAAAGAAGAACTCTTCATTTGAGCAAACAATTGCTGAAGGTTGATCTACTTTCTATTTAATTCAGTA

ACATATAAATGATGTCTCTTATATACTACTATGCATTGACAAAAATATATTAGTTTGGCCTTTAGTTCTT

GCTTCTCTTTCCC

>MSTRG.226.1 gene=MSTRG.226

TCCCTACTCTCTGCACTTTTGGCTATTTCTTTTATATTTTCTCTCAGCTTGCCTAGCGGTTATGTTCAGC

CTGGGACTTGGACAGGCACTTCAGTTACCCGCTCCGTAACTGAGGTCTCTGTGCCTCCTCAGAGATTTAC

TTATAAATGGCCTTCTCTGGACAATCTTTTCACACCGGGGAAAGCCGATCGGCATGAATATGACGGATTC

AGCTAGAGGGTCAAAGCGGGATAAAGTCTTAGATTTGACAGAGGCAAGGAAAGGTGAGGAAAATCCAGGC

ATCCAGCTGTATGCTCAAAAACAAACCTCCAGCGCAAGTGGAGACTGTTCTAAGTGACTGCTCCTCAGCA

GCAGTTCACCAACCTAAGCGCAAGAGCCATCGTTACATTCCACAGGCGGTGATTATTTTTTTCAACTTAC

TTTAAGGGGACTTGTGTGTTCATGCCTTGCTAATAATAGTTGTCATTTGTTTCTTGTATTCACATATGAA

ATGTTTTTGCAGCTCAAGGTAATACGTAGTTGCAGCTTAGTCATTCCACATGCAATTTTATTTTACAAGC

ACTTGTGATTTATGATAGAAATCTTCTCATTTATTGTGTCATTTCTCGGACTTCAATCTTCTTTTCAGGG

AGATCTTGTATGGGCAATCAGTTCAGACACGATTCATTCTCTCGTGGGACTCTCTGAACTTTTCTTTGTA

CTTCCGCTTATGAATTCAGGCCTGTGAGCTTTCAAGGATGTTCTTGCATTGGCGTCACTCGACAGGGCCA

ACAATTGAGCTGTTGAGAAACAAAAAAATAAAAGAGGCCTTCGAAGATGGTTTTAGGAGTTGCGTAAAGA

TTTTTATGGCCATTGATCCGAAATATATTGGGCAGACAAATTTGGCGAAGGGGTGGCCAAATGGATGAAA

GAATTTTCTGTTATGGAGGCCGTTGTATTACTGAGAGAAATGCTCTGTGAAAGCGAGAGATAGAGCACAC

CTTTAAAATCCAAATACAAAAGATGATGTTTGCTCAGAGAAAACAGAGGATGTACTTCAACTACTTTAAC

TTAAAAATTTATGAAATGTTTTGGGACTTCTTTTAGTGTGAGCAGCCTTTGAGGCTTTGCAAATTGTAGG

ATAATGGTGGACAAAAATTGTCTTGTCTTGATTAAGATATTGTATAAGCTGCTATTTATTTGGTTCTCAA

CTTTTAGACAAATGTGTTATCTTTAATGTAAAAGTTTAAATTTGCAA

>MSTRG.227.1 gene=MSTRG.227

TTTATTTTTAGGTATAAAATCACTCGTCGTGTAGGTTAAGGCGGCAGCTTCTCAATTACATTAGGGCATC

GCCTAGATTAGGGCATCGCCGCAACGATCTATGATAATCGTGTGTGTATGTATCTATAGTGTAAAGTGCT

GAATCGTTTCGTGATGCTATCTGTTTCGAAGTCTAAAAACAGTAAACAATGAAGAAGATGCTCCAGAAAA

ACAAAACAATGATCTGCCTGATTGGGTTGGCAAGGGAAAACAGTAGTTCTACCAAACCCGCCTCAAAGTG

GACAACGAGAAATGCCAACATTATCATTCCGGAATTCCCACTTTGCTTGATATTTGTCAGACAACAAAAA

TTATTTGTTACATGTACAAACTTCATACAGTATATGCAAAATCGTGACAGTCAATCTTTATCTCTCTAAC

AAGCTTTATATGTCCAATGTACTCTTTGTGTCCAGTGGTCATGCAGCTGATATGAGGCAAAAGAACCCAA

AGGCTGGCAACCCATTTAATGGAATGTATATAATACGTTTGTGTGATGACAGAGGATCGATTCAGCACTT

TGTAAAAAAAATTCATAGTTTACATGCTACTCACTTGACGTATATGTAATTAATATTATTTTTAAAATTT

TATACTAAAGAATGTCAAATCTTATATCTATATAAAGCAAAATCTCGGGCGTCTTACGTGGCGCTCTTTA

CGCTCTTGAATCAAAATCCCTGCTCTCTGCACTTTTGATTGTTGTTAGGAATTAATCTAATGTCCATATT

TGATGTAAATATTTTGTTACAGGCGAAAGCATTGGAAGCAGCTATGAATGACATTAATAACTCATTTGGA

AATGGAGCTGTAACAAGATTAGGCAATGCTGGTGGATCTCTTGTGTATGTTGACTCTTTTCCAAGTGGTT

GTCTGACATTGGACATTGCATTGGGTGTCGGCCTTCCTAGAGGAAATATTGTTGAGGTATGTTTTCGTAC

ACACTTTGAGGTATTGTAAAACAAACTCAGTACAATCTTTTTCACGTGAAGATTATCTCGTTTAATATCT

GGGATACGTATGGCTCAGAGATGGTGAAGTAAACAATGAAGATGATGCTCCAAAAAGATGGAAAACAGTA

ACTAAGGCTGGCTAATCTGCTACTTATCTGCTATTAAGAAGCTGCTGCTTGTCTACTTGGTTATTTCGTT

GAAGATAAAGCAACGCGACATATCGGGAACTTTTGAAATTGCAAGAACTAAGTTAAAAGCTGGCAATGTA

GATGATTTTTTTCCAAGTAGTTGAACCTACCAATGTATTCCTTACCATATTGTCTTATTCTTATTTCTAT

GTAACATCAATGCTCAAAAGACTTAAAATCTAAGTTACTCAAGTAGTTTTTATATTTGTCTGTTTATTTA

TTTACCTATATAATGTATATAATACGTTTGTGTGATGACAGAGGATCGATTCAGCACTTTGTAAAAAAAT

TCATAGTTTACATGCTACTCACTTGACGTATATGTAATTAATATTATTTTTAAAATTTTATACTAAAGAA

TGTCAAATCTTATATCTATATAAAGC

>MSTRG.227.2 gene=MSTRG.227

GTGGTCATGCAGCTGATATGAGGCAAAAGAACCCAAAGGCTGGCAACCCATTTAATGAAATTGCACCTAT

TAAATGGGAGATAATTTGTGACCACTGCGAGGGATTCCTTGTGACTCGGCGAACTGTGTGCGAGAGAACA

TCAGAAGAGCACACAATTGCTGAAGGCGAAAGCATTGGAAGCAGCTATGAATGACATTAATAACTCATTT

GGAAAAGGAGCTGTAACAAGATTAGGCAATGCTGGTGGATCTCTTGTTGACTCTTTTCCAAGTGGTTGTC

TGACATTGGACATTGCATTGGGTGTCGGCCTTCCTAGAGGAAATATTGTTGAGTAAACAATGAAGATGAT

GCTCCAAAAAGATGGAAAACAGTAACTAAGGCTGGCTAATCTGCTACTTATCTGCTATTAAGAAGCTGCT

GCTTGTCTACTTGGTTATTTCGTTGAAGATAAAGCAACGCGACATATCGGGAACTTTTGAAATTGCAAGA

ACTAAGTTAAAAGCTGGCAATGTAGATGATTTTTTTCCAAGTAGTTGAACCTACCAATGTATTCCTTACC

ATATTGTCTTATTCTTATTTCTATGTAACATCAATGCTCAAAAGACTTAAAATCTAAGTTACTCAAGTAG

TTTTTATATTTGTCTGTTTATTTATTTACCTATATAATGTATATAATACGTTTGTGTGATGACAGAGGAT

CGATTCAGC

>MSTRG.227.4 gene=MSTRG.227

TTTTAGTTAAACCAATTTATTTTTAGGTATAAAATCACTCGTCGTGTAGGTTAAGGCGGCAGCTTCTCAA

TTACATTAGGGCATCGCCTAGATTAGGGCATCGCCGCAACGATCTATGATAATCGTGTGTGTATGTATCT

AGAGTGTAAAGTGCTGAATCGTTTCGTGATGCTATCTGTTTCGAAGTCTAAAAACAGTAAACAATGAAGA

AGATGCTCCAGAAAAACAAAACAATGATCTGCCTGATTGGGTTGGCAAGGGAAAACAGTAGTTCTACCAA

ACCCGCCTCAAAGTGGACAACGAGAAATGCCAACATTATCATTCCGGAATTCCCACTTTGCTTGATATTT

GTCAGACAACAAAAATTATTTGTTACATTGGTCATGCAGCTGATATGAGGCAAAAGAACCCAAAGGCTGG

CAACCCATTTAATGGAATGCGAAAGCATTGGAAGCAGCTATGAATGACATTAATAACTCATTTGGAAATG

GAGCTGTAACAAGATTAGGCAATGCTGGTGGATCTCTTGTTGACTCTTTTCCAAGTGGTTGTCTGACATT

GGACATTGCATTGGGTGTCGGCCTTCCTAGAGGAAATATTGTTGAGGTATGTTTTCGTACACACTTTGAG

GTATTGTAAAACAAACTCAGTACAATCTTTTTCACGTGAAGATTATCTCGTTTAATATCTGGGATACGTA

TGGCTCAGAGATGGTGAAGTAAACAATGAAGATGATGCTCCAAAAAGATGGAAAACAGTAACTAAGGCTG

GCTAATCTGCTACTTATCTGCTATTAAGAAGCTGCTGCTTGTCTACTTGGTTATTTCGTTGAAGATAAAG

CAACGCGACATATCGGGAACTTTTGAAATTGCAAGAACTAAGTTAAAAGCTGGCAATGTAGATGATTTTT

TTCCAAGTAGTTGAACCTACCAATGTATTCCTTACCATATTGTCTTATTCTTATTTCTATGTAACATCAA

TGCTCAAAAGACTTAAAATCTAAGTTACTCAAGTAGTTTTTATATTTGTCTGTTTATTTATTTACCTATA

TAATGTATATAATACGTTTGTGTGATGACAGAGGATCGATTCA

>MSTRG.227.5 gene=MSTRG.227

TTTTAGTTAAACCAATTTATTTTTAGGTATAAAATCACTCGTCGTGTAGGTTAAGGCGGCAGCTTCTCAA

TTACATTAGGGCATCGCCTAGATTAGGGCATCGCCGCAACGATCTATGATAATCGTGTGTGTATGTATCT

AGAGTGTAAAGTGCTGAATCGTTTCGTGATGCTATCTGTTTCGAAGTCTAAAAACAGTAAACAATGAAGA

AGATGCTCCAGAAAAACAAAACAATGATCTGCCTGATTGGGTAAAATTCTGATCTTTGTAATGAAAGGCT

TGTTTAATGGACTAGCTAAATGCAATTTAATTGTGGTCCGTTTTGAACATATATGTGACAACTGGTAGTG

GTCGGCTATATGCTGTTGATAGATCGTGCCCTTGATCGGAATTTTTTGTTGTCTGTTTTGAACATATACG

TGGTCAGATTTATTGATGGGTTTTGAATGCTGGTAACTAACCCTATGATATATTGTGCCCTTGGTTGTGT

TGGTTAGCTGTATCGGAATATTTTGTCATAACATAAGTGGGTAATTTGTGTTAGTTTTGAGCGCCCTTGT

TCTTGTACTATAATATGGAAGGTAATTGAAAAACTAAACCACATTTCGATCTCTATCTATGAAAATACTT

TCTCTTATTGGAGAGATGATAAAGGGAAGCCATAATCTCGAAATTGAACGATTAATGGTCATTAAGAATC

CAGATATTGTTGATAAGATTTTATTGTATCTTGTTGGATTACATTTATGGCCTGTTTGGGAAATTGGATT

TCATTTGAAATTTTGGATTTGATCAAATAATCTGTTTGGGAATGTGGATTTGGATTTCATTTGAAATCCA

GACATTCAAATATTTGTATAAATTTGAGTGTTGGAGATGACAACTTAAATCTTGTCATTTGAAATGAAAT

GCATGTCTCCAAATGGCCTCTTATTTAAATTAAAACATCGAGGGAGGATCCATTTGAATCAGCGCTATCG

TGCAATTGGGATCAATTATATATGTTGGCAAGGGAAAACAGTAGTTCTACCAAACCCGCCTCAAAGTGGA

CAACGAGAAATGCCAACATTATCATTCCGGAATTCCCACTTTGCTTGATATTTGTCAGACAACAAAAATT

ATTTGTTACATGTACAAACTTCATACAGTATATGCAAAATCGTGACAGTCAATCTTTATCTCTCTAACAA

GCTTTATATGTCCAATGTACTCTTTGTGTCCAGTGGTCATGCAGCTGATATGAGGCAAAAGAACCCAAAG

GCTGGCAACCCATTTAATGGAATGTATATAATACGTTTGTGTGATGACAGAGGATCGATTCAGCACTTTG

TAAAAAAAATTCATAGTTTACATGCTACTCACTTGACGTATATGTAATTAATATTATTTTTAAAATTTTA

TACTAAAGAATGTCAAATCTTATATCTATATAAAGCAAAATCTCGGGCGTCTTACGTGGCGCTCTTTACG

CTCTTGAATCAAAATCCCTGCTCTCTGCACTTTTGATTGTTGTTAGGAATTAATCTAATGTCCATATTTG

ATGTAAATATTTTGTTACAGGCGAAAGCATTGGAAGCAGCTATGAATGACATTAATAACTCATTTGGAAA

TGGAGCTGTAACAAGATTAGGCAATGCTGGTGGATCTCTTGTGTATGTTGACTCTTTTCCAAGTGGTTGT

CTGACATTGGACATTGCATTGGGTGTCGGCCTTCCTAGAGGAAATATTGTTGAGTAAACAATGAAGATGA

TGCTCCAAAAAGATGGAAAACAGTAACTAAGGCTGGCTAATCTGCTACTTATCTGCTATTAAGAAGCTGC

TGCTTGTCTACTTGGTTATTTCGTTGAAGATAAAGCAACGCGACATATCGGGAACTTTTGAAATTGCAAG

AACTAAGTTAAAAGCTGGCAATGTAGATGATTTTTTTCCAAGTAGTTGAACCTACCAATGTATTCCTTAC

CATATTGTCTTATTCTTATTTCTATGTAACATCAATGCTCAAAAGACTTAAAATCTAAGTTACTCAAGTA

GTTTTTATATTTGTCTGTTTATTTATTTACCTATATAATGTATATAATACGTTTGTGTGATGACAGAGGA

TCGATTCA

>MSTRG.227.3 gene=MSTRG.227

AGTGTAAAGTGCTGAATCGTTTCGTGATGCTATCTGTTTCGAAGTCTAAAAACAGTAAACAATGAAGAAG

ATGCTCCAGAAAAACAAAACAATGATCTGCCTGATTGGGTTGGCAAGGGAAAACAGTAGTTCTACCAAAC

CCGCCTCAAAGTGGACAACGAGAAATGCCAACATTCTCATTGCGGAATTCCCACTTTGCTTGATATTTGT

CAGACAACAAAAATTATTTGTGGTCATGCAGCTGATATGAGGCAAAAGAACCCAAAGGCTGGCAACCCAT

TTAATGAAATTGCACCTATTAAATGGGAGATAATTTGTGACCACTGCGAGGGATTCCTTGTGACTCGGCG

AACTGTGTGCGAGAGAACATCAGAAGAGCACACAATTGCTGAAGGCGAAAGCATTGGAAGCAGCTATGAA

TGACATTAATAACTCATTTGGAAAAGGAGCTGTAACAAGATTAGGCAATGCTGGTGGATCTCTTGTTGAC

TCTTTTCCAAGTGGTTGTCTGACATTGGACATTGCATTGGGTGTCGGCCTTCCTAGAGGAAATATTGTTG

AGGTATGTTTTCGTACACACTTTGAGGTATTGTAAAACAAACTCAGTACAATCTTTTTCACGTGAAGATT

ATCTTGTTTAATATCCGGGATACGTATGGCTCAGAGATGGTGAAGTAAACAATGAAGATGATGCTCCAAA

AAGATGGAAAACAGTAACTAAGGCTGGCTAATCTGCTACTTATCTGCTATTAAGAAGCTGCTGCTTGTCT

ACTTGGTTATTTCGTTGAAGATAAAGCAACGCGACATATCGGGAACTTTTGAAATTGCAAGAACTAAGTT

AAAAGCTGGCAATGTAGATGATTTTTTTCCAAGTAGTTGAACCTACCAATGTATTCCTTACCATATTGTC

TTATTCTTATTTCTATGTAACATCAATGCTCAAAAGACTTAAAATCTAAGTTACTCAAGTAGTTTTTATA

TTTGTCTGTTTATTTATTTACCTATATAATGTATATAATACGTTTGTGTGATGACAGAGGATCGATTCA

>MSTRG.227.7 gene=MSTRG.227

CACTCGTCGTGTAGGTTAAGGCGGCAGCTTCTCAATTACATTAGGGCATCGCCTAGATTAGGGCATCGCC

GCAACGATCTATGATAATCGTGTGTGTATGTATCTATAGTGTAAAGTGCTGAATCGTTTCGTGATGCTAT

CTGTTTCGAAGTCTAAAAACAGTAAACAATGAAGAAGATGCTCCAGAAAAACAAAACAATGATCTGCCTG

ATTGGGTTGGCAAGGGAAAACAGTAGTTCTACCAAACCCGCCTCAAAGTGGACAACGAGAAATGCCAACA

TTATCATTCCGGAATTCCCACTTTGCTTGATATTTGTCAGACAACAAAAATTATTTGTTACATGTACAAA

CTTCATACAGTATATGCAAAATCGTGACAGTCAATCTTTATCTCTCTAACAAGCTTTATATGTCCAATGT

ACTCTTTGTGTCCAGTGGTCATGCAGCTGATATGAGGCAAAAGAACCCAAAGGCTGGCAACCCATTTAAT

GGAATGTATATAATACGTTTGTGTGATGACAGAGGATCGATTCAGCACTTTGTAAAAAAAATTCATAGTT

TACATGCTACTCACTTGACGTATATGTAATTAATATTATTTTTAAAATTTTATACTAAAGAATGTCAAAT

CTTATATCTATATAAAGCAAAATCTCGGGCGTCTTACGTGGCGCTCTTTACGCTCTTGAATCAAAATCCC

TGCTCTCTGCACTTTTGATTGTTGTTAGGAATTAATCTAATGTCCATATTTGATGTAAATATTTTGTTAC

AGGCGAAAGCATTGGAAGCAGCTATGAATGACATTAATAACTCATTTGGAAATGGAGCTGTAACAAGATT

AGGCAATGCTGGTGGATCTCTTGTGTATGTTGACTCTTTTCCAAGTGGTTGTCTGACATTGGACATTGCA

TTGGGTGTCGGCCTTCCTAGAGGAAATATTGTTGAGTAAACAATGAAGATGATGCTCCAAAAAGATGGAA

AACAGTAACTAAGGCTGGCTAATCTGCTACTTATCTGCTATTAAGAAGCTGCTGCTTGTCTACTTGGTTA

TTTCGTTGAAGATAAAGCAACGCGACATATCGGGAACTTTTGAAATTGCAAGAACTAAGTTAAAAGCTGG

CAATGTAGATGATTTTTTTCCAAGTAGTTGAACCTACCAATGTATTCCTTACCATATTGTCTTATTCTTA

TTTCTATGTAACATC

>MSTRG.227.6 gene=MSTRG.227

TTTATTTTTAGGTATAAAATCACTCGTCGTGTAGGTTAAGGCGGCAGCTTCTCAATTACATTAGGGCATC

GCCTAGATTAGGGCATCGCCGCAACGATCTATGATAATCGTGTGTGTATGTATCTATAGTGTAAAGTGCT

GAATCGTTTCGTGATGCTATCTGTTTCGAAGTCTAAAAACAGTAAACAATGAAGAAGATGCTCCAGAAAA

ACAAAACAATGATCTGCCTGATTGGGTTGGCAAGGGAAAACAGTAGTTCTACCAAACCCGCCTCAAAGTG

GACAACGAGAAATGCCAACATTATCATTCCGGAATTCCCACTTTGCTTGATATTTGTCAGACAACAAAAA

TTATTTGTTACATGTACAAACTTCATACAGTATATGCAAAATCGTGACAGTCAATCTTTATCTCTCTAAC

AAGCTTTATATGTCCAATGTACTCTTTGTGTCCAGTGGTCATGCAGCTGATATGAGGCAAAAGAACCCAA

AGGCTGGCAACCCATTTAATGGAATGTATATAATACGTTTGTGTGATGACAGAGGATCGATTCAGCACTT

TGTAAAAAAAATTCATAGTTTACATGCTACTCACTTGACGTATATGTAATTAATATTATTTTTAAAATTT

TATACTAAAGAATGTCAAATCTTATATCTATATAAAGCAAAATCTCGGGCGTCTTACGTGGCGCTCTTTA

CGCTCTTGAATCAAAATCCCTGCTCTCTGCACTTTTGATTGTTGTTAGGAATTAATCTAATGTCCATATT

TGATGTAAATATTTTGTTACAGGCGAAAGCATTGGAAGCAGCTATGAATGACATTAATAACTCATTTGGA

AATGGAGCTGTAACAAGATTAGGCAATGCTGGTGGATCTCTTGTGTATGTTGACTCTTTTCCAAGTGGTT

GTCTGACATTGGACATTGCATTGGGTGTCGGCCTTCCTAGAGGAAATATTGTTGAGTAAACAATGAAGAT

GATGCTCCAAAAAGATGGAAAACAGTAACTAAGGCTGGCTAATCTGCTACTTATCTGCTATTAAGAAGCT

GCTGCTTGTCTACTTGGTTATTTCGTTGAAGATAAAGCAACGCGACATATCGGGAACTTTTGAAATTGCA

AGAACTAAGTTAAAAGCTGGCAATGTAGATGATTTTTTTCCAAGTAGTTGAACCTACCAATGTATTCCTT

ACCATATTGTCTTATTCTTATTTCTATGTAACATC

>MSTRG.227.8 gene=MSTRG.227

TTTTAGTTAAACCAATTTATTTTTAGGTATAAAATCACTCGTCGTGTAGGTTAAGGCGGCAGCTTCTCAA

TTACATTAGGGCATCGCCTAGATTAGGGCATCGCCGCAACGATCTATGATAATCGTGTGTGTATGTATCT

AGAGTGTAAAGTGCTGAATCGTTTCGTGATGCTATCTGTTTCGAAGTCTAAAAACAGTAAACAATGAAGA

AGATGCTCCAGAAAAACAAAACAATGATCTGCCTGATTGGGTTGGCAAGGGAAAACAGTAGTTCTACCAA

ACCCGCCTCAAAGTGGACAACGAGAAATGCCAACATTATCATTCCGGAATTCCCACTTTGCTTGATATTT

GTCAGACAACAAAAATTATTTGTTACATGTACAAACTTCATACAGTATATGCAAAATCGTGACAGTCAAT

CTTTATCTCTCTAACAAGCTTTATATGTCCAATGTACTCTTTGTGTCCAGTGGTCATGCAGCTGATATGA

GGCAAAAGAACCCAAAGGCTGGCAACCCATTTAATGGAATGCGAAAGCATTGGAAGCAGCTATGAATGAC

ATTAATAACTCATTTGGAAATGGAGCTGTAACAAGATTAGGCAATGCTGGTGGATCTCTTGTGTATGTTG

ACTCTTTTCCAAGTGGTTGTCTGACATTGGACATTGCATTGGGTGTCGGCCTTCCTAGAGGAAATATTGT

TGAGGTATGTTTTCGTACACACTTTGAGGTATTGTAAAACAAACTCAGTACAATCTTTTTCACGTGAAGA

TTATCTCGTTTAATATCTGGGATACGTATGGCTCAGAGATGGTGAAGTAAACAATGAAGATGATGCTCCA

AAAAGATGGAAAACAGTAACTAAGGCTGGCTAATCTGCTACTTATCTGCTATTAAGAAGCTGCTGCTTGT

CTACTTGGTTATTTCGTTGAAGATAAAGCAACGCGACATATCGGGAACTTTTGAAATTGCAAGAACTAAG

TTAAAAGCTGGCAATGTAGATGATTTTTTTCCAAGTAGTTGAACCTACCAATGTATTCCTTACCATATTG

>MSTRG.227.9 gene=MSTRG.227

TTTATTTTTAGGTATAAAATCACTCGTCGTGTAGGTTAAGGCGGCAGCTTCTCAATTACATTAGGGCATC

GCCTAGATTAGGGCATCGCCGCAACGATCTATGATAATCGTGTGTGTATGTATCTATAGTGTAAAGTGCT

GAATCGTTTCGTGATGCTATCTGTTTCGAAGTCTAAAAACAGTAAACAATGAAGAAGATGCTCCAGAAAA

ACAAAACAATGATCTGCCTGATTGGGTTGGCAAGGGAAAACAGTAGTTCTACCAAACCCGCCTCAAAGTG

GACAACGAGAAATGCCAACATTCTCATTGCGGAATTCCCACTTTGCTTGATATTTGTCAGACAACAAAAA

TTATTTGTTACATTGGTCATGCAGCTGATATGAGGCAAAAGAACCCAAAGGCTGGCAACCCATTTAATGA

AATTGCACCTATTAAATGGGAGATAATTTGTGACCACTGCGAGGGATTCCTTGTGACTCGGCGAACTGTG

TGCGAGAGAACATCAGAAGAGCACACAATTGCTGAAGGCGAAAGCATTGGAAGCAGCTATGAATGACATT

AATAACTCATTTGGAAAAGGAGCTGTAACAAGATTAGGCAATGCTGGTGGATCTCTTGTGTATGTTGACT

CTTTTCCAAGTGGTTGTCTGACATTGGACATTGCATTGGGTGTCGGCCTTCCTAGAGGAAATATTGTTGA

GGTATGTTTTCGTACACACTTTGAGGTATTGTAAAACAAACTCAGTACAATCTTTTTCACGTGAAGATTA

TCTTGTTTAATATCCGGGATACGTATGGCTCAGAGATGGTGAAGTAAACAATGAAGATGATGCTCCAAAA

TGATGAAAAACAATTATCTAAGGCCCGCTAATTTGCTAGTAACCATATTGTCTTATTCTTATTCCTATGT

AAGATCAGGGCTCAAAAGACTTAAAATAGATGATGTTCAAGTAACCATTAAGTTACTTGAAGTATTTCTA

GTATTTGTCTGTTTATTTATTGACCTGTATAATGTATATAATAAGCTTGTGTGTGTGTTAAAACAGGATT

CTGCATTTTGG

>MSTRG.227.10 gene=MSTRG.227

AGTGTAAAGTGCTGAATCGTTTCGTGATGCTATCTGTTTCGAAGTCTAAAAACAGGTATATATGTTGTGT

TTTTAAGCCCTAGATTTCTCAATTCTGATTGTTTAATAGTTAATTTATCTAATTTGAAATACACTAATTA

TCAGATTTCTTCTTGCAGTAAACAATGAAGAAGATGCTCCAGAAAAACAAAACAATGATCTGCCTGATTG

GGTTGGCAAGGGAAAACAGTAGTTCTACCAAACCCGCCTCAAAGTGGACAACGAGAAATGCCAACATTCT

CATTGCGGAATTCCCACTTTGCTTGATATTTGTCAGACAACAAAAATTATTTGTGGTCATGCAGCTGATA

TGAGGCAAAAGAACCCAAAGGCTGGCAACCCATTTAATGAAATTGCACCTATTAAATGGGAGATAATTTG

TGACCACTGCGAGGGATTCCTTGTGACTCGGCGAACTGTGTGCGAGAGAACATCAGAAGAGCACACAATT

GCTGAAGGCGAAAGCATTGGAAGCAGCTATGAATGACATTAATAACTCATTTGGAAAAGGAGCTGTAACA

AGATTAGGCAATGCTGGTGGATCTCTTGTTGACTCTTTTCCAAGTGGTTGTCTGACATTGGACATTGCAT

TGGGTGTCGGCCTTCCTAGAGGAAATATTGTTGAGGTATGTTTTCGTACACACTTTGAGGTATTGTAAAA

CAAACTCAGTACAATCTTTTTCACGTGAAGATTATCTTGTTTAATATCCGGGATACGTATGGCTCAGAGA

TGGTGAAGTAAACAATGAAGATGATGC

>MSTRG.228.1 gene=MSTRG.228

GGTCTCTGTGCCTCCTCAGAGATTTACTTATAAATGGCCTTCTCTGGACAATCTTTTCACACCGGGGAAA

GCCGATCGGCATGAATATGACGGATTCAGCTAGAGGGTCAAAGCGGGATAAAGTCTTAGATTTGACAGAG

GCAAGGAAAGGTGAGGAAAATCCAGGCATCCAGCTGTATGCTCAAAAACAAACCTCCAGCGCAAGTGGAG

ACTGTTCTAAGTGACTGCTCCTCAGCAGCAGTTCACCAACCTAAGCGCAAGAGCCATCGTTACATTCCAC

AGGGAGATCTTGTATGGGCAATCAGTTCAGACACGATTCATTCTCTCGTGGGACTCTCTGAACTTTTCTT

TGTACTTCCGCTTATGAATTCAGTTCTAGGCCTGTGAGCTTTCAAGGATGTTCTTGCATTGGCGTCACTC

GACAGGGCCAACAATTGAGCTGTTGAGAAACAAAAAAATAAAAGAGGCCTTCGAAGATGGTTTTAGGAGT

TGCGTAAAGATTTTTATGGCCATTGATCCGAAATATATTGGGCAGACAAATTTGGCGAAGGGGTGGCCAA

ATGGATGAAAGAATTTTCTGTTATGGAGGCCGTTGTATTACTGAGAGAAATGCTCTGTGAAAGCGAGAGA

TAGAGCACACCTTTAAAATCCAATTACAAAAGAGGATGTTTGCTCAGAGAAAACAGAGGATGTACTTCAA

CTACTTTAACTTAAAAATTTATGAAATGTTTTGGGACTTCTTTTAGTGTGAGCAGCCTTTGAGGCTTTGC

AAATTGTAGGATAATGGTGGACAAAAATTGTCTTGTCTTGATTAAGATATTGTATAAGCTGCTATTCATT

TGGTTCTCAACTTTTAGACAAATGTGTTATCTTTAATGTAAAAGTTTAAATTTGCAATTTTAAAGTTTTT

TATTTTTTTTTCATTCCTTCTTTCACCAGCTTCACATACCTCGATCTTTGATAACCTTTTTTCTGGAGCC

ATTCAACATCAGCTTGGTAATCTTTTCAAATTACAGCAGCTTAATCTCGGCCACACTGGTATTCTACAAG

GTGGCATATTTGGTTGGTTATTGAATTTATCTACTCTGGT

>MSTRG.228.3 gene=MSTRG.228

GGTCTCTGTGCCTCCTCAGAGATTTACTTATAAATGGCCTTCTCTGGACAATCTTTTCACACCGGGGAAA

GCCGATCGGCATGAATATGACGGATTCAGCTAGAGGGTCAAAGCGGGATAAAGTCTTAGATTTGACAGAG

GCAAGGAAAGGTGAGGAAAATCCAGGCATCCAGCTGTATGCTCAAAAACAAACCTCCAGCGCAAGTGGAG

ACTGTTCTAAGTGACTGCTCCTCAGCAGCAGTTCACCAACCTAAGCGCAAGAGCCATCGTTACATTCCAC

AGGGAGATCTTGTATGGGCAATCAGTTCAGACACGATTCATTCTCTCGTGGGACTCTCTGAACTTTTCTT

TGTACTTCCGCTTATGAATTCAGGCCTGTGAGCTTTCAAGGATGTTCTTGCATTGGCGTCACTCGACAGG

GCCAACAATTGAGCTGTTGAGAAACAAAAAAATAAAAGAGGCCTTCGAAGATGGTTTTAGGAGTTGCGTA

AAGATTTTTATGGCCATTGATCCGAAATATATTGGGCAGACAAATTTGGCGAAGGGGTGGCCAAATGGAT

GAAAGAATTTTCTGTTATGGAGGCCGTTGTATTACTGAGAGAAATGCTCTGTGAAAGCGAGAGATAGAGC

ACACCTTTAAAATCCAATTACAAAAGAGGATGTTTGCTCAGAGAAAACAGAGGATGTACTTCAACTACTT

TAACTTAAAAATTTATGAAATGTTTTGGGACTTCTTTTAGTGTGAGCAGCCTTTGAGGCTTTGCAAATTG

TAGGATAATGGTGGACAAAAATTGTCTTGTCTTGATTAAGATATTGTATAAGCTGCTATTCATTTGGTTC

TCAACTTTTAGACAAATGTG

>MSTRG.228.2 gene=MSTRG.228

ACAATGAAGATGATGCTCCAAAAAGATGGAAAACAGTAACTAAGGCTGGCTAATCTGCTACTTATCTGCT

ATTAAGAAGCTGCTGCTTGTCTACTTGGTTATTTCGTTGAAGATAAAGCAACGCGACATATCGGGAACTT

TTGAAATTGCAAGAACTAAGTTAAAAGCTGGCAATGTAGATGATTTTTTTCCAAGTAGTTGAACCTACCA

ATGTATTCCTTACCATATTGTCTTATTCTTATTTCTATGTAACATCAATGCTCAAAAGACTTAAAATCTA

AGTTACTCAAGTAGTTTTTATATTTGTCTGTTTATTTATTTACCTATATAATGTATATAATACGTTTGTG

TGATGACAGAGGATCGATTCAGCACTTTGTAAAAAAAATTCATAGTTTACATGCTACTCACTTGACGTAT

ATGTAATTAATATTATTTTTAAAATTTTATACTAAAGAATGTCAAATCTTATATCTATATAAAGCAAAAT

CTCGGGCGTCTTACGTGGCGCTCTTTACGCTCTTGAATCAAAATCCCTGCTCTCTGCACTTTTGTCTATT

TCTTTTATATTTTCTCTCAGCTTGCCTAGCGGTTATGTTCAGCCTGGGACTTGGACAGGCACTTCAGTTA

CCCGCTCCGTAACCGAGGTCTCTGTGCCTCCTCAGAGATTTACTTATAAATGGCCTTCTCTGGACAATCT

TTTCACACCGGGGAAAGCCGATCGGCATGAATATGACGGATTCAGCTAGAGGGTCAAAGCGGGATAAAGT

CTTAGATTTGACAGAGGCAAGGAAAGGTGAGGAAAATCCAGGCATCCAGCTGTATGCTCAAAAACAAACC

TCCAGCGCAAGTGGAGACTGTTCTAAGTGACTGCTCCTCAGCAGCAGTTCACCAACCTAAGCGCAAGAGC

CATCGTTACATTCCACAGGGAGATCTTGTATGGGCAATCAGTTCAGACACGATTCATTCTCTCGTGGGAC

TCTCTGAACTTTTCTTTGTACTTCCGCTTATGAATTCAGGTAAGCTGAATTATTATTGCTTGATAGAATG

AATTGATAAGCTTTATGGTGCCAGCAAGTAGAAAGCCAGGTTATATTCGCACAACGTCCTTTATCTGAGG

CCACCAAGTCTCTACCTCTTTATACTGCTTTTCCCAACTGATGTTTTTAAATAAGTTGTATAACTTTGTT

GTTCATTCTATGACGATGTGCCTAAGTTTGTCTGCTTATATATTGTTAGTTCGTAATCGTAATAAATTGT

GATTACACATAGATATAGACTATTCTTGTAGAGATAAAGTTTAATGTTTTAAAGAATATCTTTTTATTTA

TAAGATTGTTTTCAGTTCTAGGCCTGTGAGCTTTCAAGGATGTTCTTGCATTGGCGTCACTCGACAGGGC

CAACAATTGAGCTGTTGAGAAACAAAAAAATAAAAGAGGCCTTCGAAGATGGTTTTAGGAGTTGCGTAAA

GATTTTTATGGCCATTGATCCGAAATATATTGGGCAGACAAATTTGGCGAAGGGGTGGCCAAATGGATGA

AAGAATTTTCTGTTATGGAGGCCGTTGTATTACTGAGAGAAATGCTCTGTGAAAGCGAGAGATAGAGCAC

ACCTTTAAAATCCAATTACAAAAGAGGATGTTTGCTCAGAGAAAACAGAGGATGTACTTCAACTACTTTA

ACTTAAAAATTTATGAAATGTTTTGGGACTTCTTTTAGTGTGAGCAGCCTTTGAGGCTTTGCAAATTGTA

GGATAATGGTGGACAAAAATTGTCTTGTCTTGATTAAGATATTGTATAAGCTGCTATTCATTTGGTTCTC

AACTTTTAGACAAATGTG

>MSTRG.229.1 gene=MSTRG.229

GATACCATCATATCTCAGTAAATCAGTGTTTGTCGAGCACAAATTGTTCTTGCCGATGATTAGTTGCCCA

AAAATACTGTAAGACATACTGTAAAAAGAATTCAAGAGGCCGATAACAGCAGCGGAGATAATAAAGGCAG

TGATTCCCAGTTGCAAGGTCTGCAATTTTTTCTGTCTTCGAGTCTTTGGAATGTTGCTGTTATGGAATGC

CCAAGGAAAATATAACTTCGGAATCAAGAAGTTTCGAAAGG

>MSTRG.230.1 gene=MSTRG.230

ATGGCGTTACTCAGCAACGCTCTCCGGCAAGCCTTCATGCCGAAGCACGAGTACGACAATCTGCGCGAAG

AAGACAAAGCGATGATCCAATTACAACGCCCCGTCTTGATTTCCCTCTTGTTGTGTATCGTGATTGTGAT

CGTTGTGTCCACCTCGATTAGCGTGAAGATTGTGTTCCCGGCGGAGGACGGGAAACGCGTGTTTTGCCGG

GATTTGAGGATTCAGCCGTTGTCGATAAATGTGAGCTCTGGTGGCGGTGGCGGCGGGGAGGATGTGTTTC

CTGGGGCGTTTTATTTGACGGATCAGCAGACGGTGGATTATTATTGGATGGTGGTGTTTATGCCGTCGGT

TTTGG

>MSTRG.231.1 gene=MSTRG.231

CAATACTCGTAAGAAAGCCCAGGCCTTTTAGTTAACCCAATTTATTTTTAGGTATAAAATCACTCGTCGT

GTAGGTTAAGGCGGCAGCTTCTCAATTACATTAGGGCATCGCCTAGATTAGGGCATCGCCGCAACGATCT

ATGATAATCGTGTGTGTATGTATCTATTTGAGTGTAAAGTGCTGAATCGTTTCGTGATGCTATCTGTTTC

GAAGTTTAAAAACAGTAAACAATGAAGAAGATGCTCCAG

>MSTRG.231.2 gene=MSTRG.231

CAATACTCGTAAGAAAGCCCAGGCCTTTTAGTTAACCCAATTTATTTTTAGGTATAAAATCACTCGTCGT

GTAGGTTAAGGCGGCAGCTTCTCAATTACATTAGGGCATCGCCTAGATTAGGGCATCGCCGCAACGATCT

ATGATAATCGTGTGTGTATGTATCTATTTGAGTGTAAAGTGCTGAATCGTTTCGTGATGCTATCTGTTTC

GAAGTTTAAAAACAGTAAACAATGAAGAAGATGCTCCAGAAAAACAAAACTTTGGGTAAAATTCTGACAA

TACAGGTATGGAAGCGTCCCGTGGACGTGATGTTAATCTTATGGCCCCTGTTATCAGCTCAGATGTCAAC

CTCGGTAATACTAACACCGAGCTTTCCCTGGCCATCCTCCCTCCACAAGCACCAGGCGGAACTATTCCAA

GGAAACGACGCAGGCCTTGTTACGGGTGGATTGATGATGAGGACGACATTGATGACATTGTGCACTTAAG

AACTTCTCCTGG

>MSTRG.231.3 gene=MSTRG.231

CCTTTTAGTTAACCCAATTTATTTTTAGGTATAAAATCACTCGTCGTGTAGGTTAAGGCGGCAGCTTCTC

AATTACATTAGGGCATCGCCTAGATTAGGGCATCGCCGCAACGATCTATGATAATCGTGTGTGTATGTAT

CTATTTGAGTGTAAAGTGCTGAATCGTTTCGTGATGCTATCTGTTTCGAAGTTTAAAAACAGTAAACAAT

GAAGAAGATGCTCCAGAAAAACAAAACAATGATCTGCCTGATTGGGTTGGCAAGGGAAAACAGTAGTTCT

ACCAAACCCGCCTCAAAGTGGACAACGAGAAATGCCAACATATTCGTTGCGGAATTCCCACTTTGCTTGA

TATTTTGGTCATGCAGCTGATATGAGGCAAAAGAACCCAAAGGCTGGCAACCCATTTAATGAAATTGCAC

CTATTAAATGGGAGATAATTTGTGACCACTGCGAGGGATTCCTTGTGACTCGGCCAACTGTGTGCGAGAG

AACATCAGAAGAGCACACAATTGCTGAAGGCGAAAGCATTGGAAGCAGCTATGAATGACATTAATAACTC

ATTTGGAAAAGGAGCTGTAACAAGATTAGGCAATGCTGGTGGATCTCTTGTTGACTCTTTTCCAAGTGGT

TGTCTGACATTGGACATTGCATTGGGTGTCGGCCTTCCTAGAGGAAATATTGTTGAGGATACGTATGGCT

CTGAGATGGTGAAGTAAACAATGAAGATGATGCTCCAAAAAGATGGAAAACAGTAACTAAGGCTGGCTAA

TCTGCTACTTATCTGCTATTAAGAAGCTGCTGCTTGTCTACTTGGTTATTTCGTTGAAGATAAAGCAACG

CGATGTATTCCTTACCATATTGTCTTATTCTTATTTCTATGTAACATCAATGCTCAAAAGACTTAAAATC

TAAGTTACTCAAGTAGTTTTTATATTTGTCTGTTTATTTATTTACCTATATAATGTATATAATACGTTTG

TGTGATGACAGAGGATCGATTCAGCACTTTGTAAAAAAAATTCATAGTTTACATGCTACTCACTTGACGT

ATATGTAATTAATATTATTTTTAAAGTTTTATACTAAAGAATGTCAAATCTTATATCTATATAAAGCAAA

ATCTCGGGCGTCTTACGTGGCGCTCTTTACGCTCTTGAATCAAAATCCCTGCTCTCTGCACTTTTGGCTA

TTTCTTTTATATTTTCTCTCAGCTTGCCTAGCGGTTATGTTCAGCCTGGGACTTGGACAGGCACTTCAGT

TACCCGCTCCGTAACTGAGGTCTCTGTGCCTCCTCAGAGATTTACTTATAAATGGCCTTCTCTGGACAAT

CTTTTCACACCGGGGAAAGCCGATCGGCATGAATATGACGGATTCAGCTAGAGGGTCAAAGCGGGATAAA

GTCTTAGATTTGACAGAGGCAAGGAAAGGTGAGGAAAATCCAGGCATCCAGCTGTATGCTCAAAAACAAA

CCTCCAGCGCAAGTGGAGACTGTTCTAAGTGACTGCTCCTCAGCAGCAGTTCACCAACCTAAGCGCAAGA

GCCATCGTTACATTCCACAGGGAGATCTTGTATGGGCAATCAGTTCAGACACGATTCATTCTCTCGTGGG

ACTCTCTGAACTTTTCTTTGTACTTCCGCTTATGAATTCAGGCCTGTGAGCTTTCAAGGATGTTCTTGCA

TTGGCGTCACTCGACAGGGCCAACAATTGAGCTGTTGAGAAACAAAAAAATAAAAGAGGCCTTCGAAGAT

GGTTTTAGGAGTTGCGTAAAGATTTTTATGGCCATTGATCCGAAATATATTGGGCAGACAAATTTGGCGA

AGGGGTGGCCAAATGGATGAAAGAATTTTCTGTTATGGAGGCCGTTGTATTACTGAGAGAAATGCTCTGT

GAAAGCGAGAGATAGAGCACACCTTTAAAATCCAAATACAAAAGATGATGTTTGCTCAGAGAAAACAGAG

GATGTACTTCAACTACTTTAACTTAAAAATTTATGAAATGTTTTGGGACTTCTTTTAGTGTGAGCAGCCT

TTGAGGCTTTGCAAATTGTAGGATAATGGTGGACAAAAATTGTCTTGTCTTGATTAAGATATTGTATAAG

CTGCTATTCATTTGGTTCTCAACTTTTAGACAAATGTGTTATCTTTAATGTAAAAGTTTAAATTTGCAAT

TTTAAAGTTTTTTATTTTTTTTTCATTCCTTCTTTCACCAGCTTCACATACCTCGATCTTTGATAACCTT

TTTTCTGGAGCCATTCAACATCAGCTTGGTAATCTTTTCAAATTACAGCAGCTTAATCTCGGCCACACTG

GTATTCTACAAGGTGGCATATTTGGTTGGTTATTGAATTTATCTACTCTGGT

>MSTRG.231.4 gene=MSTRG.231

CAGGCCTTTTAGTTAACCCAATTTATTTTTAGGTATAAAATCACTCGTCGTGTAGGTTAAGGCGGCAGCT

TCTCAATTACATTAGGGCATCGCCTAGATTAGGGCATCGCCGCAACGATCTATGATAATCGTGTGTGTAT

GTATCTATTTGAGTGTAAAGTGCTGAATCGTTTCGTGATGCTATCTGTTTCGAAGTTTAAAAACAGTAAA

CAATGAAGAAGATGCTCCAGAAAAACAAAACAATGATCTGCCTGATTGGGTTGGCAAGGGAAAACAGTAG

TTCTACCAAACCCGCCTCAAAGTGGACAACGAGAAATGCCAACATATTCGTTGCGGAATTCCCACTTTGC

TTGATATTTTGGTCATGCAGCTGATATGAGGCAAAAGAACCCAAAGGCTGGCAACCCATTTAATGAAATT

GCACCTATTAAATGGGAGATAATTTGTGACCACTGCGAGGGATTCCTTGTGACTCGGCCAACTGTGTGCG

AGAGAACATCAGAAGAGCACACAATTGCTGAAGGCGAAAGCATTGGAAGCAGCTATGAATGACATTAATA

ACTCATTTGGAAAAGGAGCTGTAACAAGATTAGGCAATGCTGGTGGATCTCTTGTGTATGTTGACTCTTT

TCCAAGTGGTTGTCTGACATTGGACATTGCATTGGGTGTCGGCCTTCCTAGAGGAAATATTGTTGAGGAT

ACGTATGGCTCTGAGATGGTGAAGTAAACAATGAAGATGATGCTCCAAAAAGATGGAAAACAGTAACTAA

GGCTGGCTAATCTGCTACTTATCTGCTATTAAGAAGCTGCTGCTTGTCTACTTGGTTATTTCGTTGAAGA

TAAAGCAACGCGACATATCGGGAACTTTTGAAATTGCAAGAACTAAGTTAAAAGCTGGCAATGTAGATGA

TTTTTTTCCAAGTAGTTGAACCTACCAATGTATTCCTTACCATATTGTCTTATTCTTATTTCTATGTAAC

ATCAATGCTCAAAAGACTTAAAATCTAAGTTACTCAAGTAGTTTTTATATTTGTCTGTTTATTTATTTAC

CTATATAATGTATATAATACGTTTGTGTGATGACAGAGGATCGATTCAGCACTTTGTAAAAAAAATTCAT

AGTTTACATGCTACTCACTTGACGTATATGTAATTAATATTATTTTTAAAGTTTTATACTAAAGAATGTC

AAATCTTATATCTATATAAAGCAAAATCTCGGGCGTCTTACGTGGCGCTCTTTACGCTCTTGAATCAAAA

TCCCTGCTCTCTGCACTTTTGGCTATTTCTTTTATATTTTCTCTCAGCTTGCCTAGCGGTTATGTTCAGC

CTGGGACTTGGACAGGCACTTCAGTTACCCGCTCCGTAACTGAGGTCTCTGTGCCTCCTCAGAGATTTAC

TTATAAATGGCCTTCTCTGGACAATCTTTTCACACCGGGGAAAGCCGATCGGCATGAATATGACGGATTC

AGCTAGAGGGTCAAAGCGGGATAAAGTCTTAGATTTGACAGAGGCAAGGAAAGGTGAGGAAAATCCAGGC

ATCCAGCTGTATGCTCAAAAACAAACCTCCAGCGCAAGTGGAGACTGTTCTAAGTGACTGCTCCTCAGCA

GCAGTTCACCAACCTAAGCGCAAGAGCCATCGTTACATTCCACAGGGAGATCTTGTATGGGCAATCAGTT

CAGACACGATTCATTCTCTCGTGGGACTCTCTGAACTTTTCTTTGTACTTCCGCTTATGAATTCAGTTCT

AGGCCTGTGAGCTTTCAAGGATGTTCTTGCATTGGCGTCACTCGACAGGGCCAACAATTGAGCTGTTGAG

AAACAAAAAAATAAAAGAGGCCTTCGAAGATGGTTTTAGGAGTTGCGTAAAGATTTTTATGGCCATTGAT

CCGAAATATATTGGGCAGACAAATTTGGCGAAGGGGTGGCCAAATGGATGAAAGAATTTTCTGTTATGGA

GGCCGTTGTATTACTGAGAGAAATGCTCTGTGAAAGCGAGAGATAGAGCACACCTTTAAAATCCAAATAC

AAAAGATGATGTTTGCTCAGAGAAAACAGAGGATGTACTTCAACTACTTTAACTTAAAAATTTATGAAAT

GTTTTGGGACTTCTTTTAGTGTGAGCAGCCTTTGAGGCTTTGCAAATTGTAGGATAATGGTGGACAAAAA

TTGTCTTGTCTTGATTAAGATATTGTATAAGCTGCTATTCATTTGGTTCTCAACTTTTAGACAAATG

>MSTRG.231.7 gene=MSTRG.231

TAGATTAGGGCATCGCCGCAACGATCTATGATAATCGTGTGTGTATGTATCTATTTGAGTGTAAAGTGCT

GAATCGTTTCGTGATGCTATCTGTTTCGAAGTTTAAAAACAGTAAACAATGAAGAAGATGCTCCAGAAAA

ACAAAACAATGATCTGCCTGATTGGGTTGGCAAGGGAAAACAGTAGTTCTACCAAACCCGCCTCAAAGTG

GACAACGAGAAATGCCAACATATTCGTTGCGGAATTCCCACTTTGCTTGATATTTTGGTCATGCAGCTGA

TATGAGGCAAAAGAACCCAAAGGCTGGCAACCCATTTAATGAAATTGCACCTATTAAATGGGAGATAATT

TGTGACCACTGCGAGGGATTCCTTGTGACTCGGCCAACTGTGTGCGAGAGAACATCAGAAGAGCACACAA

TTGCTGAAGGCGAAAGCATTGGAAGCAGCTATGAATGACATTAATAACTCATTTGGAAAAGGAGCTGTAA

CAAGATTAGGCAATGCTGGTGGATCTCTTGTTGACTCTTTTCCAAGTGGTTGTCTGACATTGGACATTGC

ATTGGGTGTCGGCCTTCCTAGAGGAAATATTGTTGAGTAAACAATGAAGATGATGCTCCAAAAAGATGGA

AAACAGTAACTAAGGCTGGCTAATCTGCTACTTATCTGCTATTAAGAAGCTGCTGCTTGTCTACTTGGTT

ATTTCGTTGAAGATAAAGCAACGCGACATATCGGGAACTTTTGAAATTGCAAGAACTAAGTTAAAAGCTG

GCAATGTAGATGATTTTTTTCCAAGTAGTTGAACCTACCAATGTATTCCTTACCATATTGTCTTATTCTT

ATTTCTATGTAACATCAATGCTCAAAAGACTTAAAATCTAAGTTACTCAAGTAGTTTTTATATTTGTCTG

TTTATTTATTTACCTATATAATGTATATAATACGTTTGTGTGATGACAGAGGATCGATTCAGCACTTTGT

AAAAAAAATTCATAGTTTACATGCTACTCACTTGACGTATATGTAATTAATATTATTTTTAAAGTTTTAT

ACTAAAGAATGTCAAATCTTATATCTATATAAAGCAAAATCTCGGGCGTCTTACGTGGCGCTCTTTACGC

TCTTGAATCAAAATCCCTGCTCTCTGCACTTTTGGCTATTTCTTTTATATTTTCTCTCAGCTTGCCTAGC

GGTTATGTTCAGCCTGGGACTTGGACAGGCACTTCAGTTACCCGCTCCGTAACTGAGGTCTCTGTGCCTC

CTCAGAGATTTACTTATAAATGGCCTTCTCTGGACAATCTTTTCACACCGGGGAAAGCCGATCGGCATGA

ATATGACGGATTCAGCTAGAGGGTCAAAGCGGGTATGTATGTTATTCTTGCTGATTTTCAGGGTTGATTT

ATGTTATATGATCATAGTCGTGGGCAGTTATCTCCTTCTGATTTATCCGGAGAGCATAGCAGCAGATTTT

TATGATAAAGTCTTAGATTTGACAGAGGCAAGGAAAGGTGAGGAAAATCCAGGCATCCAGCTGTATGCTC

AAAAACAAACCTCCAGCGCAAGTGGAGACTGTTCTAAGTGACTGCTCCTCAGCAGCAGTTCACCAACCTA

AGCGCAAGAGCCATCGTTACATTCCACAGGGAGATCTTGTATGGGCAATCAGTTCAGACACGATTCATTC

TCTCGTGGGACTCTCTGAACTTTTCTTTGTACTTCCGCTTATGAATTCAGGTAAGCTGAATTATTATTGC

TTGATAGAATGAATTGATAAGCTTTATGGTGCCAGCAAGTAGAAAGCCAGGTTATATTCGCACAACGTCC

TTTATCTGAGGCCACCAAGTCTCTACCTCTTTATACTGCTTTTCCCAACTGATGTTTTTAAATAAGTTGT

ATAACTTTGTTGTTCATTCTATGACGATGTGCCTAAGTTTGTCTGCTTATATATTGTTAGTTCGTAATCG

TAATAAATTGTGATTACACATAGATATAGACTATTCTTGTAGAGATAAAGTTTAATGTTTTAAAGAATAT

CTTTTTATTTATAAGATTGTTTTCAGTTCTAGGCCTGTGAGCTTTCAAGGATGTTCTTGCATTGGCGTCA

CTCGACAGGGCCAACAATTGAGCTGTTGAGAAACAAAAAAATAAAAGAGGCCTTCGAAGATGGTTTTAGG

AGTTGCGTAAAGATTTTTATGGCCATTGATCCGAAATATATTGGGCAGACAAATTTGGCGAAGGGGTGGC

CAAATGGATGAAAGAATTTTCTGTTATGGAGGCCGTTGTATTACTGAGAGAAATGCTCTGTGAAAGCGAG

AGATAGAGCACACCTTTAAAATCCAAATACAAAAGATGATGTTTGCTCAGAGAAAACAGAGGATGTACTT

CAACTACTTTAACTTAAAAATTTATGAAATGTTTTGGGACTTCTTTTAGTGTGAGCAGCCTTTGAGGCTT

TGCAAATTGTAGGATAATGGTGGACAAAAATTGTCTTGTCTTGATTAAGATATTGTATAAGCTGC

>MSTRG.231.8 gene=MSTRG.231

TAGATTAGGGCATCGCCGCAACGATCTATGATAATCGTGTGTGTATGTATCTATTTGAGTGTAAAGTGCT

GAATCGTTTCGTGATGCTATCTGTTTCGAAGTTTAAAAACAGTAAACAATGAAGAAGATGCTCCAGAAAA

ACAAAACAATGATCTGCCTGATTGGGTTGGCAAGGGAAAACAGTAGTTCTACCAAACCCGCCTCAAAGTG

GACAACGAGAAATGCCAACATATTCGTTGCGGAATTCCCACTTTGCTTGATATTTTGGTCATGCAGCTGA

TATGAGGCAAAAGAACCCAAAGGCTGGCAACCCATTTAATGAAATTGCACCTATTAAATGGGAGATAATT

TGTGACCACTGCGAGGGATTCCTTGTGACTCGGCCAACTGTGTGCGAGAGAACATCAGAAGAGCACACAA

TTGCTGAAGGCGAAAGCATTGGAAGCAGCTATGAATGACATTAATAACTCATTTGGAAAAGGAGCTGTAA

CAAGATTAGGCAATGCTGGTGGATCTCTTGTTGACTCTTTTCCAAGTGGTTGTCTGACATTGGACATTGC

ATTGGGTGTCGGCCTTCCTAGAGGAAATATTGTTGAGTAAACAATGAAGATGATGCTCCAAAAAGATGGA

AAACAGTAACTAAGGCTGGCTAATCTGCTACTTATCTGCTATTAAGAAGCTGCTGCTTGTCTACTTGGTT

ATTTCGTTGAAGATAAAGCAACGCGATGTATTCCTTACCATATTGTCTTATTCTTATTTCTATGTAACAT

CAATGCTCAAAAGACTTAAAATCTAAGTTACTCAAGTAGTTTTTATATTTGTCTGTTTATTTATTTACCT

ATATAATGTATATAATACGTTTGTGTGATGACAGAGGATCGATTCAGCACTTTGTAAAAAAAATTCATAG

TTTACATGCTACTCACTTGACGTATATGTAATTAATATTATTTTTAAAGTTTTATACTAAAGAATGTCAA

ATCTTATATCTATATAAAGCAAAATCTCGGGCGTCTTACGTGGCGCTCTTTACGCTCTTGAATCAAAATC

CCTGCTCTCTGCACTTTTGGCTATTTCTTTTATATTTTCTCTCAGCTTGCCTAGCGGTTATGTTCAGCCT

GGGACTTGGACAGGCACTTCAGTTACCCGCTCCGTAACTGAGGTCTCTGTGCCTCCTCAGAGATTTACTT

ATAAATGGCCTTCTCTGGACAATCTTTTCACACCGGGGAAAGCCGATCGGCATGAATATGACGGATTCAG

CTAGAGGGTCAAAGCGGGATAAAGTCTTAGATTTGACAGAGGCAAGGAAAGGTGAGGAAAATCCAGGCAT

CCAGCTGTATGCTCAAAAACAAACCTCCAGCGCAAGTGGAGACTGTTCTAAGTGACTGCTCCTCAGCAGC

AGTTCACCAACCTAAGCGCAAGAGCCATCGTTACATTCCACAGGGAGATCTTGTATGGGCAATCAGTTCA

GACACGATTCATTCTCTCGTGGGACTCTCTGAACTTTTCTTTGTACTTCCGCTTATGAATTCAGGCCTGT

GAGCTTTCAAGGATGTTCTTGCATTGGCGTCACTCGACAGGGCCAACAATTGAGCTGTTGAGAAACAAAA

AAATAAAAGAGGCCTTCGAAGATGGTTTTAGGAGTTGCGTAAAGATTTTTATGGCCATTGATCCGAAATA

TATTGGGCAGACAAATTTGGCGAAGGGGTGGCCAAATGGATGAAAGAATTTTCTGTTATGGAGGCCGTTG

TATTACTGAGAGAAATGCTCTGTGAAAGCGAGAGATAGAGCACACCTTTAAAATCCAAATACAAAAGATG

ATGTTTGCTCAGAGAAAACAGAGGATGTACTTCAACTACTTTAACTTAAAAATTTATGAAATGTTTTGGG

ACTTCTTTTAGTGTGAGCAGCCTTTGAGGCTTTGCAAATTGTAGGATAATGGTGGACAAAAATTGTCTTG

TCTTGATTAAGATATTGTATAAGCTGC

>MSTRG.231.6 gene=MSTRG.231

CAGGCCTTTTAGTTAACCCAATTTATTTTTAGGTATAAAATCACTCGTCGTGTAGGTTAAGGCGGCAGCT

TCTCAATTACATTAGGGCATCGCCTAGATTAGGGCATCGCCGCAACGATCTATGATAATCGTGTGTGTAT

GTATCTATTTGAGTGTAAAGTGCTGAATCGTTTCGTGATGCTATCTGTTTCGAAGTTTAAAAACAGTAAA

CAATGAAGAAGATGCTCCAGAAAAACAAAACAATGATCTGCCTGATTGGGTTGGCAAGGGAAAACAGTAG

TTCTACCAAACCCGCCTCAAAGTGGACAACGAGAAATGCCAACATATTCGTTGCGGAATTCCCACTTTGC

TTGATATTTTGGTCATGCAGCTGATATGAGGCAAAAGAACCCAAAGGCTGGCAACCCATTTAATGAAATT

GCACCTATTAAATGGGAGATAATTTGTGACCACTGCGAGGGATTCCTTGTGACTCGGCCAACTGTGTGCG

AGAGAACATCAGAAGAGCACACAATTGCTGAAGGCGAAAGCATTGGAAGCAGCTATGAATGACATTAATA

ACTCATTTGGAAAAGGAGCTGTAACAAGATTAGGCAATGCTGGTGGATCTCTTGTTGACTCTTTTCCAAG

TGGTTGTCTGACATTGGACATTGCATTGGGTGTCGGCCTTCCTAGAGGAAATATTGTTGAGGTATGTTTT

CGTACACACTTTGAGGTATTGTAAAACAAACTCAGTACAATCTTTTTCACGTGAAGATTATCTTGTTTAA

TATCCAGGATACGTATGGCTCTGAGATGGTGAAGTAAACAATGAAGATGATGCTCCAAAAAGATGGAAAA

CAGTAACTAAGGCTGGCTAATCTGCTACTTATCTGCTATTAAGAAGCTGCTGCTTGTCTACTTGGTTATT

TCGTTGAAGATAAAGCAACGCGACATATCGGGAACTTTTGAAATTGCAAGAACTAAGTTAAAAGCTGGCA

ATGTAGATGATTTTTTTCCAAGTAGTTGAACCTACCAATGTATTCCTTACCATATTGTCTTATTCTTATT

TCTATGTAACATCAATGCTCAAAAGACTTAAAATCTAAGTTACTCAAGTAGTTTTTATATTTGTCTGTTT

ATTTATTTACCTATATAATGTATATAATACGTTTGTGTGATGACAGAGGATCGATTCAGCACTTTGTAAA

AAAAATTCATAGTTTACATGCTACTCACTTGACGTATATGTAATTAATATTATTTTTAAAGTTTTATACT

AAAGAATGTCAAATCTTATATCTATATAAAGCAAAATCTCGGGCGTCTTACGTGGCGCTCTTTACGCTCT

TGAATCAAAATCCCTGCTCTCTGCACTTTTGGCTATTTCTTTTATATTTTCTCTCAGCTTGCCTAGCGGT

TATGTTCAGCCTGGGACTTGGACAGGCACTTCAGTTACCCGCTCCGTAACTGAGGTCTCTGTGCCTCCTC

AGAGATTTACTTATAAATGGCCTTCTCTGGACAATCTTTTCACACCGGGGAAAGCCGATCGGCATGAATA

TGACGGATTCAGCTAGAGGGTCAAAGCGGGATAAAGTCTTAGATTTGACAGAGGCAAGGAAAGGTGAGGA

AAATCCAGGCATCCAGCTGTATGCTCAAAAACAAACCTCCAGCGCAAGTGGAGACTGTTCTAAGTGACTG

CTCCTCAGCAGCAGTTCACCAACCTAAGCGCAAGAGCCATCGTTACATTCCACAGGGAGATCTTGTATGG

GCAATCAGTTCAGACACGATTCATTCTCTCGTGGGACTCTCTGAACTTTTCTTTGTACTTCCGCTTATGA

ATTCAGGTAAGCTGAATTATTATTGCTTGATAGAATGAATTGATAAGCTTTATGGTGCCAGCAAGTAGAA

AGCCAGGTTATATTCGCACAACGTCCTTTATCTGAGGCCACCAAGTCTCTACCTCTTTATACTGCTTTTC

CCAACTGATGTTTTTAAATAAGTTGTATAACTTTGTTGTTCATTCTATGACGATGTGCCTAAGTTTGTCT

GCTTATATATTGTTAGTTCGTAATCGTAATAAATTGTGATTACACATAGATATAGACTATTCTTGTAGAG

ATAAAGTTTAATGTTTTAAAGAATATCTTTTTATTTATAAGATTGTTTTCAGTTCTAGGCCTGTGAGCTT

TCAAGGATGTTCTTGCATTGGCGTCACTCGACAGGGCCAACAATTGAGCTGTTGAGAAACAAAAAAATAA

AAGAGGCCTTCGAAGATGGTTTTAGGAGTTGCGTAAAGATTTTTATGGCCATTGATCCGAAATATATTGG

GCAGACAAATTTGGCGAAGGGGTGGCCAAATGGATGAAAGAATTTTCTGTTATGGAGGCCGTTGTATTAC

TGAGAGAAATGCTCTGTGAAAGCGAGAGATAGAGCACACCTTTAAAATCCAAATACAAAAGATGATGTTT

GCTCAGAGAAAACAGAGGATGTACTTCAACTACTTTAACTTAAAAATTTATGAAATGTTTTGGGACTTCT

TTTAGTGTGAGCAGCCTTTGAGGCTTTGCAAATTGTAGGATAATGGTGGACAAAAATTGTCTTGTCTTGA

TTAAGATATTGTATAAGCTGC

>MSTRG.231.5 gene=MSTRG.231

CAATACTCGTAAGAAAGCCCAGGCCTTTTAGTTAACCCAATTTATTTTTAGGTATAAAATCACTCGTCGT

GTAGGTTAAGGCGGCAGCTTCTCAATTACATTAGGGCATCGCCTAGATTAGGGCATCGCCGCAACGATCT

ATGATAATCGTGTGTGTATGTATCTATTTGAGTGTAAAGTGCTGAATCGTTTCGTGATGCTATCTGTTTC

GAAGTTTAAAAACAGTAAACAATGAAGAAGATGCTCCAGAAAAACAAAACAATGATCTGCCTGATTGGGT

TGGCAAGGGAAAACAGTAGTTCTACCAAACCCGCCTCAAAGTGGACAACGAGAAATGCCAACATATTCGT

TGCGGAATTCCCACTTTGCTTGATATTTTGGTCATGCAGCTGATATGAGGCAAAAGAACCCAAAGGCTGG

CAACCCATTTAATGAAATTGCACCTATTAAATGGGAGATAATTTGTGACCACTGCGAGGGATTCCTTGTG

ACTCGGCCAACTGTGTGCGAGAGAACATCAGAAGAGCACACAATTGCTGAAGGCGAAAGCATTGGAAGCA

GCTATGAATGACATTAATAACTCATTTGGAAAAGGAGCTGTAACAAGATTAGGCAATGCTGGTGGATCTC

TTGTTGACTCTTTTCCAAGTGGTTGTCTGACATTGGACATTGCATTGGGTGTCGGCCTTCCTAGAGGAAA

TATTGTTGAGGATACGTATGGCTCTGAGATGGTGAAGTAAACAATGAAGATGATGCTCCAAAAAGATGGA

AAACAGTAACTAAGGCTGGCTAATCTGCTACTTATCTGCTATTAAGAAGCTGCTGCTTGTCTACTTGGTT

ATTTCGTTGAAGATAAAGCAACGCGACATATCGGGAACTTTTGAAATTGCAAGAACTAAGTTAAAAGCTG

GCAATGTAGATGATTTTTTTCCAAGTAGTTGAACCTACCAATGTATTCCTTACCATATTGTCTTATTCTT

ATTTCTATGTAACATCAATGCTCAAAAGACTTAAAATCTAAGTTACTCAAGTAGTTTTTATATTTGTCTG

TTTATTTATTTACCTATATAATGTATATAATACGTTTGTGTGATGACAGAGGATCGATTCAGCACTTTGT

AAAAAAAATTCATAGTTTACATGCTACTCACTTGACGTATATGTAATTAATATTATTTTTAAAGTTTTAT

ACTAAAGAATGTCAAATCTTATATCTATATAAAGCAAAATCTCGGGCGTCTTACGTGGCGCTCTTTACGC

TCTTGAATCAAAATCCCTGCTCTCTGCACTTTTGGCTATTTCTTTTATATTTTCTCTCAGCTTGCCTAGC

GGTTATGTTCAGCCTGGGACTTGGACAGGCACTTCAGTTACCCGCTCCGTAACTGAGGTCTCTGTGCCTC

CTCAGAGATTTACTTATAAATGGCCTTCTCTGGACAATCTTTTCACACCGGGGAAAGCCGATCGGCATGA

ATATGACGGATTCAGCTAGAGGGTCAAAGCGGGTATGTATGTTATTCTTGCTGATTTTCAGGGTTGATTT

ATGTTATATGATCATAGTCGTGGGCAGTTATCTCCTTCTGATTTATCCGGAGAGCATAGCAGCAGATTTT

TATGTATGCGCAGTTCGTTGTCTTTATCAATTTCTATGTGTATTTGAATTGGTGTCTGTGTTTCATTAGG

GTTTTGATGCTTAATTTATCGTGGATTTTGATTTTTTAATCGGTGATGATTAGGATAAAGTCTTAGATTT

GACAGAGGCAAGGAAAGGTGAGGAAAATCCAGGCATCCAGCTGTATGCTCAAAAACAAACCTCCAGCGCA

AGTGGAGACTGTTCTAAGTGACTGCTCCTCAGCAGCAGTTCACCAACCTAAGCGCAAGAGCCATCGTTAC

ATTCCACAGGGAGATCTTGTATGGGCAATCAGTTCAGACACGATTCATTCTCTCGTGGGACTCTCTGAAC

TTTTCTTTGTACTTCCGCTTATGAATTCAGGTAAGCTGAATTATTATTGCTTGATAGAATGAATTGATAA

GCTTTATGGTGCCAGCAAGTAGAAAGCCAGGTTATATTCGCACAACGTCCTTTATCTGAGGCCACCAAGT

CTCTACCTCTTTATACTGCTTTTCCCAACTGATGTTTTTAAATAAGTTGTATAACTTTGTTGTTCATTCT

ATGACGATGTGCCTAAGTTTGTCTGCTTATATATTGTTAGTTCGTAATCGTAATAAATTGTGATTACACA

TAGATATAGACTATTCTTGTAGAGATAAAGTTTAATGTTTTAAAGAATATCTTTTTATTTATAAGATTGT

TTTCAGTTCTAGGCCTGTGAGCTTTCAAGGATGTTCTTGCATTGGCGTCACTCGACAGGGCCAACAATTG

AGCTGTTGAGAAACAAAAAAATAAAAGAGGCCTTCGAAGATGGTTTTAGGAGTTGCGTAAAGATTTTTAT

GGCCATTGATCCGAAATATATTGGGCAGACAAATTTGGCGAAGGGGTGGCCAAATGGATGAAAGAATTTT

CTGTTATGGAGGCCGTTGTATTACTGAGAGAAATGCTCTGTGAAAGCGAGAGATAGAGCACACCTTTAAA

ATCCAAATACAAAAGATGATGTTTGCTCAGAGAAAACAGAGGATGTACTTCAACTACTTTAACTTAAAAA

TTTATGAAATGTTTTGGGACTTCTTTTAGTGTGAGCAGCCTTTGAGGCTTTGCAAATTGTAGGATAATGG

TGGACAAAAATTGTCTTGTCTTGATTAAGATATTGTATAAGCTGC

>MSTRG.231.9 gene=MSTRG.231

CAGGCCTTTTAGTTAACCCAATTTATTTTTAGGTATAAAATCACTCGTCGTGTAGGTTAAGGCGGCAGCT

TCTCAATTACATTAGGGCATCGCCTAGATTAGGGCATCGCCGCAACGATCTATGATAATCGTGTGTGTAT

GTATCTATTTGAGTGTAAAGTGCTGAATCGTTTCGTGATGCTATCTGTTTCGAAGTTTAAAAACAGTAAA

CAATGAAGAAGATGCTCCAGAAAAACAAAACAATGATCTGCCTGATTGGGTTGGCAAGGGAAAACAGTAG

TTCTACCAAACCCGCCTCAAAGTGGACAACGAGAAATGCCAACATATTCGTTGCGGAATTCCCACTTTGC

TTGATATTTTGGTCATGCAGCTGATATGAGGCAAAAGAACCCAAAGGCTGGCAACCCATTTAATGAAATT

GCACCTATTAAATGGGAGATAATTTGTGACCACTGCGAGGGATTCCTTGTGACTCGGCCAACTGTGTGCG

AGAGAACATCAGAAGAGCACACAATTGCTGAAGGCGAAAGCATTGGAAGCAGCTATGAATGACATTAATA

ACTCATTTGGAAAAGGAGCTGTAACAAGATTAGGCAATGCTGGTGGATCTCTTGTGTATGTTGACTCTTT

TCCAAGTGGTTGTCTGACATTGGACATTGCATTGGGTGTCGGCCTTCCTAGAGGAAATATTGTTGAGGTA

TGTTTTCGTACACACTTTGAGGTATTGTAAAACAAACTCAGTACAATCTTTTTCACGTGAAGATTATCTT

GTTTAATATCCAGGATACGTATGGCTCTGAGATGGTGAAGTAAACAATGAAGATGATGCTCCAAAAAGAT

GGAAAACAGTAACTAAGGCTGGCTAATCTGCTACTTATCTGCTATTAAGAAGCTGCTGCTTGTCTACTTG

GTTATTTCGTTGAAGATAAAGCAACGCGACATATCGGGAACTTTTGAAATTGCAAGAACTAAGTTAAAAG

CTGGCAATGTAGATGATTTTTTTCCAAGTAGTTGAACCTACCAATGTATTCCTTACCATATTGTCTTATT

CTTATTTCTATGTAACATCAATGCTCAAAAGACTTAAAATCTAAGTTACTCAAGTAGTTTTTATATTTGT

CTGTTTATTTATTTACCTATATAATGTATATAATACGTTTGTGTGATGACAGAGGATCGATTCAGCACTT

TGTAAAAAAAATTCATAGTTTACATGCTACTCACTTGACGTATATGTAATTAATATTATTTTTAAAGTTT

TATACTAAAGAATGTCAAATCTTATATCTATATAAAGCAAAATCTCGGGCGTCTTACGTGGCGCTCTTTA

CGCTCTTGAATCAAAATCCCTGCTCTCTGCACTTTTGGCTATTTCTTTTATATTTTCTCTCAGCTTGCCT

AGCGGTTATGTTCAGCCTGGGACTTGGACAGGCACTTCAGTTACCCGCTCCGTAACTGAGGTCTCTGTGC

CTCCTCAGAGATTTACTTATAAATGGCCTTCTCTGGACAATCTTTTCACACCGGGGAAAGCCGATCGGCA

TGAATATGACGGATTCAGCTAGAGGGTCAAAGCGGGTATGTATGTTATTCTTGCTGATTTTCAGGGTTGA

TTTATGTTATATGATCATAGTCGTGGGCAGTTATCTCCTTCTGATTTATCCGGAGAGCATAGCAGCAGAT

TTTTATGTATGCGCAGTTCGTTGTCTTTATCAATTTCTATGTGTATTTGAATTGGTGTCTGTGTTTCATT

AGGGTTTTGATGCTTAATTTATCGTGGATTTTGATTTTTTAATCGGTGATGATTAGGATAAAGTCTTAGA

TTTGACAGAGGCAAGGAAAGGTGAGGAAAATCCAGGCATCCAGCTGTATGCTCAAAAACAAACCTCCAGC

GCAAGTGGAGACTGTTCTAAGTGACTGCTCCTCAGCAGCAGTTCACCAACCTAAGCGCAAGAGCCATCGT

TACATTCCACAGGGAGATCTTGTATGGGCAATCAGTTCAGACACGATTCATTCTCTCGTGGGACTCTCTG

AACTTTTCTTTGTACTTCCGCTTATGAATTCAGGTAAGCTGAATTATTATTGCTTGATAGAATGAATTGA

TAAGCTTCTTTATCCTTGCTATATGCTATGGTAACTCTCTCCACATATGCACACAGCCACACACTTAACT

AACTTCGTATCAGACTCTTTGAGGCATTTAATTTTTGCATCAACTAGCTACTATTGCTCTACCTGAT

>MSTRG.231.10 gene=MSTRG.231

AAAGCCCAGGCCTTTTAGTTAACCCAATTTATTTTTAGGTATAAAATCACTCGTCGTGTAGGTTAAGGCG

GCAGCTTCTCAATTACATTAGGGCATCGCCTAGATTAGGGCATCGCCGCAACGATCTATGATAATCGTGT

GTGTATGTATCTAGAGTGTAAAGTGCTGAATCGTTTCGTGATGCTATCTGTTTCGAAGTCTAAAAACAGT

AAACAATGAAGAAGATGCTCCAGAAAAACAAAACAATGATCTGCCTGATTGGGTTGGCAAGGGAAAACAG

TAGTTCTACCAAACCCGCCTCAAAGTGGACAACGAGAAATGCCAACATATTCGTTGCGGAATTCCCACTT

TGCTTGATATTTTGGTCATGCAGCTGATATGAGGCAAAAGAACCCAAAGGCTGGCAACCCATTTAATGAA

ATTGCACCTATTAAATGGGAGATAATTTGTGACCACTGCGAGGGATTCCTTGTGACTCGGCCAACTGTGT

GCGAGAGAACATCAGAAGAGCACACAATTGCTGAAGGCGAAAGCATTGGAAGCAGCTATGAATGACATTA

ATAACTCATTTGGAAAAGGAGCTGTAACAAGATTAGGCAATGCTGGTGGATCTCTTGTTGACTCTTTTCC

AAGTGGTTGTCTGACATTGGACATTGCATTGGGTGTCGGCCTTCCTAGAGGAAATATTGTTGAGGTATGT

TTTCGTACACACTTTGAGGTATTGTAAAACAAACTCAGTACAATCTTTTTCACGTGAAGATTATCTTGTT

TAATATCCAGGATACGTATGGCTCTGAGATGGTGAAGTAAACAATGAAGATGATGCTCCAAAAAGATGGA

AAACAGTAACTAAGGCTGGCTAATCTGCTACTTATCTGCTATTAAGAAGCTGCTGCTTGTCTACTTGGTT

ATTTCGTTGAAGATAAAGCAACGCGACATATCGGGAACTTTTGAAATTGCAAGAACTAAGTTAAAAGCTG

GCAATGTAGATGATTTTTTTCCAAGTAGTTGAACCTACCAATGTATTCCTTACCATATTGTCTTATTCTT

ATTTCTATGTAACATCAATGCTCAAAAGACTTAAAATCTAAGTTACTCAAGTAGTTTTTATATTTGTCTG

TTTATTTATTTACCTATATAATGTATATAATACGTTTGTGTGATGACAGAGGATCGATTCAGCACTTTGT

AAAAAAAATTCATAGTTTAC

>MSTRG.231.12 gene=MSTRG.231

GCCTTTTAGTTAACCCAATTTATTTTTAGGTATAAAATCACTCGTCGTGTAGGTTAAGGCGGCAGCTTCT

CAATTACATTAGGGCATCGCCTAGATTAGGGCATCGCCGCAACGATCTATGATAATCGTGTGTGTATGTA

TCTAGAGTGTAAAGTGCTGAATCGTTTCGTGATGCTATCTGTTTCGAAGTCTAAAAACAGTAAACAATGA

AGAAGATGCTCCAGAAAAACAAAACAATGATCTGCCTGATTGGGTTGGCAAGGGAAAACAGTAGTTCTAC

CAAACCCGCCTCAAAGTGGACAACGAGAAATGCCAACATTTTCGTTGCGGAATTCCCACTTTGCTTGATA

TTTTGGTCATGCAGCTGATATGAGGCAAAAGAACCCAAAGGCTGGCAACCCATTTAATGAAATTGCACCT

ATTAAATGGGAGATAATTTGTGACCACTGCGAGGGATTCCTTGTGACTCGGCCAACTGTGTGCGAGAGAA

CATCAGAAGAGCACACAATTGCTGAAGGCGAAAGCATTGGAAGCAGCTATGAATGACATTAATAACTCAT

TTGGAAAAGGAGCTGTAACAAGATTAGGCAATGCTGGTGGATCTCTTGTGTATGTTGACTCTTTTCCAAG

TGGTTGTCTGACATTGGACATTGCATTGGGTGTCGGCCTTCCTAGAGGAAATATTGTTGAGGTATGTTTT

CGTACACACTTTGAGGTATTGTAAAACAAACTCAGTACAATCTTTTTCACGTGAAGATTATCTTGTTTAA

TATCCAGGATACGTATGGCTCTGAGATGGTGAAGTAAACAATGAAGATGATGCTCCAAAAAGATGGAAAA

CAGTAACTAAGGCTGGCTAATCTGCTACTTATCTGCTATTAAGAAGCTGCTGCTTGTCTACTTGGTTATT

TCGTTGAAGATAAAGCAACGCGACATATCGGGAACTTTTGAAATTGCAAGAACTAAGTTAAAAGCTGGCA

ATGTAGATGATTTTTTTCCAAGTAGTTGAACCTACCAATGTATTCCTTACCATATTGTCTTATTCTTATT

TCTATGTAACATCAATGCTCAAAAGACTTAAAATCTAAGTTACTCAAGTAGTTTTTATATTTGTCTGTTT

ATTTATTTACCTATATAATGTATATAATACGTTTGTGTGATGACAGAGGATCGATTCAGCAC

>MSTRG.231.11 gene=MSTRG.231

AGGCCTTTTAGTTAACCCAATTTATTTTTAGGTATAAAATCACTCGTCGTGTAGGTTAAGGCGGCAGCTT

CTCAATTACATTAGGGCATCGCCTAGATTAGGGCATCGCCGCAACGATCTATGATAATCGTGTGTGTATG

TATCTAGAGTGTAAAGTGCTGAATCGTTTCGTGATGCTATCTGTTTCGAAGTCTAAAAACAGTAAACAAT

GAAGAAGATGCTCCAGAAAAACAAAACAATGATCTGCCTGATTGGGTTGGCAAGGGAAAACAGTAGTTCT

ACCAAACCCGCCTCAAAGTGGACAACGAGAAATGCCAACATATTCGTTGCGGAATTCCCACTTTGCTTGA

TATTTTGGTCATGCAGCTGATATGAGGCAAAAGAACCCAAAGGCTGGCAACCCATTTAATGAAATTGCAC

CTATTAAATGGGAGATAATTTGTGACCACTGCGAGGGATTCCTTGTGACTCGGCCAACTGTGTGCGAGAG

AACATCAGAAGAGCACACAATTGCTGAAGGCGAAAGCATTGGAAGCAGCTATGAATGACATTAATAACTC

ATTTGGAAAAGGAGCTGTAACAAGATTAGGCAATGCTGGTGGATCTCTTGTGTATGTTGACTCTTTTCCA

AGTGGTTGTCTGACATTGGACATTGCATTGGGTGTCGGCCTTCCTAGAGGAAATATTGTTGAGGATACGT

ATGGCTCTGAGATGGTGAAGTAAACAATGAAGATGATGCTCCAAAAAGATGGAAAACAGTAACTAAGGCT

GGCTAATCTGCTACTTATCTGCTATTAAGAAGCTGCTGCTTGTCTACTTGGTTATTTCGTTGAAGATAAA

GCAACGCGACATATCGGGAACTTTTGAAATTGCAAGAACTAAGTTAAAAGCTGGCAATGTAGATGATTTT

TTTCCAAGTAGTTGAACCTACCAATGTATTCCTTACCATATTGTCTTATTCTTATTTCTATGTAACATCA

ATGCTCAAAAGACTTAAAATCTAAGTTACTCAAGTAGTTTTTATATTTGTCTGTTTATTTATTTACCTAT

ATAATGTATATAATACGTTTGTGTGATGACAGAGGATCGATTCAGCAC

>MSTRG.231.13 gene=MSTRG.231

CAGGCCTTTTAGTTAACCCAATTTATTTTTAGGTATAAAATCACTCGTCGTGTAGGTTAAGGCGGCAGCT

TCTCAATTACATTAGGGCATCGCCTAGATTAGGGCATCGCCGCAACGATCTATGATAATCGTGTGTGTAT

GTATCTAGAGTGTAAAGTGCTGAATCGTTTCGTGATGCTATCTGTTTCGAAGTCTAAAAACAGTAAACAA

TGAAGAAGATGCTCCAGAAAAACAAAACAATGATCTGCCTGATTGGGTTGGCAAGGGAAAACAGTAGTTC

TACCAAACCCGCCTCAAAGTGGACAACGAGAAATGCCAACATATTCGTTGCGGAATTCCCACTTTGCTTG

ATATTTTGGTCATGCAGCTGATATGAGGCAAAAGAACCCAAAGGCTGGCAACCCATTTAATGAAATTGCA

CCTATTAAATGGGAGATAATTTGTGACCACTGCGAGGGATTCCTTGTGACTCGGCCAACTGTGTGCGAGA

GAACATCAGAAGAGCACACAATTGCTGAAGGCGAAAGCATTGGAAGCAGCTATGAATGACATTAATAACT

CATTTGGAAAAGGAGCTGTAACAAGATTAGGCAATGCTGGTGGATCTCTTGTTGACTCTTTTCCAAGTGG

TTGTCTGACATTGGACATTGCATTGGGTGTCGGCCTTCCTAGAGGAAATATTGTTGAGGATACGTATGGC

TCTGAGATGGTGAAGTAAACAATGAAGATGATGCTCCAAAAAGATGGAAAACAGTAACTAAGGCTGGCTA

ATCTGCTACTTATCTGCTATTAAGAAGCTGCTGCTTGTCTACTTGGTTATTTCGTTGAAGATAAAGCAAC

GCGACATATCGGGAACTTTTGAAATTGCAAGAACTAAGTTAAAAGCTGGCAATGTAGATGATTTTTTTCC

AAGTAGTTGAACCTACCAATGTATTCCTTACCATATTGTCTTATTCTTATTTCTATGTAACATCAATGCT

CAAAAGACTTAAAATCTAAGTTACTCAAGTAGTTTTTATATTTGTCTGTTTATTTATTTACCTATATAAT

GTATATAATACGTTTGTGTGATGACAGAG

>MSTRG.231.14 gene=MSTRG.231

CCCATACCTGTATATCATTTTATTTCTAAAAAGACATGATAAGATTGTTGTTAGGAATTAATCTAATGTC

CATATTTGATGTAAATATTTTGTTACAGGCGAAAGCATTGGAAGCAGCTATGAATGACATTAATAACTCA

TTTGGAAAAGGAGCTGTAACAAGATTAGGCAATGCTGGTGGATCTCTTGTGTATGTTGACTCTTTTCCAA

GTGGTTGTCTGACATTGGACATTGCATTGGGTGTCGGCCTTCCTAGAGGAAATATTGTTGAGGATACGTA

TGGCTCTGAGATGGTGAAGTAAACAATGAAGATGATGCTCCAAAAAGATGGAAAACAGTAACTAAGGCTG

GCTAATCTGCTACTTATCTGCTATTAAGAAGCTGCTGCTTGTCTACTTGGTTATTTCGTTGAAGATAAAG

CAACGCGACATATCGGGAACTTTTGAAATTGCAAGAACTAAGTTAAAAGCTGGCAATGTAGATGATTTTT

TTCCAAGTAGTTGAACCTACCAATGTATTCCTTACCATATTGTCTTATTCTTATTTCTATGTAACATCAA

TGCTCAAAAGACTTA

>MSTRG.232.1 gene=MSTRG.232

AAGCTAAATGAGTGTCAGAGATACCATCATATCTCAGTAAATCAGTGTTTGTCGAGCACAAATTGTTCTT

GCCGATGATTAGTTGCCCAAAAATACTGTAAGACATACTGTAAAAAGAATTCAAGAGGCCGATAACAGCA

GCGGAGATAATAAAGGCCGTGATTCCCAGTTGCAAGGTCTGCAATTTTTTCTGTCTTCGAGTCTTTGGAA

TGTTGCTGTTATGGAATGCCCAAGGAAAATATAACTTCGGAATCAANCGAGTCTTTGGAATGTTGCTGTC

ATGGAATGCCCGAGGGAAATATAACTTCGGAATCAAGAAGTTTCGAAAGGTAGCTTGGCAAATCAGCCCA

ACCAAAGCTGCTCAGAAGTACAACTTAGATACAAGTTGTTTTTTCAAATATGTGTGAGTCCTTGTATTGA

TATCTATCAGGCTAACGAGTGATGCCACTATGGATATGATCTTCTAGCACGTATGTATATTAATGTGTTA

ACTGTACCAATTGTTAGTATTGGTGTCCTAGAGACAATACTATTGTGTTCTATCGTAGACATTTATGGAT

TATGTTATATTGATTAT

>MSTRG.233.1 gene=MSTRG.233

GTTCCGCATTCTATAGTAAGGTATTCATTATCGAACTGATACCCGACTTCCCAAATATTGAACTCTCAGG

TGTCAGAGATACCATCATATCTCAGTAAATCAGTGTTTGTCGAGCACAAATTGTTCTTGCCGATGATTAG

TTGCCCAAAAATACTGTAAGACATACTGTAAAAAGAATTCAAGAGGCCGATAACAGCAGCGGAGATAATA

AAGGCCGTGATTCCCAGTTGCAAGGTCTGCATTTTTTCTGTCTTCGAGTCTTTGGAATGTTGCTGTCATG

GAATGCCCGAGGGAAATATAACTTCGGAATCAAGAAGTTTCGAAAGGTAGCTTGGCAAATC

>MSTRG.232.2 gene=MSTRG.232

GTGTTGAAAAGGTGAAACCTCCAACCGGTAGCCCAATATCTATGTTAATGGCTACTCCAGATGGCTCATA

TGCATTACCAAGCGGTTGATGTTCGTAAGCTAAATGAGTGTCAGAGATACCATCATATCTCAGTAAATCA

GTGTTTGTCGAGCACAAATTGTTCTTGCCGATGATTAGTTGCCCAAAAATACTGTAAGACATACTGTAAA

AAGAATTCAAGAGGCCGATAACAGCAGCGGAGATAATAAAGGCCGTGATTCCCAGTTGCAAGGTCTGCAA

TTTTTTCTGTCTTCGAGTCTTTGGAATGTTGCTGTTATGGAATGCCCAAGGAAAATATAACTTCGGAATC

AANCGAGTCTTTGGAATGTTGCTGTCATGGAATGCCCGAGGGAAATATAACTTCGGAATCAAGAAGTTTC

GAAAGGTAGCTTGGCAAATCAGCCCAACCAAAGCTGCTCAGAAGTACAACTTAGATACAAGTTGTTTTTT

CAAATATGTGTGAGTCCTTGTATTGATATCTATCAGGCTAACGAGTGATGCCACTATGGATATGATCTTC

TAGCACGTATGTATATTAATGTGTTAACTGTACCAAATATTTATCATTTAATATTTTTATTCACGGCAAT

GTTGGTTTTAACCGATATTAAGAAAATGAAA

>MSTRG.234.1 gene=MSTRG.234

CATCCATGGCGTTACTCAGCAACGCTCTCCGGCAAGCCTTCATGCCGAAGCACGAGTACGACAATCTGCG

CGAAGAAGACAAAGCGATGATCCAATTACAACGCCCCGTCTTGATTTCCCTCTTGTTGTGTATCGTGATT

GTGATCGTTGTGTCCACCTCGATTAGCGTGAAGATTGTGTTCCCGGCGGAGGACGGGAAACGCGTGTTTT

GCCGGGATTTGAGGATTCAGCCGTTGTCGATAAATGTGAGCTCTGGTGGCGGTGGCGGCGGGGAGGATGT

GTTTCCTGGGGCGTTTTATTTGACGGATCAGCAGACGGTGGATTATTATTGGATGGTGGTGTTTATGCC

>MSTRG.235.1 gene=MSTRG.235

TTTTTTTATTCAGTGTAGATATTATTTTTTTGCATCTCAATATACGTGTTAAAAAGTCAAAGTCGGTTAA

AATGTAACCGGGACAGAGGGAATATTAAACTTTCTGTCGTAGTATATTGTATAGTTTGGTTGAAAGTTAT

TAGTTTGTGATATATCTTTACTTCTTTTTAAAGGTATGTTTTCGTACACACTTTGAGGTATTGTAAAACA

AACTCAGTACAATCTTTTTCACGTGAAGATTATCTTGTTTAATATCCAGGATACGTATGGCTCTGAGATG

GTGAAGTAAACAATGAAGATGATGCTCCAAAAAGATGGAAAACAGTAACTAAGGCTGGCTAATCTGCTAC

TTATCTGCTATTAAGAAGCTGCTGCTTGTCTACTTGGTTATTTCGTTGAAGATAAAGCAACGCGATGTAT

TCCTTACCATATTGTCTTATTCTTATTTCTATGTAACATCAATGCTCAAAAGACTTAAAATCTAAGTTAC

TCAAGTAGTTTTTATATTTGTCTGTTTATTTATTTACCTATATAATGTATATAATACGTTTGTGTGATGA

CAGAGGATCGATTCAGCACTTTGTAAAAAAAATTCATAGTTTACATGCTACTCACTTGACGTATATGTAA

TTAATATTATTTTTAAAATTTTATACTAAAGAATGTCAAATCTTATATCTATATAAAGCAAAATCTCGGG

CGTCTTACGTGGCGCTCTTTACGCTCTTGAATCAAAATCCCTGCTCTCTGCACTTTTGGCTATTTCTTTT

ATATTTTCTCTCAGCTTGCCTAGCGGTTATGTTCAGCCTGGGACTTGGACAGGCACTTCAGTTACCCGCT

CCGTAACTGAGGTCTCTGTGCCTCCTCAGAGATTTACTTATAAATGGCCTTCTCTGGACAATCTTTTCAC

ACCGGGGAAAGCCGATCGGCATGAATATGACGGATTCAGCTAGAGGGTCAAAGCGGGATAAAGTCTTAGA

TTTGACAGAGGCAAGGAAAGGTGAGGAAAATCCAGGCATCCAGCTGTATGCTCAAAAACAAACCTCCAGC

GCAAGTGGAGACTGTTCTAAGTGACTGCTCCTCAGCAGCAGTTCACCAACCTAAGCGCAAGAGCCATCGT

TACATTCCACAGGGAGATCTTGTATGGGCAATCAGTTCAGACACGATTCATTCTCTCGTGGGACTCTCTG

AACTTTTCTTTGTACT

>MSTRG.235.2 gene=MSTRG.235

CTATTATTTGGTATAAGAATTGAATTTTTTTATTCAGTGTAGATATTATTTTTTTGCATCTCAATATACG

TGTTAAAAAGTCAAAGTCGGTTAAAATGTAACCGGGACAGAGGGAATATTAAACTTTCTGTCGTAGTATA

TTGTATAGTTTGGTTGAAAGTTATTAGTTTGTGATATATCTTTACTTCTTTTTAAAGGTATGTTTTCGTA

CACACTTTGAGGTATTGTAAAACAAACTCAGTACAATCTTTTTCACGTGAAGATTATCTTGTTTAATATC

CAGGATACGTATGGCTCTGAGATGGTGAAGGTATGCTTGTTATATAGTTATAATGTACTTGAAAATCTAT

TAAGTTTTCTAGTATTCAATTCACTGGTTTTCAAATTTCTTCTTGCAGTAAACAATGAAGATGATGCTCC

AAAAAGATGGAAAACAGTAACTAAGGCTGGCTAATCTGCTACTTATCTGCTATTAAGAAGCTGCTGCTTG

TCTACTTGGTTATTTCGTTGAAGATAAAGCAACGCGACATATCGGGAACTTTTGAAATTGCAAGAACTAA

GTTAAAAGCTGGCAATGTAGATGATTTTTTTCCAAGTAGTTGAACCTACCAATGTATTCCTTACCATATT

GTCTTATTCTTATTTCTATGTAACATCAATGCTCAAAAGACTTAAAATCTAAGTTACTCAAGTAGTTTTT

ATATTTGTCTGTTTATTTATTTACCTATATAATGTATATAATACGTTTGTGTGATGACAGAGGATCGATT

CAGCACTTTGTAAAAAAAATTCATAGTTTACATGCTACTCACTTGACGTATATGTAATTAATATTATTTT

TAAAATTTTATACTAAAGAATGTCAAATCTTATATCTATATAAAGCAAAATCTCGGGCGTCTTACGTGGC

GCTCTTTACGCTCTTGAATCAAAATCCCTGCTCTCTGCACTTTTGGCTATTTCTTTTATATTTTCTCTCA

GCTTGCCTAGCGGTTATGTTCAGCCTGGGACTTGGACAGGCACTTCAGTTACCCGCTCCGTAACTGAGGT

CTCTGTGCCTCCTCAGAGATTTACTTATAAATGGCCTTCTCTGGACAATCTTTTCACACCGGGGAAAGCC

GATCGGCATGAATATGACGGATTCAGCTAGAGGGTCAAAGCGGGTATGTATGTTATTCTTGCTGATTTTC

AGGGTTGATTTATGTTATATGATCATAGTCGTGGGCAGTTATCTCCTTCTGATTTATCCGGAGAGCATAG

CAGCAGATTTTTATGTATGCGCAGTTCGTTGTCTTTATCAATTTCTATGTGTATTTGAATTGGTGTCTGT

GTTTCATTAGGGTTTTGATGCTTAATTTATCGTGGATTTTGATTTTTTAATCGGTGATGATTAGGATAAA

GTCTTAGATTTGACAGAGGCAAGGAAAGGTGAGGAAAATCCAGGCATCCAGCTGTATGCTCAAAAACAAA

CCTCCAGCGCAAGTGGAGACTGTTCTAAGTGACTGCTCCTCAGCAGCAGTTCACCAACCTAAGCGCAAGA

GCCATCGTTACATTCCACAGGGAGATCTTGTATGGGCAATCAGTTCAGACACGATTCATTCTCTCGTGGG

ACTCTCT

>MSTRG.235.3 gene=MSTRG.235

CATTACTCGTAAGAAAGCCCAGGCCTTTTAGTTAACCCAATTTATTTTTAGGTATAAAATCACTCGTCGT

GTAGGTTAAGGCGGCAGCTTCTCAATTACATTAGGGCATCGCCTAGATTAGGGCATCGCCGCAACGATCT

ATGATAATCGTGTGTGTATGTATCTAGAGTGTAAAGTGCTGAATCGTTTCGTGATGCTATCTGTTTCGAA

GTCTAAAAACAGTAAACAATGAAGAAGATGCTCCAGAAAAACAAAACAATGATCTGCCTGATTGGGTTGG

CAAGGGAAAACAGTAGTTCTACCAAACCCGCCTCAAAGTGGACAACGAGAAATGCCAACATTTTCGTTGC

GGAATTCCCACTTTGCTTGATATTTTGGTCATGCAGCTGATATGAGGCAAAAGAACCCAAAGGCTGGCAA

CCCATTTAATGAAATTGCACCTATTAAATGGGAGATAATTTGTGACCACTGCGAGGGATTCCTTGTGACT

CGGCCAACTGTGTGCGAGAGAACATCAGAAGAGCACACAATTGCTGAAGGTATGTTTTCGTACACACTTT

GAGGTATTGTAAAACAAACTCAGTACAATCTTTTTCACGTGAAGATTATCTTGTTTAATATCCAGGATAC

GTATGGCTCTGAGATGGTGAAGTAAACAATGAAGATGATGCTCCAAAAAGATGGAAAACAGTAACTAAGG

CTGGCTAATCTGCTACTTATCTGCTATTAAGAAGCTGCTGCTTGTCTACTTGGTTATTTCGTTGAAGATA

AAGCAACGCGACATATCGGGAACTTTTGAAATTGCAAGAACTAAGTTAAAAGCTGGCAATGTAGATGATT

TTTTTCCAAGTAGTTGAACCTACCAATGTATTCCTTACCATATTGTCTTATTCTTATTTCTATGTAACAT

CAATGCTCAAAAGACTTAAAATCTAAGTTACTCAAGTAGTTTTTATATTTGTCTGTTTATTTATTTACCT

ATATAATGTATATAATACGTTTGTGTGATGACAGAGGATCGATTCAG

>MSTRG.236.1 gene=MSTRG.236

CCGCACAATCCATCTTCATTCAGGTTCAGGCCCAAGCCCAGTTAGTCATTGTTTTCTCCGCTGCTTGTGT

GTATATGTACGTATACGTGTGAGATGAGGATGAATCAATGGCGATTCCAAAGATTTAAGATCAATTCTCA

CACTCTACATTGTTATGATACTTGTTTCTATCGCTTCCACTCTCAAATTTTAAACAATGAAGAAAGTGCT

TCAGGAAGCCGATAAGAATGGTCGAAGAAAGCGCTTAATTAATAAATAAGGTAGTGGCATCAATGGACTG

GCCTGAGGTCACCACATACAGAGAACTCTACACACATGGTCAGTGAGAGGAAATATTAAGGAATCTCTAC

TGCACTTGTGAAACTGAAAGGGGTGTTGTTGCTAGTGGCATGACCAGGGACTATTTGATGGCATTCTACC

AGCAAACAGAACTCCGACCTAAACGGATCATATTCTACAGGGATGGAGTCAGTGAAGGTCAGTTTAATCA

GGTCTTGCTGAAAGAGATGGATGCCATTAGGACGGTTGATGTGTTTTATTTTTTGTCTGCCAGGCGCTGC

AATCAAATTATTTACAATGGCAGGAAATCAACTGATCGGAGTGGCAACATATCACCTGGTACTGTTGTAG

ATATGACCATCTGCCATCCCACTGAGTTTGACTTTTATTTGTGTAGCCATGCTGGTATACAGGACGGACT

TTGACTTTGTTGTCGGCAATGCACAGACTAATGAGGGTGTTGAAAAGGTGAAACCTCCAACCGGTAGCCC

AATATCTATGTTAATGGCTACTCCAGATGGCTCATATGCATTACCAAGCAGTTGATGTTTTTAAGCTAAA

TGAGTGTCAGAGATACCATCATATCTCAGTAAATCAGTGTTTGTCGAGCACAAAATGTTCTTGCCGATGA

TTAGTTGCCCAAAAATATTGTAAGACATACTGTAAAAAGAATTCAAGAGGCCCATAACAGCAGCGGAGAT

AATAAAGGCCGTGATTCCCAGTTGCAAGGTCTGCATTTTTTCTGTCTTTGGAATGTTGCTGTCATGGAAT

GCCCGAGGAAAATATAACTTCGGAATCAAGAAGTTTCGAAAGGTAGCTTCGCAAATCAGCCCAACCAAAG

CTGCTCAGAAGTACAACTTAGATGCAAGTTTTTTTTTCAAATATGTGTGAGTCCTTGTATTGATATCTAT

CAGGCTAACGAGTGATGCCACTATGGATATGATCTTCTAGTGAAATTTGTATATACATTCATTTATGCAC

GTATGTATATTGTGTTAACTGTGCCAAATATTTATCATTTAATATTTTTATTCACGGCAA

>MSTRG.236.7 gene=MSTRG.236

CCGCACAATCCATCTTCATTCAGGTTCAGGCCCAAGCCCAGTTAGTCATTGTTTTCTCCGCTGCTTGTGT

GTATATGTACGTATACGTGTGAGATGAGGATGAATCAATGGCGATTCCAAAGATTTAAGATCAATTCTCA

CACTCTACATTGTTATGATACTTGTTTCTATCGCTTCCACTCTCAAATTTTAAACAATGAAGAAAGTGCT

TCAGGAAGCCGATAAGAATGGTCGAAGAAAGCGCTTAATTAATAAATAAGGTAGTGGCATCAATGGACTG

GCCTGAGGTCACCACATACAGAGAACTCTACACACATGGTCAGTGAGAGGAAATATTAAGGAATCTCTAC

TGCACTTGTGAAACTGAAAGGGGTGTTGTTGCTAGTGGCATGACCAGGGACTATTTGATGGCATTCTACC

AGCAAACAGAACTCCGACCTAAACGGATCATATTCTACAGGGATGGAGTCAGTGAAGGTCAGTTTAATCA

GGTCTTGCTGAAAGAGATGGATGCCATTAGGACGGTTGATGTGTTTTATTTTTTGTCTGCCAGGCGCTGC

AATCAAATTATTTACAATGGCAGGAAATCAACTGATCGGAGTGGCAACATATCACCTGGTTACATATTGT

TCTATTGTATCTTTAAAAATCTTTATTATTCTGATTAGCAAATAACTGGCTTTTGTATACCAGGTACTGT

TGTAGATATGACCATCTGCCATCCCACTGAGTTTGACTTTTATTTGTGTAGCCATGCTGGTATACAGGGT

GTTGAAAAGGTGAAACCTCCAACCGGTAGCCCAATATCTATGTTAATGGCTACTCCAGATGGCTCATATG

CATTACCAAGCAGTTGATGTTTTTAAGCTAAATGAGTGTCAGAGATACCATCATATCTCAGTAAATCAGT

GTTTGTCGAGCACAAAATGTTCTTGCCGATGATTAGTTGCCCAAAAATATTGTAAGACATACTGTAAAAA

GAATTCAAGAGGCCCATAACAGCAGCGGAGATAATAAAGGCCGTGATTCCCAGTTGCAAGGTCTGCATTT

TTTCTGTCTTTGGAATGTTGCTGTCATGGAATGCCCGAGGAAAATATAACTTCGGAATCAAGAAGTTTCG

AAAGGTAGCTTCGCAAATCAGCCCAACCAAAGCTGCTCAGAAGTACAACTTAGATGCAAGTTTTTTTTTC

AAATATGTGTGAGTCCTTGTATTGATATCTATCAGGCTAACGAGTGATGCCACTATGGATATGATCTTCT

AGTGAAATTTGTATATACATTCATTTATGCACGTATGTATATTGTGTTAACTGTGCCAAATATTTATCAT

TTAATATTTTTATTCACGGCAA

>MSTRG.236.2 gene=MSTRG.236

AAAATGCCATCTGTACCCGCACAGTCCATCTTCATTCAGCTTCAGGCCCAAGCCCAGTTAGTCATTGTTT

TCTCCGCTGCCTGTGTGTATATGTACGTATACGTGTGAGATGAGGAAGAATCAATGGCGATTCCAAAGAA

TTAAGATCAATTCTCACACTCTGCATTGTTATGATAGTTGTTTCTATTGCTTTTACTCTCAAATTTTAAA

CAATGAAGAAAATGCTTCAGAAAGCCGATAAGAATGGTCGAAGAAAGCGCTTAATTAATAAATAAGGTAG

TGGCATCAATGGACTGGCCCGAGGTCACCACATACAAAGAACTCTACACACATGGTCAGTGAGAGGAAAT

ATTAAGGAATCTCTACTGCACTTGTGAAACTGAAAGGGGTGTTGTTGCTAGTGGCATGACCAGGGACTAT

TTGATGGCATTCTACCAGCAAACAGAACTCCGACCTAAACGGATCATATTCTACAGGGATGGAGTCAGTG

AAGGTCAGTTTAATCAGGTCTTGCTGAAAGAGATGGATGCCATTAGGACGGTTGATGTGTTTTATTTTTT

GTCTGCCAGGCGCTGCAATCAAATTATTTACAATGGCAGGAAATCAACTGATCGGAGTGGCAACATATCA

CCTGGTTACATATTGTTCTATTGTATCTTTAAAAATCTTTATTATTCTGATTAGCAAATAACTGGCTTTT

GTATACCAGGTACTGTTGTAAATATGACCATCTGCCATCCCACTGAGTTTGACTTTTATTTGTGTAGCCA

TGCTGGTATACAGGGTGTTGAAAAGGTGAAACCTCCAACCGGTAGCCCAATATCTATGTTAATGGCTACT

CCAGATGGCTCATATGCATTACCAAGCAGTTGATGTTTTTAAGCTAAATGAGTGTCAGAGATACCATCAT

ATCTCAGTAAATCAGTGTTTGTCGAGCACAAAATGTTCTTGCCGATGATTAGTTGCCCAAAAATATTGTA

AGACATACTGTAAAAAGAATTCAAGAGGCCCATAACAGCAGCGGAGATAATAAAGGCCGTGATTCCCAGT

TGCAAGGTCTGCATTTTTTCTGTCTTTGGAATGTTGCTGTCATGGAATGCCCGAGGAAAATATAACTTCG

GAATCAAGAAGTTTCGAAAGGTAGCTTCGCAAATCAGCCCAACCAAAGCTGCTCAGAAGTACAACTTAGA

TGCAAGTTTTTTTTTCAAATATGTGTGAGTCCTTGTATTGATATCTATCAGGCTAACGAGTGATGCCACT

ATGGATATGATCTTCTAGTGAAATTTGTATATACATTCATTTATGCACGTATGTATATTGTGTTAACTGT

GCCAAATATTTATCATTTAATATTTTTATTCACGGCAA

>MSTRG.236.3 gene=MSTRG.236

AAAATGCCATCTGTACCCGCACAGTCCATCTTCATTCAGCTTCAGGCCCAAGCCCAGTTAGTCATTGTTT

TCTCCGCTGCCTGTGTGTATATGTACGTATACGTGTGAGATGAGGAAGAATCAATGGCGATTCCAAAGAA

TTAAGATCAATTCTCACACTCTGCATTGTTATGATAGTTGTTTCTATTGCTTTTACTCTCAAATTTTAAA

CAATGAAGAAAATGCTTCAGAAAGCCGATAAGAATGGTCGAAGAAAGCGCTTAATTAATAAATAAGGTAG

TGGCATCAATGGACTGGCCCGAGGTCACCACATACAAAGAACTCTACACACATGGTCAGTGAGAGGAAAT

ATTAAGGAATCTCTACTGCACTTGTGAAACTGAAAGGGGTGTTGTTGCTAGTGGCATGACCAGGGACTAT

TTGATGGCATTCTACCAGCAAACAGAACTCCGACCTAAACGGATCATATTCTACAGGGATGGAGTCAGTG

AAGGTCAGTTTAATCAGGTCTTGCTGAAAGAGATGGATGCCATTAGGACGGTTGATGTGTTTTATTTTTT

GTCTGCCAGGCGCTGCAATCAAATTATTTACAATGGCAGGAAATCAACTGATCGGAGTGGCAACATATCA

CCTGGTTACATATTGTTCTATTGTATCTTTAAAAATCTTTATTATTCTGATTAGCAAATAACTGGCTTTT

GTATACCAGGTACTGTTGTAGATATGACCATCTGCCATCCCACTGAGTTTGACTTTTATTTGTGTAGCCA

TGCTGGTATACAGGGTGTTGAAAAGGTGAAACCTCCAACCGGTAGCCCAATATCTATGTTAATGGCTACT

CCAGATGGCTCATATGCATTACCAAGCAGTTGATGTTTTTAAGCTAAATGAGTGTCAGAGATACCATCAT

ATCTCAGTAAATCAGTGTTTGTCGAGCACAAAATGTTCTTGCCGATGATTAGTTGCCCAAAAATATTGTA

AGACATACTGTAAAAAGAATTCAAGAGGCCCATAACAGCAGCGGAGATAATAAAGGCCGTGATTCCCAGT

TGCAAGGTCTGCATTTTTTCTGTCTTTGGAATGTTGCTGTCATGGAATGCCCGAGGAAAATATAACTTCG

GAATCAAGAAGTTTCGAAAGGTAGCTTCGCAAATCAGCCCAACCAAAGCTGCTCAGAAGTACAACTTAGA

TGCAAGTTTTTTTTTCAAATATGTGTGAGTCCTTGTATTGATATCTATCAGGCTAACGAGTGATGCCACT

ATGGATATGATCTTCTAGTGAAATTTGTATATACATTCATTTATGCACGTATGTATATTGTGTTAACTGT

GCCAAATATTTATCATTTAATATTTTTATTCACGGCAA

>MSTRG.236.4 gene=MSTRG.236

AAAATGCCATCTGTACCCGCACAGTCCATCTTCATTCAGCTTCAGGCCCAAGCCCAGTTAGTCATTGTTT

TCTCCGCTGCCTGTGTGTATATGTACGTATACGTGTGAGATGAGGAAGAATCAATGGCGATTCCAAAGAA

TTAAGATCAATTCTCACACTCTGCATTGTTATGATAGTTGTTTCTATTGCTTTTACTCTCAAATTTTAAA

CAATGAAGAAAATGCTTCAGAAAGCCGATAAGAATGGTCGAAGAAAGCGCTTAATTAATAAATAAGGTAG

TGGCATCAATGGACTGGCCCGAGGTCACCACATACAAAGAACTCTACACACATGGTCAGTGAGAGGAAAT

ATTAAGGAATCTCTACTGCACTTGTGAAACTGAAAGGGGTGTTGTTGCTAGTGGCATGACCAGGGACTAT

TTGATGGCATTCTACCAGCAAACAGAACTCCGACCTAAACGGATCATATTCTACAGGGATGGAGTCAGTG

AAGGTCAGTTTAATCAGGTCTTGCTGAAAGAGATGGATGCCATTAGGACGGTTGATGTGTTTTATTTTTT

GTCTGCCAGGCGCTGCAATCAAATTATTTACAATGGCAGGAAATCAACTGATCGGAGTGGCAACATATCA

CCTGGTTACATATTGTTCTATTGTATCTTTAAAAATCTTTATTATTCTGATTAGCAAATAACTGGCTTTT

GTATACCAGGTACTGTTGTAAATATGACCATCTGCCATCCCACTGAGTTTGACTTTTATTTGTGTAGCCA

TGCTGGTATACAGGGTGTTGAAAAGGTGAAACCTCCAACCGGTAGCCCAATATCTATGTTAATGGCTACT

CCAGATGGCTCATATGCATTACCAAGCAGTTGATGTTTTTAAGCTAAATGAGTGTCAGAGATACCATCAT

ATCTCAGTAAATCAGTGTTTGTCGAGCACAAAATGTTCTTGCCGATGATTAGTTGCCCAAAAATATTGTA

AGACATACTGTAAAAAGAATTCAAGAGGCCCATAACAGCAGCGGAGATAATAAAGGCCGTGATTCCCAGT

TGCAAGGTCTGCATTTTTTCTGTCTTTGGAATGTTGCTGTCATGGAATGCCCGAGGAAAATATAACTTCG

GAATCAAGAAGTTTCGAAAGGTAGCTTCGCAAATCAGCCCAACCAAAGCTGCTCAGAAGTACAACTTAGA

TGCAAGTTTTTTTTTCAAATATGTGTGAGTCCTTGTATTGATATCTATCAGGCTAACGAGTGATGCCACT

ATGGATATGATCTTCTAGTGAAATTTGTATATACATTCATTTATGCACGTATGTATATTGTGTTAACTGT

GCCAAATATTTATCATTTAATATTTTTATTCACGGCAA

>MSTRG.236.5 gene=MSTRG.236

AAAATGCCATCTGTACCCGCACAGTCCATCTTCATTCAGCTTCAGGCCCAAGCCCAGTTAGTCATTGTTT

TCTCCGCTGCCTGTGTGTATATGTACGTATACGTGTGAGATGAGGAAGAATCAATGGCGATTCCAAAGAA

TTAAGATCAATTCTCACACTCTGCATTGTTATGATAGTTGTTTCTATTGCTTTTACTCTCAAATTTTAAA

CAATGAAGAAAATGCTTCAGAAAGCCGATAAGAATGGTCGAAGAAAGCGCTTAATTAATAAATAAGGTAG

TGGCATCAATGGACTGGCCCGAGGTCACCACATACAAAGAACTCTACACACATGGTCAGTGAGAGGAAAT

ATTAAGGAATCTCTACTGCACTTGTGAAACTGAAAGGGGTGTTGTTGCTAGTGGCATGACCAGGGACTAT

TTGATGGCATTCTACCAGCAAACAGAACTCCGACCTAAACGGATCATATTCTACAGGGATGGAGTCAGTG

AAGGTCAGTTTAATCAGGTCTTGCTGAAAGAGATGGATGCCATTAGGACGGTTGATGTGTTTTATTTTTT

GTCTGCCAGGCGCTGCAATCAAATTATTTACAATGGCAGGAAATCAACTGATCGGAGTGGCAACATATCA

CCTGGTACTGTTGTAAATATGACCATCTGCCATCCCACTGAGTTTGACTTTTATTTGTGTAGCCATGCTG

GTATACAGGACGGACTTTGACTTTGTTGTCGGCAATGCACAGACTAATGAGGGTGTTGAAAAGGTGAAAC

CTCCAACCGGTAGCCCAATATCTATGTTAATGGCTACTCCAGATGGCTCATATGCATTACCAAGCAGTTG

ATGTTTTTAAGCTAAATGAGTGTCAGAGATACCATCATATCTCAGTAAATCAGTGTTTGTCGAGCACAAA

ATGTTCTTGCCGATGATTAGTTGCCCAAAAATATTGTAAGACATACTGTAAAAAGAATTCAAGAGGCCCA

TAACAGCAGCGGAGATAATAAAGGCCGTGATTCCCAGTTGCAAGGTCTGCATTTTTTCTGTCTTTGGAAT

GTTGCTGTCATGGAATGCCCGAGGAAAATATAACTTCGGAATCAAGAAGTTTCGAAAGGTAGCTTCGCAA

ATCAGCCCAACCAAAGCTGCTCAGAAGTACAACTTAGATGCAAGTTTTTTTTTCAAATATGTGTGAGTCC

TTGTATTGATATCTATCAGGCTAACGAGTGATGCCACTATGGATATGATCTTCTAGTGAAATTTGTATAT

ACATTCATTTATGCACGTATGTATATTGTGTTAACTGTGCCAAATATTTATCATTTAATATTTTTATTCA

CGGCAA

>MSTRG.236.6 gene=MSTRG.236

AAAATGCCATCTGTACCCGCACAGTCCATCTTCATTCAGCTTCAGGCCCAAGCCCAGTTAGTCATTGTTT

TCTCCGCTGCCTGTGTGTATATGTACGTATACGTGTGAGATGAGGAAGAATCAATGGCGATTCCAAAGAA

TTAAGATCAATTCTCACACTCTGCATTGTTATGATAGTTGTTTCTATTGCTTTTACTCTCAAATTTTAAA

CAATGAAGAAAATGCTTCAGAAAGCCGATAAGAATGGTCGAAGAAAGCGCTTAATTAATAAATAAGGTAG

TGGCATCAATGGACTGGCCCGAGGTCACCACATACAAAGAACTCTACACACATGGTCAGTGAGAGGAAAT

ATTAAGGAATCTCTACTGCACTTGTGAAACTGAAAGGGGTGTTGTTGCTAGTGGCATGACCAGGGACTAT

TTGATGGCATTCTACCAGCAAACAGAACTCCGACCTAAACGGATCATATTCTACAGGGATGGAGTCAGTG

AAGGTCAGTTTAATCAGGTCTTGCTGAAAGAGATGGATGCCATTAGGACGGTTGATGTGTTTTATTTTTT

GTCTGCCAGGCGCTGCAATCAAATTATTTACAATGGCAGGAAATCAACTGATCGGAGTGGCAACATATCA

CCTGGTTACATATTGTTCTATTGTATCTTTAAAAATCTTTATTATTCTGATTAGCAAATAACTGGCTTTT

GTATACCAGGTACTGTTGTAAATATGACCATCTGCCATCCCACTGAGTTTGACTTTTATTTGTGTAGCCA

TGCTGGTATACAGGACGGACTTTGACTTTGTTGTCGGCAATGCACAGACTAATGAGGGTGTTGAAAAGGT

GAAACCTCCAACCGGTAGCCCAATATCTATGTTAATGGCTACTCCAGATGGCTCATATGCATTACCAAGC

AGTTGATGTTTTTAAGCTAAATGAGTGTCAGAGATACCATCATATCTCAGTAAATCAGTGTTTGTCGAGC

ACAAAATGTTCTTGCCGATGATTAGTTGCCCAAAAATATTGTAAGACATACTGTAAAAAGAATTCAAGAG

GCCCATAACAGCAGCGGAGATAATAAAGGCCGTGATTCCCAGTTGCAAGGTCTGCATTTTTTCTGTCTTT

GGAATGTTGCTGTCATGGAATGCCCGAGGAAAATATAACTTCGGAATCAAGAAGTTTCGAAAGGTAGCTT

CGCAAATCAGCCCAACCAAAGCTGCTCAGAAGTACAACTTAGATGCAAGTTTTTTTTTCAAATATGTGTG

AGTCCTTGTATTGATATCTATCAGGCTAACGAGTGATGCCACTATGGATATGATCTTCTAGTGAAATTTG

TATATACATTCATTTATGCACGTATGTATATTGTGTTAACTGTGCCAAATATTTATCATTTAATATTTTT

ATTCACGGCAA

>MSTRG.236.9 gene=MSTRG.236

TCCATCTTCATTCAGGTTCAGGCCCAAGCCCAGTTAGTCATTGTTTTCTCCGCTGCTTGTGTGTATATGT

ACGTATACGTGTGAGATGAGGATGAATCAATGGCGATTCCAAAGATTTAAGATCAATTCTCACACTCTAC

ATTGTTATGATACTTGTTTCTATCGCTTCCACTCTCAAATTTTAAACAATGAAGAAAGTGCTTCAGGAAG

CCGATAAGAATGGTCGAAGAAAGCGCTTAATTAATAAATAAGGTAGTGGCATCAATGGACTGGCCTGAGG

TCACCACATACAGAGAACTCTACACACATGGTCAGTGAGAGGAAATATTAAGGAATCTCTACTGCACTTG

TGAAACTGAAAGGGGTGTTGTTGCTAGTGGCATGACCAGGGACTATTTGATGGCATTCTACCAGCAAACA

GAACTCCGACCTAAACGGATCATATTCTACAGGGATGGAGTCAGTGAAGGTCAGTTTAATCAGGTCTTGC

TGAAAGAGATGGATGCCATTAGGACGGTTGATGTGTTTTATTTTTTGTAAGTAATTACTACCTTCACTTG

TTAGTTGTTACCCCTATAGTTCAAGTTTGATGAATGGGAAGATATATCTGTTTTTAAGGTCTGCCAGGCG

CTGCAATCAAATTATTTACAATGGCAGGAAATCAACTGATCGGAGTGGCAACATATCACCTGGTTACATA

TTGTTCTATTGTATCTTTAAAAATCTTTATTATTCTGATTAGCAAATAACTGGCTTTTGTATACCAGGTA

CTGTTGTAGATATGACCATCTGCCATCCCACTGAGTTTGACTTTTATTTGTGTAGCCATGCTGGTATACA

GGGTGTTGAAAAGGTGAAACCTCCAACCGGTAGCCCAATATCTATGTTAATGGCTACTCCAGATGGCTCA

TATGCATTACCAAGCAGTTGATGTTTTTAAGCTAAATGAGTGTCAGAGATACCATCATATCTCAGTAAAT

CAGTGTTTGTCGAGCACAAAATGTTCTTGCCGATGATTAGTTGCCCAAAAATATTGTAAGACATACTGTA

AAAAGAATTCAAGAGGCCCATAACAGCAGCGGAGATAATAAAGGCCGTGATTCCCAGTTGCAAGGTCTGC

ATTTTTTCTGTCTTTGGAATGTTGCTGTCATGGAATGCCCGAGGAAAATATAACTTCGGAATCAAGAAGT

TTCGAAAGGTAGCTTCGCAAATCAGCCCAACCAAAGCTGCTCAGAAGTACAACTTAGATGCAAGTTTTTT

TTTCAAATATGTGTGAGTCCTTGTATTGATATCTATCAGGCTAACGAGTGATGCCACTATGGATATGATC

TTCTAGTGAAATTTGTATATACATTCATTTATGCACGTATGTATATTGTGTTAACTGTGCC

>MSTRG.236.8 gene=MSTRG.236

CTCCAAAATGCCATCTGTACCCGCACAGTCCATCTTCATTCAGCTTCAGGCCCAAGCCCAGTTAGTCATT

GTTTTCTCCGCTGCCTGTGTGTATATGTACGTATACGTGTGAGATGAGGAAGAATCAATGGCGATTCCAA

AGAATTAAGATCAATTCTCACACTCTGCATTGTTATGATAGTTGTTTCTATTGCTTTTACTCTCAAATTT

TAAACAATGAAGAAAATGCTTCAGAAAGCCGATAAGAATGGTCGAAGAAAGCGCTTAATTAATAAATAAG

GTAGTGGCATCAATGGACTGGCCCGAGGTCACCACATACAAAGAACTCTACACACATGGTCAGTGAGAGG

AAATATTAAGGAATCTCTACTGCACTTGTGAAACTGAAAGGGGTGTTGTTGCTAGTGGCATGACCAGGGA

CTATTTGATGGCATTCTACCAGCAAACAGAACTCCGACCTAAACGGATCATATTCTACAGGGATGGAGTC

AGTGAAGGTCAGTTTAATCAGGTCTTGCTGAAAGAGATGGATGCCATTAGGACGGTTGATGTGTTTTATT

TTTTGTCTGCCAGGCGCTGCAATCAAATTATTTACAATGGCAGGAAATCAACTGATCGGAGTGGCAACAT

ATCACCTGGTACTGTTGTAAATATGACCATCTGCCATCCCACTGAGTTTGACTTTTATTTGTGTAGCCAT

GCTGGTATACAGGTAATCATCTTAATACAAGACTACATGGATTTTGTACAGGCAGCAGTAATAAATGCAT

GTGTTGTGTGGCAACACAATAGAGATACAAATGTACATATTTATTAATATACAAATAGAGAAACAAATGT

AAAGATCTTAATCAAAATAAACACAAATACACTCATATTAAAAAACTAACATGAACCAATGAGATATAGA

AACCATTAAAAATCAACTAGTTTATTCAATCTACCAAAACATCTAAGTAAAGGAAATCCAGATCTGTTCA

TATTTTACCGGTAACAATTAGCTAGTTAGGATATTAAATAAGAACAGAAAATTTATGCTCACTAGTTCTC

CACGAATTTAGGAAGCTTGTTGATTGAATATCATATACTAAAACACACAATTTCTACTATACATACTGTG

ACTATTTTAATGTCATTAGCCTAATTCATAACTTACGAAAAATGCTTATGGAGTATGTAGATTCAGGATA

GGTAAATAAAAGGATCAGGGAACATGTAAGCTAACAAACACTGCATTGTATCTTGTTGAATAATGAATTT

GTTTAAAATAAAACTTGCTCTGACGCACGTGATCTACGATGGAGTTATATACGCATGCCATTGCATTTAC

ATGTATAGGTTTCCCTATGAACTTTATTTAATGTGATATCTTAATGGACAAAATTTAATATTTTAATATT

CTCTTATCTGTTAATGCATTCATAGATTATGACACTCAAAATATTTTAATGGGCAATAGGACGGACTTTG

ACTTTGTTGTCGGCAATGCACAGACTAATGAGGGTGTTGAAAAGGTGAAACCTCCAACCGGTAGCCCAAT

ATCTATGTTAATGGCTACTCCAGATGGCTCATATGCATTACCAAGCAGTTGATGTTTTTAAGCTAAATGA

GTGTCAGAGATACCATCATATCTCAGTAAATCAGTGTTTGTCGAGCACAAAATGTTCTTGCCGATGATTA

GTTGCCCAAAAATATTGTAAGACATACTGTAAAAAGAATTCAAGAGGCCCATAACAGCAGCGGAGATAAT

AAAGGCCGTGATTCCCAGTTGCAAGGTCTGCATTTTTTCTGTCTTTGGAATGTTGCTGTCATGGAATGCC

CGAGGAAAATATAACTTCGGAATCAAGAAGTTTCGAAAGGTAGCTTCGCAAATCAGCCCAACCAAAGCTG

CTCAGAAGTACAACTTAGATGCAAGTTTTTTTTTCAAATATGTGTGAGTCCTTGTATTGATATCTATCAG

GCTAACGAGTGATGCCACTATGGATATGATCTTCTAGTGAAATTTGTATATACATTCATTTATGCACGTA

TGTATATTGTGTTAACTGTGCC

>MSTRG.236.10 gene=MSTRG.236

CTCCAAAATGCCATCTGTACCCGCACAGTCCATCTTCATTCAGCTTCAGGCCCAAGCCCAGTTAGTCATT

GTTTTCTCCGCTGCCTGTGTGTATATGTACGTATACGTGTGAGATGAGGAAGAATCAATGGCGATTCCAA

AGAATTAAGATCAATTCTCACACTCTGCATTGTTATGATAGTTGTTTCTATTGCTTTTACTCTCAAATTT

TAAACAATGAAGAAAATGCTTCAGAAAGCCGATAAGAATGGTCGAAGAAAGCGCTTAATTAATAAATAAG

GTAGTGGCATCAATGGACTGGCCCGAGGTCACCACATACAAAGAACTCTACACACATGGTCAGTGAGAGG

AAATATTAAGGAATCTCTACTGCACTTGTGAAACTGAAAGGGGTGTTGTTGCTAGTGGCATGACCAGGGA

CTATTTGATGGCATTCTACCAGCAAACAGAACTCCGACCTAAACGGATCATATTCTACAGGGATGGAGTC

AGTGAAGGTCAGTTTAATCAGGTCTTGCTGAAAGAGATGGATGCCATTAGGACGGTTGATGTGTTTTATT

TTTTGTCTGCCAGGCGCTGCAATCAAATTATTTACAATGGCAGGAAATCAACTGATCGGAGTGGCAACAT

ATCACCTGGTACTGTTGTAAATATGACCATCTGCCATCCCACTGAGTTTGACTTTTATTTGTGTAGCCAT

GCTGGTATACAGGGTGTTGAAAAGGTGAAACCTCCAACCGGTAGCCCAATATCTATGTTAATGGCTACTC

CAGATGGCTCATATGCATTACCAAGCAGTTGATGTTTTTAAGCTAAATGAGTGTCAGAGATACCATCATA

TCTCAGTAAATCAGTGTTTGTCGAGCACAAAATGTTCTTGCCGATGATTAGTTGCCCAAAAATATTGTAA

GACATACTGTAAAAAGAATTCAAGAGGCCCATAACAGCAGCGGAGATAATAAAGGCCGTGATTCCCAGTT

GCAAGGTCTGCATTTTTTCTGTCTTTGGAATGTTGCTGTCATGGAATGCCCGAGGAAAATATAACTTCGG

AATCAAGAAGTTTCGAAAGGTAGCTTCGCAAATCAGCCCAACCAAAGCTGCTCAGAAGTACAACTTAGAT

GCAAGTTTTTTTTTCAAATATGTGTGAGTCCTTGTATTGATATCTATCAGGCTAACGAGTGATGCCACTA

TGGATATGATCTTCTAGTGAAATTTGTATATACATTCATTTATGCACGTATGTATATT

>MSTRG.236.11 gene=MSTRG.236

CTCCAAAATGCCATCTGTACCCGCACAGTCCATCTTCATTCAGCTTCAGGCCCAAGCCCAGTTAGTCATT

GTTTTCTCCGCTGCCTGTGTGTATATGTACGTATACGTGTGAGATGAGGAAGAATCAATGGCGATTCCAA

AGAATTAAGATCAATTCTCACACTCTGCATTGTTATGATAGTTGTTTCTATTGCTTTTACTCTCAAATTT

TAAACAATGAAGAAAATGCTTCAGAAAGCCGATAAGAATGGTCGAAGAAAGCGCTTAATTAATAAATAAG

GTAGTGGCATCAATGGACTGGCCCGAGGTCACCACATACAAAGAACTCTACACACATGGTCAGTGAGAGG

AAATATTAAGGAATCTCTACTGCACTTGTGAAACTGAAAGGGGTGTTGTTGCTAGTGGCATGACCAGGGA

CTATTTGATGGCATTCTACCAGCAAACAGAACTCCGACCTAAACGGATCATATTCTACAGGGATGGAGTC

AGTGAAGGTCAGTTTAATCAGGTCTTGCTGAAAGAGATGGATGCCATTAGGACGGTTGATGTGTTTTATT

TTTTGTCTGCCAGGCGCTGCAATCAAATTATTTACAATGGCAGGAAATCAACTGATCGGAGTGGCAACAT

ATCACCTGGTACTGTTGTAAATATGACCATCTGCCATCCCACTGAGTTTGACTTTTATTTGTGTAGCCAT

GCTGGTATACAGATTATGACACTCAAAATATTTTAATGGGCAATAGGACGGACTTTGACTTTGTTGTCGG

CAATGCACAGACTAATGAGGTTGGTTGGATTAAATCTCTTAGGTGTTTACCATATTGTTGCTTAATAATG

CATTTGAAGGATGGGAGGTGATCTTTACTTGATGTAGCAATGAAGATGAAAAACAGATCACCTATTTGTA

ATATTGATATGCGGATTATGCTTGTCTGTGCTCTATTATGCTTATTTTTGCTCTAATGTTGGATGTTAAT

ATGCTTCCATTATCCTACAAATGGAGTTAGGGTGTTGAAAAGGTGAAACCTCCAACCGGTAGCCCAATAT

CTATGTTAATGGCTACTCCAGATGGCTCATATGCATTACCAAGCAGTTGATGTTTTTAAGCTAAATGAGT

GTCAGAGATACCATCATATCTCAGTAAATCAGTGTTTGTCGAGCACAAAATGTTCTTGCCGATGATTAGT

TGCCCAAAAATATTGTAAGACATACTGTAAAAAGAATTCAAGAGGCCCATAACAGCAGCGGAGATAATAA

AGGCCGTGATTCCCAGTTGCAAGGTCTGCATTTTTTCTGTCTTTGGAATGTTGCTGTCATGGAATGCCCG

AGGAAAATATAACTTCGGAATCAAGAAGTTTCGAAAGGTAGCTTCGCAAATCAGCCCAACCAAAGCTGCT

CAGAAGTACAACTTAGATGCAAGTTTTTTTTTCAAATATGTGTGAGTCCTTGTATTGATATCTATCAGGC

TAACGAGTGATGCCACTATGGATATGATCTTCTAGTGAAATTTGTATATACATTCATTTATGCACGTATG

TATATT

>MSTRG.236.12 gene=MSTRG.236

GCCATCTGTACCCGCACAGTCCATCTTCATTCAGCTTCAGGCCCAAGCCCAGTTAGTCATTGTTTTCTCC

GCTGCCTGTGTGTATATGTACGTATACGTGTGAGATGAGGAAGAATCAATGGCGATTCCAAAGAATTAAG

ATCAATTCTCACACTCTGCATTGTTATGATAGTTGTTTCTATTGCTTTTACTCTCAAATTTTAAACAATG

AAGAAAGTGCTTCAGGAAGCCGATAAGAATGGTCGAAGAAAGCGCTTAATTAATAAATAAGGTAGTGGCA

TCAATGGACTGGCCTGAGGTCACCACATACAGAGAACTCTACACACATGGTCAGTGAGAGGAAATATTAA

GGAATCTCTACTGCACTTGTGAAACTGAAAGGGGTGTTGTTGCTAGTGGCATGACCAGGGACTATTTGAT

GGCATTCTACCAGCAAACAGAACTCCGACCTAAACGGATCATATTCTACAGGGATGGAGTCAGTGAAGGT

CAGTTTAATCAGGTCTTGCTGAAAGAGATGGATGCCATTAGGACGGTTGATGTGTTTTATTTTTTGTCTG

CCAGGCGCTGCAATCAAATTATTTACAATGGCAGGAAATCAACTGATCGGAGTGGCAACATATCACCTGG

TTACATATTGTTCTATTGTATCTTTAAAAATCTTTATTATTCTGATTAGCAAATAACTGGCTTTTGTATA

CCAGGTACTGTTGTAGATATGACCATCTGCCATCCCACTGAGTTTGACTTTTATTTGTGTAGCCATGCTG

GTATACAGGTAATCATCTTAATACAAGACCACATGGATTTTGTACAGGCAGCAGTAATAAATGCATGTGT

TGTGTGGCAACACAATAGAGATACAAATGTACATATTTATTAATATACAAATAGAGAAACAAATGTAAAG

ATCATAATCAAAATAAACACAAATACACTCATATTAAAAAACTAACATGAACCAATGAGATATAGAAACC

ATTAAACATCAACTAGTTTATTCAATCTACCAAAACATCTGAGTAAAGGAAATCCAGATCTGTTCATATT

TTACCGGTAACAATTAGCTAGTTAGGATATTAAATAAGAACAGAAAATTTATGCTCACTAGTTCTCCACG

AATTTAGGAAGCTTGTTGATTGAATATCATATACTAAAACACACAAATTCTACTATACATACTGTGACTA

TTTTAATGTCATTAGCCTAATTCATAACTTACGAAAAATGCTTATGGAGTATGTAGATTCAGGATAGGCA

AATAAAAGGATCAGGGAACATGTAAGCTAACAAACACTGCATTGTATCTTGTTGAATAATGAATTTGTTT

AAAATAAAACTTGCTCTGACGCACGTGATCTACGATGGAGTTATATATGCATGCCATTGCATTTACATGT

ATAGGTTTCCCTATGAAATTTATTCAATGTGATATCTTAATGGACAAAATTTAATATTTTAATATTCTCT

TATCTGTTAATGCATTCATAGATTATGACACTCAAAATATTTTAATGGGCAATAGGACGGACTTTGACTT

TGTTGTCGGCAATGCACAGACTAATGAGGGTGTTGAAAAGGTGAAACCTCCAACCGGTAGCCCAATATCT

ATGTTAATGGCTACTCCAGATGGCTCATATGCATTACCAAGCAGTTGATGTTTTTAAGCTAAATGAGTGT

CAGAGATACCATCATATCTCAGTAAATCAGTGTTTGTCGAGCACAAAATGTTCTTGCCGATGATTAGTTG

CCCAAAAATATTGTAAGACATACTGTAAAAAGAATTCAAGAGGCCCATAACAGCAGCGGAGATAATAAAG

GCCGTGATTCCCAGTTGCAAGGTCTGCATTTTTTCTGTCTTTGGAATGTTGCTGTCATGGAATGCCCGAG

GAAAATATAACTTCGGAATCAAGAAGTTTCGAAAGGTAGCTTCGCAAATCAGCCCAACCAAAGCTGCTCA

GAAGTACAACTTAGATGCAAGTTTTTTTTTCAAATATGTGTGAGTCCTTGTATTGATATCTATCAGGCTA

ACGAGTGATGCCACTATGGATATGATCTTCTAGTGAAATTTGTATATACATTCATTTATGCACG

>MSTRG.236.13 gene=MSTRG.236

GCCATCTGTACCCGCACAGTCCATCTTCATTCAGCTTCAGGCCCAAGCCCAGTTAGTCATTGTTTTCTCC

GCTGCCTGTGTGTATATGTACGTATACGTGTGAGATGAGGAAGAATCAATGGCGATTCCAAAGAATTAAG

ATCAATTCTCACACTCTGCATTGTTATGATAGTTGTTTCTATTGCTTTTACTCTCAAATTTGTAAGTAAT

TTCCTTCCAGATCTTCAATTAATTATTTCTCTGAATCAAAAATAAACAATGAAGAAAATGCTTCAGAAAG

CCGATAAGAATGGTCGAAGAAAGCGCTTAATTAATAAATAAGGTAGTGGCATCAATGGACTGGCCCGAGG

TCACCACATACAAAGAACTCTACACACATGGTCAGTGAGAGGAAATATTAAGGAATCTCTACTGCACTTG

TGAAACTGAAAGGGGTGTTGTTGCTAGTGGCATGACCAGGGACTATTTGATGGCATTCTACCAGCAAACA

GAACTCCGACCTAAACGGATCATATTCTACAGGGATGGAGTCAGTGAAGGTCAGTTTAATCAGGTCTTGC

TGAAAGAGATGGATGCCATTAGGACGGTTGATGTGTTTTATTTTTTGTAAGTAATTACTACCTTCACTTG

TTAGTTGTTACCCCTATAGTTCAAGTTTGATGAATGGGAAGATATATCTGTTTTTAAGGTCTGCCAGGCG

CTGCAATCAAATTATTTACAATGGCAGGAAATCAACTGATCGGAGTGGCAACATATCACCTGGTTACATA

TTGTTCTATTGTATCTTTAAAAATCTTTATTATTCTGATTAGCAAATAACTGGCTTTTGTATACCAGGTA

CTGTTGTAGATATGACCATCTGCCATCCCACTGAGTTTGACTTTTATTTGTGTAGCCATGCTGGTATACA

GGGTGTTGAAAAGGTGAAACCTCCAACCGGTAGCCCAATATCTATGTTAATGGCTACTCCAGATGGCTCA

TATGCATTACCAAGCAGTTGATGTTTTTAAGCTAAATGAGTGTCAGAGATACCATCATATCTCAGTAAAT

CAGTGTTTGTCGAGCACAAAATGTTCTTGCCGATGATTAGTTGCCCAAAAATATTGTAAGACATACTGTA

AAAAGAATTCAAGAGGCCCATAACAGCAGCGGAGATAATAAAGGCCGTGATTCCCAGTTGCAAGGTCTGC

ATTTTTTCTGTCTTTGGAATGTTGCTGTCATGGAATGCCCGAGGAAAATATAACTTCGGAATCAAGAAGT

TTCGAAAGGTAGCTTCGCAAATCAGCCCAACCAAAGCTGCTCAGAAGTACAACTTAGATGCAAGTTTTTT

TTTCAAATATGTGTGAGTCCTTGTATTGATATCTATCAGGCTAACGAGTGATGCCACTATGGATATGATC

TTCTAGTGAAATTTGTATATACATTCATTTATGCACG

>MSTRG.236.14 gene=MSTRG.236

GCACAGTCCATCTTCATTCAGCTTCAGGCCCAAGCCCAGTTAGTCATTGTTTTCTCCGCTGCCTGTGTGT

ATATGTACGTATACGTGTGAGATGAGGAAGAATCAATGGCGATTCCAAAGAATTAAGATCAATTCTCACA

CTCTGCATTGTTATGATAGTTGTTTCTATTGCTTTTACTCTCAAATTTGTAAGTAATTTCCTTCCAGATC

TTCAATTAATTATTTCTCTGAATCAAAAATAAACAATGAAGAAAATGCTTCAGAAAGCCGATAAGAATGG

TCGAAGAAAGCGCTTAATTAATAAATAAGGTAGTGGCATCAATGGACTGGCCCGAGGTCACCACATACAA

AGAACTCTACACACATGGTCAGTGAGAGGAAATATTAAGGAATCTCTACTGCACTTGTGAAACTGAAAGG

GGTGTTGTTGCTAGTGGCATGACCAGGGACTATTTGATGGCATTCTACCAGCAAACAGAACTCCGACCTA

AACGGATCATATTCTACAGGGATGGAGTCAGTGAAGGTCAGTTTAATCAGGTCTTGCTGAAAGAGATGGA

TGCCATTAGGACGGTTGATGTGTTTTATTTTTTGTCTGCCAGGCGCTGCAATCAAATTATTTACAATGGC

AGGAAATCAACTGATCGGAGTGGCAACATATCACCTGGTTACATATTGTTCTATTGTATCTTTAAAAATC

TTTATTATTCTGATTAGCAAATAACTGGCTTTTGTATACCAGGTACTGTTGTAAATATGACCATCTGCCA

TCCCACTGAGTTTGACTTTTATTTGTGTAGCCATGCTGGTATACAGGTAATCATCTTAATACAAGACTAC

ATGGATTTTGTACAGGCAGCAGTAATAAATGCATGTGTTGTGTGGCAACACAATAGAGATACAAATGTAC

ATATTTATTAATATACAAATAGAGAAACAAATGTAAAGATCTTAATCAAAATAAACACAAATACACTCAT

ATTAAAAAACTAACATGAACCAATGAGATATAGAAACCATTAAAAATCAACTAGTTTATTCAATCTACCA

AAACATCTAAGTAAAGGAAATCCAGATCTGTTCATATTTTACCGGTAACAATTAGCTAGTTAGGATATTA

AATAAGAACAGAAAATTTATGCTCACTAGTTCTCCACGAATTTAGGAAGCTTGTTGATTGAATATCATAT

ACTAAAACACACAATTTCTACTATACATACTGTGACTATTTTAATGTCATTAGCCTAATTCATAACTTAC

GAAAAATGCTTATGGAGTATGTAGATTCAGGATAGGTAAATAAAAGGATCAGGGAACATGTAAGCTAACA

AACACTGCATTGTATCTTGTTGAATAATGAATTTGTTTAAAATAAAACTTGCTCTGACGCACGTGATCTA

CGATGGAGTTATATACGCATGCCATTGCATTTACATGTATAGGTTTCCCTATGAACTTTATTTAATGTGA

TATCTTAATGGACAAAATTTAATATTTTAATATTCTCTTATCTGTTAATGCATTCATAGATTATGACACT

CAAAATATTTTAATGGGCAATAGGACGGACTTTGACTTTGTTGTCGGCAATGCACAGACTAATGAGGTTG

GTTGGATTAAATCTCTTAGGTGTTTACCATATTGTTGCTTAATAATGCATTTGAAGGATGGGAGGTGATC

TTTACTTGATGTAGCAATGAAGATGAAAAACAGATCACCTATTTGTAATATTGATATGCGGATTATGCTT

GTCTGTGCTCTATTATGCTTATTTTTGCTCTAATGTTGGATGTTAATATGCTTCCATTATCCTACAAATG

GAGTTAGGGTGTTGAAAAGGTGAAACCTCCAACCGGTAGCCCAATATCTATGTTAATGGCTACTCCAGAT

GGCTCATATGCATTACCAAGCAGTTGATGTTTTTAAGCTAAATGAGTGTCAGAGATACCATCATATCTCA

GTAAATCAGTGTTTGTCGAGCACAAAATGTTCTTGCCGATGATTAGTTGCCCAAAAATATTGTAAGACAT

ACTGTAAAAAGAATTCAAGAGGCCCATAACAGCAGCGGAGATAATAAAGGCCGTGATTCCCAGTTGCAAG

GTCTGCATTTTTTCTGTCTTTGGAATGTTGCTGTCATGGAATGCCCGAGGAAAATATAACTTCGGAATCA

AGAAGTTTCGAAAGGTAGCTTCGCAAATCAGCCCAACCAAAGCTGCTCAGAAGTACAACTTAGATGCAAG

TTTTTTTTTCAAATATGTGTGAGTCCTTGTATTGATATCTATCAGGCTAACGAGTGATGCCACTATGGAT

ATGATCTTCTAGTGAAATTTGTATATACATTCATTTATGCAC

>MSTRG.236.15 gene=MSTRG.236

GCACAGTCCATCTTCATTCAGCTTCAGGCCCAAGCCCAGTTAGTCATTGTTTTCTCCGCTGCCTGTGTGT

ATATGTACGTATACGTGTGAGATGAGGAAGAATCAATGGCGATTCCAAAGAATTAAGATCAATTCTCACA

CTCTGCATTGTTATGATAGTTGTTTCTATTGCTTTTACTCTCAAATTTTAAACAATGAAGAAAATGCTTC

AGAAAGCCGATAAGAATGGTCGAAGAAAGCGCTTAATTAATAAATAAGGTAGTGGCATCAATGGACTGGC

CCGAGGTCACCACATACAAAGAACTCTACACACATGGTCAGTGAGAGGAAATATTAAGGAATCTCTACTG

CACTTGTGAAACTGAAAGGGGTGTTGTTGCTAGTGGCATGACCAGGGACTATTTGATGGCATTCTACCAG

CAAACAGAACTCCGACCTAAACGGATCATATTCTACAGGGATGGAGTCAGTGAAGGTCAGTTTAATCAGG

TCTTGCTGAAAGAGATGGATGCCATTAGGACGGTTGATGTGTTTTATTTTTTGTCTGCCAGGCGCTGCAA

TCAAATTATTTACAATGGCAGGAAATCAACTGATCGGAGTGGCAACATATCACCTGGTACTGTTGTAAAT

ATGACCATCTGCCATCCCACTGAGTTTGACTTTTATTTGTGTAGCCATGCTGGTATACAGGGTGTTGAAA

AGGTGAAACCTCCAACCGGTAGCCCAATATCTATGTTAATGGCTACTCCAGATGGCTCATATGCATTACC

AAGCAGTTGATGTTTTTAAGCTAAATGAGTGTCAGAGATACCATCATATCTCAGTAAATCAGTGTTTGTC

GAGCACAAAATGTTCTTGCCGATGATTAGTTGCCCAAAAATATTGTAAGACATACTGTAAAAAGAATTCA

AGAGGCCCATAACAGCAGCGGAGATAATAAAGGCCGTGATTCCCAGTTGCAAGGTCTGCATTTTTTCTGT

CTTTGGAATGTTGCTGTCATGGAATGCCCGAGGAAAATATAACTTCGGAATCAAGAAGTTTCGAAAGGTA

GCTTCGCAAATCAGCCCAACCAAAGCTGCTCAGAAGTACAACTTAGATGCAAGTTTTTTTTTCAAATATG

TGTGAGTCCTTGTATTGATATCTATCAGGCTAACGAGTGATGCCACTATGGATATGATCTTCTAGTGAAA

TTTGTATATACATTCATTTATGCAC

>MSTRG.236.16 gene=MSTRG.236

GCACAGTCCATCTTCATTCAGCTTCAGGCCCAAGCCCAGTTAGTCATTGTTTTCTCCGCTGCCTGTGTGT

ATATGTACGTATACGTGTGAGATGAGGAAGAATCAATGGCGATTCCAAAGAATTAAGATCAATTCTCACA

CTCTGCATTGTTATGATAGTTGTTTCTATTGCTTTTACTCTCAAATTTTAAACAATGAAGAAAATGCTTC

AGAAAGCCGATAAGAATGGTCGAAGAAAGCGCTTAATTAATAAATAAGGTAGTGGCATCAATGGACTGGC

CCGAGGTCACCACATACAAAGAACTCTACACACATGGTCAGTGAGAGGAAATATTAAGGAATCTCTACTG

CACTTGTGAAACTGAAAGGGGTGTTGTTGCTAGTGGCATGACCAGGGACTATTTGATGGCATTCTACCAG

CAAACAGAACTCCGACCTAAACGGATCATATTCTACAGGGATGGAGTCAGTGAAGGTCAGTTTAATCAGG

TCTTGCTGAAAGAGATGGATGCCATTAGGACGGTTGATGTGTTTTATTTTTTGTCTGCCAGGCGCTGCAA

TCAAATTATTTACAATGGCAGGAAATCAACTGATCGGAGTGGCAACATATCACCTGGTTACATATTGTTC

TATTGTATCTTTAAAAATCTTTATTATTCTGATTAGCAAATAACTGGCTTTTGTATACCAGGTACTGTTG

TAAATATGACCATCTGCCATCCCACTGAGTTTGACTTTTATTTGTGTAGCCATGCTGGTATACAGATTAT

GACACTCAAAATATTTTAATGGGCAATAGGACGGACTTTGACTTTGTTGTCGGCAATGCACAGACTAATG

AGGTTGGTTGGATTAAATCTCTTAGGTGTTTACCATATTGTTGCTTAATAATGCATTTGAAGGATGGGAG

GTGATCTTTACTTGATGTAGCAATGAAGATGAAAAACAGATCACCTATTTGTAATATTGATATGCGGATT

ATGCTTGTCTGTGCTCTATTATGCTTATTTTTGCTCTAATGTTGGATGTTAATATGCTTCCATTATCCTA

CAAATGGAGTTAGGGTGTTGAAAAGGTGAAACCTCCAACCGGTAGCCCAATATCTATGTTAATGGCTACT

CCAGATGGCTCATATGCATTACCAAGCAGTTGATGTTTTTAAGCTAAATGAGTGTCAGAGATACCATCAT

ATCTCAGTAAATCAGTGTTTGTCGAGCACAAAATGTTCTTGCCGATGATTAGTTGCCCAAAAATATTGTA

AGACATACTGTAAAAAGAATTCAAGAGGCCCATAACAGCAGCGGAGATAATAAAGGCCGTGATTCCCAGT

TGCAAGGTCTGCATTTTTTCTGTCTTTGGAATGTTGCTGTCATGGAATGCCCGAGGAAAATATAACTTCG

GAATCAAGAAGTTTCGAAAGGTAGCTTCGCAAATCAGCCCAACCAAAGCTGC

>MSTRG.236.17 gene=MSTRG.236

GCACAGTCCATCTTCATTCAGCTTCAGGCCCAAGCCCAGTTAGTCATTGTTTTCTCCGCTGCCTGTGTGT

ATATGTACGTATACGTGTGAGATGAGGAAGAATCAATGGCGATTCCAAAGAATTAAGATCAATTCTCACA

CTCTGCATTGTTATGATAGTTGTTTCTATTGCTTTTACTCTCAAATTTTAAACAATGAAGAAAGTGCTTC

AGGAAGCCGATAAGAATGGTCGAAGAAAGCGCTTAATTAATAAATAAGGTAGTGGCATCAATGGACTGGC

CTGAGGTCACCACATACAGAGAACTCTACACACATGGTCAGTGAGAGGAAATATTAAGGAATCTCTACTG

CACTTGTGAAACTGAAAGGGGTGTTGTTGCTAGTGGCATGACCAGGGACTATTTGATGGCATTCTACCAG

CAAACAGAACTCCGACCTAAACGGATCATATTCTACAGGGATGGAGTCAGTGAAGGTCAGTTTAATCAGG

TCTTGCTGAAAGAGATGGATGCCATTAGGACGGTTGATGTGTTTTATTTTTTGTCTGCCAGGCGCTGCAA

TCAAATTATTTACAATGGCAGGAAATCAACTGATCGGAGTGGCAACATATCACCTGGTACTGTTGTAGAT

ATGACCATCTGCCATCCCACTGAGTTTGACTTTTATTTGTGTAGCCATGCTGGTATACAGGTAATCATCT

TAATACAAGACCACATGGATTTTGTACAGGCAGCAGTAATAAATGCATGTGTTGTGTGGCAACACAATAG

AGATACAAATGTACATATTTATTAATATACAAATAGAGAAACAAATGTAAAGATCATAATCAAAATAAAC

ACAAATACACTCATATTAAAAAACTAACATGAACCAATGAGATATAGAAACCATTAAACATCAACTAGTT

TATTCAATCTACCAAAACATCTGAGTAAAGGAAATCCAGATCTGTTCATATTTTACCGGTAACAATTAGC

TAGTTAGGATATTAAATAAGAACAGAAAATTTATGCTCACTAGTTCTCCACGAATTTAGGAAGCTTGTTG

ATTGAATATCATATACTAAAACACACAAATTCTACTATACATACTGTGACTATTTTAATGTCATTAGCCT

AATTCATAACTTACGAAAAATGCTTATGGAGTATGTAGATTCAGGATAGGCAAATAAAAGGATCAGGGAA

CATGTAAGCTAACAAACACTGCATTGTATCTTGTTGAATAATGAATTTGTTTAAAATAAAACTTGCTCTG

ACGCACGTGATCTACGATGGAGTTATATATGCATGCCATTGCATTTACATGTATAGGTTTCCCTATGAAA

TTTATTCAATGTGATATCTTAATGGACAAAATTTAATATTTTAATATTCTCTTATCTGTTAATGCATTCA

TAGATTATGACACTCAAAATATTTTAATGGGCAATAGGACGGACTTTGACTTTGTTGTCGGCAATGCACA

GACTAATGAGGGTGTTGAAAAGGTGAAACCTCCAACCGGTAGCCCAATATCTATGTTAATGGCTACTCCA

GATGGCTCATATGCATTACCAAGCAGTTGATGTTTTTAAGCTAAATGAGTGTCAGAGATACCATCATATC

TCAGTAAATCAGTGTTTGTCGAGCACAAAATGTTCTTGCCGATGATTAGTTGCCCAAAAATATTGTAAGA

CATACTGTAAAAAGAATTCAAGAGGCCCATAACAGCAGCGGAGATAATAAAGGCCGTGATTCCCAGTTGC

AAGGTCTGCATTTTTTCTGTCTTTGGAATGTTGCTGTCATGGAATGCCCGAGGAAAATATAACTTCGGAA

TCAAGAAGTTTCGAAAGGTAGCTTCGCAAATCAGCCCAACCAAAGCTGC

>MSTRG.236.18 gene=MSTRG.236

CACAGTCCATCTTCATTCAGCTTCAGGCCCAAGCCCAGTTAGTCATTGTTTTCTCCGCTGCCTGTGTGTA

TATGTACGTATACGTGTGAGATGAGGAAGAATCAATGGCGATTCCAAAGAATTAAGATCAATTCTCACAC

TCTGCATTGTTATGATAGTTGTTTCTATTGCTTTTACTCTCAAATTTTAAACAATGAAGAAAATGCTTCA

GAAAGCCGATAAGAATGGTCGAAGAAAGCGCTTAATTAATAAATAAGGTAGTGGCATCAATGGACTGGCC

CGAGGTCACCACATACAAAGAACTCTACACACATGGTCAGTGAGAGGAAATATTAAGGAATCTCTACTGC

ACTTGTGAAACTGAAAGGGGTGTTGTTGCTAGTGGCATGACCAGGGACTATTTGATGGCATTCTACCAGC

AAACAGAACTCCGACCTAAACGGATCATATTCTACAGGGATGGAGTCAGTGAAGGTCAGTTTAATCAGGT

CTTGCTGAAAGAGATGGATGCCATTAGGACGGTTGATGTGTTTTATTTTTTGTCTGCCAGGCGCTGCAAT

CAAATTATTTACAATGGCAGGAAATCAACTGATCGGAGTGGCAACATATCACCTGGTACTGTTGTAAATA

TGACCATCTGCCATCCCACTGAGTTTGACTTTTATTTGTGTAGCCATGCTGGTATACAGGGTGTTGAAAA

GGTGAAACCTCCAACCGGTAGCCCAATATCTATGTTAATGGCTACTCCAGATGGCTCATATGCATTACCA

AGCAGTTGATGTTTTTAAGCTAAATGAGTGTCAGAGATACCATCATATCTCAGTAAATCAGTGTTTGTCG

AGCACAAAATGTTCTTGCCGATGATTAGTTGCCCAAAAAT

>MSTRG.236.19 gene=MSTRG.236

CACAGTCCATCTTCATTCAGCTTCAGGCCCAAGCCCAGTTAGTCATTGTTTTCTCCGCTGCCTGTGTGTA

TATGTACGTATACGTGTGAGATGAGGAAGAATCAATGGCGATTCCAAAGAATTAAGATCAATTCTCACAC

TCTGCATTGTTATGATAGTTGTTTCTATTGCTTTTACTCTCAAATTTTAAACAATGAAGAAAATGCTTCA

GAAAGCCGATAAGAATGGTCGAAGAAAGCGCTTAATTAATAAATAAGGTAGTGGCATCAATGGACTGGCC

CGAGGTCACCACATACAAAGAACTCTACACACATGGTCAGTGAGAGGAAATATTAAGGAATCTCTACTGC

ACTTGTGAAACTGAAAGGGGTGTTGTTGCTAGTGGCATGACCAGGGACTATTTGATGGCATTCTACCAGC

AAACAGAACTCCGACCTAAACGGATCATATTCTACAGGGATGGAGTCAGTGAAGGTCAGTTTAATCAGGT

CTTGCTGAAAGAGATGGATGCCATTAGGACGGTTGATGTGTTTTATTTTTTGTCTGCCAGGCGCTGCAAT

CAAATTATTTACAATGGCAGGAAATCAACTGATCGGAGTGGCAACATATCACCTGGTTACATATTGTTCT

ATTGTATCTTTAAAAATCTTTATTATTCTGATTAGCAAATAACTGGCTTTTGTATACCAGGTACTGTTGT

AAATATGACCATCTGCCATCCCACTGAGTTTGACTTTTATTTGTGTAGCCATGCTGGTATACAGATTATG

ACACTCAAAATATTTTAATGGGCAATAGGACGGACTTTGACTTTGTTGTCGGCAATGCACAGACTAATGA

GGTTGGTTGGATTAAATCTCTTAGGTGTTTACCATATTGTTGCTTAATAATGCATTTGAAGGATGGGAGG

TGATCTTTACTTGATGTAGCAATGAAGATGAAAAACAGATCACCTATTTGTAATATTGATATGCGGATTA

TGCTTGTCTGTGCTCTATTATGCTTATTTTTGCTCTAATGTTGGATGTTAATATGCTTCCATTATCCTAC

AAATGGAGTTAGGGTGTTGAAAAGGTGAAACCTCCAACCGGTAGCCCAATATCTATGTTAATGGCTACTC

CAGATGGCTCATATGCATTACCAAGCAGTTGATGTTTTTAAGCTAAATGAGTGTCAGAGATACCATCATA

TCTCAGTAAATCAGTGTTTGTCGAGCACAAAATGTTCTTGCCGATGATTAGTTGCCCAAAAAT

>MSTRG.236.20 gene=MSTRG.236

CACAGTCCATCTTCATTCAGCTTCAGGCCCAAGCCCAGTTAGTCATTGTTTTCTCCGCTGCCTGTGTGTA

TATGTACGTATACGTGTGAGATGAGGAAGAATCAATGGCGATTCCAAAGAATTAAGATCAATTCTCACAC

TCTGCATTGTTATGATAGTTGTTTCTATTGCTTTTACTCTCAAATTTTAAACAATGAAGAAAATGCTTCA

GAAAGCCGATAAGAATGGTCGAAGAAAGCGCTTAATTAATAAATAAGGTAGTGGCATCAATGGACTGGCC

CGAGGTCACCACATACAAAGAACTCTACACACATGGTCAGTGAGAGGAAATATTAAGGAATCTCTACTGC

ACTTGTGAAACTGAAAGGGGTGTTGTTGCTAGTGGCATGACCAGGGACTATTTGATGGCATTCTACCAGC

AAACAGAACTCCGACCTAAACGGATCATATTCTACAGGGATGGAGTCAGTGAAGGTCAGTTTAATCAGGT

CTTGCTGAAAGAGATGGATGCCATTAGGACGGTTGATGTGTTTTATTTTTTGTCTGCCAGGCGCTGCAAT

CAAATTATTTACAATGGCAGGAAATCAACTGATCGGAGTGGCAACATATCACCTGGTTACATATTGTTCT

ATTGTATCTTTAAAAATCTTTATTATTCTGATTAGCAAATAACTGGCTTTTGTATACCAGGTACTGTTGT

AGATATGACCATCTGCCATCCCACTGAGTTTGACTTTTATTTGTGTAGCCATGCTGGTATACAGGGTGTT

GAAAAGGTGAAACCTCCAACCGGTAGCCCAATATCTATGTTAATGGCTACTCCAGATGGCTCATATGCAT

TACCAAGCAGTTGATGTTTTTAAGCTAAATGAGTGTCAGAGATACCATCATATCTCAGTAAATCAGTGTT

TGTCGAGCACAAAATGTTCTTGCCGATGATTAGTTGCCCAAAAAT

>MSTRG.236.21 gene=MSTRG.236

CATCTTCATTCAGCTTCAGGCCCAAGCCCAGTTAGTCATTGTTTTCTCCGCTGCCTGTGTGTATATGTAC

GTATACGTGTGAGATGAGGAAGAATCAATGGCGATTCCAAAGAATTAAGATCAATTCTCACACTCTGCAT

TGTTATGATAGTTGTTTCTATTGCTTTTACTCTCAAATTTTAAACAATGAAGAAAATGCTTCAGAAAGCC

GATAAGAATGGTCGAAGAAAGCGCTTAATTAATAAATAAGGTAGTGGCATCAATGGACTGGCCCGAGGTC

ACCACATACAAAGAACTCTACACACATGGTCAGTGAGAGGAAATATTAAGGAATCTCTACTGCACTTGTG

AAACTGAAAGGGGTGTTGTTGCTAGTGGCATGACCAGGGACTATTTGATGGCATTCTACCAGCAAACAGA

ACTCCGACCTAAACGGATCATATTCTACAGGGATGGAGTCAGTGAAGGTCAGTTTAATCAGGTCTTGCTG

AAAGAGATGGATGCCATTAGGACGGTTGATGTGTTTTACACACATGGTCAGTGAGAGGAAATATTAAGGA

ATCTCTACTGCACTTGTGAAACTGAAAGGGGTGTTGTTGCTAGTGGCATGACCAGGGACTATTTGATGGC

ATTCTACCAGCAAACAGAACTCCGACCTAAACGGATCATATTCTACAGGGATGGAGTCAGTGAAGGTCAG

TTTAATCAGGTCTTGCTGAAAGAGATGGATGCCATTAGGACGGTTGATGTGTTTTATTTTTTGTCTGCCA

GGCGCTGCAATCAAATTATTTACAATGGCAGGAAATCAACTGATCGGAGTGGCAACATATCACCTGGTTA

CATATTGTTCTATTGTATCTTTAAAAATCTTTATTATTCTGATTAGCAAATAACTGGCTTTTGTATACCA

GGTACTGTTGTAGATATGACCATCTGCCATCCCACTGAGTTTGACTTTTATTTGTGTAGCCATGCTGGTA

TACAGGGTGTTGAAAAGGTGAAACCTCCAACCGGTAGCCCAATATCTATGTTAATGGCTACTCCAGATGG

CTCATATGCATTACCAAGCAGTTGATGTTTTTAAGCTAAATGAGTGTCAGAGATACCATCATATCTCAGT

AAATCAGTGTTTGTCGAGCACAAAATGTTCTTGCCGATGATTAGTTGCCC

>MSTRG.236.22 gene=MSTRG.236

CACAGTCCATCTTCATTCAGCTTCAGGCCCAAGCCCAGTTAGTCATTGTTTTCTCCGCTGCCTGTGTGTA

TATGTACGTATACGTGTGAGATGAGGAAGAATCAATGGCGATTCCAAAGAATTAAGATCAATTCTCACAC

TCTGCATTGTTATGATAGTTGTTTCTATTGCTTTTACTCTCAAATTTTAAACAATGAAGAAAATGCTTCA

GAAAGCCGATAAGAATGGTCGAAGAAAGCGCTTAATTAATAAATAAGGTAGTGGCATCAATGGACTGGCC

CGAGGTCACCACATACAAAGAACTCTACACACATGGTCAGTGAGAGGAAATATTAAGGAATCTCTACTGC

ACTTGTGAAACTGAAAGGGGTGTTGTTGCTAGTGGCATGACCAGGGACTATTTGATGGCATTCTACCAGC

AAACAGAACTCCGACCTAAACGGATCATATTCTACAGGGATGGAGTCAGTGAAGGTCAGTTTAATCAGGT

CTTGCTGAAAGAGATGGATGCCATTAGGACGGTTGATGTGTTTTATTTTTTGTCTGCCAGGCGCTGCAAT

CAAATTATTTACAATGGCAGGAAATCAACTGATCGGAGTGGCAACATATCACCTGGTACTGTTGTAAATA

TGACCATCTGCCATCCCACTGAGTTTGACTTTTATTTGTGTAGCCATGCTGGTATACAGGGTGTTGAAAA

GGTGAAACCTCCAACCGGTAGCCCAATATCTATGTTAATGGCTACTCCAGATGGCTCATATGCATTACCA

AGCAGTTGATGTTTTTAAGCTAAATGAGTGTCAGAGATACCATCATATCTCAGTAAATCAGTGTTTGTCG

AGCACAAAATGTTCTTGCCGATGATTAGTTGCCC

>MSTRG.236.23 gene=MSTRG.236

CACAGTCCATCTTCATTCAGCTTCAGGCCCAAGCCCAGTTAGTCATTGTTTTCTCCGCTGCCTGTGTGTA

TATGTACGTATACGTGTGAGATGAGGAAGAATCAATGGCGATTCCAAAGAATTAAGATCAATTCTCACAC

TCTGCATTGTTATGATAGTTGTTTCTATTGCTTTTACTCTCAAATTTTAAACAATGAAGAAAATGCTTCA

GAAAGCCGATAAGAATGGTCGAAGAAAGCGCTTAATTAATAAATAAGGTAGTGGCATCAATGGACTGGCC

CGAGGTCACCACATACAAAGAACTCTACACACATGGTCAGTGAGAGGAAATATTAAGGAATCTCTACTGC

ACTTGTGAAACTGAAAGGGGTGTTGTTGCTAGTGGCATGACCAGGGACTATTTGATGGCATTCTACCAGC

AAACAGAACTCCGACCTAAACGGATCATATTCTACAGGGATGGAGTCAGTGAAGGTCAGTTTAATCAGGT

CTTGCTGAAAGAGATGGATGCCATTAGGACGGTTGATGTGTTTTATTTTTTGTCTGCCAGGCGCTGCAAT

CAAATTATTTACAATGGCAGGAAATCAACTGATCGGAGTGGCAACATATCACCTGGTACTGTTGTAAATA

TGACCATCTGCCATCCCACTGAGTTTGACTTTTATTTGTGTAGCCATGCTGGTATACAGGGTGTTGAAAA

GGTGAAACCTCCAACCGGTAGCCCAATATCTATGTTAATGGCTACTCCAGATGGCTCATATGCATTACCA

AGCAGTTGATGTTTTTAAGCTAAATGAGTGTCAGAGATACCATCATATCTCAGTAAATCAGTGTTTGTCG

AGCACAAAATGTTCTTGCCGATGATTAGTTGCCC

>MSTRG.237.1 gene=MSTRG.237

ATTGAGTTGTGTTTTGGAGCCATTTGATATCACGTGATGCTGTCTGATTTCTTCCTACTTCACATCACTT

TCCACCGCAGTGGTTTAGACAGTTTCGGCAAAGAGACAACAAGCTCATGGCTCCAAAATCCCCACGAGCT

GAATCTTTTCCTCTTCAATCCATTTCTTACAGGTCAGTTTGAATTTTTGTTCAATTTGAATTCAAAACC

>MSTRG.236.24 gene=MSTRG.236

GCACAGTCCATCTTCATTCAGCTTCAGGCCCAAGCCCAGTTAGTCATTGTTTTCTCCGCTGCCTGTGTGT

ATATGTACGTATACGTGTGAGATGAGGAAGAATCAATGGCGATTCCAAAGAATTAAGATCAATTCTCACA

CTCTGCATTGTTATGATAGTTGTTTCTATTGCTTTTACTCTCAAATTTTAAACAATGAAGAAAATGCTTC

AGAAAGCCGATAAGAATGGTCGAAGAAAGCGCTTAATTAATAAATAAGGTAGTGGCATCAATGGACTGGC

CCGAGGTCACCACATACAAAGAACTCTACACACATGGTCAGTGAGAGGAAATATTAAGGAATCTCTACTG

CACTTGTGAAACTGAAAGGGGTGTTGTTGCTAGTGGCATGACCAGGGACTATTTGATGGCATTCTACCAG

CAAACAGAACTCCGACCTAAACGGATCATATTCTACAGGGATGGAGTCAGTGAAGGTCAGTTTAATCAGG

TCTTGCTGAAAGAGATGGATGCCATTAGGACGGTTGATGTGTTTTATTTTTTGTCTGCCAGGCGCTGCAA

TCAAATTATTTACAATGGCAGGAAATCAACTGATCGGAGTGGCAACATATCACCTGGTACTGTTGTAAAT

ATGACCATCTGCCATCCCACTGAGTTTGACTTTTATTTGTGTAGCCATGCTGGTATACAGGGTGTTGAAA

AGGTGAAACCTCCAACCGGTAGCCCAATATCTATGTTAATGGCTACTCCAGATGGCTCATATGCATTACC

AAGCAGTTGATGTTTTTAAGCTAAATGAGTGTCAGGGATACCATCATATCTCAGTAAATCAGTGTTTGTC

GAGCACAAATTGTTCTTGCCGATGATTAGTTGCCCAAAAATACTGTAAGACATACTGTAAAAAGAATTCA

AGAGGCCGATAACAGCAGCGGAGATAATAAAGGCCGTGATTCCCAGTTGCAAGGTCTGCAATTTTTTCTG

TCTTCGAGTCTTTGGAATGTTGCTGTTATGGAATACCCCAGGAAAATATAACTTCGGAATCAAGAAGTTT

CGAAAGGTAGCTTGGCAAATCAGCCCAACCAAAGCTGCTCAGAAGTACAACTTAGATACAAGTTGTTTTT

TCAAATATGTGTGAGTCCTTGTATTGATATCTATCAGGCTAACGAGTGATGCCACTATGGATATGATCTT

GTAGTGAAATTTATATATACATTCATTTATGCACGTATGTATATTAATGTGTTAACTGTACCAAATATTT

ATCATTTAATATTTTTATTCACGGCAATGTTGGTTTTAACCGATATTAAGAAAAT

>MSTRG.236.25 gene=MSTRG.236

GCACAGTCCATCTTCATTCAGCTTCAGGCCCAAGCCCAGTTAGTCATTGTTTTCTCCGCTGCCTGTGTGT

ATATGTACGTATACGTGTGAGATGAGGAAGAATCAATGGCGATTCCAAAGAATTAAGATCAATTCTCACA

CTCTGCATTGTTATGATAGTTGTTTCTATTGCTTTTACTCTCAAATTTTAAACAATGAAGAAAATGCTTC

AGAAAGCCGATAAGAATGGTCGAAGAAAGCGCTTAATTAATAAATAAGGTAGTGGCATCAATGGACTGGC

CCGAGGTCACCACATACAAAGAACTCTACACACATGGTCAGTGAGAGGAAATATTAAGGAATCTCTACTG

CACTTGTGAAACTGAAAGGGGTGTTGTTGCTAGTGGCATGACCAGGGACTATTTGATGGCATTCTACCAG

CAAACAGAACTCCGACCTAAACGGATCATATTCTACAGGGATGGAGTCAGTGAAGGTCAGTTTAATCAGG

TCTTGCTGAAAGAGATGGATGCCATTAGGACGGTTGATGTGTTTTATTTTTTGTCTGCCAGGCGCTGCAA

TCAAATTATTTACAATGGCAGGAAATCAACTGATCGGAGTGGCAACATATCACCTGGTTACATATTGTTC

TATTGTATCTTTAAAAATCTTTATTATTCTGATTAGCAAATAACTGGCTTTTGTATACCAGGTACTGTTG

TAGATATGACCATCTGCCATCCCACTGAGTTTGACTTTTATTTGTGTAGCCATGCTGGTATACAGGGTGT

TGAAAAGGTGAAACCTCCAACCGGTAGCCCAATATCTATGTTAATGGCTACTCCAGATGGCTCATATGCA

TTACCAAGCAGTTGATGTTTTTAAGCTAAATGAGTGTCAGGGATACCATCATATCTCAGTAAATCAGTGT

TTGTCGAGCACAAATTGTTCTTGCCGATGATTAGTTGCCCAAAAATACTGTAAGACATACTGTAAAAAGA

ATTCAAGAGGCCGATAACAGCAGCGGAGATAATAAAGGCCGTGATTCCCAGTTGCAAGGTCTGCAATTTT

TTCTGTCTTCGAGTCTTTGGAATGTTGCTGTTATGGAATACCCCAGGAAAATATAACTTCGGAATCAAGA

AGTTTCGAAAGGTAGCTTGGCAAATCAGCCCAACCAAAGCTGCTCAGAAGTACAACTTAGATACAAGTTG

TTTTTTCAAATATGTGTGAGTCCTTGTATTGATATCTATCAGGCTAACGAGTGATGCCACTATGGATATG

ATCTTGTAGTGAAATTTATATATACATTCATTTATGCACGTATGTATATTAATGTGTTAACTGTACCAAA

TATTTATCATTTAATATTTTTATTCACGGC

>MSTRG.236.26 gene=MSTRG.236

AAAATGCCATCTGTACCCGCACAGTCCATCTTCATTCAGCTTCAGGCCCAAGCCCAGTTAGTCATTGTTT

TCTCCGCTGCCTGTGTGTATATGTACGTATACGTGTGAGATGAGGAAGAATCAATGGCGATTCCAAAGAA

TTAAGATCAATTCTCACACTCTGCATTGTTATGATAGTTGTTTCTATTGCTTTTACTCTCAAATTTTAAA

CAATGAAGAAAATGCTTCAGAAAGCCGATAAGAATGGTCGAAGAAAGCGCTTAATTAATAAATAAGGTAG

TGGCATCAATGGACTGGCCCGAGGTCACCACATACAAAGAACTCTACACACATGGTCAGTGAGAGGAAAT

ATTAAGGAATCTCTACTGCACTTGTGAAACTGAAAGGGGTGTTGTTGCTAGTGGCATGACCAGGGACTAT

TTGATGGCATTCTACCAGCAAACAGAACTCCGACCTAAACGGATCATATTCTACAGGGATGGAGTCAGTG

AAGGTCAGTTTAATCAGGTCTTGCTGAAAGAGATGGATGCCATTAGGACGGTTGATGTGTTTTATTTTTT

GTCTGCCAGGCGCTGCAATCAAATTATTTACAATGGCAGGAAATCAACTGATCGGAGTGGCAACATATCA

CCTGGTTACATATTGTTCTATTGTATCTTTAAAAATCTTTATTATTCTGATTAGCAAATAACTGGCTTTT

GTATACCAGGTACTGTTGTAGATATGACCATCTGCCATCCCACTGAGTTTGACTTTTATTTGTGTAGCCA

TGCTGGTATACAGATTATGACACTCAAAATATTTTAATGGGCAATAGGACGGACTTTGACTTTGTTGTCG

GCAATGCACAGACTAATGAGGGTGTTGAAAAGGTGAAACCTCCAACCGGTAGCCCAATATCTATGTTAAT

GGCTACTCCAGATGGCTCATATGCATTACCAAGCAGTTGATGTTTTTAAGCTAAATGAGTGTCAGGGATA

CCATCATATCTCAGTAAATCAGTGTTTGTCGAGCACAAATTGTTCTTGCCGATGATTAGTTGCCCAAAAA

TACTGTAAGACATACTGTAAAAAGAATTCAAGAGGCCGATAACAGCAGCGGAGATAATAAAGGCCGTGAT

TCCCAGTTGCAAGGTCTGCAATTTTTTCTGTCTTCGAGTCTTTGGAATGTTGCTGTTATGGAATACCCCA

GGAAAATATAACTTCGGAATCAAGAAGTTTCGAAAGGTAGCTTGGCAAATCAGCCCAACCAAAGCTGCTC

AGAAGTACAACTTAGATACAAGTTGTTTTTTCAAATATGTGTGAGTCCTTGTATTGATATCTATCAGGCT

AACGAGTGATGCCACTATGGATATGATCTTGTAGTGAAATTTATATATACATTCATTTATGCACGTATGT

ATATTAATGTGTTAACTGTACCAAATATTTATCATTTAATATTTTTATTCACG

>MSTRG.236.27 gene=MSTRG.236

CATCTTCATTCAGCTTCAGGCCCAAGCCCAGTTAGTCATTGTTTTCTCCGCTGCCTGTGTGTATATGTAC

GTATACGTGTGAGATGAGGAAGAATCAATGGCGATTCCAAAGAATTAAGATCAATTCTCACACTCTGCAT

TGTTATGATAGTTGTTTCTATTGCTTTTACTCTCAAATTTTAAACAATGAAGAAAATGCTTCAGAAAGCC

GATAAGAATGGTCGAAGAAAGCGCTTAATTAATAAATAAGTGGCATCAATGGACTGGCCCGAGGTCACCA

CATACAAAGAACTCTACACACATGGTCAGTGAGAGGAAATATTAAGGAATCTCTACTGCACTTGTGAAAC

TGAAAGGGGTGTTGTTGCTAGTGGCATGACCAGGGACTATTTGATGGCATTCTACCAGCAAACAGAACTC

CGACCTAAACGGATCATATTCTACAGGGATGGAGTCAGTGAAGGTCAGTTTAATCAGGTCTTGCTGAAAG

AGATGGATGCCATTAGGACGGTTGATGTGTTTTATTTTTTGTCTGCCAGGCGCTGCAATCAAATTATTTA

CAATGGCAGGAAATCAACTGATCGGAGTGGCAACATATCACCTGGTACTGTTGTAAATATGACCATCTGC

CATCCCACTGAGTTTGACTTTTATTTGTGTAGCCATGCTGGTATACAGATTATGACACTCAAAATATTTT

AATGGGCAATAGGACGGACTTTGACTTTGTTGTCGGCAATGCACAGACTAATGAGGTTGGTTGGATTAAA

TCTCTTAGGTGTTTACCATATTGTTGCTTAATAATGCATTTGAAGGATGGGAGGTGATCTTTACTTGATG

TAGCAATGAAGATGAAAAACAGATCACCTATTTGTAATATTGATATGCGGATTATGCTTGTCTGTGCTCT

ATTATGCTTATTTTTGCTCTAATGTTGGATGTTAATATGCTTCCATTATCCTACAAATGGAGTTAGGGTG

TTGAAAAGGTGAAACCTCCAACCGGTAGCCCAATATCTATGTTAATGGCTACTCCAGATGGCTCATATGC

ATTACCAAGCAGTTGATGTTTTTAAGCTAAATGAGTGTCAGGGATACCATCATATCTCAGTAAATCAGTG

TTTGTCGAGCACAAATTGTTCTTGCCGATGATTAGTTGCCCAAAAATACTGTAAGACATACTGTAAAAAG

AATTCAAGAGGCCGATAACAGCAGCGGAGATAATAAAGGCCGTGATTCCCAGTTGCAAGGTCTGCAATTT

TTTCTGTCTTCGAGTCTTTGGAATGTTGCTGTTATGGAATACCCCAGGAAAATATAACTTCGGAATCAAG

AAGTTTCGAAAGGTAGCTTGGCAAATCAGCCCAACCAAAGCTGCTCAGAAGTACAACTTAGATACAAGTT

GTTTTTTCAAATATGTGTGAGTCCTTGTATTGATATCTATCAGGCTAACGAGTGATGCCACTATGGATAT

GATCTT

>MSTRG.236.28 gene=MSTRG.236

CATCTTCATTCAGCTTCAGGCCCAAGCCCAGTTAGTCATTGTTTTCTCCGCTGCCTGTGTGTATATGTAC

GTATACGTGTGAGATGAGGAAGAATCAATGGCGATTCCAAAGAATTAAGATCAATTCTCACACTCTGCAT

TGTTATGATAGTTGTTTCTATTGCTTTTACTCTCAAATTTTAAACAATGAAGAAAATGCTTCAGAAAGCC

GATAAGAATGGTCGAAGAAAGCGCTTAATTAATAAATAAGTGGCATCAATGGACTGGCCCGAGGTCACCA

CATACAAAGAACTCTACACACATGGTCAGTGAGAGGAAATATTAAGGAATCTCTACTGCACTTGTGAAAC

TGAAAGGGGTGTTGTTGCTAGTGGCATGACCAGGGACTATTTGATGGCATTCTACCAGCAAACAGAACTC

CGACCTAAACGGATCATATTCTACAGGGATGGAGTCAGTGAAGGTCAGTTTAATCAGGTCTTGCTGAAAG

AGATGGATGCCATTAGGACGGTTGATGTGTTTTATTTTTTGTCTGCCAGGCGCTGCAATCAAATTATTTA

CAATGGCAGGAAATCAACTGATCGGAGTGGCAACATATCACCTGGTACTGTTGTAAATATGACCATCTGC

CATCCCACTGAGTTTGACTTTTATTTGTGTAGCCATGCTGGTATACAGGGTGTTGAAAAGGTGAAACCTC

CAACCGGTAGCCCAATATCTATGTTAATGGCTACTCCAGATGGCTCATATGCATTACCAAGCAGTTGATG

TTTTTAAGCTAAATGAGTGTCAGGGATACCATCATATCTCAGTAAATCAGTGTTTGTCGAGCACAAATTG

TTCTTGCCGATGATTAGTTGCCCAAAAATACTGTAAGACATACTGTAAAAAGAATTCAAGAGGCCGATAA

CAGCAGCGGAGATAATAAAGGCCGTGATTCCCAGTTGCAAGGTCTGCAATTTTTTCTGTCTTCGAGTCTT

TGGAATGTTGCTGTTATGGAATACCCCAGGAAAATATAACTTCGGAATCAAGAAGTTTCGAAAGGTAGCT

TGGCAAATCAGCCCAACCAAAGCTGCTCAGAAGTACAACTTAGATACAAGTTGTTTTTTCAAATATGTGT

GAGTCCTTGTATTGATATCTATCAGGCTAACGAGTGATGCCACTATGGATATGATCTT

>MSTRG.236.29 gene=MSTRG.236

CACAGTCCATCTTCATTCAGCTTCAGGCCCAAGCCCAGTTAGTCATTGTTTTCTCCGCTGCCTGTGTGTA

TATGTACGTATACGTGTGAGATGAGGAAGAATCAATGGCGATTCCAAAGAATTAAGATCAATTCTCACAC

TCTGCATTGTTATGATAGTTGTTTCTATTGCTTTTACTCTCAAATTTTAAACAATGAAGAAAATGCTTCA

GAAAGCCGATAAGAATGGTCGAAGAAAGCGCTTAATTAATAAATAAGGTAGTGGCATCAATGGACTGGCC

CGAGGTCACCACATACAAAGAACTCTACACACATGGTCAGTGAGAGGAAATATTAAGGAATCTCTACTGC

ACTTGTGAAACTGAAAGGGGTGTTGTTGCTAGTGGCATGACCAGGGACTATTTGATGGCATTCTACCAGC

AAACAGAACTCCGACCTAAACGGATCATATTCTACAGGGATGGAGTCAGTGAAGGTCAGTTTAATCAGGT

CTTGCTGAAAGAGATGGATGCCATTAGGACGGTTGATGTGTTTTATTTTTTGTCTGCCAGGCGCTGCAAT

CAAATTATTTACAATGGCAGGAAATCAACTGATCGGAGTGGCAACATATCACCTGGTTACATATTGTTCT

ATTGTATCTTTAAAAATCTTTATTATTCTGATTAGCAAATAACTGGCTTTTGTATACCAGGTACTGTTGT

AAATATGACCATCTGCCATCCCACTGAGTTTGACTTTTATTTGTGTAGCCATGCTGGTATACAGGGTGTT

GAAAAGGTGAAACCTCCAACCGGTAGCCCAATATCTATGTTAATGGCTACTCCAGATGGCTCATATGCAT

TACCAAGCAGTTGATGTTTTTAAGCTAAATGAGTGTCAGGGATACCATCATATCTCAGTAAATCAGTGTT

TGTCGAGCACAAATTGTTCTTGCCGATGATTAGTTGCCCAAAAATACTGTAAGACATACTGTAAAAAGAA

TTCAAGAGGCCGATAACAGCAGCGGAGATAATAAAGGCCGTGATTCCCAGTTGCAAGGTCTGCAATTTTT

TCTGTCTTCGAGTCTTTGGAATGTTGCTGTTATGGAATACCCCAGGAAAATATAACTTCGGAATCAAGAA

GTTTCGAAAGGTAGCTTGGCAAATCAGCCCAACCAAAGCTGCTCAGAAGTACAACTTAGATACAAGTTGT

TTTTTCAAATATGTGTGAGTCCTTGTATTGATATCTATCAGGCTAACGAGTGATGCCACTATGGA

>MSTRG.236.30 gene=MSTRG.236

GCACAGTCCATCTTCATTCAGCTTCAGGCCCAAGCCCAGTTAGTCATTGTTTTCTCCGCTGCCTGTGTGT

ATATGTACGTATACGTGTGAGATGAGGAAGAATCAATGGCGATTCCAAAGAATTAAGATCAATTCTCACA

CTCTGCATTGTTATGATAGTTGTTTCTATTGCTTTTACTCTCAAATTTTAAACAATGAAGAAAATGCTTC

AGAAAGCCGATAAGAATGGTCGAAGAAAGCGCTTAATTAATAAATAAGGTAGTGGCATCAATGGACTGGC

CCGAGGTCACCACATACAAAGAACTCTACACACATGGTCAGTGAGAGGAAATATTAAGGAATCTCTACTG

CACTTGTGAAACTGAAAGGGGTGTTGTTGCTAGTGGCATGACCAGGGACTATTTGATGGCATTCTACCAG

CAAACAGAACTCCGACCTAAACGGATCATATTCTACAGGGATGGAGTCAGTGAAGGTCAGTTTAATCAGG

TCTTGCTGAAAGAGATGGATGCCATTAGGACGGTTGATGTGTTTTATTTTTTGTCTGCCAGGCGCTGCAA

TCAAATTATTTACAATGGCAGGAAATCAACTGATCGGAGTGGCAACATATCACCTGGTACTGTTGTAGAT

ATGACCATCTGCCATCCCACTGAGTTTGACTTTTATTTGTGTAGCCATGCTGGTATACAGGTAATCATCT

TAATACAAGACTACATGGATTTTGTACAGGCAGCAGTAATAAATGCATGTGTTGTGTGGCAACACAATAG

AGATACAAATGTACATATTTATTAATATACAAATAGAGAAACAAATGTAAAGATCT

>MSTRG.240.1 gene=MSTRG.240

ATGTAGGAGGGCCAAAGTCCCGGCCTTACAAAATCAATCATTGTCAGTAATTTCAGTTGCAAGGAGATTG

CTGATGTTCTGGCATTTTCCAGGATTCCTTCTGCAATCAATCAATTTTATCAAGTTAGAGAGGGGTCTCC

TGTTTCTTTTGTTGCTGGTTCTTCAAACAAACATGGAAAGGGACAATCGACTTCGCCTGATTTGCCAGCG

GTGT

>MSTRG.241.1 gene=MSTRG.241

AGAAAATCCAAGACATACTAAGTTGTACAGAAATTTTAAGTAAAAGATGCCTATCCCAGAAATTTGTTTG

AGCTCCGGAGATGCGAAAGCTATGCCAATTATGGGACTGGGATTAGGAGTTCCTGATCTAACACCATAAG

TTATGAATAATACGAAGAAAGTTGTGCTTGACGCGATTGAGCTTGGTTACAGGATATCAAATCTGTTTGG

ACAGCAATGTAGGAGTGCCAAAGTCCCGGCCTTATAAAATCAATCATTCTCAGTAATTTCAGTT

>MSTRG.244.1 gene=MSTRG.244

TATGGACTTGAACTCTTCTCTGCATGCCAGCATTGTTGAAATTAAGAAAGGTCTCTTCTTGAAATTCTAC

TTAACTCTAGCATTTTGTATGCATATTCTTGACACTAGTGTAATAAATAGAACTGTTATCAGTATATTCT

TTTTATCGAAAATGATGTGATCAATTTGTTATTTATTTCATCTTTTGCTAGTTTTGTTGTTCTTCAAACC

ATAGTTTTCATGTGATTGTTTGCAAGTGATAACTAATAATCATGGTCCATGTGGCAGTAATTTTTTTAAA

TTGCCGTAGATTAATTTAGCAGTTGAAGGAGCAAGACACACAGTGTGGCATTGAATCGAGCTTGGGGGAG

GCCCTTTATCCAGGTTTAGGATGCAAGCCAAGGGAGAGAGATGCTTTATGGAGAAAGTATGGAATTGGTA

GTCTGAGTTACTGTTGGCCTTTTCATCTTGTTGGGGATTTGTGTTCAGAATGAACAAGCGGGAGTCGTGA

GATTCTAGCATGACAATAGACTACACTGATTTTCGGTATATTAAAAACATTAAGAATGTTGATAACAGAC

TAGTACAATATAAGGCATTTGCATCTGGCAAATTTACGTGAGAGTTTTTGAAGAGGTTGATGAAGTTCAC

ATTGTGAAATTGTATATGGTACTTACTAATTCATAATCTTATTTTGATGCTGATCTCATGTTAAATAATG

TGAATGCTTTTTGTTACAGATTCGGCATCTACTATATTTCATGTTGATATATGTGCCGTTTCAAAACTTC

AGGCCCCTGTGATATATATTCAGTTTAGAGTTTTAGCAGAGTTGCATTTGTAGCCAGTGGCAGTTCTCAG

AGAGGCCGTGCAGAAATGTGGTGGAAGGATGACAAATC

>MSTRG.244.2 gene=MSTRG.244

TATGGACTTGAACTCTTCTCTGCATGCCAGCATTGTTGAAATTAAGAAAGGAATTGGTAGTCTGAGTTAC

TGTTGGCCTTTTCATCTTGTTGGGGATTTGTGTTCAGAATGAACAAGCGGGAGTCGTGAGATTCTAGCAT

GACAATAGACTACACTGATTTTCGGTATATTAAAAACATTAAGAATGTTGATAACAGACTAGTACAATAT

AAGGCATTTGCATCTGGCAAATTTACGTGAGAGTTTTTGAAGAGGTTGATGAAGTTCACATTGTGAAATT

GTATATGGTACTTACTAATTCATAATCTTATTTTGATGCTGATCTCATGTTAAATAATGTGAATGCTTTT

TGTTACAGATTCGGCATCTACTATATTTCATGTTGATATATGTGCCGTTTCAAAAC

>MSTRG.256.1 gene=MSTRG.256

CTAAACATGAGAGCTGCTGTTTCAAATTGGCGCCAAAAAACATTAAACTTGTCTTGTGCTCAAACTCATC

ATCTAGCTGCTTCTGCTGATCAAACAACTCCCAAGACATTCTTCAACTGCGATTCCCATGTTAAACACTT

GTGTAATGACAGGAAATTCTCGGAAGCCATTCACATTTTGTGTAAAAACAAGCGATTATCCGAAGCAATT

GAGCTGTTAAGCCGTATTGATCGGCCTTCTCCGGCGATATATTCGACTCTCTTGCAGCTCTGTATACAGC

AGAGAGCGGTTGATTTGGGTAAAAAAGTGCATGATCATGCGGAGAATAATGTGGGTTTTAGAGCTGGGAT

TGTGATATGTAATCGTTTGTTGGATTTGTATTGTAAATGTGGGTGTGTTGGGGATGCGCGGAAGGTGTTT

GATGAAATGTCTGCGAGAGATGTGTGTTCTTGGAATATAATGGTATCTGGGTATGTGAAAGCGGGGGATT

TGGGGGGTGCTAGGGGGTTGTTTGATGAAATGCCGGAGAGGGATTATTTTTCATGGAATGCGATGGTTTC

GGGGTATGTGAGGTATGATCAGGCGGGTGAGGGGTTGAAGTTGTTTAGGGTGATGGAGAAGAATGGGGAT

TTGGGGAGTAGTAAGTTTGTGGTTTCGAGTGCTCTTTCGGCTTGTTCTGCTGCGAGATGTTTGCGTTCTG

GGAAGGAGATTCATGGGCATATAGTGAGGACGGGGTTGGATTCTGATGAAGTGGTGTGGAGTTCGTTGTC

CGATATGTATGGGAAATGTGGTAGTATAGATGATGCAAGGCATATATTTGACAAATCTTTGAATAGAGAT

GTTGTTTCGTGGACAGCAATGATTGATAGGTACTTTGAGGATGGAAGGAAGGAAGAGGGGTTTAAATTAT

TTATGGAGTTGTTAAGTTCGGGGATTAGACCAAATGAGTTTACTTTTGCTGGGGTTTTAAATGCTTGTAC

AAGTCAATCTGCAGAATATCTGGGGAAACAAGTTCATGGCCATATGATTCGGATTGGGTGTGATCCGTCA

TCATTTGCAACTAGCACACTGGTCCACATGTATTCCAAGTTGGGAAACATGGAGAATGCTTATAGGGTTT

TCAAAGACATACTGGAGCCTGACCTGGTTTCATGGACTTCTTTAATCAATGGATATGCTCAGAATGGTCA

ACCTCATGAAGCTCTTCGGTTATTCGAGTTGCTTCTTAAGTCTGGTAATAAACCCGATCACGTTACTTTC

ATTGGAGTTCTCTCTGCTTGCACCCATGCTGGTCTAGTTGACAGAGGGCTAGAGTATTTCCACTCAATAA

AAGAGAAACATGGTTTAAAATACACTGCAGATCACTATGCTTGTGTTGTTGATCTCCTGGGTCGTTCTGG

CAGATTTAAAGAGGCTGAAGATATCATTGTAAGCATGCCCATGAAACCAGATAAGTTTCTTTGGGCTTCC

TTGCTTGGTGGTTGCAGAATTCATGGAAATATAGAATTGGCAAAACGAGCAGCTGAAGCGTTATTTCAGA

TTGAACCTGATAATCCAGCTACTTATGTTACTCTTGCAAATATTTATGCCACTGCTGGTAAATGGGATGA

AGTGGCAAAACTCAGAAAGGCTATGAATGACAAAGGAGTTGTCAAGAAACCAGGTATTAGTTGGACAGAG

ATCAAAGGAAAAGTCCACACTTTCATGGTGGGAGATAAATCCCACCCATATTCTACTAAAATATACGACT

ATCTGTCAGAACTGTTCAGAAGAATGAAGGAAGAAGGGTATAATCCCGAGACAGACTATGTGCTGCATGA

TGTAGAAGACGAACAAAAGGAGCAAAATATATTCTACCATAGTGAGAAGCTCGCAGTTGCATTTGGAATT

ATTGCTACTCCACAGGGAACCACAATTAAAGTTTTTAAGAACTTAAGAACTTGTGTTGATTGCCATACCT

CATTCAAATATATCTCCAAAATTGTTAAGAGAAAAATAATAGTAAGGGATTCAACCCGATTCCATTGTTT

TGAGGGAGGAAGCTGTTCTTGCAAAGACTATTGGTGATTTAGGATATATTAACTTTGAAAGTTATTGGAT

GATTGTTGGTTGAAGTGAGCTATGAACTTATGGGACCTATGGTTTGAACTGTTTGTTGTTTTGTGGTTAC

ATTCGTTCTCTTTTACTTCCTCAAACTTTCAGTTCCTCATTCGTATTAAAAAGTTAATATCATTTATCTC

TTCATCGTCTTTGGAGAGTTCTTCTATAGAGTTTGGCCACGGGCAGCACCATCAAGTAGTTCATGCATGA

GTTAATCTCAGAGTTTTTCACTGTGAGATCAACCTATCCCCACAGCTGGTTCCTCGAGCAGAATTCTATA

CAGTTACTGATAGTATACAGGAGTAGTACAAGCAGATTAATTATTGTAGAAACATTTCTTTATAGAATTG

ATGGTGGCGAAACTATAGACTGCTTCAAGAAACTCTGTAGGTGTATGTCATCAAAAGTTTGAATTGTGTT

CTGAGGCCTGTACATTAGTAATTGTTTAT

>MSTRG.257.1 gene=MSTRG.257

TTACATATATGCATATGCTTATTCATCAACATCTGAAAAACACAGAGCTATCTTTCTTATGATATGGAGC

AACTAGATCAGAATTCAGAGCTTGTTGTTATACCAATAAATCCCTTTCCTTTCGTCTTCCTCGACATAGA

CACAGTCTTTTATAGCACTCCACAGCGATTTGTAAAATATTGTGCCATCATACCGGTAGTACTCTCCGAG

TATCGGTTTGATTGCTCTGTTGGCCTCCATTGCATGGTAATGTGGGATTGATGTGAAGAGATGATGTGCT

ACATGAGTATTGGTTGCTTGATAATAGATTGTGTCAAGGATTCCGTAGTTTCGATCAATGGTAGACAATG

ATCAATGGTAGACAAGAAGTTAGAACAATGGAGAAAATTGTGGTGCTCTTCATTCTCATCGTTACATCAA

GTGTTGTTCAGGGTTGCATGACTTGGAATGAGACGGAAATGTCGCAAACAGTGGCAAAAATAAATCAGAT

CACGGAGAGTTTTATGGTTTTTCCAAAAAGGATTATGTATGTGCATAAGGAGAAAAACACTTGTTCTGGG

TATGAAAAGACAAACTGTGATTGACGCTCATGTATTTTAATTGATAAATAATCATGAAATGCGTATTGAT

AATTAAATCACTTTAGTTTATATTAGTTTGTTTGTGT

>MSTRG.257.2 gene=MSTRG.257

TTACATATATGCATATGCTTATTCATCAACATCTGAAAAACACAGAGCTATCTTTCTTATGATATGGAGC

AACTAGATCAGAATTCAGAGCTTGTTGTTATACCAATAAATCCCTTTCCTTTCGTCTTCCTCGACATAGA

CACAGTCTTTTATAGCACTCCACAGCGATTTGTAAAATATTGTGCCATCATACCGGTAGTACTCTCCGAG

TATCGGTTTGATTGCTCTGTTGGCCTCCATTGCATGGTAATGTGGGATTGATGTGAAGAGATGATGTGCT

ACATGAGTATTGGTTGCTTGATAATAGATTGTGTCAAGGATTCCGTAGTTTCGATCAATGGTAGACAATG

ATCAATGGTAGACAAGAAGTTAGAACAATGGAGAAAATTGTGGTGCTCTTCATTCTCATCGTTACATCAA

GTGTTGTTCAGGGTTGCATGACTTGGAATGAGACGGAAATGTCGCAAACAGTGGCAAAAATAAATCAGAT

CACGGAGAGTTTTATGATTTTTCCAAAAAGGATTATGTATGTGCATAAGGAGAAAAACACTTGTTCTGGG

TATGAAAAGACAAACTGTGATTGACGCTCATGTATTTTAATTGATAAATAATCATGAAATGCGTATTGAT

AATTAAATCACTTTAGTTTATATTA

>MSTRG.258.1 gene=MSTRG.258

AATCATGTCCCACGAAGAATCTTCTCAACAAGCAGCAACACCCCAAGTGGAAAAGGGGCTCTGGCTAGCG

GATGTGTTTAACGTGCTATACAGAGGCTGTGACATGATAATGCCGGAAAAGCAGATCGGCACCATCAAAG

TCAGTGATGATCAGAATGATCATTACTTGTACAAGGTGGTAGATGCAGATATGACTAAGTCATCACTTCA

ACAGATAGATGAACTTATGTTGAATGGTCCTGAGATATCATCAATCTCAGCAGAGGATTCCCTGAAAATT

GATGTTGATCTTTTTTGTGGAGCTTTCAAGGACACTTTATATATCGATGATTGCCCCAGCGATGATTCGA

TTGAGACGGATTCTCCCCTCGAAATGAAAATCGTATCAGAAGATGGCAAAGGAGAGATTTATCTTCTGTA

TGCAATATTTGATAATGCTGTAGAAGCCCATTTGGAGGTTAAGCTGTTCGCTACTTTTGAGGTTTATGGA

TGTATTGCTGCGAGTACAAGCAAAATTGAAGTCCCCGGCTATGCACATATGCTCTTTTTAGAGAGACCTT

ACAGTAAAACAAAAGCAGGACCCCTTAATCCCCCTAATCTATTACTGCATTCCAAATCCGTAGTAGCTGT

GCCGTTGGAGTCTGAGTTGCTTGTGGACATTCACTTGATGAGGGGTGATGAGATTGATATTCATGAAGGC

ACCGTTAGTTTTGTGGCTAAACGGACAGGTATCAGTACAGAAGTCGTGAGGGGTGCAAAATGGAAAATTG

AATTGGAAATCTGCTGGAAGTGTGGGACATGAGCCGTGGCAGCCTAGAAATTATATTTAGACTAGGTGCT

TAAATTCTCCACTGTAACGAGATAGTAAGTAGGGATGCATATATTTATGCCAAGAGTTTCTAATGTTTTT

GGACTGAAGTACTGTATGGTTTTTGTTGTACCTTGTACGGATCCATCTTAAAAACTCATTAGAAATTGGA

CATGGCATAAACCAGCTTCAAATGTTAAAAGTAATATGTTAC

>MSTRG.259.1 gene=MSTRG.259

TCTTTCAGCTTCCTCAAAACCAGAGAGAGAGAGAGAGAGCGAGAGAGGTAGAAATCATAGAATCCGTGAG

GCTATGGAAGACAGGGAAACCAGAGATTTTAGGCCAAGAAAGAGAGACCGTGGAGGAGAATCGAGTCTGC

CTCCAATGGATATCTACTGCTCACGCTACGTCTCCAAACTTCTCAGCTTCACCGTCAAAAGAAAGCTAAA

AAGACTTGGTGAAGTTACTTCTGTAATTTGCTCCCCTGATCATGAATCTGTTGAGAAGCTGAGGGCTCTC

AAACAATTGGCTTCTTTGATTGTAGAAGACACAAGACTTGGTGATTATCTGGACTCGAAGGTGCTCCCGG

AACTTATTAGCTTCCTCGAGTTTAAAGAACAGAAAACCCCCCTCCAGGCCGATGGTCGGGTTAGGAAAAA

ACCCGTCTTGGAAGATTTAGTTAACAGGAGATTCCAGTACTATTCTGCAATCGTTCTCACGAATATTGTA

TCGTCCAAGAAAGGCTTTAAGTTCAAAGATAAAGCAATCCCGGCTCTTGTGAAACTTATAACTACTTGGC

ATTGGTTCGTGCAGATTCAGGTTCTGTGTGCATTGTCCAATATAGCCAATGTATTTCCTGAAACCTGTGA

TGCCATCATTAAACATGGGGCATTTGAAATCTTGGAGTCTATTATTACAAATGGGAGGAACCACTATGCT

CTGCGAAAAAGTGCCTGCGAGCTTTTGTCAGCCCTTTGTAAACTGTATGTACCCTTTGAGAAGAGAAAGA

TAGTGTTACATACCCTCAAGACAGCAATATTCTCCGAAAGTGAGTCTGTACTAGTACCAACATGCCTTGC

GCTTTCTAATTTCTCCAACAAAAGATTTGTGGATGTTGGTGCTGAAGTATACAAACGTATTCTTCATCTG

ATTGAGTATACAATACCGGAGGTAGCCCTTTCTGCCCTCAGAGTAATTGGAAATCATGTGAGATGGTGTG

ATAATGACCGGCTTCAGTCTATAATTATTGGAGGTTTACCACTTCACCTTATGGGCTTGTTACACCATAA

ATATACGGAAGTAAAGAAGGAGACTTGCTGGATAATTTCAAACATTATTGCAGCTGCAAACAATTCGCAG

TTACAAGTGGTGCTTAACTGTCAGTTGATAATACTTCTCGTCGGATTTGTTCAAAAAAACAAGGTGGAAT

TGATGGAAGTTGCATCCCGTGCAATTTCAAATGCTGTACTTATTAGTGACAAAAATCAGTTCGAATATCT

TAGAGAGAAATGCTTTGAGCCTTCGAATACCACACGATATTTTTCCAATGATGCAAGGACGATTGCTGCC

TGTTTACAAATACGTGACAATATCTTATATCGTGCCAAGGTTTTGGGATGTGCTTCTCTATCTCAAGAAA

TGGAAAGATTGAGCATCGGAGCGCATGACTATTCTAGAGCTGAATATGCTCAACATTCTAAAAGGATTGG

TAAGAAAAAGAAGACCAAAAAAAGGTAAGCAAATGTCCACTAGTCTTGCACCAAATTAACTAATGATGGA

AGATTTCGTACTAATTTATGAGAGAGATAAAATATACTCGCAATTAAAAATAG

>MSTRG.265.1 gene=MSTRG.265

ACAAATATTGTAAGTGGCTGATCGGAAAGATATCCCGAGGAAATGTTTGCATGAAAAGTTTGGCCATTGT

CATAATTGCCCGTGGAGTGGGTACTGTTATGTCCCCCACCATCGAGTCTTTGGATTGGAAGCAACTTTCC

GTCTTCTTCAACCAGCAATCCTTCTTAACCAGGTAAAGAAGCAAATTTGCAGCCAGGAGGTATCCAGGAC

CTGCAACATTGATCCATAATGATATTGACATCAAACAGAACAAGTTCCGCTTCAATAAAAAGTATGAGAT

GTCTGAGATGCTCTTGGGTGAATTATGAACCCTTCGAATATATAAATAGTTGAGAGAAGATAGACTTGGG

TTTTAATATCTGATGAAAATATAATAATGGTGGTTTATGGCATGAACTTTTAGTACTTTTCGGATACATA

GATG

>MSTRG.266.1 gene=MSTRG.266

GTCCTGGAACGTACAGGTCATGGACTAAATATTGAGAATCGTTTTCGTGATGGGACCTATACAATAATGC

GTGATATATCAAATAGAGACCTCAATCCAAACCTGGGTTCTGACAATCTCCAACTTCGCTTAAACGTTGA

TGCACCCCCTGGTAATTGTACTTTCAGCCACACTATAAAACTAGACATCTTAAATTCATACAAATTGGAC

CCATTGTGTTGTCTGGTATTATTGATCCGTTGAGCTACTTTCTATGTAAACCCAAACGTAATGGCCGGGG

TCCTAACATTGAGAAGCAATTGAGGTTTTCCATGATAGCCAGTCCACTATGTCAAAATGAAATTAGCTGT

TCTGTCGTGAGTGAATCAGTAGTAAAAAAGCGCAAAGGCCGTGGGTTTAGCACTGAGAAACGGATACAGA

TGCGGGAGAAAGAAACACCTCATGGTCAGACACACGAGAACCACTCCTCAGCCATCAACACAGATTAAAG

GCGGCAAAGCATTCACACTCCGTCTTTAAGACCTTCATCTTCGTATTCATCTGTATTGACTGATATATCT

GGCAAAGAGGGACGAACCAATCCCACTACTTGGAATTCTCCTACCGGTATCAAGAATCTGCTTGATGTGT

TTGAGGACGAAGCTGATCACCTAGGTCAAAATCAGCAGCCTCTTGATCATGCTACTAGACCTTCTCCGTT

GTGTAATGTTGATCAGAATGTATCTGTTGGCTGTGACAATCTGTTGGCTATGAGTTGGATTCTACCATCA

AAGGTACATAGATTTTGTACTAGGTTGGATTCTATTTAGATTGTTTTTAAGGTTTGATTATACTGTGATC

TCACCTTCTATTTCTCTACCTGCCAACCCATTACCACTTTTGTATAATGCTGCGCCATGTATGTTTTTGT

CTCTTTTTACTGGATTATAACTATGTACAATTTTGTACATACTTTTGGATCAAAGTAGAGGTAGATATTA

ATTTAATACAGTTTTTCTTTCAC

>MSTRG.266.2 gene=MSTRG.266

GGCAGGTAATCTGAAATTGCATTTCTCTCGCACAATCGGCGCCGATTCAGACCATGAGGTTGTTTTTTCA

AATTTGAATGACAGATAAGTGGCAATTAATAATATGTATCAAGCTTAATCTGTATCATGCTCTTCTGAAG

CGTATCATTCATAATATTCACAGGATTGCAGGCATGGAGGATCATTCGTGAATCGAGAGTCTTCGCTTCT

AATCTCGATCGCGATCTAACAAGACAATCGCTTGACACGCTTTACTGACGGGTTGTTCTCACTCATCCTA

TCAAAGTAGTGGATTCTCACCTTCCACTTACGCCTCGAAGGGCCACGACAGTGGCTCGAAGACGTGGGTT

AGAGTGTTCCGATAATACAGATAATGGATAACGTCGTGTTCATCGCAATGCTCTCCTTCACCAGCGGGTT

GCAAATATGCAGAAACATGCCAGACCCGCTGAAGACAAAAATCAGGTAGGAGAATGGTATAATGCAGGAG

CCCAAAAGGATGCAGATCAATCAGCCTCTTCAATTGATGTTAGTGGGAGCGGCATGTTTGTATCCGGGAT

CACACAGAATGATCCCGGAACAATCACTGGACAAGTTTCAACTGACATGATAGGTTATCGACACCTTTAC

AACTATTAAATTATATTGTTTATAATTAATCAATTTGGATGTACGCTGCCCTCAAATCTTACAATTCATA

CTCATAAACCAGGAAACTCTAAAAGAAGACGTGGTCCAAATGTTGAAACTCTTCTATCGGATAATGTGTT

GAAGAAGGCGCATTCTCAAGACAAGGAAAACCATGTTCCTGTTCATGTCCTGGAACGTACAGGTCATGGA

CTAAATATTGAGAATCGTTTTCGTGATGGGACCTATACAATAATGCGTGATATATCAAATAGAGACCTCA

ATCCAAACCTGGGTTCTGACAATCTCCAACTTCGCTTAAACGTTGATGCACCCCCTGTAGTAAAAAAGCG

CAAAGGCCGTGGGTTTAGCACTGAGAAACGGATACAGATGCGGGAGAAAGAAACACCTCATGGTCAGACA

CACGAGAACCACTCCTCAGCCATCAACACAGATTAAAGGCGGCAAAGCATTCACACTCCGTCTTTAAGAC

CTTCATCTTCGTATTCATCTGTATTGACTGATATATCTGGCAAAGAGGGACGAACCAATCCCACTACTTG

GAATTCTCCTACCGGTATCAAGAATCTGCTTGATGTGTTTGAGGACGAAGCTGATCACCTAGGTCAAAAT

CAGCAGCCTCTTGATCATGCTACTAGACCTTCTCCGTTGTGTAATGTTGATCAGAATGTATCTGTTGGCT

GTGACAATCTGTTGGCTATGAGTTGGATTCTACCATCAAAGGTACATAGATTTTGTACTAGGTTGGATTC

TATTTAGATTGTTTTTAAGGTTTGATTATACTGTGATCTCACCTTCTATTTCTCTACCTGCCAACCCATT

ACCACTTTTGTATAATGC

>MSTRG.266.3 gene=MSTRG.266

GTCCTGGAACGTACAGGTCATGGACTAAATATTGAGAATCGTTTTCGTGATGGGACCTATACAATAATGC

GTGATATATCAAATAGAGACCTCAATCCAAACCTGGGTTCTGACAATCTCCAACTTCGCTTAAACGTTGA

TGCACCCCCTGTAAAAAAGCGCAAAGGCCGTGGGTTTAGCACTGAGAAACGGATACAGATGCGGGAGAAA

GAAACACCTCATGGTCAGACACACGAGAACCACTCCTCAGCCATCAACACAG

>MSTRG.272.1 gene=MSTRG.272

ATTTACATATATGGACAATAGCACCAAATCTGTTATACAGCTATGATATCTAAATCTGACTAGATATGAC

TATTATACAGCTAATACTTTTAACACCCTCCCTCAAATTGATGTTGTTGGTAAATCTACAAGCATCAATT

TGCCAACAAGAAAGTTGTGGCGCTGACGTGGCATCGATTTGGTAAAAATATCAGCGATCTGTAATGTAGT

GGAGGTATGTGGGAGAGTAATGACCAAGTGGTCATAGGCCTCGCGAATAGAGTGGCAGTCAACCTCGATG

TGTTTCGTGCGTTCATGGTAAACTGGGTTAGCTGCGATTTGAATGGCGTTGGTGTTATCACCATGTAAGG

GAGTTGGTTCAGCTTGGGAAAACCCAAGTTCTGCAAGGAGCCCACGCAACCAAATAATCTCAGAACAAGC

AGCAGACATGGATCGATATTCTGCTTCCGTGGATGACTTAGAGACGCGATTCTGCTTTTTACATTTCCAT

GAAATAAGGGAGTCTCCAAGAAACATACACCAACCAGTAATAGATCTTCTAGTGTCTGGGCAACCAGCCC

AATCGGCATCACTATATGCCTGAAGCTTGAGTGAGGAACCAGAAGGGAAAAACAGGCCGCGATTAGGAGT

TCCAAGAATATATCGAATGATCCGTTTGACTGCTGATAGGTGGAAATGCCGAGGACACTGCATGAACTTA

CTTACAATATGGACAGCATAAGAGATGTCTGGTCGAGTGATTGTAAGATATATGAGGCTGCCAACTAGCT

TCCGATATAATGTAGGATCTGAAAGTAGCTCCCCATCATTTCTGCGATATTTGACATTAATTTCCATTGG

GGTTTCAACAGAATTAGTGCCATTGAGTCTTGCCAATTCAACCAGATCCTGAATATACTTGTGTTGGTTC

AAAAAGATACCTTCTGGTCGATGATGTACCTCTAATCCTAAGAAGTAGTGTAACTTTCCAAGATCTTTCA

TTTCAAAAGTAGAACACAACAGATTTTTAGGATGATTGATGGCTTGTATATCAGAACCTGTAACAAGAAT

ATCATCCACGTAGACAAGAAGCACAACTATACCCTTACTTGTCTTTAGCAGAAAAAGGGAAGAGTCATAT

TTACTTTGTATATATGAGAAACCAAGTAGAGTGGATTTGAACTTTTCAAACCAAATTCTAGGTGCCTGCT

TCAGACCATACAAAGATCGCTTGAGTTTGCACACATGATTAGGTGAAATAGGCATGCCATTGGGAAGTGT

TATGTAAACTTCTTCTTTGAGGTCACCATGTAAGAAAGCGTTTTTTACATCCATTTGAGACAAGGTCCAT

GATTGAGACGCCGCAATGGCTAGTATGGTTCGCACAATAGTCATCTTAGCTACAGGGGCAAACGTCTCTT

CGTAGTCAATGCCGTACTCTTGTTTGTTTCCAAGAACTACTAACCGAGCCTTATATCGCTCCACAGAACC

ATCAGGATGAAGTTTTACAGAGAATACAAACTTGCTGTCCAACGGTTTTGCAGATGAGGGACGCACTACA

ACATCCCATGTTTGGTTTTTTTCAAGTGCAAGTAGCTCTGTTTCAATAGCTTCCCGCCAACATTCATGCT

TCATTGCCTGTTTATATGATGATGGAATAGAAATGGGAGACACAGTAGTTGTCAATGATACCGGATTAGA

AAAACCATACCTGTCGGGAGGCTGACTAACTCGAGTACTGTGTCTTCGAGTGGCAGGTGCATTTGCTTGA

ACAAGATCAGCAGCAGGGGAATCAGTTTGAGGGGGTTGAGCAGCTTGAACATGATGAGTAGCTTGATTTG

GATCAGGTTCTGTGGTTTCTGACATGTTTAAGGTTTTCACAGCTTTCGGGCGTCGTTGATATACCAAGAG

TGGTTTAGAGGGAGGCTCTGAATTTGACGATACTGAAGTAGATAAATGTTCTGCAGAAGAGTTAGTAAAT

AATGGTACAATTGATAATGAAGGAGCATGATGATCATTGTAGGTAGAGAAAAAAGGCTGATTTTCTAAGA

AAATTACATTTCTTGAAACCCTTATACGTCGTAAATTAGGATCATAACAGAGAAATCCTTTTTGGTGCTG

AGCATATCCTAAGAAAGCACATCTTATCGATTGAGCTGTGAGTTTTGTGCGCTCATGTGGAGGTAGGTGA

ACAAAACATACACACCCAAAAGTACGAAGACCAGAATAATCAGGAGTATAACCAAAGAACCGAGTGAAAG

GTGAAACATGATTTAAGGATGAAGAGGGTAACCGATTTATGAGGTATACTGCAGCAGAAAGAGCTTCACA

CCAGAACCTAGCGGGAACAAATGATTCTAACAAAAGGGTACGTACAACATCAAGAAGATGACGGTTTTTT

CTTTCAGCTACCCCATTTTGTTGAGGAGTTGAAGGACAAGACCGTTGAAATATTATGCCAATCTTTTGTA

AAAATTCTTGAAACAAATGAGACGTATATTCACCCCCATTATCCGAGCGAAGAATTTTAATATTTGCACG

GAATTGTGTTTGAACATATGCATGAAACACCTTGAAGGTTGAAAGGACCTCAGCTTTGGAAAGAAGGAAA

TATATCCATGTAAAACGACTGTAATCGTCAATAAATGTGACAAAGTACTTATAGTTAGCATGGGAAACAA

CCGGTGACATACCCCAAACATCACTATGTATTAAATCAAAAGCTCGAGTTGCACTAGATGTATGAATGGG

AAATGGTAAAGTCTTGCTTTTTCCAAGTTTACAAGAATTGCAATCAAACTGGACAGAAGATAGCTTTTCA

TTCCCAAGTACACCAGATTTCAACAAAACACGAAGAACATTGTTATTAGGGTGTCCTAGACGCTTATGCC

ATGCATGGTAATCAACACGGGCAGAATTACAAGAAACAATGGGAAAGGAGCAGCTTGGAGATAAAGGAAA

TTAGATAGGAAACAAACGTTCAACTTTAGGCCCCCTCGTGATCACCCTCCCCGACTGTTGATCCTGCACA

ACACAACCAGGTTTTGAAAATGTCACTTTGCAATTATTTTCAACCAATTGACCAACAGAGATAAGATTAG

TAGTAAGGGCAGGAGAGACAAAGACATCGGTTAGGGAGGAAGAAATATCACCGGTTTCTGTAATAG

>MSTRG.273.1 gene=MSTRG.273

AATATATATAATGAGGACTCTCCAGCAATGGGAGATAAACTTTTCCCCACATCTTTTACTTCTAATATTC

TTGTCATGGTATCAGAGCCACAATTCTGATTCGCCTCTTGCTCAAGATCTGTTCTTGTTCTTCGTTCTGT

TCTTGTTCATAGTTCTTGTCTATTGACAAAGCTTCTTTCCTGGTTACTCTTGATCTTTTGAGTCATGCTT

TT

>MSTRG.276.1 gene=MSTRG.276

TTCAGACAAGTCAATGTCCTTGAATTTGGTCCCTGCAATAGATTCAGCCTCCCAACCAGTGCCGAACACG

AACTCCACAGGCTCATAGCCCCTACAATCAAACTGCATCATAGGAGCATATTTACCTGATTCACTTAGTT

CTTGAGTCAGTGCACTGCCTTTGCCACTGATCATAGTCACATTTCCATCCCTCCCACAAAATTTACACTT

CTGAACAAGATGAGCGGTACCCTTCCCCATAGGCACAACTTCAT

>MSTRG.280.1 gene=MSTRG.280

CGCCAGCATACATGTATGGATTGGTTGGCATCCCGAGTACACCAGGGCCGCCATGGTCGGAGTAGTATAT

GAAAATGTGATCATTTGGACCACTATCCACAACCTTCCCACTCCCTCCAGTAGTAGCAGATTTGTCGCCA

AGAATAGCAGCAAAAACATTGTTGACGGTAACATCTTCACCAGTGTAATCCTTTGGAACACCTTTATATA

CATCAGAACCATGAGGGCTGTTAATTATTACTCCAGGCCTAGGATTCTCTTCATCATATGCAATGTCATC

ATACATGAAAACAACAATATTTTCTTCTTTGAC

>MSTRG.290.1 gene=MSTRG.290

TGCTCTGTAGAGAGCACATCATAGGTAGTATGACTGCCTCCCTGCAACTCATAGCTATAAATGCTAGTCA

CACTTGAAGGAGAAGAACTCCCAGAGCTGCACACACTATGTGAATTCTCCAATATCTCCCGAAAAACCAG

CTGTAACATCTTTACTGAATCAATTCCTGGTATCCCATGATCTGGAATCTCAAGTGAATCATTCAATAAC

ATTATGTCATCTGTTGCTACTGAACTCCAGTAGTGACCAAATTTTTCGTGCTCTTGATCATATTCCACTG

CAGTTGCAGATATATATTCTATAATCACTTCTGCAACTGACAACAGATATGACAACACAGTATCCTCCCA

ATTGGACTTCTTCATCCGCTGAACTTCCTTAAACTGCTTAACCTGATCATCCACCCGTAAAGCCAACTGA

TTCGACGTGAGACTAACAGGTGATGAGATGTCAAGCGGAAGAACTCCAACCTCACATTTGACTACCAAGC

AATCATCTTTCAAATAGCCCAATTCCTCCAAATAGCTCTTGTTCATGAAAAGAGGGAATCCCCACTGAAT

ACCGCGGCTTGGACTCACAGAAACTTGCTTCTCCCTGCCCCTGCCGAGCAACCTGCAGGCCTTATGCTTC

CCCTCCCCACTCTGATCAACAAGTGTCAACTGAAACATTGCATACATATCAGTTTCTTCACTCATTAAGC

ACACATACAGCGATAAGTATCCGCTACACTGATCATTTATGCCTCTGGGGTATAAACTGATTGCCCACTT

GTGCCTACCAGCATTAAAAGTATTTGATCTCAAGTATGTTCCAGCTCTGAGACCTCGTTGTGAGTAGTGC

TGAAATGAATACGTCTGTGATCCCATCTCTATTGTAACTCCACTTAGTACTGAAGCAGATGGTAGCTGGC

AGAGAGTGGGAAATTCTAATGCTGTTGAGCTGCTTTCTCTGGATGCTGCAGACCCTTCAGGTGTCGAAGA

TGATGATGAGGCCATTGTTTCTGTGATTTCCGCGTCTTGTGAAGGCTAAGAAAATGCAATTTTCTGCATT

GGATTATGATACAATCTTGTAAAAGATTTAGCAGGTTTAGTGAAGATCTTAGAATAAGGCTTGATATTAA

AAATTATTGAGGTTATAGAAGGGCTAGGCTCATTTTAAAGATTAGTGGTGCTATGTTATCAGACGCGGTA

GCGAGGATGATTGCCATTGACTTGCTTTTCTGAGTCTTTTTATTTAACTTCCAGGGTCTTCTTGAATCTA

GCAGTGTTGTTTGTACAGTCT

>MSTRG.294.1 gene=MSTRG.294

GCCGATTGCTACTGAAGGCACAATCTTAATGTAATTAATGCTGAGGCCAGCAAATAACTGTCTCCAACCT

TGATCACGTGCAATAGTTGTAAGCCCTTCCCATGAGCTTCTGTATCTGGCTCCAGAATGTGTTAAAGATT

GCAAATGTTCTACCTGCATCTGTCTCCTTACAACATCTAATGGGTATGTAAAGGTCTGCCCAAATAAACC

AGCAAGA

>MSTRG.295.1 gene=MSTRG.295

AAAACAACACTTTTCACACTACTTTCTATCTCTTTGCAAAATTCTCAACTCCTGCGGCGTAGATCTAAGG

AGTTTGAGAAAACCCTAATCGTACTTTTCAAGAACCCACTTCACTCTTCAACTCATCTTCAACAATCATG

GCTGGAGCTGCACCAGAAGGTTCTCAATTTGATGCACGTCAATTTGACGCAAAAATGACCGAGTTACTTG

GTGCTGAGGGAGAAGATTTCTTTACATCCTATGATGAGGTTTATGACAGTTTTGATGCTATGGGATTGCA

GGAGAATCTTTTGAGAGGCATCTATGCTTACGGTTTTGAGAAACCATCTGCAATTCAGCAGAGGGGAATT

GTTCCATTCTGCAAGGGGCTAGATGTTATTCAACAGGCACAGTCTGGTACTGGGAAAACAGCAACTTTCT

GCTCTGGAGTTCTGCAGCAGCTTGATTATAGTGTTGTTGAATGTCAAGGTCTGGTTCTTGCTCCTACTCG

TGAACTAGCACAACAGATTGAGAAGGTTATGCGAGCTCTTGGTGATTATCTTGGTGTAAAGGTTCATGCT

TGTGTTGGGGGAACCAGTGTCCGTGAAGATCAGCGCATTCTGTCAAGTGGAGTTCATGTCGTGGTTGGTA

CTCCTGGCCGTGTGTTCGACATGTTGCGAAGACAATCTCTGCGCTCAGATTACATCAAGATGTTTGTTTT

GGACGAAGCTGATGAAATGCTCTCCAGAGGATTCAAGGATCAGATTTATGATATCTTCCAGCTGTTACCT

CCCAAAGTCCAGGTTGGGGTCTTCTCTGCCACCATGCCTCCTGAGGCCCTTGAGATCACAAGGAAGTTCA

TGAATAAGCCTGTGAGGATTCTGGTAAAGAGAGATGAGCTCACTCTTGAGGGTATCAAACAATTTTATGT

TAATGTTGACAAGGAGGAATGGAAACTGGAAACACTTTGTGATCTTTATGAGACCTTGGCTATTACTCAG

AGTGTCATCTTTGTTAATACCAGGCGCAAGGTTGATTGGCTGACTGACAAAATGCGCAGCCGTGATCACA

CAGTCTCTGCCACTCACGGAGACATGGATCAAAACACCAGAGATATAATTATGCGTGAATTCCGTTCTGG

TTCTTCTCGTGTGCTCATTACCACCGATCTTCTGGCTCGTGGTATAGATGTCCAACAAGTATCCCTTGTA

ATTAACTATGATTTGCCGACTCAGCCAGAGAACTACCTTCATCGTATTGGTCGTAGTGGACGTTTTGGAA

GGAAAGGTTTTGCAATCAACTTTGTGACCAAGGATGATGACAGGATGTTGGTTGACATACAGAAGTTCTA

CAATGTAGTAGTGGAGGAGCTGCCAGCCAATGTTGCCGATCTTCTTTAGGGTGGAATTCATCTGTTTATT

TAAGTGTTACTTAAAAGTTACCTTGCTGTCCAATCTTGGTAGGTTTACTATCCGAGACACATATTTTGGG

TATCGAACTCCCTTTACAACTTCTTCCTAGTTTGTTGTAATTTTGTTTTTGTTTTTTTTGCGCAAAATTT

GGTGAATTTCTGTATCCTATATTCTGATACGATTTGCAGAATGGTATGCTTGAAGTCGAACACTTGAGCT

CAAGTGCTGAATCTTAAGTTTTGATTTTTTTCTCATCTGCAAGGATTTTGTCATTTTGCTGTCCGTCTTG

TTTCAGTTGGGTTATAAAGGTTATGTGAGATGTTTGAATATTTATTTACCTTGTGATTC

>MSTRG.295.2 gene=MSTRG.295

AAAACAACACTTTTCACACTACTTTCTATCTCTTTGCAAAATTCTCAACTCCTGCGGCGTAGATCTAAGG

AGTTTGAGAAAACCCTAATCGTACTTTTCAAGAACCCACTTCACTCTTCAACTCATCTTCAACAACTTGA

TTATAGTGTTGTTGAATGTCAAGGTCTGGTTCTTGCTCCTACTCGTGAACTAGCACAACAGATTGAGAAG

GTTATGCGAGCTCTTGGTGATTATCTTGGTGTAAAGGTTCATGCTTGTGTTGGGGGAACCAGTGTCCGTG

AAGATCAGCGCATTCTGTCAAGTGGAGTTCATGTCGTGGTTGGTACTCCTGGCCGTGTGTTCGACATGTT

GCGAAGACAATCTCTGCGCTCAGATTACATCAAGATGTTTGTTTTGGACGAAGCTGATGAAATGCTCTCC

AGAGGATTCAAGGATCAGATTTATGATATCTTCCAGCTGTTACCTCCCAAAGTCCAGGTTGGGGTCTTCT

CTGCCACCATGCCTCCTGAGGCCCTTGAGATCACAAGGAAGTTCATGAATAAGCCTGTGAGGATTCTGGT

AAAGAGAGATGAGCTCACTCTTGAGGGTATCAAACAATTTTATGTTAATGTTGACAAGGAGGAATGGAAA

CTGGAAACACTTTGTGATCTTTATGAGACCTTGGCTATTACTCAGAGTGTCATCTTTGTTAATACCAGGC

GCAAGGTTGATTGGCTGACTGACAAAATGCGCAGCCGTGATCACACAGTCTCTGCCACTCACGGAGACAT

GGATCAAAACACCAGAGATATAATTATGCGTGAATTCCGTTCTGGTTCTTCTCGTGTGCTCATTACCACC

GATCTTCTGGCTCGTGGTATAGATGTCCAACAAGTATCCCTTGTAATTAACTATGATTTGCCGACTCAGC

CAGAGAACTACCTTCATCGTATTGGTCGTAGTGGACGTTTTGGAAGGAAAGGTTTTGCAATCAACTTTGT

GACCAAGGATGATGACAGGATGTTGGTTGACATACAGAAGTTCTACAATGTAGTAGTGGAGGAGCTGCCA

GCCAATGTTGCCGATCTTCTTTAGGGTGGAATTCATCTGTTTATTTAAGTGTTACTTAAAAGTTACCTTG

CTGTCCAATCTTGGTAGGTTTACTATCCGAGACACATATTTTGGGTATCGAACTCCCTTTACAACTTCTT

CCTAGTTTGTTGTAATTTTGTTTTTGTTTTTTTTGCGCAAAATTTGGTGAATTTCTGTATCCTATATTCT

GATACGATTTGCAGAATGGTATGCTTGAAGTCGAACACTTGAGCTCAAGTGCTGAATCTTAAGTTTTGAT

TTTTTTCTCATCTGCAAGGATTTTGTCATTTTGCTGTCCGTCTTGTTTCAGTTGGGTTATAAAGGTTATG

TGAGATGTTTGAATATTTATTTACCTTGTGATTC

>MSTRG.295.3 gene=MSTRG.295

CAAGTGTCTCTAGCTTGAATAAAAGCACTCTCCCTCAGCACTGCTATTACTTCTTCGGGAATATTGATGG

TCTTTTTGTCTTCGACTCATGCTGCATCTTCGCTCATTCATTGTTACCGGTAAATCCTAGAGTTGAAGAT

ACTTCTATGAACTATAGTTTGTGCTCTAGAGGTGCAATGGATTCATTTGATCTAATGACTCTAAACCAGT

CCAACTACAAAATCTGGAGAAGTTGTATGGAGTCCTATCTTGCTAGTGAAGATTTGTGGGATATTGTGTG

TGGAGCACATGGTATTTCTTTGCAAAATACCCGAGTGAATGATGATGCTCTCAAAGATTGGAAGAAAAAG

AACGGCAGAGCTGAATTTGTCTTAAAGAGATCCATCTCTGATGAACTCTTTCATCATATTCTCTATTGCA

AGTCAGCAAGTAGTATCTGGGAAACTCTGACTAGTGTATTCAATAAAACTGACAAGGAAAATTCACTAAG

GGTGTTAGAGAATGAGCTTGAAAATGCCAGACAAGGTACTTTGTCTATTTCACAGTTTTTCTTGAAAATC

AAGAACCTGTGCTCAGAGATTTCTAAACTTACCCCAGATGAGCCGGTTTCTGACGCCCGGCTGAAGTGCA

TAGTTGATCATGGCTTGAAGCCCGAATTCGTTCCCTTTGTGAAATCAATCCAAGGATGGCCTGTACAACC

ATCTTTTGAAGAATATGAGAATATATTGGTTTCACTGGAGGGCTTAGCAAAGAAAAGTGCAGAAATTACG

TCTAAGAATGAAAGGGAAAGCGATTTGAATGCAAGAAAGCCTAAAAATAATAAAGACAAAGTCATGGCTG

GTTGTGTACCAGAAGCTTCTTGCTTCAATACAAGTATGACTGAGCTGAGCAGACTTGCTGCTGACGGAGA

AAAATTCTTTACAACATGTGATGAGGTTTATGACAGCTTTGATGCAATGGGATTGGACGAAAATCTTTTG

AGAGGCATATATGCATTTGGTTTCGAGAAGCCATCTGCGATTCAGCAGCTCGGGATTGTTCCATTCTGCA

AGGGACTAGATGTTATTCAACAGGCAGAGTCTGGTATCGGAAAATCTGTAATTTTTTGCTCTGGAGTTTT

GCAGCAACTTGATTATGATGTTGTAGAGTGTCAAGCACTGGTTCTTGCTCCTACACGTGAACTCGCACAA

CAAATTGAGATGGTTATGCAAGCTCTTGGTGGTCATCTTGGCGTAAGGGTTCTTTCCTGTGGGTGGAATA

GTGCCCGTAATGATCAACGTATTCTGTCAAGAGGTTTTCATGTTGTGGTTGGTTCTCCGAGCCGTGTATT

ACACCTGTTGCGGAAACAGTCTCTTTGTTCAGATCACATTAAAATGCTTGTTTTGGACGAAGTGGATAAA

ATTCCGTCGTTAGACTTCAAAGAGATTAATGATATCCGGCAGTTGCTACCACCTAAAATTCAAGTTGGAA

TTTTCTCGGCCACCACCATACCTCCTGAGGCTCTTGAAATCACAGGAAAATTCATGAATAAGCCTGTGAG

AATTCTGGTAAAGAGAGATGGGCTCAATCTTGAGGGTATCAAACAATTTTATGTTAATGTTGACAAGGAG

GAACGGAAACTGGAAACACTTTGTGATCATTATGAGACCTTGGTTATTTTTCAGAGTGTCATCTTTGTTA

ATACTGAGTGCAAGGTTGATTGGCTGACTGACAAAATGCGCAGCCGTGATTACACAGTCTCTGCTACTCA

TGGAGACATGGATCAAAACACTAGAGATTTTATTATGCATGAATTTCATTCTGGTTCTTCTCGTGTGCTC

ATTACCACCGATCTTCTGGCTCGTGGTATAGATGTCCAACAAGTATCCCTTGTAATTAACTATGATTTGC

CGACTCAGCCAGAGAACTACGTTCATCGAATTGGCTGTATTGGACGTTTTGGAAGGAAAGGTTTTGCAAT

CAACTTTGTGACCAAGGATGATGTCAGGATGTTGGTTGACATACAGAAGTTCTACAATGTAGTAGTGGAC

GAGCTGCCAGCCACTTTTGCCGATCTTTAGTGTGGAATTCATCTGTTTATTAAATTGTTACTTGAAAGTT

ACTCAGCTGTCGAATCTTGGCAGGTTTACTATCCGAGACATATATTGGGTATTGAACTCCCTTTACAACT

TCTTCCTAGTATGTTGTAATTTTGTTTTTGTTTTATGCGCAAAATTTGGTGAATTTCTGTATCCTATATT

CCGATACAATTTGTCGAATGGTATGCTTGAAGTTGAACACTTGAGCTCAAGTGCTGAATCGTAAGTTTTG

ATTTTTTTCTTCTCTTCTCCAAGGATTTCGTCATTTTGCTGTCCTTGTTTCACCTGGGTTATAAAGGTGA

TATGAGATGTTGAATATTTATT

>MSTRG.295.4 gene=MSTRG.295

GTTCAACTCCGAGGTTTTGTATACAAAACAACACTTTTCACACTACTTTCTATCTCTTTGCAAAATTCTC

AACTCCTGCGGCGTAGATCTAAGGAGTTTGAGAAAACCCTAATCGTACTTTTCAAGAACCCACTTCACTC

TTCAACTCATCTTCAACAATCATGGCTGGAGCTGCACCAGAAGGTTCTCAATTTGTCTGGTACTGGGAAA

ACAGCAACTTTCTGCTCTGGAGTTCTGCAGCAGCTTGATTATAGTGTTGTTGAATGTCAAGGTCTGGTTC

TTGCTCCTACTCGTGAACTAGCACAACAGATTGAGAAGGTTATGCGAGCTCTTGGTGATTATCTTGGTGT

AAAGGTTCATGCTTGTGTTGGGGGAACCAGTGTCCGTGAAGATCAGCGCATTCTGTCAAGTGGAGTTCAT

GTCGTGGTTGGTACTCCTGGCCGTGTGTTCGACATGTTGCGAAGACAATCTCTGCGCTCAGATTACATCA

AGATGTTTGTTTTGGACGAAGCTGATGAAATGCTCTCCAGAGGATTCAAGGATCAGATTTATGATATCTT

CCAGCTGTTACCTCCCAAAGTCCAGGTTGGGGTCTTCTCTGCCACCATGCCTCCTGAGGCCCTTGAGATC

ACAAGGAAGTTCATGAATAAGCCTGTGAGGATTCTGGTAAAGAGAGATGAGCTCACTCTTGAGGGTATCA

AACAATTTTATGTTAATGTTGACAAGGAGGAATGGAAACTGGAAACACTTTGTGATCTTTATGAGACCTT

GGCTATTACTCAGAGTGTCATCTTTGTTAATACCAGGCGCAAGGTTGATTGGCTGACTGACAAAATGCGC

AGCCGTGATCACACAGTCTCTGCCACTCACGGAGACATGGATCAAAACACCAGAGATATAATTATGCGTG

AATTCCGTTCTGGTTCTTCTCGTGTGCTCATTACCACCGATCTTCTGGCTCGTGGTATAGATGTCCAACA

AGTATCCCTTGTAATTAACTATGATTTGCCGACTCAGCCAGAGAACTACCTTCATCGTATTGGTCGTAGT

GGACGTTTTGGAAGGAAAGGTTTTGCAATCAACTTTGTGACCAAGGATGATGACAGGATGTTGGTTGACA

TACAGAAGTTCTACAATGTAGTAGTGGAGGAGCTGCCAGCCAATGTTGCCGATCTTCTTTAGGGTGGAAT

TCATCTGTTTATTTAAGTGTTACTTAAAAGTTACTCAGCTGTCGAATCTTGGCAGGTTTACTATCCGAGA

CATATATTGGGTATTGAACTCCCTTTACAACTTCTTCCTAGTATGTTGTAATTTTGTTTTTGTTTTATGC

GCAAAATTTGGTGAATTTCTGTATCCTATATTCCGAT

>MSTRG.295.5 gene=MSTRG.295

CTACTTTCTATCTCTTTGCAAAATTCTCAACTCCTGCGGCGTAGATCTAAGGAGTTTGAGAAAACCCTAA

TCGTACTTTTCAAGAACCCACTTCACTCTTCAACTCATCTTCAACAATCATGGCTGGAGCTGCACCAGAA

GGTTCTCAATTTGATGCACGTCAATTTGACGCAAAAATGACCGAGTTACTTGGTGCTGAGGGAGAAGATT

TCTTTACATCCTATGATGAGGTTTATGACAGTTTTGATGCTATGGGATTGCAGGAGAATCTTTTGAGAAG

ATTTCTTTACATCCTATGATGAGGTTTATGACAGTTTTGATGCTATGGGATTGCAGGAGAATCTATACAG

AAGTTCTACAATGTAGTAGTGGAGGAGCTGCCAGCCAATGTTGCCGATCTTCTTTAGGGTGGAATTCATC

TGTTTATTTAAGTGTTACTTAAAAGTTACTCGGCTGTCCAATCTTGGTAGGTTTACTATCCGAGACAGAT

ATTTGGGTATTGAACTCCC

>MSTRG.295.6 gene=MSTRG.295

AAACCGTAGCAATATTGAGCAATAAAATCAAACGGCGCCGTTCAACTCCGAGGTTTTGTATACAAAACAA

CACTTTTCACACTACTTTCTATCTCTTTGCAAAATTCTCAACTCCTGCGGCGTAGATCTAAGGAGTTTGA

GAAAACCCTAATCGTACTTTTCAAGAACCCACTTCACTCTTCAACTCATCTTCAACAATCATGGCTGGAG

CTGCACCAGAAGGTTCTCAATTTGATGCACGTCAATTTGACGCAAAAATGACCGAGTTACTTGGTGCTGA

GGGAGAAGATTTCTTTACATCCTATGATGAGGTTTATGACAGTTTTGATGCTATGGGATTGCAGGAGAAT

CTTTTGAGAGGCATCTATGCTTACGGTTTTGAGAAACCATCTGCAATTCAGCAGAGGGGAATTGTTCCAT

TCTGCAAGGGGCTAGATGTTATTCAACAGGCACAGTCTGGTACTGGGAAAACAGCAACTTTCTGCTCTGG

AGTTCTGCAGCAGCTTGATTATAGTGTTGTTGAATGTCAAGGTCTGGTTCTTGCTCCTACTCGTGAACTA

GCACAACAGATTGAGAAGGTTATGCGAGCTCTTGGTGATTATCTTGGTGTAAAGGTTCATGCTTGTGTTG

GGGGAACCAGTGTCCGTGAAGATCAGCGCATTCTGTCAAGTGGAGTTCATGTCGTGGTTGGTACTCCTGG

CCGTGTGTTCGACATGTTGCGAAGACAATCTCTGCGCTCAGATTACATCAAGATGTTTGTTTTGGACGAA

GCTGATGAAATGCTCTCCAGAGGATTCAAGGATCAGATTTATGATATCTTCCAGCTGTTACCTCCCAAAG

TCCAGGTTGGGGTCTTCTCTGCCACCATGCCTCCTGAGGCCCTTGAGATCACAAGGAAGTTCATGAATAA

GCCTGTGAGGATTCTGGTAAAGAGAGATGAGCTCACTCTTGAGGGTATCAAACAATTTTATGTTAATGTT

GACAAGGAGGAATGGAAACTGGAAACACTTTGTGATCTTTATGAGACCTTGGCTATTACTCAGAGTGTCA

TCTTTGTTAATACCAGGCGCAAGGTTGATTGGCTGACTGACAAAATGCGCAGCCGTGATCACACAGTCTC

TGCCACTCACGGAGACATGGATCAAAACACCAGAGATATAATTATGCGTGAATTCCGTTCTGGTTCTTCT

CGTGTGCTCATTACCACCGATCTTCTGGCTCGTGGTATAGATGTCCAACAAGTATCCCTTGTAATTAACT

ATGATTTGCCGACTCAGCCAGAGAACTACCTTCATCGTATTGGTCGTAGTGGACGTTTTGGAAGGAAAGG

TTTTGCAATCAACTTTGTGACCAAGGATGATGACAGGATGTTGGTTGACATACAGAAGTTCTACAATGTA

GTAGTGGAGGAGCTGCCAGCCAATGTTGCCGATCTTCTTTAGGGTGGAATTCATCTGTTTATTTAAGTGT

TACTTAAAAGTTACCTTGCTGTCCAATCTTGGTAGGTTTACTATCCGAGACACATAACTATGATTTGCCG

ACTCAGCCAGAGAACTACCTTCATCGAATTGGCTGTATTGGACGTTTTGGAAGGAAAGGTTTTGCAATCA

ACTTTGTGACCAAGGATGATGACAGGATGTTGGTTGACATACAGAAGTTCTACAATGTAGTAGTGGAGGA

GCTGCCAGCCAATGTTGCCGATCTTCTTTAGGGTGGAATTCATCTGTTTATTTAAGTGTTACTTAAAAGT

TACTCGGCTGTCCAATCTTGGTAGGTTTACTATCCGAG

>MSTRG.295.7 gene=MSTRG.295

GTTCAACTCCGAGGTTTTGTATACAAAACAACACTTTTCACACTACTTTCTATCTCTTTGCAAAATTCTC

AACTCCTGCGGCGTAGATCTAAGGAGTTTGAGAAAACCCTAATCGTACTTTTCAAGAACCCACTTCACTC

TTCAACTCATCTTCAACAATCATGGCTGGAGCTGCACCAGAAGGTTCTCAATTTGATGCACGTCAATTTG

ACGCAAAAATGACCGAGTTACTTGGTGCTGAGGGAGAAGATTTCTTTACATCCTATGATGAGGTTTATGA

CAGTTTTGATGCTATGGGATTGCAGGAGAATCTTTTGAGAGGCATCTATGCTTACGGTTTTGAGAAACCA

TCTGCAATTCAGCAGAGGGGAATTGTTCCATTCTGCAAGGGGCTAGATGTTATTCAACAGGCACAGTCTG

GTACTGGGAAAACAGCA

>MSTRG.295.8 gene=MSTRG.295

CTACTTTCTATCTCTTTGCAAAATTCTCAACTCCTGCGGCGTAGATCTAAGGAGTTTGAGAAAACCCTAA

TCGTACTTTTCAAGAACCCACTTCACTCTTCAACTCATCTTCAACAATCATGGCTGGAGCTGCACCAGAA

GGTTCTCAATTTGATGCACGTCAGCAACTTTCTGCTCTGGAGTTCTGCAGCAGCTTGATTATAGTGTTGT

TGAATGTCAAGGTCTGGTTCTTGCTCCTACTCGTGAACTAGCACAACAGATTGAGAAGGTTATGCGAGCT

CTTGGTGATTATCTTGGTGTAAAGGTTCATGCTTGTGTTGGGGGAACCAGTGTCCGTGAAGATCAGCGCA

TTCTGTCAAGTGGAGTTCATGTCGTGGTTGGTACTCCTGGCCGTGTGTTCGACATGTTGCGAAGACAATC

TCTGCGCTCAGATTACATCAAGATGTTTGTTTTGGACGAAGCTGATGAAATGCTCTCCAGAGGATTCAAG

GATCAGATTTATGATATCTTCCAGCTGTTACCTCCCAAAGTCCAGGTTGGGGTCTTCTCTGCCACCATGC

CTCCTGAGGCCCTTGAGATCACAAGGAAGTTCATGAATAAGCCTGTGAGGATTCTGGTAAAGAGAGATGA

GCTCACTCTTGAGGGTATCAAACAATTTTATGTTAATGTTGACAAGGAGGAATGGAAACTGGAAACACTT

TGTGATCTTTATGAGACCTTGGCTATTACTCAGAGTGTCATCTTTGTTAATACCAGGCGCAAGGTTGATT

GGCTGACTGACAAAATGCGCAGCCGTGATCACACAGTCTCTGCCACTCACGGAGACATGGATCAAAACAC

CAGAGATATAATTATGCGTGAATTCCGTTCTGGTTCTTCTCGTGTGCTCATTACCACCGATCTTCTGGCT

CGTGGTATAGATGTCCAACAAGTATCCCTTGTAATTAACTATGATTTGCCGACTCAGCCAGAGAACTACC

TTCATCGTATTGGTCGTAGTGGACGTTTTGGAAGGAAAGGTTTTGCAATCAACTTTGTGACCAAGGATGA

TGACAGGATGTTGGTTGACATACAGAAGTTCTACAATGTAGTAGTGGAGGAGCTGCCAGCCAATGTTGCC

GATCTTCTTTAGGGTGGAATTCATCTGTTTATTTAAGTGTTACTTAAAAGTTACCTTGCTGTCCAATCTT

GGTAGGTTTACTATCCGAGACACATATTTTGGGTATCGAACTCCCTTTACAACTTCTTCCTAGTTTGTTG

TAATTTTGTTTTTGTTTTTTTTGCGCAAAATTTGGTGAATTTCTGTATCCTATATTCTGATACGATTTGC

AGAATGGTATGCTTGAAGTCGAACACTTGAGCTCAAGTGCTGAATCTTAAGTTTTGATTTTTTTCTCATC

TGCAAGGATTTTGTCATTTTGCTGTCCGTCTTGTTTCAGTTGGGTTATAAAGGTTATGTGAGATGTTTGA

ATACTCCCTCCGCCCCCTGAGTAGTATACGGGGAACGGGGACGCGGCACGGACTTTAATACTCCTGTAAA

GTGTAGTTGTG

>MSTRG.296.1 gene=MSTRG.296

TCTATGTTACCCGGACTCTCCATTTTTGTGCCTATACCCGTGTCCACCGGACATGACATGGGTGTGGGTA

TGGGATCTGTGTCGGATCCTTCTTATTACAACCGGACACCTCACCTTGTAACATCCCATGTCAAAATGAA

AACTTAGGTCTTGTTCCAACTTGTTTATAGAGATCAAGATCCATGAATGACTTGTTTCTATGTTGTTCTA

TTGATTTTATGCTTCAATTTTCAGCATATGTGATTTATGTGCAATAGACACTATTTTTATGAATGTACCA

TAAATCAGGCTATATTAAGTTTGACTTGCATGGAGCTTAAGTGCCAAGTCCCCGTACCCGAGTCTAATTC

CGGATCCACATCCACGAATCCTAATTATCCTAAAACTACGAATCTGACTCTTGGATCTGTACCCGTGTCC

GATACCCATACCCGTGTCCGGGTAACTTAG

>MSTRG.304.1 gene=MSTRG.304

TGTTTACAGACATGGAATCGAATACTGATACTATCATGGACGTGGAAAGAGGTGCAGGGAGAAACAAAAG

GAAGTGTATTTTTTGGAGCATCGGATGCAGAGAAGAAACAACTTGTGGATGCTATACTTGCCGGTGAAAT

ATAAGCATTACTGCACTTATTTATAAACTTATGAATGTTTCGAGACTTGTCGATTTTTATTTTGTACTTA

TTTGATGATGTTCAATTTAGAATTTTGTATCATTTTGCCTTTAGTTAATTGCATTTATTTTGTCGATT

>MSTRG.304.2 gene=MSTRG.304

TGTTTACAGACATGGAATCGAATACTGATACTATCATGGACGTGGAAAGAGGTGCAGGGAGAAACAAAAG

GAAGTGGTCTGAAGAAGAAGATGAGAAATTGGTCGAGTCTCTTATTGAACTTGTGAACAATGGAGCATAC

AAAGCTGACAATGGTTTCAAGCCGGGATATCTTGGATTTCTGGAAAACTCATTGAGCGTCAAACTACCAA

CTTCAGGCCTCAAGGGGAAGCCACATATCGAGTCTCGAATAAAAACTCTCAAGAAGGATTTTAATATGGT

TTATGACTTAAGATATGGAAGCACATCTGGTTTTGGTTGGAATTCTGAAAATCAACTAGTCACGGCATCA

AGAGATGTATGGGCTGAATATGCCAAGTCTCATGTTGGGGTTTTGAAATGGAGAAGCACACCATTTCCAT

TTTTCGACGATCTTGCAATCATCTTTGGCAAAGATAGAGCAACTGGTCATAATGCTGAAAGTGCAATGGA

GGCGGAAGAAAACATTAATCTAGAAGAAGCTGCTCAAGGAAAGAATGATGATTCGAGTATGCAAGCATCT

GATGAGTCTACTTCTAGAAGGAGTTCAAAGAGGAAAAAGTTTGATGCTGAACAAATGGCTGAGGTGATGT

ATAATGCTTCTAAAATGATAGCGGCTGAGTTTGCAAATTCAACTAAAGTGATGGCAGCTGAATTTGCGAG

TTCCACCAAGCTGTTGATTGCTGCAGAAACTGATAGGTTGGAGAAAAAAGAAAAGCTAATGGACGAGTTG

TCAAAGATATCTGACATTGATGTTGTGCAAAGGTTCAAGGCTGCAAAGAAGATAGCTGATAGTGAAAATT

TGATGGTGTTATTTTTTGGAGCATCGGATGCAGAGAAGAAACAACTTGTGGATGCTATACTTGCCGGTGA

AATATAAGCATTACTGCACTTATTTATAAACTTATGAATGTTTCGAGACTTGTCGATTTTTATTTTGTAC

TTATTTGATGATGTTCAATTTAGAATTTTGTATCATTTTGCCTTTAGTTAATTGCATTTATTTTGTCGA

>MSTRG.309.1 gene=MSTRG.309

GCCAAGACTGCAGGTAAAAGGTCACACCAGAGAGGCATAACTCATTACAATGAAAGCTCCAACGAGTATA

GGCCAGAGACGAGGATGGCGAAAGAAACTATAGCTTCCACTGCAATCATCATTGGCACCACTGTAACATC

CGGGTCTGGTTGACTTAGCCACACCTCCTTCAAAAGCGAAGCTGCTGTTGCCTTTGCCATTGCTGAACAG

AACACACCAAAAATACGGACCTCCTCCATCTGCCCCGCTAACACCTGCACCCACCTCAGAGTGGTTACTG

TTGTGGACAATTTCCAAGGCCTTGTCATTTGTCACTAGGATGTCTGAGAATGCTTGCTTAGGCTGGATAT

ACTTTGACTGGCATCCGAGTAGACGTCCAGTGAAAGGGGCTAGTGATGAGACCTCAACACCGCAGTTGGG

GGCAAAGGTTTCAGCAATTTCAGCATCAGGAGGCTTCTTGGCGTTTTTCCCTCCCACTGAATCACAATCT

CCTCCGTATGCCTTTATGTACTGCAGGGCTATGCATGCCAGGCCCGGATTGTTAAAGAGTGCTGATGATC

CTTTTCGTGCAGTTCTATTACTGTTAAGCTCCGCGACTAGCTTATCAGCAGGGTTATCTGTCACTTTCTC

TGAGACTTTATATCAAAGAAAATGTTCAGATTAAACGAGCCTAT

>MSTRG.310.1 gene=MSTRG.310

GTTTGAACTTAGGTTAATTTTTTTGCTAGAGAGGAAGCAAGGAAAAGAAAAGAGGAAGACGAGAAAAAGA

GGAGGAGAATGACATCGAATTTTTATTGGATTTGAAGTTGAGGTTGCAAGGCGGGCTATGGGGAGAACGC

AACTTGTGGACTTAGAAGAGTTCGCTTTATGGATTGTATCTTTTGTAGACTTCTAGAATTTTATTCGAAT

TTAGTGATTGTGGATTAAATTTGAATTTGAATATATTATTTTTAGTTTCATGGAATGTAAGTTGGATTTG

TTCGAATTGAGTTTTTTATTGATATACTATGGAC

>MSTRG.311.1 gene=MSTRG.311

GTGAAAGCATTGCCTGATGAACTGGGGAATATTGAATCTCTAAGAGAGGTCAGAATATCGGGAATAGGAG

TTTCGAAATTGCCAGACAGCATTTGCAACCTGAGATCATTGGAAATTCTTGATATATCTTCTTCTGATAC

ACTAGTCAGGTTACCTGATCAACTGTGGAAGCTTACAAGTTTATTAGAGCTAAATGCATGTTCTGCTAGT

CGTTTAGAGAAAGTTCCTGACATAGAATCAAGCCAGACTTCATTACCACTGACAAAGTTGAATTTATCTA

ATAGTAATATCACTGCCCTGCCATCCGGTATTAGTCAGCTCTCAAACCTAGAACGTCTTCATCTTTCGAA

ATGTGATTATCTGTTGTCCATAACAGAACTTCCTCCTAATCTGAAGTACATTTCTGCCATACACTGTACA

TCTTTGGAAAGAGTAAATCTATCCAATTTGAAACTGTTGCGGAGGTTGCAACTCACAAATTGCAGTGCTT

TGACAGAGATTCTAGGCCTGGAGGAACTCACTTCTTTAGAATATCTGCATTTGCTAGATTGCCGCAGCTC

TCTACTGACACATACTTTAACAAAGCCACTGTTCCAGATGTTCTCTGGTTTTGAGAAAAAAATCAATATT

TCGGTTGGCGTGGAAGACTTTCCAGATTGGATTATTCCAGATGAAGATGGAGATGCGTCATGTTCTGTGA

ACTCTTTCGAAGAAACTTCAGATGGAGACGCGTCATTATCTGTGAATTTGCGACCAAACTTGTCACACAA

TTACTTGGGGATGATTCTCTGCTTTGACCAGCGATATTACGATGCCTGTTATTCTGTTATGACTTCCACA

AGTAATATATTAGAGAGCCGTATATATGACTCAGGTATAGTAATAGTTCCGAGATCAATCTTTAGAGTCA

CAGATAGCGATCATACAATCAAACTTACATCAAAAGGAGTAAGTGGATATTGGATTCATCTACTGTACAA

AAACGAGGACAACACATTTGCACCCAGACTTTGGATGCATCTACTGTACAAAAATGAGGACACCAGCATC

ACTCACATTGTTGCAGATGCAGGA

>MSTRG.312.1 gene=MSTRG.312

AAAATGCTGGCAAGACTTTGCACTTTTTGACTAGCAACTCGTATAACAACATTCAACATTGACAGCTGTA

TTGCAAATTTGAAGCTCTTATCTTTTGTATCTAATTCTCCTCCACACAGCATCTATCTCTTCCCCTTATC

TTATCCAATGGCTTCAACAAGTTCTCAAACCCGCACTACCTGGGACGTTTTCTTGAGCTTCCGTGGCACA

GACACTCGAATTGGTTTCACGAGTCATTTATACTCTGCCCTGGATCGCAATGGTATTCGAACTTTTATGG

ATGATCCTGAGCTACGTCTTGGAGATGGGATCACGGCTGAATTGCTCAAAGCTATTCAGGAATCGGAAAT

TTACATTGTTGTCCTCTCGGTAAATTATGCTTCTTCGAGTTGGTGCCTTGATGAGCTGGTAGAGATTCTT

GATTGTAAGGAAACAATGAAAAGATTGGTTATTCCTGTGTTTTTCAACATTGATCCTTCAGTGTAGAGCT

GTCAAACGGATAATTCGGATACGGATATGCCTATATCCATATCCGTATCCGTAATGGCCGGATTCAGATA

CGGATAATATCCGCATTATTTCGGATACGGATAATATCCGCTTTTTCTCGGATACGGATGCGAATATTTC

GGACGGATGTCGGATATTTTTGATGACTCTACCGTATTAACGATGATTTTCAAACAATTTTTTGATGACT

AAGCTCA

>MSTRG.313.1 gene=MSTRG.313

GTCCAGTGAAAGGGGCTAGTGATGAGACCTCAACACCACGGTTGGGGGCAAAGGTTTCAGCAATTTCAGC

ATCAGGAGGCTTCTTGGCGTTTTTCCCTCCCACGGAATCACAATCTCCTCCGTATGCTTTTATGTACTGC

AGGGCTATGCATGCCAGGCCCGAATTGTTAAAGAGTGCTGATGATCCTTTTCGTGCAGTTCTATCACTGT

TAAGCTCAGCTAATTTAACAATGAGAATTTTAGATCTGTCAAAAGGATGTACGTTGAGGAGCTTCTCACC

TGCAACAAGTATTGAAGCAGCAACCAGTTCAGTGCCTATTTCATGTGGGCGATTGTACCTTTGCCGACTC

ATTTCATAATCTTCAACTTCTTCTATTTTCTGTTCTCTCTGTCTTTCAGTAGAACCTCATGGTATATATT

TGTCAATGAAGAAGGATACTTAATATTTGAAGCCTCTCCGAGTTACTTCCATATTCAGCAATAAAATTAA

TAATTTCAGATACTACTGTGATATCACCTCCACAAGGAAACTACACATCGTTGGCACAGGCATTCGTCAT

CAGAGAAACTACTACTTATAAGCTCGACATTTCTTGCAACTCTAGAATGCGCATACAATACTCACCATTA

TCACTTGACAAACAGACAAGAAACTACACAACTCTTGCAACTCCGAGG

>MSTRG.314.1 gene=MSTRG.314

CAAACCCACACCCACTTGCATATATAGTGTGCTATACTAGAGAAGCACATTGATACACACGCACAGATTC

AAAATTTCAAATTCAAATCACATCCTCTTCTTTTTCTCCTTTACATTTGATTATCTTCTCCTCTCTCCCT

CTGATTTGTGGCTGGAAATCAAGCATAAGAGAATCTACTCAGAAAAGGCGTGAATGAATATAAGATTCAG

GTAGCACGAATCCAACTCGAGGACTACAAGAGGAGGATTTCTTTTTTGACAATTATTTCATTGTTCTCAG

GTGATTTTCTGTTGAGATTGCAAAGTCGTTGCTTGATAACAAATGTTGTGTGATCTTATCTGATTTACAC

AATTTGGAAGCTGTAGACTCATTAACTCCCAAGTTGAGTTCAGTCAATTGGAGCCGCCGCATGGCAATTC

CTTACAAGAAGAAAACAACGATGTTCTTTTTGATGAGGATGAAGAAATTTCTGAAGTTGATTCTAAAGAT

TCAGAGGATGAATCTGACCCAGAGACTGATGATTTGGCTATAGGTTTGACTTAAAATGCGAAGCCTATAC

AACGAAAAGGGAAGCATGCAGCAGCCAGACCAAGGTGCAGATAACGCAATGAATAAGGTTGAGACAATTG

ACTCAGGCAAATATTCAAAATCTTCCAGCT

>MSTRG.314.2 gene=MSTRG.314

CAAACCCACACCCACTTGCATATATAGTGTGCTATACTAGAGAAGCACATTGATACACACGCACAGATTC

AAAATTTCAAATTCAAATCACATCCTCTTCTTTTTCTCCTTTACATTTGATTATCTTCTCCTCTCTCCCT

CTATTTGTGGCTGGAAATCAAGCATAAGAGAATCTACTCAGAAAAGGCGTGAATGAATATAAGATTCAGG

TAGCACGAATCCAACTCGAGGACTACAAGAGGAGGATTTCTTTTTTGACAATTATTTCATTGTTCTCAGG

TGATTTTCTGTTGAGATTGCAAAGTCGTTGCTTGATAACAAATGTTGTGTGATCTTATCTGATTTACACA

ATTTGGAAGCTGTAGACTCATTAACTCCCAAGTTGAGTTCAGTCAATTGGAGCCGCCGCATGGCAATTCC

TTACAAGAAGAAAACAACGATGTTCTTTTTGATGAGGATGAAGAAATTTCTGAAGTTGATTCTAAAGATT

CAGAGGATGAATCTGACCCAGAGACTGATGATTTGGCTATAGGTTTGACTTAAAATGCGAAGCCTATACA

ACGAAAAGGGAAGCATGCAGCAGCCAGACCAAGGTGCAGATAACGCAATGAATAAGGTTGAGACAATTGA

CTCAGGCAAATATTCAAAATCTTCCAGCT

>MSTRG.315.1 gene=MSTRG.315

GTGTTAGTGGGGCAGATGGAGGAGGTCCGTATTTTTGGTGTGTTCTGTTCAGCAATGGCAAAGGCCACAG

CAGCTTCGCTTTTGAAGGAGGTGTGGTGAAGTCAACCAGACCCGAATGTTACAGTGGTGCCAATGATGAT

TGCAGTGGAAGCTATAGTTTCTTTCGACATCCTCGTCTCTGGCCTATACTCGTTGGAGCTTTCATTGTAA

TGAGTTATGCCTCTCTGGTGTGACCTTTT

>MSTRG.316.3 gene=MSTRG.316

AAAATTGCAGTACTTTGACAGAGATTTTAGGCCTGGAGGAACTCACTTCTTTAGATGAACTATACTTGCG

AGGTTGCCGCTACTCTCTACTTACACATACTTTAACAAAGCCATTGTTCCAGATGTACTCTGCTCTTGAG

CAAAAAATCAATATTCAGCTTGACGTGGAAGGCTTTCCAGATTGGATTATTCCAGATGCGTCATCTTCTG

TGAACTCTTCCAAAGAAACT

>MSTRG.317.1 gene=MSTRG.317

CTAATATCTTTGCTATCCACCAGTTACAATATCTTCAAAACTTACATACAATGGCTTCAACAAGTGCTGA

AACTCCCACTAGCTGGGACGTTTTCTTGAGCTTCCGTGGCATAGACACTCGACTTGGTTTCGTCAGTCAT

TTATATTCTGCCCTGGATCGCAATGGTATTCGAACTTTTATGGATGATCCTGAGCTACGTCTTGGAGATG

GGATCTCGGCTGAATTGCTCAAAGCTATTCGGGAATCCAAAATTTACATTGTTGTCCTCTCTGTAAATTA

TGCTTCTTCGAGTTGGTGCCTTGATGAGCTGGTAGAGATTCTTGATTGTAAGGAAACAATGAAAAGATTG

GTTATTCCTGTGTTTTTCAACATTGATCCTTCAGTGGTGCGATATCAAAAAGGGAGCTTTGAAGAACATT

TTAGAGTGCATGGAGTTAGATATGCGGATAAAATGGAAAGAATGGAGAATTGGCGCCGTGCACTTAGTCA

ACTTGCTGAGAATTCAGG

>MSTRG.318.1 gene=MSTRG.318

AATCTTTTCATTGTTTCCTTACAATCAAGAATCTCTACCAGCTCATCAAGGCACCAACTCGAAGAAGCAT

AATTTACAGAGAGGACAACAATGTAAATTTTGGATTCCCGAATAGCTTTGAGCAATTCAGCCGAGATCCC

ATCTCCAAGACGTAGCTCAGGATCATCCATAAAAGTTCGAATACCATTGCGATCCAGGGCAGAATATAAA

TGACTGACGAAACCAAGTCGAGTGTCTATGCCACGGAAGCTCAAGAAAACGTCCCAGCTAGTGGGAGTTT

CAGCACTTGTTGAAGCCATT

>MSTRG.316.4 gene=MSTRG.316

TTTGAACATTCTACTAGCCACTTGTATTGCAAAACTGAAGTTCCTATCTTTTGTTTCTAATTCTCCTTCT

CCCCAGAACATCTTTTGTAAAAAAGGCTCTGCTCCCATATACAATATTTTCAAAACTTACATAAATCTCT

TCCCAATGGCTTCAACAAGTGCTGAAACTCCCACTAGCTGGGACGTTTTCTTGAGCTTCCGTGGCACAGA

CACTCGAATTGGTTTCGCCAGTCATTTATACTCTGCCCTGCATCGCAATGGTATTCGAACTTTTATGGAC

GATCCTGAGCTACGTCTTGGAGATGGGATCTCGGCTGAATTGCTCAAAGCTATTCGGGAATCCGAAATTT

ACATTATTCTCCTCTCGGTAAATTATGCTTCTTCGAATTGGTGTCTTGATGAGCTGGTAGAGATCATTAA

TTGTAAGGAAACAATGAATAGATTGGTTATTCCTGTGTTTTTCAACATTGATCCTTCAGCCGTGCGATAT

CAAAAAGGGAGCTTTGAACAACATTTCATAAGGCATGAAATTAGATATGCTGATAAAATAGAAAGAGTCA

ATAATTGGCGCCGTGCACTAGGACAAGTTGCTGAGAATTCGGGAACACATGTAGACGGAAAAAAGTCTGA

AGCTGATATTGTCAACGAAATTGTCAAGGAGATTCTGCTTCAAATAAAACCCACTTCTTTAGATGTTGCC

AGATATCCAGTTGGATTGGATCCCCGAGTTAAAGACATAACAGCAGCATTGTTGAGCAGTGGCACAAAAG

GTGTCATTAAGATCGGTATATATGGTATGGGTGGCGTAGGCAAAACAACTCTTGCCAAGGCACTTTTTAA

TAAACTCTTGCTAGGAAGCTTTGAAGGAAGCTGTTTCCTCGAAAATGTGAGGGAAATTTTGGGGACTATA

AAAGGCCTCGAGTCATTGCAACAACAACTTATAAGTGATGTTCTTAAAATTAGTAAGGACGAAGTTAAAA

TTAGCAGTGTTGGTCAAGGAACTGAGCAGATAGAACGAAGAATTTGTTCTAGAAAAATTTTGGTTGTTAT

TGATGACTTAGAGCACAAGGAAAAATTTGAGTCACTGGTGAGACCATTTGCTCCTGGGAGTGTAGTTATC

ATAACTACAAGGAATGAAGAGATACTTCGGGGAATTGAAGTAGAAACTCAGTATCGATACAAGGTTAATG

AGATGGGTGACGCTGAGGCAGAGATGCTATTTTTCAGACATGCATTTCGAGATACTAAACCAAACGATAA

TCTAATGATATTGTCTAAAGACATTCTACGTCTTGCTGGAGGACTCCCCTTGGCTCTCGAGGTTTTTGGC

TCGTACTTGCATAACACACCCGAGGTTAAATGGAAACCTTACATTGAGAAACTGCAGCGAGATCCCGACA

ACAGTATCGAGCAAAGACTTATAATTAGCTTGGATGCCTCTGGATCAGATGATCCCCTGCTAAAGAAGAT

GTTCCTTGACATTGCTTGTCTTTTCATTGGAAGGACGAAAGAATTTTTGATTAAAATATTGGATACTTAC

TATCCCCACGCGGATGATAAAATTGATATTCTCGAGAAAAGGAGTCTGTTAACATTTAATGACAGAGACG

AAGTACGAATGCATGATCTGCTTCATGATATGGGAAAGAAAATTGCGCGTAACAACTGTCCTGATGAGCC

TGGAAAGCATAGCAGATTATGGGTATCAAAAGAAATATGCGATGTGTTGGAGAAAGACAAGGGAACAGAA

GCGATTGAAGGTTTCTCGTTCGAACGCTTTATAATTAATGGAGGATCGTTTTCTATGGATAGATTGAGAA

GAATGACTAAATTGAGATTTCTTTACTTGGATGGCATTATTCTCACTGGAAAGTTTAAACTGACACTACA

AGTTTTGAGGTGGTTTTGTTGGAATGAGTGTCCTTTAAAGTGTTTACCTTTTGATTTTTGCCCTGAAAAA

CTTGTTATTCTTGAGTTGCCGAAGAGCAAACTGACAACAATGTGGGAGGTAAAGATGGTTTCAGATGTTT

TGGAGAAGCTAAAGACTATAAACATGTTTTTTTCCCAAGATTTAACTTACAGCCCAGATTTCCGAAGACC

ACCGCGTCTCTTTAGTTTAAAAACTCTAAACATGTCATTTTCTGAAGATTTGACTACCACCCCAGACTTC

AAAAGACTGCCGAATCTTGAAAATCTGAATCTTGAGGGCTGTAGAAGTTTGAAGGAGGTCCATGAATCAG

TTGGAAGTTTGGAGAGGCTTGTTTCCCTAAATTTGAAGGATTGTGCAAACCTTAGAAGTCTTCCAGATAC

TATCTGCAAGTTGGAAGCTTTGGAAGTTCTGTGTATTGATCACTGTATAAGGCTAAAAGCACTGCCAATA

GAATTAGGGAACATTAAATCCCTAAAAGAGCTCATTGCTTCGGGAACAAGTTTTCCGAAATTACCAGATT

CGATCGGAGATCTTAGTAAGCTTGTTAAGCTGAAATGGACTCGATGTTCCTTATACGACATGGAACTTGA

ATCTCTTCCAAACACCATTTGCAACTTAAGACAATTGGAAGTTTTAATTGTTACTGTGAAAGCATTGCCT

GATGAACTGGGGAACATTGAATCTCTAAGAGAGCTCAATACATCGGGAATAGCCGTTTCAAAATTGCCCG

ACACCATTTGCAACCTGAGATTATTGGAAATTCTTCATATTACAGTTTCTGAGACACTAGAGAGATTACC

TGATCAACTGTGGAAGCTTACAAGGTTGTTGGAGCTAAAAGTAAGTTGTACTAGTCATTTAAAGAAACTT

CCTGACATAGAATCAAGCCAGACTTCATTACCATTGACAGCGTTGAATTTATCTTATAGTAAAATCACTG

CCCTACCATCCGGTATTAGTCATCTCTCAAACCTAGAAAATCTACAACTTACGAGTTGTCATCATCTGTT

GTCCATAGCAGAACTTCCTCCCAATCTGAAGTACATACACGCCGATCACTGTACATCTCTGAAAAGACTG

AATCTATCCAATTTGAAACTGTTGCGGCAGTTGCACCTTAAAAATTGCAGTACTTTGACAGAGATTTTAG

GCCTGGAGGAACTCACTTCTTTAGATGAACTATACTTGCGAGGTTGCCGCTACTCTCTACTTACACATAC

TTTAACAAAGCCATTGTTCCAGATATACTCTGGTTTTGAGAAAAAAAGTCGATATTTGGCTTGAGACAGA

AAAGTTTCCAGATTGGATTATTCAATCAAGTATAGAATGCTCAGCTGAAGATGACATTTCAGATGGAGAT

GATTCTTCATCATCTTCGGTGAATTTGAAACCAAACTTGTCACACAATTACTTGGGGATGATTCTCTGCC

TTGACCTGAGTCAGAATAAGTTTGATCAACCTTTTACTGCCTCTTATTCTGTTAAGACTTCCGCCAGTAA

TATATTAGAGAGCCGTATATATGAATCAGGTATAGTAATAGTTCCGAGATCAATCTTTACAGGCACGGAT

ACCGATCATACAATCAAATTTGCATCAAGAAGAGCACGCAGACATATTGGACTACTGTACAAAAACGAGG

ACAACATCATCATTCCCAATGTTGCAGGAAGAAACTCCTCTAGAGACAAGAAAAAGAGGAAAAGGATAAA

GAGAAGGAGGAGATGGATTTAGACATGAGAGGTGAAAGGTCACACCAGAGAGGCATAACTCATTACAATG

AAAGCTCCAACGAGTATAGGCCAGAGCCGACGGTGGCAAAAGAAACTATAGCTTCCGCTGCAATCATAAT

CATTGGCACCACTGTAACATCCGGGTCTGGTTGACTTGGCCACACCTCCTTCAAAAGCGAAGCTGCTGTT

GCCTTTGCCATTGCTGAACAGAAAACACCAAAAATACGGACCTCCTCCATCTGCCCCGCTAACACCTGCA

CCCACCTCCGAGTGGTTACTGTTGTGGACAATTTCCAAGGCCTTGTCATTTGTGACTAGGATGTCTGAGA

ATGCTTGCTTAGGCTGGATATACTTTGACTGGCATCCGAGTAGACGTCGAGCGAAAGGGGCTAGTGATGA

GACCTCAACACCACAGTTGGGGGCAAAGGTTTCGTCACAGTATAATACTCTTTCCGCCCCATTCTTATAC

GTTACTTTTTGGTTCATTTTTTAAGACTTTTTAAACTATAGCTCGTAATATATTTTTAAATTATTTTTTC

GGAATAAAA

>MSTRG.320.1 gene=MSTRG.320

AGCAGCTGTGTGCCATATATTTGATTCAAGAACAAACTGGAACAGGACCTGCTAATAAGAGTTAATGAGA

CCAAGGCTTGAGGAAGTAAATTGATGGATATTAGAGAAGTCATAATGCGAAGTAGATTGTTTGTGGAAAA

CCAGACATAGCAAACAACAGAGGCTGGACAGAATTTCAGATGATGGCATCGGAGTATAAGGC

>MSTRG.325.1 gene=MSTRG.325

TTTTTCTCTCTTCTCTCTAGCCGCCTCCTCTTTTCATCTCCGACGGGGCTTCCATCGGTTTCCGGTGGAA

AGCCCCAAATCTTTTCGTTTGATTGCTAGTTTTTGATTGTTCTTGCTTTTTCATTGAGTTTCTTAATAAA

TCTCCATCTTCGGGGAAAGTTTTCTTAGATCTATATCACCGAAACTTTCGATTTTCGTTCTTATTCGCTT

CCAGAGGCCTTTTTGGATTTGTGAGTGGATCGATTATCTCCTAAGCGTCGTGGTGCAGATCTAATTGTTT

TAGTTTGTTTTGATTATAATTTGATTTCTCTCCCTGGTGAGAGTGTGTGATTTTTCTCTCCTTCTTTTGT

GAGAGTGGTTTGGATTCATAGATAAAGCCTTTCGGGAATTGCATTTGTGGTCGATAGCTCAAACGGAGTT

TTTGGAAAAGATGGTGAACACAGAGCAAGGATCTATACATGTACCATCGTCAATTTCAACATCGCTTATT

TGTCTCATCTTGGGTATGAAGACAGGCATGTTAATTTTTTCTATGTTTGTTCGACTCGATCATTCTTGTA

GATGCCGTATGGCCATTATTTATTGAATTA

>MSTRG.326.1 gene=MSTRG.326

CTGCAACAGAATAAAGGGATAGAGATGCACACTACACACACAGACGTCCTGCCCCTCACTTTCTAAATAT

AAAGCGCAAACTGATAAGCATTGCGAACGCACTCGCGCTCCTAGAAATTTCTCAAGCCTCGGTTACATTA

CAAACTGCCAAGAGCCTCGAGCGGCTCTGAGATGTATATGATCTCATATCTCTTCCTTATTTGTACTGAG

TTTGTCGAATCTGAAGGCTCCGACTGGGAGAAAGAGAGATAAATTGCGTAGAGTTTGTTCATTGTTTCTC

ATTCAATTCTCGGTTTCTGAATGACGGAGAAGGAGGAGAGTGCGGTTAGGAAGACCGGAGGTGAATCGCC

GGAGGTGAAGACGCCGGCGGCGAGCAAATCGGCGGTGGCGGCGGCGACTCCGACGGAGGTGGAGGTGGTG

AAGTGCGAGTGCTGCGAGTTTACGGAGGAGTGTACGATCGCGTATATCGCGCGTGTGCGGGAGAGGTACG

AAGGGCGGTGGATCTGCGGGCTGTGTGCGGAGGCGGTGAAGGATGAGATCGGGAGATCTGGCCGGAGGAT

CAGCGAGGAGGAGGCGCTGGAGCGGCATGCGAGTTTCTGTGAGGAGTTCAGGTCGCCGCCGGAGAAGCCG

GCGGAGGAGCTCATCTCCGCCGTGAAGCAGCTGCTGATCAGGAGCTTGGAGTCGCCGACCAGTGTCAGGT

CTAATCCGAGTAGTCCGAGGGGCAAGGACGGGTCGGATCTTGCCCGAGCTACGTTCGGGAGGTCCCAGAG

TTGTTTCCCCAGCATGGATGCCTGAAACATCACCGAACTAAACCGAACGTGCTCAGTATATCTCGTTCCC

GATCCCGAAGAAAGATGCTTCATCGAAGGCTGATGCGCCAATTGCAATGAATAAACGATTATTGTAAAAT

AAATAAATTAGTTTAGGAAAATAAAACGAAATTGAATTAGAGGAAGTGAAGGTTCTTGGCCAATGCAATA

GTTAGGTTCTGGATCTTACTAATTCAGATTGTTGAGTTGTATTTCGAATAAATTAGCCATGGAAATGAAT

AAATTTGATAAATGTAGTGTTTATGTACTTATCAAAAACATGTGGTGAAGAAGATGATGGGTCCGGAGCT

AGTTCATGTTTGTACAATTCTGTTGTGTAATTGCCTTCAGAAATCAATCTTTCACCTCACCAAGTTCAAG

AACCA

>MSTRG.328.1 gene=MSTRG.328

TTGTGATGATAGCAGAAAAGCGTTAGAAATCTTTGCCAAAATTTACAGAAGTGGGAAGAGGTTAGATGAG

ATCACATTCATTACTGTGGCTAATGCCTGCAGCCGCTTGACTGGGTTAGAACAAGGCAGACAAATTCATG

CATGTGTGGTAAAGTTCGGGATCGATTTAGATCTGACGGTCATATCAGGAATCCTTGACATGTATGTTAA

ATGTGGAGCCATGGCAGATGCACACAATGTTTTCCAAGAGATTCCTATACCTGATCAAGTTGCATGGACA

TCAATGATCTCGGGATGCCTGGAAAATGGTGACGAGGATCGTGCCCTTCTAATTTACTATCAGATGAGGA

AATCTGGTATAGCACCAGATGAATACACCTTTGCTACTCTTATCAAAGCCTGCTCTTGCTTAACTGCATT

AGAACAAGGGCGACAAATCCATGCCAACGTGATCAAGTCGTTTTATGTTTCAGATACTTTTGTCTGGACT

TCACTGATTGACATGTATGCTAAATGTGGTAATATAGAAGACTCCTATAATTTATTTAAAAGAGTAACTG

TGAGGGACATTGGCCCGTGGAATGCGATGATAGTTGGTTTAGCTCAGCATGGACATGGAAAAGAGGCTCT

TAAACTCTTTAACAGTTTGAAATTTGTTGGCATAAGACCTGACAGTGTCACCCTTCTTGGAGTTCTTTCC

GCCTGTGGTCATTCTGGCCTGACTTCTGAAGCTAATTCCTATTTTCATTCAATGTATAATGATTATGGTA

TTCAACCTGAGATTGAGCACTATTCCTGCCTCGTGGATGCCCTTGGCCGTGCAGGGCGTGTTAAAGAGGC

AGAAGAACTGATTGAATCAATGCCATACAAAGCTTCTTCTTCAATGTATACAGCTCTGCTTGCCGCTTGT

AAGAGTCAAGGCAATATAGAAACCGGAAAACGGGTAGCTGACCAGCTTTTAGCTTTGGAGCCACCTGATT

CGTCATCTACTTATGTTCTTCTATCAAACATATATGCGACTGCCAACCAATGGGATAAAGTAACTGATGC

TCGAAGAATGATGAAGGGCAAGAATGTTAAGAAGGATCCTGGGTTTAGTTGGATAGATATTAAAAATAAA

GTACATCTGTTTGTGGTGGATGATAAATCACATCCTCATGCTGATGTGATATATGACAAAGTAGAGGAAA

TGATTAGGATGATAAAAAAAGATGGATATGTGCCTGATAAGGATTATGTTTTGCTTGATGTGGAGGATGA

AGAAAAGGAACGTGCTCTCTATTACCATAGTGAGAAGCTAGCTATAGCTTTCGGGCTGCTAAGCACTCCT

CCATGTACAACAATCCGAGTAATCAAGAACCTTAGAGTTTGCGGGGATTGTCATAATGCCATTAAATATA

TATCCAAGGTGTCAAAACGTGAAATCATTCTGAGAGATGCAAATCGATTTCATCGCTTTAGCAACGGGAA

TTGTTCTTGTGGAGATTTCTGGTAACATCAAATCAATTACGAGTTGCCTAAAATTTTGTCAGAATATTGA

ATGTATAATACAGTTGCGAACAAGGTGGTATATAAGCTATTTATTGAACTCGGTCGGATTATAGCCCAGG

TTCTGTAATATATGTTATGATGTGCAAACAAAGCTCATGGTGAAGAACCGAAGACATCGCAGAAAGTAAC

CCCTTTATCTCTCTTGTACTTGTAGAATTTTGTACACACTCTTGTGTATGCCAATGTGACAATGTCTATG

CATATACAATTGTTTCATTAAATAAATCTGTGAGTCAATAGTTTTCCTATTGATGCACTAATTAGGCACA

ATCTGGGGTACTTCAAATTCAAAAGCAGCAGCAAAACACATTCTTTCTGACAAGCTGGCATTGGGAAGAT

GGTGATTATCGTCGTGGCGAAGGGCATTGTTGCATCTGTCAAACAGTATTGTGATGAAGAGATTCATTTA

TCACGCATCACCTACAGGGTGGTTTGCAGCAACGTGCTTCCTGCGTATGGTTTCTACTATTAGCACAGCT

GCTGTAAAGAAGCTTGCTATGTATCAGACCAGTTCGACTTCTGAGTGTATATAGGTTTTGAAAGCTTCAA

GCTGAAGAATATAATGGAGCCTGAAGGATATATAGGTAAACCATCAAATCCAAGGATGTAGGGAACTTTA

GATTACGACTATGAGTTGAATCTCATCGACTAGTTAGATTTATGTCTACCTTGACATGCAGCATACTTGG

AGGAACATAGCCTTCATCTCTAAAGATTAGGAGATACCATTATTTATTTTGGTATAGTTGTACTTGATCG

TTTGTTTAA

>MSTRG.336.1 gene=MSTRG.336

CTTCAATTCGCGTTGGATCCGACCGAAAGCGGGCGCTCCACATTATTTTTTGAAATCCTCGGTCGGTTAT

TTTTTCCCGTCATTTCAGGAGCTGTTAACTCCACCACTTTCCATCCATGATTTCAGCAGCCAGATACTCC

TGCAATTTGAAAACTCAACGTGCCTGTGACGACTCTTTCTACCGACGCAGAGTTTCAGATTCCGGTACAC

ACTCATCTCTCAGACAATGGAATCAGGTAAGTGTCGATCTTCGCTAGGTGTACTCGAGATTAAG

>MSTRG.338.1 gene=MSTRG.338

TTTTAGTCTGTGATGCTCGGACTCTCCAGAAAACCCAGCCGTACCCGTGTCCACAGGACACGACACTGAC

ACTGACACTGGGATCCGGATCCGAACATCATCCAGAAATCAGACTCTCCATCTTAGACAACCTATCTTTA

ATTTCAGTGACCTATCTCTAAGAGGAAGATAGGAATAAGAAAATATATGAAGAAACAGAGGTGAAATGGA

TGTAAAAGGAGAAACGAAGCAACTTTTGCCTTAATTTGTGCAATTGTTTGTTATGCAGTAT

>MSTRG.347.1 gene=MSTRG.347

GGGGAATGTGATGATGAGTACAAAAGTTTGTGGAAGAGAATCTGGGCGTTGAAATTACCTCGAAAGGTGA

CACACGTCATATGGAGGTTGTGTAAAGATTGCCTTCCTACTAATACAGCACTGTACAATAGATATGTTAA

TGTAAATCCACTATGTCTGTGGTGTCATGCTGCAGCTGAAACGAATTTGCATGCTATTTTCCTCTGTGAT

TTTGCTCTGACGATGTGGCGAACGGTGGGTTTACAATCGCTGGTGCATTGTACAGACCAGGAAGCGCCTA

AGGAGATTTTTGATCGTGTG

>MSTRG.348.1 gene=MSTRG.348

AATCATACTTGTGTGCATGCATTAGCTAAACACTGATACTAAAACGGCACTTTAACTGATCAAGAAGGCC

AGTAACAGTCGTAGCAAATCCATGCAACTCTCTACCACCATTTTCATACGGACACGCCGGAATCCACTGC

AACCTCTTCATCCCCATGTCGAACAACGCCGCCTTCCCGGACCCTCTAATCACCATCGCAACATACTGAC

CATTTCCAACGCAGTTAAACCCTCTGCCACTCTCCTCAGCCTCGAACTGTGCATACAGTTGTAGTGGAAT

TCTGTAAAGCCCAGAGCCTCAGGCTTTTCGGCACGTTGAGCTTGCTTTTCTCCAC

>MSTRG.350.6 gene=MSTRG.350

CAGAAGCTTGATTCTAGGGTATTTTGAGGAGTATAGATTTATGATTTAGCTGTTTGACTGACGATTCTCT

CTGTTGTGTGCCCCTTTTTTCTAGGATTTCAAGCGCTGCTTCGCCGCTGGGAAATCAATTATCCTATTAA

ATCTATCTTTCTCGCCGGAGCTGGTTGTGTGATTGTGTATGTTTATACTAGCAAATATTGAAGCAAGGCT

GGGTGACCTGGGTCCATGTGATCCTAGAACACCATAATTGTGCAAACCTGCCTCTTCGTACTAGCACACA

CTCTCCAACATGAACTGCTGAAGATGTTCCAAAATACACAGAACGAACTGGAAAAAGACATAAGAGCATG

GAAGAGTCATAGCCAGTGTTATGAAAAGCGCAAATCGGCTCTAGGCGGTGAAAGGACCTTCTAGCGCTTA

AGCGGTAAAGCGGACGCTTAAGC

>MSTRG.351.1 gene=MSTRG.351

GATGAGTACAAGTAATGCTACATAATCTCCTCAAGCTGATCCACTTACCTTTTTTCACTGTGGTTTCTAA

AACACACTATCCTTTCCTCTAATAGCTGCTCATCACCAGGGAGAGATGAAGATACAGTGTAACGTGTGTG

AGGCGGCGGAGGCGACGGTGCTGTGCTGCGCCGACGAGGCGGCGTTGTGTTATGGTTGCGATGAGACCGT

CCATGCAGCTAACAAGCTGGCTAGTAAACACCAGAGAGTTTCTCTCACTAACTCCTCATCTCCTATGCCC

AAATGCGATATTTGTCAGGAAACAGTTGGATATTTCTTTTGTCTGGAAGATCGTGCTTTGCTATGCAGGA

AATGTGACGTCTCCATACATTCGATCAATGCCCATGTTTCAGCTCATCAACGCTTTCTGCTAACTGGAGT

GAAGGTGGGACTTGAACCTACTGATGAACGTAGTGCGCCATCTTCTTCAGGAAAGTCTCAGTCCCCTGAC

AAAGTTACTGAAACAGAGTCAAGGCCATTGTCTAAAAAGAATAACAACGTGGCTTTGGAAGGTCAAAACA

ACAGTGGACATTTCCAAGTTGGCGAAGTTGGTGAGGTGGCACCGACCAATTCGCCTTTTGGTGGAGGGTC

TGATTGTGAAAGCTTTCAACAATGGCAATTAGATGAAATATTTGGAGTAAGCGATCTCAATCAGAATTAC

AACTATATGGACAATGGGTCATCCAAGGTTGTGTACCTGAACTATCACTTTCTGGCACTTTGTGTACCTG

AACTATCACTTTCTGGCACATTGTGTACCTAGAGTTTGAAGATTCTATCGAAAAGCATACTCCCGTTAGT

TTTATGTTAACACCGTTAGTTGAAGTTGTAATAAGACTACAAAGTGAAGGGTAGTTGAGTACTTTCCAGC

GGTGTACTACCATTACTTCTAGCAATTAAACAGAGGAAGCCAGGTCTGTGCACTGCCTCAGCAACAAATT

GTGAGGTCCTGTGTAAAAATTTAATAATAGGATCCGGAATATTTATATAAACATATTACTATTGCTCAAT

TGTTACATTAATATAA

>MSTRG.351.2 gene=MSTRG.351

GTACAAGTAATGCTACATAATCTCCTCAAGCTGATCCACTTACCTTTTTTCACTGTGGTTTCTAAAACAC

ACTATCCTTTCCTCTAATAGCTGCTCATCACCAGGGAGAGATGAAGATACAGTGTAACGTGTGTGAGGCG

GCGGAGGCGACGGTGCTGTGCTGCGCCGACGAGGCGGCGTTGTGTTATGGTTGCGATGAGACCGTCCATG

CAGCTAACAAGCTGGCTAGTAAACACCAGAGAGTTTCTCTCACTAACTCCTCATCTCCTATGCCCAAATG

CGATATTTGTCAGGAAACAGTTGGATATTTCTTTTGTCTGGAAGATCGTGCTTTGCTATGCAGGAAATGT

GACGTCTCCATACATTCGATCAATGCCCATGTTTCAGCTCATCAACGCTTTCTGCTAACTGGAGTGAAGA

GTCAAGGCCATTGTCTAAAAAGAATAACAACGTGGCTTTGGAAGGTCAAAACAACAGTGGACATTTCCAA

GTTGGCGAAGTTGGTGAGGTGGCACCGACCAATTCGCCTTTTGGTGGAGGGTCTGATTGTGAAAGCTTTC

AACAATGGCAATTAGATGAAATATTTGGAGTAAGCGATCTCAATCAGAATTACAACTATATGGACAATGG

GTCATCCAAGGTTGTGTACCTGAACTATCACTTTCTGGCACTTTGTGTACCTGAACTATCACTTTCTGGC

ACATTGTGTACCTAGAGTTTGAAGATTCTATCGAAAAGCATACTCCCGTTAGTTTTATGTTAACACCGTT

AGTTGAAGTTGTAATAAGACTACAAAGTGAAGGGTAGTTGAGTACTTTCCAGCGGTGTACTACCATTACT

TCTAGCAATTAAACAGAGGAAGCCAGGTCTGTGCACTGCCTCAGCAACAAATTGTGAGGTCCTGTGTAAA

AATTTAATAATAGGATCCGGAATATTTATATAAACATATTACTATTGCTCAA

>MSTRG.356.1 gene=MSTRG.356

AAAAGAACAACTGGAGAGCTACTAAATTTAACAGAAGGAGGTAAGAAGCTGAGCATAGAAAGAAAGCTCG

GGTGCATCTTTCTTCATTACCGAGTGTGGCTATTTATAGCCAATGTGAATTTGCAAGAAGCCAAGAACAT

TAAATGCATACACAAATTCTACTCCTATAAATCATAGCCCTACTATTTGATTCATTGACCACATGCATGG

CAATCACTATATTATAACACTCCCCTTTGATTGTCATGATAGTGTGGATCATCAATTGCCTCGTTAAAAC

CTTGTCAAAGAAAAACCCAGTGGGATAAAAACTTTAACGAAGGAAAAAGAGTACAATCTCCCCTTGGAAG

AAAACATCAATCACTGGAGTTTTGTTGCAGCATATTCTTGAGTCGACG

>MSTRG.371.1 gene=MSTRG.371

GTATTAAGGGGCACCCTCTAATTATAAGGTGCTTTTATAAATACAGCTTTTGCAAGGAGCATCTGCAAAG

ACACTCTACAGGAGAACAAGGACGGGTACAGGTAACACTAAGAAGTTGTTTTGCACTCGGAATCAAGATG

GCTCAACTGAACGGAGATGGCATACAGTTATGGGCTAAAGGCATAATGCACCAAAATGAGTTTCGTTCAG

CGGTTCTGTATGCAAGAAATCTGTCAATAGTTTGGGGTTCAGATGGTCGTTACTGGAAGTGGGAACGGAT

CAGCAACCCGATAAGCAGCGAGGATCTTGAGGTTGCTAAACTAATTGAAGTATGCTGGCTACAGATCGAT

GGAAAATGTAGGGACCAGAACCTGGCTAAGGGAGTGACATATGGAGTGTATTTCATTGTCAAGCTTGAAG

AGAATCATTCCATCACCCGTCCTGTGACTCTTAAGCTTACTTGTCCCAACGGTGACACCCAAGAGAACAA

GGTGGATTTGAAGCAGCAGCCGAAGAAGGAACTGATAGGGCTTAAGGTCGGTGAATTTACTGGCTCAGGA

ACCAACAAATTTATCCGATTTTCTTTTAATGGACGGGAGGGTACCAGTTGGAAGACCGGCCTTACTTGCT

TCGGTGCTGTAATTGTGCCAATATGATTTCTATATAACTTGAACTGCCAACACCCTCCTGTTATGCCTGT

GATATAAGAAATGTATCAGATGTACTAAATGTATTAAATAATAAATGACCTGATATGTTAACGTTCGAG

>MSTRG.372.1 gene=MSTRG.372

GCGAGGATCTTGAGATTGCTGAACTAATTGAAGTATGCTGGCTGCAGATCGATGGAAAATACAGGGACCA

GAACCTGGCTAAGGGAGTGACATATGGAGTTTATTTCATTGTTAAGCTCGAAGAGAATCATTCCATCACC

CGTCCTGTGACTCTTAAGCTTACTTGTCCCAGTGGAAACACCCAAGAGAACAAGGTGGATTTGAAGCAGC

AGCCGAAGAAAGAACTGATAGGGC

>MSTRG.376.1 gene=MSTRG.376

CAACCTTCATTTTCAAAGGCCATTCTTTCATAATATATAAGGCTTTGGATATTTATCTTGTTCATTCTTT

CTATTAAAAAGAATAAATAATCAGACCAAAATCTCAATTCTGATCATTCACTAGCAATAGTACTGGTTCA

AGAAACTATAAGCCCAGATCAAGAATGGCTTATCTTCTCCAATTTCCAAACAATTTCAGGACCTTCTCTG

TTTCAGCTTCCTCCAATGGAGCTTCTCCAAGTGTGCCTCAGACAGGAGGGCCAGTGATTCTTGAACTTCC

CCTGGATAAAATAAGAAGACCTCTTATGCGTACCCGATCGAATGATCAAGAAAAGGTTAAGGATCTTATG

GATAGCATCGCGCAAATCGGTCTTCAAGTGCCTATTGATGTGCTCGAGGTAGATGGTGTCTACTATGGTT

TCTCAGGCTGTCACAGGTATGAAGCCCATCAGCGCTTAGGACTTCCAACAATCCGCTGTAAAATTCGGCG

TGGAACGAAGGAGACTTTGAGGCATCACCTCCGCTGAGTCCAAGAGCTGGGCTCACTTACAGGGCTGAAT

GATACAATTGCTCGACATCTATCTCACGGGGGTTTCACTACAAGTATTATTATAGCTTGTATACTTGAAT

ACATGCCCGATCCTGATCCTGATCACTCAATCAGAATATCAGATTGTAACTTATGTATGTATATAGTACT

CTGCTAATTGAAGATGGAGATTTGGAACGTTGCTTAGAACTGAATCCTTAGTATTATAGTATGCATCTGC

ATTATATAAAGCATACTACTACTATACTTTTTCTGACCATATGTAAAGAAGGCACACTTCATCCTTGGAA

TGTTTTAATTTTTTTTGCATGATTGCACATTTAAAGTTGACAATTGAGCACATTTACGGAACAAGTTGTT

GAGAAGGGCTTACTGAGTAATTCATGAAATACAATGTG

>MSTRG.376.2 gene=MSTRG.376

CTTTCATAATATATAAGGCTTTGGATATTTATCTTGTTCATTCTTTCTATTAAAAAGAATAAATAATCAG

ACCAAAATCTCAATTCTGATCATTCACTAGCAATAGTACTGGTTCAAGAAACTATAAGCCCAGATCAAGA

ATGGCTTATCTTCTCCAATTTCCAAACAATTTCAGGACCTTCTCTGTTTCAGCTTCCTCCAATGGAGCTT

CTCCAAGTGTGCCTCAGACAGGAGGGCCAGTGATTCTTGAACTTCCCCTGGATAAAATAAGAAGACCTCT

TATGCGTACCCGATCGAATGATCAAGAAAAGGTTAAGGATCTTATGGATAGCATCGCGCAAATCGGTCTT

CAAGTGCCTATTGATGTGCTCGAGGTAGATGGTGTCTACTATGTTTCTTTGTTTCTCCACCTACAGGTTT

CTCAGGCTGTCACAGGTATGAAGCCCATCAGCGCTTAGGACTTCCAACAATCCGCTGTAAAATTCGGCGT

GGAACGAAGGAGACTTTGAGGCATCACCTCCGCTGAGTCCAAGAGCTGGGCTCACTTACAGGGCTGAATG

ATACAATTGCTCGACATCTATCTCACGGGGGTTTCACTACAAGTATTATTATAGCTTGTATACTTGAATA

CATGCCCGATCCTGATCCTGATCACTCAATCAGAATATCAGATTGTAACTTATGTATGTATATAGTACTC

TGCTAATTGAAGATGGAGATTTGGAACGTTGCTTAGAACTGAATCCTTAGTATTATAGTATGCATCTGCA

TTATATAAAGCATACTACTACTATACTTTTTCTGACCATATGTAAAGAAGGCACACTTCATCCTTGGAAT

GTTTTAATTTTTTTTGCATGATTGCACATTTAAAGTTGAC

>MSTRG.377.1 gene=MSTRG.377

CCTTGAGCAGCGCCTAAAATAAAAAGTAAAACGACATCGATAATAGCCCTTCAGATTTCTTCTACTATCA

ATCTATCGCACTCTAATGGGCGACCGCGACGCCTATCGTGACCGCGAGAGAGATCGAGATCGAGATCGAG

AGAGAGACACCAGAAAACGCGACCGTGACCGTGAGCGCGACCGAGACCGTAGCAAACGCTCGCACACGCC

GGACCGGGCCAGATCGCGCCACGCCCGGTCGAGAACTCGCTCGCCGGACCGGTGGCGAAGCCGCTCCCGC

TCTCCCGACCGGCACAGGTCTCGCCGCCATCGCTCTCCCTCCCCGGACCACTCACGGAAGCGACAGAAGC

GCGGAGAGAGCAGTGAGAGAGACCGCCGGAGAGACTCATCCTCCGCCGCCGCCGCCGTGTCGGAGTTTGT

GGACGGGATAGTAAAGGAGAAGAAGGAGAAGAAGGGAGAGAGTGTGGAACCGGAGATGGATAAGGATGAA

GCTGAGATGATGAAGATGATGGGGATACCGGTAGGTTTCGATTCGACGAAAGGGAAGCATGTGGATGGCA

ATGATGTTAGCGGTATTAGAGCTGTTACGAAGCGGATGCCGAGACAGTATATGAATCGTCGTGGCGGGTT

TAATCGTCCCTTGCCTGCTGAAAGGAATCGTTGAGTTTTACTTAGGCCTAAGATGGAACATGGAATCAGG

CTAATGAATCAAATCAATTAGGGAATGAGCAGCGGCTTGGTATTGGCGGAAGTGAATTTTGTAAAACACC

TTTTGTATTCTTTGAATAGATGTATTGTACCTGAATCGGTTATTACTGCTAGTCAAGATAGTAGTCAGCT

TCGCCAGTCTGTTGTAACATACAAAATCGGTCTTCACATTGTATTTCATGAATTACTCAGTAAGCCCTTC

TCAACAACTTGTTCCGTAAATGTGCTCAATTGTCAACTTTAAATGTGCAATCATGCAAAAAAAATTAAAA

CATTCCAAGGA

>MSTRG.378.1 gene=MSTRG.378

TGCTCATCTTTCTCCTACGCTGAATATCATTAAAAAAATATACATCTTTCTTTGTTTCCACATGACGAGT

CTTAACTTTCCTCAAATGACCTAACATTACACATGTTCTTGGTCTTCCCAGCACCTTCCGCTTCCACTTT

GAGATCATACTCTTCAAACCCATTTTGCTCTCTCCTTTGTTTTTTTCCCGGAAAAAGACTAACTGTGCAC

AGCTTACCAAATAATCATGGTTATAAACAAATTTCTCACTGGCAATCAGGATTCGCCTGCAATTCTTTCT

GCTTCAGTGCATATTTGATTGTCCAAATGAATTAAATTCAAATTCAAATGAATTCAAATTCGCGCTGAAG

TACTCCAGCTAATTACAAATGGTTGACGGCTTCTTTAA

>MSTRG.384.1 gene=MSTRG.384

ATTTCAACAGCCCACTTAGACATCATCAAGATTTTGTCATCTGTCCCACAATTCAACATATGATGACCCC

TTTCTATAATGAAAAATAAAAATATAGCCGGATAGCCCTCCTAATTTTTCTATTCACATCGAACACTTCC

CCTCATCCAATCCAACAGTCACCGTCAAAAGTTGCTGGTGCCGTAGTTTTCGGACTTAACTCAGCTATAG

GCTACATGCTGATGCTTGCGATCATGTCGTCTAATGGCCGCGTTTTCATCGCGATTGTCGTCGGAATCGC

GGCCGGATATTTTATGCTTCGGAGTGGTGATGAGGATGATCAGGGGGGTGTTGATAATCCTTGTGCCTGT

GTCTGATTAATCTCCGTATGGTGCACAAATAACGAATCAAGAACAGATTGTCGAGTTTCATGAAACTTTT

AGTCTTTTCGACAAGATGCGATCTTTCCAGTACATAAAATTTTATGATTATGGTTATGTGATTATTTATA

GATTCACAGTATATATTGTTATAGCGTGGCTCATAATTTCATGATTAAGTATAATGGGGTGATGTGCTTG

AATCTATTAGTGTATGTGTTTTGTATCCTATTTTGTCATTCCTTAATCTATTTTGCCTGACTTGGAGGAG

AGTGGGTATGTAATCATTTTTATATTATATGTTTAGTTGC

>MSTRG.388.1 gene=MSTRG.388

GAAAGGAAAACAGTACTAAGTTAATACTTGTTCTTTCAATCGAGATAATGGCTCAAGCAGTGAGGTGTCT

TGATGGTCAGTCACTTTTGTGTCAGGCCAAGGCCGGCCAGAACATGCAGTGTCCCAGGAACCAGAACAAT

TCGAGTGTCATTGGACATCAGGATGCAAGGGAACTGGAAATCATTTGGGGTTCTGACACTCGCTACTGGA

AGTGGACAAGCATGCAAATCGGTGGACATTGCATTGAAGTTGCTGAATTGATTGATGTCTGCTGGCTGGA

AATCAATGGGAAATACGCGATGAAAAAACTTACTCCAGGAATGGATTACCAAGTGAAGTTCGTGTTGAAG

CTTAACAACAACTTTAACATTGACAAACCAGTGACCTTTAGCCTGACTACTCCAGAGGGATGCAAGGAAG

AGCATAAGGAGAACATGATGCAAATGCCAAGAAACCAAATCATTACCATCAATGCCGGGGAGTTTCAGAC

CCCCCGGAACAACTGTGGATGTGGAGAAGTCAAGTACCAGATGATGAACACTGATGGCTTCTGGAAGCGT

GGCCTGGTTGTCATTGGAGTCGTCATCGTGCCCAAATACAATTAAATAAAACCAGTATGCTAGTGAACTA

CTCCTAAATACCATCTCTATCAATAAATGCAACTCAGTTACATGGGTGGCTGTTACTAATCAGAATGATA

TGTAATAGATAACAACTACCTACTAATGGTGGCTTG

>MSTRG.391.1 gene=MSTRG.391

TTTTATGGATGTAGGAACTTATTGCCTAACAAGGTCATATCCATGGTGGTGATGGATAGGTTGAGAAGGG

GTGGAGAAGAAGAGCTAATTTGGCGATAATTGTGTCTTAATTAAGGATATTAATTATTCTTAATTAAGGT

TTTTAGTACCATTTATGGCGACCATTTATGGCGACCATTATTGGGGCATTATTGGCCCATAGAGGCATTT

AAGTGCCATAATGTTGGCTCGTTTATGTTTTTATTTTAATTTTAATTATTTTCAAATTCTCTCTCATTCG

TTTCATTCTGTTTTGTCTTATTTTCAGGATGTTTGAAAAAGATTTTAAGAATTTATTTATTAAGGCTGCT

TCCGAGAAGACGCGAACCTTGGTGTACAAGACAGTTGAGCTACAAACATAGATGGAAAGGAATAAAGAAG

ACATCATTTTTAGTCTGTTTTGTAATCGAT

>MSTRG.392.1 gene=MSTRG.392

AATACAGCTCCTTAAGTGGGACTTTTATATGGTATTGACTCTCTTATGTACTTGGTTTGAAACATAGTAC

TGACAATCTTCATTCATTTGTCCGAAACAGGCACAGGATTTCAGGACTGCAGGCACACAGATCCGTAGAA

AAATGTGGTTTCAGAATATGAAAATCAAGCTGATAGTGTTGAGTATTCTGATTGCCTTGATCCTAATCAT

AATACTGTCTGTCTGTCATGGTTTTAACTGTTGAAATTGAGCTGCACCTCTCTGGAGAACTCATTGGAAA

AAGCAGCCAAGTACTGATTTTGTCAAAGCTTTGTATTAAATGCTTGAATATACTTGTGTTAGTGTACTTT

GTATGTCGTAAGAAACCATTCACCTAAATTTGCTTCAGACATTTGGGCTGCAAGAACTTGCTAAAACGTG

CTTAGTATAGAAAAGCTGCTCAAGTTTGAGGTACTCTGATAAAAGTTGCTTTCTTCTCTATGTTAAATCC

CTAACTGTTGGCAACATTTCGTATTGGTACATACATCCGGGGATTTATTTGTACTGTGATTGTAAAATAT

TGTTGTTTATCTGTTCATAAAAGTTATGTATGCCTGAAAATTTTATACTCTAAACTACG

>MSTRG.392.2 gene=MSTRG.392

AAACAGGCACAGGATTTCAGGACTGCAGGCACACAGATCCGTAGAAAAATGTGGTTTCAGAATATGAAAA

TCAAGCTGATAGTGTTGAGTATTCTGATTGCCTTGATCCTAATCATAATACTGTCTGTCTGTCATGGTTT

TAACTGTTGAAATTGAGCTGCACCTCTCTGGAGAACTCATTGGAAAAAGCAGCCAAGTACTGATTTTGTC

AAAGCTTTGTAGTTAAATGCTTGAATATACTTGTGTTAGTATACTTTGTATGTCGTAAGAAACCATTCAC

CTAAATTTGCTTCAGACATTTGGGCTGCAAGAACTTGCTAAAACGTGCTTAGTATAGAAAAGCTGCTCAA

GTTTGAGGTACTCTGATAAAAGTTGCTTTCTTCTCTATGTTAAATCCCTAACTGTTGGCAACATTTCGTA

TTGGTACATACATCCGGGGATTTATTTGTACTGTGATTGTAAAATATTGTTGTTTATCTGTTCATAAAAG

TTATGTATGCCTGAAAATTTTATACTCTAAACTACGGGATCTTTCAGTAA

>MSTRG.392.3 gene=MSTRG.392

CAGGCACAGGATTTCAGGACTGCAGGCACACAGATCCGTAGAAAAATGTGGTTTCAGAATATGAAAATCA

AGCTGATAGTGTTGAGTATTCTGATTGCCTTGATCCTAATCATAATACTGTCTGTCTGTCATGGTTTTAA

CTGTTGAAATTGAGCTGCACCTCTCTGGAGAACTCATTGGAAAAAGCAGCCAAGTACTGATTTTGTCAAA

GCTTTGTAGTTAAATGCTTGAATATACTTGTGTTAGTATACTTTGTATGTCGTAAGAAACCATTCACCTA

AATTTGCTTCAGACATGTAGTTAAATGCTTGAATATACTTGTGTTAGTGTACTTTGTATGTCGTAAGAAA

CCATTCACCTAAATTTGCTTCAGACATTTGGGCTGCAAGAACTTGCTAAAACGTGCTTAGTATAGAAAAG

CTGCTCAAGTTTGAGGTACTCTGATAAAAGTTGCTTTCTTCTCTATGTTAAATCCCTAACTGTTGGCAAC

ATTTCGTATTGGTACATACATCCGGGGATTTATTTGTACTGTGATTGTAAAATATTGTTGTTTATCTGTT

CATAAAAGTTATGTATGCCTGAAAATTTTATACTCTAAACTACGGGATCTTTCAGTAA

>MSTRG.393.1 gene=MSTRG.393

CCTCCATCTGCTCCCCTAACAAGTCTTTAAACATCATATTCACCAGTCTTTGATAAGTAGCACCAGCGTT

CAAAAGTCCAAAAGGCATTCCAACGTAGCAGTAGAGACCACGGTCCGTGATGAAAGAGGTGTGTTCCTGG

TCTGGCTAATGCATTGGGATCTGATTGTAGCCGGAGTAGGCATCCATAAAGCTTAACAGGGCGTGGCCAG

CAGTAGCGTCAACTAGTTGATCAATCCTCGGAAGGGGAAAGCTGTCTTTTGGACATGCCTTATTCAGATC

TGTGAAATCGATACATACCCTCCATTTTCCATTGGCCTTCTTCACCAGCACTGGGTTCGCCAGCCATTCC

GGGTAATAAGCTTCTTTAACCAAACCAACCTTCTGAAGTCTATCGACCTCCGCTTTGGGAAAGCTGTCTT

TTGGACATGCCTTATTCAGATCTGTGAAATCGATACATACCCTCCATTTTCCATTGGCCTTCTTCACCAG

TACTGGGTTCGCGAGCCATTCTGGATAATAAGCTTCTTTAACCAAACCAACCTTCTGAAGTCTATCGACC

TCCGCTTTTAGTGCTTCTGCTCGTTCACCGCTGATGGGTCTGCGTTTTTGTCTGATACCTTTCTTCCTCG

GATCAATGTTCAAGCGGTGACACATAATGTTCGGATCAATTCCTGTCATATCAGCATGGCTCCAGGCGAA

TACATCAAGATTGCTTTTCAAAAAATCCACCATCTTCTTTCTCATCAGATTGTCAAGACGAGTACCAATG

AATAAGACTTTCGTTGCATCATTCTCATCAACCAGGATCCCAACTGTGTCCTCGGCTGGTCCTACCTTCT

GCAGTGTTTCTGGTAGTCTGGGGTCGAGATCGACATCTCTCAGATCGTCATCATCATCCTTCGGATGAAC

A

>MSTRG.394.1 gene=MSTRG.394

ATATGGTATTGACTCTCTTATGTACTTGGTTTGAAACATAGTACTGACAATCTTCATTCATTTGTCCGAA

ACAGGCACAGGATTTCAGGACTGCAGGCACACAGATCCGTAGAAAAATGTGGTTTCAGAATATGAAAATC

AAGCTGATAGTGTTGAGTATTCTGATTGCCTTGATCCTAATCATAATACTGTCTGTCTGTCATGGTTTTA

ACTGTTGAAATTGAGCTGCACCTCTCTGGAGAACTCATTGGAAAAAGCAGCCAAGTACTGATTTTGTCAA

AGCTTTGTAGTTAAATGCTTGAATATACTTGTGTTAGTGTACTTTGTATGTTGTAAGAAACCATTCACCT

AAATTTGCTTCAGACATTTGGGCTGCAAGAACTTGCTAAAACGTTCT

>MSTRG.396.1 gene=MSTRG.396

GTAGAAAAATGTGGTTTCAGAATATGAAAATCAAGCTGATAGTGTTGAGTATTCTGATTGCCTTGATCCT

AATCATAATACTGTCTGTCTGTCATGGTTTTAACTGTTGAAATTGAGCTGCACCTCTCTGGAGAACTCAT

TGGAAAAAGCAGCCAAGTACTGATTTTGTCAAAGCTTTGTAGTTAAATGCTTGAATATACTTGTGTTAGT

GTACAATTTGTATGTCGTAAGAAACCATTC

>MSTRG.398.1 gene=MSTRG.398

AAAGCATGTATCTTTGTGACCTAAAGGCTAAACCTGAATATGAGACCAGTTCATTAACTATCAAACATAT

CACACCCATCGCCGCTACCTGCTACAGCGAGTCGGCAACCTGCTGCAACCACATTCATCCAGCCTCAAGC

AATCAAACCCAATCTCATCCAGCTTCCAGGTCATCCGTTTTGTTTTTTGAATCGATCTCTGGCTTTTGTT

TCTTTGCGTGCCTGGACAAGTGGGGGATTTTATGCTTCTGTTATGTGTATAAGTGTATTTCAGAACTGGT

GAATGGTGATTCTTTATTTTTAATGTGCTGCAAGAAACCCATGTACTTTATTTTTAACTAGTTTGGCAGC

AATTTTCAGATTTTCATACACTATTTAGGAATTTCGGTTTATAAATTCGGTTTTCGGTTTTAACCGAAAT

TTAAAATTCGGTTCCGTTACAAAACCGAAATAATTTCGGTTCGATTTTCGGTAGACATTTTTTGTGAATT

TCGGTTTCGGTTAACCGAAAAAAAATTCGATTTCGGTTTCGGTTAACCGAATGCACAACCCTACTCCTAA

GTCCCAACTATCTCATGTACAGTTTTACTACCGAC

>MSTRG.400.1 gene=MSTRG.400

TTAAAACACAAAACCACAAGCCCTAAAACCCCAAATCTCTCTGCTTTATTTCACTAAACTTAGAGCGCCT

AGCCCTAGATTTCATCTCCTCGTCGACAATGAAGCTCGTCAGGTTTTTGATGAAATTGAACAACGAGACT

GTCTCCATTGAGCTTAAGAATGGGACAATTGTTCACGGTACTATCACAGGTGTGGATATCAGTATGAACA

CACATTTAAAGACTGTAAAACTTACACTGAGGGGGAAGAACCCAGTGACTCTAGATCACCTGAGTGTTAG

GGGGAACAACATCTGTTACTATATCTTGCCTGATAGCTTGAATCTTGAGACCTTGTTGGTGGAAGAGACA

CCGAGGGTTAAGCCCAAGAAGCCAACTGCAGGGAGGCCTGTGGGACGTGGTAGGGGACGTGGACGTGGGC

GTGGTCGTGGGCGAGGCCGTTAATTCTGATGTACTGCTACAATTTTTCCAATATTATGTTTCATTGCATT

GTTGGAGTTTATATGGTAATGTGCCAACAGTACCTAAAATGGACGATCGGAGGTTATTAGTTTAAGTTAT

GCACCGAGTG

>MSTRG.401.1 gene=MSTRG.401

TATATAAACTCTAAACATCTTTAACTTGTGAAACTCAACAACATAGGAGACTAACTCAATTTCAAGATTC

ATAAGATTATTATGGGGAAGGGCAATTCGGTGAAGACAGCAGCAGTGGTGCTTGGAGCCCTGGCTCTAGG

ATGGGCCACAATTGAGCTGGCTTTCAAGCCATGGCTTGACCAGGCTCGAAAAGCTATGGACAAGTCTGAT

CCTGCTAAGGACCCTGATGATGCCTCTACTTCTGCTCCTGCTCCTGACCCTGATGATGCCTCTACTTCTG

CTCCTGCTCCTGTTGGTGATGAAGAGCAGCCTGCTTCTTAGTTTTATGTATATGTGTGTTATTATGTATT

GGCTTCGGCCTGGATTAGCCTCCTAGATTTCGCTAATTGTGATGATTTATCGTTGGGATGTGGTTTGGAT

TATATGTGATGGATCTTTTGTTATGGAAATTTGTTTTCTCTGTATATTTGG

>MSTRG.402.1 gene=MSTRG.402

ATTGAGTTGAGAAGATCTTATCAAGCATATAATGTTATGCCTAGTTTAAGCAAAGCTTTAAAAAGATATG

GTGTATAACATCTGGAGACAGTCTATCAATTTTGTATCAAGAAACCCTCTATAAACGGACTGATACGGAT

CATCATCACAGAGATATAACCGAAGGTGAAGAGTCCAAGTCCTGCATGTTAAGAAACTATGGGGTGTACT

TTCGCCAATAGATCCCGTTGAGTAAGGTTACACCTAGTTCCCTACCCGACACCAGGAGGCTACGAAAGGA

TGAGACCCAGTCTTGATTGTCTGTTATGCCATAATGATCTAGTCTTGCTTCAGTTAGTAACAAAGATGAG

CTCTGCATATTCTGTATAGGTAGAAATAGTATTTGTTACCCTAAATGTATTACTGTCTGTTGAATGTTAA

TTGC

>MSTRG.404.1 gene=MSTRG.404

GTCATATTTCACAAGTCACCTGACATGAGTCATTTAGCGTGACCCTTCATCTCTCTCCCATGTGAAACCG

AAATGTAAAAAACAAACGGCTTAACAATCGATTGATGATCCGAACGAAAAGAAGTGAGCGTGACGCCTTT

GTGAACGCAACCAAAAAACCCAGCTCATCACTCATGACAACATAGCCCCCGTCACCTCCACGACTTCATC

TTCTTCCAACCTATACATACATTATACTTACAACTTTTTCATCATCCTAACAACTCTCTCTCCCCATTTT

CCTCATCTCCCTTCACTTTTTATACATTTTAATCATCTGGGCTGCTCTTAAATCTTTCTAAATTATCATA

ATTCTCACTATATATTCATTCTTGATTTTTGAAAATTTTCTTGGGTTTTCTTGAATTTTTGTGATCAGCT

CTGTTTATTAGTCTAAGATTCTGCTAGTTTGATCAAATTCTGTAGATTTTGCTGTTCAATTTTATTTATT

TAGTGCAATTTGTTTCGTTTTGTGCAATGTTCTTGATCAAATGTGATGCTTGGATTGTTAGTTAATGAGG

TGTTGAAGTGAATTGACGATGTTTTCAAGAAAATTGGTTAGGCTATTCATAAAGGGTTAGCGGCTTTGCG

GCGCAGCTTGTTGCTGGTTGAATGAATTGATTGTTACTTAAAAGGAAAAAAATAGGAATTGGAAATAGAA

ATAAAGGTTAAATTGTGAATTTTGAGCAATGGAGAATTTGACTGAACCATCATCATCGATAAGTTTTAGT

TCTTCTAATGGCTCAACTGGCTACACTTTAACCGAATCATCTGTCTCGGAGAAAGTATCTAGTCTTCAAG

TTGTGAGTTTATATAAGCTTAGTGCTAGCTTAGAGCAACTTTTGATTGATAGTGTTGGTGAGTATAGTGA

TGCGGATATTGTAGTTGAAGATATTGCTGTAGGAGTCCACCGTTGTATATTAGCTGCTAGGAGTAAATTT

TTCGATGCACTTTTTAAGAAAGAGAAGAAGGGTGGTTCTGCGGGGAAGGGAAGTAGACCGAGATATTGTA

TGAGTGATATGTTGCCATTTGGTAATGTTGGATATGAGGCCTTCTTAATTGTCTTGAGTTATTTGTATAC

CGGAAAGCTGAAGCCTTCTCCAGCGGAGGTGTCAACTTGTGTTGATGATGGATGTGCTCATGATGCCTGT

CGACCTGCTATTAACTTCTCCGTGGAATTGATGTATGCATCTGCCATATTTGAGGTCCCGGAGCTGATTT

CACTTTTCCAGCGGCGTCTTTTCAACTTTGTGGAAAAGGCTCATGTTGAAGATGTCATCCCAATCATCCT

TGCCGCTTTCCACTGTCAATTAACTCAGCTTCTCGATCAATGTATTCATAGAGTCGCACGATCAGATCTC

GACAGCATTTCTCTTGAGAAGGAATTCCCGTTTGAAGTTGCAGAGAGAATTAAGTTGCTCCGTGATAAAT

TACAGGGTGTTAAGTTACAGGCAGATGATAGTGCCATGATACCCGAGGATCCGTTGCGTGAAAAGAGTAC

AAGGAGAATACACAAGGCTTTGGACTCTGATGATGTTGAACTTGTAAAACTTCTTCTGACCGAGTCTAAT

ATAAGTTTAGATGAAGCCAATGCCCTGCATTATGCTGCTGCATACTGTGATCCAAAAGTTGTATCTCAGT

TGCTTAGTCTGGATCTGGCTGATGTCAACCTTCGAAATGCACGGGGCTACACAGTTCTCCATGTTGCTGC

ATTACGTAAGGAACCATCAATAATTGTACCACTTCTAAGCAGAGGAGCTTGTGCCTTAGAAACAACACTA

GATGGACGGAGTGCTGTCAGTATCTGCAAGAGGCTGACAAGGCCAAAGGATTTTCATGCTAAAACAGAAA

AAGGCCAAAAATCAAACAAAGACAGATTATGCATTAATGTTTTAGAGCAAGAGGTGCACAGGAATCCAAT

GGCTGGAAATTCATCAAGGTCCTCTTCAAACATAACTGATGATCTGCACTTGAAGCTTTTATGCCTTGAA

AACAGAGTGGCATTTGCACGGTTACTTTTTCCGTACGAGGCCCAGCTAGCCATGGAAAATGCCAGCGCAG

ACACAACGTCTGAGTTCTCTAGCATTTTGGAATCCAAAGTTTCAAGTGGCAATTTAAGAGAGGTAGATCT

GAATGAGACACCCATAGCACAGAACAAAAGACTTCTCTCAAGGATGGAAGCCCTCTCGAAAACAGTTGAG

ACAGGTCGGCGGTATTTCCCACATTGCTCGGAAATCTTGGATAACTTTATGGTGGATGATCTACCTGATT

CGTTCTACCTTGAAGAGGGCAGTGCAGAAGAACAGGAATTCAGAAGAAAGCGTTTTATCGAGCTTAAAGA

GGATGTTCAAAAAGCATTTACCAAGGACAAAGCTGAGTTAGAGAGGGTTGGCTTGGGCTCTTCACTATCC

ACATCCTCTAATAAGAAGAGACGTAAATTTCAAAGTTGAAAATTTTATATATGTATCCAAGTTAAAAATT

TTATATATGTATCCGAATTTTGTCATTCCTGACATACTGTGTATAAACTGAGTCATCTACATGAAAATTC

AGATGTAAGATTGGCAACTTTA

>MSTRG.410.2 gene=MSTRG.410

TGAGTTTTCTCCGATTTATCTCTTCTCAATTTCTCCATTAATCTCATCTCTTTCATCAATTTCTCAAGCA

CCTTGCTTAATTAGTTCTTTAACTTGCATAAGATATATAGATGATTTATACATTCATGATCCATGTGTGT

AAATCTAATATTCTAATGTAGATGAACTTGTTTTGCAGACATGTGTGTGAATCATGACTTGGAACAGCAA

CACGATGTTTGTACACATGCAGATGCTGATCAAACACTATACGTCGATTATAAAATCTAGATGATTAACA

CTGTTCTTGTGTGTCTGTATATATCTAGACATGACTGATTACTACTGTGTGTTTCAGTTTCAGTCGGCTT

CGTAATTACATGTACATGCATCTAGGTTTTTCCCTATGATTTTTTTACAACATAGATAGATCTGTGTCTG

CATGGTGACACAGACTACTCCGAAAATAAACTTCTTGCTTTGTATATAACATTTCAGTTTTTATTCAATG

TATTATAGGTGACAGGATCTAGGATTCTTGTTCAGACAGCACTAATTCTGATTAAGATAAAAGGAAGACA

TGGGTCAGGTGAGGTGCAGCTTTAAAATATTCTGTTGCTCTTCCAATCCAAAAAAGAAAGTAAAAAGGAA

TATTACAATGCTGTATAGTGTTTGATGGAACAAATTCCCATCCCATTATCAAAAAGATGCCTGGCCTTTT

ACATTTTGTTTCTGATTCGCTTAGCCATCTTTTTTACTTTTTTGTTCTGGTATCCACTTTAGTTCGACAA

AATATGAC

>MSTRG.413.1 gene=MSTRG.413

AGGTGTTCAGTTTGTTCAAACGAGGGACTCATTTATGATTTCTTTGGCGGCAGTCCTGTAGGCATACACC

AAAAGCGGAAATTGGGTCCTCAATATATAGTTCTTGTGATTGTAGTTTTATAGCACATCTAGGCCTAGTA

TAATTTGTTTTGGAAAGAACAATTCTGGTGATCTTTTTAAACTTTGCCTATTGTTCTTTTTTATTCTCAG

AAGTAGCTCAAACCTTATCATTCTTTAAAAGTGCCTTAAATTCATCAATGAGGTGTTGTGTTTATTTTCA

A

>MSTRG.415.1 gene=MSTRG.415

TTTAAATTAGGGAAGAAATGAGGGGTGCATGGAGAGTGCTAAAGTGACTAGATTTACAGTACTTGAACTT

CTTCCGTACGCCAGTGATAGACCCGTTTACTAACCCGAATTTGCCACCATGTCCACCGGAGTTCTCCCCG

TCGATTTACTCGAGGAGATCTTCAGCAAGCTTCCAGTGAAGTCCCTAATCTCCCTCACCATCATCTGCAA

ATCCTGGCTCGCACTAATCTCATCACAATCTTTTGCAAAAACCCATCTTTTCAAGTACCCTCTTGACCCC

TCTTCTCATTTGCTTCTTCTCCACCGCCCCTGTCGCGAAACAATCACCATTTCCGAGCTCAATTTCTCGG

GATTGCCAAGAATTGTGGAAACCCCAATTCCAGAATCTGAACCCCCGCCGATTCAAGCCCCGTCGACTCC

GGTGGTCTCGATCTGTTGTGCGAGAAATGTTCAAGTTCGTCAAGAAGTTAGGAGCTGTTTGTTGAGATTG

GTGGGCTCGATTAATGGGTTGGTGTGTTTTTCTACGCCTACTCTTAGGCCTACTAATGTTGTGATTTGGA

ACCCTGCTACTCGGAGATTTAGGGATGTGGCGGTGTCGCATTTGAATTATAGTAACCCTTTGAGGATTTA

TGTGGCTTTTGGGTATGATGGTGTTGGTGATGATTATAAGGTGGTCTGTGTTTATCGGTTTCGGGTTAGG

AATAATCCGGAGATGTCGTTTAGGTTTAGGATGTTTTCGTGTAGGGATGGTTGTTGGAAGGATGTAGAGC

CGGGGTTCGGGTTTAGTTTGGGGTTTATAGTGGGGTGTGTGATCGTAAAGGGGAATCCTTTTTGGATGGG

GTTTTACAAGGATAAAGAGATTTGGCTGACGCTTGATGTTCAAACTGAGGTGATCCGGATGTTTTCTGGG

CCGAAGTATGTAAGGGGTGCATCTACTTCGATCTCAATTATGGCTTTAGGTGATAATGCTGCTCAGATTG

TGTATTCACCCGGAATTGAATCAAGTCATATGATTCATGTGTATTGTTTGGAAGAAAGTAGTGGTACGTG

GAATTTGATGTACGCCATAGAGTCGATTGGGTTACAGAGGCCTGTACACATGGAATGCTATAAGAATGGC

AAGGTTGTTACAAAGGATAGGAACGGCAAGTTGTTCACGTATGACTTGATAAGTAAAGAAATTAAGGATC

TTGGGGTTGGAGAAGGCATGGAGGATCATTACATCGCCATTAATTACATCGAAAGTCTTGTTTCCATCGA

AAGTATGGAGAAAGTAAGAGAAGAAGCTGCTGAAGAAGCAGGGCAGGAGACTGGACTTGCTTCAAATGAT

AGGGATGCTTCGGATGCTATGATGTCAGTAAGGATAGATGAATCTCTTAAAGATGTATGAGAATCAAGGT

TACTGTTGCGGGTTGTTGGCTTTCAATTGAACTGAAGAAGTATACAAGCATGGAGGGACTGAAATGCTGT

TTCATGTACAATAGACTATTGTTGTATAATCTTGAAAAAATTGTATTATGTATTACTAGCTGTTTTCTGT

ATCAGAGACCTCAGCAAGTCTGTTGTGTTTCTATTTTTTTTGACTTTAAAATACAAATTGTGTTGTCTGA

TACTATCTGAGCTTGATCTTGATTTATGTTTACTTGGTTCTGGTTGGGTTCGAAACAGATATATGGTCAT

TTTTTATCTTGCCTTTGTATGCATAATACTGGAATGTGGTG

>MSTRG.415.2 gene=MSTRG.415

TTTAAATTAGGGAAGAAATGAGGGGTGCATGGAGAGTGCTAAAGTGACTAGATTTACAGTACTTGAACTT

CTTCCGTACGCCAGTGATAGACCCGTTTACTAACCCGAATTTGCCACCATGTCCACCGGAGTTCTCCCCG

TCGATTTACTCGAGGAGATCTTCAGCAAGCTTCCAGTGAAGTCCCTAATCTCCCTCACCATCATCTGCAA

ATCCTGGCTCGCACTAATCTCATCACAATCTTTTGCAAAAACCCATCTTTTCAAGTACCCTCTTGACCCC

TCTTCTCATTTGCTTCTTCTCCACCGCCCCTGTCGCGAAACAATCACCATTTCCGAGCTCAATTTCTCGG

GATTGCCAAGAATTGTGGAAACCCCAATTCCAGAATCTGAACCCCCGCCGATTCAAGCCCCGTCGACTCC

GGTGGTCTCGATCTGTTGTGCGAGAAATGTTCAAGTTCGTCAAGAAGTTAGGAGCTGTTTGTTGAGATTG

GTGGGCTCGATTAATGGGTTGGTGTGTTTTTCTACGCCTACTCTTAGGCCTACTAATGTTGTGATTTGGA

ACCCTGCTACTCGGAGATTTAGGGATGTGGCGGTGTCGCATTTGAATTATAGTAACCCTTTGAGGATTTA

TGTGGCTTTTGGGTATGATGGTGTTGGTGATGATTATAAGGTGGTCTGTGTTTATCGGTTTCGGGTTAGG

AATAATCCGGAGATGTCGTTTAGGTTTAGGATGTTTTCGTGTAGGGATGGTTGTTGGAAGGATGTAGAGC

CGGGGTTCGGGTTTAGTTTGGGGTTTATAGTGGGGTGTGTGATCGTAAAGGGGAATCCTTTTTGGATGGG

GTTTTACAAGGATAAAGAGATTTGGCTGACGCTTGATGTTCAAACTGAGGTGATCCGGATGTTTTCTGGG

CCGAAGTATGTAAGGGGTGCATCTACTTCGATCTCAATTATGGCTTTAGGTGATAATGCTGCTCAGATTG

TGTATTCACCCGGAATTGAATCAAGTCATATGATTCATGTGTATTGTTTGGAAGAAAGTAGTGGTACGTG

GAATTTGATGTACGCCATAGAGTCGATTGGGTTACAGAGGCCTGTACACATGGAATGCTATAAGAATGGC

AAGGTTGTTACAAAGGATAGGAACGGCAAGTTGTTCACGTATGACTTGATAAGTAAAGAAATTAAGGATC

TTGGGGTTGGAGAAGGCATGGAGGATCATTACATCGCCATTAATTACATCGAAAGTCTTGTTTCCATCGA

AAGTATGGAGAAAGTAAGAGAAGAAGCTGCTGAAGAAGCAGGGCAGGAGACTGGACTTGCTTCAAATGAT

AGGTATCCCGACTTCTCTTAGGATGCTTCGGATGCTATGATGTCAGTAAGGATAGATGAATCTCTTAAAG

ATGTATGAGAATCAAGGTTACTGTTGCGGGTTGTTGGCTTTCAATTGAACTGAAGAAGTATACAAGCATG

GAGGGACTGAAATGCTGTTTCATGTACAATAGACTATTGTTGTATAATCTTGAAAAAATTGTATTATGTA

TTACTAGCTGTTTTCTGTATCAGAGACCTCAGCAAGTCTGTTGTGTTTCTATTTTTTTTGACTTTAAAAT

ACAAATTGTGTTGTCTGATACTATCTGAGCTTGATCTTGATTTATGTTTACTTGGTTCTGGTTGGGTTCG

AAACAGATATATGGTCATTTTTTATCTTGCCTTTGTATGCATAATACTGGAATGTGGTG

>MSTRG.416.1 gene=MSTRG.416

TTTCTGTATTATATGTAAAATGCTAGGATAAGTATTTTATTTAGAGAAGGGAGGGAGAAGAACAACAGTG

AAAGAGATAGGAAAATTGTGGCAAAGATTATTACTATTACATGTTCTTCTTGGGAAACTTCTTTACACTT

CTCCCAATGCAATAAAACAAGAGTGGTTTCCCATTTTCTTGGAGAAGAACAACATAATCTTGTATCCCCA

GCTTTATCAAATCCTTGGTTTTCTCACAAATTTCTTCTAGCTATGCAGAGTCAACTAGTGTGCAGGGGAT

GTAGGACGATGCTTCTGTACCCACAAGGAGCAGCAAATGTTTGTTGTGCAGTGTGCAGCGTAGTTACTAC

CGTACCTCCTCCAGGAATGGATATGTCCCAACTAATATGTGGAGGTTGCCGTACATTGCTGATGTATACT

CGTGGAGCTTCAAGTGTGAGATGCTCGTGCTGTCACACCGTGAGCCTTGTGCCAGCACCAAACCACCAAG

TTGCTCACATCAACTGTGGCAATTGCACCACAACACTTATGTATCCCTATGGATCTCAATCTGTCAGATG

TGCCATTTGTCAATACGTAACCAACATAAATATGGGTAATACAAATATCCCTGTTCCTGCAAGCAGACCC

GATGGAACAGCTTCATCTGGAACAACAACTGCTACTTCAACAACTCAAACAGTTATTGTCCAGAATCCTA

TGTCCGTTGATGAAAGTGGAAAAACAGTGAACAACGTTGTTGTTGGTGTTACAACTTGAGAAAACTGATT

TGATTGTAACAACCGAGCTTCAAGACTCTTCAAATCTGTATCTGTTGGTCAGTGACTACTTGTGTTTTCA

AATGTGACATGCATAATAGGACGGTAACATATCCTGCCATTTTTCAGGACGGGAAAATTTTCTTTTTGGA

TGTAAAATGATGTAAAAACGACTTAGAACTATAGAGGATGGTGATGGCTAGTAATTAGTTGTGTGAAAAT

CTTTTGTTTTTATCGGCAATCAAGCCAGTTGCCTGGAATATATATGCAAGTGCAGAGAAAGAGGGGCTAA

TGTGCACCTGTCTTTGTTAAATAAAATTTAGTTTATTCATTGTTGTTGTAGATTGTTGTCTGATGTATGT

TCTGTCAGCTACTGACTTCCTTGATTTTATGGGAAGAATAACTTCTCTTGATATATATAAATGAATTTTC

ATTCTA

>MSTRG.417.1 gene=MSTRG.417

TTTAAACGTATATAATAAAATATATAATTAACCCGACCCGGAAACCTTCGTAAGCCTGTGACAGGTCGAT

TTTGAGCAAAAGAAAGCAGAGCCTCCGTTTAAAGCAGGGTAAGAAAGTTTGAAGCTTGACTGTCTTTTAG

AAGTTGAGATGGCAGTTAAAGATTTTGTAGAGGTCTGGCATTTTGTGTGGTCAATGGAATTTTGGAGGAT

GGGAGTTTGTTGGACATTGTCTCTTCTTTTGGCGTATGCTCGATTGCTTGGTCAGAGGTTTTTGACTCGA

GGGAACAAATCGTATGGTCGTTGTTGGAAACACAGTGGTGCTAGAAAGCCTGTCTGCGTGATCACAGGCG

CAACGTCTGGTCTCGGTGCTGCAGCAGCAGCTGATCTGGCAAGGAATGGATTCTATGTTGTTCTTGTTGG

AAGATCATCTCATCTGCTATCGAAGACTATTTCTGAGATCAGATTGCAAAATAAAGCTGCGGATCTCAAA

GCATTTCATCTAGACTTATCATCATTTGGGTCAATTTTAAAGTTTGAAGCCAACCTTCAGCAGTGGCTTC

TGGATTCAGATTTACATTGTTCTATACAGATCTTAATTAATAATGCTGGAATACTTGCAACGTCATATAG

ATGCACTCCTGAGGGCTATGATCAGATGATGGGAACTAATTATATTGGCGCATTTGCTCTAACAAAGGTT

CTACAGCCGCTATTGGAAAAAAGTCCTACCCCTGCTCGGATTGTTAATGTTTCATCATTTACACATTGGA

ATGCACTATCTGGCATTCGCGTTGACAAGGAAACTGTATCTGGGATGTCCTTTTCAAAATTGAAATGCTA

TCCTTATGCTCATATATACGAGTATTCAAAATTATGCTTACTACTATTCACTTACGAGCTTCACCGACAG

TTTGATATCTTGCAAAAGCCACATCAGATATCTGTAGTTGCCGTAGATCCTGGAGTAGTGAAAACCAACA

TTATGCGAGAAATTCCTTCATGTCTCTCAGAGTCTGCATTTTTGGTCCTGAGAGTTCTGGGCCTGTTGCA

GCCACCTGAAGTTGGGGTAACCTCCATTGTTGATGCAGCCCTTTCCCCTCCTGAAGCATCTGGATCTTAC

TTCTTTGGTGGGAGTGGCAGAACTCTTAAATCCTCTGAACTTTCTTATGACAAAGACTTTGCAAAAGAAC

TCTGGAAAACCTCATGCGAGTTGTTCCTAGATGCACAGCAGCTATCTCAGGAGACTTCCAGATGATCTCA

AGGTGTATAGATGCGATTGAAATGTTCTTGATAGTATGTACTCTATGCTTTTGTGCTGAACGAGTATCAG

ACCGCAAATGTACATAAGCATACACCTTGTCATCATATCACATTGAACACAGTTTTCTAACACATTTATT

CATACAAAATTGTACAATTAGATTCATCTTCTGTGATTTACCTTGTAGATATTATTAACCAAC

>MSTRG.424.1 gene=MSTRG.424

TATGATTCATACTCTCTATTTCTCCAAAAAATTAGATCAAAAGATGAATCCAAACAACACTCCTCCTACA

AATTCTCAAAATTCAAACCCTCAATTTCCATATCCATATCCAAATCCTTTTTTCCCAAACACACAAAATT

TTAATTTCAATTCTCAAACCCCGAATTATAATGCATAATATGATTGTTGAAGACGAGAGAGACACTTATG

CCACGCAATTTGGTCCTTTACCAATTTATGATGATGCAACAAATGGTTTATCTCAACCAAATTTAGGTGA

AGAACCCTTTATCCCGTATGAAAGGTATATCCAAAATACGTTACAAATGCGTGATAAACGGACACATCGT

CAGCTACAAAATGACTTGGTTGAGCACATCTCGCAGTTCCATAATAGTCGTTAGTTTCTTATTGTACTAT

TTTTAATTGTTTGTTTCCTTGTGTAATGTTTTTAATTTATTGTTTTTAATTCAGTTGAATGTATTTTATT

TTAATA

>MSTRG.426.1 gene=MSTRG.426

AAGTTTTAAAATCTTAATCGTGATCAGCATGATTAAACTGAAAATGACCTGCAAAAAAAAAATTGAGACC

AGCTTATGATATTATAACCACACGTAACTATTAATCTAGCAGTATCACTTAATCTGTGCTATTAAATATT

GCTTTTTCCTCGATGAACTTATTGTTATGTCACATTCGGAATCAGAAAATACAGTATATGTTGTAATCAA

TCTGATCTGGTAACTATTCTGCAGTTTGACAAATTTTATTGAAGTGACAGATGGACATTAATGTTGCAGT

GAGGGGTTATGGCTGGCTTATGCTCTTAGATTTTGATGTTTTCGATTTTGTTACATATAAAATAAACTGG

TAACATGTTCATAAAATGATTGAGGTACAGATTAGTAAACTGATGATCATCTAGAAATAATTTGACATCA

CGGTTGTCTTGATAATTTCCAGTTGTTCTGTTCTGAATTCTGAGTGAATTGGAATACATCAAATATACAC

AACAGTTTCAGACATTCATACGTTTATTTATATGTGAATATGTACCTCATGGTAAATATGTAGATGCTCT

TCAAGATACTTGAGCACAAAGGTAATACTCTAACAGAGCTGGTAATCTGCAGTTCCATTGGCTCAGAGGG

CAGGATATCTGTTCAAGATCCAGTATATTTCCTCCGTCTCATTCAATTGTATACGTTACTTTTTGGCACG

CCTTTCAAGACTCTTGTAAGGTATAATATTATTTTTTACTGAAAAAAATTTGATGCTTAAACTTTTATTC

AAAAAAGAAAATTTTAAAAAAACATTATGGAACTATACTTTATAGGAGCATCAAACTGTGTGCAAACAGT

GAACGTATACCATTCACCGGACGGAGGGAGTATTTTGTATTGGACTCATCATGGAAAAACTAATGCAGAT

CAAAATATGCAGAAGGCCAGAGAATTTTTTGAATTCATGGCTCCATATGTGACCAGCTCTCCAAGGGAAG

CATTTTTGAACTACAGAGATCTTGACATTGGTCAGAATCCAATAGGCAATGCAAGCTATGCAACTGCC

>MSTRG.426.2 gene=MSTRG.426

AAGTTTTAAAATCTTAATCGTGATCAGCATGATTAAACTGAAAATGACCTGCAAAAAAAAAATTGAGACC

AGCTTATGATATTATAACCACACGTAACTATTAATCTAGCAGTATCACTTAATCTGTGCTATTAAATATT

GCTTTTTCCTCGATGAACTTATTGTTATGTCACATTCGGAATCAGAAAATACAGTATATGTTGTAATCAA

TCTGATCTGGTAACTATTCTGCAGTTTGACAAATTTTATTGAAGTGACAGATGGACATTAATGTTGCAGT

GAGGGGTTATGGCTGGCTTATGCTCTTAGATTTTGATGTTTTCGATTTTGTTACATATAAAATAAACTGG

TAACATGTTCATAAAATGATTGAGGTACAGATTAGTAAACTGATGATCATCTAGAAATAATTTGACATCA

CGGTTGTCTTGATAATTTCCAGTTGTTCTGTTCTGAATTCTGAGTGAATTGGAATACATCAAATATACAC

AACAGTTTCAGACATTCATACGTTTATTTATATGTGAATATGTACCTCATGGTAAATATGTAGATGCTCT

TCAAGATACTTGAGCACAAAGGTAATACTCTAACAGAGCTGGTAATCTGCAGTTCCATTGGCTCAGAGGG

CAGGATATCTGTTCAAGATCCAGTATATTTCCTCCGTCTCATTCAATTGTATACGTTACTTTTTGGCACG

CCTTTCAAGACTCTTGTAAGGAGCATCAAACTGTGTGCAAACAGTGAACGTATACCATTCACCGGACGGA

GGGAGTATTTTGTATTGGACTCATCATGGAAAAACTAATGCAGATCAAAATATGCAGAAGGCCAGAGAAT

TTTTTGAATTCATGGCTCCATATGTGACCAGCTCTCCAAGGGAAGCATTTTTGAACTACAGAGATCTTGA

CATTGGTCAGAATCCAATAGGCAATGCAAGCTATGCAACTGCC

>MSTRG.431.1 gene=MSTRG.431

CNGCAACCAAATTGGCGCCAAGTTCTGGGAAGTCATCTGCGATGAGCATGCAATCAACCAGTCAGGATTC

TACGATGGAACCACCGATCATCTTCAGCTCGAGAGGATCAACGTGTATTATAACGAAGCTAGTGGAGGGA

GGTATGTGCCTAGAGCTGTGCTGGTGGACTTGGAGCCAGGGACTATGGATGCTGTGAGGACAGGGCCTTA

TGGCCAGATTTTTAGGCCGGATAACTTCGTTTTCGGGCAGTCGGGTGCGGGGAATAACTGGGCTAAAGGG

CATTATACGGAGGGGGCAGAGCTGGTTGATTCTGTTCTTGATGTTGTTAGAAAAGAGGCTGAGAATTGTG

ATTGTCTTCAAGGATTCCAAGTATGTCATTCTCTGGGTGGTGGCACTGGATCTGGGATGGGCACTCTTCT

CATTTCTAAAATCCGAGAGGAGTATCCAGATCGGATGATGCTAACATTTTCAGTCTTCCCTTCTCCGAAA

GTATCTGACACCGTTGTTGAGCCATACAATGCCACACTTTCTGTTCATCAGCTTGTTGAGAATGCAGATC

AGTGCATGGTCTTGGATAATGAGGCACTCTATGACATCTGTTTCCGAACCCTCAAGCTTACTACCCCCAC

ATTCGGTGATCTCAACCACCTGATCTCTGCTACCATGAGTGGTGTCACATGTTGCCTTAGGTTTCCTGGT

CAGCTGAACTCTGACCTAAGGAAACTAGCAGTTAACCTCATTCCATTTCCGCGACTCCACTTCTTCATGG

TTGGTTTTGCGCCCTTGACATCAAGAGGGTCCCAGCAATATCGTGCTCTCACTGTCCCAGAACTGACCCA

ACAGATGTGGGATGCCAAGAACATGATGTGCGCTGCTGACCCACGAAATGGTCGCTACTTAACAGCGTCA

GCTATGTTTCGTGGTAAGATGAGCACAAAAGAGGTTGATGAACAGTTGATTAATGTCCAGAACAAGAACT

CATCATACTTTGTTGAGTGGATACCGAACAATGTCAAGTCTAGTGTCTGTGACATCCCGCCAAAGGGCTT

AAAAATGTCTTCAACATTTATTGGCAACTCAACCTCTATTCAGGAGATGTTCCGACGGGTTAGTGAACAA

TTCACAGCCATGTTCAGGCGAAAGGCTTTCCTGCACTGGTACACTGGTGAAGGAATGGACGAGATGGAAT

TCACAGAGGCCGAGAGTAACATGAATGACCTTGTGGCTGAATACCAACAATACCAGGATGCAACAGCTGA

GGAGTACTACGAGGATGAAGAAGAAGCAGAAGCTGAAGCTTGAGTCTAAGATTGGTGCATCTTTGGATCT

CTCTCAGCAGGTTCATATGTGCTTATCATTGTGAATGAATGAAATAGTGTGACATAGTTACTCTGTATTG

AATGATATGTCAGATTGAAGATTATGTGTTTTACATAAATGACCGGAAATGTTTGTATTTAACAGAATGC

TGCTTGGGGTTTGAAGTATACTGCAATTTCTTGCCTTA

>MSTRG.432.1 gene=MSTRG.432

GGAAAAGTGAATCTAGTTTGGACTGACAGTCCCACTAGCTAGACCCCCTTTAGGCTTTAAGCCTGTGAAT

CCTCTGGGAGGATTTTTGAACCTCACTTTGTAGTCCTTTTTCCAGAACATCAATAGCAGAAGATCAGTTG

GTTCCAAGGATGCTGAAGTTTGTAAACTGAGAACTGAGGATTTGGATCTTATGTACTAAAATAAGATCTT

TCTTGTCCATAACACAAGTGGTAGAGCATATTTTGCGACATTCTTCTAAAAAAATCACTGGATTTGAGTG

CAAAGCACCAAGAGTCAATGGATATGCTTGCTTGGACCCGAAACTTGTTAAAGCAGACCACTTCTCAGTA

GGAGGGCTGCATCTGGATGGCAGCACGTCGAATTCCCTTGGCACTTTTGTTAACAGAATCACCGTAGCTG

AATTGCCTGGACTGAATACACTAGGTCTGTCTAATGTACGTGTCAACTATGCACGACATGGTGTAGTCCC

TCCCCACATACACCCTCGTGCCACTGAGATATTAACAGTTCTAAAGGGCCGCCTTCGTGTTGGATTTGTC

ACTGGAAACCCTGAGAATAGGCTTATAACAAAGGTTCTTGGAGAAGGTGATGTGTTTGTCTTCCCTATGG

GGCTTGTTCACTTCCAACAAAATGTTGCAGAGGGAAATACTACTGTACTAGCTTTCTTAAGCGCTGCAAA

CCCTGGGGTGATCACAGTCGCCAATGCTGTTTTTGGATCAAATTCACCGATAGGTGATGATGTACTAGGA

AAGGCATTCCAGGCAGATGCTAGGACCATTAAAGCTTTCAAATCTCAGTTTTAAGTATTCATTCACAGAT

ATGTAGTTTGCTAGTATGAGTCTATATTGATTTTTTTTATTTGTTTAAGTTATAATAAAATTTGTTTCTT

TACAATGGA

>MSTRG.437.1 gene=MSTRG.437

AAGTGGGTCACCTTTTTTCCAAGCCACCACTTGAGATTTTTCAGGCTCATCGAGAGACAAAGAATGGATG

GCTGGAAAAGAAGCTAAAAAAGGCTCGTATTTGGTCAAAATGGCTTTGACCTCGTTAATTTCCTGGACAG

TGAGTGGGTCAAGTGGGTGGAGTGGGATGGCATCGGTGATCTCCTTCCATTTCTGGTTCGGGTTTCGGTT

CGGGTTCGAGTTTGATTGACGTGAGCCTGCGGGCTCGCGGCTCGAGAAGGTGCACCTGAGTGTCTTGGAT

CCGGTGCAGTCAAAGTAGCCAGGTTTACTCAAAGTATACCATGTTACTGCAGAGATGAAAAGCATGAAGA

TGAGGAAAGAGAGGCGATTTAAGGAGCAAATCTCCATAACTTAACAAGTCTTCTCTCCACAATTTGGCTC

TTCTTTTTCTGCATATACATATAATATAGAATTCTTGTACGTCAGGCTCAGCGCTCCCTCT

>MSTRG.442.1 gene=MSTRG.442

GTTTAATGTGCCCTGGAGACTCTAGTGTTACATTTTGGGTTCAGGGAGTTGAATGTTGAGAGAGAACTTT

CAGAGTTCAGAAACGACATCTCCGGCCCAGGAAAAGACTTGGGGCAACATACATTTCAGTACTTCCATTC

AGCAACGAAACCAGGGGCAAGGTATCATCTGCCTAGATAAAGACTTGGAGCAAAATACATTTTAGTACTT

CCATTCAGCGATGAAACCAGGGAC

>MSTRG.450.1 gene=MSTRG.450

CACGTTGTCAATTTTAAAAAATCAACACATACCACACGACCCCAATCGAGCTTTGATTTCTGCTCAACCA

CCCACCACAACCAAAGAAACCAATTCTGATTCAGTTCGTTTCAATAATTTGATCTTAAACTAAACCCGAA

ACTCAGAACCTCTGGATCTTATCAGCCTTAACAACCGGATTAGCAACCTTAATAGCTTCCTTTCCGACAG

TCTTGAAATCAAACTCACACTCATGCTTCTCCGGATACCTATGGACCCCACAGTACACGTCCCCACACTT

GCACTTAAACCCCAGAACTCCCACCTTCTTCCGACAGCTCTTACACTTGTTCGCCGACTTCACCTCCTCA

TTCTCCGAAACTCTCGCCGAAGATTCCGCCACATGAACCAACTGAACCGGCTCAACCACTTTCGGCTTCA

CCGAAAACCGGTCGACTATTTTATCCACCGCTTTTTTCGCCGAAGCCGCTTGCTCTTCATGAATAACAAG

GTCTCTGTGACACTTTGAACACAAATTATTCGTTGAAGCCGAGCCGAAAAAGCCGCACCCGTTGGCGCAG

AGTATCGGCTCCGACGTCTTGTAACTCGTGTCATCGTCGTTTCCGTTAGAACTCATCTCCGATCTAAAAA

AATTTAAAATTTAAGATAATAATTCAATCGACGAATAAATTGTGTTTGTGTGGAATATATAGGTGTTGAA

TAAAGTGGATGAGTATAAACAGAAGTTGATTGATTTGTACGCGGTGTGGATTAGGACTCAATATGAGAGG

CGGTAGTAAAAAACGACGTGTATATCTCATGTACATGTAAACAAAATAATATTTTATATCGATAAACG

>MSTRG.453.1 gene=MSTRG.453

CTCTTCCACTCCTCTCTTTCGTTTCTTGATTTTATTTCAAATTTGAATCTCGCTGTGTGTATCTGTCTCT

ATCTCTCCTTCTTACTCTCATATGTTTCTCTCTCATATTTTAACATATAGTCATTGATGATTGATTGTTC

TTGCAGAAGCTTGATTCTAGGGTATTTTGAGGAGTATAGATTTATGATTTAGCTGTTTGACTGACGATTC

TCTCTGTTGTGTGCCCCTTTTTTCTAGGATTTCAAGCGCTGCTTCGCCGCTGGGAAATCAATTATCCTAT

TAAATCTATCTTTCTCGCCGGAGCTGGT

>MSTRG.454.1 gene=MSTRG.454

CATGGGTCACTCACGTGCACATTCACCCTCAAGTCTCCACTCTCTCACTCTCTCAGCGTGAGTCATTCTC

TCTCAAAAACACCATAGCCCTAATTTCCTTTCCATCTCATATCGCCGCTGATTAGTTCAAGTCGCTCATT

GTCCGCCGCTCCATCTCCGACTACAATGTTCGCTTCCGCCTCTGCTCGTAAATCGTGTAGACATTCATGT

TAATTGCTAGATCTAGTATAAGATATTCAAATCATTCAGTTTATCAAACCCTTGTACTATAGTGAAATGT

ACCCCTGTTTCTTTTACTTCACTTCTGATGCCGAATAAGTGGTCCAGATCTTAGATTTTTTTTGCAAGTC

TGTAACAATAAACACAACAAAGAGTCCTGATTTAAGGTGAGTTGATTTAGGCATTCGTTTCTTGACATCT

TCATATTACAAAAACATGAGTTTGTTTAGATTATAC

>MSTRG.456.1 gene=MSTRG.456

CTTCTGGATGGATATGCATGGGGGTTTCTTCTCTTGTAATCATGACTCCCCTTCATCATTACACTTCTTG

AATCTAACAATCTCCTCTCCCTCTCTCTCAGAGCCTACAGGCCACCATGTAATATACCACATTCACCCAT

AACTAACCATAACATTCAGCTATTTATACCAATCCCTCTACAAGTTCTAATTCTACTCAATCCTTGTAAT

ATATACTTCCTATTCGATAAAATTGAACGATGGGAAGCGCAAAACAGCTTTAAAAACTGGGAAGATGTGT

ATCGGAAGTACCTCCTCCGGATGGACTTCGTCGTCCTTCTCTTTGTACAAAAGAAAAGCAAGATCCAAAA

GATACAGGATTCAAGTTCATAGGCTTAACAGGAGAAAGAGAAGTAGAGATGATATGAAAGAGAAGAACTT

GAAGCTGTACATGGAGAACATTACTATCTTTCATGAGAATGAGAAGCTGAGAATGAAAGCTAGTCTTCTA

CACCAAGAAAACCTAAGCTTGATGTCTGAGTTTCAGAAGAAAAAACCCTCTCATCAGCACCAATGCGTAT

CCTCCACCCTAAATTCCCCAACTTACCACCAAGACTGAGTCCCTAGCTAGAATCGTTTCATTGCTTTGCA

TTATTATTATTGATCTACTCATCAAGCTGCTATTCAATAATTAATGTTAATCTTAATAAGAAGATGTACT

TTAATGGGTGGGAAGAGATCAAGGTGCAGAGCCGTAGTTCATTGTGGAAGTCGCAGTGGAGGAGTACTAC

CACATATTCTGTCGTGTACATCCTTTTAAGAGTATCAAATAATATTGAGAGCTCTAGAGCTCTACAAGAC

TCACTAGTCTTTGACAAAGGCATAACTCTAGTTATTTTTTCAGAATGTGTTGTTTAAAGTAACCACCTTC

CTTGTTTAAATAATTTCTCTTCTGTAATCCGTAGACACATCATACACTAATAAACTCACATCTCTTGATC

CCAATAGTCTTTCTAAGTTGTTTTGATGATTTAAGAGATTTAATACAAGTCCTTGCTCAAATATAAAAT

>MSTRG.456.2 gene=MSTRG.456

TAATTCTACTCAATCCTTGTAATATATACTTCCTATTCGATAAAATTGAACGATGGGAAGCGCAAAACAG

CTTTAAAAACTGGGAAGATGTGTATCGGAAGTACCTCCTCCGGATGGACTTCGTCGTCCTTCTCTTTGTA

CAAAAGAAAAGCAAGATCCAAAAGATACAGGATTCAAGTTCATAGGCTTAACAGGAGAAAGAGAAGTAGA

GATGATATGAAAGAGAAGAACTTGAAGCTGTACATGGAGAACATTACTATCTTTCATGAGAATGAGAAGC

TGAGAATGAAAGCTAGTCTTCTACACCAAGAAAACCTAAGCTTGATGTCTGAGTTTCAGAAGAAAAAACC

CTCTCATCAGCACCAATGCGTATCCTCCACCCTAAATTCCCCAACTTACCACCAAGACTGAGTCCCTAGC

TAGAATCGTTTCATTGCTTTGCATTATTATTATTGATCTACTCATCAAGCTGCTATTCAATAATTAATGT

TAATCTTAATAAGAAGATGTACTTTAATGGGTGGGAAGAGATCAAGGTGCAGAGCCGTAGTTCATTGTGG

AAGTCGCAGTGGAGGAGTACTACCACATATTCTGTCGTGTACATCCTTTTAAGAGTATCAAATAATATTG

AGAGCTCTAGAGCTCTACAAGACTCACTAGTCTTTGACAAAGGCATAACTCTAGTAACCACCTTCCTTGT

TTAAATAATTTCTCTTCTGTAATCCGTAGACACATCATACACTAATAAACTCACATCTCTTGATCCCAAT

AGTCTTTCTAAGTTGTTTTGATGATTTAAGAGATTTAATACAAGTCCTTGCTCAAATATAAAA

>MSTRG.479.1 gene=MSTRG.479

GTGAAGAGGGCTAAAGTTGAGTCCAGTGACAAGGAGAAATCTGACAATGGTATAGCTACTCGGAAAATCA

TTTTGGGCAGACCTATCTCGGGTAAGGCATTCTTCAATTGCGGTGTAATTAAGCTGTTCTCTGACTTAGG

GTTTGAATCTTTGATTGTTGATTTACCGAAAATCTGCTACTCCGTGTGTTGATTTGGGTTTGTCTGCTAA

GGAGATTATGGAATTCATGGATGAACTCCGTGATAACCATAAGCAGTTGGTTGAGGGTCAGAAGTTGTTG

TCTGAGCAAATGGATGATTTGGCCAACCAATTTCAGTTCTGGAAAGATATAGTTTTTGGAGGAAAGACAG

GGAATACACCAGAGAAGTGTAGCTCTGGTAGTTTTGTTCATGAGCTTCAGAAGCGGATGTATGGGTCTGG

TGGATCTTCTGATTTCAAGTTTACTTTTACTTCTACTGATGATGCCACTGACAGTCCTAGGCCACGTACT

GCAATGGATGCTCTTAAGGAGGCTGCCGGTACTGATTCTGCTTATGCTGATGCTGGAAATTTGATGATGG

AACAGAATCGTGTTGCTGCGGAGCTTACCAAGAAGTTTGCACAGGAGAAGCCTGACAAGGATGATGAGGA

GACCTAAATTTCTAACCTTTTCGATGATTAAGGGGGA

>MSTRG.487.1 gene=MSTRG.487

CTCGTAGATTCCCATCTTCACATCTTTCTCTTCCTTTTAAATACTCCAAACTAGTAGCTCCTCAATACAT

AAACACCTACAAGATTTTTATAATCACAATAGATACACGCATATATAATCAATTTCTCCCACGCAATCAA

ATACCCAAAAACCCTTCTTTCTTGTTATTTTTTAATCAAAAAATTCCATCTATTTACCAAACAACACAAG

AATTACTAGATTATCATCTTATAGTGATTTTTTGATCTGGGTTGTTGAAAAAGGTGCTAAATTTGGTCTT

TCTTCTGCTGGGTTTTAGTCTGTCT

>MSTRG.489.1 gene=MSTRG.489

TGCTTGATTCACATGACGGATGATGACTTAGAAAAGCTTCAGCATGACTCCCATTCTTGGACAGCAAGAA

GTCTTGCCAGTCTCCTCAATCTGCCCGACAAGAATCCTGCAGTACTGCTTCATTGATGATTTTGCTTCTG

ATTTCTTTTCAACACTTCAAAAACCTTGTTACTACTTTGATGCCAACTCGAGCAACAGATAATAATCAGC

CTGGGACGATGTAACAAATATTACAGTACGAGTGATCAGTTTGTGTAACAGTGGGAATCAAGTGACACGA

ACCGAG

>MSTRG.490.1 gene=MSTRG.490

TGCTTGATTCACATGACGGATGATGACTTAGAAAAGCTTCAGCATGACTCCCATTCTTGGACAGCAAGAA

GTCTTGCCAGTCTCCTCAATCTGCCCGACAAGAATCCTGCAGTACTGCTTCATTGATGATTTTGCTTCTG

ATTTCTTTTCAACACTTCAAAAACCTTGTTACTACTTTGATGCCAACTCGAGCAACAGATAATAATCAGC

CTGGGACGATGTAACAAATATTACAGTACGAGTGATCAGTTTGTGTAACAGTGGGAATCAAGTGACACGA

ACCGAGACCAAACAATCACCTACAATTATTAATAAAAGGATCAACATATAGTAAGCCATGGACAAGCCTG

CGGATGACATTCAACACAAATTTGAAGACTCAAGTCTAGTTGAAAGAGCACCGGATGCAATGACAAGCGA

AATGGAAAGCTTTAGTCTCGAAAAGCTCCACCATACCATATCTACAGGAATGGCCGAAATCGAGTTGTAA

AATTATGCATCAAGCTCCTAATTAAATGTGTGCCATCTCAGACAGCCTTTGTTGAATCAACTTATGAAGA

AAACTGAAGCCCAAACCGGATACTTCATCTAATGGTTCAAGCCAACAAGAAAGGAAGTGCTGATTTTTGA

GCCCCCGTTTCCAAAACAACACAGATAAAGAAATTAAAGTTCTGGGAACATCTACCAACTATTTAGAAAA

CAAGCCTCATTCAGACCCTCCGAGGTCGAGAAAAAAGGAAAAAGAAGAAGAAGGGAGAAGAATGACAATA

CTGCAGCAAACTGATGGATAACTTTTGGTAGAGAAACAACACCCGACAAAACTATCTCTCCAGGATCATC

AACGACTATTCAGGCCAGATACAATATTTGTCGAGCTGCTGATAGAGCTTCGAGCATAGCATTCTTGCTG

TTATAAGATAAGGAAAAAAGATTTCAACAAGTCGGAAGCTAGAGACGGAATTCAACAGCATCAACACAAC

TTCCCAGAAAATAAGATGTGGAGCAGAACTATTGGTTGTTAACTACAATCTATAGTGAAAGAATTGTGTT

TGATAAATACCAGAATGACACATGAGTTCCAAATTAATGTCACGTCGGGTGATCTCGTAAGACATGGCCC

CTGCAATTTTCTATGGGTTATCTAGAGTTATAATAAAAGCACCATCAGAGCACAAGTCTGAAGACAGGAT

CGTGCAAATCGGAGGGCTAAAATCTCGATAAAAGTATATCAGGCGACAAAGTCATCAAAAACAACAATGT

CATGAAAGTCACAAGGTAACAAAGTCAAAAAGTCACACAAGTCATAAGGCAAAAAAAAAGCGAACTACAT

CCCTGATAACAAAGTATACACATGACGAACTACACCCGTTCCTATCAAATACCATGGGACTTCAACGCCC

ATACACAAACACAACACATGACGGACTACACCCCGTTGCTATCAAATACCATGGGACTTCAACGCCCATA

CACAAACACAACACATGACGGACTACACCCCGTTGCTATCACATACCATGGGACTTCAACGCCCATACAC

AAACACAACACATGACGGACTACACCCCGTTGCTATCACATAACATGGGACTTCAACGCCCATACACAAA

CACAACACATGACGGACTACACCCCGTTGCTATCACATAACATGGGACTTCAACGCCCATACACAAACAC

AACACATGACGGACTACACCCCGTTGTTATCACATAACATGGGACTTCAACGCCCATACACAAACACAAC

ACATGACGGACTACACCCCGTTGCTATCACATAACATGGGACTTCAACGCCCATTCACAAACACAACACA

TGACGGACTACACCCCGTTGCTATCAAATA

>MSTRG.502.1 gene=MSTRG.502

GCAAGTGTCATGATTGTGCCTAGACCAAATATGACCTCTTGCAGATATTCTATCCATATAAAATTATGCA

AGTCAACAGATTTGCAATAACAAAAAAGATCAGAGCTTCTAGTTCATCCTCTGTGCAGCTTCCTTGGCAA

GGATCCTCTCTTTGGCCCTTGTTTTGGCTTGTGACTTTGAACCCATAATACCTCCTCCCCACTTCTTACG

GTACTCCTCATATTTGTCATTAAAATTGGCCTTAATAGCTTCCAATATTTTACTGAACTCCATCTTGTCC

TCATTTTTCACAGTGGTCAGGCACAAAACAGATGCAG

>MSTRG.509.1 gene=MSTRG.509

CGTACATAGATAGAGATGATTTTAAGTATCGGGAAAGACCCGAATAAAATAAATTGAGAATGGGTTGTGC

ATTCATCAATCTCAAATCACTGCTACTTGTCATATGCCTGCTGTTGCTTTCTTCCCATGATGTTCTTGCA

AAGTCTCGTCGCCCAATTTCTGAAACTGAGATCCGAGAGAAGAAGAATCGGTGCTTTGAAGATATTGACA

ATGGTTTATGGGGTCAACAGTGCACGTCTTCGATGATAGCCAAGGAAAATTGTATGTTGCAATGTGTGTC

CCCGCCTTGTTATGAACTTGTCTATGAAGGGGATCCGTTAGAAGAAGGGGAGAAAGATTATAGCAGGAGC

CAGGAGTACAAGTACTGCATGCACAGGTTATCTCTAGGGGAGAGCATAGATGGCATTCGAGGTTCCTTTG

ACATGTAAACATGGAAACACGAAACGGTAGCTATCATGGGACACTCAGATGCATCAGCAGTGAGAGTTGC

ATATTAAACTGAAAATGACTAGGCACTGTTCGCCATTATGATTAGTAGTTAATGTGCTTTGTATGAATAG

GCATCACGTCTTATTGGTAAAACTGCAAAAGAAGAAACCGAGTAGGGCATAATGCAAGAAACATTTTCAA

CTCTTTACAACTGACCAGAAGAGATTTGCATTGTATCTGTTTATTACCAACGGATTGACAAATATGTTGT

GCTGTAGTATCATGTGGTTGCAAGATTATTGTTGTATCTTTGTCATACAACTATACAAGTGTGTTAAATG

TTTAGTTTGGAGCCTATTTTATTTTTA

>MSTRG.511.1 gene=MSTRG.511

ATGAAATGATATAAATAGTAATGAAATAAAAGAAAAAGAATTTGATTCCAGAAACCCATGTTTCATATCC

ACCTCCCCCCTTGGGTATGAAAATCTCATACCTTGTGGGTTTGAGGAATGGGGTTGATAAAAGAAAATTT

TCAACCAAACATCAGGTATGAGTTTGGAATGAACAAAACCCATACCTGATACCAGAGAAGCCCCAACCAA

ACGACCCC

>MSTRG.516.1 gene=MSTRG.516

AATTACTCCCTCCGTCCCTTCCATTTCTTTACACTTTCCTTTTTGGGGTGTCCCATCCAATTCTTTACAT

TTCAAAACTTACCCAAAATAGTCAATGGGTCCCACCACTTCTTCACTTTACTTTCCTTTTCACACTACTA

TTACTCCACTATCTTCTTTTTATACATTAAAAATCAATGGGTCCCACCACTTCACCCACTTTTCTTTCTC

TTTTCCACTACTTTATACATATTTCTTAACATCCGTGCCCAACCCAAACGATAAGAATTGGGAGGGACGG

AGGGAGTA

>MSTRG.522.1 gene=MSTRG.522

GGTGCGTCTAGTGGTAGCTCAATTGTTGCATTGTTTTGATTGGGAACTTCCAAACGGAATGCAGCCATCA

GAACTTGACATGACTGAGGAATTCGGGCTCGTTGTTGCAAGGGCTACGCACCTTATGGCCATTCCGACCT

GTCGCCTTCACCAAAGCTAGGCGCCAATCAAGTAATACCGCTTGCTATAAACATTCCATCATGTACTTGC

TTTTATGTAGTAGTGCATTTAAGTGTCTTGTACACAGGTATCGTAACCCAGAAAAACCTCTATGTACTGA

GATGAATATGCATTGAG

>MSTRG.525.1 gene=MSTRG.525

AGGAAAAAAATATGCCAACATGATCATTGTTTTAGGAAGAGGCGATTCTCAACAGATTGTAGTAGAAATA

ATATATAAAATTTTGCTAATTATATCCACAAAGGTAAGACAACTCTACTTTTTCTTTTAATTACTTTTCT

TTTATAATCAACAATATGCATTTAATTTTGATAATTATGCACACAACCAAGAGGTATTGTATGAAGCTGG

ATATGTCCCTTCAAATTCTGAAAAGCGTCCTTTGGGGGTTATCATCTCTATTATTCAAATTGCTTTCCAT

GCAACTCCAGAGTTTACTTGCTCACAAGGTGCTGTAGAGGAACTTCAACTGTGCTTTTATGAGTATTTTA

AGATATGAATAGGGTGGCGGAAATGGGGAATATTGCTCTAGGCTTGGATTCTTAATAAGGAGAAAAAAAT

TGATTCTAAGGCTCCCCAGGGTTCTCGGCAACTGCCAAGACACTCACTGCTTTTGACCTCATTGTTATTG

GTACATTAATTTGTTTAGCTCCCTTATAACAGTATATTTTCTCGTTCAGACATATGGTTTTTTGTTTTAC

ACCTTGTCAAATATATTGTGTTTACTTGTTGCAGAATGCAGATACATCAGTAAAATGTATTTTATGGTTA

GTACTTTCTTGATGACTGAAAGGGGAGCCATCTTTTGTGCACCCCTTTGGTTTTGCCCAACGGTGTTATA

ATTATATTAGATTAATATAAAGTAATCACTATTGTATACGATATGGGAATAGCCATTGGTGCAAGTATCT

ACATTTCGGTTGGAACAATTACTAGAGGGCATGCAAAACCTATTTTTGATATCTGTTGTCTTATCGGTAG

AATGTCAGCTGATTTCTTAACGTGAGCTGGCTTATCACTGCCCTATGGTCGTTTATGGTCGTAAGTGCTT

ACAGGAACGAAGGCACAGGGGACATTGATGACGACTGTTCCCCAATTTTTTTGATAATCATGTATATTTT

CTCTTAAATTATGTATATCACATGTAAAAAAGAATTAATTTATGATCCTCA

>MSTRG.526.1 gene=MSTRG.526

CTTCAATCGTCTTCAAGGGAGTCAGCGGTTACATGAACAGGCAGGCGGCAGTTCCTCCATTTTAATTGGA

TGATGGAGTCGCCGGTGACAGGAAAGCGAAGCGAGTGTCTCCGGGGGGAGTGTTATGGATTCCGATGTGA

TTGAGAGGCCTCTGCCGACGCTGATTAATCGGAGTTCGAAATCAAAAACTCTTGTAGTTAAGGAGTGTAT

TTTTTGGAATCCGTAGAACCAATGGATGATTTTCATGCATGTTTCCAGTTTTGCTTTAAG

>MSTRG.528.1 gene=MSTRG.528

TTTTTTTTTATTTTATTTTGTTTTTATTTCTCACGTTTTTAGTTCACGTTTTTTTTTATTTTATTTTGTT

TTTATTTCTCACGTTTTTTTTCTCATATTTTTTTTTTATTTTATTTTTACATGTTTAGTAGTAATTGTTG

TAGCTAGTATCACGAGCATTTGCATGAATTATGAAGTTAAGCCAAGTTAATTGCCTATGATGAGTTATGT

GCGTAGTTCTTGATTAGTAATTTCAATTGCAATGCTAGGTTAGTACTTAGAAATTGCTTGTGTTGGAAAT

TGTAATTCACATATGTTCTTGAGATAGGATTTAGACGAAATTCCATGAATTCTAGGATTGAATTGAACAC

CCTTCAGCTTAGGTCTGTAAATCTTAAGTTTGGGGGAACTGAGGAGTAGATATACCTTTCTAAAAAAAAA

AATAGATAGAAAAAAAAAAAAAAGTAGTAAATAAATTCACTAAAAAAATAAGTGGTAGAATTAACTAGGT

TGAGCTCATTAGTACTCGAGTAATTAAGTCTGAGGGGACTTTGTGCCTAACAACCTAAAGCCCTTCGTGG

TTTGGGATTTGTTGACCCAACGCTCGCTACATGGGTACTAGTGCATAAATCTTTAGGGATCTCAACCATT

GCACGGTTAAATAAACCACTAGAATAGAGTGAATAATTGGTG

>MSTRG.532.1 gene=MSTRG.532

CCAGTTTCTCTGCACGAATACCGAGAGAAACCGAGCGGTGTTCTTCGTGTTCTTCATAATCAAACGTGGA

TTTGAAGGCGTTATTGGAGTCAGAATCAAGCGTGCGAATAGTCAAAACGAAGATATCGAGGAGTAGAATC

CATATCAATGATCAAAAACAACATGAGATTCCCAGAAAGGCTAAAAGATGAGATGATTGATTGAGATTGT

GGATAAGGCTAATGCTAGGCGAGGAGACAGAATTGAGGATCCAGTGGTCGAGGAGTGTTCAGGAAAGTTG

ATGAGGGTGGTGCAAGCTAGAGCTGTGCTATAGGGATTCAAGTGAAAGTCATAGCTGGAATAGTTAACAG

TGATTCTTCTTATCGAAGATGATCTGACTTGGTAAGAAAAGTTGTGTAATATTAGTGGTGGATTCTAATT

TCAATACTGTAACCTGGAAGCGATCCTGTTGTTTTTGGGGGTTAAATGTTCTCTTAAAATTTTCTTATT

>MSTRG.533.1 gene=MSTRG.533

TGCAGTTTAATGGACTGGAGTTTGATTGATTCCTTGACGTGGCCAATTTTTACAGCCGGATATATGATGA

TGATGGGGTATACTGAGGGGCCTAAATGGAAAGGGTTTTACGTGGATGATTCTCCTGATACATCTACGTG

GCTTTACTAAGTCGACTAAAGCAGGAGTATGAGCATGATCTAATCCAAAATGACCTACCAGATGTCTTGC

AAAGGCATATGGATGGCCAATATAGCTTTGGTTGCAGTTTGTCGAGGCTGGCCTGCATTTCCTTTCATCC

AAGTCGACATAAAAGCATGGATCTGGCTT

>MSTRG.534.1 gene=MSTRG.534

TGGATGATTCTCCTGATACATCTACGTGGCTTTACTAAGTCGACTAAAGCAGGAGTATGAGCATGATCTA

ATCCAAAATGACCTACCACATGTCTTGCAAAGGCATATGGATGGCCAATATAGTTTGTCGAGGCTGACCT

GCATTTCCTTTGATCCAAGTCGACATAAAAGCATGGATCTGGCTTGGAGTTTGCCTCTTGAGCAATGAAT

ATTGATAAATTCAATCCGGAGTGAGGAAAAGCTTTATTCACAGTCAAATGTAATCTGGAGCCAAATGTAT

TTTTGGCTTTCTCCTTCCTCCATGCAATATCAGAGTTGTTAATGCCAAAAAGGTTAGAGCTTACAAGCTC

CATTAAACAGATATGTTTTATGTCATGGAACCACATTGCTTATGCTATTATGCTCTCTCAGGTTTTACAT

TTTTACATTTTTACATATATAAAAAACAATACCATGCAATGTAATCAGATTCCCTACAGGATGTGAATAA

TGGGAGATTATTTGCATAATCATATTGAATTGTATTATTTTGTTAGCATGGTGACATTTAAAACTAAATG

ATGTATAGTTTATATATAAATACTGAGTTAATCGACTTACTAGAATTTTCGTGTAGGCACTAGAATGTGA

GTCTATATAGTTATTTTAAAACAGAATATACCAGTGTGTTGCCACATGTTTCACATTAAATCAAATGATG

CAGTTTCATATCTAATATGTGGAAAAAGAAGTTTGTGTAATTTAACAGTGTAGATGTAACTACATGACTC

TTTATCTGGTGTATGTGTTTCTATTTGGTGTGTATATAGTGGAATTCTGGCATATCTATGAATGCAATGA

AGCTGATGGGTTTCACATGCTGGTGGTTTTTGGTAGTTTAATTTAGTAGGTCCTTGAACAGAGTTTAGTG

GTTTAAATTAGCAATTGAAGGTAACACAATATCTATACATAAAAAAATTATTAATTTTTTTGGTCAGGTT

AAGTGATAGTTTTACTTAGAAAACCGACTCTACTTCTTGCGCCATTGGAAATGAGTTGGAAATTGCTAGC

ATTACCTCGGATTAGATATACTGATAAAGCAGACTCATCCCAGATTAATGGTTGTATCTCACTTTCTTCT

CAAATTCTTTGAC

>MSTRG.537.1 gene=MSTRG.537

GTGTGCGAATTAATTTGTAGGCGCTGTCGGTGTTACTCTCACAGCACAGCTTCTAGTAGTTAGCATCTCT

CTCTCTCTCTCTCTCTCGTGTAGACATACATTATATAGATATACACACTCTACCATCTCTCTCTCTCTCT

CTCTCTCTCTCTCTCTCTCTCTCTCTCTCTCTCTCTCTCTCTCTCTCTCTCTCTCTCTCTCTCTCTCTCT

CTCTCTCCGTATATACGCACAACAATATCTCTATATCATCCATATTGTTGTGAGTCTTGAGCTTCAACTA

TCTTTTCTTCGCTTCTTTCCTACGAAATGCATTGACAAGATCTCCCTCCCTCCCCTTTAGCTCTCTCTCT

CTCTCTGCGTTTATACTGTCCCAATTGTCAGCTACCTACTTCATTCAATCTGTTCTCCAGATCTTCCTAG

CTTATCTCAATCCTCACTTAACTCATTCATGTGAAGCTAGCTTCGCTTTGTAACTGTATTTTGGAGGTAG

ATCTTGAGATCTAATTAGCAAAGCTGGCGGCTACAGTAGCTTGGAGAGGGGTTGCTGCCAGCAATGGCAC

AGCCCTGTCATCTAATGATCTGGAAAGAAATGGGGATGCTAAATCTCATGACCAAGAGCCTCCAACACCA

CATTCACTCTTAAAGATGAGTTCGAGGGAGCGCAGTAGCATGGAGGATCCAGATGGAACTTTAGCTAGTG

TTGCTCAATGCATTGAGCAGTTGCGCCAGAGTTCCTCATCTGCACAAGAAAAAGAGTTCTCCTTAAAGCA

ACTATTGGAGCTTATTGATACACGTGAAAATGCTTTCAGTGCTGTTGGATCACATTCCCAGGCAGTTCCA

GTGTTGGTATCCCTTCTTAGATCAGGATCACTTGGGGTAAAAATACAGGCTGCTACAGTTTTAGGGTCAC

TGTGCAAGGAGAATGAACTAAGGGTGAAGGTATTGCTGGGGGGTTGCATTCCACCACTTCTTGGTCTACT

CAGGTCTAGCTCAGCGGAAGCTCAGATCGCTGCAGCGAAGACGATATATGCTGTTTCTCAAGGTGGTGCT

AAGGATCATGTTGGCTCAAAGATATTTTCAACTGAAGGAGTTGTGCCAGTTCTGTGGGGGCAGCTTGAGA

AGGGATTGAAGGCTGGAAACGTGGTTGATAACTTACTGACTGGAGCTTTGAAAAACCTTTCAAGCAGCAC

TGAGGGTTTCTGGTTTGCAACAATACAAGCTGGGGGATTAGATACACTTGTAAAGCTGCTTGCAACTGGA

CAATCTAACACCCAAGCAAATGTTTGCTTTCTCCTTGCATGCATGATGATGGAGGATGCATCTGTTTGTC

CTAAAATTCTGGCCGCTGAGACTACCAAGCAACTCCTAAAGCTACTAGGGCCCGGAAATGAAGCCTCGGT

CAGAGCAGAAGCTGCTGGTGCTCTAAAATCTCTATCTGCTCAGTGCAAAGAAGCAAGGCGGGAAATCGCA

AGTTCTAATGGTATACCCGCTTTGATAAATGCTACAATTGCTCCTTCAAAAGAGTTTATGCAAGGTGAGT

ATGCACAGGCTTTGCAGGAGAATGCAATGTGTGCTCTTGCAAATATATCTGGTGGTTTGTCATTTGTCAT

CTCAAGTCTTGGCCAAAGTCTTGAATCCTGCACGTCGCCTGCACAGGTAGCTGACACGTTAGGGGCATTA

GCTTCTGCACTAATGATTTATGACAGCAAAGCAGAAAGTACAAGAGCTTCAGATCCTGACGATGTTGAGA

AGACTCTTGTGAAGCAGTTCAAACCTCGCTTGCCATTTCTTGTGCAGGAGCGTACAATAGAAGCTCTTGC

CAGTTTGTATGGAAATTCCACACTTTCTGGTAAACTTGCAAATTCTGATGCAAAGCGTTTGCTAGTTGGT

TTGATCACAATGGCAAGCGATGAAGTTCAGGATGAGTTGATACGATCACTTCTGGTATTGTGTAACAATG

AAGGTACACTATGGCATGCCCTTCAGGGACGTGAGGGAATTCAATTATTGATCTCTCTTCTTGGGCTGTC

ATCTGAGCAACAACAGGAGTGTGCCGTCGCATTACTTTCCCTTTTATCCGATGAGAATGATGAAAGCAAA

TGGGCCATCACAGCTGCTGGAGGTATACCTCCACTTGTTCAGATATTAGAGACAGGATCTCCAAAAGCCA

AGGAAGATTCTGCAACAATCCTCGGAAACCTCTGTAATCACAGTGAAGATATTAGAGCATGTGTTGAAAG

TGCTGATGCTGTTCCTGCTTTGCTGTGGCTATTAAAGAATGGAAGCTCCAATGGCAAAGAAATTGCTGCA

AAGACATTGAATCATCTGATCCACAAATCAGATACAGCAACCATTAGTCAGCTCACCGCATTACTTACAA

GCGATCTACCCGAATCTAAAGTGTATGTTTTAGATGCACTAAAAAGTTTGCTATCTGTGGCCCCTCTTAG

TGATATGTTGCGTGATGGAAGTGCTTCAAACGATGCTATCGAGACAATGATTAAAATATTAAGTTCTACC

AGAGATGAGACACAGTCTAAGTCTGCATCAGCTCTTGCTGGAATATTTAATCTTAGAAAGGACTTGCGTG

AAAGTCCCATAGCTGTTAAGACTGTTAGGTCGGTCATGAAGCTTTTACATGTAGAATCCGAAAGCATATT

GGCGGAGTCTACTCGTTGCCTTGCTGCAATATTTCTGTCAATTAAGGAGAACCGTGACATGGCTATGATT

GGGAGAGATTCATTGCCTACACTAGAGGTGCTTGCAACTTCGTCTTCACTGCAAGTTGCAGAGCAGGCCA

TTTGTGCTTTGGCTAACCTTCTTTTGGATAGTGAAGTTTCAGAAAAAGCCAAGCCTGAAGAAATTATTTT

GCCTTCTACTAGGGTATTGCGTGAAAGCACAGTTACTGGAAGGAGCCATGCAGCAGCAGCAATTGCCCGG

CTTCTGAGTTCTCGTAAAATTGATGCTGAGATTGCTGACTGTGTGAACCGTACAGGAACAGTGCTTGCAT

TGGTTTCGTTCCTAGAATCAGAAAATGCCGGGACTGCTGCCATATCAGAGGGGTTAGAGGCACTTGCCAT

TATATCGAGATCTGGAGGAGATAAAGGACAGAACAAACCTGCATGGGCAGTTCTGGCTGAATTCCCAGAT

AGCATAATCCCGATCGTTTCATGCATTTCTGATGCAAACCCCTTGTTGCTTGATAAGGCTATAGAAATAT

TGTCACGGATTTCTTTGGCTCAACCTGTTGTTTTGGGGAACAATATTGTGAGTGCTTCAGGGTGTATCTC

GTCAATTGCTAGACGGGTAGTTAGCTCTTCAAAGGAAGCAGTAAAGATTGGAGGCACCACCCTTCTTGTT

TGTGCTGCCAAAGTAAATCTTCAGAGAGTTGTAGATGATTTGAATGAATCATATTCTTGCGCATATCTCA

TTCAGTCCCTTGTAGGAATGCTGACCTACACAAAAAATCCTCTGGTTGGTGACCAGGGAAGTAATGAAGG

CGTAAATATTCTTGGGCATGCTGAAGAACTAAAAGCTGGTGAGAATGAGACGAGTAGATATGTCATCTAT

GGTTCAAATACCGCAATATGGCTTCTCTCTGCTCTTGCTAGTAATAACGAAAAGAGCAAAACTCAGATCA

TGGAAGCAGGTGCCATTGAAGTTCTAACTGAGAGAATCAATCAGTGTTTATCAGAATATTCTCAGGTCGA

TTCTAAAGAGGAAAGCAGCATTTGGATATGTGCTTTACTTCTAGCAATTCTATTTCAAGATAGAGATATT

ATACGTGCGCATGCAACAATGAAATCTGTACCTGTACTGGCTAATTTGTTGAAGTCAGAGGAGGCAGCAA

ACAGATATTTTGGTGCACAAGCCATTGCCAGTCTAGTATGCAATGGTAGCAGGGGGACTCTTCTATCAGT

CGCTAATTCAGGGGCTGCAGGTGGACTCATTTCACTGCTTGGGTGTGCTGATGCTGATATATGTGATATG

CTTGACTTGGCAGACGAATTTTCCTTGGTGCGGTATCCTGAACAAGTTGCTCTTGAGAGGTTGTTTAGGG

TTGAGGATATCAGAGTTGGTGCTACTTCAAGAAAAGCAATACCTGCACTAGTTGATCTACTTAAACCGAT

TCCAGATCGTCCAGGGGCACCATTTTTAACCTTAGGGCTTCTGCTTCAGCTTGGTAAAGATTGCCCTTCG

AATAAAATTCTTATGGTAGAAGCTGGTGCTCTGGAAGCATTGACCAAATACCTTTCACTTGGCTTACAAG

ATGCAACTGAAGAAGCAGCTACGGATCTCTTAGGTATGTTATTTAGTACTGCTGAGATACGTAGACATGA

AGCAGCATTTGGTGCCGTAAGCCAACTTGTAGCAGTATTGCGATTGGGTGGAAGAGGTGCAAGGTATAGT

GCTGCTTTAGCATTGGAAAGCCTTTTTTCTGCAGACCATATTAGGAATGCAGAATCTTCCAGGCAGGCTG

TACAGCCTTTAGTGGAAATCCTTAATACAGGTTTGGAGAAAGAGCAGCATGCTGCTATTGCTGCATTAGC

TAGGTTACTGAGTGAAAATCCATCAAGAGCCTTGGCAGTTGCTGATGTTGAAATGAATGCGGTAGATGTT

CTTTGCAGGATTCTTTCATCAAACTGTTCACTAGAGCTGAAGGGGGATTCGGCTGAGTTGTGCTGTGTAC

TTTTTGGAAATACAAAGATCAGATCCACCCTGGCTGCAGCGCGCTGTGTGGAACCTCTGGTTTCTTTGCT

CGTGGCTGAGTATAGTCCTACTCAGCACTCTGTTGTCCGTGCATTAGATAAACTCGTTGATGACGATCAA

TTAGCTGAGCTCATTGCTGCACATGGAGCAGTTATACCTCTTGTAGGCCTTCTGTATGGCCACAACTATG

TGCTCCATGAGGCTATTTCCAGAGCTCTTGTGAAGTTGGGAAAAGACAGGCCTTCTTGTAAGATGGAGAT

GGTGAAGGCTGGGGTGATTGAGAGTGTACTTGATATTCTCCATGATGCACCAGATTTTCTGTGTGCTGCT

TTTGCAGAATTGCTACGAATATTAACCAATAATGCTACCATTGCCAAGGGTCCATCTGCAGCGAAAGTGG

TTGAGCCTTTCTTTGTGTTGTTAACAAGAGCAGAGTTCGGACCTGATGGACAGCATAGTGCTTTACAAGT

TCTTGTGAATATCTTAGAACATCCTCAGTGCCGTGCTGATTATACCTTAACATCACATCAAGTGATCGAA

CCCATCATTCCATTACTTGATTCTCCTGCACCGCCAGTGCAGCAGCTAGCAGCTGAACTTCTTTCTCATT

TACTCCTAGAGGAACATCTTCAGAAGGATTCAGTTACACAGCAAGTAATTGGCCCACTTATGCGAGTTCT

TGGTTCTGGTATACACATTCTGCAGCAGAGAGCTGTTAAAGCTCTTGTTAATATTGCACTGATTTGGCCC

AATGAAATTGCCAAGGAGGGTGGCGTCAGTGAGCTATCCAAAGTGATATTGCTAGCTGATCCCTCTTTGC

CTCATGTCTTGTGGGAATCAGCTGCTTCTGTTTTGTCCAGTATTCTTCAGTTCAGTTCTGAGTACTATTT

GGAAGTGCCTATTGCGGTTTTGGTGAGATTGCTTCGATCTGGTTCAGAATCTACAGTAGTTGGTGCATTA

AATGCTCTTCTAGTTCTGGAAAGTGATGATGCTACAAGTGCTGTAGCAATGGCTGAAAGTGGAGCCATTG

AAGCTCTTTTAGAACTTCTTAGATGTCATCAGTGTGAGGAAACTGCTGCAAGACTTCTGGAGGTATTGCT

GAACAATATTAAGATCAGAGAAACAAAAGCCACAAAGTCTGCAATAGTACCACTATCACAGTATCTCTTG

GATCCACAAAGCCAAGCTCAACAAGCAAGATTACTAGCAACTCTTGCTCTAGGGGATTTATTTCAGAATG

AGGTTCTTGCTCGATCTGCTGATGCTGTTTTAGCTTGCCGTGCTTTAGTAAATCTGCTTGAGGATCAACC

TTCAGAAGAAATGAAAGTGGTTGCAATATGTGCATTGCAAAACCTTGTAATGTACAGTAGATCAAATAAG

AGAGCAGTTGCTGAAGCTGGAGGTGTTCAGGTCGTGCTGGATCTGATTGGTTCAAGTGAACCAGATACAT

CAGTTCAGGCTGCAATGTTTGTTAAGCTTCTCTTCTCTAACAACACAATTCAAGAGTATGCTTCCAGCGA

AACCGTTAGAGCTATAACAGCTGCAATTGAAAAAGATTTATGGGCTAATGGTGTAGTGAATGAAGAGTAT

CTTAAAGCTCTTAATGCACTCCTTGGCAACTTCCCACGTTTAAGAGCCACAGAACCTGCAACCCTGAGCA

TTCCCCATCTAGTGACAGCCCTCAAGACTGGCTCAGAGACAACTCAAGAAGCAGCATTGGATTCACTGTC

TCTTCTCAAGCAAGCTTGGGCAGCATGCCCTGCCGAGGTCTCCAGAGCGCAATCAACAGCTGCTTCTGAG

GGGATTCCCCTGTTACAGTACTTAATTATGTCTGCCCCACCTCGAGTTCAAGATAAGGCAGACCATCTTT

TGCAGTGTCTGCCGGGAACATTGACAGTGACTATAAAACGTGGAAAGAATATCAAACAGTCAGTTGGAAA

TCCAAGTGTGTTTTGCAAGCTAACACTTGGCAACACCCCCTCCAGGGAAACCAAGATTGTGTCGACCGGA

CCAAATCCCGAGTGGGATGAGCCCTTCCAGTGGCAATTTGAAAGTCCTCCAAAAGGCCAAAAGCTTCACA

TTTCTTGCAAGAACAAGAGCAAGATGGGAAAGAAATCGTTTGGAAAAGTCACTGTCCAGATTGATCGAGT

CGTTACCCAAGGAGCAGCTGCAGGAGAGTACTTCCTGCTACCTGAAAGCAAGAGTGGTAGTAAAAGGAGC

CTGGAAATAGAATTTCAATGGACTAATAGCAATAACATGCCTCAGTCTGAAGCTTAAAGAGATTTGCCTG

GTGTTAATTTTTTCTTTTTATCGAATATAATTATTGTGCTGGTGTACATAATATCTGGAAGGTTTATTTT

TGTAGTTAGTTGCTGCAATCCGGAAAGTCGGAAACTGCAATTCATTGTATTCTTTTGCAAAGTGTGGTAG

TTTTAGTCATCTTCTGTTCTTTTTGTTATAGTTGTATTGGAGTTGTATTCTGTGTTTGATATTCAGACTC

GAGAGGTTAATAAAATCTTGTGTATTGAATTCATTCTTACCTGTTTT

>MSTRG.537.2 gene=MSTRG.537

GTGTGCGAATTAATTTGTAGGCGCTGTCGGTGTTACTCTCACAGCACAGCTTCTAGTAGTTAGCATCTCT

CTCTCTCTCTCTCTCTCGTGTAGACATACATTATATAGATATACACACTCTACCATCTCTCTCTCTCTCT

CTCTCTCTCTCTGCACAACAATATCTCTATATCATCCATATTGTTGTGAGTCTTGAGCTTCAACTATCTT

TTCTTCGCTTCTTTCCTACGAAATGCATTGACAAGATCTCCCTCCCTCCCCTTTAGCTCTCTCTCTCTCT

CTGCGTTTATACTGTCCCAATTGTCAGCTACCTACTTCATTCAATCTGTTCTCCAGATCTTCCTAGCTTA

TCTCAATCCTCACTTAACTCATTCATGTGAAGCTAGCTTCGCTTTGTAACTGTATTTTGGAGGTAGATCT

TGAGATCTAATTAGCAAAGCTGGCGGCTACAGTAGCTTGGAGAGGGGTTGCTGCCAGCAATGGCACAGCC

CTGTCATCTAATGATCTGGAAAGAAATGGGGATGCTAAATCTCATGACCAAGAGCCTCCAACACCACATT

CACTCTTAAAGATGAGTTCGAGGGAGCGCAGTAGCATGGAGGATCCAGATGGAACTTTAGCTAGTGTTGC

TCAATGCATTGAGCAGTTGCGCCAGAGTTCCTCATCTGCACAAGAAAAAGAGTTCTCCTTAAAGCAACTA

TTGGAGCTTATTGATACACGTGAAAATGCTTTCAGTGCTGTTGGATCACATTCCCAGGCAGTTCCAGTGT

TGGTATCCCTTCTTAGATCAGGATCACTTGGGGTAAAAATACAGGCTGCTACAGTTTTAGGGTCACTGTG

CAAGGAGAATGAACTAAGGGTGAAGGTATTGCTGGGGGGTTGCATTCCACCACTTCTTGGTCTACTCAGG

TCTAGCTCAGCGGAAGCTCAGATCGCTGCAGCGAAGACGATATATGCTGTTTCTCAAGGTGGTGCTAAGG

ATCATGTTGGCTCAAAGATATTTTCAACTGAAGGAGTTGTGCCAGTTCTGTGGGGGCAGCTTGAGAAGGG

ATTGAAGGCTGGAAACGTGGTTGATAACTTACTGACTGGAGCTTTGAAAAACCTTTCAAGCAGCACTGAG

GGTTTCTGGTTTGCAACAATACAAGCTGGGGGATTAGATACACTTGTAAAGCTGCTTGCAACTGGACAAT

CTAACACCCAAGCAAATGTTTGCTTTCTCCTTGCATGCATGATGATGGAGGATGCATCTGTTTGTCCTAA

AATTCTGGCCGCTGAGACTACCAAGCAACTCCTAAAGCTACTAGGGCCCGGAAATGAAGCCTCGGTCAGA

GCAGAAGCTGCTGGTGCTCTAAAATCTCTATCTGCTCAGTGCAAAGAAGCAAGGCGGGAAATCGCAAGTT

CTAATGGTATACCCGCTTTGATAAATGCTACAATTGCTCCTTCAAAAGAGTTTATGCAAGGTGAGTATGC

ACAGGCTTTGCAGGAGAATGCAATGTGTGCTCTTGCAAATATATCTGGTGGTTTGTCATTTGTCATCTCA

AGTCTTGGCCAAAGTCTTGAATCCTGCACGTCGCCTGCACAGGTAGCTGACACGTTAGGGGCATTAGCTT

CTGCACTAATGATTTATGACAGCAAAGCAGAAAGTACAAGAGCTTCAGATCCTGACGATGTTGAGAAGAC

TCTTGTGAAGCAGTTCAAACCTCGCTTGCCATTTCTTGTGCAGGAGCGTACAATAGAAGCTCTTGCCAGT

TTGTATGGAAATTCCACACTTTCTGGTAAACTTGCAAATTCTGATGCAAAGCGTTTGCTAGTTGGTTTGA

TCACAATGGCAAGCGATGAAGTTCAGGATGAGTTGATACGATCACTTCTGGTATTGTGTAACAATGAAGG

TACACTATGGCATGCCCTTCAGGGACGTGAGGGAATTCAATTATTGATCTCTCTTCTTGGGCTGTCATCT

GAGCAACAACAGGAGTGTGCCGTCGCATTACTTTCCCTTTTATCCGATGAGAATGATGAAAGCAAATGGG

CCATCACAGCTGCTGGAGGTATACCTCCACTTGTTCAGATATTAGAGACAGGATCTCCAAAAGCCAAGGA

AGATTCTGCAACAATCCTCGGAAACCTCTGTAATCACAGTGAAGATATTAGAGCATGTGTTGAAAGTGCT

GATGCTGTTCCTGCTTTGCTGTGGCTATTAAAGAATGGAAGCTCCAATGGCAAAGAAATTGCTGCAAAGA

CATTGAATCATCTGATCCACAAATCAGATACAGCAACCATTAGTCAGCTCACCGCATTACTTACAAGCGA

TCTACCCGAATCTAAAGTGTATGTTTTAGATGCACTAAAAAGTTTGCTATCTGTGGCCCCTCTTAGTGAT

ATGTTGCGTGATGGAAGTGCTTCAAACGATGCTATCGAGACAATGATTAAAATATTAAGTTCTACCAGAG

ATGAGACACAGTCTAAGTCTGCATCAGCTCTTGCTGGAATATTTAATCTTAGAAAGGACTTGCGTGAAAG

TCCCATAGCTGTTAAGACTGTTAGGTCGGTCATGAAGCTTTTACATGTAGAATCCGAAAGCATATTGGCG

GAGTCTACTCGTTGCCTTGCTGCAATATTTCTGTCAATTAAGGAGAACCGTGACATGGCTATGATTGGGA

GAGATTCATTGCCTACACTAGAGGTGCTTGCAACTTCGTCTTCACTGCAAGTTGCAGAGCAGGCCATTTG

TGCTTTGGCTAACCTTCTTTTGGATAGTGAAGTTTCAGAAAAAGCCAAGCCTGAAGAAATTATTTTGCCT

TCTACTAGGGTATTGCGTGAAAGCACAGTTACTGGAAGGAGCCATGCAGCAGCAGCAATTGCCCGGCTTC

TGAGTTCTCGTAAAATTGATGCTGAGATTGCTGACTGTGTGAACCGTACAGGAACAGTGCTTGCATTGGT

TTCGTTCCTAGAATCAGAAAATGCCGGGACTGCTGCCATATCAGAGGGGTTAGAGGCACTTGCCATTATA

TCGAGATCTGGAGGAGATAAAGGACAGAACAAACCTGCATGGGCAGTTCTGGCTGAATTCCCAGATAGCA

TAATCCCGATCGTTTCATGCATTTCTGATGCAAACCCCTTGTTGCTTGATAAGGCTATAGAAATATTGTC

ACGGATTTCTTTGGCTCAACCTGTTGTTTTGGGGAACAATATTGTGAGTGCTTCAGGGTGTATCTCGTCA

ATTGCTAGACGGGTAGTTAGCTCTTCAAAGGAAGCAGTAAAGATTGGAGGCACCACCCTTCTTGTTTGTG

CTGCCAAAGTAAATCTTCAGAGAGTTGTAGATGATTTGAATGAATCATATTCTTGCGCATATCTCATTCA

GTCCCTTGTAGGAATGCTGACCTACACAAAAAATCCTCTGGTTGGTGACCAGGGAAGTAATGAAGGCGTA

AATATTCTTGGGCATGCTGAAGAACTAAAAGCTGGTGAGAATGAGACGAGTAGATATGTCATCTATGGTT

CAAATACCGCAATATGGCTTCTCTCTGCTCTTGCTAGTAATAACGAAAAGAGCAAAACTCAGATCATGGA

AGCAGGTGCCATTGAAGTTCTAACTGAGAGAATCAATCAGTGTTTATCAGAATATTCTCAGGTCGATTCT

AAAGAGGAAAGCAGCATTTGGATATGTGCTTTACTTCTAGCAATTCTATTTCAAGATAGAGATATTATAC

GTGCGCATGCAACAATGAAATCTGTACCTGTACTGGCTAATTTGTTGAAGTCAGAGGAGGCAGCAAACAG

ATATTTTGGTGCACAAGCCATTGCCAGTCTAGTATGCAATGGTAGCAGGGGGACTCTTCTATCAGTCGCT

AATTCAGGGGCTGCAGGTGGACTCATTTCACTGCTTGGGTGTGCTGATGCTGATATATGTGATATGCTTG

ACTTGGCAGACGAATTTTCCTTGGTGCGGTATCCTGAACAAGTTGCTCTTGAGAGGTTGTTTAGGGTTGA

GGATATCAGAGTTGGTGCTACTTCAAGAAAAGCAATACCTGCACTAGTTGATCTACTTAAACCGATTCCA

GATCGTCCAGGGGCACCATTTTTAACCTTAGGGCTTCTGCTTCAGCTTGGTAAAGATTGCCCTTCGAATA

AAATTCTTATGGTAGAAGCTGGTGCTCTGGAAGCATTGACCAAATACCTTTCACTTGGCTTACAAGATGC

AACTGAAGAAGCAGCTACGGATCTCTTAGGTATGTTATTTAGTACTGCTGAGATACGTAGACATGAAGCA

GCATTTGGTGCCGTAAGCCAACTTGTAGCAGTATTGCGATTGGGTGGAAGAGGTGCAAGGTATAGTGCTG

CTTTAGCATTGGAAAGCCTTTTTTCTGCAGACCATATTAGGAATGCAGAATCTTCCAGGCAGGCTGTACA

GCCTTTAGTGGAAATCCTTAATACAGGTTTGGAGAAAGAGCAGCATGCTGCTATTGCTGCATTAGCTAGG

TTACTGAGTGAAAATCCATCAAGAGCCTTGGCAGTTGCTGATGTTGAAATGAATGCGGTAGATGTTCTTT

GCAGGATTCTTTCATCAAACTGTTCACTAGAGCTGAAGGGGGATTCGGCTGAGTTGTGCTGTGTACTTTT

TGGAAATACAAAGATCAGATCCACCCTGGCTGCAGCGCGCTGTGTGGAACCTCTGGTTTCTTTGCTCGTG

GCTGAGTATAGTCCTACTCAGCACTCTGTTGTCCGTGCATTAGATAAACTCGTTGATGACGATCAATTAG

CTGAGCTCATTGCTGCACATGGAGCAGTTATACCTCTTGTAGGCCTTCTGTATGGCCACAACTATGTGCT

CCATGAGGCTATTTCCAGAGCTCTTGTGAAGTTGGGAAAAGACAGGCCTTCTTGTAAGATGGAGATGGTG

AAGGCTGGGGTGATTGAGAGTGTACTTGATATTCTCCATGATGCACCAGATTTTCTGTGTGCTGCTTTTG

CAGAATTGCTACGAATATTAACCAATAATGCTACCATTGCCAAGGGTCCATCTGCAGCGAAAGTGGTTGA

GCCTTTCTTTGTGTTGTTAACAAGAGCAGAGTTCGGACCTGATGGACAGCATAGTGCTTTACAAGTTCTT

GTGAATATCTTAGAACATCCTCAGTGCCGTGCTGATTATACCTTAACATCACATCAAGTGATCGAACCCA

TCATTCCATTACTTGATTCTCCTGCACCGCCAGTGCAGCAGCTAGCAGCTGAACTTCTTTCTCATTTACT

CCTAGAGGAACATCTTCAGAAGGATTCAGTTACACAGCAAGTAATTGGCCCACTTATGCGAGTTCTTGGT

TCTGGTATACACATTCTGCAGCAGAGAGCTGTTAAAGCTCTTGTTAATATTGCACTGATTTGGCCCAATG

AAATTGCCAAGGAGGGTGGCGTCAGTGAGCTATCCAAAGTGATATTGCTAGCTGATCCCTCTTTGCCTCA

TGTCTTGTGGGAATCAGCTGCTTCTGTTTTGTCCAGTATTCTTCAGTTCAGTTCTGAGTACTATTTGGAA

GTGCCTATTGCGGTTTTGGTGAGATTGCTTCGATCTGGTTCAGAATCTACAGTAGTTGGTGCATTAAATG

CTCTTCTAGTTCTGGAAAGTGATGATGCTACAAGTGCTGTAGCAATGGCTGAAAGTGGAGCCATTGAAGC

TCTTTTAGAACTTCTTAGATGTCATCAGTGTGAGGAAACTGCTGCAAGACTTCTGGAGGTATTGCTGAAC

AATATTAAGATCAGAGAAACAAAAGCCACAAAGTCTGCAATAGTACCACTATCACAGTATCTCTTGGATC

CACAAAGCCAAGCTCAACAAGCAAGATTACTAGCAACTCTTGCTCTAGGGGATTTATTTCAGAATGAGGT

TCTTGCTCGATCTGCTGATGCTGTTTTAGCTTGCCGTGCTTTAGTAAATCTGCTTGAGGATCAACCTTCA

GAAGAAATGAAAGTGGTTGCAATATGTGCATTGCAAAACCTTGTAATGTACAGTAGATCAAATAAGAGAG

CAGTTGCTGAAGCTGGAGGTGTTCAGGTCGTGCTGGATCTGATTGGTTCAAGTGAACCAGATACATCAGT

TCAGGCTGCAATGTTTGTTAAGCTTCTCTTCTCTAACAACACAATTCAAGAGTATGCTTCCAGCGAAACC

GTTAGAGCTATAACAGCTGCAATTGAAAAAGATTTATGGGCTAATGGTGTAGTGAATGAAGAGTATCTTA

AAGCTCTTAATGCACTCCTTGGCAACTTCCCACGTTTAAGAGCCACAGAACCTGCAACCCTGAGCATTCC

CCATCTAGTGACAGCCCTCAAGACTGGCTCAGAGACAACTCAAGAAGCAGCATTGGATTCACTGTCTCTT

CTCAAGCAAGCTTGGGCAGCATGCCCTGCCGAGGTCTCCAGAGCGCAATCAACAGCTGCTTCTGAGGGGA

TTCCCCTGTTACAGTACTTAATTATGTCTGCCCCACCTCGAGTTCAAGATAAGGCAGACCATCTTTTGCA

GTGTCTGCCGGGAACATTGACAGTGACTATAAAACGTGGAAAGAATATCAAACAGTCAGTTGGAAATCCA

AGTGTGTTTTGCAAGCTAACACTTGGCAACACCCCCTCCAGGGAAACCAAGATTGTGTCGACCGGACCAA

ATCCCGAGTGGGATGAGCCCTTCCAGTGGCAATTTGAAAGTCCTCCAAAAGGCCAAAAGCTTCACATTTC

TTGCAAGAACAAGAGCAAGATGGGAAAGAAATCGTTTGGAAAAGTCACTGTCCAGATTGATCGAGTCGTT

ACCCAAGGAGCAGCTGCAGGAGAGTACTTCCTGCTACCTGAAAGCAAGAGTGGTAGTAAAAGGAGCCTGG

AAATAGAATTTCAATGGACTAATAGCAATAACATGCCTCAGTCTGAAGCTTAAAGAGATTTGCCTGGTGT

TAATTTTTTCTTTTTATCGAATATAATTATTGTGCTGGTGTACATAATATCTGGAAGGTTTATTTTTGTA

GTTAGTTGCTGCAATCCGGAAAGTCGGAAACTGCAATTCATTGTATTCTTTTGCAAAGTGTGGTAGTTTT

AGTCATCTTCTGTTCTTTTTGTTATAGTTGTATTGGAGTTGTATTCTGTGTTTGATATTCAGACTCGAGA

GGTTAATAAAATCTTGTGTATTGAATTCATTCTTACCTGTTTT

>MSTRG.537.3 gene=MSTRG.537

CGGTGTTACTCTCACAGCACAGCTTCTAGTAGTTAGCATCTCTCTCTCTCTCTCTCTCTCTCTCTCTCTC

TCTCTCTCTCTCTCTCGTGTAGACATACATTATATAGATATACACACTCTACCATCTCTCTCTCTCTCTC

TCTCTCTCTCTGCACAACAATATCTCTATATCATCCATATTGTTGTGAGTCTTGAGCTTCAACTATCTTT

TCTTCGCTTCTTTCCTACGAAATGCATTGACAAGATCTCCCTCCCTCCCCTTTAGCTCTCTCTCTCTCTC

TGCGTTTATACTGTCCCAATTGTCAGCTACCTACTTCATTCAATCTGTTCTCCAGATCTTCCTAGCTTAT

CTCAATCCTCACTTAACTCATTCATGTGAAGCTAGCTTCGCTTTGTAACTGTATTTTGGAGGTAGATCTT

GAGATCTAATTAGCAAAGCTGGCGGCTACAGTAGCTTGGAGAGGGGTTGCTGCCAGCAATGGCACAGCCC

TGTCATCTAATGATCTGGAAAGAAATGGGGATGCTAAATCTCATGACCAAGAGCCTCCAACACCACATTC

ACTCTTAAAGATGAGTTCGAGGGAGCGCAGTAGCATGGAGGATCCAGATGGAACTTTAGCTAGTGTTGCT

CAATGCATTGAGCAGTTGCGCCAGAGTTCCTCATCTGCACAAGAAAAAGAGTTCTCCTTAAAGCAACTAT

TGGAGCTTATTGATACACGTGAAAATGCTTTCAGTGCTGTTGGATCACATTCCCAGGCAGTTCCAGTGTT

GGTATCCCTTCTTAGATCAGGATCACTTGGGGTAAAAATACAGGCTGCTACAGTTTTAGGGTCACTGTGC

AAGGAGAATGAACTAAGGGTGAAGGTATTGCTGGGGGGTTGCATTCCACCACTTCTTGGTCTACTCAGGT

CTAGCTCAGCGGAAGCTCAGATCGCTGCAGCGAAGACGATATATGCTGTTTCTCAAGGTGGTGCTAAGGA

TCATGTTGGCTCAAAGATATTTTCAACTGAAGGAGTTGTGCCAGTTCTGTGGGGGCAGCTTGAGAAGGGA

TTGAAGGCTGGAAACGTGGTTGATAACTTACTGACTGGAGCTTTGAAAAACCTTTCAAGCAGCACTGAGG

GTTTCTGGTTTGCAACAATACAAGCTGGGGGATTAGATACACTTGTAAAGCTGCTTGCAACTGGACAATC

TAACACCCAAGCAAATGTTTGCTTTCTCCTTGCATGCATGATGATGGAGGATGCATCTGTTTGTCCTAAA

ATTCTGGCCGCTGAGACTACCAAGCAACTCCTAAAGCTACTAGGGCCCGGAAATGAAGCCTCGGTCAGAG

CAGAAGCTGCTGGTGCTCTAAAATCTCTATCTGCTCAGTGCAAAGAAGCAAGGCGGGAAATCGCAAGTTC

TAATGGTATACCCGCTTTGATAAATGCTACAATTGCTCCTTCAAAAGAGTTTATGCAAGGTGAGTATGCA

CAGGCTTTGCAGGAGAATGCAATGTGTGCTCTTGCAAATATATCTGGTGGTTTGTCATTTGTCATCTCAA

GTCTTGGCCAAAGTCTTGAATCCTGCACGTCGCCTGCACAGGTAGCTGACACGTTAGGGGCATTAGCTTC

TGCACTAATGATTTATGACAGCAAAGCAGAAAGTACAAGAGCTTCAGATCCTGACGATGTTGAGAAGACT

CTTGTGAAGCAGTTCAAACCTCGCTTGCCATTTCTTGTGCAGGAGCGTACAATAGAAGCTCTTGCCAGTT

TGTATGGAAATTCCACACTTTCTGGTAAACTTGCAAATTCTGATGCAAAGCGTTTGCTAGTTGGTTTGAT

CACAATGGCAAGCGATGAAGTTCAGGATGAGTTGATACGATCACTTCTGGTATTGTGTAACAATGAAGGT

ACACTATGGCATGCCCTTCAGGGACGTGAGGGAATTCAATTATTGATCTCTCTTCTTGGGCTGTCATCTG

AGCAACAACAGGAGTGTGCCGTCGCATTACTTTCCCTTTTATCCGATGAGAATGATGAAAGCAAATGGGC

CATCACAGCTGCTGGAGGTATACCTCCACTTGTTCAGATATTAGAGACAGGATCTCCAAAAGCCAAGGAA

GATTCTGCAACAATCCTCGGAAACCTCTGTAATCACAGTGAAGATATTAGAGCATGTGTTGAAAGTGCTG

ATGCTGTTCCTGCTTTGCTGTGGCTATTAAAGAATGGAAGCTCCAATGGCAAAGAAATTGCTGCAAAGAC

ATTGAATCATCTGATCCACAAATCAGATACAGCAACCATTAGTCAGCTCACCGCATTACTTACAAGCGAT

CTACCCGAATCTAAAGTGTATGTTTTAGATGCACTAAAAAGTTTGCTATCTGTGGCCCCTCTTAGTGATA

TGTTGCGTGATGGAAGTGCTTCAAACGATGCTATCGAGACAATGATTAAAATATTAAGTTCTACCAGAGA

TGAGACACAGTCTAAGTCTGCATCAGCTCTTGCTGGAATATTTAATCTTAGAAAGGACTTGCGTGAAAGT

CCCATAGCTGTTAAGACTGTTAGGTCGGTCATGAAGCTTTTACATGTAGAATCCGAAAGCATATTGGCGG

AGTCTACTCGTTGCCTTGCTGCAATATTTCTGTCAATTAAGGAGAACCGTGACATGGCTATGATTGGGAG

AGATTCATTGCCTACACTAGAGGTGCTTGCAACTTCGTCTTCACTGCAAGTTGCAGAGCAGGCCATTTGT

GCTTTGGCTAACCTTCTTTTGGATAGTGAAGTTTCAGAAAAAGCCAAGCCTGAAGAAATTATTTTGCCTT

CTACTAGGGTATTGCGTGAAAGCACAGTTACTGGAAGGAGCCATGCAGCAGCAGCAATTGCCCGGCTTCT

GAGTTCTCGTAAAATTGATGCTGAGATTGCTGACTGTGTGAACCGTACAGGAACAGTGCTTGCATTGGTT

TCGTTCCTAGAATCAGAAAATGCCGGGACTGCTGCCATATCAGAGGGGTTAGAGGCACTTGCCATTATAT

CGAGATCTGGAGGAGATAAAGGACAGAACAAACCTGCATGGGCAGTTCTGGCTGAATTCCCAGATAGCAT

AATCCCGATCGTTTCATGCATTTCTGATGCAAACCCCTTGTTGCTTGATAAGGCTATAGAAATATTGTCA

CGGATTTCTTTGGCTCAACCTGTTGTTTTGGGGAACAATATTGTGAGTGCTTCAGGGTGTATCTCGTCAA

TTGCTAGACGGGTAGTTAGCTCTTCAAAGGAAGCAGTAAAGATTGGAGGCACCACCCTTCTTGTTTGTGC

TGCCAAAGTAAATCTTCAGAGAGTTGTAGATGATTTGAATGAATCATATTCTTGCGCATATCTCATTCAG

TCCCTTGTAGGAATGCTGACCTACACAAAAAATCCTCTGGTTGGTGACCAGGGAAGTAATGAAGGCGTAA

ATATTCTTGGGCATGCTGAAGAACTAAAAGCTGGTGAGAATGAGACGAGTAGATATGTCATCTATGGTTC

AAATACCGCAATATGGCTTCTCTCTGCTCTTGCTAGTAATAACGAAAAGAGCAAAACTCAGATCATGGAA

GCAGGTGCCATTGAAGTTCTAACTGAGAGAATCAATCAGTGTTTATCAGAATATTCTCAGGTCGATTCTA

AAGAGGAAAGCAGCATTTGGATATGTGCTTTACTTCTAGCAATTCTATTTCAAGATAGAGATATTATACG

TGCGCATGCAACAATGAAATCTGTACCTGTACTGGCTAATTTGTTGAAGTCAGAGGAGGCAGCAAACAGA

TATTTTGGTGCACAAGCCATTGCCAGTCTAGTATGCAATGGTAGCAGGGGGACTCTTCTATCAGTCGCTA

ATTCAGGGGCTGCAGGTGGACTCATTTCACTGCTTGGGTGTGCTGATGCTGATATATGTGATATGCTTGA

CTTGGCAGACGAATTTTCCTTGGTGCGGTATCCTGAACAAGTTGCTCTTGAGAGGTTGTTTAGGGTTGAG

GATATCAGAGTTGGTGCTACTTCAAGAAAAGCAATACCTGCACTAGTTGATCTACTTAAACCGATTCCAG

ATCGTCCAGGGGCACCATTTTTAACCTTAGGGCTTCTGCTTCAGCTTGGTAAAGATTGCCCTTCGAATAA

AATTCTTATGGTAGAAGCTGGTGCTCTGGAAGCATTGACCAAATACCTTTCACTTGGCTTACAAGATGCA

ACTGAAGAAGCAGCTACGGATCTCTTAGGTATGTTATTTAGTACTGCTGAGATACGTAGACATGAAGCAG

CATTTGGTGCCGTAAGCCAACTTGTAGCAGTATTGCGATTGGGTGGAAGAGGTGCAAGGTATAGTGCTGC

TTTAGCATTGGAAAGCCTTTTTTCTGCAGACCATATTAGGAATGCAGAATCTTCCAGGCAGGCTGTACAG

CCTTTAGTGGAAATCCTTAATACAGGTTTGGAGAAAGAGCAGCATGCTGCTATTGCTGCATTAGCTAGGT

TACTGAGTGAAAATCCATCAAGAGCCTTGGCAGTTGCTGATGTTGAAATGAATGCGGTAGATGTTCTTTG

CAGGATTCTTTCATCAAACTGTTCACTAGAGCTGAAGGGGGATTCGGCTGAGTTGTGCTGTGTACTTTTT

GGAAATACAAAGATCAGATCCACCCTGGCTGCAGCGCGCTGTGTGGAACCTCTGGTTTCTTTGCTCGTGG

CTGAGTATAGTCCTACTCAGCACTCTGTTGTCCGTGCATTAGATAAACTCGTTGATGACGATCAATTAGC

TGAGCTCATTGCTGCACATGGAGCAGTTATACCTCTTGTAGGCCTTCTGTATGGCCACAACTATGTGCTC

CATGAGGCTATTTCCAGAGCTCTTGTGAAGTTGGGAAAAGACAGGCCTTCTTGTAAGATGGAGATGGTGA

AGGCTGGGGTGATTGAGAGTGTACTTGATATTCTCCATGATGCACCAGATTTTCTGTGTGCTGCTTTTGC

AGAATTGCTACGAATATTAACCAATAATGCTACCATTGCCAAGGGTCCATCTGCAGCGAAAGTGGTTGAG

CCTTTCTTTGTGTTGTTAACAAGAGCAGAGTTCGGACCTGATGGACAGCATAGTGCTTTACAAGTTCTTG

TGAATATCTTAGAACATCCTCAGTGCCGTGCTGATTATACCTTAACATCACATCAAGTGATCGAACCCAT

CATTCCATTACTTGATTCTCCTGCACCGCCAGTGCAGCAGCTAGCAGCTGAACTTCTTTCTCATTTACTC

CTAGAGGAACATCTTCAGAAGGATTCAGTTACACAGCAAGTAATTGGCCCACTTATGCGAGTTCTTGGTT

CTGGTATACACATTCTGCAGCAGAGAGCTGTTAAAGCTCTTGTTAATATTGCACTGATTTGGCCCAATGA

AATTGCCAAGGAGGGTGGCGTCAGTGAGCTATCCAAAGTGATATTGCTAGCTGATCCCTCTTTGCCTCAT

GTCTTGTGGGAATCAGCTGCTTCTGTTTTGTCCAGTATTCTTCAGTTCAGTTCTGAGTACTATTTGGAAG

TGCCTATTGCGGTTTTGGTGAGATTGCTTCGATCTGGTTCAGAATCTACAGTAGTTGGTGCATTAAATGC

TCTTCTAGTTCTGGAAAGTGATGATGCTACAAGTGCTGTAGCAATGGCTGAAAGTGGAGCCATTGAAGCT

CTTTTAGAACTTCTTAGATGTCATCAGTGTGAGGAAACTGCTGCAAGACTTCTGGAGGTATTGCTGAACA

ATATTAAGATCAGAGAAACAAAAGCCACAAAGTCTGCAATAGTACCACTATCACAGTATCTCTTGGATCC

ACAAAGCCAAGCTCAACAAGCAAGATTACTAGCAACTCTTGCTCTAGGGGATTTATTTCAGAATGAGGTT

CTTGCTCGATCTGCTGATGCTGTTTTAGCTTGCCGTGCTTTAGTAAATCTGCTTGAGGATCAACCTTCAG

AAGAAATGAAAGTGGTTGCAATATGTGCATTGCAAAACCTTGTAATGTACAGTAGATCAAATAAGAGAGC

AGTTGCTGAAGCTGGAGGTGTTCAGGTCGTGCTGGATCTGATTGGTTCAAGTGAACCAGATACATCAGTT

CAGGCTGCAATGTTTGTTAAGCTTCTCTTCTCTAACAACACAATTCAAGAGTATGCTTCCAGCGAAACCG

TTAGAGCTATAACAGCTGCAATTGAAAAAGATTTATGGGCTAATGGTGTAGTGAATGAAGAGTATCTTAA

AGCTCTTAATGCACTCCTTGGCAACTTCCCACGTTTAAGAGCCACAGAACCTGCAACCCTGAGCATTCCC

CATCTAGTGACAGCCCTCAAGACTGGCTCAGAGACAACTCAAGAAGCAGCATTGGATTCACTGTCTCTTC

TCAAGCAAGCTTGGGCAGCATGCCCTGCCGAGGTCTCCAGAGCGCAATCAACAGCTGCTTCTGAGGGGAT

TCCCCTGTTACAGTACTTAATTATGTCTGCCCCACCTCGAGTTCAAGATAAGGCAGACCATCTTTTGCAG

TGTCTGCCGGGAACATTGACAGTGACTATAAAACGTGGAAAGAATATCAAACAGTCAGTTGGAAATCCAA

GTGTGTTTTGCAAGCTAACACTTGGCAACACCCCCTCCAGGGAAACCAAGATTGTGTCGACCGGACCAAA

TCCCGAGTGGGATGAGCCCTTCCAGTGGCAATTTGAAAGTCCTCCAAAAGGCCAAAAGCTTCACATTTCT

TGCAAGAACAAGAGCAAGATGGGAAAGAAATCGTTTGGAAAAGTCACTGTCCAGATTGATCGAGTCGTTA

CCCAAGGAGCAGCTGCAGGAGAGTACTTCCTGCTACCTGAAAGCAAGAGTGGTAGTAAAAGGAGCCTGGA

AATAGAATTTCAATGGACTAATAGCAATAACATGCCTCAGTCTGAAGCTTAAAGAGATTTGCCTGGTGTT

AATTTTTTCTTTTTATCGAATATAATTATTGTGCTGGTGTACATAATATCTGGAAGGTTTATTTTTGTAG

TTAGTTGCTGCAATCCGGAAAGTCGGAAACTGCAATTCATTGTATTCTTTTGCAAAGTGTGGTAGTTTTA

GTCATCTTCTGTTCTTTTTGTTATAGTTGTATTGGAGTTGTATTCTGTGTTTGATATTCAGACTCGAGAG

AAAAATTGCTGCTCCTTCCATCTTGGTGGTTTCTTTACGTACCAGCTGGACACGGAGACAAGACAAACAT

ATAATAATTTTTCTATAAATTTTTA

>MSTRG.537.4 gene=MSTRG.537

TCTCTCTCTCTCTCTCTCGTGTAGACATACATTATATAGATATACACACTCTACCATCTCTCTCTCTCTC

TCTCTCTCTCTCTCTGCACAACAATATCTCTATATCATCCATATTGTTGTGAGTCTTGAGCTTCAACTAT

CTTTTCTTCGCTTCTTTCCTACGAAATGCATTGACAAGATCTCCCTCCCTCCCCTTTAGCTCTCTCTCTC

TCTCTGCGTTTATACTGTCCCAATTGTCAGCTACCTACTTCATTCAATCTGTTCTCCAGATCTTCCTAGC

TTATCTCAATCCTCACTTAACTCATTCATGTGAAGCTAGCTTCGCTTTGTAACTGTATTTTGGAGGTAGA

TCTTGAGATCTAATTAGCAAAGCTGGCGGCTACAGTAGCTTGGAGAGGGGTTGCTGCCAGCAATGGCACA

GCCCTGTCATCTAATGATCTGGAAAGAAATGGGGATGCTAAATCTCATGACCAAGAGCCTCCAACACCAC

ATTCACTCTTAAAGATGAGTTCGAGGGAGCGCAGTAGCATGGAGGATCCAGATGGAACTTTAGCTAGTGT

TGCTCAATGCATTGAGCAGTTGCGCCAGAGTTCCTCATCTGCACAAGAAAAAGAGTTCTCCTTAAAGCAA

CTATTGGAGCTTATTGATACACGTGAAAATGCTTTCAGTGCTGTTGGATCACATTCCCAGGCAGTTCCAG

TGTTGGTATCCCTTCTTAGATCAGGATCACTTGGGGTAAAAATACAGGCTGCTACAGTTTTAGGGTCACT

GTGCAAGGAGAATGAACTAAGGGTGAAGGTATTGCTGGGGGGTTGCATTCCACCACTTCTTGGTCTACTC

AGGTCTAGCTCAGCGGAAGCTCAGATCGCTGCAGCGAAGACGATATATGCTGTTTCTCAAGGTGGTGCTA

AGGATCATGTTGGCTCAAAGATATTTTCAACTGAAGGAGTTGTGCCAGTTCTGTGGGGGCAGCTTGAGAA

GGGATTGAAGGCTGGAAACGTGGTTGATAACTTACTGACTGGAGCTTTGAAAAACCTTTCAAGCAGCACT

GAGGGTTTCTGGTTTGCAACAATACAAGCTGGGGGATTAGATACACTTGTAAAGCTGCTTGCAACTGGAC

AATCTAACACCCAAGCAAATGTTTGCTTTCTCCTTGCATGCATGATGATGGAGGATGCATCTGTTTGTCC

TAAAATTCTGGCCGCTGAGACTACCAAGCAACTCCTAAAGCTACTAGGGCCCGGAAATGAAGCCTCGGTC

AGAGCAGAAGCTGCTGGTGCTCTAAAATCTCTATCTGCTCAGTGCAAAGAAGCAAGGCGGGAAATCGCAA

GTTCTAATGGTATACCCGCTTTGATAAATGCTACAATTGCTCCTTCAAAAGAGTTTATGCAAGGTGAGTA

TGCACAGGCTTTGCAGGAGAATGCAATGTGTGCTCTTGCAAATATATCTGGTGGTTTGTCATTTGTCATC

TCAAGTCTTGGCCAAAGTCTTGAATCCTGCACGTCGCCTGCACAGGTAGCTGACACGTTAGGGGCATTAG

CTTCTGCACTAATGATTTATGACAGCAAAGCAGAAAGTACAAGAGCTTCAGATCCTGACGATGTTGAGAA

GACTCTTGTGAAGCAGTTCAAACCTCGCTTGCCATTTCTTGTGCAGGAGCGTACAATAGAAGCTCTTGCC

AGTTTGTATGGAAATTCCACACTTTCTGGTAAACTTGCAAATTCTGATGCAAAGCGTTTGCTAGTTGGTT

TGATCACAATGGCAAGCGATGAAGTTCAGGATGAGTTGATACGATCACTTCTGGTATTGTGTAACAATGA

AGGTACACTATGGCATGCCCTTCAGGGACGTGAGGGAATTCAATTATTGATCTCTCTTCTTGGGCTGTCA

TCTGAGCAACAACAGGAGTGTGCCGTCGCATTACTTTCCCTTTTATCCGATGAGAATGATGAAAGCAAAT

GGGCCATCACAGCTGCTGGAGGTATACCTCCACTTGTTCAGATATTAGAGACAGGATCTCCAAAAGCCAA

GGAAGATTCTGCAACAATCCTCGGAAACCTCTGTAATCACAGTGAAGATATTAGAGCATGTGTTGAAAGT

GCTGATGCTGTTCCTGCTTTGCTGTGGCTATTAAAGAATGGAAGCTCCAATGGCAAAGAAATTGCTGCAA

AGACATTGAATCATCTGATCCACAAATCAGATACAGCAACCATTAGTCAGCTCACCGCATTACTTACAAG

CGATCTACCCGAATCTAAAGTGTATGTTTTAGATGCACTAAAAAGTTTGCTATCTGTGGCCCCTCTTAGT

GATATGTTGCGTGATGGAAGTGCTTCAAACGATGCTATCGAGACAATGATTAAAATATTAAGTTCTACCA

GAGATGAGACACAGTCTAAGTCTGCATCAGCTCTTGCTGGAATATTTAATCTTAGAAAGGACTTGCGTGA

AAGTCCCATAGCTGTTAAGACTGTTAGGTCGGTCATGAAGCTTTTACATGTAGAATCCGAAAGCATATTG

GCGGAGTCTACTCGTTGCCTTGCTGCAATATTTCTGTCAATTAAGGAGAACCGTGACATGGCTATGATTG

GGAGAGATTCATTGCCTACACTAGAGGTGCTTGCAACTTCGTCTTCACTGCAAGTTGCAGAGCAGGCCAT

TTGTGCTTTGGCTAACCTTCTTTTGGATAGTGAAGTTTCAGAAAAAGCCAAGCCTGAAGAAATTATTTTG

CCTTCTACTAGGGTATTGCGTGAAAGCACAGTTACTGGAAGGAGCCATGCAGCAGCAGCAATTGCCCGGC

TTCTGAGTTCTCGTAAAATTGATGCTGAGATTGCTGACTGTGTGAACCGTACAGGAACAGTGCTTGCATT

GGTTTCGTTCCTAGAATCAGAAAATGCCGGGACTGCTGCCATATCAGAGGGGTTAGAGGCACTTGCCATT

ATATCGAGATCTGGAGGAGATAAAGGACAGAACAAACCTGCATGGGCAGTTCTGGCTGAATTCCCAGATA

GCATAATCCCGATCGTTTCATGCATTTCTGATGCAAACCCCTTGTTGCTTGATAAGGCTATAGAAATATT

GTCACGGATTTCTTTGGCTCAACCTGTTGTTTTGGGGAACAATATTGTGAGTGCTTCAGGGTGTATCTCG

TCAATTGCTAGACGGGTAGTTAGCTCTTCAAAGGAAGCAGTAAAGATTGGAGGCACCACCCTTCTTGTTT

GTGCTGCCAAAGTAAATCTTCAGAGAGTTGTAGATGATTTGAATGAATCATATTCTTGCGCATATCTCAT

TCAGTCCCTTGTAGGAATGCTGACCTACACAAAAAATCCTCTGGTTGGTGACCAGGGAAGTAATGAAGGC

GTAAATATTCTTGGGCATGCTGAAGAACTAAAAGCTGGTGAGAATGAGACGAGTAGATATGTCATCTATG

GTTCAAATACCGCAATATGGCTTCTCTCTGCTCTTGCTAGTAATAACGAAAAGAGCAAAACTCAGATCAT

GGAAGCAGGTGCCATTGAAGTTCTAACTGAGAGAATCAATCAGTGTTTATCAGAATATTCTCAGGTCGAT

TCTAAAGAGGAAAGCAGCATTTGGATATGTGCTTTACTTCTAGCAATTCTATTTCAAGATAGAGATATTA

TACGTGCGCATGCAACAATGAAATCTGTACCTGTACTGGCTAATTTGTTGAAGTCAGAGGAGGCAGCAAA

CAGATATTTTGGTGCACAAGCCATTGCCAGTCTAGTATGCAATGGTAGCAGGGGGACTCTTCTATCAGTC

GCTAATTCAGGGGCTGCAGGTGGACTCATTTCACTGCTTGGGTGTGCTGATGCTGATATATGTGATATGC

TTGACTTGGCAGACGAATTTTCCTTGGTGCGGTATCCTGAACAAGTTGCTCTTGAGAGGTTGTTTAGGGT

TGAGGATATCAGAGTTGGTGCTACTTCAAGAAAAGCAATACCTGCACTAGTTGATCTACTTAAACCGATT

CCAGATCGTCCAGGGGCACCATTTTTAACCTTAGGGCTTCTGCTTCAGCTTGGTAAAGATTGCCCTTCGA

ATAAAATTCTTATGGTAGAAGCTGGTGCTCTGGAAGCATTGACCAAATACCTTTCACTTGGCTTACAAGA

TGCAACTGAAGAAGCAGCTACGGATCTCTTAGGTATGTTATTTAGTACTGCTGAGATACGTAGACATGAA

GCAGCATTTGGTGCCGTAAGCCAACTTGTAGCAGTATTGCGATTGGGTGGAAGAGGTGCAAGGTATAGTG

CTGCTTTAGCATTGGAAAGCCTTTTTTCTGCAGACCATATTAGGAATGCAGAATCTTCCAGGCAGGCTGT

ACAGCCTTTAGTGGAAATCCTTAATACAGGTTTGGAGAAAGAGCAGCATGCTGCTATTGCTGCATTAGCT

AGGTTACTGAGTGAAAATCCATCAAGAGCCTTGGCAGTTGCTGATGTTGAAATGAATGCGGTAGATGTTC

TTTGCAGGATTCTTTCATCAAACTGTTCACTAGAGCTGAAGGGGGATTCGGCTGAGTTGTGCTGTGTACT

TTTTGGAAATACAAAGATCAGATCCACCCTGGCTGCAGCGCGCTGTGTGGAACCTCTGGTTTCTTTGCTC

GTGGCTGAGTATAGTCCTACTCAGCACTCTGTTGTCCGTGCATTAGATAAACTCGTTGATGACGATCAAT

TAGCTGAGCTCATTGCTGCACATGGAGCAGTTATACCTCTTGTAGGCCTTCTGTATGGCCACAACTATGT

GCTCCATGAGGCTATTTCCAGAGCTCTTGTGAAGTTGGGAAAAGACAGGCCTTCTTGTAAGATGGAGATG

GTGAAGGCTGGGGTGATTGAGAGTGTACTTGATATTCTCCATGATGCACCAGATTTTCTGTGTGCTGCTT

TTGCAGAATTGCTACGAATATTAACCAATAATGCTACCATTGCCAAGGGTCCATCTGCAGCGAAAGTGGT

TGAGCCTTTCTTTGTGTTGTTAACAAGAGCAGAGTTCGGACCTGATGGACAGCATAGTGCTTTACAAGTT

CTTGTGAATATCTTAGAACATCCTCAGTGCCGTGCTGATTATACCTTAACATCACATCAAGTGATCGAAC

CCATCATTCCATTACTTGATTCTCCTGCACCGCCAGTGCAGCAGCTAGCAGCTGAACTTCTTTCTCATTT

ACTCCTAGAGGAACATCTTCAGAAGGATTCAGTTACACAGCAAGTAATTGGCCCACTTATGCGAGTTCTT

GGTTCTGGTATACACATTCTGCAGCAGAGAGCTGTTAAAGCTCTTGTTAATATTGCACTGATTTGGCCCA

ATGAAATTGCCAAGGAGGGTGGCGTCAGTGAGCTATCCAAAGTGATATTGCTAGCTGATCCCTCTTTGCC

TCATGTCTTGTGGGAATCAGCTGCTTCTGTTTTGTCCAGTATTCTTCAGTTCAGTTCTGAGTACTATTTG

GAAGTGCCTATTGCGGTTTTGGTGAGATTGCTTCGATCTGGTTCAGAATCTACAGTAGTTGGTGCATTAA

ATGCTCTTCTAGTTCTGGAAAGTGATGATGCTACAAGTGCTGTAGCAATGGCTGAAAGTGGAGCCATTGA

AGCTCTTTTAGAACTTCTTAGATGTCATCAGTGTGAGGAAACTGCTGCAAGACTTCTGGAGGTATTGCTG

AACAATATTAAGATCAGAGAAACAAAAGCCACAAAGTCTGCAATAGTACCACTATCACAGTATCTCTTGG

ATCCACAAAGCCAAGCTCAACAAGCAAGATTACTAGCAACTCTTGCTCTAGGGGATTTATTTCAGAATGA

GGTTCTTGCTCGATCTGCTGATGCTGTTTTAGCTTGCCGTGCTTTAGTAAATCTGCTTGAGGATCAACCT

TCAGAAGAAATGAAAGTGGTTGCAATATGTGCATTGCAAAACCTTGTAATGTACAGTAGATCAAATAAGA

GAGCAGTTGCTGAAGCTGGAGGTGTTCAGGTCGTGCTGGATCTGATTGGTTCAAGTGAACCAGATACATC

AGTTCAGGCTGCAATGTTTGTTAAGCTTCTCTTCTCTAACAACACAATTCAAGAGTATGCTTCCAGCGAA

ACCGTTAGAGCTATAACAGCTGCAATTGAAAAAGATTTATGGGCTAATGGTGTAGTGAATGAAGAGTATC

TTAAAGCTCTTAATGCACTCCTTGGCAACTTCCCACGTTTAAGAGCCACAGAACCTGCAACCCTGAGCAT

TCCCCATCTAGTGACAGCCCTCAAGACTGGCTCAGAGACAACTCAAGAAGCAGCATTGGATTCACTGTCT

CTTCTCAAGCAAGCTTGGGCAGCATGCCCTGCCGAGGTCTCCAGAGCGCAATCAACAGCTGCTTCTGAGG

GGATTCCCCTGTTACAGTACTTAATTATGTCTGCCCCACCTCGAGTTCAAGATAAGGCAGACCATCTTTT

GCAGTGTCTGCCGGGAACATTGACAGTGACTATAAAACGTGGAAAGAATATCAAACAGTCAGTTGGAAAT

CCAAGTGTGTTTTGCAAGCTAACACTTGGCAACACCCCCTCCAGGGAAACCAAGATTGTGTCGACCGGAC

CAAATCCCGAGTGGGATGAGCCCTTCCAGTGGCAATTTGAAAGTCCTCCAAAAGGCCAAAAGCTTCACAT

TTCTTGCAAGAACAAGAGCAAGATGGGAAAGAAATCGTTTGGAAAAGTCACTGTCCAGATTGATCGAGTC

GTTACCCAAGGAGCAGCTGCAGGAGAGTACTTCCTGCTACCTGAAAGCAAGAGTGGTAGTAAAAGGAGCC

TGGAAATAGAATTTCAATGGACTAATAGCAATAACATGCCTCAGTCTGAAGCTTAAAGAGATTTGCCTGG

TGTTAATTTTTTCTTTTTATCGAATATAATTATTGTGCTGGTGTACATAATATCTGGAAGGTTTATTTTT

GTAGTTAGTTGCTGCAATCCGGAAAGTCGGAAACTGCAATTCATTGTATTCTTTTGCAAAGTGTGGTAGT

TTTAGTCATCTTCTGTTCTTTTTGTTATAGTTGTATTGGAGTTGTATTCTGTGTTTGATATTCAGACTCG

AGAGGTTAATAAAATCTTGTGTATTGAATTCATTCTTACCTGTTTTCTCTTCAGGGGTTAAGTAA

>MSTRG.537.5 gene=MSTRG.537

TATAGATATACACACTCTACCATCTCTCTCTCTCTCTCTCTCTCTCTGCACAACAATATCTCTATATCAT

CCATATTGTTGTGAGTCTTGAGCTTCAACTATCTTTTCTTCGCTTCTTTCCTACGAAATGCATTGACAAG

ATCTCCCTCCCTCCCCTTTAGCTCTCTCTCTCTCTCTGCGTTTATACTGTCCCAATTGTCAGCTACCTAC

TTCATTCAATCTGTTCTCCAGATCTTCCTAGCTTATCTCAATCCTCACTTAACTCATTCATGTGAAGCTA

GCTTCGCTTTGTAACTGTATTTTGGAGGTAGATCTTGAGATCTAATTAGCAAAGCTGGCGGCTACAGTAG

CTTGGAGAGGGGTTGCTGCCAGCAATGGCACAGCCCTGTCATCTAATGATCTGGAAAGAAATGGGGATGC

TAAATCTCATGACCAAGAGCCTCCAACACCACATTCACTCTTAAAGATGAGTTCGAGGGAGCGCAGTAGC

ATGGAGGATCCAGATGGAACTTTAGCTAGTGTTGCTCAATGCATTGAGCAGTTGCGCCAGAGTTCCTCAT

CTGCACAAGAAAAAGAGTTCTCCTTAAAGCAACTATTGGAGCTTATTGATACACGTGAAAATGCTTTCAG

TGCTGTTGGATCACATTCCCAGGCAGTTCCAGTGTTGGTATCCCTTCTTAGATCAGGATCACTTGGGGTA

AAAATACAGGCTGCTACAGTTTTAGGGTCACTGTGCAAGGAGAATGAACTAAGGGTGAAGGTATTGCTGG

GGGGTTGCATTCCACCACTTCTTGGTCTACTCAGGTCTAGCTCAGCGGAAGCTCAGATCGCTGCAGCGAA

GACGATATATGCTGTTTCTCAAGGTGGTGCTAAGGATCATGTTGGCTCAAAGATATTTTCAACTGAAGGA

GTTGTGCCAGTTCTGTGGGGGCAGCTTGAGAAGGGATTGAAGGCTGGAAACGTGGTTGATAACTTACTGA

CTGGAGCTTTGAAAAACCTTTCAAGCAGCACTGAGGGTTTCTGGTTTGCAACAATACAAGCTGGGGGATT

AGATACACTTGTAAAGCTGCTTGCAACTGGACAATCTAACACCCAAGCAAATGTTTGCTTTCTCCTTGCA

TGCATGATGATGGAGGATGCATCTGTTTGTCCTAAAATTCTGGCCGCTGAGACTACCAAGCAACTCCTAA

AGCTACTAGGGCCCGGAAATGAAGCCTCGGTCAGAGCAGAAGCTGCTGGTGCTCTAAAATCTCTATCTGC

TCAGTGCAAAGAAGCAAGGCGGGAAATCGCAAGTTCTAATGGTATACCCGCTTTGATAAATGCTACAATT

GCTCCTTCAAAAGAGTTTATGCAAGGTGAGTATGCACAGGCTTTGCAGGAGAATGCAATGTGTGCTCTTG

CAAATATATCTGGTGGTTTGTCATTTGTCATCTCAAGTCTTGGCCAAAGTCTTGAATCCTGCACGTCGCC

TGCACAGGTAGCTGACACGTTAGGGGCATTAGCTTCTGCACTAATGATTTATGACAGCAAAGCAGAAAGT

ACAAGAGCTTCAGATCCTGACGATGTTGAGAAGACTCTTGTGAAGCAGTTCAAACCTCGCTTGCCATTTC

TTGTGCAGGAGCGTACAATAGAAGCTCTTGCCAGTTTGTATGGAAATTCCACACTTTCTGGTAAACTTGC

AAATTCTGATGCAAAGCGTTTGCTAGTTGGTTTGATCACAATGGCAAGCGATGAAGTTCAGGATGAGTTG

ATACGATCACTTCTGGTATTGTGTAACAATGAAGGTACACTATGGCATGCCCTTCAGGGACGTGAGGGAA

TTCAATTATTGATCTCTCTTCTTGGGCTGTCATCTGAGCAACAACAGGAGTGTGCCGTCGCATTACTTTC

CCTTTTATCCGATGAGAATGATGAAAGCAAATGGGCCATCACAGCTGCTGGAGGTATACCTCCACTTGTT

CAGATATTAGAGACAGGATCTCCAAAAGCCAAGGAAGATTCTGCAACAATCCTCGGAAACCTCTGTAATC

ACAGTGAAGATATTAGAGCATGTGTTGAAAGTGCTGATGCTGTTCCTGCTTTGCTGTGGCTATTAAAGAA

TGGAAGCTCCAATGGCAAAGAAATTGCTGCAAAGACATTGAATCATCTGATCCACAAATCAGATACAGCA

ACCATTAGTCAGCTCACCGCATTACTTACAAGCGATCTACCCGAATCTAAAGTGTATGTTTTAGATGCAC

TAAAAAGTTTGCTATCTGTGGCCCCTCTTAGTGATATGTTGCGTGATGGAAGTGCTTCAAACGATGCTAT

CGAGACAATGATTAAAATATTAAGTTCTACCAGAGATGAGACACAGTCTAAGTCTGCATCAGCTCTTGCT

GGAATATTTAATCTTAGAAAGGACTTGCGTGAAAGTCCCATAGCTGTTAAGACTGTTAGGTCGGTCATGA

AGCTTTTACATGTAGAATCCGAAAGCATATTGGCGGAGTCTACTCGTTGCCTTGCTGCAATATTTCTGTC

AATTAAGGAGAACCGTGACATGGCTATGATTGGGAGAGATTCATTGCCTACACTAGAGGTGCTTGCAACT

TCGTCTTCACTGCAAGTTGCAGAGCAGGCCATTTGTGCTTTGGCTAACCTTCTTTTGGATAGTGAAGTTT

CAGAAAAAGCCAAGCCTGAAGAAATTATTTTGCCTTCTACTAGGGTATTGCGTGAAAGCACAGTTACTGG

AAGGAGCCATGCAGCAGCAGCAATTGCCCGGCTTCTGAGTTCTCGTAAAATTGATGCTGAGATTGCTGAC

TGTGTGAACCGTACAGGAACAGTGCTTGCATTGGTTTCGTTCCTAGAATCAGAAAATGCCGGGACTGCTG

CCATATCAGAGGGGTTAGAGGCACTTGCCATTATATCGAGATCTGGAGGAGATAAAGGACAGAACAAACC

TGCATGGGCAGTTCTGGCTGAATTCCCAGATAGCATAATCCCGATCGTTTCATGCATTTCTGATGCAAAC

CCCTTGTTGCTTGATAAGGCTATAGAAATATTGTCACGGATTTCTTTGGCTCAACCTGTTGTTTTGGGGA

ACAATATTGTGAGTGCTTCAGGGTGTATCTCGTCAATTGCTAGACGGGTAGTTAGCTCTTCAAAGGAAGC

AGTAAAGATTGGAGGCACCACCCTTCTTGTTTGTGCTGCCAAAGTAAATCTTCAGAGAGTTGTAGATGAT

TTGAATGAATCATATTCTTGCGCATATCTCATTCAGTCCCTTGTAGGAATGCTGACCTACACAAAAAATC

CTCTGGTTGGTGACCAGGGAAGTAATGAAGGCGTAAATATTCTTGGGCATGCTGAAGAACTAAAAGCTGG

TGAGAATGAGACGAGTAGATATGTCATCTATGGTTCAAATACCGCAATATGGCTTCTCTCTGCTCTTGCT

AGTAATAACGAAAAGAGCAAAACTCAGATCATGGAAGCAGGTGCCATTGAAGTTCTAACTGAGAGAATCA

ATCAGTGTTTATCAGAATATTCTCAGGTCGATTCTAAAGAGGAAAGCAGCATTTGGATATGTGCTTTACT

TCTAGCAATTCTATTTCAAGATAGAGATATTATACGTGCGCATGCAACAATGAAATCTGTACCTGTACTG

GCTAATTTGTTGAAGTCAGAGGAGGCAGCAAACAGATATTTTGGTGCACAAGCCATTGCCAGTCTAGTAT

GCAATGGTAGCAGGGGGACTCTTCTATCAGTCGCTAATTCAGGGGCTGCAGGTGGACTCATTTCACTGCT

TGGGTGTGCTGATGCTGATATATGTGATATGCTTGACTTGGCAGACGAATTTTCCTTGGTGCGGTATCCT

GAACAAGTTGCTCTTGAGAGGTTGTTTAGGGTTGAGGATATCAGAGTTGGTGCTACTTCAAGAAAAGCAA

TACCTGCACTAGTTGATCTACTTAAACCGATTCCAGATCGTCCAGGGGCACCATTTTTAACCTTAGGGCT

TCTGCTTCAGCTTGGTAAAGATTGCCCTTCGAATAAAATTCTTATGGTAGAAGCTGGTGCTCTGGAAGCA

TTGACCAAATACCTTTCACTTGGCTTACAAGATGCAACTGAAGAAGCAGCTACGGATCTCTTAGGTATGT

TATTTAGTACTGCTGAGATACGTAGACATGAAGCAGCATTTGGTGCCGTAAGCCAACTTGTAGCAGTATT

GCGATTGGGTGGAAGAGGTGCAAGGTATAGTGCTGCTTTAGCATTGGAAAGCCTTTTTTCTGCAGACCAT

ATTAGGAATGCAGAATCTTCCAGGCAGGCTGTACAGCCTTTAGTGGAAATCCTTAATACAGGTTTGGAGA

AAGAGCAGCATGCTGCTATTGCTGCATTAGCTAGGTTACTGAGTGAAAATCCATCAAGAGCCTTGGCAGT

TGCTGATGTTGAAATGAATGCGGTAGATGTTCTTTGCAGGATTCTTTCATCAAACTGTTCACTAGAGCTG

AAGGGGGATTCGGCTGAGTTGTGCTGTGTACTTTTTGGAAATACAAAGATCAGATCCACCCTGGCTGCAG

CGCGCTGTGTGGAACCTCTGGTTTCTTTGCTCGTGGCTGAGTATAGTCCTACTCAGCACTCTGTTGTCCG

TGCATTAGATAAACTCGTTGATGACGATCAATTAGCTGAGCTCATTGCTGCACATGGAGCAGTTATACCT

CTTGTAGGCCTTCTGTATGGCCACAACTATGTGCTCCATGAGGCTATTTCCAGAGCTCTTGTGAAGTTGG

GAAAAGACAGGCCTTCTTGTAAGATGGAGATGGTGAAGGCTGGGGTGATTGAGAGTGTACTTGATATTCT

CCATGATGCACCAGATTTTCTGTGTGCTGCTTTTGCAGAATTGCTACGAATATTAACCAATAATGCTACC

ATTGCCAAGGGTCCATCTGCAGCGAAAGTGGTTGAGCCTTTCTTTGTGTTGTTAACAAGAGCAGAGTTCG

GACCTGATGGACAGCATAGTGCTTTACAAGTTCTTGTGAATATCTTAGAACATCCTCAGTGCCGTGCTGA

TTATACCTTAACATCACATCAAGTGATCGAACCCATCATTCCATTACTTGATTCTCCTGCACCGCCAGTG

CAGCAGCTAGCAGCTGAACTTCTTTCTCATTTACTCCTAGAGGAACATCTTCAGAAGGATTCAGTTACAC

AGCAAGTAATTGGCCCACTTATGCGAGTTCTTGGTTCTGGTATACACATTCTGCAGCAGAGAGCTGTTAA

AGCTCTTGTTAATATTGCACTGATTTGGCCCAATGAAATTGCCAAGGAGGGTGGCGTCAGTGAGCTATCC

AAAGTGATATTGCTAGCTGATCCCTCTTTGCCTCATGTCTTGTGGGAATCAGCTGCTTCTGTTTTGTCCA

GTATTCTTCAGTTCAGTTCTGAGTACTATTTGGAAGTGCCTATTGCGGTTTTGGTGAGATTGCTTCGATC

TGGTTCAGAATCTACAGTAGTTGGTGCATTAAATGCTCTTCTAGTTCTGGAAAGTGATGATGCTACAAGT

GCTGTAGCAATGGCTGAAAGTGGAGCCATTGAAGCTCTTTTAGAACTTCTTAGATGTCATCAGTGTGAGG

AAACTGCTGCAAGACTTCTGGAGGTATTGCTGAACAATATTAAGATCAGAGAAACAAAAGCCACAAAGTC

TGCAATAGTACCACTATCACAGTATCTCTTGGATCCACAAAGCCAAGCTCAACAAGCAAGATTACTAGCA

ACTCTTGCTCTAGGGGATTTATTTCAGAATGAGGTTCTTGCTCGATCTGCTGATGCTGTTTTAGCTTGCC

GTGCTTTAGTAAATCTGCTTGAGGATCAACCTTCAGAAGAAATGAAAGTGGTTGCAATATGTGCATTGCA

AAACCTTGTAATGTACAGTAGATCAAATAAGAGAGCAGTTGCTGAAGCTGGAGGTGTTCAGGTCGTGCTG

GATCTGATTGGTTCAAGTGAACCAGATACATCAGTTCAGGCTGCAATGTTTGTTAAGCTTCTCTTCTCTA

ACAACACAATTCAAGAGTATGCTTCCAGCGAAACCGTTAGAGCTATAACAGCTGCAATTGAAAAAGATTT

ATGGGCTAATGGTGTAGTGAATGAAGAGTATCTTAAAGCTCTTAATGCACTCCTTGGCAACTTCCCACGT

TTAAGAGCCACAGAACCTGCAACCCTGAGCATTCCCCATCTAGTGACAGCCCTCAAGACTGGCTCAGAGA

CAACTCAAGAAGCAGCATTGGATTCACTGTCTCTTCTCAAGCAAGCTTGGGCAGCATGCCCTGCCGAGGT

CTCCAGAGCGCAATCAACAGCTGCTTCTGAGGGGATTCCCCTGTTACAGTACTTAATTATGTCTGCCCCA

CCTCGAGTTCAAGATAAGGCAGACCATCTTTTGCAGTGTCTGCCGGGAACATTGACAGTGACTATAAAAC

GTGGAAAGAATATCAAACAGTCAGTTGGAAATCCAAGTGTGTTTTGCAAGCTAACACTTGGCAACACCCC

CTCCAGGGAAACCAAGATTGTGTCGACCGGACCAAATCCCGAGTGGGATGAGCCCTTCCAGTGGCAATTT

GAAAGTCCTCCAAAAGGCCAAAAGCTTCACATTTCTTGCAAGAACAAGAGCAAGATGGGAAAGAAATCGT

TTGGAAAAGTCACTGTCCAGATTGATCGAGTCGTTACCCAAGGAGCAGCTGCAGGAGAGTACTTCCTGCT

ACCTGAAAGCAAGAGTGGTAGTAAAAGGAGCCTGGAAATAGAATTTCAATGGACTAATAGCAATAACATG

CCTCAGTCTGAAGCTTAAAGAGATTTGCCTGGTGTTAATTTTTTCTTTTTATCGAATATAATTATTGTGC

TGGTGTACATAATATCTGGAAGGTTTATTTTTGTAGTTAGTTGCTGCAATCCGGAAAGTCGGAAACTGCA

ATTCATTGTATTCTTTTGCAAAGTGTGGTAGTTTTAGTCATCTTCTGTTCTTTTTGTTATAGTTGTATTG

GAGTTGTATTCTGTGTTTGATATTCAGACTCGAGAGGTTAATAAAATCTTGTGTATTGAATTCA

>MSTRG.537.6 gene=MSTRG.537

TATAGATATACACACTCTACCATCTCTCTCTCTCTCTCTCTCTCTCTGCACAACAATATCTCTATATCAT

CCATATTGTTGTGAGTCTTGAGCTTCAACTATCTTTTCTTCGCTTCTTTCCTACGAAATGCATTGACAAG

ATCTCCCTCCCTCCCCTTTAGCTCTCTCTCTCTCTCTGCGTTTATACTGTCCCAATTGTCAGCTACCTAC

TTCATTCAATCTGTTCTCCAGATCTTCCTAGCTTATCTCAATCCTCACTTAACTCATTCATGTGAAGCTA

GCTTCGCTTTGTAACTGTATTTTGGAGGTAGATCTTGAGATCTAATTAGCAAAGCTGGCGGCTACAGTAG

CTTGGAGAGGGGTTGCTGCCAGCAATGGCACAGCCCTGTCATCTAATGATCTGGAAAGAAATGGGGATGC

TAAATCTCATGACCAAGAGCCTCCAACACCACATTCACTCTTAAAGATGAGTTCGAGGGAGCGCAGTAGC

ATGGAGGATCCAGATGGAACTTTAGCTAGTGTTGCTCAATGCATTGAGCAGTTGCGCCAGAGTTCCTCAT

CTGCACAAGAAAAAGAGTTCTCCTTAAAGCAACTATTGGAGCTTATTGATACACGTGAAAATGCTTTCAG

TGCTGTTGGATCACATTCCCAGGCAGTTCCAGTGTTGGTATCCCTTCTTAGATCAGGATCACTTGGGGTA

AAAATACAGGCTGCTACAGTTTTAGGGTCACTGTGCAAGGAGAATGAACTAAGGGTGAAGGTATTGCTGG

GGGGTTGCATTCCACCACTTCTTGGTCTACTCAGGTCTAGCTCAGCGGAAGCTCAGATCGCTGCAGCGAA

GACGATATATGCTGTTTCTCAAGGTGGTGCTAAGGATCATGTTGGCTCAAAGATATTTTCAACTGAAGGA

GTTGTGCCAGTTCTGTGGGGGCAGCTTGAGAAGGGATTGAAGGCTGGAAACGTGGTTGATAACTTACTGA

CTGGAGCTTTGAAAAACCTTTCAAGCAGCACTGAGGGTTTCTGGTTTGCAACAATACAAGCTGGGGGATT

AGATACACTTGTAAAGCTGCTTGCAACTGGACAATCTAACACCCAAGCAAATGTTTGCTTTCTCCTTGCA

TGCATGATGATGGAGGATGCATCTGTTTGTCCTAAAATTCTGGCCGCTGAGACTACCAAGCAACTCCTAA

AGCTACTAGGGCCCGGAAATGAAGCCTCGGTCAGAGCAGAAGCTGCTGGTGCTCTAAAATCTCTATCTGC

TCAGTGCAAAGAAGCAAGGCGGGAAATCGCAAGTTCTAATGGTATACCCGCTTTGATAAATGCTACAATT

GCTCCTTCAAAAGAGTTTATGCAAGGTGAGTATGCACAGGCTTTGCAGGAGAATGCAATGTGTGCTCTTG

CAAATATATCTGGTGGTTTGTCATTTGTCATCTCAAGTCTTGGCCAAAGTCTTGAATCCTGCACGTCGCC

TGCACAGGTAGCTGACACGTTAGGGGCATTAGCTTCTGCACTAATGATTTATGACAGCAAAGCAGAAAGT

ACAAGAGCTTCAGATCCTGACGATGTTGAGAAGACTCTTGTGAAGCAGTTCAAACCTCGCTTGCCAGTTT

GTATGGAAATTCCACACTTTCTGGTAAACTTGCAAATTCTGATGCAAAGCGTTTGCTAGTTGGTTTGATC

ACAATGGCAAGCGATGAAGTTCAGGATGAGTTGATACGATCACTTCTGGTATTGTGTAACAATGAAGGTA

CACTATGGCATGCCCTTCAGGGACGTGAGGGAATTCAATTATTGATCTCTCTTCTTGGGCTGTCATCTGA

GCAACAACAGGAGTGTGCCGTCGCATTACTTTCCCTTTTATCCGATGAGAATGATGAAAGCAAATGGGCC

ATCACAGCTGCTGGAGGTATACCTCCACTTGTTCAGATATTAGAGACAGGATCTCCAAAAGCCAAGGAAG

ATTCTGCAACAATCCTCGGAAACCTCTGTAATCACAGTGAAGATATTAGAGCATGTGTTGAAAGTGCTGA

TGCTGTTCCTGCTTTGCTGTGGCTATTAAAGAATGGAAGCTCCAATGGCAAAGAAATTGCTGCAAAGACA

TTGAATCATCTGATCCACAAATCAGATACAGCAACCATTAGTCAGCTCACCGCATTACTTACAAGCGATC

TACCCGAATCTAAAGTGTATGTTTTAGATGCACTAAAAAGTTTGCTATCTGTGGCCCCTCTTAGTGATAT

GTTGCGTGATGGAAGTGCTTCAAACGATGCTATCGAGACAATGATTAAAATATTAAGTTCTACCAGAGAT

GAGACACAGTCTAAGTCTGCATCAGCTCTTGCTGGAATATTTAATCTTAGAAAGGACTTGCGTGAAAGTC

CCATAGCTGTTAAGACTGTTAGGTCGGTCATGAAGCTTTTACATGTAGAATCCGAAAGCATATTGGCGGA

GTCTACTCGTTGCCTTGCTGCAATATTTCTGTCAATTAAGGAGAACCGTGACATGGCTATGATTGGGAGA

GATTCATTGCCTACACTAGAGGTGCTTGCAACTTCGTCTTCACTGCAAGTTGCAGAGCAGGCCATTTGTG

CTTTGGCTAACCTTCTTTTGGATAGTGAAGTTTCAGAAAAAGCCAAGCCTGAAGAAATTATTTTGCCTTC

TACTAGGGTATTGCGTGAAAGCACAGTTACTGGAAGGAGCCATGCAGCAGCAGCAATTGCCCGGCTTCTG

AGTTCTCGTAAAATTGATGCTGAGATTGCTGACTGTGTGAACCGTACAGGAACAGTGCTTGCATTGGTTT

CGTTCCTAGAATCAGAAAATGCCGGGACTGCTGCCATATCAGAGGGGTTAGAGGCACTTGCCATTATATC

GAGATCTGGAGGAGATAAAGGACAGAACAAACCTGCATGGGCAGTTCTGGCTGAATTCCCAGATAGCATA

ATCCCGATCGTTTCATGCATTTCTGATGCAAACCCCTTGTTGCTTGATAAGGCTATAGAAATATTGTCAC

GGATTTCTTTGGCTCAACCTGTTGTTTTGGGGAACAATATTGTGAGTGCTTCAGGGTGTATCTCGTCAAT

TGCTAGACGGGTAGTTAGCTCTTCAAAGGAAGCAGTAAAGATTGGAGGCACCACCCTTCTTGTTTGTGCT

GCCAAAGTAAATCTTCAGAGAGTTGTAGATGATTTGAATGAATCATATTCTTGCGCATATCTCATTCAGT

CCCTTGTAGGAATGCTGACCTACACAAAAAATCCTCTGGTTGGTGACCAGGGAAGTAATGAAGGCGTAAA

TATTCTTGGGCATGCTGAAGAACTAAAAGCTGGTGAGAATGAGACGAGTAGATATGTCATCTATGGTTCA

AATACCGCAATATGGCTTCTCTCTGCTCTTGCTAGTAATAACGAAAAGAGCAAAACTCAGATCATGGAAG

CAGGTGCCATTGAAGTTCTAACTGAGAGAATCAATCAGTGTTTATCAGAATATTCTCAGGTCGATTCTAA

AGAGGAAAGCAGCATTTGGATATGTGCTTTACTTCTAGCAATTCTATTTCAAGATAGAGATATTATACGT

GCGCATGCAACAATGAAATCTGTACCTGTACTGGCTAATTTGTTGAAGTCAGAGGAGGCAGCAAACAGAT

ATTTTGGTGCACAAGCCATTGCCAGTCTAGTATGCAATGGTAGCAGGGGGACTCTTCTATCAGTCGCTAA

TTCAGGGGCTGCAGGTGGACTCATTTCACTGCTTGGGTGTGCTGATGCTGATATATGTGATATGCTTGAC

TTGGCAGACGAATTTTCCTTGGTGCGGTATCCTGAACAAGTTGCTCTTGAGAGGTTGTTTAGGGTTGAGG

ATATCAGAGTTGGTGCTACTTCAAGAAAAGCAATACCTGCACTAGTTGATCTACTTAAACCGATTCCAGA

TCGTCCAGGGGCACCATTTTTAACCTTAGGGCTTCTGCTTCAGCTTGGTAAAGATTGCCCTTCGAATAAA

ATTCTTATGGTAGAAGCTGGTGCTCTGGAAGCATTGACCAAATACCTTTCACTTGGCTTACAAGATGCAA

CTGAAGAAGCAGCTACGGATCTCTTAGGTATGTTATTTAGTACTGCTGAGATACGTAGACATGAAGCAGC

ATTTGGTGCCGTAAGCCAACTTGTAGCAGTATTGCGATTGGGTGGAAGAGGTGCAAGGTATAGTGCTGCT

TTAGCATTGGAAAGCCTTTTTTCTGCAGACCATATTAGGAATGCAGAATCTTCCAGGCAGGCTGTACAGC

CTTTAGTGGAAATCCTTAATACAGGTTTGGAGAAAGAGCAGCATGCTGCTATTGCTGCATTAGCTAGGTT

ACTGAGTGAAAATCCATCAAGAGCCTTGGCAGTTGCTGATGTTGAAATGAATGCGGTAGATGTTCTTTGC

AGGATTCTTTCATCAAACTGTTCACTAGAGCTGAAGGGGGATTCGGCTGAGTTGTGCTGTGTACTTTTTG

GAAATACAAAGATCAGATCCACCCTGGCTGCAGCGCGCTGTGTGGAACCTCTGGTTTCTTTGCTCGTGGC

TGAGTATAGTCCTACTCAGCACTCTGTTGTCCGTGCATTAGATAAACTCGTTGATGACGATCAATTAGCT

GAGCTCATTGCTGCACATGGAGCAGTTATACCTCTTGTAGGCCTTCTGTATGGCCACAACTATGTGCTCC

ATGAGGCTATTTCCAGAGCTCTTGTGAAGTTGGGAAAAGACAGGCCTTCTTGTAAGATGGAGATGGTGAA

GGCTGGGGTGATTGAGAGTGTACTTGATATTCTCCATGATGCACCAGATTTTCTGTGTGCTGCTTTTGCA

GAATTGCTACGAATATTAACCAATAATGCTACCATTGCCAAGGGTCCATCTGCAGCGAAAGTGGTTGAGC

CTTTCTTTGTGTTGTTAACAAGAGCAGAGTTCGGACCTGATGGACAGCATAGTGCTTTACAAGTTCTTGT

GAATATCTTAGAACATCCTCAGTGCCGTGCTGATTATACCTTAACATCACATCAAGTGATCGAACCCATC

ATTCCATTACTTGATTCTCCTGCACCGCCAGTGCAGCAGCTAGCAGCTGAACTTCTTTCTCATTTACTCC

TAGAGGAACATCTTCAGAAGGATTCAGTTACACAGCAAGTAATTGGCCCACTTATGCGAGTTCTTGGTTC

TGGTATACACATTCTGCAGCAGAGAGCTGTTAAAGCTCTTGTTAATATTGCACTGATTTGGCCCAATGAA

ATTGCCAAGGAGGGTGGCGTCAGTGAGCTATCCAAAGTGATATTGCTAGCTGATCCCTCTTTGCCTCATG

TCTTGTGGGAATCAGCTGCTTCTGTTTTGTCCAGTATTCTTCAGTTCAGTTCTGAGTACTATTTGGAAGT

GCCTATTGCGGTTTTGGTGAGATTGCTTCGATCTGGTTCAGAATCTACAGTAGTTGGTGCATTAAATGCT

CTTCTAGTTCTGGAAAGTGATGATGCTACAAGTGCTGTAGCAATGGCTGAAAGTGGAGCCATTGAAGCTC

TTTTAGAACTTCTTAGATGTCATCAGTGTGAGGAAACTGCTGCAAGACTTCTGGAGGTATTGCTGAACAA

TATTAAGATCAGAGAAACAAAAGCCACAAAGTCTGCAATAGTACCACTATCACAGTATCTCTTGGATCCA

CAAAGCCAAGCTCAACAAGCAAGATTACTAGCAACTCTTGCTCTAGGGGATTTATTTCAGAATGAGGTTC

TTGCTCGATCTGCTGATGCTGTTTTAGCTTGCCGTGCTTTAGTAAATCTGCTTGAGGATCAACCTTCAGA

AGAAATGAAAGTGGTTGCAATATGTGCATTGCAAAACCTTGTAATGTACAGTAGATCAAATAAGAGAGCA

GTTGCTGAAGCTGGAGGTGTTCAGGTCGTGCTGGATCTGATTGGTTCAAGTGAACCAGATACATCAGTTC

AGGCTGCAATGTTTGTTAAGCTTCTCTTCTCTAACAACACAATTCAAGAGTATGCTTCCAGCGAAACCGT

TAGAGCTATAACAGCTGCAATTGAAAAAGATTTATGGGCTAATGGTGTAGTGAATGAAGAGTATCTTAAA

GCTCTTAATGCACTCCTTGGCAACTTCCCACGTTTAAGAGCCACAGAACCTGCAACCCTGAGCATTCCCC

ATCTAGTGACAGCCCTCAAGACTGGCTCAGAGACAACTCAAGAAGCAGCATTGGATTCACTGTCTCTTCT

CAAGCAAGCTTGGGCAGCATGCCCTGCCGAGGTCTCCAGAGCGCAATCAACAGCTGCTTCTGAGGGGATT

CCCCTGTTACAGTACTTAATTATGTCTGCCCCACCTCGAGTTCAAGATAAGGCAGACCATCTTTTGCAGT

GTCTGCCGGGAACATTGACAGTGACTATAAAACGTGGAAAGAATATCAAACAGTCAGTTGGAAATCCAAG

TGTGTTTTGCAAGCTAACACTTGGCAACACCCCCTCCAGGGAAACCAAGATTGTGTCGACCGGACCAAAT

CCCGAGTGGGATGAGCCCTTCCAGTGGCAATTTGAAAGTCCTCCAAAAGGCCAAAAGCTTCACATTTCTT

GCAAGAACAAGAGCAAGATGGGAAAGAAATCGTTTGGAAAAGTCACTGTCCAGATTGATCGAGTCGTTAC

CCAAGGAGCAGCTGCAGGAGAGTACTTCCTGCTACCTGAAAGCAAGAGTGGTAGTAAAAGGAGCCTGGAA

ATAGAATTTCAATGGACTAATAGCAATAACATGCCTCAGTCTGAAGCTTAAAGAGATTTGCCTGGTGTTA

ATTTTTTCTTTTTATCGAATATAATTATTGTGCTGGTGTACATAATATCTGGAAGGTTTATTTTTGTAGT

TAGTTGCTGCAATCCGGAAAGTCGGAAACTGCAATTCATTGTATTCTTTTGCAAAGTGTGGTAGTTTTAG

TCATCTTCTGTTCTTTTTGTTATAGTTGTATTGGAGTTGTATTCTGTGTTTGATATTCAGACTCGAGAGG

TTAATAAAATCTTGTGTATTGAATTCA

>MSTRG.541.1 gene=MSTRG.541

CAAGAAAAAGAAGAAAAGGGAAAAGAAAAGAAGAAAAAAAGAGAGTTGGCAACTAAGGGACACATGGAGG

GCCACAATTCTCCCAACACTCAAGGCACATGGATGGTCAAGATGGAGTCACCCTACCCCTAAACCCTAAT

TCTTTAGTTATAAATACCCCTCTTCTTTTACTTGTAATCACCAACCTAGCAACCTAGTAACCCTAGCCAC

CTACTCACCTTTTCTACCTAGTCATTTCTATAGTTAGTATCTTAGATAGATTTACATTTTGCTAGCCCCA

ATTTCCTTTCCAAGCTTGTATACACTTTTTAATTTCTA

>MSTRG.545.1 gene=MSTRG.545

TTCCCCCCTTTCTCATCTCTCACTCTTTTGTTGTGCCTTCAGACTTTTCATAAATCCATACCTACCCACA

GTGTATACACACTCCAGCTCTCATTCAAAATAATAGTTTTTCATGGGCAACTCTCATCCGGCCTCTGCCA

TTCACGACTTGGGGGTTTTGTTCCAGATCTGGTGCACTTCTTCCATTAGGGTTAAGATTGGAACCAATTT

AAATGGGGATTTATACATCGAGTAGATTAGACTTGTAATAAGGTATTATTCTCAATTTTAACTATTTTTC

ATCCTCTCTTTGTCTCTCTTCTTTTGTCA

>MSTRG.546.1 gene=MSTRG.546

CTTGAGAGGAGAAATTTGGAAGATTTCCAGGAGAATTCAAGGACGAGATTTAGAAGAAAACAACACTAGC

TAATAATGAGTTCAAATAATTAAGTAAAGACAGAAAATGGCACAAATGAGAAGGGTAATTCGGAAGCATA

TAATAGCAAAATCAATGGCAAAGCGTGAGGCTGTGGCACGAAGGGTGCGAAGACTTGTTCCAGTAACGAG

AATGTTTAATGATTATAATAATGGTAGATTGTACTCTTGAACTTATCATTGTAATGTTAGAATAGACTGC

GTGGAAGTGTAAGATCGACCCTGGACTTGTGGGGCGAGCTATGTAACGTAAGACAGGTATGCTATTACAG

TGTGACGGCCTAGATCCGGGATATACGGATCTGGGGGCGTTACAG

>MSTRG.546.2 gene=MSTRG.546

GAGAAATTTGGAAGATTTCCAGGAGAATTCAAGGACGAGATTTAGAAGAAAACAACACTAGCTAATAATG

AGTTCAAATAATTAAGAATATTTACTATCTGGAAAGGCGAGCTCAATTTGAGACAAGAATATTTTGTGTT

ATTATCAAAAGACGCCAGTAAAGACAGAAAATGGCACAAATGAGAAGGGTAATTCGGAAGCATATAATAG

CAAAATCAATGGCAAAGCGTGAGGCTGTGGCACGAAGGGTGCGAAGACTTGTTCCAGTAACGAGAATGTT

TAATGATTATAATAATGGTAGATTGTACTCTTGAACTTATCATTGTAATGTTAGAATAGACTGCGTGGAA

GTGTAAGATCGACCCTGGACTTGTGGGGCGAGCTATGTAACGTAAGACAGGTATGCTATTACAGTGTGAC

GGCCTAGATCCGGGATATACGGATCTGGGGGCGTTACAG

>MSTRG.547.1 gene=MSTRG.547

AAACAAATGGTAGACTCCTGATAGCACGGAGTTGGCTGCTTCTTCCTACAATTCTTTCATTCACGAAAAG

AGGGTCTTCTTTGGAGAAGATTTGCAGTTTATTGAGGCCTGAACCTTTGATTCCTTTCGCGGCTTTCTCG

ATTTTTGATTCTCAGATTTCTGATTCGAAGTATTCTATCCCTGTTGAATAACCTTTTAACAAAAAGGTTC

TTTATATTTAAGTCCCATATCTTCTTTCCTTGGCTGTGATGCCCTCCCTTTCTTGCCATTCTTTTCACGG

TTGTATAGCTCAC

>MSTRG.550.1 gene=MSTRG.550

TAGGGCTGTAAATGAGTTGATACGCTCGATAACCGCTCGATGTTCGGTTCGAAGAAAGCTCGGTTCGACC

TCGGTTCGATTCATAAACGAGTCGATCTTGAGCACAATTTTAAGGTTCGTTTTATAAACGAGCTGAACTT

GAGCACAGTAGAGTTCGACTCGATAGTTCGCGAACAAGTTCGAATTATGAGTTCGTGAACAGAGTTCGTG

AACATGGTTCGTGAACCTGGCTCGTG

>MSTRG.551.1 gene=MSTRG.551

GTGGTAGAATTAACTAGGTTGAGCTCATTAGTACTCGAGTAATTAAGTCTGAGGGGACTTTGTGCCTAAC

AACCTAAAGCCCTTCGTGGTTTGGGATTGTTGACCCAACGCTCGCTACATGGGTACTAGTGCATAAATCT

TTAGGGATCTCAACCATTGCACGGTTAAATAAACCACTAGAATAGAGTGAATAATTGGTGTGTGAAGTCT

TGTGGTGTTCGTAACGCATTTATACTGCGAAGCGCTCTGACCTTAGACCGAGCAATTAATCACCAATAAA

AAGAAAAAAAGAAGAAAAAAGATATAGTGAATAAAATAGAAGGGTGTGGACATTCTTTGGGACCTTGGCT

TTTAGTTGGACTTGAGTGACGGG

>MSTRG.552.1 gene=MSTRG.552

GTGCCTATTTTTTGTTTATTTTCGTAGTATTAATCGCTTATTTATTTTATTTCAGGAAATTGTAGATAAA

TAAAGAAAGAAGAGAAAATGCAAGAAAAAGAAGAAAAGGAAAAGAAAAGAAGAAAAAAAAAGAGTTGGCA

AATAAGGGACACATGGAGGGCCACAATCTTCCCAACACACAAGGCACATGGATGGTCAAGATTGAGCCAC

CCTACCCCTAAACCCTAATTCTTTAGTTATAAATACC

>MSTRG.555.1 gene=MSTRG.555

GTTGGTTTAGGGAGAACATTAGTGATACTTGTCCTATAGGGATGAACGATTGTCACACGCTAGAGAAAGT

CTTTGTGCATAAAAATTTTCGAAAATATTTTGACTCTTTCTCGAAGCTTCCTCCCATCTTTCCTTCCGGA

ATTGGTTGATTTGCTTCCGTCTTCAAACCTTCCTTCATGAAGTTCAGGTATGGGGAAGGAAATGTGTGTG

TGCGTGGGTAGTGTTAGTGTTTCGTTCGTTAATAAAAATCCCCTTCATATCCATTTCAGGTATTTTCTCT

TTACTCTCTCATTTCAATTCTCTCTTTTGCATACATTGAGGACAATGCATGATTTAGGTATGGGGAGGGC

TTTAGTGTAATTCTTAAAAAATGAAAATCCCTAAAAAATGAAAAAAT

>MSTRG.556.1 gene=MSTRG.556

AAAAAAAAGAGAAGAATAAGAGTTTGTGAAGTTTGGCTAGATATTTGTTTCGATAGAGATTGGGTCAGTT

CAAAATTGGTTGAAAAAGAGAAAATGTTATTAGACGATTAGGATCCATCAAAGGCCTTAAATTACTTACA

CTCCTTGGATTTCTAGTAAATTTTTGTGTCGATTCAAGTAGATTAGAGACAACTGTATCTTGATTATTAG

TAGATGAGTCTTTAAGTCGTATTTTCCTTAAAAACGCTTTTTGTTAACACCAACGAAATTAGAGTAGAGT

CGAGTATACACTTGCTAGAATGTCTCGAAGATTTTATGGGTATATCTTCTGTAAACCCTTAGGAGACAAC

>MSTRG.557.1 gene=MSTRG.557

TTACTCCCTCCGTCCCCCTCAATTGTTTACATTGGAGGGGGACACGGAGACCAAGACAATGTATGAAAAA

TGAGTAAAATTAGGTGAAAAGTGAGTAAAGTGGTGGGACCCATCAATATTTAATAATAGATTTGAGATAG

TGGAGGAAGGTAGTGGGTGTAATAGTAGTTTTTATTGTTAAATATGAGATAGTGGAAGAAGATAGTGGGT

GTAATGATGAAAAAACTTACTATTTATGGTAACGTAAAGAAATGAGAGGGACATCCCAAAATAGTAACGG

TAAAGAAATG

>MSTRG.559.1 gene=MSTRG.559

GAAAAATTCCTGCAAAAAGCGTTACAGTGAACACGAGATTCCTATAAGCAATTGGATCAGCCTCAGGGTC

TTGAACTTTTTGAACCATTGAAGATATCAAGAGTGAAACTACAGCCACTGGTCCGATAGCTATTTCCCGT

GAACTTCCCATGAACGCATAGATGATCGGAGGGACAACGTTCGTGTCTGGAATCAAAAAC

>MSTRG.561.1 gene=MSTRG.561

TGATAAGTGGTATTTATACACACTTATAGGCCTTCATTTCCACTTAAATTGGTTGATTGTACTTAAGTGT

TTAGTGTCTTTTGATGTGTTTTTATTGTTTTTCTGTGCAGGTCACGAGTTGAGGTGATGAAGTGATTTTC

TATCATTTTAGGTTGGTTTTGGTGCATTATTTGCAAGAATAGAGAATTGTTGATTCACCGCGACTTGGGA

TTTTGTGGGTATATCTTCTACAAACCCTCACGAGAGTACACTCGTCCACTAGAGCTATCTAGGGGTTTAA

AGGGCTTGTTGCATATGCTAAATGCAACCGTGATCACCTACGGAAGTGGTACTAGAAGCGAGGAAGGACA

TGCACG

>MSTRG.562.1 gene=MSTRG.562

TGGGTACTAGTGCATAAATCTTTAGGGATCTCAACCATTGCACGGTTAAATAAACCACTAGAATAGAGTG

AATAATTGGTGTGTGAAGTCTTGTGGTGTTCGTAACGCATTTACACTGCGAAGCGCTCTGACCTTAGACC

GAGCAATTAATCACCAATAAAAAAAAAAAAAAAATAGTGAATAAAATAGAAGGGTGTGGACATTCTTTGG

GACCTTGGCTTTTAGTTGGACTTGAGTGACGGGGTGTTAGTTTTGAACTTTTACGATTCTTGATTAGGAT

TCTGTGGGAGTAGTAGTTTTTTATTAGTTTTAAACTTTTACGATTCTTGATTAGGATTCTGTGGGAGTAG

TAGTTTTTGTCTTGTCTCAATAAAAGAGCATGCTTGCACACACTGGCACTCCACACTTGTAAATAGAAAA

GTAATTGACTTGGTAGCGAGAGAGATGAAATTCAAGAACATTAGCACAATCTGAGGTATTATAATTGACA

AGGCAATTACTTCATAGTTCACTCATTCATCATGTAGTTGCATTCATGCATTGCATTGTATTATTGTATA

GTGTCGTTGACGCTTGAGGACAAGCATCAGTTTAAGTTTGGGGGTGTGATAAGTGGTATTTATACACACT

TATAGGCCTTCATTTCCACTTAAATTGGTTGGTTGTACTTAAGTGTTTAGTGTCTTTTGATGTGTTTTTA

GTGTTTTTCTGTGCAGGTCACGAGTTG

>MSTRG.563.1 gene=MSTRG.563

TTTTAAATTGTTTGAGGGGAATAGTAACTTGTGAATATCTATCCCGGATGAGATTTTAATTCTGAAAATG

TTTAGTGATTATCATTGCTAGACCTGTGATATAATTTTTGAGACATGGTATTGATTCATGAATATCATGT

AATGGTTTTGAAAATGGGATGGATACAAATCAAAGTGAAAACTGGAAATGGATCAGATATCTCTATTTGG

AGTTGCGTAAGAGGTTCCGTTATTTGATAACGACCCAGCATGGTGGCGTAAGAGGCTTGCGTGGTTTACA

CACCCAAAACCACCTGGTCCAGCGTCTAAAGACCTAGCTAGTCTTTAGGTTCCGG

>MSTRG.564.1 gene=MSTRG.564

CATTTATTCAGTCATTTTATGATGTTTTTGTTTCGTTTTATTAAAGCAAACCCAGTACCCAATTTCCCCT

GCTTTCGTTAAACACAAAACCCCTGTTTTGTGGTTATTTAATCAAACCCATTTCCTTTTCTTTTTAAAAC

ATTCAAACCCTCTTCGTTTTATCAAAATCTCACCTCTGTTTTCTTTGCATCAGTTAGCTTGTAGCGAGTC

GAGTAGCTGGATGCCTTGATTTCTTTTAAATTTCTTGCTGTGATTTATTTTGCCATCGACGAGTTCAAGA

TGCCGAGTTGCCTGCGGAGTCTGGTCTTAGTTTGACTCCTGTAGTTTTGTTAGAGTTCGATAGCTGAGTT

GTAGCTTTCTGTTCTGAGTTTTGTTTGCGTGATTTCTCTGTTGGTTGTGTTTTCTTTTACTGAGTTTAGC

AACGAGTTGTCACTGGACTTCACTGAATAGAATCGTCCGAGTTCACCGAGTTGTTTTTCTGCTGAGTCCG

AGTTTTGTTTTGTTGAGTCACTGAGTCGATGTTGTTAGAGTTTGTTTTCGTTGAATCCGAGTATGCTGTG

ATTTGAGCTTGCTGCGTGTATGCTTTCATTTGTTCAGTTTAGTTTTTTCTTTTTGTTATTTCTGCTGTTT

GTCGAGTTGCTTTATTGCCTTGCTGCGATTTGTTTGCTTCCTGTTGAGTTCTTGAGTTTGGGTAGGTTCT

GTATCATGTTGGTGTATTGAGTTGCTACCCCTTTTTCTCTTTTGTTTAGAAACCACTCGCACCTGCTTTG

TTTTTCTTTTGTTGCGTCGCAAAAACGAGTACTTGGAATTTCCGAGTTTAGGTGCTGT

>MSTRG.565.1 gene=MSTRG.565

ATTATACTCCCTCCATCCCATTTTAAGTGTCCACTTTGCAAATTTCACACATCTTAAGAAACAATTAATG

TAATGCTTTTATTATTATCTTCACACACCTATTCACCTTGGTTTTTATAAATATTAATTACGTATAGCAC

CACTCCTAGTGTGTATTTGACCTTTCTTCATATTGGAATTAGCTGGTTGTATTTAATACTATATTGAAAC

TAGTGAAAAATTGCATGGGTTAAAGAGTATGACACATATTTTGGGAATTTTTTTTTTGGCAAAGTGGACA

CTTAAAATGGGATGGAGGGAGTA

>MSTRG.566.1 gene=MSTRG.566

CTTGTCAGATCTTCTAGATAGATCAGAAGCTTCACAATTTGCTAAAGGATCGCGCGAAGCAAGCCAGGAC

ATTTCAGGTCATATCTTTGGGATGATGTCCACTATATTGTGCGAGGTGGATGAAATGGTGCGCTCTCTGC

CTACTCGTGAGGTCACGCAGCCCGTAGTGGATCAGGAACTGAGCAAGTTGGCTGCTGCCACTTTTCCAGA

CCCAGTCCAGGAGTTATTCCGGACTGATTATGTGCGTGCAGCGAAGGGATTACTTAGTCAAATCCTCAAG

AACAATGACAAAGTCATTGTAGAGGTCAGTCAGCAGCAGCAGTTAGAGGGGAATGGGAAGGAGCAGCAGC

AGCAGCCAGAGGGGCAGCCAGAGGGGAAGGGGAAGGAGGCTGCTAGCACAGAGAAAAATCCGTATGAAGA

TGACTATGATGTAAATGCGTTTTTATACCCTCTGTTAGATTAGCAGCAGTACTGGTGGTAATTTGGAATT

TCATACTTTTGGTTGATGGTTTAGTAAGCCAGTGAATATTCTAACTTTTGGTGGATGGATGAATTTCTAA

CTTTTGGTGGAT

>MSTRG.567.1 gene=MSTRG.567

AAGGATCGCGCGAAGCAAGCCAGGACATTTCAGGTCATATCTTTGGGATGATGTCCACTATATTGTGCGA

GGTGGATGAAATGGTGCGCTCTCTGCCTACTCGTGAGGTCACGCAGCCCGTAGTGGATCAGGAACTGAGC

AAGTTGGCTGCTGCCACTTTTCCAGACCCAGTCCAGGAGTTATTCCGGACTGATTATGTGCGTGCAGCGA

AGGGATTACTTAGTCAAATCCTCAAGAACAATGACAAAGTCATTGTAGAGGTCAGTCAGCAGCAGCAGTT

AGAGGGGAATGGGAAGGAG

>MSTRG.572.1 gene=MSTRG.572

TGTACGCTCGAAGACGTCAGGAAGTCGAAATGGTATTGCAAGGCGAATAAGGGATGAATCGAGTAAGCGA

AGCGATGTGGCGAATAGAGTACGGCTAGAGATGGTCTAGAAATTATGATATGGATAGATTGTGAAAGTGG

AAAGATGAGATAGAGATGTGAGACTGGAGTCAGATTTAAGAAAGGATTAAAGATGAATGGCTAGTCTATG

ATTGCGAGTAAAGCTAAGGCTAGGCGAGGAGTGTTCAGGAAAGTTGATGAGGTTGGTGCAAGCTAGAGCT

GTGCTACAGAGATTCAAGTGCAAGTCATAGCTGGAATAGTTGACAGTGATTCTTCTTATCGAAGATGATC

TGATTTGGTAAGAGATGTTGTGTAATAATAGTGGTAGATTCTAATTTCAATACTGTAACCTGGATGCGAT

CCTGTTGTTTTAGGGGG

>MSTRG.572.2 gene=MSTRG.572

ATTGGACCCCGCTTTATTAATATAAAGCTATTAAATACGTATTTTCATGTCATAAACGATCCAAAGGGGT

ACCAAGTATTTGTAAATATAAATAGACTTTTCTATAGCTGATTTTCACCGTAGAAATCATTTTCACCAGT

TTCTTTTCACAAATACCGAGAGAAACCGAGCGGTGTTCTTCGTGTTCTTCATAATCAAACGAGGATTTGG

AGGTGTTATTGGAATCCGATGGAGGTGTACGAATAGTCAAAACGAAGCTATCGACGCGTAGAATCATGTT

TAATCACCCGTTTGTATGCAGATTTATCAGGATTTGGACGAGACGTGTATGCCCGAAGACGTTCAGAAAG

TCACGTGGTGTTACAAAGCGAATAAGGGATGAATCGAGTAAGCGAAGCGAAGATGTACGGTAGATAGAGT

ATAGCCAGAGATGGTCTAGAGACTATGAGATGCATAGATTGTGAAAGTGGAAAGATGAGATTGAGATGTG

AGACTGGAGTCAGATTTAAGAAAGGATTAAAGATGAATGGCTAGTCTATGATTGCGAGTAAAGCTAAGGC

TAGGCGAGGAGTGTTCAGGAAAGTTGATGAGGTTGGTGCAAGCTAGAGCTGTGCTACAGAGATTCAAGTG

CAAGTCATAGCTGGAATAGTTGACAGTGATTCTTCTTATCGAAGATGATCTGATTTGGTAAGAGATGTTG

TGTAATAATAGTGGTAGATTCTAATTTCAATACTGTAACCTGGATGCGATCCTGTTGTTTTAG

>MSTRG.573.1 gene=MSTRG.573

CTTGCCGGATATTTAAACTCCGGCGGCGTTCTTAGCTTTGATCTAATTTTCTGGCTTGATTAATGATTTG

TTTTGAAATCCGAGACCATATACGAGTTCCTGGGATGCTATTTAATAAATCTGGAGTTGTTTGCTGAGTT

CTGGGCGCGGTGGCAACACCGGCGACTCGCCGGCGGCGAGTCTGGGGGCGGCGACGGCGAAAATGCCGTT

TAGCCCCCAAACTTTTGGAGGTGGTGAAGATTTTGTACTCTAGTTTCTTTTCTGGCAAATTGACCCCTGA

CACTTTTATAAGTTACACTTTACACCCTTGC

>MSTRG.574.1 gene=MSTRG.574

AAAAAAAAAGAAAAAAAAATAGAAAAAGCAAAATTGAATGGCGGTCGTGACTCACTTACACTGAGAAGCG

TCCCAACCTTAGACTTAGCAAGAAAAAAAAAGAAAAAAAAATAGAAAAAAATAGGAGAGGCATAGGAATT

CTTTAGTGACCTTAAATTGTAGTTGACTCTTTAGTAAAGAAGCGTTTAAGCCTTATTCTGATTAACGGCT

CATAGTGAGGACTTTGTGAAGTTTGATGGTCTTTGTTTTGTTTCACTAGAGAGCATGCTAGCACACACGA

TGCACTTCACACTAGTAGTGGAAGAGAACATGGATAAGATGTGAAAGAGATGAATGCCGAGA

>MSTRG.577.1 gene=MSTRG.577

TATCCGACAAACTCCGTCTCACCTAATTTAACAAATCAGTAGGCTGACAGCAAAATTTGCTTATATAGTT

TGATCTAATATTGCTTTTTCTCCGTCCTCTCTGATTTGAGTCACGATTAGAGGATCTGTTAACTACCATC

TGCTATACACTACTCTTCACCTGCCCTTTGACATGAAGTTGGTATCATACTAAAGTGGGGAGGGGTTCTG

TAAGTTAATATGGGAAGTATGAAGAATTATAAAGACCGCTTATTCCGTCCTTGCACGCTTTGACTGAGCT

TGTCTCATCCAGTCGATGCACTCCTCGGCAGACCCAGAAGGGTGAGCAAGCTGCCATTCAAGCAACTTTT

CTACCTGCAATATGATGGAATAATTTTAGCATACACAGTTGCTTAGATAGAGTCATTAGCATATATAAGG

TTGGAAAGTCCGCACCCATATATAGTTAATTTAGCAGTAAAAGAGGATGGCTTACCCATTCCCTCACAAG

AGGTCCTCCTGTCTTTAGCTGCAAAACGCTCATAATTTCCTTTCCGTTAACCAATGGCTTCACTTCCCAT

ATTTTGTCAAGACCTTCATTCAGTATCACAAAGAAGAG

>MSTRG.579.1 gene=MSTRG.579

ACAAGAATGAGAAGGTTGAGGAATGATGCTGAAAATATATTAGTCAATCTGGACAGAGGTTACCTGTTCA

ATCTGGATAATAGTTATGTGCTCAATTTGGACATCAAATACAAGCTTTATCCAGGCAGCATCTGCAATCT

GGGCAACTTGTTTCCACTCAATCTGGACAACGCGTTCGAGCTCCATATGCATAACAATCAAGACAAGGTC

CTTATTTTTGATATTTTTGGTATTGTTGGTTGTTATGGTTTATAAGTAGTAATGCTATCCGATCCGGTTT

GTTCTTTTGGATATCCACATTTTGAATATGTATTATATAATGAAACGTTGGG

>MSTRG.582.1 gene=MSTRG.582

ATCTACTTTTCCATGATACTAGTATCACTCAAGTTAAAGTGCGTTTAGATGATGCACAATGTAATGAATA

TTGCAAATTTGAGTATCCGCTTGACTTGCCCACTTGTTTGTGGAATTTGCAATCAAATGGTTTAATCTGT

TTGTCTAATATGTTTAATGAAAATGTGGGTTATAATCCCGACATTTATCTCTGGAATCCTCTTGTTCAGA

AATTTAGGAATGTGCCGGACTCTCCCTTTTCGATGTTTACATTTAGGGAGACTAAGTGGAATGCTTTAGC

TTTTGGGTTTTTGCCGGAAGTTAATGATTATGTTGTGGTACATGTTATCAAACCTATTTCGACGGCTGCA

CCCCCTTGCTACTTAATTTCTAATCCTGACATAGGTTATGAAGAATACCCACACTCAGTCAAGATTGGTG

TTTATAGTCTCAACAGTAATTCTTGGAAGGAAGTATGTCAGGATAAGGTATTTGTTGATTACATGAGTAC

GGATGTATCGGTATTTGTTGATGGTACTGCATTTTGGGTAGGCTATAATAATGAGGATTTGTGCCAGTTA

GTTATGTACTTTGATACGAAGACAAATGTACTGGGACAAGTCATTTTACCTGATGACATTATATTTCGTG

CATGTCAGCTTGAGAATCCACTTATTCTTCCATTTGGTCAATCGATTGCTTACTTTGTTGAGGTCGGTGA

GGATGATGCAAGCGAGGATGATGAGGACTATGGATCTCCTCATATGGACATATGGGTATTGAACAAAGAT

ACGATGGGTGAGTTTTCTTGGGAGGAAAAGATGAGTGTTACTTTAAGTGAAAATGTTTGGCCTGAAGTCT

TGGGTATAAGGAACAATGGTGAGCCAATACAAGCAAAATTGGACAAACTGATTTCATATGATCTTGATAC

TCATGAACCATATGTTTTTGTTGAATCATGTGAACATTTGACTCCAACTCCCTATTACAAGGAGGGCTCT

AAATCACCTTTTGTCATTCGTCCTTTCTTGGAAACTCTACTTTTTCTTGATATGGATTGAGAAATTTGAA

CGAAATGTTTGTCGGACCCTTGTTATCGCAATACGAACTTGCTAGAGTTCTTTAACTCTTCATTTTTCTA

GATAGAGAATTAGAGATGATGATGTAGACGTCATTATTTGTGCATCAAAAGTGGACCTCGAACTAAGATT

TCAACAAATGATTGCTAGTGACATGGCTCGTCTGAATTTAAGGAATCCAGTGTTGAATCAGGCAAGTGCT

TCCCATTTGACAACTTCATGGCAATTCCGGTTCCTATGGTATTAGCAGAACTCGAGTATAGCAGATAGAT

GTTATGGGGAAGGTTAAAAACAGATAGTAGCTATACTACCTCTGTTGAAGTTGTAAGCAACACAAAGGGA

AGCTAATTGTCTTAATTTCTTATGTTTTTGATAAAGTTCTAATGGAAGTGTGTTTATGATCAATAGCTAA

TGTAAGTGTACCCTATATCTTTTGTCATAGTAGTTCATCCTGTTAAAGAAAGAGAGTCATAGTTGTTGTT

ATAGTTACATAGGACTGGGTTCGATTCGAAACGGGGAACGAAATCGGGTACGTGGAACATTAGTTTTAGT

GAATCTGGTACGATGGAACGTATAGGTACGATGGAACGTATGGGTACGATTG

>MSTRG.585.1 gene=MSTRG.585

CTTGTAAACATTCTCTCGACTTCACCTCACAAACACTCTGAGCGACGAAAACAAGAGAGAGATAGATGGC

GAGATAAAGAGAAATGGCGATAAGAAGAAGATGAGAGAGAGTGAAGAAAGCACCCCGTAGAGGATACAAA

AGAGATTTGCATAAATTATGGAACAGATACAGGTAAGACAATTGTGACTTTGATGGAACATAATCGAATT

ATGGAACAGATACAGGTTAGACAATTCTGACTTTGATGGGTGATCCAAATGGAGAACAGAAACAGGTGAT

ATTCGTATGAATTATACTGAATATTATTGCTTTAGGCTGTTGCTTCTAGTTCCTTTGCTTTGATGTCAGT

ATATTTTTTAATCACTCGTATTTTTTCGTTGTAATGATGGGAATGGGTTAGCATTGTACAACTTGCAATA

AACCAAGGACTTTAAGTGATGTACAATATCAACATATATTCTGCTTTTAACATTCTTCTGCTTTTGAGGA

AGCTCTTGCCCTTGTAACTACAGACAAACTATTTGAAAGACTGGTTTAATTATCATCCACTTGTCGCATA

CCATGTCTTATATGGCGGAGCATGTGGTTGGTACTGGTTTATTTGGTGTTGTATATCAGGTGCGCAAAAT

GAGGGACTTTTTCCCAGGTTTCTAATAAATGGATAAGTTGAGTTTCAGGGTTTTTTTAGTAATGTCCGCT

TGG

>MSTRG.585.5 gene=MSTRG.585

CTTCGGTCGATATTTGATTTGTTAAGGTTTTAATGGCTGTTGAAGAAAGTTGTTGATATTCTGTCCAAGC

ATCTTGATCTTTCAGACCAATGCTAAATATTGAGAAAAGATTTGAGGAGGTGAAGAAAGAAAAATCCAGT

CCACAAGCAAAATTTTAAGAGGAAACACAAAAGTCTATATTCAAAAGAATTCAGCAGGCTCTGATGGATG

GAAAACGAGCTCGATCAGTTTGAAAAGGATGTTATAAAGCACATGTGTTTACTTGCTATATCCCTATGAT

GTGTGCTTGATTAGTAATACCTTTTAGCAAATTGGATGTTGAAAAATTGGTGTATTGATATTTTGTCTGT

TTTCTTGATTGTTCTAGACGTGTAGTTGGTAATAATCGTAGTATTGTTTTAGGATGTTCTTGTAATGAAC

CTTTGAAGGTGGATTAATGTATTTTCGAAATGGTCATGTCAAATTTTTAACAAG

>MSTRG.586.1 gene=MSTRG.586

ATTTTATATTCTAAACTTATACTTCATCTCTACCCATTCCCCACCAACCACCCCCACCCCAGCCCCACTC

ACCACCATTGTCGCACATCAGCACACCACCATCACGCACACCACTGCCACCGAACATCATATCGACAGAT

CTCTCTCTCCAGCAACACGGCAGGCCGCACTCTCTCTCGCACTGCAGCAGCAAGACAGGCTCACCGGCAT

CGCAGTGGTCGCCGACATCGCTACATCACCACGCGGAAACTATTATTTCTGATGTTTCCAACAATTCCGG

CGATTTTCCACTGCCTCATTCACACCAGGCCTAGCCTTTCCTCCGCGCGCATCACCATGGCGATTATATA

CATACACATCTGTTCAATCTCTTCCATCTCTCTCTTGTTCATTTTCAGCGATTTTACTTTGTAATTAGTG

ATTTTTCAGATTTGGTGGTTTGTAGATGGTGGTGGTTGTGATTATTGTAGTGATTTATACTTAAATATGG

TGATTTTTATTATGATGTTGGTGGTCGGAGTTGATGAATGGAGCTAGTCGCCGGATCTGGTGAGCATGAA

TAG

>MSTRG.588.1 gene=MSTRG.588

ATTTTTTTGAAATCGTCGGTGGGTTATTTTTTCCCGTCATTTCAGGAACTGTTAACTCCACGACTTTCCA

ATCCTGTGAGGCTGCTGGATTAATCATACTGCAGTTCTTCTCTATACGAATATAGATCATATTGGTGGCC

ATGGAAAATGGACATCATATCACCGTCACTGTTTGTATCGTTCCATCTATGGACCACGGAGTGCATTCTA

AGTGTTAGGAATTCTACTACTGCGCCCATTATTTCGTTGGATTCGCTTTCTGCTGATTCACAGGATTCAG

CATTTAATAGTTTTCATGGGAGAAAAAATTTCAACACCCAGGTTTAAGTCTGGTGTCACACAAACCATCA

GGTGCGATTGATAATGCTTAATTTCTCTGCAACACGTTCAAGTCAATTTTGTTGTAATATTTTACGCCAT

TGCAAAGGAGGCTCCACGGAGGTGATTGTCTGTTATATTTTAAAAACATCTAATGCATACATGATCCTTA

TATATACAAGCTCTACATGTGTTGCTTTTGTCTTCACAGCTACAAGGTCACCTTTTTCAGTTGTAAGGGA

CACAGTGAGGAAGACTACATCCCTTGCAAAGTTGCCTCCATTTCAATTACAATCCTCTACAATCTCTTCC

GGTTTTAAGTCTCATTCATCCGGCGAGTGTGATATTTTAAAGGACCCTATTTTAGGACATTTATATTTTT

ACTGTTTTGTTGATCATTTTTTTGCACATATTGTTTCAATTCATATTATTTAATATTAGGTATCATTGGC

CTATTGTGAGCCACGTATAAATTAGTTGTTAATGAAGAAAAGATTTTCATAAAGAAACTCTATTTTGTTG

CTGATGTGTGTAAAAGCAAATAAGTTAATGCGGGGTTCTAGGATTTCTATTGGTCTATTTTTTTAAACGC

AGATTCTTTGCTTTCTACAGTGAAACATATAAGTCCTGAC

>MSTRG.593.1 gene=MSTRG.593

CATCATCCGCCTCTTGTGCATCTCTTTCTCTGTATAAATATCTCTTTCTTTCCCATCATTTAATTCAAGA

TCTGAATCACAGACACCCCTTTTCACATTCTCTCTGGTTTCCACTCAGATATGTGCTTTTGCAGCTAGTT

CTTGCACTTAGATCCACTAAACCCACTTGAAAGGTACAAGCTTTTTCAATACCCAGATCATAATTTGTT

>MSTRG.595.1 gene=MSTRG.595

CTCCAATCTTTTTGGCCATTGTTCTGGCCACTGGCTACCATGTTCTGACACTTTTTCTGGTACTTTGTGT

ATGCATGCTTGAAGCTGCACATTCCAGACTGCATCTGGCTCGTCAGCTTTGTTGCAGAGTGGGGGTTCAT

TTTGTGATCTTTTCTCATAGCACTCATTCGATGATGGTTTTCGGTATATTGCTGCTCCTACTTCATTCAA

GTCATCACGGCTAATTTTTACTAGTTCCCAGCACATGGACTTCGTTAGTTCAGACATAGCTTTCCATATG

T

>MSTRG.601.1 gene=MSTRG.601

TCATGAAATAGTCCAAACTCCTTCCTTGCACATCCAGCCATTCAGTACAGCAATCATTGGGCAATACTAC

TATAGCAATCATTTGTATGATTACATTGATGATCGGTTGTTGCAGTTAATCAGGCATGTCAAACTTGGAT

TAATCTATTGTCCCATTCTGGCAGCATACTCCTTAATGCATGTAGTCAGCTCCGAGGCTCTCATTGCCTC

GGAGCTTCAAGTAACCACTATCAAGTTTCCAGTATTACTTGACCACATACATCTACTACTAGTCAGGGTT

GGAACTGTCATGTTTTTGTGAGTTTCAAAGGCAAGGATACTCGCAACCCCTTTACAGGCCATCTCATTGC

AGACAGACGAGGATGAATTACCTAGAGGTGAAGAAATATCACCACGGCTATGGAAAGCAATGCACGGATC

AAGTATTTGTATCTGTATACTTGTTTTCTCAAACAACTACGCTTCATCAAGCTTGTGGAAATTCTTGATT

GCCAGAAACATTGTGTGTCCTATATCCTATCATGTGAATCCTTTGCATTTACGTTATCAGAGGGGAAGTT

TCGCAGATATTTTTGCAAGGTGATATTCATATCGACAATGGAGAAGCAAAAGATAAAAACTTGGCAATTT

GCACTATCTGAAGCAGCTGATCTATCAGGATTCAAGTTTTTGACATCTATTTATCAATCTTAATTACTTC

AAATGCTTCACCAGAATTATTTTCT

>MSTRG.604.1 gene=MSTRG.604

CTTCATTTCCACTTAAATTGGTTGGTTGTACTTAAGTGATTAGTGTCTTTTGATGTGTTTTTAGTGTTTT

TCTGTGCAGGTCACGAGTTGAGGTGATGAAGTGATTTTCTATCATTTTAGGTTGTTTTTGGTGCATTATT

TGCAAGAATAGAGAATTGTGGATTCATCGCGACTTGGGATTTTGTGGGTATATCTTCTACAAACCCTCAC

GAGAGCACACTCGTCCACTAGAGCTATCTAGGGGTTTAAAGGGCTTGTTGCATGTGCTAAATGCAACCGT

GATCACCTACGGAAGTGGTACTAGAGCGAGGAAGGACATGCACGCGAGAACGAGCTGAAAACGAAGAATC

AGGGCTGCCAGGGCAGACAGTAGCGCGCCCGCGCTAACCTCTAGCGCGCCCGCGCTACCTACTGCTGCCA

CTAGCGCGCCTGCGCTAGTTTAGCGCGGTCGCGCCCAGCAGAAGTGCCCCGGGCCGATTTTTACAGCTTT

TCTGACGGATTTTAGCAGCGGTCGAGGCCCGGTTGACTTGGTCAATTTATTTTAATTTCTCGTGTGTAAA

ACCCTAGCCTCCAAG

>MSTRG.605.1 gene=MSTRG.605

AAAATAAAATAGAAAATGCTGCATTTTATTTAAATTTAAGCTTGGTTATGGTATAAACTTGCTGGATGTT

TCTGACTATATATATGTGTTTTTGGCAGTATGCATGCATCAGCAACCATGAGCTTAAGGACAGCCTTGTT

GTTACTAGCCTTGATGGTTTCAGTAACTTCATCTGTTCCAAAAGATAGAGGTGCATACATAGTTCACATG

TACAAGTCGAAGATCAAAAGTCTGCGGACAATAAAAAAATGGCATGAATATGTCATGGACTCTATCCATA

GATTGTCAATTCAAGAAAATAATGAAGAAGAAGCATATCCACCCAATCTTCTATACGTTTACGGAACAAC

CATATTTGCTTTTTCTGCAGCACTCACTAAATCAAACCTAAGATCATTAGACAAAATCAACGGTTTTCTT

TATGCCATTCCTGATGAAATGCTAAGCATCCACACCACTCATTCACCAGGTTTTCTCAGCCTAAATCCTG

GCAGGGGACTCTGACATCCCAAGAACTTGACTTCTGATGTCATAATCGGTATTCTAGACACTGAAATATG

GCCAGGACACATCAGTTTCTTCGACACTGGCATGCCTCCAGTGCCCTACAATTGGAAAGCCAAATGCGAA

GAGGGCACAAACTTTACAATTTCAGATTGTAACAGAAAGCTAATCGGAGCAAGAGCTTTCTTCAAGGGTT

ATGAATCTGTTAGAGGAACAGATTAATCAAACAGAGGATTGCAGATCTCCCCTTGATGCAGAAGGCCATG

GTACACATGCTGCATCAACTGCAGCTGGTAATCTTGTACCAGGTGCAAGCTTTCTTGGTAATGCTAAAGG

CGCAGCTACTGGAATGATGTTCACAGCAAGAATTGCAGCATATAAAGTTTGTTATGCTTTCGGTGGTTCA

AGCTTAGATAATTTAGCTGCCATGATACAGCTATTAGTGATGGGGTCAATATATTGTCACTCTCTTTGGG

TGGCATTGCAAAGCCTTATTATGAACATAGCATAGCTATTGCATCACTTGGTGCATTTCAACAAGGGGTC

TTTGTTTTATGCTCAGCTGGACAGGAGCTTTCCAACCACATTTAAACTCAGGAATGGTCAACTATTTAAA

GGGCCTTCATTGTGTTCCGGGAAAAATATTAAAAGCTTGCAACCTGTGTATGGTGCAGCTGCAGGTATAC

AAGGGGCTCAGTACTGTTCTAATGGCTCTCTTACACCGGAGCTTGTGAAGGGAAAGATTGTAGTTTGTCA

AAGAGGAGGGAGTAGCCGAGCCAAAAAAGCAGAACAGGTACTGTTTGCAGGGGGTGCTGCAATGCTGCTT

GTAAACACTGATAATGAAGGTGAGGACATCTTTGCTGATCCACATATAATCCCTGCAACACCTTTGGGAG

CTTTAGCTGCAAGTGCTTTAGATGCTCTATGTCGGTCAAGGGGCTTATTATCTTTTTTTGATTTGTTTGA

ATATGAGGATTTTATGAAAGCGGATTTATGGACTCTACCCCACTTAACAACGATCAATATTGAATATCGT

TCTTATAAAAGACTCAAGGTTCACAAATCATGTTGGAGGCTGCCTGTCTTGACAACTCTATGGTCTTTCC

TCAGAATATGATTGACTACTTTCTCGCACTTACCGGTCTAAGGAATCTTGTCATTGATTTTTGCCAACAC

TATATCAGACGTCTTGTCATTTGGTGAACCTAAAAATTAAGGTCAGCTTATCTTCTGTACATATACTTAG

GCTCAAAAATTGTGGTATGGCCACAAAAACTGTGTAATTTTCAAGCTATTGGTTTCTATCGGGTGTCTTA

TGAAGGTTCTAAGTTGGAGATTGCAGCTATTAAATTTTGGGATCTTCTATTAAGCTCGACGGTCTGGAGA

CACTATCAAGATGTGATTGAAACTATGTTCTCCCAGCTTGGCAGCGCCAAGATTCTTAGTATTGACTTGG

TGACCATTCAGGATAAAATATCTGGAAAATTTGAATATAAAATCCCATCAAAATTAGTCAAGAATGCAAC

AAAATGTTGAGATGAATGCTGAGGTGCTTTCTGAGTATCGGGCTCTTTTTACTGACTTGCAGGGATGGGG

ATTCAAGTTAAATTGGCTGATAAACCATCTCACCTTTATTGAACCGCTCTTGTTTTCAAAGACTAAACTT

AACGAACTTCATGCAATTGACTTTCACATTAATGATGCCAAACGTAAATTACACGATTTGCAAACTCTTC

ATGTCGAGAAAATGACACATTCAGAAAGCTCTCCAAGTAAGAGTACTAGCCTTGCGGTTATTTCTGGCTA

CCTTGGAGATGTTATACTGTAAGAACAAAAAAAAAACTTGACAGAATGTTACTATATATCTGTGTTTGAT

CCTCGGTAATTTCTATGTAATTTATGATATTTATTCAAGAATTTCAAGAACACTTTTCCTTTGGAGAGGG

TAATAGTTGATTTAAGTTATGCCTCCAGTCC

>MSTRG.605.2 gene=MSTRG.605

ATAAACTTGCTGGATGTTTCTGACTATATATATGTGTTTTTGGCAGTATGCATGCATCAGCAACCATGAG

CTTAAGGACAGCCTTGTTGTTACTAGCCTTGATGGTTTCAGTAACTTCATCTGTTCCAAAAGATAGAGGT

GCATACATAGTTCACATGTACAAGTCGAAGATCAAAAGTCTGCGGACAATAAAAAAATGGCATGAATATG

TCATGGACTCTATCCATAGATTGTCAATTCAAGAAAATAATGAAGAAGAAGCATATCCACCCAATCTTCT

ATACGTTTACGGAACAACCATATTTGCTTTTTCTGCAGCACTCACTAAATCAAACCTAAGATCATTAGAC

AAAATCAACGGTTTTCTTTATGCCATTCCTGATGAAATGCTAAGCATCCACACCACTCATTCACCAGGTT

TTCTCAGCCTAAATCCTGGCAGGGGACTCTGACATCCCAAGAACTTGACTTCTGATGTCATAATCGGTAT

TCTAGACACTGAAATATGGCCAGGACACATCAGTTTCTTCGACACTGGCATGCCTCCAGTGCCCTACAAT

TGGAAAGCCAAATGCGAAGAGGGCACAAACTTTACAATTTCAGATTGTAACAGAAAGCTAATCGGAGCAA

GAGCTTTCTTCAAGGGTTATGAATCTGTTAGAGGAACAGATTAATCAAACAGAGGATTGCAGATCTCCCC

TTGATGCAGAAGGCCATGGTACACATGCTGCATCAACTGCAGCTGGTAATCTTGTACCAGGTGCAAGCTT

TCTTGGTAATGCTAAAGGCGCAGCTACTGGAATGATGTTCACAGCAAGAATTGCAGCATATAAAGTTTGT

TATGCTTTCGGTGGTTCAAGCTTAGATAATTTAGCTGCCATGATACAGCTATTAGTGATGGGGTCAATAT

ATTGTCACTCTCTTTGGGTGGCATTGCAAAGCCTTATTATGAACATAGCATAGCTATTGCATCACTTGGT

GCATTTCAACAAGGGGTCTTTGTTTTATGCTCAGCTGGACAGGAGCTTTCCAACCACATTTAAACTCAGG

AATGGTCAACTATTTAAAGGGCCTTCATTGTGTTCCGGGAAAAATATTAAAAGCTTGCAACCTGTGTATG

GTGCAGCTGCAGGTATACAAGGGGCTCAGTACTGTTCTAATGGCTCTCTTACACCGGAGCTTGTGAAGGG

AAAGATTGTAGTTTGTCAAAGAGGAGGGAGTAGCCGAGCCAAAAAAGCAGAACAGGTACTGTTTGCAGGG

GGTGCTGCAATGCTGCTTGTAAACACTGATAATGAAGGTGAGGACATCTTTGCTGATCCACATATAATCC

CTGCAACACCTTTGGGAGCTTTAGCTGCAAGTGCTTTAGATGCTCTATGTCGGTCAAGGGGCTTATTATC

TTTTTTTGATTTGGTATCGACACTATTTGTTTGAATATGAGGATTTTATGAAAGCGGATTTATGGACTCT

ACCCCACTTAACAACGATCAATATTGAATATCGTTCTTATAAAAGACTCAAGGTTCACAAATCATGTTGG

AGGCTGCCTGTCTTGACAACTCTATGGTCTTTCCTCAGAATATGATTGACTACTTTCTCGCACTTACCGG

TCTAAGGAATCTTGTCATTGATTTTTGCCAACACTATATCAGACGTCTTGTCATTTGGTGAACCTAAAAA

TTAAGGTCAGCTTATCTTCTGTACATATACTTAGGCTCAAAAATTGTGGTATGGCCACAAAAACTGTGTA

ATTTTCAAGCTATTGGTTTCTATCGGGTGTCTTATGAAGGTTCTAAGTTGGAGATTGCAGCTATTAAATT

TTGGGATCTTCTATTAAGCTCGACGGTCTGGAGACACTATCAAGATGTGATTGAAACTATGTTCTCCCAG

CTTGGCAGCGCCAAGATTCTTAGTATTGACTTGGTGACCATTCAGGATAAAATATCTGGAAAATTTGAAT

ATAAAATCCCATCAAAATTAGTCAAGAATGCAACAAAATGTTGAGATGAATGCTGAGGTGCTTTCTGAGT

ATCGGGCTCTTTTTACTGACTTGCAGGGATGGGGATTCAAGTTAAATTGGCTGATAAACCATCTCACCTT

TATTGAACCGCTCTTGTTTTCAAAGACTAAACTTAACGAACTTCATGCAATTGACTTTCACATTAATGAT

GCCAAACGTAAATTACACGATTTGCAAACTCTTCATGTCGAGAAAATGACACATTCAGAAAGCTCTCCAA

GTAAGAGTACTAGCCTTGCGGTTATTTCTGGCTACCTTGGAGATGTTATACTGTAAGAACAAAAAAAAAA

CTTGACAGAATGTTACTATATATCTGTGTTTGATCCTCGGTAATTTCTATGTAATTTATGATATTTATTC

AAGAATTTCAAGAACACTTTTCCTTTGGAGAGGGTAATAGTTGATTTAAGTTATGCCTCCAGTCCAACAA

GTTATCAGTGGC

>MSTRG.606.1 gene=MSTRG.606

TTGAATGGTGTTGTTCATTGCTTGAGGACTTCAAGAAAGCTTGTTGTTTTCAAGGGAGTCGAGTAATTAA

AACTGAAGATCGATGGTCTCCTCCAAATAATGGTAGCATAATGTTGAATGTTGATGCTAGTTTCTTAAGT

TGTAACAATTCTGGAGGAGTTGGCGTGGTGGCTCGTGATCATGATGGTAAGGTTCTCGATGCATCCACTA

AGTTTATGTCATTTGTACCTAATGTGTTGTTGGCTGAACTAAATGCAATCAAGGAAGGGGTCATCTTGGC

TACTAGTAAAGGCTGGAACAAATTTTGTATAGTGTCTGATAGCCAAAATGCAATAGCTGCACTACACAAC

TTTCCGAGGTATTGCAGTGACTTAGATCCCCTCTTGTCAAGTATTATCAATCTTCTGCCTAGTATAGGTT

TTGAGGGTTTTATATTTAAGCCGAGAAACACTAATGGTGTGGCTCATAGTTTAGCCAATTATGCTTTGCA

TGCCAAAGTTAGTGCTCTTTGGGGTGGGGTTCTTCCCTTAGTTGCCGGTCAAGCTTTAGCACAAGACTGG

CCTACCTCTTTGTAACTAATATTTGTTTAATGAAGTTAATTTCCTTTCAAAAAAAAAAAAA

>MSTRG.607.1 gene=MSTRG.607

AGTCCTGTAAGTGAGTGAGGTTTCTCTCTAAGTTTCTAGACTTTTCAACTTCTATATTTCTTTCTCTCTC

TAAAACCCTTTCTCTCTCTAATAGTTTTCTGGGTTGCATGGCTGATTTAATAGAGGAGATAGTCTTTAAG

ACTAATAATCTTGTAGTTGATAATGAAGAAGAGTGGGTGATCAATGATGATATTGAAGATCCAAGCGAAA

AAGCTCTCGTGGGAAGAATTTTAGCGAAGAAACAATGTAATAGGAAGTTCATTCACCAAGTATTTGAAGG

GATGTGGAGAACTAAGGGAAAATGGGAGGTTAAAATACTAAAGCATGACAATAGAAGTACTTATGTTGGA

TTTACTTTTCAAGAAATTGAAGATATGAAATGGGTAAAAGAAAAAATGCCATGGAACTTTGGTGGGGGAC

TGTTGATACTAGAAGAGTGGCCAAAAAGCGGTAAGTGGCAAGATGCTAAACTAGATGGTGTTCCATGTTG

GATCAGAATGCGAGGCTTTCCTTGGAAAGTATTAAATCTCACTAATGCAAGGAAAATGGGCGGTTTAGCT

GGGGAGATTCTAGATCTGAGGTGGAAAGATGGTGCTAGAAATCTTCTACACAATTATGCAAGGGTAAGAA

TCAAATTTCCAATAAACAAATCTATTTTTGTGGGAAAATATATCTTGGTGGAAGGCAAAAAAGAATGGGT

GCAATTTAAATTTGAGAGGTTGCCCCTTCTTTGTTTTAAGTGCGGAATATGGGGCCATGATAGATCTGAA

TGTAATGAGCCATCAGTTAAGGTGGAAAACATTGATGGAAAATTATATGATTTATATGGTGGATGGCTAA

AGGAAGCTGATAGTAGAGAAAATTGTTTTTCAATCAATAGTAGTGCAGAGGAAGATGGGAGCAAGGAACA

AGTTCTTGAAACAAGGGGAATGTTGAATGCTAATAAGGGAGATGTAGTGAGAAAAGAGAACCAGATAAAA

GAAAAGTCCAAGGGAAATTTAGTGTTTACCGGAGAATCGCAGCTAACTGAGGAAAATACAAAAAATCAGA

TTGTCAAAACAAAGGATAACAACATGCGTGTGATGGATGCAGAGATGATAGCTGGTTCTGTTCAATCAGG

GGGATTGGATAAAATATTTAATTGTCAAGAGAGAGAGTCATTTGGGGAAGAAGAGGTCCTAAATAATGTA

GCAGAACAAAATAGAGAAAACATTATGCCGGAGAAAGTGGAAACTTCATCCACTTCTAAAGGTAGTGAAG

GAAAAAAAAGAAAAGGTGTAGTCAATACGGAAGAAAGTAAAGACCAGCCAAACAAAGAAGATCATGAATT

AAAGAAAAAGATTAAGAGCAAGTTTGAAGATGCCGGGCAAAGTAACAAGGAGGACCAGAATAAAAGTGAA

GAGCATGCATTTGTCATTGGTAAGCATGATCCAATTTTGCAGCAGAAATGTAATATTTCGATTAAGAAAA

TGGCTAGAGGAAGAATGCAGAGTCTTGCAAGAAAAGAAAGAGATGGCTCGAAACATCAACATGCATCTGA

AGGTAATAAGCAAATGGAATTAGTAATATACCCAGTTGATGATAATTCTTATATAGGAGTGGACTTTTCT

AAAGAATTAGATGGTTCTTCTCCTTTGTCAGAAATTGTATCGGCGTCCCTTGCTAGTCAAGGGCGCCGGG

AGCCATGAATTGCTTATGTTGGAATGTCCAGGGGGTTGGGAGAACCTGGACATTCAATATGTTAAAATCC

CATATTAAGAAATTTAATCCTGATTTGGTTTTTTTATCGGAGACGAAGGCATTGTATCATCAGGCTGAAG

TTGTAGCTCGTAATTTAAATTTTAATAATTTCTGGGTTGTTGAAAGAAAGGGACTTGGAGGAGGTTTGCT

TTTGATGTGGTCTGATAAATTGGCTGTAAAGGTCCTTAACTGGAATTTAGGGAACATAACAGCAGTGGTA

GCTGGTAAAGGATTCAGACCATGGGTCTTTACTGGTTTTTATGGGAATCCAGATCCCAAACAAAGAAAAC

ACTCATGGGAGCTACTTTGCAGAATTAGAAGAAGCATTAATGGTGCTTGGTTGGTTGTAGGGGATTTCAA

TGAGATTGCAAGTGCAAATGAGAAATATGGAGGGTGTGATAGAAATCCTGCAGCAATGCGTAGATTTAGG

GAAGCTATGGATAATTGCAAGCTCATTGATTTTTCGAATATCAAGAGTGAATTTACTTGGTACAAAGGGC

ATGGTCGCCATTTGGTAATGGAAAGACTTGATAGGTGCCTCTGCGATGATGAATGGTTAAATTTATTCCC

AGAGGTATCTACTTCCTTGTTGGAGTGGGGAGGATCAGATCATAGACCTATGCTAATTAATCTCTTCTCT

CAAAAGCAGAATGTCAAAATTGGGGCTAAAAAAAGAAACACCAGGTTTCATTTTGAGGAGGCATGGTGTG

ATGAGCCTGAATGCAAAAATATTGTTGAAAATCTTTGGGAGAATAGTGGCCATTGTAGAAATAGTTGGGA

GTTAAAGAAAAAGATCAGGAAGTGCGGAGATTCTCTATCTGATTGGAACAAAAGACAAAGAGCAGAATTT

GGAAGAGAGCTGAAAGAAGCTAGGCAAAGGCTTGCTGATTTATCAAAGGAAAATGATCCAATGTTATGGG

AATGTATAAAGAAAGAAGAAGACAGAGTGAAGGGCCTTATGCAGAAGGATGAGATATATTGGAGGCAAAG

AAGCCGCTCACTATGGCTTAAGTGGGGTGATAAAAATACAAAATATTTTCACCGTAAAGCAACAACGAGA

AGAAAGAAGAATGAGGTGAGAGGGATTACGGATGAAGCTGGAATTTGGCAAGAGGATGAGGAAGTGGTTC

ATCAGTGCTTTTATAAGTATTTCTCTAAATTATTCACATCTTCGCACCCAGATATCAACTCTATTGATGA

GGCTATTGATGAGTTAGAGGAAAAAATTTCAAGAGAAGATAATATGAAGCTGCTAAGTCCTTTCACTAGT

GATGATGTGCTAAATGCGGTAAAAGATATGAACCCAACCAAAGCTCCAGGTGAAGATGGGTTGCCAGCGT

TGTTTTATCAGAAATTTTGGACCTCCCTGAAGAATGAGATTTGGAATGTCAGAAATAGCTGGGTTCATGG

AAAGGTACTGCCTATGGGATCTTCATTGGTTGAATGGTGTTGTTCATTGCTTGAGGACTTCAAGAAAGCT

TGTTGTTTTCAAGGGAGTCGAGTAATTAAAACTGAAGAACGATGGTCTCCTCCAAATAATGGTAGCATAA

TGTTGAATGTTGATGCTAGTTTCTTAAGTTGTAACAATTCTGGAGGAGTTGGCGTGGTGGCTCGTGATCA

TGATGGTAAGGTTCTCGATGCATCCACTAAGTTTATGTCATTTGTACCTAATGTGTTGTTGGCTGAACTA

AATGCAATCAAGGAAGGGGTCATCTTGGCTACTAGTAAAGGCTGGAACAAATTTTGTATAGTGTCTGATA

GCCAAAATGCAATAGCTGCACTACACAACTTTCCGAGGTATTGCAGTGACTTAGATCCCCTCTTGTCAAG

TATTATCAATCTTCTGCCTAGTATAGGTTTTGAGGGTTTTATATTTAAGCCGAGAAACACTAATGGTGTG

GCTCATAGTTTAGCCAATTATGCTTTGCATGCCAAAGTTAGTGCTCTTTGGGGTGGGGTTCTTCCCTTAG

TTGCCGGTCAAGCTTTAGCACAAGACTGGCCTACCTCTTTGTAACTAATATTTGTTTAATGAAGTTAATT

TCCTTTCAAAAAAAAAAAAA

>MSTRG.609.1 gene=MSTRG.609

GAGAAACCGAGCTGCGTTCTTCGTGTTCTTCGTAATCAAACGAGGATTTGAAGGTGTTATTGGAATCCGA

TGGAGGTGTATGAAGAGTCAAAACGAAGCTATCGACGCGTAGAATCAAGTTTAATCATCCGTTTATATGC

AGATTCAACAGGATTTGGACGAGACGTGTACGCTTGAAGACGTCAGAAAGTCGGAATGGTATTGCAAAGC

GAATAAGGGATGAATCGAGTAAGCGAAGCGATGTGGCGAATAGAGTATAGCTAGAGATGGTCTAGAGATT

ACGAGATGCATAGATTGGGAAAGTGAAGAGATGAGATAGATGTGAGA

>MSTRG.612.3 gene=MSTRG.612

TGAATATCCGCTAAACATAATAATAATATTCTTTTTAACACGTTAGCTTTTGAAACTGCTGAATATGATC

CTGCTATTTATCGGCGACGGTGCCGTGGAAAACTCGTTCTTGGGAAGCCTCGGACCTAGGCGCTCTCTTC

TGGCTTCTGAGCCTCCTTTTGCTGATATCGACTGAGGATATGCTGCTGCATAGGTAAATACTAGTCACCT

AGGGCTTTGAAAGGAAGGAAAACGGGTCCTGGGAAATTTACTCATATCCAAGTTGAATTTGACAGAAGTG

GTCTCACTGCATGGTAGGAGAGACATTGTATGAGTTGACAGTTAACCGCGGAACTGAGGCCTGATGTATC

ATTCAATTTTATGGTCACATGATTAGATTTTTGCTTCTTCTTTTCCCTTAGCCAAGCTGATTACTTAATT

TTACGACTATGTATGTGAAACATCTCTAAGAGTTTTCACAGATGCATCCTTTTAACATGTCTT

>MSTRG.615.1 gene=MSTRG.615

CTCCGTTTACTCTCTCTTTTCCCTTCCTCTCTATGTATCCCTTTTTCTCTCTCTCTACTCTATTTCCATC

TCATTGCTCTCGTTTCTCTCGCTCATTGCTCTCTTCTTCGTTCTGTCTCGTGTTCTCTCCTGCCCCCTTT

TCTTTTCTCTGATTTTCAATCTCTCCTCCTCTTACTCTCTTTATCTTTTTAAAAACAAATATAAATTTAC

ACAATCAAAACAAGAAAAACTGAGATAACATCATAAGAGACAAATGTTGCTGACTTGAATAAAACGAAGG

AGTTCGTCGATAATGGGGATCGGAGCTTCATCACCGGCGAAGAGCGGCCGCGATTGTATCGCTAGCCGGC

GGACAGGATGAAGCATGGCGACAACAAGAATGACGAGAAGAGAATGTTGATGTTGAGCCATTTCATGTTG

TAGTAGAGATTCTTTAAATTTGATTTTTAGATTTGCTTTTGGTTGGTTGCTTTTGAGTTGATTTCTAGTT

TGTTATTTTATTTACTTTTTAGGTTTCAATGTTGATATTAGTATAGACCCGGAGGGGTTTTCATTTTTTT

CATTTGCAACCAGTTGTAACTTATTATACAACTAAATGTAATTTTCAAAAAAAA

>MSTRG.616.2 gene=MSTRG.616

ATATAGATCAATCATTGCATATACATACACACCGTCGATACATGCCTACATATATAATCTTCACGACACA

CACACAATACGGTTGCGCCAAAAGCGGTTAAATCGAAATGAACGTCGGAAATATCGCCTCCAGCCCGGCC

GACCAATCGCCGGAAAAAGGCAAACCCATCGGAGTATCCTCCACTCCGGCGAACAAATCGCCGGGAAGTG

ACGCGATCAAACTACCGCCTTCGCTTGCCGCGAAGAATGACGTTCCGATAAAATCGCCGTTGCCGCGGTT

GCAGACGCAGTCGGAGGCGACGGCGTCGGCGGGAGAAGATGCTCCGGCGCTTCCGCCGCATATAAGGAAT

GCTATGTTCCCGACTGCTGATCCTGGATCCGTTGTTAGCGGGGGAGGCATAAGTTTTCTCACTGGAAACC

GAAACGCAAAGTTCAGCTATGGTTATTCTAGTTTCAAGGGTAAAAGATCTTCAATGGAGGATTTCTATGA

AGCAAGTATATCTGAAGTTGATGGTCAGATGGTTGCCTGCTTTGGCGTCTTTGATGGTCATGGTGGCTCC

AGAACTGCAGAATACCTGAAGAACAATCTTTTCAAAAATCTAAGTAGCCATCCAGATTTTATTAAAGACA

CCAAGTCAGCTATTGGTTTAACGATCTGGATTGCAGTTGATTCATTTCGCCGGACAGATGCGGATTACCT

CAATGAAGAAAAAACCCAACAAAAAGATGCAGGATCAACTGCGGCAGCTGCTGTATTGGTGGGGGATCGG

TTGCTTGTAGCTAATGTGGGAGATTCTAGAGTTGTTGCATCTAGAGCCGGCTCAGCTATACCTCTATCCG

TTGATCACAAGCCCGATAGATCTGATGAACGTGAGAGGATTGAACAGGCTGGGGGTTTCATCATTTGGGC

AGGAACTTGGAGGGTTGGCGGTGTTCTTGCTGTTTCCCGTGCATTTGGAGATAAACTACTAAAGCCATAT

GTGGTGGCTGATCCAGAAATTCAGGAAGAAGAAATTGAGGGTTTTGATTTTTTAATCATTGCAAGTGATG

GACTTTGGAATGTCTTTTCGAATAAGGATGCTGTGGCTATGGTGCAGGATATTTCTGATGCAGAAGCAGC

ATCTAGAAAACTCATAGAAGAAGCTTATGCAAAAGGGAGCTCGGACAATATAACGTGTATGGTTGTCCGA

TTTGATGGCCAGTGAGAGATGCATAAAGGTCTTGTATTGCCCACCTTGTATCTTCTTGCATTTGAGATAA

ATTACTCTTCAAACTACAAATACCAAGGATCTTGCAGGCTTGAAGAACTGGAGTCTTAAAATGGAATCTT

GTGATTATTGTAATGAAGAAGCTTCAGTACATTCCGTGTTTTTTAGGCTAACTAGTAATCAACTTCTTGG

TGTTTGTATTATCACATGTCACATTTTTTATTGTTTCTTAATAGATGTATCCACACAAATGCTAAATTAT

GTTTCTGTAGTCTATATGTATCGGACTTCTCAAAAAGGTACAAGTATGAGTTGAGGATGAGAGCAAGTTT

GGAACTTGTGGGCCAGTGTCACTGCTGGCATGTTGTGTTTGTTGTTTCTAAACACATCTGACTTGTAATA

ATGTGTGCATATTTATAAAGGCTACTTGATTTGAACAA

>MSTRG.616.1 gene=MSTRG.616

ATATAGATCAATCATTGCATATACATACACACCGTCGATACATGCCTACATATATAATCTTCACGACACA

CACACAATACGGTTGCGCCAAAAGCGGTTAAATCGAAATGAACGTCGGAAATATCGCCTCCAGCCCGGCC

GACCAATCGCCGGAAAAAGGCAAACCCATCGGAGTATCCTCCACTCCGGCGAACAAATCGCCGGGAAGTG

ACGCGATCAAACTACCGCCTTCGCTTGCCGCGAAGAATGACGTTCCGATAAAATCGCCGTTGCCGCGGTT

GCAGACGCAGTCGGAGGCGACGGCGTCGGCGGGAGAAGATGCTCCGGCGCTTCCGCCGCATATAAGGAAT

GCTATGTTCCCGACTGCTGATCCTGGATCCGTTGTTAGCGGGGGAGGCATAAGTTTTCTCACTGGAAACC

GAAACGCAAAGTTCAGCTATGGTTATTCTAGTTTCAAGGGTAAAAGATCTTCAATGGAGGATTTCTATGA

AGCAAGTATATCTGAAGTTGATGGTCAGATGGTTGCCTGCTTTGGCGTCTTTGATGGTCATGGTGGCTCC

AGAACTGCAGAATACCTGAAGAACAATCTTTTCAAAAATCTAAGTAGCCATCCAGATTTTATTAAAGACA

CCAAGTCAGCTATTGTTGATTCATTTCGCCGGACAGATGCGGATTACCTCAATGAAGAAAAAACCCAACA

AAAAGATGCAGGATCAACTGCGGCAGCTGCTGTATTGGTGGGGGATCGGTTGCTTGTAGCTAATGTGGGA

GATTCTAGAGTTGTTGCATCTAGAGCCGGCTCAGCTATACCTCTATCCGTTGATCACAAGCCCGATAGAT

CTGATGAACGTGAGAGGATTGAACAGGCTGGGGGTTTCATCATTTGGGCAGGAACTTGGAGGGTTGGCGG

TGTTCTTGCTGTTTCCCGTGCATTTGGAGATAAACTACTAAAGCCATATGTGGTGGCTGATCCAGAAATT

CAGGAAGAAGAAATTGAGGGTTTTGATTTTTTAATCATTGCAAGTGATGGACTTTGGAATGTCTTTTCGA

ATAAGGATGCTGTGGCTATGGTGCAGGATATTTCTGATGCAGAAGCAGCATCTAGAAAACTCATAGAAGA

AGCTTATGCAAAAGGGAGCTCGGACAATATAACGTGTATGGTTGTCCGATTTGATGGCCAGTGAGAGATG

CATAAAGGTCTTGTATTGCCCACCTTGTATCTTCTTGCATTTGAGATAAATTACTCTTCAAACTACAAAT

ACCAAGGATCTTGCAGGCTTGAAGAACTGGAGTCTTAAAATGGAATCTTGTGATTATTGTAATGAAGAAG

CTTCAGTACATTCCGTGTTTTTTAGGCTAACTAGTAATCAACTTCTTGGTGTTTGTATTATCACATGTCA

CATTTTTTATTGTTTCTTAATAGATGTATCCACACAAATGCTAAATTATGTTTCTGTAGTCTATATGTAT

CGGACTTCTCAAAAAGGTACAAGTATGAGTTGAGGATGAGAGCAAGTTTGGAACTTGTGGGCCAGTGTCA

CTGCTGGCATGTTGTGTTTGTTGTTTCTAAACACATCTGACTTGTAATAATGTGTGCATATTTATAAAGG

CTACTTGATTTGAACAAGGTTCGTCGTTACTTGGCTACTAGCTAAAAGTTTATGCGTGGTAGCATAG

>MSTRG.616.3 gene=MSTRG.616

ATTGCATATACATACACACCGTCGATACATGCCTACATATATAATCTTCACGACACACACACAATACGGT

TGCGCCAAAAGCGGTTAAATCGAAATGAACGTCGGAAATATCGCCTCCAGCCCGGCCGACCAATCGCCGG

AAAAAGGCAAACCCATCGGAGTATCCTCCACTCCGGCGAACAAATCGCCGGGAAGTGACGCGATCAAACT

ACCGCCTTCGCTTGCCGCGAAGAATGACGTTCCGATAAAATCGCCGTTGCCGCGGTTGCAGACGCAGTCG

GAGGCGACGGCGTCGGCGGGAGAAGATGCTCCGGCGCTTCCGCCGCATATAAGGAATGCTATGTTCCCGA

CTGCTGATCCTGGATCCGTTGTTAGCGGGGGAGGCATAAGTTTTCTCACTGGAAACCGAAACGCAAAGTT

CAGCTATGGTTATTCTAGTTTCAAGGGTAAAAGATCTTCAATGGAGGATTTCTATGAAGCAAGTATATCT

GAAGTTGATGGTCAGATGGTTGCCTGCTTTGGCGTCTTTGATGGTCATGGTGGCTCCAGAACTGCAGAAT

ACCTGAAGAACAATCTTTTCAAAAATCTAAGTAGCCATCCAGATTTTATTAAAGACACCAAGTCAGCTAT

TGTTGATTCATTTCGCCGGACAGATGCGGATTACCTCAATGAAGAAAAAACCCAACAAAAAGATGCAGGA

TCAACTGCGGCAGCTGCTGTATTGGTGGGGGATCGGTTGCTTGTAGCTAATGTGGGAGATTCTAGAGTTG

TTGCATCTAGAGCCGGCTCAGCTATACCTCTATCCGTTGATCACAAGCCCGATAGATCTGATGAACGTGA

GAGGATTGAACAGGCTGGGGGTTTCATCATTTGGGCAGGAACTTGGAGGGTTGGCGGTGTTCTTGCTGTT

TCCCGTGCATTTGGAGATAAACTACTAAAGCCATATGTGGTGGCTGATCCAGAAATTCAGGAAGAAGAAA

TTGAGGGTTTTGATTTTTTAATCATTGCAAGTGATGGACTTTGGAATGTCTTTTCGAATAAGGATGCTGT

GGCTATGGTGCAGGATATTTCTGATGCAGAAGCAGCATCTAGAAAACTCATAGAAGAAGCTTATGCAAAA

GGGAGCTCGGACAATATAACGTGTATGGTTGTCCGATTTGATGGCCAGTGAGAGATGCATAAAGGCTTGA

AGAACTGGAGTCTTAAAATGGAATCTTGTGATTATTGTAATGAAGAAGCTTCAGTACATTCCGTGTTTTT

TAGGCTAACTAGTAATCAACTTCTTGGTGTTTGTATTATCACATGTCACATTTTTTATTGTTTCTTAATA

GATGTATCCACACAAATGCTAAATTATGTTTCTGTAGTCTATATGTATCGGACTTCTCAAAAAGGTACAA

GTATGAGTTGAGGATGAGAGCAAGTTTGGAACTTGTGGGCCAGTGTCACTGCTGGCATGTTGTGTTTGTT

GTTTCTAAACACATCTGACTTGTAATAATGTGTGCATATTTATAAAGGCTACTTGATTT

>MSTRG.617.2 gene=MSTRG.617

GTTGTCATTCATGACTAAAACTAAAACAATAATTTGTGACAACAATCTCTGGATTCATTATATTACATAA

ACAAGTAACGTATCTAATCTGCACATATTCTTTATATTCTCTCCGATTATTCTCCCCTCGAAATTAATCT

CCGACTCATAAGTATAGCATGTATGATTCAAATCCGAATAGCACCTTCGATTTTCTCGGACAGACCAAAC

ACAATTGTCATGTTGGGATGGATCCTGGTGTCAGAGAATCAGAGAAGCCTAACAGATTAAGCAACATGAG

TGGTTCGGGTGTTTGTTCAGGGACGGATCTGGTAAGAGGCCCGGGGCATCAGAAACCGTCTTTCAACAAG

GGTAGGAATTCTCCTCCGTCTCTGAGGGTTCAGGCCATTGCAAGAGGCCAGAAGGAGCTTATGGAGATGG

TCAAGGCCATGCCCGAGTCGACGTACGAGCTCTCACTAAAAGACCTCGTTGATCATCACCAGAATGGAAA

TGAGTCGGTAGAAATTGATCAACAGAAGCAAGAAGAGGAAGTAAAAATCGATGAAGAAGAGAGTAATGGT

AATAAAAACAAGGGTGGGGTGAGAGTAAGACAAGGAAGTGTCAGATTAAGACAAGGGAGTGTGAGGAAGC

AAGCAAGTCTCAAGAAAGTGGAAAAGAAGATGGCAAGAAATGGGAGCAATATAAGTGAGAGTGGCAACAA

AGGCTTGTTTTTGAAGATGGTATTTCCCATGCCATTTGGAGGAGGGACCAATAAGAAGGTTGTGAAAGCT

AAGACTAGTAGTACTTCTAAGACAACAGCAAAGGAGAATCCTTATGCTAAAGTGTGTTCAAGTAGTGTTG

GGAATAAGGATCCTAGTATTGGTGATAAATCTTCCAAGAGTAGTAAGCATAAGGAGTGGTGGAAGAGGAG

GTATCCGGTGGCGTCGGACGGGAGTGAGAGCAGTGGATTGAGTAGCAATGGGAGCAGTGGGAGCAGTGGG

AGCTCGGCCAGCAATGACAGCAGTGGAAGCAGCATTAGCAACAAGAGCATCAGGTTTTTTCACTCAATTT

ACTCTGTTTTTCTTTTTCTTCTTCTTCTTTGGTGTCAAAGCTGCTAGAATTGTATTTTTCATGATTTTGA

TTCATTTTACAAGCTCTTAGATTGGTAATAGTCAGATGTTATTTAGAATTTAAGTTCTGAGATTTCACAA

AATACTGGAAAGTCCAAAGGGACGGAATCCTGATTTCTTAAAATTCTGATTTCGATTCTCCCTCGCATGC

TTTGCAATGTGAAACGGAAATCTATAGTCCACAGATATCACCCTCCGGAAGACAAGAGACGTTTAGCATG

CAAGATACTGTTAATAAATATATATTCAGATTTCAATATTGAAATGTTTTAGTTATGTAGTAATATTTTC

TCATAAAAATGACAGTGACTCGTATTCAAAACGGATGAGTTACAGATAATCATGAGTCAGAGATGCTGAG

CTTAAAAAAACAGATTTCTAGAAGGAATATCGAGTTTTAAAATTTAAAGCAAAAAAACGTTTATGTATAT

CTACACGCAAACGTTACAACCCATGCAACTTTAACACAG

>MSTRG.617.1 gene=MSTRG.617

GTTGTCATTCATGACTAAAACTAAAACAATAATTTGTGACAACAATCTCTGGATTCATTATATTACATAA

ACAAGTAACGTATCTAATCTGCACATATTCTTTATATTCTCTCCGATTATTCTCCCCTCGAAATTAATCT

CCGACTCATAAGTATAGCATGTATGATTCAAATCCGAATAGCACCTTCGATTTTCTCGGACAGACCAAAC

ACAATTGTCATGTTGGGATGGATCCTGGTGTCAGAGAATCAGAGAAGCCTAACAGATTAAGCAACATGAG

TGGTTCGGGTGTTTGTTCAGGGACGGATCTGGTAAGAGGCCCGGGGCATCAGAAACCGTCTTTCAACAAG

GGTAGGAATTCTCCTCCGTCTCTGAGGGTTCAGGCCATTGCAAGAGGCCAGAAGGAGCTTATGGAGATGG

TCAAGGCCATGCCCGAGTCGACGTACGAGCTCTCACTAAAAGACCTCGTTGATCATCACCAGAATGGAAA

TGAGTCGGTAGAAATTGATCAACAGAAGCAAGAAGAGGAAGTAAAAATCGATGAAGAAGAGAGTAATGGT

AATAAAAACAAGGGTGGGGTGAGAGTAAGACAAGGAAGTGTCAGATTAAGACAAGGGAGTGTGAGGAAGC

AAGCAAGTCTCAAGAAAGTGGAAAAGAAGATGGCAAGAAATGGGAGCAATATAAGTGAGAGTGGCAACAA

AGGCTTGTTTTTGAAGATGGTATTTCCCATGCCATTTGGAGGAGGGACCAATAAGAAGGTTGTGAAAGCT

AAGACTAGTAGTACTTCTAAGACAACAGCAAAGGAGAATCCTTATGCTAAAGTGTGTTCAAGTAGTGTTG

GGAATAAGGATCCTAGTATTGGTGATAAATCTTCCAAGAGTAGTAAGCATAAGGAGTGGTGGAAGAGGAG

GTATCCGGTGGCGTCGGACGGGAGTGAGAGCAGTGGATTGAGTAGCAATGGGAGCAGTGGGAGCAGTGGG

AGCTCGGCCAGCAATGACAGCAGTGGAAGCAGCATTAGCAACAAGAGCATCAGGAAAAAAGGTGGATTGC

TAAAGGGATGCGCACCCTTCTATTACAACAAGAACAAAACAATAGTGGAGTGAGTTGCGTGTTTGAAAGC

TGCCCAGCTCTGCAATTCATACTTTCTGTATACAGATTCAGCTTCATGTATAATGCCCCCATCTCAATTA

TATGATACATTTTTATACATGGTATTGATAGTTTCGATTTTGTGTAATTTTTGTTTTTGATCTTGTCATT

CTTTTTGTTGTGTTAGAAAATAAATTGATTTTGTTTTGTCTCTAAA

>MSTRG.621.1 gene=MSTRG.621

GTAATACTGAATGTTGTTCATATTGTCATCTCATTTAACTTTTGTTAATATTATATGGCCTTCTAGTCTT

CTGACAGCTTTCCATGGTATCGAGACTCAAGAATTATCATTTCACAGAGTCGATTGGTTATTGATTAGAG

AACAATAATAAAATTTTGGCATATCAATATGCCACTATAGGATCACTACATGACGTATTACACGGGAGGA

AGGGTGTACAAGGGGCTGAACCCGGTCCTGTTCATAGTTGGGTCGAAAGAGTTAAAATTGCTTATGGTGC

AGCGAGAGGCCTTGAGTTTCTACATGAAAAGGTTCAGCCTCCAATTGTACATCGTGATGTCAGATCCAGC

AACGTCCTTCTGTCTGATGATTTTGTACGTAGGATAATGTTGACTTCTCTATTCTGCACATAATAAAACC

ATCCCCTAATTTATTCATATGTGTTTTTGTAGAAGCCTATGATAG

>MSTRG.624.1 gene=MSTRG.624

AATCGCTTATTTATTTTATTTCAGGAAATTGTAGATAAATAAAGAAGGAAGAGAAAATGCAAGAAAAAGA

AGAAAAGGAAAAAGAAAAAGAAAAAAAAAAAGAGTTGGCAACTAAGGGACACATGGAGGGCCACAATCTT

CCCAACACACAAGGCACATGGATGGTCAAGATGGAGCCACCCTACCCCTAAACCCTAATTCTTTAGTTAT

AAATACCCCTCTTCTTTTACTTGTAATCACCAACCTACCAACCTAGTAACCTAGCCACCTACCCACCCTA

CCCCTAAACCCTAATTCTTTAGTTATAAATACCCCTCTTCTTTTACTTGTAATCACCAACCTACCAACCT

AGTAACCTAGCCACCTACCCACATTTTCTACCTAGTCATTTCTATAGTTAGTATCTTAGATAGATTTATA

TTTTGCTAGCCCC

>MSTRG.625.1 gene=MSTRG.625

GAAAAAAGCAATGGTAACTTTTTAGGTTTAATTGAAATGTTAGCTGAGTATGACCCAATTATCCAAGAAC

ATGTTAGTCGTATCACTAATGATGATATTTATATTCATTATCTAGTTAACTCAATTCAAAATGTGTTGAT

AATTTTGCTTGGTTCTGCAATTAAAAGTGAAATCATTAGAAGAGTAAAACGTGCAAAATATTTCTCTGTG

ATACTTGATTGTACACCCGATGTAAGTCACCAAGAGCAAATGTCTTTGATATTAAGATATGTGAATGTCT

CATCGAATCCTATTACTGTTGAAGAATCTTTTTTAGGATTTTTTGGATGTGAATGATACAACTGGACAAG

GACTTTTTGATGTTTTACAAAATGAATTAAAAATGCTTATACTTGATGTGTGAAGACAAGGTTATGATAA

TGGTTCAAATATGAAAGGAAAACATCAAGGAGTACAAAAAAAATTATTGGAAATTAATTCTACAGCTTTT

TATACTCCTTGAGGTTGTCATAGTCTAAACTTGACATTATGTGATACGGCTAATACATGTGGTAAAGCTA

AAGATATTTTTGGAGTTATAGAACGCATATACACAATTTTTTCATGTTCTACAAAAAGATGGTAAATTTT

GAAAGAAAATGTGGGATTGACTTTAAAGTCATTATCATCTACTCGTTGTGAGATTCGTGTTGAGAGTGTC

AAAGCTATAAGATTTCGAATGTTAGATATACGAGAAGCTTTGCTTCAAGTAGCGGATATTGATAATGATT

CAAAAGTGAAGAGTGAATCTAAATCGTTAGCAATGAATGAGCTTGGTGATTTTGAATTTTTAGTTGCAAT

TATTATTTGGTATGAAATATTACATGCTGTTAATATTGTGAGTAAATTTTTGCAATCAAAGGACATGCTT

GTTGATGTAGCTATTGAAAAAATAAAAGGGTTAGTATCATTTTTTGAGGACTATAGAGAAACTGGTTTTA

ACGATGCTTTAAATAGTGCAAAAGAACTTGCTACTGAAAGAATGTTGATCCTGTATTTCCCCAAAAGCGT

GAAATTCGAAGGAAAAGGTTCTTTGATGAAAATTTGGACTCATCATCAAGTGCTCCATTATCCGCAGAGG

AAAAATTTAGAGTTGATTATTTTCTTAACATAGTTGATCAAGCGCTTAGTTCTCTAAATAGAAGATTTGA

GGAATACAAAAATTATGAAAATATTTTTGGGTTCTTGTTTACTTATAAAAAGTTTAAGTCATTGGATGAT

AAGAGTTTGAAAAATTCATGTGTTCAAATAGAAAATGCACTAAAAAATGATGAGTTATCTGATATTGATG

GGAATGACTTATATATGGAATTGAAATTGCTTAGAGACTTTCTTCCTGCTGACATTGTAGGAGCTACTAA

TGTTTTAAAATATTTGAAAGATTTGATTGTTTCCCCAATGGGTTTATTGCTTATAGAATTTTATTGACTA

TTCTTGTGACTGTTGTATCTGCAGAGAGGAGTTTTTCAAAACTGAAGTTGCTGAAGACTTACATGCGCTC

CACAATGCTACAGGAAAGACTTAATGAATTAGCGATGATAGCAATTGAAAGTGATCTCTTACAATATGTA

GAATATGAATATTTAGTTGATGATTTTGCATCGAAAAGTTTTCGAAGGATGTCACTTTTCAAATAAAATA

TGTAAACTATTGTGAAGTTATGATCAATGTAAT

>MSTRG.628.1 gene=MSTRG.628

GATATCTGAGACGAGCATCCGTCCACTGAAAGAAGTTCATTAATATGTTCAAGCCTCAGTGAAGAATAAA

GTTCAATATGTTTCTGCTATCAAGATTGCCTGAAGATCAAGTATCAAAGACCAGAAGATTCCAGTATATT

TAATTTGATTATTTATAATCAAATTTATCGAAGCAACATCTAACCCGGAATGACTGATCAAGTATATTAT

CAAGCAAAGGATTCAAAGAATTATTTCAGTTCGAGTCGTTCGTGAACCAGACCAGTGCACAGGACGTCAG

GACATACGAAAGTATTCAGTGGATTCGATCAACGAATTAATTGATCAAGTCAATCGAGCTACTCCAAAAG

TTATTGTCAAAGAACTCATTGATATGTATATGTATATATATATATTAATTGTGAATTACCTGAATCTAAA

TAATTCAAGTAATTAAATTAACACAATTAATAATATATATATGTATATATATTAGTGCCAATGG

>MSTRG.630.1 gene=MSTRG.630

TGCAAGTCAAGTGGGTGGCGGTTATTGACGGTTTTCGGAAGTTTTATTGAAGGGAAACTGGCAAAGGTCA

AGGCTCCAATTGGCAAGCACAAGCAGGGCAGTTATGGGCAGTTGAGAGCTTCATGGGGCAGTTTTATAAC

TGCCCGGCAGTTAGAAAACTGCCAAGTTGTGGCTGATTCATATTAATGTTAGGCATTTCAGATGTAAACA

TAGAGAGAGAGAGAGAGAGAGTAGAGTTCTATATTGAGGGAATATAGACTAGAGATAGATATATTGTGGT

GTATATCTAGCCTAAGAGAAGAAATTTGGGAGAGTAAAGTGTGTAAACTCTTGCCATAACTCTTGGGTTT

TGTGGGTGTCCTTTGTAATTGTGTGATGCAATAATATAGGTTGGCTTCGGCTATAAATCTGTG

>MSTRG.634.1 gene=MSTRG.634

AAAAGTTACCTGCAGCATTGTATAGTGAGCAGCAGAGGTGATGGGCAGGGGACCAATTGTGGCAGAGTCC

ACTTCACCAGTCGGCAATGGCACTGCCGGGTTGCTGTTTATAAATGGTATGCCTCCACCGGGACTTGAGG

CCGGAAGTAGTGCAGTGGTAGGTGCAGGACCCGGGGATGTCGAATCGCGGGGACGAAAAGGCCATTGGTA

CGATGGGGAGGACGATACTGCCCCACCGGGACTCAGGGGAGGAGATGCCAAGGGTGAGGAGACTTGAGGC

TGGAAAATGGATACTTTGATGGCAAGTTTTTGGCCTTGTTCACATGGTTTTGTGGTGCCATTGTTGAAGG

AGAAATAGAAGTACCCGAGTCTAGATGGAT

>MSTRG.635.1 gene=MSTRG.635

AAGTAAAGTTTCAAGGGTTTGTCTAGCGTGTGCCCATGGGCACATGCTAAGCACTAAAATCTATGCATTT

GGAGGGTTTTGATTGGTGTGGTTGTTGTAAATTCAGGGGGGTCCACCATTATTAGAGAGTGTAAGCCAAT

CAAAAGCTTCCAAATTCATAGATTTTGGCACTTAATGTGTGCCCATGGGCACACCATAGAAAAACCGA

>MSTRG.636.1 gene=MSTRG.636

TCACTTTTGTTCTTTAATTCATCGAGGTTTAGGGCTTCTCTTAAGTGTCAATGGAACAAGGTAACGATTC

AAGTTCAACTCCGAGTGTATCATCAAGTTTGGTTGGTATGATATCAACGAAGTGTGGTGAAAATATCTCT

AAAAGCGCTGATCTGGATTGTGTAATAATTTCGCAAGAAGATAATCTTATTATTGGGCGTGATCAAGCTA

TTCAAGGAAAAGTGCAACCAATCAAAATATAGAAGACCTTGAAGTCATAAGTTTGGAATCATTTTGATCG

GATTGAAAGCGAAATAAAAAGTACAAAAAAATTTGATGAAATTTGTAAGTATTGTCACACTACATTTAAT

GGGGCAAGTGATCAGGGGACGACACATTTAAAAAATCACACAAAAACTTGGCGGATGAAATTAATAAAAG

TTCCTATATCACAAATGTTGTTGGGAAAGCAGGTTAGTGGTGAGGATTCTCGACCAAAAGTAGGAGTTCA

TAAGTTTGATAATGCTTCTAATAATTTAGGGATGCTCTTAGGTATAAAAATTTGTTCAAATGGAAAAAGA

GCACTTGTTCACAATGGTGACATGTTCCATATGCGTTGTGTGCCAAATATTCTTAATTTGATTGTTAAGA

GTGGGTTAGAGATGCTTGATCATTTAGTTGAAAAAATTCAGAAAACTGCAAAATATATAGGATATTCTTC

CCAAAGGGATGAGAAGTTTACCCCTGCTTTGTCTCAAACCAAACTTAATACTAAGAGACGAATCCCTTTT

AACCAATCAATACGACCTCGTAGAGCAGCTTTGATGTTGACACTCGTTGGAACTTAACATATGAAATGAT

TGCGGCATTAATTGAACTCCAACCAGCCATTTGATCGTCTAAATGAATTGGATCCTGATTTTAAATGTTT

CCCTTCAGAACAAGGATGGGAAAATGGAAAGAAAGTGTTTGATTGCTTAAAGATTTTTTCAGATATTACG

AAAAAACATCCTGGTGTGAAATATCCAACTGCAAATCTGTACTTTATTGATGTGATTCAAATTTGTAGAA

GTATTAAAATGTGGGCTGAATCTAGTGATGATTGGATTATTAGTGCAATGGGTAATAAAATGCAACTTAA

ATTTGATAAGTATTGGGGAAAATGCAACAAGTTGTTGACTGTCGCTGTTATTCTTGATCCTAGATATAAA

ATGTCCATCGTCTCTTATGCTTATAAAGGTATCTTTGATATTCATGCTGACTTTCAGGTTAATAAAAATA

CGTGAGTTATTGCTAGAGATTTTTAATGAATATTCTTAGAAATTTGCAAGTAGTAGAGGTTTTGTAGTAA

ATTCTGGTAATGCAAGTTTTGGTTGTTCAAGTTCAGTTGGTGGGGAGTGGTTGAGAGGTTTTCAGGATTT

CATTGCAAGTAGCAATTTGAGGGAAAATACAAGAAAAAGTGAGCTGGATGAATATTTAGAAGAAGGGTTG

TTTTCGATGGATAAAGAAGTGGAGTTTGACATTCTTCATTGGTGGAAACTTCATGGACCTATATTTCAAA

TCTTTGCCCGGATGGCACGAGATATTAGCTATACCTACTTCTTCTGTTGCCTCAGAAAATTCTTTTAGTA

AGTGTCGTAGAATTATAACGGACACTCGGTCTTCTCTAAATGCTGACTCAATTGAAACTTTGATGTGTGT

GAAGGATTGGCTCCCCGAACTCAAGGATGGTAATCAATCTGGAAAAACTTCAGAGGATGATGTTTTTAAT

GATTTGGATAATGAATGGGATTTGGACTTCTAAAAGAATCAGGACTGTGAATCTAATTTAGCAGCTGCAA

GTGGTATTTCACACTCTGTA

>MSTRG.636.2 gene=MSTRG.636

TCACTTTTGTTCTTTAATTCATCGAGGTTTAGGGCTTCTCTTAAGTGTCAATGGAACAAGGTAACGATTC

AAGTTCAACTCCGAGTGTATCATCAAGTTTGGTTGGTATGATATCAACGAAGTGTGGTGAAAATATCTCT

AAAAGCGCTGATCTGGATTGTGTAATAATTTCGCAAGAAGATAATCTTATTATTGGGCGTGATCAAGCTA

TTCAAGGAAAAGTGCAACCAATCAAAATATAGAAGACCTTGAAGTCATAAGTTTGGAATCATTTTGATCG

GATTGAAAGCGAAATAAAAAGTAATCAATCTGGAAAAACTTCAGAGGATGATGTTTTTAATGATTTGGAT

AATGAATGGGATTTGGACTTCTAAAAGAATCAGGACTGTGAATCTAATTTAGCAGCTGCAAGTGGTATTT

CA

>MSTRG.638.1 gene=MSTRG.638

CTTTTTAAGAGAATATTTCTGTTCCTATACATGCAGAATTGAAATGGCTAGTGATATGGAATCAAATAAT

TCATCAGTAGAAAGAGGCGCTGGAAAAAATAAAAGGAAATGGACTGAAGATGAAGATGGAAAACTAGTCG

AAGCTTTGATGAAGACGCTAAAAAAGGATTTCGATGTTGTGTACGACATCTGCTATGGAGCAAACAGTTC

GGGGTTTGGATGGAACGCAGAAGACAATGTTCTTACGGCACCGAGGGATGTCTGGGTTCAATATCTTAAG

GTTCATCCTGGAGCAGCTAAATGACAGAATACTGCCATGCCATCATTCAAAGAACTTTCAGTTATTTTGG

GAAAAGATAGAGCAACCGGAAATATGGTTGAAAATCTTGAAGATGTGGTAGGAGAATTGAACGCTGAAGA

TGCTGATGAATTATCTCTTTAGGAGGACCTGCAACGAAGCACACATTCAGACGAGTCAACAAGCAAGAAA

AGGAAGATAGGAAATGTAGAATCATTGTTGGAAGCTGTGTATGCTGCTTCAGACAGGATTTCCAATCAGT

TTGAGGCGTCAACCAAGTTACTTATTGCAACAGAAGAAGATATGATGCAGAAAAAGAAGCAACTAAATGA

CGAACTATCAAAAATTCCAAATCTCACAGTATTGCAGAAGCTTCAAGTTCCAAAGAAAATAGCCAAGGAC

GAAG

>MSTRG.639.1 gene=MSTRG.639

AAATCTTTGTGTCGATGCAAGTAGATTAGAGACAACTGTATCTTGATTATTAGTAGATGAGTCTTTAAGT

CGTATTTTCCTTAAAAACGCTTTTTGTTAACACCAACGAAATTAGAGTAGAGTCGAGTAGACACTTGCTA

GAATGTCTCGAAGATTTTATGGGTATATCTTCTGTAAACCCTTAGGAGACAAAACTCCTCTTGTAGAGAT

CGTCTACGAGTTTAACGGGTTGGTGAACCTAAAATCACCCGTCACGCCTACGAAAAGGAGCCTAGGAATC

GCATTTAGTTTTCTTAGTTTATTTCTTTTCTTTTGATTTTTACTCGAGGACGAGCAAAAGATAGGTATGG

GGAAGTTTGATTAGTCCATATTTTTGCATATTTAAGTGC

>MSTRG.640.1 gene=MSTRG.640

TGATTGGACTATAAATAGAAAGTGTAACATGTTGAAAAGTAGCAACACAGAAACTTAAACAGAAAAGATA

TAGGAATTAGGAAGCATGAAAGTTCTGAATCGAAACATGCTAATGCTTGTGTTAGTTTTGTCCTTGATAT

GTAATATATCTCACTTGTATGCTGCTGATCCTACTGATGGATTCACCAACATACCACTGACTGAAGCCAA

TTTCAAAATTCAGAGGCCTTATAATTTACCAGTCGAGGCACGATACAGCTTTGAAAATGGCGTTCGCAGG

ATGTGGGTGTACGCTAATGACAAGTCCCACAGCCCAAATAGCCAGACACAACCCCGCACTGAAGTTCGAA

TTATGGGACATGACTACTCATCTGGAGTATGGCAGTTTGAAGGATATGGTTTCGTCCCAAATGGAACCAC

TGGACCTTCAGTTGTGCAGATACATGGTGCCACCCACGAATCGTCTACAATAATTTTAAGGATCTATGAC

GGGGAGCTGAGGTACTATAGTGGTGATCTTATAGCCTCCAATATGTACGATAAGTGGTTCAAGGTTAACT

TAATCCATGATGTGGATCAAGGGAAAGTCACCGTTTTCATTGATAACCAACAAAAGTTTCAGACCAAGGA

CAGAGGGCCTGGAGACCTGTATTTCAAGTGTGGAGTTTATGCTGCTCCCAAAAACATCAGTTACTATATG

GAATCAAGGTGGAGAGACATCAAATTATACAAGAAGTGAACTGCTTAATAGCGAAAACTGTTGAAAATAT

AACCATTGGTGTTTGATATTTGTATCATGTATTTAATCTATCTACAAATCAAGGTTGGTGTGTGAGTATA

GACTTTCGATTTCTGATCAACTTTGGC

>MSTRG.641.1 gene=MSTRG.641

TTTTATTTTGTGCTGAAGCGTTCTCTTCTCTTTTGTATAAAGCAGAGATGGATAAAAAAAAATAAATAGT

TTAAAGTTTGGCAAAAATAGTGTTATGGTGTCCCATTTATTTTTTTGCAGATGATAGATAGCCTGATATT

TATGGAAGGTAACGAGGAGGCATGTAACAATTTCAAGAAAATTGTTGAGAAGTACACTAAAGCATCAGGG

CAATGTATAAATTTTAGCAAGTCTAAATTGTGTGCTGGAAAACTTGTCGGGATTGAAATGCAAGAAATTT

GGCTAAAATTTTGGGGGTGAGGTTAGTTAGACATTTTGAAAAATATTTAGGTTTGCCGACACATGTGGGG

AGAAGTAAGAATGATGTGTTTAGATTTATTAAAGAAAGAGTGTGGAATAAACTCAAAGGATGGAAATCTC

ATCTATTTTCTGTTGCCGGTAGAGAGATACTTATCAAGGCGATCATACAAGCTGTTCCAGTATTCACTAT

GAGCTGTTTTAGAATTCATAAAGGAATTGTTCAAAAAGTTCACAGTATGATGGCTCGATGCTGGTGGGGT

TCAAATGATAAAAAGAGAAGAATTCACTGGTGCAGATGGAATTATTTGTGTGAGCCAAAGGAAAGAGGAG

GATTAGGGTTTAGAAATCTGGAGTTTTTTTTTAATCAGTCCCTCTTGGCGAAACAAATATGGAGAGTCAT

AAGAAATCCTAATTCGCTTTGCAGCAGAGTCTTAAAGGGTTGTTATTTTCCTAACATTTCAATTCTAGAA

GCATCATGTGGTTCAAGTGCATCTTCAGTTTGGAGGAGTCTGATATGGGGGAGTGAGATTATCTTTAAGG

GTTATAGATGGAGAGTGGGTGATGGTAATGATATTAGAATCATAGAAGATTCCTGGATTCCCAGACTGAA

AACTTTCAAAATTTTTGAAAAACGTCATATTCCTCCTAATCTTTATGTAATCGATCTTAAACGAGGAGAT

GGATCATGGGATGAAGAATTTATCAGAATGAACTTTTGCAAGGATGATGCTGAGGTAATCACTCAGATTC

CATGCTGTGATAATGATCTAAAAGATAAGATTATTTGGCATTATTCCAAGGATGTGGAGTATAGTGTGAA

GAGTGGATATAAGTTGGCCATGAAATCGAACAATCAAGCTGAAGCTTCAAATATGAATGAAACTGAGAAT

TGGTGGTCTTCGGTTTGGAAAATGAAAATTCCACAGAAAATCAAATTCTTTATCTGGAAAGTTACTCACA

TGTGGTTGCCTACTAATTTTATTATTTATCGTAAAGGTATCACTAACTCTCCTGAATGTTTGCGATGTAA

CATTGGAGCTAATGAGGATATCATGCATGCATTATGGTCATGCCCCGCTACTTCTAAAATCTAGAAGATT

GTTGGATGGTGGCAGGTGATGAAAAATTATAACGCTTTAGATGTAAGTAGCTTTCTAATTAAAATGAAGC

AAAGATTGACAATGGAAGAATTTGAATTATTTTCTGTCATTAGTTGGCAAGTCTGGTACACGAGGAATAA

TGTTGTTCATTGTGGAATTTGTCCTAAACCTAATGTTCTGGTTAGCTGGTGTTGTGAATATGTCAAAAAT

TTTAAACTTTTAAATGTTTATCACATGAAGGTGGCTGATAGATCAAAATGCAAATGGAAAGTGCCTTCTC

CAGGTATGCTCAAGCTCAATGTAGACGGAAGTATAAATATGAAGAATCCAGGATGTGGTTTAGGTATTGT

GGTGTGCAATGACAAAGGAGATTGTATCGAAGCAAGGTGTGTTTATTTGAACAAATGTAGTTCTCCTTTA

GCTGCTGAATTGTTAGCTATTAAGGAGGGGTTGAAGACTGCCAG

>MSTRG.646.1 gene=MSTRG.646

ATTATTTGGCGGATTGTGGATTTTCAAATAGACTCCAGTTTTTAGCTCCCTTCCGTGGTGTTCGTTATTA

TCTTCAAGATTTTCATGGTCAAGGTCGTCATCCTGAAAATGCCAGAGAATTGTTTAATATTCATCATGAT

TCCTTGAGGAATACAGTGGAACGATTATTCAGAATATTCAAATCACGATTTACAATTTTCAAGACGGCGC

CTCCTTTTTCATTTCAAACACAAGCGGAGATGGTTTTCGTTTGTGCAGGCTTACATAATTTTCTTCGCAA

AGAATGTCGTTCAGATGGATTTCCAGTTGAAGCAGATAATGAAACTTCAGCAACAAAACAAGAAGAGTAT

TTTGAACCACTTATTCATGAAACACAGGAGCAACAACGTGAAAATGCTAATGCGTGAAGAAATGATTTTG

CGCATGAAATGTGGATGAATGTTGAAGATAGGCGGAGCTA

>MSTRG.648.1 gene=MSTRG.648

GATAGATCCGGATAGGTCAATCCGGACCCACCAACTTTCACTATCCGGGTACCTCGCCGGCTTACAACCA

AGGACGTCAACCCTGGAAGGAGACTCAACCCGGGTAGGCACCTAATCCACCCTACCCGGGTACACATGTA

CCGGCCGACAACCACTGCACAAGGCACAGACTGACACGATGAAGACAAATGTAACGGTCAACTACTGACG

ATAGAGACAAGCCAGGGAACATTCCAAAATACTACTCCTACGCTGACACCTGGACACGTGTAGGGGATCG

AACCCCTACGCGTGTAGGACCAGGATCCAGGTACTCACCCTTAAACCCTAATCCTTGGCCTATAAATAGA

CCAAGGATCAAGAGTCTAAGGGTGACTCTCTCT

>MSTRG.649.1 gene=MSTRG.649

CACAAAACCGAGAACTGCATATCCCTCAAGTATTTCATCGAAGAACAGATCAGCAAAGGAAACATGGGGC

AATATGTAGCTCGGAACACAGCCGACAAAACAGGAGGATCCGGGAAGCAGAAAAACATTGTCAACGTAGT

GCTGGGAGGATCCTGTTCCCCTCCCCCTAGCCCGGGCTCCTGTCAAGAAGTAATGTCCATCCAAGCCTAC

CCCGAAC

>MSTRG.650.1 gene=MSTRG.650

GTCTGCGAAAGATTCTTTTGCTTCTGCTCTTCGGAATTTGGGAGGCGTGCATGTGGCCCCGAAGCCTCAA

CCGAAGAAGAAATCAGGGAAGAAAGGGGACTCGAAGAAAGGGGAGTCCGGGTCGAAGAGAGGGGAGTCCG

GGTCTTCCCGTCAAGGGACGGAGGGTACTGGGCATTCCAGTGCTAACCTGGGTGCAACTCCGGATCCGGA

GTTGATCGCTGATGCTGAAGAGGTGGAGAGCCCGGATCAGGGGCCGAAAAGAAAGAGGAAGAAGACTGAT

GCTGATTCTTCTGAAAGGGAAGTCATTGATATGACAGTGGAGGGTCCGGGTAAGGCAGCTATGGAGGTTG

TGGATCTTGGAGTCGATGTTGGGGGCGGTTCCCGGCCGGCTCCTAAGTTTGGCAAGTACCCGGTTAAGAA

AGTGCTCGGGTTGATGTCCGAACTTCCCTCTGACCAAGACTGGGAAATGATGGAGGATGAGGGTCTCGCC

ACCAATTTCAAAGAAATCGGGAACTTGTGGGGTCAGCTCGGTGGTCGTCTGGCCGGGTTCAATACCCATG

CTTTGAACAGCTTGAAGCAGGAGAGGGATCTTTCCGTGAAGAGCCTCGCCCGGGTCACGAAGCTGGAGAA

GGATCTGGATGTGGAGAAGTCAGCCCGGGAGGCTTTGGAGTCCGGGGTAGCCTCTAAGATCAAAGAAGCT

GAGATCCGGAAAGAGGCTGAGCTGAATCAGAAGATTAAGGATGCTGGGACCCGGGCTGACAGCGCTGAGA

AGAAGGTTTCCGGGCTGGAGAAGGAGGTGGCTGATCTGAAGAAGCTCTTGGAAGGGAGAGAGGAACCGGA

GAAGGTTATTGCGGAATTCCAAAAGTCCTCTGCTTATGCGGATGCCTTGGCCGCCGCTGGAGCTCTTGAA

GTCGTCCGGTGTTGGCATGTTGCCGAGCGGCACATCAAGACTGACCCGGAGGCTTCCTTGCAGAGTTTTA

TAGAGCTCTATCTTGCTGCTAAGGATGAGATCAAGGCTGGGAAAGGAGAACCGGAGCCTTACGAAGGCCC

CTCTCCTAGCTTTCTCCCTCCTGCCAATCCGGGTGCTGATGCTGATCTGGGCTCTTCCTCCGATTCTGAA

CCGGTTGATGGGACTCCGGTTGAGAAGACTCCGGCTGATGAAACTCCTGCTGATGAAACTCTTGCTGACT

GATCC

>MSTRG.651.1 gene=MSTRG.651

GATAGATCCGGATAGGTCAATCCGGACCCACCAACTTTCACTATCCGGGTACCTCGCCGGCTTACAACCA

AGGACGTCAACCCTGGAAGGAGACTCAACCCGGGTAGGCACCTAATCCACCCTACCCGGGTACACATGTA

CCGGCCGACAACCACTGCACAAGGCACAGACTGACACGATGAAGACAAATGTAACGGTCAACTACTGACG

ATAGAGACAAGCCAGGGAACATTCCAAAATACTACTCCTACGCTGACACCTGGACACGTGTAGGGGATCG

AACCCCTACGCGTGTAGGACCAGGATCCAGGTACTCACCCTTAAACCCTAATCCTTGGCCTATAAATAGA

CCAAGGATCAAGAGTCTAAGGGTGACTCTCTCT

>MSTRG.652.1 gene=MSTRG.652

TTATAGACGTCGAAATTTCAAGAAGTTGAAAGACCTCATCCCAGTATTGATGGACAACCAATTTCCCCAA

CTTTCATACCAAGATCTTCTGCTTGCTACAAATGAGTTTTCTCCAGATAATATGCTCGGTAGAGGAAGAT

ATGGTTCAGTTTACAAAGGAGTTCTCAAAGCAATGGAGCAAATAGTTGCTGTGAAGGTACTAAACGTTGA

AATACATCGAGCAAAGAAAACTTTTTTGGCAGAGTGTCAAACATTGAGAGAGATTCGACACCGGAATCTT

ATCAAGATCATTACAGCATACTCTACCATTGATTTTAAGGGCAATGACTTCAAGGCATTGGTTTTTGAGT

ACATGGCTAATGGGAGTGTGGACAACTGGTTGCATCCAAGCCCTTCCCATCAGCGGAATGAAAGAAACTT

GACTCTACTACAGAGATTAAATATTTCCATCGATGTTGCAATGGGATTGGACTACCTACATCATCACAGC

CATGAAAGTATCATTCATTGTGACATAAAGCCAAGTAATATTCTTCTTGATGAGGATTTTGTTGCTCATA

TTGGGGATTTCGGTTTCGCGAGGTTTTCTCATGGTACCACAAGTGATGTCAATCAAACACAGACGAGTTC

AACAAGTGTACATGGAACATTTGGATATGTTCCTCCAGAGTATGGAATAGGTGGAGAGGTATCTACAAAG

GGTGATGTGTATAGCTATGGAATTCTTCTACTAGAAATTTTCTCAGGGAAATGCCCTACTGAAAGCAGCA

TATTAAAAGACGGTTATAGTAATCTTCATAATTATGTGAGGGCAGCACTCCCACGAAGAGTGATGGATAT

TGCTGATCCGCAGATTGTACTTGATCAAGAAGAACATGGCTTGACTGTAAATCAATCATACAGCAGGGCT

TCCATGAAAGTATGCCTCACATCAATATTTGAAGTAGGGATATTATGCTCTGAAGAGTTGCCACAAAAAC

GCATAGACATTAATGTTGCTATTAAGAGATTACAGGCAGCACAAGACACAGTTCTGCAGCACAGGCAGTA

AGTTGGGTGAGAGTGGAAATCTCTATACTTTCTAAATAAAGGGATCTATATGTTGATCCTGATGTGAAAA

TAAAGACTTTGTACCTATTTATTTCCCATATTAACAGATTACATGCGGCAGAAGACACAACTCTGCAGCG

CAGGCAGTAAGATGGATGAGAGTGGCAATCTCTATACTTAATAAATAAAGGCATCTCGATATATATGATT

CTAATGTGACAAGAAATAATTTTTACCTATGTATTTCCC

>MSTRG.653.1 gene=MSTRG.653

CCCTGGCTCGTAGAGCCGTGTTGTTGTTGATCAGCCATCGTCATGTGTTTCCTCGTAGAATCTTTAGCAA

GTTTCCCACAGACGGCGCCAATTGTTCGTGCAAGATTTTGCGGGGTTAAATGCTAAAGAATTAAACTAAG

AATAAAATGAACAAGTAACGAGACAAAGATTGGTTACGCGGAAAACCCCCGAAGAGGAAAAAACCGCGGG

AGGGTCGATGCCCTGCCAAATATGTGTTTCACTATGTAGATAATTTGCGTACAATGTTTTTGATATGAGG

CTGCTTATTGCTTGCCCTTTACAGCTTGAATCTAAGCCCTATTTATACAAGTCATATTGTAACTACTACT

CCTACTAATGCTCCACGTGCAGCTGTAACTTATCTCTCCTACTTATCCGCATATGACCACCTCAATGCTT

ATCTTCCGCCTTTGACTCGTTCTCCGCGAATTCAAACAATCGGCATGCCGTCACGTGCACACGTCCCGTC

TTGTCACCCTTCTGTGATGTCAAG

>MSTRG.654.1 gene=MSTRG.654

GATTTTGCGGGGTTAAATGCTAAAGAATTAAACTAAGAATAAAATGAACAAGTAACGAGACAAAGATTGG

TTACGCGGAAAACCCCCGAAGAGGAAAAAACCGCGGGAGGGTCGATGCCCTGCCAAATATGTGTTTCACT

ATGTAGATAATTTGCGTACAATGTTTTTGATATGAGGCTGCTTATTGCTTGCCCTTTACAGCTTGAATCT

AAGCCCTATTTATACAAGTCATATTGTAACTACTACTCCTACTAATGCTCCACGTGCAGCTGTAACTTAT

CTCTCCTACTTATCCGCATATGACCACCTCAATGCTTATCTTCCGCCTTTGACTCGTTCTCCGCGAATTC

AAACAATCGGCATGCCGTCACGTGCACACGTCCCGTCTTGTCACCCTTCTGTGATGTCAAG

>MSTRG.656.1 gene=MSTRG.656

GCCAGTTACACAATCTCATCCAAAGAACTGGACGATTAGATATCAACAAACCCTCCTGTTCGATTTTATA

CCCCCAACCAGGAATCTTACGTATTCTTGCTGCTTTGATTATATATGGACATGGGAATATAGAGTTGTCA

AAGTATCATGTCGTCAAGCTCCAGTACTATACATAATCGACTCATCTTTAACCCATAGGAGTGTGTGAAT

TTGTTTTTTAGAAGTTTTCCCAAAACACTGCAACCAGTGGCAAGTGGATGAGCACATCAGCATTAAAACT

TGATCTGTCAATCAAAAGTCCCTACAGGATCACGAACTGGAGCAGGCCAGGATGTTATTAATCTGCTGGG

CGCTAGCATGTCTGAAGTGTGTCCGAGAAAAAGAGTTATACCACAAGTAAGCTGAGAACTGGGATTAACA

AAAGTGTGTATCCTGCAATCAAAGTAG

>MSTRG.656.2 gene=MSTRG.656

CAATCTCATCCAAAGAACTGGACGATTAGATATCAACAAACCCTCCTGTTCGATTTTATACCCCCAACCA

GGAATCTTACGTATTCTTGCTGCTTTGATTATATATGGACATGGGAATATAGAGTCCCTACAGGATCACG

AACTGGAGCAGGCCAGGATGTTATTAATCTGCTGGGCGCTAGCATGTCTGAAGGCTACGACAGAAAATAT

CATTGTCGGTTCTCATGTTTGGATTGAAGATCCAGAAATAGCTTGGATCGATGGACAAGCTACAAAGATT

AATGGACAAGAAGTTCAGATAGATACTACTAGAGGAAATTCGGTGCTTCTAGCGGAGGACTCGGTGCCCA

TGTGAACTGGCATGTTTGATGCAAATCAAGTGAAAGCTGTAGGAAATGAAAGAGCAAGGTGTTTCTGATG

ATAAACTATTACTTCTCAAGGGCATGAGTGGAGTTTTCAGGTCAGGGGTTCTAACTGCTCTAATGGGTGT

TAGTGGTGCTGGACTGTTGGTAAAACAACTTTGATGGATGTATTGGCTGGTCGGAAACTCGGAATTGGTG

CTGGTTTCATTGCTGATGTTTTGGACATGAGCATTCGTGACAAAGTAATTCAAGTAGGCATTTTGGCAAA

AGCTATAATGTCGTTGACTTCAGTTAATAAATTTTATGGAGAGGCTCTTTCGAGCTTTTATGCACAATTA

TCTGGCAAGATTTTAGTCTAAATTGCATCAAGAGTATGATTACTAAGTAGGTAGTGAATGTTCTATTGAT

TTTTGAGACATCTTGAACAGG

>MSTRG.660.2 gene=MSTRG.660

ACTGTACCACGTGTATCGGTACGTGTATACCTCTGGTTTCTCTCTATCTCTTTTCTCACCCTTGACCTCG

ATCTCTCTCATTTTAATCTCGCTACCAATCTCTCCCGCACTCTCCTTCTCTTTCCCCAATGTGACTCACA

TCTCCTCCTTCAATTTCCCACCAACCGTCAACGAGCCCCATTGCTGCGGTAATCTGATTCAGCATCTTTC

TCCGCGGATTCTGCTCCGTTTCTTCTCCCTGTGTTGTGTCTCTGTGGTACCCTCTGTAGTGTTCTCGATT

CCGTTGCCCAACCACCATCCAACAGCCATTGCTGTCCACTTTTCCGGCAACCCGAGCCTTGCCTCGTCTT

GTGTCTTAATTTCAAATTGAACGAGACGTTGATCGAGGCATTGATCGAGAAGTGATTGAAATGGGTATTC

AGAAATAGCTAAGTTGTGTTGCAAGTTGAAGTTAGGCAGAGATTTGCTATAGATAAAATCTTGTCAGAAG

CTGGCAGAAAGGAATGATATGGCACTACATAGTGTTACTAGAAATCTGAAATAGACACCGG

>MSTRG.660.1 gene=MSTRG.660

ACTGTACCACGTGTATCGGTACGTGTATACCTCTGGTTTCTTCTCATTTTCTTTTCTTCTCTCTCTATCT

CTTTTCTCACCCTTGACCTCGATCTCTCTCATTTTAATCTCGCTACCAATCTCTCCCGCACTCTCCTTCT

CTTTCCCCAATGTGACTCACATCTCCTCCTTCAATTTCCCACCAACCGTCAACGAGCCCCATTGCTGCGG

TAATCTGATTCAGCATCTTTCTCCGCGGATTCTGCTCCGTTTCTTCTCCCTGTGTTGTGTCTCTGTGGTA

CCCTCTGTAGTGTTCTCGATTCCGTTGCCCAACCACCATCCAACAGCCATTGCTGTCCACTTTTCCGGCA

ACCCGAGCCTTGCCTCGTCTTGTGTCTTAATTTCAGTGGAATTGAACGAGACGTTGATCGAGGCATTGAT

CGAGAAGTGATTGAAATGGGTATTCAGAAATAGCTAAGTTGTGTTGCAAGTTGAAGTTAGGCAGAGATTT

GCTATAGATAAAATCTTGTCAGAAGCTGGCAGAAAGGAATGATATGGCACTACATAGTGTTACTAGAAAT

CTGAAATAGACACCGGAGTTCAGGCAAGTATATATCCAACCTACTTTCTCTTTAAAAGATTTTGCTCCTG

CAATGCTTTTGTTATGCTTTGAATATTGCAAAACTCCTTATAGAATTAGACTTCTGCTAATTAACTAGTA

TTCTGGAATTACCCAAGTATGAAATGGAATACCCTTTGTCACTTATATCTTGCAAATATATCAACTTAAT

TATCATCCTTATCTTCTTCAACTGTCAGTTACTTAATATTGAGCTTAGAACCTTGTTTTCTTACCTCCCT

TTTAACTGCAACAAATTGTGTTCTTGCTTATAAACTAGTTCACTTGTTTGTTCCTGACCCTGAAATGAAA

CCCCATTTCTTGAAAGACCTTAGTTATCATCATACAACCTACCCCGCCATCATTATAAAATACATTGATA

CACTTGATTCAGTAACCTATACCCTTGATAAAGAATACTTAATTAGTA

>MSTRG.660.3 gene=MSTRG.660

CGTGTATCGGTACGTGTATACCTCTGGTTTCTCTCTATCTCTTTTCTCACCCTTGACCTCGATCTCTCTC

ATTTTAATCTCGCTACCAATCTCTCCCGCACTCTCCTTCTCTTTCCCCAATGTGACTCACATCTCCTCCT

TCAATTTCCCACCAACCGTCAACGAGCCCCATTGCTGCGGTAATCTGATTCAGCATCTTTCTCCGCGGAT

TCTGCTCCGTTTCTTCTCCCTGTGTTGTGTCTCTGTGGTACCCTCTGTAGTGTTCTCGATTCCGTTGCCC

AACCACCATCCAACAGCCATTGCTGTCCACTTTTCCGGCAACCCGAGCCTTGCCTCGTCTTGTGTCTTAA

TTTCAGTGGAATTGAACGAGACGTTGATCGAGGCATTGATCGAGAAGTGATTGAAATGGGTATTCAGAAA

TAGCTAAGTTGTGTTGCAAGTTGAAGTTAGGCAGAGATTTGCTATAGATAAAATCTTGTCAGAAGCTGGC

AGAAAGGAATGATATGGCACTACATAGTGTTACTAGAAATCTGAAATAGACACCGGAGTTCAGG

>MSTRG.660.4 gene=MSTRG.660

CGTGTATCGGTACGTGTATACCTCTGGTTTCTTCTCATTTTCTTTTCTTCTCTCTCTATCTCTTTTCTCA

CCCTTGACCTCGATCTCTCTCATTTTAATCTCGCTACCAATCTCTCCCGCACTCTCCTTCTCTTTCCCCA

ATGTGACTCACATCTCCTCCTTCAATTTCCCACCAACCGTCAACGAGCCCCATTGCTGCGGTAATCTGAT

TCAGCATCTTTCTCCGCGGATTCTGCTCCGTTTCTTCTCCCTGTGTTGTGTCTCTGTGGTACCCTCTGTA

GTGTTCTCGATTCCGTTGCCCAACCACCATCCAACAGCCATTGCTGTCCACTTTTCCGGCAACCCGAGCC

TTGCCTCGTCTTGTGTCTTAATTTCAAATTGAACGAGACGTTGATCGAGGCATTGATCGAGAAGTGATTG

AAATGGGTATTCAGAAATAGCTAAGTTGTGTTGCAAGTTGAAGTTAGGCAGAGATTTGCTATAGATAAAA

TCTTGTCAGAAGCTGGCAGAAAGGAATGATATGGCACTACATAGTGTTACTAGAAATCTGAAATAGACAC

CGGAGTTCAGGCAAGTATATATCCAACCTACTTTCTCTTTAAAAGATTTTGCTCCTGCAATGCTTTTGTT

ATGCTTTGAATATTGCAAAACTCCTTATAGAATTAGACTTCTGCTAATTAACTAGTATTCTGGAATTACC

CAAGTATGAAATGGAATACCCTTTGTCACTTATATCTTGCAAATATATCAACTTAATTATCATCCTTATC

TTCTTCAACTGTCAGTTACTTAATATTGAGCTTAGAACCTTGTTTTCTTACCTCCCTTTTAACTGCAACA

AATTGTGTTCTTGCTTATAAACTAGTTCACTTGTTTGTTCCTGACCCTGAAATGAAACCCCATTTCTTGA

AAGACCTTAGTTATCATCATACAACCTAC

>MSTRG.661.1 gene=MSTRG.661

ATCTTTCCCAACATCTCACATGAGTCGGCGTCACCTACACTATACATAATTTCTATTTGTACAACAACAT

GAGTGTACATGAATCTTCCTATCCAATTAATCTACGTGATATGCATGGACTTATATGGTTGTCTTTATTT

CCACATTCTATTTAAATAGCTAGCCGGTGCTCGCATAAGTGGAAACTAGAAAACAAAGAGATTGAGGTGA

CTCGAATAAAAATTGTACGAGATCAAAGATGGACAAGAGCAGGTCATTCCCGGAGTACTCATCGTGCGGT

GAATTGGAGTCGGAGAATCGAGCGAATGCTTATTCGTTCAACGGGCCAAACGCCACACCGCCTAGTGATC

CAGAGCTCAAGAGGAGGAAGAGGATTGCTGCATATAACATGTTCACCACTGAAAGAAAACTCAAGTCTAC

TATGAGGGAGAGGCTCAAGTGGATTAAGAATAAGTTCACCGATGTTCGCTATAGTTTCTGAAGTCATTTA

TCAATCTATCTATCTATCTGAGTACAAATAGTTGTGTAACTAAAGAAGCTATATCCTCTGCAATGGATTT

CAGTTCTGCCCTTTGTCGACAAGCTTGTGCCTGTTCAGAGGGATCATTATAAATTATTAAAGTGTGTCGT

TCACGGTACATAAAAATTCCGCATCCGACCGAAGAGTTTTCGGTTTTGAATCATGGTGATCATAAAAATG

TAATCAAGCTTCGGAAATGATATATTCTAAGCGTCCTCCTATTATTTTTCTACATTTGTAAGCATTGTTG

TTCTCGTGTGTGTGTGTTTTTCGATTGTTGGCATCAAGATGCGCCCCTCGGAATCAAACAATCAAATACT

TGATAGTTGAAAACATCTTGTAATGGTTGCCGAGTCGCAATCAATATAACACAAAAGGACAAGTATAGAA

ACTAAAACAATGTTTG

>MSTRG.666.1 gene=MSTRG.666

GTCAAACCCTAATGTAAAATAACTAATCCCTTTCAAATCATTTCAGTTAAACATTCTTATTTATACACAC

ATACACATACCTGTGTCTATCTCCTTGTTTTCACACAAATTCATCGCCAAATCGCAAGAGCTCACACTAC

AAGATCCGAAATTCACACTCATTACACACTTCTAAGCTCCTCACGGAAAACTAAGAGCTAAAGACGGGAG

TTCTGTGTTGTGTTGTTAATGCTGAAAGCAGAAGTTCATTTATGATACAGAAGAGGTAGATATCCTCTTT

TGGCCGAGAATTTGATTTTGAAATAATGGCGCCATCAAGCAAGGCTGATAAAAAGGCTGCAATAGATGCA

GCTGCATGGATGTTTAATGTAGTCACTTCCGTCGGGATCATCATTGTCAATAAAGCCTTAATGGCGACAT

ATGGATTTAGTTTTGCTACAACATTAACTGGTTTGCATTTCCTTACGACAACTTTGATGACGGGTGTCCT

TAGATGGTTGGGATATATCCAGCCTTCTCATTTACCTATTTCAGAGCTGCTCAAATTTGTCCTCTTTGCA

AATTTCTCTATTGTAGGGATGAATGTTAGCTTGATGTTGAACTCAGTCGGATTCTACCAGATCGCAAAGC

TAAGTATGATTCCTGTATCCTGCTTATTGGAAGTTGTCCTAGACAAAATCAGATATTCAAGAGATACAAA

ACTGAGCATCTTAGTCGTTCTGATGGGTGTTGCTGTCTGTACTGTTACTGATGTGAGTGTTAATGGTAGA

GGATTCATTGCTGCTTTTATTGCAGTGTGGAGTACTTCTATGCAGCAATATTATGTTCATTACCTTCAGC

GGAAATACTCACTTAGTTCTTTTAATTTGTTGGGGCACACTGCGCCAGTCCAAGCTGCCTCTCTTTTGTT

AATAGGCCCCTTTCTGGATTATTTATTGACTGACAAGAAAGTATATGAATTTGACTTCAACATGGCATCT

GCGACATTCTTATCACTTTCGTGCATAATCGCAATAGGGACCAACCTGAGCCAGTTCATTTGCATTGGCA

GATTCACAGCAGTGTCATTTCAAGTACTTGGCCATATGAAAACAATCCTTGTTTTGATCTTGGGATTCTT

GTTCTTTGGAAAAGAGGGTCTTAATATACAAGTGGTCATTGGCATGCTCATAGCCATAGTTGGAATGGTC

TGGTATGGGAATGCATCTTCTAAGCCAGGCGGCAAGGAACGCCGCAGCCACTCTATGTCTAGGAGCAGCC

AACAGAAGCTTGTTCCATCTTTAGAACCAACTGAACTTGATGACAAGGTTTAATCAGATCTCAAAATTTA

ATTGGAGCATACAGAGGGTAGCTGATACAATAGGTACCAGGAAAGCTTTAGCAATCCGATTCTAGATACA

AACAGTCAAATACTGCCATATCTTTCCCGATTCTGTAATTCTCATCTTTCATTGTGGAAATTCTTGGGGG

TTTAAAACTCAACTATATAGAGCTCTATGTTTTCCCATTGTTATAATTTATAATAATTTATACTTGTGCA

TTTTTTTATACAATTAAAATATTCTTTCTTCAGGTTGACCTTATATGAGT

>MSTRG.666.2 gene=MSTRG.666

CTTATTTATACACACATACACATACCTGTGTCTATCTCCTTGTTTTCACACAAATTCATCGCCAAATCGC

AAGAGCTCACACTACAAGATCCGAAATTCACACTCATTACACACTTCTAAGCTCCTCACGGAAAACTAAG

AGCTAAAGACGGGAGTTCTGTGTTGTGTTGTTAATGCTGAAAGCAGAAGTTCATTTATGATACAGAAGAG

ATGCAGCTGCATGGATGTTTAATGTAGTCACTTCCGTCGGGATCATCATTGTCAATAAAGCCTTAATGGC

GACATATGGATTTAGTTTTGCTACAACATTAACTGGTTTGCATTTCCTTACGACAACTTTGATGACGGGT

GTCCTTAGATGGTTGGGATATATCCAGCCTTCTCATTTACCTATTTCAGAGCTGCTCAAATTTGTCCTCT

TTGCAAATTTCTCTATTGTAGGGATGAATGTTAGCTTGATGTTGAACTCAGTCGGATTCTACCAGATCGC

AAAGCTAAGTATGATTCCTGTATCCTGCTTATTGGAAGTTGTCCTAGACAAAATCAGATATTCAAGAGAT

ACAAAACTGAGCATCTTAGTCGTTCTGATGGGTGTTGCTGTCTGTACTGTTACTGATGTGAGTGTTAATG

GTAGAGGATTCATTGCTGCTTTTATTGCAGTGTGGAGTACTTCTATGCAGCAATATTATGTTCATTACCT

TCAGCGGAAATACTCACTTAGTTCTTTTAATTTGTTGGGGCACACTGCGCCAGTCCAAGCTGCCTCTCTT

TTGTTAATAGGCCCCTTTCTGGATTATTTATTGACTGACAAGAAAGTATATGAATTTGACTTCAACATGG

CATCTGCGGTAAGTACTGCATACTGTTTTGTTTATTTCCGTGAACTCCGTGAAGTAGCTGAATAGGTTCA

GCTTTAGAAGTGTTTATGTGGCATGTCATGTTGAACTCCTGCTTTTTTCCAAATACTAAGATCTCACAAT

TTTTTCTCGTGCAGACATTCTTATCACTTTCGTGCATAATCGCAATAGGGACCAACCTGAGCCAGTTCAT

TTGCATTGGCAGATTCACAGCAGTGTCATTTCAAGTACTTGGCCATATGAAAACAATCCTTGTTTTGATC

TTGGGATTCTTGTTCTTTGGAAAAGAGGGTCTTAATATACAAGTGGTCATTGGCATGCTCATAGCCATAG

TTGGAATGGTCTGGTATGGGAATGCATCTTCTAAGCCAGGCGGCAAGGAACGCCGCAGCCACTCTATGTC

TAGGAGCAGCCAACAGAAGCTTGTTCCATCTTTAGAACCAACTGAACTTGATGACAAGGTTTAATCAGAT

CTCAAAATTTAATTGGAGCATACAGAGGGTAGCTGATACAATAGGTACCAGGAAAGCTTTAGCAATCCGA

TTCTAGATACAAACAGTCAAATACTGCCATATCTTTCCCGATTCTGTAATTCTCATCTTTCATTGTGGAA

ATTCTTGGGGGTTTAAAACTCAACTATATAGAGCTCTATGTTTTCCCATTGTTATAATTTATAATAATTT

ATACTTGTGC

>MSTRG.668.1 gene=MSTRG.668

GTCAATTTAATTGTCTAGTAATGTTCTTTCTTCTGTTTCCAAGATCTGGACCTGATAGAATTTTCTTGGC

AGTGTGGATTGGACGTTGCAGAACACAGGTGTAAGCACAAGCAATAACTCCTCCGTGGTCACAAGGTCTC

CTGTTGGTGGTGTACACTCTAGCTTGGATGTACAATCAATTTGGTCATATGATTAAAAAGGATGTTAAAA

GGATAAGGAGAACTAATAGTGGACTTGTTAAATATACTTCCTGTTCAGAAGAAAGGAAAAAGGTTTCCTC

TTTTGTCTGGGATTTAGAATACTGGATCCAATATATCGTCCTTGAGTTTATCAGGGAAGGTTTACTTGCA

ATAGCATATATGACTTGCTAGTGCAAGGCATGTGCTAGGAGGCAGTAGGTAACATATCAAGCTTTTTTGG

TATAATATCTTGGTCCGTCTTCCTTGAAAATTGCGTTTATTTTACTTACAAATGAG

>MSTRG.674.1 gene=MSTRG.674

CATTATGATCCGTAGTAGTATCACAATATCATTAACGACGACAACTTAGTTGCGATACATGCATAACGTC

AAACATCTTTATCATGTACAAATTTGCCAGGAAGATGTCTTCAGGACCAATTATTTTGCACTTCTAAGGA

ACTCACATCTTCAAGTTGTGTTTGATGTTGCTGTTGCGTTTCCCTCCCCGTCTCTCTTTTAATTATTTTT

TGGATTTCTGATGCTTTGTGGGTTTATCTGACAGGTGTGCCATGGCACTTGTTCATCAGCGGATTAAGCG

TATATTTTACACTTTTCCTAATCACAATGCTGGGGCGCTTGGAAGTTTGCACAGATTGCAAGGAGAAAAA

AGCTTAAATCACCACTATGCTGTTTTCAGAGTCTTTTTACCAGAATCTATATTGGATAGAGATGATGTTG

TCACTGCTGTTTCTGGAAGTGATAAAAACAAAACACATGTTCTATAGGAGGTATGTACTTGCAATTAATT

ATGCCCCTTGTTTTTTATAGGAGGGAACACTTCCATACTACCATGGTAGCAGATCATCTCCAAGAGACTC

TCTAATTGAGCTCTTAAGTAGAAAAATAAAGAGCCATGTCCAAAATAGAGCTCCAAGAGACTCTTAGAGG

CTTTTTAAAAAGTAAAGAGCTCACTCTCTCTCCTCTCTTTTAGGAGCCGCA

>MSTRG.675.1 gene=MSTRG.675

CACTTTAACCGTACGGGTTAACGTGTAACCTAGTTCTATATGAATTCTTATTACTGTGTAACTATTATAA

TTCTTTTCAACAAAACGAGGCCCAAGAAAACATCAACAGCAGGAGTGGGAGCTTAAATAACCCACCCCCA

ATTGCTTGTATATATCTGTACGCTAAGAAAGGAATGGATATGAATGAGAAAATAGAGATATTGAAATCAG

AATTTCAAAAGCTGTCAAAGGATGGACAAGCTTGGGTGCAACATTTATCAGTGGAGATGAATAAACAAGA

GTGGTTTCAGAAGTTGCCCCCTTTTATTCAAAACATACCTCCACTTCAACTCTATGTTGCACTTGTGGTC

TTGTTCCTCACCATGTTTTTCTTCTTTATAGTTCGGTTATTCAACCGCAAGACATCTAATACCATATTAC

TCACTGGGCTAAGTGGAAGTGGAAAAACTGTCCTATTCTATCAG

>MSTRG.676.1 gene=MSTRG.676

GAGCTGTAGAAGAAGAGTGAAGAAGTAATGGAAAGAGCCAATGATGCAAGTAATGGTGGTGGTGATCTAG

GTGGAGGTCATGTGATCAAGACAGAAGACGATGAGAAGGTAACAATCAAGGTGCAAAAAGAAGGAATGGC

TGATTTCTCCTTCACTATGAACCGTGATCAACCATTAAGAAAGCTTATGATCGCGTTCTGTGAGCGTCTG

AGTCTAGGTGATTATAATAACTTGTGTTACTATTT

>MSTRG.679.1 gene=MSTRG.679

GTAGTGGTTGACAGTAGACTCAAAAAGCTCTTTCTTGCTGTGTAGTAGCTGGGTTCGATTCCTTGGAACT

ATGCAATTTTTTCGTTTCCATGTCACTCGGCTTAAATGGGCTGTACAAATGAGTCCATGAAGTTGGACCA

CTGCTAGCATTGCATATGAAGAAATTGATGAAGCGGCTTTGATAAAGCGGCTTGGATTTGATGGGCCTCC

AAATGGTTAATCTCCTTGGGCCTGGACTGGATCAGAGCAAGTGGAATGGGCTGCAAATGGAATGGGCTTA

GAGCAACTACATAGTAGCGCCTAGAATATTAGGCGGAAGCAGTAATGTGAGAAACTAGTGAAAACTATGG

ACGCAGGTTTAGCTGCTAAATAGGCTTAAGGCCCAGCCCAGCCGAAGAGGGAAACGCTTTAATGCGAGCA

GGTCAGCAACGTGAAGACAGAAGGCTAATACGCCAGTTTAATAAAAACTAGTTAAAAAA

>MSTRG.682.1 gene=MSTRG.682

CATTATAGTTATTTTCATCGAAACACAGAGACCCGTAGGTACAACAAATTTTAACACAGAGATGCTATAA

TTAACTTGTCACTTTCAAGTTTTCAGATGCGTGGTGCATTGTGTTTTACATATTTAAATAAAGATCAAGT

CTTCAGACAAGGTTACAAGTTTAGACATGGGTGCAAAGACTGATCAAAGTTCAAGATACAACAAATATGC

ATATGCTTGTCTCTTGTGTTCTATCCTGGTGAACATATTCTTTTCTTATGATCACTTGGTGAGTAACAAG

TGGAGTTTGAGTTGGAGCACAAGTGCAGCAGCAGAAGCAGAAGCTGTGGCATCGATTTCGTGTTCGGGCC

ACGGGAGAGCTTACTTGGATGGTTTAGTGGTTGATGGTAGGCCAATTTGTGAGTGTAATACTTGTTATGA

TGGCCCTGACTGCTCAAATTATATCCCTGATTGTCCTGCTGATGTAAACAGTGGGGATCCCTTGTTTTTG

GAGCCATTTTGGATGCAAAATGCAGCTAGTAGTGCACTAGTAGTGGCAGGGTGGCATCGAATGAGCTATT

CATTCGGTGACCATACATTTACCTCGCAAGAACTGGAGAAAAGCATCCGTAAATTACATTCTATAGCTGG

AAATGCAGTTGCAGAAGGAAAATATATTGTTTTTGGAACAGGCTCTACCCAATTACTCAATGCAGCAGTC

CATGCCCTCTCCTTGGATAACTCATCTACACCTTCAAAAGTTGTAGTTTCCATCCCATTCTACCCGGCTT

ATGAATTGCAGACTGAATTTTTTGCAACTGAGGACTATGAATTCTATGGCGATACATCGTTGTGCAAAAA

CAGTTCTGAAGACTCTCTGAATGTGATTGAGTTTGTAACTTCACCTAACAATCCTGATGGAAAATTGAGG

AAAGCGGTTCTTAATGGTCCATATGCCAAGTCAATCAATGACCATGCCTATAATTGGCCACATTTCACTG

CAATTCCATCTCCATCAGATGAAGACCTTATGATCTTTACACTTTCCAAGCTCACGGGTCATGCAGGCGT

TCGATTTGGGTGGGCATTGATAAAAGACAAGGATGTATATGAGAGAATGTCAACATATACGGATGACAAT

ACTTTTGGAGTCTCCAGGGACAGTCAGTTACGAGCTTTGAAGCTATTAAAGGTAGTTCTTGAAGGCGATG

GAAGATCAATCTTTAAATACGCTTATGAAACCATGAGAAAACGTTGGGAAGGCTTGAGCCAAGCAGTATC

ATTGTCCACACGCTTCACCATCCAAGAAATCGCGCCCCGCTTCTGTACTTTCTTTCAACATGTCAGAGGA

CCTTCTCCAGCTTATGCATGGCTGAAATGTGAGAGGGAAGAAGATGAACTATGTTACCAAGTCCTCCAAG

CTGGTGGCATACTTGGTCGTGATGGCACTCTATTCGGTGCTAGCAGTCGATATGTTCGCCTCAGCCTCAT

CAAGTCTCAAGATGATTTTGATCAGTTGCTACAACATTTAAACAAATTAGTTCTCCAGGAAGCTGGCAGT

ACTAAAGCAGATATCTGAAATTCTTTTGCATATCAAGGAGAAATACATATTTAATGATGTACATTTACGC

AAGTTTACGAGTCTGCTTATTGGCTCGGTCTATTAGTTCATGTAAATACAATAATAATACATTGTTGCTT

CCTTTGCAACTTGTATAAGTGTTTGTGTAGCTGTGTCTGTAATGTCAACTTGAAGTTGACATGTATTTAT

ACATCGATGATGCATTTTCCAGAGAGATCGACTTGAAG

>MSTRG.685.1 gene=MSTRG.685

GCTCCAACTGCAACTTTATACCTCGTCTCCCAATCCAGATTTCTGTTGCTCTGCAGAAGTTGTTGGAGAT

TCCCATTCGAGATATAGTTGTAGAGCAAGAGTTTGACACTCTTATTGGAACAGTACCCTAAAAGCTTTAC

TATGTTCCTATGCCGAATGTGCCCAAGGATTTGAATTTCTGCAGCAAAAGAGTCTATAGCTGCCTCTTCA

TCTCCCTTTGTTTTCCAAAGCTTCTTTACTGCTATCAACTCTCCATTGGGCATTTCTGCCTTGTAAACAA

CTCCAGAACAGCCTTTCCCGATCACATTTTCATCTTTGAGGCAATCTAGGATGTTGTCGATAGTGAAATT

TAGCTTTTGAAATGGAATGAATGTCCAGGGATAAGAAAAATCTTCTCGGCCCGATGAAGAGGCTAATGTG

CAAGATGATTTGTCTGCCATATACCTGTGATTCCTTGATACGAAAATCCAGAAGGCAACAACTGCTATTG

TCACAGAAGTTAAAATAACAGCTACCAAAGCTACAGTCTTGGCAGATTTTAATCTGTTCTTGCGAAACTG

ATGTGAGGAGCAAGTATAACCGTTAACAGATTGGCAAAGATCTTCATTTTCAAGGAATGAGTCGGAAGAT

AATGTTTTGAAAAAAGGTGTCACGGGTATAGGACCTGAAAAATTATTGTATGAAATATTTAAAGAAGTAA

GGCTTGTGAGGGAACCTAAAATTGCAATCTTTCCGTGGAGCATGTTGTGCGAAAGATCGAGGGCTTGCAA

TTGTGTCAAATCAGACATAGTCTCAGGGATCTCACCTGTAAATCCATTTGAGCTCAAGTCTAGACTAATG

GTTAAGCTTGTAACATCACCAATCTCATCTGGGATTGCACCAGAAAGACTATTGAAGCTCAAATCAAGTA

GAGTTAGTTTCTGAAGATTCCTAATGGACTTTGGTATTGAACCAGTAAGCAAATTGTTGTTGAGAATCAT

TTTATTTAAGTAACTGAAGTTCCCAAAACTCCAAGGAATTTCTCCAGTAAAACCGTTCCTGCTGAGATCA

AGCTGCTCTAAATTCACAAGCTCCCCCAACCGAGATGGTATATCACCACTGATATGGTTATTGTGCACAT

CAAAAAGCTCAAGAACCGTTATATTAGCAATCTCCGCAGGCAAACCACCAGAGAAATGATTTGCATATAG

ATCAAGAAACACAAGATTTTGCAATTGGCCCAACTCCTTTGGAATGTGTCCTGAGAGCTGGTTCTCCCCA

AGTCTTAATCTCACTAAAGACTGGCAATTAGCAACACTTGGAGGCAGCCCTCCCGACAAAGAATTCCCCA

GTAATAGTAGCTTGCTCAGTTTTTTCAAACCAAATAACTCCTCAGGTATACTGCCAGTGAGTTTGTTCTT

AGAAAGATCGACTGAATATAGTTCAGTGCAGTTCCCAAAAGAAGCTGGTATAGTTCCAGAGACCAAATTC

CCCCAAAGAAAGAAACTCTGCAAGAATTTCAGATTTCCAATTTGTGATGGAATTGGACCTGAAAGTTGAT

TCTTGTCAAGCTGAAGAGCTGTAAGACTAGTGCAGTTACTCAACTGCCATGGAATAGACCCTGTAAGAAA

ATTATCAGACAAATGCAGCTCTTCAAGAAGTACTAGCTTCCCCAAATCACCTGGAATTACACCGCTTAAC

TCATTTGCAGAAACATCAAACACTACTAGGGATGAACAATTTGATATTTCACCCGGGATAGGTCCACTAA

GAGAGTTTCCCCACAGGAGCAAGCTAGTAAGTTTCTGTAACTTGCCCAACTGTTGAGGCATTGAACCAGT

GAGCTTGTTCATATGCAAATACAAATTCCTCAGCTCTGAACACAGCCCAAGTTCAGGAGGGATTCTCCCA

GATATTTCAGTGTCGTAAATTGCCAAAGTTTGGAGATTGATCAAGTTTCCAAATGTAGGCGGTATGACAC

CAGATAGTCCAGTAGCTGCCGCGCCAAATGTTGTGAGATTAGTGAGTAATCCCAACTGCAGAGGTATTGC

TCCACTCAAATATGGATTACCCCCAATCCTAAACTGTTGTAAGGAACTTAATGAGCCTAACTGCGACGGA

ATCGACCCATTAAGTAGATTATCTTGAACACAAAACACTTGCAAAGATGAAAGATTCCCAATTTGTGGAG

GTATAGAATCTGTGAGTTGATTAGTATTCAAGAACAGAAACTGCAGTGAACTGAGCTGGCCTAGTTCAGG

AGGGATTGACCCTGACAACGAATTCGAAGACAAGTCCAAAAGATGCAGATGACTAAATGAGCCAAAATAA

GATGGAATATGACCAGAAATATTAGTAGAAGAGAGATTAAGAAGCTGCAGAGATGACAGTGAAGAAAGCT

CTAAAGGCAAACCAGAGAGATTGAGAAAAGTGTTTGGTATAGAAACAGAAATAACCCTGTCTTGTGGAGA

ACAAGTAATACCTTGCCAAGAACATGGAGTTTGGCTTGAAGGGGTCCAAGTAGACAAAACAGAAGATGAA

GATTTTGCATAATGGTCTGCAGTAGAGACCAGAGAGAGCAGAGCTTGGCCATCACTAGAGAGAGATGTCA

CAAGTTCTGGCCTTGTAAAAACTACCAAAAAACAAGAATAAACCAAGAAAAGTGAAGAGACTAAATCAAG

GGTTGCTTTATTTTTCTCCATTTTTTGAGACTTAAAATCAAGAAATGGGCTTACAAGTATGGTGGTGGTG

GTGGTGATGGTGAAAGTGAAAGTCCACTCTAGTAGACAACATGCATGTGAAGAACACAAATGTGAGATAA

TAATAAGAATGCTTAAAACTCACATTTGTGTGTGAATGTGTGTAAATGTTTCATTGCATGTGAACAAAAA

GAGAAACTTCAAAAGGAAACTAGTAGTGTTATTGTTGTTGTGTATCTTGAAGCTATCATCTTACCGGAAA

ATAAAAATGTGTTCTTGAAACTCCGGCAGGCAGGGG

>MSTRG.694.1 gene=MSTRG.694

AACCACCAACATGTTTTTCTTGACATCATGTATGATGAACACTCTGGCTTCGTACTACCTTAACTTTGTG

CTACCTTAAACACCTGATGAAATGAAATGCATCACCACTACACAGCTCTCTTCACACAATGCCTGAAAAC

ACAAAACCTTAACTTGGGAAAACAACTTCACTCCCACCTTATTAAAACATCTCTAACACTAGACACTTTC

TACACTAACCGCCTAATAGATATATATTGTAGATGCAATTCATTAGAATCTGCTCAAAAGGCATTTAATC

AACTTCCAGCAAAGAACACTCATTCATGG

>MSTRG.704.1 gene=MSTRG.704

AAAAAAACAAATTTTGGGTCACTTTTAAGGCCCAAAAGTACTGACCCATAACAAAAAAAAGAATTTTTGG

CCCAAATAAAAAGGCTATATCAGGCTCTCAGCTTTCAACAACTTTAAAAGGGGAAAACCGTAGCCGTTCA

AATAACCCCAATCCTTCAAGTTCAAAACAGAGCAAAGGTAAGAGGCCTCGACGTTTAATAGATGAGGACG

AAGAAGAAGAACGGAATTTGAATTTGAAGAGAACACTTTAATACATGAAGAAGAGTCAAATCAGGATTTT

GAAGATGACTTTGGTGAAGATGATTGATAAAATATCTACCTGTTAAGAGCTGGAAGCCTGAAATTCTGC

>MSTRG.706.1 gene=MSTRG.706

CTGAGATATTATCCTACTATCGTGTCTAATCGATCAAGACGTGTAGATCGAGAATGTTCAGAATTAATCT

GATGGTGTTGCAAGTTGGAGCAAGAATAATCTGAGAAGACAAACGTAGAAAAAGTATTGTCAGAAGCTAC

CTTGGAACATATGGTTGTGCGATATAGTGCTAATAGAATCGTGGTGAGACATCGGAGTTAAGGCA

>MSTRG.707.1 gene=MSTRG.707

AACCCTTTTCAGCTAGGGATAATGATGATTCACTTTAGACTAGGCCAGGATTGCAAGTGGTTTAGTTTAG

GCAGATGGTCAGGTAGAGTATCCAGGACGATATTATTACGACTAGATGCAGTTGGAGTTCTGCAATTCAA

TTCAGTTGGAAATTAGATTTATTTATTTTGTTATCAACTACTCTTCTTATCGAGGTTTTCTTATCGAGGT

TTAGCTGATAATGTAGTTTAATTTGTAATTAGGATTTTATAGTTTTAGTTTCTGATTTCAATGCTCTAAC

CTGTATGCAATCCTGTTTTCAGGGGGT

>MSTRG.709.2 gene=MSTRG.709

GTTTTAACACCCGGTTCTCGGCGGTTAAGAACGAATAAGAACACAACGCGAGATCTGGAAGTAATGTACA

GCTGAAAGATGTTGTCGGCGACGAGCAGAAGCTCCGCATCTTCATTACACTTGAAATCCACCAGACATTT

CCATAGGAAACCATCTCCATACACAATCACGCTTTACATAATCGCCCTATTTGCTCTCTCTATTTTCATT

TTTATTTACTACAGTAAAGATATCTTAGACATTGATGATCAGCACAAGCCCTTTTCGTCTCAATCCCACC

AGCTGTCAGATGAACAAGTCCAGGGAGCTCTCTCGAGCTATGGTTTACATCAATGCGTCGAACCAACTGC

AAAATATAAAGCGGCACTAGGATCGGATCATTATATAACTGTAAGAAGTAATGGAGGACTGAATCAAATG

CGAACTGGTATATCCGATATGGTGGCTATTGCACACATTATGAATGCAACCTTAGTAATTCCACAGCTGG

ATAAGCGCTCATTTTGGCAGGACAAAAGTAAATTTTCTGATGTATTTGACGAGCCACACTTCATTAGAAG

CTTACAAGGAGATATTCGGATTATCAGAGAGCTACCTGGGGAACTTGTATCTCTCCCTCATGCCAGAAAG

CACTTCACCTCCTGGTCTGGAATGGGATACTTTGAGGAGATGAAACAAATATGGAAGGACCACCAGGCAT

TACATGTTCCGAAGTCAGATGCTCGCCTTGCCAACAATGATCTTCCCCTTGATATCCAGAGGTTGAGATG

TAGGTCGCTTTATCATGCTCTTCGCTTCACACCTCCTATAGAGAATCTAGGGAAGAAGCTAGTGGAGCGG

CTGAGATTGCGTGCGGAAAGATATATTGCTCTTCACCTCAGATATGAGAAAGACATGTTATCTTTTACTG

GGTGTACCTATGGTCTTACTGATGCTGAATCCGAAGAGCTGAGAGTGATGAGGGAAAACACCAACCATTG

GAAGATGAAGAAGATAAATTCAACAGAGCAAAGAATTGGAGGGTTTTGTCCACTAACTCCTAAAGAGGTT

GGCATATTTCTTCAAGGTCTTGGTTATACTTCATCGACACTAATATACATTGCGGCTGGGGAGATTTATG

GTGGAAGCACTCACCTTAAAGAGCTTACATCTCGCTTTCCGAATGTTGTTTTCAAGGAAACAATTGCTAC

TGAGGAAGAGCTGAAAGAATTAACCAATCATGCATCTCAAACTGCAGCTCTTGATTATATAATATCAGTA

GAGAGTGACGTATTCATTCCATCGCATTCAGGTAACATGGCAAGAACTGTTGAGGGGCATCGCAGGTTTC

TAGGCCACCGAACAACAATTACCCCTGACAGGAAAGGTCTGGTTGACATATTTGATAAAATGGATAGTGG

ACTGCTTAATGAAAGCTCATCGTTGTCGCGCCTTATAATTAAGCTGCACAAAAACAGGCGTGGTGCTCCC

AGAAAAAGAGGCGGCCCTCCTCCAGGACTCAAAGGCCGAGCAAGGTTAAGGTTTGAAGAACCCTTCTACC

AAAATCCATACCCAGAATGTATATGTGATTCAAAAGTCACTACAAATAGTAGTCATAGTGCGTAGGTCAG

CTTGCACTTATCACTAAAAAAGATGTTCCGAGACATTAGATAATTCAAATTAGAATGACAGTGCTTTTTA

CTGCTGCAAGCCTTTTTGTAATTGAACTGGTTCCTTTTTTTGGTTCTTCGATTCTTTCTAGAATCTGTAT

TTTATGTTAAGATCCTGTACAATCAATACAAGTAGTTATC

>MSTRG.709.1 gene=MSTRG.709

GTTTTAACACCCGGTTCTCGGCGGTTAAGAACGAATAAGAACACAACGCGAGATCTGGAAGTAATGTACA

GCTGAAAGATGTTGTCGGCGACGAGCAGAAGCTCCGCATCTTCATTACACTTGAAATCCACCAGACATTT

CCATAGGAAACCATCTCCATACACAATCACGCTTTACATAATCGCCCTATTTGCTCTCTCTATTTTCATT

TTTATTTACTACAGTAAAGATATCTTAGACATTGATGATCAGCACAAGCCCTTTTCGTCTCAATCCCACC

AGCTGTCAGATGAACAAGTCCAGGGAGCTCTCTCGAGCTATGGTTTACATCAATGCGTCGAACCAACTGC

AAAATATAAAGCGGCACTAGGATCGGATCATTATATAACTGTAAGAAGTAATGGAGGACTGAATCAAATG

CGAACTGGTATATCCGATATGGTGGCTATTGCACACATTATGAATGCAACCTTAGTAATTCCACAGCTGG

ATAAGCGCTCATTTTGGCAGGACAAAAGTAAATTTTCTGATGTATTTGACGAGCCACACTTCATTAGAAG

CTTACAAGGAGATATTCGGATTATCAGAGAGCTACCTGGGGAACTTGTATCTCTCCCTCATGCCAGAAAG

CACTTCACCTCCTGGTCTGGAATGGGATACTTTGAGGAGATGAAACAAATATGGAAGGACCACCAGGCAT

TACATGTTCCGAAGTCAGATGCTCGCCTTGCCAACAATGATCTTCCCCTTGATATCCAGAGGTTGAGATG

TAGGTCGCTTTATCATGCTCTTCGCTTCACACCTCCTATAGAGAATCTAGGGAAGGTTCTGAGCATACAT

GACGGTGACACCACAGAAGCTAGTGGAGCGGCTGAGATTGCGTGCGGAAAGATATATTGCTCTTCACCTC

AGATATGAGAAAGACATGTTATCTTTTACTGGGTGTACCTATGGTCTTACTGATGCTGAATCCGAAGAGC

TGAGAGTGATGAGGGAAAACACCAACCATTGGAAGATGAAGAAGATAAATTCAACAGAGCAAAGAATTGG

AGGGTTTTGTCCACTAACTCCTAAAGAGGTTGGCATATTTCTTCAAGGTCTTGGTTATACTTCATCGACA

CTAATATACATTGCGGCTGGGGAGATTTATGGTGGAAGCACTCACCTTAAAGAGCTTACATCTCGCTTTC

CGAATGTTGTTTTCAAGGAAACAATTGCTACTGAGGAAGAGCTGAAAGAATTAACCAATCATGCATCTCA

AACTGCAGCTCTTGATTATATAATATCAGTAGAGAGTGACGTATTCATTCCATCGCATTCAGGTAACATG

GCAAGAACTGTTGAGGGGCATCGCAGGTTTCTAGGCCACCGAACAACAATTACCCCTGACAGGAAAGGTC

TGGTTGACATATTTGATAAAATGGATAGTGGACTGCTTAATGAAAGCTCATCGTTGTCGCGCCTTATAAT

TAAGCTGCACAAAAACAGGCGTGGTGCTCCCAGAAAAAGAGGCGGCCCTCCTCCAGGACTCAAAGGCCGA

GCAAGGTTAAGGTTTGAAGAACCCTTCTACCAAAATCCATACCCAGAATGTATATGTGATTCAAAAGTCA

CTACAAATAGTAGTCATAGTGCGTAGGTCAGCTTGCACTTATCACTAAAAAAGATGTTCCGAGACATTAG

ATAATTCAAATTAGAATGACAGTGCTTTTTACTGCTGCAAGCCTTTTTGTAATTGAACTGGTTCCTTTTT

TTGGTTCTTCGATTCTTTCTAGAATCTGTATTTTATGTTAAGATCCTGTACAATCAATACAAGTAGTTAT

CATATATCAATTATACATTTTACTTGTA

>MSTRG.716.1 gene=MSTRG.716

TACTTATCATAAAAAGCAAAAGCAAGTGCATCAAAACTCTTCTCACAACCACAATTCTAAACTGCATATG

CATAAAAATGTCACCTCTGCACCAAACACTCAAGCTTCTAGGCACACACACACTCCAAATGCTAAACCTA

CTGTGAATGCTCCTAAAAGTGTGTTAGTTGATCCATCCAATCTGTCTAAGCTTGATAACATGCTCTTGAA

TGTGCAACCTGGTATGATGATCTATGTTTCTCAGAATGGTGTGGTGCATAACATGTTTCTATCTGCTAAG

TGGGAAAACATG

>MSTRG.719.1 gene=MSTRG.719

TGTGTTGTTCTGGTATTGTTAATTGTTATGGATGCAGGAGCTTCTTGCATAACAAGATCGTAACATCTAT

GGTGGTGGTGGAGAGATTCAGAAGGGGCGGAGATGAAGGATGCTAATTTGGCGATAATTGTGCCTCAAGT

AAGGATATTAAATATTCTTAATTAAGGGTTTTTAGTACCATTTATGGCGACCATAATTGGGGCATTTATG

GCCCATAGAGGCATTTAAGTGCCATAATGTTGGCTCATTTATGTCTTTATTTTAATTTTTTTCAAATTTA

CTCATAATTCTTTTATTCGATTTTGTCTTATTTTCAGGATGTTTAAAAAAGATTTTATGAATTTACTTAT

GAAGGCTGCTTCTGAGAAGACGCGAACCTTGGTGTACAAGACATTTGAGCTACAAACATTGATAGCAGGG

AATAAAGAAGACATTTTTAGTCTGCTTTGTAGATGATGTAATTTTGTAGGGCACTCTTCTTTCCCCTTGA

GTTTTTTTCCACAGGGTTTTACTCTTGAAGGCTTTTAACGAGGTCTTTTTCGGTGGGTATCTCTTCTGAC

TTGAAAGGCTATGTTA

>MSTRG.720.1 gene=MSTRG.720

GGGATGAATATGCCGATGAATATGCCTTTCTAAAAAAAAAGAGAGAAAAAAAAAAGTGAAAAAAAAAGAA

TAAAAATTATCAGGTACTCCTTTGAGATCAATGGGTAAAATGCAATGTCATGTGAAAGGGACGATCCATG

TACTTGTGCTGGTATTAAGTGCAAATGTATTAGAATAAGCATATTCTTGGTGACTTTTCACAACCCTTGA

TTACTGGAGAATTACTTGATTAAAAAAAAAAAAAAAAAAAGCGAAGTTGAATGGCGGTCGTGACTCACTT

ACACTGAGAAGCGTCCCAACCTTAGACTTAGCAAGAAAAAAAAAAAAAAAAAAAAAAAAATAGGAGAGGC

ATAGGAATTCTTTAGTGACCCTAAATTGTAGTTGACTCTTTAGTAAAGAAGTGTTTAAACCTGATTCTGA

TTAACGGCTCATAGTGAGGACTCTGTTGAAGTTTGATAGTCTTTGTTTTGTTTCACTAGAGAGCATGCTA

GCACACACGATGCACTTCACACTTGTAGTGGAAGAGAACATGGCTAATATGTGAAAGAGATGAATGCCGA

GAATGTTGCATATTCTGAGGATTCTATGTTAACAAGGATTACACTTCATGATTCGAGTAGACGTAGGCAT

ATAGTTGCATTCATGCATTGCATTAGTAGTTTGTGTGTGTGTCTATGCTTGAGGACAAGCATCGGTTTAA

GTTTGGGGGTGTGATAAGTGGATTTTATATCCACTTGGAATGCCTTCGTTTCGACTTAAATTGATGATTT

GGCCTCAAGTATGTGGTGTTTTTGATGTGTTCTTGTGGTGTTGTGTAGGTTTCCTAGGAGGAGGATAAAG

GGTGGATTAAAGGCTAAAAGGTGAAAGAAACGGCGTAAGGATCGAGCGAGCAAGGAAGGAGAAAAGGGAA

GGAAGGTTCCAAGAGCGTTCGGGGCTGCCCAAATGGGGTCTCCACCCCATTTACGGGGTCTCCACCCCGT

TCTGAAGAAGAAAAATCTGCCAAGATGGGGTCTCCACCCCACTGATGGGGTCTCCACCCCACTTCAGCAG

AATAGAGGCTGCCAAGACAGGGTCTCCACCCCACTGACGGGGTCTCCACCCCACTGACGGGGTCTCCACC

CCATTCTGTTGGGAGAAAAAGCCCGAATTGCCTATTCTTGATCCGTTTGAAGGCCCGATCGCTGTGGGAC

TTAAATCAAAGCTCAAGGGAGAAGCCTAATAACCTAGAAACATTCCAAGGAGCACGTGACGGCTACG

>MSTRG.721.1 gene=MSTRG.721

ATAAAAGTTAGCATTATCAGATTTTTCATAAATATTTTTAATTTTTTATACAAACGGATCAAAAAATATA

GAAGCCGCCTTAACATAAAAGTCGGCTTCTTAGCATAAAAACACAAAACACAGAAGTCTTGTTACCCAAA

TAGGCACCGAAATTTTAAGCATCCTTTCTTAAGTGACCAAGCTCATAGAGTTGCTTAAATTTGTGGTAAA

CAAAGTGTGTATCCATGAAACCTGTGTTGTGTGATCATCAGCTACTTGTGGCGGTGTTTCTCGGTAATAT

TAGCAGCCTGGCAGTTCCTACTTACTCTAGGACAACATCGAGATCAAAGGGACGGAAAATGAAGAGGAAA

AAGAAAAAAGAGTACTAGAGGGGTGCTCAAGAACAAGTAACAAATGCTTTGTTCTTGTAACACCTTTGGC

TTCTAGACGCAACCAGTTTGTATGGCACCAATATCGATCTCCATGTCCTTCTTAATCTTGCATCTTGTTC

TGTTTTCCTTTCCACTTTCCTAGTTGCTTGCTTTCATCTTTGTGTTGCTTTCCTTGGCTTTCATGTTCTT

TTGTTTGGTTTTCATGACTCTGTTATTGCAAGAAACCGGAGTATATAAATAAATTGTGCCAGTTTCTGTT

TTTCGA

>MSTRG.725.1 gene=MSTRG.725

TCCATTGCTGCTCTCAGACAAGGGCGATTCAGAGTTCAGACTCTCGCATCGACTCCACAGATTTCAATAA

GCTCCATGTCCGACGCATCTCAAGGATTCAGTTTACCATTGCAACGATTTTGAAATCTAAAATTATCTAA

GCATACAGAATTGAAATGGCTGGTGATATGGAATCAAATAATTCATCGGTACTAGAAAGAGGCGCTGGAA

AAAATAAAAGGAAATGGACAGAAGAAGAAGATGAAAAACTAGTCGAAGCTTTGATGGAACTTGTGAATAC

CGGAAAGTTCAAAGCGGACAATGGCTTTAAACCTGGATACCTGACATTTCTTGAGAGTTCACTGCAAACC

AAGCTTCCTACAGCACGTATAAAGGGAAAGCCTCATATAGAGTCTCGAATGAAGACACTAAAAAAGGATT

TCAGTGCTGTGTACGACATCAGCTATGGAGCAAACAGTTCGGGGTTTGGATGGAACGCAGAAGACAATGT

TGTTACGGCGCCGAGGGATGTTTGGGTTCAATATCTTAAGGTTCATCCTGGAGCAGCAAAATGGCAGAAT

ACTGCCATGCCATCATTCAAAGAACTTTCAGTTATTTTTGGAAAAGATAGAGCAACCGGAAATATGGCTG

AAAATCTTGAAGATGTGGTAGAAGAATTGAACACTGAAGCTGCTGATGAATTATCTCTTCAGGAGGACCT

GCAACGAAGCACACATTCAGACGAGTCAACAAGCAAGAAAAGGAAGAAAGGAAATGTAGAATCATTGTTG

GAAGCTGTGTATGCTGCTTCGGACAGGATTGCCAATCAGTTTGAGGCGTCAACCAAGTTACTTATTGCAG

CAGAAGAAGATATGATGCAGAAAAAGAAGCAATTAAATGACGAACTATCAAAAATTCCAAATCTCACGGT

ATTGCAGAAGCTTCAAGTTGCAAAGAAAATAGCCAAGGACGAAGATCTTATGATATTGTTTTTTGCAGCA

CCGGCCGAGGAAAAAATCATCTTTGTTCATGCTGTTCTAGATAACCAGATTTGAGCAAACTTTGTTATTA

AGAACCTTTGTTTTTGTTAGGTCCAATCAGACGTAGAAGGGGGGGGTTGAATACGTCGTAC

>MSTRG.725.2 gene=MSTRG.725

ACCAGTTTCTTTTCACGAATACCGAGAGAAACCGAAGTGTGTTATTCGTGTTCTTCGTAATCAAACGAGG

ATTTGGAGGTGTTATTGGAATCCGATGGAGGTGTATGAATAGTCAAAATGAAGCTACCGACGCGTAGAAT

CTAGTTTAATCATCCGTTTGTGTGCAGATTCAACAGAATTGAAATGGCTGGTGATATGGAATCAAATAAT

TCATCGGTACTAGAAAGAGGCGCTGGAAAAAATAAAAGGAAATGGACAGAAGAAGAAGATGAAAAACTAG

TCGAAGCTTTGATGGAACTTGTGAATACCGGAAAGTTCAAAGCGGACAATGGCTTTAAACCTGGATACCT

GACATTTCTTGAGAGTTCACTGCAAACCAAGCTTCCTACAGCACGTATAAAGGGAAAGCCTCATATAGAG

TCTCGAATGAAGACACTAAAAAAGGATTTCAGTGCTGTGTACGACATCAGCTATGGAGCAAACAGTTCGG

GGTTTGGATGGAACGCAGAAGACAATGTTGTTACGGCGCCGAGGGATGTTTGGGTTCAATATCTTAAGGT

TCATCCTGGAGCAGCAAAATGGCAGAATACTGCCATGCCATCATTCAAAGAACTTTCAGTTATTTTTGGA

AAAGATAGAGCAACCGGAAATATGGCTGAAAATCTTGAAGATGTGGTAGAAGAATTGAACACTGAAGCTG

CTGATGAATTATCTCTTCAGGAGGACCTGCAACGAAGCACACATTCAGACGAGTCAACAAGCAAGAAAAG

GAAGAAAGGAAATGTAGAATCATTGTTGGAAGCTGTGTATGCTGCTTCGGACAGGATTGCCAATCAGTTT

GAGGCGTCAACCAAGTTACTTATTGCAGCAGAAGAAGATATGATGCAGAAAAAGAAGCAATTAAATGACG

AACTATCAAAAATTCCAAATCTCACGGTATTGCAGAAGCTTCAAGTTGCAAAGAAAATAGCCAAGGACGA

AGATCTTATGATATTGTTTTTTGCAGCACCGGCCGAGGAAAAAATCATCTTTGTTCATGCTGTTCTAGAT

AACCAGATTTGAGCAAACTTTGTTATTAAGAACCTTTGTTTTTGTTAGGTCCAATCAGACGTAGAAGGGG

GGGGTTGAATACGTCGTAC

>MSTRG.725.3 gene=MSTRG.725

CTTGTTCTGAGACTATTTCTGTTCTTGTATATGCAGAATTGAAATGGCTGGTGATATGGAATCAAATAAT

TCATCGGTACTAGAAAGAGGCGCTGGAAAAAATAAAAGGAAATGGACAGAAGAAGAAGATGAAAAACTAG

TCGAAGCTTTGATGGAACTTGTGAATACCGGAAAGTTCAAAGCGGACAATGGCTTTAAACCTGGATACCT

GACATTTCTTGAGAGTTCACTGCAAACCAAGCTTCCTACAGCACGTATAAAGGGAAAGCCTCATATAGAG

TCTCGAATGAAGACACTAAAAAAGGATTTCAGTGCTGTGTACGACATCAGCTATGGAGCAAACAGTTCGG

GGTTTGGATGGAACGCAGAAGACAATGTTGTTACGGCGCCGAGGGATGTTTGGGTTCAATATCTTAAGGT

AATTAATTATCACATATAATTAATAATACCAGATCTGCTTTTCCTGGCAATTTCTTACACTAACATGGTT

ATCACACTCCATGACTCCACCAAGTCTCCTTCCACCAAGACATTGCAGAATAAAAAAAACAAAAACATGT

ACTTAGCAGTTGCAATGTGCAACTAAATTCCACAATTATGAAAGGATTATATTATTAGGGAAGCCATAGT

AGGTTGTTTGGTTGAGGGAATTGAGTTTTTATTATTAAAAAGTGCTGGATGTAGTGGA

>MSTRG.725.4 gene=MSTRG.725

ACCAGTTTCTTTTCACGAATACCGAGAGAAACCGAAGTGTGTTATTCGTGTTCTTCGTAATCAAACGAGG

ATTTGGAGGTGTTATTGGAATCCGATGGAGGTGTATGAATAGTCAAAATGAAGCTACCGACGCGTAGAAT

CTAGTTTAATCATCCGTTTGTGTGCAGATTCAACAGGATTTGGACGAGACGTGTACGCTCGAAGACGTCA

GGAAGTCGAAATGGTATTGCAAAGCGAATAAGGGATGAATCGAGTAAGCGAAGCGAAGATGTACGGTAGA

TAGAGTATAGCCAGAGATGGTCTAGAGACTATGAGATGCATAGATTGTGAAAGTGGAAAGATGAGATTGA

GATGTGAGACTGGAGTCAGATTTAAGGCAAGCTTAAAGATGAGATGATTGATTGAGATTGTGAATAAGGC

TAATGCTAGGCGAGGAGACAGAGTTGAGAATCCAGTGGTCGAGGAGTGTTCAGGAAAGTTGATGAGGTTG

GTGCAAGCTGAAATAGTTGACAGTGATTCTTCTTATCGAAGATGATCTGATTTGGTAAGAGATGTTGTGT

AATTAATAGTGGTAGATTCTAATTTCAATACTGTAACCTGGATGCGATCCTGTTGCTTTTAGGGGGTTTA

AATGTTCTCTTTCG

>MSTRG.725.5 gene=MSTRG.725

ACCAGTTTCTTTTCACGAATACCGAGAGAAACCGAAGTGTGTTATTCGTGTTCTTCGTAATCAAACGAGG

ATTTGGAGGTGTTATTGGAATCCGATGGAGGTGTATGAATAGTCAAAATGAAGCTACCGACGCGTAGAAT

CTAGTTTAATCATCCGTTTGTGTGCAGATTCAACAGAAAGGCTTAAAGATGAGATGATTGATTGAGATTG

TGAATAAGGCTAATGCTAGGCGAGGAGACAGAGTTGAGAATCCAGTGGTCGAGGAGTGTTCAGGAAAGTT

GATGAGGTTGGTGCAAGCTGAAATAGTTGACAGTGATTCTTCTTATCGAAGATGATCTGATTTGGTAAGA

GATGTTGTGTAATTAATAGTGGTAGATTCTAATTTCAATACTGTAACCTGGATGCGATCCTGTTGCTTTT

AGGGGG

>MSTRG.725.6 gene=MSTRG.725

CGCGTAGAATCTAGTTTAATCATCCGTTTGTGTGCAGATTCAACAGGATTTGGACGAGACGTGTACGCTC

GAAGACGTCAGGAAGTCGAAATGGTATTGCAAAGCGAATAAGGGATGAATCGAGTAAGCGAAGCGAAGAT

GTACGGTAGATAGAGTATAGCCAGAGATGGTCTAGAGACTATGAGATGCATAGATTGTGAAAGTGGAAAG

ATGAGATTGAGATGTGAGACTGGAGTCAGATTTAAGAAAGGCTTAAAGATGAGATGATTGATTGAGATTG

TGAATAAGGCTAATGCTAGGCGAGGAGACAGAGTTGAGAATCCAGTGGTCGAGGAGTGTTCAGGAAAGTT

GATG

>MSTRG.728.1 gene=MSTRG.728

AGCAGATAATGTAACAATTGGAAGGAACAATAACTAGAAGTGTTGTGTGATTTCTGGAGCCCATGATTAA

AAGGAAATTTTATTGACGTGAATGATTAGTCCACTTAAGTGGTATTGCTGTTCAAATTCAGAAAGTGCAG

AATCTGTGTTCCCAACTTACGACTATACTTCAATGGCATTACTGGAAGGAGTCACACTATGGGATGATTA

TCACAAATTTATATTTCAAGTCGGCAGAAATCTGCTTCCTATGCAATGTTAGGATATTT

>MSTRG.737.1 gene=MSTRG.737

TACAACCACAGTTCCCAACCTAATCTCTATTCAAAATTTTTAATCAATCAAAACAAATTCACATTGCCGC

TTCTCCAATTACTTGCGCGCAGATGGAAAGAAAATTCGAGCTGGGTAAAGATGCGGTTGATCTAGGTATT

CAACCACCATTCAGTCGCAATGTTTGTAAGTTGCTCAGATTTTGATACACTCTTCTTGGTTGATTGATTC

ACAACTTTTTACATATACCATTCAGTCGAAATGTTTACAGATTTTGTTACCCTCTTTCCTCTTTTCCGAA

TTGATGGAGAACACACTTTGATAGTAAGTTTGTTTTTGTTGCAGTTTATAGTTTGATGTGAACTGTTTGA

TGAAATGGCTGAAAGAAAAGGCTATTAATTCTTGCACTCTGTATTGTTAATGGCTTGATTTTGGGATGTA

TATACAAGTTTCTTTGAATATTTATTGTTTGAAACTAACTTATAATGTGATATATTTTGATGTAGACGAA

ATCAAAGATGCAGATTGATGTGTATGGCCCTGTTGTAACTGTCTGGACTTTGCCTGAGAAGAAGGATTCT

CATTTCCAGGAGGATCACATTACGGAGCAAAATGATGTAGGGCTAGAACCTAAAGTGATGTTAGATCCAG

CAACATACTTCTGGATCGTAAGTGAAATCCACAAGTATTAGATTTTGGACTTACCAATTTGTTTGAATAA

AAGATGGTCAGAATCCAGCAGCTTGGATGTTGGATATAACTTCAGAATTGCTTGAAGCAATAAATGGAGT

CGACTTTGCTGAAGTGTACAAGACATAACTATACAGGAGAAACAAAGAATTGACAGGGGTTCTGTTTGGA

AGAATTTTTCTGCTTGGATTATGAACAAAAAGTGGATTCTTTACAGTGAAGTGTGGTGCTGAGCTGTGGA

AGGGGCAAAAGTTCGAAAAGCAGTGATAAGGAAGAAGAGAAACTGAAGGATGTGGTACATGTCAGAGTGA

GAAGGGGCCACTGGCCCTCAAATTTGCAGGCTGCAGTTGCAGAGACAGATCTGATGGTCCTCATAATATG

TGCGGAGAGTTGCTCAAAATTAGCGGCAATTTAGGAAGGTAACGTGTCTTCACTAGCTATTGTGCAGGTA

AAAACTTATGTTGGAACTGTTATAACATGGAAACTGCTCTATTTGATAGTAAAAAATGGCACTGACAATC

AAAGGAAGTATTTTCTACACATGTTACAACCTTACAGATGAATTACTACTAAATTTTACAAACTAAATAT

ATATTCAACCAAGCCAAGAG

>MSTRG.741.1 gene=MSTRG.741

CCAGACCTAACGATGTTATGCTTTGTTTCTCGCCTCTCTTACTTGGTCTGTTTCAGATGCCTTATGTCAG

GCAGGAAGAGGTATACTCGGTGCCCACCTTTTTGCTCATCGACGCTCCCTTCAATCCCTTGGTAAGCGAG

GTGGAGAGGGCAGAGCTACCCAGCGCATCCATAGATTTGTCTCTGTAGAATCAGAAGTGGTGAAAGCTCT

GTTTGAGCATCATGATTCTTGCCATTTTGCTTATATATTTCAGGTAAATGGAGCAACCTCTTGGGAAAAG

TGTTATCTTTCTCTGGTGGTTACAGTTTTATCTCATAAGGGCTGCCTAATGTTTCCAAGCTGGTATGTGT

AGGAACAATCGGACCTTGGCCCTGCGTTTTATGATATTAATTGAAAGTTCGAACAGAATATTAACCTTAA

TCGCTACGGCGCAAGCGATTCAAATCCACGTCTTCTACTGTATTCCTTGATTCTCCTTGGACGAAGTGTG

GCCTTCGATCTCCCAAGGTGTGCCTCTCCTTAAAGCTCCGAAAACCCAAAACCCTAGCTGTCTCCTTGAG

AAAAACGTCTGCCTCTCTCAAACTCTAGGATGAATTCGTTTTGGTGTGTCTTTCAGCTTTAGGAGAACGA

GAATATATATAGGCTACAGTAGGGATCATGGACCATTAGGTCAGGCCTTCTAATGACATTCCGGGCCAGG

CCCAATGTCATTAAATATTAATTCAATCCACTAAAGAATTAATATTTGCACTACCTTTCCTAATTCCGTA

AATTAATTATTTAATTCGGCTCCCATATTAATTGCTTATAAATTCCC

>MSTRG.741.2 gene=MSTRG.741

GACCTAACGATGTTATGCTTTGTTTCTCGCCTCTCTTACTTGGTCTGTTTCAGATGCCTTATGTCAGGCA

GGAAGAGGTATACTCGGTGCCCACCTTTTTGCTCATCGACGCTCCCTTCAATCCCTTGGTAAGCGAGGTG

GAGAGGGCAGAGCTACCCAGCGCATCCATAGATTTGTCTCTGTAGAATCAGAAGTGGTGAAAGCTCTGTT

TGAGCATCATGATTCTTGCCATTTTGCTTATATATTTGTAAATGGAGCAACCTCTTGGGAAAAGTGTTAT

CTTTCTCTGGTGGTTACAGTTTTATCTCATAAGGGCTGCCTAATGTTTCCAAGCTGGTATGTGTAGGAAC

AATCGGACCTTGGCCCTGCGTTTTATGATATTAATTGAAAGTTCGAACAGAATATTAACCTTAATCGCTA

CGGCGCAAGCGATTCAAATCCACGTCTTCTACTGTATTCCTTGATTCTCCTTGGACGAAGTGTGGCCTTC

GATCTCCCAAGGTGTGCCTCTCCTTAAAGCTCCGAAAACCCAAAACCCTAGCTGTCTCCTTGAGAAAAAC

GTCTGCCTCTCTCAAACTCTAGGATGAATTCGTTTTGGTGTGTCTTTCAGCTTTAGGAGAACGAGAATAT

ATATAGGCTACAGTAGGGATCATGGACCATTAGGTCAGGCCTTCTAATGACATTCCGGGCCAGGCCCAAT

GTCATTAAATATTAATTCAATCCACTAAAGAATTAATATTTGCACTACCTTTCCTAATTCCGTAAATTAA

TTATTTAATTCGGCTCCCATATTAATTGCTTATAAATTCCC

>MSTRG.741.3 gene=MSTRG.741

ACCTAACGATGTTATGCTTTGTTTCTCGCCTCTCTTACTTGGTCTGTTTCAGATGCCTTATGTCAGGCAG

GAAGAGGTATACTCGGTGCCCACCTTTTTGCTCATCGACGCTCCCTTCAATCCCTTGGTAAGCGAGGTGG

AGAGGGCAGAGCTACCCAGCGCATCCATAGATTTGTCTCTGTAGAATCAGGGCGGCACAAAATTAAATAA

GTTGTGCCTATGGTGAAGACAAAGATCTGATGACATGAAAGTATGTGACCAAAATTCTCCTAATCAACGC

ATAGGTATACAGGTGGTCCGTTGGAACCTTGGCTTGCCTTTACTCCTTGACAACTTTAATATAACATACA

ATTATATGGATTTTCAATTTACAAATTGTTGAGTTACGTAGTGAAGAAATTGTATCCTGG

>MSTRG.742.1 gene=MSTRG.742

AATGGTTTGTCTAACACCATTTAGACTGTGAACCTTTATTATATTATATAAGGAACTCGACAATCTAATC

TTCTGCTATCCCATTTGATACTAGATTGTCTACAATATATAATATACAGACATTGTGAAAACATGCATTC

AAGATTCTCAAATAATTGTTATATACTTCAAACGAATATTTCAGTATAACCCTAACAGTATGAAGATCTG

AAGCATGAGTATGATGGTCAGACTACTGGACTTATATCCATAACATGTTGGGCAATCTGGCGGGCCCGAA

TGAATTGGTATGGAGGCAACATAATCCGGTGACTACAAGAGTGGTAAATAATGCAATCACTTTTCCTGAT

ACATGGAATCTTGCTCGGGTATATTATTCTAATCATCCGGATACTGCTACAAATCATGTTGAAGTAATAG

AGTGGTGGTCTTAACCTGAAACTGGTAATATTAAGTTTAATTGTGATGGCTCTATCTTTGGCTCTGAGAA

CAAGTGTGGATTCGATTGGCTGGCTAGTGATCATGATGGTGTGTTGTTAGAGACTACTGCTAATGTTCTA

TTAGGGATAGTGAAATTGCAGCTTGCTGGGGTAATTAGTTTAAAGGGGTTGTGAGTTGAATCTAAGATCA

AGATTAATACTTACAATACTTTGAAGTCTCCATTGGTGTACATACTGATTGTTATCTGCTGGTGCAAGCA

GTAAATAGCACACATTCCTTGCTCTCTGCTCAGGTCTTGTTGCGGGATTGCAACGAATTGCTTCAAATAT

TGTCAAATGCTTCTAGATAGAATACTTCCCTAAAACAACTCGGGAACAGGGCAACCATTTGATCAGCCTA

TGTTTCCTTATTTCTGTATAATTACACCTGCCTTGTATCCACTGTACAATGTACATGACATATGGCCATT

AATCTTGCATTTGCAGTTTGCACACAGTCGAAACTATTATATATGGC

>MSTRG.746.1 gene=MSTRG.746

CACGCACTTCTCCTGAGCAAATTAATTCAATGTTAAAAGACAGTTCAACCCCTCAGGCCAAAGCTAGATA

TATTACCGAAGCAGTGTTTTTCCGCCATTCTGCCCTTCCGACCCTGTCTCCGAACTGGCTCCCGGCCTAA

CTATCATCCTCGGCGGTCATTTCTGTCGGCCCATCTCGACTCCTTCCGTCTTCTGTCCGCAGCGCTGCTT

CCGTGGTGATCAAGAGGAGCGTGGTTACCTCATTTCAGAGCTACTTTGTTGGTTGAGAGGCCTCTCGGTT

GTACTCAAGCATCTCTTCGGAGCTTCTATCTCTCCGAGGCCTTCAATCCATTGCGGCTAATTCCGGGAGG

CAAATGGTTGAACCGCGCCAGTTCTACATCTAACTCAGATTCCCCTGCTGCGAGAGAAGCTGTCTTGGTA

GGAGATCTGGTCAAAGATCAAGAGTGAATGCAATTTGGAGCGTGGAAGGTTATATTCATTGGAGGAGCAA

ATTGAACATAAGTTGGATTTCATCAAGCAGGAACTTAACAACATTGAAGTCCCTATAGTACTGGTGCGTA

TTAGTTCTTAATAAC

>MSTRG.746.2 gene=MSTRG.746

CTTCCGACCCTGTCTCCGAACTGGCTCCCGGCCTAACTATCATCCTCGGCGGTCATTTCTGTCGGCCCAT

CTCGACTCCTTCCGTCTTCTGTCCGCAGCGCTGCTTCCGTGGTGATCAAGAGGAGCGTGGTTACCTCATT

TCAGAGCTACTTTGTTGGTTGAGAGGCCTCTCGGTTGTACTCAAGCATCTCTTCGGAGCTTCTATCTCTC

CGAGGCCTTCAATCCATTGCGGCTAATTCCGGGAGGCAAATGGTTGAACCGCGCCAGTTCTACATCTAAC

TCAGATTCCCCTGCTGCGAGAGAAGCTGTCTTGGAGATCTGGTCAAAGATCAAGAGTGAATGCAATTTGG

AGCGTGGAAGGTTATATTCATTGGAGGAGCAAATTGAACATAAGTTGGATTTCATCAAGCAGGAACTTAA

CAACATTGAAGTCCCTATAGTACTGGTGCGTATTAGTTCTTAATAAC

>MSTRG.747.1 gene=MSTRG.747

TATTGAAGTAACAACTATTATATATCTATTTTCTAGCTAATTTAAATGTGTTTCCTACCTCTGCATCCCT

TTGAGACATTGTTATATAGTAAGAGGTGTAACAAATAGCATATTCCAAAAAGAGAACATTTTGAAATAGA

TCAACAAAGATTGCTAATACAGACCGAGCTTCAGAGCTTGTCATATTTGATGCTCATAGAAACTCTTCAC

AGATTATTTAAAGCGTGTCCGTTTTTTTGTTGTGTATTATTTTCGTATCAGCTGAAACGGTGAATGTTTA

GAGAGACCAGAGACCAGAGACCAGAGATGGAGCTAGATTGTATCTTGGCAGCAGCTTGGACTTTTGAGCC

TAGAATACCTCCTCATATATAATGGTATCTTCTATTTGGTTGTTTTTGGTTAATAGTTTATCTGTAATAC

TGGATGTAGACTCGAATCCAATACTATGTTCCTGTATAATGTTTGATTATAATTAAAATTTGGTTGATTT

ATGCTT

>MSTRG.750.1 gene=MSTRG.750

GTTTTTTAGTTTGTTATATAATGCCATCTCAAGGTAGCTTATTCTTCAGAGCCAAACAGCTTTCAATACT

CATGGAGAAAAGCAAATCATTCCCAGATTATTCTGGCTCCTACGTGGAAGCTCGGTTCGGGTTCGAGGAT

CGGTGCAAAGCCTTTAGCTTCAATGGCCACGGCCCAGGTGAAAATCCAGAGGCTAAGAGGAGGAAGAGAG

TGTCCAGTTATAATATGTACACTGTGGAGGGTAAGCTCAAGTCTTCCTTCTTGAACAGTTTTAAGTGGAT

CAAGAACAAGTTTGGCTCTGACGGATACTATGATTGATTCGATGAGATATTATTTAATTCTCTCATGTCC

TCCGAGCAAGTAAATTTAGGGGGTGTATTCAATTGAGATTCTGGTTTATGTGACGTTGAGGTTATGGAGA

TTATGTTTTAACCTTGGTTTATGTGAATATGTTTGCGATGAATGTGTAATTTCAAGTTTGTTAATTTATC

ACTTTCGAAA

>MSTRG.751.1 gene=MSTRG.751

TTTTAAATTATTATGAAGAATCGCTTTGTTCTGTTTATACATTTCATGAAGCTGATTAGATCAATAACAA

AATATTGAGGCAAAAACAGGGAATATGATTTTAATTAGATCTAAACTTCATTCTAATTGTTGTTGGTGCT

GGACGAGAGACAGCATTTGCAGCTGCACTATTAGGATATTATGCATTGATGAAAAAGCATGGCAGTGAAT

GCTTGCTGAAGCTGTATTACATGCCCATG

>MSTRG.758.1 gene=MSTRG.758

CTCCACCAAGTATAACAACGCTCTTAGGTAACTCTTCCAAGCTCAAAGCCTCATCAGAAGTTATGCCCAG

CTCCTGCCCAGGAATTGGTGGACGCTGAGCTCTACTACCAGTTGCAATCAGAATGTGCTTCGTCGAGTAA

CTAAGTTTAGTGCCGTCTAGTTGTGTTACCTCTACCTCATTCGGACCAACCACCTTTCCTTCACCTTCAA

ACAGTTTAACTCCCGCATTGGCAAGCATGCGCTTGTAGATTCCATTTAACCTGACAATTTCCTCCGTCTT

TTTCTGCAATAATTTTTTCCAGTTAAAATCAACATTTTCATTTAGTTCCCAC

>MSTRG.764.1 gene=MSTRG.764

CAAAATTTGAAGAGGCAAATTTTTTGAGGACATAAACCAGTATCCACACAAGTCCAGATTTTTGAACATT

CAGAGTATCAAACACGCTTTATTGACTCGTTCTCTCCCTCGCCTCGTCTCTCTTCTCAATCTCTCCGTTC

TCTCTCACTCTCTCCGTCTGTCTCTCTCTTCATCTCTTTAGATAAAAATGAGGTCGATTTCACTCGTTCT

CGTGGAATTGGAGCAAGCTCTCACAAATCACCTGGATTTTGAATTAAATTTAGGGTTTATTCAATTGATT

TGTTCTTGGATCAAAATTGGGGGTTTTCATGCCTTTGATTCAGGGGTTTTTTGGTTATATTGTTTTTTCT

CTTAGATCTTTAACTCTGTCTCTAATTTGTTTTATATCTTTGTTCTGTTTTAGGCTGGCTTTGGGTTCCT

TAGGGATTGACTGATTTAAATCTTTGATGTGCGCTTCTTCAGCTTAAGGTTAGGTTGTGGATGTTGGATT

TGCAGTTAGAAAATTCAAGAAAGGCGAAAAAGTTATGGCCATGCTTGGAACAACGACTGGAGGTGGATTA

GTTGAGTATGCAGTAGCAAAGTAAGAAGTAACAGTTCTAAGGCCACTTGAACTATCAGCTGGTGAATGTG

CCGCATTAGGCACTCAAGGGTTAACCACTTACCAATGCCACATCACAGCGGGAGTTGTTCTATGAAAAGA

CGCCAACCTATGAACATACTAACTGTTGCAGCTTGAGGCGGTGTTGGTCACTACGCAATTCAGCTCGCAA

AGCTAAGGAACACTCATGTCACTATAACAGGTGGCTCTCGCAACATTGATTGTGTGAAGAAGACCTAAAG

GATGAGTTGGATTTGTTAGGTAGTTTAGTTTGGTTGGATTTGTTTTGAACTTTGGTTTGTATGAATGATT

AGTTAATTTTGCAGTTGATATAGTAAACTTTTGCATAGTAGTTTAATGTTGCATTAACACATATATATCA

GATATTTAGTACAAATATATGTTGTAGAAAGTAGCCCTTGTGGGC

>MSTRG.764.2 gene=MSTRG.764

CAAAATTTGAAGAGGCAAATTTTTTGAGGACATAAACCAGTATCCACACAAGTCCAGATTTTTGAACATT

CAGAGTATCAAACACGCTTTATTGACTCGTTCTCTCCCTCGCCTCGTCTCTCTTCTCAATCTCTCCGTTC

TCTCTCACTCTCTCCGTCTGTCTCTCTCTTCATCTCTTTAGATAAAAATGAGGTCGATTTCACTCGTTCT

CGTGGAATTGGAGCAAGCTCTCACAAATCACCTGGATTTTGAATTAAATTTAGGGTTTATTCAATTGATT

TGTTCTTGGATCAAAATTGGGGGTTTTCATGCCTTTGATTCAGGGGTTTTTTGGTTATATTGTTTTTTCT

CTTAGATCTTTAACTCTGTCTCTAATTTGTTTTATATCTTTGTTCTGTTTTAGGCTGGCTTTGGGTTCCT

TAGGGATTGACTGATTTAAATCTTTGATGTGCGCTTCTTCAGCTTAAGCGCGCAGGTTGTAAGCAATGCT

AACAAATTATTAGGACTTGTCAATGAGAAGAAAAAGAAACAAAATTGGTTTGATTATTATCAACTGAAGT

ATGCTAGATATCAGTCAAAGAGACCGATTATTAAGGTTAGGTTGTGGATGTTGGATTTGCAGTTAGAAAA

TTCAAGAAAGGCGAAAAAGTTATGGCCATGCTTGGAACAACGACTGGAGGTGGATTAGTTGAGTATGCAG

TAGCAAAGTAAGAAGTAACAGTTCTAAGGCCACTTGAACTATCAGCTGGTGAATGTGCCGCATTAGGCAC

TCAAGGGTTAACCACTTACCAATGCCACATCACAGCGGGAGTTGTTCTATGAAAAGACGCCAACCTATGA

ACATACTAACTGTTGCAGCTTGAGGCGGTGTTGGTCACTACGCAATTCAGCTCGCAAAGCTAAGGAACAC

TCATGTCACTATAACAGGTGGCTCTCGCAACATTGATTGTGTGAAGAAGACCTAAAGGATGAGTTGGATT

TGTTAGGTAGTTTAGTTTGGTTGGATTTGTTTTGAACTTTGGTTTGTATGAATGATTAGTTAATTTTGCA

GTTGATATAGTAAACTTTTGCATAGTAGTTTAATGTTGCATTAACACATATATATCAGATATTTAGTACA

AATATATGTTGTAGAAAGTAGCCCTTGTGGGC

>MSTRG.768.1 gene=MSTRG.768

CGAAAAAAAAGCGGTAACATTCCCGCTCAATCCAAACACACTGGACCTGTCTCACAACACACACATCCCA

CGGAGAACCCACCTTACACTCTTGTCAAATCAGGGTTTGGGTGGTTCTGCTAGCTAATCAAAGGAGCTGT

GTCATGGCTGCCAAGAATCTCACATCTCTCTTACTGCTTTTAATTTGGTTTTGTGTTTTACTTGTTTTGA

TTTTTTTTGTTTCTAGCCACTGATTTCTTTCTTTTTTCTTTCAAATATACAAGTGATTAGGGCTCTTAAT

TTGCGCTTTTGGCAGGGACTTGAAAATAGGGCTTTTAGCGGGGCCATGAACTAGGCTTTGAGCTAGAAAT

TCATCAAAGGCAAAATGTCTCAGCGAGCGGTGCTCGCCCTAACTCACACGGGTCTTTCCACTATGGTCAA

ATACCAGTGACTGATACAATTGTCTTAAAAAGTGTTCCCCTGGTAATGATTGATGGGAAACACAGAGCTA

CACTAAATGGTATTTCATTCATCAATCCTGATACACCCATCAGGCTTGCAGACAAGTACAATATTAAGGG

GGATTATAAGCTCGACTTCCCCAGTAGACCACTCAACGGACCACTTCGATCGGAAAGCTCTGTCATCAAC

GCTACATACAAAAGATTTATAGAAATTTTATTGCAGAACAATGGATGGCTTATGGAGAATGGACTGAAAA

TAATCGGGGCTCCTATAATAAGTGGGATGCTATAGCTCGCTCCACCCCACAGGTACTTCCCGGAGGATGG

ACAACAATTCTAGTATCCCTAGACAATTATGGAGTATGGAACCTTAGAGCTGAGAACCTGGACAGATGGT

ATTTAGGGCAAGAGACATATATGAGAGTTTTTAATCACGAAGCTGATCCAAGCAACAAAACAGAATTCCC

TGTCCCTGATAATGCTCTCTTCTCTGATGCTCTTTCACGATTGCAGATGCCACAGAAGTCCTCAACATTA

ATCATCAATGACAGGTCAAATTCGAATTATCAGCTGCTTCTGGTGACATTATCTGCTTTCATATTCATGT

GGTTATGATCTTCATCATATTGGCGAGGCATTGGGCAGTTTTTACCGGTTTTTGAAGGACAAATTATTAA

CCAAGTAGTTTTATGCTTCGGTATATTATGATTATGTACGTAGTTATATACATTATGAATATTCGAT

>MSTRG.768.2 gene=MSTRG.768

TTCCCGCTCAATCCAAACACACTGGACCTGTCTCACAACACACACATCCCACGGAGAACCCACCTTACAC

TCTTGTCAAATCAGGGTTTGGGTGGTTCTGCTAGCTAATCAAAGGAGCTGTGTCATGGCTGCCAAGAATC

TCACATCTCTCTTACTGCTTTTAATTTGGTTTTGTGTTTTACTTGTTTTGATTTTTTTTGTTTCTAGCCA

CTGATTTCTTTCTTTTTTCTTTCAAATATACAAGTGATTAGGGCTCTTAATTTGCGCTTTTGGCAGGGAC

TTGAAAATAGGGCTTTTAGCGGGGCCATGAACTAGGCTTTGAGCTAGAAATTCATCAAAGGCAAAATGTC

TCAGCGAGCGGTGCTCGCCCTAACTCACACGGGTCTTTCCACTATGGTCAAATACCAGTGACTGATACAA

TTGTCTTAAAAAGTGTTCCCCTGGTAATGATTGATGGGAAACACAGAGCTACACTAAATGGTATTTCATT

CATCAATCCTGATACACCCATCAGGCTTGCAGACAAGTACAATATTAAGGGGGATTATAAGCTCGACTTC

CCCAGTAGACCACTCAACGGACCACTTCGATCGGAAAGCTCTGTCATCAACGCTACATACAAAAGATTTA

TAGAAATTTTATTGCAGAACAATGGTATAGTAGTTCAGAGCTTTAACATGGAAGGCTATTCCTTTTTTGT

AGTTGGGATGGCTTATGGAGAATGGACTGAAAATAATCGGGGCTCCTATAATAAGTGGGATGCTATAGCT

CGCTCCACCCCACAGGTACTTCCCGGAGGATGGACAACAATTCTAGTATCCCTAGACAATTATGGAGTAT

GGAACCTTAGAGCTGAGAACCTGGACAGATGGTATTTAGGGCAAGAGACATATATGAGAGTTTTTAATCA

CGAAGCTGATCCAAGCAACAAAACAGAATTCCCTGTCCCTGATAATGCTCTCTTCTCTGATGCTCTTTCA

CGATTGCAGATGCCACAGAAGTCCTCAACATTAATCATCAATGACAGGTCAAATTCGAATTATCAGCTGC

TTCTGGTGACATTATCTGCTTTCATATTCATGTGGTTATGATCTTCATCATATTGGCGAGGCATTGGGCA

GTTTTTACCGGTTTTTGAAGGACAAATTATTAACCAAGTAGTTTTATGCTTCGGTATATTATGATTATGT

ACGTAGTTATATACATTATGAATATTCGATAGATTTGACTCAT

>MSTRG.769.1 gene=MSTRG.769

GCTTCTTTTGCTTTTCAGTTACCAGTATTTTGTCTTTTGTTGACACTTGTTGTCTTGTAATAAGCCAAGA

CTAATTAGGGGTATTTCTGGTTTTTCTTATTCTGTTATTGTGGTCCTGTGTTGATCCCAATGCTCCCATC

TTTTAGTATATATTGATGGGACAGTAGTAGTGAGACTAAGAATGTTGTTTTACTTTTATGAGAATTAAAG

CTGAGCTCCATTTTCTCCCTGTTTCTCTGTGCATTTGTCTTTGTTTTCATGGTATCAGAGCATAAGTGTG

CAGATATTCAGGGCTTTCTTTGAGTTTCTTTCACAGCTGGTAATACTCTTTACACATTCTTCTTTTAGTT

GTGTTCATATAATACCTGTTAGAGATCTCTTTCTGTCATGGCCATGAATGGAAGGTACAACTTTGCGGAT

ATGCAAAACCCGTTGTTCCTTCATCCTTCTGATGGGCCGCTGTCCATCAGCGTTGACAAACTGCAGGGTG

CTGGTGATTATAGAAGCTGGAGGAGGTCATTTGAGATTCAGATCTCATCCAAGCGCAAACTGGGTTTCTT

GAATGGAACTGTCACTAGAAGCACTACTGATGAGACTCAAGGAGTACAATGGGACACCTGCAATGATCTG

GTCATAGCTTGGATTCATGGTAATGTTTCTGACAATATCAAACGGTCTATTTTGTTTATTCACTCGGCTC

ATGAGATCTGGAAACAACTTGAGAAACGTTTCCAACTCAGTAATGGGTCTAGAAAGTATAAACTAAACAA

GGAACTGTTTAATATGAGACAAGAAAAAATGAAAATCCATGAATATTATACTGCTGTGAGTAGTGTTTGG

GAAGAAATAGATTCTATGAATATGTTGCCATCTGTTACAAATGTGACAGAAGATGTTAGGGCTTTGCTTA

AGGCTATTGATGTGCAGAAAGAAGAGTCAAAACTGTTTGTGTTTTTGAATGGCTTGGAGGAGGCATATGG

TTCTGTTAGAAGTCAGTTGCTGATGCAACACCCCTTACCCACAGTAGAGGCTGCTTGTGCAATGCTTCAA

CAGGAGGAATCTCAAAGGGATGTGTTGCCTGGTGGTGAAATGGAACATATGGCTATGTACACCAAGGGTC

CAGCTGAGAGGAATGCTATGTCTTGCACTGCTTGTGGTGGAAGGAGCCACACCAGTGACAAGTGCTGGAG

TGTCACTGGTTATCCAAAGTGGCACTACAAGTATAGGAAGCCAGTGGCCAAGGGAACTCAGTCACCTGGA

CAGAACAGGTGGAATAATGCCAGCAATACCAGCAAGGCAAACAAACAAAAGATGGCCAACAATGCTACTG

TAAGCCAAGATGGAAGTCAAGAGGAAAATGTGATGTTTAGCTCTCAACAGCTTCAGCACTTGCTGAAATT

TATGCCTGTTGCTGGCTCTGGAAGTGCTCACAGAGGTTCTGACACAGAGGAGGAGCTGGAGATGGCCTTC

TCAGGGATGGTTGCTTGCAACCTATCTGTTACAAGTTCAAATGCATGGATTGTTGATTCAGGTGCAAGTG

ATCACATGACAGGCACTTTGGAAAAATTGATCAATATCAAGCCTGTCAATTCTGCCTTGACCATCACTCT

TCCAACAGGAGATGCTGCAAAAATAACCCATATGGGGGACATGAAGTTGAAGAATGGTCTGCTCTTGAGC

AATGTTCTTGTGGTTCCCAAATTTCATCATAATCTTCTGTCCATTCACAAATTGGCCTTGGATAATGAAT

GTGATGTGAAATTTTCTCCTGAAACTTGTGAAGTTGTGTGTACTAAAACAAGGATTCTAAAGGCAGTTGG

CAGAGTCTATAATGGTCTCTATTACTTGTTTGATAATTCTGAAGCTCCTCAATGTGCTGCTGCAACCACA

GTGGATCACAATCTGTGGCATCTGAGACTTGGACATGCCTCAGTCACGAAAATGAAACACATCCCAATGC

TGCAGAAATCACAGTTTGCCTCCACTGGACAGGTTTGTCTTACCTGTCCTATGTCAAAGTTTACAAAGTT

GCCATATGATTTGAGTGAATCACATGCTAGTAAGCCCTTTGATTTGGTGCATTTAGACACTTGGGGACCA

TATAAAGTTTGCACTAGGAAGAAATTTAAGTATTTCTTAACATTAGTAGATGATCATTCTAGAATGACTT

GGGTTTATTTGATGGAACAGAAGTCAGATTACTTTGCATGTTTTAAGGCTTTTTTCAAATATGCCAAAAA

TCAATTTGGCAAGTCTATAAAATGTATAAGGTCTGATAATGCATCAGAATTTGCAACTGATGTATGTGTT

CAATTTTATAATGATGTTGGAATTGTTCACCAAAAATCATGTGTGAATAGACCTCAGCAGAATGCTAGAG

CTGAGAGGAAACATAGACATGTTCTTGAAGTGGCTAGGTCTTTAAAATTCCAATCTGGTCTGCCTCTGTC

CTATTGGGGGGAATTTGTTTTGACAGCAGTTCATATCATAAACAAACTTCCCACACCTGTTCTGAAAAAC

AAAACTCCTTTTGAAGTTTTCTACCATAAACCCACTGAGTATGATACTCTAAAGGTCTTTGGTTGTCTTG

CCTTTGCCTCAAATCCAGCTACTGGTACTGACAAATTCAGTGCAAGGGGGGTGCCATGTGTTATGTTGGG

CTATCCACCTAATACCAAGGGATACAAACTCTTAAACCTCTCAAAAATGCAG

>MSTRG.770.4 gene=MSTRG.770

GCAATTCTAAATGATTTTGTATTTTGTAGAAGGATCTTAATTAATTTAGCAGGTGACTCGGGTATATTCA

CATGAGTCATATGTCTATTCTGGACATGGTGATAGCTTGCTTGTATAGTTGTATGTGCCCGTGGTGTTTA

TTGGTGTAATTAGATTAGTCATACAATTCGGGTCAGTATATGCAGAGAAAATGGCAAGCGTGTGGGATAT

ACTGTGAAGGATAGTGACTCTGAAGCAGACGGTATTTATACTTTGATTTTTATTTTCTTTTTTGTTTTCT

TTTTCCCCCTGTAACCCTTCTCTCAGTTTGTACCTATTTTCATCTACACTGGTAGTAGGTGATGAATATT

GGGGCTCGAGGGTTTAAACTGAGACCAATGTGTTTAGTGGGCTTTATCTGTATATCATGAATTTTCATAT

AATTTAGTAGATACGCACTTGTCCCAATGGATGGATTGATAAATGTATTTTGGGGAATTTTGCCTTTAGC

AAAATGTTATGTTCTGGGTGCTGTTGTGCCTTAATGTTATGACTTTTTTCAGCTTATAATGGAATGGGAT

ACTTGTATGTAAAAGGTTATGGAGCGGAGAAGAAGAACTTTACCAAAATCATAGAGAAAGAACTGCAAGG

GGAGGAATAAAGGCGTTGTAGGGCTTGTGTTGGCGAGGCAGAGGAAGAGACATGTGTCGAGAGAATGGAA

CCCTATGCATGGATTTAAACTTCTGATGGTTCTATACAAGAAGTGGAACGAGAGATTGCAATGTTATGCC

CCACTGCTGTTCATGAAATGGCATCAGGACTTGGATCGTCAAAGAGCAATCCTATATCACTACCTGCACG

AGTTAAACCTACTTCTTTGAGCTTAGTTATTGAATACTGCAGATTTCATCTAGCACACCGCTCTTACCAG

GACCAAGATACAAAAATCTGCAAGATCATACTCAAGATGTCTATAGTGACAAAGTTACGCTGTATCACTC

TGGATGTTACTGGCACATTAATAGCTTACAAGGGGGAGCTTGGTGAATACTATTGCATGGCAGTCAAATC

TGCAGACTTCCGTGTCCAAACTACAAGCGTGTGCATGAGGGCTGTAAAATTGCATACACAAACACGGCAA

AAGAACATCCCAGTTTTGGGTTTGCAGAGAAAATTCCCAACATTGTGTGGTGGAAAACTTGTGTTAGAGA

CTCTTTTGTAAAGGCAGATATGATTATGATGAGGAGATATTTGATAAAGTTTCAAACGCATATACTCCAC

CTTTGGCTCTGTTTCTCCTTATACTATGTTTTCAGATTCCAAGCCTTTCCTAAGATGGGCCAAGGGCGAA

CAACCACTTACTGTTTCCTCATTATTTGTTCACAAATAAGGGATGAAAGTCATTGTAGCTTTATGGGAGA

GCAAGGCAACCACCTTCCTCGACTTATTGCCAAAGGAAGGAAATGAGCCTGTTTTTGTGGTTATAACTGG

CTTAATGGCCAAAAAATACTCAGAAATTCGTAAGGGAACTCGTCAGTTTTGACAAATCTGTCTAAGATGA

TGTTGAGGGTGTATACAAGGAAAATCCGCAAGCATTTTTTTAGTAAAACTTAATGGAATATAGTTGTAAT

TATCTTATGTTATCTTACTTGTGATACTTTTGGC

>MSTRG.770.5 gene=MSTRG.770

CTGGTATATAAAACCATATTGCCAGATCTCTGATCTATATTGATTTTCTAAATTTATGCAATTCTAAATG

ATTTTGTATTTTGTAGAAGGATCTTAATTAATTTAGCAGGTGACTCGGGTATATTCACATGAGTCATATG

TCTATTCTGGACATGGTGATAGCTTGCTTGTATAGTTGTATGTGCCCGTGGTGTTTATTGGTGTAATTAG

ATTAGTCATACAATTCGGGTCAGTATATGCAGAGAAAATGGCAAGCGTGTGGGATATACTGTGAAGGATA

GTGACTCTGAAGCAGACGGTATTTATACTTTGATTTTTATTTTCTTTTTTGTTTTCTTTTTCCCCCTGTA

ACCCTTCTCTCAGTTTGTACCTATTTTCATCTACACTGGTAGTAGGTGATGAATATTGGGGCTCGAGGGT

TTAAACTGAGACCAATGTGTTTAGTGGGCTTTATCTGTATATCATGAATTTTCATATAATTTAGTAGATA

CGCACTTGTCCCAATGGATGGATTGATAAATGTATTTTGGGGAATTTTGCCTTTAGCAAAATGTTATGTT

CTGGGTGCTGTTGTGCCTTAATGTTATGACTTTTTTCAGCTTATAATGGAATGGGATACTTGTATGTAAA

AGGTTATGGAGCGGAGAAGAAGAACTTTACCAAAATCATAGAGAAAGAACTGCAAGGGGAGGAATAAAGG

CGTTGTAGGGCTTGTGTTGGCGAGGCAGAGGAAGAGACATGTGTCGAGAGAATGGAACCCTATGCATGGA

TTTAAACTTCTGATGGTTCTATACAAGAAGTGGAACGAGAGATTGCAATGTTATGCCCCACTGCTGTTCA

TGAAATGGCATCAGGACTTGGATCGTCAAAGAGCAATCCTATATCACTACCTGCACGAGTTAAACCTACT

TCTTTGAGCTTAGTTATTGAATACTGCAGATTTCATCTAGCACACCGCTCTTACCAGTATACTTCAAATG

CATTATATTATGATGCAACCCTGGAATATAACTCTGTGTATGGCAAAGTAATCTGATTGGTGCTAGATAC

GTGTACATGAAGTTGATTATAGACTGAGGAGTAAATAGAGGACCAAGATACAAAAATCTGCAAGATCATA

CTCAAGATGTCTATAGTGACAAAGTTACGCTGTATCACTCTGGATGTTACTGGCACATTAATAGCTTACA

AGGGGGAGCTTGGTGAATACTATTGCATGGCAGTCAAATCTGCAGACTTCCGTGTCCAAACTACAAGCGT

GTGCATGAGGGCTGTAAAATTGCATACACAAACACGGCAAAAGAACATCCCAGTTTTGGGTTTGCAGAGA

AAATTCCCAACATTGTGTGGTGGAAAACTTGTGTTAGAGACTCTTTTGTAAAGGCAGATATGATTATGAT

GAGGAGATATTTGATAAAGTTTCAAACGCATATACTCCACCTTTGGCTCTGTTTCTCCTTATACTATGTT

TTCAGATTCCAAGCCTTTCCTAAGATGGGCCAAGGGCGAACAACCACTTACTGTTTCCTCATTATTTGTT

CACAAATAAGGGATGAAAGTCATTGTAGCTTTATGGGAGAGCAAGGCAACCACCTTCCTCGACTTATTGC

CAAAGGAAGGAAATGAGCCTGTTTTTGTGGTTATAACTGGCTTAATGGCCAAAAAATACTCAGAAATTCG

TAAGGGAACTCGTCAGTTTTGACAAATCTGTCTAAGATGATGTTGAGGGTGTATACAAGGAAAATCCGCA

AGCATTTTTTTAGTAAAACTTAATGGAATATAGTTGTAATTATCTTATGTTATCTTACTTGTGATACTTT

TGGC

>MSTRG.774.1 gene=MSTRG.774

ATTTAGTTTTTTTTGGTAATTAAGTCAAATTTCTGACAGAGGCGGCATCCGTTCAAACATTACCACATCT

TCAGTGGCTGTGGTGTCACCCTTTGGACTCGAAAACCACTTCCCACATCGACACATAAGCTCTCTACCAA

AGTCGTCTGTTTTGATATGCCACAGTCACACCCACAGACTATTACTTACACATTTCATCCGTGTTACTGA

CACAAAATCTTGCCGACATAATCTTTGCTCCATCACTGTACCTAACCATCATAAGGTAACGATGATTACT

TAAATTCTCAATATTATATCATCTTTTATGTTTTAACAACAACCCACTTTGTAAAGCTGCGGTTTTTATT

TTATTAACACAATCTGATTGAGTT

>MSTRG.775.1 gene=MSTRG.775

TACATATTACAACTCCTGTGATTCAGGCAAAGCACTACTAGTACATTTCCAGAGAGAAAGTTTAGAAAAC

AATGGGGGCCTGTGCAACCAAACCAAAGGCAGATGCTACCGATGTTCCCGCACCCCTACCGGAGAAAGAT

GAGGTTGAGAAGAAGGAACTCACTGTTGTTGGCTCAGTTATCGAGACCAAGGAGTTTGTTGATGCTGACA

AGTCAAGGTCTCTCAGTAATCTTTTCAAACAGAGTGAAGAAGTGAAAGGCTCGACTGGAACTCACTCAAC

ACCGACAGAAATGGTGAAAGAGGAGGCATGTGAAGCTGAAGAAGGCACAGATGTTTCTGAAACTAAGACA

ATTGAAGTTATAAAATCTGAAATATCAGCAGAAACAACTATAACACCAACTGTTGTTCCTGTTTTTGAAG

CAGAAAAAGCAATTGAAGTAGAAGCAGCTACCGATAAATTTGAAATTTCATTAGAGAAAACAGCAATGCC

AATTGCCGATGCTCCTGCTTTGGTGAAACCGGAAGAATCAATAGAAGTCGAAACAGCTAAAGAAGAGACT

GAAACTTTGGAAGAAAAAACATCAACACTCACTAGTCATGTTCCTGTCGACGATGGAGCCGAGAAAGATA

TTGACGCAGCATTAGCCACTGAGAACCAAGAAACACTTGCAATTGAAGATAAAAAGATTGATGAACATGA

CAAAGGCATAGAAACTCCAATATCAGAATCAGTTGTAGAGGTTGATAGCAGTAAAACTGAGTTTGTTGAA

GACGGAACATCTTCCCCAGTTGTGTATGCTCATGTTGAAGCAGACACTAAGATTGAGGCAGCCGCTAATG

AAACCCAAATAGCAGAGGCAGTCGAGGAAAAAAAAATTGAGGAACATGAGGAAGTCGAAGAAACACTGAA

CCCTGAATCACCTAAGAACACTGGAGAAGAAAGCAAGGAAGCTACAGAACTTGAAAAACCAGCAATATTA

CCTGAAAAGGAGAACCAAGATTCTAAAATAGAAGAAGAAATAGCAGTCACTGAAAACCAGAAGCCTCCAG

AGGAGACAACAACAGAAACTGAAGTAGCTGCTGAGATTGTCAAACCATGACAGATAAAGCCTTGAAGATG

ATGTCAAATTTAATTTCTTTTGGAATATATATCATATACTTTTGCCGAACATGAGTCTGTTGAAGTTGCT

TACAAGTGCAGTTGAACTGATTGCAGGGTGCTGCTGTTGTGATACTTTTTTCCTGTTGATTGATTGCTGT

CATTTCTGTTCGCTTCCGAGTTTTTAGTAAGCTGGTAGGTTATTACGTGTTTTTGTTACATATAGTTTTC

TTAGTGCAGTAATGCGATCATGACTTGTGCATCTGATGTTTTCTTGACAATTATATTACTTGTCAATGTA

TGAAAACTAAATGAAGGTTAAGATGACTTCGTCTACCTGTTGTTTACTTGCTTAATGAATTTTGCGATGT

TGGGACTTGTAATAA

>MSTRG.777.1 gene=MSTRG.777

TATCTCCTCTTCTTTACTTGCCCTCCGCCTGGCGGCGTCTCCGTTGATCCCGGCCTTCGTACCACCATAA

ACACACTCAACACACACACACACACACAACGACCGCAGAACACACGCACGCACACATGCGACCCTCCTCT

CTCCTCTCTTTCTCCCCTCACTGCTGCCTTCTCCCTTTCTCCCGGTGGTCGCCGGCGTGAACCGCCGGAC

GACACTCCGCCCTCCCTCCTTTCGTTGTTTCCTCCCAACCAACACCCTTTTACCTTCATCAGCGGCTCGC

TTTTCAATTTATGTTGTCATCCGATGAATGTAAGCCGGCGGATTTCTATTCAAGGACGGAGATTGATTTG

TGTTATGATATCGGGTGTCGGCTCAGTCTCGTGCTGGAGAGTTAGGACCGCGGCGGTGGCCGAAGACGAT

GACGGTGACGGCCGTTGTTCCGGATCAGTCGGTTTCAGAATGGAAGGCGTTAATTGTGCCTTAATTGAGA

GCATTAATTGTCTCTTAATTTGTGCCTTAATTGAGGCTTAATTGTCTTTTAATTGAGAGTTTAATGTTTG

TTAATCGTGCCCTAATTGAGGGCTTAATTTGTCTCATTTATTGAGTTATCGTGATTTTTGTTTATTTTGA

CCCGTACAAGCATTTTAGCATATATTTTTTGTTTTAAATTAATTGTCATTTGTCGCGTGTTTGGTTTTAA

ATTATTTCTGCACTATTTCTTTCCTTAGTGCTCACTAATAATTTTTCACTTAATGTCTACAATCTTCTTA

CAGGTTTATATTTCAAGGGTTTTTGGAGGTAGCAGATGAAAGGTGAATATAAATGTACAAGCCACTGTAG

CAGATTGACGAAAGTAGAGGCTGTAGCAGATTGACGAAAGTAGGAAGATACGAGTTTTTCTTGTTATTTC

TTTGTAACTAATGTAATTGTGTTTTGTGGGTACTCTTTTTCCCCTTGAGTTTTTTTCCCATAGGGTTTTA

CTCTTGCAGGGTTTTAACGAGGCCCGGTTGGTGGGTTTTCTTTTCGGACATTAAAGGTCATATTGTCTAA

ACGTCCGGGAGTACTTGCTTACTCTTCGGTTAGATGGTATACTTTTTTGGATGAAAGAAACCTCTGTCTC

ATTAGACCTTGTATTTTCGATACTCAGTATCAATGGAAATTTATTTTCCTGTTCAAAAAAA

>MSTRG.779.1 gene=MSTRG.779

CAACTAAATCGAAAAAGAAATTCAACCTCAATACAAATAATATCGAACTTAGAAACACTTCCTGTGCACG

ATTGATGGACCAGATTCATCATATTCGCCCTTGGATATCCACATCTGTTGGAATGTGCTGAGAGATGCAA

GAATCGATCCTCCAATCCAGACACTGTATTTTCTCTCTGGTGGTGCAACAACTTTGATCTTCATGCTGCT

GGGAGCAAGAGCAGTAATTTCCTTGCTCATACGATCTGCAATACCAGGGAACATTGTTGAACCACCACTG

AGCACTATGTTTCCATACAGATCCTTCCTGATATCGACATCACACTTCATGATGGAGTTATAAGTAG

>MSTRG.782.1 gene=MSTRG.782

CATTAATGGCGCTATAATATACCTCATTAAGGCACAATAAATTTGGTATACACCAATTAATCCATTAACC

ATAAAAGGTCCAAAAATCCATCATAAAATGGCAACAAGTCATCATAAATGATCCTTCCTTTTACGGATAC

AAGAACCACCAATTATGGCACAAAGACGGAAATATAATAATACTCTATTTCAAAATATGTAACTTTCCC

>MSTRG.785.4 gene=MSTRG.785

GGTAGCAGGACCATCATTAACCAATATTATGTCCTTCATATCATTCTACGACACTGAGTCAACAGTTAAG

AAATTTCCTTGAGAATGATCTTCCAAACCCGTTTGATGTCCAAACAAGATATTATGTGTCAACTATGGAT

ATAGATTTGGGACATAATATATCAAATGCATTGAGAATGTCCGTTGAGTGTGGTCGGCAAAATAATGATC

TCATGCGTAGTTTACGGATGAACATTGAAAAATTCATCGATGGCTTGATGCCTAAAGACGTGGAAAAGGC

TCAACAGAATTTGGCACGCCTATATAGTAGACAAATGTGTACTTCAACGTACGCACAAAGTTCCACGAGA

ATTTCATCTTCTTCCGTACAACGAGGGTTGGTTGTACCATATAGATTTGAAGGAGTTTTCATCGCCAAGG

GTAAGGAGGATTTTATTTGTACCAGAAATTTGGTACCCGGTGAAGCTCTTTATGGTGAGGAATTAATATT

TGTTCAGAATGAGGGTGGAACCGAAGTTGAATACAGAGTATGGAATCCGCTCAGGTCGAATTTGGCTAAT

GCAATCATGTGCGGTGTTACCAATATATGGGTTAAACCTGGATCTCGTATTTTATACCTGGGAGATGATG

TGTGTCAAATTACACTTTCACATTTGTCTGATCTTGTTGGTCTGGATGGTTTGGTCTATGTAGTGGGGTT

GTCTGATGTTGCAGTAAAAATGGCGGAGAAGAGGCCAAATGTTATAACAGTTTTCCAAAAACCTTCATGT

ATTGGGAAGTATCGCATGGTTGTTGGCATGGTGGATGTAATTTATGCTGATATACATTCAGACGAGGTAC

ATACTATTACTAGTAATATTATATACTATCTTCGAGCTGGTGGGCATTACTTGGTCTACACACGGGTTTG

TTTCTGAACTATAATATTTATCAAGTACACTGTAAAGTAATAAAGTTTTTTGAAGTTTATTTCAAATATA

TTGATCTTTTGTGAATTATAGGCAAGCAACATCTATTCAACTACTCGAGGTAAAGATCCATTTGCTTACC

ACCATCGACAAAAGGAGTTTACACCAATTGAAGCAGTCATGTTGGAACCGATTGACAGAGAATATGCAAT

GGCTGTTGGTGGTTTCCGTATGCCGGAAGAATAGATTTACAATCACAATTGTGTTTTCGGGAACTTTCTT

TGAAACTTTGGTTTTCATTCCTTTCTTTTAACGAAATGAAGCTT

>MSTRG.788.1 gene=MSTRG.788

TAGAGTTCCAGAAGTAAATCTTACGTTGTTCAAGACTTCAATCACCGACGCAATCAAGACAAGGCCCTTC

ACAATCTAGACAGAGCATGCCCACTCCATCACGACAACTCAGTACTCAATCACGCCAACACACACCTGAA

GCTTAGTCACGACAACACACAACTGATGTCCAATCAAATCCAAACTCGCTTAATGTATTGTCTATTGTTG

GTAATGATAGTGTATGTTGGAATATTGGCTCCACAAGTGGGGATGGTAAAATATCCAAGGAACATTTTTT

TGTTTTAACAGTTCATGTAAGTTAATGCTATGCGGTTTGTTCTTTTGGATATTCCACATTTTGAATATGT

ATTATATAATGAAATACTGGAATTATTATAAGTTTTTTGTTGCTATTTAA

>MSTRG.789.1 gene=MSTRG.789

AATAGATTTGAAGGGAATAGATTATGTATTTTGACTCTTTCTGTCTCTTTTTTGCAGGCTGTAAGCTTAC

TGACAGCATCCCTGGTATCTACTTCAGTTTTAACCAATGGCCTTAAGCACCTGATACTGAGCCAACCAAA

GACAAAGAGATTAGTAAAGAAGCCGAAGGTAGATATATAATTAAGGAAAGGTTGCTTGCATGGTACCAAA

AG

>MSTRG.797.1 gene=MSTRG.797

CTTGTGTTTTGTTTGGTTTTTGATGTTTTTGGCGATTTTTCGTGTTACATGTGAGACGCTTCGTTCGCTT

CGCATACGACGAGGCAACTTGATCGATAGCATAGCGGTGGTGGCCGATGGTGGCGGTGACGACGATTTCA

GAACCGGATCCGGTTTCAGAAAGGAAAGCGCGCTAATTATGCCTTAATTGAGGGCGGTAATCGTGCATGG

TTAATTGCGCCTCTTTATTGAGGACGTAAATTTTGCCTCAATTATTGAGTGCGTAAATTGTGCCTCTTCA

TTGAGGGCGTTAATTGGGCCTTATTTAAGGGCGTTAATTGTGTCTTAATTAAGGATATTAATATTTTATT

ATTATGCCTTAATTAAGAGCTTAATTGTCTCAATCATGAGTTATCATGATTGTGGGAATATTTTGATGTA

ATTTGTTAAATATATCGACTTATATTTCTTCTTGTTTTCTTGTTTTATTTTCAGGATTATTGGAAGAAGA

CCGGTTTTCTTTTCGGACATTAAAGTTTTATTGTCTAAATTTCCCCGGTCATCTTGCATGTCCGACGGTT

AGATAATAAGCTTTCTTGAATGAAAGAATTATCACGGTGAATTGTCGATTCTCGTTGAGATTTATTCTCA

TTTTTCAAAAAAAAAAAAAAAAA

>MSTRG.808.1 gene=MSTRG.808

GTCGTAGATTTGCTGATAATGGTGAAAAACATAATTCATGTGGTGTGGAGGAGATAACAAAGATGCCTAA

TAATTGTAAACGAAGTCGTGCTATTAGTAAGGAGATTGGAGAGTCGATTAAAATTTCGAAAGAATCTATG

GAAAGGCAGAAAATTTTACGCGGGTCCCTCGAAGATTGTGAATTTCCGAAATCCACAGAATTCAAGATGC

CTAATAATTAAAAACAAAGTCGGGCCATTATTAAGGAGATTGGAGAGTCGATTAACATTTCGAAAGACTC

TAAGGAAAGTCAGAAAGTTCTACGGGGTTCCCTCGAAGAGTGTGAATTTCCGAATTCCACAGAATTCAAG

GAACTTCGTAAAGATCTAGACTACTATCAGTATCTCCAACGTCAATTGGTTGTACACGTCGTTGCAGCGT

CTAAAAACAAAGAAACAAATAGTACAGCGCAACATTTCATGAAAAGGTTCATCCAAATGAACCAGGACTA

CAACAACATCATTGGTTTTCTTGTAGACAAAAACAAGTAGATTTTTAAGGGGTTTGTTATTTTTAATTGA

CTGAAGTTTTAACGCTGTTAAGAATTTGACAAACGCACTTTTCCGCAGTGGATTGTATGTTATTTAAAAT

TATAAGATTTTTATGGATTTTTATCTCTAATGGTTTGTTATTTCCTTCATATATT

>MSTRG.810.1 gene=MSTRG.810

GTGACGCGGGTCATTTTCCTCTAGGGTCCTGACAACTGGTTGTCAGGGATTTTCAGAGGGCGTCGGATTT

ATATTTCAAGAGATTTCTCATATTAACGGAGCTGGGTGTTAAATGTTGTGGCGGACCAAAACCCTTTTCT

CTATCAACTCAGTGTTCTGATGCTGTACTTCTGTATGCTGTGTCATGTGTGCCGGTATCGAACCCAATAT

TTCGGGATATACAAGCGACTATTCAATAGCGATCACTAAACCCTACCGTATATAGAAGAAGAAAGGAAGA

AGTTTTTATGACGAGCCAAAAAAAGAAAAGTTTTGCAAGGGTATTACATTTTGAAATTATAACGACACAC

TGATACTGGGGATTGGGAGACAAACAAAGAAACCCTAGGCTCTATTTACTGATGGATTCTTGCTGAAATC

TTTGAAATAATTGTGTTTTGAGTTGAGTTTTGAGGTGGGGTGTGTTCCTTATTAATGAATGATGAGTAAT

AGTGGCACTAATAGGCCCTGTGGGGATCGGGGTGATGATGTATTGGACTACTTGAAACGAATGCAGGCGG

AGAATCCTGCATTCTGTTATGCTGTTCAGGATGAGAATCTATTCTGGGTTGACCCTGTTTCCAGAATTAA

TTATACCCATTTTGGAGATACTGTGAGATTTGATACTACATTTAGGAGCCAACATGTTGATTTACCGTTT

GTAACGTTTACTGGATTAAATCATCATGGCCAGCCTGTATTGTTTGGATGTGGATTGCTTTTTGATGAGT

CTGAGGCTTCGTTGATTTGGGTATTGCAGACTTGGCTACAGGCCATGTCTGGACAAGCTCCGATTTCTAT

CACAACTGATCCGGATAGGTTGATACAGTTGGCTGTAGCACATGTTTTTCCGGGAAGTCGTCAGCGGTTT

TGCAAGTGGTGCATCTTGAAAGAAACGCAAGAGAAATTGACTCATATTTATCAGACTCATTTAAGTTTCG

AAGCGGACTTTAGGAAGTGCATTAATAACACAGAGACAATTGAAGAATTTGAGTCGTGTTGGCAGTCGCT

TTTAGATAGATACTATCTTATGGATAATGAATGGCTTCAGTCTATGTATGGTGCTCGTCATCAATGGGTT

CCAGTCTACATGAGGGACACGTTCTTTGGAGAATTGTGTACATCAGAGAGTAGTGATGCTATAAGCTCGT

TCTTTAATGGGTATATAGATGCAGCTACCACCATTCAGGCCTTAATTACGCAGTATGAGAAAGCTGTAGC

GATTTGGCATGAAAAGGAAATGAAGGCAGACCATGATACCATGAATACTATACCAATTCTCAAAACACCG

TCTCCTATGGAAAAACAAGCCGCAAATCTTTATACGAGGAAACTTTTCCTAAAGTTCCAGGATCAACTGG

TGGAAACTATGGCCTATCCTTCAACCAAAGTTAAGGAATCAGGAGCCGTCACCCTATATCGAGTAGCAAA

ATTTGAGGAAGACCAGAAAGCGCACACTGTAACATTTAATAGCTACGAAATGAAAGCTAACTGCAGCTGC

CAAATGTTTGAGTATTCTGGCATTATTTGTAGACATATATTGTCAGTTTTTAGAGCAAAAAATGTGCTTA

CACTTCCTTCTCAATATATAATGACAAGATGGACAAGAAATGCAAAGAGTGGACAATCTGTTAGTGAAAT

ATCTCCACAATTGCCAACCGATGCAGAAGATCCCTTAGTTGTTCGCTGCAATAATCTCCGCCAAGAAGCA

ATGAAGTTTGTTGAAGAAGGGGCCAAATCTATACATATTTATAATGCAGCAATGGATGCTTTGCAAGAAG

CTTCAAAGAAGGTTGCCACTGCCAAGAACAAAAAGTTTGTAACTACAGATGGAGACAGTTTGGCAAATGG

TGGTAACCAGCACGCATGTGGTGAACATCAAATGACGCGGCATCAGTCCATGGTTGAGAAGCAAAAGAAA

ATTTCTGAATTGACTGCTGAGTTGGATAATACAAATCAGCGATGCGAAGTTTATCGTGCAAATCTACTTG

CAGTATTGAGAGATATGGAGGACCAAAAGCTAAAGCTATCTGTTAAAGTTCAGAATGTAAGGCTAAGCCT

AAAAGATTAGACGTGGATAGCTACACAGAAATGGTCATCTTTATCATCTTGCCTGCTTGTATAAAGGTAA

ACCTGTTAGATTTTTGAGCATCTGGTAAACTTTAGGAGACTTAGAATTGTAACTTGGAGTTTTTGATAAA

CATTAGGAGACTTGGAATTGTATTTGGTAAACATCGGGAGACTTAGTTTGATATCAGAAGTTTCTGTGTG

TCAAGTCTTAAAGCTTTTTTGCATGTATAAATATCTGTTGGTTGTCCAATAGTTTCTATTATCACCTTCA

CACAGAGGTCTGTGTTTCAGTGTTATTAAGGATAGTTTTATAATGTTCGCC

>MSTRG.813.1 gene=MSTRG.813

CACAATACTTCTTCCTTTTTATCTCAAACATCAGTCTCCTTCTAATCTCTCCCCCTCTCGAAATCTTCTC

TGTGTCGACAGCCTCTCACTCGAACTGTCCCTTCTCCTATTCTGTTAAACCTCCCCGCTGCCGATATGCT

CCACACAAGTCGTCGCCGCTGGTTCCGATTCCGGCCAAACCTGTTGTGGTCACCCCGCGAATTTCTTCTT

GTCCATGCCTACTGCTTCTTCCCGGTCGGAAGTTATCGATTCTTTTCTCTCCGGGTTTCTCTATCATTGA

GTCTCCAAAACCTGCATCAGCTGTGGCATATTTGTCCAACGACACAAGCTTTATTCCACCCCTCGAGTTC

AACACAGTTCGAATTGAAGACATTGACCGAGGCATTAATAGAGAAGTGATTGAAAGGGGTATTCAGAAAT

AACCAAGTAGTGTTACAAGTTGAAGTAGGGCGGAGATTTGTTATAGATAAAAGCTTGTCAGAAGCTGGCA

GAAAGGAATGATTATGCGGTATATAGTGTTACTAGGAATCTGAGCTAAACGTTGGAGTTAAGGCAAAGAT

AAAGATGATTCACTTAAGACTAGGCAAGGATTGCAAGTGGTTGAGTTTCAGGCAGATGGTCAGGTAGGGT

ATCCAGGACGATATCATTACGACTAGATGCAGTTGGAGTTCTGCAATTCAATTCAGTTGGAAATTAGATT

TGTTTATTTGGTTATCAGCTACTCTTCTTATCGAGGGTTAGCTGATAATGTAGTTTAATTTGTAATTAAG

ATTTTATAGTTTTAGTTTCTGATTTCAATGCAGTAACCTGTATGCGATTCTGTTTTCAGGGGGTGAAATT

TATTTTGAATTTAAATCATTTTAAAAAAA

>MSTRG.814.2 gene=MSTRG.814

CATTTAAGTGCCATAATGTTGGCTCGTTTATGTCTTTATTTTTATTTTTAAATTCACTCACATTCGTTTC

ATTCTGTTTTGTCTTATTTTCAGGATGTTTGAAAAAGAATTTAAGAATTTATTTATTAAGGCTGCTTCCG

AGAAGACGCGAACCTTGGTGGACAAGACAGTTGAGCTACAAACATAGATAGAAAGGAATAAAGAAGACAT

CATTTTTAGTCTATTTTGTAATCGATGTATTTTTGTAGGGTACTCTTTTTCCCCCTTGAGTTTTTTTCCC

ACAGGGTTTTGCTCTTGAAGGGTTTTAACGAGGCCCTTTTTT

>MSTRG.814.1 gene=MSTRG.814

ACCATTTATGGTGACCATTATTGGGACATTAATGGCCCATAGAGGCATTTAAGTGCCATAATGTTGGCTC

GTTTATGTCTTTATTTTTATTTTTAAATTCACTCACATTCGTTTCATTCTGTTTTGTCTTATTTTCAGGA

TGTTTGAAAAAGAATTTAAGAATTTATTTATTAAGGCTGCTTCCGAGAAGACGCGAACCTTGGTGGACAA

GACAGTTGAGCTACAAACATAGATAGAAAGGAATAAAGAAGACATCATTTTTAGTCTATTTTGTAATCGA

TGTAATTTTTGTAGGGTACTCTTTTTCCCCCTTGAGTTTTTTTCCCACAGGGTTTTGCTCTTGAATGTCT

ATTTTGTAATCGATGTAATTTTTGTAGGGTACTCTTTTTCCCCCTTGAGTTTTTTTCCCACAGGGTTTTG

CTCTTGAAGGGTTTTAACGAGGCCCTTTTTT

>MSTRG.819.1 gene=MSTRG.819

TAATACTACTCTTCTGCAAGATTATAAATTCAAGAATTCTTTGACAAAAGGATTGTTTGTGAAGCAAATG

ATGAATGGAGAAGGTACGATGAAGAATTTGGAAGAAGAATCGATTGACTGGAGAGGAAGACCCTCCAAGT

CTAACAAACATGGTGGCATGTCAGCTGCTGTTTTTGTTCTTGGGTTACAGGCATTTGAGATGATGGCAAT

TGCTGCTGTTGGGAACAATCTCATAACCTACGTTTTCAATGACATGCACTATCCCTTATCCAAGGCTGCC

AATATTGTCACAAACTTCGTCGGAACCGTTTTTCTGCTCTCCCTCCTTGGTGGCTTCCTTTCTGATTCTT

ATCTTGGGAGTTTTGCCACTATGCTCATCTTTGGATTCGTCGAGCTTTCAGGTTTTATATTACTCTCGGT

GCAAGCTCACTTTCCACAACTGAAGCCACCTAAGTGCAATATGCTATCATCAAGCGCATCAGCTGAGTGC

GCGGAAGCAAAGGGCTACAAGGCTTTAATATTCTTCGTGGCCTTATATTTGGTAGCTTTAGGAAGTGGAT

GTTTGAAACCGAACATAATCTCACTTGGAGCTGATCAATTCACAAAACAGGACTCCAAGCAGTCCAGAAA

GCTTTCCTCATACTTCAATTGTGCCTATTTTGCCTTCTGCATGGGCGAGCTTATTGCCCTTACGGTCCTC

GTTTGGGTTCAGACACATTCGGGCATGGACGTTGGTTTTGGAGTCTCCGCGGCTGCCATGACTGTCGGAC

TCATTTGCCTCGTATCAGGAACTTCTCTTTACAGAAACAATAGGCCTCGAGGAAGCATTTTCACACCAAT

TGCACAAGTTTTTGTAGCTGCAATCACAAAGAGAAAGCAGGTATGCCCTTCCAATGCGGATTTGCTCCAT

GGAAGCAAAACCATCGTGCTCCAACGCGTTTCTGCCTCATCTCCAGCCACCAGCACTCTTCTGCATACAG

AAAAGTTCAGGTTCTTAGACAAGGCCTGCATCAAAACTGAAGATGGAAATGAGATGAAGGAAAGTCCGTG

GAAATTATGCACCGTGTCACAAGTTGAACAAGTGAAGATACTTATTTCAGTTGTTCCGATATTTGCCTGC

ACCATCATCTTCAACACGATTCTGGCTCAACTGCAGACATTCTCAGTCCAACAAGGAAGTACCATGAACA

AACGTCTCACCAAAAACTTCCAAATCCCCCCTGCATCTCTTCAAGCCATTCCGTATATTATGCTCATCTT

TCTTGTGCCTCTTTATGAGACTGTTTTTGTTCCAATGGCACGACGAATAACAGGAAGAGACTCTGGAATA

ACTTCTCTACAGAGAGTTGGGATCGGACTATTTATAGCCACTTTCTCAATGGTCTCAGCTGCAATAGTAG

AGAACAAGAGACGAAAATCTGCATTGATGAATGAAACTCTAACCATCTTTTGGATTGCCCCACAGTTTCT

TATTTTCGGTTTATCTGAGATGTTCACAGCAGTTGGCCTCATTGAGTTCTTCTACAGCCAGTCTTTGGAA

GGAATGCAATCATTTCTAACAGCCATGACATACTGCTCGTACTCATTCGGGTTCTATTTGAGCTCTCTTT

TAGTCTCCTTGGTGAATAAGATAACGTCAGGCTCCTCTGGTGGTGGATGGCTCAGTGGCAATGATCTTAA

CACAGATAAGTTGGACCATTTCTACTGGCTTCTAGCTGCTCTGAGCTTCGTAAACTTCTTCAACTATCTG

TTCTGGTCGCGATGGTACTCCTACAATCCGTCGTTGTCCCTCACCAGTCCTCCCAATCAACCTTATGCAG

TACAAGAATCCAGTAAGAACTGTATCGAAACAAAACTGGCTAATTTACCATGAAATACATTTTTATGATC

ATCATCTTCGTCGTCGTCATCATCACCAGATCTAGTCTTTCCACGCATGTTGGAATTCGAGTTTGTTTAA

GGAAAAAGAAATACTGGAATCATGATCATGATAGCATATTAGCATTAGAGTATTCATGAACCTCTAGCAC

CTTTAAGCTAAGTTCTTAACTTTATTAATGTAATATTGTGATGCCTATGTCTACTTTGGTTAAGTTTTAA

GATTCCCTAGTGTCTGATCTTTGGTCTTAGAATCTGTATGTACAAGAATGTTAGGTCTAATTCCATGAAA

GACTTTCTCTGGATCTCTTTTGATTTTTTGTGCAAGTGGTGTAGGGTTTGTTAAATCTTTATGCTAAAAA

TTCACTAGTAAATTTTGGACAAAAA

>MSTRG.819.2 gene=MSTRG.819

CTACTCTTCTGCAAGATTATAAATTCAAGAATTCTTTGACAAAAGGATTGTTTGTGAAGCAAATGATGAA

TGGAGAAGGTACGATGAAGAATTTGGAAGAAGAATCGATTGACTGGAGAGGAAGACCCTCCAAGTCTAAC

AAACATGGTGGCATGTCAGCTGCTGTTTTTGTTCTTGGTACCTCCATTTCTTAACCTGTTTTTACATTTC

CATACATGGTGATCTGTTTACATACCACTTTCGGGTTCTCATTTGAGCACGACCCTGTACAGAGAGTACG

ATGTGGTAATATTTGTGGTTGTGAATGCAGGGTTACAGGCATTTGAGATGATGGCAATTGCTGCTGTTGG

GAACAATCTCATAACCTACGTTTTCAATGACATGCACTATCCCTTATCCAAGGCTGCCAATATTGTCACA

AACTTCGTCGGAACCGTTTTTCTGCTCTCCCTCCTTGGTGGCTTCCTTTCTGATTCTTATCTTGGGAGTT

TTGCCACTATGCTCATCTTTGGATTCGTCGAGCTTTCAGGTTTTATATTACTCTCGGTGCAAGCTCACTT

TCCACAACTGAAGCCACCTAAGTGCAATATGCTATCATCAAGCGCATCAGCTGAGTGCGCGGAAGCAAAG

GGCTACAAGGCTTTAATATTCTTCGTGGCCTTATATTTGGTAGCTTTAGGAAGTGGATGTTTGAAACCGA

ACATAATCTCACTTGGAGCTGATCAATTCACAAAACAGGACTCCAAGCAGTCCAGAAAGCTTTCCTCATA

CTTCAATTGTGCCTATTTTGCCTTCTGCATGGGCGAGCTTATTGCCCTTACGGTCCTCGTTTGGGTTCAG

ACACATTCGGGCATGGACGTTGGTTTTGGAGTCTCCGCGGCTGCCATGACTGTCGGACTCATTTGCCTCG

TATCAGGAACTTCTCTTTACAGAAACAATAGGCCTCGAGGAAGCATTTTCACACCAATTGCACAAGTTTT

TGTAGCTGCAATCACAAAGAGAAAGCAGGTATGCCCTTCCAATGCGGATTTGCTCCATGGAAGCAAAACC

ATCGTGCTCCAACGCGTTTCTGCCTCATCTCCAGCCACCAGCACTCTTCTGCATACAGAAAAGTTCAGTC

TGAAAAATCTGCAGGTTCTTAGACAAGGCCTGCATCAAAACTGAAGATGGAAATGAGATGAAGGAAAGTC

CGTGGAAATTATGCACCGTGTCACAAGTTGAACAAGTGAAGATACTTATTTCAGTTGTTCCGATATTTGC

CTGCACCATCATCTTCAACACGATTCTGGCTCAACTGCAGACATTCTCAGTCCAACAAGGAAGTACCATG

AACAAACGTCTCACCAAAAACTTCCAAATCCCCCCTGCATCTCTTCAAGCCATTCCGTATATTATGCTCA

TCTTTCTTGTGCCTCTTTATGAGACTGTTTTTGTTCCAATGGCACGACGAATAACAGGAAGAGACTCTGG

AATAACTTCTCTACAGAGAGTTGGGATCGGACTATTTATAGCCACTTTCTCAATGGTCTCAGCTGCAATA

GTAGAGAACAAGAGACGAAAATCTGCATTGATGAATGAAACTCTAACCATCTTTTGGATTGCCCCACAGT

TTCTTATTTTCGGTTTATCTGAGATGTTCACAGCAGTTGGCCTCATTGAGTTCTTCTACAGCCAGTCTTT

GGAAGGAATGCAATCATTTCTAACAGCCATGACATACTGCTCGTACTCATTCGGGTTCTATTTGAGCTCT

CTTTTAGTCTCCTTGGTGAATAAGATAACGTCAGGCTCCTCTGGTGGTGGATGGCTCAGTGGCAATGATC

TTAACACAGATAAGTTGGACCATTTCTACTGGCTTCTAGCTGCTCTGAGCTTCGTAAACTTCTTCAACTA

TCTGTTCTGGTCGCGATGGTACTCCTACAATCCGTCGTTGTCCCTCACCAGTCCTCCCAATCAACCTTAT

GCAGTACAAGAATCCAGTAAGAACTGTATCGAAACAAAACTGGCTAATTTACCATGAAATACATTTTTAT

GATCATCATCTTCGTCGTCGTCATCATCACCAGATCTAGTCTTTCCACGCATGTTGGAATTCGAGTTTGT

TTAAGGAAAAAGAAATACTGGAATCATGATCATGATAGCATATTAGCATTAGAGTATTCATGAACCTCTA

GCACCTTTAAGCTAAGTTCTTAACTTTATTAATGTAATATTGTGATGCCTATGTCTACTTTGGTTAAGTT

TTAAGATTCCCTAGTGTCTGATCTTTGGTCTTAGAATCTGTATGTACAAGAATGTTAGGTCTAATTCCAT

GAAAGACTTTCTCTGGATCTCTTTTGATTTTTTGTGCAAGTGGT

>MSTRG.820.1 gene=MSTRG.820

CCAAAGTAGACATAGGCATCACAATATTACATTAATAAAGTTAAGAACTTAGCTTAAAGGTGCTAGAGGT

TCATGAATACTCTAATGCTAATATGCTATCATGATCATGATTCCAGTATTTCTTTTTCCTTAAACAAACT

CGAATTCCAACATGCGTGGAAAGACTAGATCTGGTGATGATGACGACGACGAAGATGATGATCATAAAAA

TGTATTTCATGGTAAATTAGCCAGTTTTGTTTCGATACAGTTCTTACTGGATTCTTGTACTGCATAAGGT

TGATTGGGAGGACTGGTGAGGGACAACGACGGATTGTAGGAGTACCATCGCGACCAGAACAGATAGTTGA

AGAAGTTTACGAAGCTCAGAGCAGCTAGAAGCCAGTAGAAATGGTCCAACTTATCTGTGTTAAGATCATT

GCCACTGAGCCATCCACCACCAGAGGAGCCTGACGTTATCTTATTCACCAAGGAGACTAAAAGAGAGCTC

AAATAGAACCCGAATGAGTACGAGCAGTATGTCATGGCTGTTAGAAATGATTGCATTCCTTCCAAAGACT

GGCTGTAGAAGAACTCAATGAGGCCAACTGCTGTGAACATCTCAGATAAACCGAAAATAAGAAACTGTGG

GGCAATCCAAAAGATGGTTAGAGTTTCATTCATCAATGCAGATTTTCGTCTCTTGTTCTCTACTATTGCA

GCTGAGACCATTGAGAAAGTGGCTATAAATAGTCCGATCCCAACTCTCTGTAGAGAAGTTATTCCAGAGT

CTCTTCCTGTTATTCGTCGTGCCATTGGAACAAAAACAGTCTCATAAAGAGGCACAAGAAAGATGAGCAT

AATATACGGAATGGCTTGAAGAGATGCAGGGGGGATTTGGAAGTTTTTGGTGAGACGTTTGTTCATGGTA

CTTCCTTGTTGGACTGAGAATGTCTGCAGTTGAGCCAGAATCGTGTTGAAGATGATGGTGCAGGCAAATA

TCGGAACAACTGAAATAAGTATCTTCACTTGTTCAACTTGTGACACGGTGCATAATTTCCACGGACTTTC

CTTCATCTCATTTCCATCTTCAGTTTTGATGCAGGCCTTGTCTAAGAACCTGCAGATTTTTCAGACTATG

AATATTATGCTTTGACAAAATTAATGTAATTCATTAATCAAACACATTTGATCAAACCAAAATCTTGGAT

AGAATGTTAGAAAAACATATAAAGTTGTCCCAGTCCCACCATGATTATTTGTTTATTTTGTGTGTGCGTG

CAAGTTACACTCTGCCTACCTGAACTTTTCTGTATGCAGAAGAGTGCTGGTGGCTGGAGATGAGGCAGAA

ACGCGTTGGAGCACGATGGTTTTGCTTCCATGGAGCAAATCCGCATTGGAAGGGCATACCTGCTTTCTCT

TTGTGATTGCAGCTACAAAAACCTTTGAGA

>MSTRG.821.1 gene=MSTRG.821

ATACCCAACCCGATTATTTGCACACAAACACTCTATCCCTTTTCCCGGCAATTAAAACCCTCCCTGGAGC

CACTTCTTTCTTAGACACGCCCCCACAAATTTCACTCTTAATTCATAAAGGTTTAGAAGTTCTTTCTTTA

GGAACAGAAAATGAACCAAAGGCAACAGTTGGAGGAAAACAAACATGATGATGATGCTGCACTTTCCAAT

TTCTTGGCTTCCTTACTGGACTATACCCCCACTATACCAGATGAATTGGCTGAGCATTATCTGGCCAAAG

GTGGGTTTCAGTGTCCAGACGTACGACTTGTTAGGTTGGTAGCTGTAGCTACACAGAAGTTCATTGCAGA

TGTTGCTAGTGATGCTCTTCAGCACTGTAAAGCAAGACAAGCTGCGGTAGTCAAAGATAAAAGAGACAAG

CTTCAAAAGGATAAACGCCTGACCTTGGCGATGGAGGACCTTTCGAAAGCGCTGCAAGAGTATGGTGTGA

ATGTCAAGCATCAGGAGTACTTTGCTGATAGCCCTTCTGCTGGGTTGGATGCTGCCTCAAGAGACGAATG

AGAAACTCTACCTAATTTTTACATCATCTGCCTTCAGTAATTTGGTTGGAAATGAATGCGGCATCATGGG

CAGGATGCCTGAGGCTGCTGAAGCGTAACTGACTCCTCCACTGAGTCTGTCCCCGGTTGTTGCTGGATCT

GCAAAGGGAGGAGGCGCTGGGATTCCCAGCATCTCTCCTGCAACTAAATTTGTTAAGAATCTGCCAGTAG

GAAGTCCGGAGATGGAATCACAGCCATATGGATAGTAATTTGCTTTAGCAATAGAATTGATAAAATTGTT

GTTGCCAGTGTCGACAATCGAATCTCCAAACACAAACATGGCTGGAACCTCTGAAGCTTCTACAAATGAA

TCATGGATCGAGCAGCGGAGAAACAGCACCAGCA

>MSTRG.821.2 gene=MSTRG.821

CCCAACCCGATTATTTGCACACAAACACTCTATCCCTTTTCCCGGCAATTAAAACCCTCCCTGGAGCCAC

TTCTTTCTTAGACACGCCCCCACAAATTTCACTCTTAATTCATAAAGAAGTTCTTTCTTTAGGAACAGAA

AATGAACCAAAGGCAACAGTTGGAGGAAAACAAACATGATGATGATGCTGCACTTTCCAATTTCTTGGCT

TCCTTACTGGACTATACCCCCACTATACCAGATGAATTGGCTGAGCATTATCTGGCCAAAGGTGGGTTTC

AGTGTCCAGACGTACGACTTGTTAGGTTGGTAGCTGTAGCTACACAGAAGTTCATTGCAGATGTTGCTAG

TGATGCTCTTCAGCACTGTAAAGCAAGACAAGCTGCGGTAGTCAAAGATAAAAGAGACAAGCTTCAAAAG

GATAAACGCCTGACCTTGGCGATGGAGGACCTTTCGAAAGCGCTGCAAGAGTATGGTGTGAATGTCAAGC

ATCAGGAGTACTTTGCTGATAGCCCTTCTGCTGGGTTGGATGCTGCCTCAAGAGACGAATGAGAAACTCT

ACCTAATTTTTACATCATCTGCCTTCAGTAATTTGGTTGGAAATGAATGCGGCATCATGGGCAGGATGCC

TGAGGCTGCTGAAGCGTAACTGACTCCTCCACTGAGTCTGTCCCCGGTTGTTGCTGGATCTGCAAAGGGA

GGAGGCGCTGGGATTCCCAGCATCTCTCCTGCAACTAAATTTGTTAAGGTATAAATTTGTTAAGGGTTGA

GCATTTGGGACTCAGATTTCATAAAATGATAGTTCATATGATATTTGCTCATAAATATGACTATTCGCAT

GCGGCTTCATTTGAATGCTGAAGTAGCTACTAAGACTTAACATTACCCATAAAATCGACGACGGTTCTGC

CATTACAGAATCTGCCAGTAGGAAGTCCGGAGATGGAATCACAGCCATATGGATAGTAATTTGCTTTAGC

AATAGAATTGATAAAATTGTTGTTGCCAGTGTCGACAATCGAATCTCC

>MSTRG.821.3 gene=MSTRG.821

CCCAACCCGATTATTTGCACACAAACACTCTATCCCTTTTCCCGGCAATTAAAACCCTCCCTGGAGCCAC

TTCTTTCTTAGACACGCCCCCACAAATTTCACTCTTAATTCATAAAGAAAATGAACCAAAGGCAACAGTT

GGAGGAAAACAAACATGATGATGATGCTGCACTTTCCAATTTCTTGGCTTCCTTACTGGACTATACCCCC

ACTATACCAGATGAATTGGCTGAGCATTATCTGGCCAAAGGTGGGTTTCAGTGTCCAGACGTACGACTTG

TTAGGTTGGTAGCTGTAGCTACACAGAAGTTCATTGCAGATGTTGCTAGTGATGCTCTTCAGCACTGTAA

AGCAAGACAAGCTGCGGTAGTCAAAGATAAAAGAGACAAGCTTCAAAAGGATAAACGCCTGACCTTGGCG

ATGGAGGACCTTTCGAAAGCGCTGCAAGAGTATGGTGTGAATGTCAAGCATCAGGAGTACTTTGCTGATA

GCCCTTCTGCTGGGTTGGATGCTGCCTCAAGAGACGAATGAGAAACTCTACCTAATTTTTACATCATCTG

CCTTCAGTAATTTGGTTGGAAATGAATGCGGCATCATGGGCAGGATGCCTGAGGCTGCTGAAGCGTAACT

GACTCCTCCACTGAGTCTGTCCCCGGTTGTTGCTGGATCTGCAAAGGGAGGAGGCGCTGGGATTCCCAGC

ATCTCTCCTGCAACTAAATTTGTTAAGGTATAAATTTGTTAAG

>MSTRG.821.4 gene=MSTRG.821

CCCAACCCGATTATTTGCACACAAACACTCTATCCCTTTTCCCGGCAATTAAAACCCTCCCTGGAGCCAC

TTCTTTCTTAGACACGCCCCCACAAATTTCACTCTTAATTCATAAAGTTCTTTCTTTAGGAACAGAAAAT

GAACCAAAGGCAACAGTTGGAGGAAAACAAACATGATGATGATGCTGCACTTTCCAATTTCTTGGCTTCC

TTACTGGACTATACCCCCACTATACCAGATGAATTGGCTGAGCATTATCTGGCCAAAGGTGGGTTTCAGT

GTCCAGACGTACGACTTGTTAGGTTGGTAGCTGTAGCTACACAGAAGTTCATTGCAGATGTTGCTAGTGA

TGCTCTTCAGCACTGTAAAGCAAGACAAGCTGCGGTAGTCAAAGATAAAAGAGACAAGCTTCAAAAGGAT

AAACGCCTGACCTTGGCGATGGAGGACCTTTCGAAAGCGCTGCAAGAGTATGGTGTGAATGTCAAGCATC

AGGAGTACTTTGCTGATAGCCCTTCTGCTGGGTTGGATGCTGCCTCAAGAGACGAATGAGAAACTCTACC

TAATTTTTACATCATCTGCCTTCAGTAATTTGGTTGGAAATGAATGCGGCATCATGGGCAGGTACTGAAA

TACTTTGTTTTCTGGACCATGTCTGTTATCTCCAAATTGTATTTACATTTTAGTCCAATTTCTCACCCGA

ATGCAAGGAAAACTGTGAATCAAGATATAATGTTTAGTTGGGTGTTCATGGATTCGTAATTTTATGTATT

GTGCCTTATTCTTTATGTTTGGGTATATTTGCCAACTTAAAAAGTTTATCTTCTAAGC

>MSTRG.821.5 gene=MSTRG.821

CCCAACCCGATTATTTGCACACAAACACTCTATCCCTTTTCCCGGCAATTAAAACCCTCCCTGGAGCCAC

TTCTTTCTTAGACACGCCCCCACAAATTTCACTCTTAATTCATAAAGGTACGATCTTTCTTCTGTCACAT

CAAATTTACATGTTTCATTACCAAACCCTAGCCTATTTTTTATGATGCCGTTTCAGATTATCTACTTGAA

TTCTTGGTAGCCCAGTGGCTTCATTCTTATTTTGGGTTTAGAAGTTCTTTCTTTAGGAACAGAAAATGAA

CCAAAGGCAACAGTTGGAGGAAAACAAACATGATGATGATGCTGCACTTTCCAATTTCTTGGCTTCCTTA

CTGGACTATACCCCCACTATACCAGATGAATTGGCTGAGCATTATCTGGCCAAAGGTGGGTTTCAGTGTC

CAGACGTACGACTTGTTAGGTTGGTAGCTGTAGCTACACAGAAGTTCATTGCAGATGTTGCTAGTGATGC

TCTTCAGCACTGTAAAGCAAGACAAGCTGCGGTAGTCAAAGATAAAAGAGACAAGCTTCAAAAGGATAAA

CGCCTGACCTTGGCGATGGAGGACCTTTCGAAAGCGCTGCAAGAGTATGGTGTGAATGTCAAGCATCAGG

AGTACTTTGCTGATAGCCCTTCTGCTGGGTTGGATGCTGCCTCAAGAGACGAATGAGAAACTCTACCTAA

TTTTTACATCATCTGCCTTCAGTAATTTGGTTGGAAATGAATGCGGCATCATGGGCAGGTACTGAAATAC

TTTGTTTTCTGGACCATGTCTGTTATCTCCAAATTGTATTTACATTTTAGTCCAATTTCTCACCCGAATG

CAAGGAAAACTGTGAATCAAGATATAATGTTTAGTTGGGTGTTCATGGATTCGTAATTTTATGTATTGTG

CCTTATTCTTTATGTTTGGGTATATTTGCCAACTTAAAAAGTTTATCTTCTAAG

>MSTRG.821.6 gene=MSTRG.821

CTCTATCCCTTTTCCCGGCAATTAAAACCCTCCCTGGAGCCACTTCTTTCTTAGACACGCCCCCACAAAT

TTCACTCTTAATTCATAAAGGAACAGAAAATGAACCAAAGGCAACAGTTGGAGGAAAACAAACATGATGA

TGATGCTGCACTTTCCAATTTCTTGGCTTCCTTACTGGACTATACCCCCACTATACCAGATGAATTGGCT

GAGCATTATCTGGCCAAAGGTGGGTTTCAGTGTCCAGACGTACGACTTGTTAGGTTGGTAGCTGTAGCTA

CACAGAAGTTCATTGCAGATGTTGCTAGTGATGCTCTTCAGCACTGTAAAGCAAGACAAGCTGCGGTAGT

CAAAGATAAAAGAGACAAGCTTCAAAAGGATAAACGCCTGACCTTGGCGATGGAGGACCTTTCGAAAGCG

CTGCAAGAGTATGGTGTGAATGTCAAGCATCAGGAGTACTTTGCTGATAGCCCTTCTGCTGGGTTGGATG

CTGCCTCAAGAGACGAATGAGAAACTCTACCTAATTTTTACATCATCTGCCTTCAGTAATTTGGTTGGAA

ATGAATGCGGCATCATGGGCAGGTACTGAAATACTTTGTTTTCTGGACCATGTCTGTTATCTCCAAATTG

TATTTACATTTTAGTCCAATTTCTCACCCGAATGCAAGGAAAACTGTGAATCAAGATATAATGTTTAGTT

GGGTGTTCATGGATTCGTAATTTTATGTATTGTGCCTTATTCTTTATGTTTGGGTATATTTGCCAACTTA

AAAA

>MSTRG.823.1 gene=MSTRG.823

TGTAGATCTTAATTCTGATTGTATAATTTATAATTATTGCTTGCAGCCTCATCTTTCTAACAAAAGTATT

AAAACTATGTATTCTTGAGATTAAATGTTTGAACTAAAAGAATTAGCATTGACTGACTTTGTTTCCAACC

ATTCCTTCAATTTCTAGCTATCCTTATGATTGTTAATTCCATGGCCACCAATATTAGTGATCTTGAAATA

ACTGCCCCTTTTTCTTCATCATCTCCACCAAGCAGCTGGGATGTGTTCTTGAGCTTTTGTGGCCAGGATA

CTCGTGCAAACTTCATTTCACATCTTTACAAAGCTTTGTTTGAAGCTGAATTTTTAACTTTCAGAGATGA

TCCTGCACTTGAAAAGGGTGAAGAAATTTCACCTGGGCTACTCGGTGCAATTAGAGACTCAAACATGTTT

GTGGTTGTTATATCAGAGAACTATGCTCGTTCAAGTTGGTGCCTTAAAGAGCTCGTCCAAATTCTTACTT

GCAAGACTAGAGATTATCAGGTCGTCCCTGTTTTTTACTATGTTGAACCATCACATTTGCGGCACCAGGA

AGGCTGTGTTGGAGAAGCTCTTAAAGATCATAAAGACCGTTACTCTGATGATATGATTGACAAGTGGAGA

TCTGCTCTTGCTCAAATCGCGGAGCTATCCGGATACCATCTTAAGAAACATGCAAATGAGAATGAATCCG

AGACTATTCAACAAATTGTAGAGAACGTGACACGACAAGTATCCAAAAGGGCATTGCACCTTGAAAACGA

TCTACTTGGGATAGATTCTGCCATTGAAGAGATTTATCAGAAACTAAAGATGGAATTAAATCATGTTCGT

GCTCTTGGGATCTGTGGGATGGGCGGAATTGGGAAAACAACAATTGCCAAAGCATTCTACAACAAATATT

ACAACATGTTTGATATTAGCTGCTTTATTGATAAAGTTAAACAATATTCACAAGCAGGTAGTCCTCTACT

TCCCTTGCTTAACCAAGTACTGATTGGACTTCTCAGACGAAAGAATTATAAAGTCACTGATATTGAAAGT

GGTTTTGGACAGCTGAAACAAATTCTTTGTTATAAGAAAGCTCTCATTGTTTTGGATGATTTGGATCAGT

TAAGCTATTCAGAATTTCTAGTACCACTTTGCAACTTCTTTTCGGCTGGAAGTAGAATTATTGTAACGAC

AAGAGATGCAAATTTGGTAAATCAAATAAGAATTGATATACCAGAAGTAGATACATACATGGTGAAGACA

TTGGGAAAAGATGATTCATTGGAGCTCTTTAGTTATCATGCCTTCAGAACACCAAGGCCACCGGTACATT

TAAAAGTGCTCTCGGAAAGCTTTGTAACTTATGCTGGAGGTCTTCCATTAGCTCTTAAAGTGTTGGGTTC

ATCTTTACGCGGTAGAAATCAAGATGAGTCATTCTGGAAAGCTAAACTTGAGAAAGTTAAAGAAATTCCA

GAGGAAGGGATACTGGAAATCCTTCAATTGAGCTACGATGAATTAGGAAGTGATACGGAGAAGGCAATGT

TTCTTGATATTGCATTCTTCTTTGTTGGAAAATTCGAATATGAAGCTGTTGAGATATTTAAATCTTGTCA

TTTTCATCCAGAAGTTGGAATACCAATTCTAGTGGAAAGAAGTCTGCTAACAGTTGATGAAGATAATCAG

TTTCAAATGCATGATCTTATACAAGAGATGGGAAGGAGATTAGCCAAGAACACTCACTTATTCTTGCAAG

GAAATGCATGGGAAGGTTTGCAAAATCAAAAGGAAACAAATATAATTGAAGGCCTCATTTTAGATTTGAC

CGAATCAAAGAATCAAATCAGTTCTCAAATCTTTAACAAGCTGCCTAAATTGAGATTACTTGAGATACTT

GGTGCTCCTCGCATTAAGGAAGATTTCAAGAATTCATTTGGTGAATTAAGGTGCTTTTATTGGAGATACT

GTACATGGACTTGTTTACCTCCTAGTTTTCAACCCGAAAATCTTGTCGTTCTTAATATGCCATTTAGCAA

TTTCAAGATATTGTGGAGTGGTTCCATGCCTTTCAAGAGTCTGAAGATTATAGATGTGGATTACTCTGTG

AAGTTGGAAACGACTCCCGACTTTGGAAATTCAAAATTGGTTAAGAGGTTATTCTTTCGTGGTTGTGAAA

GCCTGCTAAAACTTCACCCATCAATCAGAGAGTTGATTCATTTGTATAATTTAGATTTGAGGGAATGCAC

CAATTTGAAAGAATTGGCTGAACCAGTTGGGCAATTGACTATATTGGGCTATCTGTATTGGTATGTGAAA

GGATCTCCGGAGCCTATGAGGCAGTGGGTTAATGTGGGCTATTTGCGGTTGGGTCATTGTAGCAATGTAA

GACGATTGCCAGAACAATTGGGTGACATGAAAAGTTTAACGTATCTTGATGCAAGTCACACATCAATCAC

AGAACTCCCAGATTCTATTACTCAAGTCAAGAGCTTGGTTGAGTTGAACTTATCCAGTTGCAAGAAGTTT

AGAAAGTTACCTCAAGAATTTGGGAGTATGGAAGGCTTGGAGATATTTAGTGCATGTGATACTGCAATCG

AACAATTGCCAGATTCGTTTGGAGGCCTCATTAATTTGCTCCAATTGATATTGAGTTCCTGCCAGAAGCT

TAAAAATCTACCGGATGACATATGGAAGCTCAAGTTGCTTGCAGGACTATATCTTGATGAGTGTTCAAAT

TTGGAGCGATTGCCTGAGCAATTGGAGAAAATGCAATGTTTAGAATCACTTGATGCATCTCATACAGCAA

TTGAGAAACTACCTGATTCTATTGGACAACTGAGCAAGTTAAAGTACTTGGGTTTGAGTAATTGTGAGAA

TCTTAAATATATACCGAATAGCATATGGAAGCTTACATCACTTACACACTTATATCTTTTGAAAGCATCC

ATCAGTATAACTACTTTACCCAATACAATGAAGGATACAAAATTGGTTTCCTTGGGTCTGACTTGTGATA

TAAGATTATGGTTGCCTATGATTTCAAGTTTCTCTTTGTTGTAGGAGTTATATCTTCGTGATGAAGGTGA

AAGTTTATCTTCAATGGAGCCATTTAGTCTCTCTAAGCTTCACAACTTACGGTTACTCGAACTGAATAGA

TGTACAAGTTTTGGGTCATCATTTCCTGAACTTCCTCCGAGTCTAACACACTTAAGCCTATATAACAATG

CAACATTGATAAATCTACCTGATTTATCCGGCTTGAAACAGTTGAAGAAGTTGGATATAGTAAATTACAG

CAGCCTTGAATCACTTCCTCCACTTCCTCCTCATCTTCATTCATTTATCGTTTATCATTGCCCAAGTCTA

CATGATCTACCAGATTTGTCAATATTGAAAGATTTGACATTTTTGAGTTTTGGCAAGTGGGAGCATAGTT

ACCATCAGATTGCAGAATGGTCCAGCTACAAAAACAGTGGGCGTACACTCCAATTTAATATCCCACCACT

CACGGGAGATGACTTCTTAGGTCTTGCGCTCTGGATTCTTTTTACATCCAAAAACAGTGACAATTTTAGC

ATCAGAGCTGTTGTTACTAATAAAACAAATGGTATAACAATGTACTATGAGATCCCTGTATATTGTAATA

GAGTTGGTGATGTATATAATCTGGTAAGATGCATTAATGGAGATGAAATGTTGATGAGAAGTGGAGACAG

CATTCACATTTCTTTTGAAAGGCAGTTATATAGGTTTTCTGACGGAGATGTAGAATTTCCTTACGAAGAA

GTGAAGATGAAGAAGTGTGGAGCTCATGTGATACAAAGATCACCGATCACCCCCAACTTCCCGTGTTTTT

TCAATAACTTGGTAACTGGTGTTTAGATGTTTGTACGTATAAATGTGTTTCATTCATCATATTTATATTT

CCCTTTTCTTTTAATCAATAAAAAAACTCTTATTTGTACGCCTGCGTTTGAGAGCTGTAATTGAATCCAT

ATTCAATATTGCTTGATTATCTGTAAGCACTTTCATGTTTGTAAATTCTCTGTGCCGCACACACAAACAA

TCATTCTCACATTTTGTAACTATCCTTCTTTCATGTCTTCCACGAGGTAATTGATTAATTGTGCCTTGTT

CTTCTTCCTCATTATTGTCTCCACCTTGTCGTGGTGTCTTAATGAGCTCATACTGATCCTTAGGTGCCCG

AGAATAGAGATTTGGACTATTCCTGTATTTTACTATGTTGATCCATCAGTCCTAAAAGTATAGGATCATC

ATTTTCGGAACTTCCAACTAATCTAAAACACTTAAGCAACGCTACAAGATCAGTCAGATATGTCAATGTT

GAAAGAATTGGCGTCTGTGACTTTTGGTAGG

>MSTRG.824.1 gene=MSTRG.824

GTTAAATCTGCCCTAAATTTTGATTTTTGGATCACCACCTCGAGACGCAAAACTACGCTTGAAGATGACC

TCAAGAAAAGAAAAGGCCTTGTCGAGTCGATTGCAATGGCAACAACAGTTGATGGGAACACTGAAGGTAA

ACATGCCAACGGGAATGCTTAGACTACACATATCCCTAAAGAGAGAGACAGCTCCTTGTTTCATTTTAAA

AGCGCGCAAGCAGCTGAGTGAAGTGAAAATTGCAAGCGAGAGGTCGCATTTTCAGCATATTATGGAACAA

GAAAAGAGTATCAACGAATGATTATCGGAAAAATATTAGTACCTGCTGTAACTTTAAATGTGAACCGGGG

ATATGTCCTAATGACACTCCCAATTTCATGTTTATCATTTTAAAAAATTGGAATTCAAGACGTCGACGTT

TCTGCGAGGGTAAGAGTTGGATCATTAATGGATCGATGAAGCAAATGAATTCTTTAGAGCTGTACCGGAA

CCCCCAAGTACTAAATTACCAGTTCATATCAATATCTGTTACTGTAAGCACTTTATTTTTACTGCCTTGT

GTAATTTCAATTGTTTATGTTGCTTTGCTTTATGTTCCAGGAATCCTCACCAGTTTATTTATTTAATTAA

TTATCATGACATATTTGTACTAATTTAATCATGCCAAACTCAATAGATTCTGT

>MSTRG.826.1 gene=MSTRG.826

ATTCCCTCAGACATACCTCTTGTAGCAGAACTCGACACATTGTCCGTTGCTCTTTTGCCGTCATGAAAAT

TTGTTGCCGAATAATTGATGTTTTGCTTTCCGTTCGTCGAATTCCGCACATAGTTGTTACCGTTTTCGGG

TGAAATGAGGTCCGGTAATCCGGATACTTGTGCCGTATTCATTTGACCATAGCTGAAACCAAATTCTTCT

AGGGCACCGACTCTGAGATTGGATTGTGAAGGTATTAAAAGTGAAAAATCATGTCCTTGCTTCGACGTAA

ATATCGATGAAGTGGATGATGAGGACTGGAGGAAATTAGAGTGATTAGGGTTTTGAAGTTGCTGATGTTT

CAAAAACCCTAGTTCAGAAGTGCTAGGAAAAATAAGAGGAGATCGAGTATTGTAAGATGGAGTCTGAGTG

ATATTGATGCTGGATCTGATAGAGGAAGGTGGACTAAAGAGATCAAGCCTGCTTCGAGGGAATGTCGAGG

ATTGCGAAAAAGGAGGCGCTGGGATTCCGGTGAATTCTTGGACCATGGCTCTGAAATTTGAGGTGTCTGT

GGTGAGAATAGTTGTGGGCGCGCGCCTGGATGCTCTTGATCTTTTCTTGGGGTTTCTTGTTACTTGTGTC

AGCTTATGTTGGTCAATTTTTCGCGTCGAGGGCTGAGATGATGATGCAGGTAGTACTAGGGTTGGTTGAT

TTTGTTTGAAATGGTGTGGAGGGAATGAGAGGGTTTGGGATTGAGGGGCTAAATAAGGGTTGGTGAGTTG

TTGGAGTTGAAGCTGTGGGGGTGGATTGAAAAAGTTGGAGGTATTAGTATGTGGATCATACATGGTTGGT

GGTGGTGGAGATAGATGGTTATAGATGGGGTTAAAGGGTGAATTAGCGGCGGTGCGTGGATCGTAATCCT

CCTCGCCACCGCTTGATTGCGTGCTTCCGCTGTTTCCAGAATCCATTAAAGGCTGTGTTTTATGAAAGAG

GGGAGAGAGAGTGGGGGAGGGAGAGAGATTGGGGGAGCTTTGTATCTCCAAATTTGTATGAAAAACTGAG

AGGCTTGTTTCACTTTAAAACATACTAAAAAGGGTTAGATTGGGTTTGATTTTGATGTCCACAAAGAACA

GGGAGGAAGGAGAAAACAGAAAAGAAGAGGAAACCAATAATCTATAAATCTCTGATCTTCTTTTGGGAAG

GGAGCGTGAGGCGG

>MSTRG.827.1 gene=MSTRG.827

ACGGTGAAAATGTAACAGCAAAAACAAAATAACAGCACACTCTCGTGGATAATATTCGGAACAAGAAAAA

TTTCAAATCATCACTCAAACAGCAACAACATGATCATCCCTGGCTAAACCCTTCAAGATCTTCCGGCCAC

CATGGCTACTCGTACCCGAACAATTATCTCGAATTGCGGTCAGCTCATACGGTGATCGATCAAGTGCAAT

CCATTGCTCGATGGTGAATAATTGTTGCGCACAAGGAATATGCGCGCCGGTGGTTGTCACCTCAATGTAG

TTACAATACCATCCGTGACCTGATCCTGTTCCATCAGATGTTATGTTCAGTTTACAGGCCGGTCCCGGAA

GACACGGGCCTTTTCCGCTGAAGATATCGAGATTGCCTCGTTCGAAATAGTTGTAATCAGGTCCCATGAG

TCCTCCCCAGGCTTCCAGATTGTTGATCCGGACTCCGTACCCTTCTGCGTCGTAGAGTGTCAGTGTCATT

ATCGAATCCGTTCCCCCTTTGATCACCGAGCCTGTCCGGACGTAGACCGTGTATACGCATTCATCGTCTG

ATCTGACGACGGAGGAGATGGCGAGGAGGATGAAGAGATTGAAGCTAAGATAAGAGAGTCGTCCCATGGC

CTTTGACAATCTACTTTTCTGTCTGTGTGTGATGTTGAGAGAGTGAGGGGTTTGGATTAGATCGAATAGG

AATATGCAATATTTGTAAAGGGTAAGGAGGAAATATCTAGATAGTGCTCTATTTTTCTGTTCAGCAAAGA

AAAATATATTGTATGAAATATAGTACATAAACGAAATAGTAGTAATTAATTGAAATTTG

>MSTRG.830.1 gene=MSTRG.830

GATGGCGATCAATAGAATAATCGATCGATTCTCTGGTCGATCTCCTTTTTCGTCACAATTCCGATTCGAT

GGTGATTTTTGGGCTTTATTTTAAAATCGGTAATATCATTCCTTTATTTTTTAAATCAAATCCTCTCCGT

CTCTTTAAAAGGTACGTCTTTTCCTATGTTTGTTGTGTGTTTTTCGCTTGTTTTGTTGGCTTTTGTATTT

TATTATGAATGTAGATGTACATTATTGC

>MSTRG.832.1 gene=MSTRG.832

TACACCGCGCCCAATTGAATTGAAATTAAATCACAACAAGCTTAACTATGGTGTCTTCCGATACTCAGCT

TGCTCTTCAGTCAGACGTTGAATCCCCGCTCTCAACTCTCCTCTATGACCTCTCACAACAGGTACAGGGA

GTGATGGAGAGCATGCTTAAGATGACAAATGAAATTGATCAACACTCTGCTGGCATCATGGAAGAAATAG

AAAGGTGCAAAGACACTGTATGTGAGAGAAAGAAAGCTTTAGATGAAGAAAAGGACCGTTTTCAGAAAGC

TGCTTACACAGTTCTAGACTTGCTAAATAACAGGCTCATCAGCTAAAAAAGATGTACAGGTGATCATTTT

CTCTTAAAGCATCCAAATCTTCTACTTGAGAACCATCTTCCATGAGTACTATATCTCCTTTTTTCCCAAA

CTTCTTTTCTGTATCAGCAAAAAGACAAGTAAATTACAATGCAACTCTCACTATCTATCCCTGTATACTC

GATATATAACTGATCTTAGTTCATTTCATTTCCGAAGTCCGAACTCATAACGATGCCTA

>MSTRG.832.2 gene=MSTRG.832

AGCAGGTGGAGTATATAAATAGGCTTAACCCTGTTTAAAATTTGGGAATCAAATACACCGCGCCCAATTG

AATTGAAATTAAATCACAACAAGCTTAACTATGGTGTCTTCCGATACTCAGCTTGCTCTTCAGTCAGACG

TTGAATCCCCGCTCTCAACTCTCCTCTATGACCTCTCACAACAGGTACAGGGAGTGATGGAGAGCATGCT

TAAGATGACAAATGAAATTGATCAACACTCTGCTGGCATCATGGAAGAAATAGAAAGGTGCAAAGACACT

GTATGTGAGAGAAAGAAAGCTTTAGATGAAGAAAAGGACCGTTTTCAGAAAGCTGCTTACACAGTTCTAG

ACTTGCTAAATAACAGGCTCATCAGCTAAAAGTAGCTTTGTTCTGTTCGAGGAAACTTCTTCATCCCGTT

TTGCAGGAGATACATAGTTGTACAAAAAGTTTACTGTTTACCCTCAGCTATTAAATATTTTAGTGCTACT

ATAGAAGCTGTATAGAGTTTCTGGAATCTAGACATAAACTTCATTTTCTCGCTTGATACCTTGTCTGTAT

GTGATGGCCTGCTAATGGTTTAGACAGTAGTATATTATCGAAATATTTGTATGGACCGTATACTGTGCTA

CAGTTTGTGGAATATCCTGTGTAAACCCTGAATATCTAGAATTTTATGTAAAATGCCTGAGTTGAATGCA

GTAATCCTTTAATAAACA

>MSTRG.832.3 gene=MSTRG.832

GTGGAGTATATAAATAGGCTTAACCCTGTTTAAAATTTGGGAATCAAATACACCGCGCCCAATTGAATTG

AAATTAAATCACAACAAGCTTAACTATGGTGTCTTCCGATACTCAGCTTGCTCTTCAGTCAGACGTTGAA

TCCCCGCTCTCAACTCTCCTCTATGGCTAATATTGAAATTGCATGTATTTATGTTGCAGACCTCTCACAA

CAGGTACAGGGAGTGATGGAGAGCATGCTTAAGATGACAAATGAAATTGATCAACACTCTGCTGGCATCA

TGGAAGAAATAGAAAGGTGCAAAGACACTGTATGTGAGAGAAAGAAAGCTTTAGATGAAGAAAAGGACCG

TTTTCAGAAAGCTGCTTACACAGTTCTAGACTTGCTAAATAACAGGCTCATCAGCTAAAAGTAGCTTTGT

TCTGTTCGAGGAAACTTCTTCATCCCGTTTTGCAGGAGATACATAGTTGTACAAAAAGTTTACTGTTTAC

CCTCAGCTATTAAATATTTTAGTGCTACTATAGAAGCTGTATAGAGTTTCTGGAATCTAGACATAAACTT

CATTTTCTCGCTTGATACCTTGTCTGTATGTGATGGCCTGCTAATGGTTTAGACAGTAGTATATTATCGA

AATATTTGTATGGACCGTATACTGTGCTACAGTTTGTGGAATATCCTGTGTAAACCCTGAATATCTAGAA

TTTTATGTAAAATGCCTGAGTTGAATGCAG

>MSTRG.833.1 gene=MSTRG.833

AGCAGCAATGAAGCATCAGTAGCTGCATCTCACCTACGTGTCTAAGACAGAGTTGAATAAAAAGGAAAAT

AGCATTAGGTCAGTTTATATGAGATAATCAGGAGGCGGCCGAGAGAAAACTAGGGTTCCTCCTTTTCTTT

AAAATCCATCAATTGATTTGAAGCATCTGCACAGGTGAGGATCTTGTGTATTTCTTGTGTGATTATACAT

ACATATCTGTTGTATGGACTATGTGTTTGTAGTTGGGTTGATTGTTGTTGTATTGATGGTTCGGCTGGGT

GTGGGTGGAAGTCTGGGCTGGTGTCGGAGCTGGCGGAGCGGCCAGCGACGGTGACGCGCTCGCCGGAGTT

GCACATGATAAAGCAGGGGCGGGCTGTTGGGTTGCAGCGCCGCTGCTGCGTGAGAAAGCAAGGGGAAAGA

GGTTGGTTTGTGCTGACAGGAATAGAGGTGGCTGGGACAGGGCGGCATGGCCGCGATTGGACAGTGGTAG

ACAGCGCCGCCTCGACGCCGATGAGGTGTGGCGGCGTGCTGCAGCAACGGGCCAGAGAGAGAGATATGGA

GAGAGGCCAGTGGTGGAGAAGGAATCCGGCGAGGGGAGGAGCCGCTGAGATTCCGACGAAGCTGGTGGTC

CGGGGCTGTCTGGGGTGGGCTGCCGGACAGGGCTTGATTTCAAAACTCAAAAACCAAGATGTTAATTTCA

GTTTATTATTATTATTGTTGTTGTTGTTGT

>MSTRG.837.1 gene=MSTRG.837

ATGTGTTTAAATTAGGTTACCCACGAACGAGATGGCGCGACTGAAGCGAACTCCAGAGCCAAGATCACCC

TGATAATCAGCAGACCTGATAAACTTGATTAACGAGAGTATTCGGAACTGAGTCGAGCGAGTTGAGTGAC

TCAGTATGTGAACTCAGTGAACTCAGTATGTGATTTGAGCTTTGAAGAGAGAATGAGTAAGAGGCTGTGT

TTGATTTTTCAATAGCTTCGCTGTAAATGAAGGCTGTCACGGGAGGCGATGGCTGTGAAAACGTGGAGAT

TGGAGTTTTGTGTGTGTTATACGTGGCTGATGTCTCAAAGGAGAGGAGCACGAGAGAGTTGGCTGGTGTT

TCTGACTCTTCAAAAGGAGTGTTTGTGTTACTCAGAAGCTTGTTGACAAACTGCTCGTGAAAATGACTCA

AAGAGCTTTGCTGACAAACTGTTTGTGAAAATGACTCAAAGAGTTGATAAATCAGAAACAATGAAGCATG

AAGATTATTAGCAGAAGCAATTGAGGTAAGAACCTTGCTCTGATACCATGTGAAAATATGAGTGCATATA

TGTATATATAACCAGTGTGTGTCACGAGAGAAGCAGTAGAAACAATGTAAGAGAGAAACTCTTATATTGC

TAGCAGGAAGCTTAAGAAATTACATGAGAGAATGAACATATTTATACAGAAAATAACATGAAGTAAGTCT

ATGCTAATGAATGTGGTGAGCTGCTCAACCACAATATTGTTCACTGCTCAACTGCAGAGTTTAAGAGCTG

GTGTTCTATTCTAGGAGTTGACATTTGTGTCAATGTGAACTAGAGACTTTAACACCAGTGATCCATATGA

ATCTTGGCGAGTAATTTATAATGGAACCCGTAAGATAGAGTACTCGTTTGTGTCAGAACTGTATGCTTGA

ACTAATTTTCGATGTCAATGCATGGGTTGAGGACTGATCATTGAAGTTTAGTTTTAAGCATTGCCGCAAC

AGCCTTGAAAAAAACCAGGGCTTTAACCTTGGCTGAACGCCTGAACCCCCTATCCTTCGTGGAGTAGTCG

AAGAGAAAATAATATGAACCATTGTCGAATCTGTCTTCTAGCTTTAATCCGAATGTCTGAATCAACGAAG

TTTCATCTATAAAAGCTATTTAGTAGTGAAATATTTTATCATTGATCTTGGTAAGCATAACCAGTGCAAC

TGATGTACATTGAAACACTGCAACTGCTAGGATCTAAGCCAACTCAATTGGCATGGCTCATATTTTCAAA

AAGGTGCAACTGATTGAAATAGATTTTGGC

>MSTRG.838.1 gene=MSTRG.838

CTATCTTCTCTTCTGTTGGCAATGCCCGGATTATTTTGTGGTGGGCTATGGACTTGATTTTGCTGAAATA

TACATGAACTTGCGTTTAATTGGGGTTTTAAAGCCTGAAATTTACAAGTTGTGATGTTAGAGAACCCAAT

TTGGGTATCAGAATTCTTCCAAATGGCAAAGTGTTGATAATATGTCGGTTTGGATTTTAATTGGTGGGAT

TATTTGAATGCAAGCTATCTATTATATTTAATCGAATGTTTTT

>MSTRG.841.1 gene=MSTRG.841

ATTTACCATCCGCTTTTCATACTTAGTTGGTGCAGATTTCTAATTATGTATCAAGGAAATACTTGAGGCA

AATGAGAATAAAGAGTCATGTTCTGTTTGCTGTAAAATTTTCGAAGGATTGGCCAACTGCTGAACAGTTG

GAAGCAGAGGCATAGCCAAGGACTGAAATGGTCCATTGATGTATCCAAACTCCTCCAACTTTCCTGTACA

AATGTTGTGACCTCAGTACCAGATTAAGACATGCATTTCCTCAATTCATTACCTTATTTATAATTTTTTG

TTTTATTGATCGACATGTGAATTGTTTATTTTACTTTAGGATTAGAAAAAGCTTCAAGGACAGCTACTGG

CAGGAAGTGGGATTTTGATGGGTTTAGGTTCAAGTTTTGAGATGTAAATTAGTCGATTTAATTAGTCAAG

TTTATTATGATTTATGTATGATTTATTGGATGTTGGATGTTGGATTTTGGGTATTGGATTTTAGATACTG

GATATATGAATTTTATGTATAATTTATTGTGGGAAGCATTATATATTTAAAGCATATATTCAAAAAAAAA

>MSTRG.841.2 gene=MSTRG.841

ATTTACCATCCGCTTTTCATACTTAGTTGGTGCAGATTTCTAATTATGTATCAAGGAAATACTTGAGGCA

AATGAGAATAAAGAGTCATGTTCTGTTTGCTGTAAAATTTTCGAAGGATTGGCCAACTGCTGAACAGGAT

TAGAAAAAGCTTCAAGGACAGCTACTGGCAGGAAGTGGGATTTTGATGGGTTTAGGTTCAAGTTTTGAGA

TGTAAATTAGTCGATTTAATTAGTCAAGTTTATTATGATTTATGTATGATTTATTGGATGTTGGATGTTG

GATTTTGGGTATTGGATTTTAGATACTGGATATATGAATTTTATGTATAATTTATTGTGGG

>MSTRG.841.3 gene=MSTRG.841

ATTTACCATCCGCTTTTCATACTTAGTTGGTGCAGATTTCTAATTATGTATCAAGGAAATACTTGAGGCA

AATGAGAATAAAGAGTCATGTTCTGTTTGCTGTAAAATTTTCGAAGGATTGGCCAACTGCTGAACAGTTG

GAAGCAGAGGCATAGCCAAGGACTGAAATGGTCCATTGATGTATCCAAACTCCTCCAACTTTCCTGTACA

AATGATTAGAAAAAGCTTCAAGGACAGCTACTGGCAGGAAGTGGGATTTTGATGGGTTTAGGTTCAAGTT

TTGAGATGTAAATTAGTCGATTTAATTAGTCAAGTTTATTATGATTTATGTATGATTTATTGGATGTTGG

ATGTTGGATTTTGGGTATTGGATTTTAGATACTGGATATATGAATTTTATGTATAATTTATTGTGGG

>MSTRG.845.1 gene=MSTRG.845

GTTGCTTTGGCTGCTCAAAAGATTGAGTTATTGGAGAAAAGGAAGGCTCAGAAGAGTAGAAGACCTAGAT

TGGCAAATCCTTTTGCATCAAAACAGGAACCATTGGATTTACATTTCACTCAATTTGAGGAGTATACTGA

TCACTTGAAGGAAACCAGGCCACTGTTTAGGGTCACTCGGGAAGAAGATATGTCTTCTCCAGAAAATTCA

GAACCAAGCCCAGAGGAAAAGTCTGAGATAATTTCTGCAATCTCAGAATATCAGATTGGGGATTCTGTGC

ATCATCTTGAGCTAAGACCTCCTAAGTTTGAAGGTGCTAGAGTCTTCGAGTCATCACGTGATTGGGAGCA

CCTTGTGGCTGTTGAACCTATCTATAATGATGACCCTATCTATGATGAAGATCCAATGGAGGGCGATGAG

TGCTGGCCTGAGTATGAACTCATTTCAGGAGTCCTAGGAGGAGATAACCTACACTCATCTTGTTTATCAC

ATTCAGATCTTTGCACAAGTGAAAGATTAGAGGTACAGTGGAGGGACAAATCAGATGAGGTATTTGAAGA

TGGTGACATGCCAAAGCTTGTTGATTACAGGGGTGAAGAGAAACAACATGTTGTTGTGGAGGAATCTTCA

ATCATCCTTACCAGCATGAGGGAATTGATTGAGGATGTTCCATGCATAGATAGTCCCGTAACTCCAAATT

ACCCTGGTCTAGAAGAGAAGTCTAGCCATCTTGATACACAGATTATGACCATCTTTGATCCAGGGGGTTA

TATGCAAAAAGATCATTCAGCTGTTGATATGTTATTCAAAGAAATGATCATGGCTAAGGAAGTGAAGCTA

CCAGAAGCGACACTAAGCAACTCAGTTGGTGGCCAACTTGGACAGTGGCAAGAGGGGCTTAGTTTGAAGA

TCTTGGCAATAAAGCAGCTGTTAAAGGAACACCCTAAGTTGAGGAAGAGAGGTTTTGTAGTTCAGTTTGA

GAATTCAGCTGGCATACAAGGCCAAGATATTCAAAAGGATGATGATGAGATTGACCAAGTCTTCACTGAG

ATTAATGAGGCTGATGAACTGTATTGTGGTTCTGTGAGGGCTGGGATAATGGGGACATCCTACAGGTACA

CTGAGCAGATACAAGGTCCAGGAGATATTATGAAATTTGAAGTCTCAAGGAAAAATGTTATAATATACAT

ATATGATCCAGGAGGTTTTTTAAGTTGTTTCTATGGAATTCGATATGTTGATGGTTCTGTAGGAATTCTA

AATATTCAGCAGGTGGTGCAAGTTTTCAATAAAAAAGAGTTGCATCTGGAAGGAAAAGCAAGCATAGCAG

TAAAGGCCTTACCAGAGATGTACATAACCCAGGTGGAAAGAGGTTGTGAAGTTTATGATCCACCATGAGG

ACATGGTGAAATTCAAGGAGGGTTGAATGTCAGGAATCAGCTTAAGTAGCAGTGTCATATGAAAATTGAT

AGTAGTAGTAGGAGTAATAGTAGTCAGGGTAATTGTGTCCTTTCATTACAAGTTAGTAGCACAAGTTAGT

AG

>MSTRG.846.1 gene=MSTRG.846

AAAAACCTGTTACAATCTCTGTGGATGATGTGATTCGGGCAAACATAGATGTGGCAGATCTTGGGAACTA

AAGCCCTCATTTAAGAAAGACGGGACGACGACTGCTGGTAACTAGGGGAGAGCATTTGGTTTGATTAACC

GATAACCGAACTGAATAACCGAAGAAATGAAACAACTAGATGAAGCACTAGAGAAAATGAACATGGATGC

AAGTTCTGCAGCAGCTTCAGCGACTGGAACAAGTCATCCACCGAAATTCTAGCCAGGTAAGTGTGTCACA

GAGAAGTACTGCTATGCAGAAAGAATTACCG

>MSTRG.847.1 gene=MSTRG.847

CTCCAATCATCACTAGTAGGCCTCCCATCAACATTATTCAACTTGGGGTGTTTTTTATACGTAGGATCTT

GTGTATCATACCTACCAAATGCCTTTTCATATTTTTCTGCAACATCTAGCATTTCATATGTCGAATTCCA

TCTAGTATCCACATCAAGAATCAAACATTTTTTAGTTTCAATTCTCTCATACGTTGCACATTCCATAAAC

AATGCCTCCCTAACGGGTGAATACCTCACATACTTAAACCGCATTTCTCATCCTCACAATTGACTCATTA

AAGACCCCCAATCCAACTTTGACAATCAAATTTAGAACATGAGCAACACACCTCACGTGTAAGTGTTCAC

CATTTAGCATGCTAGTACCCTAATTACCAATTTTATTTCTTAATATTTCCAATGCAACATCATTAGTACT

AGCATTATCAACCGTGATGGATAACACCCTAGTTATCCCCCACTCCACCAAACACTCCTCGATTGCCAAT

GCAATTGCCTTACCTTTATGACTAAGAATTGGACAAAAGTTTAGTATATTTTTATGCAACTTCTAATCCT

TATTAATAAAGTGTGCGGTTAAACACATGTAATTAATTCTTTGATTGGAAGTCAATGTGTCCGTAGTTAG

GCAAAGGCGTTGTGTTGGTTCAAGCAATACAGCTTTTAAATCATTTCTACAAAGAGAATACATACCAAAA

CAATCACGTGCAACCGTCCATCTTGATGGAATTTAAAACAAGGGTTGCACAGTAGCTATAAATTCTCTAA

AGCCCGAACCTTCCCCAAACTTGAATGGTAACTCATCAACAGTTATCATCTTAGTATATTTTTCTCTAGC

CAAATCTTTGTCAAAAGTCCAAGTAGTTAATGAAGTGCCATCCTTATTAACTTGAAAAGCTAATTTTGTT

TGACGACCCTCAACATTAAATGACTTGCAAGATTTCAGATGGTTGTGCAAACTACTAGTACCATTTTTTG

AGGGATTGTCACACAAATCCTTTGCACAATATTTGCACTTACCTTTCTTCACGTTGCCGTTTTCATCTCT

AACTTTGATGTAGTGATCCCATACATTGCTTCGAGATTCCATTGGCTTCCTTTTTCTAAGATTTGAAGCC

ACAGTTTCAGTTTGTGCTTCAAGTTCTTCT

>MSTRG.848.1 gene=MSTRG.848

AGAAGAACTTGAAGCACAAACTGAAACTGTGGCTTCAAATCTTAGAAAAAGGAAGCCAATGGAATCTCGA

AGCAATGTATGGGATCACTACATCAAAGTTAGAGATGAAAACGGCAACGTGAAGAAAGGTAAGTGCAAAT

ATTGTGCAAAGGATTTGTGTGACAATCCCTCAAAAAATGGTACTAGTAGTTTGCACAACCATCTGAAATC

TTGCAAGTCATTTAATGTTGA

>MSTRG.851.1 gene=MSTRG.851

CTTGCCACTTGTAAAGTTTAAAAAAATCATAAAATCAAAAACCCATTAAAACTATAAAATCAATACTCTG

CTCTCTCGTTGTTGTTCTCTCTACTCTTCATTCCATATGATATACGAACGGAGGAGGGTTGTAAGTGTTT

AAGATTCATCTGGAGGCAGTATACGGTCATTATCCCAATTTCCAGTGGACCCTGATAAATGACCCATATT

AGATATGGTGATCGACTTTGAAGATGAAGGTAAGAAGTCTGGTGCTGTGTTTAATTATGAATATGAAGTT

TCTTCATATGTTTACAAGATGGATCATGTGAAATTGTTGAATGATAAAGACTTAACGAAAATGTTTGCTA

GACTAAATGAGAAGAAGTTGATAGATATATGAGTAGGTACCATCACTACTAGAATTTAGTCCATAGTCAT

TGGTTTTTGACCAATGTCTATGCACTTCACTAACTGATGTCTAAGTTTGTGATGTTACATGAGGGCCTAA

GTCGACCGATGTCTATACAATGATGTTTATCCTACTTTTCTAGTAGTGCATGGTCAAGCCTAATCAGTGT

ATAAGTAGGTTTTACGACTTAGAGAATAAAATAAGGAGGGGGATAATGAGGAACTAAAGGGATGGATGAG

CAAGGGGATGATCACGAGGTAGAAGCTATGGAGGAGGGGGATAATCATAAGATAGATGCTATGGAGTAGG

GGGATGTAGATGGAGGTAGATTTGTGAAGGAAGTTATGTGTGCATGGTATTAGTGCCGTATACCATGAAA

ATGGGAATCTAGATGATTGTGTGCATCGGTGTTATATGAAGGACAGATAAAAAAAAGATATACAATTGTT

ACTTGCAACCAGTGAAAGGCTCCAACATGTGGCCGGAAAGAAGCATACAATCTATGGTTTTACTGTGGAG

AGAAGAATGCCAGGCAGACCAGAGAAGGCAAGGAGAAAGGAATTTGATGAGGATGGAAAAGGTTTCAATT

ACACTTCTTTAATAACAGAAACACTCATAGTCAGCAAAAAATTCTAGAAATAAACTATAGTCACTGTGGA

GTCCCTGGACACAACTAGAAGACATGTAAAATCCACCTAAAATGATGAACCCTCAACCAAAGAAGATACT

TGGCAAGCCAAGGGTGGAGAAGCCTATAACATGTGTTGCTCTCAAGAAAATACAACCAAGAAAGTTTGCA

AACATGGGATATTGCTTATGCATTGATCCAAAAACTGTGAAAACCTATTTGAATGTAAGTTTGTGTGTTT

TTTTTTTCTTCTGTCCATGTGCCTTGCATAGTTTTTCGACATGTATTATTTACATGTTTTTCAAAGAGGG

TTATTTGATTCAAAGTGGAGAAACTAACGTCGAAGAATTTCAAAC

>MSTRG.853.1 gene=MSTRG.853

GTTCTACATGCATGATAAAATTTTGGTAGTTTTCTGTTAAGGTGATCAAATTGTTATATCTCACTCTGGT

ATAGGTACACTTAAATATAGAGTCTAGTTACCTATCCTAAAGTTCTCAGTAAGGTTACCATTTTTAAAGT

CAAGAGCTGGTGCAGTTACCTGAGTGATGTAAATCGTCTGTGTGGATATATTATTGAGTTTTTGCATTTG

TGTTTCTTTTGTGCAGAGATCAAGACACTGACTGTTGATATGAGGGATGAGCCAGGAGCAAGACCTATTC

CCAAAGCAAAGGTAACTTCAATACAATTTACTGTGAGCAAATGAAGGGAAGGATAATTGCTGTGTTCACG

TACACAGCATTCCAAGAATGCTTTCCTATGTACTAGCTAATGTCCTCATTTTATTTTAATTAAAACTGTA

TTAAATATATGTTCTCGAGACAATCGGCGTATGGGATTATGTTGCATCTATTTGTCATCTATCCATTCAC

CTGTCTCAGATTTTTAATTGAGACTGTATTTACAGTTATATAGATGACATTCTCCAGTAAATTGGATTCA

AGATAAACTCAATGCTTTTATCTTGTAGATTGAAATATTGCTGTCTAAGACGGAGAAGTTTGAAGAGTTG

ATGGCCGCTGAGGCGGAGGAAAGAGAATTTGCTGCAGATGGTGAGGAACAGAACTGAGCTAGATTGCAGG

ACTGAGCCTGATGTTACTCATCCCTGATAATGAGCATAAGTCATTGATTATCACTTGCTCTGAAAAACCC

GAAAAAGAGGATAGGAAAACCAACTAAAATGTTACATAGCTTGGTACTTCGCTGCATGAATTGTTCTGTG

AGGAAAAAGTGGTTATTGTATTGGGTATCTGTTTTTGTTGTTAAGTCTTTGTTGCAAAGGGTTCCATTGG

CCATCAGTTCTGTTGCTTTAGACTTTTCACCTAGTGAGGATGCTTTAGAATTTGTATCACAAGCTTGTAA

TGAATAAACTATTAGAAAGATCATCTTTGGGAAACCAGATTCGGCCTCTCTTTCTCGCAGGCATGTTCTC

TTATATGGCTTCTTTAGTCTTTA

>MSTRG.853.2 gene=MSTRG.853

GTTCTACATGCATGATAAAATTTTGGTAGTTTTCTGTTAAGGTGATCAAATTGTTATATCTCACTCTGGT

ATAGGTACACTTAAATATAGAGTCTAGTTACCTATCCTAAAGTTCTCAGTAAGGTTACCATTTTTAAAGT

CAAGAGCTGGTGCAGTTACCTGAGTGATGTAAATCGTCTGTGTGGATATATTATTGAGTTTTTGCATTTG

TGTTTCTTTTGTGCAGAGATCAAGACACTGACTGTTGATATGAGGGATGAGCCAGGAGCAAGACCTATTC

CCAAAGCAAAGATTGAAATATTGCTGTCTAAGACGGAGAAGTTTGAAGAGTTGATGGCCGCTGAGGCGGA

GGAAAGAGAATTTGCTGCAGATGGTGAGGAACAGAACTGAGCTAGATTGCAGGACTGAGCCTGATGTTAC

TCATCCCTGATAATGAGCATAAGTCATTGATTATCACTTGCTCTGAAAAACCCGAAAAAGAGGATAGGAA

AACCAACTAAAATGTTACATAGCTTGGTACTTCGCTGCATGAATTGTTCTGTGAGGAAAAAGTGGTTATT

GTATTGGGTATCTGTTTTTGTTGTTAAGTCTTTGTTGCAAAGGGTTCCATTGGCCATCAGTTCTGTTGCT

TTAGACTTTTCACCTAGTGAGGATGCTTTAGAATTTGTATCACAAGCTTGTAATGAATAAACTATTAGAA

AGATCATCTTTGGGAAACCAGATTCGGCCTCTCTTTCTCGCAGGCATGTTCTCTTATATGGCTTCTTTAG

TCTTTA

>MSTRG.855.1 gene=MSTRG.855

GGTAGCCGTACTGGAAGGTGCGGCTGGATCACCTCCTTTTCAGGGAGAGCTAATCTTTGTTGGGTATTTT

GGTTTGATACTGCTTCACACCCAAAAAGAAGGGAGCTACGTCTGAGTGAAACTTGGAGATGGAAGTCTTC

TTTCGTTTCTCGACGGTGAAGTAAGACCAAGCTCATGAGCTTATTATCCCAGGTCGGAACAAGTTGATAG

GATCCCCTTTTTTACGTCCC

>MSTRG.863.1 gene=MSTRG.863

CATAAGCCATAAGCTTAGCCAAACATACTCTGAATCACTTTCTTCATAATCTGAAAGTTGAATATTTGTA

CAAATGTTGTCATGACAAGGATCAATTTTCTATATATACATGATGAGTTCGGGGGTGGGGGTATCAACTT

GAAATGATAACCCCTGCAATGACAGCTCCTCCGAGAGAAGTCCAAACACAAAATTCTCAGGAGACACCAA

AGGAGCCCAAGAGGCAGAAAGTGCAAGTTAAGAGAGCAAAAAAATCTGATACGAAGAACAACAAAGGTAA

CACTTTCGACAAAAGAAAGGATAAGATGCCATTTCTCTCCTTCGTTCCGAGGAAGAAGCTTCCTTTTGTG

TCAAGCAATGATGAAGGTAAGGACCATTGTTTGACAGGGAACAAGTCCAGAAGCTTGGAGGATGCACTAA

GGGGAACATTGGGTTATCAAGCTGTTTTTATGCCAGCACCTGGTCATAACCCATTTAGGCCAGCAGGAGG

AGCTCCACCACTTTCCAGTTATCCTTCTCGGTCTTCTCAGTCTGAGTCAATGAAGTCTTCTGCATGAGCC

AGAGGGAGCCCAAGGATAAGTACAAAAAAAAAATCAAATCAAGAATAGTTAAGGTGATCCGGTGAAACTT

GCACATGATTCAACTATATGATGTCTTATGAACTAGAATAGCTTTTCTCATTGTCTTTCATGAACTATTT

CTGTTGTCTTTTGTGATATCCATGGTGAACCAGTTCACAATGATAACCTTTATGATAAACTTTATGTCCA

AGCTAAACTTTTGTCAAAATGCTAGCC

>MSTRG.864.1 gene=MSTRG.864

TTAATGTAGATTGATCCGAGTACAATGGAAGGTTAAAGTTAAAACTTAAAACTGAGCTGAGTTGAAGTTG

CAGAGGAAGAAATTGCTAGAAAGTCCATGAAGAAGAAGGAGAAGCAGCACATGAAGACTTCATTGACTCA

GACTGAGAAGACCGAGAAGGATAACTGGAAAGTGGTGGAGCTCCTCCTGCTGGCCTAAATGGGTTATGAC

CAGGTGCTGGCATAAAAACAGCTTGATAACCCAATGTTCCCCTTAGTGCATCCT

>MSTRG.866.1 gene=MSTRG.866

GTGATTGACACTTTGCACCCCTTATGTTTAGGCGCTTTTTCGATTTGGTCTCAAACAATTTTTTTTGCCA

GTATGCACCCTAAAGTTCGAAAAGCGCTTCGCTTTGCACCCCTTTAACCACTTTCCGTTTAATTGACCGT

TAAAGTTAACAGATGTGCCCTTTTCACTTTGCACCCCACAATTTTAATTTTTTTTCATGACTATTTTGAA

ATATTTTTTTTCATGAATGTAAATATCAAAAATATTGAGAGACTTTAAGTATGGCAGTGGTTTCGAAAGC

TTATGCGGAAAACCTCTGGCGGCCAAGACCTTTGAAAGGCATTGCAAATGAAAGCACGAAAATTAGAAGT

TGCTCAGTAGAGTAAGGTTGCACAGAATAATGTAAAGATAATCTGAATATTACCTTCAAGGAAACACCTG

CTAAGGAGAACTTCTCAATTTTATGTAGACTACCAGGGATTATAACCACATTGAATGTTTGCATCTCCTC

AGAGCAATCTTCTAATAACATTAAAGAGTATTCTTCGAGGTTTTCTAACCCAGCTAGAGAATATTTGGAA

GATAACTCATGGTGTACTTGAATCAGACGTTTAAGATTAGGAGCACAAAAGTTGTTAGGGAAAAGTCCTC

TGCAAATTACCAAGGTCAACTTCTCAAGTACAGGGCAAAAGAAGATGCTTTGTCCAAAACTATAAGTAGC

ATCAACTAGCTCAAGATGAGTCAGATTAGCAAATCTTTCACATGAGGGTGTATAGGAAAACCAAACACTT

AAAAGCCTTAAATGAGTCAGGTCTAGAAGAGAGAAATTATGTGCATTAGACTCTTCAAGGAGATTATAAT

AGTTATCCAGTGTAGGTTGCTTGGTACCATTCTTTGATAATAATGAAATCCACTGATCAATATAATCATG

AACTATTTGAGTATCAGACTGATAACCTTGAGGTATGTCTAGTACTGAAAACTTGAGGATCGGACCTTTG

TGGAGTAGGAGAACTTTATTTATCACACTAATAAATCTATACGCCTTCAGTCTTGTATCATCATTTTCAG

TATATTTCTTCATCATGCTGTGAAATATCGATTCCAAATCAAAGATCAGATGTGAGATCCTAGTCCAACG

ATGCCTCCATGCTCTTGACAAGGCACTGGTTCTCACTGCGTCTTGTATTGGCAGAAAACATAGGGATAAT

TTCTCTTAAGTTCTGAGGCAACTCACTGATTATAACTTTCTTAACACCACCGACCTTTCTCCTACTAGAT

TGTGCCATCATCTTCTGCAATTAATAATTGCCAACCCCTTCAATTTATTAATTATTATATCCCAGTAGCA

TCATTTTAATCACACTAAATTCATGAACTAGGTTTAAGATCCACTAATATTCCATCATCAATTAGTGCAT

GCAGAAAATTCTTAAACATATTGAAAATATTTCAAAGGGCTTACATTCATGAAAAAAAATATTTCAAAAT

ACTCATGAAAAAAAATTAAAATTTTGGGGTGCAAAGTGAAAAGGGTAGATCTGTTAACATTAACGGTCAA

TTAAACGGAGAGTGGTCAAAGGGGTGCAAAGCGAAGCGCTTTTCAAACTTTAGGTTGCATATTGTCAAAA

AAAATTGTTTGAGACCAAATCGAAAAAGCGCCTAAACATAAGGGGTGCAAAGTGTCAATTACTCTTA

>MSTRG.874.1 gene=MSTRG.874

CTGCATTTATCTGTATGAAATATCGAAAAATTAAGTAGACATTTTACCTGCAAGTAGTTCAAATTTTAAA

CTATTTCCGTTCCATAAGAAAAAAACTATGGGGGTGTTTGGTTGGAGGCTTATGGGCTTGGTAAATGGGA

ATGGGCTTCCCATTCCGTAACCTTGTGTTTGGATTAAATTTTTAAGACAAGGGAATGGAAACTCGTAGAC

CTCTGGCATGTTAACACAAACCTGGCCTGTACCCCGAATTACGAAATTTGATTCTCAATCTTCTGCTCAC

TCGATATACATTTCTCTTTCATCGTTTCTTCGTTTCTCTCTATCATCATTCTTCCTGTCTTTCATCTCGC

CTCTCACCTCTCTCTCGATCTCGATCTCGCCTGCCCCTCTCTCTCTCTCTCTCTCTCTACTCTCCATTTG

CTCGATAGTCAAGGCTCTTTCCAAGGATGATAGAAGAATGGGTAGTCAAGTACCAAGCTGATGAAGTCTA

CTTGTTAGCTTCTCAAATAAAAAAGAACAAATTAGATAACTGGATTAGATTAATCAAGATTAAAGCATGT

AAACCAAACTAGATGAAACTGGAAAACAATTTGAAGATCATATTGTAAACCCCGTAATTAAACCTTCCAT

ATTGCTACATAGTCACTCTCTTTTAGCTATCAAATTCTCTTACTGGACTTGAG

>MSTRG.886.1 gene=MSTRG.886

ACTGATACCATCCTGATGGAACCTCTGGCCATCAAATCTCTGATTGCCCTCCTAGCAAGAGATCCATTGA

TCCTCAAACGATCAGAGAGGATGGAAGGAGTGATGAGCTTGAACTTAGGAGCTTCGGAAAGGAGCTTGTC

ATACGTGGCCTTGTCAAACAAAACCATGTTGTTCACCTTCTCCTTTTGCTTTCCCTTGCTCCACTTCTTC

TTCTTTTGCTTTCCACCGCCAGATTTGGCAGGCTTGGAGGAGGGAGGAGGAGC

>MSTRG.887.1 gene=MSTRG.887

ATTACACTAACAATCAGTTGTACATGAAAGTGGCATTGTCAATTTGTCATAACAAGTTGTGGCATGCAAC

TGAAAGAACTAGCATGTCTATCTAGAATTAATCCATGTATGGTGTGCCGTACCTAAAGCTATCTGCTTTC

GCCTCGAAGATTGGCCTGCCAAGAACGTGATCTATAAAATACAGATCAAGTCATGTAAAATAACATGAAT

TTGGTTTTCTTTATTGGAGTATTCTAATTTAGTGAGATTATGGCTTTTGTTATGCACAACTCTGATTCAC

AA

>MSTRG.891.1 gene=MSTRG.891

TCCAGCTTTTAATTCTTTGCCTATAAGTGAAGTGGAAATTCCAGTTGATATTCCAGAAGAGCAGTGTGTG

AAAACTGATCCTGGGATACGTAAGCCAATTTGTAAATTTCCGGTGAATCAACAAGACCGTATTCGGAAGG

AGTATATACAATTGGGTCCTTGTCAGCCTAAGTTGGAAAATTATCCACCAACATCTGATGGACGTGATAA

TCGTCATTTTCAATATAGATGGTTTTCATTATTTCCGTGGTTGAAGTATTCGGTAGCTACTGATAAGGTG

TTTTGTTTACTTTGCTTTCTTTTTGAAAAAGATCCACCAAAGTCACCTATGTTCACTACTAAAAGATGTT

GCAATTGGAGCAGAATGTTAGCCAGAAAGAAAGGAATCTGTGAACAACATGTTGGACAATTAAATTCTTA

TCATCAGAAGAATGTTCAAAATTGGGCTGCTTTGTCAAGAATTTCTCAGCAAATTGATAGAGTGATTGAT

AAGCTTCCACCAGAAATTGTAAAGCAAAACAGGTTGCGTTTAAAGACAACAATAGCAGCAGTAAAGTATC

TTGGCAAACAAGGGTTGCCATTTAGAGACCATGACGAATCTCCAAACTCATGTAGTAGAGGACATTTCAT

TGAAATAATCAAGAGTTACGCAGGTATGAATGATGATATTGCGAAAGTTGTTCTTGAAAATGCTCCTAAA

CATGCCATGTACATTGCACCATCAATTCAGAAAGAGATTCTAGATATAATTGCTACTAAGATTCGAAACA

AAATTCGTCAGGAAATTGGAGATGCCAAATTTTGTATCCTAGTTGATGAATCACTTGATATATCTCATAA

GGAGCAAATGGCAATTATTATCGGATTTGTTGATGAAGAAGGTTTTGTTCGATAAAGGTACTTTGAAATT

GTTAGTGTTAAAGACACTACATCTCTTAATCTCAAGAAACATATATGCCTTGTTCTTTCTACACACAAAC

TTCAAGTGCAGAATATCCGTGGCCAGGGGTACGACGGAGCTAGCAATATGCGAGGTCAGTTTCATGGATT

AAAGACATTTTTCTTCAAGACTGTCATTTTGCATATTATGTTCACTGTTTTGCTCACCGACTACAATTAG

CTTTGAATACATGTGCGATAGATGTCCCTGATATGCATATATTTTTTCAAATGCTAAGCTCCATTCTCAA

TTTTGTTGGATCTTTGTCAAAGCGTACCAATATGTTAAAAGATATTCAAGAAGCAAAGGGTTTTCTTAAG

GCATTAAGGACATTTTATTTTGTTTTTTGTTTGTTACTCATTAACAAAATTATAGGTATAACCGATTTAC

TTTCACAAGCTTTGCAAAGACAATCACAAGACATTGTTAATGCTATGAATTTAGTTTCAAGTACAAAGAC

ACTTCCTCAAGAATTGAGAGAACATGGTTAGGATAGTTTTTTTCTAAGTGTCCATCAATTTGGTGAAAAT

CATGGTCTTGATATGCCTAATATGGATGCAGCCTATTCAATCGGGACCGGTCGTGGTTGTCAACAACGTG

ATTTAGTTACAAATGAACATCACTATTACTTTAAAATTTTTAATGTTATCATTAATGTTCAACTAATGGA

GCTAAATGACAGGTTCACGGAGCAAAGTATAGAATTACTTGTTTTGAGTTCTGCATTGGACCCAAAAGAC

CAGTTTAAGAAGTTTGATATCGACAAGATTTGTGTACTTGCAGAGAAGTTTTATTTAGAAGATTTTGATG

CATCAGAGGTACGAGCTTTAAGGTTACAATTAGAGCATTATAGGTGTCAAGTAGTTTCGCATAAATATTT

TCAAGAATTATCTTCTCTTGCGCAATTATGTAGACAATTAATTGAAACAGGCTTAGCCAAGATTTTTCCC

TTGGTTGATATATTGATCATTTTGGTTTTGACACTTCCTGTTTCTACTGCTGATACTGAGCGAGCATTCT

CTGCGATGAAAATTATAAAAACAAGACTGAGA

>MSTRG.892.1 gene=MSTRG.892

GGGAGAGAGAGAGGGAGAGAGAGAGAGAGAGAGAGACTAACCATAGCAACCTCCATAGCCTCAACAGCAA

GCTTCTCCATAAGAGTTTGAGTCGCTACATACACAATCTCGTTCTTCAAGTCTATATATCAACAAATCAA

TCAACAAAAACACATAAACATACACACTTGTGAAATCATATACTAATACACCGATCAAACTTACTAGAAA

CAGCGTGAACAAGATGAATGGTATCAGCGAGACGACAGAGATGAACAATAGCCCAATCAAAAGCGTGTTT

ACTATTGGGACCATGATCTATGGCAATAAGTATGTCTCTGCCACGGCGGCGCTCGCCTGTTTCTCTCTCT

AGCTCCGGCTCAGCTCCGAGGGACGGGAGCTTAATCTCCGACCACTGCAACACCACTTCTTCTTCGGACA

CTATTTCCATTTTAGCTATTCTTGATAATTACTTTAGGAGCAGACAAGTGTTGTTTTAACTCTCTATCTA

TTATTGTATGTCTATCATTATTGCATTTACCCGTGTCCACGTTCCAATTCCAAATGGTTTCCACTTTTTA

CAAGTTTCCACTTATATTAAAGTT

>MSTRG.892.2 gene=MSTRG.892

AGAGAGGCGAGGGGCAGAGAGATAAGAGGAAGGGAGGGAGAGGGAGAGGGAGAGGGAGAGCGAGAGACAG

GGAGAGAGAGAGAGAGAGAGAGAGACTAACCATAGCAACCTCCATAGCCTCAACAGCAAGCTTCTCCATA

AGAGTTTGAGTCGCTACATACACAATCTCGTTCTTCAAGTCTATATATCAACAAATCAATCAACAAAAAC

ACATAAACATACACACTTGTGAAATCATATACTAATACACCGATCAAACTTACTAGAAACAGCGTGAACA

AGATGAATGGTATCAGCGAGACGACAGAGATGAACAATAGCCCAATCAAAAGCGTGTTTACTATTGGGAC

CATGATCTATGGCAATAAGTATGTCTCTGCCACGGCGGCGCTCGCCTGTTTCTCTCTCTAGCTCCGGCTC

AGCTCCGAGGGACGGGAGCTTAATCTCCGACCACTGCAACACCACTTCTTCTTCGGACACTATTTCCATT

TTAGCTATTCTTGATAATTACTTTAGGAGCAGACAAGTGTTGTTTTAACTCTCTATCTATTATTGTATGT

CTATCATTATTGCATTTACCCGTGTCCACGTTCCAA

>MSTRG.897.1 gene=MSTRG.897

TTCATTTCATTCGACTAGGTAATTAAACTCACGAGAAAGCAGTAACATAGGGGTAGTAACCAGTGGTCAA

AATTGAGCACAGTCAAACACTAACATAACCATAACATCTCAAAGAACACAGCAGTAACTTCTAAATTTAA

CTTCATTCACTCAATCCCCACGGCGGTAACTTCATAATTACCACCTGGATCCACCACCACCGCCTCCATA

GCCTCCATCCCCTCCCTCTCTTCTGCCACCATATCCACCGCCACCGCCTCCATAACCGCCACCACCGCCT

TCACGACGTCCTCCATATCCACCGCCGCCTCCGCCTTCACGACGTCCTCCATAACCGCCACCGCCTCCAT

AACCACCACCACCGCCTTCACGACGTCCACCGCCGCCACCACTTCCGCGAGACTGAGCCTCGTTGACGGT

GATATTACGGCCGTCCAACTCCTGGCCGTTCATTCCTTCGATCGCATCGCGCATCGACTTCTCGTCCTTG

AAAGTAACGAATCCGAAGCCTCTCGATCTCCCTGTCTCGCGATCGTTGATGATCTTGGAGTCAGTGATAT

CGCCGAACTGAGAGAAAGCTTGTTCAAGAGATTCATCGTTAGTGGCCCATGCGAGGCCACCAACGAAGCA

TCTGTATTCAACTTCTGCCATTTTAAGAGGGTATTGAAAAGAGGGATTGGATAAAGA

>MSTRG.909.1 gene=MSTRG.909

GGCAGGCCTAATGACACCACACTGTTCTGAGCCTTTAGTCCACCAAATGAGTTCAACCCACTAATGTACT

TCACATTAGGCCTCTTCTGGGATGCCCTTGTCGGGCACGCTACCGCCGCGGAGATGGCGGTGAATTTTGC

TGGACATAGTGTTGCTGATGCTGCCATTATAAAATTTTCAGACTCACCACCTGAATAGAAATAAAGAAAC

CTTGTTAAAGACAAGATATTATTAGAATTCTTGAGGTATGAGCTAGTAGAATCTGAG

>MSTRG.913.1 gene=MSTRG.913

TCTTCCCTCTCTCTTAGACTTTCAATGAGATGAATTTTTTATTAGAGTTTCGGATTTCTGTGAAATCGAA

CAATTGATGAGAATGATTGCTGATTGCAGTGGTTGTTGATTTTGATTTCGCAGTTGCTTAGACGGAAGAG

ACAGAGAAGAGAACTCGAGAAAGAAAGAGATGGACCGTAAGTCTTGGTTGATTACTTTGAATTGTTTTCT

TGTTTGTTTGAGTTGGTTGCTTTTGTGTTGATTTCTAGTTTGCTGTGTTAGTTGATTTTTTGGTTGCAAT

GCTTGTTGATTTTGCTTTTATGGTTGTTTTTGTG

>MSTRG.915.1 gene=MSTRG.915

CGCGGAGAGTTGAGCAAACTAACAAACACACACGTATATAAAGATATGATGTACCAGAGGTGGAAGTTCT

GCATGGAGTTGCTCAAGTTTGCGATTAAGTTTGTCATGGCGTTCTGTGAAGCTGTTAGTATTGTCATGGA

GCAGCAGTCCTCCCCGTCACTCTTTTCCATGAATGCGCAGAGCTTGTATATTGGTGTTCTTCCTTGACGT

ATATATTTCTTGCTCTCTAGAAATCCTGGTTCCGCTCCTTGACGTATATATTTCTTGTCTTTCAAAATCC

TGGTTCCGCGCCATTTTAGTAAAAATGTACATGATTCCTTTTGTTTATATTTACACATTTAAGTTTTTAC

ATAATTTAAAGGTCTAGGTTGATTGGTTTAATCTTTAATCATTAGTTCTTGGAATGAATGTGCCTGTGGT

TTAGTTTTGGTTGCTTTATTTTTCTGGAAAAATACTCTGTGGTTATTTGTGGTTACTCTGTTTCAGAATC

TGTAATTATGACATGCTTACAGCTGCAGCTGTCTCTGCAATCTTCAGTCACTGTACTGGTACAAATGGT

>MSTRG.918.1 gene=MSTRG.918

TAAAATATTGGGAAAATATATTGAAACGTCCTTGAACTATTCCACTTTTATTCGATAGGTCCCTGAACAA

AAGATTCCCCAAATCGGGTCCTTTAACTTTTCAAAATTTTGTATCAAGTCCTTCCGTTAAGTAGATTCTA

ACGGACGTTAGAATCTTGCTGATGTGGCGACGACTAATGCTGATCCATCGCTGACATGGCGCTGATTGTA

CATATATTAAAAGAAAAATGAAATTAAACTGAAATTAAATTTAAAAAACAAGTCACATGCCCTGCATGTG

AGTTTATAAACACGTTTATATCTCCCGATTCACCCATTTCTTTCAATCTACAAAAGGGTTTTAGGGTTCC

ATTAAAAGGAATTAGGGTTCTTAATCAGTAAATTAGATTGGGCAACTTCTATACACAAACGCACTACACT

AAATACATACTATCATCACAAACTATCTCGACATCTATTCACAGACATGAAGGCCGAAATCACCTTGTAC

ATTCATCACGGAAGTGAGATCGTGCAAGAGCGTGGCATCAAATACTACAATGGAGGTAAGGTCGAGTATG

TTTTTAATCAGAAATTAGATGTGTTATCTATTTTCATACTTCAAAAGTATGTTCGAAGTGATCTTGGTTA

TAAAGTGTTTAAAATGTATTGGCACAAGAGTGGAGTGAAGATTGTGCTGGAGAGAGTGAGAATGATGATG

GCAGTATGTACAAATCATCTGGTGATGTGTCGCCATGTCAAAAAAATTCTGGTGGATCAGCTGGTGACGT

GTCGCCATGTCAGCAAAGATTCTAACGTCCGTTAGAATCTACTTAACGGAAGGACTTGATACAAAATTTT

GAAAAGTTAAAGGACCCGGTTTGGGGAATCTTTTGTTCAGGGACCTATCGAATAAAAGTGGAATAGTTCA

AGGACG

>MSTRG.920.1 gene=MSTRG.920

CACGAGACTACGGGATAGAAATTAATCAATCAGTGTTTGTAATGGCTGAATCCGGAAAGAGTAATGAAGT

AATTGATCCCAAAACTGATCCTAAACATAAGGCTAAGTCAACTGATCCCGGATGGAAGTATGGATATTGG

CCTGATCCAGAAGTGAAAGGCTGTATTGAATGTATTCTTTGTGGAAAACAAGTAAAACAGGGGATCAATA

GACTCAAGCAACACTTGACTGGTAATTATGCCGATATAAGCAACATTTATTTATAGGCATGGTCGTCTTC

ATAATGCAATGAAGGTTAAAACAGGAGGCAGAGAACTTGTGAGACCTGGAGCGACTAGATTTGCCACATC

TTTTCTCACACTGCAGTGTTTATTCAAATTCCAAGATGCATTGAGGCAGCTCTTTGTAAGTGAGGACTGG

AGCAGGTCAAGTTTGGCTAAAACAGAGGCTGGAAAACAGGTTTATGACTTGATTATGTCAACTGCACTAT

GGAATAATATTGCAAATTGTATCAGGGCTGCACAACCGCTTCTTGTTGCCTTGAGAGTAGTCGATGCTGA

TGAACAACCAGCAATGCCAGAACTTGTTGTTGCTATGGAACATGCCAAGAAAACTATCAAAGATTCTTTT

GAAAACAATGCATCACTTTGCAAATTTGTTATTGACATCATTGAGAAACGCTGGGATACGCAGATGGGGG

TTCAAATACATGCTGCTGGTTTGTTTTTAAATCCTGGAAAGTTCTTTGACATACAAGAGAAAGACTATCG

TAATGCTTGCAAACTCCGGGAAGATTTCAATGATGTTTTGGAGAAGATGGTGCCTGATATTGAGACTAGG

AATTTGGTAAGTAACCAAACTGATAACTACGAGCATACGAGAGAGAACTTTGCAAGACAAATGGCAATTG

ATCAAAGAAAAACGAAGAGTCCTCTTGACTGGTGGAGTGCTTTTGGGGGTCGCGCACCTGACCTAGCAAT

GATAGCGAAACGGATTGTTGGCCTTTGCTGCTCTTCATCAGGCTGTGAGCGAAACTGGAGCACCTTTGAG

TTTATTCACACTAAAAGAAGAAATCGCTTGGAACATAAGAGGCTGAATGATCTCGTGTATATCCAGTACA

ATGAAAAGATTGCAGAGCGATTCCAAAAGCGCCGTGAGCTTGGAGATAAGTTCGATCCAATTGCGCTCGA

AGACTTTAACTGGAGTGGTGAATGGGTAGGTAATGCAGATGGTGATTCTGTTCATCCTGGAGAAGATCTG

CTATGGGAACATGTTGTTATAGCTGCTGGTCCGGCAAGTGCAGTGCGTAATCTTCGTAATAGAAATCGTG

ATAAAGGTGGCGAAACTAATACAAACACCAGGGAGATTTTAACATATAATCGACATCGTAATGGTGCAGG

TTCTTCTCGTGCGCACACTTCACGTACTTTTCACTTGCACGATTCTGAAAGTGAAGAAGAAGTCCAGGAC

AATGACATATTGTCAAATGAAAATGATGACGAAGATATTGATGATGATTACGGGGAAATACCGAGTGCAA

CTCAAAGTCGACCCGGAAATGAAGAACAGCCTTTTCTTGATGATTTTATCATCTAGTTTATCATAGTTTT

CTTGTAGAAATTACTAGATTTGTATGTTTTAGACCTGTTCCTCTTAACTTTATATAATATTTACTTTTGG

TATTTAATTATAGTTAATCTATGC

>MSTRG.921.1 gene=MSTRG.921

AAGATTTATAATGGTATCTTTGAGCCAATTCCAAACAAGGGTGATCCTCGATATCCTACAGAGCTTGCTT

TCCCGAGAGTTGTGCATGACCCCGATATGGAGAAGAAGAAGGGGCGAAGAAAAGCAACGAGGTACAAGAA

CGAGATGGATTTCCAGAGCCGGGGAAAGCATGGAGGGACTTCATCGAGAGACAGTTGATTCTCTAGTTGT

TTTTATTGTTGTAGTACTATGGTTCAAGTACAATTTGAT

>MSTRG.923.1 gene=MSTRG.923

TTTTCTGATTAAATCATGTGATCCCTGCTGAGTTTGTCTCCTGCACTGCTATACTTGAGTATGTGTCTAC

TTGCTCTCAGTCATGTGATAATAGATTAATATTAACTTAAAAGAAAGTCAATTGTATTTTGCTGATGATG

TGTCCTCTGCAACTTGAACTTGTGCTATTATTTGATGAGGTGTCCTGTGCTGGTTAATTCTGAGTTTATT

ATACATGATGTGTCCTGTGCTAGTTAACTTTGTGATTATTATATGTAACTTGTTCTGTTTTGCTTAACTA

TATGATACTTATAATGATATGAGATACTTATGCACTTATTACTACATATGTTTGTGAGTATGAGTCTGTG

ATTAGTGGATTATGTACTCTGCCATGGTAAGTCTCCATATACAAGACTCATATAGGGAGTATACATATAA

ATTAAGTAGAAATCTCCTAATATAGGACACTACACACACCATGCCCATGCATTAAGACTTGGCTTGGGGG

TCCATTGAGCCCCTGGGTATCTGGATCACTGGTGCATCTGGCCACTAGACTTAGGTTAGAAAATTAATGC

ATTAGGACTTAATCTCTGTGATGAATGACATAGCATCTGCATCATATCTGTG

>MSTRG.926.1 gene=MSTRG.926

AAAAAATCTGGTGTCTTGGACTATTATGATCACTGGGTACGGAAAGTATGGTCTTGGCATAGAAGCAGTA

GACCTTTTCAAGAGGATGCAGTTAGATGGTATTGAGCCTGATGGAGTAACTTACTTGGCTCTGCTAGCGG

CTTGTAGCCATTCCGGACTTGTTGAAGAAAGTCGAGAATATTTCTCAAGATTGTGTAGAGATCGTCATAT

CAAGCCTAAGGTTGAGCATTATGCTTGCATGGTTGATATTCTTGGCCGAGCTGGACGCTTGAATGAAGCT

AAGAATCTCATACACAATATGCCATTCAACCCGAGTACAGGTATCTGGCAGACATTGCTTAGCTCATGCA

GAGTACATGAGAACTTGGAAATGGGGAGAGAAGTAGAAAGGATTCTTCTGAATCTGGATGGTAACAATCC

AGTTAATTATGTTATGGTATCAAACTTGTATGTTGGCGCTGGCTACTGGAAAGAGAGTGAGAGACTGAGG

AAAGTGGCTAAAAGGAAAGGATTAAAGAAAGATGCTGGACGCAGTTGGATTGAGATTGACAAGGAAGTTC

ACTTTTTCTTTAACAGAGACGAGACACATCCACTAACCCGAAAAATCCATGAAGTATTGGAAGAAACTGA

GAGAAGGATGAGGCAAGAGATAGGGTATACTTCTGATGTACGGTATGCATTGCATGATGTTGAAGAGGAG

ACGAAGGAGGAGAGCTTGAGAGTGCATAGCGAGAAGTTGGCAATTGGGCTGGCACTACTTTGTGGTGGAA

TGGAGAAAGGGGGTAAGCCTATTCGAATATATAAGAATCTCAGAGTTTGTGGGGATTGTCATGAGTTTAT

CAAAGGCTTGTCGAAGATCTTGTCAAAGGTGTTTGTTGTGAGAGATGCTAGTAGATTTCACAAGTTTGAA

GATGGACGGTGTTCATGCAACGATTATTGGTGATATCTGTGTACACAGGACACTAAAGGTCATTATTATC

ACAAGTATATTATCTGCTGTCTGAAGTCATTAAGGTGATAAATTCCATTGCAATGAGTATATTTCGACCA

TTTGGATTATGCTCTGAAGATCTTTAAAAACTGGCAAATTGAAAGGGATGCATTAGAGGATTTTTTTTCT

TGTCTTATGATGTGGTTTGACTGCGACCTGCGGGACAAGTCCTTGGGCTATTCGAGGAAGAACTGCAAAT

AACCTGGCTGGAGACTTCTATAACTTGCCTTGAGTTACCTCGTTGGCACAAGAAACTTCAGAGTAAGGGT

CTTAAACAAGGAGAAGAATAAGTCCTAGGCCGTGGAAAGGAGAAAAATTAGTGAAAAAGACTCTTTACGC

AATATCACCTAAATGCTCATCAGTGATGCATCCAGGTAATGTCAACTATGTGGTCATCAGTGCAAGTATG

GAAGACTACATAGCGAAAGGAACTATACCAGAATTACCCTAGAAGCACTTCCAGACCCTAAAAAC

>MSTRG.927.1 gene=MSTRG.927

GCCAGTTTCTGAAGCCTGATGAATTTAACGTGCCATACGTTGTCATGATCCCTGAACTCGCAGAGGTGAT

GAAGTCCTATTTAATCAACTCAGATAATCCAACTGTGAGCATCAAGTTCCAAATAACTCAGTTGGGCATC

AAACCAGCTCCTCAGGTTGTAGATTTCTCGTCACGAGGGCCTGACACAAGGTCTCCTTGGATCCTTAAAC

CAGATATCTTGGCACCAGGAGTTGATATCTTGGCTGCATGGGCACCAAACAGAGGATCTGCAGTAATTGG

AGACGATGATTATCTGCTCACAGATTATGCCCTTTTATCAGGTACATCCATGGCGTCTCCCCATATAGTT

GGCATTGCTGCATTACTAAAAGCCACACACAGCGACTGGAGCTCAGCTGTCATACGATCTGCAATGATGA

CAACTGCTGACATATTTGACAACAGCAATGGTCCGATTATAGACATGACAACTGGAGTAAGTGGGACTCC

TCTTGACTATGGCGCAGGACATGTTAACCCTAACAAGGCCATGGACCCTGGTCTTGTCTATGATATAAAA

CCTGAAGACTACATCAGTTACCTTTGTGCCATGAACTACACCAGAGATGAGATCATGCTTCTTGCACGAA

GATCAGACTTTTCATGTGAATTTTCAACTCTTGATCTTAACTACCCTTCTTTCATAGTCATTCTCAACAA

TACTAATACAACTACTTACACCTTCAAGAGAGTGTTAACTAATACAGTTGATACAAGTGCTTCTTATCGA

GCTGTTGTGAAGGCACCATCTACTGGCATGCATGTGGCTGTGCAGCCATCAACAATTACATTTTCGGGAA

AGTACAGCACTGCAGAGTTTAACA

>MSTRG.934.1 gene=MSTRG.934

TTCGTTCTAAGAACCAAGTTTTTGACGTATTGACGGGGAGTTAATAGTCATCGCACCGTGAAGAAGAGGA

AAGACCCAAGACCTAATACCTAAGATCCAAGACCGGCGAAAGGTTTAGAAGTACAAAGACCACTGGAGGT

GCAACGACTTTGAGGAAGTAACAAGGTAGTTACGTTCCAAGACGAGTCTTGTAATTTACATTTATAGTGA

GGAGGTAACGAAGAGTTACGCTCCAAGATATATATTTGTAAAAGCTTCTCCGCCTACTGTAAAGGAGTGT

CTTTATAGTAGAGAAAATATCGAGTCCGGGGAGGAGCTCGGGGACTAGACTAGGGGTGAGTGGACTCCAA

GGGTTAAGTTAGCCACCGCGAACCAGGATAAAATCGCTCGTGTGATA

>MSTRG.941.1 gene=MSTRG.941

TTGTAATACTGGTGGTGGTGATGATGATGAAAAGGTTGAGGCTTGGATTGTTTTTGACGGCATGCAGTAT

AAGAATCTCATTACTTGGAATTCTATGATTGCAGGTTTTCAGCAGAGTGGACATTGGGTTGAAGCTTTGA

ATTTTTTTTCCTGGATGAGACATGATGGTGTTGGGTTTGATCGTGCAACGCTTCTTGGTGTCATATCTTC

CTTATGGGGTAAAAATTGTGATGATGCTAGTTTGGTGTTGCAATGCTGTCTTCAAGTCCACTGTCTGACG

ATAAAGACGGGATTCGTTTCTGAAATAGGAGTAACAACTGCGTTAACAAAAGCTTACTCCAATCTTGGAG

GAGATGTTGCTGAGTGTTACAATCTTTTTCTTGATAGTAGTGGAAATCGAGATGTAGTTTCTTGGACTGG

AATAATAACCACATTTTCTGAAAGGAATCCAGAAGAAGCTCTCTTCCTTTTTCAACAATTTTGTCGAGAG

GGCTTTTCTCCAGACCGTTATACATACTCCATTGTGTTAAAGGCTTGTGCTGGCTTAGTAACTGATAGAC

ATGCATCCGCTATACATTCAAAGGTTATCAAAGCAGGGTTTGAAGATGACACTGTACTAGCGAATTCATT

GATCCATGCATATGCAAGAACTGGATCGATCGTGAGGTCCAAAAGAGTATTTGATGAGATGGATTATCGA

GATACAGTTTCTTGGAATTCGATGTTAAAGGCTTATGGCTTGCATGGACAAGCTAAGCAAGCATTGTCCA

CCTTTGATCAAATGAATGTCCAACCTGATGCAACCACCTTTGTAGCCCTCCTCTCGGCCTGTAGTCATGC

TGGATTGGTAGATGAAGGTTCTAAAATTTTTAATACTATGGGCAGTAATTATGGAATAGTTCCTAAACTT

GATCACTTTGCATGTATGGTTGATATACTTGGACGGGCTGGTCGTATTCCTGAAGCCGAGAAGGTTATTA

AAGAAATGCCTATGGAAGCTGATTCTGTTGTGTGGAGCACTTTGCTTGCGGCATGTCGTAAACATGGTAA

AACCGAGCTGGCTAACATGGCTGCCTCAAAACTAATGGAATTGGATCCAAAAAGATCACTAGGATACGTT

ATGATGTCAAACTTATATTGCTTAACCGGTAGCTTTGGTGAAGCAGGCCTTGTACGAAAGGAAATGAAAG

GGCAGGGAGTAAAGAAGGAACCTGGTTTGAGCTGGACTGAGTGTGGTAGTCGAGTCCATGAATTTGCATC

TGGAGGTTTAAGACATCCCCAAGGAGATGCTATCCGCAAGAATCTTGAAGAGCTAGTGAAGCAGTTAAAA

GGTGTGGGTTATGTCCCTGAGACAGTCTTAGCACTGCATGATGTTGAAGAAGAGCACAAGGAAGAGCAGT

TGTATCATCACAGTGAGAAGCTGGCTTTGGTGTTTGCCTTGATGAATAGTGGTCATTCTCAACGCAATGA

CGGTATTATTAGAATTATCAAGAACATTAGAATTTGTGTCGATTGTCATAACTTTATGAAGTTTGCATCT

GGGTTAATCCAAATGGAGATTGTTGTGAGGGATTCCAATCGATTCCACCACTTTCGAGAGAGGGCATGCT

CCTGTAATGATTACTGGTAACATAAGTTGATCTACACATCCAAATAATTAGGGTTGTAAATGAAAACAGT

CTGAATGATTTATGATCAACCCGATTACAGGCGGCATTACCATCTAGATATCTACTAGGCAAGGGTCAAG

GGCTGTTAGCTGCCAGAAACTGTCTTGGAAGAAAATTCAGCTCTTCTTGTGTGTCCCATACTTTAATAGC

TGAGCTGGTTGAAAACGAGAGTTTTAGGTCATTTTCGCAGATAAAAATGTTTGCACAGGCGTAAGATTGT

GCAAGAGAGGCAGTTTATGTTACTGAAACTCCAAATGGTGCCATGTTTATTTGTTACATGTGCGTAATGA

ATCAACTCTTAATGGTAAATGG

>MSTRG.942.1 gene=MSTRG.942

GTGCATTTCAGTCATTCACGGGTCATTAGACTCAATAGACAATACCTCAAGTCTCACCTCTCATCTCCCC

TCTCAGCGATCAAATTAAAAACCCTAATTTTTATTTTCATCTCAGCCTCTCCGCCGCCTAGTTCAAGTCG

CCACCTGCAACTCTGTTGTCCACCTCCGTTGCTTTGTCTCTGAGTCCGCTGTCCGCCTCCAGGCTCTGCT

GTCTCCACGTCCGCTGTATATTGATGTTATTGTATGTATTTGAACTATAAAGCACTGTGTAAGCCTAGAG

TCTAATGCTTAATGTTGCTTAATTATTTAG

>MSTRG.946.1 gene=MSTRG.946

CTTATTTCCCAAGAGATGGATAATTGTTTTATAGAAAGAAACACAGAGCTACTACTTTGTATTGGCTCCT

TGGATCCTAGGAATTCATTTTCAAGTTTCGATAATTCAAAACTTGTTCGCCTTGCACAATTTTATCCAGA

GGATTTCTCTTTATTAGACCTTGAGCTGTTGCCAAATCAGTTGGATAATTTTATCTATGATGTGAGAACG

GATAAAGATTTATCTGGACTTCAAAATATAGGAGATCTTTCTGTAATGATGGTTAAGACACACAGACACA

CTGCTTATCCTTTAATTTATCTTCTAGTTGAGTTAGCTTTGGTTCCGCCTGTTGCCACTGTTACTATTGA

GAAAGTTTTTTCTGCTATGAAAATGATCAAGACTTATCAGCGCAATAGGATGGGAGATGCGTAGTTGAAT

GACAGTATGATGATATATATCGAGAAAGCTATTTTTGCAAGTGTGGATAACGAAGCTATTCTACAACACT

ATCAATATATGCAAAATCGCAGGATCCAACTATCATCCATCACTTCTTCACTAAAGAATAGTACAACTAT

TGGAAATTAAAGGTTAGTTGCATTTTCTTTTAGTATTGTATGTTTTTTCATGTTATGGATATTCCATAAT

GAAATATTTCAGCC

>MSTRG.950.1 gene=MSTRG.950

GGCTAATCAAATACAACCCCTATCTCTATCTATCTGTTATCAGCAACCCTCCCCTCTATCTATCTGCTAT

CTCTATAAATAATTTTCTGTGATTGAATTTCTGTCACTCTGTGATTGAGTGCTTTGATTGAATTTGATCA

GACCTTGTTAGTCACAGGCTCACAGGTAAGCTTTATATCAAGTGTCAACAGCAGGTAATTGAGTGTTGTG

TTTGTGTACTGTTTTGCGTCCATTCTGGAACCAGGAACCATTCTGGAATTCAGTCATTTAATGCTTTTAT

TGAAGTAAAGAAAGCCTGATTGATGCGGCAACTATCTTGTTGTTTATAAATTGTGGTCAGTTGGTATATA

TAGTGCTATTGTTCTATTCATTTATAAGCACCAAGCTTATCTAGTGAGAAATTGAATCTGATCTGGTAGC

AGGCTAGCAGCATTGAACAAACTTCGGTTTTTTTTTATAACCGAACCGAACAAACCGAAGTTTTTCGGTT

CGGTTAAGTTCGGTTAGAAACATAAGTTCGGTTCGGTTCGGTTTTCAGAAAAATGAAACTTCGGTTTTCG

GTTATTTCGGTTCGGTTCGGTTTCTGATCGAACCGACCGTTTGCTCAGCC

>MSTRG.960.2 gene=MSTRG.960

CGCGCGGACAGAGCGAGAGAAAGAGCGAGTGGAAGTGAGAGAGGAAGAGAACGAAACAGGGAGGTTATCG

TAAGCATATTTTCTCAGATCTGAACTTGTTTCCTTGATTTATCGCGAGCATCGAGCAGGACATGTTGGGA

AAAAGCTGATACTAAGGATTATATATTGATGTTCTTTGGAAGTGTTGGATCATGTATGCATGGTGCTGCG

CTTTCTGTTTTCTTCATTTTGTTTGGTCGGATGATTGACCTCCTAGGACACATGTCTTCAGATCCTCATA

GATCATCTTTGGTGGTTTCCAAGACGAGAGGTTCGGCAACTTGCATTCCTCGTTTGAATGGAATTTAGCC

TGTGACATTGGTGACATGAGTCCATTATTAACTGCATCTTGGATCCCTGAAGAATTTCCATACAATTGAT

GTCCAGCATTAGCATTTGACTACAACCATTCTCAGATTGTTGGCTTCTCAACTGATGGATTTTCGACTAC

AACTACCATGAGAAGATTGAATCTCATATGGGTGACAGCCCGAATGCACATTGAAGTCTACAAGTACCCT

GCTTGGAGTGACGTGGTTGAGATTGAGACATGGGGCCACAGCGAAGGAAGAATTGGGACCAGACGCGTCT

GGATTATCAGAGATTACTCCAATGGCGAAGTCATTGGTAGAGCAACAAGCAAGTGGGTGATGATGCATCA

AGATAGTAGAAGATTTCAGAAAGTCAGTGATGAAGTCCGCGATGAATATTTAGGTTTCCGCCCAAAAACC

CCGAGATTCGCATTTCCTGAAGAAGACAATTACATCTTAAAGAAGATATCGAAGTTAGAAGATCCTGCTA

CCTTTTCCAGTCTCGGCCTTGCGCCACGAAGAGTTGATCTGGACATGAACCAACATGTAAACAATTTCGC

TTACATTGGATGGATTCTGGAGAGCATTCCGCAGGACGTGATCAACACCCATGAACTACAAACTAGCTGC

TGCAATTGATAAAATTAAAAAAGCAACTACGAGTACTGAAGAATCTGAGAATGGGTTCATTAAAAAGCAT

AACTATACACAAGTCGGAGTCTTCTACGGAGAAAAAGGAAGGACTTGGGATTTTACCTTTGTTCATTTCA

GTTGATCTGAAAGGGATAATATCGAACAACTTAGTGAATATATTAAAGGTTTTGTACACACCTAACACCT

CACAGGAGAAGGAAGAGCCAAATTTACTTGAAAATTTAAGGCCCAAATTTAATATTCAGTTTGTTTATAG

AAGCAAATGTTCATGTAAATTTAAGGACCAAATTTCCTATAGAATTTTCTTGCAAATTTAGCATGCTTAC

AAT

>MSTRG.960.1 gene=MSTRG.960

CGCGCGGACAGAGCGAGAGAAAGAGCGAGTGGAAGTGAGAGAGGAAGAGAACGAAACAGGGAGGTTATCG

TAAGCATATTTTCTCAGATCTGAACTTGTTTCCTTGATTTATCGCGAGCATCGAGCAGGACATGTTGGGA

AAAAGACGAGAGGTTCGGCAACTTGCATTCCTCGTTTGAATGGAATTTAGCCTGTGACATTGGTGACATG

AGTCCATTATTAACTGCATCTTGGATCCCTGAAGAATTTCCATACAATTGATGTCCAGCATTAGCATTTG

ACTACAACCATTCTCAGATTGTTGGCTTCTCAACTGATGGATTTTCGACTACAACTACCATGAGAAGATT

GAATCTCATATGGGTGACAGCCCGAATGCACATTGAAGTCTACAAGTACCCTGCTTGGAGTGACGTGGTT

GAGATTGAGACATGGGGCCACAGCGAAGGAAGAATTGGGACCAGACGCGTCTGGATTATCAGAGATTACT

CCAATGGCGAAGTCATTGGTAGAGCAACAAGCAAGTGGGTGATGATGCATCAAGATAGTAGAAGATTTCA

GAAAGTCAGTGATGAAGTCCGCGATGAATATTTAGGTTTCCGCCCAAAAACCCCGAGGCAAGTCAAGCCA

GCGATTGATATCTTATTCCCAGATTAATTTCTGGTGATCATAATAAATTTTGTCTGCCCTCTTTTGTTTT

TGATGATACATTGTTTGATTACTATCTGGGCATTCTGTTGTGTTTCTCAAATTCGCATTTCCTGAAGAAG

ACAATTACATCTTAAAGAAGATATCGAAGTTAGAAGATCCTGCTACCTTTTCCAGTCTCGGCCTTGCGCC

ACGAAGAGTTGATCTGGACATGAACCAACATGTAAACAATTTCGCTTACATTGGATGGATTCTGGAGAGC

ATTCCGCAGGACGTGATCAACACCCATGAACTACAAACTAGCTGCTGCAATTGATAAAATTAAGAAAAAG

GAAGGACTTGGGATTTTACCTTTGTTCATTTCAGTTGATCTGAAAGGGATAATATCGAACAACTTAGTGA

ATATATTAAAGGTTTTGTACACACCTAACACCTCACAGGAGAAGGAAGAGCCAAATTTACTTGAAAATTT

AAGGCCCAAATTTAATATTCAGTTTGTTTATAGAAGCAAATGTTCATGTAAATTTAAGGACCAAATTTCC

TATAGAATTTTCTTGCAAATTTAGCATGCTTACAATG

>MSTRG.960.3 gene=MSTRG.960

AGCGAGAGAAAGAGCGAGTGGAAGTGAGAGAGGAAGAGAACGAAACAGGGAGGTTATCGTAAGCATATTT

TCTCAGATCTGAACTTGTTTCCTTGATTTATCGCGAGCATCGAGCAGGACATGTTGGGAAAAAGCTGATA

CTAAGGATTATATATTGATGTTCTTTGGAAGTGTTGGATCATGTATGCATGGTGCTGCGCTTTCTGTTTT

CTTCATTTTGTTTGGTCGGATGATTGACCTCCTAGGACACATGTCTTCAGATCCTCATAGATCATCTTTG

GTGGTTTCCAAGACGAGAGGTTCGGCAACTTGCATTCCTCGTTTGAATGGAATTTAGCCTGTGACATTGG

TGACATGAGTCCATTATTAACTGCATCTTGGATCCCTGAAGAATTTCCATACAATTGATGTCCAGCATTA

GCATTTGACTACAACCATTCTCAGATTGTTGGCTTCTCAACTGATGGATTTTCGACTACAACTACCATGA

GAAGATTGAATCTCATATGGGTGACAGCCCGAATGCACATTGAAGTCTACAAGTACCCTGCTTGGAGTGA

CGTGGTTGAGATTGAGACATGGGGCCACAGCGAAGGAAGAATTGGGACCAGACGCGTCTGGATTATCAGA

GATTACTCCAATGGCGAAGTCATTGGTAGAGCAACAAGCAAGTGGGTGATGATGCATCAAGATAGTAGAA

GATTTCAGAAAGTCAGTGATGAAGTCCGCGATGAATATTTAGGTTTCCGCCCAAAAACCCCGAGGCAAGT

CAAGCCAGCGATTGATATCTTATTCCCAGATTAATTTCTGGTGATCATAATAAATTTTGTCTGCCCTCTT

TTGTTTTTGATGATACATTGTTTGATTACTATCTGGGCATTCTGTTGTGTTTCTCAAATTCGCATTTCCT

GAAGAAGACAATTACATCTTAAAGAAGATATCGAAGTTAGAAGATCCTGCTACCTTTTCCAGTCTCGGCC

TTGCGCCACGAAGAGTTGATCTGGACATGAACCAACATGTAAACAATTTCGCTTACATTGGATGGATTCT

GGAGAGCATTCCGCAGGACGTGATCAACACCCATGAACTACAAACTAGCTGCTGCAATTGATAAAATTAA

AAAAGCAACTACGAGTACTGAAGAATCTGAGAATGGGTTCATTAAAAAGCATAACTATACACAAGTCGGA

GTCTTCTACGGAGAAAAAGGAAGGACTTGGGATTTTACCTTTGTTCATTTCAGTTGATCTGAAAGGGATA

ATATCGAACAACTTAGTGAATATATTAAAGGTTTTGTACACACCTAACACCTCACAGGAGAAGGAAGAGC

CAAATTTACTTGAAAATTTAAGGCCCAAATTTAATATTCAGTTTGTTTATAGAAGCAAATGTTCATGTAA

ATTTAAGGACCAAATTTCCTATAGAATTTTCTTGCAAATTTAGCATGCTTACAATG

>MSTRG.960.4 gene=MSTRG.960

AGCGAGAGAAAGAGCGAGTGGAAGTGAGAGAGGAAGAGAACGAAACAGGGAGGTTATCGTAAGCATATTT

TCTCAGATCTGAACTTGTTTCCTTGATTTATCGCGAGCATCGAGCAGGACATGTTGGGAAAAAGCTGATA

CTAAGGATTATATATTGATGTTCTTTGGAAGTGTTGGATCATGTATGCATGGTGCTGCGCTTTCTGTTTT

CTTCATTTTGTTTGGTCGGATGATTGACCTCCTAGGACACATGTCTTCAGATCCTCATAGATCATCTTTG

GTGGTTTCCAAGACGAGAGGTTCGGCAACTTGCATTCCTCGTTTGAATGGAATTTAGCCTGTGACATTGG

TGACATGAGTCCATTATTAACTGCATCTTGGATCCCTGAAGAATTTCCATACAATTGATGTCCAGCATTA

GCATTTGACTACAACCATTCTCAGATTGTTGGCTTCTCAACTGATGGATTTTCGACTACAACTACCATGA

GAAGATTGAATCTCATATGGGTGACAGCCCGAATGCACATTGAAGTCTACAAGTACCCTGCTTGGAGTGA

CGTGGTTGAGATTGAGACATGGGGCCACAGCGAAGGAAGAATTGGGACCAGACGCGTCTGGATTATCAGA

GATTACTCCAATGGCGAAGTCATTGGTAGAGCAACAAGCAAGTGGGTGATGATGCATCAAGATAGTAGAA

GATTTCAGAAAGTCAGTGATGAAGTCCGCGATGAATATTTAGGTTTCCGCCCAAAAACCCCGAGATTCGC

ATTTCCTGAAGAAGACAATTACATCTTAAAGAAGATATCGAAGTTAGAAGATCCTGCTACCTTTTCCAGT

CTCGGCCTTGCGCCACGAAGAGTTGATCTGGACATGAACCAACATGTAAACAATTTCGCTTACATTGGAT

GGATTCTGGAGAGCATTCCGCAGGACGTGATCAACACCCATGAACTACAAACTAGCTGCTGCAATTGATA

AAATTAAGAAAAAGGAAGGACTTGGGATTTTACCTTTGTTCATTTCAGTTGATCTGAAAGGGATAATATC

GAACAACTTAGTGAATATATTAAAGGTTTTGTACACACCTAACACCTCACAGGAGAAGGAAGAGCCAAAT

TTACTTGAAAATTTAAGGCCCAAATTTAATATTCAGTTTGTTTATAGAAGCAAATGTTCATGTAAATTTA

AGGACCAAATTTCCTATAGAATTTTCTTGCAAATTTAGCATGCTTACAATG

>MSTRG.963.1 gene=MSTRG.963

AGAGATGGTCTAGAGACTATGAGATGCATAGATTGTGAAAGTGGAAAGATGAGATTGAGATGTGAGACTG

GAGTCAGATTTAAGAAAAGCTTAAAGATGAGATGATTGATTGAGATTGTGGATAAGGCTAATGCTAGGCG

AGGAGACAGAGTTGAGGAGGAGACAGAGTTGAGGATCCAGTGGTCAGGGAGTGGTGAGGAAAGTTGATGA

GATTGGTGCAAGCTGGAATAGTTGACAGTGATTCTTCTTATCGAAGATGATCTGACTTGGTAAGAAATAT

TGTATAGTATTAGTGGTGGATTCTAATTTCAATACTGTAACCTGGAAGCGATCCTGTTGTTTTGGGGGTT

AAATGTTCTCTTAAAATTTTCTTATTATGGTTTTCAAGTTTGAAATTCAGGTTTTGAAATCATTTGATTT

A

>MSTRG.964.1 gene=MSTRG.964

CATAAAACACATACTTATTAAACACCAAACCCTCTTATTTCAACTAAACCCCCACAAAACAAAAATGGCC

GCCGGAGTGCTGAGAAACGCCGTGAAACACAAATTTTCCGGCCACTATATCCAGCAAAGACATTCATTTT

CCGACAGGCTCACCAAAACTCCGGGGCCCACCAACCCATCTCATCATTATTGTCAGCAAACTAAAGCTCA

TTACTCAAGGTTCCCTCAGTTTTTTTACAGAAAAGCCAAGGAGCTTGATTTCTCAGCTCCTGTTAATGTT

TTTTCATCACCCTTTTCATCAAATTCATCAGTTTCGAGAAATGGGCTTGTGGGGTGGTACTTGGAGATGA

TCAAGGCTAGGCCCATTCTTACAAAGTCCATCACTTCTGCAATTATTTACACTGTTGCTGATTTTTCTTC

TCAGAAAATTGTAAGTCCGTCTTCAGCGCCCTATGATGTAGTGCGAACACTGCGTATGGCTGGATATGGA

ATGCTAATATTAGGGCCATCTTTGCATTTCTGGTTCAATTTTGTATCTAAGGCCCTTCCAAAGCGAGATC

TTCTTACCACAATGACAAAGATAGTCATGGGGCAAACTCTATATGGACCTCTTATGACTGTAGTATTCTT

CTCAGTCAATGCAAGCTTACAAGGTGAAAATGGAGAGGAAATTTATGCTCGACTGAGGAGAGACTTGGTG

CCTACAATGATAAGTGGTGTAATGTATTGGCCATTGTGTGATTTTGTGACATTCAAGTTCATTCCTGTTC

AGTTGCAGCCTTTGGTGAGTAATTCATTTTCCTATCTTTGGACCGTGTACATGACATACATGGCAAGCTT

AGACAGAGCAGATGTTTAGTGATATGTAACTGGCTCTGCTGGAATCACCGGAATCAGCATCTTATGGTGC

CTATCCTCAAACTCGACAAGTTCTGGTGAATGTTTTTGGCACAGTACCATTCTACTGACATCAAGGAAGC

TGTAAGAAGTAATACAATTTCAATGCTACCTCAACTTGTTCTGCATTGTTCATCTATAGAGCTGGAGTTC

AAATAGTTTTGTAACTTTTAGGTGATACAGAATTATAATACCTCACAATTTGTAAAATGGATATCTTACC

AACCTTTAATCAGGTGCTTTTAAGTTCTGTCCATTAAAATGGATGAAGTCATTCAGTGTTGTCCATTGTC

GGGTTACTGCAATATGCATCAACATTAAATACTGTAACAATCTTGCAATTTTTGTTATTTTCTTTGTCGA

ATTCTGTTACAGTACTGAAAATGTTAATATTGATCTGTTCTTTCTTC

>MSTRG.966.1 gene=MSTRG.966

ACTACTTACATCAAAAAATTAACTAAGTTACATGCTTTGCTTAATAATATTACAAGCCAGCATAAAGTTA

TTATTAAAGATAACATATGTTCAATATAGATTTATGCGAGCTTAAACTTAAAATCTCAACCATGGAGAGG

GAGAGAGACGGAGAGAGATTGAGAGAGAGATATGGGGCACTCAAACATCTGGAACGCGCATCCCAAGACC

TACGGCCCTGGCTCTCGCGCCTGCCGTGTTTGTGGAAATCCTCATGGGATAATTAGAAAGTATGGGATAA

TGTGCTGCAGACAGTGCTTTAAAACCAATGCCAAGGAAATCGGTTTCATCAAGTATCGTTAAGTTGGGAC

TTGTCATCTACCCCCACAGTCCTTCACCCTGCCGAAGAAGACTTGTGGTACCTGTTTTATTGTCAATTTA

GTTTCATGAAGATGTAGTTTTGTTTTTCTGGAGACAATGGATCCATCCTGATGATGTAATAGATTGAGTA

ATTTGTAGACTATTAAAGCATTGGCAAATTTGAAGCTGCTATTGTTCTTTTTGTTTAGATAACAACATTT

TCATATTCTGCAAATCTCACTC

>MSTRG.967.1 gene=MSTRG.967

CTCCAGAAAAACAAAACTACATCTTCATGAAACTAAATTGACAATAAAACAGGTACCACAAGTCTTCTTC

GGCAGGGTGAAGGACTGTGGGGGTAGATGACAAGTCCCAACTTAACGATACTTGATGAAACCGATTTCCT

TGGCATTGGTTTTAAAGCACTGTCTGCAGCACATTATCCCATACTTTCTAATTATCCCATGAGGATTTCC

ACAAACACGGCAGGCGCGAGAGCCAGGGCCGTAGGTCTTGGGATGCGCGTTCCAGATGTTTGAGTGCCCC

ATATCTCTCTCTCAATCTCTCTCCGTCTCTCTCCCTCTCCCTCTCCCTCGTTGAGATTGATCTGGAATTA

TAAC

>MSTRG.969.1 gene=MSTRG.969

GCTTCATAGGTGAGTCTTCTGTGTTTTTGTGTTCGTTGAGTTATTTTTCTATTTTTCTATTTGGTTTTTA

TGTTGAAGCACAACTGTTCTTGCAGGTATGAAGCGAGTAAGAGATGATTTTTCTAACTCGCCGATTAAGA

AGTCCTTTGCTTCTTCTCTGCAGAATCGTATGTTTCTTCACTTTGATTATGATGGTTTTGTGTTCTTTTT

GCTTGTGAATTCTCTGCGAAATATGGGTTGTGAATAATTTTAATTAGTTTGTGCTTAGTTTTTACCTTTA

GGGATTCTATCGGTTTAGCTTAAGAATGGGTTGTTGGAGAAGCAAGGGTTGTTAATTGAGGGTTAAGTGT

AACTAAGGGTTGTTGGTTGTGTACTGATTTGGGGAATTTGTTGATGCCCTTTTCGGGAAAATGGGTTACT

TGACTTGTGTGGGTGAAGTTAGGAATATTTTCGTGTATAAAGATGTGATCTTTGTGACTAATTGGTTGGG

TGTGTGGGAAATACTATGATGGCTTTATCGGGATATAATTAGTTGATTAATGGCCATATAAAGAGTCATC

TGCCATCTAAGTCGAGGTTAAGCATGTGATGTATAGAGTAACATGTTTTAGT

>MSTRG.973.1 gene=MSTRG.973

TCCTAATGCTTTTTATTTTATTTCTCTGCTAGTCCTTTACTCCAGAAAGAGACTTCCTTCAACTCCATCC

TGAGTCTCCTGACTAAACATACACATTTTAAATAGCATTTGATGAGGTTTACAATACACTAGCAAGACTT

CTGGATTTGTTTATTGCCTGTAACTTGATTTAGTGCTAGTGCTGGTTCACAGAATGTTCTACCTAGCAGG

CAACATGCAATAGCGTGACT

>MSTRG.975.1 gene=MSTRG.975

GTTAATTTGAATCTATAAACCATAGTTACATAGGACTGGGTTCGATTCGAAACGGGGAACGAAATCGGGT

ACGTGGAACATTAGTTTTAGTGAATCTGGTACGATGGAACGTATAGGTACGATGGAACGTATGGGTACGA

TTGGTCCTTTTAAAAAGAATTCTATAATCATATATATAAATTCCATTATAATATACAATTTTAGTTGATA

ATTTATACATTTATAATTTTTTATAAAGAAAAAGATATTGAATAACAAATAAAAGTACACATTTATTGAT

AAAATAAATTACTTATCATATGACATAGACATGGATCTATTGTTCAAAACAAGAATGAAGTGTGTTCTTG

CGGGTCGTCACTTCCGGCGATCTGGGTCGCGACCCTGAACGTTTCCGTTCGCGTTTCCGACCGTATCGGG

TACGAACGATCTAGATCGCGGTTCGTGGCAGCGTACCCGTTTCCTTGTATCTATGCTATAAACTGACCTT

ATT

>MSTRG.977.1 gene=MSTRG.977

ATTGAATACTTACTATCCCTACGCGGATGATAAAATTAATATTCTTCAGAAAAGAAGTCTATTAACGTTC

AATGACAAAAACGAAGTGCGAATGCATGATCTGCTTCAGGATACGGGAAAGAAAATCGCGCGTAACAACT

CTCCTGATGAGCCTGGAAAGCATAGTAGATTATGGGTACGAAAAGAGATATGCGATCTGTTGGAGAAAGA

CAAGGGAACAGAAGCGATTGAAGGTATCTTGCTCAACCATTTTGAACCCGAGCTTCCT

>MSTRG.979.1 gene=MSTRG.979

ATACAAGTCAGGCAATCAAAAAACATCAAAGAAGCAACATAACGGGCCAGCCGAGCCGAAGATGGAAAAT

GCATTTTAAAGGGTGAAAATTGATGAGGTGGAATTTGCTGACAAGAAGCTGCAAGATAATTCTTGCTGAG

CAAAGAGCGGTGCTGAGATTTGATATCTAAAGCACAAGAAGTCTTGGGGCAGGTTAGAGGAAGGATAGGC

TGGATGGACAACTAGAGATTTTGTTTGATCTGAATGAGAACTTAGTGACGATGAAACAATAGCATTGAGT

GTATCTGCAGTAAGCAAGGATGCTAAATACTTGGCATATGCTCTCAGTTCGAGCGGAAGTGATTGGGTAA

CTATTAAAATAATGGATGGGGGTTCTGGACAAAAAAGTTCACAGTGATACTCTATCATCGAGACAGCAGC

CATGGAGATGAATGTGGGTTTAACATATCTAGAGAGAAGGAG

>MSTRG.980.1 gene=MSTRG.980

TGTAGCTCGAAGAGAATGATGTAAGAGTTCAATCTCAATAGGAAGCTGCAGAAAGCAACAATGTTGCATA

TAGAAGCCATCCAGCTGCATACCCTGATCACGGGGTTTTCCAGTCTCTAGAATGCAAAAATACAATGCAC

ATGGGCTACAATTTAGCAATGAATGATCATCAAATGGCTTCTGCATCACCGGCACAGAATACCAGTGGAG

TCATTCCAGGGTGGATGCTTTGATCATAAATTCTCGTTGGTGAATTGAAGAATTGAGCTTGAGCTAAGAA

GGTGGTTGCTTCACAAGATTGGCCAGAGATAACAAAGTATGGTGAATTGGTTTGTGCTCAACCCCATAGA

CAAGAGCTTATTCAGGATCTGTACGAGAGCTGGTAGGATCCAAATAGAGAAACTATGTCAGGGGCATGAT

CAAGGAACTCCTTATTTCATTCCGAAATCAACTGGACAGAAACCCCAGAGCATTATATTCTATAGGGATG

ATGTTGGTGAGGGGCGGTTCTATCAAGTTCTTCTGTATGAACTTGATGCAATTCGCAAGGATTGTGCTTC

ATTAGAGCCAAATTATCAGCCTGTTGTTTGCCAACAATCATCGCAACCGTAATTCAGTTGATGGGAGTGG

AAATATTTTGCCTGGGACTTGTTTGGATTCCAAAATCTGTAACCCCGCCGAGTTTGACTTCTATCTGACG

ATAAGTTACTGCATTTAACTGGCTTCATTTTTGTGTCTTGCAGTTAGGCCAGGTGCACAAGATCTGTTTC

AATAGTTCCACCAGCATATTATGCTCACTTGGCAGCATTTCAAGCTCGCTTCTATATGAAACCTGAAACA

TCTGATAGTGAGTCAATGGCAAGTGATGCTGCTGGTTGTCGTGGTCGTGGATGATTTGGTTCCGGTTGTT

TATCTTGTGGCGATTTGGATCGCACCTGCTCATGAAGGGTTGGAGCTTGGAAATTGGAATGCCAGTATTA

TTAAAGCGCTGGCTGAAGCTTGTGGATGCAATGAAGCGCGGAATAGGAAGCAGTACAAAGTAATGAGAGA

AAGTCGCTAAGGTCTTTAACACATTTCAGCTCATTGCCGAGGTGATCCGGTCTGACTTCTGCTATTTGGT

TTAATGTTGAGCTATTTTGGCATTGCAGTGATTATTACATAGAAGAAGCAATTTGGGAATGGAGTTGAAA

TTGATTAACATAATAACATTCTTAGTCAAACAAATATTGCTTAATGACAATCTATATATTATGATCATCT

ATAG

>MSTRG.981.1 gene=MSTRG.981

AAAAAAGAGATATAGGAATGGATCATTTCAGCCACGAGCATCCACTGATCCTGAATGAAGATTACATTGC

TAGAGCGAGTGATATGTGTAATGCCTGCAACGAAGAGATCGTCTCATGTAAGTCATTTGTTTTCAGTTGC

AGCCGCAGAAGCAGCATCACAGGCATCCCAGGTACTGATAATAAGGATGTCGATGAAAAATGTCTTAAAT

TCTTACTCCACAAAACATGTGCAGAGTCACCCCGTAGAATTCAAAGCCTAAGAAACCGAGAAGAGTTCCT

GGACCTCCTTATCAATCCCCGCGCTAACAAAACATTTTTACAAACTTTTTTGAACCGTCCAGGTGACATT

TATCCCTGTAAATTGTGTATGAAACAAATGCCATGGATTTACTGCAGTCCTGGTTTTCTCTTCTGTGTTT

GCCTTAAATGTGCCATGTTCCAAGTTGATTCTTTCATAAGTCGTAAATTTATTCATCAATCTCACACTCA

ACACCAACTAGCCCTGATCCAACGTCCAACTTCATTTAAATGTGATGCTTGCAACATGGAGGATAGTACT

AAAGACATGTCATATAAATGTGTGGATTGTCCGTTTTGGATGCACAAGAGCTGCGCGGATGCTCCTACCT

CTTTCCTCTTTAATTTTCACAAAAAACACCCTCTTATGCTCTCATTTTCTTTGCCACAAGTTTATCATAA

ATTTGCACAGTATTGCAGACTCTGCAATGGAGAGCTGGGTGAATTAAATTGGCTTTACTATTGTTCCAAA

TGCAGATATTTCGCCCATTTTCAATGTGCTAGATCAAGGCAGATGTTGAGCTCTTCTGAAAACGAAGAAT

ATCCCGATCTTGTGCACTTCCCAGCAGCTACTGAATCATCTGTAAATTTATTGTTTGAGCAATTTATCAA

GGAGATGAGTCGTCCAAAGAACGAGGAAGGATGTAGCCCAAGAGGTCGTCTTAAAGATTGGGCTCATGGT

GAACATCCCCTGCAGCTTGTTACAGTTGATCAGTTAAATGATGAGAAAGATGTATTATTCATATGCGATG

GCTGTGCCAAACCCATCCGAAATGATGGCCATCTTTTCTATGCTTGTGTTCCTTGTAAATATTTTCTGCA

TAAATTCTGCGGTGAATTACCCACCACCAACCCCCAAAGTCCATCTCATAGAATTCTATATAACAAGTAC

AGTCCAGCATACAAGCTTTTTCTTTGTCATGGCTGTCATTATTGGGGCAATGTTATATCTTTCCGCGGAG

ATGAGCCAAGAAACTCGTTGCCTTTTGAAGAACATATAGCATGCTCAGCCCTACCCAAAATGATCAAACA

CGAGTCCCACCCTCACCCACTTCACTTGAATAGTTATAAGCCTACAAGTTGCAAAGCATGTGATGATAAG

AGGTACTTTTATACATGGCACGGTTGCGAAGGTTGTGATTTCCATTTATGTGCAGGATGTATTATGAAGG

CAAGTAGAGTTAAACATGGGTGGGATCCTCACCCCCTCCGCTTGATTTTTGATCCTGGCATGGTCGAGGA

TCACGAACATGAGTTTCAATGTGAGTTCTGTTCTCAAGATTTGGACACAAACTTTTGGTTCTATCACTGC

GATGATTGTGATCTTTCGTTTCATCTAAGATGTTATGATAAATCATGTTATCTCAACTTCTCACATGTCA

AGTATGGTGCTACTGATATTATTAAAGACCATCTCCATCCTCACAGCCTCACATTGGTACTCAACAAGAA

AGTTCGTAGTTGCGGAAAATGTCACTGGTCTAGTATTGGCAGGGCTGTCCTTGAATGTGCACCATGCAAA

ACCATATTCTGTGCGTCCTGCTCGGGAATGTATCAATAACAGATCGCATCAATCGTCTATTATCTGGAGT

ACTTG

>MSTRG.992.1 gene=MSTRG.992

CTTGATTAAAAAAAAGAAAAAAAATAGAATAAGCGAAGTCGAATGGCGGTCGTGACTCACTTACACTGAG

AAGCGTCCCAACCTTAGACTTAGCAAGAAAAAAAATTAGAAAAAATAGGAGAGGCATAGAAATTCTTTAG

GGACCCTAAATTGTAGTTGACTCTTTAGTAAAGAAGTGTTTAAACCTGATTCTGATTAACGGCTCATAGT

GAGGAC

>MSTRG.997.1 gene=MSTRG.997

GGTGTTTTCGTGGTCGATCAAAGTCATAGCAGGCGTCGGATAAGCATATGCGCACGTTATAAAGAGCCAG

GCCCCCATCACATCCCCTGCGCAAATTAGGGTTCTCCCCCCAAATCAATTGAATTTCTTGTCTCCTTTCC

TTTCTCTTGCCCCCTATTTAATTCAACAATCTCTACAGCTACTTTTCAAATTTGCAGCCTACTGTTTAAT

TATTCTTGTTATCGTATGGTTTGTGTTTGACCAGAAGTAGGATTTGAGGGGGCTATGATGTCTGAAGAGA

CTGAAAATATTATGGTCGTTTATGATGATCACTCTGAGCAAAGGTCTCTGTCATTGGATGATACAAGCAG

CACTGAGGAATCACCAGATGAATCAAGATTCTCCTTGGAAACAAATGATGCAATTCCTTATATTGGGCAA

AGATTTACCACTCACGATGCTGCCTATGACTTCTACAGTGAATTTGCCAAAAGAAACGGATTTTCTATTC

GCCGTCATCGCACAGAAGGTAAAGATGGAGTAGGCAAAGGGCTAACGAGACGATACTTTGTTTGCCACCG

TGCAGGCAATACTCCCGTCAAAGCCCTAACTGAAAATAAGCCTCAACGAAATCGCAAGTCCTCACGATGC

AATTGCCAAGCATATATGCGGATAAGCAAGACAACTGAAATGGGAGCAGCAGAATGGCGAGTCACTGGAT

TTGGAAACAATCATAATCATGAACTTCTAGAAGCGAACCAAGTTCGTTTTCTTCCTGCATATCGAACCAT

CTCAGATACAGATAAGAGCAGGATTCTTATGTTTGCGAAAACTGGAATTTCAGTGCAACAGATGATGAGG

CTCATGGAGCTAGAGAAATGCGTGGAACCTGGATATCTGCCCTTCACAGAGAAGGATGTTCGAAATCTGC

TCCAGTCATTTAGGAAAGTAGATCCAGAGGACGAGAGCATAGATCTACTGAAAATGTGCAGAAACATTAA

GGATAAAGATCCAAATTTTCAGTTTCAGTATACACTTGATTTAAACAATAGGCTGGAGAATATTGCCTGG

TCATATGCCTCTTCAATCAGGTCATATGAGGTGTTCGGTGATGCAATTGTATTTGATACAACCCGCCGCT

TAACTGCATTTGACATGCCCCTTGGAATATGGGTTGGAATGAATAACTATGGTATGCCTTGCTTCTTTGG

TTGTGTTCTTCTAAGAGAGGAAAACTTGAGATCATTCTCCTGGGCTCTGAAGGCATTCGCTGGTTTCATG

AATGGTAAGGCTCCACAGACAATATTAACCGATCAAAATATGTGTCTGAAAGAAGCAATATCAATTGAGA

TGCCTACCACCAAGCATGCCCTTTGCATTTGGTTAATTGTGGCCAAGTTCCCGTCTTGGTTTAATGCTGT

CCTGGGAGAACGCTACAATGAGTGGAAGGTTGAGTTTTATCGATTATTTAATATGGAGTCAATCGAGGAA

TTTGAACTTGGGTGGAGGGACATGGTCGCTTCCTTTGGACTTCATACTAACAGGCATATAGCAAGTTTGT

TCGCATTAAGATCACATTGGGCATTGCCATACTTAAGAAGCTATTTCTTTGCTGGAATGACCACTGCTGG

ACACGCGAAGTCTATTAATGCTTTCATCCAACGGTTTCTCAGTGCACAAACAAGGCTTTCACATTTTGTG

GAGCAAGTTGCCACTGCTGTGGATTTTAAAGATCAAGCAGGAGAACAACAAACAATGCAACAGAATATCC

AAAACATATGCCTGAAAACTGGGGCTCCTATGGAATCGCATGCGGCCATAATATTGACCCCTTTTGCATT

CTCTAAGCTTCAAGAACAACTTGTCTTGGCTGCCCATTATGCATCATTTCAGATGGATGAAGTTTTTCTT

GTTAGGCACCACACAAAGCTTGAAGGAGGACGGAGTGTTTATTGGTCTCCACAGGAAGGAATAATTAGTT

GCAGTTGCCATCAGTTTGAGTTTTCCGGAATACTTTGTCGGCATGCCCTGCGAGTTCTCTCTACTGGAAA

TTGCTTTCAAATCCCAGAGAGGTACCTTCCTCTCCGATGGCGTAGAATTAGCACTCCATCTGCCAAGCTC

CTCCATACTAGTTCAAGCGATCATGCTGAACGAGTGCAGTTATTACAGAATATGGTGTCAACTCTTATCA

CAGAATCTGCCAAGTCAAAAGATCGTCTTAATATTGCAACTGACCAAGTTTCAATGCTTCTGTCTCGGAT

TAAAGAGCATCCTGTACCCTCATCAGGCACGAGAGAAATTGCTTCTTATAAAAGGAATCTTGATTACAGG

TGATACACCATAGAACCATACCAAGATTTTGTTGTAGAAAGCAGTGATTAACATTTAAACCACAAGGATT

TAGTGCTTCACATATGGAATGATACCTCTCTTTCTTAAAGAGTTGTAGATGCCTTGTTGCTGATACTCTG

TGCTTGACATTTTTGTAATTACAGACTGACAGTTGGTTTTACTTGTGAGGTACTATAACTTATACTAACA

TATCTATTAAAGTGGAGTATGTGCAAGTTCATATTTTTATGTGTAAAATTTT

>MSTRG.999.1 gene=MSTRG.999

ATCGGATTCGGATTTCCCTAGTATCCAAATCCAAATCCAAAATATCGGGTTCGGTAAGATCCATTCGGAT

CGAAACGGTCGGGTATTCATTCGGATCGAAACGGTCGGGTATCCATTCGGATCGGTTTTTTCTGTCTATC

CCTACTCGGACCGACTAGACGATAGAGATACGAATACTATTCAGACAAATTTGGGTCCTCGAAGAAGTTG

AGAATGGCGGCTGCGATAATGAACGGCGCCGGAATCGGAGTAATCGCCGGAAAACAGAAAACGGTGTTGA

TAACAGGAGTAAGTAGAGGCCTCGGAAAATCACTAGCTCTTGAATTAGCAAAGCTGGGGCACACCATAAT

AGGGTGTGCTCGATCACTGGAGAATCTCAATGCTCTACAATCTCAGCTCAATGACACCAAGAATAAGCAC

TTTTTCATGACTGTTGATGTGAGGTCAAATAGTAGTGTAGCAGAGTTGGCAAAGGCTGTCATGGAAAAGA

ACGGTGTTCCGGATATCATAGTGAATAATGCAGGAACTATAAATAAAAATAACAAAATCTGGGAGGTACC

AGAGGAGGAGTTTGATGCAGTCATCGACACAAACTTAAAAGGAACAGCAAATATGTTGCGTCACTTCATC

CCTCTTATGATCGAGAGCAAGCAAGGGGTCATTGTCAATATTTCATCTGGATGGGGAAGATCTGCTGCAG

CACAGGTGGCACCTTATTGTGCTTCAAAATGGGCAGTAGAAGGTCTGAGCAGATCAGTTGCAAAGGAGTT

GCCAAGTGGAGTGGCAATCGTTGCGCTAAATCCAGGTGTGGTTAACACAGATATGCTTGCTTCATGCTTT

GGTAATTCGGCCTCTTTGTACCAGCAGCCCGAGGCATGGGCTCCAAAAGCAGCAACAATGATACTTAATC

TTACTGCAGCAGACAATGGTGCATCTCTGACCATATGATGTTTGAGCTAAATGGTGCCTTGGTTATTTGT

GGATAAGACCAGAAGAGGACTGATGTTTCTATAGATATTGTAAAAATGAGTATTTTCAGACAAAATACTG

ATAAGTATGTTCGCCTTTTTTTTTTTACTTAAGTTCATATTGCATAAATTATCCAGCCTACGGTTGCAGA

CTTAACTGATAAGTTTAGATGCATAGTTAGGTATGCTACAGTTTCCTCATGCCTTATATAACAGCTAATA

TTTTTAATGCCAGTATGACTGCGAGATCAATGTTGGTAACTTAACAAGGTCCTGCCTCTGCTTAATAAAG

G

>MSTRG.999.2 gene=MSTRG.999

ATCGGATTCGGATTTCCCTAGTATCCAAATCCAAATCCAAAATATCGGGTTCGGTAAGATCCATTCGGAT

CGAACGGTCGGGTATCCATTCGGATCGGTTTTTTCTGTCTATCCCTACTCGGACCGACTAGACGATAGAG

ATACGAATACTATTCAGACAAATTTGGGTCCTCGAAGAAGTTGAGAATGGCGGCTGCGATAATGAACGGC

GCCGGAATCGGAGTAATCGCCGGAAAACAGAAAACGGTGTTGATAACAGGAGTAAGTAGAGGCCTCGGAA

AATCACTAGCTCTTGAATTAGCAAAGCTGGGGCACACCATAATAGGGTGTGCTCGATCACTGGAGAATCT

CAATGCTCTACAATCTCAGCTCAATGACACCAAGAATAAGCACTTTTTCATGACTGTTGATGTGAGGTCA

AATAGTAGTGTAGCAGAGTTGGCAAAGGCTGTCATGGAAAAGAACGGTGTTCCGGATATCATAGTGAATA

ATGCAGGAACTATAAATAAAAATAACAAAATCTGGGAGGTACCAGAGGAGGAGTTTGATGCAGTCATCGA

CACAAACTTAAAAGGAACAGCAAATATGTTGCGTCACTTCATCCCTCTTATGATCGAGAGCAAGCAAGGG

GTCATTGTCAATATTTCATCTGGATGGGGAAGATCTGCTGCAGCACAGGTGGCACCTTATTGTGCTTCAA

AATGGGCAGTAGAAGGTCTGAGCAGATCAGTTGCAAAGGAGTTGCCAAGTGGAGTGGCAATCGTTGCGCT

AAATCCAGGTGTGGTTAACACAGATATGCTTGCTTCATGCTTTGGTAATTCGGCCTCTTTGTACCAGCAG

CCCGAGGCATGGGCTCCAAAAGCAGCAACAATGATACTTAATCTTACTGCAGCAGACAATGGTGCATCTC

TGACCATATGATGTTTGAGCTAAATGGTGCCTTGGTTATTTGTGGATAAGACCAGAAGAGGACTGATGTT

TCTATAGATATTGTAAAAATGAGTATTTTCAGACAAAATACTGATAAGTATGTTCGCCTTTTTTTTTTTA

CTTAAGTTCATATTGCATAAATTATCCAGCCTACGGTTGCAGACTTAACTGATAAGTTTAGATGCATAGT

TAGGTATGCTACAGTTTCCTCATGCCTTATATAACAGCTAATATTTTTAATGCCAGTATGACTGCGAG

>MSTRG.1005.1 gene=MSTRG.1005

CTTGGATCTGTGATCCATGGATCATACGACTTTTCTTGCCAAACCACGTAGTCAACATTAATCTTTGACT

TGACCCGTAATGATCATCGATGACGTTATCCAACTCCAGTTTTCTCGGCGTTTACTTTGTCATGCTACCT

CTTGTTCAGATGGTATCTCGACCTCGACCATTATATATAGCAGGCTGTCCGTGACAATGATGAAAGAACG

CAACTTACTAGTTACTAGCAAACATCATTCTTTCTTTGCTCTCTTGTTAGCTCTTTGGTTTAATCAAAAT

GAAGGCCTGGATATACGGTGAATATGGTGTGTCAGGAGATGTTTTAAAATTCGAACCCGAATACAGTGTT

CCTGAGATTAAGGATGATCAAGTTCTTATCAAGGTTGAGGCTGCTGCTCTTAACCCCGTCGATTATAAGC

GTATGCTTGGTTGGTTCAAAGGCATCGACTCCGCTTTACCTCATGTCCCTGGATTTGATGTAGCTGGAGT

GGTGGTTAAAGTAGGAAGTAAAGTTAAGAAGCTGAAGGAAGGAGATGAAGTGTATGGTAATGTTAATGAA

GATCCTCTGAAGGATTGTAAGAGGACGGGAACGCTCGCGGAGTTCACGGCAGTTGAGGAACATATACTGG

CAATTAAGCCCAAGAATTTGAGCTTTGCTGAAGCAGCTAGCCTTCCGCTGGCTCTTTCCACTGCTTACCA

GGGTTTCGAGCTCGGGGACCTCACACCTGGTAAATCTGTCCTTATCTTGGGTGGTGCTGGCGGTGTTGGA

ACGCTAGCAATTCAGCTAGCAAAACATGTTTTTGGCGCATCAAAAATCGCGACGACTTGCAGCGCTGGCA

AAATTGAGCTGGTGACTAGTTTGGGTGCAGATGTGGCAATTGACTACGCCAAGGAACAATTCGAGGAGTT

GCCTGAGAAGTATGATCTAGTTTTCGATGCAGTTGGTCAATGCGACAAGGCCGTGAAGGCGATCAAAGAA

GGTGGAAAAGTTGTTACAATATCAGGCCAGGTGACTGCACCTGGTGTCACATTTCTAGTGAATTCAAAAG

CATCGGACTTGGAGGCAGTCAATCCCTACTTGGAGAGTGGGAAGTTGAAGGCATTGTTGGACCCCAAATC

CCCCTTCCCATTCTCTCAGACCGTCGAAGCATTCGCCTATCTTGAAACTGGCAGAGCTATCGGCAAAGTA

GTTATTCATCCTATTGCCTAATAGTCTACATTGATCAAGTTTTGTTTTGCTTTAGCTTTCTTGTTTTGTG

GTGTTATGAATAAATCTGCAGCAGATACTCTGTCCTATCTTGTTCGAAAATCGTATTGTTCTAGTACTGT

AAATGTTCCTTAATCCTCAATGTTATTTTACCAAGCTCCCTGCTCTGATTGTCAAATATCAAGAACACTG

TTCTCGTTTACATATTCAAATGTTATCAAACAGCAAATAGTATGGGC

>MSTRG.1006.2 gene=MSTRG.1006

AATTAATAAAAAGACTCTCTGTCTCTGTCTCTCAATTATTAAACAGTAAAAAACACAGAGCTTAAAACCC

AATTCTCTCTCGTTAGGGTTTCACAAATATATCGAGTTCAATTGATGCGAATTGGGTTCGAATTGGAGGC

TTCCGATCAGTGAATTGCAGTTTAATTCCGTAACAGCATACGAGATGGAAGGAGACGGTGATGAAATTAG

CGTACACGAACTCGTCTCCAATCTCTCTAATTACAAACAACAGCTTCAACAGGTTAGGAGCCTCTTGGAT

GATGATCCTGGGAACTCTGAATATGAAGACATGGAAAAAGAGCTTGCAGAGGTAATTGCTTTGACGGAAG

AACTCTTGGAATCTGCAAAGCAAAATGAGAAATCAGGCATGATTACTGGAACAGGTTCTGATGTATCTCA

TAGTTTGCACCATCCTGAAGGAGTCTCTCCACATTACCTGGAATCAGGTCTCATGTCTGATCAGTCTGAC

AGACTCCCTATTGGCACCAAAGTTCAAGCAGTTTATAGTGAAGATGGGGAGTGCTGATTTTCAGGTATGA

TGCAACGATTGATGCATTTACACCAAACGGCTATCTGGTGTCATATGAGGGATGGGGCAATAAGGAAGAG

GTGGATCCTGCAAATATTAGGCCGTTGCAAGAAGGGGCTGTAGATACAATTTTAGAAGCTGAGAAGGAAG

CTGAGGCTACAAAACTTGCTCTCAAGCGGAAAATTGCTCAGGCTGCTTCTGCTGATACTGACCTCCAGTC

CCGTAGTTTACCAGCAAAGCTACGTATTGAGCCTGAGGACAGTGAAGATGTGAAAGCTGCCAAACGTAAG

AAGATACATGCTTTCAAGTCAAAGATGCGGAAGGAGCAACTTGAGATGACACAGAATAAGAGGCAGAATG

CATGGCAGCAATTTCAGACCACCAAAGGCCGAGCTAAGAAGATTGGTTTTTTCTCTGGTCGCAAACGTGA

GAGCATTTTTAAATCTCCGGATGACCCTTTTGGAAAGGTTGGTGTAATTGGAAGCGGAAAAGGATTGACG

GACTTTCAGAAAAGGGAAAAGCACTTGCATCTGAAAGGAGCAAATGCTGATGGCACAGATGAATAGATAG

TCAAAAGATTGATGAGGGTTGTCTGAGGCGTCTTTTCTGTATGTTACTTTGAATTTGTATGCTGTTCATC

TAGCTTCAGTGCAAAATGTTCCGATATATGACAGATAAAGTAGGCTGTAACCCTAACATATACTTGTGAA

CACAAAGTTTTAAATTTGGTTAACCTGGTTTATGATATATGTGAGGTTCCTGTTGCGATTTTAGATTACA

ATATGTTGCTATGTTACTAATGTCTG

>MSTRG.1006.1 gene=MSTRG.1006

ATAAATTAATAAAAAGACTCTCTGTCTCTGTCTCTCAATTATTAAACAGTAAAAAACACAGAGCTTAAAA

CCCAATTCTCTCTCGTTAGGGTTTCACAAATATATCGAGTTCAATTGATGCGAATTGGGTTCGAATTGGA

GGCTTCCGATCAGTGAATTGCAGTTTAATTCCGTAACAGCATACGAGATGGAAGGAGACGGTGATGAAAT

TAGCGTACACGAACTCGTCTCCAATCTCTCTAATTACAAACAACAGCTTCAACAGGTTAGGAGCCTCTTG

GATGATGATCCTGGGAACTCTGAATATGAAGACATGGAAAAAGAGCTTGCAGAGGTAATTGCTTTGACGG

AAGAACTCTTGGAATCTGCAAAGCAAAATGAGAAATCAGGCATGATTACTGGAACAGGTTCTGATGTATC

TCATAGTTTGCACCATCCTGAAGGAGTCTCTCCACATTACCTGGAATCAGGTCTCATGTCTGATCAGTCT

GACAGACTCCCTATTGGCACCAAAGTTCAAGCAGTTTATAGTGAAGATGGGGAGTGGTATGATGCAACGA

TTGATGCATTTACACCAAACGGCTATCTGGTGTCATATGAGGGATGGGGCAATAAGGAAGAGGTGGATCC

TGCAAATATTAGGCCGTTGCAAGAAGGGGCTGTAGATACAATTTTAGAAGCTGAGAAGGAAGCTGAGGCT

ACAAAACTTGCTCTCAAGCGGAAAATTGCTCAGGCTGCTTCTGCTGATACTGACCTCCAGTCCCGTAGTT

TACCAGCAAAGCTACGTATTGAGCCTGAGGACAGTGAAGATGTGAAAGCTGCCAAACGTAAGAAGATACA

TGCTTTCAAGTCAAAGATGCGGAAGGAGCAACTTGAGATGACACAGAATAAGAGGCAGAATGCATGGCAG

CAATTTCAGACCACCAAAGGCCGAGCTAAGAAGATTGGTTTTTTCTCTGGTCGCAAACGTGAGAGCATTT

TTAAATCTCCGGATGACCCTTTTGGAAAGGTTGGTGTAATTGGAAGCGGAAAAGGATTGACGGACTTTCA

GAAAAGGGAAAAGCACTTGCATCTGAAAGGAGCAAATGCTGATGGCACAGATGAATAGATAGTCAAAAGA

TTGATGAGGGTTGTCTGAGGCGTCTTTTCTGTATGTTACTTTGAATTTGTATGCTGTTCATCTAGCTTCA

GTGCAAAATGTTCCGATATATGACAGATAAAGTAGGCTGTAACCCTAACATATACTTGTGAACACAAAGT

TTTAAATTTGGTTAACCTGGTTTATGATATATGTGAGGTTCCTGTTGCGATTTTAGATTACAATATGTTG

CTATGTTACTAATGTCTG

>MSTRG.1007.1 gene=MSTRG.1007

TGAGAAGTCATGTAATCTATCCAAGTGAAGTGAAGATCGATCGGTAATTATAAAGTATCAATGGCTATTT

GGAAGGAAGACTATCAACCGCTATATCAACATCATCATTATCCCATAACTCAAAGAAGACTATCAACTGC

TATTTCAGTGTATTATCAATTGATAATATTAGCAATTGATTATCAGCAATTGACAGCTACAAGTCAGGTA

AAAGACAAAGTTCACATGGGTTGATGTGACAGATGCTGCACCAGCTTTGGAAGCTAAACCAAAAGGAGTT

TAGTCAAATATTGAGAACCTCGAAGCCTACCATTTCTGGCAAAGCAACAAGAATTTCAAGCTGCCAAATG

CAGTATCAAGTCCAAGAAATTCTATATTTGTAC

>MSTRG.1010.2 gene=MSTRG.1010

ATTTTGAATTCAAACCATACGAGCCAAACCCAGTTCCTCTTCTTCGACCCAAACTCAAACCTCAAATTCT

CTCTCTGCTGTGTCCCCCCTTTTCTAGGGTTTCAAGCGCCGCTTCGCTGTTGGGAAATCAGTCCTCCAAT

TAAATCTCTCTCTCTCTCTCTCTATGGCGCTGGTAATCTCTCTACCTCTCTTCGTTTCTTAATTCGATTT

ACTTTAATTACATTTAATTAGGATTTTATAAGTCCTTTTTACTGTCAGATCAAATGCTCTTTTGATCCGA

TTCGTGCTTCTATCTTTGCTGTGTGCCCCTTTTTCTAGGGTTTCAAGCGCTGGTTCGCCGCTGGGAACCA

ATTCTCCGATTAAATATCTCTCTCTCCGGAGCTGTCCAACTTCAATGGCAATTTCAGGAAAGGAGTTGAC

TGGACTTGGTTTATGCTCCTGTTCTCTCACAGAAGCAAATAAACTGAATAGAAAATTGCGTGAGGTCTAT

AATTCCTAGGAAGCATTTGCATGGTTCAATGTTACTGCAATCTCTCTACAACACTTTCTGAGAACATGGA

TGTGCGTAATGACATGTGGATCTCTGGGTTTTGGCTAATTGTTTTATACATGTAGCTAATTATCAAACTG

CACTTCTTCAGCTATTCTTGAAAACGCAACAGACATAGTTGCATTGCTTGATTTCAAGTCACATATAAGC

GAAGACCCTTGAGGAGTTCTCAACTCTTGGAATGATCCCATATATTTTTGTCAATGGCCTGGAGTTAAAT

GTGGTGTCAAATACCAGAAAGTCATCTCCGTTGAGCTTAAAGGTCAGAGTTTGGTCGGGAACATAAAAAA

GCGGTTGTTCTTAAACAATTTGGCAATAATTTACCCGACTTAGGCCTCGAAGATCAAGAATCTGATCATG

ATTTCATGTCCATCTGCAAGGTCAGTCTTTCAGTCATCTTATATGTACATTTGTCCTTAAGTTTTGTTTG

CATGATTTGATATTATAGATTCTCTCAAGTGCACAATAATAAAGCTCTAATTCAAGCTAGAAACATATTT

CTTCTAAAAATTAGAGCTTTTCTCTTTGCTGCACTTTCACAGGTATGCCATAGCACCCCACAGGCTTGAT

TATAATTAATTTATCTATTATTTTTTTCCTGAAATACACTCGCGATGAGATCCAGCATTTTCAGGGAATA

AAGATGATCTAAAGAATTTATATCTCTTCCTAAATCATAAAGGAACAGAATCAGAAGCATATAGTAAACT

GGTGACATTTCTAAACATTTCTATTCTCTGCTGATTTAATCTATTGTACTCACCATGTATTTGATATGAG

ACCTTGATGAATTATATACGAC

>MSTRG.1010.1 gene=MSTRG.1010

ATTTTGAATTCAAACCATACGAGCCAAACCCAGTTCCTCTTCTTCGACCCAAACTCAAACCTCAAATTCT

CTCTCTGCTGTGTCCCCCCTTTTCTAGGGTTTCAAGCGCCGCTTCGCTGTTGGGAAATCAGTCCTCCAAT

TAAATCTCTCTCTCTCTCTCTCTATGGCGCTGGTAATCTCTCTACCTCTCTTCGTTTCTTAATTCGATTT

ACTTTAATTACATTTAATTAGGATTTTATAAGTCCTTTTTACTGTCAGATCAAATGCTCTTTTGATCCGA

TTCGTGCTTCTATCTTTGCTGTGTGCCCCTTTTTCTAGGGTTTCAAGCGCTGGTTCGCCGCTGGGAACCA

ATTCTCCGATTAAATATCTCTCTCTCCGGAGCTGGAAAGGAGTTGACTGGACTTGGTTTATGCTCCTGTT

CTCTCACAGAAGCAAATAAACTGAATAGAAAATTGCGTGAGGTCTATAATTCCTAGGAAGCATTTGCATG

GTTCAATGTTACTGCAATCTCTCTACAACACTTTCTGAGAACATGGATCTATTCTTGAAAACGCAACAGA

CATAGTTGCATTGCTTGATTTCAAGTCACATATAAGCGAAGACCCTTGAGGAGTTCTCAACTCTTGGAAT

GATCCCATATATTTTTGTCAATGGCCTGGAGTTAAATGTGGTGTCAAATACCAGAAAGTCATCTCCGTTG

AGCTTAAAGGTCAGAGTTTGGTCGGGAACATAAAAAAGCGGTTGTTCTTAAACAATTTGGCAATAATTTA

CCCGACTTAGGCCTCGAAGATCAAGAATCTGATCATGATTTCATGTCCATCTGCAAGGGAATAAAGATGA

TCTAAAGAATTTATATCTCTTCCTAAATCATAAAGGAACAGAATCAGAAGCATATAGTAAACTGATTCGA

CCTCCTAGAGATGCTCCTATCAGTTGGTATACAATAGAGGCTTTGCAGAGAATGAAAGAACATGTTTTAG

GGTTCGATGATTACATTACAACTTTTTCACAAGCAAGGGGTCAAGACATAGTCAGGGGTGTGAATTATGC

ATCTGTAGCTGCTGGAATCAAAGAAGAAACGGGGCAGCAGCTGGT

>MSTRG.1010.3 gene=MSTRG.1010

CAAACCCAGTTCCTCTTCTTCGACCCAAACTCAAACCTCAAATTCTCTCTCTGCTGTGTCCCCCCTTTTC

TAGGGTTTCAAGCGCCGCTTCGCTGTTGGGAAATCAGTCCTCCAATTAAATCTCTCTCTCTCTCTCTCTA

TGGCGCTGGTAATCTCTCTACCTCTCTTCGTTTCTTAATTCGATTTACTTTAATTACATTTAATTAGGAT

TTTATAAGTCCTTTTTACTGTCAGATCAAATGCTCTTTTGATCCGATTCGTGCTTCTATCTTTGCTGTGT

GCCCCTTTTTCTAGGGTTTCAAGCGCTGGTTCGCCGCTGGGAACCAATTCTCCGATTAAATATCTCTCTC

TCCGGAGCTGGTAATCTCTCTGCCCACTGCTTTTTACTGTCGAATACAATGCTCTTTTGGTCCGATTTGT

ACTTTTCGTTATTAGTTGAGGTGGAAAGGAGTTGACTGGACTTGGTTTATGCTCCTGTTCTCTCACAGAA

GCAAATAAACTGAATAGAAAATTGCGTGAGGTCTATAATTCCTAGGAAGCATTTGCATGGTTCAATGTTA

CTGCAATCTCTCTACAACACTTTCTGAGAACATGGATCTATTCTTGAAAACGCAACAGACATAGTTGCAT

TGCTTGATTTCAAGTCACATATAAGCGAAGACCCTTGAGGAGTTCTCAACTCTTGGAATGATCCCATATA

TTTTTGTCAATGGCCTGGAGTTAAATGTGGTGTCAAATACCAGAAAGTCATCTCCGTTGAGCTTAAAGGT

CAGAGTTTGGTCGGGAACATAAAAAAGCGGTTGTTCTTAAACAATTTGGCAATAATTTACCCGACTTAGG

CCTCGAAGATCAAGAATCTGATCATGATTTCATGTCCATCTGCAAGCATTTTCAGGGAATAAAGATGATC

TAAAGAATTTATATCTCTTCCTAAATCATAAAGGAACAGAATCAGAAGCATATAGTAAACTGATTCGACC

TCCTAGAGATGCTCCTATCAGTTGGTATACAATAGAGGCTTTGCAGAGAATGAAAGAACATGTTTTAGGG

TTCGATGATTACATTACAACTTTTTCACAAGCAAGGGGTCAAGACATAGTCAGGGGTGTGAATTATGCAT

CTGTAGCTGCTGGAATCAAAGAAGAAACGGGGCAGCAGCTGGT

>MSTRG.1010.5 gene=MSTRG.1010

CTTTTTCTAGGGTTTCAAGCGCTGGTTCGCCGCTGGGAACCAATTCTCCGATTAAATATCTCTCTCTCCG

GAGCTGGAAAGGAGTTGACTGGACTTGGTTTATGCTCCTGTTCTCTCACAGAAGCAAATAAACTGAATAG

AAAATTGCGTGAGGTCTATAATTCCTAGGAAGCATTTGCATGGTTCAATGTTACTGCAATCTCTCTACAA

CACTTTCTGAGAACATGGATCTATTCTTGAAAACGCAACAGACATAGTTGCATTGCTTGATTTCAAGTCA

CATATAAGCGAAGACCCTTGAGGAGTTCTCAACTCTTGGAATGATCCCATATATTTTTGTCAATGGCCTG

GAGTTAAATGTGGTGTCAAATACCAGAAAGTCATCTCCGTTGAGCTTAAAGGTCAGAGTTTGGTCGGGAA

CATAAAAAAGCGGTTGTTCTTAAACAATTTGGCAATAATTTACCCGACTTAGGCCTCGAAGATCAAGAAT

CTGATCATGATTTCATGTCCATCTGCAAGCATTTTCAGGGAATAAAGATGATCTAAAGAATTTATATCTC

TTCCTAAATCATAAAGGAACAGAATCAGAAGCATATAGTAAACTGGT

>MSTRG.1010.4 gene=MSTRG.1010

CTTTTTCTAGGGTTTCAAGCGCTGGTTCGCCGCTGGGAACCAATTCTCCGATTAAATATCTCTCTCTCCG

GAGCTGGAAAGGAGTTGACTGGACTTGGTTTATGCTCCTGTTCTCTCACAGAAGCAAATAAACTGAATAG

AAAATTGCGTGAGGTCTATAATTCCTAGGAAGCATTTGCATGGTTCAATGTTACTGCAATCTCTCTACAA

CACTTTCTGAGAACATGGATGTGCGTAATGACATGTGGATCTCTGGGTTTTGGCTAATTGTTTTATACAT

GTAGCTAATTATCAAACTGCACTTCTTCAGCTATTCTTGAAAACGCAACAGACATAGTTGCATTGCTTGA

TTTCAAGTCACATATAAGCGAAGACCCTTGAGGAGTTCTCAACTCTTGGAATGATCCCATATATTTTTGT

CAATGGCCTGGAGTTAAATGTGGTGTCAAATACCAGAAAGTCATCTCCGTTGAGCTTAAAGGTCAGAGTT

TGGTCGGGAACATAAAAAAGCGGTTGTTCTTAAACAATTTGGCAATAATTTACCCGACTTAGGCCTCGAA

GATCAAGAATCTGATCATGATTTCATGTCCATCTGCAAGCATTTTCAGGGAATAAAGATGATCTAAAGAA

TTTATATCTCTTCCTAAATCATAAAGGAACAGAATCAGAAGCATATAGTAAACTGGTGACATTTCTAAAC

ATTTCTATTCTCTGCTGATTTAATCTATTGTACTCACCATGTATTTGATATGAGACCTTGATGAATTAT

>MSTRG.1010.7 gene=MSTRG.1010

AGTTGCAGTATTGTTAGATGACATGATTGAAACTGCGGGAAAGGAGTTGACTGGACTTGGTTTATGCTCC

TGTTCTCTCACAGAAGCAAATAAACTGAATAGAAAATTGCGTGAGGTCTATAATTCCTAGGAAGCATTTG

CATGGTTCAATGTTACTGCAATCTCTCTACAACACTTTCTGAGAACATGGATGTGCGTAATGACATGTGG

ATCTCTGGGTTTTGGCTAATTGTTTTATACATGTAGCTAATTATCAAACTGCACTTCTTCAGCTATTCTT

GAAAACGCAACAGACATAGTTGCATTGCTTGATTTCAAGTCACATATAAGCGAAGACCCTTGAGGAGTTC

TCAACTCTTGGAATGATCCCATATATTTTTGTCAATGGCCTGGAGTTAAATGTGGTGTCAAATACCAGAA

AGTCATCTCCGTTGAGCTTAAAGGTCAGAGTTTGGTCGGGAACATAAAAAAGCGGTTGTTCTTAAACAAT

TTGGCAATAATTTACCCGACTTAGGCCTCGAAGATCAAGAATCTGATCATGATTTCATGTCCATCTGCAA

GCATTTTCAGGGAATAAAGATGATCTAAAGAATTTATATCTCTTCCTAAATCATAAAGGAACAGAATCAG

AAGCATATAGTAAACTGGT

>MSTRG.1010.6 gene=MSTRG.1010

AGTTGCAGTATTGTTAGATGACATGATTGAAACTGCGGGAAAGGAGTTGACTGGACTTGGTTTATGCTCC

TGTTCTCTCACAGAAGCAAATAAACTGAATAGAAAATTGCGTGAGGTCTATAATTCCTAGGAAGCATTTG

CATGGTTCAATGTTACTGCAATCTCTCTACAACACTTTCTGAGAACATGGATCTATTCTTGAAAACGCAA

CAGACATAGTTGCATTGCTTGATTTCAAGTCACATATAAGCGAAGACCCTTGAGGAGTTCTCAACTCTTG

GAATGATCCCATATATTTTTGTCAATGGCCTGGAGTTAAATGTGGTGTCAAATACCAGAAAGTCATCTCC

GTTGAGCTTAAAGGTCAGAGTTTGGTCGGGAACATAAAAAAGCGGTTGTTCTTAAACAATTTGGCAATAA

TTTACCCGACTTAGGCCTCGAAGATCAAGAATCTGATCATGATTTCATGTCCATCTGCAAGGGAATAAAG

ATGATCTAAAGAATTTATATCTCTTCCTAAATCATAAAGGAACAGAATCAGAAGCATATAGTAAACTGGT

GACATTTCTAAACATTTCTATTCTCTGCTGATTTAATCTATTGTACTCACCATGTATTTGATATGAGACC

TTGATGAATTATATACGACTTTTCATTTATTTATTGTGAATATGTCTGTAGCTTGATTTTTTTTGTATCA

TTCAAAATTGCTATGAAGATCTTTTTAACGAGTTTGACAAGTACAAAAAGATTGAATGATTGTGATACTT

TTATTATGGATAATACTTCTTGGAATGAATGGTATAGTTTATTAACTTAATGATGGGAGGAGTTGAAGGA

TAGTACTGATTAAGTGATGATATTGTTCAAAATTTCTTTAAAATTGTAAATAATATATGAGTTTAATTCA

TCTCTGCCATTATTTGTAATCAAAGCTAGAATTGTGATTTGATCTTCGAGTCTCCATCTTAGTGATGGTC

TCTTATTAGAGCTACTGACAAGAGGTCTCTTACAGACTGGGTGAACTCTCATCCAACAGTGGGTTTCTAA

CCTTGATAACTGAGTTTGATTTTCAGTATTTATAATTTTACTTTTAGGTAAATATTATGAAAACTAATCA

TGCGAAGACGTTTGAATATATATGACAAGAGAATAATGCTTGCAAATTTCAACTAGAATAATGCTTCTAA

GTATGAGAAGTTAAACAAGAAAGAATTGTAAAATTCTTTGCTCAATTTAGAGATCAGATGATCACGATGT

TGGGTTTTAGCAGGAGCATGGGAGCAACTATTTATGTAGAAATATAGAATTCTGCAATAAAAAGGCAGAA

TATATTCTTGTTTTAGTATTTACTAATACCAGAATTTAATAGTAGTTACAATAATGATTCCTCTGTTTGC

CCAATGATTTCCATGGTATATATGATGTCGAGCATTTTGTCAATGCGCTAAGGTATGATCTACAAATTAT

GAAGAGGATTCCTGAAATTCGAAAAAAATGGGAAGATGAAGAATATAAAAGCTTTCCTGGTAACAAAGCT

TTGCCTTTTGCCTCCAATATATTGCTTATAGAATATTAAACTATAGCTGTGTCATTTTGGTTTAGATTTG

ACCTCCTAGAGATGCTCCTGTCAGTTGGTAATTTGGTATACAATAGAGGCTTTGCAGAGGATGAAAGAAC

ATGGCAGTCATCATGCAATTTCAATTCATCTTATAATTCCAACATATGTAGCTTCATTCATAATCACAGC

ACGCATACTTGATTATCATAATTGAATAATCATATAATTTCTTAACTGCACTCACATACTAAAGACAAAG

AATATAAAAGCTTTCCTGGTAACAAAGCTCAGCCTTTTCCCTCCAATATATTGCTTATAGAAGCTTAAAC

TATAGTTGTGTCATTTTGCTTTAGATTCGACCTCCTAGAGATGCTCCTATCAGTTGGTATACAATAGAGG

CTTTGCAGAGAATGAAAGAACATGGTAGTCATCATGCAATTTCGATTCATCTTATAATTCCAACATATGT

AGCTTCATTTTCATAATCACAGCACGCATACTTGATTATTACAATTGAATAATCATATAATTTCTTAACT

GCATTCACGTACTAATTGTATATGAAACTGAAATTATAACTTTCTTCCTTACATGACAACTGAAGTTTTA

GGGTTCGATGATTACATTACAACTTTTTCACAAGCAAGGGGTCAAGACATAGTCAGGGGTGTGAATTATG

CATCTGTAGCTGCTGGAATCAAAGAAGAAACGGGGCAGCAGCTGGTACATCTACATTTATAGTTCTTTGT

TCTATTTCTTGGAAGTATTATGTATTATCAATCTTCACTTCCCTCTGCTAATATTTTTAATAAGAAAAAG

TATGTTGACAAACAATGATGTGGTTATTATAAAGAGCATAATATTTTGCTCTGGATATGTTGGGTACTTT

CGGAGATTTTAGTTACTTCCAAGAAGCAACGCGGTCACCTGTTTTATAAGTCCTTGGCATGTTATTTAGA

TATACCATACACGTCTCCAGAGTAATGAGGATCAAATCAAAACCTTCGTGTTCTGTTGGTTTAAGTTAGA

CTATGCATTTTTACTTAAGTTTGC

>MSTRG.1010.8 gene=MSTRG.1010

ATGTATTGAGCACTTAAGTACTGGAAAAAGACCATAGTCATTTTGTATAGCATTTGGTGTTTCTGAGCTC

TAAAGATTTCGTTATAGCAGCTATCAATATTATAACAGTCCAACTTCAATGGCAATTTCAGGAAAGGAGT

TGACTGGACTTGGTTTATGCTCCTGTTCTCTCACAGAAGCAAATAAACTGAATAGAAAATTGCGTGAGGT

CTATAATTCCTAGGAAGCATTTGCATGGTTCAATGTTACTGCAATCTCTCTACAACACTTTCTGAGAACA

TGGATACCCTTGAGGAGTTCTCAACTCTTGGAATGATCCCATATATTTTTGTCAATGGCCTGGAGTTAAA

TGTGGTGTCAAATACCAGAAAGTCATCTCCGTTGAGCTTAAAGGTCAGAGTTTGGTCGGGAACATAAAAA

AGCGGTTGTTCTTAAACAATTTGGCAATAATTTACCCGACTTAGGCCTCGAAGATCAAGAATCTGATCAT

GATTTCATGTCCATCTGCAAGGTCAGTCTTTCAGTCATCTTATATGTACATTTGTCCTTAAGTTTTGTTT

GCATGATTTGATATTATAGATTCTCTCAAGTGCACAATAATAAAGCTCTAATTCAAGCTAGAAACATATT

TCTTCTAAAAATTAGAGCTTTTCTCTTTGCTGCACTTTCACAGGTATGCCATAGCACCCCACAGGCTTGA

TTATAATTAATTTATCTATTATTTTTTTCCTGAAATACACTCGCGATGAGATCCAGCATTTTCAGGGAAT

AAAGATGATCTAAAGAATTTATATCTCTTCCTAAATCATAAAGGAACAGAATCAGAAGCATATAGTAAAC

TGATTCGACCTCCTAGAGATGCTCCTATCAGTTGGTATACAATAGAGGCTTTGCAGAGAATGAAAGAACA

TGTTTTAGGGTTCGATGATTACATTACAACTTTTTCACAAGCAAGGGGTCAAGACATAGTCAGGGGTGTG

AATTATGCATCTGTAGCTGCTGGAATCAAAGAAGAAACGGGGCAGCAGCTG

>MSTRG.1016.1 gene=MSTRG.1016

CCCGGTGAAAATTAAAGAAAAGAAAAACAGAAGCTCTCGAGTCTGAAGTTACGCTCTCCGCTCTCTTCTC

ACCTCGCTTCACTTTGACCACTTTCTCTTGCTAATTGATTGTGCATCTCAATTGTAAAATTGGGTTCATC

ACCTTAGTTGGGTTCAGCCTTCGCTCTCTTTCATTCTCGCCAACTGAAATAGCCAATTGAAAGATCGAAT

TCAGGGGTCTTCATAAGTTTCATGCGTTTCCAAGAGTTTCTTCTGATTGGTATATATTTTAATTATTTCA

TCGATTTTGTACTAACTATGATCGACAATCTACATTTTTCAGATGATTTGAGCTTTTTCTCGAAGCTTGG

AATTGGGGCTTTTACATGGAATTTCATTTGGGAATTATCCAACTGGAGTTTGTTTATGGCGCTTATCTCT

CTCATTAAGTTAATTACTGTTAATTTGCTTTTTGTTTCATTCGTTTTTAGAATTGCTTAGATATCAGTAA

TCTGAATTGGTCGATTCTGACAGTTGGGTATGTTGTGTGTGTTTGTGTACGCTTCGGTAGATTATTAATC

ATTCTGTTTGAAATTTTTTGAATGAGAATCGAATCTAGTTTTGTGTACATCATCGTGTTCTGTTTCCGAT

AGTTGCTTGATTATGAATCTGTTGGTTACAAATTTTCTGTCTAAATCCATTGATGTGTACAGCTGTGTGA

GTGTGATTTGTGCACAGATATCCTTGATTAGTGATTCTATTTACGGACTAACAAATCAGAACTAGATCTA

TGAAAGGTTTCGTGTGTATGATACGTAAGCTGAATCTATTGGATATTATCGAAATTTGTTCTCGTGTTTC

AGTGGTTAATTTCTGATAATTGTAGGATTTTGTTGAAACTGTTCGCCGCGTGCTTAGGAATTACAAAGAC

TAATTGACTATATGGAACCGAACCTAGTTTTGTGTGTAGGATTTTGATTTGATTCTACTATCTGCACAAG

TACGACTAATGATTACTAGTACTCTGTATATTTCAATGTGCTGTGTGTGATTGTTTATGATTACTAGAAC

TCAGTATATTTCAAATATTCACCTCTCTGTGT

>MSTRG.1017.1 gene=MSTRG.1017

TGTTAAGAAGAGAAAAGTAATAGATCTACTTTAACAAGAAAATGAGAGCAGGAAGAGGAAAATTCATTCA

GCAAAAAACAAGATTGTACCAAGTCTGAGAATTACAACCAGGGTGGAGGCATAAATAACAACCTAGTACT

ATTACTGACACAGGACCATACACCAACAGGCTGTGCATTGGCCCGTTAGAATTAAACAAGCAAAACGATG

TAGTTTAATAAACTAACAGATTCCCTCCACTAAGCTGCAAATGACCTCTATTGTATTCCCGCCTTTTATG

ATAGATAAATTGGTTTGGGCCAATCGTAACAGCTGCC

>MSTRG.1018.1 gene=MSTRG.1018

GAGAGGCACATCCAAAATAATAACGGTAAAGAAATGAGAGGGACATAAGAAGTATTAGTTATTTTTATGA

TAGAGATAGCGGGCGCAAATTATGGGTCCACGAGAGGAGATAAGATTGGTTAGATGAAAAGGTATCGTGG

GTGTAGTGTTCCACTATTTTGCGGGCTTTTAGTTAAATCTATTCATCATCCTTGGCATTTCTTCTCATAC

ACAGCAGGTGTTCCAACACTGTATGTGGGATTTAGGATTTCCTTATGGGCTTTTTCATGTTTCTTCTTGA

AGATACCACAAAAAATATAAAAAACAAGGGCGCACGAGTTTTGAGAATTCAACCACTTGAGCTCAATTTC

ACATTTGAGTTGGAGAAACAAAATAAGTGTGCCTTTACCCTCACAAATACAAGTGATGATGATCATGTAG

CATTTCAGGTAATCACACCATGGCCATTAACTTTTCCTGAAACAGGCATCATGAAGCCAAATTCAACATG

TAAACTCACAGTTATAATCCAAGCTCAAAGTTCATACATAGATTTTATGAGGTGCAAAGACAAAATTTTG

ATTCAGAGCAAAATTTTACCTTTTCGGACAACTCAAGATGATGTTTCACCTGAAATGTTTGCAGATGATT

ATATAAACTACACAGAAGAAAACACGCTATGGATTGTAGCTTTACCACCACTTAGCCAAGCATATTCTCC

GGGATTAAATTTAATGATCAACATAGATCGTCTACTGGGAAAGTTGGACAGTTTAGAGGCAGAGTTTAAC

ACCTCAAAGACAAAGGAAGGGGGAGGCACAACCACTCGAGTGCATGTCAAACCAATTATTGAAATGTCGA

TCTGTGAGTTCGTAAGAGTTTGGGGATGGAAGATTTTGTTTCCAGGCAGCGTAACTATCCTATTATTAAT

CTTAACCTTACTCTTCGTGATATATTATCGCGGCCGGGAGTGATAAGCTGCTGCCAGTCAGAAACTTTGT

TATTCAAAATAAATTTATCTTGACTGTAGTAGTTGAAATAAGTGTTTTACTTGCTGTTTCTCATGTGCTA

ACCATGCTACTCCACACTTTTATTCGTCTTTCCTTCCTTATTAGTCATAAATATTATGCTATCTATTTAA

TTTTACATTTTAACTTTATATCACC

>MSTRG.1018.2 gene=MSTRG.1018

GAGAGGAGATAAGATTGGTTAGATGAAAAGGTATCGTGGGTGTAGTGTTCCACTATTTTGCGGGCTTTTA

GTTAAATCTATTCATCATCCTTGGCATTTCTTCTCATACACAGCAGGTGTTCCAACACTGTATGTGGGAT

TTAGGATTTCCTTATGGGCTTTTTCATGTTTCTTCTTGAAGATACCACAAAAAATATAAAAAACAAGGGC

GCACGAGTTTTGAGAATTCAACCACTTGAGCTCAATTTCACATTTGAGTTGGAGAAACAAAATAAGTGTG

CCTTTACCCTCACAAATACAAGTGATGATGATCATGTAGCATTTCAGGTAGAGTCAATCCTCACTTTTAA

TCAAGATGAGAGTAAACTTCGGGTCTTTCTAGCGTGTGCGAATGGGCACACACTGACAACCACTTTCTCC

TTAGGTGGGAGGTTTTGATTGGTATAGATTTTTGTGTATGCAGGTAGGCCATCATTTATGAATGGGATGG

CAGCCAATAAGAATCCAACACCTATTAAAAAGTGATTGTTAGTGTGTGCTCATACGCTAATGAATACGTC

CGTGTTGCAGGTAATCACACCATGGCCATTAACTTTTCCTGAAACAGGCATCATGAAGCCAAATTCAACA

TGTAAACTCACAGTTATAATCCAAGCTCAAAGTTCATACATAGATTTTATGAGGTGCAAAGACAAAATTT

TGATTCAGAGCAAAATTTTACCTTTTCGGACAACTCAAGATGATGTTTCACCTGAAATGATGATTATATA

AACTACACAGAAGAAAACACGCTATGGATTGTAGCTTTACCACCACTTAGCCAAGCATATTCTCCGGGAT

TAAATTTAATGATCAACATAGATCGTCTACTGGGAAAGTTGGACAGTTTAGAGGCAGAGTTTAACACCTC

AAAGACAAAGGAAGGGGGAGGCACAACCACTCGAGTGCATGTCAAACCAATTATTGAAATGTCGATCTGT

GAGTTCGTAAGAGTTTGGGGATGGAAGATTTTGTTTCCAGGCAGCGTAACTATCCTATTATTAATCTTAA

CCTTACTCTTCGTGATATATTATCGCGGCCGGGAGTGATAAGCTGCTGCCAGTCAGAAACTTTGTTATTC

AAAATAAATTTATCTTGACTGTAGTAGTTGAAATAAGTGTTTTACTTGCTGTTTCTCATGTGCTAACCAT

GCTACTCCACACTTTTATTCGTCTTTCCTTCCTTATTA

>MSTRG.1018.3 gene=MSTRG.1018

ATTGGTTAGATGAAAAGGTATCGTGGGTGTAGTGTTCCACTATTTTGCGGGCTTTTAGTTAAATCTATTC

ATCATCCTTGGCATTTCTTCTCATACACAGCAGGTGTTCCAACACTGTATGTGGGATTTAGGATTTCCTT

ATGGGCTTTTTCATGTTTCTTCTTGAAGATACCACAAAAAATATAAAAAACAAGGGCGCACGAGTTTTGA

GAATTCAACCACTTGAGCTCAATTTCACATTTGAGTTGGAGAAACAAAATAAGTGTGCCTTTACCCTCAC

AAATACAAGTGATGATGATCATGTAGCATTTCAGGTAATCACACCATGGCCATTAACTTTTCCTGAAACA

GGCATCATGAAGCCAAATTCAACATGTAAACTCACAGTTATAATCCAAGCTCAAAGTTCATACATAGATT

TTATGAGGTGCAAAGACAAAATTTTGATTCAGAGCAAAATTTTACCTTTTCGGACAACTCAAGATGATGT

TTCACCTGAAATGATGATTATATAAACTACACAGAAGAAAACACGCTATGGATTGTAGCTTTACCACCAC

TTAGCCAAGCATATTCTCCGGGATTAAATTTAATGATCAACATAGATCGTCTACTGGGAAAGTTGGACAG

TTTAGAGGCAGAGTTTAACACCTCAAAGACAAAGGAAGGGGGAGGCACAACCACTCGAGTGCATGTCAAA

CCAATTATTGAAATGTCGATCTGTGAGTTCGTAAGAGTTTGGGGATGGAAGATTTTGTTTCCAGGCAGCG

TAACTATCCTATTATTAATCTTAACCTTACTCTTCGTGATATATTATCGCGGCCGGGAGTGATAAGCTGC

TGCCAGTCAGAAACTTTGTTATTCAAAATAAATTTATCTTGACTGTAGTAGTTGAAATAAGTGTTTTACT

TGCTGTTTCTCATGTGCTAACCATGCTACTCCACACTTTTATTCGTCTTTCCTTCCTTATTAGTCATAAA

TATTATGC

>MSTRG.1018.4 gene=MSTRG.1018

GTTAGATGAAAAGGTATCGTGGGTGTAGTGTTCCACTATTTTGCGGGCTTTTAGTTAAATCTATTCATCA

TCCTTGGCATTTCTTCTCATACACAGCAGGTGTTCCAACACTGTATGTGGGATTTAGGATTTCCTTATGG

GCTTTTTCATGTTTCTTCTTGAAGATACCACAAAAAATATAAAAAACAAGGGCGCACGAGTTTTGAGAAT

TCAACCACTTGAGCTCAATTTCACATTTGAGTTGGAGAAACAAAATAAGTGTGCCTTTACCCTCACAAAT

ACAAGTGATGATGATCATGTAGCATTTCAGGTAGGCCATCATTTATGAATGGGATGGCAGCCAATAAGAA

TCCAACACCTATTAAAAAGTGATTGTTAGTGTGTGCTCATACGCTAATGAATACGTCCGTGTTGCAGGTA

ATCACACCATGGCCATTAACTTTTCCTGAAACAGGCATCATGAAGCCAAATTCAACATGTAAACTCACAG

TTATAATCCAAGCTCAAAGTTCATACATAGATTTTATGAGGTGCAAAGACAAAATTTTGATTCAGAGCAA

AATTTTACCTTTTCGGACAACTCAAGATGATGTTTCACCTGAAATGTTTGCAGATGATTATATAAACTAC

ACAGAAGAAAACACGCTATGGATTGTAGCTTTACCACCACTTAGCCAAGCATATTCTCCGGGATTAAATT

TAATGATCAACATAGATCGTCTACTGGGAAAGTTGGACAGTTTAGAGGCAGAGTTTAACACCTCAAAGAC

AAAGGAAGGGGGAGGCACAACCACTCGAGTGCATGTCAAACCAATTATTGAAATGTCGATCTGTGAGTTC

GTAAGAGTTTGGGGATGGAAGATTTTGTTTCCAGGCAGCGTAACTATCCTATTATTAATCTTAACCTTAC

TCTTCGTGATATATTATCGCGGCCGGGAGTGATAAGCTGCTGCCAGTCAGAAACTTTGTTATTCAAAATA

AATTTATCTTGACTGTAGTAGTTGAAATAAGTGTTTTACTTGCTGTTTCTCATGTGCTAACCATGCTACT

CCACACTTTTATTCGTCTTTCCTTCCTTATTAGTCAT

>MSTRG.1024.1 gene=MSTRG.1024

GCTGGTGAGTAGGTGGGGGAGCATACTGAGGTGGTGGGTATTGCTGGGGATAGCCCTGAGGAGGATACCC

TCCTTGAGGATACCCCTGTGGTGGATACCCTGGTTGAGGGTATCCTTGTGGAGGGTACCCCTGTGGAGGA

TACCCCTGTGGTGGATACGCATCTTTAGCCTGATCTTGTGGAGGATAACCTTGAGGAGGAGGATTGCCAA

CAGGAGGGTAACTCATGATTTTCTTGGAAGATATTAGGTGGGTTTTGTTGGTTCTCAAAGGGTTGTGAGA

AATATTTGTGTGTGTATATAACGGGGGTTGGG

>MSTRG.1026.1 gene=MSTRG.1026

CTTTTCTTCCAACAAAAGGATACTGATAACTTTGCTTGCCTTGAGTACCAAAAAATGCAAATGTGGATTC

ATTACTCACCCGTATGTTGAGTGTGCATCTTCATCATCATGCAGAGGAGATGCCACAGTCACAGCCTGGT

CATGAATTCTGCAGTACCTTTCCGAGACTGATGAAGATCTTCTCATCGAAAACACTCTCTTCACTGCACT

TGAACCCTGTTTCTCAATCTTGAACATGTTCAATGAAGATGTGTTCACTGACACTGACCTGTTTCTGCTC

AACTTGGCCATCCCAAAGCTCCCTTCACTACTATTGTTGTTATGTTCTTCTTCATTGAGTTTCTGCAGGG

TGTTGTTGATCTGTGTTTTGGGTTCTTGGGTTTTGATCTTGGTGTGTTGAGGTTCAGTGGGGATTTTGGT

TTCGAGTGGGAAAGACTGGAGATCTGAGAAATCCATGGATCTTGATTTGGTGCTGTTGTTGACCAGCTTC

ATCTTTGAGTACCTGCTCATTTTGTTTCTTAATTTGTGGTTTTGTTTTGGAGTGGAGAATGGAGTTTGTA

TTGCACTACATATACAAGGGGATAAAAGAGTTGATGGTTAAAGACTCAAGGTATCCTTCCTATCTATAAA

TAATAGATATTGTATTGAATATTTTGATGAGTGTGTAAAAGGGAAGGGAAGTGGACTTTGGAATATATTG

GTGTTACAAGATTTATACTTGAGATTTAAATACAAAAATTTATAAAAATAGTATTCTTTTCGGGTGTAC

>MSTRG.1035.6 gene=MSTRG.1035

CCCCCTTTCGCCACACTTCAACTTTCAGCCCTTTTAGTATACAGATACTCAAGTTTAGTTTCTGGTGGGA

GCTGCAGTCTACTAGCAAAGATGAGGAGCCTTACACCACCTGTTCTATTGAGAATCCAGTGTTGTAACAT

TGGAAATTGGTTTGAATCAGTTAGGGACTTAGGGCGTAGTAGTTATTCGACTTAGCGGGAAATTACACAG

AATGACATCATATTACGTTTTGTAATATTTATGGGATCTGATACATGGAAAACGGCTGCCCAGTGACGCA

CATAATATGTGCAAAAGGCTTAGCCTAGAAAGCAGGGATTCTTCACTCACCTGCCGCACCGCACACTCGA

TTCTTGGGGATTCTTTGATACTCGGGGATTCTTGTTAATGATTAAT

>MSTRG.1039.1 gene=MSTRG.1039

CTGCCTTCTTTGTACAGGATTCTTTGAGACTCATTCTCACTTGTTTTGTGAATGTCATTATTCTAGAACT

ATTTTGAATGCATGTCCTATTGCTGTTAGCACTGTCTGGACAGATATTTGTGCCGGAGGAATTCTGGTGA

ACCAAGTGGATACTATTCGTGATAATATTGCATACCTGTATGTGGCAGTAGCTTATTATCACATTTGGGC

TGAAAGGAATCTACGTCTTCATAGTCCTGGCCAATATAATCAGCCTATTATTCTAATCAGAAGAATCAAG

GATGTTGTCAGAAGCAAGTTGGTTGTTAGTAAATTATTTACTCGGTTTGCTAGACAAGATTTTACTTTAA

ATAGTTTTCTCTAGTTCTTCTGTTATCTAGTCTTTGGCTATTTGAATCTAGGAGTGTTAGCTTTTATCTC

TTTGTGACAGTTTGAGTTTCTCCCGCGGGGTATGCCCCTAGATTTGTAATCTGT

>MSTRG.1040.1 gene=MSTRG.1040

TTAGCACACATTGAATCTCTATATAACACATGCAAAAATAACACTGCACTTCAACTCAAAAATATTTCAC

ACACTTACCTTCTGCCTTCTCCTTGATCATTTACAACTAGCTAGGCAAGAAATAGAAATGTGGAAGCTGA

TCAATCTCTCATTACTTGTCATGGTCTTAATGTTTTCGGACTTGGATAAGGCGACAGGGCGCCCGGAGCC

AAATTTTCCTACCATCCCAAATTTACCATTTATACCAAGCCTGCCACCTCTAGTTTCACAAGTTCCTGGT

CTGCCCACCATTCCTGAACCTCTGTTTCCAGGTTCTTTGAGCCCAATCACCTCCCCTGTTCGCGGTCTAC

CGGGCTTTCCCAGCTTCTTTGGTCCTCCCCAAGCCCCTCCTAGCCATGATTAACAGATGTAGCTGCTAGT

GTTTAGTTGGCTTGTTAAGTGCAGGATTTGTGTTTGTGACTTTGTGGAGCACAATGTTGTTTCTTGGTAA

TAAATTTATCTATGGATGTAGTTTGTTTTCAAA

>MSTRG.1041.1 gene=MSTRG.1041

GCAAGAAAAGGGGAGAGTTGTATAGCGGTTCTGTAGTTACGGATTGTTCGAGACGAGTAGAACCAGACGT

TGGGGATCAAGACTGAAGTTAGATAAGATTTCGTATCAAGCTGCAGCAACCGCATGAGCGCCATAACAGA

AAATCAGGAATAGTTGAGGGTGATGTCTTGAGAAAGCAGAATAAGATAAGAATGTTAGCTAAGAGTGTAT

AAGCTAAGGACCAAAGCTAAGGCTAGAGATGGATTATCTGATTAGGCTGCGCATGAGAGTGATGCTAAGC

GGCAAGGATTCTAAGTGGTTAAGTTATACATCAGAGATGAGGTTGGTTGATTCTAATTTCAATACTGTAA

CCTG

>MSTRG.1042.1 gene=MSTRG.1042

CTCGCCTGAATCTCCCTCCACTGGCCAAGTCTCTCTCTCGCCGTCTCTCTCCGCGCGGCTCTTACTCCGG

CGAACCACCACTTCCTCAGCCGTTCTTCACCTCTCCCTGGGTCTCTCTCGCTGCCGTCAGCCACCAGACT

CGGCATCTCTCTCTTGCTCTGGATCGAGCATTGAAGGCGTGATACCAAGCAAAGGCAAGAAAAGGGGAGA

GTTGTATAGCGGT

>MSTRG.1045.1 gene=MSTRG.1045

CTCACGGGAGCCAGTTTTCAGTCACAGAAGTGTGACCTGTCTTGCATAAGCCAATTGATTTCTACTTGTA

CCTTGTTATCTATGGATTTTGCAAGGCAGGACACCTTCTATGTCTGATCTCTCTCCCTCTCTCTCGCGCG

ATCTCTCTCCCTCTCTCTCTCTCGATCTCTCTCTCTGTGTATAATTCAGAGCAGGTTATCAGTTATAGAA

GTGTGACCTGTCTTGCATAAGCTAGCTGATTCCTTCGTGTGACTGTAATCTATGGTCTCTGCAAGGCAGG

ACGCCTTCTATGGTTGATCCCGTTGCCGATATGTATCTTCACGTGCTCGTGTTCTTAATTATATTTTAAA

ATAATATCAAATTCGAAAAATCAACGATTTGTTTTCGATTTTCCAAGAAACCAATATTAACTTTAAAGAA

ACGGTGATTTTATTGTAATTTATCTCGGTTATGGAATTGCATTCTGGTATCCAGTACACGTTTTGCATGT

ACAGTCTTTAAGAAAATGTGTG

>MSTRG.1048.1 gene=MSTRG.1048

GAATAGATAATTAAAAGGGAAACTCCTTAAAGTTGTACTTCAACTGCCCCTGCAACAAAGCAGGACTCAA

AGTGCGCTGATAACAACAATCTTATCAGGTTTATGTGGGGCTAAGGATATTGCAGTCTCCACCTCTGCTT

TCCGCTGTAGCTGATTTAGTTGCGCGAATACCTTTTAATTAGTGGTAGGATGATTAGTACCACTATGAGT

AGACATTTTCATGTCTCCAAAGAATAACAA

>MSTRG.1050.1 gene=MSTRG.1050

GGTATGTGGGGGTAGATGAATCAGAGGATGTGCAGTTGTTCTATTATTTTGTTGAGTCGGAGAGAAATCC

GATGGTGGATCCTCTTATTCTGTGGTTGACAGGAGGCCCTGGTTGTTCTGGCTTCACTGCTCTTGCTAAT

GGAATAGGTCCACTATATTTTAGGGAAGAGAGGTACAATGGGACTTTGCCTACCCTGTATTCAAATCATA

ACACATGGACAAAGATAGCTAGTATGATATTTTTGGACTTGCCAGTCGGTACTGGTTTCTCTTACGGAAG

GACATCTGTAGCTTCTAGCTCCAATGACAGCCAAGCTTGTGCTCAAGCTGTCCAGTTCTTAAGGAAGTGG

TTACTTAGTCATCCTAAATTCCTTTCAAATCCCTTCTACGTTGGTGGAGACTCTTACGCGGGAATATTTG

TTCCAATTGTCACTCAAATCATCTCAAATG

>MSTRG.1051.1 gene=MSTRG.1051

CAGTTTTCATCATAAATCTCAAAATCTGTTGAAAACACTAACAAATTCCCAATCAATACTTATCAAATTG

TTATTTATAGGCTGCTAATCCTGACAACTCATATCTTATAGAGTCCATAACATTCTCTGCATTCTGTGAA

ATTCTTCTGAAAATGGCAGCCTACAACAATGTGATCATCCTGATGCTGGCCTCACTCCTGTTAAGATCAG

TGGTCTCCGATTCTCCGGCCCCATCGCCTTTCTCTCCGGTGCCTCAGCCGCCTTCGAGTATCGATAGTCC

AGCCCCCTCTCCAGGTGACAAGGCTCCTGCTGAGGCTCCAGTGCCAGCAGTTAAATCTCCGCCTCCTGCA

CCAGGACCAGGCCCTGTTCAGCATCTGGCTCCAACCCCTGCTGAATCTCCGAAAGCAAGTGCACCAGGGG

CGCAGTCTCCTGAGCCTTCACCGTCGATATGGACCCCGCCTGCTGAAGCCCCGGCCCCGGAAAGCAATGA

TGGAGTGAAGAGAGTGGCTGGTGGATTGGTCTGTTTTGGGGCTTTTGTTGTTGGTTTTATGGCTTAGGTC

TGTTTTGCATGCTAG

>MSTRG.1055.1 gene=MSTRG.1055

GGGATACAGTCTAAGTTACCCGGACACGGGTATGGGTATTGGACACGGGTACAGATCCAAGAGTCAGATT

CGTAGTTTTAGGATAATTAGGATTCGTGGACGTGGATCCGGAATTAGACTCGGGTACGGGGACTTGGCAC

TTAAGCTCCACGCAAGTCAAACTTTATATACCCTGATTTATGGTACATTCATTAAAATAGTGTCTACTGC

ACATAAATCACATACGCTAAAAATTGAAG

>MSTRG.1059.1 gene=MSTRG.1059

TTTCTCTTGTCAGGAGATGGAAGTTCTTTGAATGATCAGCTGAACACCACTATAAATTGGATTCCTGGAA

TAAAAGGCATTCAAATAAAAGATTTTCCGAGTTTCCTCCAGACAGCTGATTCCAATGATCACATGCTTCA

GTTTGTCATTAGTGAAGCAGAAAGATGCAAGACAACAGCCTCAGCAATAATCATCAACTCGTTCGATACC

CTGGAGTCTGATGTTTTATGTGCCCTTTCATCTATCTGTCCACCAATTTATACCATTGGCCCTCTTCAGT

TGCTAATCAACAAACTTCCAGAAAATAAGTTAAAGTCCATTGGATCAAACTTATGGAAAGAAAATGCAGA

ATGTCTGAAGTGGCTTGATACGAAGCAAACAAATTCTGTAATTTATGTAAGTTATGGGAGCATCACAGTG

ATGACACCTGAAAAGCTAGTGGAATTTGCATGGGGGCTTGCTAGTAGCCACCAAAACTTCTTGTGGATCG

TGAGGCCTGATCTGGTTGTTGGCGAGGGTGCAATATTGCCTGCTGAATTTCTTGAGGAAACTAAAGACCG

AGGGCTTATAGCCGGATGGTGTCCACAAGAACAAGTTCTTGAACACCGAGCTGTTGGAGGATTTTTGACA

CATTGTGGATGGAATTCAATGTTGGAAAGCCTATGTAGTGGCGTCCCGATGCTTTGTTGGCCATATTTTG

CTGATCAAACGACTAACTGCAAGCTGGCTTGCAGTGAATGGGGCGTTGGCATAGAGATTGATAAGAATGT

CACTAGGGAGGAAGTGGCCATGCAAGTAAGGGAACTGATACAAGGAGAGAAAGGCAAGGAACTGAGAAAA

AACGCAACGATGTGGAAGGAAAAGGCAGAGGAGGCAGTTGGATTAGATGGTTCATCAACCTCCAATCTGG

AAAAATTAGTGAAAGAGGTTTTATTTTCTGATCCCTGAAGGTGAAGGTGTGCATGAACCTTGATTGTCCA

GGAAAATGTCACTTGCAGCTCCTGAGGGCTATTGGATTAACCAAGGCACTCCATATGGGATACTCTGAAA

GTCTTTATTTCATATCAGTCTTTAATTCATTCGGATATTTAGATATTTGTACAATTTTTATCATAATCTC

TGTTCCTTTT

>MSTRG.1061.1 gene=MSTRG.1061

CCGATGCTTCAGTTCCTCGTTACTGAAGTAGAAAGATGCACGACAGCCTCAGCAATAATCTCCCACTCTT

TTAATGCTCTAGAGTCCGATGTTCTAAGTGCCCTCTCATCTATTTGTCCACCAATCCATGCCATAGGCCC

TCTTCAGTTGCTCGTCGACCAACTTCCAGAAAATAATTTGAAGTCAATTGGATCCAACTTATGGAAAGAA

GATGCAGAATGTCTAAAGTGGCTTGATACGAAGGAAACAGATTCGGTTATTTATGTCAGTTATGGAAGCA

TCACAGTAATGACACCTGAAAAGCTAGTAGAATTTGCATGGGGGCTTGCTAGTAGCAACCAAAACTTCTT

GTGGATCATTAGGCCTGATTTGGTAGTTGGTGAGGCTGCAGTATTGCCTGCTGAATTCCTCGAGGAAACC

AAAGACCGAGGTCTCATAGCAGAATGGTGTTCACAAGAACAAGTTCTTAATCATCGAGCTGTTGGTGGTT

TTTTGACACATTGTGGATGGAATTCCATGTTTGAAAGCCTATACAGCGGCGTCCCCATGCTCTGTTGGCC

ATTTTTCGCTGATCAAACGACTAACTGCAAGCTGGCTTGCAATGACTGGGGTGTCGGTATAGAGCTTGAT

AAAAATGTTACAAGAAATGCAGTGGAGTTGCATGTCAGGGAACTGATGCAAGGAGAGAAAGGCAAGGCAA

TGAGGATCAAGGCGACGGAGTGGAAGAAAAAGGCAGAGGAGGCAGTTGCATCAGATGGTTCGTCGTCCTC

TGATCTCGAAAAATTAGTGAATAAGGTCTTATTGTCCAAACCCTGAAGCCATTTTGATTTGAAAAGACAA

CAATGCATGAGTAATTCAAATCAGACC

>MSTRG.1065.1 gene=MSTRG.1065

TGCATGTGCGCGGTTACTTGATTCATCATTCTCTCCGTTTTCTTCATGAATGATTTATGCACTAGATACT

AGAATCTATTGGCTTGAATATGGAAGTCATCAGCCTCTAGCATTACTGGGGTCTCAAACCTAATGTCAGG

GACGTAGAGAAAGGTAATGGTGTACTTGGAACTGAAATCCCATATATTTACTTCGTATATCTACTAAGAA

TACTTAGTTGTTGAAGCAAAGGGACTACTTGTTGAAATGGAAGTTACAGATTCTGTGGAATGATCTCACA

ACAGCATTATTCACAGCGGATGTTTAGAATGAGAAGATAGAAAGTTGATATGTATGTGAACTTGGTACAG

AGGAATGGTATAGTACAGAGGAATGGTTTAGTAAATTGTTGCAACCCTTGCCAGCAGGGGCGGATCCAGA

AAGCCATTCCGCGCTGCCCTAGCAGAGGCGTGGTTATATTCCGGGTGTGCCAAGCTAATTTTTTTGAATA

TACCTTTATATTTTTAAACTTCATGGAGGCATACTCTTATTTTCTGAAATTTATAATGATAAAATAATAT

TACATTTTTTCAAGAGGACACATGTTCCCGTGTCCCTTAACCCTTCTCTCATTTTCTAAAATTTTTAATG

GTGAAAAACTATTAATTTTTTTAGGGGCACACATGCTCCCGGATCTTTTACTAGGCCCGTTCCCTTAAGT

TACTCAGTTATCCACTTACTCAGAATGGAATTAGAGC

>MSTRG.1073.1 gene=MSTRG.1073

AATGAGAGTCGGCCGCCATTCATAACTATCATCCATCCTTCACCGCCCCTTTCTTCCATCAAAACCACAT

CATCCCATCCATCCTCTCATCCCATTTCCTCCATTTCTATATCTTTAATTTCTACTATAATAGTTTTCAA

TAAAATCGAGATCTAAGTCCTTTTGGGTGTGATCTTGTTACCGTACCCATCTTTTTGGGTTGTTGATTGT

CCCCAATTTTTGGGTGTATGTCTTGTTGCGTTTTTGATGCGTTGTCATGTTGCGTGAAGAATGATTTCTA

CGGTGTATTGGGAGATCTGGGCAACCCACTTCAAATCTGGAATTGTGGTTATCGCGAGAGCTCACCTCGC

ACAACAAATGATTGTTCAATATATGGGGCATCTGTACCCCCAACCTCAGTTGGTGTAACTGATGGATATG

AACACTGGTCTATCACCGGTTTTTTATGTGCGCAGGTCAATTGTGATTCTACTATTTTCAGTGATGTCGG

TGTCGACTGGATTGCTCGAGAAACTTTTGTAATCATTTTTAATATCAAGAATATTTATATTCTATCTGTT

AAGCAATCTGAAAACAAAACGGCTATTTGTTTAGCTCGATTTTTGTTTCACAATCAGGTTGTAGTTGTCA

GTTTGGGGGTTTGTCCCGACTGTGTTCTTAAGTTCTATGTTTAATAAAATCTTATTGTTCGTCAAAAAAA

AAAA

>MSTRG.1076.1 gene=MSTRG.1076

AGCTGGGAAATAATGTCCATGAATTTGTTATGGGAGATAGATCACATCCACAAAGTGATGATATATATGC

AAAGCTAGCTGAGATTACAATGTTGTTACGACTAGAAGGTTATATGCCTCGCACCACCAATGTGCTCGCC

GACATTGAAGAGGAGGAAAAGGAGACTGCTCTATCATATCACAGTGAGAAGATTGCAATTGCATTCATGC

TCATTAGTACCCCACCTTGCACCCCAATCAGTGTTGTAAAGAATCTGAGGGTTTGTGCAGACTGCCATTT

AGCAATAAAACTCATATCCAAGGTTTACAACCGAGAAATTGTTGTGAGAGATCGCAGTAGATTCCACCAT

TTTAGAAATGGTTCATGTTCTTGTAGAGATTATTGGTAAACATTTGTTTACTTTATCAAAATTTTAAAGA

AATGCTAATGATGCTCTCTGTAATTAT

>MSTRG.1079.1 gene=MSTRG.1079

TTTATAACAACACATAACAAACTCAAATGGATTCTTTCCTTAGCAAACTCTCTATTCTATTTGTTCTTCT

AGTCGCCACACCCTTAGCTCTAGCTCAGACTAGGGTCGGCTTCTATTCCACCTCTTGTCCCAGAGCGGAA

TCCATCGTTCAAGCCACCGTTAAGAAACATGTTAACTCCGATTCCACTGTTGCCCCCGGACTGCTCAGAA

TGGTCTTCCATGATTGCTTCGTTCATGGTTGCGATGCTTCGATTCTTATCGATGGCAGCAATGCTGAGAA

AACCGCTAGGCCAAATCTTCTGCTGAGAGGATATGAAGTGATTGAGGATGCCAAAACACAGCTAGAGGCA

GCATGCCCCGGAGTCGTGTCGTGTGCTGATATTCTTGCTCTTGCTGCTCGTGATGCCACAGTAGAAGTCG

GTGCACTGACTTATCGCGTGCCTACTGGCCGGAGAGACGGACGAGTGTCATTGGCGTCCGACACCGCCAA

TTTGCCGAGTTTCACAGACTCAGTCGCGGTCCAGAAGAAAAAATTTGCAGATGTTGGTCTAAGCGCTCAA

GATCTTGTTACCCTTGTTGGAGGACACACTATTGGAACAGTGGCTTGCCAATTCGTGAGCTACAGATTAT

ACAACTTCACAAACTCCGGGGGTGCTGATCCTACCATCGATCCCTCGTTCCTTCCTACACTCCAATCGCT

TTGCCCTAAAAATGGCGATGGAAATAAACGTATCGCTTTGGACTACGGGAGCGGTAACAATTTTGATCGT

TCTTTCTTCAAAAATGTTCGAGATGGACGGGGAATTCTGGAGTCAGATCAGAGCTTGTGGCAGGACAGTA

CAACGAGGAACTTTGCGCAGAGATTTATGGGTATCAGAGGCTTGGCTGGATTGACTTTGAACGTCGAATT

CGGAAAATCTATGATTAAAATGACGAATATTGGGGTGAACACTGGGACGGCCGGGGAAATTCGAAAGGTC

TGTTCAAGAGTTAATTAGAGTCGAACTTTGATATTCCAGTTCTAGAGTTATGAAAACTTGAGTAGGCGGA

TGCAAAATAAGCCTGCAGTTGTAAAAGCTTCCACGAACAGCAAAGAAAGTTCATTTACAAACAAACACGT

ATTGTTTGAGTTATATATAAAGAGTATAAAGCGTCTTCAAGAACTAAGAAACAGATGTTTTTCATGTACA

ATATATATATATTTTTATATTAGGAATGGTCACTTAATGAAATACAATTTATACTTTGCTTGCTGGGCCA

>MSTRG.1079.4 gene=MSTRG.1079

TAACAACACATAACAAACTCAAATGGATTCTTTCCTTAGCAAACTCTCTATTCTATTTGTTCTTCTAGTC

GCCACACCCTTAGCTCTAGCTCAGACTAGGGTCGGCTTCTATTCCACCTCTTGTCCCAGAGCGGAATCCA

TCGTTCAAGCCACCGTTAAGAAACATGTTAACTCCGATTCCACTGTTGCCCCCGGACTGCTCAGAATGGT

CTTCCATGATTGCTTCGTTCATGGTTGCGATGCTTCGATTCTTATCGATGGCAGCAATGCTGAGAAAACC

GCTAGGCCAAATCTTCTGCTGAGAGGATATGAAGTGATTGAGGATGCCAAAACACAGCTAGAGGCAGCAT

GCCCCGGAGTCGTGTCGTGTGCTGATATTCTTGCTCTTGCTGCTCGTGATGCCACAGTAGAAGTCGGTGC

ACTGACTTATCGCGTGCCTACTGGCCGGAGAGACGGACGAGTGTCATTGGCGTCCGACACCGCCAATTTG

CCGAGTTTCACAGACTCAGTCGCGGTCCAGAAGAAAAAATTTGCAGATGTTGGTCTAAGCGCTCAAGATC

TTGTTACCCTTGTTGGAGGACACACTATTGGAACAGTGGCTTGCCAATTCGTGAGCTACAGATTATACAA

CTTCACAAACTCCGGGGGTGCTGATCCTACCATCGATCCCTCGTTCCTTCCTACACTCCAATCGCTTTGC

CCTAAAAATGGCGATGGAAATAAACGTATCGCTTTGGACTACGGGAGCGGTAACAATTTTGATCGTTCTT

TCTTCAAAAATGTTCGAGATGGACGGGGAATTCTGGAGTCAGATCAGAGCTTGTGGCAGGACAGTACAAC

GAGGAACTTTGCGCAGAGATTTATGGGTATCAGAGGCTTGGCTGGATTGACTTTGAACGTCGAATTCGGA

AAATCTATGATTAAAATGACGAATATTGGGGTGAACACTGGGACGGCCGGGGAAATTCGAAAGGTCTGTT

CAAGAGTTAATTAGAGTCGAACTTTGATATTCCAGTTCTAGAGTTATGAAAACTTGAGTAGGCGGATGCA

AAATAAGCCTGCAGTTGTAAAAGCTTCCACGAACAGCAAAGAAAGTTCATTTACAAACAAACACGTATTG

TTTGAGTTATATATAAAGAGTATAAAGCGTCTTCAAGAACTAAGAAACAGATGTTTTTCATGTACAATAT

ATATATATTTTTATATTAGGAATGGTCACTTAATGAAATACAATTTATACTTTGCTTGCTGGGCCA

>MSTRG.1079.2 gene=MSTRG.1079

TTTATAACAACACATAACAAACTCAAATGGATTCTTTCCTTAGCAAACTCTCTATTCTATTTGTTCTTCT

AGTCGCCACACCCTTAGCTCTAGCTCAGACTAGGGTCGGCTTCTATTCCACCTCTTGTCCCAGAGCGGAA

TCCATCGTTCAAGCCACCGTTAAGAAACATGTTAACTCCGATTCCACTGTTGCCCCCGGACTGCTCAGAA

TGGTCTTCCATGATTGCTTCGTTCATGGTTGCGATGCTTCGATTCTTATCGATGGCAGCAATGCTGAGAA

AACCGCTAGGCCAAATCTTCTGCTGAGAGGATATGAAGTGATTGAGGATGCCAAAACACAGCTAGAGGCA

GCATGCCCCGGAGTCGTGTCGTGTGCTGATATTCTTGCTCTTGCTGCTCGTGATGCCACAGTAGAAGTCG

GTGCACTGACTTATCGCGTGCCTACTGGCCGGAGAGACGGACGAGTGTCATTGGCGTCCGACACCGCCAA

TTTGCCGAGTTTCACAGACTCAGTCGCGGTCCAGAAGAAAAAATTTGCAGATGTTGGTCTAAGCGCTCAA

GATCTTGTTACCCTTGTTGGAGGACACACTATTGGAACAGTGGCTTGCCAATTCGTGAGCTACAGATTAT

ACAACTTCACAAACTCCGGGGGTGCTGATCCTACCATCGATCCCTCGTTCCTTCCTACACTCCAATCGCT

TTGCCCTAAAAATGGCGATGGAAATAAACGTATCGCTTTGGACTACGGGAGCGGTAACAATTTTGATCGT

TCTTTCTTCAAAAATGTTCGAGATGGACGGGGAATTCTGGAGTCAGATCAGAGCTTGTGGCAGGACAGTA

CAACGAGGAACTTTGCGCAGAGATTTATGGGTATCAGAGGCTTGGCTGGATTGACTTTGAACGTCGAATT

CGGAAAATCTATGATTAAAATGACGAATATTGGGGTGAACACTGGGACGGCCGGGGAAATTCGAAAGGTC

TGTTCAAGAGTTAATTAGAGTCGAACTTTGATATTCCAGTTCTAGAGTTATGAAAACTTGAGTAGGCGGA

TGCAAAATAAGCCTGCAGTTGTAAAAGCTTCCACGAACAGCAAAGAAAGTTCATTTACAAACAAACACGT

ATTGTTTGAGTTATATATAAAGAGTATAAAGCGTCTTCAAGAACTAAGAAACAGATGTTTTTCATGTACA

ATATATATATATTTTTATATTAGGAATGGTCACTTAATGAAATACAATTTATACTTTGCTTGCTGGGCCA

>MSTRG.1079.3 gene=MSTRG.1079

TTTATAACAACACATAACAAACTCAAATGGATTCTTTCCTTAGCAAACTCTCTATTCTATTTGTTCTTCT

AGTCGCCACACCCTTAGCTCTAGCTCAGACTAGGGTCGGCTTCTATTCCACCTCTTGTCCCAGAGCGGAA

TCCATCGTTCAAGCCACCGTTAAGAAACATGTTAACTCCGATTCCACTGTTGCCCCCGGACTGCTCAGAA

TGGTCTTCCATGATTGCTTCGTTCATGGTTGCGATGCTTCGATTCTTATCGATGGCAGCAATGCTGAGAA

AACCGCTAGGCCAAATCTTCTGCTGAGAGGATATGAAGTGATTGAGGATGCCAAAACACAGCTAGAGGCA

GCATGCCCCGGAGTCGTGTCGTGTGCTGATATTCTTGCTCTTGCTGCTCGTGATGCCACAGTAGAAGTCG

GTGCACTGACTTATCGCGTGCCTACTGGCCGGAGAGACGGACGAGTGTCATTGGCGTCCGACACCGCCAA

TTTGCCGAGTTTCACAGACTCAGTCGCGGTCCAGAAGAAAAAATTTGCAGATGTTGGTCTAAGCGCTCAA

GATCTTGTTACCCTTGTTGGTAAGCAATAAGCCAATATTACATACGTTTATCAGACATCCGAAATTTTTC

TATCTGCATGCATGCAACATTCTAAACCATATAGTACAAAAATGTGTTGCAGGAGGACACACTATTGGAA

CAGTGGCTTGCCAATTCGTGAGCTACAGATTATACAACTTCACAAACTCCGGGGGTGCTGATCCTACCAT

CGATCCCTCGTTCCTTCCTACACTCCAATCGCTTTGCCCTAAAAATGGCGATGGAAATAAACGTATCGCT

TTGGACTACGGGAGCGGTAACAATTTTGATCGTTCTTTCTTCAAAAATGTTCGAGATGGACGGGGAATTC

TGGAGTCAGATCAGAGCTTGTGGCAGGACAGTACAACGAGGAACTTTGCGCAGAGATTTATGGGTATCAG

AGGCTTGGCTGGATTGACTTTGAACGTCGAATTCGGAAAATCTATGATTAAAATGACGAATATTGGGGTG

AACACTGGGACGGCCGGGGAAATTCGAAAGGTCTGTTCAAGAGTTAATTAGAGTCGAACTTTGATATTCC

AGTTCTAGAGTTATGAAAACTTGAGTAGGCGGATGCAAAATAAGCCTGCAGTTGTAAAAGCTTCCACGAA

CAGCAAAGAAAGTTCATTTACAAACAAACACGTATTGTTTGAGTTATATATAAAGAGTATAAAGCGTCTT

CAAGAACTAAGAAACAGATGTTTTTCATGTACAATATATATATATTTTTATATTAGGAATGGTCACTTAA

TGAAATACAATTTATACTTTGCTTGCTGGGCCA

>MSTRG.1079.5 gene=MSTRG.1079

TTTATAACAACACATAACAAACTCAAATGGATTCTTTCCTTAGCAAACTCTCTATTCTATTTGTTCTTCT

AGTCGCCACACCCTTAGCTCTAGCTCAGACTAGGGTCGGCTTCTATTCCACCTCTTGTCCCAGAGCGGAA

TCCATCGTTCAAGCCACCGTTAAGAAACATGTTAACTCCGATTCCACTGTTGCCCCCGGACTGCTCAGAA

TGGTCTTCCATGATTGCTTCGTTCATGGTTGCGATGCTTCGATTCTTATCGATGGCAGCAATGCTGAGAA

AACCGCTAGGCCAAATCTTCTGCTGAGAGGATATGAAGTGATTGAGGATGCCAAAACACAGCTAGAGGCA

GCATGCCCCGGAGTCGTGTCGTGTGCTGATATTCTTGCTCTTGCTGCTCGTGATGCCACAGTAGAAGTCG

GTGCACTGACTTATCGCGTGCCTACTGGCCGGAGAGACGGACGAGTGTCATTGGCGTCCGACACCGCCAA

TTTGCCGAGTTTCACAGACTCAGTCGCGGTCCAGAAGAAAAAATTTGCAGATGTTGGTCTAAGCGCTCAA

GATCTTGTTACCCTTGTTGGAGGACACACTATTGGAACAGTGGCTTGCCAATTCGTGAGCTACAGATTAT

ACAACTTCACAAACTCCGGGGGTGCTGATCCTACCATCGATCCCTCGTTCCTTCCTACACTCCAATCGCT

TTGCCCTAAAAATGGCGATGGAAATAAACGTATCGCTTTGGACTACGGGAGCGGTAACAATTTTGATCGT

TCTTTCTTCAAAAATGTTCGAGATGGACGGGGAATTCTGGAGTCAGATCAGAGCTTGTGGCAGGACAGTA

CAACGAGGAACTTTGCGCAGAGATTTATGGGTATCAGAGGCTTGGCTGGATTGACTTTGAACGTCGAATT

CGGAAAATCTATGATTAAAATGACGAATATTGGGGTGAACACTGGGACGGCCGGGGAAATTCGAAAGGTC

TGTTCAAGAGTTAATTAGAGTCGAATTCGGAAAATCTATGATTAAAATGACGAATATTGGGGTGAACACT

GGGACGGCCGGGGAAATTCGAAAGGTCTGTTCAAGAGTTAATTAGAGTCGAACTTTGATATTCCAGTTCT

AGAGTTATGAAAACTTGAGTAGGCGGATGCAAAATAAGCCTGCAGTTGTAAAAGCTTCCACGAACAGCAA

AGAAAGTTCATTTACAAACAAACACGTATTGTTTGAGTTATATATAAAGAGTATAAAGCGTCTTCAAGAA

CTAAGAAACAGATGTTTTTCATGTACAATATATATATATTTTTATATTAGGAATGGTCACTTAATGAAAT

ACAATTTATACTTTGCTTGCTGGGCC

>MSTRG.1079.6 gene=MSTRG.1079

TTTATAACAACACATAACAAACTCAAATGGATTCTTTCCTTAGCAAACTCTCTATTCTATTTGTTCTTCT

AGTCGCCACACCCTTAGCTCTAGCTCAGACTAGGGTCGGCTTCTATTCCACCTCTTGTCCCAGAGCGGAA

TCCATCGTTCAAGCCACCGTTAAGAAACATGTTAACTCCGATTCCACTGTTGCCCCCGGACTGCTCAGAA

TGGTCTTCCATGATTGCTTCGTTCATGGTTGCGATGCTTCGATTCTTATCGATGGCAGCAATGCTGAGAA

AACCGCTAGGCCAAATCTTCTGCTGAGAGGATATGAAGTGATTGAGGATGCCAAAACACAGCTAGAGGCA

GCATGCCCCGGAGTCGTGTCGTGTGCTGATATTCTTGCTCTTGCTGCTCGTGATGCCACAGTAGAAGTCG

GTGCACTGACTTATCGCGTGCCTACTGGCCGGAGAGACGGACGAGTGTCATTGGCGTCCGACACCGCCAA

TTTGCCGAGTTTCACAGACTCAGTCGCGGTCCAGAAGAAAAAATTTGCAGATGTTGGTCTAAGCGCTCAA

GATCTTGTTACCCTTGTTGGAGGACACACTATTGGAACAGTGGCTTGCCAATTCGTGAGCTACAGATTAT

ACAACTTCACAAACTCCGGGGGTGCTGATCCTACCATCGATCCCTCGTTCCTTCCTACACTCCAATCGCT

TTGCCCTAAAAATGGCGATGGAAATAAAC

>MSTRG.1080.1 gene=MSTRG.1080

AGTCCCGCCCGCTGCCTACTGAATCCAGTCATCGTCCAAAAAGCAGTAGCAAACTACTACCAGGTCGTCC

TCCCGTCAGCTTTGTTTCCATACATATTTATGCAATCAAATTCACCATCCTCCAACCCCTTTTCACAACC

CTCCACCATTCCAATCCCCACTCTCTTCTCCTCCCAGCAAACCCCAAGCTCTTACCTCATCAAGAAATGC

ATAGCTCTCCTCCTCTCTTGCGCTTCTTCAACCCCCAAGCTCCGCCAAGTCCACGCCTTCTCCATCCGAC

ATGGCCTCCCTCTCACCAACCCAGATATGGGCAAGCACCTC

>MSTRG.1081.1 gene=MSTRG.1081

TTTATAACAACACATAACAAACTCAAATGGATTCTTTCCTTAGCAAACTCTCTATTCTATTTGTTCTTCT

AGTCGCCACACCCTTAGCTCTAGCTCAGACTAGGGTCGGCTTCTATTCCACCTCTTGTCCCAGAGCGGAA

TCCATCGTTCAAGCCACCGTTAAGAAACATGTTAACTCCGATTCCACTGTTGCCCCCGGACTGCTCAGAA

TGGTCTTCCATGATTGCTTCGTTCATGGTTGCGATGCTTCGATTCTTATCGATGGCAGCAATGCTGAGAA

AACCGCTAGGCCAAATCTTCTGCTGAGAGGATATGAAGTGATTGAGGATGCCAAAACACAGCTAGAGGCA

GCATGCCCCGGAGTCGTGTCGTGTGCTGATATTCTTGCTCTTGCTGCTCGTGATGCCACAGTAGAAGTCG

GTGCACTGACTTATCGCGTGCCTACTGGCCGGAGAGACGGACGAGTGTCATTGGCGTCCGACACCGCCAA

TTTGCCGAGTTTCACAGACTCAGTCGCGGTCCAGAAGAAAAAATTTGCAGATGTTGGTCTAAGCGCTCAA

GATCTTGTTACCCTTGTTGGAGGACACACTATTGGAACAGTGGCTTGCCAATTCGTGAGCTACAGATTAT

ACAACTTCACAAACTCCGGGGGTGCTGATCCTACCATCGATCCCTCGTTCCTTCCTACACTCCAATCGCT

TTGCCCTAAAAATGGCGATGGAAATAAACGTATCGCTTTGGACTACGGGAGCGGTAACAATTTTGATCGT

TCTTTCTTCAAAAATGTTCGAGATGGACGGGGAATTCTGGAGTCAGATCAGAGCTTGTGGCAGGACAGTA

CAACGAGGAACTTTGCGCAGAGATTTATGGGTATCAGAGGCTTGGCTGGATTGACTTTGAACGTCGAATT

CGGAAAATCTATGATTAAAATGACGAATATTGGGGTGAACACTGGGACGGCCGGGGAAATTCGAAAGGTC

TGTTCAAGAGTTAATTAGAGTCGAACTTTGATATTCCAGTTCTAGAGTTATGAAAACTTGAGTAGGCGGA

TGCAAAATAAGCCTGCAGTTGTAAAAGCTTCCACGAACAGCAAAGAAAGTTCATTTACAAACAAACACGT

ATTGTTTGAGTTATATATAAAGAGTATAAAGCGTCTTCAAGAACTAAGAAACAGATGTTTTTCATGTACA

ATATATATATATTTTTATATTAGGAATGGTCACTTAATGAAATACAATTTATACTTTGCTTGCTGGGCCA

>MSTRG.1082.1 gene=MSTRG.1082

CTTATAACAACACATTACAAGCTCAAATGGATTCTTTCCTTAACAAATTTTCTGTTCTAGTGCTTCTTCT

ACTTGTCACGCCTTTGGCATTGGCACAAACGAGGGTAGGGTTTTATTCTAAGTCATGTCCGAGAGCGGAA

TCTATTGTTCAAGCCACCGTTAAGAAACATGTCAAGTCCAATTCTGCGGTTGCTCCGGGATTGCTTAGAA

TGGTCTTCCATGATTGTTTCGTTCATGGTTGCGATGCTTCGATTCTTATCAATGGCAGCAATGCTGAGAG

AACGGCTAGGCCAAATCTTCTGCTGAGAGGATATGATGTGATTGATGATGCCAAAACACAACTAGAGGCA

GCATGCCCCGGAGTTGTGTCGTGTGCTGATATTCTTGCTCTTGCTGCTCGCGACGCTACGGTTGAAGTCG

GTGCATTGACTTACCGCGTGCCAACTGGACGCAGAGATGGACGAGTCTCATTAGCATCTGATACCTCCAA

TTTGCCGAGTTTCACAGATTCAGTCACTGTTCAGAGGAAGAAATTCTCAGACGTAGGTCTCAGCGCTCAA

GATCTTGTCACTCTTGTTGGAGGACACACTATTGGAACCGTTGCATGCCAATTCGTAAGTTACAGACTAT

ACAACTTCAAAAATACTAGCGGTGCGGATCCAACAATCAATTCCAAATTCCTTCCTACACTCCAATCTCT

TTGCCCGCAAAATGGTGATGGAAACAAACGAATCCCGTTAGATTATGGAAGTGGCGACAATTTTGATCAT

TCGTTCTTCAAAAATGTACGAGATGGACGGGGAATTCTGGAATCAGATCAGAGCTTGTGGCAGGACGGTA

CAACAAAGAATTATGCACAGAGATTTATGGGTATTAGAGGATTGGCTGGACTGACTCTGAATGTTGAGTT

TGGAAAGGCCATGATCAAAATGACGAATGTTGGAGTGAAGACGGGGAGTGCCGGTGAAATTCGCAAAATT

TGTTCAGCAATTAATTAAAAAGGATGTAGAATTTGGGGGGTTTTCCAAAACCCTACTTTTTTTTAAACAA

TAACAATTTGAAACTTGAGTAGGGAAGGTTATGTAATATAAGCATGCAATTCCATTTATCAGTCGCAAAA

TAGTGCTCCCCTCCTTTTACATCTTATTCGAATTTGTCC

>MSTRG.1083.1 gene=MSTRG.1083

CAAAACACAAAGCAAGCAGAAATGGATTCTTTCCTTAACCAGTTATCTGTTCTAGTGCTTCTTTTACTAG

TCACCCCCTCGGCTTTGGCACAAACCAAGGTCGGCTTTTATTCCACTTCGTGTCCCAAAGCCGAGACCAT

TGTTTCAGACACTGTTAAAAAGCATGTTAAGGCCAACACTGCTGTCGCTCCGGGATTGCTCAGAATGATC

TTCCATGATTGCTTCGTTCATGGTTGCGACGCTTCCATTCTTATCGATGGCAGCGATGCGGAGAAAACCG

CGACGCCGAATCTTCTGTTGAGAGGATATGATGTCATTGAAGATGCCAAAACACAACTGGAAGCAGCATG

CCCTGGAGTGGTGTCTTGTTCCGATATTCTCGCTCTTGCTGGTCGGGATTCCACTGTTGCGGTTGGTGGA

TTGACGTACAAGGTGCCAACAGGACGGAGAGATGGACGAGTGTCGCTAGCCTCCGACACCGCCAATCTGC

CAAGTTTCAGGGATTCTGTCGATGTGCAGAAGAAGAAATTTTCGGATGTTGGTCTGAGCGCTCAAGACCT

TGTGACTCTTGTTGGAGGGCACACAATTGGAACAGTGGCTTGCCAGTTCGTAAGTTACAGACTATACAAC

TTCACAAGCTCCGGGGGTCCTGATCCCACCATCGACTCCACATTCCTTTCTACACTCCAATCTCTTTGCC

CCCAAGGCGGTGATGGAAATAAGCGTATCGCTTTAGACTATGGAAGTGGGAACAATTTCGATAACTCCTT

TTTCAAAAATGTGCAAAATAACCGCGGAATCCTGGAATCAGATCAAAGCCTGTGGCAGGACAGTACAACA

CAGAACTATGTGAAAAGGTATATTGGAATAAGAGGCTTGGCTGGATTGACTTTGAACGTTGAGTTTGCCA

AAGCCATGATCAAAATGACTAACATTGGAGTCAAAACTGGCACAGATGGTGAAATTCGTAAAGTTTGTTC

AAAAATTAATTAAGGAGATCAAAAATGCAAGTTTTGGGGTTATTTAAATTTTGCTCAACTACTGGAAGAA

TTGTCCATAAGATCAGTGTATTTGATTTACGTACAGTTTGTCAAGTTCAATGATGTAATACCATATTACT

ATATATATATCCAGC

>MSTRG.1084.1 gene=MSTRG.1084

CAAAACACAAAGCAAGCAGAAATGGATTCTTTTCTTAAACAAGTATCTGTTCTAGTGCTTCTTCTACTAG

TCACCCCCTCGGCTTTGGCACAAACCAAGGTCGGCTTTTATTCCACTTCGTGTCCTAAAGCTGAGACCAT

TGTTTCAGACACTGTTAAGAAGCATGTTAAGGCCAACACTGCCGTGGCTCCGGGATTGCTGAGAATGATC

TTCCATGATTGCTTCGTTCATGGCTGCGACGCTTCCATTCTTATCGATGGCAGCGATGCTGAGAAAACCG

CTACGCCGAATCTTCTGTTGAGAGGATATGATGTGATTGAAGATGCGAAGACACAACTGGAGGCTGCATG

CCCTGGTGTGGTGTCTTGTTCCGATATTCTCGCTCTTGCTGGCCGTGATTCTACTGTTGCGGTTGGTGGA

TTGACATACAAGGTGCCAACAGGACGGAGAGATGGACGAGTGTCGCTAGCCTCCGACACCGCCAATCTGC

CGAGTTTCAGGGATTCAGTCGATGTCCAGAAGAAGAAATTTTCAGATGTTGGTCTGAGCGCTCAGGATCT

TGTTACTCTTGTTGGAGGACACACAATTGGAACAGTGGCTTGCCAGTTCGTAAGTTACAGACTCTACAAC

TTCACGAGCTCCGGGGGTCCTGATCCCACCATCGATTCCACATTCCTCTCTACACTCCAATCTCTTTGCC

CCCAAGGCGGTGATGGAAATAAGCGTATCGCTTTGGACTATGGAAGTGGGGACGATTTCGATAACTCCTT

TTTCAAGAATGTGCAAAATAACCGCGGAATTCTGGAATCAGACCAAAGCCTGTGGCAGGACAATACAACA

CAGACTTATGTGAAGAGGTATATTGGTATAAGAGGCTTGGCTGGATTGACTTTGAACGTTGAGTTTGCTA

AATCCATGATTAAAATGACCAACGTTGGAGTGAAAACTGGCACGGATGGTGAAATTCGTAAAGTTTGTTC

AAAAATTAATTAAGGAGATCAAAATTTACAGGTTCGGG

>MSTRG.1092.1 gene=MSTRG.1092

AGGAAGGCTGGCATCGCCCAGCAAAACTTGCACAACACTTCTCATACTAGGTCTGTGATCAGGATTAGGA

TGAGCGCAAAGCAGTCCGAGACTGAGCACGAGCTCAGCTTCATATTGATTGTATTCCTCCAAATTAGGAT

CAACTGCTCGAAGAATCTCTCCTTTGGAGTGTAGTTCCCTAACCCAGTCTACTAATACCAGTTCTTCAGC

AATTCTCCGGGGTTCAATTGGCCTCCTTCCACATGCCACCTCTAGCATTAGTGCACCATAACCATACACG

TCTGTGCTTGTTGTAGCCTTGCCTGTTCTGGTCAGTTCAGGGGCCAGGTAACCCAGTGTTCCTACGATGT

GCGTTGTCTGTGGGTTCAAACCATGACCATAAGTCCTGGATAAACCAAAATCTCCAAGTTTTGGTGTGAG

ATCTGCATCTATAAGGACGTTGCTCGGCTTCACGTCTCTGTGAACTACCATCTGCTCACACTCTTCATGA

AGGTATAACAATGCGTGTGCAACACCAGTGAGTATCTTAAATCTTTCTTGCCACGTCAGTAGCTTCTTTT

CCTGCTCATGTTTGTTGAACAGTAGATCATCCAGGCTTCCATTGACCACGTAGTCATATACAAGATGCAG

CTCGTCTTGTCTTCGGCACCATCCATGTAGCTGAACAAGATTACGATGCCTAAGC

>MSTRG.1096.1 gene=MSTRG.1096

CATAAAACACATCCAAAAACAATCACAATGATACAGAACACACTCAGTTTAAAACCAAAACGACCCAACA

CACGCTCTCTGCCCAAGAACCCTAAAAATCGCCGCACATCGAATTCACCGGCGTTCATCATCTCCGTCGC

TGTTCAGGCTCTCCAAGAGCTCTACACCCTCAGTTCCGCCTTTAATTCTTCAGTACCGAGGTCTTTCCTG

TGCATCTTGCTGGTAGATGCTCTATTTCGAGTGCTCGAAGCTTGAGCTTAATCAGTCAATTGGGGGCTTA

GTTCTTGAGTTGTTACCCCTTGGAAGATCGGCATCTTTTCCTCTCTAACTATTTTCGATTTGTTCTGGGG

GCTGACACATTAACTCGGATTCTCCTTCCTAGAGGCTGCGTTAATATATAGCTGAAGATTTCAGGAATTG

GTGTGAAAGTGAATGTGTCGGACTCAAAGGGATAGAAGAATGTTCTGACAATTATAATTGGATGCTTACT

TGCTGGAATTGGTATTGGCATCTCATCAACCATTGTGCCACTGTACATATCTGAGGCATTAGAACTTTTC

CGGAAGCAGGGAGTGGATTTTTTGATTGCTCCAGATGTTGCATTTCTACCAGTGAACATTTGTATCCGTG

TGGGTGCACATTGTGGTTTGTAAAATAGTTGCCAAAGCAGATCAGATTCAGATGTATATGGCCATGCAAC

ATCAAACCTTGTAGCTGGAGTGGTTTTGAAAAGGACGTCTGGACCTAGACAGCCGAGGGAAGGCCTTTTG

TTGGTTATTTTGTCTAGTTAAAGATTTTTCCAAATTTGGGGTAGATGGTTTATGGAATGACATGAATGAG

CCTGCTGTCTTTGAGGGGAGATGCTGAACTTGGAGGATGCCAAAATCACGCACATTATCACAATGTGTAT

GGAATGTTAATGGCCAGATCAACATATGAAGGCATGAAAGTAGCTGATGAGAGTAATAATCTATCAGGGA

GCATCTATATATGAGCATATCCATGGTTCCCCAACTGTAGGCTGCAAGGAGTCGCAGGACTTATTCTAAG

ACGGACCTCGACATCATTGGTTGGAGTCGCTAAATTATATCCAGTCTATTATGTAGTTTGGATTCTCATA

AGTTAGACAGATTAATTGCATGTTATCGGATATTTTAAGATGCACATTTTAAATTTATGTTTAATTAGAT

GTTTGGAGTAGATAAATTTGAATGTTGGACTAGTTGTATTGGATGAATGTTGGATCTACATTGG

>MSTRG.1096.2 gene=MSTRG.1096

CATAAAACACATCCAAAAACAATCACAATGATACAGAACACACTCAGTTTAAAACCAAAACGACCCAACA

CACGCTCTCTGCCCAAGAACCCTAAAAATCGCCGCACATCGAATTCACCGGCGTTCATCATCTCCGTCGC

TGTTCAGGCTCTCCAAGAGCTCTACACCCTCAGTTCCGCCTTTAATTCTTCAGTACCGAGGTCTTTCCTG

TGCATCTTGCTGGTAGATGCTCTATTTCGAGTGCTCGAAGCTTGAGCTTAATCAGTCAATTGGGGGCTTA

GTTCTTGAGTTGTTACCCCTTGGAAGATCGGCATCTTTTCCTCTCTAACTATTTTCGATTTGTTCTGGGG

GCTGACACATTAACTCGGATTCTCCTTCCTAGAGGCTGCGTTAATATATAGCTGAAGATTTCAGGAATTG

GTGTGAAAGTGAATGTGTCGGACTCAAAGGGATAGAAGAATGTTCTGACAATTATAATTGGATGCTTACT

TGCTGGAATTGGTATTGGCATCTCATCAACCATTGTGCCACTGTACATATCTGAGGCATTAGAACTTTTC

CGGAAGCAGGGAGTGGATTTTTTGATTGCTCCAGATGTTGCATTTCTACCAGTGAACATTTGTATCCGTG

TGGGTGCACATTGTGGTTTGTAAAATAGTTGCCAAAGCAGATCAGATTCAGATGTATATGGCCATGCAAC

ATCAAACCTTGTAGCTGGAGTGGTTTTGAAAAGGACGTCTGGACCTAGACAGCCGAGGGAAGGCCTTTTG

TTGGTTATTTTGTCTAGTTAAAGATTTTTCCAAATTTGGGGTAGATGGTTTATGGAATGACATGAATGAG

CCTGCTGTCTTTGAGGGGAGATGCTGAACTTGGAGGATGCCAAAATCACGCACATTATCACAATGTATAT

TATATCAACATATGAAGGCATGAAAGTAGCTGATGAGAGTAATAATCTATCAGGGAGCATCTATATATGA

GCATATCCATGGTTCCCCAACTGTAGGCTGCAAGGAGTCGCAGGACTTATTCTAAGACGGACCTCGACAT

CATTGGTTGGAGTCGCTAAATTATATCCAGTCTATTATGTAGTTTGGATTCTCATAAGTTAGACAGATTA

ATTGCATGTTATCGGATATTTTAAGATGCACATTTTAAATTTATGTTTAATTAGATGTTTGGAGTAGATA

AATTTGAATGTTGGACTAGTTGTATTGGATGAATGTTGGATCTACATTGG

>MSTRG.1096.3 gene=MSTRG.1096

CATAAAACACATCCAAAAACAATCACAATGATACAGAACACACTCAGTTTAAAACCAAAACGACCCAACA

CACGCTCTCTGCCCAAGAACCCTAAAAATCGCCGCACATCGAATTCACCGGCGTTCATCATCTCCGTCGC

TGTTCAGGCTCTCCAAGAGCTCTACACCCTCAGTTCCGCCTTTAATTCTTCAGTACCGAGGTCTTTCCTG

TGCATCTTGCTGGTAGATGCTCTATTTCGAGTGCTCGAAGCTTGAGCTTAATCAGTCAATTGGGGGCTTA

GTTCTTGAGTTGTTACCCCTTGGAAGATCGGCATCTTTTCCTCTCTAACTATTTTCGATTTGTTCTGGGG

GCTGACACATTAACTCGGATTCTCCTTCCTAGAGGCTGCGTTAATATATAGCTGAAGATTTCAGGAATTG

GTGTGAAAGTGAATGTGTCGGACTCAAAGGGATAGAAGAATGTTCTGACAATTATAATTGGATGCTTACT

TGCTGGAATTGGTATTGGCATCTCATCAACCATTGTGCCACTGTACATATCTGAGGCATTAGAACTTTTC

CGGAAGCAGGGAGTGGATTTTTTGATTGCTCCAGATGTTGCATTTCTACCAGTGAACATTTGTATCCGTG

TGGGTGCACATTGTGGTTTGTAAAATAGTTGCCAAAGCAGATCAGATTCAGATGTATATGGCCATGCAAC

ATCAAACCTTGTAGCTGGAGTGGTTTTGAAAAGGACGTCTGGACCTAGACAGCCGAGGGAAGGCCTTTTG

TTGGTTATTTTGTCTAGTTAAAGATTTTTCCAAATTTGGGGTAGATGGTTTATGGAATGACATGAATGAG

CCTGCTGTCTTTGAGGGGAGATGCTGAACTTGGAGGATGCCAAAATCACGCACATTATCACAATATCAAC

ATATGAAGGCATGAAAGTAGCTGATGAGAGTAATAATCTATCAGGGAGCATCTATATATGAGCATATCCA

TGGTTCCCCAACTGTAGGCTGCAAGGAGTCGCAGGACTTATTCTAAGACGGACCTCGACATCATTGGTTG

GAGTCGCTAAATTATATCCAGTCTATTATGTAGTTTGGATTCTCATAAGTTAGACAGATTAATTGCATGT

TATCGGATATTTTAAGATGCACATTTTAAATTTATGTTTAATTAGATGTTTGGAGTAGATAAATTTGAAT

GTTGGACTAGTTGTATTGGATGAATGTTGGATCTACATTGG

>MSTRG.1096.4 gene=MSTRG.1096

ATCACAATGATACAGAACACACTCAGTTTAAAACCAAAACGACCCAACACACGCTCTCTGCCCAAGAACC

CTAAAAATCGCCGCACATCGAATTCACCGGCGTTCATCATCTCCGTCGCTGTTCAGGCTCTCCAAGAGCT

CTACACCCTCAGTTCCGCCTTTAATTCTTCAGTACCGAGGTCTTTCCTGTGCATCTTGCTGGTAGATGCT

CTATTTCGAGTGCTCGAAGCTTGAGCTTAATCAGTCAATTGGGGGCTTAGTTCTTGAGTTGTTACCCCTT

GGAAGATCGGCATCTTTTCCTCTCTAACTATTTTCGATTTGTTCTGGGGGCTGACACATTAACTCGGATT

CTCCTTCCTAGAGGCTGCGTTAATATATAGCTGAAGATTTCAGGAATTGGTGTGAAAGTGAATGTGTCGG

ACTCAAAGGGATAGAAGAATGTTCTGACAATTATAATTGGATGCTTACTTGCTGGAATTGGTATTGGCAT

CTCATCAACCATTGTGCCACTGTACATATCTGAGGCATTAGAACTTTTCCGGAAGCAGGGAGTGGATTTT

TTGATTGCTCCAGATGTTGCATTTCTACCAGTGAACATTTGTATCCGTGTGGGTGCACATTGTGGTTTGT

AAAATAGTTGCCAAAGCAGATCAGATTCAGATGTATATGGCCATGCAACATCAAACCTTGTAGCTGGAGT

GGTTTTGAAAAGGACGTCTGGACCTAGACAGCCGAGGGAAGGCCTTTTGTTGGTTATTTTGTCTAGTTAA

AGATTTTTCCAAATTTGGGGTAGATGGTTTATGGAATGACATGAATGAGCCTGCTGTCTTTGAGGTGTAT

GGAATGTTAATGGCCAGATCAACATATGAAGGCATGAAAGTAGCTGATGAGAGTAATAATCTATCAGGGA

GCATCTATATATGAGCATATCCATGGTTCCCCAACTGGCTGCAAGGAGTCGCAGGACTTATTCTAAGACG

GACCTCGACATCATTGGTTGGAGTCGCTAAATTATATCCAGTCTATTATGTAGTTTGGATTCTCATAAGT

TAGACAGATTAATTGCATGTTATCGGATATTTTAAGATGCACATTTTAAATTTATGTTTAATTAGATGTT

TGGAGTAGATAAATTTGAATGTTGGACTAGTTGTATTGGATGAATGTTGGAT

>MSTRG.1097.1 gene=MSTRG.1097

ATAAATACTCTCAATTATCGACAATATCTTCATAAGCAATTCAAAATCTCTTTATTCGACTCCTTTTCTT

GCAAATACTTTACATTTTCTGCTCTGATCTTCATGGCTTCCTTAAGTGATGTTCAAATATCAATGCCTAG

CGATAGACAAGTACCTGAAAAAGAGTACTTTGATTACTCACAAAGGGCACAATGGCTTCGAGCAGCTGTA

TTAGGAGCAACAGATGGCTTGGTTTCAGTTGCATCATTAATGATGGGTGTAGGAGCTGTTAAAAATGACG

TTGCAGTCATGATTCTTACTGGTTTTGCTGGCCTAGTAGCAGGTGCGTGCAGCATGGCTATCGGGGAGTT

TGTGTCTGTATACTCACAGCTAGATATTGAGGTAGCTCAGATGAAAAGAGACAAGAGAATGCTGGACAAT

GAGAAGAATGATGATGAAAGCGAAAAGGAACAACTGCCAAATCCCTTTCAGGCTGCATTAGCGTCAGCTC

TTGCATTTTCTTTAGGGGCTTTAGTTCCATTACTAGCTGCTGCATTTATAGGTGATCATAAGGTGAGGCT

TATTGTGGTGATTGCTGCGGTAACCTTTGCATTGCTCGTGTTTGGAGGAGTTGGAGCAGTGTTGGGAAAG

ACCCCGGCTCTGAAATCTTGCGCTAGGCTGCTGATTGGTGGCTGGATAGCAATGGCTATTACCTTTGGTT

TAACCAAATTAATTGGCTCTAGCGGTTTGTAGAGAGTGATCATTAAAGTAATATTAATATTGATATGAAT

GATAATATATTTAGTATTCTCCTGCCCCTTTGTCCTTTTCTTTCTGAATAGAAATATAATTTCTTTCCTG

TCCTTGTAAAGAAATGTTGGTAGTTCCAAAGAAAGGTCACCTCCTTTTGTTGTGGTTGAAGATCGTAAAA

GTAAAGATGGTAACTTATGTAAAGGCTTAATGCAAAAGTATATGTAAATTCTTCAGGAAAG

>MSTRG.1099.1 gene=MSTRG.1099

ACAAACTCAAGCACACTGTTGGTTCTGTCTCTAGCTATATTTCTCTCAATAATTTCCTTAATTTTCTCTG

AAACTTTGTTATCATCATGGCAGTTGCCATCAAAGTTCATGATCATCCCAACAAACAATTTGAACAACAA

TCCGCGCAAGCATTTGAAACTGAGGAGGCTTTTGATTACTCACAGAGATCACAGTGGCTCAGAGCTGGTG

TGCTGGGAGCAAACGACGGCTTGCTCTCTACTGCATCCCTCATTATGGGCGTTGGAGCTGTCAGAACTGA

CCAAAAGACCATGATCCTTAGCGGAATTGCAGGACTGGTTGCTGGAGCTTGTAGCATGGCCATCGGAGAA

TTTGTGTCAGTGTATTCTCAATATGACACGGAAGTGGAACAAATAAAGAGGGAAGCTGTGCATGGAAATA

AAGCAAAGGCCGAGTTAGAGGCGAAGAAGAAGAAATTGCCTAGCCCTGTGAAGGCCGCGCTAGTGTCAGC

CTTCATGTTTGCCATAGGAGCCCTGGTGCCCCTACTGGCTGCAGCATTTGTGAAGAATTATCATGTGAGG

TTAATAGCTGTCACAGCTGCAGTAAGCTCTGCATTGATAGGGTTCGGAGGTCTGAGTGCGATGCTGGGAA

AAGCACCTGTCATAAGGTCTATATCTCGGGTTTTAGCTGGAGGGTGGTTAGCCATGGGGATCACTTTCGG

TTTAACCAAATTAGCCGGCTCTGCTGTTTCATTTTAGGCTTATTAGATCCGCAAAATCTTCCCTTTCTGT

TAAACACAAGTTTATTCAGCTATGATCTAGCTTTCTTCCTAGAATAATATCGTTTCAAGATCAGTTCGAT

CTCTCTTTTGTCTAGAGAGCATGTGTTATGTGTGTGTTTCTACGAGGGCAGCC

>MSTRG.1101.1 gene=MSTRG.1101

CAATTTTTTAACCTTCATACTTAGAAATTGTGCCCCAAAGTGTGTCCGAGACGTATAAATAGGAACCAGA

ACTGTACGACATGCATTCTCTTACCATAACATGATTAACTTGAGACCATCGTCTCTTCGACTAGCTTCAT

CACTTCTTCACTCTCTTCCAAACCCTAACCCATTACACTCACATCATCTAGACACACGACAACTTCGCTC

TACTTGCTCGGACTCCCAAAAACACGCGCATGCTCTCAACCTGTTTGATCAAATGCTCCAGACACAACCC

CTTCCTTCTATCAAAGACTTCACTAAATTCTTAAACATACTTGTAAACTCCAGACAGTACTACACTGTCG

TCTCTTTGTTTACTGACATGTGTTTTTTGTCCTTTCCTGTCGATTTGTGTACTTATAATACTGTGATCAA

CTGTTGTTGTCATGTAGGGAGATTTGACTATGCATTTTCGCTGCTGAGTGGGGTTATTAAGCGTGGGTTT

GGCCCGGATGTTGTTACTTATACGACGCTTGTGAGGGGGTTGATTAGGGGAGATATGGGCGTGGAGGCGG

AGGGGTTGGTTAGGAAGTTAGTTAGGTGGAATGAGGTTGAGTTTGATGTCGTTATGTATAGTACGATTGT

CTATGGACTTTGTAGAGAAGGTGAGGTTTACGCGGCGATTGAGTGTCTTAGGTTTATGGAAAAGGAGAAT

TGTGAAGCTAATGCGATAACTTATAATACGATTATTGATGGTTTGTGTAAGAGGGGACTTGTTGATGGTG

CGTTGGACCTTTTTTCTGAAATGAAGAAGAAAGGTGTGGGAGTGACGGTTGTAACTTACACTTCGTTGAT

TCATGGTTTGTTCAAGTCGGGGCAATGGGAGGATGCTATCTTATTGTTTGATTCGATGGATGTTTCTCCG

GATGTGTATACTTATAGTGTATTGGTTAATGGGTATGGGAAGAGAAAGATGGTAGAAGAGGCTAGGAATA

TTGTCTTGAGAATGATTAAAATGGACGAATATCCTAATATAGTCACGTATACTACACTCATGCAAGC

>MSTRG.1102.1 gene=MSTRG.1102

CAAGCACCCAAATACACGGAACATGAATGTACATGAGACGAACGATTTGCCAGGTCTATGTGTAGCAATC

TTTCACTTTAAAATCCAAAGGGAACTTGCATTAGTGATCAGACAATCACACAATCGCAATCCGAACAATG

TAAAGCATGTCTGACATTAGAGATAATCCTCCTTGCATAAGCCATGAAATCAGCATTTAGCAACACTATC

ATCTTAAAATGGAGTAATGGACATATTGTGCTGGAAATTATAACGGATGGTTGGTGTCACCTGACTAGCA

AAATAGGGCACTTAACCCTGGCCACCTCAGAAAACCAGTGTTACAAGAGAATCCTATTTTATCTTTTTTA

GTAACTTGTACTGCATTCCTTTTCCTCCCAAGTCAATTGCTTATACTATTATACATGACACATAAGTAGC

AG

>MSTRG.1107.1 gene=MSTRG.1107

CAACACTCTCCCTCGGCTCTCTCCAGATTCTTTCCGCGATTTTTTCGATCCAACGGCCACGACTCACGCG

TCCTCCAATCGTACTGCTAGGGCTCATCTACGCAGGGTATAGCTTAAGAAGATGGCGATGAAAATGAGGC

AAACAGAAGATTTAAAGAGAAGCACAGGAGATTTTGTGTTGGACTCCAATTGGTGAAAGCAAAGAATCCA

CACGGCTTTGCCCATCCCAGTTTTGTGTCAAGTCTAGCGATGTATCTATATTCGCAACAACAGAAGGGGA

GGAAG

>MSTRG.1108.1 gene=MSTRG.1108

CATACTCAGTTTTCAGCGAGCTCATATCTGACTAGTGCCTAGGGTTTATCTTGAAAGAGAGACTGCAATC

TATCTCGGCGGTGTTTCTTCAAGATAGTCTCTTATTCACCACCACCACCACCACCACGATAACGACTAGG

TACCGTTTGTTCTTATTTTCACTTATTTATTTTCTCATCGCCAACTAGTGTTGTTCATTTTGTGTCTCTG

TTAATATTATTTCGACCCCTAATTAGCATTCTTATCTCGCACAGCAATTTTCAGAAATTTAATTATGGTT

TCGATAAAGTCGACTATGGTCGTGTTGGGTTTGATAAAATGCTAACAGGCGATTGAGGTCGTTTGCAAGA

CTGGTCACACTTCTAAAAGTTATGGATGATTTGACTTGTCGTACTTGTTAATTATTTTGTTTGGATATAT

TCTCTGATATGATAGAAAATGAGGAAATGGAGTACTGTTCCCGAGGGCAAACTAATGGAGCATTGGCTGT

GCTGAATACAACTTGATCTAATGATGCACTGAACCTTGTTAGACGGTCTTTGCTAAAATCAGTGTGTTGA

TATGACAGTGTCTTTCTTTTCGGTGATGAAATGCAATGTATTAATTCCTGAGGGTGGCACTTACAATATC

ATTATTTGTGGATGTCTTCGGAATAAGAAGGATGCTGAAGCATGTAAACTGTTAGATAATATAGTTGATT

GCGGCTTCTTAGAAGATGCATTTACAACCTCCCTTTTTATTACGACATTTACTTTTATCGGAAGTACATG

ACCCAAATTTGGTTCAACAAACTTGAAGATGCTCTTCTTGTGTTCGATAAAATCCTGTACTTGAAGATGC

TCTTGTGTTCGATAAAAATTTGGTTCAACAAACTTGAAGATGCTCTTGTTGTGTTCGATAAAATTCTCAA

CTTGCAGATGTTCTTGTTGTGTTCGATAAAAATTTGGTTCAACAAACTCGAAGATGCTCTTCTTGTGTTC

GATAAAAATTTGGTTCAACAAACTTGAAGATACTCTTGTTGTGTTCGATAAAATGCTCAACTTGAAGTTG

CTCATTGTTTTGTTCGATAAAATGCTCAGAACTCGAAGATGATTTTTTGTGTTCAATAACTTGAAGTTGC

TCTTGTCGTGTTCGGTAAAATGCTCAGAACCTCTCCCTAATCCTATTGACTTTACTCGGCTGTTGTGTGC

AATTGTTGATAGGGAACGAGACACCGCATCTGTTCCTCACTTAACAAATTTATGTGACCTAAGCAAATCT

GTCAATATTATCACCTTTAATATATCAGGAGGCTTCTTTTTCACAGATTTTATCAAATTCAAAGACTTTA

AACCTTAAAGTGCCATCATTAATGGCCTTTGTAAGACAGGTCATACTTCTATGGCCCTTTCTAAGAGGCA

TTTACCGAATGTTCGGACGTACTCTCCGGTTGTTCAGGTCCTCTGCAACTTCAATAGGAGGTTGATATGT

TACTTGAAAATATGATTGAGGTGAAAATCTCTTGAAATATGTACACTTTTGACATATTGGTCGATGCTTA

TTCCATGTTTAGGAAGTTGGACGGCGCAAAGTGGATAATTGAGATGATGAACAGACGGGGTCAATATCCT

GATATAGTCACCTACGATACTATCATGCAGGCATATTGTTTACAAGGCCAAATGAACAGAGCATTGTCTG

TGCAGAAGACCATCAGAAGTAGGAGGATAATGCCCGATTGTAGCACATATGCTGCCATTCTGTTGGATGC

ATATGCAGGCAGCTGAAATTGGACAGGCACATACCTATGGATCTTTACAGAAACATGCCTGTGATGCTTC

GAGGCCCACATATGTAACACAACATTTTCTTACACAGGCTGTCAGAAGGGTAAATCTATGGAAGTACTGA

CGTTCTTCTATAGGATTAAAGATCAAGACATAAGGTAGCTTTTGCTGCATACAGAACTTTGTTGGATGGT

CTTTGCAAGAACCATTATGTTGATAAAGCACCATCTTTATTTCACGCAGTGAAATGCATTGTATTAATTC

CCATGGGTGACTATTACAATAGTGTTGTCTGTGAATGACTTCGGAATAAGAAGTATGATGAAGCATGTAA

ATGTTTAGACAAAATGGTTTATTGTGGCTTCTTAGCATATGCATTTACAACCTCTTTACTGCAACACCTA

CCCTTATCAGGCACAGGATCTAATGTATCAGGATCTAAGGTAACTTTGGATAAGTTAAGTGGTTCATGAT

CTTTCTTATGTACCTAAAGTATGCATTGTAAAATAGTTCTTATTGTCTACTTTAAGATTACCTTTTTTTC

CCGTAAGGATTCATGTTTTGACGACTTAACTCTCATATACTTCTTACAATCTTTTAAACTCTTGTTTTTG

GTTTAGTTCTCGTCTATGTTTCCAGTTAACAGAAGCATGACTGATGCACTATGCCTTGTCCTAGAGCCAG

TGTAGTACTAAGCAGCATGCTCGCCCTCTCATTATCAAATATGCACACATCCTATATACATACACATTGG

TTGC

>MSTRG.1108.2 gene=MSTRG.1108

CTCAGTTTTCAGCGAGCTCATATCTGACTAGTGCCTAGGGTTTATCTTGAAAGAGAGACTGCAATCTATC

TCGGCGGTGTTTCTTCAAGATAGTCTCTTATTCACCACCACCACCACCACCACGATAACGACTAGGTACC

GTTTGTTCTTATTTTCACTTATTTATTTTCTCATCGCCAACTAGTGTTGTTCATTTTGTGTCTCTGTTAA

TATTATTTCGACCCCTAATTAGCATTCTTATCTCGCACAGCAATTTTCAGAAATTTAATTATGGTTTCGA

TAAAGTCGACTATGGTCGTGTTGGGTTTGATAAAATGCTAACAGGCGATTGAGGTCGTTTGCAAGACTGG

TCACACTTCTAAAAGTTATGGATGATTTGACTTGTCGTACTTGTTAATTATTTTGTTTGGATATATTCTC

TGATATGATAGAAAATGAGGAAATGGAGTACTGTTCCCGAGGGCAAACTAATGGAGCATTGGCTGTGCTG

AATACAACTTGATCTAATGATGCACTGAACCTTGTTAGACGGTCTTTGCTAAAATCAGTGTGTTGATATG

ACAGTGTCTTTCTTTTCGGTGATGAAATGCAATGTATTAATTCCTGAGGGTGGCACTTACAATATCATTA

TTTGTGGATGTCTTCGGAATAAGAAGGATGCTGAAGCATGTAAACTGTTAGATAATATAGTTGATTGCGG

CTTCTTAGAAGATGCATTTACAACCTCCCTTTTTATTACGACATTTACTTTTATCGGAAGTACATGACCC

AAATTTGGTTCAACAAACTTGAAGATGCTCTTCTTGTGTTCGATAAAATCCTGTACTTGAAGATGCTCTT

GTGTTCGATAAAAATTTGGTTCAACAAACTTGAAGATGCTCTTGTTGTGTTCGATAAAATTCTCAACTTG

CAGATGTTCTTGTTGTGTTCGATAAAAATTTGGTTCAACAAACTCGAAGATGCTCTTCTTGTGTTCGATA

AAAATTTGGTTCAACAAACTTGAAGATACTCTTGTTGTGTTCGATAAAATGCTCAACTTGAAGTTGCTCA

TTGTTTTGTTCGATAAAATGCTCAGAACTCGAAGATGATTTTTTGTGTTCAATAACTTGAAGTTGCTCTT

GTCGTGTTCGGTAAAATGCTCAGAACCTCTCCCTAATCCTATTGACTTTACTCGGCTGTTGTGTGCAATT

GTTGATAGGGAACGAGACACCGCATCTGTTCCTCACTTAACAAATTTATGTGACCTAAGCAAATCTGTCA

ATATTATCACCTTTAATATATCAGGAGGCTTCTTTTTCCACAAAAGTTTGCCGAAGGCAGACTTTTGTTC

ACAGATTTTATCAAATTCAAAGACTTTAAACCTTAAAGTGCCATCATTAATGGCCTTTGTAAGACAGGTC

ATACTTCTATGGCCCTTTCTAAGAGGCATTTACCGAATGTTCGGACGTACTCTCCGGTTGTTCAGGTCCT

CTGCAACTTCAATAGGAGGTTGATATGTTACTTGAAAATATGATTGAGGTGAAAATCTCTTGAAATATGT

ACACTTTTGACATATTGGTCGATGCTTATTCCATGTTTAGGAAGTTGGACGGCGCAAAGTGGATAATTGA

GATGATGAACAGACGGGGTCAATATCCTGATATAGTCACCTACGATACTATCATGCAGGCATATTGTTTA

CAAGGCCAAATGAACAGAGCATTGTCTGTGCAGAAGACCATCAGAAGTAGGAGGATAATGCCCGATTGTA

GCACATATGCTGCCATTCTGTTGGATGCATATGCAGGCAGCTGAAATTGGACAGGCACATACCTATGGAT

CTTTACAGAAACATGCCTGTGATGCTTCGAGGCCCACATATGTAACACAACATTTTCTTACACAGGCTGT

CAGAAGGGTAAATCTATGGAAGTACTGACGTTCTTCTATAGGATTAAAGATCAAGACATAAGGTAGCTTT

TGCTGCATACAGAACTTTGTTGGATGGTCTTTGCAAGAACCATTATGTTGATAAAGCACCATCTTTATTT

CACGCAGTGAAATGCATTGTATTAATTCCCATGGGTGACTATTACAATAGTGTTGTCTGTGAATGACTTC

GGAATAAGAAGTATGATGAAGCATGTAAATGTTTAGACAAAATGGTTTATTGTGGCTTCTTAGCATATGC

ATTTACAACCTCTTTACTGCAACACCTACCCTTATCAGGCACAGGATCTAATGTATCAGGATCTAAGGTA

ACTTTGGATAAGTTAAGTGGTTCATGATCTTTCTTATGTACCTAAAGT

>MSTRG.1108.3 gene=MSTRG.1108

CAGTTTTCAGCGAGCTCATATCTGACTAGTGCCTAGGGTTTATCTTGAAAGAGAGACTGCAATCTATCTC

GGCGGTGTTTCTTCAAGATAGTCTCTTATTCACCACCACCACCACCACCACGATAACGACTAGCAATTTT

CAGAAATTTAATTATGGTTTCGATAAAGTCGACTATGGTCGTGTTGGGTTTGATAAAATGCTAACAGGCG

ATTGAGGTCGTTTGCAAGACTGGTCACACTTCTAAAAGTTATGGATGATTTGACTTGTCGTACTTGTTAA

TTATTTTGTTTGGATATATTCTCTGATATGATAGAAAATGAGGAAATGGAGTACTGTTCCCGAGGGCAAA

CTAATGGAGCATTGGCTGTGCTGAATACAACTTGATCTAATGATGCACTGAACCTTGTTAGACGGTCTTT

GCTAAAATCAGTGTGTTGATATGACAGTGTCTTTCTTTTCGGTGATGAAATGCAATGTATTAATTCCTGA

GGGTGGCACTTACAATATCATTATTTGTGGATGTCTTCGGAATAAGAAGGATGCTGAAGCATGTAAACTG

TTAGATAATATAGTTGATTGCGGCTTCTTAGAAGATGCATTTACAACCTCCCTTTTTATTACGACATTTA

CTTTTATCGGAAGTACATGACCCAAATTTGGTTCAACAAACTTGAAGATGCTCTTCTTGTGTTCGATAAA

ATCCTGTACTTGAAGATGCTCTTGTGTTCGATAAAAATTTGGTTCAACAAACTTGAAGATGCTCTTGTTG

TGTTCGATAAAATTCTCAACTTGCAGATGTTCTTGTTGTGTTCGATAAAAATTTGGTTCAACAAACTCGA

AGATGCTCTTCTTGTGTTCGATAAAAATTTGGTTCAACAAACTTGAAGATACTCTTGTTGTGTTCGATAA

AATGCTCAACTTGAAGTTGCTCATTGTTTTGTTCGATAAAATGCTCAGAACTCGAAGATGATTTTTTGTG

TTCAATAACTTGAAGTTGCTCTTGTCGTGTTCGGTAAAATGCTCAGAACCTCTCCCTAATCCTATTGACT

TTACTCGGCTGTTGTGTGCAATTGTTGATAGGGAACGAGACACCGCATCTGTTCCTCACTTAACAAATTT

ATGTGACCTAAGCAAATCTGTCAATATTATCACCTTTAATATATCAGGAGGCTTCTTTTTCCACAAAAGT

TTGCCGAAGGCAGACTTTTGTTCACAGATTTTATCAAATTCAAAGACTTTAAACCTTAAAGTGCCATCAT

TAATGGCCTTTGTAAGACAGGTCATACTTCTATGGCCCTTTCTAAGAGGCATTTACCGAATGTTCGGACG

TACTCTCCGGTTGTTCAGGTCCTCTGCAACTTCAATAGGAGGTTGATATGTTACTTGAAAATATGATTGA

GGTGAAAATCTCTTGAAATATGTACACTTTTGACATATTGGTCGATGCTTATTCCATGTTTAGGAAGTTG

GACGGCGCAAAGTGGATAATTGAGATGATGAACAGACGGGGTCAATATCCTGATATAGTCACCTACGATA

CTATCATGCAGGCATATTGTTTACAAGGCCAAATGAACAGAGCATTGTCTGTGCAGAAGACCATCAGAAG

TAGGAGGATAATGCCCGATTGTAGCACATATGCTGCCATTCTGTTGGATGCATATGCAGGCAGCTGAAAT

TGGACAGGCACATACCTATGGATCTTTACAGAAACATGCCTGTGATGCTTCGAGGCCCACATATGTAACA

CAACATTTTCTTACACAGGCTGTCAGAAGGGTAAATCTATGGAAGTACTGACGTTCTTCTATAGGATTAA

AGATCAAGACATAAGGTAGCTTTTGCTGCATACAGAACTTTGTTGGATGGTCTTTGCAAGAACCATTATG

TTGATAAAGCACCATCTTTATTTCACGCAGTGAAATGCATTGTATTAATTCCCATGGGTGACTATTACAA

TAGTGTTGTCTGTGAATGACTTCGGAATAAGAAGTATGATGAAGCATGTAAATGTTTAGACAAAATGGTT

TATTGTGGCTTCTTAGCATATGCATTTACAACCTCTTTACTGCAACACCTACCCTTATCAGGCACAGGAT

CTAATGTATCAGGA

>MSTRG.1108.4 gene=MSTRG.1108

CTGACTAGTGCCTAGGGTTTATCTTGAAAGAGAGACTGCAATCTATCTCGGCGGTGTTTCTTCAAGATAG

TCTCTTATTCACCACCACCACCACCACCACGATAACGACTAGAAAATGAGGAAATGGAGTACTGTTCCCG

AGGGCAAACTAATGGAGCATTGGCTGTGCTGAATACAACTTGATCTAATGATGCACTGAACCTTGTTAGA

CGGTCTTTGCTAAAATCAGTGTGTTGATATGACAGTGTCTTTCTTTTCGGTGATGAAATGCAATGTATTA

ATTCCTGAGGGTGGCACTTACAATATCATTATTTGTGGATGTCTTCGGAATAAGAAGGATGCTGAAGCAT

GTAAACTGTTAGATAATATAGTTGATTGCGGCTTCTTAGAAGATGCATTTACAACCTCCCTTTTTATTAC

GACATTTACTTTTATCGGAAGTACATGACCCAAATTTGGTTCAACAAACTTGAAGATGCTCTTCTTGTGT

TCGATAAAATCCTGTACTTGAAGATGCTCTTGTGTTCGATAAAAATTTGGTTCAACAAACTTGAAGATGC

TCTTGTTGTGTTCGATAAAATTCTCAACTTGCAGATGTTCTTGTTGTGTTCGATAAAAATTTGGTTCAAC

AAACTCGAAGATGCTCTTCTTGTGTTCGATAAAAATTTGGTTCAACAAACTTGAAGATACTCTTGTTGTG

TTCGATAAAATGCTCAACTTGAAGTTGCTCATTGTTTTGTTCGATAAAATGCTCAGAACTCGAAGATGAT

TTTTTGTGTTCAATAACTTGAAGTTGCTCTTGTCGTGTTCGGTAAAATGCTCAGAACCTCTCCCTAATCC

TATTGACTTTACTCGGCTGTTGTGTGCAATTGTTGATAGGGAACGAGACACCGCATCTGTTCCTCACTTA

ACAAATTTATGTGACCTAAGCAAATCTGTCAATATTATCACCTTTAATATATCAGGAGGCTTCTTTTTCC

ACAAAAGTTTGCCGAAGGCAGACTTTTGTTCACAGATTTTATCAAATTCAAAGACTTTAAACCTTAAAGT

GCCATCATTAATGGCCTTTGTAAGACAGGTCATACTTCTATGGCCCTTTCTAAGAGGCATTTACCGAATG

TTCGGACGTACTCTCCGGTTGTTCAGGTCCTCTGCAACTTCAATAGGAGGTTGATATGTTACTTGAAAAT

ATGATTGAGGTGAAAATCTCTTGAAATATGTACACTTTTGACATATTGGTCGATGCTTATTCCATGTTTA

GGAAGTTGGACGGCGCAAAGTGGATAATTGAGATGATGAACAGACGGGGTCAATATCCTGATATAGTCAC

CTACGATACTATCATGCAGGCATATTGTTTACAAGGCCAAATGAACAGAGCATTGTCTGTGCAGAAGACC

ATCAGAAGTAGGAGGATAATGCCCGATTGTAGCACATATGCTGCCATTCTGTTGGATGCATATGCAGGCA

GCTGAAATTGGACAGGCACATACCTATGGATCTTTACAGAAACATGCCTGTGATGCTTCGAGGCCCACAT

ATGTAACACAACATTTTCTTACACAGGCTGTCAGAAGGGTAAATCTATGGAAGTACTGACGTTCTTCTAT

AGGATTAAAGATCAAGACATAAGGTAGCTTTTGCTGCATACAGAACTTTGTTGGATGGTCTTTGCAAGAA

CCATTATGTTGATAAAGCACCATCTTTATTTCACGCAGTGAAATGCATTGTATTAATTCCCATGGGTGAC

TATTACAATAGTGTTGTCTGTGAATGACTTCGGAATAAGAAGTATGATGAAGCATGTAAATGTTTAGACA

AAATGGTTTATTGTGGCTTCTTAGCATATGCATTTACAACCTCTTTACTGCAACACCTACCCTTATCAGG

CACAGGATCTAATGTATCAGGATCTAAGGTAACTTTGGATAAGTTAAGTGGTTCATGATCTTTCTTATGT

ACCTAAAGTATGCATTGTAAAATAGTTCTTATTGTCTACTTTAAGATTACCTTTTTTTC

>MSTRG.1110.2 gene=MSTRG.1110

GTGACATGATTATATTTTCAAGGATTCGTCCAGGCAAATATAAAGTCGAAAAATAGTTTTAACAGCTTGA

ATTTATATGTCTGGCCAAACAGTAGTACAGTACAGTTAACTGAGCCACCTGATCAATTCCCACCAAGAAA

ACTGTAGGAAGCAAATCATATTATGCAGAGGAACACAACGTTGCGTCGAAGAATGATCGGAGATTATTTC

TGTAGTGTGTAAAACAGCTCCTAAAAGAGTCTCGTTAGAAGACGTCATCCAGTTTGGCTTGTAGATGAAG

TTCTCATCATAATCAGACGACGCTTGCGCAAATTGCTCATGATCCAGCACTCCGCTTTGGAGCTGTTCAT

GGAGCTGTGCTGATGATGAAGTTGCTGATGGAAAGGGCACTTACTTGCAACGTAACAATGGCAACATGGT

CCTTGTCTTCCCATGGATGATAAAGTTGCTGATGGAAAGGGCACTTACTTGGAACGTAACAATGGCAACA

TGGTCCTTGTCTTCCCATGCGTGGAGGTAGTCAGGGCTCAGGAGTTTGAGATTTGGGGGTATTGAGATCT

CTTCTGAGCCGCTCTTTTGATGATTCATGTTTGTATGCTTAATATTTAATATCTCTTCGTGTCTCTGTTA

CTGTTATTGCAATATTATCAAGAAAGTGTAAGGACCAAGATTATACTGCTGACGGTAAATGAAATTTTAT

TAGGCTGGTTCTTTTTTATCCTTTGTATCTTGTATTAGTATAATTACTTGAAATTTAATTAGGCTGAGTT

TTTACCTTTCGACACAAATGATCAAGCTTAAAAGAAAAGAAACGACATACTAGATGCAGCTCTTGCATGC

TCCGGACTCTCATGAGCACTGCAATTCCTTTTCCA

>MSTRG.1110.1 gene=MSTRG.1110

GTGACATGATTATATTTTCAAGGATTCGTCCAGGCAAATATAAAGTCGAAAAATAGTTTTAACAGCTTGA

ATTTATATGTCTGGCCAAACAGTAGTACAGTACAGTTAACTGAGCCACCTGATCAATTCCCACCAAGAAA

ACTGTAGGAAGCAAATCATATTATGCAGAGGAACACAACGTTGCGTCGAAGAATGATCGGAGATTATTTC

TGTGATGATGAAGTTGCTGATGGAAAGGGCACTTACTTGCAACGTAACAATGGCAACATGGTCCTTGTCT

TCCCATGGATGATAAAGTTGCTGATGGAAAGGGCACTTACTTGGAACGTAACAATGGCAACATGGTCCTT

GTCTTCCCATGCGTGGAGGTAGTCAGGGCTCAGGAGTTTGAGATTTGGGGCAATCATCTTGCACGGCCGT

CTACAGGGGAGGAAAGCGTTTCAGCATGGTCCTTTCTTCCTCTACTTTGCGGACTTTATATTGATTTTTT

TTTATTAATTTCAAGAATCCAGAAGTTCTTCCAGATGCTGAAATCTCATTTGATAACCTAAAACGTGCAT

TTAGTGTTGCTGCAACTGGCTTCTTGTAAGCCTGTTTGTTCTGGGGGAGGTGATCTGTAAAAATAATAAA

AATGATGAGGGCCATATATGATTGAGTCGCATATGATTGAGATATATCTTCTAATATATATTTCAAAAAT

CTCTAA

>MSTRG.1110.3 gene=MSTRG.1110

TACTAATTGCCAGTTTCACTTCACTCTTTCGCTGGCAGAGTGTGTAAAACAGCTCCTAAAAGAGTCTCGT

TAGAAGACGTCATCCAGTTTGGCTTGTAGATGAAGTTCTCATCATAATCAGACGACGCTTGCGCAAATTG

CTCATGATCCAGCACTCCGCTTTGGAGCTGTTCATGGAGCTGTGCTGATGATGAAGTTGCTGATGGAAAG

GGCACTTACTTGCAACGTAACAATGGCAACATGGTCCTTGTCTTCCCATGGATGATAAAGTTGCTGATGG

AAAGGGCACTTACTTGGAACGTAACAATGGCAACATGGTCCTTGTCTTCCCATGGTACTTCTTTTTTACT

AATACTGCAAGGCAACCAAAGAATTTGTGCAGACAAAATTAAAGTTTGAAACAGCAAAAAATCATATACC

ACAATGAAACTGGAAAATAGCAGGAGTATATAAGATTTATCGTTGCTACTCAAATGGAAAATAAATATGA

GAATATTTTATATAGACTTGCATATGCCAATTAAGCACTCCAGATTTTGATTTAAAAAACTTTAATTGAA

AAGGTGGAAGGTGAACAATTTATTATTATTTTTTCTATCTTCTGATTCACATGAAATCTGCATCGTAGAT

TCCATGAATTTCCAAATCTGTGACATTTAAATCAGTACAAATCGTTGATTTGTGCCTACCGCTGATAATT

TGTAAAGGAGTAGTGATTTTAGATTTCGACTTGATGAGAGTACTCCTAATTTCAGTTCTAAAAAGAGGGT

GTGCTACCTTGATCAAACCAAGGAGCGATATTCAAGAAGAGAGCATTTAGTGTTTTTGTCAATACACCAT

GTTTGGCCTATGTAATAAACATATTGGCAAAAGTAATAAGCTCATCAATATATATCTAACACCTCAACTA

ACTCACCTACCTTGAAGTTTAAACTTAGTTAATGTTAAATATAATCATGTTTATTAATGTCATTTACAAA

AAAAATTCCCTCTCCGTCTCTTCCCTTATTCTTGCTCTTCCCCATTTTTAGTCACCCTTCCCATTATTCT

CTCTTCTCCATTTCTTAATTATCAACTAGGCCACTACTTATTTTTGTAATTGTTTTATTATCATGTTCCA

CTAATTTTCCACCTTTCTCTTCTTTTTTCTCCAAAATTATTGATCGAGACCGATATCAATATTTGATCAA

AATATATATATACCCATCCTCCAGAGTACCAAAGTTAAACTTAACTTAAGTTGGTTTAGTTATGATGAAA

AGATAGTTTAGTCTTTTTATCAGTAAACTTGTTCACTGATTCCGGCTGTATTCAAGAACTTATCCCTCTA

TTTATGTCTTTTTTTTGAAAGTGTCAATATTTGTCTCTTGAACACTTATAATAGAAACAATAATAATCAA

ATCATTGGACAACCGCTAAATAGATTTAAGAACTATACTACCCATCACTCTATCACAGCATATATATTCA

TAATAAACATGTCCGTGAGGATTGAGATCACAATTATATGGTTACAAATTTTAAGTGAGTTGCTATTAAA

TGTTATTTAGTATAATTTCCAAGTATCTGAAGAGTCTCCTTAGAACTCTCAATTAACCCTAAAATGACCA

TGGTTACATTGGTGAACGTGTTTCGAGTGCCTCCACACAACAGGCTATATAAACACCCCACTAGTTACCC

CCGAGTCACTCAGTTAAAACAAAACAACACTTGTCTTCTTCTTGTAAACACAAATGGCAGAGGTGGAAAA

TGTTGAAGCCAACTCCAGGTGGCACATGCCCCGCAAGGGTATGGTGAAGATCAATGTCCACGGTTGCTTC

TTTGACAACCCATTACCAAACGGCAATGTTTCAGGCATAGGAGCTGTCATCAGAAATAGCAGAGGTAAAA

TAGTGAGGATGCTCTCGGGTTCTTTAGGGATACAAAACAGAAGGATCAATGAATATCAAGCTCTTTTGGA

AAGTTGTAAGCGATCTTTTGTCGAAGACTAGGAGCATTTTATCTTGGAATCTGACCACCTCGACTCATTC

TAGGAATGGAAGAACTCCTCTCTTGAAGGAGTGCACCCGAATCATGCCGAGATGGTGCAGCAGCTTAACT

AGAGAAGTGATGACAGAAACATCCATATGGAAGTCACTCTACTGAGCCAAGATATATGGCTGTGCATGAA

GCTGATATTGAACTTGTTGTGCTCAATGGAGAAGAAGCTCCAGTAGCAGAAGTTGAAGATGAAGAGGCGT

CCGTGGATGCAAATGGTGACAGAGTTGAGGTGATAGAATACATGGACTAGCTTGCATGCCTCACCAAGTC

TGCTAACCTACGAAACAGGTTTCCCTATGCAGTTTTTAGTATTCCAATATATTAGTCAGTTGTTGTTTTT

GCTTTAAGTGGATGGAGTTAGTTTTGCTAGCTGATGGTTGGATTTGATTCGCCCCTTTTGTAAGGTCTAG

GTGTAGGCCTTAATCGATATGTTGGG

>MSTRG.1113.1 gene=MSTRG.1113

CCCCTCCCGCCGCTCGGAACCTGCTTAACATTTTCTCGTCAATGGCTTTCTCTCTTCGGAGATCAATGCC

ATTACGAGCTGCATTTCCATTTATCCACAACACCCCATTATCCCCATCAACATCATCCTTATCCACAACC

CGTCGCAGAGAAGTTCAAACAATCCTCCTCAAGAGCTCGAAATCGGGTTTCGAAAACCTCGGCGACGCTC

TCCACCTGTTCGACGAAATGCTCCAGTGGCGCACCCTGTTCTCCGTTGTCAACTTCACTCAATTACTATC

TGGGCTCGTTAAAATGAGACATTACTCTGTGGCCGTGTCATTGTTCCACCAAATGTGCGCTTGGAACATT

CCTGTTAACACGTATACGATCAGTACCGTGATCAACTGCTATTGTCGTCTGAATATGATCCAATGTGGGT

TCTCGATGTTGGGGTGGATGATCAAGCGTGGTTTTAGGCCTAATGTGGTTACGTACACGGGTTTGATTAA

AGGGTTGTCGTCGCCTGTGGAGGCGAGGGGATTTATTAAGAGGCTGATACGGCTGAATGAGATTGAACCG

GATGTCGTGTTGTATAGTGCTGTTATTGACAGGCTTTGTAGAAAGAATTTTGTTGATCAGGCTCTTGATG

TTTTTTCTGATATGAGGCAGGAAGGGGTGGATGCTAATGTTGTTACGTACAACTTGTTGGTTCAGGGGCT

CTGCAAGTTAAATCGGTGGGAAGAGATTGATCGGTTGTTAAGAGAGATGGATGTCATGAATGTATGTCCG

GATGTTTACACGAGCAGTATATTAGTTAACGCGTATTGCAGAGTAGGGAGGGTTGAAGATGCAGAGAATG

TGATTAAAATGACAATTGAACGAAGTGAGTGTCCTAATGTAGTAACATATAATACTCTAATGCACGGATA

TTGTCTGAGAGGAAAAATTGATGGAGCAGTGGCAGTGGTGAGTACGATGAAAAGTATGGGGGTAGTTCCT

GATATAGTGACATGCGGCATTTTGTTAGACGGCTTTTGTAAGAAAAATGGCGTCGATGAAGCAATGTCGT

TGTTTCACGAAATGGAAAGGAATGGGTTGATTCCTAATATTTACATATATGGTATCCTGATAAATGGATT

GTGCAACGACGGGCAGCTTAGTAAGGCAAAAATTCTCTTTGATAACCTTTCGTCGATTAGGTTGGAACCA

GACGTCAAGACATACACTATGATGATTCAAGGATTTTGTCAGAAAGGGCTTCTGGATGAAGCAAATGAAC

TGCTCAGGCAAATGGAAGCTGAAAATTGTTTGCCAGATAATGTCACCTACAACATTCTTATCCGCGGATG

TATCCAGAATAAGAAGTACAATGAAGCCTGTGGACTCATGGACAGAATGTGTTGGCCGTCTGGTGGTGGC

TTCTCACTGGATTCATGTACCACCTGCATGATACTGGATGTTCTGGGTACCGAAAAGGATGATCCATCTA

CTCTTGCTTTGCGCAAAAAGTTCTTGACGTAGTTACTCAGCATTCAGGGGTTTTGGTTGCAGAAAGATGA

AGAAGTTGAGTCGATCCTGGGTTTTACGTGCTCCGTGGGATCATGAAGTGTGCTTTTGTAGCTGGTCTGA

CATGTAACACTCTTTGCAAGTGCCTTCTTTCGGAGAAAAGGTTTGCAGGGGAGAGATATATGTTCCAACT

GAACCAACTGTTGTTACCTGTATGCCATTTTTGACGGACTTTACAAAACAGACAACACTTCGACAGCCAT

CAGTTTGTTCACATCTACGGTTTTTGATGAAACGGCTATCAGGCATTGTCGAGGCTTCTTAGCAGCTGGA

CTTGCGACCTCCATGTTACAAGGCAGTGTCGAGGCTTCTCAGCAGCTGGACTGCAGCTGCAGTTGCAACA

TTCATGTTTACTGCATCAGCTACGTAATGCCAGGAACTATCTGTCATTTATTGCTTCTCAGGATTGTAAA

GGTACAAAGTTTGATAGGGGTAGGGATCATTTAACTGCCAAGTTCTTCACTTTGTTCTCAAATGTTAAAG

ATTTATACAGTAATGGCACACCAATTGCATTTTATGCATTATATTTTCGTAAACTTTGGATATATTACAG

AGTGCAG

>MSTRG.1121.1 gene=MSTRG.1121

GGGAATTTTAGAAAAAATGTTACACAAACCTGGAATTCAAACAAATAGCATTTTAAGATCAAACCAGAAA

CACACAAGTATTCTCAATCTCACGCATGTGCATAAACAACCCTATTAACTCCTCCCACGCACAAAATTTA

CAGAACATGAGGATATAAACACATATATCTTACAATCATGAAACAAGCTTGAAGTTCATGGATAACGGAC

TAGTTTTTAGATATAAGTCATAGTCTCTTAGGTAAGAGGGACTTCCCCACTTTGATGAGCTTCTTTCACT

CGCTTATCATACTCATCCTCGTCCTCGTCATCTACCTCAGCTTCTAGATCTTCCTCTTCTTTCTCATCTT

CTTCCTCTCCCCATCCAAAGCCTCCAATAGAACGTGAAACTGGAGGAGGACTCCTGGCCTCTTCCACAAT

CCTCATTATGTCGTCCACAC

>MSTRG.1128.1 gene=MSTRG.1128

TGACTCTACCCAAACACATTACACCAACCCTAATTACACGCACGCTCTCCCGCCTCCCGACAATGCTCAC

CGCCGGCCGCTGACTTCAAGCATCTGTATATATAGTCAAAAATCAAATCAAAAATTCAAAGCATCAAAGG

CATCAGAAGAAATCAGCGACATCAAGCCTTCAGAACACTTTTGGCTACATATGTTGATGTTTCATATCTT

GATTTTTCATAATATTGTTTTCTTAAAATTCAAAAATCCAAAAAATTCGGTTATTTCATAACCAAAACCG

AATTAACCGAAATATTTCGGTTCGGTTACGGTTCGGTTTAATATATAATTCGGTTCGGTTCGGTTTATGA

AAATTTTGACATTTCGGATTTCGGCTATTTCGGTTCGGTTCGGTTTTGAACCGAACCGACCGAATGCTCA

GCCCTACTTCTCATGGTTGGAGGTGCTCTTATATATTAATAGTCAATGGATCTCACCACTTCACCCCCTT

TTTTTCCTTTTTTCCACTATTGTATACATATTTCTTAACCTCCGTGCCCAAACACAATGTAAACAATTGG

GTGGGACGGAGGGAGTATATAATAGAAACTATGCAATGACTACCTTTTTCTGCAAATTTCAAATGACTCT

GGTTTCTATTCTCGTGGTAGGAATATACAATGAACAAAATTATAGATTATGTTTCACTACCGGATAAATT

TGTATGAAGGTTCTTGCAAAATTGGAAACTACGAATAGCAATCATGATTTGGGGGCGGGGGGGTTAATGG

AATTGTGGAGGATCAGAAGTGTAGGAAATTACTGTTATCCTATTCCCACCTTAGAGAATTGCCCAAAGAA

GTAAATGAGATATTCCTTAATTTAACAGCCCGAGTAGAATTTGAATTCAGATAATTATGATAATGTTGTG

AAATATTGTTGATTTTCACTAACAAGTCAGAACTTGCGCGAAGAATGTTTATTCAATACAATTTGAGAAT

GAAGCTTATTTTTCAAGGCTTTGGAACAAGACCGGCTTTGATTTTCTGAATTGTGGCAACGTCAGTTTTA

AAAGAGAGAGCCAAGATGTTGTCGTCGATCCCTGTGGCAAACACAGTGCTTGGAATTGAGACAGTTCCTG

CACTACTGCTCCCGAATGCAGAAACTGCTAGAACAGGCTTCTGTGCATCCGCATTAAACTGGAAGTGAAC

AAGTCCTTTGGGAAAAACAAACAAGTCACCAGTCTGCAGGGTCTGAGTGTAGAGCTTGTTAGTCGTGTCA

ATAAGGCCAACTTCCAGAGTCCCATCAAGGACGAAAAGAAGCTCAGATGCACGAGGGTGAGTGTGAGGAG

GGTTTGTCGTGCCAGCAGGGAACTGAAGAGCAGCATACGAAACACTCTGACCATTGAGCGCTGGAAATTC

GGCCATGCTTGCTTTCAAGACCTTAAAAGCAGGTGGAGTTTGTGCACCAACAAGTACCCTGAAGCCAGTG

AATGTAAAGAAGTTCCCATCAATCGAGGTTACGTTGGATGGGACTATGAAGTCGGTGAGGATATCTGGAT

CACCAGCCATTGTGATCTCGACGACAGAGAAAGAAGCCACCAAGAAAAAGAACCAGCAGAGAAATGATCT

TGGTGATGATCCCATGGTGTATGTAGTCTACACTTAAAGTGAACTTTTGGTTTCTTTTCCTTGATGTTGA

CTAGAAACCCACAAGAGGCAAGTATTTATAGTATGGAATTATGATCTTGATCCTTGGACTCTAGTTGTTG

TTGCACCTGGAACACTTCTCATTGTGGAATGTGGACTACAATGTTTTAACTTTGTTATTGATGTTGTGTT

GCTACAGCTGTGCCAAAACCTATGAGAATATGTTGAAGATATAGTTTATATTCCATTTGGTATTATCATA

CGTTGCAGATTAGCTGCTCCCTTTGTTTTTCTTATGTACAGAAATATCTTCTTTAAGAATAAGTAACCGG

CT

>MSTRG.1128.2 gene=MSTRG.1128

ATTACACCAACCCTAATTACACGCACGCTCTCCCGCCTCCCGACAATGCTCACCGCCGGCCGCTGACTTC

AAGCATCTGTATATATAGTCAAAAATCAAATCAAAAATTCAAAGCATCAAAGGTATCTAATCGACTACTC

ACTAAGTACTATATATATCTTCTGATTTGCCTACTTTGTGTTTAGATTTTATACCCAATCAGATGTTGGC

CTGTGAGTCTGTGACTTTTGAGTACCTGCATACAGGCATCAGAAGAAATCAGCGACATCAAGCCTTCAGA

ACACTTTTGGCTACATATGTTGATGTTTCATATCTTGATTTTTCATAATATTGTTTTCTTAAAATTCAAA

AATCCAAAAAATTCGGTTATTTCATAACCAAAACCGAATTAACCGAAATATTTCGGTTCGGTTACGGTTC

GGTTTAATATATAATTCGGTTCGGTTCGGTTTATGAAAATTTTGACATTTCGGATTTCGGCTATTTCGGT

TCGGTTCGGTTTTGAACCGAACCGACCGAATGCTCAGCCCTACTTCTCATGGTTGGAGGTGCTCTTATAT

ATTAATAGTCAATGGATCTCACCACTTCACCCCCTTTTTTTCCTTTTTTCCACTATTGTATACATATTTC

TTAACCTCCGTGCCCAAACACAATGTAAACAATTGGGTGGGACGGAGGGAGTATATAATAGAAACTATGC

AATGACTACCTTTTTCTGCAAATTTCAAATGACTCTGGTTTCTATTCTCGTGGTAGGAATATACAATGAA

CAAAATTATAGATTATGTTTCACTACCGGATAAATTTGTATGAAGGTTCTTGCAAAATTGGAAACTACGA

ATAGCAATCATGATTTGGGGGCGGGGGGGTTAATGGAATTGTGGAGGATCAGAAGTGTAGGAAATTACTG

TTATCCTATTCCCACCTTAGAGAATTGCCCAAAGAAGTAAATGAGATATTCCTTAATTTAACAGCCCGAG

TAGAATTTGAATTCAGATAATTATGATAATGTTGTGAAATATTGTTGATTTTCACTAACAAGTCAGAACT

TGCGCGAAGAATGTTTATTCAATACAATTTGAGAATGAAGCTTATTTTTCAAGGCTTTGGAACAAGACCG

GCTTTGATTTTCTGAATTGTGGCAACGTCAGTTTTAAAAGAGAGAGCCAAGATGTTGTCGTCGATCCCTG

TGGCAAACACAGTGCTTGGAATTGAGACAGTTCCTGCACTACTGCTCCCGAATGCAGAAACTGCTAGAAC

AGGCTTCTGTGCATCCGCATTAAACTGGAAGTGAACAAGTCCTTTGGGAAAAACAAACAAGTCACCAGTC

TGCAGGGTCTGAGTGTAGAGCTTGTTAGTCGTGTCAATAAGGCCAACTTCCAGAGTCCCATCAAGGACGA

AAAGAAGCTCAGATGCACGAGGGTGAGTGTGAGGAGGGTTTGTCGTGCCAGCAGGGAACTGAAGAGCAGC

ATACGAAACACTCTGACCATTGAGCGCTGGAAATTCGGCCATGCTTGCTTTCAAGACCTTAAAAGCAGGT

GGAGTTTGTGCACCAACAAGTACCCTGAAGCCAGTGAATGTAAAGAAGTTCCCATCAATCGAGGTTACGT

TGGATGGGACTATGAAGTCGGTGAGGATATCTGGATCACCAGCCATTGTGATCTCGACGACAGAGAAAGA

AGCCACCAAGAAAAAGAACCAGCAGAGAAATGATCTTGGTGATGATCCCATGGTGTATGTAGTCTACACT

TAAAGTGAACTTTTGGTTTCTTTTCCTTGATGTTGACTAGAAACCCACAAGAGGCAAGTATTTATAGTAT

GGAATTATGATCTTGATCCTTGGACTCTAGTTGTTGTTGCACCTGGAACACTTCTCATTGTGGAATGTGG

ACTACAATGTTTTAACTTTGTTATTGATGTTGTGTTGCTACAGCTGTGCCAAAACCTATGAGAATATGTT

GAAGATATAGTTTATATTCCATTTGGTATTATCATACGTTGCAGATTAGCTGCTCCCTTTGTTTTTCTTA

TGTACAGAAATATCTTCTTTAAGAATAAGTAACCGG

>MSTRG.1132.1 gene=MSTRG.1132

ATTTTGTTGAGACAAGCATTTTATATTGTGGAGATTGGTTTTATTTTATTTTTGTCAATAATTATGCACT

ATTACAGTTAGTGGAGATATGCTCTGTTATTTATCATAGTCTTAATCTTTGCATTTCAGATGACCTTTTG

CATCTGTATCCCTCTACTTTTAGATCATTTTCGAGTTACATAAATCTTGCTGTATTGTGTTTTCTAAAAT

TTTACTCCTCATGTTCACTTATAGAAGCTGTTAAAGTTTGTACAAGTCATAGTTATGCATTGGACTTTAT

GATCTGGAGTTGATTAGAAATAATGTTAAGTCACTGAAAAAGCCTCTCCTTTGAGCAGTCCTGCAGGCCG

CAACCCTCCTTATGTTTCTGATCCAATTTAGCAGTCCTGCATTTGAAGCTGAGTGATTTTTCTAAGCCAA

GCCATAGTTCATCGAAAAATCCTTCACCAGTTACCCCCATCCTCCTTCTCCCAGTGAACCTGCTGGATCT

GGTTCTATTCTTCTATGCCGCGCCCAACAATATTTGATGGATCAGAACCAACAGCTACTCCGAATTCAGG

AGTTCCTCTGTCTCTTGACCTGATGCTGTTCTTCACAGCAAGTAGTCTACTTCCCATTGTGCTGAAGACA

AGGAAAACAATCTACAGCTCCACCAACTGATCAGCCAACTGTACCAGCTCAGAAAAGGGGATATGAAGCA

TGGAACCCTGAGATGGATAACATCTTAACTTCAACATTGTATGATCAAATTAATGAAGGAAATAAAGGCG

ATGGAAATTTTAAAACTCAAGCTTACCAGGCTGTGGTTGATAAGCTGCGGGTTGAAAGGAGTATATTGGC

TACTGTGGATCATGTGAAACACCGAATTAAGATTTGGAAGAAACACTATGGTGTGATTTCTGATATTCGA

AACCTTACCAAATTTAATTGGGACGAAGAAAAGAAGATGTCAGATCTTGTAGAATGGAAAGCTTAATGTG

AGACAGGGATGTTCAATTTGGATGATTGTATTGGTGGTTTGGTTCAAAAGAAGAAGAATCATCAAGCAGC

CATGTAGTCATACAGAAAGAACAGAGTCTGTAACAACTGAATGCCAGATTCTATAGTTATCATCTTTAAG

CAACAATGAGAATAAAAAGATAGATGACCGAGGAAAGCCCTCTCAAGTAGCACAAGGCCTTAGCAAAGTG

AAACTAATACACGTGCATTGTCAAGTTTTATTTAAACATCCAATTGTTTTCGATTTCTATTCGTCAATCA

TTTGATGCTTAATTTATAATCAAC

>MSTRG.1132.2 gene=MSTRG.1132

GGAAGAATATAGAGATGATCTTGCAAAGAAAATGTTGCACGGACTAGAGCTGCATAAATGATTGGTTTCT

GTTTTGACCTTTTTGTACCATTTCTGATGCAGCTGTTGGCATATATTTTGTTGAGACAAGCATTTTATAT

TGTGGAGATTGGTTTTATTTTATTTTTGTCAATAATTATGCACTATTACAGTTAGTGGAGATATGCTCTG

TTATTTATCATAGTCTTAATCTTTGCATTTCAGATGACCTTTTGCATCTGTATCCCTCTACTTTTAGATC

ATTTTCGAGTTACATAAATCTTGCTGTATTGTGTTTTCTAAAATTTTACTCCTCATGTTCACTTATAGAA

GCTGTTAAAGTTTGTACAAGTCATAGTTATGCATTGGACTTTATGATCTGGAGTTGATTAGAAATAATGT

TAAGTCACTGAAAAAGCCTCTCCTTTGAGCAGTCCTGCAGGCCGCAACCCTCCTTATGTTTCTGATCCAA

TTTAGCAGTCCTGCATTTGAAGCTGAGTGATTTTTCTAAGCCAAGCCATAGTTCATCGAAAAATCCTTCA

CCAGTTACCCCCATCCTCCTTCTCCCAGTGAACCTGCTGGATCTGGTTCTATTCTTCTATGCCGCGCCCA

ACAATATTTGATGGATCAGAACCAACAGCTACTCCGAATTCAGGAGTTCCTCTGTCTCTTGACCTGATGC

TGTTCTTCACAGCAAGTAGTCTACTTCCCATTGTAATAAATCTTTTGTAGATTTGTTCCCTTGCTAGATT

CTTGTTTTTACCTATTTTCACACATATGTAAACCATGGAAAAGTGGATAACAAGGTCATCTTCACCTTTG

ACGGTTTACTTTACTACATGCATAATAAAAATAAATAATTCAGTACTAAACATGATTGTCTTTTTAAATC

TTCTGAAGTTATCATGATTGATTTGCCTAATTCTGTGCAACCCTCTGATTTAAATCTTGTCAGAAGTGAG

ATCTGTTCTTACATTCCTGCTCGGTAGCTGTCTTTTGCTCTGTGCTCGGTATTAAGATGGCTAGTTACTT

GTGTTTTTGAATTAGTATGACTGTTTGAAGTGGATCATCCAACTGATTGTCCAGAATTAGGTGAGAAGCA

TGAGCCCTGTGGGATGTGAGTTTTTATGCGACAATGTGATAACTTTGTTAAGGCTGTTAATTAGTGCAGG

ATTGTTCTTTTCTTCTGCAAAGAAACATTTACATTCGTTTTTGTGTTTAGTGTTACTATGGATTAAAATT

TATTAATGTTGTAGGAAACTAAGGATTGTCTGTCTTTTTCGTTTTATTTACTTGTATTTTCGCAAAAGTA

TTGCATGAATCTACTCATTAATAATTATTTTTAGCATAAATATCTTAGCTATAATTCTTGGAAACAATTA

TTGAGCTCACTTTCACTTCTGAAACGAGGTTGTGCAGTGGTGTTCATAATTTATGCAACGTTTTGCTTAA

CATTCTTTGGTACTCGATCATCTATACTTTTGTTTTGCAGGTGCTGAAGACAAGGAAAACAATCTACAGC

TCCACCAACTGATCAGCCAACTGTACCAGCTCAGAAAAGGGGATATGAAGCATGGAACCCTGAGATGGAT

AACATCTTAACTTCAACATTGTATGATCAAATTAATGAAGGAAATAAAGGCGATGGAAATTTTAAAACTC

AAGCTTACCAGGCTGTGGTTGATAAGCTGCGGGTTGAAAGGAGTATATTGGCTACTGTGGATCATGTGAA

ACACCGAATTAAGATTTGGAAGAAACACTATGGTGTGATTTCTGATATTCGAAACCTTACCAAATTTAAT

TGGGACGAAGAAAAGAAGATGTCAGATCTTGTAGAATGGAAAGCTTAATGTGAGGTAAATTGTTATGTAC

TTGAAATAATCATGCATTGTTTAGCGTCTATTTTTAATTATACTAGGTCTTCTAAAAAGAATCCAACATC

TGCTGGTACTATCTCTGTCCAGTGAGCTGGAGTATTGAAAATTAAAGAATCATTTCATCTATTACATCTA

TTGTTGCGCGTAATCCAACTGAGCATCCGTGGATTGTATTCTGTAAAATGATTGGGAAGCATTTACTGTT

TTCTTAAGAAATGATAGAAGATTTAATTTAGTCTGGCTCATGCTACTACTATCTGCAATTTAGACCTATA

AACTAAGACTGATTGCACCATTATTTTACAAGTTGTCATACACATAACATACTCTGCTATGAATTGCTTA

GACAGGGATGTTCAATTTGGATGATTGTATTGGTGGTTTGGTTCAAAAGAAGAAGAATCATCAAGCAGCC

ATGTAGTCATACAGAAAGAACAGAGTCTGTAACAACTGAATGCCAGATTCTATAGTTATCATCTTTAAGC

AACAATGAGAATAAAAAGATAGATGACCGAGGAAAGCCCTCTCAAGTAGCACAAGGCCTTAGCAAAGTGA

AACTAATACACG

>MSTRG.1132.3 gene=MSTRG.1132

GTGGAGATATGCTCTGTTATTTATCATAGTCTTAATCTTTGCATTTCAGATGACCTTTTGCATCTGTATC

CCTCTACTTTTAGATCATTTTCGAGTTACATAAATCTTGCTGTATTGTGTTTTCTAAAATTTTACTCCTC

ATGTTCACTTATAGAAGCTGTTAAAGTTTGTACAAGTCATAGTTATGCATTGGACTTTATGATCTGGAGT

TGATTAGAAATAATGTTAAGTCACTGAAAAAGCCTCTCCTTTGAGCAGTCCTGCAGGCCGCAACCCTCCT

TATGTTTCTGATCCAATTTAGCAGTCCTGCATTTGAAGCTGAGTGATTTTTCTAAGCCAAGCCATAGTTC

ATCGAAAAATCCTTCACCAGTTACCCCCATCCTCCTTCTCCCAGTGAACCTGCTGGATCTGGTTCTATTC

TTCTATGCCGCGCCCAACAATATTTGATGGATCAGAACCAACAGCTACTCCGAATTCAGGAGTTCCTCTG

TCTCTTGACCTGATGCTGTTCTTCACAGCAAGTGCTGAAGACAAGGAAAACAATCTACAGCTCCACCAAC

TGATCAGCCAACTGTACCAGCTCAGAAAAGGGGATATGAAGCATGGAACCCTGAGATGGATAACATCTTA

ACTTCAACATTGTATGATCAAATTAATGAAGGAAATAAAGGCGATGGAAATTTTAAAACTCAAGCTTACC

AGGCTGTGGTTGATAAGCTGCGGGTTGAAAGGAGTATATTGGCTACTGTGGATCATGTGAAACACCGAAT

TAAGATTTGGAAGAAACACTATGGTGTGATTTCTGATATTCGAAACCTTACCAAATTTAATTGGGACGAA

GAAAAGAAGATGTCAGATCTTGTAGAATGGAAAGCTTAATGTGAGTGAGCTGGAGTATTGAAAATTAAAG

AATCATTTCATCTATTACATCTATTGTTGCGCGTAATCCAACTGAGCATCCGTGGATTGTATTCTGTAAA

ATGATTGGGAAG

>MSTRG.1134.1 gene=MSTRG.1134

TCCGGTGCTTCTCAGGCAATGCCCTGTGCTACTTCCTTAGGTAGTGCATTTTAATCAAAGTTCATGGTGC

CTCCTGACTGAACTTGAATTTTATGATTGTGCTTCATCTGCATCGGAGTCACTGCCTGAAGAACTCCCTG

AATCACTTGAAGAAGACGAGTCACTGCCAGAACTGCCAGAGCTACTCGAACTACTACTAGCATTACCATT

TCCAGTACCATTCTTAATACCATTATCCTGACCATGCCCACCATCATCCTTCTCAATTTCAACAGGTGGA

AAACTATTTTCCGGCATCTCATCCCCAATGTCAACATCCTCATCCC

>MSTRG.1135.1 gene=MSTRG.1135

CTCATATTTCGTACACACAGCTCCACATAAATTGAGAGAAAACTCAGGGAGCTAATCAAATCTCAAATCT

CATACTGTCCTTGTGAAAATTCCGGGAGAATTTCAGATCTCATCTGCCATCTGCATCCTCTTGTTCGGTC

TGTTTTATGTACAATCTATCGTTATGTGTATGTATGTTTTACCGATTTTATAAATCATGAATTTTGTGTT

TAATAGTCGACCAGACATAATGTGTAAGGGTGTTTATCGATTTTATAAGGCGTGGATTTTGTGCTCAGGC

ATGTTGGTGTGATAATGGTGGTTTTGACTTCTCCGGCCGGCAAGTAACTTGTACGATGGATCAAAGTAGA

TGAGATCTGAAATGGTAATACTCTCTGCAGGTGGTCCAGTTTTACATCTATGGAAAATTGGAAATAATAC

TAGTAAGTATAAAAAGGACGATCAAATTTAGTTTGGTGTGCATTAATGCATCATTTGGAGTGAATTTGAG

GGCATGCGAAGATTATATATGATGAATCGGAACACCAAGGATGATGCAGAGCGCTCTATAAATCATTCTT

TTAATTTTTATAATTTTTTGAGAGATTATAGGTAAGATAGC

>MSTRG.1153.1 gene=MSTRG.1153

CCCTTCTTTAATATTAATTAGTTTTGGCTCTCTACATTAGAGCTCTGAAAACTTTAATCATATACACATA

TACATTTCTAAAATGTTGATCATGTCGATATAAAAATTGTGTGGACAGCCATAAAAGAGTTTCTGCTCAG

GTTGCTTTGAGATGAGTAACAGTATACAAGTAGTTGCCGGGGTGTAATTACATTAAATAAAGGAGATGGT

ATTCTGCCTCGCTGCCAATCATCTTCAACGATTTTGGTTACAGAATTATGTCTCATTCCCATTTTAAATG

TTCTTATTTAGAAGAGTGTGAAATGATTTTGCTCAGGATTTTTTCCCCAAGGAGATCTTGCGCGGGGGAC

GACAATGCTATGGTTGCTCGTGCCAAGCTTCTTCTTGAGGAGTTTGTGCAGTTCCATCCAATTGGAATAT

ATGGTGCTGACCGCCTCATTAATGAAGGTTCTCGCGGTGAAGGAGGCATTCTTAGAAACAGTGAAGTGTT

TACCTCCATCATCTGGGAACTGTGCTCCACGCTCGGCTCAAACGCAAGAAACCGGGAGGGAGATGGTCGT

ACAATCCGTTTTCGTTACTTGGATGCTCAAAATATTGTGTTTTTTTTTAAATTTCAGTGGTTAATCGGTG

CATAGAGCTGAACTTGAAGGGTTTATATAAATTTAAGCCGAATTAATATGAAGAACGTAACTAATCAAAA

TAACTACAACCTATCAAAATATTGTGAATCGTACGACATTATTATTTTATTCTACTTA

>MSTRG.1155.1 gene=MSTRG.1155

CTCCTGCTCGTTTACGCCAACATCTCCACCTCTATCACCTTTTTCTTCAGTTTCAATATCTTCTTCTTGC

TCTTCGTTTTCATGCTCATCTTCTATTCCAGTTTCTTCTTCATGTTTGTCCTCTTCTCCATTTTCTTCTT

CATGTTTGTCCTCTTCTCCATTTTCTTCATCATGCTTGTCTTCTTCTCCAGACTCTTCTTCTTTCTTACC

TTCTTCGGTTTCTTCTTCATTTTTGTCTTCTTCTGCAGTTTCCTCTTCATGCTTCTCTTCTTCCTCGCTT

TCTTCTTCATGCTTATCTTCTTTCTCGATTTCTTCTCCATGTTTATCTTCTTCTCCAATTTCTTCTTCAT

TCTTATCTTGCTCTCCATTTTCTTCTTCACTCTTATCTTCTTCAACAATTTCTTCTTCAACTTCTTCATC

ATGACTCTCAAGATCCTTGATTGGCCCCTTTTCTTGTGAGAT

>MSTRG.1158.2 gene=MSTRG.1158

GCTAGAAGGATATATATAAGAAAGTCTTATGCTGAATCTACGTAATCAACTGCACTACCACTGGAAGTTG

CTTAAGTGAATAGGATATCTGTAGTTGAATCTATCTTAGCGACACACTGATAAAGATATCTCTGGACGTG

TGAGTTTTCGAAAAATATCATTTAACAGGTCACAATATGTGGTCACCTAGAGGAGCGTAGTGACTTTTAA

GGTTGTGTTAACAATTGTCGTATATAAAATTTTCAGTTCTGCAGCTTGTTGTTTAAATGTGATGCAATTT

TCGCGCCCTCTATGTTGATGTTTGTTTTCGAAATGGAAGTGAAGTATCTACCTTTGATTTGTGCATATAT

TATTATCTATCTAAATGATTTTTATTTTTTTTTTGTTTCTCAAGGCTGGGGATGGAGAAGATATTAGAAG

ATAAAATGGTGTGTAGATAATTATCAGTACTCTTAGTAACTGGTCCTTGCAATGTTTCCATCATGTGCTC

GAGCTCTTTTGTCCCGAAAGAGTAACTATCGCTACAGAGACTTGAACAGAATTTTAAATTTTCTGTCGAA

TTATAGCAGTATTTCACCACCCTCCTACTATTTATGCAAAGCATCCTTCATCTCATCTGCAAATACGAGA

TCTGTCCTGGATACTTTTGATATAAATGGTCATCAGACGAAAAGCTGGGTTTTAAATCAATTTAAATTGA

TTCCAGAATCATGTATTTTCCTGGCTCAATGCAAGGCTGGATTTTGTACTAGTCCAGAGATTAATGTGAA

TCTTGATTCGGAAGAAGCATCTAGTGATTCTGTTAGTAGTGCTTCTAGTTCTGATCCCAAAGTAAAGTTA

ACAAGTAATGATTCCATCGAGTTTAGTAAAGTTGATGCAAATATGCTTCCAACTGTACTGCTTATTGGTC

GGCCAAATGTTGGCAAATCTGCTCTTTTCAATCGTTTGATCCGACGAAGAGAGGCTCTTGTTTACAATAC

TCCTAATGATCATGTTACTCGAGATATACGAGAAGGGATTGCAAAACTGAGTGATTTACGGTTTAGGGTA

TTGGATTCTGCTGGTTTAGAGGCAGAGGCTTCTTCTGGGTCTGTTCTTGGAAGAACAGCAGGAATGACGG

AGAATGTGTTAAAGCGGTCTCAGCTTGCACTTTTCCTAATAGATGCAAGAGATGGGTTGCAACCAATGGA

CTTGGACGTTGGAAAATGGCTGCGCAAGAATGCACCTGAATTGAAAACTATCGTCTTAATGAATAAAGCG

GAGTCACTGGATGATGGCCTTGGTTCTCTTGCTGCAGCTTCTGGCGAGGCCCTTAGGTTGGGATTTGGAG

ATCCAATTGCCCTCTCTGCTGAGACTGGTCAGGGTATGGTGGAACTCTATGAAGCTCTTAGGCCATTGCT

TGTGGATTATATGGTTCAACGTCTGAAAGGAAATTAGCTCCTCCGAGGATGAGGACTCAAAGACCCCATT

ACAGCTGGCGATTGTGGGGCGACCAAATGTAGGAAAGTCAACCTTGTTGAATACATTACTTCAAGAGGAG

CGTGTTTTAGTGGGGCCAGAAGCTGGACTGACAAGAGATTCAGTTAGAGCTGAGTTCGAATATCATGGAA

GAACCATCTACATGGTTGACACTGCAGGTTGGCTGCAGCGGACAAATCAGGAGAAGGGACCAGCATCGCT

AAGCATTATGCAATCAAGAAAAAACCTAATGAGGGCCCATGTAGTTGCTTTAGTCCTAGATGCTGAAGAG

ATTGCTAATGCTAGGAGAACAATGAAGCATGATGAAGTAGTCATAGCAAGGAGAGCTGTGGAAGAAGGAC

GTGGTTTGATTGTTGTGGTGAACAAGATGGACCTTTTGAGAAATTCTAGGTCATATGAGAGTGTCATCAA

GGCTGTTCCTGAAGAAGTTCAGACTGTTATACCGCAGGTAACAGGAATTCCTGTTGTGTTTGTTTCAGCA

TTGGAGGGAAGGGGCCGGATAGCTGTAATGGATCAGGTGATTGATACCTATGAAAAATGGTGTTTAAGGT

TGCCAACTGCTCGTCTTAATCGATGGCTACGGAAGGTTA

>MSTRG.1165.1 gene=MSTRG.1165

CGAAAGTACATAAATTTATGAACCCCTCGGGCCCTCATTTTCCAACTTCCACTCCTCCCTTCTCTTCTCC

AACCTTCTCCTCTGCTTTACATTCTGTCTCCCTCTGCATACAATGAATGCGAATGAAATGAGTTACATTC

CAATATTCAGGCCAGAGATGGGGTCTTACTCTGAGACAGAGAAGAGAGCATTGTTTCTCAGGAGTTACCA

TTTCTCAAGAAAGCAGTCCTTTTCAGAAAGAATCAAGAAATCTTTCTTCAGGATCAAGAGGGTCATCAGG

GTCAGGCTCCGATCCGCCAAAAGGCTGAGGAAAATAGTGTGGTTCAGACTCAAGAATGGAATGTTCTGCT

CTACTAGAACCAGAACGAGGTTTCTTCGAATGCAAACTAGAGATGCACAAAAAATCTGGACCCGAAGACG

CTGCAAAGCGTATGTTAGTTCATGGCCATCTTCTTGTTTATGGTAGGAGGTAAATCAGGGTGATTATTGC

AATGTGTTTGTTGGAGGAATTGATTTAGGAAGATTTGTAGTTAATTGTATGGATTATAGTTTATGCTGTT

GGCTAGGTTTCTAATAAGAATTATTTTACCGCTGGAT

>MSTRG.1167.1 gene=MSTRG.1167

ATATATTTTAACGAGTTTGGTGGACACTTTGATCTGAATCCTTTTTACGTCTAAACAGGTGTTAAGATGA

ATGGATATGCCTGCAAAGAGAACTTCACTGCTGATGATTTCTTTTTCGCGGGATTGTCCAAGCCAGGACT

CACCAACAACACATTTGGTTCATTGGTGACCGGAGCCAATGTGCAGAAGATCCCCGGACTAAACACACTC

GGGGTCTCATTTTCACGCATAGACTATGCCCCTGGTGGCCTCAACCCTCCTCACACGCACCCTCGTGCCA

CAGAAATTGCGTTTGTGCTTGATGGAGAGCTTGAGGTTGGATTCATTACGACAACAGGTACCCTGTACTC

CAAGACCGTCAAGAAAGGTGAAATCTTCGTGTTCCCTAAGGGACTTGTCCACTTCCAGAAGAACAATGGC

AAGGTACCTGCTGCAGTCATTGTCGCGTTTAATAGTCAGTTGCCTGGCACTCAATCCATTGGCGCCACAT

TGTTTGCCGCGGTGCCCCCAGTGCCAGATTATGTGCTGTCACAAGCATTCCAAATTGGCACCAAAGATGT

TCAGAAAATTAAGTCAAGATTTGCACCAAAGTGATGTTAATGTAACATTGCATCTGCAGGTTTCTGTTGA

TTGTATTTCGGAAAAATTTGTGAGGTCAGTTGTTACAGTAGATGAGGTTTACTTTAGTCCTTTGCGCGTT

GAGAATCTTAAACTTGTTTGTAATGCTGTACTGATATTTAAAATTTTCTTCTCTGAGTACTTCTAATAAA

ATGGGAGGTTAATGTTCTTGGTGA

>MSTRG.1169.1 gene=MSTRG.1169

GGCATCATTCTCTTCTTCAACTACAGGAGGCTTGGTCTCATTCTTCACTTCCTCAACATTAGGAGTTTCG

GTTTTGTTCTTCGAACCTCCAACCACTGGATTCTCAACAATCTTCTCTTCTACAGGAGTTTCTGTCACGA

TCTTCTCGTCTACTGAAGGTTCTGTCATGATCTTCTGTTCGACTAGAGGAGCTCCAGTAGCATAATTCTC

TTCTACAACTAAAGGAGTCCCAGTCATGCTTTTCACTTCTTGAACAATAGGAGTCTCGGCACTGTTTCTT

GTTTCTTCAACTGTAAGAATCTTAACATTCTTCTCTTCAACTGGAGGCTCAGTTTCAAACTTCTCTTTCA

TTGGAGGAGGCTCTATACCGCTGTTCTCTTCTTTGGTTTCAGAAGTCTCAGTTGCGGTCTTTTCTTCATC

TATAATTTGCTCTGTCATGACCTTTGTCTCTTCAACCACAGGAGCCTGAACAAGCTTTTCTTGTCCTGCC

TCAGATAAGATTTCCTCATTTGTTACAAGTTGCTCATTGGTTTCCTCAGTTTCTTTAACGGCAAAAGTGT

GTAGAATCTTTTCTACTGGAAGAGGCTCAATCACGATCTTCTCTTCTACCACAGGAGGCTCTGCAGTGAT

CTTTGCTTCTTCAATTATAGGAGTCTCAGTTTCAGTCTTTGTTTCCAATACAAGTTTCTCAATTGTGTCG

TCTGCTACTTCAACCGCAGGAGTCTGAACAAGTTTATCTTCCACTGCAGGAGTCTCAGTTGACAACTTGT

CTTCCACCGCAAGTGCCTCAATGGTGTCCTTTTCTTCTTCAACCACAGGAGTCTGAACAAGCTTATCTTC

CACTAAAGGAGTCTGAGTTGATATCTTCTCTTCCACTACGGGAGACTCAGTGGTGTTCTTCTCTTCTTCA

GTTACAAGACTCTGAACAAGCTTTTCTTCCACTACAAAAGGCTCAGTAACAATGTTCTCTTCTACTAGAG

GAGGCTCAGCAACGATCTTTGCTTCTGCAGCTACATGAGTCTCACTTGCGACATTTAATTCAATTATGGA

AGGCTCAAAACTCTTATCTTCAACGTTTGATTCAGGTGCGATAGAAGGAAGTACTTCAGAAGCCAGAGGC

TGGACTTCTTCAGCCAAAACAACACTTGGACTAGTGAAGTCCTCAGATTTCAGCTTTTCCTCTGAAGTTG

GCTCCATTTCTACTGTTGACATTTCTTCCTCAACAGTTATCTTTTCCTGTGTAACTACAGAAGACTCGAT

ATTCTTGCCTTCAATTTTCTCTTCAGCATCAGCTACAGGTGCAATAGGAGCTTTCTCTTCCTCGTCTCTT

TCCTTTAGCTCTTCATTCGTTGGGGCCTGAATTGGCTTTGCCTCAATCTTATCTTCAGATTCAGACAAAG

GCGCCACAGGAACTGGCAACGAGTTGAGTTCTGGCTTCTCTACCTCAGCTGGAGAAATGATCTCTGGTTC

ACATGCTTTTACCTCTTCATGAACTTCCTGTGCCTGCTGCAATAATGTCAAAATCAATCATTTCCATTCA

AGGAAAAAGATTACAATGCCTC

>MSTRG.1171.1 gene=MSTRG.1171

GGCTTATCTCATTTCAAGTTTCAACGATCACTAATTTTGTCACACTTCATTCTCCTTGTAATGGCAGAGC

TGGATTATGAACTGCCTCGTTTTATAAGAGTGTACAAAGATGGTACCGTGGAACGATTAATGGGCACGGA

CACGGTTCCTGCTGGCATTGATCCCGAAACCGGTGTCGAATCCAAAGACGTGCTCGTCACACCAGAGAGC

GGAGTATCTGCTAGGCTTTATCTACCGAAAATCAGAGACCCGGCCAAAAAACTTCCGCTTGTGATTTATT

TCCACGGGGGAGGATTTTATTGTTTTACACCGTTTTCCACTTTTTACCAGCCATTTCTTAATTCTTTTGT

CGCGGAAAGCAACATTATTTTGATATCGATCGATTATAGAAGGCCACCGGAACATCCGGTCCCGGTGGCC

TACGATGATTCTTGGAATGCTATCCGATGGGTGGCTTCTCATTGCAATGGCACTGGCCCGGAGGCATGGC

TCAGCGAGTATGCCGATTTTGGACGTGTACATATATCGGGAGATAGTGCGGGTGCAAATATCGCGCATTG

CATGGCTATGCGGGCCGGGTCGGAGAAAGTTAATGGGCTGGACTTTGATGGGCTGATTCTCATGCATCCG

TATTTTTGGGGCAAGGATCCAGTGGGTGGAGAAACGGATGATACCGAAAAAAGGGCCTATATTGAGAAGT

TTTGGGTATTAGCATGCCCAACAAGTAGTGGAGCGGATGACCCGTGGATGAACCCGGGTTTGGATCCGAA

ATTGTCGAGCTTGGTGTGCAAAAGGGCTTTGGTTTTTGGTACAGAAGATGATTTTCTGAAATATAGAGCC

AAGTATTATGGTGAGGTGTTGGGTAAAAGTGGATGGGCTGGAAAGGTGGAGGTGGTGGAGACTGAAGGTG

AAGCCCATGTGTTTCATTTGTACAAGCCCAAGATTAAGAAGAGCATGGAATTGGTCCATAAAGTAATGTC

TTTTATATACGAAGAGAATGCCAATCCTATTTAAACCCGAAATTTAGTGTTACGGGG

>MSTRG.1172.1 gene=MSTRG.1172

GACTATGATTCAATCTCATGTAACTAACCCCCCTCACAAGTCACCACATGAATTTATACATACCAAACTC

AAATCACACTACAAATCCATCACACTTTTCGAGAAACAACAATGGCATCCGACGAAGAATTAGTTCACAA

CTTCCCTCCTTACTTTCAGGTCCTCAAAAACGGCGCCGTCAAGAGGTTCTTTAACGACTCTTTTTTTCCG

ACCACCGGCGATGAGCCCAACAGCGACGGCGTTTTGACCCGGGATGTTGTGGTCTCGACGGAGCCTAAAG

TGGGTGCGAGGCTTTTCTTGCCGAAGATCACTGACCCGACCCGGAAGCTGCCGCTCCTGGTTTATATCCA

TGGAGGTGGGTTTTCGATCGGGTCGGCTTTTTTTAAGCTGTTTACGGATTATGTGAGTTCCCTCGTTGCG

CTTTCTGGGGTTGTGGCGGTGTCGGTGGAGTATCGGTTGGCGCCGGAGAATCCGATTCCGGCGTGTTATG

AGGATACTTGGGAGGTTTTGAAGTGGGTTGGTGGTCACGGGTCGGGTCAAGGGTCCGACCCGTGGTTGAA

CAAGTACGCGGATTTCGGGCGGGTGTTTTTGGCGGGGGATAGTGCGGGTGCGAATATTGCACACAACATG

GCTGTTCGGGTCGGGTCGGAGGATCCGGGGGTGAAGATCTCGGGTATGGTTTTGGTGCACCCGTTTTTCG

GGAATGATGAGGTGAGTCCAATGTGGAAGGTGATATGTCCGGATACAAGTGGGCCTGATGACCCGAGGTA

TAACCCGGCTGCAAATTCGGGTTTGATGGGGAAGATTGGGTGTGGGAAGGTGTTAGTTTGTATCGCCGGG

AATGATGAGTTGAAGAGTAGGGGATTGAGTTATTACAAGGCGTTGAAGAAAAGTGAGTGGAGCGGCGAGA

CGGAGATTTTTGAGACCGAAGGTGAGGAGCATGTGTTTCATGTTTTTAATCCGGCTTGTGAGAAAGCTGC

TGTGTTGCGAGAGAAAATTGCTCGTTATTTCAACAATTGACTGGAATCAAGTAGGCTGTCGGGGATGAAT

TAGTAGATGACAATGATCAATAAAATGAGAAGATCGTGAGACTTAGGTTTATTTTATTTTGTGTAGGTTA

TCTTTGTTTAATAAATCATGTGTTATTTACTTCAAAAATGAAACGTAGTTGAGGTTTAGAGTGAAATTTT

TTCGAATGTACCTTGAAATATGTATCAGAGAATTTGAATCTTTAGGCCTTGATAATAATGGATATTGGCT

GATGAAATTT

>MSTRG.1174.1 gene=MSTRG.1174

AAATTCTGAAACTCTGAGTTACGAAATGGCTTCTGCGGTGAAGAAGAAGAAGAAAGCAACAATCACAATT

GAATCTCAAGGCGAAGACAGGCTCAGCGCACTGCCAGATGAACTGATTCACAAAATCTACTCATTTCTAG

ATACTAAAGAGGCAGTTCAAGCCAGCGTTCTCTCCAAACGATGGAAGCTTATGTGGACTACTCTCCCTTT

CATTTCTTTTGGTCGGTACCTGGAATCTATGGACATTTTCACCATACATGCTTTGTTGAATATACCCAAT

TATGCTATGTTTATTCGCCATGTATTGACTAACCGAAACCATCAATCGAATATTTCCGAATTGAGAATGT

GTGTTCCCGCGAGATTAACTGTACCGCAGAGTGTTTCGAGGAAGAGGTTTCGTGGATGTGTGATTCAGAG

GTTTATCCAGTATGCAATTGATCATAATGTGGAATGTTTAGATATTGCCTTACGTTATAACCATAAGGCT

CGTAAGTTATCTCCCTTTAGCTCCAAGGTTTTGAGAGAACTCACATTAGATATGCCTCTTGAGGTATGTC

CAGTGGAACCGGATTGTTGGGATTTGCCTGCTTTAACTACTCTTCACTTGAAAAATGGTTCCCGTTCCAT

GTTACATTACTGGGAACGGGGTGTACATCCCAAGTTACCTGAGTTATGGCTGACCTCCTTGCCTGCCTTG

AGAACGTTGCAGCTTGACTCTTGGGATATATCAGAAACGTCTTTTTCTCTGCCTAATCTAACATCCCTCC

GTCTGTGTTACTGTAAAATATCCAAAATGGTATGGGATCTCCCAGCTTTAATGAATCTACATCTGGATGA

TAGTGCTCTTCCCAAGAATCTGAGTGACATGTTCTCCAACCTTGACAGTCTAAAAAATCTCACATTAGTT

ATACTCAAGTCTATTCAAGAAGTTCATATCATATCTTGTCCTCAATTGCTGAACCTGAATATCAGGATCA

GTCGTGAACCTCCTAATGATGCTCCTCATGGTATGATTGTGGTTTCGGCTCCACAGCTCTGCAACTTCAG

TTCTTTTGGTATATTTTTAATTACGTTTAAAGCTGCTGAGGTGGAGAATATCGATGTAAAGTTAAAGGAT

TGGCTCAAATATGTGGGCCCCGGAGATATGCAAATATACCGTCAGTGGTTTACATGTATGCTTTCAGGAC

TTGGTGCTGCAAAGAATCTTTCCTTTGACTTGGAGAGCATTCAGGCACTTGCTAAGCATTACGATCTAGC

AAGTTGTTCTTCTCCATTCTATAACTTGAAGTGTGTGAAGCTACCAAAAGAATATAGAGAATCAAGTGTA

CCTGGTTTTCTACGACGCTACTTACTTAATGGCTCTCCTGGAGCAACCATTGTCACATCACTGCCTCGGA

CCCAAACACGAGCAGTTTCAGTGGCAGCTGAAAATATGGTGCTCCCAGAGCCCATGGCAGCTCCAACTAA

GGAACTTGGTGATTGTCAAAATATAAATGAAGCTGTGAGTTCTGACACTCTTGACACGGGGGTTCGAGAA

CCGTATGTGATGGAGAATTCTTCAGATAATGCCAACAGATCTAGACAAACTAATCCTGCTCTTGTGGGGG

TAAGCAACGATCGGGCAAGCTCTTCTCAGGGTAATAAAGATATCGGGTTATGGCAGGGCCATAAGGTCAA

CTCTGAGTTTGTATGTCTACTCGATCGCATTTTCCACAAATACCCGGAAACCTTTGATCAATTCACCTCA

AAAAACAAGAAATTATGTACAATGAATTTGAATATGCTGTGCACCTCTGTGAACGACTTCACCAAAACAT

TCTTGAATGAGGTTGATTCTGAAATGATTGCTGATTACAGGGATGCATTTGCTTACTTGGGGAATCGAGG

ATTGAACATAGGATGGTTGAGAAGGCGCCTGAACTATATAGAACATCTTAAGTTCTCAAAGCCCCTGATT

CCCGAGCTTCATGCAATTGACTGTCATATTGATGATGCTAAAAGGCAAAATTTGCAGAGTAAACTACATG

TTTTGCAGGCTTTCCGCAGGGCGAAGATGTCAGAGATTCAAAAAGCTTTTGGAACTATGGGTACTAGACT

TACCGCTGGTTTCATCGGAGATGATCTCCTATCTGGTCCCTAACCTTCATAAATTTGTTGTGCCATTGAC

AGTTTTCTGGATGTGTAAATAACTTTCATTACTGTCCTGGTTATTGACTTTCTATGCTGTTGTTCTTGTT

TGGTTTTCTAAGAATTAGGAACTGTTATAGAAGTCTCTGAATGCTT

>MSTRG.1178.1 gene=MSTRG.1178

CATAATTTATCTTTTATAAATCAAAGTTATCACTTCACTTGCTTCATATTTCATATCAATTTCTTTTATA

ATGGATCCTGATGATATCATAGCATATTTGGTTCAAGAAGAAGTGGAGGAAGAGGATGAAATGAATATTG

CTCTGGCTTCATTTTCGGAACTCATTCGCAGAAGTAGTTCGAGGGTAGTTCGTCATGGAGGATCAGTGTT

CAACCACCGTATGATTCGCCGTAATAGAGATTTAGGTAATCAACTTATCTATCGTGATTATTTCAGTGAT

AATCCTACATATCCGGATTATATTTTTCGTAGGAGATTTCGTATGAGAAGATCATTATTTGTTCGAATTC

AAGCTGCGATTGAGTCACATGATCCCTACTTCGTGCAAAGATATAATGCAGCAGGAATTCTTGGTCTATC

ATCACTCCAAAAGATAACTGCAGCATTAAGAATTATTGCATATGGAGTCCCTGCAGATGCAATTGATGAC

TATATTCGAATTGGTGAAAGCACCGCAATTGAAAGTTTAAAAAAATTTGTCACAGCAATTGTGCAGATTT

TTGGTGAGCAGTACTTAAGAAAGCCAAATAGTGCTGATATTAATAGATTGATGGAGGTTGCTAAGCAGCG

TGGTTTTCCGGGGATGCTAGGCAGCATAGATTGTATGCATTGGAGATGGAAAAACTGCCCTACAGCAGCG

TGCTTTCCTCCCAATTGGCCTTTCAAAGGCTGAGTTTCCTAACATTTCACAATCCGGAGTTGAGGGAGAA

GAAATTTTTGGACTGTCCTTTTGGAAAATTTTTGCTTTCTTTGTACCAAACTCAATATTCCATTTAGGCA

AATGTCTCAGTAGATTCCAGCAGTGCTCAAGAGAAAAGTTGTGTTTGCAAAGATTTTTATACATATCCTT

CGCATTATTAATCTGTTAAAAAAATCCACAACTCTTGTATCATTCATGTGAGTAGTCTATTAAATACAAA

TTACTAGGAAAAAAATTACCTTATCAGCTTCAGAGCACCCACTTTTTCCTTCTACTTGGTTATAATATCC

TTGAAATTTATTGACGGCAAGTTGTATCACGGACCAACGATGCATCAATGAACAGCTGGTACGATCAGAT

TGAAAGTCTTTATGTTTCATAAAGTAGATGTGCACTCTTTGCCAG

>MSTRG.1181.1 gene=MSTRG.1181

ACTTGCTCTAAAAGATAAAAGACAGAGAGCCAAGAAATACATATGAAAACCATAAGAAATACAAAAGGGT

AATAGTTTTGACAGATGAAATCTTTGCAATTGGAAGCTAAAATAATATCCCATAAGCTTTCTGCTTGCGA

AGCCATGGGCTGGAATACAAGAAAGAAGGTTTTAAAAGGTGTTTTTGTTACCTTGTCTGGTATGGACTAA

ATTCAGCGGCAAGGCCTAATTAATCGGACAACTCAGAAACCAGAAGCTCGACATCAACTGCAATGGACTC

CCTGGCACCCTCTGGCGGCAACGGCGTCGTTTCACCCACCGAAACAACAGCAGCAACATCCTCACAAGAT

CCTAAAAACCTAGATGAAACCATCAATTCAATTCAAAAATCCCTGGGATTGCTTCACCAGCTAAATCTCA

CCGTCTCTTCTTTCAATCTCGCTTCTCAACCGCGTCTCCTTCAACGCCTGAACAACCTTGTGTTAGAACT

AGATAACATGTCGAAATTGCCGGAGAAGTGCAACATTGATGTGCCTATGGAAGTGTTCAATTTGATTGAT

GATGGGAAGAATCCGGATGAGTTTACGAGGGATGTGTTGAATAAATGTATTGCCAAAAATCAGATTACCA

AAGGCATGGCTGATGCGTTAAAGGGTTTGAGAAAGCATCTTCTTGAAGAGCTTGACCAATCATTTCCTGA

TGAAGTGGAAACTTATCGAGAGATACGTTTTGCATCTGCTGCTGAAGCTAAATGGGTTGCCCAAGCGCAA

AGTATGTTACCAAATGGGGATGTTAAGGTTAAACCTGAGGTATAGGATACAGTCACCAAGAATTCAAATT

AGTTATTTAAGAATCACCAATCAACCATGTTTTTAAGTCAAGATTGTTGATACCATTAGTATGTCTATGT

AAATTGTAAAAGTTTCAAGCTTTACATGAACCTATAGTTGCCTGTGGCAGTGGTACAGTAAAAAATAAGT

TCTCATTATGTTTTACCTTTGGGCATGACATTCCACCAAGAAGTCTCACTTGAGATTTGCACAAGATTGT

AATGGCAGGTTAGTTGAGTGGTACTAAATGGATAGTGTAATTATATGTTGAAGGGGTAAACTAAGACGAT

AGAGTCTCTTCATTAGGAGGTTTGCATGTAACCACCCGGTGTTACCTGGGCTAACCATTGAGATTAAATA

TGTACTTTCTTAACCATCAATCTTGGTTATTTGTCGTTTGTCTTGGATATTTGAAAGAACTCATTCTTAC

AT

>MSTRG.1181.2 gene=MSTRG.1181

AAGACAGAGAGCCAAGAAATACATATGAAAACCATAAGAAATACAAAAGGGTAATAGTTTTGACAGATGA

AATCTTTGCAATTGGAAGCTAAAATAATATCCCATAAGCTTTCTGCTTGCGAAGCCATGGGCTGGAATAC

AAGAAAGAAGGTTTTAAAAGGTGTTTTTGTTACCTTGTCTGGTATGGACTAAATTCAGCGGCAAGGCCTA

ATTAATCGGACAACTCAGAAACCAGAAGCTCGACATCAACTGCAATGGACTCCCTGGCACCCTCTGGCGG

CAACGGCGTCGTTTCACCCACCGAAACAACAGCAGCAACATCCTCACAAGATCCTAAAAACCTAGATGAA

ACCATCAATTCAATTCAAAAATCCCTGGGATTGCTTCACCAGCTAAATCTCACCGTCTCTTCTTTCAATC

TCGCTTCTCAACCGCGTCTCCTTCAACGCCTGAACAACCTTGTGTTAGAACTAGATAACATGTCGAAATT

GCCGGAGAAGTGCAACATTGATGTGCCTATGGAAGTGTTCAATTTGATTGATGATGGGAAGAATCCGGAT

GAGTTTACGAGGGATGTGTTGAATAAATGTATTGCCAAAAATCAGATTACCAAAGGCATGGCTGATGCGT

TAAAGGGTTTGAGAAAGCATCTTCTTGAAGAGCTTGACCAATCATTTCCTGATGAAGTGGAAACTTATCG

AGAGATACGTTTTGCATCTGCTGCTGAAGCTAAATGGGTTGCCCAAGCGCAAAGTATGTTACCAAATGGG

GATGTTAAGGTTAAACCTGAGGTATAGGATACAGTCACCAAGAATTCAAATTAGTTATTTAAGAATCACC

AATCAACCATGTTTTTAAGTCAAGATTGTTGATACCATTAGTATGTCTATGTAAATTGTAAAAGTTTCAA

GCTTTACATGAACGGTGAACCTATAGTTGCCTGTGGCAGTGGTACAGTAAAAAATAAGTTCTCATTATGT

TTTACCTTTGGGCATGACATTCCACCAAGAAGTCTCACTTGAGATTTGCACAAGATTGTAATGGCAGGTT

AGTTGAGTGGTACTAAATGGATAGTGTAATTATATGTTGAAGGGGTAAACTAAGACGATAGAGTCTCTTC

ATTAGGAGGTTTGCATGTAACCACCCGGTGTTACCTGGGCTAACCATTGAGATTAAATATGTACTTT

>MSTRG.1181.3 gene=MSTRG.1181

AAAAGGGTAATAGTTTTGACAGATGAAATCTTTGCAATTGGAAGCTAAAATAATATCCCATAAGCTTTCT

GCTTGCGAAGCCATGGGCTGGAATACAAGAAAGAAGGTTTTAAAAGGTGTTTTTGTTACCTTGTCTGGTA

TGGACTAAATTCAGCGGCAAGGCCTAATTAATCGGACAACTCAGAAACCAGAAGCTCGACATCAACTGCA

ATGGACTCCCTGGCACCCTCTGGCGGCAACGGCGTCGTTTCACCCACCGAAACAACAGCAGCAACATCCT

CACAAGATCCTAAAAACCTAGATGAAACCATCAATTCAATTCAAAAATCCCTGGGATTGCTTCACCAGCT

AAATCTCACCGTCTCTTCTTTCAATCTCGCTTCTCAACCGCGTCTCCTTCAACGCCTGTATGTAATTGAA

CAACCTTGTGTTAGAACTAGATAACATGTCGAAATTGCCGGAGAAGTGCAACATTGATGTGCCTATGGAA

GTGTTCAATTTGATTGATGATGGGAAGAATCCGGATGAGTTTACGAGGGATGTGTTGAATAAATGTATTG

CCAAAAATCAGATTACCAAAGGCATGGCTGATGCGTTAAAGGGTTTGAGAAAGCATCTTCTTGAAGAGCT

TGACCAATCATTTCCTGATGAAGTGGAAACTTATCGAGAGATACGTTTTGCATCTGCTGCTGAAGCTAAA

TGGGTTGCCCAAGCGCAAAGTATGTTACCAAATGGGGATGTTAAGGTTAAACCTGAGGTATAGGATACAG

TCACCAAGAATTCAAATTAGTTATTTAAGAATCACCAATCAACCATGTTTTTAAGTCAAGATTGTTGATA

CCATTAGTATGTCTATGTAAATTGTAAAAGTTTCAAGCTTTACATGAACGGTGAACCTATAGTTGCCTGT

GGCAGTGGTACAGTAAAAAATAAGTTCTCATTATGTTTTACCTTTGGGCATGACATTCCACCAAGAAGTC

TCACTTGAGATTTGCACAAGATTGTAATGGCAGGTTAGTTGAGTGGTACTAAATGGATAGTGTAATTATA

TGTTGAAGGGGTAAACTAAGACGATAGAGTCTCTTCATTAGGAGGTTTGCATGTAACCACCCGGTGTTAC

CTGGGCTAACCATTGAGCACCACCAGAATTTCCCTTTAGTTTTAAATTGCCC

>MSTRG.1181.4 gene=MSTRG.1181

ACTTGCTCTAAAAGATAAAAGACAGAGAGCCAAGAAATACATATGAAAACCATAAGAAATACAAAAGGGT

AATAGTTTTGACAGATGAAATCTTTGCAATTGGAAGCTAAAATAATATCCCATAAGCTTTCTGCTTGCGA

AGCCATGGGCTGGAATACAAGAAAGAAGGTTTTAAAAGGTGTTTTTGTTACCTTGTCTGGTATGGACTAA

ATTCAGCGGCAAGGCCTAATTAATCGGACAACTCAGAAACCAGAAGCTCGACATCAACTGCAATGGACTC

CCTGGCACCCTCTGGCGGCAACGGCGTCGTTTCACCCACCGAAACAACAGCAGCAACATCCTCACAAGAT

CCTAAAAACCTAGATGAAACCATCAATTCAATTCAAAAATCCCTGGGATTGCTTCACCAGCTAAATCTCA

CCGTCTCTTCTTTCAATCTCGCTTCTCAACCGCGTCTCCTTCAACGCCTTTTGGCAGGAACAACCTTGTG

TTAGAACTAGATAACATGTCGAAATTGCCGGAGAAGTGCAACATTGATGTGCCTATGGAAGTGTTCAATT

TGATTGATGATGGGAAGAATCCGGATGAGTTTACGAGGGATGTGTTGAATAAATGTATTGCCAAAAATCA

GATTACCAAAGGCATGGCTGATGCGTTAAAGGGTTTGAGAAAGCATCTTCTTGAAGAGCTTGACCAATCA

TTTCCTGATGAAGTGGAAACTTATCGAGAGATACGTTTTGCATCTGCTGCTGAAGCTAAATGGGTTGCCC

AAGCGCAAAGTATGTTACCAAATGGGGATGTTAAGGTTAAACCTGAGGTATAGGATACAGTCACCAAGAA

TTCAAATTAGTTATTTAAGAATCACCAATCAACCATGTTTTTAAGTCAAGATTGTTGATACCATTAGTAT

GTCTATGTAAATTGTAAAAGTTTCAAGCTTTACATGAACGGTGAACCTATAGTTGCCTGTGGCAGTGGTA

CAGTAAAAAATAAGTTCTCATTATGTTTTACCTTTGG

>MSTRG.1181.5 gene=MSTRG.1181

ATAATATCCCATAAGCTTTCTGCTTGCGAAGCCATGGGCTGGAATACAAGAAAGAAGGTTTTAAAAGGTG

TTTTTGTTACCTTGTCTGGTATGGACTAAATTCAGCGGCAAGGCCTAATTAATCGGACAACTCAGAAACC

AGAAGCTCGACATCAACTGCAATGGACTCCCTGGCACCCTCTGGCGGCAACGGCGTCGTTTCACCCACCG

AAACAACAGCAGCAACATCCTCACAAGATCCTAAAAACCTAGATGAAACCATCAATTCAATTCAAAAATC

CCTGGGATTGCTTCACCAGCTAAATCTCACCGTCTCTTCTTTCAATCTCGCTTCTCAACCGCGTCTCCTT

CAACGCCTGAACAACCTTGTGTTAGAACTAGATAACATGTCGAAATTGCCGGAGAAGTGCAACATTGATG

TGCCTATGGAAGTGTTCAATTTGATTGATGATGGGAAGAATCCGGATGAGTTTACGAGGGATGTGTTGAA

TAAATGTATTGCCAAAAATCAGATTACCAAAGGCATGGCTGATGCGTTAAAGGGTTTGAGAAAGCATCTT

CTTGAAGAGCTTGACCAATCATTTCCTGATGAAGTGGAAACTTATCGAGAGATACGTTTTGCATCTGCTG

CTGAAGCTAAATGGGTTGCCCAAGCGCAAAGTATGTTACCAAATGGGGATGTTAAGGTTAAACCTGAGGT

ATAGGATACAGTCACCAAGAATTCAAATTAGTTATTTAAGAATCACCAATCAACCATGTTTTTAAGTCAA

GATTGTTGATACCATTAGTATGTCTATGTAAATTGTAAAAGTTTCAAGCTTTACATGAACGGTGAACCTA

TAGTTGCCTGTGGCAGTGGTACAGTAAAAAATAAGTTCTCATTATGTTTTACCTTTGGGCATGACATTCC

ACCAAGAAGTCTCACTTGAGATTTGCACAAGATTGTAATGGCAGGTTAGTTGAGTGGTACTAAATGGATA

GTGTAATTATATGTTGAAGGGGTAAACTAAGACGATAGAGTCTCTTCATTAGGAGGTTTGCATGTAACCA

CCCGGTGTTACCTGGGCTAACCATTGAGATTAAATATGTACTTT

>MSTRG.1184.1 gene=MSTRG.1184

CTAGAGCTGGCCAAATGAGCGGTTTCGAACCGCTCGTTCGGAACTGAACCGTCAAATTATTCGTTAAAAT

TCGACCCGTAACGAATTTGAACCGGCCCGGACCGGCTCGTTAAATATCCCGAATCGAATACGAATCAGGA

TATAAGAAATCGTAACCGGTCCGTAACCGGCCCGCGAATTACCGAGCTAAGAACCGGGCGAACGACTCGC

GGTTCGAACGAACCAACCAAACGAACGAACCGCAAGTCAACGAGCTGCCACGAGCTGTCAACGAGCTGGA

CGAATCGAACGGGCTGGACGAATCGAACGAATCAACCGGTTCGAACGAATCCAGGCC

>MSTRG.1189.1 gene=MSTRG.1189

GGACAACACAAAATCCCTGTAAATGTTTTAGACAGCATTCAAGAAATCAAAAACCTAACAAACCCATCAT

TTAAAACTCAAACCCCATCATTCTTAGCCACTAATCTCCTCAAATCATATTTTGGAAGGGGCTTAATTGA

GGAAGCACGCACACTGTTCGATGAAATGTCTGAGAGAGACGTTGTTGCCTGGACTGCCATGATTTCTGGT

TATGCTTCTTGCAATTACTATACCAATGCATGGAATGTGTTTTTGAAAATGTTGAGGGGATGTGGTGATC

CCCCGAATGCGTTTACGCTGTCTAGTGTGTTGAAGGTTTGCAAGGGGATGAATTCACTTTCTTCTGGGGC

TTTGGTTCATGGGTTGGCTCTTAAACATGGTTTGAATGGTTGTTCTTATGTTGATAACTGTTTGCTGGAT

ATGTATGCTACTTGTGGCTGTAATATGGATGCTGCATGCGCGGTTTTTCAAGATATGGAGGTTAAGATCC

CAGTGGCGTGGACTATTTTGATCACTGGGTACACTCATCGGGGTGATGGTCATGGAGCTCTACAAGTTTT

TAGACAAATGTGG

>MSTRG.1190.1 gene=MSTRG.1190

AGTCCCTGTATATATTCTCACAAATGGGAATAGAGGGGCTTAGTCCCAACAGCTTCACATATTCAAGCAT

TATAGCGGCATGTGCTAATTTAGCAGTTTTAGGTTGTGGACAGCAAGTTCATGGAGGGATTTTTCAAAGA

GGTCTCGAAGGGGATTTACCTCTGGCTAATGCCCTTATAGACATGTATGCCAAGTGTGGAAGCATAATGG

ATTCATGTAGAATTTTTAGTCAAATGAACCGTAAAGATTTGGTATCATGGACTTCTATGATGATTGGGTA

TGGGAGCCATGGTTATGGCAAAGAAGCTGTTACTTTGTTTAATGAAATGGTCAAATCAGGTATTCGGCCT

GATCGGATTGTATTTGTGGCAGTCATTGGGGCCTGTAGCCATGCTGGCATGGTTGATGAAGGCATGCGTT

ACTTCAAATCGATGACTGTTGATTACAAAGTTTCCCCCAATCAGGAGGTTTACGGATGTGTAGTTGATTT

GCTTGGTCGAAGTGGTAGAGTTGAGGAAGCTTTTAAACTGATAGAAAGTATGCCATTTAAGCCTGACGAA

TCTGTTTGGGGAGCGCTTTTAGGAGCTTGTAAGGCTCATGGAATTTCAGATCTTAGCAAACGTGCAGCTC

AGAGAGTACTAGAGTCGAAGCCTAGTGTTGCAGGAAGTTATGTGATGCTAGCAAATATTTTTGCAGCTCA

GGGTGATTGGGGGGATTTTGCAAGAATGAGAAAGTTGATGAGGGGAACCGGAAACAAGAAAGAGGCTGGT

AGGAGTTGGGTGGAGCTCAGAGACCAGGTCTATAGCTTTGTTGTTGGAGATGTAATCGGTCCTCATATTG

TATTGGCTCGTGAAGTCACTGAATTATTTAGTCAGCATATGCAAGAAACAGCTTGTTTTCGTGATCTCGA

TTTCTTGTTTTATGACTCCTTATGTGGGCTTCACCCTGTTGTAAGCCACAGTATTGATCCAAAGGCTATT

TAAGGAATATTTGTCTAGCCGAATATGTCCATTTCTTAGTCTGTACTCTATAGCCACAGGTTGATAAACA

GAAAAATCAAGCTATAGGGAATGTTCCTAGTAGCAATTGGTTTGCTTGACAGCGGGTTTTGACAAGGAAC

TTAAGTTGAAAAAGCACAGGAGTGCGGTGCTTGCCATACCTCGAAAGGATCCAAGGCTTTTCAAATTTGC

TTCAGCAATTGCAAAAATGTGCATGATATGGACAGCATAGACCGCCTCACTACGTTGTTCAGAGAAATAG

AAATGGCTGATACCTTTAGTTGATGAACTGCAAAGGTTAAAGTTTATACCAAAGAAAGGCCGAATACAAT

GCAGCTGGTTCTTGTGTAGAAGCCAAAGTTCATGACTGACACTTAAACATAAGACAGATTACATTTTTCT

GCAAAGTTGGGGGCGATTGGTACTTAAATATGAACATGGAAAGTACACTACACTCAATATGTTCAAGTGT

GTTAAATTCTCGAATTCATTCTTTAATGGCCTACATGCTGGAATTTTTATTATAACTACTGTACTATTGG

GGTCAAGCATGCGTGATTCTAGGGGTTGGATCACCGCAACCCATACCTAATTTCTGGAATAAGTTTTGTT

TCTATGGATAAGTTTCCTTGTGGAAGCATTGGTCGATCTCCTATGCTTGTCATAGACTACCTATGATTGG

GTGCTCAAGTCTCCTGGACCATCAGTGTAAAATTATGGATAGACAAGTTGTTCGGTAGTTGGCATGTGGT

TAGGTTGCCCACCTTAAATACCATTCCGGCGTTCTCACTTGGTGGAGCACAGCCTTCCTG

>MSTRG.1197.1 gene=MSTRG.1197

ATTAATTATTGTTCCTTTATTTTTCTAGCTTATTAATTGTTACAGTTATCCGTCAAAAAAAATAAAATTT

TGGTACAGTTAAAAGACCTTTTGCTTTCCATTTCAAAGCTTCCAGTTCTGATCGATCCTCAGATTCCCAG

TTTCACACAGTATTCTGAGTTTCACACAGTCTCTTCGCCATCTGCGGTGATCCAAGCTCTTGGAATTTGA

AAATTTTCGTGATCGAGTAAAATTATGAAAATTGGCTCCTCAACGTTGATCACAGCGCCACATGCGTGAT

CAGATGCAGCAGAGTAATACTGCGGCAACTGCTATGTATGATTAGCCTAGGACTACTGGTGGATCGCTGC

GCAGTGCGGGTTCTGCTACTGATGCTGGTGATGCAGTCATGGCACGGTGGCTTCATTATGCCTGGTTGCA

GCATCTGGCCTCACCCTTGGCTACTTCCGCTGCAAATGATCATCGCAATATGCTCATGCAGATACATCTG

AATTCTGAATTGAATTGTTATCCGAAAAATTGAATTTACCTTTTACGCTTTTTGTTCAGTGCTAATCATT

TCAACATATGTGCAATGTTTATGTTTTTGTTTCCTGATCAAACATGATATAGGAATTGGAATTTTTAATA

TATGTATATCATATGCATTTTACATTTTATTGGTTTAGTATAAGATTTTGTTTGGCAATAACAGTCGTTA

ATGTTATTCTTTGTCAGTTTGATTGTTCGAATTGTTAATTTTTGTTGCACCAAGGTACGAGTTAATATAA

TTTGTTTTGATAAGAAATTAAGTGTTGCCAAATGCTATAAATTTATGGAATTATTAGCTTATGATCAATT

TGGTGTTTTATTATTCCTTTTCTAGGGTTATGGAGCACAATCTGCTGAGGAGAAGCAGAGGCTTTTCAAA

CTAATGAGACACCTCAATCTTAATGGAGAGTACGGTTCAGAACCATATACTCCGACTGCACAAAATTCTG

TTGCTTTGTTCCTGTCGTGGGATCCTTGCAAGAGGCCAACTGCCTTGGAAGTTCTTCAGCATCCTTTCTT

CCAGCAGAGTTGCTATTATGTTCCTCCGATCCTCCAATAGCTGCAGTTGCAAGGACTCCTCCTGGATACA

ACCAAGTATGACAAGACTCTGAGAAATCCTGCTGTGAAGAATTTTATATAACTATTCCATCTCCCGTGAT

TTTTTTGATGCTGACTTGTTAGTTACTACATGTTTGGTCAGTATCTTCCGTAATTATTTATCTGTTGTAC

TTTTTTTTGGTGTTGCATACTTGCGAAATTGTTGTAATATTTGAGTTGTGTATATTTTCTTTCATCTAGC

CATACTTTATGCGGTCTGACACATGTAGAATGCCCTGACAGTGCTCAGTGTTCGACAGTATTGAACTTTG

C

>MSTRG.1198.1 gene=MSTRG.1198

AAAAAACAACAAAAACAAAAATCCCAACAAACAAATATAAACACACCCACAGAACTCTCTCCCTTCGACT

TTCTCAAACTCAAACTCACGTCCATCTCTCTCAACTCTCTCAAATAGTATGAACCCTAATTCCTCTCTAC

ACTCCACAACTCGGTCTCTCCGCAGCTTTCAATCTCTACAAGCTCAGCCTCATCTCTCCAAACCTCAGTC

TCTCTTCAACGAAGCTTATCAAAATCAAAACTACCCTCTCGCTCCCAAGGTTAAAGCTGCTATCTTGGTG

TACAAAGGTCGAAAGCTCAGTTATATATATTATGGGTTGAGGCTGATGACACTAGCGATCATATCGTTCC

ATTTCCTCGTGGATGTGAAGAACTACCTGCAGATGTGCCGAAGATCTTACTAAGAAGCCAAAAAATAAAG

AGATTGCCAATGGGAAAATGTCAAAGCAGAAGAAGCATGTTTCCTCGAGTGAGTTAGACGGGGTTAAACA

AGAGATATCTTTCGAATGTTATAATAATAAATTTCTTTCAGCTGCCGGAGACAGTGTAGACTCATGGAAC

AAATCCTTGGCCAAGGCTTCCAAGAATTCGAGGGGTGAAGCTGTTTACTTGTACTCTATCTATTAGTTCA

GTCAAATCATAAACTTCAGCACGGTGGTGGAGAAGCATCTGAAGCTGATTTTTTTGAGGTTTACAACGTG

GTTCTGTATCTTGGCATAGTTGGCATAATTGTGTATGTTTTGATTTCATGTTGTCGGATATTTTGGCTAG

TGTACAATTAGAATTAGAAAAGATGGATGAATATTTCTTGTAATTGATGTTTGGATTAGATGAAAATGTT

GTACTTTTTGATGTTTGGATATTGGTTGGCTGGATATGTATATAATTGATATTATGCAGTAATTTGGATC

TGGTAGCTTCTTGGTTTGGCAAAAAA

>MSTRG.1199.1 gene=MSTRG.1199

TAAACAGAGAGAAACTTAGCATGGCAATGAGAAGGGTAGGTTTTTCATTCATGTTGGTTACATTGTTTTT

TGCATGCGCAACAACAAATGACGAGAAGGACAAGGAGGAATGTACGCAACAGTTGGTGGGGTTAGCCACG

TGTTTGCAGTATGTGGGAGGAAACGCAAAATCTCCAACTCCAGATTGTTGTAATGGTCTCAAACAGTTGT

TGAAAACCAACAGAAAGTGTTTGTGTGTGATTGTTAAGGATCGGAACGACCCGGATTTGGGGCTTGACAT

TAATGTCACCCTTGCTTTAGGCCTCCCTTCTGTTTGTCATGCCCCTGCCAACGTTTCCCAGTGCCCTGCT

CTTCTACATTTGGCTCCTAATTCAGCAGATGCCCAGGTGTTCTATCAGTTTGGCCGTGCCATGGTTTCGG

GTAATGCTACTGAGTCGGGCACTAGAGGCGCAGTTAAAGCTAATGCAACACGGAGCAGCAGTAGTGAAAC

GAGCAGTGTTCCACAAAAGAACTCCGGGGGTGACAGTAAACAGGGAAGATTTGAATCGAAGTTGCAACTA

GTGGTTCTTTTTGTAATCGTCGACGTTATATTTCAAATATTTAACTAGTAATCTTCGGTGACAGAGGCAG

GTGTAAGTTATAAGATTTAATGGTGGTTGTTCATTCTCTCCTGCTGCAATCTTGTGTTTCTTCCTTTGTA

TATCGTTTCAGTTTGTTAATGGAAATTGTCAGATTTGTCGTATTTTAACATGGTAATAAATTAATAATCC

TCCTCTGCACTGGAATAAAATGCTAGAAATACCAGCACGTTCCTCTATTTAGTGACAACTTTGTCAATGA

ATAATACAAATTGATGAGGCCTAGTGAAGGAGGTGGTTTATCCATTTGTCGACGACGCTTGGAGAACCAT

ACCAGGGACGCGACTCATCATCTGAACCTACCGGAGAGTGGTAGACTCCATCCAAACTTATGTTCAGTGC

ACCTTCGAGATGGGCAGATACTTCTGGCACGACGCCATCACCCCATACGTCTGCCTGCCCGCAAACCTGA

TACATTGTTACAGCAATTAGAAAATTTAGGCTCTGAGCTTGATTATTCACTGACAATCAAGTAGACTACC

ACATGTACACTAAATGGAAAAAGAGGATGTTTACCTGTTTGTAGCCCTGGCCAACAAAACGAGCACGAAA

TGTAGTGCCACTGGATGTTGCTGTAAGACTTGTTCCATTTTCAATCATCACCTCTGAAGCTGGTTGCTGC

ACGGGAACAGGAATAGTGGATACAGAATTTGTGTCTTCATTTCCGAAGAAACGAGCACCTTGGATATACC

TGTAGGTTATGTAAACCAGCAATTTGTTTAAGGCATTAGGCTC

>MSTRG.1201.1 gene=MSTRG.1201

GGACTGTTGAACATGTCTAAAATCTGTTTAGAGTCTTGGTGATGCTTCGGAAGTTTTAAATAGATTTGTT

GTTGGTTGATTGATCAGAGTTGGATGGAAAAGGGGTGAAGGGCTGTTGGGCGGAACTCGCGTCGGCAAGG

CCGACGGAGTTTCGCGGCGGATGCAAGGGAGGGGGACCGGAGAAGAAAAGGGGGAAGGTTAGAGAAGTTG

GTTAAAACTTGATAATATAGATGTACATAGATCTTATGGAAAAAAAGATACATCTGGGCTTCGGGACTTC

GCAGGTGGCTGCTAAACTTGATATAACCAAATCAGTAGTGGGTCCATTAAATCACTTCTTTGTTCAGTTT

GTGTTTTGTTTATATTCGTATACGAGCTTTCGCTCACTACTGATCCATGTTTCATTATTTTCCTCGGGCT

GGTTATGTGGGCTACTTTCGGAAAGAGCGTGAGTTATGGTATTTGGAGTATCTAGTTGTATTTTTTCATC

GGGCTAATCGAAAGAGCACGAGTTACGATATAAAGAATATCTAGTTGTATATTTTCAAACTGCTACTCCC

GTAAGATTTTCAGATGTTGCCGCTCACTCCCTCCGTGTTCCTTTCTTTTGTGTGCGGTCAAGATTTAGTA

TTAGTTAATTACACATATTGTATCAAATTCTCTAATCTCATTAATCCATACACTCAGTTCGTTGTTGGCG

TATTAAACATAGATATCGACACAAACATTATATCTATGTTAACCAGTGCGTACCAATGGATCATCACGGA

TGTTGGTTCATCGAAGGATTTGACAAGATTTCACACGGTTTCTAAATGTTTTTGGCGCGGTACCCGTATT

CTTATGGTGGGGGTGTTCTGAATTCGTGTCACCATAGTCTACAATGCTATTACATGAGGGTGGGGGAAGG

CAAGAGTGAGGCTCCGTTGGATTGGTGGCGGTTAGTGTATTGACGGTTGTTGCTTTGGCAGTGTAGATGA

AGCAGAACCGACATGGATTGATTACACAAGAATAAATCAAGTTAATTCAAGAAAGTATGCAAATAATGGC

AGTTCTTATCCAGTTTGATATCTCCGTTTCAGATCATTGTCACCCATCATCGACAAACGGGAATTGTTTA

GGTAAGTCCCATGTGAAGAAAGCCATGGGCTAAGGTAATATATTAATATTATATCACACACTATTTATCT

CTCTAATTTAAGGGCATGGTTACGTATATCTTAGCCTCCTTAGATGATGCTTTAAGCGGGAATATGTTTG

GTTTGGTTTGGTTGGATATTTTCATTCTAAGATTGTAAGTTGTTGGAATTTATTATAATTTTAGTGGATT

CAAGTGTTTAATTCGGGCCATTAAATCATTACTAAGTTCTAGTAATCGTATATATGTG

>MSTRG.1204.1 gene=MSTRG.1204

TTACTACATCCACACATATATACACATTAAATCTTAATCTCGAATAAGGAGCATAAGTAGAAAACAAACA

AGTTCAAGATAACAAAATACACAATGAGTACCTTAACAGTCATTTTTCCACCTTTAAACTCATTTTCTTG

ATCCGGGTCAAGAGCAGGCAGCAAAGAAAACTCAAACCCATCAATATCCTCCCAATCATCTGCATGCCCA

TCAAGCGTTATGAGCCCGGGTCTAAATTCCGCCCCGACCCGTTTCTCCGGATCCGGGTCGCACCCCCAGG

GTCCAGACTCTTCATGTGCTCTCACTCGGGTTGTCCCGAATGTTATTAACCCGAAGAGGACGACCCGGAG

AAGAAGCATTGGACAGGAAGTGAAGGGGTTGTGTGAGAAGTGTCTGACCAGAGATTGAAACAATGTTTAA

GTATATAAGTTTATTGTAAGTATATATATTGTGTGTGGCCGCCATGGTCTACTGGTAAAACTCATTAAAT

TACTC

>MSTRG.1205.1 gene=MSTRG.1205

TGGCACTTCTGCAATTTTACTACACGTTCAAGGGCAACATCTTTCGTGTCATCTATAAAACTAATCAAAC

AAGCCTGCAGTTGCACACGAAAAAAGAGAGAAAAAGTAAAATGTCTGCAGTCACAGAAGAAATCAAGGCA

AGTGCAACAGAAGTGTACCATGGAAATGAAATGTGTCAAATAAAATCAAAATTTATTCTCAGAGAAATGG

GTTTACCAAATGGCCTTCTGCCACTCAAAGATATGGAGGAATGTGGGTATGTCAAAGAAACAGGCTTTGT

GTGGCTCAAACAGAAGAAAAGAACTGATCACACTTTCCAAAAAATTGGAAAGCTCACTTCTTATGCTAAT

GAGGTCACAGCGGTTATTGAGAAAGGCAAGATCAAGAAACTGACTGGGGTCAAGACTAAGGAGCTCTTGT

TTTGGATCAAACTTAGTGATATTTATGTTGATGATCCTCCTACTGGGAAGATCACTTTCAAGACTCCAAC

AGGTCTGTACCGGACCTTCCCTACGTCTGCTTTCGAGATTGAGGGTTGTGAAGAACCCAGTGGCCAAGTG

AAGGAGGTCTGATTGCCTCTGAAAATGCACAGTTTGGTTCATCAAATTTTCTGTGATTCTACTATTATTG

TTGTTAGTATTTGAAAAATGTCCTATGATTATGCTGGTAGTAATCATTTGGGCCGACAATAAATGCAATA

TTTTTGTGTAACAAAAGCGATAAGCCTCTATGTTTTACTAAAATCTACTGTCCATTTGAGTTCTGTATTC

AGATCATTCCAGATCACCGTG

>MSTRG.1208.1 gene=MSTRG.1208

ATGTATTGTAGGTGTAATTTGTTTTCTGAAGCACGTCTATGTTTTTATGGAATGATTCAGAGAGACTTAG

TTACTTGGAATATTTTTATTGCTGGATATGAAATGTTGCATCCAAATGAGTCCCTGTACATATTCTCACA

GATGGGATTAGAGGGGTATCGTCCCAACAGCTTCACATATTCAAGCATTATAGTGGCATGTGCTAA

>MSTRG.1212.1 gene=MSTRG.1212

GATTCATTCCCGTTGCCAAATTAAAACGCCGGGCCACGTAAAAACAGAGCTCTCCTCTCTTGCACAGCGA

TTCTTCTCCTGACGATATCTCTCTCTACGAACAGCGATCTCTCTCTCTCCCAAGCGCTCCAGCACTCTCT

CTCTTCCAATTAGCACTCAAGCACACACTCTCTCGGTTTCGTTACATCAGTGATCTTTTCCAGTGAAACA

ATACAGCAGCTTGTTCCTTCAGTTGCAGATGCAATATTGGTCCTCGGACAAGGAAGCATATTCGCTTATA

CAATTTCGTAGTTCCTGGGTGTTTGATCTTTCTGTAGTAAGCAGGGCTGTTTCCAAAAAGAATTATAGCC

ATGGCCCGACTTCCGTTGACAATGATTCAACATATCCGGAAGAAGAAAGCTATTTTAGTATATACACTAT

GGATGCGAATGATTTTATCGTTCAATGCTGTTTTTCATCTTATGTTTCTCTCCCAATTGACGAGAAAAAC

TTTGAGAGAGCGATACACGATTTTTAAGATGGAGCGAAATCAATATATACATAGGCTGATTCATGAAGGC

CCTGCTAGCTGTGTTGAAAATCTTCGAATGGATAAAAGATCGTTTTATAAGTTGTGTTTCTTGTTGGAGT

CAAGGGGTTATGTGCAACAAAGTAGGCATGTTAACGTGGTAGAAAAAGTTGCTATGTTTATTTATATTCT

AGGACACCACACAAAAATGCGAATTGTTCACACAGATTTTCAATGATCAAAACAATCAATTACTAGGTAT

TTTCATGCGGTACTCAATGGGGTTTTAAGGTTGCAAGGAGATTATTTTAAAAAACCTGAACCAATTCAAA

GCAACTCAACGGATAAAACATGGAAGCATTTTAAGAACTGTTTGGAAGCTTTGGATGGGACACACATTAA

GATGACCATACCCGTAGATGATAGGCCAAAGTACAGAAATAGGAGAGGTGAAATTACAACAAATGTTCTT

GGAGCTTGTTCACCTAATTTGCAATTTCAGTACGTGTTGACGGGTTGGGAAGGATCAGCCGCAGATGGAA

GAGTCTTAAGAGATGCTATATATAGACATAGATTGATTGTCCCAACCGGTACTTACTATCTAGTTGATGC

TGGTTATACTAATGGGGAAGGTTTTTTGGCACCATTTAGAGGAGAAAGATATCATTTGAATGATTGGGAT

GACGATAATCAGCCCACATCTGCCCGTGAATATTACAATATGATGCACTCGTCAGCTAGGAATGTAATTG

AGAGATGTTTTGGCATCCTAAAGAAGAGATGGGCGATCCTTAGAAGCCCATCATTTTATCCCGTGAAGAC

ACAAAATAGGATCGTCTTAGCTTCTTGTATCCTTCATAATTACATTCGGATTGAATCAGAATTTGATCAA

GCTGAAATTGAGTTTGATCAGCATGGGCCAGAGGACCATCATGAGCAAAATGAGCATGAGGCAGACGAAG

ATCACATCACAACTGTTGGCACTTCAGATGCATGGACGACATTTAGAAACAACCTAGCCGACTCGATGTT

AATTAGTTGGAATCATTAGATTTTTCTATAATCTTCCTAACTGGTGTATAAGACTATGCTATCGGTGGTT

ATATGTCATTTTTGTGCTGCCATCTTTATGTTTAAATCACTTTGCAGAAGTTTCTAACCATGTTTTACTT

CAATTTATGGTGTTTGTAGGTTTAATCAGAAAGTGAACAACTCCTGGAGTATATATATTTGGTGGCTCGC

CCAAATGGAAGTATGCAGCTCTCTCATTTTGTTGTTATATTTGGAGTAATGAGCATGGTCCTCCAGAACC

AAAGACATGAAGATTTTACATGGTCATGTAGCGAAACTTTGGTGTACAGTTGTTAAAATAGGTGTCTTGT

ATTAGTATCTTAAACTTGATGGGTGCTTGTACGAGACTTGTCGGCATTTATTTTATACTTATTTTGATGA

A

>MSTRG.1213.1 gene=MSTRG.1213

GATTCATTCCCGTTGCCAAATTAAAACGCCGGGCCACGTAAAAACAGAGCTCTCCTCTCTTGCACAGCGA

TTCTTCTCCTGACGATATCTCTCTCTACGAACAGCGATCTCTCTCTCTCCCAAGCGCTCCAGCACTCTCT

CTCTTCCAATTANAACATTGAAAAAAATGACTCCCGAATACCTTTTGGTGCTTGTCGCAAAAGGGAAGGA

TATCAAGAAAATACATCAAGGCGTAAAAGCTGGAAACATGAGGGTGCTGACTATGAGTCACGTCAATTTT

GGGGAAAGGTGAGACCACAGGGACAATATAACTGAGGTAATCTTAATTTTTAGGAATATACTACGTCGGG

CGTAGTGGTCATTTTGAATGAAGGTTACACTGAATCAGAATGTACTATGCTTTTGCTCACAATATGAGTA

TGTATCTTTTTAATATGTGAACGGACTAGAAAATGTATTCCTTGTTTGGTTTCTCTTGTAATCAGAACTT

TGTGGCTGATGTTGCTATGCCGCTCCTGTTATCTCAGAGACCTTTAATGACCCTTCTGCTATAATTTTTA

GTGTGTGCGATTATTGAATGTTAGTACCTTTCTGCATTTAGTGAACACGACATTATTGTGGATGTTTTAC

TTTTTACTTGTTGGACCTGCTTGTTAATATGGATGCAGAGAAATTTTGATGAGATGAGTAATGCATGTTG

CAGGG

>MSTRG.1217.1 gene=MSTRG.1217

CCCGATTCCCAAAGGGAAACCCCATGTCCCATGGCTTCTACTTGTTTCTCTGCTAGGAATAATGAGAGTA

ATAGTATTGACGAAGAAGAAAAGCAGGATGATTGTTTTTGTGTGTTTGCTGATACAAACATGGGTACTCA

CATTGCCATTTCTATCTCCCCACTTTCTACTGCTGCCCATTTTGCAAGTACTCTCTCTCTCTCTCTCTCT

CCCTCTCTCTC

>MSTRG.1219.1 gene=MSTRG.1219

CTGAAGTTTGTGTAAAAGTTCAGGTGGGTAGTCAAGTGCTTAAAACCAAGATAAGTCCAGCTCGGACACT

AAATCCACTCTGGAATGAAGATTTGGTCTTTGTTGCTGCTGAACCTTTTGAGGAGCAGTTAGTTCTGACA

GTTGAAGACCGAGTTCACCCTACTAGAGATGATGTTCTGGGGATGATTACCCTACCACTGACTATTTTCG

AGAAGAGGCTTGACCACAGGCCAGTTCACTCTCGCTGGTTCAACCTCGAGAAATTTGGTTTTGGTGCCTT

GGAAGTTGACAGGAGAAAGGAGCTCAAGTTCTCAAGTAGGATTCACCTGAGAGTTTGTCTCGAAGGCGGA

TACCATGTCCTGGATGAATCCACCATGTACATAAGTGATCAAAGGCCTACTGCAAGACAGCTCTGGAAAC

AGCCTATTGGAATACTAGAAGTGGGAATATTGAGTGCACAAGGACTTCTGCCAATGAAGATGAAGGATGG

TCATGGAAGTACAGATGCTTACTGTGTTGCTAAGTATGGACAGAAATGGGTTCGAACAAGAACGATTCTT

GACACATTTAGTCCTAAATGGAATGAGCAGTACACCTGGGAAGTTTATGATCCGTGCACGGTGATCACAT

TGGGAGTTTTTGACAACTGCCACCTGGGAAATGATAAACAAGGCACCACTTCAGCAAGGGATTCAAAAAT

TGGAAAGGTAAGAATTCGCTTATCAACCCTTGAAGCTCATCGAATATACAAACATTCGTATCCTCTTCTT

GTTCTTCATCCCTCTGGGCTGAAGAACATGGGTGATCTCCAGCTTGCCATTCGGTTTACCAACTTGTCTG

TAGCTAACATGATATACTTGTATGGGCATCCGTTACTGCCCAAAATGCATTACTTGAATCCTTTCACGGT

GAACCAAGTAGACAACTTAAGGTACCAAGCAATGAATATTGTCGCGGTGAGGCTGGGACGAGCAGAGCCA

CCATTAAGAAAGGAGGTGGTGGAGTACATGTTAGATGTGGATTCCCACATGTGGAGCATGAGAAGAAGCA

AAGCTAATTTTTTCAGGATAATGTCGCTCTTGTCGGGGATGTTCACTGTGAACAGATGGTTTGGGGATGT

TTGTCACTGGAAAAATCCGATCACATCAGTTTTAGTACACATACTGTTTTTGATACTAATCTGGTACCCG

GAGCTGATTCTTCCAACTCTGTTTCTATACATGTTTCTCATTGGATTGTGGAACTATAGGTTCCGGCCTA

GGCACCCGCCTCATATGGATACTAAGCTATCATGGGCAGAAGCAGTACACCCTGATGAGCTTGATGAAGA

ATTTGACACATTTCCTACATCCAGGCCACAAGATGTGGTCCGTCTGAGGTATGATAGGCTTCGAAGTGTA

GCAGGGAGGATTCAAACTGTGGTCGGAGATATTGCAACGCAAGGGGAGAGATTTCAGTCACTACTAAGCT

GGAGGGATCCTAGAGCGACGAGTTTCTTCATAATGTTCTGTCTATGTGCAGCTGTGGTGCTTTATGTGAC

TCCATTTCGTGTGGTGGCTTTAGTTGCAGGATTATATCTATTACGACATCCTAGGTTTCGAAGCAAATTG

CCTGCTGTTCCAAGCAATTTCTTCAAAAGATTGCCAGCTCAAAGTGATAGTTTGCTGTGACATAATAACT

AATAAGCTCACAACCTTTGTTTCTTTTTCTACTTGTTTGCATACTCTAATAAAGGTTTGCTGTGTTTGTA

ATTTCTTTTCTTAGTCTCTGGCTTTTACTGGCTC

>MSTRG.1225.1 gene=MSTRG.1225

CATTACTCCTATCCATGTAACTACGCGATAATCATGGAAACTGATTGAAATTTGAATTTCTGAAAAGATT

TAGTCATGGACATGTATCAATCATCCAGAGTAGTGCCTCGCAGCCGCAGACGTTCAAATCCTCTCAGGGT

ACCAAGCCTACATATTCGTCTTGCTTTCTAATATCATTGCGGAGCTCACAAAACGGCCCGCCTAGGTTCA

TACGCTGTAAATAATGCACCACTGGAAAAGGACAACACAGTAGG

>MSTRG.1226.1 gene=MSTRG.1226

TGATAAGTGCATATTTTTGTATATTTTAAATGCCTAATTATTGCTTGTTTTCACTTATTTATTTGTTTAT

TTGTCTTTTTTTAGGAAAATTGCAAAAGCTTGAAGAAATAAAAAAATAAAGCAAAAAGGAGAAAAAGATG

GGAAGAAAAGGATCATAGGGGAATATTCTCAAGGAAGCATTTAGATGGGACACCTACACCCCCCCTCTAC

CCTAATTTCCCCCTATAAATACA

>MSTRG.1230.1 gene=MSTRG.1230

TATGGAATGAGCGCGAGATGATCTTGAAGGTGAATGTAAACGAGCAGATGAATTTTGTTTCTTAAAATCC

TGAGGCACTTGGCTGCTTCCCAGGGGAGAAGCTTCTTCTTCACTGTAGTCAACAGACTCTGTTTCCATGT

TGCCTGCTGAATTTTGTTGTTCGACCAAGTTAGCCGATTGAGGTAAATGCCCCCTTAATGAACGAGCAGA

TGCAGGCCGTTGATCAACTTTAGTACTGACAGGTTTAGATGCTCGACTGCTTTCTGAGGATGAAGATTGT

TGTTCTTCAAAATTGACAGGTTTAGATATCCGGTGAGCCAATGATCGCAGAGGAGCATGTTCTGTTGAGG

ATGCACGAACTGACTAATAGAAACAACATTTCATTTTCAATTGTTACCACAATGTTTAACTTTGAAACAA

AAAATTGTGCAGAAACAAAAAAATAATTTCAGGGAGCAATATTGCAG

>MSTRG.1235.1 gene=MSTRG.1235

CAGGTTCTGGAGATTTTAAGAGGTGATAAATATGACCCCCACTGTCCAAGAGTGCAACAAAAGCGATTAC

TTCAAAGAACCTACTCAGAAGAGATTAATGATGCAGAAGAATACAACTCAACAAAATATTTACATAACGT

CGATCGACTTAAGGAAATAGCTTTGGAGACATAAAAGCGCTATAACAACTTCAGCTAATTGTAATCATTT

CTTGATGTAGAGTAACCAGATGACCTTCCACAAAAGTCATGTAAACTACTTTGGAGCTTTCCCCAGTGTA

GTTCTTTTGATCATTTCCTTAATAGCTAATATATTACTTTTATG

>MSTRG.1238.1 gene=MSTRG.1238

GCTCCTCTTGGTCCACTCTAAGGACATGCTTAGTGTCAATCATTTCAATTTCTCCAACATACTTATCAAT

CACTTTACGAACAACACCTTTTTGCTTGTAATAACCCTTCTCCGCCAAAGGCTTGCTCATAACCTTAACA

ACAATTCCCTCCCACAACCAGTAATCCTTCCTATTAGTACGCTCTTTGGCCTTCTCTTGCTCCCTCATCA

ACTCATCCAAAGCCGATTTTCCACCAGCTTTCTCATTACCATCTTTCTTTGCTCTCTCTTTCTTACTCCC

C

>MSTRG.1247.1 gene=MSTRG.1247

TGAATAAAATTCTTTTGTATATCAATTTAATGATCTAATGGGTAGCAACCGGGATAGATCTTTTGCAACT

TATTTCTGTTTCCGCAGCCTGTGCTGTTTAGTGATTCTTTCTGTGTGTTTATGACTGTTATCGCGTTTGA

CTTGTAGCGAAACACCTTCTTGTGCTAGTGAATCTTGAAGTTGGCTCTATGTTCCTGGTTTCAATAGATT

TGATACGCTCTTTGGTCCTGTTTTTAGTCAGACAGGTGTGTGATTTTCAAGCTAGTGTGAAACCAGGTAT

TCCCCATCTATAACTAAACTCCCGCAAAAGAGCTCTAGACCACCTTTGCATTTAAAAAAGAAGAAGAATG

AAAATGGTTAATAGGGATTAGGGAGGAACTTGAAGTTACTTCATTTTTGTGATAAGAGCTTGTCCATAGA

TATATTCCCTGAGAATATTTGTTAGTTGGTTAGGGTTTGGAGTAAGATTAAGTCGATTAATCTTAAATGC

GGTTTGAGCTATGAAAGCCAACCAAAATTTATTCTTATATATGCACTACGATGAGAGAAAATTTTCTCCC

ACTTTTCAGGTATGAAGTACTTTTAGTATTTTACTGTAATTCATTGGATTCCATGAAATACCGACATGGT

AAGAAGCA

>MSTRG.1253.1 gene=MSTRG.1253

CTTCCCCCCTTCATTTCATTTGAGACATTTAGACACAGACACCAAACTCCACACACTCGCACAAACTCAC

TCCACAGTCCACCCACCCCACTCACTGACCTCGGCTGAAACACCTCTCGGCTGCTCAACTTACCTCTCCG

TTCAAATACACAATCTCTACTGCTGAAATCACCAGTGATTAGGGCTTCGATTGAATTGGGGCTCCAGTGG

AAGTAGGGTTCAACAGCGTTTCACTTTAATTGCAGCTTCGATTTGCTTTGATTTAAGATACAATCTGCTG

TCATCCTTTGCTTTTCACTTTAAATGTTTGCTTTGTTTTAGATTTTATGTGTAAATGCATGTCTATCTTA

GTAGCTTTGTTGTGATTTGTGGTCATATGTTATATTACCCATCCCTCTCCCTCTCCCTCTCCCTCTCTTC

CTCTCCCTCTGTGTCCCCTCTGTCTCTCCCCCTTCTTTATATCTTGGTGAATTTGTTAAAGGTGTAGAGC

TTGTTGAGTCAAATTTAATTTTTGTTTAGTGAAGATATGTATCAAGTTATTAAACATGTTAACAGTGTTC

AGTAATCAATTTTCTATGATATTTCTTAAAGAATTGTTTCTTCAAGTTCTGTTTCTAACTTAAAATACAA

ACTACCATATAGAACCATATCCATTAATGTTGTGAGATGCATTCTCTCAAATCCACAAATATATCACTAT

GTACTTCATTTTTTTCTGGTTTTTTGTGTTCTCGTGAAGTTATTCAGGTACTGGAAAGTGTTGAAACAAC

TGCAGAGAAAGCGACCCGATAGTAGGAGCTTGGTCAGGTGTGTTCAGGAGCTGAGGATGAAGGGGGCTAG

CTTTCATTTGTTTAAAGAGCCTCTGAATGCAAAGAGAATGAAGAGTTCCAGTGTCGAGATCAAGAGAAAG

CCATTGACAGGGTGTTCTTAGAGTCTCACAACTATTTGTCTAATCTGTTTGACTGGCTTGCTCTTCCTGC

TTCCAGAATTATTCTCTGTGTTTAAAGCTACTTCCAAATGCAGCCATACTCCAGGTCCTGAGGCTTCAAT

CTTTAATTTAATAGATTTTATAGATGATGCTCAAATGGCTAAGGCCGATGACATTTCAAGGAGCTGTATG

ATGGAAGCAAGGTTCTTGAAATTTATGGCTAGAGAAACTATGTAGCTTCGTTTGATTTGGTAGTTATTTT

GATTTGGTTCGAATTATTAGGTACTTTAGTTTGTATAAAGAAACAATTTATTGGGA

>MSTRG.1253.2 gene=MSTRG.1253

CTTCCCCCCTTCATTTCATTTGAGACATTTAGACACAGACACCAAACTCCACACACTCGCACAAACTCAC

TCCACAGTCCACCCACCCCACTCACTGACCTCGGCTGAAACACCTCTCGGCTGCTCAACTTACCTCTCCG

TTCAAATACACAATCTCTACTGCTGAAATCACCAGTGATTAGGTCTTCAATCGGGGCTGGGGCTTTGATT

GAGAGCTTCTCCATTTCGCGGAATTAGGGCTTTTGGTGTATAGTTCCTTGTCTTCATTTTGCTTTTCATT

TTACTTTTCGCTATAATATTCTTAATTTTGTGTCTTTATTCTTCTTAGGGCTTCGATTGAATTGGGGCTC

CAGTGGAAGTAGGGTTCAACAGCGTTTCACTTTAATTGCAGCTTCGATTTGCTTTGATTTAAGATACAAT

CTGCTGTCATCCTTTGCTTTTCACTTTAAATGTTTGCTTTGTTTTAGATTTTATGTGTAAATGCATGTCT

ATCTTAGTAGCTTTGTTGTGATTTGTGGTCATATGTTATATTACCCATCCCTCTCCCTCTCCCTCTCCCT

CTCTTCCTCTCCCTCTGTGTCCCCTCTGTCTCTCCCCCTTCTTTATATCTTGGTGAATTTGTTAAAGGTG

TAGAGCTTGTTGAGTCAAATTTAATTTTTGTTTAGTGAAGATATGTATCAAGTTATTAAACATGTTAACA

GTGTTCAGTAATCAATTTTCTATGATATTTCTTAAAGAATTGTTTCTTCAAGTTCTGTTTCTAACTTAAA

ATACAAACTACCATATAGAACCATATCCATTAATGTTGTGAGATGCATTCTCTCAAATCCACAAATATAT

CACTATGTACTTCATTTTTTTCTGGTTTTTTGTGTTCTCGTGAAGTTATTCAGGTACTGGAAAGTGTTGA

AACAACTGCAGAGAAAGCGACCCGATAGTAGGAGCTTGGTCAGGTGTGTTCAGGAGCTGAGGATGAAGGG

GGCTAGCTTTCATTTGTTTAAAGAGCCTCTGAATGCAAAGAGAATGAAGAGTTCCAGTGTCGAGATCAAG

AGAAAGCCATTGACAGGGTGTTCTTAGAGTCTCACAACTATTTGTCTAATCTGTTTGACTGGCTTGCTCT

TCCTGCTTCCAGAATTATTCTCTGTGTTTAAAGCTACTTCCAAATGCAGCCATACTCCAGGTCCTGAGGC

TTCAATCTTTAATTTAATAGATTTTATAGATGATGCTCAAATGGCTAAGGCCGATGACATTTCAAGGAGC

TGTATGATGGAAGCAAGGTTCTTGAAATTTATGGCTAGAGAAACTATGTAGCTTCGTTTGATTTGGTAGT

TATTTTGATTTGGTTCGAATTATTAGGTACTTTAGTTTGTATAAAGAAACAATTTATTGGGA

>MSTRG.1253.3 gene=MSTRG.1253

ATTTAGACACAGACACCAAACTCCACACACTCGCACAAACTCACTCCACAGTCCACCCACCCCACTCACT

GACCTCGGCTGAAACACCTCTCGGCTGCTCAACTTACCTCTCCGTTCAAATACACAATCTCTACTGCTGA

AATCACCAGTGATTAGGTCTTCAATCGGGGCTGGGGCTTTGATTGAGAGCTTCTCCATTTCGCGGAATTA

GGGCTTTTGGTGGCTTCGATTGAATTGGGGCTCCAGTGGAAGTAGGGTTCAACAGCGTTTCACTTTAATT

GCAGCTTCGATTTGCTTTGATTTAAGATACAATCTGCTGTCATCCTTTGCTTTTCACTTTAAATGTTTGC

TTTGTTTTAGATTTTATGTGTAAATGCATGTCTATCTTAGTAGCTTTGTTGTGATTTGTGGTCATATGTT

ATATTACCCATCCCTCTCCCTCTCCCTCTCCCTCTCTTCCTCTCCCTCTGTGTCCCCTCTGTCTCTCCCC

CTTCTTTATATCTTGGTGAATTTGTTAAAGGTGTAGAGCTTGTTGAGTCAAATTTAATTTTTGTTTAGTG

AAGATATGTATCAAGTTATTAAACATGTTAACAGTGTTCAGTAATCAATTTTCTATGATATTTCTTAAAG

AATTGTTTCTTCAAGTTCTGTTTCTAACTTAAAATACAAACTACCATATAGAACCATATCCATTAATGTT

GTGAGATGCATTCTCTCAAATCCACAAATATATCACTATGTACTTCATTTTTTTCTGGTTTTTTGTGTTC

TCGTGAAGTTATTCAGGTACTGGAAAGTGTTGAAACAACTGCAGAGAAAGCGACCCGATAGTAGGAGCTT

GGTCAGGTGTGTTCAGGAGCTGAGGATGAAGGGGGCTAGCTTTCATTTGTTTAAAGAGCCTCTGAATGCA

AAGAGAATGAAGAGTTCCAGTGTCGAGATCAAGAGAAAGCCATTGACAGGGTGTTCTTAGAGTCTCACAA

CTATTTGTCTAATCTGTTTGACTGGCTTGCTCTTCCTGCTTCCAGAATTATTCTCTGTGTTTAAAGCTAC

TTCCAAATGCAGCCATACTCCAGGTCCTGAGGCTTCAATCTTTAATTTAATAGATTTTATAGATGATGCT

CAAATGGCTAAGGCCGATGACATTTCAAGGAGCTGTATGATGGAAGCAAGGTTCTTGAAATTTATGGCTA

GAGAAACTATGTAGCTTCGTTTGATTTGGTAGTTATTTTGATTTGGTTCGAATTATTAGGTACTTTAGTT

TGTATAAAGAAACAATTTATTGGGATGTTTAATATTTAAATAGTATATTTTCAA

>MSTRG.1268.1 gene=MSTRG.1268

AGTGCTCAAGTCTTCGACAATGGGTGAAATAACTATTGAAAATTGCAGGTGAAATGTACAAAACATCCTT

CTTGTTCAGGCAAGCCTGCAAGAGTTATGATTACTGTTTTTTTTCCCGGAGGTGTTTGCGCGTCCGACAG

AGCTCATTTCGATTTAAGTGGAACGGCCTTTGGCGTCATGGCCGTTCCCGGCCAAGAGAAACTCCTCCAG

AATGCTG

>MSTRG.1270.1 gene=MSTRG.1270

CAAGTAATTAGATTTTCATTTAATTACAGATGAGATGGAAATGGGCAAGACTGTTGGAGCCATCACACTT

GTACTGGCTAAGCCGTGGATGGAGCTTTCCCCAACCAAGTTCATCTTTTGGGTTGCCAAAGTGACGCAAT

GGGTAAATGAGATTAGCCGCTTTACCTTGACAGGTAGCAACAAGGTGCTGGTGTATCATGGATCCAAGAG

GGGAAGGTGCCTTGATAAATCTGAACAAGGATATGAACGTGACTGGTCAGGAGACTCTGATAAGAGGCCT

GGTGATAAGAGCAATCATGTGACAGGGAGCATGGATATAGGGAGAACCGATACCGCCAGTACAGCGAGCA

TAGGTACAGGGAGCACCGCTACAGGGACCACCAGTACAGGGAGCATGGGTACAGGGAGGAAAAGGATGAC

TATGACAATCATCGGGCTTACAAGGATGATTGTGGTCGGGGACAAATCGGCAGGATCTAAGGGCAAATCG

CGTGTTGTGGAGGAGGATCACAGATCTCGTTCAAGGGATTCTTACCATGGAAAAAGGAGACGTGGTCTAC

CGTTCATTTAAATTTTTACATTCATATGCAGTACTCTTGATTCTCTATTTATCAAAACAATTCTTCTTGC

CACATTTTTAAAGGTTCGATCAATTCTTTGAATAATAATAATTGACATGGCTTCCCGTCAAATCTTGGTT

CTGGCCCGTCGAAGCTGAATACAGGAAATATGTGTTGCCAGAGAAACAAAAATGCACGTGGTGTGGAAAG

TTATTTCATGAAGCATTGTTGGCTCATCACTGAGCTATTATTTGTTATTTGAAAACAAGCAGTCAAAGAG

AGGAAGAGTAAAAATGTTGGGGTGGATGTTACTGGGTCCGGGGATGATCGGAAGCTTGGGAAGAAGAATA

TGTGAGATGGTTAGAATTTCTGGAGGATGCAGACCCAGATATGGATAATGACCATCAAGCTTGGGAATTT

ATTCATGTTGACCTTAGACTCTTCCTTTATGAGAATGCTCAATTTACATGATGAAATATTATGCATATAG

CATTTGCTCAAGTTTTAATAAATAGTCTTTTAATTTTTATTTGGTAAATTAGTAAAATATATGGTTGATT

GTACCAGATTTTATTTTGCGGTCAGGCCGGAACACTTGTCTAAATGCCAGCTGTGGGGCGAATGTATGGA

ACTTGGTTGACTGCAACTTTTTAAGTCTGCAACTTTTATAAGATGAGGTTTATTTGATTTGTAATGTGTG

TAAACTCACAATTTGCTCTTCCATGTATTGTTTATCCATGTCTCTTGCGGTTATCCATGCCTCTTGTCAA

GATAATGGGGCTGTTTTGCTTAGTTTAAAAGACACTATTTATTACTTAAAGTCATTTGACTTGGTTTAAA

AGAAATTATTCACTCCATTCTTAAA

>MSTRG.1270.2 gene=MSTRG.1270

TGTATGGAACTTGGTTGACTGCAACTTTTTAAGTCTGCAACTTTTATAAGATGAGGTTTATTTGATTTGT

AATGTGTGTAAACTCACAATTTGCTCTTCCATGTATTGTTTATCCATGTCTCTTGCGGTTATCCATGCCT

CTTGTCAAGATAATGGGGCTGTTTTGCTTAGTTTAAAAGACACTATTTATTACTTAAAGTCATTTGACTT

GGTTTAAAAGAAATTATTCACTCC

>MSTRG.1272.1 gene=MSTRG.1272

AGAACCCTAGTGGTTCTATGGTTCTATTTTAGGGAGTGGTTCTCTCTTGAGCGCGACCCTAAAAAGTATA

TATATTCTTATTTCTACATAGAAAAGATATATTTAAGTCCATGTATATAAAGAGATATCGGGCTTGTCAA

AGCCCAGTAAACCCTAGGATGAAAATCACATATAACATATTTGATGGCGCCGTGGATAGGCATGAGTGTT

CACAAAATTTACATTAGCAGATTTGTTCAATATAGAGATTTGATCAACTATACATTGAGTTATGATCCAG

ACTTGTTTTCATCGATATTGATTTTATAATCTTCATGCTAATGATTTTATTCAGCGTTTGGGAAGGGAAT

ACACGGATTCTATCATGATCGAACTCGCTAATACATCTACAATTACTAAGCATCATTGCAGTTCTGTTCC

GAGATGAAGGAGGTTACTAGGTTTAAGGGTTCGCTTGAAGGCTGATTTGAGGAGGTTTTAGTCAAGTTAG

ATGTGACAGAGATGGAGAAAAGGGTTATATCTAAGGAGATGATTCGAATTCGTATAATAAAGCTTGGGTT

GAGCTTGAAATGAAAAGGGAGATTAATTTGGTACAAATTCCGGGTGAAGAAAGATGCAGAAAACTTGAGC

TGGGGAGGAAAAGATGGAAGAGAAGATAAAGGAACTAGAAATAAAAACAAAAAAGTCCAGCAGCATTGAG

TTCTCGATTGTAAAACCCGAGCCATGGAGCGATGATGGTTCTTATGCAGATATAAGATTTTCTGTTACGA

TGGATGGAAACAACTTGCTCTTGTATTTGGTCAACCATAAAGGGGATCTTGATTCAATGTCTGATGAAGT

TTATGAGGCACTTGGGAAATTAATGGAGCCTGGAAAACTGGTTTTAGATGCACTATAAGGCTTTTACACA

AAAAAAAAAGCCGAGGAGTTTGAAGACAATGTAGTTTGTAGAAGCAGCATTCTTTTGTTAGAGCAGTTCA

GGAGAATTTCCCCTCATATTCAGTCATACCTCCTCACTAGAGTAAAGCAACTACATCTCTACTTTGATAT

GAGAGGTACACCGATAACACAAGTTTATTGTAGATATTACGCGAGGCAACTGTCTCAAACTTGGACAATA

AAGAGGGGGAATCTACACATATGAGCATGCATTAACTAAACCTTCGGATGTTTGGAACAGGTTTTCTCCA

GTCAAGATCCTTATTTTATGCATAATACATGAACAAGCTGTTTGCACGGTACTTTTTCACTTGGATGCCA

TGCCATGTGCAGGAGCTTCGCCATTTAATCAATTGCGTCTTCGTGTGCTTCGTTTGCTTATAAAACTGGG

CATACAATGGAAGCTTCAAAGATATACAGTATTGTAATGATCATTTATACATAATGACATTACAAACGGG

AGGAGAAACCTACTTAACATAACATAACATGATCTTTTGTTTTAGAGTAAAAAAGCTTCTTAGGCGTGTC

TTCCATGTCGTGGATTTAGAGAGGATTGGCAACAAGGTGAATACAGGGTCCACAAGCTTAAATAGAGGAG

CTCTCAGACAGCTCACTTATGGGGTTCGCCTTTAGCATTTGTGTGGATAGAGCATCTCCTGTGGAAGACT

TGTAAACTTCCTCACATGGCTTCCAAGGTGATCATATTTGTAAATCTCAAGTTTTCGTGTAGAATATCAA

TGTACAAAATATAATAAAATGTAGATTGTATGTATTGTCCCTTTTTGATATGAAAGTATACACGGTGTGT

GTTGTCCACAAACACAATACACATAATAGTGTTTTGTGACACTTCTTCACCA

>MSTRG.1277.1 gene=MSTRG.1277

ATTTTTGTTCTGGGAGTAATTTAAGCTTGCTATGGGTTTCTTTGGGAATGGATGTTTTGTAGATGCAGGA

TGTATAATTATGCATAATTAACATCTTTTGTTTAGCCTTTTCAATTACCAAGGCTTTCTACTGTTTCTCA

TGTTCGATTTTCTTTTGTAGATTTTGTCTGCAACAAGCTAATTGCTTCAATTCGAGCCTAAAGTGGAAAC

TCAAAGATTCAGAGAATAAAGTGGCCGAAGAAGCTGATTTTGATGATAAATTTTGGGCTCATTTTGATAG

ACGGCCTATTAGGTCAAGGGATGTTGAGAGGCTTCATATGATTCACTTTTTTTCATGCAGTGGTGTTGTC

AACTTGGCTTTCAACAATGTCAC

>MSTRG.1278.2 gene=MSTRG.1278

AAACAGGCTAGTTTAAAGAATTAACCCAAGAGATCCGATCCAGTAATCACCGGCAGTGCTCCACAACACC

CCCTCTCTCTCTCTCTCTCTCTCTCTCTCTCTCTCTCTCTGGATTTTGGTTAGAACAGTGCCTCACTCTC

CTCAACCTTCTGCTTACTTCTGTATTCGACAGTTCGCATACTTACTGCTCAGTACTGATCTTTTGCCTCC

CACTTCTATCCCCCACCTGCAAAACTTTCATCTTGAACTGAATAGATAAAGCATTGGAAAATGAAAGCAT

TGGGATGGTGGTTAGTTATAGTTGGATGTCTCCGTTTAGCTTCCGTTTGGTTTGGTTTCTTTGACATTTG

GGCACTCCGTCTCGCCGTATTCTCTCTATCACCCATGACTGAAGTGCATGGGAGAACATTTGGAATCTGG

ACTCTTCTTACCTGCACTCTTTGCTTTCTTTGTGCTTTTAACCTCGAAAACAAGCCACTTTATACTGCTA

CATTTTTGTCATTCGTTTATGCATTTGGTCATTTTTTGACAGAGTATCTCGTATATCAAACAATGGCCAT

CGGTAATCTCACAACAGTCAGCATATTTGCAGGCACGTCAATGTTATGGATGCTGTTACAGTGGAATGCA

CATCAACCCAAGCTTCAGGTTACACTAAAGGTGGAGTGAAATCAGAATCAACTCTTGCTGTATTATTACT

TGTGCAAAGGAACAGTGTATTACTTTGGGTTTGTCATCTGTAATTGTACTATTAGCATTCAGTTTTGGCT

ATTTATTCCCGGAGAATTTAGCATTTGACCGTGATTCAGAATAGAACAATTTATGCTCAAAATTCAAATA

CACAAATTTGCAATCGTGTTGAATTTTTGCTTGGTTGTTCG

>MSTRG.1278.1 gene=MSTRG.1278

TTCTAATTAAAAAGATTATGAGAAGCACCGGTAAAGCCAGAGATGAGGCAAACAGGCTAGTTTAAAGAAT

TAACCCAAGAGATCCGATCCAGTAATCACCGGCAGTGCTCCACAACACCCCCTCTCTCTCTCTCTCTCTC

TCTCTCTCTCTCTCTCTCTGGATTTTGGTTAGAACAGTGCCTCACTCTCCTCAACCTTCTGCTTACTTCT

GTATTCGACAGATAAAGCATTGGAAAATGAAAGCATTGGGATGGTGGTTAGTTATAGTTGGATGTCTCCG

TTTAGCTTCCGTTTGGTTTGGTTTCTTTGACATTTGGGCACTCCGTCTCGCCGTATTCTCTCTATCACCC

ATGACTGAAGTGCATGGGAGAACATTTGGAATCTGGACTCTTCTTACCTGCACTCTTTGCTTTCTTTGTG

CTTTTAACCTCGAAAACAAGCCACTTTATACTGCTACATTTTTGTCATTCGTTTATGCATTTGGTCATTT

TTTGACAGAGTATCTCGTATATCAAACAATGGCCATCGGTAATCTCACAACAGTCAGCATATTTGCAGGC

ACGTCAATGTTATGGATGCTGTTACAGTGGAATGCACATCAACCCAAGCTTCAGGTTACACTAAAGGTGG

AGTGAAATCAGAATCAACTCTTGCTGTATTATTACTTGTGCAAAGGAACAGTGTATTACTTTGGGTTTGT

CATCTGTAATTGTACTATTAGCATTCAGTTTTGGCTATTTATTCCCGGAGAATTTAGCATTTGACCGTGA

TTCAGAATAGAACAATTTATGCTCAAAATTCAAATACACAAATTTGCAATCGTGTTGAATTTTTGCTTGG

TTGTTCG

>MSTRG.1278.3 gene=MSTRG.1278

CAAACAGGCTAGTTTAAAGAATTAACCCAAGAGATCCGATCCAGTAATCACCGGCAGTGCTCCACAACAC

CCCCTCTCTCTCTCTCTCTCTCTCTCTCTCTCTCTCTCTCTGGATTTTGGTTAGAACAGTGCCTCACTCT

CCTCAACCTTCTGCTTACTTCTGTATTCGACAGATAAAGCATTGGAAAATGAAAGCATTGGGATGGTGGT

TAGTTATAGTTGGATGTCTCCGTTTAGCTTCCGTTTGGTTTGGTTTCTTTGACATTTGGGCACTCCGTCT

CGCCTGACTGAAGTGCATGGGAGAACATTTGGAATCTGGACTCTTCTTACCTGCACTCTTTGCTTTCTTT

GTGCTTTTAACCTCGAAAACAAGCCACTTTATACTGCTACATTTTTGTCATTCGTTTATGCATTTGGTCA

TTTTTTGACAGAGTATCTCGTATATCAAACAATGGCCATCGGTAATCTCACAACAGTCAGCATATTTGCA

GGCACGTCAATGTTATGGATGCTGTTACAGTGGAATGCACATCAACCCAAGCTTCAGGTTACACTAAAGG

TGGAGTGAAATCAGAATCAACTCTTGCTGTATTATTACTTGTGCAAAGGAACAGTGTATTACTTTGGGTT

TGTCATCTGTAATTGTACTATTAGCATTCAGTTTTGGCTATTTATTCCCGGAGAATTTAGCATTTGACCG

TGATTCAGAATAGAACAA

>MSTRG.1278.4 gene=MSTRG.1278

TATAATTCTGTGAAATTGACCATTTATTTCAAATGTGGTATCATGGGTAAAATATATTAAAATGTGGCAA

TACTGTCAGTTTTTCTGAAATGTATATTTGTCCAGTTCGCATACTTACTGCTCAGTACTGATCTTTTGCC

TCCCACTTCTATCCCCCACCTGCAAAACTTTCATCTTGAACTGAATAGGTGTGTTACTTGTTTATTTATA

TAATAGAAATTGGGCGGGCTATACGTATGAGTTTTAAATTGATTATACTGTTGCATCTATAACAGATAAA

GCATTGGAAAATGAAAGCATTGGGATGGTGGTTAGTTATAGTTGGATGTCTCCGTTTAGCTTCCGTTTGG

TTTGGTTTCTTTGACATTTGGGCACTCCGTCTCGCCTGACTGAAGTGCATGGGAGAACATTTGGAATCTG

GACTCTTCTTACCTGCACTCTTTGCTTTCTTTGTGCTTTTAACCTCGAAAACAAGCCACTTTATACTGCT

ACATTTTTGTCATTCGTTTATGCATTTGGTCATTTTTTGACAGAGTATCTCGTATATCAAACAATGGCCA

TCGGTAATCTCACAACAGTCAGCATATTTGCAGGTATGTCAATGTTATGGATGCTGTTACAGTGGAATGC

ACATCAACCCAAGCTTCAGGTTACACTAAAGGTGGAGTGAAATCAGAATCAACTCTTGCTGTATTATTAC

TTGTGCAAAGGAACAGTGTATTACTTTGGGTTTGTCATCTGTAATTGTACTATTAGCATTCAGTTTTGGC

TATTTATTCCCGGAGAATTTAGCATTTGACCGTGATTCAGAATAG

>MSTRG.1286.1 gene=MSTRG.1286

TATATACCTATATCACTTCTTGTAGTAGATGGTATTTAAGGCTAATGGGTCCAGTTGTTTACCTTCAGTT

AATCACTGTATTAATCTCTCTCTCTCTGTCTCTCTCTCCCAACTAAACAAGACCCTCCCCTCTTGAAGCT

GCTCTTTAGTTCCAACTCCCAAGCTGTGTTTTTCTTCTTCCTTTTTATGCATAAAATTGGCCTTTTGGAG

TTGACATTTATTTGGATTTCTTTACCCTCTTTCTGGGATCAGATTTGAGAATTGGGGGTTTGATTTGAAG

AGTTCTTGATCTGGGTTTATTGTTTTGAAGATGGAGCAATCTGCTGAAAGGTCTCCAAATGGACACAGGG

GTTTTAGAGTTCAAGCTCCCTTGGTGGATTCCATTTCTTGCTACTGCAAAGTAGATGCTGGATTAAAAAC

TGTTGCAGGGGCAAGGAAATTTGTCCCAGGATCAAAACTTTGCATCCAGCCTGATATCATTCCTCGTGCA

CACAAGACAAAGAGCTCTCGCAGGGAGAGGACCAGAGTTCAGCCACCCCTCTTGCCTGGACTTCCTGATG

ATCTTGCTATTGCTTGTCTAATTAGAGTTCCGCGTGTCGAACACAACAAACTCCGTCTTGTATGCAAGAG

GTGGAACAGGCTTCTGGCAGGAAACTTTTTCTACTCTCTTAGGAAGAGTCTTGGAATGGCTGAAGAGTGG

ATTTATGTTATCAAAAGAGACCGTGATGGAAGGATCTCTTGGCATGCATTTGATCCTACCTATCAGCTAT

GGCAGCCACTGCCACCTGTTCCAGGTGAGTACAGTGAAGCTGTTGGATTTGGTTGCGCTGTTTTAAGTGG

TTGTCACCTTTACTTATTTGGAGGAAAAGATCCCATCAAGGGTTCAATGAGACGTGTCATTTTCTATAGT

GCTCGAACAAATAAATGGCATAGGGGTCCAGACATGCTCCGGAAAAGGCATTTTTTTGGTTCTTGTGTCA

TAAATAACTGTCTTTACGTGGCTGGTGGTGAATGTGAGGGAATTCAAAGGACTCTCCGTACTGCTGAAGT

TTATGATCCGAATAGGAACCGTTGGAATTCTGTTTCAGACATGAGCACTGCAATGGTTCCTTTTATTGGG

GTAGTTCATGATGGAATGTGGTTCCTAAAAGGACTTGGGTCCCACAGAGAGGTTCTTAGTGAAGCATACA

TTCCAGGAACAAATTCATGGACGCCAATCAGTGATGCGTTGGTCGCTGGGTGGCGCAATCCGAGCATCTC

TATGAATGGGAAGCTGTATGCATTAGATTGCCGTGACGGGTGCAAACTTAGGGTGTATGATGAAGCTACA

AACTCTTGGAGAAGATTTATTGACAGTAAGCTCCATCTCGGTGGCTCTCGTGCTCTCGAGGCTGCTGCTC

TTGTTCCTCTCAATGGAAAGCTCTGCATTATTCGGAACAACATGAGCCTCAGTTTGGTTGATGTATCGAG

TCCAGACAAACGTGTTGAGAGTAATCCACAGCTTTGGGAAAATATTGCTGGGAAAGGCCACTTCAGGACT

CTATTCACAAATATTTGGTCTAGTATAGCTGGTCGGAATGGACCAAAGAGTCATATTGTACATTGTCAGG

TACTTCAGGCTTGAGTTCTGACGGCATGTCATTGCTGCTGTTGTACATTCCCAAAGATCTGCTGTAGCTT

TTGCCTTCCGAAGAGACTTAAGAGAACAAAGACGCCAAATTGAATAGTTTTCCCTGAAAATTCAAAGCGA

GATTATGATTTCTAGCGGAATTAGTTTCATGTCCAGATTCAGGTAGTATTTAGAGGACGTGCATTCACAA

GGTGATGGTGAGGTTTTGCTGGTTTTGGCATATAGCTGCTGTATAATTTCTCAATATTTTCACATATGAG

CAAAAATTAGCTGCAAGCATATTTTATCATCATCAGCTGAATCTGCAGTAGTATGCCTTTTGGCATCTTA

GGCAAAAATAAAGATTCTTACAACTTTGTGAATACACAAGCGTTTAGACCATGGATTACATTTTTGTGGA

ATGGATATTGATAAAATGTCACTTTTAATTGTGTGCCCTGGCTAGAAGCTATTCAGTTTGTTCATTGTTG

TTTGGTAAATTTCATTATCACATTGTCAAGTTCAAC

>MSTRG.1286.2 gene=MSTRG.1286

GGAAGGAAGGAAATATTTGTACAAGACAAGTCTCTAGAGAGAAGGAGATAATTTATGTGGATTCCATTTC

TTGCTACTGCAAAGTAGATGCTGGATTAAAAACTGTTGCAGGGGCAAGGAAATTTGTCCCAGGATCAAAA

CTTTGCATCCAGCCTGATATCATTCCTCGTGCACACAAGACAAAGAGCTCTCGCAGGGAGAGGACCAGAG

TTCAGCCACCCCTCTTGCCTGGACTTCCTGATGATCTTGCTATTGCTTGTCTAATTAGAGTTCCGCGTGT

CGAACACAACAAACTCCGTCTTGTATGCAAGAGGTGGAACAGGCTTCTGGCAGGAAACTTTTTCTACTCT

CTTAGGAAGAGTCTTGGAATGGCTGAAGAGTGGATTTATGTTATCAAAAGAGACCGTGATGGAAGGATCT

CTTGGCATGCATTTGATCCTACCTATCAGCTATGGCAGCCACTGCCACCTGTTCCAGGTGAGTACAGTGA

AGCTGTTGGATTTGGTTGCGCTGTTTTAAGTGGTTGTCACCTTTACTTATTTGGAGGAAAAGATCCCATC

AAGGGTTCAATGAGACGTGTCATTTTCTATAGTGCTCGAACAAATAAATGGCATAGGGGTCCAGACATGC

TCCGGAAAAGGCATTTTTTTGGTTCTTGTGTCATAAATAACTGTCTTTACGTGGCTGGTGGTGAATGTGA

GGGAATTCAAAGGACTCTCCGTACTGCTGAAGTTTATGATCCGAATAGGAACCGTTGGAATTCTGTTTCA

GACATGAGCACTGCAATGGTTCCTTTTATTGGGGTAGTTCATGATGGAATGTGGTTCCTAAAAGGACTTG

GGTCCCACAGAGAGGTTCTTAGTGAAGCATACATTCCAGGAACAAATTCATGGACGCCAATCAGTGATGC

GTTGGTCGCTGGGTGGCGCAATCCGAGCATCTCTATGAATGGGAAGCTGTATGCATTAGATTGCCGTGAC

GGGTGCAAACTTAGGGTGTATGATGAAGCTACAAACTCTTGGAGAAGATTTATTGACAGTAAGCTCCATC

TCGGTGGCTCTCGTGCTCTCGAGGCTGCTGCTCTTGTTCCTCTCAATGGAAAGCTCTGCATTATTCGGAA

CAACATGAGCCTCAGTTTGGTTGATGTATCGAGTCCAGACAAACGTGTTGAGAGTAATCCACAGCTTTGG

GAAAATATTGCTGGGAAAGGCCACTTCAGGACTCTATTCACAAATATTTGGTCTAGTATAGCTGGTCGGA

ATGGACCAAAGAGTCATATTGTACATTGTCAGGTACTTCAGGCTTGAGTTCTGACGGCATGTCATTGCTG

CTGTTGTACATTCCCAAAGATCTGCTGTAGCTTTTGCCTTCCGAAGAGACTTAAGAGAACAAAGACGCCA

AATTGAATAGTTTTCCCTGAAAATTCAAAGCGAGATTATGATTTCTAGCGGAATTAGTTTCATGTCCAGA

TTCAGGTAGTATTTAGAGGACGTGCATTCACAAGGTGATGGTGAGGTTTTGCTGGTTTTGGCATATAGCT

GCTGTATAATTTCTCAATATTTTCACATATGAGCAAAAATTAGCTGCAAGCATATTTTATCATCATCAGC

TGAATCTGCAGTAGTATGCCTTTTGGCATCTTAGGCAAAAATAAAGATTCTTACAACTTTGTGAATACAC

AAGCGTTTAGACCATGGATTACATTTTTGTGGAATGGATATTGATAAAATGTCACTTTTAATTGTGTGCC

CTGGCTAGAAGCTATTCAGTTTGTTCATTGTTGTTTGGTAAATTTCATTATCACATTGTCAAGTTCAAC

>MSTRG.1292.1 gene=MSTRG.1292

AAAATTGTGAAAACTGACCCAACTGTATATAAGTAAATGGATAGGAGGGAAGGAGTACAATACAAATGCA

AACACATACAATTAAAGGCAACCCTTAATTCAATTATATCTGCAAACCTGAATGCTTTCATCGAGGGAGA

CTTATGACTGACAATGCTTTGATTGTATTTGAAGCCAATCATTACATTCACAGGAAAACACGGCGTTACT

AGAGTGGTTGGATTCAAGGTGGAAAGGGTGGATAAAAATTAATACAGATGCTGCTCCTTTCCGTGATGAG

GGTATTATAGTAAGTGCTGTTATAAGGGATTCTCATGGATGCTTCATAGGGGCTAGGTGTTGTAAGGTGG

AAGGAGCTTGAAGTCCAAGAGAAGTAGAAGCTTTGTCTTGGGTCATAAAAAGAAATTATAAACATTGTGT

GTTCGAGACTGATTCGAAGTGCCTTGTGATTGCGTGTAATGGATCTTCAAGTGAGGCTTTTTTTTAAAAA

CTTATGTTAGATTGTGTTAATTTGTTAAAGCACATGAATCTTGTTCTAGTTAAGTTTGCTTTTCAATCTG

CGAATACATTGATACATGAACTAGCAAAAGCCGCTTATTGTGCTAGTTATGTTTGTTTTTCGATCTGTGA

ATACATTGACGTATTGACGTATGAACTAGCAAAAATTGCTTGTTACTTGTTGTATGTCACATATTAGGGA

ATAGTAAGCACTAGCAAAAGTTGTCAAATATTAGGAAATAGTATGCACTAACAAAAGTTGC

>MSTRG.1295.1 gene=MSTRG.1295

CGAAGACATGTCAGGTCTGAGTCCGGAAATTGCTCAGCATTGTTTGCAGGTGGATCCGTCCGCCAAACCA

GTGAAACAGAGGAAGAGGCCTTTTGCAATGGAAAGGCAGAAGGTTCTTCATTGAAGCTTCTGTTGTAGTT

GCATGTATATGGTTAGCATTTTGGGTATTTAGTGGGATCAACTTCTTTTATTTTTCCCAGAACTTTGCTC

ATTGGAACTTTTATGTAGTCAACTCTGGATGAGATTAGACTTGAGTTGCATCTATTTCGCTGCTATAGTG

CAAGATTGTAGGGTGTATCCGGTTGATCAGATCAATGAGTTAACTAATTCAATAGCCATGAAATAGAATT

GATTTTTCAATCTATATTTTTTATATACAGTTATAATTTGTTAAA

>MSTRG.1297.1 gene=MSTRG.1297

TGAGAAGCGGCAAAATTGCCGCCTTCCACAAAAACTTACTCAAACCTCAATCCCCAGTACAAACTCTCTC

ATTCTCTCCCGACTTAGCTCCCCTCTCTTTCTTTCTCGATTTGAAGCAACTCAAGTGTTGTCAGATTAGT

AATTTCTTTCATATCGGGATTCTTCAAACTCTCAACTGATGGGATTTACAATTAGGGTTTTGATTTGGGT

TGGGGTTTTACAATTCGAGATATTACTAGTGCTTCGACATTTATTGTGACTCAGACGTTAAGATCTCTAT

CAAGATATGGAAGGACTGTCATAGCCTCTATTCATCAGCCAAGCAGTGAAGTGAAAGATTATTTATTTTG

GCAAGGCTTCACAAGCTGCTATTCAATATATGTATACATTACCATTGTGGGAACATGCAGAAGTGGCATC

CTGACAAACACAAAGGAATACTTAACAAGATTTTAAGGGATGATACTTACTTGTAATGGGCTGGGTCTTG

GTCACTCATCAATTTGTTTGGTGATTATTAGAGTCAGAAGATTTGCTGGAGCAGCTGGAGGTGCAGTTGA

AAGGACATGCTATGCAGCTTCTTCAGTATAATTGGGGATCCATCTAAAGGATTTTTTGTTATGCCCAGGG

ATATAGCACCCATTTCTTCTGCGTCGCTTCTTTTCTGTGGACTACTACAATTGCATTTACGCTTCATCGT

ACTGTTGTTAGACACAAAGCAGATGTAGAGGACTTGGAGCCTATGTTTCATCTGTATGTTTGGGCTTGGG

AATATTTATCTTTAGTAGGAACTGATGTTTACACAGTATTTCTACCACTTAAGAATGGGGGAGGTCCCAC

CTTTCTAATCAGGAGCTAACACAAAATCAGTTTCTTCTCCTGCAACAAAGGCACCTAAAGCAAGAAGCCC

AAATATTGTTAATCGTAGGGGACCTCGTACTCTGGCATCCAAGCTCTGCTCATTGGAGTCGGTGAAGAAT

CAGGCTAATTTAGACAATGAAGAGGCCCATAAACAGCTAGCTGCCTTGTCCGCGAATCTTGAGGAGAGAC

ACTTCTTTCTAAAAACATAGGGCGTAGATTACCCGTTGCTGAATACAGGGGTGAAAGCCATCAATCTCCA

CCTTGCAAAAGGCGCTTCTCCAAAGTCTCTGATCAAATCCCTTCAACAGTTTACCTGTACAAGAATGAGG

ATGAAATTACTTAGTTGTGATATTATGTAACTCTATTTTTCATTGTAACTCGTGTATAATTCATCTTAGT

TGAGTGATTTATGATTCCACACATGTACATTGTATAAATTAATTGAAAATATCAATCATCATTTTTATAG

ATACTAATAGATAATGTAATTATTAATGTTGGC

>MSTRG.1297.2 gene=MSTRG.1297

CAAACTCTCTCATTCTCTCCCGACTTAGCTCCCCTCTCTTTCTTTCTCGATTTGAAGCAACTCAAGTGTT

GTCAGATTAGTAATTTCTTTCATATCGGGATTCTTCAAACTCTCAACTGATGGGATTTACAATTAGGGTT

TTGATTTGGGTTGGGGTTTTACAATTCGAGATATTACTAGTGCTTCGACATTTATTGTGACTCAGACGTT

AAGATCTCTATCAAGATATGGAAGGACTGTCATAGCCTCTATTCATCAGCCAAGCAGTGAAGTGAAAGAT

TATTTATTTTGGCAAGGCTTCACAAGCTGCTATTCAATATATGTATACATTACCATTGTGGGAACATGCA

GAAGTGGCATCCTGACAAACACAAAGGAATACTTAACAAGATTTTAAGGGATGATACTTACTTGTAATGG

GCTGGGTCTTGGTCACTCATCAATTTGTCTTAACCATTTTCAGATGTTATGAATCAGTTTGGTGATTATT

AGAGTCAGAAGATTTGCTGGAGCAGCTGGAGGTGCAGTTGAAAGGACATGCTATGCAGCTTCTTCAGTAT

AATTGGGGATCCATCTAAAGGATTTTTTGTTATGCCCAGGGATATAGCACCCATTTCTTCTGCGTCGCTT

CTTTTCTGTGGACTACTACAATTGCATTTACGCTTCATCGTACTGTTGTTAGACACAAAGCAGATGTAGA

GGACTTGGAGCCTATGTTTCATCTGTATGTTTGGGCTTGGGAATATTTATCTTTAGTAGGAACTGATGTT

TACACAGTATTTCTACCACTTAAGAATGGGGGAGGTCCCACCTTTCTAATCAGGAGCTAACACAAAATCA

GTTTCTTCTCCTGCAACAAAGGCACCTAAAGCAAGAAGCCCAAATATTGTTAATCGTAGGGGACCTCGTA

CTCTGGCATCCAAGCTCTGCTCATTGGAGTCGGTGAAGAATCAGGCTAATTTAGACAATGAAGAGGCCCA

TAAACAGCTAGCTGCCTTGTCCGCGAATCTTGAGGAGAGACACTTCTTTCTAAAAACATAGGGCGTAGAT

TACCCGTTGCTGAATACAGGGGTGAAAGCCATCAATCTCCACCTTGCAAAAGGCGCTTCTCCAAAGTCTC

TGATCAAATCCCTTCAACAGTTTACCTGTACAAGAATGAGGATGAAATTACTTAGTTGTGATATTATGTA

ACTCTATTTTTCATTGTAACTCGTGTATAATTCATCTTAGTTGAGTGATTTATGATTCCACACATGTACA

TT

>MSTRG.1298.1 gene=MSTRG.1298

ATATTTAGTGAAGATGAGGCTCAAAACTTCTTTCAGCAACTAATCTCAGGAGTAAGCTACTGTCATTCAA

TGCAAGTATCTCATGGGGACTTGAATCTGGCCAACATTATGCTAGATGGAAGCCCTGCAAGAAGATTGTA

AATATGTGATTTTGACTTCTCAACGGTGAATCTTTTATAGACGCGGATACAATTGATGGAATGGTTGATA

CAATCTGAGTCAACCAGATAAAAATATGTAACAGAGACAAAGTCAAATGGTGCATGTTCATTTTTTAAGT

TTGTTTTTACTGGAAATCTGCAGACCTTTGATAAGATGACACTTCTTTTGTATTTGGAGCAAATTAATGC

TACTGCTGGTTTTTACAGACAATAAGGTGCATGGGAAACATGGGCAAACTGGCCATTCTGGAGAACTGGA

ACTTGGAACTCGCAAACTCCTAAAACAGTGAGTGATCCAATGTTCGACGTTAGTTTTGATTGTGACAAGG

AGAACGATGGAGTTCCAAGGGCATTTATCATTAAGAATAATCATCATCATGAATTCTACCTTAAGACCCT

TACTCTTGAAGATGTTCCTGGCCACAATCGTGTTCATTTTATTTGCTTGCCACATTTTACAAGAAAAAAT

TGAAGAAATAGTAAAACTATATGAAGGGGAGATCAAACTATATGATGGCCCTGTATTGTCAAGTGTTCGA

GATCGTATATCATCACAAATTCTCAGAAAACTACTTTGAAGTGATGGTGAGCCAGTGTTTCAAATCCCTA

TGCCACAAGTGATTAAAGATGAAGTGCGCAAAAAGGGTGGACTGAAGTGCGTGAAAAGGGTCATGGTGAC

AAGAAAGATGAACCTTGGTGGCCTAAAATGCAGACTCGAAATGAGCTAATAGAGTCATGTACTACCATAA

TTTGGCTGGCTTCTGCTCTTCATGCAGCAGTAAACTTTGGACAATATCCATATGATGGCTACCTCCCTAA

CTGTCCAGCAATGAGTAGGCGCTTCATTCCTGTTCCTGGCACATAGGAAAATTGAGTTTTGTAGAATATG

CTTCTTTTTCTTCTGATTTGCATAAAGAAAAGGAAGATGTTTAGGTTAATTCGAAAGTGCTAAGATTTTC

CCATGAAGTGTTTATGAGTTCTTATCTATTTTTGGTCATGTAAAGTTAAATGTGCT

>MSTRG.1299.1 gene=MSTRG.1299

CATAGTGCCTGTTTGTTTAGCAGAAGCAGGGGCTTCTGCTTCTGACTTCTGCTTCTTGACCCGTTTGTGT

AAAGAAGTAGAAGCACTTTTAAGAAGCTGAGAATGCTAGCTTCTCTCTCACAGCTTCTACTTTTTTTCCA

AACACTTTATTCACTTATTTACTTCTCACTTCTAATTCACTTCTTTATTTTAAACAAAGAAGCACTTCTT

TTAAGCTTAGCCAAACGGCCCCATAAACTCAAAATTCTTATCCCAGTTGCGAATCTCCCTCCCCTTTCTT

TGGTAGGGTTCTTCTTCTTCTTCTCTTTTTTAACATGACTGGGATTTTAATTATATTTCTCCGTCTCTCA

CAAATTTAAAAAATTGCTCTTCATTGTTCACTGCAGATGATTCATTAGAACTATTTTTGAACAACATAGG

TTTAATTTTGTAGCAGTGATACGCAGACTCCTCTTTGACGGAGAATGTCTCTCCTCCATGTAATGCAAAT

CTGTGAAATAAATCGGTTAAATTTTTTAGGTTGAGTAGTTGTTTCAGTTGTCTCAGGTTTCGAAGTGTCT

TTAAAAGGATGGCATATGCTCTAATTCAATGACTGATTTGAACTGACAAAGAAATGATAATCACAATATT

TTCAATCTTAGAGCAATCTCAGTCACACTTACCAACTGGTTTGCCTAAAATCAGAACTGGAATTCAAAGC

TCATCTGAGAGTTTGAAGAAAATAGAAGGAAATATTTTAGTTGGAAAAAAGGCAATAGCAGATGCTGGAT

ACTTCATATGAGAACTGACTCTTTGTTTGTGTGTAATCAGCAGGTTCTGTGAAATGTGGTGTGGAGCAGA

TCATATTCATATAGATCATAATCATACTATAAAGTAATAATTTATGTTAAAAATCTTAACTTTTTGATGT

TCAACTTCTTATGCCACTTTAAATAATCTTGAAGCACAGACTGAGAGTTTGGAAAATGAGTTTAAAAAGC

GGTCAGAAAAATTTTCAGGGTCTTTGATTACCGTAAATGAACTTGAAAATCATATAAAGATCTTACATGA

AGAGCTGGAAAAGCGGGCACAAGTTTTTGAAGGTGATTAGGAAGTACTTATAAGTACTAAAGTTGAGCAG

GAAAAGAGAGTCATAAAAGCAGAGAAAGCCCTGAGAAAAGCGAGCCATAAAAGTAGAGGAAGCCCTGAGA

AAAGCACAATGCTAGAATGTAAATACAGCCGAGAGATTGCAAAAGGAATCACTTTAGATGTGAATGATGA

GCTAGCTACAAAAGCACTGACTGAAGCAAATGAATTTCATCTAGTGAAAAACTATCTAGAGGAAATGCTC

CATCAAGCCACGAAAGAGGTTAAGTCAGTTAAGGGGTTTATGAAAGAATTTAAACCAAATACTTTCCAAG

TCAAAACAAATGGAACAGGTACCATCAGAGACATAGTGAATATGCAGGAACTTAAAGAAAGCTTGCTGAT

GAAACTCACATGCTCCTTTCAGAAGAAATCTTAATGCTGCAGTCTGAAATTGAAATACTTGCAAAAGAAA

ACAGTGTATTCTCTGAGCAGAAAAGGCAGAATGATAATTTGAAAGTTGAGCTAAATCATAAT

>MSTRG.1299.2 gene=MSTRG.1299

ATCTCCCTCCCCTTTCTTTGGTAGGGTTCTTCTTCTTCTTCTCTTTTTTAACATGACTGGGATTTTAATT

ATATTTCTCCGTCTCTCACAAATTTAAAAAATTGCTCTTCATTGTTCACTGCAGATGATTCATTAGAACT

ATTTTTGAACAACATAGGTTTAATTTTGTAGCAGTGATACGCAGACTCCTCTTTGACGGAGAATGTCTCT

CCTCCATGTAATGCAAATCTGTGAAATAAATCGGTTAAATTTTTTAGGTTGAGTAGTTGTTTCAGTTGTC

TCAGGTTTCGAAGTGTCTTTAAAAGGATGGCATATGCTCTAATTCAATGACTGATTTGAACTGACAAAGA

AATGATAATCACAATATTTTCAATCTTAGAGCAATCTCAGTCACACTTACCAACTGGTTTGCCTAAAATC

AGAACTGGAATTCAAAGCTCATCTGAGAGTTTGAAGAAAATAGAAGGAAATATTTTAGTTGGAAAAAAGG

CAATAGCAGATGCTGGATACTTCATATGAGAACTGACTCTTTGTTTGTGTGTAATCAGCAGGTTCTGTGA

AATGTGGTGTGGAGCAGATCATATTCATATAGATCATAATCATACTATAAAGGTCTTTGATTACCGTAAA

TGAACTTGAAAATCATATAAAGATCTTACATGAAGAGCTGGAAAAGCGGGCACAAGTTTTTGAAGGTGAT

TAGGAAGTACTTATAAGTACTAAAGTTGAGCAGGAAAAGAGAGTCATAAAAGCAGAGAAAGCCCTGAGAA

AAGCGAGCCATAAAAGTAGAGGAAGCCCTGAGAAAAGCACAATGCTAGAATGTAAATACAGCCGAGAGAT

TGCAAAAGGAATCACTTTAGATGTGAATGATGAGCTAGCTACAAAAGCACTGACTGAAGCAAATGAATTT

CATCTAGTGAAAAACTATCTAGAGGAAATGCTCCATCAAGCCACGAAAGAGGTTAAGTCAGTTAAGGGGT

TTATGAAAGAATTTAAACCAAATACTTTCCAAGTCAAAACAAATGGAACAGGTACCATCAGAGACATAGT

GAATATGCAGGAACTTAAAGAAAGCTTGCTGATGAAACTCACATGCTCCTTTCAGAAGAAATCTTAATGC

TGCAGTCTGAAATTGAAATACTTGCAAAAGAAAACAGTGTATTCTCTGAGCAGAAAAGGC

>MSTRG.1301.1 gene=MSTRG.1301

GGTCGGTTCAATTGGGGGCTAGGGTTCCTCTCTTTTCTTTTATCGTTTCAGTCGATTCAAGCAGGGGTTA

AGAGTAAGGTAATTTCCAGAGTTTTATCTGCGCGACTCAGAGGAGGCAAGATGCCATCCTATAGATGATC

TATTTTGGTAAGTCATAATACATATTTTGTATTAAATAGTTATGACCCTAGTATGCTTCTCATCGGAGTT

ATGCTAGTGGTATAAGATTTGCGTTCCAGAGATTATTGTATATTATGTTTTGATTTTGGATTCTTGTTTC

TAGCGCTATAGCCTAAAAAGATCTTGGATTTGGGGTCAGATTGTTTTATTTTCTGTTATTTATTGAACAG

ATTCGATTTTA

>MSTRG.1302.1 gene=MSTRG.1302

TCAGAAACATCAGATAAGATTTCAGAGGCGGTTCAATTATTGTGGGTATACTCTCTCTTTCCTTATCTTT

TTTCTCTCTCCGTAAACATCCATCCCCTCGTTTATACACATTCGATTGTATAATTATTATCTGTGATTGC

TTATGGTTTGAATCGGTTGTTGATTGGGGTTTGAAATCGGGATTTGTGTGGCTGTTGGTGGGCCCGGAGG

TTGAGATGGTGGTCGGTGGAGGCGGATATCGCCGGCGAACGGCGGGTTAAGGGAGCCGGCGAATATATGG

AGCTATTAGATGTGGCTGATTGTGTGTGGCCGGCCATTGGTGGTTGAACGGCGGCTCGATTCACAGCACC

GCGTTGGGCCGATGTTAGCAGCGGTGGCGGCTGTGATCGAGCAGTGAAAGGGAGGAATAAGGAGGCCCCG

GTGGCGGCTGGGTGCAACAGATTAGGCGGCAACGCCGCTTGTTTTGTGGCCGGGCGAGGAGGGAGAGAGA

GAGAGAGAGAGAGAGCTGGGTAGGAGGCTGCCGGAGAATTTTCCGGCGAGCGCTGACCGAGGCTGGCGGT

GGTGGTGACCCGGGGAGCCCGGGAAGATATTTCGGAGAAAATCAAAATTAATATATATATTTTTGG

>MSTRG.1303.1 gene=MSTRG.1303

GGCCGGTCCACATAGAATCCGAAAATAACTAACCCTATCCTAGCAAAAGGGGAAAACGCCGCTGTTGATA

CAAATCTGGTGTTTAAAGAGACAAGGCAGAGAGATAAGACACAGAGAGAAAGATGTGGAGATGGATGAGG

AAAGGGTCTCTGAACGAATGGAAGCCACTTTTTCGCAGATCAATGGAGAAAGCGGAGGAGATTTATGTGA

GCAATTGGTTCGATATCTTGGTCGTGAGGGGTGGCTTGGCTGTCACCATTGCCGCTTCAACTGCGAGGCA

CCGTAAGAATAAGAAGAATTCATAGATGCTGCTCGTACGATCTGCTTATATATTTATATTTATGTAAACA

CCAATGTTGTGAGTAATTATGTCTATTAGATGAACATGGCTGGAAGTAAACTACTTTTATTAGAACTCTG

TCTTTTGTGGTGTTGATCTTACAACTCTTATCATGTTTGATCTGTTGGTTGTTGTTTATCTTCAAAAAGA

AGAATATTGATTTTTAGCTCAGTAATTACTAATTTCGGATACAGTC

>MSTRG.1305.1 gene=MSTRG.1305

GTTTTATAAACCGTAACTACGTGCATTTGCTGGGCTCAAAGATAAAAATGTGGTATTTGGGCCCAGATAA

CATGGGGCAGGTAAAACCCAAGTACCTTCTGGTTCCTAACCCTAGTATAAGGGAGGCTTTGTGTCTGCAA

GAAATTATTGAGATCGAGAGGGCGCGAGAGAGAAGGATGTCGTTTCTAACACCCTTTAAGCGAGCCAACC

AGTATCTTCGTACACACAGGACAGTGGATCGAACGTTGAGCCTTATGGCTATTGTTGCTATCGGATCCGC

GTTCTGGATTTCCAGTGGGCCCGACAAGAAACCAGACAAGGATGAAGCTGTCCGAACTCGTTCCTAGACT

CACCCAACTGTTATTACTAACATCTCTGGTGCGATGTTATATATATATATATTAATCAACTCTTATCGTC

TCTTTAATGGAGCTTTGTCCACTGAATGTTGAGATCTAACTGCTTGCAGTAATTGCAATTGTTATTTATT

TACTAGACTATGTTCTCAATTTCAATTGTGGGTATTTGCGTGTAAAAAAAAAAAA

>MSTRG.1305.2 gene=MSTRG.1305

CGTGCATTTGCTGGGCTCAAAGATAAAAATGTGGTATTTGGGCCCAGATAACATGGGGCAGGTAAAACCC

AAGTACCTTCTGGTTCCTAACCCTAGTATAAGGGAGGCTTTGTGTCTGCAAGAAATTATTGAGATCGAGA

GGGCGCGAGAGAGAAGGATGTCGTTTCTAACACCCTTTAAGCGAGCCAACCAGTGGATCGAACGTTGAGC

CTTATGGCTATTGTTGCTATCGGATCCGCGTTCTGGATTTCCAGTGGGCCCGACAAGAAACCAGACAAGG

ATGAAGCTGTCCGAACTCGTTCCTAGACTCACCCAACTGTTATTACTAACATCTCTGGTGCGATGTTATA

TATATATATATTAATCAACTCTTATCGTCTCTTTAATGGAGCTTTGTCCACTGAATGTTGAGATCTAACT

GCTTGCAGTAATTGCAATTGTTATTTA

>MSTRG.1307.1 gene=MSTRG.1307

AAGGAAGTCCTCCCCAAGGGTGACTAATGAAGCTGTTCAGAAGGTTGCTTCCGCTCTAAAGGGTTCTGAT

CACCGGCGTGCTACCAACGTTGGTGCAAGACTTGATGCACAACAAAAGAAGCTTAATCTCCCAATTCTCC

CAACTACTACCATTGGATCTTTCCCTCAAACTGTTGAACTCAGAAGGGTGCGCCGTGAATACAAGGCAAA

GAAGATTTCAGAGGAAGAATATGTTAATGCCATCACAGAGGAAATTAGCAAGGTTGATTTCAGAGGAAGA

ATATGTTAATGCCATCACAGAGGAAATTAGCAAGGTTGTCAAAATACAAGAGGATCTTGATATTGATGTC

TTGGTCCATGGAGAACCATAG

>MSTRG.1307.2 gene=MSTRG.1307

AGGAAGTCCTCCCCAAGGGTGACTAATGAAGCTGTTCAGAAGGTTGCTTCCGCTCTAAAGGGTTCTGATC

ACCGGCGTGCTACCAACGTTGGTGCAAGACTTGATGCACAACAAAAGAAGCTTAATCTCCCAATTCTCCC

AACTACTACCATTGGATCTTTCCCTCAAACTGTTGAACTCAGAAGGGTGCGCCGTGAATACAAGGCAAAG

AAGATTTCAGAGGAAGAATATGTTAATGCCATCACAGAGGAAATTAGCAAGGTTGTCAAAATACAAGAGG

ATCTTGATATTGATGTCTTGGTCCATGGAGAACCATAG

>MSTRG.1307.3 gene=MSTRG.1307

AGGAAGTCCTCCCCAAGGGTGACTAATGAAGCTGTTCAGAAGGTTGCTTCCGCTCTAAAGGGTTCTGATC

ACCGGCGTGCTACCAACGTTGGTGCAAGACTTGATGCACAACAAAAGAAGCTTAATCTCCCAATTCTCCC

AACTACTACCATTGGATCTTTCCCTCAAACTGTTGAACTCAGAAGGGTGCGCCGTGAATACAAGGCAAAG

AAGATTTCAGAGGAAGAATATGTTAATGCCATCACAGAGGAAATTAGCAAGGTTGTCAAAATACAAGAGG

ATCTTGATATTGATGTCTTGGTCCATGGAGAACCATAGAGGAAGAATATGTTAATGCCATCACAGAGGAA

ATTAGCAAGGTTGTCAAAATACAAGAGGATCTTGATATTGATGTCTTGGTCCATGGAGAACCATAG

>MSTRG.1307.4 gene=MSTRG.1307

GGAAGTCCTCCCCAAGGGTGACTAATGAAGCTGTTCAGAAGGTTGCTTCCGCTCTAAAGGGTTCTGATCA

CCGGCGTGCTACCAACGTTGGTGCAAGACTTGATGCACAACAAAAGAAGCTTAATCTCCCAATTCTCCCA

ACTACTACCATTGGATCTTTCCCTCAAACTGTTGAACTCAGAAGGGTGCGCCGTGAATACAAGGCAAAGA

AGATTTCAGAGGAAGAATATGTTAATGCCATCACAGAGGAAATTAGCAAGGTTGTCAAAATACAAGAGGA

TCTTGATATTGATGTCTTGGTCCATGGAGATTTCAGAGGAAGAATATGTTAATGCCATCACAGAGGAAAT

TAGCAAGGTTGTCAAAATACAAGAGGATCTTGATATTGATGTCTTGGTCCATGGAGAACCATAG

>MSTRG.1307.5 gene=MSTRG.1307

GCTTCCGCTCTAAAGGGTTCTGATCACCGGCGTGCTACCAACGTTGGTGCAAGACTTGATGCACAACAAA

AGAAGCTTAATCTCCCAATTCTCCCAACTACTACCATTGGATCTTTCCCTCAAACTGTTGAACTCAGAAG

GGTGCGCCGTGAATACAAGGCAAAGAAGATTTCAGAGGAAGAATATGTTAATGCCATCACAGAGGAAATT

AGCAAGGTTGTCAAAATACAAGAGGATCTTGATATTGATGTCTTGGTCCATGAAGAATATGTTAATGCCA

TCACAGAGGAAATTAGCAAGGTTGTCAAAATACAAGAGGATCTTGATATTGATGTCTTGGTCCATGGAGA

ACCATAG

>MSTRG.1307.6 gene=MSTRG.1307

GATCACCGGCGTGCTACCAACGTTGGTGCAAGACTTGATGCACAACAAAAGAAGCTTAATCTCCCAATTC

TCCCAACTACTACCATTGGATCTTTCCCTCAAACTGTTGAACTCAGAAGGGTGCGCCGTGAATACAAGGC

AAAGAAGATTTCAGAGGAAGAATATGTTAATGCCATCACAGAGGAAATTAGCAAGGTTGTCAAAATACAA

GAGGATCTTGATATTGATGTCTTGGTCCATGGAGAACCATAGAGGAAATTAGCAAGGTTGTCAAAATACA

AGAGGATCTTGATATTGATGTCTTGGTCCATGGAGAACCA

>MSTRG.1311.1 gene=MSTRG.1311

CTTAAATGACTCAGTATACCAAGCAAATAACCAGATTTCAAGCGTGGAGAACAAGCTAGTAGTATAATTG

AATATCTAGCTAGTAAAACATTAATATTGAAAATTAAGCTCATTGATTTACATTCTGCCGTTTGATCGTA

ACACATTCAAACCTGGAGTACAAAATTTTGTTGGGCCAAGAAGAACACCAACACCCAAATCTAAAAGAGA

GACATTTCTATATATATGTATGTGGGTATTAAGCTTGAAGGTGTTGATGAGAATGGTTTGTAGAAGAGAC

AGGCATGGCTAAGTATCGAGGCTTCTCGTCTCCGGCCATGATGACAAGATTAATCTTCGGCTCCATTTCG

GGCTGCAATAAAATCACCGGTTTAGCCGGTTTTTCCTCGTCGGAAGTGTTTTGAGCAGACCGGTCGGAGT

ATTTTCTGTATGAGCAAGCTAAGATCAGTAAAGCCATCGCGATTAGCGCCAGCATTAAGCCCAAACCGCC

GAATAAATATGGAATCGGAGAATTCATATTTACGTGGTGGCCGCCGGCCGACGGTGATGCCGGAGTGGTG

GCGTTAACT

>MSTRG.1312.1 gene=MSTRG.1312

GAGTTAAGTGCGTAGTTCTTGATTAGTAATTTCAATTGCAATACTAGGTTAATATTTGGAAGTTACTTGT

GTTGGAAATTGTAATTCACATATGTTCTTGAGATAGGATTTAGACGAATTTCCTTGAATTCTAGGATTGA

ATTGAACATCCTTCAGCTTAGGTCTGTAAATCTTAAGTTTGGGGGAACTGAGGAGTAGATATACCTTTCT

AGAAAAATAAAAGAAAAAAAAAAGAGATAAAAGAGAAAAGAAAAAAAATAGAAAAAAAAAGAAAAAGAAA

AGAAAAAATAGTATCAATAAATTCACTAGAAAATAAGTGGTAGAATTAACTAGGTTGAGCTCATTAGTAC

TCGAGTAATTAAGTCTGAGGGGACTTTGTGCCTAACAACCTAAAGCCCTTCGTGGTTTGGGATTGTTGAC

CCAACGCTCGCTACATGGGTACTAGTGCATAAATCTTTAGGGATCTCAACCATTGCACGGTTAAATAAAC

CACTAGAATAAAGTGAATAATTGGTGGGTGAAGTCTTATGGTGTTCGTAACGCATTTACACTGCGAAGCG

CCTAACCTTAGACCGAGCTATTAATCACCAATATAAAAATTTTAATATGTAGAATAAAATAGAAGGGTGT

GGACATTCTTTGGGACCTCGGCTTTTAGTGGACTTAAGTGACGGGGTGTTAGTTTTTAAACTTTTACAGT

TCATGATTGGGATTCTGTGGGGAGTAGTAGTTTTATCTTGTCTCAATCAAAGAGCATGCTTGCACACACG

GGCACTCCACACTTGTAATAGAAGAATATTTAATTTAGTCGCGAGCGAAGTGAAAGTTGAGAACACTAGC

ACAATCTGAGGTATTATAATTAGCAAGGCAATGGCTTCATGATTCGATCATTCATCAC

>MSTRG.1320.1 gene=MSTRG.1320

CTCTGCACGAATACCGAGAGAAGACGAACGGGTGTTCTTCGTGTTCTTCATATTCAAACGAGGATTTGAA

GGCGTTATTGGAATCCGATGGAGGCGTATAAATAGTCAAAACGAAGCTCTCGACGCGTAGAATCTATTGT

AGTCATCAATATCAGCGCTGAATCGACAGAAAAGTTGAAAGATGAATTGCATGATTGAGATTGCGAGTAT

GGCTAAGGCTAGGCGAGGAGTCAGAGTATAGAGTATACAGAGGACAAGTGGAAATGTTCAGGGAGATTGT

CAAGGTTGGTTCTAGATAGAATTGTTGAGATTCAGTTGTAATCATAATTTAAATAGATTGGCAGTGATTC

TTCTCATCGAAGATGATCTGAGGTTGTAAGACATTGTTTATTTATAGTGGTTGATTCTAATTTCAATACT

GTAACCTGGATGCGATCCTGTTTTTAGG

>MSTRG.1320.2 gene=MSTRG.1320

GTCAGTTTCTCTGCACGAATACCGAGAGAAGACGAACGGGTGTTCTTCGTGTTCTTCATATTCAAACGAG

GATTTGAAGGCGTTATTGGAATCCGATGGAGGCGTATAAATAGTCAAAACGAAGCTCTCGACGCGTAGAA

TCTATTGTAGTCATCAATATCAGCGCTGAATCGACAGGGTTTGGACGAGACGGATACGATCGAAGACGTT

CAGAAGGTCAAACGATGTTACAAAGCGAATAAGGAACGAATCGAGTAAACGAAGCAGTGTGGCGAATAGA

GTATAGCCAAAGATGGTCTAGAGACTATGATATGCATAGAGTGTGAAAGTGGAAAGATGAGATTGAGATA

TGAGACTGGAATCGGATTTAAGAAAAGTTGAAAGATGAATTGCATGATTGAGATTGCGAGTATGGCTAAG

GCTAGGCGAGGAGTCAGAGTATAGAGTATACAGAGGACAAGTGGAAATGTTCAGGGAGATTGTCAAGGTT

GGTTCTAGATAGAATTGTTGAGATTCAGTTGTAATCATAATTTAAATAGATTGGCAGTGATTCTTCTCAT

CGAAGATGATCTGAGGTTGTAAGACATTGTTTATTTATAGTGGTTGATTCTAATTTCAATACTGTAACCT

GGATGCGATCCTGTTTTTAG

>MSTRG.1326.1 gene=MSTRG.1326

CAAAACTAGAAAAGAGGTTCTGAGGAGAAGAGAATCAGCAGGAGAGATTTGGAGTCGCAGAAGGAAAACA

AATCAAATACTAAGAAAATTGATTAGCTGGAAGGCGTGCTTGTTAGTCAACTTAAATTGGAACGTGGTGC

ACTGTAAGTATATACTGTACCCTTCACTGTTGATATTTTGTGATGTTTCGGTGAAAAATTTGTTGATGAT

TTTCTGATGAGAAATGAGTATTCTTGCATTGATACTTGATGAACTGATGATGATGCTTCCTTATTGGAAG

TAAAGCTAAACCAATTGTTTAGCTGGCCGGGATCCCAACTTGATGAATTGATGAACTTATCGTGCTTGAA

TACTCATACTCATACTCATCTTGGTGTGTGAAATTGTTTTGATAACTCTGATGAAATATTGATTTGGTTC

ACCAAGTTTTGTGAATAAAACTCAGTTGAAATGTAAGTATATACTGTACCCTTCACTGTTGATATTTTGT

GATGTTTCGGTGAAAAATTTGTTGATGATTTTCTGATGAGAAATGAGTATTCTTGCATTGATACTTGATG

AACTGATGATGATGCTTCCTTATTGGAAGTAAAGCTAAACCAATTGTTTAGCTGGCCGGGATCCCAACTT

GATGAATTGATGAACTTATCGTGCTTGAATACTCATACTCATACTCATCTTGGTGTGTGAAATTGTTTTG

ATAACTCTGATGAAATATTGATTTGGTTCACCAAGTTTTGTGAATAAAACTCAGTTGAAATGTACGTATA

TGCTTACTGAGCTTTATAAGCTCATCCCTTTTTATGTACTAACCTTACAGGGAAGAACTCTGGAGAAGGA

TATTCATGCAAGGGTAAAAGTGTGATGAACCGAAGTGCTAGACTTGATTGGATTTGCAAGTAGAAGTTTA

ATAGAAGTATTATGTAATCGAAAGAAATATTTGTGAACTTTTAGAAGAGTGATGTAAGACTCTGATGTAA

GGTAAAATAAGCCTTTAAGTTATGTTAAAGTTTGAGATTATGAAGATGTATTTCGGTGTGTTCAGTAGAA

TGGCACCCCCGGGTCGGGTTGCTGAGGTCATCCCGGGCCGGGGGCGTCACAACTACAGTAAATATGTGGA

TTACATGGAACTTAACTCTTTGT

>MSTRG.1333.1 gene=MSTRG.1333

CCGGATCAAAAGCTCAGTGCTCTTCTGGTACTTCCTGATCTCTCGCAACGCCACCGTCCCTGGCCTGAAT

CTGTGGGGCTTCTTCACTCCTCCGGTTGCCGGTGCCGACTTTCTCGCCGCTTTTGTCGCCAGCTGCTTCC

TCGGGGCCTTTCCTCCCGTCGATTTTCTCGCTGTTTGCTTTGTTCGTGCCATTACAATACAACTAAGATA

CAGATATATATTCGTTTGTGTGTATAGGTTGTTG

>MSTRG.1336.1 gene=MSTRG.1336

AAACACTTATGTGATATGTTTCTCACTTATGTGCTAGTAATTAACCCATTCTGATATACACCATTATCAT

CCCTACTTCATTACTATCATATCCATGTTCTGAAGCACAAGATTTAGACACAGAAAGCTTCTTGATTCAC

AACCCAAGAAAATCAGCTTATTAAGGAAAAAGAAACTTCTGCTAGCTCCACTACCTTAAAAAACCAAAAC

AAACCAAATCCATATTGCTTCCTCATCTAAAAACCAAATCAAGCTCATCAAGAAAACCTCAAATCCTGCA

ATTACTTCAACCAGTGCTTCCTTCATCTTCAACCAAAAACCAAACTAAGATTCTCAAGCCCACTACTTCA

ACTTCAGATTCTTCACTCAAAAACAAAACTCATCAAACCCAGATCAACTTCCAACTCACCCACCATCAAA

ACCCAACTAAAGAAGCTCAATTCTACCACACCAAAGCCATTATTCACAGCTAAGAAATCTTCAGATCTTA

CTACAAAGCCCAAACAGACAAAAACAGAAAAGCCCAAGCTACAAAAAAAGAAAAAGAAAAAAAGAAAGCA

GCCCGTTTGGATAGATGAAGATGAGGAGGATTTAGTTTCAGATTTCAGAGTTCTACCATCAAGATTCCAA

GAAACATTAATTCCAGATTTGGAAAGATTCTCCACATCTTCCAAAGTTTATCTCACTAGAGCCAATAAAG

AAAATATGCACCTGCCAATCGTTACACATGTACACTGTACACATGTATGGCGGCCGACTCATTATTCTGC

CTATAATTATCATGTTCATATATGATTAACTGAAAATTGTTGTTTAACCAATCTTCTGGAAGATTGTGTG

CAGCATGCATATGTGTATCAGGTTATAGTGTAGTCAAGCTTCAAACAGGAC

>MSTRG.1339.1 gene=MSTRG.1339

GATTTTTATGGAGAAATCTAAGGATTCAGCTTGCGGCCCTATAGATCGATCACTGTTGTTCATGCAGGAG

GATCATATTAGCACCTCCATCTGGGAGGGCGATGATAGGCTCAGGTTTGATGTCAGACAGTATACCACAT

GTATGGATAAGTGGGTTTTGGATGATGAGCAGAGGCATCTTCTTGATTCTTGGGGTTTTGGGGTATTTGC

TCACCCTCAGGTTGTTCGACAGAACGACATTGCCCTGATAAGAGCCCTTGTTGAGAGATGGAGGCCGGAG

ACAAACACCTTTCACTTCCCATTCGGTGAGATGACCATCACACTAGAGGATGTCTACATGCTTATGGGTC

TGCCGATCAGCGGTCGTGCTCTTACTTTCACAGATCTTGCTACTCCACAGCAGTATTGGCTTACTCAGTG

GGAGGATTTGAGGTTGGAGAAGGGGCCTGTCCGAAAGAAGAAGTCTGACAAGAAGAAGAAGAAGAAGAAG

TCTAAAGGGAAGGGGAAGTCTGAACGGGAGAAGATGTGGTCAGAAAACGCGAGTGCAGTTAGTTTGTATT

CATTGCGGGAAACATATAAGAAGAAGCCAAAGTCAAAGCGTCCTGAATATTTAGAGCAGGATACCAGGGT

GTATACCCGAGCCTATGTGTTCTGGATCATAGGGGCTATTCTTTTTCCTACATCATCCAGAGCTACTGTC

CATCCCCGATACATACAGTTGCTGCAGAAGACGGATGAAATCATTGATTATGCCTGGGGTGCTGGTGTGT

TGGCATATCTGTACATTCTCTAATCCACCGAGAGCTTCCCCCATCACTTTCAACACTTTTGAGGGTCCTA

TTCCCGATTTGTTCAACGTCTCAATTAAAGATCGGGCAACCGGACCTATATTACGCGATCTTTGCATGAA

TTGTACTTTGTCTGGGGTAACCATTGGATGATTGTGTTCTAACTCTACTGTGGTTATTTCCCATTTATCC

GTCTTCACTCTATGAACAACCGAAATTCGAGCCTTACATTTTGTCCGATCAATTGTATCACGAGTCCTTT

TTAAATTCAATTCCTCATCAACATTAGCCGAACTTGAACATACTCTCCCACCTAGCCGGC

>MSTRG.1340.1 gene=MSTRG.1340

CACATATTTATGTATATGTTTAGCATATATATTCATAAAAAAACCGCAAGTCTTGAAGCTCGGTACTAGT

CAATCTCTAACAAGATCTAAGGGGCTTCCCACAGAGACAATCATTGTTGAGAAACGACGCCGCTTCAAGA

TGATCAAACGGTGAACCAACTGGAATCCGACCACATAGATGATTGTGACTCAGATCAAGATGACCCACGT

AACTTGCAGCTGATATGGACTTTGGAATAGGACCCTTGAGCTGATTATAAGATAAATCCAGGGCTGTGAA

ATATGATCGTGGCCCGAAAACATCGGGGATGTTACCTTCAAGCGAGTTCCTGCTCAGATTTAGTATACTC

ACTCCAGAGTTGATTAATGTTGTGGGTATGCTGCCTGATAATTTGTTACCGTCGAGATTTAGTGTTGCGA

GGACTCCCATTTTGCCCAGGGATGCTGGAATTGCACCCGAAATCTGATTTAACGAGAGGTCTAAATCAGA

AAGTCGGTATATGTATGATATTGAGTTCGGAATCGGACCAGTGATCCGATTCCGACTCAATAAAGCTCGG

CTCATCATGCGGAGCTTTCCAATGTCTCGGGGAAGTGGCCCCGAGATTTGGTTGTTACGGAGATCTAAAT

GCATTAGAGCTGTTAAACCGGATATAGACCGAGGTATCCAGCCCGAAATGTTGTTGTCGGCAACGTTTAG

CACCGTGAGTCGTTTCAATCGGCCGATATCTTTCGGGAGCTGGCCTGTAAGCTTGTTTCCGATTAAATCA

AGCATTCGGAGATACGGAAGATACGAAACGCAATCTGGAATCGGTCCAGTAATGCCTTTCCAATCTGCTA

TAATGATACTTGACAGCCACGTTAACTTGCAAATAGCTGGCGAAATCGTACCGGTCATATAACCGGTACG

ACGAGCCTTCTGGAATAACGGATCCTCCGATTCGCCACGGAGAATAATATCCGCGACTCGACCATTAGTC

GAGTCGCAACTAACTCCGTACCAATTATGGCAACAATCGCTGCCCGTCCACGACTTGAAAATTCCCAAGT

ACGGTTCGTGCAGGGAAGCTTTGAACGCGAGCAAAGCTTCCCTGTCCGAAGGCGGGCAGCCAGTAACTGC

CGAAACCGTGGCTAATGCTACTATAGCTGTGATGAACAAAAAAGAAGTCATTTTAAGTTGAGTGACTAAT

GTCATGCACATTGAAGTGTGAGCAAGATAAGAGAGTTATATAGTGAAAAAATTGAATGACGGTTTTAGCC

TTCACGTGTGGTGTAGAGAGGTGAGTGTACAGGTAAACATGGGTAAGGGATAAGATGGAGACTGGGAATT

ATTGCGATGAAGGTGCAAGGAAATGTGAGGTGCTGGAGCATGTGTCATTTAGTCTCTTCATGTAATTTGT

GCATACACAGAGGTGCCTACCCCATCAAATTTATTATTTCTGTATTTTTGCATTAAATTCCGCAATGTGA

ATTAATATTCGAAAGGGAATAATATCCTTTTTCTGTATGAAAGATTG

>MSTRG.1343.1 gene=MSTRG.1343

GATATCATGAAATTTTGCCTACATTTAAGGGATTATATCGCAGAAATATATCTCCCCATAGCAAATGTCC

ATTATGTGGTTATGCTGATGATTCCAATGCACATGCGGTGTTCTGGTGCGTCTTTGCGCAAGAGGTATGG

GAGCTAATGGAATACCCCTTTCTTGTGGGACGAAAAGAGGAGATCTCCTTTAAAGATGTGTTGTTATATT

TAACAGAGCTCTTAGAGAAGGAAGATATTGTGATGGTGTTTATAGCAACATGGGGAATTTGGACAGAAAG

AAATAAATTAACCCATCAACAAAAACGGAGAACGCCA

>MSTRG.1352.1 gene=MSTRG.1352

TGGTGATTTATAAATATATGGAGGAGGTGGTGAGGGTGATGGTGGTGGTGGTGATTTATATTTATACGGA

GGTGGTGGTGATTTATATGAATAGTGTTTCGGTGGTGGTGGAGGAGATTTATAAATGTACCGAGGTGAGG

GTGAAGTGTATCCATACCGTTTTGGTGGTGGAGACGCATAAACGGAGTACCAACTGTCGTGGGCACTTAC

ACTAGCCGCCACCACACATATAGCCGAGACATATAACAGG

>MSTRG.1353.1 gene=MSTRG.1353

CATATGTCGAATTCCATCTAGTATCCACATCAAGAATCAAACATTTTTGAATTTCAATTCTCTCGCATAC

TTTGCACATTCCATAAACAATGCCTCCCTAGCGGGTGAATACCTCACATACTTAATCACAATTGACTCAT

TAAAGACCCCCAATCCAACTTTGACAATTAAATTTAGAACATGAGCAACACACCTCAAGTGTAAGTGTTC

ACCATTTAACACGCTAGTACCCCAATTACCAATTTTATCTCTTAACCTATCCAATGCAACATCATTAGTA

CTAGCATTATCAACCGTGACAGATAATAC

>MSTRG.1363.1 gene=MSTRG.1363

CTCTTCCCCACTAAACACAACCAACATACACACATACATAATGCTGTGAAAATTACACATAATTACACCC

ACCCATCTCTCTCTCTCTCTCTCCCCCTCTCTCTCCCCTCGCCCGACGCCCTCTCTCCCGTCTCCCGTCT

CCCGTCTCTCTCTCTCCCCCTCTCTCCCGTCTCTCTCTCTCGGATGGCGACGATTGGAGCATCAAACACA

GCAGCAATAAAGAAAGACCCAAATCATGAAATTCAGAGGTTTGATGGGGTGAGGAAGACGAGCTTTTGTT

CGAATTCACTTCTTTTGCGCAAAAAGATTCAATTTTTGCCAACCCCATATGCCAACAATTCTTGTTTCAC

CATCAGAGCTGTATCTACTCCAGTCAAGCCAGATACGACTGCTCAACCTAAGAGAAGCAAGGTTGAAATA

TTTAAAGAACAAAGCAACTTCATAAGATATCCTCTTAATGAAGAGTTGTTAACGGATGCCCCCAATATAA

ATGAGCCTGCAACTCAATTGATAAAGTTCCATGGAAGCTATCAACAGACTAACCGAGAGGAGCGTGGTTC

CAAGTCTTACTCATTTATGCTTCGAACTAAAAACCCTTGTGGGAAGGTTTCAAACAAACTATACTTGGCC

ATGGATGATCTTGCAGATGAGTTTGGAATAGGCACTCTTCGTTTGACAACAAGGCAAACATTTCAGCTCC

ATGGTGTTCTGAAAAAGGATCTCAAGACAGTAATGAGTACCATCATCCGATGTATGGGCTCAACGCTTGG

AGCATGTGGTGATCTCAACAGAAATGTTCTTGCTCCTGCTGCCCCAATTGTCAAGAAAGATTACCTTTTT

GCGCAACAAACTGCAGAAAATATTGCTGCATTATTAACTCCCCAGTCTGGTTTTTATTATGATATGTGGG

TGGATGGGGAGAGAATTATGTCAGCAGAACCTCCTGAAGTCACAAAAGCTCGCAATGATAATTCCCATGG

AACAAACTTCACTGATTGCCCGGAGCCTATATATGGAACACAGTTCTTACCAAGGAAATTCAAAGTTGCA

GTTACCGTCCCAACAGATAATTCGGTGGATTTATTTACTAATGATGTTGGTGTTGTTGTCATCTCTGATG

ACAATGGAGAGCCTCAGGGATTTAACATTTATGTTGGTGGTGGTATGGGAAGAACACATAGGAATGAATC

AACTTTTGCTCGTCTTGCTGAGCCATTGGGCTATGTTCCAAAAGAGGATATACTGTCTGCAGTGAAAGCT

ATTGTTGTTACGCAACGTGAGAATGGAAGAAGAGATGACCGCAAGTATAGTAGAATGAAATATCTAATAA

GTTCATGGGGAATTGAGAAGTTTAAATCTGTTTTTGAGCAGTACCATGGAAAGAAAATTGAGCCCTGCCG

TGAGTTGCCTGAGTGGGAATTTAAAAGTTACTTGGGATGGCACAAGCAGGGTGATGGTGGTTTATTTTGC

GGGCTTCATGTTGATAGTGGTCGTATCAAGGGAACAATGAAGAAGACATTAAGGGAAATAATTGAGAAGT

ACAACTTGAACGTGCGTATCACACCAAACCAGAACATCATATTATGTGATATTCGTCAGGCATGGAAGCG

CCCAATCACCACAGCTCTTGCACAATGTGGTTTACTGCATCCTAGATATGTGGACCCCCTCAATTTAACA

GCAATGGCATGCCCAGCTATGCCACTCTGCCCTCTGGCAATAACTGAAGCTGAGCGTGGGATACCAGATA

TTCTCAAGCGGGTTCGAGCTGTGTTTGAGAAGGTTGGTCTCAAGTATAATGAATCTGTTGTAATAAGAAT

AACTGGCTGCCCTAATGGCTGTGCTCGTCCCTACATGGCTGAACTCGGATTAGTTGGAGATGGTCCTAAT

AGTTATCAGATCTGGCTTGGAGGGACACCAAACCAGACATCTTTAGCAAGAACATTCATGAACAAAGTTA

AACTTCAAGACCTCGAAAAAGTATTTGAACCATTGTTTTATTACTGGAGAACGAGGAGGCAGTCTAAAGA

ATCATTTGGCGAATTCTCTAATAGAATGGGGTTTGAGAAGCTACAAGAATTGGTAGACAAATGGGAAGGT

CCACCTCAGTCGTTGGCTCAATATAATTTAAAGCTTTTCGCTGACAAAGAGACATATGAGGCTGTGGATG

AACTTGCAAAAATCCAAGACAAGAATGCTCATCAATTAGCCATGGATATAATTCGCAATTATGTTTCTTC

GCACCAAAATGGCAAAATTGAGTGAGCCATAAAGCACGTTTCACTAGGATGCTATTATTGTTGCTAGTCT

TGGGAGCCCAGAAAACTTCTTCAACATCTGCTGTTACTTTTCGGTAACAACGGATAATAATGAGTCTGTT

AGCTGCACATGCATCCAAAGTCTTTAAGTAATATCATCAACTTTCACCTTCTACAAGGTTGTTTTTATTT

GTTTTGCTTGAAATTAGTTTGAAACTTGGTTTTTGTAATATGCTAAAACTTAGTGTCTGTAAAAGTCTTC

GGGATTATTGTTTAACAAGTAGTCCAACCTTTAAGGTTTTTCTGTTGGATGACTGAAGCAATTAAATCCA

TTTTTAAGTAAC

>MSTRG.1364.1 gene=MSTRG.1364

ATTGCAGTAACGATTTGGATAAAAATTACTTTCTTCTGAACAATGCACTTTAATCACAAGACAAAGAATC

AGACATTGCCATATGAACCAAAAAACACCAGTCTCAACATACAAACAGCTCAAAAACATCAGTTGCGACA

AATACGGGCAGTTGGATTGGTAACACAAACTTGAGCATACTTCTTTTTGGCCTCGGGGCTTCCACGCTTT

GGCTCATCAGCCAACGCAATGTTGGCCACCGAGCATGCTGCTGCAGCCGCGACAGCAAACAACAATTCCC

TCCTGGCATTCTTGTTATTCACTTCAAAACTCTCCTTAGTGACTTCTGAAGCCTTTGACATGACAACATT

GCTTCTGCGGACACTTATCATTGGCTGCTTTGCGGAGGCAGCTAACATGACTGATGAGCCTGTCCCGAGG

AATGAGGATGTCATAGTTACTGCAGCCATTTTGCTTCTGGGAAATATTACCTGGTCTCTTTTCAATGTTA

ACAAGTCTGCAGATAGTTATGTGTGTTTATAGACAAATTTCATCCATTTCCTTATCTTAGTTATTGTGTT

GTCTCGAATGAGATGATGCCATGTGTCACTTTATCTATGCATAAATCCGTGGTTCAG

>MSTRG.1367.1 gene=MSTRG.1367

ATATATCCTCTGCAGTTGGTGGGTGGAAACTGGATTGGACAAGATGAGCTTTGCAAGAGACAGGTTGGTA

GAACATTATTTCTGGTGCAATGGAACAGTTCCTGATCCAGAATATAAGGCTTTCCGAGATATGGGAACCA

AAGTGATTTGCTTACTAGTAATCATTGATGATTTGTACGATATATATGGCTCATTAGAAGAGCTGGAGCT

CTTTACAGATTATGTGGACAGATGGGACATAACAGAAATTGACAGGCTCCCGAAGAAATTGAAGACAGTT

CTTCTTGCAATGTTCAATACTACAAATCAAATAGGATACTGGACACTGCAAGAACGCGACTTTAACATTA

TACCATATTTAAGTAAACAA

>MSTRG.1371.1 gene=MSTRG.1371

ATCTCATGTACTGCAGCCGCAGTAGTATCCAAGTGACAAAAGCTTTTCTTTAGTAGAAATATTTTACCTC

ACAGCTATCAAGTTTTGCAACACATGATCTAGCAACTACGGCAGAATAGAAGCAAGGATACTGGGAGTCG

ACCTGAGCAAGATTACCATTCCAGCCAAAGTGCATATGAGAGTGGCGGGGGAGATGGACCTGGTTTGGAG

CTGGCAATTTTAACAGAGGAAATAGTCCATGCCAATGGTTCAGAAAGCTAATGGACCACCTGTAATGACT

AAAAGCAATAGTCTTACAGCAACTTCAGATAGCAGGTCTTTCATAGCTGGTTCACATTGATGTTAGTTGC

GAGAACACAACGTATTAGGCACAAATGTATAGTTAACAGAACAACATAATAATGTGGATCTCAAGTAAAA

AGGTTTAGTAAGATGGTGCAAAAGCAATATAGGTCAAGGACAGATTTTGCGCAGATCTCAAGTCTCAAGT

TTCAGCTGTGTGGCTCGTGATGAGTATGAAGGACTGATTGAGGCTAAGAATTTTTGTCGAATGGGAAGAA

TTGCACCCGATGAGGCAGAAGCAACTGGTATAAAGAGGCCTTGAGTTTTATTAAACAAAAATCACGGGGA

AAAGTGGTCGTTGAAACAGACTGTCTTGTAGCTTTGAAAGCAATTAGAAGTCCCCTAGCCATGTCTTCAT

ACTTTGGTGATATCTCCAGGCCTGCAAATTACTTATGGCTGAGTTGAACAATGTTATTATTGTTTTTGTA

AGACCGTCTGCAAATGCAGTTGCCCATGCTCTAGCAAGGGAAACTCATGTAGTAGCTGATCATACATTTG

GTGGAGAAGATG

>MSTRG.1373.1 gene=MSTRG.1373

AAATATCTCCCACAAAACTAGATCTAGCCGCCCATCCAGTTCTCTCCTTTTATACTGTCCTTCCCTATCA

TTTCTTTTCTTGGGATTTGTGAATTCTTTTCTTGGGGGTTGTGAATATTGTTCATCTGTTAGAATGCAGG

AATTGAAGTTTGCTAAACAATGAAGAAAATGCTTCAAAACTAGGGGGATTTTTATGTGTATACGATTGTT

TACCAAGTACAATGAAGCCAGGAGTACCCGCTATAACATATAAAACAAAGGCAAAAAGTTCTGCTCTCCT

GCTGCCAAGAAGGCTGATGGTGTTGACGTGGAGGC

>MSTRG.1374.1 gene=MSTRG.1374

GTGAAAGTGAGTCATTTAATGTATCGATAATAACTGGCTCTTCAAGAGGAATTTTAAAAGGATCTAGCCA

TGAAAAAAATTTGAATTGTTCATCTTCAATGTCAAATGACACTACTGCTTTGGAAAGTCTATCTTCAATA

CCTGCAGAATCAGTAACCCAATGAAATGCCTATTTTAAAGGTCAGATGAAATAAGGAGCAGAGATACCAA

AAAGTATCACTGGATCATTGAAATGAGGGNCAGAATCAGTAACCCAATGAAATGCCTATTTTAAAGGTCA

GATGAAATAAGGAGCAGAGATACCAAAAAGTATCACAGGATCATTGAAATGAGGGGAACTTGATCACGCT

CATGACATGGAAGAGTTGTTACATCATGTGCAC

>MSTRG.1375.1 gene=MSTRG.1375

GCTCGACTCCTTCAGTCTTCCACGCCCACACTGAAGCACTGATTCCACGATGATTAAGAGAAGTGAATTT

ATCTGATTAACAAGTTTATGTATCTAGCTATCTGATTGTTTTAGATCTGTTAATTCTTCATTTTGTGTGC

TTTTTCTGTTTATATGTTCTGTTGTCGTCCCTAGATGTTTGGATTTTTGTTGTTTTTTTGGGTGAATTAG

GGTTTCGTGTGATATGAATGTGTTGTTTTTGTTCATACGTTTGGTTGATTTGTGCTTGTGCTTTATTCGT

GTTCTAAATGTTAATTTGAGAACCAAATGAATTAAGAATTGAAAACCTTTAGGTTTGTATAGTTGTAACT

GGTGATTGGGATGCAATTCAGTATTGATATGTATTTGAAACGAATTACTGATATGCATCTGAGCAAAGAG

TCGGGAGTGGTTGATATAAATTTGTGTTCAACCTTTTCAGGTGATCAGTAACAAGAAGTTACCCAAATGA

GGATTCATTAGACAAATCCATGGAGAGTGATAGAAGAATATGTGGCAGATTGATTTGTTCTGTAGCCAAA

GAATCTCCACAGCAAGTCGAAGAAACGACCATGGACATGCGAAAGTGAGGAACAAACAGTTGGCTGCTCC

AACCACAGTGTTTATTCGAGATTCGAAGGTATCACCAACAACTAATTCTTCTTCAGTATTTGGATATTCT

CAGATTATCTGGCTTGTCACAATCAAGAGTTTGCTATAACGAGGTCTTTACCGGTCACTAGAAATTTCTT

CACACAATGTGAAACAGCGGATTTTTGGAAAATCCTAATTGTCGTAGTATATATGATGTAAAAACTTTTC

AATATCATGAGCCTGATCAAGTGCCCCTAATTTCAATGATCCAGTGATACACTCTGGTATCTCTGCTCCT

TATTTTATCCGGCCTTTAATATAGGAATTTCATTGGGTTACTGATTCTGCAAATATTGAAGATAGACTTT

CCAAAGCAATAGTGTCATTTGACATTGAAGATGAACAATTCAATTCTTATTTACATGGATGGATTCTCCT

AAAATTCCTCTTGAAGAGCGAGTGATTATCGATACATTAAATGACTCACTTTCATTGGTTGTTCCTCGCA

AAAGTTTTCTAAACTCAAGCTTTTATATATGGGTGATGGATGACTACGGGAAATAGGTATCTTGGACTAG

GCAGTCCATCACCCATATATAAAAGCTTGAGTTTGGAAAACTTGTGTGAGGAATAATCAGTGAAAGTGAG

TCATTTAATGTATCCATAATCACGGGCTCTTCAAGAGGAATTTTAAAAAGATCTAGCCATGAAAGAAATT

TGAATTGTTCATCTTCAATGTCAAATGACACTACTGCTTTGGAAAGTCTATCTTCAATAATGCCTATTTT

AAAGGTCGGATGAAATAAGGAGCAGAGATACCAAAAAGTATCACTGGATCATTGAAATGAGGGGCACTTG

ATCACGCTCATGACATGGAAGAGTTGTTACATCATCATGTGCACTACGACAGTTAGGATATTGGACAAAT

GATCTTCCTATAGTGACAACATATGAGAGCGATTTTTTTTATGACCCTAAAATCCAAGAACAGAAGAACA

CTCCAGTTTTTTGTATTTCACAACATTCTGCATTTATGTTGAAACTTTGGAAACAGTCACCTGGGATATT

GAATATGCAGGAAGGGCTGTTCAGTGATGGCACACATCCTGTTGAGTAAAAACTAATGTAACATGTGATG

GGAGCACAACAATTTTTTGAAGTTGATTATTATTAGATGATATTGCCGAGATAATGCTCATCAGAAGAGA

GTGACTGGCCTTGCCTATTCTAGTGCATTAAATATCCTTTTATCGTCTGGATAAGATGCTCAGGTACATT

TTTAAAGTGGAAGGCTAATAGAAGATATCAGATGCATGGGCACATTTCTGTCAAGTGTCACCAGGACCAA

AGACATTTGTTGGTTGTACATGAAAGTTGTATAGCCACTCCTGATCGGGACCTAATCTCGAGCCCTGCAA

GCAGTAGATTCCCGGGGAATTGATCAGTGATGCTGCATATATATTCCTGTGATGGCCAGTCCATATATGC

CTGCTTTCTTAACAGGCACATTGTTTTCTTGGACTCGGCATCTTTGACCATAATTTTTTTTTATCAATCC

GAGTGCATATTTTACCTTCTAAGTAAAGCCGGGTGATGTATGCTTCTACTCTTGCGGCTAATCCTACAAA

TTAAAACCCAATCAGTTTGCTCTCGGGCTTAGTGATGGTGGCGTACTAGTTATAGAACTTGAGGAAGAGG

ATATTGGGGTTGGCATTGTAGCACGCCTTAGCTTGAGACCAGCAACAACTGAGAACTTGATATGAGCTTT

TGTGATAATCAAAGCCCAGTGTCTCTTGAAATCTATATGTTTTAAAACATACATCTGGGGATTGGATTAG

TGCAAGAGTTGTGAAGTTTTTTTTTTTGCTTGACTATGTTAAGTGATAAATGTTCAATTCAAATATACAC

CTAGTGCAGAATTTGAAATGAGAATTGTCAACAGAGTATGAAAATGAGAAAAGAAAGCCAATATTTTGGC

ATATACAA

>MSTRG.1375.2 gene=MSTRG.1375

TATGGGTGATGGATGACTACGGGAAATAGGTATCTTGGACTAGGCAGTCCATCACCCATATATAAAAGCT

TGAGTTTGGAAAACTTGTGTGAGGAATAATCAGTGAAAGTGAGTCATTTAATGTATCCATAATCACGGGC

TCTTCAAGAGGAATTTTAAAAAGATCTAGCCATGAAAGAAATTTGAATTGTTCATCTTCAATGTCAAATG

ACACTACTGCTTTGGAAAGTCTATCTTCAATAATGCCTATTTTAAAGGTCGGATGAAATAAGGAGCAGAG

ATACCAAAAAGTATCACTGGATCATTGAAATGAGGGGCACTTGATCACGCTCATGACATGGAAGAGTTGT

TACATCATCATGTGCACTACGACAGTTAGGATATTGGACAAATGATCTTCCTATAGTGACAACATATGAG

AGCGATTTTTTTTATGACCCTAAAATCCAAGAACAGAAGAACACTCCAGTTTTTTGTATTTCACAACATT

CTGCATTTATGTTGAAACTTTGGAAACAGTCACCTGGGATATTGAATATGCAGGAAGGGCTGTTCAGTGA

TGTACATTTTTAAAGTGGAAGGCTAATAGAAGATATCAGATGCATGGGCACATTTCTGTCAAGTGTCACC

AGGACCAAAGACATTTGTTGGTTGTACATGAAAGTTGTATAGCCACTCCTGATCGGGACCTAATCTCGAG

CCCTGCAAGCAGTAGATTCCCGGGGAATTGATCAGTGATGCTGCATATATATTCCTGTGATGGCCAGTCC

ATATATGCCTGCTTTCTTAACAGGCACATTGTTTTCTTGGACTCGGCATCTTTGACCATAATTTTTTTTT

ATCAATCCGAGTGCATATTTTACCTTCTAAGTAAAGCCGGGTGATGTATGCTTCTACTCTTGCGGCTAAT

CCTACAAATTAAAACCCAATCAGTTTGCTCTCGGGCTTAGTGATGGTGGCGTACTAGTTATAGAACTTGA

GGAAGAGGATATTGGGGTTGGCATTGTAGCACGCCTTAGCTTGAGACCAGCAACAACTGAGAACTTGATA

TGAGCTTTTGTGATAATCAAAGCCCAGTGTCTCTTGAAATCTATATGTTTTAAAACATACATCTGGGGAT

TGGATTAGTGCAAGAGTTGTGAAGTTTTTTTTTTTGCTTGACTATGTTAAGTGATAAATGTTCAATTCAA

ATATACACCTAGTGCAGAATTTGAAATGAGAATTGTCAACAGAGTATGAAAATGAGAAAAGAA

>MSTRG.1375.3 gene=MSTRG.1375

GCACACACTATTCAGTAAAACCTAATGTGGCTTTTGGGAGGTTTTGCTCTCTTTTGTATAGTAGGGCGCA

CATCCTGTTGAGTAATTTGATGCAGGGTCTTTACCAGTCATTAGGAAATTTCTGCACACAATGTGAAGCA

GCAGATTTTCGGAATATCCTAATTGTTGTAGTGTATATGATGTAACAACTCTTCATGACATGGAAGAGTT

GTTACATCATGTGCACTACGACAATTAGGATATTGGACAAATGATCTTTCTATAATGACAACATATGAGA

GCGATTTTTTTTTATGACCCTAAAATCCAAAAACGGAAGAACACTCCAGGTTTTGGTATTTCACTACATT

CTGCCATTATGTCGAAACTTTGGAAACATTCACCTAGGATATTGAATATGCACAAAGGGCTGTTCAGTGA

TGTAATTCCCTACCCTTGCTTAGTATATCTATTAACACTGCATGATATAATATCTGAAGCATTGTTGAAG

AATAATAAATTATAACATCTGGTTAAAAATTTGAAAGGAAATATATCTCTGTAATAGTTTTGAAATGTTA

CGGAAAGTTTAACACTAACTAATGTTGCTTTTGTGAGCTTTGTCTACTTTAAATTGTAGCGCACACACTG

TTCAGTTAAAGCAATGTGGCTTTCGGGAGCTTAATTTGCTCTAATTTGCTTTGTAGAGCATACAATATTC

AGTAAAACCTAATGTGGCTTTTGGGAGGTTTGCTCTCTCTTTTGTATTGTAGGGCACACATCCTGTTGAG

TAAAAACTAATGTAACATGTGATGGGAGCACAACAATTTTTTGAAGTTGATTATTATTAGGTACATTTTC

TATCAGTAACCACCACAACAATAATGTCTGC

>MSTRG.1375.4 gene=MSTRG.1375

ATTTTTTGATCTTTTTATCTTAAAAATATTTCTTTCCTTCTCTTTTAGTTTTCAAATACTTTTATTCACA

TGTGTATTGTTGATGTGATTTATCTCTCTTGTAATACTACCCATTTACAGTTTGTAAAACTAGTTATTAC

TATTCTATACTTCTACTACAATCGGACCATGGAAAAAAACTTTTCCCGGCCACCTGCTGCAACTTCAGTT

ATGTTCTATCCTGAGGAAAATAATATCATATCGTCTGCAGGGAGGATGGCTCTATACTAATCTATGGTGT

TCAGAAAGATATGGTCTTTATCTATATATACTGATATCTAAGTGTTGTACATTAAAAATGAATTATCATT

TACCGATTAAGTTCTTATTATCTATTTGGCTTATATAGATGATATTGCCGAGATAATGCTCATCAGAAGA

GAGTGACTGGCCTTGCCTATTCTAGTGCATTAAATATCCTTTTATCGTCTGGATAAGATGCTCAGGTACA

TTTTTAAAGTGGAAGGCTAATAGAAGATATCAGATGCATGGGCACATTTCTGTCAAGTGTCACCAGGACC

AAAGACATTTGTTGGTTGTACATGAAAGTTGTATAGCCACTCCTGATCGGGACCTAATCTCGAGCCCTGC

AAGCAGTAGATTCCCGGGGAATTGATCAGTGATGCTGCATATATATTCCTGTGATGGCCAGTCCATATAT

GCCTGCTTTCTTAACAGGCACATTGTTTTCTTGGACTCGGCATCTTTGACCATAATTTTTTTTTATCAAT

CCGAGTGCATATTTTACCTTCTAAGTAAAGGTACCTGCCTTCTGTATTATATCTACAGCCGGGTGATGTA

TGCTTCTACTCTTGCGGCTAATCCTACAAATTAAAACCCAATCAGTTTGCTCTCGGGCTTAGTGATGGTG

GCGTACTAGTTATAGAACTTGAGGAAGAGGATATTGGGGTTGGCATTGTAGCACGCCTTAGCTTGAGACC

AGCAACAACTGAGAACTTGATATGAGCTTTTGTGATAATCAAAGCCCAGTGTCTCTTGAAATCTATATGT

TTTAAAACATACATCTGGGGATTGGATTAGTGCAAGAGTTGTGAAGTTTTTTTTTTTGCTTGACTATGTT

AAGTGATAAATGTTCAATTCAAATATACACCTAGTGCAGAATTTGAAATGAGAATTGTCAACAGAGTATG

AAAATGAGAAAAGAAAGCCAATATTTTGGC

>MSTRG.1376.1 gene=MSTRG.1376

TGAAGTTTGCTAAACAATGAAGAAAATGCTTCAAAACTAGGGGGGTTTTTATGAGTATACAATTGTTTAC

CAAGTACAGTGAAGCCAGGGGTACCCATTAGCTCTAAGCAGTACATGTTATAACATATAAAACAAAGGCA

AAAAGTTCTGCTCTCCTGCTGCCAAGAAGGCCGATGTTGTTGACGTGGAGGCGTGAGAGTCGTTGGATGT

GATCTGTTCTGCTAGTTTTTACTGTTTTAGTTTAATGTGGTTAATGGGATGCAGGTCTAGGCTAGTAGTA

GTCTAGTCATAAAAGTTCAGTATTTTGGGTGTTTAGATGGATGGTTAAATTTGAACTTGATAGACAGCTT

TGT

>MSTRG.1377.1 gene=MSTRG.1377

CACAATCACATCACATATATATATATTTGTATTATTTCAATCGCTGTGTTCTTGTGTCTCCTCTCCTTCA

AAAAAAAATAAAATGGCGATACAACTTACAGGATTGCAAACAAGTCACGGAAGAGTCACGGTTCTTGTTT

GATTTAAGATATTACAAATTAGTANTTGCCTCTTTATTTTTCCTTCAGGATTAGGGCTTTTACTATTCAT

AAAATTGATTGTTGTAAACTGGGTTCCTAAATTCAGGTCTTAAATACAAGGTGGAAGGCTATAATCTTTG

ATATTTATACATCATAATTCAGAATTGTGCTTCCTCAGTGATCAACGATTTACTCCTGAGACTCCTAAGC

CTTGAGCTTCTGAAGAGGAAGCAGCTAAGGCATATGATTTGGCTGTAATTGGGAAAGATGCTTTGACAAA

CTTTGATATAAAGAAAACTAAAATACCATCCCATGGACCAAATTGGCATCTTGGAAGATCAGGAATATCC

AAAAAAATAAAGAAATATGACCAGCTTAATTTGGAGAATGGTCAGAGTATTGCTGATCATACTGATTTGA

AAAAGGCTATAAGGGATGAAGTTCTTGCTGAGATGGAGGAGATGATCGAGAAAAAAGTTCATGATAAGGT

GACAAAGATATTTTGCAAACTGGGAGAGGTAGATCCTTATTTCAACGACATCGATCTTGAAGCGGTGTTG

CGTACTTCTGATGATGAACTTGAACATGGTGATGAATAAAGATTTTTAGTAGTCACTTTTTATGTTTACA

TCATCTTCTCTGTGCTTTTGGATTGATTTTTGTAAGGATGATTTATAAGTTGGTACCTTTGTCCAGTTTC

TATGTATCTTTTCAACTTCATTACTGAGTTTGTAATTTGAATTCTATTTTGGTCAATTTTCAAGTTATGA

AAATTTCAGAGTAT

>MSTRG.1377.2 gene=MSTRG.1377

CACAATCACATCACATATATATATATTTGTATTATTTCAATCGCTGTGTTCTTGTGTCTCCTCTCCTTCA

AAAAAAAATAAAATGGCGATACAACTTACAGGATTGCAAACAAGTCACGGAAGAGTCACGGTTCTTGTTT

GATTTAAGATATTACAAATTAGTANTTGCCTCTTTATTTTTCCTTCAGGATTAGGGCTTTTACTATTCAT

AAAATTGATTGTTGTAAACTGGGTTCCTAAATTCAGGTCTTAAATACAAGGTGGAAGGCTATAATCTTTG

ATATTTATACATCATAATTCAGAATTGTGCTTCCTCAGTGATCAACGATTTACTCCTGAGACTCCTAAGC

CTTGAGGGTTGCGTAGGAATGGTAACAATGTAGAGTAAGGTGGATGAATCATCTCAGCCCTGATTTGAAA

GGGGAAATTTTACTACTGACGAAGATCAGATAATTATCCGTCATCAAGCTCTTAATGGCAACCGCTTCTG

AAGAGGAAGCAGCTAAGGCATATGATTTGGCTGTAATTGGGAAAGATGCTTTGACAAACTTTGATATAAA

GAAAACTAAAATACCATCCCATGGACCAAATTGGCATCTTGGAAGATCAGGAATATCCAAAAAAATAAAG

AAATATGACCAGCTTAATTTGGAGAATGGTCAGAGTATTGCTGATCATACTGATTTGAAAAAGGCTATAA

GGGATGAAGTTCTTGCTGAGATGGAGGAGATGATCGAGAAAAAAGTTCATGATAAGGTGACAAAGATATT

TTGCAAACTGGGAGAGGTAGATCCTTATTTCAACGACATCGATCTTGAAGCGGTGTTGCGTACTTCTGAT

GATGAACTTGAACATGGTGATGAATAAAGATTTTTAGTAGTCACTTTTTATGTTTACATCATCTTCTCTG

TGCTTTTGGATTGATTTTTGTAAGGATGATTTATAAGTTGGTACCTTTGTCCAGTTTCTATGTATCTTTT

CAACTTCATTACTGA

>MSTRG.1377.3 gene=MSTRG.1377

TTTCAATCGCTGTGTTCTTGTGTCTCCTCTCCTTCAAAAAAAAATAAAATGGCGATACAACTTACAGGAT

TGCAAACAAGTCACGGAAGAGTCACGGTTCTTGTTTGATTTAAGATATTACAAATTAGTANTTGCCTCTT

TATTTTTCCTTCAGGATTAGGGCTTTTACTATTCATAAAATTGATTGTTGTAAACTGGGTTCCTAAATTC

AGGTCTTAAATACAAGGTGGAAGGCTATAATCTTTGATATTTATACATCATAATTCAGAATTGTGCTTCC

TCAGTGATCAACGATTTACTCCTGAGACTCCTAAGCCTTGAGCGTTGCATAGGAGTGGTAACAGCTGTAG

AGTAAGGTGGATGAATCATCTCAGCCCTGATTTGAAAGGGGAAATTTTACTACTGACGAAGATCAGATAA

TTATCCGTCATCAAGCTCTTAATGGCAACCGGTGTTGGCTCTTTTATTTATTCATTTCTCTTGGCCTCTT

AAATAGGACAATGAAAATAAGAATTTATTAGCGTACGTGTTTAATATACAATCTTAAAGCCGC

>MSTRG.1380.4 gene=MSTRG.1380

TGCAGACTAAGAAATTTACTGAAACGGAGGAGCTCAATCGCAACCAAGAAGAGAAATTCATGGCTTTTAC

AAAAGCCTCAGATGAACTCCTAGAACAGTATAGAATTGAGAATGACAGACTTATGGCAGAAAATGATGAT

TTGAGATGCGAACTGGCTTCAACAAGATCTCGCATGGATGATAAAATCGCTGAGTATCAGAAGCTTTTGA

TGGAAGAAAATCACAAAATTCTCATTGTGTGCAAGAAGAACATAGACCGAAGCGTGAGAAGACTGATAGT

GCTCATATTTCAACTATTTCAAAGTTCAATAAGCGAAAGAGAAGTAAAGAAAGAGAAATTGCAGTGGGAA

TTTCACAACCCAAGAATGTAAAGACTAATGAAAAAGGTCGTACCTGTTTCTTTTGCAAGAAATCCGGTCA

CGTAAAAAAGAAATGTCCCGAGTTCGCTGCATGGCGCATGAAGAAAGGGTTGAATTTGGAGCTAGTCGTC

AAGTGATGTTGAAGGATTGATCTGTGTGGGGGTTGAGTCAATGACGATGCAGGACTTGTTGAACTCGTTA

GCTTTTTTTGGATTATCTTTTGATATCGGTCAGTCTATTAAATTAGGGGAGACTAATATAGTGCCATCAT

TTTGATGGAATTAGGTTTTCATTTTTGTTTTGGACAATAT

>MSTRG.1380.5 gene=MSTRG.1380

CGCTGGGCTTAAATCAAAATTGAAGTTACCATCCGCTTCAGTAGTTCGATTTGAGGGCTTTCTACTTGCT

TCTTCATCTCTTGAATTACCCCTTTTGTTCTTTTCATCTTCCTTTCAGACTTCACCATGGAAGCGCTTTA

CTCCAAGCTCCATGAAAAATACACTCATCTCAAGACTAAGAAATTTACTGAAACGGAGGAGCTCAATCGC

AACCAAGAAGAGAAATTCATGGCTTTTACAAAAGCCTCAGATGAACTCCTAGAACAGTATAGAATTGAGA

ATGACAGACTTATGGCAGAAAATGATGATTTGAGATGCGAACTGGCTTCAACAAGATCTCGCATGGATGA

TAAAATCGCTGAGTATCAGAAGCTTTTGATGGAAGAAAATCACAAAAAAGAACATAGACCGAAGCGTGAG

AAGACTGATAGTGCTCATATTTCAACTATTTCAAAGTTCAATAAGCGAAAGAGAAGTAAAGAAAGAGAAA

TTGCAGTGGGAATTTCACAACCCAAGAATGTAAAGACTAATGAAAAAGGTCGTACCTGTTTCTTTTGCAA

GAAATCCGGTCACGTAAAAAAGAAATGTCCCGAGTTCGCTGCATGGCGCATGAAGAAAGGGTTGAATTTG

GAGCTAGTCGTCAAGTGATGTTGAAGGATTGATCTGTGTGGGGGTTGAGTCAATGACGATGCAGGACTTG

TTGAACTCGTTAGCTTTTTTTGGATTATCTTTTGATATCGGTCAGTCTATTAAATTAGGGGAGACTAATA

TAGTGCCATCATTTTGATGGAATTAGGTTTTCATTTTTGTTTTGGACAATAT

>MSTRG.1385.1 gene=MSTRG.1385

AGGCGAGTGTGCCTTGCAATTACGTTATCAAGCCCTTCCTCAAAGATAAGATCAAGTGCTGCCCTCAGCC

CATATAATAGTTGGATGGAAGGAGTGTATGGCCAGTATGATCCCATGTTATAGAACTTTAAGTAGTCTTT

CCAGTCAAAGAAAACTCTGACCGACTTTGCAGTTTTGGATGCTTCAAGAGCTTTGGGGCTTGCACAAACA

ATTCCCAATCCAGTGGGAAGAGAAAGAGCTTTTTGGGAGCCAGTTAGAGCCACATCCACACCCCATTCAT

CCATACGGAAATCAAGTGCACAGATGGAGGACACGCCATCAACAAGCAAAAGAGCTGGAT

>MSTRG.1386.1 gene=MSTRG.1386

AGGAAATTCATGATAGTTGCATTTAAATTAGAGCATATGGTTATGTAATTTGACATGCATTTTTTTTAGT

GGATTTTTTTTTTCTTATTTCTTTTTAGTTTGCATTTGCATGATTCGATGAAAGTTGTAGACCAAGTTAA

TTTCCTATGATGAGTTTATTTGCATGATTTCTTGGATAGTAATCTTGATCATATGTGATGTCTTGCATGC

TTAGAAATTGCTTGTGCGGAACTTGTATTACACTATAACTGTTCTAGAGACGAGATTTAGACTAGCTTAT

TTCTTGAATCTTAGCTATGAGCCGAGCGTAGAATTTTGACTATTCCCTTTTGAGCTTAGAGTCTATAATT

CTTAAGTGAGGGGGAGCTTTGGGATATACATGCCTTTCAAAAAAAAAAAAGAGAGAGAAAATATATACAT

GTATTAGCAATAAAGTTCACGAAAAAAAAATGGTGGTATAATTGACTAGGTTGGGCTCATTAATACTCGA

GTAATTAAGTCTGAAGGGACTTTGTGCCTAGTAACCTAAAGCCTTTCATGGTTTGGGATTACTGACCCAA

CACTCGCTAAATGGGTATTAGTGCATAAATCTTTTGGGATCTCAACCATTGCACGGTCAAATAAGCCACC

TAAACCGCATGTGAACAATTTGGTTGGTGAAGTCTTATGGTGGTCGTAACTCATTTACACTGAGAAGCGT

CCTGACCTTAGACCGAGCATGTACACACCAAATAATAGATATTGAAAAAAGAAAAGAAAAAGAGAAGCAT

ATATAGGAGAGGCATGGAAGTTCCTTGGTGACCCTAAACTGTAGTTGGCTCTAATGTAACGAAGTGTTTA

AGCATTATTCTAATTAACGACTCATGGTTAAGATTCTATTGAACTTAATTTGTTGGTCTCTATTTCGTTT

CATTAGAGAGCATGTTGAGCACACACGATGCACTCCGTACTTGCAGTGAGAGTTGCATGACTAATATGTG

TGATAGATGAAATCCTTGATTATTGCATATTCTGAGGATCATATGTTGACAAGGTTGATAATTTCATGGA

TGGAATCTATGCATATTCATTTAGCTTGCATTCATATTAGTTGCATTCTTAAGACTAGACTGAAAAAAAA

AAAAAAAAAATTTTGTGGGTACATTCACTACAGGCCCTCACGAGACCTTACTCGTCCACTAGGGCTGCCT

AGAGGTTTAAAGGGCTTGTTGCATATGCCAAATGCAACCGTGATCACCTACGGAAGTGGTATATATCTAT

TAGGTGTAAGTTTTATTTATTTTATTTGCCCGAGGACGTGCAAAACACTAAGTGTGGGGATATTTGATAG

AGCATATATTTATATATGTTTTTATATGATTTTATTCCCATAATTTTAGTATTTTGTAAATAATCGGGAT

GTTACTAATTAATTTTAAGTGTTTAATGGCAGGAAAATAAATATTCCTAAGGTTGGATTAAATCAAGCTT

ATTTGTGCTTAATCATATAGAAATTCAGATTTTATGGATAGACTGCAAAGCAAATCTTGTTTGGGCCATA

ACTTGAGATCTGGACATCAGCATTGGGCGATTCAACAGCCCATACGAAGATATCGGAATTTTCTACAAGT

TTAAAGTGGTCCAGATGTCAATATCCAACTGGAGCAGCCCAAAATCAGGCCTGAAAGTCAGCCTGCGAAT

TTCTTTTCAGAATCCAATTTATTTTAGGAAACCGGATTTGTTTTCCAAGGAGCCCATTAAAAACACTATT

ATTATTAGGGTTTAGATTATTAGAGATATATCGGAGATTATTTTGTATTGAGAGAGAGAGAGAGAAACAC

TTGTAGAGGGAGATAGAGCAAAGGATTGAAGGGATTCATCCGGTGTTTTAAGGAGCTTTTAAAGTTGTAT

TCTTCTACTCTTGACTCTTGTACTCATGGTTTGGTTTACTCATACA

>MSTRG.1391.1 gene=MSTRG.1391

TAAAAGATGTCCATCCGTGATTATCATTTGGACTCCACCCAACACAAAACTAAAAGTACACACACACAAG

TCACAACACTGTTTGGACTCCCAGGCTTACTAAACAAGACTACCTCTCAGCCATCCTATACATACGATAC

TTACAACAATTTTCAGATAAACCCCATCTCTCTCTCTCTGGTCATCTTGCCTATTGAATATATTATAGAT

TAGGTGAAAAAAGCAGAAAAACCCACTCTTACCATCTTCCTAAGACACCCCATTTTCTAACTCCACCTAA

TTTAACGCCCCTTTTGGAGAGAGAAGCTGGGTTCTGGTTTTTAGAGAGAGAAAGTGCTTTTTTGGGTTTC

TTGTTTGTGAATCTGTTTGGTTTGTTCATACCCCATTGCTCAGATCTTGTTTATTTTGTGTTTTTGATGT

GGGTTTAAATTATGTGACATTCTTGAGCTCAGATTGTTGATTGCTGTCAATGCACATAGTTTGGGTGGCT

TATTGTGTGTAAATGTCTGCTCCTTTGGCTTTGGCAGAGAGGCCTCGAATCGCTGCTTCGAATACGGTTT

TCAAGAGTGGGCCACTATTCATATCCTCAAAAGGCCTCGGTTGGAAATCTTGGAAGAAGCGTTGGTTTAT

CCTTACACGCACCTCTTTAGTCTTTTTCAAAAATGATCCTAGTGCACTTCCCCAGAAAGGTGGTGAAGTA

AATTTGACGTTAGGAGGCATTGACTTGAATAATTCAGGGAGTGTTGTTGTCAGAGAAGATAAAAAGCTTC

TAACTGTTCTGTTTCCTGATGGGCGCGATGGGAGAGCATTTACTCTTAAGGCCGAATCATCAGAAGACTT

GTATGAATGGAAGGCGGCTTTGGAACTTGCTCTTGCACAAGCCCCTAGTGCAGCCCTTGTTATTGGACAT

AATGGAATTTTTCGGAATGATGCTAATGATACAATGGAAGGTTCATTTAATCAATGGAGAGAAAAGCGTC

CTGTGAAGTCTTTGGTGGTTGGAAGGCCCATTTTGCTTGCATTAGAAGATATAGATGGAGGACCATCCTT

CTTAGAGAAAGCTCTGCGATTTCTCGAGATGCATGGAACCAAAGTAGAAGGAATTCTGCGACAATCAGCA

GATGTTGAGGAAGTAGATAGCCGAGTTCAAGAATATGAACAAGGGAAGAGTGAATTTGGTGAAGACGAGG

ACGCTCATGTAATTGGTGACTGTGTTAAGCATGTTCTCCGGGAACTACCTTCATCTCCAGTTCCGGCTTC

TTGCTGCACTGCTTTGTTGGAAGCTTACAAGATTGACAAAAAGGAGGCTCGGCTGAATGCCATGCGTTCA

GCGATACTGGAAACGTTTCCCGAGCCAAACAGGCGGCTATTACAAAGGATTTTAAAAATGATGCACACAG

TTTCTATACATGCACCCGTAAATAGGATGACTCCCTCTGCTGTTGCCGCTTGTATGGCTCCGTTACTTTT

ACGACCTCTTTTGGCCGGTGAATGTGAATTGGAGGATGATTATGATATGAATGGTGATAGTTCTGCTCAG

CTTCTTGCTGCTGCAAATGCTGCGAACAATGCTCAAGCTATCATCACAACACTTTTAGAGGATTATGAAA

GCATATTTGATAATGATGCTATGCATAGATGCTCTATTTCTGCCGATTCTCAAATCGGTAACAGTGTGAG

TGACGATTCAACTGATGATGAAACCCTGGACAATAAGAAAAATGGTTACCATGATGTAGAAAATGAAGTT

CAGCAAGAGACTGATGATGATCAGGATCGTGTGCTTAGTGGAAAGTTGAGTGAAACAAGTGGTTATGCTG

GTAGCGATCTTTATGATGATAAGGCATACAGAGGTGATGATTCAGAGGTTGGATCCCCTCGAGTTGATTA

CAACTTAGGGGCAAAGTCCAGTACCCCTGTAGGCTCTCATCCACATGTGGATTCTCGAGCCAGGGAGCCA

CAGGTTCTACAACATAAGGCAGGGGAGAATGGATTGGAAACCTCACCGGTATTACGGGGTGGTGACTCTT

ATCGTTCTATGGGGCAAATTCTATCTTCAATGGATCAAGGGCTACCTTTGTCAGTCCCTGGACTGGAATC

AAACATTGAAAAGACAATCACCAAACCTACCAGCTCCAATCTTAATGCCAAACGGTCAACATTTTGGGGA

AGGAACAGTACCAGGAAGACACCCTCAATGGATTCAATTGATTCTTCTGGTGAAGAAGAGCTTGCTATCC

AAAGGCTTGAAAGTGCAAAGAATGACCTGCGACACCGAATTGCAAAGGAGGCTAGGGGGAATGCTATATT

ACAAGCTAGCTTGGAGAGAAGGAAGCAAGCTTTGCATGAGCGTCGCATGGCACTTGAGCAAGATGTTTCG

AGACTTCAGGAGCAGTTGCAAGCTGAGAGAGATCTAAGGGCTGCACTGGAAGTTGGATTAAGCATGTCCA

CCGGCCAATTTTCAGGCTCTCGTAGTATGGATACAAAGACAAGGGCGGAACTTGAGGAGATTGCTCTTGC

TGAAGCTGATGTTGCTAGATTGAAGCAAAAAGTTGCCGAACTCCATCATCAACTTAATCAGCAACGTCAG

AGTCATTATGGCTCCATCTCTGATTCTTGTGATCGCCACCAACATGTCCATAATAATCATAATACCCAAC

AGAAATATTATCAGCAAGATTTTGATACATCTCTTGCTTTTGTTAATCATGAGAGGAAACAAAGAACTGA

GGAGAATTTGTTGGGGACCAGAAATCTTAAAGGACAAGGAATAACACCTAGTAACGGTAGTAGACAACCT

GTTCGGAAGCATCTCATGGATTCAGCAAGTTTTAGTGATTCTAAAAGTACCGAGGCCTCGACAAGTATTT

CTATGGATGAATTTGGTGCTGTTGATTCTGCGTCTGTCCCTTCCTCTTCGAGAGCAGCAGAGGTGATGGA

TTATCCTAGGCATCCATCTGTAGCCTCTTCAACCCTCATCGAGTTGACATCACGTCTTGACTTCTTCAAA

GAACGCCGCTCTCAACTTATGGAACAACTCCACAACCTAGATTTGAACTATGAAACAGGACCGGCACAAG

ATTTAATGTATAAGCCATCGACACCAAACTGGAACTGATAGCAACCAGATGCTCAGGTTTATAGTTGACA

CCTCCACATCTCAGCATTTGTACATCCTTCTTCCCCGGGCACATAATCTTTTTTCCCTTGTGGTATCCCT

TCCTGAATATTTTTTTTGTGTGAAATTCCTTTTAATGTAATTTGTACCAAGAGTCTGCTTAGTCTTAGAT

GAGATTTTGTTATAAATATTAACTCGGTTCACACTAATGTATCTGTAATTTGCTAAGTGTGTTAATACTT

CCTGTATATATGTTGGTGATTTGTTTATCAGAACAAAGCTGGAGAAAACAGATGCTATTGCCTTGTGAGA

TTATTTCAGCAGCTCTGTTTCGCTTCTAATTGTTTCTGTCAATTGTCATTATTTTTCTTTTTTTTCTTTT

GCTAAATAATTGTCATTATTTTTCTTCCCTTCAAAGGTAATAATAAGC

>MSTRG.1391.2 gene=MSTRG.1391

CACAAAACTAAAAGTACACACACACAAGTCACAACACTGTTTGGACTCCCAGGCTTACTAAACAAGACTA

CCTCTCAGCCATCCTATACATACGATACTTACAACAATTTTCAGATAAACCCCATCTCTCTCTCTCTGGT

CATCTTGCCTATTGAATATATTATAGATTAGGTGAAAAAAGCAGAAAAACCCACTCTTACCATCTTCCTA

AGACACCCCATTTTCTAACTCCACCTAATTTAACGCCCCTTTTGGAGAGAGAAGCTGGGTTCTGGTTTTT

AGAGAGAGAAAGTGCTTTTTTGGGTTTCTTGTTTGTGAATCTGTTTGGTTTGTTCATACCCCATTGCTCA

GATCTTGTTTATTTTGTGTTTTTGATGTGGGTTTAAATTATGTGACATTCTTGAGCTCAGATTGTTGATT

GCTGTCAATGCACATAGTTTGGGTGGCTTATTGTGTGTAAATGTCTGCTCCTTTGGCTTTGGCAGAGAGG

CCTCGAATCGCTGCTTCGAATACGGTTTTCAAGAGTGGGCCACTATTCATATCCTCAAAAGGCCTCGGTT

GGAAATCTTGGAAGAAGCGTTGGTTTATCCTTACACGCACCTCTTTAGTCTTTTTCAAAAATGATCCTTG

CACTTCCCCAGAAAGGTGGTGAAGTAAATTTGACGTTAGGAGGCATTGACTTGAATAATTCAGGGAGTGT

TGTTGTCAGAGAAGATAAAAAGCTTCTAACTGTTCTGTTTCCTGATGGGCGCGATGGGAGAGCATTTACT

CTTAAGGCCGAATCATCAGAAGACTTGTATGAATGGAAGGCGGCTTTGGAACTTGCTCTTGCACAAGCCC

CTAGTGCAGCCCTTGTTATTGGACATAATGGAATTTTTCGGAATGATGCTAATGATACAATGGAAGGTTC

ATTTAATCAATGGAGAGAAAAGCGTCCTGTGAAGTCTTTGGTGGTTGGAAGGCCCATTTTGCTTGCATTA

GAAGATATAGATGGAGGACCATCCTTCTTAGAGAAAGCTCTGCGATTTCTCGAGATGCATGGAACCAAAG

TAGAAGGAATTCTGCGACAATCAGCAGATGTTGAGGAAGTAGATAGCCGAGTTCAAGAATATGAACAAGG

GAAGAGTGAATTTGGTGAAGACGAGGACGCTCATGTAATTGGTGACTGTGTTAAGCATGTTCTCCGGGAA

CTACCTTCATCTCCAGTTCCGGCTTCTTGCTGCACTGCTTTGTTGGAAGCTTACAAGATTGACAAAAAGG

AGGCTCGGCTGAATGCCATGCGTTCAGCGATACTGGAAACGTTTCCCGAGCCAAACAGGCGGCTATTACA

AAGGATTTTAAAAATGATGCACACAGTTTCTATACATGCACCCGTAAATAGGATGACTCCCTCTGCTGTT

GCCGCTTGTATGGCTCCGTTACTTTTACGACCTCTTTTGGCCGGTGAATGTGAATTGGAGGATGATTATG

ATATGAATGGTGATAGTTCTGCTCAGCTTCTTGCTGCTGCAAATGCTGCGAACAATGCTCAAGCTATCAT

CACAACACTTTTAGAGGATTATGAAAGCATATTTGATAATGATGCTATGCATAGATGCTCTATTTCTGCC

GATTCTCAAATCGGTAACAGTGTGAGTGACGATTCAACTGATGATGAAACCCTGGACAATAAGAAAAATG

GTTACCATGATGTAGAAAATGAAGTTCAGCAAGAGACTGATGATGATCAGGATCGTGTGCTTAGTGGAAA

GTTGAGTGAAACAAGTGGTTATGCTGGTAGCGATCTTTATGATGATAAGGCATACAGAGGTGATGATTCA

GAGGTTGGATCCCCTCGAGTTGATTACAACTTAGGGGCAAAGTCCAGTACCCCTGTAGGCTCTCATCCAC

ATGTGGATTCTCGAGCCAGGGAGCCACAGGTTCTACAACATAAGGCAGGGGAGAATGGATTGGAAACCTC

ACCGGTATTACGGGGTGGTGACTCTTATCGTTCTATGGGGCAAATTCTATCTTCAATGGATCAAGGGCTA

CCTTTGTCAGTCCCTGGACTGGAATCAAACATTGAAAAGACAATCACCAAACCTACCAGCTCCAATCTTA

ATGCCAAACGGTCAACATTTTGGGGAAGGAACAGTACCAGGAAGACACCCTCAATGGATTCAATTGATTC

TTCTGGTGAAGAAGAGCTTGCTATCCAAAGGCTTGAAAGTGCAAAGAATGACCTGCGACACCGAATTGCA

AAGGAGGCTAGGGGGAATGCTATATTACAAGCTAGCTTGGAGAGAAGGAAGCAAGCTTTGCATGAGCGTC

GCATGGCACTTGAGCAAGATGTTTCGAGACTTCAGGAGCAGTTGCAAGCTGAGAGAGATCTAAGGGCTGC

ACTGGAAGTTGGATTAAGCATGTCCACCGGCCAATTTTCAGGCTCTCGTAGTATGGATACAAAGACAAGG

GCGGAACTTGAGGAGATTGCTCTTGCTGAAGCTGATGTTGCTAGATTGAAGCAAAAAGTTGCCGAACTCC

ATCATCAACTTAATCAGCAACGTCAGAGTCATTATGGCTCCATCTCTGATTCTTGTGATCGCCACCAACA

TGTCCATAATAATCATAATACCCAACAGAAATATTATCAGCAAGATTTTGATACATCTCTTGCTTTTGTT

AATCATGAGAGGAAACAAAGAACTGAGGAGAATTTGTTGGGGACCAGAAATCTTAAAGGACAAGGAATAA

CACCTAGTAACGGTAGTAGACAACCTGTTCGGAAGCATCTCATGGATTCAGCAAGTTTTAGTGATTCTAA

AAGTACCGAGGCCTCGACAAGTATTTCTATGGATGAATTTGGTGCTGTTGATTCTGCGTCTGTCCCTTCC

TCTTCGAGAGCAGCAGAGGTGATGGATTATCCTAGGCATCCATCTGTAGCCTCTTCAACCCTCATCGAGT

TGACATCACGTCTTGACTTCTTCAAAGAACGCCGCTCTCAACTTATGGAACAACTCCACAACCTAGATTT

GAACTATGAAACAGGACCGGCACAAGATTTAATGTATAAGCCATCGACACCAAACTGGAACTGATAGCAA

CCAGATGCTCAGGTTTATAGTTGACACCTCCACATCTCAGCATTTGTACATCCTTCTTCCCCGGGCACAT

AATCTTTTTTCCCTTGTGGTATCCCTTCCTGAATATTTTTTTTGTGTGAAATTCCTTTTAATGTAATTTG

TACCAAGAGTCTGCTTAGTCTTAGATGAGATTTTGTTATAAATATTAACTCGGTTCACACTAATGTATCT

GTAATTTGCTAAGTGTGTTAATACTTCCTGTATATATGTTGGTGATTTGTTTATCAGAACAAAGCTGGAG

>MSTRG.1391.3 gene=MSTRG.1391

TAAAAGTACACACACACAAGTCACAACACTGTTTGGACTCCCAGGCTTACTAAACAAGACTACCTCTCAG

CCATCCTATACATACGATACTTACAACAATTTTCAGATAAACCCCATCTCTCTCTCTCTGGTCATCTTGC

CTATTGAATATATTATAGATTAGGTGAAAAAAGCAGAAAAACCCACTCTTACCATCTTCCTAAGACACCC

CATTTTCTAACTCCACCTAATTTAACGCCCCTTTTGGAGAGAGAAGCTGGGTTCTGGTTTTTAGAGAGAG

AAAGTGCTTTTTTGGGTTTCTTGTTTGTGAATCTGTTTGGTTTGTTCATACCCCATTGCTCAGATCTTGT

TTATTTTGTGTTTTTGATGTGGGTTTAAATTATGTGACATTCTTGAGCTCAGATTGTTGATTGCTGTCAA

TGCACATAGTTTGGGTGGCTTATTGTGTGTAAATGTCTGCTCCTTTGGCTTTGGCAGAGAGGCCTCGAAT

CGCTGCTTCGAATACGGTTTTCAAGAGTGGGCCACTATTCATATCCTCAAAAGGCCTCGGTTGGAAATCT

TGGAAGAAGCGTTGGTTTATCCTTACACGCACCTCTTTAGTCTTTTTCAAAAATGATCCTAGTGCACTTC

CCCAGAAAGGTGGTGAAGTAAATTTGACGTTAGGAGGCATTGACTTGAATAATTCAGGGAGTGTTGTTGT

CAGAGAAGATAAAAAGCTTCTAACTGTTCTGTTTCCTGATGGGCGCGATGGGAGAGCATTTACTCTTAAG

GCCGAATCATCAGAAGACTTGTATGAATGGAAGGCGGCTTTGGAACTTGCTCTTGCACAAGCCCCTAGTG

CAGCCCTTGTTATTGGACATAATGGAATTTTTCGGAATGATGCTAATGATACAATGGAAGGTTCATTTAA

TCAATGGAGAGAAAAGCGTCCTGTGAAGTCTTTGGTGGTTGGAAGGCCCATTTTGCTTGCATTAGAAGAT

ATAGATGGAGGACCATCCTTCTTAGAGAAAGCTCTGCGATTTCTCGAGATGCATGGAACCAAAGTAGAAG

GAATTCTGCGACAATCAGCAGATGTTGAGGAAGTAGATAGCCGAGTTCAAGAATATGAACAAGGGAAGAG

TGAATTTGGTGAAGACGAGGACGCTCATGTAATTGGTGACTGTGTTAAGCATGTTCTCCGGGAACTACCT

TCATCTCCAGTTCCGGCTTCTTGCTGCACTGCTTTGTTGGAAGCTTACAAGATTGACAAAAAGGAGGCTC

GGCTGAATGCCATGCGTTCAGCGATACTGGAAACGTTTCCCGAGCCAAACAGGCGGCTATTACAAAGGAT

TTTAAAAATGATGCACACAGTTTCTATACATGCACCCGTAAATAGGATGACTCCCTCTGCTGTTGCCGCT

TGTATGGCTCCGTTACTTTTACGACCTCTTTTGGCCGGTGAATGTGAATTGGAGGATGATTATGATATGA

ATGGTGATAGTTCTGCTCAGCTTCTTGCTGCTGCAAATGCTGCGAACAATGCTCAAGCTATCATCACAAC

ACTTTTAGAGGATTATGAAAGCATATTTGATAATGATGCTATGCATAGATGCTCTATTTCTGCCGATTCT

CAAATCGGTAACAGTGTGAGTGACGATTCAACTGATGATGAAACCCTGGACAATAAGAAAAATGGTTACC

ATGATGTAGAAAATGAAGTTCAGCAAGAGACTGATGATGATCAGGATCGTGTGCTTAGTGGAAAGTTGAG

TGAAACAAGTGGTTATGCTGGTAGCGATCTTTATGATGATAAGGCATACAGAGGTGATGATTCAGAGGTT

GGATCCCCTCGAGTTGATTACAACTTAGGGGCAAAGTCCAGTACCCCTGTAGGCTCTCATCCACATGTGG

ATTCTCGAGCCAGGGAGCCACAGGTTCTACAACATAAGGCAGGGGAGAATGGATTGGAAACCTCACCGGT

ATTACGGGGTGGTGACTCTTATCGTTCTATGGGGCAAATTCTATCTTCAATGGATCAAGGGCTACCTTTG

TCAGTCCCTGGACTGGAATCAAACATTGAAAAGACAATCACCAAACCTACCAGCTCCAATCTTAATGCCA

AACGGTCAACATTTTGGGGAAGGAACAGTACCAGGAAGACACCCTCAATGGATTCAATTGATTCTTCTGG

TGAAGAAGAGCTTGCTATCCAAAGGCTTGAAAGTGCAAAGAATGACCTGCGACACCGAATTGCAAAGGAG

GCTAGGGGGAATGCTATATTACAAGCTAGCTTGGAGAGAAGGAAGCAAGCTTTGCATGAGCGTCGCATGG

CACTTGAGCAAGATGTTTCGAGACTTCAGGAGCAGTTGCAAGCTGAGAGAGATCTAAGGGCTGCACTGGA

AGTTGGATTAAGCATGTCCACCGGCCAATTTTCAGGCTCTCGTAGTATGGATACAAAGACAAGGGCGGAA

CTTGAGGAGATTGCTCTTGCTGAAGCTGATGTTGCTAGATTGAAGCAAAAAGTTGCCGAACTCCATCATC

AACTTAATCAGCAACGTCAGAGTCATTATGGCTCCATCTCTGATTCTTGTGATCGCCACCAACATGTCCA

TAATAATCATAATACCCAACAGAAATATTATCAGCAAGATTTTGATACATCTCTTGCTTTTGTTAATCAT

GAGAGGAAACAAAGAACTGAGGAGAATTTGTTGGGGACCAGAAATCTTAAAGGACAAGGAATAACACCTA

GTAACGGTAGTAGACAACCTGTTCGGAAGCATCTCATGGATTCAGCAAGTTTTAGTGATTCTAAAAGTAC

CGAGGCCTCGACAAGTATTTCTATGGATGAATTTGGTGCTGTTGATTCTGCGTCTGTCCCTTCCTCTTCG

AGAGCAGCAGAGGTGATGGATTATCCTAGGCATCCATCTGTAGCCTCTTCAACCCTCATCGAGTTGACAT

CACGTCTTGACTTCTTCAAAGAACGCCGCTCTCAACTTATGGAACAACTCCACAACCTAGATTTGAACTA

TGAAACAGGACCGGCACAAGATTTAATGTATAAGCCATCGACACCAAACTGGAACTGATAGCAACCAGAT

GCTCAGGTTTATAGTTGACACCTCCACATCTCAGCATTTGTACATCCTTCTTCCCCGGGCACATAATCTT

TTTTCCCTTGTGGTATCCCTTCCTGAATATTTTTTTTGTGTGAAATTCCTTTTAATGTAATTTGTACCAA

GAGTCTGCTTAGTCTTAGATGAGATTTTGTTATAAATATTAACTCGGTTCACACTAATGTATCTGTAATT

TGCTAAGTGTGTTAATACTTCCTGTATATATGTTGGTGATTTGTTTATCAGAACAAAGCTGGAGAAAACA

GATGCTATTGCCTTGTGAGATTATTTCAGCAGCTCTGTTTCGCTTCTAATTGTTTCTGTCAATTGTCATT

ATTTTTCTTCCCTTCAAAGGTAATAATAAGC

>MSTRG.1396.1 gene=MSTRG.1396

CCAGACACACTCGTAATCCACTTATTCCGACCCTTAATTTTGCGGTAAAGGCTGAGCCAGACATGCTCCG

GACATTTTGTTCCCGCGTAATCACAGTCCGTGGACGGATAGAATTTCAAACCGCACCCACCGATCACAAC

TGCCGGCACAACTCTCTCCGAATAACACAAACTTATGTTAAGCTCCACTTTGTCCTTTGTTCTTTGTTAG

CTCAACCGCCAAATCAATATCCTTCTCTGTCAAATTATAATTTAGTTCATTCAACATCTTCCTCGCAACT

GACAAAGGGACATTCAAACCTCCTAATCTAAAGCCAACAATGTGTCATGCTACATATAATGGCTTGAATG

GTGCCCATCACTTGGTCTTGCGTGCAGCGACCACCAAGGATCCTAACTGGTTGTCGGATCAAGGGGCAAA

GGAATATTTGCTGATTAAGCTGGGGCGCCACTATTCGTGTCCTCTATGACGCCAAAATTGAAGAGTTTGA

GAAAGATTTGGCATTTGGGAGTAAGTTAGTAACAAAGCTTATGTACCAATATGAACGGTTCTCTTACTTC

TATGTGCAGCTGTAATGCTCTGATTCCCTTGCAAACTGCGTCGTTCATGTGAGTAAAGATCACATCCTAC

TTCCTCCAATCATATCATTGTTTTGGCCCTAGAACCTCTGAAAGGAACCAAGGATGAGGACAATACAATA

TACTTGGTGACAGTAAAGAAGCAGACCAACCAAATATCCGGCATGATTCAGATGGTTCACGGGAGGGTAG

GTCTTCTGACCAAGTTGACGTATTAACAGCTGTCCGGAGAGCCATCTGCCAAGTCTTAGCGACGCCAGCA

AGATATAAATGTTTCCATAAGCATACAGTTTAGTGTCCTTTTGTGATCAACTTACTATATTTTATGATCT

CTACATCAGACGATCATTATGTAAATTACATTAGTGTTGTTTTCGTGGTAACTTGTAACTAGGTATGATC

AGCCCATGAACAGTTTAGTTCTGGATTTGATTGAAGACTTCTTCAGCCTGCAGACCTTTGTAGTGGACCT

AAAAGCTTTGCAATTGATCTCCTGTCAATTCTTGCTCTAACTAGACTTGTTTCTCTAAATGAGAGATGTG

G

>MSTRG.1396.2 gene=MSTRG.1396

GGACATTCAAACCTCCTAATCTAAAGCCAACAATGTGTCATGCTACATATAATGGCTTGAATGGTGCCCA

TCACTTGGTCTTGCGTGCAGCGACCACCAAGGATCCTAACTGGTTGTCGGATCAAGGGGCAAAGGAATAT

TTGCTGATTAAGCTGGGGCGCCACTATTCGTGTCCTCTATGACGCCAAAATTGAAGAGTTTGAGAAAGAT

TTGGCATTTGGGAGTAAGTTAGTAACAAAGCTTATGTACCAATATGAACGGTTCTCTTACTTCTATGTGC

AGCTGTAATGCTCTGATTCCCTTGCAAACTGCGTCGTTCATGGTGACAGTAAAGAAGCAGACCAACCAAA

TATCCGGCATGATTCAGATGGTTCACGGGAGGGTAGGTCTTCTGACCAAGTTGACGTATTAACAGCTGTC

CGGAGAGCCATCTGCCAAGTCTTAGCGACGCCAGCAAGATATAAATGTTTCCATAAGCATACAGTTTAGT

GTCCTTTTGTGATCAACTTACTATATTTTATGATCTCTACATCAGACGATCATTATGTAAATTACATTAG

TGTTGTTTTCGTGGTAACTTGTAACTAGGTATGATCAGCCCATGAACAGTTTAGTTCTGGATTTGATTGA

AGACTTCTTCAGCCTGCAGACCTTTGTAGTG

>MSTRG.1402.1 gene=MSTRG.1402

GTTTTCCAACATATCTCAGTTATTCTTCTGCCTCTTTTCTTCTTATTTAATTTTTATATATCTATATAAA

GCAAAATTCCGGGCAGCTGACGTGGCTATTCTCTCCTTGCTCTTAAACTGAATTCTCTCTTCTCAGGTTT

TCCAACATATCTCAGTTATTCTTCTGCCTCTTTTCTTCTTATTTAATTTTTATATTTATTTTTTTATTCC

TGACTGATCGGTTTCGTAACTGAAGCCTCGTCATCTCCTCTGACAGTTTGTTATAAATGATCCTCCGGCA

ATCGTTTCTTTTGGCTTGGTTATTTCTTACTCGAAAATTAGCTTCACAGACACTATTGGACCTCTTTGTT

GGGCTGGCTTGGCTATAGCAATTGCTGCAACAGCTGTTATTCTTTTGGTGGGCGTCCTCTTCTTCATCTC

TAGCTTAGTCGTGAATCTTCTTCAGGTATTATTTTCCCAGGTTTCCACTCTTTCTTTTGTGATTTACCAA

ATGTTCCTGGGATTTTACATATTTGCCGAAAGATTAAAAGGGTGGTTGCGAAGTTGTTGTGGCTGGAACT

TTGGTTGATTGGTGGGCTGCTGTTAAGGAGTTAAAGGAAAGCGGTTGCAGATACATTTTATGGAGATGTA

ACAGTGAAAGATACCGAAGTTTTCATATCCAATATCTAGACTTTCCGGTATTTAAATATTTCTTTATTAC

TTTAGAAACAATAGCTATAAGTTGCTGGGACACGAGGGAGTATTAGTCAGGAAAGATATATTTGTTTGCA

ACTACTAAAATCAATAAATTATCAAGAATTTGCTTTATCATACTTGATTCGGACTATAAACCACTGGTGA

CAAGGCGCAGGTGCTTGATGCCACACAGTTTGGCTACTTTAAGGTGCTGGGGAAGGGCTCAGGGACGAAG

CCGATTGTAGTAAAGGCCAAGCTGGTGTCGAAGATAGCGGAGAAGAAGATTGAGATGGCCCGCGCTGTTG

TGCTTACTGCTTAGGATTGGTTGTTTCTGTTTTAATTGAAATGGACTATACTGATTCTTATGGTTATTAT

GGATTCTCACGTTGTGGAACACTGTTTTTGGATAAATTTATGGTTAATGGTACTTTTAATTTTGGATTTG

GT

>MSTRG.1402.2 gene=MSTRG.1402

TTCTTCTGCCTCTTTTCTTCTTATTTAATTTTTATATATCTATATAAAGCAAAATTCCGGGCAGCTGACG

TGGCTATTCTCTCCTTGCTCTTAAACTGAATTCTCTCTTCTCAGGTTTTCCAACATATCTCATTTGTTAT

AAATGATCCTCCGGCAATCGTTTCTTTTGGCTTGGTTATTTCTTACTCGAAAATTAGCTTCACAGACACT

ATTGGACCTCTTTGTTGGGCTGGCTTGGCTATAGCAATTGCTGCAACAGCTGTTATTCTTTTGGTGGGCG

TCCTCTTCTTCATCTCTAGCTTAGTCGTGAATCTTCTTCAGGTATTATTTTCCCAGGTTTCCACTCTTTC

TTTTGTGATTTACCAAATGTTCCTGGGATTTTACATATTTGCCGAAAGATTAAAAGGGTGGTTGCGAAGT

TGTTGTGGCTGGAACTTTGGTTGATTGGTGGGCTGCTGTTAAGGTCAGTTTCGGGTCGGAGTTAAAGGAA

AGCGGTTGCAGATACATTTTATGGAGATGTAACAGTGAAAGATACCGAAGTTTTCATATCCAATATCTAG

ACTTTCCGGTATTTAAATATTTCTTTATTACTTTAGAAACAATAGCTATAAGTTGCTGGGACACGAGGGA

GTATTAGTCAGGAAAGATATATTTGTTTGCAACTACTAAAATCAATAAATTATCAAGAATTTGCTTTATC

ATACTTGATTCGGACTATAAACCACTGGTGACAAGGCGCAGGTGCTTGATGCCACACAGTTTGGCTACTT

TAAGGTGCTGGGGAAGGGCTCAGGGACGAAGCCGATTGTAGTAAAGGCCAAGCTGGTGTCGAAGATAGCG

GAGAAGAAGATTGAGATGGCCCGCGCTGTTGTGCTTACTGCTTAGGATTGGTTGTTTCTGTTTTAATTGA

AATGGACTATACTGATTCTTATGGTTATTATGGATTCTCACGTTGTGGAACACTGTTTTTGGATAAATTT

ATGGTTAATGGTAC

>MSTRG.1402.3 gene=MSTRG.1402

ACTGAATTCTCTCTTCTCAGGTTTTCCAACATATCTCAGTTATTCTTCTGCCTCTTTTCTTCTTATTTAA

TTTTTATATTTATTTTTTTATTCCTGACTGATCGGTTTCGTAACTGAAGCCTCGTCATCTCCTCTGACAG

TTTGTTATAAATGATCCTCCGGCAATCGTTTCTTTTGGCTTGGTTATTTCTTACTCGAAAATTAGCTTCA

CAGACACTATTGGACCTCTTTGTTGGGCTGGCTTGGCTATAGCAATTGCTGCAACAGCTGTTATTCTTTT

GGTGGGCGTCCTCTTCTTCATCTCTAGCTTAGTCGTGAATCTTCTTCAGGTATTATTTTCCCAGGTTTCC

ACTCTTTCTTTTGTGATTTACCAAATGTTCCTGGGATTTTACATATTTGCCGAAAGATTAAAAGGGTGGT

TGCGAAGTTGTTGTGGCTGGAACTTTGGTTGATTGGTGGGCTGCTGTTAAGGTGCTTGATGCCACACAGT

TTGGCTACTTTAAGGTGCTGGGGAAGGGCTCAGGGACGAAGCCGATTGTAGTAAAGGCCAAGCTGGTGTC

GAAGATAGCGGAGAAGAAGATTGAGATGGCCCGCGCTGTTGTGCTTACTGCTTAGGATTGGTTGTTTCTG

TTTTAATTGAAATGGACTATACTGATTCTTATGGTTATTATGGATTCTCACGTTGTGGAACACTGTTTTT

GGATAAATTTATGGTTAATGGTACTTTTAATTTTGGATTTGGTGGTGTTTGTTTGGTTTTTATTGTGTTG

TGATTGATAATGTTTGATTGTGTATGTGGTCGGATTGGTTCTTGTAGTGATGGTTGTAGTCGAGGCTCTG

ATTGTGGTTTGAGA

>MSTRG.1405.1 gene=MSTRG.1405

ATGCTTTCCGGTAGAACTAAACCTTGCCATTGTTTTTCAACAGGGGTAGAGGATATGCAGAGGACGATGC

TAAAACCATTATTGTGCAGATACTAGGAGTGGTTGCCTTCTGTCATCTACAAGGTGTTGTGCACCGTGAT

CTGAAATCGGAGAACTTTCTTTTCGCAGCAAGAGACGGGGATGCCCCATTGAAGATCATAGATTTTGGCC

TATCTCATTTCAGCAGGCCAGATGAACACCTCAATGATATTGTTGGCAGTGCATACTATGTCGCGCCTGA

AGTGCTCCACAGATCATATAGCATTGAAGCTGATATTTGGAGTATAGGTGTCATAGCATATATGCTCTTA

TGTACCATCTGCCCAGGCAACGAGGCAGGTACTGAGTTGCTGCTTTTTAACTACAGCTGAAATGATGTGG

CTTCTCAATCTTCTTCGTGAATATGATGTTTCTTTATTACTGTATGTCATACACCTATTATATACCGTGA

TAACTTAAATGTCACCCATCTTAGTGTAAATCCTATTTTTCATTCTAGAATGAAACATTTAACACTTTCA

TCTTATTTTACCAGAGAAATTATTCGTACAAAAACTTTATCTCATCCAGGAGTGGCCCTGATTATAGACG

GATAAGACTTGAGCCTAGTGCCCTAATGTTTAAGGCCCTCAAAATTTTGAACCGTTAGTATGTAAATGCA

TGAATTATACAGAATATATGCAGACCAGGCCCC

>MSTRG.1409.1 gene=MSTRG.1409

CATGTTTTCTTGTAGAAAATAAAGAATTCTCATAATAGAAGGAATGGAGCAAAAAGTGCATTATTTGAAG

TGATTTCTAGCAAAATTCGGATTTTGGGATTAATTGTGTTGTGTTCATTGTTTTGATCATAGCCGGAGTT

CTGTAAGTCCGTTTTGGACCCGTAAACAGCCGTTGGAAAGCTTGCGAGAAGACCTTTAATTCAAGTAAGG

TTTCAGAATTTATTGACATTCCAATCAGCCCAGAATATGGCCAGAAGCAGAAAGTCCGAATTTTCTACTT

TGAACTCCTATTCTATTTGATAAACTTATTTGTAGTATTTATTTTATTCAAAATATAGTATAGGTAAAAT

AGATCAGTTATTATTTTGTTTTCTAATGGGCTTGGGAGTTACATGGGCCAAGAGACACAAAAACATCACA

GGAGAGAAGTTACAAAAATTAGACAATAGACACAGGGAGAGAGGTACAGGAAGAAGTTGCACGCAAGAGA

GCAAGGAGGAAGGAAGCAATTGGAAGAGACAAATATAGAGTTTTTCTTCTCTTCTTTACCTTTCTTAATT

GTAACATGGATTTGATGTCTTTATACTTGTGTTTAGTTCTTAGTAGTGTGATGTTGAACTAATTTCTCTT

ATAGGCCTTTAATGGATTCTTTTGATTATATATAA

>MSTRG.1410.1 gene=MSTRG.1410

TAGAAAATAAAGAATTCTCATAATAGAAGGAATGGAGCAAAAAGTGCATTATTTGAAGTGATTTCTAGCA

AAATTCGGATTTTGGGATTAATTGTGTTGTGTTCATTGTTTTGATCATAGCCGGAGTTCTGTAAGTCCGT

TTTGGACCCGTAAACAGCCGTTGGAAAGCTTGCGAGAAGACCTTTAATTCAAGTAAGGTTTCAGAATTTA

TTGACATTCCAATCAGCCCAGAATATGGCCAGAAGCAGAAAGTCCGAATTTTCTACTTTGAACTCCTATT

CTATTTGAGAAACTTATTTGTAGTATTTATTTTATTCAAAATATAGTATAGGTAAAATAGATCAGTTATT

GTTTTGTTTTCTAATGGGC

>MSTRG.1412.1 gene=MSTRG.1412

AAATCGTAAACCCGAGGCTATTTACAAAGTTTCCTGCCCAACAAAGCTTAAGCTCACAAATACATAGATT

GGTTTGTTCAGTCCAGTTCTTGAATTCTCATCAGAGCTATATACATTGTTCTGTGGCTTGCTTAGTGAGA

TGAGGCCAGAAAGAGTCGCCTCAAACATTGTTTTATTGGATCGAGTTCCATGGCGTGGCATTGTTCCTGT

CCTGTCTTGGAATTATCATCCATCAGTGGAACTGTTAGCCTGTTTGGCTCTCAGTGGAACTGTTAGCCTG

TTTGGCTCTCCCTGGTATCTAATTGTTTCAGTCAAACGAATTTATCTTGCATGTACTCGCATTAAGTTCT

GTA

>MSTRG.1423.1 gene=MSTRG.1423

GCGGAGGCAACTAAAGATGTCACTGATGGTGAGAGTTCGGAGAATGACATGATGGGGATGATGATGGTGA

TGGCGATGAGGATGATTGAAATGAATCTAACTAGCTATATGCATTATTGGTAGGGATGTTGATTTTGTTA

AAACATTGGTTTTTTAAACCGAGACCGAACTTATAACTTTTTTTATTCAGCTGTAACGTATATTTGCATG

GTTTCGGTTATTTTGTTGAAAG

>MSTRG.1424.1 gene=MSTRG.1424

TGGGATTTTGTGGGTATATCTTCTACAAACCCTCACGAGAGCACACTCGTCCACTAGAGCTATCTAGGGG

TTTAAAGGGCTTGTTGCATATGCTAAATGCAACCGTGATCACCTACGGAAGTGGTACTAGAAGCGAGGAA

GGACATGCACGCGAGAACGAGCGAGAAACGAAGAAACGGGGCTGCCATCACAGACAGTAGCGCGCCCGCG

CTATCCACTGGGACAACTAGCGCGCCTGCGCTAGTTTAGCGCGGCCGCGCCCAGCAGAAGTTCCCCGGGC

CGATTTTTACAGCTTTTTAGCGGATTTTCGCAGCGGTCGAGGCCCGGTTGACTTGGTCAATGTATTTTTA

TTTCTCGTGTGTAAAACCCTAGCCTCGGAGAAGTAGTTTTAAAAAAAAAAAAAGAAGAGAGAACAATATC

GTAGTTAAGTATTATTCATTCAAGCACATTATCAGTTTGTATTAGATCGTTCTTCGCGAGGAAGTGACAG

TTTCGAGCGAGATCGTGAACATCAAATCTGTAACGTGTGATCTTTATTATTAATTCAAGCATTTTTATTC

C

>MSTRG.1425.1 gene=MSTRG.1425

ATATGAACACTGCAGTAGCTGATGTCACTGCTGCCATGATCACCGAGTACAGGGCACTATTTTCCGACTT

GCAGAGATGGGGATTTGACATAAGTTGGCTGGTCAGACGTCTGAACTATGTGGAAGAACTCCGAAATTCA

AAACCCTTAATTAATGAGCTTCGTGCAATTTTTTCTCGTGTTCATGATATGCAGACTCTTTCTCCTGAGA

AGATGACAGAGATTCACAAGGCTTTTGGAACCATGGATGCTGGCCTTTCTCTTCACTTTATTGGCGTCAA

CTTACTTCCTAGTTCCTAGTTTTCTTATGGTCACCAAGGCAAGATGATTTTACAGATATAGGAGTTTAAG

AGTGGTGCATTAGTTTCTCAGAAAATGGATTATCTGAATTTAAATGTTTTAGTTTTCAAAATGTCATAAA

ATACATACAAAGAAACTACATTTTTTGTAGGTTATTA

>MSTRG.1426.1 gene=MSTRG.1426

AGTTGTTGCTGTGGCTAAATGTTCTCCTGCAGTTAGTGATTTTTTTGATCATCTTTGCCGTATTGTGAAT

TTGGTTGGTGCTTCTTGTAAGAGAAAGGATATGCTTCGATAAAAACAACATGAACTTACATTGAAACGTT

TAGAAAGTGGTGAGCTATTTTCTGGAAAAGGGAAGAATCAAGAAACCTCTTTAACTCGGCCAGGAGATAC

ACGGTGGGGTTCACATCATAGAACAATACTTCGTCTTTTTTTGATGTGGCCATCGGTAGTGGAAGTGCTT

GGGGATATACATCAAGATGCTACTAATCAAGAATTAAAAGGTATTGCAGAAGGCTTACTTGAAAAAATGA

AGACATTTAAATTTGTCTTTATGTTGCATTTGATGAAAGTTATATTGGC

>MSTRG.1428.1 gene=MSTRG.1428

CTCCAAGGTTAGGAGGAAGACTTTATCAGCAATACATGGTGGACGCTTTTTCCACCATTGAGCAGACTCG

GTTATGGTGGTTTCGTACTCACCAAACCATACTCCGGAATGAACTATACACGCACATTTCTGATTCTGTT

AGGAAAGGAGATGCGAATGCCACCAACGTCGGTAAAGGCATAATACTCCCAGCTGGATATGTTGGCTCCA

AAAGGTATATGCAACAGAATTTCCAGGATGCATTGGCTGTTTGTCGTCATGTTGGCCATCCTGATGTTTT

TTTGACGATGACATGTAATTCCATGTGGGATGAAATTCAAAGGATGATGAATTTTGTGCCCGGATGCATT

CCTGCTAACTGTCCGGATATAATATCTCGGGTTTTCCGATTGAAATTGGAACAGTTAACTAATGATATTA

AGAAGAAAGGTCACTTTGGGAAATGTATTGGAGTGATGTATGTTGTCGAGTTTCAAAAAAGAGGACTGCC

CCATGTACATATGTTAATCTGGCTTGATGCACCATCAAAAAAGTATCTCAAGGAAAATGTCGACAAGTTC

GTTTCAGCTGAAATTCCAGATCCTTTGATTGATCCTGAAGGTTGTGCAGCAGTCAAGGCGTATATGATGC

ATGGTCCATGTGGATTGCAAAATCCAAAATCTCCTTGCATGAAGAATTTCAAATGTATACGCCACTTTCC

CAAAAAGTACAGCTCACAGACTGTGTTTGATGACAGTGGCTTCCCACTTTATATGCGGCGCAAATCAAAC

ATCAGTGTTAATGTTGGCAAGGCTGATCTTGACAATCGATGGGTTGTCCCATATAACCGCGAGCTTTTGG

TAAAATACCAATGTCATATGAACATTGAGATATGCTGCCATGCACGTAGTATAAAATATCTATTCAAATA

TTGCCTCAAAGGTCACGACACTGCCACAGTCGAAATCACTGGACGTCGCAAGAAAAAGCAAACTGATGGT

GCTGACCAGCCTGTTGATGAAATTCAAGCATATTTTGATGGTAGATATGTATGCGGCGCTGAAGCAGCAT

ATCGAATTTTTGGATTTCCAATACATTATAGGTCTCTGTCTGTTGAGCGCCTCCCATTTCATTTGCCCGG

TCAGAAAAGCTGCACCTTTCGTGCAACTGAAAAGCTTGAGAAAGTGCTAGAGCGTGAAAATGAAAGGCTT

AGCAAGCTGGAAGCTTTTTTCTTGCTGAACAAGTTTGATTTGAAGGCACGTTCTTACACATATGATGAAA

TCCCACAGCATTATGTGTGGAACGATTTGGCGAGAAGGTGGACTATTAGGAAAAGAGGAACTCAGATTGG

GCGTCTGTCTTACACTCATCATAGCAGTGGGGAAGTGTGGTACTTGCGCATGCTATTGACAAAAGTTCGA

GGTCCGACATCATTCGAAGACATTAGAACTGTCAATGGTGTACTCTGTCAGACATTTCGGGATGCATGTA

AAGAATACGGATTGCTCGATGATGACAAAGAATGGCATGATGTTTTAGACCAATGTTCTGTTGGTGGATT

GCCTCCGCAAATACGCCAACTGTTTGTACACATTATTGTTAATTGCAAAGTGACTGATTTGGCGTCTTTG

TGGGATAAACATTGGAAGCAGATGGTAGATGACATTCTTTTGAAGCGAAGATTACTAAGTGAAAATCAGA

ATCTCATTCTTAATGATAAGCAGCTCCAGTTCTACGCCCTTGCAGAAATTGATGACCTGCTGCGTGCAAT

TGGCAAAAGTTTGAAGAATTTTTCCCAACTGCCACAACCACCTCTAAGTTACCTGCATCACGGAACTAAT

AACTTGATAATTGAAGAGACAAGCTACAACCTCTCAGAGATGAAATGTCAACACGATAGTCTAATTGCAC

AGTGCAACGAAGAACAAATTAGTGTTTATAAAGCTGTCTTATCTTCAATTGAGTCCAATCAAGGTGGATT

ATTCTTTGTGTATGGCAGCGGTGGCTGTGGAAAAACCTTTCTATGGAGAACACTAATAGCTAAACTACGA

TCTGCTGGTGACATTGTTCTTCCAGTGGCCTCATCTGGCATTGCTGCTACACTCATGCCTGGTGGTCGTA

CCGCGCACTCGCGCTTCAAAATTCCCATAGTGCTTGATGACTATTCGCTATGCAACATTGGCCATAATTC

GGACATTGCTGAGCTTATTAAGCAGACCAAGCTAATAATATGGGATGAAGCACCCATGCAACATAGGTAC

GCTTTTGAATGTTTGGACAGGTCGTTGAAGGATATAATGAAATCAGTTGATCCTGCGCGTGCTCTTTTGC

CATTTGGTGGTATAACTGTTGTTCTAGGCGGTGATTTTAGGCAGATACTACCTGTCATTTCCCAAGGAGA

TCGAGCTGAGATAGTTTCAGCATGTATCACAAGATCACGTCTGTGGTCAATTTGCAAGATTTTTCTCCTG

ACCAAAAATATGCGTCTTAATGAAGGAAAATCTGAGAAAGAAGTCGAGGAACTCCTGCAGTTTGCCAAGT

GGGTTCTCGATGCAGGTGATGGAAAGTTGGAGGCTCCCGACAAGGAATTATTTCCATCTGCCAAAGAAGA

CGACATATTGATTCCACCTCAATTCTGTGCAAAAGAACAGGAAAACTCTGTTGACAATATGATTGCAAGC

ACTTTTCCTAATTTTGTTGAAAAATGCAAAGATGCCAAATATTTGAGTGAGCGAGCCATTCTAACTCCTA

CAAATCAAACTGTAAGTAATCTTAACTCTGCCATAGTTCAGAATTTGCCAGGTGAGTCATTTTCATATTA

CAGTGTAGATAGTGCTGAGGATTTTGGTGGAACCGAAGATGATTTAAACTCGGCTTTCCCAATTGAATAT

CTCAATTCAATGAATGTTCCTGGAATGCCACCACATGATCTTAAATTGAAAGTGGGAGTCGTTGTCATGT

TGATGAGAAACTTGAACCAAACGTTAGGTCTGTGTAACGGCACACGGATGATCATTACCAAATGTCTGAA

GTTTTGCGTTGAGTGTGAAGTTATATGTGGTTCTTTCATTGGAACACGTCATTTTATCCCGCGGATGGAG

TTGTGTCCAAGCGATACTAGGCTGCCTTACAAGCTGATTCGTAAACAAATGCCACTGCAAATATGCTACG

CCATGACTATTAATAAATCCCAAGGTCAGTCACTTTTACAAGTTGGATTATATCTGCCTAAAGGCGTATT

TACGCATGGTCAGTACTATGTGGCAATTAGTAGAGTAACCTCTCCTCAAGGACTAAGAATTTTCATCGAC

GATGAGAGTGGCCAGCCTACTAATATAACTGCTAATGTTGTGTACAAAGAGGTTTTCTACGCCTTGCCAA

AGACCTAGAGAAAATCTGTTAAGTATGCGTTTGTTGTGTCCAGTGTTTTTGGTATTTGATTAATCTTTGT

TTTTGAGGACCAATATGATTCTGTTATTCTGTTGACAATGTTGTTTTGGTTACTAAGAACAATGTAAAGT

TACGCCTATCTATTATTTTACCTC

>MSTRG.1437.1 gene=MSTRG.1437

CAAAATAATTCGCTTTTCGAGAGAAACACCTCTGTTTACGATTCACACTTCAAATTCAAACTATCCCTTC

TGTTTAAGCCTAGGTTTTCCCAAGAATATGAGCACCCAGATTATTCACCGAAACATGGGCCAAGTCATCC

AAGAAGAAGAAGAAGAAGATAGAATCAGCAGATTGCCTGATGAACTGCTCTCTCACATCCTCTCCTACCT

CGATTCTGATGCCGCTGTACAGACCTCCGTCCTTTCAAAACGATGGGTAAATCTATGGACTACTCTCCCC

TCTCTCTTATTCAAAACCAACACCCAATATTTTAATTCCGATAACTATGATCTCGGTTGCAAATTTATCA

CTAATTTTCTCAATTCTCGAAACCAAGATACTGTTCTTTCCACTATTAAGGTCCATTGCATTAAAAGCAA

TGATGAATGCATTTCAGCATTGTGTTTAGACTATGCACTTGCGAACAATGTTGAGAATTTACGCTTGTGC

GGCGGAAAATTCAGGGATGCCATTTGTCTTGAGTCTGATTCCCTTAAAACATTGCATCTTACTTGGTGTG

GGAAATTTCAGCCAATTGATGATTGGGTGTTGCCCTCTTTGAATTCCTTGTATCTTCGGCATGTTTGGTT

TGGGGATCACGTTTCGGGGTTTGAGAACTTGAAGGAATTAACTCTGTCGGGATGCTCTGAATGTTGGTGG

AATTTTGAGGGGGATTTTACTATAAACTGTGTAAATCTCGAACGGCTTGCCTTGGGCCCTGATATCTCAC

ATTGTGATATGGCTGTTTTTGCGCCAAAGCTATCGTACTTTGAATTTCGTTCTGGTCATGTACCACAGTT

CTCTGCGGGGGATGGATTCCCTTGTATAAAAGAAGTGGATATTGATATTGAACTTAGGAATGATGATTTC

GGGGGTTTATCAGAAGACGATGAGTACAGGATGGCGTCTGCTATGCCAAATTTTATTTCTATGTTAGATG

CAGTCAGGGAAACTCCGGTTCTCAAACTTTCAAGTGAAACAATTCAGTTTCTTCGCAGGGTTCCAGATCT

TTCAGAATATCAGCCGTCTCCATTATGCAACCTCAAGTTTCTCTACCTGCAAGGCCTTGATGAAATTCCC

TCCAGGTCCACTGATTATGTCATTAACTACTTGCTCAGTAATTCCCCTCATGCTGAGATTCTCAAAGGTA

ATCTTGGACGGAATCAGAAATTTGTGAGGATTGATCACTATATCACAGACTTGTAAGGATGGTCGCCTTA

TCACGTCTCAGTGATATGCCATTGCCATGAAAAATGCTAGCACCAATGAATTACCAGAAGTGCAGAACTA

TCTGGCTGATTGTATATCTTTTATCTCTAAGAACCTTTTTATGGATATCTCAGAAGGCTTTGTTTTGTTC

TTTTCTATTACAGGTTGTCAATCAAAACGAATTACTAGATGTTTTATTGTTTAAGTGTGATCCACAGAGC

ATACATCTCCTTCGCACGATTTTTTGTTTGCATTTACTTTAACATTTTCTTCAAGTATCAAGTGTCAGGC

TTGTTATCAAGCCAAAATACATTTATCATTCTAATGTTCAAGCAACTAATACAATCAATC

>MSTRG.1438.1 gene=MSTRG.1438

TACTCCCTCCGTCCCAGTCATTTGTATACAAATGGCTGGGACACAGAGACCAATAAATTGTGTAAGAAAT

GAGTAAAGTTGGATGAAAAGTGGGTAGAGTGGTGGGACCCATTAATATTTAATAATAGATTTGAGATAGT

GGAGGAAAGTAGTGGGTGTAATAGTGTTTATATTATTATAGAATAGAGATAGTGGAGGAAAGTATTTGGT

>MSTRG.1443.1 gene=MSTRG.1443

CTCTCCTCACCCCCTCTGCTAATCTCTCACACTCTCTCCCGTACATATCTCCTCCATTCCTCTTCCACCT

TTGTTCCCCGTTCTTTTCTTTTCCCCCGAAACCCGCCGCCGCCGCTGTTCAAACTCCGGCCAGTCGCCGG

CACCACCACTGTGTTGTTTCTTCTGATTAATTCACCCGCGATTCTTCGAGTTCCTAGGCAATCCAGACGC

AGGCTTGCTGTTTTCGGACGAATCCAGCCCTGTTCTTCAGCCTCCAGTGTTGTGTTAGCGCGGTTCCGGT

TAGAGAGCCTAACAGTATAGTGCAGTCTTATCTCTAGCATTCTATTATCAAG

>MSTRG.1444.2 gene=MSTRG.1444

CTGAACCATTTCATTCCCCTGCATAAGTTTTCGTTTCCCCCCTTCAGCATTAACCCTAATTTTGATTTTC

CCCCCTTATCTTTCTCTCCAAACTCCAATCGGCAGGTCGACGGCCTTCCCCCTTTATGCTTCTCTCTCTC

AGGTGTCTCTCTTCTCTTCTCTTCTCTCTCTGACCATCTAGCTTTATTCTCTTCTCTCTGTCCAATATTC

TCTCTCGTTGTCTCTCTTCTCTGTCCGGCTCTCTCTTCTCTTCTCTGTAGTTGTCTCTCTTCTCTTTATC

CTCTTGTTTTCTTGTTTTTCTTCTTCACTTTGTGTTGGAGTAGCCCTGTGCCTAGAGGGAGTGGCCTTTT

TTGGCAGAAGCTTGAGGTTCATCAGAAGCACTTAAATCGACTCAACAAGCCCCCCATCAAATCCATTCAG

GTCATATCTTTACATCTACTCACATTTTTATGTATGTATTTAGCATTTGTGTCACATTTGTTTTTGCATT

TGTATAGTATTTTGATTTTGTTTATAGGAAAGTATAGTTCTTAATTGGTGTTGTTTTGAATGAGCAGAGC

CCAGATGGGGATACCATTTACTTTATTCCAATCTCACATCAGCCTGCTTTTGAACACCTTCTCCTTAAGG

ATCAGAATATCTAGGTCCCCTTTTTAATTTTTGAGAAAATGTGCTTAAAGAGATGACATTTTGAGAGCAA

GCTCAGTTAAAGATATGATAAAAAGAAGCATAGATTTCTTTATTTATTTTCATCCCCCTCATCTCTCTCT

CTCTGTTCTGTTTTTTTGTAAGGGAGGCTATTTTATTGTTTAGTTTTGACTTACCGGAGTAGCTGCCTTG

TTTGAATCTGCCGGAATGATGCTCCTCCACCAGAATCCAGGAGGAGCTCTAGATGCTGCTGCTGTTGTAT

TGCTCACTCTTTTCACTCCCTCTAGCCGCGCCTCTTCTTCATTTCTGCCGATGAACAAAAAGCTGCTCTA

TGGGGGCCTCTGGCCAACCAAGGATGGAAAGCATGTGTCGATTCAGCTACTGGCCCTAGTATTAAATGGA

GTTGTCGTCAAGAAGACAACAAATACGGAGTAGATCTTAAAGCCCTATTGTCTAACACTAGGAGTTATTA

TTTTTCTTGCAACAATTATGGAGGTCAAACTCCAATAAGCTGCTCTATCTTGCAAAGGGCTTTCTGTTGG

CTTCTATCTACTCTATATAGGCTCTTGTCAATTGTGTGAAACAAATTTTTGTATGTGATTGTACGTTGTT

ACTATAGTTGGTAATAAAAACT

>MSTRG.1444.1 gene=MSTRG.1444

CTGAACCATTTCATTCCCCTGCATAAGTTTTCGTTTCCCCCCTTCAGCATTAACCCTAATTTTGATTTTC

CCCCCTTATCTTTCTCTCCAAACTCCAATCGGCAGGTCGACGGCCTTCCCCCTTTATGCTTCTCTCTCTC

AGGTGTCTCTCTTCTCTTCTCTTCTCTCTCTGACCATCTAGCTTTATTCTCTTCTCTCTGTCCAATATTC

TCTCTCGTTGTCTCTCTTCTCTGTCCGGCTCTCTCTTCTCTTCTCTGTAGTTGTCTCTCTTCTCTTTATC

CTCTTGTTTTCTTGTTTTTCTTCTTCACTTTGTGTTGGAGTAGCCCTGTGCCTAGAGGGAGTGGCCTTTT

TTGGCAGAAGCTTGAGGTTCATCAGAAGCACTTAAATCGACTCAACAAGCCCCCCATCAAATCCATTCAG

AGCCCAGATGGGGATACCATTTACTTTATTCCAATCTCACATCAGCCTGCTTTTGAACACCTTCTCCTTA

AGGATCAGAATATCTAGGTCCCCTTTTTAATTTTTGAGAAAATGTGCTTAAAGAGATGACATTTTGAGAG

CAAGCTCAGTTAAAGATATGATAAAAAGAAGCATAGATTTCTTTATTTATTTTCATCCCCCTCATCTCTC

TCTCTCTGTTCTGTTTTTTTGTAAGGGAGGCTATTTTATTGTTTAGTTTTGACTTACCGGAGTAGCTGCC

TTGTTTGAATCTGCCGGAATGATGCTCCTCCACCAGAATCCAGGAGGAGCTCTAGATGCTGCTGCTGTTG

TATTGCTCACTCTTTTCACTCCCTCTAGCCGCGCCTCTTCTTCATTTCTGCCGATGAACAAAAAGCTGCT

CTATGGGGGCCTCTGGCCAACCAAGGATGGAAAGCATGTGTCGATTCAGCTACTGGCCCTAGTATTAAAT

GGAGTTGTCGTCAAGAAGACAACAAATACGGAGTAGATCTTAAAGCCCTATTGTCTAACACTAGGAGTTA

TTATTTTTCTTGCAACAATTATGGAGGTCAAACTCCAATAAGCTGCTCTATCTTGCAAAGGGCTTTCTGT

TGGCTTCTATCTACTCTATATAGGCTCTTGTCAATTGTGTGAAACAAATTTTTGTATGTGATTGTACGTT

GTTACTATAGTTGGTAATAAAAACT

>MSTRG.1445.1 gene=MSTRG.1445

CTTCATATCTAATTACAAGTTCTGCCACCGGAAAGCTGCACATTCTTAAGTCACGTGAATTGATTACCGT

CTTTGGCTAAGCATTTCATGAGCAGTCAATGGTGACCACCAGGTCCACCATGAGTTCCATTAGCTTCTGT

CCAAAACAATGTTGTATTTTTCCAGAAACTACTTTCTTTAGTGTATCAAGAAAAGTTACAATGACTTGAA

GCTCCATGATTTCAACTCATTCAAATATGATATTGGTTGGTGCTGGTGGTTTCAAGATCTTCACATCCCC

TAAATTTAATGCTTGGTTGATATTATTTGTTCATCAATATTTCTTTTTTACTGAGTTTTTCTAAATAATA

TTCAGCAGGTGTTTGATCAAATGGGCCACTGAACTTCTTCATTTTAAAAAATAAATTCACATGCCTTTGC

AAATCTTACTTATTTACAGTATTTGTTGTTGAATGAGAATGGTCTTCCTGATGTTATTGACAAAACCTGG

ACAACATTGAGTCAACTATTGTTATTTTAATTTTTGTCGGCTTTGTGTTATATATATCCCAAATTTTATG

TCACGACCTGAGGTGGGTTTGGTGTTGTTTACACTGGAGAATTGCATGATTGCACTCAGATTGCAGTGTA

AAGAATGGAATTTGGGTTGTGTGTACAAAAGGACTGAATGAGTTCGAAGCAGAATTGTTATTGCTGATAG

GATAGAAATTGTTTCCTGGAAGCTCAAGATGGCAAGTCAGAGTGCTAGAGGCATTTTTTGTCACACTTTA

TAAGAGAGTTCAAGGTTTGGAAAGTTATAATCCCACAAATCCAGTAGATTCAGCTGGGTATCTTGAGCAT

TGGATTAAACCACCGGTGAACTGCGTTATAGGCAACTGTGATGCTGCGATCTTTACATGAGGTGCTCGGT

CTGGAGTTGGTTGGATACCAGAGGCCCAAAGATATTAATGGGATTTTGATCAGAGCTGTTGCTCCTGAAA

TTCTTTGGGCTGAGACAATAGCTATTCGGGTGGTGCTTAGTTGGATCTAGCGGGAAATAAAGGCAAGTGA

AGCTAATATTTGTAACAATTAGTTGTCGTATGTGGGGGAATCTGACTGCCAACTTGCTATAAGGGTCATT

CTTAACAGGGAAGCAATTTACTCGCCCTTTAGTAATGCAGTTGATAAATGCAGGAACATCATAGATTCTT

TAAACAATATTAAAATTGTTTTTGTAAAATTATCAGGGCTGCAGATTGGCTTGCATGGTCATCGAGTTCT

ATGCCAGGTTGTATTGCCAGTGGGGGTTCTCAAATTTTAAGCTTCATAGCATTTTATATGCTGATTTGAG

TTAATGAAAATTTAACATTCCTGAAAAAATCTCTCATAGACATTTGAAATGTATACTTAATGGAATTTTT

TAACCTTTGTTATTCTTTTTATTGATGATTGTTGCAACATGAGTTCCATGTAACTGAGAAAACAGAAGAA

CGACCAGCAGCCACGAATGCATTTGAGTTAATATCCATGTTGTGGGGGACTCAATCTTGGAAATCTTTTT

CAATGTAGAGCAGGTATTCAATTAAATATTGGAGACCTCCCACAAATTCAAATATTATACAAGTGATTTA

AGTTCTGAAAAAGAAGCAGGTGTCTGGGAGATTGTCC

>MSTRG.1447.1 gene=MSTRG.1447

TTATAATCTATGTTACCCGGACTCTCCATTTTTGTGCCTATACCCGTGTCCACCGGACATGACATGGGTG

TGGGTATGGGATCTGTGTCGGATCCTTCTTATTACAACCGGACACCTCACCTTGTAACATCCCAGGTCAA

AATGAAAACTTAGGTCTTGTTTCAACTTGTTTAAAGAGATCAAGATCCATGAATGACTTGTTTCTATGTT

GTTCTATTGATTTTATGCTTCAATTTTCAGCGTATGTGATTTATGTGCAGTAGACACTATTTTTATGAAT

GTACAATAGACTAGTCTATATAAAGTTTTACTTGCATGGAGCTTAAGTGCCAAGTCCCCGTACCCGAGTC

TAATTCCGGATCCATGTCCACGAATCCTAATTATCCTAAAACTACGAATCTGACTCTTGGATCTGTACCC

GTGTCCGATACCCATACCCGTGTCCGGGTAACTTAGATTAT

>MSTRG.1448.1 gene=MSTRG.1448

GAGCAATCTAATGCAAAAGGTACAACTTTTACTTTTGAACAAGATGGAGAACCTATATATGTGACATACG

AAGATGTTGATCAATTTCTAAAAAGGAGCTGGTTAAATATACCCATCTTGGAAATCTTTTCAAAGTACAT

TGGGAAGTTGTGCAAAGAGCTAAACGATGATAGGTTTGCATTTATGTCACCTTCTAGACTAGTGATAAAT

CACAAGCGTGAATACAATCGTATTGAGGATGCAACTCAATATATGACACAATTTTTAGTTGCAAACAAAG

ATAAGCGTTTCATCATTGCACCATACATCCAAGATGATCATTGGATGTTACTTCTGTTTTGTTTGGATGA

GAGTGTTATTTATGTGTTTGATTCCTTGAGAAGGGAACGAGATATAAGGTTGACGACCCCGGCACGAACG

GCATTTAAACTTTATGTACCACAAGGTGGAAAGAGAAATAACAGAAAGGAATTTCTTTGGTCTCATACAG

ATGTTCAGTGTCCTCAACAAGAAGGAGGTACGGAGTGCGGATTTTTTGTTATGAGGTACATGTATGAAGT

AGTCAAGCTTTCTCAAAAAAATCCCAACATCAATTGGAAAGAGGGTCTTGGTTCAAGGAGGTATCTAAAG

AAGGAAATCAATGAGGTTCGAGAGTTGTGGGCTGAATATTTTACTGTTGAATGTCTATAATCAGACATAC

ACTCTATGAAGGCCAGTTCAAGTATTTTGGAGAAAAGCACGAGTGAAGAAAAGACTCGGTCTGATTAGTG

GATTTCCTAGACTTTTTCACTATTGATTAGTGGTATTGACATGGTGTAGTAGTACTTATATGATTGATGG

ATGTAATAGAAGTGGAT

>MSTRG.1450.1 gene=MSTRG.1450

AATACCCACTAATTTGCCTGACCCATAACCCGTTTACCCATTTACCCGTATTAACCGCCTAATACGACCC

ATTTGCCTCAGATTTTGAATTGCTCCATCCTCCAAGACGCTATCATCACGCAGACTGTTACATACCCAGT

CTATCATCCTCCAAGACTCCTCACTCTGTTCTTTATTTCAAAATCATTATCAATCTCCAGATGATACGAG

ATCAATGGCCTTGGTACCTATCAATGACCTAAACTATTCACATTCAGTTCTGGCGAATCAACTTCCCTAA

TCTACCCATGAAAGAACGGGTAATCAGGGTTTATACTTTGGTGTTTCGTCACTTGCTCTTTCAGCCACTT

CATCTCTCCGCTCTCTCTGCAAATGAAATGGTAA

>MSTRG.1451.1 gene=MSTRG.1451

AAAAAATTCCAATAGCCATGTCAATTGATGTACAATCGAAGAAGAAACTGAAATTCACCGACGAAGACAG

AATCAGCGCATTACCCGATGATCTAATTCATCAAATTCTCAAGTCTGTGGATACCAAAGAAGCAGTTCAA

ACCAGCATTCTCTCGAAACGATGGAAGTTTGTATGGACTACTCTCCCTTTCCTCAAATTCAAATGGGCTC

ACCCACGTTTCCAAGATACCAGTGTTAATAAAAGCAATGCTAAATTTACCCGTCATGTTTTGAATTATCG

AAACCATAATTCTTCGATTTCGTGTTTGGAGCTTAAGTACCTCACTCCTGGTTTATTGGATAGGTTTATT

AAGTATGCTATTGCTCATCATGTTGAGTGTCTTGATGTTCATTTACGACACAAGCATAAGCCGTATATGT

TAGGGAGTTTCAGCTCTGGTTCGATTAAGAAACTTAAAGTGAGGATGAGTTTTGAGGATATTGTGCCTGA

ATCTGATTGTTGGGATTTGCCTGCGTTAACGAGTTTACATTTGTGTCGGCTTTTTCTGTTTTTTGAAAAT

CAGAATTTGCCGGAGAGATGTATAACTTGTTTGCCCGCGTTGAGAGATTTGTGTCTTGAAGATTGGGACT

TGAGGGAATCGTCTTTATCTTTCAATTGGCCGGGTTTGACGAGTTTCTGTCTGATTAAGTGCATATTGCC

TACAAAAGTTTGGAATTTTCCGGGTTTGAAGAGTTTAGAGCTTCTCGATGTGACGTTTCCGAAAAATATG

AGGGAGATATTCACTGCACTTGTCAGTCTAAAAGATCTCATATTATCTTTTGATGAGGTTTCCCAGACAA

AATATTTCTTACATTGTCCTGAATTGGTGAACCTGGAGATCAGAACCCGGTACATAAGCCCATTCTTTGA

GATTGTTAAAAGAAACTTTGTGGTTTCTGCACCAAAACTCAAAAATTTCACTTCTGTCGGTATCTTCACG

GTCAAATTTGATGTTCCTGAGTTGGACAATGTTTATATGAAATTGCGTGGTTGGATTGATGATCAGAGTT

TTTCTCCAAACCAGTTAAAGGAGTATTATCGACCGTTTAGACAAATGCTACTGGGGCTTGGTGGTGCCAA

AATCCTTCATTTCGAGTTGGAGACAATAAAGGCACTTTCATCAATCTCTGACTTTCTTGGAAGATCTCCC

CCTCCATTCTGTAACTTGAAGTATGTCAAACTACCACATGGATCTGAAGAAGCAATCATTTCTAACAATT

TAAGAAGCTACTTACTTGGTGCCTCTCCAACAGCCTCCATTGTCACAACATTTTCACAGAGCAACATGAA

AAATTCTCATACAACAACAGCTTCTGTGACATCTCAAAATGTGGCTCTACAAAAGCCTTTGGTTTCTTCA

ACAAAGGTTCTGGTTGATTCTGGAAATGTACCTAAAACTGTGTGCATTAATACTGTGGACGTAGGGGTTC

AAGCAGAGCATGTGGTGCGGAATTCTGTACAACATACCTACAGTGACAGGGTTAGACATGTTGGTTCCCC

TGTTGACGGGACCGACAATGATCATGCGAGATCTTCTTCAGGCAATCCGGATATTAGATTATGGAGGGGT

TATGATGTTAACTCAAAATTTGTATGTATACTCAATGCCATAATGGATAAGTACCCAGAAACATTTGAGC

ACTTCGCGACCAGAAATAGAAATTTCAGTGAAATGAAGCTAAATATGTTGTGCACCTCAGTACATGACTT

TGTCAAAATGCCTGTGCTCATAGTTGATGCTGAAATACTTGCTGGGTACTGGGATGTATTTGCTGACTTG

CAGAAACTGGGATTTGATGTAAGTTGGCTGGTGAGTCGTCTGGTCATTGTTGAACAGCTATACACCGCGC

ACCCCCTACTTCTTAAGCTTCATGCAATTGACTGTGATATTGATGATGCCCAAAGGATGTTTCAAGATTT

GCAGTTTAATATGAATGAAATCAGTAGTATATTGCAAAATCTGCAGACTCTTCGTAATAAGAAGTTGCAA

GAGATTCGGAAAGTTGTTGGAACTACGGCTGCAAAACTTGCTGGTGGCTACATTGGAGATGATTTGTTGC

CTGTCACCTAATTTACTTGGTGGTATCAGTATGTTTGTTTAGATGCCTCGGTAGTCTATTGTTTTTTGGC

TTTTATTGTTGCTTAAAACTAAATTGCAAGATTTGTAGGGATGAATCCGAAACTTGCAAGATCCTCAGCC

TCTTCATGAGAGATGCAACAGATTGAAAAAGCTTTTAAAAGTATGGGTTCGAATTGTTTTAGGCTACATT

GAAGATGATCCATCGCCTGGTCCCTAATTTCCTCAGAGTTAGTCGTATGCTTGAGTATTGTTTGGCTGCC

TTGTATCTTGAATATTATCTGCTCTCATGAGTATAGTAAGTAGATTTCTGGTGCCTTGTATCTTGTATTG

ACTTTGGATTAAACTGTTAGTAGTCTTTTCCTTCCTGC

>MSTRG.1451.2 gene=MSTRG.1451

AGGCACTTTCATCAATCTCTGACTTTCTTGGAAGATCTCCCCCTCCATTCTGTAACTTGAAGTATGTCAA

ACTACCACATGGATCTGAAGAAGCAATCATTTCTAACAATTTAAGAAGCTACTTACTTGGTGCCTCTCCA

ACAGCCTCCATTGTCACAACATTTTCACAGAGCAACATGAAAAATTCTCATACAACAACAGCTTCTGTGA

CATCTCAAAATGTGGCTCTACAAAAGCCTTTGGTTTCTTCAACAAAGGTTCTGGTTGATTCTGGAAATGT

ACCTAAAACTGTGTGCATTAATACTGTGGACGTAGGGGTTCAAGCAGAGCATGTGGTGCGGAATTCTGTA

CAACATACCTACAGTGACAGGGTTAGACATGTTGGTTCCCCTGTTGACGGGACCGACAATGATCATGCGA

GATCTTCTTCAGGCAATCCGGATATTAGATTATGGAGGGGTTATGATGTTAACTCAAAATTTGTATGTAT

ACTCAATGCCATAATGGATAAGTACCCAGAAACATTTGAGCACTTCGCGACCAGAAATAGAAATTTCAGT

GAAATGAAGCTAAATATGTTGTGCACCTCAGTACATGACTTTGTCAAAATGCCTGTGCTCATAGTTGATG

CTGAAATACTTGCTGGGTACTGGGATGTATTTGCTGACTTGCAGAAACTGGGATTTGATGTAAGTTGGCT

GGTGAGTCGTCTGGTCATTGTTGAACAGCTATACACCGCGCACCCCCTACTTCTTAAGCTTCATGCAATT

GACTGTGATATTGATGATGCCCAAAGGATGTTTCAAGATTTGCAGTTTAATATGAATGAAATCAGTAGTA

TATTGCAAAATCTGCAGACTCTTCGTAATAAGAAGTTGCAAGAGATTCGGAAAGTTGTTGGAACTACGGC

TGCAAAACTTGCTGGTGGCTACATTGGAGATGATTTGTTGCCTGTCACCTAATTTACTTGGTGGTATCAG

TATGTTTGTTTAGATGCCTCGGTAGTCTATTGTTTTTTGGCTTTTATTGTTGCTTAAAACTAAATTGCAA

GATTTGTAGGGATGAATCCGAAACTTGCAAGATCCTCAGCCTCTTCATGAGAGATGCAACAGATTGAAAA

AGCTTTTAAAAGTATGGGTTCGAATTGTTTTAGGCTACATTGAAGATGATCCATCGCCTGGTCCCTAATT

TCCTCAGAGTTAGTCGTATGCTTGAGTATTGTTTGGCTGCCTTGTATCTTTAGATTTCTGGTGCCTTGTA

TCTTGTATTGACTTTGGATTAAACTGTTAGTAGTCTTTTCCTTCCTGCTG

>MSTRG.1452.1 gene=MSTRG.1452

AAAAAACATTTTCTTGCAAAGTAGGGGCTAAAGCCATCTCCAACGGGAGAACCTGTTTTCGGTCCAAATT

CAAATACCCAATTTTTATGAAATTAGGGTTCCAGAGAAGAAAGCAAAACATCCATCTGAATGGACACCCA

ATCGAAGAAGAAAGCGAAAGTCGCCGACGAAGACAGAATCAGCGCATTGCCCGATGAGCTACTCCATCAA

ATCCTCAAGTTCCTCGACACCAAATCAGCATTTCAAACAACCATTCTCTCCAAACGCTGGAAGCTTGTAT

GGACTACTCTCCCTTTCCTCAAATTCATCTGGGATTTCCCCGATAAACAAAACACAAGTAACAACAAATC

CATCGCTAAATTCACCCATCACGTATTATATCATCGAAACCGCCATTCTTCAATTTCGCATTTGGAGCTT

ATGTTCCTCACTCCTGGCTTATTGGATAGGTTTATTAAGTATGCTATTTCTCATAATGTTGAGGTTCTTG

ATGTTCATTTTCGGTTCAAGCATAAGCCCTATAATTTGTCGAATTTCACCTCTAGTTCGATTAAGAAGCT

TAAAGTGAGGATGAATTTTGAGAGGATTGTGTCGGAGTCTGATTGTTGGGATTTGCCTGCTTTAACGAGT

CTACATTTGCGGGGGCTTTTTTGGGCTTATGAACCCGAGACTTTACCGGAGATTTGTATAACTTGCTTGC

CTGCGTTGAGAAGTTTGTGTCTTGAAGATTGGGATTTCAAGGATCTGGTGAACGGAAGATCGTCTGTGTC

TTTTGATTGGCCGGCTTTGACAAGTTTTAGTCTGATTAAGTGTAAGTTGCCTACAAAAGTTTGGAATATG

CCGTATGTGAAGAGTTTAGAGCTTCACAAGGTGACGTTCCCGGAGAATATGAATGACATCTTCGCTGCAC

TTGTCAGTCTACAAAGTCTCATATTATCTTTTAAAGAGGTTTCGCATAATGGTCTTTTCCTTACTTGTCC

TGAATTGGTGAACCTGGAGATCAGAACCTGCTGTGGTGACAAGTTTTATAAAAATCTCGAAAGAAACATT

GTGGTTTTGGCTCCAAAACTTAGATATTTTACTTCTGTTGGTATCTTTACTGTCAAATTTGAAGTTCCTG

AGTTGGGCTATGTTAATATCAAATTGCGGGGCTGGATTGATGATCAGGAGTTCTCTCCAAACAAGTTAAG

GCAATATCATCGAGAGTTTAAACAGATGCTGCCGGGGCTTGGTGGTGCCAAGATTCTTAATCTTGAGTTG

GAGACTATGAAGGCACTTTTATCAATCTCTCCCTTTCTTGTAAGTTCTCCCTCTCCATTTTATAACTTGA

AGTATGTGAAACTGCCACATGGATGTGAAGAAGCAAGTATTCCTAGCAATTTAAGAAGCTACTTACTTGG

TGCCTCTCCAACAGCCACCATTGTCACAACATTTTCACAGAACATGAATTCTCCTACAACTTCTGTGACA

TCTCAAAATGTGGTGCCACAAAAGCCTTTGGTTTCTTCAACCAAGGTGCTTGTTAATTCTGGATATGAAT

CTGTGCCCATTGATACTGTGGACATGGGGGTGCAAGTAGAGCATGTGGTGCAGAATTTTGTAGAATATGC

TTCCAGTGACAGGGTTAGACATGTAGGTTCCTCAGTTGAAGGGACCGACAATGGCCATGTGAGCTCTTCT

TCGGGAAATATCGATTCTGGGTTATGGAGGGGTTATGCTGTTAACTCTGAATTTGTAATCCTACTCAATG

CCATAATGGATAAGTATCCAGAAACCTTTGAGCACTTTACTACCAAAAACAAGAAATTCGGTACAATGAA

GCTCAATATGTTCTGCACCTCAATAAATGACTTTGCCAAAATCTCTATGATCAAGGTTAATACTCAAATG

ATTACAGAGTACAGGGATGTATTTGCTGACTTAGAGAAATTGGGATTCAATGTAAGTTGGCTTTTGAGTC

GTCTGAACTTTATTGAACAGCTTCGGATTTCACAGCCCCTACTTCCTGAGCTTCATGCAATTGATGGTGA

TATTAATGATGCCAATAGTAAATTACAAGATTTGCAGATTGATATGGATAAAATGAAAACTAAATTGCAA

AATCTAAAGACTCTTCGTGAAAAGAAGATGCAAGATATTCAGGAACTTTTCGGAACTATGGGTAGAAACC

TTGCTGTTGGCTACGTTGGAGATGATCTGTTGCCTGGTTCCTTATTTACTTGAAGTTAGTTGTTTGTTTG

AATTGTTTGGATGCCTCAGTATATCATCTGCTGTCTGCTCCCGCACTTTTACTTTTTGGCCGTCATGCTT

ATTGAGTTGGTATTGAATCATTTGATACTAAACTATAAGATTTTCATGGATGAGTCCAACTTAAAATGCA

ATATCTAGATACTGATGACAAGATGCAAGAGATTGAGAAAACTTCTAGAACTGTGGGTACAAATCTTGAC

GTTGACTACAATGAAGATGATCTGTTGTCTGGTGCCTAATTTACTTGGAGTCAGTTGTATGATTGAGAAT

TTCATGGATGCCATGGTCTAGTACTATTTTGGTTTCTTGCTTATTGACTTTGTTTTGAACTGTTAGCAAA

GTTTGCTTTTCTACTGTTTAATGGAAGTGCTTGTATGCTTGTACTATGTCTTGCAAAATTCTGTTTCACT

TAACCTAATCATATTCGTAGCATGTTATCATGAG

>MSTRG.1453.1 gene=MSTRG.1453

TTTTAAGATCGGACCGAGGAGGAGAATTCACTAGTAATCAATTTCATAATTCTGCAAAGAAAATGGAATA

AAAAGGGAGTTAACAGTACGCCGTACACCTCAACAAAATGGTGTGGCGGAACGGAAGAATAGGACAATTG

TTGAGATGGCCAGAAGCATGCTCCAAGGTAAAAAATTACCTAAATATTTATGGGCGGAGGCTGTTAATAT

AGCAGTTTATATTCTGAACAGGTCTCCTACAAAAGCTGTGAGAAATAGGACGCCGTATGAAGCTTGGTTT

GAAGAAAAACCCAAGGTTGATCATTTAAAAATATTTGGCAGCATCGCCTATGCTCATGTTCCTAAAGAAA

ATCGAGACAAGCTGGATGAAAAGGGCGAAAAATGCATATTCATTAGATATAGTGATGAATCGAAAGGTTT

CAAGTTGTTTAATCCTATAACCAATCAGCTTGTAATTTCTAGAGACGTCATTTTTGACGAATTTGCAACA

TGGTGTTGGGAGAACGTTGTGGCTGATAATTATCAGAAACTTGACTTGGTTGAGCCTCTATCTACAAACG

AAGGAGCACCAGAAAATCAAGAAATGAGCATTCGACCCGAAGATGTGGATGCATATTCAGAATTTCCACT

ACAAAAAACTCGATCTTTGGAAGATATTTATCGAAACTGCAACATGGCCTTACTATCCATTGAACCTCAA

AGTTTCGAGGAAGCTTCACAAGAAAAATCATGGAGAATAGCAATGGATGAAGAACAGAAAATGATCGAGA

AAAATCAGACTTGGGAGCTCGTGAATAGGCCTGAAAACAAAGAAGTTATTGGACTAAAATTGGTCTATAA

AACAAAATATAATGAGGATGATTCAGTTCAAAAATATAAAGCTCGTTTGGTGGCGAAAGGGTACTTTCAG

AAGCCTGGAGTTGATTTTAACGAAACTTTTGCACCGGTTGTTCGAATGGAGACTATAAGAACAGTCCTTG

CTTTAGCTGCGCAACTCAAGTTAAATGTATACCAACTAGATGTAAAATCTGTTTTTCTTAATGGAGATCT

CGAAGAAGAAATTTTTGTTGAACAACCACAGGGATACGTCATCAAGGAGCATAAAAATAAAGTTTATCGA

CTCAAAAAGGCTCTTTACGGATTGAAGCAGGCACCGAGATGCTGGAATAGCAAAATTGATGAATATCTTC

TGAAAAATGGTTTCATTAAAAGTCCATCTGAACCATCACTGTTCATCACTCTATGTCAAGAAGGAGGGTA

CAGATTTTTTTTATTTTTTTGCTTGTACGTAGATGATCTGATCTACACCGGTACAAATGAATAAATGGTT

GAAGATTTCAAAAAGGCCATGATGAAGGAGTACGAGATGACTGACCTTGGTCTTATGAAATATTTTCTCG

GAATGCAAGTAAGGCAATCTCCTGGAGAAATATTTATTTCTCAAGAGAAGTATGCTGATGATATACTCAA

GAAGTTCAACATGTCAAATTTTAAGTCTCTTTCTACTCCAATGGCGGTTAATGAAAAACTATCCAAGTAT

GATGAAAGTGAGAAGGTAGATGAGAAATTGTATAGAAGCTTAGTTGGTTCTTTGATTTACCTTACGAACA

CAAGGCCAGATTTGGTACAAGCTGTTAGTGTGGTTTCGAGATTTATGAGTGAGCCAAATAAAACTCATTT

TGTTGCAGCAAAAAGGATACTCTGATATGTCAAAGGAACAAACTGTTATGGACTGCTCTACAGGTTGGAA

GGAGATAGTAAGCTAGTAGGTTATACAGATAGTGATTGGGCAGGTTGTGTGGATGATAGAAAAAGTACTA

TTGGCTATGTGTTAAGTCTCGGCACAAAAACCATCTCCTGGTCATCAAGGAAGCAAAAGACGGTAGCCTT

ATCCTCTGCAGAAGCAGAATATGTGGCAGCTACAAGTGCAGCCTGTGAAGCAATATGGTTAAAGAGAATT

CTAGCAGATATGCAGCAACAAGTGGATGGCTCTACTGATATATATTGCGACAACATGTCAGCCATTGCAA

TGACAAAAAACCCTGTTTTTCACAGTCGAACAAAACATATTGAAATACATCATCACTTCATTCGAGAACT

TGTTGAAGATGGTGAGCTGAAAATTCTGCATTGTAAAACTGGTGAACAACTAGCGGATGTCTTCACTAAG

GCGTTGCCGGCTGATAAGTTCAACTTCTTTCGAGAAAAGCTTCGAGTAAAGATTTTACCAGATTAAGGGG

AGTGTTGAAAAGTGAACTAGCGGATGTCTTCACTAAGGCGTTGCCGGCTGATAAGTTCAACTTCTTTCGA

GAAAAGCTTCGAGTAAAGATTTTACCAGATTAAGGGGAGTGTTGAAAAGTAATCTGTTAAATCTTGAAGC

TTCTAGATTTGAGGAAATGTCAAGAGTCTGGAAGTTTAGAAGTCTAAGTGAAAAGTTAAGTGCAGTTAAT

CTAGTCTGCAGATGTCATTAATTATATATCTAGAATGATCTAATGTTAGCTATAAATTCAAGTGTAC

>MSTRG.1454.1 gene=MSTRG.1454

GATTTGAAGGCGTTATTGGAGTCGGAATCAAGCGTGCGAGTAGTCAAATCGAAGATATCGAAGAGTAGAA

TCCATATCAATGATCAAAAACAACATGAGATTCACAGAAAGGCTTAAAGATGAGATGATTGGTTGAGATT

GTGAATAAGGCTAATGCTAGACGAGGAGACAGAGTTGAGGATCCAGTGGTCGAGGAGTGTTCAGGAAAGT

GATGAGGTTGGTGCAAGCTAGAACGGTGCTATAGGGATTCAAGTGTAAGTCGGAGCTGGAAGAGTTAACA

GTGATTCTTCGTATCGAAGATGATCTGATTTGGTAAGAATTGTTGTGTTAGTGGTGGATTCTAATTTCAA

TACTGTAACCTGGAAGCGATCCTGTTGTTTTTGGGGGTTTAAATGTTCTATTTCGAATCATTTATTAAAG

TTTAAAGATTTTCTGGAAATTCTTTTATG

>MSTRG.1455.1 gene=MSTRG.1455

CATGCGTACTCAGTTCAACAAAATCTGAAGAGTTTTTCAGATCAGATTTTATTATTCAATTTTACACTTA

CTCTCTACTTTCACTTTCTAACCAACACTCTTACACTCTGTTCTCTGCAAATTCAAATCTTCATCTACAC

ACTTCCTCAAACATCTTCTTCTCATAATCTCTGATCTAATGGCCACTACAGCTTTTATCTTGGCAGGTGT

TGAATTTGTTCCAAATAACTATGCAGCCATTCTTGACACTGCTGAAGCTCCAAGGGATTATCATCCCTTT

CAGCGATTCTTGGCACAGAGTGTCATCGGTACAGCTCTCACCGCTCCTGCAAGACTTTCAGGAAGCCAAA

TCATCAGCTTTTGGAGGTCGGGTACATATGACAATGGTGGTGCAGATGGATCACCATCAATTGTGTTTTC

CTATGACGGGGAGGAGTATGCAGTTACTCCAGCAACAGTTCGTCAGGCACTCAACTTGCCGGAACATCCT

GCATACATTACAAATGGAGATGCAAATCTCAGAACGATGATGCTTGATTTGGGGTACTACGAATCTCTGG

ATAAACTGGGTCAGTTGAAGCGTCC

>MSTRG.1456.1 gene=MSTRG.1456

CTGGGTTGGATCGTTTTATCGGGTCTAACCCGACCGACCCAACCCGTCTTCACCACTGCTGTCCGGGTAG

ACACCAATTAGGGTTCTTCACTCCTTTCCTAATTCAGTGTAACTAGAAATCACCTACTACTACTATCCAT

TCATTTCTGTATTGTCTGAGTCTTCATCAGTCTCTCAATGGATGGTCCTATGAAGCAGAAATCAAAAATC

ACAAGCGAAGATGAAGAGCAAGAAGACAGAATCAGCAGATTGCCAGACGATCTAATCCATCAAATCCTCA

AGTTTGTAGACACCAAATTGGGTGTTCAAACCAGCGCTATCTCGAAAAGATGGAAGCTCGTGTGGACTAC

TCTCCCATTCCTCAAATTCAGATGGGATCAATCTCGATACGATGGAAGCTCGTGTTCGTCTAAAAGCACA

ACAAATCTGGCCCGTCATGTTTTGAAGCACCGCAACCATCAAGCCCAGATATCGTATTTGGAGCTGGCGT

TCCTGCCTTCAGGTTTATTCGCAAAGTTTGTTAGTTATGCTATTGCTCACAGTGTGCAACACTTGAATGT

TCATTTTCGAGAAAAGCATAAGCCTTACAAGTTATCTAACTTTAATTGTGATTGTATTGAGAAACTCGAG

ATGAGAATGGAACTTGATGATTCTCTGCGGGAATCGGATTGTTGGGATTTACCGGCTTTAGCAAGTCTAC

GCCTGAGAGGGTACTTGGTTTCGGGTCGGCTAGATAAGTTACCGGAAATGTGTGTGACCTGCTTGCCTGC

GCTACGATCCTTGAATCTTGAAGAATGGAATTTGGAGCGTTTTTCTTTCAGTTTGCCTAATTTAACAAAT

TTCCGGCTATGGTGCTGCCGTTTGCCCCAAACAATTTGGGATTTCCCAGTTTTAAAAAGTCTAGAGTTAT

GTAGTGTGGAATTTCCCGAGAATATGAGTGACATATTTGCTGCACTTGTCAGTCTACAGAATCTCACAAT

ATATTTCTACGGGTCGCATGCATTGTCTGCAAAAGATTGTTTCATAAATTGTCCTCAATTAATCAACCTC

CAGATTATAACCAGCTACAGTGATCGTGGTAAGGCCAACATTGTGGTTGCGGCACCAAAAATTCGCAATT

TCACATCGGTTGGTATATTTACAATCACATTCGGAGCTTCTAAGTTGGATACTGTGAATATAAAGTTAAA

GGATTGGATAAATTCCGGTAGCTATAAGTGGTTGCACCAGAAGGAGTTGAAACAGGATTATCAACGGTTT

AAATTCATGCTACCAGGACTTGCCACTGCCAAGATTCTTAAACTTGACTTGGAAACCATAGAGGCACTTT

CATTGATATCCGACTTTCTTGGAAGTTCATCTCCATTCTATAACTTGAAGTATGTGAAACTTCCACATGG

ATGTAAAGAGTCAAGTATATCTACTTCTCTAAGAAAGTACTTACTCGGTGCCTCCCCAACAGCCACTATA

GTCACAACATTTCCCCAGAACCATGTGATTCCTTATACAGCGACAACTTCTCTGACAACCCAGTATGTAG

TACGAGAAGAGCCCTTGGCTGCTGTAACCAGGGTATTGACTGATGTTGAAAGAACATATAAAAAAATGCG

CAATGAAACTATGGACATGGGGGTGCAGGAAAAGCATGTTTTACATGCTGACAAGGTTAGAAATTCTGTT

TACCTCATTGAAGGTATTGGCAATGATCGTCTAACTTCTTCAAGGGGGGGAAGTAATCTTGGGTTATGGC

GAGGCCATGAAGTTAACTCTGAATTTGTATGCCTACTTAATCGGATCATGAATAAGTACCCCGAAACCTT

CGAGTCCTTCGCCATTAAAAACAAGAAGTTTTGTACAGTGATGCTGAATATGTTGTGCACCTCAGTAAAT

GACTTTATCAAATTCACGATGACTGAAGTTGATAGAAAGATGATTGCTGAGTACAGGGATGTATTTGCTG

ACCTGAAGATATTAGGATTCGATGTTAGTTGGCTCGCGGGTCGTCTGAATTATATAGAACATCTCCAATT

TTCACATCCCTTACTCACTGAGCTTTATGCACTTGACGGTCATCTTAATGGTGCTAGAAGTAAATTACAA

TATTTGCAGACTCATAGTGGTGATACCGAAATTAGATTGCAAGACCTACAAACTCTTCGTGATGAGAAGA

TGCAGGAGATTGAGAAAGCTTTTGGAACTTTGGGTACAAAACTTGTTGCTGGCTGTATCGGAGATGAGTT

ATTTATTTAGTCCCTAATCTTCTTGGAATTATTTTTGTTTAAGAGTTCTGGATGCCTTCTTTACTTGTTT

ACTCTCTGGTCCTTGAGTCTTTAGTATGTACGTGTCGCGTATATTGTCTTTGTGTTGATCTGTTAGCTGT

GTTTGGTTTGCTACTGTTTGTCAAGGTGTGAATCATGTAAGATGTTGCATTCACAAAAGGATTTTCTAGC

ATATTATATATTGAGCTATGCTATATATGGTGCTATTTTTATAAATACTTCCTCAGACATTTCTGCAGTT

CTTATTCTTGCAAGTTTGTGTTCCCCATCCTATTAATTGATGTAAAAATACTGACATGGACATTGGGAAT

CCGGC

>MSTRG.1456.2 gene=MSTRG.1456

GTTTTATCGGGTCTAACCCGACCGACCCAACCCGTCTTCACCACTGCTGTCCGGGTAGACACCAATTAGG

GTTCTTCACTCCTTTCCTAATTCAGTGTAACTAGAAATCACCTACTACTACTATCCATTCATTTCTGTAT

TGTCTGAGTCTTCATCAGTCTCTCAATGGATGGTCCTATGAAGCAGAAATCAAAAATCACAAGCGAAGAT

GAAGAGCAAGAAGACAGAATCAGCAGATTGCCAGACGATCTAATCCATCAAATCCTCAAGTTTGTAGACA

CCAAATTGGGTGTTCAAACCAGCGCTATCTCGAAAAGATGGAAGCTCGTGTGGACTACTCTCCCATTCCT

CAAATTCAGATGGGATCAATCTCGATACGATGGAAGCTCGTGTTCGTCTAAAAGCACAACAAATCTGGCC

CGTCATGTTTTGAAGCACCGCAACCATCAAGCCCAGATATCGTATTTGGAGCTGGCGTTCCTGCCTTCAG

GTTTATTCGCAAAGTTTGTTAGTTATGCTATTGCTCACAGTGTGCAACACTTGAATGTTCATTTTCGAGA

AAAGCATAAGCCTTACAAGTTATCTAACTTTAATTGTGATTGTATTGAGAAACTCGAGATGAGAATGGAA

CTTGATGATTCTCTGCGGGAATCGGATTGTTGGGATTTACCGGCTTTAGCAAGTCTACGCCTGAGAGGGT

ACTTGGTTTCGGGTCGGCTAGATAAGTTACCGGAAATGTGTGTGACCTGCTTGCCTGCGCTACGATCCTT

GAATCTTGAAGAATGGAATTTGGAGCGTTTTTCTTTCAGTTTGCCTAATTTAACAAATTTCCGGCTATGG

TGCTGCCGTTTGCCCCAAACAATTTGGGATTTCCCAGTTTTAAAAAGTCTAGAGTTATGTAGTGTGGAAT

TTCCCGAGAATATGAGTGACATATTTGCTGCACTTGTCAGTCTACAGAATCTCACAATATATTTCTACGG

GTCGCATGCATTGTCTGCAAAAGATTGTTTCATAAATTGTCCTCAATTAATCAACCTCGAGATCGAAACC

AGCTACGGTGATCGTGGTAAGGCCAACATTGTGGTTTCGGCACCAAAAATCCGCAATTTCACTTCTGTTG

GTATATTTACAATCACATTCGGAGCTTCTAAGTTGGATACTGTGAATATAAAGTTAAAGGATTGGATGGA

TTCTGGGAGCTACGAGTGGTTGCACCGGAAGGAGTTCAAACGGGATTATCAACGGTTTAAATTCATGCTA

CCAGGACTTGCCACTGCCAAGATTCTTAAACTTGACCTGGAAACCATAGAGGCACTTTCGTCGATATCCG

ACTTTCTTGGAAGTTCATCTCCATTCTTTAACTTGAAATATGTGAAACTTCCACATGGATGTAAAGAGTC

AAGTATATCTACTTCTCTAAGAAAGTACTTACTCGGTGCCTCTCCAACAGCCACTATAGTCACAACATTT

CCCCAGAACAACATCATTCCTTCTACAGCGACAAGTTCTCTGACAACCCAATATGTAGTACGAGAAGAGC

CCTTGGCTGCTCGAACCAAGGTATTGACTGATGTTGAAAGAACATATAAAAAAATGCGCAATGAAACTAT

GGACATGGGGGTGCAGGAAAAGCATGTTTTACATGCTGACAAGGTTAGAAATTCTGTTTACCTCATTGAA

GGTATTGGCAATGATCGTGTAACTTCTTCAAGGGGGAATAGTAATCTTGGGTTATGGAGAGGCCATGAAG

TTAATTCTGAATTTGTATGTCTACTTAATCGGATCATGAATAAGTACCCCGAAACCTTTGAGTCCTTCGC

CATAAAAAACAAGAAGTTTTGTACAGTGATGCTGAATATGTTGTGCACCTCAGTAAATGACTTTATCAAA

TTCACTATGACTGAAGTTGATAGAGAGATGATTGCTGAGTACAGCGATGTATTTGCTGACCTGAAGATAT

TAGGATTCAATGTTAGTTGGCTTGTGGGTCGTCTGAATTATATAGAACATCTCCAATTTTCACATCCCCT

ACTGACAGAGCTGTATGCACTGGATGGTCATCTTAATGATGCCAGAAGTAAGTTACAGTATTTGCAGACT

AGTGGTGATACCATAATCAAATTGCAAGACCTACAAACTCTTCGTGATGAGAAGATGCAGGAGATTGAGA

AAGCTTTTGGAACTTTGGGTACAAAACTTGTTGCTGGCTGTATCGGAGATGAGTTATTTATTTAGTCCCT

AATCTTCTTGGAATTATTTTTGTTTAAGAGTTCTGGATGCCTTCTTTACTTGTTTACTCTCTGGTCCTTG

AGTCTTTAGTATGTACGTGTCGCGTATATTGTCTTTGTGTTGATCTGTTAGCTGTGTTTGGTTTGCTACT

GTTTGTCAAGGTGTGAATCATGTAAGATGTTGCATTCACAAAAGGATTTTCTAGCATATTATATATTGAG

CTATGCTATATATGGTGCTATTTTTATAAATACTTCCTCAGACATTTCTGCAGTTCTTATTCTTGCAAGT

TTGTGTTCCCCATCCTATTAATTGATGTAAAAATACTGACATGGACATTGGGAATCCGGC

>MSTRG.1456.3 gene=MSTRG.1456

CCGACCGACCCAACCCGTCTTCACCACTGCTGTCCGGGTAGACACCAATTAGGGTTCTTCACTCCTTTCC

TAATTCAGTGTAACTAGAAATCACCTACTACTATCCATACATTTCTGTATTGTCTGAGTCTTCATCAGTC

TCTCAATGGATGGTCCTATGAAGCAGAAATCAAAAATCACAAGCGAAGATGAAGAGCAAGAAGACAGAAT

CAGCAGATTGCCAGACGATCTAATCCATCAAATCCTCAAGTTTGTAGACACCAAATTGGGTGTTCAAACC

AGCGCTCTCTCGAAACGATGGAAGCTTGTGTGGACTACTCTCCCATTCCTCAAATTCAGATGGGATCAAT

TTCGATACAACGGAAGCTCGTGTTCGTCGAAAAGCACAACAAATCTTGCCCGTCATGTTTTGAAGCACCG

CAACCATCAAGCCCAGATATCGTATTTGGAGCTGGCGTTCCTGCCTTCAGGTTTGTTCGCGAAGTTTGTT

GATTATGCTATTGCTCACAATGTGCAACACTTGAATGTTCATTTTCGAGAAAAGCATAAGCCTTACAAGT

TATCTAACTTTAATTGTGATTGTATTGAGAAACTCGAGATGAGAATGGAGCTTGATGATTCTCTGCGGGA

ATCGGATTGTTGGGATTTACCAGCTTTAACAAGTCTACGCTTGAGAGGGTACTTGGTTTCGGGTCGGCTA

GATAAGTTACCGGAAATGTGTTTGACCTGCTTGCCTGCGCTACGATCCTTGAATCTTGAAGAATGGAATT

TGGAGTGTTTTTCTTTCAGTTTGCCTAATTTAACAAATTTTCGGCTATGGGGCTGTCGTTTGCCCCAAAC

AATTTGGGATTTCCCAGCTTTAAAAAGTCTAGAGTTATGTGATGTGGAATTTCCCGAGAATATGAGTGAC

ATATTTGCTGCACTTGTCAGTCTACAGAATCTCACAATATATTTCTACGGGTTGTCTGCAAAAGATTGTT

TCATAAATTGTCCTCATTTAATCGACCTCGAGATCGAAACCAGCTACGGTGATCGTGGTAAGGCCAACAT

TGTGGTTTCGGCACCAAAAATCCGCAATTTCACTTCTGTTGGTATATTTACAATCACATTCGGAGCTTCT

AAGTTGGATACTGTGAATATAAAGTTAAAGGATTGGATGGATTCTGGGAGCTACGAGTGGTTGCACCGGA

AGGAGTTCAAACGGGATTATCAACGGTTTAAATTCATGCTACCAGGACTTGCCACTGCCAAGATTCTTAA

ACTTGACCTGGAAACCATAGAGGCACTTTCGTCGATATCCGACTTTCTTGGAAGTTCATCTCCATTCTTT

AACTTGAAATATGTGAAACTTCCACATGGATGTAAAGAGTCAAGTATATCTACTTCTCTAAGAAAGTACT

TACTCGGTGCCTCTCCAACAGCCACTATAGTCACAACATTTCCCCAGAACAACATCATTCCTTCTACAGC

GACAAGTTCTCTGACAACCCAATATGTAGTACGAGAAGAGCCCTTGGCTGCTCGAACCAAGGTATTGACT

GATGTTGAAAGAACATATAAAAAAATGCGCAATGAAACTATGGACATGGGGGTGCAGGAAAAGCATGTTT

TACATGCTGACAAGGTTAGAAATTCTGTTTACCTCATTGAAGGTATTGGCAATGATCGTGTAACTTCTTC

AAGGGGGAATAGTAATCTTGGGTTATGGAGAGGCCATGAAGTTAATTCTGAATTTGTATGTCTACTTAAT

CGGATCATGAATAAGTACCCCGAAACCTTTGAGTCCTTCGCCATAAAAAACAAGAAGTTTTGTACAGTGA

TGCTGAATATGTTGTGCACCTCAGTAAATGACTTTATCAAATTCACTATGACTGAAGTTGATAGAGAGAT

GATTGCTGAGTACAGCGATGTATTTGCTGACCTGAAGATATTAGGATTCAATGTTAGTTGGCTTGTGGGT

CGTCTGAATTATATAGAACATCTCCAATTTTCACATCCCCTACTGACAGAGCTGTATGCACTGGATGGTC

ATCTTAATGATGCCAGAAGTAAGTTACAGTATTTGCAGACTAGTGGTGATACCATAATCAAATTGCAAGA

CCTACAAACTCTTCGTGATGAGAAGATGCAGGAGATTGAGAAAGCTTTTGGAACTTTGGGTACAAAACTT

GTTGCTGGCTGTATCGGAGATGAGTTATTTATTTAGTCCCTAATCTTCTTGGAATTATTTTTGTTTAAGA

GTTCTGGATGCCTTCTTTACTTGTTTACTCTCTGGTCCTTGAGTCTTTAGTATGTACGTGTCGCGTATAT

TGTCTTTGTGTTGATCTGTTAGCTGTGTTTGGTTTGCTACTGTTTGTCAAGGTGT

>MSTRG.1456.4 gene=MSTRG.1456

ACTCCTTTCCTAATTCAGTGTAACTAGAAATCACCTACTACTACTATCCATTCATTTCTGTATTGTCTGA

GTCTTCATCAGTCTCTCAATGGATGGTCCTATGAAGCAGAAATCAAAAATCACAAGCGAAGATGAAGAGC

AAGAAGACAGAATCAGCAGATTGCCAGACGATCTAATCCATCAAATCCTCAAGTTTGTAGACACCAAATT

GGGTGTTCAAACCAGCGCTATCTCGAAAAGATGGAAGCTCGTGTGGACTACTCTCCCATTCCTCAAATTC

AGATGGGATCAATCTCGATACGATGGAAGCTCGTGTTCGTCTAAAAGCACAACAAATCTGGCCCGTCATG

TTTTGAAGCACCGCAACCATCAAGCCCAGATATCGTATTTGGAGCTGGCGTTCCTGCCTTCAGGTTTATT

CGCAAAGTTTGTTGATTATGCTATTGCTCACAATGTGCAACACTTGAATGTTCATTTTCGAGAAAAGCAT

AAGCCTTACAAGTTATCTAACTTTAATTGTGATTGTATTGAGAAACTCGAGATGAGAATGGAGCTTGATG

ATTCTCTGCGGGAATCGGATTGTTGGGATTTACCAGCTTTAACAAGTCTACGCTTGAGAGGGTACTTGGT

TTCGGGTCGGCTAGATAAGTTACCGGAAATGTGTTTGACCTGCTTGCCTGCGCTACGATCCTTGAATCTT

GAAGAATGGAATTTGGAGTGTTTTTCTTTCAGTTTGCCTAATTTAACAAATTTTCGGCTATGGGGCTGTC

GTTTGCCCCAAACAATTTGGGATTTCCCAGCTTTAAAAAGTCTAGAGTTATGTGATGTGGAATTTCCCGA

GAATATGAGTG

>MSTRG.1456.5 gene=MSTRG.1456

ACTCCTTTCCTAATTCAGTGTAACTAGAAATCACCTACTACTACTATCCATTCATTTCTGTATTGTCTGA

GTCTTCATCAGTCTCTCAATGGATGGTCCTATGAAGCAGAAATCAAAAATCACAAGCGAAGATGAAGAGC

AAGAAGACAGAATCAGCAGATTGCCAGACGATCTAATCCATCAAATCCTCAAGTTTGTAGACACCAAATT

GGGTGTTCAAACCAGCGCTATCTCGAAAAGATGGAAGCTCGTGTGGACTACTCTCCCATTCCTCAAATTC

AGATGGGATCAATCTCGATACGATGGAAGCTCGTGTTCGTCTAAAAGCACAACAAATCTGGCCCGTCATG

TTTTGAAGCACCGCAACCATCAAGCCCAGATATCGTATTTGGAGCTGGCGTTCCTGCCTTCAGGTTTATT

CGCAAAGTTTGTTAGTTATGCTATTGCTCACAGTGTGCAACACTTGAATGTTCATTTTCGAGAAAAGCAT

AAGCCTTACAAGTTATCTAACTTTAATTGTGATTGTATTGAGAAACTCGAGATGAGAATGGAACTTGATG

ATTCTCTGCGGGAATCGGATTGTTGGGATTTACCGGCTTTAGCAAGTCTACGCCTGAGAGGGTACTTGGT

TTCGGGTCGGCTAGATAAGTTACCGGAAATGTGTGTGACCTGCTTGCCTGCGCTACGATCCTTGAATCTT

GAAGAATGGAATTTGGAGCGTTTTTCTTTCAGTTTGCCTAATTTAACAAATTTCCGGCTATGGTGCTGCC

GTTTGCCCCAAACAATTTGGGATTTCCCAGTTTTAAAAAGTCTAGAGTTATGTAGTGTGGAATTTCCCGA

GAATATGAGTGACATATTTGCTGCACTTGTCAGTCTACAGAATCTCACAATATATTTCTACGGGTCGCAT

GCATTGTCTGCAAAAGATTGTTTCATAAATTGTCCTCAATTAATCAACCTCCAGATTATAACCAGCTACA

GTGATCGTGGTAAGGCCAACATTGTGGTTGCGGCACCAAAAATTCGCAATTTCACATCGGTTGGTATATT

TACAATCACATTCGGAGCTTCTAAGTTGGATACTGTGAATATAAAGTTAAAGGATTGGATAAATTCCGGT

AGCTATAAGTGGTTGCACCAGAAGGAGTTGAAACAGGATTATCAACGGTTTAAATTCATGCTACCAGGAC

TTGCCACTGCCAAGATTCTTAAACTTGACTTGGAAACCATAGAGGCACTTTCATTGATATCCGACTTTCT

TGGAAGTTCATCTCCATTCTATAACTTGAAGTATGTGAAACTTCCACATGGATGTAAAGAGTCAAGTATA

TCTACTTCTCTAAGAAAGTACTTACTCGGTGCCTCCCCAACAGCCACTATAGTCACAACATTTCCCCAGA

ACAACATCATTCCTTCTACAGCGACAAGTTCTCTGACAACCCAATATGTAGTACGAGAAGAGCCCTTGGC

TGCTCGAACCAAGGTATTGACTGATGTTGAAAGAACATATAAAAAAATGCGCAATGAAACTATGGACATG

GGGGTGCAGGAAAAGCATGTTTTACATGCTGACAAGGTTAGAAATTCTGTTTACCTCATTGAAGGTATTG

GCAATGATCGTGTAACTTCTTCAAGGGGGAATAGTAATCTTGGGTTATGGAGAGGCCATGAAGTTAATTC

TGAATTTGTATGTCTACTTAATCGGATCATGAATAAGTACCCCGAAACCTTTGAGTCCTTCGCCATAAAA

AACAAGAAGTTTTGTACAGTGATGCTGAATATGTTGTGCACCTCAGTAAATGACTTTATCAAATTCACTA

TGACTGAAGTTGATAGAGAGATGATTGCTGAGTACAGCGATGTATTTGCTGACCTGAAGATATTAGGATT

CAATGTTAGTTGGCTTGTGGGTCGTCTGAATTATATAGAACATCTCCAATTTTCACATCCCCTACTGACA

GAGCTGTATGCACTGGATGGTCATCTTAATGATGCCAGAAGTAAGTTACAGTATTTGCAGACTAGTGGTG

ATACCATAATCAAATTGCAAGACCTACAAACTCTTCGTGATGAGAAGATGCAGGAGATTGAGAAAGCTTT

TGGAACTTTGGGTACAAAACTTGTTGCTGGCTGTATCGGAGATGAGTTATTTATTTAGTCCCTAATCTTC

TTGGAATTATTTTTGTTTAAGAGTTCTGGATGCCTTCTTTACTTGTTTACTCTCTGGTCCTTGAGTCTTT

AGTATGTACGTGTCGCGTATATTGTCTTTGTGTTGATCTGTTAGCTGTGTTTGGTTTGCTACTGTTTGTC

AAGGTGTGAATCATGTAAGATGTTGCATTCACAAAAGGATTTT

>MSTRG.1456.6 gene=MSTRG.1456

GAAGTTTTGTACAGTGATGCTGAATATGTTGTGCACCTCAGTAAATGACTTTATCAAATTCACGATGACT

GAAGTTGATAGAAAGATGATTGCTGAGTACAGGGATGTATTTGCTGACCTGAAGATATTAGGATTCAATG

TTAGTTGGCTTGTGGGTCGTCTGAATTATATAGAACATCTCCAATTTTCACATCCCCTACTGACAGAGCT

GTATGCACTGGATGGTCATCTTAATGATGCCAGAAGTAAGTTACAGTATTTGCAGACTAGTGGTGATACC

ATAATCAAATTGCAAGACCTACAAACTCTTCGTGATGAGAAGATGCAGGAGATTGAGAAAGCTTTTGGAA

CTTTGGGTACAAAACTTGTTGCTGGCTGTATCGGAGATGAGTTATTTATTTAGTCCCTAATCTTCTTGGA

ATTATTTTTGTTTAAGAGTTCTGGATGCCTTCTTTACTTGTTTACTCTCTGGTCCTTGAGTCTTTAGTAT

GTACGTGTCGCGTATATTGTCTTTGTGTTGATCTGTTAGCTGTGTTTGGTTTGCTACTGTTTGTCAAGGT

GTGAATCATGTAAGATGTTGCATTCACAAAAGGATTTT

>MSTRG.1457.1 gene=MSTRG.1457

GACTGATAGGCTGCAACGCAGCTAATCAAGGCAACAGTGGAGGTGTAGACAGGATCAGCAGTTTGCCAGA

TCAACTACTTCATCACATCCTCTCCTTCTTGGAAACTCGATTGGCGGTTCAAACCAGCGCTCTCTCGAAA

CGATGGAAGCTTATTTGGACGACTCTGCCTATTCTCAATTTCACTTGGTCTTGGTCTAATATTAGATCCG

GTAATAAGTTCATTAACAATATCTTTGCTCGTCGAAACAATAATTCTGATGTATTGAAACTCAATCATTT

TTTCTTTTATGAATCCCCGCAATTTCATGGGGTTCAGAAGCTTGCAAATCATGCAGTTTCACATAATGTT

CGTCAAGTAAATGTAGATGTTACTGGGGCTGATTGTATTGTGACTATGAAATTAACAGGACCTAATGCCA

TCGTATTGAATTTAAAGCTACCCTTTGATCAATTCCTGAAAAAATTAGATTCTTGTTGGCGTTTACCCGC

TTTAACAACTCTACACTTGGTACGCACTGATTATTCGAAACCCTACAAGTTACCTGCATCATATTTGGTT

GACTTGCCTGCTTTGCGGACTCTGTTACTTGATGGTTTCGAGTTACCAAGTTCTGTTAGCCTGCCTGCTT

TGCATACTCTCATACTGCATATTGTCGAATTCCCTGAAAATATGAGTGAGTTCTGCCGGGTCTTCCAGGC

GCTTGGAAACCTTCAGAATCTCACATTATTTTTCTGGACCACCTTAAGGTGTGATTGTGTTATAGATTGT

CCCCAGTTGATTAACCTGGAAATCAATTACTGCACTAGCAAATCAGGAAAGAGTATAGTGTCTAGTAATT

CTGGTAAAATAATGGTTTTGACTCCGAATATCCGCAACTTCAGTTCTGTTGGCATCTTCCCAATCGAATT

TGGAGCTTTTGACTTGGAAAATGTGAATATACAATTACGGGATAGCGCTGTAATCAAGACCGTTGCACCA

TCTAAGATGAAGGAAAATTTTAAGAATCAGTTTACTGATATGTTTTGGGGACTGTCCCGCACAAAAATTC

TTACACTTGACTCGAACATCATTGAGGCACTATCTGCTCTATCGGACTTTCTTGTTCAGACATCCTCTCC

TTTTATTATGTTAGAGTATGTGAAGGTACCGCATGGATATAAAGAATCAAGTATGTCCACATACCTTAAA

TGGTTCCTACTTGGTCGCTGTCGGAAAGCCACTATTGTCACTACATTGCCTCAGAAAATCGAGATCAAAG

AGCCAGCATCTTCGGCAGCGAAGAATGTGCCACCACAAGACCCCTTGACAGCTCCAACAAAGGCACGGGC

TGGCTCTAAAAAGGTGCATTCAGAAGTAAATGCTGACAGGGCGACACAGATTGGTGTTACAGTGGAAGGG

TAGGGCAATGACGGGGTGACTTCTTCTAGAAGGGATAGTGATTTCGGGTTATGGCAGGGCCATGAGGTGA

ACTCTGAGTTTGCACGTCTACTCCATAACATTATGAATAACTACCCTGAAACCTTTGAGCACTTTACAAC

CTCAAGAAGTTATCGACTATGAAGCTGAATATGTTATGCACCTCAGTGAATGTGTTTACCAAAATCTCTA

TGACTCAGGTTAATTCTGAGATGATTGCCGAGTACAGGGATGTATTTGCCGGCTTGCAGAAATTCGGGTT

CAACGTAAGTTGGCTTGTGAGCCGTCTAAACTATATTGAAAAGCTCCAGAATTCACAACCTCTTTCCGAG

CTTCATGAGATTGACTGCCATATCAATAGTGCCAAAAGTAAGTTACAAGATCTAAAGACTCTTCGTAAGA

AGAAGATGCAAGAGATTCAGAAAGCTTTTGGAACTATGGGCACAAACCTTGCTGTTGGCTGCATTGGGAA

TGATCTCCTCCGTGCTCCTGAATTTAATGGAAATTGATGTGTGTTTGAAGGTTGGTTAAATGTTGCACCG

TCTGGTTGTCAAGTCTTTAGTTACTTCTTGACTTTAAACCGAACTGTTTTTGGTGTATGGTTTTTGCTTA

ATTGTCATGGAAGTCTGTATGGTGCTTGGTATCTTGTACAACTCCATCCCAAAAAGGGGTGCCTGCATTT

ACTTTGCCAAAGTAATGTGGTTTGGTGACAAGTTATTGTTGATTTTG

>MSTRG.1457.2 gene=MSTRG.1457

CTCGTCGAAACAATAATTCTGATGTATTGAAACTCAATCATTTTTTCTTTTATGAATCCCCGCAATTTCA

TGGGGTTCAGAAGCTTGCAAATCATGCAGTTTCACATAATGTTCGTCAAGTAAATGTAGATGTTACTGGG

GCTGATTGTATTGTGACTATGAAATTAACAGGACCTAATGCCATCGTATTGAATTTAAAGCTACCCTTTG

ATCAATTCCTGAAAAAATTAGATTCTTGTTGGCGTTTACCCGCTTTAACAACTCTACACTTGGTACGCAC

TGATTATTCGAAACCCTACAAGTTACCTGCATCATATTTGGTTGACTTGCCTGCTTTGCGGACTCTGTTA

CTTGATGGTTTCGAGTTACCAAGTTCTGTTAGCCTGCCTGCTTTGCATACTCTCATACTGCATATTGTCG

AATTCCCTGAAAATATGAGTGAGTTCTGCCGGGTCTTCCAGGCGCTTGGAAACCTTCGGAACCTCACATT

ATTTTTCTGGACAACCTTAAGGTGTGATTGTGTTATAGATTGTCCCCAGTTGGTCAATTTGGAAATCAAT

TACTGCACTAGCAAATCAGGAAAGAGTATAGTGTCTAGTAATTCTGGTAAAATACTGGTTTTGGCTCCGA

AAATCCTCAATTTCAGTTCTGTTGGCATCTTCCCAATCAAATTTGGAGCTTTTGACTTGGAGAATGTGAA

TATACAATTACGGGATAGTGCTGTAATCAAGACTGTTGCACCATCTAAGACGAAGAAAATTTTTCATAAG

CAGGTTACTCATATGTTAGTGGGACTGTCCCGCACAAAAATTCTTGCACTGGACTCGAACACCATTGAGG

CACTCTCTGCTCTTTCGGACTTTCTTGTTCAAACTTCCTCTCCTTTTTTTATGTTAGAGCATGTGAAGGT

ACCGCATGGATATGAGGAATCAAGTATGTCCACATACCTCAAATGGTTCCTACTTGGTTGTTGTCCAAAA

GCCACTATCATCACTATATTGCCTCAGAAAATCGAGATCATAGAGCCAGCATCTTTGGCAGCTCAGAATG

TGCTACCACAAGGCCCCTTGACAGATCCAACAAAGGTACTGGTTGGCTCTAAAAAGTTGCATACAAAAAT

AAATGATGACAGGGCGACACAGTTTGATGTTACAGTGGAAGGGTTGGGCAATGACGGGGTGAGCTCTTCT

AGAAGGGATAGTGATTTTGGGTTATGGCAGGGCCATGAAGTGAAGTCCGAGTTTGCACGTCTACTCCATA

ACATTATGAATAACTACCCTGAAACCTTTGAGCACTTTACCACCTCAGACAAGAAGTTATCAACTATGAA

GCTGAATATGTTATGCACCTCAGTGAATGTGTTTACCAAAATCTCTATGACTCAGGTTGATACTGAGATG

ATTGCCGAGTACAGGGATGTATTTGCTGGCTTGCAGAAATTCGGGTTCAACGTAAGTTGGCTTGTGAGCC

GTCTAAACTATATTGAAAAGCTCCGGAATTCACAACCTCTTTCTGAGCTTCATGTGATTGACTGCAATAT

CAATAGTGCCAAAAGTAAGTTACAAGATCTTGAGACTCTTCGTAAGAAGAAGATGGAAGAGATTCAGAAA

GCTTTTGGAAGTATGGGAACAAACCTTGCTGTTGGCTGCATTGGGAATGATCTCCTCCGTGCTCCTGAAT

TTAATGGAAATTGATGTGTGTTTGAAGGTTGGTTAAATGTTGCACCGTCTGGTTGTCAAGTCTTTAGTTA

CTTCTTGACTTTAAACCGAACTGTTTTTGGTGTATGGTTTTTGCTTAATTGTCATGGAAGTCTGTATGGT

GCTTGGTATCTTGTACAACTCCATCCCAAAAAGGGGTGCCTGCATTTACTTTGCCAAAGTAATGTGGTTT

GGTGACAAGTTATTGTTGATTTTGTTAGACTATGCATTACTAGCAATACTAATTATTGTTCCTTTTGTCA

GAATTATACATTAGTGTCG

>MSTRG.1457.3 gene=MSTRG.1457

TCTGCTCTATCGGACTTTCTTGTTCAGACATCCTCTCCTTTTATTATGTTAGAGTATGTGAAGGTACCGC

ATGGATATAAAGAATCAAGTATGTCCACATACCTTAAATGGTTCCTACTTGGTCGCTGTCGGAAAGCCAC

TATTGTCACTACATTGCCTCAGAAAATCGAGATCATAGAGCCAGCATCTTTGGCAGCTCAGAATGTGCTA

CCACAAGGCCCCTTGACAGATCCAACAAAGGTACTGGTTGGCTCTAAAAAGTTGCATACAAAAGTAAATG

CTGACAACAATGTTGACATGGAAGTGCAAGAAGAGCCCGGGGTGCAGAATTCTGCAGTAAATGCTGAGAG

GGTTGAACAATTAAGTGGTCCAGTTGAAGGGACTGCCATTGGCCGGGTGAGCTCTTCTGGGGGAGATGGT

GATTTTGTGTTATTGCAGGGGCATGAAGTTAACTCTGAATGACCGGGTGAGCTCTTCTAGAGAAGATAGT

GATTTCGGGTGATGGCAGAGACATGGAGTTCTCCGAGTTTACATGTCTACTCGATAATATTATGAATAAG

TACTGAAAAACCTTTGAACACTTTATGAGCCCTTTTGATTGATCGTCATATTGATGATGCCAAAAGTAAG

TTACAAGATTCACAAACTCATACTGATAATACCATAACTAGTTGCTAGAACTACAGACTCTTCTTGCGAG

GAAGATGCAAGAGATTCAAAAAGCTTTTAAAAGCTTTTGGAACTATGCGCGCAAAGCTTATATTTGGCTA

CATTAGAAATGATCGTCCATTTTGACAATATTATTTGTAAGGTTGAGACTTGTATGGATACCTTAGTATC

TTATCTGTTATTTGATTTT

>MSTRG.1457.4 gene=MSTRG.1457

TCTTCTAGAAGGGATAGTGATTTCGGGTTATGGCAGGGCCATGAGGTGAACTCCGAGTTTGCACGTCTAC

TCCATAACATTATGAATAACTACCCTGAAACCTTTGAGCACTTTACCACCTCAGACAAGAAGTTATCAAC

TATGAAGCTGAATATGTTATGCACCTCAGTGAATGTGTTTACCAAAATCTCTATGACTCAGGTTGATACT

GAGATGATTGCCGAGTACAGGGATGTATTTGCTGGCTTGCAGAAATTCGGGTTCAACGTAAGTTGGCTTG

TGAGCCGTCTAAACTATATTGAAAAGCTCCGGAATTCACAACCTCTTTCTGAGCTTCATGTGATTGACTG

CAATATCAATAGTGCCAAAAGTAAGTTACAAGATCTTGAGACTCTTCGTAAGAAGAAGATGGAAGAGATT

CAGAAAGCTTTTGGAAGTATGGGAACAAACCTTGCTGTTGGCTGCATTGGG

>MSTRG.1457.5 gene=MSTRG.1457

GGTCATCAAACTGCCAAAATCGAAACCCAAAACATACCTCTTCTCGTATTGATCGCTTGTAATTGTACTC

CTCACTAAGCATATCCAGCATGCCTATATCAAATCGAAGATGAAGAAGAAGATAGAATTACTAGATTGAT

GATCAATTACCTAGTGATGATGTAGCTTGTAAGCTTTTCTAACCTTTTTAGTAACATTTTTTTAGAATTA

TTATAAATTGTTGCCTTAATTACTCCATTTACACCTTGATCATATAGTCTCTGCAAATTGTTGTGGTTGC

TAATCGGAAAAAGGCTCGTTGAATCCCTGAGTATTCATTTGAATTCCACACAACGTTTACTGACTTTTTG

CTGATTAGTTCTCTTCGTATTTGTTATCAATCATATATTGTGATTTATTTTGATGTTCACTGCATTATAT

CGGATGATGAATTTGCAATTGAGATATGGATTAAAAAGGTAACTAGAATTACGAATGATTCTTCTGTATC

GTGTGTAGTATTACGTGATGATTTTGTTCTCAAATACCAAAGCCTGGTAAGCCGCTCTAGACGGTCCCAA

CACGAATGTGCAAATCTTGACGGTTTTGTTTCTTATGTGCGCAAGAGAGGTGAAGGTAAATTATTTTTAT

ATCTGTTTGATGTGAATGTTGTTTTTGTGTCTCTTCTTGCTAAAGCTGACTATAAGCTAGAATTTAATAA

CGGTTAAGTTTTATACAAGAGAATGGATCTTTTGTGGGCAAAGGTCAGATTTTTAATAGTTTGTTTCATT

TAATTGTTAATAAAGTTACGTCGCTTAATACTAGTTATGGCCTCTATTTGTTTGATAGAAGATGAAAGTG

TTTACCATACAAAGTTAGGCCAAACTAATAATGAATACATTAATAGCTCGGATTATTGGTTTTTGAAGGA

CATTAAGTTAATATAGAGTCAAAGTCCACTAACAGAACCTTGCAAGACAATAAAATTCGGAGAATGTAAA

TCTTCTGTTTAATTCATTCTGATTGATTGTCTAATACATCAAGAGGCTGTAAAAGTAGCAACTTTTGTGG

ATGACTTCTCTACGGATACACTTTAGCTTATTTTGTTGAGGAACAAATATGAAGCAGAAATAAATTTATA

GTTTATAAAGGTTAAGCTGAAAGTAAATTGGATAAGAAAATTATTATCTTAAAACAAATAAAGGCAGAGC

AGTTGGTTTTGATCAATTTTTTCGGGCACTGTGAGGAAAATGGTATAGCTTAAACACTCCCTCTAACGGA

GCAACAGAAAGAAGAAAAACAGGACTTGAAAGAAATGACTAATTTCTTGCTAGTTAGATCCGATGCTCTG

TTATCTCTATGAGGTGAGGCCTTCATCAGTTATTGACATATTTACTAAAGAATATAGGAAAGACAGCATA

TGAACTTTGTGAGACCTAAACACATGATCCTAATTTTTTGATAGTCTGGGGTGTCTTGCTAAGGTGTTGT

AACGAGACCTAAAAATTGGTAAATTGGTTGTAAAACTACGAATTGTTTGTTTTTCAGAAAATGGTACTGG

TTAATGCTGATTATTTTCTAGTGATTTATTAAGAATGTATTTCCTTTAAAAGTTAGAATTAGTGGATCCC

CAAATGATCTTCTAGAGACCTCTATTAGACACCTCCATTAGACAAGTAGCTAAAGATTTTTGTGATACAG

AAAGTCAGAGGATAAAATGTGCTTGGTTATTGTGGTGATTATGTTTATTTGTTCTTAGTGGATAATGATC

CGTTAACTCGATTTACAGTTAAATTTACCTAAACCATGTTTGACCACCTTCCTTTATAAATTGTCCTACT

CTTTAGGATTTACGAAAATCTTTTTTATCATTTTGTTTGCCCTGTCATTAACTCTCTGTCTAGTTTGACA

CCATGTTTCTTATTTGTTTATACCTTATAGTGATATTACCGTCAGCCGGAGAAACAGATGGTGTTCGATA

ATGTTAAATTGCTAAGTTATTTCTTAGCGTGCCTTCATTGTAAATTTAGTTATGCATTGTTAAGTAATCC

AACCTTTCAAGTCATGTTCAGTTAAACCTGTCCTATATATCAGTATCCTGTATCACTATAGGTTGTTAGA

AGTGTTTTTCAATCTTTAAAGGAAGTGAGTTATAAATTGTTTGTTATGAAGACTTGTTATACATTCACTC

TAGTAGTGAACACATAGTTATATCTTTAGCTATTGTATGTACTAAGCATCATTTGTACCTTTGAGCTTTA

ATATTCCATTCAAAGATGACAATCAAAACTTCACCGACCTTTAAGGTTAGACAATCAAAAATTCAACGAC

CTTTGAGGTTACTTGTGTGCAGTGACCCGGGGGGGGGCCCTGGCCCTATATATACCAACAAAACTCTATC

TGTCTCTCGCTTGGTATGGTTTGCGTAATGTTGATGTTGGGAAAGTGGTTACAGCAAATCATCAGTTAAC

ATCTCTTGACATTCAAAAGCAAATCGTTAGTGCTTGTGCATCCGAAACAAAGAAAGCTGTGATTCGTGAT

ATTGGAGTAGTATTGGAGTATTTCTCTTTATTACTTGGCAATGCCCGTGATAATTTTGTTAAAGAACAAA

TGGCTGTAGTTATAAGATACATAAATAATTATGGCTGTGTTTTGGAACGGTAGTTGTCCATGTTAAAGAT

ACTACTGCACTTTCTTTAGAGAGTGCTAAAATTTCTTTTCCAAACATGGATTGTCGTCTGTCGATATCAA

AAACTAGAGGCAAGGATATGACGGAGCTCCGAACATGCGGGGGGAGCTTAATGGTTTCAACTGGTATATA

TATGAAGGCCCTCCTCGGGAAAACTTCCTGCCTCCGTCACTGCTTGTGTGCTTGTTGGCTCTCCTAAATT

TGGGTTCAGTGTTTGATTTTCTTAAATTGGTCATTTGGCTAAGAGGGCAAAAGATATTATTGGCATAGAA

TATTGATATTTACTACAATTTTTACTATATATGATTAACAAAATATAAACTTTTTTTAGACTGAATTGGT

AAAAGCTATAAGTTGGCTAGGAATTGCCGTATAAAAGATTCGGATATTCGAAACAGTGTGAATCTCTTTT

CCATGCGATTCGTATACTGGGTTGGTTTTGTTATCCTTTCCCAGTACACCTCTATTAAACAGACATTGCA

TTCCATTTCATACTTTATAAATTAGGGCTAGAGACGAGTAGTGAATAAGATACAGTATGTCATTGAGTCG

CCCCTTACGGAACACAAAAGCCAGAATCACAGTTGAAGATGAAGAGGACAGAATTAGCAGGTTGCCGGAT

GAGCTAATTCATCAAATTCTCGAGTTCGTGGATACCAAATTGGCTATTCAAACCAGTCTTCTATCTAAGC

GATGGAAACTTATATGGAAGACTCTCCCTTTTCTCAAGTTTAGATGGGTGCAACAGGGTTCACCTACAAA

CACAAATAATTTTACCCGTCAAGTTTTAAGTCGCCGAAACCATCAATCCCAGATTTCCAGTCTGGAGCTT

GCTTTCCTCCCTGCAGGTTTATTTGCAAAGTTTGTTGAGTATGCTATATCTCACCATGTTCAATACTTAA

ATGTTCATTTACGTCGTGATCATAAACCTTATAAACTATCCAACTTTAGCTCTAATTCGATTCAGAAACT

TCAATTGGAGATGAAGTTTGGAGATTCTATTTCAGAATTAGATTGTTGGGATTTACCTGCATTAAACACT

CTGCACTTGACAAGTTACTATAATAATCATGTCGTAGGAATCCTATCAGAATTATGTTTAACCTGCTTGC

CTGCTTTAAGAACTCTGTGCCTTGAATATTTTGATTTGAAGTCCATGTCTATTAGTTTGTCTTCTTTAAC

AACTCTTCGGCTGTTTAAGTGCAAGTTGCCCATTATAGTATGGAAATTTCCAGCTTTGAAAAGTCTAACG

CTTCATGATGTGGTGTTTCATCGAGATATGAGTCACATGTTCGCTGCACTTGTCACTCTGCAAGATCTCA

CATTAGTTTTTGAAAAGGAGCATATACAAGATTGTGTTATACCCTCTCCTCAACTAGAGAATCTGGAGAT

CAGGTCCCGCTACGTATCCTACACTCACAGGATTGTGGGTAACATTGTGGTTGTGGCACCAAAACTCAGA

AATTTTACTTCTGTTGGTATCTTTCAAATCATATTTGGAGGTTCTAAGGTGGATAATGTGAATATAAGAT

TGCGTGGTTGTTTTGATGGCATGATCTATCCATTGTTAAAGGAATATTATCAACAGTTAACATTTATGCT

TCCAGGCTTTGCTAGTGCCAAGATTCTTAATCTTGATCTCGAGACCTTGCAGGCACTTTCTTCCATTGTG

GACCGTCTTGAAATTTCTCCGTCTCCATTCCATAACTTGGAGTATGTGAAACTACCTTGTGGATTTGTAG

AATTTGAAGAAAAACGTGTTGCTGATATCTTACGAAGATACTTACTTGGTGGCTCTCCAACAGCAACTTA

TGTCACTACATCATTCCAGAACATGATGCCTGCTACACCGGCAGCTTCTGTGACAGCTCAGAATGTGGTG

CTACAGGAGCCCTTGGCTGCTTCTACCAAGGTGTTGGTTGGTTCTAAAAATATAAATAAAAATGTATGCA

TTGACAATGTTGACATGGGGATGCACGAAGAGCCCGTACTGCAGAATTCTGCAGTAAATGCTGACAACAA

TGTTGACATGGAAGTGCAAGAAGAGCCCGGGGTGCAGAATTCTGCAGTAAATGCTGAGAGGGTTGAACAA

TTAAGTGGTCCAGTTGAAGGGACTGCCATTGGCCGGGTGAGCTCTTCTGGGGGAGATGGTGATTTTGTGT

TATTGCAGGGGCATGAAGTTAACTCTGAATGACCGGGTGAGCTCTTCTAGAGAAGATAGTGATTTCGGGT

GATGGCAGAGACATGGAGTTCTCCGAGTTTACATGTCTACTCGATAATATTATGAATAAGTACTGAAAAA

CCTTTGAACACTTTATGAGCCCTTTTGATTGATCGTCATATTGATGATGCCAAAAGTAAGTTACAAGATT

CACAAACTCATACTGATAATACCATAACTAGTTGCTAGAACTACAGACTCTTCTTGCGAGGAAGATGCAA

GAGATTCAAAAAGCTTTTAAAAGCTTTTGGAACTATGCGCGCAAAGCTTATATTTGGCTACATTAGAAAT

GATCGTCCATTTTGACAATATTATTTGTAAGGTTGAGACTTGTATGGATACCTTAGTATCTTATCTGTTA

TTTGATTTTGATCCTTGTTCTTCAAGTCTGGTAACTTTATGTTA

>MSTRG.1457.6 gene=MSTRG.1457

GGTCATCAAACTGCCAAAATCGAAACCCAAAACATACCTCTTCTCGTATTGATCGCTTGTAATTGTACTC

CTCACTAAGCATATCCAGCATGCCTATATCAAATCGAAGATGAAGAAGAAGATAGAATTACTAGATTGAT

GATCAATTACCTAGTGATGATGTAGCTTATATGAGTCACATGTTCGCTGCACTTGTCACTCTGCAAGATC

TCACATTAGTTTTTGAAAAGGAGCATATACAAGATTGTGTTATACCCTCTCCTCAACTAGAGAATCTGGA

GATCAGGTCCCGCTACGTATCCTACACTCACAGGATTGTGGGTAACATTGTGGTTGTGGCACCAAAACTC

AGAAATTTTACTTCTGTTGGTATCTTTCAAATCATATTTGGAGGTTCTAAGGTGGATAATGTGAATATAA

GATTGCGTGGTTGTTTTGATGGCATGATCTATCCATTGTTAAAGGAATATTATCAACAGTTAACATTTAT

GCTTCCAGGCTTTGCTAGTGCCAAGATTCTTAATCTTGATCTCGAGACCTTGCAGGCACTTTCTTCCATT

GTGGACCGTCTTGAAATTTCTCCGTCTCCATTCCATAACTTGGAGTATGTGAAACTACCTTGTGGATTTG

TAGAATTTGAAGAAAAACGTGTTGCTGATATCTTACGAAGATACTTACTTGGTGGCTCTCCAACAGCAAC

TTATGTCACTACATCATTCCAGAACATGATGCCTGCTACACCGGCAGCTTCTGTGACAGCTCAGAATGTG

GTGCTACAGGAGCCCTTGGCTGCTTCTACCAAGGTGTTGGTTGGTTCTAAAAATATAAATAAAAATGTAT

GCATTGACAATGTTGACATGGGGATGCACGAAGAGCCCGTACTGCAGAATTCTGCAGTAAATGCTGACAA

CAATGTTGACATGGAAGTGCAAGAAGAGCCCGGGGTGCAGAATTCTGCAGTAAATGCTGAGAGGGTTGAA

CAATTAAGTGGTCCAGTTGAAGGGACTGCCATTGGCCGGGTGAGCTCTTCTGGGGGAGATGGTGATTTTG

TGTTATTGCAGGGGCATGAAGTTAACTCTGAATGACCGGGTGAGCTCTTCTAGAGAAGATAGTGATTTCG

GGTGATGGCAGAGACATGGAGTTCTCCGAGTTTACATGTCTACTCGATAATATTATGAATAAGTACTGAA

AAACCTTTGAACACTTTATGAGCCCTTTTGATTGATCGTCATATTGATGATGCCAAAAGTAAGTTACAAG

ATTCACAAACTCATACTGATAATACCATAACTAGTTGCTAGAACTACAGACTCTTCTTGCGAGGAAGATG

CAAGAGATTCAAAAAGCTTTTAAAAGCTTTTGGAACTATGCGCGCAAAGCTTATATTTGGCTACATTAGA

AATGATCGTCCATTTTGACAATATTATTTGTAAGGTTGAGACTTGTATGGATACCTTAGTATCTTATCTG

TTATTTGATTTTGATCCTTGTTCTTCAAGTCTGGTAACTTTATGTTA

>MSTRG.1457.7 gene=MSTRG.1457

TCAAACTGCCAAAATCGAAACCCAAAACATACCTCTTCTCGTATTGATCGCTTGTAATTGTACTCCTCAC

TAAGCATATCCAGCATGCCTATATCAAATCGAAGATGAAGAAGAAGATAGAATTACTAGATTGATGATCA

ATTACCTAGTGATGATGTAGCTTTATTACGTGATGATTTTGTTCTCAAATACCAAAGCCTGGTAAGCCGC

TCTAGACGGTCCCAACACGAATGTGCAAATCTTGACGGTTTTGTTTCTTATGTGCGCAAGAGAGGTGAAG

GTAAATTATTTTTATATCTGTTTGATGTGAATGTTGTTTTTGTGTCTCTTCTTGCTAAAGCTGACTATAA

GCTAGAATTTAATAACGGTTAAGTTTTATACAAGAGAATGGATCTTTTGTGGGCAAAGGTCAGATTTTTA

ATAGTTTGTTTCATTTAATTGTTAATAAAGTTACGTCGCTTAATACTAGTTATGGCCTCTATTTGTTTGA

TAGAAGATGAAAGTGTTTACCATACAAAGTTAGGCCAAACTAATAATGAATACATTAATAGCTCGGATTA

TTGGTTTTTGAAGGACATTAAGTTAATATAGAGTCAAAGTCCACTAACAGAACCTTGCAAGACAATAAAA

TTCGGAGAATGTAAATCTTCTGTTTAATTCATTCTGATTGATTGTCTAATACATCAAGAGGCTGTAAAAG

TAGCAACTTTTGTGGATGACTTCTCTACGGATACACTTTAGCTTATTTTGTTGAGGAACAAATATGAAGC

AGAAATAAATTTATAGTTTATAAAGGTTAAGCTGAAAGTAAATTGGATAAGAAAATTATTATCTTAAAAC

AAATAAAGGCAGAGCAGTTGGTTTTGATCAATTTTTTCGGGCACTGTGAGGAAAATGGTATAGCTTAAAC

ACTCCCTCTAACGGAGCAACAGAAAGAAGAAAAACAGGACTTGAAAGAAATGACTAATTTCTTGCTAGTT

AGATCCGATGCTCTGTTATCTCTATGAGGTGAGGCCTTCATCAGTTATTGACATATTTACTAAAGAATAT

AGGAAAGACAGCATATGAACTTTGTGAGACCTAAACACATGATCCTAATTTTTTGATAGTCTGGGGTGTC

TTGCTAAGGTGTTGTAACGAGACCTAAAAATTGGTAAATTGGTTGTAAAACTACGAATTGTTTGTTTTTC

AGAAAATGGTACTGGTTAATGCTGATTATTTTCTAGTGATTTATTAAGAATGTATTTCCTTTAAAAGTTA

GAATTAGTGGATCCCCAAATGATCTTCTAGAGACCTCTATTAGACACCTCCATTAGACAAGTAGCTAAAG

ATTTTTGTGATACAGAAAGTCAGAGGATAAAATGTGCTTGGTTATTGTGGTGATTATGTTTATTTGTTCT

TAGTGGATAATGATCCGTTAACTCGATTTACAGTTAAATTTACCTAAACCATGTTTGACCACCTTCCTTT

ATAAATTGTCCTACTCTTTAGGATTTACGAAAATCTTTTTTATCATTTTGTTTGCCCTGTCATTAACTCT

CTGTCTAGTTTGACACCATGTTTCTTATTTGTTTATACCTTATAGTGATATTACCGTCAGCCGGAGAAAC

AGATGGTGTTCGATAATGTTAAATTGCTAAGTTATTTCTTAGCGTGCCTTCATTGTAAATTTAGTTATGC

ATTGTTAAGTAATCCAACCTTTCAAGTCATGTTCAGTTAAACCTGTCCTATATATCAGTATCCTGTATCA

CTATAGGTTGTTAGAAGTGTTTTTCAATCTTTAAAGGAAGTGAGTTATAAATTGTTTGTTATGAAGACTT

GTTATACATTCACTCTAGTAGTGAACACATAGTTATATCTTTAGCTATTGTATGTACTAAGCATCATTTG

TACCTTTGAGCTTTAATATTCCATTCAAAGATGACAATCAAAACTTCACCGACCTTTAAGGTTAGACAAT

CAAAAATTCAACGACCTTTGAGGTTACTTGTGTGCAGTGACCCGGGGGGGGGCCCTGGCCCTATATATAC

CAACAAAACTCTATCTGTCTCTCGCTTGGTATGGTTTGCGTAATGTTGATGTTGGGAAAGTGGTTACAGC

AAATCATCAGTTAACATCTCTTGACATTCAAAAGCAAATCGTTAGTGCTTGTGCATCCGAAACAAAGAAA

GCTGTGATTCGTGATATTGGAGTAGTATTGGAGTATTTCTCTTTATTACTTGGCAATGCCCGTGATAATT

TTGTTAAAGAACAAATGGCTGTAGTTATAAGATACATAAATAATTATGGCTGTGTTTTGGAACGGTAGTT

GTCCATGTTAAAGATACTACTGCACTTTCTTTAGAGAGTGCTAAAATTTCTTTTCCAAACATGGATTGTC

GTCTGTCGATATCAAAAACTAGAGGCAAGGATATGACGGAGCTCCGAACATGCGGGGGGAGCTTAATGGT

TTCAACTGGTATATATATGAAGGCCCTCCTCGGGAAAACTTCCTGCCTCCGTCACTGCTTGTGTGCTTGT

TGGCTCTCCTAAATTTGGGTTCAGTGTTTGATTTTCTTAAATTGGTCATTTGGCTAAGAGGGCAAAAGAT

ATTATTGGCATAGAATATTGATATTTACTACAATTTTTACTATATATGATTAACAAAATATAAACTTTTT

TTAGACTGAATTGGTAAAAGCTATAAGTTGGCTAGGAATTGCCGTATAAAAGATTCGGATATTCGAAACA

GTGTGAATCTCTTTTCCATGCGATTCGTATACTGGGTTGGTTTTGTTATCCTTTCCCAGTACACCTCTAT

TAAACAGACATTGCATTCCATTTCATACTTTATAAATTAGGGCTAGAGACGAGTAGTGAATAAGATACAG

TATGTCATTGAGTCGCCCCTTACGGAACACAAAAGCCAGAATCACAGTTGAAGATGAAGAGGACAGAATT

AGCAGGTTGCCGGATGAGCTAATTCATCAAATTCTCGAGTTCGTGGATACCAAATTGGCTATTCAAACCA

GTCTTCTATCTAAGCGATGGAAACTTATATGGAAGACTCTCCCTTTTCTCAAGTTTAGATGGGTGCAACA

GGGTTCACCTACAAACACAAATAATTTTACCCGTCAAGTTTTAAGTCGCCGAAACCATCAATCCCAGATT

TCCAGTCTGGAGCTTGCTTTCCTCCCTGCAGGTTTATTTGCAAAGTTTGTTGAGTATGCTATATCTCACC

ATGTTCAATACTTAAATGTTCATTTACGTCGTGATCATAAACCTTATAAACTATCCAACTTTAGCTCTAA

TTCGATTCAGAAACTTCAATTGGAGATGAAGTTTGGAGATTCTATTTCAGAATTAGATTGTTGGGATTTA

CCTGCATTAAACACTCTGCACTTGACAAGTTACTATAATAATCATGTCGTAGGAATCCTATCAGAATTAT

GTTTAACCTGCTTGCCTGCTTTAAGAACTCTGTGCCTTGAATATTTTGATTTGAAGTCCATGTCTATTAG

TTTGTCTTCTTTAACAACTCTTCGGCTGTTTAAGTGCAAGTTGCCCATTATAGTATGGAAATTTCCAGCT

TTGAAAAGTCTAACGCTTCATGATGTGGTGTTTCATCGAGATATGAGTCACATGTTCGCTGCACTTGTCA

CTCTGCAAGATCTCACATTAGTTTTTGAAAAGGAGCATATACAAGATTGTGTTATACCCTCTCCTCAACT

AGAGAATCTGGAGATCAGGTCCCGCTACGTATCCTACACTCACAGGATTGTGGGTAACATTGTGGTTGTG

GCACCAAAACTCAGAAATTTTACTTCTGTTGGTATCTTTCAAATCATATTTGGAGGTTCTAAGGTGGATA

ATGTGAATATAAGATTGCGTGGTTGTTTTGATGGCATGATCTATCCATTGTTAAAGGAATATTATCAACA

GTTAACATTTATGCTTCCAGGCTTTGCTAGTGCCAAGATTCTTAATCTTGATCTCGAGACCTTGCAGGCA

CTTTCTTCCATTGTGGACCGTCTTGAAATTTCTCCGTCTCCATTCCATAACTTGGAGTATGTGAAACTAC

CTTGTGGATTTGTAGAATTTGAAGAAAAACGTGTTGCTGATATCTTACGAAGATACTTACTTGGTGGCTC

TCCAACAGCAACTTATGTCACTACATCATTCCAGAACATGATGCCTGCTACACCGGCAGCTTCTGTGACA

GCTCAGAATGTGGTGCTACAGGAGCCCTTGGCTGCTTCTACCAAGGTGTTGGTTGGTTCTAAAAATATAA

ATAAAAATGTATGCATTGACAATGTTGACATGGGGATGCACGAAGAGCCCGTACTGCAGAATTCTGCAGT

AAATGCTGACAACAATGTTGACATGGAAGTGCAAGAAGAGCCCGGGGTGCAGAATTCTGCAGTAAATGCT

GAGAGGGTTGAACAATTAAGTGGTCCAGTTGAAGGGACTGCCATTGGCCGGGTGAGCTCTTCTGGGGGAG

ATGGTGATTTTGTGTTATTGCAGGGGCATGAAGTTAACTCTGAATGACCGGGTGAGCTCTTCTAGAGAAG

ATAGTGATTTCGGGTGATGGCAGAGACATGGAGTTCTCCGAGTTTACATGTCTACTCGATAATATTATGA

ATAAGTACTGAAAAACCTTTGAACACTTTATGAGCCCTTTTGATTGATCGTCATATTGATGATGCCAAAA

GTAAGTTACAAGATTCACAAACTCATACTGATAATACCATAACTAGTTGCTAGAACTACAGACTCTTCTT

GCGAGGAAGATGCAAGAGATTCAAAAAGCTTTTAAAAGCTTTTGGAACTATGCGCGCAAAGCTTATATTT

GGCTACATTAGAAATGATCGTCCATTTTGACAATATTATTTGTAAGGTTGAGACTTGTATGGATACCTTA

GTATCTTATCTGTTATTTGATTTTGATCCTTGTTCTTCAAGTCTGGTAACTTTATGTTATCTTTATGTTA

TCATGTTTATT

>MSTRG.1457.10 gene=MSTRG.1457

CAAACTGCCAAAATCGAAACCCAAAACATACCTCTTCTCGTATTGATCGCTTGTAATTGTACTCCTCACT

AAGCATATCCAGCATGCCTATATCAAATCGAAGATGAAGAAGAAGATAGAATTACTAGATTGATGATCAA

TTACCTAGTGATGATGTAGCTTTGATATTACCGTCAGCCGGAGAAACAGATGGTGTTCGATAATGTTAAA

TTGCTAAGTTATTTCTTAGCGTGCCTTCATTGTAAATTTAGTTATGCATTGTTAAGTAATCCAACCTTTC

AAGTCATGTTCAGTTAAACCTGTCCTATATATCAGTATCCTGTATCACTATAGGTTGTTAGAAGTGTTTT

TCAATCTTTAAAGGAAGTGAGTTATAAATTGTTTGTTATGAAGACTTGTTATACATTCACTCTAGTAGTG

AACACATAGTTATATCTTTAGCTATTGTATGTACTAAGCATCATTTGTACCTTTGAGCTTTAATATTCCA

TTCAAAGATGACAATCAAAACTTCACCGACCTTTAAGGTTAGACAATCAAAAATTCAACGACCTTTGAGG

TTACTTGTGTGCAGTGACCCGGGGGGGGGCCCTGGCCCTATATATACCAACAAAACTCTATCTGTCTCTC

GCTTGGTATGGTTTGCGTAATGTTGATGTTGGGAAAGTGGTTACAGCAAATCATCAGTTAACATCTCTTG

ACATTCAAAAGCAAATCGTTAGTGCTTGTGCATCCGAAACAAAGAAAGCTGTGATTCGTGATATTGGAGT

AGTATTGGAGTATTTCTCTTTATTACTTGGCAATGCCCGTGATAATTTTGTTAAAGAACAAATGGCTGTA

GTTATAAGATACATAAATAATTATGGCTGTGTTTTGGAACGGTAGTTGTCCATGTTAAAGATACTACTGC

ACTTTCTTTAGAGAGTGCTAAAATTTCTTTTCCAAACATGGATTGTCGTCTGTCGATATCAAAAACTAGA

GGCAAGGATATGACGGAGCTCCGAACATGCGGGGGGAGCTTAATGGTTTCAACTGGTATATATATGAAGG

CCCTCCTCGGGAAAACTTCCTGCCTCCGTCACTGCTTGTGTGCTTGTTGGCTCTCCTAAATTTGGGTTCA

GTGTTTGATTTTCTTAAATTGGTCATTTGGCTAAGAGGGCAAAAGATATTATTGGCATAGAATATTGATA

TTTACTACAATTTTTACTATATATGATTAACAAAATATAAACTTTTTTTAGACTGAATTGGTAAAAGCTA

TAAGTTGGCTAGGAATTGCCGTATAAAAGATTCGGATATTCGAAACAGTGTGAATCTCTTTTCCATGCGA

TTCGTATACTGGGTTGGTTTTGTTATCCTTTCCCAGTACACCTCTATTAAACAGACATTGCATTCCATTT

CATACTTTATAAATTAGGGCTAGAGACGAGTAGTGAATAAGATACAGTATGTCATTGAGTCGCCCCTTAC

GGAACACAAAAGCCAGAATCACAGTTGAAGATGAAGAGGACAGAATTAGCAGGTTGCCGGATGAGCTAAT

TCATCAAATTCTCGAGTTCGTGGATACCAAATTGGCTATTCAAACCAGTCTTCTATCTAAGCGATGGAAA

CTTATATGGAAGACTCTCCCTTTTCTCAAGTTTAGATGGGTGCAACAGGGTTCACCTACAAACACAAATA

ATTTTACCCGTCAAGTTTTAAGTCGCCGAAACCATCAATCCCAGATTTCCAGTCTGGAGCTTGCTTTCCT

CCCTGCAGGTTTATTTGCAAAGTTTGTTGAGTATGCTATATCTCACCATGTTCAATACTTAAATGTTCAT

TTACGTCGTGATCATAAACCTTATAAACTATCCAACTTTAGCTCTAATTCGATTCAGAAACTTCAATTGG

AGATGAAGTTTGGAGATTCTATTTCAGAATTAGATTGTTGGGATTTACCTGCATTAAACACTCTGCACTT

GACAAGTTACTATAATAATCATGTCGTAGGAATCCTATCAGAATTATGTTTAACCTGCTTGCCTGCTTTA

AGAACTCTGTGCCTTGAATATTTTGATTTGAAGTCCATGTCTATTAGTTTGTCTTCTTTAACAACTCTTC

GGCTGTTTAAGTGCAAGTTGCCCATTATAGTATGGAAATTTCCAGCTTTGAAAAGTCTAACGCTTCATGA

TGTGGTGTTTCATCGAGATATGAGTCACATGTTCGCTGCACTTGTCACTCTGCAAGATCTCACATTAGTT

TTTGAAAAGGAGCATATACAAGATTGTGTTATACCCTCTCCTCAACTAGAGAATCTGGAGATCAGGTCCC

GCTACGTATCCTACACTCACAGGATTGTGGGTAACATTGTGGTTGTGGCACCAAAACTCAGAAATTTTAC

TTCTGTTGGTATCTTTCAAATCATATTTGGAGGTTCTAAGGTGGATAATGTGAATATAAGATTGCGTGGT

TGTTTTGATGGCATGATCTATCCATTGTTAAAGGAATATTATCAACAGTTAACATTTATGCTTCCAGGCT

TTGCTAGTGCCAAGATTCTTAATCTTGATCTCGAGACCTTGCAGGCACTTTCTTCCATTGTGGACCGTCT

TGAAATTTCTCCGTCTCCATTCCATAACTTGGAGTATGTGAAACTACCTTGTGGATTTGTAGAATTTGAA

GAAAAACGTGTTGCTGATATCTTACGAAGATACTTACTTGGTGGCTCTCCAACAGCAACTTATGTCACTA

CATCATTCCAGAACATGATGCCTGCTACACCGGCAGCTTCTGTGACAGCTCAGAATGTGGTGCTACAGGA

GCCCTTGGCTGCTTCTACCAAGGTGTTGGTTGGTTCTAAAAATATAAATAAAAATGTATGCATTGACAAT

GTTGACATGGGGATGCACGAAGAGCCCGTACTGCAGAATTCTGCAGTAAATGCTGACAACAATGTTGACA

TGGAAGTGCAAGAAGAGCCCGGGGTGCAGAATTCTGCAGTAAATGCTGAGAGGGTTGAACAATTAAGTGG

TCCAGTTGAAGGGACTGCCATTGGCCGGGTGAGCTCTTCTGGGGGAGATGGTGATTTTGTGTTATTGCAG

GGGCATGAAGTTAACTCTGAATGACCGGGTGAGCTCTTCTAGAGAAGATAGTGATTTCGGGTGATGGCAG

AGACATGGAGTTCTCCGAGTTTACATGTCTACTCGATAATATTATGAATAAGTACTGAAAAACCTTTGAA

CACTTTATGAGCCCTTTTGATTGATCGTCATATTGATGATGCCAAAAGTAAGTTACAAGATTCACAAACT

CATACTGATAATACCATAACTAGTTGCTAGAACTACAGACTCTTCTTGCGAGGAAGATGCAAGAGATTCA

AAAAGCTTTTAAAAGCTTTTGGAACTATGCGCGCAAAGCTTATATTTGGCTACATTAGAAATGATCGTCC

ATTTTGACAATATT

>MSTRG.1457.8 gene=MSTRG.1457

CAAACTGCCAAAATCGAAACCCAAAACATACCTCTTCTCGTATTGATCGCTTGTAATTGTACTCCTCACT

AAGCATATCCAGCATGCCTATATCAAATCGAAGATGAAGAAGAAGATAGAATTACTAGATTGATGATCAA

TTACCTAGTGATGATGTAGCTTTATTACGTGATGATTTTGTTCTCAAATACCAAAGCCTGGTAAGCCGCT

CTAGACGGTCCCAACACGAATGTGCAAATCTTGACGGTTTTGTTTCTTATGTGCGCAAGAGAGATATGAG

TCACATGTTCGCTGCACTTGTCACTCTGCAAGATCTCACATTAGTTTTTGAAAAGGAGCATATACAAGAT

TGTGTTATACCCTCTCCTCAACTAGAGAATCTGGAGATCAGGTCCCGCTACGTATCCTACACTCACAGGA

TTGTGGGTAACATTGTGGTTGTGGCACCAAAACTCAGAAATTTTACTTCTGTTGGTATCTTTCAAATCAT

ATTTGGAGGTTCTAAGGTGGATAATGTGAATATAAGATTGCGTGGTTGTTTTGATGGCATGATCTATCCA

TTGTTAAAGGAATATTATCAACAGTTAACATTTATGCTTCCAGGCTTTGCTAGTGCCAAGATTCTTAATC

TTGATCTCGAGACCTTGCAGGCACTTTCTTCCATTGTGGACCGTCTTGAAATTTCTCCGTCTCCATTCCA

TAACTTGGAGTATGTGAAACTACCTTGTGGATTTGTAGAATTTGAAGAAAAACGTGTTGCTGATATCTTA

CGAAGATACTTACTTGGTGGCTCTCCAACAGCAACTTATGTCACTACATCATTCCAGAACATGATGCCTG

CTACACCGGCAGCTTCTGTGACAGCTCAGAATGTGGTGCTACAGGAGCCCTTGGCTGCTTCTACCAAGGT

GTTGGTTGGTTCTAAAAATATAAATAAAAATGTATGCATTGACAATGTTGACATGGGGATGCACGAAGAG

CCCGTACTGCAGAATTCTGCAGTAAATGCTGACAACAATGTTGACATGGAAGTGCAAGAAGAGCCCGGGG

TGCAGAATTCTGCAGTAAATGCTGAGAGGGTTGAACAATTAAGTGGTCCAGTTGAAGGGACTGCCATTGG

CCGGGTGAGCTCTTCTGGGGGAGATGGTGATTTTGTGTTATTGCAGGGGCATGAAGTTAACTCTGAATGA

CCGGGTGAGCTCTTCTAGAGAAGATAGTGATTTCGGGTGATGGCAGAGACATGGAGTTCTCCGAGTTTAC

ATGTCTACTCGATAATATTATGAATAAGTACTGAAAAACCTTTGAACACTTTATGAGCCCTTTTGATTGA

TCGTCATATTGATGATGCCAAAAGTAAGTTACAAGATTCACAAACTCATACTGATAATACCATAACTAGT

TGCTAGAACTACAGACTCTTCTTGCGAGGAAGATGCAAGAGATTCAAAAAGCTTTTAAAAGCTTTTGGAA

CTATGCGCGCAAAGCTTATATTTGGCTACATTAGAAATGATCGTCCATTTTGACAATATTATTTGTAAGG

TTGAGACTTGTATGGATACCTTAGTATCTTATCTGTTATTTGATTTT

>MSTRG.1457.9 gene=MSTRG.1457

CAAACTGCCAAAATCGAAACCCAAAACATACCTCTTCTCGTATTGATCGCTTGTAATTGTACTCCTCACT

AAGCATATCCAGCATGCCTATATCAAATCGAAGATGAAGAAGAAGATAGAATTACTAGATTGATGATCAA

TTACCTAGTGATGATGTAGCTTTATTACGTGATGATTTTGTTCTCAAATACCAAAGCCTGGTAAGCCGCT

CTAGACGGTCCCAACACGAATGTGCAAATCTTGACGGTTTTGTTTCTTATGTGCGCAAGAGAGGTGAAGC

TTTGAAAAGTCTAACGCTTCATGATGTGGTGTTTCATCGAGATATGAGTCACATGTTCGCTGCACTTGTC

ACTCTGCAAGATCTCACATTAGTTTTTGAAAAGGAGCATATACAAGATTGTGTTATACCCTCTCCTCAAC

TAGAGAATCTGGAGATCAGGTCCCGCTACGTATCCTACACTCACAGGATTGTGGGTAACATTGTGGTTGT

GGCACCAAAACTCAGAAATTTTACTTCTGTTGGTATCTTTCAAATCATATTTGGAGGTTCTAAGGTGGAT

AATGTGAATATAAGATTGCGTGGTTGTTTTGATGGCATGATCTATCCATTGTTAAAGGAATATTATCAAC

AGTTAACATTTATGCTTCCAGGCTTTGCTAGTGCCAAGATTCTTAATCTTGATCTCGAGACCTTGCAGGC

ACTTTCTTCCATTGTGGACCGTCTTGAAATTTCTCCGTCTCCATTCCATAACTTGGAGTATGTGAAACTA

CCTTGTGGATTTGTAGAATTTGAAGAAAAACGTGTTGCTGATATCTTACGAAGATACTTACTTGGTGGCT

CTCCAACAGCAACTTATGTCACTACATCATTCCAGAACATGATGCCTGCTACACCGGCAGCTTCTGTGAC

AGCTCAGAATGTGGTGCTACAGGAGCCCTTGGCTGCTTCTACCAAGGTGTTGGTTGGTTCTAAAAATATA

AATAAAAATGTATGCATTGACAATGTTGACATGGGGATGCACGAAGAGCCCGTACTGCAGAATTCTGCAG

TAAATGCTGACAACAATGTTGACATGGAAGTGCAAGAAGAGCCCGGGGTGCAGAATTCTGCAGTAAATGC

TGAGAGGGTTGAACAATTAAGTGGTCCAGTTGAAGGGACTGCCATTGGCCGGGTGAGCTCTTCTGGGGGA

GATGGTGATTTTGTGTTATTGCAGGGGCATGAAGTTAACTCTGAATGACCGGGTGAGCTCTTCTAGAGAA

GATAGTGATTTCGGGTGATGGCAGAGACATGGAGTTCTCCGAGTTTACATGTCTACTCGATAATATTATG

AATAAGTACTGAAAAACCTTTGAACACTTTATGAGCCCTTTTGATTGATCGTCATATTGATGATGCCAAA

AGTAAGTTACAAGATTCACAAACTCATACTGATAATACCATAACTAGTTGCTAGAACTACAGACTCTTCT

TGCGAGGAAGATGCAAGAGATTCAAAAAGCTTTTAAAAGCTTTTGGAACTATGCGCGCAAAGCTTATATT

TGGCTACATTAGAAATGATCGTCCATTTTGACAATATTATTTGTAAGGTTGAGACTTGTATGGATACCTT

AGTATCTTATCTGTTATTTGATTTT

>MSTRG.1457.11 gene=MSTRG.1457

AAACTGCCAAAATCGAAACCCAAAACATACCTCTTCTCGTATTGATCGCTTGTAATTGTACTCCTCACTA

AGCATATCCAGCATGCCTATATCAAATCGAAGATGAAGAAGAAGATAGAATTACTAGATTGATGATCAAT

TACCTAGTGATGATGTAGCTTTGCAAGTTGCCCATTATAGTATGGAAATTTCCAGCTTTGAAAAGTCTAA

CGCTTCATGATGTGGTGTTTCATCGAGATATGAGTCACATGTTCGCTGCACTTGTCACTCTGCAAGATCT

CACATTAGTTTTTGAAAAGGAGCATATACAAGATTGTGTTATACCCTCTCCTCAACTAGAGAATCTGGAG

ATCAGGTCCCGCTACGTATCCTACACTCACAGGATTGTGGGTAACATTGTGGTTGTGGCACCAAAACTCA

GAAATTTTACTTCTGTTGGTATCTTTCAAATCATATTTGGAGGTTCTAAGGTGGATAATGTGAATATAAG

ATTGCGTGGTTGTTTTGATGGCATGATCTATCCATTGTTAAAGGAATATTATCAACAGTTAACATTTATG

CTTCCAGGCTTTGCTAGTGCCAAGATTCTTAATCTTGATCTCGAGACCTTGCAGGCACTTTCTTCCATTG

TGGACCGTCTTGAAATTTCTCCGTCTCCATTCCATAACTTGGAGTATGTGAAACTACCTTGTGGATTTGT

AGAATTTGAAGAAAAACGTGTTGCTGATATCTTACGAAGATACTTACTTGGTGGCTCTCCAACAGCAACT

TATGTCACTACATCATTCCAGAACATGATGCCTGCTACACCGGCAGCTTCTGTGACAGCTCAGAATGTGG

TGCTACAGGAGCCCTTGGCTGCTTCTACCAAGGTGTTGGTTGGTTCTAAAAATATAAATAAAAATGTATG

CATTGACAATGTTGACATGGGGATGCACGAAGAGCCCGTACTGCAGAATTCTGCAGTAAATGCTGACAAC

AATGTTGACATGGAAGTGCAAGAAGAGCCCGGGGTGCAGAATTCTGCAGTAAATGCTGAGAGGGTTGAAC

AATTAAGTGGTCCAGTTGAAGGGACTGCCATTGGCCGGGTGAGCTCTTCTGGGGGAGATGGTGATTTTGT

GTTATTGCAGGGGCATGAAGTTAACTCTGAATGACCGGGTGAGCTCTTCTAGAGAAGATAGTGATTTCGG

GTGATGGCAGAGACATGGAGTTCTCCGAGTTTACATGTCTACTCGATAATATTATGAATAAGTACTGAAA

AACCTTTGAACACTTTATGAGCCCTTTTGATTGATCGTCATATTGATGATGCCAAAAGTAAGTTACAAGA

TTCACAAACTCATACTGATAATACCATAACTAGTTGCTAGAACTACAGACTCTTCTTGCGAGGAAGATGC

AAGAGATTCAAAAAGCTTTTAAAAGCTTTTGGAACTATGCGCGCAAAGCTTATATTTGGCTACATTAGAA

ATGATCGTCCATTTTGACAATATTATTTGTAAGGTTGAGACTTGTATGGATACCTTAGTATCTTATCTGT

TATTTGATTTTGAT

>MSTRG.1457.12 gene=MSTRG.1457

AAACTGCCAAAATCGAAACCCAAAACATACCTCTTCTCGTATTGATCGCTTGTAATTGTACTCCTCACTA

AGCATATCCAGCATGCCTATATCAAATCGAAGATGAAGAAGAAGATAGAATTACTAGATTGATGATCAAT

TACCTAGTGATGATGTAGCTTGCTTTGCTAGTGCCAAGATTCTTAATCTTGATCTCGAGACCTTGCAGGC

ACTTTCTTCCATTGTGGACCGTCTTGAAATTTCTCCGTCTCCATTCCATAACTTGGAGTATGTGAAACTA

CCTTGTGGATTTGTAGAATTTGAAGAAAAACGTGTTGCTGATATCTTACGAAGATACTTACTTGGTGGCT

CTCCAACAGCAACTTATGTCACTACATCATTCCAGAACATGATGCCTGCTACACCGGCAGCTTCTGTGAC

AGCTCAGAATGTGGTGCTACAGGAGCCCTTGGCTGCTTCTACCAAGGTGTTGGTTGGTTCTAAAAATATA

AATAAAAATGTATGCATTGACAATGTTGACATGGGGATGCACGAAGAGCCCGTACTGCAGAATTCTGCAG

TAAATGCTGACAACAATGTTGACATGGAAGTGCAAGAAGAGCCCGGGGTGCAGAATTCTGCAGTAAATGC

TGAGAGGGTTGAACAATTAAGTGGTCCAGTTGAAGGGACTGCCATTGGCCGGGTGAGCTCTTCTGGGGGA

GATGGTGATTTTGTGTTATTGCAGGGGCATGAAGTTAACTCTGAATGACCGGGTGAGCTCTTCTAGAGAA

GATAGTGATTTCGGGTGATGGCAGAGACATGGAGTTCTCCGAGTTTACATGTCTACTCGATAATATTATG

AATAAGTACTGAAAAACCTTTGAACACTTTATGAGCCCTTTTGATTGATCGTCATATTGATGATGCCAAA

AGTAAGTTACAAGATTCACAAACTCATACTGATAATACCATAACTAGTTGCTAGAACTACAGACTCTTCT

TGCGAGGAAGATGCAAGAGATTCAAAAAGCTTTTAAAAGCTTTTGGAACTATGCGCGCAAAGCTTATATT

TGGCTACATTAGAAATGATCGTCCATTTTGACAATATTATTTGTAAGGTTGAGACTTGTATGGATACCTT

AGTATCTTATCTGTTATTTGATTTTGAT

>MSTRG.1457.13 gene=MSTRG.1457

CCCAAAACATACCTCTTCTCGTATTGATCGCTTGTAATTGTACTCCTCACTAAGCATATCCAGCATGCCT

ATATCAAATCGAAGATGAAGAAGAAGATAGAATTACTAGATTGATGATCAATTACCTAGTGATGATGTAG

CTTTCTCTGCAAATTGTTGTGGTTGCTAATCGGAAAAAGGCTCGTTGAATCCCTGAGTATTCATTTGAAT

TCCACACAACGTTTACTGACTTTTTGCTGATTAGTTCTCTTCGTATTTGTTATCAATCATATATTGTGAT

TTATTTTGATGTTCACTGCATTATATCGGATGATGAATTTGCAATTGAGATATGGATTAAAAAGGTAACT

AGAATTACGAATGATTCTTCTGTATCGTGTGTAGTATTACGTGATGATTTTGTTCTCAAATACCAAAGCC

TGGTAAGCCGCTCTAGACGGTCCCAACACGAATGTGCAAATCTTGACGGTTTTGTTTCTTATGTGCGCAA

GAGAGGTGAAGGTAAATTATTTTTATATCTGTTTGATGTGAATGTTGTTTTTGTGTCTCTTCTTGCTAAA

GCTGACTATAAGCTAGAATTTAATAACGGTTAAGTTTTATACAAGAGAATGGATCTTTTGTGGGCAAAGG

TCAGATTTTTAATAGTTTGTTTCATTTAATTGTTAATAAAGTTACGTCGCTTAATACTAGTTATGGCCTC

TATTTGTTTGATAGAAGATGAAAGTGTTTACCATACAAAGTTAGGCCAAACTAATAATGAATACATTAAT

AGCTCGGATTATTGGTTTTTGAAGGACATTAAGTTAATATAGAGTCAAAGTCCACTAACAGAACCTTGCA

AGACAATAAAATTCGGAGAATGTAAATCTTCTGTTTAATTCATTCTGATTGATTGTCTAATACATCAAGA

GGCTGTAAAAGTAGCAACTTTTGTGGATGACTTCTCTACGGATACACTTTAGCTTATTTTGTTGAGGAAC

AAATATGAAGCAGAAATAAATTTATAGTTTATAAAGGTTAAGCTGAAAGTAAATTGGATAAGAAAATTAT

TATCTTAAAACAAATAAAGGCAGAGCAGTTGGTTTTGATCAATTTTTTCGGGCACTGTGAGGAAAATGGT

ATAGCTTAAACACTCCCTCTAACGGAGCAACAGAAAGAAGAAAAACAGGACTTGAAAGAAATGACTAATT

TCTTGCTAGTTAGATCCGATGCTCTGTTATCTCTATGAGGTGAGGCCTTCATCAGTTATTGACATATTTA

CTAAAGAATATAGGAAAGACAGCATATGAACTTTGTGAGACCTAAACACATGATCCTAATTTTTTGATAG

TCTGGGGTGTCTTGCTAAGGTGTTGTAACGAGACCTAAAAATTGGTAAATTGGTTGTAAAACTACGAATT

GTTTGTTTTTCAGAAAATGGTACTGGTTAATGCTGATTATTTTCTAGTGATTTATTAAGAATGTATTTCC

TTTAAAAGTTAGAATTAGTGGATCCCCAAATGATCTTCTAGAGACCTCTATTAGACACCTCCATTAGACA

AGTAGCTAAAGATTTTTGTGATACAGAAAGTCAGAGGATAAAATGTGCTTGGTTATTGTGGTGATTATGT

TTATTTGTTCTTAGTGGATAATGATCCGTTAACTCGATTTACAGTTAAATTTACCTAAACCATGTTTGAC

CACCTTCCTTTATAAATTGTCCTACTCTTTAGGATTTACGAAAATCTTTTTTATCATTTTGTTTGCCCTG

TCATTAACTCTCTGTCTAGTTTGACACCATGTTTCTTATTTGTTTATACCTTATAGTGATATTACCGTCA

GCCGGAGAAACAGATGGTGTTCGATAATGTTAAATTGCTAAGTTATTTCTTAGCGTGCCTTCATTGTAAA

TTTAGTTATGCATTGTTAAGTAATCCAACCTTTCAAGTCATGTTCAGTTAAACCTGTCCTATATATCAGT

ATCCTGTATCACTATAGGTTGTTAGAAGTGTTTTTCAATCTTTAAAGGAAGTGAGTTATAAATTGTTTGT

TATGAAGACTTGTTATACATTCACTCTAGTAGTGAACACATAGTTATATCTTTAGCTATTGTATGTACTA

AGCATCATTTGTACCTTTGAGCTTTAATATTCCATTCAAAGATGACAATCAAAACTTCACCGACCTTTAA

GGTTAGACAATCAAAAATTCAACGACCTTTGAGGTTACTTGTGTGCAGTGACCCGGGGGGGGGCCCTGGC

CCTATATATACCAACAAAACTCTATCTGTCTCTCGCTTGGTATGGTTTGCGTAATGTTGATGTTGGGAAA

GTGGTTACAGCAAATCATCAGTTAACATCTCTTGACATTCAAAAGCAAATCGTTAGTGCTTGTGCATCCG

AAACAAAGAAAGCTGTGATTCGTGATATTGGAGTAGTATTGGAGTATTTCTCTTTATTACTTGGCAATGC

CCGTGATAATTTTGTTAAAGAACAAATGGCTGTAGTTATAAGATACATAAATAATTATGGCTGTGTTTTG

GAACGGTAGTTGTCCATGTTAAAGATACTACTGCACTTTCTTTAGAGAGTGCTAAAATTTCTTTTCCAAA

CATGGATTGTCGTCTGTCGATATCAAAAACTAGAGGCAAGGATATGACGGAGCTCCGAACATGCGGGGGG

AGCTTAATGGTTTCAACTGGTATATATATGAAGGCCCTCCTCGGGAAAACTTCCTGCCTCCGTCACTGCT

TGTGTGCTTGTTGGCTCTCCTAAATTTGGGTTCAGTGTTTGATTTTCTTAAATTGGTCATTTGGCTAAGA

GGGCAAAAGATATTATTGGCATAGAATATTGATATTTACTACAATTTTTACTATATATGATTAACAAAAT

ATAAACTTTTTTTAGACTGAATTGGTAAAAGCTATAAGTTGGCTAGGAATTGCCGTATAAAAGATTCGGA

TATTCGAAACAGTGTGAATCTCTTTTCCATGCGATTCGTATACTGGGTTGGTTTTGTTATCCTTTCCCAG

TACACCTCTATTAAACAGACATTGCATTCCATTTCATACTTTATAAATTAGGGCTAGAGACGAGTAGTGA

ATAAGATACAGTATGTCATTGAGTCGCCCCTTACGGAACACAAAAGCCAGAATCACAGTTGAAGATGAAG

AGGACAGAATTAGCAGGTTGCCGGATGAGCTAATTCATCAAATTCTCGAGTTCGTGGATACCAAATTGGC

TATTCAAACCAGTCTTCTATCTAAGCGATGGAAACTTATATGGAAGACTCTCCCTTTTCTCAAGTTTAGA

TGGGTGCAACAGGGTTCACCTACAAACACAAATAATTTTACCCGTCAAGTTTTAAGTCGCCGAAACCATC

AATCCCAGATTTCCAGTCTGGAGCTTGCTTTCCTCCCTGCAGGTTTATTTGCAAAGTTTGTTGAGTATGC

TATATCTCACCATGTTCAATACTTAAATGTTCATTTACGTCGTGATCATAAACCTTATAAACTATCCAAC

TTTAGCTCTAATTCGATTCAGAAACTTCAATTGGAGATGAAGTTTGGAGATTCTATTTCAGAATTAGATT

GTTGGGATTTACCTGCATTAAACACTCTGCACTTGACAAGTTACTATAATAATCATGTCGTAGGAATCCT

ATCAGAATTATGTTTAACCTGCTTGCCTGCTTTAAGAACTCTGTGCCTTGAATATTTTGATTTGAAGTCC

ATGTCTATTAGTTTGTCTTCTTTAACAACTCTTCGGCTGTTTAAGTGCAAGTTGCCCATTATAGTATGGA

AATTTCCAGCTTTGAAAAGTCTAACGCTTCATGATGTGGTGTTTCATCGAGATATGAGTCACATGTTCGC

TGCACTTGTCACTCTGCAAGATCTCACATTAGTTTTTGAAAAGGAGCATATACAAGATTGTGTTATACCC

TCTCCTCAACTAGAGAATCTGGAGATCAGGTCCCGCTACGTATCCTACACTCACAGGATTGTGGGTAACA

TTGTGGTTGTGGCACCAAAACTCAGAAATTTTACTTCTGTTGGTATCTTTCAAATCATATTTGGAGGTTC

TAAGGTGGATAATGTGAATATAAGATTGCGTGGTTGTTTTGATGGCATGATCTATCCATTGTTAAAGGAA

TATTATCAACAGTTAACATTTATGCTTCCAGGCTTTGCTAGTGCCAAGATTCTTAATCTTGATCTCGAGA

CCTTGCAGGCACTTTCTTCCATTGTGGACCGTCTTGAAATTTCTCCGTCTCCATTCCATAACTTGGAGTA

TGTGAAACTACCTTGTGGATTTGTAGAATTTGAAGAAAAACGTGTTGCTGATATCTTACGAAGATACTTA

CTTGGTGGCTCTCCAACAGCAACTTATGTCACTACATCATTCCAGAACATGATGCCTGCTACACCGGCAG

CTTCTGTGACAGCTCAGAATGTGGTGCTACAGGAGCCCTTGGCTGCTTCTACCAAGGTGTTGGTTGGTTC

TAAAAATATAAATAAAAATGTATGCATTGACAATGTTGACATGGGGATGCACGAAGAGCCCGTACTGCAG

AATTCTGCAGTAAATGCTGACAACAATGTTGACATGGAAGTGCAAGAAGAGCCCGGGGTGCAGAATTCTG

CAGTAAATGCTGAGAGGGTTGAACAATTAAGTGGTCCAGTTGAAGGGACTGCCATTGGCCGGGTGAGCTC

TTCTGGGGGAGATGGTGATTTTGTGTTATTGCAGGGGCATGAAGTTAACTCTGAATGACCGGGTGAGCTC

TTCTAGAGAAGATAGTGATTTCGGGTGATGGCAGAGACATGGAGTTCTCCGAGTTTACATGTCTACTCGA

TAATATTATGAATAAGTACTGAAAAACCTTTGAACACTTTATGAGCCCTTTTGATTGATCGTCATATTGA

TGATGCCAAAAGTAAGTTACAAGATTCACAAACTCATACTGATAATACCATAACTAGTTGCTAGAACTAC

AGACTCTTCTTGCGAGGAAGATGCAAGAGATTCAAAAAGCTTTTAAAAGCTTTTGGAACTATGCGCGCAA

AGCTTATATTTGGCTACATTAGAAATGATCGTCCATTTTGACAATATTATTTGTAAGGTTGAGACTTGTA

TGGATACCTTAGTATCTTATCTGTTATTTGATTTTGATCCTTGTTCTTCAAGTCTGGTAACTTTATGTTA

TCTTTATGTTATCATGTTTATT

>MSTRG.1458.1 gene=MSTRG.1458

CTGAGATTCACCACTCCAACTCCCACCGTACAAGCCGAAGGCGAAGACCGCATAAGTGATTTACCCGATG

AACTCATCCATCACATTCTCTGGTTCTTAGATGTGCAAACTGCAGTACAAACCTGCATTCTCTCCAAACG

ATGGAAGCCTGTGTGGACTACTCTTCCGTTTATCAATTTCACGAGATACGATCAACTACACGGTGAATAC

ATTCGAGGCAGCACTTTCAATTCTTTTCGATGTCTCTTCTTGGATAGCTTCACTGCCGAGATGAACTTGT

TTGACAAGTTTTTCTCCCACCGAAACCATGATTCACGCATATTCAAATTGTACCTCTCGTTTTTCTTTGT

TCCGCCCTCGAATTTGGTTCACAAGTTTATCAATTACGCAATTTCGCGTCAAGTGCAGGACTTGAACGTG

CATATACCAGAATTTGTTACTGATTTATCGACCTTTAATTCTGCTTCTGTGGAGAAACTGACTCTGAGTC

TCGATTTTTGTAAGGTGTTCAAATTCTCGGATTGTTGGGATTTACCTGCTTTAACAACTCTACGCTTGAT

ACTCCCTTCGTTAAGCCCTGGCGGAATAAGTATTCCTGAATTATATTTGATCTGTTTGCCTTCTTTACAG

ACTCTGTTTCTCGATCGAATTGAGTTACCAGATTCTCTTGATTTGCCTGATTTAGTAACTCTCTATCTCG

CACGGTGCAAATTGCCCTCTGAAATTGGGGATTTCTCGGCTTTATCAACTCTCATACTTGATGATGTGGA

ATTCCAGGAAAATGAGACGAATTTATTCCCTGCACTTGTCAATCTTCGGAATCTCACCTGGTTTTTCCGC

CAAGGATTAATCAAGGCAGATTGTGTTATAAATTGTCCTCAATTGGTGAACCTGAATATCTATGCCTATA

CCTATGTGACTGCGTTTAATAATCATTCTGGTAATTTTATACTTGTGGCGCCTAAAATCCGCAATATTAG

CTGTGTTGGTTTATTCCCAATCACATTTGGACTATCTCAGTTGGAGAGTGTGAATATGAAAATAAGGGAT

AGAACTGATGGAAGAAAGATTATTCCAGGATCTAAGTTGATGAATATATATTATCGTGGATTTACTTATA

TGTTTCAGTCTCTGGGTAGTACAAAGATTCTTACTCTCGACTTGGAGACCATTGAGGTTTTATCTTCTAT

TTCAGACTCTCTCGTAAATTCTCCTCCTCCATTCTATAACTTAGAGTATGTGAAGGTGCCAGAAGGATGT

AAAGAATCGAGTCTGTCGACTGACCTTAAATGCTACTTGCTTGGTCGCTCTCCAGAAGCTACTATTGTCA

CTAAATTGCCTCAGAAAAATATTGTTCCTCATAGTGAGGCAGCTTCAGTGACAGCTCGGAATGCAGTGCT

GCATAAGTCCTTGAAATCCCCAATAAACATGTTGGTTTGTCCTGAAAACCTAAATAAAACTGTGTGCATT

GACACTTTGGACATGGGGGTGCAAGAGAAGATGGCACAGAATTATGTAGTAAGTGCTGAAAGGGTTACAC

AAATTGATGCTCCAGTTGAAGATAGTGATAGAATTAGCTCCTTCGGGTTATGGCAGGGTCACGAGGTTAA

ATCCGAGTTTGTATGTCTACTCGATCTTATAAAGACAAAGTACCCAGAAACCTTTGAGAATTTTACTACC

AACAACAAGAGAGCTAGTACAACGAAGCTAAATATGTTGTGCACCTCAGTAAATGACTTTATTAAAATCT

CTCTGAGTGATATCGATACTGAGATGATTGCTGTGTACAGGGACGTATTTTCCTGCTTAAAGCAATTTGG

ATTCAACGTAGGATGGCTTGTGAGCCGTCTGAAATATGTAGAACAGCTCCAATTTTCACAGCCTCTACTT

TCCGAGCTTCATGCAATTGACTGTCATATTGAGGATGCAAAGAGTAAGCAAGATATGCAGGTCCGTGTTA

GCAACACAGGAAGTAAATTGCAAGATCTACAGAACCTTGGTGCTGCCAAGGTGCAACAGATTCAGAATGC

TTGTGGAGCAAAAGGTAAAAACCTTGTTGTTGGCTACGTTGGAGATGATCTATTTTCTGATCCCTAATTT

TTCAAGAATGGTCTGAATCTTTTTGGCAATGTGTTGATAATGTGCTTCAGAATATATATCTTCTCAGAGT

TTGAGCTGCCCTTAGTTATATTCAGTGTCATTCAG

>MSTRG.1460.1 gene=MSTRG.1460

CAAACGGAGGAGTGGAACAACTAATATCTTTATCAAGAAATTGGACAAGTCGATCGATCACAAAGCCTTG

CAAGATACCTTTTCAAGTTTTGGAACTATTCTTTCCTGCAAGATAGCGACCGATAATTCTGGCCAGTATA

AAGGCTATAGGTTTGTCCAGTATGATAGTGAAGAAGCTGCTAATACTGCAATTGAAAAGCTAAATGGCAT

GTAGTTGAACATGATTTGCAACTGTGATATTCTGGTACTTAGAATTGTTTAGATGGGGCAGTTCATCTTT

GGTGTATTCTTTTGTGTTCGATGATTTTGATTGGAAAAGAAGTACTAAATGTACCAGGCTGTCATATATA

TAAATAGTACATTTCAAATCGGCCATTACAAAACTGTTGCAGTAGCTATAAAAAA

>MSTRG.1461.1 gene=MSTRG.1461

ACACACCACAACTAAAGCGTTGACATCTTGACTCTGCATAAAATTCACATTGACTTTTCAACCCCCACCC

CTACTCCTCTTCCTCCTCAAAAGCACTCACAGGAAAACTCCTCGAAATCCCCGTAGGGGTACCGAACGTA

ATTTTCCCAGATTCCGGGTCATCAATCCGGATATCCGAAAGACTGACCCAGATGAAAAGCTCTCTGGTCT

TGACACCTGTCAGCTTCTTCATCCGTCGGTTCTCAACGAACGCCGTCACTTCCGTATCATAACTTGTCCT

GCGCCCTATCGCACGGAACAAATGCTCAGTCTTCTTCTTCTGTTTGAGCCACACGAAGCCTGTTTCATGA

TTGTGTCCTACCTCAACGATGTCGTTTAACGGCAGCAGGCCGGTGGGCAGAGCGATGCTACGGAGAAGCT

CCTTGGATTTTTTCTTGCAGAGGGCCTCGTCGGTGATGATCTCGGCGTTCTCGCGGTGGTCTTGGATCAC

TGACGTCATTTTCCGGTGGGTTTAGAATGGTTTTTTAGTGCATGTAATAAGAGATTTTGGGCTGGTTTTT

ATAGGATGAATTTTGTGTGCGTGTTGATGTGTAGGAGGAGGAGATCGTGGAGAATAAAGGGGTGGAATAT

TTCTTTTTAATATGGAGATTTTTTGGTGTATTATTGATGATATTGGTTTTATAATGGACGTTAAGAATCA

TATGTTTGTAACTTTGCCAAGATATTTGCAGAGGTGTGTAAAGAGTTTTAATGGAAGATCACAACCCAAA

TAGGCAAATATGTGAATTTGGCAGTGGGTTTGGCTTTGATAACTTGCAATATTTTTTGTAAGACCATCTC

CAATCATAATTTTATAAATAAAGCAAC

>MSTRG.1467.1 gene=MSTRG.1467

ATCGCTGCTATATGTTTCATCCACCTGACTATCTTCATCCCCATCCTCACCCCCTGTCACAAACTCTAGC

ATTTCACCATCATCATCTTGGTCATCCATCATGAATTCACCATCATCATCTTCCTCGTCCTCCAACTCGC

TTGCACTATCATCATTACTCATATCGTCCATATCACTTCCACCATCTCCCACAATTCCCAACAATGGATC

CTCATCTATGTCTTCATCGTCATCTTCATCTTCATCCTCAGTCTCGGCATCATCGGGATCCGAGTCATCA

TCAGTTGGCCTCCGCCTACCTATCTCAAAAATCCTGGCCGAAGAATACATTTCATCCGG

>MSTRG.1469.1 gene=MSTRG.1469

TTACATTTTGCGTATCAGGTAAAGCCACGAGCCTTTCCATTTCTAGGAAGTCATCCATGAGACTGTCATC

AAGTGAAGAAGCTGTAAGGCTTTTTCCTAAAGCTTTTACGTTTCTGAAGTTAGCAGACTCTGATACTGAT

GACGTGGAAGGATTTTGTTTCCAATCATTTGGCCCCAATTCATGAATCCTATGAGTATCATTATCAACCA

GTGATAGCCTCTCTCCAATATCAGACTGGCTGTCTGTGAAGGATTCCACATAAACAGAAGATTCAGTAAT

AGACCTTTGCTCATTAAATGATGGTGCTCTGCGAGCCACAGCTTTGAGTCTACGACACTCGGTTTCAAGC

TTAGCCACCTTCTTTACACTTTCCAGATTTTGTCTGCTAGCTGATTCAGCAGCTTCAGTACTCAAGTCCC

TCTCAACAGTCCTCAATTCTAGCTCCTCTGATCTGGAAAGGAGCTCATGTTTAAGGGATGAGATCTCCTT

TTCCGCAGCTTCAAGCTTAGAGCGAAGATCAATATGCCAGGAGGTGGCACCTTCACCTTTTGCGGCTTGA

AGTTGGGACTGAAGCTCGACAAGCTGTTTCTGAAATCCAGATTTAGTAGCTTCCCATTCATTGGTTTTCT

TAACAATAGCTTCATGGATCTTTTGCTCTTGTTCTTCTCTTTCGTGCCGGAGCTGTCTCATGCATTCCCT

GAGAGCTACGTCAAGATGGATGGTACGATCTTCCAGCGATGAACTTCTCTGGGTTAAGGATTCTAATTGT

TGCTTTAGTGCAAACACTTCATTCTCAGCTTTCTCCCATCCTTGTATATTTGAATAACAAACAATTATCA

TATTAACAAGCAGTAACCAAGAATAC

>MSTRG.1476.1 gene=MSTRG.1476

CACAAATTCCGAACATGCCAAAGTGGTGATATAACCGTTACACCACAATTATTATCAAGCAATGTATGAA

CATACAAAAGGTAAGCTCCCCATATAACTCATCAATACCAAAGATCTCGCAAACCCCTTGTTAATCTCTC

TAGAAATGGCCGATTTTTCATTCCTTTCTGACAGTGATGACGATAAGAAAGTAGATGATCTCTTATCCCA

AGTCATGGATCACTCTGTTCTTGAACAAGTTTCCGCTATAAATTGCTCTGGTTTTACAGACTCTGTTCTT

CCATCTCAGCTTGAAACCAGGTTTAGTAGACTCAAGTCATTCCCTGCTGCAAAACCCATGTCTCAGATTT

CCCCAGTGAGGCCTTTTAGTGCTCGGCTCGAGCCCGACTCGAAAAAGAAAAAATCAGCTCAGGGTGATCG

GTCTTCTGTGAATGAGATGGGCTTGGATGAAACTCGAGAGAAGGATTTGAAGTCGAAATCACAGCATGGG

GTTTTGGATTTTGATGAGGAAGTGTCGGATTTTTCGATAGAGAGTGATGGTTGGGTAGAGAAAAAAGATG

GGGAGTTGAGTGAGAGGTTGAAGGTGAGGACTAGGTCGTTTCGATCACCTCTGGATTCGAGGAGTTCTTC

GGGGGGTTCTCCGTCGCCTCCGAAGGGAATTGGTTGTTTTTGGTGTTCTCCGAAAAGGGTTTTGAGTAGG

AAGAGTAAAGAGTGGAGGGCTGATCATGATCGGGGCAAGAATGAGGAGTTTTTGATGGATATGAGTGATT

TTTCGACTAAAGGGCAGGAAAAATTGTTGAAGAAACTGATGAAGGAAGAAGAGAAAATTAATCGGGAGGC

TGCGAAGATTGTGAAATTGGTAAAACAAGCGTCTGCGAGGATGGAGATTTCGGAAGATGAGCTAAGTGAT

GATGAGAGAACAAAGTGAATATATGAGCATGTTTTACTAAACTTTATACTACTAAAATGCAGCATTGTAT

AGATAAATGCTATCCGATTTTGTTTCAATTGTATGATCATAGAGTGTGTTTTGTTTCAATTGTATGAACA

TAGAGTGTGAGGTTTAATCTGCTAAGTATATGATTGTATTATTATTCTCCATAGGAAAGAGAAGATGATA

AATAATATATAGCTCTTTTCTCGGATTGTTCATATACAATCTGTTAGTGGTGTATCATGTTATGAGTGCC

TTGCACAGATTAGTAATTTCATGTTTATGCATTTGAGGGAG

>MSTRG.1479.1 gene=MSTRG.1479

ATCCGAGATACTGTATACATATTCTCTTCCCACAATCTTTTTTTTAATCACTGGCCTTAATTAATTTTCA

GTAATAAAATGTAAAGATGAATCCTCACAACTCTCTCTAATTCTTCTATAAATCCGTATTTCCGGTCTAG

TTAAGCTGCATGATCTCAACTTTGTTCTGATCACCCAAAAAAAAATGTGGGGAAGCAACCATAACTGCCA

CCACCATCATCAAGGACACCATCCAAGCCATGGAGGCTATCCTCCCCCGTGCTACCCCCCGGCAGCTGGT

TGTCCTCCACCCGCGCCTTACGGACCATCTGGATACCCTCACGGATACCCCCACCATTCAGGACATGGAT

CTGGTGTGGTAGGAATGCTTGCTGCTGCAGCATCATACGGGGCTCACCGTATAGCACACGGCCACGGTCA

CAGTCACGGCTATGCACACAGCAGCTACTATCGACATAAACACGGCAAATTCAAGCACGGGAAATTCGGA

AAACATGGAAAATTCAAGTTCAAGAAGTGGAAGTGACACTTTCAGCTGCAATCACAGAGTTCCTTTCAAG

GCTTGTATTAAATAAATTATTTTCGCCACAATTCAACTACTTCAGTGACTGCACTTGTTGCAGTTCACCT

TGGTTTCTTTCGATTTCTCATTGAGGAAGATGTGATGTCTTGTATATCTAGCTCGATATCTCGGTCACTT

AATCGAAGCAAAATAACATGAGAACTTGTGTAAATACTCCGATGTGTTCCAGACATGACAGAGTATAATT

GAAAGATATATATGTGGACTTGTGTAAATACTCCGATGTGTTCCAGACATGACAGAGTATTATTGAAAGA

TATATATGTGGACAGCCTATGTGTCATGATCTTCACAATAGAGCTCGTGAACTATAGAGATTTAATT

>MSTRG.1479.2 gene=MSTRG.1479

AACAGACGATAGCTTATTTATCAGGCTAAGAAGATCCGAGATACTGTATACATATTCTCTTCCCACAATC

TTTTTTTTAATCACTGGCCTTAATTAATTTTCAGTAATAAAATGTAAAGATGAATCCTCACAACTCTCTC

TAATTCTTCTATAAATCCGTATTTCCGGTCTAGTTAAGCTGCATGATCTCAACTTTGTTCTGATCACCCA

AAAAAAAATGTGGGGAAGCAACCATAACTGCCACCACCATCATCAAGGACACCATCCAAGCCATGGAGGC

TATCCTCCCCCGTGCTACCCCCCGGCAGCTGGTTGTCCTCCACCCGCGCCTTACGGACCATCTGGATACC

CTCACGGATACCCCCACCATTCAGGACATGGATCTGGTGTGGTAGGAATGCTTGCTGCTGCAGCATCATA

CGGGGCTCACCGTATAGCACACGGCCACGGTCACAGTCACGGCTATGCACACAGCAGCTACTATCGACAT

AAACACGGCAAATTCAAGCACGGGAAATTCGGAAAACATGGAAAATTCAAGTTCAAGAAGTGGAAGTGAC

ACTTTCAGCTGCAATCACAGAGTTCCTTTCAAGGCTTGTATTAAATAAATTATTTTCGCCACAATTCAAC

TACTTCAGTGACTGCACTTGTTGCAGTTCACCTTGGTTTCTTTCGATTTCTCATTGAGGAAGATGTGATG

TCTTGTATATCTAGCTCGATATCTCGGTCACTTAATCGAAGCAAAATAACATGAGAACTTGTGTAAATAC

TCCGATGTGTTCCAGACATGACAGAGTATAATTGAAAGATATATATGTGGACAGCCTATGTGTCATGATC

TTCACAATAGAGCTCGTGAACTATAGAGATTTAATT

>MSTRG.1479.3 gene=MSTRG.1479

TCTCTAATTCTTCTATAAATCCGTATTTCCGGTCTAGTTAAGCTGCATGATCTCAACTTTGTTCTGATCA

CCCAAAAAAAAATGTGGGGAAGCAACCATAACTGCCACCACCATCATCAAGGACACCATCCAAGCCATGG

AGGCTATCCTCCCCCGTGCTACCCCCCGGCAGCTGGTTGTCCTCCACCCGCGCCTTACGGACCATCTGGA

TACCCTCACGGATACCCCCACCATTCAGGTCCAATTTAGCCGACGATTATAGCCAACGCCGTTGGATATG

CTCTAAGTAGATAAACTAAGAATGTGGATGAATTTTTTCAGGACATGGATCTGGTGTGGTAGGAATGCTT

GCTGCTGCAGCATCATACGGGGCTCACCGTATAGCACACGGCCACGGTCACAGTCACGGCTATGCACACA

GCAGCTACTATCGACATAAACACGGCAAATTCAAGCACGGGAAATTCGGAAAACATGGAAAATTCAAGTT

CAAGAAGTGGAAGTGACACTTTCAGCTGCAATCACAGAGTTCCTTTCAAGGCTTGTATTAAATAAATTAT

TTTCGCCACAATTCAACTACTTCAGTGACTGCACTTGTTGCAGTTCACCTTGGTTTCTTTCGATTTCTCA

TTGAGGAAGATGTGATGTCTTGTATATCTAGCTCGATATCTCGGTCACTTAATCGAAGCAAAATAACATG

AGAACTTGTGTAAATACTCCGATGTGTTCCAGACATGACAGAGTATAATTGAAAGATATATATGTGGACT

TGTGTAAATACTCCGATGTGTTCCAGACATGACAGAGTATTATTGAAAGATATATATGTGGACAGCCTAT

GTGTCATGATCTTCACAATAGAGCTCGTGAACTATAG

>MSTRG.1489.1 gene=MSTRG.1489

CTGCTTATAACTGAAATCAAAAATCAATTTCCTGTAAAAAAAATGAAAGGGACGATGATTAGCAGCTCTG

CACGATACCCTCAACATGGGTCATGCTATACCATAAGTATAATGGCACTATCAGAAGGAATCATAGTGAT

CATATCAAACGAGTTCACCCTACATGTTTGATAAATTGTGCAAGTGAATTGGATCTTCTTCACATTGATT

CAATTTGAAATGTTTCACTGTACCGTGAGAATCCTCTTCCCTTAAATCAAATAGTGATTTTATGCAGAAG

CACTTGACCATAGTCACTCATCAGTCATCAGGGCAGACTGGTCGTATTTCTCCAACTAAATTGAGGATGA

AGCTTATGGATCACCAGAATTAAAAAGATGATTCTGATATTATATCAGCTCGGACGCATCCTAAGCTTCA

GGATATCGAGTTCGTCGAGAACAGATTATTAGCTTCTAATGTTGATTATGATGAGCAAGTGTGTAGTACA

GGGAATGCACCACTGAATTCCAGCTTGAGTGGGCATGGTATCAAATGATGCTGAAGCGAATCAGAGTTCT

TTCTGATAAAAGGAATTTACGTCAGGAGATCATGGCAATGCTCCGGGCCAAAATGCAGAGGGCTGACAGT

GTTAATTCAAGCTCAGTTCACCCACTGAGAACATTAGACGATACTCTACGCAGATAAGTTGCCTTTTTTA

AAACTCTCAATTCACGCGATTATCATGTGATTTGCTGTTCGGTGTAATTTTTGTCAATATATGCAGGTAC

ATAAGCGAAAATGAGTTAATTTATGTGAATCAAATTTTCAGGAACCATTGGTTTTTAGGTGCTATAATAT

TTTGAATTTTGTTCGGAATATGTTTGTATAAGTGGTGATATTTTAAATTATGCACTAATTTGCAGCTGAT

TGTATCCATCTCTTTTTGCACAGTCCGTTGGATCAGAATTCGTATAAGCTGTTGATTTTGAAAAGACGCG

CCGAGAATCGCTGTGTGAAGATTTTTAAACAGGAGCGTGTTCCAGAGACTGAGTTACAGGTCCGAGTTTT

AACCGTGTGGCGTGTTCCAGAGACTGAGTCACAGACAAAGCTCGAGCCTTTCTACCTTCAATCACCTGTG

AGTGGTTTGATATCAGATGGAGGTCTTGCATAAGACATCCAGAAAGTATCATCATTTTGACATATCCGTG

AGCTGGAGACTGCGTACTCATGAGAGCATCTAAACCTGATGCGCCACCGTACGTCGCTCGTGTGGAAAAT

TTAGAAACTGATAGAAAAGGTAGAGTTAGTGTTCGTGTGAGATGGTACTACAGGCCAGAGGAGGCGAGTT

GTGGGAGACAACCATTTCATGGAGCCAAGGAATTGCTTTTGTCAGATCACTATGACAATCAGAGTGCCAA

GACAATTGAAGGAAAGTGCATTGTGCACCCTTTTAAGGAATACACAAAGCTAGAAGATGTTGGAGCTGAG

GATTACTACTCTCGCTTCGAGTACAACGCTAGTACTAAAGAAATCACACCCGATCGTATTTCAGTATACT

GCAAGTGCGAAATGCCTTACAATCCAGATGATTTGATGATACAGTGTGATGGTTGCAAAGACTGGTTCCA

TCCAGTCTGTGTGAAAATGACTACTGCACAAATAAAGCAGTTAAGTAACTATCGGTGTGATCAATGTTCT

TCTTAACATTGTGAAGTACTCCTGGCTAGGTTGCCTGCAAATGCCAAAGACAAGACTGTTGTTCAATTGC

TATGGTCCAAGTGGCGGGGAAAAAGCAGCTGGTTTTTATTACATTTTGTTTTATTAACTGAATTACTGTA

ATAATGCACTATTTTTAGCTGCTTAGGGCCTTAGGTTGGTGTTCTTCAGCAATTACCTGACTTTGTTATG

TTAAAAGCTATCAATTGTGTAAGTGTTTTATTTAGATGGTGAGTGTTGATGGCAAGCAGTACTTTGTGCA

AAGAAAGACTCGGGGAGTTTAATTTGGAATTGTAGAAATTTTATCGGGCACAGGCTATTTTATCTTTTTT

GTTTAGCTTTGCAAG

>MSTRG.1489.2 gene=MSTRG.1489

CTGAAATCAAAAATCAATTTCCTGTAAAAAAAATGAAAGGGACGATGATTAGCAGCTCTGCACGATACCC

TCAACATGGGTCATGCTATACCATAAGTATAATGGCACTATCAGAAGGAATCATAGTGATCATATCAAAC

GAGTTCACCCTACATGTTTGATAAATTGTGCAAGTGAATTGGATCTTCTTCACATTGATTCAATTTGAAA

TGTTTCACTGTACCGTGAGAATCCTCTTCCCTTAAATCAAATAGTGATTTTATGCAGAAGCACTTGACCA

TAGTCACTCATCAGTCATCAGGGCAGACTGGTCGTATTTCTCCAACTAAATTGAGGATGAAGCTTATGGA

TCACCAGAATTAAAAAGATGATTCTGATATTATATCAGCTCGGACGCATCCTAAGCTTCAGGATATCGAG

TTCGTCGAGAACAGATTATTAGCTTCTAATGTTGATTATGATGAGCAAGTGTGTAGTACAGGGAATGCAC

CACTGAATTCCAGCTTGAGTGGGCATGGTATCAAATGATGCTGAAGCGAATCAGAGTTCTTTCTGATAAA

AGGAATTTACGTCAGGAGATCATGGCAATGCTCCGGGCCAAAATGCAGAGGGCTGACAGTGTTAATTCAA

GCTCAGTTCACCCACTGAGAACATTAGACGATACTCTACGCAGATAAGTTGCCTTTTTTAAAACTCTCAA

TTCACGCGATTATCATGTGATTTGCTGTTCGGTGTAATTTTTGTCAATATATGCAGGTACATAAGCGAAA

ATGAGTTAATTTATGTGAATCAAATTTTCAGGAACCATTGGTTTTTAGTCCGTTGGATCAGAATTCGTAT

AAGCTGTTGATTTTGAAAAGACGCGCCGAGAATCGCTGTGTGAAGATTTTTAAACAGGAGCGTGTTCCAG

AGACTGAGTTACAGGTCCGAGTTTTAACCGTGTGGCGTGTTCCAGAGACTGAGTCACAGACAAAGCTCGA

GCCTTTCTACCTTCAATCACCTGTGAGTGGTTTGATATCAGATGGAGGTCTTGCATAAGACATCCAGAAA

GTATCATCATTTTGACATATCCGTGAGCTGGAGACTGCGTACTCATGAGAGCATCTAAACCTGATGCGCC

ACCGTACGTCGCTCGTGTGGAAAATTTAGAAACTGATAGAAAAGGTAGAGTTAGTGTTCGTGTGAGATGG

TACTACAGGCCAGAGGAGGCGAGTTGTGGGAGACAACCATTTCATGGAGCCAAGGAATTGCTTTTGTCAG

ATCACTATGACAATCAGAGTGCCAAGACAATTGAAGGAAAGTGCATTGTGCACCCTTTTAAGGAATACAC

AAAGCTAGAAGATGTTGGAGCTGAGGATTACTACTCTCGCTTCGAGTACAACGCTAGTACTAAAGAAATC

ACACCCGATCGTATTTCAGTATACTGCAAGTGCGAAATGCCTTACAATCCAGATGATTTGATGATACAGT

GTGATGGTTGCAAAGACTGGTTCCATCCAGTCTGTGTGAAAATGACTACTGCACAAATAAAGCAGTTAAG

TAACTATCGGTGTGATCAATGTTCTTCTTAACATTGTGAAGTACTCCTGGCTAGGTTGCCTGCAAATGCC

AAAGACAAGACTGTTGTTCAATTGCTATGGTCCAAGTGGCGGGGAAAAAGCAGCTGGTTTTTATTACATT

TTGTTTTATTAACTGAATTACTGTAATAATGCACTATTTTTAGCTGCTTAGGGCCTTAGGTTGGTGTTCT

TCAGCAATTACCTGACTTTGTTATGTTAAAAGCTATCAATTGTGTAAGTGTTTTATTTAGATGGTGAGTG

TTGATGGCAAGCAGTACTTTGTGCAAAGAAAGACTCGGGGAGTTTAATTTGGAATTGTAGAAATTTTATC

GGGCACAGGCTATTTTATCTTTTTTGTTTAGCT

>MSTRG.1489.3 gene=MSTRG.1489

ATCAATTTCCTGTAAAAAAAATGAAAGGGACGATGATTAGCAGCTCTGCACGATACCCTCAACATGGGTC

ATGCTATACCATAAGTATAATGGCACTATCAGAAGGAATCATAGTGATCATATCAAACGAGTTCACCCTA

CATGTTTGATAAATTGTGCAAGTGAATTGGATCTTCTTCACATTGATTCAATTTGAAATGTTTCACTGTA

CCGTGAGAATCCTCTTCCCTTAAATCAAATAGTGATTTTATGCAGAAGCACTTGACCATAGTCACTCATC

AGTCATCAGGGCAGACTGGTCGTATTTCTCCAACTAAATTGAGGATGAAGCTTATGGATCACCAGAATTA

AAAAGATGATTCTGATATTATATCAGCTCGGACGCATCCTAAGCTTCAGGATATCGAGTTCGTCGAGAAC

AGATTATTAGCTTCTAATGTTGATTATGATGAGCAAGTGTGTAGTACAGGGAATGCACCACTGAATTCCA

GCTTGAGTGGGCATGGTATCAAATGATGCTGAAGCGAATCAGAGTTCTTTCTGATAAAAGGAATTTACGT

CAGGAGATCATGGCAATGCTCCGGGCCAAAATGCAGAGGGCTGACAGTGTTAATTCAAGCTCAGTTCACC

CACTGAGAACATTAGACGATACTCTACGCAGATAAGTTGCCTTTTTTAAAACTCTCAATTCACGCGATTA

TCATGTGATTTGCTGTTCGGTGTAATTTTTGTCAATATATGCAGGTACATAAGCGAAAATGAGTTAATTT

ATGTGAATCAAATTTTCAGGAACCATTGGTTTTTAGCTGATTGTATCCATCTCTTTTTGCACAGTCCGTT

GGATCAGAATTCGTATAAGCTGTTGATTTTGAAAAGACGCGCCGAGAATCGCTGTGTGAAGATTTTTAAA

CAGGAGCGTGTTCCAGAGACTGAGTTACAGGTCCGAGTTTTAACCGTGTGGCGTGTTCCAGAGACTGAGT

CACAGACAAAGCTCGAGCCTTTCTACCTTCAATCACCTGTGAGTGGTTTGATATCAGATGGAGGTCTTGC

ATAAGACATCCAGAAAGTATCATCATTTTGACATATCCGTGAGCTGGAGACTGCGTACTCATGAGAGCAT

CTAAACCTGATGCGCCACCGTACGTCGCTCGTGTGGAAAATTTAGAAACTGATAGAAAAGGTAGAGTTAG

TGTTCGTGTGAGATGGTACTACAGGCCAGAGGAGGCGAGTTGTGGGAGACAACCATTTCATGGAGCCAAG

GAATTGCTTTTGTCAGATCACTATGACAATCAGAGTGCCAAGACAATTGAAGGAAAGTGCATTGTGCACC

CTTTTAAGGAATACACAAAGCTAGAAGATGTTGGAGCTGAGGATTACTACTCTCGCTTCGAGTACAACGC

TAGTACTAAAGAAATCACACCCGATCGTATTTCAGTATACTGCAAGTGCGAAATGCCTTACAATCCAGAT

GATTTGATGATACAGTGTGATGGTTGCAAAGACTGGTTCCATCCAGTCTGTGTGAAAATGACTACTGCAC

AAATAAAGCAGTTAAGTAACTATCGGTGTGATCAATGTTCTTCTTAACATTGTGAAGTACTCCTGGCTAG

GTTGCCTGCAAATGCCAAAGACAAGACTGTTGTTCAATTGCTATGGTCCAAGTGGCGGGGAAAAAGCAGC

TGGTTTTTATTACATTTTGTTTTATTAACTGAATTACTGTAATAATGCACTATTTTTAGCTGCTTAGGGC

CTTAGGTTGGTGTTCTTCAGCAATTACCTGACTTTGTTATGTTAAAAGCTATCAATTGTGTAAGTGTTTT

ATTTAGATGGTGAGTGTTGATGGCAAGCAGTACTTTGTGCAAAGAAAGACTCGGGGAGTTTAATTTGGAA

TTGTAGAAATTTTATCGGGCACAGGC

>MSTRG.1489.4 gene=MSTRG.1489

TTTGGAAACAGTGTTACATCAGACCCGGCTTTTCAAATTCCAGCTCCCTGTACTTTTTTTTTTATCCCCA

AAAAGTCTAGGGTTTACAGAACAATCTCGAATTTTACCAAATGACAAAAGACCCATTTTCTTAATATTAT

TTTATTCCACCCCTTCTTTTATTATTCCCCATTCTCTCTATATATTCTGTAACACACAATTTACACACAC

ATTACATTACTACTGTTTTATTTTCAAGAAAACCCCGTTTTCGAGATTTCAGAGCTTGATTCCATGGCAA

AGACCAGACCAGGCAAAAGGGACTTGGAGTCTTACACCATCAGAAGCACCAACAAAGTTGTAAGAGCTGG

AGATTGTGTTCTCATGAAATCATCTGAACCTGATAAACGGCCATATGTTGCTCGTGTTGAAAAAATTGAA

TCTGATACTGGGGGTAACGTTATGGTCAAAGTCAGATGGTACTATAGGCCAGAGGAGGCAAGTGGGGGAA

GGCGACAGTTCCATGGCGCGAAGGAATTGTTTTTGTCTGACCACTATGACATTCAGAGTGCCTACACAAT

TGAAGGGAAGTGCATTGTGCACTCCTTTAAGGACTACACCAAGCTTGAAGATGTAGGTGCTGAGGATTAC

TACTCACGATTTGAGTATAAAGCTGCTACTAAAGAATTCACACCTGATCGTGTTGCAGTTTACTGCAAAT

GCGAGATGCCTTATAATCCGGATGATTTGATGATACAGTGTGATGACTGCAAAGACTGGTTTCATCCTGG

CTGTGTGAAATTGACTACTGCGCAAGCAAAGCAGTTAAGTAACTACCTGTGTGATGAATGTTCTCCTGAA

CAAAATGGGAAGGACTCTTCTGGCCAGGTAGCCAGCAAACGCCGAAGGACCTGAGCTTTGTTGGTTAGCT

GTGGTCTGCTGGTTGGAAGAAGCAGCTGGTTTATCCCATATTTTGTTTTCTCAAATGAATGACTGTAATG

GTGCACTGTTTCTAGCTGCTTAGGTTGGTGTTCTTCAGCAATTACCTGACTTTGTTATGTTAAAAGCTAT

CAATTGTGTAAGTGTTTTATTTAGATGGTGAGTGTTGATGGCAAGCAGTACTTTGTGCAAAGAAAGACTC

GGGGAGTTTAATTTGGAATTGTAGAAATTTTATCGGG

>MSTRG.1490.1 gene=MSTRG.1490

GTTGGAAGAAGTGGCTGCTTTCTTCTATGTTCTGTTTCTTAACTGAATGGCTGTAATGTTGAACCATTTC

TAGCTACTTAGGTCGGTGATTTTCAAAAATTATCAAATTGTATGTTGTTGAAACCTAGCGATTGTGAAAC

TGCTTTAACTTCAGCTGGTGACTGGTGATGGTAAGCAGTACTCTGTGCGAAGAAAGTGTCGGGAGTTTAA

TTGAAAATCTTTGAACTTGTCTCCATATCGCTTGTCACCAAGCATATTAGAGCATCTCCTACATATAACT

ATATT

>MSTRG.1491.1 gene=MSTRG.1491

GTTGGAAGAAGTGGCTGCTTTCTTCTATGTTCTGTTTCTTAACTGAATGGCTGTAATGTTGAACCATTTC

TAGCTACTTAGGTCGGTGATTTTCAAAAATTATCAAATTGTATGTTGTTGAAACCTAGCGATTGTGAAAC

TGCTTTAACTTCAGCTGGTGACTGGTGATGGTAAGCAGTACTCTGTGCGAAGAAAGTGTCGGGAGTTTAA

TTGAAAATCTTTGAACTTGTCTCCATATCGCTTGTCACCAAGCATATTAGAGCATCTCCTACATATAACT

ATATT

>MSTRG.1492.1 gene=MSTRG.1492

AGAGCACTTTCTCCCCAGTCCCAGAATCAGGATATTGTTTAACAAATATATTTTGGTCATTTTTCGCAAC

AAAAATGCTCGTGATATACCGGGTCTGTTATCGTATTTTCTGATGATTCTTATTTTGTACATACACAATC

ACTACAGCTATTTGATTTTCAAGAAAAGCCCGTGGTATGGTTCCCAAGCTTGTTTCAATGGCAAATTTCA

GACCTGGCAAACGAGGCTTGGTCTTTTACAGCATCAGAGCCACCAATAAATTTATACCAGAAAGCGGAAA

AGGGGAGCAGATATTTGCCAAATCAGGGGAACCGGCGAAATTGTTAGGGCTGGAGATTCCGTGTTCATGG

AATCATCTGAAACTGATAAACAGATCTATGTTGCTCGTGTCGAAAGATTCGAAACTGATAGCGAAGGCAA

GTTTATGGTTAAAGTGAGATGGTACTATAGCCCAGAGCAGGCTGGTGGTAGGAGACAGTTTCATGGGGCA

AAGGAATTACTTTTGTCAGATCAATATGCCACTCAGAGTGCCAACAAAATTGAAGGAAAGTGCAATGTGC

ACTCGTTCCTGAACTACACCAAGCTAGAAAATGTAGGTGCTCGGGATTACTACTGTCGCTTTTGGTACGA

AGCTGGTACTAAAGAAATTACACCCAATAAAGTTCCAGTGTACTGCATATGTGAAACACCTTACAATCCA

GATGATTTGATGTTACAGTGTGATGACTGCAAAGCCTGGTTCCATCCTGGCTGTGTGAAAATGACTATTG

CACAAGCAAAGCAGTTAAGTAGCTATCAGTGTGATCAGTGTTCTTCTTAACAAGATAGGAAGCACTCTCC

TGGCCAGGTGCCTGCAAAGTCTGAAGGACATGATCGTCACTGGAAGCTGTGGCCTAAGTGGTTGGAAGAA

GTGGCTGCTTTCTTCTATGTTCTGTTTCTTAACTGAATGGCTGTAATGTTGAACCATTTCTAGCTACTTA

GGTCGGTGATTTTCAAAAATTATCAAATTGTATGTTGTTGAAACCTAGCGATTGTGAAACTGCTTTAACT

TCAGCTGGTGACTGGTGATGGTAAGCAGTACTCTGTGCGAAGAAAGTGTCGGGAGTTTAATTGAAAATCT

TTGAACTTGTCTCCATATCGCTTGTCACCAAGCATATTAGAGCATCTCCTACATATAACTATATT

>MSTRG.1492.2 gene=MSTRG.1492

AGAGCACTTTCTCCCCAGTCCCAGAATCAGGATATTGTTTAACAAATATATTTTGGTCATTTTTCGCAAC

AAAAATGCTCGTGATATACCGGGTCTGTTATCGTATTTTCTGATGATTCTTATTTTGTACATACACAATC

ACTACAGCTATTTGATTTTCAAGAAAAGCCCGTGGTATGGTTCCCAAGCTTGTTTCAATGGCAAATTTCA

GACCTGGCAAACGAGGCTTGGTCTTTTACAGCATCAGAGCCACCAATAAATTTCTGGAGATTCCGTGTTC

ATGGAATCATCTGAAACTGATAAACAGATCTATGTTGCTCGTGTCGAAAGATTCGAAACTGATAGCGAAG

GCAAGTTTATGGTTAAAGTGAGATGGTACTATAGCCCAGAGCAGGCTGGTGGTAGGAGACAGTTTCATGG

GGCAAAGGAATTACTTTTGTCAGATCAATATGCCACTCAGAGTGCCAACAAAATTGAAGGAAAGTGCAAT

GTGCACTCGTTCCTGAACTACACCAAGCTAGAAAATGTAGGTGCTCGGGATTACTACTGTCGCTTTTGGT

ACGAAGCTGGTACTAAAGAAATTACACCCAATAAAGTTCCAGTGTACTGCATATGTGAAACACCTTACAA

TCCAGATGATTTGATGTTACAGTGTGATGACTGCAAAGCCTGGTTCCATCCTGGCTGTGTGAAAATGACT

ATTGCACAAGCAAAGCAGTTAAGTAGCTATCAGTGTGATCAGTGTTCTTCTTAACAAGATAGGAAGCACT

CTCCTGGCCAGGTGCCTGCAAAGTCTGAAGGACATGATCGTCACTGGAAGCTGTGGCCTAAGTGGTTGGA

AGAAGTGGCTGCTTTCTTCTATGTTCTGTTTCTTAACTGAATGGCTGTAATGTTGAACCATTTCTAGCTA

CTTAGGTCGGTGATTTTCAAAAATTATCAAATTGTATGTTGTTGAAACCTAGCGATTGTGAAACTGCTTT

AACTTCAGCTGGTGACTGGTGATGGTAAGCAGTACTCTGTGCGAAGAAAGTGTCGGGAGTTTAATTGAAA

ATCTTTGAACTTGTCTCCATATCGCTTGTCACCAAGCATATTAGAGCATCTCCTAC

>MSTRG.1493.1 gene=MSTRG.1493

AGTTCCTGACCACACTTTTGTAGGAGTGGAATTCGTTACCTTCATTCAAGTTTTGCATTCAGTCTCACTC

TTTGTGATTAATGCTTCCGGTTCCTGCCAGAAGATTTGGACCACCTGTTTTTTTTCATAAGGTGAAACTC

CTGCGTGTCTTCGGGGTCTTTTTGGCTATGGCAATTCAGCAATACTTGAGAGATGGTATAACCAGAGATG

AGAGAGCTAGAGGCCAAGGACACAAATGCCATGAAAGAAACTGCCACAGATGCATTTGCCATATTCGGAA

ATTTGTCTTTTCCCCAGTTAGATCTCCAGTCCTCAACTCGTGTTGCTGCTGACGATGAAGACGACATTAG

AAGATAAGCCAAAATCTGCATGCATTTTTCAGACACCATCATCAATCATTGAAATGCATTACTAGATAAA

TTATGCTCCACTGCAGAAGTTGATACCGGAGAGTCTTAAAGGAACGGATATTTATGCTGCTCTGAAGTTT

AAACAAAAACCAAAATAAAAAAAAATTTCAGGTGGATTTTTACTCAATTAATCGAAACCATCTACTTAGG

AATATCATAATATTTTATAATCTAGCTCTTGTCTCTTATGCAGTGCGTTCAGAGGGCCTCTCTTATTCTT

TTAGCTTTTTTCTCTTAGATTATGGCTAGGCATCATTTCGGCAGTAGACTTTCACTCACCTTTCAATCTA

CATAAATAGTGCTTTGAAAACGTCCTAAAGAACTGCATCTCTAATTTTATGAATTGATCAATGTCACCTC

AAGGAGTTGATTAAAATTAACCAGATAGTGATATCAGGGTACGTTTGTTGCATTCTTCTTTTACACAGGA

CTTGATCAACTTCGATAAAAAATGATGAATTCAAGCAGAATAAAGCAGAATGACAATCAAAAGTAGACGG

AAAAAGACACCTGATCGGAGGCAAAATCAACAATATTGCGGAGATGAGATGCAGGCACAAGAAGTTTATT

ACCATCAGAGGAGAATAACTGGTATGCAAGATCAAATCCTTGTAAGGCCGAGTATATAAATCCTATCACA

TTCACTGTCAAACAGTACCTGCAGCAGTTCCATATAATTTTATGAATAGAAATCAACCTAGGAAACTAAA

GAGCAAGTCTTAAACATACCTAAATTCGATATAACGATGAAATGAGTCAAGAGTCCAGCCTTTATCACGA

TCAGCAGCCATTATGGAAAATGAAATCAAGCAAAACAAAAACCCGAAAACCCGAAACCAGAAATTCACCT

TCTTAAGCACCTGATCATCACCGGTTCTTTTTGTCGACTGCTTTCGACTGATTTCTTTTTGTGGAGCTGG

CGAGGGATGTAAACTGTGTTCAGGTCCTTCTCCGTCGCGACTAAAGGATGACGGAGAAAAAGAGGATATG

GATGACTCGTATGGCGGAGGGGAGGGATTCTGCGATCTGAAGTTCTGCTGCTTCTTCTCCATGAGATACT

ATTGTTGTCAACTGACAACTAACTGTAATAAATAAATGATGAATCTTTCGTTTGGAGGGAAGTAAGCAAA

ACAGAGTAGACTAATACTGTGTCATGTTTGCTAGTTAAAATTTCTTAATATAGGTAAAAGAAATTCAGAA

TTCTCTGATTATTTGAG

>MSTRG.1496.1 gene=MSTRG.1496

TCTTTATTTATGCAAATTATGTGTTACTACATTATTGAGCAATTGTTCATGTGTAGTAGTAACACAAAAA

AGCCTCGCAGCTCATATTAGGGACTGCTCATAGATTTAGATGTCTTTAAAACTGTGACTTGGGCACAAGC

AATTAACAATCACTTTTTGGAGTTTATAGAAGAGTTAAACCAAAACTTGTTATTGACAGACAATGGAGTA

A

>MSTRG.1499.1 gene=MSTRG.1499

CAGAGTGACATCCGAATGAGAAACACTGACATCTGAATGAGAAACAGTGCAAGTAATAAAAATTTACGCG

TGTGCAAGTGAATATATAAGGCAGGGATCAAGAACTGAGCCCAATCAAACCATTTCTCTTCCCTCTCTTC

TCTCTGCAACGATTTATATATCAACAAAACTTCAATATCACTCAAATAAACAACTTTTAGTCCCCCAGAA

AAATCCATGGCAGAAGAAGCTGAAGATTATCAGAAACAAGAAGAGGGTTTCGTGAGAAAAAGCTCAACAT

TGATCTCCATCCAAGATGGATCTGATGATCTCAAGAATTTCTTCACATGGGTTTTTCTTGATCAATCAAA

TATTTGGAGAACTGGTCTTTCATGGTCCATCTTCTCTCTTCTTGCTATTGGTGTGCCACTTTTGTCTCAT

TTTGTGTTTCAGTGCAATTCTTGTGATCATAATCATAGAAGACCCTTTGATTCCATTGTTCAGTTGTCTC

TCTCTTTTGTCTCTGTTCTTTCTTTCATTAGTTTGTCTTCTTTTACAAGAAAATATGGGCTCAGAAGATT

TCTTTTTCTTGATAAATTGTATGGTGTTAGTGATATTGTTCAGAAAGGATATGCTCACCAGCTTCATATC

TCAATGAAGCTGTTGTGTTCTTTTGTTCTTCCGTGCTTCGTTGCTGATTGTATGTATAAGATATGGTGGT

TCACCTCAGGGAGAAATCAGATTCCCTATTTTTTAAATGCCTTTTTGAGCAACACAGTTGCTTGTTTTTT

GTTGCTAGCTTCGTGGTTGTACAGAACATCAATATTCTTCCTCGTATGTGTTCTTTTTCGATTGATATGC

TACCTTCAGATACTCAGGTTGGAAGACTTTGCTCAATTTTTCCACAAAGAATCCGACGTTGGTTTAATCT

TGCTTGAACACTTGACAGTAAGAAGAAATCTCCGTATTATAAGTCACAGGTTCCGAGTATTCATTCTGCT

GACTCTAATCTTGGTAACAGCAAGTCAGTTCGCTTCTCTGCTTGTTACAACGGAAACTGGCTCTGAAGTT

AATATATTTAATGCTGGAGAACTTGCGTTAACCTCAATGACCCTAGTGGCTGGGCTCTTCATATGTCTGC

GTAGTGCTGCAAAAATTACACACAGAGCACAGGCTATCACATGCCTTGCTACAAAGTGGCACGCATGCTC

CACAATCAACTCTTTTGATGACGTTGACAATGAAACTGCAAGCTCTCAGTCTCAGTCTCAGACTATCTCA

CCCAGGGTAAATTATATTGACGGTCACTGGGGATCTGACAATGACGAAGGCGATGAAGATGATATAGACA

ATACAGATATGACGCCGATTTTCACCCATACAATATCCTACCAGAAGAGACAAGCTCTTGTGACATACTT

TGAGAACAACAAAGCGGGGATTACTGTGTTCGGATTTATGCTAGACCGCACATCACTCCACACTATATTT

GCTATTGAGATGTCCCTTACCCTCTGGCTGTTGAATAAGACAATTGGTATTTCATGATCTTCTCCATACA

CGATGCTGAAAATACCTTTAGTTTTCTTGTTTAGATATGTCTATAGTCAAGATCAATTTATGGTGTAACT

GGCCTTGTGATGAGAGGCACTTAAAACCTGTGTAATTTAATGTGTGTATATATATATATCATAAAACATG

ATATGTATATGTTGTACTCTGGCAGTCTGGCCTAGTATTGTACTTAGTTGTGCATGAAGAGATAAGAGTG

TATATTATTAACAATATTGTTGAGAAGTTGAATAATCTGAGGCAGAATTCAGAAAATCGTATACGATTGA

TTCAGAGTTGAAGAGAAAAGTGTATGAATTCTTGTGGTTGTTGGCTGAATCAGTACTAAAATAACGGAAA

GTCTGGCCAGTATTTGATAAGCCAACTATGGTAACGCTTCTTGGAAAGCTTACTGTAAACAGTTTTTTTT

TTTGTAACATTAATGGTCATTGCTGGATAGTTGAGATCCTTAGGTGTGAATGTGTCTGCTGTAGCGAGCT

GGTACTGTTATCTCCTGAAATAATTCTAAGTTTGGAGTTACTGTTATCAATGCTGCATGCAGAGCATTCT

AAGATGGTCTTCTTTGGATGTTTCATACTAGTCCAGGATTTATAGCTTTTAGTGGATTAAAGTGTCCTGA

TCCATAAGCAAACTCTCTGCGTCTGGATTCTTTGAAACATCCATGCGCCACGCTAGCATGTTATGCAACA

TGAAGCATGCTGGTGGTTTCATCAACTCAGCTTACTGTGATAGTAAATTATGGCAAAAACGAAATTTTAC

TTGCAGAGTTG

>MSTRG.1504.1 gene=MSTRG.1504

CCTGCCACAATCATCCTTCAAGATGGAAATTTCTAAAATTAATTTTGTAACAATAAACAAAAACATTGAT

ATGTTCAGGCTATGGTGGATTGATCTATGGACTTGGCCAACTGACAATCAAACGAATTTTCGTATGTGAG

ACTACATTTGTTCTAAGGGTCAGGCATTTACAGGTAGAGACGAGGACGATGACTTTATGGGAATGCAAAT

TTTCAAGCGTTCCAGTATGGCTTTTCCGCTGACAAGCCCTGTGTGAAGAGTCAATGGAAGTCCTTCCTTG

ACACACTCGTCGTATTTCTCCAGAGCAGATACGATGGCGCTGAAATCTGGGCGTTTGGCTGGATTTGCAG

CCCAGCAGTGCTTTATCAGATGTGCAAGCGCGGGTAGACAGCTTGGTGGCATTGGGGGTCGCTCGTTCTG

CAGAATTTGTAAAATGTAGTAGGAAAGTAAGCATCAAGATTCAATAGTTCTGATTATTATTATTTTTATT

TTTTTGAGAATTTGAGTG

>MSTRG.1505.2 gene=MSTRG.1505

CCTCGGCCAAACGGCTCTTAACTCAGTCCATAGCCCCTTGTTAATTCAGGTGAGAGGCTTGGTTTCACTC

GGTCGGTTGCAATACGCAAAAGAAAGATGGACTCGCCGGGGCAGAACACTTCTCTCCAACGACTCCAGAA

TGTTGAAAAGAGAATAGTTCGGGTACTGGAGCTAGCAGGAGGAGTGATGGATGAATTAGCCAACCCCAGT

GGCTCCAGGAAGGATTTGGTCAGTGCCCATTGCGCCGAGTTCATGAAACTCATCAAGGACATCCAAATGA

CACTGAGGGAAGAAATTAAGAGTGCATGTGATTATCGTCCATTCGAAAAGTGTGACTACATAGCGAGAGT

TTCTAACGAGATATGCTGCAAAAAACTAGAATATGTTGCTGCTCAGTTGGATGGTATGAAACAAACAATT

GATGAATATCATAATGCAATATGAAGTTTCCCACATGCCAATTTTGGCAGTTTGTTGAATTTTCATATTG

TGAAGCTGATTCGAATTGTTAGAACGATGATTTACATTCATATGTTCAAGGATTTGCTAAAGTTTAACTA

TTGGAAATCCCAATATATTACATAGTCACCATATCATTCTCATTGTTCTAGGTGCTTATATATTTCCCCA

TCAGCTATTTTTTTGGTCATTCCACCAAAAATTATTAGTTTATTACAGACTCTGAAAAATTTCATCTGTT

AATCATATTGTCCACACGTTTATTT

>MSTRG.1505.1 gene=MSTRG.1505

CCTCGGCCAAACGGCTCTTAACTCAGTCCATAGCCCCTTGTTAATTCAGGTGAGAGGCTTGGTTTCACTC

GGTCGGTTGCAATACGCAAAAGAAAGATGGACTCGCCGGGGCAGAACACTTCTCTCCAACGACTCCAGAA

TGTTGAAAAGAGAATAGTTCGGGTACTGGAGCTAGCAGGAGGAGTGATGGATGAATTAGCCAACCCCAGT

GGCTCCAGGAAGGATTTGGTCAGTGCCCATTGCGCCGAGTTCATGAAACTCATCAAGGACATCCAAATGA

CACTGAGGGAAGAAATTAAGAGTGCATGTGATTATCGTCCATTCGAAAAGTGTGACTACATAGCGAGAGT

TTCTAACGAGATATGCTGCAAAAAACTAGAATATGTTGCTGCTCAGTTGGATGGTATGAAACAAACAATT

GATGAATATCATAATGCAATATGAAGTTTCCCACATGCCAATTTTGGCAGTTTGTTGAATTTTCATATTG

TGAAGCTGATTCGAATTGTTAGAACGATGATTTACATTCATATGTTCAAGGATTTGCTAAAGTTTAACTA

TTGGAAATCCCAATATATTACATAGTCACCATATCATTCTCATTGTTCTAGGTGCTTATATATTTCCCCA

TCAGCTATTTTTTTGGTCATTCCACCAAAAATTATTAGTTTATTACAGACTCTGAAAAATTTCATCTGTT

AATCATATTGTCCACACGTTTATTT

>MSTRG.1505.3 gene=MSTRG.1505

CCTCGGCCAAACGGCTCTTAACTCAGTCCATAGCCCCTTGTTAATTCAGGTGAGAGGCTTGGTTTCACTC

GGTCGGTTGCAATACGCAAAAGAAAGATGGACTCGCCGGGGCAGAACACTTCTCTCCAACGACTCCAGAA

TGTTGAAAAGAGAATAGTTCGGGTACTGGAGCTAGCAGGAGGAGTGATGGATGAATTAGCCAACCCCAGT

GGCTCCAGGAAGGATTTGGTCAGTGCCCATTGCGCCGAGTTCATGAAACTCATCAAGGACATCCAAATGA

CACTGAGGGAAGAAATTAAGAGTGCATGTGATTATCGTCCATTCGAAAAGTGTGACTACATAGCGAGAGT

TTCTAACGAGATATGCTGCAAAAAACTAGAATATGTTGCTGCTCAGTTGGATGGTATGAAACAAACAATT

GATGAATATCATAATGCAATATGAAGTTTCCCACATGCCAATTTTGGCAGTTTGTTGAATTTTCATATTG

TGAAGCTGATTCGAATTGTTAGAACGATGATTTACATTCATATGTTCAAGGATTTGCTAAAGTTTAACTA

TTGGAAATCCCAATATATTACATAGTCACCATATCATTCTCATTGTTCTAGGTGCTTATATATTTCCCCA

TCAGCTATTTTTTTGGTCATTCCACCAAAAATTATTAGTTTATTACAGACTCTGAAAAATTTCATCTGTT

AATCATATTGTCCACACGTTTATTT

>MSTRG.1514.1 gene=MSTRG.1514

GCATTGTTTTATATTCTTTTCCTGCTGAAAACGGTGGGATAAGCCCATTCTAATCTCTCGGCTAAAACAA

AAAAACACGCTCGAAGCATCCGATCGAGCTTCAGACTCTCGCTATTCAGGCAAGGACGATTCAAACTCTC

GCATCTTCGACTCCGCCGATTTCAATAAGCTCAAAATCCGATCGACGCATTTCAGCGATTCACTTTACCA

TTTGCAACCATTTTGAAACCTAAATCAATCTGAGCATACTGGCCATAAAGCAGAAGAAACTTTTCATGAT

AGATTACCATGATTTGCTACTACCATATGTGAAGAAAGTGAGGCAGATAAAAGGGACATCTCTATATGGA

TCAAGGGCCTTGTTTTTTCTAACTCCTATTGGCACATTGAAGCCAGTAGCCATTGAGCTTGTCCGATCTC

CAATGAATGGGAAACCTCAGTGGAAGAAAGCATATTCTCCAGGGTTTGATGCGACAAGTGTTTGGCTTGG

AAGCTCGCTAAAGCTCATTTATTGGCACATGACTCTGGTTTTCATCAGCTTGTTAGTCACTGCCAGATCT

GCATCTTTGACTGAATGCGGCTTCATCTTAAATTTCTGATATTGTTCAGAGTGCTTTATTCCCTGATGCT

GGCTTGGAAAGTGATCTTGGTCATGCTGGAGAAGTGGCAGGGCTAGTAATAGTTCAAAATCTTGACCTTT

TCTTGGAGAAATTTTGTATTTTATTTGTAAATTATGAAGGAGAATTAGTAGTGGCGAAAGTTTGGTTTTT

TGTAAATTTTTAATATTTATTATAGTTGATTTAGGTACTACTAATTTGTGTAATGAAATGTGAAGTATGC

TATTGACATGAAATTAATCCATTTGTCTATTTGATTTATAAA

>MSTRG.1514.2 gene=MSTRG.1514

GCATTGTTTTATATTCTTTTCCTGCTGAAAACGGTGGGATAAGCCCATTCTAATCTCTCGGCTAAAACAA

AAAAACACGCTCGAAGCATCCGATCGAGCTTCAGACTCTCGCTATTCAGGCAAGGACGATTCAAACTCTC

GCATCTTCGACTCCGCCGATTTCAATAAGCTCAAAATCCGATCGACGCATTTCAGCGATTCACTTTACCA

TTTGCAACCATTTTGAAACCTAAATCAATCTGAGCATACTGGTGTTCGATAAAAAGCCTCTAACAACTAT

ATGATGAATTACAGCAATGGCAAGACTTTCACTAACTAAAAGTATGGTTTTTTTTTTATGATTTTAATTG

TTACGGCCCGAGGTATTGGAGTGTTTTTTATGGTTGCACTTCAGGGGGTAATGTACTTAATAAAACCAAA

AAATCAAAAGTAGAAGTGTTGATTTAGGGATCGGAGAAGTGAATTGGTTGAGGAACAATTTTACGGGGTT

GCTTGAAGGAGTAGAGTACTGCGCCACAATCAATAGAGTGATCTGGCATTTTTTTAGGGATTTTTTTAGG

TTTGTTTGGTCATATAAACAAGGGTATAGCTTATTATTCTGAAAATGGGTGCTTCTGTTATCTTGGTAAG

ACTGCAGTGCTTTCATAATTATGTTCTCATGGGGGTTCAGTAGGATTTACGGTTAATGCATGACTGGATT

ATGGTGAATGGATGGAAGAATTTCTAGCCAGACAAGCTGGTTATTTGTTTGGTGTTTAAGAAAGAATTGT

GTTTAATGAGTCTGAAATGTTTTGTTATTATTGCTTCATTTTGGACTGTTGGGTACAAAGCTGGTCATTC

ATTTTGATGTGTCGAAACAACTTTGGTTGCCAGGTTAAAAATTAGGAGCCTAAATCTCACCATCTGTGGC

ATAAGACTCTATAGGGAGTGGTTCAAGGGTTCAAAAACCCTTATATATGATAATAATACATATCAATCAG

CTTATATATGATAATTATAAATATCAATGCAGGCCATAAAGCAGAAGAAACTTTTCATGATAGATTACCA

TGATTTGCTACTACCATATGTGAAGAAAGTGAGGCAGATAAAAGGGACATCTCTATATGGATCAAGGGCC

TTGTTTTTTCTTTGTAAATTATGAAGGAGAATTAGTAGTGGCGAAAGTTTGGTTTTTTGTAAATTTTTAA

TATTTATTATAGTTGATTTAGGTACTACTAATTTGTGTAATGAAATGTGAAGTATGC

>MSTRG.1514.3 gene=MSTRG.1514

GATAAGCCCATTCTAATCTCTCGGCTAAAACAAAAAAACACGCTCGAAGCATCCGATCGAGCTTCAGACT

CTCGCTATTCAGGCAAGGACGATTCAAACTCTCGCATCTTCGACTCCGCCGATTTCAATAAGCTCAAAAT

CCGATCGACGCATTTCAGCGATTCACTTTACCATTTGCAACCATTTTGAAACCTAAATCAATCTGAGCAT

ACTGGTGTTCGATAAAAAGCCTCTAACAACTATATGATGAATTACAGCAATGGCAAGACTTTCACTAACT

AAAAGCCATAAAGCAGAAGAAACTTTTCATGATAGATTACCATGATTTGCTACTACCATATGTGAAGAAA

GTGAGGCAGATAAAAGGGACATCTCTATATGGATCAAGGGCCTTGTTTTTTCTAACTCCTATTGGCACAT

TGAAGCCAGTAGCCATTGAGCTTGTCCGATCTCCAATGAATGGGAAACCTCAGTGGAAGAAAGCATATTC

TCCAGGGTTTGATGCGACAAGTGTTTGGCTTGGAAGCTCGCTAAAGCTCATTTATTGGCACATGACTCTG

GTTTTCATCAGCTTGTTAGTCACTGCCAGATCTGCATCTTTGACTGAATGCGGCTTCATCTTAAATTTCT

GATATTGTTCAGAGTGCTTTATTCCCTGATGCTGGCTTGGAAAGTGATCTTGGTCATGCTGGAGAAGTGG

CAGGGCTAGTAATAGTTCAAAATCTTGACCTTTTCTTGGAGAAATTTTGTATTTTATTTGTAAATTATGA

AGGAGAATTAGTAGTGG

>MSTRG.1515.2 gene=MSTRG.1515

TTGAATTTTGAGCTATGAAATCAACAATCACGAGCTTCCCAAAACCCAAATCATACATGTATTTTTTCAG

CTTGAAACATCCCAGAAATCCTTTGATTTTTTATGCTAAATCAACCCTTTTCAACAGGCTTCACACTCAC

CCTTCTTTCTCTCCCGGAGAGATTGTGGGGCCCATGTTCTCCGGCAAGACCACCACCACTGTCTCGCCAA

GACTCCCTCCTCTCACCAGGATGAGGCCAACAAACACTTCATGTATTTGGACAAAACGTTGAAAGGTTTG

ATATACTCGGGAAGGCTGGTACATGCAATTGATTTGTTGTGTCGCTCAGAGGTGGCAGTGGAATCTCAAA

CATATCGTCTTCTTTTGCAAGAATGCATATTTAAGGGAGACTATAGTAAAGGAAGAAGAATCCATGCGCA

GATGGTTGTTGTTGGATTTGTTCCTGATGAATTTCTTAAGGTCAAGTTACTTATACTGTATGCAAAATCT

GGGAATCTAGATACTGCTCATATACTCTTTGATCAATTACGTGAGAAAACTTTGATTTCATGGAATGCAA

TGATAGCAGGTTATGTTCAGAACAGTCTTGAAGTTGTGGGATTGAATCTTTATCATAGTATGAGACAGTG

TGGTTTTCTACCTGATCAGTTCACATTCTCATCAGTCTTGAGAGCCTGTGCCACTTTGACATTACTCGAA

CAAGGGAAACGAGTCCATTGTCTCTTGATAAAGAGTCAGATTACTGAAAATGTCGTAGTTAATTGTGCCC

TTATTGATATGTATTTTAAGTGCAGCTGCCCCTATGATGGTCAGTTAGTATTTGATACATCTCCTGTGAG

GAATGTTATTACATGGACTTCTTTGATATCTGGATATGGTCAGCAGGGAAGAGTAATAGAGGTTTTGGCA

TCCTTTAATAAGATGATATCTGAAAATTTTAAGCCAAATTATGTGACTTTTCTTGCAGTCCTTTGTGCGT

GTAGCCATGGAGGTTTGGTTAACGAAGGTTGGGAATATTTTTTGTCGATGACAAGAGATTTTGGCATTAA

ACCAAGAGGGAAACATTATGCAGCAGTTGTTGATATCTTAGCAAGAGCTGGAAGGCTAGAAGAAGCCTAC

ACATTTGTAAGAGACTCACCTTGTAATGAACGCTCGGTAATATGGGGTTCTCTTTTGGAGGCTTGTAAAA

TGCATGGTAACATGGACATGTTGAAGTTTGCTGCAAGTAAATATTTTGAGTTGGAACCTGATAATGCAGG

GAAGTATGTTGTCTTGTCTAATGCTTATGCAGCTGTTGGCTTATGGGACAAGGTCTCTAAAATAAGGAGT

GAAATGAAGGAAATGGGATTAAAGAAGGAGCCAAGTTATAGCATGATTGAGGACCAAAATGGGGTACACT

ACTTTTATATGGGTCGTAATCCTCACAAAAATATTGAAGGAATATACGAGGTTATTAAAGCATTGACTTG

CATCCTAAAGGATGATGGCCATGTTCCTGACATAAGCAGCGTCTGGCAAGAAGATTAATGCAGTCAGCTT

AAATAAAGTTGTAGATGAGGGGGGACCTCATAAGTTCTTCTGGGGAGGCTAATTTACTCGAAAGATCAGG

AACCTATGTGGACTTTGTTTTTTGATTTAGGTACAGCCTGGTAGGAGTTCGATGCAATCAGCAGGTACTA

CTCAAAGAAAAAGGCTGAACCATATAAGTTTCTGAGGTACTATTTGAGAGACATGGATCACCGAGAGTTT

GAAGTTGAAGATAATTTTGCAAAGTAAATAATATTAGTATGGGGCACAATGTATCCGCTAATAACATGAC

TTACTAATTTGAATGGTTCTTGGAATACATGTGTGGCCAGTGGCAAGGGCAAGCAACATCATGATGGCAT

AATTGTCTCAGAGTTTTTGGTTCGTTCAGGCCATGTTGTTGTTTTCTTAACATAAGCTGGAGAATTTTTA

GGAAATGTTTATCTTCAATTGGTGTAAGTGGCTGCTTAAGGATGTCATTTGAAGGAATTGCATAATTTTT

ACCTTCCAGGTAGCTGAGGATCGCTCTCCTGTTCCTGTTCAAACCATTTAGGTTGGCCATCTTCGTAGAA

GTCAATTTTATGCTCCACATCTCTATTTTCTGTACACCTGGTCTGAAACCTGGGATAGAAGTGTCAATTG

ATAACAGAAAAAGAATGCTTTTATTGGGCGTACTAATTTCTGCTGAACAGACTGATTCTACTGGAATGTA

AATTAAGGTATATTATTACTTTTTAATTTTGCAGGCTAGCATTTCCTGTTACACTCTAAACTAGCAGTAT

GAGCTTATAAATTAATCTGCTATCAAGTACAAACTTGTGGATTTCCAGACTACTTTCGCCATTCTGTTGC

TGCCTCCGCAGTTTCTTTATTACACTGTTGCTGCTGCTGCAGCCTGCTGGACTGGTTTTTCTGCTGCCAT

ATAAATTCATATCTCTGCTGTTTTACAGATTCCATACAGTTGTAGCCTTGCTTCTAAACTATGATCATTT

TGCAAAGCAGGTATTCACAGAGACTAGAGCACTTGTGCCACTACATTCTTTTTATTCATCAAGAGCTCTC

AAACAGCATTGCTAACGAGGGTAGTTCTGCATGGTGCCATCCTTTCTTGTCTTTGCTTACTCTGCAGTCT

GCCATGTGAAATTTACACTTTGTTTCGACACTTACAGGTGACTGAAGCTAGAGATATGCAAGAAGCAGAA

GCTTTTGGAGGATATGTATAAATTTCAAGGGTGTTGATTTTCTGATGGGGACCAACACATTTTAGAATGT

GCCTTCTTTTGTTAGGCAGACAGGAAAACTAAAGCAAAAGCCGTCGATAGCCAAGTAAGAAAACGCTACT

GACGATGCATTTAGGTTTGGTGTGGCTTATTGTTGAATGTTTATGCTTTCATGTTGCAGGTCCAGAGCTG

TTTGGTTGTGCTTTTAAGTTATATCAGATGGAGATACTATTGTACAAGTACTGGTAGACTAGATGGAGAA

AAAACAATGTGTTTCTTCACTATTTTTCAATCATCTTTAAAGACTTGATTTCATGTACCTCTTGGTTTAT

ATATGAGATTCAAATGTAGACGGCGAGTAAAGATAAAACAGCACTGCAAGATGGTTATATAGAAAGACAA

TACATAA

>MSTRG.1517.1 gene=MSTRG.1517

CGACCCGCTGCTCACTGTAGTGCAGAAGCTTTTTGCAGCAAATACAAACACTTGTTCCCAGGTTAGAGCT

CTTCATTCGGCTCTGTTTCTCATATATTTTTCAACTCTATATACACATATACATACATGTATACACACAA

TTGCTTATAACCCTGAAAAGTTTATGTATGTGTAATGCCAATCAGTTGTTTGCTTAGCTTTTGCTTGTTT

ACAATTACCCAATAAGAGTTATGCTCAAAAGATTGAATCTTGGGTGTCATTTTGTCAAAAAGGTTCAATC

TTTGTATAAAAATCCAATCTTTACATATTGTAGTTCAATTGGGGATGATGGAAATGTTGTTGGTAATTCT

GATTATAAGAATGGGTCATCCCCTGAAACAGATTGGGAAAGTTTGCTTGAACCATTTGACATTGAACAGC

TTCGAAAGTCCCTTAATCGACTTACACCGTATCAGCTTAATAAGTTGCTTCAGCTCCCCCTTGATGTCCC

TACTTCTATGGATTTGTTCCGTAAGGCGGGTACTCAAATGGGGTATACTCATACATTTGATGTGTATTGT

ACTTTGATTAATAAACTTGGAGCGGCCAGGGAGTTTAAGTTGATTGATGGATTGTTGCTGCGGATGAAAG

ATGACGGGGTAGTTTTACAGGAGTCACTTTTTATTACTATAATGAGGCATTATGGTAGAGATGGTTTACC

TGGTCAATCGACGCGGTTACTTTTTGATATGAGGGAAGTTTTTAGTTGCGAACCTAGTTTTAGGTCATAT

AATGTTGTTCTGGATATATTGGTTGCTGGGGATTGCCCGGCTGTTGCACCGAATATTTTTTATGAAATGT

TGCACAAGGGGATCTCTCCAACTGTTTTCACGTTTGGTGTGGTGATGAAAGCGCTTTGTATGGTTAATGA

GGTGGATTCTGCATGTTCACTTTTAAGAGATATGACAAAACATGGGTGTGTGCCGAATTCAGTTATTTAC

CAGACACTTATACATGCGCTTTGCAAGGTAAATAGAATAAATGACGTGATAAGATATTTGGAGGAAATGT

TTCTGATGGGTTGCACACCTGATGTTGATACTTTTAATGATGTTATTCATGGTCTTTGCCGTGCTAAGAG

GATACCTGAAGGTGCTAAATTGGTCGATAGAATGCTTCTCCGAGGCTTCACTCCTAGCAAGTTTACTTAT

GGAATTTTAATGGATGGATTATGCCGAACTGGTCAAGTTGATAAAGCTAGAGCTTTGCTGGAGAGAGTGC

CTGAGCCTAATGTTGTTTTGTTCAATACTCTGATTAATGGTTATGTGGCCAATGGACAGTTTGATGATGC

CAATGCTGTTTTGAGGCAGAGGATGCCCGCTGTTGGTTGCAATCCAGATATCTATACATATAACATAATC

ATCCGTGGTCTATGTTCGAGAAAATCTTTTTCTTTAGCACGAAATTTGATGAAAGAGATGGAAATTATGG

GTTGCAAGCCCAACCTAATAACATACACGAGTTTGATCAATGGGTTCTGCAAGGTCGGCAGATTGAAGGA

AGCAGAAGAAGCTGTCTGTGAAATGTCCGCATTGGGGATCCGCTTAAACACTGTTGGATACAACTGCCTG

ATACATGCTTTAGCTAAGGATGAGAAGGTTAAGGAGGCTCTGGAACTATACGCTGACATGGTAAACGGTG

GATGTAAACCAGATGTGTTCACTTTCAATTCTCTGATATATGGGCTCTGTAGGGTTGATCAGATAGAAGA

TGCCTTAGCATTAGTTCGAGATATGTTATTGGAAGGTGTCATTGCCAACACTGTAACTTACAATACCTTA

ATTAATTCTTTTTTGAGGAGAGGTGAATTGCAGGAAGCACTAAAACTTATAAATGATATGTTATATAGAG

GTTGTGCCCTCGATGAAATTACATACAACTCCATGATAAAGGCACTATGCATGGAGGGGTCTGTTGAGAA

AGCATTGGGACTCTTTGAGGAAATGATCCGGAAGGGACTGAATCCCAAAATCATATCATGCAACATTTTG

ATCAGTGGCTTATGTAGAATTGGAAAAATAGAAAATGCATTTGAATTTTTAAAAGATATTATACAGCGAG

GACTGAAACCTGATGTAGTCACTTATAATAGCCTCATAAATGGTTTGTGCAAGATGGGGCATGTGCAAGA

AGCTCTGAACCTCTTTGAGAAGATACAACTTGAAGGAATCCGTCCAGATGCTATCACTTATAGCACTTTG

ATCTGTTCATTTTGTAAAGCAGACATGCTTGATAATGCATATGCACTTTTATGCAGAGGCGTGAGCAATG

GATTCATACCTAATGATGTCACATGGTATTTTTTGATCAGATATTCCGTCATGAAAAAGTATGAGGAGAT

CTCAATAGTTCATCGTTTGGAACTCACTTGGAGAAGATTGGGGATCTAAAATGACATATTTATGCCCATC

AAATATATTGAGTCTCCACGAGTCCTGCCTGCTGAATTTGTTCGACAGAACTATTCAAACTGAAATCCAG

CTACCTCTCAAGCAAGAAAGAAAAGAATGGTATCTTACCTACTCTAGACAAGGAGAAGCACCGCCTCACT

GTCTAGATTTTTCTCAAGCATTGTTGAGGTAAGTTAATTAACACTGGAGTAAAATCCTGGCAGTCTTATG

ACCTCTGTTCATCAGATATTTGCAGGGTGTGTTATTCAGGTTGCATGATTTCTTGTCATGTAAGATTATT

TTATACTTTTGGCCCTATATTCTCAGGAGTCAGGCCACTTAACGGGAAAAATATAGGTAATCAATTCTAT

TAAGTCTTGATACTTGCGTTACTATCTATTCCTTTCCTTGGTCTACAATTTATGGTAAAATCTAGCAAGC

AATGAGAAAAATCAGAGATGCTTGGATTGAAGGTTAAACTGAAGACTCCCCACTAAACTGGTGCTCTAAC

CGAGCTAAGAAAGCAATTTTATCGTTGTATTGATTCATCTGTCATTTCTGTAAAACAAGGAAAAAAAGGA

TAAATACATAAATCCTCCCTTGTTTACACATGTAATTATTGAAAGAAAGATATGAGGAGGCCCCAAGAAT

CATGATTTATGACATCAGACAAAATAAATCAAACATAACATGATGCTAGGATGACCTAACAAAGAAAGAG

AGAACTCACTTAGTCCATTTTTTATATTTTAAATGCTAGATAAGTTCATCAGTAAGCTGTCTAAATTTTT

TTTATTGACAGAGTTGCTACATATTTTACAGTAAAGATTTTTGTAATGTATCTGTTTTCTTCAAATAAGT

GGTATAATAAATGGAAGGTCAAAGGCTTTTCATTGTTTTTAACATCTCAAGTTCTTAACACTTCAAGAAA

CACACACAGACACACACACAAAATGATTTATTCTCTTTGAAATTGCTGTTTGCTGCTTGTTATGTGTAGC

GGTAAACAGTTACCATCCAACATGAAGGTCAATTGAAAACATTTCTTAATGTGCAACCTTTGCGATGCAT

GAGTCATTTGCCAGTTTATATACTCTAAGTCTTTATCATACTAGACTAATCGCCAGATAAGTTGAATTTA

GGTAAAAATAGATACTACTCGGTCTTTTCAAATACTGCATAGTGCATAGCATCTTCTCAACCCCTCCCAA

GCAGTGGGCGGAGCCCGGGACTTTTAGTTTGTGCCTGTTGAAGAATGAATTGTCCATCAATGGCTGCTGA

AAGCGGAGACTTCTCTATCTTATTTAAGATGCCGTCAACTACAGTATCAAGGTCATACACGATGGCTTGT

GAAGTACTAATATACTTGCCACTCAATTTCTACATTATGCCAGAGAAGGTCTGCTCGCAGGGGCTGCTTT

TGTGTAGATGAGCTTTTGTGGGCGGTGGCGTGCTGCTGTTCTTGGGCGTGTACTTGAAGTTATGCTGTAT

AATGTTTACAATTTACAGTAAAGGAATCATTTATATGTTCAGGAAGAGAAATGAAAACAATTATTCTATG

TTTGCAATCGACAAGTCATGTTTTTGTTGTAAATTCTGTTTCAGCTCTAAACTTAAGCTTTTAGAGATTA

TTTCAGCTCCGCATACAGTGAATTAAAATATATTGAGGATAAAATTAAAGAGTATATGAAAATCTTGACA

GTTGGTTAATCAGATTCTACATGCTATGTTCGTCTCTCATCTAGTCTATCAACAGTTGTACAATGCTATC

TAAACTTTATTTGATTGCAGAGTACAAACAAGCAAAATCAGAGGATAAACATTCAACCAGAAGCCATGCA

TCTGTGATCTACGGCTGTGTTATGCACTTTTCTATCAACTACTTTTGCTTCAACGTCCTCGTCTTTCCCT

A

>MSTRG.1517.2 gene=MSTRG.1517

TTTTTGATATGAGGGAAGTTTTTAGTTGCGAACCTAGTTTTAGGTCATATAATGTTGTTCTGGATATATT

GGTTGCTGGGGATTGCCCGGCTGTTGCACCGAATATTTTTTATGAAATGTTGCACAAGGGGATCTCTCCA

ACTGTTTTCACGTTTGGTGTGGTGATGAAAGCGCTTTGTATGGTTAATGAGGTGGATTCTGCATGTTCAC

TTTTAAGAGATATGACAAAACATGGGTGTGTGCCGAATTCAGTTATTTACCAGACACTTATACATGCGCT

TTGCAAGGTAAATAGAATAAATGACGTGATAAGATATTTGGAGGAAATGTTTCTGATGGGTTGCACACCT

GATGTTGATACTTTTAATGATGTTATTCATGGTCTTTGCCGTGCTAAGAGGATACCTGAAGGTGCTAAAT

TGGTCGATAGAATGCTTCTCCGAGGCTTCACTCCTAGCAAGTTTACTTATGGAATTTTAATGGATGGATT

ATGCCGAACTGGTCAAGTTGATAAAGCTAGAGCTTTGCTGGAGAGAGTGCCTGAGCCTAATGTTGTTTTG

TTCAATACTCTGATTAATGGTTATGTGGCCAATGGACAGTTTGATGATGCCAATGCTGTTTTGAGGCAGA

GGATGCCCGCTGTTGGTTGCAATCCAGATATCTATACATATAACATAATCATCCGTGGTCTATGTTCGAG

AAAATCTTTTTCTTTAGCACGAAATTTGATGAAAGAGATGGAAATTATGGGTTGCAAGCCCAACCTAATA

ACATACACGAGTTTGATCAATGGGTTCTGCAAGGTCGGCAGATTGAAGGAAGCAGAAGAAGCTGTCTGTG

AAATGTCCGCATTGGGGATCCGCTTAAACACTGTTGGATACAACTGCCTGATACATGCTTTAGCTAAGGA

TGAGAAGGTTAAGGAGGCTCTGGAACTATACGCTGACATGGTAAACGGTGGATGTAAACCAGATGTGTTC

ACTTTCAATTCTCTGATATATGGGCTCTGTAGGGTTGATCAGATAGAAGATGCCTTAGCATTAGTTCGAG

ATATGTTATTGGAAGGTGTCATTGCCAACACTGTAACTTACAATACCTTAATTAATTCTTTTTTGAGGAG

AGGTGAATTGCAGGAAGCACTAAAACTTATAAATGATATGTTATATAGAGGTTGTGCCCTCGATGAAATT

ACATACAACTCCATGATAAAGGCACTATGCATGGAGGGGTCTGTTGAGAAAGCATTGGGACTCTTTGAGG

AAATGATCCGGAAGGGACTGAATCCCAAAATCATATCATGCAACATTTTGATCAGTGGCTTATGTAGAAT

TGGAAAAATAGAAAATGCATTTGAATTTTTAAAAGATATTATACAGCGAGGACTGAAACCTGATGTAGTC

ACTTATAATAGCCTCATAAATGGTTTGTGCAAGATGGGGCATGTGCAAGAAGCTCTGAACCTCTTTGAGA

AGATACAACTTGAAGGAATCCGTCCAGATGCTATCACTTATAGCACTTTGATCTGTTCATTTTGTAAAGC

AGACATGCTTGATAATGCATATGCACTTTTATGCAGAGGCGTGAGCAATGGATTCATACCTAATGATGTC

ACATGGTATTTTTTGATCAGATATTCCGTCATGAAAAAGTATGAGGAGATCTCAATAGTTCATCGTTTGG

AACTCACTTGGAGAAGATTGGGGATCTAAAATGACATATTTATGCCCATCAAATATATTGAGTCTCCACG

AGTCCTGCCTGCTGAATTTGTTCGACAGAACTATTCAAACTGAAATCCAGCTACCTCTCAAGCAAGAAAG

AAAAGAATGGTATCTTACCTACTCTAGACAAGGAGAAGCACCGCCTCACTGTCTAGATTTTTCTCAAGCA

TTGTTGAGGAGTCAGGCCACTTAACGGGAAAAATATAGGTAATCAATTCTATTAAGTCTTGATACTTGCG

TTACTATCTATTCCTTTCCTTGGTCTACAATTTATGGTAAAATCTAGCAAGCAATGAGAAAAATCAGAGA

TGCTTGGATTGAAGGTTAAACTGAAGACTCCCCACTAAACTGGTGCTCTAACCGAGCTAAGAAAGCAATT

TTATCGTTGTATTGATTCATCTGTCATTTCTGTAAAACAAGGAAAAAAAGGATAAATACATAAATCCTCC

CTTGTTTACACATGTAATTATTGAAAGAAAGATATGAGGAGGCCCCAAGAATCATGATTTATGACATCAG

ACAAAATAAATCAAACATAACATGATGCTAGGATGACCTAACAAAGAAAGAGAGAACTCACTTAGTCCAT

TTTTTATATTTTAAATGCTAGATAAGTTCATCAGTAAGCTGTCTAAATTTTTTTTATTGACAGAGTTGCT

ACATATTTTACAGTAAAGATTTTTGTAATGTATCTGTTTTCTTCAAATAAGTGGTATAATAAATGGAAGG

TCAAAGGCTTTTCATTGTTTTTAACATCTCAAGTTCTTAACACTTCAAGAAACACACACAGACACACACA

CAAAATGATTTATTCTCTTTGAAATTGCTGTTTGCTGCTTGTTATGTGTAGCGGTAAACAGTTACCATCC

AACATGAAGGTCAATTGAAAACATTTCTTAATGTGCAACCTTTGCGATGCATGAGTCATTTGCCAGTTTA

TATACTCTAAGTCTTTATCATACTAGACTAATCGCCAGATAAGTTGAATTTAGGTAAAAATAGATACTAC

TCGGTCTTTTCAAATACTGCATAGTGCATAGCATCTTCTCAACCCCTCCCAAGCAGTGGGCGGAGCCCGG

GACTTTTAGTTTGTGCCTGTTGAAGAATGAATTGTCCATCAATGGCTGCTGAAAGCGGAGACTTCTCTAT

CTTATTTAAGATGCCGTCAACTACAGTATCAAGGTCATACACGATGGCTTGTGAAGTACTAATATACTTG

CCACTCAATTTCTACATTATGCCAGAGAAGGTCTGCTCGCAGGGGCTGCTTTTGTGTAGATGAGCTTTTG

TGGGCGGTGGCGTGCTGCTGTTCTTGGGCGTGTACTTGAAGTTATGCTGTATAATGTTTACAATTTACAG

TAAAGGAATCATTTATATGTTCAGGAAGAGAAATGAAAACAATTATTCTATGTTTGCAATCGACAAG

>MSTRG.1517.3 gene=MSTRG.1517

GAAAATTTGACTTTGGACCCGACCCGCTGCTCACTGTAGTGCAGAAGCTTTTTGCAGCAAATACAAACAC

TTGTTCCCAGATATTCCGTCATGAAAAAGTATGAGGAGATCTCAATAGTTCATCGTTTGGAACTCACTTG

GAGAAGATTGGGGATCTAAAATGACATATTTATGCCCATCAAATATATTGAGTCTCCACGAGTCCTGCCT

GCTGAATTTGTTCGACAGAACTATTCAAACTGAAATCCAGCTACCTCTCAAGCAAGAAAGAAAAGAATGG

TATCTTACCTACTCTAGACAAGGAGAAGCACCGCCTCACTGTCTAGATTTTTCTCAAGCATTGTTGAGGT

AAGTTAATTAACACTGGAGTAAAATCCTGGCAGTCTTATGACCTCTGTTCATCAGATATTTGCAGGGTGT

GTTATTCAGGTTGCATGATTTCTTGTCATGTAAGATTATTTTATACTTTTGGCCCTATATTCTCAGGAGT

CAGGCCACTTAACGGGAAAAATATAGGTAATCAATTCTATTAAGTCTTGATACTTGCGTTACTATCTATT

CCTTTCCTTGGTCTACAATTTATGGTAAAATCTAGCAAGCAATGAGAAAAATCAGAGATGCTTGGATTGA

AGGTTAAACTGAAGACTCCCCACTAAACTGGTGCTCTAACCGAGCTAAGAAAGCAATTTTATCGTTGTAT

TGATTCATCTGTCATTTCTGTAAAACAAGGAAAAAAAGGATAAATACATAAATCCTCCCTTGTTTACACA

TGTAATTATTGAAAGAAAGATATGAGGAGGCCCCAAGAATCATGATTTATGACATCAGACAAAATAAATC

AAACATAACATGATGCTAGGATGACCTAACAAAGAAAGAGAGAACTCACTTAGTC

>MSTRG.1518.1 gene=MSTRG.1518

GAAAATATGTTTCAAATATTAAACGGGATCTTCGTGAGGAACGCCGATACATTCAACCCTGAGAACACTC

ATCTCCCGCAATCATCTTCTCCAAATCTCTGCTTATCCTCAAAACCACTTTAGATGTTACCCTAACATAG

CTAGTAAAATGTTTTCACGGGTCACTGATGAGGTCTCGTATGCCCGCCACCTGTTTGACACAAGTCCCCA

ACCAGATACTTTAGATACTGATGATTAGAGGTTTTGGGTCTGATTTGTTTGTTCATACGGCGTTGATTGA

AATGTACGGGAGATACTCTGGGCTCGTAATTGTAAACCCCAGATGTGAATCGGCAAAGCATTTACAGAAA

CTCATGGAGAGTAGGTACTAGGTAATGGATGGAGAAAATTATATATTCACTTACATAATTACTATGTAAA

CAAAGTTGAAGCCTCAGCAAGAAAAGCAGAACTTTTGTATAATTGTCTGAAATTCTCTAAGTTCTTGCTT

TAGAACACATATACTTTTTATATACTTTTGTGTTCCAACAAACTTTAAAGTCGTAAACCACTTTCTTTTT

ATCTCTTTTTAATTTGAAGTCTTTAGGAACTTTGATTTAAGAGAATGGAAAGCAGTGAAGCAAATATTGG

AGAATTCCCTGATGTGAATTACGTTATTTGGTATTCCCAATACGTGAAGAGGATTGCTGGACATTTTCCT

CTCGCGGAAGACTCGACTAGAGTCGAAGTATACTGCGGGATCTCAATGAGAACCTTCAGATACTTATGCT

CTCTTGTGATGGAGCCTATGTCAAATTCACAAATATATAAATTTAACGATAACAGGGAATTAACTGTGGT

TGATAAAGTAGCCATAGCTATGAGAAGGCTAAAATTGGGCAAGTTGCTGAATTCAATTGGTGGTGAAATT

GGGACGCACTCATTTACCGCTGCTAAGGTCACATGGTCCTTTGTGGAGGTCTTGGCAGAAAATGGCCATC

GCCACCTCCGTGGGCCTGCTTCTGCCCCTGAAGAGGGAATGAGTGCTGAGACATTTAATAGGCTAAAGAA

AGAATGGAAGGTTCTTACAGCTGGAATGAAATCCGAAAAGCTTCAAAGTATGACCAAGGCCTGCTGCATC

TTGCATAACATTGTCATTGACATGAATGATGAGAGGCAACGTGAGCTATCTCTGTTGGAAGGTCATCATC

CAGACTTTTTGCATCACAGATTTTAAAGATTAGTTTTGTAATTGTTAGAAAGCTTTGAATCATGTCTTCC

AGATATTTCTTTGGTTTCCGCTTTTTTTTCGTGGTTGTGTAATTATATGTTTTTGTGGTCACTAATATTT

AGTTGATG

>MSTRG.1518.4 gene=MSTRG.1518

ATTCACTTATGTAATTACAATGTGAACAAAGTTGAAGCCTCAGCAAGAAAAGCAGAACTTTTGTATCATT

GTCTGAAATTCTCTAAGTTCTTTATTTAGAACACAGTACTCCACAGGAAGATGATGGATAACTGGAGCTG

CTCAAGTAAACTTGGCAGGTTTACTTCAATTACTCTCTGTTTCTGTCAGGACATTTTCCTCTCGCGGAAG

ACTCGACTAGAGTCGAAGTATACTGCGGGATCTCAATGAGAACCTTCAGATACTTATGCTCTCTTGTGAT

GGAGCCTATGTCAAATTCACAAATATATAAATTTAACGATAACAGGGAATTAACTGTGGTTGATAAAGTA

GCCATAGCTATGAGAAGGCTAAAATTGGGCAAGTTGCTGAATTCAATTGGTGGTGAAATTGGGACGCACT

CATTTACCGCTGCTAAGGTCACATGGTCCTTTGTGGAGGTCTTGGCAGAAAATGGCCATCGCCACCTCCG

TGGGCCTGCTTCTGCCCCTGAAGAGGGAATGAGTGCTGAGACATTTAATAGGCTAAAGAAAGAATGGAAG

GTTCTTACAGCTGGAATGAAATCCGAAAAGCTTCAAAGTATGACCAAGGCCTGCTGCATCTTGCATAACA

TTGTCATTGACATGAATGATGAGAGGCAACGTGAGCTATCTCTGTTGGAAGGTCATCATCCAGACTTTTT

GCATCACAGATTTTAAAGATTAGTTTTGTAATTGTTAGAAAGCTTTGAATCATGTCTTCCAGATATTTCT

TTGGTTTCCGCTTTTTTTTCGTGGTTGTGTAATTATATGTTTTTGTG

>MSTRG.1518.2 gene=MSTRG.1518

CCGCAATCATCTTCTCCAAATCTCTGCTTATCCTCAAAACCACTTTAGATGTTACCCTAACATAGCTAGT

AAAATGTTTTCACGGGTCACTGATGAGGTCTCGTATGCCCGCCACCTGTTTGACACAAGTCCCCAACCAG

ATACTTTAGATACTGATGATTAGAGGTTTTGGGTCTGATTTGTTTGTTCATACGGCGTTGATTGAAATGT

ACGGGAGATACTCTGGGCTCGTAATTGTAAACCCCAGATGTGAATCGGCAAAGCATTTACAGAAACTCAT

GGAGATACTCCACAGGAAGATGATGGATAACTGGAGCTGCTCAAGTAAACTTGGCAGGTTTACTTCAATT

ACTCTCTGTTTCTGTCAGGACATTTTCCTCTCGCGGAAGACTCGACTAGAGTCGAAGTATACTGCGGGAT

CTCAATGAGAACCTTCAGATACTTATGCTCTCTTGTGATGGAGCCTATGTCAAATTCACAAATATATAAA

TTTAACGATAACAGGGAATTAACTGTGGTTGATAAAGTAGCCATAGCTATGAGAAGGCTAAAATTGGGCA

AGTTGCTGAATTCAATTGGTGGTGAAATTGGGACGCACTCATTTACCGCTGCTAAGGTCACATGGTCCTT

TGTGGAGGTCTTGGCAGAAAATGGCCATCGCCACCTCCGTGGGCCTGCTTCTGCCCCTGAAGAGGGAATG

AGTGCTGAGACATTTAATAGGCTAAAGAAAGAATGGAAGGTTCTTACAGCTGGAATGAAATCCGAAAAGC

TTCAAAGTATGACCAAGGCCTGCTGCATCTTGCATAACATTGTCATTGACATGAATGATGAGAGGCAACG

TGAGCTATCTCTGTTGGAAGGTCATCATCCAGACTTTTTGCATCACAGATTTTAAAGATTAGTTTTGTAA

TTGTTAGAAAGCTTTGAATCATGTCTTCCAGATATTTCTTTGGTTTCCGCTTTTTTTTCGTGGTTGTGTA

ATTATATGTTTTTGTG

>MSTRG.1518.3 gene=MSTRG.1518

TGCATTCAATGTGTAACACCCCAAGCCAAAGTGAAGGAGATGGTTCAACAACTCACCATAGACCAAACCA

AAAGAGCTAACGACTTTTCTACAACATTTATAAATGCGTCCACACTTACATACTATGAAATGTTACAGTT

TTTCCACATTATATAAGTTTAACGTCCTCGTCGAGCCCAAAAACTGAAGAAATCCTTTTAAGGCACTTTT

GGCAGGGTCGACCTCGAACAAAATTTGTTATAGTCCAAGCCACACATACTACATCATTCAACAACCAACT

ATAAGTTGATACAAAAGAGCTATCAATTCCGATACAATATTAGTTGTAAGTTTGTGCTAATAAAGGCATT

GATATAACTTAGTGACGTGGGATGTTGCAAATTTCTTCCGTGACACCACTGAATATGTTAGTTAGCTACT

GACTAGACACATTACGATTGTGACCACTAAATATTAGATTCCGGAAATAGTGCTATGTATATATTGTATG

GAAATTTTCATTTATATAATATACTATATAGATACTGATAATAGTAAAATGGTCAAGCTAAATTGAACAA

ATGTTTTCTGGTTTTAACAAAAAGAAGTGCAATGGGCAAAGAGAAATACATAATAATGAATAAAATACTG

GAAACTATATTTTGAATATAAATTATATTTAAATAATACTTCCTCTCCCATTTTGTTTGGCCCACTTTCT

TTTTTAGGATGTCCCGAGAAGAATGTACCTAACTTTATATAGGGGCTGTTTGGGTGAACTTAAAATAAGT

GTTTTCTGCTTAAAGTAAAAAAAGGTGAAGTAGAAGTTAGTCAAGACTTATAAGTGATTAAAGTGTTTGG

GAAAAAAGTAGAAGTCATGAAACAAAATCCTAACTTTTTTATAAGTGTTTCTCGACTTTTTACACAAACG

GTACGAATAAGTGCTTCTAACTTAAAACTCTAGAAGTGGGGCTTAAAAGCCCCGCCCAAACACCCACATA

GTATGTTTTAAGATAGCTATAACCACATGTTCACGTAAATGCTTAGTCATCATTCACTAACTCATCTTTT

GTGAATACATATCAAATAGTCATACTCAAATTTTTTAAATGTGCGTAAAATTGATTAGCTCATGCTTTAT

GGGATGGAGTGAGTATATTTTAATTACTTTATTTTACTTCATATATGAGCTACAACTTTGAAATAAAATT

TATTTTATGCCCAATTACAATTTCCTTAGGATAAATTTGTATATATGAAGCCCACCCCCAGTATTTACAA

ACATACTCCTTATACTATTCTATTTCTTTTAACCTCCATCAACGCGCTGACGTCATCCCAGACCCATCCA

TCTCACTCTTTATATACTTCTGTGTTCCAACAAACTTTAAAGTCGCAAACTTTAGAGTTATTGTACGGTG

GTAAGGAAAATCAATCTCCATTCCTCAAACCACAAAAGAATGGAAAGAAGTGAAGCTAATCGTGGCAAAT

TCCCTTTGAATTACGTTATTTGGTTTGCCGAATTCGTGAAGAGGATTGCTGGTATGTAAATCTTCAAATT

CTTAAGTTCTCATTTGGTGGTGATGAATTGTTCTTGTCGATTTCGTTATGGTTTGGTATGTACTACCTTT

TTTTTCCCGAAGTAATTTTATGTTTGCTTGTTCATCTGTGTATGATGAAATTCTAATTTGTGACCAGTTG

ATTCGTCTTTTGATTTATTGAAAATTACTTTAGATGGAGAATGAATAAATCCTCAGTCTGTGGTTCGGAG

TCTTATTTGACCTAGAATTTAATTGTTGCTTAGTACAATGAAGATATAAACATTTAAATTTTAATTCTGG

GAGTTATGTTATGTGTATATAAATTTTGCTGATGGGATTAGAGTCATTAGATTTGAAAGGAGTTGAGGAA

GGTGATGTGATTCGTATGTAGTTCTTGTGGATATTTTAGTTTCTTTGTTTATACTCTATAATATGGGAAA

GAAAGCATTTCAGATTTTGACATTGGTGCATACATAGAAGAAAATATGAGGTAAATCCAAACTATACCGA

GTATAATCATCTTCTGATTATATGATCTATAAGTCACCAGTGAGAATTCAGGACTAGCTTTTTTTCCAAA

GTAGCAAGGGATTGATTTTAGGCAAGTCGCTATTCAAAATACTTCTATTTCCAGTTCCACAAGGTCAAAC

TAGTAATTAAGGTCAAACTGCTTCCAGTTGTCTTAGTCTTTGGAACAATATCTAATCTGTTGTTTTCGAA

TATGTTGTTATTTACTTAGTTCTTGTAAGGGTTTGTTCAGTCAAACAGCTATATTATGGTAGATATAATG

TTTTCCTGGGAGCTTGTAACTTGCTATGAATACCGAAAGTATCCACTTATAATCAGTTCATATTGATTCT

GATTTGATATGAAAGCTGTAGCATTCTTTGCAGATTGGAACCTTACTGAAAGCGATATATCCCTGTGTGT

TAATCACAAGCAGTTTACTGATTTACATTTTTGACATTCTAGCTTCATTTTGTTTATACAGTTTACAGGT

CGATGTGTGTATGTATACTGAAATGAATGGAATAGTATGGTGTTATACATTAATCTAATATGTTCAAGTT

TTAAGCCTGATAAACACATTATGGCAGTGTCTTAATCGTATGTATAAAAATGCATTTTCTTCTCTTCTTT

GCAGGACATTTTCCTCTCGCGGAAGACTCGACTAGAGTCGAAGTATACTGCGGGATCTCAATGAGAACCT

TCAGATACTTATGCTCTCTTGTGATGGAGCCTATGTCAAATTCACAAATATATAAATTTAACGATAACAG

GGAATTAACTGTGGTTGATAAAGTAGCCATAGCTATGAGAAGGCTAAAATTGGGCAAGTTGCTGAATTCA

ATTGGTGGTGAAATTGGGACGCACTCATTTACCGCTGCTAAGGTCACATGGTCCTTTGTGGAGGTCTTGG

CAGAAAATGGCCATCGCCACCTCCGTGGGCCTGCTTCTGCCCCTGAAGAGGGAATGAGTGCTGAGACATT

TAATAGGCTAAAGAAAGAATGGAAGGTTCTTACAGCTGGAATGAAATCCGAAAAGCTTCAAAGTATGACC

AAGGCCTGCTGCATCTTGCATAACATTGTCATTGACATGAATGATGAGAGGCAACGTGAGCTATCTCTGT

TGGAAGGTCATCATCCAGACTTTTTGCATCACAGATTTTAAAGATTAGTTTTGTAATTGTTAGAAAGCTT

TGAATCATGTCTTCCAGATATTTCTTTGGTTTCCGCTTTTTTTTCGTGGTTGTGTAATTATATGTTTTTG

TG

>MSTRG.1518.5 gene=MSTRG.1518

GAAAATATGTTTCAAATATTAAACGGGATCTTCGTGAGGAACGCCGATACATTCAACCCTGAGAACACTC

ATCTCCCGCAATCATCTTCTCCAAATCTCTGCTTATCTGATGAGGTCTCGTATGCCCGCCACCTGTTTGA

CACAAGTCCCCAACCAGATACTTTAGATACTGATGATTAGAGGTTTTGGGTCTGATTTGTTTGTTCATAC

GGCGTTGATTGAAATGTACGGGAGATACTCTGGGCTCGTAATTGTAAACCCCAGATGTGAATCGGCAAAG

CATTTACAGAAACTCATGGAGAGTAGGTACTAGGTAATGGATGGAGAAAATTATATATTCACTTACATAA

TTACTATGTAAACAAAGTTGAAGCCTCAGCAAGAAAAGCAGAACTTTTGTATAATTGTCTGAAATTCTCT

AAGTTCTTGCTTTAGAACACATATACTTTTTATATACTTTTGTGTTCCAACAAACTTTAAAGTCGTAAAC

CACTTTCTTTTTATCTCTTTTTAATTTGAAGTCTTTAGGAACTTTGATTTAAGAGAATGGAAAGCAGTGA

AGCAAATATTGGAGAATTCCCTGATGTGAATTACGTTATTTGGTATTCCCAATACGTGAAGAGGATTGCT

GGACATTTTCCTCTCGGAAGACTCGACTGGAGTCGAAAAATACTGTGGGATCTCAATGAGAACCTTCGTA

TACTTATGCTCTCTTGTGATGGAGCCAATGTCCAATTCACACATATGCAAATTTAACGATAACAGAGAAT

TGACCGTGGTTGATAAAGTAGCCATAGCGATGAGAAGGCTTAAATTTGGCAAGTCACTGAACTCAATCGG

TAAGGAAATTGGTAAGCACTCATTTACTGCTGATAAGGTCACACGGTCCTTTGTGGAGGCCTTGGTAGAA

AATGGCCTTCGCCACCTCCATGGGCCAGCCACTGCCACTAAGGAAATAAGGACTGCTGAGGCAGTGGATA

GGCTAGAGGAAGAACGGAAGGTTCTTACAAGAGAAGCGTCAAAATATGACCACGGCTTGCCGCATCTTGC

ATAAGATTGTCATTGACATGAATGATGAGAGGCATCGTTAGCTATGTCTGTCACAAGGTCATCATCCAGG

CTTTATGACTCAGAAAAATTAGTTTTGTAATTGACAAAAAAGTTTTGAATCATCTCTTCCGAATATTTAT

TTGGTTTCTGCTTTTTTATTTGTATTTTGTGTAATTATTTATTTTTGTAGTAATTAATTTTTAGTTCATG

CGTTTGTTTA

>MSTRG.1518.6 gene=MSTRG.1518

GAAGCAAATATTGGAGAATTCCCTGATGTGAATTACGTTATTTGGTTTGCTGAATTCGTGAAGAGGATTT

CTGGACATTTTCCTCTCGGAAGACTCGACTGGAGTCGAAAAATACTGTGGGATCTCAATGAGAACCTTCG

TATACTTATGCTCTCTTGTGATGGAGCCAATGTCCAATTCACACATATGCAAATTTAACGATAACAGAGA

ATTGACCGTGGTTGATAAAGTAGCCATAGCGATGAGAAGGCTTAAATTTGGCAAGTCACTGAACTCAATC

GGTAAGGAAATTGGTAAGCACTCATTTACTGCTGATAAGGTCACACGGTCCTTTGTGGAGGCCTTGGTAG

AAAATGGCCTTCGCCACCTCCATGGGCCAGCCACTGCCACTAAGGAAATAAGGACTGCTGAGGCAGTGGA

TAGGCTAGAGGAAGAACGGAAGGTTCTTACAAGAGAAGCGTCAAAATATGACCACGGCTTGCCGCATCTT

GCATAAGATTGTCATTGACATGAATGATGAGAGGCATCGTTAGCTATGTCTGTCACAAGGTCATCATCCA

GGCTTTATGACTCAGAAAAATTAGTTTTGTAATTGACAAAAAAGTTTTGAATCATCTCTTCCGAATATTT

ATTTGGTTTCTGCTTTTTTATTTGTATTTTGTGTAATTATTTATTTTTGTAGTAATTAATTTTTAGTTCA

TGCGTTT

>MSTRG.1518.7 gene=MSTRG.1518

CCTCAACCATCTTCTCCAAATCCATGCTTTAATTCTCAAAACCGCTTTAGACATCAACGTTGTTGTCCTA

ACCAAGCTATTAGGTAAGATGTTTTCACGGGTCACCGATGAGGTCTCGTATGCCCGCCACCTGTTTGACA

CAAGTCCCCAACCAGATACTTTAGATACTGATGATTAGAGGTTTTGGGTCTGATTTGTTTGTTCATACGG

CGTTGATTGAAATGTACGGGAGATACTCTGGGCTCGTAATTGTAAACCCCAGATGTGAATCGGCAAAGCA

TTTACAGAAACTCATGGAGAGTAGGTACTAGGACATTTTCCTCTCGGAAGACTCGACTGGAGTCGAAAAA

TACTGTGGGATCTCAATGAGAACCTTCGTATACTTATGCTCTCTTGTGATGGAGCCAATGTCCAATTCAC

ACATATGCAAATTTAACGATAACAGAGAATTGACCGTGGTTGATAAAGTAGCCATAGCGATGAGAAGGCT

TAAATTTGGCAAGTCACTGAACTCAATCGGTAAGGAAATTGGTAAGCACTCATTTACTGCTGATAAGGTC

ACACGGTCCTTTGTGGAGGCCTTGGTAGAAAATGGCCTTCGCCACCTCCATGGGCCAGCCACTGCCACTA

AGGAAATAAGGACTGCTGAGGCAGTGGATAGGCTAGAGGAAGAACGGAAGGTTCTTACAAGAGAAGCGTC

AAAATATGACCACGGCTTGCCGCATCTTGCATAAGATTGTCATTGACATGAATGATGAGAGGCATCGTTA

GCTATGTCTGTCACAAGGTCATCATCCAGGCTTTATGACTCAGAAAAATTAGTTTTGTAATTGACAAAAA

AGTTTTGAATCATCTCTTCCGAATATTTATTTGGTTTCTGCTTTTTTATTTGTATTTTGTG

>MSTRG.1518.8 gene=MSTRG.1518

CACGGATTGAGGATTTATTCATTCTCTATCTAAAGTAATTTTCAGTAAATTAAAACACAAATCAACTAGT

CACAAATTAAAATTTCATTAAACACAGATAAACAAGCAAAGATAAAATTACTTCGGAAAAGAAAAGGTAA

TACATACCAAACCATAACGAAATCGACAAGAACAGTCACCACTTAATGAGAACTTAATATACATGAAGAT

ATACATACCAACAATCCTCTTCACGAATTCGGCAAACCAAATAACGTAATTCGGAGGGAATTTGCCACAA

TTAGCTTCACTTCTTTCCACTCTCTTAAAGTAAATTTCTTAAAAACTTAAAATTAAAAGGATATAAAAAG

AAAGTGCTTTGAGGAATGGAGATTGATTTTCCTTACCACCGTACAATAACTCTAAAGTTTGCGACTTTAA

AGTTTGTTGGAACACAAAAGTTTATAAAGAGTGAGATGGATGGGTCTAGGGTGACTTTCTGTTGTTATTT

GGATTTGCCTCATATTTTCTTTTATGTATGCCTCAATGCCAAAATCTGAACTACTTTCTTTCCCGTAGTG

TCACGGAGAAAGTTTGCTACTGCCACCTGACTAAGTTAGGTATGTTATATCAATGCCTTTATGAGCACAA

CTTATAACTAATACTGCATCGGGATTGATACCTCTTTTGGATCAACTTATAGTGGGTTGTTGAATGATGT

AGTATATTAGGATTGGACTGTAATGAATTTTGTCAGAGCTCGACCCTGCCAAAAGTGCTTCACAAAATTT

CTTCAGCTTTTGGGCCCCACGAGGACGTCAAACTAATTGTGTGGAAAAACTGTAACATCCTACTGTGGAC

ATTTTCCTCTCTTGGAAGACTCGACTAGAGTCGAAGAGTACTGCGGGATCTCAATGAGAACCTTCGTATA

CTTATGCTCTCTTGTGATGGAGCCAATGTCCAATTCACACATATGCAAATTTAACGATAACAGAGAATTG

ACCGTGGTTGATAAAGTAGCCATAGCGATGAGAAGGCTTAAATTTGGCAAGTCACTGAACTCAATCGGTA

AGGAAATTGGTAAGCACTCATTTACTGCTGATAAGGTCACACGGTCCTTTGTGGAGGCCTTGGTAGAAAA

TGGCCTTCGCCACCTCCATGGGCCAGCCACTGCCACTAAGGAAATAAGGACTGCTGAGGCAGTGGATAGG

CTAGAGGAAGAACGGAAGGTTCTTACAAGAGAAGCGTCAAAATATGACCACGGCTTGCCGCATCTTGCAT

AAGATTGTCATTGACATGAATGATGAGAGGCATCGTTAGCTATGTCTGTCACAAGGTCATCATCCA

>MSTRG.1518.9 gene=MSTRG.1518

CAAATATGTTTCAAATATTAAACAGGATCTTTGTGAAGAACGCCGATACAGGGTTTCAACCCTGAGAACA

CTCATCTAACCTCAATCATCTTCTCCAAATCCCTGCTTATCTCCAAAACCGCTTTAGATATCAACGTTGT

TGCCCTAACCTAGCTATTACGTAAGATGTTTTCACGGGTCACCGATGAGGTCTCGTATGCGCGCCACCTG

TTTGACACAAGTCCCCAACCAGATACTTTAGATACTGATGATTAGAGGTTTTGGGTCTGATTTGTTTGTT

CAGACGGCGTTGATTGAAAATGGGAGATACTCTGGACTTGTAATTGTAAACCCCAGATGTGAATCGGCAA

AGCATTTACAGAAACTCAGGGAGAGTAGGTACTAGGTAATGGATGGAGAAATGTATATATTCACTTGTGT

AATTACTATGTAAACAATGTTGAAGCCTCAGCAAGAAAAGCAGAACTTTTGTATCATTGTCTGAAATTCT

CTAAGTTCTTGCTTTAGAACACATATACTTTTTATATACCTTTGTGTTCCAACAAACTTTAAAGTCGCAA

ACTTTAGAGTTATTCTCCGGTGGTAAGGAAAAACAATCTTCATTCCTCGAACCACTTTTTTTTTTTATCT

CTTTCTAATTTGAAGTTTTTAAAAACTTTGATTTAAGAGAATGGAAAGCAGTGAAGCAAATATTGGAGAA

TTCCCTGATGTGAATTACGTTATTTGGTTTGCTGAATTCGTGAAGAGGATTTCTGGACATTTTCCTCTCT

TGGAAGACTCGACTAGAGTCGAAGAGTACTGCGGGATCTCAATGAGAACCTTCAGATACTTGTGCTCTCT

TGTGATGGAGCCAATGTCCAAGTCACACACATACAAATTTAACGATAATAGAGAATTGACCGTGGTTGAT

AAAGTAGCCATAGCGTTGAGAAGGCTAAAATTTGGCAAGTCACTGAACTCAATCAGTAAGGAAATTGGTA

CGCACTCTTTTACTGCTGATAAGGTTACAAGGTCCTTTGTGGAGGCCTTGGTAGAAAATGGCCTTCGCCA

CCTCCATGGGACAGCCACTGCTACTGAGGAAAGAATGACTGCTGAGACAATGGATAGGCGGCTAAAGGAA

GAATGGAAGGTTCTTACACTTGTAATGAAAGCTGAGAAGCGTCAAAGTATTAGCACAGCTTGCCGCATCT

TGCATAACATTGTCATTGAGATGAATGATGAGAGGCATGGTTAGCTATCTCTGTCACAAGGTCATCATCC

AGGCTTTATGACTCACAAATTTTAAAGATTAGTTTTGTAATTGTTAGAAAGTTTTGAATCATCTCTTCTG

AATATTTCTTTGGTTTCCGCTTTTTTATTTGTATTTTGTGTGATTATTATTTGTGGTCATTAATATTTAG

TTCATGAGTTTGTTTAGTATTTTT

>MSTRG.1518.10 gene=MSTRG.1518

GTTATCCCCAAATATGTTTCAAATATTAAACAGGATCTTTGTGAAGAACGCCGATACAGGGTTTCAACCC

TGAGAACACTCATCTAACCTCAATCATCTTCTCCAAATCCCTGCTTATCTCCAAAACCGCTTTAGATATC

AACGTTGTTGCCCTAACCTAGCTATTACGTAAGATGTTTTCACGGGTCACCGATGAGGTCTCGTATGCGC

GCCACCTGTTTGACACAAGTCCCCAACCAGATACTTTAGATACTGATGATTAGAGGTTTTGGGTCTGATT

TGTTTGTTCAGACGGCGTTGATTGAAAATGGGAGATACTCTGGACTTGTAATTGTAAACCCCAGATGTGA

ATCGGCAAAGCATTTACAGAAACTCAGGGAGAGTAGGTACTAGGACATTTTCCTCTCTTGGAAGACTCGA

CTAGAGTCGAAGAGTACTGCGGGATCTCAATGAGAACCTTCAGATACTTGTGCTCTCTTGTGATGGAGCC

AATGTCCAAGTCACACACATACAAATTTAACGATAATAGAGAATTGACCGTGGTTGATAAAGTAGCCATA

GCGTTGAGAAGGCTAAAATTTGGCAAGTCACTGAACTCAATCAGTAAGGAAATTGGTACGCACTCTTTTA

CTGCTGATAAGGTTACAAGGTCCTTTGTGGAGGCCTTGGTAGAAAATGGCCTTCGCCACCTCCATGGGAC

AGCCACTGCTACTGAGGAAAGAATGACTGCTGAGACAATGGATAGGCGGCTAAAGGAAGAATGGAAGGTT

CTTACACTTGTAATGAAAGCTGAGAAGCGTCAAAGTATTAGCACAGCTTGCCGCATCTTGCATAACATTG

TCATTGAGATGAATGATGAGAGGCATGGTTAGCTATCTCTGTCACAAGGTCATCATCCAGGCTTTATGAC

TCACAAATTTTAAAGATTAGTTTTGTAATTGTTAGAAAGTTTTGAATCATCTCTTCTGAATATTTCTTTG

GTTTCCGCTTTTTTATTTGTATTTTGTGTGATTATTATTTGTGGTCATTAATATTTAGTTCAT

>MSTRG.1519.1 gene=MSTRG.1519

GTTTGAGTAATTAACTATTTATATTGTTACACTTTGATTCTTATTGTCTGAGTGTGTGTCTGTCATTAGG

ATGGGTGGGAAAAAACATAACTACAATGCCTGGGAAGAGAGAACTATTTCTGAGGTCAATGTCGAATATC

ACTTGAGATCGTATTCAGGAATTTCACTGCTTGCTGCTGTTGGTAATGAGAACAGAGAAGGTCGGATGAT

GTATGTAGTTGCGAAGGAGTATCTTGAGGCTTTTGGGTCGAATGCAACCATTAATGCATATACACAGTGG

AGAAACCGGGACGATGTAGTGCAGTGGCTCATATCATTGGTGAGAAATGTTGACTCGCCGCCTGCTATAT

CAGATTCCGACCTTGTTCCAAGGTACAAACGTCATGGTTCAAGTATAGATGAAGAGGAGAAGCCTGCCCA

GGGAAATTCTATAAAGAGCTTTCCTAAGAAGTTAAGGAGTAGCTTAGTGCACCAGAAACAGGATAATGGC

AGGTCAAATGCAATAAATCTAGTCCCGCCAAACGAAGCAGCAGCAAGTGAACCAACCGATTCCAACCTTG

TTCCAAAGAACAATGGTCATGGTTCAAGTATAGATGAAGAGGAGAAGCCTGCCCAGGGAAATCCTATAAA

GAGCTTTCCTAAGAAGTTAAGGAGTAGCTTAGTGCACCAGAAACAGGATACTGGCAGGTCAAATGCAATA

AATCTAGTCCCACCAAAAGAAGCAGGCTGTGAACCAACCGGACAGTTTACTCCACGGGAATACTCGAACA

GAGTTGAAACATTCTGTGGGATCTCGATGAGAACCTTCAGAGACTTATGCACTCTTGTGAAGGAGCCTCT

GGGCAATTCACCGAAATACAAATTTAACGATAACAGGGAATTAACCGTGGATGCTAAAGTGGCCATAGCT

ATGAGAAGGCTTAAATTGGGGAAGTCGCTGAACTCAATCGGTGATGAAACTGGGATGCACTCATTTACTG

CTGACAAGGTCACACGGTCCTTTGTGGAGGCAGTAAATGGCCTTAGCCACCTCCATGGGCCTGGCGCTGC

CACTGAAGGGGGAACGAGTGCTGAGACATTGGATAGTGCCGTTGAAGAGGGAATGAGTGTTGAGAAATTG

GATAGGCTAAAGGAAGAATGGAAGGTTCTTACAACCGGAATGAAACCTGAGAAGCTTCAGAGTATGATCA

AGGCCTGCTGCATCTTGCATAACATTGTCACTGATATGAATGATTAGAGGCATCGTGAGCTATCTCAGTC

GCAAGGTCATCATCCGGACTTTATGACTCACAAATTTTAAAGATTAATTTTGTATTTGTTAGAAAGATTT

GCACCATCTCTTCCAAATATTTATGTGGTTTCAACTTTTTATTCGTATGTTGTGTGATTCTTTATTTTTG

TGGTCCGTGATCTTTTGTTCATGAGTTTGTTTAGTATCTTTATTTTACATGCTTTTGTTCAGATGGTTTG

>MSTRG.1520.1 gene=MSTRG.1520

GGAAAATCTGTTTCAAATATTAAACAGGATCTTCGTGAAGAACGCCGATACAGGGATTCAACCGTGAGAA

CACTCGTCTAACCTCAATCATCTTCTCCAAATCCCTGCTTATCTTCAAAACCGCTTTAGATATCAACGTT

GTTGCCCTAACCTAGCTATTACGTAAGATGTTTTCACGGGTCACCGATGAGATCTCGTATGCCCGCCACC

TGTTTGACACAAGTCCCCAACCAGATACTATAGATACTGATGATTAGAGGATTTGGGTCTGATTTGTTTG

TTCATACGGCGTTGATTGAAATGTACGGGAGATACTCTGGACTCGTAATTCTAAACCCCAGATGTGAATC

GGCAAAGCATTTACAGAAACTCATGGAGTACTAGGTGATGGATGGAGAAATTTATATATTCACTTATGTA

ATTATTATGTAAACAAAGTTGAAGCCTCAGCAAGAAAAGCAGAACTTTTGTATCATTGTCTGAAATTCTC

TAAGTTCTTGCTTTAGAACACATATACATTTTATATACTTTTGTTCCAACAAACTTTAAAGTCGCAAACT

TTAGAGTTATTGTCCAGTGGTAAGGAAAAACAATCTTCATTCCTCGAACCACTTTCTTTTTATCTCTTTC

TAATTTGAAGTTTTTAAGAACTTTGCTTTAAGAGAGAAAGCAGTGAAGCAAATATTGGAGAATTCCCTGA

TGTGAATTACATTATTTGGTATGCCCAATACGTGAAGAGGATTGCTGATGAGTAAGAAGGATGTTTTTGC

CTGGAGTGCAATGATAATGGGGCTTGCTAACCATGGGTACGGTGAGGCTGCACTTGATCTTTTTTCTAAA

ATGATATCTGAAGGAACTAAACCCAATGATATCACCTTCATAGGTGTGCTGACAGCATGTAGTCACATTG

GCTTATTTGACAAGGGGTGGACATATTTTAATGCTATGAGTGATATCTATTGTATTCCTCCCAAAATTGA

ACACTATGGAAACATGGTTGACATTCTGGGCCGATCAGGGAGACTTCAGGAAGCTAGAGAACTTATTAGA

AGCATGCCTTTCGAACCGGATGCAGTAATTTTGAGAGCCCTTCTTGGAGCGTGTAAAATTTATAGGAATG

TAGAACTGGCAGAAGAAGCTATAATAAAACTTGTAGCACTAGAACCCAATGTTGACAGGAATTATGTTCT

GTTGTCAAATATATATTCTCAAGCCAAACAGTGGGACAAGGTTGTCAATGTGAGAAGGATTATGAAAAAC

AACAAAATCCAGAGGATCCCAGGGAGCAGTTCAATTGAAATTGGGAACGAAGTCCATGAATTTATTTCTG

GTGATGAATCACACTTGCAATCTGTTGAAATCTATAAGATGTTATCTGAGATTATGAATAGATTGAAAGA

AGCAGGCTATGTACCTTTAACAACCTCTGTTCTGCAAGATATTAATGATAAAGAGAAAGAGACATTGCTT

GTTCGTCACAGCGAAAAGCTTGCAATAGCTTTTGGTATTCTTAATACAGCTCCAGGAGTCTCAATTCGAA

TTGTTAAGAATCTCCGTGTTTGTGATGACTGCCATTCAGCCATAAAAATAATCTCCATAGTTTATAATAG

GAAGATCATCATTAGGGACCGGAATCGGTTTCATCATTTTGTTGCAGGGACATGCTCATGTAAGGATTAC

TGGTGATGCTCCAAAAAAGGGGAAATTGTCTACCAATTACAAATGCAGTTACAGGTCTTATTGTATCTAA

TGATTAGTAGTAATGTTGGGTTTCATGAAATGGAGGTTAACACTGTGCAGGTTATTCAGACTCCTATACT

CAGTGG

>MSTRG.1520.2 gene=MSTRG.1520

TTATATTATGCCCAATTACAATTTTCTAAGGATAAATTTGTATATATGAAGCCCACCTCCAGTATTCACA

AACATACTCCTACTCTTACTATTCTATTTCTTTTAACCTCCATCAACACGCTGACTTCACCCAAGACCCA

TCCATCTCACTCTTTATAAACTTTTGTGTTCCAACAAACTTTAGAGTTATTGTACGGTGGTAAGGAAAAT

CAATCTCCATCTCAAACCACTTTCTTTATATCTCCTTTAATTTTAAGTTTTTAAGAACTTTGCTTCAAGA

GAATGGAAAGAAATGAAGCTAATTGTGGCAAATTTCCTTTGAATTACGTTATTTGGTTTGCCGAATTCGT

GAAGAGGATTGCTGGACATTTTCCTCTCTCGGAAGACTCGACTAGAGTCGAAGAATACTGCGGGATCTCA

ATGAGAACCTTCAGATACTTATGCTCTCTTGTGATGGAGCCTATGTCAAATTCACAAATATTTAAATTTA

ACGATAACAGGGAATTAACTGTGGTTGATAAAGTAGCCATAGCTATGAGAAGCCTAAAATTGGGCAAGTT

GCTGAATTCAATTGGTGGTGAAATTGGGACGCACTCATTTACCGCTGCTAAGGTCACATGGTCCTTTGTG

GAGGTCTTGGCAGAAAATGGCCATCGCCACCTCCGTGGGCCTGCTTCTGCCCCTGAAGAGGGAATGAGTG

CTGAGACATTTAATAGGCTAAAGAAAGAATGGAAGGTTCTTACAACTGTAATGAAACCTGAAACGCTTCG

ATGTATGACCAAGGCCTGCTGCATCTTGCATAACATTGTCATCAACATGAATGATGAGAGGCAACGTGAG

CTATCTCTGTCGGAAGGTCATCATCCAGGCTTTTTGCATCACAGATTTTAAAGATTAGTTTTGTAATTGT

TAGAAAGCTTTGAAATGACTCTTCCAAATATTTCTTTGGTTTCCTCTTTTTTTACGTGGTTGTGTAATTA

TTTGTTTTTGTGGTCACTAATATTTAGTTGATGAGTTTGTTTAGTATCTTTATTATACATGTATACTCTT

TTATATGCATATCTGAGATATTAGAAAAGGACGCCTTTGCTAATGTACTAGTTACGCAAACTGATGATCC

>MSTRG.1520.3 gene=MSTRG.1520

TTTGTGTTCCAACAAACTTTAAAGTCGCAAACTTTAGAGTTATTCTCCGGTGGTAAGGAAAAACAATCAT

CATTCCTCGAACCACTTTTTTTTATCTCTTTCTAATTTGAAGTTTTTAAGAACTTTGATTTAAGAGAATG

GAAAGCAGTGAAGCAAATATTGGAGAATTCCCTGATGTGAATTACGTTATTTGGTTTGCTGAATTCGTGA

AGAGGATTTCTGGACATTTTCCTCTCTCGGAAGACTCGACTAGAGTCGAAGAATACTGCGGGATCTCAAT

GAGAACCTTCAGATACTTGTGCTCTCTTGTGATGGAGCCAATGTCCAATTCACACACATACAAATTTAAC

GATAACAGAGAATTGACTGTGGTTGATAAAGTAGCCATAGCGTTGAGAAGGCTAAAATTTGGCAAGTCAC

TGAACTCAATCAGTAAGGAAATTGGTACGCACTCTTTTACTGCTGATAAGGTTACAAGGTCCTTTGTGGA

GGCCTTGGTAGAAAATGGCCTTCGCCACCTCCATGGGACAGCCACTGCTACTGAGGAAAGAATGACTGCT

GAGACAATGGATAGGCGGCTAAAGGAAGAATGGAAGGTTCTTACACTTGTAATGAAAGCTGAGAAGCGTC

AAAGTATTAGCACAGCTTGCCGCATCTTGCATAACATTGTCATTGAGATGAATGATGAGAGGCATGGTTA

GCTATCTCTGTCACAAGGTCATCATCCAGGCTTTATGACTCACAAATTTTAAAGATTAGTTTTGTAATTG

TTAGAAAGTTTTGAATCATCTCTTCTGAATATTTCTTTGGTTTCCGCTTTTTTATTTGTATTTTGTGTGA

TTATTATTTGTGGTCATTAATATTTAGTTCATGAGTTTGTTTAGTATTTTT

>MSTRG.1520.4 gene=MSTRG.1520

ACAGGATCTTCGTGAAGAACGCCGATACAGGGATTCAACCGTGAGAACACTCGTCTAACCTCAATCATCT

TCTCCAAATCCCTGCTTATCTTCAAAACCGCTTTAGATATCAACGTTGTTGCCCTAACCTAGCTATTACG

TAAGATGTTTTCACGGGTCACCGATGAGGTCTCGTATGCGCGCCACCTGTTTGACAGAAGTCCCCAACCA

GATACTTTAGATACTGATGATTAGAGGTTTTGGGTCTGATTTGTTTGTTCAGACGGCGTTGATTGAAAAT

GGGAGATACTCTGGACTTGTAATTGTAAACCCCAGATGTGAATCGGCAAAGCATTTACAGAAACTCAGGG

AGAGTAGGTACTAGGTAATGGATGGAGAAATTTATATATTCACTTGTGTAATTACTATGTAAACAAAGTT

GAAGCCTCAGCAAGAAAAGCAGAACTTTGGTATCATTGTCTGAAATTCTCCAAGTTCTTGCTTTAGAACA

CATATACTTTTTATATACCTTTGTGTTCCAACAAACTTTAAAGTCGCAAACTTTAGAGTTATTCTCCGGT

GGTAAGGAAAAACAATCATCATTCCTCGAACCACTTTTTTTTATCTCTTTCTAATTTGAAGTTTTTAAGA

ACTTTGATTTAAGAGAATGGAAAGCAGTGAAGCAAATATTGGAGAATTCCCTGATGTGAATTACGTTATT

TGGTTTGCTGAATTCGTGAAGAGGATTTCTGGTATGTTAATCTTTAAATTCTCATGTATATAAAGTTCTG

ATTTGGTGGTGCTCAACTGTTCCTGTCGATTTCGTTATGTTTTGGACATTTTCCTCTCTCGGAAGACTCG

ACTAGAGTCGAAGAATACTGCGGGATCTCAATGAGAACCTTCAGATACTTGTGCTCTCTTGTGATGGAGC

CAATGTCCAATTCACACACATACAAATTTAACGATAACAGAGAATTGACTGTGGTTGATAAAGTAGCCAT

AGCGTTGAGAAGGCTAAAATTTGGCAAGTCACTGAACTCAATCAGTAAGGAAATTGGTACGCACTCTTTT

ACTGCTGATAAGGTTACAAGGTCCTTTGTGGAGGCCTTGGTAGAAAATGGCCTTCGCCACCTCCATGGGA

CAGCCACTGCTACTGAGGAAAGAATGACTGCTGAGACAATGGATAGGCGGCTAAAGGAAGAATGGAAGGT

TCTTACACTTGTAATGAAAGCTGAGAAGCGTCAAAGTATTAGCACAGCTTGCCGCATCTTGCATAACATT

GTCATTGAGATGAATGATGAGAGGCATGGTTAGCTATCTCTGTCACAAGGTCATCATCCAGGCTTTATGA

CTCACAAATTTTAAAGATTAGTTTTGTAATTGTTAGAAAGTTTTGAATCATCTCTTCTGAATATTTCTTT

GGTTTCCGCTTTTTTATTTGTATTTTGTGTGATTATTATTTGTGGTCATTAATATTTAGTTC

>MSTRG.1521.1 gene=MSTRG.1521

GTCATGGTTCAAGTATAGATGAAGAGGAGAAGCCTGCCCAGGGAAATCCTATAAAGAGCTTTCCTAAGAA

ATTAAGGAGTAGCTTAGTGCACCAGAAACAGGATACTGGCAGGTCAAATGCAATAAATCTAGTCCCACCA

AAAGAAGCAGGCTGTGAACCAACCGGACAGTTTACTCCAGGGGAATACTCGAACAGAGTTGAAAAATTCT

TTGGGATGTCGATTAGAACCTTCAGAGGCTTATGCACACTTGTGAAGGAGCCTCTGTGCAATTCACCGAA

ATACAAATTTAACGATAACAGGGAATTAACCGTGGATGCTAAAGTGGCCATAGCTATGAGAAGGCTTAAA

TTGGGGAAGTCGCTGAACTCAATCGGTAACGAAATTGGGATGCACTCATTTACCGTGGATAAGGTCACAC

GGTCCTTTGTGGAGGCAGTAAATGGCCTTAGCCACCTCCATGGGCCTGGCGCTGCCACTAAAGAGGGAAT

GAGTGCTGAGACATTGGATAGTGCCGTTGAAGAGGGAATGAGTGTTGAGAAATTGGATAGTGCCGTTGAA

GAGGGAATGAGTGTTGAGAAATTGGATAGGCTAAAGGAAGAATGGAAGGTTCTTACAACCGGAATGAAAC

CTGAGAAGCTTCGGAGTATGATCAAGGCCTGCTGCATCTTGCATAACATTGTCACTGATATGAATGATTA

GAGGCATCGTGAGCTATCTCAGTCGCAAGGTCATCATCCGGACTTTATGACTCACAAATTTTAAAGATTA

ATTTTGTATTTGTTAGAAAGATTTGCACCATCTCTGCCAAATATTTATGTGGTTTCAACCTTTTTATTCG

TATGTTGTGTGATTCTTTATTTTTGTGGTCAGTGATTTTTTGTTCATGAG

>MSTRG.1522.1 gene=MSTRG.1522

GGTCAGTAAAATGGATGGGCACATTGTGCTCGAAGAGGAGCATATATGCACGCCGCCTTGTGGATTCCGA

AATAAACGTTACAGTGCATTGCATATGTTTCTCGCGAAAGTGCACTATCGCAAACATTCACCTAAGCTGT

GGCTGGTTCAGCTCCACAAGGGTGATTAACCCCAGTTTCTTTTCCGGATTTGGTATGAAGATGGCCTTGT

ATAAAAGAGCAAACCCCTGTTCCTTGGCAGTCCATTACTTTCCGGATGGGTGGGAAAAAACATAACTACC

ATGCCTGGGAAGAGAGAACTATTTCTGAGGTCAATGTCGAATATCACTTGAGATCGTATTCACGAAATTC

ACTGCTTGCTGCTGTTGGTAATGAGAACAGAGAAGGTCGGATGACGTATGTAGTTGCGAAGGAGTATCTT

GAGGCTTTTGGGTCGAATGCAACCATTAATGCATATACACAGTGGAGAAACCGGGACGATGTGGTGCAGT

GG

>MSTRG.1523.1 gene=MSTRG.1523

CAGAATGTTATCCCCAAATAAGGAAAATATGTTTCAAATATTAAACAGGATCTTCGTGAAGAACGCCGAT

ACAGGGATTCAACCGTGAGAACACTCGTCTAACCTCAATCATCTTCTCCAAATCCCTGCTTATCTTCAAA

ACCGCTTTAGATATCAACGTTGTTGCCCTAACCTAGCTATTACGTAAGATGTTTTCACGGGTCACCGATG

AGATCTCGTATGCGCGCCACCTGTTTGACAGAAGTCCCCAACCAGATACTTTAGATACTGATGATTAGAG

GTTTTGGGTCTGATTTGTTTGTTCAGACGGCGTTGATTGAAAATGGGAGATACTCTGGACTTGTAATTGT

AAACCCCAGATGTGAATCGGCAAAGCATTTACAGAAACTCAGGGAGAGTAGGTACTAGGACATTTTCCTC

TCTCGGAAGACTGACTAGAGTCGAAGAATACTGCGGGATCTCAATGGGAACCTTCAGATACTTGTGCTCT

ACTCCTTCAGAACTTCTGAATTCTTTGAGCTCTACTCCGTGAATTCCTCATGTTTATCGAACCCTATGCA

CCTAGTGTGACTATACTATCTAACTCCTCTTCTTGTTATCCGAG

>MSTRG.1523.2 gene=MSTRG.1523

CAGGATCTTCGTGAAGAACGCCGATACAGGGATTCAACCGTGAGAACACTCGTCTAACCTCAATCATCTT

CTCCAAATCCCTGCTTATCTTCAAAACCGCTTTAGATATCAACGTTGTTGCCCTAACCTAGCTATTACGT

AAGATGTTTTCACGGGTCACCGATGAGATCTCGTATGCGCGCCACCTGTTTGACAGAAGTCCCCAACCAG

ATACTTTAGATACTGATGATTAGAGGTTTTGGGTCTGATTTGTTTGTTCAGACGGCGTTGATTGAAAATG

GGAGATACTCTGGACTTGTAATTGTAAACCCCAGATGTGAATCGGCAAAGCATTTACAGAAACTCAGGGA

GAGTAGGTACTAGTGTCACGGAGAAAGTTTGCTACTGCCACCTGACTAAGTTAGGTATGTTATATCAATG

CCTTTATGAGC

>MSTRG.1524.1 gene=MSTRG.1524

TGCAGTGGCTCATATCATTGGTGAGAAATGATGACTCGCTGCCCGCTTTATCAGATTCCTACCTTGTTCC

AAGGCACAAACGTCATGGTTCAAGTATAGATGAAGAGGAGAAGCCTGCCCAGGGAAATCCTATAAAGAGC

TTTCGTAAGAAGTTAAGGAGTAGCTTAGTGCACCAGAAACAGGATACTGGCAGGTCAAATGCAATAAATC

TAGTCCCACCAAAAGATGCAGCGGCAAGTGAACCAACCGATTCCGACCTTGTTCCAAAGAACAAACGTCA

TGGTTCAAGTATAGATGAAGAGGAGAAGCCTGCCCAGAGATATCCTATAAAGAGCTTTCCTAAGAAGTTA

AGGAGTAGCTTAGTGCACCAGAAACAGGATACCGGCAGGTCAAATGCAATAAATCTAGTCCCACCAAAAG

AAGCAGCAGCGAGTGAACCAACCGGACGGTTTACTCCACGGGAATACTCGAACAGAGTTGAAATATTCTG

TGGGATCTCGATGAGAACCTTCAGAGACTTATGCACTCTTGTGAAGGAGCCTCTGTCCAATTCACCGAGA

TACAAATTTAACGATAACAGGGAATTAACCGTGGATGCTAAAGTGGCCATAGCTATGAGAAGGCTTAAAT

TGGGAAAGTCGCTGAACTCAATCGGTGATGAAACTGGGATGCACTCATTTACTGCTGACAAGGTCACACG

GTCCTTTGTGGAGGCAGTAAATGGCCTTAGCCACCTCCATGGGCCTGGCGCTGCCACTGAAGGGGGAACG

AGTGCTGAGACATTGGATAGTGCCGTTGAAGAGGGAATGAGTGTTGAGAAATTGGATAGGCTAAAGGAAG

AATGGAAGGTTCTTACAACCGGAATGAAACCTGAGAAGCTTCGTTGTATGATTAAGGCCTGCTGCATCTT

GCATAACATTGTCACTGATATGAATGATTAGAGGCATCGTGAGCTATCTCAGTCGCAAGGTCATCATCCG

GACTTTATGACTCACAAATTTTAAAGATTAATTTTGTATTTGTTAGAAAGATTTGCACCATCTCTTCCAA

ATATTTATGTGGTTTCAAC

>MSTRG.1529.1 gene=MSTRG.1529

CTACATTCAGAAATCTGTGTTGTGTATAGTTCATGATTGTTAGATGATTAGAGGGTCCTCACCAAGGTGA

CACGGAGCAGCAAAGTTATGATATGTGTTCGTGTTTTAATGAGCTCTGCAGGTGTTGGTGTTTCTTATGG

CAAGTTGACAATGTAGAGTTAGAGCATGCAGTCCCATACTCTTGAATTATAATGATGAATTCCCTCTGGT

CCAAAAAAAGTTCCATTTCCAGGGAAGTTTATCCAGCTAGCAAGGTGTGATCCCATACCACACCTTCGGC

TAGAAATGTGGGAGTGCTAGCCTAATCTGGTTCTGGCTTTTTTATGGCTTCATTATTTTCCATACTTCAT

ATTCCAAATATGTTACTGGAATGAGGCAACATCAGAAAAGATGAGGAAGAAATTGTAAGACACTTTTGAT

AACATCATCCAGAAGCATGTTAAAAGAAGTCTAATAGAAAACACCAGAGGGTCCATGACACTGACCGACT

GAAGAAGATCTAGTTAAATTTTTGTCTATGAATGAACAATGGCAAAACTCTATAATACCCATTACTTGCA

AATATAACCATGTAACTCTTGCAAGTTTACTTAATTCATTTCGTCGTATGCATGTCAGGTCTGATGTACT

GGGCAGTGAGGCGCGGTATAGTCTAAATTAGGAGGGGGGTATATCCAAATTGATAAAATTTCTT

>MSTRG.1533.1 gene=MSTRG.1533

TCATTGTCGAGATCAGCTGACTCAAGATCCACAGCTGTCAATTTTCTTTCAAGAACCCAATCCACCAGCA

ATTTTCGCCTCTCCTCACTCCAGAATTCAGCAAAGTAGACGAATAATTGTGCTAAACATATCGTACTAGT

TAGCATCCAGAAGATGGCAAATAAGCGCCCTCCTCCAGTGGAGAAGCTTTTGTCCCCATAGCCTAGAGTG

GTC

>MSTRG.1535.1 gene=MSTRG.1535

AAGCCAGTGTGTGCTTATGAAGATTAATGAATTATCAGCTATTACCTACTCATATTTCTTCTTCAACATG

AATCTTCTCTTTAGAAGATTCTATTGTTTTCTTGAATGAGTCTTCTGGTGCATCAGGTGAAAAAATGGCC

AAATTCTCTCAGGCTAGAGAAGGGTGCACCAAAATACTACGATATAGAAATAAATTTCATTATTAAACAT

AGACAGTTGATTCGGCTGAGAACATCTTGTTTGGGAGAATAGTGAAAACAATTGTTCAACACTTCTTAGA

AAGTGCAGAACAATTATTTTAAAAACTTTTCACTTTTTAAATGTTGTATTAATTTATACTAGAATAATAG

TGTTCTCAGTTCTCACATTAAAAACCGGTAGACAAGTTTATATAACTAGCCAGCCTCCTACAATTTAATC

TCGTATACTATAATTTATAACTACAACAGTTGTATTTTTAAAAATATGTGTAAATTATTTTGCTATATAG

GAGAACTAAGTATGTTATGTATTTATATTTACATTACAGGTTCTGTAAACACTTTTTCACTATTTTTTAA

CCAATTAATTAGACCATTTTATAATTATCTTTTTATTTACATAGATTATCAATCAATTTCTTAATAAATG

ATGTCATATTTTCACAAATCACAATTTATAAAGAGTAGTGCTAGGTGCACATAATTGTGTACAGAAAATT

TGTACATAATGATGTGGCAAGTGATGTGGTGGGTTTTAATTGGGGTGGTTGGTGTAAAGGGGCCATCCAA

TTAAAATTCGTCACATATGTCATTATGTAAATGTTTTTGTACACAATTATTTGCACCAAGCATTTTCTAT

TTATAAAACTAACATGCCCATGTTCTTCAAGTAGTTTTTATATTTTTTAATACATAAATGTTCTCATTAT

TATGTTTTGATTACAA

>MSTRG.1537.1 gene=MSTRG.1537

ATTAATCATATTTGCCGCTGAATTCTTTTCCCTTTCCGCTGTTTCCCTCTTAATTTTCCTAGACTCCACC

TCCAGAGCTTTTACTAATGTGTCTACCTTCTTCATGAGTACCTTAATTTCTTCATCTTTTGCACTTAAAT

TGTTATCTTTCTTCTCGCAAGACTTTCTAAGACTTATGACCTCTTTCTGCAGCTTGTCATACAGAAACCC

CGACACTGAGTCATCCGTCTGAGGATTCAAGCCTGTAATACTTTCTGAATCTTCTCTCACAGCGACATAT

GGTTCAACATTGTCTAGAACTTCACTTCCATTCTCCTTCATCTTGTCGATGTTCACGTCATTTTCTTTAC

CCGTCTTAGCTCTTTCTAGTTCATCAATATTACCAACCTTACTTCTGGATGACCACAAACCTTTTCTTAC

AAGAAGCTCTCCAAACCCATATCTCATTTTCAAGCTATTTCCCCGTTTAAGTTCCTTCTCTAAAATCGGT

TTTTGTAAATTTCTATTAATGGCGGACGCCCTTGGCTGTGATACAGACCTCATCCTTTCACCATTAGATA

CATTCTCAGTACTGCTAGAGTTTCCAGGTTTTGGGCTTCCAGAAGAAGCTTTTTGATTGACAGTTATATT

AGGTGTTTGCTTCAAACCTTCTTCCAGCGTCTTTAGTCTCAGTTTGAGTTTGTCCTGAATTCAGAAGTGA

ACGATGCTGATATATTAGCAAAACAATTTTATGTTTGTGTTATATATAAATTAAGATAAATAGACAAATA

AGTTCACCTTAAGTTGTGCTTCGGTATTAGCTCTTCTCTCTGATATGGCTAACTTGTCCCTTAGTCGTTG

CATTTCAGCCTGCAGATCACAGAGAGCTTTTATTTATATTTTTCTGAATATGTGGACATCATTGCATAAG

CGAAAACACATAATAGATACCTGCATAACCCTTCTTTCTTCTAGCCACTGCTTAACTGGCATAACTTTGT

CATTTTCATCCTTCCATTCATTAGCCACTACAGTTGCTACCCGGTTAGCTGAGACCTTGATTCTTGCCAG

CTCTCTTTCCAATGTCCGGTTCTCTTCCTGATAAGATTATGCCAAAACTAATGTAATTATCCATAAATGG

CTATTATACTTCACTAGCGAAAGGTACTCTCACAACATAAATTTTTTACATTTAGTTCAGAGATCCGTCT

CCTGTAGTCACGGATGGTGTTAGCTGCTGCTCCACCAACTAGAATAGCCTCCTCAAGTTCCTGAATGGTT

TCTCCAAGTTTTTGGACCTCAATCACCTTTTGACGATTTGTTTTCTCAAGGATTTTGTTCTCCTCCTGAA

AGTGGAGAACTCATATTTTGTATAAAAAAAAGTTCTCAAGGATTTTGTAAATTTTCAGCATACAATATCA

ATATATGTCCGGCAACGCAGAAACTCTTTTCTGGAAAGGAGGGAGTGAAAAGATCAAGTGTCGAACCTGG

CAGATCTCAATTTGTCGTTTAAGCTCATAGTTCTGGTTCTGAATCTCCTCTACAACCAGTGCCCTCTCTA

GAGCACTTTTAAGAATCTTCTCCACTTCAAGCAGTGCAGTTTCTTTCGACTTAGTGAGTCGTTCCAAAGC

CCTCTTCTCCTCCTGCAGTGTCATCACCTGGGAGAAGAAACAAAGGCGTATGAGCCTAAAAGAATCATGT

ATCCACACTTTAGCCATTTAGATATGCTATACCTCGTGTTTATACATCTTGATGTCTGCTTCTAGAGGAG

CGATGACTGATTCAATAGGAAAAGAATCGTCGTCCTTTTGATCTGCATGAACCCTTCTAACAGTTG

>MSTRG.1540.1 gene=MSTRG.1540

ATCACTGAACAATTCAGGGGCAGAAGAATACTCAGTGTAGTACTTGGGACCATAATATTCACTCCTCTTC

ATTTCCATTGCTTTATCATCTTCATCTTCCTTGGCTTCTTCCGCTTCTTTAGATTCTTCTTTCTTCTCTT

CCTCTTTCTTTTCTTCCTCCTTTTTCTCTTCGTCCTTCACCACGCTTGCTTGCTTCCTGGTTTTCTTGTA

CACATCATTCACTAACTTTTCCGGATCAAGAACACCTTTCACAGTTACTTGATCATTTGCTA

>MSTRG.1544.1 gene=MSTRG.1544

GCAGAAATAATTATATACTGATTTGTTGGAAAAAATCTTTAGTGTGCAGATTCTAAGTGGGTAAAGTTTG

TATTTTAATGTTGCGGGTGTTATGTAAATTGTAAGAACATACAGTGATGGGAAATATGAATGGTAAAGAA

GATGGTAAGGCCCTTGATAGGCCATCTGCTGGTGCTGATATGGTTGAGGAGGATGGTGGTCTTCTTTTTG

ATGGTGAATAATATCTGTTTGGGTGTATAT

>MSTRG.1557.1 gene=MSTRG.1557

CCCCACTGCCAACTATCCACTGCCCACCTCTTCCTCTTTTATCTACTGCCATCCGCCACTACCTATTGCT

GCTGCAGCGACACAACCACCAACCACCAGATCTGCTACAACGTCGTCGTAGTCGAGGCGGCGACCTCGTC

TCCAGTCACGCACACAAAAAACACAGATCTCTCTCCCTCTCCCTCCCCATCTCTCTCGCCAGAAACGCGA

CTACTGCCTCCCCTCTAAAAGCCCCGTCCTTATTCTTCCATTTTGTTTCTTCTCTATTCATCGCAGATCC

ACATCACCTCCGATTTCGATACTCCAATCTGGAGATGGTATTGGTGGAGGTTCAGTGATTTTTGTTTTTA

CGGTGGTGATTTCATTTTGTTGATGGTGTTTGTTTGTGGTAATATAGTAATTTTTGTTATGGTCGGAGTT

AATGATGGATATGGGGCAGAAATAGAGATTGGATGTAGTGAATGAGAGAGAGCGTGTGATGTAGAGAGAG

AGAGAGAGGAGATTGAGAGAGACAGTGAGGAAGCTGATGAATTGGAAATGATGATTTTCTGTTGTGTTTT

TTATTTTTTCCGAAGGTGATTTTGTTTTGTATTTGGTGATTTTTCGGATTCGGGGGTGGCGGTGGTGGTT

TGTAGCTGGTGTTGGTGTGGTGATTTTAATTTTCTGGCGTTGATTTTTGATTTTCTGGCGGTGATTTCTT

ATTTTCTGGTGGTGATTTTATGTTTCCGATGGTGATTTTATGTTTGCCGGCGGTGATTTTCTGGTGGTCG

CCGGAGCTGGTGGTGGTGGTCAGCGGTGGTTGAAGGTGATGGGTAGTTGAATTAGAAGATGGTGAAATAT

TAAAATATTAAATGAATGGTGAGAGATGAATATAATGTATTTAAATGAGTGGCTGAGATTAGATCTCATA

TCTCACTATAAGATTGGTTCTCACCGGAGCATGACTCTATATA

>MSTRG.1560.1 gene=MSTRG.1560

ATTAAAACCACACGCTCAGCCCCTTTTAAAAAACGCTCGCGCTCTCTCCCTGTCTCCCATATGTGGGGTT

CTGAGAGAGAGATTGTTTATCCTCTTCCTCATCCACAGCCCAAGTGTATCACCACTACCATATTGACATT

GCAGAGTTGGCTGGCAAGCTTGTTCAGGACATACCAATTGATCAACCAACTGACATTGTTATGACAACTG

ATCTGGACGGCTTTCCCCTTTAGAAAAGATCATATATAAATGGTGTAAAGTCAACGGATTTGAGATCACA

ATGACCTCTTCATATACGGTCTTGGGTTGATTTTCAAAAACATAGAAAGAAAGAAAAGCACTCTGTTAAT

CTGCCTAGAATTGCTAGTCCAGTAATGTCAACTGTGCTGCCTGCAGCACCACTGAACATTTTCCCACAAG

AATTATGTTGCAAATATGAGGGGACAATTTAAAGTGGTTCCTGGGAGATTTGGTCTTGGGGCAATGGAAG

TCCCCCGGCCCCCAGAGGTACTGGAAAGCGGTCACTTGAGCAGTGGTCTGCAGACTCTATATCATGAACC

AGCTGGTGCAACCGAGGCGTGATACCATCAAGCAGTCTCCTGAGTAGTGGGTGGCAGACTCTGAACTGTC

ACCCGAACAATGGCCTGCAGGCCCTGAACCAAGAAAATGCTGGTGATACTGCGATAGCGATGTTGATCTG

GGAAATCCAGTTAACATGGACAGTCACAGCTTTGAAGAAGGCACTAGTGCAAATCAAGTGCTGGACGATC

TGCCTAGTGAGGAGGAAGTTAGTAATATGATACAGGATCTTGAAGCCTATGATTTATTTGATTTGATAAA

TTAAAACCAAACAAAGGAGGAATATAGCGTAAGTTCGTTCTTTAAAGGCTTACTTGACTTGTCTGGCTGG

CCCCAAATGCTCAAGTTAAATGATTGACCTCAATCTGGGTTTTGTGGATCATCTCCCAAGTCATAATGTG

GAGTTCTTACACACATCCACAGAGCGGCTACTTGAACCTTGCTGTCAAGTTACCCGAGGGCCATCTGAAG

CCAGACAAAGGGCCAAAGCTGTATACTGCTTATGGCTTTTCCCAGCAACTCGGACGTGGTGACTGTCTCA

AAACTCAGCTATTATAATTCAGATGCGGTATGATCTAGTTTGTAAATAGAGTGAATAACTTTCACAAACA

ATGAAGAATATGCTCCAAATTTACAACCTCTCCCAAGTTAAGTTGTTTTTAATTGTGCAAACAATTTTGT

TACCTATGCAATGTATATATCAATGTATGTCATAGAATCA

>MSTRG.1560.2 gene=MSTRG.1560

CCCCTTTTAAAAAACGCTCGCGCTCTCTCCCTGTCTCCCATATGTGGGTATGATTTAGGACTACCAATTG

GTATTTCTTCAATTGACCACCATAAATTAGGGTTTTTGGTTTGAATCAGGGGTTCCTTTTTATTTCAATT

TATCAACTCAAGAGTTCAATTGCTCTGATTAGTTTGGTTTCTCTCTTTGTTTCAGGGTTCTGAGAGAGAG

ATTGTTTATCCTCTTCCTCATCCACAGCCCAAGTGTATCACCACTACCATATTGACATTGCAGAGTTGGC

TGGCAAGCTTGTTCAGGACATACCAATTGATCAACCAACTGACATTGTTATGACAACTGATCTGGACGGC

TTTCCCCTTTAGAAAAGATCATATATAAATGGTGTAAAGTCAACGGATTTGAGATCACAATGACCTCTTC

ATATACGGTCTTGGGTTGATTTTCAAAAACATAGAAAGAAAGAAAAGCACTCTGTTAATCTGCCTAGAAT

TGCTAGTCCAGTAATGTCAACTGTGCTGCCTGCAGCACCACTGAACATTTTCCCACAAGAATTATGTTGC

AAATATGAGGGGACAATTTAAAGTGGTTCCTGGGAGATTTGGTCTTGGGGCAATGGAAGTCCCCCGGCCC

CCAGAGGTACTGGAAAGCGGTCACTTGAGCAGTGGTCTGCAGACTCTATATCATGAACCAGCTGGTGCAA

CCGAGGCGTGATACCATCAAGCAGTCTCCTGAGTAGTGGGTGGCAGACTCTGAACTGTCACCCGAACAAT

GGCCTGCAGGCCCTGAACCAAGAAAATGCTGGTGATACTGCGATAGCGATGTTGATCTGGGAAATCCAGT

TAACATGGACAGTCACAGCTTTGAAGAAGGCACTAGTGCAAATCAAGTGCTGGACGATCTGCCTAGTGAG

GAGGAAGTTAGTAATATGATACAGGATCTTGAAGCCTATGATTTATTTGATTTGATAAATTAAAACCAAA

CAAAGGAGGAATATAGCGTAAGTTCGTTCTTTAAAGGCTTACTTGACTTGTCTGGCTGGCCCCAAATGCT

CAAGTTAAATGATTGACCTCAATCTGGGTTTTGTGGATCATCTCCCAAGTCATAATGTGGAGTTCTTACA

CACATCCACAGAGCGGCTACTTGAACCTTGCTGTCAAGTTACCCGAGGGCCATCTGAAGCCAGACAAAGG

GCCAAAGCTGTATACTGCTTATGGCTTTTCCCAGCAACTCGGACGTGGTGACTGTCTCAAAACTCAGCTA

TTATAATTCAGATGCGGTATGATCTAGTTTGTAAATAGAGTGAATAACTTTCACAAACAATGAAGAATAT

GCTCCAAATTTACAACCTCTCCCAAGTTAAGTTGTTTTTAATTGTGCAAACAATTTTGTTACCTATGCAA

TGTATATATCAATGTATGTCATAGAATCACCTCC

>MSTRG.1560.3 gene=MSTRG.1560

TTTCTCTCTTTGTTTCAGGGTTCTGAGAGAGAGATTGTTTATCCTCTTCCTCATCCACAGCCCAAGTGTA

TCACCACTACCATATTGGTAATATTTCTCTCCCTCCCTCCCTCTGTTCTTGTTATATGTACTTTCTAATT

TTTTTGTCTTGGAATCAGACAAGTGTTTGCTTTGTCGAAGGTGGTACAAAAACTATAATGGTTTTTTAAA

TATTTTATAGAATATGTTTAACTCGATCATATTTATTTGCTTGATGTATAATGTTTTCGTATTCAATATA

GTAATTTGTTTTATCAAGTGGAAACACTCGAATAAACATAGGAAAGCTTAAAGGTGTTTAGTCAGGAAGA

AAAAATCAGTAGAAAGATTGAGTTGGTGGAAAGAAATTGGAGAAGGGCATTTGATGAGTCAATTCAGTGC

CTCCTCCAGCGGCAAAACAAGTGAGAACTTCTTGAACTTCTTAAACTTCAAACACGAGGAATGACAGGGA

AGGCTATTTGCACCCGTCTAATAATTTTAAGGTTTAAAATTATGAAGAGGGCAATGCCATTGGATGATCA

GATGAACATATCTTCTGATGAGTCATTATCTTCTGAACCAGACATTGCAGAGTTGGCTGGCAAGCTTGTT

CAGGACATACCAATTGATCAACCAACTGACATTGTTATGACAACTGATCTGGACGGCTTTCCCCTTTAGA

AAAGATCATATATAAATGGTGTAAAGTCAACGGATTTGAGATCACAATGACCTCTTCATATACGGTCTTG

GGTTGATTTTCAAAAACATAGAAAGAAAGAAAAGCACTCTGTTAATCTGCCTAGAATTGCTAGTCCAGTA

ATGTCAACTGTGCTGCCTGCAGCACCACTGAACATTTTCCCACAAGAATTATGTTGCAAATATGAGGGGA

CAATTTAAAGTGGTTCCTGGGAGATTTGGTCTTGGGGCAATGGAAGTCCCCCGGCCCCCAGAGGTACTGG

AAAGCGGTCACTTGAGCAGTGGTCTGCAGACTCTATATCATGAACCAGCTGGTGCAACCGAGGCGTGATA

CCATCAAGCAGTCTCCTGAGTAGTGGGTGGCAGACTCTGAACTGTCACCCGAACAATGGCCTGCAGGCCC

TGAACCAAGAAAATGCTGGTGATACTGCGATAGCGATGTTGATCTGGGAAATCCAGTTAACATGGACAGT

CACAGCTTTGAAGAAGGCACTAGTGCAAATCAAGTGCTGGACGATCTGCCTAGTGAGGAGGAAGTTAGTA

ATATGATACAGGATCTTGAAGCCTATGATTTATTTGATTTGATAAATTAAAACCAAACAAAGGAGGAATA

TAGCGTAAGTTCGTTCTTTAAAGGCTTACTTGACTTGTCTGGCTGGCCCCAAATGCTCAAGTTAAATGAT

TGACCTCAATCTGGGTTTTGTGGATCATCTCCCAAGTCATAATGTGGAGTTCTTACACACATCCACAGAG

CGGCTACTTGAACCTTGCTGTCAAGTTACCCGAGGGCCATCTGAAGCCAGACAAAGGGCCAAAGCTGTAT

ACTGCTTATGGCTTTTCCCAGCAACTCGGACGTGGTGACTGTCTCAAAACTCAGCTATTATAATTCAGAT

GCGGTATGATCTAGTTTGTAAATAGAGTGAATAACTTTCACAAACAATGAAGAATATGCTCCAAATTTAC

AACCTCTCCCAAGTTAAGTTGTTTTTAATTGTGCAAACAATTTTGTTACCTATGCAATGTATATATCAAT

GTATGTCATAGAATCACCTCC

>MSTRG.1560.4 gene=MSTRG.1560

TTTCTCTCTTTGTTTCAGGGTTCTGAGAGAGAGATTGTTTATCCTCTTCCTCATCCACAGCCCAAGTGTA

TCACCACTACCATATTGGTAATATTTCTCTCCCTCCCTCCCTCTGTTCTTGTTATATGTACTTTCTAATT

TTTTTGTCTTGGAATCAGACAAGTGTTTGCTTTGTCGAAGGTGGTACAAAAACTATAATGGTTTTTTAAA

TATTTTATAGAATATGTTTAACTCGATCATATTTATTTGCTTGATGTATAATGTTTTCGTATTCAATATA

GTAATTTGTTTTATCAAGTGGAAACACTCGAATAAACATAGGAAAGCTTAAAGGTGTTTAGTCAGGAAGA

AAAAATCAGTAGAAAGATTGAGTTGGTGGAAAGAAATTGGAGAAGGGCATTTGATGAGTCAATTCAGTGC

CTCCTCCAGCGGCAAAACAAGTGAGAACTTCTTGAACTTCTTAAACTTCAAACACGAGGAATGACAGGGA

AGGCTATTTGCACCCGTCTAATAATTTTAAGGTTTAAAATTATGAAGAGGGCAATGCCATTGGATGATCA

GATGAACATATCTTCTGATGAGTCATTATCTTCTGAACCAGACATTGCAGAGTTGGCTGGCAAGCTTGTT

CAGGACATACCAATTGATCAACCAACTGACATTGTTATGACAACTGATCTGGACGGCTTTCCCCTTTAGA

AAAGATCATATATAAATGGTGTAAAGTCAACGGATTTGAGATCACAATGACCTCTTCATATACGGTCTTG

GGTTGATTTTCAAAAACATAGAAAGAAAGAAAAGCACTCTGTTAATCTGCCTAGAATTGCTAGTCCAGTA

ATGTCAACTGTGCTGCCTGCAGCACCACTGAACATTTTCCCACAAGAATTATGTTGCAAATATGAGGGGA

CAATTTAAAGTGGTTCCTGGGAGATTTGGTCTTGGGGCAATGGAAGTCCCCCGGCCCCCAGAGGTACTGG

AAAGCGGTCACTTGAGCAGTGGTCTGCAGACTCTATATCATGAACCAGCTGGTGCAACCGAGGCGTGATA

CCATCAAGCAGTCTCCTGAGTAGTGGGTGGCAGACTCTGAACTGTCACCCGAACAATGGCCTGCAGGCCC

TGAACCAAGAAAATGCTGGTGATACTGCGATAGCGATGTTGATCTGGGAAATCCAGTTAACATGGACAGT

CACAGCTTTGAAGAAGGCACTAGTGCAAATCAAGTGCTGGACGATCTGCCTAGTGAGGAGGAAGTTAGTA

ATATGATACAGGATCTTGAAGCCTATGATTTATTTGATTTGATAAATTAAAACCAAACAAAGGTATTCTC

TTTCAGTTTTTTAGCAACTATTCAGAAGTT

>MSTRG.1560.5 gene=MSTRG.1560

GAACTTCTTAAACTTCAAACACGAGGAATGACAGGGAAGGCTATTTGCACCCGTCTAATAATTTTCAGAA

ACATAGAAAGAAAGAAAAGCACTCTGTTAATCTGCCTAGAATTGCTAGTCCAGTAATGTCAACTGTGCTG

CCTGCAGCACCACTGAACATTTTCCCACAAGAATTATGTTGCAAATATGAGGGGACAATTTAAAGTGGTT

CCTGGGAGATTTGGTCTTGGGGCAATGGAAGTCCCCCGGCCCCCAGAGGTACTGGAAAGCGGTCACTTGA

GCAGTGGTCTGCAGACTCTATATCATGAACCAGCTGGTGCAACCGAGGCGTGATACCATCAAGCAGTCTC

CTGAGTAGTGGGTGGCAGACTCTGAACTGTCACCCGAACAATGGCCTGCAGGCCCTGAACCAAGAAAATG

CTGGTGATACTGCGATAGCGATGTTGATCTGGGAAATCCAGTTAACATGGACAGTCACAGCTTTGAAGAA

GGCACTAGTGCAAATCAAGTGCTGGACGATCTGCCTAGTGAGGAGGAAGTTAGTAATATGATACAGGATC

TTGAAGCCTATGATTTATTTGATTTGATAAATTAAAACCAAAC

>MSTRG.1560.6 gene=MSTRG.1560

GACGAATCCATCCCGTATCCCGTGTTGTGTTTCGTGTCGTGTCGTGTGGACACGGGTACGGTCGCTGGAG

TAGGCGAGTCGGAGTAACACAGTGTATATCAATCTTCAATATCATCGGTGGATTACTTTCACAAACAATG

AAGAAAATGCTTCAAATTTACAACACCTCCCGAGTCATCAAAGTAATTTGTCCTGAAGGTTTAAAATTAT

GAAGAGGGCAATGCCATTGGATGATCAGATGAACATATCTTCTGATGAGTCATTATCTTCTGAACCAGAC

ATTGCAGAGTTGGCTGGCAAGCTTGTTCAGGACATACCAATTGATCAACCAACTGACATTGTTATGACAA

CTGATCTGGACGGCTTTCCCCTTTAGAAAAGATCATATATAAATGGTGTAAAGTCAACGGATTTGAGATC

ACAATGACCTCTTCATATACGGTCTTGGGTTGATTTTCAAAAACATAGAAAGAAAGAAAAGCACTCTGTT

AATCTGCCTAGAATTGCTAGTCCAGTAATGTCAACTGTGCTGCCTGCAGCACCACTGAACATTTTCCCAC

AAGAATTATGTTGCAAATATGAGGGGACAATTTAAAGTGGTTCCTGGGAGATTTGGTCTTGGGGCAATGG

AAGTCCCCCGGCCCCCAGAGGTACTGGAAAGCGGTCACTTGAGCAGTGGTCTGCAGACTCTATATCATGA

ACCAGCTGGTGCAACCGAGGCGTGATACCATCAAGCAGTCTCCTGAGTAGTGGGTGGCAGACTCTGAACT

GTCACCCGAACAATGGCCTGCAGGCCCTGAACCAAGAAAATGCTGGTGATACTGCGATAGCGATGTTGAT

CTGGGAAATCCAGTTAACATGGACAGTCACAGCTTTGAAGAAGGCACTAGTGCAAATCAAGTGCTGGACG

ATCTGCCTAGTGAGGAGGAAGTTAGTAATATGATACAGGATCTTGAAGCCTATGATTTATTTGATTTGAT

AAATTAAAACCAAACAAAGG

>MSTRG.1561.1 gene=MSTRG.1561

CTGTCTTGCAGGAGGAATATAGCGCAAGTTCGTTCTTTAAAGGCTTACTTGACTTGTCTGGCCGGCCCCA

AATGCTCAAGTTAAATGATTGACCTCAATCTGGTTTTTGAGAATCATCTCCCATGTCATAATGTGGAGTT

CTTAAGTGCTTTGCCTTTCAAGGAGTACACACATCCACAGAGCGGCTACTTGAACCTTGCTGTCAAGTTA

CCCGAGGGCCATCTGAAGCGAGACAAAGGGCCAAAGCTGTATACTGCTTATGGCTTTTCCCAGCAACTCG

GACGTGGTGACTGTCTCAAAACTCAGCTATCATAATTCAGATGCGGTATGATCTAGTTTGTAAATAGAGT

GAATAACTTTCACAAACAATGAAGAATATGCTCCAAATTTACAACCTCTCCCAAGTTAAGTTGTTTTTGA

TTGTGCAAACAATTTTGTTACCCATGCAATGTATATATCAATGTATGTCATACAATCACCTCCAACCATT

TATCAATTATCGT

>MSTRG.1562.2 gene=MSTRG.1562

CAAAATATTAAAACTACACGTTCATAAAACTCTCTCTCTCGCTCTCTCCTTTCTCCCTCTCACTTAGGTT

CCTCTCTGCGTCTCTCAACCGATATGTGGGGTTCCGAGAGAGATTCTGTTTATACAAGTTACCGAACCAC

CATATTGAGGAAACACTCGAATAAACATAGGAAAGCTTAAAGGTGTTTAATCAGGAAGGAAAAAAACCAG

TAGAAAGATTGAGTTGGTGGAAAGAATTTGGAGAAAGGCATTTGATGAGTCAATTCAGTGCCTCCTCCAG

TGGCAAAACAAGTGTGAACTTGAACTTCAAACACGAGGAATGACGGGGAAGGCTATTTGCACCCTTCTAG

TAATTTTCAGGGGCTTCGACTGAATTGGGGCTTCTCTAGAACTAGAGCTTCAACAGCTTTTAACTTTAAG

TGGGGTTTCGGTTTGCTTGGATTTAAGTTCTATAGACATTCAAGCCGGGGATCGGAACTCGGAAGCATTT

TCAGACCCATTGGTCAAAAGGGGAACCAGTGATTGTTACTATTGTGCTTGACAGTACATATGGTCTAAGC

TGGGAACCAATGGTGATGTCTCTAGCAATCCATGATAAGTCCCACACTGATGTGATTTGTCTTAAATTGT

TTGAGCTGGTGCGAGGAGGAATATAGTGTATGTTCGTTCTTTAAAGGCTATACAGAGGGTATACACTTGT

CTGCCTGGCCCCAAATGCTCAAGTTAAATGACTGGCCTCAATCTGGTTCGTTTGATGATCATCTCACGTC

ATAATGTGGAGTTCTTAAGTGCTTTGCCGTTCAAGGAGTACACACATCCACAGAGCGCCTACTTGAACCT

TGCTGTCAAGTTACCCGAGGGCAATCTGAAGCCAGACACAGGGCCAAAGCTGTATACGCTTAGATAATGC

ACAAACACGCCTAACCAATAAGCATCAGAACATAAATAGTCTTGGCTAAAAGGTTGATAGTACAACACGT

AACAGAAAATCTGTAGATGTCTGTAAATAGTTAGTTAAAGCCCTGAACATGCCTCTTGGCAATAGCTACA

AGTGTAGTTTGATTTTTAACTTAAAATAGAATAAGAAAATAGCTTTTGGTTTCTATTTTTGCACATTTTC

GGAGCTCTGAGATTCATACATAATAGTTTTCCTCGAGTTGTTTAACCATATGGTTATATTGAATAGGGTT

CCTATCTGTTTTACCTTTTGCATCCGGTTTTGATGAGATTGTACAATTATATACTTGCTTATATCAGGCA

AGTCATGGGCTACCAGACTATGTACTTTTTGTAATTTGAATCTTATTTCTGTAACTTTGGCTTCATTTTA

ACTTATTGCCTCTATTTCTTGGTTTGTGTGGATGCAATATCTCTTA

>MSTRG.1562.3 gene=MSTRG.1562

AAAATATTAAAACTACACGTTCATAAAACTCTCTCTCTCGCTCTCTCCTTTCTCCCTCTCACTTAGGTTC

CTCTCTGCGTCTCTCAACCGATATGTGGGTATGATTTACGACTTGGACTTGACTCCGTCTTCTTCATTTA

GTTATACCCATTGGTATTTCTTCAGTTGACCGGTTCCGAGAGAGATTCTGTTTATACAAGTTACCGAACC

ACCATATTGAGGAAACACTCGAATAAACATAGGAAAGCTTAAAGGTGTTTAATCAGGAAGGAAAAAAACC

AGTAGAAAGATTGAGTTGGTGGAAAGAATTTGGAGAAAGGCATTTGATGAGTCAATTCAGTGCCTCCTCC

AGTGGCAAAACAAGTGTGAACTTGAACTTCAAACACGAGGAATGACGGGGAAGGCTATTTGCACCCTTCT

AGTAATTTTCAGGGGCTTCGACTGAATTGGGGCTTCTCTAGAACTAGAGCTTCAACAGCTTTTAACTTTA

AGTGGGGTTTCGGTTTGCTTGGATTTAAGTTCTATAGACATTCAAGCCGGGGATCGGAACTCGGAAGCAT

TTTCAGACCCATTGGTCAAAAGGGGAACCAGTGATTGTTACTATTGTGCTTGACAGTACATATGGTCTAA

GCTGGGAACCAATGGTGATGTCTCTAGCAATCCATGATAAGTCCCACACTGATGTGATTTGTCTTAAATT

GTTTGAGCTGGTGCGAGGAGGAATATAGTGTATGTTCGTTCTTTAAAGGCTATACAGAGGGTATACACTT

GTCTGCCTGGCCCCAAATGCTCAAGTTAAATGACTGGCCTCAATCTGGTTCGTTTGATGATCATCTCACG

TCATAATGTGGAGTTCTTAAGTGCTTTGCCGTTCAAGGAGTACACACATCCACAGAGCGCCTACTTGAAC

CTTGCTGTCAAGTTACCCGAGGGCAATCTGAAGCCAGACACAGGGCCAAAGCTGTATACGCTTAGATAAT

GCACAAACACGCCTAACCAATAAGCATCAGAACATAAATAGTCTTGGCTAAAAGGTTGATAGTACAACAC

GTAACAGAAAATCTGTAGATGTCTGTAAATAGTTAGTTAAAGCCCTGAACATGCCTCTTGGCAATAGCTA

CAAGTGTAGTTTGATTTTTAACTTAAAATAGAATAAGAAAATAGCTTTTGGTTTCTATTTTTGCACATTT

TCGGAGCTCTGAGATTCATACATAATAGTTTTCCTCGAGTTGTTTAACCATATGGTTATATTGAATAGGG

TTCCTATCTGTTTTACCTTTTGCATCCGGTTTTGATGAGATTGTACAATTATATACTTGCTTATATCAGG

CAAGTCATGGGCTACCAGACTATGTACTTTTTGTAATTTGAATCTTATTTCTGTAACTTTGGCTTCATTT

TAACTTATTGCCTCTATTTCTTGGTTTGTGTGGAT

>MSTRG.1562.4 gene=MSTRG.1562

TATAGTAAAAGCTGTGATTGGATTCTATTGGTTTTATATTATAACATTCATCTCAATTGCATTGTTGTTA

ATTTTTTGTGTCGATTTATGATATCGTAAAGGCTTGATTACTTTTCCGGAAAAACCAAAGCTATGTGTAT

TTCTGATATATTCTTGACAAAGAAGATATGGAGTAGATGAATATTTTATAGAAGTATGTTTAACTTGATC

AGATTAATGTGCTTGATGTATAATTATTTCGTATTTAATATAGTAATTTGTTTTATCAAGAGGAAACACT

CGAATAAACATAGGAAAGCTTAAAGGTGTTTAATCAGGAAGGAAAAAAACCAGTAGAAAGATTGAGTTGG

TGGAAAGAATTTGGAGAAAGGCATTTGATGAGTCAATTCAGTGCCTCCTCCAGTGGCAAAACAAGTGTGA

ACTTGAACTTCAAACACGAGGAATGACGGGGAAGGCTATTTGCACCCTTCTAGTAATTTTCAGCGAGCTG

TGATTTAGATGCTCCGACGACCATCAACTTTTCCGGCGACGGAAAAGAAGTGTAGAGATTTTCATGGACG

AATCCGTCCCGTATCCCGTGTCGTGTCGTGTCAACACGGGTACGGTCGCAGGAGTAGGCGAGTCGGAGTA

ACACAGTGTATATCAATCTTCAATATCATAGGTGGATTACTTTCACAAACAATGAAGAAAATGCTTCAAA

TTTACAATACCTCTCGAGTCATCAGACTAATTTGTCCTGTAGGGGCTTCGACTGAATTGGGGCTTCTCTA

GAACTAGAGCTTCAACAGCTTTTAACTTTAAGTGGGGTTTCGGTTTGCTTGGATTTAAGTTCTATAGACA

TTCAAGCCGGGGATCGGAACTCGGAAGCATTTTCAGACCCATTGGTCAAAAGGGGAACCAGTGATTGTTA

CTATTGTGCTTGACAGTACATATGGTCTAAGCTGGGAACCAATGGTGATGTCTCTAGCAATCCATGATAA

GTCCCACACTGATGTGATTTGTCTTAAATTGTTTGAGCTGGTGCGAGGAGGAATATAGTGTATGTTCGTT

CTTTAAAGGCTATACAGAGGGTATACACTTGTCTGCCTGGCCCCAAATGCTCAAGTTAAATGACTGGCCT

CAATCTGGTTCGTTTGATGATCATCTCACGTCATAATGTGGAGTTCTTAAGTGCTTTGCCGTTCAAGGAG

TACACACATCCACAGAGCGCCTACTTGAACCTTGCTGTCAAGTTACCCGAGGGCAATCTGAAGCCAGACA

CAGGGCCAAAGCTGTATACGCTTAGATAATGCACAAACACGCCTAACCAATAAGCATCAGAACATAAATA

GTCTTGGCTAAAAGGTTGATAGTACAACACGTAACAGAAAATCTGTAGATGTCTGTAAATAGTTAGTTAA

AGCCCTGAACATGCCTCTTGGCAATAGCTACAAGTGTAGTTTGATTTTTAACTTAAAATAGAATAAGAAA

ATAGCTTTTGGTTTCTATTTTTGCACATTTTCGGAGCTCTGAGATTCATACATAATAGTTTTCCTCGAGT

TGTTTAACCATATGGTTATATTGAATAGGGTTCCTATCTGTTTTACCTTTTGCATCCGGTTTTGATGAGA

TTGTACAATTATATACTTGCTTATATCAGGCAAGTCATGGGCTACCAGACTATGTACTTTTTGTAATTTG

AATCTTATTTCTGTAACTTTGGC

>MSTRG.1562.5 gene=MSTRG.1562

CGATGATGACGATTCATGTCTACTGCAATGCCATATGAACATCGAGTGTTGCAAAATTATGTTGTATTAG

AGGCACTAATCCATACCTACTGCAACGATAACCATACATGCCCTACTACTAAACATTGGATAGAGACAAG

TTTCCCCCATTTTTCTTCTTTCACTCCACACTTGGGGACACACCGACATTTCACTCCACAAACTCGCCTC

ACTCCACACACTCACCTCACTCTCTAAAGCTCACCTCTCTGCTCAAACTCACTAGCGATCAGGTAAAATC

GGAGAACCTCTCCCCTCAAACTTACCTCTCAATTGATTAGGTTTGGTTTGAGGTCTTTAATTGGGGGTTT

GATTGGCGGCTTCTTCATTCCGCGGAACTAGGGCTTCCGGTTTATACTTCTTCATTTTTTTTTCACTATA

ATGTTTTTAGTTGTGTCTTTATTTTTCTCAGGGGCTTCGACTGAATTGGGGCTTCTCTAGAACTAGAGCT

TCAACAGCTTTTAACTTTAAGTGGGGTTTCGGTTTGCTTGGATTTAAGTTCTATAGACATTCAAGCCGGG

GATCGGAACTCGGAAGCATTTTCAGACCCATTGGTCAAAAGGGGAACCAGTGATTGTTACTATTGTGCTT

GACAGTACATATGGTCTAAGCTGGGAACCAATGGTGATGTCTCTAGCAATCCATGATAAGTCCCACACTG

ATGTGATTTGTCTTAAATTGTTTGAGCTGGTGCGAGGAGGAATATAGTGTATGTTCGTTCTTTAAAGGCT

ATACAGAGGGTATACACTTGTCTGCCTGGCCCCAAATGCTCAAGTTAAATGACTGGCCTCAATCTGGTTC

GTTTGATGATCATCTCACGTCATAATGTGGAGTTCTTAAGTGCTTTGCCGTTCAAGGAGTACACACATCC

ACAGAGCGCCTACTTGAACCTTGCTGTCAAGTTACCCGAGGGCAATCTGAAGCCAGACACAGGGCCAAAG

CTGTATACGCTTAGATAATGCACAAACACGCCTAACCAATAAGCATCAGAACATAAATAGTCTTGGCTAA

AAGGTTGATAGTACAACACGTAACAGAAAATCTGTAGATGTCTGTAAATAGTTAGTTAAAGCCCTGAACA

TGCCTCTTGGCAATAGCTACAAGTGTAGTTTGATTTTTAACTTAAAATAGAATAAGAAAATAGCTTTTGG

TTTCTATTTTTGCACATTTTCGGAGCTCTGAGATTCATACATAATAGTTTTCCTCGAGTTGTTTAACCAT

ATGGTTATATTGAATAGGGTTCCTATCTGTTTTACCTTTTGCATCCGGTTTTGATGAGATTGTACAATTA

TATACTTGCTTATATCAGGCAAGTCATGGGCTACCAGACTATGTACTTTTTGTAATTTGAATCTTATTTC

TGTAA

>MSTRG.1563.1 gene=MSTRG.1563

CTTTTGTACCTCATATAAGCTTTTGTCCGGGACTCATACATACAGATCTATATATGTTCACATCATATTG

CCATATCTTTCTTCTCTATTTCAGAAACAAACGAAAGTGCAGCGGCAGTACTGTCTGCTCTACTTCCCCA

TTTGATTTGTTGTGATTTCAAATTCAAAATTCAAATTTATATCCTCTGATTAGATTCAAGCTCATTTAAA

TCTCATTTCAAGCTCTCTAACCTCGCCTCTCCTGCTCCAAGCGATTTTACCAATATCCATGAAGTGTTTG

ATAAAATGTGAAGATTTGGGGGACTTTCTGTCTTCATGTTTGGTGTGAGTGGAATTGTGGAGGAAGGAAA

AGACGAGGAGGGAACGGAAGGCAAAAAGATGAGGTAGAATTAAGCTGCTGTAGCAGTGGAACTCTTGATA

TATTGTGTCCCTCTATCCTCTGGGAATGGCAACAGCATTTAAAGTGAAGAACAATGCTCGCAAAGCTATT

AAACTAACCGGTGCAATACCAAGTCATTTCAAATTCAAGAAGCGAAGTGGTACTGGGATTTAGGGCCTGA

GAAACTGCTCTTACTGCTCTTAATTGTAATTACATAAGTTATTAGGAAGTTCAATAAAATGAATGCAGGA

ACCATTTGATTAAAAAATTTAAACACGCCGTTATCTTATTATTTGGG

>MSTRG.1563.2 gene=MSTRG.1563

CTTTTGTACCTCATATAAGCTTTTGTCCGGGACTCATACATACAGATCTATATATGTTCACATCATATTG

CCATATCTTTCTTCTCTATTTCAGAAACAAACGAAAGTGCAGCGGCAGTACTGTCTGCTCTACTTCCCCA

TTTGATTTGTTGTGATTTCAAATTCAAAATTCAAATTTATATCCTCTGATTAGATTCAAGCTCATTTAAA

TCTCATTTCAAGCTCTCTAACCTCGCCTCTCCTGCTCCAAGCGATTTTACCAATATCCATGAAGTGTTTG

ATAAAATGTGAAGATTTGGGGGACTTTCTGTCTTCATGTTTGGTGTGAGTGGAATTGTGGAGGAAGGAAA

AGACGAGGAGGGAACGGAAGGCAAAAAGATGAGTGCAGGTAGAATTAAGCTGCTGTAGCAGTGGAACTCT

TGATATATTGTGTCCCTCTATCCTCTGGGAATGGCAACAGCATTTAAAGTGAAGAACAATGCTCGCAAAG

CTATTAAACTAACCGGTGCAATACCAAGTCATTTCAAATTCAAGAAGCGAAGTGGTACTGGGATTTAGGG

CCTGAGAAACTGCTCTTACTGCTCTTAATTGTAATTACATAAGTTATTAGGAAGTTCAATAAAATGAATG

CAGGAACCATTTGATTAAAAAATTTAAACACGCCGTTATCTTATTATTTGGG

>MSTRG.1563.3 gene=MSTRG.1563

CTTTTGTACCTCATATAAGCTTTTGTCCGGGACTCATACATACAGATCTATATATGTTCACATCATATTG

CCATATCTTTCTTCTCTATTTCAGAAACAAACGAAAGTGCAGCGGCAGTACTGTCTGCTCTACTTCCCCA

TTTGATTTGTTGTGATTTCAAATTCAAAATTCAAATTTATATCCTCTGATTAGATTCAAGCTCATTTAAA

TCTCATTTCAAGCTCTCTAACCTCGCCTCTCCTGCTCCAAGCGATTTTACCAATATCCATGAAGTGTTTG

ATAAAATGTGAAGATTTGGGGGACTTTCTGTCTTCATGTTTGGTGTGAGTGGAATTGTGGAGGAAGGAAA

AGACGAGGAGGGAACGGAAGGCAAAAAGATGAGGTATTTAATTTGCAGAAAATGGGGTATTTGAGACTTG

CAAGCTGCGGATTACTGTCAAATATGATACTGTATTTTTATAATGGATTACAGGATGAGCTTGACAAAAG

GCCTTGGGTTCATCATTGCTTCAAGTGTCTGTCATTGCATAATTACTGTGTAGTTATTTCTTTGCAGGTA

GAATTAAGCTGCTGTAGCAGTGGAACTCTTGATATATTGTGTCCCTCTATCCTCTGGGAATGGCAACAGC

ATTTAAAGTGAAGAACAATGCTCGCAAAGCTATTAAACTAACCGGTGCAATACCAAGTCATTTCAAATTC

AAGAAGCGAAGTGGTACTGGGATTTAGGGCCTGAGAAACTGCTCTTACTGCTCTTAATTGTAATTACATA

AGTTATTAGGAAGTTCAATAAAATGAATGCAGGAACCATTTGATTAAAAAATTTAAACACGCCGTTATCT

TATTATTTGGG

>MSTRG.1563.4 gene=MSTRG.1563

GCCATATCTTTCTTCTCTATTTCAGAAACAAACGAAAGTGCAGCGGCAGTACTGTCTGCTCTACTTCCCC

ATTTGATTTGTTGTGATTTCAAATTCAAAATTCAAATTTATATCCTCTGATTAGATTCAAGCTCATTTAA

ATCTCATTTCAAGCTCTCTAACCTCGCCTCTCCTGCTCCAATGCAGGTAGAATTAAGCTGCTGTAGCAGT

GGAACTCTTGATATATTGTGTCCCTCTATCCTCTGGGAATGGCAACAGCATTTAAAGTGAAGAACAATGC

TCGCAAAGCTATTAAACTAACCGGTGCAATACCAAGTCATTTCAAATTCAAGAAGCGAAGTGGTACTGGG

ATTTAGGGCCTGAGAAACTGCTCTTACTGCTCTTAATTGTAATTACATAAGTTATTAGGAAGTTCAATAA

AATGAATGCAGGAACCATTTGATTAAAAAATTTAAACACGCCGTTATCTTATTATTTG

>MSTRG.1563.5 gene=MSTRG.1563

AACGAAAGTGCAGCGGCAGTACTGTCTGCTCTACTTCCCCATTTGATTTGTTGTGATTTCAAATTCAAAA

TTCAAATTTATATCCTCTGATTAGATTCAAGCTCATTTAAATCTCATTTCAAGCTCTCTAACCTCGCCTC

TCCTGCTCCAAGTAGAATTAAGCTGCTGTAGCAGTGGAACTCTTGATATATTGTGTCCCTCTATCCTCTG

GGAATGGCAACAGCATTTAAAGTGAAGAACAATGCTCGCAAAGCTATTAAACTAACCGGTGCAATACCAA

GTCATTTCAAATTCAAGAAGCGAAGTGGTACTGGGATTTAGGGCCTGAGAAACTGCTCTTACTGCTCTTA

ATTGTAATTACATAAGTTATTAGGAAGTTCAATAAAATGAATGCAGGAACCATTTGATTAAAAAATTTAA

ACACGCCGTTATCTTATTATTTGG

>MSTRG.1564.1 gene=MSTRG.1564

ACAGTTGATCAAAATATTAACCACACACGCTCAGCCCCATTTAGAAAACTCTCGCAGTCTCTCCTTTTCT

CCCTCCCTCACTTAGGGTTTCTCTTTCTGTCTCACAACCCATATGTGGGTATAATTGACCACCGTAAATT

AGGGTTATTGGGTTTGAAACTTTGAATCAGGGGTTCTTTTTTGTTTCATTTTTTCAACTCAAGAGTTCAG

TTGCTTTGATTAATTTGGTTTCTCTCTTTGTTTCAGGGTTATGAGAGAGTTTCTTATAGTGTGTTAAAAA

CTCTAAGAAGAGTTAGCTTATTGATTCTGTTTGTGACCGTGTGCACAGGCTTCAGTGCTCTGCAACTGTT

TATACATGTTGCAGAACCACCATATTGTGGAAACACTCGGATAAGCATAGGAAAGCTTAAAGGTGTTTAG

TCAGGAAGAAAAAATCAGTAGAAAGATTGAGTTGGTGGAAATAAATTGGAGAAAGGCATTTGATGAGTCA

ATTCAGTATCTCCTCCAGTGGCAAAACAAGTGAGAACTTCTTGAACTTCAAACACGAGGAATGACGGGGA

AGGCTATTTGCACCCGTCTAGTAATTTTCAGGGCTTGGATTGAATTGGGGCTTCACTGGAACTAGAGCTT

CAACAGCTCTTAACTTTAAGAGGCGCTTCGGTTTTCTTTGATTTAAGCAACTATTACGGAACTTCTATTG

CTGACTACCACCGGAGCTGCACAAAATGTCCGTATGATCTATGCATCGCTTGTTGTCGGGACTTCCGTGC

TGGCTCTTTGCATGCAGGTCATATGAGAAGCATTAACCAATATAAAAACCTCAGGGTGGGCTATTTACAT

GGTGAAACATGTAAACTTTGTAGCAATAGAACTGTGGGCACATCTACTGAAACATCCCCTGAAGGCAAGA

CCTGAATGGAGACCTAACAAGGATGGTAGCATCTCTTGTCCACCTAAGAGTATTGGTGGGTGTGGTGGTA

AAGGGATTCTAAAGCTGAACCAAGTCTTGCGACTTTCAAACAAGATATTGAAAGCTGAAGACTTATATGA

ACTGTATAATCTGGATGGCATGCCAGAAGATCCCATACACTGGTGCTCGTGTTATAATTTGGCACGTGAT

AACACGGCCACAAAGAAAATAAGGAAAGCCACTTCTCGAGAAAACTCTGATGACAACACTTCTACTCTCC

AAGTTCTATAGACATTCAAGCCGGGGATCGGAAGCATTTTCAGACCCATTGGTCAAAAGGGGAACCAGTG

ATTGTTTCTAGTGTGCTTGACAGTACATATGGTTAAGCTGGGAACCAATGGTGATGTCTCGAGCAATCCA

TGATAAGTCCCACACTGATGTGACTGTCTTAAATTGTTTGAGCTGGTGCGAAGAGGAATATAGCGTAAGT

TCGTTCTTTAAAGGCTATACAGAGGGTATACTTGTCTGGCTGGCCCCAAATGCTCAAGTTAAATGATTGG

CCTCAATCTGGTTTTTGAGGATCATCTCCCATGTCATAATGTGGAGTTCTTAAGCGCTTTGCCTTTGAAG

GAGTACACACATCCACAGAGCGGCTACTTGAACCTTGCTGTCAAGTTACCCGAGGGCCATCTGAAGCGAG

ACAAAGGGCCAAAGCTGTATACTGCTTATGGCTTTTCCCAGCAACTCGGACGTGGTGACTCTGTCTCAAA

ACTCAGCTATCATGATTCAGATGCGGTATGATCTAGTTTGTAAATACAGTGAATAACTTTCACAAACAAT

GAAGAAAATGCTCCAAATTTACAACCCCACCCGAGTCATCATACTAATAGTCCATCCTGAGATCATAATC

ATTTTTGGTAATTTTATTCGGCAAGGTTTCTC

>MSTRG.1564.2 gene=MSTRG.1564

ATTAACCACACACGCTCAGCCCCATTTAGAAAACTCTCGCAGTCTCTCCTTTTCTCCCTCCCTCACTTAG

GGTTTCTCTTTCTGTCTCACAACCCATATGTGGGTATAATTGACCACCGTAAATTAGGGTTATTGGGTTT

GAAACTTTGAATCAGGGGTTCTTTTTTGTTTCATTTTTTCAACTCAAGAGTTCAGTTGCTTTGATTAATT

TGGTTTCTCTCTTTGTTTCAGGGTTATGAGAGAGTTTCTTATAGTGTGTTAAAAACTCTAAGAAGAGTTA

GCTTATTGATTCTGTTTGTGACCGTGTGCACAGGCTTCAGTGCTCTGCAACTGTTTATACATGTTGCAGA

ACCACCATATTGTGGAAACACTCGGATAAGCATAGGAAAGCTTAAAGGTGTTTAGTCAGGAAGAAAAAAT

CAGTAGAAAGATTGAGTTGGTGGAAATAAATTGGAGAAAGGCATTTGATGAGTCAATTCAGTATCTCCTC

CAGTGGCAAAACAAGTGAGAACTTCTTGAACTTCAAACACGAGGAATGACGGGGAAGGCTATTTGCACCC

GTCTAGTAATTTTCAGGGCTTGGATTGAATTGGGGCTTCACTGGAACTAGAGCTTCAACAGCTCTTAACT

TTAAGAGGCGCTTCGGTTTTCTTTGATTTAAGCAACTATTACGGAACTTCTATTGCTGACTACCACCGGA

GCTGCACAAAATGTCCGTATGATCTATGCATCGCTTGTTGTCGGGACTTCCGTGCTGGCTCTTTGCATGC

AGGTCATATGAGAAGCATTAACCAATATAAAAACCTCAGGGTGGGCTATTTACATGGTGAAACATGTAAA

CTTTGTAGCAATAGAACTGTGGGCACATCTACTGAAACATCCCCTGAAGGCAAGACCTGAATGGAGACCT

AACAAGGATGGTAGCATCTCTTGTCCACCTAAGAGTATTGGTGGGTGTGGTGGTAAAGGGATTCTAAAGC

TGAACCAAGTCTTGCCAGACAGTTGACTTTCAAACAAGATATTGAAAGCTGAAGACTTGTATGAACTGTA

TAATCTGGATGGCATGCCAGAAGATCCCATACACTGGTGCTCGTGTTATAATTTGGCACGTGATAACACG

GCCACAAAGAAAATAAGGAAAGCCACTTCTCGAGAAAACTCTGATGACAACACTTCTACTCTCCAAGTTC

TATAGACATTCAAGCTGGGGATCAGAAGCATTTTCAGACCCATTGGTCAAAAGGGGAACCAGTGATTGTT

ACTAGTGTGCTTGACAGTACATATGGTTAAGCTGCGAACCAATGGTGATGTCTCGAGCAATCCATGATAA

GTCCCACACCGATGTGACTTTCTTAAATTGTTTGAGCTGGTGCGAAGAGGAATATAGCCCAAGTTCGTTC

TTTAAAGGCTATACAGAGGGTATACTTGACTTGTCTGGCTGGCCCCAAATGCTCAAGTTAAATGATTGGC

CTCAATCTGGTTCGTGTGAGGATCATCTCCCACGTCATAATGTGGAGTTCTTAAGTGCTTTGCTGTTCAA

GGAGTACACACATCCACAGAGCGCCTACTTGAACCTTGCTGTCAAGTTACCCGAGGCCAATCTGAAGCCA

GACACAGGGCCAAAGCTGTATACGCTTAGATAATGCACAAACCACGCCTCTTGGCAGTAGTATGTGTAGT

TTGATTTTTAACTTACAATAGAATTAAAAAATAGCTTTTGATTTCTATATTTGCACATTTTTGGAGCTGA

AAGATTCATACATACTCCCTCTGTCCCTCTCATTTCTTTACACTTTTTTTACATTGTTCGATACGCATTT

TAAGGCGTATATAAATATAGTTTCATAACTTTTTTCCAAAATTTTTCTTTTTTGTATAAAAATTTAGATA

GTAAACATTTTTTCAGAAGAAAAGAATTCAAAATAATTTGTCAAGCTATATTTTATAAGAGCATTGAATG

CGTGCCGAGCCCCCGTCCCCCAATGTAAACAATTGAGGGCGATGGAGGGAGTAATAGCTTTCCTCAGTTG

TTTAAACATATACTTGCTTCTATCAGGCAATGGCTACCAGACTCTGTACTTTTTGTAATTGGAATCTTGT

TTCTGTGAGTTTGGCTTGATTTTAACTTATGTTATGGCCTCTATTTCTTGGTTTGTTTGGACTTTGG

>MSTRG.1564.3 gene=MSTRG.1564

GAAAACTCTCGCAGTCTCTCCTTTTCTCCCTCCCTCACTTAGGGTTTCTCTTTCTGTCTCACAACCCATA

TGTGGGTATAATTGACCACCGTAAATTAGGGTTATTGGGTTTGAAACTTTGAATCAGGGGTTCTTTTTTG

TTTCATTTTTTCAACTCAAGAGTTCAGTTGCTTTGATTAATTTGGTTTCTCTCTTTGTTTCAGGGTTATG

AGAGAGTTTCTTATAGTGTGTTAAAAACTCTAAGAAGAGTTAGCTTATTGATTCTGTTTGTGACCGTGTG

CACAGGCTTCAGTGCTCTGCAACTGTTTATACATGTTGCAGAACCACCATATTGTGGAAACACTCGGATA

AGCATAGGAAAGCTTAAAGGTGTTTAGTCAGGAAGAAAAAATCAGTAGAAAGATTGAGTTGGTGGAAATA

AATTGGAGAAAGGCATTTGATGAGTCAATTCAGTATCTCCTCCAGTGGCAAAACAAGTGAGAACTTCTTG

AACTTCAAACACGAGGAATGACGGGGAAGGCTATTTGCACCCGTCTAGTAATTTTCAGGGCTTGGATTGA

ATTGGGGCTTTACTGGAACTAGAGCTTCAACAGCTCTTAACTTTAAGAGGCGCTTCGGTTTTCTTTGATC

TAAGCAACTATTACGGAACTTCTATTGCTGACTACCACCGGAGCTGCACAAAATGTCCGTATGATCTATG

CATCGCTTGTTGTCGGGACTTGCGTGCTGGCTCTTTGCATGCAGGTCATCTGGGAAGCATTAACCAATAT

AAAAACCTCAGGGTGGGCTATTTACATGGTGAAACATGTAAACCTCGTAGCAATAGAACTGTGGGCACAT

CTACTGAAACATCCCCTGAAGGCAAGACCTGAATGGAGACCTAACAAGGATGGTAGCATCTCTTGTCCAC

CTAAGAGTATTGGTGGGTGTGGTGGTAAAGGGATTCTAAAGCTGAACCAAGTCTTGCCAGACGGTTGACT

TTCAAACAAGATATTGAAAGCTGAAGACTTATATGAACTGTATAATCTGGATGGCATGCCAGAAGATCCC

ATACACTGGTGCTCGTGTTATAATTTGGCACGTGATAACACGGCCACAAAGAAAATAAGGAAAGCCACTT

CTCGAGAAAACTCTGATGACAACACTTCTACTCTCCAAGTTCTATAGACATTCAAGCCGGGGATCGGAAG

CATTTTCAGACCCATTGGTCAAAAGGGGAACCAGTGATTGTTTCTAGTGTGCTTGACAGTACATATGGTT

AAGCTGGGAACCAATGGTGATGTCTCGAGCAATCCATGATAAGTCCCACACTGATGTGACTGTCTTAAAT

TGTTTGAGCTGGTGCGAAGAGGAATATAGCGTAAGTTCGTTCTTTAAAGGCTATACAGAGGGTATACTTG

TCTGGCTGGCCCCAAATGCTCAAGTTAAATGATTGGCCTCAATCTGGTTTTTGAGGATCATCTCCCATGT

CATAATGTGGAGTTCTTAAGCGCTTTGCCTTTGAAGGAGTACACACATCCACAGAGCGGCTACTTGAACC

TTGCTGTCAAGTTACCCGAGGGCCATCTGAAGCGAGACAAAGGGCCAAAGCTGTATACTGCTTATGGCTT

TTCCCAGCAACTCGGACGTGGTGACTCTGTCTCAAAACTCAGCTATCATGATTCAGATGCGGTATGATCT

AGTTTGTAAATACAGTGAATAACTTTCACAAACAATGAAGAAAATGCTCCAAATTTACAACCCCACCCGA

GTCATCATACTAATAGTCCATCCTGAGATCATAATCATTTTTGGTAATTTTATTCGGC

>MSTRG.1564.4 gene=MSTRG.1564

GAAAACTCTCGCAGTCTCTCCTTTTCTCCCTCCCTCACTTAGGGTTTCTCTTTCTGTCTCACAACCCATA

TGTGGGTATAATTGACCACCGTAAATTAGGGTTATTGGGTTTGAAACTTTGAATCAGGGGTTCTTTTTTG

TTTCATTTTTTCAACTCAAGAGTTCAGTTGCTTTGATTAATTTGGTTTCTCTCTTTGTTTCAGGGTTATG

AGAGAGTTTCTTATAGTGTGTTAAAAACTCTAAGAAGAGTTAGCTTATTGATTCTGTTTGTGACCGTGTG

CACAGGCTTCAGTGCTCTGCAACTGTTTATACATGTTGCAGAACCACCATATTGTGGAAACACTCGGATA

AGCATAGGAAAGCTTAAAGGTGTTTAGTCAGGAAGAAAAAATCAGTAGAAAGATTGAGTTGGTGGAAATA

AATTGGAGAAAGGCATTTGATGAGTCAATTCAGTATCTCCTCCAGTGGCAAAACAAGTGAGAACTTCTTG

AACTTCAAACACGAGGAATGACGGGGAAGGCTATTTGCACCCGTCTAGTAATTTTCAGGGCTTGGATTGA

ATTGGGGCTTCACTGGAACTAGAGCTTCAACAGCTCTTAACTTTAAGAGGCGCTTCGGTTTTCTTTGATT

TAAGCAACTATTACGGAACTTCTATTGCTGACTACCACCGGAGCTGCACAAAATGTCCGTATGATCTATG

CATCGCTTGTTGTCGGGACTTGCGTGCTGGCTCTTTGCATGCAGGTCATCTGGGAAGCATTAACCAATAT

AAAAACCTCAGGGTGGGCTATTTACATGGTGAAACATGTAAACCTCGTAGCAATAGAACTGTGGGCACAT

CTACTGAAACATCCCCTGAAGGCAAGACCTGAATGGAGACCTAACAAGGATGGTAGCATCTCTTGTCCAC

CTAAGAGTATTGGTGGGTGTGGTGGTAAAGGGATTCTAAAGCTGAACCAAGTCTTGCCAGACGGTTGACT

TTCAAACAAGATATTGAAAGCTGAAGACTTATATGAACTGTATAATCTGGATGGCATGCCAGAAGATCCC

ATACACTGGTGCTCGTGTTATAATTTGGCACGTGATAACACGGCCACAAAGAAAATAAGGAAAGCCACTT

CTCGAGAAAACTCTGATGACAACACTTCTACTCTCCAAGTTCTATAGACATTCAAGCCGGGGATCGGAAG

CATTTTCAGACCCATTGGTCAAAAGGGGAACCAGTGATTGTTTCTAGTGTGCTTGACAGTACATATGGTT

AAGCTGGGAACCAATGGTGATGTCTCGAGCAATCCATGATAAGTCCCACACTGATGTGACTGTCTTAAAT

TGTTTGAGCTGGTGCGAAGAGGAATATAGCGTAAGTTCGTTCTTTAAAGGCTATACAGAGGGTATACTTG

TCTGGCTGGCCCCAAATGCTCAAGTTAAATGATTGGCCTCAATCTGGTTTTTGAGGATCATCTCCCATGT

CATAATGTGGAGTTCTTAAGCGCTTTGCCTTTGAAGGAGTACACACATCCACAGAGCGGCTACTTGAACC

TTGCTGTCAAGTTACCCGAGGGCCATCTGAAGCGAGACAAAGGGCCAAAGCTGTATACTGCTTATGGCTT

TTCCCAGCAACTCGGACGTGGTGACTCTGTCTCAAAACTCAGCTATCATGATTCAGATGCGGTATGATCT

AGTTTGTAAATACAGTGAATAACTTTCACAAACAATGAAGAAAATGCTCCAAATTTACAACCCCACCCGA

GTCATCATACTAATAGTCCATCCTGAGATCATAATCATTTTTGGTAATTTTATTCGGCAAGGTTTCTCA

>MSTRG.1564.5 gene=MSTRG.1564

AAGAGTTAGCTTATTGATTCTGTTTGTGACCGTGTGCACAGGCTTCAGTGCTCTGCAACTGTTTATACAT

GTTGCAGAACCACCATATTGTGATTTGTTTTATCAAGTGGAAACACTCGGATAAGCATAGGAAAGCTTAA

AGGTGTTTAGTCAGGAAGAAAAAATCAGTAGAAAGATTGAGTTGGTGGAAATAAATTGGAGAAAGGCATT

TGATGAGTCAATTCAGTATCTCCTCCAGTGGCAAAACAAGTGAGAACTTCTTGAACTTCAAACACGAGGA

ATGACGGGGAAGGCTATTTGCACCCGTCTAGTAATTTTCAGGGCTTGGATTGAATTGGGGCTTCACTGGA

ACTAGAGCTTCAACAGCTCTTAACTTTAAGAGGCGCTTCGGTTTTCTTTGATTTAAGCAACTATTACGGA

ACTTCTATTGCTGACTACCACCGGAGCTGCACAAAATGTCCGTATGATCTATGCATCGCTTGTTGTCGGG

ACTTCCGTGCTGGCTCTTTGCATGCAGGTCATATGAGAAGCATTAACCAATATAAAAACCTCAGGGTGGG

CTATTTACATGGTGAAACATGTAAACTTTGTAGCAATAGAACTGTGGGCACATCTACTGAAACATCCCCT

GAAGGCAAGACCTGAATGGAGACCTAACAAGGATGGTAGCATCTCTTGTCCACCTAAGAGTATTGGTGGG

TGTGGTGGTAAAGGGATTCTAAAGCTGAACCAAGTCTTGCGACTTTCAAACAAGATATTGAAAGCTGAAG

ACTTATATGAACTGTATAATCTGGATGGCATGCCAGAAGATCCCATACACTGGTGCTCGTGTTATAATTT

GGCACGTGATAACACGGCCACAAAGAAAATAAGGAAAGCCACTTCTCGAGAAAACTCTGATGACAACACT

TCTACTCTCCAAGTTCTATAGACATTCAAGCCGGGGATCGGAAGCATTTTCAGACCCATTGGTCAAAAGG

GGAACCAGTGATTGTTTCTAGTGTGCTTGACAGTACATATGGTTAAGCTGGGAACCAATGGTGATGTCTC

GAGCAATCCATGATAAGTCCCACACTGATGTGACTGTCTTAAATTGTTTGAGCTGGTGCGAAG

>MSTRG.1564.6 gene=MSTRG.1564

GTAACATTCTCGAAATTTTATAATTATATAGATAATTTCATTTTTGTCCAAAAATCGCGCGAGTAATCGC

GCTTCGCGAGTTGCGCTTCAAAATTGGATCTTGCGCTTTTTTTATTTTGCGTGTTTCACAACACTGACCA

TATTGTGGAAACACTCGGATAAGCATAGGAAAGCTTAAAGGTGTTTAGTCAGGAAGAAAAAATCAGTAGA

AAGATTGAGTTGGTGGAAATAAATTGGAGAAAGGCATTTGATGAGTCAATTCAGTATCTCCTCCAGTGGC

AAAACAAGTGAGAACTTCTTGAACTTCAAACACGAGGAATGACGGGGAAGGCTATTTGCACCCGTCTAGT

AATTTTCAGGGCTTGGATTGAATTGGGGCTTCACTGGAACTAGAGCTTCAACAGCTCTTAACTTTAAGAG

GCGCTTCGGTTTTCTTTGATTTAAGCAACTATTACGGAACTTCTATTGCTGACTACCACCGGAGCTGCAC

AAAATGTCCGTATGATCTATGCATCGCTTGTTGTCGGGACTTCCGTGCTGGCTCTTTGCATGCAGGTCAT

ATGAGAAGCATTAACCAATATAAAAACCTCAGGGTGGGCTATTTACATGGTGAAACATGTAAACTTTGTA

GCAATAGAACTGTGGGCACATCTACTGAAACATCCCCTGAAGGCAAGACCTGAATGGAGACCTAACAAGG

ATGGTAGCATCTCTTGTCCACCTAAGAGTATTGGTGGGTGTGGTGGTAAAGGGATTCTAAAGCTGAACCA

AGTCTTGCCAGACAGTTGACTTTCAAACAAGATATTGAAAGCTGAAGACTTGTATGAACTGTATAATCTG

GATGGCATGCCAGAAGATCCCATACACTGGTGCTCGTGTTATAATTTGGCACGTGATAACACGGCCACAA

AGAAAATAAGGAAAGCCACTTCTCGAGAAAACTCTGATGACAACACTTCTACTCTCCAAGTTCTATAGAC

ATTCAAGCTGGGGATCAGAAGCATTTTCAGACCCATTGGTCAAAAGGGGAACCAGTGATTGTTACTAGTG

TGCTTGACAGTACATATGGTTAAGCTGCGAACCAATGGTGATGTCTCGAGCAATCCATGATAAGTCCCAC

ACCGATGTGACTTTCTTAAATTGTTTGAGCTGGTGCGAAGAGGAATATAGCCCAAGTTCGTTCTTTAAAG

GCTATACAGAGGGTATACTTGACTTGTCTGGCTGGCCCCAAATGCTCAAGTTAAATGATTGGCCTCAATC

TGGTTCGTGTGAGGATCATCTCCCACGTCATAATGTGGAGTTCTTAAGTGCTTTGCTGTTCAAGGAGTAC

ACACATCCACAGAGCGCCTACTTGAACCTTGCTGTCAAGTTACCCGAGGCCAATCTGAAGCCAGACACAG

GGCCAAAGCTGTATACGCTTAGATAATGCACAAACCACGCCTCTTGGCAGTAGTATGTGTAGTTTGATTT

TTAACTTACAATAGAATTAAAAAATAGCTTTTGATTTCTATATTTGCACATTTTTGGAGCTGAAAGATTC

ATACATACTCCCTCTGTCCCT

>MSTRG.1564.7 gene=MSTRG.1564

TCCACCTAACCACGAATGAATGTGGACCACATGATGTCCAAGTACAGTCCTTTCTTATGCCTTATCATCT

TCTTTTCCTTTTTCTCTTTCATCTGCCTCATCATCCTCATATCATCTATCTATTTCATCGTCTTCTTCTG

TGATCATCTTCTTCCATTTTTATTCTTCATCTTCTCTCTTCTTCTTCCCTTCTTATTCTTCTGACACACC

GCCTTTAAATTCTCCGGCGACCTTCATATTTTTCCGGCGCCAGATAAGAAGTGTCGGTTTTGCATATGGT

GAATCCACCCCGTATCCCGCGTCGTGTCGGGTCGACACGAGTACGGCCGCCGGAAGTGCCGAGTCGGAGT

AACACAGTATGTATATCAATCTTCAATATCATAGGTGGATTACTTTCACAAACAATGAAGAAAATGCTTC

AAATTTACAACACTTCCCGAGTCATCAGTCTAATTTGGTTTGGCTGGAAGGGCTTGGATTGAATTGGGGC

TTCACTGGAACTAGAGCTTCAACAGCTCTTAACTTTAAGAGGCGCTTCGGTTTTCTTTGATTTAAGCAAC

TATTACGGAACTTCTATTGCTGACTACCACCGGAGCTGCACAAAATGTCCGTATGATCTATGCATCGCTT

GTTGTCGGGACTTCCGTGCTGGCTCTTTGCATGCAGGTCATATGAGAAGCATTAACCAATATAAAAACCT

CAGGGTGGGCTATTTACATGGTGAAACATGTAAACTTTGTAGCAATAGAACTGTGGGCACATCTACTGAA

ACATCCCCTGAAGGCAAGACCTGAATGGAGACCTAACAAGGATGGTAGCATCTCTTGTCCACCTAAGAGT

ATTGGTGGGTGTGGTGGTAAAGGGATTCTAAAGCTGAACCAAGTCTTGCCAGACAGTTGACTTTCAAACA

AGATATTGAAAGCTGAAGACTTGTATGAACTGTATAATCTGGATGGCATGCCAGAAGATCCCATACACTG

GTGCTCGTGTTATAATTTGGCACGTGATAACACGGCCACAAAGAAAATAAGGAAAGCCACTTCTCGAGAA

AACTCTGATGACAACACTTCTACTCTCCAAGTTCTATAGACATTCAAGCTGGGGATCAGAAGCATTTTCA

GACCCATTGGTCAAAAGGGGAACCAGTGATTGTTACTAGTGTGCTTGACAGTACATATGGTTAAGCTGCG

AACCAATGGTGATGTCTCGAGCAATCCATGATAAGTCCCACACCGATGTGACTTTCTTAAATTGTTTGAG

CTGGTGCGAAGAGGAATATAGCCCAAGTTCGTTCTTTAAAGGCTATACAGAGGGTATACTTGACTTGTCT

GGCTGGCCCCAAATGCTCAAGTTAAATGATTGGCCTCAATCTGGTTCGTGTGAGGATCATCTCCCACGTC

ATAATGTGGAGTTCTTAAGTGCTTTGCTGTTCAAGGAGTACACACATCCACAGAGCGCCTACTTGAACCT

TGCTGTCAAGTTACCCGAGGCCAATCTGAAGCCAGACACAGGGCCAAAGCTGTATACGCTTAGATAATGC

ACAAACCACGCCTCTTGGCAGTAGTATGTGTAGTTTGATTTTTAACTTACAATAGAATTAAAAAATAGCT

TTTGATTTCTATATTTGCACATTTTTGGAGCTGAAAGATTCATACATACTCCCTCTGTCCCTCTCATTTC

TTTACACTTTTTTTACATTGTTCGATACGCATTTTAAGGCGTATATAAATATAGTTTCATAACTTTTTTC

CAAAATTTTTCTTTTTTGTATAAAAATTTAGATAGTAAACATTTTTTCAGAAGAAAAGAATTCAAAATAA

TTTGTCAAGCTATATTTTATAAGAGCATTGAATGCGTGCCGAGCCCCCGTCCCCCAATGTAAACAATTGA

GGGCGATGGAGGGAGTAATAGCTTTCCTCAGTTGTTTAAACATATACTTGCTTCTATCAGGCAATGGCTA

CCAGACTCTGTACTTTTTGTAATTGGAATCTTGTTTCTGTGAGTTTGGCTTGATTTTAACTTATGTTATG

GCCTCTATTTCTTGGTTTGTTTGGACTTTGG

>MSTRG.1565.1 gene=MSTRG.1565

CTTTCTTTAATTTATAGTAGAGACTTGGTTTTATTATAGTTTCTATAGCACACAAAACTTACCTAGCACT

TTTTGGATTTTCATAAATGATTCATATTGGTCCTGATTCCTGAAGCTCCAAGGAATTGTCTGATGGAAGC

GAGGTTCTTGAAATTTATGGCTAGAGCAACTACATGGCTTTGTTTGGCAGCTTTGACTTAATTGTTGATG

GTCGTTCATTCAACAATCTGCTTTAGTACAAGCAGATTGTTTAATATAAGGGAAACGTTAAGATGGTTTT

GGTTGAATATATTAGTGAAATGTTGC

>MSTRG.1564.8 gene=MSTRG.1564

AAGTATGGGTTTGCCTGATTCCTCGTACCACCAAACGACCCCTTGCGGGGAGTATGGGTATAATTGACCA

CCGGTCCACCGTAAATTAGTGTTATTGGATTTTAAACTTTGAATCAGGGGTCCTTTTTGTTCATTTTTTC

AACTCAAGAGTTCAATTGCTTTGATTAGTTTGGTTTCTCTCTTTGTTTCAGGGTTCTGAGAGAGATTCTT

GTAGTGTGTTATAAACTCTCAAGCATGGATTTCGTTTTGCTGCTGATATCTCAATGATGTACATTAAGAC

GGTTTCTTGATTAGCAGAGTTAGCTTATTGATTCTGTTTGTGACCGTGTTCACAGGCTCCAGTGCTCTGC

AACTGTTTATACAAGTTGCAGAACCACCCTCTTGGCTTGATTACTTTTCCGGAAAAACCAAAGCCTATTT

CTGATATATTCTTGACAAAGAAGATATGGAGGAGATGAATATTTTATAGAATATTTTTAACTTGATCAGA

TTTATGGGCTTGATGTATAATGTTTTCGTATTCTATATAGTAATTTGTTTTATCAAGTGGAAACACTCGA

ATAAACATAGGAAAGCTTAAAGGTGTTTAGTCAGGAAGAAAAAATCAGTAGAAAGATTGAGTTGGTTGAA

AGAAATTGGAGAAGGGTATTTGATGAGTCAATTCAGTGTCTCCTCCAGCGGCAAAACAAGTGAGAACTTC

TTGAACTTCTTGAACTTCAAACACGAGGAATGACGGGGAAGGCTATTTGCGCCCGTCTAGTAATTTTCAG

GGCTTGGATTGAATTGGGGCTTTACTGGAACTAGAGCTTCAACAGCTCTTAACTTTAAGAGGCGCTTCGG

TTTTCTTTGATCTAAGCAACTATTACGGAACTTCTATTGCTGACTACCACCGGAGCTGCACAAAATGTCC

GTATGATCTATGCATCGCTTGTTGTCGGGACTTGCGTGCTGGCTCTTTGCATGCAGGTCATCTGGGAAGC

ATTAACCAATATAAAAACCTCAGGGTGGGCTATTTACATGGTGAAACATGTAAACCTCGTAGCAATAGAA

CTGTGGGCACATCTACTGAAACATCCCCTGAAGGCAAGACCTGAATGGAGACCTAACAAGGATGGTAGCA

TCTCTTGTCCACCTAAGAGTATTGGTGGGTGTGGTGGTAAAGGGATTCTAAAGCTGAACCAAGTCTTGCC

AGACGGTTGACTTTCAAACAAGATATTGAAAGCTGAAGACTTATATGAACTGTATAATCTGGATGGCATG

CCAGAAGATCCCATACACTGGTGCTCGTGTTATAATTTGGCACGTGATAACACGGCCACAAAGAAAATAA

GGAAAGCCACTTCTCGAGAAAACTCTGATGACAACACTTCTACTCTCCAAGTTCTATAGACATTCAAGCC

GGGGATCGGAAGCATTTTCAGACCCATTGGTCAAAAGGGGAACCAGTGATTGTTTCTAGTGTGCTTGACA

GTACATATGGTTAAGCTGGGAACCAATGGTGATGTCTCGAGCAATCCATGATAAGTCCCACACTGATGTG

ACTGTCTTAAATTGTTTGAGCTGGTGCGAAGAGGAATATAGCGTAAGTTCGTTCTTTAAAGGCTATACAG

AGGGTATACTTGTCTGGCTGGCCCCAAATGCTCAAGTTAAATGATTGGCCTCAATCTGGTTTTTGAGGAT

CATCTCCCATGTCATAATGTGGAGTTCTTAAGCGCTTTGCCTTTGAAGGAGTACACACATCCACAGAGCG

GCTACTTGAACCTTGCTGTCAAGTTACCCGAGGGCCATCTGAAGCGAGACAAAGGGCCAAAGCTGTATAC

TGCTTATGGCTTTTCCCAGCAACTCGGACGTGGTGACTCTGTCTCAAAACTCAGCTATCATGATTCAGAT

GCGGTATGATCTAGTTTGTAAATACAGTGAATAACTTTCACAAACAATGAAGAAAATGCTCCAAATTTAC

AACCCCACCCGAGTCATCATACTAATAGTCCATCCTGAGATCATAATCATTTTTGGTAATTTTATTCGGC

AAGGTTTCTCATGCTTTATACAATTGTCCATGTGATTTAAGCGATCTAGTTAGTCTTAC

>MSTRG.1564.9 gene=MSTRG.1564

ACCACCAAACGACCCCTTGCGGGGAGTATGGGTATAATTGACCACCGGTCCACCGTAAATTAGTGTTATT

GGATTTTAAACTTTGAATCAGGGGTCCTTTTTGTTCATTTTTTCAACTCAAGAGTTCAATTGCTTTGATT

AGTTTGGTTTCTCTCTTTGTTTCAGGGTTCTGAGAGAGATTCTTGTAGTGTGTTATAAACTCTCAAGCAT

GGATTTCGTTTTGCTGCTGATATCTCAATGATGTACATTAAGACGGTTTCTTGATTAGCAGAGTTAGCTT

ATTGATTCTGTTTGTGACCGTGTTCACAGGCTCCAGTGCTCTGCAACTGTTTATACAAGTTGCAGAACCA

CCCTCTTGTAATTTGTTTTATCAAGTGGAAACACTCGAATAAACATAGGAAAGCTTAAAGGTGTTTAGTC

AGGAAGAAAAAATCAGTAGAAAGATTGAGTTGGTTGAAAGAAATTGGAGAAGGGTATTTGATGAGTCAAT

TCAGTGTCTCCTCCAGCGGCAAAACAAGTGAGAACTTCTTGAACTTCTTGAACTTCAAACACGAGGAATG

ACGGGGAAGGCTATTTGCGCCCGTCTAGTAATTTTCAGGGCTTGGATTGAATTGGGGCTTTACTGGAACT

AGAGCTTCAACAGCTCTTAACTTTAAGAGGCGCTTCGGTTTTCTTTGATCTAAGCAACTATTACGGAACT

TCTATTGCTGACTACCACCGGAGCTGCACAAAATGTCCGTATGATCTATGCATCGCTTGTTGTCGGGACT

TGCGTGCTGGCTCTTTGCATGCAGGTCATCTGGGAAGCATTAACCAATATAAAAACCTCAGGGTGGGCTA

TTTACATGGTGAAACATGTAAACCTCGTAGCAATAGAACTGTGGGCACATCTACTGAAACATCCCCTGAA

GGCAAGACCTGAATGGAGACCTAACAAGGATGGTAGCATCTCTTGTCCACCTAAGAGTATTGGTGGGTGT

GGTGGTAAAGGGATTCTAAAGCTGAACCAAGTCTTGCCAGACGGTTGACTTTCAAACAAGATATTGAAAG

CTGAAGACTTATATGAACTGTATAATCTGGATGGCATGCCAGAAGATCCCATACACTGGTGCTCGTGTTA

TAATTTGGCACGTGATAACACGGCCACAAAGAAAATAAGGAAAGCCACTTCTCGAGAAAACTCTGATGAC

AACACTTCTACTCTCCAAGTTCTATAGACATTCAAGCCGGGGATCGGAAGCATTTTCAGACCCATTGGTC

AAAAGGGGAACCAGTGATTGTTTCTAGTGTGCTTGACAGTACATATGGTTAAGCTGGGAACCAATGGTGA

TGTCTCGAGCAATCCATGATAAGTCCCACACTGATGTGACTGTCTTAAATTGTTTGAGCTGGTGCGAAGA

GGAATATAGCGTAAGTTCGTTCTTTAAAGGCTATACAGAGGGTATACTTGTCTGGCTGGCCCCAAATGCT

CAAGTTAAATGATTGGCCTCAATCTGGTTTTTGAGGATCATCTCCCATGTCATAATGTGGAGTTCTTAAG

CGCTTTGCCTTTGAAGGAGTACACACATCCACAGAGCGGCTACTTGAACCTTGCTGTCAAGTTACCCGAG

GGCCATCTGAAGCGAGACAAAGGGCCAAAGCTGTATACTGCTTATGGCTTTTCCCAGCAACTCGGACGTG

GTGACTCTGTCTCAAAACTCAGCTATCATGATTCAGATGCGGTATGATCTAGTTTGTAAATACAGTGAAT

AACTTTCACAAACAATGAAGAAAATGCTCCAAATTTACAACCCCACCCGAGTCATCATACTAATAGTCCA

TCCTGAGATCATAATCATTTTTGGTAATTTTATTCGGCAAGGTTTCTC

>MSTRG.1564.10 gene=MSTRG.1564

TTATCATCTTCTTTTCTTTTTTCTCTTTCATCTGCCTCATCGTCCTCATATCATCTATCTATTTCATCGT

CTTCTTCTATAATCATCTTCTTCCCTTTTTATTCTTCATCTTCTCTCTTCTTCCCTTTTTATTCTTCTGT

TACACCCCCTTTAAATTTTCCGGCGACCTTCATATTTTTCCGGCGCTAGAAAAGAAGTGTCGGTTTTGCA

TATGGTGAATCCACCCTGTATCCTGCGTCGTGTCGGGTCGACACGGATACGGCCGCCGGAAGTGCCGAGT

CGGAGTAACACAGTATGTATATCAATCTTCAATATCAAAGGTGGATTGCTTTCACAAACAATGAAGAAAA

TGCTTCAAATTTACAACACCTCCCGAGTCATCAGACTAATTTGGCTGGAAGGGCTTGGATTGAATTGGGG

CTTTACTGGAACTAGAGCTTCAACAGCTCTTAACTTTAAGAGGCGCTTCGGTTTTCTTTGATCTAAGCAA

CTATTACGGAACTTCTATTGCTGACTACCACCGGAGCTGCACAAAATGTCCGTATGATCTATGCATCGCT

TGTTGTCGGGACTTGCGTGCTGGCTCTTTGCATGCAGGTCATCTGGGAAGCATTAACCAATATAAAAACC

TCAGGGTGGGCTATTTACATGGTGAAACATGTAAACCTCGTAGCAATAGAACTGTGGGCACATCTACTGA

AACATCCCCTGAAGGCAAGACCTGAATGGAGACCTAACAAGGATGGTAGCATCTCTTGTCCACCTAAGAG

TATTGGTGGGTGTGGTGGTAAAGGGATTCTAAAGCTGAACCAAGTCTTGCCAGACGGTTGACTTTCAAAC

AAGATATTGAAAGCTGAAGACTTATATGAACTGTATAATCTGGATGGCATGCCAGAAGATCCCATACACT

GGTGCTCGTGTTATAATTTGGCACGTGATAACACGGCCACAAAGAAAATAAGGAAAGCCACTTCTCGAGA

AAACTCTGATGACAACACTTCTACTCTCCAAGTTCTATAGACATTCAAGCCGGGGATCGGAAGCATTTTC

AGACCCATTGGTCAAAAGGGGAACCAGTGATTGTTTCTAGTGTGCTTGACAGTACATATGGTTAAGCTGG

GAACCAATGGTGATGTCTCGAGCAATCCATGATAAGTCCCACACTGATGTGACTGTCTTAAATTGTTTGA

GCTGGTGCGAAGAGGAATATAGCGTAAGTTCGTTCTTTAAAGGCTATACAGAGGGTATACTTGTCTGGCT

GGCCCCAAATGCTCAAGTTAAATGATTGGCCTCAATCTGGTTTTTGAGGATCATCTCCCATGTCATAATG

TGGAGTTCTTAAGCGCTTTGCCTTTGAAGGAGTACACACATCCACAGAGCGGCTACTTGAACCTTGCTGT

CAAGTTACCCGAGGGCCATCTGAAGCGAGACAAAGGGCCAAAGCTGTATACTGCTTATGGCTTTTCCCAG

CAACTCGGACGTGGTGACTCTGTCTCAAAACTCAGCTATCATGATTCAGATGCGGTATGATCTAGTTTGT

AAATACAGTGAATAACTTTCACAAACAATGAAGAAAATGCTCCAAATTTACAACCCCACCCGAGTCATCA

TACTAATAGTCCATCCTGAGATCATAATCATTTTTGGTAATTTTATTCGGCAAGGTTTCTCATGCTTTAT

ACAATTGTCCATGTGATTTAAGCGATCTAGTTAGTCTTAC

>MSTRG.1564.11 gene=MSTRG.1564

GATTGACAGCTTCTTCATTCCGCGGAACTAGGGCTTCCGGTTTATACTTCTTCATTTTTTTTCACTATAA

TGTTTTTAGTTGTGTCTTTATTTTTCTCAGGGCTTGGATTGAATTGGGGCTTTACTGGAACTAGAGCTTC

AACAGCTCTTAACTTTAAGAGGCGCTTCGGTTTTCTTTGATCTAAGCAACTATTACGGAACTTCTATTGC

TGACTACCACCGGAGCTGCACAAAATGTCCGTATGATCTATGCATCGCTTGTTGTCGGGACTTGCGTGCT

GGCTCTTTGCATGCAGGTCATCTGGGAAGCATTAACCAATATAAAAACCTCAGGGTGGGCTATTTACATG

GTGAAACATGTAAACCTCGTAGCAATAGAACTGTGGGCACATCTACTGAAACATCCCCTGAAGGCAAGAC

CTGAATGGAGACCTAACAAGGATGGTAGCATCTCTTGTCCACCTAAGAGTATTGGTGGGTGTGGTGGTAA

AGGGATTCTAAAGCTGAACCAAGTCTTGCCAGACGGTTGACTTTCAAACAAGATATTGAAAGCTGAAGAC

TTATATGAACTGTATAATCTGGATGGCATGCCAGAAGATCCCATACACTGGTGCTCGTGTTATAATTTGG

CACGTGATAACACGGCCACAAAGAAAATAAGGAAAGCCACTTCTCGAGAAAACTCTGATGACAACACTTC

TACTCTCCAAGTTCTATAGACATTCAAGCCGGGGATCGGAAGCATTTTCAGACCCATTGGTCAAAAGGGG

AACCAGTGATTGTTTCTAGTGTGCTTGACAGTACATATGGTTAAGCTGGGAACCAATGGTGATGTCTCGA

GCAATCCATGATAAGTCCCACACTGATGTGACTGTCTTAAATTGTTTGAGCTGGTGCGAAGAGGAATATA

GCGTAAGTTCGTTCTTTAAAGGCTATACAGAGGGTATACTTGTCTGGCTGGCCCCAAATGCTCAAGTTAA

ATGATTGGCCTCAATCTGGTTTTTGAGGATCATCTCCCATGTCATAATGTGGAGTTCTTAAGCGCTTTGC

CTTTGAAGGAGTACACACATCCACAGAGCGGCTACTTGAACCTTGCTGTCAAGTTACCCGAGGGCCATCT

GAAGCGAGACAAAGGGCCAAAGCTGTATACTGCTTATGGCTTTTCCCAGCAACTCGGACGTGGTGACTCT

GTCTCAAAACTCAGCTATCATGATTCAGATGCGGTATGATCTAGTTTGTAAATACAGTGAATAACTTTCA

CAAACAATGAAGAAAATGCTCCAAATTTACAACCCCACCCGAGTCATCATACTAATAGTCCATCCTGAGA

TCATAATCATTTTTGGTAATTTTATTCGGC

>MSTRG.1567.1 gene=MSTRG.1567

AGAAAACTTGAAAAGCCCAAGCTATAAACTCATTCTACTCTCTTACAAATACCCAGACATGTACAACAGC

AGCAGCTTTTGTTAACTCGTTGCATCTCCCAGCTCTCATCTTCAGTCTCGCTTCTCAGGCAATTCAGTAG

GCATATTTGTGGTTTGTTGATATTTATGGAGGTCGACGAGATCGACGTGCTTGGATCTCAGGCCCTCGCT

ACTGCTGATGCCTGCTTTCAGGCTGCTCTCCACCAAACTCGAGCTTGGAAGAAACAAGCAAGGGAGCACA

AGAAGGCTACCAAGGCCTTAAAGGCGAGATTGAAGGACCTCCGGATCAAGCACATGGAGCAGGAGGAGCT

CCTAAGGAAGACTCAAGCGGAGTTGGTTGAGGCGAGGGAAGATAGGGAAACTATCATAGATAAATACATG

GATTCAGAGGCCTTTCAACAGCTCTTGGTCCAGCATGACGACCTCCTCTATCGGAAGTACTATAACGATG

GGTGGAATGATGCACTTAAGGAGGTGAACAGGCGACATCCAGGATTATTCAATCCTAAGGTTGACTTTCC

TTGTCCTTTAGATCTTATTCCCGCCACTCAGCCGACTTCGGGGGGTGGTCAGTCCCCGCGCTTTGCTGCT

GATCCGAGGATCCTTAGCTCGACTCGTTCCCTGCTTCGAGGTGAGCCTTCGGAGGCTTCAGATGATAAGA

CCTCATTTTAGATCAGATTTCTCCATGGCCCTGCATGGCCTTATTTATTTAGACTTTTGTTTGTAAAATC

TTTGACTTTAACATTAATCGTATTTTATTTGGTTTCCTCATTCCTCTTT

>MSTRG.1577.1 gene=MSTRG.1577

CCCATCCGGCACTTATAAATCGAAGAAATTAGGGCTTGAAAATGGCGTAAAAGAAATCCCCAATTCTCGT

TTGGCGGGAAGAAACAATTGGACAATCATACTTTTAAACATCCACCTTTCTTCTTTCGATGTCGTTTCTC

GAAGAATTTCAGGCCAATTTGGAATCATTGCCAAATATCATACAGAGGAAGTATGCCATACTGCGGGATT

TAGATAAAAGTTTACAAGTGATTCAGCAGCAAAATGAACAACGTTGTGGACAAGAACTTGAAAATTTTAA

GCGAGGAGTTAGATCTGGTGCTATTATGCCTGACTCTTCACTTGCCAAATACTCAGATGAGGCACTTGAT

GAGCAAAAACATGCCGTTAGACTTGCTGACGAGAAGGTTGCATTGGCTGTGCAAGTATATGATTTGGTAG

ATTCTCATATACAACAACTTGATCAGCAAATGAAAAAGTTCAATGAAGATCTTCAGCGTGAAAGAGATAG

CAATGGTGCAACTGGTTCACTTCCCTCAAACAATGATGGCACTGCAAAAACTGGAAGGAGCGGAAGTGGT

AGAGGAGGTCGCAAGAAGACAGGCCTTGCTGCAGCAGCGGTGGCATCAGCAGCACCAGCAGTGCCTGCAA

ACCCAACAAGCATGGATTTAGATCTACCAGTGGATCCAAATGAACCCACATATTGTCTTTGCAACCAAGT

GAGCTATGGTGACATGGTAGCATGCGACAACCCTGATTGCAAGATAGAATGGTTTCATTTTGGTTGTGTT

GGTGTGAAAGAACAACCCAGGGGAAAGTGGTATTGCTCTGACTGTGCCGGACTGCAGAAGCGCAGGAAAG

GTAAGTAATGCTTGAGATATTCTGGGAGTTTGTCCAGATCCAGTGTGTTTGTTCTTCGAGGAACAAGCAC

TTGATGAAAAGATTACTTACTGTATAATGCCCAAATCATGTTTAGTTCAGTCATACTCTGCGCTTGGATT

CGTCTTTGTGACATAATTTCACTTTTTGCAATTTCTTTGCTCTCATGTTATAGTTTAGATTGTTTTGAAC

CTTTCTCGATCCAAAGGTAGATAAAAAAATTAGAATTCATGAATATATAACAAATGTTTTTTTAAGCATC

TGTTTGCTTGTAATTAATATACATGTATATTATGACTTCCA

>MSTRG.1580.1 gene=MSTRG.1580

CAGGTAAGGTATGGTACAGTAAGATCAAAGCTTGATTTGTTCATTTGGTGAGGGTCTTACAGAAGGGCTT

TGGGTCTGCTGGTGCAGGGTTTTTAAATGTGGAAATTTCTGTGTTGCTTCCTAAGAAATTTGCTTGTAAG

GTCTGATCTCGGAGATTGTTGTTTGTCTAGATGCTGAGGTTGTTTTGATGTTTATAAATTTCAGAGGCAT

TATTTTCTAGTTACACAGAT

>MSTRG.1583.1 gene=MSTRG.1583

ACCACCACTATCGTAACCACCGCCACCGACACCATAAAAACACATATCTGTATAAAGGAGAGACCCTTAC

ACTATCATCACCTACACTTGCATCTGCCGCCACCATCATATCCCCACAGCATCAACACCATCACAATCAC

CGCCACGACACCTTAATCGCCACATCTACATCACAAATCTGAATAAAATTGTACAAAAAGAAGGGGGGG

>MSTRG.1584.1 gene=MSTRG.1584

ATCATTAATTCTCGTCAGTACCGATGGCTATCAATACATCTCGTATAAAAATCACCGCCTTCAAACATTA

CTTATAATTCCTAGATTCAGCCACAAATCACCAATTCATATTACAACATCACTAATGTGTGCTGCAAAAA

TCACCAATTTACATGGTACCAATCACTAAAAAAATCACTATATCATTGTATATATCATCTTCATTACCCG

CAACCTCACACATGCTATCACAATCTTATCGCCTCAGTACCACCACTA

>MSTRG.1602.1 gene=MSTRG.1602

GGTGATTATCACATTCATCTCCGTCAAATACGTCACCACGGCGAACATCTCATCAATTCAACATTATCAC

GAATAACGCAAATGCATCTGGACCGTCCATTTGCCGCTGCCTTTGATTCTAAATATTCACTGCCTAGTTG

GGACAAACGTGCCAATGAAACCAAATGCTACTGTGAAGAACGCCCCGGATTGCAAGAACAAATATACAAA

CATGTGATCATTGCCGTGCAACATGTTGTATACAGCACAATGAAGCATGAATATCCCCATTATGAACTCC

AGCTTGTGAATCCTGTTCAAATTCAGCGTATGAGTTACAAACAAGAAAATGATGAATGTTTCCTATTGTT

AGCCTAATTTTTACAAAACATTAGTAGAGCAAGAGTAATTAAATATAACCTCTCTCCACATTGGAATTTG

GATTTAAAAAAAGAAACTCTAGCTTGGTTATTTCTTCTATTGCCAAGCTTTTCCGTCACAACCCATTCAT

TGACTCGACTGGCTTCCAATAGCCCGATTATTGATGCCTTAGTTCGGTGAAGGGACATGACATTTTCAAA

CAGTATCCAGAATACGAGAAGATGCAAGGATCTGCTTGCCAGCATATCAATTCAGGTTAATCAATATCTT

TTTTGGAAGTTATGATATATCTCTTAAATGGAATGTAATACAGTTATTAGAATTACTACCTAGGAGTGCC

ACCAGTGTTCAGAATTGTGATGATTGCTGGAATATATATGGCTAATGGCTTTGGAAGATAAACCTCAGGA

AGCAAAACGCATGCAGGAATAACCACACAGTAGAAGAAGAAGGTTACCCAATGTGCAACGATCTTCCTAA

CAAAGAAGAATGCGTATATGAGGTGAATCTTCTTCCATACAGACACGCGCTGCAAAATAAAACAAGGTTG

GCTGGACCACATGACCAGCGGTGCTGCTGGAATCGAATAAAATTACAGGGTGAGATAAAATTACCGACAC

ATCTCCTACAAAGACAAACTTCCAACCCTGCAGACTGGCTCTGACTGCAAGATCCATGTCCTCCACTGTG

GTTCGCTCTTTCCATCCTCCTGCATCCTTAATTGCTTGAATTCGCCATACACCAGCGGTATCTGGATCAA

GACTGTCGGATGCTCTCTATTCCGTTCCATTCTTTCTTTCACAGAATCCAGATTGTATTGTGTGTATCTC

TTTTTCCCCAATGCTTTAACATATAGGATCACGGCTGTCATGTACAGCCGTTCAATGAAAAGCATAAGAG

ACATAATCACACATACAAATAGTGCAAAATACAAGAGTGGCACAATTACTGATGCTCTGAGTTTCTTCTG

AGCATCATTGATGCTCATTGTTATCTCATCAGGCTCTTGGAAGACTGAATTTCTCATATTGTAAGGTGTG

GGGATAAAACTGAAGTGCGAAGCAAAAAAGGAAATTGATGTGGGAAAAAGCTTAGATGCAGAGCTTTAGA

AGAAAGATAAAGATGAGGATAGCTAGGCTTTAGCTGGAAGCATTCTACAAGAGGAGTGAACGATTGAAAT

TGAAGTCTTTCGTTAGCAAGGACACCCGGAGAACATGCACGCAATGATGCTGCTACGTAGCTACTTGCTA

GCTGGAACCGCCAACAATAAAAGGAAGTCATCAAATTGTTGATCGAAGATCCTTACACCACCCTCTCTTC

TGAATTTGTTTATATCCTTGTCTTCTGATAACGTTTGAGAATGTCATACCTTCCATCCCTCTATGGCAGC

AATAGAAGCTATCCAAGTCAACGAATGAATTGTGACTATTAAGATCCAGAGTTCGGCGCATTCTTGGCTT

TATCATTGGGTTTGTTGGCACCTTTGTCCCGAGTATGCAGTGAAGACAAGGGCACTTCCTTGACAGGGTC

TTGTTTTGGTTATAGATATTTTTGGTGCTACACAGAAGATTGGATTAGAGAGATTCTGAACAGTTTATTC

AAGAGGGTCTGAAGAATCTAATACTGATTGAGTGAAAGGAAATTGAAAAGCTTCCTTTTACTATATGTTG

GCCTATCGTATGATCCGCATTGCCATATTGGTTACCAAATGCAATCACGTTGATCGCCGTGCTGCAACAT

ATTTAAAGCGAGTGTCTCTAGAGATTCATCTGTTGAGGTTAGTGAGGAGAATATGAATTCGGCTCAAGAG

GTCAAAGAGAAGGCTATGGGAGCGATATCCGAAGGTCCTTGTGGATTTGAGAGAAAAGCTTTCCACCGCT

CTTTCACAATGATTCCTCAATTAATATCAATTTGTTATGATTAGATCATGATAGAAGAAGTTCCTGCTGA

CACGCAACCAAAAGAGAAAACAAGTTTTTAAGCTCTGTTCCACAATCCTGGATTGGCTTCATTTTTCTAG

GTAGGAACACCAAATTGAATGAGTTAATAAGATCAGTATGGAGGACAGACATGCAGTTGCTGAAGCTCAG

TTTCAAACTATATTTATATGAAGATTTAGTCATGAAAGTCATTAAATCTCAGCCTGAGGTTTCACGGAGT

TTTCTGTTTTGTTGAGGAATTAGTAAGAATCCGGTTACTTGTTAGAAAAAGCTAAACTTGAATTTTCTTC

AGTTTCCTTTGACCACCTTTCTTTAACATGTGCCAATGGGTAAAAAAAAAAAAAAAAAACAGATTTGAAG

TAGAACCCCTAGAAATCATAGGCCCATCAGTCTCAATTCCTGATATCTAATACATTATGTAGATTTTGGC

ATTTGCAAATCCTTTCTTAATTCTAATTTAGTGATGCAGGATTTAACTTTCTATAGTAACAAGCTTAAAT

TATATCATCATAATTCAAAGTATACAAGTTAATCTGCAACGGAGATATACAATGTACTGATCAGTCAAAG

AATAAGATACATGTAAATGAAC

>MSTRG.1602.2 gene=MSTRG.1602

GAAGATTGAAACATTAGTGAGGGTCTGTATTGTTGTTTTGGATATAGGAATTGTACTCTTTATACGTATC

GGTTTGATTCTGTTAAGTTTGATTGAGTTAAGCCGAGCTAGAGAGAGGCTTTGTCTCGTAAGAATCTTTC

GAGTTTGTGCCTGATTTCTACAGTGTTATAGCATTTTAAAATCGAATGAATTCCACCACTATGTGATTTC

GTGTTGAAGTTAGTTTTTGTAAACCAACAAAAGTCGAAGGGTATGGTAAAAACCCCAATTGACCACCAAC

ATAATCTTGTAAAGACCTTGAATCTTGCTTGAATATGACAAGAATTACAAATTCTCGATATCATCTAGAG

TCAGATGTACCTCCCTAGTGATGAACACTAGTAAATAATTTCAAGCATTCTATGGATTTTATAGGAGAAG

CACTAACAATATACATTATGGACAATCAAGGGATTCAAGTTAATGTTTTTGACAGGGATAAGATGATAAA

TAAGTGTGTATCCTATTCTTTTATTACAGACATATACTAGATTTATAATATGATCTACGTTGGTATGCTA

ATAAAATCGAAATCAGATTTCAGAACAGTCCTAATACCTACTCTTCCCTAGATGCTGCAGGATTTATGCA

CCCTTAATTTTTTATTTGGAGGATAATTAAATGGTACAATTTATAGGGATGCAGGTTAAAATGACTTTTG

ATTGTTGTCTATCACATATTACCAAGTCAGAAGTCCAATTGTTGAAAATGCAGTAAAAGGTTTGAGTACA

GTATCTTTGAAACCGTTTTAAACACACAGGGGACAATCTTTATTTTAAGGTAGATGCATATAACCATACA

TACATTACACTGCAACTAAGCACCTCTCTTCGCACCATTACCAGAAGTCACCCCTGACAAGATACAAACT

AGTTTTGTACAAGTGGGGTGATATTCATAAACATTAACATTCACCAACTTGTCATTGTCTTCCTCATTGC

TTATTCTCAGATATTCACTGCCTAGTTGGGACAAACGTGCCAATGAAACCAAATGCTACTGTGAAGAACG

CCCCGGATTGCAAGAACAAATATACAAACATGTGATCATTGCCGTGCAACATGTTGTATACAGCACAATG

AAGCATGAATATCCCCATTATGAACTCCAGCTTGTGAATCCTGTTCAAATTCAGCGTATGAGTTACAAAC

AAGAAAATGATGAATGTTTCCTATTGTTAGCCTAATTTTTACAAAACATTAGTAGAGCAAGAGTAATTAA

ATATAACCTCTCTCCACATTGGAATTTGGATTTAAAAAAAGAAACTCTAGCTTGGTTATTTCTTCTATTG

CCAAGCTTTTCCGTCACAACCCATTCATTGACTCGACTGGCTTCCAATAGCCCGATTATTGATGCCTTAG

TTCGGTGAAGGGACATGACATTTTCAAACAGTATCCAGAATACGAGAAGATGCAAGGATCTGCTTGCCAG

CATATCAATTCAGGTTAATCAATATCTTTTTTGGAAGTTATGATATATCTCTTAAATGGAATGTAATACA

GTTATTAGAATTACTACCTAGGAGTGCCACCAGTGTTCAGAATTGTGATGATTGCTGGAATATATATGGC

TAATGGCTTTGGAAGATAAACCTCAGGAAGCAAAACGCATGCAGGAATAACCACACAGTAGAAGAAGAAG

GTTACCCAATGTGCAACGATCTTCCTAACAAAGAAGAATGCGTATATGAGGTGAATCTTCTTCCATACAG

ACACGCGCTGCAAAATAAAACAAGGTTGGCTGGACCACATGACCAGCGGTGCTGCTGGAATCGAATAAAA

TTACAGGGTGAGATAAAATTACCGACACATCTCCTACAAAGACAAACTTCCAACCCTGCAGACTGGCTCT

GACTGCAAGATCCATGTCCTCCACTGTGGTTCGCTCTTTCCATCCTCCTGCATCCTTAATTGCTTGAATT

CGCCATACACCAGCGGTATCTGGATCAAGACTGTCGGATGCTCTCTATTCCGTTCCATTCTTTCTTTCAC

AGAATCCAGATTGTATTGTGTGTATCTCTTTTTCCCCAATGCTTTAACATATAGGATCACGGCTGTCATG

TACAGCCGTTCAATGAAAAGCATAAGAGACATAATCACACATACAAATAGTGCAAAATACAAGAGTGGCA

CAATTACTGATGCTCTGAGTTTCTTCTGAGCATCATTGATGCTCATTGTTATCTCATCAGGCTCTTGGAA

GACTGAATTTCTCATATTGTAAGGTGTGGGGATAAAACTGAAGTGCGAAGCAAAAAAGGAAATTGATGTG

GGAAAAAGCTTAGATGCAGAGCTTTAGAAGAAAGTAAGGAAGGGGATGATTTTTATAGAGGGGGAACCGG

GGAAGGGATAACTATGGAGTGCATCTGATGGATGCTGTGCTTTGGCAATTACGTAGTATATGCTTAACAT

AGAAAAATTTCAGGATAAAGATGAGGATAGCTAGGCTTTAGCTGGAAGCATTCTACAAGAGGAGTGAACG

ATTGAAATTGAAGTCTTTCGTTAGCAAGGACACCCGGAGAACATGCACGCAATGATGCTGCTACGTAGCT

ACTTGCTAGCTGGAACCGCCAACAATAAAAGGAAGTCATCAAATTGTTGATCGAAGATCCTTACACCACC

CTCTCTTCTGAATTTGTTTATATCCTTGTCTTCTGATAACGTTTGAGAATGTCATACCTTCCATCCCTCT

ATGGCAGCAATAGAAGCTATCCAAGTCAACGAATGAATTGTGACTATTAAGATCCAGAGTTCGGCGCATT

CTTGGCTTTATCATTGGGTTTGTTGGCACCTTTGTCCCGAGTATGCAGTGAAGACAAGGGCACTTCCTTG

ACAGAAGATTGGATTAGAGAGATTCTGAACAGTTTATTCAAGAGGGTGAGCGGCTTGTGTTGAAGTTATT

TCTATCTTCGATTCTCCGCATTGCCATATTGGTTACCAAATGCAATCACGTTGATCGCCGTGCTGCAACA

TATTTAAAGCGAGTGTCTCTAGAGATTCATCTGTTGAGGTTAGTGAGGAGAATATGAATTCGGCTCAAGA

GGTCAAAGAGAAGGCTATGGGAGCGATATCCGAAGGTCCTTGTGGATTTGAGAGAAAAGCTTTCCACCGC

TCTTTCACAATGATTCCTCAATTAATATCAATTTGTTATGATTAGATCATGATAGAAGAAGTTCCTGCTG

ACACGCAACCAAAAGAGAAAACAAGTTTTTAAGCTCTGTTCCACAATCCTGGATTGGGTAATTGATAAAA

TTTATTGAACTTCATCATATTATATGAACTGAATGCTTGTGTAGTGTA

>MSTRG.1602.4 gene=MSTRG.1602

TCCTAATACCTACTCTTCCCTAGATGCTGCAGGATTTATGCACCCTTAATTTTTTATTTGGAGGATAATT

AAATGGTACAATTTATAGGGATGCAGGTTAAAATGACTTTTGATTGTTGTCTATCACATATTACCAAGTC

AGAAGTCCAATTGTTGAAAATGCAGTAAAAGGTTTGAGTACAGTATCTTTGAAACCGTTTTAAACACACA

GGGGACAATCTTTATTTTAAGGTAGATGCATATAACCATACATACATTACACTGCAACTAAGCACCTCTC

TTCGCACCATTACCAGAAGTCACCCCTGACAAGATACAAACTAGTTTTGTACAAGTGGGGTGATATTCAT

AAACATTAACATTCACCAACTTGTCATTGTCTTCCTCATTGCTTATTCTCAGATATTCACTGCCTAGTTG

GGACAAACGTGCCAATGAAACCAAATGCTACTGTGAAGAACGCCCCGGATTGCAAGAACAAATATACAAA

CATGTGATCATTGCCGTGCAACATGTTGTATACAGCACAATGAAGCATGAATATCCCCATTATGAACTCC

AGCTTGTGAATCCTGTTCAAATTCAGCGTATGAGTTACAAACAAGAAAATGATGAATGTTTCCTATTGTT

AGCCTAATTTTTACAAAACATTAGTAGAGCAAGAGTAATTAAATATAACCTCTCTCCACATTGGAATTTG

GATTTAAAAAAAGAAACTCTAGCTTGGTTATTTCTTCTATTGCCAAGCTTTTCCGTCACAACCCATTCAT

TGACTCGACTGGCTTCCAATAGCCCGATTATTGATGCCTTAGTTCGGTGAAGGGACATGACATTTTCAAA

CAGTATCCAGAATACGAGAAGATGCAAGGATCTGCTTGCCAGCATATCAATTCAGGTTAATCAATATCTT

TTTTGGAAGTTATGATATATCTCTTAAATGGAATGTAATACAGTTATTAGAATTACTACCTAGGAGTGCC

ACCAGTGTTCAGAATTGTGATGATTGCTGGAATATATATGGCTAATGGCTTTGGAAGATAAACCTCAGGA

AGCAAAACGCATGCAGGAATAACCACACAGTAGAAGAAGAAGGTTACCCAATGTGCAACGATCTTCCTAA

CAAAGAAGAATGCGTATATGAGGTGAATCTTCTTCCATACAGACACGCGCTGCAAAATAAAACAAGACTC

CGGAGTTGTCTTACTCTGCAAAGCATGATTTCCTTTGTCATTTTTTTGAAAAGGTTGGCTGGACCACATG

ACCAGCGGTGCTGCTGGAATCGGGTGAGATAAAATTACCGACACATCTCCTACAAAGACAAACTTCCAAC

CCTGCAGACTGGCTCTGACTGCAAGATCCATGTCCTCCACTGTGGTTCGCTCTTTCCATCCTCCTGCATC

CTTAATTGCTTGAATTCGCCATACACCAGCGGTATCTGGATCAAGACTGTCGGATGCTCTCTATTCCGTT

CCATTCTTTCTTTCACAGAATCCAGATTGTATTGTGTGTATCTCTTTTTCCCCAATGCTTTAACATATAG

GATCACGGCTGTCATGTACAGCCGTTCAATGAAAAGCATAAGAGACATAATCACACATACAAATAGTGCA

AAATACAAGAGTGGCACAATTACTGATGCTCTGAGTTTCTTCTGAGCATCATTGATGCTCATTGTTATCT

CATCAGGCTCTTGGAAGACTGAATTTCTCATATTGTAAGGTGTGGGGATAAAACTGAAGTGCGAAGCAAA

AAAGGAAATTGATGTGGGAAAAAGCTTAGATGCAGAGCTTTAGAAGAAAGTAAGGAAGGGGATGATTTTT

ATAGAGGGGGAACCGGGGAAGGGATAACTATGGAGTGCATCTGATGGATGCTGTGCTTTGGCAATTACGT

AGTATATGCTTAACATAGAAAAATTTCAGGATAAAGATGAGGATAGCTAGGCTTTAGCTGGAAGCATTCT

ACAAGAGGAGTGAACGATTGAAATTGAAGTCTTTCGTTAGCAAGGACACCCGGAGAACATGCACGCAATG

ATGCTGCTACGTAGCTACTTGCTAGCTGGAACCGCCAACAATAAAAGGAAGTCATCAAATTGTTGATCGA

AGATCCTTACACCACCCTCTCTTCTGAATTTGTTTATATCCTTGTCTTCTGATAACGTTTGAGAATGTCA

TACCTTCCATCCCTCTATGGCAGCAATAGAAGCTATCCAAGTCAACGAATGAATTGTGACTATTAAGATC

CAGAGTTCGGCGCATTCTTGGCTTTATCATTGGGTTTGTTGGCACCTTTGTCCCGAGTATGCAGTGAAGA

CAAGGGCACTTCCTTGACAGGGTCTTGTTTTGGTTATAGATATTTTTGGTGCTACACAGAAGATTGGATT

AGAGAGATTCTGAACAGTTTATTCAAGAGGGTCTGAAGAATCTAATACTGATTGAGTGAAAGGAAATTGA

AAAGCTTCCTTTTACTATATGTTGGCCTATCGTATGATCCGCATTGCCATATTGGTTACCAAATGCAATC

ACGTTGATCGCCGTGCTGCAACATATTTAAAGCGAGTGTCTCTAGAGATTCATCTGTTGAGGTTAGTGAG

GAGAATATGAATTCGGCTCAAGAGGTCAAAGAGAAGGCTATGGGAGCGATATCCGAAGGTCCTTGTGGAT

TTGAGAGAAAAGCTTTCCACCGCTCTTTCACAATGATTCCTCAATTAATATCAATTTGTTATGATTAGAT

CATGATAGAAGAAGTTCCTGCTGACACGCAACCAAAAGAGAAAACAAGTTTTTAAGCTCTGTTCCACAAT

CCTGGATTGGCTTCATTTTTCTAGGTAGGAACACCAAATTGAATGAGTTAATAAGATCAGTATGGAGGAC

AGACATGCAGTTGCTGAAGCTCAGTTTCAAACTATATTTATATGAAGATTTAGTCATGAAAGTCATTAAA

TCTCAGCCTGAGGTTTCACGGAGTTTTCTGTTTTGTTGAGGAATTAGTAAGAATCCGGTTACTTGTTAGA

AAAAGCTAAACTTGAATTTTCTTCAGTTTCCTTTGACCACCTTTCTTTAACATGTGCCAATGGGTAAAAA

AAAAAAAAAAAAACAGATTTGAAGTAGAACCCCTAGAAATCATAGGCCCATCAGTCTCAATTCCTGATAT

CTAATACATTATGTAGATTTTGGC

>MSTRG.1602.3 gene=MSTRG.1602

TCCTAATACCTACTCTTCCCTAGATGCTGCAGGATTTATGCACCCTTAATTTTTTATTTGGAGGATAATT

AAATGGTACAATTTATAGGGATGCAGGTTAAAATGACTTTTGATTGTTGTCTATCACATATTACCAAGTC

AGAAGTCCAATTGTTGAAAATGCAGTAAAAGGTTTGAGTACAGTATCTTTGAAACCGTTTTAAACACACA

GGGGACAATCTTTATTTTAAGGTAGATGCATATAACCATACATACATTACACTGCAACTAAGCACCTCTC

TTCGCACCATTACCAGAAGTCACCCCTGACAAGATACAAACTAGTTTTGTACAAGTGGGGTGATATTCAT

AAACATTAACATTCACCAACTTGTCATTGTCTTCCTCATTGCTTATTCTCAGATATTCACTGCCTAGTTG

GGACAAACGTGCCAATGAAACCAAATGCTACTGTGAAGAACGCCCCGGATTGCAAGAACAAATATACAAA

CATGTGATCATTGCCGTGCAACATGTTGTATACAGCACAATGAAGCATGAATATCCCCATTATGAACTCC

AGCTTGTGAATCCTGTTCAAATTCAGCGTATGAGTTACAAACAAGAAAATGATGAATGTTTCCTATTGTT

AGCCTAATTTTTACAAAACATTAGTAGAGCAAGAGTAATTAAATATAACCTCTCTCCACATTGGAATTTG

GATTTAAAAAAAGAAACTCTAGCTTGGTTATTTCTTCTATTGCCAAGCTTTTCCGTCACAACCCATTCAT

TGACTCGACTGGCTTCCAATAGCCCGATTATTGATGCCTTAGTTCGGTGAAGGGACATGACATTTTCAAA

CAGTATCCAGAATACGAGAAGATGCAAGGATCTGCTTGCCAGCATATCAATTCAGGTTAATCAATATCTT

TTTTGGAAGTTATGATATATCTCTTAAATGGAATGTAATACAGTTATTAGAATTACTACCTAGGAGTGCC

ACCAGTGTTCAGAATTGTGATGATTGCTGGAATATATATGGCTAATGGCTTTGGAAGATAAACCTCAGGA

AGCAAAACGCATGCAGGAATAACCACACAGTAGAAGAAGAAGGTTACCCAATGTGCAACGATCTTCCTAA

CAAAGAAGAATGCGTATATGAGGTGAATCTTCTTCCATACAGACACGCGCTGCAAAATAAAACAAGACTC

CGGAGTTGTCTTACTCTGCAAAGCATGATTTCCTTTGTCATTTTTTTGAAAAGGTTGGCTGGACCACATG

ACCAGCGGTGCTGCTGGAATCGAATAAAATTACAGGGTGAGATAAAATTACCGACACATCTCCTACAAAG

ACAAACTTCCAACCCTGCAGACTGGCTCTGACTGCAAGATCCATGTCCTCCACTGTGGTTCGCTCTTTCC

ATCCTCCTGCATCCTTAATTGCTTGAATTCGCCATACACCAGCGGTATCTGGATCAAGACTGTCGGATGC

TCTCTATTCCGTTCCATTCTTTCTTTCACAGAATCCAGATTGTATTGTGTGTATCTCTTTTTCCCCAATG

CTTTAACATATAGGATCACGGCTGTCATGTACAGCCGTTCAATGAAAAGCATAAGAGACATAATCACACA

TACAAATAGTGCAAAATACAAGAGTGGCACAATTACTGATGCTCTGAGTTTCTTCTGAGCATCATTGATG

CTCATTGTTATCTCATCAGGCTCTTGGAAGACTGAATTTCTCATATTGTAAGGTGTGGGGATAAAACTGA

AGTGCGAAGCAAAAAAGGAAATTGATGTGGGAAAAAGCTTAGATGCAGAGCTTTAGAAGAAAGTAAGGAA

GGGGATGATTTTTATAGAGGGGGAACCGGGGAAGGGATAACTATGGAGTGCATCTGATGGATGCTGTGCT

TTGGCAATTACGTAGTATATGCTTAACATAGAAAAATTTCAGGATAAAGATGAGGATAGCTAGGCTTTAG

CTGGAAGCATTCTACAAGAGGAGTGAACGATTGAAATTGAAGTCTTTCGTTAGCAAGGACACCCGGAGAA

CATGCACGCAATGATGCTGCTACGTAGCTACTTGCTAGCTGGAACCGCCAACAATAAAAGGAAGTCATCA

AATTGTTGATCGAAGATCCTTACACCACCCTCTCTTCTGAATTTGTTTATATCCTTGTCTTCTGATAACG

TTTGAGAATGTCATACCTTCCATCCCTCTATGGCAGCAATAGAAGCTATCCAAGTCAACGAATGAATTGT

GACTATTAAGATCCAGAGTTCGGCGCATTCTTGGCTTTATCATTGGGTTTGTTGGCACCTTTGTCCCGAG

TATGCAGTGAAGACAAGGGCACTTCCTTGACAGGGTCTTGTTTTGGTTATAGATATTTTTGGTGCTACAC

AGAAGATTGGATTAGAGAGATTCTGAACAGTTTATTCAAGAGGGTCTGAAGAATCTAATACTGATTGAGT

GAAAGGAAATTGAAAAGCTTCCTTTTACTATATGTTGGCCTATCGTATGATCCGCATTGCCATATTGGTT

ACCAAATGCAATCACGTTGATCGCCGTGCTGCAACATATTTAAAGCGAGTGTCTCTAGAGATTCATCTGT

TGAGGTTAGTGAGGAGAATATGAATTCGGCTCAAGAGGTCAAAGAGAAGGCTATGGGAGCGATATCCGAA

GGTCCTTGTGGATTTGAGAGAAAAGCTTTCCACCGCTCTTTCACAATGATTCCTCAATTAATATCAATTT

GTTATGATTAGATCATGATAGAAGAAGTTCCTGCTGACACGCAACCAAAAGAGAAAACAAGTTTTTAAGC

TCTGTTCCACAATCCTGGATTGGCTTCATTTTTCTAGGTAGGAACACCAAATTGAATGAGTTAATAAGAT

CAGTATGGAGGACAGACATGCAGTTGCTGAAGCTCAGTTTCAAACTATATTTATATGAAGATTTAGTCAT

GAAAGTCATTAAATCTCAGCCTGAGGTTTCACGGAGTTTTCTGTTTTGTTGAGGAATTAGTAAGAATCCG

GTTACTTGTTAGAAAAAGCTAAACTTGAATTTTCTTCAGTTTCCTTTGACCACCTTTCTTTAACATGTGC

CAATGGGTAAAAAAAAAAAAAAAAAACAGATTTGAAGTAGAACCCCTAGAAATCATAGGCCCATCAGTCT

CAATTCCTGATATCTAATACATTATGTAGATTTTGGCATTTGCAAATCCTTTCTTAATTCTAATTTAGTG

ATGCAG

>MSTRG.1602.5 gene=MSTRG.1602

AGGGACATGACATTTTCAAACAGTATCCAGAATACGAGAAGATGCAAGGATCTGCTTGCCAGCATATCAA

TTCAGGAGTGCCACCAGTGTTCAGAATTGTGATGATTGCTGGAATATATATGGCTAATGGCTTTGGAAGA

TAAACCTCAGGAAGCAAAACGCATGCAGGAATAACCACACAGTAGAAGAAGAAGGTTACCCAATGTGCAA

CGATCTTCCTAACAAAGAAGAATGCGTATATGAGGTGAATCTTCTTCCATACAGACACGCGCTGCAAAAT

AAAACAAGGTTGGCTGGACCACATGACCAGCGGTGCTGCTGGAATCGAATAAAATTACAGGGTGAGATAA

AATTACCGACACATCTCCTACAAAGACAAACTTCCAACCCTGCAGACTGGCTCTGACTGCAAGATCCATG

TCCTCCACTGTGGTTCGCTCTTTCCATCCTCCTGCATCCTTAATTGCTTGAATTCGCCATACACCAGCGG

TACCTATGAACATAATGATTAAGAATTATATCGAAATTGTTTTGTGGAGTTGTTTTTCACTCGTTATGCT

TATATTGATTTATACTGGGGTTTTGATTTACCATTGAATCCAAAAAATTGGCATGTTGAGGAGCCTACTT

CTTGCTCAA

>MSTRG.1604.1 gene=MSTRG.1604

TGATGAAGACCTAAGGATACAAGCAAAACATGGGAAGGAGGATTAAAGTGGATGGATGGTTTTTATCTAG

AAAGTTTAAGAGAAAGATGGGAACATAGTTTCCTACAAAAGGAGAAAAAAAAAGGTACTATTCCTTCCCC

TCAACAGGAGGACTGTGTGGCATTAGAGGAGATGGAGTGATGGAATTGTGAAGAGTGAAGGTACTTGGTT

CGGGCTTTTGTGGAAATAAAAGAGGTTTGACAGTAAGATTGATATGATAGCTAGATAGTGACACCC

>MSTRG.1611.1 gene=MSTRG.1611

CTCTTCAAATAAAATTTATTTCTTTTAGATATGGCTTCAAGTGTGGGACGTAATTCAGCAGGTTTTCATC

CAAGCATTTGGGGGGACAAATTCATCCCCTCCAGTGACTCAGCTCTTAAGAAAACTGATGTGGACTGCCA

AGAGGAAGAGAAGCTCCAACTGCTAAAACAGGAAGTGAAGAAGATGCTGACTGGTGCAGAGATTCCTCTG

CAAGAGTTGATATGCTTAATTGATGACATTCAACGTCTGGGGCTGTCTTACCACTTTGAAGCCGAGATTG

ATAAATTATTACACTGCGTCAAGGATAGCTTTATCGACTATTATGGCACTAAGAATAATGATAATTTACA

TGATGTTGCACTAAGTTTTCGCCTACTTAGACAAGAAGGGCATAATGTTTCATCTGACGTATTCTCCAGA

TTCCAGGACAGCAACGGGAAGTTTATGGAGGAATTGGCTAAGGATGTGAGAGGAATGCTAAGTTTGTTTG

AAGCATCACACATGAGAGTGCATGGAGAAAGTATACTGGAAGATTCTCTTGAATTTACCACCTCTCACCT

TAATTCATATCTGGATTCCAACCCGAATGTTCCACTAGCAGACCTTGTTCGTCGTTCATTGAAGTACCCC

CTGCGTAAGAGTTTTAATAGGATGGTGGCAAGGTTTTACATTTCCATCTACCAAAAACTTGAGTGGCATA

AGCAAGTGCTGTTGGAATTGGCAAAATGTGATTTTAACCTAGTGCAAAAGGTTCATCAAAATGAGCTAGG

TTATATCACAAGGTGGTGGAAAGATTTAGATTTCAAAAACAGAACTCCTTTTGCAAGAGATAGGGTAGTA

GAATGCTACTTCTGGATAACAGGGGTGTACTTTGAGCCCCGGTATGCAGCAGCCCGAAAATTTCTAACCA

AAACTATCTCCTTAACATCCATAATTGATGACATATATGATGTGTATGGAACCCCAGAAGAGCTTGTGCA

ACTCACTGATGCGATTGATAAATGGGATATGAGTATCTTAGATGAGCTGCCGGAGTACATGAGATACGCT

TATAAGCCTCTATTGGATGTTTTTGCTGAAGCTGAAAAGGAGATAGCAAAGGAAGGGCTGCCGACGTTTG

GAGTCGATTATGCTAAAGATGCATTCAAGAAGTTGACAGTAACATATCTCCACGAAGCTAAATGGCTCCA

GACCCAGTACTTCCCATCGTTTGAGGAGTACATGAGAGTTGCCCTTGTTTCTGGGGCGGTCAAAATGCTC

TCAGTCGCAAGCTTCGTGCGAATGGGTGACATTGCAACCAGAGAAGCATTCGAGTGGCTGTCTAAAGATC

CCTTAGTTGTCAATGGTTTATCCGTGATTTGCAGACTCAGCGACGACATTGTGGGACATGAGTTCGAGAA

TCAGAGACCCCATATACCATCAGCTGTGGAGTGCTACATGAAATCACGTGGTGTTACAAAGGAAACTGCT

TATGCTGAATTACGGAAACCCATCATAGAAGCATGGAAAGACATGAACGAGGAATGCCTCCTTCCAGAAG

CCCCTCCTAAGCCGTTGCTCGAAAGGGTCTTCAATCTGGCTCGTGTCATAAACTTTCTTTATGACGGCCA

TGATGGATACACACATTCCTCAACCAGGACAAAAGATATGATAACCTCGGTTCTCATAAATCCCATTCCA

GCATAAGCCTCCAACATTATTAATTATAAATCCCATTCCAAAAGATATATATGTAATATGTGGCTTTTGG

TACTGTTCGTTGTTTCAATAATCTATAAGACCTGTGAGCTGGGCGGCAGGAGCTTTTCTGTATGTACTAC

TCCTAATTAATGAATAAGACCTGTAAGGTTGGAGCTATGGTCTCTGTAATAAGCATGCATGTAGCAGTAC

TGTTGTACTCTTTCATATTATAATATTATTAATTACTCCTTCCGTTTCAAA

>MSTRG.1616.1 gene=MSTRG.1616

AATAATTGTATATTTGATTTTAGGGGTAATTTTACCACTGATAATAATCAGATAAAGACCCAGTTTTTCG

ACGTTTTTACGTATTGAGTCTATATCTAAGAAGATAAAGAAGGGACGTTTATATTGGTATGGAGATGTTC

GACGCAAATGTAACTCAGCACCGGTTAGGAGAGTCGAACACATATCGGTTCGGGGCAAGAGAAAGAAAGT

CGGCCTTGTCGGATATGAGCTGATCAATTAAGTTTAGACATGGAAGCCCTGAATCTCACATGTGATATGA

CGTTAGATAAAAATGCTTGGAGACAGCGTATTAGAGTAGTTGAGACTCGATCTCGTGGTTTAAAGAGATT

ATGAGTATTATGGCGTTATGCTTTAAGGATAAGGTTCAGCTCAGACATTGATAGATGCATTAGCGACAGG

TGGTTTGGAATATTGTGGAGCTCTGTTTTGAAGACAAAGCTCGGATCATTCATTGCCAGCTATATTCACG

ATAAAGGTAGGTCGCATGTGTCAGAAATCGTGGGCCAATGTACTATTTACTTAATTATATCTCACATAAT

ACTTTGTATTGCACTAAACTGGGTTCTTCACAGAAATAGCCTCCCTACTCTTCGGCGTAGGGACATCGTC

TGCGTATATCTTACTCTCCTCAACCCTGCTCTAAGTGAGATATACAAGATATGTTTTGTTTGGTTTGGTT

TATAAGACCCAGCTTGTGTACATAGCCGATTTATAATTTCTAAAAGAATCGTCACACGATAATAAAATTG

TGACACATAAATAATAAATGGGCCGTGTGCTTTTAAGTGTC

>MSTRG.1618.1 gene=MSTRG.1618

CGGAGTTGAAATTGAAGAACAAGAGCCATGCTTTGATCAAAACCCACTGGATTTGGAACTACCCAGATAT

TTCCTTGCATCATCTGGACTAGTTTCCTGCAAATGCCAAAGCTCAAGCCTTCCTCAATTCTCTTGTTCCA

ATATCTTTCCTCACCAGAACTATTCATCAATGCATCATTATTCTTGATACCAATCTCAAATTTTACATGA

ACCTGCCCATCGAACGAATTTGTTCTCCATGTTCCCCATCGTTGATCATTTCTTCCCTGGCTTCCCCTCT

CTGAGAATACCCTTATGGTTAGACGCCCTTGTTTGTTGGTACCCTTTAGGAGATTTCCTACCATATGTAA

AATAACCTGGAAAACTCTCCTCTCATCCCCCATAACATAATCTGG

>MSTRG.1619.1 gene=MSTRG.1619

AGAGTGGCCTCCCTCTCCTCAACTCAGTCCTAAATCCTCTCAAGGATAAGATGAAAAGCAAGATGGATGA

TGTTCAATCCCCTGCTCAGAGGGTGTTTGAGGAAAGCCCTAAACCAGTAGAGAAAAGTGAGGAGAAAAAA

CCACCTGGGTCAGAAGCTAAAAGCGAGGAGAGTAAGCCGAAATCTTGGGCTAATGTGGTTAAAAAAGATG

TTCCTGAGCTGAAATTTAACTATTATCCGAAAGCTAAAGGTGTTTCACTCCTTGAACCTCCGGATGAGGT

TTTGAAGAAAGGTAATGACAAATATAAATTTTGTGTTGTGGGCACTTTCTCGAAAGGTACTTTGGCATAT

AAATTGGTTTCGGAGTTTGCTTTTAAGTTCTGGAAATCATGGGGTCTGTTGTCAGTGCACCAGAAGGACA

AATCCACGTTTATGTTTCGCTTTTCTGACCTTATTGGTGTCAACAATCTATTATCTCGGGGAACCTGGTA

TATCGAGCGTAGGCCAATGATCATGACAGCCTGGGGACAAAAACCAGGGAGTTCGATCATCACTAGCATG

CCGCTTTGGGTCAAGTTCTATAATCTTCCTGATAGTTACTGGACAGAGGATGGTCTTGCTCACGTTGCTA

GTGTAATTGGCAAACCGATTGGCGCGGACCAACTCACTGCAAAATTGGAAATTCTCCCATTCGCCAAAAT

GCAAGTTAACTACACATTAGGTGACCCCTTACCTAATGACATCAATGTCACTGTTATTGACCCGGTAACT

GAAGAACGCTCTATTGCTAAAGTCTTGGTCTCTTACCCGGTAAGACCACTTTTTTGTACTGGATGTAACT

CTTTGGGCCATTCGCCCTCTGCTTGTCCGAAAATTAAAAGGGTGTGGAAGCAAAAGGATGTGAGTTCTGC

GGAGAGTCCTCCTGGTGATGCCCAGAACCCGCATACATCTTATACAGCTGCTGAAGTTAATAATCCAAAT

GGGGATAGCCCTGCTGAGCACTCTCATGCTACCCCGTTTAAGAGTTCCAATCCTGACGATGATTGGACAG

AAGTTAAAAGGAAAAAGCCACAATCACCTATGGAATCTGAGTATTCCCCTACACCTCCTGTCAAATTCAA

AAATCTCAAGAAAATTGATGAAATTGAGGGTAAAAAGGCTCATCCAGTACCACCGGGCAGCCCCAAGCGT

CTAACGAAGTCTCAAAAGAAGAAGCTGAAAACTTCAATGGGGAGCAGTCCCCAAGCCCAAGCCTAATATT

TATGATAATCGATTTTTGTTCCTACAATATTCGAGGGTTGCATAATAAAGTATCTTTTGTGAAGGATTTT

TTGAGCGTCAACAAGATCTCCATGGCAGCCCTCCTGGAAACTCACGTTAAAAGGGAAGACGCATTGTCCA

TTTCTTCTATTATTGCTCCGAGATTTAAATGGATTTTTAATTACGATTATCATGATAATGGCAGAATTTG

GTTGGGCTGGGATGAGAATATTTGGAATGTGAATTTTGTATGCTCCTCGGCTCAGCACATTACTTGTGAT

GTTAAACGTATTGATGGGTCTTGCTCTTGTCTCATATCTTTTGTATATGCTTCGAATCAAGTGTATACAC

GTAAATTACTATGGTCAGATCTTATTAATATGCAGGACAGGCTCTCGATTACTGAGCAGCAGAGGCCTTG

GTGCCTCCTTGGAGATTTTAACTCCTTTCTTTTTGATTTTGAAACTAACAGGAC

>MSTRG.1620.1 gene=MSTRG.1620

CAAGTTTCTTTTGGCAGGTTAAAACAAATGGCATTTACAAGATTGATCCGGTCAAGATTGATCCAGTCAG

TAGTTTTTCAAAGGCAGCTCCATGTCATCTCAAGATCATCCACCAACATCAAACCAGCTTCTCCAACTCC

CTCGAAGCTCAACCGATACAACATTCCTTTATATGATCGCATGGCTGCAGATGCTCATATGCCGTTGCTT

TTATTGTATCCCAGCTACAACTCTGATCATAGTCTCAAACTGATTCCAGATGATGCATCATTGTCAGATC

TTCTGAAGAAGTCTTTGTCAGAGACACTGTCTAAGTATTACCCGTTTGCTGGAAGGCTACGCTCTGGATC

CTACGTTGAGTGCAACGATGAAGGGGTTCATTTTGTTGAAGCTGAAATTGGATGCAAGTTGTCAGAAGTT

CTGGAAAAAGCTCCTGTTAAGGAAGAAGAAGAAGGCTTAGGCCATCTCTTTCCGGCTTGCACAATTTGGA

ATCATTCATCTAAGATGTATCCTGGTGTCGTAATGCACGTTCAGCTTAGCCATTTTACTTGTGGAGGAGT

AGCCATTGCAGTTACCCTTCATCACCATCTCGGAGATGCTCTTACTCTGTGCTCATTTCTAAGGTACTGG

GCAAATTTGTCACTCCACTCCGGTGACCACAAGAAGTTATTACATCTCTGTCCGCGTTTGGTGAATGAGC

TGCTGCCTCCCTCTGATGATGATTCAATCAAAAATATTTCATATCCTGATAAGAATTGGACCACTAAGGA

GGTGGTATTTCCCAACAGAAAGCTAGCAGAACTGAAGGTGGTTGTAGGAAATGAAGACAAGCTCGATGGC

ATTGCAGAAGATCAAAAATACACCCGCAATGAGCTCCTGACTGCACTCCTTTACAGATGTCTTGTAGCAG

CGGTAGCTGAGACTAATACAGGAGCGGACAAAGGATCAGTTCTGATTAGAGGTGTAAACGTAAGGCCTAT

GCTTGATCCGCCATTGCCAGAAACAAGTGTGGGGAATATTGTAACTGTAAACTACATTCCAACAAGCACA

GAGAGTGAGACCAAGTACAGAACGCTGGTGGCGCGGATGAGGGAAGAGAAGAGGCGGCTCAGAGGAATCA

AAAATTTAGATGGACATGAACTGGGACCAAAGTTAGTTGAGTTCAGTAAAAATAACTACAGGTTCTTCAT

TATCAGCAGCATATGTAATTTTCCAATATATGAGGCGACGGATTTCGGGTGGGGAAAGCCGATCAAAGCC

ATTCTTGTTGACACGGGGATGGTCGATATCATTTCTTTGATGGATACTGCGAATGACGAAATTAGAGCTG

TTGTAGGATTGGGAGAGCAGGATATGAAAAATTTTCTAGCACAGAAAGAGCTGCTTACTTATGGTTCTTT

TAAATAGTTTGTTGATGCCCCATTACTCTCAAAATAAGAAAGGAGCTGTTTGCTTATGCTTCTTATCATA

TCCAACTCTGTTTCTGAAGTGTTTAC

>MSTRG.1630.1 gene=MSTRG.1630

CCCTAACCCCACTTTTCTTCTAAAATGGCCGCAAACAACATAATTTAGGAGTTTGGGAAGTTGCTTAATT

TCTGATGGATTGAATGGGGGAAGTGAGTTGGGGTCGTAGGGTGTTGACAACAGGAGGTGGCCGGAGTAGC

GAAACGCCGTCACGGCTGTGATGTCCGGCGGCGGAGGTGATTTCGCTGCTGTTGCAGTGGCGCCGGAGCA

AGTATGGAGAGGAAAGGTTTTTCCTGGGGGCTGTGTAAATTGAATAGTGTATTGTTTTTGTGTGTAGGCA

GCGAGGGAGAGAGACGGTGAGAACTGGAGAGAGGGAGAGGCGAGGATGGAGAGAAACTAGAGAGAAGGAA

GCGATGAACGAAGGAGAAGAAAGAGAGAAGTGAGGGTGGAGGTCGCCAGATTTTGGCCGGAAAATGGGGT

AGAGGCCGAGTTTATTAAATCCGTCTTGGCGGATGTACAGATCGGGCCTCTACGAGTCCAGCCATGGTGG

TTATGCAAGCATTGGGTGTCGCCGGTATTTGACTGGACGATGGTGGCCACCGGCGGCGGTCATTGAAGGT

GAAGCTGAGTGAGGAAGATGGTGAAGGAAGGAGATGAGG

>MSTRG.1631.1 gene=MSTRG.1631

CGCTACGTTGCCTCTTTCAAAATCAAAACTAGGGTTAGAGGCTTAGAGCTGCGGCGCCCCCAAACCCCAT

TATCACACCAAGAATCCAAGATCTGGAGTATCACCGAGAATCACCGAATCAGAATTAAATCTATTTGCTG

TCATGTCGACTTCACAATCGGCTCCTAACTCTCAGACGGCTCCTAATTCTCAGTCCATTCCAGAACAACA

ATCTGAACCATCTTCAACACTGGAGCTACAATCAAATTATTAATTTAATTTTGTTTTTTCGGATTTTCGG

GTATCGGATTTCGGGTTTCGGGTTTTGGTTATCGGATTTCAGTTATATCCAAATCAAATCCAAATAAAAT

CGGGTTTCGGGTATACCCGAATCCGATCCGAAATCCGTTGGATCGGATTTGGATATCAGTTTTCGGGTAA

TTTTCGGGTTCGGATCGGGTTCGGGTACAGTATAAAATTTCGGGTTTGGATATGGATATAGTCATATCCG

ATCCAATCCGACCCGATTGCCATCCCTATTGATGGGTGTATTCATTCATCACCATATATTTATGGACACA

GAATCTGTGTACATATGTACAGATATACAGGTATTATTTAATGGTGGTGACAGATGAATGTAGCTAAGAG

GTTCTTGAATAAAGAGAATTAATGGGGTTTGGAATTGTTTAGTGGGTGAAGAGTGGTCAGAAGCACATGC

ATGTCGATCCCCATATGCTTGTGTTGGGTAGGACTTGAGACTAGTTGGGAGTAGAAACATTATATTATGA

GACAGATGATTCACAATCTCACATGCTCATTCATCTTTTTTGAAACAGCACTGGCTCCATCTCCAGCACC

CAGTGACACATTTATGAAATATGTAGGTCATGTATATGCAAAATCTACAGCTCTTCTTTTTAATTTTAGA

ACAGTGATTTAAGAATTTATGAAATATGTAGGTCATGTATATGCAAAATCTACAGCTCTTCTTTTTAATT

TTAGAACAGTG

>MSTRG.1639.1 gene=MSTRG.1639

CATCTTCTCACACCCTATCATGCATGCATTCTTATATCCCAACTCAAATCCCTAAGCAATAATAGTGCAA

CACTACTGTATTTAAACTTTTTAGCATTTCTAACGCTATATTTTCCTAGCCACGCTATAAATATCCATGC

ATGACCTCTGCTAAAATCACCATATAATTAACCCCTTCCTCTGTTCTGGTTATTTTTGTTGTCAAACGCC

TCATCACAGATCCACAGGCTTTGAAAGATGAGTTCAAAGCCGAGTCTTTCACAAGTTCTTTTCATATTTC

TTAACCTTCTCTTTCTTGCACTAGTAGTCAATTGCACTCGCAATTGTAACAATCCTCCAGACCATAAACA

TAAACATAAACATAAGCATCATAAACACAAGTCAAGCTCGCCTTCCTCTCCTACTCCCCCTGCCCCAGAG

AGTCCTAAGCTGCCCCCTGTTTTTCCCGGGACTCCGAGACTACCTCCAGCCTTTCCACTGCCTTCAAGTC

CTCCATATGCATTCCCGGGGACTCCAAGACTCCCAGGCTTTCCCCTGCCTTCAAGCCCCCCTTTCGCCTT

TCCCCCGGGGACTCCTCCAGTCCCAGGCTTTCCCCTGCCTACAAGCCCCCCTTTCGCGTTCCCTCCCCGG

ACTCCACGAGTCCCAGGCTTTCCACTGCCCCGAATAAGCCCCCCATTACCAGGCTCCCCAAAAACAGAGA

CTTGCCCGAGAGATGCATTCAGATTCGGGGCATGCGCCAAGGTGCTTGGTGGCCTAGTGGGCGGTGAAGT

GGGAAAAGCTCCGAAACAGCCATGCTGTAGTCTCTTTGGAGGCCTTGTGGAGGCAGAGGCAGCTGTTTGT

CTCTGCACTGCAATAAAAGCCAACTTTCTGGGATTCAATCTCAATATTCCAGTCGCAATCAGCATGGTTC

TTAACGTTTGTGAAATGAGGACTCCACCGAATTTCCAATGCTCATAAAAATAAATCTTAATTTCCTTTCC

TTTAATTTCTATTACGTGGCTTTTGCCTCGTGTGCATTATATTTGCAGCTGCCATTATATTTGCATTGCA

ATTGTATCATCTTCAGCTTGTGGATTACTGTAATCATTGCATCATCATTTGTTGAAATAATAATTACTAA

GTTTCCTTTATGGAGAAAATAGAAAA

>MSTRG.1647.1 gene=MSTRG.1647

CTCTTCAAAAAAGTTTTTTATCTTCTCCTCATAATTTGGTAACTTTTCTGGGCACACCTCACAGAAGTCC

GTGTAAGAGTATCCACGTGATTCACGAATTTCCTTCAACTTCTCATCTGTTTCATAATTATCTGCATCTA

GTCGCCAGCTAAGCACCCCAAGTTCGGCAAGTTTCTCTAGCGAGACAAATTCCTTTGGATTTCGGTGATG

GGGTAATCGTTGGTCTTCATCACTATCATCCATGTACCATGCCTGTATAACTTCCTCCCTCTCATCCTTG

CTGATAGACCCCATTTGCGCCTTGTATCAAAAAGTTTTATTAAAGATTTAAGTTCTC

>MSTRG.1655.1 gene=MSTRG.1655

AGTCAATGAGTGATGGTGGCTTTTGTTTTTTTTTGATAATTCACTCCACAACTTGTATGATTTCAACTTC

ATACTATCACACACAGAACAATTCTTGAAATTGCCTAATAACATAATCATCAGGAAACATAAGCCTAGAA

CTTTAAATGACTTTTACACGAGAAATCCTTAAGACCTTGGATTTACAAGGGGTTTCTTGAAGGACTTTTA

CACTAGAAATCCTTAAAACCTTGAATCTACACCGACTACACGGGTTTTCTTGAACTCTGAGTGATCTCAC

TAGAAAGAATACTAAAAATCACCATATCTCTAGTCCTGCCCTTCAAAACAACATACTTCCTCAACACCCC

TTCCCTCATAAACCCCGCCTTCTCCATCACCTTCTGCGATCCCACGTTGTCCACATCAACCAGAGCCTCC

AGCCTCTCCAAATGAGGCCACTCAGCAAATACCATCTGCACCACCATCTTGACAGCCCTGGTCACAATCC

CTCTGCCCCAGTAGCTGGAAGCTAAAACATACCCAAGCTCCCCTCTACAAACATCATTCCCTTGGAATGG

AGTCACGGAAATGGATCCTATGGCGCGGTTGTTGACACATATGGCTCTCCACCAGGGATGGGGAGCCACA

GTGTTGGCCATGTAGTCTATGGCTTCTTGTCTTGAGGTGTAAGTTGACCATGAACAGAATTTGCTCACCT

TATCATCTGTAGCCCATTTCATCATATCATCTATATCATTCTCGTCCTGCAGTCTCAGACTTATGTCTGA

ACAGTCCTCATTTTCTGCTAGTGTTTCTTTTAAGTTCAGCTGAAGCTCCTCCATGAGGAAGTTGATGTTT

TTGTTGAGGATATTTTGTGTATGTTTTCTTATGAGGATTATGCTATCAGTGAGGCCAAGATATAGGGCCA

AAGTAATTGCCAGGAAAGTTTATACTACTTGATTAATGATTAGTGATTTATTTGCTATGAAAGCTCTTTT

TATGTCC

>MSTRG.1657.2 gene=MSTRG.1657

CCCGCAACCCACTCTCCACAGAACAGTGGTTACAAACTAAACATCTATCCATTAACACGAGGATCATCGG

AGCACAATTCTTGCAAAACGAGTCGGCGGAAATCTCTACAAAGTTCTTCGCTTGCAGCACCAACACCTTC

ATCCAAGATCAGCAGTAGAGATGTTCAGAGGTAGTAAGTGTTAGAGTCCAGCTTCAAAAAGACGAGTTGT

GGATCATAGCAGTTGTTTAGCAGGGGTTGAGGGCATTAGAGGTGAATGGAAACTTGCAGGAATCGGGGGC

AATAATGCCTCAAGCGATGAAGACAATGATTTTGTAATCCTGCAGAATACTTCTCGAAAAGCAATAATAT

AATCCACAAGAGAAATGGTAGTGCAGTAGAGGAAAAAGACGGTTCGAAACACATTGTTCAGAGTGGGACG

AAAAGATGAGCAGCAATTTAAAAGCACCAAGAGAACAAGGGAGAACGAGGTACAAAGGCAAGGTCAAGAA

GACGATAACATATGAAAACTACATACAAAACAAGTTTTCCCCTTCAATAATGTCTGATGTTCTTGGAAAT

TTGCCGATGAACAAAAAAATTGGGTGACAGCAACCGAGTTTGGTGGTCTTTTGAGTTTTAGGATGGTGTG

GTACACACACTCATTGGGGTACAATGCAGGATTTTGTTCTTTTGTTCTGCAGATTTTGTAGATTCTACCA

TCTTTTAATTGCTTCATCATTGTCTACTTTAACATTATTATATTCTCA

>MSTRG.1657.1 gene=MSTRG.1657

GTTGTCTTAACGCATTATACGGAATCATAATCCCGCAACCCACTCTCCACAGAACAGTGGTTACAAACTA

AACATCTATCCATTAACACGAGGATCATCGGAGCACAATTCTTGCAAAACGAGTCGGCGGAAATCTCTAC

AAAGTTCTTCGCTTGCAGCACCAACACCTTCATCCAAGATCAGCAGTAGAGATGTTCAGAGGTATGTTGC

AGTACTCTGTATTACTCTAATTGAAAAATCGTCTATGAATTAGTGAATAATCGTGGGTTGATTATGAGAA

GGTAGTAAGTGTTAGAGTCCAGCTTCAAAAAGACGAGTTGTGGATCATAGCAGTTGTTTAGCAGGGGTTG

AGGGCATTAGAGGTGAATGGAAACTTGCAGGAATCGGGGGCAATAATGCCTCAAGCGATGAAGACAATGA

TTTTGTAATCCTGCAGAATACTTCTCGAAAAGCAATAATATAATCCACAAGAGAAATGGTAGTGCAGTAG

AGGAAAAAGACGGTTCGAAACACATTGTTCAGAGTGGGACGAAAAGATGAGCAGCAATTTAAAAGCACCA

AGAGAACAAGGGAGAACGAGGTACAAAGGCAAGGTCAAGAAGACGATAACATATGAAAACTACATACAAA

ACAAGTTTTCCCCTTCAATAATGTCTGATGTTCTTGGAAATTTGCCGATGAACAAAAAAATTGGGTGACA

GCAACCGAGTTTGGTGGTCTTTTGAGTTTTAGGATGGTGTGGTACACACACTCATTGGGGTACAATGCAG

GATTTTGTTCTTTTGTTCTGCAGATTTTGTAGATTCTACCATCTTTTAATTGCTTCATCATTGTCTACTT

TAACATTATTATATTCTCA

>MSTRG.1663.1 gene=MSTRG.1663

CACCAGTAGTACTCACCACTTTACTAAAAGCAGGCCATTCAATCTCCCCAATCTCTTCACCCACCCCTTT

CCAAAAATCATCACTTTCTTGTTTACTCTCGTCTCTCTCTTTCATGGCCTTCTTGATTTCTGCTCCTATT

TCAGAAAGTTGGTCACTTTCTGGTTCTTGGGGTGCGGGTGTTTCTGTTTCTGAAGTGGCTTCTTGAGACA

CGCTGCTGCTGTTGGGGTCTTGAATGGATTTGATTATGGGGAAGTGTGATTTTTTGAGGGGTTTTGAAGT

TGAA

>MSTRG.1669.1 gene=MSTRG.1669

GAATATGAGTAACTATCTAAGGACCATAGTCCTAACTGTTGAGTAATTTTACTTTTAATAAAAAGCAAGG

GGGGCACGCCTCTACTGAGGAAGGTGCTGAACCACTTGCTCATGCAATCATCATCTCTCCAGTATATATA

AACACTAGCCTCTGACCACATTCTTGCATTGCCTTCTCATATTTCCACACACAAATCAGAAGAAAGGTGT

GATGGGTCTGGAGAGAGCTGATCTGGGTGTGTTGGTGATGGTGGTGATATGTTGTTGCATGCAAATAGCA

GGAGGTACAGTTTACAAGGTGGGGGACTCAGCTGGCTGGACAACCATTGGGAAAGTTGATTACAAGCAGT

GGGCTGCTACTAAAACCTTCAAACTTAATGATGTTATCTTGTTTGAGTACAGCTCTCAGTTCCACAATGT

GATGCAAGTGAAACATGTACACTACAAGTCATGCAATGCATCTTTCCCAATTGCCACTCACACATCAGGC

AATGACTCCATCACCATCACCAAGCACGGCCACCATTTCTTTCTCTGCGGGGTTCCTGGTCACTGCCAAG

CTGGTCAGAAAGTCGACATCAACGTGCTTCGCCTTTCTGATTCACTTCCTCCCTCCTCCCCTCCTGCTGC

ATTGCCCACTTCTTCACCCATTGTGCCTGCACCTTCTCCGAGCAAGGCAGCTCCATTCTCTGTTTTGAGC

TGGTTTTCGTGTTACCTTTGGTTGGTTATGCCTGTTATTGTAGCCTTGTTTTGAGATTTTGATCAATCAG

GAAGATATGTATTTACTCTCTGGGGAGTCTTTGTAAACTGAACCAATATTTGATGGAATTAATGTCATAG

GGTCTTAATAATACTGCTTGATTTTGTATTTGCTACAAAGTTTTTCATTTCTATGATAGAATTAATGTTG

CATTCTTT

>MSTRG.1672.1 gene=MSTRG.1672

CTTATTTGTCATTGATATTCAAGAAAACTTTTATCTAGATTCTAACAGAAATGGTTTCTAGCAACAACAA

AGAGGTCGTCTCGGAGCTCTCACCCATCCTCAGGGTCTACAAAGACGGCACGGTGGAACGGTTCTTCAAT

TCACCGCACGCTCCGCCATCACCCCTAGGCAACACTACTGGCGTTCTGTCGAAAGATATCGCCATAACAC

CCGATATCTCAGCTCGATTATACCTCCCTAAACTCGCATCTTCAGCCAACGAAAAACTCCCGATTTTAGT

CTACTTTCACGGAGGAGGCTTTTGCATCGAATCGCCTTTTTCGATACTCTGTCACCGATACATAAATTCC

ATAGTTTCGCAATCAAATGTTATCGCGGTCTCGGTTGATTACAGGTTAGCTCCAGAGCATCCTTTGCCTA

TAGCTTATCAAGACAGCTGGGCTGCTCTGCAATGGGTCGCATCTCATAATACCACCACCAAGACCACCAA

AAAGGATCCTTGGTTGATTCGTCACGGTGATTTTAATCGAGTTTATTTAGGCGGCGATAGCTCGGGTGCT

AATATAGCTCATAACATGGCCATGCAGGCCGGTATCGAAACATTGCATGGTGACATGAAGATTTACGGTT

TAGTTCTGTGTCAGCCTTATTTCTGCGGTTCAAAATTATTTGGAAACGAGAGTTTGGTCGAAAAAAGATT

ATTACACGAGACATGGATGTTTGTGTATCCTGATGCGAAGGGAGGGATTGATAATCCCATGATTAATCCG

TTTGCTTATGGTTCTCCGAGGTTGTCTCAGATTCAATGTCACAGGTTGTTTGTCTGTGTGGCGTCGAAGG

ATGAACTGAGGGAAAGAAGAGTTTACTATTGTAATGCAGTGAAAGAAAGTGGATGGAAAGGGGAAGTGGA

CTTGTATGAAGTGGAAGGTGAGGGCCATGCCTTTCATTTTTTTGACTTTGACGGCGAGAAAGCACAAGAC

ATGGTCAAACGTTTGGCGGCTTTTCTTTGTTAGTTTTGGTCATTCTATGTTTTGTGATTTTGTGTTTATC

TTCTTTTGAAGGGATTTTGTAATTTTGTGTTTATGTGAATGCTTAGTTGATGTATTGATGATTTGGACTG

CAAGAATGAAGT

>MSTRG.1673.1 gene=MSTRG.1673

ACGAGCACCACGACCTCCGAACTTCTTCGGCTCGCACCTTCTGGGATCAGCAACAAGCAAAGTCCTATCA

TACCTCACCAAAATATCCTTAATCTCCTTCTTCGACTCTTCGTCGACGAACTTCTGGTAATACGCAACCA

GAGCCTTCGCAATTGACTGACGAATAGCGTAGATCTGCGAGGTGTGACCGCCTCCTTTGACGCGGATCCT

CATGTCGACGCCGGCGAAGCGGTGGCGACCGAGGAGGAGGATTGGCTCGAAGGCCTTGTAACGGAGGATC

TCCGGCTGAACGAGCTCGATTGGAACGCCGTTGATTTTGATCAAGCCGCGGCCGCGCTTGCAGTGAGTTA

CGGCGACGGCGGTCTTCTTCCTGCCGAAGCATTGAACTGATTCCGGTGGTGTTGCCGCCATGATTCTCTC

TCTGTATCTCTCTCCCTCTGTGTCTCTCCCTCTCTCTCTGGAATGCGGAG

>MSTRG.1678.1 gene=MSTRG.1678

GTTTACAGATCAGTTGTGGCTTTCATTGCCCGATGTGGACATCTTACAAGCTGTATAGCAGAGGAAGGAT

CCGAATTGGATGCCAAATAAGTGGATTAGAAGCTTTGGAGAAACTGAAGATTAGTAGTATGGAGATGAAG

ATTAGTACTATGGAGATGAAGTTGAAGAACAAAGGAACTGGAATTTTTTTCTAGAGTTGTAATCAAGTTG

TGTTGCTTGTTCTAGTTGTAATCTTCATTTGTGATTTAAAGTTTATTGTAATCAGAATTTGTGAATTAAA

ATGATTGCATTTTTTTTATCTTAATTGGC

>MSTRG.1679.1 gene=MSTRG.1679

GTTTGATCAACTGAAACTGAGAGCCCTTAAAGGACAGCAGACCCGCACCCAAGATGCAAGGTTATGGTGC

AATCTGCTGAATCTGATGACGAAGAAACAGATGAGGGTAAAAAGAAACCCAAGAAAACATTTTTTGGTGC

ACTTAATACCATATGGAAAACAGAAGGCCTACTAGGATTTTTTAAAGGTCTGGATGCCCAGATCCTGAAG

ACAGTGCTGAGCTCTGCATTGCTCCTAATGATCAAGGAGAAGATAACGAAAACCACATGGGTCTTGCTAC

TAGCAGTCCGGAGGACTTTGTTCTTAACCCGGACCAAGTTGAAGAGCTCGTAACATGAAATGTACAAGTA

TAGCTCATGGATTTACTTATACATAAGGATAACCTACCATAATGTCGGGGAGTTCCTTTTGGTGATGTTT

CAATTATTTTGTGAAGGATGGTAGAAGATATTCTTTGTATCATTTATAAACTTCTTACCATATATGTTTT

CCCCTTCACAGGGTTGATTTCATTCTTCCCTCCCTGTTGAAGTTTTCGCAGGGTCTAATAAATGACATAA

ATTTGTTCCCTGGCTGGTTCTTTAATCATTTTCAAGCTCTAGCTTTGATTCATTACTCATTGTGAAGCCG

TATCGATTTACTATATATGCAATTACGCATATGATTTGTGC

>MSTRG.1680.1 gene=MSTRG.1680

TATTAAGAGTATTTACTTCTGTCTATGACAAGAAGTTGGTTCTGGTTTTCTTCTTCTTGTCATTGTAATT

TTTCTAGCAGTCATTTGTGATTTTGATTGGGAAATTTAATATTCATCTCTCAGGCAATAGAAATGCAAAG

AGATCAAGATACAGTTTTAGTGTACTTTCAGTTAATAGTCATAAAGTACAAGCTGCCGGTAGTCAGACCT

TCACACCAGTATATCAACCGGAGCCACCAACACCTAGGAAAGAGGAAGAAGCTCTGAAATCACCAAAATA

TAAGATTTTTACATTCAATGATCTTGCAGTAGCTACCAGAAAATTCCATCATCACAGTCTTTTAGGGGAA

GGTGGTTTTGGTTGTGTTTATAAGGGTTGGATTGATCAAAACACACTTGCTGCTACGAAATGGGGTACTG

GTCTGGCTATTGCAGTAAAGAGATTAAATGTAGAAGGATTTCAAGGTTGCGCGGAGTGGATGACAGAAAT

CAATCATCTGGATACACTTTGTCATCCTAATATTATAAAATTGATTGGGTACTGCTCCGAAGAGGAGCAT

CGGCTTCTAGTATATGAGTTCATGCCTCAGGGCAGCGTGGAAAAGCTTCTATTTAGGAGGGATTCAAACT

TTAAACCGCTCACTTGGAAGCTTCGTATGAGTATTGCACTTGGTGTAGCAAAAGGGCTTGCATATCTTCA

CAGTTCAGAAGCAAGAGTGATCTACCGCGATCTCAAAGCATCCAATATACTGATTGATTCTAATTACAAT

GCAAAGATCTCTGATTCTGAGTTGGCAACAAACGGTCCAGATGCTGGAGAAAGCCATGTCTCCACCAGAG

TTATGGGAACATACGGTTATGCAGCTCCGGAATATATAACTACAGGTCATCTGTATTTGAAAAGCGATGT

CTTCAGCTTTGGAGTTCTTCTCCTAGAACTTTTGGCAGGACGAAGAGTAATTGATTTGACCCTTCCAACA

ACAGAACATAACCTTGTTAATTGGGCTAAGCCTTGCCTTACCAGCAAGCGTAAGATTTTGCATGTCATAG

ATGCAAGCATGGAAGGCCAATATACAGTTGGAGCAGCTTTAGCAGCTTCTTTACTCGCACTTCAATGCCT

CTCAGAGGATCACAACTTAAGACCTGATGCGAACCAGGTTGTAAAAGCACTAGAGCAGCTTCAGGAATTT

GTGAACTCAGGATAATATCACAAGTGCAGCCCTTGACAAGGTCTAGAAGCATCCCTGTTCTTGAACTAGA

AGCTCACATACAGGTAAAAACACTTCAATCATCGACATCAGGACTTGCAGACGGAGAGTTTATAGAACTG

AACGTGTACCAGGGGTACTTCACTTTAAATTTGAAATTGCTCGTAAATTTTATGGTTGGATGGCTACATG

GTGTGTAAAAATTACAGTACTTGCTGAATAAATCTATAACATAATGCAATATATGAAAATGAAA

>MSTRG.1681.1 gene=MSTRG.1681

GCTCTCTGATTTTGGGATTGCAAAAGACGGACCAGATCACGGAAATACCCACGTGTCCACAATGATTTTG

GGAACACGCGGTTATGCAGCTCCAGAATATGTGGCTACAGGTCATCTAACTTATAAAAGCGATATCTATG

GCTTTGGAGTGGTTCTCCTGGAAATTATAACTGGACGACGCATAGTTGACAGAAAACGTCCTCCAGAAGA

ACGTAGCCTTGTAGATTGGGCTAAGCCTTACCTGACCAGCAAGCACAAGGTTTTGCATGTCATGGATCCA

GACATCAAAGGCCAATATACTGTCAAAGAAGCATTCAGAGCCTCTTCCCTTGCTCTTAAATGCCTCTCAG

GAGCTACCAGATCAAGACCTGACGCAAACCAGTTGGTAAAAGAATTAGAGCAGCTCCAGGATGTGTAATT

TTCGGGGAACAATTGCAAGTGAATCCTTTCGAAACCCCCCCAAAGAAC

>MSTRG.1682.1 gene=MSTRG.1682

GTTCAACATCCAATATTGAGGTAGGGCCTCATTTTCAACTTTTGCTTAGGCCCCAAATTTGGTTGAGCCG

GCCCTGATTCCACTCATCTTGTTACTGAAAGCTAGTCTATCTTTTCTAAATATATTATTTTTGAATTTCA

CACTAATCTGTTGAAGAAGTTCTTGTATAACATTGATTTAACTCGGGTCCAATGACGGAGAAGTCGAATG

GGGTGGTGATTCTCATCACTCTATTGAATATCTTGCTCTATAAGCTCCAGAAACTCCAAGAAGCTCAACA

AAAAAGGCGCTTACCGTTGTCTGTTCGACAATTGTGTCGTCGTTTCTCATTTGCAGACATCAAGTCGGCA

ACAGAGAACTTTAAGGAAGATTTGGTGATTGGAAAAGGTGGATTTGGAATTGTGTATAAAGGTTTTTTGG

ACTTCAAAGAGAGGATTGTTGCTATAAAGAGATCGAAAGCCATGTCAAAGCAAGGATGCAAGGAGTTTCT

TACAGAGATTAAGATGCTTTCACAGTTTCAACACAGTCATTTAGTCTCTCTCATTGGTTATTGTGATGAC

TTTGAGGAGATGATTCTAGTTTATGATTTTATGCCTTCTGGGAGCCTTGCTGATCATCTGCATAAAAGGA

CCAGAAAGGGCAATTCTTCTTTGCCTTCTTTGTCGTGGGTGCAGCGCCTCAAGATCTGCATTGGGGCAGC

CCATGGACTAGACTACCTTCACACTGGTACAGGCATAGATGATAGAGTTATACACAGAGATGTCAAGACT

TCAAACATTCTTCTGGATGACAATTTGGCAGCTAAAATTTCAGACTTTGGATTGTCCAGAACAGGTCCAG

CAAATCAGACTTGTACATATGTAAGCACTCGGGTCAAAGGGACACACGGTTATCTAGATCCTTACTATGT

TGCAACGCACAGACTAACTAGGAAATCTGATGTCTATGCATTTGGTGTAGTGTTATTTGAGGTCTTGTGT

GGAAGACCAGCAGTTGATACAGGTCTGGACGAAGAGCAAATTGGTTTAGCGGGATGGGCACAACACTGCT

TCAAAGAGGGACTCTTGGAACAAATTATTGATCCCAGTATTAAGGCGGACATTTCTTCTGATTCCCAAAC

CACATTTGTTGACATTGCTATCAAGTGTTTACATAATCAACCCAAATTGCGTCCAACTATGGCTGAGGTT

GTGGTTGGTCTTGAGTCTGCACTTGCATTGCAAGAAAAGAGTACATATTATATCTTACAGGAGATAATGC

CTCCTGATTATGCAACCCAAGAAGATGAAGATTCTTCGACACTTGAGGTTAAAAATACTAATCACAAGCG

AGAACATGAATTTGCAGGAGGAGAAGTTGTAAATGTTTCTAAAGATTGGCATCACGATAAGCAGTCTTTT

CCTAGAATGAAATTCACAAAAAGATTAAGTGGTCTTCTTTCAGTTACAGCTCGGGTGTTCTCAGGGTATG

TGTCAGGCAATAGACCTGCAAAGACATCAAAATACAGGACTGGCATACTATCAGTTGTAAGAAACAAATT

TCGGTCCGTCAATACTAAGACCTTTGTGCCAGTATATCCAGCGATGCCACCACCACCTCGGAGAGAGACA

CAATTTCTGCAATCACTAGAGTTGAAGTGTTTTACTTTCAATGATCTTGAGAAAGCTACTAGGGCTTTCC

GTCCTGACAGTATCTTAGGAGAAGGTGGGTTTGGTCGTGTTTATAAGGGTTGGATTGATGAGAACACATT

TGCAGCTGCAAATCCCGGTTCTGGTCTGGTCGTTGCAGTAAAGAGATTAAATTCAGAAGGACTTCAAGGT

CACCGTGAGTGGTTGGCAGAAATCAATTATCTAGGGTCACTTTGTCATCCTAATCTTACGAAACTGATCG

GGTACTGCGTAGAAGATGACCACCGACTTATAGTATATGAGTTCATGCCTAAAGGCAGCTTGGAAAATCA

TCTATTTAGAAGGGGTCCTCACATACAACCGCTCTCTTGGAATATTCGTTTGAGGATAGCTCTTGATGCA

GCAAAGGGACTTGCATATCTTCACAGTCCAGAAGCAAGAGTGATATATCGTGATTTCAAGACATCTAATA

TATTGCTTGATTCTGAATACAATGCAAAGCTCTCAGATTTCGGTTTGGCAAAAGACCAACCACAACATGA

CACCCACGTGTCCACAAGAGTCATGGGAACATATGGTTATGCAGCTCCGGAATATGTGGCTACAGGTCAT

CTATCAACGAAAAGCGATGTATATGGCTTTGGAGTCGTTCTCCTGGAAATTTTGACAGGACGAAGAGTAA

TAAACAAGAACCTTCCACAACCTCAACATAACCTTGTTGATTGGGCTAAGCCTTATCTTACCAGCAAGGA

TAAGGTTTTGCATGTCATGGATGCGAAAATTAAAGGCCAATATACTGTTAGTGCAGCTTTAGGTGTTTCT

TCCCTCGCATTTAAATGCCTGTGTACAGAGCCTAGGTTCAGACCTGACGCAAACCAGGTGGTGATAGCAT

TAGAAGAGCAGATTCTGAATCTCAAACTTTCAAAGAGAAGGCCACGAAAAGGTTTATAAGCATGAGTACC

GTTCTTTCAAGACTTTTAAAAGCAGCTCTTTGGCTCTATAGGGATTGGGGAGCAAAGGGAGAAAAGGAGA

AACAGAAACTGTATTGAATTTTCAGATGAATACATCTGAATGTGAAGCTCTATATGTTACGTGTAAAAAT

CTGTCAACCGAACACGAAAACAGAGCAATAGACTACTTAATTATAGAGATTCATAGTGCAAGTATATGGA

CACTGTTCTTAGCTAAAGATTGTGGCAGAATATCCGGA

>MSTRG.1687.1 gene=MSTRG.1687

CTACAACTAAATAGACACGTCAGATAGGTAAAATTACAGTGCACCTCAGAGCAGCAGATTGGAAACTTCA

AGATTGAAGGGCATTCTTAAACACAGCTAAACTACACCTTTTCAACATCTGTTATAGTTTCTTTCTCATC

AAGTTCAACTAGAATTGAATGGATGCTGGGTTTTTCATCTGTAACTAGAAGGCAACAACACTTGATTAGG

TGTTTGGGAGGTTGTAATAACCACTATAGATACACAACAATTCTACATAATCACAATCAGCAGCAGAAAA

AAATGCAGTCTGTGACACCTATTTTAGAAGCTGATGCTGTCAGTATAATTAAGTCAATTACTCCAGCTCT

TGATCATTCTAGGCACAAGGGCCAAGCAGGAAAGATAGCTGTTGTAGGTGGATGCCGCGAATATACAGGT

GCCCCATACTTCTCTGCTATTTCGGCTCTGAAAATTGGTGCAGATTTGTCACACGTCTTCTGTACTAAAG

ATGCAGCCCCTGTTATAAAATGTTACAGCCCTGAGTTGATAGTACACCCTGTTTTAGAAGAATCATACAG

TGTCAGTGATGATGAAAAAAGCTCAAGAGCAGACAAGGTTCTCGCTGAGGTTGATAAGTGGATGGAAAGG

TTTGACTGCCTTGTTATTGGCCCGGGTCTTGGAAGAGATCCATATCTTCTGGATTGTGTTAGTGATATCA

TTAAGCATGCAAGGCAGTCCAAAGTTCCAATGGTCATAGATGGGGACGGACTATTTCTTGTTACAAACTG

TCCTGATCTTGTAAGTGGGTATCCCTTAGCAGTCCTGACTCCAAACGTGAATGAATACAAACGACTGGTT

CAGAAAGTACTAAATGGTGAAGTAAATGATCAAAATGGACCTCAACAGTTAATTAATCTTTCCAAGGGCA

TGGGTGGTGTAACTATACTAAGGAAAGGAAAATCTGATCTCATCAGTGATGGTGAAACAGTGAGATCAGT

AAGCATCTATGGTTCTCCAAGGCGATGTGGTGGTCAGGGTGATATACTTTCTGGAAGTGTTGCATTATTT

CTTTCCTGGGCTCGTCAACAAGATGGGGAATCTAAACTAAACATGAATCCTACAATGTTGGGTTGCATTG

CTGGGTCAATCTTAATGCGGAAGGCAGCATCATTTGCTTTTGAACACAAAAAAAGGTCAACACTTACCAG

TGATATCATTGAGTGCTTAGGAAGAAGTATGGAAGATATTTCCCCAGTTAACTGATCGTTCAGCAGCGTA

ACAGTACTAGTTAAACCACTGCTCTATATGTCATTACCTTCTACAGTTATAATTTCTTTTGCTGTATGTG

CTTCTTTCTCCTTTGGTATGTGTTGTTTTAGACCTTAAATGTTCGAAAGCTTAAAAGATTATACATCTTG

AGCAATGTTACACACACCAGATGGATTTATATGTCTTTACTTTTTGACAGTAAAGTTTGGATGTTGGGAA

AATCCACTTTTTCATAATACGAAGTACTTTGAACTTACAAAAGTGCAAGCTCTATAAAATTCACGAAACA

ATGATTAAGACTGATCACGAATGAAGACAGACAGAATACTTTGCACCGTCTAAGCTTCGGACATGAGCCT

GTAGTTTGAGCATGTTGTTTGTCTAAACAGAAGTGAAGAACTGAAC

>MSTRG.1687.2 gene=MSTRG.1687

GTTTCTTTCTCATCAAGTTCAACTAGAATTGAATGGATGCTGGGTTTTTCATCTGTAACTAGAAGGCAAC

AACACTTGATTAGGTGTTTGGGAGGTTGTAATAACCACTATAGATACACAACAATTCTACATAATCACAA

TCAGCAGCAGAAAAAAATGCAGTCTGTGACACCTATTTTAGAAGCTGATGCTGTCAGTATAATTAAGTCA

ATTACTCCAGCTCTTGATCATTCTAGGCACAAGGGCCAAGCAGGAAAGATAGCTGTTGTAGGTGGATGCC

GCGAATATACAGGGTGCAGATTTGTCACACGTCTTCTGTACTAAAGATGCAGCCCCTGTTATAAAATGTT

ACAGCCCTGAGTTGATAGTACACCCTGTTTTAGAAGAATCATACAGTGTCAGTGATGATGAAAAAAGCTC

AAGAGCAGACAAGGTTCTCGCTGAGGTTGATAAGTGGATGGAAAGGTTTGACTGCCTTGTTATTGGCCCG

GGTCTTGGAAGAGATCCATATCTTCTGGATTGTGTTAGTGATATCATTAAGCATGCAAGGCAGTCCAAAG

TTCCAATGGTCATAGATGGGGACGGACTATTTCTTGTTACAAACTGTCCTGATCTTGTAAGTGGGTATCC

CTTAGCAGTCCTGACTCCAAACGTGAATGAATACAAACGACTGGTTCAGAAAGTACTAAATGGTGAAGTA

AATGATCAAAATGGACCTCAACAGTTAATTAATCTTTCCAAGGGCATGGGTGGTGTAACTATACTAAGGA

AAGGAAAATCTGATCTCATCAGTGATGGTGAAACAGTGAGATCAGTAAGCATCTATGGTTCTCCAAGGCG

ATGTGGTGGTCAGGGTGATATACTTTCTGGAAGTGTTGCATTATTTCTTTCCTGGGCTCGTCAACAAGAT

GGGGAATCTAAACTAAACATGAATCCTACAATGTTGGGTTGCATTGCTGGGTCAATCTTAATGCGGAAGG

CAGCATCATTTGCTTTTGAACACAAAAAAAGGTCAACACTTACCAGTGATATCATTGAGTGCTTAGGAAG

AAGTATGGAAGATATTTCCCCAGTTAACTGATCGTTCAGCAGCGTAACAGTACTAGTTAAACCACTGCTC

TATATGTCATTACCTTCTACAGTTATAATTTCTTTTGCTGTATGTGCTTCTTTCTCCTTTGGTATGTGTT

GTTTTAGACCTTAAATGTTCGAAAGCTTAAAAGATTATACATCTTGAGCAATGTTACACACACCAGATGG

ATTTATATGTCTTTACTTTTTGACAGTAAAGTTTGGATGTTGGGAAAATCCACTTTTTCATAATACGAAG

TACTTTGAACTTACAAAAGTGCAAGCTCTATAAAATTCACGAAACAATGATTAAGACTGATCACGAATGA

AGACAGACAGAATACTTTGCACCGTCTAAGCTTCGGACATGAGCCTGTAGTTTGAGCATGTTGTTTGTCT

AAACAGAAGTGAAGAACTGAAC

>MSTRG.1691.1 gene=MSTRG.1691

AGAGATGCTTACCAACAGATTGCTCAACGCAATCTCTCCTCCTCTTCTTCCACTGCCACTGCTCCCCCAA

TTCACAAGGTCTCTCTCTCTCTCTCTCTCTCTCTCTCTCTCTCAGCAGACTTTCTCCCTCCCTCTCTCTC

TCTCCTTCTCTCTCTCCCTCCCTCACTCTCTCCCTCTCTCTGTCTGTGTTTCCGTCTCTCCCACCCTTCT

CTCTCTCCGTGTGTGTATTTTGTGTGACCCCATACATACATTTTACAAACATCTTAATGTTGTTTAGAAA

TTTGGTGAAAATATATTTTAGTTGATGAAATTTGGTTTTTGGTGTTTTGGGCGATTTTCATGTTATATGT

GATACGTTTCGTTGTTCGCTTCGCATACGACGAGACAACTTGATCGATAGCATAGCGGTGGTGGCCGATG

GTGGCGGTGACGGCGGTTTCAGGACGGGATCCGGTTTCAGAAAGGAAAGCGCGCTAATTGTGCCTTAATT

GAGGGCGGTAATTGTGCCTCGTTAATTATGCCTCTTTATTGAGGGCATAAATTGTGCCTCAATTATTGAG

GGCGTAAATTATGCCTCTTTATTGAGGGCGTTAATTGGGCCTTATTTGAGGGCGTTAATTGTGTCTTAAT

TAAGGATATTAATATTTTATTATTATGCCTTAATTAAGAGCTTATGTTAAATATATC

>MSTRG.1694.1 gene=MSTRG.1694

GTCACTTGTGATCTGATACGGCTGGTCACAACAATCTATTAATGTGCACATATATATCTAAACCTGTTGT

ATATGAGTGATGAATGAGCAGTTTGGTACAAGAGTACAGTGTATAGACATATCACAGATAGTTGTAGTTG

AGGATAAGGAGGGTGTCGGCTTGCATGACAGAGGAGGCATGGGAGGAAGGTCAGCAGAGAGATTTGCAAG

GTGGAAGAAGATTGGGAGACAAGTGAGAGCAAAGGTTTACATAATCAGGCTTTGCATTGTTATGCTTCTT

TGTTGGGACAACTCTAACTAAGTACACACAACATTGTTTGCTCAGCCTCAGCTTCATCCGCTTTTCATCA

TTTTCGTAAATTCTTATGTAACCTTAATTGCTATCTTTTTTATATATAACCTAAACTCAGCTCGGTTGCA

TTTGTAATTTAGGCTATGTGAAATCAGGTGAAAAATAACGTAATCTGTCTTTAGGAA

>MSTRG.1698.1 gene=MSTRG.1698

ACACCGTCTAAAGTCTGAAGAGTCGAGAGTCCATTTAAGTGAGCAACAAATACACACTCATACAGAATCA

TAGATAGACGCCGAAGCACACACTCTGACACTACCCTAATTCCTAACTCAGCTGCTCACTGCGACAAATT

CAGCGGCGAGCCCACCACAACGGCTGCTCAGCTCCTCTCTAACTCTCTCCCTCCCTCCCGGCTCCAGTGC

CCACAGCTGCAACCCCCAATTTCTAGTTCAGAGTGACCGCGGATTGTTTCTTGCAAGAGCTGTAATGCAG

ACAGATGGGTTTGATTACTTAAAGCAGAGCTGCCCTCTGTCATCTCTGAATTGTTGGAGTGTGTGGCGAG

GATCAGTGAGCATCCTGTTATTTCATTTGGTTACAGGAATGAACCTTCCCTTGACGGCGCTGATATGAAT

GGCAGGCGAGTGAAACAAAGAATTTACTGAGCGAAAATCATTAAGCATTCTGTTATTTCATGTAATTACA

GGAATAGACCTTCCTCTGATGGTGCTGAGTGCTGACATGAATGGCAGGCGAGGGAAACAAAGAATTTACT

GAGCAAAGAAACAGCGTAGTCGGCTCAAAATTGTTATCCAGTAATGAATGTAATCTCTAGAACTTTTAGT

AACCTTTGGATTGGATAATATCCTCCAAGTCGTGTATATATTAGCGTTATGGCTTTTAGCCAAATCCTTT

AATTGGCATGTCTATATTAAGGGCCAGTTAGTAACTTTTGGATTGGATAATATCCTTTAAGGAATAGTCG

CGTGTCTCGACTGACCTAGACTACCGACTAAAAGTAGTCGAGCAAACAGAGCAAAAAGTCGAGAGAAAAG

CACTTTGAGAGCATTACAACCATAGCAATCAAATGGGAGAAAAGGCCAAGTATAACATGAACTGAAACAT

GGCAGAGAAAACTAGAAACACTTGCGCAGTGCCACCACCATTTTCATTTCAGGAATGGCTTTGCCAACTA

CATGTATAAAATGTAATTATACATCTACGATAGCATGAAATGTCATTAAAAACTTACTCATTACATACAT

GATTGTTGGCACTACTCATCAGAAACGCAAAATGGTCTTCAAATTTTTTTTTTTGACAGAAGCAGAGTTG

GTAAAAAACCAACT

>MSTRG.1698.2 gene=MSTRG.1698

ACACCGTCTAAAGTCTGAAGAGTCGAGAGTCCATTTAAGTGAGCAACAAATACACACTCATACAGAATCA

TAGATAGACGCCGAAGCACACACTCTGACACTACCCTAATTCCTAACTCAGCTGCTCACTGCGACAAATT

CAGCGGCGAGCCCACCACAACGGCTGCTCAGCTCCTCTCTAACTCTCTCCCTCCCTCCCGGCTCCAGTGC

CCACAGCTGCAACCCCCAATTTCTAGCTGTAATGCAGACAGATGGGTTTGATTACTTAAAGCAGAGCTGC

CCTCTGTCATCTCTGAATTGTTGGAGTGTGTGGCGAGGATCAGTGAGCATCCTGTTATTTCATTTGGTTA

CAGGAATGAACCTTCCCTTGACGGCGCTGATATGAATGGCAGGCGAGTGAAACAAAGAATTTACTGAGCG

AAAATCATTAAGCATTCTGTTATTTCATGTAATTACAGGAATAGACCTTCCTCTGATGGTGCTGAGTGCT

GACATGAATGGCAGGCGAGGGAAACAAAGAATTTACTGAGCAAAGAAACAGCGTAGTCGGCTCAAAATTG

TTATCCAGTAATGAATGTAATCTCTAGAACTTTTAGTAACCTTTGGATTGGATAATATCCTCCAAGTCGT

GTATATATTAGCGTTATGGCTTTTAGCCAAATCCTTTAATTGGCATGTCTATATTAAGGGCCAGTTAGTA

ACTTTTGGATTGGATAATATCCTTTAAGGAATAGTCGCGTGTCTCGACTGACCTAGACTACCGACTAAAA

GTAGTCGAGCAAACAGAGCAAAAAGTCGAGAGAAAAGCACTTTGAGAGCATTACAACCATAGCAATCAAA

TGGGAGAAAAGGCCAAGTATAACATGAACTGAAACATGGCAGAGAAAACTAGAAACACTTGCGCAGTGCC

ACCACCATTTTCATTTCAGGAATGGCTTTGCCAACTACATGTATAAAATGTAATTATACATCTACGATAG

CATGAAATGTCATTAAAAACTTACTCATTACATACATGATTGTTGGCACTACTCATCAGAAACGCAAAAT

GGTCTTCAAATTTTTTTTTTTGACAGAAGCAGAGTTGGTAAAAAACCAACT

>MSTRG.1700.1 gene=MSTRG.1700

TGTTATTAAAAGCGCGCATATGCGCACGCATATGCGATATGCGATGCGATGCCCGCCTGAAGCGCGTCGC

TTCGGAGATGCGATGCGCCTTCAATGATGCGCTCGCTTTTATGTGCAGCGAGTGCTTATGCGCTGTTCAA

AAGCGCAAAATAAGCGTGCTTCTTTTAAAATGAGTCATTTTTAAAGGAAATGGGTTCAGAAAAAACCCTG

TGGACCTTATAGTAACTAAAACAAATAGATATAGACCAGATTTACATGCAAGAACAAGAGATATCGAACC

GGAGACTAAAATTAAACTCATCTTCTTTTATGGTTATTGCATATAGATACCCATATATATAAGATACACA

TATCTTGTATATACTATTATATATTTATACATACAAGTGTGGTATATACACATATGACTTGAGTATGATG

AAGAAGACTTTGTTAAGGATCCGAATTATATTAGTAAAGATGATGATTTCCTGAATATTGATGACGAAGA

AGTGGAAGAATGGACTAAGTTTGGAATTCTTTGCTACCTTGACTTTAGAAGAATTTGGTTGCTTTATCAA

TAAATGTCATCTTATGTGTAAATTAACAGTTTTCTGCTTATATATCATTTATATATGTATATTTATTTGT

CTTCGATGAATCCGCGCATTTCGCGTTGCGCTTATGCTCTAGGGGGCCTTTGCGCTTCTGTGCGCATATC

GCGTTTAATAACAC

>MSTRG.1709.1 gene=MSTRG.1709

GTTGAAACAACTCAACAAACCTTGTCTTCATCTCCAACATTGCCACCCTGTATCTTCTAATCAATCTCCC

ATAATCACCATCCACATGGTTTTATACCCCACCCATGAATCTTTTTCTGACCATAACTTTTTCATATTCT

AAAATCAAAATCATAACTCTGGTCTCTGCCCCTTAAAAATCCCACTTTCTCACACCCCCTCCTCATTACA

CATTTATTGTAGTGTTTGTTGCACTTGGAAAAACATATCATTCTTTCTTTTTTTATAAATGGCATCTCTG

GAGGCAATGCTTATGGTAGTAACACTCACTACCATGTCTTTTATGATCTCACCTTCATTTTCTTTAACTT

CTCATCAGCTCAAAATAGATACAGTACCAGCAGCACCAGCATTGTTGCCTAATACTGTCCCTGTATCTCC

ATCTACTAAGATATCACCCGATATTTCGCCACTTCTGCCATCTTCTGGTGGCGTCATGCCTTCAACTGAC

TCTGCTTCTATCCCAATTATCCCTTCAACCCCGAGCCAGAACCCTGATGAAGTAGCCGCTTTTAGCCCGG

ATTCTGCAGTTTCACCATCCGGGCACTTAAGTGACTCGTCTGCAGTGACACAGAGCATGATCAGGTCTCT

AAAGTTCGCAGTTTTTATGGTCTTGTTTTCATGTTGGACAATTCACAATTTTGCAGTTTAAGAAGGTTTC

TGTTGGCTTATATGTTAGTTTATGCAAAAAATTATTCTATGTATAGTGGATATTACAATAATGCACTACA

GTTCAAGGTGAGTGGCATGTTAAGTAGCAGTTGAGTTGATTACTTGATTGGGGTGGGGGAAATTCCAAAA

GTAAATCTATCTGGTCAGAAGATTAATCATTTATATTACTTTTGTCATTGTTTTAATTTACA

>MSTRG.1711.2 gene=MSTRG.1711

ATCAGGTCCATTGTTGCATATTTCTTTCTCATCAGAGGCGTGTTGTCCTTACCTCGACCGGGGTCTCTAT

CTCATTCGCAGTGTGATCAAGTGGGCAATGCCCTCGCAGCCTCGCTTCGCTGACAAGAGGGAGATCTTGT

AGCACTTACGGATTAGGCCCGGTGTGGCTCCAACAGAGACCGAGCCTGACATCAGCCTAAGTCATTGTGT

AATATTGCTTGTGAATGACATCAACTTACAGTGCAGGGTTTCATTTTCCATTAAGGTGCCAAC

>MSTRG.1720.1 gene=MSTRG.1720

CATTTTCCTATAACAAATCATATTTTCACTAGCGTCTCCTTTTTATTAAAAATCTCTTTTTCACTTGTCT

CTCCTCTACGCTTCGAAAACCCTAGATTTTGAAATCGAAGAGTCATGGCGAAGCTTTCGGAACCAAGTGA

CTCGACATCGGAGGAAGAGCAAGTCAATGACGCTCAGATGAATGAGGAGGAAGACGAGGAGGAGCTTCAA

GCCGTCGCCAGAACCGCCGCCGATTCCGACGACGATGACGGCGAAGCTGCCGGAGATGACCAGGAAGACG

ACGATGAGGAAGGTGATGGGGTTGGTAATGGTGAAATTAGCAGGCGTGAGAAAGCAAGGTTAAGAGAGTT

GCAGAGAAAAAAGAAGCAAAAGATTCAAGAGCTCTTGGATGCCCAGAATGCTGCCATAGATGCTGATATG

AACAACAAGGGGAAGGGCCGTTTGAAGTATCTCCTGCAACAAACAGAGATCTTTGCTCACTTTGCCAAGA

GTGACCAGGCGGCACAGAAGAAGACAAAGGGAAAGGGTCGTCATGCTTCAAAAGTTACTGAAGAAGAAGA

AGACGAGGAATATCTTAAGGAAGAAGATGGACTTACTGCGGGAGGAAGCACAAGGCTACTCGTTCAACCC

TCTTGTATTCAGGGAAAGATGAGGGATTACCAACTTGCTGGTTTAAACTGGCTTATACGTTTATACGAGA

ATGGTATAAATGGAATTCTTGCAGATGAGATGGGGCTTGGTAAAACATTGCAAACCATCTCATTGCTTGG

CTATTTGCATGAATTCAGAGGCATCACTGGTCCCCACATGGTTGTTGCTCCAAAGTCAACTCTAGGCAAT

TGGATGAATGAGATTCGACGTTTTTGCCCTACACTCCGTGCAGTCAAATTTCTAGGAAATCCTGATGAGC

GAAAATACATACGTGAAGAATTACTAGCTGCGGGAAAGTTTGATGTTTGTGTGACCAGTTTTGAAATGGC

AATTAAAGAGAAGAATGCCTTACGGCGCTTTAGTTGGCGCTACATCATAATTGATGAAGCTCACAGGATC

AAGAATGAAAACTCACTGCTTTCCAAAACCATGAGGCTCTTTAGCACTAATTATCGCCTTCTTATCACCG

GCACTCCACTGCAGAATAATCTCCATGAACTCTGGTCCCTTTTAAATTTTCTACTACCAGAGATTTTCAG

CTCCGCGGAAACATTTGATGAATGGTTCGCAATATCTGGTGAAAATGATCAAGAGGAGGTTGTTCAACAG

CTTCACAAGGTCCTCCGGCCTTTTCTTCTGCGAAGGTTGAAATCAGATGTGGAGAAAGGTCTGCCCCCGA

AAAAGGAAACTATACTTAAGGTTGGGATGTCCCAACTGCAGAAGCAGTATTACAAGGCTTTATTGCAGAA

AGATCTCGAGGTTGTAAATGCTGGCGGTGAGCGCAAGCGCCTACTTAATATAGCAATGCAGCTGCGTAAA

TGCTGTAATCACCCTTATCTTTTTCAAGGTGCCGAACCTGGTCCTCCATATACTACAGGAGATCACCTTG

TAACAAATGCCGGAAAAATGGTTCTGCTAGATAAGTTGCTCCCGAAGCTTAAGGAACGTGATTCAAGGGT

TCTTATATTTTCACAGATGACAAGACTGTTGGACATCCTGGAAGACTATTTGATGTATCGTGGATATCTT

TACTGTCGAATTGATGGAAATACAAGTGGTGAAGATCGTGATGCTTCTATCGATGCCTTCAATAAGCCAG

AAAGTGAAAAGTTTGTCTTCTTATTATCAACTAGAGCTGGAGGCCTTGGAATTAATCTTGCAACAGCTGA

TATAGTCATTTTATATGACAGTGACTGGAACCCCCAAGTTGATTTGCAGGCTCAGGACCGTGCCCACAGG

ATTGGACAAAAGAAAGAAGTGCAAGTCTTCCGTTTTTGCACTGAGTATACTATTGAGGAGAAGGTTATTG

AAAGAGCTTATAAAAAACTTGCGCTTGATGCATTGGTTATCCAACAAGGGCGATTAGCTGAGCAAAAGAC

TGTTAACAAAGATGAGCTCTTGCAAATGGTTAGATTTGGTGCTGAAATGGTTTTTAGTTCCAAGGACAGC

ACAATTACAGATGAAGATATAGATAGAATCATTGCTAAAGGAGAAGAAGCAACAGCAGAGCTTGATGCAA

AAATGAAGAAATTCACAGAAGATGCAATCCAGTTTAAAATGGATGATACTGCTGATTTGTATGATTTTGA

AGATGACAAGGAAGAGAACAAGGTTGATTTCAAGAAGATTGTTAGTGATAATTGGGTTGAACCATCAAGA

AGAGAAAGGAAGCGCAATTACTCGGAATCAGAATACTTCAAGCAAACAATGCGGCAAAATGGTCCTGCAA

GACCAAAAGAGCCTCGAATTCCTCGCATGCCTCAGTTGCATGACTTCCAATTTTTCAATACCCAAAGGCT

CACGGAGCTCTATGAAAAAGAAGTTCGTTTTCTCATGCAAGCTCATCAAAAGAATCAGGCAAAAGACTCT

GTAGAAGCTGATGAATCTGAAGATGGTGGTGATCCATTGACTGCCGAAGAGCAAGAAGAAAAGGAGCAGT

TGCTGGAAGAGGGTTTTTCAACATGGAGTAAGAGAGACTTTAACACTTTCACCAGGGCTTGTGAGAAGTT

TGGTCGAAATGATGTGAAAGGTATTGCCTCTGAAATGGATGGTAAATCAGAGGAAGAAGTTGAGAGATAT

GCAGAAGTTTTTAAAAAGAGATACAAAGAAATAAGTGATTATGACAAGATCATAAAGAATATTGAAAGAG

GAGAGGCTAGAATTTCCCGTAAGGACGAGATCATGAAAGCAATTGGAAAGAAATTGGATCGCTACAAGAA

TCCTTGGTTGGAACTGAAGATCCAGTATGGTCAGAACAAAGGAAAATTATACAATGAAGAATGTGATCGC

TTTATTTTGTGCATGACTCACAAAGTTGGTTATGGGAATTGGGATGAGCTTAAAGCAGCATTCCGGACTT

CGCCCTTGTTTCGCTTTGATTGGTTTGTCAAATCGCGTACTACTCTAGAGCTTGCGAGGAGATGTGATGC

ACTCATTCGATTGGTGGAGAGGGAAAACCAAGAATTTGATGAGCGGGAGAGGCAGGCTCGTAAAGAGAAG

AAACTTGTCAAGAACACGACACCTTCCAAGAAGTCTGGGGGAAGGCAGTCTGCTGAGAGCCCACCATCTA

GTATGAAGAAGAGGAAGCAACTGTCGATGGATGATTATGTCACCTCGGAAAAGAGGAGAAAATGATGCTG

GAACAAATTTTCAAGTTAATGAATGCCCTTTAGTCAGGGTGTTATTTTTTTGCTATACCTTTTGCTGGAG

GGTTTAAGCATCTGAACAGTGATAGAATTAGGCAGTGATGAGCTATGTATGCAATGACTAGGTGTTGAAT

TATAAACGTATCTACTTAAAGTTGTCTATTGTACAACTTATCATATTTTAACTTAAATTTAGAAGTTTTT

CAATAGCTA

>MSTRG.1725.1 gene=MSTRG.1725

CAAGGTCGGGCACATCGACGATGTGCAGGAACTGAGGAAGGCTAAACAATCAAACGTTCCTCAACGATTT

GTCCGTGACATGGCTGAAAGGCCAGAACTTGCCACTACAGTCACTCACCCTAGTGACATTCCTGTGATTG

ATTTGTCGAACCTTGTTAAGGGTGGCAAGGCTCGGTTTCAAGCAGAAATTTTGAAACTCACTGCTGCTTG

TGAGGAGTGGGGATTTTTTCAGGTGATAAATCATGGAATTGATGTGGAAGTTCTGTCAGACATAGAGAAG

ATGGCCATGGAATTTTTCATGCTCCCTTTAGAGGAAAAACAGAAATATCCAATGGCGCCAGGGACAGTTC

AGGGCTACGGACAAGCATTCGTGTTTTCAGAAGATCAAAAGCTGGACTGGTGTAACATGTTTGCTCTTGG

TTTAGAGCCACACTACATGAGAAACCTTAGTTTATGGCCCAAAAATCCTCCTACTTTTAGTGAAAGTATA

GAAAATTATTCAAGGGAGATCAGGAAACTGTGCAAGAGTCTTATGAGTTTTATAGCAGTGAGCCTTGGGC

TAAACAAAGACGTGTTTGAAGATATGTTCGGGGTAGCTGTGCAAGCTGTGAGAATGAACTACTATCCAGC

ATGCCCAAGGCCAGACCTTGTTATGGGATTAAGCCCTCACTCTGATGGGAGTGCTCTCACAGTTCTGCAG

CAAGGAAAGGGTAGCTCGGTCGGACTTCAAATCCTCAAGAACAACACTTGGGTCCCAGTGCAGCCCATTC

CAAATGCCCTAGTCATCAACATTGGTGACACCATAGAGGTACTAACCAATGGGAGATACAAAAGTGTGGA

ACATAGAGCAGTGACTAACAAGGAAAAAGATAGGTTGTCCATAGTTACATTTTATGCACCAAGCTATGAC

ATTGAGCTCGGACCAATGCCAGAACTGATGGATGAAAAGAACCCTACCAAGTTCCGAAGATACAACCATG

GAGAGTATAGCAAACACTATGTAACCAACAAACTCCAGGGAAAGAAAACACTGGAATTTGCAAAGATTCA

CCCCCAAGCTTCATTCTAAAAAAAAGGGCCAGGCACAGTTTCACAACTTGGAGCAGCAATTAACTATGAA

CAGAAACTATAACATAATTGTAATCTAAAATTTGGCTTGAACTATTTATCAGTATCTGCAAGTTGGAAGG

TGGAGAATCATAAAATATGGTTCTTTAGACTGTACATGTATCTCAGTATCTGTAACTCAAAATATCTAGT

GATGCATATTATCAATAATATATTATGGGCC

>MSTRG.1727.1 gene=MSTRG.1727

TTTTATTTTATTATTCTCAATTTTCTCTCTCTCTAAAACTGCGACCAGCGAGAGAGAGATAGCGAGAAAG

AGAAAGAGAGAGAGAGTCTTGCATATTTCCTCTCTCTAAAACACCTCGGTCTCGTCACGAATTAACCCCA

ATGAAACCTTAGCTTCTTCACACAAGATGGAGACGGAAAAATTGGACCAGAATGGGGATTGTAGTATGGA

CTTGTCTGATGACCAGTTGCTCAATACAGAATCTTCTGATGCATGCGGCGTGTTTGATGAACCAGAGTTG

GTTCCTCGTGTTGGCGATGAATTCCAGATTGAAATTCCAGCCCTTGTCACAGGACCTGAGTATTATTACT

ATTTGAAGTCCCCAGTTGATGCTGAAAACAGGGGTCATGTTCCCTCTGATTTTGGGCTGGGATTGTCAGT

ACCAGTCACGTGGAACAAGAATGTTAACAAGGTTATGAAAAAGGGAGAGCAGGACTTTTCTGGTCATTCA

ACTGAAAGTGTTGAAGAGACAGTAGACGGATACCAAGGGTCAAGCAATTTTTTGGTTCCTGGGGTTTGTG

CCGACGTATGGAATGATATCGAAGAGGAGAGCTTTCTCCTAGGTTTGTACATCTTCGGGAAGGAGTTTTC

CCAGTTGAAGAGGTTTGTTGGAAGCAAGAAGATGGAAGATATACTTTTCTTATATTATGGGAAGTTTTAT

CGGTCTGTTCAGTTTACTAGATGGACAAAAATCACAAAACGCCGCAAAGTGAAAACCAAAAAGTGTGCAT

ATGGCCAACGGATGTTTAGTGGTTTGGGGCTACGGGAACTGCTATCTAGGATTCTTCCCCATGTGTC

>MSTRG.1731.1 gene=MSTRG.1731

GTTGGATCAGTTGCACCTTCCTTGTAATAAGCAAAGACTAAGCTACCATCATCATGCATGCTCTCCCCAA

CGAAAAATTGAAGGTCCTTAATCTTGCCGAGCAGGTATTTGGTTGCCCCTTCAATACCCTTCTTGAATTC

CTCTTTCTTGTCATCCTCTAACTTGGCTGTCAAGTTTTTAATGTACTTCTTGATATAAGCCACGAACTGC

TTCTTGTCAAAAGGAGGTTGCTCCTGGAGCCTAAAGGTGTCCACAATATCGACCACCTTCTGGGCCTGAT

CATCAACACCTTCATCTTCAC

>MSTRG.1733.1 gene=MSTRG.1733

AGAGATGAAGGAGCGGGAATCTGTAAATTCATCATGACAAGCTCCCTTCTGCAACTTGAAGGTTAGTTAT

CTTACAACCCCATTAAGATTGGCGATAGCACGTTTGGTTTTCATTTTTTGCAGCTCTCTTATGACTTATG

CATCAGGTTTAGAATTATTGATATATGACTCAAGTATATAGCAACAAAAACAATGTCAACTAAAGAAATA

GGGTGGTGTTACTTGTTAGATCTGTCGGTGGCTTTCTATGTTCAAATTAAATTATACAGCGACGAGCTCA

CAAACTTCCCTCTCTCTGTTGCTTGAGGATGTTGATACCTGAAAGCAATGTCTTATACATAGGAAGTTCG

TTGAATCCAGTAAGTTCGTTGCATAACAATTATTTGAAAAAATTTGAAGATACTATTACAGCGGAAGAAG

GAAGAAATTTTTCAGAATGCGTTAAAGCCAATACATTAGCATTAGAACATTCTTCATTGTTCACTGCTCC

TTCCTGGGAAACTAATGTAAAGAAAGTGAATATGGATTTAATCTCTGTATATGTAGATTTACACTTGTAG

AATCTCTAAATAGAGAGGTTTAAGCTGTAGCGATACGTAGTAATATATATAGAGAGATATAGACTCATGA

TCATTCGTCTGAATCATCATCACCACCACCAAAAATGGAAATCCTGTTGTACAGCAAGAAAAGAACCAGA

ACAAGAAACAGAGCAACTCCGATGGGCGATCCACTGGCCCGGTGTATGGATTCAGGTCCTCCAGAAGAGA

AAAGACTTGATACAAAAGAGCCACTTTCAGATGAGAGAAACTGTATAGTAAGTAGCAGTATAATGGGCAG

TAGAAAAAGACCAACTGGACTTAAGAGCTCAGCTATAAACTCCGTCAAGGCCTCACCTTGCTCGCCAAGA

ATTGAGGGAACAACCACCAGCAGTACTACCACTACCACTAGCAGAATACCGTAGGGGGCGGAGCCACGAT

GGTCATCGACGGCCATGTGGTTTCCACTAGGGTACATTGTTACTTGACAGGAAGTTGTAGTGGTGTTTTT

TGTTCTGCTTTTGAAACTTCTTTTTTGTGTAAGGTGAGAATGTGAGATGTATGTTGAGTGAATGAGAAGA

CTAGGGTGGGTGCTCTAGTAGGGTTGGTATTTTTGAAATAGGTCACTCAATCTTTTGCTTTTAATAATGC

ACACCAAAAACATAAGCAACTTTCCATGGC

>MSTRG.1737.1 gene=MSTRG.1737

CTCAAATCATCAACAGGACTAGATTCTACCAAATTTGTGCCACGCATGACAGTAGATTTACTGCGGTTAA

AAGTGACTGCATTGGATGATGATGAAGGTGGGTGATTCTGATGCTTGTTTTTTGGGGTTGGGAGAAATGG

GTTTCTCGATTTCTTGAAATGATGATACGGGTTGTTGTTCGCATTGATGGGGAGTACAGATGAAGAGTGA

TGTGATGTTGAAGAAGAGGAAGCAGTAGCTTTAGTAGTGACAAGAAATGGGGAGGAGACTAGTTTGCCAA

GCTTTTCTTGGAGACAGAAGGCACATATACCACCACTAGGGCTGTTGTTTTTGTATGGATGATATGTGCA

TTGCATTGAAGCATCGCCATAATCATCATCAACTACATTTACCATGGTTTTATGGTGAGATGGAAATGAG

TTGTTCTTGGTGGTGTTCAAGACACTTTGGTGGTGGTGTGGAGTGAGGAGATTTTGTGGAGGAGTGTGTG

TCTTGTAGCTTGTGTGTGTGTTGGAGTTTAGTAGAGAATGGGGAGACTGGTAGTCCCAAGTAGCACGTTA

TTCCTCGTTTTAAGGAGAACATGACTTGTGGTTTGCACCGCCACGCTCATTCTCGAACACTATTTTAC

>MSTRG.1739.1 gene=MSTRG.1739

CTCTGCTACATCTTTCTCCACGCTGTGCTTCTCAAATGCACCGATAGCAATGTTGATTGCTTCAGTTTGC

AAATCGTCCTTCATATCCGCGCTTTTGATGATTACTTTCTTGCCGGTCGCGGATGAAGAGGCCGCCGGCC

GGGAATTCCGGTCATCTCCGTTTCGCTTTGCTGCTTCACTCATTGAGTTGGAGATTTTTG

>MSTRG.1743.1 gene=MSTRG.1743

CGGCAAATAATGAATACTCAAGCTCACCAAAGCCTTCACTTTATCCGGCCTAAACAAACTCAAATGCCAC

GCAATATACGCACCCCAGTCATGCGCCACCACAAACACCTTCTCCTTCGGCGCCACCACGTCCAGCAACG

CCACCAGGTCGCCGACGACATACGGCACGCTGAACTTGGACGGGTCATCGAGTGGGGCCCCCGTGGTGTC

TCCGTAGCCACGGAGATCAGGCGCCACCGCGCGGTAGCCGTGGGAGGCGAGGAAATGGATCTGGTGGCGC

CAGGAGTACCAGAGCTTGGGGAAGCCGTGGATGAAGAGGACGGTGGGGCCCTGGCCCTTTTCGGCTACGT

GGATGTTGAGGCCATTGGCGCTGATCGTTCTGTGCTCAATGCCTTCCATGTTTGACTTAAAAGTGAGCTT

TTTTAGTATTATTTTGTGGGCGCAAGGCAGTGGTGAACAAGAGATAACAGGTGGTTAATTATTTATAGAT

GATGGAGCCTCTGCCGCTCTGCGTATTGTAACGTATGGATAAGTTCTAGAAGTTGGGCCGGTTGTGCTAC

ATTCGGAGTTCAGCCCAGAAAGTTTTTATGGAATTAAGGAGCACAAATGGTTTCTGTTTTTATGAATGAT

TATATTTGGTGAATCATGTAAATCTGTAATAAATGTCACCGTCCTTCATTACCAGAAAAAATACATTC

>MSTRG.1747.1 gene=MSTRG.1747

CTGACAATAAAAACACGCACCGACATTATTTTTATGATCTTTATATACAATCACTCAAAGCATCATTTTT

TTGTTATTTCACTCAAACTCTTGCATTAGATGATTAGCAATCTGGTCCAGGTTATCCATCATCATCCACC

GATAAACATCAGATTCAGTCACGCCAATCTGGCTCATGATCTCAGCCTCCCATCCCAACAAGCTTGATTC

CATGTTCGAAACCTGGTGATCATCCTTCAGCTCCAACTCATCAGCACCATGACTTGCACAGTAGTTAGGA

GGCTGAAATGTTGTCGAAAACTCCATTCCCATCAGAGGGCTGCTAAAATTCGGAGAATTTATTTCGGTAA

TTGCCTTAGGCGTGCACTGTTTCTTGGCTAGAGGTGGTAGTGGTACTGGTTCAGGTGGGTGGTCATCTGG

AAAGTTGAGCCGGGCCTTGTCGCCGCGGATACGCTTGGCGGCCTCATCATAAGCCCTGGCTGCTTCTTCC

GGGGTGCTGAAGGTGCCCAGCCAGACTCGGACACCTTTCTGAGGATCACGAATTTCGGAAGCCCATTTGC

CCCAGGGTCTCTGCCTGATCCCTCTGTACTTGTTCTTTCGACTTCTCTTATTGCCCTCTTGGCTCAGCTT

CTCACCCTCCTCACTCACCTCCTCTTTGGTGATGATCTTCTTGGGCTTAAGAGGTCGACCATGAAGAGGA

AGTTGTGGGAGTTTTGCATCGAAGTGATCAGTCCAGCTGCCAGATAGTTGTGGCAAGGTAGAAGCAAAGC

CTGGTTGATCCCAAAGGTCATGTTTGTTCACTTTCCGGCTGCCAGTTGTCAGCGGAGGACGTTCAAAGTC

AGAAATAATTGCACCACCGCACATCTTGGTGGACTAGCTACGTCTCGAGAGAGAGAGAGAGGTGGCTTGA

ATAAGAGAGGAACGAACAATTGTTAAAAACAGG

>MSTRG.1752.1 gene=MSTRG.1752

TACTCCCTCCGTCCCATCAGGTTCTTTACGTTACTTTTCGGCACGCATTTTGAGACTCCTATAAAATATA

GTTCCATAATGTTTTTTATTTTAAAAAAAATCCTGAAAAAAAGTTAGACTTTTAAACTTTTATTCAGAAA

AAAAATAAACTTAAAAAAGGTTATTGAACTATATTTTATAGGAGCCTCAAAATGCGTGCAAACAGTGAAC

GTAAACATCCTGGTAGGACGGAGGGAGTACT

>MSTRG.1753.1 gene=MSTRG.1753

TAAATAGCGGCATCACTTCTGCTATTTTTACACACTAAAACCTCACTACACTCTCTCATTGCCTCATCAA

GTTTTGCAACTTGAATCCTTCTAAAACAACATGGGTGTCCAAAAGACTGAGGTTGAGGCTCCTTCCACCG

TGTCAGCAGAGAAAATGTATCAGGGATTTCTCCTTGATATGGACACTGTTTTCCCCAAGGTTCTTCCTCA

GCTTATCAAGAGTGTCGAGATTCTTGAGGGCGATGGTGGTGTTGGAACCGTCAGACTCGTCCATCTTGGC

GAAGCAACTGAATATACCACAATGAAGCAAAAGGTAGACGTAATCGACAAGGCCGGATTAGGCTACACCT

ACACCACCATCGGAGGTGACATCCTCGTGGAAGGACTCGAATCCGTGGTGAATCAATTCGTTGTTGTGCC

AACAGATGGAGGCTGCATTGTGAAGAACACCACCATTTACAACACCAAAGGCGATGCTGTGCTCCCCGAA

GACAAGGTGAAGGAAGCCACCGAAAAGTCTGCCCTGGCCTTCAAAGCTGTTGAGGCTTACCTCCTAGCAA

ACTAAACCGGCCAAGTTGTCTTGCACTTTCTGTTTCATAGTATCACCAGGGCGTACTGTGCTATTATGAA

TCGAGTTTGCTTGTTTGTTTACGAGTATGTATGAGTTGAACTGAGTGTAAACTGCTGGGATATGTTATAT

GTACACCTCGGGTATATATATCTGAATATCATCAATAAAACATCGATCGTGCTCTATGCCCTCTTTGCCC

TTTCTGAGAAATTGAATTTCACGCTG

>MSTRG.1754.1 gene=MSTRG.1754

CTCAGTTCAACTCATACATACTCGTAAACAAACAAGCAAACTCGATTCATAATAGCACAGTACGCCCTGG

TGATACTATGAAACAGAAAGTGCAAGACAACTTGGCCGGTTTAGTTTGCTAGGAGGTAAGCCTCAACAGC

TTTGAAGGCCAGGGCAGACTTTTCGGTGGCTTCCTTCACCTTGTCTTCGGGGAGCACAGCATCGCCTTTG

GTGTTGTAAATGGTGGTGTTCTTCACAATGCAGCCTCCATCTGTTGGCACAACAACGAATTGATTCACCA

CGGATTCGAGTCCTTCCACGAGGATGTCACCTCCGATGGTGGTGTAGGTGTAGCCTAATCCGGCCTTGTC

GATTACGTCTACCTTTTGCTTCATTGTGGTATATTCAGTTGCTTCGCCAAGATGGACGAGTCTGACGGTT

CCAACACCACCATCGCCCTCAAGAATCTCGACACTCTTGATAAGCTGAGGAAGAACCTTGGGGAAAACAG

TGTCCATATCAAGGAGAAATCCCTGATACATTTTCTCTGCTGACACGGTGGAAGGAGCCTCAACCTCAGT

CTTTTGGACACCCATGTTGTTTTAGAAGGATTCAAGTTGCAAAACTTGATGAGGCAATGAGAGAGTGTAG

T

>MSTRG.1755.1 gene=MSTRG.1755

GTTAAAATATAGATGTTTCTTTGAAAAAAGTGAAAATTTCATTGATTCTTCTTTGAGTATTTTTATATTG

CTTGTATTATCTCTTCTCTTGTGATGAATTATGTTTGCATATCTTTCTTCACTTTTCCTACATCAATAAA

GGCTGTCTATCCAGTTCGTGGCTTGCTTGTCAAGTTGTTATCTTCTAAAGTATAGCAATAGCTTTTTTTT

TTTTTAATATTACAGTGAATACCTACCACTCATCAAATATAATCTGACAGAAAGCTTGGTGAAGAGAGGG

ATGGGGTGTCTTCATCTCATGGCAGTAGCGACCATTGTGGCAGATGTAGGTTAGAGAATGGTATCAACTA

TGTTCAGGTAGATAGGAACTTGCAGAATCAAGAGAATGCTGCCAAAGAATGCTCAAGTAGGTTATCGTTG

AGAAACAATCTTGTAACAGGATCAGCAGTTAATGAAAGTGAGATTAGTAATCCTCCTGGTCTTCTTCTGT

TTGAGTATTTCGCACATGATCTGCCATATGCTCGTGAACCTTTGGCTGACAAGATTTTAAAGCTTGCATC

TCAATTTCCGGAACTGAGGACATGCAGGAGCTGTGATTTAACACCTTCAAGCTGGATTTCTGTTGCCTGG

TATCCAATATATAGGATACCTGTAGGTACAACTCTGAACAATATAGATGCCTGCTTTTTGACTTTTCATT

CATTGTCAACGCTATATGGGAATACTGGTACTAATTTGCCGCAAAACCAAAGTTCAACAGCTAAGCAATT

TGGTAATGGTGGCATGTCCTTGCAGTTGTCCCTACCAATATTTGGGCTTGCTCCATACAAGTTTAAAGTC

TCTGATTGGAGTCCAGAAGGGGAATATGATTGCCATAAGATAAACTCTTTATTCCAAGCAGCTGATAATT

GGCTGCGGCTTTTGCAAGTTGATCACCCAGATTATAGGTTCTTTAAGGAAAGGGGAAAAGGGACGTACTC

CCTTGTGCCGATCGAAGAGGGTAAGATGGGCGCGGCCTTACATCTACCCAAGAGAGAGTAGAGAGTCCGT

TTCCAGAAAAATGACCCCTCCGGCATGCTGGAGAAGAGAGTCTAACCCATGACCACAATTAGATTATAGG

TTCTTTCTATCCACAATTCGTTATGGAGGTGATGAGAACGTGGATAAACGTGTAAAGTCTGAATTGGCAA

AGAACATAACAACATGTTCCGGGTCCATCTGCAGTCTCTATCCTTGTGTATTTTTTAGGTCCCTGCAATA

CTAACATCTCCTCAATTGTCCTGGGAAGGAATCAGAATTCAACTGTCGTAAGAAAGAAAGTTTGAAGAGG

ACGAGGAACAACATGGCGACTAGCAAAGATTCTAGTACAGTAGTATGGTAATACAGAGTTCCATGTTTTC

TTTACATGCTAAAATTTGGAACTCAGAAAGGAGTTTCTACTTTTGAAATCTAGGTGGGAATTGCTACTAT

GATGACGACGAAAATATTAGATCATATTTGTAAATCTGTTTTAGTTGCTAAAATGTATAAGCAATTGTTT

TTAGTTTGTACAAGTACAAGGATTTTGTTTAACTGTGTATTAGTTATTTTATCAGGGTATAACTTGTTGA

TAAGTAGAGAGCACCTGTAACATTGACCTTTGTGAATTTCTCCACTAACCCCTTGCCATTTTTA

>MSTRG.1755.2 gene=MSTRG.1755

TTGAGTATTTTTATATTGCTTGTATTATCTCTTCTCTTGTGATGAATTATGTTTGCATATCTTTCTTCAC

TTTTCCTACATCAATAAAGGCTGTCTATCCAGTTCCAATAGCTTTTTTTTTTTTTAATATTACAGTGAAT

ACCTACCACTCATCAAATATAATCTGACAGAAAGCTTGGTGAAGAGAGGGATGGGGTGTCTTCATCTCAT

GGCAGTAGCGACCATTGTGGCAGATGTAGGTTAGAGAATGGTATCAACTATGTTCAGGTAGATAGGAACT

TGCAGAATCAAGAGAATGCTGCCAAAGAATGCTCAAGTAGGTTATCGTTGAGAAACAATCTTGTAACAGG

ATCAGCAGTTAATGAAAGTGAGATTAGTAATCCTCCTGGTCTTCTTCTGTTTGAGTATTTCGCACATGAT

CTGCCATATGCTCGTGAACCTTTGGCTGACAAGATTTTAAAGCTTGCATCTCAATTTCCGGAACTGAGGA

CATGCAGGAGCTGTGATTTAACACCTTCAAGCTGGATTTCTGTTGCCTGGTATCCAATATATAGGATACC

TGTAGGTACAACTCTGAACAATATAGATGCCTGCTTTTTGACTTTTCATTCATTGTCAACGCTATATGGG

AATACTGGTACTAATTTGCCGCAAAACCAAAGTTCAACAGCTAAGCAATTTGGTAATGGTGGCATGTCCT

TGCAGTTGTCCCTACCAATATTTGGGCTTGCTCCATACAAGTTTAAAGTCTCTGATTGGAGTCCAGAAGG

GGAATATGATTGCCATAAGATAAACTCTTTATTCCAAGCAGCTGATAATTGGCTGCGGCTTTTGCAAGTT

GATCACCCAGATTATAGGTTCTTTAAGGAAAGGGGAAAAGGGACGTACTCCCTTGTGCCGATCGAAGAGG

GTAAGATGGGCGCGGCCTTACATCTACCCAAGAGAGAGTAGAGAGTCCGTTTCCAGAAAAATGACCCCTC

CGGCATGCTGGAGAAGAGAGTCTAACCCATGACCACAATTAGATTATAGGTTCTTTCTATCCACAATTCG

TTATGGAGGTGATGAGAACGTGGATAAACGTGTAAAGTCTGAATTGGCAAAGAACATAACAACATGTTCC

GGGTCCATCTGCAGTCTCTATCCTTGTGTATTTTTTAGGTCCCTGCAATACTAACATCTCCTCAATTGTC

CTGGGAAGGAATCAGAATTCAACTGTCGTAAGAAAGAAAGTTTGAAGAGGACGAGGAACAACATGGCGAC

TAGCAAAGATTCTAGTACAGTAGTATGGTAATACAGAGTTCCATGTTTTCTTTACATGCTAAAATTTGGA

ACTCAGAAAGGAGTTTCTACTTTTGAAATCTAGGTGGGAATTGCTACTATGATGACGACGAAAATATTAG

ATCATATTTGTAAATCTGTTTTAGTTGCTAAAATGTATAAGCAATTGTTTTTAGTTTGTACAAGTACAAG

GATTTTGTTTAACTGTGTATTAGTTATTTTATCAG

>MSTRG.1758.1 gene=MSTRG.1758

GGTGATTGCTTAGATGCCATTTGAAGTTGAAGTAGATGTTCTTATATATATTTTTAAGAGTAATAACATT

GGTTTAATATGTAAGAAATGTGTGGCAGGATGGTTGGAACTAGACCCTGCATCATGATACAAACGACACC

AAAATCTAGTGCTTATTATTACCTTTTAGGTACACGAACAGGGAATACACAATGAGATTTAGTGTCGTGT

CTGGGTACACAACACGGCACGGAAGTTCATAAAACAAAAAAGCTCCGCAGCCTGTGCTTTCGGACAAGGA

CCATGAAGAGAGAGCTAGGAAATGTGAGTTTGTTCAAGGGCTGCTGTTGTCAACTTTTCTCGATGATGGC

TTATAATTAAAGGATAACGCTTATATTCGAAGGATAAAATGTAGCAGCAACAAATTTGGATGATATAGTC

ATATTATAGTATAAAGAAAGGGGAGGGACATTAGCGAGTGTCTTTAAGACTCTGAGGGAGAGCCGGAGAG

GAACCATGGATTCAGTATACTGTGTGTTCAGAGTTGTATGTGCATAATGAAGTAGTATGAAGTATGAACT

TCCTTAATTTTGTACTTTTCTGTTTTCTGCTGCGTTTTATTTTCGCTTTCAGTAAAAATTAAACAATTTT

TATTATTTATATATCGTTATCAGTGAAA

>MSTRG.1768.1 gene=MSTRG.1768

TAAAGCTGAAACCACAAAGCCCACAACCCAGACCTAAAACGACGTCGTTAACACAATTCAACCCCCTTAT

TGCTAAACCCTGAATCACTTCGCCTCTTTAACCCCGACCTATCAACGGCAGGTTCTTGAAGCTTCTTTCT

GGGTTTATATCGGCCGAGTCAAGATGAGATTGGAGAAGTGCTGGTTTTGTTCTTCAACCGTGTATCCTGG

TCACGGTATTCAGTTTGTGCGTAACGATGCCAAGATATTCCGATTTTGTAGGTCCAAATGCCACAAGAAT

TTTAAGATGAAGAGAAATCCCCGTAAAGTTAAATGGACCAAGGCATACAGGCGGTTACACGGGAAGGACA

TGACTCAGGATTCAACATTTGAATTTGAAAGGAAGCGTAACAGGCCAGAGAGATACGACAGGAATCTCGC

TGAGAATACCCTCAAGGCCATTAAGAAGATTGTGAAAGTTAGAGCTGATAGGCAGAGCAAGCACATTTTG

AATAGAATGAAAGGCAACAAATCTGCGGAGCGCAAACGGGATGCAAAGGAGCTTGAACAGAGCATTCACA

TGGTCAAAGCTCCTGGTGCCCTTCAAAAAGATCCATCCCTTACATTGGTCAAGGTTACCCGGCGTTCAGA

AGAAAATGGCATGAAAGAATAGTCTGGTTTTACAAAATTCTTTTGCAGTTTCATTGTGTTGATTGCTGTA

GCATCAGAGTATTGATGAGACGAAGACACTTAATATTTGAAGTACTTTTGGTATCTTGTGACTATGGAAA

CGTCATAAACAAGTTTGTTTTAATGCACATCGAATCTTAATGGTAATTTTATTTTGTTTACTCATTGCTT

CAGTAATTGCCAATTTGCCATCTTGAACTGGATAC

>MSTRG.1769.1 gene=MSTRG.1769

ATTTGGCACTAGATCTGACAAAGAATTCGAAATGGCTGAAGCAGTAGTGTCGATTGTTGTGGCAAGGGTC

ACTGATTTATTGATTCAAGAACCCCTGGCTCTTCATGGACTCAAGGATGAAATTCAACAAGTGGTGACCA

AGCTCCAGTTGATCAAGACATTCTTACGAGATGCTGATTCGAGGATAAGTGAGTGTCAAGTCCGGACTTT

GGTTGCTGACATACGAGCACTTGCTTATGATGCTGAACATGTTGTAGAAAGCTTTATTGTTAAAGCTTCA

TCCTCTGCAAGAAACAGAAGGAAACAAGCAATCAAGATCAAAGATATAGAAAGCAAGATGTCTCTGCTCT

CTGATCGCATCCGTGATAATAATATTAAATCAACATCAGAATCATCAAATTCATTGTCGGAAGCACCTGG

AAAGCTAAAGCGATTTCACTCTTTCACAACCGTCGAGCCAGAGATATTTGTGGGATTTCATGGAGCCGTT

GATCGACTGGTGGGACATTTGGTGAAGAAAAGTGATGATTGTTATCCGCTTATCTCTATTTGTGGAATGG

GCGGTCTAGGTAAGACAACTCTCGCTCAAAAAATATACAATCATCCAACCATTAAAGCATCCTTTGCTGG

TTTAGCCTGGGTTTCCATTTCGCAGAAGTGGCAAACGAAACTTGTGTTGCAACGAATACTGATATGTCTC

ATCCATGAAAAGAAAGAGGAAATTCTTACTTGGGACGATGACAAGTTAGTGGAGAATCTGCTAGAAATCC

AGCAGAGGAAAAAATGCTTAATAGTCCTTGATGACATATGGACTACTGATGCTTGGGATTCTATAAAATC

GGCCTTCACAGCTCAAACATCTGTAAGCAAATTAATGCTTACCAGCCGTAATGTGGAGGTTGCTGTGCAT

GTAAATCGAGAAGGATTCATATACCAACCAGAATGCCTAAATGAAGAACAAAGTTGGGAGCTACTTAAGT

TGAAAGCAGTTCCCAGAGGAGATCATCTAGATTCCAGAGATATTTTTACTGTGATGGAAAAATTAGGAAG

AGAAATGGTTAAAAAATGTGCCGGCCTTCCACTAGCTATAGTCATTTTAGGCGGAATTCTTGTAACAAAA

CCTTCACTGATGGAGTGGGAGAAGGTATATGCTGATAGTTTAGGGTCCCTGGAGAAGGGAAAAGGATTGG

GAGAAGATCAGCAAAGAGAATTATTTCATGTTCTAGTCTGGAGTTACAATGATTTACCCCCACAATTAAA

GCCATGTTTTCTGTATTTGGGTAAATTTGGTGAAGATGAACAGATAGAAGTAGAGACTCTATATCAGTTA

TGGATTGCCGAAGGGTTGGTACTGTCGACTGATAAAAGGGAAGGAGAAACAATGATGCAAGTAGCTGAAT

CATATATGGGGGAACTGATCCATAGGAGTATGGTTCAAGTGACATTTGAAGATATGGAAAACTCGCGTAC

AAAGATAAAATCTTCCTCACTTCATGACCTTATGAGAGACCTATCTCTCTTCCAAGCAAAAGTAGAAGAT

TTTTCTGAAGTTATTGATCTTCGAGAAGGAAATGATTTTCAGTTCAATCCTTCAGCAGATTTAAGGTCCA

TTGACAGTCGGCAACTTGTAGTTTATTATGATGAGGAACATATAAGCAAACAAAGTAAATCCTATTTTGT

CAAGAGGTCCAATGAACAACAATATCGTTCAATGTTACTGTTCAATGACGGTGTATTCACTTGTTTGCCG

AAAGTAGTGGGGTCACATGTTGCCAATTTCAGGTTCTTAAGAGTTTTTTGCATGGAAAATGCTGAAGTAG

ATGGTATACTTTATCCTAGAAGTTTTGTGAGAGCGTTAGGAAGCCTTGTGTACTTGAGATATCTTAGTGT

AAGGAATGTTTCCTTGTCATTGATTCCATCTATACAGAACATGGTGCTACTCCAGACTCTCAAATTAGAT

CTATTCAGACCTATATCTATTCCATCATGGCTTTCAGGAAATATTCTGGGCAAGCTGGGACATTTGCGGC

ATTTGTGTTTACCTCTTAGTGATTCGACAACTCCAAAGAAACAAAAATTAAGGTTTGAGGGATTGGGCAA

GTTAGAGACACTGGAAAACTTCAGTAATGTGTGGTGTGAGGCCAAGGATATTCCGCAACTAATCAATCTT

CAAAAACTAACATTAGAAGTAGATAGTGGTTGTGATGATGTGAGACAGACCTTGAAAAATCTGTCATCAT

CTTCAAATCTCGGGTACTTGGGAATTTCTCTTATTTTACATGATCAACCAGGAGAGAACGTGAGAGAGTT

GTTTTGGGATTATAATTCCATCCTTGAGAAATTAAGCATAACTGGTCGGCTTGCAGAATTGGCTCAATTA

TTTGAGAACCCACCGCGGAATATTTTAAATAATCATGTTAAAGAATCTTTAATTCGTATCACTTCCTTGA

CATTAAAGTCATCGCTCCTGGTAGATGACCCAATGCCAGTACTGGAGAAGATTCCTACGCTGCGTAATCT

CGATTTATCCTACAACTCATTTGTGGGGAAGGAGATGGTCTGCTCGTCCATGGGTTTCCCGCAGCTCACC

AGTCTTGTACTAGACTACCTCCCCAACTTGGTGAAATGGAATATAGAGGAAGGTAGCATGTCCGTTCTCA

CCAAGTTAGAGATTTTTGATTGTTCGATGTTAGAGGAGCTTCCGGAAGGCCTGATATTTCTCACTTCCCT

TCGGAATTTATTTTTATACCGTATGCCTAAAGATTTCATACACAAACTCCGCACGGTAAATGGCAAACGG

GGACAGGACTTTTACAAAGTTGCTCACGTTCCCCTCCTTGCTATTTTCAATGGGAGTAAGTGGCAATACG

TGGAATAAGGAGCTGATCACTTCTAGAATCTCACAACCTCTTAATTTCCTTTTTTCAATACTCATGAATG

GGTTTTCTTGTTTAATATTGTTTTTCTTTTATCAATATAGTCTTTAACCCGTGCCAAGCACGGACGAGTA

TATAA

>MSTRG.1774.1 gene=MSTRG.1774

GTCATTCTTCAACCAAAGAAACCACATCTATCTCTTGGATGGTCTCACATCATCCTCTCCACCATCATCA

GAATCATCAGAATCTTCTTTGTCTGATCTCTTCCTCTTTGCTGGAGCCTCAACCTTCCCACCAGTGTCAT

CATCTTCATCAAATTCATCTTCCTCACCATTTTCTTCTGGTTCAAAATCGCTAGCCTCTTCTTCCTCTTC

GGCACCAACCACAGGCCTAATCAGGTACTCAGTTCCCATATCCTCCTCTTCAGCTTCCTCATCATCGCCT

TCGTCATCATCGTCATCGTCGTCGTCATCGTCATCGCCATCATCATCCTCATCATCCTCCCCCTCCTCAT

CTTCATCTTCATCATCTACCGACTGAACCGGCGGTCCACCAGCGGTGTGCAACACCTCCTGCTCATCCAC

CTCATCATCGTCATCATCATCATCCTCATCGTCATCGTCGTCATCATCATCTTCATCATCATCTCCATCT

TCCTCCACTTCACTGTCGCCTTTCTCGGGATCATCAACGACGTCATCCTCGCCGTCCAATTCCACCACCT

CGTTCTCCTCCACAAACTCCTTGTCAATCACAGCCATCGAAATTCGAATAGAATTTTAAATTAGAAATCG

AATTGTTTTGAGATTTTTGGGGATTGAAAGTGTGAGAAAAGGGGGATTTTTTGAGACTGAGGTTCTTCGT

TACGCAGAATAGGTTCGGGGGATATTTGATCTGGACCGTTGATGACGCGGATCGATGACGCGGAAGGAAA

ACGGACGGTTCAGATTGGACTATTTGTGTTTCTAGGTTCTTGCGGCGCCTCTGTGTTGTTTTTGCTTCTG

GAGTTTACAAGTCACGAGACATGCACGTGAGGTGTTTAGGGTTTGGGGATTTGTTGGATTCTTTTCAGTT

TTCTCCCCCTACTTGTCTCTCTTCAACTTTGGTCATCCAAGAAGTCAATACTCAGTCTTGCCCTCTAACC

AATTTTCTATTTCATTTTGTCGTTATCATTTGAGTATCATTTTATCACCGAATAATTAATTCTTTGTCAA

TGACTTCGACACAAATTACGAATTTTTTAAGATATAAT

>MSTRG.1779.1 gene=MSTRG.1779

GTTTCTCCCCAGCCTCTATCTATCTCCTCTCGCGCACCCTTCTTTCTCTCTTCCTCTCCTTTTCAAATTA

CAACATCTCCTTCTCATCTCGATCTACATCTTTTATCTTCTATACACATGTACCTACATAATCATAAGAT

ATTTTGAGTGTGATTAGACTTGTGAATAAAATGGGAAATAGATCTGCGTAAGATGTTGGATCGACGCTTT

GAGTGTGGTTGTTGGTTGGAGAAACAGGTTTGGGTGTTGAATCAGAGGTGTTGGTGATGTCTTTGTCGGA

AGTGCGAGTGTGATGGCGTGGGTTGAAATAGGATGTGTAACGGAGTTTCGATGAAGATGAATTGCCGGAG

TTTATGGTGGTTGAAGCTAGGTGGATTGGATTTTGTAGTGATTTTTGATTTGCTTTTAGTTGATTTGATG

GTTGCTTTTGTGATATTAGTTTTAACCCCG

>MSTRG.1783.1 gene=MSTRG.1783

GGAAGATGGAAAACCAAAGTACAGAGGAGAAAAAGGGTGGGAAATGTGGAGGAGATTATAAACACTTGGT

GCTAGGAAAGTTTACACAAGAAACAACTCAACAAGAGATAGATGATCTCATCAAGGGTTATGCCAATCTC

GTCTCTCTCGTCCCTCCCATGATATCATTTCGCTGGGGCAAGGATGTGAGCCAGGAGAATTTGAACCAAG

GTTATACCTATATCTTCGAGTCGACCTTTGACAGCACAGAAGATATTGCAGAATATGTAGCTCATCCTGC

TCATGTCGAATATGCAAATAAATTACTCCCTAAATTGGAGAAAGTGATCATCGTCGATTACAAACCTGTT

GCTGTTCATCCCTGAGCTGAATGCAGAACAGAAATCAATTATCTTTCCACTCCTGTTCTTGTATTTTCAT

TTATGTGTGAACCAGACTGTTTGCCTCAGCAACTTAATAAGCTGCAGCTGGATTAAATAATGCACTTTGT

GGCTCATTTGCTTTACCCCATCTACTATGCAGATTATGTTTTTATTTTA

>MSTRG.1783.2 gene=MSTRG.1783

GGAAGATGGAAAACCAAAGTACAGAGGAGAAAAAGGGTGGGAAATGTGGAGGAGATTATAAACACTTGGT

GCTAGGAAAGTTTACACAAGAAACAACTCAACAAGAGATAGATGATCTCATCAAGGGTTATGCCAATCTC

GTCTCTCTCGTCCCTCCCATGATATCATTTCGCTGGGGCAAGGATGTGAGCCAGGAGAATTTGAACCAAG

GTTATACCTATATCTTCGAGTCGACCTTTGACAGCACAGAAGATATTGCAGAATATGTAGCTCATCCTGC

TCATGTCGAATATGCAAATAAATTACTCCCTAAATTGGAGAAAGTGATCATCGTCGATTACAAACCTGTT

GCTGTTCATCCCTGAGCTGAATGCAGAACAGAAATCAATTATCTTTCCACTCCTGTTCTTGTATTTTCAT

TTATGTGTGAACCAGACTGTTTGCCTCAGCAACTTAATAAGCTGCAGCTGGATTAAATAATGCACTTTGT

GGCTCATTTGCTTTACCCCATCTACTATGCAGATTATGTTTTTATTTTA

>MSTRG.1783.3 gene=MSTRG.1783

GGAAGATGGAAAACCAAAGTACAGAGGAGAAAAAGGGTGGGAAATGTGGAGGAGATTATAAACACTTGGT

GCTAGGAAAGTTTACACAAGAAACAACTCAACAAGAGATAGATGATCTCATCAAGGGTTATGCCAATCTC

GTCTCTCTCGTCCCTCCCATGATATCATTTCGCTGGGGCAAGGATGTGAGCCAGGAGAATTTGAACCAAG

GTTATACCTATATCTTCGAGTCGACCTTTGACAGCACAGAAGATATTGCAGAATATGTAGCTCATCCTGC

TCATGTCGAATATGCAAATAAATTACTCCCTAAATTGGAGAAAGTGATCATCGTCGATTACAAACCTGTT

GCTGTTCATCCCTGAGCTGAATGCAGAACAGAAATCAATTATCTTTCCACTCCTGTTCTTGTATTTTCAT

TTATGTGTGAACCAGACTGTTTGCCTCAGCAACTTAATAAGCTGCAGCTGGATTAAATAATGCACTTTGT

GGCTCATTTGCTTTACCCCATCTACTATGCAGATTATGTTTTTATTTTA

>MSTRG.1787.1 gene=MSTRG.1787

CTCGACTTAAGATATGGATAAGGGTAAACGTAAATTTGAAGAAAGCAATCGTATTACTAGATACTTTTCT

AGAGCTGACAAGCAACAAGCCAATCCTGCTTCCAATAATGAGTCTGGTATCAATGTGGAGGGAGTTTATG

AATCAATGGGAAGGTTTGTTTGCTCAGCAGAAATTTCTTTTTCATCTATAAAAAAGTTGACAATGATGAT

GAACCCAAGCTTTCCTTTTAGTTATCTTGAACTAAAACGTAGTTGCTTGAAGGTTTATCAAGAAGAGAAA

GCAAAAGTTGTGCAGACTATGGAAAAATTGGACGGGTTGATTGCCCTGTCGATGGACATACTCAGGCGAG

ACACTTATGATTATTATAGTCTTGAGCTTCCTGAAGGACGAACAGTTTTTGACTATTTGTGCTTAAGAGC

TCACTTTATTGATGATAACTGGGAACCAAAGTCATGGGTTATTTGTTATATTGATACAGACCACATCTTT

AAAAATGGCGCTGAGAAGACCATTTTGAAATGCCTTTCAGATTTTCACATTGAGAAAAAGATATCCACCA

TAACACCCAGAGACAGATCTGAATATGATGACATGATCGAGACTTTGAAGAATCTCCCTAATGAGAAGAA

AAAGCTACAAATTAAGTGCCAGCAATTTGGAATATATTGTTGTTCGAATCTTTTATGTCAAATGGTGGAA

GTGGCTTTTGAAGAAATTGAAGATATCATTGATAGATTGTCAGTACGTTCCTCTCATTGGCATATTACAT

TAAGCAGTCTACAAGAAGCAATAGATTTGGAGGCTAAAGGAAAATTTTTGGAGCAATATCATTCCGAGCA

TGAGGACTTACCCGATGAGAAAGAATGGAAGAAACTTAGATATGTTTGTAGACTGGTGGCATACATATAT

AATGCAGCAGAAGTATTGTTCTTCACAAAAAATCCTACAGCCAGCCTATATCTTCATAATCTGCATCAGC

TTCAGGCAAGCTTGAGGAAAGAATCTATGAGTACTGACAGGTTGGATCTAGCTACAGATCTGCTACAGAA

GTTTGATGAGTACTGGAATGACATGTTTTTGTTATTGGCTATCGCTACTGTGATGGATCCGCGCTGTAAG

ATGAAATATATAGAATTTTCCTTTTTGAAGTATGATGATAACTCCGGAAACTCAAAAGTCACAACTATTC

TGGAGGCCATTCGGGGGATATATGATGATTATAAGATGCATAGTACTGAAGCACTGAAATCTTCGAAACT

TTCTGAATCAAAGCCTTCTGAGTTAAAGCCCCCTGATCGAGATTCTGAAGAAGAAACTTTGGAAGAAGAA

GAAGAAGATCTCCCTATGCATACACTTGAGAGACTAAAAAACTGCAACTTTGGGTTTAATTGTATGGATG

AGTATAATGAGTTTATTAAACCAGGTGATCAACCCCCAAAGTCTGAACTGGAATGGTACTTTGATGAGCC

TGTCTTGCCATGGACGAAGGATTTCGACTTAATGAGTTGGTGGAGAACCGAAAGCCCCAAGTATCCTGTA

CTGTCCAAGATGGCGCGCGACCTTTTAGCAATACCATTCTCTGTAGCCAGTTCATATGAAGCTTTTGTTT

ACCAGGATTATCGAATGGCTGATAAATCTCTTGATTCGTTAGGACCAGTTCTAATTAATGCCTTGGCGTG

CACTCGGAGCTATTCTCCAAAGTACTGAGATCTGTTGACTTAGTGAGGAAAAGCAAGTCATTTGGCGGGA

CTCTGGAGGATGGATCAAGAACTAGTTATTATTATGTGTGACTAAATCAGAACTCGGGTGTTTTGTCGTG

AATTTTATGTCTATTAGTCTGCATGGTATTTAGTTTTCTTTGGCCTGATAAACCTGGAGTATGGTATTTT

ATTTGCTTTGAGTGTCTTGGCACTTAACAGACATCTTAAGCTTATGCATCTTGGTTTCAAGGTCTTG

>MSTRG.1788.1 gene=MSTRG.1788

TGGATTACATATATAATGCAGCAGAGATACTGTACTTCACAAAAAATCCTACTGCTAGCCTATATCTTCA

TAATCTTCTGGAGCTTCAGGCAAGCTTGAGGAAAGAGTCTATCAGTCCTGACAGGTTAACTATAGTCACA

GATCTTCTAAAGAAGTTTGATGAGTACTGGAATGACATGTTCTTGGTGCTGGCTATTGCTACTGTGCTGG

ATCCACGCTGCAAGATGAAGTATATAGAATTTTCATCTCTGAAGTATGAGGATAATTCAGGAAACACGCA

AGTCATATCTGTTCTTGAGGCAATTCGGGGGATATATGATGATTATAAGATGCATAGTCTTGAATCACAG

ACTTCCAAGCCTTCTGATCCAAAGCCCTCTGGTTTAAAACCTTCTAAATCAGATTCAGACTCGGACGGGG

ATGAACTTTGTTCTGACTGCAAAGGAATTCCAAAGCATACACTAGAGAGACTCGAAAATTGTAACTTTGG

GTTTAATTGTTTAGATGAGTATAATGAGTTTCTTAAACCAAGCAATGGACCCCCAAAATCTGAATTGGAG

TTGTACTTTGATGAGCCTGTCATGCCATGGAGCAAGGATTTTGACTTAGTGAGTTGGTGGAGAACTGAAA

GCCCCAAATATCCTATATTATCCAAGATGGCCCGCGACTTTTTGGCAATACCATTTTCACTAGTCAGTTC

ATATGAAGCTTATTATTACAGAGATTCTCGAGCGGCTGATAAAAGTCTTGCTTTTCTGGGACGAGATTTA

GTTAATGCATTGGTGTGCAGTCGGAGCTATTTTAAAGAGGACTGAAGTTCTTCATCGTATTTAGGGAATA

GCAAGTCATCTGGAAGGGGTTCAGGAATCAGGAGGCGTGGATCCTGCATCATCGGATCAAAGCATTGGTG

GTTTTGTCATGGATTTTGTTTGTTGGCTTCGTTCGTGAGTAGTTTGCATTGCATTAGATCTAGTTTCCTT

TTGGCAGAAAACGCTGACTTCAAGTATTATGTGTTTGCATTGCATTAGATCTAGTTTCCTTTTGGCAGAA

AACGCTAACTTCTAGTATTGTGCGATTTGTTTGAAGTATTTTAGTAG

>MSTRG.1788.2 gene=MSTRG.1788

AGTAGACTACAGTCTCTTTAGACTCATTTTACCCTAGAGTCTTTTCGCATTCTCCGTCGTTTCGTCTTCT

AATCAGACATGGATAAGGGTAAACGTAATCCGGGAGTGAACCCGGGCTTTTCTTTTAGTTATGGTGAATT

TAAAGGCAGTTGCTTAAAAGTTTATCAGGAAGGGGTAGCAAAAGTTATTAAGATTTTGGAAAATTTAGAT

GGACAGATTGCCCTCTCGATTGAGTTATTGAGGCTGGACACTTATGGTCTTCCTCGTCCCAAAGATTGTG

ATGAGCCAGAGATTATTGACTATTTGTGTGTCAGAGCTCATTTTATTGATGATAATTGGAAACCAAAGTC

GTGGGTTCTTTACTATGTTGATGCAGACCACATTATGGAAACTGGCTATGAGAAGACAATTCTGAAATGT

ATTTTGGATATGCGCATTGAGAATAAGATATTCACTGTGACACGTGCATATGATAATATGTATTTTGAAC

TGATTGAGGTTGTAGAAAACCAGGCTAAGGAGAAGAAAACACTGCAATCAAATTGCCAGCCATTTAAAAT

ATATTGTTGTGCTAACCTTTTAGGCAGAATGGTGGAAGCAGCTTTTAAGGACATTTCTGACATTATCTTG

AAAATAACACAAAAAAATTTTATGTACTGGCACCTTACATTCAGTACACTACAAAATGCGATGAAGGCGG

AGGCTGATGGAGAATTTTCAGAGCTGTTTGGTTGTGATTATTTTGATCCACCTACTGAGGAAGAATGGAC

TAAACTTAAATATGTTTGCAGACTGGTGGATTACATATATAATGCAGCAGAGATACTGTACTTCACAAAA

AATCCTACTGCTAGCCTATATCTTCATAATCTTCTGGAGCTTCAGGCAAGCTTGAGGAAAGAGTCTATCA

GTCCTGACAGGTTAACTATAGTCACAGATCTTCTAAAGAAGTTTGATGAGTACTGGAATGACATGTTCTT

GGTGCTGGCTATTGCTACTGTGCTGGATCCACGCTGCAAGATGAAGTATATAGAATTTTCATCTCTGAAG

TATGAGGATAATTCAGGAAACACGCAAGTCATATCTGTTCTTGAGGCAATTCGGGGGATATATGATGATT

ATAAGATGCATAGTCTTGAATCACAGACTTCCAAGCCTTCTGATCCAAAGCCCTCTGGTTTAAAACCTTC

TAAATCAGATTCAGACTCGGACGGGGATGAACTTTGTTCTGACTGCAAAGGAATTCCAAAGCATACACTA

GAGAGACTCGAAAATTGTAACTTTGGGTTTAATTGTTTAGATGAGTATAATGAGTTTCTTAAACCAAGCA

ATGGACCCCCAAAATCTGAATTGGAGTTGTACTTTGATGAGCCTGTCATGCCATGGAGCAAGGATTTTGA

CTTAGTGAGTTGGTGGAGAACTGAAAGCCCCAAATATCCTATATTATCCAAGATGGCCCGCGACTTTTTG

GCAATACCATTTTCACTAGTCAGTTCATATGAAGCTTATTATTACAGAGATTCTCGAGCGGCTGATAAAA

GTCTTGCTTTTCTGGGACGAGATTTAGTTAATGCATTGGTGTGCAGTCGGAGCTATTTTAAAGAGGACTG

AAGTTCTTCATCGTATTTAGGATTGGAGAGAGTCAAGTCCCTGAGCAAATTTTGTTGGGAATAGCAAGTC

ATCTGGAAGGGGTTCAGGAATCAGGAGGCGTGGATCCTGCATCATCGGATCAAAGCATTGGTGGTTTTGT

CATGGATTTTGTTTGTTGGCTTCGTTCGTGAGTAGTTTGCATTGCATTAGATCTAGTTTCCTTTTGGCAG

AAAACGCTGACTTCAAGTATTATGTGTTTGCATTGCATTAGATCTAGTTTCCTTTTGGCAGAAAACGCTA

ACTTCTAGTATTGTGCG

>MSTRG.1788.3 gene=MSTRG.1788

TCAGATATGGATAAGGGTGGCTTTTCTTTTAGTTCTGATGAACTTAAAGGTAGTTGCTTAGAAATTTATC

AGGAAGAGAGAGCAAAAGTTATTAAGATTTTTGAAAAGTTAGATGGACGGATTGCCCTCTCGATTGAGTT

ATTGAGGCTGGACACTTATGGTCTTCCTCGTCCCAAAGATTGTGATGAGCCAGAGATTATTGACTATTTG

TGTGTCAGAGCTCATTTTATTGATGATAATTGGAAACCAAAGTCGTGGGTTCTTTACTATGTTGATGCAG

ACCACATTATGGAAACTGGCTATGAGAAGACAATTCTGAAATGTATTTTGGATATGCGCATTGAGAATAA

GATATTCACTGTGACACGTGCATATGATAATATGTATTTTGAACTGATTGAGGTTGTAGAAAACCAGGCT

AAGGAGAAGAAAACACTGCAATCAAATTGCCAGCCATTTAAAATATATTGTTGTGCTAACCTTTTAGGCA

GAATGGTGGAAGCAGCTTTTAAGGAC

>MSTRG.1788.4 gene=MSTRG.1788

TCAGATATGGATAAGGGTGGCTTTTCTTTTAGTTCTGATGAACTTAAAGGTAGTTGCTTAGAAATTTATC

AGGAAGAGAGAGCAAAAGTTATTAAGATTTTTGAAAAGTTAGATGGACGGGTTGCCCTCTCGATTGACTT

GTTGAGGCTGGACACTTATGGCCTTCCTCGTCCCAAAGATTGTGATGAGCCAGAGATTATTGACTATTTG

TGTGTCAGGGCTCATTTTATTGATGATAATTGGAAACCAAAGTCGTGGGTTCTTTACTATGTTGATGCAC

TCCACATCATGAAAACTGGCTACGAGAAGACAATTCTGAAATGTATTTCGGATATGCGCATTGAGAATAA

GATATTCACTGTAACTCGTGCGTATGATATTATGTATTCTGAACTGATTAAGGTTGTAGAAAACCAGGCT

AAGGAGAAGAAAACACTGCAATCAAATTGCCAG

>MSTRG.1788.5 gene=MSTRG.1788

CTTTGGACTCATTTTACCCTAGAGTCTTTTCGCATTCCCTGTCGTTTCGTCTCCGATCAGATATGGATAA

GGGTGGCTTTTCTTTTAGTTCTGATGAACTTAAAGGTAGTTGCTTAGAAATTTATCAGGAAGAGAGAGCA

AAAGTTATTAAGATTTTTGAAAAGTTAGATGGACGGGTTGCCCTCTCGATTGACTTGTTGAGGCTGGACA

CTTATGGCCTTCCTCGTCCCAAAGATTGTGATGAGCCAGAGATTATTGACTATTTGTGTGTCAGGGCTCA

TTTTATTGATGATAATTGGAAACCAAAGTCGTGGGTTCTTTACTATGTTGATGCACTCCACATCATGAAA

ACTGGCTACGAGAAGACAATTCTGAAATGTATTTCGGATATGCGCATTGAGAATAAGATATTCACTGTAA

CTCGTGCGTATGATATTATGTATTCTGAACTGATTAAGGTCGTAGAAAACCAGGCCAAGGAGAAGAAAAC

ACTTCAATCAAAGTGCCAGCCATTTAAAATATATTGTTGTGCTAACCTTTTAGGCAGAATGGTGGAAGCA

GCTTTTGAGGACATTTCTGATGTAATCTGGAAAATATCGCAAAAAAATTATATGTACTGGCACCTTACAT

TCAGTACACTACAAGAGGCGATAAAGGCGGAGGCTGATGGAAAATTTTCAGAGCTGTTTGGTTGTGATTA

TTTTGATCCACCTACTGAGGAAGAATGGACTAAACTTAAATATGTTTGTAGACTGGTGGATTACATGTAT

AATGCGGCAGAGATACTGTTCTTCACAAAAAATCCTACTGCTAGCCTGTATCTTCATAATCTTCGGGAGC

TTCAGGCAAGCTTGAAGAAAGAGTCTATCAGTCCTGACAGGTTAACTATAGTCACAGATCTGCTAAAGAA

GTTTGATGAGTACTGGAATGACATGTTCTTGGTGTTGGCTATTGCTACTGTGCTGGATCCACGCTGTAAG

ATGAAGTATATAGAATTTTCATATCTGAAGTATGAGGGTAATTTGGGAAACATGCAAGTCACATCTGTTC

TGGAGGCAATTCGTGGGATATATGATGGTTATAAGATGCATAGTCTTGAACCGAAGACTTCCAAGCCTTC

TGATTCGAAGTCCTCTGATTTAGAAGCTTCTGAATCAGACTCAAACTCAGACTGGGAAATTTTTTCTGAT

TGCGAAGAAATTCCAAAGCATACACTCGAGAGACTCCAAAATTGCAACTTTGGGTTTAATTGTCTAGATG

AGTATAATGAGTTTCTTAAACCAAGCAATGGACCCCCAAAGTCTGAATTGGAGTTGTACTTTGATGAGCC

TGTCATGCCATGGAGCAAGGATTTTGACTTGGTGAGTTGGTGGAGAACTGAAAGCCCCAAATATCCTATA

CTATCCAAGATGGCCCATGACTTTTTGGCAATACCATTTGCACTTGTCAGTTCAAATGAAGCTTATTACT

ACAGAGATTCTCGAGCGGCCAGTAAAAGTCTTGCTTTCCTGGGACGTGATCTCGTTAATGCATTGGTGTG

CACTAGGAGCTATATTAAAGAGGATTGAAGTTCTTTGTCGTATTTAGGAATAGCAAGTCATCTGGCAGGG

GTTCAGGAGCGTGGATTGTGCATCATCAGATCAAAGCTTTGGTGGTTATGTCGTGGATTTTGTTTGTTGG

GTTCGTTCATGAGTAGCTTGCCTTGCAATAGATCTAGTTTCCTTTTGGAAGAAAACGCTGACTTCAAGTA

TTATGGGATTTGTTTGAAGTATTTGAGTAGACAATACAATGCTTTCTATTAAATTCCTATTTGCGT

>MSTRG.1794.1 gene=MSTRG.1794

CCGCAACCGAATTATTTTAAAGAGTCATGACATAACACAATTCAAAACATATTCTAACCGAATTCTCGTA

ATGAAATGAGTAAACAAACAATAACGATTAGTTAATTACTCGACTAATCACCACCGGCTTTCTGATAAAC

ATAAGTAAGGCCACACTTGCCGCAATAATGGCGATCAAAATGATTCGCCATGAACGTGCCGGCCCCACAC

TCAGCATTAGGACACTCCTTCCTCAACCTCTGCACCTTCCCACTCTCATCCACCTTATAGAACTGAAGCA

CGGCGAGCTTGACTTTCTTATGCTTGTGCTTGATCTTCTTAGGCTTGGTGTAGGTCTTCTTCTTGCGTTT

CTTGCCGCCGCCGCGGAGGCGGAGGACGAGGTGGAGAGTGGATTCCTTCTGGATGTTGTAGTCGGCTAGG

GTTCTGCCGTCCTCGAGCTGTTTTCCGGCGAAGATGAGGCGTTGCTGGTCGGGAGGGATGCCTTCTTTGT

CTTGGATTTTGGCTTTGACATTGTCGATGGTGTCGGAGGATTCCACCTCCAGGGTGATGGTTTTGCCGGT

GAGGGTTTTTACGAAGATCTGCATTGTGGGGGAGTG

>MSTRG.1801.1 gene=MSTRG.1801

GAGACAGTAGTGCTCTATGCCTATATCTACACAACTGCTGACACTACTATTCGACTTTAATTTCCATTTG

TTTCATGCCCTTCTTGAAATGGCCGCCAAGAAGCTGTGCGAGTCATTCTCCAAGAGCCTGTTGGATGAAG

TTCATCGATGCGGGTGCATGAAGCAGACCGGGGTGAGTCTCCGGTACATGATGGAATTTGGGTCCAAGCC

CACTCTCAGAAATCTTTTGATTTCAGCTCAGTTTCTTCACAAAGAGCTCCCTATCAGGATTGCTCGCAGA

GCTATTGAGCTTCAGAAACTACCCTATGGTTTATCTGATAAGCCTGCTGTTCTTAAGGTGCGCGATTGGT

ATTTGGATTCATTTCGTGACCTTAAATCCTTCCCTGACATCAAGGACAATAATGATGAGTTAGAATTCAC

AAAAATGATTAAGATGATCAAGGTGAGACACAATAATGTTGTGCCCACAATGGCCTTGGGAGTCCAACAA

TTGAAGAAAGCTCTGCATCCAAAGGTTCATAATGACGGTCTAGAAGAAATTCACCAGTTTCTGGATCGCT

TCTACATGTCTAGGATTGGAATACGTATGCTTATAGGGCAACATGTGGAATTACACAATCCCAATCCTCC

TCCAGATGTTGTAGGCTACATACATACTAAAATGTCTCCTCTGGAGGTAGCAAGAAGTGCCAGTGAGGAT

GCTCGTGGCATTTGTCTGCGTGAATATGGCAGTGCACCTGATGTTAATATCTACGGGGATTCTAATTTTA

CTTTCCCGTATGTTCCAAATCACTTGCATATGATGGTTTTTGAGTTGGTTAAGAACTCATTGCGAGCTGT

CCAAGAGAGGTTTATGGACTTGGATAAAATTGCACCTCCTGTTCGAATTATAGTGGCTGACGGGTTAGAA

GATGTTACCATCAAGATTTCTGATGAAGGTGGCGGCATAGCAAGAAGTGGTCTGCCTAAAATTTTTACAT

ATCTCTACAGCACTGCAAAAAACCCATTGTGTGAGCAGCCAGATCTCGAAACAGCTAATGTGGCGACAAT

GGCTGGGTATGGATACGGGCTCCCGATTAGCCGTTTGTATGCTCGGTATTTTGGAGGGGATCTGCAAATT

ATCTCTATGGAAGGATATGGGACTGATGCTTATCTTCATTTGTCTCGCTTGGGAGATTCACAAGAGCCCC

TGCCATGATACCTCCATTCTCAATCAAGATTCAACAAGCTAGCAATACTTTCACAGTACAATGCACATGT

AAATACTCCGAACATATAAAATTGCAGCATATCTTCAAATATATACTTGATGTAAATATGCAAAGCTTTA

CTTTGTATAAAAGTTCATATGTGCGTCTTGTATATGGCTATATGATCCTTTACCATTGTTAGACACTCAC

AACATGATTGTAGAGCTGAAGTGTTATTTCCCGATTTTTCTGGTATCCCATCTGATGATGCAGTACAATT

TGAAACTCTTCCTGGG

>MSTRG.1824.1 gene=MSTRG.1824

ATATGATAGCCTAGGGGGACTCTAATATAAAGGAACTTTTACATATAGGAGAAAGAGGCTGAGGACATGG

CAAACTTGACTCTAACAGTTTTTGGAAGAATTTATGTAAACTACAGTATTATTTGGTAGCATGGGAGGAA

TAATTTTCTGCGTGTTTCCATAACTTGGTGAAAAGGAAATTGCAGTATTATATTTAGTTTGTTTATTTAG

TAGGTGGACTATATTGCAAGAGGGTCTTTGTTTTGTCAGATGCCCCTACAACATAAACACATAGTACTTT

TATATAACCAGAAGGTCTAGGATTGGATACTTGTATGGAAGTTATGTTTTTATCTTCAGCTGTACGCATT

GTATGCAACACTTGCAGTAGTAGTTATATATTTCGTTAAATTTCAGCAATTAGAAGTTACTGCAGCAGTA

AACTAATAACTGGAGTACAATACAATTGTGAAGTATAGGCACTAAATTGTTTATTTTGTTCTATGAATAG

AGTTCTTGCCTGATCCCTCATTGTAAAACTTAATTTTTACGAGCAAAAGGAAGAGAATTCTCAGATGCTT

GACCAAATCATCAAGCTGTGCAGTTTTGAAAGTATGTGCAGTCTTGTGAGGTTAATAAGACTGGGAATTT

ACGTGGAAGTAAAAATCGATATATATTTGTGTGGTGGCATGGTTGGCGAATGGAAGAAATATTTAACTGC

GGAGATGGATTCTAAGCTGGATCAGATTACTGAAGAAAGTTTTGTGGTTCGTGATTGTATATGGCATGAT

GCTGACACCTGTATATATTTGACAAGTACTGGCCTTGCTTTTCTTATTCTCGTTCCATTTCAGACTACTC

TGTGTGTGTATGTCTTCTCTGTGTTTAACCCCATGTTCAGTTATGTCAAACTTTATAAATTGCTTGGAAT

AGAA

>MSTRG.1829.1 gene=MSTRG.1829

AATCATTTCACCAGTTTCTCTGCACGAATACCGAGAGAAACCGAGCGGTGTTCTTCGTGTTCTTCATAAT

CAAACGAAGATTTGAAGGTGTTATTGGAATCCGATGGAGGCGTATGAATAGTCAAAACGAAGCTATCGAC

GCGTAGAATCTAGTTTAATCATCCGTTTGTGTGCAGATTCAACAGGATTTGGACGAGAAGTGTATGCCCG

AAGACGTTCAGAAAGTCACGTGGTGTTACAAAGCGAATAAGGGATGAATCGAGTAAGCGAAGCGAAGATG

TACGGTAGATAGAGTATAGCCAGAGATGGTCTAGAGACTATGAGAGGCATAGATTGTGAAAGTGGAAAGA

TGAGATTGAGATGTGAGACTGGAGTCAGATTTAAGGCAAGCTTAAAGATGAGATGATTGATTGAGATTGT

GAGTAAGGCTAATGCTAGGCGAGGAGACAGAGTTGAGGATCCAGTGGTCGAGGAGTGTTCAGGAAAGTTG

ATGAGGTTGGTGCAAGCTGGAATAGTTGACAGTGATTCTTCTTATCGAAGATGATCTGATTTGGAGCATC

TAGGCATATTTGCTCCGACAAGGCGACTTTCTCTAGCCTTGTAGCTTCTGATGCTGGTGAGAAGCTCTAC

ATGGGAAACTCTGCGAC

>MSTRG.1830.2 gene=MSTRG.1830

CCCCAACCATCTTACTTTTATGATGCCCCGACTCCTCTCAAAACTCTTGCGTTCTCTGCACACAAGTCCG

ATGGGTTCACTAATCGATTCACTGGGTTCTTGATTTGTCACTACGCTTGAGCCGAGCCTGCAGAATGTTT

TCTTAATTAAGCTGGTCTTCTGACTAGATGATCTGGTGTATCATACTCTTGTTCGTTGAAGAATAATTAG

AGGTTTTATAAAGCATCAGCCTCTTGGACAATATTTCATATTCAACAGGTCCACCTTTTGTTGTTTAAGC

TGCCTGTAAAAGAGCAAATAATGCAACCTAGTATCGACCGTTACTTCTGGATCCAGTTTTTATAAAAGTT

TAGACGGATAGTTTCCGCAAGTTTATGGTCATCTTTTTGATGTTATGCAAACTTCTGGTGCACTTACAAA

CCTATTTGTTGCAGCAGTCCTAGCACTACATGTAAGTTCTCCAAACTGGTTTCAGGAATGAAAGATGACA

GCTCAATGCCTACTTTCTATCCAAGGATGTTATCATTTGGAAAATAAAGATCCCACATATTTCTACTTAC

AAGAGTTACCGTGCTTTCAAGTTTATTTACATGCTTTTTGAAGGATAGATCGATGGACGTTTTATCTTTG

ACATTTTTGTTGAAGGTTCTTCACCGTTATAAATTTAGAATGTTCGCATGTAAGAATGTTCTTTAGTTAT

CACGTGTCTTTGTTGAATTTATAAATTATGTAAAATTTTC

>MSTRG.1830.1 gene=MSTRG.1830

CCCCAACCATCTTACTTTTATGATGCCCCGACTCCTCTCAAAACTCTTGCGTTCTCTGCACACAAGTCCG

ATGGGTTCACTAATCGATTCACTGGGTTCTTGATTTGTCACTACGCTTGATGATCTGGTGTATCATACTC

TTGTTCGTTGAAGAATAATTAGAGGTTTTATAAAGCATCAGCCTCTTGGACAATATTTCATATTCAACAG

GTCCACCTTTTGTTGTTTAAGCTGCCTGTAAAAGAGCAAATAATGCAACCTAGTATCGACCGTTACTTCT

GGATCCAGGTTTCAGGAATGAAAGATGACAGCTCAATGCCTACTTTCTATCCAAGGATGTTATCATTTGG

AAAATAAAGATCCCACATATTTCTACTTACAAGAGTTACCGTGCTTTCAAGTTTATTTACATGCTTTTTG

AAGGATAGATCGATGGACGTTTTATCTTTGACATTTTTGTTGAAGGTTCTTCACCGTTATAAATTTAGAA

TGTTCGCATGTAAGAATGTTCTTTAGTTATCACGTGTCTTTGTTGAATTTATAAATTATGTAAAATTTTC

TCCAAGTTTAAATAATATGGATTATTAGC

>MSTRG.1830.4 gene=MSTRG.1830

CCCAACCATCTTACTTTTATGATGCCCCGACTCCTCTCAAAACTCTTGCGTTCTCTGCACACAAGTCCGA

TGGGTTCACTAATCGATTCACTGGGTTCTTGATTTGTCACTACGCTTGATGATCTGGTGTATCATACTCT

TGTTCGTTGAAGAATAATTAGAGGTTTTATAAAGCATCAGCCTCTTGGACAATATTTCATATTCAACAGG

TCCACCTTTTGTTGTTTAAGCTGCCTGTAAAAGAGCAAATAATGCAACCTAGTATCGACCGTTACTTCTG

GATCCAGGTTTTGTTCCAATCCTGACTTGCAACTTTCAGAATACAGATTAGGGGAACGGGAATCTAATTT

TCTATCTGCATAAAAGGACACGCTGTTAACAAATCAATTGAAGTAAATGAATGTCAGGGGAGTACAACAT

CCATGTTCCTGTCTGCATTCTCCGGGTCTACTTTTACAGTTTTTATAAAAGTTTAGACGGATAGTTTCCG

CAAGTTTATGGTCATCTTTTTGATGTTATGCAAACTTCTGGTGCACTTACAAACCTATTTGTTGCAGCAG

TCCTAGCACTACATGTTTCAGGAATGAAAGATGACAGCTCAATGCCTACTTTCTATCCAAGGATGTTATC

ATTTGGAAAATAAAGATCCCACATATTTCTACTTACAAGAGTTACCGTGCTTTCAAGTTTATTTACATGC

TTTTTGAAGGATAGATCGATG

>MSTRG.1830.3 gene=MSTRG.1830

CCCAACCATCTTACTTTTATGATGCCCCGACTCCTCTCAAAACTCTTGCGTTCTCTGCACACAAGTCCGA

TGGGTTCACTAATCGATTCACTGGGTTCTTGATTTGTCACTACGCTTGATGATCTGGTGTATCATACTCT

TGTTCGTTGAAGAATAATTAGAGGTTTTATAAAGCATCAGCCTCTTGGACAATATTTCATATTCAACAGG

TCCACCTTTTGTTGTTTAAGCTGCCTGTAAAAGAGCAAATAATGCAACCTAGTATCGACCGTTACTTCTG

GATCCAGGACACGCTGTTAACAAATCAATTGAAGTAAATGAATGTCAGGGGAGTACAACATCCATGTTCC

TGTCTGCATTCTCCGGGTCTACTTTTACAGTTTTTATAAAAGTTTAGACGGATAGTTTCCGCAAGTTTAT

GGTCATCTTTTTGATGTTATGCAAACTTCTGGTGCACTTACAAACCTATTTGTTGCAGCAGTCCTAGCAC

TACATGTTTCAGGAATGAAAGATGACAGCTCAATGCCTACTTTCTATCCAAGGATGTTATCATTTGGAAA

ATAAAGATCCCACATATTTCTACTTACAAGAGTTACCGTGCTTTCAAGTTTATTTACATGCTTTTTGAAG

GATAGATCGATGGACGTTTTATCTTTGACATTTTTGTTGAAGGTTCTT

>MSTRG.1830.5 gene=MSTRG.1830

CAACCATCTTACTTTTATGATGCCCCGACTCCTCTCAAAACTCTTGCGTTCTCTGCACACAAGTCCGATG

GGTTCACTAATCGATTCACTGGGTTCTTGATTTGTCACTACGCTTGATGATCTGGTGTATCATACTCTTG

TTCGTTGAAGAATAATTAGAGGTTTTATAAAGCATCAGCCTCTTGGACAATATTTCATATTCAACAGGTC

CACCTTTTGTTGTTTAAGCTGCCTGTAAAAGAGCAAATAATGCAACCTAGTATCGACCGTTACTTCTGGA

TCCAGTTTTTATAAAAGTTTAGACGGATAGTTTCCGCAAGTTTATGGTCATCTTTTTGATGTTATGCAAA

CTTCTGGTGCACTTACAAACCTATTTGTTGCAGCAGTCCTAGCACTACATGTTTCAGGAATGAAAGATGA

CAGCTCAATGCCTACTTTCTATCCAAGGATGTTATCATTTGGAAAATAAAGATCCCACATATTTCTACTT

ACAAGAGTTACCGTGCTTTCAAGTTTATTTACATGCTTTTTGAAGGATAGATCGATG

>MSTRG.1830.6 gene=MSTRG.1830

CAACCATCTTACTTTTATGATGCCCCGACTCCTCTCAAAACTCTTGCGTTCTCTGCACACAAGTCCGATG

GGTTCACTAATCGATTCACTGGGTTCTTGATTTGTCACTACGCTTGAGCCGAGCCTGCAGAATGTTTTCT

TAATTAAGCTGGTCTTCTGACTAGATGATCTGGTGTATCATACTCTTGTTCGTTGAAGAATAATTAGAGG

TTTTATAAAGCATCAGCCTCTTGGACAATATTTCATATTCAACAGGTCCACCTTTTGTTGTTTAAGCTGC

CTGTAAAAGAGCAAATAATGCAACCTAGTATCGACCGTTACTTCTGGATCCAGGTTTTGTTCCAATCCTG

ACTTGCAACTTTCAGAATACAGATTAGGGGAACGGGAATCTAATTTTCTATCTGCATAAAAGTTTTTATA

AAAGTTTAGACGGATAGTTTCCGCAAGTTTATGGTCATCTTTTTGATGTTATGCAAACTTCTGGTGCACT

TACAAACCTATTTGTTGCAGCAGTCCTAGCACTACATGTTTCAGGAATGAAAGATGACAGCTCAATGCCT

ACTTTCTATCCAAGGATGTTATCATTTGGAAAATAAAGATCCCACATATTTCTACTTACAAGAGTTACCG

TGCTTTCAAGTTTATTTACATGCTTTTTGAAGGATAGATCGATG

>MSTRG.1830.7 gene=MSTRG.1830

CCCGACTCCTCTCAAAACTCTTGCGTTCTCTGCACACAAGTCCGATGGGTTCACTAATCGATTCACTGGG

TTCTTGATTTGTCACTACGCTTGATGATCTGGTGTATCATACTCTTGTTCGTTGAAGAATAATTAGAGGT

TTTATAAAGCATCAGCCTCTTGGACAATATTTCATATTCAACAGGTCCACCTTTTGTTGTTTAAGCTGCC

TGTAAAAGAGCAAATAATGCAACCTAGTATCGACCGTTACTTCTGGATCCAGTTTTTATAAAAGTTTAGA

CGGATAGTTTCCGCAAGTTTATGGTCATCTTTTTGATGTTATGCAAACTTCTGGTGCACTTACAAACCTA

TTTGTTGCAGCAGTCCTAGCACTACATGTAAGTTCTCCAAACTGGTTTCAGGAATGAAAGATGACAGCTC

AATGCCTACTTTCTATCCAAGGATGTTATCATTTGGAAAATAAAGATCCCACATATTTCTACTTACAAGA

GTTACCGTGCTTTCAAGTTTATTTACATGCTTTTTGAAGGATAGATCGATGGACGTTTTATCTTTGACAT

TTTTGTTGAAGGTTCTTCACCGTTATAAATTTAGAATGTTCGCATGTAAGAATGTTCTTTAGTTATCA

>MSTRG.1830.8 gene=MSTRG.1830

CGACTCCTCTCAAAACTCTTGCGTTCTCTGCACACAAGTCCGATGGGTTCACTAATCGATTCACTGGGTT

CTTGATTTGTCACTACGCTTGATGATCTGGTGTATCATACTCTTGTTCGTTGAAGAATAATTAGAGGTTT

TATAAAGCATCAGCCTCTTGGACAATGTCCACCTTTTGTTGTTTAAGCTGCCTGTAAAAGAGCAAATAAT

GCAACCTAGTATCGACCGTTACTTCTGGATCCAGTTTTTATAAAAGTTTAGACGGATAGTTTCCGCAAGT

TTATGGTCATCTTTTTGATGTTATGCAAACTTCTGGTGCACTTACAAACCTATTTGTTGCAGCAGTCCTA

GCACTACATGTAAGTTCTCCAAACTGGTTTCAGGAATGAAAGATGACAGCTCAATGCCTACTTTCTATCC

AAGGATGTTATCATTTGGAAAATAAAGATCCCACATATTTCTACTTACAAGAGTTACCGTGCTTTCAAGT

TTATTTACATGCTTTTTGAAGGATAGATCGATGGACGTTTTATCTTTGACATTTTTGTTGAAGGTTCTTC

ACCGTTATAAATTTAGAATGTTCGCATGTAAGAATGTTCTTTAGTTATCA

>MSTRG.1842.1 gene=MSTRG.1842

GCTAGCAATATTCTATCTACCCGATTATCTACTCTATCTTCAACACAAACCTTTTCACGGCAATCTCTCA

GATATGAAGCACCAAGTTCAAAAAAATGTTGTTGGGGTTCCTGTAAGCTCATCTGTGTATTCAAATAAGA

GATCATCAGGCAACCTCGTAACAGACCATAATCTTGCTATCAAGTGCTACAATCCGCCTGCAAAGACAAA

TTCCAAATTTAGTCACAATAAACTAGTATCAACAAATTATAGGACAAACAAGCTTGTCGGAAGAACAGAA

AGTTTTGCACAAAGAGTCAGGGAGCACGTGAGATTAGGCCCCAAAATTACTGAAACCCTGAAGGGCAAGC

TGAGTTTAGGAGCGAAGATTCTCCGTTTAGGCGGAGTTAGCAAAGTTTTCAACAAGAATTTCAACACAAG

AACAGGAGAGAAGTTACTAAAAGCTTCCCAATGCTATCTGTCAACAACAGCAGGTCCAATCGCAGGACTC

CTCTTCATTTCCACCGACAAGATCGCCTTCTGCAGTGAGAGGACAATGAAAGTATCTTCTCCAACCGGAG

AAATATTAAGAGTACATTATAAGGTGATGATCCCTCTAGGGAAGATTAAGAGAGCCAACAAGAGCACAAA

TATCGAAATTCCATCACAGAAGTATATAGAGATAGTGACCGGTGATGATTTTGATTTTTGGTTCATGGGA

TTCATTAATTATCAGAGAACTTTCAACTATCTAGAACAAGCAATCTCTCAAATGCAATGAAGAAAGAAAT

GTTTTTATCCTTTCTGCCTTGCTGCATGCAACGCACGTATATAACTTGAGCTATAGATTACACATATGGA

AGTTGGAAGGTGCATTCTCTGAAAATATTTGGATGTAAAACAAAGTGTAGATACATGAATATTTCAGTTG

GATTAGATATTAAAAGTTCATGATTCTGCCTTATCTGTGTGGCTTATATGTTTTAATTTAAGTGCA

>MSTRG.1843.1 gene=MSTRG.1843

GTTGATTTAATTCATCAGACAATATACATCTGCAAGAGAAGAATATTCAATCAGGCTTTAGAGACAGAAT

CAAGCATTTTACAGTTCATCAGACATGAAGAACCAGATTCTAGGACAGGTTATTGGAATTCCAGTCAGTT

CAGCAGCATATCCAGCTGAGATGCCTCAACAAGGAATTTTATGTATCACTTCCGATCAATACTACGCATC

TCCAAAATTTAGACAGAGTAAGATGGACACTGTTTTCGATAGGATGAACCAATTGGCAAAAGGCATCCGT

GAACATGTGAAAATTGGGCCTAAGTTGAGTGAAACAGTGAAGGGAAAGTTGAGATTAGGAGCTAGAATTC

TCCAAGTAGGGGGTGTCAAGAAAGTGTTTAAACAAAACTTCAGTTTCAGTGAAGGAGAAAAACTGTTGAA

GGCTTCACAATGCTATTTATATACAACTGCAGGTCCAATAGCCGGATTACTATTTATCTCAACCGACAGA

ATTGCTTTCTGTAGTGAAAGATCAATGAAGGTCTCTTCTACAACAGGGGATTTGCTTCGGATTCATTACA

AGGTTTCAATCCCATTAAACAAAATAAAGAGAGCAAATGAAAGTGAAAATATGAAGGATCCATCAAGAAA

GTACATTGATATAGTTACAAAGGATGACTTTGAGTTTTGGTTCATGGGATTTCTTAACCATCAAAGAACT

TACAAGTTTCTTCAGCAAGCAGTGTATGAATCTCAATAAACACTATAAAAAAAAACCTGTTCATTCTTCC

CTTGTTCCATAACATAATTTGTGTTATGGGATTAATATGTAAGTATAGTACTAATGTCTCTGTAATCTAG

AATTCATTTTCTTAATGAAAATTCACTTGTTTCCAGATATGACAGCAACACTGCACACTCTCTGCTAC

>MSTRG.1844.1 gene=MSTRG.1844

TATTTTTGCACTCAAAAAACACTCCTTATATATAAATACTTCTCTTCAAACATGGAGTAAAAGTTGCAGA

TTCTGAGGATGATTTTAGGGAATCTGCTATGTTTATGGTTTGATAAATTTGTTCCTTGATTATTTGATTT

CAACTAACATGGTATCAAAAGTCGGTTTACTTGCATCAGAAAGCAGGCAGGCCATCTCAGAAAGTTCTCT

TCAAAATGCTTACACAGTCAGCCTAAAACAAACAAGGTAGCCACACTGAAGAAGGCTTGGCCCTTTGAAC

AAGCTCCTTTTGCAGCTAGCAGACATGCCATCTTCTATATACATTTTCTATTATTCAACTTTATCCATTC

ACCTTTATTGATATTAATGATGGAATGACGAAAAGATGACTCAGGCAACTGAAAGTAAAGATCCAAATAC

CGACCTCATTCTATAAATACCGTTTGAATAAACAATATTCAAATCCATCTTACACTAAACCAGTCATTAC

CTCAGAGTTTAAAGGGTATTACTATTTACCAGAGAGTTGGTGATCCAGATAAAACGCGAAGATTCTACAA

TTTGCTCACAATGAAGAACCAGGCTGGGAAACATATTATTCCATATTCACCAGGCTCTGCAATTCCTGTT

GAGAGGCCTCTAAGAGGAGTGCTAACTAGCCATTGCCATGCATCACCAAAATTCAGATGGGGTAGATTTG

ATTCAGTATTTAGTAGGATGAACAGCTTCGGTGAAAAAATAGATACTTTAGCAGAAGGCATCAGAGAGCA

TGTGAGAATAAGGTCCATAACTGAAACTATGAAAGGGAAGTTGAGCTTAGGAGCAAAAATATTACAAGTA

GGTGGAGTGAAGAAAGTGTTCAGACGAAATTTTAGTGCGAAAGAAGGGGAGAAGCTATTGAAAGTTTGTC

AATGCTATTTGTCGACCTCAGCAGGTCCATTGGCAGGCCTACTCTTTATTTCCACTGATAAGGTTGCCTT

CTGCAGTGAAAGATCGATCAAAGTTTCTTTCCCTACTGGAGAGTTTCACAGAATACGTTACAGGGTCGTC

ATCCCACTTGGAAAGATCAAGAAAGTGAATGAGAGCGAAAATATGAAGCAGCCTTCAAGGAAGTACATGC

GGATAGTTACAAGAGATGATTTTGAATTCTGGTTTATGGGACTCTTGAATTATGCAAGAACAATCATATC

TATTCAGAATGCCGTCTCTCATGCACAATGAATTGAACTCGGATCAAAAATCAGCAAAATGTATTATACC

CTGAATCTCTGATGCAACTGGAAATACACTAAGCTATGTAGTTAGCAGTGCACTAGTAGTAAGTAGTAAC

CTAGCTTAGGGTGTACATATTTTGTTTCTTAAAATTCATCTGCCTTGATGAAAAGATTTGCTTGTAAATG

AAGATGTCAAAAT

>MSTRG.1844.2 gene=MSTRG.1844

TATTTTTGCACTCAAAAAACACTCCTTATATATAAATACTTCTCTTCAAACATGGAGTAAAAGTTGCAGA

TTCTGAGGATGATTTTAGGGAATCTGCTATGTTTATGGTTTGATAAATTTGTTCCTTGATTATTTGATTT

CAACTAACATGGTATCAAAAGTCGGTTTACTTGCATCAGAAAGCAGGCAGGCCATCTCAGAAAGTTCTCT

TCAAAATGCTTACACAGTCAGCCTAAAACAAACAAGGTAGCCACACTGAAGAAGGCTTGGCCCTTTGAAC

AAGCTCCTTTTGCAGCTAGCAGACATGCCATCTTCTATATACATTTTCTATTATTCAACTTTATCCATTC

ACCTTTATTGATATTAATGGTGGAATGACGAAAAGATGACCAACTAAAAGTAAAGATCCAGATGCCGACC

TTATTCTATAAATACCGGTTGAATAAACATATTTAAATCCATCTTACACTAAACCAGTCATTGCCTCAGA

GTTGTAAGGGTATTATATTACCTCACAATGAAGAACCAGGTTGGAAAGAATGTTATTCCATATTCACGAA

GCTCTGCAATTCCTGTTGAGAGGTCTCTGAGAGGAGTTCTAGCCGTCCCTGTTAGCCGTTGCCATGCATC

ACCAAAATTCAGATGGGGTAGATTTGATTCAGTATTTAGTCGGATGACCAGCTTTGGTGAAAAAATTGAT

TCTTTAGCAGAAGGCATCAAAGAGCATGTGAGAATAAGGTCCGTAACTGAAACTATGAAAGGGAAGTTGA

GTTTAGGGGCAAAAATCTTACAAGTAGGGGGAGTGAAGAAAGTGTTCAGACGAAATTTTAGTGTGAAAGA

AGGGGAGAAGCTACTGAAAGTTTGTCAATGCTATTTGTCGACCTCAGCAGGTCCGTTAGCAGGCCTACTC

TTTATTTCCACTGATAAGGTTGCCTTCTGCAGCGAACGATCAATCAAAGTTTCTTGCCCTACTGGAGAGT

TTCAAAGAGTACATTACAGGGTCGTGATCCCACTAGGAAAGATAAAGAAAGTGAATGAGAGTGAAAATCT

GAAGCAGCCTTCAAGGAAGTACATGCGGATAGTTACTAGAGATGATTTTGAATTCTGGTTTATGGGACTC

TTAAATTATGAAAGAACGATCATATCTATTCAGAATGCCCTCTCTCATGCACAATGAATTGAACTCGGAT

CAAGAATCAGCGGTTTTACCCCTGAATCTCTGATGCAGCTGGAAATACACTGAGCTATGCTATGTTACCC

GGACTCGGCTGAAAGTGTCCAAACTGGATACGTGTCCGAGTGTCGGACTCGGCAATATTTTGAAAATTTC

ATATGTTTTTGGCCTAAAATAAGTGTCGGAGTGTCCATACCCATGTCCGAGTGTCGAGTGTCCGACATGG

GTACTCGAGGCAAAATGAAGAGTCCGGGTAACGTAGGAGCTATGTAGTTAGTAGTGCACTAGTACTAACC

TAGCTTAGAGTGTACATATTTTGTTTCTTAGAATTCATCTGCCTTGATGAAAAGATTTGCTTGTAATTTG

AAAATGAAGATGTCAATTAAACTTGTTTATATAATGTGTTTCACCA

>MSTRG.1849.1 gene=MSTRG.1849

CTACACTACAAACAACTGTAGTCTACGTATAGTTACCACCAGCTGAAGGGGATATTACTATTTCTAGGGC

TTTTTTCTGCGTATTCAAGAAAGTCTAAACACAAACACACTCTCAAATTCGCTGCTTATAGATTTCAGGA

GGTTTCATGGTATTTTTAAATAGGGTGTCTCGGATAAAAAGATAGTCTTTTGTTACGGTTGCAATGGAAA

CTATAGATATTGTGTCATCAGATAGTAGTGATAGTGATTGGGATCTGGATGAGGTTAAGGCACTGATTGA

TTCTGTTCCGTCAGAGTCTACTGCTCATCCAATTTCAACAGTTCAAGGTCAATCGTCCCAAGTATCTACT

TCAAATGGAATCTCGGCAAAACCAAGACCAGGATTGGCTGCCAATCAAAGGGCCCTTACATCCAGGGGAA

ATACCTCTGGTAGCATTTTGACCGTTGATAATAGACCTTCACGTAAACTGCCTTCACCAGCAAGTGTGAA

TGCTTACAACGGAAGCTCATTACACCGTAGCCAGCAAATACAGGGTCATATTCATAAGTCTCCAGGCTCA

ACTAGCAGTTTGGGAGTTTCAAACCTGAGAATAACCCAAGATAATAATCCTACGAGTTCAGCTCATAAAG

ATAACAATCAATATACACCTTTACAAGGCTCTAAAAGGGCACTTCCACCATCTATACATCGTCAAGTACC

TTTAGTGGACAATGTGGGTAGCAGTCAGATGCGTGAATCTTATAGAAAAACTCTTTTTTCTCAACAAAAT

TCAGCATATGGCATGAGCCATATGAAAAATCAAACTGGCAGGGGAAGTGATGATGAAATAGTAATGTTCG

ATAGCAGCGGGTGTAGGATTCAACCTCCATCACATGCAAAGTCTACTTTAACATCACAGCATACCAGTTC

GACAGATACAGTTTTTCGCTCTGCAGTTGGCCAAGACAGAGTAGCTGAACCTGATGAGAGGCTAATTTTC

CAAGCAGCATTGCAGGATCTAAGTCAACCTAAAGTTGAAGAAGATCTACCTAATGGCCTCATGACAGTCC

CTCTTCTTAGACACCAGAAAATTGCACTAGCCTGGATGTTTCAAAAGGAAACTAAAAGCACTTATTGTTC

TGGTGGAATTCTTGCTGATGATCAGGGCCTTGGTAAAACCATTTCAATGATTGCCATTATACTGAAGCAG

AGATCTTTACAGTCGAAATCCACACCAGAAGATCAAAACCAGCTGACAACTGAAGCCCTAAATTTAGATG

ATGATGATGATGATGATAAAATTACTATTCTTGATGATGACAAAAAAGATGGGGAATCTAATGACCTTAA

ATTGCTACCAGAAGCAAGTACTTCTAGACGGGAATTTAAGAGTTCAAGGCCTCAAGCAGGGACATTGGTT

GTGTGTCCAGCAAGTGTGCTTCGTCAATGGGCCAGGGAGCTGGCTGAGAAAGTTGCAAAAGAAGCTCCAC

TTAAAGTTCTAATTTATCATGGAGGCACTAGAACAAAGGATCCTAGTCAACTAGCAGCATATGATGTAGT

TCTCACAACATATGCTCTTGTGGCGAATGAGGTCCCAAAACAACCTGTGGTTGATGAGGATGAAGATGAT

ATGAGAAATGCGGAAAGGCATAGATTTTCCGCTGAGTTCTCTACTAACAAGAAACAAAAGAAAATGAATG

GTGCTGGTAAGAGACGAAAGAAAGGTAATAAAGGGATTGACTTTTCTGATATTGACTTGGATTCTGGTGC

TCTTGCAAGAGTGCGTTGGTTTAGGGTCATATTGGATGAAGCACAGACAATAAAGAACCACAGAACGCAG

GTGGCCAGGGCCTGCTGTAGTCTAAAAGCAAAAGTAAGATGGTGTTTATCTGGAACACCCATACAAAATT

CAATAGATGAATTGTTCAGCTACTTCAGATTTTTGCGATATGATCCATATAGTGCCTACAAACCTTTTAT

TAGTATGCTAAAGAACCCCATTTCCAGAAATTCTGTCCAAGGATATAAAAAGCTTTATGCTGTGCTCAGG

ACCGTAATGCTTCGTCGAACTAAAGACACATTCATTGATGGAGAGCCTATTATTACATTACCATCAAAAA

CAATAAATCTAATTAAGGTGGACTTCTCTGCCGAGGAACGGGCTTACTATCAAAAACTTGAAGCTGATTC

ACGCTCACAGTTCAAAGCCTATGCTGCTGCTGGCACTTTGGGTCAAAATTATGCAAATATACTGCTCTTG

CTCTTGCGCCTTCGGCAAGCTTGTGACCACCCGCTGCTAGTAAAAGGCTTCACTTCGGATTCTGTTGGAA

GATATTCCTCGCAGATGGCAACAACTCTTCCTAGGGATATGCTAATAAACTTGTTGAAGCAACTACAGAC

GACCTCAGCCATTTGTGGTTGGTGCAAAGATCCACCCGAAGATGCAATTGTCACCATGTGCGGACATGTT

TTCTGCTATCAGTGTGTATCAGAATTCTTAAGGGGAGATGAAAATACATGTCCTTCACCTGAATGCAAAG

AACAACTTGGTCCTGATCTTGTGTTTTCGAAAGCTACCTTGAGAAGTTGTCTATCTGATGATCAAGATGA

TAATCCCTCTAGTTCATCCCAATCTGAAAAATCTTCCGTAGTTTTGCAGAGGGACTATATTTCTTCAAAA

ATTAAAGCTGCTTTGGCAATTCTTATGACTCATTGCAAATCAAACAGTCAAATGTTGCAGTTGCATGGCG

TGGTTTCTTCAGATCGAGAGGAAGAACTATTAGAGTCAGGTGGTTCATGTTCGGATGCACAAATTGGAGC

TCCAAGAAAGGCTATTGTTTTCTCTCAGTGGATTGGCATGTTGGATTTATTTGAAATGGCACTGAAACAG

TCTGGTCTAGAGTATAGGACACTTGATGGTACAATGACATTATCTGCAAGAGACAAGGCTGTCAGGGAAT

TTAACACTGACCCCGAGGTAACAGTTATGCTCATGTCGCTAAAAGCTGGTAACCTTGGTCTGAATATGGT

TGCTGCATCGCATGTGATCCTTTTGGATCTTTGGTGGAATCCAACTACTGAAGATCAGGCTATTGACAGA

GCTCATCGAATCGGACAGACTCGTCCTGTCACTGTATCAAGGCTCACTATCAGAGATACTGTTGAAGATA

GGATATTGGCTCTTCAGGATGAGAAAAGGAAAATGGTTGCATCTGCATTCGGTGAGGATCAAAGTGGGGG

CACAGGAAGTCGATTAACTGTGGAAGACCTGAAATATCTATTTATGGGTAACAATAACATGGTTTGACAG

GTGAATCTAGCATGCACAAGTTTTGCCATAAAAAAGAAAATGTAGTATGTAATGAATTTCTAAATACTTG

TAGGAAACAAAACATTAGCATTTAAGTGTAGAATTAACCATCGGACCTAGTAGGTTTGCGCTCTGCATAA

TCTGGTGCTGCACCAGTTTTTGGGTCAGCAAATGGAGCAATTTTGTTGTGATAATATATGTGGATACTAG

ATACTATGTATGTAGGGGATGTAAATATAAGAACCAAAAATTTGTACTTGATCAAACGCAAAATCCGAGA

AACAGCGGGAGCAAGCCATTTGGTCGGGACTAATGCCCAGATCATTAGTTTTGTAATTATGGTAAATTGG

ATAAAACGTATTGTTCATTGTTGCGATTAAAGACAGAAAATTGTTAATGTTAGTAGTTTGGGCAATGTTG

TTGGAATCAACAAGAGCAGGGCGTTGTCAGGTCATATTGAAGAATGAAAGAGTTATCGAGAAATGAATTG

ATTAAACGTTTACAGCAGCACGAATTAGATCTTGGACAGTAGGCTCAAAATCTAGTACCA

>MSTRG.1856.1 gene=MSTRG.1856

TAACAAATGACATTAAGAAAAAATCTCATTTTGGTGTTTGTGTTGGAGTGATGTATGTTGTGGAGTTTCA

AAAGCGTGGCCTCCCCCATGTTCACATGCTCATTTGGCTTGACAGCGCTTCAAAGATTTATCTAAAGAAT

AATGTAGACAAGTTTGTCTCTGCTGAAATACCCGATCCTAAAAAAGACCCTGTGGGATATGCTGCGGTGA

AGGCTTTTATGATACATGGACCGTGTGGACTCCAGAATACGAAATCTCCCTGCATGAAAGGTCTAAAGTG

CATACGTCATTTTCCGAAAAAGTACTGTCCGCGCACAATGTTTGATGAAAGCGGATTTCCAATGTACAAA

CGGCGTAGACAGAACATAACAGTCCATGTTAGGAATGCTGATTTGGATAATCAGTGGGTAGTTCCTTATA

ATCGGGACTTATTGGTCAAATACCAATGTCACATGAACATTGAGATATGTTGTCACGCTCGCAGTCTCAA

GTATTTGTTCAAATACTGTTTAAAGGGTCATGATACTGCCACGATACATGTTACTGGGAGGAAGAAGAGA

ACAGCCAACTCTAACGGTGATGAGCCTATCGATGAAATTGATGCCTACTTTGATGGTAGGTACATATGTG

GAGCAGAATCGGCTTATAGGATTTTTGGTTTTCCTATACACCACCGTACAATATCTGTTGAAAGGTTACC

ATTCCATCTACCAGGACAAAAGAACTGCACATTTCACTCAAACCAGCCTTTGGATAAGGTAGCTGAAAGG

GAAAAGGACAGACTTAGTAAACTTGAAGCGTTCTTCCTTTTATGTAATACTGATGCTACAGCTCAGCAAT

ATACCTATCAGGAGATCCCAAAGCATTATGTATGGAATGATGTTGAAAGGAAATGGAATCGAAGAAAAAG

AGGATATCAGATAGGACGACTATCTTATACTCATCACAGCAGCGGCGAAGTATGGTTTTTGAGATTATTG

CTAACCAAGGTGCGCGGCCCAACTTCCTTTGAGAAACTCAGAACTGTCAATGGAATATGCTATGAGTCCT

TTCGTGATGCTTGCAAAGAGTATGGTTTGCTGGATGACGACAAGGAATGGCATGAAGTTCTTGATCAATG

CTCCTCTGGTGGCTTGCCTCCTCAGATTCGTCAACTTTTTGTTCACATAATAGTGAATTGCAAGGTAACA

GATTTGTATAATTTGTGGAATGCTCACTGGAAGCAAATGGTTGATGATATTATTTTAAAGAGGCGACGCA

TAAATAAAGATGAAAGCCTAATGCTGGATGATAAGCAGCTACAATTTTATGCTCTGGCAGAAATTGATGA

TTTGCTTAGATCAATTGGGAAATCTTTAAAGAATTTTCAGCAGCTGCCACAACCTCCTGAAATGTATCTG

AATCATG

>MSTRG.1872.1 gene=MSTRG.1872

AACAAGAAGAGCTTCTGAATCTTCAACATTCCTCCGGAGGAGCGAACTCGGTTATGGCTTCTTCTGCGTC

AGCTGCTGCTCCTCAGAGAAAAAAGAGAAACCAGCCCGGAACTCCTAGTCCTGATGCTGAGGTGATAGCT

CTATCTCCGGATACACTAATGGCTACAAATCGATTTGTTTGTGAGGTATGCAACAAGGGTTTTCAGAGGG

AGCAAAACTTGCAGCTGCACAGAAGAGGACACAATCTTCCGTGGAAGCTTAAGCAAAAGACAAACCGGGA

CGTGAGGAAGAAGGTCTACCTTTGCCCTGAACCTGGCTGCGTCCATCACGATCCGTCAAGGGCTCTCGGG

GACCTCACTGGAATTAAGAAACACTTCTCCCGCAAGCATGGTGAGAAGAAATTTAGGTGCGAAAAGTGTA

ACAAAAAATATGCAGTGCACTCCGACTGGAAGGCTCACTCTAAGACTTGCGGCACTAAAGAGTACAGATG

TGAATGTGGCACTACCTTCTCAAGGCGTGATAGCTTCGTGACTCATCGAGCCTTCTGTGATGCGCTTGCT

CAAGAGACGGCACGTCATACACTGCTCCCCGCATCACTCAGCAACATGGCTGCTACAATTTACGCACACG

CTAGTGGCGTTATGAGCAGCTTGGGCTCGTCCGGGCCTCTCGTTTCAGCTGACCGTCAGGAGCCATCGTC

TCCTGTATTCGGGCTTGGTGCTCCAGCAGCTGCGCCAAGAAGTTCATTCACTGGCTTCCTATCAGGCCAG

TCTGCTGGTGGATCTGCTTTTCGACCCATCCAGTCGCTCACAATGCCTGTTCAGCCACCCGAAGCTAAGC

ATAATTTCTCAAGATTTTCTGATTTTTCGGGCCTTCAGGGCTTTTCAAACAACACTACTTCTGCTGGCCC

TATGTTCAACTTGGGATTTAATGGCAACGCTGATGTGGACAGTCCTCATCACGAGGTTGATCTTAACATG

AACAGCCCCGGTCTTCTTACTGGTTTGATGATGAATGCGGATCCTCCTCCGGTTTATAGTCATATTAATA

CTATTCAGTCTCCGCAGCTATCAGCTACAGCTCTCCTTCAAAAGGCGGCGCTCATGGGAAGCTCTACAAG

CAGCGCTGCCTTGATGAAGGGCTTGGGAAGCTCTTCGTCTAGCGGAGTAAAGTTTAGTAACTATGGTGGA

GCTTTCGGAGCTGGAATATTGAGTGGAGGTGGCGGGGCTAGTGCAAGTATATTTCAAGATATGTACAGCT

CAATCAATGCAGGTGGTGGGGGTTATGGGAATGGAGGGTTTGGAGGCGGAATGGGATTTAGCGGAAACAA

CCCATACGGAGGAGTAATGTACAATGTCAAGAATGATCATGAGGGGATGGAATTGGATGAAAAGGAGAAT

AGCATGGCTATGGAGCCGCAAAGTGTTCAAAGCGGGTACTGTGACAAGCTTACTAGGGATTTTCTCGGCA

TGGGAAAGAGGAGGACAAGCTTTAGTGGTGGAATGGAGGATGATCACAAAGGGAATGGAATAGCATCGAA

CTCGCCGGATAGTGAAGCAACCACCTCAGCTGAGCCTAACCCCTTTGTCCTGAGCCGCCGCGGCTTTTGA

TCCAGCAACAATTCAGGCCGGAGACTCAAGATGATCAAGGCATGCAGACAGACGCTTAATGTGATTTCAG

TGACCTTGCATCTTTTACAAGAATCAATGAGAACAATCAGCAAAAAACTCATCTTTCTCTTTTTTCTTGG

AACAAAACTTTTGATGGAGATACTTCGGTTTAGTTTTTATGTTTTGCTTAAACTTTAAAAATTACAGAGA

GACGCCCCCCGCAGGCTTTGAACTTAGCTAGAGGTCGGTTTTTATTTGTTTTTGTGGTAAAAGTTTCGGA

TTGTGCTTGTTAAGTTTGAATCACC

>MSTRG.1875.1 gene=MSTRG.1875

CTCACACTTCCATGGCAGATAAAACTCAGATTGGCGACACTTGTTGAGGGGAATGAGGAGATGGGCACAC

TGGTCTCTGTATGGGATTGGGACGCGGTTATCCACCATTTCTTGTTGGGTTGCTATCATCTTCTTCGATG

ATCCAAGGGGCACTTCTTCCATTTTTCTGTGTAGACTTTTTTGTCGTGATTGCCCTCAATTTAG

>MSTRG.1877.1 gene=MSTRG.1877

GTATTTCATATATGCCTTCATGGATTTTTGTTTACCACGGGAGATAATGGCATGATAGCTTTTGATCTAA

ACAAGGAGGTGTTTATTTGTGATATCAAACTTCCCGGATGCTCCAATGAACCTCGTGAAGAATGTTATGA

CGCTCGAATTGTTGACTTCAAGGATTCTATTGCTGTCATTACCTCTGATGATAAGGAGCTCAGTGGAAAA

CTTAACTTGTGGACACTAGATGACGAGGCATGCCTTCGTGGTGGTAGAGTTAAGGCATTATGGACTTTCA

TGCTTAGTATTGATGTAGTATTCAAGTATACAGAGAACCAGCAGATGCTTTTAGTCACGAAATCAAAACT

AGTCAATTGGTCTGATGATGAAGAAGATTTTAGTGAGGAAGTTAGCATTAATTTGGAAAATGTCTATGAT

GATTTTATTGTTATTTCAGATGATAGTGAGGAAAGTAATTAAGAAGATGATAATGAAGAAAGAACCGCTG

ATTTCGAAGACAATGAGAGGAGTAATATTATCTCTAAAGATGATAATGAGGAAACAAACAGAGATTCTGA

AGATGATAATGAGGAAAATACCTCAGATTCTGGCAGTTAATGATGGCATTACTCATCTCAGTCAGTGTTT

TTTTCCTTTCATTTCCCTCAGTTTATTGCACATTTTCGAAATTTTATGTGGAATAATTTGATGTTTCTTT

CCATGAGTTGGACATGATAAATATCTGGTATAAGTCTTGCCTGTTCTGTTCAGTTCTTGGAAATTTTTTT

GGGCTATGTAGATGCTACTGGTGTGTGC

>MSTRG.1878.1 gene=MSTRG.1878

CAGCCCCCATTCCCAACAATCAGTAGCGGAAAACCTAACCCCTGAGATGGCGAAGCGGACTGAGGGGATG

GAGATGTTAACAGACGAAGATCTTTTAACCGAGGTGCTCTCTAGACTCCCTGTAAAATACTTACTCCAAT

GCAAAGCCGTGTCAAAACCCTGGTACTCTCTCATTTCCAGCCCTAGTTTCATCAAATCTCATTATCAACG

TCATGCTATTACAATCAACCGCGCTGTAGATGATAATAATGAACAAGGTCCTACTCTCATTGTACGCCTT

TTCAAAGGAGAAGATCTTGATGACCATCCAGAAGAAGACGGCTCCTATTCACTATTCCGGCTCGGTACTA

CTGATGATTATATAGCCAAGCTTAAGTTTCCTTTCTCCAGAGGTGATTATCCCTTCACGCCTCCCTCTAT

TCTCGTTGGTTGTGTTTGCGGAATTGTTTGTGTTTGTGTTGATCCCTCTGCTCCTATTGATGCTAATTAT

TGCTTGCCTACTCCTGATAAAGCTACTGATATATACATTTGGAATCCCGCAACTAGGCAGTCCAAACTCA

TTCCCTCGCATGCTCTTTACGATCGTGATGGTATGAATAAAGTTGCTTTAGGCTTCGGGTTTGATCCCGT

TGATTTTGATTTTAAGGTTGTTAGGGTTGTATCTCGTGCTATTTTTCCTCAGGTCTATTCTGCAAATAGA

GATGCTTGGCAAAACATTGAGACTAACTTGACTGATTTTCCCGAAGAATTTATATTTCATATATGCCTCC

ATGGATTTTTGTTTACCGCGGGACAGAATGGTATGATAGCTTTCGATCTAAACAAGGAGGTGTTTATTTG

TGATATCAAACTTCCTGAATGCTCCATTGACCCGGGTGAAGAATGTTATCATGCTCGAATTGCTGACTTC

AAGGATTCTATTGCTGTCATTATCTCTCATGACGAGGAGCTCAGTGGAAAAATTAACTTGTGGACACTAG

ATGACGAGGCATGCCTTCGTGGTGGTAGAGTTAAGGCATCATGGACTTTCATGCTTAGTATTGATGTAGG

TGTGTATTTCATGTATGTGCAGGGCCTTTTCAGTAGTGTTGAATTCTTACTGGTAGCTGATCATGAATGG

TTCTCCTATAATTCTGACAAGAAAGAGGCCAGAAGGATTGATACTGATCCTTTTTTCGATTTCGATGCAG

TATTCAAGTATACAGAGAACCAGCAAATGCTTTTAGTCACGAAATCAAAACTAGTCAGTTGGTCTGATGA

TGAAGAAGATTTTGGTGAGGAAAGTAGCATTAATTTTGAAAATGTCTATGATGATTTTATTGGTATTTCA

GATGATAGTGAGGAAAGTAATTAAGAAGATGATAATGAAGAAAGAACCGCTGATTTCGAAGACAATGAGA

GGAGTAATATTATCTCTTAAGATGATAATGAGGAAACAAACGGAGATTCTGAAGATGATAATGCAGAAAA

TACCTAAGATCCTGGCAGTTAATGATGGCATTACTCATCTTAGTCAGTGCTTTTTTCCGTTCATTTCCCT

CAGTTTATTGCACATTTTCGAAATTATATGTGGAATAATTTGATGTTTCTTTTCATGAGTTGGAC

>MSTRG.1881.1 gene=MSTRG.1881

CCAAAATGGGGAAAAAATATAAGGGCCAAGAATACACCCCCAAATTTGAGGTCTCACTGTACACCCCGCA

TCTCACCTACCCGCCATTTTTTCCTACCTTTTTCTTCTCTCTTCTCTGTAGATACAACCTCCAGTTCACT

TCAAGTCTAGCTTCCATGGAATAGCTTGTTATAATCGATTCGCCGTCTCCGAAGGTTCATC

>MSTRG.1882.1 gene=MSTRG.1882

CTTCAGTTTCCCTTCTTTAGTTTTGCCTCTTCAGGGGCCTTAATGTTTCCCTTTCAACCGTTGATCCATT

ACAAATTCATCACACAAAATCACCCGTGGTTGAAGAATGCAGTATAGATGCTTCAGTGTAGAAGTTGGGC

TCATGCGATTTATGTGAGATTGCATGTAATTCATTCTACCAATCAGGTTTCTCACATATTTTAAAGGTAC

AGTTGTTAAGGAGCTTTTCTAAAATTCTTTATGTATATTAAATCTCTTCTGATATGGTCGCCATGCTACA

GTAGGATCGACAGAAATAGTTTCTCGGATGGGAAAGGGGGTTGTGCTGTCATGCTAAAGGGGTTGTGATA

TCATGTTTATGTTTCTGGAAATAGTATTTGAACTACAATGATGCTTTCTGTTTCTTCAATTTTTGTTCTG

ATATTTGAAGAGCTCTACCAAATTATATTCCTTTTTATTTATTGTCCCACTGGATTGGAAAGGAGTGCTA

CAAGAGAGGACCATATGGTAATGATATTTACAGGAAGAATGTGCCTCATATCAGGCTGGAATTTTGTGGC

ACGGTATGCTGTTTACTTTTAGAAAGTTCTATTTTGCTTCACATACTTAATTATGCACCTCTTCTCTACA

AATTTTCACTTTCACTGATAGCCACTCAGTTGCATCAAATGAAAACTCTGATCTATATGTTGAAGAGTCT

TAGGTATGTAAGGGCACCAAAAAAGAAACAACATGTTTTAATCAACATGAATATTTAAAGACCACATTAA

TATTTCTGCTACAGACATGGAGAGTGGAGATGCCACAGGTTTACCTTGGCAAAACGAGTTTTCCAAAAAA

CATTGAAACCTAGCCAGCTTGATGGATGTGCTTTTCTGTACCTAATTATATAAGAGTATATTTCTCAAGG

TCTTTTTGATCAGCTCATTCTACCAATATATGGCATCAAGTTTTCTCGGACATGGCTCTGAAAGCTAAAG

GTTATGCATAGTCTGTTTTTAACTGAAACTTGCCTTAAATTTTGATTTACAAGTACTATAATCTATAATT

ACTTTAGTATTATTTTTTTCCTTCTTATGTTTTTACAGTTCCAGACTTTAGAAGGAAAAACGCTAATACA

TATATTCATGTGTACAATGGCTATAATGGTTGCAGTCCGTAACAAGGTGTGAGTACCCTGAAATTTGAGA

TATGGATTATAGGCACAAAAGGCATTTAAGCTTCAGGCGAAAAAGTTTGTTTCAGGTTTCAGATTGCGAT

TTGGAGTGATATCATAACATATATTTTGTAAATCAAATATTTTTTATTTTTGCCAGAGATTTGATACTGC

ACCTGGATGACTAGATGAGTTGGTTTGAAGTTTAGGAAACACTAAGATAATTCTGTAGTTAATCAGTAAC

TTAGCTTATCAGTACTAGCAATATATCATTAAATTGCTCCGGCAGACTTGCAGTGTGCTCCTTCGTCCAA

GTTCTTTCATTCTTTTTTATAGGTAGTGAGTTAAAGTATAGCTTCTAATGGAAATTGTAAAGTCAAAAGG

GAAATTTGCTGCTCAGGTTGCTTTCAGATTGGGCTTATGATGCTATGAGCAAGGCGCTGTTATTATGTTC

GAGAGCTTCAATAAAGAAAGACTAAGCAGTATCTACGGATTTTTGACTAGGAGTTGAGCGATGAAGAGTC

CGAAAAGGTTACTGAAATTCCTCATAGCAAGGCAAACCTTGATGAGATGTTAATCTTTGAAACCAATCAT

GTGATTGAGAGAGTATATCAAACATGTAACCATCCTTTTCAATATTTATTCAACTTCACTCTGTAGTGAG

GTTTCTTTAAGACCTTTGCAACTTCTTTCGTCTATATTTTCAGCTTCACTATGTAGTGAGGTTCCTTCA

>MSTRG.1884.1 gene=MSTRG.1884

CCCTCAATATTATCTCTTGATAGTGATGCTTTGCAAAACCTTGAGTTATTCAAAGGATTGGTGTAAAAGC

AGCTAGAGGCTGCTGAAATGAAGGTTATGGTTGCTCCTTTGCTCCAAAGCGAATCTGGTCAATTTTCTAT

TGCTGATATGGTTCTTCACACACTGCTGGACTGGCAGCCAGAAGTCTGGGATCTACCACCGTTCGAAGAC

TATGTGTGCCAATTAATCCACTTATAGCATATATAAGTGTAGAACGTACATCTTTATAATTTTGTTCAAT

TGACTGGTAGTGAGTAGCAGGATGTACTTTGATTTGGTAGTTACTTTGGTTTTTAGCAACTTAGATTTGG

TTGGAATTTTGGGATGTTTAATATTTAAAGAGTATATTTTTAATTTCTTTCAGTAAATAATGTTGCCTTT

GATTAATAAAATGTTGCAGCTCA

>MSTRG.1884.2 gene=MSTRG.1884

CCTATCCACTCTGGACATTTCTCTTTGCCTCTCTTCTAGATCTAGCCAGTTCTCTCTCTCTCTCAGATTT

CTTTCAACTGAGCTGTAGATAAAAGTCAATTGGCTTCAATTGAGAGTTAAACTTTCACAGATCAATGGAA

ATGGGGACTCCTTGAACCGATTCGAGCTTCAAACAAAACTAGGGTTTCATTGCTAATTTGGGGTTGAATA

AGTTGGGACTTAGCTTAAGCTTCTACTTCAAGCTAATGAAAGGGCAACTGCTTGAAAAATTGCTAAAGAG

TCGATGGAATTGATTGATGATGAACGTGTGGAACTCATGGATATTGCAGCCTCTTGGAAAAGGGTTGCCC

TCAATATTATCTCTTGATAGTGATGCTTTGCAAAACCTTGAGTTATTCAAAGGATTGGTGTAAAAGCAGC

TAGAGGCTGCTGAAATGAAGGTTATGGTTGCTCCTTTGCTCCAAAGCGAATCTGGTCAATTTTCTATTGC

TGATATGGTTCTTCACACACTGCTGGACTGGCAGCCAGAAGTCTGGGATCTACCACCGTTCGAAGACTAT

GTGTGCCAATTAATCCACTTATAGCATATATAAGTGTAGAACGTACATCTTTATAATTTTGTTCAATTGA

CTGGTAGTGAGTAGCAGGATGTACTTTGATTTGGTAGTTACTTTGGTTTTTAGCAACTTAGATTTGGTTG

GAATTTTGGGATGTTTAATATTTAAAGAGTATATTTTTAATTTCTTTCAGTAAATAATGTTGCCTTTGAT

TAATAAAATGTTGCAGCTCA

>MSTRG.1884.3 gene=MSTRG.1884

CCTATCCACTCTGGACATTTCTCTTTGCCTCTCTTCTAGATCTAGCCAGTTCTCTCTCTCTCTCAGATTT

CTTTCAACTGAGCTGTAGATAAAAGTCAATTGGCTTCAATTGAGAGTTAAACTTTCACAGATCAATGGAA

ATGGGGACTCCTTGAACCGATTCGAGCTTCAAACAAAACTAGGGTTTCATGTTTGAATTGGGGGTGTTTG

GTTTTTGGTTCTGTTTTTGTGAGACTCACTCTGTTTTGGTACTAATTTTGTTTTATCTCTTTGTTCCAGT

GCTAATTTGGGGTTGAATAAGTTGGGACTTAGCTTAAGCTTCTACTTCAAGGGAAGTCTCTCTTCTTGGA

AATGTAAAGGTAGCCTGGACATGTTCAGGTTCATACAGAATTATCTCCATACTGGCATGGAAATATATAT

GTAGGTGACACCTCCTTGACGCCTCCTTTATTTATTAAGTGTTTCCATTTTATCTGTGTTTTAAAATTAG

CTAATGAAAGGGCAACTGCTTGAAAAATTGCTAAAGAGTCGATGGAATTGATTGATGATGAACGTGTGGA

ACTCATGGATATTGCAGCCTCTTGGAAAAGGGTTGCCCTCAATATTATCTCTTGATAGTGATGCTTTGCA

AAACCTTGAGTTATTCAAAGGATTGGTGTAAAAGCAGCTAGAGGCTGCTGAAATGAAGGTTATGGTTGCT

CCTTTGCTCCAAAGCGAATCTGGTCAATTTTCTATTGCTGATATGGTTCTTCACACACTGCTGGACTGGC

AGCCAGAAGTCTGGGATCTACCACCGTTCGAAGACTATGTGTGCCAATTAATCCACTTATAGCATATATA

AGTGTAGAACGTACATCTTTATAATTTTGTTCAATTGACTGGTAGTGAGTAGCAGGATGTACTTTGATTT

GGTAGTTACTTTGGTTTTTAGCAACTTAGATTTGGTTGGAATTTTGGGATGTTTAATATTTAAAGAGTAT

ATTTTTAATTTCTTTCAGTAAATAATGTTGCCTTTGATTAATAAAATGTTGCAGCTCA

>MSTRG.1884.4 gene=MSTRG.1884

ATGCAACGTACCCAGATTAACTTACCCAGTTAATCATAAACCAGGATTAACTTCTCTATATCCTTGTTAG

TCAGTCTATGGTCTCGAGTACTCTGGACAGAGTTCAGGGCTTTGTGTAATTAGTACTTTATATTTTGGCA

AGTTGTCTTTAGTAAAATGTTGATTTCTTATAATTAATCTAATTGTTGTGAATTCTTATAATTAATCAGG

GAAGTCTCTCTTCTTGGAAATGTAAAGGTAGCCTGGACATGTTCAGGTTCATACAGAATTATCTCCATAC

TGGCATGGAAATATATATAGTCGATGGAATTGATTGATGATGAACGTGTGGAACTCATGGATATTGCAGC

CTCTTGGAAAAGGGTTGCCCTCAATATTATCTCTTGATAGTGATGCTTTGCAAAACCTTGAGTTATTCAA

AGGATTGGTGTAAAAGCAGCTAGAGGCTGCTGAAATGAAGGTTATGGTTGCTCCTTTGCTCCAAAGCGAA

TCTGGTCAATTTTCTATTGCTGATATGGTTCTTCACACACTGCTGGACTGGCAGCCAGAAGTCTGGGATC

TACCACCGTTCGAAGACTATGTGTGCCAATTAATCCACTTATAGCATATATAAGTGTAGAACGTACATCT

TTATAATTTTGTTCAATTGACTGGTAGTGAGTAGCAGGATGTACTTTGATTTGGTAGTTACTTTGGTTTT

TAGCAACTTAGATTTGGTTGGAATTTTGGGATGTTTAATATTTAAAGAGTATATTTTTAATTTCTTTCAG

TAAATAATGTTGCCTTTGATTAATAAAA

>MSTRG.1884.5 gene=MSTRG.1884

CTCTGGACATTTCTCTTTGCCTCTCTTCTAGATCTAGCCAGTTCTCTCTCTCTCTCAGATTTCTTTCAAC

TGAGCTGTAGATAAAAGTCAATTGGCTTCAATTGAGAGTTAAACTTTCACAGATCAATGGAAATGGGGAC

TCCTTGAACCGATTCGAGCTTCAAACAAAACTAGGGTTTCATTGCTAATTTGGGGTTGAATAAGTTGGGA

CTTAGCTTAAGCTTCTACTTCAAGGGAAGTCTCTCTTCTTGGAAATGTAAAGGTAGCCTGGACATGTTCA

GGTGCAAGTGTTTGTCGAGGTTCATACAGAATTATCTCCATACTGGCATGGAAATATATATGTAGGTGAC

ACCTCCTTGACGCCTCCTTTATTTATTAAGTGTTTCCATTTTATCTGTGTTTTAAAATTAGCTAATGAAA

GGGCAACTGCTTGAAAAATTGCTAAAGAGTCGATGGAATTGATTGATGATGAACGTGTGGAACTCATGGA

TATTGCAGCCTCTTGGAAAAGGGTTGCCCTCAATATTATCTCTTGATAGTGATGCTTTGCAAAACCTTGA

GTTATTCAAAGGATTGGTGTAAAAGCAGCTAGAGGCTGCTGAAATGAAGGTTATGGTTGCTCCTTTGCTC

CAAAGCGAATCTGGTCAATTTTCTATTGCTGATATGGTTCTTCACACACTGCTGGACTGGCAGCCAGAAG

TCTGGGATCTACCACCGTTCGAAGACTATGTGTGCCAATTAATCCACTTATAGCATATATAAGTGTAGAA

CGTACATCTTTATAATTTTGTTCAATTGACTGGTAGTGAGTAGCAGGATGTACTTTGATTTGGTAGTTAC

TTTGGTTTTTAGCAACTTAGATTTGGTTGGAATTTTGGGATGTTTAATATTTAAAGAGTATATTTTTAAT

TTCTTTCAGTAAATAATGTTGCCTTTGATTAATAAAA

>MSTRG.1885.1 gene=MSTRG.1885

GGAGAGAAAAGTACCCAAAGGGCCAAAGTGATCAAGAAAAGGAGAAATATGCGAAAAGGAAGCAGAAGAG

TCACATGGGTTTCCAATACACAGCGACCAAACGAGCCTAAAAGCTTTGAATTACTCAACCAAAACCCTAA

TGCCCGGGCGAAGAAACTCTCATCGGACACGTTAACAGAAGCTTTAGGCTTTGGTTTTGATTGCTTACAT

TATGATTTCAAGGTTGTTAGACTTGTATCGTGCTCTGTTCCTGTTGAAATTTATTCTGCAATTAGGAATT

CCTGGGAAATCATTAGGCATAATCCCACGGATACTCCTTATGCCTATGAAGAGGTATGATGGCTTTTGAT

CTACACAAGGAGGCCTTTATTTGTAACATTGGGCTGCCTATCAGCTCTCAAATTGCTGTTATTGTTGACT

TGGAGAATTCTGTTGCTGTTTGTAACCTTGATCAATTGCATAATCAGTTTAGCTTGTGGACATTAGACGA

TGAGGCATGCCTTAGTGGTGGAGGTGGAAGTAAGGCATCATGGACGACGAAGTTCATTGTTGATTTAGGC

GTGCCACTAGATCTTGGGGAAGGTGTTTTCAACATTGTTGAGTTCTTGATACAACCTCAAGAACTTGGAC

GTCTATGGTTTTCATACAACTCTGAGAAGAAAGTGGGCAAAAATGTCAATGTCAATTCTTATTCTGAGAA

GTATACAGAGAGCTTGCTTTCTGTTGCAGGACCCGAACTAATCAGGATAGCGAGGATCATCTTCCGGACT

TGTAGTTGGCTCTCCAAAAATTCTCAACAAATCTGGCGAGCAATATGAAGTACTAAAGCCTGGTGGATAA

TCAAGGCTGTTGTGTTGATACAAGTCTGTATGAATAGATATGAAAGATTCTTCGCAAGCCAGGATCTATA

CCAGTCAACTGGCCTGCTGATAAAGAAGATAATTATGAAAGTGAAGTGGATCTGGAAGATTGTAATTATG

ATAGCGAAGTGGATCTGGAAGATTGTAATTATGATAGCGAAGTGGATCTGGAAGCTGGTAATTATGAAAG

AGAAGTGGATCTGGAAGCTGGTAATTGTGAAAGCGAAGTGTATCTGGAAGTTGATACCGAGGAAAGCAAT

GTTGCCTCTGAAGATGATAAGGAAAGCAGTGAAGATTCCGAATATGATCCTGAGGAGAGCATTACTGATT

CCGAAGATGATAATGTGGAGAGCTACGAAGACGATATTGAGGAAAGCTCTGAAGCTGATAATGGGGAATG

CAGTGATGATTCTGCTAGTGAGTAATGAGAATAATTGTCACAATTATTTTATTTCTGTAGCCTTCAATTT

TCTCAATCAAATTTACTATCACAAAATCATGTGCATTGACATTGTTAAACTTTTACTATATATTAAAGTT

GTCAAACTTTTGTTTT

>MSTRG.1886.1 gene=MSTRG.1886

TCCTAATGCTTTTTATTTTATTTCTCTGCTAGTCCTTTACTCCAGAAAGAGACTTCCTTCAACTCCATCC

TGAGTCTCCTGACTAAACATACACATTTTAAATAGCATTTGATGAGGTTTACAATACACTAGCAAGACTT

CTGGATTTGTTTATTGCCTGTAACTTGATTTAGTGCTAGTGCTGGTTCACAGAATGTTCTACCTAGCAGG

CAACATGCAATAGCGTGACT

>MSTRG.1890.1 gene=MSTRG.1890

ATTAAACCCTGAACTAACCTCAGCCAGCACACGCGGATCACTATGATTCCTAACAGCCTGCACAATCGCC

CTCACCTTCTTATACGGATCCCCACATCCAAAAACCTCCAGTCCAACAAACACACCATCACACCCCAATT

GCATCATCAACGCTGCATCAGCCGGCATCACAATCCCCCCACACGCAAAATGCACAACCGGAAGCCTCCC

CATTTGTTTCGTCTGCATCACTATATCATAAGGAGCACCAATCCTCTTTGAAAACGTGAAAACCTCATCA

TCATCCATATTTGACAAAATCCTAATATCCCCCATCACTCTCCTGACACTCCCAACCGTCTCAACAACAT

CACCTAACTTCATCCCATCCCCTTGTGTCCTAATCATTGCCGCGCCTTCTCTAACCCGCCTCAACGCCTC

TCCCAAATCCTTACATCCACAAACAAACGGGACACGGAAATTATGCTTATTAATAAAATGGTCCTCATCA

GCAATACCAATCACTTCACTCTCATCAATATAATCAGCCTCCACCGCCTCCAGAATCTGACCCTCGACGA

AATGCCCAACCCGGGCTTTCGCCATTACAGGAATCGAAATAGCCCTTTTAACTTCCTTCAATAATCCCGG

GTCAGGCATCCGTGAAATGCCACTTTCAACACTCTCAACAACAACTACACACTGATTCAGCGATCTTGGC

TTGCTCAGCATTATTAACCTCGATAATCACGCCGCCCCTCAGCATCTGGGCAAGACCCACCTTCATTGAT

AACGAACTGTTCTTGTTTGCGTCGGTTATTGCGGTGGTGGTGTAGATTGTAACGGCGTTTTCGTCGTCTG

CCATTGTTGCACACAAGGTGTTTGATTTTTCGCTGAGTTTGAGCTTGTAACGAGGGTTTTTTTCAGATGG

GTTTGAAGATATAAGGGGGAGTATGGCGGTGCCGGTGGTTGATTATGGGTTCCAGAAGTTTCTATAAGAT

TTCGGGAAAATTGTAGAAAGTTAAGTTCAGATGATACAAAACTGGTTTGGTTTTGTATTG

>MSTRG.1901.1 gene=MSTRG.1901

TTGCAACTTAATATTACGGCTCACAGTTGCCATGTTGCTATATATATACACAACTTCTGACAGCTTATTT

TGCTGTAGGCACGTTTGGTATCCATACTGGATGATTACAAACTTACAAGACAACAGAAAGAGGAGGCAAA

GAAGCGATACAAGGACCAAAAGAAGCTCCAAGACTTGCTATTAACAGAAAAAGAATCCATTTATGGATCA

AAGCCTAGTCCGAGAAGAAGCCCCAGTTTCAGGAAGACAAATGGCTATCATGCTAACGGAAATGGATCGA

TGACTCCCACACCTCGCAGGAACTCAGTTGGATGTGCGACTCCAGAGCTCATGACACCACGTTCATATTC

TGGGCGTCAAAATGGGCATTTCAAGGAGATGAGAAGACTGTCTACTGCACCACTGAATTTTGTAGCCATG

GGAAAAGAAGATATCATATCTTTTTCTTCTGTTTGTGGTTCTGAGCCAGGGTCTCCTCCTCATAGTTGAA

TTTGGAACTTAGGCTGAACCAGCAACTCTACATTATGGATTTCAATAAACAATGCTGGTTTTGATGTAGC

AAAAGGCTTGTGAATATAAAGGGGGAATAATCTACTCTAGTGTATGATGGCTAGTATGTTAGTAACTACA

TAGACAGCGATGACTGACATATGAGGACCTGTGTTGCTCTCTTCGAAGAAATATCAAGATTTGGGAAATG

ATAGAATGTTGCGGATATGCAACTGCCAGTTTTTGTTTTTGTTTGTTACTAAAATATGTGTGTTGAGAAA

GATGCAACCAATTGGCCATGTGTTTGTTGATGAT

>MSTRG.1902.1 gene=MSTRG.1902

CTATAATTCTCGTAAAACAGAAAAGGGTGAAAATTTTGGGCATTAGGTTTCATAGTGAGAGTTTGGATTG

AAGAAAACAGCCTAATTTCAAACATAAACCCTCTTCAGGTTTTCCATGGCACCCCAGTTGGAAATTTCAG

GTAAATTTGAATTGGGTTTTGTCTGAATAAATCATTCATCTTCTATCTCACTTATCTCACTTGTATCTTT

TTATTTGTTTTTGTATAGTTAGAGAACCTGGGGGCTTAACTATATGGACTGGACCACCTTTCAACAATTC

AATACCAAGTGTGAAGCTTGAAAAAATCCCTTGTAGCAGTGCAAGTTTTTGTGAAAATGGAGCGAGAATT

ATGGTGATAAACCCTGGTTCTGGTATATGTATTTATGATGCTACAAGTTTTGAGCAAGTCAGGACTTTTG

ATGTGGTGGATACACTTGCAGCTACATTGTCTCCTTGTGGGACTTATCTGCAGACTTTTCAGAAACCCGT

TAATCCGAAAAATGTTACTTTGTGGGATATTGGGTCAGGGGATGCTGTTTATCAGTTGTTCCAGAAGAAT

ATG

>MSTRG.1902.2 gene=MSTRG.1902

AATTCTCGTAAAACAGAAAAGGGTGAAAATTTTGGGCATTAGGTTTCATAGTGAGAGTTTGGATTGAAGA

AAACAGCCTAATTTCAAACATAAACCCTCTTCAGGTTTTCCATGGCACCCCAGTTGGAAATTTCAGAGAA

CCTGGGGGCTTAACTATATGGACTGGACCACCTTTCAACAATTCAATACCAAGTGTGAAGCTTGAAAAAA

TCCCTTGTAGCAGTGCAAGTTTTTGTGAAAATGGAGCGAGAATTATGGTGATAAACCCTGGTTCTGGTAT

ATGTATTTATGATGCTACAAGTTTTGAGCAAGTCAGGACTTTTGATGTGGTGGATACACTTGCAGCTACA

TTGTCTCCTTGTGGGACTTATCTGCAGACTTTTCAGAAACCCGTTAATCCGAAAAATGTTACTTTGTGGG

ATATTGGGTCAGGGGATGCTGTTTATCAGTTGTTCCAGAAGAATAT

>MSTRG.1914.1 gene=MSTRG.1914

TATTTCCAACCAACAATCATATTTATATACTATCATTGGCAAGAGAGGAGAGACTGTCAAGAAGCTGTGT

GTCGAGACTGGTGCTCGTGTCAATGTTCTTCCTGGGCTTGTCACTTTCCTCATCGCGTTGTTCAAAAAAT

AGGTATGTGGATTGTGCTCGGGAGGTGTTTGATGGAATGACGGAGAGAAATGATGTTTGCTGTATGGCAA

TCCATCTATAGCCAAGTCAACCAAGCTCGTCCAAATCAGGAGGGGAGTAGAATCAATTTCATATAGCGCA

CAGCAGCGTGCGGAGATGACTCATGGCAGGGGGAGAAGCAGTGAAGAGGGAACTCTCTCTTACAATGTCA

AAGAACGACTACAGCAAATGTCATGTTTAACTCTCTGAAGGGTCAGATACAACAGTAAGACGAGGGATGA

GAAAGAAATAGAGTGGATCTA

>MSTRG.1918.1 gene=MSTRG.1918

ATGAATTAGTTTTGTTGGGGGTTGCGAATGTGGTCTAGTTCACAACTCTGTCTGTTAACTGGAAAATGGC

GATGTTATTGATGCTTGGATTGATCAATTGGGTGGCGCTGGCTTCTCCTGGTCATAATTTGTATGCATTT

AATCAGGTTATCTAGTTAAATTTTTATTGTTTTAACCTCTCAGCAACGTTCGTACTTAATATCTGTAAAA

TGGTCTACTACAGCCTACATATGTGTTTTGTTATGGTAATGTTTGTGTTTAATTTCATATGGTTTGTTCG

CCTTGATGTATGTTGAACGCTGTAATTTTGGAGATCTATTATCTATTTTCGAAGTTTATAGTATCTAAAA

CTTGGTAAGCCTTTGTTTGAGGAGTG

>MSTRG.1921.1 gene=MSTRG.1921

CTTCATTTCCATCATCTCATCATTCCTCATACTTGCAATCACACACAATCTAACACAACCCTTCAACTCT

TTATCTCCAACACAGCCTTGAGAAAACATACAAGTTCTCGTATCACACAAATTTTAAAGACTTTCTACAT

AAACCAAAGAATTAGCTAGTAATGGAGCAAGAAAGCCATGTTATGAAAGCAACTGAGCTCAGGTTAGGGT

TACCAGGATCCGATGAGCCCGTGAAGAAGAGTAACAACAAGCGAGCTTCATCAGAGATGAATAATAATTT

GTCGAATGCTGAAAAGGAAGATCATCAAGAATCACCTGTCCCCCCAACCAAGACACAAGTCGTTGGTTGG

CCACCAGTCAGAGCTTACCGCAAAAATGTTTTGCGACAAAAGAAGACGGAGGCAGATAGCAGCGGGATGT

ATGTTAAAGTGAGCATGGATGGAGCTCCGTTCCTCAGAAAGATCGATCTTAAGGTGTACAACAACTACAC

TGAGCTCTTAAAAGCCTTCGAAAACATGTTCAAGTGCACTATAGGTGTGTATTGTGAAAAAGAAGGATAT

AATGGATCTGAGGATGCCCTAACTTACGAAGATAAAGACGGGGACTGGATGCTAGCAGGAGATGTTCCAT

GGGAGATGTTTATTGGTTCCTGCAGAAGGATGAGGATTATGAAAGCCTCCGAAGCTAAAGGATTGGGATG

TCTGTAATGTAACTTGCATACACGCAACTATACACACACTCCTGAGCCTGGGTTCTTCAAGAATATCGAT

GCTAAAAGAGTATACATTCCACCCAACTCGAAGCTGGTTGTTTTTTTGGTCACGAACCAGTAAATCATGT

TCTGAAAAGTACATGACATACTCCCAGAAACAGAGGCAATTGTTTGAGACTTTGTAAAAATTATCTTGCG

AGTCTTTTCTTTTGGGCGTTTGCCCCTTAAGATGTAATTATGTTCGGCTTTTCCATGGAAATACAGTATG

TTTACTGCAAGTTTTA

>MSTRG.1926.1 gene=MSTRG.1926

TTTGTTTGCTTCTTGGAAAATCACCAATATCATGGAAAAGTAAAAAGTAAACAGTGGTGGCTAGGAGTTC

AGCTGAAGCAGAGTATAGAGCAATAGCTCATACTGCAGGTGAGCTCACTTGGATGGCTGCATTGCCGAAA

GATATGGGACTGCATCTTCCTCCTGCTGTTCTTCATTATGATAATGTGATTTCTATTTCAACTGCAGCAA

ATCCTGTGCAACATGAACAAAC

>MSTRG.1930.1 gene=MSTRG.1930

TTGTGACAAAAGACACCGGTACTCGTATAGTAGTGGCATACTTCATCGCGGAAGAGTAATCTAAACATAG

CAGTGTAGCACATCGGAAGCTGGAGATTTTCTGTTCTGCAGCCCCCTGACGAGCGTAGATGTCAAGTGTT

GAACAGCTATTGCAGTTACATATTCAGAGATCAAGTGATGGTCTTTAAACATAACCAGTTGGCCACAATG

AAGCTGCAATAACAATACAATCTCCTCTTCGTTCTGGGAATCAGTTTAAATAACTTTCATTCTTCGTGTG

TCATGCTGAAATTCTGAGTAATTTGTGGACAGGCAGACTTCATTCTCATCTTCTAAGATAGCTGAGAAAT

GGTCTTTTTGCTACAGGTATGAACTTTACGAAGTAACCAATTGGCCAAGATAAAGCCGCAATGGCAATAC

AAGCACCCCATTGCCCCCAATTCAATTTTTCTGTGTCCGCAAATTTCTTTAGGAACTCTACCATTACAAT

CTGAAGAATGATTGTTATTGCAATGATCCCTAGAAACAAATGGTTTTTAAGTATACCTGCAAACACATTC

TTCTTCTCCAATTTCCTTGCATTGAATTCATTGAAGACTTGGCAAAGGACAAAAGTGTTAAAGATTAGAG

TATTCTTGACCGTCTCATTCACATTAAAGATTGACTTACCCTTGAATTGCAAAGTTAGGAGGATTGTTAT

CTGGTATATTGCTTGTGCTAACAGGTTTCTCCACATGATATTTGTAATGAGGGGTTCAGTTCGACCCACC

GGTGGATTATGCATAAGCTCATTTGTAGGCCGCTCTGTAGCAAGTGCTAAAGCACCTAGTGTGTCCATAA

TGAGGTTGACCCACAATAGTTGGACTGTTGTTAGAGGAACTTCACCAGCTGAAACTGATGCAACGAAGTT

GATGACAAGAGCTGCAACATTAACAGTGAGCTGAAATTGGATAAATTTCTGGATATTGTTGTATACACAT

CGCCCCCATCTCAAAACTGTGGCCACAGAGGCGAAATCATCATCCAGTATAACGATGTCTGAGCTTTCTT

TGGCTACCTCCGTGCCTTGAATTCCCATGGAAAGTCCAATATCAGCTTCTTTCAAGGCTGGTGCATCATT

TGTTCCATCTCCTGTGACAGCAACCACATGGCCTTTTTGTTTCAAGCACTGTACCATCAGCAGCTTGTCG

AATGGAGACGACCTTGCCATGACACGAATATTCTCAACCTTCTCCATTCTTTCTTCAGATGTGTAATTCC

GGAACTCGATCCCTTCTACCACTTCCCCACTATTAACTTGTTGGTCAGGTCCAAGTATTCCGCATTCTGT

AGCAATAGCCTTTGCTGTGAAAACATTGTCTCCAGTGATCATTTTAATATCAACCCCAGCAGATCTGCAT

GTTTGAATGGCTTTTTTTGCTCCTGGTCTGCAAGGATCCTTTAGACCAACAATTCCTAGCAAGGTTAGCC

CATCTTCTTTTAGCTTTTTGTTGCTTGTCCCTTCGCCTTTATGTTGCATATCTTCTTCCTGCATTTTACT

ATGAGCAAAAGCAATGCAACGAAGACTGCTTGCTGCCATTCCTTCGATGATCTTCTCGAACTGTGTTCTT

TCATATTCACTTATCAAGTTTACCACTCCATTGCTCTGATAATAATGGGAACACATTGCTAGGACCATCT

CAGCTGCTCCTTTCCAATGTACATGGATGGTGTTATCAGCTCTCCTTCTGGCCAAAACACCACTTCGTTT

CTTCTCCGAGTTAAATGTTTCAATATGAATAATATTAGAGTTTTGCTTCAGTTCTTCCATATTCATTCCC

AATTTCAGGACAGCCCAAGAGAGGATCGCCTTTTCAGTTGGACTACCAGAATACTCTGGCATATTTCCTG

AGACTGGTTTGTAGACGCTGCCTGTGGTGTTCAAACCAACTCCTTGGTGAAACAACTCAGCAACTTCTGG

TGAAATTGCATCAGAATAATCTTCTCCCATGTTTTCTTGTCCAAGCCAAAACTTTGTTACCGTCATTTGG

TTCATTGTCAATGTGCCTGTTTTGTCGGTGCAAATAACTGTGGCCGACCCCATGGTTTCACAAGCTGATA

GCTTCCTGACCATTGCCTGATCAGCCATCATTCTCTTCATGGAGTATGCTAGGGTGAGCGTTACAGCCAA

TGGAAGCCCTTCAGGAATTGCAACCACTACTATAGTGACTGCAGTAGATACAATACGTAGTACTGAATTA

ATTATATCATTTACTTCTGTTTTGCCACCATGGTACTCCTGATTCCCGTCATCGTCTTCTGTATTTCCGG

TAAAGTAGCGAGCCAACATAACTGTAAGTACTAGAAAAGCGACTACAAGGCCGACTTTCCCAATTGATGA

AGTCAATTTGTCAAGTCGAGCTTGTAATGGTGTCTGTTCATTGGAATCATGGTTTATTGAGCTCATCATT

TGGCCCCAAGCAGTATTCGTCCCAACTGATACAACTAGCATTCGAGCATGGCCATCTGCTACTTTTGAAC

CTGAAATCAGGAACGGGTTTTGCACAAGGTCCACTTCTACATGGTCGCTCTCTCCTGTCATGCTTGACTC

GTCCACTTGTAGAGAATGTCCATTCAAGAACACTCCATCTGCAGGTATTTGATCTCCGATTCCCAGGAAC

ACGACATCCCCGACTACGATGTCAAAGATGGATATCTTTTGCCTCCTTCCGCCTCTCATAACATCTAGCA

TAATATTATCACTTATTTTTGACAATTTTTGGAATTGTCTCTCTTGTCTGAAGTTACTAGCAGCTGATAC

AGCAATTACTAAAAAGACTGCTACAAATATACTACCTCCCTCATACCAACCTTCCTTCAAACCATCTTCT

CTGATGCCAAAGGCTAGAGAGAGAGCAGCACAGACCAGCAAGATGAGAATAGTGGTGTCTTTGAAAGCCT

CTACGATATGAAAAAGTAGCCCTTTCGGTGGTGGCTTCTGGTAAGTATTTGGACCGAATGTGTTGCGTCT

TTGGATAACATCTTCTTCAGTGCCCTGGATGCCATTGTCAGGATCAGTTTGAAGAGCCCTGGTTAAGCCC

TCAACATCCCCTACTCTGCGGAGCTCTTCCAAATTTTTGTTCCTCATCATATCCGTAAGTTTTGTTTTGT

CAATGGTGTCAAAGCTAGAACATCCAGGCAGATTTTCTGTTTTATCAGGATGCACACTATCATTCTCTTC

TCGGACATTGGGTTCAATGTTGAGAACAGTATACGAAGGGGTGGATATAATTTCTGAAATGGGAGAAAAG

TTTGTTTTCTTCGGGACAATATCTCTGGCTACAGAGATCATGCCCCTGGAGAAATAGATGGCTACAAATG

CAACTCGCCATCTTTTTTGTGCTTTGGTGAGGGTTTGGGTGCTTATGTTAAGTATAAGAGTGGCATAATC

TTTGGGTAAAATGACTGTGCTCACCATATTCTTGATTTCAAGATATTTTTTCTTCAAATTCTGCCTCAGA

AAGGTTTGAACTTCTTTTAGCTACACGGAGCATGCTATCATTTATACTAAACAAGTGAGCTCAGTCGGCT

GTAAAGGAAATTTCGCGGCAATTTCGCTCCGCTCATATTCAAAACTCTAATACTCTAATAAAGTACTTTC

GTACATAATTGTACTTTGTGAAATTTAGTTTTCTATGTTTTGAACGAGTCTTTTGTGCTACTCTTAAGTC

TTTGCGCACGATGGAGACTTGCAGAACTA

>MSTRG.1930.2 gene=MSTRG.1930

AAGACACCGGTACTCGTATAGTAGTGGCATACTTCATCGCGGAAGAGTAATCTAAACATAGCAGTGTAGC

ACATCGGAAGCTGGAGATTTTCTGATTTTTTTGGTGCATAGTTCTGCAGCCCCCTGACGAGCGTAGATGT

CAAGTGTTGAACAGCTATTGCAGTTACATATTCAGAGATCAAGTGATGGTCTTTAAACATAACCAGTTGG

CCACAATGAAGCTGCAATAACAATACAATCTCCTCTTCGTTCTGGGAATCAGTTTAAATAACTTTCATTC

TTCGTGTGTCATGCTGAAATTCTGAGTAATTTGTGGACAGGCAGACTTCATTCTCATCTTCTAAGATAGC

TGAGAAATGGTCTTTTTGCTACAGGTATGAACTTTACGAAGTAACCAATTGGCCAAGATAAAGCCGCAAT

GGCAATACAAGCACCCCATTGCCCCCAATTCAATTTTTCTGTGTCCGCAAATTTCTTTAGGAACTCTACC

ATTACAATCTGAAGAATGATTGTTATTGCAATGATCCCTAGAAACAAATGGTTTTTAAGTATACCTGCAA

ACACATTCTTCTTCTCCAATTTCCTTGCATTGAATTCATTGAAGACTTGGCAAAGGACAAAAGTGTTAAA

GATTAGAGTATTCTTGACCGTCTCATTCACATTAAAGATTGACTTACCCTTGAATTGCAAAGTTAGGAGG

ATTGTTATCTGGTATATTGCTTGTGCTAACAGGTTTCTCCACATGATATTTGTAATGAGGGGTTCAGTTC

GACCCACCGGTGGATTATGCATAAGCTCATTTGTAGGCCGCTCTGTAGCAAGTGCTAAAGCACCTAGTGT

GTCCATAATGAGGTTGACCCACAATAGTTGGACTGTTGTTAGAGGAACTTCACCAGCTGAAACTGATGCA

ACGAAGTTGATGACAAGAGCTGCAACATTAACAGTGAGCTGAAATTGGATAAATTTCTGGATATTGTTGT

ATACACATCGCCCCCATCTCAAAACTGTGGCCACAGAGGCGAAATCATCATCCAGTATAACGATGTCTGA

GCTTTCTTTGGCTACCTCCGTGCCTTGAATTCCCATGGAAAGTCCAATATCAGCTTCTTTCAAGGCTGGT

GCATCATTTGTTCCATCTCCTGTGACAGCAACCACATGGCCTTTTTGTTTCAAGCACTGTACCATCAGCA

GCTTGTCGAATGGAGACGACCTTGCCATGACACGAATATTCTCAACCTTCTCCATTCTTTCTTCAGATGT

GTAATTCCGGAACTCGATCCCTTCTACCACTTCCCCACTATTAACTTGTTGGTCAGGTCCAAGTATTCCG

CATTCTGTAGCAATAGCCTTTGCTGTGAAAACATTGTCTCCAGTGATCATTTTAATATCAACCCCAGCAG

ATCTGCATGTTTGAATGGCTTTTTTTGCTCCTGGTCTGCAAGGATCCTTTAGACCAACAATTCCTAGCAA

GGTTAGCCCATCTTCTTTTAGCTTTTTGTTGCTTGTCCCTTCGCCTTTATGTTGCATATCTTCTTCCTGC

ATTTTACTATGAGCAAAAGCAATGCAACGAAGACTGCTTGCTGCCATTCCTTCGATGATCTTCTCGAACT

GTGTTCTTTCATATTCACTTATCAAGTTTACCACTCCATTGCTCTGATAATAATGGGAACACATTGCTAG

GACCATCTCAGCTGCTCCTTTCCAATGTACATGGATGGTGTTATCAGCTCTCCTTCTGGCCAAAACACCA

CTTCGTTTCTTCTCCGAGTTAAATGTTTCAATATGAATAATATTAGAGTTTTGCTTCAGTTCTTCCATAT

TCATTCCCAATTTCAGGACAGCCCAAGAGAGGATCGCCTTTTCAGTTGGACTACCAGAATACTCTGGCAT

ATTTCCTGAGACTGGTTTGTAGACGCTGCCTGTGGTGTTCAAACCAACTCCTTGGTGAAACAACTCAGCA

ACTTCTGGTGAAATTGCATCAGAATAATCTTCTCCCATGTTTTCTTGTCCAAGCCAAAACTTTGTTACCG

TCATTTGGTTCATTGTCAATGTGCCTGTTTTGTCGGTGCAAATAACTGTGGCCGACCCCATGGTTTCACA

AGCTGATAGCTTCCTGACCATTGCCTGATCAGCCATCATTCTCTTCATGGAGTATGCTAGGGTGAGCGTT

ACAGCCAATGGAAGCCCTTCAGGAATTGCAACCACTACTATAGTGACTGCAGTAGATACAATACGTAGTA

CTGAATTAATTATATCATTTACTTCTGTTTTGCCACCATGGTACTCCTGATTCCCGTCATCGTCTTCTGT

ATTTCCGGTAAAGTAGCGAGCCAACATAACTGTAAGTACTAGAAAAGCGACTACAAGGCCGACTTTCCCA

ATTGATGAAGTCAATTTGTCAAGTCGAGCTTGTAATGGTGTCTGTTCATTGGAATCATGGTTTATTGAGC

TCATCATTTGGCCCCAAGCAGTATTCGTCCCAACTGATACAACTAGCATTCGAGCATGGCCATCTGCTAC

TTTTGAACCTGAAATCAGGAACGGGTTTTGCACAAGGTCCACTTCTACATGGTCGCTCTCTCCTGTCATG

CTTGACTCGTCCACTTGTAGAGAATGTCCATTCAAGAACACTCCATCTGCAGGTATTTGATCTCCGATTC

CCAGGAACACGACATCCCCGACTACGATGTCAAAGATGGATATCTTTTGCCTCCTTCCGCCTCTCATAAC

ATCTAGCATAATATTATCACTTATTTTTGACAATTTTTGGAATTGTCTCTCTTGTCTGAAGTTACTAGCA

GCTGATACAGCAATTACTAAAAAGACTGCTACAAATATACTACCTCCCTCATACCAACCTTCCTTCAAAC

CATCTTCTCTGATGCCAAAGGCTAGAGAGAGAGCAGCACAGACCAGCAAGATGAGAATAGTGGTGTCTTT

GAAAGCCTCTACGATATGAAAAAGTAGCCCTTTCGGTGGTGGCTTCTGGTAAGTATTTGGACCGAATGTG

TTGCGTCTTTGGATAACATCTTCTTCAGTGCCCTGGATGCCATTGTCAGGATCAGTTTGAAGAGCCCTGG

TTAAGCCCTCAACATCCCCTACTCTGCGGAGCTCTTCCAAATTTTTGTTCCTCATCATATCCGTAAGTTT

TGTTTTGTCAATGGTGTCAAAGCTAGAACATCCAGGCAGATTTTCTGTTTTATCAGGATGCACACTATCA

TTCTCTTCTCGGACATTGGGTTCAATGTTGAGAACAGTATACGAAGGGGTGGATATAATTTCTGAAATGG

GAGAAAAGTTTGTTTTCTTCGGGACAATATCTCTGGCTACAGAGATCATGCCCCTGGAGAAATAGATGGC

TACAAATGCAACTCGCCATCTTTTTTGTGCTTTGGTGAGGGTTTGGGTGCTTATGTTAAGTATAAGAGTG

GCATAATCTTTGGGTAAAATGACTGTGCTCACCATATTCTTGATTTCAAGATATTTTTTCTTCAAATTCT

GCCTCAGAAAGGTTTGAACTTCTTTTAGCTACACGGAGCATGCTATCATTTATACTAAACAAGTGAGCTC

AGTCGGCTGTAAAGGAAATTTCGCGGCAATTTCGCTCCGCTCATATTCAAAACTCTAATACTCTAATAAA

GTACTTTCGTACATAATTGTACTTTGTGAAATTTAGTTTTCTATGTTTTGAACGAGTCTTTTG

>MSTRG.1930.3 gene=MSTRG.1930

CACCGGTACTCGTATAGTAGTGGCATACTTCATCGCGGAAGAGTAATCTAAACATAGCAGTGTAGCACAT

CGGAAGCTGGAGATTTTCTGATTTTTTTGGTGCATAGTTCTGCAGCCCCCTGACGAGCGTAGATGTCAAG

TGTTGAACAGCTATTGCAGTTACATATTCAGAGATCAAGTGATGGTCTTTAAACATAACCAGTTGGCCAC

AATGAAGCTGCAATAACAATACAATCTCCTCTTCGTTCTGGGAATCAGTTTAAATAACTTTCATTCTTCG

TGTGTCATGCTGAAATTCTGAGTAATTTGTGGACAGGCAGACTTCATTCTCATCTTCTAAGATAGCTGAG

AAATGGTCTTTTTGCTACAGGTTTCTCCACATGATATTTGTAATGAGGGGTTCAGTTCGACCCACCGGTG

GATTATGCATAAGCTCATTTGTAGGCCGCTCTGTAGCAAGTGCTAAAGCACCTAGTGTGTCCATAATGAG

GTTGACCCACAATAGTTGGACTGTTGTTAGAGGAACTTCACCAGCTGAAACTGATGCAACGAAGTTGATG

ACAAGAGCTGCAACATTAACAGTGAGCTGAAATTGGATAAATTTCTGGATATTGTTGTATACACATCGCC

CCCATCTCAAAACTGTGGCCACAGAGGCGAAATCATCATCCAGTATAACGATGTCTGAGCTTTCTTTGGC

TACCTCCGTGCCTTGAATTCCCATGGAAAGTCCAATATCAGCTTCTTTCAAGGCTGGTGCATCATTTGTT

CCATCTCCTGTGACAGCAACCACATGGCCTTTTTGTTTCAAGCACTGTACCATCAGCAGCTTGTCGAATG

GAGACGACCTTGCCATGACACGAATATTCTCAACCTTCTCCATTCTTTCTTCAGATGTGTAATTCCGGAA

CTCGATCCCTTCTACCACTTCCCCACTATTAACTTGTTGGTCAGGTCCAAGTATTCCGCATTCTGTAGCA

ATAGCCTTTGCTGTGAAAACATTGTCTCCAGTGATCATTTTAATATCAACCCCAGCAGATCTGCATGTTT

GAATGGCTTTTTTTGCTCCTGGTCTGCAAGGATCCTTTAGACCAACAATTCCTAGCAAGGTTAGCCCATC

TTCTTTTAGCTTTTTGTTGCTTGTCCCTTCGCCTTTATGTTGCATATCTTCTTCCTGCATTTTACTATGA

GCAAAAGCAATGCAACGAAGACTGCTTGCTGCCATTCCTTCGATGATCTTCTCGAACTGTGTTCTTTCAT

ATTCACTTATCAAGTTTACCACTCCATTGCTCTGATAATAATGGGAACACATTGCTAGGACCATCTCAGC

TGCTCCTTTCCAATGTACATGGATGGTGTTATCAGCTCTCCTTCTGGCCAAAACACCACTTCGTTTCTTC

TCCGAGTTAAATGTTTCAATATGAATAATATTAGAGTTTTGCTTCAGTTCTTCCATATTCATTCCCAATT

TCAGGACAGCCCAAGAGAGGATCGCCTTTTCAGTTGGACTACCAGAATACTCTGGCATATTTCCTGAGAC

TGGTTTGTAGACGCTGCCTGTGGTGTTCAAACCAACTCCTTGGTGAAACAACTCAGCAACTTCTGGTGAA

ATTGCATCAGAATAATCTTCTCCCATGTTTTCTTGTCCAAGCCAAAACTTTGTTACCGTCATTTGGTTCA

TTGTCAATGTGCCTGTTTTGTCGGTGCAAATAACTGTGGCCGACCCCATGGTTTCACAAGCTGATAGCTT

CCTGACCATTGCCTGATCAGCCATCATTCTCTTCATGGAGTATGCTAGGGTGAGCGTTACAGCCAATGGA

AGCCCTTCAGGAATTGCAACCACTACTATAGTGACTGCAGTAGATACAATACGTAGTACTGAATTAATTA

TATCATTTACTTCTGTTTTGCCACCATGGTACTCCTGATTCCCGTCATCGTCTTCTGTATTTCCGGTAAA

GTAGCGAGCCAACATAACTGTAAGTACTAGAAAAGCGACTACAAGGCCGACTTTCCCAATTGATGAAGTC

AATTTGTCAAGTCGAGCTTGTAATGGTGTCTGTTCATTGGAATCATGGTTTATTGAGCTCATCATTTGGC

CCCAAGCAGTATTCGTCCCAACTGATACAACTAGCATTCGAGCATGGCCATCTGCTACTTTTGAACCTGA

AATCAGGAACGGGTTTTGCACAAGGTCCACTTCTACATGGTCGCTCTCTCCTGTCATGCTTGACTCGTCC

ACTTGTAGAGAATGTCCATTCAAGAACACTCCATCTGCAGGTATTTGATCTCCGATTCCCAGGAACACGA

CATCCCCGACTACGATGTCAAAGATGGATATCTTTTGCCTCCTTCCGCCTCTCATAACATCTAGCATAAT

ATTATCACTTATTTTTGACAATTTTTGGAATTGTCTCTCTTGTCTGAAGTTACTAGCAGCTGATACAGCA

ATTACTAAAAAGACTGCTACAAATATACTACCTCCCTCATACCAACCTTCCTTCAAACCATCTTCTCTGA

TGCCAAAGGCTAGAGAGAGAGCAGCACAGACCAGCAAGATGAGAATAGTGGTGTCTTTGAAAGCCTCTAC

GATATGAAAAAGTAGCCCTTTCGGTGGTGGCTTCTGGTAAGTATTTGGACCGAATGTGTTGCGTCTTTGG

ATAACATCTTCTTCAGTGCCCTGGATGCCATTGTCAGGATCAGTTTGAAGAGCCCTGGTTAAGCCCTCAA

CATCCCCTACTCTGCGGAGCTCTTCCAAATTTTTGTTCCTCATCATATCCGTAAGTTTTGTTTTGTCAAT

GGTGTCAAAGCTAGAACATCCAGGCAGATTTTCTGTTTTATCAGGATGCACACTATCATTCTCTTCTCGG

ACATTGGGTTCAATGTTGAGAACAGTATACGAAGGGGTGGATATAATTTCTGAAATGGGAGAAAAGTTTG

TTTTCTTCGGGACAATATCTCTGGCTACAGAGATCATGCCCCTGGAGAAATAGATGGCTACAAATGCAAC

TCGCCATCTTTTTTGTGCTTTGGTGAGGGTTTGGGTGCTTATGTTAAGTATAAGAGTGGCATAATCTTTG

GGTAAAATGACTGTGCTCACCATATTCTTGATTTCAAGATATTTTTTCTTCAAATTCTGCCTCAGAAAGG

TTTGAACTTCTTTTAGCTACACGGAGCATGCTATCATTTATACTAAACAAGTGAGCTCAGTCGGCTGTAA

AGGAAATTTCGCGGCAATTTCGCTCCGCTCATATTCAAAACTCTAATACTCTAATAAAGTACTTTCGTAC

>MSTRG.1938.1 gene=MSTRG.1938

ATATTTTCCAAAAATATTATTGTTTGCAGGGCTGGCATCAAGGCCACTCACGTTCGCATATACCGTCAAC

TAAAAGCATATACTTGTGCACTAATTCTGTGGACCTAACAAACGAAATATACACTGAAAAATAAAAATTG

TATTATCCTCTGAGAGCTTTGATCACAATTTTCTGCACCATCACACTGCAAAACTTGTGATGGCCATCTT

TCTCCCTGTGTTGGGCCTGTTTCTTGTATTCATTGTGCAAGATTTGACGGGCTTGCTGTGCTTTTGTTTC

TGGATGTAGCTTCCATCTTGAATTTCTTCTGGCCTGGAACCAAGGCCAATCAACTTCGTAACTACCGTCA

ACCAGCATATACTAGACCAAACAAATATGTGATGAACTTTTGCTCTGACCAAAAAAAATTTACTGTCCTT

ACAGAGCTCTGATCTCAATTTTCTGGCAACTCTTAAGATAGATTATGATCTGTGGTTGTTGAATCAAAGC

TGTAAAGGATCTTCGGGCTGCAAGTCATTCTAGTTCTGAACACCATCACACTGCAGGACTTGTGATGGCC

ATCTTTCTCCCTGCTCTGGGGCTGTTTCTTGTATTATTTGTGCAAGATATGACGGGCTTATTGTGCTTTT

GTTTGTGGATGTAGCTTCGATCCGATCATGTGTAGCAGTTGAGGGGAAATCTTGAAATTTCTGGGCTTGA

ATCCTAGCATTTTTCATCAAAAAGTTGGAGGCATGGTTAAAGACTTAAAAATGAGTAAGACGGATCACAA

TGTTTTTGTACTGATCAACTTTAAGGTGTCGAGTGAGGGATTGTTCTATCTCTTATGAAAGGGGCCCTGG

CGCCTTGTTATTAGAGAACGGGGAGTACTATCTCGATTATTGTCCCATTTCACTCGATTATTGCGCCCAA

GATCAAATGATCTGCCATTTTGCATGCTTATTGTACAAAGTGTGCTAAACTTATCTTCAACTTTCTTGTA

TGGCGCCTTCTGGCAAATGAAATAACATGAAGCAGACTCTTAAAAACACGTGTTTTGAAGGTTCTGTTTT

CGCTGATCTACTTGTTAATTGTTATGTGGTGCATCTGCACTTTGGTACGACCCAGCACAAAACAAATATT

CGTATGAGCCGTATTGTCCCAGCTAAGACATCTTAGTAGTTGAGATGATATCCTCCTTTGCTAAAAATCT

TGGTATTAGCCTGGAGTATTTGTTGCGACCACGAAGTTGTTTACGTGGTCATGCCAAGATCCAGAAACTC

GTAGAATATCTGGATGACCCTCCTATTATTTGCCGCTTCATGTTTGGATGTGTTGATAATTCCCGAATCT

TTATCTGATGGAGTTGTGTTTATTTTTCCTCCATTCTTTTGTAAATGGTGTGTGTTCCTGTATCTTATTG

TGTGAGTTTCAGCTGGGTGAGCAACTGTTCCAACCGGCTCGAACCGGACCTGAAAAAATCGTTGGAATTT

TAGAAACCGATTCAGTTCTTAATTAACCTTTAAGGACCCCTAAATTTCGATTTTGTTTCGAAC

>MSTRG.1938.2 gene=MSTRG.1938

CAAGGCCAATCAACTTCGTAACTACCGTCAACCAGCATATACTAGACCAAACAAATATGTGATGAACTTT

TGCTCTGACCAAAAAAAATTTACTGTCCTTACAGAGCTCTGATCTCAATTTTCTGGCAACTCTTAAGATA

GATTATGATCTGTGGTTGTTGAATCAAAGCTGTAAAGGATCTTCGGGCTGCAAGTCATTCTAGTTCTGAA

CACCATCACACTGCAGGACTTGTGATGGCCATCTTTCTCCCTGCTCTGGGGCTGTTTCTTGTATTATTTG

TGCAAGATATGACGGGCTTATTGTGCTTTTGTTTGTGGATGTAGCTTCGATCCGATCATGTGTAGCAGTT

GAGTCGAACTTATCAGGGGAAATCTTGAAATTTCTGGGCTTGAATCCTAGCATTTTTCATCAAAAAGTTG

GAGGCATGGTTAAAGACTTAAAAATGAGTAAGACGGATCACAATGTTTTTGTACTGATCAACTTTAAGGT

GTCGAGTGAGGGATTGTTCTATCTCTTATGAAAGGGGCCCTGGCGCCTTGTTATTAGAGAACGGGGAGTA

CTATCTCGATTATTGTCCCATTTCACTCGATTATTGCGCCCAAGATCAAATGATCTGCCATTTTGCATGC

TTATTGTACAAAGTGTGCTAAACTTATCTTCAACTTTCTTGTATGGCGCCTTCTGGCAAATGAAATAACA

TGAAGCAGACTCTTAAAAACACGTGTTTTGAAGGTTCTGTTTTCGCTGATCTACTTGTTAATTGTTATGT

GGTGCATCTGCACTTTGGTACGACCCAGCACAAAACAAATATTCGTATGAGCCGTATTGTCCCAGCTA

>MSTRG.1940.1 gene=MSTRG.1940

ACCTATGTTACCCGGACTCTCCATTTTTGTGCCTATACCCGTGTCCACCGGACATGACATGGGTGTGGGT

ATGGGATCTGAGTCGGATCCTTCGTATTACAACCGGACACTTCACCTTGTAACATCCCAGGTCAAAATGA

GAACTTAGGTCTTGTTTCAACTTGTTTATAGAGATCAAGATCCACGAATGACTTGTTTCTATGTTGTTCT

GTTGATTTTATGCTTCAATTTTCAGTGTATGTGATTTATGTGCAGTAGACACTATTTTTATGAATGTATA

ATAAATCAGTCTATATAAAGTTTTACTTGCGTGGAACTTAAGTGCCAAGTCCCCGTACCCGAGTCTAATT

CCGGATCCATGTCCACG

>MSTRG.1942.1 gene=MSTRG.1942

CTTCCATTAACACGCTGAGAATGCATGTGAACGTGTGGTCCCTCGTTGTTGCTCCGTGCGAGTGAAGAAA

GCGGAGGAGGGAAAGATCGGTGCTTTTGCATCATCTTTTTATCAGCAGCGCCAGAGTAACTATATTTTGC

AACTACAGGAAACTCATCTCCCAAGTGATTCTTGATATCTTCATGTCGAGATTCCTCTTCTTTGTCGAAG

TCACTCGCCTCCGAGTTAGCCTCAGAAGAGGAATAAGACGAAAACCCGTCAGACCCCGTCTCAGATCCTA

GGTTTTCAGTACATATCTCAAGGCTCTTCTCGCTCAAAGTGCTGCTACTTTTCTTCAAAAGAGGATGAAC

ATAAGGAGGTGGAAGTTCAGCCTCATCACTCTTTTTCTGCGATATAATTGTGCCCCAAACATCATCTTGC

TTCTGCACACGAACTCTGCCTTCTCCTTCGGACAAAGAATCAAGGGCTGAGATGATGGCAAGCTGGCCCG

AGGAAGCAATTTTCTTCATGGTCGGATGCTTGAAACCATTCTGCGCCAACCATGTTTTCGAGGACATGTC

AGCCGAGAGTGTTCTGCGAAAGGAGGTCCCTTCAGCAGGCTTTAGACCTTGCGAATCATCGCAACGAAGA

ACAGAAGTGATCCCTTGCTTCTCCATCATGGTTGGCGACAGCTCCTCAAGAAAGCAGTTGTTGTTTGGTG

CTGAGCTAGACTTGTTGACATACATCTCTTCCATTTGGAGAAAATCTTCTCTGCAAACCAAAGGAGTGAT

TAAGTATGAGAAGACTAATGGAGATCAAAACTAGAGATCAAGAAAGCTGCTAGCTTTTTACATGCAGTAC

TTAAAAATACCAAAAAAACAAAGAGAACTACGTCCATACGCAAATACTATGATAGCATGTGCCCACAGAC

ACACACCGGAATGAACCGATATAATAATATAAGAGAGAAATGAAATGTAGTTGCTTACCTTGAGAAGCAA

GAGAGTTATCCCCTGGAGAGTGAGAAGAGAGTAGAGAGTGAAGTTGATCCTTGGGGGAGTGGGAGGAGGC

AAGGGAGGTGAGTGTTTGGGATGTGGCTATGAATGCAATTTACAATGGGTGGAAATTGGCGAGTATTTGT

AGGCGCTCAGAATTGCCTAGGGAATCGGGTACTTGGGACCCGCCTGCATTGCCCATTTCATCTGTTTTTC

TCTATACGACTCTGATTTTCTTGATGTAAGAACCATCTTTGTAATTGTGAACCAGCTCACAGATCATAAC

AAGGATTGCAAGGATTTGAATTCAAACTCAATATTTTGAAAATTTTGGGTTGAATTATCTCCATCTGATG

TGATCCACAGAGGAGAATAGTGACCTCAAAG

>MSTRG.1943.1 gene=MSTRG.1943

ACAGAGCGCATGACAGAAATAAGATCAGTGGAACGAGTGAAATGTTCCAAGCCAAGAATCCCTTTGAGTT

GCTGCAGGCACACCACAGTGGCGGCTCCTGCCATGAATCCAACAATTGTTGCATGCGACAGAAAGTCCAC

TATGAATCCCAGCCTGAATATGCCCAGAGAGGCTTGTAACACCCCGGCAAAGAAACAGGCTGTGAAAGCC

AGGTGAAGGTAAAGACTAGGGTTCTCTGTTGCATCCACTTCTTTTCCCAGCATTGAAGCTGTCAGTAGTG

AGGCCACGGCCACCGTCCCGACGGCCAAATCTCGCGAACTTCCCATCAAGGCGTAGACCAATGGCGGTAT

GAAACTCGAATCTGGTTTATATTCACAAATTTAGTAACCAGTTAGTAAATCAGGTTCGTAGGATTAAAGT

TGTTACTGACAAGGTTCGGAGATGATACTATATTATTCAATGCAGATAAATTATCAATCAGGTTCGTAAT

ATTAAAATTGTTACTGTATTATTCGAAACAGATAAACTATCGTATGAAAAGTGTAATAATATTGATAATA

TAGGATAATATGTAAAACTCACATAAGCCGAGAATAGGGGGCAAATTTGCTAGCTTGGCATAACTGATTC

CCTGAGGAATGGCCAAGCTAGCAATAGTGATTCCGGCTATGAGGTCGGATTTGAAGAAACTCAAGCTGTA

ACGCGGCCCCCATTCGAAAATCGGGAACACATATTGTACACCTAGTTTGAGTTTGGTGGATAATGGTTGG

TTTTTGAATTGGCGTAGAGGGTCATCAGGGAAGAAGGTTTCTTTGAGATTATTTGCAAACGACTTGGTGA

ATGGTTGCGGTGGCGGAATCGCGACACGGTGCGAACACTCGACATTCTCCGACGGGTACGAGTAGTCTGC

GTTACCCATTTCACGGCAAATAAATATTTTCGAACTCTGTATGAAAATTACTAAGTTGAATATGTGGAGA

AAGTATGAAGTGTTGAGGAGCAGAGTATACAACATTGGAGGTGTGCTAGGAGCTTTATATAGTGGAAGTG

TGAAGAAATTGTGAGTGGACGTTTGGGTCCCACACATGTTCAGGGGTGCCACGTGGCTCAGAGGTCTTCA

AACTTATACTGCTGCCAACCTTGGGCCTGGTTATCATTTAGATTTCCCTGACGAGAGAAAGCAAAGTGTT

AGTGGCTCACGTATATGTGTGCACAATGCCGTGTATACTATAGAGTCCCTTAATTTTCCGTCCCACAATA

AGATTTCTTCGACTCTGACACTTTATTTAGTCAGAATTATGATTTTGTAATCCGGATCAAAAATGTATAA

TAATCCTGATTCCGTCACTGTCTACGATTTTATCGGACTCGAATTATATACATGATAAATGATATGGATA

TGATCTGA

>MSTRG.1945.1 gene=MSTRG.1945

GAAACACAACTCTGATTAAAAATACAGACACGAACGTGTGAAATAGCGAAATTAGCTCTCTCTCTCTCTC

TCTACCTGCACACAGTGTGTTTGAAATCGATGCAATTACCTGAGCGCCAGTACGGCGAGCAGCATCAATG

ATAGAATCAGCTTTAAGATAACTCAACCGAGCTGCTGCAGGACCGATATGGATCGCTTCATCTGCGGATT

TCACGTGCAGGCTGTGCTTATCAGCATCGCTATAAACGGCGACAGTTCGAATCCCTAATCTCTTCGCCGT

TCTCATGATTCTACACGCAATTTCGCCTCTATTAGCTATCAATATCTTCTCTATCTTTACATTTTTAACA

CTTCTAGTTGAGTAAAGTCTCGCCGGAATTACAAAGTGATTGCTGCTCCGGTTAAATTTCCGGCGAATTA

TTGCCGCTAGTAATGACATTGGAG

>MSTRG.1949.1 gene=MSTRG.1949

CTTATTATCACTTCTGCATTTTGATACACCTTAATTTACGTCGGTTGTTATGTTTCAACTCTTTGATTGC

AGCAAAATACAGGAGTGCCAGATGACGACTTTGTTTCATACGTGAACAAATGGTGTGAAATAGGAGCTTC

TCTTGTAGGAGGTTGCTGCAGAACTACACCCTACACTATCAGAGGCATTTATAGGACTCTAGCTTCGAGA

TCGGTCTCGTGAACGTGATCAAGCAGGGAAAACAACCAAAAGTTGATTAAGAGGTCTAGTTTCAGCTTAT

AGTCTTATACTGTTCAGATCCCCTTTATGGGATTTAAATTCTATATTTGGTTGCCAAGAAATATTTTTTT

TTTCTCACCAATTAAATTGGGAGGATAGAAGGTTGATCCAATATTTGGATGCAGAATATGATATGTTGAA

TTGTTGAGGGTCTGATAGTATGTACTATGTACTATGAGAAATTCAAATTTTAGTAATATGATCTTTTTTA

TGTTTGTTCGAGTCGA

>MSTRG.1952.1 gene=MSTRG.1952

CACCTCTTCTCCGTGAGGAGTAGTCGCACTTTCCTCCAACAAGGCGCGCCCTCCTAAAGGGGTACGCCAT

ATCCAACCAGTATTCTATCAAGAGAACGAATTGCGCGCCTCCCTCTGACAGGGCGCGCCCTCTGGTATGT

CTGATGCATTAATTGCTTCATATGATATTTTGTGAAGACAAACATACTCAGGAAGGAGCTCACTAGTCTT

GAAGAGGTC

>MSTRG.1954.1 gene=MSTRG.1954

TAGTTATTATCATGCTGTATATTGTGAAACGGACAATCTATTGTATCTGTACTAATATTTATTGACATGT

GATCTGTGCACAGCAGTTTCTTCTTAGTCATGTCTAATAAGGTCTTCCTTGTTGTCAGTTGGAAAATCCT

TAAGATTTATTGTGTATCCTGTTGGCCATGGCAGAGCCATAGTATATAATTTTACTGTCGCTGTTTGAGT

TTTTCATAAGACATTAAAAGTATCTGATTTGGTGGTGATGTATGGACTTTTAATGCAG

>MSTRG.1955.1 gene=MSTRG.1955

CTGGTCACTACAATCAACACTAACGTTCCAATGCCGATAAAAAGACCGCGAATCGGTTATTCAACCAGCT

CCGGCGAGAACGGCCGGAGGTCTCACCGCCGATCAATTCTCCCACGGAACGCCGCTGATGCTTTTTGTAC

GTATAGTATCATTGGGCACTATCACATTTAGCCATGCGAGAACAGAGCTTTATTTATGAAGGTAGCATAC

AGCCTGACAAGACCTGAAAGAACAAGGTGTAAATAAAAACTGTTACATTTTTCCTTTATGAAAGACTCTG

TCTTCTATTTCCTTAAATGTTCTATACTTCCCTGTATATTGCTAGAGACCAGAGCTCTTTAGTAATTGCG

AGATGCTTAAGATATTTATCAGTGAAGTTTAGTTTTTGTTTACTTTTCCATTTCTAAAGAGGCCAGCAGG

ACTCTGCTATATCTCAGGTGCTGGATGGAGAGAAAAGCTCTGTTAGATTTGACGTAACCTTCTTTTTTTG

GTTTAATGGAGTATGTATTACTATAGAACACTAA

>MSTRG.1962.1 gene=MSTRG.1962

ACAAACTGCCCACTCCTCTCTCCCCACCTCTCCTCCACCTCCTCCACCTCCTCTCCCTTCGGCACCACCA

CCACCAGCTCGCCGTCGCTGAACGCCGCGCTCGCCATCTCCGGCAAAGTCGCCTCCGGCAACCGGAATCT

CCACACATCAAGCTCCAGCTCATCCACCGACAAGTCCTCCACGTCTTCACCCCCTCCTCGTACGACAATC

TTGGTGATTCCCGGGTGAATATTCACAGAGTGGGCCCTGATGTCGTCCCCGATGGCGGCGTCCTCGGTTG

TGATAACGAAGCGGAGTGAATCGGAAGTCTCGTGGACCAAAACGTCGGCGTTTGAGTGGAAGGGGAGTTC

GAGAACTTTCGCGAAGATGTGGGGGAGTCTGCGGAGCTGCTTCTTGTTGTAGCGGCATCTCGCGTTTCGG

TGTTCTTGAATCGGAGCTGGGTGAACCCTCATCGTCTGTATTTTATTTTCGAATTATTTTTTGTATCGGT

TTGTTTGGTTGGTTCAAGTAGAGATGAATAATGGAATGTGGGAGAGAATGCGGAGGTAAAGAGATGAGGG

GATTTGTGGGTGAGTTATATAAGAATCGGAGTGTCATTTTTTGATTTGGACGTG

>MSTRG.1966.1 gene=MSTRG.1966

TAAGAAAATTATAATTTACCTGGGTTTTATGTTTCTTGCAGGATAAGTTGATTCCTCTAATGTTGCCTGT

TTGCCGGTCCAGTTAGAGGTTATTGCCATGAAAGCCTTTGGCAACTCCAGTTGCAGAGACAAAAGGAATT

TACAAGATCCGCATGCTCTTCGAATGAGGATGCTGACAGAATTTCATATGGGAATTGGAACTATGGATAC

TATTCTCGATGTTGGCTAAATTGGAGATGATTTAATTTCTGGTCCTTAATATGTAGTCGTATTTCTTCAT

TTCGTCAGTGTTCTTGATTGTCAGTCTAATTCATACTGTAACTATTAATTGTATACTTGGTAACTGGTTT

GAATGCCTTGGTATCTTGTGATCTCTTTCGACTATAATATTATGTTCTGGACTTGTGAATGT

>MSTRG.1979.1 gene=MSTRG.1979

CTCCAGTAAACTCTCATCCTATTAGCTTTCTGTTACGCAACCTAATAAAATGTGTTGGTGTTCCGACTAT

GCAGATGAAATGACTGAGCATGACCGAATAAGTACACTAGTCGACTAAGACAGGACGGTTCACTAGATGC

ATCATGGACCGAGTGCTTCGGATTAATGGCAATGGAGGTGAGGTCTTGCATTTTGCAGGAGACAAATGAT

AAGATATAAGTGGAGAGGCCACTGAGAATGCATAAGCAGGATATCTTATGGTAAGAGTACAATGAGACCT

CTTTCGACTGATGTGATCACTTGTTTCCATTACTTGTTGTCAGCCTCATTTTTCATTTCTCA

>MSTRG.1981.1 gene=MSTRG.1981

TGTATGCTGAAACTAAATTTTCTTCTAGTTGCTCATATGCTGTAGCTTTATCTATTTAATGTTTCATTGA

ATATGAACTATCTAAAGCTTGATATTTTTTCACTCATGCGTTACCTTCTATACAACCAGCGAGTTTTCCA

AACTTGGATGCTGTAGATTCGTGTAGCTCATCACTAGCCCACATGGGAAAGATCAGTAAGTTGGTGAATA

GGCAGACGAAGAAACCCATACCAATTGTAGAGAGTCGTTCACGGGCCAATGCCATGACTTTATCAGCTCG

AACACCAGACACAACCACCAGACTGAAGG

>MSTRG.1982.1 gene=MSTRG.1982

ATTCAAGTATCTATTATTACGGTCACAGATGATGTATCATCTCATGTATATCTTTTAAGGCCATCCTCAC

TTCTTTCTGTTTTAGGAAAGGAACAGGGAACAAGTGAGGATATATAGTTTCTTCTTACATATATACAGGC

TAATCATGCCAAAATTGACTGCCAAGGTTATATGTTCATGCATATTAGAGTTTATATATACAGCACCGAC

TGCAATTCTTATTTGATGCTGAAATTAGCCATCTTTCCTAGTTCTTCCACCTCTTTCGCTAATACTTCCA

CTTTGTCCACTATCTCCATCAACAAAAACATAAAACTTGCAATGGCGATACTTTCACCTGTATCTAGTTC

CTCTAGCTTTGGAGATTTTAGGAGGCTTAATTCCATTCTCAGGGTCTGCAGCTTTGGCAACATCAAGATT

GTTGCATGGCACCTCTTCATTTTAATGATGCTTTCACCCAGCTCTCTCAAAATCTGCACAAGTTTTGATC

CAACAATCTTGCATGGCTCTTTAAGTTCTTGTCTTGTTGTCGATGATGGCTGTGGAAAATTTGAGATCAA

CTATTTAAGTTGCACTAGATATGAAGATATCGTTCTGGACGGTGTAAAAGTTTATAATTGTTTATATATA

GAGTTATGCCACTATCATTTCAGCGATAAGGGATTTGAGTATCACTGACCTGTTTGGGAGATTCAACACA

TCCTTTCAAACAAATAATGGTCGCAGAAAGATCTCTAAGAGCTTCTCCTATGCGCAGATACTTTTCCCAA

GGATAATGAAAGCCAAATTTTCCATGCCTCGGTTCCCACTTTGCAAAATTGGCCTGAAATAGGTTGTAAA

CTAATTCGTAAGCTATAAGAGTTTTATTTGTTAACACATTTGGTGAATATTTCACTTCATTGATATA

>MSTRG.1989.1 gene=MSTRG.1989

CTCAAAAATTTATCCACAAACTTATCAACATCAACACAACTCATCTCAACTACATCATCCTCACTTGAAT

AACACGCTTTTTCCCTCTTTTCACCCTTTTTAAATCCCCCCTCAAATCTTCTCGATTTCGAACTCTCACC

CTTCTTGACCACACTCGGCCCTTCATCATCTTCATCAGAAACCAAATCCACGTGCTCCACCTTAATCTCC

GGTCCGGCAGCTCCGGCGCCGGAAGTTCTGCCCTTCCTGGCCCTCTCCTTCCTTAAAGTCTCATGCAAAA

TCTTATCAAAACCCAACCTCGTGCGCTTAGCTACCGGCAACTCTCCAGAATCCATAACTGCAAATAAGTT

GCTGTGTGTTCTAAGAGAGAGA

>MSTRG.1991.1 gene=MSTRG.1991

GAACAATAGAAATACTATTTGACTCGTGAAGTGTCTTCATAAATGTTCTTTCTTTACTGGTGCCTTCATA

AATGTTGTGAGGACTCTGTTTGTCCTATTCCTATGAGGTACTCACACTTTTTCCGAATTCCGATTCACTT

GTCAACGGAGGGAGTACTTGATTTCTATATTCCCATCGTCAAGATGATGCTTTTATGCATGCGAGGACGG

AGGATCTTGCAACTCGCCGCACAGTAAACGATCTTTATATTGGAATCTTAGAGTTGTAATCCGTTGGAAT

TTTAATACTAG

>MSTRG.2003.1 gene=MSTRG.2003

ATCGTAAAGCCCAAGACCAGTTTAGTATAAATCTATTCGAACCCCGAAATGTGAGTCAGAAACCCTAAGA

TAAACAAGCCGCAAGGTACTGCACAACGAAGATTGGAGTTTGGAGAAGGAAGCTTATGTCATCTTTCAGT

GTGCTTGCCCGACGAACGTGACAGAGTAGCAGGCTAGCAGCAGCTCCAGTGTCGCTTCACATTGAACAAT

TTGCAATTAAGTGTTTATAAGGAGAATAGGCGCTGGGAGAAAAGGTGTGTTCCCCTAATTTGTCTAATAA

CATTCACCATCTTTTCTATATCGTTGTTGTGGACTTTAATTTTACTGCTACATGTAATCTTAACGTCAGT

TCAGTCATTTCTTTGTTAATAGCCATTGTTGCTTAAAATCCAGTCATTTTCTCTTGTTTTACTGCTCCTC

TTTCGTTTTGTTTTCAGTTCTGTTGTGCCTGTCTGTCTTATTGTTTCTAAGGGGC

>MSTRG.2003.2 gene=MSTRG.2003

ATCGTAAAGCCCAAGACCAGTTTAGTATAAATCTATTCGAACCCCGAAATGTGAGTCAGAAACCCTAAGA

TAAACAAGCCGCAAGGTACTGCACAACGAAGATTGGAGTTTGGAGAAGGAAGCTTATGTCATCTTTCAGT

GTGCTTGCCCGACGAACGTGACAGAGCAGGCTAGCAGCAGCTCCAGTGTCGCTTCACATTGAACAATTTG

CAATTAAGTGTTTATAAGGAGAATAGGCGCTGGGAGAAAAGGTGTGTTCCCCTAATTTGTCTAATAACAT

TCACCATCTTTTCTATATCGTTGTTGTGGACTTTAATTTTACTGCTACATGTAATCTTAACGTCAGTTCA

GTCATTTCTTTGTTAATAGCCATTGTTGCTTAAAATCCAGTCATTTTCTCTTGTTTTACTGCTCCTCTTT

CGTTTTGTTTTCAGTTCTGTTGTGCCTGTCTGTCTTATTGTTTCTAAGGGGC

>MSTRG.2003.3 gene=MSTRG.2003

CCCAAGACCAGTTTAGTATAAATCTATTCGAACCCCGAAATGTGAGTCAGAAACCCTAAGATAAACAAGC

CGCAAGGTACTGCACAACGAAGATTGGAGTTTGGAGAAGGAAGCTTATGTCATCTTTCAGTGTGCTTGCC

CGACGAACGTGACAGAGGCTAGCAGCAGCTCCAGTGTCGCTTCACATTGAACAATTTGCAATTAAGTGTT

TATAAGGAGAATAGGCGCTGGGAGAAAAGGTGTGTTCCCCTAATTTGTCTAATAACATTCACCATCTTTT

CTATATCGTTGTTGTGGACTTTAATTTTACTGCTACATGTAATCTTAACGTCAGTTCAGTCATTTCTTTG

TTAATAGCCATTGTTGCTTAAAATCCAGTCATTTTCTCTTGTTTTAC

>MSTRG.2012.1 gene=MSTRG.2012

TGTTACTAGTGGTTCGTCGCATGACCTGTCTGATGATGATGATCTAGATATTGAAGCTGGTCCATGTGAG

CAGAGTGACCAGGTTGATGTAAAACGTATTAAAATAATGGTCTCTAACAGAGAGTCTGCGAGGCGTTCCA

GAAGAAGAAAATAGGCTCATTTAGCAGATCTCGAGCAACAGGTGGATCAGCTTCGCGGAGAAAACTCAAC

TCTGTTCAACAATCTGACAAATGCTAACCAGCAATTCAAAGATGCAAATATGAATAATCAAGTTCTGAAG

TCAGATGTGGAAGATTTAAGAGCCAAGGTACCAAAGATTTTCAACTTTCAACAGCACAAATGTTAATATT

ATTCCAATCTGCAAC

>MSTRG.2013.1 gene=MSTRG.2013

GTTCCACCGGTGACAATCTCAGGACAGCAGGTACCATATGCAGATTGGGTTGTTAGTGTTGGCGATGGGG

TCGTGCCAACAGTACCATCTGTTGAGGGTAATGAACCATGCTGGATAGAGATTCCTTCAGAATTGAACCT

TGATCCCGGAGATGATGGGAAAAAGGCTATTATTGATGCTGTGTATGATGAATTATGCAACAGCTCTAAT

GATTCAGGATATTTCAGGGACCGTGCTATCTTGACTCCATTGAATGAAGACGTTGATCTTATTAACAAAG

AGGTCATGAAAAGATTTACAGGCCAGTCCAAAATATATCGAAGCGTGGATTCAGTTTGCAAAAGCACAGT

TAACTATGAATCAATGGAAAGCATGTACACACCAGAATATTTGAATTCTTTAAGAATAGCAGGAGTACC

>MSTRG.2014.1 gene=MSTRG.2014

AGTAATCCTAAAAGAACTATGTTCCAAAATAACATCCGGATGTACAACAACGCATTCGGGTTTACTTCAG

TTGGAGCTAATATAGATAAAAGTATTAATAATGGGCGTGGACCTTTTGTTTATCGTATTCATGGTGTAAT

TTACCATCAAATAGGTTCACTGTTTGCTGAAGAATCCAAAAAGCCAGTTTTCTCGCAAATATATATGTAC

GATAATCTAGAACAGGTTAACAAGCGTATGAATTTTCCCAATTCTGAAGAACCCCTTGATAGTGAAATAA

CCGAATTGTTATCTGTGATGTTGCACCGAGTGAATGCTTTGGTGGATATATATAGACAGGTTCGAGATAG

ATACAAAGAATCAGAAGTAATTCCAGGAAAGTTGCGTTTAATTGCAAATAGGGACACAGATGGTCGAGAA

AGTAATGTTCCAACCAATGCCTATGACTTTGCAGGATTAGTTGCAAACAGTAATCTAGCAGATGATAGGG

ATATACTTGTCCACCCTCATGATGGAGTTCTGCATAGGATTTCAGTGCTCCATCCATGTTATATGTCCCT

TCAGTACCCGCTTCTATTCCCGTGGGGTGAAGATGGTTTCAGAATTGATATCTTACATTCAGGGATAAGT

GAAAAATCAGGAAATAAAAGAGATGTGGTTACCATCAGAGAATATTATTGTTACAGACTTCAGTACAGAG

ATGCGGAAGGGCATACTTTAATTGGCGGGGGGCGGTTATTCTTACAGTTTGTAGTTGATGCATGGGCATG

TATTGAGCATACTAGACTTACATGGGTCTTAACACACCAAACAATATTGAGGTCTGATCTATATAACAAT

GTTGTTGATTCTGTTAATAAAGGAGATACTAATGCTTCTACAATTGGAAAACGGATTGTATTGCCATCCT

CATTTACAGGTAGCCCACGTTACATGCAACAAAATTATCAAGATTGTATGGCTATTTGCAGAAAATTCGG

AAGCCCGGATCTGTTCATTACATTCACATGCAATCCACAGTGGCCAGAAATCAAGGAGTATTGTGATAAG

GTGCCACACTGTATCCCAGCCGATAGGCCCGATGTTATGGCTAGAGTGTTCAAGATTAAGCTTGAATTGC

TTCTAGAAGATTTGACAGATAACCATGTTCTTGGAAAAGTCATTGGAGTTGCATATACCATTGAATTTCA

GAAAAGGGGATTACCCCATGCACATATAATTGTGTGGCTTGAGGAATCTGACAAGTGTCACTCAACTGAT

GACATTGACCAGTTAATATGTGCTGAAATTCCTGACAAAGAAAGTGA

>MSTRG.2019.3 gene=MSTRG.2019

GCCTGATCGGGTCACGGATTTCTGTAAGAGGAACAAGTTACAGATGATCATAAGAGCTCATGAATGTGTT

ATGGATGGGTTTGAACGGTTTGCTCAGGGACAGTTGATCACTCTTTTCTCTGCGACGAACTATTGCGGGA

CTGCCAATAATGCTGGAGCTATACTGGTAGTTGGTAGAGGGTTGATGATTGTGCCTAAATTGATACATCC

ATTGCCACCACCCATGCATTCACCAGAGACATCTCCGGAAAGGGGAATGGATGACACATGGATGCAGGTA

AAGAACTAAACATTCAAAGACCGCCAACTCCTACCAGAGGGCGTCCACAGCCTGATCTTGATCGGAATTC

ACTTGCATACATCTAAATTTGGTCAATGTTCGAGAACAATGTTTACGTGAAACCTGCAGTCCTCTGGCTA

CAATATAGTTCATTTTTGCTGTCATCGAGACTACATACTACTTTTCTGTCCTTTAATGACATTCAAGAGA

AGCTTGTGCATACAGAAGGGGAGCCAGCAGTAGCCATTGGAGGTGTGTGATGACTGTTATATGTTGTGTT

TTCAGATTACTTGTATAATAGAGCCTGTGGTTGAGAGTCAGTTTAGTGATGATTTGTGTAGTCAGAGGGG

AGGGTGGTCCCTTGAGCTGTGTGTATTGTAATATAGATAGTGAATAATTTA

>MSTRG.2029.1 gene=MSTRG.2029

AACTTTTCTTTTCCTTTCACATCTCCTTTTTGAACCCAATATCAATAATCATCCCACAATCACACACTTG

CATGTTTTTCTGCAATTCCTCATCTAAAAGATAAAACACATTGCCCAACTAAAAGTTCCTCACATTCATT

CCCTCATTATTACACAATGCCTGCACAATCTTCTCTTTTCTTGCCACTGATTATACTTTCTATGGCCTCA

CTAGCCTCATCTCAGCTGCCAATATCTGTTATTTCAGCAGCACCTGCACTTCTTCCATTTGAGCCTCCTC

TTTCCTCTCCTCCAGCCTTAAGCCCCGCGCTCTCACCGGACATGCTGCCTGTATTTCCTTCTGAGACGGC

TCCTTCTCCATCCGAATCGACGTTTCCAGTCATACCATCCAGTCCAAGCCCTCCAAATCCTGATGAAATG

TTTGCTCCAGGCCCAGTTTTGCCTTTTGCACCATCAGGCTTGCCTGAGTCCTCAGCATTGAGTCTGAGCA

CAGCTGGCTTTTCGAGTTCAGTGATGCTGATTCTCGGCATTCTTGCAGTTTTTGCTTGAGGAAACTTGGC

ATAAGTACTCTTCAAGGTTCATTATTGGTTATGAATGAGGTATTATTAAGGGTGGTTACAATCATTTGTG

GTTTATAATTTACTTAATTTTGTAAACATTTACTACTAGTTTCTTGTTGCTTTAAGTATATAATTGATAC

TTGTTTAATTTAATGAATTTGAAGAAATATGGTAATATTTACTTCAAATCTTTATCAACC

>MSTRG.2048.1 gene=MSTRG.2048

CTCTATTTAGACACCTTTTTCTTCCCAGCAATTGCAATCTTCTTTCCTTTCAATGTCCGGCAGTTGTTCT

TTGTCCGTTGCCCCCTGCAAGGCAATCCCTGAATATGTCGTATACCTCGGTAGCATTGAATCTCCTTCAA

CCTCCTAATATTTAGTGCATTGAATCTCCTAAGGTCTCCTTCAATCATGTACTTGGAAACTTCATCACGG

AGAGTAATGAGCTCTTCTTCAGACAAGTCCTTAGTGATCTTATTCTCCA

>MSTRG.2056.1 gene=MSTRG.2056

GGTTCCGGTTAGAGAGCCTAACAGTATAGTGCAGTCTTATCTCTAGCATTCTATTATCAAGATGCCAAGG

ATTAGAGCAGCTGCTTATAAGGTTTCCCAGAGTTATCTTTTGTGCTAGGAAAATCGCAGGATGATGCTAG

GAGTCTTGGACTAAGTGTGGATTAGTTAGTACGACTTGTTTCTTTGAAGTTGTATTATAGTGTCAGTATG

GTTGTAGTGGAAAACTCATGTAGTAAGGAACCTATGTCGTTGTTCGTTTGCAGTAACAGTTTAAACCTTT

CGCAGACTTAAACCTGGCAAGATCTTTGGTCAGGGGTGTAATGGTATATATTTATTATGTTAATCTATTC

AGCTGTCATTAGTTTATTTTATTGTTTAGTTTATTCTTTTTAGTGACACCGAATCTGCACCCCAGATTCT

GGGCTGTCACAGTTGGTATCAGAGCTACAGGTTTAAGTCACTGTGTTAGGTTAGGTTAGGTAGAGAGTCT

ACCCTTATTAGTGATATTTGGACTAGGT

>MSTRG.2057.1 gene=MSTRG.2057

GGTTCCGGTTAGAGAGCCTAACAGTATAGTGCAGTCTTATCTCTAGCATTCTATTATCAAGATGCCAAGG

ATTAGAGCAGCTGCTTATAAGGTTTCCCAGAGTTATCTTTTGTGCTAGGAAAATCGCAGGATGATGCTAG

GAGTCTTGGACTAAGTGTGGATTAGTTAGTACGACTTGTTTCTTTGAAGTTGTATTATAGTGTCAGTATG

GTTGTAGTGGAAAACTCATGTAGTAAGGAACCTATGTCGTTGTTCGTTTGCAGTAACAGTTTAAACCTTT

CGCAGACTTAAACCTGGCAAGATCTTTGGTCAGGGGTGTAATGGTATATATTTATTATGTTAATCTATTC

AGCTGTCATTAGTTTATTTTATTGTTTAGTTTATTCTTTTTAGTGAC

>MSTRG.2067.1 gene=MSTRG.2067

GGCTCATGACTCTGCTGGCTGGCTGGCTGGCTGCTGCTGTACCAGGTAGGGGGTAGGAAGGAAAAGATAG

TTCTATCGAACAATTTTGGCCTCCCTTGCGGTATAAAGGTATATATGTTCCGTGTTTAGGCTGTTGTGGC

ACCATCTTCTTTTTGATCACCGAAGATACGTTGCCTGAAGTCAGATACCATCAGAGGCAGACCCTTGCGA

AGAGCTACCTTGGGCTCCCAACCAAGAAGATC

>MSTRG.2068.1 gene=MSTRG.2068

TGTTTTAGCTGAATCTAGTTTAGAAAAGTACCTTTAATTTAGTTTTCGTTTAGTTTGTTATAATTCTTTA

TCATTACCTCTGTTTTGTGGCCATCCTTTGACATGTTTGTCCACACTTGGAGACACACTTGCAATTCCAT

ATTCTTTATTGTACTCTGCACAACGATACACAATATAAGCTATTCTTATGTATGTCTAGTTCGTGCAAAA

GGGGTGTG

>MSTRG.2072.1 gene=MSTRG.2072

CTTCTACATCAAAAAAAAGAGAGAGACAACAGAACCAAATCAAACAATTTAATCCTCCTCTTCTCCTTCG

TTTTCAGCAATGTTAAAGTATCGCAACTCGTAAACATTTCGGTCTTTGTTGGAGGCAATAACCCTCAACC

AATCCCTCACATTGTTCTTCTTCAGGTACTTCTTTGTCAAGTATTTCAAATACCTCTTGGAGAAGGTACT

GTCGGAGGTGACAGTGATCTTGCTCTTCTCACGCACGACGGTGACGGAGTCTCCGAGAGCTCCGGCTTTC

CCGCCGACCTTAATTCTCTCCTGCAGAAACTTCTCGAGAGAAGCGATGTCCATGATCTTATCTTCCACAG

GCTTTCCGCAGTCGATCACGAACGTCGCTCCCTTCTTCTTCCCCCCCTTTCCGGCGCCTCCGGTGACTTT

GCTCATCTTTACTCTTCTCTG

>MSTRG.2073.1 gene=MSTRG.2073

CGTAAACATTTCGGTCTTTGTTGGAGGCAATAACCCTCAACCAATCCCTCACATTGTTCTTCTTCAGGTA

CTTCTTTGTCAAGTATTTCAAATACCTCTTGGAGAAGGTACTGTCGGAGGTGACAGTGATCTTGCTCTTC

TCACGCACGACGGTGACGGAGTCTCCGAGAGCTCCGGCTTTCCCGCCGACCTTAATTCTCTCCTGCAGAA

ACTTCTCGAGAGAAGCGATGTCCATGATCTTATCTTCCACCGGCTTTCCGCAGTCGATCACGAACGTCGC

TCCCTTCTTCTTTCCCCCCTTTCCGGCGCCTCCGGTGACTTTACTCATCTTTACTCTTCTCTGGTGGAAA

TGTGAAGCGGCCGTCTGCAACTCGATACGAG

>MSTRG.2075.1 gene=MSTRG.2075

CCCACTTCTTGTCTGGCCAAAATATCCATTCCCAGATTCTCCGCTTCGGGTTTGGTTCTGACCCTTTTGT

TCAAACTTGTTTGATTAACATGTACTGTTTTTGTGGTCATTTGGGTTGTGCACACCAAGTGTTTGATGAA

ATTGATGAACCGGATTTACTGTCGTGGAATTCGATGGTGAATGCGAATGTGAAGATGGGGTTGGTTGATG

TGGCGCGGGGGTTGTTTGATGCAATGCCGGTTAAGAATGTTGTTTCTTGGAGTTGTATGCTTAATGGGTA

TGTTAAGTTTGGGAAGTATAAGGAGTGTTTGGCGTTGTTTAGGGAAATGCAGAAGTTGGGGGTGGATTGT

GTGAGGCCTAATGAGTATACGATGTCGGTCTTGTTGTCTGCGTGTGGGAAATTGGGGGCGCTTGAACATG

GGAATTGGGTGCATGCTTATATTGAGAGATGTGGGATGGTGATTGATGATGTCTTAGGGACTTGTTTGAT

TAATATGTATGCGAAATGTGGGAGGGTTGATAGGGCGAGATGGGTTTTTGATAATTTGGGTTCGAGTAAG

GATGTGATCACGTGGAGTGCCATGATCTCTGCCTTTGCTATGCATGGGCACTCTGCTGAGTCTCTTGCCT

TTTTTTCAAGGATGATGTGTAGTGGGATTAAACCTAATGCTGTTACGTTCTTGGGTGTACTTTATGCGTG

TGTGCATGGTGGATTAGTGAATGAAGGGGAGGCATATTTTGAGAAAATGACTAGGGAGTTTGGTATTGTC

CCTTCAATCAATCACTATGGTTGTATGGTTGATCTTTACGGAAGAGCTGGTCTTATTGACAAAGCATGGA

ATGTTGTTAATTCAATGCCAATGGAGCCAGATGTTCTTGTATGGGGTGCTCTTCTAAGTGGGTCTAGAAT

AGCTGGGGACATTAAGACATGTGAGATTGCAGGCAAGAAATTGATTGAACTAGAACCTACAAATAGCTCA

GCCTATGTACTTCTGTCAAATGTGCATGCCAAGTTGGGTAATTGGAAGGATGTCCGGAATGTACGAGATC

TTATGGAGACAAAGGGAGTCAAGAAAGTTCCTGGCTGCAGCTTGGTGGAGCTGGATGGTGTAATACATGA

GTTTTTTGTTGGAGATGAGTCTCATCCAGAGACGAGGGAAATATATATGATGCTTCAGGAGATGATGAGC

AGATTAAAACTAGAAGGATATGTGAGCAACACCGGAGAAGTATTACTTGATTTGGATGAAGAAGCAAAAG

AGAAGGCACTTTGGCTTCACAGTGAAAAATTGGCAATTGCATTCGCCATATTAAAGACAAGTGCTGGTAC

ACCTATCCGCATTGTAAAAAACCTAAGGATATGTAGAGATTGTCATGTTGCCATAAAGATGATATCTAAT

ATATATGGTCGAGAGATTGTTATAAGAGATTGTAACCGCTTTCACCATTTTACAGAAGGAATATGCTCCT

GTGGTGATTATTGGTAAAGTTAAAATTAGAAGGTAGTCTTCAAGTTACAAAAATACCCCTGCAGATGACT

TTATAAAAAACTATAAAATTTA

>MSTRG.2079.1 gene=MSTRG.2079

CCGGACTTTCAGCCAGGCGATAAAGTTCGAGAATAGTACATAACAATTTGTGAACTCTAGTGACGAAATC

TGAGACATGGTCTTTCAATTTTGACAAAAACAAAAGAAAGCTTGTGAAACAATGGCGGAGATGAATGTAG

GCAGCGAAGTTCCCGTCAACAATACAATTTACATCAACAATATCAACGAAAAAATCAAACTTGACGAGTT

GAAGAAATCACTGCACGCTATATTCAGTCAATTTGGGAATATACTGGAGATTCTAGCTTTCAAAACCCTA

AAACATAAGGGCCAAGCTTGGGTTGTTTTCGAAGATGCTTCGTCGGCTTCTCGAGCCATTCAGAAGATGC

AGAATTTCCCCTTCCACGATAAGCCGATGAGGATACAATATGCAAGGACGAAATCTGATATTATTGCTAA

GAAGGATGGAACTTTTGTGCCACGTGAAAAAAAGAAGAGGCACGAGGAGAAAGGAAAGAAGCGGAAAGAA

CATGATGCAAATCAAGCTGCAATGGGATTGAATCCTGCTTACAATGGTGCTTATGGTGCAGCTCCTCAGT

TGTCACAGATACCATACATGGCTAAGTCTGCTGTTCCTGAGGCTCCTGCTCCACCAAATAGCATTTTGTT

TATCGAGAATCTTCCCCATCAGACAACTCCAATGATGCTTCAAATGCTTTTCTGCCAGTATCCTGGGTTT

AAGGAAGTCAGAATGGTGGAAGTCAGACCAGGGATAGCATTTGTTGAGTATGAAAATGAGGCTCAGTCAA

CTCAAGCTATGATTGGACTCCAAGGATTCAAGATAGTGCCCGAGAACCCAATGTCGATCACCTACGCCAA

GAAATAGAAGGGTGTTTATTTAGCTGTGGTTTTATTAATTGTAGTAATCTTGGGAGTTCCCGGGGAAGGG

AAGTGTTTACTTGAGCTATGGGGCATTTAAATGGTCTTTTATCAATCCAAGTAGATGATGTAATGGTTTG

TGACTTTCTTTAATATTGTTAGTTCTGCAGAATTACTGTAATTATAGCACTGGATTTAGCTGAGTTGATA

TTTTAGTTCATAAACTGGCATTTCCAAGCTTTGCTAATTTTCATCGAAATGCCAGACTTGACTTGCTACT

ACAAATTAAATCAGATGTTGTGATGGTTACAGAA

>MSTRG.2079.2 gene=MSTRG.2079

GAGTTGAAGAAATCACTGCACGCTATATTCAGTCAATTTGGGAATATACTGGAGATTCTAGCTTTCAAAA

CCCTAAAACATAAGGGCCAAGCTTGGGTTGTTTTCGAAGATGCTTCGTCGGCTTCTCGAGCCATTCAGAA

GATGCAGAATTTCCCCTTCCACGATAAGCCGATGAGGATACAATATGCAAGGACGAAATCTGATATTATT

GCTAAGAAGGATGGAACTTTTGTGCCACGTGAAAAAAAGAAGAGGCACGAGGAGAAAGGAAAGAAGCGGA

AAGAACATGATGCAAATCAAGCTGCAATGGGATTGAATCCTGCTTACAATGGTGCTTATGGTGCAGCTCC

TCAGATACCATACATGGCTAAGTCTGCTGTTCCTGAGGCTCCTGCTCCACCAAATAGCATTTTGTTTATC

GAGAATCTTCCCCATCAGACAACTCCAATGATGCTTCAAATGCTTTTCTGCCAGTATCCTGGGTTTAAGG

AAGTCAGAATGGTGGAAGTCAGACCAGGGATAGCATTTGTTGAGTATGAAAATGAGGCTCAGTCAACTCA

AGCTATGATTGGACTCCAAGGATTCAAGATAGTGCCCGAGAACCCAATGTCGATCACCTACGCCAAGAAA

TAGAAGGGTGTTTATTTAGCTGTGGTTTTATTAATTGTAGTAATCTTGGGAGTTCCCGGGGAAGGGAAGT

GTTTACTTGAGCTATGGGGCATTTAAATGGTCTTTTATCAATCCAAGTAGATGATGTAATGGTTTGTGAC

TTTCTTTAATATTGTTAGTTCTGCAGAATTACTGTAATTATAGCACTGGATTTAGCTGAGTTGATATTTT

AGTTCATAAAC

>MSTRG.2083.1 gene=MSTRG.2083

CAATCCAGCACCGCCATTACAATCAAAGCCACACCCACACTGCGGACTCTCGACAAAATTATCGATATAA

AACCCCGCCGACAAAACCCCCACCGAAAACTCCAGATCATCTCCCTCCCGGCTCACTTCCCCGATATTCA

GCCACAAGAACAAAATCTTCACGCTTATCCCGCTCAGATTCTTCAGCTTCCGATCCGATATCGTCCCCGA

TATCGTTGACTTGTATCTCAGCCTGTAGTCCTGCACGAAGAACTCGCAGGTTTTTTCTAGGTAGACTTTG

AACTTGCCGGTGCTCCGGTCGAGCTCGTAGTTGGTGATGCCGACGGGGAGCATGCCGACCGGGAAGTTGT

ATTGTTGGAGGACTTCGTAGACGGTGGGGGTGGCGGCGGCGGCG

>MSTRG.2087.1 gene=MSTRG.2087

CACAATATATAAATCAAACCCTAAAGCCCCTGCTGTTTCTTTTTCCTCTGGCCTTTTCAAATTTCCTTCC

CTTTGATCGAACATATGGCTTGCTGTGACTGTGTGGCACACCAGGAGCCTTGCCAAAGTGCTTAACCGCT

TCACGGGAATTCTTAGGGCCTCTAAGAAGTACCGTGTTCTGCCCAAGAGGGGCCCTAAGAGCGAGCTGGT

CAAATGTCAAGCATTCACCTCCTG

>MSTRG.2089.1 gene=MSTRG.2089

GTGTACTTGAGATCAAGATGAGCACTGAGGTTCAGCAGGCTTCTAGAGTTGATCATGGAATTCAGACCAA

TATGACGTCCACCTCATCTGCAGCCCCAACAATTAGTCCAAAAATTTCAAGGTTTGCAGCAAAGTCTGGA

TTTGTGATACCAAAAAATAAGTTGTTAGGCTCGCTGGTACCTGCATATCGGGGTAGTAAAAAGCCGGAAG

AAACTGCTCCTATTAGTGACGAAAGTAGTAAACAGACTCGAAGGAAAACAAAATGGGGGCCTGATCAGAC

ACAAGATGCTAATGTTTCCTTGATTTTAGTTAGCGAAATATGGATTTGGTAGTTGCTTGGACTTGTGTAG

CTAATTTGTTGTGTAGCTATGTGGAATTGGTTGGATTTGTTGTGATGTTTGTATTTAAATAATATATTTT

TAATTCGTTTTAGTAAGTAACGTTGCCA

>MSTRG.2093.1 gene=MSTRG.2093

CTAGTGCTAAACAAGTACTGATACAGAATTCCCCTTCAAGCAGCAACAGGAAGTTCCTCACTTCTTTGAT

TTCTTCGATGGCTGAGATTGAGCTGGAGCAGCTGCTGCGGCAGGCTCTTGTGGTGCTTTCTCTCCAGGTC

CCTGAGCAAGACGTTCCTCCCTCCTAGCATGTTTCCTTTCCCTGCTTGCCTTATTCTTTGCTCTCTTAGC

CTCAAACTGGTCAGAGAGGGTCTTCTCACGAGCCTTTTCAGCCTTGGATTTGTGGATACTCTCCATGAGT

ACTCTCTTATTCTTGTAGACATTACCTTTCACTTTCATGTACATATCATGGTACATGTGCTTATCAATCT

TCTTTGCTTCACGGTATTTGCGGAGAAGGCGCCTGAGCACTCTCATCCTGCGCATCCATAGAACTTTTGT

GGGCAACCTAGCCTCCCTGGTTCCCCTTCGCTTACCATATCCAGCATGACGACCCTTCCTCTTTGCTTCC

TTCATCCTGCGTGCACGGGAACGGGAGTGAATCTTGGTTGGCTTCCTAATGATAAACCCATCCTTCACTA

GCTTGCGGATGTTCTGACGAGAATTGGCCATGGAGATCTCATTGCTTTCATTTGGATCAAGCCACACCTT

TCCCTTTCCGCACTTGAGGACGCTGGAGGCCAGACGCTTCTGCAACTTGAGCGACACCATCTTTTTTCTC

CGCAGCTGCGCTCA

>MSTRG.2095.1 gene=MSTRG.2095

GAAAATTTAGTAATATGCATCTTGACTTTGGGCTGATTTGTAAAGTTTTTTCGCCTGGTTGAATATTATT

AGCTCTAGTATTTTTAGACATATTAAATCAAAAGAGTTATTGCTTTGCACTATATTTTTTAGTTAAGGAA

GGGGAAATTTCTTAATCTGATTTTTTGTGGAAATTGGCACAATGAGATTGTTGTCACAAAAGGGAAAGAA

TTTCCTTGAGATCAAACTCTGATATGTTCTCATCATATATGTTGAAATAATAGTGATTTCATGCTGGGTG

ATTGCTATCTTTTGTTCTTTTTAGTCTGAATAACTCTGAGATAGTTATAGACTTATAGGAACTTTGTATG

CATAAGTGGTTTTCCTTGAGTACTATTTCCCAGTACAATCGAATTGCTTTGGCTGTGTCATATTGCTCAT

CTAGACATGTCTATAATCTTACACTATCACATAAATACCTTATAATTTAGTTTTAG

>MSTRG.2096.1 gene=MSTRG.2096

ATTGGACAGCCCAACAAAATTGGAGCCAATCCACCGGAAACTAAACACACTTCCCGGTACTCATTTTTGG

CAATTCCTGGCGGCTTTGAGGTACGCTACCTTACTTTCTAGATCTTTCCATTTTCTCGCATTTGATCGTT

CGTACAAATTTGATTTTGATCTTTTTGAGTGTGAGTGCAGGAGTTGAAGTCTCGCAACATGTCGATTTCT

GTAGCGTATCGGCGATTAACTAGCTGCGGATCCACGTCGTCTGGTCTTGCAAACGCCAGTACTCTCAGAT

TTTTCTCTTCTTTTCGAGAGGAACGAGATACTTTTGGTCCAATTCAAGTTCCTTCTGACAAGTTATGGGG

TGCCCAAACCCAAAGATCACTACAAAATTTTGAAATTGGTGGTGAGCGTGAGAGAATGCCCGAGCCCATT

GTTCGTGCTTTTGGCATCCTTAAAAAATGTGCTGCCAAGGTTAACATTGAGTATGGGCTTGACCCATCCA

TAGGTAAAGCAATAATGCAAGCTGCTCAGGAAGTGGCTGAAGGGAAATTGAATGATCACTTCCCCCTGGT

GATATGGCAGACTGGCAGCGGAACTCAAAGCAACATGAACGCCAATGAGGTAATTGCAAATAGAGCTGCA

GAAATTCTTGGTCACCAAAGAGGTGAAAAGTTTGTGCACCCAAATGATCATGTGAACAGATCCCAATCTT

CTAATGATACATTTCCAACTGTAATGCACATTGCAGCTGCAATGGAAATCAATTCACGATTTATACCGAA

GTTGAAACAGTTGCATACTACTCTAAACTCGAAGTCGGTTGAGTTTAAAGATATTGTTAAGATAGGACGA

ACTCACACACAAGATGCAACACCTTTGACACTAGGGCAAGAGTTCAGTGGTTATACAACCCAAGTGAAGT

ATGGAATTGATAGAGTGATGTGCACTCTCCCTCGCATGTATCAGCTTGCACAGGGCGGCACTGCTGTGGG

AACTGGATTGAACACAAAGAAAGGGTTTGATGTTAAAATTGCTGCAGCAGTAGCTGAGGAGACTAAGCTA

CCATTCATAACAGCGGAAAACAAGTTTGAAGCGCTGGCTGCTCATGACGCTTTTGTTGAAACTAGTGGGG

CTTTAAATACAGTTGCTGCATCTCTTATGAAGATTGGAAATGACATACGGCTATTGGGAAGTGGGCCACG

GTGCGGTCTTGGTGAACTTATACTACCTGAGAATGAACCAGGCAGCAGTATTATGCCTGGGAAGGTAAAT

CCTACTCAATGCGAGGCTCTCACGATGGTCTGTGCTCAGGTCATGGGAAATCACGTGGGAATTACTGTGG

GTGGTGCAAATGGACATTTTGAGCTCAATGTTTACAAGCCAATGATCGCTAGTAGTCTCCTACATTCCGT

GAGATTGCTAGCAGATGCTTCTGCTTCTTTTGAAAAGAATTGTGTGAGGGGAATTGAGGCCAACAGGGGA

AGAATCTCAAAATTGCTACACGAGTCACTCATGCTCGTCACCTCCTTGAACCCTAAAATTGGCTATGACA

ATGCCGCAGCAGTTGCAAAGAAAGCTCACAAGGAGGGAACTACTTTAAAGGAAGCTGCTTTAAACTTAAA

AGTGCTCACCAGTGAGGAGTTTGACGAACTTGTAGTGCCTGAAAAAATGATTGGTCCCTCAGATTGATTC

ATAGTGAGTGTAAAAGTTTCAAGAATTCGCACCCACAACACAGCATAGAAATAAAATCCAAAATGTGATT

ATCCTTTACTGGAACAATATTATTCTTGAGGTTCTCTCCTGTGTAGAATTCATTTTTGTTTTTTCTGGAG

ATTCCAGATGTGACACTTGAAATGGGGGAACCTCCGCATTTTTTTGCGTAATGCAACATAACAGTTTACG

GATAAACGCAGTTTATAGAGAATGATTAACGGATGATCAAACCAAGTTTTTTTTGCTAATTTTACTTTTG

GTCACATCTGGTCTTAACATACAGTTTACATGTACCATGCTATGCATATACC

>MSTRG.2096.2 gene=MSTRG.2096

ATTGGACAGCCCAACAAAATTGGAGCCAATCCACCGGAAACTAAACACACTTCCCGGTACTCATTTTTGG

CAATTCCTGGCGGCTTTGAGGAGTTGAAGTCTCGCAACATGTCGATTTCTGTAGCGTATCGGCGATTAAC

TAGCTGCGGATCCACGTCGTCTGGTCTTGCAAACGCCAGTACTCTCAGATTTTTCTCTTCTTTTCGAGAG

GAACGAGATACTTTTGGTCCAATTCAAGTTCCTTCTGACAAGTTATGGGGTGCCCAAACCCAAAGATCAC

TACAAAATTTTGAAATTGGTGGTGAGCGTGAGAGAATGCCCGAGCCCATTGTTCGTGCTTTTGGCATCCT

TAAAAAATGTGCTGCCAAGGTTAACATTGAGTATGGGCTTGACCCATCCATAGGTAAAGCAATAATGCAA

GCTGCTCAGGAAGTGGCTGAAGGGAAATTGAATGATCACTTCCCCCTGGTGATATGGCAGACTGGCAGCG

GAACTCAAAGCAACATGAACGCCAATGAGGTAATTGCAAATAGAGCTGCAGAAATTCTTGGTCACCAAAG

AGGTGAAAAGTTTGTGCACCCAAATGATCATGTGAACAGATCCCAATCTTCTAATGATACATTTCCAACT

GTAATGCACATTGCAGCTGCAATGGAAATCAATTCACGATTTATACCGAAGTTGAAACAGTTGCATACTA

CTCTAAACTCGAAGTCGGTTGAGTTTAAAGATATTGTTAAGATAGGACGAACTCACACACAAGATGCAAC

ACCTTTGACACTAGGGCAAGAGTTCAGTGGTTATACAACCCAAGTGAAGTATGGAATTGATAGAGTGATG

TGCACTCTCCCTCGCATGTATCAGCTTGCACAGGGCGGCACTGCTGTGGGAACTGGATTGAACACAAAGA

AAGGGTTTGATGTTAAAATTGCTGCAGCAGTAGCTGAGGAGACTAAGCTACCATTCATAACAGCGGAAAA

CAAGTTTGAAGCGCTGGCTGCTCATGACGCTTTTGTTGAAACTAGTGGGGCTTTAAATACAGTTGCTGCA

TCTCTTATGAAGATTGGAAATGACATACGGCTATTGGGAAGTGGGCCACGGTGCGGTCTTGGTGAACTTA

TACTACCTGAGAATGAACCAGGCAGCAGTATTATGCCTGGGAAGGTAAATCCTACTCAATGCGAGGCTCT

CACGATGGTCTGTGCTCAGGTCATGGGAAATCACGTGGGAATTACTGTGGGTGGTGCAAATGGACATTTT

GAGCTCAATGTTTACAAGCCAATGATCGCTAGTAGTCTCCTACATTCCGTGAGATTGCTAGCAGATGCTT

CTGCTTCTTTTGAAAAGAATTGTGTGAGGGGAATTGAGGCCAACAGGGGAAGAATCTCAAAATTGCTACA

CGAGTCACTCATGCTCGTCACCTCCTTGAACCCTAAAATTGGCTATGACAATGCCGCAGCAGTTGCAAAG

AAAGCTCACAAGGAGGGAACTACTTTAAAGGAAGCTGCTTTAAACTTAAAAGTGCTCACCAGTGAGGAGT

TTGACGAACTTGTAGTGCCTGAAAAAATGATTGGTCCCTCAGATTGATTCATAGTGAGTGTAAAAGTTTC

AAGAATTCGCACCCACAACACAGCATAGAAATAAAATCCAAAATGTGATTATCCTTTACTGGAACAATAT

TATTCTTGAGGTTCTCTCCTGTGTAGAATTCATTTTTGTTTTTTCTGGAGATTCCAGATGTGACACTTGA

AATGGGGGAACCTCCGCATTTTTTTGCGTAATGCAACATAACAGTTTACGGATAAACGCAGTTTATAGAG

AATGATTAACGGATGATCAAACCAAGTTTTTTTTGCTAATTTTACTTTTGGTCACATCTGG

>MSTRG.2099.1 gene=MSTRG.2099

ATTTAATTGCTCTGCAAAAAGAAAAATAACCTTTATTCCTGCAGTTACTCATCTCCCTAATCTCAAGGGC

TTCCATTCTGAAGCTGGTGCTTCTTGCTAACGCATGGTACGAAAACGGGTGTTTGACATTCAAAATTACA

TTTTCTGCTTTAGGAAGGATTACTCTATGTTACCTGGACTCTTCATTTTGCATGGATTACCTGTATCGGA

CATTTGGCACCCAGACATAGATATGGACAACTAGGACACTTATTTTAAGCTATTTATATGTGAAATATAC

AAAATTTTGGGAGTCCAACAGTTTGACACGGTATCCATGTGGACACTTTTAGCTGAGTCCAGATAACAAA

GGGATTACTCATGAAAGGTCATCTGTGATCTACACAGGCTGTTCAACAAATTTTCCGGTTTGATCAATGA

TTTTATGCCAGTTTGTGCTTTTATTGACTACATAGTTATCTATAAAATTAATGATTTTCAGTGTTTAATA

GGCTATACAGTATGGTGTCTCGTCCCTGCATTGGCTGGTATAGTCATGTTGCTGAACAAGGTATATCTTT

CACCTGGCTGTTCAAACACTCAAATCGATAATTGAGACCAACTTCATCACTGAGTGATGCAGTTGATATC

TTTTATCGACTTGAGTGATGAATATGAAAGTTAACCCGTTTACAAAGTAAAGTGTGCAAATACTTAAACT

TACCTTTTCTTGGTACTGCTAATTATAATACAAGATATCACTACAATAGACAAACTTCGTGCAAGCGATG

TGGATGTTGCCTCATCACTTTCAAGCATTATCACATTAAGATTGACTTGTCCGCATAAGGTATCAAGACA

AGAAAACAAAGTTGATTCAAGATCATCTTGATATATATGATCTAAGAGTATATGCTCTCTAATAATGTTT

TTTCTCTTATTTATTGTTAAAAAGTGGAAACAGCAGAAATATAATTGAGGACTTCTGTGTTATATTGCAA

TTCCAGAGAGAAAGAGCGGGAGGGCGGTTTGTGTTTTATGGGCATTTAGAAAAATTAATGGTGGTAACAA

AGTTTTCAATCGAAAGCATGGCGAATGTATTTGTTATTAATATCCTCTCATTGTCATTAGATGGATGATA

TAAGGTTTCCTTCATATAGATGCACTTATCCCTCTCCCCCTTCATGCATTTTCTTCCCATGCATCTTGGT

CCATATTTTTCAATAGCAATTATCCTGTCAGCGTGATTTGTTCATTCCTTGACTCCCTTTGCCTTAATAA

TTTGATAACGTTTTATTTATTTTATGAGAACTTTGATAGAATATAATTCTGAACATACTGGGCTTACTAC

CAGTACTTATATGATATGGTCACTTTTCTGCAGCACTGTTTACCAGTTTATTGTTTTTGAAATGTATACA

CAGATCATGGGTCCAAGTCTCAGGAACAAGGGAGCTATTCTAGGATGTTTACCTTGGTTTAGTTGAATGC

AATAGAAAGGAAAGCTCCTTTGTTTGGTTGTTCAACACGAAAAAGCAGTACAATCCCTACATTTCCTTCA

GTAGGGTTGAGTGACCCTGAAAGATGGTTAGATGGCGATTATGTTACAGTACACGACAATGTTGAGAAAG

TGTTCAGTGTCTTTGACTGGCTCTGTTTTGTGGATATCATTTTCTTTCCCTTGTCGCTCAAGCCTGATAC

ATAATAGTGTTCGAATGTCAATTGTTTCTGGACTGACATCACTAATATTTAAGGCCCTCTCTCACGTCGG

CAAAAATATCTCGAGTTGGGAGGAATTGCTAATTTGCTGAGACGAAGAACCTACTTTTTCCTTTTGTGCA

TGTGCTTATGGTATTCGTATGTGGAAACGATATAGTTTATCGGGTGATATCAATTCCAAATTTTGCCAAT

GTAATTGTTGATCGAATAATATCATTTCTAAATTATTTTATTGTAGTCAGTAACCCGTCGGTATTCTGAA

TTTCTGAGTAATCAGTTGCCGAATCGGAGATTATTTAGTTATTCA

>MSTRG.2112.1 gene=MSTRG.2112

CTCAAATTAGCCATCCTCTTTGTAAAACTTTCCCTCATCTCCCTAACAGCCTCACGATGCTTGTAGTTTT

CTTCATCTTCTTCCTTGTCACGAATTCTACCAAGATCTGTGGCATCCCTATTATATTCCAACTCTAATTC

TTCTAACGTCCGGGGAAACATAGGATAATCCATCTTATTGCTAGTTTTCAAGTCAGATCCACGTCTCTCT

GTATCCCACCCATCTGTTTGGTATCCTCCCC

>MSTRG.2115.1 gene=MSTRG.2115

ATTTCGTACAAATCCTTGACCGCCACCTGTCAGTCAACACTGTCTATATTCACCTCATCCTCCTTCCTTC

ACTGTCCCCTCTCTCTCCCTACTCTCTCTCCTCCCTCTCTCTCTTCCTCAATCTCTCTCTCTCTCTCTCT

CTCTCTCTCTCTCTCCCTCTCTCCCTCAATCTCTCTCATTCTTACAACATGGAGAGGAAGCAGACTGTGA

AACAAGAGGAGGCTCAGAAGCAGAAGAGTTTTGTAGTGGATGATCAAACTGATAACAACAACATCAGTAG

CAGCATCAGCAGCCCATTAATGTTATCTGATATGTTAGGGTTTGGTGATCAGAAGAGCTCAGTGGGCTTC

ATGGAGCTCCTGGGCCTTCAAGATTTTAACAACACTTGTAGCAACAGTTCCATCTTTGATTTATTAACAG

CAGAAGAGCTTGTAGTGGGCCGGAGCCCAGTTCAGAATCAAACTCAACTCAATCCTTATCAAGAATCATC

AGAGGTTTTGAACAATCCCCCAACTCCCAACTCCTCTTCACTCTCTTCTGTTTCAAGTGGTGAAGAAAAG

ACTAAAGTTGTGGCTGATGAACAAGAACAACAGCAGCAGCACAAAGTCACTAAACCACAGTTGAAAGGCA

AGAAGAGTGGGAGTCAGAAGAAAAAGAAACAGACAGAGCCAAGATTTGCATTCATGACAAAGAGTGAGGT

TGATCATCTGGAAGATGGCTACAGATGGAGAAAGTACGGCCAAAAAGCTGTCAAAAACAGCCCTTTTCCT

AGGAGTTACTATAGGTGCACTAGTACCTCATGTAATGTGAAGAAAAGAGTGGAGCGTTCTTTCAATGATC

CAAGCATAGTTGTGACAACATACGAAGGCCAGCATACACATCAGAGTCCGGTGATGCCTCGGGGCATTCA

TGCTGGAGTTTCATCGGTTTCTGGTATCTCTCCCTTGAATTTCCCGGCCAACATTCATATGAACAACCTC

ATGATGCAACAGCAGCAGAACTCAAACAATTTTGCATTGCCTTCAAATTGTAGCCAGAATACAATGATGA

ATAATTATCCTAACGGTTATACAATTGCAGACGGGGGAGGTCTCGGGTTCGGGCCCTCCTCAGCTATGGC

GTTCTTGAGAGACAATGGCCTGCTTCAAGATGTGGTGCCTTCCATGATAAGAACAGAAGATCAGTAGATG

CATAACATCAAGTTTCATATAGTACGACTTATAAGTTATAAATATGTACTTATTCAGTAATTCTGTTTAG

TGACTGACTAGAACTTGTAGCTTAATCTGTAGATGGTCCATCATAACTTATGCAGTTTATTTCTTTTGTT

GATCAGCAAGTAGCTTCTTGCAATCTCTGATGTTTCTCAGTCTGACATGCTTCTTTGATCCAATGAAGAT

TCTGGTTTCTTACTTTCTTCAAATGTTGTATATTCAATGTCCATTCAACTTCAACCTTGTTATTTTTGCC

AGTTTTACTTTGTCCTTTGTCATGTGTACATAGGGGTCCAATTAAC

>MSTRG.2116.1 gene=MSTRG.2116

AGATGATCAACCTCACTCTTTGTCATGAATGCAAATCTTGGCTCTGTCTGTTTCTTTTTCTTCTGACTCC

CACTCTTCTTGCCTTTCAACCTAAACTCCCCCGAAACAAAACCAAATCAACACTCAACAAGTCTTCACTT

TTCCACAAATGCCTTTTACTTTTTTACTTTTTAGTTTTTAGAAAAACCAATACAAGAGTAGTAAATGAAA

ACTTACTGTGGTTTAGTGACTTTGTGCTGCTGCTGTTGTTCTTGTTCATCAGCCACAACTTTAGTCTTTT

CTTCACCACTTGAAACAGAAGAGAGTGAAGAGGAGTTGGGAGTTGGGGGATTGTTCAAAACCTCTGATGA

TTCTTGATAAGGATTGAGTTGAGTTTGATTCTGAACTGGGCTCCGGCCCACTACAAGCTCTTCTGCTGTT

AATAAATCAAAGATGGAACTGTTGCTACAAGTGTTGTTAAAATCTTGAAGGCCCAGGAGCTCCATGAAGC

CCACTGAGCTCTTCTGATCACCAAACCCTAACATATCAGATAACATTAATGGGCTGCTGATGCTGCTACT

GATGTTGTTGTTATCAGTTTGATCATCCACTACAAAACTCTTCTGCTTCTGAGCCTCCTCTTGTTTCACA

GTCTGCTTCCTCTCCATGTTGTAAGAATGAGAGAGATTGAGGGAGAGAGGGAGAGAGAGAGAGAGAGAGA

GAGAGAGAGAGAGATTGAGGAAGAGAGAGAGGGAGGAGAGAGAGTAGGGAGAGAGAGGGGACAGTGAAGG

AAGGAGGATGAGGTGAATATAGACAGTGTTGACTGACAGGTGGCGGTCAAGGATTTGTACGAAATATACC

AAACAAACAAAAAGAGGAGGGGCGGGTGAGAGTGCATTAAATGCTATAAGTAAGAGTGGTAGTGGTCTAC

GCTTTCATACGTGGTTTTGACCGGTAATCTGGTTACTGTGTGAACGCGATCCATGTGTACGTGGCTTATT

ACGGCTTGTACAAAGTTTGAGTTTGGCTTTAGTAATCTAGTACGTATGTTATGTGCATGTGTGTCGACTA

GATTATTCTCCTTTAGTTAAAATAAATGTAACAATTTTTTAGTGACAAATGGTTCGGATATTCGGAACAT

GTGTTCTAGTCTTCTGCCTTAAATCTCGCCATACATCAGTCGTCAGTGTGTCTTAGTTTGAATGAAAGGG

TTCCAGAGGTGATCT

>MSTRG.2120.1 gene=MSTRG.2120

CCTCAAGCTTGTCAAGGCAATCATTAGTCATCCCTGTAGCCCAATCGCTGAAGATTTTAGCTTCAAGATC

TTCTCCTACCATCTCCCTGTAACATACGTTGCAGGTGTCGATATTTGCAGAGATTGAAGTAAGATCCATA

TTGACCTTGTAATAGTTATCTGATTCCAGGTCTTCAATGGTGTTCTTCATGTCATCAAGGGCAGCTTCGT

ACACATCTTGGCATGTCTCTAAGCAATCATACGTTGGTTTTGAGGTAGCTGGATGCTTCACCTGCGTCTC

AAGGCCCTGAAGATGTTCTCTAGTTTTCTGCATTCCAAGAATCATGGCAGAATTGATGAATCGGGTCATC

TGGGTCTTGGAATCAACAGGTGCAAGTGCCTCAGGGACGGGGGTCGATTCTGCTTGAGGAAATAAGGCGA

GTAGGAGCGAGGCGATGGACATGACAAGGACCACCTGGCCAATGTTGGAAGCCATGGTCTCTTCTATCTA

TTTAAGAAGAGAGCAGGCTAACCAACAGAGGGAGGGATTGAAAGAGAGCCTGGTGAAATATAAAAGACCG

AATCTCGAAAGGCCTGAGATTTAAGGGTGGTTGGATATTTGCCCAGTTATCAAGGATTTAAGAGTGGCCA

AGCTTGAAATTAGGAATCTTATGAAATATATTTGGGATACTTAAGAAAGTATAGAATATTATATGTACAA

GAAATCTG

>MSTRG.2144.1 gene=MSTRG.2144

GCACTTGCCTGGTATTTTGGTATGGCTCTGTTTCATGTTCCTTGTTATGGACTATATGCACTCCTCGTAA

AAGCAAAAGACTTTGTTTTCTCAAGGATCTTCCCCCATGCATTTTTAAGGGTTACAATGGAGAAGAAGCA

ACCTTGAAATCTCCACGTTCCCCCACCTATTGTTAATGCTCATTTCTGATTGTATTAAACATTGAACGAT

TAGTCAGGTGAAGTAGTCAGGTACACTTCACCATTTGTAATGGATGTCAGATTCTATAATTTATGCACAT

ATATCGTACGAAGTATAGTTTAGTTTTACTCGTTTGTATTGTTTATCAGTTGTCGGAGGTCAAAACGGGG

TAACATAAAATGCATGGTGCATGGCTTAGGCAAGAACAGAAGAATGCAACTTAAGTCTGAACACAGGGAA

AGAATGTATAATCACAGGAATTATGTCTCAATCTTTACTTGAACAGTCGTGTCTGAATCTGTAGATATAT

AGAAGGTTCAGCTCTCAATCTTTTATTCCAAGAAAGGATAAGGAGTTTTACTACATGGAAGATATATACT

TG

>MSTRG.2149.1 gene=MSTRG.2149

TTACAGCTGAGAAGAATACACACACGGATACACGCAACTTAGAGAGCTCTCGAAATTGAAGATCAAAACC

TCATTCTTCAAGAACAAATCGAGCTTGCTGCAAGAACAAATCGAAGCTTTGAAGGACAAAATGAAGCTAT

TAGAAGGTTGATTTATAGGAGGAATCAAAGATTATGCTAAGACGAAAAGGGATGTTGGCCGCATTCTCGT

TCTCCTAAGTCTTTATGCTATTTACTATTGCGGCTTCTCCTTGAGTTTTTCGAGATTGTCGACATTTGCA

TGCTATTGGCAGAGGTGTAAAAGGGATGAAGTTAGATTCGTGGAGATGATTTTAATGGTTTATGCGCTTT

TGGGGAGATTCCCTTGCAGATGGAATATTGTAGCTTTTAGTAAAATTAATTTCATAATTCTTTATGCTTG

TAATTTTGTTTTAACTTGTTATTTTATTTCCTAGTTTTTTGTTGGTGGTTTCTTGGAGAGTGTTCCTTGA

AACTTAAGTTAGAGTTGAGAGCTTTTTTCTCATGTTTGTATATTTTTTTTAATCTATAAAATTCATACGA

G

>MSTRG.2161.1 gene=MSTRG.2161

CTGGCACTCTGAAATACTCCCAAGTCCCAATACAAAACTCAGTGGATACTTGGTGTATAATTAATCTTAG

GTAATGTACTCCACATGGTCGTCGGCAGTTCTAGCCGTTGGTTTGACTCCGTCCTTCCACCGCCACATTT

CACCGGCCGGCGAAAAACTCCGTCAGTTATGTAAATCATCATATTTTTCAGAGTCTTCAACAAAAAGACG

ACGATCAGTTGCGTTATTGGCGAGTAGTTCGAGTGGCAGAATCTTGAAAATAGGCGAGAATTCAGAGGCT

GATAAGAGATTGAGCCTGAAAGATTACTTTGAACAGTCGGAGTATTTATTCCGATCAGACGGTGGACCTC

CGCGGTGGTTCTCCCCGTTAGAGTGCGCTTCTCAGTTGGATGACTCGCCTCTCCTTCTGTCTCTTCCAGC

AATTGATGGTCTCGGATCCATGCTTACCCTGCATTTCCAATCACTTGGAAAGATATTCGACATTTGGTGC

TTGCATATTCCTATTGACGACCGAACACCATTTCCGGAGCTGGTGAAGGGGGTTGTAGAGACGATTAGAT

CAGAGCATTGCCATTCACCAAAGCGGCCCATATATCTTGTTGCAGAATCATTTGGGGGTTGCTTAGCACT

AGCTGTTGCTTCCTGTTGTCCTGACATTGATCTTTTATTAGTTCTGGCTAATCCAGCCACATCTTTCAGC

AAGTCCCAGCTACAACCTCTTATACCCCTGCTAGAAATCATGCCTGATATAAGCCGCTTTAGCGTGCCTT

ATATGCTGAGCTTCCTGGCAGATATCCCTTCGAAGATGGTGATGGGAACCCTGGAGGAGAGACTTCCTGT

TCAACAAACAATTGGAGATATAACTGATGCTTCCATAGCATTGTCATCATATCTTTCTGTGATGGCAGAT

GTTCTATCAACGGAATCACTTCTGTGGAAGCTGAAGATGGTCAAATCAGCTAGTGCATATGCTAATTCAC

GTCTTCATGCCGTGAAATCTGAAACACTAGTCCTTTCCAGTGGTAGAGATCTGTTGCTTCCAAGTGTTGA

AGAAGGAGAAAGACTACAGAAAGTTCTCCCAAAATGCAAGGTTCGGACATTTAATGACAGTGGCCATGCT

TTCTTTTTGGAACGTACTCATGATCTGGTTAGTATCTTAATGGGTACCAGTTTCTATCGCCGAGGAGCAT

ATCATAACTATGCATCAGATTATATTCCGCCAAGTCCTTCTGAATTCAGAAAAATATATGAGTCAACAAG

CTGGTTAGAGAATGCAACAAGTCCTGTGGTTATGTCGACAATAGAAAGTGGGAAGATTGTGAGGGGCCTT

GCAGGGATTCCTAAAGAAGGCCCAGTCTTATATGTAGGCAATCACATGATGTTGGGACTTGAATTAGCGC

CTCTAGTAACCCGCTTTATGGTGGAGCGAGATATACTTCTGCGAGGAATGGCACATCCACTGATGTTTAC

AAAGTTAAAAGAAGGAAATTTGCCTGATGTATCATCATTCGATTATGTTAGAGTTATGGGTGCTGTTCCT

GTATCAGCAACTAACTTTTACAGACTCTTCTCTGTTAATTCACATATCTTGCTGTACCCCGGAGGTGTGC

GTGAGGCTCTTCATCGAAAGGGAGAGGAATACAAACTATTCTGGCCGGAACAGTCTGAGTTTATTAGGAT

GGCAGCTAGATTTGGAGCAAAGATAATACCATTTGGCGCTGTTGGAGAAGATGATATATTTCAATTGCTT

CTCGACTACAATGACCAAGCAAAGATTCCCTTTCTTAAATCTTTTTACGAGGAACTAAGTAAAGAAGCTG

TGAACTTGAGGAGTGACAGCACAGGCGAGGTAACAAATCAAGAACAGCACATGCCAGTATTGCTTCCTAA

ACTTCCAGGTCGCTTTTATTATCTTTTTGGAAAGCCAATTGGAACTGCAGGTAGGAAGCAGGAACTAAAG

AGCAAAGAAAAATCACATGAACTGTATCTAGAAGTACAGACTGAGGTCGAGAGATGCCTTGGATATCTGA

AGGAAAAACGAAAATATGATCCATATAGGAATATAGTACAGCGGTTGGCCTTTCAGGCGAGGCATGGCTT

GGAATCTGAAATTCCTACATTTGAGCATTGATAATTTGTTGTTATAAGGGTCCTGATCCTCTCAACTGTT

TGCAAGGAACATGTTAACAATGAGGAGTTATCCAAGAGAAAGAGGATGAAAACAGGTTTCTTACAGGTTT

GTAGAGGTTCAACAAATATAACCAACATCAGGAGGTTGACTATATTTATGGACAAGGGGAATGTATCATT

GGAGCTTAAGTTTTTTTTTACATAATCTCTATATTTTTTGTTTATAGTATGTATTAATGATATTTAGCTA

AGGATTTTACATGATTAGAGAATAGAGATGCTCTAAAGACAACTAAAATAGCAAATTAACTAAATTAGGT

TAAAATTGATTCATAAGTATATATTGTTGAAACTAGTACAAAATATGGCAAGATGGATGATTGCGTATTG

CAAGTTAATCTGTTGTTTTGGAAAGTTTCCAAGAGTGTGAGCAATTGGTTGAGCTAGAAAGAAGAAACAA

CTACTATTCTTTCATGCCTTGTGAATGAAAGCAACAACTTGTTACAGTTGATCACTCGAATATTTATTTA

CATTATATATATCAGTAGCATTTTACAAGTTGGTGATTTAACGATTATCAGCAAATATGAGAGGCAGTAG

CAAAGCAAGAACTGTAGCCATTTTACTAATTATGTATACGTATACAGTCGGATTAGTGTTTCAGTGTGAT

GGAGCGTCAGAGCAGCGGGAGGCTGCGGGGCAAAAATGCATCAAGGATTGTGGTTTTAGAGTGGTGAAAT

GTGGCATCGCGTGTAACAAATATGGAGAATACAATATTTACTCTTGTGTTGTAGAATGTGGTCGCTCAAA

TATGATTTGTTTGGCCTCCTGCGTCAGATTACCGCCCCCGCCCCCGCCTCCCCTGATGTTTCTAAATTCG

TGAC

>MSTRG.2161.2 gene=MSTRG.2161

ATACTCCCAAGTCCCAATACAAAACTCAGTGGATACTTGGTGTATAATTAATCTTAGGTAATGTACTCCA

CATGGTCGTCGGCAGTTCTAGCCGTTGGTTTGACTCCGTCCTTCCACCGCCACATTTCACCGGCCGGCGA

AAAACTCCGTCAGTTATGTAAATCATCATATTTTTCAGAGTCTTCAACAAAAAGACGACGATCAGTTGCG

TTATTGGCGAGTAGTTCGAGTGGCAGAATCTTGAAAATAGGCGAGAATTCAGAGGCTGATAAGAGATTGA

GCCTGAAAGATTACTTTGAACAGTCGGAGTATTTATTCCGATCAGACGGTGGACCTCCGCGGTGGTTCTC

CCCGTTAGAGTGCGCTTCTCAGTTGGATGACTCGCCTCTCCTTCTGTCTCTTCCAGCAATTGATGGTCTC

GGATCCATGCTTACCCTGCATTTCCAATCACTTGGAAAGATATTCGACATTTGGTGCTTGCATATTCCTA

TTGACGACCGAACACCATTTCCGGAGCTGGTGAAGGGGGTTGTAGAGACGATTAGATCAGAGCATTGCCA

TTCACCAAAGCGGCCCATATATCTTGTTGCAGAATCATTTGGGGGTTGCTTAGCACTAGCTGTTGCTTCC

TGTTGTCCTGACATTGATCTTTTATTAGTTCTGGCTAATCCAGCCACATCTTTCAGCAAGTCCCAGCTAC

AACCTCTTATACCCCTGCTAGAAATCATGCCTGATATAAGCCGCTTTAGCGTGCCTTATATGCTGAGCTT

CCTGGCAGATATCCCTTCGAAGATGGTGATGGGAACCCTGGAGGAGAGACTTCCTGTTCAACAAACAATT

GGAGATATAACTGATGCTTCCATAGCATTGTCATCATATCTTTCTGTGATGGCAGATGTTCTATCAACGG

AATCACTTCTGTGGAAGCTGAAGATGGTCAAATCAGCTAGTGCATATGCTAATTCACGTCTTCATGCCGT

GAAATCTGAAACACTAGTCCTTTCCAGTGGTAGAGATCTGTTGCTTCCAAGTGTTGAAGAAGGAGAAAGA

CTACAGAAAGTTCTCCCAAAATGCAAGGTTCGGACATTTAATGACAGTGGCCATGCTTTCTTTTTGGAAC

GTACTCATGATCTGGTTAGTATCTTAATGGGTACCAGTTTCTATCGCCGAGGAGCATATCATAACTATGC

ATCAGATTATATTCCGCCAAGTCCTTCTGAATTCAGAAAAATATATGAGTCAACAAGCTGGTTAGAGAAT

GCAACAAGTCCTGTGGTTATGTCGACAATAGAAAGTGGGAAGATTGTGAGGGGCCTTGCAGGGATTCCTA

AAGAAGGCCCAGTCTTATATGTAGGCAATCACATGATGTTGGGACTTGAATTAGCGCCTCTAGTAACCCG

CTTTATGGTGGAGCGAGATATACTTCTGCGAGGAATGGCACATCCACTGATGTTTACAAAGTTAAAAGAA

GGAAATTTGCCTGATGTATCATCATTCGATTATGTTAGAGTTATGGGTGCTGTTCCTGTATCAGCAACTA

ACTTTTACAGACTCTTCTCTGTTAATTCACATATCTTGCTGTACCCCGGAGGTGTGCGTGAGGCTCTTCA

TCGAAAGGGAGAGGAATACAAACTATTCTGGCCGGAACAGTCTGAGTTTATTAGGATGGCAGCTAGATTT

GGAGCAAAGATAATACCATTTGGCGCTGTTGGAGAAGATGATATATTTCAATTGCTTCTCGACTACAATG

ACCAAGCAAAGATTCCCTTTCTTAAATCTTTTTACGAGGAACTAAGTAAAGAAGCTGTGAACTTGAGGAG

TGACAGCACAGGCGAGGTAACAAATCAAGAACAGCACATGCCAGTATTGCTTCCTAAACTTCCAGGTCGC

TTTTATTATCTTTTTGGAAAGCCAATTGGAACTGCAGGTAGGAAGCAGGAACTAAAGAGCAAAGAAAAAT

CACATGAACTGTATCTAGAAGTACAGACTGAGGTCGAGAGATGCCTTGGATATCTGAAGGAAAAACGAAA

ATATGATCCATATAGGAATATAGTACAGCGGTTGGCCTTTCAGGCGAGGCATGGCTTGGAATCTGAAATT

CCTACATTTGAGCATTGATAATTTGTTGTTATAAGGTTTCTTACAGGTTTGTAGAGGTTCAACAAATATA

ACCAACATCAGGAGGTTGACTATATTTATGGACAAGGGGAATGTATCATTGGAGCTTAAGTTTTTTTTTA

CATAATCTCTATATTTTTTGTTTATAGTATGTATTAATGATATTTAGCTAAGGATTTTACATGATTAGAG

AATAGAGATGCTCTAAAGACAACTAAAATAGCAAATTAACTAAATTAGGTTAAAATTGATTCATAAGTAT

ATATTGTTGAAACTAGTACAAAATATGGCAAGATGGATGATTGCGTATTGCAAGTTAATCTGTTGTTTTG

GAAAGTTTCCAAGAGTGTGAGCAATTGGTTGAGCTAGAAAGAAGAAACAACTACTATTCTTTCATGCCTT

GTGAATGAAAGCAACAACTTGTTACAGTTGATCACTCGAATATTTATTTACATTATATATATCAGTAGCA

TTTTACAAGTTGGTGATTTAACGATTATCAGCAAATATGAGAGGCAGTAGCAAAGCAAGAACTGTAGCCA

TTTTACTAATTATGTATACGTATACAGTCGGATTAGTGTTTCAGTGTGATGGAGCGTCAGAGCAGCGGGA

GGCTGCGGGGCAAAAATGCATCAAGGATTGTGGTTTTAGAGTGGTGAAATGTGGCATCGCGTGTAACAAA

TATGGAGAATACAATATTTACTCTTGTGTTGTAGAATGTGGTCGCTCAAATATGATTTGTTTGGCCTCCT

GCGTCAGATTACCGCCCCCGCCCCCGCCTCCCCTGATGTTTCTAAATTCGTGAC

>MSTRG.2161.3 gene=MSTRG.2161

TTGCAGCAATTGATGGTCTCGGATCCATGCTTACCCTGCATTTCCAATCACTTGGAAAGATATTCGACAT

TTGGTGCTTGCATATTCCTATTGACGACCGAACACCATTTCCGGAGCTGGTGAAGGGGGTTGTAGAGACG

ATTAGATCAGAGCATTGCCATTCACCAAAGCGGCCCATATATCTTGTTGCAGAATCATTTGGGGGTTGCT

TAGCACTAGCTGTTGCTTCCTGTTGTCCTGACATTGATCTTTTATTAGTTCTGGCTAATCCAGCCACATC

TTTCAGCAAGTCCCAGCTACAACCTCTTATACCCCTGCTAGAAATCATGCCTGATATAAGCCGCTTTAGC

GTGCCTTATATGCTGAGCTTCCTGGCAGATATCCCTTCGAAGATGGTGATGGGAACCCTGGAGGAGAGAC

TTCCTGTTCAACAAACAATTGGAGATATAACTGATGCTTCCATAGCATTGTCATCATATCTTTCTGTGAT

GGCAGATGTTCTATCAACGGAATCACTTCTGTGGAAGCTGAAGATGGTCAAATCAGCTAGTGCATATGCT

AATTCACGTCTTCATGCCGTGAAATCTGAAACACTAGTCCTTTCCAGTGGTAGAGATCTGTTGCTTCCAA

GTGTTGAAGAAGGAGAAAGACTACAGAAAGTTCTCCCAAAATGCAAGGTTCGGACATTTAATGACAGTGG

CCATGCTTTCTTTTTGGAACGTACTCATGATCTGGTTAGTATCTTAATGGGTACCAGTTTCTATCGCCGA

GGAGCATATCATAACTATGCATCAGATTATATTCCGCCAAGTCCTTCTGAATTCAGAAAAATATATGAGT

CAACAAGCTGGTTAGAGAATGCAACAAGTCCTGTGGTTATGTCGACAATAGAAAGTGGGAAGATTGTGAG

GGGCCTTGCAGGGATTCCTAAAGAAGGCCCAGTCTTATATGTAGGCAATCACATGATGTTGGGACTTGAA

TTAGCGCCTCTAGTAACCCGCTTTATGGTGGAGCGAGATATACTTCTGCGAGGAATGGCACATCCACTGA

TGTTTACAAAGTTAAAAGAAGGAAATTTGCCTGATGTATCATCATTCGATTATGTTAGAGTTATGGGTGC

TGTTCCTGTATCAGCAACTAACTTTTACAGACTCTTCTCTGTTAATTCACATATCTTGCTGTACCCCGGA

GGTGTGCGTGAGGCTCTTCATCGAAAGGGAGAGGAATACAAACTATTCTGGCCGGAACAGTCTGAGTTTA

TTAGGATGGCAGCTAGATTTGGAGCAAAGATAATACCATTTGGCGCTGTTGGAGAAGATGATATATTTCA

ATTGCTTCTCGACTACAATGACCAAGCAAAGATTCCCTTTCTTAAATCTTTTTACGAGGAACTAAGTAAA

GAAGCTGTGAACTTGAGGAGTGACAGCACAGGCGAGGTAACAAATCAAGAACAGCACATGCCAGTATTGC

TTCCTAAACTTCCAGGTCGCTTTTATTATCTTTTTGGAAAGCCAATTGGAACTGCAGGTAGGAAGCAGGA

ACTAAAGAGCAAAGAAAAATCACATGAACTGTATCTAGAAGTACAGACTGAGGTCGAGAGATGCCTTGGA

TATCTGAAGGAAAAACGAAAATATGATCCATATAGGAATATAGTACAGCGGTTGGCCTTTCAGGCGAGGC

ATGGCTTGGAATCTGAAATTCCTACATTTGAGCATTGATAATTTGTTGTTATAAGGGTCCTGATCCTCTC

AACTGTTTGCAAGGAACATGTTAACAATGAGGAGTTATCCAAGAGAAAGAGGATGAAAACAGGTTAGAGA

AAGTTTCTTACAGGTTTGTAGAGGTTCAACAAATATAACCAACATCAGGAGGTTGACTATATTTATGGAC

AAGGGGAATGTATCATTGGAGCTTAAGTTTTTTTTTACATAATCTCTATATTTTTTGTTTATAGTATGTA

TTAATGATATTTAGCTAAGGATTTTACATGATTAGAGAATAGAGATGCTCTAAAGACAACTAAAATAGCA

AATTAACTAAATTAGGTTAAAATTGATTCATAAGTATATATTGTTGAAACTAGTACAAAATATGGCAAGA

TGGATGATTGCGTATTGCAAGTTAATCTGTTGTTTTGGAAAGTTTCCAAGAGTGTGAGCAATTGGTTGAG

CTAGAAAGAAGAAACAACTACTATTCTTTCATGCCTTGTGAATGAAAGCAACAACTTGTTACAGTTGATC

ACTCGAATATTTATTTACATTATATATATCAGTAGCATTTTACAA

>MSTRG.2164.1 gene=MSTRG.2164

TCCTCATCTTCTTCCTTCTTGCTGCTCTTAGCAGTATCCACCCTCTCTGGAGGCCCAGCTCGAAATGGGG

TACGCCTTCGTCGATTGCGCTTAGCAGCTTCTCGGGTCTTTCGCTTCTTCTCATCCTGTTTATTCTCATG

ATACTTCCTCCTCTTACACTCCTGAATGATCCCAGCCCTCATGACTTCCCTCCTGAACCTACCCAAAAGC

TTCTCTTCAGG

>MSTRG.2170.1 gene=MSTRG.2170

TGGGTATCGATCATCCACTCCTCAACACTTTCTACCACATTCTAGACGATCATCCACTCCTCAACACTTT

CTACCACATTCTAGACGATCAGCACGAGAGCAACAACAAGAACAAGTCTGAACAGGCGAGGAGCTATGTT

CGCGACGCGAAAGCCATGGCGACGACTCCAGCTGATGTTAAAGAGTATCCGAATGCCTATGTGTTTGTGG

TGGATATGCCTG

>MSTRG.2176.1 gene=MSTRG.2176

ATCCATTCCTTTGGTATCTGTTGCAAGAATTCGTATCTTTCCTTTCTATATTTAAAAGTGCATATGTTCG

AATCTATTCTTTCTATTCTTTAAGAATCTAAAGAATTTCCGTGTTATTAATTAATTCGAAGTTCAGATTG

TCATTGAAGCATGTGTTGCTCAGTGATAGTTGGAGCTTTGAATGAATTGGGATCTTCTTGATTATTCAAC

CCGCCATTGTCATGCTGGTTTAGCAGGCTAAATTCAGTTGCTTAGCCGATAATGGAGGATACTGAATAGT

TGAGGATTTCCTGAACTGTGATTTATGAAATGATGGATCATGATTTATGAATAGATTATTATCAAAGTTT

GAATTGAAATTATTCTGTTGTTGATGATATTTATGATGATGTTCTGTTAATTTACATTGGCTTCTTGAAT

TGAATTAGAGAATTG

>MSTRG.2177.1 gene=MSTRG.2177

CAGGAGAGGTGACTGTTACTTGAATAAATGCCTTTAAGCTTTGCTTTTGGGACTCTGCACAATAGAACTA

TACTCACATTTTGAAGCTCTGTCCGAGTTCGTTGAAATTAGGGGAGAAAGAATGTTCTTTCTACATGAGA

ACTGGTTCATGCAAGTATGGGAAAAGATACAGGTTTCATCATCCGAATCCTGCAACACCGGAAGGAGGTG

ACACTATTTCACCAGGCTCCTGCTTACTGAACATCACAAGGCATTCTGCCTATTCCCCAGCACTTTCGAT

GATCATTTCAGCAATTAGATCCAGCTTCTATGCACATCCGCAGGGCCAGGTGAAAGTAGATGAATATCCT

GAATGACTTGGACAGCCCAATTGCAGTTTCTAACGGAGACGGCTATCAATCTTAGTGACAAAGGACTGCC

TTTGCAACCTG

>MSTRG.2178.1 gene=MSTRG.2178

GGCCAAAGTGTCTCCTCACACTTTAGCCGTTTGGGTCGACCTGTAAGTGTGATCATTCAGTTGGTGGCAC

TGCAACTTCTCAGATGGGGATGATCAAGGCCGTATCCCAGAGTCTACTTGCTTTGCAGATGAACATGAGG

AGTGGGAAACGGAGGCGATGGGGTCTGCTTGAGTATCATACAGTAGTTGCAAAAATCGTTTCTCGAAAGT

ATTTTTTTTCTGGGAACATGGTCAACCATAAATTCTTATGTTCTCGATTCTTCATCCACCGATGCAGAAA

ATGCTCCAGTTTTCCATTTTCATATGAGGCCGGAAAGTTTATTGGCAACAAGTCGGCACGAATAGTAACT

GATTACAAAACTCTGTGAAATTTACAAAGT

>MSTRG.2183.1 gene=MSTRG.2183

GGAACCTCAAAAGTGGGAGCCAAATTCATGCCTCTGTAGCTCTTCTTCTCCTCATCAGTACAGCTGAGGA

GCCGGGTAATTGGGCCGGGCTGTTGGGCCTTCTTGTAGCTCCGCATCAGACCCACTGTTTGCCAGTCGAA

GTAGTTCTGGCACTCCACAGAAAACAATGTATGGATCCTCCTCCCTTTCTCCTCAGCCCAGCCCTTTTTC

ACCATTAAAACCAACACAAACACGCCTACTAGAAACAAGACGCCTAATTTACCCATCACTCAATGAATCT

GATCACTTTTCGATCTCGATTTCATCCGCTGTGTTCCGAGGAGAGAAATGTTCAAGAAATGGTGATTAAA

TTATTGATGCAGAGAAAAGTCAAAATCGAAGTGTCGTGTGTGCATTCTTGTACATGTTTTTTGTGTCAAA

TGGACTTGGAATCGTGATCGGAGGCTGGTAAAAGGCTTGCTGCGTTCTAGTTTTAGATTATACCAATTCT

TTTCGGGTTGTTCAGTTATTTTATTATTATTATTATTATTACTTTGCCG

>MSTRG.2188.1 gene=MSTRG.2188

CTTAATTGGGTTCTGTAGCTGGGTAAGTTCATGTGAGACCCGACCATGGCTCAGGTGAGGGGTTCCAATA

ACCCTAAGTGGTTTGTTACATTATTAATTCGATTGAATCTAGTATTGATCCCAAGTTTGTTGAATTCGAG

TTATTTGATTTAATTTTATTTGTCGTCGTTTTTTTATATTTTGATATTCATCCATTTTATATATTGTGTT

GTTCTGGTATTGCTAATTTTATGGATGTAGGAACTTTTTGCCTCACAAGGCCATATCCATGGTGGTGATG

GATAGGTCCAGAAAGGGTGGAGAAGAAGAGCTATTTTTGGCGATAATTGTGCCTTAATTAAGGATACTAA

TTATTCTTAATTAAGGTTTTAGTACCGTTTATGGCGACCATTTATGGCGACCATTATTGGGGCATTATTG

GCCCATAAGAGGCATTTAAGTGCCATAATGTTGGCTCGTTTATGTCTTTTATTTTAATTTTAAATTTGCT

CACATTCGTTCATTCTGTTTTGTCTTATTTTCAGGATGTTTGAAAAAGCATTTAAGAATTTATCTATTAA

GGCGTCTTCCGAGAAGACGCGAATTTTGGTGGACAAGACAGTTGAGCTACAAACAAAGATAGAAAGGAAT

AAAGAAGACTTCATTTTTAGTCTATTTTGTAATCGATGTATTTTTTTGTAGGGTACTCTTTTTCCCCCTT

GAGTTTTTTTCCCACAGGGTTTTGCTCTTGAGGGGTTTTAACGAGGCCCTTTTTTTGTGGGTTTTCTCTT

CGGACTTGAAAGGTTATGTTATCTAAGCGTCCGGAGTATTTACATACTTTCGGTTAGATGATATACCTTC

TTGGATGAAGGAAACTCTGTCCATCCGGATTGTAATTTTTTTCTTGGTTTAATACAACCGTTTATTGTGA

CAAAAAAAAAAAAAAA

>MSTRG.2195.1 gene=MSTRG.2195

TTCCAATATAAAACAATTTGCACACAGCAGCAGAGATGGGTAACGAGGTTTTGTGGTTGGCAAGGCGAGT

TGTGTACAAAATAAAATCAGGGAAAACGAAAGAAACAAAAAAGCAGAAGATGAGTGGAAAGTAAAGGAAA

GTGTTTGAATCAAGGAGAAATGAGAACTTGTCCAAGTTCATGGTTGGATTTGCCGCCCTTACTATACGGA

ACGATCAGGAGAGGTGCAAATCTGTTGCATTTACCAACGTCCTCCTCAACCATTAGACCAAAATCATCAA

CAAGTTTCAGTAAACTCATCCACCACAAACCCACCATTTTAAACAGCAGCAGCAGCACTAATGAAGTGGA

GGTGAAGGTGGAGGTGGACACAGAAACCCGAAAACATAATTTGCTGAAAGCAATTCAAGATACGCAGAGG

GGAATTTCCGCAACTCCCCTCCAGCGTTCTTGTATCGAGGAGGCTCTCGTTAGCCTAGAAGCTCTTGATG

CTGCTTTACCGTTGGACCTTGATTTGTTGAATGGCAATTGGCGGTTGCAGTATACTTCTGCTCCTGATGT

TCTTATCCTTTTCGAATCCGCTAACAGACTTCCCTTTTTCCAGGTTGGCCAAATTTATCAAAGCTTTGAA

TGCCAAGCTGAATCCAAGGGAGGCTACGTATGCAATGTCGTTCGCTGGAGTATCCCCAAACTCTTAGAGG

AAGAAGAAGGTGCCACTCTCCTTGTTTCTGCAAAATTCTCTGTAGTTTCTTCACGTAATATCTTTCTCCA

GTTTGAAGAGATAACTATTAAGGACATATACATTAGTGAAGAGGTGCAAGCTTTAATAGCTCCAGCGATT

CTTCCACGGACGTTTTTTAGTTTGCAGATCTTGCAGTTCATCCGCACATTTGAAGCACAATATCCGGTGG

AAAATCCAGGGAGGCAATCTGTCGGAGGACTGTACTATCTCTCATATCTGGATAACAATATTCTGTTGGG

TCGTGCTGTTGGAGGTGGAGGGGTATTTGTATTTACTAGAGCGCCTACTCTTGTGTGACTTACGTGTTTG

TTAAATATTTTATTTTTATGTACCCTGGATGGACATAGTCATTTGAGACGTTAAAATTATATTCGTGCAA

TTAATTTGTTATTGATTCCTAAATTGCCTTGCGGGTCAGTTTTTTGTTTTGCGTTGCACAAACACCTGTA

GATGCTATGTA

>MSTRG.2205.1 gene=MSTRG.2205

CTATTGCTACTGTGTTCAAGGGTTCCAAGCTTAATGGAAGGTTATGAGATCCACGGCCAGCACTCATTGT

TCTTGACATAGATATGTGGACACCAGGATCATGGTGAGTGCAAGTAGCATTGAAGAGGATGGAATTGTTC

ACTGCCTGTACATTGCTGTTCACATAAGGAAGTGACAATGCCTTTGTCATCGCCTCCTTAGGATTCCCAT

CCTTTTTCGATTTACCTTCTTCACTGCTGCTGCTGCTACTGCTACCGCTTCTGGATTTCTCTCCTTCACC

CCAAGTCCGAAACTTTCCTTGGAAGTTGACAGGGTTGGTGCCCAAGTCCTTGTTCTTATGAGGCCTCAAC

TCCATTATTGCTCCTTTGTTGTCACCTGCAACTGTTATGATCCTCATCCCGGAACTATTAGAATTTGCAG

TGTTTTTGTGGCTACCTTTGTCTTTGTGATGAGTCTTTTTATTTGTTTCTCGATCTCCTTTATGGGGATT

GTCAGTGCTACGAGGTTCAGTGTTGTACAAGTTTGGCTTTTCTTTGGTTTCCTGCACAAGGACAGCTTTC

TGCTCGAATTCCAGAGGATGTTTTAGTTCTCGCTCGTTGTCAGACCTTAACTGTGCTGGAGGGAGAGTTA

GTGGGGATGGAGGATGAACAATGCGCCGATTGATAGGAGATGCTGATGGAGTATTTGGCTGAGAGCTTAC

GGGTTTGGGGAAGGATGTTGAGTTTGTTGCAGGAACATTGATAGATCCGGCTGCATAGGGTGATGGTTTT

GGAGCAGAGGTTTGAGCTTGGGATACTGTTTTGGGTGCAGGAGAATATATTGTGCCACGTGACGGAGATG

GATCTTTGGGCTTAGGCGACTGAGGAAGGGAAGAAGAGGAGGGCAGTGGAATCACGGGAGCCAGTGTTGC

TTTTGGAGCAGAAGACACTGGAGATGATCTGTTCGCGGTGGATGGGGACTGTGTTACAATTGGTGAAGGT

GGAGCAACAGTAGTTATTTTTGATGGTGGTGTGGCTGGAACTGGGGCAGGAGCAGCAGGAGGAGACCTTG

TCACTGGTGCAACAGAAGCTGGTGGCACTTGAGTATTGGCCACAGGTGGAGGTGGAGCAGAAACAGGTGC

TTGTGGTGCAGGTTCTTGAGTTTGAGTAGGCGGTGCCCTAAATGCAGGTCGAATAGCACTTGGCCTTGGC

TGGGGAACTGGATCAGGCTGTGGGGGAGGGGCCGACCGCGCAGCCATAGTAGCTAAACGAAACCATGGAC

GTGCTGGAGGTTGGCTAGCCATTAATTAGAGAACTTCAAAGAAAGTGATACCTTTCTGGTTGATTTTCAA

GTACTCTGGAGGATGGTTAATGATAAGCACAAATATTTGGGATGACAATGCTCCTACATATATAGTTTGG

TGACCCTGGAGTAAGCATGAAACGTAGCCCCCTCCATTTTCTCAAGCAGTGAGGCAAGGTATCAGATTCA

GAAAGCACTTATACAGCTCACAAATTTCCCTACCATGGTTTCGTATCAAATGTCAGCTGGGGTGCATTTC

AAGAAAGGTTACGGAGATTGCCTGCGTTGGGCCCCATCCGGGTAAGTTAACGGTCATATAAATCACGTAG

CCTCTGTTTTAAAAGTCGGTGAAGTTTTATTAACCGTTCACAAGAATTCCCCCATTATATAGTGCGCTGA

CTACTGTGCTAATTGCAGCTGGACATTCTAAGCACATGCAGAATCATTGGCTTATAGCAAGTCTGATTCT

GTCTTAACTAAAGCTCAAAAAGTATAGGCCATTCATTATACATGCAGAAACATGTCAAGTACCAAGATTC

TAAATTTGATATTCACAAAAGATTTTCGAGTACTCACTTTAGATATTGTCTGTGCCTTGCTTGATGATAA

TGTAATCCTCCTGCTGCCTCTTACCTTTTGGATGGCTTCAACTGAATGTCTTTCAAATAAAGGCATCTAT

TCTTTCATCTATTCAGTACAAGTATTACTGAATAACGGAGTCCTCCGG

>MSTRG.2224.1 gene=MSTRG.2224

AAAGTAGATAAATAGGATTGAAATAAATTTTATATATTTGATCTCTGATCAATCTCGAAACCCCTGTTCC

CTAAATCTGGAAATGGATCCACCAACTTCTTGGGACTCCTTGAGGAAACAGGCAAGGAGGCTTGAAGCTC

AGTTGGATGATCAAATGCATTTATATCGTAAGTTAGTATCAAAAAATGTGGGCAATGGTAATGAGAATGA

CCTGGAAGCGGAAATAGATCAACTGCTCAAACAACTTCAATTAGTAAATTCACAGATGCAAGCGTGGGTG

TCGTCCGGAGGATCAGAAATCTTTTCCCATACATTGACACGTCATCAGGAAATTCTTCATGACCTGAATC

AGGAGTTTATGCGACTTCGTTCTAGCCTTAGAGCCAATAAAGAACATGCTTCACTGCTTGAGGACTTCAG

GGAGTTCGACCGTGCAAGATTAGATCTGGAAGAAGGTGGATCTTATGAGCAGGCATCTCTACTCAAGGAG

CGAGCAGCTATCAGCAGAGGTACAGGACAGATGGACTCGGTAATTTCTCAAGCACAAGAGACTCTTGGTA

CACTTGTGTTTCAACGTTCAACATTTGGAGGCATCAACTCAAAGCTTAGTAATGTTAGCAGTCGTCTTCC

AACGGTGAACAGTATTCTCTCTGCAATAAAGAAGAAAAAGTCCATGGATACTATTATCCTTTCGTTGGTT

GCATCCATCTGCACGTTTCTCATTTTAATTTATTGGTTGACCAAATAGAAAGTACATGTAATTTCTGATG

GGAGCCCCTCCTGCTGTACTGTTTGTAAACATCCTTTTTTGGTTTTTTGTTAAGTTGTGCTATTCTTTCT

ATTCGCCATTACATTAGATGGCATTGTATACACCCTGTTGGTTTGTTGCACTAGTTACCATGCCTTGTAT

AAAAAAACAACCCAGAAATCAAGATTACAGATTATATTAGTTATTTTGGTCTTGCTAAAATGAAGCCCCC

>MSTRG.2236.1 gene=MSTRG.2236

TATGTGTCGGTACTCGGGTCCTAGTTTTAGCTGTAAATCATAAGTTTGGAAGAAGTAAAGCTTTGTATCT

ACAGTGCGCCCACAGAGAAAACACAAGTGAAGTCCGAGCAGGAAGTCAATTCACTTGTGTGCTCTCGAAT

CATGGCAAATCGGCACACCATTATTCTTATGCAGACTTCCCAAAGCAGAGCTACTCGGACATTTATGGAT

TATGATTCAATTAGTCAAGCAATAAATGGTGTATGTGGGCTTTACGAAAGGAAACTTAAGGATTTAAATC

CAGCAGTAAGAAACATCACTTATGACATTGCCGATCTTTACAATTTTATCGATGGGCTTGCTGATATGAG

TGCATTAGTTTATGACCATTCAATTCAGGCATACCTCCCGTATGATCGGCAGTGGATCAAACAGCGTACA

TTTCATCATTTGACGAAATTGGCCAATTAAATTATATTGTATCATTGTCTTCTATATATTTTCTGAGGTG

GCCACTATATGGTTCTAGTCCTATAAGACATCTGGAGTAGAATTTGTAGTTGGGGCCTTGGAAGTCTAGT

TGCAAAGATGCTTTGTAGCTTATGTAGCACCTGAGAATTACACCTCGGTTGCCTGATTTACTGTATGTTA

TAGCTGTATGTCTTTTACTTTTTTGGATGAAAGTATCTGAATATATATGGTCATTTTTATGGAAACTTTA

AACTGCTAAACTTTTTTGAAAGTCTTGTGGTACAATAGGTCTGATTCACCCG

>MSTRG.2243.1 gene=MSTRG.2243

GTTTGCCATGGATGTGATTGTTTCACCTGGTCAAAAGCATGCTGTCCAACTGCTGTAGTTTCATCAAAAA

GTGAGTTATCCATCTGTTTTGATTCCAAGTTGGAGTAAATTACAGGGGTGTGACTACAGTTTACACTGAT

TTGCAAAAATGTGGCTGAATTTTAAAATTAGCAGAAAGGTGGCTGACATGCGACTCCATCTTTTAAAAAG

TTGGCTGCCGTTAAGGGCGTTAGTTTTCAACCATTAACTTAATTAAAAAACATAAAGTGAAGGGGTATTT

ACGTCTCTTCCTCTACACAACCCTCATTCACCCTGCAAACAAGAAAAATAATCTACTGTTTAGAGATTGT

ATTAAGTCATTAAGGATCATCATCAGCATCGATTTTCTTGGACGTTATATTCTTTTTCCGCTGATTCTAT

TGTCTCCATTTTAGAGTGAGGTTTCGAGGCCATTGAGGGGGCAACTGTGGTATCCCAAACCATCTCTTAC

CTTTTTGTGCTTTTGTGTGTGTGATCGTGGAGGGACTGAGAGGGAGAGTGAAAGTTTGAGTTTTTTCACA

GAGGGAGAACTTTTAAAAATATATATAATTATACCCTCTCAACCTCTCAACCTCTCTTCTATTACTCGGT

TAAGATCTATTTAATGGTAGCCAATTTCTTGAAAAGCGGAGAAAAGTGCATTTTGTCTGTGTATATTTTT

ACATGTTGTTTACCTATATTTTGGGCTTTACGAC

>MSTRG.2244.1 gene=MSTRG.2244

AAACTTCTTTCACCACATTTTCTCTCTAATGACATCCGAAAATAGCTAAAATATCACCTCAATTCCTCCC

ATTTGCATAGAAGATTTACTCCTTTCATATAAATAAGGTTGTCTTGAAAAATGAATAATAACCGATCTTG

GATGTATGAGAGAATTGATGATAGAGGATTTTTAAATGCTCTATTTAATTCCGGTGTGGAGGAATTTATG

AATTATGCTATATCTCAACCAACGTCTATGGGCGGGACAAGCATACAATGTCCTTGTTCAAAGTGTAAGA

ATCGAAAATATTGGAATGGAGATACTGTGAAGCTACATTTATTGAAAAATGGATTTGTGAAAGATTATTA

TGTATGGAGTCGACATGGGGAGCCATACATTAAAGATGGACATGTCGAACATTCTTCAACAAATTATTCA

AATATTTCACGGGGCACAGATGAGAACAACTTAATGTACAATATGATGATTGATGTAGCTGGCCCGTTCA

CCGCCGCGAAGGCTCAACGTAAAGATCCAACTCCGGCCGATGTTTATCTTGTTACACATACGGTGAAGCA

TGACAAAAAGATTTTTATTACCAAGAAAGCGGAACAAGTATATAACAGGGTTGAGACACTACGTGAGGAA

CGTTCGACACCCATTGAAGGTTCTGATGAACATCGAGTTGTTGATGAAGACCAATTGTTTCTTGAAGCTG

ATGGCGGACTAGACAAAAAAAACAAAGTTTATGGCATGAGTTCATTACAAAGTGTTATGTATGGACTAGA

AACTGAGGGTAGTCGTTACAGGGGCTCCAACTTCAACAATGAAAAATATAAGCAAATGCAGGGGGAGTTG

CAGGACATGAAGAACCAAGTAAATGAATTACAGGAGATGAGAAACAAGGAATTAGAAGAGATCAGAAAGC

AGATGGAAGAGATGAAGAGCCAACTTGCTATAGTGTTTAACAATCGGAATGGAAATTAAACATTAGTGTC

TGACTGGCTGCATTTTATTTATGTATTAAGTTTGTATGACTTCTGAATTATAACTTGGATGTTGCTTTTA

CTTATTAATTTATGTCATTTTCCCTTTATTTAATTAAACAA

>MSTRG.2244.2 gene=MSTRG.2244

TTGTTAAAGAATTCTTATGTAGGTGGGTCTTGAAAAATGAATAATAACCGATCTTGGATGTATGAGAGAA

TTGATGATAGAGGATTTTTAAATGCTCTATTTAATTCCGGTGTGGAGGAATTTATGAATTATGCTATATC

TCAACCAACGTCTATGGGCGGGACAAGCATACAATGTCCTTGTTCAAAGTGTAAGAATCGAAAATATTGG

AATGGAGATAC

>MSTRG.2256.2 gene=MSTRG.2256

CACAGACAAATTTTCTCTCTCTAGAAACTAGTAATAAAGAAACAAAGAAAAGAAGAAAAGCAGCGAGAAG

CACGCAGTAGCTACACAGTGTATGTTGAGTATTCTTCTTCTTTCTCCTTGTTTACTCTTTCTTTCATCAT

GCCTTGTGCAGTTGTTTCATTCATTTCTGGTTTGATTTGATCATCTCATTTTGTTTGAAAACCTAATTAG

AGCTACCGCAATTATTAAATCCACTAGTGGTGTGTATTTGTTGTTGGTGATTGCTATGGGATACTTGAAT

TCTGTTGTGCCATCGGCTAGCCAGGTTCCTGGTCAATCTGATTCTCCTGTTTCCGGTGGCGGTCTCAGTC

AAAATGGAAAGTTTAGCTATGGTTATGCAAGCTCGCCTGGGAAGCGATCTTCCATGGAGGATTTTTATGA

GACAAGGATAGATGGTGTTAACGGAGAGATAATTGGTCTGTTCGGAGTTTTTGATGGTCATGGTGGTGCT

CGTGCTGCAGAATATGTTAAAGAAAATCTCTTTAATTATCTCATCAGTCATCCAAAATTTATATCTGATA

CAAAATTGGCAATAGCGGATGCATATAACCATACAGACTCCGAGTTTCTAAAATCAGAAAATAATCAACA

CAGAGATGCTGGATCCACAGCTTCCACAGCAATCCTTGTCGGTGATCGCTTACTTGTAGCAAATGTTGGG

GATTCCAGAGCTGTAATCAGCAGGGGTGGCAATGCTATTGCTGTATCTCGTGACCACAAGCCAGACCAAA

CAGATGAGCGCCAACGGATTGAAGATGCTGGAGGATTTGTTATGTGGGCAGGAACTTGGAGAGTGGGAGG

CGTGCTTGCTGTTTCTCGTGCTTTTGGTGATAGACTGTTGAAGCAGTATGTTGTTGCTGATCCTGAAATT

CAGGAAGAGAAGGTTGATAGCTCACTTGAATTTCTTATCCTCGCTAGTGATGGGCTATGGGATGTTGTCA

CAAACGAGGAAGCTGTTGGGATGGTCAAGCCAATTCAGGATCCGGAAGAGGCAGCAAAAAGATTAATGCA

AGAAGCTTACCAGAGGGGCAGTGCTGATAATATTACTACTGTTGTTGTTCGATTTCTGGAAGATCAAGTT

GGAACCTCTATCAGCGGCTCTGGTTAGGTTAATTTCTATTAGAATATTTAGACTTTCTCAACACAGATGG

ATGAATGAAAAACCCCTGAAGACGAGTTTAGTGTGTTTTAAGTATTGGAAACATTAGTTGCGTCAAATAT

AGGTTATGTACCTTCAAACACCGGAAGTGGTTTCTTACCTTAGAAATTTGCATCTTGCTGCTGGTATGTT

CTTTATATGGTTGGTCTTGAGTTATACTCCGTTTAGTTTTATAGGTCCCAAGGTGCTTTGACCGCATAGT

TCACATTATCCTTAAGTTACTATTTTGTAAGTTTTTTTATGAATAGTTCTTTTATGA

>MSTRG.2256.1 gene=MSTRG.2256

CTGAAATAAGAAGAGGAAGAAATCTCAAGAAAACAATCAATTTCATCATCTACTCACCAAACTCGCAACA

GAAAATAATGAATGATAGATGATGTAGTATTACAATATATGCCACCTCATCCTCTATGCCTGTCTATAAA

TACACACAGACAAATTTTCTCTCTCTAGAAACTAGTAATAAAGAAACAAAGAAAAGAAGAAAAGCAGCGA

GAAGCACGCAGTAGCTACACAGTGTTTGATTTGATCATCTCATTTTGTTTGAAAACCTAATTAGAGCTAC

CGCAATTATTAAATCCACTAGTGGTGTGTATTTGTTGTTGGTGATTGCTATGGGATACTTGAATTCTGTT

GTGCCATCGGCTAGCCAGGTTCCTGGTCAATCTGATTCTCCTGTTTCCGGTGGCGGTCTCAGTCAAAATG

GAAAGTTTAGCTATGGTTATGCAAGCTCGCCTGGGAAGCGATCTTCCATGGAGGATTTTTATGAGACAAG

GATAGATGGTGTTAACGGAGAGATAATTGGTCTGTTCGGAGTTTTTGATGGTCATGGTGGTGCTCGTGCT

GCAGAATATGTTAAAGAAAATCTCTTTAATTATCTCATCAGTCATCCAAAATTTATATCTGATACAAAAT

TGGCAATAGCGGATGCATATAACCATACAGACTCCGAGTTTCTAAAATCAGAAAATAATCAACACAGAGA

TGCTGGATCCACAGCTTCCACAGCAATCCTTGTCGGTGATCGCTTACTTGTAGCAAATGTTGGGGATTCC

AGAGCTGTAATCAGCAGGGGTGGCAATGCTATTGCTGTATCTCGTGACCACAAGCCAGACCAAACAGATG

AGCGCCAACGGATTGAAGATGCTGGAGGATTTGTTATGTGGGCAGGAACTTGGAGAGTGGGAGGCGTGCT

TGCTGTTTCTCGTGCTTTTGGTGATAGACTGTTGAAGCAGTATGTTGTTGCTGATCCTGAAATTCAGGAA

GAGAAGGTTGATAGCTCACTTGAATTTCTTATCCTCGCTAGTGATGGGCTATGGGATGTTGTCACAAACG

AGGAAGCTGTTGGGATGGTCAAGCCAATTCAGGATCCGGAAGAGGCAGCAAAAAGATTAATGCAAGAAGC

TTACCAGAGGGGCAGTGCTGATAATATTACTACTGTTGTTGTTCGATTTCTGGAAGATCAAGTTGGAACC

TCTATCAGCGGCTCTGGTTAGGTTAATTTCTATTAGAATATTTAGACTTTCTCAACACAGATGGATGAAT

GAAAAACCCCTGAAGACGAGTTTAGTGTGTTTTAAGTATTGGAAACATTAGTTGCGTCAAATATAGGTTA

TGTACCTTCAAACACCGGAAGTGGTTTCTTACCTTAGAAATTTGCATCTTGCTGCTGGTATGTTCTTTAT

ATGGTTGGTCTTGAGTTATACTCCGTTTAGTTTTATAGGTCCCAAGGTGCTTTGACCGCATAGTTCACAT

TATCCTTAAGTTACTATTTTGTAAGTTTTTTTATGAATAGTTCTTTTATGA

>MSTRG.2256.3 gene=MSTRG.2256

AGAGGGACGGAGGGAGTAATAAAGAAACAAAGAAAAGAAGAAAAGCAGCGAGAAGCACGCAGTAGCTACA

CAGTGTTTGATTTGATCATCTCATTTTGTTTGAAAACCTAATTAGAGCTACCGCAATTATTAAATCCACT

AGTGGTGTGTATTTGTTGTTGGTGATTGCTATGGGATACTTGAATTCTGTTGTGCCATCGGCTAGCCAGG

TTCCTGGTCAATCTGATTCTCCTGTTTCCGGTGGCGGTCTCAGTCAAAATGGAAAGTTTAGCTATGGTTA

TGCAAGCTCGCCTGGGAAGCGATCTTCCATGGAGGATTTTTATGAGACAAGGATAGATGGTGTTAACGGA

GAGATAATTGGTCTGTTCGGAGTTTTTGATGGTCATGGTGGTGCTCGTGCTGCAGAATATGTTAAAGAAA

ATCTCTTTAATTATCTCATCAGTCATCCAAAATTTATATCTGATACAAAATTGGCAATAGACTCCGAGTT

TCTAAAATCAGAAAATAATCAACACAGAGATGCTGGATCCACAGCTTCCACAGCAATCCTTGTCGGTGAT

CGCTTACTTGTAGCAAATGTTGGGGATTCCAGAGCTGTAATCAGCAGGGGTGGCAATGCTATTGCTGTAT

CTCGTGACCACAAGCCAGACCAAACAGATGAGCGCCAACGGATTGAAGATGCTGGAGGATTTGTTATGTG

GGCAGGAACTTGGAGAGTGGGAGGCGTGCTTGCTGTTTCTCGTGCTTTTGGTGATAGACTGTTGAAGCAG

TATGTTGTTGCTGATCCTGAAATTCAGGAAGAGAAGGTTGATAGCTCACTTGAATTTCTTATCCTCGCTA

GTGATGGGCTATGGGATGTTGTCACAAACGAGGAAGCTGTTGGGATGGTCAAGCCAATTCAGGATCCGGA

AGAGGCAGCAAAAAGATTAATGCAAGAAGCTTACCAGAGGGGCAGTGCTGATAATATTACTACTGTTGTT

GTTCGATTTCTGGAAGATCAAGTTGGAACCTCTATCAGCGGCTCTGGTTAGGTTAATTTCTATTAGAATA

TTTAGACTTTCTCAACACAGATGGATGAATGAAAAACCCCTGAAGACGAGTTTAGTGTGTTTTAAGTATT

GGAAACATTAGTTGCGTCAAATATAGGTTATGTACCTTCAAACACCGGAAGTGGTTTCTTACCTTAGAAA

TTTGCATCTTGCTGCTGGTATGTTCTTTATATGGTTGGTCTTGAGTTATACTCCGTTTAGTTTTATAGGT

CCCAAGGTGC

>MSTRG.2256.4 gene=MSTRG.2256

AGAGGGACGGAGGGAGTAATAAAGAAACAAAGAAAAGAAGAAAAGCAGCGAGAAGCACGCAGTAGCTACA

CAGTGTATGGTTTGATTTGATCATCTCATTTTGTTTGAAAACCTAATTAGAGCTACCGCAATTATTAAAT

CCACTAGTGGTGTGTATTTGTTGTTGGTGATTGCTATGGGATACTTGAATTCTGTTGTGCCATCGGCTAG

CCAGGTTCCTGGTCAATCTGATTCTCCTGTTTCCGGTGGCGGTCTCAGTCAAAATGGAAAGTTTAGCTAT

GGTTATGCAAGCTCGCCTGGGAAGCGATCTTCCATGGAGGATTTTTATGAGACAAGGATAGATGGTGTTA

ACGGAGAGATAATTGGTCTGTTCGGAGTTTTTGATGGTCATGGTGGTGCTCGTGCTGCAGAATATGTTAA

AGAAAATCTCTTTAATTATCTCATCAGTCATCCAAAATTTATATCTGATACAAAATTGGCAATAGCGGAT

GCATATAACCATACAGACTCCGAGTTTCTAAAATCAGAAAATAATCAACACAGAGATGCTGGATCCACAG

CTTCCACAGCAATCCTTGTCGGTGATCGCTTACTTGTAGCAAATGTTGGGGATTCCAGAGCTGTAATCAG

CAGGGGTGGCAATGCTATTGCTGTATCTCGTGACCACAAGCCAGACCAAACAGATGAGCGCCAACGGATT

GAAGATGCTGGAGGATTTGTTATGTGGGCAGGAACTTGGAGAGTGGGAGGCGTGCTTGCTGTTTCTCGTG

CTTTTGGTGATAGACTGTTGAAGCAGTATGTTGTTGCTGATCCTGAAATTCAGGAAGAGAAGGTTGATAG

CTCACTTGAATTTCTTATCCTCGCTAGTGATGGGCTATGGGATGTTGTCACAAACGAGGAAGCTGTTGGG

ATGGTCAAGCCAATTCAGGATCCGGAAGAGGCAGCAAAAAGATTAATGCAAGAAGCTTACCAGAGGGGCA

GTGCTGATAATATTACTACTGTTGTTGTTCGATTTCTGGAAGATCAAGTTGGAACCTCTATCAGCGGCTC

TGGTTAGGTTAATTTCTATTAGAATATTTAGACTTTCTCAACACAGATGGATGAATGAAAAACCCCTGAA

GACGAGTTTAGTGTGTTTTAAGTATTGGAAACATTAGTTGCGTCAAATATAGGTTATGTACCTTCAAACA

CCGGAAGTGGTTTCTTACCTTAGAAATTTGCATCTTGCTGCTGGTATGTTCTTTATATGGTTGGTCTTGA

GTTATACTCCGTTTAGTTTTATAGGTCCCAAGGTGC

>MSTRG.2259.1 gene=MSTRG.2259

TCCAGCTTGTGAAATCAAAATTAAAAATACTCACTATATACAAGACATATTGCTAGTAGTTGCAGGCAGC

TGCAAGTCTGCAACGCCAATGGCAACCATGGAAAAGTGCTCCACCTCTTAATAAGAGGGCACTACGCCGT

AAGTGGCCGTTGAAATGAGAGCCTCCACATCATCAATTTCTGAGCGCTTAGCGAATCTACCCTTGATTCT

GGGCCTGGTCTCTGCATAAGCCTTTCTTGAAGCATATCTGATTGTTTTCTCAAATTTCCGGTTCTTTCTT

TTCTCTCTGTACCTCATAACTCTAGCTTCTCTATCGAGTCCGGAGACAGGATTAGGAAGGGGTTCTGAGT

TGGGATGTTGGGTGATATTGTTTGATACATCTGTCATGGCATTGTGATCAGGAACTAGGCCAACATCCAG

TGATGAGGATGATACACTTTGGCTGATGGACTGAGAGTTGAAG

>MSTRG.2261.1 gene=MSTRG.2261

TTCAAATATGTGAGAGCAGGATACGTAGGATTGTAGAGTTGTCTCCGGTGAAGGCGCAAGTTTTGTGGCG

CCAAAGGCCAAACAGAAACAGATTATTCGTTTATATTACAAAACGCTTCAGAGAGACATTTGGGTGCTGT

GATGCCAACATAACAAATGAGTAGTATTCGAAAGGATGATGACATTTGGCTTGATCCATAGCAGTAAGGA

TTATTTCGTGTTCCTTGCTATCCTTGGCTAAAGTGGGGTCGATCTCCTGAACGCTCACTAGTAGCTGAAG

AATACTTTTCTCTTAGGACAAATGCAAGATAGTATATTTGTGGACTGATGATTCACACTAGCGAAAGGAG

TAGTGTCATATTGTATCTGTTTGTTCAATGTGTACTATTCAACTATATTTGGCTCTTCTATAATTGACCT

GCACTTTTAGTTTATAATCAAAATGAGTTTGAGAACGACAAAACTCCAACTCTTCTTGAAATTTCAGGGT

AGGTTCTATACTCTAGAATAAACTTGCACACAGACTCT

>MSTRG.2263.1 gene=MSTRG.2263

CAAACAATCAAACCCTAATAATCAGACATCCGCCTCTTTCATTCTAATTTTCCACTCTCAATTTCCACGC

GGCTGTACTAGGGGCGGCAATCAGACCTCCGCCTCTTCATTATTTCGCATCCACAAATCGAATCTAGTAT

CAAATTTGCAAGCCAAGCGCATCAAATCCTCAAATCGACTCAAATCCAAGCCAAGCGCATCAAATCTTGA

AAGATTAATGGAGAGTGAAAGCGAAAATCAAAACGAAAATGTTGATGAGCAAGAGCAGCCCGAGATTGAG

CCTCCTGCTGAACAAAAAACTGATGACGATGTTGTAGAGGTCGAGATGTCTAACGCATCAGCAATGTCAA

AGAGGCGACTAAGTGCCGATTTTTGGAAAGAGTTTAATATGCTGCCTCCGGGGCCGGATAAGAAGCAATG

GTGTTCGTGTAAAAGATGTGGTGCCACATATGCTTGTCCGAGTACCTATGGGACAGGTAATTTGCATAAA

CACTTAGAGAGGTGTAAGCGGAGAGATACGATAGATATTGGGCAGTTGCTTATAAGTTGTGATCATGGGT

CTGTTTTTTTAGGAACTAAGAAGTTTGATCAACAGAAATTTAGGGAGTTATTAGTTGTTGCTATAATTAT

GCATTGCCAACCATATCAATTCGTAGAATATGAGGGTTTTCGAGCTGTGTTTCGTTATTTAAATGAGAAT

GTCCAGACAATCTCAAGAAATACTTGCAAATCTAATGTGAATAAGTTGTATGCTAGAGAGAAAGAAAGGC

TAAAATCTTTGTTGCAATCATCTCCGGGTAGAATTTTTTTGACATCAGATTTATGGTCTTCTATAATTTT

AGATGGTTATTTGTCTTTAACTGCACATTTTGTTGACAAAGATTGAAATTTGCAAAAAAGTGTACTAAAT

TACACTCTAATGTCACATCCTCATACGGGTGTGAATTTAAGTGAAAAGATTTACTCATTACTTTGTGAGT

GGGGTATTGAGCATAAAGTGTTTGCAATGACTTTGGATAATGCGAGTGCAAATATTTGCTTTATTAATTT

GCTGATTTCTCAATTAAATATAAGAAATTTGTTGGTGAGTGATGGTGATTATGTGCATATACGTTGTTGT

GCACATATCTTGAATCTAGTTGTACAAGATGGTTTGAAGGTGATTGGTGAATGCCTTGAAAATATTAGAG

AGACTATTAAGTATTTGAAAGGGTCACAAGTGACATAAACAAAGTTTATTGAATGTGTTAAAAATATGAC

AATTGAGCTTAAAAAGGGGTTAAGACAAGATTGTTCAACAAGATGAAATTCCACTTATGTGATGCTTGAT

GAGGCTTTTATATTATCGTCGTGCTTTTTGCCATTTGGAGCTTTGTGATTCAAGTTACACATATTGCCCT

TAGAAGGATGAGTGGGACAAAGCGGAAAAGATTTGTAAGTTTTTTAGTGTCTTTTCTGAGATAACTCAGT

TATTTTCTGGAAGTAAGTATCCTACTTCGAATTTGTACTTCCCGTTGGTTGCAATTGCTCATTTAACTTT

AAAGGATAAGTTGAATAGTGAGGATTATTTTATGAGTTCCATGGCAGCTAAGATGTATGAAAAATTTTCT

AAATATTGGGCTGATTTTAGCACCATCTTGGCTATTACTTGCATCCTTGATCCTCGATACAAGCTGAGTG

ATTTCTTTTATAAGAAATTTTATGGTGATGATTCCTTGCAATTCAGGGATCTTAAAAACAAGCTCTTTTC

ACTTTTTAACAAGTATTTTGTTAAGGTGTCTTCGGGTGCAAGTGCAGGGACCTCAACTCAAAGACGAGGT

GGCCTGAATATGTGTGAAAATATTTTTGCTAACGAGGCAGAAGATGTAATGAAGACTTATTCCAGGATCC

ACCCAATTCATCCCAGGCGTGATTGAGTCCCACCAAAGCTACAAACAAGTATCGTGATGTCGTGCATGCT

CCAAATAGGCCCAAGAGCTCCCCATAGACCTCAAACGATCGCACCAGGTGCCACTCTAACTCCAGTGAGG

CTTGCCCCCGTTTCAAAAGACCCATCTCCAGATTCATTGCTAATCCCGCAATCCTTTTCATTCTTTTAAT

AGCATTAGATTAGCATTGTTTGTCATTTGATTCCTACAATATTCACTTTCTGTGTATCCATCACTAAAAG

TATGCTTACAAATAACTTTAAGATTTTATTTGACCAATTTTTATAGCTATA

>MSTRG.2277.1 gene=MSTRG.2277

TTAGTATGTCGATTACGTGGTTTAATGGCTTCTTCACGTTTTGAGATCATCATCGCCGTCCAGATTCGCC

ATTACAAGTGAGATGGATAACACCGTTTTTTGGTTTGTATTGGGGATAACAAAGCTACCTAGGATTTAAG

CTATTTGCTGCCGTTGATGGAACTAATCGGATATGAGCCAATCACCACCGCAACCATTCAGGATAATTTG

GAAGCGAGCTGGTTCGTGGCTTATGGGTGTTCTTTTCTCATTCATTTAAGCTTTATTGTCTAAGCGTCCG

GAGTTGTGCTTGCATGACTTTCGGTTAGATGATAAACTTTCTTGAAAGAAAGATATCCCGGATGCGGTTT

ATTGTATTTCGATACAATTCATCTACGTATATGTAGACCTTACAAAAAAAAAAAAAAAAAACTTTC

>MSTRG.2278.1 gene=MSTRG.2278

GGGATATGGGCTTAGCAAGGCAATGGTCCAGAATGTTCTGCACGTGACTACTAGGAAAGAAAGAGCTATA

AATACGAGGAGGTGTGTTCCATATCCTAGTACTATTATCGTTCCCTGAACTCCATACATATTCATCAGTA

TAAGGAATACATAATTTTTGACAAATGGGCAAGAGGAAGTCGAGGGCCAAGCCAGCACCCAAGAAAAGGA

TGGACAAACTGGATACAGTATTTAGCTGCCCATTCTGTAATCATGGAAGTAGCGTGGAATGCCGTATCGA

TATGAAGAATCTGATTGGTGAGGCTTCATGCAGGATCTGTCAAGAAAGTTTTAGCACAACTGTTACTGCT

CTGACCGAAGCAATCGATATATACAGTGAGTGGATCGATGAATGTGAACGTGTGAACAACCCTGAGGAAG

ATGGTGCAAAAAATCTTGAAGATGATGATGAGGAGGAGGCTTATGAGAATGTTTAGCTGCTGTCTGATGA

TATCAAGTTGTAATTTCCAAATGATCCTAATTTCCAAATGATCCTAACAGCGGTTGGGCATGTCAGTCTT

TGTTCTCTTAGATGTCATAGTAAAGACATAGGCAAATATACTTGTGTATGTTTTCTGACTTTTTATGGAT

GGTTGTGTTTCTGACATAAATCTATCTCCTAACCATTCTACATGTTGCTGTTAATTTGTTGCATGCTGCT

TGTGAAAATGTCTTGGAGTGCTATACATCCTTGGAACTCTCAGCAATAGTTTTTTTTTTTTTAAAGTTTT

CCTTCTTAAAAGGCTTATCCTTCTCCCCCTGCTA

>MSTRG.2279.1 gene=MSTRG.2279

GCCAAGTCCATCTCGAACAGATAAAATTCCTTGACATCCATCGTGGCCTCATACAAACTATTATATCCTT

TATCACTCGACTTGTCTTTCTTCTCACCGTTGCTCAACGATTCATATTTCTCGCCCAATTTGTCCTTTTC

ACTATTTCCCCACAATCCAAGTCTCTTCTTCTCATCACACTTTTTTACTGACTGTGAAATCACGCGATTC

GATTCAATAAACAGGTCAGGATCATCCCTAAACAAATCCTTACAAATCTGATCCACCTGACAATAACTAA

TCCTCTTCTCATAATACGCCAAAACACTTTCAGTAAACTCAATCAAAACACGAAACGCAACATCATTCTT

CTCCACACACCGATCACGAATCCTATCGAACAATTCGCTAGAAGATTCACATAAAAACCATGACTTATCA

GCCGTGCAGAGCCTCTTGCTGGGAGGATGATCTTCATCACGAATCACGTGATCATCGTCGTCGTCGTTCT

CGTTGTTGTTGGCAGCAGAGAATTCAGACGTGTTGGGCAAGAAAAGCTGGAGAGCGTGGAGAAGGTCAGG

ATGATCGTCGGCGAAAAGAGAGTTGAGTTTGGAAATGAGACGAGTATGATCTTGTTGCAGCTTCAGACTA

GCGAGAACTTGGGCAAACTCGTTGAATTTAGAAGGTTGGTTAGTGAATCTCCGTTTCACAACATAAACAA

AATCGATGGCTTCTTGTTGAAGGTCTTTCATGGTGGTCTATGTCGTAAGTTAACGCTGAGTATGTGTTGG

TATATATACATAACTCTGGCTACTCCCATCAAACCTAGGAGTGGAGTTTGCTTTCTCACTTCGAATCATA

TTGGGATTTGTAAATTGTGATACAACGGCTCCTTTTCTCTGTTTTTTTTGAGTTTACCCAGAAGCATGAA

TATGTTCAAGTTAATATGACGTTCCCATCAGTTTTTTTTTTAA

>MSTRG.2285.2 gene=MSTRG.2285

GTTGGTTCCTCGTTCCTCGTATTCAAAGGGGAACTTGAGCTTCACTTGCAGGCCGCAGCAGATCTACATC

ACTGCACCATTTGTCTCTGTACCAGGTTAATGATATAAAACAAAATTCAAATTTAATGTACAATATGGAA

TATGCACTTGGCAATTTATGAAGTTTCTGCAACACGCTTAGGAATAGTTTCTCCTCAGACTCCAATTACT

GTTTTGCTGATTAGTTCACTAACACAAAATGGCACTGAGAATGGTTGCAAGTAATATGAGAACTACGCTT

TTGCCAAGTCATCGACATTCAGAAAAGAAATGCTATCCACACTCAGACCAAGACAGAACTTTCACTTGTT

CGAAGAGCAATTAAGGTGAAGCAGGATCGGGAACGACTCACTCCGGAATATATGTTTCACATCATGCTTC

AAGTTTTGGTGAGTTTCTTACATTTAAAGTCGTAGTTTTTACTTCTCATATAATATGTCAAATTCAGAAC

TTAATGTAATCCTTTATTCTTTTAGATACTTGTAGAAAATTATAGCTAGAGAGAGATCGATTCAGAGCGA

GAGATAAAATTGTATTAACAAAGCTAAAAAATCACAGACTTTACACCTTTGTATAAGAAAGTGTGGCCAA

GTATATCATAGTAAATAACTAAACTAACCCTTTCCTTTAACTAACTTGACACCTAATAGTAGAATTATTC

AGGAGCTTTATGCTTAATAGCCTCCTTTAAAATGGAGGGTACAGACTCCCAATTTGCCAAGTAAAATTAA

TAAAATGAGCTGCGGGTAAGGATTTGGTAAACCAGTCTGCAATTGGAGCTTGGTTGATAGGTAAGTTGGT

GTAATGAATCCAGACTTGACCTCACCACGAACTAGGTGACAATCTAACTCGATATGCTTGGTATGGTCAC

GAAGAACAGAATTTTATGCATTGTGGATTGCAGAAGTATCATCACAAAATAAAGTTAATGGTGTACGAAT

TGTTATTGATAATTCCTGCAAGAGATTGAATAGCCAAGTTGCTTTACAGACGGCATTAGCCATGGAGCGA

TATTCAGCATTTCAGTCTCAGCTGCAGATCATGCTACAATGTTCTGTTTCTTGTATATCTAGGACAATAA

AGAATTGCCCATCATAGCCAGTAACTAGAAATGGATTGTTGAGTCATGTGGCTTTTTCCCAGTCATCATC

ACTGTAAATAGACAAGCAAAGAGTATTGGGGGCCCTTGGTCTGGAGATTTCTTGATATAATGTAGCAGTT

TTAAAACAGCATTCAAGTGAACTGTAGTCGGATTGGCCATATCTTGACTCAAAATGTGTGTTGCAACATC

TGGTCTAGTAATCATGAGATAAATACCATCAGTCTGTAAGAACTACCATCGCTCAAAGGAGTTTCAGCAG

TATTTGTAAAAGATTGTTTAGATTTCATAGGTATTTTACCTGGTTTTGAGGCTAATAAGCCAACAGAAGG

AGGTTTAGAGCATACTTCCTTTGATGGATGTAAACAGACCGAGTGATTTCCAGTCCCAGCAAGTAATAAA

GATGCCCTAGTAAGAATTATGTCATCGACCTAGACTAACAATAATATGAAGGAAGTAGCAGTTTAATGTA

TGAACATGAACTTTCACACCTAGAGTGATAGAAACCATAAGATATTACCTCAGTTTTGAATTTTGTAAAC

CACTCTCTTGGTGATGGTTTTAAATCGTAAATTGATCTAACAAGTTTGCAGACAGGGTGATGTCCTTTAA

ATCCAGCTGGAACAGACACTTCAGGTGGCAACCTTTAAAAACCTCCTCATGTAAATCACCATATTTGAAC

GCATTGTTTACATCCAGTTGTGTGATGCGCCCACCTTTTATAGCAGCAAGAGACAAAATTTGATGAACCG

TGGTCATTTTGCCCGTCGTTGCAAACGTTTGAAAGTAGTCTATATACCAGCTATTTGTGTAAATCCACGA

GCAACAAGACGAGCTTTATAATGATCTAAAGACCTGCCTGCCTTGTATCTCTCCTTAAACAACCACTTAC

ATCTCACTACTTTCTTCTTCCTTGTCAGGAGGCAAATCTGTGATGATCAAGGTGTTGCTAGCCTCCATTA

CTGCCAACTCTGTATGCATGGGTTTGCAACAATGAGGATCTTGTACTGCTTCATGATAGTATGTAGGTTC

AAAGATTTTGTTGGAATTGGCTATAAATTTTTGGTAGTGATCAGAAAATATGACATGAATCTACGTAAGT

GAGATAACCAAAGTACCAAACACAACATTAGAATTTGTAAGCCAATTGTACCGTAATCCTGTGTTTGTCA

TTGCTTCTCAACTACTAGGTCAGCCTCAGCTCAAATTCTGGCATCTAGTGTTAAGAATATAGGTGTCCCG

CCTCCACCAAGTAGAATTCTGTGGTGGTACACCCTGCACTGTTATAGTGCAGCTTTCCGAGTACATTGCA

AGTATTTGCCAATAATGTTCTATATATACTCTGTAGAATATTCTTCTAGTTGCTTCTCCCATTTCCAAGT

GTCTTATATAGAGATCTGCTACATTCCATCATATTCGAAAATGCCCTAGACAGATCTGGACATCCCTAGA

ATTTAGGTAGAAAATAACCTGTTCAATAAAAAGGAGGGGTGCCCTTATCAGCCCTTGCACTCGTAGATGG

TTCTGGTAGCCGGGACCCGGTGTCAGATGCCTGGCACCGTAGTGCCTTGAGGCTTTATAGGCTTTGCGTC

CCTGACACGCAGCCATGTTAGGCCATAGGTTAGTGATGTAGGGCATGCATTTTTTTGTTAACTGTTATAT

TACTCGATATTATTAATTTGTTTGTGTCAGGGTGGTTTGAATGGGGAAAAGATTTCTGACTTGAATTGAG

CCATGTGGAATATCCAGGTTTCGAGCACTGATATGATTATTACTCATTCTCTGTCGTCCAACCATAGGGG

AGCTAGTTGCAGGAGATCAGAATTCTAGCCAGTCCTTGGTCCGCAAAATTTCCTCACAGAATATCTTTCG

TTCAGTGATTGCAGAAGCTGGATCTTTAAATGTGAAATATGAAAATCTTGTTGGAGTGGTAGTTGCAGAT

CATTCTGCATTGAAGTTGGTAGAGGAGAAATTGCAAAGTTTATATGGGTTATATGATTCAACGATTGCAG

ATGCTCAACTTCAGAAAGTAGGATATGGAAATTCATTGCCATTCTTGGTTCAAGTTTGAATGCCTTAAGA

ACATACTTGATGTTGTATCAGTAGGAGATGGAAACACAATCTCTAGCTCTCCACAAAAAATTCTTAGGGT

TCGGCTTGTTGTGGGATGATCATTCCTTCTATATACAGCATTCGCGATGATTCTCATATCAAAAACAACT

GCTCCTTTATCGCGTCGGTCACTTTCTATATCTTTCTCCCATGCGGCTCCTGCGATGTTCTTTTGATGCG

CATTTGGTATTTGAAACTTGAAACTGATACTCCCTCTTTCAGTTTTTGCCAATACAAAAGGATTCACATA

ATTGGTGTCCAGAAGATTATCTTCATAGACCAATGCAGCTTCAACGTACTCATATTGAACTTCATACCGC

TCGCTTGGATTTCTGACACTCATGTTTGCGCTCAATGTCGCGTTTATCCCGGTGTTATCAAGTCTGAAAA

CTGTCATGTTTAATGATTGTGCGTAAAATACAGGAATATCAGAGCCATAAATAAGCCAGGCTAGGAGGCT

TAGTACTGTTACTCCAAAAAATATCATGATCAAGACCACAAGAAACACACGAGCAAATTTGTAACCCGGG

CTTCTATGAGGAGGTCCGATTAATACAAAGTTTTGTGCAGTGGTGGGGTAATAGCCGGGTGGTGTAGGTT

TCACGGGACCATAATGGTAATCAGATTTAGGGTAGACAGGATATGGAGGATGGGCTTGATATGGTGGTGG

AAAAGGATATCCCTGAGGTGGTGGTGGTGGTGGTGGTTGTGACAGCTCTGTTACAAATTTTTCAGACTTG

GGATCACCTTCCGAGGAGGAATCATCCTTAGTTTCTCCCTCTCCCATTCCATAGGCATCAAGAGAGAGAA

TAAAGATGAGAGATGAATCAATGTTATGTGAAGGTTGGAATTGTAGAAAGAAAGAGAGGAAGATGAGGTG

AAAATGTTAAGACAATTATGCGATGATATGCGTTTTTGGATAGTTGAGCTTTGTTTTGTTGAGCCATGCA

TGAAATACATTTACAAATCATTGGTCGAAAATGAGATGTAATTATGTAAAGAATGGTGTACATAGAATAG

AAGCAGGAGGACACATGGAGGTTTGGGGTATGTTCAGGTTCAAGTCACAAATACTTTTGTTCATAATTTC

AACAATTTCATCTCATAATTTTTATCTTAACCACAAAATGCATCTATTAGATCATATTCCATGCATCTTT

TTGGATCATACTCGGTCTAGGCCTCAAGGGTAGTTCGGATGGGATATGCTTGTGTTGAAGCTTGAGATTT

GTAAAAGAACTGTATTTTAAAAATTGAAATTTG

>MSTRG.2285.1 gene=MSTRG.2285

GGGAAGTGTGTGTTGGTTCCTCGTTCCTCGTATTCAAAGGGGAACTTGAGCTTCACTTGCAGGCCGCAGC

AGATCTACATCACTGCACCATTTGTCTCTGTACCAGAACACAAAATGGCACTGAGAATGGTTGCAAGTAA

TATGAGAACTACGCTTTTGCCAAGTCATCGACATTCAGAAAAGAAATGCTATCCACACTCAGACCAAGAC

AGAACTTTCACTTGTTCGAAGAGCAATTAAGGTGAAGCAGGATCGGGAACGACTCACTCCGGAATATATG

TTTCACATCATGCTTCAAGTTTTGGAGGCAAATCTGTGATGATCAAGGTGTTGCTAGCCTCCATTACTGC

CAACTCTGTATGCATGGGTTTGCAACAATGAGGATCTTGTACTGCTTCATGATAGTATGTAGGTTCAAAG

ATTTTGTTGGAATTGGCTATAAATTTTTGGTAGTGATCAGAAAATATGACATGAATCTACGTAAGTGAGA

TAACCAAAGTACCAAACACAACATTAGAATTTGTAAGCCAATTGTACCGTAATCCTGTGTTTGTCATTGC

TTCTCAACTACTAGGTCAGCCTCAGCTCAAATTCTGGCATCTAGTGTTAAGAATATAGGTGTCCCGCCTC

CACCAAGTAGAATTCTGTGGTGGTACACCCTGCACTGTTATAGTGCAGCTTTCCGAGTACATTGCAAGTA

TTTGCCAATAATGTTCTATATATACTCTGTAGAATATTCTTCTAGTTGCTTCTCCCATTTCCAAGTGTCT

TATATAGAGATCTGCTACATTCCATCATATTCGAAAATGCCCTAGACAGATCTGGACATCCCTAGAATTT

AGGTAGAAAATAACCTGTTCAATAAAAAGGAGGGGTGCCCTTATCAGCCCTTGCACTCGTAGATGGTTCT

GGTAGCCGGGACCCGGTGTCAGATGCCTGGCACCGTAGTGCCTTGAGGCTTTATAGGCTTTGCGTCCCTG

ACACGCAGCCATGTTAGGCCATAGGTTAGTGATGTAGGGCATGCATTTTTTTGTTAACTGTTATATTACT

CGATATTATTAATTTGTTTGTGTCAGGGTGGTTTGAATGGGGAAAAGATTTCTGACTTGAATTGAGCCAT

GTGGAATATCCAGGTTTCGAGCACTGATATGATTATTACTCATTCTCTGTCGTCCAACCATAGGGGAGCT

AGTTGCAGGAGATCAGAATTCTAGCCAGTCCTTGGTCCGCAAAATTTCCTCACAGAATATCTTTCGTTCA

GTGATTGCAGAAGCTGGATCTTTAAATGTGAAATATGAAAATCTTGTTGGAGTGGTAGTTGCAGATCATT

CTGCATTGAAGTTGGTAGAGGAGAAATTGCAAAGTTTATATGGGTTATATGATTCAACGATTGCAGATGC

TCAACTTCAGAAAGTAGGATATGGAAATTCATTGCCATTCTTGGTTCAAGTTTGAATGCCTTAAGAACAT

ACTTGATGGAGATGGAAACACAATCTCTAGCTCTCCACAAAAAATTCTTAGGGTTCGGCTTGTTGTGGGA

TGATCATTCCTTCTATATACAGCATTCGCGATGATTCTCATATCAAAAACAACTGCTCCTTTATCGCGTC

GGTCACTTTCTATATCTTTCTCCCATGCGGCTCCTGCGATGTTCTTTTGATGCGCATTTGGTATTTGAAA

CTTGAAACTGATACTCCCTCTTTCAGTTTTTGCCAATACAAAAGGATTCACATAATTGGTGTCCAGAAGA

TTATCTTCATAGACCAATGCAGCTTCAACGTACTCATATTGAACTTCATACCGCTCGCTTGGATTTCTGA

CACTCATGTTTGCGCTCAATGTCGCGTTTATCCCGGTGTTATCAAGTCTGAAAACTGTCATGTTTAATGA

TTGTGCGTAAAATACAGGAATATCAGAGCCATAAATAAGCCAGGCTAGGAGGCTTAGTACTGTTACTCCA

AAAAATATCATGATCAAGACCACAAGAAACACACGAGCAAATTTGTAACCCGGGCTTCTATGAGGAGGTC

CGATTAATACAAAGTTTTGTGCAGTGGTGGGGTAATAGCCGGGTGGTGTAGGTTTCACGGGACCATAATG

GTAATCAGATTTAGGGTAGACAGGATATGGAGGATGGGCTTGATATGGTGGTGGAAAAGGATATCCCTGA

GGTGGTGGTGGTGGTGGTGGTTGTGACAGCTCTGTTACAAATTTTTCAGACTTGGGATCACCTTCCGAGG

AGGAATCATCCTTAGTTTCTCCCTCTCCCATTCCATAGGCATCAAGAGAGAGAATAAAGATGAGAGATGA

ATCAATGTTATGTGAAGGTTGGAATTGTAGAAAGAAAGAGAGGAAGATGAGGTGAAAATGTTAAGACAAT

TATGCGATGATATGCGTTTTTGGATAGTTGAGCTTTGTTTTGTTGAGCCATGCATGAAATACATTTACAA

ATCATTGGTCGAAAATGAGATGTAATTATGTAAAGAATGGTGTACATAGAATAGAAGCAGGAGGACACAT

GGAGGTTTGGGGTATGTTCAGGTTCAAGTCACAAATACTTTTGTTCATAATTTCAACAATTTCATCTCAT

AATTTTTATCTTAACCACAAAATGCATCTATTAGATCATATTCCATGCATCTTTTTGGATCATACTCGGT

CTAGGCCTCAAGGGTAGTTCGGATGGGATATGCTTGTGTTGAAGCTTGAGATTTGTAAAAGAACTGTATT

TTAAAAATTGAAATTTG

>MSTRG.2285.3 gene=MSTRG.2285

GGGAAGTGTGTGTTGGTTCCTCGTTCCTCGTATTCAAAGGGGAACTTGAGCTTCACTTGCAGGCCGCAGC

AGATCTACATCACTGCACCATTTGTCTCTGTACCAGAACACAAAATGGCACTGAGAATGGTTGCAAGTAA

TATGAGAACTACGCTTTTGCCAAGTCATCGACATTCAGAAAAGAAATGCTATCCACACTCAGACCAAGAC

AGAACTTTCACTTGTTCGAAGAGCAATTAAGGTGAAGCAGGATCGGGAACGACTCACTCCGGAATATATG

TTTCACATCATGCTTCAAGTTTTGGGTGGTTTGAATGGGGAAAAGATTTCTGACTTGAATTGAGCCATGT

GGAATATCCAGGTTTCGAGCACTGATATGATTATTACTCATTCTCTGTCGTCCAACCATAGGGGAGCTAG

TTGCAGGAGATCAGAATTCTAGCCAGTCCTTGGTCCGCAAAATTTCCTCACAGAATATCTTTCGTTCAGT

GATTGCAGAAGCTGGATCTTTAAATGTGAAATATGAAAATCTTGTTGGAGTGGTAGTTGCAGATCATTCT

GCATTGAAGTTGGTAGAGGAGAAATTGCAAAGTTTATATGGGTTATATGATTCAACGATTGCAGATGCTC

AACTTCAGAAAGTAGGATATGGAAATTCATTGCCATTCTTGGTTCAAGTTTGAATGCCTTAAGAACATAC

TTGATGTTGTATCAGTAGGAGATGGAAACACAATCTCTAGCTCTCCACAAAAAATTCTTAGGGTTCGGCT

TGTTGTGGGATGATCATTCCTTCTATATACAGCATTCGCGATGATTCTCATATCAAAAACAACTGCTCCT

TTATCGCGTCGGTCACTTTCTATATCTTTCTCCCATGCGGCTCCTGCGATGTTCTTTTGATGCGCATTTG

GTATTTGAAACTTGAAACTGATACTCCCTCTTTCAGTTTTTGCCAATACAAAAGGATTCACATAATTGGT

GTCCAGAAGATTATCTTCATAGACCAATGCAGCTTCAACGTACTCATATTGAACTTCATACCGCTCGCTT

GGATTTCTGACACTCATGTTTGCGCTCAATGTCGCGTTTATCCCGGTGTTATCAAGTCTGAAAACTGTCA

TGTTTAATGATTGTGCGTAAAATACAGGAATATCAGAGCCATAAATAAGCCAGGCTAGGAGGCTTAGTAC

TGTTACTCCAAAAAATATCATGATCAAGACCACAAGAAACACACGAGCAAATTTGTAACCCGGGCTTCTA

TGAGGAGGTCCGATTAATACAAAGTTTTGTGCAGTGGTGGGGTAATAGCCGGGTGGTGTAGGTTTCACGG

GACCATAATGGTAATCAGATTTAGGGTAGACAGGATATGGAGGATGGGCTTGATATGGTGGTGGAAAAGG

ATATCCCTGAGGTGGTGGTGGTGGTGGTGGTTGTGACAGCTCTGTTACAAATTTTTCAGACTTGGGATCA

CCTTCCGAGGAGGAATCATCCTTAGTTTCTCCCTCTCCCATTCCATAGGCATCAAGAGAGAGAATAAAGA

TGAGAGATGAATCAATGTTATGTGAAGGTTGGAATTGTAGAAAGAAAGAGAGGAAGATGAGGTGAAAATG

TTAAGACAATTATGCGATGATATGCGTTTTTGGATAGTTGAGCTTTGTTTTGTTGAGCCATGCATGAAAT

ACATTTACAAATCATTGGTCGAAAATGAGATGTAATTATGTAAAGAATGGTGTACATAGAATAGAAGCAG

GAGGACACATGGAGGTTTGGGGTATGTTCAGGTTCAAGTCACAAATACTTTTGTTCATAATTTCAACAAT

TTCATCTCATAATTTTTATCTTAACCACAAAATGCATCTATTAGATCATATTCCATGCATCTTTTTGGAT

CATACTCGGTCTAGGCCTCAAGGGTAGTTCGGATGGGATATGCTTGTGTTGAAGCTTGAGATTTGTAAAA

G

>MSTRG.2285.4 gene=MSTRG.2285

CACCTTTGTATAAGAAAGTGTGGCCAAGTATATCATAGTAAATAACTAAACTAACCCTTTCCTTTAACTA

ACTTGACACCTAATAGTAGAATTATTCAGGAGCTTTATGCTTAATAGCCTCCTTTAAAATGGAGGGTACA

GACTCCCAATTTGCCAAGTAAAATTAATAAAATGAGCTGCGGGTAAGGATTTGGTAAACCAGTCTGCAAT

TGGAGCTTGGTTGATAGGTAAGTTGGTGTAATGAATCCAGACTTGACCTCACCACGAACTAGGTGACAAT

CTAACTCGATATGCTTGGTATGGTCACGAAGAACAGAATTTTATGCATTGTGGATTGCAGAAGTATCATC

ACAAAATAAAGTTAATGGTGTACGAATTGTTATTGATAATTCCTGCAAGAGATTGAATAGCCAAGTTGCT

TTACAGACGGCATTAGCCATGGAGCGATATTCAGCATTTCAGTCTCAGCTGCAGATCATGCTACAATGTT

CTGTTTCTTGTATATCTAGGACAATAAAGAATTGCCCATCATAGCCAGTAACTAGAAATGGATTGTTGAG

TCATGTGGCTTTTTCCCAGTCATCATCACTGTAAATAGACAAGCAAAGAGTATTGGGGGCCCTTGGTCTG

GAGATTTCTTGATATAATGTAGCAGTTTTAAAACAGCATTCAAGTGAACTGTAGTCGGATTGGCCATATC

TTGACTCAAAATGTGTGTTGCAACATCTGGTCTAGTAATCATGAGATAAATACCATCAGTCTGTAAGAAC

TACCATCGCTCAAAGGAGTTTCAGCAGTATTTGTAAAAGATTGTTTAGATTTCATAGGTATTTTACCTGG

TTTTGAGGCTAATAAGCCAACAGAAGGAGGTTTAGAGCATACTTCCTTTGATGGATGTAAACAGACCGAG

TGATTTCCAGTCCCAGCAAGTAATAAAGATGCCCTAGTAAGAATTATGTCATCGACCTAGACTAACAATA

ATATGAAGGAAGTAGCAGTTTAATGTATGAACATGAACTTTCACACCTAGAGTGATAGAAACCATAAGAT

ATTACCTCAGTTTTGAATTTTGTAAACCACTCTCTTGGTGATGGTTTTAAATCGTAAATTGATCTAACAA

GTTTGCAGACAGGGTGATGTCCTTTAAATCCAGCTGGAACAGACACTTCAGGTGGCAACCTTTAAAAACC

TCCTCATGTAAATCACCATATTTGAACGCATTGTTTACATCCAGTTGTGTGATGCGCCCACCTTTTATAG

CAGCAAGAGACAAAATTTGATGAACCGTGGTCATTTTGCCCGTCGTTGCAAACGTTTGAAAGTAGTCTAT

ATACCAGCTATTTGTGTAAATCCACGAGCAACAAGACGAGCTTTATAATGATCTAAAGACCTGCCTGCCT

TGTATCTCTCCTTAAACAACCACTTACATCTCACTACTTTCTTCTTCCTTGTCAGGAGGCAAATCTGTGA

TGATCAAGGTGTTGCTAGCCTCCATTACTGCCAACTCTGTATGCATGGGTTTGCAACAATGAGGATCTTG

TACTGCTTCATGATAGTATGTAGGTTCAAAGATTTTGTTGGAATTGGCTATAAATTTTTGGTAGTGATCA

GAAAATATGACATGAATCTACGTAAGTGAGATAACCAAAGTACCAAACACAACATTAGAATTTGTAAGCC

AATTGTACCGTAATCCTGTGTTTGTCATTGCTTCTCAACTACTAGGTCAGCCTCAGCTCAAATTCTGGCA

TCTAGTGTTAAGAATATAGGTGTCCCGCCTCCACCAAGTAGAATTCTGTGGTGGTACACCCTGCACTGTT

ATAGTGCAGCTTTCCGAGTACATTGCAAGTATTTGCCAATAATGTTCTATATATACTCTGTAGAATATTC

TTCTAGTTGCTTCTCCCATTTCCAAGTGTCTTATATAGAGATCTGCTACATTCCATCATATTCGAAAATG

CCCTAGACAGATCTGGACATCCCTAGAATTTAGGTAGAAAATAACCTGTTCAATAAAAAGGAGGGGTGCC

CTTATCAGCCCTTGCACTCGTAGATGGTTCTGGTAGCCGGGACCCGGTGTCAGATGCCTGGCACCGTAGT

GCCTTGAGGCTTTATAGGCTTTGCGTCCCTGACACGCAGCCATGTTAGGCCATAGGTTAGTGATGTAGGG

CATGCATTTTTTTGTTAACTGTTATATTACTCGATATTATTAATTTGTTTGTGTCAGGGTGGTTTGAATG

GGGAAAAGATTTCTGACTTGAATTGAGCCATGTGGAATATCCAGGTTTCGAGCACTGATATGATTATTAC

TCATTCTCTGTCGTCCAACCATAGGGGAGCTAGTTGCAGGAGATCAGAATTCTAGCCAGTCCTTGGTCCG

CAAAATTTCCTCACAGAATATCTTTCGTTCAGTGATTGCAGAAGCTGGATCTTTAAATGTGAAATATGAA

AATCTTGTTGGAGTGGTAGTTGCAGATCATTCTGCATTGAAGTTGGTAGAGGAGAAATTGCAAAGTTTAT

ATGTAAATTTCTAATATTTCCTATATTTAAGGGGTTATATGATTCAACGATTGCAGATGCTCAACTTCAG

AAAGTAGGATATGGAAATTCATTGCCATTCTTGGTTCAAGTTTGAATGCCTTAAGAACATACTTGATGGA

GATGGAAACACAATCTCTAGCTCTCCACAAAAAATTCTTAGGGTTCGGCTTGTTGTGGGATGATCATTCC

TTCTATATACAGCATTCGCGATGATTCTCATATCAAAAACAACTGCTCCTTTATCGCGTCGGTCACTTTC

TATATCTTTCTCCCATGCGGCTCCTGCGATGTTCTTTTGATGCGCATTTGGTATTTGAAACTTGAAACTG

ATACTCCCTCTTTCAGTTTTTGCCAATACAAAAGGATTCACATAATTGGTGTCCAGAAGATTATCTTCAT

AGACCAATGCAGCTTCAACGTACTCATATTGAACTTCATACCGCTCGCTTGGATTTCTGACACTCATGTT

TGCGCTCAATGTCGCGTTTATCCCGGTGTTATCAAGTCTGAAAACTGTCATGTTTAATGATTGTGCGTAA

AATACAGGAATATCAGAGCCATAAATAAGCCAGGCTAGGAGGCTTAGTACTGTTACTCCAAAAAATATCA

TGATCAAGACCACAAGAAACACACGAGCAAATTTGTAACCCGGGCTTCTATGAGGAGGTCCGATTAATAC

AAAGTTTTGTGCAGTGGTGGGGTAATAGCCGGGTGGTGTAGGTTTCACGGGACCATAATGGTAATCAGAT

TTAGGGTAGACAGGATATGGAGGATGGGCTTGATATGGTGGTGGAAAAGGATATCCCTGAGGTGGTGGTG

GTGGTGGTGGTTGTGACAGCTCTGTTACAAATTTTTCAGACTTGGGATCACCTTCCGAGGAGGAATCATC

CTTAGTTTCTCCCTCTCCCATTCCATAGGCATCAAGAGAGAGAATAAAGATGAGAGATGAATCAATGTTA

TGTGAAGGTTGGAATTGTAGAAAGAAAGAGAGGAAGATGAGGTGAAAATGTTAAGACAATTATGCGATGA

TATGCGTTTTTGGATAGTTGAGCTTTGTTTTGTTGAGCCATGCATGAAATACATTTACAAATCATTGGTC

GAAAATGAGATGTAATTATGTAAAGAATGGTGTACATAGAATAGAAGCAGGAGGACACATGGAGGTTTGG

GGTATGTTCAGGTTCAAGTCACAAATACTTTTGTTCATAATTTCAACAATTTCATCTCATAATTTTTATC

TTAACCACAAAATGCATCTATTAGATCATATTCCATGCATCTTTTTGGATCATACTCGGTCTAGGCCTCA

AGGGTAGTTCGGAT

>MSTRG.2285.5 gene=MSTRG.2285

GTGTGTTGGTTCCTCGTTCCTCGTATTCAAAGGGGAACTTGAGCTTCACTTGCAGGCCGCAGCAGATCTA

CATCACTGCACCATTTGTCTCTGTACCAGAACACAAAATGGCACTGAGAATGGTTGCAAGTAATATGAGA

ACTACGCTTTTGCCAAGTCATCGACATTCAGAAAAGAAATGCTATCCACACTCAGACCAAGACAGAACTT

TCACTTGTTCGAAGAGCAATTAAGGTGAAGCAGGATCGGGAACGACTCACTCCGGAATATATGTTTCACA

TCATGCTTCAAGTTTTGGAGGCAAATCTGTGATGATCAAGGTGTTGCTAGCCTCCATTACTGCCAACTCT

GTATGCATGGGTTTGCAACAATGAGGATCTTGTACTGCTTCATGATAGTATGTAGGTTCAAAGATTTTGT

TGGAATTGGCTATAAATTTTTGGTAGTGATCAGAAAATATGACATGAATCTACGTAAGTGAGATAACCAA

AGTACCAAACACAACATTAGAATTTGTAAGCCAATTGTACCGTAATCCTGTGTTTGTCATTGCTTCTCAA

CTACTAGGTCAGCCTCAGCTCAAATTCTGGCATCTAGTGTTAAGAATATAGGTGTCCCGCCTCCACCAAG

TAGAATTCTGTGGTGGTACACCCTGCACTGTTATAGTGCAGCTTTCCGAGTACATTGCAAGTATTTGCCA

ATAATGTTCTATATATACTCTGTAGAATATTCTTCTAGTTGCTTCTCCCATTTCCAAGTGTCTTATATAG

AGATCTGCTACATTCCATCATATTCGAAAATGCCCTAGACAGATCTGGACATCCCTAGAATTTAGGTAGA

AAATAACCTGTTCAATAAAAAGGAGGGGTGCCCTTATCAGCCCTTGCACTCGTAGATGGTTCTGGTAGCC

GGGACCCGGTGTCAGATGCCTGGCACCGTAGTGCCTTGAGGCTTTATAGGCTTTGCGTCCCTGACACGCA

GCCATGTTAGGCCATAGGTTAGTGATGTAGGGCATGCATTTTTTTGTTAACTGTTATATTACTCGATATT

ATTAATTTGTTTGTGTCAGGGTGGTTTGAATGGGGAAAAGATTTCTGACTTGAATTGAGCCATGTGGAAT

ATCCAGGTTTCGAGCACTGATATGATTATTACTCATTCTCTGTCGTCCAACCATAGGGGAGCTAGTTGCA

GGAGATCAGAATTCTAGCCAGTCCTTGGTCCGCAAAATTTCCTCACAGAATATCTTTCGTTCAGTGATTG

CAGAAGCTGGATCTTTAAATGTGAAATATGAAAATCTTGTTGGAGTGGTAGTTGCAGATCATTCTGCATT

GAAGTTGGTAGAGGAGAAATTGCAAAGTTTATATGGGTTATATGATTCAACGATTGCAGATGCTCAACTT

CAGAAAGTAGGATATGGAAATTCATTGCCATTCTTGGTTCAAGTTTGAATGCCTTAAGAACATACTTGAT

GTAGGAGATGGAAACACAATCTCTAGCTCTCCACAAAAAATTCTTAGGGTTCGGCTTGTTGTGGGATGAT

CATTCCTTCTATATACAGCATTCGCGATGATTCTCATATCAAAAACAACTGCTCCTTTATCGCGTCGGTC

ACTTTCTATATCTTTCTCCCATGCGGCTCCTGCGATGTTCTTTTGATGCGCATTTGGTATTTGAAACTTG

AAACTGATACTCCCTCTTTCAGTTTTTGCCAATACAAAAGGATTCACATAATTGGTGTCCAGAAGATTAT

CTTCATAGACCAATGCAGCTTCAACGTACTCATATTGAACTTCATACCGCTCGCTTGGATTTCTGACACT

CATGTTTGCGCTCAATGTCGCGTTTATCCCGGTGTTATCAAGTCTGAAAACTGTCATGTTTAATGATTGT

GCGTAAAATACAGGAATATCAGAGCCATAAATAAGCCAGGCTAGGAGGCTTAGTACTGTTACTCCAAAAA

ATATCATGATCAAGACCACAAGAAACACACGAGCAAATTTGTAACCCGGGCTTCTATGAGGAGGTCCGAT

TAATACAAAGTTTTGTGCAGTGGTGGGGTAATAGCCGGGTGGTGTAGGTTTCACGGGACCATAATGGTAA

TCAGATTTAGGGTAGACAGGATATGGAGGATGGGCTTGATATGGTGGTGGAAAAGGATATCCCTGAGGTG

GTGGTGGTGGTGGTGGTTGTGACAGCTCTGTTACAAATTTTTCAGACTTGGGATCACCTTCCGAGGAGGA

ATCATCCTTAGTTTCTCCCTCTCCCATTCCATAGGCATCAAGAGAGAGAATAAAGATGAGAGATGAATCA

ATGTTATGTGAAGGTTGGAATTGTAGAAAGAAAGAGAGGAAGATGAGGTGAAAATGTTAAGACAATTATG

CGATGATATGCGTTTTTGGATAGTTGAGCTTTGTTTTGTTGAGCCATGCATGAAATACATTTACAAATCA

TTGGTCGAAAATGAGATGTAATTATGTAAAGAATGGTGTACATAGAATAGAAGCAGGAGGACACATGGAG

GTTTGGGGTATGTTCAGGTTCAAGTCAC

>MSTRG.2285.6 gene=MSTRG.2285

CGTATTCAAAGGGGAACTTGAGCTTCACTTGCAGGCCGCAGCAGATCTACATCACTGCACCATTTGTCTC

TGTACCAGAACACAAAATGGCACTGAGAATGGTTGCAAGTAATATGAGAACTACGCTTTTGCCAAGTCAT

CGACATTCAGAAAAGAAATGCTATCCACACTCAGACCAAGACAGAACTTTCACTTGTTCGAAGAGCAATT

AAGGTGAAGCAGGATCGGGAACGACTCACTCCGGAATATATGTTTCACATCATGCTTCAAGTTTTGGGTG

GTTTGAATGGGGAAAAGATTTCTGACTTGAATTGAGCCATGTGGAATATCCAGGTTTCGAGCACTGATAT

GATTATTACTCATTCTCTGTCGTCCAACCATAGGGGAGCTAGTTGCAGGAGATCAGAATTCTAGCCAGTC

CTTGGTCCGCAAAATTTCCTCACAGAATATCTTTCGTTCAGTGATTGCAGAAGCTGGATCTTTAAATGTG

AAATATGAAAATCTTGTTGGAGTGGTAGTTGCAGATCATTCTGCATTGAAGTTGGTAGAGGAGAAATTGC

AAAGTTTATATGGGTTATATGATTCAACGATTGCAGATGCTCAACTTCAGAAAGTAGGATATGGAAATTC

ATTGCCATTCTTGGTTCAAGTTTGAATGCCTTAAGAACATACTTGATGGAGATGGAAACACAATCTCTAG

CTCTCCACAAAAAATTCTTAGGGTTCGGCTTGTTGTGGGATGATCATTCCTTCTATATACAGCATTCGCG

ATGATTCTCATATCAAAAACAACTGCTCCTTTATCGCGTCGGTCACTTTCTATATCTTTCTCCCATGCGG

CTCCTGCGATGTTCTTTTGATGCGCATTTGGTATTTGAAACTTGAAACTGATACTCCCTCTTTCAGTTTT

TGCCAATACAAAAGGATTCACATAATTGGTGTCCAGAAGATTATCTTCATAGACCAATGCAGCTTCAACG

TACTCATATTGAACTTCATACCGCTCGCTTGGATTTCTGACACTCATGTTTGCGCTCAATGTCGCGTTTA

TCCCGGTGTTATCAAGTCTGAAAACTGTCATGTTTAATGATTGTGCGTAAAATACAGGAATATCAGAGCC

ATAAATAAGCCAGGCTAGGAGGCTTAGTACTGTTACTCCAAAAAATATCATGATCAAGACCACAAGAAAC

ACACGAGCAAATTTGTAACCCGGGCTTCTATGAGGAGGTCCGATTAATACAAAGTTTTGTGCAGTGGTGG

GGTAATAGCCGGGTGGTGTAGGTTTCACGGGACCATAATGGTAATCAGATTTAGGGTAGACAGGATATGG

AGGATGGGCTTGATATGGTGGTGGAAAAGGATATCCCTGAGGTGGTGGTGGTGGTGGTGGTTGTGACAGC

TCTGTTACAAATTTTTCAGACTTGGGATCACCTTCCGAGGAGGAATCATCCTTAGTTTCTCCCTCTCCCA

TTCCATAGGCATCAAGAGAGAGAATAAAGATGAGAGATGAATCAATGTTATGTGAAGGTTGGAATTGTAG

AAAGAAAGAGAGGAAGATGAGGTGAAAATGTTAAGACAATTATGCGATGATATGCGTTTTTGGATAGTTG

AGCTTTGTTTTGTTGAGCCATGCATGAAATACATTTACAAATCATTGGTCGAAAATGAGATGTAATTATG

TAAAGAATGGTGTACATAGAATAGAAGCAGGAGGACACATGGAGGTTTGGGGTATGTTCAGGTT

>MSTRG.2292.1 gene=MSTRG.2292

GACTACAGGAAGCTAAATTCAATGACTATAAAGAATAAGTTTCCAATTCCTCTAATTGAAGAACTGTTAG

ATGAGCTTCATGGGGCTAAGTACTTCACTAAGCTAGATCTAAGAGCTGGCTACCACCAAGTCAGGATGAA

AGAAAGGGATGTAGAAAAAACAGCTTTTAGAACTCATTTGGGGCACTATGAATTTACTGTCATGCCATTT

GGCCTCACCAATGCCCCTGCCACTTTTCAGAGTTTGATGAATGAAGTCTTCAGGCCCTATTTGAGGAGTT

TCACTCTGGTTTTTTTTTATGACATTTTGGTCTACTCTAAGACCAGGGAAGATCATCTGGTACACTTAAG

GAAGGTCTTTCAGCTAATGAGGCAACACACTTTATTTGCTAAGCTTTCTAAATGTGAATTTATGACTGCT

CAAGTTGCTTACTTAGGACATGTCATATCACAACAAGGGGTAGCTGTAGACCAAGCTAAAGTTGCTGACA

TGGCTTCCTGGCCTATACCTCAGAATATAAAGGCACTAAGGGGGTTTCTGGGCTTAACTGGATATTATAG

GAAATTTGTGAAGGATTATGGTACTATAGCCAGGCCTCTGACAGACTTACTGCAGAAAGACAATTTCCAC

TGGAATAAAGAAGCTACTGAGGCTTTCAGGGCTCTTAAGGAGGCTATGGCTAGTACTCCAGTACTGAGGC

TGCCAGACTTTAGTAAGGAGTTTACTATTGAGTCTGATGCTTGTAACTCTGGTATTGGTGCTGTGTTGAC

TCAAGAAGGACAGCCTCTGGCTTATTTCAGCAAGGCATTGGGTGCAAGGGGACAAGCAATGTCTACCTAT

GAAAAAGAACTAATGGCAGTAGTAGCAGCTGTGAAGAAGTGGTCTGCCTATCTCATGGACAAGCATTTTA

CTATCAAGACTGATCATTGGTCCTTGAAGTATTTGACTGAGCAAAAGATCAGTAATTTGCTGCAGCAAAA

GTGGATTAGCAAGTTATTAGGCTATGACTACACAATTGTGTATAAAAAAGGCAAAGACAACACTGTGGCT

GATGCTTTGTCCAGGAAATTTGAAGATTCTGCAAATGGAAGTGCTGGCAACCATGAGGACATGGCTGAAT

TTAAGGGGGAAGGAATGTCAGGATCAGAATGTCAGGATCAGAGCAATACTAGTAATAGCAAGTAGGATAG

GGGTAACTAAGTAAATTTCCATTCTGTTATGACAGCTGTGTTAGCAAGAGTCTATAAATAGTCAAGGTGT

GCTGTGAATTGCTATTATGCCTAATATGAAATACAACTTCTTCTCCAAGTTTCTCTCTCTCTCTTCTCTC

TAACTGATTTTCCCTCTCTTTCTCTCTCTAAACTCTGTCTCTCTCTCCTGATCTGATCTCTTTCTCTCTC

ATTTCTACATCTCTCTGCCCTAATTCATCTCTATCTAGCTATAATCTACTGTTCTGTTACTTGTTTTGCT

GCTATTTCAGTTCATAAACTCAGAAAACAC

>MSTRG.2293.1 gene=MSTRG.2293

ATCCCTCCATGTCCATTTTCACCATCTCTATCACCAACAAATCCATCCCACCTCCATCGCTCCATCCTGC

ATCGTCTGCACCTCTGCCGTCTCCAAAATATCTATTTAATAATTATTCAGTTTTATCCACTCCCCGTCTC

ACCTCCGTCTTTCTCGATCACTGTCATCATCTTTGATTACCACTACTATGCTGAACGCGAATTTCGAGAC

AAAAGCTTCGAGCAAATTGGACGATTTATCTGGATGCTAATCAATCTGGGTGCTTAGCTGCGAGAGGGGT

TTGGAGTTTTGAAATTTTAGAGAAGGGAATAAGGAAGACAGCGGAAAAGAAAAAGTGGCGGGGACAGCGT

GGTGTTAATATTGGTGGTTGGCAGCAATATGGGCTGATGAGTAGCGTGGGTGAGCGGTTACAGCTAAACG

TGAGCGGGGAGAGTGTGGTGTTGTTGGTGGCTGTAGTGAGGTGGCTGGCGCGGCGGCATCATGGGTCGAT

GAGTAGCATGGGTGAGTGGTTAGCTAGACGTGAGGTGCTATCAGATTGGAGTTGTAAAACAGAAGTTTTC

GCTTTCCAACAATTGGCTGGATGGTGCTGGTTCATTCTTCACCTCAACTGCATCTTTTGGTCCCAAAAAA

CTTGGCTGGTTTGTTGAAGAAAGAAGTCTTGCCCACATCCTCTTTGAGCAAGGAGTGTGCAATACACATT

GTCAGAGGCATTACTCGGAATAGCACG

>MSTRG.2293.2 gene=MSTRG.2293

CTCCAATTTCGGCAATTCCACGTCCGACATCCCTCCATGTCCATTTTCACCATCTCTATCACCAACAAAT

CCATCCCACCTCCATCGCTCCATCCTGCATCGTCTGCACCTCTGCCGTCTCCAAAATATCTATTTAATAA

TTATTCAGTTTTATCCACTCCCCGTCTCACCTCCGTCTTTCTCGATCACTGTCATCATCTTTGATTACCA

CTACTATGCTGAACGCGAATTTCGAGACAAAAGCTTCGAGCAAATTGGACGATTTATCTGGATGCTAATC

AATCTGGGTGCTTAGCTGCGAGAGGGGTTTGGAGTTTTGAAATTTTAGAGAAGGGAATAAGGAAGACAGC

GGAAAAGAAAAAGTGGCGGGGACAGCGTGGTGTTAATATTGGTGGTTGGCAGCAATATGGGCTGATGAGT

AGCGTGGGTGAGCGGTTACAGCTAAACGTGAGCGGGGAGAGTGTGGTGTTGTTGGTGGCTGTAGTGAGGT

GGCTGGCGCGGCGGCATCATGGGTCGATGAGTAGCATGGGTGAGTGGTTAGCTAGACGTGAGGTGCTATC

AGATTGGAGTTGTAAAACAGAAGTTTTCGCTTTCCAACAATTGGCTGGATGGTGCTGGTTCATTCTTCAC

CTCAACTGCATCTTTTGGTCCCAAAAAACTTGGCTGGTTTGTTGAAGAAAGAAGTCTTGCCCACATCCTG

TCAGTTATTACAACCTTATTCTGCATCTCGGTGATACGCTGCACATGCACTATCTGATTACAAATCTGGC

AATGCTGATTGTAAAAATTAAAAAAAAAAACTACGCAAATTTTGTCACTCCTATTTACTACCTTTCCCCT

TTCCTGACAACTAAAGAAAGAAATGTGACACAGGAAAGAGTAATGGTCTACAAGAAAGAAAAAACAATCT

TACACCTATATTAATTGAAGCAACTTGTGAAAATTAGTTACATGTGTAGCAAGAAAGTTGGAGCAGAATT

GTCATTAAAC

>MSTRG.2330.1 gene=MSTRG.2330

AATAATTTGGGGATAAACAAACCGGTTCGATAGTCATTCTTGTTCTTTTCAATTATTCTCATCATGGCTA

GTAATCTAGTCCAAGCAATGAATAACATCTCTTTAGAGAACGAGGAAGGAGGAAGAGAGATTAATGATGA

TGTTCATAATCAACCCTTTAACGCAAATCTTTTTGTTGTTGGGCGATTTATTAACAAGGGACGAGTCAAT

TTTGAAGCAATGCAGCACACTCTAGCCATTCTATGGAAGCCAGGTAAAGGTGTCTCCATAAAGGAATTGG

ATTCAAATTTATATCTATTTCAATTTTATCATGAGATTGACGTTAAAAGAGTTATGGAGGGCTGCCCTTG

GTCATTTAACAGGAAAGCTTTGGTGATGTCGAGATTAAAACAAGGACAGAATCCTCGTTATGTTGATCTC

AACACCATGGATCTCTGGGTACAGATTCGTGATTTAAAGGTCGGATTTATGTCGGAAACAGTCCTTAAAG

GAATTGGGAATTATGTGGGGAATTATGTTCGATCTTGCCCATCAAACTTCACGGGTGTTTGGAGAGAATA

TATGAGGATCAGGGTCTCAATTGATCTGGCAAAACCATTAAAGAGACGGATGAAAATCAAGATGGCGGGA

GATGCGTGGTTCTGGATTAATTTTAAATATGAAAATGTTCCAAGTTTTTGTTTCATTTGTGGCATTGTAG

GTCATTCGGAACGATTTTGTAGTCAGTTGTTTGAAAAAGCGGAGAGTGAAATAGTTAAACCTTATGGTGA

GTGGATGAGAGCTCCTCTTCGAAGACAAGTGAAACCAATAGGAGCCAAATGGTTACGAAATGGTGAAAGT

GGTAGTAAATATGCAGCAGATTTGCAGATTCAGAGCAAACGGAACGAAGACGATGAGAGCAATCAAGACT

CAGAAAAAATAGGCTTGTATAAAGAAGTTTTAATGGACTCGGCTGATATTACTGAGACTAAAATGGCTGA

TGAAAATTCAGGTAACAGTTCGATTTCAAAAAACGTATATGGCGCGAGTACTCATGGGAGTGCTCGCCTA

TCATTATGAGTTTAATAAGTTGGAATTGCCATGGGATTGGGACTCCGTGGCCTTTTCAATTCCTTAAGGA

GATCACTCCCCAAAAGAGACCCGGTTTTATTTTCCTGTGTGAAATTATTAGTAAGAAAGCTACTGTAGAA

AGGTTTCAACGAGCAGTTGGCTTTAAGGGCTGTAGTAGTAGAATCCCATGGTCACAATGGTGGTTTTGCT

TTACTCCAGCGTATAAGGATGAAGTCCTTTCACGTTCCTACTTGATTGACAGGAATTTACGACGAGCCTG

ACAGAAATAAAAGGTATGAAACATGGCAACTCATTCGCCATCTGTCTGTAAATAATTCCTTAACTTGGGT

GTTAATAGGTGATATGAATAATGTCTGCTCGCAAGAAGATTAGAGAGGTGGTCGTCCATATCCTCAAACT

CTGATCAGAGGTTTTCTTGATGTTCTATAAGACTGCAATCTCTCAGACATGATGAATCTTATTGGCCAGC

CATATACATGGGAAAGAGGTGCTGGCAATGATTATTGGATCGAACTAAGGCTGGATCGTGCCTTGTCAAC

AGCTGATTCTCAAACATGTTCAAGGATGCTACTCTCACAAATTTGGAAGTCCCCACGTCAGACCATTGCC

CTTTGTTTCTGGAGTTTTATAAAGCTCAACAGATAGTCAAAACTCGAATCTTTTGCTTTGAAAATGCATG

CCTCCGCGAGCCAATGTACCAACAGCTAGTGGCGGATGTTTGGGATAGAAATCAAGGAAGATCTTTTTTT

GACAAAATCACTGAATGTACTCAAATCTTATCCGAATGGGGCCAAGAAATCACAGGTAGCTTCAAACGTC

GAATCCAAGCATACAAAAGAGTGATTAAAGCTCTAAAAGGTCGAAGAGATGACTTCTCAGTTAAAGTAGT

AAAGGAGAATCAGAAGCAGCTTGCTGAGGTTTACCTCAACAGGAAATTTTTTGGAAACAACGGTCCAAAC

AGTTGTGGTTAAAAAAGGGGACCAAAACACCAAATTCTTTCATGCAGCGGCTAAGAATAGAAGGACTATA

AACCAGATCAAAAATCTCAAGAATAAAGATGGCATTACAGTTGATTGGAATTCGGGTTTAGAAGATGTGA

TAGTGAGCTTTTTCACAGAATTGTTTAAGGCGTCAGATACTGAATGGAGCTTAATTACATATACTGAATG

AGTTATGGCCTATTTTCTATGTCGTCTGTAACTGTAGTATGTACTCAGATCGTTCCATTAATATATATTC

CCTTTTGGCC

>MSTRG.2333.1 gene=MSTRG.2333

CAAATCACACAGCTAATCGACCTTCTCCGGCTACTTTTCTTCCATGCTGTCATACATTCATCATGTATCA

AATTATTACATGTGGCGCATGCTATCAACTTCTCTTGCAGTGTCATTTCCTCCAAACAAACCGGACATAA

CGTCCCCTCTTCGATTTTAAAAGATTGTTGGTTAGTATGACTATTACTGCTACTACATTGTCTTTCTTGG

AAAAACAGCTGATGAAATCGCTCACGAATGCTAGCTCCGGCCATGGCTTGGCTCGACATGGGGGCAGAGA

GTAGGCGGTGTAGCTGACAGGGACGCAGAGTTCTTCTCCAGAGACATGCGTCGTCGAGGGAGATGTTGAG

TACTTGAATGTACACGAAGAGTATGTGTTTGCATGGAGTGGTACGGTCAGGGCAGGAACATGCAGGGGTA

GACGATAAAGTGACCGTGTACACATTTGCAGTGGCTCCAAGGATGTAAAAGGTGGCAGCGGAGCGGTGGA

GGAGACGGAGGCCGTGGCGGAGTGCACGGATTACGCGTTCGGCTAGAGGTTGCGTTGGGTAGAAACGGGG

GCGTTGAGGTTGAGGGCGGGTGTGGCTGGAAGAGGATGCGTTGGA

>MSTRG.2342.1 gene=MSTRG.2342

TTCATTTTTAGTTGATCAAATTCAACACTCGACTGTGGGTAAAATCATGGATATAGATACTTTGTACGCC

GCTGCCATTGCTGGGGACTCTGATGCCATACTTAAACTACAGACTGAAGCTGAGAAACTGAGCACACATA

ACAGAACCATCCTTCACAACGAAACCAGGTACGGAAACACAGTGACAGAATTTGCCAACAAAAACCTGTT

GATCAAGCTGGAGTCACGGGGGAGTAACACTGCACTTCACCTTGCAGCAGAGCATGGACACGCTCAAGTG

GCCGAGCTCCTCATTGATGCAGCCCGACACTTGCCTTGTTCTTCAGCTAAATACCGTGATGATTGTGTTA

CTTGTTTTCAAGCTTTTCTTCGGCAAACTAATTATAGTCAGAGCACTGCCTTGCACATAGCAGCGCGCAG

AGGTCACGCGGCTATTGTGAAGATGTTAGTAGAGGCGGATCCGAGTGATAGACATGTTCAAAATAGTGTC

GGTGAAACTCCAATCTACATTGCTGCTAATTTGGGGCATTTTGATATAGTCAAGATGATTTGTACAGCTT

GCTCAGCTCCGGCTTTGGACGGCCCTGATGGGACCACTGCTTTCCATGCCGCTGTTGAGAAGCTCCATCA

AGCAAAGAGAGAAGATAGAGATGTGATAAAGGTATATATTAATGCAGCCAAACGCTTCATCAGTTATGAA

TCTTTGTTTCTTCCAAATCGGAAGGGTGAGACTGCATTAGAGCTTGCAGTGCGGGTAAATCACTTAGAGG

TGGTTGAGCTGATATTAGCAGAAGATCCAGCATATAAACGTGACAATGTAGATAAAAATGATGAGCTCCT

AAGTCTAATCTACATAGCTGCCGAACAGAGGTACAAGGATATGGTCAAATTACTCTGTCAAACTTATGAA

AGTCGAAATAGCCTTTGTCATAAAGGTCATATTGCTTTCCTTGCTGCTATTAGGGGGCGCGATAAAGAAT

CTGCACTACGTCTTTTGGGAGAGGACAGACGTCTTGTAACCTATGCTGACGATAAAGGCTGGACAGCACT

TCACTATGCTGCAGATTGTGAATTTGATTCAATACTTAATGCCATAGTAGAATCACACGTGCATCCTGGG

TACCAATCTGTTTCTGGAGAGATAGTACCAATGCCACTTACTCTTGCAGCCAAGAAAGGATATACATCTA

CCGTAATACAACTTCTGAAATTATTGCCAGCTTCTTCATGTGTAGATGTTAACAACGAGGGTCGAAACAT

ACTTCATCTTGCAGCCATTCAAAGCAATATAGATATGATACAAAGTATACTAAAATATTGTCCAGAAGAG

TATACAAACAAAATTTTGAATAACAAGGATGTCAATGGTAATACACCTCTGCATCTGCTTATACGTGAGG

GCTGCTTCGTTCCTCAACTCATAAAGCATAAAGGAGTTGATACAACAGCAGAAAACTACAAAAATTGGAC

TCCATTAGACATGTTATATTTCAGTGATGAAATCATAGCCGACCAGGTACGTATTAAAGAAGCAATAGAC

ACTCAAAATAATGAGCCTTGGGGGTTTTGGCGTAGAAGCATCAAGGGGAAATCCAATCTGCAGGAAAGTT

TAGTGCCCCCAAGTAAACGGTCAAAAAAAGATGCAGCATTTGAGAAAGCTAAGAATTTGTTGAGAAATGA

AAAGCGCAAATGGATGAAAGAAGAGCTTGAACGATATAAAAAAAGAACCAACACCCAGATAATTGTTACT

GCGCTCATTACTACAGTAACTTTTACTGTTGGATTTACGTTGCCTGGAGGATACCATCAGAGTGGAGAAT

CTGATGAAGGGTTGGCTCTTCTTTCCCGAAAGACAGCTTTTAAAATATTTATGGTAGCAGATGCAATAGC

TCTCCTACTATCAACATCTTCTTTGTTTCTATACTTTATTGAATCAATGTATGAAGATCCCCAAAAAGTT

GCAAACATTAATGCTGCATCAACTGGGCTCAGCATTGTTTCTGTTATAGCAATGATGTTGACTTTCATCA

CGGGAACATACGTTGTGCTATCCCGCTCACCAGTCCTTGCGATTAATGTCTGTGTCATTGGCTCCTTGTT

CTTTGTTTTTGTCATGGTACTGTTGATCAAATTAATGTATGATTATAAACTGGAAAGACATGTAGTGTGA

GTATGTCCA

>MSTRG.2343.1 gene=MSTRG.2343

TTCACCGACACTATTTTGAACATGTCTATCACTCGGATCCGCCTCTACTAACATCTTCACAATAGCCGCG

TGACCTCTGCGCGCTGCTATGTGCAAGGCAGTGCTCTGACTATAATTAGTTTGCCGAAGAAAAGCTTGAA

AACAAGTAACACAATCATCACGGTATTTAGCTGAAGAACAAGGCAAGTGTCGGGCTGCATCAATGAGGAG

CTCGGCCACTTGAGCGTGTCCATGCTCTGCTGCAAGGTGAAGTGCAGTGTTACTCCCCCGTGACTCCAGC

TTGATCAACAGGTTTTTGTTGGCAAATTCTGTCACTGTGTTTCCGTACCTG

>MSTRG.2345.2 gene=MSTRG.2345

TATCTATCTATATTATAAAGGCTGTTGTTTTTTTACCCTTGAAATGACCATTCCACCCATGTGTATTTGT

TGTATTTCCAGAAACACTTATGTACAATACCCGTATGTATACGTTGGTGTCATCCACACCTTCCTTTCCC

TTACCAAAGCTCCACGTGTTGCCCATCCCTCTCTCTACGATTCTCTCCTCCAAAATCATCTTACTACATT

CAAAATTTTCAAATTTAACTCCTTCCGCAATTTTTCAGTTGTAATACATGGATTCTTTAGAAAGAAGCCC

TCTCAAGTCGCCGGCATAGGAGTTGGTTTCTGGGAAATCATCCAATACATGTCAAACTTTCAAAATTAAT

TGTAATTGTGGCAGATAGTACAAAAATGGGAACAAAGAAATCTTGGCTTCCAAAACAAAGAATTGACGAG

CTGTATTACTTGATTTCACGAAGGTCAAGAGATGGTTGTGGGAGAAAGAGGCTGAACAGGCTAGG

>MSTRG.2345.1 gene=MSTRG.2345

TATCTATCTATATTATAAAGGCTGTTGTTTTTTTACCCTTGAAATGACCATTCCACCCATGTGTATTTGT

TGTATTTCCAGAAACACTTATGTACAATACCCGTATGTATACGTTGGTGTCATCCACACCTTCCTTTCCC

TTACCAAAGCTCCACGTGTTGCCCATCCCTCTCTCTACGATTCTCTCCTCCAAAATCATCTTACTACATT

CAAAATTTTCAAATTTAACTCCTTCCGCAAGTAAAGCTTCTAATCCCATTTTGTTTTTCGCCGCTTTTCA

TTGTAATCCAATTGTCTCCATTCATTGTATCACTTCTTCGTTTATATAGTTTTTCAGTTGTAATACATGG

ATTCTTTAGAAAGAAGCCCTCTCAAGTCGCCGGCATAGGAGTTGGTTTCTGGGAAATCATCCAATACATG

TCAAACTTTCAAAATTAATTGTAATTGTGGCAGATAGTACAAAAATGGGAACAAAGAAATCTTGGCTTCC

AAAACAAAGAATTGACGAGCTGTATTACTTGATTTCACGAAGGTCAAGAGATGGTTGTGGGAGAAAGAGG

CTGAACAGGCTAGGTAAGTTCCACCAATTTTGAAATGGCTGATGCTTCTGCAGTCGAGAAGCAATGGCGA

AATCAATTCAGTTTCTTAAATGTCTCAGTCCCAAAACAATTAGAAGATATAGTGGTCCTGGCACACAAAA

GATATATCATACAAAGGCACAAGCCGGGGACCTTCGAGTTTATTTTTGGATGAATTTTATGAGGGTAATA

TATTAAGGTTTCCATTTAAGGTTTTTTTTTTGGATTATTTGTAAATTAATTACAAAAATAATTTTACAGT

ATATTCACTTTAATTATATTAATTCCAGTCCTTTTTAATTCCTTATCATGTTAGGAAAGTGTTCAAGGAT

TTCCACTCAGAAAAATGTTAGGAAAATGTTAAAGGATTTTCTTTACAGAATTTGCCCCAATTTAATAACA

TGAAAACTACTGGAAACCTACAATTTTGTGCTATCTGTATTTCCCTGGAGTTGTGAATTCACAGTATCTG

CACAAATAAGTTTGCATAGTAGATATTGGTTGATTAGGTATGTTTTTCCGTATGAAATAAATGGAAGAAT

ATGCTAATTGCATATAAATTTCCATTTTTTTCTTTTAGTATGCTCAAAGGTATAGAATGCAAAATATCCT

TCACTCGACTGAACCATTTCGAACTTAACATATATACTTTCTAAATGCTTACACTATAAAAGCTTCATTA

GAAAATTGCATAAGGCTCAACACCAGTTTTGGTCTGAATGTAGTGGCATTTAACAAGTAAAAAGACATTG

CACCTACACTATAAAAGTTGTAGATGACCCGAGGACCTTGAACAAAATTCTCTATGTGAGGCCTCCCCAC

AACACATTTTTTTTCAATGATTTTCTGTCTTTTTAGGAGAAAAAGATGGGCAAAACTCTTGAGAAGATAT

ATGTTCCAGAGGAGCAAGTCTTGAAGAATATTAAAGATGAGGAATTTTTCAAATTTTTTTGCTTGACATT

ATTATAATAATTGACCAGAGCAAGTCTTGAAGAACATTGAAGGTGAGGGGTTTTTTAAATTTTTTTTGCT

TCACAATGTTATAATAATTGATCATTATCTTATGATCCTGGCAAAGTTGTTTACAAAGTTTTTTTTGTCC

TTTATGTTAAACTCTATGAGGAATCCCCTATTCGGATCAATGTGCTGCTATTTATCGGTCATTCTACATT

TGTGAAGGGAGATTAAACCAGTTTTGAGATTGAACCTTCATTTGGTGTGGAGGCTTCAGCACTTTATCCT

GATGTCAAATACAAAACTGAAGAGGAGTATATCAACAAATTTGTCTAAACCAAATCTATTTTGTCTTTAA

AATATTTTCTTTTTTACAATTCATATAATATAATTTTTCCAAATTCAAAAACAACAATCTATTCACTAAA

CAAAAATCATCCGCG

>MSTRG.2349.1 gene=MSTRG.2349

GTTCCTACAATTTACAAAACTACCAAGGTAGTCCATTACGGGCCCAAAATGAAACGCCGCCGTTTAAGTG

TGATTAGGCTTTTTAGGTTGTGATCCACATTATTGTTTCTCAACTCCGCGGCGTAAATCCTCGGAGTTTG

AGAAAACCCTAACCCTAATTTTCAAATCGCAATCTTCTACTTTCAGTCATGGCCGGAGCTGCTCCCGAAG

GTTCTCAATTTGACGCTCGTCAGTTTGATGCAAAAATGACTGAGTTACTTGGGGCTGATGGACAAGAATT

CTTCACCTCATATGACGAGGTCCACGATAGTTTCGATGCAATGGGATTGCAAGAAAACCTTTTAAGAGGC

ATATATGCATACGGTTTTGAGAAGCCTTCTGCCATACAGCAAAGGGGTATTGTTCCGTTTTGCAAGGGAC

TGGATGTTATTCAACAAGCACAATCTGGGACTGGAAAAACAGCAACTTTTTGCTCTGGAATTCTTCAGCA

ACTTGATTATGCTGTAGTTGAATGTCAGGCACTGGTCCTGGCACCTACTCGTGAACTTGCACAACAGATT

GAGAAGGTTATGCGAGCTCTTGGTGACTATCTTGGTGTAAAGGTTCATGCTTGTGTAGGAGGGACTAGTG

TTCGAGAAGATCAGCGCATCCTCTCAAGTGGGGTTCATGTTGTTGTTGGTACCCCTGGCCGAGTGTTTGA

CATGTTGCGAAGGCAGTCTCTTCGTGCAGATCACATCAAAATGTTTGTACTTGATGAAGCTGATGAAATG

CTGTCAAGAGGTTTCAAGGATCAGATCTATGATATCTTCCAGTTGCTACCCCCAAAAATTCAGGTTGGGG

TTTTTTCTGCAACCATGCCTCCAGAGGCACTTGAGATCACCAGGAAATTCATGAACAAGCCTGTAAGGAT

TCTTGTGAAACGTGATGAGCTTACTCTTGAGGGTATTAAACAGTTTTATGTTAATGTTGACAAAGAGGAG

TGGAAATTGGAAACACTTTGTGATTTGTACGAGACTTTAGCCATTACGCAAAGTGTCATATTTGTTAATA

CTAGGCGAAAGGTTGATTGGCTCACTGACAAAATGCGAAGCCGTGACCACACAGTTTCTGCTACTCATGG

AGACATGGATCAGAATACTAGAGACATTATCATGAGGGAATTTAGATCTGGATCATCCCGTGTGCTTATC

ACCACTGACCTTCTGGCCCGTGGTATTGATGTGCAGCAAGTGTCACTTGTTATCAACTATGATCTCCCAA

CTCAACCAGAGAATTACCTTCATCGTATTGGGCGTAGTGGACGATTTGGAAGGAAAGGTGTTGCAATTAA

TTTTGTGACCAAGGATGATGATAGGATGCTGTTTGACATACAGAAGTTTTACAATGTAGTAGTTGAGGAG

CTGCCAGCCAATGTTGCTGATCTCCTTTAGTAAGGTTTTGATGCGATGAGGTAAAATTGCTTGTTATGCA

AAAGCCTTGGTTTTTCATACTATTTACTAGTAGAATATTTATTGGACTTCCATCTGGGAGATGTGGATCT

CATCAATTGCATTGGGACCTCTCATTTCCAGTCGGCTTGTAATCTGAATATGTGTTTGCAATTTTTGGGG

ATTTTTTTGTTTTTTTTATATAGCTTGCAAGTTAAATTGACTATTATCTACTCTTTTTTTAAAAAATTTA

TCTGTAG

>MSTRG.2350.1 gene=MSTRG.2350

CATGGAAGCATCGTGATTTTAGAGATTTTAGGGTAGTGGATGCTATGATCTGAATCTCCTAATTAAGGTA

ACTGAATAAGATTTGATTAGCTGGTTAATTACAATACGTATGGAAGCTGTATATTTACACATCATTACAA

ATCGATAAAACTACTTTAGATCGAGGGAACAAATCCAGCCATGATGAAGAAGAATTTCGATGTCAATCAC

GATCCATATGACGTTAATAGAGAGATAGACAGAGACTTCTGCATTCTCATATACCATTGACAATCACATG

AAGATAATGTCAATCTTGATTGCTGGTAGTGTTAACTATAGAAAGATATATATGAAGATGTCAACCAAGT

CATGTCACCATTTTAGTCTCAGAGGAATTCAATCAAAGTATTGGTTGATGTGGCCATCTTTCAAGAACAT

AATGCAATTGCGCTGGGTATCTTAGCTAGAAACTCAGACAGAGATATGACTGAAGCCATGGCAATCAAGG

AGGCTTTAAGTTGGTGTAAGGAGACTAATACAACTGACCAGGAGGTCATGATTGAATCTGATTGCCTGGT

GATAGTCCTGGCAATACGAATTGCCTGGTGATAGTCCTGGCAATACGAAGCAGTGTTCCAATGTTCTGGC

TAGCAGGGTGGCAATTGGCATGCCCATAGTTTAGCTAGAGCTTGGTATTCCTCACTATTCCATCCGTGGT

GTTTGGTAGAGTTGATGTACCTCAGAAAGTTCTCAATTGTATTATGAACGATTCTCTTTGCGTTAAAAAA

AAAAAAGATCTAGTTTGTCCA

>MSTRG.2350.2 gene=MSTRG.2350

CATGGAAGCATCGTGATTTTAGAGATTTTAGGGTAGTGGATGCTATGATCTGAATCTCCTAATTAAGGTA

ACTGAATAAGATTTGATTAGCTGGTTAATTACAATACGTATGGAAGCTGTATATTTACACATCATTACAA

ATCGATAAAACTACTTTAGATCGAGGGAACAAATCCAGCCATGATGAAGAAGAATTTCGATGTCAATCAC

GATCAAATTCATCTACTCAATCGTTGTAAAGAAGCAAACTTAATCATTCAATAAGGTTGAACATAACCTT

ATAAAATGATGTGGTAGCATATTAGTCTTAGGTTTTGAAAAATGGTGATTGGAACGGGTTTTGAGAATTG

GTTTTCGTATTCTTTAACCAAGATTTGTTAAGATGTTCTGTTGAGATCCATATGACGTTAATAGAGAGAT

AGACAGAGACTTCTGCATTCTCATATACCATTGACAATCACATGAAGATAATGTCAATCTTGATTGCTGG

TAGTGTTAACTATAGAAAGATATATATGAAGATGTCAACCAAGTCATGTCACCATTTTAGTCTCAGAGGA

ATTCAATCAAAGTATTGGTTGATGTGGCCATCTTTCAAGAACATAATGCAATTGCGCTGGGTATCTTAGC

TAGAAACTCAGACAGAGATATGACTGAAGCCATGGCAATCAAGGAGGCTTTAAGTTGGTGTAAGGAGACT

AATACAACTGACCAGGAGGTCATGATTGAATCTGATTGCCTGGTGATAGTCCTGGCAATACGAATTGCCT

GGTGATAGTCCTGGCAATACGAAGCAGTGTTCCAATGTTCTGGCTAGCAGGGTGGCAATTGGCATGCCCA

TAGTTTAGCTAGAGCTTGGTATTCCTCACTATTCCATCCGTGGTGTTTGGTAGAGTTGATGTACCTCAGA

AAGTTCTCAATTGTATTATGAACGATTCTCTTTGCGTTAAAAAAAAAAAA

>MSTRG.2350.3 gene=MSTRG.2350

CATGGAAGCATCGTGATTTTAGAGATTTTAGGGTAGTGGATGCTATGATCTGAATCTCCTAATTAAGGTA

ACTGAATAAGATTTGATTAGCTGGTTAATTACAATACGTATGGAAGCTGTATATTTACACATCATTACAA

ATCGATAAAACTACTTTAGATCGAGGGAACAAATCCAGCCATGATGAAGAAGAATTTCGATGTCAATCAC

GATCAAATTCATCTACTCAATCGTTGTAAAGAAGCAAACTTAATCATTCAATAAGGTTGAACATAACCTT

ATAAAATGATGTGGATGGTGATTGGAACGGGTTTTGAGAATTGGTTTTCGTATTCTTTAACCAAGATTTG

TTAAGATGTTCTGTTGAGATCCATATGACGTTAATAGAGAGATAGACAGAGACTTCTGCATTCTCATATA

CCATTGACAATCACATGAAGATAATGTCAATCTTGATTGCTGGTAGTGTTAACTATAGAAAGATATATAT

GAAGATGTCAACCAAGTCATGTCACCATTTTAGTCTCAGAGGAATTCAATCAAAGTATTGGTTGATGTGG

CCATCTTTCAAGAACATAATGCAATTGCGCTGGGTATCTTAGCTAGAAACTCAGACAGAGATATGACTGA

AGCCATGGCAATCAAGGAGGCTTTAAGTTGGTGTAAGGAGACTAATACAACTGACCAGGAGGTCATGATT

GAATCTGATTGCCTGGTGATAGTCCTGGCAATACGAATTGCCTGGTGATAGTCCTGGCAATACGAAGCAG

TGTTCCAATGTTCTGGCTAGCAGGGTGGCAATTGGCATGCCCATAGTTTAGCTAGAGCTTGGTATTCCTC

ACTATTCCATCCGTGGTGTTTGGTAGAGTTGATGTACCTCAGAAAGTTCTCAATTGTATTATGAACGATT

CTCTTTGCGTTAAAAAAAAA

>MSTRG.2354.1 gene=MSTRG.2354

CATTTGTTAACCAAACGTTAACTTCTGCTCAATTAAACCTTATATAAAATTGAAAAAAATTTAAAACATT

ATCAACATCAGGCTTCTGGCCTACATCTCGAACCAAATCCTAAATTCCTCCACCAGAACCGGATAACTCC

GAGTTTTAACAACCGAGTGAACAGCCGAGCCAAGATCATCCTTCTTGTAACCGATCCGCAATCCAGCAGC

TCGCTTCTTCTTCCTCGTATTCGAAACGTGTCGAGTAATCGAGTTAGTAATAATCGGCGTACGTGGCAAC

AATCGGCGAAGGTTGCGCTAGGGTTTGCTGAATCTGCGGAGTAACAAGAGTGAGATCAGAGAATAAATTC

TCATCGAGAGGCTGAGAAGCGAACTCCACCGCTGGAGATTTGGAGGCCAGCGCCGAAGGATCCGTCGATC

CGGTTGATGATTCCGGTGGTGTTTGGAATCATGAGAATTGTTATCGATAGGAGTAGATTGTTGAGTTGAT

AATGGTGGGAGTTAGGGTTGAGAGGGATCGAATAGCACCATTCCTTCGAGCAATGAGTCCATGTTGTCCG

ACTTTTAACCCACATCGGCTGTAATTTCTTCTGTTTTTAAGCTTTTTTAAAAAAAATTTATTTAAGGTTT

AATTGAGGGGAAGTTAACATGTGGCTAACAGATGTTGACGAGAAGGGTATGGCACATAACGTTAGAATAA

TA

>MSTRG.2355.1 gene=MSTRG.2355

AAAAAAGCTTAAAAACAGAAGAAATTACAGCCGATGTGGGTTAAAAGTCGGACAACATGGACTCATTGCT

CGAAGGAATGGTGCTATTCGATCCCTCTCAACCCTAACTCCCACCATTATCAACTCAACAATCTACTCCT

ATCGATAACAATTCTCATGATTCCAAACACCACCGGAATCATCAACCGGATCGACGGATCCTTCGGCGCT

GGCCTCCAAATCTCCAGCGGTGGAGTTCGCTTCTCAGCCTCTCGATGAGAATTTATTCTCTGATCTCACT

CTTGTTACTCCGCAGATTCAGCAAACCCTAGCGCAACCTTCGCCGATTGTTGCCACGTACGCCGATTATT

ACTAACTCGATTACTCGACACGTTTCGAATACGAGGAAGAAGAAGCGAGCTGCTGGATTGCGGATCGGTT

ACAAGAAGGATGATCTTGGCTCGGCTGTTCACTCGGTTGTTAAAACTCGGAGTTATCCGGTTCTGGTGGA

GGAATTTAGGATTTGGTTCGAGATGTAGGCCAGAAGCCTGATGTTGATAATGTTTTAAATTTTTTTCAAT

TTTATATAAGGTTTAATTGAGCAGAAGTTAACGTTTGGTTAACAAATGTTGACGGGAGGGGTATGGCACA

CAACGTCAGAATTGTATAAGTTCAAGAAGC

>MSTRG.2361.1 gene=MSTRG.2361

ATAAAGAGCGCGGGAGCGCGGAGGAGGACTGGACTCCTACTTGTCGGATCTGACAACATTATAAACACTC

AGAGATCAGTCAAAACCGAATACATATACAGCAACTCACCGGATGCTCGGAGGAAGACGGATTTTTCTTT

TCAGACATTTAAGTTTTATTGTCTAAATTTCCCCTGTCATCTTGCATGTCCGACGGTTAGATAATAAGCT

TTCTTGAATGAAAGAATTATCACGGTGAAT

>MSTRG.2362.1 gene=MSTRG.2362

GGGGCCAAGGCCACCCCTTACCCCCCCTTCCCTCCGTCCCTGCTCGATCAGCAAGGTACATAGCAGTCAA

TCCGGTATTTCATGAACGCGCTAAGCACATCGACATCAATTGTCACCTGGTCATGGAAAAATTGAAGAAA

GGTCTGATCACTACCACCTACATCTCATCCAAGGAACAGCCTGCTCATTTGGTAACTAGTAATCTGCACA

TAACTAGTAGAGGGGCTAGATCAGTTTGGAAAGAAATGTCGAAGGATATTACACCAGCCCTCTAGTTGAT

TAGGTTGCACTATAAAAAACATTTCACTGAACTCTAATTTCAAGGTTTAGGAACTTGGGTAAAACATGCT

TGTTGGCT

>MSTRG.2366.1 gene=MSTRG.2366

GTATAAATAAAGACACAGGAGCAAAGAAAGAGGCACAATACTACATTTGGGGGTTTAGGGTTTTTATTAT

CAATCTCGTGTTGTGTCTGTTTTAAGTTGTATCAAAAGATGGGTGAAGATGGCAGCCGTTTGAAGGGTTC

AGTCAAGTGGTTCAATGACACCAAGGGCTTTGGATTCATCACCCCTGACGAAGGCGGTGAGGATCTGTTT

GTTCATCAATCTTCCATCAAGTCTGATGGGTTTCGCAGCCTCGCTGAAGGGGAGTCGGTTGAGTTTGTCA

TCGAGAATGGCAGTGATGGCAGGACCAAGGCCGTTGATGTCACTGGCCCAGATGGTGCTTCTGTCCAGGG

AAGTGGAAGGGGAGGCGGCGGCGGCGGTGGTAGAGGCGGCGGCGGCTATGGTGGTGGCGGAGGCGGCTAC

GGTGGTGGCGGCGGCGGATATGGTGGTGGTGGCGGTTATGGCGGCGGTGGCGGCGGAGGCTGCTACAAGT

GTGGGGAGAATGGTCATATGGCGAGGGATTGCGACAATGGCGGAGGCGGTGGAGGCTATGGAGGTGGTGG

CGGCGGCTATGGCGGAGGCGGTGGCTATGGAGGTGGTCGTGGTGGAGGACGTGGTGGGGGCCGTGGCGGC

GGCGGCGGAGGTGGCTGCTATCAGTGTGGTGAGGACGGCCATTTCGCCAGGGACTGTCCTAATTCCAACC

GTTAGGTTTCGGGGTTGTGCTGCTGCGATGGGGTAATATGTGCTTCCGTGTTGATGCATTTTGCTCTCTC

GCTTTCTTTACTGTTAATTAATCTTAATCCTCCTATCCTGGTGCGACTCTTGGTCTTTTGATTAATTAAG

GATTATTATTATTATGTAGGTG

>MSTRG.2366.2 gene=MSTRG.2366

AGGAGCAAAGAAAGAGGCACAATACTACATTTGGGGGTTTAGGGTTTTTATTATCAATCTCGTGTTGTGT

CTGTTTTAAGTTGTATCAAAAGATGGGTGAAGATGGCAGCCGTTTGAAGGGTTCAGTCAAGTGGTTCAAT

GACACCAAGGGCTTTGGATTCATCACCCCTGACGAAGGCGGTGAGGATCTGTTTGTTCATCAATCTTCCA

TCAAGTCTGATGGGTTTCGCAGCCTCGCTGAAGGGGAGTCGGTTGAGTTTGTCATCGAGAATGGCAGTGA

TGGCAGGACCAAGGCCGTTGATGTCACTGGCCCAGATGGTGCTTCTGTCCAGGGAAGTGGAAGGGGAGGC

GGCGGCGGCGGTGGTAGAGGCGGCGGCGGCTATGGTGGTGGCGGAGGCGGCTACGGTGGTGGCGGCGGCG

GATATGGTGGTGGTGGCGGTTATGGCGGCGGTGGCGGCGGAGGCTGCTACAAGTGTGGGGAGAATGGTCA

TATGGCGAGGGATTGCGACAATGGCGGAGGCGGTGGAGGCTGCTACAAGTGTGGGGAGAATGGTCATATG

GCGAGGGATTGCGACAATGGCGGAGGCGGTGGAGGCTATGGAGGTGGTGGCGGCGGCTATGGCGGAGGCG

GTGGCTATGGAGGTGGTCGTGGTGGAGGACGTGGTGGGGGCCGTGGCGGCGGCGGCGGAGGTGGCTGCTA

TCAGTGTGGTGAGGACGGCCATTTCGCCAGGGACTGTCCTAATTCCAACCGTTAGGTTTCGGGGTTGTGC

TGCTGCGATGGGGTAATATGTGCTTCCGTGTTGATGCATTTTGCTCTCTCGCTTTCTTTACTGTTAATTA

ATCTTAATCCTCCTATCCTGGTGCGACTCTTGGTCTTTTGATTAATTAAGGATTATTATTATGTAGGTTT

AAACTTTTTTAAGGATTTCAGATGGTTTTTGGTGGTTTGGTTTATCAATCAACTGTGTGATGCTTGTTCT

CTTTCGGATTTACTCAGTGTTGTCTATTATTATGCTTTTCGTTTCTCATTATTGTTAATATGCTCATGAA

TGAATGGATGTGTGGAATTAAGCTGATGACAACACTCAGTGGTGCTCTTTATAACGTGCTCTCTTTTCTT

TTTTCATTA

>MSTRG.2366.3 gene=MSTRG.2366

GAAAGAGGCACAATACTACATTTGGGGGTTTAGGGTTTTTATTATCAATCTCGTGTTGTGTCTGTTTTAA

GTTGTATCAAAAGATGGGTGAAGATGGCAGCCGTTTGAAGGGTTCAGTCAAGTGGTTCAATGACACCAAG

GGCTTTGGATTCATCACCCCTGACGAAGGCGGTGAGGATCTGTTTGTTCATCAATCTTCCATCAAGTCTG

ATGGGTTTCGCAGCCTCGCTGAAGGGGAGTCGGTTGAGTTTGTCATCGAGAATGGCAGTGATGGCAGGAC

CAAGGCCGTTGATGTCACTGGCCCAGATGGTGCTTCTGTCCAGGGAAGTGGAAGGGGAGGCGGCGGCGGC

GGTGGTAGAGGCGGCGGCGGCTATGGTGGTGGCGGAGGCGGCTACGGTGGTGGCGGCGGCGGATATGGTG

GTGGTGGCGGTTATGGCGGCGGCGGTGGCTATGGAGGTGGTCGTGGTGGAGGACGTGGTGGGGGCCGTGG

CGGCGGCGGCGGAGGTGGCTGCTATCAGTGTGGTGAGGACGGCCATTTCGCCAGGGACTGTCCTAATTCC

AACCGTTAGGTTTCGGGGTTGTGCTGCTGCGATGGGGTAATATGTGCTTCCGTGTTGATGCATTTTGCTC

TCTCGCTTTCTTTACTGTTAATTAATCTTAATCCTCCTATCCTGGTGCGACTCTTGGTCTTTTGATTAAT

TAAGGATTATTATTATGTAGGTTTAAACTTTTTTAAGGATTTCAGATGGTTTTTGGTGGTTTGGTTTATC

AATCAACTGTGTGATGCTTGTTCTCTTTCGGATTTACTCAGTGTTGTCTATTATTATGCTTTTCGTTTCT

CATTATTGTTAATATGCTCATGAATGAATGGATGTGTGGAATTAAGCTGATGACAACACTCAGTGGTGCT

CTTTATAACGTGCTCTCTTTTCTTTTTTCA

>MSTRG.2366.4 gene=MSTRG.2366

AAGAGGCACAATACTACATTTGGGGGTTTAGGGTTTTTATTATCAATCTCGTGTTGTGTCTGTTTTAAGT

TGTATCAAAAGATGGGTGAAGATGGCAGCCGTTTGAAGGGTTCAGTCAAGTGGTTCAATGACACCAAGGG

CTTTGGATTCATCACCCCTGACGAAGGCGGTGAGGATCTGTTTGTTCATCAATCTTCCATCAAGTCTGAT

GGGTTTCGCAGCCTCGCTGAAGGGGAGTCGGTTGAGTTTGTCATCGAGAATGGCAGTGATGGCAGGACCA

AGGCCGTTGATGTCACTGGCCCAGATGGTGCTTCTGTCCAGGGAAGTGGAAGGGGAGGCGGCGGCGGCGG

TGGTAGAGGCGGCGGCGGCTATGGTGGTGGCGGAGGCGGCTACGGTGGTGGCGGCGGCGGATATGGTGGT

GGTCGTGGTGGAGGACGTGGTGGGGGCCGTGGCGGCGGCGGCGGAGGTGGCTGCTATCAGTGTGGTGAGG

ACGGCCATTTCGCCAGGGACTGTCCTAATTCCAACCGTTAGGTTTCGGGGTTGTGCTGCTGCGATGGGGT

AATATGTGCTTCCGTGTTGATGCATTTTGCTCTCTCGCTTTCTTTACTGTTAATTAATCTTAATCCTCCT

ATCCTGGTGCGACTCTTGGTCTTTTGATTAATTAAGGATTATTATTATGTAGGTTTAAACTTTTTTAAGG

ATTTCAGATGGTTTTTGGTGGTTTGGTTTATCAATCAACTGTGTGATGCTTGTTCTCTTTCGGATTTACT

CAGTGTTGTCTATTATTATGCTTTTCGTTTCTCATTATTGTTAATATGCTCATGAATGAATGGATGTGTG

GAATTAAGCTGATGACAACACTCAGTGGTGCTCTTTATAACGTGC

>MSTRG.2366.5 gene=MSTRG.2366

AAGAGGCACAATACTACATTTGGGGGTTTAGGGTTTTTATTATCAATCTCGTGTTGTGTCTGTTTTAAGT

TGTATCAAAAGATGGGTGAAGATGGCAGCCGTTTGAAGGGTTCAGTCAAGTGGTTCAATGACACCAAGGG

CTTTGGATTCATCACCCCTGACGAAGGCGGTGAGGATCTGTTTGTTCATCAATCTTCCATCAAGTCTGAT

GGGTTTCGCAGCCTCGCTGAAGGGGAGTCGGTTGAGTTTGTCATCGAGAATGGCAGTGATGGCAGGACCA

AGGCCGTTGATGTCACTGGCCCAGATGGTGCTTCTGTCCAGGGAAGTGGAAGGGGAGGCGGCGGCGGCGG

TGGTAGAGGCGGCGGCGGCTATGGTGGTGGCGGAGGCGGCTACGGTGGTGGTGGCGGCGGCTATGGCGGA

GGCGGTGGCTATGGAGGTGGTCGTGGTGGAGGACGTGGTGGGGGCCGTGGCGGCGGCGGCGGAGGTGGCT

GCTATCAGTGTGGTGAGGACGGCCATTTCGCCAGGGACTGTCCTAATTCCAACCGTTAGGTTTCGGGGTT

GTGCTGCTGCGATGGGGTAATATGTGCTTCCGTGTTGATGCATTTTGCTCTCTCGCTTTCTTTACTGTTA

ATTAATCTTAATCCTCCTATCCTGGTGCGACTCTTGGTCTTTTGATTAATTAAGGATTATTATTATGTAG

GTTTAAACTTTTTTAAGGATTTCAGATGGTTTTTGGTGGTTTGGTTTATCAATCAACTGTGTGATGCTTG

TTCTCTTTCGGATTTACTCAGTGTTGTCTATTATTATGCTTTTCGTTTCTCATTATTGTTAATATGCTCA

TGAATGAATGGATGTGTGGAATTAAGCTGATGACAACACTCAGTGGTGCTCTTTATAACGTGC

>MSTRG.2366.6 gene=MSTRG.2366

AAGAGGCACAATACTACATTTGGGGGTTTAGGGTTTTTATTATCAATCTCGTGTTGTGTCTGTTTTAAGT

TGTATCAAAAGATGGGTGAAGATGGCAGCCGTTTGAAGGGTTCAGTCAAGTGGTTCAATGACACCAAGGG

CTTTGGATTCATCACCCCTGACGAAGGCGGTGAGGATCTGTTTGTTCATCAATCTTCCATCAAGTCTGAT

GGGTTTCGCAGCCTCGCTGAAGGGGAGTCGGTTGAGTTTGTCATCGAGAATGGCAGTGATGGCAGGACCA

AGGCCGTTGATGTCACTGGCCCAGATGGTGCTTCTGTCCAGGGAAGTGGAAGGGGAGGCGGCGGCGGCGG

TGGTAGAGGCGGCGGCGGCTATGGTGGTGGCGGAGGCGGCTACGGTGGTGGTCGTGGTGGAGGACGTGGT

GGGGGCCGTGGCGGCGGCGGCGGAGGTGGCTGCTATCAGTGTGGTGAGGACGGCCATTTCGCCAGGGACT

GTCCTAATTCCAACCGTTAGGTTTCGGGGTTGTGCTGCTGCGATGGGGTAATATGTGCTTCCGTGTTGAT

GCATTTTGCTCTCTCGCTTTCTTTACTGTTAATTAATCTTAATCCTCCTATCCTGGTGCGACTCTTGGTC

TTTTGATTAATTAAGGATTATTATTATGTAGGTTTAAACTTTTTTAAGGATTTCAGATGGTTTTTGGTGG

TTTGGTTTATCAATCAACTGTGTGATGCTTGTTCTCTTTCGGATTTACTCAGTGTTGTCTATTATTATGC

TTTTCGTTTCTCATTATTGTTAATATGCTCATGAATGAATGGATGTGTGGAATTAAGCTGATGACAACAC

TCAGTGGTGCTCTTTATAACGTGC

>MSTRG.2366.7 gene=MSTRG.2366

CACAATACTACATTTGGGGGTTTAGGGTTTTTATTATCAATCTCGTGTTGTGTCTGTTTTAAGTTGTATC

AAAAGATGGGTGAAGATGGCAGCCGTTTGAAGGGTTCAGTCAAGTGGTTCAATGACACCAAGGGCTTTGG

ATTCATCACCCCTGACGAAGGCGGTGAGGATCTGTTTGTTCATCAATCTTCCATCAAGTCTGATGGGTTT

CGCAGCCTCGCTGAAGGGGAGTCGGTTGAGTTTGTCATCGAGAATGGCAGTGATGGCAGGACCAAGGCCG

TTGATGTCACTGGCCCAGATGGTGCTTCTGTCCAGGGAAGTGGAAGGGGAGGCGGCGGCGGCGGTGGTAG

AGGCGGCGGCGGCTATGGTGGTGGTCGTGGTGGAGGACGTGGTGGGGGCCGTGGCGGCGGCGGCGGAGGT

GGCTGCTATCAGTGTGGTGAGGACGGCCATTTCGCCAGGGACTGTCCTAATTCCAACCGTTAGGTTTCGG

GGTTGTGCTGCTGCGATGGGGTAATATGTGCTTCCGTGTTGATGCATTTTGCTCTCTCGCTTTCTTTACT

GTTAATTAATCTTAATCCTCCTATCCTGGTGCGACTCTTGGTCTTTTGATTAATTAAGGATTATTATTAT

GTAGGTTTAAACTTTTTTAAGGATTTCAGATGGTTTTTGGTGGTTTGGTTTATCAATCAACTGTGTGATG

CTTGTTCTCTTTCGGATTTACTCAGTGTTGTCTATTATTATGCTTTTCGTTTCTCATTATTGTTAATATG

CTCATGAATGAATGGATGTGTGGAATTAAGCTGATGACAACACTCAGTGGTGCT

>MSTRG.2370.1 gene=MSTRG.2370

TGATATATCTTAATAAATCGGAAAATTTTAGACATTGTTGCGGTATCATAATAAAAATGTGGTACAATAA

TGTGTAGTAATGTAAGGTACCTGGTCATGGAGGAGTGAGGCAATGGTGAGGTACCAAGCAGTGTAAGTGG

TCATGATGAGGCCTAAGAATGACCAGATTCTGTAGTTGTGGAATGAGGGTATGAAAACTGTGGTAGCACA

GCAAGCTCCAAATATGTAAGTCCAAGTTCTCTTGTCCAGATTATCATTTATGTAGTATATGTTGCTGCAA

CAAGAATCTTGATGTTAACAGAC

>MSTRG.2371.1 gene=MSTRG.2371

AAGCAAGTAGACCACATGATTGCAGCCAAACGGGCGTGGTCTGTTTTTTATGAAGTAGAGAAACCACTTT

AATCACATGGCCAATCCCAAATCCCCTCCCTGGACTCGAGAACTCATCTTGACCCCTCAGATTAAAGCCC

ACAACCCTAAACCACTCTTATTTCCATGCTCACCTTTCTTTTGTTTTACATACACATGTACATTCTTGAC

ATGCTAGGCCGGAAACACGGCTAACCTACGCTGCAGTCCCGACTAAACTTCAAAATTTTATTGAATTTTT

CTTAAGCCCGG

>MSTRG.2372.3 gene=MSTRG.2372

AAAAATAACACTCTGTTTGGTTGGCCTGAATGGAATCATTTTAGCCCAGAATAGAATGGGCCATTCCATT

CCAAAAATAAAAATTTCTGTTTGGTTGGCCCAAGGAATGGAATAGCCCATTCTTTTATTCATTCCAATCC

AATCACTAATCAGACAATCGATTCATTTTTGGAAGTGCGGGTTAAGCCCATTCCATTCACTCACGAGTCA

CGCCCGCCTCTCTCACGCAGCTCTCCCTCGCCTCCCTCACGCAGCTTTCTCTCGCCTCTCTCCGCCCCTC

TCTCTCGCCTCTCTCTCTTCCTCGCATCTCTCTCTCATCACTCCTTCCTCGCATCTCTCTCTTCTTTTTC

TGCCTCTCCCCAGTCGCCTCCATTGCTGCTCTCAGACAAGGGCGATTCAGAGTTCAGACTCTCGCATCTT

CGACTCCGCAGATTTCAATAAGCTCCATGTCCGACGCATCTCAAGCATTCACTTTACCATTTGCAACGAT

TTTGAAATCTAAAATTATCTAAGCATACAGTAAGTTAGGTGTGTGTCTTTCAGACGTTCAAAAATATCTG

CTAAAACAGGAGAAGTTTCGGCTGCAATCATAGTTACAAGGTAAAGGTGGAAGATATTGTTCAGAGTGCT

GGCTCTAATTTTTTGTGAAGATATTTTTATTCCCTGAGACCGGCTTGGAAAGTGGAGTTGGTCATGCTGG

AGGAGTGGGAGGCTAGTAATAGTTTGAAATCTTGACGACGAGTTAAGTGCCATGTCGGTTATGTGCCATC

TGGACAACGAGTGGTAGGCCTAAGCAAGCTCTCTTGGGTTGCTGATGTTTTTGCAAGAAGGTTTGAAGAT

CCATCACTGACAGATGATGTCTGTTCAGCTCTACATTATAGAAAAATGCCTATAGGTCTTGCGGGTCTTC

GGTGTTCTCATTTTCACTTCACAAATTTTGCATTTTACATGTAAATTATGGACAACTAGTAGCGGCCGAA

GTTTGGATTTTATTTGTAAATTTTTAATATTTATTATAGTTGATTTAGGTAC

>MSTRG.2372.1 gene=MSTRG.2372

AAAAATAACACTCTGTTTGGTTGGCCTGAATGGAATCATTTTAGCCCAGAATAGAATGGGCCATTCCATT

CCAAAAATAAAAATTTCTGTTTGGTTGGCCCAAGGAATGGAATAGCCCATTCTTTTATTCATTCCAATCC

AATCACTAATCAGACAATCGATTCATTTTTGGAAGTGCGGGTTAAGCCCATTCCATTCACTCACGAGTCA

CGCCCGCCTCTCTCACGCAGCTCTCCCTCGCCTCCCTCACGCAGCTTTCTCTCGCCTCTCTCCGCCCCTC

TCTCTCGCCTCTCTCTCTTCCTCGCATCTCTCTCTCATCACTCCTTCCTCGCATCTCTCTCTTCTTTTTC

TGCCTCTCCCCAGTCGCCTCCATTGCTGCTCTCAGACAAGGGCGATTCAGAGTTCAGACTCTCGCATCTT

CGACTCCGCAGATTTCAATAAGCTCCATGTCCGACGCATCTCAAGCATTCACTTTACCATTTGCAACGAT

TTTGAAATCTAAAATTATCTAAGCATACAGTTTTATTGAATCATATATCTTTGCTGTTGGGGTTTCAACT

CTTCTGAACTGGGGAATTTTCTTCAAGGAGATAATTTTAGGTTTGGCTCTGGGCATTCACAAGCTTTTAA

ATTGGCAGTCTCAAGCTTATTGGTGTAATTTTTGGTCAGTCTGAAAAGCCATATAAGTTAGGTGTGTGTC

TTTCAGACGTTCAAAAATATCTGCTAAAACAGGAGAAGTTTCGGCTGCAATCATAGTTACAAGGTAAAGG

TGGAAGATATTGTTCAGAGTGCTGGCTCTAATTTTTTGTGAAGATATTTTTATTCCCTGAGACCGGCTTG

GAAAGTGGAGTTGGTCATGCTGGAGGAGTGGGAGGCTAGTAATAGTTTGAAATCTTGACGACGAGTTAAG

TGCCATGTCGGTTATGTGCCATCTGGACAACGAGTGGTAGGCCTAAGCAAGCTCTCTTGGGTTGCTGATG

TTTTTGCAAGAAGGTTTGAAGATCCATCACTGACAGATGATGTCTGTTCAGCTCTACATTATAGAAAAAT

GCCTATAGGTCTTGCGGGTCTTCGGTGTTCTCATTTTCACTTCACAAATTTTGCATTTTACATGTAAATT

ATGGACAACTAGTAGCGGCCGAAGTTTGGATTTTATTTGTAAATTTTTAATATTTATTATAGTTGATTTA

GGTACTACTATTTTGTGTAATGAAATGGAAAGTATGCTATTGAC

>MSTRG.2372.2 gene=MSTRG.2372

AAAAATAACACTCTGTTTGGTTGGCCTGAATGGAATCATTTTAGCCCAGAATAGAATGGGCCATTCCATT

CCAAAAATAAAAATTTCTGTTTGGTTGGCCCAAGGAATGGAATAGCCCATTCTTTTATTCATTCCAATCC

AATCACTAATCAGACAATCGATTCATTTTTGGAAGTGCGGGTTAAGCCCATTCCATTCACTCACGAGTCA

CGCCCGCCTCTCTCACGCAGCTCTCCCTCGCCTCCCTCACGCAGCTTTCTCTCGCCTCTCTCCGCCCCTC

TCTCTCGCCTCTCTCTCTTCCTCGCATCTCTCTCTCATCACTCCTTCCTCGCATCTCTCTCTTCTTTTTC

TGCCTCTCCCCAGTCGCCTCCATTGCTGCTCTCAGACAAGGGCGATTCAGAGTTCAGACTCTCGCATCTT

CGACTCCGCAGATTTCAATAAGCTCCATGTCCGACGCATCTCAAGCATTCACTTTACCATTTGCAACGAT

TTTGAAATCTAAAATTATCTAAGCATACAGTTTTATTGAATCATATATCTTTGCTGTTGGGGTTTCAACT

CTTCTGAACTGGGGAATTTTCTTCAAGGAGATAATTTTAGGTTTGGCTCTGGGCATTCACAAGCTTTTAA

ATTGGCAGTCTCAAGCTTATTGGTGTAATTTTTGTAAGTTAGGTGTGTGTCTTTCAGACGTTCAAAAATA

TCTGCTAAAACAGGAGAAGTTTCGGCTGCAATCATAGTTACAAGGTAAAGGTGGAAGATATTGTTCAGAG

TGCTGGCTCTAATTTTTTGTGAAGATATTTTTATTCCCTGAGACCGGCTTGGAAAGTGGAGTTGGTCATG

CTGGAGGAGTGGGAGGCTAGTAATAGTTTGAAATCTTGACGACGAGTTAAGTGCCATGTCGGTTATGTGC

CATCTGGACAACGAGTGGTAGGCCTAAGCAAGCTCTCTTGGGTTGCTGATGTTTTTGCAAGAAGGTTTGA

AGATCCATCACTGACAGATGATGTCTGTTCAGCTCTACATTATAGAAAAATGCCTATAGGTCTTGCGGGT

CTTCGGTGTTCTCATTTTCACTTCACAAATTTTGCATTTTACATGTAAATTATGGACAACTAGTAGCGGC

CGAAGTTTGGATTTTATTTGTAAATTTTTAATATTTATTATAGTTGATTTAGGTACTACTATTTTGTGTA

ATGAAATGGAAAGTATGCTATTGAC

>MSTRG.2378.1 gene=MSTRG.2378

CGTGGCAAAAACAAAAAAAGAAAAAAAAAAATGCAATGCACTAGATCGAGGGTAGTATTGTCAGCTACAC

TAACAAAAACAGAAACAGTACTTAGTCAAACGTTAGTGGTATACTTCTTAACCAAATCTCCCCGAGATCT

TATATATAGTCCAATTCTTCCCACTACATTTTTCATAGCTCCACTTTCATACAAAATCAACTTTCACCAT

CTACCTCTCTTTCTAGCTTCAACAGATATACTAACCTGCTCCACCACCATGGCTATGGCTTTCATGATGG

CGACAAAGGGTATGTGCGTGACAGAGGAATGCAAGAAATCGTTCATGGAAATGAAGTGGAAGAAGGTGCA

CAGATACATAGTCTACAAGATTGACGAGAAATCGAAGCTAGTGACCGTGGATAAGGTGGGCAGCTCCGGC

GAAAGCTACGATGACTTGGCCGCATCTTTGCCGGAGGACGACTGTCGATACGCGGTGTTCGATTTCGATT

TTGTGACAGTTGACAATTGCCGGAAAAGCAAGATCTTCTTTATTGCATGG

>MSTRG.2378.2 gene=MSTRG.2378

GTCCAATTCTTCCCACTACATTTTTCATAGCTCCACTTTCATACAAAATCAACTTTCACCATCTACCTCT

CTTTCTAGCTTCAACAGATATACTAACCTGCTCCACCACCATGGCTATGGCTTTCATGATGGTAAGCATT

CAAACATATGTTTCTTAAATTTAAGTGTGTTACACTATTTTTCTATTTTACGTGCATATATTTATATTAG

TTCCGACTGTTTCAAAATTCTGATTCTTTTTGTTACATGCACACACCGACAAGACACATATCCATCCATG

TAGTCGATAAATTTTTGGTTTTTTGTTTCCGGTTCGATCTGGACCTAGATCTCGTGTAAATAAAATATAA

TCGATAAATTTGCATGAATAATTGACAGGCGACAAAGGGTATGTGCGTGACAGAGGAATGCAAGAAATCG

TTCATGGAAATGAAGTGGAAGAAGGTGCACAGATACATAGTCTACAAGATTGACGAGAAATCGAAGCTAG

TGACCGTGGATAAGGTGGGCAGCTCCGGCGAAAGCTACGATGACTTGGCCGCATCTTTGCCGGAGGACGA

CTGTCGATACGCGGTGTTCGATTTCGATTTTGTGACAGTTGACAATTGCCGGAAAAGCAAGATCTTCTTT

ATTGCATGGCATGGATGGTTGTTTTAGGGCTCCCACGGCATCAAGAATCAGAGCAAAAATTCTGTATGCA

ACTTCGAAAGACGGATTAAGAAGAGCATTAGATGGAATTAGCTATGAAGTACAGGCAACTGATGCTGCGG

AGATGGGTTTCGACGTCATTACTGATCGTGCTAAGTAATGATTAAGCAAGGCGATTAGGATTAACTAAAG

TTTGTTTTGGAATCTTTATAATTTTGAATAATGCTGCTGCTGCTTGTTGCTTTTCTTTACTCGTTAGTTG

TTAACATCAACTTTTTTTCTAATCTCCATGTACATAATGCCCCAATAACTAATCTATGATTAACACATGC

TGTGTTTTTGTTAATTAATGGAAAATACCCTTATTGTTGCCTTGCT

>MSTRG.2381.1 gene=MSTRG.2381

GAAAAGAAGTAGTGGGACCCATTAAGCTTAACCGCCGGCTCATCCTTCGCCAACGGCCACTGAACTTCAT

CTCCGACCCTAGCAAGAACCGCTACGACATTATCACCCTGTCGAAGCCTGACAATGTTTAATTCACCAGA

GGCAAGTTGAACACTCCGCTTCTTATCAATCAAATGAACAATCGCACCGGGGATTTTAATCAGAATCTCC

TCTGACGACTCAAACGCCGGCGGGTTCTCGCCGGAATCGCGATGAATGGGGACGTCGGAATCGGTTTCCG

GGAACAGTTTTTCGGCAACTTCTTGAGAGTCTATAGACGGGTACAAAGAAGACATGGTGTTGTGCGTATG

TGAAGATTGTAATTTGATTAATTAAATATGTGTGTATTGGTTAGA

>MSTRG.2393.1 gene=MSTRG.2393

CTGCTTTTTGTGGCCAATGACAGTGGCTACAGTTATGTAAAGAAACAGGAGAGTGGCTATGAACTCAGCT

ATAAGAGCTCTGTAAAAAGACCAGAGCTTGAGTTCACCCATGTCAAGAAGAGGAGCTGGCGGTGGGTCCA

CATAGTCTTTTCCATGGTGATGAGCTTGTCCTTCCTCACTCACTTCTTTCATTTTGTTTCTCTCTCTACA

ATTCTTTCTTTCTTCTTTTTGTGTTTCTCTCTCTAGAGCTTTGTAACTGCC

>MSTRG.2399.1 gene=MSTRG.2399

ATTTAGGTTCGGAGTCCGGAATTTAGCGGATTATTTACGGCTTCTCGTGAGTGTTCGATTAGCTATTTTC

GAGGTACTTAGGGGGATTCGGGACTGTGTGATGGAGAGCAGCCCATTCTTTTCATTGTTTAGGACTTTAT

GGACTTCTAGCATTTATTCGGATTTTATTGTTTAGAGATTCTATTTATTCGCATTATTCGGATTAAATTA

TTTTCGAGGATTTATATCATTCGAGTACTTATTAGAACTGTTTATCATCGTTTG

>MSTRG.2399.3 gene=MSTRG.2399

ATTTAGGTTCGGAGTCCGGAATTTAGCGGATTATTTACGGCTTCTCGTGAGTGTTCGATTAGCTATTTTC

GAGATGCCTCCTAGACGTGAGAATCCTAGGCGCGATAACGTTAACATTGGTGCAGCTGAGTTAGCTCAGT

TGATAGCTCAAGCAGTGACTCAGGCTTTGCAACAAGCTGGGCAAAATCAGGGAAACCCAGGAAATAATGA

GGAGCAACAAAATCAGCCTGATGCACTAGCTTGGTTCGATAGGTTTGTTAAGCAGAAACCGGATTCTTTT

CATTCAGCACCACAGCCTATTGATGCTGAAAATTGGATTGTTCATCTTGAGAAGATTTTTGATGCATTGG

GATGTGATGACGCTACAAAGGTGAGGTTGGCTGTGTATAAGTTAGAGGGGGATGCTCAGAGGTGGTGGAG

AGGAGTGAAGACTATCAGGGGTGATGCATTTGTTGAGGCATTGACTTGGCAGGGGTTCACAGATTTGTTT

TATGAGCAGTACTTCTCAAAGGCAGAGAAGGATGGTTATATGAGAGAGTTTAATTCTATTGAGCAGAAGC

ATGATGAGAGTATCACTGAGTATCTTGCGAGATTTATCAGGTTGGCGGGATTTGCTGGGACTTCTGCAGG

GACTGCAGCACAGCAGGCAGAAAAATTTAAATGGGGG

>MSTRG.2399.2 gene=MSTRG.2399

ATTTAGGTTCGGAGTCCGGAATTTAGCGGATTATTTACGGCTTCTCGTGAGTGTTCGATTAGCTATTTTC

GAGGTACTTAGGGGGATTCGGGACTGTGTGATGGAGAGCAGCCCATTCTTTTCATTGTTTAGGACTTTAT

GGACTTCTAGCATTTATTCGGATTTTATTGTTTAGAGATTCTATTTATTCGCATTATTCGGATTAAATTA

TTTTCGAGGATTTATATCATTCGAGTACTTATTAGAACTGTTTATCATCGTTTG

>MSTRG.2399.4 gene=MSTRG.2399

AGCAGAGGTGGCTGATGCAGCCAAGGATGTTGAGAAGGAGCGTATAGACTTCCGGACTAACAGATCTGAC

AGTGGTCGTAAGAGGGGTAGAGATGATCAGCTAGTCACAGCACAGGGTAGACAGGGGTATGGTGGTCAGA

ACAGTCGGTATGGACAGTGGCGTGGACAGAACCAGAATAGGGGAGGTCATACAGCTCATGGTCAGAGACA

GAATCAGTATAGTGGTCAGGGGCAGGCTCAGCAGGGGCAGACTCAGCAGTTTCAGCAGCAGCCTAGACAG

TGGCAGAATCGTCAAAGGGGACAGGGACGTTATCCAGCGTATGGGGGTAACCCCAATATGATTCCGGTGG

CTCCTTGTGCTACATGTGGTAGACACCATCCGGGCAGAGCTTGTTATAAACAGACTGGGGCATGTTTCTC

GTGTGGTTCGATGGATCATAGGTTAAAGGACTGTCCACGTAATAATGGTGATGGAGGTAGTGGCAGTGGA

GGTCAGCAGAATCAGCAGAATCCTACAGGCAGAGTCTTCGCATTGACTTCTTCTCAGGCAGCAACTAATC

CAGGTTCGGAGTCGGGAGTTTAGCGAATTATTTGCGGAATCTCGTGAGTGTTCGATTAGCTATTTTCGAG

GTACTTAGGGGGATTCGGGACTGTGTGATGGAGAGCAGCCCATTCTTTTCATTGTTTAGGACTTTATGGA

CTTCTAGCATTTATTCGGATTTTATTGTTTAGAGATTCTATTTATTCGCATTATTCGGATTAAATTATTT

TCGAGGATTTATATCATTCGAGTACTTATTAGAACTGTTTATCATCGTTTG

>MSTRG.2405.1 gene=MSTRG.2405

AGGTACAAATCAGCTGATGATGTGTCGCCATGTCAGCATGATTCAGCGGGTGATGTGTCACCACATCAGC

AAGATTCTAACTCTGTTAGTATCTACTTAACGGAAGGACTAGATTCCAGCGTTTGGAAAAGTTTAATGAC

CCGGTTTGAGGAATCTTTTGTTTAGGGACCTATCGAATAAAAGTGGATTAGTTTAAGGACGTATGAATAT

ATTATCCCT

>MSTRG.2408.1 gene=MSTRG.2408

TTGTAAATTCAGGTAACTACTATACATCTTGTAGTACTTTGTCTTGTAGTTTGTTTTTTCTCCGGCAGAA

TGCTACTGTTAAATTAGTAATTATTAATTTAATCAGGGCTTAAATGATTTTCCATCTCATATTTTCTAGG

ATTATTATATTTTATGAGTTGCATTGGGTAGGATGATGATAATGTTCACTTATGTGAAACCTTTATTTGT

AATTTGTAGCTCTGTTTTATCAATTTAATGTTTTGGTTTAGCTTCTTGATTGCAATTTTATGATTTGATT

TTAGCTTCTTGGTTGCAATTGATATGGCGTCTACTGTTGTATTAAATATGTGGTGAATGATGTGGCTTGA

TTATATTTTTTATAGTGGTGATAAGAATTATGCTACTAGTAGTAATAGTCTCCAGTGTTGTTGCTTCCTA

TTTTTTGGCTGGGAGTATTTTCTTGACCTCATCTAATGCACTCAGGGCGGCGGCTAAGTGGAGTATTAAT

TCTGCAAGGGGTCTGGGATTTGCAGAGTGATTTGGGGTATGCTGATTCAAGGAGTTTATTAGGAAATTTA

CTATGCTGCTGCCTGATAGTTGTGTTGCTTTAATTTTCAGAGGCACGATTGTCTGGTTACATAGATTTGA

TTATAGTTTTAAGTTATGTGTGCATCTCAACAGATCCTAACTCCAGAAGTCCGACTGACTGAGTGAGACA

TACCAATTACAGTATATGTGTTAATGTGCTGCTGCAGCACAGAAAAATAAATTTTGTTGTGGGAAAAATA

TATTCGTGAAAGAAATGTCTGGCTTCGATGATTAATGTCATCAGAGATTTTATTTTGCGACCATAAAAGT

TAAGAAATTTGACTAGTGCTTGTCATATAGTACTGCAGAAAGTGTTATATAGTTAGTATTGGAACAAGTT

GTCCACCTAATTCAAGTG

>MSTRG.2410.1 gene=MSTRG.2410

TTGCAATATACTCAGTATCAAGAATGAAACTTTATATCAAAGTGATAACAAAACCTAAACAAACAAAATC

TCACGACTCTGCTACAACCTCAAACTCTTTAATCAAATCAGAATCAACATTCTCCGCAGGCTCCCAGGTG

GCCTCATCAATATCCATCCATTTCACCAAATACTCTCTCACTCCACTCTCACCTTCTCTGCACTCTAATA

TACACTCCACCACACCATACTCCAACCCTTGCTCAAAATCCCTTACCAAATCCTCACTAATCATCTCCCC

TTTCACCCACTCATTATCTCCACCGTCCATCCATTTCACCAGATACTCCACCTTATCATCTCTGCCTCTC

TTCTCTATGATCTCCTGCACCTCCGCGTATTCGAATATCGCCCCTTCGAGTACCTTAATCACGTTCTCTA

ACCCTAATCTTCGCGCAAATTGCATGGGCGGAGTTGCTTTTAAAACCTCTCGCGCCAAATCCAACGGCGT

ACGTTTCCTGTCATCTTCGACCTCGGGATCCGCGCCATAATCGATTAGTAATTTGGCCACGTTTGGCTTC

ACGTACCCGGCTGCCATATGTAGAGCCGTGAAGCCGCCGCTTGTGTCCTTGCGGTCCAAGTCGGCTCCGG

CTTCGGCTAATATCTTCACGCACGGCTCCGCGCCGAGCCCCGAGACGAATATCAAAGCGGTACGACCGTT

AACATCAATCGCGTCGACATCTCTCTCGTCCTCGGCTTCGATGAGCCGAGCCAGAGCCGAGTCATCGCCT

TTCTTGGCGGCTTCCCACCAGGGAGAATCGAACTCGGCGACCACATCTTTCGCGATGAATTCGGACGGGA

CCCAGGTAGGGGCATGATCGTCTTTCCACTCGATTAGGTACTCCATAGTGTTCGACACGGCTCGGCTCCC

GATGATTTTTTTGACTTCCCCGTAGGTTTCTTCCTCATTGTAGTCTTGGAAGGTGAGTTGAGAGGAGGGG

GATTGGTTATCGACAGAGAGGCTGAG

>MSTRG.2411.1 gene=MSTRG.2411

AGAACACCGCGTCTTGCTTAATTTAAATACTTCTCCGCACTTGAAATACTCTTTCTGCTACCCACTGACA

TCACAAACCCCAGTTTCTGACTCATCTTTTAACCAATTCACATTCACATTCTTCCTTCATTCCAATCTTC

ATAATCATTTTCTTCCTCTAAAATACTCTCGATTGGGAACAAAATAATAATTCTAAAGAAATGCATGCGA

AATCAGACTCCGACGTGACGAGCGTCGAGGCCTCATCGCCGGCACGGCCACTCTACTACGTGCAAAGCCC

ATCGAATCATGACCCGGACAAGATGTCTTACGGGTCGAGTCCATTCGGGTCACCGGGTCACCACTACCAC

TGCTCCCCCATCCACCATTCCCGAGAATCGTCGACGTCGAGATTCTCGGCTTCGTTGAAAACCACGGCGA

GGAACGCCGGGTATTGGAAGAAACTGAACCGCTCCGGCGACGGAATCGAGGAGGGTGACGAGGATGACGA

GTACGATGAAGAGGAAGAGAGAGAGAAGAAAGCGATAAAGAGGTTTTATGTTGTGTGTTTTGTGGTGTCG

TTTGTTGTTTTGTTCTCTATCTTTTCGTTGATCTTGTGGGCCGCGAGCACTTCTTACCCTCCTGAAGTTT

ACGTCAAGAGGTTGGTGTTTGAGAGGGTGAATGTGCAATCGGGTATGGATTTATCGGGGATCGCAACGGA

TATGTTGACTTTGAATTCAACGGTCAAGATTTTCTATCGGAACACTGGTACATTTTTTGGAGTTGATGTT

AAAGTTGCGCCCATTCGACTTTTGTATTACAAGCGTAGCCTGGCCAGTGGTCCTGAGGTGAAGTTCCATC

AACATAGAAAGACTCAACGGACTATAATAACCACCATGGGAGCACATCAACTTCCACTATATGGTGGAGT

GCCTGGTTTAACTGCTAGGAATGGCGAAATTGACAGCATATCTGTACCACTGAACCTTACTATCGGCATA

AGGTCGAGGGCTTATGTTTTGGGAAAACTGGTGAAGCCCAAGTTTTACCGGACCATCAGATGTGAAGTTA

CTTTACATGGCAGCAAGCTTGGGAAGCCCGTCAATCTTACGAATTCTTGTTTTCATCATAACTAATCATC

CAAATAACCCCAACAATCGCTTAAACTTGCTCAGGTTTTTCTGGTGCTTTATCATTAGGATGTTTATCAT

TAGGATGTTTTTGTGGAAATTCGTCATGCATCATTATTGTACTTTATTGTGCCATTATTGTACTGTAATT

TGTAGTCATTTTGATGTATTTTTTAGCTTATACCATATCTGACCCAGTGTATATAGGAAAGATGTTGGAG

AAATTGTAGATCATCTGCATATATATTGATACAGACAATAGGAACTTTTCTTGAA

>MSTRG.2413.1 gene=MSTRG.2413

GGATCATATAGGTACAGGCGAGGATCGAGGCCCAGAGTCTTCTGCTACTGCTGCTACAGCTACTGTTGCT

GATGCTTCTGCTGCTGCTGCCACATCTACTGCTACTGTAGCCGAGGCTTCTACTCGCCCTGCTGCTGCTG

CTGCTGCAGCTACTACTACCGGTGCCGGTGACTGGTGACCTAGTTTTTCTCTAGGCCTAAGTTTGCCTTC

ACAGCCAGTACCTGAGCAGCAGACTTCTCAGATGCGAGATACCAGTACAGATCAGCCTCCTCCTGTTGTG

CAGCGTCAGCACAGGATTCGGCCTTGGCATGGTACACCAGAGGTACCCCCTTTTGACTTGGGCAGCTCTT

CACAGTCTACACAGTCTACGCAGCCTAGTCAGCCTGCTACGTCTACGCAGCCTAGTCAGCCTGCTACGCA

GCCTAGTCAGCCTGCTACGCAGCCTAGTGTACCCCCTTTTGACGTCTTTTATGTTCGACGGTCAAAGAGG

GCTAAGAAGGTTCCTGACTGTGGTACTGGTTCACATAGGGGATGATGATATTTGTAGGTTTTTGATGACA

TTCTAGTTATAGGATGCTGAACTTGTGATATATGATAGCACGGATCTTGTAAATATTTTAATTTCATGTC

GTATGTTTCTGTTTTA

>MSTRG.2417.1 gene=MSTRG.2417

AAAAGAGAGAGAGAGAGAGATTCGGCGTGAATGAAAGAGGCCACTGAGCTCCTATAATGTCTCACAACCT

TGTAAAACCCTGAAAGGAAGGCCAAAAAGACACAATCCCAGAGTCTTGTCCATTCACTTTACAGAAGGCT

TGATGGATCAAACAAACCATCCAAGTGTCATACAGAAAATAAATTTTGGTGTCCAGCACCCTTCTCTGAA

CCAGAGAAAATGCAAAGGTGGAAATTATAGCATTGCTGGTTTGCAACATCCAATGGTGCAGCCAAATCGA

TCCAGACATGATATATCAATTACATCGTCCGCTGTGTCACCAGTTTTTGTCCAAGCACCTCAGGAGAAAG

GCATTAAAGGGTTTCTCACTGATTTTCTCATGGGAGGAGTCTCTGCAGCTGTATCCAAAACTGCTGCTGC

TCCAATTGAACGCGTGAAGCTTCTGATCCAGAACCAGGATGAGATGCTCAAATCTGGGAGGCTTTCTGAG

CCATACAAAGGTATAGGTGATTGTTTCAGCCGAACAATGAAGGATGAAGGTGTTGTATCGTTATGGAGAG

GAAATACTGCCAACGTCATTCGTTACTTCCCAACTCAAGCCTTGAACTTTGCTTTTAAAGACTACTTCAA

GAGCCTTTTCAACTTCAAGAAAGACAGGGATGGATACTGGAAATGGTTTGCTGGTAACTTGGGCTCTGGA

GGTGCTGCTGGTGCTTCATCCCTCCTCTTTGTTTACTCGCTGGACTATGCTAGAACCCGTCTAGCAAATG

ATGCAAAGGCAGCAAAGAAAGGAGGAGGACGGCAATTTAATGGCTTAGTTGACGTCTACAAGAAAACATT

GGCATCAGATGGTGTTGCAGGGCTTTATCGTGGATTTACCATTTCATGTGTTGGTATCATAGTTTATCGT

GGTCTCTACTTTGGAATGTATGATTCCTTGAAGCCAGTCCTCCTCACTGGGAAGATGCAGGATAGTTTCT

TTGCTAGCTTTGCCCTTGGTTGGGTTATCACTAATGGTGCTGGATTGGCTTCCTATCCCATTGACACTGT

CCGCAGAAGAATGATGATGACTTCTGGTGAAGCAGTGAAGTACAAAGGCTCAATGGATGCACTTTCTCAG

ATAATGAAGAATGAAGGTACTAAATCGCTCTTCAAGGGTGCTGGTGCCAACATTCTACGTGCTATTGCAG

GTGCTGGTGTTCTTGCTGGTTATGACAAGCTGCAAGTGCTAGTTCTCGGAAAGAAGTACGGATCAGGTGG

TGCATAAAAAACACAAATGCCCCTCGGCCCTTCAACATTAGATCATGATGAAAATAATTCAAATTTTTTA

GCTATCGCACCAACAATCTTTTTTCAATAAATCAGTTTGATCATGGGAGTGAGTACTCCTGAACAGCTGA

GCCCCACAATCACCAGACAAGTATCATTTCAGTTTTGTAAGAACACAATATTTACGGTAGTTTTAGATAT

ATTATGTGGTAATTTCTATTGTGAAATCCTTTTTATTGCTAGGGGATAGTTCGTATCTCTCTGCAATTAC

ATTACACTTTGTACCAAGTCCGAAGTTCCAGTGGAACAAAGTCATGGATACAGAGTAAACCCTTACCCTT

ACTGTATTGAAACCAGAAGTTATATCAAGTCTGGAACAATCACGAGTGTCAGTTTCTGAGTGTCGGGTCT

TGTCACTGGACATTTCAATTTCCATCCTCGCTTCTTAATACTTGTTTCATTAGTTTCTATGACTGAAG

>MSTRG.2417.2 gene=MSTRG.2417

AGAGAGAGAGATTCGGCGTGAATGAAAGAGGCCACTGAGCTCCTATAATGTCTCACAACCTTGTAAAACC

CTGAAAGGAAGGCCAAAAAGACACGTACACGACCCCTTCTCTTTCTAACAATTCACCACCTATATTTCTC

ATCTACATAATTTATTTTCTTGTATTTTATCTTGCTCTACATTGTTGCATCCTTTCTTGCGAATCCCAGA

GTCTTGTCCATTCACTTTACAGAAGGCTTGATGGATCAAACAAACCATCCAAGTGTCATACAGAAAATAA

ATTTTGGTGTCCAGCACCCTTCTCTGAACCAGAGAAAATGCAAAGGTGGAAATTATAGCATTGCTGGTTT

GCAACATCCAATGGTGCAGCCAAATCGATCCAGACATGATATATCAATTACATCGTCCGCTGTGTCACCA

GTTTTTGTCCAAGCACCTCAGGAGAAAGGCATTAAAGGGTTTCTCACTGATTTTCTCATGGGAGGAGTCT

CTGCAGCTGTATCCAAAACTGCTGCTGCTCCAATTGAACGCGTGAAGCTTCTGATCCAGAACCAGGATGA

GATGCTCAAATCTGGGAGGCTTTCTGAGCCATACAAAGGTATAGGTGATTGTTTCAGCCGAACAATGAAG

GATGAAGGTGTTGTATCGTTATGGAGAGGAAATACTGCCAACGTCATTCGTTACTTCCCAACTCAAGCCT

TGAACTTTGCTTTTAAAGACTACTTCAAGAGCCTTTTCAACTTCAAGAAAGACAGGGATGGATACTGGAA

ATGGTTTGCTGGTAACTTGGGCTCTGGAGGTGCTGCTGGTGCTTCATCCCTCCTCTTTGTTTACTCGCTG

GACTATGCTAGAACCCGTCTAGCAAATGATGCAAAGGCAGCAAAGAAAGGAGGAGGACGGCAATTTAATG

GCTTAGTTGACGTCTACAAGAAAACATTGGCATCAGATGGTGTTGCAGGGCTTTATCGTGGATTTACCAT

TTCATGTGTTGGTATCATAGTTTATCGTGGTCTCTACTTTGGAATGTATGATTCCTTGAAGCCAGTCCTC

CTCACTGGGAAGATGCAGGATAGTTTCTTTGCTAGCTTTGCCCTTGGTTGGGTTATCACTAATGGTGCTG

GATTGGCTTCCTATCCCATTGACACTGTCCGCAGAAGAATGATGATGACTTCTGGTGAAGCAGTGAAGTA

CAAAGGCTCAATGGATGCACTTTCTCAGATAATGAAGAATGAAGGTACTAAATCGCTCTTCAAGGGTGCT

GGTGCCAACATTCTACGTGCTATTGCAGGTGCTGGTGTTCTTGCTGGTTATGACAAGCTGCAAGTGCTAG

TTCTCGGAAAGAAGTACGGATCAGGTGGTGCATAAAAAACACAAATGCCCCTCGGCCCTTCAACATTAGA

TCATGATGAAAATAATTCAAATTTTTTAGCTATCGCACCAACAATCTTTTTTCAATAAATCAGTTTGATC

ATGGGAGTGAGTACTCCTGAACAGCTGAGCCCCACAATCACCAGACAAGTATCATTTCAGTTTTGTAAGA

ACACAATATTTACGGTAGTTTTAGATATATTATGTGGTAATTTCTATTGTGAAATCCTTTTTATTGCTAG

GGGATAGTTCGTATCTCTCTGCAATTACATTACACTTTGTACCAAGTCCGAAGTTCCAGTGGAACAAAGT

CATGGATACAGAGTAAACCCTTACCCTTACTGTATTGAAACCAGAAGTTATATCAAGTCTGGAACAATCA

CGAGTGTCAGTTTCTGAGTGTCGGGTCTTGTCACTGGACATTTCAATTTCCATCCTCGCTTCTTAATACT

TGTTTCATTAGTTTCTATGACTGAAGGCAGTAACTACATAAGCTGTATATTTGAGACTTCCTC

>MSTRG.2418.1 gene=MSTRG.2418

ATATATCTGCTTAAAGAAAATAGTTGTCGCAGAATCCAAAGTGCCTTGTTCCCTATCCAAAACTCCCTGG

TATCATCTTCTAATTCAAAAGAAAGATGTTGGAGATGTTTTTTGTAGTAGCTTACGCTGCGGCGCCATGG

ACTCTGTATATTCCTCCGGTGAAAAACTTGAGCTTATTTGTGGAGACCATGGAAGAGCTATACAGGGAGT

CCAGTTATTATGCTAACAGGCTCTACCCACGCCTTCGATATGCTTGCTCTAGGATCTTGAACTGCCTTCT

TTGCAATGATACAAGGTAGATTGTTGTTATATACTTAGCCTAGTTATGTTGAAAGAGCTTGTACTTTGGT

TATGAGGTTGTAATTATTTCGGTGTTCTTTGTTGTGAAAATTCACATGAGGAGTTCTTTTTTCATCTATC

AATAAATGGTGGAAATTGTTTGACTTTTCATGATTTCATAAGTTGATTCTATTTGTAAGTAAGC

>MSTRG.2423.1 gene=MSTRG.2423

TTCTTTCCAGTGAAAGCGGAGAAGGTTGGTAGCGATGACCCTCAGGCTAGTGGTAAGGCCAGGATTCCAG

GTTGAGAGACGGAGGAAGCTATCAGCAGCACTTGGCTTGTGATATATGTACTAGTTTGAGTTTCGAGATA

CGAATATTATGTAAGATTGTAAGAATTGAATTTATATTTGGGATTTGGGTTTGTAATAATTAAAGTTAGA

TTGTGGCTTGTTTTATACTTTAACCTGTTGCGATCCATGGACAGTTAAGGGTCACTGCATATATTATATT

ATTAATTACAGGTTTATGTTATGGTGTGGACCCCAAACTTCTGACCCGGGTTTGGAGGGCGCCACA

>MSTRG.2429.1 gene=MSTRG.2429

CTTGTCATCTAACTCTTATCCCCCGTCTATTTTCTCTATTTTTAGTATTTACATATAAATCTTTTCCTAA

CTATCTCTTTATAATAATCCACTACATCAGTATATACTTCACACCACTGGTACTACTTTTCTCTCACATT

GATCCAATCTCTACTTTCCAGGGCTTTCAGATTCTATCTTAAAGGGCAGGCAGGTATGGCGAATGGTGGA

GCTTTGGATCTTACTTCTCATCTTGGTGGAAAGATTCAGAAAGATGATGTTCTTGATGCTGTTGACAAGT

ATGAGAAGTATCATGTTGCTTTTGGAGGGGAAGAAGAAGGGAGAAAAGCAAACTACACGGACATGGTTAA

TAAATACTATGATCTAGTCACAAGCTTTTACGAATATGGTTGGGGAGAATCATTTCATTTTGCACCCAGA

TGGAAAACAGAATCTCTTCGAGAAAGCATCAAGCGGCATGAACATTTCCTTGCTCTACAACTGGGTCTTA

AGCCTGGGCAGAAGGTTCTGGATGTAGGATGCGGAATTGGTGGACCACTGAGGGAAATAGCTAGGTTCAG

TTCCACTTCCGTTACAGGGCTGAACAACAATGAATATCAGATAACAAGGGGAAAGAAACTAAATCACATG

GCAGAACTAGACCAGACCTGTAATTTTGTGAAGGGAGATTTCATGAAAATGCCATTTCAAGATAATAGTT

TTGATGCAGTCTATGCAATTGAAGCTACTTGCCATGCCCCTGATGCGGTCGGGTGCTACAAAGAGATTTA

TAGAGTGTTAAAGCCCGGTCAATGTTTTGCTGCATATGAATGGTGCATGACTGATTCCTTTGATCCCAGC

AATGCAGATCACCAAAGAATCAAGGCTGAAATTGAGCTTGGTGATGGCCTCCCTGACATCAGGTTGACTA

CACAATGTATTGAAGCTTTGAAACAAGCAGGTTTTGAAGTCCTATTCGAAAAAGATCTGACGATTGGCTC

ACCCTTACCTTGGTATTTACCTCTGGATAAAAGTCATTTCTCGCTAAGCAGTTTCAGGTTAACAGCACTT

GGACGTTTTATAACAAAAAACATGGTTGTTGCCTTAGAATATGTTGGAATTGCACCAAAGGGTAGCCAAA

GAGTTCAAGCTTTCTTGGAGAAAGCTGCAGAGGGACTTGTTGCTGGTGGAAAGAAAGAGATATTCACACC

ATTGTACTTCTTCTTGGCTCGGAAGCCTCTTTCAAGTTAAATCGGTGTGCCTCTATTGAATTCTTTGTGA

GGATTTCAAAGGGATAACTAGCTATGAAATTGGTGCTAGACCACTAATAGAACATCAGTGGTAGATGGTT

TCAGTTTGTGAGTTGTGTTTCTTTTTTCCCGTAATAGTGTTTCATTGGTTTTCCGGGGGATGATCTGTGC

ATTACTTAGTTCATGTACAACACTTAAAAGTTGTTTGAGCTTTCAAGAGATGTTAGTTTTCTAAATTAAT

CTTTCCTGGCTTGCAAATTTTAAGATCATCGATTTTTCCTATGGGAAGCAACAGTTAATGTTTGACCAAT

ACCAAGTATGCTATTGTGG

>MSTRG.2430.1 gene=MSTRG.2430

ATAAAATTGGAGTTCCATTGATGCAAATGATTTTCTCAAATATAATAATAATAATAAGCTTGCGCCGTTG

GGTGAAAACAAATCACCCAACAATAGAATAAAGAATGACGTCAGCAGTATCCATGCTCACGCCAGCCTTG

GACCCGCCCTGCTTGACCCGACCCGAACCCCCGCCGCCGTGAAGGTGGGTGTCATGGTGGTGGTGATTCT

CCGCCGGTGGGTGGTAGTACATACGCGCCGTCTGGGGGCAGTGAGAGTAGGTTCTTGCCACGATTCGAAC

ACCTGCATCAAGCAACTCCATGTACTTGGCC

>MSTRG.2433.1 gene=MSTRG.2433

CTTCACTGTTGTCCCAAGACCACTCTTCTTCTTACCCTTCTTTTCAGCTGACTCTTCTGCGGATACAGGC

AGTTGACTTTTTCCCTGTTCATCACCATCTGCCTTCTTTTTCGTTTTCTTTGCTGGCTTCTCCACATTCT

TATCAGTAGTGACCTGTCTTTCATTCACGTCAGCCTTCTTATTCGTTTTCTTTGCTGGCTTCTCCTTCTC

CACATTCTTAACTGTGGCAACCTGTCCTCCAGCATCAGCCTTCTTTCTCTTCTTCTTAAGCTCAACTTCT

TTGACATCCAAATCGTGCTTATCATCAGTTTCTTTAATCACATTTTCCTCAATAGCACCAACTCTTCCTC

CACGAGCAGCTTTTTTAGTCTTCTTGCCTGACTTATCTTCCCCAATAGCCTTTTCTTTCGCCGAGCCCCT

CTTCTTTGTCTTCTCCTCCAGTTCACTCTCATCATTCACCTTCTCACTCTCTTTCTCAC

>MSTRG.2449.1 gene=MSTRG.2449

GCTGAATTTCAATAAATGACTTGGCTGCAGGAAGTTGCTCTTTTTCTTCCATACCATCTTTGGCATCAGC

TATAGGCTGTTTCTGCTCACAGTCTTTCAGTGATTCAGATGTTCGAATTCCATGAAGCAACTCTGTGACA

AGAACTTCTTTTTCCTCTATTCCATCTCTGGCCTCAACAATAGAAATCTTCTGGTCATGCTCTTCCTGGC

CTGCTTCTTCATTTTTAG

>MSTRG.2452.1 gene=MSTRG.2452

GAATATCATCATCCGTGACTTTAAGAGCGTCTCTCATCCATTGCAACGAAGGAACAAACTGCCGGAAATA

ATATTGACGCCGATCGAGAACGGGGATATTAACGTAAAGATTATCTTCTTCCTGGAGGGAGATACGTCGA

GCATAATAGATATCAGCGTTGGAACGGTGTCGTTTAAAGATTGAGAAAAGAGTCACGAGTATAAGAGCTA

CTCCTATGTTGATTCCTGCCGATGCTAGTAGGCTTTCCCCTGTTATCATCCTCCTTAAATCTCCT

>MSTRG.2456.1 gene=MSTRG.2456

CAGTTTGCTTCTCCCCTCTCTTTCTCGCTTACTGTATAAACAACTTGATGCTTTCTTTCGCTGTCAATGG

GTCTAGTTTTTATGTAAAGAAGCTTAAGAATATCTTCGCTCTTCACTCTCAGTTACTCACTTACTAACAG

CATATATTTTAGCCTCGTGTATCAGGATCTTGAGTTTGAGTTTGTCATTTAGTATCGATGTCTTATGTAC

CATCATCTTTAAGGGTTATGTTTTAAGGATCAGTTTGGCAGGGTGTTGCTAGTTCTTCATAATATATGTT

TAAAGTGACTCCAGGGGGTTTATAAACAATGAGATGGCCTCTCTCAAGTTGTACGCCGCAGAATTCAATC

GTAGTTAGTATTTCAGCGGAGGTATTGCTTTAGTATATATCCAGCGTGTTGGAACTTGGAAAGGATTAGA

TGGTCTTTTTGAAGCATGTTTAATTAATTTGAATTTGAATATAATTTATA

>MSTRG.2457.1 gene=MSTRG.2457

ATAGCCTAGTGGTTAAGGGCTTATCCTCTGTCGTCCCAGGTCTTGGGTTCGACCCTGGCTCATGTCGAGA

ATATCTTAGAACAGGTCTTGGGTTCGACCCTGGCTCATGTCGAGAATATCTTAGAACAGATACTCATTTG

TAAGGCTATAAGCCATATTTGTCAAAAAGAAAAAAAAAGTGGTTGTTTGGTTTGTTAGATTTT

>MSTRG.2465.1 gene=MSTRG.2465

TTTAGAATCTTCTTCATGTATATACATTTTCTGTCAGAGATCCAAACACAAACTTCAGCATCTTCTGTCT

TAAGTTTTGGTTCATAATTATCACACACACGCACTGATTTTACGGGCATTACACGTTTTTAATCTTCTTG

ATCTCTCAATTACGCGCTGTCTCTCCCGAGATTCTGCTGCTCTGCACACCCCCTTTACACTCTTTTATCT

TTCTCCTTCAAATTTGTATTCTTTTCTTCAGGTTTGGATGATCGTTTGCCTCTTTAAGGTT

>MSTRG.2467.1 gene=MSTRG.2467

AAAATTAAAAAGAAAAGTACACTAAATTTCAGCATGCCACAGCAACCAACAAAAACAAAACAAGCTTAAA

GAAGATCACCCAAATTAAGTGGCAGATAATTATTGAAATTTAACACGGAATTTTAATTAATCTAAGTGCA

TGTCTCCCGGGCTACTCCAAGCCTTGAATTTGGAACATCGAAGAGCACTCTATGGTTCTGTTGTTGCAAG

CTGGCAATGACATTGAGGACAGAGTTCACGTTGTCTGGTGCCCCTGCCATGGCGAGGCAAGTGGTGCTGC

CAGAGCTGCTGTGGATCAGGAAATTGTCTTGTGGCAGGGTCACGTTCATGCCGGAGAACATGAATGTGAT

CGAGGGGATGGTGATCGGGACATTATAACACGTGTCGAATCCGCCTAGCGATGATACGGTTGCGGTTTTG

CCCATACGTCTGCGGAATTCGTTTCTCACTGGAATGTAGGCTGATTGCACAAGCCGAGTGAAGGTGGTGC

CTGAATCAAAAACGGTTCCTGCTCCGGTGGTGGGATTGAATGCGAATGCGCTAGCCGGAATGTCAACAAT

TTTCTTGCCGACTTTGATGGCCACTAAGTTGACATAGTAAAGTGATGATCTTCTCGGGTTCTTTAGCAAT

TGGGTGTACTTGATCCGAGTGGGTTGAGCGGTGGGACCCAACCGGAGCGAACCGGAGAAGTTGGGTGACT

TGTAGCTAGGCAGGCAATAAGAGAATGTGGATTTGTAGAGGCTCTGAGTTTGTGACAGTAGTGACAATGG

ACCCCTGCCGAGGCCTAGAAGGCCTTGTGGTGGCACCGAGTTGCCTGTGGTCTTCTGGATGCACCCGAAA

GTGTAGCTTGGGATGACGTCTAATGCGAGTTTTAAGGAATCTTGTGAGAGGTTTGCCGCCACCGTGGAGC

CTCCGTAGGTTGTGTTGAAGCTGCAAGCAATGGCGGTGCAAGTGGTGGGGTTGAGGACCTAATTGGTTCA

TCATTGAGGATCATGTTAGTAATGGACATGGGGAATGAGTGAAAAAATTGAAAAAGTTACTGTATTTTAC

CTGTTTGCATTGGGGAGCTTGACAGCCAAGAGTCTTGTAAGATGTCGATTTTTCGGAAGCGAAAGTGGAA

GAAGGGCAGCCAATGCAGCCAGAGCAGGGCACCCAAGCAGCATCACTACTGGTATCCACAGCCATAAGGA

ATGTTTGTGGAGGTGTTCCAATCATAGCCCTCACCACATAAGTTGGGCTTTGTATAATCCCTCTCCCTGA

GGCAATAGGCACCACAGATTTCTTCGCGACAGCGAGGCTTGACAAGTACTGGAGCCTGGACTTGTCTTCT

GACTGCATTTGAAGCACACTTTCCTCCCATGAGACGTGGTTCTGCGACCGGAATGGGGAGCATGGGCTGT

TGACATGGAAGACTTGGAGGGTGGAGGCTTGCTGTGGCTTGTCACATGTTTGGCCTTGGGCTGTAGAAAT

GAAGACAAGTGCAAATGCAAAGAAGATCATGTTGGAGAGGGTTAAGGGGTTTGAGGCCTTCATAGCTGAA

AAAGAGTGATGGTCTTATTAGTAAGAGCTTAAGGTGTGTGGATATATATAGGAAGAGATATTGTTTAAGT

GAGGGGTGTGGTGAATTAGTTTATGAGTCTTGAGTCTTGACTCCCCAGTCTCTAGTATAGTGATTTATGC

TCTGTATTCCAGTAATTTGTGTTTGTTTGATAGCCAACTCGTGCCTCTAGTTTCCCCAAAGGCCACCCAT

TCATAATGCAATGCAAGAAAATTTCACACTACCCCATGTACCCATGTCTACAAAATAAGAGGACCAATGG

TGAAAAG

>MSTRG.2469.1 gene=MSTRG.2469

TGAAAACCTCAGCCGCCCTCACCCCCGTTCCATGCCTCATTTTAAGCAATAATATTAATTTCATAACCTA

GCCAATCTCAGGATTGAACGTAATGAATACACATTCTTAGAATGGTATGCACACAGAAGTACCATAAATA

ATGAAGAAGATGCTCCACAAAATTATTTTGCTTTTTTGGATTTTCTGCATCGCGGGGATGATTAAATCGA

AGGAGGGTGAGCGACCCATACATATCTCTAAACGTCCGAGGTGTATCTGATGATTTTTCGGGCACCGTCA

GGATTGTAAAGTGAGGTTCCTACGTGTGATGTACTTAGAGTGACCTATTTGTGCACTTGTTGGTTTTTAT

TAGTATTTGAGGGAAGTGAACACTAGCCAAGGTGCTTGCTAATTGTAATGGAGATGACAAG

>MSTRG.2476.1 gene=MSTRG.2476

CTTGAAATCCCCAAGAATATTATGCGTAACAAGTACTGCTCTTTGGTATCAGCTGAAAATGGTGATGTGC

TTAAAAAGTACAAGATTCCTTTTCAAACTAAGAATGGTTCTTTGCAATTATGTGGGGCTTTTAATGGGAT

TCTTTGTTTGAATGCACTTGAGTCAGACCCTAATGTTGATATTGAGTATCAGGGAATGTATCTGTGGAAT

CCTTGTGTTGGGAAATACAGAGCCCTTTTCTCGACTTGTTTCAAGAAACGGGGATTTTGTGTTTATTCAC

TTGGGTTGGGGGTTTATGAGCCTACGTATGATTTTCGGGTTGTTAGAATTGTGTATCCCGCGGATGACAG

GGGTTATGCGCTTGGGAAGGTGCCTCCTAAGGCCGAGGTTTATAGCTTGAAGAGGAATACTTGGAGGAGG

ATTAAGGACCCTGGTATTGGTTTAGTTGGTTTTCATACTGGGGTCACTGTTGGTAATAATATGACATACT

GGCTAAATACAAAAGCTCCTCGAGGTTATAAGGAAGAGGCGTGGGTGTTGTCGTTTAATTTCAATGATGA

GGTGTTTGGGATGTTTAAATTGCCGGATCATGTTCGTTATTGTTTGGGAGTGAAGGCAGAGTTTAGACTC

TTGAAGGTTGAGGGTAAACTTGCTGTTTTTGTTCTAAGTGAAAAAAAAGGAAGTAATGGAAATTGCGGCC

AGCCTTGTTATATATGGCTCATGAGCCATGAAGATCATGAGGTGTCGTGGAATTTGCGTTTTAAAATTGT

ACTCAAGAAACGTGAGTGGCCTTTGGGTGTTTCAAGGGGTGGAACACTTTTATTGGTGTCACCAGGTGGG

TTCAGTACAGTTTTAGCTTCATTTAATCTCATGGGTGGAGATCTTCAACAAAGTAAGCCACTGAACTGTG

GACCTGGTGCTGTAGACACTCTTTTCATAGAGAGTTTGCTCATGCTTGAGGGACGGGATGAACTGTTGAA

ATCTGCATAGAGAGCTTTCCTGAATCCTTCTGAAACTCCTGGCAATTTAATTGCTAATTTTTCTTCAGGA

GGGAGACGACTGCAGTAGTTGCATATATATGGACTGATGGAGTAGTTATATGGAGATATTATTGAACTCT

AAGTGGTATAATCTATTTATGATTAGACATGTCCCAAGTTTATGTAAAACTTATGCCTATCAGACCATCC

ACGTGTTACTTTACTGTCTTTTGCTACTTATATCATCTGCAGTTCTGCACAATCATAATGTGTTTGGT

>MSTRG.2485.1 gene=MSTRG.2485

GGTCCACAGCAGTCGAACCTAGGCCACCTGTCAAAGCAAGCTGATAGAACCGAGAGAAACAGAAACGCTG

CTCTCTCTGCAGCCGCTTTCTCTTCGTTAATTCCAATGCACTCCCCTCCCTCCATATAAATACAATCCCC

CATCTTTTCCCTCTTTTTCACCACCACCACCCCCATCTTCTCTTTCATTTTGATTCCTCTCCAATTAATT

ATCATCACTATGAAGCTCAAACCCTCACCGACCTCGCCATCCCAAACCCTAACCAACGCGCCGTCGCAAG

ATGCGCGTCTCCTCGTGCGCGAGACTCTCCGCATCAGCGCCAATCTCGCCTCCGCTCCGCCCGCCGAGAA

GCCCGCGCCGCCGCCGCAGCTGCCTCCTGTAGAGCCGGGATTTCAGGACCAGTTTGTCGATTCGAGCTTC

AGGTTGATTTGTAGCGAGGAGATCGACGGACGCCGCTGGAATTATTTTGCTCAGGATATCGGTTCTGTTA

AGTTTAAGAAAGGCTCGATTCGCGCCGTCAGCTTGCAGTCTCCGAAAGCTCCTGTTGATGAGTTGGTGTC

ATTTGTTAGATCATATGTAGTTCCAGAAGGTTTTCCTGATAGTGTGATACCCTCTTATGTTCCATACATG

ACATGGAGAGCTTTAAAGCATTTCTTTGGGGGTGCAATGAGTGTTTTTACCACTCAGACGCTTCTGAATT

CAGTTGGGGTCTCTAGAAATCGATCTACACCTGGGGCTGTTGCTATTAATTGGATTCTCAAGGATGGAGC

AGGCCGTGTTGGGAAGATGCTTTATTCTCGTCAAGGGAAAAAATTTGACTATGATCTGAAACAGCTGCGG

TTTGCTGGTGATCTCTTGATGGAGTTGGGAGCTGCTGTGGAATTAGCAACTGCAGCTGTACCACACCTTT

TCCTTCCTTTAGCATGTGCTGCCAATGTAGCCAAGAATGTTGGTGCGGTCACATCAACATCTACTCGTAC

CCCAATATACAAAGCCTTTGCAAGAGGAGAAAACATCGGTGATGTTACTGCAAAAGGAGAATGTGTTGGA

AATATTGCAGATCTGTTGGGTACTGGATTTAGCATCATGGTTGCAAAAAGAAATCCATCTTTGGTTACCA

CCTTCGCTTTACTCTCTTGTGGATACGTATTCAGCTCCTATCAAGAGGTCAAATCTGTTGTTCTGCATAC

ACTAAATCGTGCAAGATTCACTGTTGCCGTAGATTCTTTCCTTAAGACTGGGCAAGTACCAACATTGCAG

GAGGGAAATGCAATGGAAAGTGTATTTAATTTACCATGGTCAAACCACAGCCCCATTGTTCTTGGATCAA

GATTCAAAGATGCGTTCCAGGATGCAAACTCTTACCTTGCCATAGAGCCTGTTTTTGAGAGAGAAAGATA

TGTAGTTACATATAATCCTTCTAAGGGTAATATATATGCGCTGTTCAAGGATCAAGCAAAACCCGATGAC

ATATTGAAAGCTACATTCCATGCTAATGTACTATTGCATATCATTCGTTCATCAAACGGGAATAGGTTTC

CCAGGAGAGAAAATGAAACTGATCCTTCAGTTTTGCTGCCCTCTTCAACCAATCTTGAAGCTCATATTGC

AGACTCTTATAAGATGGTTTCAGCTTTGTATGCGCCATTTAAGAGCAAAGCTAAAGAACAGGGCTGGGTG

ATGTCAGAATCTCATCTAAATCCTGGCCGAGCTCGGCTCTGTGGATTGACCAAGTGAATGTGTCAAATCA

ATGACATCTTCATCCGTCCAGAGCATCCAATTTTCAAGAGATTAGATAGCTTATTTAAACTCGGAGAGAG

GAGCTACTGCAGTTCGTGCATCAACCGAATTGATCATCTTTAAGAGTATCAGATACCCTAAGTAAGAAGA

TGTTCCCTATTGTATATCAACCATGACTTGTACAACCAGCAATGTGTGTTATGCCCATATTTACGTTTGT

AGGAGAAATCAGTTCTAGTGGATAGATAAAATGTATTAGTGCTTGTATATGAAATATAAATAAACCAGTT

TTGTCTTATGTTGTAATCTCTATCGATAAAAGGTCTTTTTCCCATGTGGGGCCTTCTGTTATTCTGTGGA

GGTAAGATGCATCCATATCATAGCGCATTACTGGCCGTCTTAAGGAACAAGGCCAACATATTATCTATGC

TCTTTCTAAAGCTATATTTAGGAGCTTTGGGTTGTTATGTTACCTACATATAGTGGATTATACAGATTGT

CAATCTAATTAGTTGCTGTGC

>MSTRG.2485.2 gene=MSTRG.2485

CCTAGGCCACCTGTCAAAGCAAGCTGATAGAACCGAGAGAAACAGAAACGCTGCTCTCTCTGCAGCCGCT

TTCTCTTCGTTAATTCCAATGCACTCCCCTCCCTCCATATAAATACAATCCCCCATCTTTTCCCTCTTTT

TCACCACCACCACCCCCATCTTCTCTTTCATTTTGATTCCTTGTAGCATTTCTTTGGGGGTGCAATGAGT

GTTTTTACCACTCAGACGCTTCTGAATTCAGTTGGGGTCTCTAGAAATCGATCTACACCTGGGGCTGTTG

CTATTAATTGGATTCTCAAGGATGGAGCAGGCCGTGTTGGGAAGATGCTTTATTCTCGTCAAGGGAAAAA

ATTTGACTATGATCTGAAACAGCTGCGGTTTGCTGGTGATCTCTTGATGGAGTTGGGAGCTGCTGTGGAA

TTAGCAACTGCAGCTGTACCACACCTTTTCCTTCCTTTAGCATGTGCTGCCAATGTAGCCAAGAATGTTG

GTGCGGTCACATCAACATCTACTCGTACCCCAATATACAAAGCCTTTGCAAGAGGAGAAAACATCGGTGA

TGTTACTGCAAAAGGAGAATGTGTTGGAAATATTGCAGATCTGTTGGGTACTGGATTTAGCATCATGGTT

GCAAAAAGAAATCCATCTTTGGTTACCACCTTCGCTTTACTCTCTTGTGGATACGTATTCAGCTCCTATC

AAGAGGTCAAATCTGTTGTTCTGCATACACTAAATCGTGCAAGATTCACTGTTGCCGTAGATTCTTTCCT

TAAGACTGGGCAAGTACCAACATTGCAGGAGGGAAATGCAATGGAAAGTGTATTTAATTTACCATGGTCA

AACCACAGCCCCATTGTTCTTGGATCAAGATTCAAAGATGCGTTCCAGGATGCAAACTCTTACCTTGCCA

TAGAGCCTGTTTTTGAGAGAGAAAGATATGTAGTTACATATAATCCTTCTAAGGGTAATATATATGCGCT

GTTCAAGGATCAAGCAAAACCCGATGACATATTGAAAGCTACATTCCATGCTAATGTACTATTGCATATC

ATTCGTTCATCAAACGGGAATAGGTTTCCCAGGAGAGAAAATGAAACTGATCCTTCAGTTTTGCTGCCCT

CTTCAACCAATCTTGAAGCTCATATTGCAGACTCTTATAAGATGGTTTCAGCTTTGTATGCGCCATTTAA

GAGCAAAGCTAAAGAACAGGGCTGGGTGATGTCAGAATCTCATCTAAATCCTGGCCGAGCTCGGCTCTGT

GGATTGACCAAGTGAATGTGTCAAATCAATGACATCTTCATCCGTCCAGAGCATCCAATTTTCAAGAGAT

TAGATAGCTTATTTAAACTCGGAGAGAGGAGCTACTGCAGTTCGTGCATCAACCGAATTGATCATCTTTA

AGAGTATCAGATACCCTAAGTAAGAAGATGTTCCCTATTGTATATCAACCATGACTTGTACAACCAGCAA

TGTGTGTTATGCCCATATTTACGTTTGTAGGAGAAATCAGTTCTAGTGGATAGATAAAATGTATTAGTGC

TTGTATATGAAATATAAATAAACCAGTTTTGTCTTATGTTGTAATCTCTATCGATAAAAGG

>MSTRG.2493.1 gene=MSTRG.2493

CAAAGGGAGTCTTGTTCGGTTTGAACTCCGTGCCCGTTGCTCGTTGGTTTAATCAGAAATATTTCTAATT

ATGACTCATCTCTAATTCTCTATCGTCAAATCTCATAGATTCTGCCTCTACTCTCTCTCTCTCTCTCTCT

CTCTCTCTCTCCCTCCCTCTCTCTCATATGCATATACAGTCGCCACCTCTTGCTCTACAAATCCTTCTCG

CCTTCCCCGGATCACCCTCAGATAAGCACTCAGAAAACTTATATTTACAACTG

>MSTRG.2503.1 gene=MSTRG.2503

TGTGCAATAAAATTCAAGCTCACCTGTCTGTCTATCATGGATGTGTATGTAGAGTACTATTTTCAAGGCG

CATATATAGAGGCACACTCATAGCACCACTTTCGTGTCCGCTCTCCACGTTGGCCAGTTACAAAACACTC

ATCATCTTCCTGAGATGCTATTCAAGTAATGATAAAAGAGCAGCAGCATGTTAAAGCTGTTAGATATGCT

TGTGCTTTTGGGCTTCGTGAAAGGTTTCAGCCTGCTTCACTCCTGAAAGACTTCCTGAAGAATGCTGCAG

AGGTGTCAAAGACTTTAGGTGATAATATAAACGGCCCAGTTAAAAATAAAGACAAGGACATAGACAACAT

TGTGGCTTCGCTAAAGGAGGCACTCGTTTGCATTTCTCATTACAAACTTGAATTAGAATATTCTCCAGAA

TGCATTGAGATTTTTATTCAACAGCTTGTACAGCAGAAGGAAGATGGAAAAGTTTGCTCATCAACCTCCA

AAAATGAAGCAGACATACAACATGAGGCCGAGAAAAATTGTAAATTAGCCGAAGTTGTACCTGACCCAAC

TGAAACATGTGCCATTCAGCCTCAGAAAATATCAAGGAAACGACCTGCTGCAGGCTCTGATCTTGGCCCG

AAAGCTCAACTCCGACGGAGCAACAGGAAACTTAGACGTACTGAACCACTAGTAGAAACTTCCTCAAATG

TTCCGCCTCCTGCTTTTCACTCGGCGGCAAATGTTATCACCTGAGAGGAGACCTATATGCAGAAAATGAT

CATGCCTTTTGTATTCAATCTGCAGAATACACTATATGGCTGACTTGGTTTCTGTTTGTGTCTTGTATAG

CATAATTATTTGTTTGGTGTCAAAGAATGGCAACAAAATGTTGCACAACATGGTCTAGATCACTGAGCTT

TTTCAGAGTTCAGGAGTTCGGTCATCCGGGGAGCTGCAAAAGCCTCTTACTTCAAGATGAATGAAGACTT

TGTCACTGCATTTGTGCAGTGATGATTTCTGATGTCTTGCTACGTCCATGAGGGCCTAAATCTTCATAGC

GTCTAATAATCTGGCTCGAGTCTTGTTAATAAAGCATTAGAAGTTCATCAACAGACTCAAAAGTCGAGTC

TTTGGGGAAAATTTTACGGTACTGC

>MSTRG.2508.1 gene=MSTRG.2508

CTTTTACACTTCTAACACCTTATTTTCTCTCTAATAGCATCCAAAAATAGCTGAAAATTCACCTTAATCC

TTCCCATTTGCTAAGAAGCTTTACTCCATTCTTGTAAAATAAGGTGGTCTCGAAAAATGAATAATAATCG

ATCTTGGATGTATGAGAGGATTGATGATAGAGGATTTTTGAATTCTCTATTTATTTCCGGGGTGGAGGAT

TTTATGAATTATGCTATATCTCAGCCAACGTCTATGGGTGGGACAAGTATCCAATGTCCTTGTTCAAAGT

GTAAGAATCGAAAATTTTTGAATGGAGATATGGTGAAGCTGCATCTATTGAAGAATGGATTTGTGAAAGA

TTATTATGTATGGAGTCGACATGGAGAGTCATCCATTTATAATGAAAATGACGAACATCCTTCAACAAAT

CATTCCAATATTTCACGGGGCACGGATGGGAGCAACCTAATGTACAATATGGTGATTAATGCAGCTGGTC

CAAGTTTTGATCCACATGATTCAGAAGAGATGCCGAATGCAGCAGCACAAAATATATACAACATGTTAAA

GTCTTCAGAACGAGAGTTATATGATGGTTGTGAATCTTCACAGTTGGCTGCCATGGCCCAAATGTTGAGT

TTGAAATCTGATCATCACTGGTCGGAGGCATGTTATGATCAGACATCACAATTTGTCAAAGGTATCTTGC

CTGAAGACAATACATTTCTTGATAGTTTCTATAGCACAAAGAAACATATGGAAGGATTGGGTCTACCTTC

CATTCATATCGACTGTTGTGTTAATGGTTGCATGATATATTGGAACGAAGACATTGACATGGAGTCATGC

AAATTTTGTTCTCAGCCGAGATACAAGATTAGAGTTCACAGATCCACAAGAGAGAGAAAGAAAGTAGCTG

TTCAAAAGATGATATATTTTCCACTGGCCCCGAGATTGCAAAGGTTATATGCCTCACCGACTACTGCAGC

TCACATGAGATGGCATGCCGATCACTATAAGGAAGATGGTGTGATGCGTCATTGTTCAGATTCAGAGGAG

TGGAAGCAATTTGATAGAGCGCATCCATTATTTTCCTCAGAGATAAGAAATGTGAGACTTGGACTTTCTG

CTGATGGATTTCAGCCATTTGGTAGTTCCGGCAAGCAATATTCTTCTTGGCCAATTATAGTGACTCCATA

TAACTTACCCCCCTGGATGTGTTCAAAAGAAGAGTACATGTTTCTATCCATCCTTGTACCTGGTCCGAGA

AATCCAAAGCAGAAGATAGATGTTTTTCTTCAACCATTGGTTTCCGAGTTAAAAATGCTATGGGAAGTTG

GTGTAGAGACATGGGATACTTCTTTGAAACAAAATTTTCAGATGCGAGCGGCTTTGATGTGGACTATAAG

TGACTTTCCTGCTTATTAAATGGAATGGGGATGGAAAACTGCTGGTCATTTAGCTTGTCCTCATTGTGCT

CATGATCATGATGCTTATAATCTCAAACATGGAGGAAAGACAACATGGTTCGATAATCATCGAAAATTTT

TACCTAGAAATCATCCATTTCGAAAGAATAAAAATTGGTTTACCAAGGGGAAGGTTGTGTCTGAATTTCC

TCCACCTATTCGGACAGGTGAAGATGTCTTACAAGAAATTGAGTCACTTGGTTTGATGAAAATCACTGAA

CTGGGAAGTGAGGAACATAATGCAAGAATCATCAAAACTTATAATTGTGGATGGAAAAAGCGAAGCATCT

TTTGGGATTTTCCATATTGGAGGACGTTATCAATTAGACATAATCTTGATGTTATGCACATTGAAAAAAA

TGTGTTTGAGAATCTTTTTAACACCATAATGGCCATTGAAGGTAAAACCAAAGATAATGCTAAAGCTAGA

GCTGATATTGCACTAATCTGTCGAAGGCCTGAGTTAGCAATCAATGAGTCAACACAAAAATATCCAAAAG

CATGTTACAGCTTGGATAAAAAGGGTAAAGAAGCAGTTTGCAAATGGTTACAAGATCTTAAATTTCCGGA

TGGATATGTTTCAAATATGGGACGTTGCATCGACATGAAAAAATACAAGTTATTTGGAATGAAAAGTCAT

GACTGTCATGTGTTTATGCAAAGACTTCTGCCAATTGCTTTTCGAGAATTTCTTCCTAATAATGTTTGGG

AAGCTGTGACAGAGTTGAGCCTTTTTTTCAGGGATCTAGCTTCATCCACTTTGAAGGTTAGTGATATGTG

TAGACTTGAAGAACAGATTCCAGTGATACTTTGCAAGTTGGAACGCATATTTCCACCAGCTTTGTTTGAC

TCAATGGAACATCTTTTGGTGCATTTACCGTATGAGGCACGTATAGCTGGTCCTGTACAGTATCGTTGGA

TGAGTGCACGAAGACTTCGACTAACCAGTAGTCCTAATCCTGGGAACTCTACAGAACACACTCCAATCCG

ATCACAGCGATCACAGACGGAACACACACCTGCACAGTCAAGGGTTCTATCTTCAGCATCTGCGAATGAA

CCTGAACATGCCAGTGAATCGGAAGATCCATATTGGGAAGTTGGCTCCATGCATGATGACGGGCGTATGC

GGGTCGAAGTAATTAGTGGATTGTTAGAGCCTTCAGGTAGATGCTCGCGAGCTATCACAGATTCTATATC

TGAGAGGCAGGATCCTGATGGATTCTCCTGGAAGGCAGTGTCCAAGGAGGTGAAAGATTTTTATTTCGAC

GAGTTCAAGAAAACATGTGTTTGGAGACAAGACGACAGAATAATACATAAAGCATGGGCAAAGAAAGCGA

GTGTTCGTTATACAAACTTTTGCAGTGATGCCCGGGCAAGATGGGAAAAAGGTGGACAGGATAGTAGAGT

TGCTATGCATGTATGGTTGCGTTGGGTTGAGTTTTGGAAAACCCCGAATTTTCAAGTTAAATCAAATACT

CAAAAGAAGAATCGCAAGGGTGGAGCTGACAAATACCCTCCAACCCACACAGGTGGTTCAGCCTCTTTAA

GAACCCATGCCGCTGTGCTGGCGAAGACTCAAGGTAAAGATCCAACTCCGGCCGATGTTTATCTTATTAC

ACATACGGTGAAGCATGATAAAAAGACTTTTGTTACCAAGAAAGCAGAACAAGTTTATAAAAGGGTTGAG

ACACTACGTGAGGAACTCTCAGCACCCATTGAAGGTTCTGATGAACCTCAAGTTGTTGATGAAGACCAAT

TGTTCCTTAAAGCTGCTGGTGGACTTGACAAAAAAAGCAGAGTTTATGGCATGGGTTCATTACAGAGCGT

TATATATGGACCGGAAACTGGAGGTAGTCGTCATAGGGGCTCCAATTTCAGACATGAAGAATATAAGCAA

ATGCAAGTGGAGTTGCAGGACATGAAGAACCAAGTGAAGGAGTTACAAGAAATGAGAAACCAGGAACTAG

AAGAGATGAGAAAGCAGATGGAAGAAATGAAGAACCAACTTGCATTGGTGTTTAACCATCAGAATTAAGT

TTGCCTGCCTGCCTGGCCTGACTTTTATTTATGTTAAGTTTGTATGACTTCTGAAATATTGTAGTTTGGA

TGCTGCTTTTGTTTATAACTTGGGCTGTTGCTTTTGCTACTAAATCTATGACTGTTCCTTTATTTAATTA

AGCAGTTTATTTTATTAATTGTATAATGCATG

>MSTRG.2508.2 gene=MSTRG.2508

CTTTTACACTTCTAACACCTTATTTTCTCTCTAATAGCATCCAAAAATAGCTGAAAATTCACCTTAATCC

TTCCCATTTGCTAAGAAGCTTTACTCCATTCTTGTAAAATAAGGTTAGAGCCTTCAGGTAGATGCTCGCG

AGCTATCACAGATTCTATATCTGAGAGGCAGGATCCTGATGGATTCTCCTGGAAGGCAGTGTCCAAGGAG

GTGAAAGATTTTTATTTCGACGAGTTCAAGAAAACATGTGTTTGGAGACAAGACGACAGAATAATACATA

AAGCATGGGCAAAGAAAGCGAGTGTTCGTTATACAAACTTTTGCAGTGATGCCCGGGCAAGATGGGAAAA

AGGTGGACAGGATAGTAGAGTTGCTATGCATGTATGGTTGCGTTGGGTTGAGTTTTGGAAAACCCCGAAT

TTTCAAGTTAAATCAAATACTCAAAAGAAGAATCGCAAGGGTGGAGCTGACAAATACCCTCCAACCCACA

CAGGTGGTTCAGCCTCTTTAAGAACCCATGCCGCTGTGCTGGCGAAGACTCAAGGTAAAGATCCAACTCC

GGCCGATGTTTATCTTATTACACATACGGTGAAGCATGATAAAAAGACTTTTGTTACCAAGAAAGCAGAA

CAAGTTTATAAAAGGGTTGAGACACTACGTGAGGAACTCTCAGCACCCATTGAAGGTTCTGATGAACCTC

AAGTTGTTGATGAAGACCAATTGTTCCTTAAAGCTGCTGGTGGACTTGACAAAAAAAGCAGAGTTTATGG

CATGGGTTCATTACAGAGCGTTATATATGGACCGGAAACTGGAGGTAGTCGTCATAGGGGCTCCAATTTC

AGACATGAAGAATATAAGCAAATGCAAGTGGAGTTGCAGGACATGAAGAACCAAGTGAAGGAGTTACAAG

AAATGAGAAACCAGGAACTAGAAGAGATGAGAAAGCAGATGGAAGAAATGAAGAACCAACTTGCATTGGT

GTTTAACCATCAGAATTAAGTTTGCCTGCCTGCCTGGCCTGACTTTTATTTATGTTAAGTTTGTATGACT

TCTGAAATATTGTAGTTTGGATGCTGCTTTTGTTTATAACTTGGGCTGTTGCTTTTGCTACTAAATCTAT

GACTGTTCCTTTATTTAATTAAGCAGTTTATTTTATTAATTGTATAATGCATG

>MSTRG.2512.3 gene=MSTRG.2512

AGAAATTATATGATGAAGGAGCAAGAAACTTGTGGATCCATAACACAGGTCCTCTTGGGTGCTTACCGCA

GAATATTGCCAAATTTGGGACAGATACCTCAAAGCTTGATGAGCTTGGATGTGTAAGCACACATAACCAA

GCTTCAAAACTCTTGAACTTGCAGCTTCATGCACTCACCACAAAGCTAAAGGCTCAGTTGTCAGAGGCAA

ATATCACCTATGTCGACATTTACACAATAAAATCCAATCTCATTGGAAACTACTCAAGATATGGATTTGA

ACAGCCCTTGGCTGCTTGCTGTGGCTACGGAGGATTGCCTTTGAACTACGACAGCAGAGTTGCTTGTGGA

CAAACTAAAGTGATCAACGGAAGCACGATTACTGCAAATGCATGCAGTGATAGTACTGAGTATGTCAACT

GGGATGGAATTCATTACACTGAAGCTGCGAACCAGCATGTTTCATCACAAATACTCACTGGAAAATATTC

CGACCCGCCATTTGCTGATAAGATGCCATTTCTTCTTAAGCTCAAGTTCTGAACCCAAGATATACTAATC

AACATTCTTTGTATTTCTCATATGTTATTCCACAATGATATAGTAACTAACTAATGGAAGTAATGAAAAT

CATGTCATTTTATGTCGATGAAAGAAA

>MSTRG.2532.1 gene=MSTRG.2532

CATGGCTAAAGGCATAACTCCAACCCTCCCTGCCTTCTTCATCGCAGTTTTTTTAGTCATCACAACAACC

GCTACCACCTACACAACCACCACCATCACCACCACCACAAACAACGAAACACCGATCAACAGCAAATGCA

AGATAGAGAGGGGAGAGATCAAAGACTGCTTGATTTATTTATCACCGGAAGGTGATTTCTTTTTGGACGA

GTGCTGCTCACAGTTAGAAGAATTCAAGTCCGAGTGCCAATGCAAGGCACTGAAAGAAGTTGTGTGGCAA

ATAGAACACCAGAGGAAAAGTGATGGATATGGAATGACGCCTGAAGAAAAGAAGCAGATACAGGAAATTG

CACAGGATCTTCCCAGGCGGTGCTACTTGGATTTTTCTGGTTCCTGCACGATTATGCCAGATCAACCTGC

AGCTACTTATTAAATAAATATTCTGAAATGATGCTTGTAGATCGATGCAGAATTCATGTAAAGCAAAAGC

ATTTAATAAAACTAGTTTGTTAAGCCTCAAATGAAGGAG

>MSTRG.2536.1 gene=MSTRG.2536

CTCAAGATTGTTGCATTATAAAGATTTATCACCAAACATGTTTTTGTTTTAACAGAGAAAGTGGAAACAA

CTGTCTGTGCTTTAGTCTGAAAACATGAAATTAGGTACTTTTGCTAAACTAAACAAGGAATATACAACTC

ACACTTATATATGTTTATATAAACTTGCACAATTACGGCCGACGGTGGTCCTGCTTGCCAAACATAAATT

TATCCAGAGGGCCCTCCCAGAAACCCTTTTTAATATTTCACGAAACAGTGTGGACTGATATATGTGATTA

AATTGAATATATATAGTGAAATATTGATAGTTGGTGGCTTGTTGCATTATTGAAGATACCGGAGGTAGAT

AAATGTTGAAGAGGAAATCCATTCTATTGTGCATGGCATGTGAACATCTTTAATATGTGGTTTCATGTTC

TTGTGAATTACAGGGTGTGACAACACAACAAGTAATATTATGTGTACAAGGCATCTGTTGTGTATAAGGC

ATCTAGGTTTATAGCTAGCTAGTTTTGAGGATATGATAGAAGGATCCTAACTTTCATCTAGAGTTTAAAG

TACTCGACATTATTCGATTACACAAAAATTAGTGAATCCATAAAGATCTTTCAATACTGTCCTCGAAGAT

GTG

>MSTRG.2537.1 gene=MSTRG.2537

GCTTACTTGATCACTAAATAAATAAATAATTAACTCAATTCAAAATGATTTCGAATTAATTTCGAATCCC

AATAACTCTCCTGTTCATCATCATCATCTTCACAACAATCGCTTAAACGACACAGTTTCAGCCCAACAAA

CTCCGATACGCTGCCGTTTGTAAATATCCTTGATTCTCGGACTCCCATGGCTACTCCTGCACAATTGGTG

CTGCAGTGTTGGAGCTTGTGGTTGCACAAGTAATGTATTTGCAATGGATGGATCTGAAAGATCCTATCTA

CCTCTATATGAAATCTAGAGGAACAGCTTGATGTGATCGTGAAATGCCTAATGTTTTTAACCATTTCTAG

TTTTTACTTGCC

>MSTRG.2538.3 gene=MSTRG.2538

TAAATCTAAAATGGAAAATTATCTAATTAGATTTGTTGGGCGTCCATCCAATACCGTTATGCCACCTTGC

AAAACCCAATTTATGTCGCAATCCATCCATGCCTCCCTCACGCGCCTCCTAGAAACTTCTATATTCTTAG

TGACGAAGTATTCAACTACCAAAGCTTAACCGACGTCAACGTTCATCTACACGCTTTCACCCCATTTCTC

GACTTCCGTGCCGGTTATCTACATCTACAACGCCTGCCTATATATACGAGATTATTTCGTTGAAGATTTG

CGATTATGGCCCAAATTAACGACTCAAATCAACAAAAGGTACTGTATTTATTTATAATTTTTATCTTTTT

AGGTTCGATTGATCAGCTAGGGTTTAGCTTTGTGATTGTTTAATTATTTATTGATGTATTCGTTGTTTGC

TGATGAATTTCTGTTAAATGTACAAATGATAAGTAATGTTCTTGATGATAAAAAACTGAACCCGTAAATT

TGATTGGAGCGTTCCTGCCGTGAAGAGGAGTTTAACAGCAATTAGGAAAAAGTCTTTACCCTTATTGTGG

TATATATACA

>MSTRG.2543.1 gene=MSTRG.2543

AAAATTCAGCAGCCGCGTGCGTATACACAACAGATATATACAGTCCTCCTCTTCATCTCTTTCTTCTTCC

TCCTTTAATCATCTCTTTCTTCTTCTTTCTTTAATCAGTATGTTGTACAGTACAACATGAAGTAACACCG

TGGATTTTTTGTTTCGTCGGAATTAGATGTCGTCTCGCCGGAGTTTGTGTAGATCGCCGAAATTGACGAA

CCAAACACGAATTAGGGCGATCTGGGACGGAAACGACCCGTGTTCCGAGTCGGACGAACCGGTTCGGGAT

ACGTGGCAGCGTACCCGTTTCCGTGCAACTATG

>MSTRG.2548.1 gene=MSTRG.2548

ATGCTGCCGAAAGCCGTGAAATCCTCCTTTTCCTGGAACAAAGTTAAAGGCAGACAACGGATGTCTCGTT

TCCCAAGAAACCGATAATTTCGCGTGACTCTCCTCTCACTCCATATCTACACTACACACGCCGTATCTAC

ATACACAACTCAACCGCCTTTCCGATCGCCGCCTCGGCGGCGATCAGCGCCGTCCGGTCATGCCTCGGGA

ATCGTGCCAAATCGACGAGTGTACAATCGTTGAATTCTGGTTCGATAATCACATCTGAACGGTTCAGCTT

CATGCCACAGTTTACTGGAGATATGCAGGAAGGGTGATGCTGACATTACTTATAAGGATTTAAAAACATT

CGATCAGAATTTCTCAGACCAGAGGCTATGCTTGGAATCCTAGTTACTTTCAGAGGGGGGAGGGGGGCGG

GCAGTACTGCAGTAGTAGACATACTTAAAATCTTAAATGGGAGCCCATGGATGAATCATTTATATCTTTC

TGACATTAAAACATTATTCTGCAATCTGATAGCAACAATAAAAGATAGTTTGGCACAAAGACTTATAACA

GAACAGGTTCATTAAAAGTCTAGATGCGACAAGGTTGATGCTTGCTTCACTTCTTTGTTTCTGTTTTCTT

GGAGTTGCAATCAACAGCAGCTTTGTCTCCAGCAGAGATGGAAGCAGCAAGTCTATGCTTCAACTCAGTA

AACAGAGCCCTATTCTCCTGTTTCAGCTCCGCAGCTTGCTTCCTAAGCCTCTCATTCTCTTGCATGATTT

TGCAGTTCTTCAAGTAAAGTTCTGAGTTGTTCTTGTCCATGCTGGGTGCTTGCTTTTGCCTGCAACAAAA

ATAAACAAGGAAAACAATTGTCAGACAACCTGTTCTCTTTATCAGTAAAACATATTATCTAATCTAGTAG

TGAAAAAGTGTGTGTGTGTGTGTGTGTGTGTGATGGACAAATATGTGTTAAATTTTGGATAGCTAAAACT

TGTAAACCTTGTTGAGATTACTGATTATCTGTATGTTGTTTTGAGTTGAGATGTCTAGGAATGCATGTGA

ATGTCTCTATTTATAGGCACACCAACTACACATATATGTCTGGATGAAAAAAGTTTATTATAGCTCATGA

TTGTAATATGCAACAAAAACATGAACAAACCATCATGATTTATTCCAATACCAGAAAATGTTTATCAGGA

AAACCCCCTTTTTAAAAAAAACTTGCTCCGAACACGAGACTTTAACCGTTTCGGCAAGGTAAGGCTTGCG

CCTCGTCGTAGCTCATGAAATCCAG

>MSTRG.2548.2 gene=MSTRG.2548

ATGCTGCCGAAAGCCGTGAAATCCTCCTTTTCCTGGAACAAAGTTAAAGGCAGACAACGGATGTCTCGTT

TCCCAAGAAACCGATAATTTCGCGTGACTCTCCTCTCACTCCATATCTACACTACACACGCCGTATCTAC

ATACACAACTCAACCGCCTTTCCGATCGCCGCCTCGGCGGCGATCAGCGCCGTCCGGTCATGCCTCGGGA

ATCGTGCCAAATCGACGAGTGTACAATCGTTGAATTCTGGTTCGATAATCACATCTGAACGGTTCAGCTT

CATGCCACAGTGTAACTGTCTTTTTGCTGCAGTTTACTGGAGATATGCAGGAAGGGTGATGCTGACATTA

CTTATAAGGATTTAAAAACATTCGATCAGAATTTCTCAGACCAGAGGCTATGCTTGGAATCCTAGTTACT

TTCAGAGGGGGGAGGGGGGCGGGCAGTACTGCAGTAGTAGACATACTTAAAATCTTAAATGGGAGCCCAT

GGATGAATCATTTATATCTTTCTGACATTAAAACATTATTCTGCAATCTGATAGCAACAATAAAAGATAG

TTTGGCACAAAGACTTATAACAGAACAGGTTCATTAAAAGTCTAGATGCGACAAGGTTGATGCTTGCTTC

ACTTCTTTGTTTCTGTTTTCTTGGAGTTGCAATCAACAGCAGCTTTGTCTCCAGCAGAGATGGAAGCAGC

AAGTCTATGCTTCAACTCAGTAAACAGAGCCCTATTCTCCTGTTTCAGCTCCGCAGCTTGCTTCCTAAGC

CTCTCATTCTCTTGCATGATTTTGCAGTTCTTCAAGTAAAGTTCTGAGTTGTTCTTGTCCATGCTGGGTG

CTTGCTTTTGCCTGCAACAAAAATAAACAAGGAAAACAATTGTCAGACAACCTGTTCTCTTTATCAGTAA

AACATATTATCTAATCTAGTAGTGAAAAAGTGTGTGTGTGTGTGTGTGTGTGTGATGGACAAATATGTGT

TAAATTTTGGATAGCTAAAACTTGTAAACCTTGTTGAGATTACTGATTATCTGTATGTTGTTTTGAGTTG

AGATGTCTAGGAATGCATGTGAATGTCTCTATTTATAGGCACACCAACTACACATATATGTCTGGATGAA

AAAAGTTTATTATAGCTCATGATTGTAATATGCAACAAAAACATGAACAAACCATCATGATTTATTCCAA

TACCAGAAAATGTTTATCAGGAAAACCCCCTTTTTAAAAAAAACTTGCTCCGAACACGAGACTTTAACCG

TTTCGGCAAGGTAAGGCTTGCGCCTCGTCGTAGCTCATGAAATCCAG

>MSTRG.2548.3 gene=MSTRG.2548

TGCTGCCGAAAGCCGTGAAATCCTCCTTTTCCTGGAACAAAGTTAAAGGCAGACAACGGATGTCTCGTTT

CCCAAGAAACCGATAATTTCGCGTGACTCTCCTCTCACTCCATATCTACACTACACACGCCGTATCTACA

TACACAACTCAACCGCCTTTCCGATCGCCGTCCGGTCATGCCTCGGGAATCGTGCCAAATCGACGAGTGT

ACAATCGTTGAATTCTGGTTCGATAATCACATCTGAACGGTTCAGCTTCATGCCACAGTTTACTGGAGAT

ATGCAGGAAGGGTGATGCTGACATTACTTATAAGGATTTAAAAACATTCGATCAGAATTTCTCAGACCAG

AGGCTATGCTTGGAATCCTAGTTACTTTCAGAGGGGGGAGGGGGGCGGGCAGTACTGCAGTAGTAGACAT

ACTTAAAATCTTAAATGGGAGCCCATGGATGAATCATTTATATCTTTCTGACATTAAAACATTATTCTGC

AATCTGATAGCAACAATAAAAGATAGTTTGGCACAAAGACTTATAACAGAACAGGTTCATTAAAAGTCTA

GATGCGACAAGGTTGATGCTTGCTTCACTTCTTTGTTTCTGTTTTCTTGGAGTTGCAATCAACAGCAGCT

TTGTCTCCAGCAGAGATGGAAGCAGCAAGTCTATGCTTCAACTCAGTAAACAGAGCCCTATTCTCCTGTT

TCAGCTCCGCAGCTTGCTTCCTAAGCCTCTCATTCTCTTGCATGATTTTGCAGTTCTTCAAGTAAAGTTC

TGAGTTGTTCTTGTCCATGCTGGGTGCTTGCTTTTGCCTGCAACAAAAATAAACAAGGAAAACAATTGTC

AGACAACCTGTTCTCTTTATCAGTAAAACATATTATCTAATCTAGTAGTGAAAAAGTGTGTGTGTGTGTG

TGTGTGTGTGATGGACAAATATGTGTTAAATTTTGGATAGCTAAAACTTGTAAACCTTGTTGAGATTACT

GATTATCTGTATGTTGTTTTGAGTTGAGATGT

>MSTRG.2553.1 gene=MSTRG.2553

GTTCACTTCTCCTCTTTTCTTCCACTTCCTGCTCCCTTCGCCTCTGCTTCTCTGCAATTTCATCCATCTT

TGCCCGGTGCTCAGCCTCCTCTTTCTTGCGTCTTTCAGCCGCTTCACGCTTGCGAGCTTCTTCCTCTTCT

TGAAGCTTTTTCAGTCTTTCTTCCTCAACACTCAAATAAAACAACATTTTTCTCTTACTTTCTCTCTCCT

CCTTCCGTGCCTGCAGGATTTGTTCAATTCGTTCATGTCTTTCCTTTCTCAACCTCTCATACTCAGCTTC

CCGATGTTTCAC

>MSTRG.2558.1 gene=MSTRG.2558

AGCAAGTGGAACGAAAGTATGTGAGCTGTTCATTCCTGGTGAGAAGCAGTGGGATATTTCTAAGGTTCAT

AATTTAGTTACAAATTATGACGCCAAAATTTTTTTGGCTATACCGATCCCAAGAAACCAGGTTCCAGATC

GCATTGCTTGGAATTTCACCAATGATGGCAAGTATTCAGTGAAATCTGGGTATAAATTTTGGCATGATCA

AAATAGTGAGCGTCAAAGAGTACCATGTAGTCCAGGATGGGTGAAGCTATGGAAGTTGGCGATTCCTCAC

AAATTTCGAGTCTTCTTGTGGCGTATATGCAGAAGTAATCTGCCTGTCCGTTACCAGCTGAGAGGAAAGG

GTGTTCAAACGGCTATCATATGTCCACTGTGCTTGAATGATATCGAGCATCTTTTACATATTTTCTTGGA

TTGTTCTTTTGCTAAGAGGTGCTGGAATTTACTGGGTTTAGAGTTTGATACATCATCAGCAGAATCATGT

TCTGAATGGTTGCTGCAAAGATTAGCTGAAGAGGATGGTGAGAAAATGGTGCAAGTAGCAATAGTGCTAT

GGGGGATTTGGTCGGCGAGGAACATGTTAGTCTGGGAAAATAAAAAAGTAACTCCGGAGAATGCAATGCA

ATGGAGCATAAACCAGGTGAGGCAATGGCAGGAAGTGCAGGTTGTGAGGTCGAGTAACGATGTCATAGGT

ATGCCAAGTCATCGACCAGAATTAGCGCTATGGAATGCTCCAGCTACAGGAAAGCTTAAAGTTAACGTGG

ATGCATCAGTGGTCTCAGGAGCTTCTAGCTTTGCCATTGGAATGGTTCTGCGAGATCATTTAGGTGGGTT

CTGTAACGCACGCAATCTTCGAAGAGAAGGAGAGATTTCAGTTTTCGAAGCAGAAGCTTACGGTGTACTG

GAGGCTATTAGGTGGGTGCATGATCTGGGAATTTCAAATGTGGAAATTGAAAGTGATTCGCTACTCACAG

TTCAGGCCATTCTTCAGGGTACAACAAACTATCATGAGGTGGGAAATCTGCTATACGAAAGTCGGATACT

AATCAGTGATCGCCCGGATATCTCTATAGCGTTTATTAAAAAGCAAGCCAACAAAGTAGCTCATATGTTA

GCAAGG

>MSTRG.2559.1 gene=MSTRG.2559

TGAAAAAGAAGCGAGTACCCGAGAAGGCGATGCCAGAAGTAGCCACAACGCAGGTCTGAATAACAGTATT

CTCCTGCCTCGTAAAGGGCTGCTTGAGCAGCCCACTCTTGCTCAAAAACGAGGTCCACGTCTTCACAAAG

AAGAAGCCCAGCAGGCCCGCCGAGACATTCAACGAAGGTATAATCCCCGTAGTCAAGTTCAGCTTCATGA

CAATGAAAGTGAACAATATTCCCAGGATGAAGCTGACGACGAAGGCTCGGAGCGTGAGCTGTTTCTGCCA

GGGCGGCACTTGTTTGGACTCGAACATCATTTCTACGGACTCGGCGAACTCGTCGTCGTTGTCGGCTGTG

GTGTACTTGGGTTTGTTTGGTTCCTCGGAGGGGTCGTTGCGGCGTGTCGCCATGGCGGGGAGAGTGTGGG

GTGAGGGAGGTGTGGTTTTCGAGGAGGGTGGTGTCCTCGAAGTGATGAATGGTAATTATGGGAGGGAGAT

GAGATGTTGGGCTAGTTTTGGTTCAAGAATTTAAAAGTTGATAGGATTTTTTCTCTCTCGAATCTTAAGT

TTGAGCAGGGTTACAGAAATCGGAAATCGGACCTAATTGGTTGAACTGCCGATTCAGGGATTAATCGGAA

ATCGGG

>MSTRG.2563.1 gene=MSTRG.2563

CAGGAACACCAGAAGAAGCTCAGAGTAAGTCTGGAAAACTAGTTTATTGCTAATCTTACATAGCGAAGCG

CACAGCAAATATATACAAAAACGATGGTTTTACTTATTCCAAAGACACCATTTACATTGATTAAGCTTAA

GTAGCAAAAAACACCAAATCAACCGCACTGCGTACTCTGCTCTCTCGCTTCGATCTCTTAGTGATGATTC

TTGCTCTTAGTCTCCCCACTGTAAAGCCAGTCATAAACAGTAGGAGTGTTGGGACTAGGCTTGTCAAAGA

CCTGAGATCCCATTCTCCTGCTAGCAAGGTTGCTCCCTGGATTAAACACACTCCTCCACACGTTGTCCTT

GCGCGGCGCCATCGCCGACGGAGTTGTTGGCGTTGTCGGCGTTGCTGGTGCCTCCGGAGTCGACGGCATA

GACAGCGACCTCACATATTTCCCACTATCATCCCCGGCCGTATCGGTAGAAGTAGTGACATAGAGCTTCC

GGAACTTGCCGAGGCCTCGTTCAGGCTGAGGTCCTGCCATAACATCGTCCCATATTTTATCCAACATCTT

ATCTCTCTATTTCTTTCTCTCAG

>MSTRG.2565.1 gene=MSTRG.2565

CCGCACTTATCAATCGCCTTCCGTCTTCCTCCTCTTCTTCTTACTCAATCCACTCTGCTCCGCAGACTCC

CCCGAAAAATCGCCCTCAATCTCCTCGCCTTTCCGTTTCTTCTTCTTCGTGACCTCACTCTTTTCCACCA

GAATTCCAGCATTTCCCTCGATCTCCTCCACTTTATCCCCCTCCCCCTTAATCTCCGTCTCGTCCTGTCC

GAAATCAAATGCGATGATTTGAGAGTGTTTTTTGAGGGACTTAGAAGAGGAGGATGATTTGAGGTCCTTG

TGGAACTGATTCAATTCGGCGAAAGAGGTGGCAGCGCGGCGGAGATAGACGGAGACGGCGTGAGAAGCTC

CGTTCTCCATAGAGACGAAATTGGAGAGGATTCTCGAAGCTGTGGCTAAAGAGACTGGCTTTGAGGATAC

AACGTGTGCTGATACTGACTTCATCGTGATCTTGAATGAGCAATTGAATTTTCTAGGGTTTTGGAGACTC

AATTGAGAGTTGAAGTTGGGTCTAGGGTTTATA

>MSTRG.2574.1 gene=MSTRG.2574

CATATTAAAGCTTACATAATTCAGCACGTAACAAACTGGTATCAGAGCCATGGCTGGGCCATCGCTCCAA

AAACATGAACAGTCCATTAAGTTGATCGAGGAGGCGCAAGCTGCAAGCACCCAGAAACATGATGAAACCC

TTCTCCAGATTGCAAATCTAGCTACTCAAATGGCTAATATGGGTACTCAGTTTAAGGGTTTGGCCACGCA

GATGCAAGCTGATCGTGACACGATGTCACGACAGATTCATGGTTTGACGGAGCAGTATCAAGCACAGGTG

GCATCAATTCAACCCCTGCTGCACCAACAACCCAACTTCAGCAAGGCAACCGCCTCTCCTTCCGCAAACG

CATCTTCCTCGAACTCCCCGATCTCGTCAGCTCTCAAAATTGGTAATGTTTCTGCCCCAACTTCACTTTC

CCCATTCGTCACTCACACTCTCACAAACCCTCTCCCAAGGCCCCTAAACTCCTCCCTTAATCCACTACGC

CCACCCATTGACCCACCCGTCACCCACAGTTTCATCACCTCCGCCTTAACTGCACCTCCCTTCATCCCTA

ACCAGCATATTAATTTTCGAGCACAGCACACGAATTTCATACCACATACTACCCCACCAGTAAGTTTCCC

TCAAACTCTCACCACCGGGGTTCTCAGTAATGGGGTTTTCAACCCTCCTCAAGGCTGGATTCCAAACTTG

GATAGCTCTCTGCCCAGGCCCCCTTATCATACCCAGTTTAAGTTTCCTAAGATGGACTTTCCACGTTTTG

ATGGAAAAAATGTACGTGGCTGGATAAGTAAAGCTGAGAAGTTTTTCCTCCTCAACCCTGTAATGGATTC

CACCACCAAGGTCATCTACGCAGCTTTGTATTTGGACGCAGAAGCTGACTACTGGTACCAAACCATACAA

GCCCAGTACCCAGGGTTAGGCTGGGATGCTTTCGTTTCTTTTTTACTCCGGAGATTCAATACAGGTAATC

AGGGCAATCTCATCGGTAAGTTTAACAAACTGACTCAGACTAGTACTGTTGATGCCTATGTAACTCAGTT

TGAGGAATTGCGTGGTTATTTGATGAGTACGTATTCGTTTCATACTGATGAGTTTTACCTGTCTAGCTTT

CTTAGTGGATTACGCCCCGACATTCAGCAGGCACTATATATCCACAACCCTTGTACCTTACAGGAAGCTA

TCGATAAAGCTCGGGAGCAAGAAATCTTTATTGACATAATTGAGAAAAAGTTGAAGGGCCCAAAAACTTC

ATACTCACCCAACCCATTTAAATCTCCGGCAGATGCCACAAAACCCAACTCCACCAGTTCACCTAACCCT

GTACCCAAACCCCTCACTGAAACAGGACACTCTAAACCTATTACTACCTTTCCAGGCCCCAAACCGAACC

CTGTAATTAAACGTATTTCTCCTGCAGAGATGGCACGACGCCGGGAACGGGGATTATGTTACAATTGTGA

TGATGTTTACTCTCCAGGGCATAAATGTGCTCGCCCTCAGCTTTATTTGATGGTTGGGGAAGGTGATTCG

GAGGAATCCGACACCGATGAAACCGATACCCCTGAGATGGAGACTCAACCTGAGTTTGGTATCTCTATCC

ATGCTTTACAAGGCACTCAAGGGTTACATACCTTGAAGGTGGCAGGGCAAATTCAGGGTCACCCTATCCA

GTTGCTTATCGATACAGGCTCCACACATAATTTTGTCTCCCCAGAATTCATTCAGCGCCTTGGCATCAAG

GCATCTTCCTGCAATTCACTCAATGTTCAACTAGCTAATGGCACTACAGCTAGCTGTGGAGGCAAGGTTG

AAAACCTCAGCTGGACAATGGGTACTGTCACATTTGTCTCTGACTTTTATGCCATCCCACTGGGAGGTTA

TGACGGCATCTTGGGGGTACAATGGCTCTCACAGGTGAGCCCGGTCAGCTTTGACTTCACTGCTCAACAG

ATTGAAATTCACTGGAAGGGTAACCGGATAGCACTTAAACAAGAAGTTTCCAACCCAGCCAGTATTAAAG

TCCTTCACAACAAGAAACATCTGCCAAAGTCACCTGGCCATGCTTGTTTCTTGGTCCAATTGACGGCCAT

AGAGGCCCCGAAGGACGTGGAACCTCCCAATCCACCAATTGTGCAAACGGTGATTGATAAATATGATGAT

ATCTTCCAAGTCCCCAAATCACTACCACCAGCACGCAAAGCTGATCATCTCATTCCTATCATGGAAGGCA

GTAAACCAGTGAATCTTAATCCTTATAAATGCCCCTACCTTCAGCGTGTGGAGATCGAGAAAATGGTAAA

AGAGATGCTAGACAGTGGCATCATCCGCCACAGCCAGTCACCTTATTCTTCCCCTGTTTTGCTCGTTAAA

AAACGTGATAACAGTTGGCGATTCTGCGTTGACTATCGGGCCCTTAACGCCATCACAATCAAAAACCGTT

TTCCAATTCCTTTAATTGAGGAAATGATGGATGAACTCCATGGCAGCAGGATCTTTAGCAAGCTCGATCT

TCGCAGCGGCTATCACCAAATCAGGGTTCAAGAACAAGACATTCATAAAACTGCCTTTAAAACTCAGCTA

GGCCATTATGAATTTGTGGTGATGCCATTTGGCTTGACAAACGCCCCGGCAACGTTTCAAGGGGTAATGA

ACGAGATTTTCAGGGATCTCATTGGCAAGTTCGTACTCGTCTTCTTCGACGATATACTTATATATAGTAC

AACAATGGAGGACCATTGTGAGCACCTGGAACTTGTCTTTCAGCATCTACGAGATCACCAATTATTTGTT

AAGCGGAGTAAGTGCGCCTTCGCCAGGACTCACATCGAGTACCTGGGGCATATAATCAGCCATGACGGGG

TTGCAGCAGACTCAAACAAAATCGTGGCCATGCAACAATGGCCTTCCCCGAAAAATCTCAAATCTCTACG

TGGCTTCTTGGGCTTAACAGGGTATTACAGACGCTTTGTGAAGAATTATGGCGCCTTAAGTAGGCCTCTG

ACTGAGTTACTTAAAAAGGGTGCTTTTAGGTGGTCACCTCAGGCTGAGGATGCCTTCCAACTTCTCAAAA

CAGCCATGTGTAACACTCCTGTCCTCGCCCTTCCAGATTTTTCACAGCCATTTGTGGTTGAAACCGACGC

TTGTAAATCAGGAGTGGGTGCTGTCCTAATGCAATCTGGAAGGCCACTTGCTTTCTTTAGCAAGGCCATT

TCACAACGGGATATGGGTCTGTCCACCTATGAAAAAGAGTTGTTGGCTGTTGTTTTGGCAGTACAGAAAT

GGCGTGGCTACTTGCTTAGCCAACCTTTCACTATCCGCACGGACCAAGAAGCCCTCAAACATCTCCTCAA

TCAGAAAATCACTACTCTTGTTCAGCAGAAATGGCTCACCAAACTGCTCGGATTTGACTACAAGATTGAG

TATAAAAGGGGTCATGACAATGTTGTAGCAGACCCTCTTTCACGTCTCCATGATGCGGAAGAAGATAGTA

CTGGAGAACTTAAGGCTACATCTGTGGTTATACCACAATGGAAAACAGACCTCAAATTATCTTGGGAACA

AGATCCAGCGTTACAATCTACTTTGGCACAACTAGCAATTAGTCCCACCTCGGATCCCAACTATACCTTG

GATGGGGGGGAACTTCGCTACAGGGGACAACTGGTTGTGGGGGCAAGTGGCACACTTCGGCATGACATTA

TTCATAATCTGCACAGCAGCCAGGAAGGTGGTCATTCAGGAATTAATGCCACGATAAAGAGGATCACCTC

ACTATTTTGGTGGCCGAGCATCACTAAAGAAGTTACTTCCTTTGTTCAGGCTTGTGATATTCGTCAGCGT

TACAAAGTCGAGCATGTTCCATCTCCGGGACTTCTGCAGCCTATTCCAATCCCGGCTCAAGCTTGGGAAA

CAGTTACAATGGACTTCATCGAAAGTCTCCCAAAATCGAAGGGGTATGATACTATCCTGGTAGTAATTGA

TAAGTTCACAAAGTATGCTCATCTTCTTCCCCTCAAACATCCGTTCACTGCCACACAAGTTGCCCAGGAG

TTCCTCAATCATGTTGTTAAGCTGCATGGTCCACCCAAAGTTCTCATTTCGGACAGAGACAAGATTTTTA

CGAGCCAATTTTGGTCCCACCTCTTTGCAGCGATGGGAACATCTTCTCATCTCACCACAGCGTACCATCC

CCAGTCTGATGGCCAGAGTGAACGCTTAAATCAGTGCGTAGAAATGTATTTGCGCTGCCTCTCAAGCCAA

AAACCAACAGACTGGGTTCACTGGCTTCCAATGGCTGAGTGGTGGTACAATTCCTCCCACCATTCTGCTA

TCAAGATGACTCCCTATAAGGCTCTTTACGGCCAGGACCCCCCCAGCACCAATTATCATCAGGCCCAACG

AACGAACATTGCTTCTGTGGATCAGTTTATTTCTCAACGTGCAGAACTCCAACACCTTCTCAAGTCCAAC

CTCACACTCGCCCAGGAAAGAATGGTCTTATATGCTAATAAGAACAGAACAGAGCGCCACTTCAATGTCG

GAGATTTTGTCTTCTTGAAAGTTCAGGCCTTTCGACAGATTTCTCTCAAGGGAACCAGAGATAACAAGTT

CTCTGCCCGATACTATGGACCTTACAAAATCATCCACAAGATAGGTTCTGTTGCTTATCGCTTGGAATTA

CCAACTGCAGCCCGTATTCACAACACTTTCCACGTCTCCCTCCTCAAGAAACATATCGGATCACACCACA

GTGTCTCTCAAACCCTGCCAACAACTGTTGCCCCAGCTGACAATCCAGAACCTGCAGCTATATTAGACCG

TCGTCTTCTCAGAAAGGGAAATAAGCCAGGAGTCTCCTTGCTTATTCACTGGAAGGATCAGTTACCCGAA

GATGCCACTTGGGAAGACTATGATTACATTCAGTCCAAATTTCCAACTTTTCTCGGCGCGGTCGCCGCTT

TTCAAGAACAGGGCAATGTCACGAGTGGAAAAGAAATTAGTCACACTTGATTTTATTTAGCTGACAAACA

ATATCATGTGCTTAATGGGCTGAATTTATTATTTGGGCTGGCGTAACAGTAGAAGGATCTAGATGGAGAG

AAACGACGTAGTTTTGTTATATCTGTTATTTGTCTTTTGTTATAAGTGGGGGCTGTGTAACAACTTCTTC

TTCAAGCAATGAAATGAATTATTCCTTCTCTCTCTAAA

>MSTRG.2576.5 gene=MSTRG.2576

GTAGAACATTCAAAATTGAACCTCTAAACTTGTTCATTCACTCAAAAACAAATGGTGGTCAAAGTGTTTG

GTTCTCTTAGAGGAGCTTGTCCTCAGAGAGTCCTGGCTTGTCTTTTGGAGTTGGATTTGGATTACGAGTT

TGTTCATGTCGACCTCGACTCTGCCTTTCGGACAAGTTCCTGCTATAGAAGAAGGGGACTTCAAGCTGTT

TGGTAAGAGCAAGATCACTTGAAAATTGAAATCTCTACTTAAAAGTTGATAAATTTACGTTTTGGACAAT

TATCAGTGAAGGACTCGTTTATTTTCAGAGTCTAGAGCCATCATAAGGTACTATGCAGCAAAGTACGCGG

AAAAAGGTGTCAAGCTTATTGGGAATACTTTAGAGGAACGAGCCCTCGTTGAACAATGGCTGGAAGTTGA

AGCACACAATTTCAATCCCATGCTTTACACACTGGTACTTCAGCTCCTGGTGTTCCCAAAAATGGGAGGA

AAAACGGACCATGTTTTAGTTCAAGAATGCGAAGAAAAACTGGAGAAAGTGCTTGATGTGTACGAGCAGA

GGCTGTCAGAGAGTAAATATCTGGGCGGTGATTATTTTTCACTGGCTGACCTGAGCCATCTCCCGGGAAT

CCGATACTTGATGAATGAAATTGGATTGGCACACTTGGTGACAGATAGAAAG

>MSTRG.2576.6 gene=MSTRG.2576

GTAGAACATTCAAAATTGAACCTCTAAACTTGTTCATTCACTCAAAAACAAATGGTGGTCAAAGTGTTTG

GTTCTCTTAGAGGAGCTTGTCCTCAGAGAGTCCTGGCTTGTCTTTTGGAGTTGGATTTGGATTACGAGTT

TGTTCATGTCGACCTCGACTCTGCCTTTCGGACAAGTTCCTGCTATAGAAGAAGGGGACTTCAAGCTGTT

TGAGTCTAGAGCCATCATAAGGTACTATGCAGCAAAGTACGCGGAAAAAGGTGTCAAGCTTATTGGGAAT

ACTTTAGAGGAACGAGCCCTCGTTGAACAATGGCTGGAAGTTGAAGCACACAATTTCAATCCCATGCTTT

ACACACTGGTACTTCAGCTCCTGGTGTTCCCAAAAATGGGAGGAAAAACGGACCATGTTTTAGTTCAAGA

ATGCGAAGAAAAACTGGAGAAAGTGCTTGATGTGTACGAGCAGAGGCTGTCAGAGAGTAAATATCTGGGC

GGTGATTATTTTTCACTGGCTGACCTGAGCCATCTCCCGGGAATCCGATACTTGATGAATGAAATTGGAT

TGGCACACTTGGTGACAGATAGAAAG

>MSTRG.2576.7 gene=MSTRG.2576

AGAGAACACATCAATATCATTAATCACTTGCATTGTGTAGAACATTCAAAATTGAACCTCTAAACTTGTT

CATTCACTCAAAAACAAATGGTGGTCAAAGTGTTTGGTTCTCTTAGAGGAGCTTGTCCTCAGAGAGTCCT

GGCTTGTCTTTTGGAGTTGGATTTGGATTACGAGTTTGTTCATGTCGACCTCGACTCTGGTGAGCACAAA

CAACCTCCCTTCTTACTCCGCCAGGTAACATTTCTACCATGCCTAAATATTTTGCGTAGTTCTTGGCATT

CTAGAATCTAGATTAGCTTCCTTGTTTGATAATTTGTCTGATAAGGTCTCATTGTGACTTGTATATTCTT

GGCAGCCTTTCGGACAAGTTCCTGCTATAGAAGAAGGGGACTTCAAGCTGTTTGGACTCGTTTATTTTCA

GAGTCTAGAGCCATCATAAGGTACTATGCAGCAAAGTACGCGGAAAAAGGTGTCAAGCTTATTGGGAATA

CTTTAGAGGAACGAGCCCTCGTTGAACAATGGCTGGAAGTTGAAGCACACAATTTCAATCCCATGCTTTA

CACACTGGTACTTCAGCTCCTGGTGTTCCCAAAAATGGGAGGAAAAACGGACCATGTTTTAGTTCAAGAA

TGCGAAGAAAAACTGGAGAAAGTGCTTGATGTGTACGAGCAGAGGCTGTCAGAGAGTAAATATCTGGGCG

GTGATTATTTTTCACTGGCTGACCTGAGCCATCTCCCGGGAATCCGATACTTGATGAATGAAATTGGATT

GGCACACTTGGTGACAGATAGAAAG

>MSTRG.2576.4 gene=MSTRG.2576

CACTTACCGGTTGTTGGCATGTGACAACCCCCAATCCAGCATGAACTGGCTACTGAGCCACCACTTCAAT

CCACTGCTCTTGCAAGTAACACCATTGTTCAATTTCTTGTGTACCTCTATATAAAGAGAACACATCAATA

TCATTAATCACTTGCATTGTGTAGAACATTCAAAATTGAACCTCTAAACTTGTTCATTCACTCAAAAACA

AATGGTGGTCAAAGTGTTTGGTTCTCTTAGAGGAGCTTGTCCTCAGAGAGTCCTGGCTTGTCTTTTGGAG

TTGGATTTGGATTACGAGTTTGTTCATGTCGACCTCGACTCTGGTGAGCACAAACAACCTCCCTTCTTAC

TCCGCCAGCCTTTCGGACAAGTTCCTGCTATAGAAGAAGGGGACTTCAAGCTGTTTGAGTCTAGAGCCAT

CATAAGGTACTATGCAGCAAAGTACGCGGAAAAAGGTGTCAAGCTTATTGGGAATACTTTAGAGGAACGA

GCCCTCGTTGAACAATGGCTGGAAGTTGAAGCACACAATTTCAATCCCATGCTTTACACACTGGTACTTC

AGCTCCTGGTGTTCCCAAAAATGGGAGGAAAAACGGACCATGTTTTAGTTCAAGAATGCGAAGAAAAACT

GGAGAAAGTGCTTGATGTGTACGAGCAGAGGCTGTCAGAGAGTAAATATCTGGGCGGTGATTATTTTTCA

CTGGCTGACCTGAGCCATCTCCCGGGAATCCGATACTTGATGAATGAAATTGGATTGGCACACTTGGTGA

CAGATAGAAAG

>MSTRG.2578.1 gene=MSTRG.2578

AAGAAGAAGATAACTCCCCCTCTCCATTATTTTTCATAATATCCAAACTGTTACAATTACAAACCCTTCA

AGACCAGAGAGAGAGAGAGGAAGAGAGAGAGAGTTGTTTCCTCATCTCATCTACAAAAACACACACAAAT

TCCCTAGATAAACACACACACAAAACCTGCATATGCAAAATTTGACGTTTTGTGTTTAGCTAATAACTGA

TCCGAACAACGAACCACCG

>MSTRG.2590.1 gene=MSTRG.2590

GGTACATTTCCTTGAGCTCTGGTCCACGATCTAACTTCCTTCCTACCACCCACTTCACCTTTGTAAATCC

GATGCTGTCTATAAGTGGCGCATAAACCTCCTCCAGCTGCTCACCGACGCAGAAGAAATGGTCCAGCCAA

AACAAACCACCGGGCCTCAGCACCCTGTAGATGTCAAAGAACATGAAGTGCAGCAATGTTGTTGGTATCC

AGTTACTCATAACGTGCATGGAGTGGACGATATCCAATGTGTTGTCGAAGAATGGTAGCCTCTGGGAAAT

ACTGAGGTACAAAGGCACAACCCCTCTTGCAGCTATAAAATTATTGAAAGGACCGTTCAAGTTCATTGAT

GTTGTAACAATGGTTACATTTCTTGCCTGCATTCTGATTGCAAATGTGGCCACTCCGCCTCCAATGTCAA

GCCCGATTCGGATTGTTCCTGGCTTTTTCACTGCCAGGACTTCGTCTATAGAGAAATCAACTCCACCACC

ACTGGTTGGTTTAGTCCAACGACTCTTCTCTACTCCTTTAAGATCAAAACAGTCTTTGCAGTCATCAAAG

CCTTTCTGATTACGCTTGCGGTTAATTAGACATGTGTAGTTCTTGCAGGTATATGCAGTCCAGACAACTG

ATGAATCTGATGGGGTGGACCAGAAGCTAGCTGGAAGAGGATATGGCTCCACGTAGTCCTGTGGAGCAGC

TGCACGGCATCTTCTACGAGGTAGTGGCTCACATCCTTTAAGGAGGAGCTTTTGTGCTAATAGCTCATCA

TCAGGGCATGATCCATTGACCTTATATGTCATGTATTTACGGAGAATGTCAGGAAAGAGGGTACAGGGGC

GGCCAACTGGAGGGACGATTTTGTCAGAATCAAAATTCGTGTTGAATCCGAATGGAAGTTCATGTGGAGA

TGTGAACGCGATGAATTCAGATGGGAGGTTAGAACTTGCCTGGGTGACATGATCCTGTATAGCTGTTGGA

GTGTTTCCGGAAAAATTTAGAGAGCATCCAGAGGAATTAAAGGTGGTCGATATGAACAAGGTAAGAAGAT

TTGTCGATAGAAGAAGGAAAATATAGTAGAATTTGGTAAACTGAGAGTGGGATTTCTTAGGAGAATCCAT

TATATGATTGATAGTAACTTTGAGCTGAAGAAGAGTAATGATCAATATATAAGATATTGCAGGTCAAAAT

ATGTGTTTGAAATGTAAACTAAGATACAGATTGACTAGATTTGTTTTGTTGTTAGTTTAGTATAGCTGTA

GCATTGAAGAATGAGATGATGGAAATTTTATGCATAGATTGCATGTGCAGGTG

>MSTRG.2592.1 gene=MSTRG.2592

CTATTTCACTTTTTTTACTATATACTCATGATCTGCTACAATCAGCCCTACCGAATCCAACTCTTCTGTT

GCTCACATCAAACTCCACCCATAGATTTTGTTGATGGAAATTCCCTATTATATTACTTGGGATGCCAAGC

GTTTCTGACCGTCCGATCCCTAGACAGTGGACCCCGCTTCCCACGTCGTCTAAAATTCTACTCTTCTCGA

TCAGAACCTCTGCGCCTTTCTCGAATTCGAATACCATGTCGCCTAGTAGCCGTCCGATCATGATCGAATC

TCCGCCGTTGAAACACATTTCGAGTGTGTTTTCGTAGACGTAATTTTTACGCATTTTGGGGCCCACTAGT

CTTAAGACCTCTTCTCTTATTTTTGTGTATGCTTCGTCGACTAGGTAGGTGTACTCCGTGCCGGAGTCGA

TCATCGTCTGGCCGGAGCCACCGAGGTTTGGCCGGAAAACGGCCGCTGATATGTTGAGTTTTCTCCCGCC

CATTCGTATTCCGACCATGGGGAGCGTGTAGGCTGACGGGTCGAGATTCGGCATGCGTTGACTTTTTGGG

AAAGTCAAGATGTTGACGTAAGCGAATGTGTTTGAGTTCGGATTCTGGCCGAGGTAAAAAGAGCCTGTCG

GCGAAGTTTTCGCGAGTCGCTGACGTGTGGGGACACAGTAAGAAAATTTATTTAATTTCGCTTGGGAAGG

AAACGAGAATCTCCCGAGGTTCATTCCTAAAATGCCCTCGGCCTCGGCTGACTTGGCAGCGCAACCGAGG

ATCAGAGGAGGGGTACTTTGGGAAGGCGAAAATGTTAGTTTTTCCTTGACTAAATTGCCCTCGGCGTAGG

TCCCATCAGCATAGAAGTATGAATAGTGGCAGTGTCTATTTTGGTCGCATGAGGTGGGAAGGGTATAATC

GGGAATAGCGGGTTTACAAATAGGGTGATTGCAAGGGAGAATGGAGAAAGTTGATGAGAGCGCAGGATCA

AACGACGTCGGATCAGGTTTTTTATTACACTGAATCCATGACAACTGACTTCCTGTGTCAAGAACCATGT

TTTGAGTTTGTGGTGGGGTCCCTATTGGAAGAGAAACAATCAGAGCCATTGAATACTTGAAAGACGATCT

GACATCATAGTTTAGGTATGAGTTTGGAGAGATTTGTGTTTTGGGTCTTTGTGAGGAAGATGATTTGATG

AGTGAAGCTCTGAGAGTAGAGTGAGGAGAGAGAGGGAGAGATGATAGAGAGAAGGAGAGAGATAGAGATT

TGTTATGTAAGGAAGATGATTTGATGAGTGAAGCTCTGAGAGTAGAGTGAGGAGAGAGAGGGAGAGATGA

TAGAGAGAAGGAGAGAGATAGAGATTTGTTATGTAAGGAAGATGGGTTGTGAGTGTTTGGATTATAGTTT

GATTTAGCTGAGGAGAAGCATATGGAGATGAGAAAAATAAGTGCTTTCAGAAGGGCTTTGGACTGAGCCA

TGGAGGTAATTGATATTTGAGAGGGCGGGGGAGCAGAGGCTTATGTATGGGGGAGAGTTGGAGTTTGAAT

GTTTGATATAAAATTCTAGTGTGGTGTGAAGATGATGGATACTGTATTTGTAATTGTGTGTGCATGTTTA

TATATATACCATTAAGGTGGGAGGAGAGGCACAGGGAGTGTAAGGGTGAGCTGAGACCATTTCTTTTAAC

AGCGGTTTGGGCCAGATGTGTGTTTCATGGTTAATGACGTCTGTTTCATGATTAATGAATCGAATAATTA

AGAATTGAAATCTCGAATTCATGAGTAGAAATTTGAATTATTCATTTTGTGATATGAAATGAGGATTTCA

TTCCTATTTTAC

>MSTRG.2592.2 gene=MSTRG.2592

CTATTTCACTTTTTTTACTATATACTCATGATCTGCTACAATCAGCCCTACCGAATCCAACTCTTCTGTT

GCTCACATCAAACTCCACCCATAGATTTTGTTGATGGAAATTCCCTATTATATTACTTGGGATGCCAAGC

GTTTCTGACCGTCCGATCCCTAGACAGTGGACCCCGCTTCCCACGTCGTCTAAAATTCTACTCTTCTCGA

TCAGAACCTCTGCGCCTTTCTCGAATTCGAATACCATGTCGCCTAGTAGCCGTCCGATCATGATCGAATC

TCCGCCGTTGAAACACATTTCGAGTGTGTTTTCGTAGACGTAATTTTTACGCATTTTGGGGCCCACTAGT

CTTAAGACCTCTTCTCTTATTTTTGTGTATGCTTCGTCGACTAGGTAGGTGTACTCCGTGCCGGAGTCGA

TCATCGTCTGGCCGGAGCCACCGAGGTTTGGCCGGAAAACGGCCGCTGATATGTTGAGTTTTCTCCCGCC

CATTCGTATTCCGACCATGGGGAGCGTGTAGGCTGACGGGTCGAGATTCGGCATGCGTTGACTTTTTGGG

AAAGTCAAGATGTTGACGTAAGCGAATGTGTTTGAGTTCGGATTCTGGCCGAGGTAAAAAGAGCCTGTCG

GCGAAGTTTTCGCGAGTCGCTGACGTGTGGGGACACAGTAAGAAAATTTATTTAATTTCGCTTGGGAAGG

AAACGAGAATCTCCCGAGGTTCATTCCTAAAATGCCCTCGGCCTCGGCTGACTTGGCAGCGCAACCGAGG

ATCAGAGGAGGGGTACTTTGGGAAGGCGAAAATGTTAGTTTTTCCTTGACTAAATTGCCCTCGGCGTAGG

TCCCATCAGCATAGAAGTATGAATAGTGGCAGTGTCTATTTTGGTCGCATGAGGTGGGAAGGGTATAATC

GGGAATAGCGGGTTTACAAATAGGGTGATTGCAAGGGAGAATGGAGAAAGTTGATGAGAGCGCAGGATCA

AACGACGTCGGATCAGGTTTTTTATTACACTGAATCCATGACAACTGACTTCCTGTGTCAAGAACCATGT

TTTGAGTTTGTGGTGGGGTCCCTATTGGAAGAGAAACAATCAGAGCCATTGAATACTTGAAAGACGATCT

GACATCATAGTTTAGGTATGAGTTTGGAGAGATTTGTGTTTTGGGTCTTTGTGAGGAAGATGATTTGATG

AGTGAAGCTCTGAGAGTAGAGTGAGGAGAGAGAGGGAGAGATGATAGAGAGAAGGAGAGAGATAGAGATT

TGTTATGTAAGGAAGATGGGTTGTGAGTGTTTGGATTATAGTTTGATTTAGCTGAGGAGAAGCATATGGA

GATGAGAAAAATAAGTGCTTTCAGAAGGGCTTTGGACTGAGCCATGGAGGTAATTGATATTTGAGAGGGC

GGGGGAGCAGAGGCTTATGTATGGGGGAGAGTTGGAGTTTGAATGTTTGATATAAAATTATAGTTTGATT

TAGCTGAGGAGAAGCATATGGAGATGAGAAAAATAAGTGCTTTCAGAAGGGCTTTGGACTGAGCCATGGA

GGTAATTGATATTTGAGAGGGCGGGGGAGCAGAGGCTTATGTATGGGGGAGAGTTGGAGTTTGAATGTTT

GATATAAAATTCTAGTGTGGTGTGAAGATGATGGATACTGTATTTGTAATTGTGTGTGCATGTTTATATA

TATACCATTAAGGTGGGAGGAGAGGCACAGGGAGTGTAAGGGTGAGCTGAGACCATTTCTTTTAACAGCG

GTTTGGGCCAGATGTGTGTTTCATGGTTAATGACGTCTGTTTCATGATTAATGAATCGAATAATTAAGAA

TTGAAATCTCGAATTCAT

>MSTRG.2592.3 gene=MSTRG.2592

CACATTTCTGGTGTGTTTTCGTAGACGTAATTTTTACGCATTTTGGGGCCCACTAGTCTTAAGACCTCTT

CTCTTATTTTTGTGTATGCTTCGTCGACTAGGTAGGTGTACTCCGTGCCGGAGTCGATCATCGTCTGGCC

GGAGCCACCGAGGTTTGGCCGGAAAACGGCCGCTGATATGTTGAGTTTTCTCCCGCCCATTCGTATTCCG

ACCATGGGGAGCGTGTAGGCTGACGGGTCGAGATTCGGCATGCGTTGACTTTTTGGGAAAGTCAAGATGT

TGACGTAAGCGAATGTGTTTGAGTTCGGATTCTGGCCGAGGTAAAAAGAGCCTGTCGGCGAAGTTTTCGC

GAGTCGCTGACGTGTGGGGACACAGTAAGAAAATTTATTTAATTTCGCTTGGGAAGGAAACGAGAATCTC

CCGAGGTTCATTCCTAAAATGCCCTCGGCCTCGGCTGACTTGGCAGCGCAACCGAGGATCAGAGGAGGGG

TACTTTGGGAAGGCGAAAATGTTAGTTTTTCCTTGACTAAATTGCCCTCGGCGTAGGTCCCATCAGCATA

GAAGTATGAATAGTGGCAGTGTCTATTTTGGTCGCATGAGGTGGGAAGGGTATAATCGGGAATAGCGGGT

TTACAAATAGGGTGATTGCAAGGGAGAATGGAGAAAGTTGATGAGAGCGCAGGATCAAACGACGTCGGAT

CAGGTTTTTTATTACACTGAATCCATGACAACTGACTTCCTGTGTCAAGAACCATGTTTTGAGTTTGTGG

TGGGGTCCCTATTGGAAGAGAAACAATCAGAGCCATTGAATACTTGAAAGACGATCTGACATCATAGTTT

AGGTATGAGTTTGGAGAGATTTGTGTTTTGGGTCTTTGTGAGGAAGATGATTTGATGAGTGAAGCTCTGA

GAGTAGAGTGAGGAGAGAGAGGGAGAGATGATAGAGAGAAGGAGAGAGATAGAGATTTGTTATGTAAGGA

AGATGGGTTGTGAGTGTTTGGATTATAGTTTGATTTAGCGAGGAGAGAGAGGGAGAGATGATAGAGAGAA

GGAGAGAGATAGAGATTTGTTATGTAAGGAAGATGGGTTGTGAGTGTTTGGATTATAGTTTGATTTAGCT

GAGGAGAAGCATATGGAGATGAGAAAAATAAGTGCTTTCAGAAGGGCTTTGGACTGAGCCATGGAGGTAA

TTGATATTTGAGAGGGCGGGGGAGCAGAGGCTTATGTATGGGGGAGAGTTGGAGTTTGAATGTTTGATAT

AAAATTCTAGTGTGGTGTGAAGATGATGGATACTGTATTTGTAATTGTGTGTGCATGTTTATATATATAC

CATTAAGGTGGGAGGAGAGGCACAGGGAGTGTAAGGGTGAGCTGAGACCATTTCTTTT

>MSTRG.2594.1 gene=MSTRG.2594

AATTTTTCGATGATATGATATATTTTACAGGGTAATTTATGCTGAATTCTAACTGGATTATGTTTTATTT

ACTTGAATTTCGATCATTAAATTATGCGTAGAGATTAAATGTGGTGAGTAATATTTGGATATAGATATGT

GGTCCAGTTTGAAAGTGGTTCTGTATTTGATTGGGACTTTCAAGTTGAGAAGAAGGGTGATAAGATTATG

ATTGCCGAAAGTAAATGATGAACAAGGTTGTCTTAATCTTCTTTTATCTATGGTAGGTCAATACACAAGC

ATCTCGAAGTTCTTAACTGGAGAAGGTTGTTGGGTTGTAAT

>MSTRG.2595.1 gene=MSTRG.2595

GTGATCGATTTGTTTGAAATGTTAATATTATGCATCGAGGAGAGCCGAATCGGGGTTAATATGGGCCCCA

TATGATGGGAACTCAATTTTCTCAAAGCCTCAAAGGGGGTAGCACACCGGAATCTGAGAAAAATTGAGTC

CATATGATGAATATTTTTACATAGTCGTACGTTCCAAGAAGACGGCAGTGGTATAGTTCTTCGAGCGGTG

TAGAATCCTCTGATTGTATGAGCATTCAGGAAACATGTGTGCATTTGTTTCTTTTCTATTGATGCAGAAA

TAGCAGGAGTCCACAGTGGTTACTCCATAATCAACAAGATTATCTGCATTATTTACTCTATCATGCATGA

TCAGCCAATGCAAAGAGGAGTATCCAGGAACACTCAATTTATGCCAAACGCATTAACCCAGGAAACAGGG

GGACTGGAGTGTCTAATGGAGTTCCAGAAATTTGTTACCTTAACTTACCAGAGAATATTGGTCATCCCTT

ATGTTTAAGTCATAGGAGAAATTTTGCTTTTAAGTCATGGTGATTCGAGGCAGGGAGGGACCATCCAGTA

GGGTTAAAATATGTGCCACAGTAGCATCATTACTCAACCCGGATCGGCTGATGAGTTGGTGAAAAGGATA

AGTGCATACAGGTCTTTATTAAACCAAGGATCAAACCAGAGTTATATGTTGCTGCCAGTACTAATTTTAT

ACAAAACATGCTGCAAAGCAATTGATCGTAAGGAGAAACATTCTTGAGAATCCAAGAAGCATTGTAAGGA

CAATTCATTGTCCAGAAGGAGTGTTTTGTGACTTTTTTCCTCAGCCAATCAGCCTAAAGTGAGGTAACTC

TCTAGGACCAATGGCGCAAATCTTTTATACCAAGGCCTCCCTCGTGCTTAGGAAGAGTGACAATTTTCCA

GGAGACTTTAGCTTTAGTTCGGTTCTGGTATGACCATTCCAAAGGAAATGAGCAAGTTTCTTTTGATAGT

TTTCAAGATGGTAATAGGAAGGATGAAATGAGAACTCCAGTATGCTTGAACAACAAATAAAACCGAAGAT

ATGAGTTGTAGTCTTCCAGCAAAGGACAAGCATTTTGCCACCCAAGAGTCAATACGAGAGGTGATTTTAG

TAATAAGAGTCTGACAGTGATAGGCATGAAGCCCTTGGCAATGAGGGGAACTCCTAAAAAAATTGCAGGC

AGAATCCCATTCCTGATACCATACTCATGTTCAATCCACGTTTTAAGGCAAGGGGTCGCATTCGAAAGGA

AGTAATTACTTTTGAGAATGTTATTTTCATAGTTACCAAAGACATGAAGTGCCTGCAAATCCAGATGTTG

GGATTCCAGCCTGACCAAATTCTGCCTCTCGGAGAAAAATGGTAGTTATCATCCCAGCGCCAGTGATTAG

CCGTGCACTATACTTGAGTGGATATGTTGAAATATAAGTATAAGATCCATAAAGGGCCTCCTCTAACACC

TTGGGAATGGTTCTCTACATGGTATCACAGCTCTGGTTTGCGGGAGGTCTCGGGTTCGAACCCCTGCCAC

CCCCCAATATTCTCACAATTTAAAATGGACATGAAGGCAATTTATACCACTCTTCAACTCAAATAATGAG

ATTTCGAGTGGGGGGAACACCTTAAGATTATGTAAGTGTTTGGAAATTAGCTATATCAAAGCCTGAAATT

CGTATAATGTTGTGAGTTTTCGCAAATTTAAGGGTTTTAGGGATTTATGG

>MSTRG.2596.1 gene=MSTRG.2596

ATGGAACACAATCATGGTACTCTTTCACCGCAAAGTATCTGCAGCAAATTCTCCCTCAAGGCTTCACAGA

ATTGTTGTTTCTGCGATCATGGAAGTGACGACATATGGATCCATGTTGGAAGCAGGCCTACGGTCCTCAA

AGTAACCCTTGCCTTCTTTTTCAGTGTCTCTACCCACCCGGACTGATGCTCCACGGTTTGCAACACCCCA

TGAGAACTTGTGAATACTGGCAGTCTCGTGCTTTCCAGTGAGACGACGTTCGTTGCCTTCACCATATGCA

GCAATGTGTTGTGCATGCTTAGTCTCAAGTTTGGCAATAGCCTTCTTGATGATCTCAAATCCTCCCTCAT

TTCTCATGGACTTGGTGCTGTAATTCGTGTGGGCACCTGCACCATTCCAATCACCAGGGATAGGCTTAGG

ATCCAATGAAACAACCACCCCAGCAATTTCAGTTATCCTCTCAAGAATGTAACGGGCCACCCACAGTTCA

TCACCTGCTGAGATACCAACAGCTGGTCCCACTTGGAATTCCCATTGTCCAGGCATAACCTCCCCATTTA

TGCCACTTATGTTAATACCAGCATACAGACATGCTTTGTAATGAGCATCAACAATGTCCCGTCCAAAGGC

TTTATCAGCACCAATACCACAG

>MSTRG.2599.1 gene=MSTRG.2599

TGAAAATACATAAATTACATATACATTTTTGTAAGTATCATTAATTTCATCTGGGCTTTTCTTCAATTAT

GGGTTTGTGGGATTCTCTTCTCAACTGGCTCCGGAGTTTATTTTATAAACAGGAAATGGAACTTTCCTTG

GTAGGTCTTCAGAATGCAGGAAAGACATCTCTTGTTAATGCCATTGCTACAGGTGGTTACAGTGAAGACA

TGATTCCAACTGTTGGCTTTAACATGCGGAAAGTAACAAAAGGCAATGTAACTATCAAACTTTGGGATCT

TGGAGGTCAACGGAGGTTTCGCACTATGTGGGAGCGTTATTGCCGGGGCGTTTCGTCGATTCTGTATGTA

GTTGATGCAGCTGACAGAGATAGCGTTCCTATATCACGGGCAGAGCTGCATGAGCTCTTGAGTAAACCTT

CGCTTAGCGGAATCCCATTGCTTGTTTTAGGAAACAAAATAGACAAGTCAGAAGCTCTCTCAAAGCAAGT

CTTGATGGAGCAACTAGGTCTTGATACAATCACAGACAGAGAAGTCAGTTGCTACATGATCTCTTGCAAG

GACTCCGTGAACATTGATGCTGTTTTGGACTGGCTAATCAAACACTCTAAAACAGCAAAATGAAAACTTT

TTGCAACTTGGTCCGTGTTCATATGTAAGTTTGTTTGCAAGTTTGATGTTTATATATTACTCGAAAATCC

TTGTAATAAGTGAGTGATATTGCTGAAAATTCGTTTGAAAAATATACGTGGATTTTCTGGATAATTCCAT

TTTTTTTAACAAAAAAA

>MSTRG.2606.1 gene=MSTRG.2606

TTCTGATGCCAATAACATTTAACTAATCACGCAGGCAGTCTAGGTAATTCAGAAAATAAAGTTTTTCACA

AGCGATCTTCAAGTTTGATGTTGAAGTAGAAAAATCGCGAGAATTGAACAAGGATCATGGTCAAGAGCGA

CTGCATTTGGGATTTTTTTTTTTAAATCGGCACTTGCTTTACACTCCCCAGTAAGTAGAGTTAGATATTA

ATTTAATTATGCGTCCAAAGCACTTTAAAGTGTGGATTTTTTTTACTTTCTACAGTATGTATTAAAAAAA

TAATAAGAACGTGGTATTGTGGTAGATATTATCTATTTTTGGAAATTATATGGACATGGTTTGAACCCCA

CTCCCTTCTCATTCAAGTCCAAAATTTAACGCAATAGGACTAAAGGATAATTTTATTACCTGGAAGGTGT

TTTACATTATTAATGGTTTGATTTTCGTTCAACATATAAGTTATTGGAGTG

>MSTRG.2615.1 gene=MSTRG.2615

CTCATTTCTCATCCCTTTAATCACAAAACACTGGCCCAACAAACAAAAGGCCTTCCCATTATCCTCACTC

AACCTCACGGCCTCCAACAAATCAGCCACCGCCGAGTCAACTCGCCGCCGCCGATTCACCGCCATCCTCA

ACTCGGCCCTCGTAAACAACGCATCCCCTCTCTCCTTCTCCGACAAGGCCTTCACCGCCGGGAAAGTCAA

CGCCTGGTCAACGCACTTAAGTGCCGCCGACTTGTGGCCCATCAGATCCAGGGCCAGGCCTTTGAGGATC

AACGTGGCGCCATCTCTTGGCTCGAGCGAAAGGGCTTGGTCAGCGAGATTCACGGCGTCTTTAGCGAGGT

TGAGTGAGGTAGTGCGGTTGGCGGTGGATCGGGCTCGAGATAAGAGGTGGGCCCCTTGGATGAAGTGACG

GTGGGCTTGGGCTTTGAACCGGTTTTTAACACGGATGTTTGTAAGGGTTTGTTTGGAGAATTTGTTGACG

GCGTAGAATATGCAAAGTGTTAATACGAGGATAGATAGTTGCATTTGTTTGGAGAATTTGTTGACGGCGT

AGAATATGCAAAGTGTTAATACGAGGATAGATAGTTGCACGATTATGTCCGAAGTCATTTTTCGTAGATA

CTGTAACGAGACTCAGG

>MSTRG.2624.1 gene=MSTRG.2624

CTTTAAGCAATTTCCTGACTTGTTCCAGTGTGACAAAGAGCACCACAGTGAATGGACCCTGCCTCGATAT

AGTCGGAATAAACCCCTTGTACAAAGCCATCACGCCCTCTGATCGGACGGTCTTCATTGCACAATCTATC

GCACCACTATAAGGCGGCGCATTCCCTGGCTCCACTTTCATGTTCATAACTCGGGTCTTGATCACGTCCA

CGGGATTGGTCACCACCGAGGCTACGAACCCGGCCGCGAAGCTCGCGGTCACGTGGGTCCCGAGCCCGTC

TTTCATTAAATTGTTATCGAGGATCGATTCTTTGATTTGATCGTAAGAGGCGAGCTGTGATGCGGTGACT

AGCATGGCGCGGTTCACCGTAAGGGATGAGCCGCGCCACAGGGAAGTGATGCCCTCGGATTTGCGCATTT

GGGTGATGGCGTCGATGACGGACTTGTAGTTGCGTCGTTGCGACGCGGGGAGCCTTCCGTCGGCCTGCAT

ACGGACCATGGCCACGTCGGCGGGGTTGCCGACGGCGGCGCCGATGCCGCCGGCGATGAGTCCGGCGCCG

ATCTTTGTCATTAACGGCATGCTGCCGGTTTCCGGGTCGGTCCATTTTTGCTTCATGACGTCGTACAAGC

CCATACGGGTTGTTGAGTAGAGGGTCTGCCGGAGCATGGTCGCCGAGACGCCGGAGAAGAGCGCTTTGAC

CCCGTCTTGCTGGATGATCTTGACGCCGACGGCGATGGGTCCCACGCGCGGGGGAGGGAGGTTGATGCGC

GTGGAGGATGTCTGGAAAGCCAGAGCGGGTCGGAGCGGCTGAACAGACCCGGATGATTCGCCCTGCAGCT

GCATTCGGACCTTGATTAGATCAAGTGGGTGGGTCGAACATCCAGCTATGATTGATGCTATCCCTCCTTC

AACAAATCCCTTAACACCCATCTCTCTTAAATTTTTCTCAAAAAAGATTTATATAAATTCAAAAATTCAA

GATTTCGAGAAACGAGAATCAAATAATGAATGTGTGTATATATGTATGTATAGGTGTTGTTATTGCCAAG

GGGAGGTGGGGTTCATTTCACTGGAGATGAAGAGATGAGAATGGAGGCTAAAGGAGATAGAGGAGGAGTG

GGGGACGGTAGGTCTTTGAGACATAAGCCGTGGGGGAGATGAATTTATATGTGGAGAAATAATGAAGGGT

TGGGTTTTGTGGGAGAGATTTGGGAGAGATTATATAGGGTGGGAGAGAGAGGGTGGAAAGAGAGAAGAAG

GAAAGAGGGAACATTGACATACAGAAACTCGTAGCTGTCTTTAATTGGAGTTTGAATGTGTGTAGCAACT

TTGCACGTGATGTATGGTAGTTCACATTTCTGCAATCCGCGTATTTGGGAATATTAGTATTACGATTTCT

CGGAAGATTATTATTCCCCTGTCT

>MSTRG.2632.1 gene=MSTRG.2632

GTAAAAGAAACCCGAAATACACATTAATTCATGATTCCGTCGAGTCCAACAGATTAAAAGAATTTCTTCT

GTTCAAACCAAACCTCGACGACGAAAACTTCACCGAGTCACTCGACGTCAGAACTCGCTCCCCAAAACGA

GCCGACTCAGTATACCTCTCCTTATTTTCCTTCACCAAAAATTTACCCTCATCTTTCATCCCAACCCTAC

TCTGTAACTCCTCACTCAAACCCCATAACGTCCCACCCATTGATTGCTCCGTTACAATAAATGAACCTAA

CAACCTCCCCTTGCATTCATTCAATCTCTTCATGATGCAAACCAACTCGAACGATTCACCGAAGCTTAGG

TGACTCAGTCTCTCCTTGAACTCACTGAGTCGAAGTATCAACTCGTTGACGCATGATAAATAATCATCTT

CTACTAAAGCTATTAGCTCATGAATAAGCTTGTTGGGTTGGAGTGAGACTGCTCCATCGGGTACTTTGCA

GATTTGCTCGAGAACTGCGATGAGTGAATCAACGGCTCTGATTAAATCGTGGTTCAAGGTTGATGAAACG

ACGTCGACTTGTTTTTCTTTCTCGATTTTACTTAGAGATGAACCGAGAAAGAACCCTAGAGTTCTGGAGG

TTGAGATTAGGGTTTCGAGGTAGAGCGCGTACCACCGGACGTAAGCGGAGAGGAGCCAGGTGGTTGCGGT

GGTGCCGTCGCGGAATCCGGAGAGTTTGAGGTAGTTGTGGCCGCCGGTGGCCGGGAAAATCGAGAGTTGA

TCTTGGAGGATGAAGGATCCGCGTTTGATTATGTGGTGGATGGTGAGGAGGCACTTGAGAGCAACGGTGG

AGTTTCCGGTGCGGTGGAGGCGGTCCATGAGGGATTCGATAAGGGTGGACGCGGTGGCGCGTGATCCATC

ACCGGCGGATAAGAGAATGGAGAGATGGTGGTCTTGGGGGATGGTGGACGGCGCGTGGGAAGTGGCGCGT

AAGACAGCGAGGTGGAGAGTTGATTTGGAGAGTATGGTTGCTTTGGAAAGTGATGCCTTGTCTTTTATAA

CTCCGATCATATCCCGGAATTTCGTTATCTTCCCCATCTAAAATATATTAACAACACAAGTTCAGTTGTG

TTTTGTTTTTCTAGGAGAGAAAGAGGTGTTGATGTTCAAATAAGCTTTGTTCCGAGCCTGTTGACAATAT

TTTTTTGAAATGATTGTTTTGACGATATTGAC

>MSTRG.2634.1 gene=MSTRG.2634

CTCTTTTTCTAATCAGCCGATTCCAATGTCCATGATAAAAAAGATTTGAAGATATGATATGTTATTATAT

TTGTTTTGTCTGTGGTTGTAAATGGTCTGTGCTCACTAATCTTCTGCTGCCACCTCCAAGATCTGGTTCC

TCTCATGTACAAATCATTTCTCTGTTTCAGGGTTCATGTGCTAGCATGCCTCATGTTTTTTGTTCTGTTG

GCAATGGATAAAATACTTGGATGTGTTAGTTCAGCATATGCCTTTCTGCTGCATTTGGTCCATCTCCTAA

TACGTGTCGGATCTTCGAGGTTAAATAAATTGCAGATTATAACCCTCCGTCTATCTGTAGTTGAACTCCT

GTTGATTCTAAACCACGGTTCAAATCGCGATCTACTATAGATTGGCTGTGGCAACTAGCTGGCTACTTGT

GTCCTAAAGGCGAAGGCACTCGGCCATGTTTGTTCGGAGGAATTATAATCT

>MSTRG.2636.1 gene=MSTRG.2636

CCCCATTAGTAGCATATGAAGAAATGGCTGCCTTCGGGCTGTTATTGTCTTCGTTTAGGATTGCACTTGA

ATCGCTATCATCTGATGACCCGTCTTTGAAGTCCGGGAAAAATGTAGTTAGATCAGCAGCCATTGTGATC

CCATTGTTGCTTTGATGATCAAAACTTTCAAAATTAACCTCTGTATTAATATCTTCACATTCGATCGATG

GACTTTTCAGTACTACATCAGTAGCTTTCTCACTGTCTTCAGTCACCATCATCTCTTCTTTCACTGAATG

TTCACTGGTTTTATCAGTGTTATCCTCTCCCAACTTCGAATTCAGCTCTCGTATCTCTTTGAGCAGAGAT

TGATTTTCTTGTTGAAGAGTATCATTCTTAAGCTTGAGGCTGTCATAATTGGCTTTAAGAACACCATAGT

CTCTCTCCAATTGTTTGGTTTTCCATCGGGCACGGCGATTTTGGAACCAAACCGCAACTTGTCTAGGCTG

CAAACCAAGTTCTTGAGCAAGCTTCACTTTTCTTTCTGGTTCAAGCTTGTTTTCAACTTCAAAATTCTTC

TCCAATGCCTTGACTTGATTTATGCTCAGTCTCCGCTTTTTCTCCGAGATCTGCCCCATTTCTTCTACAC

AACCTTCTTCATCTAACCCCTCCAACATAGACTGAAATTCCCTGCTGTAAACATGGTTGTTTCCTGGACT

CCCTTCTTCTGTTGTTGGACAGATGGACATCAAAGCACCC

>MSTRG.2641.1 gene=MSTRG.2641

CCAGGTCTGTATCTGTGAGGCTTCTTGACTCCTCCAGTAGTGGGAGCAGACTTGCGAGCAGCCTTGGTTG

CAAGCTGCTTCCTGGGGGCCTTCCCTCCGGTTGACTTACGGGCGGTTTGTTTTGTACGGGCCATCTCTTA

ATTCTCCTGTGAGTTGTTTGGCGACTTGTTCTTCTTTTCTTGCTCCTCGAAAGATGAAAGTTTG

>MSTRG.2644.1 gene=MSTRG.2644

GGTTCTTCATTTTTTCAAGTCTTTTGAAGCCTATTTTGGTGTTCGATCCTCAAGATTTGGTGTTTGTTCT

TCGCACCTTCAAAGATTCATCAAAATCTCAAGGTTTTGGAGGCCTATTTTGGTGTTTGATCTTCAAGATT

TGGTGTTTGTTCTCCAAAGTTTCATCAAGATTTCAAGCTTTCGGAGGCATCTTTTGGTGTTTGATCTTCA

AGATTTTGGTGTTTGTCAATGGCCGAAAATCACTCTATTGCTGAGGAGAGGGCAAAAAGGTTGGCTGCTC

GAGCTTCAAAAAGAGCTGTTGAGATTATCGATGATGACCAGGGTCGTGGCGATAAAGTCACGAAGAGGAT

GATGAAGAAGCTTGCTGAGAGAGCCAACCCTCCTGAGCCTACCACTCCTGCTTTCCTCCAGCACAGTCAG

CAAGCTGGGAAGAGGTTTAGAGATGCCGATGGTCACATGCAGACTGCTTTCATGCCGGCTTGGGGTATTT

GTGATCAGCATTCGGTGTCTGGGAGTTCTCTCCTAGCTAAGGACTGGTCTAGGGAGTGCATAACTCCTCC

TGATCGGACCAACATTGTGGAGAGAACGAGCCTTGACGAGGCGGAGCAGCTTGGTGCTAGTGCGGCCTAT

AATCTCAACTCCTACTTCCAGGCGGCTATTTATCAAGCTAATGCCTTCAAGGAGTCTGAGAGGGCTCTTC

AGAAAGAAAAGGTTAGCTTGACTAGAGCGATGAACGATTGGGAGAGCCGTGCTCATTCGGCTGAGGACAT

GGTCACATCTCTTCAAAACAGGATGAAGACTCTCGAGGAAGGTTTGGAGAAGAACAAGGGGATGATTATC

GACGAGTACATGGACTCTGACCCCTTCCTCCAGTTCATGGACGACCATGATGATAGGGTGCGTCCCTCGG

TTTTCACTTCTGGCTGGGACAAGGCCCTCGAGACCGTTGTTGCTGATCACCCGGGAATGTTTGATCCTTC

TCGGTATCCTAGCCCTTATCTACCAGCTCAGTCTTTGGCCACTGCATCTTCCTCCCATGTTCCCGCCGTT

GTTGATGTTGAGGAGGTGGAAGGAGCAGGGGTCGCAGTTGGAGATCAGACTCCTCACAGTGCGTGGCCTA

GCTGGGAGTAGTTCCAGCTCCTCTTACTAGTCTTACGAGACTTTATCTATGCGTATTTTATTATCTATCT

AGGACTTGTTTATTCGGGCCCTGCGTGGCTAAAGCTTTGTGCTTGCTTGACTTCTATTTGGGAATGTAAT

ATGACTTTAACTTATCTTGTAGCCTTGCATGGCTTTTATGTACTCGACTTATCGTATCTTTTATAAGGGT

TGTACCACAACTCTTAAGTTATTAATCTAACTATTTCGGATTTG

>MSTRG.2647.1 gene=MSTRG.2647

GGCTTTGGCAATTCTTTGCTTCTTCTTAGCAATTCTAGCTCTCTTCCTTTGCAGAGTCAATGGAGTCACT

AGCCTCTGAATTTTGGGAGCTTTACTGATCTTCTTCCCATTCTTGCTTGTAAATGTACGACGATAGGTGT

TCACATACTTCCTGACATCATCATCCTTTGTCAGGTTAAAGAGCTTCCTGATCTTGGAAGCTCTCTTGGG

ACCTCTCATCCTTGGTTTCTCAACATCAGTCAGGCCAGGAAGATCATTCTC

>MSTRG.2652.1 gene=MSTRG.2652

ATTTATGTGGTTTTTTCAGGGTACTCGACAGAAGTTAAGGACGGAAAGCCGAGCAGGACCTTGTGGGTGT

TTTCTTATCGAACATTAAAGCTTTATTGTCTAAGCGTCCCCGGTCATCTCTGCATGTCCGAGGGTTAGAT

GGTAAACTTTCTTGAACGAAAGGAACTCCTGGGAGCGGCTTTTGTATTCTGATACAAATCGTTTATACGA

GAAAGTATCTTGTGACAAAAAAAAAAAAAAAAAAAAAAAAAA

>MSTRG.2653.1 gene=MSTRG.2653

TCTTCCCCGTAACTCTGGGTTATTTAATTGCGGGACAAGTAATCCAGGTTGGATACTCAGTTGTTTTTAA

CATCTCTGAAATGCATATTGAAGATACCATCAGTGTGCGTCAAGAATAAGTGCCAATAACTACAGCTACT

AATTGCAGAATATGGGAACAATTCCTACATTGTACAACTTTGCTGCAATCATCCCAGTAGTAATGATAGC

AATAATGATCACAATGTTGCTTCCCAAATGGCACAACAGAAAGAATTTAATCTCAAGAACCAATCTGATT

CACATTATGATGTATTGTTGCATGGTATATTTTACTATGTATTAAGTCCTAAAGTATCTAACCAAAGAGA

TTTCATTATCTCTTTGGTGTAGATGTTTTTT

>MSTRG.2654.1 gene=MSTRG.2654

CTAGTAGAATCACCTGCTAAAGTATCAAGAAGAGTAGATTTGCCACAGCCAGAAGGACCCATGATGGCCA

AACACTCACCAGGCCTTACATAGCCCGTGAGCCCCTGCAGAATTGGGCGGCGACCACCATTCTTCCCTGG

CACAGTCACCCACAACTCACTCCAGGTGACATAAATTCCAGAAGCTTGACTATCATCCTGTTGTTCTTCC

CTTCTTGAAAACCTTGATTCAGTAACTTCACTTTTCACTGATCTTGATCTCCTGCTACCGGAGCCATAGA

GCCGAATAGAAGGAGTTTGAAGGCAGCCGACATGAGGGGGTGACCGTAGAGGAGTGTTTGTCGAGGAGGC

TGAGTTATAGAAGGGCACATCATCAGCTTCAAGTTCAGGAATGTAGTAATTGTGGTGTTGCTCAATGTTA

GTGTTAATTTCATCTTCATCTGTCGGAGGTGTTGGAGTTTCAGATTGATCATTTATATGCTTTGCATTGA

ATTTCAGAAGAAGTGTTCTTGTTGGATGAGCTAGGGGAGTCAATCTAGACAAGTTATTTGGAGAATCCAT

GTGAAATGTGTATGACTAGATTGTCTTTAGTAAGAGCTAGCTAGCAGCAACATTGAAGGAGAGTCTTATA

CATATTATCATTACTCTTACTTTCATGTACATGCTCCTTTTTCACCACAATTATATTAGAAAAATCCAGA

ACAGTATATAAATGTTATTTAAGAAGTTAATTAGCAAAATAAAAGAAGTTGGCTGATTTTAGC

>MSTRG.2660.2 gene=MSTRG.2660

ATTTTGGCAGTTTTACTCTCTGATTAACTGCACTTGTCTGGTCACCTGCTAGTTCACTTAGTCTGTTGGT

TCATGTAAATGTATAAAATGTCAACACCTACTTCCATAAATTGGTTGATTATGATTTCGAATGTTCCTTC

TGTATTCACTGTAGCCTGTAGGTACTTTTTCACAGAAAAATGAAATATGGTCTGTATATAGGGGGTGTTT

GGTTGGGGGTTAATAGGCTTGGGAAACAGGAATGGGAAGCCCATTCCATAACATAGTGTTTGGTTCAGAT

TTTTGACAAAGGGGAATGGGAACCTGTATCACTCTAGCCAGTTAACACAAACCTCACCTGCAGCCCGGGT

TACTAAAATGCATTCCCGACCAAGCTAATATATCTGTCTATCATCTCCTCCCTTTTACTTCATCCCTCTC

GATCTCTCTCCCTCTCTCCCTGACTCTCTCGCCCTCCGCCTCTTTAGATATCGCGCCTATCTCTCTCTGT

GAATCTATCCCCCTCCATCTCCTTCGATCCCCCCTCTCTCTGTCTCACTCTCTTCCAATCTCTCTTCCCA

TCTCTCTCAGCCATATAACTCTGTAATCACGGCTCTTTTGAAGACCCTTGGACACTGCCACGGTTCGGCT

TCAAATCCAAGAGAAAGATTCTGCAGCAATATCTCAATTTGAATTCTTCACCTGCCCAACTTGAAACAAT

TAAGGAACTGAAATGTGGAGATTTGGATGAGCCAGGAGTCTTGAGTGAGCAGCAGCAGCCAGCAGCAGGC

CCAACACTCAGCAGACTGCAGACTAGGACTAGGACTTTAACAAGAATTCATTCACAACCAAACTTTGTGT

TCTGGCTACTGTTATTTTAAATATTTGAATGTTCAAACTAATTCTGATGGAGACTAGCAGATCTTTGAAT

TGTG

>MSTRG.2660.3 gene=MSTRG.2660

GTACTTTTTCACAGAAAAATGAAATATGGTCTGTATATAGGGGGTGTTTGGTTGGGGGTTAATAGGCTTG

GGAAACAGGAATGGGAAGCCCATTCCATAACATAGTGTTTGGTTCAGATTTTTGACAAAGGGGAATGGGA

ACCTGTATCACTCTAGCCAGTTAACACAAACCTCACCTGCAGCCCGGGTTACTAAAATGCATTCCCGACC

AAGCTAATATATCTGTCTATCATCTCCTCCCTTTTACTTCATCCCTCTCGATCTCTCTCCCTCTCTCCCT

GACTCTCTCGCCCTCCGCCTCTTTAGATATCGCGCCTATCTCTCTCTGTGAATCTATCCCCCTCCATCTC

CTTCGATCCCCCCTCTCTCTGTCTCACTCTCTTCCAATCTCTCTTCCCATCTCTCTCAGCCATATAACTC

TGTAATCACGGCTCTTTTGAAGACCCTTGGACACTGCCACGGTTCGGCTTCAAATCCAAGAGAAAGATTC

TGCAGCAATATCTCAATTTGAATTCTTCACCTGCCCAACTTGAAACAATTAAGTAATGGCTCGTTTGGGA

GTGACACGAAGAAGAAGAATATTGTTAAAGCTTCAAGTTAACATAATGATAATCAAGTTTTTCGTTATTT

TTGTTCCTTGGTATTTGATGGTGATAACACTATTGAGTGAATCTTATTTTCGTTTGGGAGGTGGAGGATG

ATGATGATCTCGAAGATGATCCTCGATATATTGAAGTTGTTAGAACAATTCAGACATCTCATTTTTGGAC

CAACTATACAAATCAACTTGCTTTGGGGATGTACAATGAGTGGAGAAATAGAAGTTAGAAGTCAGAAGTC

AGGAACTGAAATGTGGAGATTTGGATGAGCCAGGAGTCTTGAGTGAGCAGCAGCAGCCAGCAGCAGGCCC

AACACTCAGCAGACTGCAGACTAGGACTAGGACTTTAACAAGAATTCATTCACAACCAAACTTTGTGTTC

TGGCTACTGTTATTTTAAATATTTGAATGTTCAAACTAATTCTGATGGAGACTAGCAGATCTTTGAATTG

TGACAATTTTTTTTCGAATGTTTTAATGTCATTCTTGTAAGACTGCAAGTTACAAGTTCTTAAATTTTCA

AA

>MSTRG.2668.1 gene=MSTRG.2668

TTGTGTCTCTAGCTAATTAGCAAACTACAACTGATCGCTTCCCCTCTTCTAACAAGAGAACTCCAAAGCA

CCATCCAGTGCAAAAAGTTCACAATTTAAACAACTTTTACACTGACATCCATCACCGCAACTCTTCCTCT

GTTTCACAGCCTTTGTTGCAACTGGTATAACAGCTCCGTCAACTTTCATTCCAATCTGCACAGCAGGAGG

CTTCACACCATCAGGCAACCTCTTCCCCGTTTCCATCACAACCCATTTCTCAACATCCATCGTCGACGAT

TCCAAAACAGTTTTCCAAGGAACTTCGGCCTTCCCCAAGAGCTGTGACTTTCGGGTTCTTCCAAAGAATG

AAATTCCGCTCCTGCATCTAAGTTCAAAAATCACAGTTCCTTCTAATAGCATGCTCCTCATAGACTCTTT

GGTTCCAAAACAATCTAAGGAGAAAGACTGGTTCCACACCATATTTGAAGATGACACTTCTCGGCTCTGG

AATCGGACTCTGTTCTTGTTTCCTGCTGACATGTAGCATCTTACGAAAAAGCTGCTTCCTGTGGGAATAA

GATCGGCATTTCTAGCTCTGATGACACTTAGCTCACAGCTAATAGCTGCTTGATCTTGAAATTTATCCAT

GCAAGATTATATAAACTGGTTAATAATTTTTGCTTATGCTTTTGTACGAGTGAGTGAAAGAGAGTTATGA

ATTTGATGAAGTTGAGGGAGTACTTTGTAGTATATATAGAGAGATTACAGTTAGTAATTAGCCACTTGAT

TAG

>MSTRG.2672.1 gene=MSTRG.2672

CCTACACACTTCACAGACAGTAGGCTCTCATCAGCCTTAACTTAACCGCCTCTCCAGAACTTTCCTTACC

GCCTAGGCCTATACATACACCAGCCAGCCAGGGTTTATACAACATACAGTGTAACTAAAGAGGAAAATCA

TGTTCAAAAAGTTGGTAGAGGCCAAAACTCATCAGCGACTATCCGGCGCAGACCGTAAAAAACTTAAACG

AACGATTAGAGATCGTTTTCCTAATGCTTCTGATGCTCACATTGACATCTTAATCCNC

>MSTRG.2685.1 gene=MSTRG.2685

GAGAGAGAGATGGGTCACCTGGTAAAGGATTGTAGTAACCAAGAACTTCCAAGTGCTTGCCATCACGAGG

AGATCTGCTATCAGCTGCCATGACTCTGTAAAAAGGCTTGTTCTTGCATCCCAATCTTGATAATCTTAAT

CTCACCACCATCTTTCCTCTTTCTTTTTCTTCAATTCAGGGTAGAGAGAGTGAAATCCAATGAGATGGAA

AAAGCACACTGAAGTGAAGAAAGGGTTTTTCCCATTTCAGGGGTTTTATTTATTTAGCATAACTCCTGGT

TCTATT

>MSTRG.2686.1 gene=MSTRG.2686

GGCCTTGCTCCCGAGTTTTTTTAGGAGAGAAAGCAAGTGAATGCAAGTGAAAGCAAGTGTGCTTTCACTT

TCATCCCACTTTTGGAGTGAAAGTTTTGGAGTGAAACTATGCTATATTTGCACTCACTTTCACTCACTTT

CATCTCTTTAAAAAACTCGGGAGCACACAAAGGAGTGAAAGTTAGATAACTTTACTTCCTCTTCACTTGC

TTTCCTCCCATTTTAAAACTCGGGAGCAGGGTGT

>MSTRG.2690.1 gene=MSTRG.2690

TGCTTTTACTTTATGTCTGAGCCAGATATGATATTCTCACTATGAGTCCATTATGAACATGCTACAAATC

TAAACTTGAAACGGGATGTAACTTGTAAGATCAAATTTTCCATATATCATGCATGTGTTAGTTCTGTTGG

TAGAAACATTATGTGGCCAATCCTATCTGCAAGAAGAACTCTCAAATGTGCACCACTTTGTATTGTCAAG

TTTTAAGCTGCCTTGAAACTTTTGTCCCTGATATCATTTTTTTCTGGGGAAACCTCCGTATAGTGTATCA

TAAATTGGATGGATGGCCTCTGTATAAGAAGAATAACAATCCAACTTGCTTTAGCTTTTGTTGCTTATTA

GGTTAATCTGATTATTACTTAGCAAAAGAAAAAAAATGAAAGATTGACAACAAGATGTCACACCAGCTAG

GATAGTTATCTACTACAAATCAGGAATTGTTAGGAATACAATATAACTCTGGTAAGCCCTCTTCCTGACA

ACTTGAGATTTAGGTGAACTGAGAACCTAACGGTAATCCATTTTAACTATTGTCACACTTTACGAGATTT

ATGTTAACTAGGATATAATGTTAAACTACACTCTACTAGGATAACAGCATATCAGACTTTGATGGGAGTC

TTTTGAAGAAATTGACAGGAACAGAAGGCATCTTGTTTCTGAACTTCGGATGGCGCATCCAGTACAGCCC

GTGAAGCACTGCAATGACCTGAAAGGGAGTGAGGTAAAACAAAGTTGCCCACATGAGAGCAAATATAATG

AAGATTGCGGTTGCTCTTGAATCTCGCCAGCTAAGTATTGCCAAAGCTCTCTCCGCTTGTGTAGCTAAAT

CTCCGAGGACTGTTTGCACTCTTCCTGCCACACTACGCATCCGATCATACCTCATCCTAATTAGGTCTGT

AGGTCGCGAAGATGGAAATGTGTCAAATTCCTCGTCTAGCTCATCTGGATGCACGTTCTCAGCCTGTGAG

ATTCGTGCATCCATATGTGGAGGCTTTCGGGGTCTGAACTGGTAGTTCCATAAACCAATCACACACAAGT

AGAGAAATATTGTAGGGAGGATTAGTTCAGGGTAGCAAACCAGGATCAAGAAAAGTACGTGCACAAGGCA

TGTTGTCAGAGGGTTTCTCCAGTTGCAAATTCCATCCAACCATTTCCAGATAGCTGAAAATCCTGAGAGG

AGTGAAATTATTCGGTAAAAATTTGCTTTGCTTCTCCTGAGGCTCCACATGTGATAGTCCACATCAAGCA

TGTACTCAACTGTTTCGCGTCTTAGAGGTGGTTCTGCCCTGCTTAACCGTGCTGCAACTATCTGCATTGC

TTGGTGTCGTAGCCAGTCAATGTGTCGGACAGAAATAGGCTGGACATAGTGCATTTTGGGGAGTAATGGC

TTGCTATATTGGGTCAACATATTAACCCAAGCTACACAAGTGAAGCGTATGGCTAAATGAAGCTCTCCGT

GTTTCTTCAAACCAGAGGGTTGCAGAACCAGCAAGGGATAGTTATGTGTATAAATTCTATCAGTTTCAAG

AGTTGATAGACGAATCCTTACTTTCCCGATCCTCTGATCTCTTGCATCCTCTTTGTTTCCATTAACATGA

TGATTGTCGAATACACCAATAGTAATTACAGTGCAGGGATCATAAACTTCCCACGTATACTGCTCGTTCC

ACCTGGGAGACAGTGTGTTAAGAAGAGTTCTAGTACGTACCCATTTATTTCCGTACTTGGCTACACAATA

TGCATCTGTACTTCCACCTTCTCGGATTTTCATTGGTAGCAGATTTTGAGCACTCAATATCCCTACTTCA

AGAATTCCTATTCTTTGTTTCCTCAAGTACTTGGATGATGGCTGAAGATCACTACTGAAATGAGTCGTTT

CGTCTAAGACATGGTACCCGACGTCTAAGCAGAGGCAGAGGCGGATTTTGCTAGAAAACTTGACTTCTTT

CTCGCCATCTTCATGCACAGAGTGAGAAGGTCTCTGAAGATTTAACCAGCGAGGATCCGGGAGCTTACTT

GAATCCACCCTTGGTGGAATCTCTCTGACCGGTACCATTGCTCTTCCCATATTATCCTCCCTTCCCCCTT

CACCACGGTCTTCAACACTCACAATAAGGACGTCTTCAAAAGGCTCAGAAGCTACAAACATAAGCTCTTC

ATTCCAAGCAGCATTTATGTTCCTCGTTTGGGAAGGCCGGGTGACCCTGATCTGATGTCCTAGCTGTATT

CTCACATATGGCTGTGGTAGTCTTGTTCTATCAGAAGGCACAAGATCTTGGGCTTCAATGACATGGACGC

GAAGATAGTACAACTTAGGCGAGAAATAAACCTTTGAGCGTGTACTCGCTAGGTTATGATGATTAATATC

ATGAGCATCACTATGCCATGCCTCAGGAAAAGCCTCATCAGCCTGGGTTCCAAGCCAAACTGCAAGCATA

ATCTCTCCCTGACCAGCCAATCTATACCACTGGGGAGCCAGAGGACTATCAGGTGGCACCCGAAGAGGCA

CTTCCACTACATCGAAAACCACCTTCCCAACATAATCATCCTTCCCCAAATCTTTAGCCCTCACTATCAC

CTCAATCACATTTGACTGCAGCCTTTCTTTAGAGAACGCAAACACACTGTTCCACATCGGATCCTGGTTA

TGCTCCAAGTGTTTTGTCAAGCCCCTGTAGTTTCCCACCTTTACTTCAACATAAGGGTCAAGACTTCCAT

TCACATCCAAAGCAGGAAGATCTCTAGCTTTCACCACATTGATGTACAAGTAATGCACTTTCTCCACCAA

ATCATAAGTACTCGCCATCTTGTCACCTCCCAAATACCCCATCCTTGCTGCCACTGGTGGCCGCGTCTCA

ACCAACCCGAATTCAGGCTTCTGCATGGGAATTTGATTTTGCATAACCATAGGTGGCCTTGGTGGGATAA

ACTGAGGCCTGGTTTCAAGAACTACTGGCTTCTGCATTGGAGGCATAGGACCAGTTGGGCCACCAGTTCC

AACAGAGTAGAAAGTCCTAATCTCTTTCTCCTTCTTCCTTTTTATCTTCTGTTTGCTAAAATCATTAAAC

TCATCATCCTGAGCCTTCTTATCATACCTTTTAACATTTTCTTGCACCGGAGTTTCTCTACTTTCCATGC

CTTTCTTCGTCTTTTCTTTTGGTAATTCAGCTTGATCTGATGAAGCGCCATCCTCATCTTTGCCATCAGC

AAGATACAGCCTGAGCGCGATATCACCACGAATATGAGAAAATATACCTCGTTTCTCAAGAGGATATCGT

TGCACCGTTGCTTCAGACTCGGACTTAGGAACTGACAGACCAGATAACCGAACTCGACCAAGAAAATTGT

GGTTGCCATGACTAGTATTTTCATTGTAGATGACAACTTCAATAGTTTTCCTAGAAAGATCCTGCAAGTC

ATCGAATTTAAAGACAAATTTTTCGTGCCACGAGGGGTTAAGATCCTTAATCCTTGTCTGTGTCCTTTGA

TGCTGATCATCTATATCTACCTCAACAAAGGGACTTGCCGAGCCTTGTCCATCTTTCGGCATCAGGTCAT

TGGCCTCAAGGACTTCAACTACAAGCCTCCCCATGATAGCTGTGCACTAAATTTTCTCAACAAAGATGGT

GAAAAGTTGCAGGGGTTTTAGACTGAAGTACTTTTATTGTTACTGAAATGTATCATCAACGATGGTCAAG

TGTATACATGCTATCCGCTGAGCAAAAGTCGAAACTCGAAAAAACAGCTATTTTCAGAAGGGTGTGTGCA

TACATGTTGCAGAGGAGGTGTAAAGTGGAGAAAGTTATGATGGTGTGAGAACTAAAGAAATCAGAAACTG

CATTATTACCAGCATTATTAGTGTGTGATCTAGAGGAACTAATGTGAGTTTCACACTTTTGGTCACATTT

TGACTTGAATCACTTTTTTCAAGTTTTACTTGTGTTAGTAACATG

>MSTRG.2694.1 gene=MSTRG.2694

ATATTTGGCACAAGTTCTTGAAGTTTTCGTATACGGTCACTTATTCGCGTCCTCCGAACCCTCTCGGCTA

TACTGCGAGGATGAGTAGCACAGCCTCGCTTTGCCCGAATTCTACAAGGAACCGAATCTTCGAAAAGCTT

CTGCATATCAGCATCAACACCATTGAACTCCCCACTCTTTTCCCCTTTCTCTCGAGGCCGTTTAGTCGAC

GAAAGCGAGTCATAATTCGCCGGAACTCCAAAATTCGAGAAGTACGCATCGGAGCCACCGGCCGGACTAA

TCTGGCCGAGAAACTCCGCCGGCGAGCTGTTCTGCCGGAGAAACGGCGACGATCGGAGAAGCTCCGGATC

TAGAAATCCAGGTTTGGCCATAGTCTCCGGCGGGGAGAAGTAGTGAGTGGCCGGCGGCGGCGGCTTCGCC

GGCGAGGGATGGAAGATGCTCTCCTTGGCGTCTTCTTCTTCTTCGGGCTCTTCTTCGGATTCGAGAAGAG

CTTCGAGCCATGTGGCCGGAGCTGAGCGGAAGCGGGCGAGTCCGCGACTCAGTCCTGAGTTAGCCGAGTT

GCCTGTCGGTTGCATAATGAAAGTGTGTGAGAGTGTGTGTATTTTGTCTAGATTTCTGCTGCGGGGCTTT

AATTTCAGACACTTTTTGTTGGGGGCAGTGGGAATCCAATTTACTATAAATAGAAACATACATCTACTTA

CATTATCTGTATGTATAGCTTTTGCCTAACACAATCTACTGC

>MSTRG.2697.1 gene=MSTRG.2697

TAATAGGCATACCAAGACATAAATAAATGAAGCGACGACGATTGTTCGATATCGACCTAAATACGAATCA

GCGATAATTGCTCCCAACAGCGGCGTGATCAGGGAGAAGCCAAACCAAGCATTCACATTAGCCGCAGCAG

CCGCCGTGGATTGACCCAAGTCTCCGGTAAGATATGTTATTAAATTCAAGCTGATTCCGCAATAAGCGAT

TCTCTCGGAAATTTCAGCACCTACGAAACCAATTGATTTTTTATATTACTTTGGGATATATTAATAAATT

TCAACACATAGTATATAGTGACTGGTGAGTGGTGAGTGATACCTATAATGAGGAGTGCCGATCGCCATCG

GCCGGAAGTTGATCGATACACTGGCCGGTTTTTGTAGTCAACGGCGAATTTTACGACGTCGTCGGATAGG

AGTGGAGTGATGATGGAAGAAGCGGCGGTCATTTGATTTATTTCACTCTCTGGACTTGTTTGTCATGTTT

AGGTCGTTGATCTTTCGAGCATCATCTTGTCAAAATTAAGTGGGTGTTTGGCTGGCACTTATAAGTCCAG

CTTCTGAATTA

>MSTRG.2698.1 gene=MSTRG.2698

AATATTTTGGTAGTTGTTGGGAGAAATTTTATTGCAAGTAATCTCCGTAAATTAGTTGAAAGCGGGTAGA

AGTTAATTGCTTACTCAGTTTAATAAAATAGTTGGCAGCTTAAAATTTATCCGCGTGTCTCCACACTTCT

CTTCCTTCAGCCACGTATACACCCCTGTGAACCCTTTTCCTTTTATTCCCTGTGAAACCTCATCTCTCTC

GTTAAATCACAGTAATCGGTTCTCTCGCAAATATCTCCCCAATCTCACGCCATTCCTTCGGTCTCTCGCT

ACCATCTCCCTATACACTTCCCTCCATCTCTCGCCTTTGTACGTACGGTCGCCATATTCCATCGGGATTC

TCTGCCGCCGCTTCCTTTTCCGGCCAGAACCAACACCGCCTATTTCTTCCTCTTCCGGCGACCACCATAG

CCCTGCACCAGTTATAAATCAGTGGTTAACTCCGTCTGCGTCGACAACTGAAGTACCGTGTTACTCTTGG

AAAATCAACCAAGACTTGTATATCAAGGACATTCAGAAGTAGTTGAATAGCTTTGCGAACTGAATCAGGT

ATAGTTCCTTAAGACGAAGCTGTGCTGTGAGATAGCCCGGTAGAATTAAATCGGTATGAACATCAGCAGA

ACTGAGTTAACTCCGTCTGCGTCGACAACTGAAGTACCGTGTTACTCTTGGAAAATCAACCAAGACTTGT

ATATCAAGAAGAGTAATGCTAGGCAAGGATTCCAAGTGATAAGTTGTGACTGAATGATCAGGTAAAGTAT

TCAGGAAGATGTCTGCGAGATTAGTGGTAATTAGAGTTGGCTTAATTTTGCTTCAAATGTAAAATAAATT

TATTAAATATTATCAGCTACTCTGATAATTTAGATTGGGTTGTAATAAGATTTATTAGTTTTGATTCTGA

TTTCAATACTATAGCCTGTTTGCGATCCTGTTTAAAGGGG

>MSTRG.2699.1 gene=MSTRG.2699

GACATATGATCCTGAATCCTACTATGAATTATCATCTTTCTTACATATAAATTGCAATTGAAAGAAAATT

CATAAAATGATCCTGGTTATAGTCAAGTTCCTCAACTTACAATTGCCCAAATCTAAAACAACAAACAGTC

AATAAACCAACAATATTTACAAATTCCCCGAAGGACCTCTCACAATTTCTGACTACTCTGCATCACAAAC

GGATGAGACAACAGCTCCTCAGCCGTCCACCTCTTACTACTCTCCTTCTGCAAACAACACTCAATAAAAC

TCCTAAACTCCCCCGACACACTCTCCGGCAGACTCGGAGACTCCCCGAAGCAAATAGCGCACATAAGCGT

CGCCCAATCAGGCCTCTGCCCCGGCGACAACAGCGGAAAATGCCCCAAATATAATTCCAGCAATGTCAAG

CCTAAACTCCAAATATCAGCCGCGTATCCGTTGTAATCACGCCCGTAGGTCTCGGGATCGAACCGCTCCG

GGCTCATGTACGCGCAAGTACCTACATATGAGTTGCATGGGTCGAGGGTTCGGCACATGAGTTTGCTCAC

CCCGAAATCGGAGATTTTGATCTCGAATTTTGAGTTTACGAGTAAATTTGCTGGCTTGATGTCGCGGTGG

ATGATTTTGTTTGTGTGGATGTAGTTGAGGCCGTTGAGGACTTGAAAGGCGACGTTGGATAAGACCGGTT

CGGAGAAGGTGCCGGTTTTTTTGAGGAGCGTGTCGAGATTGCCGGCGTCCATGAACTCCATGAGGATTTG

GATGTCGCCGTCTTGGCGTTCGGTGATGTTGTGGCAGTTGATGACGTAGGGGGAGTCGATGCGGCGGAGG

ATGTCGACTTCGCGGAGGATCTGGCGGCGCGTGGTGGAGTCGGTGTTGGCGTGGATGTGTTTGAGCGCGT

AGGTTTTGGAGGTTTTGTTGTTGTGGACTTTGTAGACGGTGCCGCCGTTGCCGTGGCCCAGGACCTGGAG

CTTTTCGAGGTCGGGGGAGATGGTGGTTGTAGCCGTTGTGGTGGTTGTCGGTGGCAGAGGGAGGGGGCAG

CGTGGGAGGTGGAGGCTGAGCTGGCGGCGTTTCTGAACAAGAGCCATGGAATTTGTGTGCAAGAAGTTGA

GGACTCAGACTCAGAGACTCAGTTGTTGTGTTGTGGAAGGAGGGATATTTATATAAGTGGGGAGATTAAA

GCACGTTGGATTTGATTTGGAAACGAGAGGGAAGGCCACATGAGTGTGCGAGGGAGGTGGGAATATGGTT

ATCACAATGAGGACCTCCATTTCTATTTTATTAGCATTATCCAAAAATGTGTGATTAATTATCGACTAAT

TCATGTTTTAATCAATTATTTGCTTAATTTTAGTTTCAGAAAAATAAGTACTTGACTAGTTTCAAATCAA

>MSTRG.2702.1 gene=MSTRG.2702

GGCAGGATCAGTTGGAAACCATGCTCTCACTTTCTTTCACAAATCCTTCACCTGTCCAGTCTTTGTCTGA

TGCCAGTCCTTTTTTCATTGCCTTTTTCTTCTGTACCGCTTCTTTTTGCATCCTGACCAATTCTGTGTGG

GTTGTAAAATACTGAAGAGACATCCCTCTGAATTCTTCTATGGATGAAAAGTTGTGCATTTTCATAAAAT

CTTTC

>MSTRG.2713.1 gene=MSTRG.2713

AAAAACATCTCTCTTTCTCTAATCTCTACCCGTTACAATTCTTCAGGCGAAGTCCCAAATTTGTAAAACA

TGACACTCTAAACCCTAACGACTATGAGGGTCTCGTAGCAGAGAAACATATAATCCTTTCCTTTTGTTTG

GTATGCTAATCATATTCATGTCATGGGATTCGTTTTTCATCCTGGAAATCATAATTACATCCGGCTGGTT

TTGTAGCCAAAGGCTGCTCTATAGGCTTTTAGGTGCTGGATTCTTGGTGGAGGCTTTAAAGGATGGGGTT

AGAGACCGTCACCAAGTTTGGGATGAGGATGGATAGCCGTCGTGTATGTACGGTTACCAGACTAGCTTGA

TTGTAGATTTGTAGTGATATGTTCGATCAGCGGCACATTGACACTTGGGTGGCAATTCAAGTCACGGTTG

AATGACTGAACCAAGAGCTGGTTTTCAGATGAAAGACTAGCTGCAGTTTTGTTGTTGGCTTGCTGCAATA

TGGTGCGCCTTCTGGTTTTTCCATGAATATGGGAAAAGTGATGGCCAAAGATGGACGTTACCAGCGTCTG

AGATTTTATGTGATTGTAGGCTTTAAAGGGTGTGACTAGAGACAAAGGTGAATAGGCAACAAAGTTTCGG

AACAGGTTGGATAACCAAATCAGGGAACCTTTTTAAGCTAGCTTGAGTGTATTGACTTGTTCTATCAGCT

GTGCATTTACACTTGGGTGGCAAGTTAAGTCACGGTTTGGGTTCGAAAGCTTATTTTGGAAGTGGTTGCA

GTGTGGCAGTGGTTACTGGTCAAATAATATGCAGATGCGGTTTTAAAAACATTTATAGATGTGCTACTAG

CCTAGAACTATAAATATACTTGATTAAGGTCTAGTTGTTTAAATTGAAGTACTTTATATTCTCAACTAGC

AATTTTACTATATTGTTCAAACTTTAAACATTGTGTGTAG

>MSTRG.2713.2 gene=MSTRG.2713

CTTTTATTCTGAATCTTCTTAATTATTATTTAACTGTGTTACTTTTTACAAGTATACTGAGATATGAAAT

GATTATTATTCAATCATTTATGCTTTACTTTCAAATACGTATCCATATTTGTTGTTATGGACACCCCAAC

ATTTATATAAAAGAAATTACAGAGATAAGAGCTTTCGACCTATCTTAATTTTTAGATCTCCCCCAAAATT

AGCAGATTACAAAAATAAGTGCCGCACATTCTCGTGCAATTTTTCACCCAGTCACATAATGGCAGTGCGT

GAGAAATTAAAGAGGAAGAAGACCGTGGCGGCGGACCTACCAGGAGAAGTAGTCTACTGGGAGATACTCA

CCCGACTCCCAATCAAATCTGCGGTACGTTTTAAATGTGTATCTAAATCATGGCTATCTCAAATCTCTAG

TCCCAAATTTGCAAAAAGACACCTCACTTGCAACGCCACTCAAAACCCTAGTGACTATGACCGTCTCATT

GCTCAAAAAGAAAACAAAATTGTCATTCTTTCTCACTACAATGAAACATTTGCACTCCCATTTTCTTGTC

GCGGTTTAGTTGGTTCTGTTCGTGGCCTAGTTTGTCTTAGTCATGACGAGAAGTTGTCTTTGTGGAACCC

TGCAATACATCAATCCAAGGAATTCACTCTTCCACCAAACTGTTTTGATTATATTGAATATATGGGATTT

GGTTTTGATCCCGTAAGTAATAACTTCAAGGTTGTGCTTTTGTCAAGTGATTTTAGGTCTGCTAGTGTAT

ACAACTCTGACTCCGATTCTTGGACTAATGTATTTGTTCCTGATAATGTGCATCTTCTTATGAAAAAAGA

AAGGACCTTCTTGACAGATTCTATGACTATTGTAAAGGACTGTCCGTATTGGAGATTTATTACATTTGCT

AAAGCGCCTGCTACAGATACGACAGATACGCTCACTCCAACTTCTTTGACACCGATGAAGTTTGATGCTG

GAAGTAACGATTTCATTTTGTTGCCTGAATTTCATTTTGATACGAGTGTACGTGCAGAGAGTGATGCATT

TAGTTTCCAACTGGTGGATATGAATGATCATCTTACTTTGATATTTTATGATCAAAACTCTTCAAATCGT

ATGCTGGATATATATTCTTTGAAGGGTGAGGAGGGAAGTTGTGTTTGGAGTCGGATATACAGTATTGGAC

CGCTTGATTTTCATAAACGGCATTGGTTTGTGTGTCAGGGATTTTATGGTGGCGAGATTGTAATTGAGGA

TAATGGAATGTTTTCATGTTTTGATCAGAAAAGGGGTACAATCAAGCGCCTTAGAACAACTGTTACAACT

GTAACTGTTACCTCTTTAAGCTGCTTCAGATATACGCCTAGCATGTTTTTTCTTGAAGGAATGAAGTCTG

TTTATCTAAGAACCCAAACTCGCCGTACAACTGGTCTATGCTCTAGAGTCCCTCGGCGATTAGTCGATTC

ATTTAAAAATTGATCATTTTCTTAAGGTTTGGGTTCGAAAGCTTATTTTGGAAGTGGTTGCAGTGTGGCA

GTGGTTACTGGTCAAATAATATGCAGATGCGGTTTTAAAAACATTTATAGATGTGCTACTAGCCTAGAAC

TATAAATATACTTGATTAAGGTCTAGTTGTTTAAATTGAAGTACTTTATATTCTCAACTAGCAATTTTAC

TATATTGTTCAAACTTTAAACATTGTGTGTAGTCTGCTTCTTGTTTAGAATTTCTCGCTATCTAGTACAT

GTTTAGTTTTGTGGCTAATGGGTGATTGAATGATTGCTGTATCATCTATCTAATTGTATTCTATTATCTC

ATAATCTTTATCAGTTTATGTATATCATGGATTATGGAGTCGAGCTTAAGGAGCTTAACAAGTTGAATTG

AGTAATCGACTCATTAAGTTTATCTGACAAAATTTTATATTTGAACCCGAGCTCAAATTCTTATCAAACC

AG

>MSTRG.2714.1 gene=MSTRG.2714

GGCTACAAATTACAATACAGTAATAATACAGCACACAATAGTATACAAACCGCAAGGCTTAAATTCCGAA

ATTTCCAGCTCAAAACATCCATCATCGACATCTCCTGACAAGATAAAATCAAATGATTCATTATCTTGTT

CCCAAAATAAAGATGCAGAGATTATTGGGAGCCGGAAAATCTGTAGCAGCAGCAGCTGCGGGAGCTTTTA

CGAGGAAGCACCGTGCTCTCTCTTCATCTTCTATGCTCTTCGATGATACTCAGATTCAGTTTAAGGAAAG

CGTTGCACAGTTTGCTCAAGAAAATATAGCACCTCATGCATCAAAAATTGATAAAGCAAATTCTTTTCCG

AAGGAGGTCAACTTATGGAAACTAATGGGAGATTTTAATCTTCTTGGAATAACTGCTCCAGAGGATTATG

GGGGACTTGGTCTTGGTTATTTGTATCATTGCATAGCAATGGAAGAAATTAGCCGAGCTTCTGGATCGGT

GGGCCTTTCCTTTGGTGCACATACCAATCTGTGCATTAATCAGTTGGTGAGGAATGGAAACGCTACACAG

AAGCAAAAATACTTGCCTAAGCTTATCAGCGGAGAGCATGTGGGAGCACTTGCAATGAGTGAACCTAATG

CTGGTTCGGATGTTGTTAGCATGAAATGCAAGGCTGATAGTGTTGATGGGGGTTATATTTTGAATGGAAA

CAAGATGTGGTGCACGAATGGCCCAACTGCTCAGACACTGGTTGTATATGCAAAAACGGATATTGCTGCC

AAGTCAAAAGGAATTACAGCATTCATTATCGAGAAGGGAATGCCTGGATTTAGTACTGCACAGAAGTTGG

ACAAACTTGGGATGCGAGGCAGTGACACATGCGAGCTTGTGTTCGAGAATTGTTTTGTCCCTAGAGAAAA

TATTCTTGGGCAGGAAGGAAAAGGAGTTTATGTTATGATGTCTGGGCTGGATTTGGAGAGGCTTGTTCTG

GCTGCTGGACCAGTGGGCCTCATGCAGGCATGCCTTGATATTGTTATTCCGTATGTTCGGCAGAGAGAAC

AGTTTGGCAGCCCGATTGGACAGTTCCAGTTTATACAGGGGAAAGTTGCTGACATGTATACTTCCTTACA

ATCTTCAAGATCGTATGTTTACTCAGTCGCAAGGGACTGTGACAATGGAAGAGCGGATCCAAAGGATTGT

GCTGGTGTAATACTCACAGCAGCTGAGAGAGCCACCCAAGTCGCATTGCAGGCTATTCAGTGTTTAGGTG

GCAATGGATATGTAAACGAGTATGATACTGGGCGTCTTCTTCGCGACGCCAAATTATACGAAATTGGAGC

AGGAACTAGCGAGATTAGAAGAATGATTATTGGCCGGGAGCTGTTTAAGGAAAGCTGATAAAGAAAAATT

ACAGGAAGAACTACTTGAAGTTGCATGTTTCACTTCTTTGTGGTTTACAAACTGTTTTACTCTCACAATT

TAAATGTAAAAGTTTCTCTGGACCCTAACAAAAGGTAGGGTATATGCAACGTTTACATGAACTCTTATCT

TAACTTTATTTGTTGTTCTGACACTGATCTTATAAATTACCAAAATCTTTGTGCATCTTCGGTCTATTCT

TTGAGGGGGCCACAATAAATGAATTCCCG

>MSTRG.2718.1 gene=MSTRG.2718

AAAACATAATTATTGGAGTTAAATAGTCTTAGAAATGATGTGCGTGTGATGAACGCAGAATTGCAAAAGC

ACCACCACCAAACTCATAATATGGAGAAGAAGCCACACTTTAAACCGGAGCTTCTTGTTGTGGTAGTTTT

TGGAATACTACAAAACCTTGTCCAGCAAGCAAGTGCAGATTTCCATTCACCACTCATCAAGTATCTCCCC

GGTTTTCAAGGACCGCTCCCTTTTCGACTCACAACTGGGTACGTTGGCGTGGATGAATCAGATGATGTGC

AGTTGTTTTATTATTTTGTCGAGTCTCAGACAAATCCGAGAGATGATCCTCTAATACTTTGGATGACAGG

AGGTCCTGGTTGTTCTACTATCACCGCCTTCGCTTATGAATTCGGTCCAGTATATTTTGAGCAAGTGGAG

TATAACGGCACTTTGCCAACCTTAATTTTAAATCCCCACACATGGACAAAGGTAGCAAATATACTATTTT

TGGACTTGCCTGTTGGTACCGGTTTTTCCTACGCCAAGACTTCTATTGCTTCTCAATCTAATGACACTCA

AGCATGTGCTCAGGCTCTACAATTCTTAAAAAAGTGGTTTATGAGTCATCCTCAATTCCTTGCGAATCCG

TTTTATGTTGGTGGAGACTCTTACGCTGGAATCTTTGTTCCGGTTATTACACAGATGATTTCACATGTAA

ATGAGGCTAAAATAGGTCCTGAGCAACCCATTAATCTGAAGGGTTACTTACTTGGAAATCCTACGGCATT

TCCTAGTGAGATAAACTTTCGCTACACATTTGCTCATGGAATGGGAATTATTTCAGATGAACTATATGAG

TCATTGAGAAGAAATTGTGGGCTTGATTATCAAAAATCAGATATTGATAATGCAGAGTGTTCACAAAATA

AAGCGGCTTATGAAGAGTGTAGGAGTGGACTCTTTAAACCACAAATATTGGAGAAAAATTGTGTTGAACC

GCTGCGCAGGTCATTATCTTATGAAAAAACAGAGGTCATGGATAAATTTGGAGGCCATAATAATCCTAAA

TTACCACTATCTGCATTTGAATGTCGTAATGATGGATATCCGCTTTCTAGTTACTGGAGTAATGATGAGA

ATGTGCAGAAAGCGTTGCATATCAGAAAGGGGTCCATAGGAACATGGGTAAGATGCAAAAAAGAGTTGTT

ATTTGAAAAGGTGATCTTAGATACGAGATCATATCATGCAAATCTCAGCAAAAAAGGTTATAGATCTCTC

ATATACAGTGGAGACCATGACATGGTGGTGCCATTTCAATCAACTCAAGCTTGGATAAGAGGTCTAAACT

ATTCCATCATTGATGATTGGCGACCATGGATTGTCGAAGGCCAATATGCCGGTTACACAAGGACGTACTC

TAACAAAATGACTTTTGCTACCGTAAAGGGGGCAGGCCATACAGCACCAGAATACAAGCCTGCAGAATGT

TATGCTATGTTCAATAGATGGTTATTAGAGGAACCTTTATAAAACTACGTTTTGAGCATGTACTATGACG

GAAATTATATTAATAAAAATATTTGTTTGTATATGAATTGTTGTGCATTTTGACAACTCAAACTTCATCA

TATTATATTATC

>MSTRG.2718.2 gene=MSTRG.2718

AAAACATAATTATTGGAGTTAAATAGTCTTAGAAATGATGTGCGTGTGATGAACGCAGAATTGCAAAAGC

ACCACCACCAAACTCATAATATGGAGAAGAAGCCACACTTTAAACCGGAGCTTCTTGTTGTGGTAGTTTT

TGGAATACTACAAAACCTTGTCCAGCAAGCAAGTGCAGATTTCCATTCACCACTCATCAAGTATCTCCCC

GGTTTTCAAGGACCGCTCCCTTTTCGACTCACAACTGGGTACGTTGGCGTGGATGAATCAGATGATGTGC

AGTTGTTTTATTATTTTGTCGAGTCTCAGACAAATCCGAGAGATGATCCTCTAATACTTTGGATGACAGG

AGGTCCTGGTTGTTCTACTATCACCGCCTTCGCTTATGAATTCGGTCCAGTATATTTTGAGCAAGTGGAG

TATAACGGCACTTTGCCAACCTTAATTTTAAATCCCCACACATGGACAAAGGTAGCAAATATACTATTTT

TGGACTTGCCTGTTGGTACCGGTTTTTCCTACGCCAAGACTTCTATTGCTTCTCAATCTAATGACACTCA

AGCATGTGCTCAGGCTCTACAATTCTTAAAAAAGTGGTTTATGAGTCATCCTCAATTCCTTGCGAATCCG

TTTTATGTTGGTGGAGACTCTTACGCTGGAATCTTTGTTCCGGTTATTACACAGATGATTTCACATGTAA

ATGAGGCTAAAATAGGTCCTGAGCAACCCATTAATCTGAAGGGTTACTTACTTGGAAATCCTACGGCATT

TCCTAGTGAGATAAACTTTCGCTACACATTTGCTCATGGAATGGGAATTATTTCAGATGAACTATATGAG

TCATTGAGAAGAAATTGTGGGCTTGATTATCAAAAATCAGATATTGATAATGCAGAGTGTTCACAAAATA

AAGCGGCTTATGAAGAGTGTAGGAGTGGACTCTTTAAACCACAAATATTGGAGAAAAATTGTGTTGAACC

GCTGCGCAGGTCATTATCTTATGAAAAAACAGAGGTCATGGATAAATTTGGAGGCCATAATAATCCTAAA

TTACCACTATCTGCATTTGAATGTCGTAATGATGGATATCCGCTTTCTAGTTACTGGAGTAATGATGAGA

ATGTGCAGAAAGCGTTGCATATCAGAAAGGGGTCCATAGGAACATGGGTAAGATGCAAAAAAGAGTTGTT

ATTTGAAAAGGTGATCTTAGATACGAGATCATATCATGCAAATCTCAGCAAAAAAGGTTATAGATCTCTC

ATATACAGTGGAGACCATGACATGGTGGTGCCATTTCAATCAACTCAAGCTTGGATAAGAGGTCTAAACT

ATTCCATCATTGATGATTGGCGACCATGGATTGTCGAAGGCCAATATGCCGGTTACACAAGGACGTACTC

TAACAAAATGACTTTTGCTACCGTAAAGGGGGCAGGCCATACAGCACCAGAATACAAGCCTGCAGAATGT

TATGCTATGTTCAATAGATGGTTATTAGAGGAACCTTTATAAAACTACGTTTTGAGCATGTACTATGACG

GAAATTATATTAATAAAAATATTTGTTTGTATATGAATTGTTGTGCATTTTGACAACTCAAACTTCATCA

TATTATATTATC

>MSTRG.2718.7 gene=MSTRG.2718

AGGGGTCTATAGGAACATGGATAAGATGCAAAAAAAGAGTTGTTATTTGAAAAGGTGATCTTAGATACGA

GATCATATCATGCAAATCTCAGCAAAAAAGGTTATAGATCTCTCATATACAGTGGAGACCATGACATGGT

GGTGCCATTTCAATCAACTCAAGCTTGGATAAGAGGTCTAAACTATTCCATCATTGATGATTGGCGACCA

TGGATTGTCGAAGGCCAATATGCCGGTTACACAAGGACGTACTCTAACAAAATGACTTTTGCTACCGTAA

AGGGGGCAGGCCATACAGCACCAGAATACAAGCCTGCAGAATGTTATGCTATGTTCAATAGATGGTTATT

AGAGGAACCTTTATAAAACTACGTTTTGAGCATGTACTATGACGGAAATTATATTAATAAAAATATTTGT

TTGTATATGAATTGTTGTGCATTTTGACAACTCAAACTTCATCATATTATATTATC

>MSTRG.2718.5 gene=MSTRG.2718

TACAATGGGACTTTGCCCACACTAATATTAAATCCCCACACATGGACAAAGGTAGCAAATATACTATTTT

TGGACTTGCCTGTTGGTACCGGTTTTTCCTACGCCAAGACTTCTATTGCTTCTCAATCTAATGACACTCA

AGCATGTGCTCAGGCTCTACAATTCTTAAAAAAGTGGTTTATGAGTCATCCTCAATTCCTTGCGAATCCG

TTTTATGTTGGTGGAGACTCTTACGCTGGAATCTTTGTTCCGGTTATTACACAGATGATTTCACATGTAA

ATGAGGCTAAAATAGGTCCTGAGCAACCCATTAATCTGAAGGGTTACTTACTTGGAAATCCTACGGCATT

TCCTAGTGAGATAAACTTTCGCTACACATTTGCTCATGGAATGGGAATTATTTCAGATGAACTATATGAG

GTTTGTTTGTTTTTAGAAGTATTTATTTTTTAATGATTTTGATTTATCAAATCCACTTATAACTTTCTTG

ATTATAGTCATTGAGAAGAAATTGTGGGCTTGATTATCAAAAATCAGATATTGATAATGCAGAGTGTTCA

CAAAATAAAGCGGCTTATGAAGAGTGTAGGAGTGGACTCTTTAAACCACAAATATTGGAGAAAAATTGTG

TTGAACCGCTGCGCAGGTCATTATCTTATGAAAAAACAGAGGTCATGGATAAATTTGGAGGCCATAATAA

TCCTAAATTACCACTATCTGCATTTGAATGTCGTAATGATGGATATCCGCTTTCTAGTTACTGGAGTAAT

GATGAGAATGTGCAGAAAGCGTTGCATATCAGAAAGGGGTCCATAGGAACATGGGTAAGATGCAAAAAAG

AGTTGTTATTTGAAAAGGTGATCTTAGATACGAGATCATATCATGCAAATCTCAGCAAAAAAGGTTATAG

ATCTCTCATATACAGTGGAGACCATGACATGGTGGTGCCATTTCAATCAACTCAAGCTTGGATAAGAGGT

CTAAACTATTCCATCATTGATGATTGGCGACCATGGATTGTCGAAGGCCAATATGCCGGTTACACAAGGA

CGTACTCTAACAAAATGACTTTTGCTACCGTAAAGGGGGCAGGCCATACAGCACCAGAATACAAGCCTGC

AGAATGTTATGCTATGTTCAATAGATGGTTATTAGAGGAACCTTTATAAAACTACGTTTTGAGCATGTAC

TATGACGGAAATTATATTAATAAAAATATTTGTTTGTATATGAATTGTTGTGCATTTTGACAACTCAAAC

TTCATCATATTATATTATC

>MSTRG.2718.4 gene=MSTRG.2718

AGCAAGCAAGTGCAGACTTCCATTCACCACTCATCAAGTATCTCCCCGGTTTTCAAGGACCGCTCCCTTT

TCGGTTCACAACTGGGTACGTTGGCGTGGATGAATCAGATGATGTGCAGTTGTTTTATTATTTTGTCGAG

TCTCAGACAAATCCGAGAGATGATCCTCTGATACTCTGGATGACAGGAGGTCCTGGTTGTTCTACTATCA

CCGCCTTCGCTTATGAATTCGGTCCAGTATATTTTGAGCAAGTGGAGTATAACGGCACTTTGCCAACCTT

AATTTTAAATCCCCACACATGGACAAAGGTAGCAAATATACTATTTTTGGACTTGCCTGTTGGTACCGGT

TTTTCCTACGCCAAGACTTCTATTGCTTCTCAATCTAATGACACTCAAGCATGTGCTCAGGCTCTACAAT

TCTTAAAAAAGTGGTTTATGAGTCATCCTCAATTCCTTGCGAATCCGTTTTATGTTGGTGGAGACTCTTA

CGCTGGAATCTTTGTTCCGGTTATTACACAGATGATTTCACATGTAAATGAGGCTAAAATAGGTCCTGAG

CAACCCATTAATCTGAAGTCATTGAGAAGAAATTGTGGGCTTGATTATCAAAAATCAGATATTGATAATG

CAGAGTGTTCACAAAATAAAGCGGCTTATGAAGAGTGTAGGAGTGGACTCTTTAAACCACAAATATTGGA

GAAAAATTGTGTTGAACCGCTGCGCAGGTCATTATCTTATGAAAAAACAGAGGTCATGGATAAATTTGGA

GGCCATAATAATCCTAAATTACCACTATCTGCATTTGAATGTCGTAATGATGGATATCCGCTTTCTAGTT

ACTGGAGTAATGATGAGAATGTGCAGAAAGCGTTGCATATCAGAAAGGGGTCCATAGGAACATGGGTAAG

ATGCAAAAAAGAGTTGTTATTTGAAAAGGTGATCTTAGATACGAGATCATATCATGCAAATCTCAGCAAA

AAAGGTTATAGATCTCTCATATACAGTGGAGACCATGACATGGTGGTGCCATTTCAATCAACTCAAGCTT

GGATAAGAGGTCTAAACTATTCCATCATTGATGATTGGCGACCATGGATTGTCGAAGGCCAATATGCCGG

TTACACAAGGACGTACTCTAACAAAATGACTTTTGCTACCGTAAAGGGGGCAGGCCATACAGCACCAGAA

TACAAGCCTGCAGAATGTTATGCTATGTTCAATAGATGGTTATTAGAGGAACCTTTATAAAACTACGTTT

TGAGCATGTACTATGACGGAAATTATATTAATAAAAATATTTGTTTGTATATGAATTGTTGTGCATTTTG

ACAACTCAAACTTCATCATATTATATTATC

>MSTRG.2718.6 gene=MSTRG.2718

AGCAAGCAAGTGCAGACTTCCATTCACCACTCATCAAGTATCTCCCCGGTTTTCAAGGACCGCTCCCTTT

TCGGTTCACAACTGGGTACGTTGGCGTGGATGAATCAGATGATGTGCAGTTGTTTTATTATTTTGTCGAG

TCTCAGACAAATCCGAGAGATGATCCTCTGATACTCTGGATGACAGGAGGTCCTGGTTGTTCTACTATCA

CCGCCTTCGCTTATGAATTCGGTCCAGTATATTTTGAGCAAGTGGAGTATAACGGCACTTTGCCAACCTT

AATTTTAAATCCCCACACATGGACAAAGGTAGCAAATATACTATTTTTGGACTTGCCTGTTGGTACCGGT

TTTTCCTACGCCAAGACTTCTATTGCTTCTCAATCTAATGACACTCAAGCATGTGCTCAGGCTCTACAAT

TCTTAAAAAAGTGGTTTATGAGTCATCCTCAATTCCTTGCGAATCCGTTTTATGTTGGTGGAGACTCTTA

CGCTGGAATCTTTGTTCCGGTTATTACACAGATGATTTCACATGTAAATGAGGCTAAAATAGGTCCTGAG

CAACCCATTAATCTGAAGTCATTGAGAAGAAATTGTGGGCTTGATTATCAAAAATCAGATATTGATAATG

CAGAGTGTTCACAAAATAAAGCGGCTTATGAAGAGTGTAGGAGTGGACTCTTTAAACCACAAATATTGGA

GAAAAATTGTGTTGAACCGCTGCGCAGGTCATTATCTTATGAAAAAACAGAGGTCATGGATAAATTTGGA

GGCCATAATAATCCTAAATTACCACTATCTGCATTTGAATGTCGTAATGATGGATATCCGCTTTCTAGTT

ACTGGAGTAATGATGAGAATGTGCAGAAAGCGTTGCATATCAGAAAGGGGTCCATAGGAACATGGGTAAG

ATGCAAAAAAGAGTTGTTATTTGAAAAGGTGATCTTAGATACGAGATCATATCATGCAAATCTCAGCAAA

AAAGGTTATAGATCTCTCATATACAGTGGAGACCATGACATGGTGGTGCCATTTCAATCAACTCAAGCTT

GGATAAGAGGTCTAAACTATTCCATCATTGATGATTGGCGACCATGGATTGTCGAAGGCCAATATGCCGG

TTACACAAGGACGTACTCTAACAAAATGACTTTTGCTACCGTAAAGGGGGCAGGCCATACAGCACCAGAA

TACAAGCCTGCAGAATGTTATGCTATGTTCAATAGATGGTTATTAGAGGAACCTTTATAAAACTACGTTT

TGAGCATGTACTATGACGGAAATTATATTAATAAAAATATTTGTTTGTATATGAATTGTTGTGCATTTTG

ACAACTCAAACTTCATCATATTATATTATC

>MSTRG.2718.3 gene=MSTRG.2718

AACGCAGAACTGAAAAGGCACCAGCATCAAACTCATAAAATGGAGAAAAAGCCATATTTAAAAGTGCTCA

GTGTCTGCAACATACAGCTTCTTGTCGTAGTTACTGCAATACTACAAAACCTTGTGCAGCAAGCAAGTGC

AGACTTCCATTCACCACTCATCAAGTATCTCCCCGGTTTTCAAGGACCGCTCCCTTTTCGGTTCACAACT

GGGTACGTTGGCGTGGATGAATCAGATGATGTGCAGTTGTTTTATTATTTTGTCGAGTCTCAGACAAATC

CGAGAGATGATCCTCTGATACTCTGGATGACAGGAGGTCCTGGTTGTTCTACTATCACCGCCTTCGCTTA

TGAATTCGGTCCAGTATATTTTGAGCAAGTGGAGTATAACGGCACTTTGCCAACCTTAATTTTAAATCCC

CACACATGGACAAAGGTAGCAAATATACTATTTTTGGACTTGCCTGTTGGTACCGGTTTTTCCTACGCCA

AGACTTCTATTGCTTCTCAATCTAATGACACTCAAGCATGTGCTCAGGCTCTACAATTCTTAAAAAAGTG

GTTTATGAGTCATCCTCAATTCCTTGCGAATCCGTTTTATGTTGGTGGAGACTCTTACGCTGGAATCTTT

GTTCCGGTTATTACACAGATGATTTCACATGTAAATGAGGCTAAAATAGGTCCTGAGCAACCCATTAATC

TGAAGGGTTACTTACTTGGAAATCCTACGGCATTTCCTAGTGAGATAAACTTTCGCTACACATTTGCTCA

TGGAATGGGAATTATTTCAGATGAACTATATGAGTCATTGAGAAGAAATTGTGGGCTTGATTATCAAAAA

TCAGATATTGATAATGCAGAGTGTTCACAAAATAAAGCGGCTTATGAAGAGTGTAGGAGTGGACTCTTTA

AACCACAAATATTGGAGAAAAATTGTGTTGAACCGCTGCGCAGGTCATTATCTTATGAAAAAACAGAGGT

CATGGATAAATTTGGAGGCCATAATAATCCTAAATTACCACTATCTGCATTTGAATGTCGTAATGATGGA

TATCCGCTTTCTAGTTACTGGAGTAATGATGAGAATGTGCAGAAAGCGTTGCATATCAGAAAGGGGTCCA

TAGGAACATGGGTAAGATGCAAAAAAGAGTTGTTATTTGAAAAGGTGATCTTAGATACGAGATCATATCA

TGCAAATCTCAGCAAAAAAGGTTATAGATCTCTCATATACAGTGGAGACCATGACATGGTGGTGCCATTT

CAATCAACTCAAGCTTGGATAAGAGGTCTAAACTATTCCATCATTGATGATTGGCGACCATGGATTGTCG

AAGGCCAATATGCCGGTTACACAAGGACGTACTCTAACAAAATGACTTTTGCTACCGTAAAGGGGGCAGG

CCATACAGCACCAGAATACAAGCCTGCAGAATGTTATGCTATGTTCAATAGATGGTTATTAGAGGAACCT

TTATAAAACTACGTTTTGAGCATGTACTATGACGGAAATTATATTAATAAAAATATTTGTTTGTATATGA

ATTGTTGTGCATTTTGACAACTCAAACTTCATCATATTATATTATC

>MSTRG.2718.9 gene=MSTRG.2718

AGAGACTATGACAGTCCTAAATTAGGATCATCAGCATTCGATTGTCGTAATAATGGATATCCTCTTTCTA

GTTACTGGAGTAATGATGAGAATGTGCAAAAAGCGTTACATATAAGAAAGGGGTCTATAGGAACATGGAT

AAGATGCAAAGATGAATTATTATTTGAAAAAGTGATCACGGATACTAGACCATACCATGCAAATCTTAGC

AGAAAAGGTTATAGATCTCTCATATACAGTGGAGACCATGACATGGTGGTGCCATTTCAATCAACTCAAG

CTTGGATAAGAGGTCTAAACTATTCCATCATTGATGATTGGCGACCATGGATTGTCGAAGGCCAATATGC

CGGTTACACAAGGACGTACTCTAACAAAATGACTTTTGCTACCGTAAAGGGGGCAGGCCATACAGCACCA

GAATACAAGCCTGCAGAATGTTATGCTATGTTCAATAGATGGTTATTAGAGGAACCTTTATAAAACTACG

TTTTGAGCATGTACTATGACGGAAATTATATTAATAAAAATATTTGTTTGTATATGAATTGTTGTGCATT

TTGACAACTCAAACTTCATCATATTATATTA

>MSTRG.2718.8 gene=MSTRG.2718

GTGTGATGAACGCAGAACTGAAAAGGCACCAGCATCAAACTCATAAAATGGAGAAAAAGCCATATTTAAA

AGTGCTCAGTGTCTGCAACATACAGCTTCTTGTCGTAGTTACTGCAATACTACAAAACCTTGTGCAGCAA

GCAAGTGCAGACTTCCATTCACCACTCATCAAGTATCTCCCCGGTTTTCAAGGACCGCTCCCTTTTCGGT

TCACAACTGGGTACGTTGGCGTGGATGAATCAGATGATGTGCAGTTGTTTTATTATTTTGTCGAGTCTCA

GACAAATCCGAGAGATGATCCTCTGATACTCTGGATGACAGGAGGTCCTGGTTGTTCTACTATCACCGCC

TTCGCTTATGAATTCGGTCCAGTATATTTTGAGCAAGTGGAGTATAACGGCACTTTGCCAACCTTAATTT

TAAATCCCCACACATGGACAAAGGTAGCAAATATACTATTTTTGGACTTGCCTGTTGGTACCGGTTTTTC

CTACGCCAAGACTTCTATTGCTTCTCAATCTAATGACACTCAAGCATGTGCTCAGGCTCTACAATTCTTA

AAAAAGTGGTTTATGAGTCATCCTCAATTCCTTGCGAATCCGTTTTATGTTGGTGGAGACTCTTACGCTG

GAATCTTTGTTCCGGTTATTACACAGATGATTTCACATGTAAATGAGGCTAAAATAGGTCCTGAGCAACC

CATTAATCTGAAGGGTTACTTACTTGGAAATCCTACGGCATTTCCTAGTGAGATAAACTTTCGCTACACA

TTTGCTCATGGAATGGGAATTATTTCAGATGAACTATATGAGTCATTGAGAAGAAATTGTGGGCTTGATT

ATCAAAAATCAGATATTGATAATGCAGAGTGTTCACAAAATAAAGCGGCTTATGAAGAGTGTAGGAGTGG

ACTCTTTAAACCACAAATATTGGAGAAAAATTGTGTTGAACCGCTGCGCAGGTCATTATCTTATGAAAAA

ACAGAGGTCATGGATAAATTTGGAGGCCATAATAATCCTAAATTACCACTATCTGCATTTGAATGTCGTA

ATGATGGATATCCGCTTTCTAGTTACTGGAGTAATGATGAGAATGTGCAGAAAGCGTTGCATATCAGAAA

GGGGTCCATAGGAACATGGGTAAGATGCAAAAAAGAGTTGTTATTTGAAAAGGTGATCTTAGATACGAGA

TCATATCATGCAAATCTCAGCAAAAAAGGTTATAGATCTCTCATATACAGTGGAGACCATGACATGGTGG

TGCCATTTCAATCAACTCAAGCTTGGATAAGAGGTCTAAACTATTCCATCATTGATGATTGGCGACCATG

GATTGTCGAAGGCCAATATGCCGGTTACACAAGGACGTACTCTAACAAAATGACTTTTGCTACCGTAAAG

GGGGCAGGCCATACAGCACCAGAATACAAGCCTGCAGAATGTTATGCTATGTTCAATAGATGGTTATTAG

AGGAACCTTTATAAAACTACGTTTTGAGCATGTACTATGACGGAAATTATATTAATAAAAATATTTGTTT

GTATATGAATTGTTGTGCATTTTGACAACTCAAACTTCATCATATTATATTA

>MSTRG.2718.10 gene=MSTRG.2718

TGCAATACTACAAAACCTTGTGCAGCAAGCAAGTGCAGACTTCCATTCACCACTCATCAAGTATCTCCCC

GGTTTTCAAGGACCGCTCCCTTTTCGGTTCACAACTGGGTACGTTGGCGTGGATGAATCAGATGATGTGC

AGTTGTTTTATTATTTTGTCGAGTCTCAGACAAATCCGAGAGATGATCCTCTGATACTCTGGATGACAGG

AGGTCCCGGGTGTTCTACGATCACTGCCTTCGCTTATGAATTCGGTCCAGTATATTTTGAGCAAGTGGAG

TATAACGGCACTTTGCCAACCTTAATTTTAAATCCCCACACATGGACAAAGGTAGCAAATATACTATTTT

TGGACTTGCCTGTTGGTACCGGTTTTTCCTACGCCAAGACTTCTATTGCTTCTCAATCTAATGACACTCA

AGCATGTGCTCAGGCTCTACAATTCTTAAAAAAGTGGTTTATGAGTCATCCTCAATTCCTTGCGAATCCG

TTTTATGTTGGTGGAGACTCTTACGCTGGAATCTTTGTTCCGGTTATTACACAGATGATTTCACATGTAA

ATGAGGCTAAAATAGGTCCTGAGCAACCCATTAATCTGAAGGGTTACTTACTTGGAAATCCTACGGCATT

TCCTAGTGAGATAAACTTTCGCTACACATTTGCTCATGGAATGGGAATTATTTCAGATGAACTATATGAG

TCATTGAGAAGAAATTGTGGGCTTGATTATCAAAAATCAGATATTGATAATGCAGAGTGTTCACAAAATA

AAGCGGCTTATGAAGAGTGTAGGAGTGGACTCTTTAAACCACAAATATTGGAGAAAAATTGTGTTGAACC

GCTGCGCAGGTCATTATCTTATGAAAAAACAGAGGTCATGGATAAATTTGGAGGCCATAATAATCCTAAA

TTACCACTATCTGCATTTGAATGTCGTAATGATGGATATCCGCTTTCTAGTTACTGGAGTAATGATGAGA

ATGTGCAGAAAGCGTTGCATATCAGAAAGGGGTCCATAGGAACATGGGTAAGATGCAAAAAAGAGTTGTT

ATTTGAAAAGGTGATCTTAGATACGAGATCATATCATGCAAATCTCAGCAAAAAAGGTTATAGATCTCTC

ATATACAGTGGAGACCATGACATGGTGGTGCCATTTCAATCAACTCAAGCTTGGATAAGAGGTCTAAACT

ATTCCATCATTGATGATTGGCGACCATGGATTGTCGAAGGCCAATATGCCGGTTACACAAGGACGTACTC

TAACAAAATGACTTTTGCTACCGTAAAGGGGGCAGGCCATACAGCACCAGAATACAAGCCTGCAGAATGT

TATGCTATGTTCAATAGATGGTTATTAGAGGAACCTTTATAAAACTACGTTTTGAGCATGTACTATGACG

GAAATTATATTAATAAAAATATTTGTTTGTATATGAATTGTTG

>MSTRG.2718.11 gene=MSTRG.2718

TGCAATACTACAAAACCTTGTGCAGCAAGCAAGTGCAGACTTCCATTCACCACTCATCAAGTATCTCCCC

GGTTTTCAAGGACCGCTCCCTTTTCGGTTCACAACTGGGTACGTTGGCGTGGATGAATCAGATGATGTGC

AGTTGTTTTATTATTTTGTCGAGTCTCAGACAAATCCGAGAGATGATCCTCTGATACTCTGGATGACAGG

AGGTCCCGGGTGTTCTACGATCACTGCCTTCGCTTATGAATTCGGTCCAGTATATTTTGAGCAAGTGGAG

TATAACGGCACTTTGCCAACCTTAATTTTAAATCCCCACACATGGACAAAGGTAGCAAATATACTATTTT

TGGACTTGCCTGTTGGTACCGGTTTTTCCTACGCCAAGACTTCTATTGCTTCTCAATCTAATGACACTCA

AGCATGTGCTCAGGCTCTACAATTCTTAAAAAAGTGGTTTATGAGTCATCCTCAATTCCTTGCGAATCCG

TTTTATGTTGGTGGAGACTCTTACGCTGGAATCTTTGTTCCGGTTATTACACAGATGATTTCACATGTAA

ATGAGGCTAAAATAGGTCCTGAGCAACCCATTAATCTGAAGGGTTACTTACTTGGAAATCCTACGGCATT

TCCTAGTGAGATAAACTTTCGCTACACATTTGCTCATGGAATGGGAATTATTTCAGATGAACTATATGAG

TCATTGAGAAGAAATTGTGGGCTTGATTATCAAAAATCAGATATTGATAATGCAGAGTGTTCACAAAATA

AAGCGGCTTATGAAGAGTGTAGGAGTGGACTCTTTAAACCACAAATATTGGAGAAAAATTGTGTTGAACC

GCTGCGCAGGTCATTATCTTATGAAAAAACAGAGGTCATGGATAAATTTGGAGGCCATAATAATCCTAAA

TTACCACTATCTGCATTTGAATGTCGTAATGATGGATATCCGCTTTCTAGTTACTGGAGTAATGATGAGA

ATGTGCAGAAAGCGTTGCATATCAGAAAGGGGTCCATAGGAACATGGGTAAGATGCAAAAAAGAGTTGTT

ATTTGAAAAGGTGATCTTAGATACGAGATCATATCATGCAAATCTCAGCAAAAAAGGTTATAGATCTCTC

ATATACAGTGGAGACCATGACATGGTGGTGCCATTTCAATCAACTCAAGCTTGGATAAGAGGTCTAAACT

ATTCCATCATTGATGATTGGCGACCATGGATTGTCGAAGGCCAATATGCCGGTTACACAAGGACGTACTC

TAACAAAATGACTTTTGCTACCGTAAAGGGGGCAGGCCATACAGCACCAGAATACAAGCCTGCAGAATGT

TATGCTATGTTCAATAGATGGTTATTAGAGGAACCTTTATAAAACTACGTTTTGAGCATGTACTATGACG

GAAATTATATTAATAAAAATATTTGTTTGTATATGAATTGTTG

>MSTRG.2718.12 gene=MSTRG.2718

CCACCACCAAACTCATAATATGGAGAAGAAGCCACACTTTAAACCGGAGCTTCTTGTTGTGGTAGTTTTT

GGAATACTACAAAACCTTGTCCAGCAAGCAAGTGCAGATTTCCATTCACCACTCATCAAGTATCTCCCCG

GTTTTCAAGGACCGCTCCCTTTTCGACTCACAACTGGGTACGTTGGCGTGGATGAATCAGATGATGTGCA

GTTGTTTTATTATTTTGTCGAGTCTCAGACAAATCCGAGAGATGATCCTCTAATACTTTGGATGACAGGA

GGTCCTGGTTGTTCTACTATCACCGCCTTCGCTTATGAATTCGGTCCAGTATATTTTGAGCAAGTGGAGT

ATAACGGCACTTTGCCAACCTTAATTTTAAATCCCCACACATGGACAAAGGTAGCAAATATACTATTTTT

GGACTTGCCTGTTGGTACCGGTTTTTCCTACGCCAAGACTTCTATTGCTTCTCAATCTAATGACACTCAA

GCATGTGCTCAGGCTCTACAATTCTTAAAAAAGTGGTTTATGAGTCATCCTCAATTCCTTGCGAATCCGT

TTTATGTTGGTGGAGACTCTTACGCTGGAATCTTTGTTCCGGTTATTACACAGATGATTTCACATGTAAA

TGAGGCTAAAATAGGTCCTGAGCAACCCATTAATCTGAAGTCATTGAGAAGAAATTGTGGGCTTGATTAT

CAAAAATCAGATATTGATAATGCAGAGTGTTCACAAAATAAAGCGGCTTATGAAGAGTGTAGGAGTGGAC

TCTTTAAACCACAAATATTGGAGAAAAATTGTGTTGAACCGCTGCGCAGGTCATTATCTTATGAAAAAAC

AGAGGTCATGGATAAATTTGGAGGCCATAATAATCCTAAATTACCACTATCTGCATTTGAATGTCGTAAT

GATGGATATCCGCTTTCTAGTTACTGGAGTAATGATGAGAATGTGCAGAAAGCGTTGCATATCAGAAAGG

GGTCCATAGGAACATGGGTAAGATGCAAAAAAGAGTTGTTATTTGAAAAGGTGATCTTAGATACGAGATC

ATATCATGCAAATCTCAGCAAAAAAGGTTATAGATCTCTCATATACAGTGGAGACCATGACATGGTGGTG

CCATTTCAATCAACTCAAGCTTGGATAAGAGGTCTAAACTATTCCATCATTGATGATTGGCGACCATGGA

TTGTCGAAGGCCAATATGCCGGTTACACAAGGACGTACTCTAACAAAATGACTTTTGCTACCGTAAAGGG

GGCAGGCCATACAGCACCAGAATACAAGCCTGCAGAATGTTATGCTATGTTCAATAGATGGTTATTAGAG

GAACCTTTATAAAACTACGTTTTGAGCATGTACTATGACGGAAATTATATTAATAAAAATATTTGTTTGT

ATA

>MSTRG.2718.13 gene=MSTRG.2718

TACAATGGGACTTTGCCCACACTAATATTAAATCCCCACACATGGACAAAGGTAGCAAATATACTATTTT

TGGACTTGCCTGTTGGTACCGGTTTTTCCTACGCCAAGACTTCTATTGCTTCTCAATCTAATGACACTCA

AGCATGTGCTCAGGCTCTACAATTCTTAAAAAAGTGGTTTATGAGTCATCCTCAATTCCTTGCGAATCCG

TTTTATGTTGGTGGAGACTCTTACGCTGGAATCTTTGTTCCGGTTATTACACAGATGATTTCACATGTAA

ATGAGGCTAAAATAGGTCCTGAGCAACCCATTAATCTGAAGGGTTACTTACTTGGAAATCCTACGGCATT

TCCTAGTGAGATAAACTTTCGCTACACATTTGCTCATGGAATGGGAATTATTTCAGATGAACTATATGAG

TCATTGAGAAGAAATTGTGGGCTTGATTATCAAAAATCAGATATTGATAATGCAGAGTGTTCACAAAATA

AAGCGGCTTATGAAGAGTGTAGGAGTGGACTCTTTAAACCACAAATATTGGAGAAAAATTGTGTTGAACC

GCTGCGCAGGTCATTATCTTATGAAAAAACAGAGGTCATGGATAAATTTGGAGGCCATAATAATCCTAAA

TTACCACTATCTGCATTTGAATGTCGTAATGATGGATATCCGCTTTCTAGTTACTGGAGTAATGATGAGA

ATGTGCAGAAAGCGTTGCATATCAGAAAGGGGTCCATAGGAACATGGGTAAGATGCAAAAAAGAGTTGTT

ATTTGAAAAGGTGATCTTAGATACGAGATCATATCATGCAAATCTCAGCAAAAAAGGTTATAGATCTCTC

ATATACAGTGGAGACCATGACATGGTGGTGCCATTTCAATCAACTCAAGCTTGGATAAGAGGTCTAAACT

ATTCCATCATTGATGATTGGCGACCATGGATTGTCGAAGGCCAATATGCCGGTTACACAAGGACGTACTC

TAACAAAATGACTTTTGCTACCGTAAAGGGGGCAGGCCATACAGCACCAGAATACAAGCCTGCAGAATGT

TATGCTATGTTCAATAGATGGTTATTAGAGGAACCTTTATAAAACTACGTTTTGAGCATGTACTATGACG

GAAATTATATTAATAAAAATATTTGTTTGTATA

>MSTRG.2719.1 gene=MSTRG.2719

ATGATGTGCGTGTGATGAACGCAGAATTGCAAAAGCACCACCACCAAACTCATAATATGGAGAAGAAGCC

ACACTTTAAACCGGAGCTTCTTGTTGTGGTAGTTTTTGGAATACTACAAAACCTTGTCCAGCAAGCAAGT

GCAGATTTCCATTCACCACTCATCAAGTATCTCCCCGGTTTTCAAGGACCGCTCCCTTTTCGACTCACAA

CTGGGTACGTTGGCGTGGATGAATCAGATGATGTGCAGTTGTTTAATGTTTAGAAAAATTATCATAATAG

TGATGACCAATCCTAATAGTGTAATACATTTTCAACATGTGTCAATAGTTATTATGGTTATGGAGGATTC

AAGGCATTCAAAGACGCACCCACTGAAATATCACATTATATATATTTCAAATATTTTCGACACATGCTAA

GCTTACGAGACCACTAAAAGGATAATATTAAGATTTTCTCGTACTAACCAATGATTTTTCTTCCCCAAAA

AGAATCTTAATTCATTACCACAAAATGCATGCAGGTTTTTGTATGTGTGTCCCCCTACACACACACAACA

CAAATTATAAAGAAGGTTCGTGTGTGTGTACACACATACACGATACACATATTTATAGTGTGTGGTTGAT

CACTTTAAAATGTGCCAAAATTATAAGTATGACATCTTGACAATATATTTTTAAAAAAATGACATAAAAC

GTCAACACAATAGTGAATACTCATCTATAGTAACTAGTTATTACCTAGATTTTGACATCAATTGCCTGCC

CTTGCATTTTTTATAATTACCTCCAATATCAATTTTCATAGTTTTAAAGTTCACATTGTTCCC

>MSTRG.2723.1 gene=MSTRG.2723

CAACCTCACCTGAACCCACCAAGATACCCAGTGTCTAATGTCCTAAACTGTCTAATGTCTAATACTGTGT

AGGTTTTTCATGAATCATGATCCATCAACTCTGTTAAATATTATCGGCAATAGCCAATCACACATAATAA

CTCGCCGTCTTCTCGTCCTTTCGATCACAGTTGTTAAATTTTACCTTAATAATTGGCACCCTGTACAAGG

GTCCAAATCATCCACAAATCAACTACTACCGGGTAATGAATCAGATAAACAAAAGTTGCAGGTTTCTGCA

ATTGGATGTCCTAAACAAGTTTAATCTATGGTCTTATTGTATCAGTTACACACAAACCTGTGTTATTGAG

TTGCAAGACCTCTACCCGCTCCATTCGCTACGCGATACTTAACATAGCATTGTGGAAGAACGATTGAGCA

GCCTTTAGGATTTATGTTTTTATGCATGCGTGGCGTAAGAATCGATATGAGGATGCAATGCCTGGATACA

AAATTCTTTACCTCTTCAACTTCTCTGGCGCCAATACAACTTCCTTCCCGAATGGAAACTCATCTTTGGT

ATGTTATACCTAATGAAGTTAAGAGTGAACAACTCTTAAACAAATACTTGGAATTTCTACCAAAATCTGA

AAAAGAACATGTTTTTAGCTTGCGTGGCGATGAATTGCAAAAAGGAGCTCTGCTTGCTCGTGCTCTTGTT

CGTACTACCATTGCAAGATATCAGATAAATCAAGTTAGTCCAAGGTCTTTGAAGTTTAGGAAGAACATCC

ATGGGAAGCCTGAGGTAGACTGGCAACAGATTGAGGATTGGCATCCTCAACAATTGCATTTCAATATTTC

ACATACCAAATCTCTAGTAGCATGTGGAGTGTCGACGCATAATCCAATTGGCATAGACATAGAAGAAAAG

GAACGGAAGATTAAACACAGTATCTCATCATTTGCTAAAAGATACTTTTCCCAGCAAGAGGTGCAACTAT

TATGTTCTATTTCAGACCCTGAAATTCAGCGCCAGGAGTTCATAAAATTATGGACACTCAAGGAGGCATA

TGTTAAAGCATTAGGAAGGGGATTTTCTGGATCACCATTCCAAACTCTTACCATCCGGTCTGTAGCTGAT

AGCAAAGAAAGCTTTCATCTTTCTGAAAGTTCCAGTTTTGAGAATTCTGACATAGTGGTCGAGTCTTTAG

AGGATTCTGCCAACCTCACAGGCGATTGGCAATTTGCACTCTTGGAATTGGTTGGGTCTCATTATGCTGC

CATTTGCTTAAGAAAGGACTGTGCCAATAAATATAAAAGCCATCCTATGAAAATGAGTGTGTGGAGGACA

ATTCCACTTGTTGAAGATGAATATGTTTCCGGGACTGATTCTGTCATAGCTGTTGGGGGGCTAATGTAAG

CAGCAATTTTTGTGCAAGTTAACTAACCGGTTATAGTTGTTGCTGGCATGATGTAATCATGATTTCAGAT

TGGTATTGCAAGTATTTAACTAATGGTATTGAATGTGTTTTCGAGATAAATTGGTCCTCTTGATCTTGTA

TCTAGGGATTACAACTTACAATTGTGTAGACTAGCTAGAGAGGTTAATGAGGTATGACATTTGATCTAAC

TGTTCTGGATTGTATTGCAATTGTAATGCTTACAATTATTTATATGAGAAGCAAAGATTGCTGTTGCATA

TGCTGTACACTCTTT

>MSTRG.2723.2 gene=MSTRG.2723

AAATATTATCAGCAATAGCCAATCACACATAATAACTCGCCGTCTTCTCGTCCTTTCGATCACAGTTGTT

AAATTTTACCTTAATAATTGGCACCCTGTACAAGGGTCCAAATCATCCACAAATCAACTACTACCGGGTA

ATGAATCAGATAAACAAAAGTTGCAGGTTTCTGCAATTGGATGTCCTAAACAAGTTTAATCTATGGTCTT

ATTGTATCAGTTACACACAAACCTGTGTTATTGAGTTGCAAGACCTCTACCCGCTCCATTCGCTACGCGA

TACTTAACATAGCATTGTGGAAGAACGATTGAGCAGCCTTTAGGATTTATGTTTTTATGCATGCGTGGCG

TAAGAATCGATATGAGGATGCAATGCCTGGATACAAAATTCTTTACCTCTTCAACTTCTCTGGCGCCAAT

ACAACTTCCTTCCCGAATGGAAACTCATCTTTGGTATGTTATACCTAATGAAGTTAAGAGTGAACAACTC

TTAAACAAATACTTGGAATTTCTACCAAAATCTGAAAAAGAACATGTTTTTAGCTTGCGTGGCGATGAAT

TGCAAAAAGGAGCTCTGCTTGCTCGTGCTCTTGTTCGTACTACCATTGCAAGATATAAATCAAGTTAGTC

CAAGGTCTTTGAAGTTTAGGAAGAACATCCATGGGAAGCCTGAGGTAGACTGGCAACAGATTGAGGATTG

GCATCCTCAACAATTGCATTTCAATATTTCACATACCAAATCTCTAGTAGCATGTGGAGTGTCGACGCAT

AATCCAATTGGCATAGACATAGAAGAAAAGGAACGGAAGATTAAACACAGTATCTCATCATTTGCTAAAA

GATACTTTTCCCAGCAAGAGGTGCAACTATTATGTTCTATTTCAGACCCTGAAATTCAGCGCCAGGAGTT

CATAAAATTATGGACACTCAAGGAGGCATATGTTAAAGCATTAGGAAGGGGATTTTCTGGATCACCATTC

CAAACTCTTACCATCCGGTCTGTAGCTGATAGCAAAGAAAGCTTTCATCTTTCTGAAAGTTCCAGTTTTG

AGAATTCTGACATAGTGGTCGAGTCTTTAGAGGATTCTGCCAACCTCACAGGCGATTGGCAATTTGCACT

CTTGGAATTGGTTGGGTCTCATTATGCTGCCATTTGCTTAAGAAAGGACTGTGCCAATAAATATAAAAGC

CATCCTATGAAAATGAGTGTGTGGAGGACAATTCCACTTGTTGAAGATGAATATGTTTCCGGGACTGATT

CTGTCATAGCTGTTGGGGGGCTAATGTAAGCAGCAATTTTTGTGCAAGTTAACTAACCGGTTATAGTTGT

TGCTGGCATGATGTAATCATGATTTCAGATTGGTATTGCAAGTATTTAACTAATGGTATTGAATGTGTTT

TCGAGATAAATTGGTCCTCTTGATCTTGTATCTAGGGATTACAACTTACAATTGTGTAGACTAGCTAGAG

AGGTTAATGAGGTATGACATTTGATCTAACTGTTCTGGATTGTATTGCAATTGTAATGCTTACAATTATT

TATATGAG

>MSTRG.2723.3 gene=MSTRG.2723

GATTATGCAGAACATAGCATTGTGGAAGAACGATTGAGCAGCCTTTAGGATTTATGTTTTTATGCATGCG

TGGCGTAAGAATCGATATGAGGATGCAATGCCTGGATACAAAATTCTTTACCTCTTCAACTTCTCTGGCG

CCAATACAACTTCCTTCCCGAATATCAGATAAATCAAGTTAGTCCAAGGTCTTTGAAGTTTAGGAAGAAC

ATCCATGGGAAGCCTGAGGTAGACTGGCAACAGATTGAGGATTGGCATCCTCAACAATTGCATTTCAATA

TTTCACATACCAAATCTCTAGTAGCATGTGGAGTGTCGACGCATAATCCAATTGGCATAGACATAGAAGA

AAAGGAACGGAAGATTAAACACAGTATCTCATCATTTGCTAAAAGATACTTTTCCCAGCAAGAGGTGCAA

CTATTATGTTCTATTTCAGACCCTGAAATTCAGCGCCAGGAGTTCATAAAATTATGGACACTCAAGGAGG

CATATGTTAAAGCATTAGGAAGGGGATTTTCTGGATCACCATTCCAAACTCTTACCATCCGGTCTGTAGC

TGATAGCAAAGAAAGCTTTCATCTTTCTGAAAGTTCCAGTTTTGAGAATTCTGACATAGTGGTCGAGTCT

TTAGAGGATTCTGCCAACCTCACAGGCGATTGGCAATTTGCACTCTTGGAATTGGTTGGGTCTCATTATG

CTGCCATTTGCTTAAGAAAGGACTGTGCCAATAAATATAAAAGCCATCCTATGAAAATGAGTGTGTGGAG

GACAATTCCACTTGTTGAAGATGAATATGTTTCCGGGACTGATTCTGTCATAGCTGTTGGGGGGCTAATG

TAAGCAGCAATTTTTGTGCAAGTTAACTAACCGGTTATAGTTGTTGCTGGCATGATGTAATCATGATTTC

AGATTGGTATTGCAAGTATTTAACTAATGGTATTGAATGTGTTTTCGAGATAAATTGGTCCTCTTGATCT

TGTATCTAGGGATTACAACTTACAATTGTGTAGACT

>MSTRG.2723.4 gene=MSTRG.2723

TAATAGAGATAGTTTACTAGACATAAGTCAACAAAGTTCAAAAACTTGATTTTTACAAACCGATTAGCTG

GATTCCATGGCTCCAAATTTTCTCTATTTTGGATGTTCGTCCCAAGAGTTTAAGTTTTAAGAAAAAGCTC

AATTAGTTTGAGTCTGGGGTGGATGGTTATTCATCCAGCGCCTCGTATGAAAAAGCAGTTCTTCCCCTAC

AGAAGTAGTACATACCAGTGTACCCTTCGATGACACGTGTGGCTCACGACGCGCGGCTTGCATCTCAAGC

GAGGCGACTTGTGAGATGCAAATGGAGAGAGGACAGTATATTTGAACCTAAAAGCAGCCCAGTGTTTGAA

ATGGAAAGATGAAACAGTCCTAAATGTAAGATGCGGCTTTTAATCTGTTATATACCAGTAGCTCTTTTGA

GAAAACTGCAACTCTGCCACAACTTGCTTAATCTTCTCTACCCTTCATCTTATTTTGTTTCCCCCTTCTC

TTCGACTCTTCCATGTACAGAATTTATGATGGCTTGGTGATCAGTCTATGATTTGCTTTTAGTTTATGAG

GTGGACTGTGGTGATTTATTCGAAAATTGTGTAGCTTAAAACTGAACTTATGCTTCTTTAATAGAGGATG

ACTCTTTATGAGGAGGGTCCTGAGCCTCACCGCGCCTCTCGCCTTTAATAAACATTACCGTATGTACGTA

TCTATCCTGATGAAAGTGAGGGAGGCAAGATAGAGTGAGGCAGACGAGCTTATGCTTCTGTTCTGAGAAA

TTTAAACAAGAGGCATATGTTAAAGCATTAGGAAGGGGATTTTCTGGATCACCATTCCAAACTCTTACCA

TCCGGTCTGTAGCTGATAGCAAAGAAAGCTTTCATCTTTCTGAAAGTTCCAGTTTTGAGAATTCTGACAT

AGTGGTCGAGTCTTTAGAGGATTCTGCCAACCTCACAGGCGATTGGCAATTTGCACTCTTGGAATTGGTT

GGGTCTCATTATGCTGCCATTTGCTTAAGAAAGGACTGTGCCAATAAATATAAAAGCCATCCTATGAAAA

TGAGTGTGTGGAGGACAATTCCACTTGTTGAAGATGAATATGTTTCCGGGACTGATTCTGTCATAGCTGT

TGGGGGGCTAATGTAAGCAGCAATTTTTGTGCAAGTTAACTAACCGGTTATAGTTGTTGCTGGCATGATG

TAATCATGATTTCAGATTGGTATTGCAAGTATTTAACTAATGGTATTGAATGTGTTTTCGAGATAAATTG

GTCCTCTTGATCTTGTATCTAGGGATTACAACTTACAATTGTGTAGACTAGCTAGAGAGGTTAATGAGGT

ATGACATTTGATCTAACTGTTCTGGATTGTATTGCAATTGTAATGCTTACAATTATTTATATGA

>MSTRG.2725.1 gene=MSTRG.2725

TAGATCATCTAACTCAAAAAACTTTCATTCACATCACAGCTAAAAGTCTACAAATAGAGAAACAGAGTAA

ATCAAGAAATCGAAGAGCGAAGAAGATGGCTTCAAACATAGTATGGAACTCCAAAGACAAGAAGTTTGAG

ACACAAGACAAAGAAGCTTACTTAGTTTATGAGCTCAGAAATGGTGGCAAAGTCATTGATATTCTCCACA

CTTTTGTTCCCCCTTCTAAAAGAGGACTTGGTCTCGCATCTCATCTATGCGTTTCAGCTTTTGATCATGC

ACAGTCTCATTCTTTGTCTGTTATTCCCTCTTGCTCCTATGTCTCTGATACTTTTCTTCCTCGAAATCCA

TCTTGGAACCATCTTGTGTACAAGGAGGGGGACACCAAATCAAATATGTGAAACTGTTATCGTGCATTGT

TCTTCTGGAATTGTGTTTGGAGTTCCAAGCATTAGAAAACTCTTTTATGAGTTCTTAATAAATTTATAAG

TACGCATGTTGTAAGATATATCCTTACACAGATATCTGTATCTGTGGTGAATGGTTGGTGCTGTTAAAAA

TGTTGGATTTAAATGAAGTTGCTTAGATTGTTTGAGATTTGCAAAATAAGTTGATTAAATATCTACTGGC

ACTTTTTTCCTGTATAATCTCCAAGTTTTTCAATCCTTCTGATAAACTATGAAAATTCACAGGCAAAGTT

TGATACGGATGTATCCGACTGCCAGAAAATGTCAGGTTTCTCATTTCTGGAGATGACTGGATCATCCAAG

TACACCTCTTAAGGTCATGAACTCAAAACGCACAGCTCCAGCAAAATCATATTCTCATGGAGAATTATGA

ATTCCAAGTGAAGATCTCCAGAAAAACAAATAACCTCCTCTCCTTCCTGTCCTGCAGCATGGTATTACAA

GTCACCACCACTCACCGCTACTTCATCAAGGGGCTGGTTTGGAAAAGCCAAGAAACCAATGTACTGGTGA

TCAAATATTAGAGATCAATTTGATAATTAAGCGGTGGTTACAAAAACGAGC

>MSTRG.2727.1 gene=MSTRG.2727

AGTGATCCATTCGTATGGAGGGGACCCCGGCTCCCCCCTCCCGAAAACAACGTTGGTGTAAATGGCCATA

GCACCAACGGAGACCGTCCCGGAGCGGCCAAGCGGGAGGGGAAACTTCCCCCCTCCCGCTGGGAAGATTT

TTAAGAGACCAGATGGTAGTGTTCCCAAAGACATTCATGTGGGGTGCTTGGCGTAGACTTGATGGTATTT

TGCCCAAAGACACTCCTATGGGGCGTTTGG

>MSTRG.2733.1 gene=MSTRG.2733

TCATGAATTACCCATTTAATAAAATTGTCTCTTTAGCAATACCCAAACCCTAGTTTTCTATTTTGCGTGA

GCCCAATTCATTTTCTTGAAATAGGGTTTTGTAGTTCTTGGGGGTTCTTGAGAGTGAGGTGTTGTAATTT

GGGGGGTTTTGGGTCGAATTTGAAGAAATATGTCGACCCAGAAAAGAATTTCGTCGGGAAAATCGATGGT

GGTGAAGAGGCAATCGGAGAATGTTGGAAAGGCTTCTGGGATTGTGCCTAAGAAAAGGCCTGCATTAGCT

AATATCACTAAT

>MSTRG.2735.1 gene=MSTRG.2735

AGTACTATTCTCCGCCCCTGTCAATTTTCAGTCTCTTCTTCCACAAGAAAACGCGTCTCCTGCATATCAG

TAAACCCTTCTCCTCTCTCGCTTCATTCCAACCACATACATATACAATCTCTACTTAAACTATTGTGTGT

GTGATTACACACACTTACTTGACCTTCCGAATACTTTTTATGTTGATAATAATATTTGGTAGCACAGCTA

GAGCATGAGCTTCTTGAGACCATCAGCACTGTACCAGCTGTTGATCACCTTTAATTCATTGAAATGGATG

CCTTGTATGTTCTTCGGATATCCAAATTTTCAAGGGATCATTAGGCTTGTGATCATGTTTCTTCTCTGGT

CAACGTTCACTGAAATGTGTCACATTTCTTCTTCCTCCATGTCTCCTACTCTTCGTGTTGGCGATCGCAT

CTTTGTCGAGAAGGCTTCATATTTCTTCAGACGTCCAGCCTTACATGATATTGTAATATTTCGAGCACAT

CTGAATCAACCTAGTTTCAGAGAAGGAGATATGCTCATTAAAAGAATTGTAGCACGAGCTGGAGATTTGA

TAGAGGTTCATGATGGGAGACTGTACGTAAATGGGCTTGCTATGAAGGAAGATTTTGTAACAGAGAAACC

AAAATATGCAATACATCCAACGTATGTACCCAGAGGTCATGTATATGTGCTGGGTGACAATCGAAATAAC

AGTTGCGACTCTCATATATGGGGTTCGCTTCCTATAAAAAATGTTGTGGGAAGATGTGCAATGCGTTATT

ACAGAAGACCAGTCATCTGATATGCTTTGGAGATAAGAATTATGGAAATACAGAT

>MSTRG.2735.2 gene=MSTRG.2735

ATTCTCCGCCCCTGTCAATTTTCAGTCTCTTCTTCCACAAGAAAACGCGTCTCCTGCATATCAGTAAACC

CTTCTCCTCTCTCGCTTCATTCCAACCACATACATATACAATCTCTACTTAAACTATTGTGTGTGTGATT

ACACACACTTACTTGACCTTCCGAATACTTTTTATGTTGATAATAATATTTGGTAGCACAGCTAGAGCAT

GAGCTTCTTGAGACCATCAGCACTGTACCAGCTGTTGATCACCTTTAATTCATTGAAATGGATGCCTTGT

ATGTTCTTCGGATATCCAAATTTTCAAGGGATCATTAGGCTTGTGATCATGTTTCTTCTCTGGTCAACGT

TCACTGAAATGTGTCACATTTCTTCTTCCTCCATGTCTCCTACTCTTCGTGTTGGCGATCGCATCTTTGT

CGAGAAGGCTTCATATTTCTTCAGACGTCCAGCCTTACATGATATTGTAATATTTCGAGCACATCTGAAT

CAACCTAGTTTCAGAGAAGGAGATATGCTCATTAAAAGAATTGTAGCACGAGCTGGAGATTTGATAGAGG

TTCATGATGGGAGACTGTACGTAAATGGGCTTGCTATGAAGGAAGATTTTGTAACAGAGAAACCAAAATA

TGCAATACATCCAACGTATGTACCCAGAGGTCATGTATATGTGCTGGGTGACAATCGAAATAACAGTTGC

GACTCTCATATATGATTTGAGGAGGAATTTGTTTATTATCTTGGCACGATCCCCTCTTTCTCTGCCTGGA

AATTGCAATAACCTGGAAAGATTGATCTCAGCCACCAGTTTACGTCTCATCTTGACAAGGCAAACAAAGT

CATCTTGAAAGGTTTAGCGTCAAGGAATGCAACATTACATAAAGTCGCGCACATACTGACATTAGGACAA

TACCCATCACATGATACTCTTCAAGTCGCATTGCATACTATTCTGAGCATTTGATCAAAATAGGGGTTTA

TTTTCAAATTCAGGACTGGATGGTAAAACACTTTGCTGGCTGGTTATATATGCACAAATTAAGTAGCATC

GGAGCGAGTCCCCTCCAGGACCACTATAAATGGGGATCCATTGTAAGTGTTTGAGGCATCCTAGCCTTAG

CATATTTTCCAAGTAATAAACTTGTGTGCTTTCTTAAATATTGTCTTCCTCTCTATATCCTTTCCATATT

CTAGTTTTCTTGTCGAGTAATCTTAATCTTTCTCTTTGTAACCCTTCTCAGTGATTGTAC

>MSTRG.2737.1 gene=MSTRG.2737

CATCCAGTTTTCCTCTTCTTGCTTAAATCCTTCTACTGAAATGAAGCTCTCAGTGTGCTTGAGAACTCTG

GCCACAAAGCTTTGAGATGGAAGTTCGATACACCGACTCTCTTTCTTCCCATGATCATACAACATTACTC

CTAATTTGTTTGTCACAACAAATTGTTTTGCACCCAAATACACAAACACCCAATCCACCTCAAAACCAGA

GTCAAATACAAACATTTTGTTCCAAAACACCTCATCCGTAACTCTATCCATAGTCCACAGACTAAACTCC

TTACCAACTAAACCATCACCAACAGACTCAACAATAACCGCAACAGAACCATCAAAATCCAAAACATTCG

ACTTCCTCATTTTCATAAATCCCGGGAACTCAATCAGCCCAAACACTTCACAACTCAAATCAAACGTCAC

AATTTCATCCTCACCTTCCAAATACAACACCCCATCCAAAACCGGACCCGATATCAAAACACTACAATAC

GGATATGAAACAAAAGACCGCAACATCACACAACCAGCCTCAATCTCCCTCCACGAATCAGCATTCGAAG

AATACAACTCAACCACCGACACCCTCACACGCTTACTATCACCCCCTTCTTTCCGATACTTAATCCTCAA

AACCTTAGAACCACCCGAAACATAATCAAACCCGAACACGACATCAACCTTAAACGGGGCTTCCTCTCGA

TTATAAGCATCCCCCAAACACACACGAGGCAATTGTTTACACGTCCTTATCACCGGATTAGACAAATAAA

TAACATCCCCAATAGCATCAGAAACGCAGACAAAGCCGTTACAAGAACCGATAAATCGCTTGGCCTCGAA

GAAGTGGGGCAGAGGGAGATTGAATTTAGAGCAATCAGAGTAGACGATAGTGGCGGCGAGACGGTCATCG

AGCTTGTCTGGACCGTAGCCAGTTTCTGAAGTGATCAAGAGAAGAGAGGGGTGAATGAGTGAATTGGGGT

GATGGGCTGCCGTGAATTTGGGCTGGGAGATGATGGACCGCCATGACTTGCAAACGGATTTGCAGCTGAG

CAGTGATGTTGCTTTTGTCCAGGAGAGAATCTGAGTTATCACTTCCTCCGGCATGTATTGGTCGGCCCGC

GAAACTTTCGCCATTTTCCGATCGAAACCCTAATTTAAGTCTGAAGATATCGATTTTGTATGCACTCAAT

AATAATTAGGTTGG

>MSTRG.2744.1 gene=MSTRG.2744

CTTCCATATCCTCAGCAGCTCTTACGCAGTTGAGCTTCTCCTTCATCGGAAGTTGAAAGAAGTCTCTGCC

AATACTGCGAACTTGGTCAAGAAATGAACTTGTCATTCCATGATTTATTGCCTGAAAAAAGGTCAACCAC

ACCAGAATCATGATACCATATAACAAATTCAAGAGCACAGGTCCTTTTAGTTGACTTAAAG

>MSTRG.2753.1 gene=MSTRG.2753

GTTCTTTAATTTGTTTCAGGGGGAATGTTAGGGAGCTCTCGTTGTGTTCCGTCAGTTCAGGAACTTGTCA

AGGACAGAAAGTCTACCATTCCAGCTTGATATGTATTCGAAAACCTGTAGTTTCCAGTTTAATAAAATCA

CTTCAAGTCGCTGGTATTGTTATGAAGATGTTACTTGACGCCCACTTCAAGTCCCTGTTATTGAGTCCGC

GTCAAGTAACGTGTTCATATCTAGAACAGGGACTCGAAGTGATTTTACTGAACTGGAAACTACAGGTTCG

AGATCTTTATGTACATATCGAGCTGGGATGGCAGACTTACTGTCCTTGACGAGTTCCTGGACTGACAGAA

CATAACGAGAGCTCCCTAGCATTTTCCGCTCTTCTTGGAAACAAATTAAAGAATTATGCTTGCAAGTTAT

GCATTTCATATGTTGTGCCCCATAGTTTATATAGCCCGAATTATGTTTCATTAACTCGTTTTTCAGGTTA

ATTGAGTTCAAGGCCTCCAGGTGAAAAAAGGATTGCACAGATATTTCTTGCTAAGACTGGATACGGTATA

GTGTGCATGATTTCTACTCCAGATTCTCATCAGATCTAATGGTCAATCAGGTCTATCTTCCTCATCCGCA

ACCTGAACAATTTACAACATTCTGTATTTGATGTAGCGACTTCTGTATTTTTTTAAGAAAACAATGTTTT

GGCTGAAATCACGACTTCCAGAGAATTTCATAGCTTAAATACAATCATATATCCTTGTTTACGGACAATA

GTACAATTTAGAAGTAACCATATATAATGCAACCGCTGAGGTAGCACAGTGATAAAACTATATTACAGGC

TAGATAGAGATCCTAAAGCAGC

>MSTRG.2761.1 gene=MSTRG.2761

CTCCAAGAGCTTATCATTTGCCTCAAACCAAAGTATTACATATCACTATATCACTCCACACTCCACTAAA

TGGGTTTAATTTCAGGCATGTTGAGTTGCTTCAAATTCAATCCCTCGTCATCTGGTCGAGTAAATGATAA

ACAAAATGTGAAAGAGCTGGAGACTTCTTCATCTTTAAACCCCAAGTTGAAAAGCAAGTCAAGTGCAAGT

AAAGCTCCCATTCCTGTTTCATATTATCCTATTAATTCTCGTCTCTCTTATTTGTAGTCAACTAGACCTA

TTGTGTACGAGC

>MSTRG.2769.1 gene=MSTRG.2769

TTTTTTTTATTATCCACAAGTACAAAATACTTTATACAGAAAGTTTGAAACATGTCCGTGGTGATAAGAA

ATAACATTATTACACCTTTTTTGTAATTTAAATAGAAAAAAAGACATAATTGAAGAATATAAATACGAGG

TCAATGCCTTTCTTAGGATTACATGCTAGTAGTTAACAGTTCAGTTACAAAATGTTAACTACTCCAGACT

ACACGAAATAACTGTCAGCCACAACAGCTGTCTGAAGGATTGTATCGCGATGACTAACTTAATTCATCAG

CCGCGTATTCATCTGTCTATCTCCAGACTCAGCAGTGGCATCATGGTCAAATACACTGGCATCAAACTTA

AGTGTGTGACACCAAAGTCCCAAACTGTGCCAAAGTAAATATATCACCAAACTTTGTACAGTGCCAATCC

GGCAGTCTGCTAAAACGCCAAAGACTCATGATTACTAGTTCAGAGCATCGTCAAATTGCTATCGCCAAAG

TAAAGTGCCGAAGAAGTCATATACAGAATCATGTATTAATATTCACATGGTACTACAACAAACGATAGGC

ATTTATCTGCAAGATGTCATTTGCTGCGAGTTTCTAATGGAGGAGTTGGAGTTCAGTGGAGATATTAGGA

TCTATATTTGTTAAAACAAGGTATTTGCCAAGAAAAGTACCCCAAATAACCACAACAGTTTAGTAGGTCT

GCATGTCACATCAGCAGAATTGGTGTTATTTGCCATCTTTATAATTCGGGCTCCAGGTGGAGGACGACAG

TCTCCGACTGTTGAGTTTTCCTGTACATGGCAGTATAGAGATGTTGAACTAATTCTTCATATGTACTGCT

TTAATTTTTAG

>MSTRG.2770.1 gene=MSTRG.2770

GGAGAAATAAAGTATTGTTATATCAAAGAATAACATACAGAGAGGGATCACTCATCCCTCTTCATCTGTT

CTTCATCTTTTTAGAAACACAGATAGAAAAAATAATACACATACCTGCTGTTTTTATATAATTATATGAT

AAATCTGTCCTTAGGATATGATCAGAATGGCATGCACTATTTGGATACATGTGAAATGAACAAAAATTAT

GATCTGCACACATTTAAATATGAAATTGTTCTTCATAGCCTTTTTTGTTGCGATTTCATTGGTCTCTCAT

GCTTCGGCTCTAAAGCAAGGACAAACGTGTTTGTTTGATAAAAACTGCAATGATGGTCTACACTGTGAAA

ATTGTTTGATAAGTGGCAATATTAGGACTCGCTGTACTCGAAAGAAACCTGTCATTCCAACTACAGTGGT

GCTAGGACTACCGTTTGATCGGTACTCGTGGCTAACGACACACAATTCATATGCTAAGACAGGTTCAAAT

TCAAAGACTAGAACTACGATTTTAGCTCCAGAGTGCCAGCAGGATAGCATTACCAATCAACTCCAAAATG

GTGTTCGAGGATTGATGCTTGATTTGTATGACTTTGAAAATGATGTCTGGTTATGTCACTCCTTCGGTGG

TCAATGCTTTGAATACACAGCTTTTCAACCAGCAATCAATGTTCTGAAAGAAGTTCAAGCATTTTTGGTA

TCAGACCCTTCTGCAATAGTCACCCTCATTATTGAAGATTATGTTGCATCTCCAAATGGGCTAACCAAGT

TATTTGATGCTGCTGGCTTAAGGCCCTTCTGGTTTCCAGTATCGCAAATGCCAAAAGATGGTGAAGATTG

GCCGTTAATAGATGACATGATCAAGCAAAATCAACGTTTAGTAGTTTTCACGTCAAGACCTGATAAAGAA

GCAACTGAAGGCATTGCTTATGAGTGGAGTTACTTGGTAGAAAATCAATATGGTGATCGTGGTATGAATG

CTGGTATGTGTCCCAACCGTGCAGAATCTCTTCCCATGAACAACAGAACAAGATCACTGGTGCTGGTGAA

CTACTTCCCAGATACACCAAGTAGTACAGAAGCTTGCAAGTCTAACTCTGCTCCTCTGACTGACATGGTG

AACACTTGTCGCGTGGTAGCTGGTAATAGATGGCCAAACTATATAGCTGTTGATTTCTACATGAGAAGTG

ATGGAGGGGGGTCCCCTGCTGCTGTAGATATGGCTAATGGAGAGCTTATTTGTGGATGCCAATCGATTCT

AGATTGCAAGGAAAACTCAACAGTCGGAGACTGTCGTCCTCCACCTGGAGCCCGAATTATAAAGATGGCA

AATAACACCAATTCTGCTGATGTGACATGCAGACCTACTAAACTGTTGTGGTTATTTGGGGTACTTTTCT

TGGCAAATACCTTGTTTTAACAAATATAGATCCTAATATCTCCACTGAACTCCAACTCCTCCATTAGAAA

CTCGCAGCAAATGACATCTTGCAGATAAATGCCTATCGTTTGTTGTAGTACCATGTGAATATTAATACAT

GATTCTGTATATGACTTCTTCGGCACTTTACTTTGGCGATAGCAATTTGACGATGCTCTGAACTAGTAAT

CATGAGTCTTTGGCGTTTTAGCAGACTGCCGGATTGGCACTGTACAAAGTTTGGTGATATATTTACTTTG

GCACAGTTTGGGACTTTGGTGTCACACACTTAAGTTTGATGCCAGTGTATTTGACCATGATGCCACTGCT

GAGTCTGGAGATAGACAGATGAATACGCGGCTGATGAATTAAGTTAGTCATCGCGATACAATCCTTCAGA

CAGCTGTTGTGGCTGACAGTTATTTCGTGTAGTCTGGAGTAGTTAACATTTTGTACTTGGTTTGTTTAAC

GTCTAACTGAACTGTTAACTACTAGCATGTAATCCTAAGAAAGGCATTGACCTCGTATTTATATTCTTCA

ATTATGTCTTTTTTTCTATTTAAATTACAAAAAAGGTG

>MSTRG.2770.2 gene=MSTRG.2770

GGAGAAATAAAGTATTGTTATATCAAAGAATAACATACAGAGAGGGATCACTCATCCCTCTTCATCTGTT

CTTCATCTTTTTAGAAACACAGATAGAAAAAATAATACACATACCTGCTGTTTTTATATAATTATATGAT

AAATCTGTCCTTAGGATATGATCAGAATGGCATGCACTATTTGGATACATGTGAAATGAACAAAAATTAT

GATCTGCACACATTTAAATATGAAATTGTTCTTCATAGCCTTTTTTGTTGCGATTTCATTGGTCTCTCAT

GCTTCGGCTCTAAAGCAAGGACAAACGTGTTTGTTTGATAAAAACTGCAATGATGGTCTACACTGTGAAA

ATTGTTTGATAAGTGGCAATATTAGGACTCGCTGTACTCGAAAGAAACCTGTCATTCCAACTACAGTGGT

GCTAGGACTACCGTTTGATCGGTACTCGTGGCTAACGACACACAATTCATATGCTAAGACAGGTTCAAAT

TCAAAGACTAGAACTACGATTTTAGCTCCAGAGTGCCAGCAGGATAGCATTACCAATCAACTCCAATTAT

TTAGATGAAACATTTAAGAGAGTCCTATCTTGGAATCGGTTACCCTGCAGAATGGTGTTCGAGGATTGAT

GCTTGATTTGTATGACTTTGAAAATGATGTCTGGTTATGTCACTCCTTCGGTGGTCAATGCTTTGAATAC

ACAGCTTTTCAACCAGCAATCAATGTTCTGAAAGAAGTTCAAGCATTTTTGGTATCAGACCCTTCTGCAA

TAGTCACCCTCATTATTGAAGATTATGTTGCATCTCCAAATGGGCTAACCAAGTTATTTGATGCTGCTGG

CTTAAGGCCCTTCTGGTTTCCAGTATCGCAAATGCCAAAAGATGGTGAAGATTGGCCGTTAATAGATGAC

ATGATCAAGCAAAATCAACGTTTAGTAGTTTTCACGTCAAGACCTGATAAAGAAGCAACTGAAGGCATTG

CTTATGAGTGGAGTTACTTGGTAGAAAATCAATATGGTGATCGTGGTATGAATGCTGGTATGTGTCCCAA

CCGTGCAGAATCTCTTCCCATGAACAACAGAACAAGATCACTGGTGCTGGTGAACTACTTCCCAGATACA

CCAAGTAGTACAGAAGCTTGCAAGTCTAACTCTGCTCCTCTGACTGACATGGTGAACACTTGTCGCGTGG

TAGCTGGTAATAGATGGCCAAACTATATAGCTGTTGATTTCTACATGAGAAGTGATGGAGGGGGGTCCCC

TGCTGCTGTAGATATGGCTAATGGAGAGCTTATTTGTGGATGCCAATCGATTCTAGATTGCAAGGAAAAC

TCAACAGTCGGAGACTGTCGTCCTCCACCTGGAGCCCGAATTATAAAGATGGCAAATAACACCAATTCTG

CTGATGTGACATGCAGACCTACTAAACTGTTGTGGTTATTTGGGGTACTTTTCTTGGCAAATACCTTGTT

TTAACAAATATAGATCCTAATATCTCCACTGAACTCCAACTCCTCCATTAGAAACTCGCAGCAAATGACA

TCTTGCAGATAAATGCCTATCGTTTGTTGTAGTACCATGTGAATATTAATACATGATTCTGTATATGACT

TCTTCGGCACTTTACTTTGGCGATAGCAATTTGACGATGCTCTGAACTAGTAATCATGAGTCTTTGGCGT

TTTAGCAGACTGCCGGATTGGCACTGTACAAAGTTTGGTGATATATTTACTTTGGCACAGTTTGGGACTT

TGGTGTCACACACTTAAGTTTGATGCCAGTGTATTTGACCATGATGCCACTGCTGAGTCTGGAGATAGAC

AGATGAATACGCGGCTGATGAATTAAGTTAGTCATCGCGATACAATCCTTCAGACAGCTGTTGTGGCTGA

CAGTTATTTCGTGTAGTCTGGAGTAGTTAACATTTTGTACTTGGTTTGTTTAACGTCTAACTGAACTGTT

AACTACTAGCATGTAATCCTAAGAAAGGCATTGACCTCGTATTTATATTCTTCAATTATGTCTTTTTTT

>MSTRG.2773.1 gene=MSTRG.2773

CCGGGTAGTCCTTGAGTAATAAAATCTTCTTCCATTGTCCCGGCATTTGAAGAAACTCTACTTGTTTCAT

CAAACATCATGGTGGTCCATTTGTCACTGGAACCATCACGAGACTGCATTTTGGAACGTCGTAAGGCTGC

CAGTAGCCCGACTTGGTCATTCTGGGAACCATGACAGTGGCCATGGAGCATCCAAATGAGGAAGATGAGC

AAGTTTATCCTATAATACATAATGATTACTCAGGATTTGCAGGAAATAATGTTAGTTAGAAGTTAGAACT

TTCATTGTGTGATTATTTATATACACT

>MSTRG.2775.1 gene=MSTRG.2775

AAATGGTGTCCTAGAAATGACTGAAATTGAGAGAGAACTTCAGCAATGCAAATAAGTTTCAGAAGATGAC

TTCTGTCACTCTGTATATTTCTCTCATGGAAAGTCCGCGATCATTACAAGGGAAACGTATACAAACAATC

ATCACATTGTTCGAAGACTTCTGATATAGAATCAACTATATTGTTATTTGTCACGGAATCTCAACTTTAT

TGAAGGCCTGCTATGAACCTCTTTAATAGAGAAAAAGATCTGTGCGGCTGAACTCGAGGGACTTCATGGC

CAGCAGCCCGAACTGTTGAGAATGTCAATCCTTGATAAACAATCTGATATCCACCAACCTGCAGAATGGA

ATAGGTCAACTGATGAAGATGTAGTGCTTAATTTACACAATTTTGATATTTTGAAGAGAGAATAACTTAA

AAGCCCCTAATGGACTTACGTCTCCAGTATCATCCAACCAGAAACTCCAAGGTTTTAGGATCTTAAGATC

CAAGGCAGCAAGGGAATATCGTGTACTGGTCACTGGTACTACTGCATCCATATCACCACTGTCCGTCAAC

ATGTTCAACTCAAAAGGTTTGCTGTACTTGCGGAAGATTGAGATAGTTTCTGGCATACGAAAGTTCACAA

GGATCATAAACACCATTGTGTTTAGTCTTCCTCGATGCATTAGAGGAATGAGAGCAAACAGGACCATAGA

TATTGTAAAAATCTATATTCCCAGCCTCATCATCAACTTTAAATTGCAAGGTGTTACATTTTGATCTGTT

ATAGTGTGGACTAGTGCAATATTGCTGAAGTCCTCGATAAGTTTCATCAGAAATCATGGCATGGCTCCAC

ATATAATCAAATGTTCCTTTATTGTCTGTAGCATCATTCATTACCCCATTTCCTACCTATCAAAAACCAG

ATTAAAATTTGGAAAAAAAACCAATGTAGACAGAACAAAACCTTCAACTTAACTGTTGTGCACACGAAAG

TTCAG

>MSTRG.2789.1 gene=MSTRG.2789

AGTCGAGGGTCGACATATTCTAAAATACCTAATCTGATCTTAACCCGCCGCAGCCATGCTCTTCTTTTGT

TCTAACCCTAAATCCCTCTCTCAGAAGTCAGATCTTACAATCGATCGATTCGATTAGTTCTTATAAGTTC

TTACGAGATTTGATCTAATTGTAGTCGAGTTTTTCAAATCATTCACCTTAGTTAAATAGTTGAATAATGG

AACAAGGTGTTGTATTTCATGTGTCCAGATTTCAGCCAAGCATTGTTAGGGATCTGCTCACTTGTTCTGT

TTGTAACTTTGTTTGCATATCAGGTCGCAGTCATAAGTAAAGTGAAACTATATTTGAGAATGTGTATTTG

ATTTTTGGTAGTATTTGAAAAAACGGATCCGGATATTATCCGTATCCGAATAATATCCGGCGGATACGGA

TACGGATATTTTTATTCTAAAATTAGCGGATACGGATACGGATTCGGATACGGTTTTTAGTTGGCGGATT

CGGATACGGATATGGGTACATCCGTATCCGTTTTATCCGTTTGACACCCCTAT

>MSTRG.2790.1 gene=MSTRG.2790

GAGAAGGGGAGAGAGAGAGAGAGAGAGACCTTTGAGACGAGGTGGGGAGGAGGGGGAGAGAAGAAGAATC

CGTGAGGAAAGAAAAAAGCGAAAATGCAACCGAGAAGGATGCCGATGAGAACAGCGATGGAGATTCTGGA

TCCGCGGACGGATCTGACTAATGAACTAGTGTTCCTAATTAATTGACCACTCCCGCTCTCTCGACGCCCA

TTGCCCATTTAGCCAGGACCTCCGATCTCACTGTTTGCGCCTGCTTTTGAACTTTCTTCTTCTTTTGTTT

CGCATCTCTCTGACTCTCCCTCACTAGTCTTGCTATAAAATACTCCGACTGTTTCATG

>MSTRG.2794.2 gene=MSTRG.2794

AGGGAAGTAACTTTGGCAACGAGGTCATTTGACACCCAGCCAGATCAATATACAGTCAGTGACGGAGAAG

GAAAAACTTGATGTCGGCGAGCGACGGAATGGAGGGCTCCGCCGAGGTGAAACGTAAAAATCCCAAGCCG

AAGCGGACCTTTAAGTCAGAAAAGGAGTTCCTAGAGTTTACTTTAAAGTATCAGCAAGTTATTGCTGAAA

GAGATTCCGCTATATCAGTTCGAGATAAGCTTGAGTCATTGTGTCGGGAATTACAACGTCAAAACAAAAT

ATTAATGGATGAATGCAAACGAGTGTCAACGGAAGGTCAAAGCTTAAGATTAGATATGTCAACCAAGTTT

CAAGATGCAATTAAGGAAATAGGCCATAAGCTGGAAGAGCAAAAAGATGAATCGCTGACTCAGCTCAAGG

AAAATGAGATGT

>MSTRG.2808.1 gene=MSTRG.2808

CCCATATCTCACTACTTATTGCTCAACACCAAATTACTCTCATACAAATTATTTTAAATTATAATTATCC

TCTCCATAGCAACCTGATGGTCCATGTTGAGATCAGTACTGTAACCCATCTGCAGATGCCATTATAGGAT

GCTGGTTTGCATACAAATAATTCATTGGCATGTTCATGTTGATGAACATGTTATCACAAGATGGCTCATC

ACCAAAAACACTGTAAGCATGGTCTTGAACACTCCCAAGATTTGCATTCCTTGCATCCATGAAGCTGATG

ATCTCATTGAGTGACTGCAGTCTGTTTGAAAGCTCAGCCACTTGGGCTGCCAAGACTGAGTTCTCAGACT

CTATGTTGAGGAAATGTTGAGTAGTGACACTCATACTTGTCATGATATGCTCATTTTCTGTTTTGAGCTG

GTTGAGCTGAGCCATCAGATCATCCAAATGCTTCTGCTTCCTCATTCTTGATCTTCTTGCTGATTCACGG

TTCGATATCATGCGCTTTCTTTTCCTCTCATCCATTAGTACCTGCAGATCTTTCTCAGAGCCTGAGTTCT

GAAGCAGAGATGACCCTGAGGAAGTTGTTGTTGCTGTTCCACTAGAGGAAGTCATGGTTTATTGAAAAAT

AAAAATACCCAGATAGTTAAATATA

>MSTRG.2810.1 gene=MSTRG.2810

CTAGATCGGATCAGTTTACATTTAGTGCACAAAAGATAAAGAGGAGATGAAGAAGAGCAAAACCCTAACT

AAAAGTATGAACCATTGTTCGTGGGCAGATCTTGATCCGGATTTGTTGGGAGAAATAAAGAAGAGGCTTT

ACTGGGCGGATCATGCTCGATTCAGTGCTGTTTGCAAGACCTGGCTAGCGGCTGAGCACGCAAAAAGAGC

CGGTGATGTATTGCCATGGTGTTTGTTGCTGTATCGAGAGAAGCTTGACAGAATCACATACCGCCTCTAT

GAACCCCTTAATCTGGACCAACCACTTATTACTCAAACTATTCATCTGGATGAGTTTTTTGACCCATCCA

CTATTCAAATGGCTGCACCACTTGTTTATCTCCATGGATGCTTGTTATTTTCTCTGCATCATATTGATCA

TACTAGCACCAATTTCTTATTAGTCTCGGTTCCTACCCGAAGAACCATTAGACTCCCTCAGTTTCACCAC

CCCTCAGTATCAAATTCTGAATTTTTAAAAGCTGTTTCCACTTACCCTTATTCTCGAGATTGTGTCTTTT

TAGCTCTTCATGTTGCTAATTCTCAACGTTGGACTGTTGGTATTTTTCGCCACGGTGATACACAGTGGAC

TACTACTGAATTTAATCTGGGAGAATTTTCCAGTCTTCAACGGAGTCAGGGTGCTGTGTTCATTCGAGGA

GTTTTTTATATACTCTGTTACAATCAAAAACTTGCATCCTATGACATTGCTTCCGGGGAATTGAAATTTG

ACTCTTTTCAGTTTCCAACTGATTGCAGGAAAAATTGTCATTTGCTAAAATTCTTCGCGTTGAATGGGGA

GCTCATGATTTTGTATCATCACTCAAAAGTTCGCAAATATATTCTTACAAGCTATGATTGGTCGCATAAA

CGTTGGGTTCCTCTGAAAAGTTTAGGGGTTAGTTCTTTGTTCATTAGCGACTGTTCTGTTTATGCAGATT

CCATAAATTATTATGGGGTGTCCTCAAACAAGATATATTACCAACAAAATGGAACCTGCCGTGTTTATTC

CTTTGAGAATGGTGAGCTGTTGGAATCGACTTCATCCGGACTTAGAAACTGGGATGGTCTCGATTATTCT

GGATCGCGTTCAGTGTGGGTTGAACCTCCTGATCTCTTGTTGAAATAATGTACCACCCAAGAGAGCTAAG

ATGACCATACAGCGGTCGCATGCCACTTGAAGAGTTGCTTAATAAGAGATTCAAGTGATCCCAGGTTCAT

GGAAGTATGTGTTAGTGATGGTTAAAATTCTGCATTGTTCTATTGACATTGGTATATTCCTTGTAATTTA

AAATTTGTGCCTTTTATTCCTGGTTATAGATGTGGTGCCTGTCCATCTCTGATCTTAGGTCTGGTAAGCC

TTGTGAGTTGTCATTGCCTCTATGTTAACAGGGATTCAACTAAAACAGTGATGTTGCTTTACTCCAC

>MSTRG.2813.4 gene=MSTRG.2813

ACTTGAACTAATTATCTAGGCTTTGCCCAACAAAGCCACTGTACCCTTGTTTTCACTCGTTGATTACTCT

ACGATTACTCTACAAGATGATATAATACATACAGTTATGTGTATATCTTGTTTCGCATTTCTCAAGTATG

TTTTGTGCATGTGAAACACTTGATAGAATTCCGGTCATTGTTGGTAATGAAGGTGATGTAAAAGAGAAGT

TATGAGTCGTATGATAGAGGTTGCTCAAAGACGAATCTACAGAGAGCAGACACAGTATTTTGTGTGGCAA

AGGGTATAAGGTTGCATTTACTTAGATGGAATGGAACGAGGGGAGAAGAGAATGAATAATTATAAAGGGG

TTTTACGGTGAAAGAAGAATGTATGAGAAAATAGTGATTATATATGATTGATACTTAATTTTTTTATGAT

GAAAATTGTAAGAAAACATGATCAAGAAGTGGAAGGAGAATTCTTTCCAAAACATGTGGGTTAGATTTCT

ATATCAAATAATTTGAAATGGAATGATTGAAGGAATGGAAATTTTATACATTTTCTTCGAAGATATGAAC

ATCAGTGGTGGTGTAAGTATGTTTGTAGAGCTTATTCGTGTGTTGTGAAGCCTGATTATGACTCCCATTT

ATAAGTATGTTGTCCC

>MSTRG.2815.1 gene=MSTRG.2815

CTTGGCTATATCTTTGCATCGATCTCTACTAGGCCTAAGCTATGAACCACCGTGGTTTCAATCTCCTCTG

GAAGTGCTTCCGCCGCTCCCTGAAGGCTTCTTCCCCTTGTACTTCTGCTTTCCGATGTTGAAAGGGACAT

ATTTGTACTTAATCATGTGCATAATTTGGCGCTTCATTGTAAGGAGAAACAGCACCAGCCAGTAAAAT

>MSTRG.2819.1 gene=MSTRG.2819

CCAAAGTTGTGGAAGAGTGTACTAGTAAGCAGTAAGGATAGTCTTATACTCTTATTGCTTTATATGTATG

AGTCAGGGATGTCATCGTTGAGCATAAGTACTGTGACTTGAAGGCCTGAACAACATTCCAAGCCATATTG

CTGGCAAGGTGAGCTGTTTTAGATGATATAGTGTTTTTTCCTGAGGAAGAGCAGGCCGTATTCTGTACTC

TTTATCATCAGGACCCCATGAGTTTT

>MSTRG.2822.1 gene=MSTRG.2822

CATACATTAACATACTGACTTATGCATACATAAAACATACACAAAGATCTATAAACTTAGATTAACATGT

TCATTATTGTTGATTTTGAGCAAGTTTATAATTATTTATTTTTTTCATTTTAATTTTTGATTGAATTTTG

CACTTGTCGACCGACTAGTCGACCAATCGATCGATTAGTCGACCATGTCGACCGACTAGTCGACAATTTG

CAAGTCGACCTCTCCAGTCGATTTTCTTTACCCGACTAGTCGACTAGTCGTCGACTAGTCGCGACTAGTC

GAGCGCCGTGACAACAATGC

>MSTRG.2826.1 gene=MSTRG.2826

CACATTGTCATCATTTGCTCTGTTCTGCAAACAACTAGTGATAATAGTGTATCGATATCTGCAAGAAGAT

GAGGAGGAGACAAGAAGATGTTTTTGAGGCAATGTGCATGACATACGAGGAGGAAGAAATGATTGTTTAG

ATAATGTGTTGAAGAATTTATTAGGGTTTTGGAATGCAGCAGAAGTAGAAGAACTGGTGACAAGGGTTTT

TAGATTCATGTCTGATTGTTTGTGTTGGTTGAGTTGGACAAGAGGAACGATGACAGATATGGATTCTGTA

ACATAGAACAGAGTCAAGGGGACTCTTATCTTATGTCATGTACGTTGGGGTAAAATTTAGAGGCCTAAAA

TCAATCTTCC

>MSTRG.2835.1 gene=MSTRG.2835

GTGTTGCTGAGTGCGAAGGCAACTAAGACTGATGATCTGGGGCGGACGCCATCAAGCAGTGCCATACAAA

AATGATCAACAGGGGAATATGCTTAACGACATGAGGAAAACTGGCAACGATTTGAATGAAACCTAATTTT

ACTCAAACCGTATGTGTTGCTGAGTGATCTTTCTCAGAAAGATAAGGTCTTCTGTGTGGAGGATAAGGCC

TGTGGTGACTATCATACTTGGC

>MSTRG.2836.1 gene=MSTRG.2836

TTCTAATCTCACCGGAGGGAATAACTTTCCGGCATTCTCTTTATATCTCTCAGTATAAATTTTTGTCAAG

TATCTGAGGGAATACTTGAATGCAATTCAAGTAATCCACGAGGGTTTCGTTTCACCTTTATTCCGTGTTT

TTTGTGTGCTTCTTTTGACGGGGGTGGCCGGGGCTCTAGCCTCGGCAGATTCCAAAGCCCCCATCTTTTA

CAACCACATACACAATTACACACACACACACACACACTTCTGTATCTATATCTCTTTATCTTTTTTTTTT

GTTTGTTTTCATCTTGATACAATATGCCTGGTTTAGCTTGTGTTGACGCCGCCATTGCACCTCCGCCGTG

CTACGCTTTTACATCCTGGGAAGCGTGTCTTCCGGCGCCGGAGATGAATTCCGGCGGAGTGGCCGATTTG

CAGTGGTCGCCGGCGCTCTCGGCCGCGTTGTACAGGATTGACAAATGGGGTGGTCCTTATTTCGGCGTGA

ATGAAAACGGCAACGTTTCGATTTACCCTCATGGTGCGAAAACGCTGAATCATCAAGAGATTGATTTGCT

TAAGGTTGTGAAGAAGGCTTCGGATGCGAAAGATGAGGGTGGGCTTGGTTTGCAGCTGCCGATTGTTATA

AGGCTGCCTGATGTGTTGAAGAATAGGGTTGAATTGATTCACTCAGCGTTCGAGTCTGCGATTGAGGCGC

AAGGGTATAATGGTCATTATCAGGGTGTTTTCCCTGTCAAGTGTAATCAGGACAGGTTTGTTGTGGATGA

CATTGTGAGATTCGGGTCCAAGTTGCGGTTCGGGCTCGAGGCGGGGTCCAAACCGGAGCTTTTACTAGCT

ATGAGTTGTTTGTGTCGGGGGAATCCGGAGGCGTTTCTTGTTTGTAATGGATTTAAGGATGTGGAGTTTA

TAACGCTTGCTTTGGTGGGGAGGAAGATGAATTTTAATACTGTGATTGTGCTTGAATTGGAACAAGAGCT

TGATATGGTAATTGAGTTGAGTCGAAAGCTAGGTGTTAGGCCTCTTATTGGGATGAGGGCGAAGCTGAGG

GCGAAGCATGAGGGACATTTTGGTGCTACTTCTGGGGAGAGAGGTAAGTTTGGACTTACAACTACTCAGA

TATTACGCGTTGTTAGGAAATTGGATGAGTGTGGTATGCTTGATTGTTTGCAATTGCTGCATTTTCATAT

TGGTTCTCAAATTCCCTCTACTGCTCTGTTGGCGGATGGGGTAGGGGAAGCAGCTCAGGTGTACTGTGAA

CTAGTTCGTCTTGGTGCTCGTATGGGTGTTATTGATATCGGGGGTGGTTTGGGAATTGATTATGATGGTT

CCAAATCACCTAATTCGGATGTTTCAGTTGATTACACTCTGGAGGAATATGCAGCTGCTGTAGTTGGCGT

AATTAAGTCTGCATGTGACCGTAAGATGGTTAAGCATCCAGTGATTTGTAGTGAAAGTGGTAGAGCAATT

GTCTCTCATCATTCAGTATTGGTATTTGAGGCTGTATCTTCAAGTAAGAATAGTGTACAACAAATGACCC

CAGTTGACGTCAGTTATTTAGTTGAGAGACTCCCTGAGGATGCTATTGTCGATTACCGGAAATTGTCTGC

AGCTGCTACTGGTGGTGAGTACGAAACTTGTTTGCTTTATTCTGATCAACTGAAGCAAAAATGTGTAGAG

CAGTTCAAGGAAGGGTCTTTTGATCTTGAACAGCTTGCTGCTGTTGATGGGTTGTGTGACTTGGTGTACA

AGGCAATCGGCGCTTCTGATCCTGTGAGGACTTACCATGTTAACTTGTCCGTTTTTACATCCATACCTGA

TTTCTGGGGTATTGGACAGCTGTTTCCGATTGTGCCTATTCATCGCCTTGAGCAAAAGCCAGTGGTGAAG

GGAACTCTCTCCGACATAACATGTGACAGTGATGGAAAGATTGGGAAGTTCATAGGGGGTGAAGAGAGCC

TACTGCTTCATGAATTGGGAGGAGGTGGTGGAGGTAGTTACTATCTAGGGATGTTTCTTGGTGGGGCTTA

TGAAGAGGCGCTCGGGGGTATTCACAATCTTTTTGGTGGGCCGAGCGTTGTTCGTGTCTCACAGAGTGAT

GGTCCTCACAGTTTTGCAGTGACAGGAGCTTTTCCTGGGCCTTCCTGTGCTGATGTTCTCCGGACAATGC

AGCATGAGCCTGAAATGATGTTTCAGACGCTCAAGCACCGTGCTGAGGAGTATGTATTCGATGAGGGTGC

TCGTGGTTTTGACCATGATGCTATAGCAACTGGGCTTGCCAGTTCTTTCCAGAACATGCCTTATCTTGTT

AAAGATGATGATGATAATGGCTACTATTATTCTGATGTTATTAACAATGGCACTGTTGTGGATTCTGATG

CACCTGAGGATGAGGAGTGGTCCGACTGCTGTGCTTAAAATTTTGTCTGGGCATCTGCAAACTGGAAGCA

TCTCTTTGTATTTGGTAGTTTGTCCTTTCAGGTCGTCTGTGTTTTAAGTTTGTGCTCTTCTTTATTTAAT

GTATGTTTACAATAATATAAGACCGAGCGAATCATTTGCATCCGGCCATCCGCAAATAAATAAGTGCCAA

GTTGGTCGGATCAGTGTATGATTGGCTTCCTGAAGCCTCTTGTCTGAGTTCTTTTATGTTGCTGTGACAA

AGTTTTGGTAAATTATTCTGCTTGTTTGTAATTTACAATTACTCTTTCCTTTCCTGACTGCCTTCTGTCA

TTCTGTGTGTAATGAGATTACAAGCAAGCTAATGATGATTTATCAGTGGTGGC

>MSTRG.2838.1 gene=MSTRG.2838

ATATTGGGCCGCTCTCAGATTCAGTTATCCCAAGTGAAAAAGATCAAAGCCTTCTCCCAAATCAGTCGCT

CAATTTCTTGAATTTTAGGTCTTAGTTGAATCTCTCTCTGTCTCCCACTCCCTTCTCTTTCTCGATCTCT

CTCTCGACTCTAGCATCAATGGCGCCGCCACCAGGACCTTATTCAGGAACCAGCACACTTGCTCTGGTGG

CTAGAGTATCCGCTTTTTCGTTTGGACTGGTTTATGGAAGCATGAAGCTCAAGTATCTTAAGGCAAAAGC

CAAATCTCAGCAGAAAGCTGCAGCCAAGGCAGGCCACTGAAAGATGCGAGTTGTTTTCACAGAGTCTTCT

AGAGGAATAACTTGTAATGATCGAAAACTATTATCATTTTGTGCGAATGTCGTACGCTGTGAGTTTTTAT

AAGTTTAGTTTTGCATCATTGTTTTGGCTTTAACCATTATGTCTATGGTTCAAATTCCTCTTGGTGGATA

TTGAGAAGTTTGAATGATGCACAATCTGAAAAAAAATAATCTGTCATGTTCACATGCATGAGTACATGCT

TTTGGCTGTGATTCTGTTTCATTGGTTGCCTCCATATGTTTGGTTTGCCATACTTGCAAGCTTTTTAATT

GTCTTGGTAGTTCAAACTTGCAGAGCCTACTTAAATTAACTGTATTTACTTTTTATTGCTGGTTTATATA

CATATATATATT

>MSTRG.2840.1 gene=MSTRG.2840

TAATCGGTCCTACCACATGTGTGTCATAACATCAGAATGCAGAAGCGCCTCTTGTAAACCCAAGGAAAGA

TAAAGACAAGTCAAAAGTATAAAGATTCGTCCACAATATCCCGCACAAACGGACCCCCTCAACAGCCCGT

GGCATGTCCTTTGCTCTGTCCCCCTCCCCACACTTTGCCTCTTCCTAATCTCTTTCATCTTCATAAATAT

TTTTGGAATCCCTCACTTTTCTGCTCTATATATATATTCCTCCCTTAGATTTTCATTCAAAACCGTACAC

ACTTCATTTGTCGCAAGCTACATGGCAATCAAACAAATCCTTAAGAAATGCTCGAGCCTAGGGAAGAAAC

AAAACGCCCACAACCGCCCAACGGATGTACCAAAAGGCCACTTTGCAGTTTATGTGGGAGAAAATCGAAC

TAGACACGTAGTTCCTATTTCATTCTTGAGGCATCCGGAGTTTCAAGGCCTGCTTCGTCGCGCTGAAGAG

GAGTTTGGATTTGATCATGACAGGGGACTCACCATTCCATGCGAAGAAAGAGTTTTCGAGTCACTTACCT

CGTTGATAATAGTTGGTTGAGTTTTAAGTTTTACGAGAGATGATGGCTGAATCTTCTTTATGTTTGATTT

TATTTTTTTAAAATATGTTTCTACGTGATGCCTTCTATCAGTCATGTTCATGCATTATTCTTGTTCACAA

GCATAGATAGGAGAAGTTCTAGTCCATGGCCAACAAGCTTTCGTTTGATTTGCATCTTAGACTAACTATT

GTACATAGTCTCACA

>MSTRG.2842.1 gene=MSTRG.2842

AAGAGTCCTTAAGTTTTGTTTAGACATGGGTATCGGATCAGTTAAGATAGCAGACGTTAAGCAGAAGCTC

CAAAGGAGTCTGCATGTCTCCAAGGATCCGAGGCACGACAGATGTTCCTAAAGGCCATTTTGCGGTTTAT

GTGGGGGAAACTCAGAGAAGGTTTGTGATTCCGATATCATACTTGAGCCATCCTTTGTTTCAAGAATTAT

TACATTGGACGGAAGAAGAGTTTGGATTTGAGCATCCTATGGGAGGATTGACAATTCCTTGTCGTGAAGA

TTATTTCAGAAGTATCACCTCTTTACTGAGTTCATCATAAAAGATTAAATCTGCTAGACTGCTTAGTTTC

TATTTAGTCATTTGCCATGTTCTACAGCGTTGAAACAACGAATATCATTTTTTTAGTAGTGAGATTTTTT

ACATATTGATTTTGTATCAACTTTTGTTGGATAATTGAAG

>MSTRG.2846.1 gene=MSTRG.2846

GTAAGAATCTTTGATAGTACAGAAGCAAAGTAAATTTCATAGCTCACAAAATGAACTGACAACAACTGAT

CATCATTTTAAGTAACTTTCGAAGCAAGGCTACCTGATCTTTGAGGCAGTATCAAACAAACCAACTTTCA

AGAAAGCTAGATTGATATTGCAGCAGCACAAGCACCAGCACAATCATCGCGGCCACACCCCATGGAAAAC

CTCCTTCAGACGAACCAGCATAATTCATCCTGCGACGCCTGTCCTGCCCAAAAAAGCTCTCTGGATTCTC

TACAGAAGACAGCCATTTCACAGCAAACACTAGAACTACTGGTGTGGCCATCAGTATCCAGCCAAAGTTT

TCTTCCGCGCTCTCCACCATAGACTCATACGACAGAAACCATTGAATACCAAGAAAGATAAAGATCACAG

CTAATATCAGAAGCACTGGATATGGCATGGGATTCAGTGAGAACACTTGTATCACCTGGGATGTCCTGCT

TCTCTTATGAGACATCTTTTTCTTTCTTAGAGTAGAACTGGAAGTTCAAG

>MSTRG.2852.1 gene=MSTRG.2852

CTACTTTTGGATTTTATTTTAAGAATTTCAGAAATTTGAAATTTTGTTTGGATATTTAACTGAATGATAA

CACCTCCGCCCTCATGTTTATAGTCCTTTTAGATCATGGCAACTCATCCACAAACTTTAATGCATCAGAC

CCTTTTTTTCTTAATTCTCCATATGTTGAACAAATTAGCAGTTAATAGAAAGAAAACGAAAATTAAATTT

TTGTGTAGTTACCAAGAACATTAACATATTCTTATAAATGATTTAGTCACTTATAAAACTGCAGGAGAAA

GTAAGGTCATATACATATGTGTTGCAGCTGTTTATTTATATGTAATTATATGATTTATTTATTTATTTAT

TTCTTGCTAGATGACCCCAACATGTGTTGCAGCTGCCTCTGACCACAAGCCTGGGTTTGTTGTATAGATA

ATAACTAGTGATGAGGTAATAGGCTTTTAGTATACTCTTAAGTTTTATGAAGCAGGTGGAGATGGCTTGT

TGACTGATGGAATTGGTGGAGGTTTTGGTATACTAAAACTTGGTACATGTGTAAAACTTGGTACATTGGG

TGGAAATGGTATACTGTCAGCCAAAGGAGACGGTGATGGGCTCAACGGCTCATCGGGCAGGGGGGATGCT

GGAGTGTCAAAACTGGGACTTGGAGATGGAGACAAGTCTACATTGTCATTGTTTGATGGGCCTAAAGGCG

AAGGCTCATCTGGTAGTGGGGATGCTGGAGTGTCAATACTGGGAGCTGGAGAAAAGCCTTCGTTGGCATC

GGGTGATGGGCCTAATGGCAATGGTTCATCTGGCAATGGGGATGCTGCAGGAGTGTCGAAACCTGAAGAT

GGACCTGATGAAAAATCTACATTGCCATCAGGTGATGGGCCTAACGGTGATGACTCAGCAGGTAACGGGG

ATGGCGGGTCAAAGCTTAAAGCTGGAGCCGGTGAGAAGTCTTTATGTTTTGACGGTGGCTGTTTTGATTT

TGGACTTGAAGGAGTACTAATTAATGCTGGAGCAGATGCTGGTGTGTCAAGTGGCAATTTGTGCTTAGAT

GAGGCTGGAGGATTGTTTTTTGCAGGATTAACAGCTGGAGGCAGGTCTTCTGGTTGTGGAGGAGAAAGTG

TTGAAACAGGTTCGTCTGAAGGCGAAGAAGATGGTGTAACTGGAGTCTCTTGTGCTAATAGATTGTTTAT

TACTTTTGGTTTTTGGAAAGGCCACAAGTAACTAGAAGTGCAACTTACTGGTGAAAAGGTAAGTTTTCCA

GAGTGTCGAATTGTATTTTTCTCTTCACTTTTGTATTGGAAGATTATCTTAAAGGGC

>MSTRG.2855.1 gene=MSTRG.2855

AAAATCCAAAGTTACAAAAAAGCTGAGAAATTTCTTCAAAAAGTTTGAAATCTAGACTAATCAGCAGCAG

CAATTACACGCCGCTACTATACATACATAGTACAGGCCGGCGATGTTCGCCGGATCTAATTCACAAAACT

GATAAGCAACTAAATGAATCTTAAAAAACCCATCTGACATGGTGTCATGCAATGCCCAAATTCTCTAAGC

GCCCATGTATTGTTCAATCTATCATCAATCTATGCTCTCAAGGTCTCTTCAAACAAGCCATTCATTCCCT

TGATATTATTGCTCGTAAAGGTACTCGTTTAGACTCGAATATACTTGCTTTTCTTCTGCGTGAGTGTGCT

GATTCCAGGTCTTTAAAAGAGGGCAGATGGGTCCATCTTCATTTGAAGCTCACTGGGTTCAAACACCCCA

GTACGTTTTTATTGAATCATTTGATTAATATGTACTGTAAATGTGGTGATCATGTTGTTGCACGCCAACT

GTTTGATAAAATGTCGGATAGGAATTTGTATTCGTGGAATAATATGGTTTCTGGGTATGTTAAGTTGGGG

ATGGTGGGTGCTGCTAGGAGGTTGTTTGATAGAATGCCGGTGAGGGATGTTGTGTCGTGGAATAGCATGG

TTGTTGGGTATGCCAAGAGTGGGGAGTTTGGTGAGTGTGTGAGGTTTTTTAAGGAGTTGAGGAGGTCGTC

GATTGGGTTAAATGAGTTCAGCTTTTCTGGGGTTTTGACTGTTTGTGTGAAAGGTAAGGATTTTGGGTTT

ACTAAGCAGGTTCATTGTCGGGTTTTGGTTGATGGGTTTTTGTCAAATGTGGTAATAGTGAGTTCTATCG

TTGATGCATATGCTGCTTGCGGGAAGTTAGATGACTCGAGAAGATTGTTTGATGAGATGAAAATTAGGGA

TGTTCTTGCTTGGACCACCTTGGTATCTGGATATGCCAAGTGGGGAAAGATGAAATTAGCTAGGGAATTG

TTTGATGCCATGCCTGAGAAAAATTCTGTCTCTTGGACTGCTTTAGTCTCAGGATATGCTAGAAAGAATA

TGGGAGATGAGGCCCTTGAATTATTTACAACAATGATAAGGTCTCGTTTCAAACCAGATCAATTTACTTT

TAGTAGTTGCCTGTGTGCTTGTGCTAGTATACCATCACTTAAGCATGGTAAAGAAATTCATGCATTTTTG

ATGAGGAGTAATTTCAGACCTAATACAATTGTTGTGAGCTCTCTCATCGACATGTATTCAAAGTGTGGAA

AGATACGAACAGGAGAAAGGGTTTTTGATGTTTCGGGTGATAAAAATGATATTATACTGTGGAACACAAT

TATATCTGCCTTGGCACAACATGGTCTTGGTGAAGAAGCAATACAAATGTTTAGATACATGGTACGCAGA

GGCTTGCAACCCGATAGAATAACTCTTGTTGTCATTTTAAATGCTTGCAGCCACTCGGGGCTTGTACAGG

AGGGCCTTTGCATTTTTGAATCCATTACAAGTTACTATGGTGTCTCTCTGGATCAAGAACATTATGCATG

CTTAATTGATCTTCTGGGTCGAGCGGGTAAGTTTGATGAGGTGCTAAAGCAGCTCAAAAGGATCCCCTTT

GAACCTGATAGTCGTGTTTGGAATGCCTTACTAGGTGTTTCTAGGATTCATGGTAACATCGAAATGGGAA

GAATGGCAGCTGAACACCTTGTTAAATTGGACCCTTTGTCCTATGCGGGTTATGTTGCGCTGTCAAGTAT

ATACGCTGCACTTGGGAAATGGGAATCTGCGGAGAACATAAGACAGATCATGAATGATAGAAATTTGAAG

AAAGACCAAGCACTTAGCTGGTGAAGAGCTAGATATAGGCTCCATGCTTCATTAGCATTTCCGTATAGGT

ATAACTTTTTCAAGGTTAAATATTTAATTTTGGAACTGCCAATCAGATGGAAGATGTGCTGCTTGATACT

TGTAGGTATGTCTTGCAAGTTTTAACTCTTTGCGAAGAGGAATCAGTAGACACGAGAGTTACTTGATCGG

ATGGCTTAGCTTGTACCATTTTTGTTAGTTTGTCTAGTAGGCGGGCTGCTGACTCTAGATTGTTAAATTT

TGACTCCATAGAACTACATTTTTACAGGACATAGGTGCTACACAATCTTTTCTGTTATTAAGTAAGAGAG

AGTTGTGTGAGCTTTAGATAAGCTCTGGGGGGGTGTTTTATTGTTGCTTGTATTTTCTTAACATCCAGAG

AGAGGTTTCTTTTTCGAAATTCTTATCAAATAAAACAAAATTTATATTCTCATTTCCAGATGATGAGAAG

TCTGTTGAGC

>MSTRG.2858.1 gene=MSTRG.2858

CTTCTTAAGCCGTCTCACTTTGCTCTCATTACACACTTTCTCTCACTACACATTCCCCCCTCTTCTCTCT

CTACAATTTCTAAACACAAATTTACATAAACACAATGCATAGCTGGGATAAATCTACGAGGCAGAGGAAA

AATCAAGCTAACCAATCATTTTCCTCGAGTCTTCTTGATGAGATCTACAGGGAGCTCGATGGTGGCGTTG

ATGAGAGGCGCATGGAAATTATAAAGCAGAATAAAGAAAGGAATGTGAAGAAATTGAGTAGTAGTGGTGG

TTTTGGAGGCAGAGGGAAGGGGAGAAGTAGCGTTGAAAATGAAGAGATGAGTAGTTTGAAAAGAGCTTGT

TTGTTGGAGAAATGGATGGAACAGAAGGTGAATGAAAAGGCTTTGGCTAGAGGGAGAGGGTCATTGGTGC

ATGATAATGATCCTTTGTTTTTTAGTTCTAGTTCTTGTTCTTCGGATTCTAGTTTCGGAGGATTCTCATC

CTCCGAGACAGAATCTTTTGGTGTGGCAAAAATAAAGAGGCCTCCCATTCCCAAACCAATTCGGACTGGT

AGTCGAATTGTTGGGAACTCTGTCGCTCAGAATGAGCGACAAAGTGAGTTCTATTTGTTTGATGATTATC

GTAATCATCAAACAAATAAAATAAACGAAGATGGCAGTCGTTTGGTAAAGTCGAAGGCCCGTGCTCAAAA

GATTTACGCAAATTTGAAGAAGATCAAACAACCTATCTCGCCAGGTGGCCGGCTCACGAGTTTTATAAAT

TCTATCTTTACTAATAGTAACCCCAAGAAAAGCAGAGAATCACACCGCAATGCCGGTAACTGTAGTGAAG

TGAGTGGTGATTGGAAGTCTAGGTCTGCTCAGACATCTACATGTTCATCAGCCTCTTCGTTTTCAAGATC

ATGTTTGAGCAAGAATTCCCCAAATTCGAAGCAGAAACGCAACAATGGGGTCCAAAGATCGGTCAGATTC

TACCCGGTGAGCGTCATAGTTGATGAAGACAGTCGTCCTTGCGGCCACAAATGCCTTTACGAAGAAGAGG

TTGAGAAATTTAAGAATCCAATTCCGTGTAGCCAGACCAATCTGAAACAGCCAATGCCGCCTGTTGTGGA

TCACAGGAGCAATGACGTTTCAAGGAATTATGAAAAATCGAAGAAGATTGATTTTGCTGCTTCAATTAAT

CAGGAAGAGGATTATGATTTTGAAGATGATTGTGCTAGTGATTTGAGCTCGGACTTATTTGAGCTTGATC

ATTTAGCATCATTTAAGGATGATAGGTTCTGCGAGGATCTTCCGGTTTATGAAACTACTCATTTAGGGCT

CAACCGTGCGATAGCAAATGGTTTGATTCGTTAAGGTTTTTGTTTAGAAATTTTCTGTTATTTTAGGGGA

TAAGCTATCAATTGGTATATCTTTTATTGTGGTTATGATTCGGTTTTTAGAAGTGAAATGGTGATATTGT

TGCATAACTAGCGTGATCAACTCAAATTCATTGTGAAATCTAACCCCTTTAAAAGTTAAGAATTTCAAAA

TTTTC

>MSTRG.2864.2 gene=MSTRG.2864

GTGATATAAGAATTTTACAGGAAGCATCAGCCTCCTGGGGATTATCTGTAAGCTTCATCATCATCATTCT

ATTGCCAATTGTCAGGAATGCAACTGAACATGCTGGAGCAGTAATTTTTGCTTTCAACAACAAGCTGGAT

ATAACTTGATCAACTCGGTTGCCTGTCCTGTGCATATTAGCCCTTGAATAATTGCTGTGTTGTTAGCACC

GAGACTCCTGGACCAACTTTGTTAATATACCTTAAAAGCTCTGTTGACATTATTATACCACGTGTTTACA

TATGAGTTATGAGAAAATATTAATGATCAAGCTGTTTTATAACAGACATCGAGGTCCGCTGTAAATGGTG

GCTGGATTTTGAAAAGAATTTCTATGTTTAAGCTTATTTTAGTTTGTCAGTTGAGGCTACTTCAATTTTC

AGTTCATTCATGGATTTCGTGTCGCCACCTAAAGTGAAGTCAGAACAATGTTGTTATTTAACTGATTTCC

AACATCTTTGAGAGTACATTCAGTAGATGAGAAGTATATGTAACACTTCAGTCCCATATCGAGGACTGTG

ACTCGGGATTTGACATGGGGATGTTAGGAATTTAAGTGGGCAAGTTTGTAACTCCCCAGTCCTACATCTG

AGGAGTATGGAGAGTTCAGTAGTCAACAGTGATCTTTTGTTAATCCATATGCTCAGGAGTCAGGTGTTTA

TCTCTTCCAATTGATACAAGTGACTTCTTAGCTCTAGCTGAGTTGATAATCGCAGTTGTTCAGTTATTCT

ATATGCTGCTACTGTTTTTACTATTTGGTGCTATTTAACAGATTTTGTTTTATTCAAGTCACCGTG

>MSTRG.2865.1 gene=MSTRG.2865

GATAAATGGTCTGCCTTTTTCTTGAATCATACACTATTTACTTATGTTTTGCTTTGACAAATTGATCATT

CCATAAGTTATGTTTAAGGTGTGGAGCATGTACCCTGTTGCAGAGAGCATTGCCAAGAAGACAGATATGT

TGGTTAGCTCAATTCTTTGGCAGCTTAAACTTAATAAACCCTGTCAGTCGTTTGTTATCCACTTATTAGC

GAAATAGTTGAAGATCCAATGAAGGCTCCTGGAGCCAGCACTCAAGAGAAATGCCGATGATGCCTGCCAA

AAGTATCTGATTCAACTTGATTAACAACCCTATTCACATTTACCAATCAGCCCCTTAATTGAGACGGTTA

AACGACTCTAGCAAATTGTTAGAGATCCTCGGGCAGGGTATGAACAGAGCTTGTCAGTCCTAAAACACGC

AAAGACCGACAAGGAGGGTATGATAACAAAAACCTCTATAATGTTGGGACTTGGAGAATCTGAGGATGAG

CTTAAGGAAGCCATGGCTGACTTAAGGGGATGTTGATATTTTAACTCTTGGGCAGTACTTATAAATATTT

ATGAAAAGATTGGATTCGAGATTTGCAGAGCAGTTTATAGGAGAGAATAAAAGGCTTGTGTGGGACAAAA

GATAAGATTAGAAGTACAACCTATAACGAGGAGAGATTGTTAAACGAAGAGCTTGTAAAAGATGGTAAAA

GAAGCTGATGTACAGAGCAGAAGCTAAAAGTGAACAAGAAAGGGACAGAGAAAAAGAATGCTTTTACTTC

AGATTTTTTCCTTTCTGCAAATTACTCACTCTGACAGAGAAACTCTTTATTTTTATGTGAGTTTCCATTT

GCACTATTATGCTTGATTTTACGGAATGGCCGGAAAATGAGG

>MSTRG.2876.1 gene=MSTRG.2876

GTAAATTTTTACACAGACAAATCCCAGTTCAATAATGATTTCTTGGCTTAAGGCTTCGGGGGCTCTGTTG

GTGCACATTCTTCCGCCTTTGGTGGTTCTGCAGGAGCTGCTTCGGGTGTTTTCTCAGGCTCTGCCTCAGG

CTCAGCAGCCACTTCTTTCTTCTCTTCTTCCACTGCTATCTCCTTGTCCTTCTCTGTTCCGCTTGTTTCT

TCAGTTGTCGTTACAGCTGGAGGTGCCTCGGCTGCTGGAACCTCCTCTTCCTTCTTCTCTTCTACCACAA

CATAAGTTGATACCTTCTCA

>MSTRG.2878.1 gene=MSTRG.2878

GAAAATCACAAATTGCCTTATCTTGATCACATTTGATTCTTTTTCACACAACCTGTTAAAAAGCCATCAA

ATAATGGAGAAGAGGTGGTGGTCACAAAGCAAAGTGAATGGCATATGCACCAAACACATGCTTCTGTCCC

AGTTTTTTCTGCCCACCTGCACATGCTCTTCTATCTTAAACTCCTTCTAAAAATCACTGCTGTTGAGTAT

AATATCTTACAACAACATATAATAAAACATCACACCTTCACACAACCTAGACATTATCTTCAAGTTCCTA

GCTGGAAAATTACCTAACAATTAAGAAACTATTAGTGTTATAATCTTGTGATTACTAAAATTAATAGCCA

TTCTTGTTTGCCATGTGAATGACAACTTCATAGAGCTGGTGCCACGGTGGCGCGTCTCTGAGCCAGCGCA

GTGCATTGAAGGTGGCGCGGGCAGGTGATGAGGTGGTCGATGGTGAGTCCAGCTGCCTGCATGAAGGAGT

GTAGAACAGTGGGTCCGACTAATCGGAATCCTCTCTTGACCATGTCTTTGCTAATGCCCTCTGATTTTGA

AGTTTTGACAGGCATTTTGTTGCATAATTTGTATTGTGTGACAATAGGGTTGTTGTTGACAAATCCCCAT

AGATATTTGTCAAATGACCCAAATTCCTTCTTAATCTGAAAACCAAATTTGTAAGCCTAGTGTTAATAGA

TTTCTTAAATTGTTATGTTGTTTAATAAACATAAAAAGATAAAAGTGTTGCAGTGATTACCTGAAGAATT

CTGTTAGAATTGTCTACAACTCCTCTGACTAGCCTTAACTCTATGTTATACTCAGC

>MSTRG.2882.1 gene=MSTRG.2882

CTTGTATATAAATATGTAAAACGTGCCGCGTTTATACAAGTTAGAGCACGTAGACTGATTTTGACTAAAA

TCTCGTGAATATCAACCCATCTTGTGAATAAATACATGATCGATTGTAACACAACATCCCATAACGCCAA

AATGTCATTTATCTACTTGAGCCTGCTACTCATTTTCGGATCAGCGGTCACTAATTTCCCCTCTGCTCTT

CGGTATCCAAATCACAGTCCCGTGTCAGTGTCAGAGTACAAGCCAATACCACCTCCCAAGCCTTCACCAG

AAGACAAACAGCTGTTCCATAAACCACCGATCCTGGTCATTTTACGACATCCAGAAACATCAAAGAAAGC

ATCAGCCGGGAGCCACAGGCCAAGTCCTGCCTCCTCACGCCCTGTGTCTTTCAACAAGCCACTACTGCCA

CCAGCACGTCCTGAAGACCCCCCGTCTCCTGCTAACAACCCTCCACCTCCTACTCGCGAGGCCTATGTTG

GACAGCCTCGGGGAAGGAATACAGGGAACTAGATTAATCTCTTCTGAATAATTGTATTTTGATTCATCAG

TGTGTGCTATTCTAGTTGCAGAAAACTACTTTATCATATTACATGATTTAAGTTTTAACGAAATCAATAA

ACTTGGCCCAGTGCTTCTGAAGTCCAATGTTCCAGCTTCAATCAACTTCAAGACAACCACATCCTGTTTA

TCAATGTCAAATGTCAACTAATACTGACAGTTTTGCGCCTGTGATTTCCACAACCTCCAGGTCCGAAAGC

GGAATTATTGTGAATCCTGTATTCTTGGCCTTGAGAAAAAAAAAAAACTCTTATAAGAACCCGTCTTCTC

TATATGATCATACACCATCCGAAACCAAACTGCACCACATGTCTGTATACTACTTGCACCTACTACTCGT

TTTAGGAGCCGTGGTTCTCACTGCTCCGTCCCTGGCTCTCCGCTATCCAAATCATAAGCCCCAATCGAAG

TACACGCCAGTATCAACAGAGTCTGGACCGCCAATTTTCAAGCCTCCACCAGGATACGAAATGTTGTTTT

ATAAAATACCAGTTCCGTTTAGTCCTCAACATCCAGAAACTTCAAAGAAGAAACCATCACCCGGGAATCA

CCGGCCAAGTCCTGCTTCTGCACCTCCTGTGTTTCCAAGCGAGGCGCCACCAAGCCCCCCTCCGCCTTTG

CGGTCAGCACCAAGCCCACCACCGCCTTCGCGATCTGACCACATGTATGTCCACCACCCTTGACAGGATC

CGCGGAGAATAAACTACACCAGTAATATGTAACCTTTCTATACATAGGATTTGATTTCTTGTAAACTTTT

GATTAAACTTTAAGAAGCCGTCTTCTTGATTGAATAATGATGTTTAAATTTTGTCATCAAATATATATAC

AATTTAAATT

>MSTRG.2886.1 gene=MSTRG.2886

CAAATCTCCCATTCACACTCACCAAAACGACACAAAAGAGAACACCCAGCAAGCTTCTTGACATGGTTGT

GTGAGAAGAGAAGATGGGGTGTGAAGTTTGTGAACCGGGAGAGATTAAGAAGAGAGAGAGAGAGAGAGAG

AAAAGTCAAGAGTGGACTTGGAAGAGAGGAGGCAGAGGAGACTTGTTTGTGGAATACTATATAGATACAG

AGATAGATACAGGGAAGAATGGTAGGAGGGAGAAATGGGTGTTGACGTGAAAGAAGGGGTGTATCACTAC

CAACTTCAGCTTTTTTCTATGTGCAAATTCTTCTAAATTCACCTAGTATTCAATTGTAATGAAATTAGAG

TTGATTTAGATTTTAAGTAAATGTTGTTTGGTGCTACGTATTACAATTTGCTTGGTCAATGCTGATCGAA

ATACGTCGATATCGACATAAATTTCAAGGAGAGTTTAAGAATGATTCAAAG

>MSTRG.2894.1 gene=MSTRG.2894

AAGAAATGGTAATACGTGCACAGTAATCTAAAGACACACATGATTCAAGTGCAATCAACCACACTCTTGC

CTGTAAAATAAGTTCAGAGAAAAATGGATATTTAAATCCGGTGGGCGCCCAAAATCCATTATATATCTGA

AGCTAACTTGGGAGCTTGGCGACATCAACGAGGTACCCACTTGAAGATTCTCTGTATTTTCACTTTCTTT

GTTTTTTCTTTACATTGCTTTTTACTTAAATGGACTTGCTCTTTTGCCTTTAATCTGTTGTTCTTTATAC

ATTTGAGGTAAATTCAACTGTTAATGCATTTACTGCATTTTTTTTTTCATTGAAATCTGGCATTTTTAAT

CTCAATAGAACATAGTAGTGGTAATTTTTTTATCCTGTATGTTATCTAGATAGTGTTATTTCTTTTCTGG

GTTTTTTTTCACTTTGAGATGGTGATAAGCTTTATGGTTTTTTAATTCAGAGTGTTTATGAGTTGATAAC

TCTAGCATTCACTGTAATTAGCTGCAGTTGTTATAATTAGGTATCATCTTATGTGGTTAATTGATGCAAG

TTTTTGAATTCATATCTATAGTGCAAGAATGTGAATATGTGATTGTGTTGAATAGCCAAACATGAACTGC

TATCCAATGTACTGCATATATGAAGAATCTAAGTAATAGTGTACGCCTCGCGGAAGTGTTTATGCACTTG

CTCAGAAATTTCACTTTCAATTAGATATTTCCTAAGATTGGTTGTTTATTATTATCAACAGTTGCCTGTA

T

>MSTRG.2911.1 gene=MSTRG.2911

TATAAACATGACTTAATATATAACATCGGAACAGGAAACTGAAAGGAAATAAAAGAGGCGACGCAACTTA

GTCCCGGCACAAGGCCTATCAAAGACATCAAGTTTGCCAGAATACATATGGTGAAACAAGGTCCCGGGAT

CAAATTCCATCGCCTTTTCTTCGCCACTACAAGTGACAGGGACTTTAGCTAGAGGATCAAGATCTCCCTA

CAACAAGTCAGCTAGCGACCCGGCATGAGACCATCCTTAGAGCCCTTGTCATACCATTCTTCCCGCGCAT

GCAACTCAGCATGCTTGGGAATCTCCAAGTGATCCTTGGTTGTTGCCTTGAGCCTCTTCGACACACGGGC

TTCCCTGGTCATGTCACCAAAGTTTCTATTACGGCACACAGGAATCTCAGTAATTGGTACATGATTCTCA

GCTCCACGAGGGGCCAACACAACTGCCTTATCAACCAATGCCTTGACACTGGCATAAGCAACTGCAGAAT

GTTCCCCACTAAGTTTCGCAACAATCTTTCTTGAAACAAGAACTTGAAATTTCAAGTGCAAGAAACAAGC

AAACTATGAAGTAAAATATGAAACAAAGACTTACTTGTGAAAAGCCCAGATGAAGACCAACTCGATCTGC

TGAGAATATTATGCAAATCTCTCTGTTGATAATATGTAATCTCTCTGTTGATAATATGTTGTTTTTCGCC

AGTTTCTCAGAAAGCTCGAATAGGACTAGTATGGGCGTTGAGTGTTATTTACACTGAAATATACGGCCAG

TGTGAAAAAAAGAGTAAAGCACACTTGCCTTATCTGTGTATTTCAGACTTTCAGTCGCATCAGTTAAGCA

TCCACAACAGTGAAG

>MSTRG.2915.1 gene=MSTRG.2915

GATGGTTTCAAAATACCAATTAGGCCAAGATCTGAAATCATCTATTTGCATCTAAGTTCAACGCCGTTAG

CAAGACTTGACAAAGGTGCACAAATTGAGGGGAATCTTCAAGAGTAGTATTGGGTCCGTAGGTTTTCAAA

ATCCAGTATTGCAAATGTGATAAAGCCCAAATTGTAGATACACAAAATGCACTAGATAATGTTAAATATA

AAAAAATTGGAGTTCAATTTTGGTGAGAAGAAAGGCAAACGGCAAGAGAAAGCTGAGCCTGTTGACACTG

GGATCGCTTCAGAGCAGTTCCACCACTTCCACAGAGACTCTTTACTGCCTCTGATTCGTCAGGGCAGATT

GTAGCCACCAAACAATTTGCGAGTGATTGGTTCAGGTGCCGGCACGCGGCATCCCTCCCTGGCCCTGCTG

GGATACGACGGTGGCACTGCAGCAGTGACTTGTACTGGCGAACACATTCTGCTGGAACAGCACCTGATTC

AACTTCTTCCGTCATTGTCAACTGTTGGTATTTACGACTTCACTTCTTTCTTCACCTGATTCAGAACAAA

ATAACATTAAATTAGTACAATTAGAGTTATAACTTATAACAAAGACATGAATTAGATCCAACTCATTCTA

CTT

>MSTRG.2916.1 gene=MSTRG.2916

GTTTTGGTCTGACCAATCAATGTTTCTTATTATATTCTTGTGATGCAATGCTTTGGTCTGATAATATATT

GAAATGTTTCTTACTTGTGATGCTTTGGTCTGTGCAATATAAACTTGTACCAGACATGTATTTCAGATTA

CTAAGTTCACATTTTTCTAAGATAATTAATAATTAATACTGATCATCATTTTGCTCTTCACATTTCAGAT

TACTAAGCATTTCAAAGTGTATGTTAAGTAGTGTGTATGGTGCGTGGTCAGTGTCTTACTTGTGTGTCTG

ACTAGATTCTGTATTTGTTGAATATATTGATGAGAAGTGGAGTGGTGATAGCTCCTCGATTCTCCCCTCT

GATCCTTATGATCGTGTTGTTGCGCGATTCTGGGCTGCGTATTTTGATGACAAGGTGGTTCCAGTCCAGA

GAGCGAAAGGGTGGCGTTGATAGAGCAGCTAGCGGAAGGATTGGTGCCGCTGGAGGAGGCTTTCAAGACA

TGTAGCAAAGGGAAGAGCTACTTCGGGGGAGATAACATTGGTTACACTGATATTGCTGTTGGAAGCTTCA

TTGGATGGATCAAGGCAATCGAGAAAATGAGTGGCATCAAGGTTCTGGCTGAGGCAAAAACACCGGGCCT

CGTCGGATGGGTGACCAAGTTTCTGTCGACTGATGCTGCTAAGAAATTTGTTCCGGAGCCTGAAGTGTAT

GTTGAGCTTCTTAAGAAGATGCAGGCTTGAGCCTGAGCTGCATAAAGGAACTGGGGGTGTGTCTGTTTGA

TCCAATAAGTGACTTGTGTTTTCTAGCAAGTTTATGTAGTATTAAAATCATAAGATAAGTACAGATCGCT

TGTAATATTTGAATTATAAGTGTGAAGTACATGTTATAATAAAGTATTTACTCATTTTA

>MSTRG.2927.1 gene=MSTRG.2927

AGATGGAAACTTGGTGATGTGCCACGCACGGCACACTTCCTCTTTCATCTCGCCACGTACCTTTTCCCCC

GAATGCTGCTCTTCCTTTTATAGTCCCTAATTTCAAAACCCCAACCCTCGTTTATGGTTTTCAATCTCAC

ATCGCTTTCTTCCTTTGACAAATTAACCCCTCGTAGCATGTATCCTTTCACCTCCACTCTATTTCCAACG

TCGTTTTCCGTTCCCACTGTTCATTCTCGCCGCCACTGCTCTCCCTGTTGCTCCGTCTCTGCTGCCACGG

TACACCGCCGCCGTTTTGCCGTCTCCAACGACCAACTCCATCCTAGAATCTGTTGATTCGAACTACAGTG

GGAATAGATCAAGACGTGTATATCGAGAATGTTCAGAATTAATCCGATGGTGTTGCAAGTTGAAGCAAGA

ATAATCTGAGATGACAGACGTAGAAAGAGTATTGTCAGAAGCTGCTTAGAAAGATATGATTGTGCGGTAT

AGTGCTATTAGAATCGTGACTAGACATCGGAGTTAAGCTAGGGATAAAGATGATTCACTTAAGGCTAGGC

AAGGATTGCAAGTGGTTGAGTTTCAGGTAGATGGTCAGGTAGAGTATCCAGGACGATGTCATTAAGACTA

GATGCAGCTGGAATCTGCAATTCAGTTCAGTTGGAAATTAGATGTGTTATTTGGTTATCAGCTACCCTTC

TTATCGAGGTTTAGCTGATAATGCAGTTTAATTTGTAATTAAATTTAATAATTTTAGTTTCTGATTTCAA

TGCTGTAACCTGTATGCGATCCTGTTTTCAGGGGGTTAAACTTTGTTCTTTTAAAATCTTTTATTAAAGC

TTTTAGGTTTTAAATTCTTTGGTTTTTATAAGCTTTTCGAAAATACAGGTTTTAAATTATTTTTATTTAC

ACGTGGCACCAGATCCAGACCCCCGGATTCCTGGGCTGTCACAGTTGGTATCAGAGCTACAGGTTATAAG

TTATTGAAATCAG

>MSTRG.2927.2 gene=MSTRG.2927

TGGAAACTTGGTGATGTGCCACGCACGGCACACTTCCTCTTTCATCTCGCCACGTACCTTTTCCCCCGAA

TGCTGCTCTTCCTTTTATAGTCCCTAATTTCAAAACCCCAACCCTCGTTTATGGTTTTCAATCTCACATC

GCTTTCTTCCTTTGACAAATTAACCCCTCGTAGCATGTATCCTTTCACCTCCACTCTATTTCCAACGTCG

TTTTCCGTTCCCACTGTTCATTCTCGCCGCCACTGCTCTCCCTGTTGCTCCGTCTCTGCTGCCACGGTAC

ACCGCCGCCGTTTTGCCGTCTCCAACGACCAACTCCATCCTAGAATCTGTTGATTCGAACTACAGTGGGA

TCAAGACGTGTATATCGAGAATGTTCAGAATTAATCCGATGGTGTTGCAAGTTGAAGCAAGAATAATCTG

AGATGACAGACGTAGAAAGAGTATTGTCAGAAGCTGCTTAGAAAGATATGATTGTGCGGTATAGTGCTAT

TAGAATCGTGACTAGACATCGGAGTTAAGCTAGGGATAAAGATGATTCACTTAAGGCTAGGCAAGGATTG

CAAGTGGTTGAGTTTCAGGTAGATGGTCAGGTAGAGTATCCAGGACGATGTCATTAAGACTAGATGCAGC

TGGAATCTGCAATTCAGTTCAGTTGGAAATTAGATGTGTTATTTGGTTATCAGCTACCCTTCTTATCGAG

GTTTAGCTGATAATGCAGTTTAATTTGTAATTAAATTTAATAATTTTAGTTTCTGATTTCAATGCTGTAA

CCTGTATGCGATCCTGTTTTCAGGGGGTTAAACTTTGTTCTTTTAAAATCTTTTATTAAAGCTTTTAGGT

TTTAAA

>MSTRG.2927.3 gene=MSTRG.2927

CGGCACACTTCCTCTTTCATCTCGCCACGTACCTTTTCCCCCGAATGCTGCTCTTCCTTTTATAGTCCCT

AATTTCAAAACCCCAACCCTCGTTTATGGTTTTCAATCTCACATCGCTTTCTTCCTTTGACAAATTAACC

CCTCGTAGCATGTATCCTTTCACCTCCACTCTATTTCCAACGTCGTTTTCCGTTCCCACTGTTCATTCTC

GCCGCCACTGCTCTCCCTGTTGCTCCGTCTCTGCTGCCACGGTACACCGCCGCCGTTTTGCCGTCTCCAA

CGACCAACTCCATCCTAGAATCTGTTGATTCGAACTACAGTGGGAATAGATCAAGACGTGTATATCGAGA

ATGTTCAGAATTAATCCGATGGTGTTGCAAGTTGAAGCAAGAATAATCTGAGATGACAGACGTAGAAAGA

GTATTGTCAGAAGCTGCTTAGAAAGATATGATTGTGCGGTATAGTGCTATTAGAATCGTGACTAGACATC

GGAGTTAAGGCAAGGATAAAGATGATTCACTTAAG

>MSTRG.2933.1 gene=MSTRG.2933

ATTGCTCAACAGAATTGTGTCCTGGCTTTCGTAGAGTCCAAACAGTACATATATTAATGTTCTACTACAA

TTTTAAAGGTACAGATTACACAAATTTAAAAAAAAAAATTCCATCACTCCGCTTCCATGAAAAGTTGATG

TACATTTCTTGTAGCATTACATTAAGAATAAACAGAGGACTGATAGATAAAAAGACACACACAGGTTTAG

ACAGAGGGGTCAATCAGATCTCCATTGAGGTAAGCTCTGTAAGCAGCCACAGGCCAGCTGAAGCCTCTGA

AGAGGAGCTTTGATGCCAACGGAACATCGATGATGATCTCCTTCTGTGTCGGCTTCTTCTCATCTCTGAT

TGCAATCAGGTAGCTCCTTCCGACCCATCCAATCCATCCGGCAATATACAAGAACAAAATCCCTGGAGTA

ATGAACTCTCCCCAGTGTCTTTGATCACCACTCACTATCAGATGTGGTAATCCATCTGATCCACATAACA

AACCTTGCTTTCCATAGTTATCAAACCTGCATATATACAATTTTCATAGTTATAGCCAATTACAACATTG

CTTATATTGATTATGGTCCATAATATTGTTAACTACCTTCTCTTGGTCTTCTCCATGGTTGCTTTGATGG

CAAGGGCAGGAGCACTATCTGGTGCATAAATTTTGAGTGAGGACTCGAGCTTCTTGAGTTGTTGCTTTTC

CCTCTTAGCAAACTGTTTAGAATCCTTGCATGGGGTGAGCCCTGCAATGTCTGCGGATGCCGGGAGCACC

GGAGCCGAGAGCAGAATTGAGGACAGTGCTAGAGCAGCTGAGAATGCCTTGAGTGATGATGCTGAGCTCT

CCTGGTCTGCAGAAGTGGAGATATTATTGTTCTGTGGTGCAGCAGAGCAGACAATATTAGCTAGCTTTGG

CTTTGCTGTGAACTGAGAGTTGAGCCTGGGGTTTGAAATTGGCTTGTAGAGGTTTGTGGGAATTGAGAGA

GCCATGGCTAGACACTGATTGTGAGCTGTTTGTTGTGTGTGTTTCTTGGGGCTGGAGTGTGGAGAGGATG

AGATGGTGTTTTGTTGTTTAGATTTAGAGAGAAGGGATTGGTGGCTGCAGAGAGGATAAGATGTGGAATT

GAATAAGTGGAAGATGTCTGAGTTGTTTGGTTCTAATTGGCTGTTGGTATTTGGTTTTTGAGTTTGTGGA

TTTTGCAATGTCTCTGCCTGATTCTGGCTTTGCTTGACTAGGATATGTGGAGCAGAGGATAACTTGGTAG

CTATTTCATCATATTTTGCTGTGTTTAAATAAATCTGATCCCTTCTCTTTGTAAGTAAATTAGAAAAGGA

CAAAGATTTTATAATATCAA

>MSTRG.2933.2 gene=MSTRG.2933

ATTTCTTGTAGCATTACATTAAGAATAAACAGAGGACTGATAGATAAAAAGACACACACAGGTTTAGACA

GAGGGGTCAATCAGATCTCCATTGAGGTAAGCTCTGTAAGCAGCCACAGGCCAGCTGAAGCCTCTGAAGA

GGAGCTTTGATGCCAACGGAACATCGATGATGATCTCCTTCTGTGTCGGCTTCTTCTCATCTCTGATTGC

AATCAGGTAGCTCCTTCCGACCCATCCAATCCATCCGGCAATATACAAGAACAAAATCCCTGGAGTAATG

AACTCTCCCCAGTGTCTTTGATCACCACTCACTATCAGATGTGGTAATCCATCTGATCCACATAACAAAC

CTTGCTTTCCATAGTTATCAAACCTTCTCTTGGTCTTCTCCATGGTTGCTTTGATGGCAAGGGCAGGAGC

ACTATCTGGTGCATAAATTTTGAGTGAGGACTCGAGCTTCTTGAGTTGTTGCTTTTCCCTCTTAGCAAAC

TGTTTAGAATCCTTGCATGGGGTGAGCCCTGCAATGTCTGCGGATGCCGGGAGCACCGGAGCCGAGAGCA

GAATTGAGGACAGTGCTAGAGCAGCTGAGAATGCCTTGAGTGATGATGCTGAGCTCTCCTGGTCTGCAGA

AGTGGAGATATTATTGTTCTGTGGTGCAGCAGAGCAGACAATATTAGCTAGCTTTGGCTTTGCTGTGAAC

TGAGAGTTGAGCCTGGGGTTTGAAATTGGCTTGTAGAGGTTTGTGGGAATTGAGAGAGCCATGGCTAGAC

ACTGATTGTGAGCTGTTTGTTGTGTGTGTTTCTTGGGGCTGGAGTGTGGAGAGGATGAGATGGTGTTTTG

TTGTTTAGATTTAGAGAGAAGGGATTGGTGGCTGCAGAGAGGATAAGATGTGGAATTGAATAAGTGGAAG

ATGTCTGAGTTGTTTGGTTCTAATTGGCTGTTGGTATTTGGTTTTTGAGTTTGTGGATTTTGCAATGTCT

CTGCCTGATTCTGGCTTTGCTTGACTAGG

>MSTRG.2934.1 gene=MSTRG.2934

AAATACCAACAGCCAATTAGAACCAAACAACTCAGACATCTTCCACTTATTCAATTCCACATCTTATCCT

CTCTGCAGCCACCAATCCCTTCTCTCTAAATCTAAACAACAAAACACCATCTCATCCTCTCCACACTCCA

GCCCCAAGAAACACACACAACAAACAGCTCACAATCAGTGTCTAGCCATGGCTCTCTCAATTCCCACAAA

CCTCTACAAGCCAATTTCAAACCCCAGGCTCAACTCTCAGTTCACAGCAAAGCCAAAGCTAGCTAATATT

GTCTGCTCTGCTGCACCACAGAACAATAATATCTCCACTTCTGCAGACCAGGAGAGCTCAGCATCATCAC

TCAAGGCATTCTCAGCTGCTCTAGCACTGTCCTCAATTCTGCTCTCGGCTCCGGTGCTCCCGGCATCCGC

AGACATTGCAGGGCTCACCCCATGCAAGGATTCTAAACAGTTTGCTAAGAGGGAAAAGCAACAACTCAAG

AAGCTCGAGTCCTCACTCAAAATTTATGCACCAGATAGTGCTCCTGCCCTTGCCATCAAAGCAACCATGG

AGAAGACCAAGAGAAGGTTTGATAACTATGGAAAGCAAGGTTTGTTATGTGGATCAGATGGATTACCACA

TCTGATAGTGAGTGGTGATCAAAGACACTGGGGAGAGTTCATTACTCCAGGGATTTTGTTCTTGTATATT

GCCGGATGGATTGGATGGGTCGGAAGGAGCTACCTGATTGCAATCAGAGATGAGAAGAAGCCGACACAGA

AGGAGATCATCATCGATGTTCCGTTGGCATCAAAGCTCCTCTTCAGAGGCTTCAGCTGGCCTGTGGCTGC

TTACAGAGCTTACCTCAATGGAGATCTGATTGACCCCTCTGTCTAAACCTGTGTGTGTCTTTTTATCTAT

CAGTCCTCTGTTTATTCTTAATGTAATGCTACAAGAAATGTACATCAACTTTTCATGGAAGCGGAGTGAT

GGAATTTTTTTTTTTAAATTTGTGTAATCTGTACCTTTAAAATTGTAGTAGAACATTAATATATGTACTG

TTTGGACTCTACGAAAGCCAG

>MSTRG.2938.2 gene=MSTRG.2938

CAGAACATTAGCAATATAAAACTGTTAATCTTAAATGAAAACGGAATGATGATACCCCTCATTCATCTCC

ACCCCATCAAGAGAAGGAGAAATAGGGGCATGACCAGAAATTCCAAAGGCCTTGATATGATCTTTCAGTG

ACCTTTTATGCTTGAAATCTGAACCGCATGTACAGAGCCAACGTTTCCCACAATTCTTTTCATGGGTTCG

CCAGTCCCCTTTTACGGCCAGACATTTGCCACACTTTCGACAACAGAATTGCTTCGTTCCATGCTTCCTC

TTGTAATGTGTCTGCAGTGTTCGGAAATCCTTCAATGGCCTGGCCCTCGGGTGATTCAGATTATTCTTGC

ACCCTTCTTCACAGCAGTAGCAGGGGATTCCTGACATTGCTCGTGGCTGCACTCCCTTCAGCGATTCAGG

TCCTTTGCGATATTGTGAGCCATGCCCCCACATGTGCATCTTCCACCAATGGTATATATAACATGCAAGC

AAAAACTATTAATGAGGAAGAACTGCAAATTCGAACATATTATAATTTGCATGAAGATCTCTAATCTGAA

CTAACCTGAAGATTGTTGTGGCGATTGAATGTTTTGTTACAGATGTGGCATGAGAAATTTGTGAAACCGA

CAAGAATTTGGGCAGGACTTGGTATCCAGTATTTGGTTATTGGTTGATGATGACTAGAGTATCTAGAGCT

ACTTTCAGAGGTGCTCATTGCAAGCCCGATATGTAAGGTGACATTCAAGTCCTCTTTATCCCCATGACAC

GACGAGTAAGAATCTTGATCTGGCATCACGGGTTTTGGGGCTTCGGATTGAAGAGAGCTCAGAATTGTGG

ATGGCAAGGATTCAAGAAAATTTCGAGTTTCCTTGGGAAAGAAGTCAACAAGTTGAGTATGATTATGGTC

TAATGTGAACGAACCATCGGAGAGCTCCATAGTGATTTTAACTGCAAAATAACTTGGAAGATACTATATA

GGCGCTTATAATCTGGACTCTGGCCATTTGTTGTCACTGACACTATGATTCTGAGACGCGTTTTGCTGAA

AATGGATATCTGCATAATCAGCGTCCAAGAATTCACAACTTTGTACTTAGAATGGTATTCAGACCGGAGA

AATTTTAAGGAGTTCTGTGATTCTGAAGGAAATGGCCTTTTTATCTTTGATTCTTGTATCAAAACGAACC

TGATGTATCAGACTCTGTAAGCGGAGATGGCAAGGTGTACGAAAACAGACCATGCCTCTACTCAAAGCAG

TGAAGATGCTGCTATATCTGTTTGAACACGAGCAAAGGTGTTCTTTGATCAATCTCTATTATGTGTATTC

TCAGCTTTACAATGAAGCTGAAAAACAGCACTATGTTGTCAGCTACTTATGCACTTGACTAGTTAGAGAT

TTCATATTCTTCTGGACCTTTGGATGCAATCCTATCATTACTTATTATC

>MSTRG.2940.1 gene=MSTRG.2940

TCTTTTTCTTCTTCAAGATAAGGTATATGCCCGCAGCACAAACTAGAACAAACCCCCCTACCGCCAATCC

AACCACCAGACCCATCTTCTTATTCTTTTTATTAGACAATATTACACCCTCTGCTGGACCCTCTGCTGCC

GGATCACTTTGACTCGGAACGTTTGTTACAGGATGAGTTGGACCTGCTTTATCTTCATCAGACTCCAGAC

TGGAATTGAATTCCCATGACCTGATTCTATTGTAAGACAATGACTTTCCTGTTGCAGCTGAAAAGCCAAC

ACTGACCCAATCTGGCAAATGATCTCTTAGATCAACCACAAAGTTGAAACTTTGTTGTGAAACACTACCA

TCGCCATCTAAGCCTGTACGGAATGAAACACTAAGATTCTTGGAACTAGAAGCATAACTTATCCGAGCTT

CATTAGTCCACCCATTCTTAATACTATCGCTCCCATTAACCCAAGGCACAACCGCCACAGATATCATAGA

GTTGATATCAACACCTGCGTGCTCACTCATATTCTCCGGGTCACAATAGTTGCTATAAATATCAAATTCA

ACAGCAACAAATTTATTATTAGCAGAGGCGTTCACAGGCTGGCTATCATTCGTAAGTCCTAATCCTACAC

CTCCTGCACCCTCTAACGGGTTCTTAGTCCCATTCGGGACCAGAAAGAATGCGATCCCATCACCATACTT

GGCCCTCTTCTCCGAATCAATCACAAAAGTGAAACGGGTAGCGAAGTCGGTAAGGTTTCCTGAGGCCTTG

TCCCATAGATGCAATGGCTGTTTATAAGTAACTCGTCCAATGCTTGAAAGAGCTGATTGTCGATTCAAGC

TTTTAGTAATGTGGATGTCTCTGCTGGATGTGAAGGCATCTCCATCATGTTCTATCTCAGGATCATCAGG

AGTGAAGCTAATATAGCTAAAATTTAAAGTTGCAGCAAAGTGGATCATACTCAGCAGTATCAGAGATGTA

ATGACAACAGAGCATCCTGCTAGGAGATGTCTGCGAACCATGCTCGAATTGATTTTATAAAATGCGTAAT

GTAGCTGATCTTGGATCTGATGTAATGATCGGAATGTATGTTACCACAAAACGTCTGTGGAACGTTGTCT

TCAGGACTACTGGTCGTTTATTAGTTATTCGATAATTTCAAGGGTGGGTGAGTTTTATTAATCATCATAG

ACGTCATTAGTATATTAATCACCATAGACGGCTAGACGTCATTAGCTTAGAGGGTAAAGCCAACCTGTTG

CAGACAATTATTTGTGCTCGAAGTGTTTTTAACTTTATTGGGAGTTTTCTAGTTTACCGTCAGATGAATG

AAC

>MSTRG.2945.1 gene=MSTRG.2945

ATGCAAAATGCCAGATATGCTGGGGTTGTCCAGCTAAGATTATGAATGGCCACGATAGAAAATTGATAGA

ATATGATTGAAATTGAAATTGGGCAACCGAAGTTTCCAATTATGTTCTTCTTGGTTGAAACATTCTAGAT

TTCTCTCTTCCTTTCTATTTATACACACGCACACACTACCATTATTATAATGTTTTTATTCTCTTGACAT

CCAAAAGTAAAGAATCTTGCAAAAGGAAGGAAGAAATAATTCAAACAAGAACAACATAATCCAAGCTTAT

CATTCTCTTCATTTTTGATATCTTGTTCTATATTCATTGTCAAAATCCAATCTTGGGAGGTGGGTCTGTG

ATTATATCTCTTGTTTCTTTGTATTTTCAAGAAAGTGTTGTTATTTCTTCTCAATCAGATTTGTTCAAGA

TCTTTCTCATAGCTATGCAGAGTCAGCTAGTGTGCAGTGGGTGTAGGACCATTCTTCTCTATCCAAGAGG

AGCCGCAAACGTATGTTGTGCAGTTTGCAATGTAGTTACTACGGTACCACCACCTGGATTGGAAATGTCC

CAACTAATATGTGGAGGTTGCCGTACATTGCTAATGTATACTCGTGGAGCTTCGAGTGTGAGATGCTCGT

GCTGTCACACAGTGAACCTTGTGCCAGCATCGGGTCAGGTTGCTCATGTCAATTGTGGCAATTGCCGTAC

AACACTGATGTATCCATATGGAGCTCCATCAGTCAAATGCGCGGTTTGTCATTATGTGACAAATGTCAAT

ATGGGTAATGCAAGGGTCCCAATTCCTGCACTGAGACCGAGCGGGGTGGCAACATCTGCATCAACAGCCT

CTACTTCACCGGCAATACCACGTCCTCAAAGTCAAACCGTTGTCGTTCAGAACCCAATGTCAGTTGATGA

AAGTGGAAAACTGGTGAGCAATGTTGTTGTTGGAGTTACAACTTAGTCAAAAGACTCAAGACTGCACAGC

TTAGTTCCAAGACAACACAATTTCAAGATTCTGTTTAGTTTGTTAAACTGGACTGAGAGATGTGTTGTTT

GTGCCTATTATTTGTAGGAAAAACTTGATGTACATTGACAGAATAGTGATTGGAAGTTTGTCACTCTTGG

TTGTGATTAATGTATTAAATAAATATTTGCCGAGTAGCAAATTATTGCGTGTAATACAGAACGTGACAAA

TGTCCATATGATATCAAGACTCTGTATGATATGCAGATTTTTGTGTATTAATATATTGGGGTCACAGATT

CAGTCAATTTTTATATTAATCCAGAGTAAGGAATTCAGTTCTATTAAA

>MSTRG.2945.2 gene=MSTRG.2945

ATGCAAAATGCCAGATATGCTGGGGTTGTCCAGCTAAGATTATGAATGGCCACGATAGAAAATTGATAGA

ATATGATTGAAATTGAAATTGGGCAACCGAAGTTTCCAATTATGTTCTTCTTGGTTGAAACATTCTAGAT

TTCTCTCTTCCTTTCTATTTATACACACGCACACACTACCATTATTATAATGTTTTTATTCTCTTGACAT

CCAAAAGTAAAGAATCTTGCAAAAGGAAGGAAGAAATAATTCAAACAAGAACAACATAATCCAAGCTTAT

CATTCTCTTCATTTTTGATATCTTGTTCTATATTCATTGTCAAAATCCAATCTTGGGAGGTGGGTCTGTG

ATTATATCTCTTGTTTCTTTGTATTTTCAAGAAAGTGTTGTTATTTCTTCTCAATCAGATTTGTTCAAGA

TCTTTCTCATAGGTAAGAAAGTACCCTTTTCTTGTTTTGATCGTAAATTTGAATCTTTTTTATTTTGGTT

GAACTTATAATTTGGGGTTGACTTTGTTGATATTTTGTTTTAGATGTATAGTGGGTGATTATAACTGTAT

TGGGTAATTTTGATGATATTTATAGGATGAGCCGATGAGGAGATGATGAATTTATGAGTTCTGTTGTTGG

TTTCGAGTTGCAGGAATGTTAATAGTTTTGTACTATGTGTGGTAGTGTCTGAAAATATAGTTGATAGAAT

TGGTGTTTTGTTGTATGAGATTGATAAGGATTATATGCATATGCTCACGATAATAAGCCCATAAGACCAA

TCTTGGAATCTTGGATGAATAGATTATTTACAGAGCGCTTTCATATGTTTCCTCGGTGTTGTAGATAGCT

CTGTATGTTGCAATAAATACAGAAATGCTGGGTGAGCAGGATTGGGTAATGCTTGACCACAGGGCTTGTT

TGTTTTGTTGAGTCTCAAGTTACGTAGTGAGCTAAAACTTCCATGGGAAATTTACATCGTACAGATCCGA

AGTAACTAAAATATAGGTGTTTGATAGATCTTCTAATACCTGATATTAAAGATTTTTCAAGCCTATTTGT

ATTTCAGATTTGCTCTTGCCATGTTGTGTGGCGTTTTTCTCTTATAACAGGGTTCTGTTATTTGATACAT

AGTAAATTAGTGGCTTGTACAGTCATGTAGAACGATTTAAGATGTATTCCCTCAAAATTTCAGGGTATTA

ATTTGGTTATTTAATTGCAGAAATAAGCTTTAGCTATTTAACTCTCTAGTTGATCAGAATTTGTATACTT

ACAAATTCAAAACTTCCATGTCCAGATGAGTCTAGGAGAGAACTAGGAACCTCAATTTCAATTAAATGGA

ACTGCTTTTACTTGAATATTATTTTCAGAGTATATTGAATAAATGAAAGAATCTGAGGATACACTTTCAA

ATTGTTGAGAGCATGAGTCTTCTAGTACCAACTGCTTAGCTCTAGTTGTTAAAGTCATAAAATTTTCCGT

TCAAGCATTTTGTGTTGGTAGGAATTCCCAAAAAGGATAGATTACTTTTTACCATACCTCCTGCTTTTCT

TTGACAAGTGTTCTGGTTACCTAGAGTGTACCAAACGGTTTATATAGAATTGGGTTTGTTGGACCTGTAG

GCTAACAATTTTTGTTCAAGGAATCAGTGACCAAGACTCCAAGAGTTTTCTCTCTCTCTCTCTCTCTCTC

TCTCTCTCTCTCTCTCTCTCTCCCTCCCTCCCTCCCTCCCTCACTTACTCACTCATATTACTGTACTGTA

GTATAATCTCAAAATACAATTGAAATTATGTAAAGTAAAGAGCTTTAGTAGTGATTATCTATAGTAAATT

ATTCAATGGTTATTCTTCAAACTTTGAAGGATCAGAGTATACCCTTCTTCTGCAAGTCCTAATCTATGAT

CCTTAGCAATAGAGACAACAATGCATTATGTTATACAAAACATAATTGGCTCTTGCGTTGTGATTCCCAC

TGGGTTGGGTTGGTGGAGTGGGCATTGCTTCATGCGGCAGCTCAATTTGCATTTGGTGTATGTATCTTCC

AGAAGACATCTAATAGTTGCGAATGAAGAAATAGTTTTTGTGGATACATGTCTCTCTAGAATGGTCATTT

TTACAAAGTTGTGTACTTATTACACACCTGGACTTCTGTTTATAAATTGAATCCCTGTAGTTAACAATGC

CAATAGTCACCCGCAATGTATTCTATTGAAATACTTCTCGCCTGAGGTCATGTTTAGACTACAGTTCTCT

TAAATGTTACTTGTGTGGTTATAATAATTTAAAGATTTTGAAGCGAAATGTTTTCTGTGTAAAACTTGCA

AGATATTATTCCACGAAGTTAAAATTCCAAAGGAGGGTGATATAACTACATTAATGCTCTTAGTACAGTT

TAGATTAATGTGTATGATGTTTTTTTGCATTACAGCTATGCAGAGTCAGCTAGTGTGCAGTGGGTGTAGG

ACCATTCTTCTCTATCCAAGAGGAGCCGCAAACGTATGTTGTGCAGTTTGCAATGTAGTTACTACGGTAC

CACCACCTGGATTGGAAATGTCCCAACTAATATGTGGAGGTTGCCGTACATTGCTAATGTATACTCGTGG

AGCTTCGAGTGTGAGATGCTCGTGCTGTCACACAGTGAACCTTGTGCCAGCATCGGGTCAGGTTGCTCAT

GTCAATTGTGGCAATTGCCGTACAACACTGATGTATCCATATGGAGCTCCATCAGTCAAATGCGCGGTTT

GTCATTATGTGACAAATGTCAATATGGGTAATGCAAGGGTCCCAATTCCTGCACTGAGACCGAGCGGGGT

GGCAACATCTGCATCAACAGCCTCTACTTCACCGGCAATACCACGTCCTCAAAGTCAAACCGTTGTCGTT

CAGAACCCAATGTCAGTTGATGAAAGTGGAAAACTGGTGAGATATTACCACCCCTCCTTTTACTTGAATA

AAATGCCGAACTTCCCAAGGTGATCACTGACAATATGTAAATTTTGTAGGTGAGCAATGTTGTTGTTGGA

GTTACAACTTAGTCAAAAGACTCAAGACTGCACAGCTTAGTTCCAAGACAACACAATTTCAAGATTCTGT

TTAGTTTGTTAAACTGGACTGAGAGATGTGTTGTTTGTGCCTATTATTTGTAGGAAAAACTTGATGTACA

TTGACAGAATAGTGATTGGAAGTTTGTCACTCTTGGTTGTGATTAATGTATTAAATAAATATTTGCCGAG

TAGCAAATTATTGCGTGTAATACAGAACGTGACAAATGTCCATATGATATCAAGACTCTGTATGATATGC

AGATTTTTGTGTATTAATATATTGGGGTCACAGATTCAGTCAATTTTTATATTAATCCAGAGTAAGGAAT

TCAGTTCTATTAAA

>MSTRG.2945.3 gene=MSTRG.2945

ATGCAAAATGCCAGATATGCTGGGGTTGTCCAGCTAAGATTATGAATGGCCACGATAGAAAATTGATAGA

ATATGATTGAAATTGAAATTGGGCAACCGAAGTTTCCAATTATGTTCTTCTTGGTTGAAACATTCTAGAT

TTCTCTCTTCCTTTCTATTTATACACACGCACACACTACCATTATTATAATGTTTTTATTCTCTTGACAT

CCAAAAGTAAAGAATCTTGCAAAAGGAAGGAAGAAATAATTCAAACAAGAACAACATAATCCAAGCTTAT

CATTCTCTTCATTTTTGATATCTTGTTCTATATTCATTGTCAAAATCCAATCTTGGGAGCTATGCAGAGT

CAGCTAGTGTGCAGTGGGTGTAGGACCATTCTTCTCTATCCAAGAGGAGCCGCAAACGTATGTTGTGCAG

TTTGCAATGTAGTTACTACGGTACCACCACCTGGATTGGAAATGTCCCAACTAATATGTGGAGGTTGCCG

TACATTGCTAATGTATACTCGTGGAGCTTCGAGTGTGAGATGCTCGTGCTGTCACACAGTGAACCTTGTG

CCAGCATCGGGTCAGGTTGCTCATGTCAATTGTGGCAATTGCCGTACAACACTGATGTATCCATATGGAG

CTCCATCAGTCAAATGCGCGGTTTGTCATTATGTGACAAATGTCAATATGGGTAATGCAAGGGTCCCAAT

TCCTGCACTGAGACCGAGCGGGGTGGCAACATCTGCATCAACAGCCTCTACTTCACCGGCAATACCACGT

CCTCAAAGTCAAACCGTTGTCGTTCAGAACCCAATGTCAGTTGATGAAAGTGGAAAACTGGTGAGCAATG

TTGTTGTTGGAGTTACAACTTAGTCAAAAGACTCAAGACTGCACAGCTTAGTTCCAAGACAACACAATTT

CAAGATTCTGTTTAGTTTGTTAAACTGGACTGAGAGATGTGTTGTTTGTGCCTATTATTTGTAGGAAAAA

CTTGATGTACATTGACAGAATAGTGATTGGAAGTTTGTCACTCTTGGTTGTGATTAATGTATTAAATAAA

TATTTGCCGAGTAGCAAATTATTGCGTGTAATACAGAACGTGACAAATGTCCATATGATATCAAGACTCT

GTATGATATGCAGATTTTTGTGTATTAATATATTGGGGTCACAGATTCAGTCAATTTTTATATTAATCCA

GA

>MSTRG.2947.1 gene=MSTRG.2947

CAAAATTTGAAGAGGCAAATTTTTTGAGGACATAAACCAGTATCCACACAAGTCCAGATTTTTGAACATT

CAGAGTATCAAACACGCTTTATTGACTCGTTCTCTCCCTCGCCTCGTCTCTCTTCTCAATCTCTCCGTTC

TCTCTCACTCTCTCCGTCTGTCTCTCTCTTCATCTCTTTAGATAAAAATGAGGTCGATTTCACTCGTTCT

CGTGGAATTGGAGCAAGCTCTCACAAATCACCTGGATTTTGAATTAAATTTAGGGTTTATTCAATTGATT

TGTTCTTGGATCAAAATTGGGGGTTTTCATGCCTTTGATTCAGGGGTTTTTTGGTTATATTGTTTTTTCT

CTTAGATCTTTAACTCTGTCTCTAATTTGTTTTATATCTTTGTTCTGTTTTAGGCTGGCTTTGGGTTCCT

TAGGGATTGACTGATTTAAATCTTTGATGTGCGCTTCTTCAGCTTAAGGTTAGGTTGTGGATGTTGGATT

TGCAGTTAGAAAATTCAAGAAAGGCGAAAAAGTTATGGCCATGCTTGGAACAACGACTGGAGGTGGATTA

GTTGAGTATGCAGTAGCAAAGTAAGAAGTAACAGTTCTAAGGCCACTTGAACTATCAGCTGGTGAATGTG

CCGCATTAGGCACTCAAGGGTTAACCACTTACCAATGCCACATCACAGCGGGAGTTGTTCTATGAAAAGA

CGCCAACCTATGAACATACTAACTGTTGCAGCTTGAGGCGGTGTTGGTCACTACGCAATTCAGCTCGCAA

AGCTAAGGAACACTCATGTCACTATAACAGGTGGCTCTCGCAACATTGATTGTGTGAAGAAGACCTAAAG

GATGAGTTGGATTTGTTAGGTAGTTTAGTTTGGTTGGATTTGTTTTGAACTTTGGTTTGTATGAATGATT

AGTTAATTTTGCAGTTGATATAGTAAACTTTTGCATAGTAGTTTAATGTTGCATTAACACATATATATCA

GATATTTAGTACAAATATATGTTGTAGAAAGTAGCCCTTGTGGGC

>MSTRG.2947.2 gene=MSTRG.2947

CACAAGTCCAGATTTTTGAACATTCAGAGTATCAAACACGCTTTATTGACTCGTTCTCTCCCTCGCCTCG

TCTCTCTTCTCAATCTCTCCGTTCTCTCTCACTCTCTCCGTCTGTCTCTCTCTTCATCTCTTTAGATAAA

AATGAGGCTGGCTTTGGGTTCCTTAGGGATTGACTGATTTAAATCTTTGATGTGCGCTTCTTCAGCTTAA

GGTTAGGTTGTGGATGTTGGATTTGCAGTTAGAAAATTCAAGAAAGGCGAAAAAGTTATGGCCATGCTTG

GAACAACGACTGGAGGTGGATTAGTTGAGTATGCAGTAGCAAAGTAAGAAGTAACAGTTCTAAGGCCACT

TGAACTATCAGCTGGTGAATGTGCCGCATTAGGCACTCAAGGGTTAACCACTTACCAATGCCACATCACA

GCGGGAGTTGTTCTATGAAAAGACGCCAACCTATGAACATACTAACTGTTGCAGCTTGAGGCGGTGTTGG

TCACTACGCAATTCAGCTCGCAAAGCTAAGGAACACTCATGTCACTATAACAGGTGGCTCTCGCAACATT

GATTGTGTGAAGAAGACCTAAAGGATGAGTTGGATTTGTTAGGTAGTTTAGTTTGGTTGGATTTGTTTTG

AACTTTGGTTTG

>MSTRG.2950.1 gene=MSTRG.2950

GAGAGTAAAATCAAAGCAACACTAATCCAATCGGCAATTCGGCATTATATCCCTTACCAAACGCGGCCTT

ATTTAACTCACCCAGTCGAATTGTTGCGTCGAGGACGAAGATTGAAGTGATCAGCTATGGCCAGTTCCGA

TCCCGACAAGCTCATCACTAAAGCCGATAAACTGACAAAGCTCAGCATGACAAGATGGAGTGCTGATTGG

AGGGGCGCTACTAGTTTATATGAACAGGCAGCTCTTGGGTTTAGACTTGCCAAAAAGTATGAGAAAGCAA

AAGTAGCATTTGAGAAAGCTTCAAAAGGACAGGAAATGCTGTCATCACCTTGGGATGCTGCTAAGCATAT

GGAGTCTGCTGCTGCTCTGGCAAAGGAACTGGGCAACTGGAGAGAAGTTGCTGATTTCTATAGACAGGCA

TCTATGTTATACAGTGAGTGTGGGAGGGCACAACCTGCCTCGGATGCACTTGCAAAAGGTGCTCGTGCAC

TCGAAGAAGCTGCACCTGATGAAGCTGTTACTCTGTATACGGAGGCTTGTGATACCCTTGAAGAGGATGG

CAAAGAACAAATGGCCTTTGATCTATATCGTGCTGCTACCAGTGTTTATATAAAGCTTGAGAAGTACTTG

GATGCTGCAACTTTTCTTATAAGGTGGGGCGTGTCAGCTGACAAGTGCAATGCAACACATAGCCAATGCA

AGGCATATCTCAGTGCAATTATTGTCTACCTGTACGCTAATGACTTCGATCAAGCGCAGAAGTGCTACAA

TGATTGTAGTCAGATTGATGCTTTCTTGAGTAGTGATCAAAGTCGATGTGCAGCTAAACTTCTTTCTGCA

TACACTGATGGTGATGTTGAAGAAATTAAGCGCGTAGCTCAGTCGAGCACTATATCCAATCTTGACCATG

CTATCATTAGGCTTGCAAGAAAGCTACCAACAGGCGACGTGAGCGGACTCAAAAGTGAGACCGGAAAGGA

GCAAGAAGAGCCACTAGATGAAGACGATCTCACTTAACTTCTCATGTTGGTAGTATTGACAACCAACCTT

TTTCTGGAGAAATCTGTTTACTTGGACGAAATACAAAATATGAAGGCGTGTGTACAAAAATGCTAGTTGA

CAGAATTGCATACTCAATGAGTTAATGGCAATATTGTTTAACATTTATATGCAGTTGCAAGTTTGACATG

AAAGGCTTTGCATAAATTAATAAATTGTTACACATTTATTTATTTGTTCATGCT

>MSTRG.2954.1 gene=MSTRG.2954

CCGAAGAGTCCAATTTTTCCAAAGGTTACAACAGGGCTGTTAGCTGTGGCATTAATAGAAATGGTCAAGA

CTTGTCTCTCTCCGCTAGCAATGAAAAACTGCATTGGTTCTGCCTTCAAAGAAGCTCCCAAGGGAGCAGT

CCATCCCACTCTGTACGTCTCATTCCTGGCGATGTTCATTACCGTTCTTTGGACTGTTCTTGACTGATTA

AGTTTTGCAATTGTGATAGAAGGCAAGTTTAGGTCTGCGCCATTAATGGTAGATGTTTTGCAGCTCATAC

CAGTGAAATTTAGGACTACTGGTGATGACCCGTTTATGCTGCAGAGGAATGACATGTAGTCATCATAAGC

TGCAATACAGAAAGCAGGTCCAGAGTTGGTTAACGTGCCAGCAGATACATCATACAAGTGACCGAGTGAC

AGTAATTTTATATACTTGAGCACAAATTTTAGAGTCCCGTTCAAATACTTTATTTGCATGATCACATCCC

AGTTAGCCAATGCATTTTTCATGTATCTATCACTACGTTTTACTTTTATCATGATATTCAACTTTCGTTT

GGTATAGTATCTATCACTAGGTTTTCCTTGTTGAGTAAGGCCTCAAAAGCCATAAAACATTCTTCATGTA

CTAGCTAACATGTG

>MSTRG.2958.1 gene=MSTRG.2958

CTTTGGTTTACCGCCTCATTGAGCTAGCCTTGATTTTACCTGTCGCGACAGCTACTGTTGAGAGAGAGTT

TTCAGCAATACTTAAGACTTATTTGCGTAACAGGATGGGGGATGAGTGGTTGAACGATAGCATGGTGGTT

TATATCGAGAAGGCCATATTTAAAGACATAGAAGATGAAGTTATCTTGCAACGCTATCAAAGTATGCAGA

ATCGAAGAATTCAGTTGTCTTCCATTCCTTCCTCCAAAAAAGGCGGTAAAAATGTCACTGGTTAACCAAT

AAAGGTATGCATATTTACTTTTCAATATTAATCCTGGTTCCGCCACTG

>MSTRG.2961.1 gene=MSTRG.2961

TCATAACCACTACCACCTCTATGCCGCCCGCCCCCTCCATAACCACCACCTCCATCTCCACCTCCACCTC

CATTATTTCTCTAACAACACAACCTCCACCTCCATCTTCACTAGCCCCATCTTAATCGTTGTTTTAACCT

CCTTCAGCCCCGTTCATACTTCTTTCTCCACCTCGACTCAACTTCAAACGCGAAGCCGCCACTATTCCGG

CAACGCTCCAACCCCCTACTTCAAGCCGCCCAAACCTCCGCTACAATCATCTTTCCTTCATCTCCACCTT

CACCTCCACCATCTCCACCACTATCACCATCATCTGAATCGAAAACGTGAACAGATCTGGAGATTCTAAG

CTGGAACATATCTGGAGATTAAGTTTTCACTAATTCGTTCAACACAGATCACTAAATCCTAAAGAGAATA

TCACTAAATTGTACTGAGAATATCACTAACTTGTATTGGTTTCTAGTTTTTATTTTAAAATTGGTTTCTA

TTTGATCATGAG

>MSTRG.2963.1 gene=MSTRG.2963

AGATGCTCTTATGCTCCTCTTCTTATCTTCACTCGAGAAGCAGCAAGCTGAGCACTTAGAATCATAATCT

TTTCTTCAACAATAAAGTTCCCATCGGCTCGAAAAGCCCTCAAGATGTCGAATGCCAACGTGAATAGCTG

GAGTTGTACATGTTGCTTCAACTATATCTGCGAAACTTGCTGTACTTGTTGCGCAATCTGTTCAGCAGAA

GCATGCCTCAGCTGTTTCTAGTAGTGAACCAGTTGAGACATGGAACTGAAACTTTAAGAACTGGAGTTAT

AACATCAGAGCAACAGGGACACAAGATGAGCTCTAAATTTGTTCATGTTCTGAGACTGTTATCAGATTTA

TCTTGTATAGTCCTGGCTTAATATCTATGCCTGATTGTATTTGATATTTATATATGTTCGCGTTTTTAAT

TACATTGATGTTTAGATTTTGGGAGCAAAATCAAACAAATTTTAGAGAAATATCTGAGAGAGTTGTATCC

TAGCCATCCGAGAGGGAAACCTCTCCCAGACTTGATATTCCTCACAATCTAAAATATTTTCATGTTCTCT

AAGATAACTTCAAACATCTCAAATATTTTTCTCCGTTTCTAATATCCGACCTAATTTTTGGTGCTTTGAC

TGTGTAGTTAAAATTGTTATTTTTGAAGATTTGTTTGTCATTTACAGGGACAGAGTGATAGACATAGGCT

GTCTGTTGCAAAAGACGGTGGGAGTAAATTGTTATCAAGGCCTTGCCAATACAATACTACAAAACATGTC

ATATAAGCCCTCTTCCTGATACCCGGAGATTTTAAAAAAATTGAACACTTAACATGGTGGTATCAGACGG

ACGAGTTCGATGCACCTCGAGATTTTGAGAGAATCGATCACCTAATAATAAACGACTTGGAATAATGTTG

AGCTAGATAGCTTGCACTAGAATAATTTCCTTTTGAAAACAACATTAAGCAGCCTTGTGTCAAATCCCAG

GCCAGTCAGTCACTGGTCATACATAATACTGGTAGCAGAGCAAGATATATAGGATTTTGTGAACTTGGTA

ACACTAGACACTAGCTAGATTCTTTTTTGGGAGGCCTTCCTCTCCCTCTTTTCATATCAACTTGAATAAT

ACCACTGCTACCTTCTCTTCTCACATATGTGAAAATACAGGTGTCCCCCCTTTTGAGTCTGTGTGCCACA

AAGAGACGAGTCCATCCAACTGTCAATGCTGCTTTCT

>MSTRG.2964.1 gene=MSTRG.2964

CAAACCAATATTTGCAAACGTGGATACTGCATATCCAATCACCTTAGCTATACAAGTCCCACACAGTTGG

CACTGGTTAGAACAATTTATTGATGTCATCAATGGCCTTAATTTTATGCTTCATTGTGTGGGAATCTCTT

GCCTAAATTATTTGACTTTATCATATATGCTTAGAGCATCTCCAACCCTAAGCACCTCTTAGGTATAAAA

TCTACGTGGATATATATTATACTTAACATCTAAATTTTAGACTACTCCAACGGAGTGTCCTCTTAGCTAT

AAATTTAGCTTCCGATAAACAAGTGCGCTACATTTGCAGATGATTTTGCTGTTATCTAAAAAGCCCACGT

CATCTCCCGCTTGAAATATTCCAGATGTAACCCTAATATTGGTCCAAAACATAATATCTTTTATTTGTAA

TATATAAC

>MSTRG.2967.1 gene=MSTRG.2967

ATTTTCACCAGTTTCTTTTCACAAATACCGAGAGAAACCGAGCGGTGTTCTTCGTGTTCTTCATAATCAA

ACGAGGATTTGGAGGTGTTATTGGAATCCGATGGAGGTGTACGAATAGTCAAAACGAAGCTATCGACGCG

TAGAATCATGTTTAATCACCCGTTTGTATGCAGATTTATCAGAAAGGCTTAAAGATGAGATGATTGATTG

AGATTGTGAGTAAGGCTAATGCTAGGCGAGGAGACAGAGTTGAGGATCCAGTGGTCGAGGAGTGTTCAGA

AAAGTTGATGAGATTGGTGCAAGCTGGA

>MSTRG.2974.1 gene=MSTRG.2974

CAAAGTTTTGAAGACAATATAAAAATGGAGGAAAAAATTGTTGTAAATTAAATAGAGCGATGATCGAAAA

AGAAGTATGATAACGACACAATGAAAATGACGGATTTGATAAGATATTCATATGCCAACGACTTGCTAAT

TTCCCATTACTCGCCTGCACAGCAGATTGGTCACACACAACCCATAATCAAATCCAAGAAAATACACACA

AACACTCCAATATTCAGTGAATTGGGTCAGTAAGTGATTGATGCAATGAATGATAAGCGAAGCAATGGGG

GAATTACAAATGGGGTTGATGATGAATTGGGTGTGATTTTGGGTTTAGATGGTGGCACTACCTCTACTAT

TTGTGTTTGTATTCCTTTTTCCGGTCAACTCGCCGACCCAGTTCCGGTTCTTGCCCGGGCAGCTGCTGGC

TGCTCTAATCACAACAGTGTTGGAGAAACAGCAGCAAGAGATACATTGGTACAAGTTATGAGTGAGGCCC

TTTTGAAATCAGGATCAAAGAAATCAGCTGTTCGAGCTGTTTGTTTAGGTGTGTCTGGTGTCAATCATCC

AGACGATCAAGAAAGAATATTAAATTGGTTGAGGAACATATTTCCAAGCCATGTTAAGCTATATGTTCAG

AATGATGCCGTGGCAGCTCTTGCAAGCGGGACTATGGGAAAACTTCATGGCTGCGTTCTTATAGCAGGCA

CAGGGACCATTTCTTATGGATTCACAGAAGATGGCCGAGAAGCTCGTGCTGCTGGTGCGGGCCCTATATT

GGGTGATTGGGGCAGTGGATATGGAATTGCGGCAAAAGCCTTGACTGCAGTTATTAGGGCTCATGATGGC

CGTGATTCACAGACAATGCTCACTGATAGGATTTTACAGACTCTTGGTCTTCCTTCTCCAGATGAACTCA

TAGGGTGGACCTATGCAGATCCATCTTGGGCAAGAATTGCAGCGCTTGTTCCAGTAGTTGTATCTTGCGC

TGAAGCTGGTGATGAGATTGCAAATCAAATTCTAAATGACGCAGTGCAAGAACTGGCTTCAAGCGTTATT

GCTGTTGTCAAGCGACTTGGCATGGGCGGGCAAGATGGGAATTCGTATTTTCCTATTGTAATGGTTGGCG

GTGTCCTGGAAGCTAACAAAAGGTGGGACATAGGTAAAGAAGTTATCAACTGTGTATTGAGGACCTACCC

TGGTGCAACTCCAGTTAGACCAAAGGTTGATCCTGCTGTTGGGGCAGCTCTACTGGCTTGGAATTACTTA

ACGAAAGAAGCAAATGGGGAATACCATAGACAGCCATAGTGGAATACGATAATAGTGTATAGTTACATCG

GACGAAATTTAAGATTGATCTGCAGTACAGATAATGTTAGGTGCTTGCGGAATATACTCAAAATGTGTAT

TTGGTGTGTAAGCACAGGGAAGTAATACACAAGCCAAAACGTGGTGTTTTGAGGATTAGTTGAAGGCAAC

ATCTATCAACTTGTTTGTGAATCTCTCGGTGCATAACTTTTTGGGCAGGTATCATCACTGGTAACTTGTA

AGCTTATGAGCAGTAACACAGCCATAATGCTACACTCATCTAACATCACAGTGCATTGCAATTCCATTTC

CGCGCTGCTCATGATTCCTCAGTGTTTTCCATTCCTAAACCTATTATCCCATGTTGCATGATAGTGGTTG

GTTTTTCTTTCCCAGTATCTGCTAGTTTATTGACCACTAAAATTAATGCAGATCCATGAAAAACAATTAA

ATTCTGGAGCGTCTGGTATTTTCTGTCCAGTTTTTTTAATTCAGGGGAATAAGCTACTAGTATGGGAC

>MSTRG.2975.1 gene=MSTRG.2975

CACATACCAGGTGACACAGAGAGTGTTGTGAACTGTACCCTTGTGGGCTTGTGGCTAGCCGAACGCGAGT

ACCTTCTGAACAACCTTGCAAACAGCAAATCAGATTTGGCCAGATGCTGATGAGTAGGCATTTTTGTTAT

TAGCAGATCTAGAATTAATACAATTTGGAGGTGTATTATATTGGTATAGCTTATAGGCAGATCTTTCATT

TTTTTTAGTTTTCAGACAGCTAAGCAGCACAAGTCATATTTAGCGATATAGTTCTCCATTTTAATGCCTG

TTTATGTGTCTCTCTGTAGCCTTCAATGTAAAGTTACTCCTCAGTGCAGTGA

>MSTRG.2977.1 gene=MSTRG.2977

GTAGAAAACACACAAAATATAATCTCAATGCAATGTATTGATAATCAAGCAACCGACAGAGAGATACTCG

TAAATATATTTCGTTACAAAATCAGATTACAATATTTATATACAACAGTCGTCTCAAAATTAATGGATGC

AATCATCTGGATTAATCGGAAATTAATTAGAAGACGGAGATTGTTTCGATTGCATTTGAAACCTAAAAAT

GAAAACAAAAGCAGCAACAGATAACAAAGCAATACCCGTAACTCCACAGACTAAACTCGCCACCAGCCAC

GCCGTGTTGACTCTTCCACCGCCGCTTCCACTCCTTGCGCCGTCGCTTCTCCGCCACGTCACGTCCGTCA

AGTTCTTCGCCTCCGGCAACGGCACCTTCGACTTAGTTCCTTCAAAAAACGACGTCGCATCAGGAGACTC

CGTCACAGTCGATGTGTTCTGGTAGGTGAGGCCATGATCCGACGGCCGGAGCTCTCTGCCGGCCGATTGT

ACGGACAAGATGAACAGGAGTATCAGCACTGCGGTGGCGCAGGTATAGTGAAAGTTCATCTTTGTTTCGA

GATTTTTGTGTTGTTTTTAGAGTGAGTTGTGAATATATACGTACAGTTTTTTGCGTGTATGGATGGAAAA

GATGCAGAGATGAGAGTGCATGCGTATATATAGTGGTATAGAGACAGTGTTTTATGGGTGTGATTGTGTG

AAGTGATAATTGAGGTTGAAGTTCTGTTAGAAAAAAAA

>MSTRG.2980.1 gene=MSTRG.2980

TGTAACATCAATTGATACAGACACATTGATAAATAAAAAATCAATAAACTTCTACATGCCATTGTTTGTT

CTTTTTCAGCTGCGAGAGCTTTGCATCCCAGCTCTCCCCTCTCTGTTCTGCGACCCTCTTCAGGGTGTCT

TGAATGCCAGGCATCATCCCCTTCAGCCCACAAAAATAAATGTGTGCCCCTCCATCCAAAAGTTTAAAGA

TTTCATCGCTGTATTCTTCTATCTTATCCTGAACATACATCTTTCCACCATTCTTGTTCTTTTGTTCTCG

GC

>MSTRG.2981.1 gene=MSTRG.2981

TTCTTTTCCTGTCTCAGGATCATAATAAACAGCTCGTCGTACACACAAACTGGCAGTCTTTCCATCAAAA

GAGTCTCCATACCTTGAGGAAGCAATTGAATAGAGACGTACATTCTGGGGATTCCCTGGCTTTTTTGGGT

TGTCACCTGGAGGAATGACACCATAGCTTTGTCCTTCCCAATAGGGTACATTGCCACCATGATCTATTAC

AATGTGACAAGTCTCTCCTGGAGCT

>MSTRG.2987.1 gene=MSTRG.2987

TAAACATAGTTTAAGTATCTACGTCGACCATCCTGACTATCCATCGTTGCTTAACATGATCTTGTATACT

TGTATCCTTGCTTAGTTGCACATTGTGCCCTGGCTCTTTATTACCTTGCTTCATTAAGCAAAAATGCCTA

TAATCATAATGGAGATATCAAAATCAAACTATGGGTTCTCCTGCCTTGGTAATTCTAAGAGATGG

>MSTRG.2988.1 gene=MSTRG.2988

AGATATTATTCTTCATTTCAAAACCTGACGTCTTCAACAGCTAAATTCTGAAACATATATCATATATTTG

GGGAGGCAAAGATAGAGGATTTGAGCTCTTAGCTACAAACACAAGCTGCCCAACAGTTCTAGAAAGACCT

GAGGTTCGCAAGTCTGCTCCTACTCCCGCGAGATTTCGTGATCAGCTTTCCATCTCAAGGGCACAATGAC

GAGAAGAATGGCAGGAGAACAATTGTAGAGAATGCC

>MSTRG.2993.2 gene=MSTRG.2993

AAATTCTGTATAATTATTGTCCAAACACTTAAAAAACGAAGGATGTGTAGATTTTTTTTACTCCGTATGC

ATTCATGGGAGAGGAGAGGCAGTGTTTAACCTGATTGCGAACACCTTAATAGGGTGCAAGAGTGGGTATT

TAAATTGTTGGAGGGATTACTTTGTATTAGTACGGCTTTTCAAGGGTCATGTATCAGAAACTTCTACCCT

CTATAGCTTTTTGGGCTACTTGTCACTTCTTGGAGTAGTGGGAATGGTCATTGCCTTCTCATTAGGAATG

GGGGCTATCTCATGGGTTATAAATTATAATGTTTGAGATTTTTCCAGTCAATAATAAAGGCCTTGCTGGA

AGTATACCAACTCTAGCCAACTGTTTCATTGCATGGTTAGTTACCCGGAGTGTGCCGTTGCTGTTGTCTT

GGACTTGGAGTACCGGAGGTTTTACTTATTTTCTTCACCTTTGAAGATGACAAATGGTTATTATGTATCT

TAGTTTTGTTCTTCTTGTCATTATTTTGTTACCTATTGCAATGTTTGGCATCTTCTACAGGAACTTTTCT

ATACTTGCTTATGTGTGCTTTCACCTTGGCATTTGCCACCGGTTTGGTTCCTGAAACTAAGGGGGAAAAC

TTTGGTGGAGATACAAGGGTCCTTCAGTTGAGATTCTTTTCTCTCTATTGCATATCTTTATCCTTAAATT

ATATTAATTGTATCTATAATATTTGCTGTGATTAAAATTATTGTATTATGTATTGAATTATATGTATAGA

AACAGATGTGATCTTTGCTATGTATTATTCGCTTACAATTATCGGATGTATTGAACAATATGTCCAGAAA

CATTGTAATATTTGCTGTG

>MSTRG.2993.1 gene=MSTRG.2993

AAATTCTGTATAATTATTGTCCAAACACTTAAAAAACGAAGGATGTGTAGATTTTTTTTACTCCGTATGC

ATTCATGGGAGAGGAGAGGCAGTGTTTAACCTGATTGCGAACACCTTAATAGGGTGCAAGAGTGGGTATT

TAAATTGTTGGAGGGATTACTTTGTATTAGTACGGCTTTTCAAGGTAGTTCAAGGGCGATCAATCCATAT

CTACATCAATTTCTTAGGCATTTGGGGTCATGTATCAGAAACTTCTACCCTCTATAGCTTTTTGGGCTAC

TTGTCACTTCTTGGAGTAGTGGGAATGGTCATTGCCTTCTCATTAGGAATGGGGGCTATCTCATGGGTTA

TAAATTATAATGTTTGAGATTTTTCCAGTCAATAATAAAGGCCTTGCTGGAAGTATACCAACTCTAGCCA

ACTGTTTCATTGCATGGTTAGTTACCCGGAGTGTGCCGTTGCTGTTGTCTTGGACTTGGAGTACCGGAGG

AACTTTTCTATACTTGCTTATGTGTGCTTTCACCTTGGCATTTGCCACCGGTTTGGTTCCTGAAACTAAG

GGGGAAAACTTTGGTGGAGATACAAGGGTCCTTCACTGACATCTTGAATGTGAAGCAAACACTGAGGGAT

GATGGCAACAGCAGTCTTGTTTCATCTGGTGGTGATTTCGAGCTTGGATTCTTCAGCCCCGGAAGCTCCA

AGAATCGATATCTTGGCATATGGTTCAAGAAGGTCTCCAACAGGACTGTTGTTTGGGTTGCTAACAGAGA

GGCTCCACTTCTTGACACTTCCGGGGTGCTCAGGATTGGTGCCAATGGAAATTTGGAACTTGTGAATGGT

AGAAACAGCACAATGTGGTTTTCTAATTCGACAAAATCTGCGAAGAATGAGCCTGTGGTGGCACAACTAA

TGGATTCCTCAAACCTTGTGATCAGGCATGAAAATGACACTGACCCTGAGAAGTATCTGTGGCAAAGTGC

TGATCACCCTACTGATCATATTTTGTCAGGCATGAAACTTGGATGGGATTTGAGTAGAGGGCTGGAACGA

TATCTTACATCGTGGAAAAGTGTTAGTGATCCTTCACCTGGGAATTACACTTTCAGAATGTACCTTGAAG

GATATCCACAGATGAAAGTGGCTGATGGTTCTAGTTCTACCACAAGTTTCAGATTTGGACCATGGGATGG

CATTCAGTTTAGTGGTGTGTCGTTTAATATCCCGGATCTCAATTTTGACTTGAAACTTATTGTGACGGAT

ACTGAAATTTATTCCAAGTTTGATGTGAACAATAATTCCAGGCCTGTGAGGTTTGATCTGAATCCTGAAG

GACTAATTGAGCTTCTAACTTGGTTTGACAATTTTAAGAATTGGACATTCAGTCAGTACATTAGTGTGCT

TTATGATCAATGTGACCCTTATGGACTTTGTGGGGCATTTGGTAGTTGTGACATTACTCGAGCTCAATCA

GGACAAAGTGCTTGTGAATGTTTAGGAGGATTCGTGCCTCGGTCTTTGGAAAAATGGAAACTTGCAGATT

GGTCCGGTGGATGTGTTCGTTCAGCTCCATTAACTTGTGGAAGTGGAGAAGAATTCCAGAAGATTCCGGG

CCTTAAATTGCCTGATACTCAGACTTCGTGGTTCAACCGAAGCATGAGCCTTGCTGATTGTGAGACCGAG

TGCAGAAAGAACTGTTCTTGTACAGCTTACTCAAATGTGGATATTAGACAAGGCGGCTCTGGATGTATGC

TATGGTTTGGTGATCTCATTGATATGAGAAATTACGCAGACAATGGCCTTCCTCTATATGTGAGAACTGC

TTCTCAACCAGCTGTGAATGGGAGCTCAAGTAAAAAGGGAGGAGTGAAGATCATTGCCATTATCATTCCT

ACTATAATAGGATTGTTTGTACTGCTAGGCCTGATTCTGATATATGTATACAGGAAGAGAAAGCTGAGTA

GTAAAGGATTAGCAACTCTGAAACCTACTAATGATGGTGGAAATGAAGATCAGGGTCAAGACCTGGAATT

GCCATTGGTTGAATTTGAAGAAATTGCCAAGGCTACGAATAACTTTTCCCTAAACAATAAGCTCGGAGAG

GGTGGTTTTGGCCCTGTCTACAAGGGAACTTTGGTTGATGGACAGGAAATAGCTGTAAAGAGACTCTCCA

AGACTTCAAACCAAGGACTCCATGAGTTTAAGAATGAAGTGAAATGCATTGCCGAACTGCAACACAGAAA

TCTTGTGAGGCTTTTAGGATGTAGCATTGAGGATGGAGAAATGTTGTTGATCTACGAATACATGTCCAAC

AAAAGTTTAGATGTCTTCATTTTCGACAAAGAGCTAAGCATGTCACTGGATTGGCCAAAGCGCTACAATA

TCATTAACGGCATTGCCAAGGGCCTACTGTATCTCCACCAGGACTCGAGACTAAGAATCATTCACAGAGA

TCTCAAAGCCGGTAACATTCTCCTCGATCATGAGATGAGTCCTAAAATTTCAGATTTTGGCCTGGCAAGA

AGTTTTGCAGACAACGAAAGTGCAGCCAACACCATTCGCGTAGTTGGAACATACGGCTACATGGCCCCAG

AGTATGCTATCGAGGGACTATTCTCTGTCAAATCCGATTCCTACAGTTTTGGCGTCCTGGTCTTGGAGAT

AGTAACCGGGAAAAAAATAAGACAATTTTACAATTCCAAGGACAACCTTAACCTCCTCGGCCATGCATGG

AAACTTTTCAATGATGGCGAATCCTTAGACCTTGTCGACGAGGCAATGTTGGGATCATGCAGTTCTTCAG

AAGTTTTGCGAGCTATTCACATCGGATTGTTGTGTGTGCAGCCATATCCGCAAGACAGGCCAAACATGAC

ATACGTTGTGTCGATGTTGGATAGTGAAAATGAGCTTCCTAAGCCAAAACAGCCAGGGTTTTTCACTGAT

AACAGAGTGCAACAGGGCTCAGAGTATCCGTCTCACCATAGTGAGTCAAGTGTAACAAATAATGATCTGA

GTATTACAGTGTTGGAGCCTAGGTAGTAATTGTACATTGTTTGGCACCATTTTTTTAAGCATATTATTTT

CCTTCTGAGATCCTGTTATGATCCGATATTTCTAAAATATTGGATGCTCCACGAGGATGAAAATTTTGTA

TGACATAATAATTTTTGAGATCAAATACA

>MSTRG.2993.3 gene=MSTRG.2993

AAACGAAGGATGTGTAGATTTTTTTTACTCCGTATGCATTCATGGGAGAGGAGAGGCAGTGTTTAACCTG

ATTGCGAACACCTTAATAGGGTGCAAGAGTGGGTATTTAAATTGTTGGAGGGATTACTTTGTATTAGTAC

GGCTTTTCAAGGTAGTTCAAGGGCGATCAATCCATATCTACATCAATTTCTTAGGCATTTGGGGTCATGT

ATCAGAAACTTCTACCCTCTATAGCTTTTTGGGCTACTTGTCACTTCTTGGAGTAGTGGGAATGGTCATT

GCCTTCTCATTAGGAATGGGGGCTATCTCATGGGTTATAAATTATAATGTTTGAGATTTTTCCAGTCAAT

AATAAAGGCCTTGCTGGAAGTATACCAACTCTAGCCAACTGTTTCATTGCATGGTTAGTTACCCGGAGTG

TGCCGTTGCTGTTGTCTTGGACTTGGAGTACCGGAGGTTTTACTTATTTTCTTCACCTTTGAAGATGACA

AATGGTTATTATGTATCTTAGTTTTGTTCTTCTTGTCATTATTTTGTTACCTATTGCAATGTTTGGCATC

TTCTACAGGAACTTTTCTATACTTGCTTATGTGTGCTTTCACCTTGGCATTTGCCACCGGTTTGGTTCCT

GAAACTAAGGGGGAAAACTTTGGTGGAGATACAAGGGTCCTTCACTGACATCTTGAATGTGAAGCAAACA

CTGAGGGATGATGGCAACAGCAGTCTTGTTTCATCTGGTGGTGATTTCGAGCTTGGATTCTTCAGCCCCG

GAAGCTCCAAGAATCGATATCTTGGCATATGGTTCAAGAAGGTCTCCAACAGGACTGTTGTTTGGGTTGC

TAACAGAGAGGCTCCACTTCTTGACACTTCCGGGGTGCTCAGGATTGGTGCCAATGGAAATTTGGAACTT

GTGAATGGTAGAAACAGCACAATGTGGTTTTCTAATTCGACAAAATCTGCGAAGAATGAGCCTGTGGTGG

CACAACTAATGGATTCCTCAAACCTTGTGATCAGGCATGAAAATGACACTGACCCTGAGAAGTATCTGTG

GCAAAGTGCTGATCACCCTACTGATCATATTTTGTCAGGCATGAAACTTGGATGGGATTTGAGTAGAGGG

CTGGAACGATATCTTACATCGTGGAAAAGTGTTAGTGATCCTTCACCTGGGAATTACACTTTCAGAATGT

ACCTTGAAGGATATCCACAGATGAAAGTGGCTGATGGTTCTAGTTCTACCACAAGTTTCAGATTTGGACC

ATGGGATGGCATTCAGTTTAGTGGTGTGTCGTTTAATATCCCGGATCTCAATTTTGACTTGAAACTTATT

GTGACGGATACTGAAATTTATTCCAAGTTTGATGTGAACAATAATTCCAGGCCTGTGAGGTTTGATCTGA

ATCCTGAAGGACTAATTGAGCTTCTAACTTGGTTTGACAATTTTAAGAATTGGACATTCAGTCAGTACAT

TAGTGTGCTTTATGATCAATGTGACCCTTATGGACTTTGTGGGGCATTTGGTAGTTGTGACATTACTCGA

GCTCAATCAGGACAAAGTGCTTGTGAATGTTTAGGAGGATTCGTGCCTCGGTCTTTGGAAAAATGGAAAC

TTGCAGATTGGTCCGGTGGATGTGTTCGTTCAGCTCCATTAACTTGTGGAAGTGGAGAAGAATTCCAGAA

GATTCCGGGCCTTAAATTGCCTGATACTCAGACTTCGTGGTTCAACCGAAGCATGAGCCTTGCTGATTGT

GAGACCGAGTGCAGAAAGAACTGTTCTTGTACAGCTTACTCAAATGTGGATATTAGACAAGGCGGCTCTG

GATGTATGCTATGGTTTGGTGATCTCATTGATATGAGAAATTACGCAGACAATGGCCTTCCTCTATATGT

GAGAACTGCTTCTCAACCAGCTGTGAATGGGAGCTCAAGTAAAAAGGGAGGAGTGAAGATCATTGCCATT

ATCATTCCTACTATAATAGGATTGTTTGTACTGCTAGGCCTGATTCTGATATATGTATACAGGAAGAGAA

AGCTGAGTAGTAAAGGATTAGCAACTCTGAAACCTACTAATGATGGTGGAAATGAAGATCAGGGTCAAGA

CCTGGAATTGCCATTGGTTGAATTTGAAGAAATTGCCAAGGCTACGAATAACTTTTCCCTAAACAATAAG

CTCGGAGAGGGTGGTTTTGGCCCTGTCTACAAGGGAACTTTGGTTGATGGACAGGAAATAGCTGTAAAGA

GACTCTCCAAGACTTCAAACCAAGGACTCCATGAGTTTAAGAATGAAGTGAAATGCATTGCCGAACTGCA

ACACAGAAATCTTGTGAGGCTTTTAGGATGTAGCATTGAGGATGGAGAAATGTTGTTGATCTACGAATAC

ATGTCCAACAAAAGTTTAGATGTCTTCATTTTCGGTAAGACTGTTGTTGATATTATTAGCTAAGCATGTC

ACTGGATTGGCCAAAGCGCTACAATATCATTAACGGCATTGCCAAGGGCCTACTGTATCTCCACCAGGAC

TCGAGACTAAGAATCATTCACAGAGATCTCAAAGCCGGTAACATTCTCCTCGATCATGAGATGAGTCCTA

AAATTTCAGATTTTGGCCTGGCAAGAAGTTTTGCAGACAACGAAAGTGCAGCCAACACCATTCGCGTAGT

TGGAACATACGGCTACATGGCCCCAGAGTATGCTATCGAGGGACTATTCTCTGTCAAATCCGATTCCTAC

AGTTTTGGCGTCCTGGTCTTGGAGATAGTAACCGGGAAAAAAATAAGACAATTTTACAATTCCAAGGACA

ACCTTAACCTCCTCGGCCATGCATGGAAACTTTTCAATGATGGCGAATCCTTAGACCTTGTCGACGAGGC

AATGTTGGGATCATGCAGTTCTTCAGAAGTTTTGCGAGCTATTCACATCGGATTGTTGTGTGTGCAGCCA

TATCCGCAAGACAGGCCAAACATGACATACGTTGTGTCGATGTTGGATAGTGAAAATGAGCTTCCTAAGC

CAAAACAGCCAGGGTTTTTCACTGATAACAGAGTGCAACAGGGCTCAGAGTATCCGTCTCACCATAGTGA

GTCAAGTGTAACAAATAATGATCTGAGTATTACAGTGTTGGAGCCTAGGTAGTAATTGTACATTGTTTGG

CACCATTTTTTTAAGCATATTATTTTCCTTCTGAGATCCTGTTATGATCCGATATTTCTAAAATATTGGA

TGCTCCACGAGGATGAAAATTTTGTATGACATAATAATTTTTGAGATCAAAT

>MSTRG.2993.4 gene=MSTRG.2993

ATAACATCGAAAATTTACTTCAACACTTGGCAAAATAAAATAAAAAGAGTACTAGGATGGATGATTTATT

GTGCTCCAGTTTTGTTATTTTCTTGATTTTTATAAACACTTGTTTTGCAGCTGACATCTTGAATGTGAAG

CAAACACTGAGGGATGATGGCAACAGCAGTCTTGTTTCATCTGGTGGTGATTTCGAGCTTGGATTCTTCA

GCCCCGGAAGCTCCAAGAATCGATATCTTGGCATATGGTTCAAGAAGGTCTCCAACAGGACTGTTGTTTG

GGTTGCTAACAGAGAGGCTCCACTTCTTGACACTTCCGGGGTGCTCAGGATTGGTGCCAATGGAAATTTG

GAACTTGTGAATGGTAGAAACAGCACAATGTGGTTTTCTAATTCGACAAAATCTGCGAAGAATGAGCCTG

TGGTGGCACAACTAATGGATTCCTCAAACCTTGTGATCAGGCATGAAAATGACACTGACCCTGAGAAGTA

TCTGTGGCAAAGTGCTGATCACCCTACTGATCATATTTTGTCAGGCATGAAACTTGGATGGGATTTGAGT

AGAGGGCTGGAACGATATCTTACATCGTGGAAAAGTGTTAGTGATCCTTCACCTGGGAATTACACTTTCA

GAATGTACCTTGAAGGATATCCACAGATGAAAGTGGCTGATGGTTCTAGTTCTACCACAAGTTTCAGATT

TGGACCATGGGATGGCATTCAGTTTAGTGGTGTGTCGTTTAATATCCCGGATCTCAATTTTGACTTGAAA

CTTATTGTGACGGATACTGAAATTTATTCCAAGTTTGATGTGAACAATAATTCCAGGCCTGTGAGGTTTG

ATCTGAATCCTGAAGGACTAATTGAGCTTCTAACTTGGTTTGACAATTTTAAGAATTGGACATTCAGTCA

GTACATTAGTGTGCTTTATGATCAATGTGACCCTTATGGACTTTGTGGGGCATTTGGTAGTTGTGACATT

ACTCGAGCTCAATCAGGACAAAGTGCTTGTGAATGTTTAGGAGGATTCGTGCCTCGGTCTTTGGAAAAAT

GGAAACTTGCAGATTGGTCCGGTGGATGTGTTCGTTCAGCTCCATTAACTTGTGGAAGTGGAGAAGAATT

CCAGAAGATTCCGGGCCTTAAATTGCCTGATACTCAGACTTCGTGGTTCAACCGAAGCATGAGCCTTGCT

GATTGTGAGACCGAGTGCAGAAAGAACTGTTCTTGTACAGCTTACTCAAATGTGGATATTAGACAAGGCG

GCTCTGGATGTATGCTATGGTTTGGTGATCTCATTGATATGAGAAATTACGCAGACAATGGCCTTCCTCT

ATATGTGAGAACTGCTTCTCAACCAGCTGTGAATGGGAGCTCAAGTAAAAAGGGAGGAGTGAAGATCATT

GCCATTATCATTCCTACTATAATAGGATTGTTTGTACTGCTAGGCCTGATTCTGATATATGTATACAGGA

AGAGAAAGCTGAGTAGTAAAGGATTAGCAACTCTGAAACCTACTAATGATGGTGGAAATGAAGATCAGGG

TCAAGACCTGGAATTGCCATTGGTTGAATTTGAAGAAATTGCCAAGGCTACGAATAACTTTTCCCTAAAC

AATAAGCTCGGAGAGGGTGGTTTTGGCCCTGTCTACAAGGGAACTTTGGTTGATGGACAGGAAATAGCTG

TAAAGAGACTCTCCAAGACTTCAAACCAAGGACTCCATGAGTTTAAGAATGAAGTGAAATGCATTGCCGA

ACTGCAACACAGAAATCTTGTGAGGCTTTTAGGATGTAGCATTGAGGATGGAGAAATGTTGTTGATCTAC

GAATACATGTCCAACAAAAGTTTAGATGTCTTCATTTTCGGTAAGACTGTTGTTGATATTATTACAAAGA

GCTAAGCATGTCACTGGATTGGCCAAAGCGCTACAATATCATTAACGGCATTGCCAAGGGCCTACTGTAT

CTCCACCAGGACTCGAGACTAAGAATCATTCACAGAGATCTCAAAGCCGGTAACATTCTCCTCGATCATG

AGATGAGTCCTAAAATTTCAGATTTTGGCCTGGCAAGAAGTTTTGCAGACAACGAAAGTGCAGCCAACAC

CATTCGCGTAGTTGGAACATACGGCTACATGGCCCCAGAGTATGCTATCGAGGGACTATTCTCTGTCAAA

TCCGATTCCTACAGTTTTGGCGTCCTGGTCTTGGAGATAGTAACCGGGAAAAAAATAAGACAATTTTACA

ATTCCAAGGACAACCTTAACCTCCTCGGCCATGCATGGAAACTTTTCAATGATGGCGAATCCTTAGACCT

TGTCGACGAGGCAATGTTGGGATCATGCAGTTCTTCAGAAGTTTTGCGAGCTATTCACATCGGATTGTTG

TGTGTGCAGCCATATCCGCAAGACAGGCCAAACATGACATACGTTGTGTCGATGTTGGATAGTGAAAATG

AGCTTCCTAAGCCAAAACAGCCAGGGTTTTTCACTGATAACAGAGTGCAACAGGGCTCAGAGTATCCGTC

TCACCATAGTGAGTCAAGTGTAACAAATAATGATCTGAGTATTACAGTGTTGGAGCCTAGGTAGTAATTG

TACATTGTTTGGCACCATTTTTTTAAGCATATTATTTTCCTTCTGAGATCCTGTTATGATCCGATATTTC

TAAAATATTGGATGCTCCACGAGGATGAAAATTTTGTATGACA

>MSTRG.2994.1 gene=MSTRG.2994

CGGCCACTCTTTTTCAATACAATGCACACATGATGCAACTAGCAACTTGCATCTTTCTTGTCTTCTTTGT

TTACAAATAATTTTATACATTGAATGATAAGGGATAAAAAAAAACAATGAACGAAGCAATGGTAATGATA

AGTTGCACTAGCTGCATTTTTTTCCTTTTTAGTGGTTTCGTATTCTTCATCACCTTTATTTCTGTAGCAG

ATGCAGCAGACACCTTGACTGGGAATCAAGCCCTTGTGGATGATGGAAAGAGCAGCATTGAATCCTCTGG

TGGGGTTTTTGAGCTGGGATTCTTCAGCCCCGGAAAGTCCACGAATCGATACGTGGGGATTTGGTACAAG

AAGATATCAATTAGGACTGTGATATGGACTGCTAATAGACAGGCTCCAATTCTTGATAGTAATTCAGGAG

TTTTGAGGATTAGGAATGCAAGATTAGAGATTTCTGATGGTGCTGATGTCATTTGGTCATCCAATTCTTC

GTCGTCCGATAGTAATAATGCTTCTGTGATTGTTCAGCTTTTGGATTCAGGCAATTTGGTGATCAGCAAT

GGGAATGCCAGTAGGTTCATTTGGCAGAGTTTTGATTTTCCTAATGACAATCTTCTGCAAGGAATGAAAA

TTGGATGGGACTTGGAGAGAGGACTAGAAAGATCTTTGGTGTCGTGGAAAAGTACTGATGATCCTTCTGT

AGGTGATTTTACACAGAGGGTAAGTAGAATTGGTTACCCTCAGATGCTTGTGTCGAAACATTCAGTTGTG

CAGTTCAGATTTGGACCATTTGATGGGATTCAGTTCAGTGGTGTTACCTATACCAGACAAGAATTGGATT

TTCAGTTGTCACTGGTTCGTGATGACAGGGAAATTTATTCGAGTTTTGGGATTAATGGGAATTCTGATCC

TGTAGTGAGGTATATGTTGCTTTCTGATGGATCGATACAAGTTCTGTCTTGGGATGATGAGAGCAAAAAA

TGGGAGGATTATTTGAGTGTTCAGAGGGATGCTTGTGATCAGTATGCAGTCTGTGGTGCCTATGGTAATT

GTAACAATACGAAATTGAGAGATGAGGCAGATTGTGGATGTTTAGGGGGGTATGTTCCGAAATCTCCAGA

AAAATGGAAATTATCAGATTGGTCTGATGGATGTGTTCCAAGAACTCAATTTGATTGTAGAAATGGAACA

GATATATTTGTTGAGTATTCAGCTATGAAATTGCCAGACACTCGGACCGCTTGGTTTAACCAGACTATGG

GGCTTGTTGATTGTAAAAAGGAGTGCTTAAAAAACTGCTCTTGTACAGCTTACTCAAACATAGATATTAG

ACAAGGTGGAACTGGCTGCTTATTATGGTTCAATGAACTTCTTGATATCAGAGATTTTAAGGACACTGGA

CTTAAATTCTATGTAAGAACTTCTTCGTTAAATACAGAGGCAAATAGAAGTTCAACAGAAAAGAAGGGAG

GAAGCAGCATTGTTATAATCATTGCTGTGTTGGTAGTGGTGGCAGTACTACTAGTCCTGATTGTGATTTT

CCTCTGCAGAAGGAGGAATATCCGTAGAAGGAAGAGAAGGAGGCGAGGAAGATTAGCATTGAATCCACAA

ACTGAAACCGGGAATGACAATGAGAGCAAAGATCTGGAGTTACCATTGTTTGATTTCCAAACCATTGCTA

ATGCTACAAACAACTTTTCCTTGAACAATAAGCTGGGCGAGGGTGGTTTTGGGCCTGTCTACAAGGGTAT

TTTGGGCCACGAACAAGAAATAGCAGTCAAAAGACTTTCAAAGACATCCAGACAAGGGCTGGATGAGTTC

AAGAACGAGGTTTCATGCATAGCAAAACTGCAGCACCGAAATCTTGTGAGACTTCTGGGATGCTGCATTG

AGCACAATGAAATGATGTTGATATATGAGTACATGCCAAACAAGAGTTTAGATACATTCATTTTCGACAA

AGAATTAAGCAAGGTACTTGATTGGCCCCAGCGCTACAACATTATAAAAGGCATCGCTAAGGGCCTTTTA

TATCTTCACCAAGACTCTAGGCTAAGAATCATACACAGAGATCTCAAAACCAGCAATGTTTTACTTGATC

ATGAAATGAGTCCGAAAATTTCAGATTTTGGCTTGGCAAGAACTTTTGGAGGGAATGAAACCGGAGTAAA

TACAGCGAAAGTAGTTGGCACATACGGCTACATGTCACCAGAGTACGCCATTGATGGGTTTTTCTCAGTT

AAATCAGACGCTTATAGCTTTGGTGTCGTCGTGCTAGAAATAGTAAGTGGAAACAAAATCAGAGGCTTCT

ACAATGCAGATAACAATCTCAATCTTCTTGGTCAGGCATGGAAAGTGTACAAAGAAGGCAACATTCTGGA

CCTAGTTGATGAGGCATTCAAGGAATCATGCAATAATTCAGAAGTTTCGCGAGCTGTACAAATCGGGTTA

TTATGTGTGCAACCATACCCGAAAGACAGACCGAATATGAGATTTGTTGTGTCCATGCTAGGCAATGACA

ATGAGCTGCCACAGCCAAAACAACAACCTGGTGTTTTTCATGACAGGAGGATGCAAGACTCTGATACTTC

ATCGAGTATGCAGGGATCTTCTACATCTAATGAATTGTCCATAACAGTTCTTAATCCTAGATAGTAAAAG

AAAAATTTGTTCTCTATATATTTTTGCTCAGTGATTTGTACAATCAATTTTTTGATACATTTGTGCTAGA

TAATTGATGGTATTCTGTTGAGGTGTGCTCTTAATCATTAATGCTTTTATCCAACACAATCTTGATAAGA

CTCTTGAATGGCCACTGCCATGCCCATACAAAGTGAGTAACTGCTTAGTCAAAGG

>MSTRG.2994.2 gene=MSTRG.2994

CACTCTTTTTCAATACAATGCACACATGATGCAACTAGCAACTTGCATCTTTCTTGTCTTCTTTGTTTAC

AAATAATTTTATACATTGAATGATAAGGGATAAAAAAAAACAATGAACGAAGCAATGATGCAGCAGACAC

CTTGACTGGGAATCAAGCCCTTGTGGATGATGGAAAGAGCAGCATTGAATCCTCTGGTGGGGTTTTTGAG

CTGGGATTCTTCAGCCCCGGAAAGTCCACGAATCGATACGTGGGGATTTGGTACAAGAAGATATCAATTA

GGACTGTGATATGGACTGCTAATAGACAGGCTCCAATTCTTGATAGTAATTCAGGAGTTTTGAGGATTAG

GAATGCAAGATTAGAGATTTCTGATGGTGCTGATGTCATTTGGTCATCCAATTCTTCGTCGTCCGATAGT

AATAATGCTTCTGTGATTGTTCAGCTTTTGGATTCAGGCAATTTGGTGATCAGCAATGGGAATGCCAGTA

GGTTCATTTGGCAGAGTTTTGATTTTCCTAATGACAATCTTCTGCAAGGAATGAAAATTGGATGGGACTT

GGAGAGAGGACTAGAAAGATCTTTGGTGTCGTGGAAAAGTACTGATGATCCTTCTGTAGGTGATTTTACA

CAGAGGGTAAGTAGAATTGGTTACCCTCAGATGCTTGTGTCGAAACATTCAGTTGTGCAGTTCAGATTTG

GACCATTTGATGGGATTCAGTTCAGTGGTGTTACCTATACCAGACAAGAATTGGATTTTCAGTTGTCACT

GGTTCGTGATGACAGGGAAATTTATTCGAGTTTTGGGATTAATGGGAATTCTGATCCTGTAGTGAGGTAT

ATGTTGCTTTCTGATGGATCGATACAAGTTCTGTCTTGGGATGATGAGAGCAAAAAATGGGAGGATTATT

TGAGTGTTCAGAGGGATGCTTGTGATCAGTATGCAGTCTGTGGTGCCTATGGTAATTGTAACAATACGAA

ATTGAGAGATGAGGCAGATTGTGGATGTTTAGGGGGGTATGTTCCGAAATCTCCAGAAAAATGGAAATTA

TCAGATTGGTCTGATGGATGTGTTCCAAGAACTCAATTTGATTGTAGAAATGGAACAGATATATTTGTTG

AGTATTCAGCTATGAAATTGCCAGACACTCGGACCGCTTGGTTTAACCAGACTATGGGGCTTGTTGATTG

TAAAAAGGAGTGCTTAAAAAACTGCTCTTGTACAGCTTACTCAAACATAGATATTAGACAAGGTGGAACT

GGCTGCTTATTATGGTTCAATGAACTTCTTGATATCAGAGATTTTAAGGACACTGGACTTAAATTCTATG

TAAGAACTTCTTCGTTAAATACAGAGGCAAATAGAAGTTCAACAGAAAAGAAGGGAGGAAGCAGCATTGT

TATAATCATTGCTGTGTTGGTAGTGGTGGCAGTACTACTAGTCCTGATTGTGATTTTCCTCTGCAGAAGG

AGGAATATCCGTAGAAGGAAGAGAAGGAGGCGAGGAAGATTAGCATTGAATCCACAAACTGAAACCGGGA

ATGACAATGAGAGCAAAGATCTGGAGTTACCATTGTTTGATTTCCAAACCATTGCTAATGCTACAAACAA

CTTTTCCTTGAACAATAAGCTGGGCGAGGGTGGTTTTGGGCCTGTCTACAAGGGTATTTTGGGCCACGAA

CAAGAAATAGCAGTCAAAAGACTTTCAAAGACATCCAGACAAGGGCTGGATGAGTTCAAGAACGAGGTTT

CATGCATAGCAAAACTGCAGCACCGAAATCTTGTGAGACTTCTGGGATGCTGCATTGAGCACAATGAAAT

GATGTTGATATATGAGTACATGCCAAACAAGAGTTTAGATACATTCATTTTCGACAAAGAATTAAGCAAG

GTACTTGATTGGCCCCAGCGCTACAACATTATAAAAGGCATCGCTAAGGGCCTTTTATATCTTCACCAAG

ACTCTAGGCTAAGAATCATACACAGAGATCTCAAAACCAGCAATGTTTTACTTGATCATGAAATGAGTCC

GAAAATTTCAGATTTTGGCTTGGCAAGAACTTTTGGAGGGAATGAAACCGGAGTAAATACAGCGAAAGTA

GTTGGCACATACGGCTACATGTCACCAGAGTACGCCATTGATGGGTTTTTCTCAGTTAAATCAGACGCTT

ATAGCTTTGGTGTCGTCGTGCTAGAAATAGTAAGTGGAAACAAAATCAGAGGCTTCTACAATGCAGATAA

CAATCTCAATCTTCTTGGTCAGGCATGGAAAGTGTACAAAGAAGGCAACATTCTGGACCTAGTTGATGAG

GCATTCAAGGAATCATGCAATAATTCAGAAGTTTCGCGAGCTGTACAAATCGGGTTATTATGTGTGCAAC

CATACCCGAAAGACAGACCGAATATGAGATTTGTTGTGTCCATGCTAGGCAATGACAATGAGCTGCCACA

GCCAAAACAACAACCTGGTGTTTTTCATGACAGGAGGATGCAAGACTCTGATACTTCATCGAGTATGCAG

GGATCTTCTACATCTAATGAATTGTCCATAACAGTTCTTAATCCTAGATAGTAAAAGAAAAATTTGTTCT

CTATATATTTTTGCTCAGTGATTTGTACAATCAATTTTTTGATACATTTGTGCTAGATAATTGATGGTAT

TCTGTTGAGGTGTGCTCTTAATCATTAATGCTTTTATCCAACACAATCTTGATAAGACTCTTGAATGGCC

ACTGCCATGCCCATACAAAGTGAGTAACTGCTTAGTCAAAGG

>MSTRG.2994.3 gene=MSTRG.2994

CTTGCATCTTTCTTGTCTTCTTTGTTTACAAATAATTTTATACATTGAATGATAAGGGATAAAAAAAAAC

AATGAACGAAGCAATGCAGATGCAGCAGACACCTTGACTGGGAATCAAGCCCTTGTGGATGATGGAAAGA

GCAGCATTGAATCCTCTGGTGGGGTTTTTGAGCTGGGATTCTTCAGCCCCGGAAAGTCCACGAATCGATA

CGTGGGGATTTGGTACAAGAAGATATCAATTAGGACTGTGATATGGACTGCTAATAGACAGGCTCCAATT

CTTGATAGTAATTCAGGAGTTTTGAGGATTAGGAATGCAAGATTAGAGATTTCTGATGGTGCTGATGTCA

TTTGGTCATCCAATTCTTCGTCGTCCGATAGTAATAATGCTTCTGTGATTGTTCAGCTTTTGGATTCAGG

CAATTTGGTGATCAGCAATGGGAATGCCAGTAGGTTCATTTGGCAGAGTTTTGATTTTCCTAATGACAAT

CTTCTGCAAGGAATGAAAATTGGATGGGACTTGGAGAGAGGACTAGAAAGATCTTTGGTGTCGTGGAAAA

GTACTGATGATCCTTCTGTAGGTGATTTTACACAGAGGGTAAGTAGAATTGGTTACCCTCAGATGCTTGT

GTCGAAACATTCAGTTGTGCAGTTCAGATTTGGACCATTTGATGGGATTCAGTTCAGTGGTGTTACCTAT

ACCAGACAAGAATTGGATTTTCAGTTGTCACTGGTTCGTGATGACAGGGAAATTTATTCGAGTTTTGGGA

TTAATGGGAATTCTGATCCTGTAGTGAGGTATATGTTGCTTTCTGATGGATCGATACAAGTTCTGTCTTG

GGATGATGAGAGCAAAAAATGGGAGGATTATTTGAGTGTTCAGAGGGATGCTTGTGATCAGTATGCAGTC

TGTGGTGCCTATGGTAATTGTAACAATACGAAATTGAGAGATGAGGCAGATTGTGGATGTTTAGGGGGGT

ATGTTCCGAAATCTCCAGAAAAATGGAAATTATCAGATTGGTCTGATGGATGTGTTCCAAGAACTCAATT

TGATTGTAGAAATGGAACAGATATATTTGTTGAGTATTCAGCTATGAAATTGCCAGACACTCGGACCGCT

TGGTTTAACCAGACTATGGGGCTTGTTGATTGTAAAAAGGAGTGCTTAAAAAACTGCTCTTGTACAGCTT

ACTCAAACATAGATATTAGACAAGGTGGAACTGGCTGCTTATTATGGTTCAATGAACTTCTTGATATCAG

AGATTTTAAGGACACTGGACTTAAATTCTATGTAAGAACTTCTTCGTTAAATACAGAGGCAAATAGAAGT

TCAACAGAAAAGAAGGGAGGAAGCAGCATTGTTATAATCATTGCTGTGTTGGTAGTGGTGGCAGTACTAC

TAGTCCTGATTGTGATTTTCCTCTGCAGAAGGAGGAATATCCGTAGAAGGAAGAGAAGGAGGCGAGGAAG

ATTAGCATTGAATCCACAAACTGAAACCGGGAATGACAATGAGAGCAAAGATCTGGAGTTACCATTGTTT

GATTTCCAAACCATTGCTAATGCTACAAACAACTTTTCCTTGAACAATAAGCTGGGCGAGGGTGGTTTTG

GGCCTGTCTACAAGGGTATTTTGGGCCACGAACAAGAAATAGCAGTCAAAAGACTTTCAAAGACATCCAG

ACAAGGGCTGGATGAGTTCAAGAACGAGGTTTCATGCATAGCAAAACTGCAGCACCGAAATCTTGTGAGA

CTTCTGGGATGCTGCATTGAGCACAATGAAATGATGTTGATATATGAGTACATGCCAAACAAGAGTTTAG

ATACATTCATTTTCGACAAAGAATTAAGCAAGGTACTTGATTGGCCCCAGCGCTACAACATTATAAAAGG

CATCGCTAAGGGCCTTTTATATCTTCACCAAGACTCTAGGCTAAGAATCATACACAGAGATCTCAAAACC

AGCAATGTTTTACTTGATCATGAAATGAGTCCGAAAATTTCAGATTTTGGCTTGGCAAGAACTTTTGGAG

GGAATGAAACCGGAGTAAATACAGCGAAAGTAGTTGGCACATACGGCTACATGTCACCAGAGTACGCCAT

TGATGGGTTTTTCTCAGTTAAATCAGACGCTTATAGCTTTGGTGTCGTCGTGCTAGAAATAGTAAGTGGA

AACAAAATCAGAGGCTTCTACAATGCAGATAACAATCTCAATCTTCTTGGTCAGGCATGGAAAGTGTACA

AAGAAGGCAACATTCTGGACCTAGTTGATGAGGCATTCAAGGAATCATGCAATAATTCAGAAGTTTCGCG

AGCTGTACAAATCGGGTTATTATGTGTGCAACCATACCCGAAAGACAGACCGAATATGAGATTTGTTGTG

TCCATGCTAGGCAATGACAATGAGCTGCCACAGCCAAAACAACAACCTGGTGTTTTTCATGACAGGAGGA

TGCAAGACTCTGAT

>MSTRG.2994.4 gene=MSTRG.2994

AACGAAGCAATGGTAATGATAAGTTGCACTAGCTGCATTTTTTTCCTTTTTAGTGGTTTCGTATTCTTCA

TCACCTTTATTTCTGTAGCAGATGCAGCAGACACCTTGACTGGGAATCAAGCCCTTGTGGATGATGGAAA

GAGCAGCATTGAATCCTCTGGTGGGGTTTTTGAGCTGGGATTCTTCAGCCCCGGAAAGTCCACGAATCGA

TACGTGGGGATTTGGTACAAGAAGATATCAATTAGGACTGTGATATGGACTGCTAATAGACAGGCTCCAA

TTCTTGATAGTAATTCAGGAGTTTTGAGGATTAGGAATGCAAGATTAGAGATTTCTGATGGTGCTGATGT

CATTTGGTCATCCAATTCTTCGTCGTCCGATAGTAATAATGCTTCTGTGATTGTTCAGCTTTTGGATTCA

GGCAATTTGGTGATCAGCAATGGGAATGCCAGTAGGTTCATTTGGCAGAGTTTTGATTTTCCTAATGACA

ATCTTCTGCAAGGAATGAAAATTGGATGGGACTTGGAGAGAGGACTAGAAAGATCTTTGGTGTCGTGGAA

AAGTACTGATGATCCTTCTGTAGGTGATTTTACACAGAGGGTAAGTAGAATTGGTTACCCTCAGATGCTT

GTGTCGAAACATTCAGTTGTGCAGTTCAGATTTGGACCATTTGATGGGATTCAGTTCAGTGGTGTTACCT

ATACCAGACAAGAATTGGATTTTCAGTTGTCACTGGTTCGTGATGACAGGGAAATTTATTCGAGTTTTGG

GATTAATGGGAATTCTGATCCTGTAGTGAGGTATATGTTGCTTTCTGATGGATCGATACAAGTTCTGTCT

TGGGATGATGAGAGCAAAAAATGGGAGGATTATTTGAGTGTTCAGAGGGATGCTTGTGATCAGTATGCAG

TCTGTGGTGCCTATGGTAATTGTAACAATACGAAATTGAGAGATGAGGCAGATTGTGGATGTTTAGGGGG

GTATGTTCCGAAATCTCCAGAAAAATGGAAATTATCAGATTGGTCTGATGGATGTGTTCCAAGAACTCAA

TTTGATTGTAGAAATGGAACAGATATATTTGTTGAGTATTCAGCTATGAAATTGCCAGACACTCGGACCG

CTTGGTTTAACCAGACTATGGGGCTTGTTGATTGTAAAAAGGAGTGCTTAAAAAACTGCTCTTGTACAGC

TTACTCAAACATAGATATTAGACAAGGTGGAACTGGCTGCTTATTATGGTTCAATGAACTTCTTGATATC

AGAGATTTTAAGGACACTGGACTTAAATTCTATAGGCAAATAGAAGTTCAACAGAAAAGAAGGGAGGAAG

CAGCATTGTTATAATCATTGCTGTGTTGGTAGTGGTGGCAGTACTACTAGTCCTGATTGTGATTTTCCTC

TGCAGAAGGAGGAATATCCGTAGAAGGAAGAGAAGGAGGCGAGGTAATCATCTCAATTTTCCAAGTATGT

TTACCTTAATGACGAATTGTGCATTATCTATATGCTTTACACCTCTCGGTGTGCAGGAAGATTAGCATTG

AATCCACAAACTGAAACCGGGAATGACAATGAGAGCAAAGATCTGGAGTTACCATTGTTTGATTTCCAAA

CCATTGCTAATGCTACAAACAACTTTTCCTTGAACAATAAGCTGGGCGAGGGTGGTTTTGGGCCTGTCTA

CAAGGGTATTTTGGGCCACGAACAAGAAATAGCAGTCAAAAGACTTTCAAAGACATCCAGACAAGGGCTG

GATGAGTTCAAGAACGAGGTTTCATGCATAGCAAAACTGCAGCACCGAAATCTTGTGAGACTTCTGGGAT

GCTGCATTGAGCACAATGAAATGATGTTGATATATGAGTACATGCCAAACAAGAGTTTAGATACATTCAT

TTTCGACAAAGAATTAAGCAAGGTACTTGATTGGCCCCAGCGCTACAACATTATAAAAGGCATCGCTAAG

GGCCTTTTATATCTTCACCAAGACTCTAGGCTAAGAATCATACACAGAGATCTCAAAACCAGCAATGTTT

TACTTGATCATGAAATGAGTCCGAAAATTTCAGATTTTGGCTTGGCAAGAACTTTTGGAGGGAATGAAAC

CGGAGTAAATACAGCGAAAGTAGTTGGCACATACGGCTACATGTCACCAGAGTACGCCATTGATGGGTTT

TTCTCAGTTAAATCAGACGCTTATAGCTTTGGTGTCGTCGTGCTAGAAATAGTAAGTGGAAACAAAATCA

GAGGCTTCTACAATGCAGATAACAATCTCAATCTTCTTGGTCAGGCATGGAAAGTGTACAAAGAAGGCAA

CATTCTGGACCTAGTTGATGAGGCATTCAAGGAATCATGCAATAATTCAGAAGTTTCGCGAGCTGTACAA

ATCGGGTTATTATGTGTGCAACCATACCCGAAAGACAGACCGAATATGAGATTTGTTGTGTCCATGCTAG

GCAATGACAATGAGCTGCCACAGCCAAAACAACAACCTGGTGTTTTTCATGACAGGAGGATGCAAGACTC

TGATACTTCATCGAGTATGCAGGGATCTTCTACATCTAATGAATTGTCCATAACAGTTCTTAATCCTAGA

TAGTAAAAGAAAAATTTGTTCTCTATATATTTTTGCTCAGTGATTTGTACAATCAATTTTTTGATACATT

TCTAATACACGATTGTTCTTGTGAGAG

>MSTRG.2998.1 gene=MSTRG.2998

ACGGTGTGTTCTAAAGAGGAGAGACAAAGAGATGGGTGATGACAACCGTGATTTATTCGAGAAATTACCT

CAAGCATTTAGACAAAAATTCACACAAGATGTATTATCACGCACAAGCCCAGCAGATGTGGTAAGAATTT

CACTTGTGTCCAAAAGTTTCCAGTCGGCGGCCAATTCTGATGTTGTTTGGGACACATTTATTCCACCTCA

TCATCGAAAACTTGTTCTTGATCGCTACTACAAATCCACCTTTGAATCTAGCAAAGACGTCTTTCTCTTC

CTCTGTGAACATATCGCTGAGAGCCATACCGACCCTCTGAAGTTTTGGTTAGATAAATGGAGTGGGAAGA

TGTGGCTCAAAATGACATATGAAAACCTCTCAATCACACAGCTTGCAACAGATTACTGGACGAGAAGCAA

AGAACAGTATCCGGTGCTAGGTCATGTGATTTGGCTTCAAGTGTATGGCAAAATACCCACTTCAAGATTG

TCACCAGAGACAGCCTATGAAGCTTATCTTGTGTTTAGACTCTCGGATTACATTTGCTATGGACTTCACA

TACCTATAGAGGCTTCTGTTGGGATCCCAGGGGAAGAAATCACAAAGGAGCTTATATATTTGGACCCTCG

AATAGCCAAGTCTGGGAGTTTTAACTATTATCCAAACACATACCCGGGGGAAGGAAATCCCTTAATAGAA

GTTAAGATGGGTGAATACTTTAACAAAGAAGGGGAGAACAGGGATGTGGAAGTAACTTTGAAAGAAGTGA

AAAGTGGAAGGCCAAAGTGTGGTGTCCATATTGCTGGGATGAAGATGCTACCTAAGCGCAATATAAACCT

TTAATAATCGTGCTCGCTCTTGTTTGGGATAGAGATCTGGACGTTAAAAATTGTTATCTGTTCATATCTT

TGACGCTTTGCGTGTGATGATGAAGACCTGTAATCGTTAGGCAACTAATATTAGCTAAGCAGTACTACTG

CAACTCTCAACATGTTTCCCGTGCTTTATTCTAAATTATTACAATGATATACTTTACTCATGAATGGTTT

ACCAAGAAACAGAACTATATGGCCATACCCTTCGTGAAACTCATGTCTGTAATGAAAGCCGGAGCCTAAA

GACAG

>MSTRG.2998.2 gene=MSTRG.2998

GTGTGTTCTAAAGAGGAGAGACAAAGAGATGGGTGATGACAACCGTGATTTATTCGAGAAATTACCTCAA

GCATTTAGACAAAAATTCACACAAGATGTATTATCACGCACAAGCCCAGCAGATGTGGTAAGAATTTCAC

TTGTGTCCAAAAGTTTCCAGTCGGCGGCCAATTCTGATGTTGTTTGGGACACATTTATTCCACCTCATCA

TCGAAAACTTGTTCTTGATCGCTACTACAAATCCACCTTTGAATCTAGCAAAGACGTCTTTCTCTTCCTC

TGTGAACATATCGCTGAGAGCCATACCGACCCTCTGTTTTGGTTAGATAAATGGAGTGGGAAGATGTGGC

TCAAAATGACATATGAAAACCTCTCAATCACACAGCTTGCAACAGATTACTGGACGAGAAGCAAAGAACA

GTATCCGGTGCTAGGTCATGTGATTTGGCTTCAAGTGTATGGCAAAATACCCACTTCAAGATTGTCACCA

GAGACAGCCTATGAAGCTTATCTTGTGTTTAGACTCTCGGATTACATTTGCTATGGACTTCACATACCTA

TAGAGGCTTCTGTTGGGATCCCAGGGGAAGAAATCACAAAGGAGCTTATATATTTGGACCCTCGAATAGC

CAAGTCTGGGAGTTTTAACTATTATCCAAACACATACCCGGGGGAAGGAAATCCCTTAATAGAAGTTAAG

ATGGGTGAATACTTTAACAAAGAAGGGGAGAACAGGGATGTGGAAGTAACTTTGAAAGAAGTGAAAAGTG

GAAGGCCAAAGTGTGGTGTCCATATTGCTGGGATGAAGATGCTACCTAAGCGCAATATAAACCTTTAATA

ATCGTGCTCGCTCTTGTTTGGGATAGAGATCTGGACGTTAAAAATTGTTATCTGTTCATATCTTTGACGC

TTTGCGTGTGATGATGAAGACCTGTAATCGTTAGGCAACTAATATTAGCTAAGCAGTACTACTGCAACTC

TCAACATGTTTCCCGTGCTTTATTCTAAATTATTA

>MSTRG.2999.1 gene=MSTRG.2999

GTTAGCATTAAAAAAGCGCTCTCACTCATCTAATTTAGATGAGCGGATATAAACCAAGTCTTTATAACAA

AGGCGTCCTCTCACGTATAAACACCGATGGCCGACGCTGATTTATTCGGAAAGCTACCTGAGGAAATCAT

AGAAGAAGTTGTTTCGCTTATTGGGCCTATGGAGGCATGCAGGCTCTCTGTGGTGTGTAAAAGGTTTCTC

TCGGCGGCGGAATCAAACACCGTGTGGGAGAGATTTTTGCCCTCTGACTATCAACAATTTCTTGATCGGG

CGGATTCTTTACCCAACATCAAATTTGCTCACAAGAAAGATGTCTTTCTCTTTCTCATTGGTAATCATGT

CATCTTTGACGGGAATACTCTGTGCTTCTGGTTACACAAGAGGAGCGGTAAGAAGTGTTTCCTTGTAGCT

GCTTCATTTACAGGTGGTGTTGAAGAATATTTCTTGAATCCAGGCTGGTGTACAAGATCTGAAAAATCAA

GATTTCAAAATCAAATTACTGAATGCAACAATCTTTTATTGTTTGAAATTCGCGGAAAGATAAGCACCTG

CATGTTATCACCAGATACGACCTATGCAATCTATCTTGTCTTTCATTCCCCAATTATTGATGAATATTTT

AATGAACCTGTTGAAGTTTCCATAGGGATTGATGGTGTGGAAAGCATTAAACAGAATGTTTGTTTGAACC

CGCATAAGATGAGGTCAGAAGGGGATGATGATCTCCAATATCCGCAACAGAGAAAAGATGACAGGTTTGA

AGTTAGATTAGGAGAATACTTTAACAGAGGAGGGGAGGAGATGAATTTGGAGATCACTATGACAAAAAAC

AGTGAGATTCCTAAGACTGGACCTGGTGTTGAAGGCTTTGAATTTCGACCAAAGGGCAGCTAATGCAGGT

TCAGGAGGCTCATACTTTTGGTTTCTGATTCCTGTTTCAAATTTGAAGTACTTGTTCAAACACTCTAATT

ATTTAGAGATAAAGGAAACTGAACTAGATAGTACAATCAGACCTGTAAATGTTCAACCAAAACACTTGTT

GCTATGTATGTGGCAATTAATTTCTAATTTGACTGTCTTTAGGCTCCGG

>MSTRG.3000.1 gene=MSTRG.3000

CAAAGAAACCTCCTTAAAATGTATTCTCGAAGAAGTTTATCGTTAAAAATATGAAAAGGATATATCAAAA

CGTAAACAAAAAAATATTTAATAGGAGAAATACTGACCCTTTTTTTTTTCTTGCTAGCTGACTTTGCAGA

ACCTGGAGTGTGAAGAATCACCCAACTGAAAGCTAGAATCCTGGGATAGTTAAGAATTGTTATAGGTCTC

ATGCTTGAAGAAATGGATAAATATTTCAGACCTCTAAGTAAAACTTACAAAATCAACTATATATTTATGT

TCTTTAGGCATGAACTTACATCTGCAGGCTGCAGCTGATATGGTATCCATGTTTGATTCCGTGAATGTAG

TAGATGTTATTCCAGAAGAAGAAGTCCCTGATTTATAAGCATCTGTAGCAACAAAAAGCTTTCTTTTGGA

AACCACATTGTCCTTCTTAATTTCTATCAGCAGCCGTACTTCTTTCCAGGTTAGTTCAGTAACTTCCTTT

GGAAATGCATTTGGTGTCTTCTTGAACCCAAGAAACAAAGAAAATTGAAAGTGTTGGTTGTTGTAATGAA

AATGCTTGTGGTATGTGTGTTGAAGTTATACATACCTTGAGGAAGTCTGCAATCAATTTTGTTGCAGTCG

TGCCTTCTCTTTGTTGCAC

>MSTRG.3001.1 gene=MSTRG.3001

GGACGAATATTCATTCACGGTATACACTACACGCCCTTCTCTTCTGTCGAGCCTGCAACTCTGCGCACAC

ACAAACACACACAGATGAAGACTGGAGAAAGAGATAGCAAAAGAATGAAGAAAGAGCAGGAGGAGAACAA

GCAGAGTGTGCCAGTGCTGCCATGGATGAGAAGCCCAATACAAGTTGATTCTTTTTCATACTGCCCACTT

GCTCAAGTCCCTAT

>MSTRG.3004.1 gene=MSTRG.3004

AGTACTTGAAGCTGTGATCCATTCCCCAACCAGGGAGGCTCTCTTAGGACCCAAAAGAAGGGCAGCTCAG

AGCGCTCCAATGCATAAGCCAACTGACTCAATTCCTCCTGGCTCAGCACTGCCTCACTCCCTAGTGCAAC

ATATAAAACACAATTGACCTTCTGCTTGTCCAGCCATTCCTTAATATAACCACACTCATCATCATCATAA

TCGAATTCATTACTAGATGGTGGCAAAAGTCCCACTGGTACAACTGGTTTCTTGTAAAGCTCAGAAACCA

GATGCAACCATTCAGGCTCAAACTCAATACTAGTTCTAATCAATACCAAGTCACATCCAGCTATGGAAGC

GCCAAATCTTACTGAATCAGATGTACGTGATTCATTCCCAGATTCACCATCAAAATTCTTCATGACCTCA

TGGAGGCGATAAGCAACATCAGATTCAAATGGAACCCATTTAGGCACCACAGTATAATCTTCACCTTTTA

ATCTTCCATCCTCCCCACTCATCAAAAATGAAGGTGGTCCTAGAAAACCCATAAAAGCAGCACTGAATAG

ACACAGATAAGCTGTTGAAATACCAAGCTCAGCTGCAAGTGAAGGTAACCAAAAAGAAGCATAGTCATAT

AGTATCCAATCTGGCTTTGGCCTTGAATTTTCAAGAAAGGTTAATAAATCTGATTTTAGTAAATCAAAGG

CTATTTTCAAGAACTGGGCTTTCGAGTTTGGTATGTCCATCGAAGATTCTTCATGTGGAGATAAGTTCTC

AACTTGCGGTAATGGAAAGGTGACTAAATCAATAAGATGGACTAAATCTGAAGGTATTTTTGGGAGCCTT

TGGATGTTTCTTGGGGTGGAAATAAATGAAATTTTATGACCTTTTCGCGCTAGTATCTTGGCCAGGCGAA

GAGAAGGTCCAAGATGTCCTATTGCTAGCCATGGAAACATTACTATATGCAGAACATTGTTTTTCTCCAT

GAATACAGAAAGACAGAGATATTCAAGTTACTTGAATCTGTTACAAAATTTATACAATTCTAGGATAAGA

TAATGAGTGCTACTATTATTGCAATTTGAATAGGATGAATGTCTTAATGTAGTTGGTCAATATGGGGGGC

TGAATTATTTTATCCTGTCCTTGTCTAATCTTAGCTGACTATATATCTATGTTCATATGCTGCAATTCTT

TTGTCTTCCTCTGGTGC

>MSTRG.3005.1 gene=MSTRG.3005

ACTACATTAAGACATTCATCCTATTCAAATTGCAATAATAGTAGCACTCATTATCTTATCCTAGAATTGT

ATAAATTTTGTAACAGATTCAAGTAACTTGAATATCTCTGTCTTTCTGTATTCATGGAGAAAAACAATGT

TCTGCATATAGTAATGTTTCCATGGCTAGCAATAGGACATCTTGGACCTTCTCTTCGCCTGGCCAAGATA

CTAGCGCGAAAAGGTCATAAAATTTCATTTATTTCCACCCCAAGAAACATCCAAAGGCTCCCAAAAATAC

CTTCAGATTTAGTCCATCTTATTGATTTAGTCACCTTTCCATTACCGCAAGTTGAGAACTTATCTCCACA

TGAAGAATCTTCGATGGACATACCAAACTCGAAAGCCCAGTTCTTGAAAATAGCCTTTGATTTACTAAAA

TCAGATTTATTAACCTTTCTTGAAAATTCAAGGCCAAAGCCAGATTGGATACTATATGACTATGCTTCTT

TTTGGTTACCTTCACTTGCAGCTGAGCTTGGTATTTCAACAGCTTATCTGTGTCTATTCAGTGCTGCTTT

TATGGGTTTTCTAGGACCACCTTCATTTTTGATGAGTGGGGAGGATGGAAGATTAAAAGGTGAAGATTAT

ACTGTGGTGCCTAAATGGGTTCCATTTGAATCTGATGTTGCTTATCGCCTCCATGAGGTCATGAAGAATT

TTGATGGTGAATCTGGGAATGAATCACGTACATCTGATTCAGTAAGATTTGGCGCTTCCATAGCTGGATG

TGACTTGGTATTGATTAGAACTAGTATTGAGTTTGAGCCTGAATGGTTGCATCTGGTTTCTGAGCTTTAC

AAGAAACCAGTTGTACCAGTGGGACTTTTGCCACCATCTAGTAATGAATTCGATTATGATGATGATGAGT

GTGGTTATATTAAGGAATGGCTGGACAAGCAGAAGGTCAATTGTGTTTTATATGTTGCACTAGGGAGTGA

GGCAGTGCTGAGCCAGGAGGAATTGAGTCAGTTGGCTTATGCATTGGAGCGCTCTGAGCTGCCCTTCTTT

TGGGTCCTAAGAGAGCCTCCCTGGTTGGGGAATGGATCACAGCTTCAAGTACTTCCTGAAGGGTTCATTC

AACGAGTCAGAGGTCGAGGGATCGTTCATGTTGGATGGGTTCCACAGGTGAAGATACTTAGTCATTCAGC

TGTTGGAGGTTTCTTGACTCATTGTGGTTGGAACTCAGTCATTGAGGCACTTGGATATGGCCGAGTTCTA

ATACTTTTTCCAGTAATGAATGACCAGGGAATAAATGCTAGGCTTTTGAACAGAAAAAAAGTTGGTTTTG

AAATTCCTAGGAACGAGAAAGATGGAACATTCACAATTGACTCAGTTGCAGATTTGTTGAGAATGGCTAT

GGTGAGTCAAGAGGGCGAATCAGTGAGGAACAATGCAAAAGAGATCAAGGACTTATTTGCAGAAGGGGTC

GAGAATGACAAATACATAGCAAATTTTGTATCACAACTTGAAAGTATTAGGGCCTCCCATCTAATGTAAT

GAACAAATATAATGAAAGTAAAGCAAATACATGTACGCTTAAGAAACTATACACTTGTAAATGTGCTCTG

TACTGTTAGCCATGCGCAAACTACAAACCACTATAATCTTTATTATTATGTTCATCAAGCTAAGGAAGCA

CTCCTGGAAATTTGTGACAAAGCATCTTCAAACTGATCATCATCTGGCTTCTCTGCAGCCAAAGGGGGCG

GAATGAATGATTTGAATACGAATTCCATGTCAGGTTCTTCCTTCAGATCTTCCCGAGCAACCTCCAGTGC

TGTGTCGTGGAATGATCTATATAATTTCTTGGAGCAAACATATGCAAAAATGAAGGAAAAGATTGGTAGC

GGTATTAGAATTGGCGTGTAATAGAACTTGTTCACTCCAAAGTAACCAAACATGGTGAGTTGGAACAGGA

TCAAGGCTGCAACAATGCGAGTATGTATGTGAGGCCACATCTGTCCGTAGCTTTCATACGAGGGAAC

>MSTRG.3007.1 gene=MSTRG.3007

CCTAAAGGTTGCATCTTTCAAAAGAAAAGAAAACACAATAATATATTTTCAGCAAAATAAATATGAATAA

ACACCAGCAGAAATCATACCAGTAGCCAAAAAGACAAAGTAGACAGCAGAATCAACTCCAGAAATACAAA

TAACATCTTTTTCAGTAGACGAAAAAGCTTCCTGGATCCAGGTAAACGGGTTCCGGGTCTTGTAGCCTGC

ATCATACGGATCCAGCCCCTTCAAGATCCGGTTAGGATAATAAACCACATTATTCCCAGGTTTCTTTGAC

AACCAAGAAAACACCAGCATGAGAAGTAGAAAAATGATGAACGACGTCGCCAAGGACGTCAAGAAGGAGC

TCAAATCCATCTGGGCATTCTTCTACAACGCAAATGCGGGGGTGTAAACGTGGTTTGTGAAGATGTGCAT

GTAATAAAATAAACAAGAAGAGGTTAAAAGAATAAGGGTGTTGGGAAGCTTAAACAATGTGCAAAGTCGG

AGACAAAAGTAATGAATATGTGTTGATATATATATGTGTGTGCTGTGATGGTTGTTAGAAGTTATATATA

TTCGTTAAGTGTATTGCTAGCTTGGGTGATGATGACAAGAAGAAAAGAGAAAGACAACAAGTTGTTAGAG

CTGGAAAAAGTTATACAAATGCAACTCGGAGGGAACTATGTGAGAAAATGATTTACAAAATTATGGTTTT

TGTTCTTTTTTCCTTTTTTTTGAAAGGAAATATTATAGTTTTTGTTCAAACTTGACAAAATTCAAACATT

GTTTCTATCATGTATCACGTTATAATCGG

>MSTRG.3029.1 gene=MSTRG.3029

TGCAGTGAACTCCACTAGTAACCCAGGTGGCACAACAGGAACCACTCATTTGATTACTTATGTTAGACGT

GGAGGCAAGGGCAAAAAGGGGAATTCAAGTAGGAGCAACTAGTAATGCTGTGAAGGGATATAAGGACAAG

GCATTAGAGAGAATGAATTTATTGAGAATTGTAGCCTGTTTTATTTACTTTCCTTTTGGGAGTGAGGGCC

ACTCGAAGAGCCTGTGTTATTCTTGTTTCCGTTGTAATCAGAAACCCTTATTATCTAGCCCTATTTTTAT

TAAGCAATTGAGTTTCATTCTAGTCTATACCAAAGTGGTATCAGAGCATTCGATCATGGCTCCACCACCT

GAGACACGACAAACCCAACGACTTGAATTAGTAGAAGAAGCAGTAGCTACGTTGCAGAGTACACTTTCTG

CGGAGGTGTCTGCGGCGGTGACTCGAGCCGCGACGGAAATGCAACAAACCCTCGTATCTCAGCTCACCAC

CAGCCTCGACCAAGTCACTCAAAAACTTCAGATCCGAATGGACCGCGTCCGAGAAAACAACGAAACCCTA

ATTGCTGAAGTGACGCGCCGCCAAGAGGAATTTCAGAACGAGATGAGATCTTCCCTTTCAACTCTCACAC

AAGGATTGGGTACCGGATCTGGACCCAAACCTTTTGCTGATATGGGATCTCGAATTCATGATATTGGGGG

TACTGGGTTTCCTTCCGGCAGCGGTGTCTTTGGAGGTTTCAGTAGCCAGAAGACTCCGGAGGACCCAGGA

GGCGGCAACCCTGCAGTGACTGGGGGTAATTGGCGGTATCGAAAGCTTGACTTACCAGTATTTGATGGGA

CAAATCCCGATGGATGGATTCTCCGGGCGGAGAGGTATTTTTCTTTTTATCGGTTGAATGAGGAGGAGAA

ACTGGAGGCGGCAGTGGTGGGTTTTGATGGGGATGCCCTCCTTTGGTACCAATGGGAAAATCGGAGACGT

CCTATTCGGCGGTGGGATGAGCTCCGTAATTTGATGTTACAGCAGTTTCGCCCAATGCATTCTGGTTCGT

TGTGTGAGCAATGGTTGGCAGTGACACAGACGGGTACGGTGGCTGAATTTAAGCGCAGATTCATTGAGAT

GGCAGCACCCCTTGAGAGGATTCCAAGAATTTATACTAATGGGCCATTTTATTAATGGGTTGAGGGAGGA

CATCAGGGCAGAAGTAAGGATGCTCGGGCTTCATAGTCTGGAACAAGCTATGGATTTAGCTTTAAAGGTG

GAGGAGAAAAATCGGGCCAGGTTAAGCACCTTGCGGGAAGGTAAACTCCAACACTCCCAGTTCCTCAAGT

TTGGGTCAAATTCACAATATACAAGTAATAGTAAACAGTCTGACAACTCTAATATGAGCCCACTTGCATT

GGTGAACCCTTCCAAATCTGTGTTCTCGACGACCGGCTCGACTGGGTCTAGTGGACAGTGGTCTCCATCA

GTGGCCAGCCCAAAATCTTCTTCTGGAGCGAGCCAGTCCAGCTCCTCTCTGCCCGTGGCACGGCCTGTTG

GTGAGGTCCGGCGGTTGACCGATAAGGAGCTGCAATATAAGAGGGACAGAGGCCTTTGTTTTCGCTGTGA

CGACAAGTGGTCAGCAGGGCATCGCTGTAAACGTAAAGAGTTGTGTGTGTTGGTGACACAAGAGCCCGAA

GATGAGGAAGATAGGGAGGTTGAAGATTGTCCGGTGACAGAAGCTGTGATTGATTGTTCAAAGCAGATTT

CCTTGAATTCGGTAGTGGGCATATCATCTCCGAAAACTATGAAGTTAAAAGGGACCCTTGCGGGACAGGA

TGTGGTAATCATGATTGATCCTGGGGCTACACATAATTTCTTGGCTTTGAATACTATCAAACGGTTAAAG

ATTCCGGTGGCAGAAACTAAAGAGTTTGGGGTAACACTTGGTAACGGGGAGGCTGTAAAGGGGATTGGCG

AGTGCAAGGGGGTGATTTTACAAATGGGGACTATAGAGGTAGTTGAAGATTTTTTGCCGTTCTCTTTGGG

AAACTCGGATGTAATTCTTGGAATCCAATGGTTGGAAAAGCTAGGGGCAGTGACAACTAACTGGAAGTTG

CAGGAGATGAAGTTTAAGATAGGCCAGGAGACTGTTTTGCTCAAAGGAGATCCTTCACTTGAACACTCGA

AGGTTTCTCTAAAGTCAATGATGCGCATCATCAAGAAAGAAGGCGGGGGGTTGCTGGTTGAGATCCATCA

GTTAGAATCCCTAGCAAGAGGGCAGAGTGTAGATGAGCAGCAACAGCCCAACCCGGAGTTTGTACAACCC

TTGTTAACAAAGTACGAGGCAGTGTTTCAGACACCACAAGGGCTACCTCCCAAGCGTAATCACGAGCATG

CTATTCACTTGAAGGCCGGGGCAAATCCAGTGAGTGTGCGGCCCTATCGATATCCCCAATTTCAAAAGGA

TGAAATTGAGCGGCTGATTAAGGAGATGCTTGAGGCAGGTATTATCAGACCCTCTTTTAGTCCATTCTCA

AGTCCTGTTTTGTTGGTTAAGAAGAAGGACGGATCATGGCGGTTTTGTGTGGATTACCGCGCATTGAACA

AGGAAACCGTTCCCGATAAATACCCAATTCCCATTATCGATGAGTTGTTGGATGAGCTAGGAGGGACAAA

GGTCTTCTCAAAACTAGACCTTAAATCTGGATATCATCAAATTCGGATTAAAGAGGAGGACATCCATAAA

ACGGCTTTTCGGACTCATGAGGGCCATTATGAATTTTTAGTAATGCCGTTTGGCCTCACGAACGCCCCTG

CAACTTTTCAATCCCTAATGAATGAGGTATTTCGTCCCCATCTTCGCCGATTTGTTTTAGTCTTTTTTGA

TGATATTCTTGTTTATAGCAAGAATGAGCTGGAACACCTCCAACATTTAGAAGTTGTGCTTAATCTGTTA

AGTACTCATTGCTTGGTTGCTAATTTTAAAAAGTGTGATTTCGGAAGGAGAGAAGTAGCCTATCTGGGGT

ATGTGGTATCTGGGGAAGGGGTGGCAGCAGATATGAGTAAGGTGAAGGCCATGGTAGAGTGGCCAGAACC

GAAGAATTTAAGGGAATTGAGGGGATTCTTGGGGCTAACAGGATATTACCGTAAATTTGTGGCAGGGTAC

GCCTCAATCGCACAGCCACTCACGGAGCAGCTTAAGAAGGACAGTTTTTCATGGAATTCAGCTGCAACAG

TGGCCTTTAACAGTCTTAAACAGCGGATGGTTAGCTCTCCTGTTTTGAGGATGCCAGATTTCAGTTTACC

ATTCGTAGTTGAGTCAGATGCATCTGGGTTTGGTATTGGTGCAGTATTGTCTCAGTTGGACAGACCAGTT

GCATTTTTCAGTAAGATTCTTGGGGTAAGAGCTCGAACGAAATCGGTGTATGAGAAAGAATTGATGGCCA

TTTGTCTAGCAGTCCAAAAATGGAAGCACTACCTTCTAGGAAGGCACTTCATCATTCGCACGGACCAGCA

GAGTTTGCGGTTCATAATGCATCAAAGGGAGTTGGGGGCTGATTTTCAGAAATGGGTGAGTAAATTAATT

GGGTTCGATTTTGAGATTGTGTATCGCACTGGTGCCTCGAACAAAGTAGCTGACGCCCTTTCCAGGCAAA

ATGAAGGAAGGGTAATGTTGTGTGCCCTTGCCAGTGCGAGTGTCATTGATTGGTCCGGATTGCAGGCCGA

AGCCCAAAGAGATCCTTTCATTTCGAGGACCATTGCTGAGATTCAAGATGAAGGCAGAGTATGTCCTGGA

TTCACTGTTGTTGATGGAAAATTATTTTACAAGGAGAGGATGGTCGTAGCTCGTAATTCAGTATACATTC

CTGAACTACTTAAAGCATATCATGACTCCCCGGTAGGAGGCCATTCAGGGGAGTTAAAAACTTATCTAAG

ACTGGCTAAGGAGTGGTTTTGGCAGGGAATGCGGGCTGCTGTGACTTCGTATGTGCGACAGTGCTTGATA

TGTCAGCAAAATAAATCTCTTACTCAGAGTCCAGCTGGGTTGTTACAACCCTTGCCAGTACCCATGCAGG

TTTGGGAAGACTTGACCCTCGATTTTATAGAAGGACTACCGGTATCATCAGGATTTGATACCATCTTAGT

TGTGGTTGACAGACTCACGAAATATGCTCACTTCCTGAAGCTTAAACATCCTTTCACCGCTTCTTCAGTG

GCGGCCGTGTTTGTGTCAGAGATCATCCGTCTACATGGGTTCCCTGCATCCATCATCTCTGACCGAGACC

GTGTTTTCATGAGTATATTCTGGACAGAAATGTTCCGCCTTCAGGGAACACAACTGAAGAGAAGCACAGC

CTACCACCCACAGACGGACGGACAGTCTGAGGTGGTGAACAAATGTTTGGAAGCATATTTGCGTTGTTTT

GCATCAAGCCACCCTCGTACATGGGCGAAATGGATGAGCTGGGCCGAGTATTGGTATAACACAGCTCCAC

ACACCTCCATCAAAATGTCTCCGTTCAAGGCCCTTTATGGCAGAGAACCACCCCAGTTGCTGCGTTGGGG

AAGCAATCAGACCCCTGTGGATAAACTGGAGGAAAACTTACAGGAACGAGACTACATACTAGATGAATTA

AGAGCAGTGCTGCTTAGCACCCAACAACGAATGAAGGCACAGGCTGATGCACACCGTCGTGATGAGTCTT

TTGAAATTGGTGATCGGGTTTTTCTGAAATTACAACCATACCGCCAACAGTCATTGGCCCGCAGACCCTA

TGAAAAACTAGCAGCTCGATTCTATGGGCCCTTTGCAGTTTTGAAGCGAATTGGTAAGGTGGCTTATCAC

CTTGATCTTCCTGCAGATAGCAAAATCCACCCGATCTTTCATATTTCTCAGCTCAAGCGAGCTATAGGCA

ATGTTTCTGCTCAGCCTACGGTACCACCGCAGTTATCAGTAGATATGGAGTTACTGGCCCAACCAGAAGC

GTTACTGAATGTGAGACAGCAGCAAGATGGCCAAACCATGTTTTTGGAGGTGTTGATTCAATGGATGGGA

CTTCCTCTGTCTGAAGCAACATGGGAACAACTGGATGAAATGCAATTGCGATTTCCGGAGTTCCACCTTG

AGGACAAGGTGCGTGTTTGGGCAGCGGGTTCTGCAGTGAACTCCACTAGTAACCCAGGTGGCACAACAGG

AACCACTCATTTGATTACTTATGTTAGACGTGGAGGCAAGGGCAAAAAGGGGAATTCAAGTAGGAGCAAC

TAGTAATGCTGTGAAGGGATATAAGGACAAGGCATTAGAGAGAATGAATTTATTGAGAATTGTAGCCTGT

TTTATTTACTTTCCTTTTGGGAGTGAGGGCCACTCGAAGAGCCTGTGTTATTCTTGTTTCCGTTGTAATC

AGAAACCCTTATTATCTAGCCCTATTTTTATTAAGCAATTGAGTTTCATTCTAGTCTATACCAC

>MSTRG.3038.1 gene=MSTRG.3038

GTGTTGTAAAAAGCGGGAATCGGACTTAATCGGTGAAGTCACGGAACAGGGATTAATCGGTGATTAATTG

AATTGATCAGGGATTAATCGGATTAATCAGTGATTAATCGGAGATTTAATCAGTTCGGCGAGTTACGGAA

CATGGATTAATCGGTGACTAATCGTGATTAATCACCGACTTTTAGAACAGAGGCAATCACACGA

>MSTRG.3042.1 gene=MSTRG.3042

CAAGGCCTTTTAATCTGGCATCGTATCCCCACTTCATCAAATCTCCTGAACGGTTATAAACTCTCTGCAA

CCCATTACTTTTCCACGTCTTTCTTCTTTTTTGACTTGGCCATCTCAACGCGCAACAGTTAATTTTATTC

TCTTCGTGTACATATTTATAGCAAAACCATACTCGTTTCATTCACTCACAAACCTACACAAAGTACACAA

TTATTTGTACATTTTTTAGAGAGTTATGGAGAACAATGTTGGCAAAGAAGGAAGCGTTGGTTTCATGCAT

CTTCTCAAGAGGGTTGGGAAGAAGAGTGGCGGTTCGTTCCGGAGATTCGGGCGGTGGAAGAAGAGATGGT

CTTTTTCTAGGTCGTCTTCGTCGTCGTCGAGTTCGTATTCGAATTCGAGGTTGCCTAGGTGGTTTGTGAA

TGATGTTTTGTTTAAAATTGTTTCGGTTTTTGAGGGGGTGGTTTTGGTTTCTACTCTCTGCATTTTCTTC

CTCTGTTGTGGATGTCACTTTTGAATTTTGGTTTGTAAATTTTATGTACTTTCTTTAACTTGATAGTTGT

TGTTTTAGTAGTTTTTAAGTGAATTATCTGAGCAAATGGCTCTTGTTGTATGTAGATTTTAAGGTGGGAT

TTACATCTGTCCAGTTCAATAAAATTAAATTAATTCTTTCTACTTTATGTTCTTTCAAGTGCTTTGGGTT

GTTGATCTGGGTTTCTTCTGTTCTGTCAATCTGAATGCTGCTGGTTCTTTCCACACTGTAACATTGTCAG

TGGAGAAACGTCTCGTTAAGTTCTGC

>MSTRG.3045.2 gene=MSTRG.3045

CATCAAACCCCATCACCAAGAACTCGCCATTTCTCTCTCTCACTCACAGATCTACCCATTTTATCTTTAA

AGTTCTAATCTTGAACCATAAAGTTTTTATCTTGATCACGAATTCTCGATTTTGAAGTTGGACTGGGTTT

ATACAACTGATAATTTGCATTGGCGAAGGATTCACAATGCCTTGGATATATTGTGTTCGCTTGGTGGTGT

TCTTTCGTGTTATACTGTAATTGATTATCTAGGTCAGCTTACAACTTGAGAATCGAGATACATCCTATAT

ATTTTGGTTGGGGTGTTGATAATGTACATTGGAATCACGTCGAGGAGATACAAAGCAGTGAAAGAAGTTG

GGAAAATTAAAATGAGTGTTGGGGTTGTTGCTTACTATCAACGGATGCAAATTTGTCAGGCTGAATATTT

CCGCCAATTGCTAAAACCTGTGACGTAGGATAGTGCTATACCATTCATTTTCTCTCAGTTTGCATACAAG

ACACCTTGAGTTGCTTAGTGCTATAGCATATAGGTTGGGTTGGATGATCACCTGTACTATAATGGAAAGC

TTGTGCGTGCTTTTGTGATCTGTTGTAGTCGTAGGAACTTCCATCCAGTTGAATTCTAATATTTTCCCTT

TGAAGCTGCAAAATTCCAGTTTAGTAAAACATCTAGTCAGTTGAAGTAGTAATGCACGGGGAACAGCAGT

TTGCCCCTCAAAAGCCAAAAGGCCACTATACAGACCAATTGGTTGGTGGTTACATGTGTAATGCTCCCAT

TTCACCCTTTTTTGGTGGAACAGTTTCCCCACCAACTCAAAATCACCTGATGCCTTTCCATGGTGTCAAG

TTTCAACCTTCTGAAGCATGCCCAAGAAATTTCATTATATGTGATCAGACTGATAACCGAAGCCAGATAA

TGTTCCATCCAACAATTTGTGGCAAATTGTGTCATCCTGGCCTGAATTTCTGTCTCCCTTCCTTTGAAGA

AGATAGAGCTAAAGTGAATACAAATATTGATGAGGGAGAAGATTCTCCGTTGAAAGAAGACTCCGATGAT

ATTGATGCATTGTTGAACTATGACGAGGAAGAGGAGGAAGAAAGTGATGATGAGGTAGTTAGCACAGCTC

GTACTTGTGGAAATTATCCAAGTGATTTCCTTGATTCTTGTTCCTCGCATGGTTCAAGACCCAAGAAGGG

CAGGTTGCCTACTTTTTCACAGGAGTCGTATGGAAATGACAGCAACTGCAGCGGCAGAAAGCGTCAGAAA

GTGAAGAAGATGGTTAAGGCATTAAAAAGAATTGTGCCTGGGGGAAACCAGATGGACACTGTCACTGTCC

TTGATGAAGCTGTGAGGTACCTCAAGTCTCTTAAAGTAGAAGTTGAGACACTGGGTGCTGGAAACATAAA

GACGTAATTCAGACTCTACTGCTTACGCAATAAATTACAAAATGATATTGCTCCCAGCGAATGGGATTAA

GATGGTCCGTGTATGTGTTAATAATGTCATGCATACTGTTGTGGTGGATGTACAGATTGCATGTACTGCT

CATGCGGACTTTTGCTGTCTAGACACCAAACAATGCTGCTATTATATTTCCTGATGCAACTCAGAGTAAG

AGTCCTTTCTGTATTATGCAAACTTCTGGTCAGAAATATGTAAACTATATTTGAATCTTGACTTTTCTGC

ATTTCGATTCTTCGTATTGGTATTATTATACAGACTAGAGTTGATTTTTATTATTAAA

>MSTRG.3045.1 gene=MSTRG.3045

CATCAAACCCCATCACCAAGAACTCGCCATTTCTCTCTCTCACTCACAGATCTACCCATTTTATCTTTAA

AGTTCTAATCTTGAACCATAAAGTTTTTATCTTGATCACGAATTCTCGATTTTGAAGTTGGACTGGGTTT

ATACAACTGATAATTTGCATTGGCGAAGGATTCACAATGCCTTGGATATATTGTGTTCGCTTGGTGGTGT

TCTTTCGTGTTATACTGTAATTGATTATCTAGCTTACAACTTGAGAATCGAGATACATCCTATATATTTT

GGTTGGGGTGTTGATAATGTACATTGGAATCACGTCGAGGAGATACAAAGCAGTGAAAGAAGTTGGGAAA

ATTAAAATGAGTGTTGGGGTTGTTGCTTACTATCAACGGATGCAAATTTGTCAGGCTGAATATTTCCGCC

AATTGCTAAAACCTGTGACGTAGGATAGTGCTATACCATTCATTTTCTCTCAGTTTGCATACAAGACACC

TTGAGTTGCTTAGTGCTATAGCATATAGGTTGGGTTGGATGATCACCTGTACTATAATGGAAAGCTTGTG

CGTGCTTTTGTGATCTGTTGTAGTCGTAGGAACTTCCATCCAGTTGAATTCTAATATTTTCCCTTTGAAG

CTGCAAAATTCCAGTTTAGTAAAACATCTAGTCAGTTGAAGTAGTAATGCACGGGGAACAGCAGTTTGCC

CCTCAAAAGCCAAAAGGCCACTATACAGACCAATTGGTTGGTGGTTACATGTGTAATGCTCCCATTTCAC

CCTTTTTTGGTGGAACAGTTTCCCCACCAACTCAAAATCACCTGATGCCTTTCCATGGTGTCAAGTTTCA

ACCTTCTGAAGCATGCCCAAGAAATTTCATTATATGTGATCAGACTGATAACCGAAGCCAGATAATGTTC

CATCCAACAATTTGTGGCAAATTGTGTCATCCTGGCCTGAATTTCTGTCTCCCTTCCTTTGAAGAAGATA

GAGCTAAAGTGAATACAAATATTGATGAGGGAGAAGATTCTCCGTTGAAAGAAGACTCCGATGATATTGA

TGCATTGTTGAACTATGACGAGGAAGAGGAGGAAGAAAGTGATGATGAGGTAGTTAGCACAGCTCGTACT

TGTGGAAATTATCCAAGTGATTTCCTTGATTCTTGTTCCTCGCATGGTTCAAGACCCAAGAAGGGCAGGT

TGCCTACTTTTTCACAGGAGTCGTATGGAAATGACAGCAACTGCAGCGGCAGAAAGCGTCAGAAAGTGAA

GAAGATGGTTAAGGCATTAAAAAGAATTGTGCCTGGGGGAAACCAGATGGACACTGTCACTGTCCTTGAT

GAAGCTGTGAGGTACCTCAAGTCTCTTAAAGTAGAAGTTGAGACACTGGGTGCTGGAAACATAAAGACGT

AATTCAGACTCTACTGCTTACGCAATAAATTACAAAATGATATTGCTCCCAGCGAATGGGATTAAGATGG

TCCGTGTATGTGTTAATAATGTCATGCATACTGTTGTGGTGGATGTACAGATTGCATGTACTGCTCATGC

GGACTTTTGCTGTCTAGACACCAAACAATGCTGCTATTATATTTCCTGATGCAACTCAGAGTAAGAGTCC

TTTCTGTATTATGCAAACTTCTGGTCAGAAATATGTAAACTATATTTGAATCTTGACTTTTCTGCATTTC

GATTCTTCGTATTGGTATTATTATACAGACTAGAGTTGATTTTTATTATTAAAGCTTTTTCCTGTAATAT

TCTGTGTTTAATTTCTCTTCAAATCTCTATTGGCC

>MSTRG.3045.3 gene=MSTRG.3045

ACCCCATCACCAAGAACTCGCCATTTCTCTCTCTCACTCACAGATCTACCCATTTTATCTTTAAAGTTCT

AATCTTGAACCATAAAGTTTTTATCTTGATCACGAATTCTCGATTTTGTTGGACTGGGTTTATACAACTG

ATAATTTGCATTGGCGAAGGATTCACAATGCCTTGGATATATTGTGTTCGCTTGGTGGTGTTCTTTCGTG

TTATACTGTAATTGATTATCTAGCTTACAACTTGAGAATCGAGATACATCCTATATATTTTGGTTGGGGT

GTTGATAATGTACATTGGAATCACGTCGAGGAGATACAAAGCAGTGAAAGAAGTTGGGAAAATTAAAATG

AGTGTTGGGGTTGTTGCTTACTATCAACGGATGCAAATTTGTCAGGCTGAATATTTCCGCCAATTGCTAA

AACCTGTGACGTAGGATAGTGCTATACCATTCATTTTCTCTCAGTTTGCATACAAGACACCTTGAGTTGC

TTAGTGCTATAGCATATAGGTTGGGTTGGATGATCACCTGTACTATAATGGAAAGCTTGTGCGTGCTTTT

GTGATCTGTTGTAGTCGTAGGAACTTCCATCCAGTTGAATTCTAATATTTTCCCTTTGAAGCTGCAAAAT

TCCAGTTTAGTAAAACATCTAGTCAGTTGAAGTAGTAATGCACGGGGAACAGCAGTTTGCCCCTCAAAAG

CCAAAAGGCCACTATACAGACCAATTGGTTGGTGGTTACATGTGTAATGCTCCCATTTCACCCTTTTTTG

GTGGAACAGTTTCCCCACCAACTCAAAATCACCTGATGCCTTTCCATGGTGTCAAGTTTCAACCTTCTGA

AGCATGCCCAAGAAATTTCATTATATGTGATCAGACTGATAACCGAAGCCAGATAATGTTCCATCCAACA

ATTTGTGGCAAATTGTGTCATCCTGGCCTGAATTTCTGTCTCCCTTCCTTTGAAGAAGATAGAGCTAAAG

TGAATACAAATATTGATGAGGGAGAAGATTCTCCGTTGAAAGAAGACTCCGATGATATTGATGCATTGTT

GAACTATGACGAGGAAGAGGAGGAAGAAAGTGATGATGAGGTAGTTAGCACAGCTCGTACTTGTGGAAAT

TATCCAAGTGATTTCCTTGATTCTTGTTCCTCGCATGGTTCAAGACCCAAGAAGGGCAGGTTGCCTACTT

TTTCACAGGAGTCGTATGGAAATGACAGCAACTGCAGCGGCAGAAAGCGTCAGAAAGTGAAGAAGATGGT

TAAGGCATTAAAAAGAATTGTGCCTGGGGGAAACCAGATGGACACTGTCACTGTCCTTGATGAAGCTGTG

AGGTACCTCAAGTCTCTTAAAGTAGAAGTTGAGACACTGGGTGCTGGAAACATAAAGACGTAATTCAGAC

TCTACTGCTTACGCAATAAATTACAAAATGATATTGCTCCCAGCGAATGGGATTAAGATGGTCCGTGTAT

GTGTTAATAATGTCATGCATACTGTTGTGGTGGATGTACAGATTGCATGTACTGCTCATGCGGACTTTTG

CTGTCTAGACACCAAACAATGCTGCTATTATATTTCCTGATGCAACTCAGAGTAAGAGTCCTTTCTGTAT

TATGCAAACTTCTGGTCAGAAATATGTAAACTATATTTGAATCTTGACTTTTCTGCATTTCGATTCTTCG

TATTGGTATTATTATACAGACTAGAGTTGATTTTTATTATTAAAGCTTTTTCCTGTAATATTCTGTGTTT

AATTTCTCTTCAAATCTCTATTGGCC

>MSTRG.3045.4 gene=MSTRG.3045

AACTCGCCATTTCTCTCTCTCACTCACAGATCTACCCATTTTATCTTTAAAGTTCTAATCTTGAACCATA

AAGTTTTTATCTTGATCACGAATTCTCGATTTTGGATTCACAATGCCTTGGATATATTGTGTTCGCTTGG

TGGTGTTCTTTCGTGTTATACTGTAATTGATTATCTAGCTTACAACTTGAGAATCGAGATACATCCTATA

TATTTTGGTTGGGGTGTTGATAATGTACATTGGAATCACGTCGAGGAGATACAAAGCAGTGAAAGAAGTT

GGGAAAATTAAAATGAGTGTTGGGGTTGTTGCTTACTATCAACGGATGCAAATTTGTCAGGCTGAATATT

TCCGCCAATTGCTAAAACCTGTGACGTAGGATAGTGCTATACCATTCATTTTCTCTCAGTTTGCATACAA

GACACCTTGAGTTGCTTAGTGCTATAGCATATAGGTTGGGTTGGATGATCACCTGTACTATAATGGAAAG

CTTGTGCGTGCTTTTGTGATCTGTTGTAGTCGTAGGAACTTCCATCCAGTTGAATTCTAATATTTTCCCT

TTGAAGCTGCAAAATTCCAGTTTAGTAAAACATCTAGTCAGTTGAAGTAGTAATGCACGGGGAACAGCAG

TTTGCCCCTCAAAAGCCAAAAGGCCACTATACAGACCAATTGGTTGGTGGTTACATGTGTAATGCTCCCA

TTTCACCCTTTTTTGGTGGAACAGTTTCCCCACCAACTCAAAATCACCTGATGCCTTTCCATGGTGTCAA

GTTTCAACCTTCTGAAGCATGCCCAAGAAATTTCATTATATGTGATCAGACTGATAACCGAAGCCAGATA

ATGTTCCATCCAACAATTTGTGGCAAATTGTGTCATCCTGGCCTGAATTTCTGTCTCCCTTCCTTTGAAG

AAGATAGAGCTAAAGTGAATACAAATATTGATGAGGGAGAAGATTCTCCGTTGAAAGAAGACTCCGATGA

TATTGATGCATTGTTGAACTATGACGAGGAAGAGGAGGAAGAAAGTGATGATGAGGTAGTTAGCACAGCT

CGTACTTGTGGAAATTATCCAAGTGATTTCCTTGATTCTTGTTCCTCGCATGGTTCAAGACCCAAGAAGG

GCAGGTTGCCTACTTTTTCACAGGAGTCGTATGGAAATGACAGCAACTGCAGCGGCAGAAAGCGTCAGAA

AGTGAAGAAGATGGTTAAGGCATTAAAAAGAATTGTGCCTGGGGGAAACCAGATGGACACTGTCACTGTC

CTTGATGAAGCTGTGAGGTACCTCAAGTCTCTTAAAGTAGAAGTTGAGACACTGGGTGCTGGAAACATAA

AGACGTAATTCAGACTCTACTGCTTACGCAATAAATTACAAAATGATATTGCTCCCAGCGAATGGGATTA

AGATGGTCCGTGTATGTGTTAATAATGTCATGCATACTGTTGTGGTGGATGTACAGATTGCATGTACTGC

TCATGCGGACTTTTGCTGTCTAGACACCAAACAATGCTGCTATTATATTTCCTGATGCAACTCAGAGTAA

GAGTCCTTTCTGTATTATGCAAACTTCTGGTCAGAAATATGTAAACTATATTTGAATCTTGACTTTTCTG

CATTTCGATTCTTCGTATTGGTATTATTATACAGACTAGAGTTGATTTTTATTATTAAAGCTTTTT

>MSTRG.3045.5 gene=MSTRG.3045

TTTTTTGGTGTTTCAGAAGTTGGACTGGGTTTATACAACTGATAATTTGCATTGGCGAAGGATTCACAAT

GCCTTGGATATATTGTGTTCGCTTGGTGGTGTTCTTTCGTGTTATACTGTAATTGATTATCTAGGAGGTC

AGCTTACAACTTGAGAATCGAGATACATCCTATATATTTTGGTTGGGGTGTTGATAATGTACATTGGAAT

CACGTCGAGGAGATACAAAGCAGTGAAAGAAGTTGGGAAAATTAAAATGAGTGTTGGGGTTGTTGCTTAC

TATCAACGGATGCAAATTTGTCAGGCTGAATATTTCCGCCAATTGCTAAAACCTGTGACGTAGGATAGTG

CTATACCATTCATTTTCTCTCAGTTTGCATACAAGACACCTTGAGTTGCTTAGTGCTATAGCATATAGGT

TGGGTTGGATGATCACCTGTACTATAATGGAAAGCTTGTGCGTGCTTTTGTGATCTGTTGTAGTCGTAGG

AACTTCCATCCAGTTGAATTCTAATATTTTCCCTTTGAAGCTGCAAAATTCCAGTTTAGTAAAACATCTA

GTCAGTTGAAGTAGTAATGCACGGGGAACAGCAGTTTGCCCCTCAAAAGCCAAAAGGCCACTATACAGAC

CAATTGGTTGGTGGTTACATGTGTAATGCTCCCATTTCACCCTTTTTTGGTGGAACAGTTTCCCCACCAA

CTCAAAATCACCTGATGCCTTTCCATGGTGTCAAGTTTCAACCTTCTGAAGCATGCCCAAGAAATTTCAT

TATATGTGATCAGACTGATAACCGAAGCCAGATAATGTTCCATCCAACAATTTGTGGCAAATTGTGTCAT

CCTGGCCTGAATTTCTGTCTCCCTTCCTTTGAAGAAGATAGAGCTAAAGTGAATACAAATATTGATGAGG

GAGAAGATTCTCCGTTGAAAGAAGACTCCGATGATATTGATGCATTGTTGAACTATGACGAGGAAGAGGA

GGAAGAAAGTGATGATGAGGTAGTTAGCACAGCTCGTACTTGTGGAAATTATCCAAGTGATTTCCTTGAT

TCTTGTTCCTCGCATGGTTCAAGACCCAAGAAGGGCAGGTTGCCTACTTTTTCACAGGAGTCGTATGGAA

ATGACAGCAACTGCAGCGGCAGAAAGCGTCAGAAAGTGAAGAAGATGGTTAAGGCATTAAAAAGAATTGT

GCCTGGGGGAAACCAGATGGACACTGTCACTGTCCTTGATGAAGCTGTGAGGTACCTCAAGTCTCTTAAA

GTAGAAGTTGAGACACTGGGTGCTGGAAACATAAAGACGTAATTCAGACTCTACTGCTTACGCAATAAAT

TACAAAATGATATTGCTCCCAGCGAATGGGATTAAGATGGTCCGTGTATGTGTTAATAATGTCATGCATA

CTGTTGTGGTGGATGTACAGATTGCATGTACTGCTCATGCGGACTTTTGCTGTCTAGACACCAAACAATG

CTGCTATTATATTTCCTGATGCAACTCAGAGTAAGAGTCCTTTCTGTATTATGCAAACTTCTGGTCAGAA

ATATGTAAACTATATTTGAATCTTGACTTTTCTGCATTTCGATTCTTCGTATTGGTATTATTATACAGAC

TAGAG

>MSTRG.3045.6 gene=MSTRG.3045

TTTGATGCTTTAAGAATTGTATAATTTGTATATCATATTTTTGAACTGTCTGTGGAAAGTACCTGTAAAG

TCATCACTATAAAAAATCAAATTGGGGCGTCTTGCTTATTTTGTGTTGATAGATTATATTCTATTTAGGA

CTTTGGCTGATTTCCAATGGGACTATGTGTCACCTTAATTCCCGTATTTGGAAAATACAAATAGTGTTAT

CTCTCCAACAGCAAATTGTATGTTTTCCAATAATATAAACTCTATATTATATCACATTTTATAATGTTTT

ATGTATTCTTTTGAAGGATACTCATTTGCAGAGTTGAGACTTACTCACTTGTTTTATGATATGACACATA

TAATTAAGTTTTAGGTCTCAATACTTCATTACAAAAATTAATTCTTCTTGTCCTTTGGTTCATATTTTAT

ATTCAATTAAGGTAGTGGTGACTAATAAGTCTGATAATATGGCTACTGAAGTTACCTTCAATTTGTTTAG

AAATTAATTGCAAAATAAAGTTTGTTCACTAATCTGATTATTGACTATTTATAATTGCTCCTTGTTTTGT

TTCATGGAGGTGTAAAGAGCTGAGAGCTACTCGCTTGTCTTTTATGATAGTGTTTCAAACTTATACTGCC

TTAATGTTACATTGATAAATGATAACTGAGGGAAATCTTTCATTCATTTTTTTGATTGATATTGGCGCAA

GCGATAATGTTAAGAATCTGTTTGATAATTTTTTTTTATCTATGTTATATATCTTATTATTTTACAAGAA

ATTAATACAATAGGAGATTGAATACCATTTTAATATATCTAATGAAGTTTCATAGTTTGTAAATTATTGG

TGTATTCTTTACCCTTGTCTGTTGTAACAGTGAAATTGATAACTGTTAGATAGGAGATCAAATTTTATAA

TGGTCTTTTTTTATGGAGCACTGTGTGGTAACCTTAGAGTTGATGATGTAGCCTCAATGTTGTGTGCTTT

GTTTTACTTCGTGTGTTCAAATATGTATTATATTCCATAGCGCTGTTTGATAATCAAGTAATCAACCTTT

GCTCCATAGCATAATTTACAAATTTATGCTGACCAATGCTTGTGATTTTGATTCTTCTTTTGTTGGCAGG

CTGAATATTTCCGCCAATTGCTAAAACCTGTGACGTAGGATAGTGCTATACCATTCATTTTCTCTCAGTT

TGCATACAAGACACCTTGAGTTGCTTAGTGCTATAGCATATAGGTTGGGTTGGATGATCACCTGTACTAT

AATGGAAAGCTTGTGCGTGCTTTTGTGATCTGTTGTAGTCGTAGGAACTTCCATCCAGTTGAATTCTAAT

ATTTTCCCTTTGAAGCTGCAAAATTCCAGTTTAGTAAAACATCTAGTCAGTTGAAGTAGTAATGCACGGG

GAACAGCAGTTTGCCCCTCAAAAGCCAAAAGGCCACTATACAGACCAATTGGTTGGTGGTTACATGTGTA

ATGCTCCCATTTCACCCTTTTTTGGTGGAACAGTTTCCCCACCAACTCAAAATCACCTGATGCCTTTCCA

TGGTGTCAAGTTTCAACCTTCTGAAGCATGCCCAAGAAATTTCATTATATGTGATCAGACTGATAACCGA

AGCCAGATAATGTTCCATCCAACAATTTGTGGCAAATTGTGTCATCCTGGCCTGAATTTCTGTCTCCCTT

CCTTTGAAGAAGATAGAGCTAAAGTGAATACAAATATTGATGAGGGAGAAGATTCTCCGTTGAAAGAAGA

CTCCGATGATATTGATGCATTGTTGAACTATGACGAGGAAGAGGAGGAAGAAAGTGATGATGAGGTAGTT

AGCACAGCTCGTACTTGTGGAAATTATCCAAGTGATTTCCTTGATTCTTGTTCCTCGCATGGTTCAAGAC

CCAAGAAGGGCAGGTTGCCTACTTTTTCACAGGAGTCGTATGGAAATGACAGCAACTGCAGCGGCAGAAA

GCGTCAGAAAGTGAAGAAGATGGTTAAGGCATTAAAAAGAATTGTGCCTGGGGGAAACCAGATGGACACT

GTCACTGTCCTTGATGAAGCTGTGAGGTACCTCAAGTCTCTTAAAGTAGAAGTTGAGACACTGGGTGCTG

GAAACATAAAGACGTAATTCAGACTCTACTGCTTACGCAATAAATTACAAAATGATATTGCTCCCAGCGA

ATGGGATTAAGATGGTCCGTGTATGTGTTAATAATGTCATGCATACTGTTGTGGTGGATGTACAGATTGC

ATGTACTGCTCATGCGGACTTTTGCTGTCTAGACACCAAACAATGCTGCTATTATATTTCCTGATGCAAC

TCAGAGTAAGAGTCCTTTCTGTATTATGCAAACTTCTGGTCAGAAATATGTAAACTATATTTGAATCTTG

ACTTTTCTGCATTTCGATTCTTCGTATTGGTATTATTATACAGACTAG

>MSTRG.3046.4 gene=MSTRG.3046

GCATAAGCCTAGTTGTGAAGGAATGTGACCAGTGAATGAGTTTGAAGTTAGATCAAGAACCTGGAGACTA

GTAAGATTTCCAAGGAATGGAGTGACTGAGCCCTCAATTTGCATCTCAACAAGGGAAATTGAGACGACTC

GTTTCGTTGAAGGTTGACACTTGATTCCAGTCCAGTTGCAGTGATGGCTAGAGTCATTCCAATCCTGAAG

AGAACCAAGCGGGTCCTTCGTAACTGAATTCTTGAATGCTTTCAGTGCTTGAATCTCTACTTCCAAATTT

AGTCCTGCACTGACTACAGTCTCAACTAGTACTGAAAAAACAAGAAGTGCTATAAAGCTAGCTTTTTCTA

GGAGAGTTGCCATTGCTCTACTTTGTGTTGAAATCAGCTGGTGATGAGGGAAGGCTGTGCAAATAGCTCC

ATCTTCATCTGAGCTCTTTGGGAATGGAAGAGTTTCCATGAGGAAGACAGTTAAAGCACCAGTCTCCGAC

AGCCCCTGGTACGGCCCAGACAGAGTCAAGTACTTGGGCCCATTCTCTGGTGAAGCGCCATCTTACCTTA

CTGGTGAATTCCCTGGTGACTATGGCTGGGACACTGCTGGACTTTCTGCTGATCCCGAGACCTTTGCCAA

GAACCGTGAGCTTGAAGTTATTCACTCCAGATGGGCCATGCTTGGAGCTCTCGGGTGTGTCTTCCCTGAG

CTATTGGCCCGTAATGGTGTCAAGTTTGGCGAAGCTGTGTGGTTCAAGGCTGGTTCCCAGATTTTCAGTG

AAGGTGGACTCGACTACTTAGGCAACCCAAGCTTGGTCCATGCGCAAAGCATCTTGGCCATCTGGGCTAC

CCAAGTTATCTTGATGGGTGCTGTTGAGGGCTACCGTGTTGCAGGAGGGCCACTAGGTGAGATTGTTGAC

CCACTTTACCCTGGAGGAAGTTTTGACCCATTAGGCCTTGCTGAAGACCCTGAAGCATTTGCTGAACTTA

AGGTGAAGGAGCTCAAGAATGGAAGACTTGCCATGTTTTCGATGTTCGGGTTCTTTGTCCAGGCCATTGT

GACTGGAAAGGGTCCATTGGAGAACCTTGCTGACCACCTTGCTGATCCAGTTAACAACAATGCCTGGGCA

TTCGCCACCAACTTTGTTCCCGGAAAATGAGAAGTGCAATTTTTAGTTTCTAAATCATTATGAGAGAGCT

GGATTATGTAAATACAAGCAATTAAATTCTACTGAATGCCAAATTTTATGTCATTTTGTCACTACTAGTA

CTAAACTAAGGACTTCTCTTGAAGCAACTAAAATTTATTTATCA

>MSTRG.3047.1 gene=MSTRG.3047

CCTTAGTTTAGTACTAGTAGTGACAAAATGACATAAAATTTGGCATTCAGTAGAATTTAATTGCTTGTAT

TTACATAATCCAGCTCTCTCATAATGATTTAGAAACTAAAAATTGCACTTCTCATTTTCCGGGAACAAAG

TTGGTGGCGAATGCCCAGGCATTGTTGTTAACTGGATCAGCAAGGTGGTCAGCAAGGTTCTCCAATGGAC

CCTTTCCAGTCACAATGGCCTGGACAAAGAACCCGAACATCGAAAACATGGCAAGTCTTCCATTCTTGAG

CTCCTTCACCTTAAGTTCAGCAAATGCTTCAGGGTCTTCAGCAAGGCCTAATGGGTCAAAA

>MSTRG.3046.7 gene=MSTRG.3046

GCTCCATCTTCATCTGAGCTCTTTGGGAATGGAAGAGTTTCCATGAGGAAGACAGTTAAAGCACCAGTCT

CCGACAGCCCCTGGTACGGCCCAGACAGAGTCAAGTACTTGGGCCCGTTGAGGGCTACCGTGTTGCAGGA

GGGCCACTAGGTGAGATTGTTGACCCACTTTACCCTGGAGGAAGTTTTGACCCATTAGGCCTTGCTGAAG

ACCCTGAAGCATTTGCTGAACTTAAGGTGAAGGAGCTCAAGAATGGAAGACTTGCCATGTTTTCGATGTT

CGGGTTCTTTGTCCAGGCCATTGTGACTGGAAAGGGTCCATTGGAGAACCTTGCTGACCACCTTGCTGAT

CCAGTTAACAACAATGCCTGGGCATTCGCCACCAACTTTGTTCCCGGAAAATGAGAAGTGCAATTTTTAG

TTTCTAAATCATTATGAGAGAGCTGGATTATGTAAATACAAGCAATTAAATTCTACTGAATGCCAAATTT

TATGTCAT

>MSTRG.3046.8 gene=MSTRG.3046

CTCTACTTTGTGTTGAAATCAGCTGGTGATGAGGGAAGGCTGTGCAAATAGCTCCATCTTCATCTGAGCT

CTTTGGGAATGGAAGAGTTTCCATGAGGAAGACAGTTAAAGCACCAGTCTCCGACAGCCCCTGGTACGGC

CCAGACAGAGTCAAGTACTTGGGCCCATTCTCTGGTGAAGCGCCATCTTACCTTACTGGTGAATTCCCTG

GTGACTATGGCTGGGACACTGCTGGACTTTCTGCTGATCCCGAGACCTTCGCCAAGAATCGTGAGCTTGA

AGTCATTCACTCCAGATGGGCTATGCTTGGAGCCCTTGGATGTGTTTTCCCCGAGCTCTTGGCCCGAAAT

GGTGTCAAGTTTGGGGAAGCTGTGTGGTTCAAGGCTGGCTCACAGATTTTCAGTGAGGGAGGGCTTGACT

ACTTGGGCAACCCGAGCTTGGTCCATGCACAGAGCATCTTGGCCATCTGGGCTACCCAGGTAATTCTCAT

GGGTGCTGTGGAGGGCTACCGTGTTGCAGGAGGGCCACTAGGTGAGATTGTTGACCCGCTTTACCCCGGA

GGAAGCTTCGACCCATTAGGCCTTGCGGAAGACCCTGAAGCATTTGCTGAGCTTAAGGTGAAGGAGCTCA

AGAATGGAAGACTTGCCATGTTTTCGATGTTCGGGTTCTTTGTCCAGGCCATTGTGACTGGAAAGGGTCC

ATTGGAGAACCTTGCTGACCACCTTGCTGATCCAGTTAACAACAATGCCTGGGCATTCGCCACCAACTTT

GTTCCCGGAAAATGAGAAGTGCAATTTTTAGTTTCTAAATCATTATGAGAGAGCTGGATTATGTAAATAC

AAGCAATTAAATTCTACTGAATGCCAAATTTTATGTCAT

>MSTRG.3048.1 gene=MSTRG.3048

CACTCTTAAAAACTACTAGCTTTATAACACTTCTTGTTTTTTCAGTTTTAGTTGGTTTTGTGTTAAGTGC

AGAAACAAGCTTGGAGGTCGAGATTGAAGCACTGAAAGCTTTCAAGAATTCAGTTACAAATGACCCTTTC

GGGGCTCTTCAGGACTGGAATGACTCCAGCCATCATTGCAACTGGACTGGAATCAAGTGTCAGCCTTCAT

CGAAACGTGTGGTCTCGATTTCCCTGGTTGAAATGCAGCTTGAAGGGTTGATTACTCCATTCCTTGGGAA

TCTTTCTAATCTCCAGGTTCTTGATCTAACTTTAAACTCATTTAGTGGTCGCATTCCTCCTCAACTAGGC

TCTTGTTCACATCTTACTGATCTTATGTTGTATCGAAATTCACTTTCGGATTCAATCCCACCAGAGCTAG

GAAACCTCAAAAATCTGCAGTATGCGGATTTTGGAAACAATTCATTTAGTGGAACCATTCCAGAAAGTTT

AACCAACTGCACTGCACTGTTAGAAATTGCCTTCATTTTCAACAATCTCACCGGTGAAATCCCATTTAGC

CTCGGTGAGTTGGTTAATTTACAGATTTTTTTGGCTCATGATAACAGTTTGACAGGCTCAATACCTGTTT

CAATTGTTAACCTTATAGATATTCAAACTCTTGATCTGGCTGAGAACAAGTTAACTGGTGAGATTCCATT

CAACATAGGAAAATTGGTTAATCTACAGTTTTTCTTAGCGCAACATAATGCTTTAACAGGCTCCATACCT

GCTTCCATTGGTAGCCTCAGAGAAATTCAGAATCTCGATCTAAATGGGAATAAATTATCAGGTCCAATTC

CCCGGGAAATAGGAAACTTGTCAAAGTTACAGATACTTCAGTTATACGACAATTCTCTCACTGGGGAAAT

TCCAGAGACATTAGGTCAGTGTATAGAACTAGAATTGCTAAACATATATAGCAATCACTTCACTGGAGCT

ATTCCGTCTGAGCTTGGAAACTTGACAAGTTTACAAGGCTTACGACTGTATGATAATAAGCTGAATTCTA

CATTCCCACTGTCGTTTCTCCAGCTAAGATCATTAATCAATCTGGAACTTTCAAATAATGAATTAGTTGG

AAGCATACCTTCCAGGATTGATTCTTTGGCATCATTGCAGATACTGATGCTACATAACAATAGGTTTACT

GGTCAGATCCCATCATCCCTAACAAATCTGACAAATCTAACCTACGTGTCCATGAGCTTCAATCTCCTCA

CAGGATCAATACCATCAAACATAGGGGCACTGCACAATTTACAGAATTTAACCTTGAATGATAATTATCT

TGGTGGAATTATACCTCCTAGTATCACTAAGTGCACACAACTTCTTAACATAGGTTTGACTAAGAATGAA

CTGACAGGGAAAATTCCTTCTGGCTTTGGGAAGTTGCAAAATCTTAGTTTTCTGAGTATAGGAAATAGTC

AATTGTTTGGAGACATTCCTGATGACCTCTTCAATTGTATTGGTCTGAGTTCGCTACTTTTAAGCGGGAA

TGATTTTACTGGATTGCTGAAACCAAGCATTGGAAGCCTCACCAATCTCAAAGTTCTGCGTCTTAGTACA

AATTCATTTTCGGGAAGGATTCCTAAAGAAATTGGGAACCTTAGTAAACTCTTTACATTATCCCTCCACA

CAAACAAATTCTCAGGCATAGTTCCGCCTGAACTGTCAAAGCTCACCTTTCTCCAGGGCTTAGCACTGCA

AAACAATCTTCTTGAAGGTTTGCTTCCAGGCAGCATTTTTGAGCTCAAAAACCTTTATGTGCTATTGCTA

AACAATAACAGTTTCACTGGACCTATCCCAGACTCTGTTTCGAAAGTCAAGCAGCTGTCACACTTGGATC

TCAGTGAAAACAGGCTTTCTGGATTAATTCCCAAGAGTATGGCAACACTGAACAGGATGATGACCTTGGA

TCTCTCACATAATAATCTCACAGGATCCATCCCAGGAACAGTTATTTCAAGCATGAGAATGATGTCAACA

TATCTCAACCTTTCATACAATTCATTTTCTGGAAGACTTCCACCTGAGATGGGAATGCTACAAGCAGTTC

AAGCCATCGAAATCTCGTCTACCAATCTCTCTGGAAGTATTTCTACACTCAAAGGCTGCATAAATTTGTT

CTCACTTGATCTGTCAAATAATAAATTATCTGGTCAAATTCCAACAGACACTTTCACTCAGCTCACTGTG

CTTACCAACCTGAATCTCTCAAGAAACCAGTTAGATGGACCACTGCCTGAAAGTTTGGCAAGCTTAAGGA

ATCTTAGCTTCCTTGATCTTTCTCAGAATAAGTTCACAGGTATCATATCTGACAAATTTTCCACTATATC

TACCTTAAAAAATCTCAACCTTTCTTTCAATCAATTTGAAGGACTTGTTCCAGAGACTGGTGTTTTTAGA

AACAAAAGTGTAACCAGTTTGCAAGGAAATCCATCCCTTTGCGGCGGCAAAGCTCTGCAGCCATGCAATG

ACCAGAAAAAATCAAAAGGCGGCTTATCCCCAAAAACTGTGGTAATTATTGCTTCATTAGCATCTTTTGG

TGTAATTATGCTTTTACTATTAGCAATTACAGTGTTCTGTAGATGCATGAAGAACAACGAAAGAAAGGAG

ATGGAGAGTGCAGCACCAGATTACACTTCAAAATCAATTCTCAAGAGGTTTCATCCAAAGGAATTAGAAG

ATGCTACTAATCAGTTCGATGAGGATAACATTTTGGGATCAAGCAGCATGAGCACTGTATACAAGGGTAT

ACTAGAAGATGGACAGGTGATAGCCGTAAAGATTTTGAAGGTGCATCTGTTTGCTGAGACAACTGAAAAA

AGCTTCAACAGAGAACTTAAAACTCTTGGTAAGCTAAAACACAGAAATTTGGTGAAGCTATTTGGTTACG

CATGGGAGAGCTCAAAGCTGAAGGGGATAGTTCTGGAATACATGGAAAATGGTAGCTTAGAGACCATTAT

ACATGATTCAGGGGTGGACCAAGCACGGTGGACACTGTCTGAGAGGATTGATGTTTTAGTCTCAGTTGCA

AGGGGTTTGGTTTACTTGCATTCAGGTTATGATTTTCCGATAGTCCACTGTGATCTAAAGCCATCTAATG

TTCTTTTTGATGGGGAGTGGGATGTTCATGTGAGTGATTTTGGGACAGCCAGAATTCTTGATATTGATAT

GAACAGCAGCAGCTCCTCCTCGGCATCAGCCTTTGAAGGAACCATTGGTTACTTGGCACCAGAATATGCA

TTTATGAGGAAAGTCACAACAAAAGTAGATGTTTTTAGCTTTGGCGTGATTGTTATGGAATTCCTTACCA

GACGACGTCCTACTGGTCTTTCTCAAGATAACGGACAGCCAATCACTTTGCCACAACTAGTTAAGAAAGC

ATTGGCAGATGGAACCGACAGATTACTTAGTGTAGTAGATCCTCAGTTAGTTTCAGATATCTCCAGGAAA

GAGCAACTAGAAAGTATCTTTACATTAGCCTTATTATGCACGTGCCAAGAGCCAGAGGGCCGTCCTGACA

TGGATGAGGTGCTATCTTCACTTTCCAAGATTAGCAAATTGGAAACGAAGACTGGGACCGTGAAAGATGA

CAGATCAAAAACAGGTCAAAGCAGCAATGCTGCAACTCAAATTCAGGGGACCGTTTCTTAGTAGCTATCA

ATGTGCAGTCAGCCAATGTGAAGGCTTAGCGAGTTTTTATCAATGTGCAGAAATTAGGGAAGGTTCATAC

TCCAAAGTTCACAGTAATTAATTTAGAGCAGCTTCAGCAACAACATAAACGAAGCAAACATCCGGATTAA

GGTTTTTCCCCGAGTATATTTATCTTCTTCTTACTATACAAATTTATTTCTCACTTATCCAGCCACAGAT

AATAGGAATGCTACAGTGAATATCCA

>MSTRG.3049.1 gene=MSTRG.3049

TTCCCTGCAAAAGAAGGAGAAGAGAGGGCCATTGCTGAAGCAGCCATTGCACAAATGAGTTAAAATAGTA

CTACTTCTAGAGAAGGTAGTGTATAGAGGTTGTTGTTAGCTTGAGTACTTGAATGCTTTGTAAGAAGGCA

TATTTATATGCATTGGAGTACATGGGTCATTATCTTCGGAGATCTATAGCTGTTTTCTCATTGGTTGATG

ATATGTTTAGCGACTGCATTCCACTTGTGGATTCATTATCTGTAGCATTGTGTGACTATGGAGTACTCAG

AGAAAAGTATGAATTATGACCACTAATGTGACTGGTTGCAAATTCATGGCTTTCACCATCAAATGATTCT

TTTATCTTTAAGAAAACAGTACGAAATGTTACATTTGAAATAATCAAATCAAAATGTTGCAGCCTTATCT

TTATATAGTTTAGTCATGTAAAACAAGTTATTGGACTAAAGTAGTGTAACATGTCCACAGTTGTAGTGAG

CCAAGTGCTA

>MSTRG.3051.1 gene=MSTRG.3051

TTCTACCATCACCATAAGTTTCCAAACAGCCAAGAAAGCAAAACTAACCTAACTCTGCGGCGGATCATTC

TGCAAAAACTCATCTACAGTTGAAGGAAACTCGAACCCTTCAACCCCATCAGAAACCAAAGCAGCTTTAG

CCAAAAGCAAATGCCCCAATATAAATGGCTCATTTTTCTCACCACGATCCATATAATACTTAATCCACCC

TTCCAATCTCTCAACTTCTTTCTTACATTTAGGTCCTTTCACTTTAACAACATTACCAAAAAACCCATCA

TCACAAACACTATTCAAATCCAACCCAACACTTTCTTGATCACCATCATCAAAACCCAGAAGCTGCATCA

AATTCTCAATATCATCAACCCCACCAGTCTCAGAAAGATCAAAATTTTCTGGGTTTTGCTTAGAAAGTGA

TTCTTGAAAAGGGTCTTTTTGGGTTTCTGGAAAATCGAGAGATTTGGTCAAGAATTCTCGAGGGGGCTCA

CTTTCTTGAAAAGTTGAAGTATTTGAAGTGGGTTGCTCAAGATGAATGTATATAGGAGAGCTAAGTGATT

GAGGGTTGGTTTGTGATGATGATGTTGAGAACTCAAGCACCTACTGGAAAACAAGCACCATGAAGGCTTC

CATAAGAGTCTCACTATTGTCGAGGGACGATAGAGGTTATACTGAACTTGTTGCAAGATGAAAAATAAGG

ATTGCTCAACAATGGGTCTCTGTCCCGCAATTTTCTTTCCATGAAGAAACCAGGTTTCTTCGGATAAGCC

AGTTTCGAATCACTGCTCAACATAAGAACTACTGCAGACATATCTGGCCTGTCGACAGGATCTTCTTGAA

CACACAATAGTCCAATATGGATAACTCGGAAAACTTTTATCAGCAACCAACCAGTGCCACAAGATTTCTG

TGCTGAAGTTTCGCAATGAAAGAAACTTCATTTATGAACTCATCGAGTCCTTGATCAGAATTTTTTGAAA

GCCTTTTCACAGCTATCTCTTGTCCATCCTCCAATATTCCCTGAAGGTAATCACCAGATTTATCATTGTG

ATGATAACACATTCTGAACTGTAGTAGATAATATGTTTTTTACATTATAGTAAGAATATGTAATTTTGTA

ATGATCTGTTTCCATAAAATGCACATGTTGAATATACGTATAATCACCTTGTACACAAGCCCGAAGCCAC

CCTCTCCAAGCTTGTTATGCAGGGAGAAGTTATCTGTAGCGTTAACAATTTTTTTGAAGTCGACTAATG

>MSTRG.3051.3 gene=MSTRG.3051

CCATCACCATAAGTTTCCAAACAGCCAAGAAAGCAAAACTAACCTAACTCTGCGGCGGATCATTCTGCAA

AAACTCATCTACAGTTGAAGGAAACTCGAACCCTTCAACCCCATCAGAAACCAAAGCAGCTTTAGCCAAA

AGCAAATGCCCCAATATAAATGGCTCATTTTTCTCACCACGATCCATATAATACTTAATCCACCCTTCCA

ATCTCTCAACTTCTTTCTTACATTTAGGTCCTTTCACTTTAACAACATTACCAAAAAACCCATCATCACA

AACACTATTCAAATCCAACCCAACACTTTCTTGATCACCATCATCAAAACCCAGAAGCTGCATCAAATTC

TCAATATCATCAACCCCACCAGTCTCAGAAAGATCAAAATTTTCTGGGTTTTGCTTAGAAAGTGATTCTT

GAAAAGGGTCTTTTTGGGTTTCTGGAAAATCGAGAGATTTGGTCAAGAATTCTCGAGGGGGCTCACTTTC

TTGAAAAGTTGAAGTATTTGAAGTGGGTTGCTCAAGATGAATGTATATAGGAGAGCTAAGTGATTGAGGG

TTGGTTTGTGATGATGATGTTGAGGTATTATTTTCAAGTTTTAATCTTTTCTCTACTTGTTTGATTGAAG

AGAAGAAGTGGGCATGTGGGTTATGTGATGATGATCGCATTTTGATAGTTGGGTGATTTTCTTGAATTCT

AATGGTGCCCTAAATATATGATCTTGTGTTGAATTGGGCAAATGCGGTACTCAATAATTTTAAATTTTCT

TTGATAGAGGTTATACTGAACTTGTTGCAAGATGAAAAATAAGGATTGCTCAACAATGGGTCTCTGTCCC

GCAATTTTCTTTCCATGAAGAAACCAGGTTTCTTCGGATAAGCCAGTTTCGAATCACTGCTCAACATAAG

AACTACTGCAGACATATCTGGCCTGTCGACAGGATCTTCTTGAACACACAATAGTCCAATATGGATAACT

CGGAAAACTTTTATCAGTGACTATCTCTATGAGTAAAACACCGAAGCTATATACATCAGATTTAACGGAG

AACTGTCCATCAATGGCATACTCCGGAGACATGTATCCACTGCAGCAGATCAAAGATAAAGATAT

>MSTRG.3051.2 gene=MSTRG.3051

CCATCACCATAAGTTTCCAAACAGCCAAGAAAGCAAAACTAACCTAACTCTGCGGCGGATCATTCTGCAA

AAACTCATCTACAGTTGAAGGAAACTCGAACCCTTCAACCCCATCAGAAACCAAAGCAGCTTTAGCCAAA

AGCAAATGCCCCAATATAAATGGCTCATTTTTCTCACCACGATCCATATAATACTTAATCCACCCTTCCA

ATCTCTCAACTTCTTTCTTACATTTAGGTCCTTTCACTTTAACAACATTACCAAAAAACCCATCATCACA

AACACTATTCAAATCCAACCCAACACTTTCTTGATCACCATCATCAAAACCCAGAAGCTGCATCAAATTC

TCAATATCATCAACCCCACCAGTCTCAGAAAGATCAAAATTTTCTGGGTTTTGCTTAGAAAGTGATTCTT

GAAAAGGGTCTTTTTGGGTTTCTGGAAAATCGAGAGATTTGGTCAAGAATTCTCGAGGGGGCTCACTTTC

TTGAAAAGTTGAAGTATTTGAAGTGGGTTGCTCAAGATGAATGTATATAGGAGAGCTAAGTGATTGAGGG

TTGGTTTGTGATGATGATGTTGAGGTATTATTTTCAAGTTTTAATCTTTTCTCTACTTGTTTGATTGAAG

AGAAGAAGTGGGCATGTGGGTTATGTGATGATGATCGCATTTTGATAGTTGGGTGATTTTCTTGAATTCT

AATGGTGCCCTAAATATATGATCTTGTGTTGAATTGGGCAAATGCGGTACTCAATAATTTTAAATTTTCT

TTGATAGAGGTTATACTGAACTTGTTGCAAGATGAAAAATAAGGATTGCTCAACAATGGGTCTCTGTCCC

GCAATTTTCTTTCCATGAAGAAACCAGGTTTCTTCGGATAAGCCAGTTTCGAATCACTGCTCAACATAAG

AACTACTGCAGACATATCTGGCCTGTCGACAGGATCTTCTTGAACACACAATAGTCCAATATGGATAACT

CGGAAAACTTTTATCAGCAACCAACTAGTGCCACAAGA

>MSTRG.3051.4 gene=MSTRG.3051

AGAAAGCAAAACTAACCTAACTCTGCGGCGGATCATTCTGCAAAAACTCATCTACAGTTGAAGGAAACTC

GAACCCTTCAACCCCATCAGAAACCAAAGCAGCTTTAGCCAAAAGCAAATGCCCCAATATAAATGGCTCA

TTTTTCTCACCACGATCCATATAATACTTAATCCACCCTTCCAATCTCTCAACTTCTTTCTTACATTTAG

GTCCTTTCACTTTAACAACATTACCAAAAAACCCATCATCACAAACACTATTCAAATCCAACCCAACACT

TTCTTGATCACCATCATCAAAACCCAGAAGCTGCATCAAATTCTCAATATCATCAACCCCACCAGTCTCA

GAAAGATCAAAATTTTCTGGGTTTTGCTTAGAAAGTGATTCTTGAAAAGGGTCTTTTTGGGTTTCTGGAA

AATCGAGAGATTTGGTCAAGAATTCTCGAGGGGGCTCACTTTCTTGAAAAGTTGAAGTATTTGAAGTGGG

TTGCTCAAGATGAATGTATATAGGAGAGCTAAGTGATTGAGGGTTGGTTTGTGATGATGATGTTGAGAAC

TCAAGCACCTACTGGAAAACAAGCACCATGAAGGCTTCCATAAGAGTCTCACTATTGTCGAGGGACGATA

GAGGTTATACTGAACTTGTTGCAAGATGAAAAATAAGGATTGCTCAACAATGGGTCTCTGTCCCGCAATT

TTCTTTCCATGAAGAAACCAGGTTTCTTCGGATAAGCCAGTTTCGAATCACTGCTCAACATAAGAACTAC

TGCAGACATATCTGGCCTGTCGACAGGATCTTCTTGAACACACAATAGTCCAATATGGATAACTCGGAAA

ACTTTTATCAGCAACCAACTAGTGCCACAAGATTTCTGTGC

>MSTRG.3053.2 gene=MSTRG.3053

TATCTATCTATATTATAAAGGCTGTTGTTTTTTTACCCTTGAAATGACCATTCCACCCATGTGTATTTGT

TGTATTTCCAGAAACACTTATGTACAATACCCGTATGTATACGTTGGTGTCATCCACACCTTCCTTTCCC

TTACCAAAGCTCCACGTGTTGCCCATCCCTCTCTCTACGATTCTCTCCTCCAAAATCATCTTACTACATT

CAAAATTTTCAAATTTAACTCCTTCCGCAATTTTTCAGTTGTAATACATGGATTCTTTAGAAAGAAGCCC

TCTCAAGTCGCCGGCATAGGAGTTGGTTTCTGGGAAATCATCCAATACATGTCAAACTTTCAAAATTAAT

TGTAATTGTGGCAGATAGTACAAAAATGGGAACAAAGAAATCTTGGCTTCCAAAACAAAGAATTGACGAG

CTGTATTACTTGATTTCACGAAGGTCAAGAGATGGTTGTGGGAGAAAGAGGCTGAACAGGCTAGG

>MSTRG.3053.1 gene=MSTRG.3053

TATCTATCTATATTATAAAGGCTGTTGTTTTTTTACCCTTGAAATGACCATTCCACCCATGTGTATTTGT

TGTATTTCCAGAAACACTTATGTACAATACCCGTATGTATACGTTGGTGTCATCCACACCTTCCTTTCCC

TTACCAAAGCTCCACGTGTTGCCCATCCCTCTCTCTACGATTCTCTCCTCCAAAATCATCTTACTACATT

CAAAATTTTCAAATTTAACTCCTTCCGCAAGTAAAGCTTCTAATCCCATTTTGTTTTTCGCCGCTTTTCA

TTGTAATCCAATTGTCTCCATTCATTGTATCACTTCTTCGTTTATATAGTTTTTCAGTTGTAATACATGG

ATTCTTTAGAAAGAAGCCCTCTCAAGTCGCCGGCATAGGAGTTGGTTTCTGGGAAATCATCCAATACATG

TCAAACTTTCAAAATTAATTGTAATTGTGGCAGATAGTACAAAAATGGGAACAAAGAAATCTTGGCTTCC

AAAACAAAGAATTGACGAGCTGTATTACTTGATTTCACGAAGGTCAAGAGATGGTTGTGGGAGAAAGAGG

CTGAACAGGCTAGGTAAGTTCCACCAATTTTGAAATGGCTGATGCTTCTGCAGTCGAGAAGCAATGGCGA

AATCAATTCAGTTTCTTAAATGTCTCAGTCCCAAAACAATTAGAAGATATAGTGGTCCTGGCACACAAAA

GATATATCATACAAAGGCACAAGCCGGGGACCTTCGAGTTTATTTTTGGATGAATTTTATGAGGGTAATA

TATTAAGGTTTCCATTTAAGGTTTTTTTTTTGGATTATTTGTAAATTAATTACAAAAATAATTTTACAGT

ATATTCACTTTAATTATATTAATTCCAGTCCTTTTTAATTCCTTATCATGTTAGGAAAGTGTTCAAGGAT

TTCCACTCAGAAAAATGTTAGGAAAATGTTAAAGGATTTTCTTTACAGAATTTGCCCCAATTTAATAACA

TGAAAACTACTGGAAACCTACAATTTTGTGCTATCTGTATTTCCCTGGAGTTGTGAATTCACAGTATCTG

CACAAATAAGTTTGCATAGTAGATATTGGTTGATTAGGTATGTTTTTCCGTATGAAATAAATGGAAGAAT

ATGCTAATTGCATATAAATTTCCATTTTTTTCTTTTAGTATGCTCAAAGGTATAGAATGCAAAATATCCT

TCACTCGACTGAACCATTTCGAACTTAACATATATACTTTCTAAATGCTTACACTATAAAAGCTTCATTA

GAAAATTGCATAAGGCTCAACACCAGTTTTGGTCTGAATGTAGTGGCATTTAACAAGTAAAAAGACATTG

CACCTACACTATAAAAGTTGTAGATGACCCGAGGACCTTGAACAAAATTCTCTATGTGAGGCCTCCCCAC

AACACATTTTTTTTCAATGATTTTCTGTCTTTTTAGGAGAAAAAGATGGGCAAAACTCTTGAGAAGATAT

ATGTTCCAGAGGAGCAAGTCTTGAAGAATATTAAAGATGAGGAATTTTTCAAATTTTTTTGCTTGACATT

ATTATAATAATTGACCAGAGCAAGTCTTGAAGAACATTGAAGGTGAGGGGTTTTTTAAATTTTTTTTGCT

TCACAATGTTATAATAATTGATCATTATCTTATGATCCTGGCAAAGTTGTTTACAAAGTTTTTTTTGTCC

TTTATGTTAAACTCTATGAGGAATCCCCTATTCGGATCAATGTGCTGCTATTTATCGGTCATTCTACATT

TGTGAAGGGAGATTAAACCAGTTTTGAGATTGAACCTTCATTTGGTGTGGAGGCTTCAGCACTTTATCCT

GATGTCAAATACAAAACTGAAGAGGAGTATATCAACAAATTTGTCTAAACCAAATCTATTTTGTCTTTAA

AATATTTTCTTTTTTACAATTCATATAATATAATTTTTCCAAATTCAAAAACAACAATCTATTCACTAAA

CAAAAATCATCCGCG

>MSTRG.3055.2 gene=MSTRG.3055

CTTTATAATCTTGATTTCTTTGGATGTTGTAGCGATGCATAGTCTATAAACAGGGAAGAAAAAATGGAGG

GACCAAGTTAGAAACCTTCCTTTTTAAACACAACTAACATCATAATCCTTGGATTGGTGATTGAGATTTC

TTTCGAATTGTTTCTTACTGACCAGAAGTGGACGGAAAATAATTTTTTCCTAAATGATTCTCTATAGTCC

TGCGTTGTGCACACATTAGAAACAACCCTTACTCGAATAGAAAAGCAAGTACATTTTTGATCCTACAATC

TTGCATCTCATCACAGATTTTCTCGGAAGGTGTACAGGTGCTCCGCGATTGGTTGCTTCTGTTGCCAGTG

CATAAGCACAATAATCTGGCTAGCAACTTGAGAAATCGGATCGAAGTTTGTCACCCTGGCCGCGATGGTA

TGGTGGTAGCCAGAATCATAAAAGAATACACAAAAAAGTGGTGGAAATCCCACCTATATATATGTGTACC

TACTAAAGACTAAAGACTAGAAGCTACAAAAAGATGCAAGATATTCAAATTGGGAATGAGTTTTGTTCAT

TATCACCCAGTTGAAGAGACGGAAGATGTACATATCTCTGACTGCCCTGTCTTTATAATATCTGGAAACT

CTTCGTGCATCGCAACAAAGGACATGTGGAACGAAATCAAAGAGGGAATATAGTGTACAGACTACCTTTT

GTCGTGATCTCAGGCCTGGACTCCATATCGGCGTTTTCTTCCTCGGACAAATCTGCCGCCCCTTCTACTG

CAGCGCGGCTGTTTGAATGCATGCAAGGGGAGAGAAGTGTTTTGCTTTAGCTTTTTTGGACAATTATTAG

GCTCCTCTTTTTAAGTTGTGCCTTATGTAAAGACCTTTTTAGATGTGTCTAATTGATATTTCGTCTCTAT

TTAATTTATTTAGGAGTTAATATGGATAACTAACTAGAAACTAATTTATATTAATTTTGGG

>MSTRG.3055.1 gene=MSTRG.3055

TGATTCTTTATAATCTTGATTTCTTTGGATGTTGTAGCGATGCATAGTCTATAAACAGGGAAGAAAAAAT

GGAGGGACCAAGTTAGAAACCTTCCTTTTTAAACACAACTAACATCATAATCCTTGGATTGGTGATTGAG

ATTTCTTTCGAATTGTTTCTTACTGACCAGAAGTGGACGGAAAATAATTTTTTCCTAAATGATTCTCTAT

AGTCCTGCGTTGTGCACACATTAGAAACAACCCTTACTCGAATAGAAAAGCAAGTACATTTTTGATCCTA

CAATCTTGCATCTCATCACAGATTTTCTCGGAAGTGGCCAGAATTGTCTAAGGTGTACAGGTGCTCCGCG

ATTGGTTGCTTCTGTTGCCAGTGCATAAGCACAATAATCTGGCTAGCAACTTGAGAAATCGGATCGAAGT

TTGTCACCCTGGCCGCGATGGTATGGTGGTAGCCAGAATCATAAAAGAATACACAAAAAAGTGGTGGAAA

TCCCACCTATATATATGTGTACCTACTAAAGACTAAAGACTAGAAGCTACAAAAAGATGCAAGATATTCA

AATTGGGAATGAGTTTTGTTCATTATCACCCAGTTGAAGAGACGGAAGATGTACATATCTCTGACTGCCC

TGTCTTTATAATATCTGGAAACTCTTCGTGCATCGCAACAAAGGACATGTGGAACGAAATCAAAGAGGGA

ATATAGTGTACAGACTACCTTTTGTCGTGATCTCAGGCCTGGACTCCATATCGGCGTTTTCTTCCTCGGA

CAAATCTGCCGCCCCTTCTACTGCAGCGCGGCTGTTTGAATGCATGCAAGGGGAGAGAAGTGTTTTGCTT

TAGCTTTTTTGGACAATTATTAGGCTCCTCTTTTTAAGTTGTGCCTTATGTAAAGACCTTTTTAGATGTG

TCTAATTGATATTTCGTCTCTATTTAATTTATTTAGGAGTTAATATGGATAACTAACTAGAAACTAATTT

ATATTAATTTTGGG

>MSTRG.3061.1 gene=MSTRG.3061

TGTAAACAGGTTCGAGATTTTGGTTTTAAAGGAAGGGGGACTTCCTATTTGAATTGAAAAGCTTTTAGAG

GAAGGGTTACTTCCAAATTGATTTGAAAAGCTTTTGATTTAGAAGCGGAGGCTTCAACTTGATTTGAAAA

GCAGATAGTAGTTGATTTTAAACTATTTTTGAACTGTTACGTTTTACTATGCTGCTTCCTTACTCGTTTT

ACTATACAGCTTTACTTACTGATTTGTTTACATCACATTTTCTTACTGTTTACCGTACATTTGAT

>MSTRG.3063.1 gene=MSTRG.3063

CAGCGGTCTTTTCCAAACCTTTTACTTTCAAATTGGACCAAAACTAAATAGATCCAACAGATCTCCCGGT

GAAGTGGGTAGGCCCAAAATTACCTTGCCCGATAATTAGGAAAACCTCACACTGTGTTCATCAAGCTTTG

GCAACTAAAGTGTAGAAGGGGCGTGCAGGAGAGAGTCTCTGATTTTAAGGCAGCAGAAGCTTTGGAATTT

CGAAAAGAAATATTAATCACAAGGCTGCTACTCTGGCAATTTCACTAGAGGCTATGTCAAATCGGCGAAA

ACTAGTGATAGAGCATAAAAGGCAGCATCTGCAGATACCTCTCTTGAAGGTGCACAAAGTGCTTAGAAAC

ATGAGGAAAGAAGCTCACACAATATGTAAGCAAGTTGTTGCTGTTGAAACTGAATATTGCATAATAGGGA

TCTTCAGAACCCTGAAATGCAGATATCTAGACGAATCGGAAGTGGAACAACTGAAGTTATTGTTCTTGTT

AAGTTCCATAAACCATGGAAGTTTATCTGAATTAAACCGAGCCTGCGTATATGAGTTACTCCGAGTTGGA

TAGTAAAAAACTTGGAAGTCATGATACTTGTATAGAATGACTGGTTGAGCAACATATGGTATGCTGGAGG

AGTTCAGAACCCACATGGAATTCTTCTAGACCACCAATTTTGGACTGATCTTCTATCCATTCAGACACTG

CAATTCCACTGCAGGCTTTTCGCCACACAGGGGAGATAGATCTCAGTTCAAGATGTCAGTTCTCAGAACT

TTCTTGAGTAAGTTCCTTGAGGACTTTTCACTCTTCACAGGAAAAGGCCAATGTTGTGATGAATATCTGG

GTATATGTAGACACAAGTCTGTCCGCAAAGGCTGGCATGTGCACAGTTCTACATTTTCGAGCACAGCTAT

CCCTTAACCAGATCACCTAGGGTCGTTTTTATATTTGGAATTCATGACAGGGCAGGGCAGTCTAAGTTGC

GAATTGTAATGATATTTTTTAATATTCCCTTTTAAGAACTCTGTAGTACCATTGCTCTTCTATGGCTAAC

TTCAACTGTGTGGTCTGTGTACAAAAA

>MSTRG.3063.2 gene=MSTRG.3063

AAACTAAATAGATCCAACAGATCTCCCGGTGAAGTGGGTAGGCCCAAAATTACCTTGCCCGATAATTAGG

AAAACCTCACACTGTGTTCATCAAGCTTTGGCAACTAAAGTGTAGAAGGGGCGTGCAGGAGAGAGTCTCT

GATTTTAAGGGATATACATGCTTCATACATGTGCAATTATGTGTAATGTCTATTGGTTGTTAAGATTACC

AATATAAATTAGTTAGTTTTATTTGTGTATAATTGTGAATTATCTTATAAATATCCACCCATTTGTGTTA

GTTTGTTAGTATAGTGAGGAAGATGATGATGTTTTATCGGTCTTATGTGCATTTCTTAATGCAATATGGG

GATTGATTTATTTATCTGATTCCTCTTATGTAATCCTTTTCGTGTGTGTTCTAATTTAATTTGTTTCTTG

ATAGATGCATGATTTTAGATGGATATTCATGAGCTTCTTTGATACCAGGATAGCTATATGATAAACTTTG

TCGATGTTGTTTCCCTTTCATTAGCTCTAATCGAAAATTGAGTTCGCATAAAAGGGAGTACCCTTTATCT

CTACTATTTGATTTGTAAGTAAGCCTTTGTATGTGTAGCTTCGTTTTGTCATCCTCGTGATTATGAGGAA

CCAACTAGATATTATTGATTGCAAATTTTTTTTAGCATTTGTTGTGCTGCATGATTTCCTCTAAAATTTT

ATCTGCAGAGCCAGATAGTTGCGGATGTGATGGGTTTGCATTAGTGAAACTGAAAACTATTAAGTAAATT

AGTTGCCTGTAACTTTTGGTGCTCGGCATTAAAGGTTTTTTTTGAAATGGATTCGATTTTGAAATGGGTA

TTTTGGACCGGGTGTTGGTTGGGTTTGTGCAGGAAGCAAACACTTTTGGTTTTGGTTACCAATTAGTATA

TATAACACAGTTTAGATAAATTGCCATCTCACAGGTGATATGATATTAGATAGGGGCTGATCGGCAACGT

CGTATTAGAGTGGTTGAGTTTGGGTCCTTGGGGTATACGGTGGGTTAGGCCTTACAGTTTTTTCATATAC

ACTAGTGTATCTTTATGTGTGTTTGTTTATATCTATATCACTCCCTGTGTTTGTCATGCACTAAGCGGGG

GGCCTTTACTAAAACAACCTACTTACTCTTTTAGAGTAGGGGAATGGTCTGGGTGCATCTTACCCTCCCA

GACCATGCTCTAAGCTGTATATATTGGGTATGTAGGCAGCAGAAGCTTTGGAATTTCGAAAAGAAATATT

AATCACAAGGCTGCTACTCTGGCAATTTCACTAGAGGCTATGTCAAATCGGCGAAAACTAGTGATAGAGC

ATAAAAGGCAGCATCTGCAGATACCTCTCTTGAAGGTGCACAAAGTGCTTAGAAACATGAGGAAAGAAGC

TCACACAATATGTAAGCAAGTTGTTGCTGTTGAAACTGAATATTGCATAATAGGGATCTTCAGAACCCTG

AAATGCAGATATCTAGACGAATCGGAAGTGGAACAACTGAAGTTATTGTTCTTGTTAAGTTCCATAAACC

ATGGAAGTTTATCTGAATTAAACCGAGCCTGCGTATATGAGTTACTCCGAGTTGGATAGTAAAAAACTTG

GAAGTCATGATACTTGTATAGAATGACTGGTTGAGCAACATATGGTATGCTGGAGGAGTTCAGAACCCAC

ATGGAATTCTTCTAGACCACCAATTTTGGACTGATCTTCTATCCATTCAGACACTGCAATTCCACTGCAG

GCTTTTCGCCACACAGGGGAGATAGATCTCAGTTCAAGATGTCAGTTCTCAGAACTTTCTTGAGTAAGTT

CCTTGAGGACTTTTCACTCTTCACAGGAAAAGGCCAATGTTGTGATGAATATCTGGGTATATGTAGACAC

AAGTCTGTCCGCAAAGGCTGGCATGTGCACAGTTCTACATTTTCGAGCACAGCTATCCCTTAACCAGATC

ACCTAGGGTCGTTTTTATATTTGGAATTCATGACAGGGCAGGGCAGTCTAAGTTGCGAATTGTAATGATA

TTTTTTAATATTCCCTTTTAAGAACTCTGTAGTACCATTGCTCTTCTATGGCTAACTTCAACTGTGTG

>MSTRG.3066.1 gene=MSTRG.3066

CTCTTTATGTCCAGTCTCCCTCACATAAAAACCCTCTTTATGTAACAAAGGATCAGCTCCTAGTTGCATA

TTTGTCCCTCCTCCCAGTCTGGCGAAAATAGACCATCTTCCTCGTCTTCATAATCCATGTAATCATTCTT

TATCGAAAGGCCAATCCTTTCCATAATGTTTCCAAATCGATCACCCTGACTTGCTTCAAAGAGTACCATC

TCAAGTTCCCTCAGAACCTTCATATTGTCC

>MSTRG.3076.1 gene=MSTRG.3076

TCTCAGCTTTTTGAAAATACTTTCGTTTCTTTACACAAATGTGTCAAGAAAAATAGAAACTAGAATCAGT

TTTTGCTTTTTGCAAACAAACGCCACCTGAGACTTGAGATGAGAGAATGAGTGGTGAAGAAGAATATGAG

GTAAGGACTGGAGGGAGCTATAGTAGTAGCCGGCTAGCCACTGTTGCTGCAAATACAAACGACTATGTAG

CCGTTGGTACTGCAAAAGCACTGAGCGGAGGTCATAGATCCGTAATTGGGAGGGCAGCTAGCCGTTGCGG

TGCAAAGAACAGCAGGCGCGTGACCCGAGTTGATTCGCGGGTCGAACAATTCACAAAAATTAGCGGCCCG

GTTCTAGTTTTCAACTTGACGACTCAGATTCAATTCATTATTTTTAACGATTTCGGCCGAATCTATTCAA

ACCTGCTCCTTCATCGAATTTCAAGGCGGTCCGGTTCAAAACCACTCATTTGGCCAGCACTGGTTATAAA

TATTGAGAAGAAGATTACAATAGATCTGATGGAATGGCACAATACAAAGGGTAAAGTATAGAGAAGATCA

ATTTTAAGGAGAGAATAGCATGGTTGCAAGTTGTTCCACACGTGAGAGGCATGATGGATACTGGAGTTCA

CTGAAACATTCAAATACTTGGAAGGACTCGGAAAGATAAATACAATACCAAGAGAAGCAGAAGGAGAAAT

GAAAAGCAGAGAATGAATGAAGAAATTAAAGAACTAACCTTCTGCATAAATTGTATGGTTAAAATAGAGG

AAGAAGCATGGCGATTCGTGTCACCATCTTTTACAGAGAGAAGAATGGGAATGGGATGGAGAATATGAAG

AATGAGAAGTGATGTATTTATACATGATCTGCAAAGAAAGAAAGCCAAGAATGGGGATTTGATAAGAGGG

GTTGGAGGAAAGTTGAGATTGTTTTGACCTTTGCAAATAAATGAGAAAGATATGTCATCATTTGACTACC

TCCTCTTCTTGTGTAATCAATTCTTACTCATCCAGCACTGTTCCTCTCTCTAATATTAATGTTTGGAATC

AATTCATCATAATCATCTTGATGTGTACCACATATGAGTTCACAGCTTTTAGCTGAAAAGCTACTTTTAG

AAAAAGTAGGTC

>MSTRG.3076.2 gene=MSTRG.3076

GCTTTTTGAAAATACTTTCGTTTCTTTACACAAATGTGTCAAGAAAAATAGAAACTAGAATCAGTTTTTG

CTTTTTGCAAACAAACGCCACCTGAGACTTGAGATGAGAGAATGAGTGGTGAAGAAGAATATGAGGTAAG

GACTGGAGGGAGCTATAGTAGTAGCCGGCTAGCCACTGTTGCTGCAAATACAAACGACTATGTAGCCGTT

GGTACTGCAAAAGCACTGAGCGGAGGTCATAGATCCGTAATTGGGAGGGCAGCTAGCCGTTGCGGTGCAA

AGAACAGCAGGCGCGTGACCCGAGTTGATTCGCGGGTCGAACAATTCACAAAAATTAGCGGCCCGGTTCT

AGTTTTCAACTTGACGACTCAGATTCAATTCATTATTTTTAACGATTTCGGCCGAATCTATTCAAACCTG

CTCCTTCATCGAATTTCAAGGCGGTCCGGTTCAAAACCACTCATTTGGCCAGCACTGGTTATAAATATTG

AGAAGAAGATTACAATAGATCTGATGGAATGGCACAATACAAAGGGTAAAGTATAGAGAAGATCAATTTT

AAGGAGAGAATAGCATGGTTGCAAGTTGTTCCACACGTACACCATTTTGTAAATATATACTTCGACAGTG

AGAGGCATGATGGATACTGGAGTTCACTGAAACATTCAAATACTTGGAAGGACTCGGAAAGATAAATACA

ATACCAAGAGAAGCAGAAGGAGAAATGAAAAGCAGAGAATGAATGAAGAAATTAAAGAACTAACCTTCTG

CATAAATTGTATGGTTAAAATAGAGGAAGAAGCATGGCGATTCGTGTCACCATCTTTTACAGGTAGAGAG

AGTCCCTCATGGTGTCCATTTGTGCTTCCTTCTGCCATCGCTCTTGTTTCTCTTTGTTAAATTTTGTTTG

TGTGTGTGTGTGTTTGTTTACATAGAGAGAAGAATGGGAATGGGATGGAGAATATGAAGAATGAGAAGTG

ATGTATTTATACATGATCTGCAAAGAAAGAAAGCCAAGAATGGGGATTTGATAAGAGGGGTTGGAGGAAA

GTTGAGATTGTTTTGACCTTTGCAAATAAATGAGAAAGATATGTCATCATTTGACTACCTCCTCTTCTTG

TGTAATCAATTCTTACTCATCCAGCACTGTTCCTCTCTCTAATATTAATGTTTG

>MSTRG.3076.3 gene=MSTRG.3076

TTTCGTTTCTTTACACAAATGTGTCAAGAAAAATAGAAACTAGAATCAGTTTTTGCTTTTTGCAAACAAA

CGCCACCTGAGACTTGAGATGAGAGAATGAGTGGTGAAGAAGAATATGAGGTAAGGACTGGAGGGAGCTA

TAGTAGTAGCCGGCTAGCCACTGTTGCTGCAAATACAAACGACTATGTAGCCGTTGGTACTGCAAAAGCA

CTGAGCGGAGGTCATAGATCCGTAATTGGGAGGGCAGCTAGCCGTTGCGGTGCAAAGAACAGCAGGCGCG

TGACCCGAGTTGATTCGCGGGTCGAACAATTCACAAAAATTAGCGGCCCGGTTCTAGTTTTCAACTTGAC

GACTCAGATTCAATTCATTATTTTTAACGATTTCGGCCGAATCTATTCAAACCTGCTCCTTCATCGAATT

TCAAGGCGGTCCGGTTCAAAACCACTCATTTGGCCAGCACTGGTTATAAATATTGAGAAGAAGATTACAA

TAGATCTGATGGAATGGCACAATACAAAGGGTAAAGTATAGAGAAGATCAATTTTAAGGAGAGAATAGCA

TGGTTGCAAGTTGTTCCACACGTACACCATTTTGTAAATATATACTTCGACAGTGAGAGGCATGATGGAT

ACTGGAGTTCACTGAAACATTCAAATACTTGGAAGGACTCGGAAAGATAAATACAATACCAAGAGAAGCA

GAAGGAGAAATGAAAAGCAGAGAATGAATGAAGAAATTAAAGAACTAACCTTCTGCATAAATTGTATGGT

TAAAATAGAGGAAGAAGCATGGCGATTCGTGTCACCATCTTTTACAGAGAGAAGAATGGGAATGGGATGG

AGAATATGAAGAATGAGAAGTGATGTATTTATACATGATCTGCAAAGAAAGAAAGCCAAGAATGGGGATT

TGATAAGAGGGGTTGGAGGAAAGTTGAGATTGTTTTGACCTTTGCAAATAAATGAGAAAGATATGTCATC

ATTTGACTACCTCCTCTTCTTGTGTAATCAATTCTTACTCATCCAGCACTGTTCCTCTC

>MSTRG.3086.1 gene=MSTRG.3086

CTTAAAACGGGGGACAAACGGGCTCTGGCCTCTGGGTACAGGGCTCTGGTCCTGGGTACAGATTTGGTTC

ACCTTCGACTTGTACACGACACGATGGACGGGCCTGGTAGAAATCGTAAAGAAGATCAGGACCTTTGGTT

TTGAAGATTGACATGAAACCCTCCTTAATTTCTACGACATTTTGCTAAAGGAGCCTTTCCTCAAAAGGTT

TGGAACCTGTGTTCAGGATGTACACAAGGAAGATCCATGGGCTTTTCCTAGGGGGAATACCTGACGAAGC

AAATGAAATATTTGTGAGAATGGAAGCCAGTGATTGCTTGTCAAATACTTTTACCATCCTTATGCTGGGC

TGTATTCATAATAAGAAGTGCAATGTAGCAGGTCTACTCATAAATGAGATGTGTGCTCAAGGTTTCTCAG

TCGATGCATCTACCACCTTCTTGTTACTTGACTAACTTGGATTAAAGGACCAGGATCTCGCTCTTCTTGC

TTTGCCTGTCTTGTTTGGAGTCTCTTGGTGCTGCAACCTTTGATTTGGACGCGCATGGAACATAGCATTT

GGATGTGCCTGAGTCGCTTGACTAGCATTTGTTTTCCCATGGTGCTTCTTCAGAGAATGAAGTAATCTTT

CTGTAATAATTACTCTTGAGCATTTTCTTGGTTTCTTGCGGAGGTGATGATTAGGTAGAGTATTGGTCGC

ATGTGGGATGAAAACTCCAGTCCCGCCTGCTGACCCGATTCTTGACCCACTCTCTCCCAGAAATACAACT

CGTATATACCAGGTTCTCTGCTACAATAGTGCAGATTGGCTTATATAGCAACATCTTAACAATAATAAGG

ATATTATCCCAACAAGGAAACTGATATGATAATTTAACCATCTCATAATACTATACTAGTATAATATTTT

TTTATGTGTTAACATTCTGGACATTTCTCGTGGACGTCGTAGATGTGGCTCGGATCATATTGATCAGATT

TTATTCTGAGTGTTTTTGTTTGGCTTTTAGAAACGGGAGGTTTGATGTACATATTCGATTGAC

>MSTRG.3097.1 gene=MSTRG.3097

ACAGTTTGTTCCTTAATTTATGCCGTTCAATTCTCTTCCTCTGAATTCAAATTTCCAATCATGTGGATTA

ACCTCTTGGATTATGTGGCATCCTCTTGGATTCATACGGGCTTCAACCTGTGATATCAAAAATCATAACC

GTGTCGCCGGTGTGTATTATCTTCGTGTTTCATCAAGTTTGATTTAAAGGTCGGAAACGAATTACTCTCA

GATGCCGTCTCTCCCTTCGATAAAAGATGCGATGCATAGTCTATAAACAGGGAAGAAAAAATGGAGGGAC

CAAGTTAGAAACCTTCCTTTTTAAACACAACTAACATCATAATCCTTGGATTGGTGATTGAGATTTCTTT

CGAATTGTTTCTTACTGACCAGAAGTGGACGGAAAATAATTTTTTCCTAAATGATTCTCTATAGTCCTGC

GTTGTGCACACATTAGAAACAACCCTTACTCGAATAGAAAAGCAAGTACATTTTTGATCCTACAATCTTG

CATCTCATCACAGATTTTCTCGGAAGTGGCCAGAATTGTCTAAGGTGTACAGGTGCTCCGCGATTGGTTG

CTTCTGTTGCCAGTGCATAAGCACAATAATCTGGCTAGCAACTTGAGAAATCGGATCGAAGTTTGTCACC

CTGGCCGCGATGGTATGGTGGTAGCCAGAATCATAAAAGAATACACAAAAAAGTGGTGGAAATCCCACCT

ATATATATGTGTACCTACTAAAGACTAAAGACTAGAAGCTACAAAAAGATGCAAGATATTCAAATTGGGA

ATGAGTTTTGTTCATTATCACCCAGTTGAAGAGACGGAAGATGTACATATCTCTGACTGCCCTGTCTTTA

TAATATCTGGAAACTCTTCGTGCATCGCAACAAAGGACATGTGGAACGAAATCAAAGAGGGAATATAGTG

TACAGACTACCTTTTGTCGTGATCTCAGGCCTGGACTCCATATCGGCGTTTTCTTCCTCGGACAAATCTG

CCGCCCCTTCTACTGCAGCGCGGCTGTTTGAATGCATGCAAGGGGAGAGAAGTGTTTTGCTTTAGCTTTT

TTGGACAATTATTAGGCTCCTCTTTTTAAGTTGTGCCTTATGTAAAGACCTTTTTAGATGTGTCTAATTG

ATATTTCGTCTCTATTTAATTTATTTAGGAGTTAATATGGATAACTAACTAGAAACTAATTTATATTAAT

TTTGGG

>MSTRG.3097.2 gene=MSTRG.3097

ACAGTTTGTTCCTTAATTTATGCCGTTCAATTCTCTTCCTCTGAATTCAAATTTCCAATCATGTGGATTA

ACCTCTTGGATTATGTGGCATCCTCTTGGATTCATACGGGCTTCAACCTGTGATATCAAAAATCATAACC

GTGTCGCCGGTGTGTATTATCTTCGTGTTTCATCAAGTTTGATTTAAAGGTCGGAAACGAATTACTCTCA

GATGCCGTCTCTCCCTTCGATAAAAGATGGTCAAAAGCCAGTGTCACTAAGTGTCCACTCTCCTTCGTCG

TCAGATAAGGCGATGCATAGTCTATAAACAGGGAAGAAAAAATGGAGGGACCAAGTTAGAAACCTTCCTT

TTTAAACACAACTAACATCATAATCCTTGGATTGGTGATTGAGATTTCTTTCGAATTGTTTCTTACTGAC

CAGAAGTGGACGGAAAATAATTTTTTCCTAAATGATTCTCTATAGTCCTGCGTTGTGCACACATTAGAAA

CAACCCTTACTCGAATAGAAAAGCAAGTACATTTTTGATCCTACAATCTTGCATCTCATCACAGATTTTC

TCGGAAGGTGTACAGGTGCTCCGCGATTGGTTGCTTCTGTTGCCAGTGCATAAGCACAATAATCTGGCTA

GCAACTTGAGAAATCGGATCGAAGTTTGTCACCCTGGCCGCGATGGTATGGTGGTAGCCAGAATCATAAA

AGAATACACAAAAAAGTGGTGGAAATCCCACCTATATATATGTGTACCTACTAAAGACTAAAGACTAGAA

GCTACAAAAAGATGCAAGATATTCAAATTGGGAATGAGTTTTGTTCATTATCACCCAGTTGAAGAGACGG

AAGATGTACATATCTCTGACTGCCCTGTCTTTATAATATCTGGAAACTCTTCGTGCATCGCAACAAAGGA

CATGTGGAACGAAATCAAAGAGGGAATATAGTGTACAGACTACCTTTTGTCGTGATCTCAGGCCTGGACT

CCATATCGGCGTTTTCTTCCTCGGACAAATCTGCCGCCCCTTCTACTGCAGCGCGGCTGTTTGAATGCAT

GCAAGGGGAGAGAAGTGTTTTGCTTTAGCTTTTTTGGACAATTATTAGGCTCCTCTTTTTAAGTTGTGCC

TTATGTAAAGACCTTTTTAGATGTGTCTAATTGATATTTCGTCTCTATTTAATTTATTTAGGAGTTAATA

TGGATAACTAACTAGAAACTAATTTATATTAATTTTGGG

>MSTRG.3097.3 gene=MSTRG.3097

AAATAATCCTAATTTTGCCCGTTTTATTTGCCTGCTCCATCCACCCGAGACCCGTTAACCCATCTGCCCA

GCACAGTTTGTTCCTTAATTTATGCCGTTCAATTCTCTTCCTCTGAATTCAAATTTCCAATCATGTGGAT

TAACCTCTTGGATTATGTGGCATCCTCTTGGATTCATACGGGCTTCAACCTGTGATATCAAAAATCATAA

CCGTGTCGCCGGTGTGTATTATCTTCGTGTTTCATCAAGTTTGATTTAAAGGTCGGAAACGAATTACTCT

CAGATGCCGTCTCTCCCTTCGATAAAAGATGGTCAAAAGCCAGTGTCACTAAGTGTCCACTCTCCTTCGT

CGTCAGATAAGGCGATGCATAGTCTATAAACAGGGAAGAAAAAATGGAGGGACCAAGTTAGAAACCTTCC

TTTTTAAACACAACTAACATCATAATCCTTGGATTGGTGATTGAGATTTCTTTCGAATTGTTTCTTACTG

ACCAGAAGTGGACGGAAAATAATTTTTTCCTAAATGATTCTCTATAGTCCTGCGTTGTGCACACATTAGA

AACAACCCTTACTCGAATAGAAAAGCAAGTACATTTTTGATCCTACAATCTTGCATCTCATCACAGATTT

TCTCGGAAGTGGCCAGAATTGTCTAAGGTGTACAGGTGCTCCGCGATTGGTTGCTTCTGTTGCCAGTGCA

TAAGCACAATAATCTGGCTAGCAACTTGAGAAATCGGATCGAAGTTTGTCACCCTGGCCGCGATGGTATG

GTGGTAGCCAGAATCATAAAAGAATACACAAAAAAGTGGTGGAAATCCCACCTATATATATGTGTACCTA

CTAAAGACTAAAGACTAGAAGCTACAAAAAGATGCAAGATATTCAAATTGGGAATGAGTTTTGTTCATTA

TCACCCAGTTGAAGAGACGGAAGATGTACATATCTCTGACTGCCCTGTCTTTATAATATCTGGAAACTCT

TCGTGCATCGCAACAAAGGACATGTGGAACGAAATCAAAGAGGGAATATAGTGTACAGACTACCTTTTGT

CGTGATCTCAGGCCTGGACTCCATATCGGCGTTTTCTTCCTCGGACAAATCTGCCGCCCCTTCTACTGCA

GCGCGGCTGTTTGAATGCATGCAAGGGGAGAGAAGTGTTTTGCTTTAGCTTTTTTGGACAATTATTAGGC

TCCTCTTTTTAAGTTGTGCCTTATGTAAAGACCTTTTTAGATGTGTCTAATTGATATTTCGTCTCTATTT

AATTTATTTAGGAG

>MSTRG.3098.2 gene=MSTRG.3098

TATAATTTGAACTTATATATATTTATGGAATGAATAGATATTCTACTATTGTTTGTAGTGCCACGTAATG

AACCTTAGACCGTTTTTCATGTCGCAATCGGTCTCCTCTTTTACTTTTGGAAATACTGTGGGTGGACTTT

ATATCTAAAATCTACTTTTTTATATTGAAAATCAACAAAACCTCAGAAAAGTGCCGAGGGACATCCTCGG

CATCATTTTTCAAAGAAAAATTGACCGCTGACCGGATATCTATTATATGCTTACTAGATATGGACCATAG

GTATATAATAATCAGAACCAGTGCCATATTGAAACTGATTTATTTGCAGAACCACGAGTGTCTTTCTGTT

CTGTTCAATAGCATGTACAGGAAATTAGACAGACTGGTGTAATCTTGGATCCTACCAAAATTTTGGACTT

TTGCGTGTGTGCACGTCTGTGTCTAAACTTTTATCTGCAATATTCAGAATTTAGCAGTAAATGACTTCCA

TCTGACATTTAGTTCTATAATTGAGGGTCCCATCCGGCCTTCCCGAGCCTGTGTTCTACTAGTTATTTAC

TGATCAGAGTTGAACTTGTGATTCTACTGTGATGACCGACCAATGTAACACCAGAGTACAAGCTTTCCTC

CCTTTTTTTGGTCATTGATTTGTAGGGATCTAATATATTTGGGTTATCCTTTGCGATGAATTAAGTTTCT

TAGAAGCCTTGAGAATCAAAGGAGTGGCCAAGCGGTGGCAAAGGAAGTGCTCGAGATGTGAGATGCAGCT

TCAGAATCCAATATTCCTACAAAAATATAAGCATCAGTCAGTTCCTGATGCAAGACAGAGACCGTGAAAC

TGCGAAAGGCATTTACCCTTTCACCACATTTCTAGAAACAGCTGTACAAAATACCATGTATGAAAGATGT

ATATCTAAGTAGAATACACGTAGTCACGTACTCATACTCTTCTTCTATGTGATAGGTAGTTATGAGAACC

ATACCTCTTGATATGCATATATGAGATGTGTATGTTTCAGATGAAATATCTCAGATCTAAAGTATATCAG

AGATGTGTATCTTCCAAATATGCATTCTAAGTATGTGCGTCTTTATGATACACATTTTAATCATGTTACA

GATTAGTTGGAACAATTCTCTGCTGGATTTTAATACTATAAGTAGAAAACTGAAGTTGCAGGGTTTGGAT

GAAATGAGGGAGCTTTGTGAATAATCTTGCAGCTTTAATTAGCGGTGGATCTGGTTCATTCTTGTATGAA

GTG

>MSTRG.3101.1 gene=MSTRG.3101

GTTAGGATCGGGTCGGTAAAAGTGGTGGGTTTGGGTCTTGATTACTCTCCGTAGAAGTGAGGCAAAAGGA

GGGGGGAGATGAGGAATGAGGGAGAGAGCAGAGAGGGGAGGGAGGGAGACTGGGAGAGAAAGAGCGAGAG

AGGATGGGTGGGAGAATATTGAACCAAGCGTTGAGGAAGATGGTGAATGCTGAGAAGAGGGGTTTTGCAA

ATACCCTTCTGTTCTTCCCTCAACTTCGTCACTTTTCATTCCTCACTCTTTGCTCTTTGAATTCATAAAC

GGGGACTCGCATCACACACACGTTACCATCAAAGAAACGTATAGTTCACCTGTTAATGGGGACTCTGAAG

ATGTTCGTTGTGTTGGTGCCAAAATAAAGCAATTCATAAAAGCACTTTTGAGATATTTTATTTTCCCTGC

TGAGGGCAGTACAGAATCAGCCACACTGGATGCTAGCAAAAATTAAATGAGGCGACAAGTCAAATTCCCT

CGAGGCCTTCTAGATCCTTGGGCTGCAGTATATTTTGTGTTGTCAGGAGAGGTCGATGCAATTTCATACT

CTCATTTAAATTCATATACTTGCTTTTGAGTGATAATGAATTGTGTTTCCCAGTTGATACTTACGAAAAG

AAACTACTATATGTTTCCAAAAGTTATACTTTTGCACATTTAACTTATCTTGATATGTAATGATGGGTGC

GAAAAACTAGGCTTATTTTAGACGAAATTACTCCACCTGGCATGGAGTTCTATTGCCTGTGCTGTTGCAG

ACAGCTTGTGCTTGTATTGTGCATTAATATGAATCTTGACAATAATATATTGCCTCAATAAATATACTGG

ACTGCTGATGGAGAAGACCTTTTCCAAACATCTGAAGGTTTGCTTTATGCAGACTAGTATATGTTGAATC

CACCTCTTTCGTGTTGAAGGTTTAGATGCTTCCCTGTCATTAAATCTGAAAGGATTTGTAGCAGTAGTCT

GTTTTTCTTTCTGTTTATTTCGTTAGCAGTATGTCCTCTGTTTTGTTATCTTTGTACCTCTTGAATCCGA

GGAGAAAGTTAGTTGCTCATTACTGCCTTTAAATTTAGGTCTAATGTTGTTTAACAATAGTGTTGATGAG

AAGAGCCAGAAGCTTCTTGTGTTTAGTTACACTGTTCTGATGATTATCGACTCTTTTTGTTTTGGG

>MSTRG.3101.2 gene=MSTRG.3101

CGGGTCGGTAAAAGTGGTGGGTTTGGGTCTTGATTACTCTCCGTAGAAGTGAGGCAAAAGGAGGGGGGAG

ATGAGGAATGAGGGAGAGAGCAGAGAGGGGAGGGAGGGAGACTGGGAGAGAAAGAGCGAGAGAGGATGGG

TGGGAGAATATTGAACCAAGCGTTGAGGAAGATGGTGAATGCTGAGAAGAGGGGTTTTGCAAATACCCTT

CTGTTCTTCCCTCAACTTCGTCACTTTTCATTCCTCACTCTTTGCTCTTTGAATTCGTAAGGGGGACTCG

CATCACACACACGTTACCATCAAAGAAACGTATAGTTCACCTGTTAATGGGGACTCTGAAGATGTTCGTT

GTGTTGGTGCCAAAATAAAGCAATTCATAAAAGCACTTTTGAGTTAAATTGATGAAATTATTCAATTGTC

ACCTGCGGCTCTGTGAATTATATGAAAATGATTGCAGCTCTGATAAATTTGATTGTTGTTTATGTGGTTA

TAGTCGGCGGGTGGCAAGTGGTAACTAATTATTACAGTACAGAATCAGCCACACTGGATGCTAGCAAAAA

TTAAATGAGGTATCCATGACCATTAATATTGCTAAGTAGAAGTAAAACTGATTCACTTTGAAATATTGTA

TTCTATTTATATGTATTTGGACATTTGCTACTGCTGCCTCCCGTAGAAGGCTGGGCATAGTAGTTGTGTA

TTAATTGAACTAAACTGTCTGTGACTGTTTCTGAATAGTCATATAGCATATGCTTCCCTGTTCTTTAATA

TTATTGCGGGGTTGTCAAAA

>MSTRG.3101.3 gene=MSTRG.3101

GGGTCGGTAAAAGTGGTGGGTTTGGGTCTTGATTACTCTCCGTAGAAGTGAGGCAAAAGGAGGGGGGAGA

TGAGGAATGAGGGAGAGAGCAGAGAGGGGAGGGAGGGAGACTGGGAGAGAAAGAGCGAGAGAGGATGGGT

GGGAGAATATTGAACCAAGCGTTGAGGAAGATGGTGAATGCTGAGAAGAGGGGTTTTGCAAATACCCTTC

TGTTCTTCCCTCAACTTCGTCACTTTTCATTCCTCACTCTTTGCTCTTTGAATTCGTAAGGGGGACTCGC

ATCACACACACGTTACCATCAAAGAAACGTATAGTTCACCTGTTAATGGGGACTCTGAAGATGTTCGTTG

TGTTGGTGCCAAAATAAAGCAATTCATAAAAGCACTTTTGAGTTAAATTGATGAAATTATTCAATTGTCA

CCTGCGGCTCTGTGAATTATATGAAAATGATTGCAGCTCTGATAAATTTGATTGTTGTTTATGTGGTTAT

AGTCGGCGGGTGGCAAGTGGTAACTAATTATTACAGAATCAGCCACACTGGATGCTAGCAAAAATTAAAT

GAGGTATCCATGACCATTAATATTGCTAAGTAGAAGTAAAACTGATTCACTTTGAAATATTGTATTCTAT

TTATATGTATTTGGACATTTGCTACTGCTGCCTCCCGTAGAAGGCTGGGCATAGTAGTTGTGTATTAATT

GAACTAAACTGTCTGTGACTGTTTCTGAATAGTCATATAGCATATGCTTCCCTGTTCTTTAATATTATTG

CGGGGTTGTCAAAATTACATGTTTTGCTTAAAATTTCTGTTTTCTGATATGCTTCAGCAGGCGACAAGTC

AAATTCCCTCGAGGCCTTCTAGATCCTTGGGCTGCAGTATATTTTGTGTTGTCAGGAGAGGTCGATGCAA

TTTCATACTCTCATTTAAATTCATATACTTGCTTTTGAGTGATAATGAATTGTGTTTCCCAGTTGATACT

TACGAAAAGA

>MSTRG.3101.4 gene=MSTRG.3101

GTAAAAGTGGTGGGTTTGGGTCTTGATTACTCTCCGTAGAAGTGAGGCAAAAGGAGGGGGGAGATGAGGA

ATGAGGGAGAGAGCAGAGAGGGGAGGGAGGGAGACTGGGAGAGAAAGAGCGAGAGAGGATGGGTGGGAGA

ATATTGAACCAAGCGTTGAGGAAGATGGTGAATGCTGAGAAGAGGGGTTTTGCAAATACCCTTCTGTTCT

TCCCTCAACTTCGTCACTTTTCATTCCTCACTCTTTGCTCTTTGAATTCGTAAGGGGGACTCGCATCACA

CACACGTTACCATCAAAGAAACGTATAGTTCACCTGTTAATGGGGACTCTGAAGATGTTCGTTGTGTTGG

TGCCAAAATAAAGCAATTCATAAAAGCACTTTTGAGCTCTGATAAATTTGATTGTTGTTTATGTGGTTAT

AGTCGGCGGGTGGCAAGTGGTAACTAATTATTACAGTACAGAATCAGCCACACTGGATGCTAGCAAAAAT

TAAATGAGGTATCCATGACCATTAATATTGCTAAGTAGAAGTAAAACTGATTCACTTTGAAATATTGTAT

TCTATTTATATGTATTTGGACATTTGCTACTGCTGCCTCCCGTAGAAGGCTGGGCATAGTAGTTGTGTAT

TAATTGAACTAAACTGTCTGTGACTGTTTCTGAATAGTCATATAGCATATGCTTCCCTGTTCTTTAATAT

TATTGCGG

>MSTRG.3101.5 gene=MSTRG.3101

ATAAACGGGGACTCGCATCACACACACGTTACCATCAAAGAAACGTATAGTTCACCTGTTAATGGGGACT

CTGAAGATGTTCGTTGTGTTGGTGCCAAAATAAAGCAATTCATAAAAGCACTTTTGAGACAGAAACAGAT

GCCTACTACTGTAAAATCTTAACCATAATTTGTTTAGGGAAGATTTGATTAGTGATTACATGGAGTTATC

TTAAAGAGGATATCAAAGCTCAATGAATAAGTTTTCTCAACATCTTTATATATTCCATCTTAGGTTCTCT

CCATCATTATGAATCACATTACCCCAAAGAACACTTCAATTATATTTTGGGAAGCATGTTCTCTGATAAG

TTTATGTTTACTACTCTTCCATATTAGTAAAAAGTGTATTCTAATTGGTGGTGATTTTGTTGGCAAAATA

TGAACACAGTCATGTACTATGACACAGAGATGTAATTTTTTTATAAAAAAAAGAGCTGTTGAATTGTTCC

TTTAAGTATATGTTACATCAGTGTTTTTGGTTGCATTTTGATGTTTATTAGATTTAGATGTCAATAGGGT

ACAATTATAGATAATGATGTAATACTTTATGTCTTGCCATGATATATTTGATCACAGAGCTATAAGTATT

TGATTAAGAAGCAATATTACTGATATTTTCATATATATATAT

>MSTRG.3105.1 gene=MSTRG.3105

AAATATTCAAACATACCAATGAAGTCAATGATGAGGCGACCGTCGGAGCACCGTCCGGTGGGGGACTTGA

AGTAAGTTTCGCCGTAAGGCGGCGAGGCGGAGGCCGGTGGGTTGTTTGTGTTTGAGAGATGAAGCTTGTT

GCCTGTGTCAGCCAGGGAGTCACCGAAGCTTATTATCGATGTGAACGTACGTTCACATGAAGGTGATTCG

CATGCAATTCTTAGAATCGGAGTTAGTGATATTAATAGAATGAGATTTATGGCCGTAATTTCCTTCGATG

TAAAAGATGAAGAGGCCATGAGTGAATATGTTACAACTACGTACTTAAGGTTGTAGAAGATATATCCGAG

>MSTRG.3109.1 gene=MSTRG.3109

CTACAAACAAACACACAAATCAAAACACACATCTCTTCTTCATCTCCACATTCTCAATTCCAATGGCGCG

TACCAAGCAGACCGCTCGCAAGTCCACCGGAGGAAAGGCACCCAGGAAGCAGCTCGCCACCAAAGCTGCT

CGCAAATCAGCTCCGGCCACCGGAGGAGTGAAGAAGCCGCACAGATTCAGGCCAGGAACCGTCGCTCTCC

GAGAAATCCGCAAGTACCAGAAATCCACAGAGCTCCTCATCAGGAAGCTCCCATTCCAGCGCCTGGTCCG

TGAAATCGCCCAGGATTTCAAGACCGATCTGCGTTTCCAGAGCTCTGCTGTGGCGGCTCTTCAGGAGGCA

GCTGAGGCTTATCTGGTCGGTCTGTTCGAGGACACTAATTTGTGCGCTATTCACGCCAAGAGAGTAACCA

TCATGCCCAAGGACATCCAGTTGGCAAGACGCATTCGTGGAGAACGTGCTTAATTTTATGCCTAATTCGA

TGAATTGGAAGGACATGTGATGTAAAGAAAGGTTGTTTCTAGGGTTTGTTAAGTAGCTTTTGATGTTCAA

GATGTAGTCCAGTAATAGTCGTGAAAGCATTTTAATCGATTATGAAGTTATCTTATTTAATGGCCTATCT

TTCTAACTGTTGAACAATGAGC

>MSTRG.3116.1 gene=MSTRG.3116

CTACAAACAAATCAAAACACAAATCTCTTCTTCATCTCCACATTCTCCATTCCAATGGCGCGTACCAAAC

AGACCGCTCGCAAATCCACCGGAGGAAAGGCCCCCAGGAAGCAGCTCGCCACCAAAGCCGCCCGCAAATC

AGCCCCGGCCACCGGAGGAGTGAAGAAGCCCCACAGATTCAGGCCAGGAACCGTCGCTCTCCGAGAAATC

CGCAAGTACCAGAAGTCCACAGAGCTCCTCATCAGGAAGCTCCCATTCCAGCGCCTGGTCCGTGAAATCG

CCCAGGATTTCAAGACCGATCTGCGTTTCCAGAGCTCTGCTGTGGCGGCTCTTCAGGAGGCAGCGGAGGC

TTATTTGGTCGGTTTGTTCGAGGACACCAACTTGTGCGCTATTCATGCCAAGAGAGTTACCATCATGCCC

AAGGACATCCAGTTGGCAAGACGCATTCGTGGAGAGCGTGCTTAATTTCATGACTAAAAAATTGTTTAGA

AGAATATCTAGGGTTTGTTATGTAAAGTAATGCTGTTTGGTTAGGGTTTGAGATTTGAAGTTGTAGCTCT

GATGAATTTTAATGAATTATCATATTATCTCATTCAATTTTGTCCCGTA

>MSTRG.3118.1 gene=MSTRG.3118

CTTATAGTCATATAGACATCTTCAAGATTGATTGATATTTCCATGACTACTCATGATGTTGCTTATGATA

TTTCCCTTTCATTTTTGATTTTGTCATTCGTTGTTCTTGTAATCATCCTGATCATCATCCTATGCAACAA

AAAGCAGGTGAAAGGCCAAGAAATTCTTCCTACCAGGAAGTCTGCACATATATATTCGTTGATCGATGTC

GATGAAGCTACTGAAGGCTTCAGCACTAGAAGGTTAGTGGGGCAAGGCCGGCTCGGAACTGTGTATACAG

GTGTGATGTCAAGAGGAGATATGGTGATTGTTAAAAGGATTCATCCGCGGTTAGTTTTGAGCAATGCTGG

TAATTCTACGTTCTCATCAGTGCTAAAGTCTCTGTGTGCAGTTCAACATCCTAATATAGTATCTATACTA

GGGTTCTCGGAGGCACCCGGGGAAAGGATCATTGTTTCGGAGTTCGTGGGGATGATCAGTTTAGAGTTTT

ATCTACATGGGAATTCTGAGGGAGTGGCTCTTTTGGACTGGGGAAATCGGTTAAAGGTTGCAGCTGGTCT

GGCTCGAGGGATCGAACATCTGCATGAAGTGGTGATGCCGCCAATAGTGCATGGTTGTGTTAAGTCTTCA

AATGTTTTGATTGATTTAAAATTTGTTGCTAAAATTTGTGATTATGGTTTATCATTTTTGACACCTCAAG

AGAAGGAAGGCCTTGTAGGGTATGTTGATGATGAGTTTTGGATTAACAAGAAGGGTGCTTCTAAGGAGAG

CGATGTTTATGGATTAGGTGTAGTTTTGTTAGAGATTTTAAGTGGAAGGAGAAGTGGGAAAGGGTTTAAT

ATTGTTGAATGGGCTTTGCCTTTGATTAAAGAAACGAAGATTAGTCAACTTTTGGATACAAGAATTGTGA

TTCCATCTGATATGAAGCCTCTTGTCAGATTGGCCAAAGTTGCTGCAGCCTGCGTCGGTAATTCTAGGAA

AAATAGACCCTCTATTGTACAGGTTGTAGCCATTTTAAACAATTTGGAAGAGGGGGCTCTGTTTATTAGT

TTTGGGTCTTTGTAAATACAGCTTCATTATATCGTTCAGACATTAGATTCTTCGTTGAAAGGAGATTATT

AATTCTATTGTTTTAATCGGTTGTATAGCTTGGATGATGTATTCATAATGTTCATGCATTTCTAGCGTCA

>MSTRG.3120.2 gene=MSTRG.3120

CATGCGTGAAAGGAAACTGTATACTATAATTGATGGGGACGGGGGAGGATGTATGTATATCTTATATTCA

TCTTCGGTTCTATATTCCAAAACTGGAATTGCAGAATACTGAGGTCACAAAATCTGCCGGGAAAGCTCCC

ACCAGAATTTGCCAAGCTACCTTACCTCCAAGTTATAGATTTGAGCATCAACTACATTGAGGGTACTCTA

CCTCTACAATGGGCCTCCTTGAGACTTCGTCAGATGATTTATACGGTATTTTGGGAGATCGGTAAGACTC

GCATCGATTTCGGAACGATGGTGGATACCGCAAGAGCTCACCTTTCACAACAAATAATTGTTCATGTTTT

GGGGACGTTTGGTGCTCCAGGATCAGTTGATTTGACTGATAGTTCTGAACATTGGGCTACAGATGATGCT

TACGTGAAGGTCAATTGCGAGACTACTAGTTTCATAACTGATGTAGGTTGGATTGCTCGAGATATCGTTG

TAACACTTTTTAGATCAAAGAATATGAATATTCTATTTGTTAAGCGATCTGAAAACAAAACGACTATTTG

TTTAGTTCGTTTTTTGTTTCACAATCAGATTGTAGTTGACAGTTCGGGGGTTTGTCCCGGCTGGGTTTAC

GATTCATATTAATAAATCCTTTGTTTGTCAAAAAAAAAAAA

>MSTRG.3120.6 gene=MSTRG.3120

CAGGACTACAGGGTCTAGATTTGCACACTGGCTCATTTACCTTACAGCAGATTCGAGCTGCAACTAAAAA

TTTTGATAGTGCTAATAAGATTGGTGAAGGTGGCTTTGGTCCTGTTTACAAGGGGGTTCTATTAGACGGA

ACCGTGATCGCTGTGAAGCAGCTTTCTCACAAATCAACGCAAGGGACTCGCCCAGAAAGAGATCGTTTGG

AATTGAATTGGTCAACGAGGTACAAGGTTTGCATTGGCATAGCAAGAGGTTTGGCCTATCTCCATGAAGA

ATCGAGGTTGAAGATTGTCCACAGAGATATTAAAGCTACCAATGTGCTGCTTGATAAAGATTTAAATCCT

AAGATATCTGACTTTGGTTTAGCTAAACTTGCTGAGGAGGACAAGACACATATAAGCACCCGAATTGCTG

GAACTTTTGGATATATGGCACCTGAATATGCAATGAGGGGTTACCTTACAGACAAAGCTGATGTTTATAG

TTTCGGAATTGTTGCCCTGGAAATTGTAAGCGGGAGGAGCAACTCTAACAGTGGGCCAAAGCAGGAGCCC

ATTTACCTTCTTGATTGGGCTTATGAACTTGAAGCTGAAGGGAAGTTAATGGAGCTAGTCGATGAGAGTT

TAGGGTCGGAATATGAGGAGGAGGTACTGTTGATGATTCATGTTGGTCTTCTATGTACTAATGCGACATC

AGCAGAGCGACCTGCTATGTCGTCGGTGGTTAGTATACTTGAAGGCAGTGTTAAAAAAAAGTGTGGATGA

ATAGAGAACTATACCATAACAGTGACTTCACCCTGATAACTAACAGTAATGGATCATGTGTCTACTCAAT

ACATATATGTAGTATCGTGTGCATACCTGAGCTGAGAGATGACATCTTGTATAGGTTACAACTGTGGGTG

TATCCTCCTGATGCAACCTTGTAAGTATAAGTCTTCTGTTGTCTCTCTCAGCAATAAGGATGGAAATTAT

ATTTGTAAGGTGTTAACACTGAAAAGACAACAGGGACCTAAACCCTTATTTATATGCCTCTTAG

>MSTRG.3121.1 gene=MSTRG.3121

TTTAGACATTTATATTCTTAACAATCATAATTATGTGGTTACGTGTGTTTTATGTTTATGTTGTTATATT

AGTAGATCTTAATTGTAATGTAATATTTTGTTTATATATTATGCCAAGCATGATTATGGAGATGTGGAGG

CATACTCTTGGTGATAACATGATGGTGAAATGAGGCCTGTGCATATTAGGATAGGGGTTGAGTAGTTCGA

TTTGATTTAATTTTTAACTCCCTATCTGCTTTAGCATAGTCGGGTTAGGAAGTTAGGATTTGTTGTATTG

CGCTAGACTGTTATGGCTTAGTTAACCATAATTATTTCCTTTTCTCGAATTTTTGTTAGGGGATTTTGAT

TTTAGTGTCCGTGAGTATGCAATACAATTAGCCGACAAGAAACCCTGGAGCATGTAGTTTGGTTTTCCCC

GTCTATTACACTTTACAATATTTTTATTGTGATGAGGACAATTATGGAGTTTCTGAATGAAGGATGGAGG

TTTTGTCAGATTTCATTCTAATTTCTTCTTTACCCTGGGTAATACTGAGTAATTGAAGTTATATGCTAGC

AGGTTTGATAAATTTTCAGTGACAAGTTCAGCAGGGTTGAAATAGAGAACCAGTGATGGTCCATTGCTTT

TGGCTCATGATTTCAAAACCTTATATGCCCATTTTCTAACATTCTTGATTCAGCCTGGAGTTCTGAGTTA

AACTTATTTTCTCGATGCAATATACGTGATGCAGGACAGGAGCACAACTTCAGTTACAAGGTAAACCGAA

TTGCTTTCTATTGTCTGCAGTACTGTTAGAGGATATATTGTTCAATAGCTGTTGCAATTACTTGGTAAGA

ATTGTAGCTTGGTTGAGAATGCAGTTACACATTACTATGTATTATTTCACCACTCCTAGCTTAAGTTTGA

AAAAATTGGTATAGTTTTCTGTTTTTGCATGTGATATCGGAGTAAAATTTAGATAGCATTACATATGTTG

GTCAGTTAGAATGGACAGATGGCTGATATAATGAAAGACAAGGCCCAAGCTGCCAAGGACAAGGCCTCTG

AAATAGTTGGATCTGCCAAGGACCGAACAGTGGAGTCCAAGGATCAAACAGGCAGCTATGTTTCGCACAA

GGCAGGCGCGGTGAAGGACATGACATGCGAGACGGCTCAGGCGGCTAAGCAATTTCGACTGAGAGAACCT

TTTATTCCGCGAGGAATAGGATGATGGACTCTGAGAGGAATGAGCTGCTATATGTTTCAAACTTGCTATA

TGTCTGTAAACATCGTAAGATAATTTAATGTCAGTTTTATTTATTATACAAAACCCCTGCACCGCAGGCA

TAAATACTAGTTGTTGATTGTTCCTTTTACTCCTTTTAGTCAATGAGCATAATTGTGTACTAAATTTTTT

AATTGATTCTGCCCCAAAATCAAGATAACTTTACAATTGTATATTTTTAGAGGTGATGTTGATTACTTTT

CAGTTTTCAGCCATTGCAGTGGTCGCAGCCTATAGTAATCTTTGTTATAGCTTGTGTTAGACTTTAGTCA

CTGACTCACTGCTACGGTGCCCCTAGTACGCCCTG

>MSTRG.3121.2 gene=MSTRG.3121

AACTAATGTTTCGCGTATCCGATTTCGTCCCCCGTATCGAATCGAACCTCGACCCATGTAACTATGGGTT

TTCCCCGTCTATTACACTTTACAATATTTTTATTGTGATGAGGACAATTATGGAGTTTCTGAATGAAGGA

TGGAGGTTTTGTCAGATTTCATTCTAATTTCTTCTTTACCCTGGGTAATACTGAGTAATTGAAGTTATAT

GCTAGCAGGTTTGATAAATTTTCAGTGACAAGTTCAGCAGGGTTGAAATAGAGAACCAGTGATGGTCCAT

TGCTTTTGGCTCATGATTTCAAAACCTTATATGCCCATTTTCTAACATTCTTGATTCAGCCTGGAGTTCT

GAGTTAAACTTATTTTCTCGATGCAATATACGTGATGCAGGACAGGAGCACAACTTCAGTTACAAGTTAG

AATGGACAGATGGCTGATATAATGAAAGACAAGGCCCAAGCTGCCAAGGACAAGGCCTCTGAAATAGTTG

GATCTGCCAAGGACCGAACAGTGGAGTCCAAGGATCAAACAGGCAGCTATGTTTCGCACAAGGCAGGCGC

GGTGAAGGACATGACATGCGAGACGGCTCAGGCGGCTAAGCAATTTCGACTGAGAGAACCTTTTATTCCG

CGAGGAATAGGATGATGGACTCTGAGAGGAATGAGCTGCTATATGTTTCAAACTTGCTATATGTCTGTAA

ACATCGTAAGATAATTTAATGTCAGTTTTATTTATTATACAAAACCCCTGCACCGCAGGCATAAATACTA

GTTGTTGATTGTTCCTTTTACTCCTTTTAGTCAATGAGCATAATTGTGTACTAAATTTTTTAATTGATTC

TGCCCCAAAATCAAG

>MSTRG.3122.1 gene=MSTRG.3122

GGATGATCAAGATTCCACCAGACCATGGCAGTTGTTTACTTCCGAAGATATAATCCTAGGTTACAAACCA

AAGAGTGTTAGAAAGCGCAAGAAATATGGAGGGAAAAAGAATATCTCTGGAAATTCGGCTAATAGATTTA

GTTATCCTCTTTCTCCGCTATCTCCTCCTAACAGTGCAAATACATCATACCCAACACATGCAATTCCTGG

GACTGTTAATCAAGGCATCTCCAGTTTATCAGTTGGAGGTACACATATATGTGGCACTAATTTTGTGAAA

GCACAATGTTCATTTCATTCTAAACTGAACGATTTTGTCTGTTTGTGGCCTTGAGAAGAAGAGTTCTTTT

ACCAGGACTCCACCAGAACATGCAAGTCCTGGGACTTTTAATCAACGCATCTCCAGTTTATCAGATGAAG

ATGGGAGATCTTTCTGTTGAGGTTAATTAGTCTAGGAGATTCGCATTGCTGCTTAGGGGGACAAGGGGAA

GGTTGTAGAAGCAATTTCTGAAGGTATTCAGCTTTCGATTTTGTGTTCGCGTGTGAGTCTGTTGTGTGGA

GTGGAGTGTTTCACATGAGAATTTTATGAGCAGAGCAGATTCAGATGCCGTATGAGAATTATTTCTGCTA

TATGAGAATTTAATTCGTTAATTGAAAAATGAAAGAGACCACACAAGATCCCATTAACTACTTGATAATT

TTAGTATTGAATTAAGCTTTGTTATATTATCAGGGAATTAATAAGC

>MSTRG.3129.2 gene=MSTRG.3129

AAAAAAAGAATTGCAAAACCCCATCAAAACCCTGAATTGCTTTCTTCTTCACAAATCGACTTTTTCACAG

AGAAGAACAAACATGGCCGCCATTGCTGCTAGATCACTCCTACGTTCCGCGCGTTTGGGAAGCACAAGAT

TCGCCAAACCCACCAAGTCCCCTTCTTCCCCTTTTCATCTCCCCAAACAAAAACCCCCCTCTTCATTCAT

TTCAAGGATGCCAGTTGAGATGAGTTGTGCTATGGATTCGATGTTACCGTTTCATACAGCTACAGCTTCT

GCGTTGCTCACTTCGAAGCTTTCTGTCACTCCCAGGATCAATTGGGCCATTGATGGTTATATGGATTCAT

AAAGGTTGATTTGGATTGGAAGGATTTTTTGCATTGGATTATCTTCTTTAGACAATTTGTTTATAAGCAG

CGCCTTTGGTTGCTAAGCTTAAATATTGTAATAGTTACTTTGTAAGAATTTCGGACTAAAAAACTGTTTT

GTATATTAGTTTGATTACATTAGCATTTAGCCGTCTATAAAAGTGTGGCATGTTATATTATTAGAGATCT

AAATTACTTTTCACTTGACTGCCACTTAGCTTGACGCGTGGGTAGTAACTAGTGAGTTTTTGAGATTGAC

ATGTCATGACTTTTTCTTGATATAGGAAAATTTGGAGTTCTAATACTGTAACATGCCACACGTTTAATAT

ATTGTTGCTATATCGTATTGTTCAAAATTATATCATCTCTAGACACATTACTTCACCCCACTAATTGGAT

TGGATAGTAGCTTGCTGAGTTTTGCAACTCTTTACAAATTTCAGGTTTTATGGATTCGTAGAGGTCGATT

TGGAATGGAAGGATTTTTGCATTGAATTGTCATCTTTAGACCTTTTCTTTTTAGCTGAGCCTTTATATTG

TGAGCTGACAATTTATAGAAGTTACTTCTGTAAGATAATATTGGACTTATAAATTGGGTTGTATTTTAGT

TGATCAAATTAACAGTCTGAAAGGGTGCGGGTTGTTACAGTTTTCTTGAACTGAATTTAGAATTAATCTC

CGTTGAATTAAGGGCAAGTTTAGGGGT

>MSTRG.3129.1 gene=MSTRG.3129

AAAAAAAGAATTGCAAAACCCCATCAAAACCCTGAATTGCTTTCTTCTTCACAAATCGACTTTTTCACAG

AGAAGAACAAACATGGCCGCCATTGCTGCTAGATCACTCCTACGTTCCGCGCGTTTGGGAAGCACAAGAT

TCGCCAAACCCACCAAGTCCCCTTCTTCCCCTTTTCATCTCCCCAAACAAAAACCCCCCTCTTCATTCAT

TTCAAGGATGCCAGTTGAGATGAGTTGTGCTATGGATTCGATGTTACCGTTTCATACAGCTACAGCTTCT

GCGTTGCTCACTTCGAAGCTTTCTGTCACTCCCAGGATCAATTGGGCCATTGATGGTTTTATGGATTCGT

AGAGGTCGATTTGGAATGGAAGGATTTTTGCATTGAATTGTCATCTTTAGACCTTTTCTTTTTAGCTGAG

CCTTTATATTATGATTGATTGTGACTCTGTGAGTTGTGGGGATGTGCTGCATCTGGAGTATGTTACAACT

TACAAGTATAAGAAGTGAATAGAATTGGTCCCTTGAGTTAAGAACACAAATTTCTTATATGTTTCTACTA

GCATATAAGTAGTATGTTGAAGGCTAGGGAAATGTTCCTGACAAAAAATAAGGCTAGGGAAATGTTTACA

CCGGAGAAAGAGC

>MSTRG.3129.3 gene=MSTRG.3129

AAAAAAGAATTGCAAAACCCCATCAAAACCCTGAATTGCTTTCTTCTTCACAAATCGACTTTTTCACAGA

GAAGAACAAACATGGCCGCCATTGCTGCTAGATCACTCCTACGTTCCGCGCGTTTGGGAAGCACAAGATT

CGCCAAACCCACCAAGTCCCCTTCTTCCCCTTTTCATCTCCCCAAACAAAAACCCCCCTCTTCATTCATT

TCAAGGATGCCAGTTGAGATGAGTTGTGCTATGGATTCGATGTTACCGTTTCATACAGCTACAGCTTCTG

CGTTGCTCACTTCGAAGCTTTCTGTCACTCCCAGGATCAATTGGGCCATTGATGGTTATATGGATTCATA

AAGGTTGATTTGGATTGGAAGGATTTTTTGCATTGGATTATCTTCTTTAGACAATTTGTTTATAAGCAGC

GCCTTTGGTTGCTAAGCTTAAATATTGTAATAGTTACTTTGTTTTATGGATTCGTAGAGGTCGATTTGGA

ATGGAAGGATTTTTGCATTGAATTGTCATCTTTAGACCTTTTCTTTTTAGCTGAGCCTTTATATTGTGAG

CTGACAATTTATAGAAGTTACTTCTGTAAGATAATATTGGACTTATAAATTGGGTTGTATTTTAGTTGAT

CAAATTAACAGTCTGAAAGGGTGCGGGTTGTTACAGTTTTCTTGAACTGAATTTAGAATTAATCTCCGTT

GAATTAAGGGCAAGTTTAGGGGTTTTGAAGTCAATTTGTGAATGATCTGTGAAATATTAATTCTGTTAAT

TGAAGTCAGTTTGGG

>MSTRG.3129.4 gene=MSTRG.3129

AAAAAAGAATTGCAAAACCCCATCAAAACCCTGAATTGCTTTCTTCTTCACAAATCGACTTTTTCACAGA

GAAGAACAAACATGGCCGCCATTGCTGCTAGATCACTCCTACGTTCCGCGCGTTTGGGAAGCACAAGATT

CGCCAAACCCACCAAGTCCCCTTCTTCCCCTTTTCATCTCCCCAAACAAAAACCCCCCTCTTCATTCATT

TCAAGGTCACTTTTAACCATATAAATCTACACACATCTATGTATTATATACACACTTATATGATTATGAT

GTAAATTTTGAATTTTACAGGATGCCAGTTGAGATGAGTTGTGCTATGGATTCGATGTTACCGTTTCATA

CAGCTACAGCTTCTGCGTTGCTCACTTCGAAGCTTTCTGTCACTCCCAGGATCAATTGGGCCATTGATGA

TGATTGATTGTGACTCTGTGAGTTGTGGGGATGTGCTGCATCTGGAGTATGTTACAACTTACAAGTATAA

GAAGTGAATAGAATTGGTCCCTTGAGTTAAGAACACAAATTTCTTATATGTTTCTACTAGCATATAAGTA

GTATGTTGAAGGCTAGGGAAATGTTCCTGACAAAAAATAAGGCTAGGGAAATGTTTACACCGGAGAAAGA

GCAATTCAGATGTTTTATTTGGTTTGAATAGAATGCA

>MSTRG.3129.5 gene=MSTRG.3129

AAAAGAATTGCAAAACCCCATCAAAACCCTGAATTGCTTTCTTCTTCACAAATCGACTTTTTCACAGAGA

AGAACAAACATGGCCGCCATTGCTGCTAGATCACTCCTACGTTCCGCGCGTTTGGGAAGCACAAGATTCG

CCAAACCCACCAAGTCCCCTTCTTCCCCTTTTCATCTCCCCAAACAAAAACCCCCCTCTTCATTCATTTC

AAGGATGCCAGTTGAGATGAGTTGTGCTATGGATTCGATGTTACCGTTTCATACAGCTACAGCTTCTGCG

TTGCTCACTTCGAAGCTTTCTGTCACTCCCAGGATCAATTGGGCCATTGATGATGATTGATTGTGACTCT

GTGAGTTGTGGGGATGTGCTGCATCTGGAGTATGTTACAACTTACAAGTATAAGAAGTGAATAGAATTGG

TCCCTTGAGTTAAGAACACAAATTTCTTATATGTTTCTACTAGCATATAAGTAGTATGTTGAAGGCTAGG

GAAATGTTCCTGACAAAAAATAAGGCTAGGGAAATGTTTACACCGGAGAAAGAGC

>MSTRG.3131.1 gene=MSTRG.3131

AATGGTTTTTAATAGTGTTTGGCTGTTAAAAATTGTTAGAAATCGAAGGAAAGTTGATGTAGAGTGCTGA

TGTGTTAATATGTTCTAACTTATAAAGTAGATAGCATGACAGAGAAAGTTGCAAGAGATTACTAAATCAC

CATCAAACAGGAGACCAACTTTCAACTGTGAAAGTATGAAACCATTAAATGGTTTTTAATAGGTTTTGGC

TGTTAAAAAATGTTAGAAATCGAAGGAAAGTTGATGTAGAGTGCTGATGTGTTAATATGTTCTAAAGTAG

TAGCATGACAGAGAAAGTTGCAAGAGATTAGGGGGAATAGGATCATGCTCTGTTGTATCAACAGAAGAAG

ATTGAAAACAGACCACCCTTGTTTCTTGAGTTACGGAGAAGGTAGAGGAAAAGATTATGGTGCGACTCCA

GGAGTTAGTTGGGAGCAAGATCTGCATCCGTCCTCATAGTTTTTGTATCTTTCGTGTATGTTATCTTTCA

TAACAATCACATAAGTACCAAAATTAGGAACATCGGCTGATTTAATTTGTGCTCTGTAGTACATCGAATA

GGATGGTGTAGGCATGTCATTTGTGTGGTTCCTGTTGCTTTTGACTGGTAGATGGGGTTGATATTTTTGT

GGAAGGATGACCTTTTTTTATTAACAGATTACTTCCAGGCTGTTCCAGCAAGTAAATGGAAACGAAATTT

TTCTTGCATTTAATTAATTAACCGACTACAGGTTTTATCTGATTGAGTTGTCAATATACAAGCTTTTGAG

TGCATACCTGCAAACTGTTCCCCTGTTCAATGCTGGTTTATCCCTCAACCCCCCAAAAAATGTCTGACGT

TTAGAAATTGACAAGCAAAAAGTTCAGACTTTTAATGGAAGAATTAAAGGCTAATGAAGGTGTAAAATGG

AGTGGATTGTTATTTACAGGCAATACTTTGAGCTGGCTCCAATACTGAATATAGTCATACGAGCAAGTGC

TTGTTTTATTTTCGTAAGATAAGATTTGCAAGTTCAATAAAGCTTAAAACTTCCTAGTTCTTATGACAAC

GTATAGAATATATTGGAATTCTCATCGTAAACAATTGATTCAACACACAGCTAACTAATCTCCGTGTAGG

TTTTCGCGTTCAATAGAGTTCATGAGTCCATGTCTTGTTAATTATCTCTATGATAAGTCATTAGATGATA

TGATCTAGTCAATTTATATTCTAGAGGTGTTAAAGAAGTATCCTTTCAAGGAAAAACGCCCTTTTTTTTA

TATCAAAATTGGGGATATACCAAATGGTTACAATATATAAATGATTTGTGTTATTCACCCAAGCTCTTAA

CTTTTACTTGTATAGTTAGAGCATGCCATACAGTTTTATGTGTTTATGTCCACATTTGTTTTTATTGTTT

TCTTTATCGATTTTTTCAGGGTTGTGAGCACTAATTTTGTCGCACGTCAACCTGAAGAGTCTTCAGTCTA

TGTAATTACATAATCTATTTACTTTGTGCTCTTAATTTTTCTTATCATGTATATGTGAAACCGTAATAGA

CCGCTGTAGACACTGATTGCTCAAGTCGAAGAGGGTACTGGTTAAGTTGTAAGGTTCATTAAGTGGGTGT

CCTACCATACGACCTGTCTGTCATGTGATCGTCCAATACACTCAGAAATTTCTTTCAGCTTAAACCACTT

AATTGATAAATGATAAACCATGATTGTTATGGATAGATACTACTGTCACATTTACATTTCAATAAACATA

TATCTTTGTTAAAATCTTGACCAATACACTCCTCTTTTCCTTTCTTTTTTGGTTTAGGGCTATAATACTG

AACATGGCTGAAATTGACTTGGAGAGTGTTCATTATATTCCATGTTTTGTACCGTCACTTTCGGAAGTTT

ATAGACCCAGTGGCTGATGATCCAAGAAGAAAAGAATTTCTTTTTTTATTATTATCATTTTGCAATATTT

GTTCTCTGGCCTGCAGCTTGGAACTATATAATCTACCAACAGCATTTACTTTGTCATAAATTTATACTGT

TGGTTTTTTTGTTTCCTTGTGAATTGTAGATGCATGACTATGAGTTGTGGGGATGTGCTGCATTTGGAGG

CAATTCAGTGGATGGTGGGATAACAATGTAATAATACAACTTTAGGCAGTTCAACGGTTAACTGTGAAAT

CAAATTGTTCAACTGTTCCTTTTTCTGGGAAGTGCAAGTAATCATATTTAAATCAGGGACGCATGCATGT

AGTATTAAATTGCTG

>MSTRG.3131.2 gene=MSTRG.3131

GTGTTTATGTCCACATTTGTTTTTATTGTTTTCTTTATCGATTTTTTCAGGGTTGTGAGCACTAATTTTG

TCGCACGTCAACCTGAAGAGTCTTCAGTCTATATGCATGACTATGAGTTGTGGGGATGTGCTGCATTTGG

AGGCAATTCAGTGGATGGTGGGATAACAATGTAATAATACAACTTTAGGCAGTTCAACGGTTAACTGTGA

AATCAAATTGTTCAACTGTTCCTTTTTCTGGGAAGTGCAAGTAATCATATTTAAATCAGGGACGCATGCA

TGTAGTATTAAATTGCTGGCACAATTACTAAATGTTGTTTAATTTCAAAATTCTTTAAAGTGGTTATGCT

TCTGGTGCAAGTCGAAAACACTTGGTTCTCCCAAAATTGCATGTAGGCATACGTCGTGGCCAGTAAGGGC

AACTGTGAACGTTCACTTTTTTAGCACTCAGATAAATGTTGCACTGTCTTTCCTGC

>MSTRG.3131.3 gene=MSTRG.3131

GTGTTTATGTCCACATTTGTTTTTATTGTTTTCTTTATCGATTTTTTCAGGGTTGTGAGCACTAATTTTG

TCGCACGTCAACCTGAAGAGTCTTCAGTCTATGGCTATAATACTGAACATGGCTGAAATTGACTTGGAGA

GTGTTCATTATATTCCATGTTTTGTACCGTCACTTTCGGAAGTTTATAGACCCAGTGGCTGATGATCCAA

GAAGAAAAGAATTTCTTTTTTTATTATTATCATTTTGCAATATTTGTTCTCTGGCCTGCAGCTTGGAACT

ATATAATCTACCAACAGCATTTACTTTGTCATAAATTTATACTGTTGGTTTTTTTGTTTCCTTGTGAATT

GTAGATGCATGACTATGAGTTGTGGGGATGTGCTGCATTTGGAGGCAATTCAGTGGATGGTGGGATAACA

ATGTAATAATACAACTTTAGGCAGTTCAACGGTTAACTGTGAAATCAAATTGTTCAACTGTTCCTTTTTC

TGGGAAGTGCAAGTAATCATATTTAAATCAGGGACGCATGCATGTAGTATTAAATTGCTGGCACAATTAC

TAAATGTTGTTTAATTTCAAAATTCTTTAAAGTGGTTATGCTTCTGGTGCAAGTCGAAAACACTTGGTTC

TCCCAAAATTGCATGTAGGCATACGTCGTGGCCAGTAAGGGCAACTGTGAACGTTCACTTTTTTAGCACT

CAGATAAATGTTGCACTGTCTTTCCTGC

>MSTRG.3135.1 gene=MSTRG.3135

CATCATGCAAGCCAAAGAGGAGGTGGCTCATTCAAATCTATCCCCAACCCACGTCGCACTATCCCCAGAT

ACGCCGCCGTGTTCGAACTCTCCGGCACCGGAGCTCCCCCACTCCCCACCACCCCACGGTGGCTCCTACT

CCCCCCTCCCATACTCTCTCCCAGAAAATCACGCATCATTATCCTCCCGGAGCTCCCTCCCCACCGCTGA

TCCGACGGCGCGTTCAGGTCCAGACCCAACCCATTTTCCGGCGCCGGAAAATTGGTCTTCGCTTTCGGCC

CTCTCAGCGACCTGGCTGCGCCGTCATACGCTAACGCAGCCTCCTCTGGCGTGTCGAACGTGCCGAGCCA

CACCCGTGTTTTCTTCCACGGGTCGCGTATCTCGGCGGCGTAACGGCCCCATGGCCGCTTTCTCACTCCT

CTATAGTGACCTTCTTTAGTGGACGCCATGGGGGTTTTAGAGAGAGAGTAAGGTGGGCCCACTGAGTTAA

CGTTTGATCGTCGGCTGAGTATTAGATGGAACAAGTGCATTATATAGAAGGGGCTTGAAGATGTTATTGT

CTGCAGCTGAGGAGTTTGGGGAGGTAGTGAGATGTGAGCCGTTGGATTTCATCGGGGGATTTTTTGTGTA

TTTATGGGAGTGTGTTGTGTGTATTATTGGGCCGTGTTTATAAACGTGTGGTATTGTTTGTTTCGTTTTT

TTCTAAAGTTGTGAAATTGATA

>MSTRG.3137.1 gene=MSTRG.3137

CATGTACCTCTCCTTCTTTGATGGTTGTGCCCACATCACCAAATCCTTTTCTCCCCCTCTTTGAGCTATG

ATTCAGGTGCTTCACATGGGTTATTTGGGTTCTTTCACAGAAATGGGTTTCAAGGCCCCTTCGCCCGTGG

AAAATCAATTCTCCGATTAAGTCTGTCTATCTCGCCGGACCTGACCCAGATGGGTGTTTGTTGCCGGAGA

CAAGTCTCGGGATCTTGCCGGAGAGTTGCTTGCCTGCTTGGCCTCTGTTCTTGAGTACATAGACCCACTT

GGTCAAAAATATATAAGGTTTCCAGAATTTATAAGCTCAAGTTTGTTCTTTAGGAGGGGCAAACACATTG

GGTCGTGCAAAAAAGCAGTAGAGGCTAATGGATGTAGAAGGAATGATGAAAGTTGTGGATAAGAGCAGGA

GAGGGGTTGGGTTGTACACAGAAGGGGTCCTGTTGGGTTGTACACAAAAGTGGATAATCACGTTGTTGCT

CAAGAGGCCAAGGGGAAGGCTGTAGAAGCTATATCGGTTTTTTGTTTGTGCTGTGCTGATGCAATTAAAG

TTCGGTGCTAGGTCGAATGAATTTAAGTATCTAAATATGGTACCGTGTTGGAAAGCTCTTCAGGATTCAA

TAACAAAACTTTCATCTATCCGGAAATTAAGCATTTGTGATTGTAAGGAGTTGGAGAGTTTGCCCACGTG

TGAAGAATCACATCCTTCAAACACTGAATATATTTAGGTGTCCTGTTCTGAGACAAAGATGCAGTAAGGG

TCAAGGGCCAGAATGGTTCAATGCGTTCCAAGTATATTTTGGGATTATAAGAGGTGGCAGTGAACTTTTT

AGCTTGGTGATTGTCTCTACTCCATCTTCCCTGCCTTGACTGTTTTGTATGTAGTAAATTTAAAGAATAG

AAAGTTATCCCGAGTTCAATAGCAGGCCTTCTATCGCTCAAGCAGTTATTTGGAATTCTGAGAACTAGGA

GAGTTTGGGAAATGGTAAAGACCTGGTGAATTATGACAATAAGCATCTAGAATTTCTCAACTCTATTTGT

TTATTTTTGAAATACCTATCTTCTGGAGCTTTGTATACAGAAGGGGTTGGGTTGGACATAATCTTTACAG

ATCAGTGAAAAAAATGCTCTAGACACCATAATATGTCTACTTGCCTATGATTTTCTTTAATTTTGTGAAA

ATTTTGGTTAATGGTTTGGTAGGCTTTGTCCTTATGATTTGACAATGAGCATTCCTGCTTCAACTCCCCC

TCAGGTTTCAAATATTTCAGAAGCATAAGTACTTTCTAAAACGATGGTTTGGAAGTGATAGGGTG

>MSTRG.3137.3 gene=MSTRG.3137

GTTTTTTTTTTGACAAAATCATGTATGAAAATATAACTTATGTTGTTTGTGTTTGTGATGTTTTTTGTTT

CGAGATGATTTTAAGTTATAGCATTCCTATTGTTTGATAAAAGCTCTAAGAGATTTTGTAATCACTTTTG

AATTGCATTTATAGCAGTAGAGGCTAATGGATGTAGAAGGAATGATGAAAGTTGTGGATAAGAGCAGGAG

AGGTATACGATAAGTGAATTTGGAGAAATTTTTTTATGTTCAAAGAAGGGGTCCTGTTGGGTTGTACACA

AAAGTGGATAATCACGTTGTTGCTCAAGAGGCCAAGGGGAAGGCTGTAGAAGCTATATCGGTTTTTTGTT

TGTGCTGTGCTGATGCAATTAAAGTTCGGTGCTAGGTCGAATGAATTTAAGTATCTAAATATGGTACCGT

GTTGGAAAGCTCTTCAGGATTCAATAACAAAACTTTCATCTATCCGGAAATTAAGCATTTGTGATTGTAA

GGAGTTGGAGAGTTTGCCCACGTGTGAAGAATCACATCCTTCAAACACTGAATATATTTAGGTGTCCTGT

TCTGAGACAAAGATGCAGTAAGGGTCAAGGGCCAGAATGGTTCAATGCGTTCCAAGTATATTTTGGGATT

ATAAGAGGTGGCAGTGAACTTTTTAGCTTGGTGATTGTCTCTACTCCATCTTCCCTGCCTTGACTGTTTT

GTATGTAGTAAATTTAAAGAATAGAAAGTTATCCCGAGTTCAATAGCAGGCCTTCTATCGCTCAAGCAGT

TATTTGGAATTCTGAGAACTAGGAGAGTTTGGGAAATGGTAAAGACCTGGTGAATTATGACAATAAGCAT

CTAGAATTTCTCAACTCTATTTGTTTATTTTTGAAATACCTATCTTCTGGAGCTTTGTATACAGAAGGGG

TTGGGTTGGACATAATCTTTACAGATCAGTGAAAAAAATGCTCTAGACACCATAA

>MSTRG.3137.4 gene=MSTRG.3137

GGTCAAAAATATATAAGGTTTCCAGAATTTATAAGCTCAAGTTTGTTCTTTAGGAGGGGCAAACACATTG

GGTCGTGCAAAAAAGTAGAGGCTAATGGATGTAGAAGGAATGATGAAAGTTGTGGATAAGAGCAGGAGAG

GGGTTGGGTTGTACACAGAAGGGGTCCTGTTGGGTTGTACACAAAAGTGGATAATCACGTTGTTGCTCAA

GAGGCCAAGGGGAAGGCTGTAGAAGCTATATCGGTTTTTTGTTTGTGCTGTGCTGATGCAATTAAAGTTC

GGTGCTAGGTCGAATGAATTTAAGTATCTAAATATGGTACCGTGTTGGAAAGCTCTTCAGGATTCAATAA

CAAAACTTTCATCTATCCGGAAATTAAGCATTTGTGATTGTAAGGAGTTGGAGAGTTTGCCCACGTGTGA

AGAATCACATCCTTCAAACACTGAATATATTTAGGTGTCCTGTTCTGAGACAAAGATGCAGTAAGGGTCA

AGGGCCAGAATGGTTCAATGCGTTCCAAGTATATTTTGGGATTATAAGAGGTGGCAGTGAACTTTTTAGC

TTGGTGATTGTCTCTACTCCATCTTCCCTGCCTTGACTGTTTTGTATGTAGTAAATTTAAAGAATAGAAA

GTTATCCCGAGTTCAATAGCAGGCCTTCTATCGCTCAAGCAGTTATTTGGAATTCTGAGAACTAGGAGAG

TTTGGGAAATGGTAAAGACCTGGTGAATTATGACAATAAGCATCTAGAATTTCTCAACTCTATTTGTTTA

TTTTTGAAATACCTATCTTCTGGAGCTTTGTATACAGAAGGGGTTGGGTTGGACATAATCTTTACAGATC

AGTGAAAAAAATGCTCTAGACACCATAA

>MSTRG.3137.2 gene=MSTRG.3137

CTCTCCTTCTTTGATGGTTGTGCCCACATCACCAAATCCTTTTCTCCCCCTCTTTGAGCTATGATTCAGG

TGCTTCACATGGGTTATTTGGGTTCTTTCACAGAAATGGGTTTCAAGGCCCCTTCGCCCGTGGAAAATCA

ATTCTCCGATTAAGTCTGTCTATCTCGCCGGACCTGACCCAGATGGGTGTTTGTTGCCGGAGACAAGTCT

CGGGATCTTGCCGGAGAGTTGCTTGCCTGCTTGGCCTCTGTTCTTGAGTACATAGACCCACTTGGTCAAA

AATATATAAGGTTTCCAGAATTTATAAGCTCAAGTTTGTTCTTTAGGAGGGGCAAACACATTGGGTCGTG

CAAAAAAGTAGAGGCTAATGGATGTAGAAGGAATGATGAAAGTTGTGGATAAGAGCAGGAGAGGTATACG

ATAAGTGAATTTGGAGAAATTTTTTTATGTTCAAAGGTACTCTCTAATTGTTACTGATCATGTATATTTG

CAAGTGTACACTATGTTTTTATGTTGAAATGTGGCAGGGGTTGGGTTGTACACAGAAGGGGTCCTGTTGG

GTTGTACACAAAAGTGGATAATCACGTTGTTGCTCAAGAGGCCAAGGGGAAGGCTGTAGAAGCTATATCG

GTTTTTTGTTTGTGCTGTGCTGATGCAATTAAAGTTCGGTGCTAGGTCGAATGAATTTAAGTATCTAAAT

ATGGTACCGTGTTGGAAAGCTCTTCAGGATTCAATAACAAAACTTTCATCTATCCGGAAATTAAGCATTT

GTGATTGTAAGGAGTTGGAGAGTTTGCCCACGTGTGAAGAATCACATCCTTCAAACACTGAATATATTTA

GGTGTCCTGTTCTGAGACAAAGATGCAGTAAGGGTCAAGGGCCAGAATGGTTCAATGCGTTCCAAGTATA

TTTTGGGATTATAAGAGGTGGCAGTGAACTTTTTAGCTTGGTGATTGTCTCTACTCCATCTTCCCTGCCT

TGACTGTTTTGTATGTAGTAAATTTAAAGAATAGAAAGTTATCCCGAGTTCAATAGCAGGCCTTCTATCG

CTCAAGCAGTTATTTGGAATTCTGAGAACTAGGAGAGTTTGGGAAATGGTAAAGACCTGGTGAATTATGA

CAATAAGCATCTAGAATTTCTCAACTCTATTTGTTTATTTTTGAAATACCTATCTTCTGGAGCTTTGTAT

ACAGAAGGGGTTGGGTTGGACATAATCTTTACAGATCAGTGAAAAAAATGCTCTAGACACCATAA

>MSTRG.3137.5 gene=MSTRG.3137

CTCTCCTTCTTTGATGGTTGTGCCCACATCACCAAATCCTTTTCTCCCCCTCTTTGAGCTATGATTCAGG

TGCTTCACATGGGTTATTTGGGTTCTTTCACAGAAATGGTAGGTGGTTATTTGTTTTCAAGCTCTGACGC

TTGGGTTCCTGCACAAGTAACAATCCATCTCAATTTGTATCTTGCTGCATACTCCAATTCAAATTTGAAT

TCAAACACCTTCTTTTCTCACACACACATCAGATGCACGTCATTATTATCTTCTTCAAATTTCAAACCAC

CGTTGAGATTCTCTCCCCATAAAAACTCAAATTCTCTCTGCCGTGTGCCTCTTTTCTAGGGTTTCAAGGC

CCCTTCGCCCGTGGAAAATCAATTCTCCGATTAAGTCTGTCTATCTCGCCGGACCTGACCCAGATGGGTG

TTTGTTGCCGGAGACAAGTCTCGGGATCTTGCCGGAGAGTTGCTTGCCTGCTTGGCCTCTGTTCTTGAGT

ACATAGACCCACTTGGTCAAAAATATATAAGGTTTCCAGAATTTATAAGCTCAAGTTTGTTCTTTAGGAG

GGGCAAACACATTGGGTCGTGCAAAAAAGCAGTAGAGGCTAATGGATGTAGAAGGAATGATGAAAGTTGT

GGATAAGAGCAGGAGAGGTATACGATAAGTGAATTTGGAGAAATTTTTTTATGTTCAAAGGGGTTGGGTT

GTACACAGAAGGGGTCCTGTTGGGTTGTACACAAAAGTGGATAATCACGTTGTTGCTCAAGAGGCCAAGG

GGAAGGCTGTAGAAGCTATATCGGTTTTTTGTTTGTGCTGTGCTGATGCAATTAAAGTTCGGTGCTAGGT

CGAATGAATTTAAGTATCTAAATATGGTACCGTGTTGGAAAGCTCTTCAGGATTCAATAACAAAACTTTC

ATCTATCCGGAAATTAAGCATTTGTGATTGTAAGGAGTTGGAGAGTTTGCCCACGTGTGAAGAATCACAT

CCTTCAAACACTGAATATATTTAGGTGTCCTGTTCTGAGACAAAGATGCAGTAAGGGTCAAGGGCCAGAA

TGGTTCAATGCGTTCCAAGTATATTTTGGGATTATAAGAGGTGGCAGTGAACTTTTTAGCTTGGTGATTG

TCTCTACTCCATCTTCCCTGCCTTGACTGTTTTGTATGTAGTAAATTTAAAGAATAGAAAGTTATCCCGA

GTTCAATAGCAGGCCTTCTATCGCTCAAGCAGTTATTTGGAATTCTGAGAACTAGGAGAGTTTGGGAAAT

GGTAAAGACCTGGTGAATTATGACAATAAGCATCTAGAATTTCTCAACTCTATTTGTTTATTTTTGAAAT

ACCTATCTTCTGGAGCTTTGTATACAGAAGGGGTTGGGTTGGACATAATCTTTACAGATCAGTGAAAAAA

ATGCTCTAGACACCATAA

>MSTRG.3140.1 gene=MSTRG.3140

ATCATGTTAATAAACACACTTTCTTTCCAGCATCTTAAACCACATTCTTAACTCTCAGCTTTCTCCACAA

CACTGTCCTTAATCTTCTCCGCAGTATCCTTGGCCGCGTCCAACGCTTCTCCGAGCTTCTGATTGGACAC

CTCCGCCACATCCTTGGCCTCACTCGCTGCCTTGGCCACTCTATCAGCCACCGTGTCTCCAGTCTCCTTC

CACTTCTCCGCTACCACGCCTCCGGTCTCTTTCAACTTATCAGCCGCCACAACTGCAGCGTCCAAAGTCT

TCACTTTCAATTCATAAGCTAAGTCAGCCGCTTTATCTGCGTCTTCTTTGGCCTTGTCAACCAAGTAATC

TGCGTTCTTCTTGCCTGCCTCTACTGCCTCGTGAGCTTCGGCTTGGAGTTTTTCATTCAAGTTCTTGACC

TTTTGTCTCGTGTCCTCGAAGCTTCCCTTTAACTCTTCTCCTGTTTTTTGTGCGAACTCTTCGAATTCTT

TGCCCAACTTACTGGGATCTTCGCTGGTGGATCCTATTCGACCCTTGGAGTATTTGGATTTGGAAATGAA

GCTAACTGTGGATACTTTAGGACTGAAAGAGGGCTGAGGATAAGCCTTTATGATGTTCA

>MSTRG.3143.1 gene=MSTRG.3143

CGGATGAAATTAGAAGAACTGTGAAAATGAGAAGGGTCACCTGCATATTCCACGGAGACTGAAACAGAAT

GGGACTGAAACAGGAATGGGGCGAGTGATGCTAATTCAAAAATTACCAACTGCTTCTCAGCTTTTCAAGG

CTTTCTTCAGCAAGAGGACCCTCAATTCTAGGCCTCTGTTTTTCTCGTCTCTCAATTGTGAAGACTATTT

GATTGAAGATAATGGCATCTGGGGATTCTAAAAAGATGCTGAACCACTGCATATACAATTATATGAAAGA

AAAGGGACTTCACGTTGCTGCTGACGTATTCGCTGTAGAAGCCAACCTTGACTTGGGTGAATGGAGATGT

CTTTTGGAAAAGGCTGGGATGAAAGGACATGGACTTTGGGAATAGCTGGGATCATGGAAAAATAAGAAGA

AAAACTTGAGCTGTTTGATTGCAAGGGATTATTTCTTCACATCGTGAATTTAACTTCTAGATATCTCTGT

GGCAAAGGGTAGCATTCCTCCTCTTGGTTTGTTGGTATTGACTCGGTGTGAAAAGATAAAAATTTGCTAG

CTCAATGTGGGAGTCATTCTGGCCTCGTTTCTGCTCTGCAAAGCCAACTATAGATGATGCCATGGCCACG

CTAGCTTCAGTGAAAGAACAAGGGGCTACTAGTTCCCAACGAACACCTCCCGAAACCAGCCAGTGGAAAG

ATGGTAATCAACCCATGAATGTAACACTGAAGATATTTCCCTTGTGCCAGCTGCATTTGCATTGGTCATA

GCGGAAGCATGGACACGTAAAATTTCTGTTCAAGTAAGGGCCTGTCCACTTATTCAGATCACCAGCATCT

ATATACATGGGCGACAAGGCCTGTAAGGGTCTATGCCATCACTAGTTAAGTCAATCACCAGTTGTTGCTG

ATATCATTTCTGTTAATAACAAGCTCACACATTGTTAATCTGTTCACCTGTCGCATACTACATGCAAGTG

ATACAGCATGTGCTTATTGTGACACTGTTTTAATTTGTACACCTCATTTGTAGGGATGGGCTTCAGATAT

ATCATTTGTGCATCGTATTTAAGGTGGTAATGTAAAGGTTTACCAGCTTTCTGTGAATAACAGTCCTTAT

CTTGATTTTTTGTAGTCATTTCTCCT

>MSTRG.3147.1 gene=MSTRG.3147

CTCAAAATCTTTAACCTTGCAAATTTTGATGACACAAAAACACACCCCTTTGTCTATTCACTCTAGTGAC

TGCATGGCTTCTTGGTAGAAGGCTCAAAAGCAAATGGACCCACACTGAACATTTCAAAGGTAGCAGGAGT

CTTTGTAGACACTGGTGGCTTTGTTGATTTCTCCAAGATCGCACCCTTCACTCCGTAACGCAGATTGGTG

GGCTTGTCACACTTCTTCTCAGGCGAAGAAACCACAAACACTCTGCACTTGTGGACTCCATAGGTGGTGA

TTGTCTTGGGGGCATTAAGTGAGAAGTAGCCATTCTTGTCTGTCTTGCCTTGGACTACTAGTGGGTACTT

TGTGTTGTTACATTGTAGCTTCACCACAGCACCTGCCATACAAAATATATTAAATGTCGGGTTCGTGTCG

AGTTTCGGAAGAATTAAAATTTATCGTGCAAAAGATAAAAAATAGGTGGAGCTTACCGAGAAGGGGAGTG

GCTCCCAAGAGAGTTTCGACGCCAGTGTAGTTGCAGGGCTTGCAGTAGACAACGCCTTGAACAGCTACGA

GCTTCCTGGCTGGGAGGGGAGTGGGGGCGTGAGATGGTGCTTTGACTGGTGGTTTGACTGGCTGAACTGG

AGGTTTGACTGGGGGCTTGAGAGGAGAAGGAGCGGGGGCGTGGGTGGGTGGTTTAGCGAGGGGTGCGGGG

GCTGGGGTGGGTGCGTGTTTGTGGTGGTGGTGGTGACCTCCGGGCTTGTGGTGGTGAGGAGCTGGGGCTG

GTGCAAGGGCGGTGGGTGGCTGGGCTATTGTGGCTGTGCCGAGAGCAAAGACAAAGGTGGAGAAGAGGAG

TGCGAGGTGGAGGAGGCCCATGGGTTTTGGAGAGAGAGCAGTCATGTTTTTAGAGAGAGAGAGAGAGAGG

AGAAATAATGTAATGTTGCAGAGGTGTGATAAAAGTG

>MSTRG.3150.1 gene=MSTRG.3150

TCAACTAATGTGAGATATGAGATCTAATCTCAGCCCCACATTTTTATCCCTAAATCAAATCACAACCACG

CATTTATATTTTAACATTTTCTTTTATTACACTCTCCCTCTTCCCTCCCCCCCTCCACTGCCCACCTCCA

CCCTCTACTGCCATCCGTCACTACCCGTTATTGCTGCTGCTATGCAACCACCAGATCTACAACATCGTCG

TAGTCGAGGCGACGACCTCGTCTCCAGCCACGCACACAAAAAACACAGATCTCTCTCTCCAGAAACACGA

CTACTTCCTCCCCTCTAAAAGCCCCATCCTTATTCTTTCATTTTGTTTCTTCTCTATTCATCGTCGATCC

ACATCATCTCCGATTTCGATACTCCGATCTAGAGATGGTATTGGCGGAGGTTCGGTGATTTTTGTTTTTA

CGGTGGTGATTTCATTTTGTCGATGGTGTTTGTTTGTGGTGCTTTACAATTAATATAGTAATTTTTGTTA

TGGTCGGAGTTAATGATGGATATGGGCAGAAATAGAGGTTGGATGTAGTGAATGAGAGAGAGCGTGTGGA

TGTGGAGAGAGAGAGAGGAGGTTGAGAGAGATAGTGAGGAAGCTGATGAATTGAAAATGATGATTTTCAG

TTGTGTTTTTTATTTTTTCTGATGGTCATTTTGTTTTGTATTTGGTGATTTTTCGGATTTGGGGGGTGGC

GGTGGTGGTTTGTAGCTGATGGTGGTGTGGTGATTTTAATTTTCTGGCGTTGATTTTTGATTTTTTGGCG

GTGATTTTTGATTTTTTGGTGGTGATTTTATGTTTGCCGGCAGTGATTTTCTGGTGGTCGCCGGAGCTGG

TGGTGGTGGTCAACGGTGGTTGAAGGTGATAGGTAGTTGAATTAGAAGATGGTGAAATATTAAAATATTA

AATGAATGGTAAGAGATGAATATAATGTATTTAAATGAGTGGCTGAGATTAGATCTCATATCTCATCATA

AGATTGGTTCTCACTGGAG

>MSTRG.3159.1 gene=MSTRG.3159

GCAACATAGCACTATCCACTAAATCTTCTTTTTTCTCTTTGCCAAGTAACATCAGATAAATCTCACCCTC

ATTCAACACATCCAAAAACCAATCCAAATCCAATCCTAAGCAAGATCAACACATACATCACAAGCAAACA

AACCAACCACATGAACACTCTACTCGAAAACATACATCAAGTTCCAAACTTTTTCCTCAAAACCCATCAA

GAATCATCACCAACATCACAAACCAAACAGTTACTCATAACCAACACATTATACAACAACCACTACACAC

AATGAACAAAAGATACAAACTTTGCATTAACCTAAGCAAACAAACAATGCAGGCTCAAATGGGTAATCTC

AAAAACAACCCATTACAAGAATCAAGCAGAAAAATCAAGAAAAGACCAATCTAGGATCTGGGTAAGAAGC

AGACCTTAAGAAAAGCCCAAGGATCAGGCCCAAAAAGATCTTTAATTTGAGGATACTTGGAAAGGATGAG

TCTCCTACGAGAAGCATGAGGCTCATCAGTGTGTGACCAGTAAAAATCTTGACCCATCTCAACCATCTTC

TCTTCTGTTTCCCTGTATAAAAACTTCACATATATTTTTCACACACACAGCACACAGCTGTTGAGTTTCA

GCTGTCTATTTTTATTCTTTTGTTTTTCTTGTATTATCCAGGAAATAGTCCTAGGAGTCCAAACATATTT

GAAGTCCCCTATTTTTGGAATCCACTTTATTTTCTTCTTGTAAGTTCTTATGGAATTCGAAGTTAGTTTT

CGAAAAATAAATAAAC

>MSTRG.3161.1 gene=MSTRG.3161

TCTAGAACAAAAGGCACCTCCTCATCGGTATCTCAAAATGGTAAAAGAAAGAGAACTAATTTGCCTACTA

TAACCACATACTATGAGTCTATAACGGTGGAGGGGAGTGAGGAAATTGGAGCAGCAGCGGAGATAAATCC

ACCTCTGAAGAAAATGGCTGCAGATGTTGTAAAACCTGCAAGAAGGCTATTCAGTGTAGATCATGGAACA

CCTCGAGTAACTAGGAATAGGGTACCGGTAGTGGAACTCACTTCTTCGGAGGAAGAATATCCAAAATGTG

ACCTCCCTCCAGTCGAACCTTCACTGGAACAAAATTCTCAAGAAGTCTTGCCTGATGATCGTGAACAACT

CATGAATGCAGCATCTGTATTAAAATGTCAGTTGGTGGCCCAGCTCACAAGTTCTGCTGAGAAATTGAGG

ACCGAGGACATGAATGATCTTGCTGATAGGTGTTACAATACACTGAGGGGATTAGGAGATGATTATACAT

CTATTAGAAGGAAAGTTCATAAACTCATTGGACAACACCAGAAGCTAGAAGATGCTGCTAGAAACAAGGA

CAACTGGAATGACCAGGACATAAAGGCTCGTTATAGTCAGCAAGTTGTGTCGCTATCTAATATGACAAAG

AAGTTATCAGGCACTGAAGAGAAATTGTCAAGGGCTAAGAATAAAAAAGAAGAGATAAAAGCTGCATTAC

TGAGGCTTAGAGAGGAGTTGTCTGACGAAGAAGAGCGAGTTAAGACACTGACAGAAGAGAGGGATATATG

CAAAGCGGATCATTCTGATGTTGAAGCTAAATTTGAGAAATTGGTCGCAGAGAAGAAGGAGGCTTACAAG

GAATTTGAGGCAATCAATGGGCAATATAATAATGCCAAGAAAGAATACGAGAGAATGAGTAATCAGTTGA

TGCAGTCTGTTGGGAAGTGACGAGGTCTCTTTTGGAGTCCTGCCCCCCTGTGAAGATCAGGATTCTGGAC

CGCACTAGGCGTCTCTTCTGAAATGGCCTTTTCATCTGAAGTACCTTCCTGGTGTTGTTGGCCATCATCT

TAGTATGTTCTTACTACATAGCAACTTGTCCACCGCAGAATTTCCTAGTAACGTAAAAGAAAGAAGATGT

TTTGCACTTTATATCGGCTTTTAGTTGAATTACTTAGACTTAATGTGTGAAGTTCAGTACTGCATTGCTT

AATATGCATTGGGTATTTTACATTGAGTAAATGTGAGGTTTATCTGGTGACATTGATCTTTGGCCAAGCC

TTCACCACTTTAAGAAATCAGG

>MSTRG.3162.3 gene=MSTRG.3162

AGATTGTCCTTGTTAGGAAATTGATGTCAGGAGAAGAAATAGACCCTCCGGAAGAAATTGATTATGTGGG

TGTGCGATTCCTCCCCATATTTCCCTTTAGTATGGAAAAGCTCTCTGATTTCGATGAGAAAGAACCACAA

TCAGCTAATGTTCTCCACTTCGTACGTATTGTGAACCCAAAGTACCATATCCCGATTGGCTGTCACCGAC

ATCCTTTGATCTGCCCCTCAATCTCTGTGAGTTTGTTCTCCACCAGTATAACATTGCTCTCCACCGGTTT

GACAGTGATAAGGATGAAGATGATGAAATATATGACAAAGTTAAAGTTCTCCGTGTGATTGATGTATTGA

GAACATTGGATGCTGAAAACGGGACAACTTACTATGTCACATTTGAAGCCTCCTTACTTGATGATGAAGG

GAATCCTCAAGTCTTCGAAACAAAATTATTTATGTCTCCTCTGATCCCAACTACAAAGATAAGGATTAAA

TTTGTCAGGGTCAAGAGCAACTGACTAAAGGGTGTTTTTAACAAAGAAGAGGAGTATTAGGAGTAAAGTG

TTAGGGACTTGTTTTTTTTCGTGGGGTTGATTGTTTCAAGGTGCATCTTTGAATAATTGTTGTAAATCAT

TTGCTCCTGAGGTTAATTGTGAATAAAAGCCGGTGGAAAGTTTACCGATGCCCAGTGAAGGGAGGTCAAG

GTATAATGATAAGCAGGAAATCACTCCTATCACTGTGCATGCTAGGAATATTGAGAATGAAAGGAATCAG

GCACAGTGTGTTTTTCATGGGTACCATTGTAGCTTCGGAGGAATTATTAAAAGTAGTAAAGCAGGG

>MSTRG.3162.1 gene=MSTRG.3162

CCTATTCATCCTTTAAGATTGTCCTTGTTAGGAAATTGATGTCAGGAGAAGAAATAGACCCTCCGGAAGA

AATTGATTATGTGGGTGTGCGATTCCTCCCCGTATTTCCCTTTAGTATGGAAAAGCTCTCCGATTTTGAT

GAGAAAGAACCACAATCAGCTAATGTTCTCCACTTTTATTGTGAACCCAAAGTACCATATCCCGATTGGC

TGTCACTGATATCCGTTGATCTGCCCCTCAATCTCTGTGAGTTTGTTCTCCACCAGTATAACATTGCTCT

CCACCGGTTTGACAGTGATAAGGATGAAGATGATGAAATATATGACAAAGTTAAAGTTCTCCGTGTGATT

GATGTATTGAGAACATTGGATGCTGAAAACGGGACAACTTACTATGTCACATTTGAAGCCTCCTTACTTG

ATGATGAAGGGAATCCTCAAGTCTTCGAAACAAAATTATTTATGTCTCCTCTGATCCCAACTACAAAGAT

AAGGATTAAATTTGTCAGGGTCAAGAGCAACTGACTAAAGGGTGTTTTTAACAAAGAAGAGGAGTATTAG

GAGTAAAGTGTTAGGGACTTGTTTTTTTTCGTGGGGTTGATTGTTTCAAGGTGCATCTTTGAATAATTGT

TGTAAATCATTTGCTCCTGAGGTTAATTGTGAATAAAAGCCGGTGGAAAGTTTACCGATGCCCAGTGAAG

GGAGGTCAAGGTATAATGATAAGCAGGAAATCACTCCTATCACTGTGCATGCTAGGAATATTGAGAATGA

AAGGAATCAGGCACAGTGTGTTTTTCATGGGTACCATTGTAGCTTCGGAGGAATTATTAAAAGTAGTAAA

GCAGGG

>MSTRG.3162.2 gene=MSTRG.3162

CCTATTCATCCTTTAAGATTGTCCTTGTTAGGAAATTGATGTCAGGAGAAGAAATAGACCCTCCGGAAGA

AATTGATTATGTGGGTGTGCGATTCCTCCCCGTATTTCCCTTTAGTATGGAAAAGCTCTCCGATTTTGAT

GAGAAAGAACCACAATCAGCTAATGTTCTCCACTTTTATTGTGAACCCAAAGTACCATATCCCGATTGGC

TGTCACTGATATCCGTTGATCTGCCCCTCAATCTCTGTGAGTTTGTTCTCCACCAGTATAACATTGCTCT

CCACCGGTTTGACAGTGATAAGGATGAAGATGATGAAATATATGACAAAGTTAAAGTTCTCCGTGTGATT

GATGTATTGAGAACATTGGATGCTGAAAACGGGACAACTTACTATGTCACATTTGAAGCCTCCTTACTTG

ATGACGAAGGGAATCCTCAAGTCTTCGAAACAAAATTATTTATGTCTCCCCTGATCCCAACTACAAAGAT

AAGGATTAAATTTGTCAGGGTCAAGAGCAACTGACTAAAGGGTGTTTTTAACAAAGAAGAGGAGTATTAG

GAGTAAAGTGTTAGGGACTTGTTTTTTTTCGTGGGGTTGATTGTTTCAAGGTGCATCTTTGAATAATTGT

TGTAAATCATTTGCTCCTGAGGTTAATTGTGAATAAAAGCCGGTGGAAAGTTTACCGATGCCCAGTGAAG

GGAGGTCAAGGTATAATGATAAGCAGGAAATCACTCCTATCACTGTGCATGCTAGGAATATTGAGAATGA

AAGGAATCAGGCACAGTGTGTTTTTCATGGGTACCATTGTAGCTTCGGAGGAATTATTAAAAGTAGTAAA

GCAGGG

>MSTRG.3162.4 gene=MSTRG.3162

CCTATTCATCCTTTAAGATTGTCCTTGTTAGGAAATTGATGTCAGGAGAAGAAATAGACCCTCCGGAAGA

AATTGATTATGTGGGTGTGCGATTCCTCCCCATATTTCCCTTTAGTATGGAAAAGCTCTCTGATTTCGAT

GAGAAAGAACCACAATCAGCTAATGTTCTCCACTTCTATTGTGAACCCAAAGTACCATATCCCGATTGGC

TGTCACCGACATCCTTTGATCTGCCCCTCAATCTCTGTGAGTTTGTTCTCCACCAGTATAACATTGCTCT

CCACCGGTTTGACAGTGATAAGGATGAAGATGATGAAATATATGACAAAGTTAAAGTTCTCCGTGTGATT

GATGTATTGAGAACATTGGATGCTGAAAACGGGACAACTTACTATGTCACATTTGAAGCCTCCTTACTTG

ATGATGAAGGGAATCCTCAAGTCTTCGAAACAAAATTATTTATGTCTCCTCTGATCCCAACTACAAAGAT

AAGGATTAAATTTGTCAGGGTCAAGAGCAACTGACTAAAGGTAATTCAATCATG

>MSTRG.3163.1 gene=MSTRG.3163

GCAGGAAAGCAATTAGGGTTCGACAATGGCGAGCGGTGGCGAGAAGAGAATAAGGAAGATTAGTGAGCTG

ACGGACGAAGAACTCGAAGTGGAAATTTGGGGAAAGACGGGAACGAGGCCGCGCATCGACAGAGCAGCCA

TTGAATAAGGAAGATTAGCGAGCTGACGGACGAAGAAGTCGAAGTGAAAATTTGGGGAAAGACGGGAACG

AGGCCGCGCATCGACAGAGCAGCCATTGATTACAGGCGTAACTTTCCATTCGCCAAGGACTTCTTGGAAG

AAAAAGGAAACCCCCCTGATCATGTACATGTTCCCCCGAGTCGCCAGATGTGGAGGAAGGACTTGTATCC

CCAATTTCTCAGAGCAGTTAAGGTTCAAGAGGAAACATACGAGGAA

>MSTRG.3163.2 gene=MSTRG.3163

AATTAGGGTTCGACAATGGCGAGCGGTGGCGAGAAGAGAATAAGGAAGATTAGTGAGCTGACGGACGAAG

AACTCGAAGTGGAAATTTGGGGAAAGACGGGAACGAGGCCGCGCATCGACAGAGCAGCCATTGATTACAG

GCGTAACTTTCCATTCGCCAAGGACTTCTTGGAAGAAAAAGGAAACCCCCCTGATCATGTACATGTTCCC

CCGAGTCGCCAGATGTGGAGGAAGGACTTGTATCCCCAATTTCTCAGAGCAGTTAAGGTTCAAGAGGAAA

CATACGAGGAAAGATACAAATTTACTGGAGATTTTGAAAAAGATGATCAAGAAGCTCTTGTGTACCGCAG

TATTGATGATTATATCAAACTTAGCGAGGACGAAAAAATCGCTACGAGAGAATATCAAAAAGATATGAGA

CGATACCACAGAGACGAAGCGAAAAGCCGTGGTTTTGATGTTGGGTCGTATCCATACAGAGATTATTTTG

GGCTCACTGGTATTGGTTACTTATACTGTTATTACAAACCACCCGAGGTGACGATCCCTAAAGCTCGTTT

GATTACTCTGCTTCATCTCTCTCGGTGGGCTGTTGCGGTGTACAACAGTTATCAGGGAAAGCGTACGTGC

TATGGTAATGTACGAGTGGTGAAGGCGATGAGTTTTGGTTGTTTTGGTACAGATTTGAATATCAGTTTTC

AG

>MSTRG.3165.1 gene=MSTRG.3165

TGTGTGCCTGTGCTTATCAAAATAAGCATATCTCTCTCTCTCTCTCTCTGCCTCTCTCTCCCTCCCTGGT

TTTTAGAAAAATGGCTTGGAACAGTACGACATCGTTTGATGATCTCTCTCAGGAAACTGCCACTAACATC

CTCTCCAGGCTCCCAGTAAAAGCTCTAATCAGGTCAACAACAGTGAACAAGAAATGGTACTCTCTAATAA

CAAACCCTCATTTCATTTCTGCTCAAATCAAGCATGCTATTTCACAATCTGATGATAATGTTGTGTTAGT

AATCCCTCCTGTTATCTCTCAAGAGAAGCATTGTTCGTTAGTCTCCATTGAAACTTCGAAAGTTCTCGAA

AAATATGAGCTCCCCTTTACTACCAGGACTAAGAGCTTGAAACTCATTGATTCACTTGATGGGCTTATGT

TGTTGACTGATTTACATGCCTTTTACGCTTCTCGGCATTTGTATCTGTGGAACCCTTGTGTTAAAACTCA

TCGACCCCTTGTTTCGTCTTGTTTCAAGAAACTTTTTGAGGACCCCTATATTAGCTATTTTATATCTGGG

TTGGGTTTTAATCAAGCTAGTAATGATTATAGAGTGGTGAGGGTTGTTTATGATGTGATAAACAAGATGA

ATCCCGCTAAGGCGGAGGTTTATAGTCTGAGTAAACAGACGTGGAGGAAGATTAAAGACCCTGTGGTTCC

AAGAATTGGTCAGAGTGATGGAGTGTATGTTGATGGTAGATTCTACTGGTTGGAAACGACACCACCTGAC

ACTCGGGAAGATGGTCGTATTCATAAAATTAAAGACCTGTGGATTTTGTCATTTGACTTTGAAAATGAGG

TGTTTGGGGAACTCAAAGTGCCGAAAATAGTTTCAAAGTGTATAGGAACAACTGCCTTGTCTAAGCTTGT

GCAGTTCGAGGGTTCACTCGCTCTCTGTGTTTTTGATGCACAGAGCACTGACAGAGGGGTCACCTATCCT

CTCCGTATATGGTTGTTGAGGAAGGAAAATGGTGTTGTCTCTTGGATTTTGCGTTTCACAGCTGCATTAA

GAGAAGTTGGGTGGCCAATCAATATTACAAACAAGGGGACAATTCTAATAGAGACGCGTCCCTCAACGAG

CCGTATTGATGTTACAAGCATAGTATCATGTAATCTGAAGTCCATGCATTACAAAAATCTTGGATTTGGT

AAGCCTAAGGGTGTAGAGGATCCCATGTTAATTCCTGAGCCTTGTACTGTAGACACTTCCTTCACGGAGA

GTTTGGTCATGTACGAGGGAGGGAAGTCCATTCTGAAGTTTTCAAAGTGATGAATCTTGTGTTGTGCAGG

TACTTGCATTGCAGATGTTGCCGATCTTGCTTTGGAAGCCGCGGCAATTCTATTACTATTAGGTGCAGAA

TATATATGAACTTTTTTTGAAAGTGCTATCTTTTGCTCTGAGGAATTCATCGAGTGACTTTTGAGCTCCA

CATAATGTAAACTCGAAGTAGTTATGTCTGGCAGATTATCTGCAGTATGTGTTCTTTGGATGGTTGCAAT

TATCATTCTCAATACCAAATTATTTACAGAAAGCTCAGGAACACGGGACATCTTACAAAGCAGAAAAAGA

AAACCTTTATAAAAAAACAATAAGAACATAAGAGAAAGAAAAACAG

>MSTRG.3172.1 gene=MSTRG.3172

AGTCACAGTAATTATTTACGCATTCATAACATAATGGCATCTATAATTTTCCCAGTTTCAACTTTGATCA

ACTTTCAGAGATTCACCACATGCAAGCCCAAAGCTGCGACAGCCTGCAAGTCTGTTATCACATCTGCCGC

TGAAACAACTAAGCTTCCTGTTCCAGCAGAGCCTATTGTCAGGCGATCAGCAAACTACAAACCTTGTCTG

TATGACAACAATTTCCTCCAGTCTATGAAAACTGAATTCAAGGGTGAAGCTGTCGATGCACGAGCTTCTG

AATTGAAGGAAGAGGTGAGGATGATATTTAATAATGTGGCCGGGCCGTTGGAGCAGCTTGAGCTCATTGA

TCAGTTGCAGAGAGTTGGGGTAGATTATCATTATCGTGATGAAATCAACCGTGCATTGAAGAGCATTCAT

CAGAATGCTGAGACTTGGGAAAAGGATTTGCATGCAACTGCTCTTGAATTTAGGCTTCTAAGACAACATG

GGCATTATATATCCCCGGAGGGATTCAACAAATTTACTGAAAACGGGAGCTTCAACAAAAGTATTCATTC

AGATGTCCGGGGACTGTTAAGTTTATATGAAGCCTCATACTTTTCTATTCAAGGAGAGTCCCTGATGGAG

GAGGCTTGGTCCTTTACAAGTAAATTACTTAAAGACTGCCTCAAAAATATTAATGATTTGAATCTCCAGA

TGCAAGTGAGACATGCTTTGGAGCTTCCACTACAGTGGAGGATCCCCAGATTTGACGTGAAATGGAATAT

AGATATGTATCAAAGAAGTGGTCACGTTGTCGTCCCTGCAGTACTTCTGGAATTTGCGAAGTTGGACTTC

AACATTAGGCAAGCGTCGAACCAAGAAGAGCTTAAGGATTTATCAAGGTGGTGGAGTCGATTAGACATGG

GAGAGAAACTTCCCTTTGCCAGAGATAGGTTGGTGATGTCTTATCTCTGGAGTTTGGCAATTGCAGGCGA

GCCTCATCACAAATATGGCAGAGAAGTTCTAACTAAAATAATCGAGCTTATTGGTGTATATGATGACGTT

TATGATATTTATGGTACCCTGGATGAACTTGAACTCTTTACAGATGTTACCAAGAGGTGGGATGTAAATG

CATTGAACGAGCTCCCAGACTATATGAAGTTGTGCTTTCTGTCACTGAACAACCTGGTGAATGATACGAC

TTACGACATCCTCAAAGACCAGAACGTTGATGTCCTAACAAATCAAAGAAAATGGTACAATGATTTATTC

GAGCGTTACATAGTGGAGGCCAGGTGGTATTACACTGGATACCAGCCAACACTAGAGGAGTATCTGAGAA

ATGGATTTGTGTCAATCGGAGGTCCTGCTGTAATGCTCTATGCTTACATCGGCACCGCGGATCCGATCAG

GAAAGAAGAATTAGAGTTCATCGAGGACTTTCCTGACATAGTACGATTGGCATGTGAAGTTTTTCGATTA

AGTGATGATTATGGAACAACTTCGGATGAGCAAGCGAGAGGAGATGTTCCGTCATCAGTAATGTGCTACA

TGTCGGATGCTGGTGTTTCAGAAGAAGTTTCTCGTGAGCACATTATGAATTTAATCAGGGAGAAATGGTC

GCAAATTAACAAACTCCGATTTTCGGAGAAGGGTAATGGTCCTTTATCTTGGCGTTTTGTTGATACTATG

CTGAATCTAGTCAGGTCAGGCCATTGTTTGTATAATGCCGGAGAGGATGGTTTTAGTGTTGAAGATGTTG

TTGCTAAAGAATCATTGGTTTCTCTTCTTGTTGAGCCCATCCCTCTCTGAGAAATTAGCGCTTATATAAG

CCAAGACAGGAGAGCTTGCCATTATCTATTTATAGTTCTGGGTTTTCCTTGTGTGCTTCTAAATAGGTGT

GCCTAAGAGGGAACACCATAGATAAGTTTGTCTCCATGGAAAATGTGAGTTTTGATTTTCGTAAGAGTGC

TTAGGCACCTTGAGCGGTGCCATCTTTTTTACTAATGTCGTTGCCTTACAATAAATATGAAAAATAAACA

ATTTTATGTTA

>MSTRG.3174.1 gene=MSTRG.3174

AATAATATTGATAGTAAATGGACGGACTTGCCCTGAAAGACACTTGGTTGTAAACAGGATATGCGAGCTT

GAGCCTTCAAGAATCAAGATAGAACTGTCAATCACACCACCATCTGCTTAAATCCAGGATTTTAGAGAAA

TGGCAACGGAGAAAACGACGTCGTTTGATGACCTTCCTCAAGAACTTGCCGATAACATCCTCTTAAGACT

CCCGGTCAAAGCCCTGATCAGGTCAACATCAGTTAACAAAAAACGGTATAGTTGCTGATCTTACTTTGAA

AGTCCTGCTTAGGTGCTGCATATGTATATGAAGTCCTTTAAAGACGCCAGTTTTTGCTGAGGGAATTCTG

TGGAAGATCTTTGAACTTGACATAATGTAATCTAATCAAGCAATCGAACAATAGTTTTCAGTATTGATTT

GTTATGGATACATCTGTTTACTAACCTATAAATGCACTACTTTGTAAATTATTAAACAGCTTGCAACTTG

CAACAGTCAATTTTATATTTTGTAAGCTCGTCACACAACTGATAAAATGAAGTGCATTTTATATATTTGA

TTGGG

>MSTRG.3174.2 gene=MSTRG.3174

ACTTGCCCTGAAAGACACTTGGTTGTAAACAGGATATGCGAGCTTGAGCCTTCAAGAATCAAGATAGAAC

TGTCAATCACACCACCATCTGCTTAAATCCAGGATTTTAGAGAAATGGCAACGGAGAAAACGACGTCGTT

TGATGACCTTCCTCAAGAACTTGCCGATAACATCCTCTTAAGACTCCCGGTCAAAGCCCTGATCAGGTCA

ACATCAGTTAACAAAAAACGGTACTCTCTAATCACAAACCCTAATTTCATTTCTGCTCAAATCAAGAATG

CTATCTCACATTGCGATGATAATTTAGTGTTAATTGTCCCCTGCCTTATCTCTGAAGAGAAACATTGTTC

CTTAATCTCTGTTGAAACTTCGGAACTTGTCGAAAAATTTGAGGTTCCCTTTACTACTAGGAGTGACTAC

TGGATACTAGTTAATTCAGTAGATGGACTTATGTTATTGACTGGTATTACTGTGTTTAATCGTACTACTA

CACATTTGTATTTGTGGAACCCTTGTCTTAAAACTTACCGAATCCTTGTTTCTTCCTGTTTCGAGAAAGT

TCTTGGTGACAGGGAAAGATTTTATTTGGTTGGGTTGGGGTTTAATAAAGTTAGTAATGATTATAGAGTT

GTTAGGATTGTTTATTTTTAGGACAAAATGGATTTGCCTAAGGTCGAGGTTTATAGTTTGAGGGAACGGA

CGTGGAAGGAGATTAAAGACCCTGTAGTTCCTAGGAGGAGTTTGCGTGACTGTGAAGGGGTTTATTTAAA

TGGGAGAATTTACTGGATGGAAGTGGAAAGGTCCTTCAGTGTGGAATCTTCTCCTTATGTTAGTAAAGAT

TTACGGATGTTGTCGTTTGACTTTGATACTGAGGTGTTTGGGGAAGTGAAAGTTCCAGAGGAGGTTTCCA

ATTGTTTAGGAGTAGGCGCCGGTTTTAAGCTGATGGAGTTTGATGGTTCTCTTGCTCTTTGTGTTAGTAA

CATACAGGTTAGGAATGGATTGTTCATCATACCTTATCGTATATGGTTGATGAGGTTAGACAATGGTGTT

ATTTCTTGGATTTTACGTTTTACAAATGAACTCAAGGAACCTGGGCTAGTAATGAATATTACAAAGACCG

GAACAATTTTAGTACAGAGACTTCCCTCGCTGCCTGTTCATGCCATTACACGCATATTCTCATGCAATCT

CAATACCATGCATTATAAAGATCTTGGACTTGGTGGGCGTTGGGGTGAAGAGGATTTCCCGACAATGCCG

CATTGTATTGTAGACACTTCTTTCATAGAGAGTTTGGTCATGTACGAAGGAGGAAAGTCTCTGTTAAAAT

TTTCAAAGTGAGCAATCATGAGTCTTTGGGTGGTATAGTTGCTGATCTTACTTTGAAAGTCCTGCTTAGG

TGCTGCATATGTATATGAAGTCCTTTAAAGACGCCAGTTTTTGCTGAGGGAATTCTGTGGAAGATCTTTG

AACTTGACATAATGTAATCTAATCAAGCAATCGAACAATAGTTTTCAGTATTGATTTGTTATGGATACAT

CTGTTTACTAACCTATAAATGCACTACTTTGTAAATTATTAAACAGCTTGCAACTTGCAACAGTCAATTT

TATATTTTGTAAGCTCGTCACACAACTGATAAAA

>MSTRG.3180.1 gene=MSTRG.3180

GTCCTTGTCCTGCATCTTTCGTTTCCTCTTCTTTGCATTCTCGGCCACCATGTCACTGAAATCTTCCCGT

TGACTACGGAGTTTCTCTTCCTCTCGGGCTTCCTCGTACTTGGCAGGCAGAACATTGTCCATAACTTCCA

ATTCTTCTGGATTTAGTGACACTTCAACTCTATCTGATTTTTGACCTCTAAGCAAATCAACCCTCTTCGC

TGCTGGCTTGTCTTGG

>MSTRG.3184.1 gene=MSTRG.3184

CCAAGCACCACCACTTCCATGGTAGACTACTTGCAGAAGCTCACAACTCAATACCTTTATACATGTTTCT

TTCCCCGGCGACTCGCCGGAAAAACACCGGCCGTGTCGAGCTCTTCGCCGTAAAAACACCGGCGGTGTCG

AGCTCTTCACCGGCATCATCAACACCTATCCCTAACCTTCATGATTAAAAGAGAATGACATGAAACATGA

GAGGATGGACTTGAATGTGAAAGTCTGAGAAGTAGAAGCACCAAGTTAAAAGTGTATAACTGTTTAAACA

AAACTGGACTAGTGGACTGGTATGCAGATGCTTGTTTGATTGAATGTTTTTGATGTTGAATGCAAACTTA

AATTTCAATGTAATAATGCAATTTAAACTTGAGTAACATTGTTGGCTTGTTTTTGCTACACAAATGTTTA

TTTTAGCAAAGTTTCACTGTTTTTGATACATTAAGG

>MSTRG.3184.2 gene=MSTRG.3184

GTTTTGCCGTCAAAACATTTTGGTTGGCTTTGGGATGAGTCTTGTGATTGATTTCTGTTATTGGATCCGA

GAATGAGTTAGTGGTTGGTTCTGAGATCGGTAATAGGAAAAAGGAAGCGACTAATATATATATGGTTGTG

TAATATAATGATACGCATATTGATTTTGCGTACGTATCAAATAAGGCAGCAAAGGCCTCATTTCCGCATT

TAGGTATTTTAATAGTTAAACTTCAATGTTAGGTTATTTTTCAATAGTATGGCTGTGGGTAAAAAATTGA

GAAAAAGGAAGACTACGGCGGCGGATCTGCCAGAGGAAGTAATTCAGAATGAGATCCTTACGCGGCTCCC

TATCAAATCAGTGGTACGCTTCAAGTCTGTATCTAAATCATGGCTTTCTTTGTTCTCTGATCCCCTATTT

GTAAAAGAACACCACCGTGCTGCTCAAAACCCTGAAGATTATGATTGTCTCATAGCTCAAAAAAACACCA

GAATTGCCATTCTTTCTCGATACAAAGAAACATTTCTACTCCCTTTAGCTAATTCCTTTTTAATTGGTTC

GATTAACGGTTTAGTTTGTCTCAGAAATGGAAAGAAGTTATTGTTATGGAACCCTGCTATACATCAATTT

AAGGAATTGACTATTCCACCCCACAAGTATTCTGTTCTGAGTAACGTCGGATTCGGTTATAATTCAGCAA

GTACTGATTACAAGGTTGTTGTTCTGTCAGTGGATAAAAGGTCTGCTACTCTTTACTGCTCTAGTTCCAA

TTCTTGGATTGATATATCTGTTCCTGATAATGTCTTCCCTAAAGATTGCAATTATCAATTAACTGGTCCA

GCTATCATCGCAAAGGATTGTCCCTACTGGACATTTATCAGATATAGTCGGTTTAAGGTTTTTATTGCTC

TGAAGTTTGATGCTGGAAGTAACAAGTTCAAGTTGTTGCCTGGACTTCACATTGATGGAGACTTAAGTAG

ATCTTGTTTTGAGTTTGTTGATATGAAAGATAGCCTTACTTTGATAATGTTTCAAGTTAAATCTTCTATG

GTGGGTTTATATTCTTTGGACGAGGAAGATGGATGTTGTGTCTGGAGTAAGATGTACAATGTGGGACCTC

TCAATTTTCACCCACTTGTGTCACCGACACAAGGATTTAAATATGGTGATGAGCTTGTATTTCGATATTA

CGGGGTGTTTTTCTGCTATGATCACGAAACACTTACAAGCAAACGTCTTGTCGGTACAAGTTCTACAAAC

ATACCCCCCTTCGTAAGGTGCTTTAGATATATGCCTAGTTTGGTTTTTCTTAAAGGAATGAAGTCTGCTC

ACTCAACTACCCAAACTCGTAGACGTGGTCTTTGCTCTAGGACTCCTTTGCGATTACTCAACTCATTACG

CGAGTAATCACTTGTTCAAGGCCATTGGTCAGTTTAGACCCTAATGACATGAAACATGAGAGGATGGACT

TGAATGTGAAAGTCTGAGAAGTAGAAGCACCAAGTTAAAAGTGTATAACTGTTTAAACAAAACTGGACTA

GTGGACTGGTATGCAGATGCTTGTTTGATTGAATGTTTTTGATGTTGAATGCAAACTTAAATTTCAATGT

AATAATGCAATTTAAACTTGAGTAACATTGTTGGCTTGTTTTTGCTACACAAATGTTTATTTTAGCAAAG

TTTCACTGTTTTTGATACA

>MSTRG.3186.1 gene=MSTRG.3186

GTTTGAGTTTGATGCAAGTGAACAGAATTGGGAACGTGGCAAGCTTTATGCACTAGTCACTATGAATGAT

TGTCTTACTTTGATGACGTATAACACGTCTAAAGATAGTGTTGTGGATGTTTATTCATTGGACGAGGAGG

GTTCGTGTGTTTGGAGTAAAATGTATAGTGTTGGACCTCTTGATCTTTATACCCATTATTTGCACTTGTT

ACAGGGTTTTAGGTATGGGGGAGAGTTTGGATGCCATAATTACGGAACTTTTGTCTGCTGTGATCCCGAA

AGAAATACAGTCAAGCGACTTGCTGGTACTACATCTCATTTGAATCTGATAAGCTGCTTTCCGTACGTGC

CTACTTTGGTTTACCTTCAAGGAATGAAGAGTGTTTATATGCAAACCCAAGCTCGTATATCTGGTAACAC

TTTCAGAACTCCTCGACGATTGATTAATTCACTAAGGGGATGATATTTTGCTCAAGGTCGACCAGCAATT

TGGACAATTTTAATGTCTGACTAAATGTTTAGTTAGACAATTAATCATTCAGTAAACAGACAGATCTTTT

GCTTGACCTCAACTGGCCGTTTAGAC

>MSTRG.3188.1 gene=MSTRG.3188

ATCTATGGCCGGCGTGTGTGTAAAAAAGACAATGAAGTCTCAAACAGAGCTGCCGGAACAAGTCCTCCAC

GACGAGATCCTCACGCGACTCCCCCTAAAATCAGTGGTACGATTTAAATCCGTTTGTAAATCCTGGTCAT

CTCTCTTCACTGAGCCCAAATTCGTAAAACAACACCTCACTCGCAGCGCCACTCAAAACCCTCACGATTA

TGATCGTCTCGTAGCTAATAATGGCCGTGCTATTGTCATTATCTCACGCTATGAAGAAACATATCTACTC

CCCCGCGACTTTAATTGCAGTTACAAATTGCTTGGTTCGGTTAATGGTTTAATTTGTCTTTATTCTGCCA

AGAAATTTTTGTTGTGGAACCCAGCCATTCTTCAATTTAAAGTTATCACCCTTCCATCAAGTTTTATGTC

TAGTTTTATATATAGAATTGGTCTCGGTTTTGATCATGTCAGTAATAATTATAAGGTTGTGGTGATCAAG

CATATCGGCGAACCGGCCTGTGTTTACTCCTCTAATTCTGATTCATGGAATGATTTACCTAAATATGAGT

TTAGGGAATGGGACTGCTATAGTCAAGAATTGTCCGTACTGGACATATGTTTCACTTTCGCGGAAACATG

ATGGGACGTGGTTACGCAGATGTGGTGGGATAAAGTTTGATGTCGGAAGTGATCAATTCAAGTTGTTGCC

TCAGTTTGATGCAGTGATGGTGTTTCGCCCTTCAAGTGATAAATTTAGTATAAAATTTGTGAATGTAAGG

GACTGTCTTACATTGATGTTGTCTGAACGATCTCATAGCAGGATGGTGGAATTATATTCTCTAGACGAGG

AGGGATGCGGTGTTTGGAGTAAGATGTACAGTGTACTGCTATACCGGGCAGATCTGTCACAAGGTTTTAA

GAACAGTGATGAGATTGTATTTAAGAAAAATGGGATGATGTACTGCTATGATCAAAAAAGCAATACATCA

AAGCGTATTCTCTCTGCAGGTTGCGGTGACATAAGCAGCTGCTTTACATATACGCCTAGTTTAGCTTTCC

TTCAAGGAATGCAATCTGTTCATTTACAAACCGAGACCCAAACTCGTGAACTTGGTTACAACTACACCAC

TCCTTGGAGGTTAATCAGTTCATTAATTAAAGAACCATTTACTGATGGTCATATTGCAGATTAGACGGTG

TTTGATTGAATGTGTGGGTAGTATTATAAGAGGGCACAAGATCATGTATAATGCTTTGGTGGCTGCGCTC

TGGAATTTCGTTCTGTTTTCTTCAATTTCTACTCAGGTTTTGTTATCCTCTTCTCATAATTTCAAATCTG

GGAAGTCCATGTAGCTTATCAATTTCTCCCATGTTAAATTTCTGTTTTGATTCGCAAATATGTTCCGACA

AATCAGATATTTACAATATTCATTGAATTGTCTCTTCTTAATATTCTAGGCTTTGGCGCTATTAGATGC

>MSTRG.3190.1 gene=MSTRG.3190

ACAGACCCGAGGAGAGCGAAGAAATCTAGAAGAAAATACTACAGTCGCAGGTCAAAAAATTTCATGCGAT

TGAAACAAGTGGCACTGCATTTACATTGCCGTCATGACTATTGTGTGTGTGTCTCTGAACAATTCTAACA

GCAATTAGCTTACGTTTTGTCCGGTCCCGTTGGTGGGGAAATGAATAGAATAAAATGGAAAGAA

>MSTRG.3191.1 gene=MSTRG.3191

GCTTCGGGATTTGATTGATCACTGTCTGCCCCGGGTCATCTCAGGATTACCGAAGTTGGATTTGCCTCCT

TCACACGAGCTTGATGGACCAGGATCTGTTACATATCTGTATTCATACAACTCATCAAAAGTTTCGGATA

CTAGGGCTGGGAGTTCAGAAAGTTCTTAAACTCTGGCCTGAAGTCTGTCATATTATCTGAAGGTGAAAAT

CAGTATTATTCTTCACAAAATGATCCTCCCATGTTTAGCCTATCTGGAAGTCATGTAGACTGGAACACTC

AGAGCCTATTTTGGTGAATTTTCGCCAAGATATTAATATTTGGTTCAAGAGAACCCTGATATTATTCCAG

CTTCATGGGCTGATTTGGCAGTCAATTGCTCACAATTTCTAGGAGCACCACGTATGAATGCTGCATTATT

CTCACAAAGTGCATACTTTACTGAATTCTCTGAGAAAAATATTCTTGGCACTGCATATGGTTCTTCTATA

GACATGGCTGACATGAAATTATGCAACATTGCTGATAAGGTTAAGGGTCTGAGAATTTTACTGGTAGGTT

GGATCAGCTGGGTGCAACGCCCAGGTGATCACCCAGGATGAAATACATGACGTACATCTTGCATCTAGTA

ACTGGTACAGGGATCTCATGTTAATGATGACCCAATATACGTATTCTTGAAGAGTTGAGTGAACCTGCAC

ACCGGCATCACTTGCCAACAACTGTTGAGCTGAATATTACATCATGAATTTCAACATCTTTTAAGCCCCA

CAATCACATTGATGTAGGCAGTGTGAGACTTGCGATAAAAGATGAGAGGTTCTTCCGAGCTGCAGTGCAG

GTGAGCTCAGGTGATTTGTGTGCCAACCATTCATTAATAACTGTGTGAATGTACATAGCTCTGACGTTAA

TTCCTACCA

>MSTRG.3194.1 gene=MSTRG.3194

AACATACTTACCATATCACTTTTTTACTTACAATAACATGACAGACAACAGTGGGACTTCGGGGTGACAA

TTTGTGTCAAGTTACCATTCTACCCCTTTGTTTCTACAGAAAAATGAAACATTCCAAGGACACGAGTGGG

TTTTCGATCAACTCATTGCTTGTCGGTGGCCACTTGCCTTCTTTATTACTTCCCTCCCTTTTTCTTGAAT

TCTCGGATAAGTCGGATCTGACCCGACCCGGTGACTTTCCTGTTCTTCTCACAGATGGGCTCCGACCCAG

ATTCTGTTTTGACCCGGCATTGTCGTTTCGGATGACGGGCGACCTTGACCTCCGGCGAGAGCTTTCTCCG

GCGTCTCTAACGGGCCTCATTTCCTTCACATGACTCTGTTCGACCCGTTTGTCTCTTCCGGGTACGATCC

GAACCCGATTCGGAGACGGTTGCTCGGGCCTCCTCGCCGGAGATTTCGCCACTCCCCTCTCTCTCCTCCC

GTGTAATTCACCGGAATACGACCGACGCTTGAACTTCTCAGGCGACTCAACTTTCCGGCGAATTTCACCG

ATTTCTTCATTTTCTCTAGAGAAGCTCTCACTCATAGAACTATAAATCTCTGAAATTTCAGAAACCTCTT

CTGGGTTTTCAAGCTTATGAATCTTTTTGAAATGGGTCTCGTTTATTCCCTCAAAATCTTGATCATTTTG

ATGTAAAGATTGAGTCTTTTTTGGAGGCTTCAAGATTCTTGGAGTTGAGGGTCTTAGCTTTGGAGTCTCG

GAGAGGACTTCTTTGACTGATTCTTCTAGTAAGTGTGGTGGTGGTGATCTTGGCAATGGTGCTTGGTTCG

ATGGTAAGTTTGATCTGTGTTTGCTTACAAAACAACCCATTGAGTGAATATGTTTGTGTTGATTTGTTTA

TCAGGAGACCAAGGGGAGCTTTGTCATGGTGATGTGTTGGAAGTTGTGATTGGGGGATTCTAGGCCGAAG

AGGATGTCTTA

>MSTRG.3195.1 gene=MSTRG.3195

CATCCTCCTTGGATTGCTCGCTTCGTGGCTGTTTCTTGTACCTGTTCAGGCTGTCGGATCAGCCGCGTGT

AGTGTTTAACCGTACGTTTCTCCGATCGCGAGACGCTAGGGCTTTTAATCCTGTCAACGATCATTGTGGT

GTTCCTGACTAGCGATGGATCGATTTTGATCTCTGCGTTGATGATTGGGGTGGCGATCGTGTGTGCGCAC

GGTGCGTTTTGAGCGCCCGATGATTTGTTTTTGGATGAGCAGGGTG

>MSTRG.3200.1 gene=MSTRG.3200

TACACAAGTTAATCACAGAAATGTGGTTAAACTTTTGGGATGTTGTTTGGAATGTGAGGTACCACTCTTA

GTATATGAACTTGTATCCAATGGTTCTTTATTGGATCATGTCCACAATATTGTTGGTGGCGCATCCTGGT

TATCACTGGAGAATCGGTTAAGAATTGCTGCTGAATCTTCTGGCGCCCTTGCATATCTTCACTCTGCAGC

TTCCATACCTATAATTCACAGAGATGTTAAATTAGCCAACATATTACTGGATGATAATAATGT

>MSTRG.3202.1 gene=MSTRG.3202

AGTCCTCATAGCGCACACAAAATGAACCTCTCCAAAATCTTCTATATTTCCTCTCTCTTGCTGCTCCTCT

GCCTCTCTGTCTCAACCCTACCAGGATCAGCTTCCAGAGCCCCGCCAAAGGGCAGTAGCAGCAGCAGTGG

TGGTGGTAACTTCCCTGGTAGCGGGGGTTACGGATATCCTGGTAACATCCCACCAGCATATCGTGGAATC

ATTGGTGGTGGCGGGTCGGGAGGATGGGGTGGTGGATATGGTGGTCCTAGTGGCGGCTCAGGGAGTGGTG

GAGTGGTGAGGCCTGTTGTGACATGCAAAGTTAAGGGACCTTGTTATGGCAAAAAACTGAGGTGTCCTTC

CAAGTGCTTCACTTCTTGGAGTCATTCTGGCAAGGGCTACGGTGGCGGAGGCGGAGGAGGTGGATGTAAC

ATGGATTGCAAGAAGAAGTGTACTGCTTCTTGCTCAACTTAGTAAGCTCTGGACACCAACCTACAGCATA

CAATGGTACCACTGAGATGCTCATATAGGATATGAGATGCTTCAGTTCATGTCATGAGTAACATATATTT

TGGTGTATGAGTATTACCTAAATAAATTACTGCCTTGAGTTTATCCAGCTAACACCCCCATAATTCGGTT

TCACAATACCTGATATAAATTCGGTTCGGTTTTCACCAAATGGTCGCCC

>MSTRG.3204.1 gene=MSTRG.3204

CCGCAAACCCTAATTTCCCCTCCTCTGTCCCTTTCTTTGCTTAATAGCAGCGGGGGCTTCCACCGGATCT

CTACCGGTGGAAAGCCCTAAAATTTTTTATTTCTTTTGTGTTTGCTTTTTTCTTCCAATAAAACTTCTTT

GGCATTTGTTTTTCCGGATCTAAGGATTCAAAACCTTTTCCGATTTGGCCTTCTTTTCTTAGATCTCAAT

TCCACGGAACTACGGATTTCTCGTTCTTCCCGGTGCTTTTATCCTTGAAAGTGGCCCGAGTTATCGTGTT

TCTGTGAGTTTTGTTCCCCAGATCTAAGATTGACTTGGTTTAGCCAATATTTCATTTACGCATTTGGTTG

TTAGGTTATCACGTTTGGTTTGGTTTTTTGTTGTCTCACCGTTTGTGTGAGTTTTGTTATTTTCTCACCG

TTTGTGTGAGTATGATGTTTGATTTAGTAATAGAAATTGTCAAGGGATTGCATTCTCAGTCGATAGCTCA

AACTGTTTGTGACGGAAGCGAAAGTGGCGAAGTTTGTGTAGATATCATCTTCAACAAATCACGTGATTTT

ATCTCCAAGATGAATGGATTAATCAGTGATTGTTTCTGTATTTTGTTTGTTCGACGCGACTGTTTATGTA

GATG

>MSTRG.3205.1 gene=MSTRG.3205

GTCTAAACACTAAATCCTTATCTCATCATACAACCCGCGTCCACCCTAACCCCACCCATATCTCCCCATT

AATACATCTCTATAAATCCATCTTTTAATACATCCTCCTCATTTCATCACCACTCTAAACTCATCTCTTC

TTCTTCCCCAATTCAAACCAAATCAATATCTTAATCATGGGCTCATCTACTAGTATGATATTTTCTGTTA

AGGCGCTGTTGATTTCAACTGGCGTTGTGTCTATAGCCTTGTTTTTAAAACTATCTCTTCCGGTTCTTAT

AGATGAAATCCCTGTCGTTTGGTCCTCGATTCTCTCGTGGCTTCAGCCTCCTTATTTGTACTTTGTTCTT

AACGCTATTATCATCATCATTGCCGCTTCATCGAAGTTTCACACAAAGATCGATGACTCCGATCATCAAC

ACCAGAATCAAATTAAAGTGCCGTTACCGTCTCCGGTCCCTATGGATCTGCCGGCTGTTTATGCTCAAAA

TGAGGTCTATCACAAAATCGAAGCTGATGATGATATTTATCGCAAAATTGAGGAACCGGAGGAGGCGTAT

CGGAAGATTGAGGATCCTGTGGAGGAATTGAAGCCGGTGGTGGAGATTAAACCGGAAGCTGTGGCGGAGA

AGGGCGAGGATAATGAGTTTGTGATATCGAGGTCCACGTGGACGCCGCCGCGGAAGGTGGAGAATGAGGA

GTTTCGAATTCCGGCGAAGGAGAAACCTCTGGTTTTGTCCAGGTTTAGCAACCGGAAGCCTGCAAAAGCT

AGTCCTGAAGGTGGAAAGAGGGCGCTGAGGGTGGCGAAGCCGAAGCGGCAAGAGACGTTGGAGAGCACCT

GGAAAGCCATAACGGACGGGCGTCACGTGCCACTCACGAGGCACCTCAAGAAGTCCGATACATTTGAGAA

TCACAACAGCCACGTGGCAGCCGATTCGCCCACGTGTTCTCCGCGGCGCGTGATCAAATCGGAGACGTTT

AGAGATCGGACGAATTACGATTCGCCGTCGCCGGTGTCGGGAGGGAAGCTGAGGAAGGAAGCGTCGCCGA

GTCAGGACGAGTTGAACCGGCGAGTGGAAGCGTTTATCAACAAGTTTAACCAGGAAATGAGGCTGCAGAG

ACAGGAGTCTCTCAACCAGTACATGGAGATGGTTAATCGAGGGTCACATTAGAAATTTTATTTTAATTTT

TAAAATTCACATTTCGTATTGAATTATAACTTTAAATGTTGCTATTATGTATACTCGCATATTGGTTTTT

GTTTTTTGGACCTGGTTGTGGTCTTTGAAAGAAGGCTCTCAAGAGGGAGTTGCAAAGTTTGAATTTTTTG

GTTTGATTATTGGTTTATGAAAAGTTGTGGTTATAGAATAGGAAGGAGGATATGATGGGGTTTGTGAAAC

TAGAGCCACCCCACTTGAATTTATTTGTATAAATCTGAAAAGTGGTCCTCAAGGAAGACCATTTTTTTTG

TTTATGGAATAATGGTTGCAAAATGGACCAAAGAGTTGTGGAAGCATGTGGGGAAATGGATGAGATGGTG

ATTTATACAGTTTCCTAGTACATACATTAACAGAAGCTTTTGGCCTCTCAAAAAAATTGGGTCCATCAAG

TAGAGCATTTGTCACATCAAGCAAAGCTAAAAATGTCAGCTAGCTCACCGTCCACAGACACTTTCGGATT

TACCCGCAACAAAACCAGACTTTGGTTCCTCTTGATTTAGTAAGAGGTACTCGTACTCTGTTGTATTAAT

CGATCTACACAAATTAACTCAACAGTAGTTAAACTTCTAGTCCCTATGTATTAAACTATAGTTGGATTCC

GTTAATGTATAATCGATCAAAATTTACGTATAGCTCAACAGAACACAGTGG

>MSTRG.3217.1 gene=MSTRG.3217

ACCTTGGCATTTATGACAGATACTCCGTAGTTGCCTTCCTCGCGCGCAATTAGTGCAAAATCAGATCATA

TGATTATTCTCCTGCGGTGCTGTAAATTGGTTAGTCGCTATGTCTTTCGAATTTTGAAATCATAACATCT

CAATGCTTTAATTTCTGGCCACTCTTCGTTGACTGATGTAGGATGGATTCATTCTCGCACTCTCCGCTGA

AGAACTCGCAGAATAGAGATGGTGAGAATTCAGAGCATGTTGCTTCTGATCTATAGAATGGATATTTATC

TTTGGGAACTCCTACTGTTATATGTGGCAAATGTGGTGCACTAATGTGGAACATGGAGAGAAACAACAAG

TCAAATATGAATTCCACTCCGACCTTTTCAATCTGCTGCAAAGATGGAGCTGTTGTGCTACCTAGGGAGG

AGAATGCACCAGAGCCTCTATCATCTCTCCTACACGGCAGTGAGCGTTCTGCAGGATTCAGGCGAAACAT

TCGAGTTTATAACAGTATGTTTGCAACGTGTTCTAGCGGTGGTAAGGTTGATCACAAGATCAGTTCCCAT

AACAAGTTTAAGCTATATTGTTATCTTTGTTTCTAAATTATGTTTCTCAGGTTTGATATTTTAATGCAGG

ATGGCTTCAGGCCCTCCAAACTTATTTATTCTCTACCATTGACAACTACACTATAAAGCAACTACTCATC

CAGGTAGGTTCCCTACAGTTTTTTTATTTTTGCGAATAAAAACCTCTTTCATATGCTTTGCATTGTATGC

CACCAAGAACTCCTCAGAGATATGTTATGCAAGTACGTCTGAATATCGAAGCTCAAGTTAAATTTCTTTT

CCAACTTTAAGAAGTGTAAAACCACCAGTCAATTGGTATGCAATGATTTTGTTTACTTCAGATTGGTTTT

AACATGATATATTACATTGTACCCTACAAACTTCCATATTAACATAGTTAATTCCACCAGTTTGGTGCAT

TAACAGCCTCTTATTGCAAACACAATAAGCTATGAACTTGACGAAACTACTTAAGTAGAAATATACCATT

ATGCCTTTTAATGCACGAAAAAGAGGCTGTTGATATCTTTTTCGTGGTTATCTCAAAAATCAATTCATAT

TTTTTAATTATAAGATATTGTATTTTCAAGTAGCAGTCATCCCATCTCAACATGTGATTTGCATAATTAT

AGATTATGAACTGTAAATCTCATATCATGGTTAAGTTGTCATTAGGATAGACTGGCAGTTGTTTACCGAT

ACTCAAAGTACCCAACTGTCACTGCTGCCGAAGTTCTCAATCTCTCAAGAAAGAGTTGTCTGTTGTTTAT

AGATCGTGTTTATCTGCAGATGTTGTGTAAAACCATGTTGCTGTCGAGTCTTGCTTATGCATTTTCTTGC

AGCATCGATTTGGCTTGGTAAATACTGAACTCTCCTTTATTCTTGAAGGCTGCTACAAATAAATTTTTTA

TGACTGGTATCTGCCCACTTGGCAATTTAGTAGAGCTCCTGCCACTGGTGCCTCTTCAAGGTCATCAATT

TCAGGTGGATCATATGCCGCTTGCTGATTGACCGGGAATTCTGCTTCCTCGAGAATCCGAGCTGAGCTGG

CACAACCAATCGTAATGGCTAGCAACAAAAGGACAGATGAAGTAGAGTTGTTTGCTATGACTACTCAGAA

TGCATACAAGCAATTACATGACAACGAAGCTCATTCAAAACCGATAACAGTATGACAGTGAACTGCATAT

ATATTATTTTTAATACTTCTGACTTGAGTTTCAGTTTCTTATTGTCCTTCGAGCATATGCATAAACTGAA

GAAAGTTGCACAAGGCAGAGTTTGGACCGACAATGATGCTGCTTTGCGAGATTTAGTTGATGCTATTGGT

GGTCTCTCCTCTCCTGTGTTGCTGCTATAGCAAAACAGAAGGTGAAGATACTCCTGGACAAAAAGGACCA

CATGTTGCAAGCTGCCTGTTGCAGTAGACATTTTATATTCAGAGTTTTGGAAGGGAACAGAAGATAACAC

AGGTTATAAGGCAGAAATGGCAATGAAAAATGGAACCAAACTGGAGAATAAAAGAGCAAGTCTTCAGAAT

AACAATGGCTTCATTGTGAATCCTTTGGCAGAAGTCCATAGCAATTGCCAGGGGATATCAAAGATTGTCT

AACTATGAACTTATCGTAGTTGGTTTTGCTCTTAGATTCCATCCTAAAGCCTTAAGATCAACATTGTAGA

ATTTTGATATTTTAGCTTCAA

>MSTRG.3217.2 gene=MSTRG.3217

ACCTTGGCATTTATGACAGATACTCCGTAGTTGCCTTCCTCGCGCGCAATTAGTGCAAAATCAGATCATA

TGATTATTCTCCTGCGGTGCTGTAAATTGGTTAGTCGCTATGTCTTTCGAATTTTGAAATCATAACATCT

CAATGCTTTAATTTCTGGCCACTCTTCGTTGACTGATGTAGGATGGATTCATTCTCGCACTCTCCGCTGA

AGAACTCGCAGAATAGAGATGGTGAGAATTCAGAGCATGTTGCTTCTGATCTATAGAATGGATATTTATC

TTTGGGAACTCCTACTGTTATATGTGGCAAATGTGGTGCACTAATGTGGAACATGGAGAGAAACAACAAG

TCAAATATGAATTCCACTCCGACCTTTTCAATCTGCTGCAAAGATGGAGCTGTTGTGCTACCTAGGGAGG

AGAATGCACCAGAGCCTCTATCATCTCTCCTACACGGCAGTGAGCGTTCTGCAGGATTCAGGCGAAACAT

TCGAGTTTATAACAGTATGTTTGCAACGTGTTCTAGCGGTGGTAAGGTTGATCACAAGATCAGTTCCCAT

AACAAGTTTAAGCTATATTGTTATCTTTGTTTCTAAATTATGTTTCTCAGGTTTGATATTTTAATGCAGG

ATGGCTTCAGGCCCTCCAAACTTATTTATTCTCTACCATTGACAACTACACTATAAAGCAACTACTCATC

CAGGTAGGTTCCCTACAGTTTTTTTATTTTTGCGAATAAAAACCTCTTTCATATGCTTTGCATTGTATGC

CACCAAGAACTCCTCAGAGATATGTTATGCAAGTACGTCTGAATATCGAAGCTCAAGTTAAATTTCTTTT

CCAACTTTAAGAAGTGTAAAACCACCAGTCAATTGGTATGCAATGATTTTGTTTACTTCAGATTGGTTTT

AACATGATATATTACATTGTACCCTACAAACTTCCATATTAACATAGTTAATTCCACCAGTTTGGTGCAT

TAACAGCCTCTTATTGCAAACACAATAAGCTATGAACTTGACGAAACTACTTAAGTAGAAATATACCATT

ATGCCTTTTAATGCACGAAAAAGAGGCTGTTGATATCTTTTTCGTGGTTATCTCAAAAATCAATTCATAT

TTTTTAATTATAAGATATTGTATTTTCAAGTAGCAGTCATCCCATCTCAACATGTGATTTGCATAATTAT

AGATTATGAACTGTAAATCTCATATCATGGTTAAGTTGTCATTAGGATAGACTGGCAGTTGTTTACCGAT

ACTCAAAGTACCCAACTGTCACTGCTGCCGAAGTTCTCAATCTCTCAAGAAAGAGTTGTCTGTTGTTTAT

AGATCGTGTTTATCTGCAGATGTTGTGTAAAACCATGTTGCTGTCGAGTCTTGCTTATGCATTTTCTTGC

AGCATCGATTTGGCTTGTAGAGCTCCTGCCACTGGTGCCTCTTCAAGGTCATCAATTTCAGGTGGATCAT

ATGCCGCTTGCTGATTGACCGGGAATTCTGCTTCCTCGAGAATCCGAGCTGAGCTGGCACAACCAATCGT

AATGGCTAGCAACAAAAGGACAGATGAAGTAGAGTTGTTTGCTATGACTACTCAGAATGCATACAAGCAA

TTACATGACAACGAAGCTCATTCAAAACCGATAACAGTATGACAGTGAACTGCATATATATTATTTTTAA

TACTTCTGACTTGAGTTTCAGTTTCTTATTGTCCTTCGAGCATATGCATAAACTGAAGAAAGTTGCACAA

GGCAGAGTTTGGACCGACAATGATGCTGCTTTGCGAGATTTAGTTGATGCTATTGGTGGTCTCTCCTCTC

CTGTGTTGCTGCTATAGCAAAACAGAAGGTGAAGATACTCCTGGACAAAAAGGACCACATGTTGCAAGCT

GCCTGTTGCAGTAGACATTTTATATTCAGAGTTTTGGAAGGGAACAGAAGATAACACAGGTTATAAGGCA

GAAATGGCAATGAAAAATGGAACCAAACTGGAGAATAAAAGAGCAAGTCTTCAGAATAACAATGGCTTCA

TTGTGAATCCTTTGGCAGAAGTCCATAGCAATTGCCAGGGGATATCAAAGATTGTCTAACTATGAACTTA

TCGTAGTTGGTTTTGCTCTTAGATTCCATCCTAAAGCCTTAAGATCAACATTGTAGAATTTTGATATTTT

AGCTTCAACTCCTTTTTGGACTTCATTTTGGCAATTACTTGCAGGCCATATTTTTTCGTGTTGGACTTAG

TAG

>MSTRG.3217.3 gene=MSTRG.3217

CATTTATGACAGATACTCCGTAGTTGCCTTCCTCGCGCGCAATTAGTGCAAAATCAGATCATATGATTAT

TCTCCTGCGGTGCTGTAAATTGGTTAGTCGCTATGTCTTTCGAATTTTGAAATCATAACATCTCAATGCT

TTAATTTCTGGCCACTCTTCGTTGACTGATGTAGGATGGATTCATTCTCGCACTCTCCGCTGAAGAACTC

GCAGAATAGAGATGGTGAGAATTCAGAGCATGTTGCTTCTGATCTATAGAATGGATATTTATCTTTGGGA

ACTCCTACTGTTATATGTGGCAAATGTGGTGCACTAATGTGGAACATGGAGAGAAACAACAAGTCAAATA

TGAATTCCACTCCGACCTTTTCAATCTGCTGCAAAGATGGAGCTGTTGTGCTACCTAGGGAGGAGAATGC

ACCAGAGCCTCTATCATCTCTCCTACACGGCAGTGAGCGTTCTGCAGGATTCAGGCGAAACATTCGAGTT

TATAACAGTATGTTTGCAACGTGTTCTAGCGGTGGTAAGGTTGATCACAAGATCAGTTCCCATAACAAGT

TTAAGCTATATTGTTATCTTTGTTTCTAAATTATGTTTCTCAGGTTTGATATTTTAATGCAGGATGGCTT

CAGGCCCTCCAAACTTATTTATTCTCTACCATTGACAACTACACTATAAAGCAACTACTCATCCAGGCTG

CTACAAATAAATTTTTTATGACTGGTATCTGCCCACTTGGCAATTTAGTAGAGCTCCTGCCACTGGTGCC

TCTTCAAGGTCATCAATTTCAGGTGGATCATATGCCGCTTGCTGATTGACCGGGAATTCTGCTTCCTCGA

GAATCCGAGCTGAGCTGGCACAACCAATCGTAATGGCTAGCAACAAAAGGACAGATGAAGTAGAGTTGTT

TGCTATGACTACTCAGAATGCATACAAGCAATTACATGACAACGAAGCTCATTCAAAACCGATAACAGTA

TGACAGTGAACTGCATATATATTATTTTTAATACTTCTGACTTGAGTTTCAGTTTCTTATTGTCCTTCGA

GCATATGCATAAACTGAAGAAAGTTGCACAAGGCAGAGTTTGGACCGACAATGATGCTGCTTTGCGAGAT

TTAGTTGATGCTATTGGTGGTCTCTCCTCTCCTGTGTTGCTGCTATAGCAAAACAGAAGGTGAAGATACT

CCTGGACAAAAAGGACCACATGTTGCAAGCTGCCTGTTGCAGTAGACATTTTATATTCAGAGTTTTGGAA

GGGAACAGAAGATAACACAGGTTATAAGGCAGAAATGGCAATGAAAAATGGAACCAAACTGGAGAATAAA

AGAGCAAGTCTTCAGAATAACAATGGCTTCATTGTGAATCCTTTGGCAGAAGTCCATAGCAATTGCCAGG

GGATATCAAAGATTGTCTAACTATGAACTTATCGTAGTTGGTTTTGCTCTTAGATTCCATCCTAAAGCCT

TAAGATCAACATTGTAGAATTTTGATATTTTAGCTTCAACTCCTTTTTGGACTTCATTTTGGCAATTACT

TGCAGGCCATATTTTTTCGTGTTGGACTTAGTAG

>MSTRG.3217.4 gene=MSTRG.3217

CATTTATGACAGATACTCCGTAGTTGCCTTCCTCGCGCGCAATTAGTGCAAAATCAGATCATATGATTAT

TCTCCTGCGGTGCTGTAAATTGGTTAGTCGCTATGTCTTTCGAATTTTGAAATCATAACATCTCAATGCT

TTAATTTCTGGCCACTCTTCGTTGACTGATGTAGGATGGATTCATTCTCGCACTCTCCGCTGAAGAACTC

GCAGAATAGAGATGGTGAGAATTCAGAGCATGTTGCTTCTGATCTATAGAATGGATATTTATCTTTGGGA

ACTCCTACTGTTATATGTGGCAAATGTGGTGCACTAATGTGGAACATGGAGAGAAACAACAAGTCAAATA

TGAATTCCACTCCGACCTTTTCAATCTGCTGCAAAGATGGAGCTGTTGTGCTACCTAGGGAGGAGAATGC

ACCAGAGCCTCTATCATCTCTCCTACACGGCAGTGAGCGTTCTGCAGGATTCAGGCGAAACATTCGAGTT

TATAACAGTATGTTTGCAACGTGTTCTAGCGGTGGTAAGGTTGATCACAAGATCAGTTCCCATAACAAGT

TTAAGCTATATTGTTATCTTTGTTTCTAAATTATGTTTCTCAGGTTTGATATTTTAATGCAGGATGGCTT

CAGGCCCTCCAAACTTATTTATTCTCTACCATTGACAACTACACTATAAAGCAACTACTCATCCAGGTAG

GTTCCCTACAGTTTTTTTATTTTTGCGAATAAAAACCTCTTTCATATGCTTTGCATTGTATGCCACCAAG

AACTCCTCAGAGATATGTTATGCAAGTACGTCTGAATATCGAAGCTCAAGTTAAATTTCTTTTCCAACTT

TAAGAAGTGTAAAACCACCAGTCAATTGGTATGCAATGATTTTGTTTACTTCAGATTGGTTTTAACATGA

TATATTACATTGTACCCTACAAACTTCCATATTAACATAGTTAATTCCACCAGTTTGGTGCATTAACAGC

CTCTTATTGCAAACACAATAAGCTATGAACTTGACGAAACTACTTAAGTAGAAATATACCATTATGCCTT

TTAATGCACGAAAAAGAGGCTGTTGATATCTTTTTCGTGGTTATCTCAAAAATCAATTCATATTTTTTAA

TTATAAGATATTGTATTTTCAAGTAGCAGTCATCCCATCTCAACATGTGATTTGCATAATTATAGATTAT

GAACTGTAAATCTCATATCATGGTTAAGTTGTCATTAGGATAGACTGGCAGTTGTTTACCGATACTCAAA

GTACCCAACTGTCACTGCTGCCGAAGTTCTCAATCTCTCAAGAAAGAGTTGTCTGTTGTTTATAGATCGT

GTTTATCTGCAGATGTTGTGTAAAACCATGTTGCTGTCGAGTCTTGCTTATGCATTTTCTTGCAGCATCG

ATTTGGCTTGGTAAATACTGAACTCTCCTTTATTCTTGAAGGCTGCTACAAATAAATTTTTTATGACTGG

TATCTGCCCACTTGGCAATTTAGTAGAGCTCCTGCCACTGGTGCCTCTTCAAGGTCATCAATTTCAGGTG

GATCATATGCCGCTTGCTGATTGACCGGGAATTCTGCTTCCTCGAGAATCCGAGCTGAGCTGGCACAACC

AATCGTAATGGCTAGCAACAAAAGAAAAGGACCTGATCTCATGGAGTTATTGCTGCATCATTCACCCAGT

ATTGCAATGTGTGATATTGGGAAAATACCAATTTGATAATAAAAAATTATTAGTTATGGAATTCATTCAT

GTAGGAATATTTAATCCGGGGAAACTATATGAGAAGATTGGTTTTGATAGAAAAATTATATCAAGAGGAA

ATTTGCAGAGCTTACACCTGATAAGCAACATTGATATACCAAGTTCAGGTTAGAAGTTAGCTCACCTATA

AGGTTTTTAAGTTGCCCAAGTTGGTTCTTGAACTTCTCTTTCCACTTGATTAAATTACAATTGACAAATA

TTTAGCTTGCAGGACAGATGAAGTAGAGTTGTTTGCTATGACTACTCAGAATGCATACAAGCAATTACAT

GACAACGAAGCTCATTCAAAACCGATAACAGTATGACAGTGAACTGCATATATATTATTTTTAATACTTC

TGACTTGAGTTTCAGTTTCTTATTGTCCTTCGAGCATATGCATAAACTGAAGAAAGTTGCACAAGGCAGA

GTTTGGACCGACAATGATGCTGCTTTGCGAGATTTAGTTGATGCTATTGGTGGTCTCTCCTCTCCTGTGT

TGCTGCTATAGCAAAACAGAAGGTGAAGATACTCCTGGACAAAAACCTAGAGCTGTCCTTAAAAGAGATC

TCACTTGATAAGGATATCAGGTACTCGCTCTGCAGGGACCACATGTTGCAAGCTGCCTGTTGCAGTAGAC

ATTTTATATTCAGAGTTTTGGAAGGGAACAGAAGATAACACAGGTTATAAGGCAGAAATGGCAATGAAAA

ATGGAACCAAACTGGAGAATAAAAGAGCAAGTCTTCAGAATAACAATGGCTTCATTGTGAATCCTTTGGC

AGAAGTCCATAGCAATTGCCAGGGGATATCAAAGATTGTCTAACTATGAACTTATCGTAGTTGGTTTTGC

TCTTAGATTCCATCCTAAAGCCTTAAGATCAACATTGTAGAATTTTGATATTTTAGCTTCAACTCCTTTT

TGGACTTCATTTTGGCAATTACTTGCAGGCCATATTTTTTCGTGTTGGACTTAGTAG

>MSTRG.3217.5 gene=MSTRG.3217

CATTTATGACAGATACTCCGTAGTTGCCTTCCTCGCGCGCAATTAGTGCAAAATCAGATCATATGATTAT

TCTCCTGCGGTGCTGTAAATTGGTTAGTCGCTATGTCTTTCGAATTTTGAAATCATAACATCTCAATGCT

TTAATTTCTGGCCACTCTTCGTTGACTGATGTAGGATGGATTCATTCTCGCACTCTCCGCTGAAGAACTC

GCAGAATAGAGATGGTGAGAATTCAGAGCATGTTGCTTCTGATCTATAGAATGGATATTTATCTTTGGGA

ACTCCTACTGTTATATGTGGCAAATGTGGTGCACTAATGTGGAACATGGAGAGAAACAACAAGTCAAATA

TGAATTCCACTCCGACCTTTTCAATCTGCTGCAAAGATGGAGCTGTTGTGCTACCTAGGGAGGAGAATGC

ACCAGAGCCTCTATCATCTCTCCTACACGGCAGTGAGCGTTCTGCAGGATTCAGGCGAAACATTCGAGTT

TATAACAGTATGTTTGCAACGTGTTCTAGCGGTGGTAAGGTTGATCACAAGATCAGTTCCCATAACAAGT

TTAAGCTATATTGTTATCTTTGTTTCTAAATTATGTTTCTCAGGTTTGATATTTTAATGCAGGATGGCTT

CAGGCCCTCCAAACTTATTTATTCTCTACCATTGACAACTACACTATAAAGCAACTACTCATCCAGGTAG

GTTCCCTACAGTTTTTTTATTTTTGCGAATAAAAACCTCTTTCATATGCTTTGCATTGTATGCCACCAAG

AACTCCTCAGAGATATGTTATGCAAGTACGTCTGAATATCGAAGCTCAAGTTAAATTTCTTTTCCAACTT

TAAGAAGTGTAAAACCACCAGTCAATTGGTATGCAATGATTTTGTTTACTTCAGATTGGTTTTAACATGA

TATATTACATTGTACCCTACAAACTTCCATATTAACATAGTTAATTCCACCAGTTTGGTGCATTAACAGC

CTCTTATTGCAAACACAATAAGCTATGAACTTGACGAAACTACTTAAGTAGAAATATACCATTATGCCTT

TTAATGCACGAAAAAGAGGCTGTTGATATCTTTTTCGTGGTTATCTCAAAAATCAATTCATATTTTTTAA

TTATAAGATATTGTATTTTCAAGTAGCAGTCATCCCATCTCAACATGTGATTTGCATAATTATAGATTAT

GAACTGTAAATCTCATATCATGGTTAAGTTGTCATTAGGATAGACTGGCAGTTGTTTACCGATACTCAAA

GTACCCAACTGTCACTGCTGCCGAAGTTCTCAATCTCTCAAGAAAGAGTTGTCTGTTGTTTATAGATCGT

GTTTATCTGCAGATGTTGTGTAAAACCATGTTGCTGTCGAGTCTTGCTTATGCATTTTCTTGCAGCATCG

ATTTGGCTTGGTAAATACTGAACTCTCCTTTATTCTTGAAGGCTGCTACAAATAAATTTTTTATGACTGG

TATCTGCCCACTTGGCAATTTAGTAGAGCTCCTGCCACTGGTGCCTCTTCAAGGTCATCAATTTCAGGTG

GATCATATGCCGCTTGCTGATTGACCGGGAATTCTGCTTCCTCGAGAATCCGAGCTGAGCTGGCACAACC

AATCGTAATGGCTAGCAACAAAAGAAAAGGACCTGATCTCATGGAGTTATTGCTGCATCATTCACCCAGT

ATTGCAATGTGTGATATTGGGAAAATACCAATTTGATAATAAAAAATTATTAGTTATGGAATTCATTCAT

GTAGGAATATTTAATCCGGGGAAACTATATGAGAAGATTGGTTTTGATAGAAAAATTATATCAAGAGGAA

ATTTGCAGAGCTTACACCTGATAAGCAACATTGATATACCAAGTTCAGGTTAGAAGTTAGCTCACCTATA

AGGTTTTTAAGTTGCCCAAGTTGGTTCTTGAACTTCTCTTTCCACTTGATTAAATTACAATTGACAAATA

TTTAGCTTGCAGGACAGATGAAGTAGAGTTGTTTGCTATGACTACTCAGAATGCATACAAGCAATTACAT

GACAACGAAGCTCATTCAAAACCGATAACAGTATGACAGTGAACTGCATATATATTATTTTTAATACTTC

TGACTTGAGTTTCAGTTTCTTATTGTCCTTCGAGCATATGCATAAACTGAAGAAAGTTGCACAAGGCAGA

GTTTGGACCGACAATGATGCTGCTTTGCGAGATTTAGTTGATGCTATTGGTGGTCTCTCCTCTCCTGTGT

TGCTGCTATAGCAAAACAGAAGGTGAAGATACTCCTGGACAAAAAGGACCACATGTTGCAAGCTGCCTGT

TGCAGTAGACATTTTATATTCAGAGTTTTGGAAGGGAACAGAAGATAACACAGGTTATAAGGCAGAAATG

GCAATGAAAAATGGAACCAAACTGGAGAATAAAAGAGCAAGTCTTCAGAATAACAATGGCTTCATTGTGA

ATCCTTTGGCAGAAGTCCATAGCAATTGCCAGGGGATATCAAAGATTGTCTAACTATGAACTTATCGTAG

TTGGTTTTGCTCTTAGATTCCATCCTAAAGCCTTAAGATCAACATTGTAGAATTTTGATATTTTAGCTTC

AACTCCTTTTTGGACTTCATTTTGGCAATTACTTGCAGGCCATATTTTTTCGTGTTGGACTTAGTAG

>MSTRG.3217.7 gene=MSTRG.3217

AGGATGGCTTCAGGCCCTCCAAACTTATTTATTCTCTACCATTGACAACTACACTATAAAGCAACTACTC

ATCCAGATGTTGTGTAAAACCATGTTGCTGTCGAGTCTTGCTTATGCATTTTCTTGCAGCATCGATTTGG

CTTGGTAAATACTGAACTCTCCTTTATTCTTGAAGGCTGCTACAAATAAATTTTTTATGACTGGTATCTG

CCCACTTGGCAATTTAGTAGAGCTCCTGCCACTGGTGCCTCTTCAAGGTCATCAATTTCAGGTGGATCAT

ATGCCGCTTGCTGATTGACCGGGAATTCTGCTTCCTCGAGAATCCGAGCTGAGCTGGCACAACCAATCGT

AATGGCTAGCAACAAAAGATGAAGTAGAGTTGTTTGCTATGACTACTCAGAATGCATACAAGCAATTACA

TGACAACGAAGCTCATTCAAAACCGATAACAGTATGACAGTGAACTGCATATATATTATTTTTAATACTT

CTGACTTGAGTTTCAGTTTCTTATTGTCCTTCGAGCATATGCATAAACTGAAGAAAGTTGCACAAGGCAG

AGTTTGGACCGACAATGATGCTGCTTTGCGAGATTTAGTTGATGCTATTGGTGGTCTCTCCTCTCCTGTG

TTGCTGCTATAGCAAAACAGAAGGTGAAGATACTCCTGGACAAAAAGATTTGATTGGGTCATCATAAAAT

AATGTGGTCTTGGAATTATGTGTTTATAGCCTAGAGCTGTCCTTAAAAGAGATCTCACTTGATAAGGATA

TCAGGTACTCGCTCTGCAGGGACCACATGTTGCAAGCTGCCTGTTGCAGTAGACATTTTATATTCAGAGT

TTTGGAAGGGAACAGAAGATAACACAGGTTATAAGGCAGAAATGGCAATGAAAAATGGAACCAAACTGGA

GAATAAAAGAGCAAGTCTTCAGAATAACAATGGCTTCATTGTGAATCCTTTGGCAGAAGTCCATAGCAAT

TGCCAGGGGATATCAAAGATTGTCTAACTATGAACTT

>MSTRG.3217.8 gene=MSTRG.3217

AGGATGGCTTCAGGCCCTCCAAACTTATTTATTCTCTACCATTGACAACTACACTATAAAGCAACTACTC

ATCCAGATGTTGTGTAAAACCATGTTGCTGTCGAGTCTTGCTTATGCATTTTCTTGCAGCATCGATTTGG

CTTGTAGAGCTCCTGCCACTGGTGCCTCTTCAAGGTCATCAATTTCAGGTGGATCATATGCCGCTTGCTG

ATTGACCGGGAATTCTGCTTCCTCGAGAATCCGAGCTGAGCTGGCACAACCAATCGTAATGGCTAGCAAC

AAAAGATGCATAAACTGAAGAAAGTTGCACAAGGCAGAGTTTGGACCGACAATGATGCTGCTTTGCGAGA

TTTAGTTGATGCTATTGGTGGTCTCTCCTCTCCTGTGTTGCTGCTATAGCAAAACAGAAGGTGAAGATAC

TCCTGGACAAAAAGGACCACATGTTGCAAGCTGCCTGTTGCAGTAGACATTTTATATTCAGAGTTTTGGA

AGGGAACAGAAGATAACACAGGTTATAAGGCAGAAATGGCAATGAAAAATGGAACCAAACTGGAGAATAA

AAGAGCAAGTCTTCAGAATAACAATGGCTTCATTGTGAATCCTTTGGCAGAAGTCCATAGCAATTGCCAG

GGGATATCAAAGATTGTCTAACTATGAACTT

>MSTRG.3217.6 gene=MSTRG.3217

AGGATGGCTTCAGGCCCTCCAAACTTATTTATTCTCTACCATTGACAACTACACTATAAAGCAACTACTC

ATCCAGATGTTGTGTAAAACCATGTTGCTGTCGAGTCTTGCTTATGCATTTTCTTGCAGCATCGATTTGG

CTTGGTAAATACTGAACTCTCCTTTATTCTTGAAGGCTGCTACAAATAAATTTTTTATGACTGGTATCTG

CCCACTTGGCAATTTAGTAGAGCTCCTGCCACTGGTGCCTCTTCAAGGTCATCAATTTCAGGTGGATCAT

ATGCCGCTTGCTGATTGACCGGGAATTCTGCTTCCTCGAGAATCCGAGCTGAGCTGGCACAACCAATCGT

AATGGCTAGCAACAAAAGGACAGATGAAGTAGAGTTGTTTGCTATGACTACTCAGAATGCATACAAGCAA

TTACATGACAACGAAGCTCATTCAAAACCGATAACAGTATGACAGTGAACTGCATATATATTATTTTTAA

TACTTCTGACTTGAGTTTCAGTTTCTTATTGTCCTTCGAGCATATGCATAAACTGAAGAAAGTTGCACAA

GGCAGAGTTTGGACCGACAATGATGCTGCTTTGCGAGATTTAGTTGATGCTATTGGTGGTCTCTCCTCTC

CTGTGTTGCTGCTATAGCAAAACAGAAGGTGAAGATACTCCTGGACAAAAAGGACCACATGTTGCAAGCT

GCCTGTTGCAGTAGACATTTTATATTCAGAGTTTTGGAAGGGAACAGAAGATAACACAGGTTATAAGGCA

GAAATGGCAATGAAAAATGGAACCAAACTGGAGAATAAAAGAGCAAGTCTTCAGAATAACAATGGCTTCA

TTGTGAATCCTTTGGCAGAAGTCCATAGCAATTGCCAGGGGATATCAAAGATTGTCTAACTATGAACTTA

TCGTAGTTGGTTTTGCTCTTAGATTCCATCCTAAAGCCTTAAGATCAACATTGTAGAATTTTGATATTTT

AGCTTCAACTCCTTTTTGGACTTCATTTTGGCAATTACTTGCAGGCCATATTTTTTCGTGTTGGACTTAG

TAGTGAT

>MSTRG.3217.9 gene=MSTRG.3217

ATGACTGGTATCTGCCCACTTGGCAATTTAGTAGAGCTCCTGCCACTGGTGCCTCTTCAAGGTCATCAAT

TTCAGGTGGATCATATGCCGCTTGCTGATTGACCGGGAATTCTGCTTCCTCGAGAATCCGAGCTGAGCTG

GCACAACCAATCGTAATGGCTAGCAACAAAAGGACAGATGAAGTAGAGTTGTTTGCTATGACTACTCAGA

ATGCATACAAGCAATTACATGACAACGAAGCTCATTCAAAACCGATAACAGTATGACATTTCTTATTGTC

CTTCGAGCATATGCATAAACTGAAGAAAGTTGCACAAGGCAGAGTTTGGACCGACAATGATGCTGCTTTG

CGAGATTTAGTTGATGCTATTGGTGGTCTCTCCTCTCCTGTGTTGCTGCTATAGCAAAACAGAAGGTGAA

GATACTCCTGGACAAAAAGGACCACATGTTGCAAGCTGCCTGTTGCAGTAGACATTTTATATTCAGAGTT

TTGGAAGGGAACAGAAGATAACACAGGTTATAAGGCAGAAATGGCAATGAAAAATGGAACCAAACTGGAG

AATAAAAGAGCAAGTCTTCAGAATAACAATGGCTTCATTGTGAATCCTTTGGCAGAAGTCCATAGCAATT

GCCAGGGGATATCAAAGATTGTCTAACTATGAACTT

>MSTRG.3217.10 gene=MSTRG.3217

AAAATACCAATTTGATAATAAAAAATTATTAGTTATGGAATTCATTCATGTAGGAATATTTAATCCGGGG

AAACTATATGAGAAGATTGGTTTTGATAGAAAAATTATATCAAGAGGAAATTTGCAGAGCTTACACCTGA

TAAGCAACATTGATATACCAAGTTCAGGTTAGAAGTTAGCTCACCTATAAGGTTTTTAAGTTGCCCAAGT

TGGTTCTTGAACTTCTCTTTCCACTTGATTAAATTACAATTGACAAATATTTAGCTTGCAGGACAGATGA

AGTAGAGTTGTTTGCTATGACTACTCAGAATGCATACAAGCAATTACATGACAACGAAGCTCATTCAAAA

CCGATAACAGTATGACAGTGAACTGCATATATATTATTTTTAATACTTCTGACTTGAGTTTCAGTTTCTT

ATTGTCCTTCGAGCATATGCATAAACTGAAGAAAGTTGCACAAGGCAGAGTTTGGACCGACAATGATGCT

GCTTTGCGAGATTTAGTTGATGCTATTGGTGGTCTCTCCTCTCCTGTGTTGCTGCTATAGCAAAACAGAA

GGTACTCGCTCTGCAGGGACCACATGTTGCAAGCTGCCTGTTGCAGTAGACATTTTATATTCAGAGTTTT

GGAAGGGAACAGAAGATAACACAGGTTATAAGGCAGAAATGGCAATGAAAAAT

>MSTRG.3217.11 gene=MSTRG.3217

TGAGATTCTTTTTGTATGTTAGACGGGAACTTTTCAGTAGGAGCGCTGAGAAAAATACACACAATTTTGA

TGGGCCTAAAGCTGGCTCAAGTTTGCATCATTGTGATAAAGTGTCATATTATGTGGTAAAAGCTAATAAT

ATCGCTGGAAATCATCATGTGAATCATAAAGAGAGTGAGGATTTACTCTTAATATGCAAGCAGCTTCTTT

ATTGAAACTAGCAATCCCCTTTGTTTGATTTACTTTAGGGACCACATGTTGCAAGCTGCCTGTTGCAGTA

GACATTTTATATTCAGAGTTTTGGAAGGGAACAGAAGATAACACAGGTTATAAGGCAGAAATGGCAATGA

AAAATGGAACCAAACTGGAGAATAAAAGAGCAAGTCTTCAGAATAACAATGGCTTCATTGTGAATCCTTT

GGCAGAAGTCCATAGCAATTGCCAGGGGATATCAAAGATTGTCTAACTATGAACTTATCGATCGATGGTC

TCAATTAAAATGTTTCTTCCTTTGTTTTTTAGATACTCCTTCCATCCCAAAATAATAGTCCTTTTATATT

TTATCTACAGGATCGATTTTGTTAATTTTAAGATATTCAATCAGAATAG

>MSTRG.3223.1 gene=MSTRG.3223

GATATATGTAGCACTCAAGAACCCATGAAGAAATTTAGACTCTTTCAGAAAATTTCCTCTGTATTTTTCT

TATTACACACTAAAATGCAAATATATTAGAAAAGAATATTGGGAAAGCTGAGATATATTTGAGAAGAAGA

AACAAGCTAGGGAGAGAATCATGAAGAAATGTGAGTTGTGCAACTCCATTGCAAGAATGTTTTGTGAATC

AGATCAAGCAAGCTTGTGTTGGCAGTGTGATGCCAAGGTTCATTCTGCTAACTTTTTAGTGGCTAAGCAC

TCTAGAACTCTTCTTTGTCAATCTTGTCAAGCTCTTACTCCATGGACTGGCTCTGGTCCTAAGCTTGGAC

CCACTCTTTCATTTTGTGAATCTTGTGTTCATGAAAATACTAATTACAATAATGATACTGATGAGGATGA

TGAGGATGAAGAGGATGATGAAGATGATGAAGAAGATGAGGAGGAAGATGACGGTGAGAATCAAGTGGTT

CCTCTTTCATCTTCATCTCTGGCTCCTCCACAAGTTACAAGCTCTTCTAGCAGTGAAGAGTCTTCTGCTA

GATTGTTTAATGGTGCAGAAAGTAGCAGAAAATCAAGATTCTCCATCTCATGGAGTAGAACTAGATGATT

TTTTAGAAAAGGGAAATGAAATGGAAGATGAAGAGGAGTGATCCAATAACACTGCTGCTCACTTTAGCAT

GTTGCTTGATCAACTTCAACTTAATGTTAATTTGTAACATTACCTCTAACTGTGACCTCCTTGTAATTAA

ATCGGAGATTAGTAACGTGTTCTATATCATCGTACAGTCGTGTGTTATATTTACCTCGTAGCATTGTTTC

TCTCGTGTACTGCATTTGAAGCAAAAATTAAAATTGGAACAAGTTATAATGTATAA

>MSTRG.3223.2 gene=MSTRG.3223

AGAAATTTAGACTCTTTCAGAAAATTTCCTCTGTATTTTTCTTATTACACACTAAAATGCAAATATATTA

GAAAAGAATATTGGGAAAGCTGAGATATATTTGAGAAGAAGAAACAAGCTAGGGAGAGAATCATGAAGAA

ATGTGAGTTGTGCAACTCCATTGCAAGAATGTTTTGTGAATCAGATCAAGCAAGCTTGTGTTGGCAGTGT

GATGCCAAGGTTCATTCTGCTAACTTTTTAGTGGCTAAGCACTCTAGAACTCTTCTTTGTCAATCTTGTC

AAGCTCTTACTCCATGGACTGGCTCTGGTCCTAAGCTTGGACCCACTCTTTCATTTTGTGAATCTTGTGT

TCATGAAAATACTAATTACAATAATGATACTGATGAGGATGATGAGGATGAAGAGGATGATGAAGATGAT

GAAGAAGATGAGGAGGAAGATGACGGTGAGAATCAAGTGGTTCCTCTTTCATCTTCATCTCTGGCTCCTC

CACAAGTTACAAGCTCTTCTAGCAGTGAAGAGTCTTCTGCTAGATTGTTTAATGGTGCAGAAAGTAGCAG

AAAATCAAGATTCTCCATCTCATGGAGTAGAACTAGATGATTTTTTAGAAAAGGTAATAAACAGTATCCT

TTTTTTTTATATAGTAGAATTTTAATTCGAATCAGCCTATTTCAGGGAAATGAAATGGAAGATGAAGAGG

AGTGATCCAATAACACTGCTGCTCACTTTAGCATGTTGCTTGATCAACTTCAACTTAATGTTAATTTGTA

ACATTACCTCTAACTGTGACCTCCTTGTAATTAAATCGGAGATTAGTAACGTGTTCTATATCATCGTACA

GTCGTGTGTTATATTTACCTCGTAGCATTGTTTCTCTCGTGTAC

>MSTRG.3223.3 gene=MSTRG.3223

GATATATGTAGCACTCAAGAACCCATGAAGAAATTTAGACTCTTTCAGAAAATTTCCTCTGTATTTTTCT

TATTACACACTAAAATGCAAATATATTAGAAAAGAATATTGGGAAAGCTGAGATATATTTGAGAAGAAGA

AACAAGCTAGGGAGAGAATCATGAAGAAATGTGAGTTGTGCAACTCCATTGCAAGAATGTTTTGTGAATC

AGATCAAGCAAGCTTGTGTTGGCAGTGTGATGCCAAGGTTCATTCTGCTAACTTTTTAGTGGCTAAGCAC

TCTAGAACTCTTCTTTGTCAATCTTGTCAAGCTCTTACTCCATGGACTGGCTCTGGTCCTAAGCTTGGAC

CCACTCTTTCATTTTGTGAATCTTGTGTTCATGAAAATACTAATTACAATAATGATACTGATGAGGATGA

TGAGGATGAAGAGGATGATGAAGATGATGAAGAAGATGAGGAGGAAGATGACGGTGAGAATCAAGTGGTT

CCTCTTTCATCTTCATCTCTGGCTCCTCCACAAGTTACAAGCTCTTCTAGCAGTGAAGAGTCTTCTGCTA

GATTGTTTAATGGTGCAGAAAGTAGCAGAAAATCAAGATTCTCCATCTCATGGAGTAGAACTAGATGATT

TTTTAGAAAAGAATAAACAGTATCCTTTTTTTTTATATAGTAGAATTTTAATTCGAATCAGCCTATTTCA

GGGAAATGAAATGGAAGATGAAGAGGAGTGATCCAATAACACTGCTGCTCACTTTAGCATGTTGCTTGAT

CAACTTCAACTTAATGTTAATTTGTAACATTACCTCTAACTGTGACCTCCTTGTAATTAAATCGGAGATT

AGTAACGTGTTCTATATCATCGTACAGTCGTGTGTTATATTTA

>MSTRG.3224.1 gene=MSTRG.3224

CATAAATCATTTTAATAAAATTTAACACATGTCATTTTATATTATTCAATTTTATGAACGAAATATTTAC

TTCGAAAAATTAGTGTATATCAACCAACATGTAAAGAAAAGGAGACAGAGGAGACAAATGAAAGCACACC

TTTTCTAAAAAATCATCTAGTTCTACTCCATGAGATGGAGAATCTTGATTTTCTGCTACTTTCTGCACCA

TTAAACAATCTAGCAGAAGACTCTTCACTGCTAGAAGAGCTTGTAACTTGTGGAGGAGCCAGAGATGAAG

ATGAAAGAGGAACCACTTGATTCTCACCGTCATCTTCCTCCTCATCTTCTTCATCATCTTCATCATCCTC

TTCATCCTCATCATCCTCATCAGTATCATTATTGTAATTAGTATTTTCATGAACACAAGATTCACAAAAT

GAAAGAGTGGGTCCAAGCTTAGGACCAGAGCCAGTCCATGGAGTAAGAGCTTGACAAGATTGACAAAGAA

GAGTTCTAGAGTGCTTAGCCACTAAAAAGTTAGCAGAATGAACCTTGGCATCACACTGCCAACACAAGCT

TGCTTGATCTGATTCACAAAACATTCTTGCAATGGAGTTGCACAACTCACATTTCTTCATGATTCTCTCC

CTAGCTTGTTTCTTCTTCTCAAATATATCTCAGCTTTCCCAATATTCTTTTCTAATATATTTGCATTTTA

GTGTGTAATAAGAAAAATACAGAGGAAATTTTCTGAAAGAGTCTAAATTTCTTCATGGGTTCTTGAGTGC

TACATATATCCT

>MSTRG.3229.1 gene=MSTRG.3229

GCCATCTCTATCACTTGCTTCTCCACATCCTACTGAACTTCCAAAAATGTACTCTGCAGCTGTTTTGCAC

AAATCAGTTTCCTTGGGCAGGTTGGACCCCCCTTTTGTGTTTCCATTTTCTTGAGCCAATTCATTTGTCT

TCTTGTTTTTCTTGGCCAAACAAGGCCAATAACCAAGTATACTTTCACAATAATTCATCACCC

>MSTRG.3233.1 gene=MSTRG.3233

CTTCTTTTCTCTAATACTCCCTCCGTCCCAGCCAATTCTTTACGTTTGGTTTGGGCACGGAGGTTAAGAA

ATATGTATAAAGTAGTGGAAAAGAGGAAGAAAAGTGGATGAAGTGGTGGGACCCATCAATTTTTAATGAA

TAAAAGAGAGATAAGGGAGTAAAAGTAGTGTGAAAAGGGAGGAAAAGTTGGGAAGTGGTGGGACCCATTG

ACTATTTTTGGTAAGTTTTGGAATGTAAAGAATTGGATGGGACATCCAAAAAAGGAAACTCTAAAGAACC

TG

>MSTRG.3237.1 gene=MSTRG.3237

GGGTGCTGGAGGGGGTCCAACCTTGATTGAAGTAGCTGCTGCAAGCGCGTCTTTTGCAAGAGAGGCAGAG

GCCAAACCAGCAGCAACAAGACCAAGCATGGCGCGGCGGCTTGTCTCAGGGTTCTCAGACGAAACCTGTT

GGGCACGAACAACGAGCCCTGCAGGTCGAGCCATGACAATTCTTTTGTTGCTGCTGGTGCTCAAGCGGTT

TGAGCCACTGATCTGCATGGTACCTTCCAAAACAGCCTGAGAAGAACCATGCAGACCAGTCATTGAAGCC

ATTGCTTGAGCCATGTATGATCTTTCTTATTGAAAAACAAAAGTTGTGGCAATTTTATATGATGGAGAG

>MSTRG.3251.1 gene=MSTRG.3251

CAACATGTACGTCACTCCAACCACTATTCCAAAACCTGCCGTGTAGCAACTTCGATTAATATCAGCCGTC

CACGTGTCAACATCACAACCATCGAAACAAACCACCTCTCCTTCTCCGTAGCCGTCTCGAAGACGCATTC

ACCCCTCTTTGCACTTATATATACAGCCCCTCATCTCCTCTTCTCCTCACAATTATCTTTCTCTTCTCAT

CATATACATACACTTTTCTGTCTCTATATTTTCATCAATTTCAAGATGCAAATCTTTGTGAAAACCCTCA

CCGGAAAGACCATCACCCTTGAGGTCGAAAGCTCCGACACAATCGACAACGTCAAGGCCAAAATCCAGGA

TAAGGAAGGCATTCCCCCAGACCAGCAGAGGTTGATCTTCGCTGGCAAGCAACTTGAAGACGGCAGGACC

CTTGCCGACTACAATATCCAGAAGGAGTCAACTCTCCATTTGGTCCTCCGTCTTCGTGGTGGTATGCAGA

TCTTCGTCAAGACTTTGACAGGGAAGACCATCACCCTTGAAGTCGAGAGCTCTGACACGATCGACAATGT

CAAGGCCAAGATCCAGGACAAGGAGGGTATTCCCCCAGACCAGCAGAGGTTGATCTTTGCTGGTAAGCAG

CTTGAGGATGGCAGGACCCTTGCCGACTACAACATCCAGAAGGAGTCAACTCTCCACTTGGTCCTCAGGC

TCCGTGGTGGTATGCAGATCTTCGTCAAGACCTTGACCGGAAAGACTATTACTCTTGAGGTTGAGAGCTC

TGACACCATTGACAATGTCAAGGCTAAGATTCAGGACAAGGAAGGGATCCCTCCAGACCAGCAGAGGTTG

ATTTTTGCTGGCAAGCAGTTGGAAGATGGCAGGACCCTAGCTGATTACAATATCCAGAAGGAATCAACCC

TTCATCTTGTGCTCCGTCTCCGTGGTGGTATGCAGATCTTTGTCAAGACTCTGACTGGAAAGACTATCAC

CTTGGAAGTGGAGAGCTCCGACACCATTGATAATGTGAAGGCCAAGATTCAGGATAAGGAAGGTATTCCT

CCAGATCAGCAGAGGTTGATCTTTGCTGGTAAGCAGCTTGAGGATGGAAGGACCCTTGCAGATTACAATA

TCCAGAAGGAGTCTACCCTTCACTTGGTGTTGCGTCTTCGTGGTGGTGATCTCTAAGCCTGTTGGAAGCG

CCCGTGGCTGTGGTTTCTGAAAAGAAGTTTGTGTTTCTGTTGAACTTGTTTGTTTTAAATCAAAGTTTGT

TTTAGTGTTTTCCCCTGTGGGGGAAAGAACAATGTGTTGTGGTGGCTTTAAGTCTTAATAAAAGATCAGT

TTAATAAGTATTTTGCTGTGGTCGTGTGATAGTTTCTGGTTTAAT

>MSTRG.3251.5 gene=MSTRG.3251

ATCAGCCGTCCACGTGTCAACATCACAACCATCGAAACAAACCACCTCTCCTTCTCCGTAGCCGTCTCGA

AGACGCATTCACCCCTCTTTGCACTTATATATACAGCCCCTCATCTCCTCTTCTCCTCACAATTATCTTT

CTCTTCTCATCATATACATACACTTTTCTGTCTCTATATTTTCATCAATTTCAAGATGCAAATCTTTGTG

AAAACCCTCACCGGAAAGACCATCACCCTTGAGGTCGAAAGCTCCGACACAATCGACAACGTCAAGGCCA

AAATCCAGGATAAGGAAGGCATTCCCCCAGACCAGCAGAGGTTGATCTTCGCTGGCAAGCAACTTGAAGA

CGGCAGGACCCTTGCCGACTACAATATCCAGAAGGAGTCAACTCTCCATTTGGTCCTCCGTCTTCGTGGT

GGTATGCAGATCTTCGTCAAGACTTTGACAGGGAAGACCATCACCCTTGAAGTCGAGAGCTCTGACACGA

TCGACAATGTCAAGGCCAAGATCCAGGACAAGGAGGGTATTCCCCCAGACCAGCAGAGGTTGATCTTTGC

TGGTAAGCAGCTTGAGGATGGCAGGACCCTTGCCGACTACAACATCCAGAAGGAGTCAACTCTCCACTTG

GTCCTCAGGCTCCGTGGTGGTATGCAGATCTTCGTCAAGACCTTGACAGGAAAGACAATCACCCTTGAGG

TCGAGAGCTCTGATACGATCGACAATGTCAAGGCCAAGATCCAGGACAAGGAAGGTATTCCCCCAGACCA

GCAGAGGTTGATTTTTGCTGGCAAGCAGTTGGAAGATGGCAGGACCCTAGCTGATTACAATATCCAGAAG

GAATCAACCCTTCATCTTGTGCTCCGTCTCCGTGGTGGTATGCAGATCTTTGTCAAGACCTTGACCGGAA

AGACTATTACTCTTGAGGTTGAGAGCTCTGACACCATTGACAATGTCAAGGCTAAGATTCAGGACAAGGA

AGGAATCCCACCAGATCAGCAGCGTTTGATCTTTGCTGGAAAGCAATTGGAAGACGGCAGGACTCTTGCT

GATTACAACATTCAGAAGGAGTCTACACTTCACCTTGTGCTTCGTCTGCGTGGTGGTATGCAGATCTTTG

TTAAGACTCTGACTGGAAAGACTATCACCTTGGAAGTGGAGAGCTCCGACACCATTGATAATGTGAAGGC

CAAGATTCAGGATAAGGAAGGTATTCCTCCAGATCAGCAGAGGTTGATCTTTGCTGGTAAGCAGCTTGAG

GATGGAAGGACCCTTGCAGATTACAATATCCAGAAGGAGTCTACCCTTCACTTGGTGTTGCGTCTTCGTG

GTGGTGATCTCTAAGCCTGTTGGAAGCGCCCGTGGCTGTGGTTTCTGAAAAGAAGTTTGTGTTTCTGTTG

AACTTGTTTGTTTTAAATCAAAGTTTGTTTTAGTGTTTTCCCCTGTGGGGGAAAGAACAATGTGTTGTGG

TGGCTTTAAGTCTTAATAAAAGATCAGTTTAATAAGTATTTTGCTGTGGTCGTGTGATAGTTTCTGGTTT

>MSTRG.3251.3 gene=MSTRG.3251

ATCAGCCGTCCACGTGTCAACATCACAACCATCGAAACAAACCACCTCTCCTTCTCCGTAGCCGTCTCGA

AGACGCATTCACCCCTCTTTGCACTTATATATACAGCCCCTCATCTCCTCTTCTCCTCACAATTATCTTT

CTCTTCTCATCATATACATACACTTTTCTGTCTCTATATTTTCATCAATTTCAAGATGCAAATCTTTGTG

AAAACCCTCACCGGAAAGACCATCACCCTTGAGGTCGAAAGCTCCGACACAATCGACAACGTCAAGGCCA

AAATCCAGGATAAGGAAGGCATTCCCCCAGACCAGCAGAGGTTGATCTTCGCTGGCAAGCAACTTGAAGA

CGGCAGGACCCTTGCCGACTACAATATCCAGAAGGAGTCAACTCTCCATTTGGTCCTCCGTCTTCGTGGT

GGTATGCAGATCTTCGTCAAGACTTTGACAGGGAAGACCATCACCCTTGAAGTCGAGAGCTCTGACACGA

TCGACAATGTCAAGGCCAAGATCCAGGACAAGGAGGGTATTCCCCCAGACCAGCAGAGGTTGATCTTTGC

TGGTAAGCAGCTTGAGGATGGCAGGACCCTTGCCGACTACAACATCCAGAAGGAGTCAACTCTCCACTTG

GTCCTCAGGCTCCGTGGTGGTATGCAGATCTTCGTCAAGACCTTGACCGGAAAGACTATTACTCTTGAGG

TTGAGAGCTCTGACACCATTGACAATGTCAAGGCTAAGATTCAGGACAAGGAAGGGATCCCTCCAGACCA

GCAGAGGTTGATTTTTGCTGGCAAGCAGTTGGAAGATGGCAGGACCCTAGCTGATTACAATATCCAGAAG

GAATCAACCCTTCATCTTGTGCTCCGTCTCCGTGGTGGTATGCAGATCTTTGTCAAGACCTTGACCGGAA

AGACTATTACTCTTGAGGTTGAGAGCTCTGACACCATTGACAATGTCAAGGCTAAGATTCAGGACAAGGA

AGGAATCCCACCAGATCAGCAGCGTTTGATCTTTGCTGGAAAGCAATTGGAAGACGGCAGGACTCTTGCT

GATTACAACATTCAGAAGGAGTCTACACTTCACCTTGTGCTTCGTCTGCGTGGTGGTATGCAGATCTTTG

TTAAGACTCTGACTGGAAAGACTATCACCTTGGAAGTGGAGAGCTCCGACACCATTGATAATGTGAAGGC

CAAGATTCAGGATAAGGAAGGTATTCCTCCAGATCAGCAGAGGTTGATCTTTGCTGGTAAGCAGCTTGAG

GATGGAAGGACCCTTGCAGATTACAATATCCAGAAGGAGTCTACCCTTCACTTGGTGTTGCGTCTTCGTG

GTGGTGATCTCTAAGCCTGTTGGAAGCGCCCGTGGCTGTGGTTTCTGAAAAGAAGTTTGTGTTTCTGTTG

AACTTGTTTGTTTTAAATCAAAGTTTGTTTTAGTGTTTTCCCCTGTGGGGGAAAGAACAATGTGTTGTGG

TGGCTTTAAGTCTTAATAAAAGATCAGTTTAATAAGTATTTTGCTGTGGTCGTGTGATAGTTTCTGGTTT

AAT

>MSTRG.3251.4 gene=MSTRG.3251

ATCAGCCGTCCACGTGTCAACATCACAACCATCGAAACAAACCACCTCTCCTTCTCCGTAGCCGTCTCGA

AGACGCATTCACCCCTCTTTGCACTTATATATACAGCCCCTCATCTCCTCTTCTCCTCACAATTATCTTT

CTCTTCTCATCATATACATACACTTTTCTGTCTCTATATTTTCATCAATTTCAAGATGCAAATCTTTGTG

AAAACCCTCACCGGAAAGACCATCACCCTTGAGGTCGAAAGCTCCGACACAATCGACAACGTCAAGGCCA

AAATCCAGGATAAGGAAGGCATTCCCCCAGACCAGCAGAGGTTGATCTTCGCTGGCAAGCAACTTGAAGA

CGGCAGGACCCTTGCCGACTACAATATCCAGAAGGAGTCAACTCTCCATTTGGTCCTCCGTCTTCGTGGT

GGTATGCAGATCTTCGTCAAGACTTTGACAGGGAAGACCATCACCCTTGAAGTCGAGAGCTCTGACACGA

TCGACAATGTCAAGGCCAAGATCCAGGACAAGGAGGGTATTCCCCCAGACCAGCAGAGGTTGATCTTTGC

TGGTAAGCAGCTTGAGGATGGCAGGACCCTTGCCGACTACAACATCCAGAAGGAGTCAACTCTCCACTTG

GTCCTCAGGCTCCGTGGTGGTATGCAGATCTTCGTCAAGACCTTGACAGGAAAGACAATCACCCTTGAGG

TCGAGAGCTCTGATACGATCGACAATGTCAAGGCCAAGATCCAGGACAAGGAAGGAATCCCACCAGATCA

GCAGCGTTTGATCTTTGCTGGAAAGCAATTGGAAGACGGCAGGACTCTTGCTGATTACAACATTCAGAAG

GAGTCTACACTTCACCTTGTGCTTCGTCTGCGTGGTGGTATGCAGATCTTTGTTAAGACTCTGACTGGAA

AGACTATCACCTTGGAAGTGGAGAGCTCCGACACCATTGATAATGTGAAGGCCAAGATTCAGGATAAGGA

AGGTATTCCTCCAGATCAGCAGAGGTTGATCTTTGCTGGTAAGCAGCTTGAGGATGGAAGGACCCTTGCA

GATTACAATATCCAGAAGGAGTCTACCCTTCACTTGGTGTTGCGTCTTCGTGGTGGTGATCTCTAAGCCT

GTTGGAAGCGCCCGTGGCTGTGGTTTCTGAAAAGAAGTTTGTGTTTCTGTTGAACTTGTTTGTTTTAAAT

CAAAGTTTGTTTTAGTGTTTTCCCCTGTGGGGGAAAGAACAATGTGTTGTGGTGGCTTTAAGTCTTAATA

AAAGATCAGTTTAATAAGTATTTTGCTGTGGTCGTGTGATAGTTTCTGGTTTAAT

>MSTRG.3251.2 gene=MSTRG.3251

ATCAGCCGTCCACGTGTCAACATCACAACCATCGAAACAAACCACCTCTCCTTCTCCGTAGCCGTCTCGA

AGACGCATTCACCCCTCTTTGCACTTATATATACAGCCCCTCATCTCCTCTTCTCCTCACAATTATCTTT

CTCTTCTCATCATATACATACACTTTTCTGTCTCTATATTTTCATCAATTTCAAGATGCAAATCTTTGTG

AAAACCCTCACCGGAAAGACCATCACCCTTGAGGTCGAAAGCTCCGACACAATCGACAACGTCAAGGCCA

AAATCCAGGATAAGGAAGGCATTCCCCCAGACCAGCAGAGGTTGATCTTCGCTGGCAAGCAACTTGAAGA

CGGCAGGACCCTTGCCGACTACAATATCCAGAAGGAGTCAACTCTCCATTTGGTCCTCCGTCTTCGTGGT

GGTATGCAGATCTTCGTCAAGACTTTGACAGGGAAGACCATCACCCTTGAAGTCGAGAGCTCTGACACGA

TCGACAATGTCAAGGCCAAGATCCAGGACAAGGAGGGTATTCCCCCAGACCAGCAGAGGTTGATCTTTGC

TGGTAAGCAGCTTGAGGATGGCAGGACCCTTGCCGACTACAACATCCAGAAGGAGTCAACTCTCCACTTG

GTCCTCAGGCTCCGTGGTGGTATGCAGATCTTCGTCAAGACCTTGACAGGAAAGACAATCACCCTTGAGG

TCGAGAGCTCTGATACGATCGACAATGTCAAGGCCAAGATCCAGGACAAGGAAGGTATTCCCCCAGACCA

GCAGAGGTTGATCTTTGCTGGTAAGCAGCTGGAGGATGGAAGGACCCTTGCTGACTATAACATCCAGAAG

GAGTCCACTCTCCACTTGGTCCTCAGGCTCCGTGGTGGTATGCAGATCTTTGTCAAGACCTTGACCGGAA

AGACTATTACTCTTGAGGTTGAGAGCTCTGACACCATTGACAATGTCAAGGCTAAGATTCAGGACAAGGA

AGGGATCCCTCCAGACCAGCAGAGGTTGATTTTTGCTGGCAAGCAGTTGGAAGATGGCAGGACCCTAGCT

GATTACAATATCCAGAAGGAATCAACCCTTCATCTTGTGCTCCGTCTCCGTGGTGGTATGCAGATCTTTG

TCAAGACCTTGACCGGAAAGACTATTACTCTTGAGGTTGAGAGCTCTGACACCATTGACAATGTCAAGGC

TAAGATTCAGGACAAGGAAGGAATCCCACCAGATCAGCAGCGTTTGATCTTTGCTGGAAAGCAATTGGAA

GACGGCAGGACTCTTGCTGATTACAACATTCAGAAGGAGTCTACACTTCACCTTGTGCTTCGTCTGCGTG

GTGGTATGCAGATCTTTGTTAAGACTCTGACTGGAAAGACTATCACCTTGGAAGTGGAGAGCTCCGACAC

CATTGATAATGTGAAGGCCAAGATTCAGGATAAGGAAGGTATTCCTCCAGATCAGCAGAGGTTGATCTTT

GCTGGTAAGCAGCTTGAGGATGGAAGGACCCTTGCAGATTACAATATCCAGAAGGAGTCTACCCTTCACT

TGGTGTTGCGTCTTCGTGGTGGTGATCTCTAAGCCTGTTGGAAGCGCCCGTGGCTGTGGTTTCTGAAAAG

AAGTTTGTGTTTCTGTTGAACTTGTTTGTTTTAAATCAAAGTTTGTTTTAGTGTTTTCCCCTGTGGGGGA

AAGAACAATGTGTTGTGGTGGCTTTAAGTCTTAATAAAAGATCAGTTTAATAAGTATTTTGCTGTGGTCG

TGTGATAGTTTCTGGTTTAATCTTGTTTTATTTTAAAA

>MSTRG.3252.1 gene=MSTRG.3252

CTCTGCTGGTCTGGGGGAATACCTTCCTTGTCCTGGATCTTGGCCTTGACATTGTCGATCGTATCAGAGC

TCTCGACCTCAAGGGTGATTGTCTTTCCTGTCAAGGTCTTGACGAAGATCTGCATACCACCACGGAGCCT

GAGGACCAAGTGGAGAGTTGACTCCTTCTGGATGTTGTAGTCGGCAAGGGTCCTGCCATCCTCAAGCTGC

TTACCAGCAAAGATCAACCTCTGCTGGTCTGGGGGAATACCCTCCTTGTCCTGGATCTTGGCCTTGACAT

TGTCGATCGTGTCAGAGCTCTCGACTTCAAGGGTGATGGTCTTCCCTGTCAAAGTCTTGACGAAGATCTG

CATACCACCACGAAGACGGAGGACCAAATGGAGAGTTGACTCCTTCTGGATATTGTAGTCGGCAAGGGTC

CTGCCGTCTTCAAGTTGCTTGCCAGCGAAGATCAACCTCTGCTGGTCTGGGGGAATGCCTTCCTTATCCT

GGATTTTGGCCTTGACGTTGTCGATTGTGTCGGAGCTTTCGACCTCAAGGGTGATGGTCTTTCCGGTGAG

GGTTTTCACAAAGATTTGCATCTTGAAATTGATGAAAAT

>MSTRG.3253.1 gene=MSTRG.3253

CTTCCAACAGGCTTAGAGATCACCACCACGAAGACGCAACACCAAGTGAAGGGTAGACTCCTTCTGGATA

TTGTAATCTGCAAGGGTCCTTCCATCCTCAAGCTGCTTACCAGCAAAGATCAACCTCTGCTGATCTGGAG

GAATACCTTCCTTATCCTGAATCTTGGCCTTCACATTATCAATGGTGTCGGAGCTCTCCACTTCCAAGGT

GATAGTCTTTCCAGTCAGAGTCTTAACAAAGATCTGCATACCACCACGCAGACGAAGCACAAGGTGAAGT

GTAGACTCCTTCTGAATGTTGTAATCAGCAAGAGTCCTGCCGTCTTCCAATTGCTTTCCAGCAAAGATCA

AACGCTGCTGATCTGGTGGGATTCCTTCCTTGTCCTGAATCTTAGCCTTGACATTGTCAATGGTGTCAGA

GCTCTCAACCTCAAG

>MSTRG.3256.1 gene=MSTRG.3256

ATAACATTCACGTACTACTATTAGATTGATCAAGCTTCCACAATCTCATCCGCAGCATCTGCAATTACAA

ACTCTGGTTGGCTCTCTTCCAGAGCATTTGGTAGACCTAATGCCCTCAATGCCAGATGTGCTGCTTTCTG

TCCTGACATCATCATCGCCCCAAATGTCGGTCCCATTCTTGGGGATCCATCAATCTCAGCCACTTCCATG

CCAGTAACAATCATCCCGGGCACAATTTCTCTAGTAAGCCTAACAATCGCGTCCTCAGCAGCATTCATAT

CAAGAGCCTTCATTCCAGGAACGCTCTCAATCATCCCGATACTCCTCAACCTCTTGACACCAGTGGCTCC

AAAAGGACCATCATGCCCACACGAGCTAACCACAATCTTAGCTTCCATGACATTCGGGTCCATGCAAGAT

TGTGTGTCATGGTTCATGGACACAAGAGCCCAATTTGTCACAACTCCAGCAACTCTGTCTCCCTTGACAA

TCAAATCCTCAGCCGCCACTGCATTGAACAGCTTCACATTAGGCCTAGCCAAAAGCTTGCTCATAATAGT

CGACGTAAAAAGTGCTGCATGCTTGATGACCACGTAGTCATCTTGCTCATCGTACTCCACCTCGAGCTCG

TCAAGGAACTTGTGAGCTGGTTTACGAACAACCATAGCTGAGAAAAGCTGTCCACCTAGCCACGCACCGC

CACCAGGGCTCACGGATTGTTCAATAATTGCTATGTTGATGTCAGGGTTCTTGCTGAGCTCATAGGCACA

TGATAGGCCAGCTGATCCTGCACCTACAACAACAACATCAGTGTCTGCATACGTGATCATGTCGGTCATG

TATCTCCGTGTCATCTCTCGGGACACGATGGACTCCTTGATAGGGTTGAATTTGAAGTTGTCTAGGTCAT

AGTTAGGAGTTGCAACCGCAGAGATGGTAGATGAGTGGTTCTTAGGGGCAGATTTGATGGATTGCACCCG

GGAGAACACGGGTGAGCCATTGAAGGAGGTTTTGTTATCGAAGATTTTCGAGGTGAGTGAAGAGGCGAGA

GTTGCAGCCATGGTTGCCATGAAGATAATATCAAAGGGAAAGTTTAGTAACAATACTTGTTAGTGAGTT

>MSTRG.3268.1 gene=MSTRG.3268

ATTTTTTTCATCAGTCTTGACTCTCTCCCTCACCATTCTATCCCGTCTCTCCTCTCTCGATCACCCCTCC

TTTCTTCTCTACTTTTCTGCTGTGTTACAAGCCACCGCGACGCCGCCCTCTGCCGCCGCGATTCTCCGGC

TAGTGTTGTCGCGGTTTCAATCAAGCCGCCTGTGTATTTCCGGCGAGCTCTCCACCCCGCACCTCCAGTT

CAATTAGTGAATTGAAGACATTGACCGAGGCATTGATCGAGAATCGATTGCAATGGGTATTCAGAAATAG

CCAAGTAGTGTTGCAAGTTGAAGTAGAGCGAAGATTTGTTATAGATAAAAGCTTGTCAGAAGTTGACAAA

AAGGAATGATATGGCGGTATATAGTATTATTAGAAATCTGAGCTAGACACTGGAGTTAAGCCAAGGACAA

AGATGTTTCACCCGATTAGGCTGCGTGTGCGTGTAAGAGTAATGATAGGCAAGGATTGCAAGTGGTTGAG

TTTCAGGCATATGGTCAGGTAGAGTATCCAGAAAGATGTCATTAAGACTAGTTGCAGTTGCAGTTGTGCA

ATTTAATTCAGTTGGGAATTAGATTTGTTTATTTGGTTATCAACTACTCTTCATATCGAGGTTTAGTTGA

TAATTTAGTTGAAGTTGTATAAAATTTGTAGTTTTGTTTTCTGATTTCAATACTATAGCCTGTATCGATC

CTGTTTTCAGGGGG

>MSTRG.3268.2 gene=MSTRG.3268

CTTGACTCTCTCCCTCACCATTCTATCCCGTCTCTCCTCTCTCGATCACCCCTCCTTTCTTCTCTACTTT

TCTGCTGTGTTACAAGCCACCGCGACGCCGCCCTCTGCCGCCGCGATTCTCCGGCTAGTGTTGTCGCGGT

TTCAATCAAGCCGCCTGTGTATTTCCGGCGAGCTCTCCACCCCGCACCTCCAGTTCAATTAGTGAATTGA

AGACATTGACCGAGGCATTGATCGAGAATCGATTGCAATGGGTATTCAGAAATAGCCAAGTAGTGTTGCA

AGTTGAAGTAGAGCGAAGATTTGTTATAGATAAAAGCTTGTCAGAAGTTGACAAAAAGGAATGATATGGC

GGTATATAGTATTATTAGAAATCTGAGCTAGACACTGGAGTTAAGGCAAGTACTCTAAC

>MSTRG.3268.3 gene=MSTRG.3268

GAGCATTTTTGGCTCATTCTTGCTTTTCAACCCTAACCTTTCAGCCAAGGACAAAGATGTTTCACCCGAT

TAGGCTGCGTGTAAGAGTAATGATAGGCAAGGATTGCAAGTGGTTGAGTTTCAGGCATATGGTCAGGTAG

AGTATCCAGAAAGATGTCATTAAGACTAGTTGCAGTTGCAGTTGTGCAATTTAATTCAGTTGGGAATTAG

ATTTGTTTATTTGGTTATCAACTACTCTTCATATCGAGGTTTAGTTGATAATTTAGTTGAAGTTGTATAA

AATTTGTAGTTTTGTTTTCTGATTTCAATACTATAGCCTGTATCGATCCTGTTTTCAGGGGGTGAAATTG

ATTTGTAATTTAA

>MSTRG.3268.4 gene=MSTRG.3268

GTAAAATAGTTAAAGCTGGTATAAGATTAGTGATTAGCCAGCTCGTTAAGGACACGTGACAGCACAAGGC

TCAGCCACGTGCCCCTATTTTCGCCACCGCAAACCCTCTTTTCTTTTCTTCTCCATACTTTTCTTTCACT

CCATTTTTTTCATCAGTCTTGACTCTCTCCCTCACCATTCTATCCCGTCTCTCCTCTCTCGATCACCCCT

CCTTTCTTCTCTACTTTTCTGCTGTGTTACAAGCCACCGCGACGCCGCCCTCTGCCGCCGCGATTCTCCG

GCTAGTGTTGTCGCGGTTTCAATCAAGCCGCCTGTGTATTTCCGGCGAGCTCTCCACCCCGCACCTCCAG

TTCAATTAGTGAATTGAAGACATTGACCGAGGCATTGATCGAGAATCGATTGCAATGGGTATTCAGAAAT

AGCCAAGTAGTGTTGCAAGTTGAAGTAGAGCGAAGATTTGTTATAGATAAAAGCTTGTCAGAAGTTGACA

AAAAGGAATGATATGGCGGTATATAGTATTATTAGAAATCTGAGCTAGACACTGGAGTTAAGCCAAGGAC

AAAGATGTTTCACCCGATTAGGCTGCGTGTGCGTGTAAGAGTAATGATAGGCAAGGATTGCAAGTGGTTG

AGTTTCAGGCATATGGTCAGGTAGAGTATCCAGAAAGATGTCATTAAGACTAGTTGCAGTTGCAGTTGTG

CAATTTAATTCAGTTGGGAATTAGATTTGTTTATTTGGTTATCAACTACTCTTCATATCGAGGTTTAGTT

GATAATTTAGTTGAAGTTGTATAAAATTTGTAGTTTTGTTTTCTGATTTCAATACTATAGCCTGTATCGA

TCCTGTTTTCAGGGGG

>MSTRG.3273.1 gene=MSTRG.3273

GTTTGACTCCAAACCGCGTCTCGAACCCGGCTCATATCCCTACCTTTAAGTGTTCGCCCCACGCCTTCGT

AAACCGAGTTGGCAGCCGAGTTGCGACTCCGAGCCAAGTACTCGTGTGGAGGAACCATATCAAAGTCACG

CTCCTCCCAATCGTCCGAGTCATGCATGGACTCAACCGACTCAACTCGGTAAATCTTTGACCAATCCGGC

ACGTTGATGGGCGCTGACGTGGCCATCTGGCGTCCACGTGGCGACGCCACGTAATCCTGACCACGGATTT

GGTGCACGATCCGCGGAGAGGAAACCCGGCCCGGATCATCAAAGGCCAATGACAACCCGCCCACGTGGCC

TTCCTCATTTCCACCTCCCCGGTGACTCCTCCTACTCACGCGCACATTTGTCTCCACGTACGCACGTGGA

GTCCACTCAGCGTTGGAACTATGATACGTGTGCTCCTCGCTGTCGACACTAACCGTGTCGACAGCGGACC

ACACGTCATCTTCCGCAAACTCCAATTCGCCGTTGACCGGACCCTGAGTTTGACGATAATTGTAGCTCCC

TAACAAACGCTCGCTTCGACTTATGGTCAGTCCTCGACCTTTGGCCATGTATCATCACAACCACCGGCCG

AAATCCGAATGAATCTTAAATTCAAGCTACCTAGTAAAAAATATTGTTTATGTGTACAAGTACATATGTT

ATATGTATGTTGTGTATGAATATTAATGTGTGTAAATGTGAGCTAGCTGATGCTATAACTCCATTAGAAA

AAAACATA

>MSTRG.3275.1 gene=MSTRG.3275

GTCACCATCTTGGATTACCATGCACACATGGCAGGCAAAAGTAGAGGTTTTGGCTTCTTTTCATATACAA

AGCTCATATGCTAGCAACACTTGCTTAGCTAAGACTACATTCAATGATGCTTGTTAGCAACATTGCATTG

CAACCTAATCTCACCCTCAGTGCCAGTGAGAGGATTGTTCTCAGTGAGCTTAGTGATGGCTCTTGAGTAC

TCCTTGAAGAAATAATCTTGGCTCTTGGCCATTTTCTTGACATATGGCTTGGTCCTCTTGTCTGTGGCTA

GTTGATGATCCACTATCAAGAGTCCCTTGTTGTCCAATATGTTTCTGTAGTAGTTGTTGTCTAGAATCAT

GGGGGTGCCTCTGTCATTTCTCACATATTGCACTGCCTTTGGGTCTGGGATCTGGTCAGGGCACTTGTGA

AGCATGTGCTCAACATGACCAGGGTTCAAGGCTGGGTCAACCTCAGGGTATAATCTGTGCACCAGCTTCA

CACAATGGGTTCTGCCAACACTGTGAGATCCTAGCAAGGCAACAAGTCCAGGGGTGTCAATTCCCATGGC

AGCAAACCTTTCAAGAACAACAGTCATGCTTTCATTGTGGTCAGGGAGGTATTGCTCTAGTATATCTGCT

CTACTTTTTCTACCATCTCTTCTTCCTGTTTTAAGAGGGATGTGTGGGCCTCCAAGAGCAACAATTCCAT

CTCTACCAGATAAAACAAGGATATCAGCACAGGAAACCACTCCGGGGCACTCTCTCTCCACAGCTTCTTT

AATAGTCTCAATATATCTGAAGTTTCTGAGGCCAAAGCTCCTGTCTGTTTCTTTCTCAGAAAGAGTCCTC

CTTGTTGAGTCCAGCAACAATGAAGCATCACATCTCTGAACAGCACAGTCATGGAAGATGTTTCTGAGCC

AAGAGAAAGCAGTGTTCTTGTGGCGCTTGTAGAGAAGCTGGACTTGTTCTTTGATGATGTCTTCAGCCTG

TGGGCATGAGTCTTTGTAAAAGTTCATGACAAGGCCTGGACCAGCTGAAGCTGTCTCTTCAGCTGCATCT

GCAAAAGCAGAGATTGCAGACAAGGATAAGAGAGCAATGAAGAAGAGAGTTTTGGAACCCATTTTGTGTG

AGATGTTATGAGTTAGCTTGCTCACTCCCAGCTACTTAAATAAGACAAGTGAGTACCTCAGCGGTGAAGA

AAAAGAGGATATGGTCAAGTCGGTCAGAAATAC

>MSTRG.3279.1 gene=MSTRG.3279

ATTAGAAACCACTGACTCTGATCAAGTTTCACAATTTTATATATAAGCAGAACTGAACGAGCATCGAAAG

AAATTATACTTGTTATATTAATGCATCTGCTCCAGCAACAATCTCCAAAATCTCTCCAGTAATCTTGGCC

TGGCGCTTCCGATTGTACACCTGAGTGTAATTCTTCTTCAGCTCCTTGGCATTGTCTGTGGCACTGCTCA

TTGCTGTCATTCTTGCAGCAAGCTCACTAGCCAGTGATTCTTGCAATGATCTCAATATTTGACTGTTGAG

GTAAAGTGGCAGCAATGCATCAAGAATCTGAACAGGGTCTTGCTCGAATTGCAGAACAGGTGAGAAATCG

ATGGTTTCAGTCCTGACCACGTCGCGTTCCACTGTCAATTTCCCTTCTTTTGTTGTTAGCCTGAAGAATT

CATCGTCTGCTGCATCGACACATGTTCCATTTATGTCACAAATTTCTCCCTTTGGTGATAATGGGAGTAG

TGTGTGGATCACCGGACTAGACTTTACCAACGACACGAATTTCGTGTACAAAAGCTCTACTTTATCAACT

TCCTCACTCACAAAAAGTGAAAACACATCATCTGCAATTGCTTGTGCTTCTTTTGCGGTTGGAAGATTAC

CCCCTTCAAGAAATTTGTCTACTGGTATATAAGGCCTTCGCAGAAAATACGAATTTCCC

>MSTRG.3280.1 gene=MSTRG.3280

ATTTTAGTTTCTCTGCTGAATAATTTGGGAGGTCTTATAATTCTGATCCCCCGAGGGCTTCTTGCCCACC

CTCAGAAACGCAGATTTCGTCTTCATTTTTCCTCCCGTGATATAGATTTGTAGTCCTTGTTTCTTGAAGT

AAATCAGGCAGCTAGAGTTGTATATAGATTCTACGTGAACTGCAATTGGAACTTGGGATTTTTAGTTCTG

AAAAATATATTTTTTTCGAGAACCAGAAGAGCTTCTAACTGCCAGGCTGAAGGCTCCACCTCAAAGGGAT

TTGGTCGAGAACCTCAGGTTTTAACAAGTAGACAAACACAATTTTTGAGAAGACAATGAATGAGAGAGTT

TTTGAGGCTAAGCTTAACGCTGAGGCGAAGGTTCTCATATTGATGATATTTGGGAAAGGTGAAAAAATCA

TCTCATGTTGCTTGTCTGGCCAAATCAACAAAAATAGAAGGATCAATTCACTTGATGAGGTGAGAAGCGA

GTTATCATTTATAGGAACCATAGCAAAGAGTGATGAGTCCGTTCAGGTGTTTGATAGAGCAGGCCTTGAA

TTTAAGTAGATGTTTGAGACAAATGGTATGGCAGAAACGACTTCCCCAATAGTGATGATAATTTATTTAA

GTGCCGTGAAAAGGAATTGTTCTCTTCATATAAGAGAACAATGTGTTCAGCTATATCCCACAGAGGTAAA

ACTCAATCAATTGAAGCGTTTTTAGTTACATTAGACACGGAATCAGATCTCATAAACATCAGCGCATTTA

TTGGCAAGAAGTCAAGAACAATGTGTTCAGCTATATCCCGCGGAGGTGAAATTTCAACGGAACTCATCTC

AATTGGGATCATTTGTCTTGCCAGGTGGAATTGATATGCTTGAATTTGAGGACTTATTTAGTGAGGAGAA

ACAGGGAATTCGAGCAAGCTAGAAAGTGAAGGCATGGGAAAGAACAGAGACGAATAGAAGCAGAGAAGGT

GGCACGTGAAGCGGACAGAGTACGTGCAAGAGGGAACCGCGAAAGGAGAATCAAAATGTTGCAAGGAGTC

AAATAGCTGAAAGAGTTCAAAGGGCTTTGACAGCTCCGACAGAAAGGATGGTGACTTGTCATATACTGAT

GTTATGCATTATTGATTTATTGCTGACGTGAAACTGAAATCAAGGAAGGGGGCACTCTAGGAGCATATTT

TGTACTGCCCCCTTTGGAGTTCAAACTTTGTGAACCTGCTGGATTCTCCGCAGTGAGAGATTGTTTCGTA

GTCCCTTATGGATAGGGCATTGATGTAGGTGTGTTGTTTTTGGATAACAGAGCTTTTATTGCTTTTATTG

TTTGCAAGTGCATGCAGAGATCCCCTCTTTCAGGAAAATGAGCATGACTGTGGCCCAAAAATTTACCCAT

ATCTGCAAGCATTTCACAAGGAAGCTGTTTTTATAAGAAGCTATTCATTGTCATCTCTGGCTCTTCTACA

GACACAGCTACACTAGTAGGCTTCTCTTGTGACTGATATTGATGAAATTTGCTTTATCGTGAATGGCATT

GAGGTGACCAACAATAATAATAGTGCAAGTCACAATTTTTGAGAGATGAGCTTGTGTGGTGGAAGTTGGG

TGTTAATTGGTGGGATTTGGGGACCATATTTTTTTTGATGGACAACCTGAGGATCAAAATTTAAAGACCC

TACCCATCTGTAACCTCAAAGATTAAGATTAAGGAAAGAAAAAAGAACTATAGAATGTTAAGAGTTAATT

TACTTAGAAATTGATTTTAACCAAAAAATCTGAATCTGATTTGCATTTTGGGTTCCAGACTCTCTGTGGT

ATTAGATACTATGTAATATTCATTGCAGGAACAATGAACAGAAACAGGAGATTTGGAATAATAAAATTAC

AGTGAAACAAAATATTAATCAGATCACAGTTCATTTTACTACAACAGAGATCTGCTATTACAGCAGTAAA

TATATTTTTGTTACTTGTTAG

>MSTRG.3280.2 gene=MSTRG.3280

AGGAGCATATTTTGTACTGCCCCCTTTGGAGTTCAAACTTTGTGAACCTGCTGGATTCTCCGCAGTGAGA

GATTGTTTCGTAGAGATCCCCTCTTTCAGGAAAATGAGCATGACTGTGGCCCAAAAATTTACCCATATCT

GCAAGCATTTCACAAGGAAGCTGTTTTTATAAGAAGCTATTCATTGTCATCTCTGGCTCTTCTACAGACA

CAGCTACACTAGTAGGCTTCTCTTGTGACTGATATTGATGAAATTTGCTTTATCGTGAATGGCATTGAGG

TGACCAACAATAATAATAGTGCAAGTCACAATTTTTGAGAGATGAGCTTGTGTGGTGGAAGTTGGGTGTT

AATTGGTGGGATTTGGGGACCATATTTTTTTTGATGGACAACCTGAGGATCAAAATTTAAAGACCCTACC

CATCTGTAACCTCAAAGATTAAGATTAAGGAAAGAAAAAAGAACTATAGAATGTTAAGAGTTAATTTACT

TAGAAATTGATTTTAACCAAAAAATCTGAATCTGATTTGCATTTTGGGTTCCAGACTCTCTGTGGTATTA

GATACTATGTAATATTCATTGCAGGAACAATGAACAGAAACAGGAGATTTGGAATAATAAAATT

>MSTRG.3281.1 gene=MSTRG.3281

CGAACCTGAAAGGCCGCCTTGTTGAAGCCACAAGCGCCTTTCCTCTCCGATATTCTCTCTTTCCTATATA

TTCACAACACAAACACACACACTCCATAAATCATCACTCATTTCTTACATATAAATCCTCAACCAAGCTT

CAAGATCATACAAACCTATACACACAGATACATACATTAACATACAATGGCCTTTGCTTCTTCAACTTCT

ATATCAACTTCTTTTTCTTCTTTCGAACAAACAAAAGTTTCGCAGATAGGGTTATATCAGCCGTTGGATC

GGGCTCATTCTTTATCGACGGTTTTAAATGTATCTAGAAAACGAGTGGTGGTTAAGGCGCTTAATGCTGA

GCCGAAGAGAAGTGAGTCAGTTGTTGTCTCTGCTGCAACTGTAGTGGCTCCTGAAGTGGTGGAGAAAGTG

GAAGTGGAGGTAGAGGATTATGAGAAACTGGCGAAGGAGCTTGAAAATGCATCACCACTTGAAATTATGG

ACAAGGCCCTAGAGAAGTTTGGGAATGATATTGCAATTGCTTTCAGTGGTGCTGAAGATGTTGCTTTGAT

TGAGTATGCTCATTTGACTGGTCGTCCATTCCGAGTGTTTAGCCTGGACACTGGAAGACTTAATCCTGAA

ACATACAAATTATTTGATGCTGTTGAGAAGCAGTATGGCATTCACATCGAGTACATGTTTCCAGATGCTG

TTGAAGTTCAGGACTTGGTAAGGACCAAGGGTTTGTTCTCTTTCTATGAAGATGGTCATCAGGAGTGCTG

CCGTGTGAGGAAAGTTAGACCTTTGAGGAGGGCTCTTAAAGGCTTGCGAGCATGGGTTACTGGACAAAGA

AAAGACCAGTCACCAGGAACTAGATCTGAAATCCCAGTTGTCCAAGTTGATCCTGTTTTTGAAGGGCTCG

AGGGTGGACCTGGTAGTTTGGTGAAATGGAACCCAGTGGCAAACGTCCAAGGTACTGACATATGGAAATT

TCTCCGTACCATGAATGTGCCTGTCAACTCATTGCACGCACAGGGATACATCTCTATTGGTTGCGAGCCC

TGCACAAGGCCAGTTCTGCCTGGTCAACACGAGAGAGAAGGAAGGTGGTGGTGGGAGGATGCTACTGCCA

AGGAATGTGGACTACATAAAGGCAATATCAAGGAAGAGAATGTAAATGGCAATGGAACTGGAGCAGTAAA

AGCCAACGGCGAAGCCACTGTCACTGACATTTTCAACAGCCAGCACGTGGTGAGCTTGAGCAGGCAAGGA

ATTGAAAATTTGATCAAACTAGAAGATAGAAAAGAGACATGGCTTGTTGTTCTTTACGCACCCTGGTGCC

AATTTTGCCAGGCAATGGAAGGATCATACATGGAATTGGCTGAGAAGTTAGCTGGAACAGGAGTGAAAGT

TGCAAAGTTCCAGGCAGATGGGGACCAAAAGGCGTATGCGCAGAAAGAGTTACAACTGGGAAGTTTCCCC

ACTATACTTTTCTATCCTAAACATTCTTCTCGTGCTGTGAAGTACCCGTCTGAAAAGAGGGATGTTGAGT

CCTTGTTAGCATTCGTGAATGCTCTTAGATGATCAACACCAAACTCTGCCGATATAGTATCAGGTCTCTG

CTGATTTATACAAGTTGAAGTAGCAGCTGTTATGGAGGAGTCCACCTCGTGATGAATGTATCATTGCCGG

AATCAACTCTGAGATGGGATGACCCTGTATATTTTCACAAAAATATATTTCTATGTGGGTTTGAGGGCAA

TGATTGTTGCTTTTGTACTCAGTTTTGCCTTTCAGTTCTAGTTCAGACATTTTGTGAGTTTTTCTATATG

ATGAAACTCAATGTTGCATTGCTCCTGATAGTGGAGATTATTCTTCAACTTCTTTTTACCAAGATTTATA

TCAAAGTGTGTTATGATCTTTTAGCAGATAAAA

>MSTRG.3281.2 gene=MSTRG.3281

TGAAGCCACAAGCGCCTTTCCTCTCCGATATTCTCTCTTTCCTATATATTCACAACACAAACACACACAC

TCCATAAATCATCACTCATTTCTTACATATAAATCCTCAACCAAGCTTCAAGATCATACAAACCTATACA

CACAGATACATACATTAACATACAATGGCCTTTGCTTCTTCAACTTCTATATCAACTTCTTTTTCTTCTT

TCGAACAAACAAAAGTTTCGCAGATAGGGTTATATCAGCCGTTGGATCGGGCTCATTCTTTATCGACGGT

TTTAAATGTATCTAGAAAACGAGTGGTGGTTAAGGCGCTTAATGCTGAGCCGAAGAGAAGTGAGTCAGTT

GTTGTCTCTGCTGCAACTGTAGTGGCTCCTGAAGTGGTGGAGAAAGTGGAAGTGGAGGTAGAGGATTATG

AGAAACTGGCGAAGGAGCTTGAAAATGCATCACCACTTGAAATTATGGACAAGGCCCTAGAGAAGTTTGG

GAATGATATTGCAATTGCTTTCAGTGGTGCTGAAGATGTTGCTTTGATTGAGTATGCTCATTTGACTGGT

CGTCCATTCCGAGTGTTTAGCCTGGACACTGGAAGACTTAATCCTGAAACATACAAATTATTTGATGCTG

TTGAGAAGCAGTATGGCATTCACATCGAGTACATGTTTCCAGATGCTGTTGAAGTTCAGGACTTGGTAAG

GACCAAGGGTTTGTTCTCTTTCTATGAAGATGGTCATCAGGAGTGCTGCCGTGTGAGGAAAGTTAGACCT

TTGAGGAGGGCTCTTAAAGGCTTGCGAGCATGGGTTACTGGACAAAGAAAAGACCAGTCACCAGGAACTA

GATCTGAAATCCCAGTTGTCCAAGTTGATCCTGTTTTTGAAGGGCTCGAGGGTGGACCTGGTAGTTTGGT

GAAATGGAACCCAGTGGCAAACGTCCAAGGTACTGACATATGGAAATTTCTCCGTACCATGAATGTGCCT

GTCAACTCATTGCACGCACAGGGATACATCTCTATTGGTTGCGAGCCCTGCACAAGGCCAGTTCTGCCTG

GTCAACACGAGAGAGAAGGAAGGTGGTGGTGGGAGGATGCTACTGCCAAGGAATGTGGACTACATAAAGG

CAATATCAAGGAAGAGAATGTAAATGGCAATGGAACTGGAGCAGTAAAAGCCAACGGCGAAGCCACTGTC

ACTGACATTTTCAACAGCCAGCACGTGGTGAGCTTGAGCAGGCAAGGAATTGAAAATTTGATCAAACTAG

AAGATAGAAAAGAGACATGGCTTGTTGTTCTTTACGCACCCTGGTGCCAATTTTGCCAGGCAATGGAAGG

ATCATACATGGAATTGGCTGAGAAGTTAGCTGGAACAGGAGTGAAAGTTGCAAAGTTCCAGGCAGATGGG

GACCAAAAGGCGTATGCGCAGAAAGAGTTACAACTGGGAAGTTTCCCCACTATACTTTTCTATCCTAAAC

ATTCTTCTCGTGCTGTGAAGTACCCGTCTGAAAAGAGGGATGTTGAGTCCTTGTTAGCATTCGTGAATGC

TCTTAGATGATCAACACCAAACTCTGCCGATATAGTATCAGGTCTCTGCTGATTTATACAAGTTGAAGTA

GCAGCTGTTATGGAGGAGTCCACCTCGTGATGAATGTATATTTTCACAAAAATATATTTCTATGTGGGTT

TGAGGGCAATGATTGTTGCTTTTGTACTCAGTTTTGCCTTTCAGTTCTAGTTCAGACATTTTGTGAGTTT

TTCTATATGATGAAACTCAATGTTGCATTGCTCCTGATAGTGGAGATTATTCTTCAACTTCTTTTTACCA

AGATTTATATCAAAGTGTGTTATGATCTTTTAGCAGATAAAA

>MSTRG.3284.1 gene=MSTRG.3284

GTAACTGCTTTGGTAGTTTACCTATTTCATGTACATATCTTCACTTTGTTCTTAAGTCTTAACTCCCGGC

CTCCTATTTTTACATTTATCAGACATTAGTATTCGATGATCTACAACAAAAAATTTGATATACTAAGATG

CAACTGTATTACATGTCTCTAAGCTTGACATAATATCTTCACCAGGCTTCTACGTCCCAACGGTTGCCCA

AAACAATACCCATCTTTCTAAGAGGTCTCAGTCCCAACCATGAGTATCATCCCCTTTTATATGTGTAATA

GTCTAACTGCACAGATGCCTCTAGATATATCTATATCTGTATCTCCTAAGAGTTGTATCTTTACCACTAA

ACTGGCATTTTAATAGAAAAATGATTCTGCTATACTAAAAGAAAGGCTAACATAGTAACATGTCAATTAA

GCTATAGAGTAATAAACAAGTAGAACTGGAATATCCTCCAAGCTAAAGAAGATATGCATGATTACCACGA

CATTGAAGAACACAATACAGCCGCTATCAGTCTAGAGTTAAACTGAAATTGTTCATTGTTGCAACACAGA

CATCCAGTTGTAATATTGATCAGAGTCTGAGCAATTATAAGCGCCACCGCTGCTGTCAATCC

>MSTRG.3288.1 gene=MSTRG.3288

CTAGTTTTCTTAGTTTATTTTCTTTTCTTTTGATTTTTACTCGAGGACGAGCAAAAGATAGGTATGGGGA

AGTTTGATTAGTCCATATTTTTGCATATTTAAGTGCCTATTTTTTGTTTATTTTCGTAGTATTAATCGCT

TATTTATTTTATTTCAGGAAATTGTAGATAAATAAAGAAAGAAGAGAAAATGCAAGAAAAAGAAGAAAAG

GGAAAAGAAAAGAAGAAAAAAAGAGTTGGCAACTAAGGGACACATGGAGGGCCACAATCTTCCCAACACA

CAAGGCACATGGATGGTCAAGATTGAGCCACCCTACCCCTAAACCCTAATTCTTTAGTTATAAATACCCC

TCTTCTTTTACTTGTAATCACCAACCTACCAACCTAGAAAACCTAGCCACCTACTCTCCTTTTCTACCTA

GTCATTTCTATAGTTAGTATCTTAGATAGATTTACATTTTGCTAGCCCCAATTTCCTTTCCAA

>MSTRG.3290.2 gene=MSTRG.3290

CTCCAGCTCCTCCTTCCCGATCCACTCTGCTGCTCGCTCCTCTTCTCCCCTGCTTTCACGCTGCTGCTCG

CTCCGGCCATCCTTCCGTCTCCGAAGAACTGCTAGATGGCAGGTGATTTGAATTCTATCACAAAGACCTA

ATCTGTTGCCTATACAGTAGACGAGAATACTATTACTTGTCGAATCTCGATCAGAATATTGCTTCAATAA

TTCAGACAGTGTATTGTTTTAAGCCTGATGGTGACAAGAAGAACACAGAAGGTGTTGGACTTGTTATAAG

AGAATCTGAGATGGTTTACAAAGCATTCAACTTCGATCAAGAGAAGCTAATGAGGTTAAAGAAGTTGATC

ATGGAAGACGGGACCATCAAGAACTGCTCCACATTTGTGGCCCTCACGGCCTTGGTGTGGCGGGCAAGAA

CTGAAGCTCTTCAGATGAAACCAGACCAACAACTCAAGCTATTATTTGTTGTTGACGGAAGATCAAGATT

GGTGAATCCTCCCTTGCCGAAGGGCTATTTTGGGAACGGGATTGTGCTAGCGTGTAGTGTGAGTAATGCC

GGAGATTTGGTGAACAAGCCATTGTGCTAGCGTGTAGTGTGAGTAATGCCGGATTGTGCTAGCATTTCAG

ATATAATTTCAGGTTTCTGCAGTATAGTTATCAGTGCTTGATTAGCATTTTTGTGTTAAGG

>MSTRG.3294.1 gene=MSTRG.3294

CCTGGGGATCATTACAAATAATACATTATGTGATATGGAAAAGCTTTCTTAATTTTAAAGTATAAATAAA

CTATATTTCTTTGTAAATGTTACGGCACCTTTCTTCTCTTCTCTATAGTTGATGCAGCCATTCTTCCATG

ATATTTATCAGTATCAATTGTATTTTGTTTGTTGGTTTATCGAGAGCTTTGTTTAGAGTACCACAGCTTC

TTCTGGCAAATCAACACCTTCTCCTTTTCTACTCCGTACTCGTTGTCTTCTTGACGACAGCTCCATTTCA

TGCCCTTGGCCATGGTTGTAATTGCATCTGTGTAGTAGACTGACTCACGGATTATTGCATATCCGTTAGG

TCGAAGAATTC

>MSTRG.3297.1 gene=MSTRG.3297

GAAATTGCTCATCTTGTAAACATTAGTGTTGCTATTAAGCAGCTACAAGCGGCGAGAGACAAACTGCTTG

AGCACAGGCAGTAATGTGGGTGAGAGTGGCTATCTCTATAATTGCACAGCAAAAATAAGGCATCTGGCTA

TTTATTTTTACCGATGTGACAAGGAAGAATGTGTAGTTTCCATATTTTTGTATTGCTATATCACTGTCAA

GTTGCTTTAGAACTTTACGTTGCTTAATTTGGATCTGCATTGCTTACTTCGTACCTGTAATACTATTGCT

TGAAATTATGCAGTCGTCAAACTTCTTTGTGTATCCAG

>MSTRG.3298.1 gene=MSTRG.3298

AAGGCATTGATAACAAATGATCCATTGAGTGTGCTGAGTTCATGGAACAACTCCATCCACTTCTGTCACT

GGAATGGCATTAGATGCAGCCGGAGACGACAGAGAGTAACAGCGCTAAACCTCACCTCACAACACTTGGG

AGGCAAATTGTCTCCTCATATTGGAAACCTCTCCTTTCTCAGGGAAATTTACCTCTCTGAAAACAACTTT

CATGGCTCAATACCAAATGAAATTGGATGGCTCTTTCGCCTACAATATCTTGATCTCGGAAGCAATTCAT

TTCAAGGTAGATTCCCTGCGAATTTGAGCCATTGCGTAGATATCAGAAACATCAGTTTGGCTGAAAATGA

ACTTGAGGGAAAACTACCTACTGAGTTTTCAACTTGGCCTAAGCTTTACTTGTTGAATCTTGGATCCAAT

CATTTTACTGGATCAATTCCACCTTCAATAGGAAA

>MSTRG.3309.1 gene=MSTRG.3309

CAGGTAGTAAAATTAAACTAAAAGATATTGAATATGTTCCTCCAAGAAATGGTGCCACCCTTTGGGAAAT

TGGCATTCCTGACCGCACTGCTGCTGAGTTTTACGTGCCTCCGCCGAACCCCATGCTTCTGAATCAGTTA

TATGTCCAAAACAATGCACAAGGGTTCAGGCAGTATGGACTGTGGGATCGGTACACAGATGAATATCCTG

ATCAAGATTTAGTCTACACCATCGGGGTTAGCAACTATCAGATAGACTGGTTCTTTGCTCATGTTAACAG

GAATATAGGAAACAAAACATATGTACCCACTACCTGGCAAATTTTATTTGATCTCGAAGAAGTTGATCAA

AACGGAAATTATACATTGCATCTGGCATTGGCTTCTGCCACCGCAGCTGAATTACAAGTTCGGATAAATG

ACTACAATCCTACCAGTCCTCACTTCACGACAGGGCTCATAGGGCAGGACAATGCCATAGCAAGGCATGG

GATTCATGGGTTATACAGGCTATATAGTGTTAATGTTTGGGGATCTTTGCTACTAGAAGGAAGAAATGTA

ATATATTTAACACAAGCAGTAGGTGTTAGGGCTTTCAGAGGAATCATG

>MSTRG.3311.2 gene=MSTRG.3311

CTTCTTGAATAGAAATGACCCGAAACAAGCATTGTATCAGAGCAGGCTTCAACAATGTGGAAGAGAGCAG

CTGCCAAATCTATACTACAGCCATGGCGCATGTTAGCATCACGGCCGATGGCGTCGAAGTCAAAGGCGAA

TCTCCCCACGGCGGTCAACTCAATGATTCTGCGCTCACTGAAAGAACACTACGTCGAAGTCAAAGGCGAA

TCTCCCCACGGCGGTCAACTCAATGATTCTGCGCTCACTGAAAGAACACTATGTTGAAGTCTCCAAAATG

ACCCCTCCTCCTAAAGTGAACCCTCCATCACCATTCACGGTTGTGAAAGGGTCTCTTGAAAATTATGGTC

CTGTTTTGACAAGGACTTACGGGAACGAGGAAATTAATCTATCCGTGATGAGGTTAACAAACATAATTCC

TGCCGGTGAGGATGATGAAGAGGATGGAATGAACCAGTTATTTCTTCATGTTGATGTGTCTAAGCCAGGG

CAGAGCAAGTCTTTGCTTTTCCTTTGTGGACTGTATCCTGATGCAGTAGGAATTCACTCAATTTCGATGA

GGGCCAAGAGCGAGGATTCGGGTTTTGTCATGGACCCAACCAAATATAATGGACCTAGATTCCAGGATTT

TGATGATAAGATGAAGGATGCCATTCATGGTTACATTGAGGAGCGAGGCATAACTGAAAGCCTCTTTCCA

TTTCTTCAAGCTTGGCTTTATGTGAAGGATCACCGGAAACTGATGGGATGGTTCAGATCAGTAGGTAGTT

ACATTACTGAAAAGAAAGAAGCTTAGATGCTCAGTTATCTTTATGTTTAATATGTCAAATGATCTCAAGT

ATTCTGGATAAGTAGGATTATATGCTTAGTTTCCATATCTTTTATAAGGGTGGTGTGTACGGATGATCCT

AATTTATTTGTGGCCATTATTTTCAAGCCTTTTGATGTTTTCACTTGTAATTTTTATGAAACTAGTCAAT

ATTGCATGTCTGCATGATAGTCTAGTACATCTTTAAGCTGCATTTTATTTGTTGAAGACATAGTTTCTCT

AGCCATCTTTGATTGTTTTTATATTAAATGCCAAC

>MSTRG.3311.1 gene=MSTRG.3311

TATCACAGTCCAATAGCTTCTTGAATAGAAATGACCCGAAACAAGCATTGTATCAGAGCAGGCTTCAACA

ATGTGGAAGAGAGCAGCTGCCAAATCTATACTACAGCCATGGCGCATGTTAGCATCACGGCCGATGGCGT

CGAAGTCAAAGGCGAATCTCCCCACGGCGGTCAACTCAATGATTCTGCGCTCACTGAAAGAACACTATGT

TGAAGTCTCCAAAATGACCCCTCCTCCTAAAGTGAACCCTCCATCACCATTCACGGTTGTGAAAGGGTCT

CTTGAAAATTATGGTCCTGTTTTGACAAGGACTTACGGGAACGAGGAAATTAATCTATCCGTGATGAGGT

TAACAAACATAATTCCTGCCGGTGAGGATGATGAAGAGGATGGAATGAACCAGTTATTTCTTCATGTTGA

TGTGTCTAAGCCAGGGCAGAGCAAGTCTTTGCTTTTCCTTTGTGGACTGTATCCTGATGCAGTAGGAATT

CACTCAATTTCGATGAGGGCCAAGAGCGAGGATTCGGGTTTTGTCATGGACCCAACCAAATATAATGGAC

CTAGATTCCAGGATTTTGATGATAAGATGAAGGATGCCATTCATGGTTACATTGAGGAGCGAGGCATAAC

TGAAAGCCTCTTTCCATTTCTTCAAGCTTGGCTTTATGTGAAGGATCACCGGAAACTGATGGGATGGTTC

AGATCAGTAGGTAGTTACATTACTGAAAAGAAAGAAGCTTAGATGCTCAGTTATCTTTATGTTTAATATG

TCAAATGATCTCAAGTATTCTGGATAAGTAGGATTATATGCTTAGTTTCCATATCTTTTATAAGGGTGGT

GTGTACGGATGATCCTAATTTATTTGTGGCCATTATTTTCAAGCCTTTTGATGTTTTCACTTGTAATTTT

TATGAAACTAGTCAATATTGCATGTCTGCATGATAGTCTAGTACATCTTTAAGCTGCATTTTATTTGTTG

AAGACATAACATAGTTTCTCTAGCCATCTTTGATTGTTTTTATATTAAATGCCAAC

>MSTRG.3311.3 gene=MSTRG.3311

TATCACAGTCCAATAGCTTCTTGAATAGAAATGACCCGAAACAAGCATTGTATCAGAGCAGGCTTCAACA

ATGTGGAAGAGAGCAGCTGCCAAATCTATACTACAGCCATGGCGCATGTTAGCATCACGGCCGATGGCGT

CGAAGTCAAAGGCGAATCTCCCCACGGCGGTCAACTCAATGATTCTGCGCTCACTGAAAGAACACTATGT

TGAAGTCTCCAAAATGACCCCTCCTCCTAAAGTGAACCCTCCATCACCATTCACGGTTGTGAAAGGGTCT

CTTGAAAATTATGGTCCTGTTTTGACAAGGACTTACGGGAACGAGGAAATTAATCTATCCGTGATGAGGT

TAACAAACATAATTCCTGCCGGTGAGGATGATGAAGAGGATGGAATGAACCAGTTATTTCTTCATGTTGA

TGTGTCTAAGCCAGGGCAGAGCAAGTCTTTGCTTTTCCTTTGTGGACTGTATCCTGATGCAGTAGGAATT

CACTCAATTTCGATGAGGGCCAAGAGCGAGGATTCGGGTTTTGTCATGGACCCAACCAAATATAATGGAC

CTAGATTCCAGGATTTTGATGATAAGATGAAGGATGCCATTCATGGTTACATTGAGGAGCGAGGCATAAC

TGAAAGCCTCTTTCCATTTCTTCAAGCTTGGCTTTATGTGAAGGATCACCGGAAACTGATGGGATGGTTC

AGATCAGTAGGTAGTTACATTACTGAAAAGAAAGAAGCTTAGATGCTCAGTTATCTTTATGTTTAATATG

TCAAATGATCTCAAGTATTCTGGATAAGTAGGATTATATGCTTAGTTTCCATATCTTTTATAAGGGTGGT

GTGTACGGATGATCCTAATTTATTTGTGGCCATTATTTTCAAGCCTTTTGATGTTTTCACTTGTAATTTT

TATGAAACTAGTCAATATTGCATGTCTGCATGATAGTCTAGTACATCTTTAAGCTGCATTTTATTTGTTG

AAGACATAGTTTCTCTAGCCATCTTTGATTGTTTTTATATTAAATGCCAAC

>MSTRG.3311.4 gene=MSTRG.3311

ATCAGAGCAGGCTTCAACAATGTGGAAGAGAGCAGCTGCCAAATCTATACTACAGCCATGGCGCATGTTA

GCATCACGGCCGATGGCGTCGAAGTCAAAGGCGAATCTCCCCACGGCGGTCAACTCAATGATTCTGCGCT

CACTGAAAGAACACTATGTTGAAGTCTCCAAAATGACCCCTCCTCCTAAAGTGAACCCTCCATCACCATT

CACGGTTGTGAAAGGGTCTCTTGAAAATTATGGTCCTGTTTTGACAAGGACTTACGGGAACGAGGAAATT

AATCTATCCGTGATGAGGTTAACAAACATAATTCCTGCCGGTGAGGATGATGAAGAGGATGGAATGAACC

AGTTATTTCTTCATGTTGATGTGTCTAAGCCAGGGCAGAGCAAGTCTTTGCTTTTCCTTTGTGGACTGTA

TCCTGATGCAGTAGGAATTCACTCAATTTCGATGAGGGCCAAGAGCGAGGATTCGGGTTTTGTCATGGAC

CCAACCAAATATAATGGACCTAGATTCCAGGATTTTGATGATAAGATGAAGGATGCCATTCATGGTTACA

TTGAGGAGCGAGGCATAACTGAAAGCCTCTTTCCATTTCTTCAAGCTTGGCTTTATGTGAAGGATCACCG

GAAACTGATGGGATGGTTCAGATCAGTAGGTAGTTACATTACTGAAAAGAAAGAAGCTTAGATGCTCAGT

TATCTTTATGTTTAATATGTCAAATGATCTCAAGTATTCTGGATAAGTAGGATTATATGCTTAGTTTCCA

TATCTTTTATAAGGGTGGTGTGTACGGATGATCCTAATTTATTTGTGGCCATTATTTTCAAGCCTTTTGA

TGTTTTCACTTGTAATTTTTATGAAACTAGTCAATTCTGCATGATAGTCTAGTACATCTTTAAGCTGCAT

TTTATTTGTTGAAGACATAGTTTCTCTAGCCATCTTTGATTGTTTTTATATTAAAT

>MSTRG.3312.1 gene=MSTRG.3312

CAGGTAGTAAAATTAAACTAAAAGATATTGAATATGTTCCTCCAAGAAATGGTGCCACCCTTTGGGAAAT

TGGCATTCCTGACCGCACTGCTGCTGAGTTTTACGTGCCTCCGCCGAACCCCATGCTTCTGAATCAGTTA

TATGTCCAAAACAATGCACAAGGGTTCAGGCAGTATGGACTGTGGGATCGGTACACAGATGAATATCCTG

ATCAAGATTTAGTCTACACCATCGGGGTTAGCAACTATCAGATAGACTGGTTCTTTGCTCATGTTAACAG

GAATATAGGAAACAAAACATATGTACCCACTACCTGGCAAATTTTATTTGATCTCGAAGAAGTTGATCAA

AACGGAAATTATACATTGCATCTGGCATTGGCTTCTGCCACCGCAGCTGAATTACAAGTTCGGATAAATG

ACTACAATCCTACCAGTCCTCACTTCACGACAGGGCTCATAGGGCAGGACAATGCCATAGCAAGGCATGG

GATTCATGGGTTATACAGGCTATATAGTGTTAATGTTTGGGGATCTTTGCTACTAGAAGGAAGAAATGTA

ATATATTTAACACAAGCAGTAGGTGTTAGGGCTTTCAGAGGAATCATG

>MSTRG.3314.1 gene=MSTRG.3314

ATAACAATCTAACTGGTGACATACCTCGGGAAGTGGCTCATTTTGCAAAATTGGAGGATCTTGATTTGTC

AGCAAACAGATTGTCGGGTATGATTCCCCGACCACTTTTTAACATATCGTCTCTCTATATTGTTGGTCTT

GCCGTTAACATGTTAAAAGGAACACTTCCATCGGATTTGGGCTTCACCCTTCCTAAGCTGCAAGAATTTT

ATGTTGGTGAGAACAAACTTTCAGGACCATTTCCACCGTCAGTTGTTAATGCGTCTAATCTTGTTTTCTT

TGATATCAATAAAAACAGTATTTTTGGGCCTATACCAAACAATTTGGGCAGCCTTCTGAATCTTGATTGG

CTAGGTCTGGGTCAGAATCCACTTGGAGAGAATATGCGGTACAATGATTGGAGCTTTCTTGATTCCTTGG

TTAATTGTACCCATCTAAAGAGATTGAGCTTGGGCGAGACTAGTTTAAGAGGGGAGCTCCCTGATTCCAT

TGCGAATCTCTCCACCACTTTGGAAGACCTGTATATGTATGGAAACTATATATATGGAAGCATACCTCAT

GGAATTGGAAAACTGGAGAAAATGATACATCTTAATTTGGGAGACAACTTCTTAACGGGGAGCATTCCAC

AATCAATTGGAAGACTATCAAAATTAGGCCTGCTAGCTTTGGCTGGAAACAACATTTCAGGTGCAATTCC

AACTTCCATTAGCAACATTACTCATTTAGTTAAACTCTATTTATATGATAATGTGCTGCAAGGAAGCATA

CCTGCTGAATTGTTTAACATCTCAACTCTAGAGGTAGTGTCCATTGCTAACAACCGGCTTGGAGGTGCAA

TACCTAATGAAATTCTGTTCTTGTCGCATTGTATTCGACTTAGTTTGTCCCAAAATTTATTTAGCGGTCC

ACTTCCATCCAACATTGGAAGCTTGAAACAACTGGTTAAACTCGGCGTTTCAAACAACAAACTCGGAGGA

GATATACCAGCTACTCTTGGTGATTGTGTAATGTTGGAGATGTTATATATGGATGGAAACCTTTTTCAAG

GTGAAATTCCATCTTCTTTTAAAGCATTGAAAAATCTTGCTTTTCTAGATCTTTCCGACAATAACATCTC

TGGGAGTATTCCGAGTTTTTTGAATGGCTTTCAATTTATCAAATTTCTCAACTTGTCACACAACAAGCTT

GAAGGTGAAGTGCCTACAGAAGGTCAGTTTTCAAATATGAGTGCCTTTTCAGTTGCTGGAAATTTGGAGC

TTTGTGGAGGCATTCCAGCATTGCATTTGGTTGCCTGTCCTCGAAAAGTCTCAAGACATAAGAAAGTAGC

ATTTGCCCCAAGGATAATACTAATTGTAGTTCTTGTACCCCTTGGTGTATTGTTTTTTTGTCTTGCGTTG

ATTTCTTATCAACGTAGAAACTCCAAGAAGTTGAATGACCCCGTCCCAGTATTGAAGGATGATCAATATC

CCAAGCTTTCATACCAAGACCTTCTGCTGGCCACAAATGAATTTTCTCCAGACAATTTGCTCGGTCAAGG

AAGATACAGTTCTGTTTACAAAGGAGTTCTCGAATCAGTGGAACACATACTTGCTGTCAAGGTACTAAAC

ATTGAAATACATGGAGCTAACAAGACTTTTCTGGCAGAGTGTGAAACATTGAGGCACATTCGTCATCGAA

ATCTTATCAAGATCATCACAGCATGCTCTAGCACTGATTTTAAGGGCAATGACTTTAAGGCATTGGTTTT

TGAGTTCATGACTAATGGGAGTGTGGACAACTGGTTGCATCCAAGTCCTTCTGATGAAGGGAATGAAAGA

AACTTGACTCTACTTCAGAGAGTAAATATTTCCATCGACGTTGCACTAGGAATGGATTACTTACATCATC

ACTCTCATGCAAGTATCATTCATTGCGACCTAAAGCCAAGTAATGTTCTTCTCGATGATGATCTTGTTGC

TCGTATCAGTGATTTTGGTTTGGCAAGATTCTACTTTGCTACTACAAGTGACATCAATCAATCACAAATG

AGTTCTACCGGTGTACGTGGAACAATTGGATATGTGCCTCCAGAGTATGGAATGGGTGGGGAGATATCTG

CTGAAGGAGATGTATATAGCTACGGAATCTTTCTACTAGAGATGTTTACAGGAAAACGACCTACATGTAG

CAGCACGTTGCTGGACAATAATAATAGTCTTCATGACTATGTGAGGAAAGCCCTCCCTTCTGGGGTGATG

AACATTCTTGACCCGCGGATCATAATAGATCTAGACGATCATGGTTTGACAACAAATCAGTCATACAAGA

AGGCTGCCTTAGAGGCATGCCTGGCATCAATATTTCAAGTGGGGATACTATGTTCACTAGAAATGCCCAA

AGAGCGCATTGACATTGGTATTGCTATCAAGCAGCTTCATGCAGCAAGAGACAATCTTCTACAGCACAGC

TAGTAACCCATGTCTATGTTTCTCTGCTTTTCCTATGATTCTCTGATCTTGTATTGAACTAAAACATGGC

TTTTAATAATATGTTTACACATTTTGCTTAATTCCA

>MSTRG.3314.2 gene=MSTRG.3314

GAGTTCTCGAATCAGTGGAACACATACTTGCTGTCAAGGTACTAAACATTGAAATACATGGAGCTAACAA

GATTTTCTGGCAGAGTGTGAAACATTGAGGCACATTCGTCATCGAAATCTTATCAAGATCATCACAGCAT

GCTCTAGCACTGATTTTAAGGGCAATGACTTTAAGGCATTGGTTTTTGAGTTCATGACTAATGGGAGTGT

GGACAACTGGTTGCATCCAAGTCCTTCTGATGAAGGGAATGAAAGAAACTTGACTCTACTTCAGAGAGTA

AATATTTCCATCGACGTTGCACTAGGAATGGATTACTTACATCATCACTCTCATGCAAGTATCATTCATT

GCGACCTAAAGCCAAGTAATGTTCTTCTCGATGATGATCTTGTTGCTCGTATCAGTGATTTTGGTTTGGC

AAGATTCTACTTTGCTACTACAAGTGACATCAATCAATCACAAATGAGTTCTACCGGTGTACGTGGAACA

ATTGGATATGTGCCTCCAGAGTATGGAATGGGTGGGGAGATATCTGCTGAAGGAGATGTATATAGCTACG

GAATCTTTCTACTAGAGATGTTTACAGGAAAACGACCTACATGTAGCAGCACGTTGCTGGACAATAATAA

TAGTCTTCATGACTATGTGAGGAAAGCCCTCCCTTCTGGGGTGATGAACATTCTTGACCCGCGGATCATA

ATAGATCTAGACGATCATGGTTTGACAACAAATCAGTCATACAAGAAGGCTGCCTTAGAGGCATGCCTGG

CATCAATATTTCAAGTGGGGATACTATGTTCACTAGAAATGCCCAAAGAGCGCATTGACATTGGTATTGC

TATCAAGCAGCTTCATGCAGCAAGAGACAATCTTCTACAGCACAGCTAGTAACCCATGTCTATGTTTCTC

TGCTTTTCCTATGATTCTCTGATCTTGTATTGAACTAAAACATGGCTTTTAATAATATGTTTACACATTT

TGCTTAATTCCACTTTAATTTTTTTTTCAAAGTCAATAACAACACATTTTGTTGC

>MSTRG.3314.3 gene=MSTRG.3314

ACTAAATGAAACCACTTATAATCTTGTGGCCAAGATTTCCTCAGAGGCCAAGATTTTCACTATTGATAGT

GATAAGCTAATCTTGTGCAGCATTTTCACTTGTCTAGTGTACAAGTACAACCATGGTTTTCAGCTTTTTC

CCATTCAGCATTCTCTTCTTTCTTGTATTGGTAACCTCAGCAAGGTTGGTATATTCGTTGTCAAACAATT

TTACTGATGAGCAAGCTTTGCTTTCATTCAAGTCCTCAATAACAGATGACCCCTCGGGTGTGCTGGAGTC

GTGGAACAATTCCATCCACTTCTGTCATTGGACAGGCGTTACATGCAGCCGCAGAAGACAGAGGGTAACT

GTACTCGACCTCTCATCACTAGGCTTGGCGGGTACATTGTCTCCTCATATTGGAAACCTCTCGTTCCTGA

GGACAATTTACCTCTTTGAAAACCGCTTTCATGGCTTGATTCCAAATGACATTGGTCGACTGTTTCGCCT

AAAAACTCTTTCTATGGGAAACAACTCTTTTCAAGGAGGTTTTCCAGCCGACTTGCATCGTTGTACAAAT

ATCAGATTCATCAATTTGCACAAAAACAATCTTCAAGGAGAATTAGCTACTGACTTTGCTTCTTGGTCTC

AGCTTTATTATTTTTCTGTTGGAAACAATCAGTTTACAGGATCAATTCCACCTTCAATAGGTAATACATC

ATCACTTCGTACTCTTTTTTTATACAAAAACAATTTTGAAGGAAATCTACCCACTGAGTTTGCCTCTTGG

TCTAAGCTTGATTTCATCGATCTTAGCAACAATCATATTAGTGGATCAATTCCACCGTCGATAGGAAATG

TATCATCACTCCGTTCGATTCATCTCGCTTGTAACAATTTAACTGGTGACATACCTCGGGAAGTGGCTCA

TTTTGCAAAATTGGAGGATCTTGATTTGTCAGCAAACAGATTGTCGGGTATGATTCCCCGACCACTTTTT

AACATATCGTCTCTCTATATTGTTGGTCTTGCCGTTAACATGTTAAAAGGAACACTTCCATCGGATTTGG

GCTTCACCCTTCCTAAGCTGCAAGAATTTTATGTTGTATTTTTGGGCCTATACCAAACAATTTGGGCAGC

CTTCTGAATCTTGATTGGCTAGGTCTGGGTCAGAATCCACTTGGAGAGAATATGCGGTACAATGATTGGA

GCTTTCTTGATTCCTTGGTTAATTGTACCCATCTAAAGAGATTGAGCTTGGGCGAGACTAGTTTAAGAGG

GGAGCTCCCTGATTCCATTGCGAATCTCTCCACCACTTTGGAAGACCTGTATATGTATGGAAACTATATA

TATGGAAGCATACCTCATGGAATTGGAAAACTGGAGAAAATGATACATCTTAATTTGGGAGACAACTTCT

TAACGGGGAGCATTCCACAATCAATTGGAAGACTATCAAAATTAGGCCTGCTAGCTTTGGCTGGAAACAA

CATTTCAGGTGCAATTCCAACTTCCATTAGCAACATTACTCATTTAGTTAAACTCTATTTATATGATAAT

GTGCTGCAAGGAAGCATACCTGCTGAATTGTTTAACATCTCAACTCTAGAGGTAGTGTCCATTGCTAACA

ACCGGCTTGGAGGTGCAATACCTAATGAAATTCTGTTCTTGTCGCATTGTATTCGACTTAGTTTGTCCCA

AAATTTATTTAGCGGTCCACTTCCATCCAACATTGGAAGCTTGAAACAACTGGTTAAACTCGGCGTTTCA

AACAACAAACTCGGAGGAGATATACCAGCTACTCTTGGTGATTGTGTAATGTTGGAGATGTTATATATGG

ATGGAAACCTTTTTCAAGGTGAAATTCCATCTTCTTTTAAAGCATTGAAAAATCTTGCTTTTCTAGATCT

TTCCGACAATAACATCTCTGGGAGTATTCCGAGTTTTTTGAATGGCTTTCAATTTATCAAATTTCTCAAC

TTGTCACACAACAAGCTTGAAGGTGAAGTGCCTACAGAAGGTCAGTTTTCAAATATGAGTGCCTTTTCAG

TTGCTGGAAATTTGGAGCTTTGTGGAGGCATTCCAGCATTGCATTTGGTTGCCTGTCCTCGAAAAGTCTC

AAGACATAAGAAAGTAGCATTTGCCCCAAGGATAATACTAATTGTAGTTCTTGTACCCCTTGGTGTATTG

TTTTTTTGTCTTGCGTTGATTTCTTATCAACGTAGAAACTCCAAGAAGTTGAATGACCCCGTCCCAGTAT

TGAAGGATGATCAATATCCCAAGCTTTCATACCAAGACCTTCTGCTGGCCACAAATGAATTTTCTCCAGA

CAATTTGCTCGGTCAAGGAAGATACAGTTCTGTTTACAAAGGAGTTCTCGAATCAGTGGAACACATACTT

GCTGTCAAGGTACTAAACATTGAAATACATGGAGCTAACAAGACTTTTCTGGCAGAGTGTGAAACATTGA

GGCACATTCGTCATCGAAATCTTATCAAGATCATCACAGCATGCTCTAGCACTGATTTTAAGGGCAATGA

CTTTAAGGCATTGGTTTTTGAGTTCATGACTAATGGGAGTGTGGACAACTGGTTGCATCCAAGTCCTTCT

GATGAAGGGAATGAAAGAAACTTGACTCTACTTCAGAGAGTAAATATTTCCATCGACGTTGCACTAGGAA

TGGATTACTTACATCATCACTCTCATGCAAGTATCATTCATTGCGACCTAAAGCCAAGTAATGTTCTTCT

CGATGATGATCTTGTTGCTCGTATCAGTGATTTTGGTTTGGCAAGATTCTACTTTGCTACTACAAGTGAC

ATCAATCAATCACAAATGAGTTCTACCGGTGTACGTGGAACAATTGGATATGTGCCTCCAGAGTATGGAA

TGGGTGGGGAGATATCTGCTGAAGGAGATGTATATAGCTACGGAATCTTTCTACTAGAGATGTTTACAGG

AAAACGACCTACATGTAGCAGCACGTTGCTGGACAATAATAATAGTCTTCATGACTATGTGAGGAAAGCC

CTCCCTTCTGGGGTGATGAACATTCTTGACCCGCGGATCATAATAGATCTAGACGATCATGGTTTGACAA

CAAATCAGTCATACAAGAAGGCTGCCTTAGAGGCATGCCTGGCATCAATATTTCAAGTGGGGATACTATG

TTCACTAGAAATGCCCAAAGAGCGCATTGACATTGGTATTGCTATCAAGCAGCTTCATGCAGCAAGAGAC

AATCTTCTACAGCACAGCTAGTAACCCATGTCTATGTTTCTCTGCTTTTCCTATGATTCTCTGATCTTGT

ATTGAACTAAAACATGGCTTTTAATAATATGTTTACACATTTTGCTTAATTCCAC

>MSTRG.3315.1 gene=MSTRG.3315

GCCATTAAATTGATTCAAGAAAATATAAAATTCCTAGTACATGCAGCTTCAGAGAATCAAATAAATTGTC

ATTAATATATGCCTTGGCAATTGAAATTTACTGTTTGGTCGTTCACAGGATATGCCTTTAATCAAGTCAT

TACCTTGTTCTATTTTATAACATCAAGCCTCAGGTTAAGAGATTTACTTTGTGGTTATTGCATATGATTT

CTGAAAAACATTGAGCTCCAGTGGATTCATGTACACCGGAATTAAATGGTAGCATCTTTCTTAACTTGCT

GGACTGTAAAACGGAGGCAGTGAACGCTGCTGAAGAGGACTGTATCAAACTGGATGGTACAACTGAGCTG

AGGGGCGGCAGGCAAGACCATGGCTCTGGTCCTGCAAATTGGATTGGGCATGACTGATTATGAATTACTT

AAATTTTTTATGTTGTTTCTGGTACACAATTTCCCACAGGCTTGGACATAGCTCAAGTCTTTACGGGAAG

CAAGACGGGTCTGTCTGAAACATCAACATGTGCAACTTGCAGGCAACAAAATGTGTTGTTA

>MSTRG.3319.1 gene=MSTRG.3319

CTTCTCACTTCTTTTATGATAAAGTCACACACTTTCCTCCAAAAGATTTTTCTTTTTTTAAATATTATCT

TTTTCTTTAATTGGTTGAAGCATGTTTATTACCACTGTCCAATCTTTATAGTCTTTTGTTTTTGTTTGTT

TGGCTTTCTTTGACTCCCCATGCTTGCTGATTTCAGCTCCATATCAACTCCTTTTTACTTGTTATGGAGC

AGCCCTCTTCAGGATCAGTTTTCTTGATGTGGGGTTTTGAATTTGGGGCATTTAAAGGCTGTAGATTGTA

ATTTATTGCTGTGAAGTGAGAGTGTATAGTTAAAGAATTTGTACTTCTTGAATTTTTTGATTGAGATGGA

GAGAAATGGGGAAGAGGATTGGAGGGAGATAGTTAGAAAGATGCTTCCTCCTGGGGCTCCCATACCTGAA

AATGAGTCTGATCTTGATTATTCGATAGCGATGGAGTATAGTGGTCCTCCAGTGTCTTATGAGCTTCCTA

AGGTAGAACCCCTTGATGTGAAGTCGAGTTCGATTGCGACAGCTTCTATGGCAGATGATTCTTTCTCTGA

TGCTAAAAGATCGGTTAGTCGAGATGTTCCACCAGTTGTAGAACCAATTTCTTTGCATGTGTCTTATATT

GCTGGTAGTGTGACTAGTCCAGCTAACCATAGTCCTAGAGTATCAGGAAGTTCTGAGTCTGAGGTGTCTG

TGTTGCACAACCGGGATTCTTCTTCGGGTTCCTTGTCAGGTTCGCCGACTTCTGTAAACAATCAACCTAA

TAATGTTCATCAACAATCAGTTAGAAGAGCCCCTGTTGTTACATTCAATACAGTTGATAGAACTGAAAGT

AAAGAGGTGGATAAGGAGGAGCCAAGATATCCTGAATATGTTGGAGTTACCAAGGAGAAAAAGAAGAAAA

AGAAAGCAAAAAGAAGAGTGTGCTATCGCTGTGGAAAAGGGAAGTGGGAAACAAAGGAGTCATGTTTGGT

TTGCGATGCAAAGTATTGTAGCAATTGTGTGCTCAGGGCTATGGGTTCAATGCCTGAAGGCCGTAAATGT

GTTACTTGCATTGGGGAGCCAATTGATGAATCAAATAGGTTGAAATTGGGTAGGCACTCCAGGTTACTGT

CCAGATTACTCAGTCCCTTGGAAGTCAAGCAGATAATGAAGGCGGAGAAAGAATGTTCAGCCAATCAACT

TCGGCCGGAGCAACTGATTGTGAATGGATTTCCACTGAAGCCAGAAGAGATGTCTGAGCTCCTTGGGTGC

CCTTTACCTCCTCGAAAATTGAATCCTGGTAGATATTGGTATGACAAGGAATCTGGCATGTGGGGAAAGG

AAGGCGAAAAACCAGATAGAGTTATAAGTTCAAATTTAAGTTTTTCAGGGAAACTCAGCCCAAATGCTAG

CAATGGAAATACTGAAGTTTACATAAATGGTCGAGAAATTACAAAACTTGAATTAAGAGTGCTAAGGATG

GCTAACGTTCAATGTCCCCGTGATACACATTTCTGGGTGTATGACGATGGTCGTTATGAGGAAGAGGGTC

AGAATAATATTAGGGGAAATATTTGGGAAAAGGCATTAACTCGTTTTGTGTGCGCATTTTTCTCCTTGCC

TGTGCCACGTGGTCAGCCTCATGGACAAAGAGATGAGCCTAGCAATTATACTACTGTCCCAAACTATCTG

GAGCAGAAAAAAGTCCAGAAGATTTTTCTAATGGGACTTGAAGGTTCCGGAACTAGTACCATCTTCAAGC

AGGCAAAATTTTTGTATGGGAACAGATTTACATTTGAGGAATTACAAGACATTAAGCTTATGATCCAGAG

CAACATGTACAAATATCTCAGTATTTTACTTGATGGACGGGAGCGTTTTGAGGAGGAGACCATGCTCAAG

AAAAATTCTCTCGGTGGAGAGGTTGATACTGATGAGAATAGTCAATGTATCTATTCTATCAACCCGAGAT

TAAAGCATTTTTCTGACTGGCTCTTAGACATTATAGCCACTGGTGATTTGGATGCATTCTTCCCTGCAGC

AACTCGTGAATATGCTCCACTGGTTGAAGAAGTGTGGAAAGACCCTGCTATACAGGAAACATACAAGAGA

AAAGATGAGCTTCACTTCCTTCCTGATGTGGCAGAATACTTCCTAGACCGGGCTGTCGAAGTGTCTAGTA

ATGAATATGAACCTTCAGAAAAGGACATACTATATGCAGAAGGAGTCACTCAGGGAAATGGTCTAGCTTT

CATAGAGTTCTCGCTGGATGATCGTAGCCCGATGTCTGAAACCTACACAGACATGGAAGCTCAACTCCCA

CCTTTGACTAGGTACCAATTAATTAGGGTAAATGCTAAAGGCATGACCGATGGTTGCAAGTGGGTGGAAA

TGTTTGAAGATGTTCGTGTAGTTGTCTTTTGTGTTGCCCTGACTGACTATGAGCAGATGTCGGTTTCCGC

TGAGAATGGCGGAAGCGGTGTGCTTCTTGTAAACAAGATGATGGAAAGCAAAGAACTATTTGAAAAGATG

ATCAGGCATCCATGTTTTAAAGACACTCCCTTTGTTTTGATTTTAAACAAGTATGATCTTTTCGAGGAGA

AGGTGAACCGGGTCCCACTAAGTACTTGTGATTGGTTCTCCGACTTCAGCCCAGTACAACCCCATCACAA

CAACCAAACTCTAGCTAATCAAGCTTATTACTATGTCGCGATGAAGTTCAAAGACCTCTACTCTTCTCTA

ACTTCTCGAAAGCTTTTTGTTTGGCAAGCTAGGGCAAGGGATCGAGTTACAATAGACGAAGCATTTAAGT

ACATAAGGGAGGTTGTTAAGTGGGATGATGAGAAGGAAGATAACTACTTTGGTATTGAGGATTCATTTTA

CAGTACAACAGATGTGAGTTCCTCCCCATTTATTAGGCAAGAGTGATTTACTGTACATGTTCTTCACGCA

TTAGATGTGTAAAGAGTAGGAAGAGATGCAGAGGCAAATAAGTACGGATGGTTTATTTTTTTTCATCCTC

AAACCTCGTGGATTGGGATTTGGGTGATTGTATTTCATGTAAAATCCGTCCAGCGGAATTGTCAAAACCT

TGTAATGATATTGTAGCTACTATGCATTCCGTCAGCCTGCTGGTAAAGATTGATGAAAATCGCTTTCACC

TGCGAGCTCTGAAAGTAATTGCATCACCATGTATTTTCTTCTAACCAGAAAATAGTTTAGGGGCAAGTTG

GATATCATGTACCTTGTATGCTGTTGTATGAACTATGAACTCTTTTAGTGTAATTGATATGCAGGTGAAG

CTCTTATTATGTT

>MSTRG.3319.2 gene=MSTRG.3319

CTTCTCACTTCTTTTATGATAAAGTCACACACTTTCCTCCAAAAGATTTTTCTTTTTTTAAATATTATCT

TTTTCTTTAATTGGTTGAAGCATGTTTATTACCACTGTCCAATCTTTATAGTCTTTTGTTTTTGTTTGTT

TGGCTTTCTTTGACTCCCCATGCTTGCTGATTTCAGCTCCATATCAACTCCTTTTTACTTGTGGCTTGTT

TTTCAAAGTTATGGAGCAGCCCTCTTCAGGATCAGTTTTCTTGATGTGGGGTTTTGAATTTGGGGCATTT

AAAGGCTGTAGATTGTAATTTATTGCTGTGAAGTGAGAGTGTATAGTTAAAGAATTTGTACTTCTTGAAT

TTTTTGATTGAGATGGAGAGAAATGGGGAAGAGGATTGGAGGGAGATAGTTAGAAAGATGCTTCCTCCTG

GGGCTCCCATACCTGAAAATGAGTCTGATCTTGATTATTCGATAGCGATGGAGTATAGTGGTCCTCCAGT

GTCTTATGAGCTTCCTAAGGTAGAACCCCTTGATGTGAAGTCGAGTTCGATTGCGACAGCTTCTATGGCA

GATGATTCTTTCTCTGATGCTAAAAGATCGGTTAGTCGAGATGTTCCACCAGTTGTAGAACCAATTTCTT

TGCATGTGTCTTATATTGCTGGTAGTGTGACTAGTCCAGCTAACCATAGTCCTAGAGTATCAGGAAGTTC

TGAGTCTGAGGTGTCTGTGTTGCACAACCGGGATTCTTCTTCGGGTTCCTTGTCAGGTTCGCCGACTTCT

GTAAACAATCAACCTAATAATGTTCATCAACAATCAGTTAGAAGAGCCCCTGTTGTTACATTCAATACAG

TTGATAGAACTGAAAGTAAAGAGGTGGATAAGGAGGAGCCAAGATATCCTGAATATGTTGGAGTTACCAA

GGAGAAAAAGAAGAAAAAGAAAGCAAAAAGAAGAGTGTGCTATCGCTGTGGAAAAGGGAAGTGGGAAACA

AAGGAGTCATGTTTGGTTTGCGATGCAAAGTATTGTAGCAATTGTGTGCTCAGGGCTATGGGTTCAATGC

CTGAAGGCCGTAAATGTGTTACTTGCATTGGGGAGCCAATTGATGAATCAAATAGGTTGAAATTGGGTAG

GCACTCCAGGTTACTGTCCAGATTACTCAGTCCCTTGGAAGTCAAGCAGATAATGAAGGCGGAGAAAGAA

TGTTCAGCCAATCAACTTCGGCCGGAGCAACTGATTGTGAATGGATTTCCACTGAAGCCAGAAGAGATGT

CTGAGCTCCTTGGGTGCCCTTTACCTCCTCGAAAATTGAATCCTGGTAGATATTGGTATGACAAGGAATC

TGGCATGTGGGGAAAGGAAGGCGAAAAACCAGATAGAGTTATAAGTTCAAATTTAAGTTTTTCAGGGAAA

CTCAGCCCAAATGCTAGCAATGGAAATACTGAAGTTTACATAAATGGTCGAGAAATTACAAAACTTGAAT

TAAGAGTGCTAAGGATGGCTAACGTTCAATGTCCCCGTGATACACATTTCTGGGTGTATGACGATGGTCG

TTATGAGGAAGAGGGTCAGAATAATATTAGGGGAAATATTTGGGAAAAGGCATTAACTCGTTTTGTGTGC

GCATTTTTCTCCTTGCCTGTGCCACGTGGTCAGCCTCATGGACAAAGAGATGAGCCTAGCAATTATACTA

CTGTCCCAAACTATCTGGAGCAGAAAAAAGTCCAGAAGATTTTTCTAATGGGACTTGAAGGTTCCGGAAC

TAGTACCATCTTCAAGCAGGCAAAATTTTTGTATGGGAACAGATTTACATTTGAGGAATTACAAGACATT

AAGCTTATGATCCAGAGCAACATGTACAAATATCTCAGTATTTTACTTGATGGACGGGAGCGTTTTGAGG

AGGAGACCATGCTCAAGAAAAATTCTCTCGGTGGAGAGGTTGATACTGATGAGAATAGTCAATGTATCTA

TTCTATCAACCCGAGATTAAAGCATTTTTCTGACTGGCTCTTAGACATTATAGCCACTGGTGATTTGGAT

GCATTCTTCCCTGCAGCAACTCGTGAATATGCTCCACTGGTTGAAGAAGTGTGGAAAGACCCTGCTATAC

AGGAAACATACAAGAGAAAAGATGAGCTTCACTTCCTTCCTGATGTGGCAGAATACTTCCTAGACCGGGC

TGTCGAAGTGTCTAGTAATGAATATGAACCTTCAGAAAAGGACATACTATATGCAGAAGGAGTCACTCAG

GGAAATGGTCTAGCTTTCATAGAGTTCTCGCTGGATGATCGTAGCCCGATGTCTGAAACCTACACAGACA

TGGAAGCTCAACTCCCACCTTTGACTAGGTACCAATTAATTAGGGTAAATGCTAAAGGCATGACCGATGG

TTGCAAGTGGGTGGAAATGTTTGAAGATGTTCGTGTAGTTGTCTTTTGTGTTGCCCTGACTGACTATGAG

CAGATGTCGGTTTCCGCTGAGAATGGCGGAAGCGGTGTGCTTCTTGTAAACAAGATGATGGAAAGCAAAG

AACTATTTGAAAAGATGATCAGGCATCCATGTTTTAAAGACACTCCCTTTGTTTTGATTTTAAACAAGTA

TGATCTTTTCGAGGAGAAGGTGAACCGGGTCCCACTAAGTACTTGTGATTGGTTCTCCGACTTCAGCCCA

GTACAACCCCATCACAACAACCAAACTCTAGCTAATCAAGCTTATTACTATGTCGCGATGAAGTTCAAAG

ACCTCTACTCTTCTCTAACTTCTCGAAAGCTTTTTGTTTGGCAAGCTAGGGCAAGGGATCGAGTTACAAT

AGACGAAGCATTTAAGTACATAAGGGAGGTTGTTAAGTGGGATGATGAGAAGGAAGATAACTACTTTGGT

ATTGAGGATTCATTTTACAGTACAACAGATGTGAGTTCCTCCCCATTTATTAGGCAAGAGTGATTTACTG

TACATGTTCTTCACGCATTAGATGTGTAAAGAGTAGGAAGAGATGCAGAGGCAAATAAGTACGGATGGTT

TATTTTTTTTCATCCTCAAACCTCGTGGATTGGGATTTGGGTGATTGTATTTCATGTAAAATCCGTCCAG

CGGAATTGTCAAAACCTTGTAATGATATTGTAGCTACTATGCATTCCGTCAGCCTGCTGGTAAAGATTGA

TGAAAATCGCTTTCACCTGCGAGCTCTGAAAGTAATTGCATCACCATGTATTTTCTTCTAACCAGAAAAT

AGTTTAGGGGCAAGTTGGATATCATGTACCTTGTATGCTGTTGTATGAACTATGAACTCTTTTAGTGTAA

TTGATATGCAGGTGAAG

>MSTRG.3320.1 gene=MSTRG.3320

CTCCATTGGGTCCGATTCATATTCTCCAAAACGTTATCCTATCTTCAATATCTAAATGTCCCATGACATT

TCTCATCTTTTACTGGCTCACAATATTTCTTATGATCTTCACAGCTCAAGGCCTCAAAAACTTTGTAGAC

CATCAGCTCACAATTCTAAACATGTAACTTCTGTTTAGTATACCACTCTTTCCATTTTCTGATATTGCAG

AGTTCTAAATTTGGATATGATGGTGTAGCTGAGAAGATGATGGTGAAGACCTCAGTGCTTCATCAACCTC

ATTTGCTTGTTCATCATGAAGATCACTCACAAGGCCAAGAATCAAGAAGTTTGAAGTTGAGAGAGCAAGA

ATGCATGTCATTGGCCAAGAAATGCAGGAACTTGAGGGAATTCAAGCAAGTGCATTGTCATGTAGTGAAG

TTTGGCTTGTTTTTCAGCTCTCTTTGCTCAAGTAATCTTCTTTTAACTTGTGCTCAATCATCTTGGGGTA

GTATGGACTATGCATGCTCAATCTTTAATCAAATTGATGATCCGGGTTCTTTTGAGTTTAATGTTATGAT

AAGAGGCTATGTTAAAGATTTGAACTTTGATGAGGCTTTGTTTATGTATCATGAAATGCTTGATATGGAA

GTTGAGCCGGATAATTTTACATTTCCATTTCTTTTGAAGGCATGTGCGCGATTGCGTGATATTAGGAAAG

GGGTACAGATTCATGGGCATGTGTTTAAGTTTGGATTTGAAGATGATGTGTTTGTACAGAATAGTTTGAT

TAACTTTTATGGGAAATGTGAGGAATTGAGGAAGTGTTGTTGTGTTTTTGAGAAAATGGAGCAAAGAAGT

GTTGCTTCTTGGAGTGCTCTTATTGGAGCTCATGCAAGAAATGGTTTGTGGGCTGAGTGTGTTGGTATTT

TCGGGGACATGATCAGGGAAGGATGTTTGAGACCTGAGGAGAGTATATTGGTTAGTGTGTTATCTGCTTG

TACCAAATTGGGAAACTATAATTTGGGGCAATGTATTCACGGGAATTTAATTAGGAATATGATTGGGCGC

AATGTTATTGTTGAGACTTCACTAGTAGACATGTATCTGAAATGTGGCTATCCAGAAAAAGGGTTGTATT

TGTTTGATAGTATGGTTGAAAAGAACCATTTGACATACAGTGTAATGATCTCTGGTCTAGCCGCTCATGG

ATGTGGTGCAGAAGCTCTAATGGTTTTCTCTAATATGCTCGAAGAAGGTCTGGAACCGGATGATGTCATA

TATGTGAGTGTTTTAAGCGCGTGCGGTAATGCTGGCCTTGTCGAGGAAGGCCTACAGCTTTTCAATAGGA

TGAAAAACGATCATAACATAGAACCAACTATTCAACACTATGGGTGTTTGGTTGATCTTTTGGGAAGAGC

TGGGAGACTAGATGAAGCTTTTGAGGTAATCACGACCATGCCAATGGAGCCAAATGATGTTGTATGGCGT

AGCCTTCTTGGTGCTTGCAAGACTTATTCAAATATCCAGATGGGGGAAATTGTAGCTGAAAAGCTGATGC

AAATGAAGTCGAGAAATGCTAGCGATTACTTGGTTCTATCACATATATATGCAGAAGCTCAGAGGTGGAA

AGATGTAGCTATTACTCGCACAAAAATGGCTGAAAGTGGATTGAGTCAGACACCTGGATTTAGCTTCGTT

GAGGTGAAGAAGAATATCTACAAATTTGTTTCACAAGACATGACACGACCACAATGTAACGGTGTCTATG

ATATGCTACACCAGATGGAGTGGCAGTTAAAATTTGAAGGGTATTCCGCAGACACTTCTCAGGTGTTGCT

TGATGTAGATGAAGAGGAGAAAACGGAGAGATTAAGTAGACATAGTCAGAAGTTGGCAATTGCCTTTGCA

TTGATACACACATCAGAAGGATCTCCTATAAGAATATTAAGAAACTTAAAAATGTGCAGCGATTGTCACA

CATACACCAAGTTCATATCCAGAGTTTTTGAACGAAGGATTATAGTTAGAGATAGGAATTTGTTCCACCA

TTTTGAAGATGGCAGCTGCTCATGCAGAGATTACTGGTAAATGAAATTTTTTATACATATCTTTTTTACA

TTT

>MSTRG.3321.2 gene=MSTRG.3321

CATTCCTCATTTTCCTTATAATTTTTGTATTAGAAAACTCAACAACACATGTAGATGCATTTGCAAACAA

TGTCACTGATCACCAAGCTTTGCTTTCTTTCAAGGCTTCCATAACAGATGATCCATTGGGTGCACTAGAC

TCATGGAACAACTCCATCCACTTCTGTGAGTGGAACGGCGTTACATGCAGTAGCAGACGACAGAGGGTAA

CAGCATTGAATCTCACCTATCAACACATGACGGGTACCTTGTCTCCTAATATTGGTGAAGGAAGATATGG

TTCAGTTTACAAAGGAGCTCGCGAATCGGTGGAGCACATACTTGCTGTGAAGGTACTAAACGTTGAAGTA

CATGGAGCAAACAAGAGTTTCTTGGCTGAGTGTGAAACGTTGAGGAATATTCGTCACCGGAATCTTATTA

AGATCATCACAGCATGCTCTAGCATAGATTTCAAGGGCAACGACTTCAAAGCGTTGGTTTTTGAATTCAT

GACTAATGGAAGTCTAGATAGCTGGTTGCATCCAAGTCCTTCCTATCAAGGAAATGAAAGAAACTTGACT

CTATTTCAGAGACTAAATATTGCCATTGATGTCGCACAGGGAGTGGATTACCTACATCATCACAGTCACA

CAAGTATCATTCATTCTGACATAAAGCCACAAAATATTCTTCTTAATGAGGATTTTGTTGCTCATATAGG

TGATTTTGGTTTGGCGAGGTTTACCTTTGCTACTACAAGTGATATCAATCAAACACAAATGAGCTCAACT

GGTGTCCATGGAACAGTTGGATATATTCCTCCAGAGTATGGAATGTTTGGAGAGATATCTATAGAGGGAG

ATGTATACAGCTATGGAATTCTTTTACTAGAAATTTTCTCAGGGAAAAGGCCAACAGAAAACAGCATATT

AATAGATGGTGTTGGTAATCTTCATGACTACGTGAAGAAGGCACTCCCGCAAAGAGTGATGGACATTGCA

GATCCGCGTACATTACTAGATCATGAAGAACATGGATCAACTGTCAGTCAAACACACAGCAGGGCTATGA

TGGAGGTATGCCTGGAATCAATATTTGAAGTGGGGTTATTATGTTCGGAAGAGATGCCACAAAACCGCAT

TGACATTTAGTGTTGCTATTAGGAAGCTACATATAGCAAGAGACAAACTTCAGCAGCGCAGAC

>MSTRG.3321.3 gene=MSTRG.3321

AGAGTTGATACCCACATTATTTTCTATTCTTTTTCTCATAGTGCAATCATGGCTTTCTGCTTTCGTCCAT

TCCTCATTTTCCTTATAATTTTTGTATTAGAAAACTCAACAACACATGTAGATGCATTTGCAAACAATGT

CACTGATCACCAAGCTTTGCTTTCTTTCAAGGCTTCCATAACAGATGATCCATTGGGTGCACTAGACTCA

TGGAACAACTCCATCCACTTCTGTGAGTGGAACGGCGTTACATGCAGTAGCAGACGACAGAGGGTAACAG

CATTGAATCTCACCTATCAACACATGACGGGTACCTTGTCTCCTAATATTGGTGATTTTGGTTTGGCGAG

GTTTACCTTTGCTACTACAAGTGATATCAATCAAACACAAATGAGCTCAACTGGTGTCCATGGAACAGTT

GGATATATTCCTCCAGAGTATGGAATGTTTGGAGAGATATCTATAGAGGGAGATGTATACAGCTATGGAA

TTCTTTTACTAGAAATTTTCTCAGGGAAAAGGCCAACAGAAAACAGCATATTAATAGATGGTGTTGGTAA

TCTTCATGACTACGTGAAGAAGGCACTCCCGCAAAGAGTGATGGACATTGCAGATCCGCGTACATTACTA

GATCATGAAGAACATGGATCAACTGTCAGTCAAACACACAGCAGGGCTATGATGGAGGTATGCCTGGAAT

CAATATTTGAAGTGGGGTTATTATGTTCGGAAGAGATGCCACAAAACCGCATTGACATTTAGTGTTGCTA

TTAGGAAGCTACATATAGCAAGAGACAAACTTCAGCAGCGCAGAC

>MSTRG.3321.1 gene=MSTRG.3321

CAATTTGATCGGTGAAGGAAGATATGCTTCCGTTTACAAAGGACATCTTGAATCAGTGGAACATTTAGTT

GCTGTGAAGGTACTAAACAGTGAAATACGTGGAGCCAACAAGACATTTTTGGCCGAGTGTGAAACACTGA

GGAACATTCGTCATCGTAATCTTATCAAGATCATCACTGCATGCTCTAGCATTGATTTTAAGGGCAATGA

CTTCAAGGCATTGGTTTTTGAATTCATGACTAATGGAAGTCTAGATAGCTGGTTGCATCCAAGTCCTTCC

TATCAAGGAAATGAAAGAAACTTGACTCTATTTCAGAGACTAAATATTGCCATTGATGTCGCACAGGGAG

TGGATTACCTACATCATCACAGTCACACAAGTATCATTCATTCTGACATAAAGCCACAAAATATTCTTCT

TAATGAGGATTTTGTTGCTCATATAGGTGATTTTGGTTTGGCGAGGTTTACCTTTGCTACTACAAGTGAT

ATCAATCAAACACAAATGAGCTCAACTGGTGTCCATGGAACAGTTGGATATATTCCTCCAGAGTATGGAA

TGTTTGGAGAGATATCTATAGAGGGAGATGTATACAGCTATGGAATTCTTTTACTAGAAATTTTCTCAGG

GAAAAGGCCAACAGAAAACAGCATATTAATAGATGGTGTTGGTAATCTTCATGACTACGTGAAGAAGGCA

CTCCCGCAAAGAGTGATGGACATTGCAGATCCGCGTACATTACTAGATCATGAAGAACATGGATCAACTG

TCAGTCAAACACACAGCAGGGCTATGATGGAGGTATGCCTGGAATCAATATTTGAAGTGGGGTTATTATG

TTCGGAAGAGATGCCACAAAACCGCATTGACATTTAGTGTTGCTATTAGGAAGCTACATATAGCAAGAGA

CAAACTTCAGCAGCGCAGAC

>MSTRG.3322.1 gene=MSTRG.3322

CCGAATACGATCTTTCACGTCTTAACTCCACCCAGTCCACTCCAAATCTCCAATCTCACAGTTCTCCAGA

ACTCACAAACACGAATGAAGTAACAGAACGACGTCGTTCCTCGTTCATCTCCATGGCGTCCATTAGACCT

AATCCTACACTTTCGGAAGATCTAATCCATGAGATTCTACTTCGCGTGCCGGTGAAATCATTATTCCGCT

GCCAATCAGTCTGCAAGAATTGGTTATTCCCTCATCAAACACCCTGATTTCGCAAAAGCGCAGCTCCGTC

ACGCAATCGCAATCGAAACCGATGATACTCTCATTATCAACCCCTGGCCACGCGAAAGCATAGCACCAAA

CAAAATTTTACTATTCCATTTCAATTCCCTCGAAATTGTCGCGGATCTTGAGCTTTCGTATCCCCCAGGC

GAGTTCGATTCTGTGACTGATATAACAGTTGTTGGTTCTGCTAACGGCATTGTTTGTGTTGCTGTTGCTT

ATATTGAAACTGATCCTGAAATTAATCTATATCTTTGGAATCCTGCTATTAGGCAATCGGAAGTTATTCC

GTTGTTTGGTCATGAGCTTGGTGCACAGGCTTTCGGGTTTGGTTATGACCCGATAGATGATGATTTTAAG

GTTGTTGTGGTTGTAACGGTGCCTGCTCTTTATGCTGAGGTGTATTCGGCTGATAGAGATTTTTGGCTTA

ATGTGCCTGATGCAGTTGATGCTCCTGTGAATGGTGATTTTGATGTGTGTGTTGAGGGATTTCTGTGTGG

TGTTGGAGAGTTTGGTGTGATGACGTTTGATTTGAACAAGGAGGTGTTGAATTGTGCTATTAAGCTCCCG

GTTATTTTTGCTGATGGTGAAACTCGTGTAATCGAGTCTAGTGAGGCTCGTCTTATCGAGTTTAATAAGT

CTATTGCTGTTATTATAGCGTGGGATGATGGATTGAATGATGATAGGAAGATTAACATGTGGACGATGGA

TGATGATGTGTGCCTTGGTGGCGGTGGAGTTGAGGCATCATGGACGCTAATGTTTAGTATTGATTTCGGT

ATGCGGTCAGATATTGTTCATGGCTACTTTACCAACAGTGATCTCCTAGTAATATTGGATAATGCTCATA

CGTGGATTTCATGTAATACCGACAAGAAAGAGGCCAACATTGTCCCACTTTCGGTTGACATGGAGAAATA

TCTTCACACTCGTCATATCTACAAGTATACAGAGAGCCTAGTTTCAGTCAATGGATTCAAGCAAGTCGAC

TGCAATGCTGGGGAAGATGATGATTAGGAAAGCAGCGCGGAGTCTGGTATTTTGGTGCCAGCAATATCTT

CGTTGAAAGAGTATGGAATGGGTGCTGAGATATCTGTAGAGGGAGATGTATATAGCTATGGGATCTTTCT

ACTAGAAATGTTTACGGGCAAACGACCTACATGTAGCAGCGCATTGCTGGACAATGATAATAATCTTCAT

GACTATGTGAGGAATGCACTCCCACAAAGGGTTATGAACATTGTGGATCCACGGATCATAATAGATCAAG

AAGATCATGGTTTAACAACAAATCATTCATACAGCAAGGCTATCTTAGAGGTATGCCTGACATCAATATT

TAAAGTGGGGATACTATGTTCAGATGAAATGCCCAAAGAGCGCATTGACATTG

>MSTRG.3322.2 gene=MSTRG.3322

CTTTAAGGCATTGGTTTTTGAGTTTATGACTAATGGGAGTGTGGACAAGTGGTTGCATCCAAGTCCTGAT

CAAGGGAATGAAAGATACTTGACTCTACTTCAAAGACTAAGTATTTCCATTGATGTTGCTCTAGGAGTGG

ATTACCTACATCATCACGCCCATGATAGTATTATTCATTGCGACCTAAAGCCAAGTAATGTTCTTCTTGA

TCAGGATTTTGTTGCTCGTATTGGTGATTTTGGTTTGGCGAGGTTCCGCTTCGCTGCTACTACAAGTGAC

ATCAATCAAGCACATTTGAGTTCTGCTGGTGTACGCGGAACAGTTGGATATGTGCCTCCAGAGTATGGAA

TGGGTGCTGAGATATCTGTAGAGGGAGATGTATATAGCTATGGGATCTTTCTACTAGAAATGTTTACGGG

CAAACGACCTACATGTAGCAGCGCATTGCTGGACAATGATAATAATCTTCATGACTATGTGAGGAATGCA

CTCCCACAAAGGGTTATGAACATTGTGGATCCACGGATCATAATAGATCAAGAAGATCATGGTTTAACAA

CAAATCATTCATACAGCAAGGCTATCTTAGAGGTATGCCTGACATCAATATTTAAAGTGGGG

>MSTRG.3322.3 gene=MSTRG.3322

ATTAGACCTAATCCTACACTTTCGGAAGATCTAATCCATGAGATTCTACTTCGCGTGCCGGTGAAATCAT

TATTCCGCTGCCAATCAGTCTGCAAGAATTGGTTATTCCCTCATCAAACACCCTGATTTCGCAAAAGCGC

AGCTCCGTCACGCAATCGCAATCGAAACCGATGATACTCTCATTATCAACCCCTGGCCACGCGAAAGCAT

AGCACCAAACAAAATTTTACTATTCCATTTCAATTCCCTCGAAATTGTCGCGGATCTTGAGCTTTCGTAT

CCCCCAGGCGAGTTCGATTCTGTGACTGATATAACAGTTGTTGGTTCTGCTAACGGCATTGTTTGTGTTG

CTGTTGCTTATATTGAAACTGATCCTGAAATTAATCTATATCTTTGGAATCCTGCTATTAGGCAATCGGA

AGTTATTCCGTTGTTTGGTCATGAGCTTGGTGCACAGGCTTTCGGGTTTGGTTATGACCCGATAGATGAT

GATTTTAAGGTTGTTGTGGTTGTAACGGTGCCTGCTCTTTATGCTGAGGTGTATTCGGCTGATAGAGATT

TTTGGCTTAATGTGCCTGATGCAGTTGATGCTCCTGTGAATGGTGATTTTGATGTGTGTGTTGAGGGATT

TCTGTGTGGTGTTGGAGAGTTTGGTGTGATGACGTTTGATTTGAACAAGGAGGTGTTGAATTGTGCTATT

AAGCTCCCGGTTATTTTTGCTGATGGTGAAACTCGTGTAATCGAGTCTAGTGAGGCTCGTCTTATCGAGT

TTAATAAGTCTATTGCTGTTATTATAGCGTGGGATGATGGATTGAATGATGATAGGAAGATTAACATGTG

GACGATGGATGATGATGTGTGCCTTGGTGGCGGTGGAGTTGAGGCATCATGGACGCTAATGTTTAGTATT

GATTTCGGTATGCGGTCAGATATTGTTCATGGCTACTTTACCAACAGTGATCTCCTAGTAATATTGGATA

ATGCTCATACGTGGATTTCATGTAATACCGACAAGAAAGAGGCCAACATTGTCCCACTTTCGGTTGACAT

GGAGAAATATCTTCACACTCGTCATATCTACAAGTATACAGAGAGCCTAGTTTCAGTCAATGGATTCAAG

CAAGTCGACTGCAATGCTGGGGAAGATGATGATTAGGAAAGCAGCGCGGAGTCTGGTATTTTGGTGCCAG

CAATATCTTCGTTGAAAGTCTCCAATTTTTATATATGTTGCTTGATGAATGATGGCTGATTGGCTGATCT

GCTGATGCCTGCTAACTGCTTAGTTGAATTTTCAGTCGGTTATGTTTGAGAATTGAGACTTTGAAGTAGT

GATCTATATTTTGTAATGCTATAACTATTGCTTGTTCAAATTTTCTAATTTGAAATGCTCAAGGATTGTT

TAAAATTACATGTTTTGCGTAATTTAGGTATAAAGTTAATGCTACTACTTACCATTAGTAGTTTCAAAAT

TCGTGTTCTGGTAAACCTAAACCCAAGCCCAGTCCGAAATCCGTTGTTAGTTGATCTAAGTATTATAGAT

CCTTTAACATATTTGAATCCATTACTCGATACTTTAATAATTTTAGTTGACATCTACTGCTGTAAGTGGG

CATCAAGCTGTAAATGTCAAAATGTATGGCAAGGCATCCGTGGGAGCAGCTTCTCTTGTTCAAAAGGTAG

TTGCATACCCAGGATTTAGAAGTACCAGTCTGTGATCCTGACTCCTAGACCTAATTACACTAGTTTCTTG

TAGATATTAAAGTTTTTGAGAGACTGGAATGATGTTTCTGTTGTTTAATTGGATAAATATTTAATTTCCA

TCACCTTCAACATAACATTCCCAGTTCTGATTTTTTTCACAACCGTTCACATAAAATTTGCATCTCGTTA

TGTTTTTCCGCTTTTCTCTCTCCAACCAAACAGAACGTTATAGTCTCAGTTAAAATTTTGGAGGACCAGT

TTCTTTCCAGTGCCATTAGCCAAACAAAAATTTCCAAGCCTGTATAGCTGGACTTACAAAGTCGTCAGTT

AACCCAATCAAATAGCTAGAAAGTGGAGACAGACTATATTGGCTAATTCATGGCAAGCAGAAGCAGTGGA

GCACCGAGAATGCATATAATATACACTGTGCATGGAAAGAAATTGGGGATTCACAAGCACTGTGAATATA

ATGTCACTACCAAGGTTGTATATAATGTCCATAAAAGTTCTACAATCAATACACCACCCCCTTGACAATT

TCTCTTATTAATTGTTCAAAACCATGATTTTCCGGTTTTGCATTCTCTTTACTCTTCTGGTATTAGCCAG

GCCACAAAAATATGTTTTTGCATTTTCGAG

>MSTRG.3330.3 gene=MSTRG.3330

CATATCCAAAGTCTGTCCCCGTAGCATATCCCGTTCATCAGGATCTAAGTGAACATTTAAATTCATATAT

GTGGGATCCTTCACTGCGAGATCCCTCGCCCGGCAACAGTATGCAGTGTTTGTATTTATGCAATAATGAT

GATCTTATTAATTTTAGCTGGACTGGTTTTAGGGGACACTTACAATGCTAACATGTAAGGTTATACAATC

ATGCAGCAGCTCAGCGCATCTGTCAGAAATCAAATTTGAAGAAAAGCGCTTTTTATTCCAGCAACTCGAT

ACTGGATATTGGAAAACAGGATCATAGCTGTGGTTTTTGTGGAGCCCGAGTTTGGACTGCTGAATTTACC

GGAAGGGCCACTGGCAATGGACCTAGAGGATATTCGATTTGTTGTGGGAAGGGTAAAGTACATCTACCTC

TATTGCGCGAGGCTCCGCCGGAATTAGCTGCGTTGTTATCTTCTACATCAAGGATGTCTAAAGAGTTTCT

CGCCAAAATTCGCATTTACAACAACTTATTTGCATTCTGTTCTTTTGGAGGGAAGGTGGATCACTCAGTA

AACGAAGGTAAAGGTCCATTTGTGTTCCGTGTATCCGGGGAAATATATCATGCCTACAGTTCTTTGATCC

CACCAGAAGGGCGTACCCCAAAGTATGCACAGTTGTACATGTATGACGCGCAACAGTCCATTACTCATCG

CCTCAACTTCCCGGGCTCTGGTAGTGAGGCTGATCCTTTGGTTGTTTCAGCCTTGGAATGCATGCTTAGG

CGTGAAAATGCCCTGGTTGGGATTTTTAGACAAGCTCGCGAAAGATTTCAAGGGGTTGATCAGATGCCGC

TTCAGCTTAGATTGTTAGAGAGGAGGACAACTGATGGTCGGTATGAGAATCTGCCCACCGAAAACGATTA

TGAATTTGCAGGCCTGGTGGTGGATGATGACTTTACGAATTCTAGGGACATAGTTGCCCACGATAAGATA

TCCGGATTGAAACATATCAGCGACCTGCATCCATGTTTTATGTCTATGCAATATCCTGTACTCTTTCCGT

ATGGGGAGGATGGCTATAGAATTGATATAAAACATCGTAATGTCACCGGATCTGAACCCAGAAAGATGAG

TACTGTGAGCCAACGTGAGTATTATGCTTTCAGATTGCAATACCGACCCGCTGAGGGGCAAACAGTTTTG

CGTGCTGGCCGGCTGTTCTTGCAATTTATAGTTGACTCGTGGTGCTCCGTTGAGCGTGGTCGATTGCAGT

GGGTTCGAACACACCAGTCTATAATCAGATCGGACCTGTATAATAATATTGTTGATAGCCTTACACGTGG

TGATTTTGTGGCCACTGATGTTGGGAAGCGAGTAATTCTGCCATCTAGCTTCACAGGTGGCTATAGATAC

ATGCAACAAAATTTTCAGGACTCCCTAGCCGTTTGTAAAGAATACGGGCACCCGGACCTTTTCATCACAT

TCACCTGCAATCCCAAGTGGGATGAAATCTCAGAGGCTGTAAAATCTTCTGGAGCTCAGGATGCCTCCGT

CCGACCGGATATTGTTGCGCGTGTGTTCAAGATGAAGCTTGACGCCATGATAGCGGACTTCACAAAAAAT

GAGGTGTTAGGCCGTGTGCTTGCAGGTTACTATGCAGAATAGCATGAAATCATGCCGGGATTGGCCAAAC

CTGTTGACCTTGGTATTTACATGATGGTCTTGCTGTTCATGTCCTCTTTGTTTCTTACGATCTAGATATT

TCTCCATAATATATTTAGCTCCATGTTCACTGCTTCCACCCTTGATTGTCA

>MSTRG.3331.1 gene=MSTRG.3331

TGTCCAAGTAAGCCATTATTTATTCCACATGGCGTGTCTCTCGTGAGGATTAATACTCTAACCTGGTTAA

AGATATCAGCAATCAGACAAGTCTTGTCTCTTTTCCTCTGGGTTCGATTCCACAAATGGAAGTTTTGCTT

TGTGTGAGTGTTACCATTATGCTGCGTGGCATTTCCGGGAGGGCATGGAGGCGTCGGACCAGAACAGGGT

ATTGCATCAGATCAGGCCTGCATGTTGATTGACATCTTTTGGTTCTTTGGGTTTACGTGACTCATATTGG

TTTTAGTAATATGGTCTTAATTGTGAACTTCCAAATGGTATCCATCCATTCATTGCTATTTAAAATTTAC

AATTTGGGTTCGGGTGAATGAATTTGGAGGAACTCCCTTCCTCCTAACAAGTCCTTCAAGCATCTCCGCG

GCAGATTAGCTATGTGATATCCTGAGCTATAATTTTAGGGAATATGCACAAACATTTACTTTAGCAGAGT

CCCTGTTCATTTCAAGGATCCATTTTCATTCCTCGTAAACAGGGTGGTGCTAGGGGAACTCTGGCCCTGA

TGGTGGTTCTGCAGATGATCTGGCACATAGAAGGCCAGTGTTGTCAGAAATCTGGGTTCTCTCTCATGCT

TGAGGCAATAGCAGAAACTTTCTCGTGCTGAAAATACATTCATTTGAGGTATTCCAAGTTATTAATTGCA

CTTTCAAATTAAGATGATTAGCATAATTAGTGTTTATGATATATATTTATATGCTAGTAATTGTTACTCT

CGTAAAAATAGAACATAAACTTATATATACCTCCGCTCTTAGCCGAGCATTCTCGAGCTGAAGCTGATGT

GTTTCAGAATCCACCTTTTTCAGATCTCCTGAACAGCGAATGCATTCAATGTTTCTCAGTGCG

>MSTRG.3332.1 gene=MSTRG.3332

AGTAAAACTACAATTTATTGATATATTAAGTACATCTAATACTTTCAATAATAATTTTTAATATCTATTC

ACAGGTTTTTCCAAATACGTCAACACCCTGGTCACGAAGAAAGACGTCGGCTTAGTCATGATTTGGGTCT

CGAACCGAGTCAAGTCAAATTCTGGTTTCAGAACAAGCGAACACAGATGAAGGTTCTGCAGGATAAAAAT

GACAGTGGTTCACTGGAGAGTGAAAATCAGAGGTTGACAGAATTGAAAAACAAAATGCTAGCCGCACTGA

GAAACATTGAATGCATTCGCTGTTCAGGAGATCTGAAAAAGGTGGATTCTGAAACACATCAGCTTCAGCT

CGAGAATGCTCGGCTAAGAGCGGAGCACGAGAAAGTTTCTGCTATTGCCTCAAGCATGAGAGAGAACCCA

GATTTCTGACAACACTGGCCTTCTATGTGCCAGATCATCTGCAGAACCACCATCAGGGCCAGAGTTCCCC

TAGCACCACCCTGCAGGCCGGGAAAAATAGCTAATTAATTAGTCAATACTGGGAGGTCCTCTTGGGCACA

ACATTGG

>MSTRG.3333.1 gene=MSTRG.3333

TTTAAAGTGAAGGGTGACGTTTTTATTGTAAATTATGGGTAGCAACAATGAACATTTGAGATAAGATGGT

GAGCATATTACTGAACTCCGATTGTTACGATCATTCGGATTGAGCTCGGCTCATTGTTACCATCACTCTG

AATTGGAACCTGAGTAGTGATCAGCTGGAAACGTTGATACTAAGAGCATCTCCAATTGGG

>MSTRG.3340.1 gene=MSTRG.3340

ACATTTTCCCCCTTTTTCCCCTAAATTGACAAACCCTTTTCTCAGCAGCGGCGACTAGAGCTAGGGTTTC

GCACGATTTCACACCGTCGGCGCAGTTTGGCAACGATTTCTCACCGATGTCTTCACAGATTATAGGAGCA

GAGGCTCGTAAGCGAAGGCGTGGTAAAGGCAGTGGCGGGGGAGGGCGTAGTAAAGGCAGTGGCGAGGGAG

CGAGGGGGGCACGTAAAATCATCAAAGAGAGTAATAAAAAGGATGACAAGGGGGCAGATAATTTGAGTGA

TGGAGAAAGTTTCATGGAAGCTTTCAAGATGGGCGGGACTAGCAAAAAAATAACATTCGAGAGGAAGAAA

CGCGATTGTAACCAAGGTGAATATGGAAGCCAACCACCCTTTAATGCAAGGCCAATCTTGTATTTTAATG

AGATGAGAACAGGCTTAATTGGCTTTGAAAAGAAGAACACTTTGGGCTATATTGTCAGAGAACTATGGAA

TGAGCGGACGGTTAACAGCAGGGGAGCCGAGCGTGAAAGTTTTTATGATGATGTTATCAAAGAGTTTAAA

CTTTATTACAAGTATCAGGATGGTTGTACAGTCGAGGAGGGGGACAGAATTGTGAGAATGCATTTGCAGA

GGAATTTCAAGTCGATTCTTAACAAGATAAAGAGCCGACTTGAGGATGACGTTGATGATCTTAGGATAGC

TGGGTACACTGACGAAGAAATCCCGTGGGGGAATTTTAAGCCTCATTACTTTACTTCCGAGGTATGGGAA

TCCGTTCTCAAATATTGGAGTTTGCCCGATACAAAGAAAAGGTCAGAAAATGGGAAGGCAGCACGCAAGA

TGTTACAATGTTACTCGCTCACCGGGGCGTTGTCTTATGATGAGAGACGGGAGAAACTTGATAAGGAAAG

AGAGGCGAGAGGGGAACCGCGGCTTACAGATCTTGAGTTCATAGATTATGTGTACAATCCTGAAGAACCG

GCAGTGAAAACCCTAAAGGTTCAAGATAAGCAAGCGAAGTCTTCTCAGCCTGCTTTGCCTGAAGATGCCG

CTTTGACTGCTGGTTTGTCTGCTGCTGACGCCGATAAAATGATAAAAAAACTGAAGCAGAGGCAAGATAT

ATTGGCGACACTGAAGGCCAAACCTCCGAGAAAAGGGAAAGCAATTCTTCGCCCGCGACAAAGCTTGTCA

GATCTTCTAGATAGATCAGAAGCTTCACAATTTGCTGAAGGATCGCGCGAAGCAAGCCAGGACATTTCAG

GTCATATCTTTGGGATGATGTCCACTATATTGTGCGAGGTGGATGAAATGGTGCGCTCTCTGCCTACTCG

TGAGGTCACGCAGCCTGTAGTGGATCAGAAACTGAGCAAGTTGGCTGCTGCCACTTTTCCAGACCCAGTC

CAGGAGTTATTCCGGACTGATTATGTGTGTGCAGCGAAGGGATTACTTAGTCAAATCCTCAAGAACAATG

ACAAAGTCATTGTAGAGGTCAGTCAGCAGCAGCAGTTAGAGGGGAATGGGAAGGAGCAGCAGCAGCCAGA

GGGGAAGCCAGAGGGGAAGGGGAAGGAGGCTGCTAGCACAGAGGAAAATCCGTATGATGATGACTATGAT

GCAAATGCGTTTTTATACCCTCTGTTAGATTAGCAGCAGTACTGGTGGTAATTTGGAATTTCATACTTTT

GGTTGATGGTTTAGTAAGCCAGTGAATATTCTAACTTTTGGTGGATGGATGAATTTCTAACTTTTGGTGG

ATGGTTTAAATATTTAAGTATATTATCTGTAGTTAAGTTGATTAGTTTGGTATTTTTTTGTTG

>MSTRG.3342.1 gene=MSTRG.3342

GCCAAAATAAGAAACTTGATCAAAAAATTTTAACCTCTGTTGAACTGTAAAAAAGCTTCCCATTTTATCA

ACAACATGGATTAATCTAATCCAATCCATCAATATTTGTCAACCCCATCTCCAAAATCAAATCTTTATGC

TTTCTTGTAAGTTGATTTTCACTATCTTCAAGTTTTTTACTGTTTGTATTAGTGAATTTCTTTACTCAGT

TGTTGCTTGAGTTGTTTAATTGAATGTTGTGCTCTTATTTGAAACTCTGGATCTCAGCTCAGTCCTTATT

TGAATTAATTCTTTGATGTGTTATATGTAATATGGTGTTGTATATTACTGTTCTTGTTCTTTGAAATAGT

TTAATTTTTCATTGTTCTGATGCAAATTTTGAAATTTTTTTGTTTTTATGGGGCATTAGTATGTTATTG

>MSTRG.3348.1 gene=MSTRG.3348

TTGGTTACATCAAATTCCTGCTTCAGGTCATGGGAATGAAAGAAACTTGACTCTACTTCAAAGACTAAAT

ATCTCGATTGATGTTGCCCATGGAGATGGAGTGGATTACCTACATAACAACAGTCACACAAGTATGGTTC

ATTGCGACTTAAAGCCCAGTAATATTCTTCTTGACGAAGAATTCGTTGCTCACGTTGGTGATTTGGGTTT

GGCAAGGTTCTCTGCTGCAAGTATAGATGAAATCAACCAAGAACAAACAGGTTCAACTGGTATCCAGGGC

ACTGTTGGATATATTCCCCCAGAAACGACCAGAAAGGTTCCGACTGAGACCTCAATTACTAAGCCTGCAC

CTAGAAGGTCCTCAGTTGAGGCGACAGACAATAAGGATGATAATGACAACAAAAACATTGCATCTCAAGC

TTTCACTTTCCGTGAGCTAGCAACTGCAACAAGAAATTTCCGACAGGAATGCCTTATAGGAGGAGGA

>MSTRG.3349.1 gene=MSTRG.3349

AGGTTGTAGCGATAAAGCAACTAGATCGTAATGGATTACAAGGAAACAGAGAGTTTCTGGTAGAAGTCCT

GATGCTGAGCCTCCTCCATCATCAACATTTGGTTAATCTGATCGGATATTGTGCTGATGGAGAACAAAGG

CTTCTTTTTTGGATGAAGATTTCAGTGCGAAGCTATCAGATTTTGGATTAAGCTTGGACCTGTTGGGGAT

AAATCTCATGTATCATCCAGATTCCAGAGTAATGGAAACATATGGATATTGCGCCCCTGAGTATCAGAGA

ACGGGTCAATTAACAGTCAAGTCTGTCTGACGTAAACAGCTTCGGGGTTGTGTTGTTGGAGATGATTACA

AGAAGAAGAGCCATTGATACCTCAAGCAGAAATGAAGAGCAAACACTAGTTGCCTGGGTAAGCTCTTTCC

CAATTAAGCCATCGAATAATTAATAATGTTATTTTGCCGCGGCCAGTGATAATTCATAAGTTGTACTAAA

ACCTTTGAGCAGGCAGGACCAATGTTCAGAAAACAAAGCAGATACTCAGAACTAACTGATCCCCTTCTAT

AAGGAGAGTATCCGGAAAAGAGCGTTAATCAAGCAATTGCAGTTGCTGCAATGTGTCTGCATGATGATGC

ATCAGTCCGTCCACTGATGATTGAGGTGGTCACTGCTCTTAGTTTCCTTGGCGACGCAGCTGAACCTATT

GTTGCTTCACCTATCACACTGCCTTCTCCACAAGCTGTGAGGCCTGTCAGCGGAAACCATTCTGATGAGA

TCAATAAAACAGAACGCTAGCTAGCAATAGCAGAAGCCAAGGAATGGTGCACAAAATCAAGAGACCAACC

GGTTTCACAAGCTTCTGTTTACCGCTGATATCCATTTTTATTTTGCACGCATGTACTATTGAACTTTAGA

TCTGAGAATCACAAAACTCAAAGACAAGGTCTTATACATTTACGCGGAGATTTTTTTAAAAGAGATGATC

CAAAAATATAGCTGCATTTCGTAGTGAAGCCTGAATCTCAAATGCATAACGTTCTAC

>MSTRG.3350.1 gene=MSTRG.3350

GTAAGGTTGGGTTATCTAATTTTTTTTTAATTAATGGGTTAAATTTTTGTTAACCCGAACTAATTGGGTT

GGGTCGGATTTTTCTAAAAACCCGCCCCAACCCGACCCATATTCACCCCTGCCCAAGAGTAGATTTGCAG

CTCCAGGACTATCAAACTGATTCCAGTCTCTCTCTAGCAAAATCAGGCATTTGGGTTAGGGTTTCACCAC

GCGATTCGAGAAGAATGTCTCGGCTAACCAGGGGAAAACGCAAAAGGGTGTCGCCGGCGCTTAAATCGGG

ACGGCCTGGAAAAAAGATGAAACCCATTCCGGAGCTTCCACAAGAATTATTCCGAGATGAAATCCTGTCA

CGCCTCCCAATCGAATCCGTGTCACGTTTCAAGTCTGTATGTAAGTCATGGCTATCTCTATTTTCCGATC

CTCACTTCGTGAAACAACATAATGATCGTCTAAACCCTAAGGAAAATGACTTGCTGGTTATTAGAAAATA

TGGCCACATTGCTATTATCTCCCGCTACCAAGAGCAATTTCTCCTCAGAAACGATGATGATTGGAATCTG

ATCGGTTCTGCCAAGGGCTTAGTCTCTCTGTGTTGTAGAAATGTAGTGTCCTTGTGGAACCCCGCAATTC

ATCAATCTAAGGAATTTCACCTTCCACCGCTAGTTTATGGTTGTCTGTGTAGAGCGGGACTCGGTTATGA

TCCTTTAAGTAAGGACTACAAGGTGGGGATATGCTATCGGTCGCTTGATAAGTTACGTCTTTGTTATGCT

GTTTACTCCTCGAGTTCAGCTTCATGGATTCATTATCAGTTTGATGATCGTGATGTTGAGATTGTTATTT

CTGATTTTGAGAATAAATCTCTAACTCCGACTACCATGGTTAGGGATTCTCCTTATTGGACTACTTCTTC

TACTACTATCTCGTCCTCGAATGAGGAGAAGGCCATTCTATGTTTGGATGCATTCAAGTTTGATGCCACA

AGTAACAGGTTCAAGTTGCTGCCTCGGTACTCCTCTGAATTCAGCAAGGATAAGAAATTTAAAGTCGTCT

GTATGAGGGACCTTCTTAGTTTCATGGCATATGAAAGATGTCCAGTTTGTAAGCTTGAGGTGTATTCTTT

GGACGAGGACAAGGGATGCAAGGGTGGTGTGTGGAGTAAGATGTACAGTTTCGGACCATTTCTACGTTTT

CGACGTTTTGACTGCCTGCAACAGGGTTTCAAGTATGGCGACGAGATTGTAATTAATGAATATGGTGACC

TTACCCGCTTTAATCACAAACTGGGGACTTTTGAGCGCATCCCGGGTGTAATGGCTCCTGATACGGGTGA

CTGCTTTACATATACACCTAGTTTGGTTTCCCTTCCAGGAATGACCTCTATTTATTTAGAAAACCATACT

CCTCTACCTTGTCATCACCCTACAATTCGTCCTCTGCGTTTAGTGAATTCACTAAGAGATCATTTGCCAT

TAGAGAAGACGACTCAAGTGTTCAATTGAATGCCAGGTTTGAAGGTCATTTTAACTGAATCTGCGCTTTA

CAGCACTCTTATTTTATGTGGGTTTTGCTATCTTTCATATTTTGGTATCTGGGAAGTTTTAAGTAGTTAT

GTTTGTTTTTCCCCTTTATATTTGTTTCCTTTAAATGTATGAAACATGAAAAAGGAAAAATGTTAGTGGT

TTGTAATGTGAAAGTAAGATGGAAGTAGTATTAAGTAATGTGAAAGACAAAAAACGAATGCCTTAGATGT

TATAGACAGTGGAATTGCTGAATGTATTTCTGCTTCCTTGTAACCCGGGGCAGTAGTTTTGTTCTGGTTT

TCTCTCTTACAATGATGATAATGATTGAGAAATGTATGGAAAGTTATTGTTAGCTGATGCAATTAAATTT

AGATGTCACAGACGGTGGAATTGCCGAATGTGTTCCTGCTAGGTTCTAATCTCTCGTTCTCAAGTCTGAC

TTAATAACATGATGGTTATGTCTACC

>MSTRG.3352.1 gene=MSTRG.3352

CCTGTACTTTAGCCGTCTTTGACGATTGCCTAAACGGACTCGGGAAATGCCACCCTTTCGTCTTCACCGC

CTTGCTGTCGGCGGCGCGTGAACTCAACGCCACGCTCACCGACCTCGACCTCTTCATCATCAACATGAAA

TCGCTGCTCTTTTTACTCTCCTCATCCTTCGGTTTATTCTCAACCTCTTCCACAACTCTCTTACTCTTCT

TAAACATCGACAGCAACGACGTCGTCTTGCCTCTGCTACTCCCGCCACTCCCCGCCGCCGGAGCATTCTT

ACTTGCTGTCACCGTCGTCGCCGCCGTCGGCAACGGCGGCTTTTCGGCGGTGAACTCGAGGTTCGGCGCG

TACTTCGAACGCGACCGGAGAAACGGTATCGCCAGCGACCTCGACTTCCGCATGGTCGGCTCGGTGTCAC

CGGAGGCCACGCCGGAGAAGAACGAGAAAGTCGAAGATGATGACGTGGACGCTGACGTGGCGGCAGGGCA

GCACTCGCAGGGGCGTGTGTTAGCGCAGTTAGGGCACAAAGTGTTGAGACGTTCACGCAAACAAGTGGGG

CAAATCCCAGTACGACGCCGTTTTGAAGGGTGTGTCGGACATTTGAAAATCTCCTCTTCATCTATATAAC

AAGCCATTAACGCAGCTTCTCGATTGTTCTTAACTGTGTGTTTTTGTGTGTGTTTTTTTTAGCTCTCTCT

CAAAGTGTATCTTGGAGTTGTGTGTTAATGTTAGGTATTGAAATTTTATGGGAGGAGTTTGTATAAATAT

GGAGAGATGGAGGGTTAGTTGCGTAAATCATGGTCAAAATAATATTGGCGACTCGGAGTAGAACGAGGGA

TTATTTAATCGTTTTG

>MSTRG.3356.1 gene=MSTRG.3356

ATTCGATCACTAATCAGTCCATCCCTTTCTCTTAATTCTCTGCATTCAACTTACTTGCCTTTCCAACTCA

CACATAATGGGTTCAGCTAGAATTGTTTTCAGTTTCATGTTACTGATAAGTTTGGTCATGACCAGCACTT

TTGTATCAGCTGCCCGTCAATTGAATGAAACTCCAAAGCTCAGGGACCAAAATGCAAATTCTGAAAAGAT

GAGTGGTGAAATTGGAAGCGAAGCAAAAGATAAAACTTTGGGTGATAAGAAATTCTTTCTTCCATTGCCA

TCACCAGATTCACCACTCCCAGAACTCCCTCCATTTCCGTTTCCATTTCAAAATCTTCCGGGGCTTCCAC

CAATGTCACTCCCCGCTCTACCAAATTTCTTCACGCCGCCTCCACAAAACTAGAAGACAGTCCACTCAGA

CACTTAGTTGTGCTTCTTATTGAACTTTCATTTCATTTGCAATTAGCATATCGTATTTCTGTTTGTTCTA

GTTACATGTTTGAAAATTGTAGTTTGTTGTGATCAATAAAAGTCGAACTGTAAGACGACATTTCTAAGTT

ATGTAGGTCATTCATTACTGTTGCATCCGATCAGTTGTTAAAATGTACTCTGAGCAATGTAATTTGCCTT

GTACACATGTAATATCAACATATCATCAATTATAATTTGATTGCTA

>MSTRG.3362.2 gene=MSTRG.3362

TTACAGTAGTTTCAATTGCGATAGCATGATCATGACCTATTGTGGATGATATTTGACAAGAACTTACCCA

TCACTTGGAATGCTATCTTCTTTTTGTCGATTTCTTGCTGTTTTGATATAAATTCTGTTAAAATTTTGCT

CCCAGGACTAATGCTTTTTTGTTGCTAGTATATTGACGAATTTTTTGACCTTGTCTCTTGCAGGGATGGT

TTTCTTTCCAAGGGATTCAATTTGAGGCTCTAAGATACCTGGAGATGGTTTATTATTACTTTGTAATGCA

TGAATGTATCAGATATGGGAGATAAGCAGACCTGTAGATCTTTTGGAGCAGAAGAGAGTTGACCCAGGGG

GCAGATATATTGATGTGCTTGGCATAATTTATTGTCCCCTGTGCTTTTGTTTATTGTTTATTTAAACAAA

ACACTGTTAGAATTTTTGAATCAGGAAGTAACTGATGTTTATATCTGATGTTATCTTCTGTTAGACTAAT

ATTTCTGCTGGTTTGTTTGTTGAAAACAATTACTCCCATTGGTATATATTTTGAATATGGACAGAGTTAT

GTTGTTCATCTAAGAGAGTTCATTCCAATTTGGTGGGCAAAGTTATATAACAGCGTTATGCTGTGTAATT

TAGTGCACTTTAACGTATGGAAAAACTCTC

>MSTRG.3364.1 gene=MSTRG.3364

ACACATTTTAGCATCAAGTCAGCACTATATCTTGTTTCTCATTTCCCGACAGACTAGTGTAGTGAAGTTA

AATTAGGGCACAACTCATCTGAGGAACAAGTTCAACACAACACATCTAAGCTTTACAATTTCAAGATGGA

ACCTCCTACTGATTTGTTGGATGAGATACTCTGCCGCCTCCCAGTCAAGTATCTTCTTCGGTGCAGATGT

GTGTCGAAAGGATGGTGCTCCCTAATTGATAGCACCCCCTTTGTTAAGAAGCATCTCAACAAGGCCCTTG

AGTGCAATGCTGGTGGTCTTATTATTAATGAAGGCGGGAAGTTTTACTTGGCTGAGGATTTTAACTCTGA

TTTGGATGGTAGTGATGATAATGTTGATGTGGTCGCGGCAGAGATAACTCGTCCACTCAAGACTCGTGTC

TCTGGTGCTAATTTTGTGGGTTCTGCTAATGGTTTAGTTTGTGTGTCCAAGAATAAGATGAATGAGTTCT

TTATATTTAATCCAGCAACCAGGAAGTCGAGAAAAATACCCAGTGCTCCACGTGAGTTTCCACTTTCGTT

TCATATGACGGAGACTTCCTTGTGTGGATTTGGATATGATCAAGTAAATGATGACTATAAAGTTGTGAAG

ATTGCTGAGTGTTATCTTCAGTTTCGCGGCATAATGGCCTTTGTCTACAGTCTCAAAACAGACTCTTGGA

AACGGATTCAGAATGTTCCTAGTAACACTCGGTTCACAGGAAATTGGGGAATGTTTGGAAATGGAGCTCT

GCACTGGTTGGCTATCAAGAATCCAGCAAACTGCATAGAAATTGTTGTTGGTTTTGATCTCGGGCTAGAA

CAGTTCAAGGAGGTCCCTTTTCCTGTTATTGAGGGCCCTTTTGTCAATTTTAATACCAGGAGTCTGGTAT

CGGATGGAAGTTCCCTTTGTGTTCTTGATATGTACCCTAATTCTCGTATAGATGTGTGGGTGATGATGAA

TAATTCCGGGGAGGAGAACCTTTGGTCCAAAGCACTTTCTGTGGATAAACACGGGATGCCTGGGTTTTCA

AGGTTTGTTAGACCTGTCCATATTTCATGGAGTAGCCAGAGTGTACTCTTAGAGATAGACAGCTCAAGAC

TTGTGTGGTATGACCTTGAAAGGAAAATGGTCAAGAATGTTAAGGTTCGTGGGATCACAAACCAGTTTGA

TTCACATATGTATACTCAAAGTCTCATACAGCTCGATGAGGACAATCAGCTGCAGAAGCCTTCACATGAC

AAGCCACCGAAGGAACAACAAAAGAGAAGGGATGATTTCCTCTCCAGGGGATTCAAACTGAGTATCTGAG

ATCAAGGACTGCACTAGATGAGAGAAGTAGAGTTGTAGATCGTGTTTGGAACAGAAGGGAGTTAACCTAA

GCAATGCATCAGTTGGCGTTCTTGTCATTGTTCATTTCTCCTCCCCTTTTGTTTGTTAACTTTAGTTGTT

GGCATTTGTTTGAGTAAACCAAGTACAATGTTAGTTTGAGTATTGAGTTGCTTTAGTTGCTATGATTATG

ATGTTGAATGATACTGTTACTGGCTTGTTTGATTAATATGACTACCGTGTTTCTTATATGTGCTGAATTT

GGA

>MSTRG.3364.2 gene=MSTRG.3364

ATTTTAGCATCAAGTCAGCACTATATCTTGTTTCTCATTTCCCGACAGACTAGTGTAGTGAAGTTAAATT

AGGGCACAACTCATCTGAGGAACAAGTTCAACACAACACATCTAAGTTTAGCTTTACAATTTCAAGATGG

AACCTCCTACTGATTTGTTGGATGAGATACTCTGCCGCCTCCCAGTCAAGTATCTTCTTCGGTGCAGATG

TGTGTCGAAAGGATGGTGCTCCCTAATTGATAGCACCCCCTTTGTTAAGAAGCATCTCAACAAGGCCCTT

GAGTGCAATGCTGGTGGTCTTATTATTAATGAAGGCGGGAAGTTTTACTTGGCTGAGGATTTTAACTCTG

ATTTGGATGGTAGTGATGATAATGTTGATGTGGTCGCGGCAGAGATAACTCGTCCACTCAAGACTCGTGT

CTCTGGTGCTAATTTTGTGGGTTCTGCTAATGGTTTAGTTTGTGTGTCCAAGAATAAGATGAATGAGTTC

TTTATATTTAATCCAGCAACCAGGAAGTCGAGAAAAATACCCAGTGCTCCACGTGAGTTTCCACTTTCGT

TTCATATGACGGAGACTTCCTTGTGTGGATTTGGATATGATCAAGTAAATGATGACTATAAAGTTGTGAA

GATTGCTGAGTGTTATCTTCAGTTTCGCGGCATAATGGCCTTTGTCTACAGTCTCAAAACAGACTCTTGG

AAACGGATTCAGAATGTTCCTAGTAACACTCGGTTCACAGGAAATTGGGGAATGTTTGGAAATGGAGCTC

TGCACTGGTTGGCTATCAAGAATCCAGCAAACTGCATAGAAATTGTTGTTGGTTTTGATCTCGGGCTAGA

ACAGTTCAAGGAGGTCCCTTTTCCTGTTATTGAGGGCCCTTTTGTCAATTTTAATACCAGGAGTCTGGTA

TCGGATGGAAGTTCCCTTTGTGTTCTTGATATGTACCCTAATTCTCGTATAGATGTGTGGGTGATGATGA

ATAATTCCGGGGAGGAGAACCTTTGGTCCAAAGCACTTTCTGTGGATAAACACGGGATGCCTGGGTTTTC

AAGGTTTGTTAGACCTGTCCATATTTCATGGAGTAGCCAGAGTGTACTCTTAGAGATAGACAGCTCAAGA

CTTGTGTGGTATGACCTTGAAAGGAAAATGGTCAAGAATGTTAAGGTTCGTGGGATCACAAACCAGTTTG

ATTCACATATGTATACTCAAAGTCTCATACAGCTCGATGAGGACAATCAGCTGCAGAAGCCTTCACATGA

CAAGCCACCGAAGGAACAACAAAAGAGAAGGGATGATTTCCTCTCCAGGGGATTCAAACTGAGTATCTGA

GATCAAGGACTGCACTAGATGAGAGAAGTAGAGTTGTAGATCGTGTTTGGAACAGAAGGGAGTTAACCTA

AGCAATGCATCAGTTGGCGTTCTTGTCATTGTTCATTTCTCCTCCCCTTTTGTTTGTTAACTTTAGTTGT

TGGCATTTGTTTGAGTAAACCAAGTACAATGTTAGTTTGAGTATTGAGTTGCTTTAGTTGCTATGATTAT

GATGTTGAATGATACTGTTACTGGCTTGTTTGATTAATATGACTACCGTGTTTCTTATATGTGCTGAATT

TGGA

>MSTRG.3367.1 gene=MSTRG.3367

GTTCCTTTTCTAAAGAAAAACATAACACCCACCTGCCATGTTCTTTCCATTTTGATTTTACTGTGAACTT

CGTTTGAGTCGCAAACCGAGTTTATCATTTTCGATTCCCCTGCTTCGAGTCTTACATCGTTTTGATGATT

TTAGAGATTGTTTTTGCTGTGTTGTACTGCGTATCGAGTCGCCGACTCCTGGGTTTTGTTTAGAGCGGAG

TCATGGCTGCGTTTCTCTTTTGGCTGAGTTTAAGTCACTGAACTGGGTTTTCTGCTTCGTGTGTTGCTGT

TATTTGATGAATGCTGAGTCGTTAATTAGTGTTTTAGGTCCGAGTTCGATTGTGAATGTTCTTCGTTAGC

TCGAGTCTAGTTTTCAGCTGGTCGAGTTGTAGCCGAGTCACTGCATTTTCTTTCTTAAACATTTAGATTC

TGAGTTATATTTGGTAGTTTTTGTCGAGTTTGCATCCTAGTCCGAGTCGCAGCTCTTGCTTCGAGTTCTG

GGTTTGTTATCGAGTTTATTCTTGCTGCGTTTTCTTGTTGAGCTGCGTTGGTTTTGAGTCGTAGCTGATG

AGTCTGAGTTTATTGTATGCGAAGTC

>MSTRG.3373.1 gene=MSTRG.3373

TAATGGGGCCGTTTGGGTTAGCTTAAAAGAAGTGACTTCTTGCTTATAGTAAAGAAGTGGAGTAGAAGTG

AGAAGTAAATAAGTTAATAAAGTGTTTGGAAAAGAAGCAGAAGCTGTGAGAGAGAAGCTAGCATTCTCAA

CTTCTTAAAAGTGCTTCTACTTTTTTGCACAAACGGGTCAAGAGAAGCAGAAGTCAGAAGTAGATTTTGA

TTCTCTTGGCCAAACAGGCCCAATGTT

>MSTRG.3388.1 gene=MSTRG.3388

GTTTACCATCCAAGTATGCTATCAAGATGATGAAAAGCGATGGTGATTCTGCTGCAGATACAAAAACCAC

TACCCATATCGAAGATATAGAAAGATTTTCCTTCTTCGACTATATTTTAGGATTGCCTTGGATTCAACCA

ATAGATGATGCAAATCAGAAGAACAAGCTTGCAGTAGCACTTGCGCAACAATTGCTAGAGAAGGAGACAG

ATTGGAGTAACTACACAGATCCAAAACATAAGTATGACGATGAGTCACCATCACGAGGAAAAAAGAAAGG

AGAAGTACTAAACCCG

>MSTRG.3389.1 gene=MSTRG.3389

ATGATTGTTCTCCACATTTGCTGCATATGCGAAATAAAGAAGGAAATACAGCAGAAGATCTACTTTTAAA

AGATTACAAAAGCAAACGAGAAGCAGCTGAGAGTGCAGTTAAAGAAATGAACCAAGGCCTAATGGTTGTG

GCAGCATTGATTGCCACTGTGAGTTTTGCAGCAGTTTTTACTATTCCTGGAGGTTTTGATCAAACTAAAG

GGAATCCTATCTTCTACAACAAACCAAGTGATCCGAAACATAGTCATAAAAATGATTTAAGTCTGTTTCT

AGGTTATGTTGGGCTTACCTTCTTTGCTTCTTTACTCGCACTGGCAACTCTTCTGTCTATTCAGCTGTCC

AGATTTAGTATGGATGATTTGTACTTCTCATTGCCCCTAAAGTACTCGATTGCCATTTCTTTTCTCTTCC

TCTCCTCCTGCTCTACAATCACCACATTTCTTCAGGCCTTAATTCTTGAGGATTACCTTTCACCGTATTA

CCTCACTTGCCTCATTATTGGTTGCGTCGTCATGGGTCTTGTTTATGTGGACTCTATATACGACGTCTTG

AGTTACTTAGTGGAGGTTTTACGGCATTCACGCACTTACAGGACTCAAGAAGGGTAATAGCTAGGCAGCT

AACTAGGGTCAGATAACATGTCTGAATAATATTAAAAACCAGTGCCAGGTATTGGTAAGAGACACATTGT

ACTCAACATCAGACCCTGTGTTTTTCTTGTTCTTTCTGATATCCCTTTCATTCCAC

>MSTRG.3390.2 gene=MSTRG.3390

TAGGACCGTACGCAACGCTATAAATAGAGCTCAAGGAAGTGAGGATTGGCATCGACAGCACAAGAATTAC

ATAAGAACTTCTTATTCTCAACTAGCAAGGAGGATGGCTCAAGTGCAGCGTGTGCGTGATGACATCGTGT

CATTGTGGACTCAAGGTGTCCAGGCTGCGAAAGACAAGAATTACATAAGAACTTCTTATTCTCAACTAGC

AAGGAGGATGGCTCAAGTGCAGCGTGTGCGTGATGACATCGTGTCATTGTGGACTCAAGGTGTCCAGGCT

GCGAAAGACAAGAGTTCCTTCGCTGCGGTGATCAGTGCAAAGGAACTCTCAATAGTTTGGGGTTCAGACA

GCCGCTACTGGAAATGGGTTTCTAAGACATACCCAATCAGCTGCCAGCCTATGGAGGTTGCTGAACTGAT

CACAGTATGCTGGCTACAAATTGACGGAAAGTACAATGCCAACAACCTCACCAAGGGAGTGAAATACGCA

GTGTATCTTGCTGTGGAGCTTAGCGACAATCTTTGCATGAATGGTCCAGTGACCCTTAAGCTTACTCGTC

CCAACGGCACTACTGAAGAGCACAAGGAAGATTTGAAGACCAAGCCAACGAATGCATTGGTGGGACTTAA

GGTTGGTGAATTCTGCAACACAGGCGCCTGCGGCTGTGAGAATACTGTCAAGTTCTCGATGAATGGATGT

GATGGAACCACCTGGAAGACTGGCCTTACTGTCGTTGGTGCTGTCATTGCCCCAGTTAACTGCTAAACAT

GAATCTGATTATCATCATGTACTAAATAATTTCCAATTGCCAACATCAACTATCTGGTGTGGCTGATATG

GAATGTGTGAAATAATCGCCAATATGTTGGTGCTTTTAATATATAAAAAGTTAAAGTAAGGAGTGTTCCA

ACTTAAAAATGACTTAGAGCCCCGGCTTTAATCACCGTCAAACAGCCTCTAAAATTGTATATGAAGGACT

CTGAGTTCATGAAGGAAAAATAAAGAAGCACCGAAC

>MSTRG.3390.3 gene=MSTRG.3390

TTTAAGTGTAACTATCAATAAGTGCACAAATAGGACCGTACGCAACGCTATAAATAGAGCTCAAGGAAGT

GAGGATTGGCATCGACAGCACAAGAATTACATAAGAACTTCTTATTCTCAACTAGCAAGGAGGATGGCTC

AAGTGCAGCGTGTGCGTGATGACATCGTGTCATTGTGGACTCAAGGTGTCCAGGCTGCGAAAGACAAGAG

TTCCTTCGCTGCGGTGATCAGTGCAAAGGAACTCTCAATAGTTTGGGGTTCAGACAGCCGCTACTGGAAA

TGGGTTTCTAAGACATACCCAATCAGCTGCCAGCCTATGGAGGTTGCTGAACTGATCACAGTATGCTGGC

TACAAATTGACGGAAAGTACAATGCCAACAACCTCACCAAGGGAGTGAAATACGCAGTGTATCTTGCTGT

GGAGCTTAGCGACAATCTTTGCATGAATGGTCCAGTGACCCTTAAGCTTACTCGTCCCAACGGCACTACT

GAAGAGCACAAGGAAGATTTGAAGACCAAGCCAACGAATGCATTGGTGGGACTTAAGGTTGGTGAATTCT

GCAACACAGGCGCCTGCGGCTGTGAGAATACTGTCAAGTTCTCGATGAATGGATGTGATGGAACCACCTG

GAAGACTGGCCTTACTGTCGTTGGTGCTGTCATTGCCCCAGTTAACTGCTAAACATGAATCTGATTATCA

TCATGTACTAAATAATTTCCAATTGCCAACATCAACTATCTGGTGTGGCTGATATGGAATGTGTGAAATA

ATCGCCAATATGTTGGTGCTTTTAATATATAAAAAGTTAAAGTAAGGAGTGTTCCAACTTAAAAATGACT

TAGAGCCCCGGCTTTAATCACCGTCAAACAGCCTCTAAAATTGTATATGAAGGACTCTGAGTTCATGAAG

GAAAAATAAAGAAGCACCGAAC

>MSTRG.3390.4 gene=MSTRG.3390

AGGATTGGCATCGACAGCACAAGAATTACACAAGTACTGCTTATTCTCAACTAGCAAGGAAGATGGCTCA

ATTGCAGCGTGATGACATCGTGTCGTTGTGGACTCAAGGTGTCCAGGCTGTGAAAGACAAGAGTTCCTTC

GCTGCGGTTATCAGTGCAAAGGAACTCTCAATAGTTTGGGGTTCAGACAGCCGCTACTGGAAATGGGTTT

CTAAGACATACCCAATCAGCTGCCAGCCTATGGAGGTTGCTGAACTGATCACAGTATGCTGGCTACAAAT

TGACGGAAAGTACAATGCCAACAACCTCACCAAGGGAGTGAAATACGCAGTGTATCTTGCTGTGGAGCTT

AGCGACAATCTTTGCATGAATGGTCCAGTGACCCTTAAGCTTACTCGTCCCAACGGCACTACTGAAGAGC

ACAAGGAAGATTTGAAGACCAAGCCAACGAATGCATTGGTGGGACTTAAGGTTGGTGAATTCTGCAACAC

AGGCGCCTGCGGCTGTGAGAATACTGTCAAGTTCTCGATGAATGGATGTGATGGAACCACCTGGAAGACT

GGCCTTACTGTCGTTGGTGCTGTCATTGCCCCAGTTAACTGCTAAACATGAATCTGATTATCATCATGTA

CTAAATAATTTCCAATTGCCAACATCAACTATCTGGTGTGGCTGATATGGAATGTGTGAAATAATCGCCA

ATATGTTGGTGCTTTTAATATATAAAAAGTTAAAGTAAGGAGTGTTCCAACTTAAAAATGACTTAGAGCC

CCGGCTTTAATCACCGTCAAACAGCCTCTAAAATTGTATATGAAGGACTCTGAGTTCATGAAGGAAAAAT

AAAGAAGCACCGAAC

>MSTRG.3390.1 gene=MSTRG.3390

ATAAGGTTAAGTGTAACTATCAATAAGTGCACAAATAGGACCGTCTGAAACGCTATAAATACAGCTCAAG

GAAGTGAGGATTGGCATAGACAGCACAAGAATTACACAAGAACTTCTTATTCTCAACTAGCAAGGAAGAT

GGCTCAATTGCAGCGTGATGACATCGTGTCGTTGTGGACTCAAGGTGTCCAGGCTGTGAAAGACAAGAGT

TCCTTCGCAGCGGTTATCAGTGCAAAGGAGCTCTCAATAGTTTGGGGTTCAGACAGCCGCTACTGGAAAT

GGGTTTCTAAGACCTACCCAATCAGCTGCCAGTCTCTGGAGGTTGCTGAACTGATCACAGTATGCTGGCT

ACAAATTGACGGAAAGTACAATGCCAACAACCTCATCAAGGGAGTGAAATACGGAGTGTATTTGGCTGTG

GAGCTTAGCGACAATCTTTGCATGAACGGTCCAGTGACACTTAAGCTTACTCGTCCCAACGGCACTACTG

AAGAGCACAAGGAAGATTTGAAGACCAAGCCAAAGAATACTTTGGTGGGACTTAAGGTTGGTGAATTCTG

CAACACAGGCGCCTGTGGCTGTGAGAATACTGTCAAGTTCTCGATGAATGGATGTGATGGAACCACCTGG

AAGACGGGCCTTACTGTCGTTGGTGCTGTCATTGCCCCAGTTAACTGAAGAATTACATAAGAACTTCTTA

TTCTCAACTAGCAAGGAGGATGGCTCAAGTGCAGCGTGTGCGTGATGACATCGTGTCATTGTGGACTCAA

GGTGTCCAGGCTGCGAAAGACAAGAGTTCCTTCGCTGCGGTTATCAGTGCAAAGGAACTCTCAATAGTTT

GGGGTTCAGACAGCCGCTACTGGAAATGGGTTTCTAAGACATACCCAATCAGCTGCCAGCCTATGGAGGT

TGCTGAACTGATCACAGTATGCTGGCTACAAATTGACGGAAAGTACAATGCCAACAACCTCACCAAGGGA

GTGAAATACGCAGTGTATCTTGCTGTGGAGCTTAGCGACAATCTTTGCATGAATGGACCAGTGACCCTTA

AGCTTACTCGTCCCAACGGCACTACTGAAGAGCACAAGGAAGATTTGAAGACAAAGCCAACGAATACTTT

GGTGGGACTTAAGGTTGGTGAATTCTGCAACACAGGCGCCTGCGGCTGTGAGAATACTGTCAAGTTCTCG

ATGAATGGATGTGATGGAACCACCTGGAAGACTGGCCTTACTGTCGTTGGTGCTGTCATTGCCCCAGTTA

ACTGCTAAACATGAATCTGATTATCATCATGTACTAAATAATTTCCAATTGCCAACATCAACTATCTGGT

GTGGCTGATATGGAATGTGTGAAATAATCGCCAATATGTTGGTGCTTTTAATATATAAAAAGTTAAAGTA

AGGAGTGTTCCAACTTAAAAATGACTTAGAGCCCCGGCTTTAATCACCGTCAAACAGCCTCTAAAATTGT

ATATGAAGGACTCTGAGTTCATGAAGGAAAAATAAAGAAGCACCGAAC

>MSTRG.3390.6 gene=MSTRG.3390

ATAAGGTTAAGTGTAACTATCAATAAGTGCACAAATAGGACCGTCTGAAACGCTATAAATACAGCTCAAG

GAAGTGAGGATTGGCATAGACAGCACAAGAATTACACAAGAACTTCTTATTCTCAACTAGCAAGGAAGAT

GGCTCAATTGCAGCGTGATGACATCGTGTCGTTGTGGACTCAAGGTGTCCAGGCTGCGAAAGACAAGAGT

TCCTTCGCTGCGGTTATCAGTGCAAAGGAACTCTCAATAGTTTGGGGTTCAGACAGCCGCTACTGGAAAT

GGGTTTCTAAGACATACCCAATCAGCTGCCAGCCTATGGAGGTTGCTGAACTGATCACAGTATGCTGGCT

ACAAATTGACGGAAAGTACAATGCCAACAACCTCACCAAGGGAGTGAAATACGCAGTGTATCTTGCTGTG

GAGCTTAGCGACAATCTTTGCATGAATGGACCAGTGACCCTTAAGCTTACTCGTCCCAACGGCACTACTG

AAGAGCACAAGGAAGATTTGAAGACAAAGCCAACGAATACTTTGGTGGGACTTAAGGTTGGTGAATTCTG

CAACACAGGCGCCTGTGGCTGTGAGAATACTGATGTCAAGTTCTCGATGAATGGATGTGATGGAACCACC

TGGAAGACGGGCCTTACTGTCGTTGGTGCTGTCATTGCCCCAGTTAACTGCTAAACATGAATCTCATTAT

AATCATCATGTACTAAATAATTTCCAGTTAATTGCCAGCATCACCTGTCTGGTATGGAATGTGTGAAATA

ATCGCCAATATGTTGGTGCTTTTAATATATAAAAAGTTAAAGTAAGGAGTGTTCCAACTTAAAAATGACT

TAGAGCCCCGGCTTTAATCACCGTCAAACAGCCTCTAAAATTGTATATGAAGGACTCTGAGTTCATGAAG

GAAAAATAAAGAAGCACCGAAC

>MSTRG.3390.5 gene=MSTRG.3390

ATAAGGTTAAGTGTAACTATCAATAAGTGGACAAATAGGACCGTCTGAAACGCTATAAATACAGCTCAAG

GAAGTGAGGATTGGCATCGACAGCACAAGAATTACACAACAACTTCTTATTCTCAACTCGCAAGGCAGAT

GGCTCAATTGCAGCGTGATGACATCGTGTCGTTGTGGACTCAAGGTGTCCAGGCTGCGAAAGACAAGAGT

TCCTTCGCAGCGGTTATCAGTGCAAAGGAGCTCTCAATAGTTTGGGGTTCAGACAGCCGCTACTGGAAAT

GGGTTTCTAAGACCTACCCAATCAGCTGCCAGTCTCTGGAGGTTGCTGAACTGATCACAGTATGCTGGCT

ACAAATTGACGGAAAGTACAATGCCAACAACCTCATCAAGGGAGTGAAATACGGAGTGTATTTGGCTGTG

GAGCTTAGCGACAATCTTTGCATGAACGGTCCAGTGACACTTAAGCTTACTCGTCCCAACGGCACTACTG

AAGAGCACAAGGAAGATTTGAAGACCAAGCCAAAGAATACTTTGGTGGGACTTAAGGTTGGTGAATTCTG

CAACACAGGCGCCTGTGGCTGTGAGAATACTGTCAAGTTCTCGATGAATGGATGTGATGGAACCACCTGG

AAGACGGGCCTTACTGTCGTTGGTGCTGTCATTGCCCCAGTTAACTGAAGAATTACATAAGAACTTCTTA

TTCTCAACTAGCAAGGAGGATGGCTCAAGTGCAGCGTGTGCGTGATGACATCGTGTCATTGTGGACTCAA

GGTGTCCAGGCTGCGAAAGACAAGAGTTCCTTCGCTGCGGTTATCAGTGCAAAGGAACTCTCAATAGTTT

GGGGTTCAGACAGCCGCTACTGGAAATGGGTTTCTAAGACATACCCAATCAGCTGCCAGCCTATGGAGGT

TGCTGAACTGATCACAGTATGCTGGCTACAAATTGACGGAAAGTACAATGCCAACAACCTCACCAAGGGA

GTGAAATACGCAGTGTATCTTGCTGTGGAGCTTAGCGACAATCTTTGCATGAATGGTCCAGTGACCCTTA

AGCTTACTCGTCCCAACGGCACTACTGAAGAGCACAAGGAAGATTTGAAGACCAAGCCAACGAATGCATT

GGTGGGACTTAAGGTTGGTGAATTCTGCAACACAGGCGCCTGCGGCTGTGAGAATACTGTCAAGTTCTCG

ATGAATGGATGTGATGGAACCACCTGGAAGACTGGCCTTACTGTCGTTGGTGCTGTCATTGCCCCAGTTA

ACTGCTAAACATGAATCTGATTATCATCATGTACTAAATAATTTCCAATTGCCAACATCAACTATCTGGT

GTGGCTGATATGGAATGTGTGAAATAATCGCCAATATGTTGGTGCTTTTAATATATAAAAAGTTAAAGTA

AGGAGTGTTCCAACTTAAAAATGACTTAGAGCCCCGGCTTTAATCACCGTCAAACAGCCTCTAAAATTGT

ATATGAAGGACTCTGAGTTCATGAAGGAAAAATAAAGAAGCACCGAAC

>MSTRG.3390.7 gene=MSTRG.3390

GTGAGGATTGGCATCGACAGCACAAGAATTACACAAGTACTGCTTATTCTCAACTAGCAAGGAAGATGGC

TCAATTGCAGCGTGATGACATCGTGTCGTTGTGGACTCAAGGTGTCCAGGCTGCGAAAGACAAGAGTTCC

TTCGCTGCGGTTATCAGTGCAAAGGAACTCTCAATAGTTTGGGGTTCAGACAGCCGCTACTGGAAATGGG

TTTCTAAGACATACCCAATCAGCTGCCAGCCTATGGAGGTTGCTGAACTGATCACAGTATGCTGGCTACA

AATTGACGGAAAGTACAATGCCAACAACCTCACCAAGGGAGTGAAATACGCAGTGTATCTTGCTGTGGAG

CTTAGCGACAATCTTTGCATGAATGGACCAGTGACCCTTAAGCTTACTCGTCCCAACGGCACTACTGAAG

AGCACAAGGAAGATTTGAAGACAAAGCCAACGAATACTTTGGTGGGACTTAAGGTTGGTGAATTCTGCAA

CACAGGCGCCTGTGGCTGTGAGAATACTGATGTCAAGTTCTCGATGAATGGATGTGATGGAACCACCTGG

AAGACTGGCCTTACTGTCGTTGGTGCTGTCATTGCCCCAGTTAACTGCTAAACATGAATCTGATTATCAT

CATGTACTAAATAATTTCCAATTGCCAACATCAACTATCTGGTGTGGCTGATATGGAATGTGTGAAATAA

TCGCCAATATGTTGGTGCTTTTAATATATAAAAAGTTAAAGTAAGGAG

>MSTRG.3390.8 gene=MSTRG.3390

ATAAGGTTAAGTGTAACTATCAATAAGTGGACAAATAGGACCGTCTGAAACGCTATAAATACAGCTCAAG

GAAGTGAGGATTGGCATCGACAGCACAAGAATTACACAACAACTTCTTATTCTCAACTCGCAAGGCAGAT

GGCTCAATTGCAGCGTGATGACATCGTGTCGTTGTGGACTCAAGGTGTCCAGGCTGCGAAAGACAAGAGT

TCCTTCGCAGCGGTTATCAGTGCAAAGGAGCTCTCAATAGTTTGGGGTTCAGACAGCCGCTACTGGAAAT

GGGTTTCTAAGACCTACCCAATCAGCTGCCAGTCTCTGGAGGTTGCTGAACTGATCACAGTATGCTGGCT

ACAAATTGACGGAAAGTACAATGCCAACAACCTCATCAAGGGAGTGAAATACGGAGTGTATTTGGCTGTG

GAGCTTAGCGACAATCTTTGCATGAACGGTCCAGTGACACTTAAGCTTACTCGTCCCAACGGCACTACTG

AAGAGCACAAGGAAGATTTGAAGACCAAGCCAAAGAATACTTTGGTGGGACTTAAGGTTGGTGAATTCTG

CAACACAGGCGCCTGTGGCTGTGAGAATACTGTCAAGTTCTCGATGAATGGATGTGATGGAACCACCTGG

AAGACGGGCCTTACTGTCGTTGGTGCTGTCATTGCCCCAGTTAACTGAAGAATTACATAAGAACTTCTTA

TTCTCAACTAGCAAGGAGGATGGCTCAAGTGCAGCGTGTGCGTGATGACATCGTGTCATTGTGGACTCAA

GGTGTCCAGGCTGCGAAAGACAAGAGTTCCTTCGCTGCGGTTATCAGTGCAAAGGAACTCTCAATAGTTT

GGGGTTCAGACAGCCGCTACTGGAAATGGGTTTCTAAGACATACCCAATCAGCTGCCAGCCTATGGAGGT

TGCTGAACTGATCACAGTATGCTGGCTACAAATTGACGGAAAGTACAATGCCAACAACCTCACCAAGGGA

GTGAAATACGCAGTGTATCTTGCTGTGGAGCTTAGCGACAATCTTTGCATGAATGGACCAGTGACCCTTA

AGCTTACTCGTCCCAACGGCACTACTGAAGAGCACAAGGAAGATTTGAAGACAAAGCCAACGAATACTTT

GGTGGGACTTAAGGTTGGTGAATTCTGCAACACAGGCGCCTGTGGCTGTGAGAATACTGATGTCAAGTTC

TCGATGAATGGATGTGATGGAACCACCTGGAAGACTGGCCTTACTGTCGTTGGTGCTGTCATTGCCCCAG

TTAACTGCTAAACATGAATCTGATTATCATCATGTACTAAATAATTTCCAATTGCCAACATCAACTATCT

GGTGTGGCTGATATGGAATGTGTGAAATAATCGCCAATATGTTGGTGCTTTTAATATATAAAAAGTTAAA

G

>MSTRG.3390.9 gene=MSTRG.3390

AGTGCACAAATAGGACCGTCTGAAACGCTATAAATACAGCTCAAGGAAGTGAGGATTGGCATAGACAGCA

CAAGAATTACACAAGAACTTCTTATTCTCAACTAGCAAGGAAGATGGCTCAATTGCAGCGTGATGACATC

GTGTCGTTGTGGACTCAAGGTGTCCAGGCTGTGAAAGACAAGAGTTCCTTCGCTGCGGTTATCAGTGCAA

AGGAACTCTCAATAGTTTGGGGTTCAGACAGCCGCTACTGGAAATGGGTTTCTAAGACATACCCAATCAG

CTGCCAGCCTATGGAGGTTGCTGAACTGATCACAGTATGCTGGCTACAAATTGACGGAAAGTACAATGCC

AACAACCTCACCAAGGGAGTGAAATACGCAGTGTATCTTGCTGTGGAGCTTAGCGACAATCTTTGCATGA

ATGGACCAGTGACCCTTAAGCTTACTCGTCCCAACGGCACTACTGAAGAGCACAAGGAAGATTTGAAGAC

AAAGCCAACGAATACTTTGGTGGGACTTAAGGTTGGTGAATTCTGCAACACAGGCGCCTGTGGCTGTGAG

AATACTGATGTCAAGTTCTCGATGAATGGATGTGATGGAACCACCTGGAAGACGGGCCTTACTGTCGTTG

GTGCTGTCATTGCCCCAGTTAACTGCTAAACATGAATCTCATTATAATCATCATGTACTAAATAATTTCC

AGTTAATTGCCAGCATCACCTGTCTGGTACGTGTGGCTGATATGGAATGTGTGAAATAATCGCCAATATG

TTGGTGTTATAATATATAAAAAGTTAAAGTACGGAGTGTTCAAGCTATTATAACGTTTAGCTCGTAATTG

CTTCAAGGTCTCTACTGTCACTTTAGCACTTTCAACTTAAAGATCTGACTAAAAGATAAGTTAAACATAA

>MSTRG.3390.10 gene=MSTRG.3390

GTGAGGATTGGCATCGACAGCACAAGAATTACACAAGTACTGCTTATTCTCAACTAGCAAGGAAGATGGC

TCAATTGCAGCGTGATGACATCGTGTCGTTGTGGACTCAAGGTGTCCAGGCTGTGAAAGACAAGAGTTCC

TTCGCTGCGGTTATCAGTGCAAAGGAACTCTCAATAGTTTGGGGTTCAGACAGCCGCTACTGGAAATGGG

TTTCTAAGACATACCCAATCAGCTGCCAGCCTATGGAGGTTGCTGAACTGATCACAGTATGCTGGCTACA

AATTGACGGAAAGTACAATGCCAACAACCTCACCAAGGGAGTGAAATACGCAGTGTATCTTGCTGTGGAG

CTTAGCGACAATCTTTGCATGAATGGACCAGTGACCCTTAAGCTTACTCGTCCCAACGGCACTACTGAAG

AGCACAAGGAAGATTTGAAGACAAAGCCAACGAATACTTTGGTGGGACTTAAGGTTGGTGAATTCTGCAA

CACAGGCGCCTGTGGCTGTGAGAATACTGATGTCAAGTTCTCGATGAATGGATGTGATGGAACCACCTGG

AAGACGGGCCTTACTGTCGTTGGTGCTGTCATTGCCCCAGTTAACTGCTAAACATGAATCTCATTATAAT

CATCATGTACTAAATAATTTCCAGTTAATTGCCAGCATCACCTGTCTGGTACGTGTGGCTGATATGGAAT

GTGTGAAATAATCGCCAATATGTTGGTGTTATAATATATAAAAAGTTAAAGTACGGAGTGTTCAAGCTAT

TATAACGTTTAGCTCGTAATTGCTTCAAGG

>MSTRG.3390.11 gene=MSTRG.3390

GTGAGGATTGGCATCGACAGCACAAGAATTACACAAGTACTGCTTATTCTCAACTAGCAAGGAAGATGGC

TCAATTGCAGCGTGATGACATCGTGTCGTTGTGGACTCAAGGTGTCCAGGCTGTGAAAGACAAGAGTTCC

TGCGCTGCGGTTATCACCGCAAAGGAACTCTCAATAGTCTGGGGTTCAGACAGCCGCTACTGGAAATGGG

TTTCTAAGACCTACCCAATCAGCTGCGAGGCTCTGGAGGTTGCTAAACTAATTACAGTATGCTGGCTCCA

AATTGACGGAGAGTACGATACCAGCAACCTGAAAAAGGGAGTGAAATACGGAGTGTATTTTGTTGTGGAG

CTTAGCGACAATCTTTGCATGAACGGTCCAGTGACCCTTAAGCTTACTCGTCCCAACCGCACTACTGAAG

AGCACAAGGAAGATTTGGAGACCAAGCCAAAGAATACATTGGTGGGACTTAAGGTTGGTGAATTCATCAA

CTCAGCATGCGGCTGTGAGAATACTGTCAAGTTCTCGATGAATGGATGTGATGGAACCACCTGGAAGACT

GGCCTTACTGTCATTGGTGCTGTCATTGCCCCAGTCTGGTAAACATGCATCTGATAAACTAAATAATTTC

CAATTGCCAACATCACCTATCTGGTTTGGCTGATGTGGAATGTGTGAAATAATCGCCAATATGTTGGTGT

TTAAATATATAAGTGATACGCCATAAACAAGTACCTGGGACTTTGAGAACGGATTTTATTAAAGATGATG

AACATCCAATGCATTAGACCCTAGAATCTCGAACTTGTCATGACCCAAATTGTAAAAATAAAAAATCACC

AAACAAAAA

>MSTRG.3390.12 gene=MSTRG.3390

GAGCATCCTACTACCGTAGATAAGGTTAAGTGTAACTATCAATAAGTGGACAAATAGGACCGTCTGAAAC

GCTATAAATACAGCTCAAGGAAGTGAGGATTGGCATCGACAGCACAAGAATTACACAACAACTTCTTATT

CTCAACTCGCAAGGCAGATGGCTCAATTGCAGCGTGATGACATCGTGTCGTTGTGGACTCAAGGTGTCCA

GGCTGCGAAAGACAAGAGTTCCTTCGCTGCGGTTATCTGTGCAAAGGAACTCTCAATAGTTTGGGGTTCA

GACTGCCGCTACTGGAAATGGGTTTCTAAGAAATACCCAATCAGTCCCCAGCCTCTGGAGGTTGCTGAAC

TCATCACAGTATGCTGGCTACAAATTGACGGAAAGTACAGTGCCACCAACCTCATCAAGGGAGTGAAATA

CGGAGTGTATCTTGCTGTGGAGCTTAGCGACAATCTTTGCATGAACGGTCCAGTGACCCTTAAGCTTACT

CGTCCCAACGGCACTACTGAAGAGCACAAGGAAGATTTGACGACCAAGCCAAAGAATACATTGGTGGGGC

TTAAGGTTGGTGAATTCTGCAACACAGCCCCCTGCGGCTGTGAGAATATTGTCAAGTTCTCGATGAATGG

ATGTGATGGAACCACCTGGAAGACTGGCCTTACTGTCGTTGGTGCTGTCATTGCCCCAGTTAACTGCTAA

ACATGAATCTGATTATCATCATGTACTAAATAATTTCCAATTGCCAACATCAACTATCTGGTGTGGCTGA

TATGGAATGTGTGAAATAATCGCCAATATGTTGGTGTTTTAATATATAAAAAGTTAAAGTACGGAGTGTT

CAGGCTATTATAACGTTTAGCTCGTAATTGCTTCAAGGTCTCTACTGTTACTTTAGCACTTTTAACTTAA

AGGTCTTACTAAAAGATTAGTTAAACATAATGAGGAACTGTATATTGTTTTGTGCTGATGTTTGTTTAAT

TAAGCACACTAAAGTTGCGTTCACTTGGAATGAAAAACTACAAAAACAGAAAGGAGAATGTATGAGACAA

TAGTGAGTGAGG

>MSTRG.3390.13 gene=MSTRG.3390

AGTGCACAAATAGGACCGTCTGAAACGCTATAAATACAGCTCAAGGAAGTGAGGATTGGCATAGACAGCA

CAAGAATTACACAAGAACTTCTTATTCTCAACTAGCAAGGAAGATGGCTCAATTGCAGCGTGATGACATC

GTGTCGTTGTGGACTCAAGGTGTCCAGGCTGTGAAAGACAAGAGTTCCTTCGCAGCGGTTATCAGTGCAA

AGGAGCTCTCAATAGTTTGGGGTTCAGACAGCCGCTACTGGAAATGGGTTTCTAAGACCTACCCAATCAG

CTGCCAGTCTCTGGAGGTTGCTGAACTGATCACAGTATGCTGGCTACAAATTGACGGAAAGTACAATGCC

AACAACCTCATCAAGGGAGTGAAATACGGAGTGTATTTGGCTGTGGAGCTTAGCGACAATCTTTGCATGA

ACGGTCCAGTGACACTTAAGCTTACTCGTCCCAACGGCACTACTGAAGAGCACAAGGAAGATTTGAAGAC

CAAGCCAAAGAATACTTTGGTGGGACTTAAGGTTGGTGAATTCTGCAACACAGGCGCCTGTGGCTGTGAG

AATACTGTCAAGTTCTCGATGAATGGATGTGATGGAACCACCTGGAAGACGGGCCTTACTGTCGTTGGTG

CTGTCATTGCCCCAGTTAACTGCTAAACATGAATCTCACAATCATCATGTACTAAATAATTTCCAATTAA

TTGCCAACATTACCTGTCTGGTACGTGTGGCTGATATGGAATGTGTGAAATAATCGCCAATATCTTGGTG

TTTTAATATATAAAAAGTTAAAGTACGGAGTGTTCAGGCTATTATAACGTTTAGCTCGTAATTGCTTCAA

GGTCTCTACTGTTACTTTAGCACTTTTAACTTAAAG

>MSTRG.3390.14 gene=MSTRG.3390

CACAAGAATTACACAACAACTTCTTATTCTCAACTCGCAAGGCAGATGGCTCAATTGCAGCGTGATGACA

TCGTGTCGTTGTGGACTCAAGGTGTCCAGGCTGCGAAAGACAAGAGTTCCTTCGCTGCGGTTATCTGTGC

AAAGGAACTCTCAATAGTTTGGGGTTCAGACTGCCGCTACTGGAAATGGGTTTCTAAGAAATACCCAATC

AGTCCCCAGCCTCTGGAGGTTGCTGAACTCATCACAGTATGCTGGCTACAAATTGACGGAAAGTACAGTG

CCACCAACCTCATCAAGGGAGTGAAATACGGAGTGTATCTTGCTGTGGAGCTTAGCGACAATCTTTGCAT

GAACGGTCCAGTGACCCTTAAGCTTACTCGTCCCAACGGCACTACTGAAGAGCACAAGGAAGATTTGACG

ACCAAGCCAAAGAATACATTGGTGGGGCTTAAGGTTGGTGAATTCTGCAACACAGCCCCCTGCGGCTGTG

AGAATATTGTCAAGTTCTCGATGAATGGATGTGATGGAACCACCTGGAAGACTGGCCTTACTGTCGTTGG

TGCTGTCATTGCCCCAGTTAACTGCTAAACATGAATCTCACAATCATCATGTACTAAATAATTTCCAATT

AATTGCCAACATTACCTGTCTGGTACGTGTGGCTGATATGGAATGTGTGAAATAATCGCCAATATCTTGG

TGTTTTAATATATAAAAAGTTAAAGTACGGAGTGTTCAGGCTATTATAACGTTTAGCTCGTAATTGCTTC

AAGGTCTCTACTGTTACTTTAGCACTTTTAACTTAAAG

>MSTRG.3390.15 gene=MSTRG.3390

AATAGGACCGTCTGAAACGCTATAAATACAGCTCAAGGAAGTGAGGATTGGCATCGACAGCACAAGAATT

ACACAACAACTTCTTATTCTCAACTCGCAAGGCAGATGGCTCAATTGCAGCGTGATGACATCGTGTCGTT

GTGGACTCAAGGTGTCCAGGCTGCGAAAGACAAGAGTTCCTTCGCAGCGGTTATCAGTGCAAAGGAGCTC

TCAATAGTTTGGGGTTCAGACAGCCGCTACTGGAAATGGGTTTCTAAGACCTACCCAATCAGGTATGTAT

AGCACATGTTGGAAAGCTTTTGGTGTTTTTGTACTACAAAATTTAATGACTGGACATTATTGACTTCCAT

TTTTCGAGACCAAAGATTTGAGATAAAGTTTTACATAATATTTGGTTTTTCGGACTTTTGTACACACAGC

TGCCAGTCTCTGGAGGTTGCTGAACTGATCACAGTATGCTGGCTACAAATTGACGGAAAGTACAATGCCA

ACAACCTCATCAAGGGAGTGAAATACGGAGTGTATTTGGCTGTGGAGCTTAGCGACAATCTTTGCATGAA

CGGTCCAGTGACACTTAAGCTTACTCGTCCCAACGGCACTACTGAAGAGCACAAGGAAGATTTGAAGACC

AAGCCAAAGAATACTTTGGTGGGACTTAAGGTTGGTGAATTCTGCAACACAGGCGCCTGTGGCTGTGAGA

ATACTGTCAAGTTCTCGATGAATGGATGTGATGGAACCACCTGGAAGACGGGCCTTACTGTCGTTGGTGC

TGTCATTGCCCCAGTTAACTGCTAAACATGAATCTCACAATCATCATGTACTAAATAATTTCCAATTAAT

TGCCAACATTACCTGTCTGGTACGTGTGGCTGATATGGAATGTGTGAAATAATCGCCAATATCTTGGTGT

TTTAATATATAAAAAGTTAAAGTACGGAGTGTTCAGGCTATTATAACGTTTAGCTCGTAATTGCTTCAAG

GTCTCTACTGTTACTTTAGCA

>MSTRG.3390.16 gene=MSTRG.3390

CAAGAATTACACAACAACTTCTTATTCTCAGCTAGCAAGGAAGATGGCTCAAGTGCAGCGTGATGACATC

GTGTCGTTGTGGACTCAAGGTGTCCAGGCTGTGAAAGACAAGAGTTCCTGCGCTGCAATTATCACTGCAA

AGGAACTCTCGATAGTCTGGGGTTCAGACAGCCGCTACTGGAAATGGGTTTCTAAGAAATACCCAATCAG

TCCCCAGCCTCTGGAGGTTGCTGAACTCATCACAGTATGCTGGCTACAAATTGACGGAAAGTACAGTGCC

ACCAACCTCATCAAGGGAGTGAAATACGGAGTGTATCTTGCTGTGGAGCTTAGCGACAATCTTTGCATGA

ACGGTCCAGTGACCCTTAAGCTTACTCGTCCCAACGGCACTACTGAAGAGCACAAGGAAGATTTGACGAC

CAAGCCAAAGAATACATTGGTGGGGCTTAAGGTTGGTGAATTCTGCAACACAGCCCCCTGCGGCTGTGAG

AATATTGTCAAGTTCTCGATGAATGGATGTGATGGAACCACCTGGAAGACTGGCCTTACTGTCGTTGGTG

CTGTCATTGCCCCAGTTAACTGCTAAACATGAATCTCACAATCATCATGTACTAAATAATTTCCAATTAA

TTGCCAACATTACCTGTCTGGTACGTGTGGCTGATATGGAATGTGTGAAATAATCGCCAATATCTTGGTG

TTTTAATATATAAAAAGTTAAAGTACGGAGTGTTCAGGCTATTATAACGTTTAGCTCGTAATTGCTTCAA

G

>MSTRG.3391.1 gene=MSTRG.3391

GGGCAACATACATACTAAGTAAGAGATAAGGGAGAGAGACAAATAGAGAGAGAGCGACTGAGAGATAGTA

ACAGTCAGAGGGAGGGACAGAGTGAGAGAGGGAGACACATAGAGAGATCGGGAAATATATTCTAAGAGAG

TGGAGATGGAGGGAAGGAGGGAGAGAGAGATTTTTAGGACATTCACTGAGAAAGCAATGTCTAAAATGTG

AGTATTAATTTATGATGGGTCCTCATTAATGTCTGGGCTAATACATTTTGCCTTTAGAAAGCCTTTCCCC

TTGCTGATTTATCTTTATTTGTATAACCCTGCAGTCATAATCTTTATTTTGGTGCGCATTATTAATTTTG

GAGTTGCTTTCTGCGCTGACATTTTTATGATTGCAGTCAGAAACTCAATTTTCCTACTAGAGCTATGTTC

TGTTTCATGACACAAAATAGCTTACAATTAGTGTATGAGGGCCTTCGATTTGGGTCCTTCGATTTGCAGT

CTAGAGGGCCTGGTTGGCAAGTTGAGACTGGTAGAATGGACAACTTAACTACTAGCAAAACTACAACAAA

CAATAACATTCCACCCCAACTTCCGATTATATCAGCACACGTATATGCCTATGTATTATTTTACCATAAT

GGCGAGTTCTCTCGATATTGCTTGATTAATTTGCATAACTTGTCTTTGCTGCAGGTATTACAATCTCGCA

CATGCAGTGAATGAAAGAGTTTTAAGGCAGCCCTCAATGCTACAAGCTGGAATCTTGAGAGATTATCATA

TTATAAGGACAACTACGTCAACTACCACAAGAGATTAATTGATTTAGTAAATATCAGACGGTTATGAGGA

CGACAACACCATGGGTAGTGCCCAGCCAGCTTATACTTCAATGAGAAAGTATGTAAGGTACTATAGAGAC

TTTGTGATTGGGCTTAACAAATCCTTGGTGGATGATTCCAGTATTGAGATATGCATGCTTGTACAAGGTT

ATTTGTAGCCATTTGAAATGTTATTAATTTAGTTTGTACAAATTATTTTAATCTTTTGTTAGGCAAAGAT

TTGACTTTCAGTTACTTGTATAAATTGTATAAGACAATAGTTTGATTTCTAA

>MSTRG.3390.17 gene=MSTRG.3390

GTTAAGTGTAACTATCAATAAGTGGACAAATAGGACCGTCTGAAACGCTATAAATACAGCTCAAGGAAGT

GAGGATTGGCATCCTCAGCACAAGAATTACACAACAACTTCTTATTCTCAGCTAGCAAGGAAGATGGCTC

AAGTGCAGCGTGATGACATCGTGTCGTTGTGGACTCAAGGTGTCCAGGCTGTGAAAGACAAGAGTTCCTG

CGCTGCAATTATCACTGCAAAGGAACTCTCGATAGTCTGGGGTTCAGACAGCCGCTACTGGAAATGGGTT

TCTAAGAAATACCCAATCAGTTGCAAGGCTCTGGAGGTTGCTCAACTAATTAAAGTATGCTGGCTAGAGA

TCGACGGAAAGTACAGTGCCATCAACCTCATCAAGGGAGTGAATTACGGAGTGTATTTTGTTGTGGAGCT

TAGCGACAATCTCTGCATGAACGGTCCAGTGACCCTTAAGCTGACTTGTCCCAACGGCACTACTGAAGAG

CACAAGGAATATTTGGAGACCATGCCAAAGAATACTTTGGTGGGACTTAGGGTTGGTGAATTCTGCAACA

CAGCCGCCTGTGGCTGTGAGAATACTGTCAAGTTCTCGATGAATGGATGTGATGGAACCACCTGGAAGAC

TGGCCTTACTGTCATTGGTGCTGTCATTGCCCCAGTTATCTGCTAAACATGCATCTGACCATCATCATGT

ACTAAATAATTTCCAATTCGCCAACATCACCTATCTGGTTTGGCTAATTAATATGGAATGTGTGAAATAA

TTGCCAATATGTTGGTATCTTAATATATAAAAAGTTAAAGTAAGGAGTGTTCAAGCTATTATAACGTTTA

GCTCGTAATTGCTTCAAGGTCTCTACTGTCACTTTAGCACTTTCAACTTAAA

>MSTRG.3394.1 gene=MSTRG.3394

GAGATGGAGCTAGTAGAGCAAGGTCCAGATTTCATAGCACTGTTCCTCCAGTTCAAAGCTTCTGCGATGG

ACCTTATCTCTGTTCTTGAGCAGAAACAAACGACTGGAGGGAGGACAAAAGTTTCCAGTTCACAAGCTGG

CAGATATTTGTGTTGCTAGTTTCGAGATAAACTTTGAAGAACAGATTTCTATGTTTAGACTCGGTTTATG

TGAAAGTTAGACTTTAAAAGCTACTGAACTAGTTAATTGGCATTTGCAGGTAATATATGGATATAGCACA

TGAAATGAATAACTA

>MSTRG.3395.1 gene=MSTRG.3395

AAATAAGGATCCTGAAAATCCTAAATCAAAAGTAAAGAAGAAAGTTAGGTTTTCTAATGATTTAGAGGTT

TTTCCATCTAATAGCCTAGTTCAAGGTAAGAGATTCACTCCGGAAGAAGATGAGAAGTTGAGAGCAGCTG

TTAAAGAGTATATACAGTCACATTGCTTAGGTGAAAAAGGTGTGGAAATGGTCATGAATTGTATTTCTCA

CCGACAAGTAAGGAACTGTTGGAATGAAATAAATAAAGCGTTACCATATAGGCCTAAATCAGCGATATAT

TGCCGTGCTCATACATTGTTTGAGAGAGGTGAGACACATGAATGGACTGAAGAAGAAAAAGAAATGCTTC

TGGAACAGTATAAAAAGCATGGTAACAATTGGAAGCTGATGGCTAAAGAATTTAAGAGACATAGACTTCA

GGTGAAGGATACATATCGTAGAATAAAAAGGGAACGAAATAGAGGACATTGGTCCCAAGAAGAATACCAA

ACCTTGTTTGACAACGTGAATATTGATTTGCGAGCTAAGCTTGATGAAGAGAAGAAATCGAAGCATGGAA

TGTTACGAGATAATATATGTTGGACTGCAATCAGTGACAAATTGTCAACTAGAGCCGACTCACTTTGCTG

TATGAAATGGTACAATCAGTTAACATCTCCTATGGTTGCTCAGGGTATATGGGCTGATTCTGATGATTAC

CGTCTGCTTGATGCACTTTTTAACTTAGATGCATGCTGCATAGAGGATGTAGATTGGGATGACCTGATGG

ATCATAGGTCTGGTGATGTATGTAGGAAGCGTTGGGACCAAATGGTGCTTCACATAGGCTTACATGGGGT

CAAGTCTTTTTCTGAGCAAGTTGAAGTTCTGGCAAAGCGATACCGTCCAGAACTAACTGAAGCACGGGAG

GCTTGGGACAGCAAACCTTTGGTCCCATGACATAGCAGACAGTTATATACTGCACAAATTACTTCTCTGG

ATATGACAAAGCCTGAGATGGAGCTAGTAGAGCAAGGTCCAGATTTCATAGCACTGTTCCTCCAGTTCAA

AGCTTCTGCGATGGACCTTATCTCTGTTCTTGAGCAGAAACAAACGACTGGAGGGAGGACAAAAGTTTCC

AGTTCACAAGCTGGCAGATATTTGTGTTGCTAGTTTCGAGATAAACTTTGAAGAACAGATTTCTATGTTT

AGACTCGGTTTATGTGAAAGTTAGACTTTAAAAGCTACTGAACTAGTTAATTGGCATTTGCAGGTAATAT

ATGGATATAGCACATGAAATGAATAACTAAATTTCCTTGTATTGAGCCTATATGGCTTAAGTTATCTTTT

CCTTGTTTCAGTCGATCTGTGTTGCAGAGGAGATTATGGAAAGAAAGGATATTTGGCCCTGAAACAGCTT

ATCTTGCTGCCCTTTCTCTTCTCCCTATATCCCTACCCTTTTTTGAATTTCTTTATTGACTTTTTTTGTT

TTTCAGAAATATATGCAACAAGCAGATTGATTGAAATGATTATGTTCAACTTGTTTGATTTCTCG

>MSTRG.3398.6 gene=MSTRG.3398

TAAACTTTTCGCTTCTGCAGAGAGTGGCACAGACATATCGAACACAAAGTTCAGACAGTCATCATCGAAG

CTATTGACCGTTAAACCCCGTTGTTAAGTGCGGCCAACACTGCTTCGCCATTCGACTAATCACACACCAC

CGCTGTTCTTTCCATCTTCCTGGTCGCTGCGTAGCTGTTCTCGAAGCCAATCATTGGCTTTCAATTCTTC

GGCCAATTAACTCCGTTTTCTGGCCAGCAGTCCACCCCATCTTTGATTATTACGATTGGTTAACTCCGTC

TGCGTTGACAACTGAAGTACTGTGTTACTCTTGGAGAATCAACCGAGACTTGTCTATCAAGGACAGATCA

AGGACATTCAGAAATAGTTAAATAT

>MSTRG.3408.1 gene=MSTRG.3408

ATTATCATATATCGTCTTGTCAAAATCCCTAGCCACAACTGCCGATCCCCAAATTTTCATTGTTCATATC

AAAATCCCTAGCCACAGCTGCCGATCCGCAAATTCTCATTGTTCTTGTCAAAAATTCTAGCCACGACTAT

CCCTCGGCAAAGTCCTCGTTCTTCAGTTTCATTTTTGAGGATAGTCATTTCATTGATATATGTCTCTATG

TCTACTGACCTCCCACTTATGATTTTGGTCATTGTGTTCATGTGTGACATGGATTCTATCATGATTGAAC

TCGCTAAATATTTATTGGTACGTGATATCTTGAATTTAAGTATATATGTTTTGATGATCATCTATTCATT

AATTTCATTTCATATGTGAATAATTATGCATAGACATTCAAATTCAAGGTTTTCGATGAAAATTGTATAG

CAATGAAATGATTGTTTTTTTATTTGTTTTGATTTATTGAATCTACTTATTTCTTTTAATGGATTTGTTT

TGAAATTATAAAAGTTGTTTTAGAATAAACTAGGAGAGCTCTTAGTTAGTAGAATTTGCTAGCCTATACT

TAATTGCTGAAATATGCTTTAGTCCAATTTAAGTTTTACTTTTCTCAGTCCACTAAAAAGTGTTGTTTTT

GTTGGATAATTAGTAGTTCACTTTGTTTATTGCTTTTACTTTGTCTAGATGGAGTTAGGTACCATATCAG

ATGGTTCATGGAAAGCTTTTCGGTAGATGGAGATTGCCTGCAGCCCATGCTACTGAATTCAACTTCTATA

AACACTGATATGTACTTTATAGAAGGAATGAATTACAACCTGTTGGTCATCTTTATCTCTTTCCTTCAAG

CTCTTCTATTGCTTCAGCAAATGAACCATATATAGCAGCACACAATTTGCAGTTGGTAGAGTATCAATTT

TGGCAATTGGGCCATGGATGCTTTACATCTGGTAGTAGGAATCGCAGCCGATCCCTTATTCAGTGCATTT

GCTACTGCTGCATTTTTGAAGTTCAGGCCTCATGTCTTGGTATCCTGTGTTGTTGTGCTTGCAATATAAA

CTGAGGTAGTGTTTGTCTTGATTTTATACGGATGAAGAATTTGACCAAGAGTTTGTACACATACTTCGTC

GACAGAGCTGCATAGTGTTTTCCAGAGGAAGTTCTGATTTTATACAGATGAAGAATTTGACCAAGAGTTT

GTACACATACTTCGTCGACAGAGCTGCATAGTGTTTTCCAGAGGAAGTTCTGATTTTATACAGATGAAGA

ATTTGACCAAGAGTTTGTACACATACTTTGTCGCATAGTGTTTTTCAGAGGAAGTTCGAGATATCGAGGA

GTTACTCTAAAAAATTGCTATATGTTATTATCTTTCTAGTTCTTGTACGCTTTTAGTTATTTGCAAAACT

TGTAGGCTTTAAGTTATTTGCAAGACCTGTATGCATTAAATAATTTTTGTAGCTCATGAATTTTGATGGT

GCTTGTGATTTTAAGACTGATTGATTGTAATATATAAATTTGATGGGAAATGCAATATTTAAAAGGTTTT

TTTAAAAAAAAAA

>MSTRG.3411.1 gene=MSTRG.3411

CGACAACAACCTCAGTACTCATCTCCGATCTCCGATCACTCTCTCCTTGTCTAGACTTGACATACAATCC

ATACACATAAGAACAAAACCCCCACCCACACAACGCCGCCGACACCGCCTTCCCACCGTCGAATCTATCG

CCATAAACCACCACTCCTCCCACCACATTCGCCCCCGTCAACGCCGTCATGCATATCGCTCCGGTGAGCC

CCGAGGTGAGAAACACCATCCCCGCGGTCCCCATGAAGCATAACTGCCACGTCACCACGTTTCCCATCAC

TGTCAGCCAGTAATATTTGTGCCCTAGATCAAACCCTAGTTCACTCTCGCGTCTCATCTCCGAAAAGCCG

CCGTTGACCGCCATGCCCGCGGTGGCTAGCACCGTGGCGGAGGCTTCCATGATCAGCTGCATTTCCATCA

CCATGGAATAACAGTAGACTTGCTTGTAAATTATTTCCATTAATGGCAAGTAGAGTGCAAATAACAAGCC

TGCGCCAATGGTGCAAAAATAGCCTATGAAATATTTCTCACGGGCCTGGTCATTGGGCTCGTCGTCGTGA

GATGAGCCCAGGCCCAGGCCCAATAAGACTGAAGCTAGGGACAAGAGGACAACACAGTTGAGATTAGAGA

ATGTGATTCTTTGTTTCACAATGATGACTGAGAAAATGAGGTTGAAAGTGAGCTGAGAAGAGAGGAGAAG

TGAGGAGGTGGAGACAGGGAGATAAGAGTTTCCCCAAGAGAAGAGGAGGTTGTTGAGGCCCAAAAGAAGC

CCAATGAGGAGAGAGTAGAGAAAGAGGTTTTGGGAAAGATGAGAGAAGGGTTTTCTTTGGGTGGATTTGA

AGAGGTGAAAAGGGAGGAAAATGGGGAGAATGAGGAGAGGGAAGCCAGCACATTGGACCCAAGTGGAGAC

CCATCTACTAGAGCCATTGTGATTGAAATAAAATTTGGAGAGTAAGCTTGATGAAACCGAGCCTAAGAAT

AGGAACATGTAGTTAAATACTAAGAGTAACATGTAGGTTTTGTTAGGCGTATTCGTCGCGATGATGGTGG

TGTTCTTGTCGACATCTCCACCGCGTAATGGTGGTGATAAGGGGTGGTTGTTATATTTTATCTGGGGGTG

GTGGTGTATTTGGGATGAGGTTTCCATTTGATAGAGGTGGAGGTAGGTTAGAAGTTTAAAAGGTGTGTGG

CTATTTATATGTGTGTTGGAGGGGGAGATGCATGTTTTGATCAAAATAACTATGTGATTGTGGCTTAAAA

GGAGGTATA

>MSTRG.3414.1 gene=MSTRG.3414

TTTGAGTTTATTATCATTCTTCGAGCAGGTGGATAAAATGGAGAAAGTCGCGTAAGGCAGAGTTTGGATC

GGAAATGATGTTGCTTCACGAGGTGCACGCGGATGAACCAGTATTGCAAGCATTTAAACTGATGAGGTAA

GAGGGATTCGGTAGGTTACCAGTGGTTGAAAGTAATGGCAATATGGTAATATAAGCATCAGAGACATTCA

GTTCCTACCACTTGCACCTGCTGTTTACAAGAAATAAAGCCTAAAGGTATCGAAGTTGGGTTATTTATAT

TTCAGCAGCTTGATCACTTGGATGCTAATCATGGTAAGATAGTTCCAGATCGCTATTATTACTCAGCTGG

TAATGTATTCACTTAAGGTTATTTGACTGCATCTAATTATGTTTTTCATTCTTTTACTTTTCATGATTCC

ATATCATCGTAGTGGAGTCGAGGATATACAAATTCAAAGTACCACACGTTAACGGTTAACTATCTTTTAT

GCCTGTAAGAGCACTATTATACTGTTTCTGGAATCCCC

>MSTRG.3415.1 gene=MSTRG.3415

ACTAGCTTTATGCCCGTGCTTCGCACACGGATACTGTTAAAATCTTGATCAATAGAACTCATGAGAGACT

ACTATCTGCATCCTTGCAAATTAATCTTGAGGCCTTGGCATGCTCCTTTCAGCTGGCTCGTATTATGAAT

ATACCTACCTTCCTTCTTGTTGGACCAAGTGATAAAAGCAAATCTCATAGCCTTCCTTCTTGTTGGACCA

AGTGATAAAAGCAAATCTCATTAAAGCTACGAAGAAGATTTGAAAGTACTTTGAAGTTTATATGTATTAA

CCTAACAATGAATCACT

>MSTRG.3421.1 gene=MSTRG.3421

GAAATTTAAAAAATGAGGTAGGGACATCCGGGGACTTATAATTGGATTCGGTCCGCACAAGGAAACAAGT

TCCAAGTACCACTGTGACTGTGTTCTTGCATAGTTATATGTGAGTGTCTGAGAGAGAGAGACAGACAGAG

AGGTTGAGAGAGATAGAGACAGGCAGAGAGACATTGAGAGACACACAGAGAGAGAGATTCAGAGAGAGAA

AGGGGGGTAATGAAATGCCAGTTGGGCGGCAGCAGCAAATCCTCAGTCAAGATTCAGAGGAATCAATTTA

TTTCTTTTCCTTTGATCTTGCAACTTCTATCCCCCCCCCCCCCCCCCCCCCCCCCCCCACCCCACCAACC

CAACAGTTAATTGTTAGAATGGCAATCAAAATTGAAGACTTGCATTTCAGTGAGTTTGCTTAATTGGAGA

GAAATGACCGAAAGAAGGGGATTGGTGGAAGCTAGTCTTGCATCAGAGTTTGTCCAATGCTGTGTTTTGA

AGTAATCCAAAAACCCACCCACTCCTTCGTGGTCAGGTTCTCAGTCTTGTTCTCTCTCTACCCATTTCTC

TTGGACCTTGCCCTCTTTGATGCACTACTCCTATTATTTTAATTTTATGTTTGTGTATAAAATATTGATT

CTGTGCATATCAGTCCTACCATGCATTGTCATGTGAGTTGTGACTATGCTGTGTAATGTTATTGCGAGGA

AGATTATACGCTATCCTCTTTCCTATTTATGTTTTGGTGAAATTATTGTACTCAAGTAACTTTATTCAAG

GTAGGAAGGAATTAAATTTCTAGGATTGGAGGTTGTTAGGGAAATTTGAGGGAGATAAGTGGCGATGAGA

GTGGACAAGATGATCAATTGTTGATGACATGATCAAATTTGTGAACATGTCATTTTCATGTGAAGACACG

ACAATTGTCAAAATCATTTGAGCTAAATAAGTTAATGGATATGTTATTGGATGTTAATTAAGTGGTCTAG

TATTTTGTTGAATTCCATTTCATCCAATTCAGTGTTAATGGATATTGTGATTATAATACCATTACTTTTT

GTTATTGTGATTATAATACCATTACTTTTTGTGAATGTGAATATGTGATTGTTATGCTGCTCCATTCGTG

AGAGTGAGTGTGTGTGTGTGTGAGAGGGGGTAATGAAAGGCCAGTTGGGCCGCAATAGCAATTTCTTAGT

AAAGATTCAGGACAACTTTTTTCCACTTAATCTTGAACAACTTGTTGCTTCAATTTATGTTAATGGGGTT

CACATACTGAGTTTGCTTAACATGGTAATTTGCTATTGGTTCATCCCATCTCTCCTACATATTTTTCCTG

TAAAATTTTGATTTTTATATGTGTGTGTTAATTGTTTTACGGAGAGGAATCTCTGAAAGAAGGGGATTGG

TGGAAGCAAGTCTTGCCCTCTAAGTTTGGTCAATGCTGTGTTTTGATGTTATCCAGAGCTCCCCCCTTCT

TAATCAGGATGGACATTGGAGAGATTGAAGAAATGATTAACGAAGCAATGGATGGAGAATGGCTGGGATG

AGAAGGAATGATTATTATTACCATAATAATCGAAGTGAAAGGCCTCTCTTTGCTGTTTGAGACTTCAGCC

TCTGTTAAAAAGGCTTGCTACTCTGTTCGAGTTAGACCCGTTGTAAAAAAAATAAATCAAAAGTTGAAGT

CCCTCTTTGTTATGAGTTTGAGTTTTAAATTCCAAATGTTGTGTAATGTTCTACTTAAATTTGTCAACAT

GTTTTAGTTTATGATTTTACTTAGTGTAGTCAGACAATGGTTTACCATAAATTTTGTTTAATGTAATACC

AGTTACAAGTTTAGTCAGACAATACTTAGAAATGCACAAAGG

>MSTRG.3421.2 gene=MSTRG.3421

CCACCAACCCAACAGTTAATTGTTAGAATGGCAATCAAAATTGAAGACTTGCATTTCAAGAGAAATGACC

GAAAGAAGGGGATTGGTGGAAGCTAGTCTTGCATCAGAGTTTGTCCAATGCTGTGTTTTGAAGTAATCCA

AAAACCCACCCACTCCTTCGTGGTCAGGTTCTCAGTCTTGTTCTCTCTCTACCCATTTCTCTTGGACCTT

GCCCTCTTTGATGCACTACTCCTATTATTTTAATTTTATGTTTGTGTATAAAATATTGATTCTGTGCATA

TCAGTCCTACCATGCATTGTCATGTGAGTTGTGACTATGCTGTGTAATGTTATTGCGAGGAAGATTATAC

GCTATCCTCTTTCCTATTTATGTTTTGGTGAAATTATTGTACTCAAGTAACTTTATTCAAGGTAGGAAGG

AATTAAATTTCTAGGATTGGAGGTTGTTAGGGAAATTTGAGGGAGATAAGTGGCGATGAGAGTGGACAAG

ATGATCAATTGTTGATGACATGATCAAATTTGTGAACATGTCATTTTCATGTGAAGACACGACAATTGTC

AAAATCATTTGAGCTAAATAAGTTAATGGATATGTTATTGGATGTTAATTAAGTGGTCTAGTATTTTGTT

GAATTCCATTTCATCCAATTCAGTGTTAATGGATATTGTGATTATAATACCATTACTTTTTGTTATTGTG

ATTATAATACCATTACTTTTTGTGAATGTGAATATGTGATTGTTATGCTGCTCCATTCGTGAGAGTGAGT

GTGTGTGTGTGTGAGAGGGGGTAATGAAAGGCCAGTTGGGCCGCAATAGCAATTTCTTAGTAAAGATTCA

GGACAACTTTTTTCCACTTAATCTTGAACAACTTGTTGCTTCAATTTATGTTAATGGGGTTCACATACTG

AGTTTGCTTAACATGGTAATTTGCTATTGGTTCATCCCATCTCTCCTACATATTTTTCCTGTAAAATTTT

GATTTTTATATGTGTGTGTTAATTGTTTTACGGAGAGGAATCTCTGAAAGAAGGGGATTGGTGGAAGCAA

GTCTTGCCCTCTAAGTTTGGTCAATGCTGTGTTTTGATGTTATCCAGAGCTCCCCCCTTTTATGTAGGAT

GGACATTGGAGAGATTGAAGAAATGATTAACGAAGCAATGGATGGAGAATGGCTGGGATGAGAAGGAATG

ATTATTATTACCATAATAATCGAAGTGAAAGGCCTCTCTTTGCTGTTTGAGACTTCAGCCTCTGTTAAAA

AGGCTTGCTACTCTGTTCGAGTTAGACCCGTTGTAAAAAAAATAAATCAAAAGTTGAAGTCCCTCTTTGT

TATGAGTTTGAGTTTTAAATTCC

>MSTRG.3421.3 gene=MSTRG.3421

CCACCAACCCAACAGTTAATTGTTAGAATGGCAATCAAAATTGAAGACTTGCATTTCAAGAGAAATGACC

GAAAGAAGGGGATTGGTGGAAGCTAGTCTTGCATCAGAGTTTGTCCAATGCTGTGTTTTGAAGTAATCCA

AAAACCCACCCACTCCTTCGTGGTCAGGTTCTCAGTCTTGTTCTCTCTCTACCCATTTCTCTTGGACCTT

GCCCTCTTTGATGCACTACTCCTATTATTTTAATTTTATGTTTGTGTATAAAATATTGATTCTGTGCATA

TCAGTCCTACCATGCATTGTCATGTGAGTTGTGACTATGCTGTGTAATGTTATTGCGAGGAAGATTATAC

GCTATCCTCTTTCCTATTTATGTTTTGGTGAAATTATTGTACTCAAGTAACTTTATTCAAGGTAGGAAGG

AATTAAATTTCTAGGATTGGAGGTTGTTAGGGAAATTTGAGGGAGATAAGTGGCGATGAGAGTGGACAAG

ATGATCAATTGTTGATGACATGATCAAATTTGTGAACATGTCATTTTCATGTGAAGACACGACAATTGTC

AAAATCATTTGAGCTAAATAAGTTAATGGATATGTTATTGGATGTTAATTAAGTGGTCTAGTATTTTGTT

GAATTCCATTTCATCCAATTCAGTGTTAATGGATATTGTGATTATAATACCATTACTTTTTGTTATTGTG

ATTATAATACCATTACTTTTTGTGAATGTGAATATGTGATTGTTATGCTGCTCCATTCGTGAGAGTGAGT

GTGTGTGTGTGTGAGAGGGGGTAATGAAAGGCCAGTTGGGCCGCAATAGCAATTTCTTAGTAAAGATTCA

GGACAACTTTTTTCCACTTAATCTTGAACAACTTGTTGCTTCAATTTATGTTAATGGGGTTCACATACTG

AGTTTGCTTAACATGGTAATTTGCTATTGGTTCATCCCATCTCTCCTACATATTTTTCCTGTAAAATTTT

GATTTTTATATGTGTGTGTTAATTGTTTTACGGAGAGGAATCTCTGAAAGAAGGGGATTGGTGGAAGCAA

GTCTTGCCCTCTAAGTTTGGTCAATGCTGTGTTTTGATGTTATCCAGAGCTCCCCCCTTCTTAATCAGGA

TGGACATTGGAGAGATTGAAGAAATGATTAACGAAGCAATGGATGGAGAATGGCTGGGATGAGAAGGAAT

GATTATTATTACCATAATAATCGAAGTGAAAGGCCTCTCTTTGCTGTTTGAGACTTCAGCCTCTGTTAAA

AAGGCTTGCTACTCTGTTCGAGTTAGACCCGTTGTAAAAAAAATAAATCAAAAGTTGAAGTCCCTCTTTG

TTATGAGTTTGAGTTTTAAATTCCAAATGTTGTG

>MSTRG.3423.1 gene=MSTRG.3423

CCCGGGTCATTGAGAGAACCGGATGTGTCTAGGCGTTCAAAGAGCCGAATGTCCAAGTCGCAGACAACAC

TGATGTCCATTCCAAACCCTGGTTTTTTGTTGAACAAAAGGAAATGGATTTTTGATACCTAAACAGAAGT

GGGTACCAGAGAGAGCTTCCACGGTGATGACAGTCGCTATAAGTTTACAAGTCATTTGTATGCTGCTCTA

GATCGCCATGGAGTCCGAACTTATAAGGATGATCCAGAACTACGCAGTGGAGAAGTAATCTCTGACGCAT

TGTTCCAAGCGATTCAGGGATCCAAGACTTACATCACTGTCCTGTCTGAAAACTATGCCTTCTCGCCTTG

GTGCCTTGACGAGCTCGTTGAAATTCTAAATTGTCATAAAACTAGGCAAAGGTTGATCATTCCTGTGTTT

TACAACATTGATCCATCAATTGTGAGACACCAACTTGGAAGTTTTAGTAAGGCTTTTGAGAAGCATCAAA

TTCGATTTGCTGGTAAAATGGAGAAAGTGACTGAGTGGCGGCTTGCACTAGCAGAGATTGCACAGTTTTC

AGGAAAACATATACATGCAAAAAGGTCTGAAGCTGATTTAGTAGATGAAATTATTGATGAGATACTTCTC

AAAATAAATCCCAAGATTTCTGATGTTGGAAAATATCCAGTGGGGTTGGATTCTCGTATCAGAGACATAA

CACTACTGTTGAACAGCCACACAGAAGGCATGATTAAGTTCGGTATATATGGAATGGGTGGAGTGGGCAA

AACAACTTTTGCCAAAGCTCTTTATAAAGAACTCTTTCCCGGAAGTTTTAAGGGAAGTTGCTTCCTTGAA

GATGTGAGTGAAGTTGCAGGAAGAATAAATGGCCTAGTATCTTTACAACAAAGACTGATCAACGATGTTC

TTAGAAGTAAGAAAATTTTGAAGGTTGACTCTGTTAACCATGGAATTAGTTCAATAGGAGCTAGAATTTG

TTCAAACAAAATCCTGGTTGTTATTGACGATTTAGACCACGTCACACAATTTCAGTCTTTAGTTGGACCA

TTTGCTTCTGGGAGTGTAGTCATCATAACTACCAGGGATGAACAAATACTGGATACAATTGAAGTCGAAC

CAAGATACAGATACATGGTGCAGGAGTTGGATGCTGCTGACTCAATGGCTCTTTTTACCCGACATGCATT

TGGAAATTCTACACCAAACAACACTCTAATGGAGTTGTCTAATGATATTCTACAGCATGCTGGTGGGCTT

CCCTTGGCTCTTAAGGTTTTCGGGGCGAACTTGTTTAACCAATCTGAGGAAGGATGGAGATGGTTTATAG

ACAAAATAAGAAGACTACCCAATAGCGATGTTGAGAGAAAACTTGTGATTAGCTTTGATGCATTGAAGAT

GGTTGATCCCACGCTGCAGGATATTTTCCTTGATATTGCTTGTTTTTTCTGTGGACGGAAGAAAGAAGTG

GCTGTGAAAATACTGGAAGGTTGTTATACGTTCATCAACCATAAAATCGACATTCTAAAGAAAAGATGTT

TATTGACTATAAATGATAGTGATGAGCTAGGGATGCATGACCTGCTTTTGGACATGGGAAGGAAAATTTC

GCGCAACAACTCTCCTGATGAGCATGGAAAGCACAGTAGATTGTGGGAATCAAAAGATATATTGGGCGTG

TTAAAGAACTGTAAGGTAACAGAGGCTATTGAGGTAATCATGGATCTCAACTTTTACAAAGAGGAATCAC

TTGAGGGAGTATCAGTCACTACAGAAACATTCAGAAGGATGTGGAAATTAAGAATTCTTCTCCTTGCGAC

TGTAAATATTACTGGAAGCTTTGAACGGACATTTGAAAATTTGAGAGTACTCTCTTGGTACCGTTGTCCA

TTGAAGTTTTTACCTTATGAATTTTTCCCACAAAAACTTGTGTCTCTTGATTTGCCTAATAGCCAACTGA

GAACAATGTGGGAGCCAAATATGGTTCCACGTGTATTTGTGAACTTGAAGTCTTTGTGGATGGGGAGTTC

TCTACATTTAACTACTATGCCAGACTTTAGTAAATTACCTTGTCTTGAAGAGTTAGATTTTAGGGGATGT

ATACAGTTGGAGGATGTCCACACAACCATCGGTAGCTTGGTCAGGCTTGTTTCCTTGAATTTGGAAAACT

GTGCAAACCTTACAAGTCTGGGAGAGAGCATTTGCAACTTGAGAGCTTTGAAAATTTTAAACATTGGGGG

TTGTAGTAGTCTGAGAGCATTGCCTACGAAATTAGGTGACCTTGAATCCCTAACAAAGCTTGTTGCTAGT

GGACTAACTGTTCTGACATTACCATATTCAATGGCACGGCTTAGTAAGCTTGTTCAGCTGGAATTAAATT

TGAACAGGAAACTTAAAAATCTTCCAGATCCCATTGGCAACCTGAGATCACTGGAGATTTTAAATATATC

TGAATGTGAAAAACTAGAAATATTACCGGATCAATTGTGGAAGATTACAAGGTTAAGGGAACTACATGCA

GGGCGTGCTAGGATGTTAACAACACTTCCTTGCATAAATTCAGGCCAGGTGATATTATCATTGAAAAATT

TAGATTTATCTAAGAGTTCCCTCACAACCCTGCCATCTGGTATTTGTCAGCTGTCAAACCTGGAAACTCT

TAATCTTCGCAAATGTCTTCATCTGTTATATATTGCTGAACTTCCTCTTAATCTGAAATGTGTAAGTGCA

GAAGGATGCAAGACTTTGCAATTTTTAACAAATCTATCCAATTTAAAACGGTTGGAGTTTTTGGACCTTA

CTGATTGTATCCGATTAACGGAGATACAAGGCTTGAAGGGGCTCAGTTCCATTCGAGAACTTCTTTTTGG

AGGTTGCCACTCAGCTCTTCTAGCTCAAACCTTCACGGAGGATTTGTTTCAGATATATTCTGGATTCGGT

CATCAAATCAAAATATACACCAAGCCAGAAGAGTTTCAGGATTGGATTTGTCAGTTACCTGATTGGATTA

TTCAATCACGTGATCTAGGATCAAAGGTGTCTTTGGATTTGCTGCCAAATTCATCAGACAATTACTTGGC

AATGATTCTGTGTTTGCACAAATGGATGAATTTTCTGGACTATTCTGTTACGACTACCACTAGTAATTTT

GAATGGACGGAAGACTATCGTACCCATTATTACCATAGCTTATCAAAATGGTCAGAACACTTACAAGGAT

CGCTGATGATAATAGTGCCGAGATCAATTTTCTTAGTCAGAGATGGTGATGACAGTATTGAAGTATCATC

AAATGTAAAAATTCTGGGGATACATCTGTTATATGGTTCAGAGATAGCAACAAGTTCCGTGGCACCACTT

TCAGGTAGAATTTGAAACAAGGTAGTGGTTCTCATTCTCCTTGGACGAATGAGTGAGCTGCAGAGTACTT

CAGATTTATAGGTGCCTGGACTGGCCATCATGTTGGAGGCTCTCGAGAACTGCGATCAGAGGTCTCAGAA

GGCTTGTGTCAAAATCATCTTGACACTTTGACATTGAGGTGGTCGGACAGCCTGTGAAAATCGGCCGTAT

AAGGGTCACAAGATTGGTGGTTACTGGATTGAGGGGAGATTGTTGGGTGGTACTAATAAAGATGAGGAAT

ATTTGTCTTGAATATTAATGATTAGTGCGGCCAAAGGAGGAGCCCAAAAAATAAATTCGGGCTTGGTCAT

GACACAGAGCTAGAGGCTCTCTATACGAGCAAGACAACGGCTGAGAGAAAGGGCCTTGATTTCTCTGAGC

TTCATCCGGGCTACTTCAAGTTGTGACATACTTATCCAGGAAGCAGTGGTGTTATTTGACAGATCAGCAA

ATCCCATGATAAGAAAATGAGGTAGCAAAAGCTTGTGTCATTTTAGTTACACTTGGGAGATGTTATGAGC

CCCTAATCTGGGTAGCATCAGTACTTGAGGGTCTGCAGATGATTGTTGTGAAGGTCATCTGATGATTGCG

CAAGTATAACATGACGTTTATGACTATGAAAATAATTTCATTTTGGTAATTATTTGTTTAAATATTTATT

TGAAATATGGATTTTTTATAAATTAGTGAATATTTTT

>MSTRG.3423.2 gene=MSTRG.3423

AGGTCTGAAGCTGATTTAGTAGATGAAATTATTGATGAGATACTTCTCAAAATAAATCCCAAGATTTCTG

ATGTTGGAAAATATCCAGTGGGGTTGGATTCTCGTATCAGAGACATAACACTACTGTTGAACAGCCACAC

AGAAGGCATGATTAAGTTCGGTATATATGGAATGGGTGGAGTGGGCAAAACAACTTTTGCCAAAGCTCTT

TATAAAGAACTCTTTCCCGGAAGTTTTAAGGGAAGTTGCTTCCTTGAAGATGTGAGTGAAGTTGCAGGAA

GAATAAATGGCCTAGTATCTTTACAACAAAGACTGATCAACGATGTTCTTAGAATTGGACCATTTGCTTC

TGGGAGTGTAGTCATCATAACTACCAGGGATGAACAAATACTGGATACAATTGAAGTCGAACCAAGATAC

AGATACATGGTGCAGGAGTTGGATGCTGCTGACTCAATGGCTCTTTTTACCCGACATGCATTTGGAAATT

CTACACCAAACAACACTCTAATGGAGTTGTCTAATGATATTCTACAGCATGCTGGTGGGCTTCCCTTGGC

TCTTAAGGTTTTCGGGGCGAACTTGTTTAACCAATCTGAGGAAGGATGGAGATGGTTTATAGACAAAATA

AGAAGACTACCCAATAGCGATGTTGAGAGAAAACTTGTGATTAGCTTTGATGCATTGAAGATGGTTGATC

CCACGCTGCAGGATATTTTCCTTGATATTGCTTGTTTTTTCTGTGGACGGAAGAAAGAAGTGGCTGTGAA

AATACTGGAAGGTTGTTATACGTTCATCAACCATAAAATCGACATTCTAAAGAAAAGATGTTTATTGACT

ATAAATGATAGTGATGAGCTAGGGATGCATGACCTGCTTTTGGACATGGGAAGGAAAATTTCGCGCAACA

ACTCTCCTGATGAGCATGGAAAGCACAGTAGATTGTGGGAATCAAAAGATATATTGGGCGTGTTAAAGAA

CTGTAAGGTAACAGAGGCTATTGAGGTAATCATGGATCTCAACTTTTACAAAGAGGAATCACTTGAGGGA

GTATCAGTCACTACAGAAACATTCAGAAGGATGTGGAAATTAAGAATTCTTCTCCTTGCGACTGTAAATA

TTACTGGAAGCTTTGAACGGACATTTGAAAATTTGAGAGTACTCTCTTGGTACCGTTGTCCATTGAAGTT

TTTACCTTATGAATTTTTCCCACAAAAACTTGTGTCTCTTGATTTGCCTAATAGCCAACTGAGAACAATG

TGGGAGCCAAATATGGTTCCACGTGTATTTGTGAACTTGAAGTCTTTGTGGATGGGGAGTTCTCTACATT

TAACTACTATGCCAGACTTTAGTAAATTACCTTGTCTTGAAGAGTTAGATTTTAGGGGATGTATACAGTT

GGAGGATGTCCACACAACCATCGGTAGCTTGGTCAGGCTTGTTTCCTTGAATTTGGAAAACTGTGCAAAC

CTTACAAGTCTGGGAGAGAGCATTTGCAACTTGAGAGCTTTGAAAATTTTAAACATTGGGGGTTGTAGTA

GTCTGAGAGCATTGCCTACGAAATTAGGTGACCTTGAATCCCTAACAAAGCTTGTTGCTAGTGGACTAAC

TGTTCTGACATTACCATATTCAATGGCACGGCTTAGTAAGCTTGTTCAGCTGGAATTAAATTTGAACAGG

AAACTTAAAAATCTTCCAGATCCCATTGGCAACCTGAGATCACTGGAGATTTTAAATATATCTGAATGTG

AAAAACTAGAAATATTACCGGATCAATTGTGGAAGATTACAAGGTTAAGGGAACTACATGCAGGGCGTGC

TAGGATGTTAACAACACTTCCTTGCATAAATTCAGGCCAGGTGATATTATCATTGAAAAATTTAGATTTA

TCTAAGAGTTCCCTCACAACCCTGCCATCTGGTATTTGTCAGCTGTCAAACCTGGAAACTCTTAATCTTC

GCAAATGTCTTCATCTGTTATATATTGCTGAACTTCCTCTTAATCTGAAATGTGTAAGTGCAGAAGGATG

CAAGACTTTGCAATTTTTAACAAATCTATCCAATTTAAAACGGTTGGAGTTTTTGGACCTTACTGATTGT

ATCCGATTAACGGAGATACAAGGCTTGAAGGGGCTCAGTTCCATTCGAGAACTTCTTTTTGGAGGTTGCC

ACTCAGCTCTTCTAGCTCAAACCTTCACGGAGGATTTGTTTCAGATATATTCTGGATTCGGTCATCAAAT

CAAAATATACACCAAGCCAGAAGAGTTTCAGGATTGGATTTGTCAGTTACCTGATTGGATTATTCAATCA

CGTGATCTAGGATCAAAGGTGTCTTTGGATTTGCTGCCAAATTCATCAGACAATTACTTGGCAATGATTC

TGTGTTTGCACAAATGGATGAATTTTCTGGACTATTCTGTTACGACTACCACTAGTAATTTTGAATGGAC

GGAAGACTATCGTACCCATTATTACCATAGCTTATCAAAATGGTCAGAACACTTACAAGGATCGCTGATG

ATAATAGTGCCGAGATCAATTTTCTTAGTCAGAGATGGTGATGACAGTATTGAAGTATCATCAAATGTAA

AAATTCTGGGGATACATCTGTTATATGGTTCAGAGATAGCAACAAGTTCCGTGGCACCACTTTCAGGTAG

AATTTGAAACAAGGTAGTGGTTCTCATTCTCCTTGGACGAATGAGTGAGCTGCAGAGTACTTCAGATTTA

TAGGTGCCTGGACTGGCCATCATGTTGGAGGCTCTCGAGAACTGCGATCAGAGGTCTCAGAAGGCTTGTG

TCAAAATCATCTTGACACTTTGACATTGAGGTGGTCGGACAGCCTGTGAAAATCGGCCGTATAAGGGTCA

CAAGATTGGTGGTTACTGGATTGAGGGGAGATTGTTGGGTGGTACTAATAAAGATGAGGAATATTTGTCT

TGAATATTAATGATTAGTGCGGCCAAAGGAGGAGCCCAAAAAATAAATTCGGGCTTGGTCATGACACAGA

GCTAGAGGCTCTCTATACGAGCAAGACAACGGCTGAGAGAAAGGGCCTTGATTTCTCTGAGCTTCATCCG

GGCTACTTCAAGTTGTGACATACTTATCCAGGAAGCAGTGGTGTTATTTGACAGATCAGCAAATCCCATG

ATAAGAAAATGAGGTAGCAAAAGCTTGTGTCATTTTAGTTACACTTGGGAGATGTTATGAGCCCCTAATC

TGGGTAGCATCAGTACTTGAGGGTCTGCAGATGATTGTTGTGAAGGTCATCTGATGATTGCGCAAGTATA

ACATGACGTTTAT

>MSTRG.3425.1 gene=MSTRG.3425

AAAAAACACAAAGACAAGCCTGTAGAAACTAAAAGGATAGATACAAACTCTACATGTACACATAAACAAC

ATAGGAGAGTACATTAAACATAAACATACATAGACATCTCTACATACACCTATACATACAGCACAGCCTA

GAGCTTCAACATCCTTGAAAAATCTCATCTTTTTACAATCAGATTCTTGTAAGTATTATTTTTCATCTTT

ACCCAGTTGATGAATCAATTTGCTATGGTTTCTTCAATCATTCAGCTTCGTAAAGGTATAAAGTTTCAAT

CTTTTGCTGGCCCAGATCTGATAAGTTCTTAATTGTTGTGTTTAGTTTGTTTTTTGTGTTGTTTTTGTGT

TTTCTTTGCGCAGATCTCTGTAAAATTTGAGTCTTTTTGTGTTTAGTTGGTTGTGTTTGACTTATTTGTT

GTGGGTTGGATTTGTTTGCTTTTTTGGTGATTTGGATTTAGTGAATTTTATGTGGGTTGCTGGATGATTG

TGTTTGTAGTTGATTGTGTGTATTTTGATGTGGTGGTAGTTGGATACTTGATTTGAGGGTGTAAAGGTTT

GAACTTTATGTTAAGAGATGCTGAAATTTACTTGATTTTGATGTGTTTTTATGGTAGGATTGTGTTTTGT

GTGTGTGAATTCAGTTCTGTATTTGTATGGCTGGTCAATGAAAATTTTGGTTGAGAT

>MSTRG.3426.1 gene=MSTRG.3426

CTTGTAAGCATCTCCCACCCATACATTCTATCTGTATCTTTCTACGTCAACATCTTCCCCCAACCACCAT

ATTTCAACTCTCCCAACCAATCTCTCTCTCATCTCTCTCTCTCTCTCTCTCCCATCTCTCCCTCTCTCTC

TCCCATCTCTCTCTCTCAAGATGATGCACATGACATTCTACTGGGGAACAAACGTCACCATCCTATTCGA

TGCATGGACTACCAGCTCCTACACTTCCTACGCAATCTCTCTCCTCCTCTGCTTTCTCGCCGCCGTCCTC

TACCAGTTCATCGAGGCCTGGCGGATCAAACTCAAGCTCCTCGCTTCATCTCCCCCGCAATCCGGCCTAA

ACGTCCCCCTCATCCAATCTAACATCGCCGGTGCCGGAAAGTCACCGTCGAAAGTCGCTGGCGCCGTAGT

TTTCGGAATTAACTCGGCTATAGGCTACATGCTGATGCTTGCGATCATGTCGTTTAACGGCGGCGTTTTT

GTCGCGATTGTCGTCGGAATCGCGGCCGGATATTTTGTGTTTCGGAGTGGTGATGAGGATGATCAGGTGG

TTGTTGATAATCCTTGTGCTTGTGCGTGATTAATTTCCGTCCTGATTGTTTGTATAATAAATTTATATTA

TTGATCACTTTTAGTTTTTAACGTTTTCGATATTAAGGTCTAATTTTTGATGCTTTTAA

>MSTRG.3428.1 gene=MSTRG.3428

GTTGCTGCTTAAAATTGAAACAAACCCTCCTGATCTCTCATTTCTTCAAGCCCCCTCTCAGTTTCTCACC

CAAAAGATCGACTCCCCTGAGCTCAAAAATGGTGAAAAATGGTAAAACAAAGAGGAAATATCACAAGCCC

ACTAAACCCCAGCATCACTCAAACCAGCCAGGAGCTAAAGAGGACAAGCCCACCATTGTAAAGACACTAC

CAGCAGATCTTCTACTTGAAATACTCTACAGGACTCCTGTTAAGTCCCTGGTAAGGTGTAAATGTGTTTG

TAAATCCTGGCTAGCTCTCTTGCATCATCCACATTCATACAAATGCACCTTGATTTCAATACTTCAAGAA

ATAACAAACAGATTTTATGCAATACCACTGATATTGCTAATTACGGTCGAAAAGTTGTGGCTTATTTACG

CGTAAATGGCCCTCCAATTCCTGTGCTCCACATTGATACAGAGTTCACTGTGAGGCCTAATTACCCTAGG

GTTATCAATTTTGCTGATTTCTCCAAGAATATGTCTCTTGGGGGTTCTGTTAATGGCATTGTTTGTCTAG

CTAATTATGCCGAAATGTTTGGGCAGTTTGTTGCGTTATGGAACCCTGCTATTAATTACTGGAAGCCCAT

TGAGTTTTCAAAAATGTGGAGTAGATATGATAGGTTGGAGGACATGTCGGTTGGGTTGGGGTTTGATGGT

GTGGCGGATGATTTCAAGATCATTCGCATTGTTCCTGTTGCTCTCCCGCCTTACTTTGAGAAGTTTCGTT

GGTCTCGAGTTGAAATGTACTCTGCTAACCAGGAAATTTGGGAAGATGTTAGTAAGGGGGGAGTTATTCC

CTTCCAGCCCAAAGTAAGACATTGTAATTTTGTTGTGAAGGGTGTTCCGTATTGGATAGGGACTGATGCA

CGACAGGAAGATATGGAGTCATTTATTCCTGATCAGCTAGAGATTTTAGGAAGAATTGATCCTTTTACGG

GGCTGTTTAAGAAGATCCTGTATCCAGAACACATTAAGAATGGGCATACTTTGGTGCATCCTGTGAACTT

GAGGGATTCAGTTGCTGCTCTAATTCAATCACCTGGTGAGCTTCCAAACCAAACGTTTGACTTGTATGTG

CGCGATGAAGATACTGCGAAGTGGACAAAGATGTATAGTATTGGCCCGTTTGATTTTGAACGAATGCGGA

TTCCTCCTCAGTGCTTTAGCACTGGTGAGATTGTACTGGAGACGTGGACAGGGGAAAATATTCATGATTC

TATAGACCGGACCACTCATATATGTGACCCTAAAAATAATATTGTTTTCCGTAACAATGAGATTGACGCG

CTACAGCCATTTTGGTTTCTTTCCTACAGTCACGTGGAGAGCCTGGTATGTGTCAAGGGAATGGTTCAGA

TTGGGAAGGAGCGCAGAGACAAGAAGACTAACCCTAAGATGAAGAAATGGATTGAATTCTTGTCTAAAGA

TTTTGAGTCGGCGTTGAATCTTTAACTGAATCTGCTGCTAGATGGGTAGTGCAGACAGGAGTAAAGCAAG

TTTGTTAAACATTATATGCCTGATGCTTCTTAATGATGTTACCTCAGTCCCTTGTGTAATGACATTGGTG

ATGAACTCTTGTTTAAATTTCAATGCAAGTCAAATTTCTAGCTTTGTATGCACTATGCAACTTTCCGCTT

TCTGTGGATAAATGTTGGATCACTGTAAACAATACAATGTTGTCGCTTGTTTCTTTACCCTTGTTCCAGT

TTCTCAGGATATATCATATTTTGCATTCAAGAACTACAGATCACATGTTGTATTAACCTACTTGCT

>MSTRG.3437.1 gene=MSTRG.3437

CTTGAAGCTTAGCTTTGGAGTTTGAGGTGACAGACACAGTCTTTTTGGACAGTTTTAGTGGGTTCAAGAA

GGGACCAGAAGCTGGTGCTTGAAGGGTCTTGTGGCTTGCAGTTGTTAGTGGCATTGCCATTGAAACAGAA

GCCATCTTTTTGGTTGATTGCTAGAGCTTGTGCTTTGTTGGTTTGGTTGTGTTTGTGAGTTCTGGTTGCT

>MSTRG.3440.1 gene=MSTRG.3440

CTTCAAGTTTCCTCCTAACAAGGCGTGAAAGATTGCTCTCATTGCACTCTCTCTCTAAAAAAAACAATTC

TCTCCTTCTTTTCAATCGTCTCTATACACCCTTTTCCTCATTTTTAGTATACTTTCACTACCCAAGAATT

TAAACACAAAGCATACACACACAGAGAAAATGTATGCTTGGGAGAGAACAGAGAAGGTGAGGAAAAATCC

GAAAAGCCCTTCATTTTCTTCGAGCCTTCTCGATGAAATCTACCGTGAAATTGACGGTAGTGATGATAGT

TACAAGGGTTTTGAGCAGCAGAAAGAGAAGAATGTGAAGAATTTGCATTGCAGCAAGAATTATGTCAATG

GTGGTGGTGACAGAGCAAAGGCTAGTGGCAGGACTGTTGAGGATGAAGAAATGACAAGCCTTCGACGGGC

TTGTTTGATCGAGAAATGGATGGAAAAGAAGGTTAATGACAAGGTCTCGGCTAGAAAAGGATCATTGGTG

CCTGAATTTGACGAGAAGTACTTAATTCAGGACAATGACCCTCTTTTTTTTAGCTCGAGTTCTTGTTCTT

CGGATTCTAGTTTCGGAGGATTTTCATCCTCCGAAACCGAATCTTTTGGAGTGGGGAACTGGAAATCTAG

TTCACTCCCTGCTCCAAAACCAATTAGGACTAATGTCCAGAAACACGATCAGTTTTATTTGTTTGATGAT

TACCGTAATCATCAAACAAATAAAACAAAAGACGGACTGATCAAGTCAAAAACCAGAGCCGTCAAGATTT

ACGAAAATCTGAAGAAAATGAAGCAGCCTATCTCACCGGGTGGTCGGCTCACTGGCTTCATCAACTCCCT

CTTCACACATGGTGATCCAAAGAAAACCAAGAATTCACATCAAAATCACAATGAGGCTTATGACTTGAAC

GCGGTTTACGACTTGAGCTCTCACAGGACACGAAATTCATCCAAATGTTCATCAGCTTCTTCTTTTTCAA

GGTCTTGTTTAAGCAAGGATTCAGCAAGGTCCAAGGAGAGACTCAACAATGGGATCAAAAGGACAGTAAG

TTTCTTTCCAGTGAGTGTCATAGTTGATGAACATTCTCGTCCTTGCGGTAATAAATACTTCGAAAAAGGC

GGTGAGAAATCGCAACGAAATCAGAACGCAAACGATTTTGTTTCCACAAAATATTGCAAAGAAAATGATG

CGGATGCTGATGATGATTATGAAGACGATGCGGCTAGTGATTCGAGTTCGGATTTATTCGAACTCGATCA

TTTGTCACTTTTTAAGACGGATACATTTTGTGACGATCTTCCTGTGTATGAAACTACCTACATGGAAAAG

AACCGTGCCATAGCAGGCAAATTCATCAAATGAGTTAACTTTTATTATTAATAGAGTTTCTAGTTCTCTT

TTTCTCATTAGCTAATAGTATTTGATCTCTGTTTTATCGATCGTGTCTGTTTTATTCCTTGAAATTTTGT

GAACAAATGCTAACTGG

>MSTRG.3445.2 gene=MSTRG.3445

GTTTGCAGTTTGCATTGTATTGTTTGTGTTATAACGAGATTGAAAGATGTTCAAAATTGAAATCTTGTTC

AGGAGATATGGATAATTTCAAGTGGCCAAGAGCTTTCAATCTCCGAAAGTGATCGGATGGTTCAGAGCGT

GAGATGATAACCTAGGGAAGTGATTGATGAGTCAAATTTAGTAAGGCAGCTTAGCTACGGGGTTACATGC

AAGCATATCAATATTCTGTAACAAATGGTGCAACAGGTCGTGAGCTGTGGGAGAAACTTGAAAATTTGGC

AGTGCACCGTTTTGGTGGACTATCAAAGCTTGGTTTATCAGCTTCAGCTTATTTCTAGTAAAGAGGCACA

AGGATATAAAGCAGTGTAGGCGTCTTTCGGTTCTCTTATGATTGTAACAGGTCAGGTCAGCTGTCAGCAT

TCAGAAACGAGTGCTGTAAGAATCGAGGGTGTTCAAATGTGAATTTATTTCACGTGGGATGCGAATGTAG

CATATTATTACTCTTCCTTGATAAGCATTATATTTGACCCTTGGCGCTTTGTATCATACTATCATGGGAT

TCCAAACTGATTAATGGGGATGAGAATGAGATTTCAGTACAAATTATGTATTGAATTCTCATCTCCATTC

CCATTCCCGTCGGCCAGTTGAAATCTTATGACTCCAACAGCAACAGACTCTGGACTCTGTAATTGTGGAT

TTCATAGTTTAGTGTGTAATAATAACCACAGTAAATATTAATTGTTGTGTTGAATCTTTGATGAGGTACT

TAGTAAGTTACTAGTATATTTAACTTTTCTTTTTAAACTGATTATATGATTCGCAATTGTTCA

>MSTRG.3445.1 gene=MSTRG.3445

TGTGAGGTTTGCAGTTTGCATTGTATTGTTTGTGTTATAACGAGATTGAAAGATGTTCAAAATTGAAATC

TTGTTCAGGAGATATGGATAATTTCAAGTGGCCAAGAGCTTTCAATCTCCGAAAGTGATCGGATGGTTCA

GAGCGTGAGATGATAACCTAGGGAAGTGATTGATGAGTCAAATTTAGTAAGGCAGCTTAGCTACGGGGTT

ACATGCAAGCATATCAATATTCTGTAACAAATGGTGCAACAGGTCGTGAGCTGTGGGAGAAACTTGAAAA

TTTGGCAGTGCACCGTTTTGGTGGACTATCAAAGCTTGGTTTATCAGCTTCAGCTTATTTCTAGTAAAGA

GGCACAAGGATATAAAGCAGTGTAGGCGTCTTTCGGTTCTCTTATGATTGTAACAGTACTTTGCATGAAT

TTTGCATATGTTCTGCATAATTCTGTTTTACCCTTCACTTAAACCCTCTGCACATGACTGGTTAATCTTA

TGCAGTCAATGTAATGAAGATATGCTCATTGCAATTGTTTGGTTATAGGTCAGGTCAGCTGTCAGCATTC

AGAAACGAGTGCTGTAAGAATCGAGGGTGTTCAAATGTGAATTTATTTCACGTGGGATGCGAATGTAGCA

TATTATTACTCTTCCTTGATAAGCATTATATTTGACCCTTGGCGCTTTGTATCATACTATCATGGGATTC

CAAACTGATTAATGGGGATGAGAATGAGATTTCAGTACAAATTATGTATTGAATTCTCATCTCCATTCCC

ATTCCCGTCGGCCAGTTGAAATCTTATGACTCCAACAGCAACAGACTCTGGACTCTGTAATTGTGGATTT

CATAGTTTAGTGTGTAATAATAACCACAGTAAATATTAATTGTTGTGTTGAATCTTTGATGAGGTACTTA

GTAAGTTACTAGTATATTTAACTTTTCTTTTTAAACTGATTATATGATTCGCAATTGTTCA

>MSTRG.3445.4 gene=MSTRG.3445

TATTGTTTGTGTTATAACGAGATTGAAAGATGTTCAAAATTGAAATCTTGTTCAGGAGATATGGATAATT

TCAAGTGGCCAAGAGCTTTCAATCTCCGAAAGTGATCGGATGGTTCAGAGCGTGAGATGATAACCTAGGG

AAGTGATTGATGAGTCAAATTTAGTAAGGCAGCTTAGCTACGGGGTTACATGCAAGCATATCAATATTCT

GTAACAAATGGTGCAACAGGTCGTGAGCTGTGGGAGAAACTTGAAAATTTGGCAGTGCACCGTGTCAGGT

CAGCTGTCAGCATTCAGAAACGAGTGCTGTAAGAATCGAGGGTGTTCAAATGTGAATTTATTTCACGTGG

GATGCGAATGTAGCATATTATTACTCTTCCTTGATAAGCATTATATTTGACCCTTGGCGCTTTGTATCAT

ACTATCATGGGATTCCAAACTGATTAATGGGGATGAGAATGAGATTTCAGTACAAATTATGTATTGAATT

CTCATCTCCATTCCCATTCCCGTCGGCCAGTTGAAATCTTATGACTCCAACAGCAACAGACTCTGGACTC

TGTAATTGTGGATTTCATAGTTTAGTGTGTAATAATAACCACAGTAAATATTAATTGTTGTGTTGAATCT

TTGATGAGGTACTTAGTAAGTTACTAGTATATTTAACTTTTCTTTTTAAACTGAT

>MSTRG.3445.3 gene=MSTRG.3445

GGTTTGCAGTTTGCATTGTATTGTTTGTGTTATAACGAGATTGAAAGATGTTCAAAATTGAAATCTTGTT

CAGGAGATATGGATAATTTCAAGTGGCCAAGAGCTTTCAATCTCCGAAAGTGATCGGATGGTTCAGAGCG

TGAGATGATAACCTAGGGTGCAACAGGTCGTGAGCTGTGGGAGAAACTTGAAAATTTGGCAGTGCACCGT

GTCAGGTCAGCTGTCAGCATTCAGAAACGAGTGCTGTAAGAATCGAGGGTGTTCAAATGTGAATTTATTT

CACGTGGGATGCGAATGTAGCATATTATTACTCTTCCTTGATAAGCATTATATTTGACCCTTGGCGCTTT

GTATCATACTATCATGGGATTCCAAACTGATTAATGGGGATGAGAATGAGATTTCAGTACAAATTATGTA

TTGAATTCTCATCTCCATTCCCATTCCCGTCGGCCAGTTGAAATCTTATGACTCCAACAGCAACAGACTC

TGGACTCTGTAATTGTGGATTTCATAGTTTAGTGTGTAATAATAACCACAGTAAATATTAATTGTTGTGT

TGAATCTTTGATGAGGTACTTAGTAAGTTACTAGTATATTTAACTTTTCTTTTTAAACTGAT

>MSTRG.3445.5 gene=MSTRG.3445

GGTTTGCAGTTTGCATTGTATTGTTTGTGTTATAACGAGATTGAAAGATGTTCAAAATTGAAATCTTGTT

CAGGAGATATGGATAATTTCAAGTGGCCAAGAGCTTTCAATCTCCGAAAGTGATCGGATGGTTCAGAGCG

TGAGATGATAACCTAGGGTGCAACAGGTCGTGAGCTGTGGGAGAAACTTGAAAATTTGGCAGTGCACCGT

GTGAGTTCATTCTTCTTCCTTTTCTTTCAAGTTCGTCCATATTGAAATCGATGTTCTTCAGGCCAAGGTT

TTCATGGCTTTTTATATAGTCGGGTCAGGTCAGCTGTCAGCATTCAGAAACGAGTGCTGTAAGAATCGAG

GGTGTTCAAATGTGAATTTATTTCACGTGGGATGCGAATGTAGCATATTATTACTCTTCCTTGATAAGCA

TTATATTTGACCCTTGGCGCTTTGTATCATACTATCATGGGATTCCAAACTGATTAATGGGGATGAGAAT

GAGATTTCAGTACAAATTATGTATTGAATTCTCATCTCCATTCCCATTCCCGTCGGCCAGTTGAAATCTT

ATGACTCCAACAGCAACAGACTCTGGACTCTGTAATTGTGGATTTCATAGTTTAGTGTGTAATAATAACC

ACAGTAAATATTAATTGTTGTGTTGAATCTTTGATGAGGTACTTAGTAAGTTACTAGTATATTTAACTTT

TCTTTTTAAACTG

>MSTRG.3445.6 gene=MSTRG.3445

GATGTAATATTTATGCAAGTTATGATAAAAAGTACAGAGAAGTTGTTTCTTGCTCCGCTTACCGGAGGTC

GAGGGTTCAAATTTTGTCAAAGACAAGATGAGCATGTGAGTGTGTTTAATTATCAAGAAAAAAATGAATT

TGTTTTTTAATAAATATTTTGTATGCCATATATATTTAGGTGGAGGGATACAAAAAGCATTATTTATCAT

AGAAGAAAAATCATATCGGTCAGGTCAGCTGTCAGCATTCAGAAACGAGTGCTGTAAGAATCGAGGGTGT

TCAAATGTGAATTTATTTCACGTGGGATGCGAATGTAGCATATTATTACTCTTCCTTGATAAGCATTATA

TTTGACCCTTGGCGCTTTGTATCATACTATCATGGGATTCCAAACTGATTAATGGGGATGAGAATGAGAT

TTCAGTACAAATTATGTATTGAATTCTCATCTCCATTCCCATTCCCGTCGGCCAGTTGAAATCTTATGAC

TCCAACAGCAACAGACTCTGGACTCTGTAATTGTGGATTTCATAGTTTAGTGTGTAATAATAACCACAGT

AAATATTAATTGTTGTGTTGAATCTTTGATGAGGTACTTAGTAAGTTACTAGTATATTTAACTTTTCTTT

TTAAACT

>MSTRG.3445.7 gene=MSTRG.3445

GTTATAACGAGATTGAAAGATGTTCAAAATTGAAATCTTGTTCAGGAGATATGGATAATTTCAAGTGGCC

AAGAGCTTTCAATCTCCGAAAGTGATCGGATGGTTCAGAGCGTGAGATGATAACCTAGGGAAGTGATTGA

TGAGTCAAATTTAGTAAGGCAGCTTAGCTACGGGGTTACATGCAAGCATATCAATATTCTGTAACAAATG

CTGTGGGAGAAACTTGAAAATTTGGCAGTGCACCGTTTTGGTGGACTATCAAAGCTTGGTTTATCAGCTT

CAGCTTATTTCTAGTAAAGAGGCACAAGGATATAAAGGTACAAAAGGGATCCCCTATTAACATCTCCATT

AATAAAGCACGTTTGTCATAATTAGTACTCTCTAATAGATTCTTCCCTATATGCAGTCCTTGTCCTAACC

ATCACAGTGAATCCAAC

>MSTRG.3447.1 gene=MSTRG.3447

CAATATTTTTGGGTATTCAACACATACAAAGTAGTAAAACATATGGACAGGGAACTATATTTAGCTCTCA

CTGAAGATGGCATTGATGTCCTAAAGGAGAAAGACTGGCTTCAAGCACGTGATCAAATGACACCTACAAA

TAATTCGGTGCTTCATGTTGCATGCCAACACGGGAGTCACAAATGCGTGCAAGAGATACTTGAACGTGCG

GATGAATCACTACTATTACAGACCAACTCGAGAGGTGAGACAGCTCTTCATCTAGCAGCAAGACAGGGGC

ACTTTGACTCTGTTGAAGCACTTATTAGTAAGGCCAAGTCTTCGGACCACCAACCTAATGATCTGGAAAA

CACAACTACGCTTCTGCAAAAGCTGATTAGAGCTTCGAATGTGGAATTTGAAACTGCCTTGCATGCGGCT

GTTCGATACAATCAGAAGAAAGTAGTTAGACTGTTAGTAAAAGAAAATTCAGGTTATTCGTATTCTCGGA

ACAAGTCTGATGAAACTCCACTTTACTTGGCTGCCGCTAGGGGTTACGGGGGTGTCACCAAAGTAATTCT

TAAGAAGTGTGAATCACCAACATTCGGGGGTCCCCACTTTACGGGGTTTGTGACTTGGCTGGCTTACATA

TAAGCACCTCTGTTTAGATGGTAATTATCAATTTTGTGATGATATGACAAAATGCATGAGATTTCTATTG

AGAAGCAACTCAGACCTCATCAAGGTATTAGATGATAATGGAAATGGACAGCGTTTCATTTTGCGGCAGA

CAACAATCTTCGGCGTTCTATTGGATTTTTATTGGATGCCAACAGAAGTGTGGCATATATAGCAGATAAA

AAAGATAAGATGACGCCTCTTCATGTAGCAGCATATAAAGGATACACGAAGGTGATAAAGAAACTTTTAA

AGTATTTGCCGGATATTGTGGATAGTGTAGATGGAAGTGGTCAAAATATTTTCCACATAGCCCTGAAGCA

AAACCAAGACTCCCTTTTGAAATTCATTTTATCAAAAACTTGGAAATTTGATACTTTAAATACACTCATC

ATACAAAAAGACAAGGAAGGGAATACGCCACTTCACCTAGTTGGCAAGCTTGGATATCATATTCCTGGAT

TGCCGCAATGGATACAGAAGATTGACCAAGAAGTAGTAGACGACACCAATTTGACTCCAGCAGAAGCGCT

TTATCAGTATTATTCTCCAATCTTAAGGGATGATGAGGTACGTGTGACAGATTGAATTTAAATTTCATAT

GGTTTTAATTGGTAACATGATAAAGTTTGTTGTTAAATTTATCGATTAGTACTCGCCATTGTTTATATGA

CCATGTCCAGGAGAACGGTGGATCACATCCAAGTAGGGACACTTTTTATAATTTTTGGGACAATTGAACT

GGCGGTAAGATACCAAAGATGGATCGATCTGATCTAAGAGGTTTCAAAGAAAGGATCGAGATACAAAGAC

AAAGGGCGAACACACATATGGTAGTGGCTGCACTTGTGACCACAGTAGCCTTAACAGCAGGCTTCGCCAT

GCCAGGTGGTTTTAACGGCAATCTGGGACCCGATCAAGGGTCCCCATTACTTCTAAGAAAGCCCGCCTTT

AATATATTTGTGGTTGCAGATACGGTTGCTCTCTTATTCTCGATTTCCTCTCTGTTTCTTTACTTCTCAT

TATCATTCCATCCGAAGAAGAAGGCCTTTTTTACGTTGTTCGCATTGGCTGTAGTGTTCAACATTGCATC

AATTGTAGCAATGATGGTGGCTTTTATTGCAGGGACTTATGGTGTACTGTCACACTCCTTAGTTCTGGCC

ATTTGTGTTTGTAGCCTCTCTTCTTTATTCTTTCTCCTTGTCTTTCCTCTTAGCTTTAGAACTTCAATAA

ACATGTTAACAGTTCCTATAGTTCTATCACGTTCGTGGCTTCGCCAGTTGTATCGCAATGTTAGAAATCG

GTACCGAAATTCGGGTATTGTGTGGAGGTGGCGACACTGGGGTATTGTCAGGAAACCATGCCAACCAGCG

AGCCCTGTTTGTACAAACTTAGTCTGAACTGTAAAATAATGCTTTCATGAATTTTATTCTGTCAAGCACA

TTTTAATAATATTACAAGATGATCCTTTGTGTGCGGCAATTAAATGAAATCTACTACCCCATTTTTGTAT

AT

>MSTRG.3447.2 gene=MSTRG.3447

TGAACGTTCAATATTTTTGGGTATTCAACACATACAAAGTAGTAAAACATATGGACAGGGAACTATATTT

AGCTCTCACTGAAGATGGCATTGATGTCCTAAAGGAGAAAGACTGGCTTCAAGCACGTGATCAAATGACA

CCTACAAATAATTCGGTGCTTCATGTTGCATGCCAACACGGGAGTCACAAATGCGTGCAAGAGATACTTG

AACGTGCGGATGAATCACTACTATTACAGACCAACTCGAGAGGTGAGACAGCTCTTCATCTAGCAGCAAG

ACAGGGGCACTTTGACTCTGTTGAAGCACTTATTAGTAAGGCCAAGTCTTCGGACCACCAACCTAATGAT

CTGGAAAACACAACTACGCTTCTGCAAAAGCTGATTAGAGCTTCGAATGTGGAATTTGAAACTGCCTTGC

ATGCGGCTGTTCGATACAATCAGAAGAAAGTAGTTAGACTGTTAGTAAAAGAAAATTCAGGTTATTCGTA

TTCTCGGAACAAGTCTGATGAAACTCCACTTTACTTGGCTGCCGCTAGGGGTTACGGGGGTGTCACCAAA

GTAATTCTTAAGAAGTGTGAATCACCAACATTCGGGGGTCCCCACTTTACGGGGTTTGTGACTTGGCTGG

CTTACATATAAGCACCTCTGTTTAGATGGTAATTATCAATTTTGTGATGATATGACAAAATGCATGAGAT

TTCTATTGAGAAGCAACTCAGACCTCATCAAGGTATTAGATGATAATGGAAATGGACAGCGTTTCATTTT

GCGGCAGACAACAATCTTCGGCGTTCTATTGGATTTTTATTGGATGCCAACAGAAGTGTGGCATATATAG

CAGATAAAAAAGATAAGATGACGCCTCTTCATGTAGCAGCATATAAAGGATACACGAAGGTGATAAAGAA

ACTTTTAAAGTATTTGCCGGATATTGTGGATAGTGTAGATGGAAGTGGTCAAAATATTTTCCACATAGCC

CTGAAGCAAAACCAAGACTCCCTTTTGAAATTCATTTTATCAAAAACTTGGAAATTTGATACTTTAAATA

CACTCATCATACAAAAAGACAAGGAAGGGAATACGCCACTTCACCTAGTTGGCAAGCTTGGATATCATAT

TCCTGGATTGCCGCAATGGATACAGAAGATTGACCAAGAAGTAGTAGACGACACCAATTTGACTCCAGCA

GAAGCGCTTTATCAGTATTATTCTCCAATCTTAAGGGATGATGAGGAGAACGGTGGATCACATCCAAGTA

GGGACACTTTTTATAATTTTTGGGACAATTGAACTGGCGGTAAGATACCAAAGATGGATCGATCTGATCT

AAGAGGTTTCAAAGAAAGGATCGAGATACAAAGACAAAGGGCGAACACACATATGGTAGTGGCTGCACTT

GTGACCACAGTAGCCTTAACAGCAGGCTTCGCCATGCCAGGTGGTTTTAACGGCAATCTGGGACCCGATC

AAGGGTCCCCATTACTTCTAAGAAAGCCCGCCTTTAATATATTTGTGGTTGCAGATACGGTTGCTCTCTT

ATTCTCGATTTCCTCTCTGTTTCTTTACTTCTCATTATCATTCCATCCGAAGAAGAAGGCCTTTTTTACG

TTGTTCGCATTGGCTGTAGTGTTCAACATTGCATCAATTGTAGCAATGATGGTGGCTTTTATTGCAGGGA

CTTATGGTGTACTGTCACACTCCTTAGTTCTGGCCATTTGTGTTTGTAGCCTCTCTTCTTTATTCTTTCT

CCTTGTCTTTCCTCTTAGCTTTAGAACTTCAATAAACATGTTAACAGTTCCTATAGTTCTATCACGTTCG

TGGCTTCGCCAGTTGTATCGCAATGTTAGAAATCGGTACCGAAATTCGGGTATTGTGTGGAGGTGGCGAC

ACTGGGGTATTGTCAGGAAACCATGCCAACCAGCGAGCCCTGTTTGTACAAACTTAGTCTGAACTGTAAA

ATAATGCTTTCATGAATTTTATTCTGTCAAGCACATTTTAATAATATTACAAGATGATCCTTTGTGTGCG

GCAATTAAATGAAATCTACTACCCCATTTTTGTATAT

>MSTRG.3448.1 gene=MSTRG.3448

GTGATCAACTCACACCTACAAATAATTCGGTGCTTCATGTTGCATGCCAACACGGGAGTCACAAATGCGT

GCAAGAGATACTTGAACGTGCGGATGAATCACTACTATTTAAGCCCAACTCGAGAGGCGAGACAGCTCTT

CATCTAGCAGCAAGACAGGGGCACTACGTTGCGGTTAAACCACTTATTAGTAAAGCCAAGTCTTCGGACC

AGCAATCTGATGATCTCGAAAACACAACTACGGCGCTGCAAAATCTGATTAGAGCACCTAATGTGGAATT

TGAAACTGCCTTGCATGCGGCAGTTCGATACAATCACGAGAAAGTAGTTGGAGTGTTGGTAAAAGCGGAT

CCAGGTTATTCGTATCCTCGGAACAAGTCTGATGAAACTCCACTTTACCTGGCTGCCGCTAGGGGTTATG

CTGGTGTGACCAAAGTAATTCT

>MSTRG.3449.1 gene=MSTRG.3449

CAAATAATTCGGTGCTTCATGTTGCATGCCAACACGGGAGTAGGAAATGCGTGCAAGAGATACTAGGGCG

TGCTGATGAATCACTACTATTAAAGATCAACTCGAGAGGTGAGACAGCTCTTCATCAAGCAGCAAGACAG

GGGCACTTTGACGCGGTTGAAGCACTTATTACTAAGGCCAAGTCTTCGGACCGGCAACCTGTTGATCTTG

AAAACACAACTACGGTTCTGCAAAATCTGATTAGAACATCGAATCTGGAATTTGAAACTGCCTTGCATGC

GGCAGTTCGATACAATCACCAGAAAGTAGTTGGACTGTTAGTAAAAGAAGATCCAGGTTTCTCGTATCCT

CGGAACAGGT

>MSTRG.3450.1 gene=MSTRG.3450

TGGGTATTCAATACATACACAGTAAAACATATGGACAGGGAACTATATTTAGCTCTCACTGAAGATGGCA

TTGATGTCCTAAAGGGGAGAGACTGGCTTCAAGTACGTGATCAAATGACACCTACAAATAATTCGGTGCT

TCATGTTGCATGCCAGCACGGGAGTATATCATGCGTTACAGAGATACTACGTGCCGATGAATCAGTACTA

TTAAAGACCAACTCGAGAGGCGAGACAGCTCTTCATCTAGCAGCAAGACATGCAGGGGCACTTTGACTCT

GTTGAAGCACTTATTAGTAAGGCCAAGTCTTGGGAGCAGCAACCTAATGATCTCGAAAACACAACTACGC

TTCTGCAAAAGCTGATTAGAGCTTCGAATGTGGAATTTGAGACTGCGTTGCATGCGGCTGTTCGATACAA

TCACAAGAAAGTAGTTAGAGTGTTAGTAAAAGAAGATCCAGGTTATTCGTATCCTCGGAACAAGTCTGAC

GAAACTCCACTTTACTTGGCTGCCGCTAGGGGTTACGGGGGTGTGACCAAAGTAATTCTTAAGAAGTGTG

AATCACCAACATTCGGGGGTCCCCATGGACGAACTGCTTTGCATGGTGCTCTGATGAGTTCAAGAGGATA

CGAATGCATGAGATTTCTGTTGAGAAGCAACGCAGACCTCATTGAGGTATTAGATGATAATGGATGGACA

GCGTTTCATTTTGTGGCAGACAACAATCTTCGGCGTTCTATTGGATATTTATTGGATGCCAACAGAAGTG

TGGCATATATAGCAGATAAAAAAGATAAGATGACGGCTCTTCATGTAGCAGCATATAAAGGATACACGAA

AGTGATAAAGAAACTTTTAAAGTATTTGCCGGATATTGTGGATAGTGTAGATGGAAATGGTCAAAATATG

TTCCATATAGCCTTGAAGCAAAACCAGGACTCCCTTTTGAAATTCATTTTATCAAAAACTTGGAAATTTG

ATACTTTAAATACACTTATCATACAAAAAGACAAGGAAGGGAATACACCACTTCACCTAATTGGCAAGCT

TGGATATCATATTCCTGGATTGCCGCAATGGATACAGAAGATTGACCAGGAAGTAGTAGACGACACAAAT

TTGACTCCCGCAGAAGCGCTTTATCAGTATTATTCTCCAATCTTAAGGGATGATGAGGTACGTGTGACAA

ATTGAATTTCAATTTCATATGGTTTTAATTGGTAACATGATAAAGTTTGTTGTTAAATTTATCGATTAGT

ACTCGCCATTGTTTATATGACCATGTCCAGGAGAACGGTGGATCACATCCAATTAGAGACATTTTTTATA

GTTTTTGGGACAATTGGACTGGCGGTAAGATGCCAAAGATGGATCGATCTGATCTAAGAGGTTCCAAAGA

AAGGATCGAGATACAAAGACAAAGGGCGAACACACATATGGTAGTGGCTGCACTTGTGACCACAGTCGCC

TTAACAGCAGGCTTCGCCATGCCAGGTGGTTTTAACGGCAATCTGGGACCCGATCAAGGGTCCCCATTAC

TTCTAAGAAAGCCCGCCTTTAATATATTTGTGGTTGCAGATACGGTTGCTCTCTTATTCTCAATTTCCTC

TCTGTTTCTTTACTTCTCGTTATCATTCCATCCCAAGAAGAAAGCCTTTTTTACGTTGTTCGCATTGGCT

GTAGTGCTCAACATTGCATCAATTGTAGCAATGATGGTGGCTTTCATTGCAGGGACTTATGCGGTGCTGG

CACACTCATTAGTCCTGGCCATTGGTGTTTCTACCCTCTCTTCTTTATTCCTTCTCATTGTCTTTCCT

>MSTRG.3450.3 gene=MSTRG.3450

ATCTTCCTTTCCTAATCCTGGTGTACAATATTTTTTGGGTGGCTAGCTATTCAATCCATACACAGTAAAA

CATATGGACAGCGAACTATATGTAGCTCTCACTGAAGATGACTTTGGTGTCCTAAGGGGAAAGGACTGGC

CTGAAGTCCGTGATCAACTCACACCTGCAAATAATACGGTGCTTCATGTTGCATGTCAGCACGGGAGTAT

ATCATGCGTTACAGAGATACTACGTGCCGATGAATCAGTACTATTAAAGACCAACTCGAGAGGTGAGACA

GCTCTTCATCTGGCAGCTAGACAGGGGCACTATGACACGGTTTCTGTACTTATCAGGATGGCCAAGATTT

TCGATAAGCGTCCTAATAATCTCGGAAACACAACTACGGCGCTGCAAAATCTGATTAGAGCATCCAATGT

GGAATTCGAAACTGCCTTGCATGCGGCAGTTCGATACAATCACCAGAAAGTAGTTGAGCTGTTAGTAAGA

AAAGATCCAGGTTTCGCGTATCCTCGGAACAAGTTTGACGAGACTCCACTTTACTTGGCTGCGGTCAGGG

GTTATGATGATGTAACTCAGGTCATTCTTGATAATTGTGATTCGCCAACGTTTGGCGGTCCTCATGGACG

AACTGCCTTGCATGCTGCAGCAATGAGTTATACAGGACATGGATGTATGAAACTCCTGATGAAAAGAAAC

GAAGACCTAATCAATGTAGCAGACAATGATGGATGGACGGCATTTCATTTCGCCGCAGATAACAACCAGT

GCTACTCAATTGAACTTTTACTGGGAGTCAACAAATATGTGGCTCATCTACCAGATCAAAAATATAAGAA

GACGGCACTTCATATAGCAGCATATAAAGGATACCACAAGGTGGTAGAGCAACTTTTGAAATATTTGCCT

GATGCGTGGGAGAGTGTAGATGGAAATGGTCAAAATATATTGCACATTGCCGTGCAACAATACAAGAAGC

CCCTGCTCAAATTCATATTATCAAAAACCCAAAAATTTGATATTTTGAATACTCTTATTATACAAAGAGA

CCAGGAAGGGAATACCCCTCTTCACTTAGCTGCCAAGTTTGGATATAATATTCCTGAGATCGGGGAGTGG

CTACAGAAGATTGATCATTATATAGTAGACAACACCAATTTAGCTCCATCAGATGTGATTTACCGGAATT

TTTATAACGACATGATGAATGAGGAAGTGGTTGGAGTATATTCAAGTAAAGACCAGATTCATGATATCTG

GAACGCTTGGACTCAAGGTAGAGTGCAAGCAACAGATAGATATGAACCGATACCTGCAAAAGAAAAGATC

ATGATGCGTCAACAAATGGCGAACAGCCATATGATAGTGGCTGCACTAGTGACCACCGTGGCCTTGACAG

CCGGTTTCACTGTGCCTGGTGGTTTTGACAGCAACCGAGGATCCCCGCTGCTTCTAAGAAAGCCGGCCTT

CAAGATATTTATGATAGCAGATACATTGGCCCTCCTATTCTCAATTTCCGCACTGTTTCTTTACTTCTCA

ATATCATTCAAACACACGCGACTGAGTGTCACTTTTTTGTTGCTCACCAGCGCTGTAGTGCTCAACGTTA

TTTCTATTGCAGCGATGATGGTGGCTTTTATTGCAGGCACTTTTGCCGTGCTCTATCATTCATTAGCTCT

GGCCATTATTGTTTCTACGCTCTCTTCTTTATTCCTTCTTCTCATCTTTTCTTTTAGCGTCCCTTTTATC

AGAAGCTACATCCCTGTGATATTGCAATTATACGGTGACCTTCGTGGGATGTATAGTTCGTTTAAATACA

CACTCGGCACCTTTTTGAGGAGAATATTGAACACCCGTTTCCGGATAAATTCGAGACCAGCAGCTGGGAC

AAATACTCGTGACTTTTCCGCAGAAAATATTGATAGAGACGTCTTAAGACAAATGGTCATGGTTGCTTCG

AGGGCGTATACGGAACAGATCAATGAAACAGCTGAAGGTGACGAGTCAACAATGTGAGCTGGCAAATTAA

AACCGCTGGTTTGTTACCAATTGTTTAGATACCGTCGGTGTGTGTGAAGGTTTAATTGGTGATAAAAATT

CAAGTTCCATTACTGCAATAGCAGCTGAGATGTAATTATAAGATACAGTATATCACAATGTTTTATTTCT

ACATGATGATTCCTTTTTTATTTTGCTA

>MSTRG.3450.2 gene=MSTRG.3450

ATCTTCCTTTCCTAATCCTGGTGTACAATATTTTTTGGGTGGCTAGCTATTCAATCCATACACAGTAAAA

CATATGGACAGCGAACTATATGTAGCTCTCACTGAAGATGACTTTGGTGTCCTAAGGGGAAAGGACTGGC

CTGAAGTCCGTGATCAACTCACACCTGCAAATAATACGGTGCTTCATGTTGCATGTCAGCACGGGAGTAT

ATCATGCGTTACAGAGATACTACGTGCCGATGAATCAGTACTATTAAAGACCAACTCGAGAGGTGAGACA

GCTCTTCATCTGGCAGCTAGACAGGGGCACTATGACACGGTTTCTGTACTTATCAGGATGGCCAAGATTT

TCGATAAGCGTCCTAATAATCTCGGAAACACAACTACGGCGCTGCAAAATCTGATTAGAGCTTCGAATGT

GGAATTTGAGACTGCGTTGCATGCGGCTGTTCGATACAATCACAAGAAAGTAGTTAGAGTGTTAGTAAAA

GAAGATCCAGGTTATTCGTATCCTCGGAACAAGTCTGACGAAACTCCACTTTACTTGGCTGCCGCTAGGG

GTTACGGGGGTGTGACCAAAGTAATTCTTAAGAAGTGTGAATCACCAACATTCGGGGGTCCCCATGGACG

AACTGCTTTGCATGGTGCTCTGATGAGTTCAAGAGGATACGAATGCATGAGATTTCTGTTGAGAAGCAAC

GCAGACCTCATTGAGGTATTAGATGATAATGGATGGACAGCGTTTCATTTTGTGGCAGACAACAATCTTC

GGCGTTCTATTGGATATTTATTGGATGCCAACAGAAGTGTGGCATATATAGCAGATAAAAAAGATAAGAT

GACGGCTCTTCATGTAGCAGCATATAAAGGATACACGAAAGTGATAAAGAAACTTTTAAAGTATTTGCCG

GATATTGTGGATAGTGTAGATGGAAATGGTCAAAATATGTTCCATATAGCCTTGAAGCAAAACCAGGACT

CCCTTTTGAAATTCATTTTATCAAAAACTTGGAAATTTGATACTTTAAATACACTTATCATACAAAAAGA

CAAGGAAGGGAATACACCACTTCACCTAATTGGCAAGCTTGGATATCATATTCCTGGATTGCCGCAATGG

ATACAGAAGATTGACCAGGAAGTAGTAGACGACACAAATTTGACTCCCGCAGAAGCGCTTTATCAGTATT

ATTCTCCAATCTTAAGGGATGATGAGGAGAACGGTGGATCACATCCAATTAGAGACATTTTTTATAGTTT

TTGGGACAATTGGACTGGCGGTAAGATGCCAAAGATGGATCGATCTGATCTAAGAGGTTCCAAAGAAAGG

ATCGAGATACAAAGACAAAGGGCGAACACACATATGGTAGTGGCTGCACTTGTGACCACAGTCGCCTTAA

CAGCAGGCTTCGCCATGCCAGGTGGTTTTAACGGCAATCTGGGACCCGATCAAGGGTCCCCATTACTTCT

AAGAAAGCCCGCCTTTAATATATTTGTGGTTGCAGATACGGTTGCTCTCTTATTCTCAATTTCCTCTCTG

TTTCTTTACTTCTCGTTATCATTCCATCCCAAGAAGAAAGCCTTTTTTACGTTGTTCGCATTGGCTGTAG

TGCTCAACATTGCATCAATTGCAGCAATGATGGTGGCTTTCATTGCAGGGACTTACGGTGTGCTGTCACA

CTCATTAGTTCTGGCCATTGCTGTTTCTACCCTCTCTTCTTTATTCTTTCTCCTTGTCTTTCCTCTTAGC

TTTAGAACTTCAATAAACATGTTAACAGTTCCTATAGTTCTATCACGTTCGTGGCTTCGCCAGTTGTATC

GCAATATTAGAAATCAGTACCGAAATTCAGGTGTTGTGAGGAGGTGGCGACACTGGGGTATTGTCAGGAA

ACCATGCCAACCAGCGAGCCCTGTTTGTACAAACTTAGTCTGAACTGTAAAATAATGCTTTCATGAATTT

TATTCTGTCAAGCACATTTTAATAATATTACAAGATGATCCTTTGTGTGCGGCAATTAAATGAAATCTAC

TACCCCATTTTTGTATATTTACAGTGATGAATTGTA

>MSTRG.3450.4 gene=MSTRG.3450

TGGGTATTCAATACATACACAGTAAAACATATGGACAGGGAACTATATTTAGCTCTCACTGAAGATGGCA

TTGATGTCCTAAAGGGGAGAGACTGGCTTCAAGTACGTGATCAAATGACACCTACAAATAATTCGGTGCT

TCATGTTGCATGCCAACACGGGAGTCACAAATGCGTGCAAGAGATACTTGAACGTGCCGATGAATCACTA

CTACTTAAGCCCAACTCGAGAGGCGAGACAGCTCTTCATCTAGCAGCAAGACATGCAGGGGCACTTTGAC

TCTGTTGAAGCACTTATTAGTAAGGCCAAGTCTTGGGAGCAGCAACCTAATGATCTCGAAAACACAACTA

CGCTTCTGCAAAAGCTGATTAGAGCTTCGAATGTGGAATTTGAGACTGCGTTGCATGCGGCTGTTCGATA

CAATCACAAGAAAGTAGTTAGAGTGTTAGTAAAAGAAGATCCAGGTTATTCGTATCCTCGGAACAAGTCT

GACGAAACTCCACTTTACTTGGCTGCCGCTAGGGGTTACGGGGGTGTGACCAAAGTAATTCTTAAGAAGT

GTGAATCACCAACATTCGGGGGTCCCCATGGACGAACTGCTTTGCATGGTGCTCTGATGAGTTCAAGAGG

ATACGTTCGTGACTTGGCTGGCTTACATATAAGCACCTCTATTTAGATGGTAATTATCAATTTTGTGATG

ATATGACAGAATGCATGAGATTTCTGTTGAGAAGCAACGCAGACCTCATTGAGGTATTAGATGATAATGG

ATGGACAGCGTTTCATTTTGTGGCAGACAACAATCTTCGGCGTTCTATTGGATATTTATTGGATGCCAAC

AGAAGTGTGGCATATATAGCAGATAAAAAAGATAAGATGACGGCTCTTCATGTAGCAGCATATAAAGGAT

ACACGAAAGTGATAAAGAAACTTTTAAAGTATTTGCCGGATATTGTGGATAGTGTAGATGGAAATGGTCA

AAATATGTTCCATATAGCCTTGAAGCAAAACCAGGACTCCCTTTTGAAATTCATTTTATCAAAAACTTGG

AAATTTGATACTTTAAATACACTTATCATACAAAAAGACAAGGAAGGGAATACACCACTTCACCTAATTG

GCAAGCTTGGATATCATATTCCTGGATTGCCGCAATGGATACAGAAGATTGACCAGGAAGTAGTAGACGA

CACAAATTTGACTCCCGCAGAAGCGCTTTATCAGTATTATTCTCCAATCTTAAGGGATGATGAGGTACGT

GTGACAAATTGAATTTCAATTTCATATGGTTTTAATTGGTAACATGATAAAGTTTGTTGTTAAATTTATC

GATTAGTACTCGCCATTGTTTATATGACCATGTCCAGGAGAACGGTGGATCACATCCAATTAGAGACATT

TTTTATAGTTTTTGGGACAATTGGACTGGCGGTAAGATGCCAAAGATGGATCGATCTGATCTAAGAGGTT

CCAAAGAAAGGATCGAGATACAAAGACAAAGGGCGAACACACATATGGTAGTGGCTGCACTTGTGACCAC

AGTCGCCTTAACAGCAGGCTTCGCCATGCCAGGTGGTTTTAACGGCAATCTGGGACCCGATCAAGGGTCC

CCATTACTTCTAAGAAAGCCCGCCTTTAATATATTTGTGGTTGCAGATACGGTTGCTCTCTTATTCTCAA

TTTCCTCTCTGTTTCTTTACTTCTCGTTATCATTCCATCCCAAGAAGAAAGCCTTTTTTACGTTGTTCGC

ATTGGCTGTAGTGCTCAACATTGCATCAATTGCAGCAATGATGGTGGCTTTCATTGCAGGGACTTACGGT

GTGCTGTCACACTCATTAGTTCTGGCCATTGCTGTTTCTACCCTCTCTTCTTTATTCTTTCTCCTTGTCT

TTCCTCTTAGCTTTAGAACTTCAATAAACATGTTAACAGTTCCTATAGTTCTATCACGTTCGTGGCTTCG

CCAGTTGTATCGCAATATTAGAAATCAGTACCGAAATTCAGGTGTTGTGAGGAGGTGGCGACACTGGGGT

ATTGTCAGGAAACCATGCCAACCAGCGAGCCCTGTTTGTACAAACTTAGTCTGAACTGTAAAATAATGCT

TTCATGAATTTTATTCTGTCAAGCACATTTTAATAATATTACAAGATGATCCTTTGTGTGCGGCAATTAA

ATGAAATCTACTACCCCATTTTTGTATATTTACAGT

>MSTRG.3450.5 gene=MSTRG.3450

CTCTCTTTCTATATATATTTTTAACTCTATATCTGTCTAAATATTACTATTTCATTTCTTTCTGTGAAGT

TCGTGACTTGGCTGGCTTACATATAAGCACCTCTATTTAGATGGTAATTATCAATTTTGTGATGATATGA

CAGAATGCATGAGATTTCTGTTGAGAAGCAACGCAGACCTCATTGAGGTATTAGATGATAATGGATGGAC

AGCGTTTCATTTTGTGGCAGACAACAATCTTCGGCGTTCTATTGGATATTTATTGGATGCCAACAGAAGT

GTGGCATATATAGCAGATAAAAAAGATAAGATGACGGCTCTTCATGTAGCAGCATATAAAGGATACACGA

AAGTGATAAAGAAACTTTTAAAGTATTTGCCGGATATTGTGGATAGTGTAGATGGAAATGGTCAAAATAT

GTTCCATATAGCCTTGAAGCAAAACCAGGACTCCCTTTTGAAATTCATTTTATCAAAAACTTGGAAATTT

GATACTTTAAATACACTTATCATACAAAAAGACAAGGAAGGGAATACACCACTTCACCTAATTGGCAAGC

TTGGATATCATATTCCTGGATTGCCGCAATGGATACAGAAGATTGACCAGGAAGTAGTAGACGACACAAA

TTTGACTCCCGCAGAAGCGCTTTATCAGTATTATTCTCCAATCTTAAGGGATGATGAGGTACGTGTGACA

AATTGAATTTCAATTTCATATGGTTTTAATTGGTAACATGATAAAGTTTGTTGTTAAATTTATCGATTAG

TACTCGCCATTGTTTATATGACCATGTCCAGGAGAACGGTGGATCACATCCAATTAGAGACATTTTTTAT

AGTTTTTGGGACAATTGGACTGGCGGTAAGATGCCAAAGATGGATCGATCTGATCTAAGAGGTTCCAAAG

AAAGGATCGAGATACAAAGACAAAGGGCGAACACACATATGGTAGTGGCTGCACTTGTGACCACAGTCGC

CTTAACAGCAGGCTTCGCCATGCCAGGTGGTTTTAACGGCAATCTGGGACCCGATCAAGGGTCCCCATTA

CTTCTAAGAAAGCCCGCCTTTAATATATTTGTGGTTGCAGATACGGTTGCTCTCTTATTCTCAATTTCCT

CTCTGTTTCTTTACTTCTCGTTATCATTCCATCCCAAGAAGAAAGCCTTTTTTACGTTGTTCGCATTGGC

TGTAGTGCTCAACATTGCATCAATTGCAGCAATGATGGTGGCTTTCATTGCAGGGACTTACGGTGTGCTG

TCACACTCATTAGTTCTGGCCATTGCTGTTTCTACCCTCTCTTCTTTATTCTTTCTCCTTGTCTTTCCTC

TTAGCTTTAGAACTTCAATAAACATGTTAACAGTTCCTATAGTTCTATCACGTTCGTGGCTTCGCCAGTT

GTATCGCAATATTAGAAATCAGTACCGAAATTCAGGTGTTGTGAGGAGGTGGCGACACTGGGGTATTGTC

AGGAAACCATGCCAACCAGCGAGCCCTGTTTGTACAAACTTAGTCTGAACTGTAAAATAATGCTTTCATG

AATTTTATTCTGTCAAGCACATTTTAATAATATTACAAGATGATCCTTTGTGTGCGGCAATTAAATGAAA

TCTACTACCCCATTTTTGTATATTTACAGTGATGAATTGTA

>MSTRG.3450.6 gene=MSTRG.3450

TCTCTTTCTATATATATTTTTAACTCTATATCTGTCTAAATATTACTATTTCATTTCTTTCTGTGAAGTT

CGTGACTTGGCTGGCTTACATATAAGCACCTCTATTTAGATGGTAATTATCAATTTTGTGATGATATGAC

AGAATGCATGAGATTTCTGTTGAGAAGCAACGCAGACCTCATTGAGGTATTAGATGATAATGGATGGACA

GCGTTTCATTTTGTGGCAGACAACAATCTTCGGCGTTCTATTGGATATTTATTGGATGCCAACAGAAGTG

TGGCATATATAGCAGATAAAAAAGATAAGATGACGGCTCTTCATGTAGCAGCATATAAAGGATACACGAA

AGTGATAAAGAAACTTTTAAAGTATTTGCCGGATATTGTGGATAGTGTAGATGGAAATGGTCAAAATATG

TTCCATATAGCCTTGAAGCAAAACCAGGACTCCCTTTTGAAATTCATTTTATCAAAAACTTGGAAATTTG

ATACTTTAAATACACTTATCATACAAAAAGACAAGGAAGGGAATACACCACTTCACCTAATTGGCAAGCT

TGGATATCATATTCCTGGATTGCCGCAATGGATACAGAAGATTGACCAGGAAGTAGTAGACGACACAAAT

TTGACTCCCGCAGAAGCGCTTTATCAGTATTATTCTCCAATCTTAAGGGATGATGAGGTACGTGAGAACG

GTGGATCACATCCAATTAGAGACATTTTTTATAGTTTTTGGGACAATTGGACTGGCGGTAAGATGCCAAA

GATGGATCGATCTGATCTAAGAGGTTCCAAAGAAAGGATCGAGATACAAAGACAAAGGGCGAACACACAT

ATGGTAGTGGCTGCACTTGTGACCACAGTCGCCTTAACAGCAGGCTTCGCCATGCCAGGTGGTTTTAACG

GCAATCTGGGACCCGATCAAGGGTCCCCATTACTTCTAAGAAAGCCCGCCTTTAATATATTTGTGGTTGC

AGATACGGTTGCTCTCTTATTCTCAATTTCCTCTCTGTTTCTTTACTTCTCGTTATCATTCCATCCCAAG

AAGAAAGCCTTTTTTACGTTGTTCGCATTGGCTGTAGTGCTCAACATTGCATCAATTGCAGCAATGATGG

TGGCTTTCATTGCAGGGACTTACGGTGTGCTGTCACACTCATTAGTTCTGGCCATTGCTGTTTCTACCCT

CTCTTCTTTATTCTTTCTCCTTGTCTTTCCTCTTAGCTTTAGAACTTCAATAAACATGTTAACAGTTCCT

ATAGTTCTATCACGTTCGTGGCTTCGCCAGTTGTATCGCAATATTAGAAATCAGTACCGAAATTCAGGTG

TTGTGAGGAGGTGGCGACACTGGGGTATTGTCAGGAAACCATGCCAACCAGCGAGCCCTGTTTGTACAAA

CTTAGTCTGAACTGTAAAATAATGCTTTCATGAATTTTATTCTGTCAAGCACATTTTAATAATATTACAA

GATGATCCTTTGTGTGCGGCAATTAAATGAAATCTACTACCCCATTTTTGTATATTTACA

>MSTRG.3452.1 gene=MSTRG.3452

GAAACTCCTTTGCTGCCCTGTCCCTGGCGGCCCTTGACCTGCAACATAGTTTGGTGGGACCGGAGGCGAA

ACTGCATAAGGATTATTGCTCCATGCAGCAGCATATGGTGGTGGTGGTGGCGGAAATGCTGGTAAAAGAG

GTGCTGGTTGATCAAAAATGTAAGGTTCTGGTGGCGAGAAATTGATTGGAGGCAGAGGTGGTGGTTGATC

ATAATAGTAAGGTTCTGGTGGCGAGAAAGTGATACGAGGCAAAGGTGGTGGCTGATCAAAGTAGTTTGAT

TCTGGTGGGGAGAGGTTGATGTGAGGCAAAGGTGGTGGTTGGTCAAAATTGTAAGGTTCTGGTGGTGAGA

GACTGATTGGAGGCAAAGGCAAATGTCGCGGTGATTTAGAAGGCATTGCTGGTGGAGCTGGTGGAGGCGG

TGAGCAGATAAAAGGGCAAGTTGCACAATTACTATAGCATAGTAGACTTGAACTTACCGGAGGTGCTTTG

AAGATATGATCTTGAGATCTTGCAGTAGAGAAGTTAACCATTTCGAAAATCCCATGCATGAAGAGAAAAA

CCAAGTAGAATCTTGGCATTATTACTTTACTCATGGGATTTTTCATTTGGGTGATTATTTACACCTATTG

TCAGTGTGTATGTTCCAGCTATACCTGGCTGGGATCCACAACTAGTACTCTTTTTGTGTGTGTGCACGCG

CGTGTGCTAATAAGGCCTTTAATTTTATCTGCTCATAGGCTGTCTGGGACAAAAGATGGGTGGTGAGGAA

ATTAGGTTTAGAGGAATAAAGATGGTAACTTTTGAGTTGAGAAGAAATGATGAATTGAAGGAGATCTTAA

TAGTGATTTTGACTTCAAATTTCAAATATTTGCTTTCGGTGGTGATTGATGCACAGATGGATTCAGTATA

GTGTAGTAAAAAGTGATCTGGAGTTTTACCAAGGGGCAATATCATGAAGTTGAATGTCTGTGACACAAAT

TTGGCGACAATGGTGAAGAATGGGACCTTAAGAGCAATGGCAG

>MSTRG.3455.1 gene=MSTRG.3455

AAAAAGGTGTGTATTTGCTATTTTCCTTGTATTATTTTTCAGAAAATCCTAAATTCTTTCGATGGAATAT

TCTTTTTATATAATATTTAGTATTGAGTATAAGAAATTGGAAAAGCAGAACATGAAGTATACAAGTCACA

CTTTCTCTGCATAAAACAAAACTTGCACAAACTAGTTTGATCAACATTTCTCAACATCTCAAAAAAAAAA

TTCTAACCTTACTTTGAAATCTTTAATCAAGAGACTTTCTATCTGCAATCATAATCATGGCTGGGGCTGC

AGAAGGTTCTCAATTTGATGCACGCCAATTTGACGCAAAAATGACTGAGATACTTAATACTGAGGGAGAA

GAATTTTTCACATCATATGATGAAGTTCATGACAGTTTTGATGCTATGGGATTGCACGAAAACCTTTTGA

GGGGCATCTATGCTTACGGTTTTGAAAAGCCATCCGCAATTCAGCAGCGAGGGATTGTTCCATTCTGCAA

GGGGCTAGATGTTATTCAACAGGCACAATCTGGTACAGGGAAAACAGCAACTTTCTGCTCTGGAGTTCTG

CAGCAGCTTGATTATGGTGTTATTGAATGTCAAGCATTGGTTCTTGCTCCCACTCGTGAACTAGCACAAC

AAATTGAAAAGGTTATGCGAGCACTTGGTGACTATCTCGGTGTAAGGGTTCATACTTGTGTGGGTGGGAC

TGTTGTCCGCGAAGATCAACGAATTCTGTCAAGTGGAGTTCATGTCGTGGTTGGTACTCCTGGCCGTGTA

TTCGACATGTTGCGTAGACAATCACTTCGCTCAGATTACATCAAGATGTTTGTTCTCGATGAAGCAGATG

AAATGCTCTCAAGAGGTTTCAAGGATCAGATTTATGATATATTTCAGTTGCTACCATCGAAAGTTCAGGT

TGGAGTTTTTTCCGCTACTATGCCTCCCGAGGCTCTTGAAATTACTAGAAAGTTCCTGAATAAGCCTGTG

AGGATTCTCGTAAAAAGAGATGAGCTTACTCTTGAGGGTATTAAACAGTTTTATGTGAATGTTGACAAGG

AGGAATGGAAACTGGAAACACTATGTGATTTATACGAGACTTTGGCCATTACCCAGAGTGTGATTTTCAT

CAATACCAGGCGCAAGGTTGATTGGCTGACTGACCAAATGCGCAGCCGTGATCACACAGTGTCTGCCACT

CATGGTGACATGGATCAGAATACGAGAGACATAATTATGCGTGAGTTTCGCTCTGGTTCTTCCCGCGTGC

TCATTACAACTGATCTCCTGGCACGTGGTATAGATGTCCAGCAAGTGTCCCTTGTGATAAACTATGACCT

GCCGACTCAGCCAGAGAACTACCTTCATCGGATTGGACGCAGTGGCCGTTTTGGAAGGAAGGGTGTTGCG

ATCAACTTTGTTACCAGGGATGATGACAGGATGTTGGTCGACATACAGAAGTTCTATAATGTGTTGGTCG

AGGAGCTGCCAGCAAATGTTGCCGATCTTCTTTAGTGTGGAACTGATGTGACAAGACTGAAAAAAACAAA

ATAGCATTGGTCCCCAATATCCTGCATTTTCTCATGGCTCGTATGCAAATCGTTTATTTCCAGCTCTAGT

AATCTATAATCAGGTTAGTGTGGAAAATTATATCTTGTTCAAGGAGATTTGTTGTCCTTTTGATTAATAC

CAAGTGTTTTTGTCCTGTTTATAATTGAGTTGTTGAACAATAAGCCCTTAACAAACTAAGCCACAATTGA

ATCCTCTGAAATTTACATAGATGTAGGCATTGATTATATGTGTTATGTTTGCATTCCTTCGCTAGCTGTA

TGTTTACTTGACAGGAAGACTTGCTTCGAAGGGTCAAAAGATCAGTGAGAACTTCTATTTCAGAAGCGAT

TGCTCTGTAAGTTGTGATTGCCTAATTGGCATGTACAGTTTTACTGTAAGTGTAAATAAACTCTTCTAAG

CCACATTTTTACCAAAATTTATGTATAATCCCAGCCGCCTTCTACTTTTCTGAATAGTTCTTGGGAAATT

TTGTCGAGAAAATAGATTTGATACCAAATAGCAGTGCTGCAAACAAATGTATCCGTAAGAGAAATTGTAA

TGAATAGCTTAGTCTTATTCAGTATCTGAATCTGCTTTTGATCGTAG

>MSTRG.3456.1 gene=MSTRG.3456

GTTTTCTTACAACCTATATATGTAGAAGATAAGTCATGGGAAGTAGATTCAATTTGTTTTAAGCACGTTG

TTTTGAATTCTTGGATCATATAAGTATTTCTAAGTTCTTTTCAGGTTTCTGCTGCTTGTGCGGTGTGACT

CATATAATGCGCTTCTTCAAGTGCTTTCTTTTTTACATTTATGTGGTAGCTTCCAACATTTAAAATATTG

TCTTCAGTTTCTTCTTTCTCTTACTGTTACATCAAAGCTCCCGCTGAGAAGCGTGATGCTTTAAATTTTT

ATTTGTTCCTGCTGTTGGCCTGCTGTAGAAGCTCGACATGGACAGACCTTAAAGTTCTACAGTGGTTGAT

TCATTCTTTTCTTATTTACTGTCGCCAAAATATTCAATCCCATTTCTTTGTTACATCTGTGTGGCATCTA

CAACATTGCCAGAATTTAGCATAATATTGTTTGATATCTGAAAAATATCTGTAATGTTGTTTTACTATGG

ATTCTGATTGAGTAAGTGCATGTTTTCTGTTGTCTGAGTGTAACTGTCTTCTTTACAGTGAATGGGTTCG

AGCCACTTAAACAGTTTGAGCTTTTCATTCATGATCGAGACTGAAGTCAAGCTACAACTCCCTTTGAAGG

AGAGTTCCAAGTCTGTTAACAGAGTGATAGAGCCGGCAACTCAGTCTGAAATTGAATATCTGAAGGAATA

GTTGAAATATTTGATTATTGTGGCAAATGTAAGAGACTGGAGTTTCAAGATAAAAGCACTTCCTAAGGAG

AACCAGCACACATGCAAATCACCTGGGCTAATGTTAAGGGAATCAGCTCTTTAGAGCATGTGGGTCCTTT

TTAATGTTACATAAGCATCATCCAGTGGATTTTAAAAAGACCCCGGTCACACTGCTTCAATGCACGAGCC

CATTTCCACTTACAGATGAGTACATCAAATCTTATCTTTCTTTATATAATATTGTTTTAATTTATAATAT

AATTCTCAAGCATTTGGAACTTGAGAATTTGGATTTGAAGAATAAGAATGTAAAAGTAGATGCACATTTA

TAAAGATTGTACAAATGAATGCATAAGGACCAAAAATAAGCACAGGTAGCAGGAGTTTACCGGGCTATGC

CCGGTTTTTGCTTTTACTCCTGTGGGTTGGGATGCTCTTAGGCTCAACGTCCTTGCAGAAAAATCATCAG

GAGGATTCTGTAAAATGATTTCTCAATAGTTTCTTGACCTTGTGGTGACAGCTGAGAAGAGAAGGCACTC

CAAGTAATGTCCGAAACTATTGGTTCAAATCTACATTTCTCTTCACTTGTATATTTTCTCAAAAAAAGAA

CTTGTATTATGAAATTG

>MSTRG.3466.1 gene=MSTRG.3466

GTTTTTCTTTATCTGGGTGATGGGGAATTTGTAGTGAAAAGTGATGGTGAAAAAATCTTTTATGACTACA

AAAAGAAAGATGCCAAGAAGCTTCTTTCACCCGCGCCTGTTGGAATGTTGGATTCAGTTGTCAAGTACAC

CGAGACCCTTGTTTCACCCAAAGATTTCAAGGATTTGAGCTTCTAAGATAATTCAATTGGCGAGCTGGCA

TTACTGTGGATTGTTTTTGCGCATTTGTTCTTAGTATGTGTCAAGTTTGCCTGAGAGTCTTGTGTAAAAC

TAATTAGCTATGTTTGTTTTTGAATCAAGTTTTTCATGAATACATATGGTTACATAGGTATTACTTGCGA

T

>MSTRG.3467.1 gene=MSTRG.3467

GCAGCAAGCATCTAGCCATCCCATCCCTCGTGTATAATCCTCGTTTATATTTTCACATTTTCATTCACTC

ATATTACACACACTTCCATCACATTATCACCCTTCTTCCCAAACCATGACTGACAGAGTCTACCCCTCTG

CCACCAAACCCAACGGCACCGCCACCACCACCGCTCCCGCCGCCACCAACCACCACAACCATCACCAACC

CCTCCCCGTCCCCNNNNNNNNNNNNNNNNNNNNNNNNNNNNCCGCCCCCCGCCACCGCCGCTCCCGCCGC

ACCCACTTCTGCTGCTGCTTCTGCTGGTCCATCTTCATCCTCCTCCTCCTCCTCACCATCGCCGCCATCG

CCGCCGCCATCGCCTACTCCATCTACCGACCCCAACGCCCCACATTCGCCTTCACCTCCCTCAAACTCTC

CCAATTCAACCTCACCACCGCCGCCGACGACTCCACCACCCTCACCACCAAGCTCAACCTCACTCTCTCC

TCCACCAACCCCAACAAAAAACTCACCTTCTCCTACTTCCCCATCCAAATCTCCGCCATCACCACCTCCG

AAGTCCAAATCGGCCAATCCTCCTTCACCAATTTCACCTCCTCCCCCAAAAACATCACCTACATCCACAC

ATCTCTCCGCAGCACATCTCAAGTCCTCGACGCCGACTCCGTCGCTTCACTCCGATCAGATCTGAAGAAA

AAATCCGGGCTACCACTAAAAGTACTTCTCGACACAAAAGTCGTTATAAAAGCCGATAATCTCAAGACCA

AAAAGCTGGGGATCAGAATTAAGTGCGAAGGGGTGCATGTCACAGCTCCCAAAGGTAAGTCTGCATCAGT

GGCTTCAGTAGGTAAAGCAAAATGTAAAGTTGATCTGAGAGTCAAGATCTGGAAATGGACATTTTAATTT

TAGTTTTTTTTTATTTACACCGATTAATTTGCTGTAATCTTTATTAATTTATTTTTTGTTTGCGGGTCCA

TGGAACTCTGGAATTCACAAAAGAAATACGATATATATTTATATATTGAGTTTGGTTATTTTAATGTTTC

GAGTAGATTGGTGGTGTAGAATGTAGATGGGTATGGATGTTTCAACAGGGTGCGTAGAGTTGTGATGTGT

AATGATAGTCTGTTAACTGGATATATATGAACCACAGTTTCTGCTATCATCATCTCGGGCCACTTGTTTT

CTGGGTGTTTTCCTACCTTTTTTCTCGTTTTTATTACATCTAATGTTCAGG

>MSTRG.3468.1 gene=MSTRG.3468

GCTGTGACATGCACCCCTTCGCACTTAATTCTGATCCCCAGCTTTTTGGTCTTGAGATTATCGGCTTTTA

TAACGACTTTTGTGTCGAGAAGTACTTTTAGTGGTAGCCCGGATTTTTTCTTCAGATCTGATCGGAGTGA

AGCGACGGAGTCGGCGTCGAGGACTTGAGATGTGCTGCGGAGAGATGTGTGGATGTAGGTGATGTTTTTG

GGGGAGGAGGTGAAATTGGTGAAGGAGGATTGGCCGATTTGGACTTCGGAGGTGGTGATGGCGGAG

>MSTRG.3470.1 gene=MSTRG.3470

TTATCATATTCCATAAGAAAAAAAACTTACCCTCCTCCATTAATCTTTCATTACACTTTGCATAATCAAA

AGGACGAAAATGCCCTTTAATGATTGTTTTCCACTTTTCAACAACAGCCCACCAAGAACCAGATCTAAAC

ATCTCATCCAATGTTTTACGAAAGAACCACTATACTTTCCCCACTTTCCAGAACAACCCCCCAACTTACA

AAAGTTTCATAAACCGCCCCAAACTTCCCATTTCTTTCAGTTAGCACCTCTGTTTATCACTTCTTTGTAC

CTCATTATCGAATCAATCCTCTGTAACTTCAGCTGCTGCTTGAATTTGTTGATGAAATCATCAGCTTTTG

CATCGACTTCTTCGTCAACGGCCGTCACTTTAGCACCCCGTTCCCTCACAGTCACCGGCCGGCGAGCCTC

CACAATGTCGTCCTCCTCGAAATGCGCAAAAGGGGACTTCATGCTCGCCGACTTCTTCATCTTCGCCGAC

AACTTCGCTGGAACCTCGCCGGAGGCCGGTTTTGTATCGGAAGTAGTTCTGCTAAAATGACTGTCGGTGA

GCTTCTCAAAATGACTGTACACTTCATCTAAGCTCTGTGGGACCACATCTTCCACTTCCGTTGATTCCTC

TTCATGGGGTTCTTGGATCTGAGATTCTGATTCATGGGGTTCTCTTAATGGGCCTGATTCTTGATGGGTT

TGTTCAAAACTGAACTGGGCTTGGACTGCCATTGTTTCATGGTCATCAAACAAGTAATGTGTTTGTGTCT

CTATAGCTGACTGCAGTTCTTGATTCAAATGTTGTTCAGTTGTGTCTGAATCTAATCCTTCAGTAAAACT

CTTGAAATTGATAGATTTGATGCGTTGGAGCACCGATGGGGATTTTTGAAGCACAGGGGTATTTTGGTCA

TTCTGGTGGTTCTGAGTGTTGGATTCTTGTTTTTCTTGTTTGTGTTCTTGGTTTTCTTGGTTTTGTTGTT

GTTGTTGAGGGGAGGTTGATAAGTTGGAGCTGAAGAGAATGGTGCCTATCATGACATTGAGAAGAACAAA

GAGGACAGTTGGAGTGAACCAGCTGTTCATTGATGCCCAGATTGAAGAAGGGTTAGACACAGATTCTTCA

AACATTGTGTGTGTGTGTGTGTGTGTGTGTGTGTGTGTTTTGTGGGGAGAGAAGGTGAGAAAGTGTGTGT

TTTGTAGGGGGGAGAGAGAGGAGTGGGAGAGTGTGGTTGATGTTATAAGGTATAGTGGTG

>MSTRG.3476.1 gene=MSTRG.3476

GCATAGTTACCCGGTTCGCGGGTCAACCTAGGGATCCGCGAACTGGTACGCGCGTTCCGAAACGCGGGTC

GAAGTCGACCCGGAATGCTCGGGATCGCGTTCCAGTTCGTCATTTCCGGCGAATCCTTTTTACTTACCGG

CGACTATAAAACGAGGAAGAAAGGAAGAAGATAAAGATGGGTTACATGACCTGTGATATATATATATATA

TATATTTATGTGGATATATGTATGTAAATTGCAGCATGACAGAAGAGGATGTGTGGTGAAGAAGATGAAG

CAGGGAGAAGATGAAGTGGGGAGAAACCGAGGAACACGTGGATGGTGGATATTAATATGAAAGTCAATAC

AGTCAACACATATTAGACAACTGGACAGTCAGTACTCTTGTTTTTTCTCTTTCTATTAGTCTTATTCACG

CAACATCAACTATGTAAC

>MSTRG.3481.2 gene=MSTRG.3481

GTAATAGAGGGGGTTCTTTATTAAAAAAAAAAAGTAATAGAGGGGTTGGAGGTAACCGCGAGAGCAGGCA

GGAACGATTCGGTGGAAGAAACAGGAAGAGAAAGAGATTGTGCAGGAGAAAACATAAGCCTGAGAAGTTA

CCCTTCTGAGGTTTTGGTTGGTATTGGCGATGAAAAGGATATAATAGTTCTAGACTGCTGGTGCATTTCA

ACAAGAAACAAAGGATCACTTGACCCTTTTCAAAAAAAAAGTCTACCATCTTACATCCTCCGAGGCTCCG

ACCACCGGAAGATAAGAGAATAGTGACCCGACCCCGTTCAGCTAGGTCGTGGAGAATCCAATTAAAACCA

TTCTCGTCGAAGCTAGTTTGGTCATTTAATCAGTCACGAGATCTGCATACCTCCAATGGGGCTCCTCAGT

ATTATTCGCAAGATCAAGAAAAAAGAAAAAGAAATGCGTATCCTCATGGTAGGGCTAGATAACTCAGGGA

AAACAACGATTGTTCTGAAGCTTAATGGTGAAGACACAAGTGTTATTAGCCCTACTCTTGGGTTCAATAT

CAAGACTATTACTTACAGCAAGTATGTGCTTAATATATGGGATGTTGGGGGGCAGAAGACTATACGGTCT

TACTGGAGGAATTACTTTGAGCAGACGGATGGCTTGGTGTGGGTTGTTGATAGTTCGGATCTTAGACGAC

TTGATGATTGTAAATTTGAACTCGACAATCTGTTGAAGGAGGAGAGGCTATCTGGATCATCACTACTCAT

TTTGGCGAATAAGCAGGACATACAAGGTGCTCTGTCCCCAGAAGAAATAGCTAAAGTCCTTAATTTGGAA

GCCATGGATAAGACCAGACACTGGAAAATTGTAGGATGTAGTGCTTACACTGGAATGGGCCTGCTGGAAG

GTTTTGATTGGTTGGTTCAGGACATCGCTTCTCGAATTTATGTACTCGACTGAATCTTTTTCAAATAGGT

ATTATACTTGCTCATGTACAGACATGTATATTAACTTATTCGAGATTGAAGAAAATTTGCTTGTTGTGGT

TAGGCTCCCTTTTACTTCGGTTGGAGGTATTTGGGATGCCTTGACAAGGGAGATAAGTTGTATGACTTAC

AAATTTTGTTTGCATGGCTTTCCAAAATTCATATATTGAGCAAGGAATGGATACAAATTTATTTATGCTA

TTTGTTTAATCTATTTTATGTAAACAAGCTGAAGCCTATGTAATTCCTGGAGGTGATTGGTTTAATCAAG

TCTCATCTCCTCACTATCTGGCAGAAATTGTGAGTTTTTGCGGAAATGTACTTTTCATCAACTCTCTTTG

GACTCGTATATTATGTCTGGTTCTCAGTATTATTACTTTTATGCCAATTAATACAAGTTTTTTGGGACCT

GTGTGAAATTTCATTTTCTTTTCCTACCCTTTTAAACCAAATCTATGTGATCTAGGTTATATATGGTGGT

CTTCTGGTTGCTAGTGGATGTGCAGAACCCACAATCTAGTTACTTTTTTCATTTGTGGTTGCAAATCTTT

TACTTGCAG

>MSTRG.3481.1 gene=MSTRG.3481

TAAATATTAGATGCAATAATAAGTAATAGAGGGGGTTCTTTATTAAAAAAAAAAAGTAATAGAGGGGTTG

GAGGTAACCGCGAGAGCAGGCAGGAACGATTCGGTGGAAGAAACAGGAAGAGAAAGAGATTGTGCAGGAG

AAAACATAAGCCTGAGAAGTTACCCTTCTGAGGTTTTGGTTGGTATTGGCGATGAAAAGGATATAATAGT

TCTAGACTGCTGGTGCATTTCAACAAGAAACAAAGGATCACTTGACCCTTTTCAAGTCAGTCACTGCCCT

ACTTATATGTATATATACTGCAAACAATGTAGATCCCATTACTCATGTTTTGGGCACATGCGTTTCATCT

AGGTTTGCATCATTTGTTTTATTTCTCTCTGTGGCCTTTTCTATTGAAGTGTTGCAAACTTTTGTCAAGT

ATTTGGTATATAATTGTCATCCTACAACACCCTTTATCATTATTGTTCAAGTTTAAAAGACACTCAAACA

TCTACCAAAATTATATTCTAATCTTTTTATATTAATGAATGTGTGTGATATAGTTGGATATATCAGATAT

GCTTTCAACCCTGTAGTTTTGTTCTGTTCCATTATAAAAATCAGTTCATCAGTTTTGTATCATTACTCAT

GCCTTAAGTCCTAACTCCTAATATGCAAACACGCAATCGACTATTTCCTGCAAAGTTGCCCTAATATGGA

TACATAATTATTGTAAATAAAAAAATTTCTACTGAAATAGTCAAACGTGGAGGCAACCTGCCATGAACTG

TGTCTTTGTTTGCGATATTATTTTTACTAACTTGATAATTTTTTTGGTGTTATGATGTCAAAGTGATTTA

CAAGTATGTTATTGATATTACACAAGATATAGTCGACTCTCAATGCTAATGAAAACTGAAACATAAGAAA

CCCCCCGCCTAGACTTTTATTTTGCCTTCTTTTGAATATTTATCACCCTTTTTCATCAACCAACTTCATG

ATTAAATGATAAATTAAGGACTTTCATTTTGCCTCCTTTTTATATCTTTCTGAAATTAAGCACTCCCATT

ATATTAAAAGTTTTAGATAACTGATACCAATATCACATAGCCAAAAATACAAATATCTGCAAGCAATGTC

AATAACACAGTAGAGGGTCCGATGCTAGACCAATACCCGTGCCTTGCACGGGTTATTATGCTAGTTTAAA

TTTCTTAAAACTTACTACCAGTATACCAACTTACATAGAAAAAAAGTCTACCATCTTACATCCTCCGAGG

CTCCGACCACCGGAAGATAAGAGAATAGTGACCCGACCCCGTTCAGCTAGGTCGTGGAGAATCCAATTAA

AACCATTCTCGTCGAAGCTAGTTTGGTCATTTAATCAGTCACGAGATCTGCATACCTCCAATGGGGCTCC

TCAGTATTATTCGCAAGATCAAGAAAAAAGAAAAAGAAATGCGTATCCTCATGGTAGGGCTAGATAACTC

AGGGAAAACAACGATTGTTCTGAAGCTTAATGGTGAAGACACAAGTGTTATTAGCCCTACTCTTGGGTTC

AATATCAAGACTATTACTTACAGCAAGTATGTGCTTAATATATGGGATGTTGGGGGGCAGAAGACTATAC

GGTCTTACTGGAGGAATTACTTTGAGCAGACGGATGGCTTGGTGTGGGTTGTTGATAGTTCGGATCTTAG

ACGACTTGATGATTGTAAATTTGAACTCGACAATCTGTTGAAGGAGGAGAGGCTATCTGGATCATCACTA

CTCATTTTGGCGAATAAGCAGGACATACAAGGTGCTCTGTCCCCAGAAGAAATAGCTAAAGTCCTTAATT

TGGAAGCCATGGATAAGACCAGACACTGGAAAATTGTAGGATGTAGTGCTTACACTGGAATGGGCCTGCT

GGAAGGTTTTGATTGGTTGGTTCAGGACATCGCTTCTCGAATTTATGTACTCGACTGAATCTTTTTCAAA

TAGGTATTATACTTGCTCATGTACAGACATGTATATTAACTTATTCGAGATTGAAGAAAATTTGCTTGTT

GTGGTTAGGCTCCCTTTTACTTCGGTTGGAGGTATTTGGGATGCCTTGACAAGGGAGATAAGTTGTATGA

CTTACAAATTTTGTTTGCATGGCTTTCCAAAATTCATATATTGAGCAAGGAATGGATACAAATTTATTTA

TGCTATTTGTTTAATCTATTTTATGTAAACAAGCTGAAGCCTATGTAATTCCTGGAGGTGATTGGTTTAA

TCAAGTCTCATCTCCTCACTATCTGGCAGAAATTGTGAGTTTTTGCGGAAATGTACTTTTCATCAACTCT

CTTTGGACTCGTATATTATGTCTGGTTCTCAGTATTATTACTTTTATGCCAATTAATACAAGTTTTTTGG

GACCTGTGTGAAATTTCATTTTCTTTTCCTACCCTTTTAAACCAAATCTATGTGATCTAGGTTATATATG

GTGGTCTTCTGGTTGCTAGTGGATGTGCAGAACCCACAATCTAGTTACTTTTTTCATTTGTGGTTGCAAA

TCTTTTACTTGCAG

>MSTRG.3481.4 gene=MSTRG.3481

TAAATATTAGATGCAATAATAAGTAATAGAGGGGGTTCTTTATTAAAAAAAAAAAGTAATAGAGGGGTTG

GAGGTAACCGCGAGAGCAGGCAGGAACGATTCGGTGGAAGAAACAGGAAGAGAAAGAGATTGTGCAGGAG

AAAACATAAGCCTGAGAAGTTACCCTTCTGAGGTTTTGGTTGGTATTGGCGATGAAAAGGATATAATAGT

TCTAGACTGCTGGTGCATTTCAACAAGAAACAAAGGATCACTTGACCCTTTTCAAGTCAGTCACTGCCCT

ACTTATATGTATATATACTGCAAACAATGTAGATCCCATTACTCATGTTTTGGGCACATGCGTTTCATCT

AGGTTTGCATCATTTGTTTTATTTCTCTCTGTGGCCTTTTCTATTGAAGTGTTGCAAACTTTTGTCAAGT

ATTTGGTATATAATTGTCATCCTACAACACCCTTTATCATTATTGTTCAAGTTTAAAAGACACTCAAACA

TCTACCAAAATTATATTCTAATCTTTTTATATTAATGAATGTGTGTGATATAGTTGGATATATCAGATAT

GCTTTCAACCCTGTAGAGGGTCCGATGCTAGACCAATACCCGTGCCTTGCACGGGTTATTATGCTAGTTT

AAATTTCTTAAAACTTACTACCAGTATACCAACTTACATAGAAAAAAAGTCTACCATCTTACATCCTCCG

AGGCTCCGACCACCGGAAGATAAGAGAATAGTGACCCGACCCCGTTCAGCTAGGTCGTGGAGAATCCAAT

TAAAACCATTCTCGTCGAAGCTAGTTTGGTCATTTAATCAGTCACGAGATCTGCATACCTCCAATGGGGC

TCCTCAGTATTATTCGCAAGATCAAGAAAAAAGAAAAAGAAATGCGTATCCTCATGGTAGGGCTAGATAA

CTCAGGGAAAACAACGATTGTTCTGAAGCTTAATGGTGAAGACACAAGTGTTATTAGCCCTACTCTTGGG

TTCAATATCAAGACTATTACTTACAGCAAGTATGTGCTTAATATATGGGATGTTGGGGGGCAGAAGACTA

TACGGTCTTACTGGAGGAATTACTTTGAGCAGACGGATGGCTTGGTGTGGGTTGTTGATAGTTCGGATCT

TAGACGACTTGATGATTGTAAATTTGAACTCGACAATCTGTTGAAGGAGGAGAGGCTATCTGGATCATCA

CTACTCATTTTGGCGAATAAGCAGGACATACAAGGTGCTCTGTCCCCAGAAGAAATAGCTAAAGTCCTTA

ATTTGGAAGCCATGGATAAGACCAGACACTGGAAAATTGTAGGATGTAGTGCTTACACTGGAATGGGCCT

GCTGGAAGGTTTTGATTGGTTGGTTCAGGACATCGCTTCTCGAATTTATGTTATATATGGTGGTCTTCTG

GTTGCTAGTGGATGTGCAGAACCCACAATCTAGTTACTTTTTTCATTTGT

>MSTRG.3481.3 gene=MSTRG.3481

TGTTAATAGATAAATTATGATAACTGTAACGTCTAATTGATTAAATAAAAAATATTTAAGCAGAATTAAA

TATTAGATGCAATAATAAGTAATAGAGGGGGTTCTTTATTAAAAAAAAAAAGTAATAGAGGGGTTGGAGG

TAACCGCGAGAGCAGGCAGGAACGATTCGGTGGAAGAAACAGGAAGAGAAAGAGATTGTGCAGGAGAAAA

CATAAGCCTGAGAAGTTACCCTTCTGAGGTTTTGGTTGGTATTGGCGATGAAAAGGATATAATAGTTCTA

GACTGCTGGTGCATTTCAACAAGAAACAAAGGATCACTTGACCCTTTTCAAAAAAAAAGTCTACCATCTT

ACATCCTCCGAGGCTCCGACCACCGGAAGATAAGAGAATAGTGACCCGACCCCGTTCAGCTAGGTCGTGG

AGAATCCAATTAAAACCATTCTCGTCGAAGCTAGTTTGGTCATTTAATCAGTCACGAGATCTGCATACCT

CCAATGGGGCTCCTCAGTATTATTCGCAAGATCAAGAAAAAAGAAAAAGAAATGCGTATCCTCATGGTAG

GGCTAGATAACTCAGGGAAAACAACGATTGTTCTGAAGCTTAATGGTGAAGACACAAGTGTTATTAGCCC

TACTCTTGGGTTCAATATCAAGACTATTACTTACAGCAAGTATGTGCTTAATATATGGGATGTTGGGGGG

CAGAAGACTATACGGTCTTACTGGAGGAATTACTTTGAGCAGACGGATGGCTTGGTGTGGGTTGTTGATA

GTTCGGATCTTAGACGACTTGATGATTGTAAATTTGAACTCGACAATCTGTTGAAGGAGGAGAGGCTATC

TGGATCATCACTACTCATTTTGGCGAATAAGCAGGACATACAAGGTGCTCTGTCCCCAGAAGAAATAGCT

AAAGTCCTTAATTTGGAAGCCATGGATAAGACCAGACACTGGAAAATTGTAGGATGTAGTGCTTACACTG

GAATGGGCCTGCTGGAAGGTTTTGATTGGTTGGTTCAGGACATCGCTTCTCGAATTTATGTACTCGACTG

AATCTTTTTCAAATAGGTATTATACTTGCTCATGTACAGACATGTATATTAACTTATTCGAGATTGAAGA

AAATTTGCTTGTTGTGGTTAGGCTCCCTTTTACTTCGGTTGGAGGTATTTGGGATGCCTTGACAAGGGAG

ATAAGTTGTATGACTTACAAATTTTGTTTGCATGGCTTTCCAAAATTCATATATTGAGCAAGGAATGGAT

ACAAATTTATTTATGCTATTTGTTTAATCTATTTTATGTAAACAAGCTGAAGCCTATGTAATTCCTGGAG

GTGATTGGTTTAATCAAGTCTCATCTCCTCACTATCTGGCAGAAATTGTTATATATGGTGGTCTTCTGGT

TGCTAGTGGATGTGCAGAACCCACAATCTAGTTACTTTTTTCATTTGT

>MSTRG.3484.1 gene=MSTRG.3484

CAATAATTCTCTAAATAAACTTCTAAACTCATAACAGACACGAGTAAACAAGCATATTAATGGATCGAGT

TCGGATCCTATGAATCTATATTACCGGGTCAAAATCTCCAGCAAACCTAAAAACCCCGGTCACTGCCTCA

CTGGAGTAAGCTGAAACGTCTTTCGTGCATCGTATTTGGTAATCAATCGCTCTTCCTTCCCATCCTCATC

CAGCTTTTGATCAAGTTTTTATGTAATTCATTGGTAAATTTTATGGATTCGACTTCAAAAAAATTAGCGA

ATATTTGTGCGGTAGATAGAATCAGTGATTTACCTGACTCACTCCTCATCGCTATACTTGCATTGCTTCC

AATTAAACCTGCTGTACGCACTTGTATCTTGTCGAAAAAATGGAGGCCTCTCTGCGAGTCACTCCCGAAT

CTTGATTTTGTTGATGTTAGCAATTCAGGCATAAATTTCACCAACTTTGTTGATAGGTTTTTGATGCGTA

GGCCGAATAACTTGAAAATCGCTAAATTCCGGTTAAATTGTTATCGTGGAGACTACTACCGGGACCGCGT

TAATGAGTGGATTATAAATGCGCTTGGACGTGATTTGAAAGAGATAAACCTTTGTCTTTCGTTTCGAGAC

TTGTATAATTTGGTCCAAGATTTTTTTTACATGTCTGCTAGTGTTGAAGTTGTTCGACTCAGTGGGAAAA

TATCAGTTGAGATTCCGGAGAATGTTACGCTTCCGAGGCTTCGGCTGCTTAGATTCGACAGAGTTATGAT

TTCGAGCTGTGAGTCAGTTGGGAAGCTTTTGTTAAACTGTCCTGTGCTTGAAGATTTGGAAATTGTCCAC

TGTGAATGGCTTACGGGCAACTCTTTCAATATTTGTGGATCTGTATTGAAGAGGTTGTCATATTATCTTG

GTTCAGATGATGCAGAATTTGTGATTCGGACTATGATTGATACACCGAGATTAGAGAATCTCCAAATTTG

TTCTTACGCAGGCGGTGATAATGATATTTTGTTCAAGGAGAACTTGCCATTCCTCAAGATAGCTGTAATC

GACATCTTTGATGATGAGATGCCACAAGGAGTTTATTACGTGTTAGGGGTGCTTAAAAAGATTAATCATG

TCAAATTTTTGACTTTGTCAGATAACACTGTGGAGGTTAGTGTCATGCCTAAATCTTAATGTAAATGAAC

TAATTTTTATCTTTGCTGCAAATTTATGTGACTGTCACGGCTTGCCTTCTTATTTTGAAGATCCTCAGCC

TGGCATATGATAATGAGTTAAGATCAGGTCATTATGAGTTTCCTCCATTTCATAACTTGACTGAGTTGGT

AATTGATGTTGACGATTTTTTTCACAAAACTTTACTGGATGACTTCCTTCAAAGCTCACCGAACCTTGAA

AGTCTTCAATTTCCGCAGGGTCTTGGTTGTAGTTCCCTTCATGAATCTTTTCGCGGCTCATGGGCCTGGT

CACAAGCACGTGTTCCAGAATGCTTATCCACACATCTTAAAACAGTATATATTAGAACATTCGGCGAAGT

AGATGAGGTATTGGCATTTGTTAAATGCCTGCTAGCTTATGGGAGTGCTTTGCGAAATGTATCAGTTGAA

ATTTCAAATTTGTCAAAAGATGCAAACGCCCGGCACGAACTGTTAAACTTACAGAGTGAATCAACAAGTT

GCAAGCTTAATATTATAGGATGAAAATGAGACATGCTATTTGTAATCACAAAATGTATGAGGTACATTTT

CTGTAATAAGCTCTGTTTTTCTTCTAGTTTTTATTAGTTTTTGCTACAATGTTTTTTTCCCCAGCATTGT

TATTTTTGCGTTTCACTGTGGCATGAAATATTAACAAGATGCATTGTTTAAGAATGATAGTTTATTGGTT

G

>MSTRG.3484.2 gene=MSTRG.3484

CAATAATTCTCTAAATAAACTTCTAAACTCATAACAGACACGAGTAAACAAGCATATTAATGGATCGAGT

TCGGATCCTATGAATCTATATTACCGGGTCAAAATCTCCAGCAAACCTAAAAACCCCGGTCACTGCCTCA

CTGGAGTAAGCTGAAACGTCTTTCGTGCATCGTATTTGGTAATCAATCGCTCTTCCTTCCCATCCTCATC

CAGCTTTTGATCAAGTTTTTATGTAATTCATTGGTAAATTTTATGGATTCGACTTCAAAAAAATTAGCGA

ATATTTGTGCGGTAGATAGAATCAGTGATTTACCTGACTCACTCCTCATCGCTATACTTGCATTGCTTCC

AATTAAACCTGCTGTACGCACTTGTATCTTGTCGAAAAAATGGAGGCCTCTCTGCGAGTCACTCCCGAAT

CTTGATTTTGTTGATGTTAGCAATTCAGGCATAAATTTCACCAACTTTGTTGATAGGTTTTTGATGCGTA

GGCCGAATAACTTGAAAATCGCTAAATTCCGGTTAAATTGTTATCGTGGAGACTACTACCGGGACCGCGT

TAATGAGTGGATTATAAATGCGCTTGGACGTGATTTGAAAGAGATAAACCTTTGTCTTTCGTTTCGAGAC

TTGTATAATTTGGTCCAAGATTTTTTTTACATGTCTGCTAGTGTTGAAGTTGTTCGACTCAGTGGGAAAA

TATCAGTTGAGATTCCGGAGAATGTTACGCTTCCGAGGCTTCGGCTGCTTAGATTCGACAGAGTTATGAT

TTCGAGCTGTGAGTCAGTTGGGAAGCTTTTGTTAAACTGTCCTGTGCTTGAAGATTTGGAAATTGTCCAC

TGTGAATGGCTTACGGGCAACTCTTTCAATATTTGTGGATCTGTATTGAAGAGGTTGTCATATTATCTTG

GTTCAGATGATGCAGAATTTGTGATTCGGACTATGATTGATACACCGAGATTAGAGAATCTCCAAATTTG

TTCTTACGCAGGCGGTGATAATGATATTTTGTTCAAGGAGAACTTGCCATTCCTCAAGATAGCTGTAATC

GACATCTTTGATGATGAGATGCCACAAGGAGTTTATTACGTGTTAGGGGTGCTTAAAAAGATTAATCATG

TCAAATTTTTGACTTTGTCAGATAACACTGTGGAGATCCTCAGCCTGGCATATGATAATGAGTTAAGATC

AGGTCATTATGAGTTTCCTCCATTTCATAACTTGACTGAGTTGGTAATTGATGTTGACGATTTTTTTCAC

AAAACTTTACTGGATGACTTCCTTCAAAGCTCACCGAACCTTGAAAGTCTTCAATTTCCGCAGGGTCTTG

GTTGTAGTTCCCTTCATGAATCTTTTCGCGGCTCATGGGCCTGGTCACAAGCACGTGTTCCAGAATGCTT

ATCCACACATCTTAAAACAGTATATATTAGAACATTCGGCGAAGTAGATGAGGTATTGGCATTTGTTAAA

TGCCTGCTAGCTTATGGGAGTGCTTTGCGAAATGTATCAGTTGAAATTTCAAATTTGTCAAAAGATGCAA

ACGCCCGGCACGAACTGTTAAACTTACAGAGTGAATCAACAAGTTGCAAGCTTAATATTATAGGATGAAA

ATGAGACATGCTATTTGTAATCACAAAATGTATGAGGTACATTTTCTGTAATAAGCTCTGTTTTTCTTCT

AGTTTTTATTAGTTTTTGCTACAATGTTTTTTTCCCCAGCATTGTTATTTTTGCGTTTCACTGTGGCATG

AAATATTAACAAGATGCATTGTTTAAGAATGATAGTTTATTGGTTG

>MSTRG.3485.1 gene=MSTRG.3485

TGAAGCTGGATAAAAAAGAGCTGAGCCCAGCCTAGCTTGTTGCGCTGGACAAGTTCACGGCCCCGTCGAA

AAAAAATACATGACAACAATCTTTCTGGGCTGTCAAAAAAAACAAGAAAACGGGGGAGGCAATCTTAAAC

CCCTTTGTCGAAGCATCACCCTTCCGGTACCATTAGAAAGCTTTACAACTTGAAGCATCATCTTTAATTC

TATTCATTCAATTCTCGCGCATCGATTTTGGTAATTAATTCTCTTCGTTTCGTTCTGTAATCCACTTTTT

GCCTAATTCATTCGTCTCCTATCATGGTCCATGGATTCAACTTCAAGAAAATTAACGAATATCTGTGACG

TAGATAGAATCAGTGATTTACCTGACTCACTACTCATCGCTATACTTGCATTGCTTCCAATTAAACCTGC

TGTACGCACTTGTATCTTGTCGAAAAGATGGAGGCCTCTCTGCGAGTCACTCCCGAATCTTGATTTTGTT

GATGTTAGGAATTCAGGCATAAATTTCACCAACTTTGTTGATAAGTTTTTGATGCGTAGGCTGAATGATT

TGAAAATCACAAAATTCCGGTTAATTTGTTCTGGTGGAGATTACTACAGGGACCGCGTTGATGAGTGGAT

TATGAATGCTGTTGGGCGTGATTTGAAAGCGATAGTCCTTGTTATTTCATTTCCAGACTTGTGTGAATTG

GTTCAAGACTTTTTTTACATGTGTACTGGTGTTGAAGTTCTTCTACTCCGTGGGAAAATATTACTTGAGA

TTCCGGAGAATGTTAAGTTTTCGAGGCTTCGGGTACTTAGCTTCCACAACGTTAGAATTTCGAGCTATGA

GTCAGTTGGGAAGCTTTTGTTAAACTGTCCTGTGCTTGAAGATTTGGATATTGTCCACTGTAAATGGCTT

ACGGGCAACTCTTTCAGTATTTGTGGATCTGCATTGAAGCGGTTGTCATATTGTCTTAGTTCAGATGATG

AAGAATTTGTGCTTCGGACTTTGATTGATACACCGAGATTAGAGAGTCTTAAAATTTGTTCTTACACATG

CGGTGATAATGATATTTTGTTCAAGGGCAACTTGCCATTCCTCAAGATAGCTGTAATCGACATCTATAAT

AATGAGATACCACGAGGAGTTGATTGCGTGTTTGGGCTGCTTAAAAAGATTAATCACGTCAAAATTTTGA

ATTTGTCAGATCACACTGTGCAGATCGTCAGCCTTGCATATGATAATGAGTTAAGAACAGGTCACTATGA

GTTTCCTCCGTTTCATAACTTGACTAAGTTGTTAATTAATGTTAAAGATTATTGTCATGAAACTTTACTG

GATGACTTCCTTCAAAACTCAGCGAAGCTTGAAAGTCTTGAATTTCCGCAGGGTCTTGTTTGTACTTCCC

TTCATGAATCTTTTCGCCGCTCATGGGGCTGGTCACAATCACGGGTTCCAAAATGCTTATGCGCACATCT

TAAAACAGTATATATTAGAAAATTCCATGGAATAAATGAGGAGTTGGCATTTGTTAAATACCTGCTAGCT

TATGGGAGTGCTTTGCGGAGTGTATCAATTGAAATTTCAAATTTGTCAAAAGATGCAGAGGCCTGGCAAG

AACTGTTAAACTTACAGACCGAATCAACAACTTGCAAGCTTAATCTTATAGATGAAAATGGGAAATGCTT

CTTATAATCACAAAGTGTATGAGAGGTTCAGAATGTTGGAGAGTTTGTCTGAACGTTGGAGAGTTTGTCT

CGACTCTATCATGCATGAATGTAGCAGAGTTTGATGGTTAGATAATGAATGCAATATTTGGTGGCACTTT

TCCTAATACACCCACCAGATAGTAGGAATTTAATGAGCCTGCAAGTCCATTGCTATTGTTGGATTAGATG

GGTGTTTGGCGGACTCTGCTTTTTGGCTTTTTTCCTGAGATGATATTTGCTGTTGTGGTTTGTTTGGTAA

AATAACAGGCATGTACTTATTTTAAATTTTAAGAGCAGAAGTTGAAAGCTACTGCAAATGTAGGATCCGT

TAAGCGGAACATAAAAGAAGGGCTTAATATTTATAAATGA

>MSTRG.3485.2 gene=MSTRG.3485

GACGGAGGGTGTAAGTTTACGAACATCTCACAGAAAACACAATCAGATGAAGCTGGATAAAAAAGAGCTG

AGCCCAGCCTAGCTTGTTGCGCTGGACAAGTTCACGGCCCCGTCGAAAAAAAATACATGACAACAATCTT

TCTGGGCTGTCAAAAAAAACAAGAAAACGGGGGAGGCAATCTTAAACCCCTTTGTCGAAGCATCACCCTT

CCGGTACCATTAGAAAGCTTTACAACTTGAAGCATCATCTTTAATTCTATTCATTCAATTCTCGCGCATC

GATTTTGGTAATTAATTCTCTTCGTTTCGTTCTGTAATCCACTTTTTGCCTAATTCATTCGTCTCCTATC

ATGGTCCATGGATTCAACTTCAAGAAAATTAACGAATATCTGTGACGTAGATAGAATCAGTGATTTACCT

GACTCACTACTCATCGCTATACTTGCATTGCTTCCAATTAAACCTGCTGTACGCACTTGTATCTTGTCGA

AAAGATGGAGGCCTCTCTGCGAGTCACTCCCGAATCTTGATTTTGTTGATGTTAGGAATTCAGGCATAAA

TTTCACCAACTTTGTTGATAAGTTTTTGATGCGTAGGCTGAATGATTTGAAAATCACAAAATTCCGGTTA

ATTTGTTCTGGTGGAGATTACTACAGGGACCGCGTTGATGAGTGGATTATGAATGCTGTTGGGCGTGATT

TGAAAGCGATAGTCCTTGTTATTTCATTTCCAGACTTGTGTGAATTGGTTCAAGACTTTTTTTACATGTG

TACTGGTGTTGAAGTTCTTCTACTCCGTGGGAAAATATTACTTGAGATTCCGGAGAATGTTAAGTTTTCG

AGGCTTCGGGTACTTAGCTTCCACAACGTTAGAATTTCGAGCTATGAGTCAGTTGGGAAGCTTTTGTTAA

ACTGTCCTGTGCTTGAAGATTTGGATATTGTCCACTGTAAATGGCTTACGGGCAACTCTTTCAGTATTTG

TGGATCTGCATTGAAGCGGTTGTCATATTGTCTTAGTTCAGATGATGAAGAATTTGTGCTTCGGACTTTG

ATTGATACACCGAGATTAGAGAGTCTTAAAATTTGTTCTTACACATGCGGTGATAATGATATTTTGTTCA

AGGGCAACTTGCCATTCCTCAAGATAGCTGTAATCGACATCTATAATAATGAGATACCACGAGGAGTTGA

TTGCGTGTTTGGGCTGCTTAAAAAGATTAATCACGTCAAAATTTTGAATTTGTCAGATCACACTGTGCAG

ATCGTCAGCCTTGCATATGATAATGAGTTAAGAACAGGTCACTATGAGTTTCCTCCGTTTCATAACTTGA

CTAAGTTGTTAATTAATGTTAAAGATTATTGTCATGAAACTTTACTGGATGACTTCCTTCAAAACTCAGC

GAAGCTTGAAAGTCTTGAATTTCCGCAGGGTCTTGTTTGTACTTCCCTTCATGAATCTTTTCGCCGCTCA

TGGGGCTGGTCACAATCACGGGTTCCAAAATGCTTATGCGCACATCTTAAAACAGTATATATTAGAAAAT

TCCATGGAATAAATGAGGAGTTGGCATTTGTTAAATACCTGCTAGCTTATGGGAGTGCTTTGCGGAGTGT

ATCAATTGAAATTTCAAATTTGTCAAAAGATGCAGAGGCCTGGCAAGAACTGTTAAACTTACAGACCGAA

TCAACAACTTGCAAGCTTAATCTTATAGATGAAAATGGGAAATGCTTCTTATAATCACAAAGTGTATGAG

CTGAATACAGCTGGTTCGATCAACTGCCTGATTCGGGATATCATAGTTTTCAAGTTAAACGGAGTCCTTG

CTATAGTTCATCCTGTATATTGGCTTTGGCACGACCATGTAAGAGGGACCATAAGACCTATTAATACATA

CCTTGTATAAACAAGTTTGATTTGCACATCAAAGCTTGGATTCCCCTATGGAGTTTGATCAAGCACTATA

TGTTGGCGACAACATTTATTCCATTGGAAAAGTTGGGATGGAACAAGCAAGAAATATTAGAGTATTAGTG

TATTATCAAATGTTATTCAATATTATTTTGATCTTGGATGCATTAGTTAATGCTTTCAGAATGGTGTCCT

AGTTCAATTGAAGTAAAATGGAGCTTCCAGAAACAGTGTGAAGAATGGCTATTTAATTCAGTTGCACTAT

AATTCTGTGATTAGTGATAAAAAGAACTTGTATTTTTGGTCGGAACCTCCTGTTGTACAATGAAACAGAA

ACAGGGGATAAATGTTGAGTTATCTCTTTAGTTAATCTTTACATATCTTGAAACAGGAATATTCACTTAT

TGATTTCAGCTTAGAAATATATCTATTTTTTCTACCTGTATCTTAGATAATTTAAAATTTTCGTGTCATC

TATGTATTCTTCAAGCACTCGAATTTCGCTCTCCCTACTAAAATTTGAGGCTGAATCTGATTTTCAGAGG

TTCAGAATGTTGGAGAGTTTGTCTGAACGTTGGAGAGTTTGTCTCGACTCTATCATGCATGAATGTAGCA

GAGTTTGATGGTTAGATAATGAATGCAATATTTGGTGGCACTTTTCCTAATACACCCACCAGATAGTAGG

AATTTAATGAGCCTGCAAGTCCATTGCTATTGTTGGATTAGATGGGTGTTTGGCGGACTCTGCTTTTTGG

CTTTTTTCCTGAGATGATATTTGCTGTTGTGGTTTGTTTGGTAAAATAACAGGCATGTACTTATTTTAAA

TTTTAAGAGCAGAAGTTGAAAGCTACTGCAAATGTAGGATCCGTTAAGCGGAACATAAAAGAAGGGC

>MSTRG.3487.1 gene=MSTRG.3487

GGTGGGTAGTGGTGGCCAAACGGGCGCCCCGTGACGAGACGAGACGAGTCGGAGGCGAGACGAGTTGATG

AATGGTGTACTCGTGGACGTCTCGTGTAAGGGTAGCGAGACGAGACGATACGAGACGAGTCGAAGGATAT

TAAACTCGTGTCCAACTCGCGAGTTTGGCGAGTTGGACGAGTTATGTATATTTTTTACTTTGCAGTTTGC

TTTGGATTAGTTGGCGAGCTGGACGTTGAACTCGTCACGTTCAACTCGTGGACCAACTCGTGAGTCATGA

GTCATGATACAGTCTCTGCAGACCAACTTGTACAGAGACGTGCGAGACGAGTCCGAGACGTGCGAGACGA

GTCGTGGACGTGCGAGACGGCCGAGACGAGTTGGCGCGAGTCGTGCGTGCGTGCCGATACGTACGAGCCG

AGACGAGGCGTCCACGTTTTCTCAAACCCAGGCTCGTGATCGTCTCGTACAGATTAACGAGATGAGGCGA

GTTTTAACTCGTCTCGCGGCGTACCGGTATTCAACGAGTTAAAAGGCGAGCCGAG

>MSTRG.3491.7 gene=MSTRG.3491

AGATGAGCAGCACTGCCGCATCATCAATGGTCTTTAACTCAGAAACTTATTACTTTAGATGTTATGTACC

TCCATGACAAACCTGGATCTGGAATCTGTCGTCAGAACTCAGAAGGTGTATGATGCACTGTGGATGCCGT

TGAACAATTCAGTGAACACGAGGGCTGCATCTTGGAAATTGTGGCCAAGGGTTTCCACATATACTCCTCC

GTCACATGGGTGTGCTATATTGCACCTCTAGCAACTTTGCATTCAAGATGATATCGCTGTTGATATTACT

ACATGTGGTGAAAGTGAAACTGTTCCAGTTTCAAGCCAATGAGTAAGGGGTGCTGAGAAAGAGCAGGCCC

GTCTTCATAATGTAGCTGATCTTGAGCTGGAAATTGCCAATTGCGAGGCTACTGATGATTTGAGGCGGCT

GCTTTGCCTATAGTAGGGCAAAGGTTCTTACTCAGAGAATCCACGAGAAAGCGCCATTCTTCTTGGCGCG

CCATTAGCGAAGATGTTTTTACTGCCTTGAACGAAATGGCAGTTGTCCAGATCAGATGTGTCTCACAAAT

GCTTTGCTTTGATTCTTTAAAGTTTTAACCATCTGTTTGTAATGTTAAACTTCTCATACCACTTTATATC

CCTTACTACTTTAATTCAAGTTCTGAAACGTACCTTTGTGATAATCAGCTAGAATTGATCTAATTTTCTC

ATTAATTCCATCAAATAATTTTACAA

>MSTRG.3493.1 gene=MSTRG.3493

TAAATACACAAAAATAGCTCAAACCCCTCCCACCTCACTCCTCTCCCTCACTCTCTCTCAAACTCACAGA

TCTCTCTCAACAATGGCGCTCTCTCACTCCACAATCTCTCTCATCGCGCTCATCGCCGTCGTCTTCTCTC

TCATCGGATCCACCGTCGCCGACGAGTCTCCCGCACCGGCGCCATCTCCGATGTCCTCCGCCGGATCTCT

CTCGCCGTCGCTTGCCGCTGTTGGTTGCGCCGTCGCCGCTCTTCTCTTCGGTTCTTTTCTCCAGATCTAA

AGAGTAGAAGTATTATATGATATGCTAGTATTATAGAGTGAAAGTGAGAGAGAGTGATTATGTTTTTGAT

TGAAGATTTTGATGTGGTGTTAGTGAGATTTATGTAGCTTTTATTACTTAATTAACTGATTCAATTATTG

ATTTTATTAATATTATTATTGTGTTTGTTTAATTGTTA

>MSTRG.3500.1 gene=MSTRG.3500

CCTACACATAGTCCCATCTTTACACGGCTGAGTCCGATACCGAGCCGGGTGTAGCCAACACTCGAAAACC

CCATGCGCGTGCTCACACGCGTCGCCCCTAACACACGCGCCTTTCCTGAAATCCGGGCACGCATTGCCGG

AATAATGGAACTTCCGGGGATCCCTCCGGCGAGCCTTCTCTCCCGGATGAGCGTACGGACACTCCGTCCA

GTCATGTGACCTGGCACGTGCGCAGCGCCTGACTTTAAACTCGTACATGCGGAAATGATCGCATGTGAAC

GCGTCCAGGGGAATATCTATGTCCTCAACTGACGGCAAGTAACGGTGTAACGCCGTTAACGTGCAGTGAG

GTGAGTGATGGTCATTTGGAGTGACTGAAAATGGGGAGTGAATCTCGGCCGTTGGATCATCGTACAGGGG

GATTTGGACGGTCGGATTTGGACGATGAAGAGGTTCTGATTCTCCGAGCATCATTGTTTGTGTGTCTGAA

TGTGTTTGTGTCTTGCAAGGTGAGACATGGTATTGGAATGGCTTCAGTTTGATGAGAATTTATACA

>MSTRG.3506.1 gene=MSTRG.3506

GTTGTTTCCATGAATTACACACCTGATAGGATTCCATCACTGGAATTACAGACCCTGGTCAATGAGGTAG

TCGCCCAGCCTTTCAACAACAATGTTGCACTGACCTGGAGTCATAGGCTCCTCGTATATGCCGATCAGTA

AAGCTTGATTGCTCTTCTTGATAGTGATCCCCCCAGATCCCTTCTTTCCCCTTATGACAGCATTAGGTTC

ACCTTGAATCACCATATACTTTGTGCCACCTAGGTATAGGCCAGTAGGTGCAAGAGATCCAGGTTCATTG

AAGTCATTCATTATTGCCGTAATTTCTTCTGGTTTAAACTGAGGAAAGG

>MSTRG.3508.1 gene=MSTRG.3508

CGATTAGCAGTACTATATATATATATGGTACTTCCAAATCTTTTCTCTCCTCGAGTGTCTTTTTATCATC

TTTAAATCTCATTTCATCTATCAAATATCTGCAACCACTTCATAATTCCTGCAGTATAATTAGGTTACAG

AGAAAACGATGGAAAGATATCCTGTAAGAGATATTAACAGCAACAAAAGAAGCCACGAAACAGGTCATGT

TTCTCAGGATAAAAGCCTGGAACTCAAACTCGCGCCTCCCGGAGAAAACAAGCCTCTTATTTCTCTGCAT

TACATATCTCATAAGTCTCATAGCAATGACAAGAACAAAAATTTGATGAGTTCTGGAACTAAAAGAGATT

TCCTTGACACGATAATCGGGGAGAAAGGCCGGGAGAGGAACTGGTTGACAGGCAGTGGTGATCAAGGATA

TCAGTCAGGTGTTAATCTTGGTGCTCCCAACAGCTCTAAGAGGATGGATGTACGTCCGATTGTGGGGTGG

CCTCCAGTTCGCTCATCAAGAAGAAATCTTCTGAGCAGCAGCAATTCCAGCAAGTTAGAGACGGATTCCT

CCAAGAACGGTGGGAAGAAAGGCGAAAAACCCGAAATCACTGAGGATAGTTTGTTTGTGAAGATAAAAAT

GGATGGAGTCCCTATCGGAAGAAAAGTTGATCTGAAATCCTATTCTGACTACGACAATCTTTCCTCCGCT

GTGGATGAACTCTTCCGAGGACTTTTAGCAGCTCAAGGGGAGATATCGAGTTCTGAGAAGGAGAAGATGG

AGAAGACGAAGACGATAACGGGATTATTAGATGGAAGAGGTGATTATACTCTTGTCTACGAGGATAGTGA

AGGAGACAGGATGCTTGTTGGTGATGTTCCTTGGCAAATGTTTATATCTTCGGCGAAAAGGCTTCGGGTT

TTGAAAGAGATCTCCTCTTCCTACCTCAGAAAAATCTAGGAACAAGGCGATGCGGCGTTTGTTGAGTCGA

AATGAAGAGGGACATGTATGGATATAAATAAGCTGAAAGATCTTCACAGCCTACTGCTTGGCCAAGTACT

CTAGTTCAATAAGCAAAACAATTATAGTAACTGATTGTTTTCCATTTTTTCCTTCAATGAAGAGGCAAAC

TGAGGGAAGTAAAATAGCTAAAGCCTAAATGTGATGTTTGCAATACACTGCCAAAGTACGGCATCTTGTT

TCTGAAAGATTATATCAGTGTGGTTATTGTAGATATGTGTGATCGTGTATGATAGAGTTTAGAGTTTATA

TGAACCCTGAGCTTCACACTGCGGAACAATACTGTAAGATCTCAGTGTACACGGCCTCTCTTTTTACCTG

TTTTCGTAAAGCCTGCATTAAAATACATACTC

>MSTRG.3511.1 gene=MSTRG.3511

GATTGATTGAGATTGTGAATAAGGCTAATGCTAGGCGAGGAGACAGAGTTGAGGATCCAGTGGTCGAGGA

GTGTTCAGGAAAGTTGATGAGGTTGGTGCAAGCTAGAGCTGTGCTATAGGGATTCAAGTGTAAGTCATAG

CTGGAATAGTTGACAGTGATTCTTCTTATCGAAGATGATCTGATTTGGTAAGAAATGTTGTGTAATATTA

GTGGTGGATTCTAATTTCAATACTGTAACCTGGATGCGATCCTGTTGCTTTTAGGGGGTTGAAATGTTCT

CTTTCG

>MSTRG.3512.1 gene=MSTRG.3512

CAAAGCATGATTACGTGTTAAGTGATATGTGATAAGTTTAAAGGGGTATATAAGTGGGTATCGATATACG

TGATATGTATATGCGATATATTGTTTAGTGACTATAGTATTATCTTATTCTCGTAAAGGATTTGGACGAG

ACGTGTACGCTCGAAGACGTCAGAAAGTCGGAATGGTATTGCAAAGCGAATAAGGGATGAATCGAGTAAG

CGAAGCGATGTGGCGAATAGAGTATAGCTAGAGATGGTCTAGAGATTACGAGATGCATAGATTGTGAAAG

TTGAAAGATGAGAATGAGCTGTGAAACTGGAGTCAGATTTAAGGCAAGCTTAAAGATGAGATGATTGATT

GAGATTGTGAATAAGGCTAATGCTAGGCGAGGAGACAGAGTTGAGGATCCAGTGGTCGAGGAGTGTTCAG

GAAAGTTGATGAGGTTGGTGCAAGCTAGAGCTGTGCTATAGGGATTCAAGTGTAAGTCATAGCTGGAATA

GTTGACAGTGATTCTTCTTATCGAAGATGATCTGATTTGGTAAGAAATGTTGTGTAATATTAGTGGTGGA

TTCTAATTTCAATACTGTAACCTGGATGCGATCCTGTTGCTTTTAGGGGGTTG

>MSTRG.3517.1 gene=MSTRG.3517

TATTGAAAATGATGCTGCTGACATCTTTACGCGTGCAATTTTTTACAAGCTTCAGGAAGAAATTTTAGCA

TCGTGCTTGGAAATGCAGATTAAGAAGATGAGTGAAGATGTTAATGGAGTCATTCATCTTGAAATTAGAG

ATGTGAAGGTTAAGGACAAAATTTTTAAGGTTTCCGTTAGTAAAGCTCATGCTGTATGTTCATGTAAGAA

ATTTGTTATGTGTGGAATAGTATGTAGGCATGCATTCTGTGGCTTGAAGCAAATAGGCGTTACAAAATTT

CCTAGAAGCCTCGTGCTTAACAGATGGATGAAAATCGCTGAGTGTGGCAGTGGATCAGATGTTGATCTGG

TATCTAATGACTATTTCAAGATGGAGAGAGTGTCTTTAAAGTTGTCAAACATATGGTTTGATTTTCGTCA

AACGGTTAACAATGCTGGTATTGATTTAGAGAAACTTGAATATGTGCACAAAGCTATAAAGGAAATAAAT

GCTGAGCTTGAAAAATATGGGGGTGATGCTGCTGAGTTTAGTAAGAAAGATCATATTGCATCTATTCTTG

GTGAACAGCCAGTTGGAGAACTTAATATTCTTGTACCAAACATTTCAAAGAACAAAGGTAATCATTTCAA

GAGGTTGATCAGTGACCGGGAGAAAGCATTGAATAAATCGAAGAAAAGGATTAGAAGATGCAAAGAATGC

TCTGCGACGACTCATGATTCTAGAACTTGTCCAAAGAAAAAAGGAGGAATAGAAACAAATGTGTGTTAAG

TGGCTTGAAAGTATTTGACAGTTTGAAGATATTATATAAACACTTTTTAAATGATGCTCAGGAAATTTGT

ACTTTAAGACAGTTCATAAGTTGACAGTGGCTTGAATGTATATTTTGATTCTTGTAGTTTATGCATAACT

CTATCTATGCTTTTAAGGCTGGGGTCTGGAAGTAGACGTTTTCGTACTCTGTAGCTAGAGAGGTACAATG

TTTTGCAAAAAAATGAGCTGTGTAATGAAATTTGTGAACCTATTGATTCATGGTTGAAATGTTCTTCCCA

GTGATAAGAGG

>MSTRG.3530.1 gene=MSTRG.3530

AGATTGTGTATGATTTATATTTACTCTTGTGATTTTTGGTTGCAGAAGCACAGATTGATTTGAACAAAAG

CGCAATCGATAAAGAAGTGGTAGAGAAGGTTTGGGACAGAATCGCTCCAGGACTCGCCTCAGAGTTTGAT

GCACCCTACACCGTTCCTCTTATTGCACCACGCCCTCTCTTGCTTCTAAATGGGGAAGAAGATCCACGTT

GCCCGCTTGCGGGTCTTGACATTCCCAAGCAAAAAGTAGCCGAGGCATATGAAAATGCTGGTTGTAAAGA

TAAATTTAAGGTGATTGCGGAACAAGGAGTTGGGCACCAAATGACAGCTTCAATGGTAAAGGAAGCAAGT

GACTGGTTTGACGAATATCTTTAATTGTAGCTTTTTTCTTGACAGTTTGAGAATACTGTCATGTGTCTTG

TCCTATCCCTTCCCTGATTTCAGTAATTGTATTTTATGTTGCAACTTCAGCATAGTATTGTAAACTTTGT

ATTTAATCAACTTGTTCCGTTCTTGCTCACCCATGTATCCATTAATTCGATCTGATCTATTTTTTATCCA

TTTGTGTTTTTTTGTTAA

>MSTRG.3531.1 gene=MSTRG.3531

TTACAATTTAAAGCAAGCAAAGTTGTCTAAGCAGGATAGGACCCCCTTCTGTACCAGACCAGACAGCAGA

GTTTATGATTGAAGTGCTGTCTTCAGTGTGTCAAAGCAAAAAAACAATGGAGGTGTTGGAAGCCATTGAA

GCCGATAGGCTCCGGTGTCAATTTCTTCATGTTTTGCACTGTCGTCGGTCCGCTCAAGTTCCACTTACTG

TGGAACCTGGAAGACCTGTGTTGGAACCCATGTATCAAGAAACTCCTAGGCCGCCAGATGGAGAGGAGAT

GCAATCTTGCCCCAAGGAAAATCACTGCGATTTCAAGGAGCTACTGAAGGAAGAGAACTTATGTCTGATC

ACTGAGGAAGGAGAACAAGGACGGGTGCCTGTCCTAGTGTTGAGCATGAAAGAAAGCAATCCTCAGAAAA

GGCCAGCTATTGTATTTCTGCATCGGACAAATACATGTAAAGAATGGTTACGTCCTTTACTTGAGGCATA

TGCCTCTCGTGGCTACATAGCTGTTGCCATCGATTCTCGTTACCATGGGGAGCGTGCTAGCAGTCCCACT

GCCTACCAAGATGCTTTAGTATCAGCATGGAAAACAGGTGATACGATGCCATTCATCTATGACACGGTTT

GGGACTTGATAAAAGTGGCTGACTATCTTACAGAGAGGGATGATATAGACCCTTCAAGGATCGGAATCAC

CGGTGAATCGCTCGGAGGAATGCATGCATGGTTTGCTGCTGTTGCTGACACTCGCTACTCAGTGGTTGTT

CCTATAATTGGTGTGCAGGGATTTCGATGGGCCATAGCAAATGGTAAATGGCAGGCTCGAGTAAACAGTA

TTAAGACCATATTTATAGAAGCACAGAAGGAACTAAACAAGAGTGAAATAGACGAAGAGGTAGTGGAGAA

GGTATGGGACAAGATAGCTCCAGGTTTAGCCTCACAATTCGATGCACCTTACACCATTCCAATCATAGCA

CCACGACCTCTGCTTATTTTGACCGGGGAGAAAGATGCTCGTTGCCCCCTACAAGGCCTTCGAGTACCTC

AACTCAGAGCAGTCCAGGCTTATGAAAATGCCGGTTGTCCAGATAAGTTCAAGATGATCGCAGAGCGCGG

AATATGGCATAGGATGACACCTTCAATGGTTAAAGAAACAAGCGACTGGTTTGACAGATATCTCAGATAA

TTCTTAGCACTATCGAGTATCGTTAATTACGACTTGTATTAACGTTTCTTTGTCGAGCAGAACATCTGCT

CTGTGTGAGCTATCTTTTTCGAGTCCTTCAGAACACTGGGGAGTGTAAATGTATGATCTATCATCTATGC

CTATGGTATGTACATGTTTGTGATTCGTTTCCTGGACTGGGAAACCTAGCAGGAAAGATGTGCTAAGTAT

GGAAGTTGCATGTACTTGTTCATCCGAAAGGGGCGGAGCTAATAAAAGACACGGAAACTCATGTTCCTC

>MSTRG.3531.2 gene=MSTRG.3531

AGCAAGCAAAGTTGTCTAAGCAGGATAGGACCCCCTTCTGTACCAGACCAGACAGCAGAGTTTATGATTG

AAGTGCTGTCTTCAGTGTGTCAAAGCAAAAAAACAATGGAGGTGTTGGAAGCCATTGAAGCCGATAGGCT

CCGGTGTCAATTTCTTCATGTTTTGCACTGTCGTCGGTCCGCTCAAGTTCCACTTACTGTGGAACCTGGA

AGACCTGTGTTGGAACCCATGTATCAAGAAACTCCTAGGCCGCCAGATGGAGAGGAGATGCAATCTTGCC

CCAAGGAAAATCACTGCGATTTCAAGGAGCTACTGAAGGAAGAGAACTTATGTCTGATCACTGAGGAAGG

AGAACAAGGACGGGTGCCTGTCCTAGTGTTGAGCATGAAAGAAAGCAATCCTCAGAAAAGGCCAGCTATT

GTATTTCTGCATCGGACAAATACATGTAAAGAATGGTTACGTCCTTTACTTGAGGCATATGCCTCTCGTG

GCTACATAGCTGTTGCCATCGATTCTCGTTACCATGGGGAGCGTGCTAGCAGTCCCACTGCCTACCAAGA

TGCTTTAGTATCAGCATGGAAAACAGGTGATACGATGCCATTCATCTATGACACGGTTTGGGACTTGATA

AAAGTGGCTGACTATCTTACAGAGAGGGATGATATAGACCCTTCAAGGATCGGAATCACCGGTGAATCGC

TCGGAGGAATGCATGCATGGTTTGCTGCTGTTGCTGACACTCGCTACTCAGTGGTTGTTCCTATAATTGG

TGTGCAGAAGCACAGAAGGAACTAAACAAGAGTGAAATAGACGAAGAGGTAGTGGAGAAGGTATGGGACA

AGATAGCTCCAGGTTTAGCCTCACAATTCGATGCACCTTACACCATTCCAATCATAGCACCACGACCTCT

GCTTATTTTGACCGGGGAGAAAGATGCTCGTTGCCCCCTACAAGGCCTTCGAGTACCTCAACTCAGAGCA

GTCCAGGCTTATGAAAATGCCGGTTGTCCAGATAAGTTCAAGATGATCGCAGAGCGCGGAATATGGCATA

GGATGACACCTTCAATGGTTAAAGAAACAAGCGACTGGTTTGACAGATATCTCAGATAATTCTTAGCACT

ATCGAGTATCGTTAATTACGACTTGTATTAACGTTTCTTTGTCGAGCAGAACATCTGCTCTGTGTGAGCT

ATCTTTTTCGAGTCCTTCAGAACACTGGGGAGTGTAAATGTATGATCTATCATCTATGCCTATGGTATGT

ACATGTTTGTGATTCGTTTCCTGGACTGGGAAACCTAGCAGGAAAGATGTGCTAAGTATGGAAGTTGCAT

GTACTTGTTCATCCGAAAGGGGCGGAGCTAATAAAAGACACGGAAACTCATGTTCC

>MSTRG.3537.1 gene=MSTRG.3537

CTTATATATCTCAGAATTTCTCTTACAATATCTCTCTCGCTCTGTGTCTCAAAACCTCTCAGAAAAAACA

CATTCTCCATCCCGCCTATGGCACCCGCAGATGTCACCCCGCATCGACCTCCGCCTCCTCTCCCAATCCG

AGCTCTTCTCTCTCCCTATGCTCTTCCTCTGCCCTTAAAACTCACTAAAACGAAATCGTCATCCCCAAAA

TCGACGGTCCCGTTTTTAATGAATCCTCCACGGCTGCGATTTTAGTTTGTAGAAGTAGGGTTTGAGATAG

GTACAGCCGCAGATTGTTTTCAGTGAGCAAGTTATAGGCGGCCCTGGTGCGCACGCAACCTGTTCGATGT

TTTGTCTCAGAGAATCAGGGGTTTAAATTGGGGCGTTTTCAGAATTAGGGTTTTACTTCGCTTTTATCTG

GTTTGAACTTCAAGAAGGTCTATGTTTAATGTTGAGTTCTTGAGGGGCTGCTTAATTCTCTTGACTTTAT

AATTAACACTGCATCCGGAGATATTCCTTTCGATCAGTGCTTATCTCTTCTCAAGAGAGGGTGTCCAGCA

TTAAGTTGTCTAAAGCTATTAAGCTAATAATAAAGCAAAGGAGGAAACTGTGTTATTACTGTGAATTATT

TACGGATGTATGACAACTTCCCCTCTCCCAGTATATGGATACATAGCTTATTTGAATGGTTGGATGATTT

GTCGAGTTGTGCACTAAAATAAACAGGGGTAGCA

>MSTRG.3537.2 gene=MSTRG.3537

CTCTTACAATATCTCTCTCGCTCTGTGTCTCAAAACCTCTCAGAAAAAACACATTCTCCATCCCGCCTAT

GGCACCCGCAGATGTCACCCCGCATCGACCTCCGCCTCCTCTCCCAATCCGAGCTTCACTAAAACGAAAT

CGTCATCCCCAAAATCGACGGTCCCGTTTTTAATGAATCCTCCACGGCTGCGATTTTAGTTTGTAGAAGT

AGGGTTTGAGATAGGTACAGCCGCAGATTGTTTTCAGTGAGCAAGTTATAGGCGGCCCTGGTGCGCACGC

AACCTGTTCGATGTTTTGTCTCAGAGAATCAGGGGTTTAAATTGGGGCGTTTTCAGAATTAGGGTTTTAC

TTCGCTTTTATCTGGTTTGAACTTCAAGAAGGTCTATGTTTAATGTTGAGTTCTTGAGGGGCTGCTTAAT

TCTCTTGACTTTATAATTAACACTGCATCCGGAGATATTCCTTTCGATCAGTGCTTATCTCTTCTCAAGA

GAGGGTGTCCAGCATTAAGTTGTCTAAAGCTATTAAGCTAATAATAAAGCAAAGGAGGAAACTGTGTTAT

TACTGTGAATTATTTACGGATGTATGACAACTTCCCCTCTCCCAGTATATGGATACATAGCTTATTTGAA

TGGTTGGATGATTTGTCGAGTTGTGCACTAAAATAAACAGGGGTAGCAAC

>MSTRG.3538.1 gene=MSTRG.3538

GAAGCATATACAGTAAGAAGAAAATGAGTGAAACAGTTTTAGAGTTGCCAAGGTTTGTTGAGTTCCGGTC

ACAATATAGCAATAACTTCTGTGGACGAAAATCCTAAAACAATCCGTCTTACCAATGTTCATGTCAAAAG

CAATTTGTGCTTGTGGCGAGCAGAGGGACCGTATTATGAATGCGTGGCTGCAGGCTACTCAGTTCGGGAT

TCTGCAAGCTGCGATGTGTTTAATGTCATCGATTGGCAGTCACTGGTGCTAAGGGTGACAATGGTAAGTT

TCTTCGGGCACTGTGGTTTCAAAAGCATAACTATCTTCAGTTTTTGGTGACTAATGTGGAAGATAATCCG

ATGCTAGTGGGGAATGAGATCTATACGGACATGCTTCACTGTAATTTCCCCTCTTTCAATACACTCGCGA

GAGAAATGAAACACTATATCAAAGTGTTTGCTCCTCTTGACATGCTGCTCTGGTTGCTGCCATAAACGCC

TTACATGAATATGGTGCGACATTTGTGAAGCCCAAGATATTAAGCATGCCATCCGTGCTCCTTTTATAAT

TACCAAACACAGTTAAGTCACTCTACGAATAACCAAATTCAAAGTTAATTGTGAGGATCGCATCCACTAT

AGATTGATGTTAATCAAACACAATAACAAGCACACAAGTAAAGAACACATACAATAAACGTGGAAACCTC

TCGAGGGAGGGTAAAATCACAAACCACTAAATAAGAATGTATTACAGGATTGGAGTCGCCCCCTCACTCA

CACTCTAAAATACAAGCACCACACATGATTTCAATCCCCTGACAACTTGGAGAACCGTCATTGTCACTTA

GCAATTCAAGTATCAACTTGGAGGAGGGTTTTTGCTTCATGATCATCAATCATGATTTGGTGATATATCT

CTTGCAACTTCCATTACGCTACTAGGTTGTCTTTTCTGGGATTCGATGCAGTGAGCTTTGACAAAATCAG

GGTCTGATATGATTGCTAACCATGTTTTGCAAACTGATTTGCATCGAAGGAGTGTTGCTACAGGTAACAA

ATTAAATGTTTTGTGAGAAGACATGATTAGGAGTGATTCGAATAATGTAACTATATATGGTAGATAAGTA

GCGGATCTTACAGCATCTTCGAGGGGAAGAACATGCTCAGTAGGCGTATCAGTGTTGCATTCTGTCATTT

TTCCAATGAGAGCCTCTTCAATCTCCTCATTCTGGGAATCCTGATTTTGTTGAAGAAGGCAAGTAGAATG

AGAATGGTGAAGGCAATGGAGTTAACGAAGCAGATTTCCATCTTCGTATGCGCATGAGGCATATGCAACA

CCATACCTATGACTAGCAGAGCAAAATATGAGTTCATTAGATAAATATAGCTTCATCAAAACTGATAATG

AAAATCTAGGTTTATAATTCTAGTGAAAATAAAGGTTTACACTAAAACAACACTACTAGAGTCCTGCAAA

TGTTTATTAACTTCCTGCAAATGAGAATTGAATGTGAAACCTAAATTAAAACAAGTAATACAATGTTCTG

CTAAAGCACTACAATATTCACATGTT

>MSTRG.3539.1 gene=MSTRG.3539

TTGTCTTTTCTGGGATTCGATGCAGTGAGCTTTGACAAAATCAGGGTCTGATATGATTGCTAACCATGTT

TTGCAAACTGATTTGCATCGAAGGAGTGTTGCTACAGGTAACAAATAAAATGTTTTGTGAGAAGACATGA

TTAGGAGTGATTCGAATAATGTAACTATATATGGTAGATAAGTAGCGGATCTTACAGCATCTTCGAGGGG

AAGAACATGCTCAGTAGGCGTATCAGTGTTGCATTCTGTCATTTTTCCAATGAGAGCCTCTTCAATCTCC

TCATTCTGGGAATCCTGATTTTGTTGAAGAAGGCAAGTAGAATGAGAATGGTGAAGGCAATGGAGTTAAC

GAAGCAGATTTCCATCTTCGTATGCGCATGAGGCATATGCAACACCATACCTATGACTAGCAGAGCAAAA

TATGAGTTCATTAGATAAATATAGCTTCATCAAAACTGATAATGAAAATCTAGGTTTATAATTCTAGTGA

AAATAAAGGTTTACACTAAAACAACACTACTAGAGTCCTGCAAATGTTTATTAACTTCCTGCAAATGAGA

ATTGAATGTGAAACCTAAATT

>MSTRG.3540.1 gene=MSTRG.3540

AGCAAATACAGTGAGAAGAAAATGAGTGAAACAGTTTTAGAGTTGCCACAAGACTCACACTATTCGAGCC

AGTTAAGTACAGCGACGACACCATCGCTCTCAGAAACGTTGCAAACAAGTTATTCTGCACTAGGCTGACT

ACCCAAGGCAAAGAAAGCTGTCTTAATGCCGGTTCTCCTACAATCAATAGCTGGGCCAAATTTAAGGTTT

CAGAGCTTGTCATTTCAAGAATGGTCAAAAATATAAAATTCCGGCTCATGGATGCAAGGATCTACAATGA

TAACATAAATGTGATGGTCAGTTCATCACACACTAATTTAGGCAATACAACCCAGGAACAGACATTTGAA

CTCTCATATACACATAAGAAGTCAAGTACCTGGAATTGCAGTATGTCAATGAAGTCAAGTGTCGAATCAT

CTCTTAAGGTACGCATCCCAAAAATCATCCAAGGCGATGTTAAACTCGGGTTTGAGATTCCAAGCACAGA

AACTTCTCCCAAAATCCAAAGTTGCAATGCCAACCACTTTGACTTCCAGTAAACTCAGCATGGACCGCAT

CCTTTAAAAATTTACTCTGGCAGTGCCTTAGTGAGTATATCAGCTCGTTGCTCTCAAAGATCTATACGGA

CATGCTTCACTGTAATTTCCCCTCTTTCAATACACTCGCGAGAGAAATAATGATACACTATATCAATGTG

TTTGCTCCTCTTGACATGCTGCTGTGGTTGCTGCCATAAACGCCTTACATGAATATGGTGCAACATTTGT

GAAGCCCAAGATATTAAGCATGCCATCCGTGCTCCTTTTATAATTACCAAACACAGTTAAGTCACTCTAC

GAATAACCAAATTCAAAGTTAATTGTGTTAGGATCGCATCCACTGTAGATTGATGTCAATCAAACACAAT

AACAAGCACGCAAGTAAAGAGCACATACAATAAAAGTGGAAACCTCTCGAGGGAGGGTAAAATCACAAAC

CACTAAATAAGAATGTATTACAAGATTGGAGTCACCCCTTACTCACACTCTAAACTTACAAGCACGACAC

ATGATTTCAGTCCCATGACAACTTGGAGAACCGTCATTGTCACTTGACAATTCAAGTATCAACTTGGAGG

AGGGTTTTTCTTTCATGATCATCAATCATGATTTGGTGATATATCTCTCGCAACTAGGTTGTCTTTTCTG

GGATTCGATGCAGTGAGCTTTGACAAAATCAGGGTCTGATATGATTGCTAACCATGTTTTGCAAACTGAT

TTGCATCGAAGGAGTGTTGCTACAGGTAACAAATAAAATGTTTTGTGAGAAGACATGATTAGGAGGGATT

TGAATAATGTAACCATATATGGTAGATAAATAGCGGATCTTACAGCATCTTCGAGGGGAAGAACATGCTC

AGTGGGCGTATCAGTGTTGCATTCTGTCATTTTTCCAATGGGAGCCTCTTCAATCTCCTCATTCTGGGAA

TCTTGATTTTTTTGAAGAAGGCAAGTAGAATGACAAAGGTGAAGGCAATGGAGTTGACGAAGCAGATTTC

CGTCTTCGTATGCGCCTGAGGCATATGCAACACCATACCTCTGACTAGCAGAGCA

>MSTRG.3541.1 gene=MSTRG.3541

ATAATCCAGTCCGGATACAAACCGTCCCGACCGTTGCCATCCCTGTGTAACATGTTTCCGTTGAACACCT

TCGTTCCCCAAATAGGGTTCCATATTCTCCCACCCAGTTCAAACCTTCACCCAGCTCATTGTGGGATCTG

GTGGAGCCAACAAGCTGCGACGACACACTTGACCTCGAAGACTCTGCTGTTACACCAGGGATTCGCGTCT

TCAAAGGGCAAACTAGGTGTTGAGGACTTGTTTAAGACACTTATTTTGAAGAAGTTTAAACACTCAGTGA

TGACACATACTGTATCTCCCGATCAATCTCAGCTCTCGCAACCTACTTCAAGTTCCGTTAAAAAGGCAAA

TAATGAAGAAAATGCTCCAAGATCAATGGTACATGACAACATGTTATGCTGGAAGGAGAAGTTGAAAGGT

TTCTTCGAAGACTTCCCAGATACAGGTTTCAAAGAAATGGTGTTTTTTGTGAGAAACAAAACGAGGATTC

TCAGAAAAATGACCGAATGCAACACTGATACGCCCACTGAGCATGTTCTTCCCCTCGAAGATGCTGATGT

GAGGAAACCTCCATGCTCCAGAGATAATTTGGTAGCTTTTGTGGGTGCAACTCTCGGTGCTTAAGACCCA

GCTGATCCTTCTAATCATGTCTTCTCACAAAACATTTTATTTGTTACCTGTAGCAACACTCCTTCGCTGC

AAATCAGTTTGCAAAACATGGTTTCCAATCATATCAGACCCTGATTTTGTCAAAGCTCACTGCATCGAAT

CCCAGAAAAAGACAACCTAGTAGCGTAATGGAAGTTGCGAGAGATATATCACCAAATCATGATTGATGAT

CATGAAACAAAAACCCTCCTCCAAGTTGATACTTGAATTGCTAAGTGACAATGACGGTTCTCCAAGTTGT

CAGGGGACTGAAATCATGTGTGGTGCTTGTATTTTAAAGTGTGAGTGATGGGGTGACTCCAATCTTGTAA

AAAATTCTTATTTAGTGGTTTGTGATTTTACCCTCCCTCGAGAGGTTTCCACGTTTATTGTATGTGATCT

TTACTTGCGTGCTTTGTTATTGTGTTTGATTAACGTCAATCTATAGTGGATGCGATCCTAACACAATTTA

ACTTTGAATTTGGTTATTCG

>MSTRG.3541.2 gene=MSTRG.3541

ACCGTCCCGACCGTTGCCATCCCTGTGTAACATGTTTCCGTTGAACACCTTCGTTCCCCAAATAGGGTTC

CATATTCTCCCACCCAGTTCAAACCTTCACCCAGCTCATTGTGGGATCTGGTGGAGCCAACAAGCTGCGA

CGACACACTTGACCTCGAAGACTCTGCTGTTACACCAGGGATTCGCGTCTTCAAAGGGCAAACTAGGTGT

TGAGGACTTGTTTAAGACACTTCAAATAATGAAGAAAATGCTCCAAGATCAATGGTACATGACAACATGT

TATGCTGGAAGGAGAAGTTGAAAGGTTTCTTCGAAGACTTCCCAGATACAGGTTTCAAAGAAATGGTGTT

TTTTGTGAGAAACAAAACGAGGATTCTCAGAAAAATGACCGAATGCAACACTGATACGCCCACTGAGCAT

GTTCTTCCCCTCGAAGATGCTGATGTGAGGAAACCTCCATGCTCCAGAGATAATTTGGTAGCTTTTGTGG

GTGCAACTCTCGGTGCTTAAGACCCAGCTGATCCTTCTAATCATGTCTTCTCACAAAACATTTTATTTGT

TACCTGTAGCAACACTCCTTCGCTGCAAATCAGTTTGCAAAACATGGTTTCCAATCATATCAGACCCTGA

TTTTGTCAAAGCTCACTGCATCGAATCCCAGAAAAAGACAACCTAGTAGCGTAATGGAAGTTGCGAGAGA

TATATCACCAAATCATGATTGATGATCATGAAACAAAAACCCTCCTCCAAGTTGATACTTGAATTGCTAA

GTGACAATGACGGTTCTCCAAGTTGTCAGGGGACTGAAATCATGTGTGGTGCTTGTATTTTAAAGTGTGA

GTGATGGGGTGACTCCAATCTTGTAAAAAATTCTTATTTAGTGGTTTGTGATTTTACCCTCCCTCGAGAG

GTTTCCACGTTTATTGTATGTGATCTTTACTTGCGTGCTTTGTTATTGTGTTTGATTAACGTCAATCTAT

AGTGG

>MSTRG.3542.1 gene=MSTRG.3542

ATCACAATATGAAGCTTAAGGGCATTTATAATTGAAACATATACACTCTGATACAGAAAATGAGTCAAGT

TGTTGTATTGCCCAGGCTTGTTGAGTTCCGATCAAAATATGCCTACAGTAAGTACTTATTGCGATATGTA

AAGGAAGAGGGGCAGAAGAAGGGGTTAGTCCAGTTCTCTGGTGAAGATGTAGGCAGTGCAGACGCGAAGT

TCGAAGTCGAAAAATCAGAAAGTGGGGATGGATCGGTGCACATAAGAAGCTGTTACAACAACAAATACCT

GCAGCTGGTGGACTACGACAACTGGATTGCTCCCAGAGCAGAAAAGCCCGACGAAGACCGATCCAAGCGT

TCGTGTACTCTGTTCAGGCCTACTTCTGTGGGCGGGGATTCCAGAACAATCAGACTTACCAATGTTTATC

TTGGAACTTATCTGTTTATGTGGCGAACAGCTTCGCCCTATTATGGATGCGTGTACTCATGCAACTCCAA

TCCAGAAGCCGATTCCTGCGACGTGTTTAGTGTCATTGATTGGCAGTCACTGGTGCTACTTCCCAAATAT

GTGGCCTTTAAGGGTGATAACGATAAATTTCTCAGTCCCATATGGTATGAAGGTCATAGTTATCTTCAGT

TTTCGGAAAGTAAAGCTGATGATTTAATGGTGGTGGGGAACGAGATAGTCCGGATCACTTCTGATGGTAA

CATTCGCATTCGTAACATGTTCAGGGACAGATACTGGTGTTTAAGCCCAAACTGGATTTGGGCGGATTCA

ACAAATTCGTCGAATAATGACACACTATTTGAGCCAATTAAATGCAGTAATGACACAATCGCTCTGAGGA

ACATGGCAAACAAGTTATTCTGCATCCGTCTGAGCACCGAAGGAAAAATTAACTGCCTTAATGCCGGCTC

TTCTACAATCAATATTAGCACGGCCAAATTTAAGGTCTCGGAGCTTATCATTTCAAGGAAGATTGATAAT

ATCAGGTTCCGGCTCATGGATTCGAGGGTATACGACGAGAAGATGCTGGTGATGGTCAGTTCCACCTACA

CGAACCGAGGCAATTCGACTCAGGAGCAGACAATTCAACTCTCATACACAAATAAGAGGTCAAGCTCATG

GAACAATAGTCTCTCCTTGACCACCAGTGTCGAGGCCACGATCACCGCAGGCATACCATTCATCGAAGAA

GGCAAAGTTACCGTTGGTGCTGAGAGCTCTGCACAGCATGCCTGGGGAGAAATTGTGGAATCCGAGGAAA

ATGTGACCATTAGCTACAAGGTTTCGATACCACCAAGGACAAAGGTGACTGTGAGAACTGTGACAACCCG

CGGTGTTTGTGATGTCCCCTTTAATTATGACCAGCGTGACGTCCTCTACACGGGGAAAACAGTTGTGACC

AACATGGATGACGGAGTTTTCACGGGCGTGAATAACTATTCTTTCTACACTGAAGAGAAAGAGGAGCCCC

TGGATAATAATAATCGAGGTAGAACTGCTACAGGATTCAGGAGGGTTGCTAAAGTGTCACGCACAGGTGT

GATAACTCGTCTCAAATGATCTGAATCAAGGTGGCTTTCGTGATGTTGGATCAGTACTATTACTCTAAGC

TATGCTTTTTATTACTACTAAATAAACAAAGACTGTATGTCAGGATTGCTATCCTATGAAATATAGCTAT

AAGTTGTGCTGTTGGGGCCAGGACCGTCTGGTATGGTAATATCTTTTAGATGGTATGTGCATAAATAAGA

ATGAGTCCTCCAAGATTATTAATCAATATATACCAGTAGTTTTTATTTAATGCACACCTTTCTATTTGTT

ATATTTTATTTGACTT

>MSTRG.3543.1 gene=MSTRG.3543

CTCGACTCGAATTACAGGCCTAATTTCCTCTATTATAAAGAGCCATCAGGATATGAAGATTACAGGCATT

CTACAACTGAAACATATACACTACTTTAATATTTATACAGCAAGAAGAAAATGAGTCAACAAGTTGTAGC

TTTGCCGAGGTTTGTTGAGTTCCGGTCACAAAATAGCAATAAGTTATTGCGATATGTGAAGGAAGATGGG

GCGGTGAAGGGGTATGTCCAGTTCTATGGTGATTTTGTAGGGAGCGCAGACGCAAAGTTCGAAGTTCACC

GGTCAACAACTGTGAATGGACTGGTCCACATAAGAAGCTGTTACAACAACAAGTTCCTGCAGGTGTTGGA

CAAAGCGCCCAACTTGCTTGCTGCCAAGGCTGATAAGCCTGACGAAGACCAATCGAGTCGTTCTTGTACA

CTGTTGAGGCCTACTTCTGTGAATGGGAATCCTAAAACAATCCGTCTTACCCATGTTAATCTTAAACTTA

ATCTATGCTTGTGGCGTGGTACTGACCCTGCCTATTATGCATGCGTAGCTGCATACTACTCAGATCCAGA

TGCCGGATCCTGTGATGTGTTTAATGTCATCGATTGGCAGTCACTTGTGATACTTCCCAAATATATTGCC

TTAAAGGGTGATAACAATAAATTTGTTACTACCATATGGTATTATGGTAAGTGCTATCTTCAGTTTTCAG

CAACTAATCTGGATGATGATCCACTCAGTGCCGGGAATGAGATTGTGCAGACTTCAAACGGTAACATTAG

CATTCGTAACTTGAACTGGGATAAGTACTGGAGGTCAGACCCGCATATCATTTATGCCGATTCAACAGTT

CCCACAAGTAATGACGCATTATTTCAGCCAGTTAAATGCAGCGAGGACACCATAGCTCTTCGAAGCATAG

CAAACAATTATTTCTGCACCATGATGACCAACGAGATCATGAAAGATTGTCTTACTGTCGGCTCGACTGC

AATCACTAGCTTTGCCAAATTCAAGGTGTCAGAGTTTGTCGTTTCAAAGACTATCGAAAATATAAACTTC

CGGCTCATGGATGCAAGGATCTACGATGAGAACATGCTCGTGATGGTCAGTTCAACTTACGCCAACCAAG

GCAATTCAACCCAGGAGCAGACATTTGAACTCTCGTACAGAAACAAGAGGTCAAGTACTTGGAACTGTAA

TTTGTCAGTGAAGGGGAGTTTCGAGGCATCTTTTAAAGCAGGCATCCCATTCATCGAGGACGGTAAAATT

ACATTCAAGTTGGAGGCTTCTGAACAATATACCTGGGGAAAAACTAAGGAAGCAGAAGAAAAAATATCCA

CTTCCTACAAGGTAACAATAGCACCAAAGACAAAAGTGACCGTGCGAATTGTGGCAACCCGTGGTATTTG

TGATGTCCCCTTTTGTTATGATCAGCGCGACGTTCTTTATACCGGGGAAACAGTCATCACTCAAATGGAC

GACGGGGTTTACACTGGCGTCAATAACTATTCTATCTACACCGAAGAAAAAGAGGAACCTCTGAATCAGA

GTACTCGGATAACAGCAGCTGCAGGTTTCAAGAGGGTTGCTAAGGTGTCACAGTCAGGTGTGATAACCCG

TCTTAAACATCCCATAATCAGCTACGGACGAGGCTATTAGTCGATCAGTACTCTAAGCTGTAACGATCAG

GTGTGATACCCGTCTTAAACATCCCATAATCAGCTACGGGTGAGGCTATTAGTCGATCAGTACTCTAAGC

TACAAATGCTTTTTATAACTAAATAAATAAAGACTAATAAGTCTATGAGGAAAACTATCCTGAATACTTT

CTATGTTGAAGCTGTGATTTTCAAGTTGTGCTGTTGGGACCAGGACTGTCTAATAACTTGTAGATGGTCT

ATGTTGAGAAATAAAACAGAGTCCTGAGATTATAATTGGTG

>MSTRG.3544.1 gene=MSTRG.3544

ATCACTTGTGATACTTCCCAAATATGTTACGTTTAAGGGTGCTAACGATAAATTTCTTAGTACCCTCTTG

TTTCATAGTAATACGTACCTTTCGTTTTCAACAACTGATCTGGACCGTGATCCACTCAGTGCCGGGAATG

AGATTGTGCAGACTTCAAACGGTAACATCAGCATTCGTAACTTGAAAGATAAGTACTGGATGTCAAGCAC

GGAATGGATTTTTGCCGATTCAACAGATCCCACAAGTAGAAACTCAATATTTCAGCCAGTTAAATGCAGC

GACGGCACAATAGCGCTTCGAAACCTAGCAAATAATTACTTCAGCACCATGATGACCAGCGG

>MSTRG.3550.1 gene=MSTRG.3550

ACACATAAACGGCGTCGGCCTGCGATTCCCGCCGTGAGTACCGCCGAGATGATCGAGCTCCATCGCCTTG

TCCACCAGCGTCTCCGCCGTGAGACCAAAACACTGCTCCTTCCAGTAGGTGTTCTGGTAAATCTTGTTTC

TGAGGATCTTCTCAATCAGATTCTGGGGATTCGTGCCCTTAATGTTCTTCGCCGCCGGGTCCGTACGGTT

CGCCATGGTTGAGATTGGGATGTTTCAAACCCTAGA

>MSTRG.3551.1 gene=MSTRG.3551

TGGATGACAAAAGCACAACTGTTGCCGTTGGAGAGGGCAATGTAGGTGTCAACACGGGCAGTAGTGGCGG

CCCAGGAACCAATGTACAAGTCGGAAAAGGCTCGGGTGTGGAAGTCAGCACAGGAACTCCTGGAGAAGAG

ACTGATGTTGGTGTTGGCAAAAATGGTGTGGTAGTCAAGTCAGGCAACAAAGGGAAGCCTGTTTACATTG

GAATCACTACAGGCCCGAGCCCGTTCGACTACAACTATGCTGCCTCTGCAGCTCAGCTTCATGATGATCC

CAATGTAGCCCTCTTTTTCTTAGAAAAAGATTTACACCAGGGAACTAGCATGAATCTACATTTCGTGAAG

TCCACTACTACGACAACTTTTCTGCCTCGTAATGTTGCTGATTCCATTCCATTCTCTTCGGATAAACTGC

CAGAAATATTTGACAGATTCTCTGTAAAACAGAACACAGTGGAATCTGAAATAATAAAGAACACACTGAG

TGAATGTGAAAGGCCAAGTATCAAGGGCGAAGAGAAGTACTGCGCAACATCACTAGAGAATATGATTGAT

TTCACGACTTCTAAATTGGGGAAAAAAGTGATTGCAGTGTCGACAGTGGTTGAAAAGAGAAGAGTAATGC

AGAGGTTCAGAGTTATTGGATTGAAAAAATTGGGGAAAAAGGCCGTGATCTGCCACAAGCAAAACTATGC

ATATGCTGTGTTTTATTGCCACAAGACAAACCATGTCAAGGCTTACACAGTTTCACTTGTTGGAAATGAT

GGAACAAAAGCCAAGGCAGCAGCAATTTGTCACACAGATACTTCGTCTTGGAATCCAAAACACTTGGCGT

TTCAGGCGCTCAATGTTAAGCCTGGAAGTGCTCCTATTTGCCATTTTCTTCCTGAGGATCACATCGTTTG

GGTTCCATATTAACTGTTTCTTCTTGATTACGATCTTGAGGAATGATTTGCCAGCTGCTGGTTGAAGGAA

TGATGTGTGTTACAAGCTTTGCTTTTTAATGTATAAGGATTGTACTTATAAGAAGGGATCCATGATCCTT

TCGCGGGTTTTAATAAAATTGCTACATGCC

>MSTRG.3555.1 gene=MSTRG.3555

CCTCTCTCTCAAATCTCTCTCTCCCTCCCTCTCTCAAATTTCTCTCCCAGTCTCTCCCTCCCTCCCTCTC

TCAAATCTCTCTCAATCTCTCTGCACATACTTTGTCTATTAAACCCCTCTGTGTCTATCTACCTTCCACT

TCTCTCTTAAAACCTCTGCTGCATTCTATGCATTAATATCACTCTCTGAACACTACACAAG

>MSTRG.3557.1 gene=MSTRG.3557

CTCGACCCTTGTCCCTGCCATCATCCCTTTCCCTGCTTCCACTCTTCTCCCGATCCCTTCGTCTTTCACT

TTCCAACTCATGGGCTCTGTCCCGGTGTCTTTCTTTATCTCTGTCACCCTCATTTTCCCTTTCCTTGTCT

CTACTTGATTTTCGCTGAGGGCTTCTGCTTCTGCTATGACTGCGAGTACGCTTTCTTGATGATTCACGAG

ACTCATCTCGTCTCTCCCTTCTA

>MSTRG.3563.1 gene=MSTRG.3563

GTGATGTTAGTTCTCGAGTTGTTTCGTGTTCGGTTTGGTAAAATATGATATGATCTGCTCAGGAAATTTG

GTTTAGTTATTTGATTAGTGTTTTGCTATGCTTTTAGCGGTGTATGTGATGTTAGTTTTCGATTGGTGTT

GTGATCAGTTTGGTAATATATGATATAATCTGCTCAAGGAAATGTGATTTCCTTGTTTGATTAGTGTTTT

GCTATGCTTTTAGGAGTGTATGTGATGTTAGTTTGCGGTTAGTGTTGTGATTGGTTTGGTAATATATGAT

ATGACCCGCTCAAGG

>MSTRG.3566.1 gene=MSTRG.3566

CCTCCTGTCACCACCACGGCCTCTATCTTCACGCCCATCACCACCCCGGCCTCTATCGTCACGCCCATCA

CCACCACGGCCTCTATCGTCACGTCTGTCACCACCACGGCCTCTATCTTCACGCCTATCATCACCATGCC

TATCATCCCGTTTCTTGTTGTCATGTGACCTCTCAATAGATCGAGAGTCATCATGTCTAGCTCTCTTGGA

TTGATGACTTGGTGGCCTCTCCTCCTGGAGACTTCTAGTATCATGTCTTTCCTCCTGCTCTCTTTCATGC

CTCCCCCGATAATCCCCCTTGCCAGCGTCATACCTCTCCTCTTCTTCCTCATCTCTTTTGTGTTTCTTTC

CATGTCTATCTTCTTCATCCTTAATTTCTTTCCTTAATTTATCATCTTCACCTCTCCTGTGCTTTTTCTC

TCTCTCATCATCTTCATCTATATTGTACTTTCTTTTTCTTTCCCTCTCATCAACTTCAACTGTATTGTGC

TTTCTTTCCTTGTAATCATTATCATCCTTCATTTTTCTTCTATCCCCACCAGAAATAGCATCACTNCCTG

CATTCCTCGACTCCACTTCTTTCCCTTTATAATCCTTATAACCTGCATTCCTCGACTCCAGTTGCTTCTT

TTTAGTGTCCTCCCTGCTATCAGATTCAGAAGATTCACTCTCTTCCCTCCTGGAAACCAATTTCGCCTTG

TGATGCACGCGTTCGGGTGTTTCATCACGACGATAACTCTCACTATCTAAACC

>MSTRG.3570.1 gene=MSTRG.3570

AACTCCTTCACCTCCCCTTTCTACTTCTCTCGCTCTTTTATACAGATTTGCGGCTTCTCGGCACCAGACG

TCTCCATACAAATTGCAGTTACGATTATCAATATTTTGATACATAGCTTTTTAATTATTCGTTCCATGCT

CGTGCGTTGAAGACTCTAGCGTCTGATCCGGATTCGAGGAAAGAAACAGGTTGCCGAACGAACAATTACA

GATTTTTTTGCGGTATTGGTCCTCTATAACAGGCCTATGTTCAGCAACTCCAAACAAGTCGTCCAAGGTT

GGCACAACCGGTGCTGGAACTTGATAGAACTAAACAGATTTTAGTACCCCAACTTGAGCCATTGATTGAA

CAACTTGTAAGGGTGAATAAGATGCGCCATTCATGCCAGCAAGCTGAGGATGCATTGTTACAAGGAATAA

AAAAACTTGAGCAGACCCTAGCTTAATGTGTTGCACATGATAACATAGGCAGGGGAAGTTATGGTTCTCG

GATGGGCACACAATGGAGAGGCCAGAAACTTTAGAGTGCTTGAACCAGGCAGATCATCTTCAACAGTAAA

CACTGCAACAGATGATTCGTAATCTTGACCACTAGCCAAGCAGCTGTGAATTGCTTGCATTTGGGGAATA

CTTTCGTGCATTTAAATTCTCTCTCGGCAGTTCATCCTCATTTAAAAGTTTTATTTCCCTACTCCCATTA

GTATCCCTAACTCAATCAAGCTAGAAAACAAAGTGTGTACAACTGTTGGCTTTTCTATATATGGTCGCTT

TATTGATTATTCTTGCTGTTTTGAGCCCTTGGTTTCGCAAGTGTATGCATGATATTTGAACTTGAATTAG

TATTGGTACATCTCATATTCCCGATTTTAGAAGTTCGTACTTGGTCAGAGTCCAATCAACACTGCCTATC

ACTGCCTGAATCAAACTCTTTAAAATGCCAGGAAGAAATTTGATTTTTGCTATAAACTATGAATATTCAG

AAGATTTGACTTGTACATTATGATCAATTTAAGGTTTATGGTAGTTTGTTCATTTTTGGTATTAATGGCT

TTTTTAGTTAATTTGAGTTGCTATATGGTTTATTTTTTTAGGAATTTACATAGGTCATTTTTATTAATTC

TATTTGTGGATTTATTTTGGAAAATTGCATGTAAATGGTAGAAAATTATTATATATTAGGCTAAACAAAA

ATACTCTGATTGCGCTTCCAAAAATGTTATTTAAGTAATAATTTCTATTGAAGCACTATGTAAGTTTTGT

GTTGCATTTGAAAAAAGCTTACTTACATAGCACTTGCGGGAACGCTATTGAAGTACGCTACTAACATAGC

GCTTACACAAGTGTTATATAAGTACATTTTACTTGCATAGTGCTGGCTTAGATAACACTTAGAAACGCTA

TGTAAGCTTAAAAATAAGCGCTATGTAAGTTAAAATCTGTAGTAATGTATGTTACAGTCAATGAGAAGTC

AGGGAGAACTTTGTTTTATTGGTTGACTGAAGCTGTTGATGATCATGCTGACAAGCCTCTTGTTCTCTGG

CTCAATGGAGGTTTTTCTTGATACCATTCAATTTATAGGTTTTGGGTTTGATTTCTTCTGGCTTGTCTTT

GTGTTATTGCTCTTGGAGAGTGTGTTGATTGGTTTCGAACCTGTTACTCTCGATGTTCAGAATAATGTGG

ATGGTAGGATTGATATTGCCTTGAAAATTGTATCTGTAATCCTGGAAGACGACGTTTGTTTTTTGACAAT

GGTGTAATCCTAATTTAAATAAATGTGTTAAATATTAATCAGGTGATGTTTTTTCAGGCCCTGGGTGTTA

TCGATTACTTTTGGCATGGCCGCGAAGATGGGACCTTTCCATGTTAATAAAGATGGGAAGACTCTCTATT

CTAATCCATTCTCTTGGAATCAAGGTAAGTTTTAGCCCTTAAAATATAATAGTTTTTTTAATTTAAACAC

CCTCTTATTTTGGTAATATGCTACACATCTGCTGAGCTGAATTGCAAGATATTTTGTACAAAACTTTTGA

ATATACTCGGCGGCAATAGAAATGGTTGTGCTGAATTGATGATCTCCTCAAGGCAACCCCATGTACATAA

TCTAACATTATATTGACCTTTGAGTTCAGGATATCAGGTTCGTAAAAAATTTGAGAGACCAGCAACTTAA

CTTGGGCTGGCACTTGTGTATTATAGAATGTACAACATAATGTAACATGAAGGAAAAGGACAACAAGTTT

TATTTTTAACTCATTTCAGTGCAGCCTATATTCTGTACATAATACTGCAACAGGAGTTTGGTGATCCAAA

AAATTCGGTTTAGTCTTTTACACCTCCATCTCCTTCTTCATATGTGTGTTTTTTGGGTGCAGCCAAGGCT

CAGAATCAATATGCACCAATAACTTCTTCTCCACAGCTATAGTGGTTCTGAGCCATTTTATTCCATGTCA

GCGCCACATCATTGCCTTGTCAGATAGGTTATACACATAATTCACGCCTCTCTGTCCCCCTCACAGTAAC

TCCTTCGCTTGGTTCTTAACCCATGAATGTTGTCGCTCTACGTTATCTATTTTAGGTCTACTCGCCTAAA

TCCTTTCGCTGACCTGATATATCTTTCGTCTCTAGCACATATAGAGTTGTAAGGTGTCGAGTTCAAAGCA

CAAAGGAGGCCACAAGGCTGTTGTTGACACCATCCAGATGCTGCAGCCCAGACAACTTGTCAGATTGCAA

ATGATAAGCTGGTGTTAGCTTCACTTGAGCTCATGACCTTGCATAAAAGACTTGCATCTAGGATGAGGTA

CATCAAATATCTTGCAGTCCAAGAAGTTGAAAGGATCTTATGTCATGTTGCAAGCGAAAAAGTTTGGAAT

TTGCTTCAGAAAATATTCACTTCTAAGTTATAATCAACAATATGGCTTTTTCACCAAAAAATTGTGCAAG

TTACCTCCTACCCAAATTCAAAAATTATGTCCATGCACCCAATGGTGGAAGTACTCATGTGCTGATATGT

GCAAGCTCGAGAAGATTTTGAAATTTAGTGTACTACGCATATATTATATTTTTAGAGTTAAAAGATGTGA

CAAGCCTGATTAAGTATAATTCGGCCATACAGAATTGC

>MSTRG.3570.2 gene=MSTRG.3570

CTCCATACAAATTGCAGTTACGATTATCAATATTTTGATACATAGCTTTTTAATTATTCGTTCCATGCTC

GTGCGTTGAAGACTCTAGCGTCTGATCCGGATTCGAGGAAAGAAACAGGTTGCCGAACGAACAATTACAG

ATTTTTTTGCGGTATTGGTCCTCTATAACAGGCCTATGTTCAGCAACTCCAAACAAGTCGTCCAAGGTTG

GCACAACCGGTGCTGGAACTTGATAGAACTAAACAGGTTAGATCATTTTGGTTGTAGAACGAGTGTTTAT

ATTCACATAGGCATGTCTGTCTATTGCTGGTGAAAATGTATGATTTCCACGTAAGCGTCTCTTGCTATGG

ATTGGAGGATTCAAGCCAGGGGCGGAGGGACTAAATGCCGAGGGTATGCCCGGCCTACCCTCAAATTCTG

GTGAAAAGGATTCCCCCCATTATGGTCATATAACATATTTTAGTACCCCAACTTGAGCCATTGATTGAAC

AACTTGTAAGGGTGAATAAGATGCGCCATTCATGCCAGCAAGCTGAGGATGCATTGTTACAAGGAATAAA

AAAACTTGAGCAGACCCTAGCTTAATGTGTTGCACATGATAACATAGGCAGGGGAAGTTATGGTTCTCGG

ATGGGCACACAATGGAGAGGCCAGAAACTTTAGAGTGCTTGAACCAGGTGACTATAACTTTCTACCAAGA

GTTCCACCTATTAAGTATCAATGCTTAAATTTTTGTAGACTTGCCATGATTTGACTGTTTTTAATATTGT

TGCTCTAATAATTACTCGCTGAATGGTTAGCTTGACTGTGTCTTTTTATATTAATAGGCAGATCATCTTC

AACAGTAAACACTGCAACAGATGATTCGTAATCTTGACCACTAGCCAAGCAGCTGTGAATTGCTTGCATT

TGGGGAATACTTTCGTGCATTTAAATTCTCTCTCGGCAGTTCATCCTCATTTAAAAGTTTTATTTCCCTA

CTCCCATTAGTATCCCTAACTCAATCAAGCTAGAAAACAAAGTGTGTACAACTGTTGGCTTTTCTATATA

TGGTCGCTTTATTGATTATTCTTGCTGTTTTGAGCCCTTGGTTTCGCAAGTGTATGCATGATATTTGAAC

TTGAATTAGTATTGGTACATCTCATATTCCCGATTTTAGAAGTTCGTACTTGGTCAGAGTCCAATCAACA

CTGCCTATCACTGCCTGAATCAAACTCTTTAAAATGCCAGGAAGAAATTTGATTTTTGCTATAAACTATG

AATATTCAGAAGATTTGACTTGTACATTATGATCAATTTAAGGTTTATGGTAGTTTGTTCATTTTTGGTA

TTAATGGCTTTTTTAGTTAATTTGAGTT

>MSTRG.3570.3 gene=MSTRG.3570

GCTTACTTACATAGCACTTGCGGGAACGCTATTGAAGTACGCTACTAACATAGCGCTTACACAAGTGTTA

TATAAGTACATTTTACTTGCATAGTGCTGGCTTAGATAACACTTAGAAACGCTATGTAAGCTTAAAAATA

AGCGCTATGTAAGTTAAAATCTGTAGTAATGTATGTTACAGTCAATGAGAAGTCAGGGAGAACTTTGTTT

TATTGGTTGACTGAAGCTGTTGATGATCATGCTGACAAGCCTCTTGTTCTCTGGCTCAATGGAGGTTTTT

CTTGATACCATTCAATTTATAGGTTTTGGGTTTGATTTCTTCTGGCTTGTCTTTGTGTTATTGCTCTTGG

AGAGTGTGTTGATTGGTTTCGAACCTGTTACTCTCGATGTTCAGAATAATGTGGATGGTAGGATTGATAT

TGCCTTGAAAATTGTATCTGTAATCCTGGAAGACGACGTTTGTTTTTTGACAATGGTGTAATCCTAATTT

AAATAAATGTGTTAAATATTAATCAGGTGATGTTTTTTCAGGCCCTGGGTGTTATCGATTACTTTTGGCA

TGGCCGCGAAGATGGGACCTTTCCATGTTAATAAAGATGGGAAGACTCTCTATTCTAATCCATTCTCTTG

GAATCAAGAGTTGTAAGGTGTCGAGTTCAAAGCACAAAGGAGGCCACAAGGCTGTTGTTGACACCATCCA

GATGCTGCAGCCCAGACAACTTGTCAGATTGCAAATGATAAGCTGGTGTTAGCTTCACTTGAGCTCATGA

CCTTGCATAAAAGACTTGCATCTAGGATGAGGTACATCAAATATCTTGCAGTCCAAGAAGTTGAAAGGAT

CTTATGTCATGTTGCAAGCGAAAAAGTTTGGAATTTGCTTCAGAAAATATTCACTTCTAAGTTATAATCA

ACAATATGGCTTTTTCACCAAAAAATTGTGCAAGTTACCTCCTACCCAAATTCAAAAATTATGTCCATGC

ACCCAATGGTGGAAGTACTCATGTGCTGATATGTGCAAGCTCGAGAAGATTTTGAAATTTAG

>MSTRG.3572.1 gene=MSTRG.3572

TAGAAAGTTAGTATAAAAACCCAACCAAAGCCTTGCTTTATTCTTTGGTTCATTCTTCTGTCAAAACTCA

AAAGAGAAAAGGGAAGTCAGGAGCTTTAACTTGGAAGTCTATTAAACAAGAATTGGTGTTTCGTCACCAT

TATTACAGCAAGGGATTTGCTTGCAACAGTGCCTGTGGAGGAGCACAAGGGAACAAAGAAAGTAGTTGCT

TAATAAAGTCATCTGCATGTATATTATGTTTATCAGTTTATGAATTGGTCATTGTTTTGACCTTTTGAAA

CAAATGGTTAGATTTCTGTCCAAGCTCGATGTGTTTAGTTTGTGTGTGTGAATTGTTCTTGTTGATTGTG

TGGATTGATAAGCGGACTTGAAAAGCTTTTCCTTTTCTTATGGTTCAAATCATTCTAAGCCAAGACTTTG

TACTATGGTAAGTAATCTGTGGTACTGTTAACTGTTGAGCAATGGTGTAATATGATGTTTCCTCAATTGG

TTGGTATGATAAGA

>MSTRG.3581.1 gene=MSTRG.3581

AACGCTTGAATTAACAGCCTGGGCAGAAGAAATACAGCGACCCCGTGAACAAAAATGCGAGTCTCTGGAG

CTGTCTGCTTATTCATCCTGTGACCTGGAACTCATAACCACTGTACATCAAGAACAAAACACACACACAC

AGTACACGCAATCACACACACAAGCTCTCATCATTCTCAACTCGAAGCTCGATTCTCAGTTTCAGAAACA

TCACGCACTGGACATTTGTAACATGATATTGGGTTTGCTGGTCAGATTTGGGTGGGGAAATGAAAATGAA

GTTGTTGCATCAAGCTTGGATGCATGGATGGTATTAGAGAGATGATGGATATTATGTACTGACTACTGAG

ACAGTCTGGATTTTATCTGATCCTTGACACAGCTGAGTTGAAGTGTCCGGATAGGGGTTGATTGCACCAA

AGTCCAAAAGCAAAGAATATGTAGGTTGTTTCAGATCCGTTGATTAAGATTTTAATCATAATTTGAAATT

TTAAATTTTTTGCCTCACTTGATAAGTGGTTCTGTATCAGTATGTAGTATGTGCATTATTACAATATGTA

TAAGATTATGACTGGGAATGTCTAGAGATGATGGAGGTAAAATTTGTGGTTCCTTTTTGTTGTGATTAAT

ACAAAATCTTGGGTTGTCCCTGCTCGTGGTTACTTAATAATCTCTGCTGTGTGAATTATATCTGGGACTC

CCAGGTATCATGTTAAGAGAATTCGGCAGAGTTGCTTCTGCATTGGACGAAATCAGGTACTACTGGTACT

TAAAGATTGCTGCCAAAGCGTGCTCTACTGAACTGGTTTTCTGGATGTGAATCGGAATAATTGAACCAAA

ACCCTGTTTTTTTTGTTTCTGTAATGTTTAAGCTATAATTAACAATTTTAGTTGATTAGTTGAGTTGAAT

TGCTTTTTGAGTGAGTTATGACTTATGAGCAGCTGCAAGGCTTATATATACAACTGAATAATGGTGTGTG

TATATGGTTTGGTGTAAAGTAGGAAGATATAGATACAATAATAGAAACAGAGAGGGAATAACATTGTAAG

ATTTTATACTTTTCCTTCAGCTCTAGTAAAGTTATATACCAAATCCTTTCTTAACAACTACCAAAGAAAC

ACAGGTTCATAAAAGCTCC

>MSTRG.3581.2 gene=MSTRG.3581

CTCCAACGTTTTTTTATAACGCTTGAATTAACAGCCTGGGCAGAAGAAATACAGCGACCCCGTGAACAAA

AATGCGAGTCTCTGGAGCTGTCTGCTTATTCATCCTGTGACCTGGAACTCATAACCACTGTACATCAAGA

ACAAAACACACACACACAGTACACGCAATCACACACACAAGCTCTCATCATTCTCAACTCGAAGCTCGAT

TCTCAGTTTCAGAAACATCACGCACTGGACATTTGTAACATGATATTGGGTTTGCTGGTCAGATTTGGGT

GGGGAAATGAAAATGAAGTTGTTGCATCAAGCTTGGATGCATGGATGGTATTAGAGAGATGATGGATATT

ATGTACTGACTACTGAGACAGTCTGGATTTTATCTGATCCTTGACACAGCTGAGTTGAAGTGTCCGGATA

GGTATCATGTTAAGAGAATTCGGCAGAGTTGCTTCTGCATTGGACGAAATCAGGTACTACTGGTACTTAA

AGATTGCTGCCAAAGCGTGCTCTACTGAACTGGTTTTCTGGATGTGAATCGGAATAATTGAACCAAAACC

CTGTTTTTTTTGTTTCTGTAATGTTTAAGCTATAATTAACAATTTTAGTTGATTAGTTGAGTTGAATTGC

TTTTTGAGTGAGTTATGACTTATGAGCAGCTGCAAGGCTTATATATACAACTGAATAATGGTGTGTGTAT

ATGGTTTGGTGTAAAGTAGGAAGATATAGATACAATAATAGAAACAGAG

>MSTRG.3581.3 gene=MSTRG.3581

CTCCAACGTTTTTTTATAACGCTTGAATTAACAGCCTGGGCAGAAGAAATACAGCGACCCCGTGAACAAA

AATGCGAGTCTCTGGAGCTGTCTGCTTATTCATCCTGTGACCTGGAACTCATAACCACTGTACATCAAGA

ACAAAACACACACACACAGTACACGCAATCACACACACAAGCTCTCATCATTCTCAACTCGAAGCTCGAT

TCTCAGTTTCAGAAACATCACGCACTGGTATTAGAGAGATGATGGATATTATGTACTGACTACTGAGACA

GTCTGGATTTTATCTGATCCTTGACACAGCTGAGTTGAAGTGTCCGGATAGGTACATGATCACTGATTCA

CAATCTTAGAATTTGGGACTGAATGTCTGATTCAGTGGTTATCACTTAGTCGGTTATCTTATTCCAGGTG

TATATGTTATATACAAAAAACAAGTGCAGTTTTCCGTGAGTAACTTTGCTAAAAACATAGATTCCCTTGT

GCTACATGTCCGATTCTGTCACAGCTATTATAATCTTTCTTAAGCTATGGTACTGAAAACGGTTCAGTAT

TGAAGACGCCTGATTTTGGGAGTGTCTGCTAGAAATCAGCAACAAACTTAAGGAAGTTTATCTGATTTTT

ATTTTGTTATTGATATCATATTGATAACTTGGTTTATAACATGCACATTGTCCTGGATTGTAGCCTAATT

TCATAGATTAATAGTTGAGAAGTGTTTTGTGCTTTTCACTTCCTCCTGTGTGTATGAGTTGGCAATGGCG

CTGTGAACTTTTTACCTATCTATAATAACTGAGTTATCTGTCTTGCTAGCCAAATGATTGGAATGAAGAA

CTTGTAAATATACAGTTTTAATCCACCCTTTCCCTCTGTATATTTGGCTTCTTATTTCTTTGATAGTTGA

TGACAAATACAGGCTTTTTTTGTTTTTAAGCTAGCTTCTGCTTTTGCATGACCGTTTGGCAAATAAGTTT

CCAGGGCTTATTTTGGGTTCTGGGGAAAGAAGTTAGAAGCTCCAATGTGTTGCTTTTTTCTCTTGCCACA

GTTGCTTATTTGTTTAATTATCCAAACACTTAATCAACTTATAAGTTCATGACTACTTATCACTTATAAT

TTGCTTCTTTATTTTAAGCAATAAGTAACTTCTTTTAAGCTAAGCCAAACGGCATATATACGTGGTGTTT

GGCAACGGCTTATAAGCCCGACTTCTGGGTTTATAAGTTAGAAGCACTTATTTGTACCGTTTGTGTAACA

AGTCAAGAAGTACTTAAAAGAAGTTAGGATTACTAGCTTTTGTTTTAGTACTTCTACTTATTTCCCAAAC

AGTTTAATCACTTATAAG

>MSTRG.3585.1 gene=MSTRG.3585

AAAAGAGAAGGAAAAGAAAAAGTAGACTACGTGGCAAGTTCTAGTGAAGTGGTGCTCACTTCTTGCTTCT

TGTGGCTGTCCGAACTCTCCTTCAAGAACCTCCCCACTTCCCAGTTCTTACATACGACAGCATATATGTA

TATGGCAAGTACAAGTAACCCAAGTGTGTCACTGCCTGAACTCCCTCAAAGATTTGTAAGAAAGATTTCA

TTTTCAACATCTTTTGATAAAACCCATGTTTTTAGAAAGTCAAGAAATGTTGTGAGGCCAAGAATTGTTG

ATGGGTGTGAAGCAAATTTTAGGTTGAGGTGTTCTGTTGCAGATGTTGATTCTACTAAAGAGAAAAACAC

TCCTGCCCATGTTGAAGTTAAGACTTGTATTTGGAAATGGAAGGGTTATTCGATTCGCTATCAGTGTGCT

GGGAGCAGTGGACCAGCATTGGTTTTAGTTCACGGGTTTGGAGCAAACAGTGATCACTGGAGGAAAAATA

TACCGGTTCTTGCACAGTCACAGAGAGTATATGCAATTGATCTTATTGGTTATGGGTATTCAGAGAAACC

CAATCCTAGTGATTTTGGAAGTGACCCGTTCTATACCTTTGAGACATGGGGTGCACAATTGAATGATTTT

TGCACTGATGTGGTTAAAGATGAAGCATTCTTCATATGCAATTCTATCGGAGGACTTGTTGGCCTTCAAG

CAGCGATTATGGAGCCACGGATCTGTAAAGGCATTCTCCTGCTTAATATATCTCTTCGAATGCTTCATAT

AACTAAGCAGCCATGGTTTGGAAGGCCTCTGATTCGAGCATTTCAGAATCTCCTAAGAAATACTGCATTG

GGAAAATACTTCTTCAAATCTGTTGCCACCCCAGATTCTGTGAAGAATATCCTTTGCCAGTGTTACCATG

ACACTTCTCAAGTGACAGACGAATTAGTCCAGGCCATACTTCTTCCTGGACTTGAGCCAGGTGCTGTGGA

CGTTTTTCTCGAGTTCATCTGTTATTCAGATGGCCCTCTTCCTGAGGAGCTCCTGCCTCAAGTGAAGTGT

CCTGTTCTGGTTGCATGGGGTGACAAAGATCCATGGGAACCAATCCAACTTGGAAGAGCCTATGGAGATT

TTGATACTGTCGAAGATTTTGTTGTTCTACCTAATGTTGGCCACTGCCCTCAGGATGAAGCACCACAGCT

TGTAAACCCACTTGTGGAGTCTTTCGTTGCTCGCCATGCTACACCAACATACTAGTTTTCCCTTGAAGGC

TATATGTCCAAGGGAAGGTCAGTTCACTACTCGTCTCCATTGACAATGAACATACAAGTTCCATGTCAGA

TCCGTGGATTTTGGAATTGACAGTGTACATTTTTGCTGTGCAATCAGATGACTAATACATAGTTTCAGCA

TGTTGCTACCTTTTTATCATGTAAATGAATCCATTACTATCACTTTGCTTTAGTAGTGCTACAGGTACTC

ACCCTTAACGCGTCCCTCTGTCTTGAACTTATTTATATATTGAAATTCCATGCATGCAGATTAAACTTTG

CTAATTTGGTATCAGCTTGTGCCTGAATTATTACATACTAACTAACCAAATCAAGTTGTGTATGCCTGTA

TGTCTTATGTCTTACTTTTGGAGACGGTAATAAGGTGCACCATTAATATAGAGATGTTTCAGTATATCTC

AAAGGCTATTCCATGAAATATCCCAG

>MSTRG.3589.1 gene=MSTRG.3589

ACCAGCATGTTCGTTTTTATATGACTATTGAATCAAACAATACAAATGTGTTGCTGAGTCTCATTTTGCA

AACTGGGGGTTGTATTATCTGTAGTTTGTGTTTGCAAAGCAAGCAACTCTGATGCAAGCCAACAAAGTGG

CAATTTTTTCTTGTTTGGCAGGTGGTCAGAGTATTTCTGCAGTCTAAACTGTGAAAACCGAACATGCATT

TTGTTAATT

>MSTRG.3597.1 gene=MSTRG.3597

TTATTACAAGATATAGATAGATGCCGTTAGCTCCTCGTACTCTTCGAGCTCCCACGCGCATATGATAACC

GCCGCCCAATTGTATGACGGTCTTCGCGAGTTCTTCGCCGAACATTTCTCCTTCATCTCCTTCAGGACGA

GCATGTAAATACACTGGCCAATCCCTGAGAGTGTTGCCAGTCAGGGTTTTAACAGTGGGAAAGAATAGGT

CAGCTGGTGTGCAGCTGATAGTTGATGAGTATGTTGATAAGATCAGCAAGTATTGCAGGATTGAGGATGT

GCAGATTAGGTCTAATCCGAAGAATGCTCGTGATGTCATGGCACAAATTGAACAGGAAGACATGGCTGTA

ATGAAAACTATCAAGCATGGCGATTGGGTAAAGCCTTGAAAACCATTGACATGGATCATGCTGTGATTTC

AGTCATTTGGTTGTGATGCTGGATGAACATGGGCTTGATGTAGGGTCTGAGCAAGTTGCTAACTTGATCG

GGGATGCCGGAAACACGGGTGCTTCGAATTTACTGTTTTGCATTGGTGGACCATATGGTCATGGAAAGCA

ATTGCGAGAACGGGCTGATGTATCAATTAAGTTGTCATCGTTGGTCTTAAATCATCAGATTGCATTGATC

GTACTCATGGAACAGCTTTACAGGGCATGGACTATACTTAAAGGACAAAAGTACCACCATTAGATGCTTT

AAGTTTTTCTTACCTGCGATCGCAATCAGGAAGTATTACATCTCGTGATTTGGTACTGGGATCATCTGCT

GGGACTTGCATTAAGCAAATTTTTTGCTTATCGCGACCGTCTCCTCTGCAGTTCCATGATCAGAAGACTT

ATTAGAGTTCATTTCTTATTCTTCCTAGAGAGGATTTCTTGTAATGTACTCCTGGAGCTTAACTAGTAAA

GTTCAGAGTATATATCTATCAATTCCCAGATAATATTACAGGGAATTTTAGACAGTACTATTACATGTGA

AAATTTTTAAGAACAGAGATATACATGCACAAGTTTGCACTACGAAGCATATGATAATTCTTTACATGAT

TTACCTTCAGG

>MSTRG.3597.2 gene=MSTRG.3597

CATTATTACAAGATATAGATAGATGCCGTTAGCTCCTCGTACTCTTCGAGCTCCCACGCGCATATGATAA

CCGCCGCCCAATTGTATGACGGTCTTCGCGAGTTCTTCGCCGAACATTTCTCCTTCATCTCCTTCAGGAC

GAGCATGTAAATACACTGGCCAATCCCTGAGAGTGTTGCCAGTCAGGGTTTTAACAGTGGGAAAGAATAG

GTCAGCTGGTGTGCAGCTGATAGTTGATGAGTATGTTGATAAGATCAGCAAGTATTGCAGGATTGAGGAT

GTGCAGATTAGGTCTAATCCGAAGAATGCTCGTGATGTCATGGCACAAATTGAACAGGAAGACATGGCTG

TAATGAAAACTATCAAGCATGGCGATTGGGTTGTGATGCTGGATGAACATGGGCTTGATGTAGGGTCTGA

GCAAGTTGCTAACTTGATCGGGGATGCCGGAAACACGGGTGCTTCGAATTTACTGTTTTGCATTGGTGGA

CCATATGGTCATGGAAAGCAATTGCGAGAACGGGCTGATGTATCAATTAAGTTGTCATCGTTGGTCTTAA

ATCATCAGATTGCATTGATCGTACTCATGGAACAGCTTTACAGGGCATGGACTATACTTAAAGGACAAAA

GTACCACCATTAGATGCTTTAAGTTTTTCTTACCTGCGATCGCAATCAGGAAGTATTACATCTCGTGATT

TGGTACTGGGATCATCTGCTGGGACTTGCATTAAGCAAATTTTTTGCTTATCGCGACCGTCTCCTCTGCA

GTTCCATGATCAGAAGACTTATTAGAGTTCATTTCTTATTCTTCCTAGAGAGGATTTCTTGTAATGTACT

CCTGGAGCTTAACTAGTAAAGTTCAGAGTATATATCTATCAATTCCCAGATAATATTACAGGGAATTTTA

GACAGTACTATTACATGTGAAAATTTTTAAGAACAGAGATATACATGCACAAGTTTGCACTACGAAGC

>MSTRG.3597.3 gene=MSTRG.3597

AGAGGCTTCTGGTCTGGGTCCTATATAATCTGTGCGCCAGAGGCTCCTGGCTTGTGCCATTTAGATGCCT

GGAACCTTATGTGCCTTGGTATGAGCTGAAAAGTTCTACCTTGAACCGTGTCTTTCCTCAAAAGATGCCA

TCACAACTATTGCTAGTATCTGACAGTAAATTTTCTCCTGTCACCTAAACTTTGATTCCAAATTATGATA

TCATGTGCAGGTTGTGATGCTGGATGAACATGGGCTTGATGTAGGGTCTGAGCAAGTTGCTAACTTGATC

GGGGATGCCGGAAACACGGGTGCTTCGAATTTACTGTTTTGCATTGGTGGACCATATGGTCATGGAAAGC

AATTGCGAGAACGGGCTGATGTATCAATTAAGTTGTCATCGTTGGTCTTAAATCATCAGATTGCATTGAT

CGTACTCATGGAACAGCTTTACAGGGCATGGACTATACTTAAAGGACAAAAGTACCACCATTAGATGCTT

TAAGTTTTTCTTACCTGCGATCGCAATCAGGTA

>MSTRG.3598.1 gene=MSTRG.3598

AGGGGTCTTTGAGGTTATTACTGCACGTTATCGAACAACGAATGGATTTTGGAAAGGAGGAAACAAACAC

ACACTAAATTTGTTTCAAGGTTTCTGTTCTTGTGGAAAATAGAGTCGGCATCACTCTCCATGTTCACATA

TAGTTGCTGCGTGTTTAGTATGTAATCTCGATTGGAAGCAATACATCGAAGGTTATCACAATCTAACAAT

ATTGTATGAAATGTGGAAGTACGAATTTTATCCACTACCCAATCAAGCTTATTGGACGTTTCCACTAGCA

TACAATTGGGAGATGTATGGTACCCTTATTGCTGACGAAAATCTGAGGAAGAAGAGGAAGAAACGTGGAC

AAAGAGGACAAAGTCAACGTATTCGAACGGAAATGGATAATTCTCAAAGAGTTAAAACTTGTGGGAGATG

CGGTCAAGAAGGTCATACAAGACGGAGTCGCCGATGTCCACATCACGGACATTAGTTTTAAAAATATTAT

GTTGTTTTAAAAATATTATTTTGTTGTATTTTAATTATCAAGTTTAATTTTATTTTAAACTTCTTTGATA

AAA

>MSTRG.3601.15 gene=MSTRG.3601

CGGAACAGTAGAGCGAGAGAGCTGAGTGGCACAGTAACTGAAGACGATGATGATGCAGGCTCGCGTATCT

CTCTGCACCAGCTCGAAGCTTCACTCTCTATTTACGCTTACTTGTAAACTTGGAATTGTATGCGTTTTCA

TCCTCATCCCCTCTCTCTCTCTCTCTCTCTCTCTCTCTCTCTCCCCCTCTCTCCCTCTCTCTTAATTTAA

ACTTTATTTCTCTTTCACTTCTTCAATCATCTCTCTCTCTCTCTCTCAGGTACTCTCTCTCTATAAATAT

TAATTTCCTCATTATTATTCATTCACTTCAATAATCAAACACTTGCTCATTACATTTGCCCACACACAAG

AACATCCCCTGTTTTTTATCGGTTAGCCAAGAATGGGGGCGAATGATGTTCTTGCGAGGCTGAACAGCTC

TGGTTTTTTACAGACTCGAGGCTTTATTGGCGGAAAATGGGTGGATGCTTATGATGGGAAGACTATACAA

GTGAACAACCCGGCTACTGGACAGGTTATCGCAGATGTTGCATGCATGGGTACTAAAGAGGCAAATGTTG

CAATTGCTTCTGCATACGAGGCATTTCAAACTTGGGGCAAGACTACTGCTGCTGAAAGGAGCAAGTTGTT

GAAGAAGTGGTATGATTTAATAATTGCCAACAAGGAGCAACTAGGTGAATTGATAACACTGGAGCAGGGA

AAGCCTCTAAATGAAGCCATTGGGGAGGTTTCTCTAGGGGCTAGTTATATCGAGTTCTATGCAGAGGAGG

CTAAACGAATCTATGGTGATATAGTTCCAGCTAATCTCCCAAATCGCCGCCTTTTTCTAATAAAGCAGCC

TGTGGGTGTCGTCGGAGCTATAACTCCCTGGAACTTCCCTTTCGCTATGATCGCTCGGAAGGTTGGTCCT

GCCCTTGCTGCTGGCTGTACTGTGGTCATAAAGCCCTCTGAATTGACTCCCTTATGCGCTTTGGCTGCAG

CTGAGCTTTCTCGTCGAGCTGGAATTCCACCGGGTGTAGTGAATGTGGTTATGGGAATTGCACCTGAAAT

TGGGGATGCTTTACTTGCAAGTACTCAGGTAAGAAAGATCACATTCACAGGATCAACAGCTGTCGGCAAA

AAGCTGATGGCAGGTGCTGCTGGGACTGTTAAGAAGGTGTCCCTTGAACTTGGAGGAAATGCACCTTGCA

TAATTTTTGATGACGCAGATCTTGAAGTGGCCATTAAAGGAACTCTGGGGGCTAAGTTCCGTAATACTGG

GCAAACATGCATCTGTGCAAATAGAATACTTGTGCAAGAAGGTATTTATGACAAGTTTGCCAGAGCTTTG

TCTGATGCTATCAAAAATTTTAAAGTCGGGGGTGGTTTCAATGAAGGAGTCGTCCTGGGACCTCTTATAA

ATGAAGCTGCAGTGCAGAAGGTTGATACATTGCTGCAAGATGCTGTCTCAAAGGGAGCGGAAGTTCTTGT

TGGCGGTAAGAGACATGGCCTTGGATTTACATTTTATGAGCCAACAGTGATAAGTAATGTTAATAAGGAG

ATGCGCATCTCAAGTGAAGAAGTGTTTGGACCAGTTGCACCTCTGTTGCGGTTTAAGACGGAGGAAGAAG

CTATCTCCATTGCTAATGACACCGATGCAGGCCTGGCTGCTTACATATTCAGTACGAATGCTCAAAGAAC

TTGGCGAGTTGCAGAAGCTCTGGAATGTGGAATTGTTGGAATTAATGAAGGACTCGTTTCATCAGAGGTG

ACTCCACATGGAGGGGTTAAGCAGTCTGGGCTCGGCAGGGAAGGCTCTAAATATGGGATGGATGAGTATT

TGGAAATTAAGTATCTGTGTCTGGGCAATCTGTAATCGGAAACTCGAGGCACCTCCGTAAAAAGCTACTG

GTTACAAAATCAGTGCTGCATTGCACATTTTCTAGTACTGTGGTTTGGTATCTTTTGTATGTTGTAACCT

TAAACTCTCCTCTGTTCGTCGTGCTCTGTTCCTTCTGTTTCTTGTAATTGTGTCTCGTATTTCAGTTCTT

TTGTGTATATTAAAAGATGTTGCAGAAATGTTTCAAGATCCATGGCTCAATTCTTTAAACCCAACGCAAA

AAATCATCTTCAAAATAA

>MSTRG.3601.14 gene=MSTRG.3601

ATCGCTTCTCCGGGTAGAGATAAAAAAAGTGTTAGTTCGAATCGGAGGCGATATATCCGAGCGGAGTAAA

ATAAAGCGATTCATGCATAAACGCACGTGTCTAACCACAAAGCTCACGGAACAGTAGAGCGAGAGAGCTG

AGTGGCACAGTAACTGAAGACGATGATGATGCAGGCTCGCGTATCTCTCTGCACCAGCTCGAAGCTTCAC

TCTCTATTTACGCTTACTTGTAAACTTGGAATTGTATGCGTTTTCATCCTCATCCCCTCTCTCTCTCTCT

CTCTCTCTCTCTCTCTCTCCCCCTCTCTCCCTCTCTCTTAATTTAAACTTTATTTCTCTTTCACTTCTTC

AATCATCTCTCTCTCTCTCTCTCTCTCTCTCTCTCTCTCTCTCTCTCTCTCAGGTACTCTCTCTCTATAA

ATATTAATTTCCTCATTATTATTCATTCACTTCAATAATCAAACACTTGCTCATTACATTTGCCCACACA

CAAGAACATCCCCTGTTTTTTATCGGTTAGCCAAGAATGGGGGCGAATGATGTTCTTGCGAGGCTGAACA

GCTCTGGTTTTTTACAGACTCGAGGCTTTATTGGCGGAAAATGGGTGGATGCTTATGATGGGAAGACTAT

ACAAGTGAACAACCCGGCTACTGGACAGGTTATCGCAGATGTTGCATGCATGGGTACTAAAGAGGCAAAT

GTTGCAATTGCTTCTGCATACGAGGCATTTCAAACTTGGGGCAAGACTACTGCTGCTGAAAGGAGCAAGT

TGTTGAAGAAGTGGTATGATTTAATAATTGCCAACAAGGAGCAACTAGGTGAATTGATAACACTGGAGCA

GGGAAAGCCTCTAAATGAAGCCATTGGGGAGGTTTCTCTAGGGGCTAGTTATATCGAGTTCTATGCAGAG

GAGGCTAAACGAATCTATGGTGATATAGTTCCAGCTAATCTCCCAAATCGCCGCCTTTTTCTAATAAAGC

AGCCTGTGGGTGTCGTCGGAGCTATAACTCCCTGGAACTTCCCTTTCGCTATGATCGCTCGGAAGGTTGG

TCCTGCCCTTGCTGCTGGCTGTACTGTGGTCATAAAGCCCTCTGAATTGACTCCCTTATGCGCTTTGGCT

GCAGCTGAGCTTTCTCGTCGAGCTGGAATTCCACCGGGTGTAGTGAATGTGGTTATGGGAATTGCACCTG

AAATTGGGGATGCTTTACTTGCAAGTACTCAGGTAAGAAAGATCACATTCACAGGATCAACAGCTGTCGG

CAAAAAGCTGATGGCAGGTGCTGCTGGGACTGTTAAGAAGGTGTCCCTTGAACTTGGAGGAAATGCACCT

TGCATAATTTTTGATGACGCAGATCTTGAAGTGGCCATTAAAGGAACTCTGGGGGCTAAGTTCCGTAATA

CTGGGCAAACATGCATCTGTGCAAATAGAATACTTGTGCAAGAAGGTATTTATGACAAGTTTGCCAGAGC

TTTGTCTGATGCTATCAAAAATTTTAAAGTCGGGGGTGGTTTCAATGAAGGAGTCGTCCTGGGACCTCTT

ATAAATGAAGCTGCAGTGCAGAAGGTTGATACATTGCTGCAAGATGCTGTCTCAAAGGGAGCGGAAGTTC

TTGTTGGCGGTAAGAGACATGGCCTTGGATTTACATTTTATGAGCCAACAGTGATAAGTAATGTTAATAA

GGAGATGCGCATCTCAAGTGAAGAAGTGTTTGGACCAGTTGCACCTCTGTTGCGGTTTAAGACGGAGGAA

GAAGCTATCTCCATTGCTAATGACACCGATGCAGGCCTGGCTGCTTACATATTCAGTACGAATGCTCAAA

GAACTTGGCGAGTTGCAGAAGCTCTGGAATGTGGAATTGTTGGAATTAATGAAGGACTCGTTTCATCAGA

GGTGACTCCACATGGAGGGGTTAAGCAGTCTGGGCTCGGCAGGGAAGGCTCTAAATATGGGATGGATGAG

TATTTGGAAATTAAGTATCTGTGTCTGGGCAATCTGTAATCGGAAACTCGAGGCACCTCCGTAAAAAGCT

ACTGGTTACAAAATCAGTGCTGCATTGCACATTTTCTAGTACTGTGGTTTGGTATCTTTTGTATGTTGTA

ACCTTAAACTCTCCTCTGTTCGTCGTGCTCTGTTCCTTCTGTTTCTTGTAATTGTGTCTCGTATTTCAGT

TCTTTTGTGTATATTAAAAGATGTTGCAGAAATGTTTCAAGATCCATGGCTCAATTCTTTAAACCCAACG

CAAAAAATCATCTTCAAAATAA

>MSTRG.3602.1 gene=MSTRG.3602

CAATAATAAAACAACAACTAGTTAAAATGGCTTCTCTTTAAACCTCAGCGGAGTCAACATCACTCTCTAG

TCAACTTTAATGGCGGATCAGCCCAAAACGACGTCGAATCGTCCATGGATTCTCGAGCTCGTCCCTCTTC

TCGTGGTCATTCTTATAGTAGCTCACGTCCTCGCTCTGGTTTACTGGGTATACAGATTAGCTACAGAGAG

ACAACCCCAGAGGAGAAAGAAGCACTAGAATTACTTCAAGATTAAAATTACCCATATGAGAATACATGAT

CCAAAATGGATTTTATATCATCAGCAGAAGCCAAGTTTCTAATACACAATCGTAAGATAGCAATAGATAT

GCAGTCTAAATTGTATTCTGAAAGGACATCATGAAATATTGCTGTGCCGATCTGGGTGTATCTTCCTGGT

AAGTACAGTTTGGACTTGGTGCTGTAGATTTTAAGGCTAGTCATATTAAAATTTATTCATGAAACACTTT

TAGTTGTGTGTACCGCCTGAAGTACTAATCCTGTAATGATGTACAATTAAAATCTCACCTACTGATTTTT

AATTAGTACCAACTACTAAGCCTGTATTGATGTACAACAAAATTTCATGTTGTTCTCATATTGCTCAGTG

TTTTATTTTGTTTATTTCTTGATTTGAGACATGCGAAGTCAAGTGGAAACGTGTTTTATTTTGTTTATTT

CTTGATTTGAGACATGCGAAGTCAAGTGGAAACGTTATTATCCTAAGCTAAGAGGAAAAAATTGACCAAG

AAATATATATCATCAATAAGTGCAGATTAAATTATTGCA

>MSTRG.3602.2 gene=MSTRG.3602

GTTAAAATGGCTTCTCTTTAAACCTCAGCGGAGTCAACATCACTCTCTAGTCAACTTTAATGGCGGATCA

GCCCAAAACGACGTCGAATCGTCCATGGATTCTCGAGCTCGTCCCTCTTCTCGTGGTCATTCTTATAGTA

GCTCACGTCCTCGCTCTGGTTTACTGGGTATACAGATTAGCTACAGAGAGACAACCCCAGAGGAGAAAGA

AGCACTAGAATTACTTCAAGATTAAAATTACCCATATGAATACATGATCCAAAATGGATTTTATATCATC

AGCAGAAGCCAAGTTTCTAATACACAATCGTAAGATAGCAATAGATATGCAGTCTAAATTGTATTCTGAA

AGGACATCATGAAATATTGCTGTGCCGATCTGGGTGTATCTTCCTGGTAAGTACAGTTTGGACTTGGTGC

TGTAGATTTTAAGGCTAGTCATATTAAAATTTATTCATGAAACACTTTTAGTTGTGTGTACCGCCTGAAG

TACTAATCCTGTAATGATGTACAATTAAAATCTCACCTACTGATTTTTAATTAGTACCAACTACTAAGCC

TGTATTGATGTACAACAAAATTTCATGTT

>MSTRG.3602.3 gene=MSTRG.3602

AATGGCTTCTCTTTAAACCTCAGCGGAGTCAACATCACTCTCTAGTCAACTTTAATGGCGGATCAGCCCA

AAACGACGTCGAATCGTCCATGGATTCTCGAGCTCGTCCCTCTTCTCGTGGTCATTCTTATAGTAGCTCA

CGTCCTCGCTCTGGTTTACTGGGTATACAGATTAGCTACAGAGAGACAACCCCAGAGGAGAAAGAAGCAC

TAGAATTACTTCAAGATTAAAATTACCCATATGCTCCTCTGACATTATTTCTCTTCTTTGCTTCGGCCCC

TTGGGCATAAAAGTTAGTGTGAGACAGTGTGTGATTTTTAACATACACCTACATTCCATTATCTCTTTAA

TTTTCTCTCTTTAAATATTTGTATTTTATCACGCGGGTCCTGATCATCATATTCGTTTTCTTATCTACAG

AGAATACATGATCCAAAATGGATTTTATATCATCAGCAGAAGCCAAGTTTCTAATACACAATCGTAAGAT

AGCAATAGATATGCAGTCTAAATTGTATTCTGAAAGGACATCATGAAATATTGCTGTGCCGATCTGGGTG

TATCTTCCTGGTAAGTACAGTTTGGACTTGGTGCTGTAGATTTTAAGGCTAGTCATATTAAAATTTATTC

ATGAAACACTTTTAGTTGTGTGTACCGCCTGAAGTACTAATCCTGTAATGATGTACAATTAAAATCTCAC

CTACTGATTTTTAATTAGTACCAA

>MSTRG.3604.6 gene=MSTRG.3604

GTTTAGCACAAGCTGATTCTGCGGACACCTCTGACTCCTCATATGATGCACCAAACGTGGTATACATTAT

AAGATTTAAGGTTTTGTGGTTTTATGTAACTGATGTAATATGTACCGAGGAGCTACTTATGATGTAAACT

ATATGAAAATTTTAACTTTTGTTGCAGGTCAACTTGATTATACAAAAGTTTTTTATGTAGTAAACTTGCA

ATAATCATAGTGGTAACCAAAAGAAAAATAGATAGGCATAACTCAAAAATTTGTGACAGATATCAGGCAA

ATAAAATCTATTTATTGTACTTACCCGAAAAAAGACTTGATGCCCATTCATCAACAATCGCCTGCAAATT

AAACACCAATAACTTGATTGCTGTAATCAGAACTCTTAGTAATAAGTTGTAGGCTTTATGATAGAAATTA

AGCATTTTACTCTTTTTTCGCTTCTTCAAGTACACATCGCAACATTACCTGCAGATCAATATTAATTTTT

CCAATTCTTCTAATGAGTCCTGCAAGATAACGCAAGACCAATAGTCCAATATATCTACTGATATAGCAGC

TTGAACTCCTGGCCTTTGCACTTTTACTACCACTAACTGGCCAGTAGACTGAAGCCTAGCTTGATAGACC

TCAGGAAATTTGTTCTATGAGATATAACTGCATGTAAAAATATACCCTGATAAACACAAAGTAGGACCAA

AAGAACATGACCAATTAAAATACCTGTCCTAAGGATGCTGCTGCAACAGGATCAGGCGAGATCTCAGAGA

AGGGCACATTAATTGGCAATCCGAGCTATTTTTCAATTGTTGATTGTCTTTTGATCAATAATTCTCGGTG

ATTGTTTAAACTGGATACAAGACACAATTAACGTAAACTCAATACCGGGTCTCTCAATTTATTGGACCGT

CATACCCCGGAAAATACTACAAGTATGTCTAGCCCTGATTCGGAAATCACTGTCGTGGTTCCTGAGCTGG

AAAAAATCGGTGTTAGGCGTCCAGCCTGGCTTTCCGTCAATTTGTGAGTGGCTAAGAAGAATGACAGGGC

ACTTACTGATGGTGGGTATGATCAATGAGGGGCCCCTTGATCAGCAAAAGCTCAAGCGTCTGATCAAGCC

TATTGATGGAAGGTGAGATCGTATTGGTTGCTATTGAGAAGCTTAATAAGGACGGATTTCAGGTAGTGTG

TTGACGGGACGTGACCAACAGAGTAATGTGAAAAACACCCACCTTCCTCTTGGTAGCTTGCAAGGAACTT

TGGATTCAAGCAGAACTCTTGGCAATTTAAGCACAACCTAATTTGATTTTCTGTTGAACTTATTTGCAGA

ACTCTATACAAGCAGGATTGGCCTCAACCCAGCAAAGTTATGTCTTGCAATCCACAAGTCGGGGTGCAGC

AACACCGGCTCCGAGACGGTCATGTTGCAGTTCATGAGATGTATGCTTCAAGTGAGAACGTGAATCCAAA

TACAACATCCAGAATAAAGCTTGCGTTTGGTACATCATCTCATCTTCAGTGCAAGAAATATGGGAGTCAA

CACCATGCATCAGGTTCACGTCTGGAAACTCTAACTCTCGCCGATAATTAATGCCAGATGCCATCGCATC

CTACCTAAGCATCAGGTTCCAATATTCCAAATCGAACTCAATCCAATCATCCATGTCATGGGTTTCAAAA

ATTAACTCAAGCCTCACGATATGATGCTGCATCATTGTCTAATCATGACGTTTAGTCCAAATTTGTATCT

TTATTTTTATACATGTGTATTCTGAGTATTGCCTGGTATTGTAGGTTCCCAAAAATGTTCCTGTAATACT

TTCATGTAGCTCAGAAATGTTCCGGTAGCTTTCATATATTGTATTAAGATTGCTATAGAATGTGACTTTT

TCTTACTCAATA

>MSTRG.3608.1 gene=MSTRG.3608

AGAATCATGAAAACGTAAATATAAACAATCATACTACACTCAATTATCAAAAAAACACAACTACCTTCTT

TAGTAACATAACCATTCATAATTCAATCAAAAAAAGATCGATATTTCTGGGGACTTACAAATTCGACAGC

ATCTCTAGCAGCCAAAACAAGTATATCAGCACAAGAAACAGTATCCGGACAGAACATCTCCACCACATTT

TTAGCCACTTCAATCACCTCGAATCCTCCCAACGAAGCATTCGCAGGATCAGACCTCTCCGTATCATTCC

CCTGCACCAGAACCGAACCATCACACCCTTCCACAAAGCAATCATGAAACAACAGCCTCAGAAGCCTCCC

TGGCACTCTCGGATCATTCGACGAAGCTGCTCTCACCGTGTCCTTCACCATGGTTTCCAGCATTGGACAC

GAAGCAGCATAAAAATCGAAGGCAAGCTCCGAAGAATGAGCTGAGTAACTCAGACACAGCAGCATTATAT

ATGTTAACATCATCTTCAGATCAAGGCCTAATGAAATACATGTAATTTTTAAGTGTTTGTCATGTTTTGT

CTTAGGACATGACTATTGACAAGGGGGAGGTTCGTGTAATAAAGGCAGCATTTATATAGTTTGGTGTTGT

CAGAGGCTTTTGCCTCCAGAGTTGCCATGACCAAATAAGTTCAGCTTGTCAGAGACTTTTT

>MSTRG.3610.3 gene=MSTRG.3610

TTTAGGGCTAGCATGCATGGGTAAAATATATTGATTACTTCAAGGCAGATGCTTCAAAAACAAAGCTGCA

GATCTACAAGCTTAGACAGAGTGGCAAGGGAATTTGGAATTTTGCCAGTGAGACCATTGTTTCCCAGATA

GACTCTTCTCATCTGATTCATACCCTTAAAAGCATCACCAGGAATCTCTCCAGAAAAGTTGTTATTTGCC

AGGTAGAGTGTTCGAAGAGCGCGGATTTTGTTGATGTTACTAGGAATTGGTCCGTCAAAGTTGTTGTTCA

TGAAGCTCACAGAACGCAGAGTTGGCAACTTTGACAGGGAATCAAAATTTATGGTGCCTGAAAGCCCCAT

GGATTCAAGTTGCAACCCATTTAAGACGTTTGCTTTGCAAAGCAAGCCATTCCAGTTCGCGTTGTTACCA

TTACACGTGCCTGAGAATAGAGTTATGTTTGTGTGTAAGAAATGCAATAACGAAATGTGCTCGGGGAGGG

ACAAATAAGAAGGATGGTAATTGGTATTGACGAGGTCTTTCCTGCAAGCTATGAGCTGTTTAAATCAAGC

CGGCATCGCTCAGTTTCAACACAGTGTTGGCCAGTTTGGCTTGATGGCATGCTTAAGATAATCTGGTATT

TGTAGTTTCCTTGTTTACTATGATGCTGACATATAATACTACTAAATTATATGCATAATCTTGTTTGCTC

TTGTTATTGATGATTAAGTTTTTTAATCTGTATTGTAATCTCAGCCGTCTCACTACTACTTACAAC

>MSTRG.3614.1 gene=MSTRG.3614

TACATTTATAATTTTTTATAAAGAAAAAGATATTGAATACCAAATAAAAGTACACATTTATTGATAAAAT

AAATTACTTATCATATGACATAGACATGGATCTATTGTTCAAAACAAGAATGAAGTGTGTTCTTGCGGGT

CGTCACTTCCGGCGATCTGGGTCGCGACCCTGAACGTTTCCGTTCGCGTTTCCGACCGTATCGGGTACGA

ACGATCTAGATCGCGGTTCGTGGCAGCGTACCCGTTTCCTTGTATCTATGATTTTGACA

>MSTRG.3619.1 gene=MSTRG.3619

TTTGAGTGGAAACAACATTTCAGGAGTCATTCCCACTTCCATTGGAAATTTAACTCAATTAATCGGTCTG

GCTGTATATGATAATATGCTCGAGGGAAGCATTCCCACTCGGTTGTTTAACATCTCTACTATGGAGAATG

TGGCCCTGGCTAACAACAGGCTTGGAGGTGAAATTCCTGAGCAGATTGGTTTTTGGTCCCGGTGCCTTAG

CCTTGATCTGTCCCAAAATCTATTCACCGGGGTGCTTCCATCTCGCGTTGGAAGGTTGAAACAATTAGTT

CAACTCGATGTTTCGTACAACAAATTAACAGGAGGTATACCCGTGACTCTGGGTGAATGTGTGATGCTTG

AGGAGCTATACATGGAAGCAAATCTTTTTCAAGGTAAAATTCCGTCTTCCTTCAAAGCATTAAAAAATCT

TGCAGTTCTAGATCTATCAACAAATAATCTCTCTGGGGATATCCCGAGATTTTTTGATAATTTTCATATG

ATATTTCTCAACTTGTCACACAACAAGCTTGGAGGTGAAGTGTCAAGAAAAGGGCTTTTTTCAAACGCAA

GTGCATTTTCGGTCGTTGGAAATCTTGAACTTTGTGGGGGTATTCAAGCATTGCAGTTTCCTGCTTGTCC

TGTAAAAGTCTTGGGAAGTAGGAAAAAAACATTTCCCCTGAGAATGATACTACTCATAGTTCTTGTACCC

CTCGGCGTCTTGTTAGCATGCCTTGCCTTTATTTGTCTTCGTCGTCGAAATTCCAAGAAGTTGGATGACC

CTGCCTTGGTGTTGAGGGGAGACCAATTTCCTAAACTTTCATACCAAGATCTTCGCCTAGCTACAAATGA

GTTTTCCGTGGATAATTTTTTAGGTCAAGGAAGATATGGTTCAGTTTACAAAGGAGTTCTCGAATCAGGG

GAAGACATAGTTGCTGTGAAGGTACTGAATATTGAAGTACGTGGAGCCAACAAGAGTTTTTTGGCAGAGT

GTGAAACATTGAGGAATATTCGTCATCGGAATCTTCTCAAGATCATCACAGCATGCTCTAGCACTGATTT

TAAGGGCAATGACTTCAAGGCATTGGTTTTTGAGTTCATGACTAATGGGAGTGTGGACGACTGGTTGCAT

CCGACGCCTTCCTATCAAGGGAATGAAAGAAACTTAAATCTACTCCGGAGATTAAGTATTGTCATTGATG

TTGCACTTGGAGTCGATTACCTACATCACCACAGTCATGCAAGTATCATACATTCTGACCTAAAGCCAAG

TAATATTCTTCTTGATGAGGATTTTATCGCTCATATTGGTGATTTTGGTTTAGCGAGGATTTCTTTTGCT

ACTACAAATGAAATCAATCAAACACAAATGAGTTCAACTGGTATACGTGGAACAATTGGTTATGTTCCCC

CAGAGTATGGAATGGGTGGCGAGATATCCATAAAAGGAGATGTATATAGCTATGGAATTCTTCTTTTAGA

AATTTTCTCGGGGAAACGACCTACTGAAAGGAGCATATTCATTGATGGTGATAGTAATCTTCATGACTAT

GTGAGAAGGGCACTCCCACATAGTGTGATGGACATTGCTGACCCCCGAATTGTACTAGAGCCAGAAGAAA

ATGGCTTGACACTAAATCAATCATACAGCAGGTCTGTCATGGAAGTATGCCTGGCATCAATATTTGAAGT

GGGGATATTATGTTCTGCAGAGGAGCCTCAAAAGCGCATTGACATTAGTGTTGCTATCAAGCTGCTACAT

GCGGCAAGAGACAAGCTTCTGCAGCGCAGACCATAACCTGGGTGAGAGTTGCAATGTCTGTAATTATATA

CCAGATATAGGCATCTTGATATTTTAAACTCCTAATCTGAAAAGGAAGAATTTTACCTATTTATTTTTCG

CATTTCTATCCTTTTTATATTGATAACGTGCTGAGAACTCTGTATTTGCTCAAAACCCAGAAGCTTGCTG

GCTCCTTCTTGCTGAAGATGTTGCCCCGGG

>MSTRG.3620.1 gene=MSTRG.3620

ACATTGCAACTCTCACCCAGGTTATGGTCTGCGCTGCAGAAGCTTGTCTCTTGCCGCATGTAGCAGCTTG

ATAGCAACACTAATGTCAATGCGCTTTTGAGGCTCCTCTGCAGAACATAATATCCCCACTTCAAATATTG

ATGCCAGGCATACTTCCATGACAGACCTGCTGTATGATTGATTTAGTGTCAAGCCATTTTCTTCTGGCTC

TAGTACAATTCGGG

>MSTRG.3622.6 gene=MSTRG.3622

GGCATGGACAGACGTCAAATTAACTTCTCCTCAAATGTCAAGGGAAACTACATATAAGATGTCACTAAAG

CATCTGCTGTTGCAGTGTACTGCTTCGAGCGACTCAAGGAACCCCTTTGTAGAGTATGCATTGCAATATT

CTGTGGCAGCAGCATATGCGAATTTCAACAGTGATAAGAAGAATGTTTTGCACAAACTACTTTTAAAAGG

>MSTRG.3628.1 gene=MSTRG.3628

AGGATGTGCCCTTGCTTTCTTTAACAAGAAAAGCACTCGGGAGAATGAATATGAGCATCATTATTTGTGA

CTCACTTGATGATGCTCGTAACATAATGAAAGGCATCTCCAAAATGCACAGTATGGGAGGAGCTTTTCTT

ACCCTACAAAGGGTCACATTCGTGATGCTATACGGCACATCATCGAAAATGCCATTCAACTCAATGCCAT

TCAACTCCGGTTGTGCGGCACTGTTGCTGATTTGAAAAGCAAAAACATGGTTGAGGTTTCTTATCTGCCT

AAGTACACTAGCCCGCCTATTAAATTCACTAGTTACCATGATTTCCGATCCAAAATAGAAATTCAAAATA

AAGATATTTGTGACTGTGTTGA

>MSTRG.3631.1 gene=MSTRG.3631

CCCTCGCCTTCAAATTGCCAACCATGGTATAGACATTCCAGTCTCCCATCAACCAATTGACCTTCAGAGA

GTTTTGCCAACCTGTGGGGACACCTATCTTCATAACACTGAAGCTCCCCATTCCCATCTTTATACAACAC

AACTTGCTTATCAAACACAGTGAGCCCCAAAGGAGAATCATCTGGAATGTTCTTGGTCAAATACAAAGGA

TACCATTCTTGAGTCCAATCATACTCAACCACTTCTCTCACACTCCCCAGCCCTTCATTCACATCAATCA

CAACATCTTGATCACCACTTTTATCACTTTCCACGGAAACCCCACCACTTATATCAGTTGCAACTGCACA

AGAACTCAAGAATTTGGCCTTAAAATGGTGGGTATTATGTGGTTGGAGTATAGATACAGAAGGGTTTGGT

TTCTTTGTGTGTATAGATGGTCTTGGCTTGTGT

>MSTRG.3633.1 gene=MSTRG.3633

CTTGACAGATGTTCATCGTCTTGGATAATACAGGACAATTTGTCGGTTTTTTGCTGGAGCAAACATATTG

ATGGAGTGTGTGCTTTCAAATGCTGCGGTTGCAAGAAGTTTTACTGAGTACTTATGTTGTGCTTTGAGGA

AAATGATCCAAATTCATGGAGATTTCAGGTGGATGGACTGGTAAAGGGTTACAACATGTTGGATGGTATA

CCAGGAAGGAAAAGTTCTATGCTTAAAAGCGTTCTGAATTTATAAGTTAGAAGTACTTATTCGTACTGTT

TGTGTAAAAAGTCAAGAAGTATTTATAAGGAGTTGAGAATATTGGTGGTGTTTGTTTACAAGAAGCTGAA

GCTGCTTGTGACTTCTGCTTTTCTTGACCCGTTTGTGTAAAGAAGTAGAAGC

>MSTRG.3641.1 gene=MSTRG.3641

ATCACTTACTGGTCACTCTGATGTAACATATGTTTCTTATAGCTGTCACTTAGTAGATGGGCAATGACTA

TAGTAGCCTCATGGGCACCACTTCATCTACCAACCACATGGTGAAGAGAAGTCACACTCAACCTAGGAAC

ACAATTCTCCCCATTACCAAAACAAAAAAGAGTGTCTTGGTGCATGTGGTTCCATTACTTTGAACCTTTC

ACGTGTTCCGTGAATTTAACTCGACAGAATCATTTATGTGTAAACACCTAGGTCCTCACCAAGCCTGACC

TTCTAGCGTTTATAAACTGAACGCAAAGCACTCCCCTTCCTCACAAATCTTCCCCCATCCCGAAACAATC

ATCGTTCATTCATTCTTTAACTCCTCTTGTTCCTTCAAACTTCTCTTCACACTCTCAAAAGACAGGGAGA

GTCATGGCACTCTCACGTGCCATATTACTCATGCTTACCTTGTCATTCATGCTCGCTGTCTTGTCTTCTG

GTCAAGCTCCAGCCATGTCCCCCTCCACCACCATGGCACCGCTACCTCTGGTGATGTTACCTACTCCGAC

CATACCACCTCCTACAATGCTGGCACCATCACCAATGGCTAGTCCGCCTAACACGGTGACTCCTCCAGCC

ATGAGCCCTGAGTCGATCAATTCACCACCTGCGCCAATGGCACCTGCAATGGGACCCACGGCACCGACAC

CTTCCATGACGGGCACACCACCCATCGCATCCACTCCAGGCTCTGCAGCTTTCAAGCATAAGAGCAGCCT

GTCCATGCTTGCATTGTTAGGAGCAGTAGTAATTCTAGTTTAAGTTCTGTTGACGGTCTTACTTACCTCT

AGTTAACGAGTAGTATATATGTATTATCTTGTGCTTTTGTTATTTCACTTATCGGGTTGTAACTTGTGCA

TTTCTCCTAGAAATGAACTGACTGTTATATATGTTACAAATATCTCGCATAATTTTGAGTTAAGGC

>MSTRG.3642.2 gene=MSTRG.3642

TACAGATTAGGGTTCCTGTCATCAGAAGTATCACTGGCACAGAAGTCGAGAAGTATGACGTTACCGTCTA

CAACAAAAACGGAAACGACGCCGGCATCACCGCCGTCGCAGAAGTTGGGCGTAAAAAGGATCCGATCATC

TTCCCCGAGAAAGAGAAAGAGAAATCGAGATTTGGATGCAAAGATAAGTGAAGCCACGAGTCCACAGCTC

CCGGAAGAATTGATCGAATCGGAGATACTGACACGCCTCCCAATCAAATCAGCGGTACGTTTCCAATGCG

TGTGCAAGTCATGGCGATCTCTATTCTCCGATCCACAGTTTGTAAATCAGCACAGCACTCGCCAAAACCC

TAACCACCGCGACTGCGTCGTGGCTGATCGAGCAGGCATTATCAGTATTCAGTCCCGCACCGAAGCGATT

TTGTCCTTTCCTAGGGATCGGGGGTCCTGTATGATCGGTTCTGCCAATGGATTAGTTTGCATCCGTTTTC

GCAACATTCTGTCCTTGTGGAACCCCGCAATTCACAAATCCAAGGAAATTCACATTCCGCCCCTAGGAGT

AGGAGTTCGCGTGGATATGGTTGGCCTCGGCTTTGATCCGATAAGTAATGATTACAAGGTGGCTGTCTCG

TACAAGCCTCGAGTGTCAGGTTGTTCTTCAGGTTATGGATATGCTGTTTACTCTGTTAATTCTGTTTCAT

GGTTACATTACCGCTTTGATGATCCTGTTTACGAGATGCTTATTAGTAAGTCATGGGATTCACTCAATCC

AACTACCGTGGTCAAGGATTCTCCTTATTGGACTTGTTCTGCCAAGACTTGTTCTGCTCGTAGGAACTAT

GCTTTGTGTTTTTCATTGAACGCACTCAAGTTTGATGCTACGAGTAACAGGTTCAAGCTGTTGCCTCGCT

TCTCCTCTGATTATGATTTTAATTTTAATTTTAGAAAGATATTTAAAGTCTTGAATATGAGGGATTTTCT

TACTTTAATGGCCTATGAACCATGTATAACTGGTAGAGCGGATGTGTTACATGTGTATTCTTTGGATGAG

GACAAGGTATGTGGTGGTTTTTGGAGTAAGATATACCGTTTGGGGCCATTTGATAATTTTTGGCGTCCTC

GGAACTTGGAACAAGGATTTAAGTATGGTGATGAGGTTGTATTACATGGTTACGGTAAGATTTTCCGCTA

TGACCAGAAAACAGATAAGTTGGAGTTCATTCCGGGTACTGCACATCGTACCTGCAGGCACTGCTATACT

TATACGCCTAGTCTGGTTTTCCTTCAAGGAATGACGTCTATTTATTCAGAGAACCAGAGTCCTCCGCTTG

GTCACCACTCTACAACTCCTCAGCGATTAATTGGTTCATTATTAAGAGAGAAATGATTTGCCAGTTTAGA

AGACAATTAAGGCCCATGGCCAGTTAAAAGAAGTGTAAGGCAAGAAAGCCACTATTACTGATTTGGGATC

TATGCTTTAGATCTCTTATGCTCATACTCAGACTTATGCTCGTATTTTGAAATTGGAAAATTATTTATAG

CTACGTTAGTTTCTTTCATGCCATTTAACATGGGTACTTTTTTATCATAAGCCTTCCTGCTACTGATTTG

CTTGCTAATATCTACATAGATCATCAGAAAACCAACCAATAAACCGTTGCGAAATCTTTCGTATTTTATC

AATTAGCTGTGTAGTTAACAAGGCTAAAGATCCCACCGAGTTTATGGACTAGACGAGACAAGCTGTAAGA

GTTCAATCAGTATATTGAAAGGTTGATTGATAAATTCTTTAGCACATCATTCCATGGCCCTTGCTCTGTA

TTTCAGAATCATTGAATGCTAAATAACCACACCTTTAATGTTATGCAAGATATCTTTTTTTTTTTTAAGT

ATCATTACTAATGACTTTATGTTGCTTTTGTAGGCCGAAAACTTGTGTGGAAGGAAAATTTGAACATTCT

TTTTATTCAGGTTTTAAAGCACCAACAAGTTATTGACGAGCTGATGGCAGAAAACAAGAAGCTCCGAGAA

ATTTTGGTAGACGACCTCAAAATTGCACCCAGCAAACTACACGGTAGCTACTCAAGTAGAAACTCTTCGC

CTTGTACTGATTGTTTTGAATGTCCTAGAAAACAAAGAAGATAGTTCAAACTTGAAAGTTAATAGTAATT

TATCAAGGTCAATCCATTGAATCTTTTGCTATATACATAATCAAGCTACCATTTTTTTAGGCAAGTTTTG

GACTCAGATGAAGGGAATATCTCAGATCATAATCAATCTATATTATACCTGGTAAAACCTTGTCAAAAGA

TTACTTTAGAAGGTCAAAGTACATGCTCTTGTAGTTACGGAGGTCACACTATATAGACCGATGCAACGAA

GGTTTGTTCTTTACTTCTATGGATGGCCTCAAGAGCCCTGTCAGAGAAGTTCAGAATTTTCCGTCATTAT

AATTGAATGGTGCATCTTCCGTCATTGATATCTATTGTGGATAGACTCCTCTATAAGCTTGAAGTTACAG

CTTACAAAGAAGTGAAAGGCAGGATTTTACACCAAGTGAAGAATCGGAAGAAAGGAGCCAATAAGGAGGC

TCTTGGAGATGCTCTTAGG

>MSTRG.3642.3 gene=MSTRG.3642

TTTTTTCTTTCCACAACTACAGATTAGGGTTCCTGTCATCAGAAGTATCACTGGCACAGAAGTCGAGAAG

TATGACGTTACCGTCTACAACAAAAACGGAAACGACGCCGGCATCACCGCCGTCGCAGAAGTTGGGCGTA

AAAAGGATCCGATCATCTTCCCCGAGAAAGAGAAAGAGAAATCGAGATTTGGATGCAAAGATAAGTGAAG

CCACGAGTCCACAGCTCCCGGAAGAATTGATCGAATCGGAGATACTGACACGCCTCCCAATCAAATCAGC

GGTACGTTTCCAATGCGTGTGCAAGTCATGGCGATCTCTATTCTCCGATCCACAGTTTGTAAATCAGCAC

AGCACTCGCCAAAACCCTAACCACCGCGACTGCGTCGTGGCTGATCGAGCAGGCATTATCAGTATTCAGT

CCCGCACCGAAGCGATTTTGTCCTTTCCTAGGGATCGGGGGTCCTGTATGATCGGTTCTGCCAATGGATT

AGTTTGCATCCGTTTTCGCAACATTCTGTCCTTGTGGAACCCCGCAATTCACAAATCCAAGGAAATTCAC

ATTCCGCCCCTAGGAGTAGGAGTTCGCGTGGATATGGTTGGCCTCGGCTTTGATCCGATAAGTAATGATT

ACAAGGTGGCTGTCTCGTACAAGCCTCGAGTGTCAGGTTGTTCTTCAGGTTATGGATATGCTGTTTACTC

TGTTAATTCTGTTTCATGGTTACATTACCGCTTTGATGATCCTGTTTACGAGATGCTTATTAGTAAGTCA

TGGGATTCACTCAATCCAACTACCGTGGTCAAGGATTCTCCTTATTGGACTTGTTCTGCCAAGACTTGTT

CTGCTCGTAGGAACTATGCTTTGTGTTTTTCATTGAACGCACTCAAGTTTGATGCTACGAGTAACAGGTT

CAAGCTGTTGCCTCGCTTCTCCTCTGATTATGATTTTAATTTTAATTTTAGAAAGATATTTAAAGTCTTG

AATATGAGGGATTTTCTTACTTTAATGGCCTATGAACCATGTATAACTGGTAGAGCGGATGTGTTACATG

TGTATTCTTTGGATGAGGACAAGGTATGTGGTGGTTTTTGGAGTAAGATATACCGTTTGGGGCCATTTGA

TAATTTTTGGCGTCCTCGGAACTTGGAACAAGGATTTAAGTATGGTGATGAGGTTGTATTACATGGTTAC

GGTAAGATTTTCCGCTATGACCAGAAAACAGATAAGTTGGAGTTCATTCCGGGTACTGCACATCGTACCT

GCAGGCACTGCTATACTTATACGCCTAGTCTGGTTTTCCTTCAAGGAATGACGTCTATTTATTCAGAGAA

CCAGAGTCCTCCGCTTGGTCACCACTCTACAACTCCTCAGCGATTAATTGGTTCATTATTAAGAGAGAAA

TGATTTGCCAGTTTAGAAGACAATTAAGGCCCATGGCCAGTTAAAAGAAGTGTAAGGCAAGAAAGCCACT

ATTACTGATTTGGGATCTATGCTTTAGATCTCTTATGCTCATACTCAGACTTATGCTCGTATTTTGAAAT

TGGAAAATTATTTATAGCTACGTATTTTATCAATTAGCTGTGTAGTTAACAAGGCTAAAGATCCCACCGA

GTTTATGGACTAGACGAGACAAGCTGTAAGAGTTCAATCAGTATATTGAAAGGTTGATTGATAAATTCTT

TAGCACATCATTCCATGGCCCTTGCTCTGTATTTCAGAATCATTGAATGCTAAATAACCACACCTTTAAT

GTTATGCAAGATATCTTTTTTTTTTTTAAGTATCATTACTAATGACTTTATGTTGCTTTTGTAGGCCGAA

AACTTGTGTGGAAGGAAAATTTGAACATTCTTTTTATTCAGGTTTTAAAGCACCAACAAGTTATTGACGA

GCTGATGGCAGAAAACAAGAAGCTCCGAGAAATTTTGGTAGACGACCTCAAAATTGCACCCAGCAAACTA

CACGGTAGCTACTCAAGTAGAAACTCTTCGCCTTGTACTGATTGTTTTGAATGTCCTAGAAAACAAAGAA

GATAGTTCAAACTTGAAAGTTAATAGTAATTTATCAAGGTCAATCCATTGAATCTTTTGCTATATACATA

ATCAAGCTACCATTTTTTTAGGCAAGTTTTGGACTCAGATGAAGGGAATATCTCAGATCATAATCAATCT

ATATTATACCTGGTAAAACCTTGTCAAAAGATTACTTTAGAAGGTCAAAGTACATGCTCTTGTAGTTACG

GAGGTCACACTATATAGACCGATGCAACGAAGGTTTGTTCTTTACTTCTATGGATGGCCTCAAGAGCCCT

GTCAGAGAAGTTCAGAATTTTCCGTCATTATAATTGAATGGTGCATCTTCCGTCATTGATATCTATTGTG

GATAGACTCCTCTATAAGCTTGAAGTTACAGCTTACAAAGAAGTGAAAGGCAGGATTTTACACCAAGTGA

AGAATCGGAAGAAAGGTACAAATTCTGATTCCATTACTGTTATCTTGCAGGTGGAACCCAAAGCTTTTAT

ATCTCCTTCAATATAAAGACACCTCTGCACATTCTTCCCCACCCCCACTCCCTCTTTAACCCCACACTCA

CCCAACAACCCACACAAACAATGACACACGCAGAGGCTTCTTTCTTGATTCTTCTCTTTTACAACAATAA

TAGCTACTAGCAACTCGTGGTAGCCTATTTCAGAGCACTACATGTCTGCAGGAC

>MSTRG.3642.4 gene=MSTRG.3642

ATTAGGGTTCCTGTCATCAGAAGTATCACTGGCACAGAAGTCGAGAAGTATGACGTTACCGTCTACAACA

AAAACGGAAACGACGCCGGCATCACCGCCGTCGCAGAAGTTGGGCGTAAAAAGGATCCGATCATCTTCCC

CGAGAAAGAGAAAGAGAAATCGAGATTTGGATGCAAAGATAAGTGAAGCCACGAGTCCACAGCTCCCGGA

AGAATTGATCGAATCGGAGATACTGACACGCCTCCCAATCAAATCAGCGGTACGTTTCCAATGCGTGTGC

AAGTCATGGCGATCTCTATTCTCCGATCCACAGTTTGTAAATCAGCACAGCACTCGCCAAAACCCTAACC

ACCGCGACTGCGTCGTGGCTGATCGAGCAGGCATTATCAGTATTCAGTCCCGCACCGAAGCGATTTTGTC

CTTTCCTAGGGATCGGGGGTCCTGTATGATCGGTTCTGCCAATGGATTAGTTTGCATCCGTTTTCGCAAC

ATTCTGTCCTTGTGGAACCCCGCAATTCACAAATCCAAGGAAATTCACATTCCGCCCCTAGGAGTAGGAG

TTCGCGTGGATATGGTTGGCCTCGGCTTTGATCCGATAAGTAATGATTACAAGGTGGCTGTCTCGTACAA

GCCTCGAGTGTCAGGTTGTTCTTCAGGTTATGGATATGCTGTTTACTCTGTTAATTCTGTTTCATGGTTA

CATTACCGCTTTGATGATCCTGTTTACGAGATGCTTATTAGTAAGTCATGGGATTCACTCAATCCAACTA

CCGTGGTCAAGGATTCTCCTTATTGGACTTGTTCTGCCAAGACTTGTTCTGCTCGTAGGAACTATGCTTT

GTGTTTTTCATTGAACGCACTCAAGTTTGATGCTACGAGTAACAGGTTCAAGCTGTTGCCTCGCTTCTCC

TCTGATTATGATTTTAATTTTAATTTTAGAAAGATATTTAAAGTCTTGAATATGAGGGATTTTCTTACTT

TAATGGCCTATGAACCATGTATAACTGGTAGAGCGGATGTGTTACATGTGTATTCTTTGGATGAGGACAA

GGTATGTGGTGGTTTTTGGAGTAAGATATACCGTTTGGGGCCATTTGATAATTTTTGGCGTCCTCGGAAC

TTGGAACAAGGATTTAAGTATGGTGATGAGGTTGTATTACATGGTTACGGTAAGATTTTCCGCTATGACC

AGAAAACAGATAAGTTGGAGTTCATTCCGGGTACTGCACATCGTACCTGCAGGCACTGCTATACTTATAC

GCCTAGTCTGGTTTTCCTTCAAGGAATGACGTCTATTTATTCAGAGAACCAGAGTCCTCCGCTTGGTCAC

CACTCTACAACTCCTCAGCGATTAATTGGTTCATTATTAAGAGAGAAATGATTTGCCAGTTTAGAAGACA

ATTAAGGCCCATGGCCAGTTAAAAGAAGTGTAAGGCCGAAAACTTGTGTGGAAGGAAAATTTGAACATTC

TTTTTATTCAGGTTTTAAAGCACCAACAAGTTATTGACGAGCTGATGGCAGAAAACAAGAAGCTCCGAGA

AATTTTGGTAGACGACCTCAAAATTGCACCCAGCAAACTACACGGTAGCTACTCAAGTAGAAACTCTTCG

CCTTGTACTGATTGTTTTGAATGTCCTAGAAAACAAAGAAGATAGTTCAAACTTGAAAGTTAATAGTAAT

TTATCAAGGTCAATCCATTGAATCTTTTGCTATATACATAATCAAGCTACCATTTTTTTAGGCAAGTTTT

GGACTCAGATGAAGGGAATATCTCAGATCATAATCAATCTATATTATACCTGGTAAAACCTTGTCAAAAG

ATTACTTTAGAAGGTCAAAGTACATGCTCTTGTAGTTACGGAGGTCACACTATATAGACCGATGCAACGA

AGGTTTGTTCTTTACTTCTATGGATGGCCTCAAGAGCCCTGTCAGAGAAGTTCAGAATTTTCCGTCATTA

TAATTGAATGGTGCATCTTCCGTCATTGATATCTATTGTGGATAGACTCCTCTATAAGCTTGAAGTTACA

GCTTACAAAGAAGTGAAAGGCAGGATTTTACACCAAGTGAAGAATCGGAAGAAAGGTACAAATTCTGATT

CCATTACTGTTATCTTGCAGGTGGAACCCAAAGCTTTTATATCTCCTTCAATATAAAGACACCTCTGCAC

ATTCTTCCCCACCCCCACTCCCTCTTTAACCCCACACTCACCCAACAACCCACACAAACAATGACACACG

CAGAGGCTTCTTTCTTGATTCTTCTCTTTTACAACAATAATAGCTACTAGCAACTCGTGGTAGCCTATTT

CAG

>MSTRG.3642.5 gene=MSTRG.3642

ATTAGGGTTCCTGTCATCAGAAGTATCACTGGCACAGAAGTCGAGAAGTATGACGTTACCGTCTACAACA

AAAACGGAAACGACGCCGGCATCACCGCCGTCGCAGAAGTTGGGCGTAAAAAGGATCCGATCATCTTCCC

CGAGAAAGAGAAAGAGAAATCGAGATTTGGATGCAAAGATAAGTGAAGCCACGAGTCCACAGCTCCCGGA

AGAATTGATCGAATCGGAGATACTGACACGCCTCCCAATCAAATCAGCGGTACGTTTCCAATGCGTGTGC

AAGTCATGGCGATCTCTATTCTCCGATCCACAGTTTGTAAATCAGCACAGCACTCGCCAAAACCCTAACC

ACCGCGACTGCGTCGTGGCTGATCGAGCAGGCATTATCAGTATTCAGTCCCGCACCGAAGCGATTTTGTC

CTTTCCTAGGGATCGGGGGTCCTGTATGATCGGTTCTGCCAATGGATTAGTTTGCATCCGTTTTCGCAAC

ATTCTGTCCTTGTGGAACCCCGCAATTCACAAATCCAAGGAAATTCACATTCCGCCCCTAGGAGTAGGAG

TTCGCGTGGATATGGTTGGCCTCGGCTTTGATCCGATAAGTAATGATTACAAGGTGGCTGTCTCGTACAA

GCCTCGAGTGTCAGGTTGTTCTTCAGGTTATGGATATGCTGTTTACTCTGTTAATTCTGTTTCATGGTTA

CATTACCGCTTTGATGATCCTGTTTACGAGATGCTTATTAGTAAGTCATGGGATTCACTCAATCCAACTA

CCGTGGTCAAGGATTCTCCTTATTGGACTTGTTCTGCCAAGACTTGTTCTGCTCGTAGGAACTATGCTTT

GTGTTTTTCATTGAACGCACTCAAGTTTGATGCTACGAGTAACAGGTTCAAGCTGTTGCCTCGCTTCTCC

TCTGATTATGATTTTAATTTTAATTTTAGAAAGATATTTAAAGTCTTGAATATGAGGGATTTTCTTACTT

TAATGGCCTATGAACCATGTATAACTGGTAGAGCGGATGTGTTACATGTGTATTCTTTGGATGAGGACAA

GGTATGTGGTGGTTTTTGGAGTAAGATATACCGTTTGGGGCCATTTGATAATTTTTGGCGTCCTCGGAAC

TTGGAACAAGGATTTAAGTATGGTGATGAGGTTGTATTACATGGTTACGGTAAGATTTTCCGCTATGACC

AGAAAACAGATAAGTTGGAGTTCATTCCGGGTACTGCACATCGTACCTGCAGGCACTGCTATACTTATAC

GCCTAGTCTGGTTTTCCTTCAAGGAATGACGTCTATTTATTCAGAGAACCAGAGTCCTCCGCTTGGTCAC

CACTCTACAACTCCTCAGCGATTAATTGGTTCATTATTAAGAGAGAAATGATTTGCCAGTTTAGAAGACA

ATTAAGGCCCATGGCCAGTTAAAAGAAGTGCCGAAAACTTGTGTGGAAGGAAAATTTGAACATTCTTTTT

ATTCAGGTTTTAAAGCACCAACAAGTTATTGACGAGCTGATGGCAGAAAACAAGAAGCTCCGAGAAATTT

TGGTAGACGACCTCAAAATTGCACCCAGCAAACTACACGGTAGCTACTCAAGTAGAAACTCTTCGCCTTG

TACTGATTGTTTTGAATGTCCTAGAAAACAAAGAAGATAGTTCAAACTTGAAAGTTAATAGTAATTTATC

AAGGTCAATCCATTGAATCTTTTGCTATATACATAATCAAGCTACCATTTTTTTAGGCAAGTTTTGGACT

CAGATGAAGGGAATATCTCAGATCATAATCAATCTATATTATACCTGGTAAAACCTTGTCAAAAGATTAC

TTTAGAAGGTCAAAGTACATGCTCTTGTAGTTACGGAGGTCACACTATATAGACCGATGCAACGAAGGTT

TGTTCTTTACTTCTATGGATGGCCTCAAGAGCCCTGTCAGAGAAGTTCAGAATTTTCCGTCATTATAATT

GAATGGTGCATCTTCCGTCATTGATATCTATTGTGGATAGACTCCTCTATAAGCTTGAAGTTACAGCTTA

CAAAGAAGTGAAAGGCAGGATTTTACACCAAGTGAAGAATCGGAAGAAAGGTACAAATTCTGATTCCATT

ACTGTTATCTTGCAGGTGGAACCCAAAGCTTTTATATCTCCTTCAATATAAAGACACCTCTGCACATTCT

TCCCCACCCCCACTCCCTCTTTAACCCCACACTCACCCAACAACCCACACAAACAATGACACACGCAGAG

GCTTCTTTCTTGATTCTTCTCTTTTACAACAATAATAGCTACTAGCAACTCGTGGTAGCCTATTTCAG

>MSTRG.3642.6 gene=MSTRG.3642

TACAGATTAGGGTTCCTGTCATCAGAAGTATCACTGGCACAGAAGTCGAGAAGTATGACGTTACCGTCTA

CAACAAAAACGGAAACGACGCCGGCATCACCGCCGTCGCAGAAGTTGGGCGTAAAAAGGATCCGATCATC

TTCCCCGAGAAAGAGAAAGAGAAATCGAGATTTGGATGCAAAGATAAGTGAAGCCACGAGTCCACAGCTC

CCGGAAGAATTGATCGAATCGGAGATACTGACACGCCTCCCAATCAAATCAGCGGTACGTTTCCAATGCG

TGTGCAAGTCATGGCGATCTCTATTCTCCGATCCACAGTTTGTAAATCAGCACAGCACTCGCCAAAACCC

TAACCACCGCGACTGCGTCGTGGCTGATCGAGCAGGCATTATCAGTATTCAGTCCCGCACCGAAGCGATT

TTGTCCTTTCCTAGGGATCGGGGGTCCTGTATGATCGGTTCTGCCAATGGATTAGTTTGCATCCGTTTTC

GCAACATTCTGTCCTTGTGGAACCCCGCAATTCACAAATCCAAGGAAATTCACATTCCGCCCCTAGGAGT

AGGAGTTCGCGTGGATATGGTTGGCCTCGGCTTTGATCCGATAAGTAATGATTACAAGGTGGCTGTCTCG

TACAAGCCTCGAGTGTCAGGTTGTTCTTCAGGTTATGGATATGCTGTTTACTCTGTTAATTCTGTTTCAT

GGTTACATTACCGCTTTGATGATCCTGTTTACGAGATGCTTATTAGTAAGTCATGGGATTCACTCAATCC

AACTACCGTGGTCAAGGATTCTCCTTATTGGACTTGTTCTGCCAAGACTTGTTCTGCTCGTAGGAACTAT

GCTTTGTGTTTTTCATTGAACGCACTCAAGTTTGATGCTACGAGTAACAGGTTCAAGCTGTTGCCTCGCT

TCTCCTCTGATTATGATTTTAATTTTAATTTTAGAAAGATATTTAAAGTCTTGAATATGAGGGATTTTCT

TACTTTAATGGCCTATGAACCATGTATAACTGGTAGAGCGGATGTGTTACATGTGTATTCTTTGGATGAG

GACAAGGTATGTGGTGGTTTTTGGAGTAAGATATACCGTTTGGGGCCATTTGATAATTTTTGGCGTCCTC

GGAACTTGGAACAAGGATTTAAGTATGGTGATGAGGTTGTATTACATGGTTACGGTAAGATTTTCCGCTA

TGACCAGAAAACAGATAAGTTGGAGTTCATTCCGGGTACTGCACATCGTACCTGCAGGCACTGCTATACT

TATACGCCTAGTCTGGTTTTCCTTCAAGGAATGACGTCTATTTATTCAGAGAACCAGAGTCCTCCGCTTG

GTCACCACTCTACAACTCCTCAGCGATTAATTGGTTCATTATTAAGAGAGAAATGATTTGCCAGTTTAGA

AGACAATTAAGGCCCATGGCCAGTTAAAAGAAGTGTAAGGCCGAAAACTTGTGTGGAAGGAAAATTTGAA

CATTCTTTTTATTCAGGTTTTAAAGCACCAACAAGTTATTGACGAGCTGATGGCAGAAAACAAGAAGCTC

CGAGAAATTTTGGTAGACGACCTCAAAATTGCACCCAGCAAACTACACGGCAAGTTTTGGACTCAGATGA

AGGGAATATCTCAGATCATAATCAATCTATATTATACCTGGTAAAACCTTGTCAAAAGATTACTTTAGAA

GGTCAAAGTACATGCTCTTGTAGTTACGGAGGTCACACTATATAGACCGATGCAACGAAGGTTTGTTCTT

TACTTCTATGGATGGCCTCAAGAGCCCTGTCAGAGAAGTTCAGAATTTTCCGTCATTATAATTGAATGGT

GCATCTTCCGTCATTGATATCTATTGTGGATAGACTCCTCTATAAGCTTGAAGTTACAGCTTACAAAGAA

GTGAAAGGCAGGATTTTACACCAAGTGAAGAATCGGAAGAAAGGTACAAATTCTGATTCCATTACTGTTA

TCTTGCAGGTGGAACCCAAAGCTTTTATATCTCCTTCAATATAAAGACACCTCTGCACATTCTTCCCCAC

CCCCACTCCCTCTTTAACCCCACACTCACCCAACAACCCACACAAACAATGAC

>MSTRG.3643.1 gene=MSTRG.3643

TGCAGACAAGATTTCCATTCACTGTCAAAATTTAGGGAGGTCTATTCCTTTTCCTCTCCTGTAGCCCTTT

GCAGATGGTATTAGGTTTGACTAGGGATTTGAATCCAGAACACATACTGCGTTAAAATTTAATACTGGAG

AAGAAGAGCATTTCATCTATCTTCTCTCTCCATGAAAGAGGTTAATTATAATCATCTAAATCTTGACAAC

TCTCCATTCTTAAACAGTGATGGTTCTGGTGATTTCACGTCCCTCTGCTCAATTAAGCAGGTAAAAACCT

ATGTTTGTTAAAGTGCTAATGATGTGTTTGATCTAACATGTCTGAAAAAGGTTCCACATGAAAAAGAAGA

AGGCAGATGGTTTATAGCAATACAGAAAACACTGAACCAAATGAAGGTCAGCATCATAATCCTCATAATC

CTGGTTTGTTGTCTTCCACTAATCCAGATTGTTCTGTTACTCCTAGACGTCCTTTGTGTCTTCCATCAAT

TTAGAGTCCACACACTAATGCTAATCTTGGTGGTTTACGCGGAAGGCATGGGAAATCACTTCTCTCTGAG

TCATTGGTTTGGAAAACACCAATGAAGGGTTCTGTACCCAGCATGAATACACAAGCTTCTCACCAATCAG

AGTTTAAGATATTCCGGGGTTTTACAGTCATCTCAATCTGGATAGTGTACCTACGTTGAGGGGGATTATT

CTGGTGATTTCATGGCTTGTGTTTTATGTTTCGTGACTAAATCCATCTCAATCTACTTCTGACTATTTTC

AATTTTTTATGAGAGACTTAGGTTCCAAATAAAAACGAAGAAGGCAAAATGTGAAAAAAGATAAAGAAAA

TATCAAACCATCTGAAGAGGGGAGTCCCAGCACTGGTGAGTTTTCTTAATGGCTGTGTGATAAAAGTGTT

AACCTCATAGAAGTTTTCTTCACTTCAGCTTTATGATTATAATCCAAGATCAGTGAGCCTGCAAACAAAC

ATAAATTAGACATTTGTATGTAAGTGTTACCCTCATTTTGTGTATTGTATGTGTTTTTCATTGTTTTGCT

TTTTTAGGTGGTCTCCAGAGCTTTTAGTAATTGTTTTCACTAACATTCTATGTTTCAAGGTCAATCTTTA

AACTTAGTATTATTGTTT

>MSTRG.3643.2 gene=MSTRG.3643

GGTTTGTTGTCTTCCACTAATCCAGATTGTTCTGTTACTCCTAGACGTCCTTTGTGTCTTCCATCAATTT

AGAGTCCACACACTAATGCTAATCTTGGTGGTTTACGCGGAAGGCATGGGAAATCACTTCTCTCTGAGTC

ATTGGTTTGGAAAACACCAATGAAGGATGGATTTTAGGGGTTTTACAGTCATCTCAATCTGGATAGTGTA

CCTACGTTGAGGGGGATTATTCTGGT

>MSTRG.3643.3 gene=MSTRG.3643

GGTTCTGTACCCAGCATGAATACACAAGCTTCTCACCAATCAGAGTTTAAGATATTCCGGTTCTGTATTC

AACATGAATACACAACTGTCTCCCAATTCTCGCTCTCCACTGTCAAATTTGACCAATCACAGTTTAAGAT

CATCTGGTGTTCCTTCCAAATTTGAAGTGATGTGTAATTATTTCAGACAACATTTGCATTTATAACCAAA

TTTAAATGCAAACTATGTTTCTTTGTTGTTAGGTGAAGGTATCAAGATGAATGCACA

>MSTRG.3643.4 gene=MSTRG.3643

TGCAGACAAGATTTCCATTCACTGTCAAAATTTAGGGAGGTCTATTCCTTTTCCTCTCCTGTAGCCCTTT

GCAGATGGTATTAGGTTTGACTAGGGATTTGAATCCAGAACACATACTGCGTTAAAATTTAATACTGGAG

AAGAAGAGCATTTCATCTATCTTCTCTCTCCATGAAAGAGTGATGGTTCTGGTGATTTCACGTCCCTCTG

CTCAATTAAGCAGGTAAAAACCTATGTTTGTTAAAGTGCTAATGATGTGTTTGATCTAACATGTCTGAAA

AAGGTTCCACATGAAAAAGAAGAAGGCAGATGGTTTATAGCAATACAGAAAACACTGAACCAAATGAAGG

TCAGCATCATAATCCTCATAATCCTGGTTTGTTGTCTTCCACTAATCCAGATTGTTCTGTTACTCCTAGA

CGTCCTTTGTGTCTTCCATCAATTTAGAGTATGGTCCATAAAGTTGCATGTTCTCACTTTAAAAATCCAT

TTCATTTATAATGATGATATAATGTCTAAATTTGACCAGTAATTCCAAACTAATAAAGTGAAATCTGGAC

TTCTTCCAGGTCCACACACTAATGCTAATCTTGGTGGTTTACGCGGAAGGCATGGGAAATCACTTCTCTC

TGAGTCATTGGTTTGGAAAACACCAATGAAGGGT

>MSTRG.3644.1 gene=MSTRG.3644

TTCGATTTTAACTGATACAAAGTTACAAACCAGCTCTAAGTGGTGAAGAATCGAAGAAAGAATTATCTGA

TGGCTTTTCAAATTAGGGTTCCGTCATCGGAAGTATCACTGGCACAGAAGTTGAGAAGTATGACATTACC

GCCTAAAACAAAAACGAAAACGACGTGGTCATCGCCGCCGCCGCCGCAGAGGTTGGGTGTAAAAAGGATC

CGATCATCTTCCCCGAGAAAGAGAAAGAGAAATCGAGATTTGGATGCAAAGATAAGCGAAGCCACGAGTC

CGCAGCTCCCGAAAGAATTGATCGAAGCGGAGATACTGACACGCCTCCCAATCAAATCAGCGGTACGTTT

CCAATGCGTGTGCAAGTCATGGCGATCTCTATTCTCCGATCCACAGTTTGTAAAGCAACACAGCACTCAC

CAAAACCCTAACCACCGCGACTGCCTCGTGGCTGATCGATCAGGCGTTATCAGTATTCAATCCCGCAGCG

AAGAGATTTTCTCCTTCCGTAGGGACCGAGAGTACTATATGATCGGTTCTGCCAATGGATTAGTTTGCAT

CAATTACCACAACGTTCTGTCCTTGTGGAACCCCGCAATTCGCAAATCCAAGGAATTTCACATTCCGCCT

CTAGGAGTAGGAGGTCTTGTGGACATGCTTGGCCTCGGCTTTGATCCGATAAGTAATGATTTCAAGGTGG

CCATCTCCTACAAGCCTCGGACGTTTGATCCGTCAGGTTATGGGTATGCTGTTTACTCCTCTAATTCTGT

TTCATGGATACACTACCGCTTTGATGATCCTGTTTACGAGATGCTTATTAGTAAGTCATGGGATTCACTC

AATCCAACTACCATGGTCAAGGATTCTCCTTATTGGACTTGTTCTGCCATTGTCCCTGCTCGTAGGAACT

ATGTTGTGTTTACATTGAACGTGCTCAAGTTTGATGCTACGAGTAACAGGTTCAAGCTGTTGCCTCGCTT

CTCCTCTGATTTTGATGTTGGAAAGATATTTAAAGTCTTGAATATGAGGGATTTTCTTACTTTAATGGCC

TATGAACCATGTATAACTGGTAGAGTGGATGTGTATTCTTTGGACGAGGACAAGGGATGTGGTGGTTTTT

GGAGTAAGATATACCGTTTTGGGCCATTTGATGATTTTTGGCGTCTTCGGGACTTGGAACAAGGATTCAA

GTATGGTGATGAGGTTGTATTACATGGTTACAGTAAGATTTACCGCTATGACCAGAAAACAGATACGTTG

AAGTGCATTCCGGGTACTAAACCTCTTACCTACGGGGACTGCTATACTTATACGCCTAGTCTGGTTTTCC

TTCAAGGAATGACGTCTATTTATTCAGAGAACCAGAGTCCCCCGCTTGGTCACCACGCTACAACTCCTCA

GCGATTAATTAGTTCATTATTAAGACAGAAATGATTTGCCAGTTCAGAAGACAATTAAGTGCCCGTGGCC

AGTTAAAAGAAGTGTAAGGCAAGAAAGCCACTATTTATGATTTGGGATCTATGCTTTAGATCTCTTATGC

TCAGTACTCAGACTTGTGCTCGTATTTTGAAATTGGAAACTTATTTAATAGCTATGTTAGTTTCTTTCAT

GTTTATTTTCTAGTATCTGTTCGAAGTTTTATATACCATGAAAGATGAGGGATTTGAAGAATAAACATGG

ACAGAAACAGAGACTACCAGTGATGGCAGGTAATATACATAATTGTATCCTTCTGGGCTGTATTTGGTGA

AAGTTCTATCAT

>MSTRG.3652.1 gene=MSTRG.3652

TCATAATCTGACTTTTAATTTACATTGAGTTTCTAAACAACGCAAATAATCACTTCAAAACCCCAACCCG

ACCAATCACCGGATTACCCGACCCGACCCGAAACACGCCTGAGACGAAACACACAATACCACGCTCAATT

TCCGGCAAGAACATCGCAGCTGAGGAGAGAAACCGGGTCGGGTCCTCAGCATTGTGTTTGAGCAAGAAGA

TGGTCAACGTGCAGGTTTATTTGCAAGCTGGTACAGACGCAGTACATATTGTGACATTCATCAGGGCTCC

ATGACATTCGTGGTTAAAGTTGAAGTTCCATGCTCACAGGAGATGAATCAGATGATGTATCTCCTGGGCT

GAAGAGAGGGCTTACTTAAACACATCAAGAGAAATGTCAAATACTTGACCGGCAGATGTGATCCCTCCTT

GTATTGAGGTGGGTCTCAGTGAACTGGATGTGGGGCAAAAACTAGTAATGGGGGAGCTTGAAGTCCATCC

CGCTTTAAAACTTGTTTAGTCGAAGGATGAAGCTGTTGTCAAGATCGTGGGAGCAAGTGTTTCGGATCAA

CAAAAGAACGATAAAGCTTCAGAAATTAAAAAAAATAGAATAGAACATATGATGCTTATTTCATTTTTTG

GCCCAGCCTATTTCGCCAGGATTAATGCTAATGGAAGGTAGAAATTATGTTTGCTTCATATACCCTGAAT

ATATTTGATAAGATTCAGAACAGTTTAGCTGCTATACAGCAAAGCTTCCCTTGTCAATATGATTTCCATG

TTATCAACAGTGATAATGGTCAAATTGGAGAACAGATTTGCCATCTATGTTATCCGGACTCTTCATTTTG

CCTCGAAGCACCCGTGTTGGACACTCGACACATG

>MSTRG.3659.1 gene=MSTRG.3659

GATATACATGTGAGTATGTGACCATGTGTGCACGACTTGTTTATACTAAATTTTATTCAAGCTAAAAAAA

GCCATTGGTGACACAAGCTTTCTGCAAAGGAGTTTGCTACCTGGCAGCTAACTAATGCATCTCTCTCTGT

CTCTCTCCATCTCTCCCTCTCTCTCTCTTTCATTCATACATCATCATAACTACTTTGTAATCTATTTTGC

TTTGTACATTCATTAGCTTCCTTAAATCTTCATTTGTAACTTCCCATAAGGTACCCATCCTTTTCTTTCT

GCTAAAATCTATGAAAAGATTGAATCTTTACATCAAGAACCAAAATTTATAACAGCCCTTTTACAAGTTT

TTGAAGTTTGATTTGGTACCATGAAAATTACTATGAAAGATTCAGAAAGAATGATATGGGATCAGATCAA

AAGCCCCCCAGGAAATGCAAGAGCTTTACCAAAACTAATGCTTTGCCTGGTCTTGATTGTTTCTGCAACT

TGTGTAGTGTTTACTGTCAAGTTTGCTGCAAATTCCCCTTCTTGTGGTGGTGAAAAATTGTACAATTCAG

TTCAAAGGAGCCTCTCTGGTCAAGATGTATCAAGGTTTTTGAGTCAGAACAAAACCCTTTTGGTTAAACC

CCCAGAAAAAACTGAGCTCAAACACATTGTTTTTGGCATTGCAGCTTCAGCTAAGCTGTGGGAGAAGAGA

AAGAATTACATTAAGTTATGGTGGAGAGGTGAAAAAATGAGAGGGATTGTTTGGTTAGATAGTCATGTTG

ATTCGAAGAAAGATGAGGCCTTGAGCTTGCCTGAGGTGAGGATCTCAGGCAAGACGGATCAATTCGCCTA

TAAGCACCCGAAGGGGCATAGGTCTGCTATTCGAATTTCAAGGATTGTATCAGAGACTTTGAAGCTGGGA

ATGCAGAATGTGAGGTGGTTTGTGATGGGAGATGATGATACATTTTTCTTGACAGAAAATTTAGTGAGGG

TTTTGAATAAGTATGATCATAATCAGTATTATTACATTGGGAGCTTATCCGAGAGTCATTTGCAGAATAT

TTATTTTTCGTATGGAATGGCTTATGGAGGGGGAGGTTTCGCAATTAGTTACCCATTGGCAAAAGCTCTG

GAGAAAATGCAAGATAAGTGCATTCAGAGATATCCCAATTTGTATGGTTCTGATGAGAGAATTCAGGCTT

GTATGGCTGAACTCGGTGTTCCACTTACTAAAGAACTCGGATTTCATCAGTATGATGTTTACGGGAACTT

ATTTGGTCTTCTTGCTGCACACCCTGTGACGCCTTTAGTGTCGTTGCATCACCTTGACGTGGTGGAGCCT

ATATTCCCAAACGCGACTCGGGTTGAAGCCTTACAACGGCTGATGATTCCAATGAAGCTTGATTCAGCAG

GTCTCATACAACAATCCATTTGTTATGAAACGAAAAAGACTTGGACCATTTCTGTGTCCTGGGGTTTTGC

TGTTCAAATCTTTCGTGGAGTCTTGTCTCCTCGCGAGGTAGAAATGCCTTCTAGGACATTTTTGAATTGG

TATAAGCGAGCAGATTACACTGGTTATGCTTTCAACACACGTCCAGTTGCACGAATCCCTTGCCAGAAGC

CCTTCGTGTTTTACTTGTCAATGGCCAGATTAAATTCCACCACCAATGAAACAGTGAGCGTATATAATCG

TCATCAAGTTCCTCATCCTGATTGCAAATGGAGGAAGATGGTATCTCCTGCAAACATCGACAAAATAGTG

GTTTACAAGAGACCTGATCTGCATCTATGGGATAGGTCTCCAAGGAGAAACTGTTGCCGGATTTTAGAGT

CGAAAAGAAAAAGCCTGGTGTTGGATGTTGGTGTTTGCAGGGAAGGCGAGATTGCTGAAATATAAGGACC

AAGCACAGAACAAGTTTTGTCTCTCTTCTTGCAATCCACATTTGTTTATTAAGTTCTTTATTTTTCAATT

CTTCTGCGTCCAATTTTCGCCAATGTACCTATTCCAGTCTCAATGTATGTAAAACGTTGTAGGACAAACC

ATACAATAAATTTCATGTTTAATGATTTTCCACAA

>MSTRG.3661.1 gene=MSTRG.3661

CCCATTTGGAAACCATACTCCATACTTCACCTTTACTCCCTCTAGTCCTCTACAACAAAACCCTAACTCT

CTCACTACGAGATGCCTCCCACCGGAAGAGGCGGCCGGCGAAACCACCTCCGGCGAACCAAACCCATTTT

ACCTTCTCACAATTTAGTGGACCCCACCACCTCAGAACCCCCAAAACCCCACCTCACAGCAAGAACACTA

CCTCCAGAGCTCCTGCTACAAATCCTCTACCGCACTGAAGTGAAGGCTCTGGTGAGATTCAAGTGTGTTT

CCAGGTCATGGTATGCTCTAATCAACCACCCCATGTTTGTCAAAAAGCATCTCTCTGTTAGTACTAGTAA

CAAGAGCAATAGTCATGTTATAGTCAATGCTAATAATGCTGGTGGTTGTGGAAAGTCTATCGTCTTGTTG

CGTGTTGATAAGGAGCCTGTTAGGTTGCTTGATATTGATACTAGTGCTAAAGATAGGGATGGGTATCCTA

GGAAGATTAATTTTGTGGATTTTTCGAAGAAGATGCATGTTGCGGGCTCGGTTAATGGGGTTGTGTTGGT

TATGCATATTGATGAGATGATGGGGAGGTTTGTTGCGTTGTGGAACCCGGGGATTAACCAGTGGAAGGCT

GTTAAGCTTGATGATTTAGGGGGTGATGATATTGAGCGGTCGGCTTTTGGGTTGGGCTATGATGAGGGGA

ATGATGATATTAAGATTGTTAGGGTTGTTACTAGGTCTAGTTGTATGAATGAGTTTATTGCGGGTTTTAA

GTGGGCTCGGGTTGAGATTTACTCAGCAAATCGGGGTTGTTGGAAAGGTGTTGATCATGAGGGTGAACTA

TTGTTTTGGCCTCGTCAGTATCATTGCAACTTTATTGTAAAAGGTGCTCCTTATTTCGTTGGAATGGATG

TCATGCCGGATAGTGCGGTAGCATTTCAGCATGAGATTTTGGCACGATTTGATCCATGTACTGGGTTTTA

TAAGAAGGTCGAGTATCCAGAACATTTTAGGGAGATGAGTACTTGGGTGCAACCTTTTGAATATCAGGAC

TCTGTTGCTGTAATGGTTCAATCTACTCCTGGATTACTTGATGAACCAATGATTGATTTGTATGTGTTGG

ATGACGACACTGCCAGTTGGACGAAGATGTACGCTATTGGGCCTTTCCCTTATAAAGAATTTCATTATCC

TCGACAGTCCTTTAAAACTGGTGAGATAGTAATTGAGGAGCAGTTACATACTCGGGACTTGTACCTCTAT

GATCCTGAAGCTAGTTATGTGAGCAGCCTCATTGGAATCGACAGTTTGAAACCACTTTGGATGCAGTCTT

ACAGTCACACAGAGAGCCTAGTTTCTGTTAAAGGGATGGAGTTGATTGGGAGGGAAAACAAAAACAAGAA

AAACAAACCCAAAAAGATTAACAGGATGGAAGTGTTGTCTAAGGAATTTGAATCAGTGTTGCATGTTTGA

CATGGGGCAGGAAGGATTGAAGCCTTACCGGTCATTTGGAAGGTCTTATGGGTTAGTAGAAGTGAGGGTT

TCCAGCGATTAAGCTTGATGTTTAAAAGATCTGGTATGTATTGGTC

>MSTRG.3661.2 gene=MSTRG.3661

CATACTCCATACTTCACCTTTACTCCCTCTAGTCCTCTACAACAAAACCCTAACTCTCTCACTACGAGAT

GCCTCCCACCGGAAGAGGCGGCCGGCGAAACCACCTCCGGCGAACCAAACCCATTTTACCTTCTCACAAT

TTAGTGGACCCCACCACCTCAGAACCCCCAAAACCCCACCTCACAGCAAGAACACTACCTCCAGAGCTCC

TGCTACAAATCCTCTACCGCACTGAAGTGAAGGCTCTGGTGAGATTCAAGTGTGTTTCCAGGTCATGGTA

TGCTCTAATCAACCACCCCATGTTTGTCAAAAAGCATCTCTCTGTTAGTACTAGTAACAAGAGCAATAGT

CATGTTATAGTCAATGCTAATAATGCTGGTGGTTGTGGAAAGTCTATCGTCTTGTTGCGTGTTGATAAGG

AGCCTGTTAGGTTGCTTGATATTGATACTAGTGCTAAAGATAGGGATGGGTATCCTAGGAAGATTAATTT

TGTGGATTTTTCGAAGAAGATGCATGTTGCGGGCTCGGTTAATGGGGTTGTGTTGGTTATGCATATTGAT

GAGATGATGGGGAGGTTTGTTGCGTTGTGGAACCCGGGGATTAACCAGTGGAAGGCTGTTAAGCTTGATG

ATTTAGGGGGTGATGATATTGAGCGGTCGGCTTTTGGGTTGGGCTATGATGAGGGGAATGATGATATTAA

GATTGTTAGGGTTGTTACTAGGTCTAGTTGTATGAATGAGTTTATTGCGGGTTTTAAGTGGGCTCGGGTT

GAGATTTACTCAGCAAATCGGGGTTGTTGGAAAGGTGTTGATCATGAGGGTGAACTATTGTTTTGGCCTC

GTCAGTATCATTGCAACTTTATTGTAAAAGGTGCTCCTTATTTCGTTGGAATGGATGTCATGCCGGATAG

TGCGGTAGCATTTCAGCATGAGATTTTGGCACGATTTGATCCATGTACTGGGTTTTATAAGAAGGTCGAG

TATCCAGAACATTTTAGGGAGATGAGTACTTGGGTGCAACCTTTTGAATATCAGGACTCTGTTGCTGTAA

TGGTTCAATCTACTCCTGGATTACTTGATGAACCAATGATTGATTTGTATGTGTTGGATGACGACACTGC

CAGTTGGACGAAGATGTACGCTATTGGGCCTTTCCCTTATAAAGAATTTCATTATCCTCGACAGTCCTTT

AAAACTGGTGAGATAGTAATTGAGGAGCAGTTACATACTCGGGACTTGTACCTCTATGATCCTGAAGCTA

GTTATGTGAGCAGCCTCATTGGAATCGACAGTTTGAAACCACTTTGGATGCAGTCTTACAGTCACACAGA

GAGCCTAGTTTCTGTTAAAGGGATGGAGTTGATTGGGAGGGAAAACAAAAACAAGAAAAACAAACCCAAA

AAGATTAACAGGTATCTTCTTTAACTATAATTTTCCCCCCTAATTTCATGTTTTAAATCGTTACAGATCC

TAGTAGACTAACTTGTGGAGTCCATTATTATATGATCCTTACGAAATAGAAGCACCAATAACTTTCCAGA

ACAGACTCTTACTTTGACAATTGATGCTACTTAAGAAGATGGAAGTGTTGTCTAAGGAATTTGAATCAGT

GTTGCATGTTTGACATGGGGCAGGAAGGATTGAAGCCTTACCGGTCATTTGGAAGGTGTGGGACTAATGG

AGGATTTTAATTATGATACTATTATCAAATGTGTAGTTACTCTTAAGTTATATGTTTCTTCTGAATCTGT

AAAAGCTTTCTAGCAAATCTCTTTGAAGGAGGGTAACCATTTATGCTTATTAAAC

>MSTRG.3673.1 gene=MSTRG.3673

TAATATCTTCCCGTTCGCTGTCTCAATTCACTTCCCAATTCGTTTTCACCCACAGGTTCTTTTCCTTTTC

TTGATTATTCCCAATCTTTATGACACCACATATTCCATCTAATTTTTAGATTTTCATTTTGTATATAAAT

TTTGATTGATGTAATAAATTATGCAATTAAAGCTGTTAAGTTTGATGGGTTTGCATAAAGTTGTGTTCTT

TATCATCTTTTCACCCAATTAAGGTTGTTGATTTTGTTGGCTTTTGCATCAAGTTGTGATCTTTATTATC

TTTTCACCCAATTTAAGTTGTTAATCTTGTTGGGTTTTGCATAAAGTTGTGATCTTTATCATAATCGCTG

CAAATTTAATTGTTAATTTTGTTGGTTTT

>MSTRG.3674.1 gene=MSTRG.3674

AGCAAAACATATATCATTATAAGTAACTTCTTAGAGATCTGAATATAGGAAATAGACCCATCTATTATTC

CACTATACAAAAATCAAAACCAAAAGAAAAAAGAGATACAGGAATGGATCATTTCAGCCACGAGCATCCA

CTGATCCTGAATGAAGATTACGTTGCTAGAGGGGGTGATATGTGTAATGCCTGCAACGAAGAGATCGTCT

CATGTAAGTCATTTGTTTTCAGTTGCAGCCGCAGCAGCAGCATCACGGGCATCACAAGTACTGATCATAA

GGATGTTGATGGAAAATGTAATAAATTCTTACTCCACAAAACATGTGCAGAGTCACCCCGTAGAATTCAA

AGCCTAAGAAACCGAGAAGAGTTCCTTGACCTCCGTGTCAATATCCATCGTCAACAAACATGGGAGCAAT

TTTTGTCCAAACGTCCAGCTGAATTTTATACCTGTAACATGTGTATGAAACAAATGCCATGGATTTACTG

CAGTCCTGGTTTTGTATTCTGTGTTTGTCTTAAATGTGCCATGTTCCAAGTTGATTATTTCATAAGCCGT

AAAATTAAGCATCAATCTCACACTCAACACCAACTAGCCCTAATCCAACGTCCATCTTCATTCAAATGTG

ATGCTTGCAATGCGGAGGATAGTATTAAAGACATGTCATATAAATGTGTGGATTGTCCGTTTTGGATTCA

CAAGAGCTGCGCGGATGCTCCTACCTCTTTCCTCTTTCATTTCCACAAAAAACACCCTCTTATGCTCTCA

TTTTCTTTGCCACAAATTCATCACAAATTTGCACAGCATTGCAGACTCTGCAATGGAAAGCTGGGTGAAT

TAAATTGGCTGTACTATTGTTCCAAATGCAGATATTTCGCCCATTTTCAATGTGCTAGATCAAGGCAGAT

GTTGAGCTCTTCTGAAAACGAAGAATATCCCGAACTTGTGCACCTCCCAGCAGCTACTGAATCCTCTGTA

AATTTATTGTTTGAGCAATTTATCAAGGAGATGAATGGTCCAAAGAACGAGGAAGGATGTAGCACTCGTC

TTAAAGATTGGGCTCATGTGACACATCCTCTGCAGCTTGTTACACACGATGAGTTACACGATGAGAAAGA

TTTATTATTGATGTGCGATGGCTGTGCCAGACCCATCCGAAATGATGGCCATGTTTTCTATGCTTGTGTT

CCTTGCCAATATTTTCTGCATAAATTCTGCGGTGAATTACCCACCACAGACCACCAAATAATTCCATCAT

ACGAGCTTTTTCGTTGTGATTGGTGTCAATATTTGGGCAATGGTATATATTTCCGCAGAGATCTGCCAAG

AAACTCGTGGTTATGGGATCTTGGACCTATTAAACAACATATAGCATGCTCAGCCCTACCCAAAATGATC

AAACACGACTCCCACCCTCACCCACTTCACTTCAATAGTCATTATACTACTTATTGCAAAGCTTGTGGAG

GGCTTTTAATTCCAAGGCACGGTTGCGAAGGTTGTGATTTCCATTTATGTGCAGGATGTATTATGAAGCC

AAGTCGAGTGAAACATCGGTGGGATCCTCACCCCCTCCGCTTGATTTATGATCCTGGCATGGTTGAGGAT

CACGAACATGAGTTTCAATGTGAGTTCTGTTCTCAAGATTTGGACACGAACCTTTGGTTCTATCACTGCG

ATGATTGTGATCTTTCGTTTCATCTCAGATGTTATGATAAATCATGTTATCTCAACTTCTCACATGTCAA

GTTTGGTGCTACTGATATTATTAAAGACCATCTCCATCATCACAGCCTCACATTTGTACTCAATAAGAAG

TTTCGTCGTTGCGGAAAATGTCACCACGGTAATATTGGCGGGCCTGTCCTTGAATGTGCACCATGCAAAA

CCACATTCTGTGGGATCTGCTC

>MSTRG.3684.1 gene=MSTRG.3684

ATCTGAGATAGGAGCGTCATGCAGTGAAAACAGAGCAGATGTCTGATGAAATATCTCGGGACCTAGCCTA

AAAAAAGAGGGAGGTGATGAATTGTGCTATTAAGCTTCCTGTTGATGGTGGTCGTGATGATGGTGACGAC

GATGATGATGACGACGAAGATGATTATGATGATGAATATGATGATGAATATGGTTTCGACTATGACAGCG

ATGGGTCTTATGAAGCTGAAGCTAATACTCGTATAATCGAGTTTAATAAGTCTGTTGGTGTTATTGTACT

GTGGGCTAATGCATTGGATTATGAATTGTCTGATTGGAGATGGAACAAGAAGGTTAAATTGTGGAAGTTG

GATGATGAGGCATGCCTTCGAGGTGGTGGAGACGAGGCATCATGGACGCTAATGTTTAGTCTTAAATTCG

ATAGACCAGCACTACTCCGTAATGGCTATTTCAGCAATGGGAATCTCTTATTATCAGTACGAAGTGATAG

TTACATGTGGATTTCATGTGATGTCGATAAGAAAGATGCCGAGATTGTCCCCCTTTCAGATGAGATGGCT

GACTGTTGTTACTGTTATGATATGTGCAAGTATACAGAGAGCTTAGTTTCACTTGCAGGATTCAAACAAA

TTAGCTGGAAAGCTGGC

>MSTRG.3685.1 gene=MSTRG.3685

TATCCTGGTGGCGGTCGTCAACTCAACCCTAATCAGGAATGGTCCTTGGATGTTGATCCAGGCACCCAAT

ACGCTCGTATTTGGGGTCGAACAAATTGTAACTCTGATGCCTCTGTTTGTGAAACCGGCGACTGTGGCAG

GCTTGACTGCACAGCATACGGCAAAACACCAAACACAGTGGCCGAATATGCATTAAACCAATTTCAAAAC

TATGATTACTACGACATTTCACTAATCGACGGTTTCAATATTCCTATGGAATTTGGTCCCACTGCAGAAG

GTGCAGATGGATGCAAACTCAGTAGATGTACAGCCGATATCACCGGGCAATGTCCTGATCCGTTGAAGTT

CCCTGGCGAGTGTAATAACCCCTGCACCGTTTATAAGACTAACGAATATTGTTGCACTCAGGTGTCGGAA

CCATGTAATCCGACAGATTATTCGAGGTTTTTTAAGGACAGGTGTCCTACTTCTCTCAGTTAT

>MSTRG.3686.1 gene=MSTRG.3686

AGCTATTATAATTTGAATATTATGAAATTTTACGTTTTACAAAATCTTAATTATATATTATTAAAATTCT

GAGAATATTTTCACAGATAAGAAAACGCGTAAAATCTCGATTGAATACACCGCATTCCCACTCTCTTTCA

ACTTCAATCGAAACCCTCGTTAGACCTTCAGTTCTCTCCCCATCATCACCACCGAGTCGTCTCAGCGACT

CAACAATGGCGCGTGTGAGGCCCAATCCGACGCTTTCAAGCGATCTACTCACCGAGATTCTGAAACGCCT

GCCAGTGAAACCCCTAGTTCAATACCAATTAGTCTCCAAGACGTGGTTATCGCTAATCAAAGACCCTGCT

TTTGCCGAAGCTCAGCTCCGTCACGCAATCTCCACCGGAACCGATGAAACCCTTATCATCACCCGTTATC

TCGATCTCCGCGGTGGTCTTAGACAGAAATTCTCACTCTTTAACATAGACTCTTGTGAAATAATGCGTGA

TGTTAAGTATCCGTGTCCTCGAAATGAGCCTATAAGATCTGAGCGTTATTTTAGACTTGTTGGTTCTGCT

AATGGCATTGTATGTGTTGCTAATTATGATTTTCACCCTGATGTTCGAGAGTTTTGTGACCCTGTGTTTC

TTTGGAATCCCGCAATTAGACAAATTAAAATTCTTCCGCCACTTCCGGTTGGTGAATTGTATGATTTGGG

GTTCGGTTATGATCCGGTGGATCGGGATTATAAGGTTGTTTTTGTTGTTAGAAAGCCTTGTCTTTCGGTG

GAGGTGTATTCGGCTAATAGTAATGTATGGCGAAAAGTGGCTTCTCCGATTCCAACAGCCCTTATTGATC

GTTTTTGCTGTACGTTCCATGTATGTGTTAATGGATTTTTATGTGGTGCGGGGGATTGGGGGATGATTGG

TATGATGGTGGTGTTTGATTTGAACAGGGAGGTGATGAATTGTGCTATTAAGCTTCCTGTTGATGGTGGT

CGTGATGATGGTGACGACGATGATGATGATGACAACGAAGATGATTATGATGATGAATATGATGATGAAT

ATGGTTTCCACTATGACAGCGATGGGTCTTATGAAGCTGAAGCTAATACTCGTATAATCGAGTTTAATAA

GTCTCTTGGTGTTATTGTACTGTGGGCTAATGCATTGGATTATGAATTGTCTGATTGGAGATGGAACAAG

AAGGTTGAATTGTGGAAGTTGGATGATGAGGCATGCCTTCGAGGTGGTGGAGACGAGGCATCATGGACGC

TAATGTTTAGTCTTAAATTCGATAGACCAGCACTACTCCGTAATGGCTATTTCAGCAATGGGAATCTCTT

ATTATCAGTACGAAGTGATAGTTACATGTGGATTTCATGTGATGTCGATAAGAAAGATGCCAAGATTGTC

CCCCTTTCAGATGAGATGGCTGACTGTTGTTACTGTTATGATATTTGCAAGTATACAGAGAGCCTAGTTT

CACTTGCAGGATTCAAACAAATTAGCTGGAATGCTGGCGAAGATGATAATTAAGAAGACACTGCTGATTC

TGGTATCGATGTGCTAGATATTGATTGCAGCAACTCTGTTCTTGTAGTCTTCTTTGGTCTGGTTTTGTAG

TCAGTTCTATTTGCAATGTCAATTTGCACTGTGTTGTCAATTTTACATCCGTATTACTGGATTCAATCCA

GTCAACTATGTTGGTAATAATATTTTTGCGAACTCATAGACTTGCATAATGATGTCTTGTGTTGCACTCA

GGCATAGGAACTCCAACAGATTATTGAAGTTTTTTTTGATGATATGTGTCCTACTTCTCTCAGTTATAAA

GCTGATCAAACAGGTTTCGAAGTTGTCCAAGTGGTACTAACTACCAAGTTCTGTTCTGTAATGAGGT

>MSTRG.3687.1 gene=MSTRG.3687

TGCTGAATTGTGCTATTAAGCTGCCTGTTATTTATGGTGATGATAATGATGAGAATGATGATGATGCTTA

TGGTTATGATGTTGGTGGTGATAATGATGTAAATGATGATAATGTTTATGATGTTGCGGAAGATGAGGGT

GATGATGATGAAGAACCTGGTGTTATTAAGTTTTATGATCCTCGTGTTATCGAGTCTAATGATGTTCGTA

TTATTGAGTTTAACAAGTCTATTGCTGTTATTATAGTGTGGGATGGTGGATTGAATGATGATAAAAAGAT

TAATTTGTGGACATTGGATGACGAAGCATGCCTCCGTGGCGGTGGAGTTGAGGCATCATGGACGTTAATG

TTTAGTATTGATTTAGGTATGCCGTTGGATTTTATTCATGGCTACACCAGCAATGGGGATATCGTG

>MSTRG.3688.1 gene=MSTRG.3688

TAATGCACTGGATTATGAAATGTCTGATTGGAGATGGAACAAGAAGGTTAAATTGTGGAAGTTGGATGAT

GAGGCATGCCTTCGAGGTGGTGGAGACGAGGCATCATGGACGCTAATGTTTAGTCTTAAATTCGATAGAC

CATCACTACTCCGTAATGGCTATTTCAGCAATGGGAATCTCTTATTATCAGTACAAAGTGATAATAACAT

GTGGATTTCATGTGATGTCGATAAGAAAGATGCCGAGATTGTCCCCCTTTCAGATGAGATGGCTGACTGT

TGTTATTGTTATGATATGTGCAAGTATACAGAGAGCCTAGTTTCACTTGCAGGATTCAAACAAATTAGCT

GGAAAGCTGGCGAAGATGATAATTAAGAAGACACTGCTG

>MSTRG.3689.1 gene=MSTRG.3689

CGTTGATTCTCGTCAAATTGTGTCTGATCTGAAGTATCCGTACACTCAAGGTGAGTCTCAACAAGTAGCT

CCTGCTTTTAGAATCGTCGGTTCTGTTAACGGCATTGTTTGTGTTGTCTTTCACCACAATCTAAGCGAAC

ATAGTTTCTATCTGTGGAATCCTGCTACTGGACAATCAAGAGCTGTTCCGGCATGTGCCTTGAGGACTTC

CCGCGCTTTGGGTTTTGGTTATGATCCGGTAGATAATGATTATAAGATTGTTAGTGTTGTGTCGAGGCCT

TCGCTTCCTACTGAGGTGTATTCGGCTAATAGGAATGTATGGCGAAAAGTTCCTAGTGAGGATTTTATGT

TTGGCATTAATGGTGGCTTCATTCTAGGCTATTTTGATGTATCTGTTAATGGATTTTTATTTGGTGTCGA

GGAGGATGCTATGATTGTGTTTGATTTGAACACGGAGGTGTTGAATTGTGCTATTAAGCTCCCTCCTGCT

ATCGCTTCTGCTATTGAAGCTGCTGATGCTGGTCTTGCTGTTATTGCTGCTGTTGGTGGTCATGACGCCC

TCATTTCTTATGCAGCTCGACCTAAAACTCGTATTATCGAGTATAATAAGTCTATTGCTGTTATAACACT

GTGGGCTAATGCACTGAATGATGATTGGACTGATCATAAGAGATTCAACAAGGCGATTAACTTGTGGAAG

TTGGAAGATGATTCATGCCTTCGAGGTGATGGAGTTGAGGCATCATGGACGCTAATGTTTAGTATTGATT

TACCGATGAGGCCAGCACATCTTCTTAATGGCTATTACAGCAATGGGGATCTCGTACTACTGGTAGCAAA

TGTTGATGACAGGTGGATATGGTGTAATGCCAGCAAGAAAGAGGCTAAGATTGTTCGAGTTGATATGGTT

GATCATTGTCACACTCATTATCTTTACAGGTACACGGAGAGCCTAGTTTCACTTTCAGGATTCAAACAAA

TCAGCTGGAATCGTGGCTATGGTGATAATTAAGGAAACACCGCTATTCTGGTATCTCTATGCTATATGTC

TTGTTAGTACCTTTGGTCTGTTATTTTAGTGACTTTTGAGAGATGTCTGCAGTATT

>MSTRG.3690.1 gene=MSTRG.3690

TAATGCACTGGATTATGAAATGTCTGATTGGAGATGGAACAAGAAGGTTAAATTGTGGAAGTTGGATGAT

GAGGCATGCCTTCGAGGTGGTGGAGACGAGGCATCATGGACGCTAATGTTTAGTCTTAAATTCGATAGAC

CATCACTACTCCGTAATGGCTATTTCAGCAATGGGAATCTCTTATTATCAGTACAAAGTGATAATAACAT

GTGGATTTCATGTGATGTCGATAAGAAAGATGCCGAGATTGTCCCCCTTTCAGATGAGATGGCTGACTGT

TGTTATTGTTATGATATGTGCAAGTATACAGAGAGCCTAGTTTCACTTGCAGGATTCAAACAAATTAGCT

GGAAAGCTGGCGAAGATGATAATTAAGAAGACACTGCTG

>MSTRG.3691.1 gene=MSTRG.3691

GTAACATTAACATGTACTCAAGCACCAAAATGAGCTACTTAGCAGCACTTTCCCTGATCATCATTTCCAC

CCTCCTAATCACCACAAGAGCTGCCAACTTCAATATCCGAAACAACTGTCCTTACGTCGTATGGGCCGGT

GCCTATCCTGGTGGCGGTCGTCAACTCAACCCTAATGACCAATGGTCATTGGATGTTGCTCCTGGCACCC

AATATGCTCGTATTTGGGGTCGAAAAAATTGCAACTCTGATGCCTCTGTTTGTGAAACCGGAGATTGTGG

TAGGCTTGACTGCACAGCATACGGCAAAACACCAAACACAGTGGCCGAATATGCACTAAACCAATTTCAA

AACTATGATTACTACGACATTTCACTAATCGACGGTTTCAATATTCCTATGGAATTCGGTCCCACTGCAG

AAGGTGCAGATGGATGCAAACTCAGTAGTTGTACAGCCGATATCATCGCGCAATGTCCTGATCCGTTGAA

GTTCCCTGGCGGGTGTAATAACCCCTGCACGGTTTATAAGACTAACAAATATTGTTGCACTCAGGCGTCG

GAACCATGTAATCCGACAGATTATTCGAGGTTTTTTAAGGACAGGTGTCCTACTTCTCTCAGTTATAAAG

CTGATCAAACAGGTAATCGAAGCTGTCCGAGTGGTACTAACTACAAGGTTGTGTTCTGTCCATGAGGATC

TTATTATTGTAGTTCTTGCTGTGAAGTTGTTGTCCATGAGATCTTATTATTGTAGTGCTTGCTATAAAGT

TGTATTTGCACATCACAAATTTTGTTGTTGTGCTCGTTCTTCTAGATAAACTTTGTTTTGATCTAAATAA

ACTTATGTACAGGGATACACAAGTCATTCTGACTTGAGCAATTTGTTATAATAAT

>MSTRG.3692.1 gene=MSTRG.3692

TAATGCACTGGATTATGAAATGTCTGATTGGAGATGGAACAAGAAGGTTAAATTGTGGAAGTTGGATGAT

GAGGCATGCCTTCGAGGTGGTGGAGACGAGGCATCATGGACGCTAATGTTTAGTCTTAAATTCGATAGAC

CATCACTACTCCGTAATGGCTATTTCAGCAATGGGAATCTCTTATTATCAGTACAAAGTGATAATAACAT

GTGGATTTCATGTGATGTCGATAAGAAAGATGCCGAGATTGTCCCCCTTTCAGATGAGATGGCTGACTGT

TGTTATTGTTATGATATGTGCAAGTATACAGAGAGCCTAGTTTCACTTGCAGGATTCAAACAAATTAGCT

GGAAAGCTGGCGAAGATGATAATTAAGAAGACACTGCTG

>MSTRG.3693.2 gene=MSTRG.3693

CACTTAGCAAGTCAACCTACGAGAAGGAATTGTTAGCTTTGGTCTCTGCTATCCACCATTGGAGGCCTTA

TTTGTTGGGTCGACAATTCATAGTTCGCACTGATCATAAGAGTCTTAAACACTTGTTAAGTCAACGTATA

TCAACTCCAGATCAGGAAAAATGGCTATCTCGCTTATTGGGCTATACATTTGAGATTGTTTATAAACCGG

GAAATGATAACAGTGCGGCTGATGCTTTGTCACGTCGTAATGAGGTGGAGGATTTGACAATTCAAGTAGC

TGATGTAACAGAATTATCAGCCATATCCAAACCTCAGTGGAAAGAGGCAGATGCCTTAGCGAAAGCCATC

AGATCGGATGAGTCCTTGTCCAAGGTTGTGACGGCTTTGGAAACTAATCAACCTACACTCTCAGGTTATG

CCCTCTCTCATGGTCAACTACTCTATAAAGGACGTCTGGTAATTCCAGCAACATCAGAATGGGTGACACG

ATTTCTTCACGAATTTCATTCAACACCCGTAGGTGGACATCCAGGCTTCCTACGAACTTACAAACGCATT

TCTAGCAACCTTTTCTGGACCGGCATGAAGACTGATATTATGAAGTATGTTGCAGAATGTAGTGTTTGTC

AACAACACAAATATCAGGCCACTACACCAGCAGGTCTGCTTCAGCCCCTTGCTTTACCAACTGCTGTTTG

GGAAGATGTCTCGTTAGATTTTGTGTCAGGCCTGCCACGGTCACAAGGTTATGACACACTATTAGTTGTG

ATCGATCGTTTCTCAAAATACAGTCATTTTTTGCTATTGAAGCATCCTCGTGTTATCGAGTCTAATGATG

TTCGTATTATTGAGTTTAACAAGTCTATTGCTGTTATTATAGTGTGGGATGATGGATTGAATGATGATAA

GAAGATTAACTTGTGGACATTGGATGACGAAGCATGCCTCCGTGGCGGTGGAGTTGAGGCATCATGGACG

TTAATGTTTAGTATTGATTTAGGTATGCCATTGGATTTTATTCATGGCTACTCCAGCAATAGGGATATCG

TGATAATACTCGACGATGCTTATGTGTGGATTTCGTGTAATGTTGACAAGAAAGAGGCCAAGATTCTCAT

ACTTTCAACTGGTATGGCTAAACATCTTTACACCCGGCATGTTTATAAGTATACAGAGAGCCTAGTTTCA

ATTGCCGGATTTGAACAAGTCAACTGGAATGCTCATGAGTCTAGATGATGATTGGGAAAGCAGCGCTGAG

TTTGGAATTTTGGTGCTAGCTTATGTTGACAGTGTCTTATTTGAAGGGGTATTCAGAACTTCCATTATAA

TCTGAGATAGGAGCGTCATGCAGTGAAAACAGAGCAGATGTCTGATGAAATTTCTCGGGACCTAGCCTAA

AAAAAGAGGGAGGTGATGAATTGTGCTATTAAGCTTCCTGTTGATGGTGGTCGTGATGATGGTGACGACG

ATGATGATGATGACGACGACGAAGATGATTATGATGATGAATATGGTTTCGACTATGACAGCGATGGGTC

TTATGAAGCTGAAGCTAATACTCGTATAATCGAGTTTAATAAGTCTATTGGTGTTATTGTACTGTGGGCT

AATGCATTGGATTATGAATTGTCTGATTGGAGATGGAACAAGAAGGTTGAATTGTGGAAGTTGGATGATG

AGGCATGCCTTCGAGGTGGTGGAGACGAGGCATCATGGACGCTAATGTTTAGTCTTAAATTCGATAGACC

AGCACTACTCCGTAATGGCTATTTCAGCAATGGGAATCTCTTATTATCAGTACGAAGTGATAGTTACATG

TGGATTTCATGTGATGTCGATAAGAAAGATGCCAAGATTGTCCCCCTTTCAGATGAGATGGCTGACTGTT

GTTACTGTTATGATATTTGCAAGTATACAGAGAGCCTAGTTTCACTTGCAGGATTCAAACAAATTAGCTG

GAATGCTGGCGAAGATGATAATTAAGAAGACACTGCTGATTCTGGTATCGATGTGCTAGATATTGATTGC

AGCAACTCTGTTCTTGTAGTCTTCTTTGGTCTGGTTTTGTAGTCAGTTCTATTTGCAATGTCAATTTGCA

CTGTGTTGTCAATTTTACATCCGTATTACTGGATTCAATCCAGTCAACTATGTTGGTAATAATATTTTTG

CGAACTCATAGACTTGCATAATGATGTCTTGTGTTGCACTCAGGCATAGGAACTCCAACAGATTATTGAA

GTTTTTTTTTGAGGATATGTGTCCTACTTCTCTCAGTTATAAAGCTGATCAAACAGGTTTCGAAGTTGTC

CAAGTGGTACTAACTA

>MSTRG.3693.4 gene=MSTRG.3693

TTTTATATGGAGTCTAGTAAATTTTAAACAAAAAATCATAAAACTAATGTAGAACACTTAAAATTTAAGC

ATAATACAAAAAAGCTATCATAATTTGAATATTATGAAATTGTACGTTTTACAAAATCTTAATTATATAT

TATTAAAATTCTGAGAATATTTTCACAGATAAGAAAACGCTTAAAATCTCGAATACACCGCATCCCCCCT

CTCTTTCAACTTCAATCGAAACCCTCGCTAGACCTTCAGTTCTCTCCCCATCATCACCACCGAGTTGTCT

CAGCGACTCAACAATGGCGCCTGTGAGGCCCAATCCGACGCTTTCAAGCGATCTACTCACCGAGATTCTG

AAACGCCTGCCAGTGAAACCCTTAGTTCAATACCAATTAGTCTCCAAGACGTGGTTATCGCTAATCAAAG

ACCCTGCTTTTGCCGAAGCTCAGCTCCGTCACGCAATCTCCACCGGAACCGATGAAACCCTTATCATCAC

CCGTTATCTCGATCTCCGTGGTGGTCTTAGACAGAAATTCTCACTCTTTAACATAGACTCTTGTGAAATA

ATGCGTGATGTTAAGTATCCGTGTCCTCGAAATGAGCCTATAAGATCTGAGCGTTATTTTAGACTTGTTG

GTTCTGCTAATGGCATTGTATGTGTTGCTAATTATGATTTTCACCCTGATGTTCAAGAGTTTTGTGACCC

TGTGTTTCTTTGGAATCCCGCAGTTAGACAAACTAAAATTCTTCCGCCACTTCCGGTTGGTGAATTGTAT

GATTTGGGGTTCGGTTATGATCCGGTGGATCGGGATTATAAGGTTGTTTTTGTTGCTAGAAATCCTTGTC

TTTCGGTGGAGGTGTATTCGGCTAATAGTAATGTATGGCGAAAAGTGGCTTCTCCGATTCCAACAGCCCT

TATTGATCGTTTTTGCTGTACGTTCCATGTATGTGTTAATGGATTTTTATGTGGTGCGGGGGATTGGGGG

ATGATTGGTATGATGGTGGTGTTTGATTTGAACAGGGAGGTGATGAATTGTGCTATTAAGCTTCCTGTTG

ATGGTGGTCGTGATGATGGTGACGACGATGATGATGATGACGACGACGAAGATGATTATGATGATGAATA

TGGTTTCGACTATGACAGCGATGGGTCTTATGAAGCTGAAGCTAATACTCGTATAATCGAGTTTAATAAG

TCTATTGGTGTTATTGTACTGTGGGCTAATGCATTGGATTATGAATTGTCTGATTGGAGATGGAACAAGA

AGGTTGAATTGTGGAAGTTGGATGATGAGGCATGCCTTCGAGGTGGTGGAGACGAGGCATCATGGACGCT

AATGTTTAGTCTTAAATTCGATAGACCAGCACTACTCCGTAATGGCTATTTCAGCAATGGGAATCTCTTA

TTATCAGTACGAAGTGATAGTTACATGTGGATTTCATGTGATGTCGATAAGAAAGATGCCAAGATTGTCC

CCCTTTCAGATGAGATGGCTGACTGTTGTTACTGTTATGATATTTGCAAGTATACAGAGAGCCTAGTTTC

ACTTGCAGGATTCAAACAAATTAGCTGGAATGCTGGCGAAGATGATAATTAAGAAGACACTGCTGATTCT

GGTATCGATGTGCTAGATATTGATTGCAGCAACTCTGTTCTTGTAGTCTTCTTTGGTCTGGTTTTGTAGT

CAGTTCTATTTGCAATGTCAATTTGCACTGTGTTGTCAATTTTACATCCGTATTACTGGATTCAATCCAG

TCAACTATGTTGGTAATAATATTTTTGCGAACTCATAGACTTGCATAATGATGTCTTGTGTTGCACTCAG

GCATAGGAACTCCAACAGATTATTGAAGTTTTTTTTTGAGGATATGTGTCCTACTTCTCTCAGTTATAAA

ACTGATCAAACAGGTTTCGAAGTTGTCCAAGTGGTACTAACTACCAAGTTCTGTTCTGTAATGAGGTATT

CTTATAATATTTGCTATGAAGTTGTTGTCCAAGAGGA

>MSTRG.3693.8 gene=MSTRG.3693

AAGATGTAGTAGCAAATTATGAGGAAGGAGAAGACATGGAGATGCCGGTAGCCAATGTGGAATGTCAAAC

AATGGACCTGCATATATACTCTACAGCAGGCATTACACGACCAAGCACTATGAAGATTCAAGGATTGTTA

CGAAAACATCCGATTGTGATCCTGATTGATAGTGGTGCCAGGCACAATTTCATCTCAGCTGATCTGGTAC

AGAAAATGGGTTTAACGGTGACTCCAACTAAGGAATTAGGGGTTAAATTGGGTGATGGACATACAGTGCC

TTGTAAGGGCTTATGTAGAGCTGTTGATATATCTGCAGGAGAGCTAAAGATTACAGCGGATTGTTACCCG

TTCAAACTCGGAGGGGTGGATCTGATTTTGGGCATTGCTTGGCCAGAAACTTTGGGAGAAGTAAGAGTAA

ATTGGCAAAAAATGACAATGACATTTGCTACAGGTTTTTTTTTACGGAAATTACAAGGAGATCCAGCTTT

ATCTACATCTCCAGTATCTTTAAAGGTAATTCTTAACACTTCAGATGTGGATATTTGTTTCATGTTATGG

GCCACTTCTGTTGTAGAATCAGAATTGTCAAAAGGTGAGCATTTGACGGAAGAACAAAAACACTCACTGA

ATCGGGTGTTGGAATGTTTTGGTGATGTATTTCAAGAACCAACAGGTTTACCTCCTCAAAGAAATTGATC

ACATGCCATCATATTGAAGCCTGGTGTTACAGCAGTGAACGTGCGTCCTTATAGATATGCTCACAGCCAG

AAAAATGAGATTGAACGTTTAGTGCAGGAGATGTTAAGTGCTGGAATTATTCAACCAAGTAGCAGCCCTT

TCTCTAGCCCTGTTATTTTGGTCAAGAAAAAGGACGGGAGTTGGCGATTCTGCATCGATTATCGTGCTCT

CAACGATGCAACCATACCAGATAAGTATCCAATTCCAGTTATTGATGAACTACTAGATGAGCTGTGCGGT

GCACAATATTTTTCAAAGATTGATTTGAAATCAGGGTATCATCAAATTCGTATACATTCCAACGACATCC

CTAAAACTGCATTCCGAACTCATGACGGCCATTATGAATTTTTGGTGCTACCTTTTGGTCTTACAAATGG

CCCGGCTACATTTCAGTCCATAATGAATGATTTATTTCGGCCATACTTACGCAAATTTGTGTTGGTTTTT

TTTTGATGATATCCTGATATATAGTAGGGACTGGGTGTCACACGTTGAACATCTGAAGACAGTGCTCCAA

GTTCTGCGATCCCAACAATTTTTTGCAAATCTGAAAAAATGCACTTTTGGTCAGCAACAAGTTGAGTACT

TAGGCCACATAATATCTGAGAAAGGGGTTGCAATGGATCCAGGAAAGTTAAAAGATGTGCAACAGTGGCC

AGTACCACGTTCGGTAAAGGCTGTTCGGGGATTTCTTGGTCTTACAGGCTATTATCGCAAATTCATCCGA

GATTATGGTAAAATTGCTCGTCCCCTGACTAATTTACTTAAGAAAGACTCATTTATTTGGAATACAGAGA

CACAAGAGGCGTTTGAGAACTTAAAACAAGCGTTAGTCACAGGGCCAGTTCTAGCCATGCCCGACTTTTC

TAAGCCATTCCTAGTGGAGTGTGATGCCTCAGGAGTGGGAATAGGAGCGGTTTTAATGCAAGAAAGTAGC

CCAATAGCTTATTTCAGTAAGGCTATTTCTGAGCGGGCACTTAGCAAGTCAACCTACGAGAAGGAATTGT

TAGCTTTGGTCTCTGCTATCCACCATTGGAGGCCTTATTTGTTGGGTCGACAATTCATAGTTCGCACTGA

TCATAAGAGTCTTAAACACTTGTTAAGTCAACGTATATCAACTCCAGATCAGGAAAAATGGCTATCTCGC

TTATTGGGCTATACATTTGAGATTGTTTATAAACCGGGAAATGATAACAGTGCGGCTGATGCTTTGTCAC

GTCGTAATGAGGTGGAGGATTTGACAATTCAAGTAGCTGATGTAACAGAATTATCAGCCATATCCAAACC

TCAGTGGAAAGAGGCAGATGCCTTAGCGAAAGCCATCAGATCGGATGAGTCCTTGTCCAAGGTTGTGACG

GCTTTGGAAACTAATCAACCTACACTCTCAGGTTATGCCCTCTCTCATGGTCAACTACTCTATAAAGGAC

GTCTGGTAATTCCAGCAACATCAGAATGGGTGACACGATTTCTTCACGAATTTCATTCAACACCCGTAGG

TGGACATCCAGGCTTCCTACGAACTTACAAACGCATTTCTAGCAACCTTTTCTGGACCGGCATGAAGACT

GATATTATGAAGTATGTTGCAGAATGTAGTGTTTGTCAACAACACAAATATCAGGCCACTACACCAGCAG

TGTGGGATGATGGATTGAATGATGATAAGAAGATTAACTTGTGGACATTGGATGACGAAGCATGCCTCCG

TGGCGGTGGAGTTGAGGCATCATGGACGTTAATGTTTAGTATTGATTTAGGTATGCCATTGGATTTTATT

CATGGCTACTCCAGCAATAGGGATATCGTGATAATACTCGACGATGCTTATGTGTGGATTTCGTGTAATG

TTGACAAGAAAGAGGCCAAGATTCTCATACTTTCAACTGGTATGGCTAAACATCTTTACACCCGGCATGT

TTATAAGTATACAGAGAGCCTAGTTTCAATTGCCGGATTTGAACAAGTCAACTGGAATGCTCATGAGTCT

AGATGATGATTGGGAAAGCAGCGCTGAGTTTGGAATTTTGGTGCTAGCTTATGTTGACAGTGTCTTATTT

GAAGGGGTATTCAGAACTTCCATTATAATCTGAGATGGGAGCATCATGCAGTGAAAACAGAGCAGATGTC

TGATGAAATTTCTCGGGACCTAGCCTAAAAAAAGAGGGGTTTTTGGTATGAGTGGTATGATGGTGGTGTT

TGATTTGAACTGGGAGGTGATGAATTATTCTATTAAGCTTCCTGTTGATGGTGGTCGTGATGATGGTGAC

GATGATGATGATGATGACGACGACGAAGATGATTATGATGATGAATATGGTTTCGACTATGACAGCGATG

GGTCTTATGAAGCTGAAGCTAATACTCGTATAATCGAGTTTAATAAGTCTGTTGGTGTTATTGTACTGTG

GGCTAATGCATTGGATAATGAAATGGCTGATTGGAGATGGAACAAGAAGGTTAAATTGTGGAAGTTGGAT

GATGAGGCATGCCTTCGAGGTGGTGGAGACGAGGCATCATGGACGCTAATGTTTAGTCTTAAATTCGATA

GACCATCACTACTCCGTAATGGCTATTTTAGCAATGGGAATCTCTTATTATCACTACAAAGTGATAATAA

CATGTGGATTTCATGTGATGTCGATAAGAAAGATGCCGAGATTGTCCCCCTTTCAGATGAGATGGCTGAC

TGTTGTTACTGTTATGATATTTGCAAGTATACAGAGAGCCTAGTTTCACTTGCAGGATTCAAACAAATTA

GCTGGAATGCTGGCGAAGATGATAATTAAGAAGACACTGCTGATTCTGGTATCGTTGTGCTAGATATTGA

TTGCAGCAACTCTGTTCTTGTAGTCTTCTTTGGTCTGGTTTTGTAGTCAGTTCTATTTGCAATGTCAATT

TGCACTGTGTTGTCAATTTTACATCCGTATTACTGGATTCGATCCAGTCAACTATGTTGGTAATAATATT

TTTGCGAACTCATAGACTTGCATAATGATGCCTTGTGTTGCTCTCAGGCATCGGAACTCCAACAGATTAT

TCAAGGTTTTTTTTTTAAGGATATGTGTCCTACTTCTCTCAGTTGTGAAGCTGATCAAACAG

>MSTRG.3696.1 gene=MSTRG.3696

CTTTAGTTATGATTCCTGGTGTTTTGGTTTTACATTTTATATATAGTTCTGTTGATCTTTTAAATGCAAT

CAAAGAAAATCCAAGACATACTAAGTTGTACAGAAATTTTAAGCGTATTTGGGCCTGCTGTCATAATATC

ATACCTCATTTCAGGTTTGTCTGCTCTGCTTTCTGTCCTTTGTTATACCGAATTTTCTGTGGAAATTCCT

GTTGCCGGAGTCTCATTTTTATTTTTTTTGTGCTAAATCTGGAGGCTCATTTGTGTATCTAAGAGTAGAG

CGTGGTGATTTTATTGCTTTTCTGGCTGCTGGAAATATACTTTTTGAGTATATTGTAGCTGGGGCAAGTG

TGCCTAGCTCATGGACTTCATATTTTGCAACATTGTGTAGCAGGCGTGGCACGCTCATGGACTAATTTAA

TTCATACTTTTTGAGAAGCCCTGAGGTTATTATTGTCCCAAAATATGTTTAATTTTTCATTGGTAGGGGA

AGGCGACGATGAAGTGGTAGATCAGCTTAAACCCCTTATTGAATCATGTAGTCCCGCGCGTTGAAAGAAG

CAAGTCTTGGAAGAGAACTCCCTGGGCTGCAGAAAGAAGTTACAGAGTATTTAAAATCCAAGAGCAGAAT

AATACATGTTTTGTTCAGGGATGAATATCGGGTCATTTCAGGGTTGGGGAGTCGCTATATCGCCCCCCAT

CCCCGAATCTGATATCGATTCCAGAGGCCGTCCAAATGCTATGGATACATTTGTCGGAAAGTAATTAACA

ATGAAATTCTGCAAATTCATTTCCACATTAGCAGCAACATGAAGTTGCGACAGCAAAAAGAGGAACTTTT

CTTTGGACTGTTAAACTAGGCCTTTCGCGCATGTCTAAATCCTCCCCGTTTTTCCCCAGGCAACTTCCAC

TCAAATTTATGCAGAAGATTTGCCAACACAAATTCATTGGTGGCCATAGCAAAAGCAATTCC

>MSTRG.3697.1 gene=MSTRG.3697

CAAAACCCTAACTCTGTATTTTGTCCTGTTTTCTTATAATCAATGGCGCCTGTTAGACGTAATCCGACAC

TACCAGACGAACTAATAAGCCAGATTCTAATACGCGTGCCGGTGAAATCATTACTTCAATACCAATCAGT

CTCCAAGACATGGTTATCGATGATTAAAAACCCTAGTTTCGTCAAATCTCAGCTCCGTCACGCCCTCACT

ACCCAAACTGATGATCAAACTCTTCTCAAAGCTGACCGTTCCTCAGTACTCCATCTCGATTCTGGCCGGA

ACGTGGCTTATCTCAAGTTTCCATTTCCTCGACGTGATGATGTAATGTATCTTAATCTTGTTGGTTCTGC

TAATGGTATTGTTTGTGTTTCTGTTGTGTTTGGTAGTCATGATGATTATAGAAACACTAGTATTTGCCTA

TGGAATCCTGCTACTACACAATCCAAATTTGTTCCGTCATCGTTTAATGCTAGTGTGAGAGTTTTAGGGT

TTGGTTATGATGGAGTAGATGATGATTATAAGATTGTTGTTTATGTAGACAATTATGTGGTGTATTCGGA

GAATAGGAATGCTTGGAGAATTGTACCTGAGACGATACATTCTCCTGGCATGGATGGTTTTGATGTATGC

ATTAATGGATTCCTGTGTGGTTGTTGTGGTGGTACTTATGCGATGATGGCTTTTGATCTGAACCGGGAGG

TGCTGAATTGTCACATTAACCTCCCTGTTACCGATGGTGATGTCGATGAGGCTCATGCTCAGCACCTGCT

GGATTGTACTATTAAGCTTCCTGTTATGAATGTTATTAATGATGCTCATGATTATGATGCTGATGATAAT

TTTGCTCGTATTATTGAGTTTAATAACTCTGTTGCTGTTATTATGATGATTGGTTCGAGTAGGAACATCA

ACACGTGGACATTGGATGATGATGCATGCCTTCGGGTTGGTGGAGTTCGAGTTGGTGGAGTTGAGGCATC

TTGGACGCCAGTGCTTAGCATTGATACACGTATGAGACCCGATCAGTTTTATGGCTACTTCAGCAATGGG

AATCTCATATTACTATTAACAAATTATGTAAGTGATGTTAAAATGTGGATTTCGTGTAATGTTGACAAGA

AAGAGGCCAAGTTTGCTCCACCTTTGCTTGATATGGCTGAACATTGTCAAGATAGCGTAATTTTCAAGTA

CAAGGAGAGCCTGATCTCACTCGCAGGATTCAGACATGTCAACTAGAATGTTGATGATTCTGGGATTATT

ATGTTTACTAATGATGGCAGTATCTCTTTTATCGTAGTCACTTATGATTGAAGGCATTTTAATTTTCCTT

TCGCTGTAAACTTTTCATCTCATT

>MSTRG.3699.1 gene=MSTRG.3699

GATAATTTGAAACCATTCTCTCTCTAATATTACTTGCTCACTCAATCAACTTCACTTTCTTGATTTGCTC

AATTCCAGATCATCCCTATCTTCTGTAATTAAGACATGGGTTCTTTGCAGAAGAGTGTTCTTGTGACTGG

AGGGGCTGGTTTTATTGGCACACACACTGTACTTGTGCTGCTTAATCAGGGCTATAAGGTCTCTATTATC

GATAATCTTGACAATGCTGTTGAAGATGCTGTCTCTAGAGTCAGGGATTTAGTGGGTTCTCAGCTTTCTC

AGAATCTTGAATTTCATTTGGGAGATATCAGAAATAAGGATGATTTGGAGAAGCTATTTTCGAAAAATAA

GTTTGATGCTGTCATCCATTTTGCGGGGCTTAAGGCTGTGGGGGAGAGTGTTGCTCATCCTTTCCGTTAC

TTCAGTAACAATTTGGTCGGATCAATCACTTTGTATGAGACAATGGCCAAGTATAACTGCAAGAAGTTGG

TTTTTTCATCCTCCGCAACAGTTTATGGGCAACCTGAGAAAATTCCATGCGTTGAGGATTTCGAGTTACA

AGCCATGAATCCATATGGGCGCACTAAGCTTTTCCTTGAAGAAATTGCTCGGGACATTCAGAAGGCGGAT

TCAGATTGGAAAATCATTCTGCTAAGGTACTTTAATCCTGTTGGGGCCCATGAGAGTGGCAAAATTGGTG

AAGATCCAAAGGGCATTCCAAACAATCTTTTGCCTTACATACAACAAGTAGCTGTGGGCAGATTACCTGA

GTTAAATGTATATGGTCATGACTATCCCACACCAGATGGTACTGCGATCCGCGATTACATCCATGTTATG

GATTTAGCAGATGGTCATGTTGAGGCACTTAAGAAGCTCACATCAGATGATATAGGTTGTATTGCCTACA

ATCTGGGGACTGGCCGTGGTACATCTGTGCTTGAAATGGTTGCGGCATTTGAGAAAGCATCTGGAAAGAA

AATCCCTTTAAAGCTTTGTGCAAAGAGGCCTGGAGATGCTACCGCTGTTTATGCATCAACTGAGAAAGCA

GAGAAAGAGCTTGGTTGGAAGGCAAAGTATGGTGTGGAGGAGATGTGCCGTGATCAATGGAAATGGGCAA

GCCAGAACCCGTATGGATACAGATCCAAGCCTGAATAGCAACACCAAAAGAAATACACAGCGTCTGCAAA

ATGGTTACCGAACCCCTTACATACTAGAAAGGGTTTGCTAAGCTTCTCCAGATATGCACTATACTGTCAA

TTTCTTTTCTGTGAATCAAAATGGTTTACAAGAAATTTGTGAATGCAAATAGAATAATACCAGTGAACTG

ATGTGAAGTTTGTGATAAACAATGTTTTACCAGTTAATCATGGATCCTTTGAATATGCTGTTGTAAAGTA

GTCCTCTTCCTCTACTCCAAAAGATAATAGTTACAAGAGTGGCAGATTGCTTGGAATTGCTTTTGGATCG

TTCGTTTTATTATCTCTGTAGGTTTTCTATAGATTGTTTCTTGTAGATATGCGGAAATTAACTGCAGTTG

TTTGCAGTACTAATATTTTCTTGCACTATACACTATTGCTTATGAATGGCTGTCTGGAACTGGCCTTGAA

ACTTACAGAGCTTCCAGCTAGATACGATATGAG

>MSTRG.3700.1 gene=MSTRG.3700

ATTCATAGAATGCAGGAAGATTATCAAATCGAGTGGCCCTTTGTGACGATGGATGTTACTATTGTATCGG

CTCAAGGCCTCAGAAGTCGAACAAATCTTTTCTCCAGCCGCGGAATTAGACCATTCACCACTCTCAGCAC

CAACATCAACGGGGTTAAACACCTTCATGTTTACAGAACAAAGGTGGATAACTATGGTGGTGCAAACCCT

GCCTGGGGCGACAAGTTCCAGCTGCCGGTTGATGAAGCCACCTTCAACTACAGTAATGTAATATCGTCTT

ACATATATTTGGAGGTTTTTACCAAACGTCTAGTTGCAGGACAGTCGACACTGGGTTGGTGCCAGATTCC

GGTGACTGATATCGTGAATGAATTTTTTCCCAAAGACGGTGTTCGCCAGCTGAGTTACCAACTGAGAAAC

AGAGATGGCTCGAGGGGTCAAGGAGTTGTTAATATTGAAGTCAAGTTGACAGGCTTATTCCATGAAAGAT

TTCAGAGGCTTCAACGGCCGCTGATTTGCAGTGGGGGATTGCCACAGCAATACGTGCCGGAGACGGTGAT

CGGCATTCCTGTTAACCAAAGCGACGGCAGAGAGTTCCTGCCCATCATCAACATGCATAAGCAGCACCCA

GTATAATTGCATAAGCACCCATCCAATCTATTTCGGGACTATATGTTACTCAGTCCGTTTCAAAATAATT

ATCTGAAAAAATGTAATTACCGTATCCCGTCAATACATTAAAATACATCGCGACAATTATTTTGAAACGG

AATGAGTAAATTTTGGTTGCCAAGGACAGTGTGTAAGCTGTAATGATTGAAGAGTAGCAGATAAGTTAAA

TGTTTTGGAATTGATTAGGAGACCATT

>MSTRG.3705.3 gene=MSTRG.3705

GTACTATATGCTACGGACTTGCAAATTTTGCCAAGAATCAAATCAAACGCGATAATTTGGCCACCGGGTC

ATTGTTAAGGTTGATAGTGTGTCGTGTTTGCGAGGAGGAAGACTGATTGGTTTGTTGAACACGTTCTTTT

ACTTTTGGACAGTAGCTGCGAGTGTCTCCGGAAGGAACTTGAGAGACATTTGGCATCTAAAGAAGCTTCT

CTCCCGCAAGAACCATCACCAGATGATGATAATGCCGTTAACCTGCTTGTGCGGATGCCAGATGGCGGCC

GGCAAGGCCGCAGATTCCTCAAGTCCCACAAGCTTCAGCATCTTTTTGACTTCATCGATGTTGGTCGGTT

GGTCAGACCTGGTACTTACAGACTGGTAAGGCCATTTCCAAGACGGGCTTTTGGTGTTGGTGAGAGCGCA

TCAACTTTTGATGAACTTGGCCTGTCCGGCAGACAGGAGGCCTTGTTTCTGGAGTTAATATAGCAGATAT

AGTTATCGTAAAATTGTAATAGGATCTTGAAGAGTTTACCCAGATGGACCAACCTTGTTTGTTAGTTGGA

TGTAAAAGTTGTTTGGAGACGGGTATTTTGTCGAAGTTTGTATCTATCGGGTATTTGGTCTTGT

>MSTRG.3706.1 gene=MSTRG.3706

CTCTTGGGGTACTACTCCGTGTATATCAGTATACGACAATTGAAGAACTCTTAAAGAAGTAGAGATATTC

ACGGGCAAAACAGAGGAGATGTTAACTCCGTGAAGATTAAGCACTCCGAGATGCGTTAAATTCTGCAACA

GTGATTTAAACACTTCTTCTTCAAGTTCAGCCCCATCGTTGTAACTGAGATCAAGTGAGACCAGTTTCTG

CAAATGAGAGATTCGAGAGGGAACTCTCCCTGATAATCCAGTATAAGAGAGGTTGAGATGCGTCAAACCT

TTTGCAAAGAAACCAAATTCCGGAGGGAACTGACTGGAGAGGGAAAAGTCATTTTGAGAGAGATTGAGGA

AGCGGAG

>MSTRG.3707.1 gene=MSTRG.3707

GCATCCTCGAGAGCATCAGGCTCTGAAGAAGTTCTATAATTTCATTCCAATGCACCAGATGACTGCAGCT

TAAATCCAGTCCCCTTAGGTCCCCTGTATTCTCGTCGCAGCTAACTCCATCCCACATGCAGTAATCAGAA

TTCATGCTCCAATTCTTTGTCTTTGGGGGTGAAGGGATATCATGAACATCATCTTCTTCGTA

>MSTRG.3708.1 gene=MSTRG.3708

AAAAAAATTTAACGGAAAAAGAAAATGAGGAATTCCATCCTCTGTTTTTCCACCATCTGTTTGTTGCTCA

TCTTCCTGCAGTTGCAGCACAACCTAGTGGCTTGTTCATTAAGCCCCCACTTGCAGAAACTTGCCTTGTT

CCATTTTAACCAAACCTTAACAATCAGTACTACTAGCTGGGATTGCGAATACGTTTCCTACGAAGAGAAT

GCAGTGTATGAAATTCCTTCACGGGCAAAGACAAAGAATTGGAGCATGAGTTCTGATCACTGCACATGGG

ATGGAGTTAGCTGCGACGACAAGACAGGGGACGTTATCGGACTGGATTTAAGTTGCAGTCAGCTGGAAGG

AGCCATTCTTCCTAACAGCGCTCTCTTCCA

>MSTRG.3709.1 gene=MSTRG.3709

TCATCAGTCTTGTATTTAGCTTGATCGAACATAATTGTTCTCTTCAATTTCTTTGCCTACTTGTCGACGT

GTTTATGTAGCGTGTCTCTACTCTGTAGCTTACACATCATTGATTCATGTTATTTTCCCCTTAGGAACTT

GAGAGACATTTGGCATCTAAAGAAGCTTCTCTCCCGCAAGAACCATCACCAGATGATGATAATGCCGTTA

ACCTGCTTGTGCGGATGCCGGATGGCGGCCGGCAAGGCCGCAGATTCCTCAAGTCCCACAAGCTTCAGCA

TCTTTTTGACTTCATCGATGTTGGTCGGTTGGTCAGACCTGGTACTTACAGACTGGCAAGGCCGTATCCA

AGACGGGCTTTTGGTGTTGGTGAGAGCTCAACAACTTTTGATGAACTTGGCCTGTCTGGCAGACAGGAGG

CCTTGTTTCTGGAGTTAATATAGCAGATATAGTTATCGTAAAATTGTAATAGGATCTCGAAGAGTTTACC

CAGATGGACCAACTTTGTTTGTTAGTTG

>MSTRG.3709.2 gene=MSTRG.3709

AATATAAATCATGCTAACAGGAGCAGATGATGAACTTGATTGCAATGAAGAACTGAAGTTCCACAACGAG

ATGAAGTCGACGAAGGAAGGAACTTGAGAGACATTTGGCATCTAAAGAAGCTTCTCTCCCGCAAGAACCA

TCACCAGATGATGATAATGCCGTTAACCTGCTTGTGCGGATGCCGGATGGCGGCCGGCAAGGCCGCAGAT

TCCTCAAGTCCCACAAGCTTCAGCATCTTTTTGACTTCATCGATGTTGGTCGGTTGGTCAGACCTGGTAC

TTACAGACTGGCAAGGCCGTATCCAAGACGGGCTTTTGGTGTTGGTGAGAGCTCAACAACTTTTGATGAA

CTTGGCCTGTCTGGCAGACAGGAGGCCTTGTTTCTGGAGTTAATATAGCAGATATAGTTATCGTAAAATT

GTAATAGGATCTCGAAGAGTTTACCCAGATGGACCAACTTTGTTTGTTAGTTG

>MSTRG.3710.1 gene=MSTRG.3710

CAGCTGGAAGGAGCCATTCTTCCTAACAGCACTCTCTTCCAACTCTCTCATCTCCGCTTCCTCAATCTTT

CTCATAACAACTTTTCCCTCTCCAGTCAGTTCCCTCCGGAATTTGGTTTCTTTGCAAAAGGTCTGACGCA

TCTCAACCTAATAGGCGGTCTTGAGCAACTGCCTTGGGGTTATATTAATTATCTTGATCTCCAAGACAAT

AAGCTGAATGGATCGTTGCCCACCTTGATGTGTAATTCCAGCTCTCTTGAGATTCTAAATCTCTCTCACA

ACAATTTGAGTGGTGTACTCCCCATTTGTCATACAAATTTGACTCAGCTTTCGGTGTTGGATCTGCGGAG

GAATAAAATTCAAGGAAACATTCCAGCAACCTTGTCAAATTTCCATAATCTGGAAACCATTAATATGAAT

GGAAACCAATTACAAGGAAGCATTCCTTCTTCATTTGCTGAATTTGATTCTTTAGCCGTTCTTGATCTTG

GAAATAATCAGATAAATGATACATTCCCCCGGTGTTTGGAAGCTCTTCCAAATCTCCAAGTTCTTGTTCT

CAAATCTAACAAGTTTTGTGGTCTTATAAACACGAGTTCTAAGATTGTACACCCATTTCCTAGCTTGAGA

ATCATTGATCTGTCGGACAATGAGTTTTCAGGTCCACTGCCAGCAAAATACTTCAGAAACTTCAAGGCTA

TGATGGATAGTGATGTAAGCAAAATAGATCGTAGTTATATGAATTATGGTTCCTATTACAGTGATTCCAC

GACTCTGGTGATTAAAGGAGTAGACATGGAGTTGGTAAAGATATTAACTATTTTTACAACCATGGATGTA

TCAAGGAACCATTTTGAAGGGGAGATTGTAGAATCAATCGGAAATCTAGTGTCACTTAGAGATCTCAATT

TATCTTACAATCATCTCACAGGCCACATACCACGCTCGATTGGAAAATTGACTGTGTTGGAGTCACTAGA

CCTCTCATCCAATGAACTTGAAGGAGAAATTCCTTCGCAACTCACTGGTTTATATTCACTTGCAGTTCTA

AATCTGTCTTGCAACCAACTCAGCGGTCATATTCCACAAGGTTCTCAGTTTAATACATTTGATGCTGATA

GCTATGGTGGTAACTTCGGGCTGTGTGGAAATCCATTGTCCAAAACGTGTGAACAAGATATTGTGGAGCA

AGAAGACGAAGAAGATGAGGATGAATGTCACTTTTTCTGTGGTTTTACTTGGGAAGCGGTGGTGATTGGA

TATGGGTGTGGAGTAGTGCCTGCATTTATCAGTGGGTACCTGATGTTGCTGGCACGGAAACCAAAATGGT

TTGCTGGAATCATTGGCAGGGAATTGGGGCTCAAGATCAGGAGGATGGAAATTAGAATGCGATACTAAAG

ATTGGAGAGGCATGAACATTTTGGCATGTAAGTTGATGGGACAAGCAGATTCAGATGCACTGGAATCAAC

CACTACTTCAGCTGTTTTGCATCCTTTGTCATTTTAAATTTTACATTTTATAATAAAAGTCAAAAAGATA

CAAGTTCCATGGACAAGTTCTGTATTGTGTAGTTCCAG

>MSTRG.3710.2 gene=MSTRG.3710

CTTGCCCACTCCAGTAAACTAGTTGTGTTGCATATTGAGAGTAATAGCTTTACTGGATCCCTACCTCGTT

CGTTGTTTCACCACTCCAGTAAACTAGTTGAGTTGTATCTTCATAATAATAGCTTTACTGGATCCCTACC

TCGTTCGTTGTTTCACCATCCATCCTTAGAGATCTTGAATCTGGCATATAATGCATTCACGGGTCCACTA

CATGAATTTCATTCTTCTTTGTCACGATTGCGATCTTTTTATTGTTCCAGCAATTTACTCAATGGATCCA

TTCCACATTCATTTTCTCAACTTGTGAATCTCACTGAACTAGATCTTTCATCAAACAATTTCACCGGGGT

TTTGGATTTGGAAATGTTTTCAGGCCTCAAATCTCTAGGAATTCTTCATCTTTCTTATAATAGTCTTTCA

GTGAGAACTACAACTATGGCCACTCTCCCTCCTACGCTTAACGACTTGGCCTTGTCATCTTGCAACATGA

AAGAGTTTCCTCACTTTTCAAGAGATGCAAACATCTTCCTCGATTTTGTAGACCTATCAAACAACAAAAT

TGAAGGGGAAATTCCACACTGGATTGGGTCGGTGGGGAGGTACCCTTTCTATCCAATCCATTATCTGAAT

CTTTCTCACAACGGACTAATATGCGGTCTTGAGCAACTGCCTTGGAGTCATGTTGAATTTCTTGATCTCC

AATACAATAAGCTGAATGGATCATTGCCCACCTTGATGTGTAATTCCAGCTCTCTCCGGGTTCTAAATCT

CTCTCACAACAAGTTGAGTGGTGTACTCCCCATTTGCCATACAAATTTGACTTCGCTTCGCGTGTTGGAT

CTGCGGAGGAATAAAATTCAAGGAAACATTCCAGCAACCTTGTCAAATTTCCGTAACCTGGAAACTGTTA

ATTTGAATGGAAACCGATTAGAAGGAAGCATTCCTTCTTCTTTTGCTGGATTTGATTCTTTAGCCGTTCT

TGATCTTGGAAATAATCAGATAAATGATACATTCCCCCAGTGTTTGGAAGCTCTTCCCAATCTCCAAGTT

CTTGTTCTCAAATCAAACAAGTTTTATGGTCTTATAAACACGAGTTCGAAGATTGTACACCCATTTCCAA

GCTTGAGAATCATTGATCTGTCGAAAAATGAGTTTTCAGGTCCACTGCCAGCAAAATACTTCAGAAACTT

CAAGGCTATGATGGATAGTGATGTAAGCAAAATAGATCGTAGTTATATGAATTATGGTTCCTATTACAGT

GATTCCACGACTCTGGTGATTAAAGGAGTAGACATGGAGTTGGTAAAGATATTAACTATTTTTACAACCA

TGGATGTATCAAGGAACCATTTTGAAGGGGAGATTGTAGAATCAATCGGAAATCTAGTGTCACTTAGAGA

TCTCAATTTATCTTACAATCATCTCACAGGCCACATACCACGCTCGATTGGAAAATTGACTGTGTTGGAG

TCACTAGACCTCTCATCCAATGAACTTGAAGGAGAAATTCCTTCGCAACTCACTGGTTTATATTCACTTG

CAGTTCTAAATCTGTCTTGCAACCAACTCAGCGGTCATATTCCACAAGGTTCTCAGTTTAATACATTTGA

TGCTGATAGCTATGGTGGTAACTTCGGGCTGTGTGGAAATCCATTGTCCAAAACGTGTGAACAAGATATT

GTGGAGCAAGAAGACGAAGAAGATGAGGATGAATGTCACTTTTTCTGTGGTTTTACTTGGGAAGCGGTGG

TGATTGGATATGGGTGTGGAGTAGTGCCTGCATTTATCAGTGGGTACCTGATGTTGCTGGCACGGAAACC

AAAATGGTTTGCTGGAATCATTGGCAGGGAATTGGGGCTCAAGATCAGGAGGATGGAAATTAGAATGCGA

TACTAAAGATTGGAGAGGCATGAACATTTTGGCATGTAAGTTGATGGGACAAGCAGATTCAGATGCACTG

GAATCAACCACTACTTCAGCTGTTTTGCATCCTTTGTCATTTTAAATTTTACATTTTTACAATAAAAG

>MSTRG.3710.3 gene=MSTRG.3710

GGTCGCTGGGGGTGTCCTATCTGAATCTTTCTCACAACAAACTAATAGGCGGTCTTGAGCAACTGCCTTG

GGGTTATATTAATTATCTTGATCTCCAAGACAATAAGCTGAATGGATCGTTGCCCACCTTGATGTGTAAT

TCCAGCTCTCTTGAGATTCTAAATCTCTCTCACAACAATTTGAGTGGTGTACTCCCCATTTGTCATACAA

ATTTGACTCAGCTTTCGGTGTTGGATCTGCGGAGGAATAAAATTCAAGGAAACATTCCAGCAACCTTGTC

AAATTTCCATAATCTGGAAACTGTTAATTTGAATGGAAACCGATTAGAAGGAAGCATTCCTTCTTCTTTT

GCTGGATTTGATTCTTTAGCCGTTCTTGATCTTGGAAATAATCAGATAAATGATACATTCCCCCAGTGTT

TGGAAGCTCTTCCCAATCTCCAAGTTCTTGTTCTCAAATCAAACAAGTTTTATGGTCTTATAAACACGAG

TTCGAAGATTGTACACCCATTTCCAAGCTTGAGAATCATTGATCTGTCG

>MSTRG.3710.4 gene=MSTRG.3710

CAAACTAATAGGCGGTCTTGAGCAACTGCCTTGGGGTTATATTAATTATCTTGATCTCCAAGACAATAAG

CTGAATGGATCGTTGCCCACCTTGATGTGTAATTCCAGCTCTCTTGAGATTCTAAATCTCTCTCACAACA

ATTTGAGTGGTGTACTCCCCATTTGTCATACAAATTTGACTCAGCTTTCGGTGTTGGATCTGCGGAGGAA

TAAAATTCAAGGAAACATTCCAGCAACCTTGTCAAATTTCCGTAACCTGGAAACTGTTAATTTGAATGGA

AACCGATTAGAAGGAAGCATTCCTTCTTCTTTTGCTGGATTTGATTCTTTAGCCGTTCTTGATCTTGGAA

ATAATCAGATAAATGATACATTCCCCCAGTGTTTGGAAGCTCTTCCCAATCTCCAAGTTCTTGTTCTCAA

ATCAAACAAGTTTTATGGTCTTATAAACACGAGTTCGAAGATTGTACACCCATTTCCAAGCTTGAGAATC

ATTGATCTGTCGAAAAATGAGTTTTCAGGTCCACTGCCAGCAAAATACTTCAGAAACTTCAAGGCTATGA

TGGATAGTGATGTAAGCAAAATAGATCGTAGTTATATGAATTATGGTTCCTATTACAGTGATTCCACGAC

TCTGGTGATTAAAGGAGTAGACATGGAGTTGGTAAAGATATTAACTATTTTTACAACCATGGATGTATCA

AGGAACCATTTTGAAGGGGAGATTGTAGAATCAATCGGAAATCTAGTGTCACTTAGAGATCTCAATTTAT

CTTACAATCATCTCACAGGCCACATACCACGCTCGATTGGAAAATTGACTGTGTTGGAGTCACTAGACCT

CTCATCCAATGAACTTGAAGGAGAAATTCCTTCGCAACTCACTGGTTTATATTCACTTGCAGTTCTAAAT

CTGTCTTGCAACCAACTCAGCGGTCATATTCCACAAGGTTCTCAGTTTAATACATTTGATGCTGATAGCT

ATGGTGGTAACTTCGGGCTGTGTGGAAATCCATTGTCCAAAACGTGTGAACAAGATATTGTGGAGCAAGA

AGACGAAGAAGATGAGGATGAATGTCACTTTTTCTGTGGTTTTACTTGGGAAGCGGTGGTGATTGGATAT

GGGTGTGGAGTAGTGCCTGCATTTATCAGTGGGTACCTGATGTTGCTGGCACGGAAACCAAAATGGTTTG

CTGGAATCATTGGCAGGGAATTGGGGCTCAAGATCAGGAGGATGGAAATTAGAATGCGATACTAAAGGTT

TGCTGTAATCTGTATCACATTCCCAGGTCTTGAATATTCTGTAATAATATAGTAATATCTGCTATCATAT

AATTCTATATTTGCAGATTGGAGAGGCATGAACATTTTGGCATGTAAGTTGATGGGACAAGCAGATTCAG

ATGCACTGGAATCAACCACTACTTCAGCTGTTTTGCATCCTTTGTCATTTTAAATTTTACATTTTATAAT

AAAAGTCAAAAAGATACAAGTTCCATGGACAAGTTCTGTATTGTGTAGTTCCAGGGCTCCTTTCAATTTC

TAACCACCCTGGACAACTTCTTTACAAAACTAACCACCCTTTTTTTCACTTCACAAAACTGAC

>MSTRG.3711.1 gene=MSTRG.3711

TCTCCCTTGACCGTAACAATTTTACTGGTCCGTTTCCCTCTTTTCTTGCCCACTCCAGTAAACTAGTTGA

GTTGTATCTTGATAGTAATAGCTTTACTGGATCCATACCTCGTTCGTTGTTTCACCATCCATCCTTAGAG

TACTTGTTTCTGGATAGTAATGCATTCACGGGTCCACTACATGAATTTGATTCTTCTTTCTCACGATTGC

GATCTTTTTCTTGTTCCCACAATTCACTCTATGGATCCCTTCCACATTCATTTTCTCAACTTGTCAATCT

CACTAGCCTAGATTTATCATCAAACAATTTCAGCGGGGTTTTGGATTTGGAAATGTTTTCAGGCCTCGAA

TCTCTACAATATCTTGGACTTTCTCATAACAGTCTTTCAGTGACAAGTACAATCAGGGGCACTCTCCCTC

CTACGCTTAACTACTTGGGCTTGTCATCTTGCAACATGAAAGAGTTTCCTCACTTTTCAAGAGATGCAAA

CATCTCCCTGGAGTTTGTAGACCTATCAAACAATAACATTGAAGGGGAAATTCCACACTGGATTGGGTCA

GTGTGGTACTCTGAGTATTATAACTTGCCCTATCTGAATCTTTCTCACAACAAACTAATAGGCGGTCTTG

AGCAACTGCCTTGGGGTTATCTTACTTATCTTGATCTCCAAGACAATAAGCTGAATGGATCGTTGCCCAC

CTTGATGTGTAATTCCAGCTCTCTTGAGATTCTAAATCTCTCTCACAACAAGTTGAGTGGTGTACTACCC

ATTTGTCATACAAATTTGACTCCGCTTTCGGTTTTGGATCTGCGGAGGAATAACATTCAAGGAAACATTC

CAGCAACCTTGTCAAATTTCTGTAATCTGGAAACTATTAATTTGAATGGAAACCAATTAGAAGGAAGCAT

TCCTTCTTCTTTTGCTCAATTTGATTCTTTAGCCGTTCTTGATCTTGGAAATAATCAGATAAATGATACA

TTCCCCCGGTGTTTGGAAGCTCTTCCAAATCTCCAAGTTCTTGTTCTCAAATCAAACAAGTTTTATGGTC

TTATAAACACGAGTTCTAAGATTGTACACCCATTTCCTAGCTTGAGAATCATTGATCTGTCGGAAAATGA

GTTTTCAGGTCCACTGCCAGCAAAATACTTCAGAAACTTCAAGGCTATGATGAATAGTGAAGTGAATGAG

GTAAAGCGTAGTTATATGGAGCATCAGTATTACAGTGATTCCACAGCTCTAGTGATTAAAGGCATGGAGA

TTGATTTAGAAAGAATTTTGACTGTGTTCACAACTATTGATGTATCGAAGAACAATTTTGATGGGGAGAT

AGCTGAATACTTTGGAAATCTAGTATCACTCAGATTTCTTAATCTTTCTCACAATCATCTCACAGGCCAC

ATACCACGCTCGATTGGAAAATTGACTGTGCTGGAGTCACTAGACCTCTCATCCAACCAACTTGAAGGAG

AAATTCCTCAGCAGCTCACTAGTGTATATTCACTTGCAGTTCTAAATCTTTCTTGCAACCAACTTAGAGG

CCGTATACCAGAAGGGGCACAATTCAATACGTTTGAAAACAGCAGCTACGCTGGTAACTTTGGGCTGTGC

GGACATCCCTTGTCGAAGAAGTGTGAATCTTATGTTAGGACACAAGAAGAAGATGACGAAGATGAGGATG

AATGTCACTTTTTCTGTGGTTTTACTTGGGAAGCGGTGGTGATTGGATATGGGTGTGGAGTGGTGCCTGC

ATTTATCACTGGGTACCTGATGTTGCTGGCAAGGAAACCCAAATGGTTTGCTGGAATCATTGGCAGGGAA

TTGGGGCTCAAGATCAGGAGGATGGAGATTAGAATGCGATGACTAGAGATTGGAGAGGCATGAACATTTT

GGCATGTAAGTTGATGGGACAAGCAGGATTCAGATGCACTGGAATCAACCACTACTTCAGCTGTTTAATT

TGCATCTTTTATCATTTTAAATTTTACATTTTACAATAAAAGTCAAAAAGATACAAGTTCCATAGACAAG

TTCTGTTTTGTTTAGTTCCAGCCTCAGGCTGTTCATTACTAAAACATGTTTGCTGATGCTTTATAATGTA

CAATTTAGTAATTTACAATATATCACAGTTTGTAATATACACTCCTAGATGATTTTATGGCCTGTTTATC

CTTAATCGCTTTCATGAACTAATGGAAATTTGTAAAAGCGTGTCACATTTGCAGTGATGCCGCTATTTTT

CCTGAATTTCCCTTGCTGTGAAGATTGTCTGAATCATGTTAAAGTCGCTGCCATCAGTATTTGCATTCAA

CGGCTTACTGATACTGGTAGAGATACTAAATTTCACAAGTTATTGATAGAATAGAAGATGAC

>MSTRG.3711.2 gene=MSTRG.3711

AAGAGTTTCCTCACTTTTCAAGAGATGCAAACATCTCCCTGGAGTTTGTAGACCTATCAAACAATAACAT

TGAAGGGGAAATTCCACACTGGATTGGGTCAGTGTGGTACTCTGAGTATTATAACTTGCCCTATCTGAAT

CTTTCTCACAACAAACTAATAGGCGGTCTTGAGCAACTGCCTTGGGGTTATCTTACTTATCTTGATCTCC

AAGACAATAAGCTGAATGGATCGTTGCCCACCTTGATGTGTAATTCCAGCTCTCTTGAGATTCTAAATCT

CTCTCACAACAAGTTGAGTGGTGTACTACCCATTTGTCATACAAATTTGACTCCGCTTTCGGTTTTGGAT

CTGCGGAGGAATAACATTCAAGGAAACATTCCAGCAACCTTGTCAAATTTCTGTAATCTGGAAACTATTA

ATTTGAATGGAAACCAATTAGAAGGAAGCATTCCTTCTTCTTTTGCTCAATTTGATTCTTTAGCCGTTCT

TGATCTTGGAAATAATCAGATAAATGATACATTCCCCCGGTGTTTGGAAGCTCTTCCAAATCTCCAAGTT

CTTGTTCTCAAATCAAACAAGTTTTATGGTCTTATAAACACGAGTTCTAAGATTGTACACCCATTTCCTA

GCTTGAGAATCATTGATCTGTCGGAAAATGAGTTTTCAGGTCCACTGCCAGCAAAATACTTCAGAAACTT

CAAGGCTATGATGAATAGTGAAGTGAATGAGGTAAAGCGTAGTTATATGGAGCATCAGTATTACAGTGAT

TCCACAGCTCTAGTGATTAAAGGCATGGAGATTGATTTAGAAAGAATTTTGACTGTGTTCACAACTATTG

ATGTATCGAAGAACAATTTTGATGGGGAGATAGCTGAATACTTTGGAAATCTAGTATCACTCAGATTTCT

TAATCTTTCTCACAATCATCTCACAGGCCACATACCACGCTCGATTGGAAAATTGACTGTGCTGGAGTCA

CTAGACCTCTCATCCAACCAACTTGAAGGAGAAATTCCTCAGCAGCTCACTAGTGTATATTCACTTGCAG

TTCTAAATCTTTCTTGCAACCAACTTAGAGGCCGTATACCAGAAGGGGCACAATTCAATACGTTTGAAAA

CAGCAGCTACGCTGGTAACTTTGGGCTGTGCGGACATCCCTTGTCGAAGAAGTGTGAATCTTATGTTAGG

ACACAAGAAGAAGATGACGAAGATGAGGATGAATGTCACTTTTTCTGTGGTTTTACTTGGGAAGCGGTGG

TGATTGGATATGGGTGTGGAGTGGTGCCTGCATTTATCACTGGGTACCTGATGTTGCTGGCAAGGAAACC

CAAATGGTTTGCTGGAATCATTGGCAGGGAATTGGGGCTCAAGATCAGGAGGATGGAGATTAGAATGCGA

TGACTAGAGGTTTGTTGTAATCTATATCACATTCCCAGGTCTTGAATAATCTGTAATAATATAGTAATAT

ATGTCTATGCGTCAGTTTAAGGATCGAGATGTGTAGCTGTGACTTGTATTTTAGAATTGCACAAGATGCT

ATACAATGGTAGCAGAGGAGGGCTGGGGATTTATTGGGTCAGTTCGAAGCACTCAAGTAGAATTGGTTGT

AGTACGACCATAAGAGTTTGAATTCTGAGGCTAATTTGACTCAACTTGTAGCCTTCCACTAGACTGCAAG

GTGAGCAAGATTGAGTCGATGTTTGGACATGAGATTATGTATACTATAACATCTGAGGCCCGTGGCAGAT

CGATGAGGCATGAAAGTTGGGAACAGGGGACGGCTGGTGCATTGGAATGAAAGTAGAAAACTATTTATAA

CGGAAGGGACCAGATTAGTTTAACTGCATAATGCATTCAGTTCGCAGCACAGAAGATTTTGGTGAGAGGG

TTTGAAGGAAAGGACATGATTCATATCATAGATTTAGATAATTAAATATAAGGCAAGGCTTTCCATTCCA

TGGCCAGTCTTTTGCTGGAAGGCTGACCTGTAGGAACAATCGTCCCAGTCATTCTTATACGCTGGTACTC

CGCTAGAAAGCCTGGTTCTGTGGACTTGAGAGTTGATATATCCTAGACTCTGAAGCCCGTAACAATCTAA

AAGATGACATGTAAGTCGAAGGAACGAGGAGGATTCGGAGCATTGGATTCAATAACAGAAAACTTCTTCA

GAGCCAGAGTCTCTTCCAGACGCTCTCTTCTGATATATATCTGATGAAGTAAAAGCATTAGTCATCACTA

CCTTGCATATTTTTTGATACAAAACATCAACTCATGTTTCTCTGGTACGCGCAAAAAATATG

>MSTRG.3716.1 gene=MSTRG.3716

TGTAATCTAACAGTTCCATCTTCTTCTTTAGGTCTCCTGCTGTTTCTACCCTTGTACTTCGAGAAAATAA

CTTCAGTTCATCCATTTTTAATTGGCTTCTCCAATTCAATGGCAGTCTGGTAGAACTTGACCTGGGTATT

AACGCGTTAGAAGGTAGAATTCCAGAGTCATTTGGTCATATGACTGCTCTGACTCATCTGAGTCTTGGCC

ACAACCAATGGTCCAATTCCAGAGTCATTTGGTCATCTGAATGATCTGAGCTATATGGATCTATCTGCAT

ATCAACTTTCTGGCGTGCTTCCGAATTCCATTCATAAATTATCCAATCTGAAAGTGAATATACGAGGTCA

CATCCCGCCTTCATCCCCTGAGTATGAAGCTGATGAAGACGATAGTGAATATACAAGGTGGCTCTATGTA

AGTTTCAGCACTAGTTTTTGGTGGATCATTGGAACTTTAATGCTCAACCGTCGTTGGAGACAAGCATATT

TCCTGTTTCTCTACGATCTACTCCCTCCATCCCAAATTAGATGAGCCTTTTGGCAAAAAAATTTATCCCA

AAATAGTTGAGCTTCTCCCTTTTCAATGCATATTTATCAACCATCTTCCATCCTTACCCTCTCTTATTTA

TCATTCCAATGAACCACTAGCTAATATACAATAAGAAGTCAACACAATAAATATGGGCAAAGATGGAAAA

AAAACAACCATTTAATATTACTTAATATGTGTGAAATGAGCAAAAGGGTCATATAAATTAGGATGGAGGG

AGTAAAAGACAATGTTATCGTGTCTCTGCGCATTGCCAAATTGCGCAGGAATGTTTAAGGCTTGAAGTCT

TGCATGATGATGCAAAATACCACTGATCTTTCTGATCAATGTTTATCAGTATCATTGCTTTATGCATTTG

TTTTTCCATCCACTCTTTTGAATCATCAATTTCAGTCTTGTCTGTTGTTGCATAGAATCTAATTCTTAC

>MSTRG.3726.1 gene=MSTRG.3726

TTGTGGTACTTATCGCGAGAGCTCACCTCGCATAACAAATGATTGTTCAATATATGGGGCATCTGTACCC

CCAACCTCAGTTGGTGCAGATGGATATGAACAATGTTCTATCACCGGTTTTTTATGTGCGCAGGTCAATT

GTGATTCCACTATTTTCAGTGATGTTGGTGTCGACTGGATTGCTCGAGAAACTTTTGTAATCATTTTTAT

TATCAAGAATATTTATATTCTATCTGTTAAGCAATCTGAAAACAAAGCGGC

>MSTRG.3730.1 gene=MSTRG.3730

TCCTGGATTTGAGCATCTAGACCACCTATATCAGCATATGATTCTAAAGGAGCTTTCTCAACCTTCATTA

CAGACACCATTGGGTCAACATCATCTTGAAGCAGACCAACCACAGAAAGAACCTTATTATGCATCAAGAT

AGCACAACCAGGCTCCAACTGATCCTTATCAACAAATGACAAAATCCCCACATAATACTCCGGTC

>MSTRG.3739.2 gene=MSTRG.3739

CCTTATTGGACTACATGTTCTGTCACTATGCCGTTCTCGAATAAGCACAGGGTTAGTCTGTGTGTGGATG

CACTCAAGTTTGATGCTACAAGTAACAAGTTCAAGTTGTTGCCTAGGTACTCATCTGAATTCAGCAGGAA

TAAGAAATTCAAAGTCGTCAATATGAGGGACCTTCTTAGTTTAATGGCGTATGAATTCAACTTTCTTGAG

GTGTATTCTTTGGACGAGGACAAGGGATGCGAGGGTGGTGTGTGGAGTAAGATGTACAATTTCGGACTAT

TTATGCGATTTGGGCGTTTTGAGTGCCTGCAACAGGGTTTCAAGTATGGCGATGAGATTGTAACTAAAGA

ATTTGGTAGGATTGCCTGCTTTAATCACAAATTGGGGACTATTAAGCACATCCCGTGGCCGTGTGTAGAC

TCTCTTGACACGCGTTTCTGCTTTCCATATACGCCTAGTTTGGTTTCACTTCCAGGAATGACCTCTATTT

ATTTAGAAAAGCAGACTCCTCTACTTGGTCATCTCCCTACAATTCCTCCTCTGCGTTGAGTGGACTCACG

AAGAGCTCATTTGCCATAATAGAAGACAACTCAAGTGTTCAATTGAATGCCAGGGGTTTGAAGGTCATTT

TAACTGATTATGTATCTGCGCTTTACAGCAATCTTATTTCATGAGGGTTTTGTTATCTTTCATATTTGGT

ATGTGGGAAGTTTTAAGTAGTTATGTTTTTTTTTTCCCATTTATATTATTTCGTTAGTCATTTGTAGCGT

GAAAGTGGGATGGAAGTAGTATTATGTAATGTGAAAGAAAAAAACAAACACTTTACATGTTATAGACAGT

GGTACTGCTGAATGTTTTTCTGCTTGCTTGTAATCTGGGGAAGTAGGTTTCATTCT

>MSTRG.3739.1 gene=MSTRG.3739

CGCGTTTCAAATCTGTATGTAAATCATGGCGCTCTCTATTCTCCGATCCACAGTTCGTTAAACAACATAG

TGATCGTCAAAACCCGAAAAACAATGACTTTCTGGTGATCACAAGCCGCAACTATATATCAAGACACATC

GCTATTATCTCCCGCTACCAAGTGGAATTTCTCCTCAACAACGATGGTTACAATCTCATCGGTTCTGCCA

AGGGCTTAGTAGGTCTGCGTAGTCGCAATGTAGTGTTCTTGTGGAACCCTGCTATTCGTCAAGCGAAAGA

ATTTCACCTTCCACCACTAGGTTTTTACTCTCTGAAGAGAGTCGGACTCGGTTATGATCCTCTCAGTAAT

GACTACAAAGTAGGAATGTGCTATCGTTCGCTTGATAAGTCTCTTAGTTATGCCGTTTACTCCTCGAGTT

CTGGTTCATGGATTCGTAATCAGTTTGATGATCGTGATGAGAAGATTCTTATTTCTATATTGGAGAATTC

TCAAATTACTGTGGTTAGGGATTCTCCTTATTGGACTACATGTTCTGTCACTATGCCGTTCTCGAATAAG

CACAGGGTTAGTCTGTGTGTGGATGCACTCAAGTTTGATGCTACAAGTAACAAGTTCAAGTTGTTGCCTA

GGTACTCATCTGAATTCAGCAGGAATAAGAAATTCAAAGTCGTCAATATGAGGGACCTTCTTAGTTTAAT

GGCGTATGAATTCAACTTTCTTGAGGTGTATTCTTTGGACGAGGACAAGGGATGCGAGGGTGGTGTGTGG

AGTAAGATGTACAATTTCGGACTATTTATGCGATTTGGGCGTTTTGAGTGCCTGCAACAGGGTTTCAAGT

ATGGCGATGAGATTGTAACTAAAGAATTTGGTAGGATTGCCTGCTTTAATCACAAATTGGGGACTATTAA

GCACATCCCGTGGCCGTGTGTAGACTCTCTTGACACGCGTTTCTGCTTTCCATATACGCCTAGTTTGGTT

TCACTTCCAGGAATGACCTCTATTTATTTAGAAAAGCAGACTCCTCTACTTGGTCATCTCCCTACAATTC

CTCCTCTGCGTTGAGTGGACTCACGAAGAGCTCATTTGCCATAATAGAAGACAACTCAAGTGTTCAATTG

AATGCCAGGGGTTTGAAGGTCATTTTAACTGATTATGTATCTGCGCTTTACAGCAATCTTATTTCATGAG

GGTTTTGTTATCTTTCATATTTGGTATGTGGGAAGTTTTAAGTAGTTATGTTTTTTTTTTCCCATTTATA

TTATTTCCTTATATGTTTGAAACATGAAAACAGAAAAACGTTAGTCATTTGTAGCGTGAAAGTGGGATGG

AAGTAGTATTATGTAATGTGAAAGAAAAAAACAAACACTTTACATGTTATAGACAGTGGTACTGCTGAAT

GTTTTTCTGCTTGCTTGTAATCTGGGGAAGTAGGTTTCATTCT

>MSTRG.3742.1 gene=MSTRG.3742

AATTTTTCAGGTGAAATTCCTCAAAGCATGTCTGGACTAAATTTTCTTGCATATTTAGATCTGTGAGACA

ATAACTTCTCGGGAAGAATACCATCTGGTACTCAACTCCAGGGATTTGATATTTCCACGTATGAGGGGAA

CGCTGGACTCTGTGGTAAACCACTCACAAACATTGGTCCTGGAGATGAACCAGTTGATCACATCCCGCCT

TCATTCCCTGAGTATGAAGCTGAT

>MSTRG.3744.1 gene=MSTRG.3744

TAGTTACACTGATGATGTTTCGGCCAGATGAAAAGGGCAGGAGCTTTCTTATGGAAGAAATTTTGCATAT

CTGAAGATGATCGATCTGTCAACCAATCAACTTACTGGAGAAATTCCCAGAGGGATAGGCAGACTTCAAG

AACTCAGGGGATTGAACTTGTCAAGAAACAGGTTTTACGGGAATGTTCCACTGGAAATTGGCCAACTGAA

AGTTTTAGAATTCATAGACTTGTCCACAACCACAAACAACTTTTCAGGTGAAATTCCTCAAAGCATGTCT

GGACTAAATTTTCTTGCATATTTAGATCTGTGAGACAATAACTTCTCGGGAAGAATACCATCTGGTACTC

AACTCCAGGGATTTGATATTTCCACGTATGAGGGGAACGCTGGACTCTGTGGTAAACCACTCACAAACAT

TGGTCCTGGAGATGAACCAGTTGATCACATCCCGCCTTCATTCCCTGAGTATGAAGCTGAT

>MSTRG.3745.1 gene=MSTRG.3745

AGAAAATAAGACCGAGAAGAATGCATAAATTATTCTTATTTGTAATTCTCTTTATAATTGCAGCTCTCAT

ATGTATTTGTATTAGTTTTCCTGTAAGAGTAGAAATATGTTTTATCGAATATTTGTGTGCGAGTTAATTT

GTAATGATAAACATAATGTGATGTCACCTATCCTTTATAGAAATATTGCTCAATTCTCTTACAATCACTT

GCAGGCACTTTATCATGCCCTCCCAATGGACCACATTACTGGTTCCAAGCTTCACAACAAGCTCGAAGGA

GAAATCAACTAGACCGCTGTGCCTAAGCTAATTGATAAGATGACAAAAGAAGGCTTCGTTGAAGCTCAAA

GCAATAAAAGACTTGATTTTCAAATTATGTCGATAGGAGCTGGATGAGCCGCAAAGGAAGCTGAACCACT

CAGAACTTCAGACAATTGGCAAGTCTGATGCCTATCAAAACGGATCCGTCAAGAGTGACCATACTGTTGT

AAAGATGAAAGAGCATGGAATCCCCAAAAGCAGAGCTCAGGCAGCAGCTTCTAAAAGAGTTTTGCACAAG

TAATAGAATTTGGCAGAACCAATGGAAATCGTGACAATTGCAGTTAGTTTGATGCAATCATGTGTAGCAA

GTCGTCGTCTCAAAGCAAGCGGTCCAGGAAAGTGCTGTCAAGGATCCTATCCTTTAGTACCTAAGAATGA

GAAGTCTCAAGTCGTCTGAAAAGAATCGGATCGTATGGCCTGATAATTTCCAGAGTTTCTTTCCACTAAT

TCCTATTTCATGAGACATAGACTTGTCTACTGCATGTATCACTTGAATTAGCTTTTTTCCTTTCTTTGTC

GATCAGTATTTTTACGCAGTATACATTCTCTCCCATTTTATACGACACCCCGTCTACTGAGCATTTTTGT

GTCTCTTTGTAAAATTAAGTGATTCATAGTTTCATACCAGGCCCGTGC

>MSTRG.3750.1 gene=MSTRG.3750

CAGTAACCCACCAAGCACTAACCCTTCTGAAATTTGAAGAGCCCCTCTTAAGTCTCTTACACGACTTACA

GCCGTTAGCCCCTTGCAGCTTGTACCATCTTCTGGACTCCATCAACAATTAGCTTCTAATTAGTAAATTA

ACCAGATCTTCTCTAGATTCCAGGACAACGGAGACGTTGTTCAGAAAACGAGCATTCCTGTCCTGATGCT

GTTGAAAATGTGGACCCGAATACAACCTCCGGAACAGGTACTACCACTTTTGGTTGTAGAGATGGATTTT

TGACGCCATCACAGAGACGGCTACACGGCGTTGATCTATCTAAAGCATCAACTCCCGCACCAGGACTATT

TCTTCCAAAATTATGTGGATCTCGAGGAATCAAAATCTGATGCCAATTGACCAAGCTGATGAGTAGCTCT

GGTGGGTGGAACACGTAAAAATACTACCATCACAAATGTTTCCATGACCAGTCCCATTACAAATCAAGTT

TGCAGGCTGGTGCAAAGGAGTAGAGTATCGGCCTAAAAAGATATGTAAATTGAGAAGGATGATGCAACTA

CCATATCCTGTTGCACTATAGATTAATTTTTGTACAAAAACACCAATGCAGGATATCAGTATAACAAAGC

ATGAAAAACTTTTGCTATTGGCTCTGTACCTTGCTTTTCACCCAGTAGCTGACAGATTGAACCTCCAGTT

TTGCTAGAATAGGGATATTGTTGTGGTTGTTTCTTTTATTTTTTTTAGAATTTGCTTTAGAAACATAGAG

GTATAATTTAATATGAAATGATATTTGTTTGTGTTGCTTCCTTTAAATGTTTACTTTAATATATGCC

>MSTRG.3753.1 gene=MSTRG.3753

CTAAAGGGGCCGGTACGGGTGGACTTGATCGTCATTTGAAAAAACATCACGGAATAACGAAAGAAAATCA

CGAAAGTGGAGCGGCACGTGGAGGTACATCGCAACAAACTCAAATTGGTGGTTTTATGACTTCGTCTCCC

GGTGGAGGTATGCCTTTCGCTTATAATCGAGATAGAATGATCGAACAATTTGCCACTTATGTTACGTTAG

ATGAGTTACCTTTTTCACATGGAGAAAGTGATAATTTAGAACACATGATAAAAACTAGCATAAATCCCGC

ATTTAGAAGAATTCCTAGGAACACACTTAAACGTCACACACAAAAACAATATTATGGTCAACGAGCACAA

TTAATAGAATATTTTCATGATTTTGATGGTATGGTTTCTTTAACTAGTGATTGTTGGAGTTCGAGTCAAG

GTGAGCCATTTATTTGTGTTACCGTGCATTGGATTGATTCTGATTATTATTTACAAAAACGAATAATTGC

ATTTGATGTTATGGATGAGAGACACACCGGTTACAACATTAAACAAAGAATTTTAGAAACAATAAATGAA

TTTAATTTGCTTCACAAGGTGTTTACGATATCCTTAGATAATGCCTCTTCGAACAATAAAGCTATGGAAT

ATATTAGGAATGATATTCCCACTATAATAGATGGCGTTTTTTTGCATGTTAGATGTTGTGCACATATTAT

TAATTTATCGGCCCAAAAAGGAATTGAACAAATAACTTTATTGTTGGAACCCATTAGAAAAATAGTTAAG

TATTTACGTTTAGCACACTCCACAAGATCTAAATATAAAAAATTGTGTAGAGAAAACGGGTTAAGGCCGA

AGAAGTGGGCTATTGATTGTCCAACGAGATGGAGTTCGACTTATAAATTACTAAAAGAGGCAATTAAATA

TAAGGTAGTTATCACTGAGTTGTATAATTCCGACCCGAGAAACCAAAATGAAGATGGATTGATTACTGAA

ACGCAGTGGGAGCTAGCAACTAGCGCACGTGATATACTTGAATGTTATGCACATGCTACTAATGTTTTTT

CTTATGTTTATGAACCAAATGTGCATCATGTTATTATTGAATGTGTTAGTATTGTTACTACTTTACGTGA

ATATGAAAAAAATCGTCATTTTGAAAATATTATTTTTGACATGAAAGTTAAGTGGATTGAATATTTTTCG

GAGTTCCCTTATATTTATGGTGTTGCATGTCTTCTTGATCCCGGTGTTAGAATGGAAGGTTTGGAAAATA

TGTTAGAACATTATTACGAAGTTTTAGGTGTTCAATATGATCATACCCTTTATGTAAG

>MSTRG.3759.1 gene=MSTRG.3759

AATATCATAGTGCGCGGGGGCAATATTGGGCTGGACCGGTTCTATATAGCATTGCTTTTAGGCTTTCACT

AGATCTAGCTAGGAGCCCACAATTTAGCGGAGTTAACCACCTCATCTCAAACTTCACCAAACCCTAATTC

TACTCTCTCTCGCTCCCCCTCTCCCCCTCCCTCCTTTCTCTCTGTGCTCCCATTATACTCTATAGTGTCA

CTAGGCTGCTATGGGCTATTGATGGTTGGAGTCGGTCTAATGCAGTTTCCAACTTGTCCTCAAGAGTCCC

TCTTGTTACAGAAGGACATAATTGAGGCCAAAGAATTCCTGAAGCAACGAGGAGTAGATGTGGGTTCTGA

TTAACGAGGATTGCTAGCATTCTTTCGGAAGCAACAAAGTTCACAACTTAATGACAGATTATAGTGATTT

CAACTTCTAAGTACCTTGTTTTCTCCCACAGCCAGAATTCACATAGTAGTTGAGGAATTGTGATTGACTT

GTATCAAAATGGGTCTCGTTGTTCTTGAAGTTGCTTACTTCTGACATTTATGTGTAAGTATGCTATACAA

AGAAGAGGATTCACCAAAGAAGAGGATTTTTACTGTATATGGTTTCACCTTACAGAAATGGAACTGTTTG

ATATTAATCTGGTCACTCTTTTTTTTTTACCTGTTTGTCGAATATGAATAATCTTTTTTACCA

>MSTRG.3759.2 gene=MSTRG.3759

AATATCATAGTGCGCGGGGGCAATATTGGGCTGGACCGGTTCTATATAGCATTGCTTTTAGGCTTTCACT

AGATCTAGCTAGGAGCCCACAATTTAGCGGAGTTAACCACCTCATCTCAAACTTCACCAAACCCTAATTC

TACTCTCTCTCGCTCCCCCTCTCCCCCTCCCTCCTTTCTCTCTGTGATACAATGAAGCATATTGTGAAAA

TTATGTCATTTTTGGTGTTGATGTCTGCTGTCTGGGTAGGATTATTGCAGGCCTCAATTGTTCCACAGAG

TTATACTTGGTTGCTCCCATTATACTCTATAGTGTCACTAGGCTGCTATGGGCTATTGATGGTTGGAGTC

GGTCTAATGCAGTTTCCAACTTGTCCTCAAGAGTCCCTCTTGTTACAGAAGGACATAATTGAGGCCAAAG

AATTCCTGAAGCAACGAGGAGTAGATGTGGGTTCTGATTAACGAGGATTGCTAGCATTCTTTCGGAAGCA

ACAAAGTTCACAACTTAATGACAGATTATAGTGATTTCAACTTCTAAGTACCTTGTTTTCTCCCACAGCC

AGAATTCACATAGTAGTTGAGGAATTGTGATTGACTTGTATCAAAATGGGTCTCGTTGTTCTTGAAGTTG

CTTACTTCTGACATTTATGTGTAAGTATGCTATACAAAGAAGAGGATTCACCAAAGAAGAGGATTTTTAC

TGTATATGGTTTCACCTTACAGAAATGGAACTGTTTGATATTAATCTGGTCACTCTTTTTTTTTTACCTG

TTTGTCG

>MSTRG.3759.3 gene=MSTRG.3759

AATATCATAGTGCGCGGGGGCAATATTGGGCTGGACCGGTTCTATATAGCATTGCTTTTAGGCTTTCACT

AGATCTAGCTAGGAGCCCACAATTTAGCGGAGTTAACCACCTCATCTCAAACTTCACCAAACCCTAATTC

TACTCTCTCTCGCTCCCCCTCTCCCCCTCCCTCCTTTCTCTCTGTGATACAATGAAGCATATTGTGAAAA

TTATGTCATTTTTGGTGTTGATGTCTGCTGTCTGGGTAGGATTATTGCAGGCCTCAATTGTTCCACAGAG

TTATACTTGCTCCCATTATACTCTATAGTGTCACTAGGCTGCTATGGGCTATTGATGGTTGGAGTCGGTC

TAATGCAGTTTCCAACTTGTCCTCAAGAGTCCCTCTTGTTACAGAAGGACATAATTGAGGCCAAAGAATT

CCTGAAGCAACGAGGAGTAGATGTGGGTTCTGATTAACGAGGATTGCTAGCATTCTTTCGGAAGCAACAA

AGTTCACAACTTAATGACAGATTATAGTGATTTCAACTTCTAAGTACCTTGTTTTCTCCCACAGCCAGAA

TTCACATAGTAGTTGAGGAATTGTGATTGACTTGTATCAAAATGGGTCTCGTTGTTCTTGAAGTTGCTTA

CTTCTGACATTTATGTGTAAGTATGCTATACAAAGAAGAGGATTCACCAAAGAAGAGGATTTTTACTGTA

TATGGTTTCACCTTACAGAAATGGAACTGTTTGATATTAATCTGGTCACTCTTTTTTTTTTACCTGTTTG

TCG

>MSTRG.3759.4 gene=MSTRG.3759

AGCTAGGAGCCCACAATTTAGCGGAGATACAATGAAGCATATTGTGAAAATTATGTCATTTTTGGTGTTG

ATGTCTGCTGTCTGGGTAGGATTATTGCAGGCCTCAATTGTTCCACAGAGTTATACTTGGTTGCTCCCAT

TATACTCTATAGTGTCACTAGGCTGCTATGGGCTATTGATGGTTGGAGTCGGTCTAATGCAGTTTCCAAC

TTGTCCTCAAGAGTCCCTCTTGTTACAGAAGGACATAATTGAGGCCAAAGAATTCCTGAAGCAACGAGGA

GTAGATGTGGGTTCTGATTAACGAGGATTGCTAGCATTCTTTCGGAAGCAACAAAGTTCACAACTTAATG

ACAGATTATAGTGATTTCAACTTCTAAGTACCTTGTTTTCTCCCACAGCCAGAATTCACATAGTAGTTGA

GGAATTGTGATTGACTTGTATCAAAATGGGTCTCGTTGTTCTTGAAGTTGCTTACTTCTGACATTTATGT

GTAAGTATGCTATACAAAGAAGAGGATTCACCAAAGAAGAGGATTTTTACTGTATATGGTTTCACCTTAC

AGAAATGGAACTGTTTGATATTAATCTGGT

>MSTRG.3760.1 gene=MSTRG.3760

CTTTCTTGAGGTAATTATAATCTGCTCCTTTTGTTGTCACTGTTGATCAAGTAGTTAATCAGTCTTCTCT

TTCACAGTCCTTTATTCACGCCACTTTCTCCCGCAAATCATACCGCATTTTCTCTCCTAAGCCCAAACAA

AAAGTCTTGATCATCTTGTTCTCGAGAAGAAAATTTTTGTCAGTACGAAAATTGAAGATGGCTGAGCTGT

GTTTGTCAAGAAGCTGCATTTCACAGATTTATCGCGTCGTTTGCACTGATTTTATTCTGTTGTTTTCTTT

CGTTTTCTGTCACCCTCTTTACTTGTCATACTTCGTTTTCTTCTCACCGTATCTTTACAAACTAGTGTCG

TTTCTTTCTCCTCTTTTCGTTACAATCTCTCTGGTTCTCTTGGCTCTCATTTTAGATAATTTTACGCCTA

AGTTGTTTCAGCCTGGCAGCAGTTTTCTGCTCTCGGCATTCTCTGCAGTTTTCGACGTATTGAGCTCAAA

ATGTGCCGAGGATTTTCAGGATTTCGAGGAGCTTGAAGTGTATAAAATCGTGTTTTTGTTGGATGAGAGT

CATGAGACCGAGACGTCTGGTTCTGATCTAGTGGAAAGCTCAGAAGCTGAAGTATCCGAGCCTGAGTGCG

AAGAAAAATACTGTCTCGAAGATGTTCATCACATTCTTGCTCTCAGTCCAGGCTCCGAGACTCGGTCAAG

TACGTTATACTTAGCCGAGGCTACTGAATTCCCATTGATTGAGGAAAAGAAAGAAATGTTGCATTCCACT

TTGGAAAAAGAAGTTGATTGTCAAGCCGCGATTTTCAATAAAGTTTTTGACGTGGAGCAAGAATCCGAGG

CTGTGGATATTAACATCGACTGTATAAACGTCGCAAAAGAACACGCAGGTACTGTAAAATCGCGGATAGA

CTCGTACAACAAAATTGTCAAAAAAGCCGTTACAAATGAACGGCGTGCTCTGAGTTTCGACCACGAGGGT

TCAAATCTCGCGAGCTACGGGTCCATGAGAAAGGAAAAGGAATGGAAGAGAACGTTAGCGTGCAAATTAT

TCGAGGAGCGACATTCTAATGTGGTGGGAGGAGGCGAAGGAATGGACTCGCTTTGGGAGACATATGAGTC

CGATTCGACCAAGCCTACACGAAAAGATGTAAAAAACAAGAACACGAAGAAAAAAACTTATGATCTCGAC

TATAATTGCAATGCAGATGAAGACGATGATGAAATCGAGATGACGAACGGGCATCTTTGCTGTTTGCAAG

CGTTCAAATTCTCTGCAAGGAAGATGAATATGGGAATGGGAAGGAGTAACATTGTTAAAATTACCAAGGC

CCTTAAAGGGATCGGTTGGTTGCATAATGTTAAGAGGCATGACAAAAAGGAGAAACCTTGAATTCGTTGT

CTATTGCTGTTGTCGCCTTAAATCCGTGTTAGTCTAGTGTATAAATATATTACTTATGTTTTTTCTTTCC

GAATGTCGGATTAATTAATAATCAGAATTTAAAATTTGTT

>MSTRG.3762.1 gene=MSTRG.3762

AAAACATAAATACATAATTGAAAAAATATTTTATTAGTATTGGTGACTTAGTTAGCATAAATACATATTA

CTTGCTATATCAGAGATTTAGCATATGAAATTTTTCTTCAAAAAGCATATGAAATTTCAGTAGTTCAGTT

TTTAGTAGTCGTCAACGGCCCTAATCGAATGTCTGATTCGAGGAGTTTTGAATGCTGTAGCCATGAATCG

AATCCATCAAGAGGTGTGGGGATTGGTAGCTCTTTTGGGTGCGCAATGGGTGCTTTAGTTGATGGGCATT

TTGATGAGGAGCTAGAGAATGGATTGGTTCTTGGGGTTTCTTGTGAGCCAGGAATTGAAAATGAACCAGA

GACCAAGGTAATTCAAGATGCTGTCAAAGTGTTGTTGCTTGGTCTAGGTGAGGATATTAATCGAGAAGGT

CTGCGAAAGACCCCCTTTCGTGTTGCCAAGGCTCTTCGAGAAGGAACAAGAGGTTACAAGCTAAAAGTCA

AGGATATAGTTCAAAGTGCTTTATTTCCTGAAGCAGGCTTGGACAATGGGGTTGGTCATGCTGGTGGAGT

GGGTGGGCTTGTTATAGTTCGAGATCTTGACCTCTTCACATACTGCGAGTTATGTTTGCTGCCATTCCAG

GTCAAGTGCCACGTCGGTTATGTCCCATCTGGACAACGAGTTGTAGGCCTAAGCAAACTCTCTAGAGTTG

CTGATGTTTTTGCAAAAAGGCTCCAAGATCCACAACGATTGGCAGATGAAGTATGTTCAGCTCTGCAGAA

TGGAATTATGCCAACAGGTGTTGCGGTCGTTCTCCAATGTTCGCATATTCACTTCCCAAATTTCGCGTCA

GCATTTCTCAACTCAAATCACCAGGGATGGTTCAAGACAGTGGTTACTTCTGGTTCAGGGGTTTTTGAAA

ATGAAAAGGACATTATTTGGGGTGAGTTTCTGAGTCTCTTAAAATTCAGAGGAATAGATATAGATGCCTG

TTCGAGAGTATCTGGTGATGGTTGCTGGTGTCCATCTCAGAATATCTCCACAGTGGAACCAGCAGATTCA

GAAATGGTTACTGCTGTAACTTCCATCATTCAGTCTCTGGGGGAAGACCCTTTAAGAAAAGAACTTCTGG

GAACTCCTACCCGTTTTGTGAGATGGCTGATGAACTTCCGAAACACAGATTTCAAGATGAAGCTAAATGG

GATTGATCGCTATAGAATGAGTCCTTTTGAATCCAATGGTGATGTTGAAGATTCTATACAATCTGAGTTA

AACTTGTCACTCTGGTCCCAATGTGAACACCATTTACTTCCATTTCATGGTGTTGTGCACATTGGTTATT

TTGGTTCTGAGATTGATGTGATTGGAAAATCCCTTTTCCAGTCTATAGTACATTTCCATGGTTTCAAGCT

TCAAGTGCAGGAAAGGCTCACCAGGCAGATTGCTGAAACTGTTTCGTTGTTGGTGGGAAAAGATGTAATG

GTAGTGGTAGAGTGTAACCACACTTGTATGATATCTAGAGGGATTGAGAAGTTTGGTAGTAGTACAGCTA

CAATGGCCGTCTTGGGTCGCTTTTCAACAGAACCTGCTGCAAGGGCAAAATTTTTGCAGAGCATACCTCC

AACCACTGGAGAATGATATGAACATTTACACACTGATGAAATTTTATACATATTTAAAAAGAGTCTGAAC

TACAGCTGTTTGACTGGCTGCTGTTCCGCCACCGATGGCATGCCTTCATAAGCAGAGGAGCAGCAGCAGC

ATATTATTCAAAGAAGCTCGGTATTGCCAGCAACTTCATTGAAATTTTGTAATGCTCTGTTGAATCGAGT

AATTGCCTTTTCTTTCGCACTCTGCTTATAGTCTGTAGAACGAAGATATGTTTCGCCACAGAAATATTTG

AGTTAGTTGTTAAAGGTTATCGGTTTCATATCATACATAGCTTATGGCACGAGTGTAGCCCACTGCTCAT

GGCAATTTTCATTTCTGTAACATTCTTGGTAGATGATGTCTTGATCTTGTTGATTTATTTGTTCCTGCAG

TATCTTTTAATCTTATTTCTTTTATAAACATTATGGTTGTCTATTTAGATGAGGGGGAAGACCCGGGTGC

TTTTCCTTTCTAGTTTTAAA

>MSTRG.3771.1 gene=MSTRG.3771

GGGCCAAAATTTCCTAACTAGATTCGAGATCAAAAATCTATTGATGTTCTTGATATTTCTAACAGCCAAA

TTTCAGAAAGATTTAAATCTCTCTTCCAACAATATTAGAGGCAAGATTTCTTCCAAACATTTTTATGCTC

AAGACATAGATTTGAGTTCAAATTATTTTGAAGGGCCACTACCACTAATTCCTGAAGATTGTGTAAGAAT

AAATTCATCTCAGAACAAGTTTTCAGGAACACTCCTTTCCTTGTCTAATTCCTGAAGGTTTCTTGATTTC

TCACATAACCAATTGTCTGGTGCACTTCCTGATGATTGGACGCATGTTAAAGGTCTGCGTCAATAGGGCA

ACTAGTTTCTCTTAATTCACAGATTTTGAGGAACAATATGCTCTACGGTGAGTTTCCTGCATCTCTGACA

AACTGCAGCTACCTTGAGTTTGTAGATTTTGCGTTGAATAAGCTATCGGGAAAACTGCCACCATGGATAG

GGGAAGGCCTCCCACAGTTGTATGCTCTGATTTTAAAATCTAATAGATTTTCCGGAAGCATACCTTATGA

AATATGTCGTCTATCCAATCTTTATTTCCTAGACTTGTCCATATGAAATATGGACTTATTCCTCCATATT

TTATGAAATTATGACTTATGACTTATGAAATATGAAATTATTTCATATTAAATTATGAAATTATGAAGCA

TACCTTATGACTTATGAAATATGAAATTATTAAAGCATGGAGGAATTATTCCTCCATGCTTTAATAATTT

GACTGCAATGATCCTGAAAGGAACTGAAGTTGATGAGCATATCTATTCTTATGAAGATCCAGAGTTATTA

AGTCCAGATTCAATTTATGATAGTTACACTGTTGATGTTTCGGCAAGATGAAAAGGGCAGGAGCTTTCTT

ATGGAAGAAATTTTGCATATCTGAAGATGATCTATCTGTCAACCAGTCAACTTACTGGAAAAATTCCCAG

ATGGATAGGCAGACTTCAAGAACTCAGGGGATTGAACTTGTCAAGAAACAGGTTTTACGGGCATGTTCCA

CTTGAAATTGGCCAACTGAAAGTTTTAGAATTCATAGACTTGTCCACAAATTTTTCAGGTGAAATTCCTC

AAAGCATGTCTGGACTAAATTTTCTTGCATATTTAGATCTCTCGGACAATAACTTCTCAGGAAGAATACC

ATCTGGTACTCAACTCCAGGGATTTGATATTTCCACGTATGAGGGGAACGCTGGACTCTGTGGTAAACCA

CTCACAAACATTGGTCCTGGAGATGAACCAGTTGATCACATCCCGCCTTCATTCCCTGAGTATGAAGCTG

AT

>MSTRG.3773.1 gene=MSTRG.3773

CCTGGCACAAGATCTTGAAGAAGCTTCATTCTTTCACTAATCTTCTCTCTTCTAACTCTTTCTGCTAAAC

TATGACTGTCAGTGGCTTGACCTCTTCTTGCTCTGACATGAATATAATCCTTCGGTGGCTCTGGAGGTTT

GTTATCCTGTTTCTTATCCTCTTCTGTAGCTATATTTGTTTCTTCTTCTTTTTTAACCCCACCATTAACA

TTTTCACTTGTCATCTTCGATCTCTTCATAACATTTGAATTCT

>MSTRG.3777.1 gene=MSTRG.3777

AATTCCCAGATTCAGCCACCAATCACTATATCATATCGCAAACATCACTAATATATGCTGCAAAAATCAC

CAATTTATACGACATTGATCACTAAAACCTATCAATATATCATTGTATATATCATTAACATTACAAGCTA

CCACACCTATGCTACCAGAATCTTATCGCCTCAGTTTTTGCCACCATCATCTTAACACCACAGCATCATC

ACCATCTCAATCACCATCACGGCACCTCATTTATACTGATTATCAGCATATATATGTCAGATGCAAATCT

CGAGGCGGAGCTCCGGCAAATTGGACGGCTAATCTAGCCGTTATCCTGACGATCATGACTTTTCC

>MSTRG.3778.1 gene=MSTRG.3778

GGGCCTCCTGACCAAAAAGGGGAATTGACCAGTGTGTTCGCTTTGGGAAGGAGCCATCCACAAACAGCAA

AATTAGGGCACAAGCGGGACGTTCTTTCTCGCCCAGTGATAATGGAATTGGCAGAGGTGTACAGGGTACA

ATATCAAGGCCGTGATGATATCTTGCGAGGTATAATTGTGTGCAAGGATAGTGAAACTGGTTACCAATGC

TTATTCAAGGATCAAGGCCTAGAAGATGAAGTCATCAACAACACACTGCTCATGTTAAATAATGACGACA

ATAACCCAGACTGTGCGCATCTTTGCCTCCAACAAAGGAGTAAGCAGACTTCACATGTACCAATTAAATC

TTTGGATCATATAATACCGAATAGTTGTATTACATCATCAATTCGTGCCAAAGATGGATTTTTAGTCGGT

TTTTATCTGTTTGATTTCACGTATGAGAAGTTGACTTTTGAATATCCGTATCACGATCCCGAGAAGTTTT

CTGTCCCTCTCAGAAAAAGCTTTGTGTCAGAAGATGGATTTAGACATTTTAACATTCTGTATGCAATATT

TAATAATGCGGTGGAAGCATGTTTGAAGGTTACGTTAACCAATTGTGTTAATCCATTTGTTCTTTGCGGA

GTTATAGCTGCTAGGACAAGCGCGATTGAACGTCCTGCTTATTCAAGTATCATCTTCAATGAAGAGTGTG

GAATTTTCATAGAGTTGGGGAGCGGCGGTGACATAGTAATTCCTTTGCTGAGAGATATTGTAGCTGTGCC

ATTGGGACATCAGCTAATTCTAGAAATCAACCTGAGTGGAAAAGATAATGATGGCAAAAAAGATTCTGAT

AAGATGGTTGAAGACTATCTAGTTTTCGATGCTATTGATGATGGGAAAAGTACCAAATCTATTAGCGGCA

GCAAAGGTGAGCTCAGAGTGGAAGTCAGTTGGAGATCTAGGGAGAAATCCCATTGAAGCTGAAGCGCAAG

TATGGGGATGTACTATATTTATTAGCCAATGTTTGTTATATCAGCCTTGTCTGACTGGGCATAAATGCTT

TTGAGTTGGGCTATTTAATTTGAAGTACATTTAGAAGATACAATTGTATTCTGTGCAATCTATTTCACTT

GAAATGTAGTAAATTTTTACATCAAGTTTCTTGTTTCATGTTCCAGCACCCTTTATCATCATAAGATTTT

TTTTTTTTTTTTTGTTTTTACATCAAGTTTCCTGTTCCAGC

>MSTRG.3779.2 gene=MSTRG.3779

CAAAATTTTTAACGCAAACAAAAACTCACCGACTGATCATTCTCAACGCAGACCCAGGCCTTCTTCTCTC

CTTGTTTTAGGGTTTATTGAGTATTTTGTTTACAGCTCTACCTTTAATCCTACCCTGATCCTCGATTTGG

TGGGACGATGGAATGGAAGAAAAGGAATCATATTCATCTATGAGGATAACAAGGAATCGGCCTGATATTC

TCCGAAATTATATTCCGAAATCCTCTGATGATCCCCCTGCCATCGCCCAATTACTTGAATTCTCTGTCAA

AAGAAAGAGACACACATTCATTCACTTAATTCGTGAAATCAAGGATGGATTCCATAATGGCCAGATGGAG

AAGATTGAAAAGTGCACTGACGAGTTAAGAAAAATGATTACAGAAGGTATAAGAATTTTTACTTTTTTAT

GTGTTGGTTGGCAGTACATTGCAAATAGATTTTTTTCACTTGATTTTTTTTCACAAACAGATCCGAGACT

GGCTCTTTTTCTTGATGTGGAGTTTCTTGAACTCCTTGTTGAGTTGCTTAAGTGTGAACAATATTCAAAC

CTCCAGGTACACGCTGCCACCATCTTTGAACCAGTAGTTACCTTTTTCCTCTGCAGAAAGGATGTTAAGG

ATCTTGCTGTTATAATAGAACGTATAATTCCAATTGTTTCGAGACTTATATGCTTTGGAGTTTATTTGGT

GCAGGTGAAGGCTCTTTGGTTATTGCGCTCTCTTGTTCCTGTAATTTCAAAATATGAAGTTGAAGAAATT

TTATCCCCAGTGGCATCTTTGATTATTGAGGATTCTACTAGTTTTAATGTGTTACATGCCGCCTCTGAGA

CTTTGCTAGTTGCTTGTGAGAAACATCCAAAAATTTCTCCTGAAAAGTTGAAGTTGGCCCTTCCGGCTTT

GATGAAGCTTATTGAGTCTAATATATTGATGTTACTGACATCTGGAAGTTGGGGACTTGCATTTTTGTGC

GACGGAAGGGGAGAGATGATAGTTGAAGATAAATACCTTGAATGTCTCATAACTCGTCTTGTTGATGTCA

TGAATGATGAAGAACGTGTTATGATCCTCTCGGCACTTCATGCAATGGGAAGTGTTGTGAAATGGGGTAG

TGATGATGTTATTGAGATTATAATCAAGGAGGACGGTGCACTGCTCATCTTGTTAGTCTTGTTAGAGGAA

GATGATTTGTACATTGTGAAACAAACTTGTTGGATACTCTCAAACATTACCGCAAGAAAAGAAAAACATA

TACGGGCTATAATTGATGGTGTCCCAGAAGGCAAGTTAATTGACCCCCTTGTCAAAATTATGCTCAATTG

CAATTTGAATGATGTAAGAAAGGAGGCGGCTTGGGCAATTTCTAATGCTATGTGTGGTGTTGGTATTGAT

CAGATTGAATGCTTGAGGAATAGCTGCTCTAATATCATTTGGGCTGATCTGTTAAATTCATTCTCCAGTG

ATAGATTATTTGTCTTGGTTACTTTAGAAGGTTTACTCAATATTAGAGTAGCGGAAGTCACTTGGAAGGG

GAACCCAGTTGATGCTGTGACGTTGCACTCGTTTTTTGAAAGCATAGTCGAAGGCATAGATCGTAGGGAT

TCCTTTATTGTACCTGAGTTCGTCAATGAGAGTGATGGGTGGACTGTATCTGGGCATTCCTTATTTCCTG

AGTTCCTCATGTCCGACGACTATGCTCTAAACTTTAAGTTTGAAGATCAGCGGTGTTCGAACATGTTTAT

AGATTATAATGGTTCGATGAAATTTTGTTAGGAGATAAAGTGAAGTCTTGTTTATT

>MSTRG.3779.1 gene=MSTRG.3779

CAAAATTTTTAACGCAAACAAAAACTCACCGACTGATCATTCTCAACGCAGACCCAGGCCTTCTTCTCTC

CTTGTTTTAGGGTTTATTGAGTATCTCTACCTTTAATCCTACCCTGATCCTCGATTTGGTGGGACGATGG

AATGGAAGAAAAGGAATCATATTCATCTATGAGGATAACAAGGAATCGGCCTGATATTCTCCGAAATTAT

ATTCCGAAATCCTCTGATGATCCCCCTGCCATCGCCCAATTACTTGAATTCTCTGTCAAAAGAAAGAGAC

ACACATTCATTCACTTAATTCGTGAAATCAAGGATGGATTCCATAATGGCCAGATGGAGAAGATTGAAAA

GTGCACTGACGAGTTAAGAAAAATGATTACAGAAGATCCGAGACTGGCTCTTTTTCTTGATGTGGAGTTT

CTTGAACTCCTTGTTGAGTTGCTTAAGTGTGAACAATATTCAAACCTCCAGGTACACGCTGCCACCATCT

TTGAACCAGTAGTTACCTTTTTCCTCTGCAGAAAGGATGTTAAGGATCTTGCTGTTATAATAGAACGTAT

AATTCCAATTGTTTCGAGACTTATATGCTTTGGAGTTTATTTGGTGCAGGTGAAGGCTCTTTGGTTATTG

CGCTCTCTTGTTCCTGTAATTTCAAAATATGAAGTTGAAGAAATTTTATCCCCAGTGGCATCTTTGATTA

TTGAGGATTCTACTAGTTTTAATGTGTTACATGCCGCCTCTGAGACTTTGCTAGTTGCTTGTGAGAAACA

TCCAAAAATTTCTCCTGAAAAGTTGAAGTTGGCCCTTCCGGCTTTGATGAAGCTTATTGAGTCTAATATA

TTGATGTTACTGACATCTGGAAGTTGGGGACTTGCATTTTTGTGCGACGGAAGGGGAGAGATGATAGTTG

AAGATAAATACCTTGAATGTCTCATAACTCGTCTTGTTGATGTCATGAATGATGAAGAACGTGTTATGAT

CCTCTCGGCACTTCATGCAATGGGAAGTGTTGTGAAATGGGGTAGTGATGATGTTATTGAGATTATAATC

AAGGAGGACGGTGCACTGCTCATCTTGTTAGTCTTGTTAGAGGAAGATGATTTGTACATTGTGAAACAAA

CTTGTTGGATACTCTCAAACATTACCGCAAGAAAAGAAAAACATATACGGGCTATAATTGATGGTGTCCC

AGAAGGCAAGTTAATTGACCCCCTTGTCAAAATTATGCTCAATTGCAATTTGAATGATGTAAGAAAGGAG

GCGGCTTGGGCAATTTCTAATGCTATGTGTGGTGTTGGTATTGATCAGATTGAATGCTTGAGGAATAGCT

GCTCTAATATCATTTGGGCTGATCTGTTAAATTCATTCTCCAGTGATAGATTATTTGTCTTGGTTACTTT

AGAAGGTTTACTCAATATTAGAGTAGCGGAAGTCACTTGGAAGGGGAACCCAGTTGATGCTGTGACGTTG

CACTCGTTTTTTGAAAGCATAGTCGAAGGCATAGATCGTAGGGATTCCTTTATTGTACCTGAGTTCGTCA

ATGAGAGTGATGGGTGGACTGTATCTGGGCATTCCTTATTTCCTGAGTTCCTCATGTCCGACGACTATGC

TCTAAACTTTAAGTTTGAAGATCAGCGGTGTTCGAACATGTTTATAGATTATAATGGTTCGATGAAATTT

TGTTAGGAGATAAAGTGAAGTCTTGTTTATTTGATGCTGCTGCTAAATCAATCTATTCAATCATTTAATT

ATTATATTTTGTTTCA

>MSTRG.3779.3 gene=MSTRG.3779

AAATTTTTAACGCAAACAAAAACTCACCGACTGATCATTCTCAACGCAGACCCAGGCCTTCTTCTCTCCT

TGTTTTAGGGTTTATTGAGTATTGCAGTTTGTTTACAGCTCTACCTTTAATCCTACCCTGATCCTCGATT

TGGTGGGACGATGGAATGGAAGAAAAGGAATCATATTCATCTATGAGGATAACAAGGAATCGGCCTGATA

TTCTCCGAAATTATATTCCGAAATCCTCTGATGATCCCCCTGCCATCGCCCAATTACTTGAATTCTCTGT

CAAAAGAAAGAGACACACATTCATTCACTTAATTCGTGAAATCAAGGATGGATTCCATAATGGCCAGATG

GAGAAGATTGAAAAGTGCACTGACGAGTTAAGAAAAATGATTACAGAAGATCCGAGACTGGCTCTTTTTC

TTGATGTGGAGTTTCTTGAACTCCTTGTTGAGTTGCTTAAGTGTGAACAATATTCAAACCTCCAGGTACA

CGCTGCCACCATCTTTGAACCAGTAGTTACCTTTTTCCTCTGCAGAAAGGATGTTAAGGATCTTGCTGTT

ATAATAGAACGTATAATTCCAATTGTTTCGAGACTTATATGCTTTGGAGTTTATTTGGTGCAGGTGAAGG

CTCTTTGGTTATTGCGCTCTCTTGTTCCTGTAATTTCAAAATATGAAGTTGAAGAAATTTTATCCCCAGT

GGCATCTTTGATTATTGAGGATTCTACTAGTTTTAATGTGTTACATGCCGCCTCTGAGACTTTGCTAGTT

GCTTGTGAGAAACATCCAAAAATTTCTCCTGAAAAGTTGAAGTTGGCCCTTCCGGCTTTGATGAAGCTTA

TTGAGTCTAATATATTGATGTTACTGACATCTGGAAGTTGGGGACTTGCATTTTTGTGCGACGGAAGGGG

AGAGATGATAGTTGAAGATAAATACCTTGAATGTCTCATAACTCGTCTTGTTGATGTCATGAATGATGAA

GAACGTGTTATGATCCTCTCGGCACTTCATGCAATGGGAAGTGTTGTGAAATGGGGTAGTGATGATGTTA

TTGAGATTATAATCAAGGAGGACGGTGCACTGCTCATCTTGTTAGTCTTGTTAGAGGAAGATGATTTGTA

CATTGTGAAACAAACTTGTTGGATACTCTCAAACATTACCGCAAGAAAAGAAAAACATATACGGG

>MSTRG.3781.1 gene=MSTRG.3781

AAAATTGCAGAAAGATTTCTACCGAGTTGTTACGACGCGTCCCGAAATATAAAAGGTACAATATTCCGAG

TTACGAATACATCACACACGTCTTGGTGTTGGGCGAAAAGTATACGAGTAGTGCAGTCTGAGGGAAATTT

AAATGCTACATACAACAATTACCCAGAAAGTTCTGAAACTCGGAGGCCGGAGAGTCGTAGAAAAATAAAC

ACACACAACACAAGATTCACTTAGTATTATACACACACTAGTCTTAACATAATCGAGATGGGGCACAGCG

TGTCGACTACATCGGCCCAACGGGCCGAGAAGCTCCGTCAAGACGGCAATCTCTACTTCTCCAAGCTCAA

ATTCGCCGCCGCTATTGATGCCTATACTGAGGCTATTACTCTGTGCCCTAATGTTCCTGTTTATTGGAGT

AATCGTGCTCTTTGTCATCGCAAGCGCAATGATTGGGACAAGGTCGAAGAGGATTGTAGGCGAGTTGTTC

AGCTTGATCGTACTTCTGTCAAGGGTCATTATATGTTGGGTCTTGCATTACTACAGAAGAAAGAGTATGC

AGAAGGGGTCAAGGAACTGGAAAAAGCTCTGGATCTGGGCAGAGGTGCAGATCCTAAGGGATACATGGTT

GAAGAAATATGGCAAGAGCTTGCCAAAGCAAGATACCTTGAGTGGGAGCAAGAATCTACCATGAGAGCAT

GGGAGCTACAAAACTTGAAAGAAGCTTGTGAGACTGCTCTCAAGGAGAAACATGAGCGGGATCTTTCTAA

TACCGGAGGGCTTGTTAATCATTCTAATAGTGTCTATCCAGAGCAGCTAAAGGCTCTGAAGAATGTAATT

GACAAAGCTATTGAAGATGACAACCAAACTGAGGTGCCGGACTATTTGTGCTGTAAAATTACACTCGACA

TCTATCGTGACCCTGTCATTACTCCTAGTGGAATCACCTATGAACGTTCAGCAATTTTTGAGCATCTCAA

CAAGGTGGGAAAATTTGATCCAATCACACGTGAATTGCTTAATCAGTCCCAGCTGGTTCCAAATCTGGCT

GTAAAGGAAGCAGTAAATGCATTTTTGCGAAAAAATGGCTGGGCGTATAAGATGTACTGATTTTCAACAG

GAATATATCACAACTGCTTTGCTTGGTGTTACTATCAGTGACAACTTTGTCCGTAGGTGATGTATAGTTT

CATACTTCTGGTCAGGCATTGATGGCCAGTATCATGCTCGTGTGTACAGCTTTAAAGTCTCCGCTTTTCA

AGCCCTCCTTTGTCCTGAATTATAAACTATTAACCGATCTTGGGTAGTTACAGCTGATAGTGAACTCTGC

TTCAGTTAGCCACGGTGACAGAATTTTTGCTTGTAAATACAATAGATCTCATTAACTCTTATAGTATTCC

AGCTTTCTGTTTTCAACCAGTCTACTGTGTGATTTCAGAATCTGAAAGTATTTCCTTTTCTTGGTG

>MSTRG.3787.1 gene=MSTRG.3787

GAGAGAGAGAGAGAGAGAGAGAGAGAGAGAGAGAGAGATTACTATTGAGCTCATCAAGCCATCGTCTGAT

ACTCTCAAAAGTGGGTCTCCTCGTAATATCATAAACCAAAAGAGCACCAAAAGCACCTCTATAATAAGCA

GAAGTAACCGCTCTAAAACGCTCCTGGCCAGCAGTATCCCACACTTGGGCCTTGACTTCCTTGCCATCAA

CTTGAACAACCTGGGTCTGAAACTCAACCCCAATAGTAGCCTTGGAGTTCTGATCAAACTCATCTCTGGC

AAATCTCGAGAGCAAATTAGATTTGCCCACAGCAGAGTCCCCTATCACCACTATCTTGAACAGATACTCC

TCTCCTTCCCCTGTTTCATCCATTGTGAGATTTGTTGGGAAGTGTGAAGATGGATTGGGGAATGTATATA

TACAAGTTTATATAAGTATGTAATGATTTGGGGAGGTTTCTCTTTGCAGCTGCAGAGACTAACTGCGCAA

TTACCGGCCTATTTTACATACTCATAACATGTCTGCATGTCTTTTTAGGCCCATTTTTCTAATGTAAATC

ATCATAATTTCGT

>MSTRG.3796.1 gene=MSTRG.3796

ATGTATCAGCAGCTTGCGTCGTTGCCGTCAACACTTCCCTTGAGAACCTCCTAAGCTTGTATGTTGTATC

TATCTATTATATTCTGTATCTATTTCTCATTCAGCGGCAGCAAGTCAGCTTCTTGACTGGAATTCGGCTA

AGAGCCCACGGTTCAGTCAATTATTGTAGATTCTACAACTTGCGAGAAGAATAACAGATTTGTTAGTTAG

TATCACTCGCCTGAATATGATCTGCATCTGATTATTCAGGCCCTTATTATACTTTCACACCTTAAAACGC

TTTTTTCTTTATCTGAATGGTTAAGCAAGTCAGCTTCTTGACTGAATCTCGATTCATAGACTACGGTTCA

GTCAATTATTGTACTTGTACAGCTTGCGATAACGATATAATCTAGTCAGTATCACTCGCCTGAATATGAT

TTTTTGCTGATTATTCAGGCCCTTATTATACTGTCACAAATTCTATTTTGTGTAAAATAATTGATTTGGA

TTGTGTGACTGCAAACTCACTCTGTCCCATAACAAATCATGTTGTGAGGCAATTTCTTGGTTTGCTATAT

TGATTTTTATCAATCCGATTTGACTCGTCTAAGTGATCTTCTCTCGGTGATCATGTAAGACACATAATTG

GATTACATTTCAGTGTTTAATCGTTTAAATCTCGTTATGAAATTGTTTTTATCAACTTTGTAAATGTTTT

TAAAAAAAATTCAATCTCTCTAATTTCGTTTCGTGATCAATTTGATTGGTATTCATGACATCTGAAGTTG

AAATTGAAACAGTAATTGTTATCTGAGATATTTGTTTCTAACAAAGAACTGATAGTTTCGAGAACCAAGC

TCACTGGGATATTCCGAAAAAACGATCAAGTGCTTTGTCTAACACTCCAACAAGAGACCTAGGTTGTGCC

TTGAGAAATGAGAAGGGATAAGACTTAAGAGTCGGAGGAAGTTTGTTCAGGACAAAACAGGAAAGAAATA

CTTCCTCCATCCCCCTGAGTTGCCTGAGTTGTATACATTGGGGGACGGGGACGCGGCAC

>MSTRG.3800.1 gene=MSTRG.3800

TTGAAGTTAGACCGAGTAACTAAAATTAAAAAGAAAAGAAAACGGAAGAAAGTAAAAGCAGTTAAAAATG

AAAAAAGGGGATGGGATAGACTTCTAGACTGAATGAATATCTTATCGTTTGTTTCTGGTTACCAAAACCC

AGACCAGCTATCTTGGGGTCTTAGGGAATGGAGTCTTCTTCTTTCTCCGCATGCCTCTCATCATCATCAT

CTTCTCCGCCATTCCGGAATCACTTTAAAACCCACAATTCTCTTCTTCTTTTCTCAAATTACAACTTTTC

CACCTCCAATTTCAATCCCATTTCTTCTTCCATAGCCAAGAAAAAAATCACACTCAAAGCTCAAACATTT

CACCAAGTTCCCAGAACAAATCTACCAATCAAGAACAACCCACTGCAAATCTTTGTCAACACCCATGTCT

CATTCCTTTCTCTAGCTTTCTCTCTACCTCTTTCTAGCTCTGCTTCTGACACAACATTGCCAGCTGAGCA

AGTCTCCAACAAGATTAACATAGAGGCTATCTTGGTCTCCATTGATGACTTTTTTAATCGATACCCTTTC

TTTGTTGCCGGAGTTACTTTCATTTGGCTTGTTGTTATCCCGCTCACCGAAGATTACTTGCAGAAGTTCA

AATTTATCTCTGCTATCAATGCTTTTCGCAAGCTTCGTGATGACCCAAGTGTTCAGCTTTTAGATATTAG

GGATCAAAAATCCTTGAGGACTTTGTCTTCTCCCATTTTGAAAATTTTGAATAAGAATGTGTGCCAGGTT

GAGTATATTGAAAAGGATGAAGATGGGTTTGTGAAGAAAGTGCTGGACTGCTTTGGGGATCCAGCAAATA

CCACAATTTGCATTCTCGACAGTTTTGATGATAACTCCTTGAAAGTTGCTGAGTTGTTGTTCAAGAAAGG

TTTCAGAGAGGCCTATGCAATTCGGGGTGGCGTTAAAGGGAAGAATGGGTGGCTGTCAATACAAGGGACA

CTTCTCCCTCCATCTGTGCACATTTTCCCGAAGAAAAAGGGAAAAAAGTTGCCACAGACTGAATTGGTTC

ACGGCCAAGGAATTAATCAACTAAGTGAAAATGCAGATCAGTCTTCCTCTGCGAGTAGCAGCGCCCAGAT

GACTAATGGATATGTAAGCAAGACTGAACCCACTGCAGAAACAAAAATTAGTGCAAGATCTTTATCTCCT

TATCCTAAGTACCCAGATCTGAAACCACCATCTTCTCCAACTCCATCGAAGCCAAAAAGCTGATACCAAG

TCCTTCAGATTTAATTTGACTCCTGCTGTTGCTTGGATCTCTTCTAGCTACCATGGAACGTAGCAACACA

TTATCGGATAAAGTGTCTCTTTCGGCATGATAGTCATAATTTGTTCTTCACATATGTTTTCAATTTTCCT

TAGTTTTTTGTTGTATTATCATCCTATTTTCCTTAGCTTTTGTTGTATTATCATCCTATTTTCCAGGATT

GGATTTCATTTGCAGCCTATTCAAATTTGTTCCAGGATGCAGCGAGTTTTTGTACACATAAATTTATCAG

AAATCAAATCTACTCGATATTTTTATTCTA

>MSTRG.3800.2 gene=MSTRG.3800

TTGAAGTTAGACCGAGTAACTAAAATTAAAAAGAAAAGAAAACGGAAGAAAGTAAAAGCAGTTAAAAATG

AAAAAAGGGGATGGGATAGACTTCTAGACTGAATGAATATCTTATCGTTTGTTTCTGGTTACCAAAACCC

AGACCAGCTATCTTGGGGTCTTAGGGAATGGAGTCTTCTTCTTTCTCCGCATGCCTCTCATCATCATCAT

CTTCTCCGCCATTCCGGAATCACTTTAAAACCCACAATTCTCTTCTTCTTTTCTCAAATTACAACTTTTC

CACCTCCAATTTCAATCCCATTTCTTCTTCCATAGCCAAGAAAAAAATCACACTCAAAGCTCAAACATTT

CACCAAGTTCCCAGAACAAATCTACCAATCAAGAACAACCCACTGCAAATCTTTGTCAACACCCATGTCT

CATTCCTTTCTCTAGCTTTCTCTCTACCTCTTTCTAGCTCTGCTTCTGACACAACATTGCCAGCTGAGCA

AGTCTCCAACAAGATTAACATAGAGGCTATCTTGGTCTCCATTGATGACTTTTTTAATCGATACCCTTTC

TTTGTTGCCGGAGTTACTTTCATTTGGCTTGTTGTTATCCCGCTCACCGAAGATTACTTGCAGAAGTTCA

AATTTATCTCTGCTATCAATGCTTTTCGCAAGCTTCGTGATGACCCAAGTGTTCAGCTTTTAGATATTAG

GGATCAAAAATCCTTGAGGACTTTGTCTTCTCCCATTTTGAAAATTTTGAATAAGAATGTGTGCCAGGTT

GAGTATATTGAAAAGGATGAAGATGGGTTTGTGAAGAAAGTGCTGGACTGCTTTGGGGATCCAGCAAATA

CCACAATTTGCATTCTCGACAGGTAGTTTTGATGATAACTCCTTGAAAGTTGCTGAGTTGTTGTTCAAGA

AAGGTTTCAGAGAGGCCTATGCAATTCGGGGTGGCGTTAAAGGGAAGAATGGGTGGCTGTCAATACAAGG

GACACTTCTCCCTCCATCTGTGCACATTTTCCCGAAGAAAAAGGGAAAAAAGTTGCCACAGACTGAATTG

GTTCACGGCCAAGGAATTAATCAACTAAGTGAAAATGCAGATCAGTCTTCCTCTGCGAGTAGCAGCGCCC

AGATGACTAATGGATATGTAAGCAAGACTGAACCCACTGCAGAAACAAAAATTAGTGCAAGATCTTTATC

TCCTTATCCTAAGTACCCAGATCTGAAACCACCATCTTCTCCAACTCCATCGAAGCCAAAAAGCTGATAC

CAAGTCCTTCAGATTTAATTTGACTCCTGCTGTTGCTTGGATCTCTTCTAGCTACCATGGAACGTAGCAA

CACATTATCGGATAAAGTGTCTCTTTCGGCATGATAGTCATAATTTGTTCTTCACATATGTTTTCAATTT

TCCTTAGTTTTTTGTTGTATTATCATCCTATTTTCCTTAGCTTTTGTTGTATTATCATCCTATTTTCCAG

GATTGGATTTCATTTGCAGCCTATTCAAATTTGTTCCAGGATGCAGCGAGTTTTTGTACACATAAATTTA

TCAGAAATCAAATCTACTCGATATTTTTATTCTA

>MSTRG.3803.1 gene=MSTRG.3803

TTTGAAGAGCTATAAGAGCCTGTTGGTCAATAGAAGAGCCCATTGGGCCGAACAAAAACTCGTTTCAGAT

AATTTTAACGAATAATTTGACGGTCCGGTGTTCGAACCCGCTCATTTGGCCAGTTCGACTCGTATCATAC

AACATCTCTCACCTGAAGTTGAATTCCAGCGCTTTACACTTGCAACAGCAAAGGATGGCGGATCAGAAAC

AAGGTTACCTCTGCATGATTACCAGACTCCCGAAATCAGCAACAACAGAGCACGTCAAGGCTAATTTTAG

CCATTTTGGCAAAATTCTTGATGCACACGTTTTCGTGGACAACTCTAATGACGTGTCAATGGGTAAAGCA

TACGTTATTTTTGAAGAAGAAAGTGGAAGGAGCGCTGCAATGAAAATGAGTGGGAAATTGTTTTGCGGAG

AGATAATTGGGGTTAAGGATGACAAATATGGTTTTGGTAGGAATGCATATGAGTTTCATGACTCATTATT

TGATTCTTACATTTCCATTGGTGAAACTCGTAAAAGAGGACCCAAGGGGGGGCATTCTATCAGCAGAAGT

AAATATGTAGGCAGGGGAAGCGGCAGGTTTGGGCCTGGGCCTCATCGTGGCGAAAGCCAGTAGTCGCGTG

AAGGATTTGCCTTGATGACTGTCCTTGATTGGCTGGATGCTTTTGATATATATATATATATATGTCCAGG

GTTATTGCAAATTCAATAAATTATATAATTAGAATCATGCTTATGCTTAGTTATAAGGTCAATTTATATC

CAAATCGTCGCCATCGTTAGCATCAGAGGATTTGACGTATATAATATCTGACATGCATTGTCAGCTACTA

TGTAAACTGTGGTGTGTGTTTCATGATTTCTATATATTTCTAATCCAAAACTAGGGATGCATGAAGGAGT

TGATTGCCCGTATCTGTAACTCCTGCAATACACCAATGAGTACAATCCATGCCTTATACTTATAGTAACT

GCTGGGATGTCCATCTTTTCTCAGCTGGAAAAGAGTCGATGTCGATTTGTCACTCCTCGCTCGTTACATT

CGGTGCCACTGGTACATTACAAGATTATATCATTAGCGCAATAGCATATATATGAAGGCGAAGTATCTTT

AAACAACATTATACATCCTCCGTCTCACATACTCCAAAAATGTGGACTACTTCGATAATTCTGTTATCTA

TAATTTCCGGAATTCGAGAGAAGGCTTACTTGTAATGTGAGGAGGAGATTCCTTGGAAGAAAACTTTGGT

TGTCTCTGTGTGTGGAGACATCTGAAGTTGTCTGTGTGTTGACATCTAAATCATCCAGTTAGCCCATGTT

AAAGCTTTACAGAACGCAACCATTCGACCCATGTCTTTCATTATTTTTTTGTCATCTTTAATAAAATCTC

ACCTGCA

>MSTRG.3805.1 gene=MSTRG.3805

AATTTTTGGTCAAAACAATTTGACCAATAAAAATAAAAATGTGACAACTATTTTGGTAAGGAGGTGAGGG

AGGGAGTAATTTGGAAAATAGCAAGTAAAAATGAAGGTGGGGGCAGAAGTGGCAATAAAACTGGAATTTG

GTAGCGAGGAGTCCCAGAGGAGGAGTGTGGGAGAAAGGGGGTGATGTTAAAGCAACTTTCATCATCATAT

TGTTCAAAAACAATACCAAGTTGGGCAAGTTTTAATAGAATTGCAAGTATTCGCATCTGCTGCAGTGGGA

TTAGTAATCATCATACACATTCAACTTGTATGGAGGCGGAGTATGTGTTCGGTGCGTACAAGATTCACCC

CAAGGAAGTGTTTTACTCTTCCCATCTTTCTTATGCCATGGTCAATCTCCGCCCTCTCCTTGCTGGTAAT

GTGCTCGTCTGCCCAAGACGCGAAGTAAAGCGCTTTGTTGATCTTACTGGTGATGAGACTAGTGACCTGT

GGATTGTGGCACAGAAAGTTGGTAAACAGCTTGAGTACTATCACAAAGCATCCTCCCTGACATTTGCAAT

CCAGGATGGTCCACAAGCAGGACAGACAGTTCCTCATGTACACATCCACATCATCCCACGAAAAGGCGGT

GACTTTGAGAGAAATGATGAAATATATGAAGCGATTGATGAGAAAGAGAAGGAATTGAAGCTGGACTTGG

ACAAGGAAAGGAAAGACAGAACCCTGGAGGAAATGGCTCAAGAAGCAGAAGAATACAGAAAGCTTTTTGT

GTAAACAAAGTGAAGCATGACATTAGTAATATTTGCTGCTTGTGTAAACAAAGTGCAGCATGCGGCGATC

CGTATTTAAAGTACTGTTGAAGATTTTTCCCCTTTTTCTTGAATAACATTCAAAAGCACAAAATTGCTTC

CTAGTTGCTAGCGATTCTGGTTATTTTAGCACAAAAAGTTACTTGCGACTTATAAAGTTGACAAGTTTAC

TAGTATTTGACCAGTAGTATGTAATAGGTGAAAAAGGTTTTGTGAGTAAATTGTAAGAGTAATTTTTCTG

AACTTGCTCCGTACTGTGCAATCAATGTCAATGTCCATCATATATTGATATTAAACATTAATACCAGATT

TTCCCTTATTGGAATGTGAAGTTGCAAAACTTAATGAAA

>MSTRG.3805.2 gene=MSTRG.3805

AATTTTTGGTCAAAACAATTTGACCAATAAAAATAAAAATGTGACAACTATTTTGGTAAGGAGGTGAGGG

AGGGAGTAATTTGGAAAATAGCAAGTAAAAATGAAGGTGGGGGCAGAAGTGGCAATAAAACTGGAATTTG

GTAGCGAGGAGTCCCAGAGGAGGAGTGTGGGAGAAAGGGGGTGATGTTAAAGCAACTTTCATCATCATAT

TGTTCAAAAACAATACCAAGTTGGGCAAGTTTTAATAGAATTGCAAGTATTCGCATCTGCTGCAGTGGGA

TTAGTAATCATCATACACATTCAACTTGTATGGAGGCGGAGTATGTGTTCGGTGCGTACAAGATTCACCC

CAAGGAAGTGTTTTACTCTTCCCATCTTTCTTATGCCATGGTCAATCTCCGCCCTCTCCTTGCTGGTCAT

ATCCTGGTTTCAACTGCTTTGCTTCTTTCTTATATGAATTTAGGCTGGTGCTTACACACACATGCATACA

CCCGTCTTCGTTCTTTTTCTATTCTCCCAACTCAGGAAGAGTTTGAGGATCACATTCAGAAGACTGATGT

TTACTCTTTTACCTAAAACTCCTTCTTAGGAGTGCCATATATTTTTCTTTCAAGTGGGCAAAGCTTGCTT

TCTTATGTGTTATGTGATGTTAATGTCTCTTACTTAACTGCTAGGAAGCACCGACTCGGATACGGCGACA

CGGGACACAGGACACGGGGATTCTCCAATTCTCAAAGAGTCTTGGATACGGGACACGGCAAAAAAAAAGT

CATAAAAATTATTATTTTTTTATATGTCAAATATTGGATTATATTAATAATAATAGAGTTTCTTTCAAAA

TGAGACACACATCTATTCTGCATTTTCATTCGAATGATCATCTTTTGTAAAAAGAACAGCCTCTATCTCC

GGTTCATCAAGGAAGAGATTTGCAATTTCTAAAACACGTGCACCCTCCTTTTTCTTGATTCCAATCCAGA

GCTTGCATTTCCTAGTGTCCCAGTCATTTTCTAGGGTTTTTTTAAGGGGTCGAGTCGGTCGCGTGTCGGA

CACGTGTCGGTCGCGTGTCGGGCGTGTGTCGGACGTGTGTCATGTGTCGACACGCCACATGGCGAGTCGG

ACACGTACTCAGCCGAGTCGGTGCCGAGTCGCCTTGTCGGCCGTGTCGGACACGGACACGGACACGGCAG

TCTTTTTGGAGTGTCGGTGCTTCCTAGCTTAACTGGCTACGCATGATCCTATTACTATGATCGAATGTTC

AAATACTAACCCTAATCAAATTAATGATTTGTCTTTTATCTGTTTCTTTCATAACATATATTTTGCGTTT

GCTCGTTTGGTGAACTTGACATATGGTACATGTGCTCGTCTGCCCAAGACGCGAAGTAAAGCGCTTTGTT

GATCTTACTGGTGATGAGACTAGTGACCTGTGGATTGTGGCACAGAAAGTTGGTAAACAGCTTGAGTACT

ATCACAAAGCATCCTCCCTGACATTTGCAATCCAGGATGGTCCACAAGCAGGACAGACAGTTCCTCATGT

ACACATCCACATCATCCCACGAAAAGGCGGTGACTTTGAGAGAAATGATGAAATATATGAAGCGATTGAT

GAGAAAGAGAAGGAATTGAAGCTGGACTTGGACAAGGAAAGGAAAGACAGAACCCTGGAGGAAATGGCTC

AAGAAGCAGAAGAATACAGAAAGCTTTTTGTGTAAACAAAGTGAAGCATGACATTAGTAATATTTGCTGC

TTGTGTAAACAAAGTGCAGCATGCGGCGATCCGTATTTAAAGTACTGTTGAAGATTTTTCCCCTTTTTCT

TGAATAACATTCAAAAGCACAAAATTGCTTCCTAGTTGCTAGCGATTCTGGTTATTTTAGCACAAAAAGT

TACTTGCGACTTATAAAGTTGACAAGTTTACTAGTATTTGACCAGTAGTATGTAATAGGTGAAAAAGGTT

TTGTGAGTAAATTGTAAGAGTAATTTTTCTGAACTTGCTCCGTACTGTGCAATCAATGTCAATGTCCATC

ATATATTGATATTAAACATTAATACCAGATTTTCCCTTATTGGAATGTGAAGTTGCAAAACTTAATGAAA

>MSTRG.3807.1 gene=MSTRG.3807

CAACTCTAATGACGTGTCAATGGGTAAAGCATACGTTATTTTTGAAGAAGAAAGTGGAAGGAGCGCTGCA

ATGAAAATGAGTGGGAAATTGTTTTGCGGAGAGATAATTGGGGTTAAGGATGACAAATATGGTTTTGGTA

GGAATGCATATGAGTTTCATGACTCATTATTTGATTCTTACATTTCCATTGGTGAAACTCGTAAAAGAGG

ACCCAAGGGGGGGCATTCTATCAGCAGAAGTAAATATGTAGGCAGGGGAAGCGGCAGGTTTGGGCCTGGG

CCTCATCGTGGCGAAAGCCAGTGGTCGCGTGAATGGATTTGCCTTGATGACTGTCCTTGATTGGCT

>MSTRG.3811.1 gene=MSTRG.3811

CAAGAAGCAAATAATTTTAATATCTTGTATAAAAAATAAAGGAAAAGATCACCTCTCTCTCTCTCTCTCT

CGATGGAGAACCCGAACCGACCTCAACAGTCAGGTACCAGTGCTCCTCCAAAACCTTGGGAGCGATCAGG

GTCTTCATCTGGTCCTGCGCCTTTTAAGCCCCCATCTTCTGGGAGCACAAGTGATATTGTTGAAGCCTCT

GGAACAGCAAGGCCGGGTGAAATTGTTACAACAGCGGATGCAAATACAGTTGCAAACAGAAATGCACTTG

GGAGACCTCTACCATCTAGGCCTTGGGAGCAGCAAACATATAACAACACCACTGCATTAGGAGGAGGTGT

TGGTTACGGTTCCGGTTTAAACTATAATTCTGGTTATGGTTCTGGAACTTACGGATCATATGGGGCTGGA

GGGTCTTATGGAGGAGGAATGTATGGAAATAGCATGTATCGAGGAGGCATGTATGGTGGCACTGGAAGTC

TCTATGGAGGAGGAATGTATGGTGGGGGAATGTATAACAGTAGTTTTGGAGGTGGAATGGGTAGTTATGG

AGGAATGGGGATGGGCATGGGAATGGGTATGGGTATGGGTGGTCCTCCATATGGCGATCAAGACCCAAAC

AATCCTTTTGCTGCTCCATCATCTCCTCCAGGTTTTTGGGTTTCTTTCATGCATGTGATGCAAGGTGTGG

TAACCTTTTTTGGCCGTGTTGCTATGTTAATAGACCAGAATACTCAGGCTTTTCACATGTTCATGTCTGC

ACTTTTACAGTTGTTTGATCGATCAGGCATGCTGTATGGAGAGCTGGCTAGGTTTGTATTTAGATTGCTT

GGAATCAGGACGAAGCCCCAGAAAGTCCAGCCCCCTGGACCTAATGGACTTCCAGGTCCACATAATTTCC

CAGGTGGTCCAAATTCCTTGGAAGGTCCCAAGGTTGCTCCATCTGGTGCATGGGATGGTGTATGGGGAAA

TGAACCAAGTAACTGAGAATACCACATGATATGCTCATTTAAATTGTCTGCATTGAGTTGAAGGAGATCA

AATGAATTGTGAAAGTGATGCCATGGCCTGTATAGATCAGATCTCCCTTTACCGCCTTGTCAAGAAATTA

TAGATTGTATTCTCCTCTCGTTATGTGTATGCTACTCAACTTATTATTGTTATTTGTGTGGATCTGTAAA

ATTTGAAGACTTGTATTTGATGTAAAATCAGTATACCTTTTCCAAGAAAGCGAGTACACATTACAATTGG

TGGTTGGGAGTTATGAATGACAAGATGTCAGGACTTTTGAAATAATAAAAAAAAAA

>MSTRG.3811.2 gene=MSTRG.3811

GGAAAAGATCACCTCTCTCTCTCTCTCTCTCGATGGAGAACCCGAACCGACCTCAACAGTCAGGTACCAG

TGCTCCTCCAAAACCTTGGGAGCGATCAGGGTCTTCATCTGGTCCTGCGCCTTTTAAGCCCCCATCTTCT

GGGAGCACAAGTGATATTGTTGAAGCCTCTGGAACAGCAAGGCCGGGTGAAATTGTTACAACAGCGGATG

CAAATACAGTTGCAAACAGAAATGCACTTGGGAGACCTCTACCATCTAGGCCTTGGGAGCAGCAAACATA

TAACAACACCACTGCATTAGGAGGAGGTGTTGGTTACGGTTCCGGTTTAAACTATAATTCTGGTTATGGT

TCTGGAACTTACGGATCATATGGGGCTGGAGGGTCTTATGGAGGAGGAATGTATGGAAATAGCATGTATC

GAGGAGGCATGTATGGTGGCACTGGAAGTCTCTATGGAGGAGGAATGTATGGTGGGGGAATGTATAACAG

TAGTTTTGGAGGAATGGGGATGGGCATGGGAATGGGTATGGGTATGGGTGGTCCTCCATATGGCGATCAA

GACCCAAACAATCCTTTTGCTGCTCCATCATCTCCTCCAGGTTTTTGGGTTTCTTTCATGCATGTGATGC

AAGGTGTGGTAACCTTTTTTGGCCGTGTTGCTATGTTAATAGACCAGAATACTCAGGCTTTTCACATGTT

CATGTCTGCACTTTTACAGTTGTTTGATCGATCAGGCATGCTGTATGGAGAGCTGGCTAGGTTTGTATTT

AGATTGCTTGGAATCAGGACGAAGCCCCAGAAAGTCCAGCCCCCTGGACCTAATGGACTTCCAGGTCCAC

ATAATTTCCCAGGTGGTCCAAATTCCTTGGAAGGTCCCAAGGTTGCTCCATCTGGTGCATGGGATGGTGT

ATGGGGAAATGAACCAAGTAACTGAGAATACCACATGATATGCTCATTTAAATTGTCTGCATTGAGTTGA

AGGAGATCAAATGAATTGTGAAAGTGATGCCATGGCCTGTATAGATCAGATCTCCCTTTACCGCCTTGTC

AAGAAATTATAGATTGTATTCTCCTCTCGTTATGTGTATGCTACTCAACTTATTATTGTTATTTGTGTGG

ATCTGTAAAATTTGAAGACTTGTATTTGATGTAAAATCAGTATACCTTTTCCAAGAAAGCGAGTACACAT

TACAATTGGTGGTTGGGAGTTATGAATGACAAGATGTCAGGACTTTTGAAATATAAATTTCTTTTTCCTG

AAAA

>MSTRG.3814.1 gene=MSTRG.3814

GCAAGTAAAAATGAAGGTGGGGGCAGAAGTGGCAATAAAACTGGAATTTGGTAGCGAGGAGTCCCAGAGG

AGGAGTGTGGGAGAAAGGGGGTGATGTTAAAGCAACTTTCATCATCATATTGTTCAAAAACAATACCAAG

TTGGGCAAGTTTTAATAGAATTGCAAGTATTCGCATCTGCTGCAGTGGGATTAGTAATCATCATACACAT

TCAACTTGTATGGAGGCGGAGTATGTGTTCGGTGCGTACAAGATTCACCCCAAGGAAGTGTTTTACTCTT

CCCATCTTTCTTATGCCATGGTCAATCTCCGCCCTCTCCTTGCTGGTCATATCCTGGTTTCAACTGCTTT

GCTTCTTTCTTATATGAATTTAGGCTGGTGCTTACACACACATGCATACACCCGTCTTCGTTCTTTTTCT

ATTCTCCCAACTCAGGAAGAGTTTGAGGATCACATTCAGAAGACTGATGTTTACTCTTTTACCTAAAACT

CCTTCTTAGGAGTGCCATATATTTTTCTTTCAAGTGGGCAAAGCTTGCTTTCTTATGTGTTATGTGATGT

TAATGTCTCTTACTTAACTGCTAGGAAGCACCGACTCGGATACGGCGACACGGGACACAGGACACGGGGA

TTCTCCAATTCTCAAAGAGTCTTGGATACGGGACACGGCAAAAAAAAAGTCATAAAAATTATTATTTTTT

TATATGTCAAATATTGGATTATATTAATAATAATAGAGTTTCTTTCAAAATGAGACACACATCTATTCTG

CATTTTCATTCGAATGATCATCTTTTGTAAAAAGAACAGCCTCTATCTCCGGTTCATCAAGGAAGAGATT

TGCAATTTCTAAAACACGTGCACCCTCCTTTTTCTTGATTCCAATCCAGAGCTTGCATTTCCTAGTGTCC

CAGTCATTTTCTAGGGTTTTTTTAAGGGGTCGAGTCGGTCGCGTGTCGGACACGTGTCGGTCGCGTGTCG

GGCGTGTGTCGGACGTGTGTCATGTGTCGACACGCCACATGGCGAGTCGGACACGTACTCAGCCGAGTCG

GTGCCTAGTCGCCGTGTCGGACACGGACACGGCAGCCTTTTTGGAGTGTCGGTGCTTC

>MSTRG.3825.1 gene=MSTRG.3825

TTTCCTTTAGCACCAAACTATCAGCACTTAACTACCACCACTAATCTGATTACTATACACCACTATATAG

TTAAACAATCGTCGTTCTGAAACGCCGCGAGAGATCGTAAAGCCCTCTCGCCGACTTAACCTCCATCGTC

GACGCATCTTTGCTCACTCTGAGAATGTGGTACAGACTCGGAGGCCTTGTGATCGACAACACCGAATGCT

CTAGTT

>MSTRG.3826.1 gene=MSTRG.3826

AGTAGTTGTCCGTATGCGTATTCTGCGGGTGCATCAAATCCCGCAACCCACTCCTGAAAGCTGCAGTTAC

ATGTAGTGCAGGATTTTCTTCTCCAAGATTGGTGTTTGAAAAGATATCGACTTTTAGTTGCAAGCGTAGG

ATAAGTACGGAAGAAAAGCAAAGAACTAATCGGAAGATGAGTGGAGGTAGGAGGCGAAGTAGCCGTTTAA

TTGAAAGCCCCTGGGTTTTCAATTTGCCAAAGAAGAGATCAGTGTTTGTAAATTTGGAGGATGATGTTGT

GACAGACAAGACAGACGTAGTTGTTGGTGATAAGAGGTCCTTGAAGAGAGCAAGACAAAAGGTAACAACC

GTTAATAATTCTGAATAACGGACGATGAATTCCAGCAGTATAGATATTCGCAGTGAAAGGATACACGGTA

ACAGGCTGAAAAGAATGGAGTTAAGCAAGGAAGAAGATGAATTTGTTACTCCAGGGGAGAATTTTGGCAG

CAGTGGCAAGAAAGGGCAATACCGA

>MSTRG.3827.2 gene=MSTRG.3827

AAATTGATTAAACTAGCATTCTTTTTCAAATTTGGGGCTACAGGTTTTAAAAGTATAAATATAGAGAGAG

TTTATTTTTCTTTCTATTCACACACATAAGCACGCGAGTAATAGATTGGGGTTTAGGGATTGATTGCAGT

CGGTGAGTTGAGCAAACCACAGAGGTAATCAATTGTCGATTTCTTTTAGCTTCTTCCTGTATATATCCTC

GTGTGTAATTGCAAATATGTGTGTATACATATATCTATACTTGCTTGCTTGTGATTGGGGGCTATGAGGA

GAGAAACAGAGGTGGTAAAAGGGAGGTCGAGGCAGAGGATTATAGCGGCCGTTGAAGATGGTTGCAAGTC

GGGCAGGTTGTTGTAGCTGCTGTTGTTGCAGCGCAGAGAGAGATGATGGAGGCGAGCAGTTGTAGAGCTA

TTTGGTGGTGGCTGAAGTCGACTGGGATGTTCAGATATAGTGAATAGTATCATGGAGCGAGGCTGGCATA

AACCAATGAAGTGAAGCTGTGCTATGGATATTTTATTAAAGTCTGTCAGAAACTAAGCAGAAGGGAACAG

TTCAGTAGATAGTACTATTAGAAATCTGAGTTAGACACCAGAGTTAAGGCAATGATTAAGATGATTCGTT

TGTCAAGCTGCGCGTAGAAGTAATGCTAGGCGAAATTGCCAGTGAGATGTCTGCAAGAATAGTATGGGTT

AGAATTGTTCTATTTCAGTTTATTGTATAATAGATTTATTTATTGGTTATCAGTTACTCTTCTTATCGAG

GTTTAACTGATAATGTAGTTTAAGTTGTAATAAGATTTGATAGTTTTTGGTTTCTGATTTCAGTGCTGTA

ACCTATTAGCGATCCTGTTTTTAGGGGGTCAAAGTGGTTTTGATTTAAATCTCTTTTAAAATGTCCAGAT

ATTTTCAAATATCTGTTTTACCTTCTTCCGCAAAATACAGGTTTTTAAATGATTTTGTTTTAGTGGCACC

AAATCCAGACCCCCGGATTTTGGGGATGTCACAATTGATATGACATTTTGTAGAATGTTAATAAATTTTT

TAATAG

>MSTRG.3827.1 gene=MSTRG.3827

AAATTGATTAAACTAGCATTCTTTTTCAAATTTGGGGCTACAGGTTTTAAAAGTATAAATATAGAGAGAG

TTTATTTTTCTTTCTATTCACACACATAAGCACGCGAGTAATAGATTGGGGTTTAGGGATTGATTGCAGT

CGGTGAGTTGAGCAAACCACAGAGAGTCGACTGGGATGTTCAGATATAGTGAATAGTATCATGGAGCGAG

GCTGGCATAAACCAATGAAGTGAAGCTGTGCTATGGATATTTTATTAAAGTCTGTCAGAAACTAAGCAGA

AGGGAACAGTTCAGTAGATAGTACTATTAGAAATCTGAGTTAGACACCAGAGTTAAGGCAATGATTAAGA

TGATTCGTTTGTCAAGCTGCGCGTAGAAGTAATGCTAGGCGAAATTGCCAGTGAGATGTCTGCAAGAATA

GTATGGGTTAGAATTGTTCTATTTCAGTTTATTGTATAATAGATTTATTTATTGGTTATCAGTTACTCTT

CTTATCGAGGTTTAACTGATAATGTAGTTTAAGTTGTAATAAGATTTGATAGTTTTTGGTTTCTGATTTC

AGTGCTGTAACCTATTAGCGATCCTGTTTTTAGGGGGTCAAAGTGGTTTTGATTTAAATCTCTTTTAAAA

TGTCCAGATATTTTCAAATATCTGTTTTACCTTCTTCCGCAAAATACAGGTTTTTAAATGATTTTGTTTT

AGTGGCACCAAATCCAGACCCCCGGATTTTGGGGATGTCACAATTGATATGACATTTTGTAGAATGTTAA

TAAATTTTTTAATAGAATCTCAATGTATTTATTAGTATGACGTTTAGTAG

>MSTRG.3827.3 gene=MSTRG.3827

TTTTCTTTCTATTCACACACATAAGCACGCGAGTAATAGATTGGGGTTTAGGGATTGATTGCAGTCGGTG

AGTTGAGCAAACCACAGAGGTAATCAATTGTCGATTTCTTTTAGCTTCTTCCTGTATATATCCTCGTGTG

TAATTGCAAATATGTGTGTATACATATATCTATACTTGCTTGCTTGTGATTGGGGGCTATGAGGAGAGAA

ACAGAGGTGGTAAAAGGGAGGTCGAGGCAGAGGATTATAGCGGCCGTTGAAGATGGTTGCAAGTCGGGCA

GGTTGTTGTAGCTGCTGTTGTTGCAGCGCAGAGAGAGATGATGGAGGCGAGCAGTTGTAGAGCTATTTGG

TGGTGGCTGAGTATGTGTATTTCGAGTCGACTGGGATGTTCAGATATAGTGAATAGTATCATGGAGCGAG

GCTGGCATAAACCAATGAAGTGAAGCTGTGCTATGGATATTTTATTAAAGTCTGTCAGAAACTAAGCAGA

AGGGAACAGTTCAGTAGATAGTACTATTAGAAATCTGAGTTAGACACCAGAGTTAAGGCAAGTATTCTTA

TTCAACTTCTATATTTAAAAGCAAGTTCTTAGTTTCTTCTATTTACTCAAGAATGATTTTGAAACTTTTT

GGAGTTC

>MSTRG.3828.1 gene=MSTRG.3828

GATCTGCTTCTTACTGATAATTCAAGGTGGCATCGCACTGTTCAAAATGTATACAAGTTGAAGGCAGACG

TGTGGAATGGGGATCGTCCGAAAAGGGAGTCTCCTGCTGTTATGCAATCCATATTGTCGGCAGTCTTGGA

TTATCATTGCAGAACTGGTCGCCCTCTTGGCAATTTGGGTGACATAGTAAGATTGTTCTTTCTGAGTGCA

CCCACAGTATTTTGGGCTTCCAACAAATCCGGTTTCGAATCACATTCGCAGAGCCATTGGAACTTGATGA

CAATTTGGTCAACTTGGCTAAAGACATGGAAGAAGTTCGAAAGATATTTCAGTTGATTCTTCGTGGGAGA

GAGTTGAACCCTGAGCTGGATTATTTGCTTTCAGTTCTGTTAACTACTTACCCCTATCCCAAACATATGA

ACTGGAAAACTCTGGTTCTTCAAAATTACACAGGTCTGTGCATAGCAAGTTACGAAGATGTTTCATGATC

ACTCTCTTCTATACTTTGGTTTCTGAAATGCAAGAGAACGAAACGCGATCCATAACCAGTAAAATGAATA

ATTGCTCTGTCCGTTTTAAGAAAAAAAAATTCAAAGGTTACAAATGGCTAAAAAGGATTCCCGAGGATGA

TTCCATCTTTTACCATATTCTTACCTACGTGCCGCCCTGCAAACAAGATCCCGTAGGTCCTGAATCCTCG

GAACCTCCTCCTCCAAAACCTCAATATAAGGATTATAGGGCGGATTCTGCTATAAAGTATTTTAGGGATT

GCCTGCATCACAACAAAGATGCCCAATGCTCATTGAGTTCGAATGACCTTGAAGTTTGTTTGACAAATCT

TTTTCCGGGTCTCGTGTTTGGGTCATATCAATGCATGTGCGAGACAAGAAAATTTCTAGGCTGGTGATGT

CATTTTCCACTTGATA

>MSTRG.3829.1 gene=MSTRG.3829

AAGCTGCAGTTGGGTGCTCGGTACTTATCTCCTCCCACGTTCTCTTGCACGGGTGATCGTGATGACTAAG

ATGCTGATGATTATGATCATGATGCTGATATTGGAGGATCTTAGTGTATCAACAAAGTGATGGTCGTAAG

GGTGATCATGCCGATTAGGATGCTGATGTTGGAGGATTTTCGAGTATTTAGGTGTTAGTATTGTACTGCT

TGGATGATATTTGGAACATATGGTTGTGTTGCTGAAATTTTAAGGATATTGATGTAAATTTTGATGTTAA

TATTGTTGGGTTATTCAA

>MSTRG.3832.1 gene=MSTRG.3832

TAATAGATTGCATTAATAAATCCGAAGAAGAATGCTCTTAATGACAAAAGAAGTTAGCAGGCTGGTTAAT

GCACTTAAAGAAGCCGAGGATGAAGTATTTGCAACAAAGGAAGAAGAAGCTTGACTTAAGAATACCCTAG

AGGAAGCTGAATCTGAGATGCACTATTTAAAGGAAGTTCTTGGGGAAGCAAAATCTGAAAGCATGAAACT

GCAGGAAAGCTTAACGGACAAAGGAAACTGAGTGTTATTCAGGAGAATAAAGAGCTTCAAAATACAGAAT

CAGCTTCTCTGATAAAGGTCAAGGAATTGTCGAAGTTGAAAAGCTCAAGGCTCTCCATCTAATTTTCCTA

TTTTTCTAGGGCTTCTATGTTTT

>MSTRG.3834.1 gene=MSTRG.3834

CAAAATAAAGAGAAACCTGATTCTCTTTGTTTCGAGAAACCAGAGGCACAGTAAAGATGGCAAGATATCG

TTTCATTCGTCAGCTCATTAGATTGGCGTTTTGATTGTTGCTTGTCCACAAGTTCCTTTTAAATTTATAT

TTGTTTTATAAATTTACGAGTGTGCAGATGATGATTAAAAATTCTTCACTTTGAACCTACTGGTGTGATT

GATTTGCTTCAATCATCCCCTGAGGATCATTTATTGTTGTCTGATGATGCACTCTTCGAATACATGGAAT

TATCGGAATAAGTGCATTGCACAGGTTATAAGCTAGTTTCTGATATTTTAGCGTAGGCTCTATACATAGA

GTCAACAGCAGGCGTTTATTTGGTAACGGTCTGACTGCCGCTTGGAGCCTAGATTAATATACCGGCTTAA

GACCCATTGAATATTTTTGTTGTTCTGTGACACAATGTGCTGGTTATATTTTTTTTTTTTTTGATAATTT

AGGTTTATATGCCTTAGGAATGAAAATATGTTTAAATACTT

>MSTRG.3835.1 gene=MSTRG.3835

TTTTCATGTTCATAGTTCCTTCTTCTATGGTTTCCAACTTTTTTCCCCAGATAGCACTTTTCTTGGCAAT

TTCATATAATTCAGCCATGGCTTCAGCTAGGTAGAGGGTCATGAAAGTGTAAATCACACAGCCTCTCAAT

CTGCAAAAAGTATCAATGTTTTGCACTTGATGTCTAAACATTGTTGGGTGCTGATCCGAATCATGGTGAT

GCTTTGGAAATTTCTCAGTGGCTGACGATGTTTCTAGGGACTTGGCAAGAGGGTCAGCGGGATCTTCAGA

GCCGCCCATCGCTTGCGGATCTTTGCAACACGGCCAATGCCAAAGAAGTAAGCTACAGAAACAGCTGATC

TTCTTTAGCGGTTATTAAGTCTGATAATTTATCTTTAGTGCAGCCTCAAATCCAATTTATGGCTCCTCTA

AATCCAGTGTTAATCTGGTTGGTAATAACCAGAAGGAATCGATATCATTATCATCCGTATCCTAGACAAC

ATATGGGAAACCTTCTAGTGCATTTGGAGACTACAATTGAAACTTATTTATGGTCTTGACAGCCTGTGGG

AAAAAGAGCCTGAAAAATAATAGTGACTCCTTTGTACTGTGAGATTAGTTCAGAACAAGAGCCAGAGGAG

GAATAGAGTGAGTCTTCACTGCGTGATACCTTCTTTGCAACTGTGGAAGCTGAGACTGATAGTAAAACTA

GAAAAGGCTGTTCCTGAAAAGAAGCTATCACTATCCAAAGGGCAAGGGAGTGGAGATAAAAGATCGGCTC

TTTGAGCTTGCTCGTTTGTTCTGGAGATCCTGGTGATTATATTGACCTTCATGAGCTTGGAATGGATCTA

TGTGCTGAGGCCTTGAGGAAACAGTGACAAGTGAAGAAGAACAAAATATGATAAGGCTGCTTTGAAATTT

TGAGAAGTGGCTGCTTTAGGTTTACCTCAACTTGAGATATGTTCATGTTTGTGAAGCGATGGAGAATCTA

CCAATCGATGATTTTTCTGCAAAGAATATGGTTGCAACACACTTCCAACTGCTTTAGGATACAGATTCTT

TAGCTAGCCAGAAGTATGAAGATGCACTTTTGATTAAACCAGACTTTTTTGAGTAAGTTGGTGATCAGGA

ATAAGTTCAAATGACAAAACTTCACAGGTCATCTTTTCTTACGAAAAAGAAGACCTCTTAAACTGGGATC

CAACAGAAACTAAAAATGTTTAATAACAGAACGGAGAAAATGCAATCTGCCACTGAATGTGGGAGAAGCC

GGAGATGCAAAGAGGCAAAGATTTTCAAAGAATCCTCGTTTCGGCTTGAAAAAAATTTGATAAAAAGAAA

GAGGAAGCAAGTAAGCAGTAGTGAAGGTGATCGGTGAGCCCTAAAAGGAAAATTTCATCCGAAGCTGCAG

AACAAGCACCATTGATACGATCACAAATACATCTATTTTGGGGTAACATGCTTTTTAAGCTATCTCCGGT

TGAATGCAAATTAGGTATGATGGATTGGAATAAAAATTTGGGCCCTGCTCTTGAGCGATTAAAACTTGAT

GGAACTTCTAAGGCTAAAATTTCAATCTTGCTGAAATATCAATGCTTGAATTTGGATGCTGTAGAAGGTA

ACGGGAAGAAATACATTGAAACTAAAGGTACAAACTAAAAAAGATACTCAAGCTTTTTGTCATTTCATCT

TGATGTTAACTAGGGAACAAGGTAGCTTTACTTCATTTGAAGAAGTATGAGTCCTTGAGCCTTCTTGCTA

TCAAGGAAGAAACACTAAAAAGATGACTACGTACCTCAAGCTGGTCCCCGATGTTGATAACAATGGAGTG

GCGCATAGGTGGAACATCAATCCATTCTCCATCCTTGAGGAGCTGGAGGCCGCTGACTTGGTCATCCTGG

AAGAGAAGGATGATACCTCCAGCGTCGGTGTGGGCACGAAGGCCTTTAATAAGGTCTGGCTTAGGACATG

GAGGATAGTTGCTCACCTTTGTTCCAAAAGTAGGACCCTTGGATCCATAAAATGCTTTCTTCAGAATTAA

GGAACAGGACTAGTGGAACAAGTATGTGCTCACATTTCCATGGACCCCTGCAAAATTTTATGAGGCTTAC

ACTATTCCAGGTATGTCTACACCTGAGTTGAAAAAAAAGTGCAAGAAAAAGTTGGTCACACATCATTATC

TGGTGAAGCATGGCAAAAGAATAGGGTGGTGCCAAACATTCTCACAAAAGTTGGTAAAAGCCAATGGGCA

GTTGGTATTCACTTAACGCCCCATTTGGCTAAATGTAAAATAAGAATTTCTGGCAAAAAATTTATATTTA

TACATTGTTGTTTGTAAATTAAGTAATTGTAAAAGTTTAATATATCTGG

>MSTRG.3835.2 gene=MSTRG.3835

CTTTTTTCCCCAGATAGCACTTTTCTTGGCAATTTCATATAATTCAGCCATGGCTTCAGCTAGGTAGAGG

GTCATGAAAGTGTAAATCACACAGCCTCTCAATCTGCAAAAAGTATCAATGTTTTGCACTTGATGTCTAA

ACATTGTTGGGTGCTGATCCGAATCATGGTGATGCTTTGGAAATTTCTCAGTGGCTGACGATGTTTCTAG

GGACTTGGCAAGAGGGTCAGCGGGATCTTCAGAGCCGCCCATCGCTTGCGGATCTTTGCAACACGGCCAA

TGCCAAAGAAGTAAGCTACAGAAACAGCTGATCTTCTTTAGCGGTTATTAAGTCTGATAATTTATCTTTA

GTGCAGCCTCAAATCCAATTTATGGCTCCTCTAAATCCAGTGTTAATCTGGTTGGTAATAACCAGAAGGA

ATCGATATCATTATCATCCGTATCCTAGACAACATATGGGAAACCTTCTAGTGCATTTGGAGACTACAAT

TGAAACTTATTTATGGTCTTGACAGCCTGTGGGAAAAAGAGCCTGAAAAATAATAGTGACTCCTTTGTAC

TGTGAGATTAGTTCAGAACAAGAGCCAGAGGAGGAATAGAGTGAGTCTTCACTGCGTGATACCTTCTTTG

CAACTGTGGAAGCTGAGACTGATAGTAAAACTAGAAAAGGCTGTTCCTGAAAAGAAGCTATCACTATCCA

AAGGGCAAGGGAGTGGAGATAAAAGATCGGCTCTTTGAGCTTGCTCGTTTGTTCTGGAGATCCTGGTGAT

TATATTGACCTTCATGAGCTTGGAATGGATCTATGTGCTGAGGCCTTGAGGAAACAGTGACAAGTGAAGA

AGAACAAAATATGATAAGGCTGCTTTGAAATTTTGAGAAGTGGCTGCTTTAGGTTTACCTCAACTTGAGA

TATGTTCATGTTTGTGAAGCGATGGAGAATCTACCAATCGATGATTTTTCTGCAAAGAATATGGTTGCAA

CACACTTCCAACTGCTTTAGGATACAGATTCTTTAGCTAGCCAGAAGTATGAAGATGCACTTTTGATTAA

ACCAGACTTTTTTGAGTAAGTTGGTGATCAGGAATAAGTTCAAATGACAAAACTTCACAGGTCATCTTTT

CTTACGAAAAAGAAGACCTCTTAAACTGGGATCCAACAGAAACTAAAAATGTTTAATAACAGAACGGAGA

AAATGCAATCTGCCACTGAATGTGGGAGAAGCCGGAGATGCAAAGAGGCAAAGATTTTCAAAGAATCCTC

GTTTCGGCTTGAAAAAAATTTGATAAAAAGAAAGAGGAAGCAAGTAAGCAGTAGTGAAGGTGATCGGTGA

GCCCTAAAAGGAAAATTTCATCCGAAGCTGCAGAACAAGCACCATTGATACGATCACAAATACATCTATT

TTGGGGTAACATGCTTTTTAAGCTATCTCCGGTTGAATGCAAATTAGGTATGATGGATTGGAATAAAAAT

TTGGGCCCTGCTCTTGAGCGATTAAAACTTGATGGAACTTCTAAGGCTAAAATTTCAATCTTGCTGAAAT

ATCAATGCTTGAATTTGGATGCTGTAGAAGGTAACGGGAAGAAATACATTGAAACTAAAGATGACTACGT

ACCTCAAGCTGGTCCCCGATGTTGATAACAATGGAGTGGCGCATAGGTGGAACATCAATCCATTCTCCAT

CCTTGAGGAGCTGGAGGCCGCTGACTTGGTCATCCTGGAAGAGAAGGATGATACCTCCAGCGTCGGTGTG

GGCACGAAGGCCTTTAATAAGGTCTGGCTTAGGACATGGAGGATAGTTGCTCACCTTTGTTCCAAAAGTA

GGACCCTTGGATCCATAAAATGCTTTCTTCAGAATTAAGGAACAGGACTAGTGGAACAAGTATGTGCTCA

CATTTCCATGGACCCCTGCAAAATTTTATGAGGCTTACACTATTCCAGGTATGTCTACACCTGAGTTGAA

AAAAAAGTGCAAGAAAAAGTTGGTCACACATCATTATCTGGTGAAGCATGGCAAAAGAATAGGGTGGTGC

CAAACATTCTCACAAAAGTTGGTAAAAGCCAATGGGCAGTTGGTATTCACTTAACGCCCCATTTGGCTAA

ATGTAAAATAAGAATTTCTGGCAAAAAATTTATATTTATACATTGTTGTTTGTAAATTAAGTAATTGTAA

AAGTTTAATATATCTGG

>MSTRG.3835.3 gene=MSTRG.3835

CTTTTTTCCCCAGATAGCACTTTTCTTGGCAATTTCATATAATTCAGCCATGGCTTCAGCTAGGTAGAGG

GTCATGAAAGTGTAAATCACACAGCCTCTCAATCTGCAAAAAGTATCAATGTTTTGCACTTGATGTCTAA

ACATTGTTGGGTGCTGATCCGAATCATGGTGATGCTTTGGAAATTTCTCAGTGGCTGACGATGTTTCTAG

GGACTTGGCAAGAGGGTCAGCGGGATCTTCAGAGCCGCCCATCGCTTGCGGATCTTTGCAACACGGCCAA

TGCCAAAGAAGTAAGCTACAGAAACAGCTGATCTTCTTTAGCGGTTATTAAGTCTGATAATTTATCTTTA

GTGCAGCCTCAAATCCAATTTATGGCTCCTCTAAATCCAGTGTTAATCTGGTTGGTAATAACCAGAAGGA

ATCGATATCATTATCATCCGTATCCTAGACAACATATGGGAAACCTTCTAGTGCATTTGGAGACTACAAT

TGAAACTTATTTATGGTCTTGACAGCCTGTGGGAAAAAGAGCCTGAAAAATAATAGTGACTCCTTTGTAC

TGTGAGATTAGTTCAGAACAAGAGCCAGAGGAGGAATAGAGTGAGTCTTCACTGCGTGATACCTTCTTTG

CAACTGTGGAAGCTGAGACTGATAGTAAAACTAGAAAAGGCTGTTCCTGAAAAGAAGCTATCACTATCCA

AAGGGCAAGGGAGTGGAGATAAAAGATCGGCTCTTTGAGCTTGCTCGTTTGTTCTGGAGATCCTGGTGAT

TATATTGACCTTCATGAGCTTGGAATGGATCTATGTGCTGAGGCCTTGAGGAAACAGTGACAAGTGAAGA

AGAACAAAATATGATAAGGCTGCTTTGAAATTTTGAGAAGTGGCTGCTTTAGGTTTACCTCAACTTGAGA

TATGTTCATGTTTGTGAAGCGATGGAGAATCTACCAATCGATGATTTTTCTGCAAAGAATATGGTTGCAA

CACACTTCCAACTGCTTTAGGATACAGATTCTTTAGCTAGCCAGAAGTATGAAGATGCACTTTTGATTAA

ACCAGACTTTTTTGAGTAAGTTGGTGATCAGGAATAAGTTCAAATGACAAAACTTCACAGGTCATCTTTT

CTTACGAAAAAGAAGACCTCTTAAACTGGGATCCAACAGAAACTAAAAATGTTTAATAACAGAACGGAGA

AAATGCAATCTGCCACTGAATGTGGGAGAAGCCGGAGATGCAAAGAGGCAAAGATTTTCAAAGAATCCTC

GTTTCGGCTTGAAAAAAATTTGATAAAAAGAAAGAGGAAGCAAGTAAGCAGTAGTGAAGGTGATCGGTGA

GCCCTAAAAGGAAAATTTCATCCGAAGCTGCAGAACAAGCACCATTGATACGATCACAAATACATCTATT

TTGGGGTAACATGCTTTTTAAGCTATCTCCGGTTGAATGCAAATTAGGTATGATGGATTGGAATAAAAAT

TTGGGCCCTGCTCTTGAGCGATTAAAACTTGATGGAACTTCTAAGGCTAAAATTTCAATCTTGCTGAAAT

ATCAATGCTTGAATTTGGATGCTGTAGAAGGTAACGGGAAGAAATACATTGAAACTAAAGGTACAAACTA

AAAAAGATACTCAAGCTTTTTGTCATTTCATCTTGATGTTAACTAGGGAACAAGGTAGCTTTACTTCATT

TGAAGAAGTATGAGTCCTTGAGCCTTCTTGCTATCAAGGAAGAAACACTAAAAAGATGACTACGTACCTC

AAGCTGGTCCCCGATGTTGATAACAATGGAGTGGCGCATAGGTGGAACATCAATCCATTCTCCATCCTTG

AGGAGCTGGAGGCCGCTGACTTGGTCATCCTGGAAGAGAAGGATGATACCTCCAGCGTCGGTGTGGGCAC

GAAGGCCTTTAATAAGGTCTGGCTTAGGACATGGAGGATAGTTGCTCACCTTTGTTCCAAAAGTAGGACC

CTTGGATCCATAAAATGCTTTCTTCAGGACTAGTGGAACAAGTATGTGCTCACATTTCCATGGACCCCTG

CAAAATTTTATGAGGCTTACACTATTCCAGGTATGTCTACACCTGAGTTGAAAAAAAAGTGCAAGAAAAA

GTTGGTCACACATCATTATCTGGTGAAGCATGGCAAAAGAATAGGGTGGTGCCAAACATTCTCACAAAAG

TTGGTAAAAGCCAATGGGCAGTTGGTATTCACTTAACGCCCCATTTGGCTAAATGTAAAATAAGAATTTC

TGGCAAAAAATTTATATTTATACATTGTTGTTTGTAAATTAAGTAATTGTAAAAGTTTAATATATCTGGT

>MSTRG.3837.1 gene=MSTRG.3837

CAAGAATGGGCAAAATGGCTTTCTCTCGCTGAATTCTGGTATAATACTACTTATCACTCAGCCACAAAAA

TGACACCATACGAAGTAATGTATGGCCAACCTCCTCCCATATATAAACCTTACATAGCAGGCTCATGTGT

GGTGGATACAATTGATAGGAGTCTTCAACAGAGGGAGAGGCTGATCGCAACTTTGAAAGAAAATCTGGCC

AGGGCCCAGAACAGAATGACTCAACTGGCGAACAAGAAAAGATCGGACAGACAATGAAGAGCTGACAGGT

GGAGCTGGGACAGGTGAAATGGCAAAAATAAAGGTGGTAGAAGAGACAAATGGAGGCAACGGCACTGGCA

GTCACACAAGGAATCTGGGAGCACATGGACCATTGGAAAACGACACCTGTCCTTTATTCATATAATATCA

GTGTAATAGAGTTTGTTAGGAAATGATTGGTGTTTTCTATAAAGCCAAGAAGAATGTATGGAAGAGACAG

CTT

>MSTRG.3842.1 gene=MSTRG.3842

TGGAAACATATGTCAAGACCAAACTACAGAAAAAAATTGTAACAGGATGCAATCTCAGAGCACAACAGTT

TAATCAATGTCGACAACTATGGAATACTTGGCACCATTTACCACTTTAAGCCTTTCATAGAAATCTGGGG

GCATCACTTCAACTCGTAATTCTTCCAGGGAGTGGAGAAACTTGATTCCTTTTGGAAGCTCCTTAAGCTT

CAGACAGTTTCTAATCATCAAATGTGACAGAACAGGCATGCTGTGCTCCTCCACCCTCCATTTCACCAAG

TTGGGAAGATTCCTCAGTACAAGACGGGTAAGTTTTGGGAAACCCGTGGCTGAACACACCAGTTCACGCC

CCATGAATGCTTTTCTGTGTATTAGTAGCTCCCGCAATGTATGAATCTTCTCCAGCACTGGCATTGGATC

TTCTTCTAGGAATGACAGCATCAGATGTAAGCGCGTGATATGAATTAAGGACAGATCAATAGCAGCGTCG

TGAAGTTGTTGCTGCTCGAATAATTGAGCCAGCTCTGGCAGTTGGCCAATTACTTTTAATTCCTGAAGGT

TGTATCTATGATCTGACAACAACTTCCTCAAGATGTCTGGACCATTTCGCGACCCTAGATCACATTGACT

GATAGTAAGGACCAAGTACTGGAGACATGAAGTCGATGACAAGGCTATAGCTGTTAAGCATTTCATCACC

TCCTCCACATCATCATAACTATCATCTACTCTTAAGGTCAGTTTCTGAAGACCGGTTAATTTGGGCAGAT

CCTTAACCTCGCACCATTGAGTATTCACGTTCTCCAGTGTCTCCAACTTGTTCAAGCCATGAAAGCGCAT

CTTTGAATTTTTCCCCAGGCTGTGAACTTTCCAATCTGGTAGATACAAATGGCGCAGACGTCCCAGCTTG

CCCATAATATTTCTTGACAGCCATGGTGGGAAAAGTATATCATTCCGTACGTCTAATTTGAGTGTTTGGA

GGAGCACCAGATTCTGTATCGATGGCAAAAGCAATAAATTAGTCTTCCTTACACTCAGGTACCGCAAGTA

AACAAGGCTGCCTAATACTCTGCCCAGATTAATACGGAAAAGGGTACCAGAAACAGTTTGTTTATCAAAC

TGGACTTCTTCCAAAGCAAACACTCTTAGTAGCCTAAAATTAGCAACATCTGACGACAACACTGGTGGCA

CACTTCTGGTCTTAAACTCGTTGAATAGTAACATTGAACGATATCGTTGCTGATTGGCTTTCTTGCCAAA

GGAGGGATTTGCTATTGTTTGCTTGTATGTTCC

>MSTRG.3843.1 gene=MSTRG.3843

CACTACTAGAAAAACAGGATTAGACATCACATTTTAGACATCTGTTAGAAAAACCACCGATGTTAAAACC

CTATTTTACATCGAATATTTTGAAACCGATTTCTATTAAAAATCTAGACATCGCCTAAGTGCAAAACCGA

TGTGTAAAACCTGTTTTTAAAAAAATTGACCGAGCTCCCCCGCTGCTTTTCCCCCCTTAGCCAATTGTTT

TACAAACAAGATTCCCCACAGAATATTTCCTCTTATTACACAAATATTCCCCCTTTACCAAAATTTTATA

TCCCACCTTCACCAAAATGTTATTTCCCCCGTTTCTTCTTTTCATTCTCGACTATGTCTCTCGACACTCT

CTCTCTCTCGACTCTCTCGTCTCTTTCTCTTCCCATCTCTGCTCTCCCTATCTGGTCTCGCTCACACCCT

CTATTTCAAGCTCCGACATTCAAATAGTAGTTAGGGTTCGATTGGGTAACCCCTAATTTGTTCTATCGGA

GACACAGCTCGAAGAAGAAACTTGAATAAATTAGTAGATAGAACTCATTTCGGAGAAACGCAAATTGCAG

ATTAGTTCGATTGGGTTTAAGAGTTTGATTGGTTCAATTGGGTTTTAGGGTTCTTATCAATTCTCACTTC

AAATCCTGCCCTAATTGAAGACATTCGCGAGTGGGAACAAGAATATCAGTGCTGAACTGCTTGTTGGCCA

AGTTGTCTCCCGTATTGCTTCTGCTCTGTTTGAATGGTTGTGGTCACTGAAGACAGATGAGCCCATTGGT

GGAAGATATGAGGCTGATTTGTGGTTTTCTCCAGGAACTTGTGCAAAGTTAAATTTAGGAGAGGAACACG

AATAGAAATCTGTTTGACTTCCCAGGGTGATTGGAACGGTGCAGGTGCAAATGAATACGAATAGAAATCT

GTTTGACTTCTCAATATGAATTTTGATTATATGTTTAAGATATTGATCCATCTTGAATATTCTTAGCTTA

ATTTTTGTATGAACTTTTCAATACTTTATATATACCACAACACCAAATGAATACGATAAGATGGTAATGA

AAGAAGGTTAACAGGAAAGCATGAGACATCTAACAAGCACACTTTCTCATGGGTAAGTTGCTTATATATA

TAAAATCTTAGTTGTAACCTGATTTTGATGCAAAAATATCGAGACTAAGACTTACCTTCAATTCAAGTAG

TTTAAAAACCTTGCAAGTAACCAAATCATTTTACTTTCTTAGTTGCCTGGTGAAACATGTGGTATTTTTT

TCTTCAAAAAGAACCTTGTTCTTGGTGTTTATTTAGAATCTTCTATCAACAATACGTTTTTTTAAGGTTT

ACAGGTAATTCTCTGTTCTTGCTCTATTACCAT

>MSTRG.3844.2 gene=MSTRG.3844

GGGAAATACAGAACGTGTGAGATTTATTTTGAGGGAATTTGCACACAAAAATCTTTTAACTAAGCTTAGT

AGATATAAGCATTCAGCACTTCACTTGGCTATATATCGAGGAGGACACACCCAAGTGGCTCAGGTCCTCA

TTGATGCAGCTCGCCATTTGCCTCCTCCTCCAGATAATGATGATAATTCAGTTACTTCTTTTCAAGCTTT

TCTCAGGCAAGGTGATAAAGATATGGACACCGCTTTACATGCAGCAGTCAAGTATGGTCACCTGGATATT

GTTAAGCTACTAGTAGAGGCTGACCCGACTGATACACATATTCAAAACTATGAGGGCAAAACACCGATGT

ACATAGCAGTGGAGAAAGGGTTCAATGATATAGCAGAGATCATCTCTACAACTTGCACTGCTCCATCTTT

GCTCGGACCTGATTGTAGTAGTGTTGTGCGTA

>MSTRG.3846.1 gene=MSTRG.3846

CTATTGCATAGTTGTTTATGGTAGTGAAAGTGTAACTTCCCACCATTTGAACTGTTCTGTAATTGGAAAC

ATATGTCAAGACCAAACTACAGAAAAAAATTGTAACAGGATGCAATCTCAGAGCACAACAGTTTAATCAA

TGTCGACAACTATGGAATACTTGGCACCATTTACCACTTTAAGCCTTTCATAGAAATCTGGGGGCATCAC

TTCAACTCGTAATTCTTCCAGGGAGTGGAGAAACTTGATTCCTTCTGGAAGCTCCTTCAGCTTCAGACAG

TTCCTAATCATCAAATGTGACAGAACAGGCATGCTGTGCTCCTCCACCCTCCATTTCACCAAGTTGGGAA

GATTCCTCAGTACAAGACGGGTAAGTTTTGGGAATCCCGTGGCTGAACACACCAGTTCGCGCCCCATGAA

TGCTTTTCTGTGTATTAACAGCTCGCGCAGTGTATGAATTTTCTCCAGTACTGGCATTGGATCTTCCTCT

AGGAATGACAGCATCAGATGTAAGCGCGTGATATGAATTAAGGACAGATCAATAGGAGCGTCGTGAAGTT

GTTGCTGCTCGAATAATTGAGCCAGCTCTGGCAGTTGGCCAATTACTTTTAATTCCTGAAGGTTGTATCT

ATGATCTGACAACAACTTCCTCAAGATGTCTGGACCATTTCGCGACCCTAGATCACATTGACTGATAGTA

AGGACCAAGTACTGGAGACATGAAGTCGATGACAAGGCTATAGCTGTTAAGCATTTCATCACCTCCTCCA

CATCATCATAACTATCATCTACTCTTAAGGTCAGTTTCTGAAGACCGGTTAATTTGGGCAGATCCTTAAC

CTCGCACCATTGAGTATTCACGTTCTCCAGTGTCTCCAACTTGTTCAAGCCATGAAAGCGCATCTTTGAA

TTTTTCCCCAGGCTGTGAACTTTCCAATCTGGTAGATACAAATGGCGCAGACGTCCCAGCTTGCCCATAA

TATTTCTTGACAGCCATGGTGGAAAAAGTACATCATTTCGTACGTCTAATTTGAGTGTTTGGAGGAGCAC

CAGATTTTGTATCGATGGCAAAAGCAATAAATTAGTCTTCCTTACACTCAGGTACCGCAAGTAAACAAGG

CTGCCTAATACTCTGCCCAGATTAATACGGAAAAGGGTACCAGAAACAGTTTGTTTATCAAACTGGACTT

CTTCCAAAGCAAACACTCTTAGTAGCCTAAAATTAGCAACATCTGACGACAACACTGGTGGCACACTTCT

GGTCTTAAACTCGTTGAATAGTAACATTGAACGATATCGTTGCTGGTTGGCTTTCTTGCCAAAGGAGGGA

TTTGCTTGTTTGCTTGTATATTCCTCGTCAAAATAAACAACAAGCTGTCTAGTGGCAGCAGCCCTCGAGC

CCACAGAAGCATTCAAATGAGAATGATTTTTTTCTCGAAGATCAATAACTTTGTATAAATCCTCCTCTTT

TGCTTGGCTTAGGGAGAGGTCTCTCATAAGATCATGAAGAGAACAAGACTTAAACTTTGTAAGCGATGAC

TCGATATCCTTATATTTCACTTGAACCATACTTCTATGGACCAGTTCCCCCAAGTACGATTCAGCCACTT

GCATCACTGTTTCTCCATTCTTTTTGTCACTTGACAGTACCATTCCTTCTGCGATCCATAACTGATATAA

AGTCTCTACTTCTATCCATTCATCTTCGTTAAATTTACCTAAATAAAGAAAGCACGGCTTCAGTTGAGGA

GGCAAGTCATTGTAACTCCAAACTAAAACCTCGTGTAATTGATGTTGATAGACTTCTCCCAGTTCTTTGC

CCCTTTCCAGAGACAACATCGTATCACGATACACCTTCTCCCACTCTATCAAGGAAGGTTTTGTCACGAG

AATTCCCCCCAACATCACTATAGCTAGTGGAAGACCCGCACAATATTCAACCATTTCTCTCCCCATTTCT

TCCATCCTTTTAACGTCTCTAGTAATATCTGTAACAAAAATGGAATCATAAACTATAATTTCAAAAGAAT

TAGATATCATGCTAAATTTTACA

>MSTRG.3850.1 gene=MSTRG.3850

GAAAAAAGAAGACTCATAAACAGACAGTCGACTAAAAGCGGGCAATGGAGAGAGAGAAAACAGAAGGTAA

CAGAGATAAAAGATCGAAGAAGGCGAGAGAGAGAGAGAGAGAGTGTGATCGAAGAAAGCTGGTAATGGAG

GAATCAATCTTTTTTTCCTAATAACTTGAAGGAGGAATTCCACCTATACGGATGGAAATTAATCAAGTGC

AAGCAGATAGAAGATCTCCTTAATTGGATTACTGGCACAAGGGACCCTACATGAACATCACTTTTACCAG

GAACTCCCCGTTATGACTTCCAGATTGGTTCTGGCAGTCACCCACCTCATCCCCAACCAGTAATATACCC

ACTCCTTCACAGTAAAATGTTAATGAACAATCTTTAAACAGAATACTTAGTTAATTCTGCTGTTTACCTC

CACTCTCTCTGCTAACTCTAGTTTTGGAAATCTTGCAGCTAAGTCCTGGTATGTGATATGATGTGTGGTT

TATAATTTTATTTCACTAGATATGGCCTGATCTTTTTGAGAAGGTGAAACTTGGAGGCATAAATTTTATT

CAAAC

>MSTRG.3850.2 gene=MSTRG.3850

GAAAAAAGAAGACTCATAAACAGACAGTCGACTAAAAGCGGGCAATGGAGAGAGAGAAAACAGAAGGTAA

CAGAGATAAAAGATCGAAGAAGGCGAGAGAGAGAGAGAGAGAGTGTGATCGAAGAAAGCTGGGAGGAATT

CCACCTATACGGATGGAAATTAATCAAGTGCAAGCAGATAGAAGATCTCCTTAATTGGATTACTGGCACA

AGGGACCCTACATGAACATCACTTTTACCAGGAACTCCCCGTTATGACTTCCAGATTGGTTCTGGCAGTC

ACCCACCTCATCCCCAACCAGTAATATACCCACTCCTTCACAGTAAAATGTTAATGAACAATCTTTAAAC

AGAATACTTAGTTAATTCTGCTGTTTACCTCCACTCTCTCTGCTAACTCTAGTTTTGGAAATCTTGCAGC

TAAGTCCTGGTATGTGATATGATGTGTGGTTTATAATTTTATTTCACTAGATATGGCCTGATCTTTTTGA

GAAGGTGAAACTTGGAG

>MSTRG.3853.1 gene=MSTRG.3853

ACAATATCTATTGGACTTTATGTGTGATATGACTACAGTCAAAAGAAAAGAAAAATGAGAAATGAAATAA

AGAAAAAGAATACAGCTGTCTAATCTGCACATACGCGTCTATTTCACGCTTCATCTTCTTCACCAGTTGA

CACAAACACATATACATACAAGCATATGTATGTATATCACACATCATCAAAACAATACAACCCACGTCTA

CTTTCTTCCTTTCTTCTCGTTTTACAGTCATCAGAGCAAGCATCTGCGGTTCGCCGGAATGACGAACTGG

AACGCGATCCCGGCCGATCCGGGTCGACTTCGACCCGTGTTCCGGTACGAGCGAACCAGTTCGCGGATCC

ATAGGCTGTACCGCGAACCGGGTAACTATGG

>MSTRG.3852.1 gene=MSTRG.3852

CCATAGTTACCCGGTTCGCGGTACAGCCTATGGATCCGCGAACTGGTTCGCTCGTACCGGAACACGGGTC

GAAGTCGACCCGGATCGGCCGGGATCGCGTTCCAGTTCGTCATTCCGGCGAACCGCAGATGCTTGCTCTG

ATGACTGTAAAACGAGAAGAAAGGAAGAAAGTAGACGTGGGTTGTATTGTTTTGATGATGTGTGATATAC

ATACATATGCTTGTATGTATATGTGTTTGTGTCAACTGGTGAAGAAGATGAAGCGTGAAATAGACGCGTA

TGTGCAGATTAGACAGCTGTATTCTTTTTCTCATTTCTCATTTTTCTTTTCTTTTGACTGTAGTCATATC

ACACATAAAGTCCAATAGATATTGT

>MSTRG.3854.1 gene=MSTRG.3854

ACAATATCTATTGGACTTTATGTGTGATATGACTACAGTCAAAAGAAAAGAAAAATGAGAAATGAAATAA

AGAAAAAGAATACAGCTGTCTAATCTGCACATACGCGTCTATTTCACGCTTCATCTTCTTCACCAGTTGA

CACAAACACATATACATACAAGCATATGTATGTATATCACACATCATCAAAACAATACAACCCACGTCTA

CTTTCTTCCTTTCTTCTCGTTTTACAGTCATCAGAGCAAGCATCTGCGGTTCGCCGGAATGACGAACTGG

AACGCGATCCCGGCCGATCCGGGTCGACTTCGACCCGTGTTCCGGTACGAGCGAACCAGTTCGCGGATCC

ATAGGCTGTACCGCGAACCGGGTAACTATGG

>MSTRG.3858.1 gene=MSTRG.3858

TACTCCCTCTGTCCCTCTCATTTCTTTACAGTTACTATTTTGGGATGTCCCTCTCATTTCTTTACGTTAC

TATAAATAGTAAGTTTTTCTCATCATTACATCCACCATCTTCCTCCACTATCTCATATTTAACAATAAAA

ACTACTATTACACCCACTACTTTCCTCCACTATCTCAAATCTATTATTAAATATTGGTAGGTCCCACCAC

TTTACCCACTTTTCATCTAACTTTACTCATTTTTCATACATTGTCTTGGTCTCCGTGTCCCTCTCCAATG

TAAACAATTGAGGGGGACGGAGGGAGTA

>MSTRG.3859.1 gene=MSTRG.3859

CAGTGGCGGAACCAGGATGAAAATTCAGGGGGGGCTCAAATATTTAACTATAGAATATCCACAATATGAA

AAAACTTACAATAATAAAAGAAAATGCAACTAACCTTTAATTTCCAATAGTTGTACTATTCTTTTCTGAA

GAAGTGATGGATGATAGTTGGATCCTGCGATTTTGCATATTTTGATAGTGTTGTAGAATAGCTTCGTTAT

CCACACT

>MSTRG.3867.1 gene=MSTRG.3867

TAGGGTAGAGCGGATGCGGGTTCGGTGCGGTTTTTGGATAAAACCGCAACCGCAACCGCATGCCCTCGGG

TTTAGAAATCAAAAACCTCAACCGCAACCTCATCCTCGGTTTCGGTTTCGGTTCCAAAACCTCGGATTCG

GTTTCTATCGGTTCGGTTTCGGTTATAAAACCGCATGAAAGAAAAAAAAAAGAATTCTACATGAATAATT

AAACTTAAGTAAAGTTATGTATCCATCCAAAGTTTAAACAAATAGAAAACATAAAAATCCATAAGGTTCT

ACATAAATACTAATACAAATTTAGTTTAGTCATGCTATTGCCTGCTAAGTAGCCTATTACATTCTGTGAA

CCAGCACTTCAAGCAAGTTACTGATTAGTCTTCATCAATGTTTAGATGTTGAATGGAAGCTCCAACACCA

ATTAGC

>MSTRG.3870.1 gene=MSTRG.3870

ATTATCTCATTCTCTCTCTCCCACTTTCTCACTCTCATTCTCTCACACTCTCGACCTACATTTTTCCGAT

GATAAAAAGCAATTTTTCAACAAGCTTAAAACAGTAATTCGGTGAGCTTGATTTTAAAAATAGCCAAAGG

TACGGCAACATAATCACCACAAGGTTCAAGCCAAGCACCTCAATGCTACCACCATGACCGAACAATATTA

TGGAACTCGGTAAATTTTAAGATTATTTTTTTTGTTCTTTTAAATATATTTTTGTGTTTTTTTTATAACT

ATGAGTTAAATGAGAATTTGTCAATACTAACGTACATGAGAATCAGTTTTTAGTTTTAGCTAAAATTACA

ATGGACATTTTAGTATTTCCTACTTCTACAATTGCATCCGAGTCCTTTTTTAGGGTCGTAATAAAAGATC

AAGTCTTGCCCCAGACATCGTAAATATATTAGTTTGAAATAAGATTGGGATCAAGTTACAAAAAGACAAC

AAAAAAGAAAAGAAGATGACTCGGAAGAAAAGGAAGATGTTTGGATGATAATGGATATTTCATCGGAATT

GGGCTGCCTGAACAAGAATATTTTTGTTTTGTATTTGTATAACTCGAATAGTGGATAGATAGAATATTTT

TGTTTTGTATTTTTAAATTTTGTTTGATTGATATTTTGTTTTAAAGTATATGCGTTTTCTTCCATCTTTC

AGAGAGGAGTTCTCTCAAATTATTTGTAATTTTTCCAAAATTAATAGAGG

>MSTRG.3874.1 gene=MSTRG.3874

GAAGTACATGGGTGGAGAGTCGGGGAAAAGAACGAGCAGAGAAGCTGAGAAAAGAGATTGGGGAAAAAGG

GGGTAAGCTAGGGTTTGTCAATTGATAGTGAGGAAAGGAGCAATCGGAGGGTCTGGTGTGGTCGGAATGA

GGCCGCGGTTGTCTTGCCATCGTTGAGGTCATTTGCTGCGGCGGTAGTGAGGGATAACAGGTTGCAACCG

GAAGCGGCAGTGTTGAGTTCCACGATGGGTGAGGGGTTCACGGCGAGGAAGTCGCGGCGATGGCTGGGTC

TACGTATATATATGTGTGTATGTACATGTCTGTGGATGGAGAGGATGTTAGCAGTGGTGGTTCGAGACCA

TGGAAAGCCGAATACTAGATACAAAGATGGTTCGCCTGTTTGGGTTGCGCTTAAGAATAACGCTAGGCAA

GGATTCCAAGTGAGTATGTTATACCTCGATGGTCAGGATAATTATTTTGGAAGATGTCAAG

>MSTRG.3874.2 gene=MSTRG.3874

GCGGTTGTCTTGCCATCGTTGAGGTCATTTGCTGCGGCGGTAGTGAGGGATAACAGGTTGCAACCGGAAG

CGGCAGTGTTGAGTTCCACGATGGGTGAGGGGTTCACGGCGAGGAAGTCGCGGCGATGGCTGGGTCTACG

TATATATATGTGTGTATGTACATGTCTGTGGATGGAGAGGATGTTAGCAGTGGTGGTTCGAGACCATGGA

AAGCCGAATACTATTCATGGAGTCCAGGGAGG

>MSTRG.3875.1 gene=MSTRG.3875

GTTGATATATACCACATGTTTATAGCATGGTTTCATTTTGTCCATAACTCTTTGTGGTTTTCAATCTTAA

AGGCGTGATTGGCCAGAGAGTGCTAACACCTGGAACCGGTTGAGCATCTTGAGGATGTTGTAGATGCTTT

TGATTAGAGGTTTTCATTGTTCCCATAATTAGCATACTCTGAATGCGAGAAAGAAGAGAGCTACACATTT

AGGATCATTGGATGAGATACCAACGATTCCAACTTGCTCCATGTCCCACTTTCTCATCTGCAGACAAATG

TTTTCAACTTTAGGCTAGTTATTATAAGGTTCTACTCCTGGCTATCTTGGCTCTCATGGGAAAATGAAAA

GAACTTTTGGGACGTGCAAGAAGCAATTGTTTTTTTTTGTTGATGAGGGGCATTAGGCCCAAATATAAGA

GGCACTTGTGGTTATGCAGGAGCAAGTTTTCATATACAAGTTGCTAGTTTATCTTTCACATTATAATATT

TACACTATAAACTGTTATTGCGTCAAATTTTGTGACTTAATTTTTTTCAAATATATTGTGAGAATATTTT

C

>MSTRG.3876.1 gene=MSTRG.3876

CTCACCTCTGTGAATCTCTCTCCTCTCAATCGCTCTCTCTCAAAATCTCTCCCTCACCATCTCTCTTCAC

TCTCCTTCATCTCTCTTTCCCGCCGCTCTCTCTCTCGGCTTTCCCTCTCCCTCGGTCGTCTCTCCCCGTC

GACCCTCTCTCCATGCCGCCTCTCTCCCCGAACAACACCGCCCGTCGCCGCCTCCTTTTCCGGCCGTTCA

GCCGCCGCCGCCGGCCGATCGAGCATTGAGGGCGTGATACCAAACAGAGGCAAGAAAAGGTGAGAGTTGT

ATAGCGGTTCAGTAGTTACGGATTGTTCGAGACGAGTAGAACCAGACATTGGGGATCAAGACTGAACTTA

GATAAGATTCGTATCAGGCTGCAGTAACCGCATGAGCGACATAGCAAAGAATCAGGAATAGTTGATGGTG

ATGTCTTGAGAAAGCAGAATAAGATAAGGATGTT

>MSTRG.3877.1 gene=MSTRG.3877

GTTATACTCTCCTGTCTTGAACTCTGTCACTAATTTTCCCAGAATTTGCTTATCAGAATTTCCTTTCCTG

TTGGATAATTCTTTGAGTTTATTCTATCAATCTCCATCTCATTCTAGAGATCAGGTTGGACTAGTATTTA

TATTTAGAAACCACAACATTGGTACCCAGAGCCAGTAGAACCTTGGACGATGGGACATGGCCGCAACACC

AGATCTTAGACTTCGGATGATGAAGGAGGTGACAACCTACTTCCAGATCTTGCAAAAATGGAGGTAAATT

TCCGCACCTTACTTGAGGACCAGAATTAACGCTTACTTAACACTTGCAACACGAATAATGTCAACATAGA

ACAAAATGCTAAGTCTATTGCTCATCTGAATGATCTTATTTTGGGTCTGTCTATGCAAGTCTCTAAATTA

GTTTCTTCTGAGGTTAAAGACTCGGATAATGTTGGCAACAATCATGAAAGAGTGAGGGGGAGTTCCAGTA

CGGACAATAATAATAAGTTTACTTCATATGCAGCTAGAATGACAAAAGTTGATTTCCCAAAATTCGACGG

CACAGATTTGAGATCTTGGTTGTTTAAATGTAATCAGTTTTTTCAACTTGATGAAACAACTGACTGAAGG

TAAGGCTAGCTGCTATTCACTTGGAGGGAAAGGCTTAATGTGGCACCAAACTTATATGCAGAGATGCAAT

AATGTTTTACCCACTTGGGCACAGTACACTGCTGATATTACTGTGAGGTTTGGTAACCTATATGATGATC

CCATGGCAGATTTAAAGGCTCTCAAACAGCAACACTCAGTATAGGAATATCATGATGAATTCGATGCTCT

TACTAGTCGGCTAAATTTACCTAAAGAATATCTTTTAAGCTGCTATTTAGGCGGACTGGAGGAGGAAATC

CAGTTATATGTCCGGATGTTTACACCTAAAACTATACAACAGGCTTTGTGCTTAGCAAAACTACAGGAAG

CCTCAAGTAAAGCCAAGCTCACCAAACAAACACTAAAAACACCTCTCTCACCCAACCCCAATAAATCAGC

CTTCCTCACAAACGTCTCTACTAAACCATCATCTCCATCATTCACCAAACCACCATACACCAAACAAACT

CCTGCCCACCTCACAACACCCTCCACTAATAAACGTACCTTAACACCTGCCGAGCTTGCTGACAAACGAG

CTAAAAACCTCTGTTTTTGGTGTGATGAGAAGTATGTTCCTGGACACAAATGTAAGGGGAAGAAGCCTCA

ATTCTTCCATATTGAAGTGGAAGATGAAGACGACAAAATTGCGCAAGATGAGCTAAACACGGAGTTGGCA

GCACCTGAAGTTGAGAATGTTTGTGCCCAAATCTCCCTTGAAGCAATTGAAGGTGTCTCTACCTTTCAAA

CTATGAGAGTAACGGGCCATGCGGGCAAGAAAGAACTACACATTTTGTTGGACACTGATAGTACACACAA

TTTTATTGATGTGAACAAGGCCTTAAAACTACAGTGTAAGGTAGAAAGCATCACACCAAGGGCGGTAAGA

GTAGCGGATAGGGGCCAACTTATATGTGATATAATGATTCGTGGCTTTATTTGGAAGATACAAGGGGTCT

CTTTCAGGACTGATGTGTTATTATTGCCGTTAACAGGCAGCGATATGGTGTTGGGTATCCACTGGTTTTC

AAGCTTGGGACCTGTTCTTTGGGATTTCTCTAAATTAACCATGCAATTTAAATATGATGGTCAGAAGGTT

AGTTTGCGAGAGGCATGGAAAATAAAAGATTGAAACACATCACATCAACTAAATTGGACAAGTTAGTGCA

CTCTGCTGGGGAGCTTTCAATGTTGCAAATTATGCCTATAGCCCCAGGCTACAGTCCCCAAGTGACAACG

ACTTTGCTCAACTGTGAAGTGAACCATGATAATGACTTAGCAGCTGTGTTAAAACAGTATTCCACAGTTT

TCAAGGACATCCAA

>MSTRG.3882.1 gene=MSTRG.3882

AAAGTAGAGACCTGACACACCCATAGAATCCAAGTTTCACTGATCCCGAAGAAACCGCTGTAAGCAGCGA

AACCCTAATTTCTCCGCACAAAATCTCTCCATCTCGGCCCCACCTCCTCGTTCTTCACACACCTCTCTCT

CCATCCACTCCACACCCTCGTCCCACTTGAGCCACTCGTCCCAGTTCTCGACTCGACAACCCGACACGGC

ATCGAGAATCTCCTGTAGCCCTACGATCCGACGGCTCAGATTTCTCCTCAGCTCCCGGATGTTCGGGTCC

CACCCCGGCACGGAGTCGAGAGTCAGCAGCAACCCCATGAGAGCCTCGTTGATTTCGAGGCGTCTCCGGT

CGTCGGAGCGGACGGCGTCGACGGTTTCTTGGCGCTGGATTAGGGTTTGGAAGGCGGCGGCGGAGGAGTG

GACGGCGGAGAGTTTTTTGAAGAGGGTTTGGGTTAGGTGGGCCCGGTAGGTGGATTGGATTTTGGCGGCG

GCGGAATCGACGGTGGGGATTGGGATGGGGTTAGGGTTTGAGGGAGGGGTAGAATGGTCATTTTTGTAGG

TGATGGGGATGGAAGTGGAGGTGTGAGAATGGTGTGTTGGTTTCATAGATGAAGAAAGTGGAAAGAGGGT

GTTGCGGATGGGGATAGAGAGGGTGAAGGTGGACTGGTGTGGATGATTGGATTGGGATTGTTTGTTGAGA

GGGGCATAATTGTAATATAGATCTTTTCAGGATTTTCTAGAGCTATTTAGAAGAAGAAGAATTTCTAATG

TATGACTAAATCAAAATTAAAAATAAAGAAAAATAGATGAGTGCGCCGGGTT

>MSTRG.3882.2 gene=MSTRG.3882

CTGACACACCCATAGAATCCAAGTTTCACTGATCCCGAAGAAACCGCTGTAAGCAGCGAAACCCTAATTT

CTCCGCACAAAATCTCTCCATCTCGGCCCCACCTCCTCGTTCTTCACACACCTCTCTCTCCATCCACTCC

ACACCCTCGTCCCACTTGAGCCACTCGTCCCAGTTCTCGACTCGACAACCCGACACGGCATCGAGAATCT

CCTGTAGCCCTACCCCGGCACGGAGTCGAGAGTCAGCAGCAACCCCATGAGAGCCTCGTTGATTTCGAGG

CGTCTCCGGTCGTCGGAGCGGACGGCGTCGACGGTTTCTTGGCGCTGGATTAGGGTTTGGAAGGCGGCGG

CGGAGGAGTGGACGGCGGAGAGTTTTTTGAAGAGGGTTTGGGTTAGGTGGGCCCGGTAGGTGGATTGGAT

TTTGGCGGCGG

>MSTRG.3890.1 gene=MSTRG.3890

AAAATAAATCACAAAACCAAACCCCCCATAAGAAAATCCTCTCCCTTCTCTGTCTTTCCTCTTTTCTCTC

AAACTGCATATATTCAATATCAATGAAGATGGTTCATGGCTTTCAAGACTAATGTTAGATCATTACGGAA

TTTCCAAATCAATGTTTCAAATGCTTGTGCCTTGGCATTGCGCAAATTTTGTGAAGAAGCAACCATTGTA

ATGTTTGAGCCTTCAGACCTGGAAATATTGATCTGGATAGGAGAGGAAATAGAGAAGAGTCATATATATT

AAGAGGACGATGAAGAATTATTTAATGCAATTGCTGTGGTACTTGCCTCCGTTTCTAAAAGATTTGTTAG

AGGAATTCAACCTGGACAACAGGCAAGATTCAATATTGATGATTCTGACCTCCATGAGATGGAAAAATGT

AAAGGAGTGGTGGTTGCATCACCTATATTTGGATTTTGCACTATGGATTGTGAGTGATATGTGGAGGATA

TACAAAAGTCGGGTGAAAAAAGATCATTATCTTCGATATGATAATGTTGTGGAAGGGCTGAAAAACAATC

CTTTTAGATTGAATATTGTAAGGATGGTTTATAAGTTAGTACCTTAGTCTCGTTTCTTTGTAGCTTTCAA

CTTGATGACTCAGTTTGTACTTTGAATTTTATTTTTGTCAATTTTCATGTAAG

>MSTRG.3890.2 gene=MSTRG.3890

CATAAGAAAATCCTCTCCCTTCTCTGTCTTTCCTCTTTTCTCTCAAACTGCATATATTCAATATCAATGA

AGGTTTTTCGCAGTGATTATGTAATTGGGTTTGGAGGGAGTGAGATGTTAGAACTTGGGTCTGCATTGGG

AAAGGGATTTGTGGAGTTTAGTGAGTGGAATTTGAGACTTGGGTGTTTGTTTGTGGTGGATTTTCAATAG

GAATTCGGAAGAACTCAAGGGCTTTATGTGCATTATGGTTCATGGCTTTCAAGACTAATGTTAGATCATT

ACGGAATTTCCAAATCAATGTTTCAAATGCTTGTGCCTTGGCATTGCGCAAATTTTGTGAAGAAGCAACC

ATTGTAATGTTTGAGCCTTCAGACCTGGAAATATTGATCTGGATAGGAGAGGAAATAGAGAAGAGTCATA

TATATTAAGAGGACGATGAAGAATTATTTAATGCAATTGCTGTGGTACTTGCCTCCGTTTCTAAAAGCTT

GTTATCTTCCTCTTGATTATGCTATTTTCCGGATTCGGACGGCCACATCTTTTTCTCCTGCAATTCAAAA

CATATCCTATATACATGAAGACTCTCTAAGAAGAAATGAGTTTGGATGTTCATAATTTGGTGGATATCCG

TCACTAGAGCATAGAAATGATTGTTATAATGTAAGAGAATCAATAAGGGCACATTGCGGATTTGTTAGAG

GAATTCAACCTGGACAACAGGCAAGATTCAATATTGATGATTCTGACCTCCATGAGATGGAAAAATGTAA

AGGAGTGGTGGTTGCATCACCTATATTTGGATTTTGCACTATGGATTGTGAGTGATATGTGGAGGATATA

CAAAAGTCGGGTGAAAAAAGATCATTATCTTCGATATGATAATGTTGTGGAAGGGCTGAAAAACAATCCT

TTTAGATTGAATATTGTAAGGATGGTTTATAAGTTAGTACCTTAGTCTCGTTT

>MSTRG.3895.1 gene=MSTRG.3895

GGGTTTGGTAGCTAATTCACCCAATTTGAGGTCCATTACCGAGTCGGTTAAGTTTTCGGAAACATAAGTT

GAAGGGGGTGAATTAGGCCTGGAAATGAAGGATTGTTCATCAGATTGGTAAAGTGTAGATCTAATTGTTC

TCATGGAACGACCCAATGGGCGTTGTAGCTTGGATGATGATGGTGGTGGTGTGTAGTAATAGGGTCGGGT

TGAGGGAAAGCGGGTCATACTTGAGTGTGAATCTCCTACAGAAACCATCTTGAATACAAAAATCTCTCTC

CCTGTTCTCTCACTCCCTGTGTTCTCTCTCCCTGTTCTCTCTCTCCCTGTTATCTCACTCCCTGTTCTCT

CTCTCCCGGTCTCTCAGTCCCTGTGTTCTCTCTCTG

>MSTRG.3896.5 gene=MSTRG.3896

TAATTAGCGTATCTTTATACAATCCTGGTCTATATCCAAGCCTCTGTTCTTACTCTCTCAGCAAAATAAA

ATTACTTTGGAACACTTGCAGGCTATTTTAGTTTTTCAGTCGACGAAACTTATCCTCATAGTTTAGTTCT

ATCAGATAAATAACACACCATTCAACACTTCTGCAGGGCTGTGCTGGGAAGTGTGAGTGATTACTGTGCT

CATCGTGCTCATTGGATGGTGACAATCGTCAGGAAGCCTAAAACCAAGTACTAAGGTTACTTGATCCGAG

CTGATGAAGAAACATCCTGGCATGACAGAAAACTGAAAGAAAGGTGCTTGCACATTATGTGTCT

>MSTRG.3904.1 gene=MSTRG.3904

TGATGATCTTTAGTTTGCCAAACCATGTAGACTAGACACTGATATAGAGAACTTAGAAGAAACATAATAT

TCAAAATTAAGACAACATATGACCACACCATATCATGCACGTATAGCGTACATTGTTCAACACTCTATTC

CTCAGTCTTGGTCTCAGCCGGAACTTCAGTGGTGGTTTCAGCCACAGCCTCAGCTGGCGTCTCCTCCACG

GGAGCTGCAGCTGCTTCCTTGGCTTCCACCGGGGCAACTGGCGCGGCTTCCACTGGAGTTTCCTCAACTA

CTGCCTCGACAACAGTCTCTACCTCAACTTCTGGAGTATCTTCCTCTGGTGCCTCTTCTACAGCTTCGGG

TGCCTCGATAAGTTCCTCAGCTTCAACTACAACCGGTTCTTCAACTTCAACTGGCTCTGCTACTGGCTCC

TCAGCTGGTATGACAGCTTCAGTAGTTTCCTCAGCCAGCTTCTCCACCTCTGGCTCGGCTGCAGCTGCTG

GTGCTACTTCTACCTCTTCCTTGATTGCTTCCGGGGCCGCGATCACCTCAGGCACAGGTGTTGTTGCTTC

TGGCAATGTTTGTGGAACTGATTCAACCTCAATAGCGGCCATTGCTAATGAATGGGAAATTGGCTGAATG

AGCTTTGATCTAAAAGTGAGGAAAATAATAGAGCGAG

>MSTRG.3906.1 gene=MSTRG.3906

CTCTCAGCAAACCTGTTAACAAGTGCAAGTGCATTACTAGTAACTTGAGCAACTGTAACAATCCTTCTCT

TGACAACAGCCTTGACATTTCCTGCCATGAAATTCCCCGAAAAGCCATCGCTACAAGTGTTCTCATTAGT

CAGAGCAGCACTCACCCATGTCTGCACATTGCTCATGTGCCACATGAAGTCCTGACCCCCATGAACCCGG

TTCATGTGGTTCAGCTCCTCCAGAGACCTCGAAAGCTGATCAACTGTGTCGCCCAAGTTATCGATACAAT

CCTGCACGGCCAGAAATTCCCTGGTTCTTATTCCTGACACTTTTGTCAGTTTTGACACGAACATTGTGGT

GAATCTGGCCTTTGCCAGGCTTACTGATAGTGCAGCTTTCGCGAGCTGGCGTTCGTTTTGTTGGATTTTG

GCCGCGTAAATTGAGAGGGACTGGTAGCATAAGGCAGGGTACAGAGTTGCTCTGCATGAAATTCTGATGA

AATTTGTGGCACCGTTGTATGAGGAGGTTGTCACAAGAGAAGACTCTGCAGTGTTAGAACTCATGAAATG

GAATGACAAGACAACTAGAAACAAGGAGTGAAGACTGAGATTTGCCATTGAAAGTGAGGTGAAGAACTTG

GAAGAACAAAACAAGAGCTTCAAGATTTATATAGAGGAGATACTCGGGTCCTGTCAAACGATTATTATAG

TCCAACACAATAAAGAAACGGCATAATAATAAATGGATCCCTCCCATATTTACAATGTTAATAATAATCG

TTTGACAGAAACTCAGACTTAATATTATATGTCTAG

>MSTRG.3909.1 gene=MSTRG.3909

CAAATAATTCGCTAAACTCCCGACTCCGAACCTAAATATAAAATTATAATCTATTATTAACGATTTCTAA

TAATCGACTGAGATATTTACAGACTCGATAATATACCAACATAATCTCAAATCCAGTGATTAATAGCGTA

CACATTAAATCAATACTCTAGTCAGAACAGGCTGATATTTGTTGCGGAAGCATATCGTAACTCATTTCCT

TTCATAATTACTAAGGAGTAAAAACTATACTCTTGTGAGCCAATATCCCCTGTACCCAATTCACCGTGCA

TGATTGGTAGCCAATTAATAAACTAACTAATACAGCATGAGTGCATGACGACATAACAAGGCAAAACATT

ATTCAATATCAAATGGCATGATACCTTTCTGCCTATAATTTCCGAGACAAAGCTCTCCCCTGGTACGTAT

TCTCCGTTCTCTCCTTCTCCTTCACTCTGTTTTTCACCCTTTTTCCTTTACTCTGCGGCTACTTCTAGGG

TTATTAATAACTGATATGTTTTTATATATAGGCCTCCAAGTATCCTATAACTAATATAACTCTAATTATT

AATATTAATCTATTT

>MSTRG.3912.1 gene=MSTRG.3912

TTAAACAAAAAAAAGTGTGAATATTCAGATATCTTGCAATGAGCAGCCAGTCTCGGCCATTCGCAGAGTA

TGGTCCAGATATTTGGGATGAGAAATTCACTTCTAATCACAGAGACTTTAAGTTGTGGAAAGCATGTACA

GAGGAACGCGATGAGCTTGAAAATGAAGTAAGGAGCATGCTGTTGTCTGCTGGTTTAAACTGGACTGAGA

AGTTGATTCTGATCAATACTATTGAACGCTTAGGAGTTGGCTACCACTTTGCTGAATATATTGAAGTCAT

GTTGGCTGAAATGCATAATGCTCATGCCTGTACTATTGAAAACTTCAATCTCTCCACTACAGCACTTTAC

TTCCGAATCTTAAGACAGCATGGTTATAACGTCTCGTCAGATATATTCAGCAAATACAAAGATATTGATG

GAAATTTCTCTGAAGCTATCACAACAGACCCGGAGGGGTTACTAAACTTGTACGAAGCAGCACATTTGAG

AACACATGGGGAAGCTATGTTAGACGAGGCACTTGTTTTTACAACATCTCACCTAACATCTATTGCAGAG

TCTTTGGATTCTGGCCCATTAGCTAAACAGGTGCGGCGTTCCTTGGAGCAGCCCTTACATAAGGGAATCC

AGAGAATCGAGGCTAAGCATTCTCTCCTTTTGTATGAGCAGAGTCGATCGAGGAATGAAGTGCTACTCAA

GTTTGCGAAGCTGGATTTCAATCTAGTTCAAATGTTGTATAAGCAGGAATTATCTCACTTAATTAGGTGG

TGGGCTGATATAGACTTTAAATCCAAATTTCCTCAGTTTCGAAGCAGGGTAGTTGAAGGTTACTTATGGG

CTGTAGCAAATGCATTTGAACCTTGCGATGCAACCGCTCGTATCATGTACACGAAGATGCTTTGTGCCCT

TTCAGTTACTGATGACCTCTATGATGCATATGGTACAATGGATGAACTAAATATGTATACTAAGGCCATT

GAGCGGTTAGACGCTGGCAGCATCGATGGACTTCCAGACCACTCTAAATTGTGCTACATCACTTCTTTAA

ATGTCTTCAGAGAATTTGATGAAGCAATGGTTAAGAAAGGAGTTTACTGTGATCTTTCTTACCTCAAAGA

AGCATTCAAAGCAAACTTGTTGGCTTTTCATAAAGAGTCTACGTGGCGGGACAAGGGCTATGTTCCACCA

CTTGAAGAATATCTCATGAACTCAACAACATCCTCTTGCATATGCATGCTTGGATTGTGTATTATTTTGG

GGAGGGGACATGGTGACACCATTGGGGCATGCAGGTGGGCAACAAAAAAACCAAAAGCTCTTGTTGCGGC

TGAAAAAATGGGGCGAATAATAAATGACATAGTGGGCTATGAGGAGGAGCACCGCCGTCCACATGTTGCT

ACATCCATCGATTGTTACATGAAGGAGTACGGGGTTTCAAAGGAAGAAGCGATTGTCAAGTTATATGAAA

TGATCGAAGACACATGGAAGGACATAAACGAGGAATGCTTGAGGCCGACACCCGTTGCAAGAAGTTGTAT

CACCACATTTCTCGAGCTGATGAGAGTAAATCATGTTACCTATAAGGATAATGATCGATACACCCATCCT

GAAGGACTGAAGCATGAAATTACCCTTATCTTGGTGGATCCGATACCGATATGAACTACAATATCCTGCA

GTTCTTGAACGGTTCTATATATAATTTCCTTTTTTGAACCTGGGAAATGTGGCGTTTATCTATGTACAAA

TATTGTTTTCCATCTATTTATGTCTTTATACACTACAAATAATCAAGCAACAAACTTGTCTAATTTGTGT

GTAACCTGAATAATATTAATTTTATGCAATATTAAT

>MSTRG.3912.2 gene=MSTRG.3912

TGTGAATATTCAGATATCTTGCAATGAGCAGCCAGTCTCGGCCATTCGCAGAGTATGGTCCAGATATTTG

GGATGAGAAATTCACTTCTAATCACAGAGACTTTAAGTAATACATAAACAGTTGTGGAAAGCATGTACAG

AGGAACGCGATGAGCTTGAAAATGAAGAGTTGGCTACCACTTTGCTGAATATATTGAAGTCATGTTGGCT

GAAATGCATAATGCTCATGCCTGTACTATTGAAAACTTCAATCTCTCCACTACAGCACTTTACTTCCGAA

TCTTAAGACAGCATGGTTATAACGTCTCGTCAGATATATTCAGCAAATACAAAGATATTGATGGAAATTT

CTCTGAAGCTATCACAACAGACCCGGAGGGGTTACTAAACTTGTACGAAGCAGCACATTTGAGAACACAT

GGGGAAGCTATGTTAGACGAGGCACTTGTTTTTACAACATCTCACCTAACATCTATTGCAGAGTCTTTGG

ATTCTGGCCCATTAGCTAAACAGGTGCGGCGTTCCTTGGAGCAGCCCTTACATAAGGGAATCCAGAGAAT

CGAGGCTAAGCATTCTCTCCTTTTGTATGAGCAGAGTCGATCGAGGAATGAAGTGCTACTCAAGTTTGCG

AAGCTGGATTTCAATCTAGTTCAAATGTTGTATAAGCAGGAATTATCTCACTTAATTAGGTGGTGGGCTG

ATATAGACTTTAAATCCAAATTTCCTCAGTTTCGAAGCAGGGTAGTTGAAGGTTACTTATGGGCTGTAGC

AAATGCATTTGAACCTTGCGATGCAACCGCTCGTATCATGTACACGAAGATGCTTTGTGCCCTTTCAGTT

ACTGATGACCTCTATGATGCATATGGTACAATGGATGAACTAAATATGTATACTAAGGCCATTGAGCGGT

AAAGCATAAATTACCCCTTTTTGTGCAGAGCCAAATTATATAATATATACCGCCCTTCTCTAACACCAAG

TACAGCCTGTTTTATTGAACATATGGGGTTTAATTATACAGGTTAGACGCTGGCAGCATCGATGGACTTC

CAGACCACTCTAAATTGTGCTACATCACTTCTTTAAATGTCTTCAGAGAATTTGATGAAGCAATGGTTAA

GAAAGGAGTTTACTGTGATCTTTCTTACCTCAAAGAAGCATTCAAAGCAAACTTGTTGGCTTTTCATAAA

GAGTCTACGTGGCGGGACAAGGGCTATGTTCCACCACTTGAAGAATATCTCATGAACTCAACAACATCCT

CTTGCATATGCATGCTTGGATTGTGTATTATTTTGGGGAGGGGACATGGTGACACCATTGGGGCATGCAG

GTGGGCAACAAAAAAACCAAAAGCTCTTGTTGCGGCTGAAAAAATGGGGCGAATAATAAATGACATAGTG

GGCTATGAGGTTCGGGGTTAATCATGACCTAATAAATATACATAAAACCTTAAATATTTTGCTTTAGTAA

GTTGTAAGACGCTAATTAGTTATTGTCTTTGTTAAATTTGAAGGAGGAGCACCGCCGTCCACATGTTGCT

ACATCCATCGATTGTTACATGAAGGAGTACGGGGTTTCAAAGGAAGAAGCGATTGTCAAGTTATATGAAA

TGATCGAAGACACATGGAAGGACATAAACGAGGAATGCTTGAGGCCGACACCCGTTGCAAGAAGTTGTAT

CACCACATTTCTCGAGCTGATGAGAGTAAATCATGTTACCTATAAGGATAATGATCGATACACCCATCCT

GAAGGACTGAAGCATGAAATTACCCTTATCTTGGTGGATCCGATACCGATATGAACTACAATATCCTGCA

GTTCTTGAACGGTTCTATATATAATTTCCTTTTTTGAACCTGGGAAATGTGGCGTTTATCTATGTACAAA

TATTGTTTTCCATCTATTTATGTCTTTATACACTACAAATAATCAAGCAACAAACTTGTCTAATTTGTGT

GTAACCTGAATAATATTAATTTTATGCAATATTAAT

>MSTRG.3913.1 gene=MSTRG.3913

TATAATCATATTATTGTTGTTGTTAATTGATTCTTATGTAGGTGGTCTTGAAATATGAATAATGACCGAT

CTTGGATGTATGAGAGGATTGATGATGGAGGATTTTGAAATTCTTTATTTATTTTCGGTGTGGAGGAATT

TATGAATTATGCTATATCTCAGCCAACGTCTATAGGAGGGACAAGTATACAATGTCCTTATTCAAAGTGT

AAGAATCGAAAATTATGGAATGGAGATATGGTGAAGCTACATATTTTGAAGAATGGATTCGTGAAAGATT

ATTATGCATGGAGTCGACATGGGGAGTCATACATTTATGATGGAAATGACGAACATCCCTTAGCAAATTA

TACAAATATTTCACAGGGCACAGATGGGAACAACTTAATGTACAATATGGTTATTGATGCAGCTGGCCCC

AGCTTTGATCTGCATAGTACAGAAGAGATCCCAAATGCAGCAGCACAAAATATATACACCATGTTAAAGT

CTTCAGAACGAGACTTATATGATGGCCGTAAAACTTCACAGTTGGCTGCCATGGCTCAAATGTTGAGTTT

GAAATCTGATCATCACTGGTCGGAGGCATGTTATGATCAAACATCACGATTTGTCAAAGGTATCATGCCT

GAAGACAACATATTCCTTGATAGTTTCTGTAGCACAAAGAAACATATGGAAGGACTGTGTGTACCTTCCA

TTCATATTGACTGTTGTGTTAATGGGTGCATGATATATTGGAACGTAGACATTGACATGGACTCATGCAA

ATTTTGTTCTCAGCCAAGATACAAGATTAGAGTTAATAGATCTAAGAGAGAGAGA

>MSTRG.3915.1 gene=MSTRG.3915

ATTAGATGTCAGTTTGTTTCTGTATATAGCCTTGCCTTTGAATAAGATACATTGACCTCATATCTTGTTT

CCTAGCTATAGTGAATGCCTTGAGCTTTTCTCTGTGTCACTCTAGTTCTCCCCGGACAGAACTAGGCAAT

GCTACAATGAATCCAGTCGAAGTCTCTAAGACATAAAGGAACTAGGTACTGCAGATAATGAAGGAATTAT

ATTCTTACTTCCATACATCACTTTGACAACTTAAGGTGGACATGGAGGTGCTCTCGGCTGAAGCTACATG

ATCTGGCATCCTCAAATCATTGCATAGCATGTTTATCTGTTCCAACACTTGATCATTTGAACATCTTGTT

TTATTACCATCAGTGTCTTGATTATCTTGACCGAGTTGAGAAGCAACAAAGTTGTCAAGTACTTGCTAAT

CAGTCAATTGACTGCAATGCTGCAATTTATAATCTTCTTGTGTGAGCATTAAGGACTGCTCAAATCCATA

TGGAACTCCTGAACTGAAGCTCACACTTGCTGCAGAATGCAGGACTTCCTGTCCTTCTAACTGAGGAGGT

TGCAGATATCGTTCATGGGAACGAAAATGCTGCAACTGACCAAGCTCTTGTTTACAAGTATAGTTCTGAT

CATTATGGAATGGTGTATAAGGATTATCAGTTATCTGCCTTCGTGATTCAAAGTCAGGCAGCAACCGGAT

TTGGTCATCATACCAGTTAGACAGTGGCTCATGCTCGTCCATTTTCCTCACCGTTGTTAATCTTTTCTTA

AAAACCCTACACACAACCCATCCCTCTTCCTGAAAGACAATATTGTTAGCAAAATTTTATA

>MSTRG.3917.1 gene=MSTRG.3917

GCATTCGCCATCAACGCCACCGTCAGCGACTTCCCAGTCGGCAAATTTGCCTTAATCCTGCTACACTCGA

CTCTAATCCCAACCTTCTTGCTCCTCACTTTCACTTTAGTATCCAGCTCGATATGCAACGGCAAGGTCGG

AGCTCGTGGACCGCTCGGTATTCTCGAAGCCGGACTCGATCTCCGACACCAGTCGTGTTCACTCGGATCC

GCAAGAACACCTCTGGTCGCGCAGAAATTCGGGTTCGGAGAAAACCCGAACCCAATCCGAAATCCGTTGG

ATCGGGTTCGGGTTTTGTTTTTCAAATCCAAACGGGTTTGGATCGGGTTCGGGTTTGGTGAAAATTTTCG

GGTTCGGGTTTGGATATCACAGATATCCGAACCGATCCGACCCGTTGCCATCCCTA

>MSTRG.3921.1 gene=MSTRG.3921

CAGAAATTGGTCGAAAGAGACCCAGACAATCTTGAAAGACTGGTAGCAATATTTGAAAGCCTGGAAACTT

TAGGGCCCTTAAGTCCCTACTTTCTCCAGATTCTTTCTCTTATTCCTCCTATTCCTGGGGTTGCTTGAGG

GTTGAGCAATCGACATGATATATTTGCTCACATTCCAGGCGAAGGTCATAATTTGCAGGAACATTCTATG

GTCTTAATAAGAGGAGGTAGAGTGAAAGATTCGCCAGGTGTGAAATCCCGTTCTGACTAGTTCCGTTTGG

TCGATTCCGGCATACATAAAGGTAGAGCTGGGCAGCGGCTTTACTTTTATGTTGTAAGAATTCAATGTAG

CCTTTAGCCTAGGCGAGGCCCCTTGTCGCTAGGGGGGGAAAGAAGAGTGGGATAGTGGGCTTCTTTCGCT

TTACTTTTT

>MSTRG.3925.1 gene=MSTRG.3925

ATTTAGACACAGACACCAAACTCCACACACTCGCACAAACTCACTCCACAGTCCACCCACCCCACTCACT

GACCTCGGCTGAAACACCTCTCGGCTGCTCAACTTACCTCTCCGTTCAAATACACAATCTCTACTGCTGA

AATCACCAGTGATTAGGTCTTCAATCGGGGCTGGGGCTTTGATTGAGAGCTTCTCCATTTCGCGGAATTA

GGGCTTTTGGTGGCTTCGATTGAATTGGGGCTCCAGTGGAAGTAGGGTTCAACAGCGTTTCACTTTAATT

GCAGCTTCGATTTGCTTTGATTTAAGATACAATCTGCTGTCATCCTTTGCTTTTCACTTTAAATGTTTGC

TTTGTTTTAGATTTTATGTGTAAATGCATGTCTATCTTAGTAGCTTTGTTGTGATTTGTGGTCATATGTT

ATATTACCCATCCCTCTCCCTCTCCCTCTCCCTCTCTTCCTCTCCCTCTGTGTCCCCTCTGTCTCTCCCC

CTTCTTTATATCTTGGTGAATTTGTTAAAGGTGTAGAGCTTGTTGAGTCAAATTTAATTTTTGTTTAGTG

AAGATATGTATCAAGTTATTAAACATGTTAACAGTGTTCAGTAATCAATTTTCTATGATATTTCTTAAAG

AATTGTTTCTTCAAGTTCTGTTTCTAACTTAAAATACAAACTACCATATAGAACCATATCCATTAATGTT

GTGAGATGCATTCTCTCAAATCCACAAATATATCACTATGTACTTCATTTTTTTCTGGTTTTTTGTGTTC

TCGTGAAGTTATTCAGGTACTGGAAAGTGTTGAAACAACTGCAGAGAAAGCGACCCGATAGTAGGAGCTT

GGTCAGGTGTGTTCAGGAGCTGAGGATGAAGGGGGCTAGCTTTCATTTGTTTAAAGAGCCTCTGAATGCA

AAGAGAATGAAGAGTTCCAGTGTCGAGATCAAGAGAAAGCCATTGACAGGGTGTTCTTAGAGTCTCACAA

CTATTTGTCTAATCTGTTTGACTGGCTTGCTCTTCCTGCTTCCAGAATTATTCTCTGTGTTTAAAGCTAC

TTCCAAATGCAGCCATACTCCAGGTCCTGAGGCTTCAATCTTTAATTTAATAGATTTTATAGATGATGCT

CAAATGGCTAAGGCCGATGACATTTCAAGGAGCTGTATGATGGAAGCAAGGTTCTTGAAATTTATGGCTA

GAGAAACTATGTAGCTTCGTTTGATTTGGTAGTTATTTTGATTTGGTTCGAATTATTAGGTACTTTAGTT

TGTATAAAGAAACAATTTATTGGGATGTTTAATATTTAAATAGTATATTTTCAA

>MSTRG.3925.2 gene=MSTRG.3925

CTTCCCCCCTTCATTTCATTTGAGACATTTAGACACAGACACCAAACTCCACACACTCGCACAAACTCAC

TCCACAGTCCACCCACCCCACTCACTGACCTCGGCTGAAACACCTCTCGGCTGCTCAACTTACCTCTCCG

TTCAAATACACAATCTCTACTGCTGAAATCACCAGTGATTAGGGCTTCGATTGAATTGGGGCTCCAGTGG

AAGTAGGGTTCAACAGCGTTTCACTTTAATTGCAGCTTCGATTTGCTTTGATTTAAGATACAATCTGCTG

TCATCCTTTGCTTTTCACTTTAAATGTTTGCTTTGTTTTAGATTTTATGTGTAAATGCATGTCTATCTTA

GTAGCTTTGTTGTGATTTGTGGTCATATGTTATATTACCCATCCCTCTCCCTCTCCCTCTCCCTCTCTTC

CTCTCCCTCTGTGTCCCCTCTGTCTCTCCCCCTTCTTTATATCTTGGTGAATTTGTTAAAGGTGTAGAGC

TTGTTGAGTCAAATTTAATTTTTGTTTAGTGAAGATATGTATCAAGTTATTAAACATGTTAACAGTGTTC

AGTAATCAATTTTCTATGATATTTCTTAAAGAATTGTTTCTTCAAGTTCTGTTTCTAACTTAAAATACAA

ACTACCATATAGAACCATATCCATTAATGTTGTGAGATGCATTCTCTCAAATCCACAAATATATCACTAT

GTACTTCATTTTTTTCTGGTTTTTTGTGTTCTCGTGAAGTTATTCAGGTACTGGAAAGTGTTGAAACAAC

TGCAGAGAAAGCGACCCGATAGTAGGAGCTTGGTCAGGTGTGTTCAGGAGCTGAGGATGAAGGGGGCTAG

CTTTCATTTGTTTAAAGAGCCTCTGAATGCAAAGAGAATGAAGAGTTCCAGTGTCGAGATCAAGAGAAAG

CCATTGACAGGGTGTTCTTAGAGTCTCACAACTATTTGTCTAATCTGTTTGACTGGCTTGCTCTTCCTGC

TTCCAGAATTATTCTCTGTGTTTAAAGCTACTTCCAAATGCAGCCATACTCCAGGTCCTGAGGCTTCAAT

CTTTAATTTAATAGATTTTATAGATGATGCTCAAATGGCTAAGGCCGATGACATTTCAAGGAGCTGTATG

ATGGAAGCAAGGTTCTTGAAATTTATGGCTAGAGAAACTATGTAGCTTCGTTTGATTTGGTAGTTATTTT

GATTTGGTTCGAATTATTAGGTACTTTAGTTTGTATAAAGAAACAATTTATTGGGA

>MSTRG.3925.3 gene=MSTRG.3925

CTTCCCCCCTTCATTTCATTTGAGACATTTAGACACAGACACCAAACTCCACACACTCGCACAAACTCAC

TCCACAGTCCACCCACCCCACTCACTGACCTCGGCTGAAACACCTCTCGGCTGCTCAACTTACCTCTCCG

TTCAAATACACAATCTCTACTGCTGAAATCACCAGTGATTAGGTCTTCAATCGGGGCTGGGGCTTTGATT

GAGAGCTTCTCCATTTCGCGGAATTAGGGCTTTTGGTGTATAGTTCCTTGTCTTCATTTTGCTTTTCATT

TTACTTTTCGCTATAATATTCTTAATTTTGTGTCTTTATTCTTCTTAGGGCTTCGATTGAATTGGGGCTC

CAGTGGAAGTAGGGTTCAACAGCGTTTCACTTTAATTGCAGCTTCGATTTGCTTTGATTTAAGATACAAT

CTGCTGTCATCCTTTGCTTTTCACTTTAAATGTTTGCTTTGTTTTAGATTTTATGTGTAAATGCATGTCT

ATCTTAGTAGCTTTGTTGTGATTTGTGGTCATATGTTATATTACCCATCCCTCTCCCTCTCCCTCTCCCT

CTCTTCCTCTCCCTCTGTGTCCCCTCTGTCTCTCCCCCTTCTTTATATCTTGGTGAATTTGTTAAAGGTG

TAGAGCTTGTTGAGTCAAATTTAATTTTTGTTTAGTGAAGATATGTATCAAGTTATTAAACATGTTAACA

GTGTTCAGTAATCAATTTTCTATGATATTTCTTAAAGAATTGTTTCTTCAAGTTCTGTTTCTAACTTAAA

ATACAAACTACCATATAGAACCATATCCATTAATGTTGTGAGATGCATTCTCTCAAATCCACAAATATAT

CACTATGTACTTCATTTTTTTCTGGTTTTTTGTGTTCTCGTGAAGTTATTCAGGTACTGGAAAGTGTTGA

AACAACTGCAGAGAAAGCGACCCGATAGTAGGAGCTTGGTCAGGTGTGTTCAGGAGCTGAGGATGAAGGG

GGCTAGCTTTCATTTGTTTAAAGAGCCTCTGAATGCAAAGAGAATGAAGAGTTCCAGTGTCGAGATCAAG

AGAAAGCCATTGACAGGGTGTTCTTAGAGTCTCACAACTATTTGTCTAATCTGTTTGACTGGCTTGCTCT

TCCTGCTTCCAGAATTATTCTCTGTGTTTAAAGCTACTTCCAAATGCAGCCATACTCCAGGTCCTGAGGC

TTCAATCTTTAATTTAATAGATTTTATAGATGATGCTCAAATGGCTAAGGCCGATGACATTTCAAGGAGC

TGTATGATGGAAGCAAGGTTCTTGAAATTTATGGCTAGAGAAACTATGTAGCTTCGTTTGATTTGGTAGT

TATTTTGATTTGGTTCGAATTATTAGGTACTTTAGTTTGTATAAAGAAACAATTTATTGGGA

>MSTRG.3929.1 gene=MSTRG.3929

CTAAAATGTAAAAATAGATAAAGTTCAGTCATCACTTTGCATTTTACTTTTTTTCCATATCTCAGTAGAG

TGAAATAAATGAAACTGAACTTTACTACACCAAAACATAAAAGAAGGCTAGGTACCTTTCTTTAACAAGA

ATTATTATTACAATTTTAAGCACAAAGTGAAATATAAGCGAATCAGATAGAAAAGAGGACTATAGCAACA

CAAAAGTAAAACTCAAGGAAATTTTTCGAGAAAAAAAAAAACTTTGGTTTAATTACACGCATTTGAAGCC

CTTGGGGACCTTCTTGCCACAGTAGTTAATCAACAAGCTCAAATCAACCGGAACATTGAGGTTAATACCC

AAGATATTGGCTTTAATCGCGGTGCAGAGACAAATAGCAGCTTCAAGATCGACAAGGCCTTCGAGGAGAC

TACAGCATGGCAGCTTAGGTGGAGTTCCAATGACCGTCCCGATCAAGCCACCTAGAACCTTGGCACACAC

TCCCAACTTGAGTGCATCTCTAGGGCAGCTTGCTGGTGGAGTGATGGGGTGGTGGTGAGGTGGTGGCGGT

GGGCAAAGTGGCTTCGGGGGTGTTGGTGTAGGTGTTGGAGTTGGTGGTGGTGGTGGAGGGCTCTTCGGCT

TCGGTGTAGGTGGTGGTGGTGGAGGGCTTTTCGGCTTCGGTGTAGGTGGTGGAGGGCAGCTAGGTTTCGG

CATTGGGGGTGGGGGGCATTTAGGTTTCGGCATTGGTGGTGGTGGGCATGGAGTGTGATGAGTTGAAGTT

ACTAAAGTGAGGAAGAGAAGGTTAAGGACTAGGAAGAAAGATGTGGATGCTAATGCCTTTGAAGCCATGC

TGCAGAGACTTCTTTGCTATGTTAGGAGAGGAGAGATTCTTGGGTTGTGAGATTAGTGAAATAGGTTATG

CAATTTATAGATGAGAAGATTGAGATTTAAATCGTTTATTAAGAAATTATTGTAACCATGTGAAATTGTT

AAACTGGCTGTGTAAGTTTGCAACTTGAAGGCTGCCTACACATCAAAAATATATCTAGGGCCGTAAATGA

A

>MSTRG.3931.1 gene=MSTRG.3931

CCGGATTCTTATTAACGGCTCATAGTGAGGACTTTATTGAAGTTTGATAGTCTTTGTTTTGTTTCACTAG

AGAGCATGCTAGCACACACGATGCACTTCACACTAGTAGTGGAAGAGAACATGGCTAATGTGTGAAAGAG

ATGAATGCCGAGAATGTTGCATATTTTGAGGATGCTATGTTAACAAGGATTACACTTCATGATTCGCTAG

ATGTAGGC

>MSTRG.3932.1 gene=MSTRG.3932

AGTTTATGTGCATAATTTCTTGGCTAGTAGTCTTTATCATATGTGATGCCTTGTTACTTGGAAACTGCTT

GTGTGAAAATTGTAATTACACTTATGCTCTAGAGATAAGACTTAGACTAACTTATTTCTTGAGTCCTCGC

GTTGAGTCGGGCGTAGAGTTTGGAGTATTCCCTTTTTGAACTTAGAGTTTGTAAATCTTAAGTTTGGGGG

AGTTAAGGGA

>MSTRG.3934.12 gene=MSTRG.3934

CTTATTTTCCTATTCCTATCGAACACCAAAAAGATATATAGACACCCATAATTTTCATATTGTGCCTGTC

TATTTTCTGATTAATGGCGATAAGTGTTCTTAAGAAGCCCAATTCATTCTTGTACAGCTTAATAGTATCA

TCTCAAGATTCAATCTTTTTACTCTCAAACTCAAAACCCGTCTCGTTTTCAACCGAATCCAATGACCCAA

TTCATAGTTCTCAGCAACTCTCACAGAAATCACATTTACATGTGTTTCAAAGGTATGGGTTTCAAGCTTC

TGAGCTCACTGATTTTCTGAAAAAGAATCATTTTTTATTGAATTACAGCTCTTCTGATATTGAGAAGTCA

CTGAAAATATTGTTATCTTTGAAGTCATGTCCCAAGTTTGTTGCTTCAGTTGTATATAGTTGTCCTAATG

TTCTTGAACTTGAGTTTCTTGAGAGATGGAAAATGGGGTTTAAGGAAAGTGGGGTTTTTGATTTTTCTTC

ATTGGCTGTTAAGAATGTTGTTGCGATTTCTAGGAAATTTGAGCTAAGCCCATGTGATTTTTTTCGTTGC

GTGGAAAGTTTTAGGAATTTCGGGTTTGGTAGTGATACTGTTGAGAAGGTTTTGGAGGTGAATCCGATGG

TGATTATGGCAGGTGAGGAGAGCATTTTGGATAAAGTTAGGTTCTTGGAAAGAATGGGACTCGAGAGGAG

GGGTATTGATTGGGTTTTAGGGAAGTTTCCCGAGTTTCTGTCGTTTGGGGTGGAAAATAGGTTGATACCG

TTGATGAGTGAATTTAGGGATTTGGGTTATAGTCGGGGTGAGGTTAAGAAAGGGATTGTGAGAGACCCGA

GAGTTATTGGGTTAGAAGTAGGAGAGCTTTCTCAATGTTTGAGGATGCTGAGGAGTTTAAAGTGTCGACT

TTCAGTAAAAGAGGAGATTTTGAAGGAAGGAACTTTTAGGGCCGGGTATGAAGTGAAACTGAGGGTTGAT

TGCCTATGCAAATATGGATTGATACGCAGAGAGGCTTTAAGTATTATTTGGAGGGAACCAAGGATGATTT

TATATGATTTAAAGGAAATTGAGAGGAAGATTGATTTTTTGTTAAATGAGATGAAACTTGATGTTCAGTG

GCTGCTGGATGTTCCAGAGTATCTGGGAATTAGTTTTGAGAAACAGATTGTTCCACGGCACAATGTCATT

GAGTATTTGAGATCAATAGGCGGGCTTGGTGAAGAGGTGGGATTGAAGTCTTTAATTAAGCTTACTAGAT

TAAAGTTTCACAATTTATATGTGAAGCCATACCCTGAGTGTGAAAAAATATATGGAAGATTTGCAGAACA

AGTTAACCTAAAAAATCAGCATCCTGTCGGGTTGTCGAAACTTTTTAAACCACAGAAGTATCCACAGTCT

AAGGAAGTTGCAAAAAACATAAGGTCTTTCATGGAACCACTAGGAAAACTGGAAAAGTAGTCTCTCCAAA

TTCCAGGGACAAATGCCTCATGGAAGCTTCAGTTCAACATTTTGAAGGGGAGTTTACAGCATGAAGTGGC

TGGGAAAATGCAGCTTTGCCACCGCTACTATTCTGCTAATTAGTGGTGACGGTTAGCATATATTGCAAGT

GATCATTATTAGCATGCTAACAAAATTCATCATGAAATCCAGTACTCTCGACAGCATAGTGCATATCTTT

AGTTCAGTACTAAACAAATGCAACCTATCCATAGTTTTTATTTGGTGATGAAGCAAGCAAGCTGTAAAGA

AAGATTATCGATGGAGGAGGAAATTTCTCTTTAGAGGATTACAGATGACTTTGTGATTGTTCTCCCAAGC

TGGGTGATTCAAGCTAACAATTGACAGGAGAAGTATGTGGAAGCCTGTTTGAGACATTCCTTTGATGTAG

TCACAATAGCTCTTCTTAGTCATATATGTATACGGTGTTATCTAGGCTATTTTAGTATTAGTCTTACTTT

GTTTATGGTTTTAGGATAGTTGGTGCGGACCATTTCAAATTCCCATGAACATATGCTAGTCCAGAAGCTA

TTAAGGTTCCATTCAGTTCCCTGACTCGAGGACAGAGATATATGAATACTCAAGGTTGCTTGGGAACTCC

CAATTCACACTGCTACCATTTCAACCATATAAGCTTGTATATGCACACATGGTGCCCGAAGTGGGAAGAG

TATCAGATTCATTGAAGTACTGTCAAGCAGTACTGAAATCTCTGAAAACTGGTTGAGCACCTGAAGTGGA

AACGTGGAGACAGTTGGCATCATCTCTTGAGGAGAGGATAAAGACTCACCAACAGGGTGGTTTCTCAACA

AACTTGGCTCCTGCAAAGTTAGTAGGTAAATTGCTTAATTTATTTGATAGCACGGCTTCTCGTGTTGTTG

GAACCCTACCACCTCCTGCATCATCGACATCTGGTAGCAATTACGGCAGCAGAGGAAGCTGCACTACCAC

CTCCACCTACCACTGCAGCCTTCCCAAATGGAACATCAGATTACAACCTGAATAGTGCCTTGAAAAATGA

AGGACCCCGCAGTAATGGAAGCCCAGACCAATGAGGCCCTACTTCTTTGGTACATAGTTCAGGAATTCCA

CCAATTCCACCCACCTCAAACCAGTTTTCAGCTCGCGGACGGATGGGAGTTAGATCAAGGTATGTTTACA

CCTTTAACAAAGGTGGTGGAAATCAAACAAACATGTTCCAGTCACCATCTGTGCCTTCAGTTAAGGCCAA

AAGCAACGCTAGC

>MSTRG.3935.1 gene=MSTRG.3935

ATAGCAATCTTTCACTGTAGTCAGTTCATACAGTGGTTCCTTATGGATGTTCTCACACTTTGCTATCTAA

ACCCTCAATATATTGTCACACTTTGCTATCTAAACCCTCAATATAGTTTTCTAACATATCCTTCAATTTT

TATTGCAGAGGGAACCTACCATGATCCTTTAAATTTAGTAGGACCAGTGTTCAAGAAAGCAGAAAAGGTG

AATCAGATCTAAAATGCTTCGAATATCCAGTTATCGCTTCATTGGACTGTTAAGCTGTTAAGAAACCTAT

TAAGCTAATTAAGCAGTCAAAAAAATCTTAAGCAGCACCGACCGCTATTTTTTTACATCTGAAATATAAT

AGCGATAAAGAACTGTAAATCGGAACGACCTACATTTGTTAAGAAACTGTGTGTTCAGATTGTTGTTAAT

TACAATTTAACTTTGCTCTTACATAGATGGTATTATATTTGAAAATTTCCAAGTATGTAGTGATGATCAG

ATCAAAAATCTGCTTAAGCACTCAAAGGCTCTCCGCTGCTTAGCGCTTTTTAGAGCACTGAGTATGACCA

AGTCAAATTCAAAGGCAAGTCGGACATCCTGAAGACAGATGATATCAACTTACAGACGGGAAGAACCTAT

ACAGAGGAAACGGTTGTATACTAGTAAGAGAAAGGAGGATATATTGAGTCAGGGAACTGAATGGAACCTG

AATAGCTTCTGGACTAGCATATGTTCGTGGGAATTTGAAATGGTCTGCACCAACAAGACAAAGTCGTGCA

CTGTCTGAATATGGCTCAAAGTTTGCTTCTGCAACTAAGTAACATATATGGGCAGCGACTATCTAAGCCA

TAATATTAACGGAAACCAATGTCAAGACTACAAAAAATTCGAAATATGAAATATCTTAGAAAGATTATAG

TGCTAAACATACTTCGCTTCTTTCTTTCCACAAGCAATCTCCAAGATGAATAAGGAC

>MSTRG.3935.2 gene=MSTRG.3935

GATCAGATCAAAAATCTGCTTAAGCACTCAAAGGCTCTCCGCTGCTTAGCTCTTTTTAGAGCATTGAGTA

TGACCAAGTCAAATTCAAAGGCAAGTCAGAAATCCTGATGACAGATGATATCAACTTACAGACGGGAAGA

ACCTAACAGAGGAAACTGCTGTATACTAGTAAGAGAAAGGAAGATATACCGAGTCAGGGAACTGAATGGA

ACCTGAATAGCTTCTGGACTAGCATATGTTCGTGGGAATTTGAAATGGTCTGCACCAACAAGACAAAGTC

GTGCACTGTCTGAATATGGCTCAAAGTTTGCTTCTGCAACTAAGTAACATATATGGGCAGCGACTATCTA

AGCCATAATATTAACGGAAACCAATGTCAAGACTACAAAAAATTCGAAATATGAAATATCTTAGAAAGAT

TATAGTGCTAAACATACTTCGCTTCTTTCTTTCCACAAGCAATCTCCAAGATGAATAAGGAC

>MSTRG.3936.3 gene=MSTRG.3936

GCACACCCTTCTTCCCAATGGATAGTTTCCTAAAAAAGCGTGGCCGGCCGAAGCTGTGCCTCACCAAAGA

GGTTGTGGAGGAGCGTAGGGCGCTGAAGCGATTGCAGAATGCCGCTCATCGTGGGTCAGGTTCTAAGAGA

GTCCTTGCTTTTTGTGATTCCTGCATTTGGAGGTGTGGTCTCTTGGGAGGGTGAAGGAGCTCCATATAAG

GAAACGGATCAGGGCATCACTTACCAGATAGAAGTGCTATGAAAAGCAGGCAGAAGATCGTAAAGAAAAT

TCCCTTCATTGAGTCAAGAATGGTGTCAAAGCCGGAGGGAATTTGATGCTGAAAGATCTGTGATTGATGA

TTGTTCCATTGCTAACCAAATTGAAAAAGTCATACCTGATGCGCCCAGTCCTGATGGTGCTGATGCATCA

AAATGGAAGGCCAATGTCTCCCGGAACACTGGCTTTGATGTGTGATGAACAAGAGATTGTGTTTGCTGCT

TCCCAAAGTGGGTGTTTGGGCAATGGTGGTAATGCATCCTCTCAGTTGCCACAAGGCGAAAGCGTAACAG

AAACCTATGTTGAATAAGAAAGAATTATCTTCACAAAGTTTCGAGATTGTTTGAACAAACTCATCACCTT

AGGGTAAATAAAAGGCCATCTGTATAAAGCGTAAGAAAGCTTATCACCATTCTTTGGGGTCCTTGGAAGG

CAAGCATCTAGAAATACATGTGTTTCCTTTGACAGCAGCAGGTGGTAAAGCCAATCATGATCAGTGGATT

AACACAATGATGAAATTTGAGGCAGCAAGTGGAAATTATATTGACAAAGAGGCAGCTAAGAACTTTGTAG

CAGAACATCTGTCAAACCAACAGGTTGCCCGTGACGTGGGAACTGATGTTGCAGTAGATTCTTCTTGAAG

AGAAAACTGTAGTATACAATATCTTTTCTGTTAGTGACATTGACGATAAGGTTACTAGGACTTTTGAAAA

TGAAGTACTTCGAGTGCCAGGAACTTTGGGGAGAAAACATTGCTTCTCATTCTGCTTGACACTGGACGAA

GGCAAAGACCAACATGTTTCTAATCATAATCACGAAGTTTATCATACTACAGACCTTAATTTCCTTCATT

TTGCTGACTGAATTGACAGAGGGATACCCATAAAGGAGGTTTCACGGGAGGAAGATTACTGGAAATTTAT

TTCACTCCTTGTCAGTTGGGCGGAAAATTAACAAGCGCTGCTATCTCGACTAACCACATTTAACCGAATT

GATGTTCCAGCTTCATCTGATCCTCTAAATGGTAGTGCAATTTATTGATAGTATATTGGCTCAAATGGTT

ACCTCGCTACGGAAGTCATTCAACTAAAAAAATTGTGTGGCCCTTGGCATGAAGTTGATAGCATAAAGGA

AGTTTCAGAGCCTGAGTTGTGCGAATATAATGAAGCTGTGAACTTGACTGGTGATATTGATATGCCAGCT

GGTCAGGCAATTCTTCACACCTCTTTGTCATCGTTTTTTGTTCTATCATCAAGTGCCATCTTGCGTTTTC

ACTAGTTCTTTAAGCTCGGAAATTTCCGCCTCATGCGTACGTGCTGTGCCATTGATCAAAGAAACTGCAA

CGTTAAAGAGTCTGGTTCGGCAGCTTGTGACTTCTTCTGCAGTCTTGCATGCGTTCATTAGGCGTAGAAT

GCCCTTTTTGTGAAATAAACTCGATATGAGGTGTGCCATGTCAGCGGTATCAGCCGGAAAGAACGGCCTG

AATGGACACTCGGTCTTACAAACGCGGCTGGCCCCTTTACAGGCTCCACATTTGTCCACTGCCACTGCAG

CAGGATCAGTGAGATCAGCTCCGGTGGTGTCTGTTGTCATCTCTACTGCTGTGCACAACAACTGCAAGTT

ATTTGGGGAGTTGGCACCCATTTAACAATCCAAGGGAATTAGACTGATGCACATAATTAATTACAGTAGT

TAAAGTCTTGAAAATACACAGAAATCACCATTCGTGCATGTAAACTTATTGTATCCATTACCATTAGCAA

TCCAACAATATA

>MSTRG.3937.1 gene=MSTRG.3937

TCACTCTCCCATTAGGGCTGCGATGAAGGAGGTTCTCACGACGAGCTTTTGTTCTTATGTTTCTTATTTC

TATTTTTTCTTACTACTCGGTTTGTAATAATGATTTGTTTTTTGATATTTCTTCTGTTTATCTGTTTCTT

TCATGGATTTTAGGGGTTTTTCAATCATCTCAATCTGGATAGTTTTCCTTCATCGAAAGGTTTCAAATGG

AAAAGAAGAAGGCAAAATGTGGAAACCAATAAAGAAAACATAAAACCATCTGAAGAGGGAAATCCTAGCA

CTGGTAGTCTGTTAAGCACAACCGAACAGAATTATCAAACTCCTACAAGGCC

>MSTRG.3938.1 gene=MSTRG.3938

ATTATCAATCTGAAAAGGATTTATACGGTATTTTGGGAGATCGGTAAGACTCGCATCGATTTCGGAACGA

TGGTGGATACCGCAAGAGCTCACCTTTCACAACAAATAATTGTTCATGTTTTGGGGACGTTTGGTGCTCC

AGGATCAGTTGATTTGACTGATAGTTCTGAACATTGGGCTACAGATGATGCTTACGTGAAGGTCAATTGC

GAGACTACTAGTTTCATAACTGATGTAGGTTGGATTGCTCGAGATATCATTGTAATACTTTTTTAGATCA

AAGAATATGAATATTCTATTTGTTAAGCGAT

>MSTRG.3939.1 gene=MSTRG.3939

GTTGTTTACCTGAGCAAATTAATTCTCTGGCTCGGTCATTCCGGACCTACTTCTCAAATCTGCTGATTAA

AACCCCCTGATCCTCGTTCATTGAAATAAATTTTATCAACAGAGAAAGACTGTCGTGCCCTTTCCGAACC

TGATTCAAAACCTGCTTCCCCATACAAGGTTACGGGAAATACAAATCTGACTGTGCCGAAACCCTTTACA

ATTGAGAAACGCCTCTTCACTTCTGTTCGAGCAGTGGAATGCACTTTAAACGGTGATTGAGACGCGCTCG

GTGCATCAATAGACTTTCTGTCTTTGAGCAAGGGTATATGTGATGTGATGATTGAGACGCGCAATCAG

>MSTRG.3940.1 gene=MSTRG.3940

AAAAAAAACACCCTACTCAGAGGCACTCCCTCAACCTCCTCTTACACACACACTCGACTGAAGAGAAAGA

AAGGGATAGAGAGAGATCGAGAGAGGAAGAGCCCGAGAAAGTTGATCAAAACATATTTGGGCTACATTTT

ACAGATCAATCAAGATGGGGACTTCAATCGAAATTGGGGTTTTTTTAGTTTAATTGGGGTTTTTTTGGTT

AATCCTCTGCAAGAGTTTAATTGAAGTTCATTTTTTGTTTCAGGGCTAATTTGGTTGATATGGGTTTTAA

TTGAATTTTGCATCTTTCGCTTCTATAGTTCAAGGATATCAATGGATTTGAAGATCTCTAAAAAAGCTTC

TGATATGGAGGACAAACTTAACAAAAACTAGAGACAATCCTCGGACATTGGCAGAGAAGAATCTCTCACT

ATATATTAATATCGAAGAACTATGTGCTAGTGGTCATAGTTAACAAGAGTCGAATCCCGATAGCAATGAA

GCTGATTTTGTTCCAGATTTGGCTCATTTTAATGCCAACTATTTGATGTACTTCACCTATTTTCGGCTAT

TTCAATTAGTGTTCTAATCTCAGAATTGTTTTTTATTTAGTAAGTATGGTATTGATCTTTGTTGTGGTTG

TGTAGGACTTGTTTGTGGTTGTCTAGGAATGACTATACTTGTGGTTAGTAACAGCCTTTGTACTATTTTT

ATTAATTTGTTG

>MSTRG.3942.1 gene=MSTRG.3942

GTGCAGAGATGGTTACGCGCGCGAAAACAAGTAATCAACTCTCGATTCTTGTCAACTTCTTTTGCTATCT

CCCTATCTATTTATATAAATACACCTATATACTTGCTTGTGTTGGTTCGTTTGGGCTTGTAATCAGGGGT

GGTGTGGTTGAATCGCTGATGGCGGTGATTTGTTTCGCCGCTGATAGAGGTGAGCGGAGGGAGCTGTTGC

GTAGGGTGAGCTTGGGGTGTATAGAGGGGAATAGATGGAGCTATTACAGAGGGCTGCGGAAGATTGTGTG

CGTGTGTCCTGTTGTGTGGGTGGTGTGTATAACTGCCCTGTACAGGTGTACTAGTTTATGTGTGTAAATA

GTTACTTTGTGTGTAAACAAAGCAGG

>MSTRG.3943.1 gene=MSTRG.3943

ATGAGTTCTAATTTTCATGAAATCCTTTCTGGATCTGGGAAAGATGTTGAAGGAAAGCATATTGACGTGG

ACTTTCTGGAGGCTCCTTGGGTCCGTCCCAGTGCTTTAGGGGGTATTCCAGTGGTTGAACGTGTAGAGAT

CGAAGATGTGGAAGATGGTTTCAGCATTGATGGGGTTGCTTCGCCTAAATCAGTGAGTGGCAATCACGAG

ATGAGCAGCACTGTTCAGACCATTCTTCGCACTTCAAAGCATGAAAGAGATGAGCTGTATTCTGCTCGTA

GAAAGCTAGGAGATGTGTTTGGTTTTCTCCGCAAGCAAGGCTTTAGCGAGGAGCAAGTTTTGGCTGATTT

GAAGAGGGAAGGGTTTGGAACTACTCTTCCAAGTAGAGACGAGAATGGCCTCCCTCTTCTCAACCCAGTT

CCAAATCCTCTCAAGGATAAGATGAAAATCAAGATGGAT

>MSTRG.3948.1 gene=MSTRG.3948

ACCAGTTTCTTTGCACGAATACCGAGAGAAACCGAGCGGTGTTCTTCGTGTTCTTCATAATCAAACGAGG

ATTTGAAGGTGTTATTGGAATCCGATGGAGGCGTATGAATAGTCAAAACGAAGCTCTCGACGCGTAGATT

ATAGTTTAATCATCAAAACAAGCGCAGAAAGGCTTAAAGATGAGATGATTGATTGAGATTGTGAATAAGG

CTAATGCTAGGCGAGGAGACAGAGTTGAGGATCCAATGGTCGAGGAGTGTTCAGGAAAGTTGATGAGGTT

GATGCAAGCTAGAGCTGTGCTATAAGGATTCA

>MSTRG.3950.1 gene=MSTRG.3950

CTGGAATCACGAATCAGATCGTCACTGTACTCAAGGTTGCAAGGCTTCGCATACATCTGGATTCCTCAAA

GGTTCAAAAAAAATTGACAAAATATATTTCCACTGGTAAAGGTTTTATTGGGACATCACATAATAGTATG

TGCTATTTGGAATCCTGTGTTCATGGCCTAGAGCTGTCCTTGGAAGAGATCTTGCTTGATTTGGCACCGT

CAACCGGAAGGATATCAGGTCTTAATATTATGTATGCAGGTCTGGACATTATTGTTGGGATACCAT

>MSTRG.3950.2 gene=MSTRG.3950

GTCATTGTTCGATTCGCTTATTCGTATATTTGTTAGCTTCTAATACTGATTGGGTCTCTACTTGACAGGA

TTCCTCAAAGGTTCAAAAAAAATTGACAAAATATATTTCCACTGGTAAAGGTTTTATTGGGACATCACAT

AATAGTATGTGCTATTTGGAATCCTGTGTTCATGGCCTAGAGCTGTCCTTGGAAGAGATCTTGCTTGATT

TGGCACCGTCAACCGGAAGGATATCAGGTCTGGACATTATTGTTGGGATACCATTGA

>MSTRG.3951.1 gene=MSTRG.3951

TATGAGCTACTAGCGCGTTTGGTATATAAAACAGGTCTGAGTTTTGAGCTTTATATATCCATGATTTCCA

CCTCTTGTACTGTGATTCTGGAGTTCTCTTGCTTTATCTAAGTTATTTGTCTTGTTCATGGTCGTAATCT

TTGGCAGGGTGTGGTTTCTTGGAGCTCTTGTCAGCCTTATTACTTTCGTGTTGCATTTCACTTCAACTTG

TTGGAAGTTTCAGACTCACATCTTTACTCTTGCTGTGCTATTGTTACTATTGCAGAAATCTTTAAGTCAA

TCCGGTTGATTTCTTGCTCCAGCTGGAAGTCTTTTGAGGGCTTTTCAAGTCTGAATGGTGCTAAATGCTA

ATATTTCTCATTGACAGTTGCCTCCATTGAATTATTTTGAGACTTATATTTCAATTTCAGGGAACAGATG

CTAACATTTCTGTCTGTTTTATTTTGATATAACCGTTGATTACTGCTTTTAAATGTTTTGTTCAATTCAT

AATAATCTTGGTTTGTAAACTTTTATTTGTTTCTTTTAACGCAAGACATAAATTGGTTCTTCTCTGTGTT

TCTTTTTTTGTACTGCTATACTTGGCAAGGATTCAGTGTGAACCTCACTTTGGGGTGGATGTACATTCAA

GGTAGTACAGTTATTGATCATTTTGTTGCATCTCATCTGCCTTGTGAGGGTGTTGAATCTCATCTGCCTC

CCAAGGGTGTTGAGATCCCAAATACTATTTCGCAGCTGGTAACTCTGGTAAGTAGAAGACTTCTTTTTAT

TTTGACAACTAAAAAGAACTTACCGTTCTTTTAGTAGGAATCAATCTCTATCTCCTTGATGAAATAGTAG

AGTTTGAAGCTACAAAAATTAGGTTTGTGAATTAAGTTTGTACTTCCATTTGTCAAATATCTGAACATGG

AACTGCAGGGTTGTCCTAAGTCTAACTAAGCTAAATCTGATAGTTGACAAGTAGAGCATAACGTGTTATA

CCAATTTTATGGCATATTGTACCCGTCAAATTATTACTTTAATAGATTATAATTTGACTTGAAATTATCC

AGCAAGTCACCAGTTATGGTAATGCCACTGACAACCTGTAGAAGATCTACATTATGATTGAATAATCATT

TGTAAGAAGGTAGACACGTAATAAATAAATTGCTTGATTAATATTATATATAATAACGGGCTAGCTTATT

AGCTCAAGACACTTAAACAGATTTAGTTCGGGTGGCATTTTTGTCTTTTTGTGCTGTAATTTCTTCTTCC

CTCTTCTCTCTTCACATGTATTGAATTTTAAAACAGTCATTTTGATTATTTTAGATCAAATTAATCTGAG

TTGGTTTATAGACATATGCTGCTAGTTGCCAAATGCTTGACTAATTGCTAAAGGAGACAACAAAAGTATC

TTCATCTACTAAATCTATAGGAAGGATATTCCTTTTATCTGCCTAGATGATTTTTACAAGATTTGGAAAC

TTAAAAATTTTAATCCATCCCACTAGAGCTGTCCATGAAGCTTCTTTTGTCCCATGGCATAGATGCAGGT

TGGTGTTAGGAATCTGTTTCAGAAGTTATTGCAGAGGATTTGGATTTATTGATATGCTTAGACGGCTTGA

CCTGTCCGAAAAATAGTTCACCCACTTGAGAAGTGAAAACATGCCTTCACATTAGTCTTTTAATTATTGC

CAGTTACAGCCTGAAAAGTAATGAAGCGTTTTCACTACTGACATGCTGTATTTTGTGGTAGGTAACCGAT

TTTTTTGGCCAAAAACATATGACCTGTATTTTCCTGCAGTTTAAGGTTACATTGTGGTCATTTCAGTTAG

TTTGAACTTTAGATGGCATCAGATCGGTAAGTGATACAATCTTTAGCAGTCATTAAAGGACTGTGCAACC

CATTTTTGGAAGTAAAGCTAGCAGTCAAGGTCCTCAAGGATGAGTTCAAGCCAGATTGGTCGTATACCTC

CCACTTGCTGTTTGGAAGTAGGAAGGAAATTAGGAGGCAGCTTTGCTCTTCATTGTGGTAGCTAAATAGT

AGCAGCACTTGTATCTTATTTGTGAATGTTCCCATATCCTATCTGGTGTATATACTTTTTAAATATATTG

GACTCTTCTCCTCTTGAAGGATAAGCTCTCGTCATTAGTCCAGTGTTCTAGTTACCTGTTGGCATGGTGT

GGATAATAGACTTTCCTATAGTATGATAATTGGTTTTTTGCAGACAATATTGGTTTCTATTGTTTCTATA

CCATTGTGCTGCCAGAGTAGTAGTGTGCTGCTTTACAATATTGCCTGTTTATTATATATCAAGAAACCTA

ATTTATCCGGCTCAGTGTGTATTTCAGCAGCTTTTTAGTTCGTGTTGCGAGTTGTTAATGACTTGACCAC

TTGTGTTCCATCATCGAGCTTCTTTTTTTTTTTTCTGGTGTGTTTGGCCCTTTTTTTCTTGTTGCCTAGT

TTCTCTGAAGATATCTTGTGCATCAATTAGTGGTTGGCTGTTTTCCCTAAATTCTTGTGGGATCGTGATT

TTCTTTCTCGTCATCTGTATATTACCGGAAATTGTTGTGGTGATCCAATGTAGCTGAAATTCGTATGAAC

CCTTCATTTTAATTTTCAGTGGATCGAGGAAATTATGTTTTAGGAGTAGAATAACGCTTTGATGCATATA

TTAAGACCATATGAATCAAGTATA

>MSTRG.3954.1 gene=MSTRG.3954

AAGGGATGATGGTGAAGAGGGACTCGACTTTCATAAAATTTCTCATGTCGATAGACTTATTATCGACGAT

CAGGAATACAATTGAATGTTCCCTCTTTTTTTCTGCTAAATCAGTTGTTTGTAAGTAGTATTTAGAGGTA

AGATGATTATGTGACTTATCTGCAATTTGTTAAATTGTTGGATTCTGTTACTTAGCGGACCCCTTTTTTT

GTATCTCATTTTT

>MSTRG.3956.1 gene=MSTRG.3956

ATCTTTTTTTGTATATCTTTCATCTTCCTTGAAAACTTGACTCTGTTCATCCACTGAAATGGTTTTTTAG

TGGATGAGGCTTTAATAACAAAACTTTCGACAGCATGTTCAGCATCATAGGCAAGCTCTCGTACCTCCCG

GACCAAAATGCGGATTGTTTCTACATCAATCCTTGAGTCAGCATCTGGTAAGAATGTCTTTATCCGCATA

AGCTCAGTCACGGCTAGTTGACTTTCATCTCTCACTACATGAAGGATTTGAGCTTTTTCAATCAACAAAT

CAGTGAGCCTTCCCACAACAATCGACACAATTGCTTCGGTCTTATTAATTTGTTTAACTTATTCTTCTGC

AGAAACCAAACAAGTTTATATAAATTTGAATAGGCTTAAATGTGTCTTCAGCACATTGGATTCTCTGTTG

CTTCTCTGGTCATAGTTAAATTCCATGTTACTGTACAGCAGCATTTGTTTATGATGTTGCTATGGGTTCT

GTGACACATGGTGAGGTCGATCTTAAGTTCAAATTGCATAACCTTGCAATGCGAAGCTGATGTTATTTTG

TTGGATTGTATTTGCTTTCGGTCGCTTCTATCATTGTTATTTATTGTTGCAATGCGATGTTGGTAACGTT

TTTGTTAAAAAGCATTTGCTTGCGGTCTATCATTGATCTTACAGCTTATGACTGTTGTTAATTGAATAAA

CTTG

>MSTRG.3958.1 gene=MSTRG.3958

CCCTTTGATAGGCTTCTCCCAGTCCTTTCCCCGCCTTTAGGGATGATTTACTATCACGATATACCTTCTC

CCACTCTATCAACGACGGTTTTGTTACCAAAATTCCACCCAAAATGACTATAGCTAGCGGAAGACCGGCA

CAATGTTCAACCATTTCCCTTCCCAGTTCTTCCATCCTTTTTACATCTCTGGTAATGTCTGCAGCACACA

AGAAA

>MSTRG.3960.1 gene=MSTRG.3960

CAGAGTACATTGTTGCCCTCAGGTTGCAACAAAACACAAAACTAGGACACTTTTAACTCTCGACTCCGCC

CTTTGTTCAAACTTTTTCTTTGTGATCTTTCTTTTCTTTATAGTTAAACTACAATATATACAGCGTTTAG

GACTGGGAAACTGCTCTCTTCAAGACGCTGAAGGCAACGAGCATAAGCAGTGATATCATAATCTTCTTTG

AAGCAAGAATCCGCAAGTTCACCTTCATGCCATGGTCTCTCATGAGCTTCACTGTTGCCCTAATTACTGA

AATAAGCCAGAATACAACGGACAAGAACACCACTTTCATTGTGTCTGCAGGTCCATTCACACTTCTTGTG

CCTCCCGACACCTCGACATAGGACCAAGCAGCGCCTTGATTATCGGTAGTTCCACTCTGATTGAGGCCAA

GTTTCTCGTTCAACCAACATGATAAAGTCACAAGCAGCTTTGCAAGCCATGCAATCTCATCATTGGAGAC

TGGTCGCTTCAACCAGTCACCCCTG

>MSTRG.3962.1 gene=MSTRG.3962

TAGAGTTAATTGCAGAGTGTGTACCTCTAGTTTGAGGCTTGTGCACTTGGTGTACCTCAGTTTTGGTGTC

TAGCAATGTGTGTACCTCAGCTTTGAAGATGCTTGCACAGTGTGTACTTTCGTGAGTTTGCAGCTTAACG

GCGTTAGATTGGAGGGACAGCGGGATAATTTGACAAAGAAAAAGGGAGGGGCAACGGCTATATGTTTGTA

TGTGTTATATGCACTCTACTATATGGTCTG

>MSTRG.3963.1 gene=MSTRG.3963

CATAAATTAAAAGAAGCTCAAAGAAAATAATTAGCTTTTTGTTGTGGTTCATTCAGATGTATCCATCAAG

CATTTTCCAGGACCACTGAAGCAATCTATGCAAAACCATGGTTGAACATCACTAACTACACAAGTAGCCT

TCCTCTTGTGTTCTGCTTATCCAGACGCAGCAAACCGATGATTCCAAGTGTCTGTTAGCAGAAGGGGAGA

GTGAAGAAGAATAGAGAGTATGAAGTGATTTGCTTTGCTATGAGGAGGTTATGCAGGAAGCGGAACCAAG

AGCGGTGATGTTTGCAGAAACACAAGAGAAAGACTTCAAATGGCATCTTGAAAAGGAATGAGAGCTGGCA

ATTGGAGCAAACACAGCAACGACAACAAAAGTAATCAAGGAAGATGAGTAGGATAGGCTTGTGAAACTGA

GCTCCTGAATTTCTGTTCTGCTTAGGATTTTCGACATAATAGAGTTATGAATAGTAATTACAAAATTGAA

AAAGTTGAACCATAATATCAAACACAGAAGGGAAAAAAATG

>MSTRG.3965.1 gene=MSTRG.3965

GATTCTGAAGTTGGTCTGATATTATAATCGGAAAATAAATTGAAGAGGAGGGACATCTTTTTATGTATAT

CTTTCATCTTTCTTGAAAATCTGACTCTGTTCATCCACTGAAATGGTTTTTTAGTGGATGAGGCTTTAAT

AACAAAACTTTCGACAGCATGTTCAGCATCATAGGCAAGCTCTCGTACCTCCCCGAGCAAAATGCGGATT

CTTTCTACATCAATCCTTGAGTCAGCATCAGGTAAGAATGTCTTTATCCGCATGAGCTCCGTC

>MSTRG.3967.1 gene=MSTRG.3967

GCTGAAGGCAACGAGCATAAGCAGTGATATCATGATCTTCTTTGAAGCAAGTATCCGCAAGTTCACCTTC

ATGCCATGGTCTCTCATGAGCTTCACTGTTGCCCTAATTACTGAAATAAGCCAGAATACAACGGACAAGA

ACACCACTTTCATTGTGTCTGCAGGTCCATTCACACTTCTTGTGCCTCCCGACACCTCGACATAGGACCA

AGCAGCGCCTTGATTATCGGTAGTTCCACTCTGATTGAGGCCAAGTTTCTCGTTCAACCAACATGATAAA

GTCACAAGCAGCTTTGCAAGCCATGCAATCTCATCA

>MSTRG.3967.2 gene=MSTRG.3967

AAGAACTACTCCTATAGAAAACGGCTCAGCGTCCTCAGTTAAACTACAATATATACAGCGTTTAGGATGG

GGAAACTGCTTTCTTCAAGATGCTGAAGGCAACGAGCATAAGCAGTGATATCATGATCTTCTTTGAAGCA

AGAATCCGCAAGTTCACCTTCATGCCATGGTCTCTCATGAGCTTCACTGTTGCCCTAATTACTGAAATAA

GCCAGAATACAACGGACAAGAACACCACTTTCATTGTGTCTGCAGGTCCATTCACACTTCTTGTGCCTCC

CGACACCTCGACATAGGACCAAGCAGCGCCTTGATTATCGGTAGTTCCACTCTGATTGAGGCCAAGTTTC

TCGTTCAACCAACATGATAAAGTCACAAGCAGCTTTGCAAGCCATGCAATCTCATCA

>MSTRG.3967.3 gene=MSTRG.3967

AAGAACTACTCCTATAGAAAACGGCTCAGCGTCCTCAGCGTTTAGGATGGGGAAACTGCTTTCTTCAAGA

TGCTGAAGGCAACGAGCATAAGCAGTGATATCATGATCTTCTTTGAAGCAAGAATCCGCAAGTTCACCTT

CATGCCATGGTCTCTCATGAGCTTCACTGTTGCCCTAATTACTGAAATAAGCCAGAATACAACGGACAAG

AACACCACTTTCATTGTGTCTGCAGGTCCATTCACACTTCTTGTGCCTCCCGACACCTCGACATAGGACC

AAGCAGCGCCTTGATTATCGGTAGTTCCACTCTGATTGAGGCCAAGTTTCTCGTTCAACCAACATGATAA

AGTCACAAGCAGCTTTGCAAGCCATGCAATCTCATCA

>MSTRG.3972.1 gene=MSTRG.3972

CGAAGCATGCCTCTGGGTCAGACTTGATATGTCTCTCGGCCACAGTCCAACAATGTATGATCTCGGTTGC

CCCAGCATCGGCCAAAGCACGGTCATACTCATCAGACTCCTGGAACTCGGAAATCACCACTGCAGGCTCT

TTCCTCTCAGCAAGCTGTGGCTTCAGACTCTCGATCTCGGTCTCAAACTCGGACATCCGAGTCTCAGCTG

CCTTCAACTTCTCGCCCAAATCGTCAAC

>MSTRG.3978.1 gene=MSTRG.3978

TTTACTTTTATCCCGAAAACACAATTCACTCCCTCCTACTTTCTCTCTCTCTCTCCTCTCTACTATGTCG

ATCTGACCTGCCTCTCGGCTCTCACCTCTCCCGTCTCTACTCTAACATCTCCACTATGCGTTTTCACATC

TCAGATAACTGAATAATCAGAATAGGGTTCATGGGTTTTCTACTAATTGTTACTATGTTGGACTATTGCC

ATCTAATTTTATTATTTCTACTGATTGGGTTTAATGTCTTTGTTTTGCAGGTGATTAGGGTTTTTCAATT

GGGGCTTTCAATCAAGGATTTACTTGTACTTGCCTAGATTCTGAATTAGCGGGATGAAAAAAATTATTGC

ATCTGCTTGCTTTGGAAAGAGTCTAGTTTATCAAGTACAGTTGTGTTGGCGAGTACTCGAGAGAGCAATG

ATTTGTACAAGGAAAAGATGTAGAATGGTGTTATAAAGCTCCGTGATCAGCAGGCTGAAGGGTAATTGTG

CTCCTGTGTATGGTCAAATGAATGAACCACCTGGTTCTGATGCTCTGGTTGGATTTACTAAGCTAGCAGT

AGCTGAGCACTTCCGAGATGCTGAGGGGCAAGATGTCCTTTTATTAGTTGACAACGATAAGCTGCTCAAT

CTGGACATCAAANTATTATTTTTCGTATATTTGATTAACAATAAGTCCAGAAATGGATATTATGATTACA

TGTAGAAATAGCAGAATCCTTACTTTTTGCTCAGGTGGGCACCTGAGGATTTACTTGTACTTGCCTAGAT

TCTGAATTAGCGGGATGAAAAAAATTATTGCATCTGCTTGCTTTGGAAAGAGTCTAGTTTATCAAGTACA

GTTGTGTTGGCGAGTACTCGAGAGAGCAATGATTTGTACAAGGAAAAGATGTAGAATGGTGTTATAAAGC

TCCGTGATCAGCAGGCTGAAGGGTAATTGTGCTCCTGTGTATGGTCAAATGAATGAACCACCTGGTTCTG

ATGCTCTGGTTGGATTTACTAAGCTAGCAGTAGCTGAGCACTTCCGAGATGCTGAGGGGCAAGATGTCCT

TTTATTAGTTGACAACGATAAGCTGCTCAATCTGGACATCAAATACAAGCTCAATCTGGGCAGCATCTTT

TTGAATATCCACATTTTGAAT

>MSTRG.3978.2 gene=MSTRG.3978

TTTACTTTTATCCCGAAAACACAATTCACTCCCTCCTACTTTCTCTCTCTCTCTCCTCTCTACTATGTCG

ATCTGACCTGCCTCTCGGCTCTCACCTCTCCCGTCTCTACTCTAACATCTCCACTATGCGTTTTCACATC

TCAGATAACTGAATAATCAGAATAGGGTTCATGGGTTTTCTACTAATTGTTACTATGTTGGACTATTGCC

ATCTAATTTTATTATTTCTACTGATTGGGTTTAATGTCTTTGTTTTGCAGGTGATTAGGGTTTTTCAATT

GGGGCTTTCAATCAAGGATTTACTTGTACTTGCCTAGATTCTGAATTAGCGGGATGAAAAAAATTATTGC

ATCTGCTTGCTTTGGAAAGAGTCTAGTTTATCAAGTACAGTTGTGTTGGCGAGTACTCGAGAGAGCAATG

ATTTGTACAAGGAAAAGATGTAGAATGGTGTTATAAAGCTCCGTGATCAGCAGGCTGAAGGGTAATTGTG

CTCCTGTGTATGGTCAAATGAATGAACCACCTGGTTCTGATGCTCTGGTTGGATTTACTAAGCTAGCAGT

AGCTGAGCACTTCCGAGATGCTGAGGGGCAAGATGTCCTTTTATTAGTTGACAACGATAAGCTGCTCAAT

CTGGACATCAAATACAAGCTCAATCTGGGCAGCATCTTTTTGAATATCCACATTTTGAAT

>MSTRG.3978.3 gene=MSTRG.3978

TTTACTTTTATCCCGAAAACACAATTCACTCCCTCCTACTTTCTCTCTCTCTCTCCTCTCTACTATGTCG

ATCTGACCTGCCTCTCGGCTCTCACCTCTCCCGTCTCTACTCTAACATCTCCACTATGCGTTTTCACATC

TCAGATAACTGAATAATCAGAATAGGGTTCATGGGTTTTCTACTAATTGTTACTATGTTGGACTATTGCC

ATCTAATTTTATTATTTCTACTGATTGGGTTTAATGTCTTTGTTTTGCAGGTGATTAGGGTTTTTCAATT

GGGGCTTTCAATCAAGGATTTACTTGTACTTGCCTAGATTCTGAATTAGCGGGATGAAAAAAATTATTGC

ATCTGCTTGCTTTGGAAAGAGTCTAGTTTATCAAGTACAGTTGTGTTGGCGAGTACTCGAGAGAGCAATG

ATTTGTACAAGGAAAAGATGTAGAATGGTGTTATAAAGCTCCGTGATCAGCAGGCTGAAGGGTAATTGTG

CTCCTGTGTATGGTCAAATGAATGAACCACCTGGTTCTGATGCTCTGGTTGGATTTACTAAGCTAGCAGT

AGCTGAGCACTTCCGAGATGCTGAGGGGCAAGATGTCCTTTTATTAGTTGACAACGATAAGCTGCTCAAT

CTGGACATCAAANTATTATTTTTCGTATATTTGATTAACAATAAGTCCAGAAATGGATATTATGATTACA

TGTAGAAATAGCAGAATCCTTACTTTTTGCTCAGGTGGGCACCTGAGGATTTACTTGTACTTGCCTAGAT

TCTGAATTAGCGGGATGAAAAAAATTATTGCATCTGCTTGCTTTGGAAAGAGTCTAGTTTATCAAGTACA

GTTGTGTTGGCGAGTACTCGAGAGAGCAATGATTTGTACAAGGAAAAGATGTAGAATGGTGTTATAAAGC

TCCGTGATCAGCAGGCTGAAGGGTAATTGTGCTCCTGTGTATGGTCAAATGAATGAACCACCTGGTTCTG

ATGCTCTGGTTGGATTTACTAAGCTAGCAGTAGCTGAGCACTTCCGAGATGCTGAGGGGCAAGATGTCCT

TTTATTAGTTGACAACGATAAGCTGCTCAATCTGGACATCAAATACAAGCTCAATCTGGGCAGCATCTTT

TTGAATATCCACATTTTGAAT

>MSTRG.3978.4 gene=MSTRG.3978

CTTTTATCCCGAAAACACAATTCACTCCCTCCTACTTTCTCTCTCTCTCTCCTCTCTACTATGTCGATCT

GACCTGCCTCTCGGCTCTCACCTCTCCCGTCTCTACTCTAACATCTCCACTATGCGTTTTCACATCTCAG

ATAACTGAATAATCAGAATAGGGTTCATGGGTTTTCTACTAATTGTTACTATGTTGGACTATTGCCATCT

AATTTTATTATTTCTACTGATTGGGTTTAATGTCTTTGTTTTGCAGGTGATTAGGGTTTTTCAATTGGGG

CTTTCAATCAAGGATTTACTTGTACTTGCCTAGATTCTGAATTAGCGGGATGAAAAAAATTATTGCATCT

GCTTGCTTTGGAAAGAGTCTAGTTTATCAAGTACAGTTGTGTTGGCGAGTACTCGAGAGAGCAATGATTT

GTACAAGGAAAAGATGTAGAATGGTGTTATAAAGCTCCGTGATCAGCAGGCTGAAGGGTAATTGTGCTCC

TGTGTATGGTCAAATGAATGAACCACCTGGTTCTGATGCTCTGGTTGGATTTACTAAGCTAGCAGTAGCT

GAGCACTTCCGAGATGCTGAGGGGCAAGATGTCCTTTTATTAGTTGACAACGATAAGCTGCTCAATCTGG

ACATCAAANTATTATTTTTCGTATATTTGATTAACAATAAGTCCAGAAATGGATATTATGATTACATGTA

GAAATAGCAGAATCCTTACTTTTTGCTCAGGTGGGCACCTGAGGTAAAGCCTATTAATTTGAAATAATTT

AATAGCTAAAACTTGTTCTAGTTTTCATGTGGATTTTCATGTGGACATGCGTATGTGTTTTCTCAGGATT

TACTTGTACTTGCCTAGATTCTGAATTAGCGGGATGAAAAAAATTATTGCATCTGCTTGCTTTGGAAAGA

GTCTAGTTTATCAAGTACAGTTGTGTTGGCGAGTACTCGAGAGAGCAATGATTTGTACAAGGAAAAGATG

TAGAATGGTGTTATAAAGCTCCGTGATCAGCAGGCTGAAGGGTAATTGTGCTCCTGTGTATGGTCAAATG

AATGAACCACCTGGTTCTGATGCTCTGGTTGGATTTACTAAGCTAGCAGTAGCTGAGCACTTCCGAGATG

CTGAGGGGCAAGATGTCCTTTTATTAGTTGACAACGATAAGCTGCTCAATCTGGACATCAAATACAAGCT

CAATCTGGGCAGCATCTTTTTGAATATCCAC

>MSTRG.3978.5 gene=MSTRG.3978

CTTTTATCCCGAAAACACAATTCACTCCCTCCTACTTTCTCTCTCTCTCTCCTCTCTACTATGTCGATCT

GACCTGCCTCTCGGCTCTCACCTCTCCCGTCTCTACTCTAACATCTCCACTATGCGTTTTCACATCTCAG

ATAACTGAATAATCAGAATAGGGTTCATGGGTTTTCTACTAATTGTTACTATGTTGGACTATTGCCATCT

AATTTTATTATTTCTACTGATTGGGTTTAATGTCTTTGTTTTGCAGGTGATTAGGGTTTTTCAATTGGGG

CTTTCAATCAAGGATTTACTTGTACTTGCCTAGATTCTGAATTAGCGGGATGAAAAAAATTATTGCATCT

GCTTGCTTTGGAAAGAGTCTAGTTTATCAAGTACAGTTGTGTTGGCGAGTACTCGAGAGAGCAATGATTT

GTACAAGGAAAAGATGTAGAATGGTGTTATAAAGCTCCGTGATCAGCAGGCTGAAGGGATCCATGACTGG

ATTTGTCTTGACCCGCTTGTAGGATTCTGGCAAATAACACCCTCTAATGAGTTCTGAACATATGGACCTC

TCAAGCAGGACTTAACCTCTCATGCCGGCCCCGTGAATCTTGTTGATTTACTTGTACTTGCCTAGATTCT

GAATTAGCGGGATGAAAAAAATTATTGCATCTGCTTGCTTTGGAAAGAGTCTAGTTTATCAAGTACAGTT

GTGTTGGCGAGTACTCGAGAGAGCAATGATTTGTACAAGGAAAAGATGTAGAATGGTGTTATAAAGCTCC

GTGATCAGCAGGCTGAAGGGTAATTGTGCTCCTGTGTATGGTCAAATGAATGAACCACCTGGTTCTGATG

CTCTGGTTGGATTTACTAAGCTAGCAGTAGCTGAGCACTTCCGAGATGCTGAGGGGCAAGATGTCCTTTT

ATTAGTTGACAACGATAAGCTGCTCAATCTGGACATCAAANTATTATTTTTCGTATATTTGATTAACAAT

AAGTCCAGAAATGGATATTATGATTACATGTAGAAATAGCAGAATCCTTACTTTTTGCTCAGGTGGGCAC

CTGAGGATTTACTTGTACTTGCCTAGATTCTGAATTAGCGGGATGAAAAAAATTATTGCATCTGCTTGCT

TTGGAAAGAGTCTAGTTTATCAAGTACAGTTGTGTTGGCGAGTACTCGAGAGAGCAATGATTTGTACAAG

GAAAAGATGTAGAATGGTGTTATAAAGCTCCGTGATCAGCAGGCTGAAGGGTAATTGTGCTCCTGTGTAT

GGTCAAATGAATGAACCACCTGGTTCTGATGCTCTGGTTGGATTTACTAAGCTAGCAGTAGCTGAGCACT

TCCGAGATGCTGAGGGGCAAGATGTCCTTTTATTAGTTGACAACGATAAGCTGCTCAATCTGGACATCAA

ATACAAGCTCAATCTGGGCAGCATCTTTTTGAATATCCAC

>MSTRG.3978.6 gene=MSTRG.3978

CACAATTCACTCCCTCCTACTTTCTCTCTCTCTCTCCTCTCTACTATGTCGATCTGACCTGCCTCTCGGC

TCTCACCTCTCCCGTCTCTACTCTAACATCTCCACTATGCGTTTTCACATCTCAGATAACTGAATAATCA

GAATAGGGTTCATGGGTTTTCTACTAATTGTTACTATGTTGGACTATTGCCATCTAATTTTATTATTTCT

ACTGATTGGGTTTAATGTCTTTGTTTTGCAGGTGATTAGGGTTTTTCAATTGGGGCTTTCAATCAAGGAT

TTACTTGTACTTGCCTAGATTCTGAATTAGCGGGATGAAAAAAATTATTGCATCTGCTTGCTTTGGAAAG

AGTCTAGTTTATCAAGTACAGTTGTGTTGGCGAGTACTCGAGAGAGCAATGATTTGTACAAGGAAAAGAT

GTAGAATGGTGTTATAAAGCTCCGTGATCAGCAGGCTGAAGGGATCCATGACTGGATTTGTCTTGACCCG

CTTGTAGGATTCTGGCAAATAACACCCTCTAATGAGTTCTGAACATATGGACCTCTCAAGCAGGACTTAA

CCTCTCATGCCGGCCCCGTGAATCTTGTTAAATGGATATTATGATTACATGTAGAAATAGCAGAATCCTT

ACTTTTTGCTCAGGTGGGCACCTGAGGTAAAGCCTATTAATTTGAAATAATTTAATAGCTAAAACTTGTT

CTAGTTTTCATGTGGATTTTCATGTGGACATGCGTATGTGTTTTCTCAGGATTTACTTGTACTTGCCTAG

ATTCTGAATTAGCGGGATGAAAAAAATTATTGCATCTGCTTGCTTTGGAAAGAGTCTAGTTTATCAAGTA

CAGTTGTGTTGGCGAGTACTCGAGAGAGCAATGATTTGTACAAGGAAAAGATGTAGAATGGTGTTATAAA

GCTCCGTGATCAGCAGGCTGAAGGGTAATTGTGCTCCTGTGTATGGTCAAATGAATGAACCACCTGGTTC

TGATGCTCTGGTTGGATTTACTAAGCTAGCAGTAGCTGAGCACTTCCGAGATGCTGAGGGGCAAGATGTC

CTTTTATTAGTTGACAACGATAAGCTGCTCAATCTGGACATCAAANTATTATTTTTCGTATATTTGATTA

ACAATAAGTCCAGAAATGGATATTATGATTACATGTAGAAATAGCAGAATCCTTACTTTTTGCTCAGGTG

GGCACCTGAGGTAAAGCCTATTAATTTGAAATAATTTAATAGCTAAAACTTGTTCTAGTTTTCATGTGGA

TTTTCATGTGGACATGCGTATGTGTTTTCTCAGGATTTACTTGTACTTGCCTAGATTCTGAATTAGCGGG

ATGAAAAAAATTATTGCATCTGCTTGCTTTGGAAAGAGTCTAGTTTATCAAGTACAGTTGTGTTGGCGAG

TACTCGAGAGAGCAATGATTTGTACAAGGAAAAGATGTAGAATGGTGTTATAAAGCTCCGTGATCAGCAG

GCTGAAGGGTAATTGTGCTCCTGTGTATGGTCAAATGAATGAACCACCTGGTTCTGATGCTCTGGTTGGA

TTTACTAAGCTAGCAGTAGCTGAGCACTTCCGAGATGCTGAGGGGCAAGATGTCCTTTTATTAGTTGACA

ACGATAAGCTGCTCAATCTGGACATCAAATACAAGCTCAATCTGGGCAGCATCTTTTTGAATATCCACAT

TTTGAATATGTATTAAATAA

>MSTRG.3978.7 gene=MSTRG.3978

CTCCTCTCTACTATGTCGATCTGACCTGCCTCTCGGCTCTCACCTCTCCCGTCTCTACTCTAACATCTCC

ACTATGCGTTTTCACATCTCAGATAACTGAATAATCAGAATAGGGTTCATGGGTTTTCTACTAATTGTTA

CTATGTTGGACTATTGCCATCTAATTTTATTATTTCTACTGATTGGGTTTAATGTCTTTGTTTTGCAGGT

GATTAGGGTTTTTCAATTGGGGCTTTCAATCAAGGATTCTGGCAAATAACACCCTCTAATGAGTTCTGAA

CATATGGACCTCTCAAGCAGGACTTAACCTCTCATGCCGGCCCCGTGAATCTTGTTAAATGGATATTATG

ATTACATGTAGAAATAGCAGAATCCTTACTTTTTGCTCAGGTGGGCACCTGAGGATTTACTTGTACTTGC

CTAGATTCTGAATTAGCGGGATGAAAAAAATTATTGCATCTGCTTGCTTTGGAAAGAGTCTAGTTTATCA

AGTACAGTTGTGTTGGCGAGTACTCGAGAGAGCAATGATTTGTACAAGGAAAAGATGTAGAATGGTGTTA

TAAAGCTCCGTGATCAGCAGGTTTGTAAGTTGTAGATGTAACCGATTGATTTGTGATGGAAATTGCTCTT

TAAAATGGCTTTAAACAATTGTCTGCTTGTTTACCATTCTTAAACAATTGTCATTAACTCTTAGGCTGAA

GGGATCCATGACTGGATTTGTCTTGACCCGCTTGTAGGATTCTGGCAAATAACACCCTCTAATGAGTTCT

GAACATATGGACCTCTCAAGCAGGACTTAACCTCTCATGCCGGCCCCGTGAATCTTGTTGTATGTAACTA

GTTACTCAAATTGCTTCCACATATCTATCAATGATCGCTTCCAACTTAAACTCTAATGATTTTTTATTAT

TTTTCGTATATTTGATTAACAATAAGTCCAGAAATGGATATTATGATTACATGTAGAAATAGCAGAATCC

TTACTTTTTGCTCAGGTGGGCACCTGAGGATTTACTTGTACTTGCCTAGATTCTGAATTAGCGGGATGAA

AAAAATTATTGCATCTGCTTGCTTTGGAAAGAGTCTAGTTTATCAAGTACAGTTGTGTTGGCGAGTACTC

GAGAGAGCAATGATTTGTACAAGGAAAAGATGTAGAATGGTGTTATAAAGCTCCGTGATCAGCAGGCTGA

AGGGTAATTGTGCTCCTGTGTATGGTCAAATGAATGAACCACCTGGTTCTGATGCTCTGGTTGGATTTAC

TAAGCTAGCAGTAGCTGAGCACTTCCGAGATGCTGAGGGGCAAGATGTCCTTTTATTAGTTGACAACGAT

AAGCTGCTCAATCTGGACATCAAANTATTATTTTTCGTATATTTGATTAACAATAAGTCCAGAAATGGAT

ATTATGATTACATGTAGAAATAGCAGAATCCTTACTTTTTGCTCAGGTGGGCACCTGAGGATTTACTTGT

ACTTGCCTAGATTCTGAATTAGCGGGATGAAAAAAATTATTGCATCTGCTTGCTTTGGAAAGAGTCTAGT

TTATCAAGTACAGTTGTGTTGGCGAGTACTCGAGAGAGCAATGATTTGTACAAGGAAAAGATGTAGAATG

GTGTTATAAAGCTCCGTGATCAGCAGGCTGAAGGGTAATTGTGCTCCTGTGTATGGTCAAATGAATGAAC

CACCTGGTTCTGATGCTCTGGTTGGATTTACTAAGCTAGCAGTAGCTGAGCACTTCCGAGATGCTGAGGG

GCAAGATGTCCTTTTATTAGTTGACAACGATAAGCTGCTCAATCTGGACATCAAATACAAGCTCAATCTG

GGCAGCATCTTTTTGAATATCCAC

>MSTRG.3978.10 gene=MSTRG.3978

CTCTCGGCTCTCACCTCTCCCGTCTCTACTCTAACATCTCCACTATGCGTTTTCACATCTCAGATAACTG

AATAATCAGAATAGGGTTCATGGGTTTTCTACTAATTGTTACTATGTTGGACTATTGCCATCTAATTTTA

TTATTTCTACTGATTGGGTTTAATGTCTTTGTTTTGCAGGTGATTAGGGTTTTTCAATTGGGGCTTTCAA

TCAAGGATTCTGGCAAATAACACCCTCTAATGAGTTCTGAACATATGGACCTCTCAAGCAGGACTTAACC

TCTCATGCCGGCCCCGTGAATCTTGTTAAATGGATATTATGATTACATGTAGAAATAGCAGAATCCTTAC

TTTTTGCTCAGGTGGGCACCTGAGGTAAAGCCTATTAATTTGAAATAATTTAATAGCTAAAACTTGTTCT

AGTTTTCATGTGGATTTTCATGTGGACATGCGTATGTGTTTTCTCAGGATTTACTTGTACTTGCCTAGAT

TCTGAATTAGCGGGATGAAAAAAATTATTGCATCTGCTTGCTTTGGAAAGAGTCTAGTTTATCAAGTACA

GTTGTGTTGGCGAGTACTCGAGAGAGCAATGATTTGTACAAGGAAAAGATGTAGAATGGTGTTATAAAGC

TCCGTGATCAGCAGGCTGAAGGGTAATTGTGCTCCTGTGTATGGTCAAATGAATGAACCACCTGGTTCTG

ATGCTCTGGTTGGATTTACTAAGCTAGCAGTAGCTGAGCACTTCCGAGATGCTGAGGGGCAAGATGTCCT

TTTATTAGTTGACAACGATAAGCTGCTCAATCTGGACATCAAATACAAGCTCAATCTGGGCAGCATCTTT

TTGAATATCCAC

>MSTRG.3978.8 gene=MSTRG.3978

CTCTCGGCTCTCACCTCTCCCGTCTCTACTCTAACATCTCCACTATGCGTTTTCACATCTCAGATAACTG

AATAATCAGAATAGGGTTCATGGGTTTTCTACTAATTGTTACTATGTTGGACTATTGCCATCTAATTTTA

TTATTTCTACTGATTGGGTTTAATGTCTTTGTTTTGCAGGTGATTAGGGTTTTTCAATTGGGGCTTTCAA

TCAAGGATTCTGGCAAATAACACCCTCTAATGAGTTCTGAACATATGGACCTCTCAAGCAGGACTTAACC

TCTCATGCCGGCCCCGTGAATCTTGTTGTATGTAACTAGTTACTCAAATTGCTTCCACATATCTATCAAT

GATCGCTTCCAACTTAAACTCTAATGATTTTTTATTATTTTTCGTATATTTGATTAACAATAAGTCCAGA

AATGGATATTATGATTACATGTAGAAATAGCAGAATCCTTACTTTTTGCTCAGGTGGGCACCTGAGGTAA

AGCCTATTAATTTGAAATAATTTAATAGCTAAAACTTGTTCTAGTTTTCATGTGGATTTTCATGTGGACA

TGCGTATGTGTTTTCTCAGGATTTACTTGTACTTGCCTAGATTCTGAATTAGCGGGATGAAAAAAATTAT

TGCATCTGCTTGCTTTGGAAAGAGTCTAGTTTATCAAGTACAGTTGTGTTGGCGAGTACTCGAGAGAGCA

ATGATTTGTACAAGGAAAAGATGTAGAATGGTGTTATAAAGCTCCGTGATCAGCAGGCTGAAGGGTAATT

GTGCTCCTGTGTATGGTCAAATGAATGAACCACCTGGTTCTGATGCTCTGGTTGGATTTACTAAGCTAGC

AGTAGCTGAGCACTTCCGAGATGCTGAGGGGCAAGATGTCCTTTTATTAGTTGACAACGATAAGCTGCTC

AATCTGGACATCAAANTATTATTTTTCGTATATTTGATTAACAATAAGTCCAGAAATGGATATTATGATT

ACATGTAGAAATAGCAGAATCCTTACTTTTTGCTCAGGTGGGCACCTGAGGTAAAGCCTATTAATTTGAA

ATAATTTAATAGCTAAAACTTGTTCTAGTTTTCATGTGGATTTTCATGTGGACATGCGTATGTGTTTTCT

CAGGATTTACTTGTACTTGCCTAGATTCTGAATTAGCGGGATGAAAAAAATTATTGCATCTGCTTGCTTT

GGAAAGAGTCTAGTTTATCAAGTACAGTTGTGTTGGCGAGTACTCGAGAGAGCAATGATTTGTACAAGGA

AAAGATGTAGAATGGTGTTATAAAGCTCCGTGATCAGCAGGCTGAAGGGTAATTGTGCTCCTGTGTATGG

TCAAATGAATGAACCACCTGGTTCTGATGCTCTGGTTGGATTTACTAAGCTAGCAGTAGCTGAGCACTTC

CGAGATGCTGAGGGGCAAGATGTCCTTTTATTAGTTGACAACGATAAGCTGCTCAATCTGGACATCAAAT

ACAAGCTCAATCTGGGCAGCATCTTTTTGAATATCCAC

>MSTRG.3978.9 gene=MSTRG.3978

CTCTCGGCTCTCACCTCTCCCGTCTCTACTCTAACATCTCCACTATGCGTTTTCACATCTCAGATAACTG

AATAATCAGAATAGGGTTCATGGGTTTTCTACTAATTGTTACTATGTTGGACTATTGCCATCTAATTTTA

TTATTTCTACTGATTGGGTTTAATGTCTTTGTTTTGCAGGTGATTAGGGTTTTTCAATTGGGGCTTTCAA

TCAAGGATTTACTTGTACTTGCCTAGATTCTGAATTAGCGGGATGAAAAAAATTATTGCATCTGCTTGCT

TTGGAAAGAGTCTAGTTTATCAAGTACAGTTGTGTTGGCGAGTACTCGAGAGAGCAATGATTTGTACAAG

GAAAAGATGTAGAATGGTGTTATAAAGCTCCGTGATCAGCAGGCTGAAGGGTAATTGTGCTCCTGTGTAT

GGTCAAATGAATGAACCACCTGGTTCTGATGCTCTGGTTGGATTTACTAAGCTAGCAGTAGCTGAGCACT

TCCGAGATGCTGAGGGGCAAGATGTCCTTTTATTAGTTGACAACGATAAGCTGCTCAATCTGGACATCAA

ATACAAGCTCAATCTGGGCAGCATCTTTTTGAATATCCAC

>MSTRG.3978.11 gene=MSTRG.3978

GTATTCTTTGTGGATTATTTAGATTTTGTTTTATGTGGAGAGAGAGACGGGAACAGAGAGGGAAAGAGGG

TGTGGGTTATAGCTAGGGAGAGAACTGAGAGATGGAGAGAAGAAAGAATAGATAAATTGAGAGATGGACA

AAGGAAGAAAGAGTTACAGAGATAGAGAGAGGGGAAGATAGAAAAAGAGTTAATTGGGTCTGAATTTGTT

GTTGAAAACAAATCTCTTAATTTAAGATTTGTTGAAAAAGTAATAGGCTTAAGGCTGCTGAGATAGCATT

CTAGGTTTCACTACATTCTTACATTCTAGGTAGCCACATATATGATTATCTAAGTTTTAATTTATTCATG

AGTGGATGACAAGTACCAATACATGTGTGAAAAAAAACACAAGATCCATGACTGGATTTGTCTTGACCCG

CTTGTAGGATTCTGGCAAATAACACCCTCTAATGAGTTCTGAACATATGGACCTCTCAAGCAGGACTTAA

CCTCTCATGCCGGCCCCGTGAATCTTGTTAAATGGATATTATGATTACATGTAGAAATAGCAGAATCCTT

ACTTTTTGCTCAGGTGGGCACCTGAGGTAAAGCCTATTAATTTGAAATAATTTAATAGCTAAAACTTGTT

CTAGTTTTCATGTGGACATGCGTATGTGTTTTCTCAGGATTTACTTGTACTTGCCTAGATTCTGAATTAG

CGGGATGAAAAAAATTATTGCATCTGCTTGCTTTGGAAAGAGTCTAGTTTATCAAGTACAGTTGTGTTGG

CGAGTACTCGAGAGAGCAATGATTTGTACAAGGAAAAGATGTAGAATGGTGTTATAAAGCTCCGTGATCA

GCAGGCTGAAGGGTAATTGTGCTCCTGTGTATGGTCAAATGAATGAACCACCTGGTTCTGATGCTCTGGT

TGGATTTACTAAGCTAGCAGTAGCTGAGCACTTCCGAGATGCTGAGGGGCAAGATGTCCTTTTATTAGTT

GACAACGATAAGCTGCTCAATCTGGACATCAAATACAAGCTCAATCTGGGCAGCATCTTTTTGAATATCC

AC

>MSTRG.3978.12 gene=MSTRG.3978

AGGATTCTGGCAAATAACACCCTCTAATGAGTTCTGAACATATGGACCTCTCAAGCAGGACTTAACCTCT

CATGCCGGCCCCGTGAATCTTGTTAAATGGATATTATGATTACATGTAGAAATAGCAGAATCCTTACTTT

TTGCTCAGGTGGGCACCTGAGGTAAAGCCTATTAATTTGAAATAATTTAATAGCTAAAACTTGTTCTAGT

TTTCATGTGGATTTTCATGTGGACATGCGTATGTGTTTTCTCAGGATTTACTTGTACTTGCCTAGATTCT

GAATTAGCGGGATGAAAAAAATTATTGCATCTGCTTGCTTTGGAAAGAGTCTAGTTTATCAAGTACAGTT

GTGTTGGCGAGTACTCGAGAGAGCAATGATTTGTACAAGGAAAAGATGTAGAATGGTGTTATAAAGCTCC

GTGATCAGCAGGCTGAAGGGTAATTGTGCTCCTGTGTATGGTCAAATGAATGAACCACCTGGTTCTGATG

CTCTGGTTGGATTTACTAAGCTAGCAGTAGCTGAGCACTTCCGAGATGCTGAGGGGCAAGATGTCCTTTT

ATTAGTTGACAACGATAAGCTGCTCAATCTGGACATCAAATACAAGCTCAATCTGGGCAGCATCTTTTTG

AATATCCAC

>MSTRG.3979.1 gene=MSTRG.3979

GGCTGCTGCATCCAAGGGAGTCCATTATTCAGCGGTTCCCGCCTTGCATTTTCTTCTTCGTTTGGTTGTT

CGCTGCCGTCTTTGTCTGCGCCATTGTTTGCACCAGAAGACTCTCCTTCAGGAAATCTTCAAGGTTGGGT

TATTTGTTCCTCTACTAAATCTGTGTCAAAATGCTGAGTTCGATGTAAAGAAAGAGGCAGTATGTGCTAT

TTCAAATGCAACTTCTGGTGGTAGCGAAGCTCAAATCAGGTACCTGGTGCGTCAGAATTGCATCAAGCCT

TTGTGTGATTTACTAGACTGCCCTGAATCTAGGATTGTTACTGCGTGTTTGGAAGGACTGGCGAATATCT

TAGAGGTAGGAGAAGCTGAAAAGAATTTGAAGAGGAACGTTAACTATTATGCTCAGCTAATCGATGAGGC

TAAGGGACTCGGGAAAATAAAGAACTTGCAGAATTACGACAACAATGAAATTCGCGAGAAGGCTGTTGAG

ATTGTTGAGACGTACTGGTTACGATACAGTGATGGGACTTGGAGTAGGTTCAGCTTCTCGAAAGCTGGTG

GCCACATACTTCAGAGCACACTTCAGAGCAAATTAAAATCTCTGATGGTAGATGATCTGGTGATGATAAA

TGAGTGAATTCTAGGATGTGGTGTTCATTTGTTTCAGTACTTTGAATTTGAAATGTGCATTCTGTTGTTT

TGTCAGGATTGTGAGGATTGTCTGCATTCATGTGAAAGCTGATGCCTAAAGTTTATGCTCCCGAAAATAT

ATATTTAATGTTCGGTTTCTTTTATTATGGCTCTTCTTTTGCCAAAATTAAGTTAGTACAAAACAAAAGT

TATTGAAGG

>MSTRG.3980.1 gene=MSTRG.3980

CTTGTAGTTAGTATATTTACTTAGGGGTTAAATATCTAACCACAAGGTGATTCTTTACAAAATTAGTGTT

AATAATGGCCTAGGTGCAAATGGGTTGGTAAAGAAAATTCTGATGAGTGGGCATGTTGTATTTGAGGAGA

GGAATTGTTCACAAGAGTCAGAGTATATGAGAGACCTAGAAGAGCTGGTCGGTAATGACAAGGTAAGGTT

CCCAATGGTGATTGTGAATGGGAAGGACTTGTGCGGAGCAGAAGAGGTGGAGGGTTTAGATTATTTTGAG

AACAAGCAAAAGCTTATATCAGTACTGAAACATCTCTATCTTTCTGCAAGAACATTAGAAAACCTTCATG

ACATGTGGATCAATGGGAG

>MSTRG.3981.1 gene=MSTRG.3981

AAAGAAAAGAGAGAAAGTTAGGCAAACAGGATAGTAAATCAGTGAGCTAATTTGGAACGGAGAAGTTTTG

GACGTTTTAGAGCACTATAATAAAAGGAGAGGAAAGGCTGGTCGGGACAAAAAGGGAGGCGGAGCTTCCG

TCGGCACTGCCAACGGAGCTTCGCGACGGGGCTGGAAAAGGGAAGAGGGATGATGGGAAGTGTGTGGTGG

TTCAGAAGGAGAAACGGTGGACTTGTCGCCGTGACAGAGCTCAGAGAGAGAGAGAGGGGGGGAGGGGACG

GCGGCGATAGGTCGCCGGAGCCGTGCTCCGACAAGCCAAGGAGCAGGTGAGGGAGATGAGGGAGAAAGAT

GGAGGTGGTAGGAGGGCAGTTTGGTCATTTTATAATTGAGAATATGAAGAAATAATAAAAAAGGGAATTG

AGAGTGTTGGCTTTTAAAGATAAAATAAATAAGAATTCAAGATTTGTTTGAAATAAAGTTTGGACAATTT

ATGGTGGTTATTGTAAGGTTGTGGTCTAGAAAATTATTATTATAGTGCCGTTAAATTAATAATAATTTTG

GATATAAATAAAGTTTGGACAATTTATGGTGGTTATTGTAAGGTTGTGGTCTAGAAAATTATTATTATAG

TGCCGTTAAATTAATAATAATTTTGGATATAAATAAATATGTGACTTAATTTGAGAAATGTTTTAGTTAA

AGACTTTGA

>MSTRG.3981.3 gene=MSTRG.3981

AAAGAAAAGAGAGAAAGTTAGGCAAACAGGATAGTAAATCAGTGAGCTAATTTGGAACGGAGAAGTTTTG

GACGTTTTAGAGCACTATAATAAAAGGAGAGGAAAGGCTGGTCGGGACAAAAAGGGAGGCGGAGCTTCCG

TCGGCACTGCCAACGGAGCTTCGCGACGGGGCTGGAAAAGGGAAGAGGGATGATGGGAAGTGTGTGGTGG

TTCAGAAGGAGAAACGGTGGACTTGTCGCCGTGACAGAGCTCAGAGAGAGAGAGAGGGGGGGAGGGGACG

GCGGCGATAGGTCGCCGGAGCCGTGCTCCGACAAGCCAAGGAGCAGGTGAGGGAGATGAGGGAGAAAGAT

GGAGGTGGTAGGAGGGCAGTTTGGTCATTTTATAATTGAGAATATGAAGAAATAATAAAAAAGGGAATTG

AGAGTGTTGGCTTTTAAAGATAAAATAAATAAGAATTCAAGATTTGTTTGAAATAAAGTTTGGACAATTT

ATGGTGGTTATTGTAAGGTTGTGGTCTAGAAAATTATTATTATAGTGCCGTTAAATTAATAATAATTTTG

GATATAAATAAATATGTGACTTAATTTGAGAAATGTTTTAGTTAAAG

>MSTRG.3981.2 gene=MSTRG.3981

AGAAAAGAGAGAAAGTTAGGCAAACAGGATAGTAAATCAGTGAGCTAATTTGGAACGGAGAAGTTTTGGA

CGTTTTAGAGCACTATAATAAAAGGAGAGGAAAGGCTGGTCGGGACAAAAAGGGAGGCGGAGCTTCCGTC

GGCACTGCCAACGGAGCTTCGCGACGGGGCTGGAAAAGGGAAGAGGGATGATGGGAAGTGTGTGGTGGTT

CAGAAGGAGAAACGGTGGACTTGTCGCCGTGACAGAGCTCAGAGAGAGAGAGAGGGGGGGAGGGGACGGC

GGCGATAGGTCGCCGGAGCCGTGCTCCGACAAGCCAAGGAGCAGGTGAGGGAGATGAGGGAGAAAGATGG

AGGTGGTAGGAGGGCAGTTTGGTCATTTTATAATTGAGAATATGAAGAAATAATAAAAAAGGGAATTGAG

AGTGTTGGCTTTTAAAGATAAAATAAATAAGAATTCAAGATTTGTTTGAAATAAAGTTTGGACAATTTAT

GGTGGTTATTGTAAGGTTGTGGTCTAGAAAATTATTATTATAGTGCCGTTAAATTAATAATAATTTTGGA

TATAAATAAATATGTGACTTAATTTGAGAAATGTTTTAGTTAAAG

>MSTRG.3982.1 gene=MSTRG.3982

GTAGGCAACAAAGCACTTTCATCCAGAATACTATGAAATTGTTTTGGGAAACACTGGCTGCCAAGTTCCC

AGAATGCCGGCTCCCAGAGCCTGAAAGATCAGTTTTACGAGGCAGTTCTTCATCCAACATCAAGGTAATT

CTTTACACAACGAGTGCTAACAATGACCTAGACGCAAATGGGTGGGCAAAGAAGGTTCTGTTGAGTGGAA

AAGTTGTATTTGAGGAGAGAAACTGTTCAAAGGAGCCTGTATATTTGAAAGAGC

>MSTRG.3984.1 gene=MSTRG.3984

GTTCCTTACCCACAAAGACTCATTCGCCCTAAACAATCGGCCCAAATGGAACAAATTCTAGAGGTTTTCA

AACAAGTCAAGATAAACATTCCTCTTTTAGATGCCATTCAACAAGTCCCTTCTTATGCCAAATGTCTAAA

AGATTTGTGTACCCATAAGAGAACCACTCATGTTCCAAAGAAAGCCTTTTTGACCTCTCAAGTTAGCTCG

ATTCTCTCGAATCAAATCCCCGTGAAATATAAGGACCCCGGTTGTCCTACAATTTCTTGTGTCATAGGCG

ATAAATTTGTGGATAAAGCTTTACTTGATTTAGGAGCGAGTGTGAATCTTCTTCCTTTCTCGGTCTACCA

AGCCCTAGGTTTAGGGGATCTCAAGACCACCAATATGACCCTTCAACTTGCCGATCG

>MSTRG.3986.2 gene=MSTRG.3986

CTAGTTTAGATTGGTACTAAACACACGCACGCCAGAGACTCCGATTGAAATATTTACGGTCATACACTTG

GTTCTTTCGATCCCCAAAATCCCAATCTACTTCCCCTTTGCTCGTCTCTCTCCCCCAAGCTTTCATCTCC

CTGATGTGAACTCCAGCGTAATTGCATCATTGAAAAAGGTTCTGGCACATGTGAGAGTGAGAATAAGTCA

AGTCTTACAAACTACAAAGTTGTGCTTCATAATAGAATCTACTTTTGCAACCACTGCCCTAGAATTGTCC

CTCATCCTGACACAAAAATTCGAAATTGTTGTTGCAGCTTCAGATCCAACAGGTGAAATACAAATTATTT

TGGAGGAAACTGAGGTTCGTACTTTGATTGGAAAAAGAGCACGATAACTACTTGCTGAGGTAACTTTATT

AGTGGAACCATATTGAAAGAGACATCATACAATATACCTATTTCAAAGAGATATATTTGTCTTATTCGTC

AAAAAGGTGAGCAGTAATTGCTCAGAGTATGCAAATAAAACAGTCTCCTTCCAAAGCCGCAAGATGCTGA

ATGGTGGAGAGATGCTGACATTCAAGTGGTCCCAAGCATGCTCATTATTGATAGTACGTTTCTCAAAGAA

GTTAATTTTAAGGTTTTTAGCTTGCGGGATAGTTACCCTTATCTTTACGGTTGAAGAAGTAATCATTCTT

CAACAGACTTTCATCGAAGAAATGGAAATTCTTCAGCCATTCATCCTCACGAGTGAAGCCCTCCTCAGAC

AGCAGCACATCTTTGTAATAGAATTGACAGTCTACATCATGCTGTTACAATAATACAATACCACATCTGT

GAGAATTGAGGACAGCAGATTCAAGGAAATTTTAAATA

>MSTRG.3986.1 gene=MSTRG.3986

CTAGTTTAGATTGGTACTAAACACACGCACGCCAGAGACTCCGATTGAAATATTTACGGTCATACACTTG

GTTCTTTCGATCCCCAAAATCCCAATCTACTTCCCCTTTGCTCGTCTCTCTCCCCCAAGCTTTCATCTCC

CTGATGTGAACTCCAGCGTAATTGCATCATTGATTAATTGATTATTGCAACGAACGCTTCCTCGGAGTTC

TGAGCCTTGATGATTTTGATCTGAAGAACCATTATGTGTGTAAAAGAAAAAGGTTCTGGCACATGTGAGA

GTGAGAATAAGTCAAGTCTTACAAACTACAAAGTTGTGCTTCATAATAGAATCTACTTTTGCAACCACTG

CCCTAGAATTGTCCCTCATCCTGACACAAAAATTCGAAATTGTTGTTGCAGCTTCAGATCCAACAGGTGA

AATACAAATTATTTTGGAGGAAACTGAGGTTCGTACTTTGATTGGAAAAAGAGCACGATAACTACTTGCT

GAGGTAACTTTATTAGTGGAACCATATTGAAAGAGACATCATACAATATACCTATTTCAAAGAGATATAT

TTGTCTTATTCGTCAAAAAGGTGAGCAGTAATTGCTCAGAGTATGCAAATAAAACAGTCTCCTTCCAAAG

CCGCAAGATGCTGAATGGTGGAGAGATGCTGACATTCAAGTGGTCCCAAGCATGCTCATTATTGATAGTA

CGTTTCTCAAAGAAGTTAATTTTAAGGTTTTTAGCTTGCGGGATAGTTACCCTTATCTTTACGGTTGAAG

AAGTAATCATTCTTCAACAGACTTTCATCGAAGAAATGGAAATTCTTCAGCCATTCATCCTCACGAGTGA

AGCCCTCCTCAGACAGCAGCACATCTTTGTAATAGAATTGACAGTCTACATCATGCTGTTACAATAATAC

AATACCACATCTGTGAGAATTGAGGACAGCAGATTCAAGGAAATTTTAAATAGAGTATACAATAAACATC

ACAGTGGC

>MSTRG.3986.3 gene=MSTRG.3986

GTTTAGATTGGTACTAAACACACGCACGCCAGAGACTCCGATTGAAATATTTACGGTCATACACTTGGTT

CTTTCGATCCCCAAAATCCCAATCTACTTCCCCTTTGCTCGTCTCTCTCCCCCAAGCTTTCATCTCCCTG

ATGTGAACTCCAGCGTAATTGCATCATTGATTAATTGATTATTGCAACGAACGCTTCCTCGGAGTTCTGA

GCCTTGATGATTTTGATCTGAAGAACCATTATGTGTGTAAAAGAAAAAGGTTCTGGCACATGTGAGAGTG

AGAATAAGTCAAGTCTTACAAACTACAAAGTTGTGCTTCATAATAGAATCTACTTTTGCAACCACTGCCC

TAGAATTGTCCCTCATCCTGACACAAAAATTCGAAATTGTTGTTGCAGCTTCAGATCCAACAGGTGAAAT

ACAAATTATTTTGGAGGAAACTGAGGTTCGTACTTTGATTGGAAAAAGAGCACGATAACTACTTGCTGAG

GTCAGTTTCTTATTATGATTTCTTAACTTTTTTACTTTCTCAAGTTATCAGTAACCAATTAGCTGAAACA

TGAACAAATATTACCCGAGTATTTTTAATATACTTGCAGACAAAGATTACACAATCATGCTAAAAAATAT

ATGAAAGCAACTCATTTTTCATCCTCATCTTAGCATGTAAAAATAAATATTTGAACTTTTACATGTCTTT

CTTATAATGCTAAACCCCAGCATGGGCGGGCATAAATATGTATGTATGTATGCATGCATGCATGTATATA

ACGTACTTAATGTTTTCATTCATGGACAGCAAGTGGGAAGATCTATATGAATAGAGTAAATGTGGCAATT

TTGAAACATTTGGATGGTATGATCCTTGATTTACATTGACCTGTTTAGTATTGCATCTCTAGCAACTTTC

CATGCTCATTGGAATTCTAAAGAAAGGAAAGGAACTTCACAATCATACCGATCATCAGCACATCCTCCTT

TGAGTTGCTCCATTCAGGTAACTTTATTAGTGGAACCATATTGAAAGAGACATCATACAATATACCTATT

TCAAAGAGATATATTTGTCTTATTCGTCAAAAAGGTGAGCAGTAATTGCTCAGAGTATGCAAATAAAACA

GTCTCCTTCCAAAGCCGCAAGATGCTGAATGGTGGAGAGATGCTGACATTCAAGTGGTCCCAAGCATGCT

CATTATTGATAGTACGTTTCTCAAAGAAGTTAATTTTAAGGTTTTTAGCTTGCGGGATAGTTACCCTTAT

CTTTACGGTTGAAGAAGTAATCATTCTTCAACAGACTTTCATCGAAGAAATGGAAATTCTTCAGCCATTC

ATCCTCACGAGTGAAGCCCTCCTCAGACAGCAGCACATCTTTGTAATAGAATTGACAGTCTACATCATGC

TGTTACAATAATACAATACCACATCTGTGAGAATTGAGGACAGCAGATTCAAGGAAATTTTAAATAGAGT

ATACAATAAACATCACAGTGGC

>MSTRG.3986.5 gene=MSTRG.3986

CTAGTTTAGATTGGTACTAAACACACGCACGCCAGAGACTCCGATTGAAATATTTACGGTCATACACTTG

GTTCTTTCGATCCCCAAAATCCCAATCTACTTCCCCTTTGCTCGTCTCTCTCCCCCAAGCTTTCATCTCC

CTGATGTGAACTCCAGCGTAATTGCATCATTGAAAAAGGTTCTGGCACATGTGAGAGTGAGAATAAGTCA

AGTCTTACAAACTACAAAGTTGTGCTTCATAATAGAATCTACTTTTGCAACCACTGCCCTAGAATTGTCC

CTCATCCTGACACAAAAATTCGAAATTGTTGTTGCAGCTTCAGATCCAACAGGTGAAATACAAATTATTT

TGGAGGAAACTGAGGTTCGTACTTTGATTGGAAAAAGAGCACGATAACTACTTGCTGAGCAACTTTCCAT

GCTCATTGGAATTCTAAAGAAAGGAAAGGAACTTCACAATCATACCGATCATCAGCACATCCTCCTTTGA

GTTGCTCCATTCAGGTAACTTTATTAGTGGAACCATATTGAAAGAGACATCATACAATATACCTATTTCA

AAGAGATATATTTGTCTTATTCGTCAAAAAGGTGAGCAGTAATTGCTCAGAGTATGCAAATAAAACAGTC

TCCTTCCAAAGCCGCAAGATGCTGAATGGTGGAGAGATGCTGACATTCAAGTGGTCCCAAGCATGCTCAT

TATTGATAGTACGTTTCTCAAAGAAGTTAATTTTAAGGTTTTTAGCTTGCGGGATAGTTACCCTTATCTT

TACGGTTGAAGAAGTAATCATTCTTCAACAGACTTTCATCGAAGAAATGGAAATTCTTCAGCCATTCATC

CTCACGAGTGAAGCCCTCCTCAGACAGCAGCACATCTTTGTAATAGAATTGACAGTCTACATCATGCTGT

TACAATAATACAATACCACATCTGTGAGAATTGAGGACAGCAGATTCAAGGAAATTTTAAATA

>MSTRG.3986.4 gene=MSTRG.3986

CTAGTTTAGATTGGTACTAAACACACGCACGCCAGAGACTCCGATTGAAATATTTACGGTCATACACTTG

GTTCTTTCGATCCCCAAAATCCCAATCTACTTCCCCTTTGCTCGTCTCTCTCCCCCAAGCTTTCATCTCC

CTGATGTGAACTCCAGCGTAATTGCATCATTGATTAATTGATTATTGCAACGAACGCTTCCTCGGAGTTC

TGAGCCTTGATGATTTTGATCTGAAGAACCATTATGTGTGTAAAAGAAAAAGGTTCTGGCACATGTGAGA

GTGAGAATAAGTCAAGTCTTACAAACTACAAAGTTGTGCTTCATAATAGAATCTACTTTTGCAACCACTG

CCCTAGAATTGTCCCTCATCCTGACACAAAAATTCGAAATTGTTGTTGCAGCTTCAGATCCAACAGGTGA

AATACAAATTATTTTGGAGGAAACTGAGGTTCGTACTTTGATTGGAAAAAGAGCACGATAACTACTTGCT

GAGCAAGTGGGAAGATCTATATGAATAGAGTAAATGTGGCAATTTTGAAACATTTGGATGGTATGATCCT

TGATTTACATTGACCTGTTTAGTATTGCATCTCTAGCAACTTTCCATGCTCATTGGAATTCTAAAGAAAG

GAAAGGAACTTCACAATCATACCGATCATCAGCACATCCTCCTTTGAGTTGCTCCATTCAGGTAACTTTA

TTAGTGGAACCATATTGAAAGAGACATCATACAATATACCTATTTCAAAGAGATATATTTGTCTTATTCG

TCAAAAAGGTGAGCAGTAATTGCTCAGAGTATGCAAATAAAACAGTCTCCTTCCAAAGCCGCAAGATGCT

GAATGGTGGAGAGATGCTGACATTCAAGTGGTCCCAAGCATGCTCATTATTGATAGTACGTTTCTCAAAG

AAGTTAATTTTAAGGTTTTTAGCTTGCGGGATAGTTACCCTTATCTTTACGGTTGAAGAAGTAATCATTC

TTCAACAGACTTTCATCGAAGAAATGGAAATTCTTCAGCCATTCATCCTCACGAGTGAAGCCCTCCTCAG

ACAGCAGCACATCTTTGTAATAGAATTGACAGTCTACATCATGCTGTTACAATAATACAATACCACATCT

GTGAGAATTGAGGACAGCAGATTCAAGGAAATTTTAAATAGAGTATACAATAAACATCACAGTGGC

>MSTRG.3986.6 gene=MSTRG.3986

GTTTAGATTGGTACTAAACACACGCACGCCAGAGACTCCGATTGAAATATTTACGGTCATACACTTGGTT

CTTTCGATCCCCAAAATCCCAATCTACTTCCCCTTTGCTCGTCTCTCTCCCCCAAGCTTTCATCTCCCTG

ATGTGAACTCCAGCGTAATTGCATCATTGATTAATTGATTATTGCAACGAACGCTTCCTCGGAGTTCTGA

GCCTTGATGATTTTGATCTGAAGAACCATTATGTGTGTAAAAGAAAAAGGTTCTGGCACATGTGAGAGTG

AGAATAAGTCAAGTCTTACAAACTACAAAGTTGTGCTTCATAATAGAATCTACTTTTGCAACCACTGCCC

TAGAATTGTCCCTCATCCTGACACAAAAATTCGAAATTGTTGTTGCAGCTTCAGATCCAACAGGTGAAAT

ACAAATTATTTTGGAGGAAACTGAGGTTCGTACTTTGATTGGAAAAAGAGCACGATAACTACTTGCTGAG

CAACTTTCCATGCTCATTGGAATTCTAAAGAAAGGAAAGGAACTTCACAATCATACCGATCATCAGCACA

TCCTCCTTTGAGTTGCTCCATTCAGGTAACTTTATTAGTGGAACCATATTGAAAGAGACATCATACAATA

TACCTATTTCAAAGAGATATATTTGTCTTATTCGTCAAAAAGGTGAGCAGTAATTGCTCAGAGTATGCAA

ATAAAACAGTCTCCTTCCAAAGCCGCAAGATGCTGAATGGTGGAGAGATGCTGACATTCAAGTGGTCCCA

AGCATGCTCATTATTGATAGTACGTTTCTCAAAGAAGTTAATTTTAAGGTTTTTAGCTTGCGGGATAGTT

ACCCTTATCTTTACGGTTGAAGAAGTAATCATTCTTCAACAGACTTTCATCGAAGAAATGGAAATTCTTC

AGCCATTCATCCTCACGAGTGAAGCCCTCCTCAGACAGCAGCACATCTTTGTAATAGAATTGACAGTCTA

CATCATGCTGTTACAATAATACAATACCACATCTGTGAGAATTGAGGACAGCAGATTCAAGGAAATTTTA

AATAGAGTATACAATAAACATCACAGTGGC

>MSTRG.3987.1 gene=MSTRG.3987

CAAATTTTCAGCCAAGTTTGAAATTTTCACACATACACTATACACGCTCGAGTCTCCTTGAATACCCTCA

ACACTCACTACACTCTCTCCCTCGCTCAACTCTCTCTCCCTTCTCTACTCTCTCTCTCTTGTTAATCTCT

CTGAAGCTAATCTCTCTCTGGCCACCATTTTCACTCAATCTCGCCATCACTTTTAATTAACTTGGATTTG

CATTTATAGTTGGAGCACAGTTGATTTGTGGGTCGAGGTTCCAATTCTTCAATTAGGGTTTTGGCTTTTC

AATCAGGGTGTTCTTAGTTTGGGTTTGCACTCCAAGACTTTATTATAAGTACAACACAGTTTTGTATTTG

GTGTGAGTGAGCAGTAAAGAATTCTATGGAAGATGCTAGTGACTACCGGTGGTATATAACCAGGAGAAGC

TTAAGCCGTTAATGGATCAACTTAACAAAGTCAAAGACTTGACTGTTCCAGAATTTGGAAATTTCCGAAC

TTGAGAAGTGCGAGCAAATGCAAGAATATGTTTATATATAATAGTGTAGTGTAGTGGGATGTTTGGTTGG

ATTCGTTTAATACTTTGATTTGGTTGGATTTGTTAGGTAGTTTTGTTTGGTTGGATTTGTTATAAACTTT

GATTTGTATGAAAGAATAGTTTAATGTTTCAGTGGATATAGTGAAATATTGCAAAGTTGTTAATTTA

>MSTRG.3988.1 gene=MSTRG.3988

TTTAGTGACTAGATTCTAGCCGAACTCGAGCCTATCTAACTGTTTACGAGCCGAATTTCGAATTTGAAAT

TGAAAGGCCGTCGAACTCGAATCCAAGCCCCGAACTTTTCGATTGAGCTCGAGCCGGAGCAGTCCGGCAG

TATTCGACTCGGCATTACACCCCTACAGTGGTCTAACAGAGAACCATCGATCCGATGGCCAATATCAAGA

GATATATTGTTGAAGTTGAGCCGGGCAAGCCGGCGATGGGACCGGTTTACCGGAGCACCTTCGGCGTCGA

GCCTCATATTCCTGGAATGGAGAGCTGCTGGGACATTTTTCG

>MSTRG.3993.1 gene=MSTRG.3993

GGTGGTTAGTAGTAATTATGTTTTGTTCGTAAGGTTGAACCATATATCTTTTCTGTCTGAAACTTCTTTG

TGCAATCTGTATGGAGCTGTACTATTAGGATGCAAATTGTTTGTTTTGCTCATTGACTTTATCAATCTTT

GTGAATGCAATATACACAAAGAAGTTGATAGGTGGTTGCTGAGGAAACTGGTTTCACCAGGGATGAGATG

ATTAGCAGCTAGGGATTCAAATTTTTTGAATACTCATGAAGTTGGTGGGCATGAAAAGTTGCAAAGTGCA

TGTGGGAGTTGTTCTG

>MSTRG.3994.1 gene=MSTRG.3994

TATTAGTAAAAATAAGTTCTCAGAATTTTTTCAGCTTAGTTTATTAAAAATTGTTTTTAGCAGGTTTTAG

CTACTTGAGTCCGCTTACAAGACTTCTCCAGCAAAGATCAGTTGGTAGCAGCAACGTCAATTGTTTCCCG

CTACCAGCATCTAGGATAAATCGGAATGATAGTGGTTGCAAGGTGACACCTGATAGGCATCCCATGAATT

CCTTAAATCGGAATGATAGTGGTTACATGGAAAGGAATTGATTTGGTTAGTACTCTTAATGTGTTCATGT

GAAGGATTGCAATGTTCTCTTGCAAAGATTTCAGTTTGAAATTCAAAATTTAAAGAATCGAAGTCCTAAT

CATCTTCTTTGTTGGTATGTATTCAGTTTTTTCTCCATTTTCATATAATTTCTTCTCCATTCTTATTTGA

TTAATTGTGCAAGGATCCACGGGTTCCTGATTAC

>MSTRG.3996.1 gene=MSTRG.3996

GCTGCTGTATTTTGCTGTTTTGAGCTTTGTCTCTGCGTTTAGTCTAGCTTGTCGAGTGGCTGGGTCATTC

TGTTTTGTCGAGTCTCAACACTGAGTCACAGCCTTCGTTTATTTAACTATTAGATTCTGTTTTTGTGATT

AGCAATTTGGTGGATTCATTTGACTTGGTAATTCATTTTTGCCGTGTGTAAGTTAGGAAGGACTCCAGTG

AACCATTGTCTAGTTTA

>MSTRG.3999.1 gene=MSTRG.3999

CTTTGCTTTGCGCTTATAAACTATATTGTAAAATAAAATTCTACTTATCTTGCACAAATCGCGTTCATCC

GTGTATGCATCAGGTAGTATTCGTCAAAGTTCTCTTGCGACGATGCTTCGACAAAGGATGGCGTCTCGAA

ATCAAAAAGAGAATAACACCATTGGCCATGCAATAGGTATCATAATTAGTACATGATTTTGTATTGAACG

TGTTTGATAATTTACTACAGAGAAAAGCTGAGAGATCGGTCCTCACGATTTCTATTGATTTCATGCTCCA

GGAATTTGAGAAAAGCCACCGGGTGAAAGTTTGGAAAGTTAGAGAGGCTTTTATGAAAGTATACGACCTA

TATATATGGGATTATGACTTAGCATTGGAATATGTTCTCTTTTTTGTATTTCTGTTCCGGTTTTTGAGTT

GTTTTATATTGTATAGATGACAAGATAATTTGTAATTACTTTATTTACATTATTTTTGCATAATTTCCTT

TCGTATTGATAGTGATTATGATGCATATATGTCCTCTAATGAATGTATTGAGCCTCTTTTTGTTA

>MSTRG.3999.2 gene=MSTRG.3999

AGGTATCATAATTAGTACATGATTTTGTATTGAACGTGTTTGATAATTTACTACAGAGAAAAGCTGAGAG

ATCGGTCCTCACGATTTCTATTGATTTCATGCTCCAGGAATTTGAGAAAAGGCAGCCACCGGGTGAAAGT

TTGGAAAGTTAGAGAGGCTTTTATGAAAGTATACGACCTATATATATGGGATTATGACTTAGCATTGGAA

TATGTTCTCTTTTTTGTATTTCTGTTCCGGTTTTTGAGTTGTTTTATATTGTATAGATGACAAGATAATT

TGTAATTACTTTATTTACATTATTTTTGCATAATTTCCTTTCGTATTGATAGTGATTATGATGCATATAT

GTCCTCTAATGAATGTATTGAGCCTCTTTTTGTTA

>MSTRG.4000.1 gene=MSTRG.4000

CTGGACTCTGGAGAGAGCAGACTTGGAAGTGAGCAGAGCACCGAACAGACTACGGAGAAGCTGGCTTTGT

ACAACTTGATTTGGGCATCCAAATTCACATCCAAACACAGATATTCATCATTGGATTAAGGTTTTAACTT

TTAAGGATGTGTGATCAACAACCCAAGAAACGAATGAAGTCACTCTTTTCTTTCTACAAGAAAGGTAGTG

ATATACAAGCACCTGAAAGGTAGTCACTCTTTTTAAGAATGAGGTTTTGACACTTCCTGTTTCTACTGCT

ACTACTGAGCGGGCCTTCTCTGCGATGAAAATTATAAAAACAAGACTGAGAAATAAGATGGAAGACGGAT

ATCTTTCGAGTTGTATGATGCTTCACATTGAAAAAGAATATGTCGATGATATTGATTCAGAGGCTGTAAT

TGATCACTTTGAGTCCACAGGAGATCGTAGAGCTCAGTTCAAATGAGAATCATGGTATACATTTTATGCA

AGTTTCGTCAAAATCTCCGAGACCCTATATATTACAATGAAATCTCTGAGACCTTATATCAAGGTGTACT

CTCCTAACTTCGTTACAAAATTTATTTGCCTGGTAAATTACATCATGTTGTACTCAGAAGTGATGATCTC

TTCAGACTTAGCAAGCCGAGTTCATCGTCATCAACTGATAAAGAACGAGGATTATCATGCTCGTGAAGGC

GATCTCTGTTATGGCTGCGGTCTACTCCTGCATGAATCATCGTCTATTTATCATTGTACCTACTTCCGTA

CCAATGCTGAAGAAGATGATTGTGCTCATGTTTTAATTCACAAAACCTGTGCACATTTACCCACAAAGCT

TAAGCATCCAAGCCACAAACATGAGCTAGAATTATCTGAACGTCCACTATCCTTTCCTATTTCCAGAGAC

AAATGTGATGTTTGTGGCATCACGTTGAAATGGTTCACTTACTATTGCGGTTTCTATAAGTCTGAATTTC

AAATTTGTCTGGCATGTATTATACCTTTTAATAAAATTACTATACACCCTCCGTATCATCCTCACAAGCT

AACCTTTCTAGCTAACCTAGCAACTTTCCGATGTGATGGCTGCCATGAGGTGGATACTGACTTTTCTTAT

ATGTGCTACACATGCCCCTTCTGGATCCACCTCAAGTGCAGTAATCTACCACCATTGTTAGCCTGTGATA

TTCACCATAAGCATCCTCTTAGGCTAACATATTCTTTACCAGAAAATTACCGTAGATTTCAACAGTCTTG

CAGAATCTGTAGACAACATTTGCGTGACCCTAGAGATTGGCTCTACCACTGTCCAGACTGCAGATTCTTT

GCGCACATCAAATGTGCTACATCACCTCCAAAAAGCCATGGTGACACCAAGTATGCCAATCAAGATTATT

CAATACATAAATCATTATCAGGTCTTGTGCAACTCCCACTGCCAGAGCATGAATCGTTGTTGCGGCATTG

TATCAAGGGAAGTACATTATGGCCGAAGGATGCTGCCCTGACACCTCCTGATCAGATCAACCACTGGAGC

CACGAACATAAACTTTTACTAAAAAATCCAACTGTTGCAAGTACTACTTCTGAAGAAGAAATATGGCCTA

TCTGTAACGGGTGCACTGAATCTATCACCATAAAAGATGGTGCATTTTATGAATGTGGAGAATGTGATTA

CATTCTCCACGGATACTGTGCCCACTTTCCACAAGAGATGTCTGACCGTGCAGGATATAGACGCAAGATA

GTGGCAATTCAACCCAGGGTCCATGATATCTGGTCTTGCGCTCGTTGTAAAGGTTTTAGAAATGGGACAC

AAATGCGATACACTTATGAAGGCGATTATGCACAGAAAAGTACAATTGAGCGCAATTTTGACAGTGCTTG

TGTTATTCTACCTGAACGCATCGTACACAAAGCTCATCAACACAAAAGTCGTCCTAACTTCTTTTTTCGT

GCATCAGCTGGTGAAAACACTACTTGTTCAGCGTGCAACCAAACACTAGACGGGTTTGACATCTGTTATG

CATGTAATGGTTTTGAATGTTTTAGATGCATACATATCGAATGTGCTCTTAAGCCACTTAGAGTGACACA

TCGATGGGATCCTCACCCCCTTCACTTGATAACTTCCGCAGAAGAAGTAATAGATCATCCTCATGAATTC

CAATGTGAATTTTGCTCAGAAGAGATTGACACGAACTATTGGTTCTACCATTGCAATGTTTGTGATTTGT

CCTTTCATTTTAAAGGATGCATCGATCCGTTTCCCTACTCAAAGGTCAAGTTTGGGGCTACCAATATTAT

GCTTGACGCTCACCAGCACGGCCTCACCTTTGTTCTTAATAAAAAGAAACACCAACCATGTCATAGATGC

CATAAAGATACATTCGGTAGGCCGGTCCTTGAGTGCATACCATGTAAATTCGTGCAACACGCCCAGCCTG

AGTTATGCACTTGATGTTAATTTGTAAATGTTCTGATTGTTTGCGTGTATTTCACATTTGTACCTGAATG

AAAAATTTGCTTGC

>MSTRG.4001.1 gene=MSTRG.4001

GTCTGACCGTGCAGGATACAGACGCAAGATAGTGGCGATTCAACCCAGGGTCCATGATATCTGGTCTTGC

GCTCGTTGTAAAGGTTTTAGAAATGGGACACAAATGCGATACACTTATGAAGGCGATTATGCACAGAAAA

GTACAATTGAGCGCAATTTTGACAGTGCTTGTGTTATTCTACCTGAATGCATCGTACACAAAGCTCATCA

ACACA

>MSTRG.4008.1 gene=MSTRG.4008

TGATAACTACGAGCATACGAGAGAGAACTTTGCAAGACAAATGGCAATTGATCAAAGAAAAACGAAGAGT

CCTCTTGACTGGTGGAGTGCTTTTGGGGGTCGCGCACCTGACCTAGCAATGATAGCGAAACGGATTGTTG

GCCTTTGCTGCTCTTCATCAGGCTGTGAGCGAAACTGGAGCACCTTTGAGTTTATTCACACTAAAAGAAG

AAATCGCTTGGAACATAAGAGGCTGAATGATCTCGTGTATATCCAGTACAATGAAAAGATTGCAGAGCGA

TTCCAAAAGCGCCGTGAGCTTGGAGATAAGTTCGATCCAATTGCGCTCGAAGACTTTAACTGGAGTGGTG

AATGGGTAAATAATGCAGATGGTGATTCTGTTCATCCTGGAGAAGATCTGCTATGGGAACATGTTGTTAT

AGCTGCTGGTCCGGCAAGTGCAGTGCGTAATCTTCGTAATAGAAATCGTGATAAAGGTGGCGAAACTAAT

ACAAACACCAGGGAGATTTTAACATATAATCGACATCGTAATGGTGCAGGTTCTTCTC

>MSTRG.4009.1 gene=MSTRG.4009

CTTCTGCAAAAAGAAAAAGAGCAGGAACTCTCTATGAGTTTGCAAAACCTCAAGCTCCACCTCCAAAGCC

TACAAAAACCATTGCATCTATCTTAAGGAAGACTCCTGAAGAAGTTGTTCAGGATAGGCATCGCAAGGAT

TTAAAACAAACTACTTTGGACAATATTCGGAGGACACCTGAAGAAAAGGAAAGGGTGCATATACACATTG

CAAAGTTCTTCTACGCATGTGGAATACCGTTTAATGCTGCAAACTCGAGACATTATGAAGTGATGATTGA

GTCAGTGGGCCAATATAGACCTGGTTTAGTTCCTCCATCTTATCACCAACTAAGAGTCCCTTTGCTTGAA

AAGGTGAAGAAACAAACTGACCAGCTACGATTGAAACATGAAAGTGCTTGGAAAAAATATGGATGTACCC

TCATGTCAGATGGATGGACAGATAGAAGAGGCAGGCACCTGATTAATTTCCTTGCAAATAGCCCAGATGG

AACATTTTTTATGGGTTCCATGGATGCTTCGAGTGAGTCACACGATCATTTGATGTTGGCAGACTTGTTA

CAAGCCAAAATTGAAGAGATTGGAAAGGAGTATGTGGTTCAAATCGTAACTGATAATGGTGCTAACTACA

AGAAGGCAGGTCAATTACTAACAACGGAAAGATTCCCACCAATGTTCTGGACTCCTTGCGCTGCTCACTG

CATAGATTTGATGTTGGAGGATGTCGGTAGCTACAAAGTATTTGATGCCACAATCAAGAACGCAAAAAGG

ATAACAACATTTATTTATAGGCATGGTCGTCTTCATAATGCAATGAAGGTTAAAACAGGAGGCAGAGAAC

TTGTGAGACCTGGAGCGACTAGATTTGCCACATCTTTTCTCACACTGCAGTGTTTATTCAAATTCCAAGA

TGCATTGAGGCAGCTCTTTGTAAGTGAGGACTGGAGCAGGTCAAGTTTGGCTAAAACAGAGGCTGGAAAA

CAGGTTTATGACTTGATTATGTCAACTGCACTATGGAATAATATTGCAAATTGTATCAGGACTGCACAAC

CGCTTCTTGTTGCCTTGAGAGTAGTCGATGCTGATGAACAACCAGCAATGCCAGAACTTGTTGTTGCTAT

GGAACATGCCAAGAAAACTATCAAAGATTCTTTTGAAAACAATGCATCACTTTGCAAATTTGTTATTGAC

ATCATTGAGAAACGCTGGGATACGCAGATGGGGGTTCAAATACATGCTGCTGGT

>MSTRG.4013.1 gene=MSTRG.4013

AAAAAAAAGAAAAAAAAATAGAAAAAAATAGGAAAGGCATAGGAATTCTTTAGTGACCCTAAATTGTAGT

TGACTCTTTAGTAAAGAAGTGTTTAAGCTTGATTCTGATTAACGGCTCATAGTGAGGACTCTGTTGAAGT

TTGATAGTCTTTGTTTTGTTTCACTAGAGAGCATGCTAGCACACACGATGCACTTCACACTTGTAGTGGA

AGAGAACATGGCTAATATGTGATAGAGATGAATGCCGAGAATGTTGCATATTCTGAGGATTCTATGTTAA

CAAGGATTACACTTCATGATTCAAGTAGACATATAGTTGCATTCATGCATTGCATTAGTAGTTTGTGTGT

TTGTCTATGCTTGAGGACAAGCATCGGTTTAAGTTTGGGGGTGTGATAAGTGGATTTTATATCCACTTGG

AATGCCTTCGTTTCGACTTAAATTGATGATTTGGCCTCAAGTATGTGGTGTTTTTGATGTGTTCTTGTGG

TGTTGTGTAGGTTTCCTAGGAGGAGGATAAAGGGTGGATTAAAGGCTAAAAGGTGAAAGAAACGGCGTAA

GGATCGAGCGAGCAAGGAAGGAGAAAAGGGAAGGAAGGTTCCAAGAGCGTTCGGGGCTGCCCAAATGGGG

TCTCCACCCCATTTATGGGGTCTCCACCCCGTTCTGAAGGAGAAAAATCTGCCAAGATGGGGTCTCCACC

CCACTGATGGGGTCTCCACCCTACTTCAGCAGAATAGAGGCTGCCAAGACGGGGTCTCCACCCCACTGAC

GGGGTCTCCACCCCACTGACGGGGTCTCCACCCCATTCTGTTGGGAGAAAAAGCCCGAATTGCCTATTCT

TGATCCGTTTGAAGGCCCGATCGCTGTGGGACTTAAATCAAAGCTCAATGGAGAAGTCTAATAACCTAGA

ATCATTCCAAGGAGCACGTGACGGC

>MSTRG.4019.1 gene=MSTRG.4019

AGATCAAGTATCAAAGACCAGAAGATTCCAGTATATTTAATTTGATTATTTATAATCAAATTTATCGAAG

CAACACCTAACCCGGAATGACTGATCAAGTATATTATCAAGCAAAGGATTCAAAGAATCATTTCAGTTCG

AGTCGTTCGTGAACCAGACCAGTGCACAGGACGTCAGGACGTACGAAAGTATTCAGTGGATTCGATCAAC

GAATTAATTGATCAAGTCAATCGAGCTGCTCCAGAATATGGCCAAG

>MSTRG.4022.1 gene=MSTRG.4022

GCTGTTGAGATTATCGATGATGACCAGGGTCGTGGCGATAAAGTCACGAAGAGGATGATGAAGAAGCTTG

CTGAGAGAGCCAACCCTCCTGAGCCTACCACTCCTGCTTTCCTCCAGCACAGTCAGCAAGCTGGGAAGAG

GTTTAGAGATGCCGATGGTCACATGCAGACTGCTTTCATGCCGGCTTGGGGTATTTGTGATCAGCATTCG

GTGTCTGGGAGTTCTCTCCTAGCTAAGGACTGG

>MSTRG.4023.1 gene=MSTRG.4023

TGGGGATTCTGTGGGAGTAGTAGTTTTTGTCTTGTCTCAATAAAAGAGCATGCTTGCACACACTGGCACT

CCACACTTGTAAATAGAAGAGTAATTGACTTAGTAGCGAGAGAGATGAAATTCGAGAACATTAGCACAAT

CTGAGGTATTATAATTGGCAAGGCAATTACTTCATAGTTTACTCATTCATCACGTAGTTGCATTCATGCA

TTGCATTGTATTATTGTATAGTGTCGTTGACGCTTGAGGACAAGCATCAGTTTAAGTTTGGGGGTGTGAT

AAGTGGTATTTATACACACTTATAGGCCTTCATTTCCACTTAAATTGGTTGGTTGTACTTAAGTGTTTAG

TGTCTTTTGATGTGTTTTTAGTGTTTTTCTGTGCAGGTCACGAGTTGAGGTGATGAAGTGATTTTCTATC

ATTTTAGGTTGTTTTTG

>MSTRG.4025.1 gene=MSTRG.4025

TTCAGATAAAAACATATCTGACAGTGTGTATCAAGCTTCAAGAACCACAAGTATTCATAAAAGCTCTGCT

TAGGTCTTGCAAGTATCCACTTTAAATAATGAAATCCTCCATGTTCTGCCATTTTTTATTCACAGGGAAA

AATATATTTTTTGAAACTTGTTGATCTTGCTTAGCTGGGTTCTGAGGTATAAAGGATTTTGGTTGGTAAT

CTATACTTGAAGCTCACCTGGTCTTAAAGTTATCACGGGGCCAAGTACCCCCACTGCCCCGAGCTGCCTC

CGTCCCTGCCTGTGGCAAGAGCATCAGGTTACTCGCTTGTATTAGCTATCAAAATTGGCGACGGTCCTGC

AGCTGAATATTTTTAACCTTGCCAAGCAATTTTCCCAACAAGTAGCCCTCTGTTTTAGTTGGCAGATGGT

ATTGATTATTTCTAGTGGTCATGTCATTCGAGTGATAGCTGCGCATAGAGCTGTAAACGAACCGAACCCG

GACGAACTCGGACGAACTCTTACCGAACTCGAATAAAATCTTAACGAACCGAACAATGTTCGATAATTTT

ATCGAACAAATTTTTCTGTCCGAACTCGAGTTCGGTAACGAAACGAACCGAACCGAACACGAACTATTCG

CGAACATGTCGAACCAAACCGAACACGGACCGAAAACGAACAGTTCGCGAACAATGTAGATTTTTAAATA

ATATTTAAGCT

>MSTRG.4027.1 gene=MSTRG.4027

CGCGTGAATTGGGAGTGCAGTATGTCCGTATCTATTGGTGGGATCAATCCTGCATTCAACTCTAGTAAAC

TAAATTGTGGTTATAACGACTGCAAGATTGACGCTAAGAACAAACCTGCGTGCTCAAAATTGCATCTGGG

TAGGCTGAAGGAGGAAAAGGAGTGAGGGACAGAAAGGTTTCAATTAACGATGAATTCAGGACACAAGTAT

TAAGAGGAATGTTGCAACGGCTAAGAAACCTGAAAAACCGAGAGTAGGTTTCGGGACAAATAAAACATCA

GAGGCACGTAAGTGTGGAGTTCGAAGTAGTGCAAAAAGGGGTCAGTTAGGAGTCCACAGGACACTGCCTA

AGCAGGACAAGATAAAGAAGCACAGTCAGAGTGGGTAAAAAGGGCATGATCTGAGTTGGATTCCGTCTGT

TTAGTTACCCACACAAGCTTGGGTACAAAATAGTGGTTTTTTAGTATATGTGTTACACTATATTAGAGTA

ACACATCTTGAAGTCCAACAGTCCTCTCCTGCTACCTAAGCTGTAACACGTGGAGGTAGTAGTAGGTCAG

TTAATGTACTGCTAAACTGTAATCACGATTGATTAAATTAAAGGCATAATTGGCTTTACGAG

>MSTRG.4027.2 gene=MSTRG.4027

CGTGAATTGGGAGTGCAGTATGTCCGTATCTATTGGTGGGATCAATCCTGCATTCAACTCTAGTAAACTA

AATTGTGGTTATAACGACTGCAAGATTGACGCTAAGAACAAACCTGCGTGCTCAAAATTGCATCTGGGTA

GGCTGAAGGAGGAAAAGGAGTGAGGGACAGAAAGGTTTCAATTAACGATGAATTCAGGACACAAGTATTA

AGAGGAATGTTGCAACGGCTAAGAAACCTGAAAAACCGAGAGTAGGTTTCGGGACAAATAAAACATCAGA

GGCACGTAAGTGTGGAGTTCGAAGTAGTGCAAAAAGGGGTCAGTTAGGAGTCCACAGGACACTGCCTAAG

CAGGACAAGATAAAGAAGGTGAGGACTTATGACTGATAAATTCAAAAGAAGTTTTCGCCGTCGATTGTGA

CAGATGTCTTGCTCAATATGTCCGCAGCACAGTCAGAGTGGGTAAAAAGGGCATGATCTGAGTTGGATTC

CGTCTGTTTAGTTACCCACACAAGCTTGGGTACAAAATAGTGGTTTTTTAGTATATGTGTTACACTATAT

TAGAGTAACACATCTTGAAGTCCAACAGTCCTCTCCTGCTACCTAAGCTGTAACACGTG

>MSTRG.4033.1 gene=MSTRG.4033

AACAATTATCACTTTTCAAACTAAACTCGGGAAAATCGTAGTGGAGATCAACATCCTACTTAAACGGAAG

GAGATTACCTTCAAATTCTTCAAATAGGTACCAAATCCTCTGTTCGACCTTGCTGGTATAATGTTTGGAC

AATTTGGAATTCCTTTCTGGGAGTTTTTTCTGGCTACATTGGTTGGAAAAGCAATTATTAAAACTCACTT

ACAGTTTTGTGAAGGTCTTTTCTTCACAAAAGTTTATGCTCAGCTGGAATTTGATGCTAGCTATGTTTAT

GAGGATGTTAATCGCTCAATTCAGTATGTTCATCGAAGTGGTTTGGTTCATAGAGGAATTCTATATGATA

CTTAAAGATACCTTGTAAAAAATGTTAGTGATAGTAAATTTGTTTATAGACTTAAATAAGTTACTCATGC

AAGTGTCTCTAACTCATTCTTTGTTATCTTTATTTAGCAGTGTCTTATGTAGTGTCACGTGTGTAACACG

CATGGCCAGCTTCTGCAATTATTAAGAACAGAGTTCCCCAACAGCATAGAGCTTGGGATATTCTAAAGGA

GCAGGGAAAGAAGCTTTTCTTGATGACTAACTCGCTATATTATTTTGGTGATGGAGGAATGCGCTTTATG

TTGGAGGTAATTAGTGTAATATTGCATCTTGCAAATTTACCATGGATGCCTCAAAAGTTTCTTACAAATC

ACAAAGTGGAAAGGTCCTGAGGTACTGACTATGTTAACCTGTAGAATTTACATGTTTAAGTTCTATATTG

TTGCAACAGATTTTAGATTCATCATACTTAACAATGGCAAAATTGTAATTTCTTCCTTTAAATTTCTCTA

TATATCCTCTTTTACAGT

>MSTRG.4033.2 gene=MSTRG.4033

AACAATTATCACTTTTCAAACTGTTCATCTCTCTCTCAAACACAAGATATAATCACCAAAAGCTCTACTC

ACTTGCTTGTATTCTTCGATTTCTCGCCGTTCCTGCAACTTTTACAAGCTAAACTCCTTCGTTTTGAACA

AATTTTGCAGAAACTCGGGAAAATCGTAGTGGAGATCAACATCCTACTTAAACGGAAGGAGATTACCTTC

AAATTCTTCAAATAGGTACCAAATCCTCTGTTCGACCTTGCTGGTATAATGTTTGGACAATTTGGAATTC

CTTTCTGGGAGTTTTTTCTGGCTACATTGGTTGGAAAAGCAATTATTAAAACTCACTTACAGGTTTGTGA

AGGTCTTTTCTTCACAAAAGATAGACTCCTGATGTACCTGCAATTACTACCAAAGTAGTTGGCTTGAGAA

ATGTTGAGTCAACCCTTGGCTAGTAACTTACACTAAAATCGAATATTTATTCAAAGTTAGCTAGCTACTG

AGAATAAATACAGTTGAGTCGAGCAAATTACCAGCTGAATTGTTAACGGGGAGCACTAAAATCATGATTA

TTATAGTATAGTATAGTCAATCCAAAACTTATTTTGTATAATCCATATGCTAGAGAAATTCAAAATAACT

AAATATGTGCTCAAAATGATAAAGTAGATTTGAGATTTGACATTTAACTAAACAGGGCTTACATACTTCA

>MSTRG.4034.1 gene=MSTRG.4034

AAAGAAGCTGCGCTTGGAAACATTCTTACAATTGCCTTCTGTGACACGGATTCGCCTATGCGATATGTAG

ACATTGGTATCCCTGCAAATAACAAAGGGAAACACAACATTGGTTGCCTCTTCTAGATATTAGCAAGGAT

GGTCTTACAGATGCGTGGTGTCATCAATCAAGGACCCAAATGGGAGGTTATGGTTGACTTGTTTTTCTAC

AGAGAGCCTGAAGAGGCAAAGGAAGAAGAAGAGGAAGCGGCATTAGCTGATTATGCAGATTACTCCGCTC

CTGCTCTTGGCGGAACTGATCATTGGAGTAGCCAATTTCCCGATGCTCAGTGGAAGAATGAAG

>MSTRG.4035.1 gene=MSTRG.4035

ATTTATATTATCAATTTGGGCAAGACATGGGATAAGCTTCAGATGGCAGCCAGAGTCATTGTTGCCATCG

AGAAGCCCCAGGACATTATTGTCGAGTCTGCCAGGCCTTATGGTCAGAGAGCGGTGTTGAAGTTTGCTCA

GTACACGGGTGCCAATGCTATTGCTGGCCGTCACACTCCTGGTACTTTTACTAATTAGTTGCAGACGTCG

TTTAGTGAGCCCCGTCTCCTCATTTTGACCGATCCAAGAACTGATCACCAG

>MSTRG.4036.1 gene=MSTRG.4036

AAAGCGAACATATTACTCTGGGCAAGCCTCCAAAACAATCCAAACACCTTTTAAATAATTAAACACTGCA

TTGCTATCTTGTCAACTTATTCACCTATTATTTGTATTTAGTTGACGCAAATAACATCATCTTCTTCCAG

GTTCAGATCAAGTTAGATATTCCTCATCATGATCCCCTGCGTTCAATGAAGATGGATTTGCTCTGCCAAC

ACTCCACACCAGTCCTTGAAGATGATAATGCCTTCAGCTCTTGTGGAAATTCATTTATGATCAAACAAGT

AAAGTTGGGTAGCAGGAAAGGGAGAGGTATCCCTCAGGCACTTCGTGCATTTGCTCGTGTTATATGTTCT

ACATCTTCACAAGAACTGAGTGATTTGGCCATTGAAGCTGCAAAAACTGACGGTCGGCTAGCTCGACAAC

CCTTACAGGACAAAGACAAAGAGTTACAAGCTCATCAGTTCTTGCTTTCAAGGATCAATCAACTAATTGA

AAATCATGATGTTGCTATGGAGTCACTTGCAGTTCCCAGTTCTCCGTGTATGTCTAACAAACATTCTCTA

AGAAGGAAACTGGCTCGAGATCTCCTTAACGGTGAGCTAGTGGTTCTTAAATCTGCTGGTGCATGGCTAA

AGAACTACTGCGATTCCTTGTAACAGAACAGAGAAGCAGCGGAACTCCCGAGTCAAGAAGAGGGTCTGCT

CTGTTGTAGTTTACAGAAAAGATAAATAAATAAATTATCTGTGTATGTGCAGTAACTTCTTTTCTCCAGT

ATCAATTAGTTCAGTCCCACCTCCCAAGCCCCCAACTGCAATAAATTAGAGCATTTTCACGTGAATGTCA

AATTTGTAAACGTTGCAACAGACATTGTACGTTTATTAACTATCTATTATTCCAACTTTGATTTCATTGA

GGGTTGCCAGTTTTTATCATG

>MSTRG.4036.2 gene=MSTRG.4036

CAGGTTCAGATCAAGTTAGATATTCCTCATCATGATCCCCTGCGTTCAATGAAGATGGATTTGCTCTGCC

AACACTCCACACCAGTCCTTGAAGATGATAATGCCTTCAGCTCTTGTGGAAATTCATTTATGATCAACGT

GTTGGTTAATAGTTTTGCATCATGTATATACTGATGATGCAGACAAGTAAAGTTGGGTAGCAGGAAAGGG

AGAGGTATCCCTCAGGCACTTCGTGCATTTGCTCGTGTTATATGTTCTACATCTTCACAAGAACTGAGTG

ATTTGGCCATTGAAGCTGCAAAAACTGACGGTCGGCTAGCTCGACAACCCTTACAGGACAAAGACAAAGA

GTTACAAGCTCATCAGTTCTTGCTTTCAAGGATCAATCAACTAATTGAAAATCATGATGTTGCTATGGAG

TCACTTGCAGTTCCCAGTTCTCCGTGTATGTCTAACAAACATTCTCTAAGAAGGAAACTGGCTCGAGATC

TCCTTAACGGTGAGCTAGTGGTTCTTAAATCTGCTGGTGCATGGCTAAAGAACTACTGCGATTCCTTGTA

ACAGAACAGAGAAGCAGCGGAACTCCCGAGTCAAGAAGAGGGTCTGCTCTGTTGTAGTTTACAGAAAAGA

TAAATAAATAAATTATCTGTGTATGTGCAGTAACTTCTTTTCTCCAGTATCAATTAGTTCAGTCCCACCT

CCCAAGCCCCCAACTGCAATAAATTAGAGCATTTTCACGTGAATGTCAAATTTGTAAACGTTGCAACAGA

CATTGTACGTTTATTAACTATCTATTATTCCAACTTTGATTTCATTGAGGGTTGCCAGTTTTT

>MSTRG.4037.1 gene=MSTRG.4037

ATTGAATCCGCTGGTCAAAATATCAGCTACGATTCAAATCTCGAATTCTGATCGCCAAATAAATTACTTG

TTGGTTCAGGGATAATGCAGCAGAGATACAATTCAAGAGCACCGGAAAATCAAGGGGTGAGTTAGAGCTG

ATGACTGGGTGAGTAACTATGATAGGTCGATGCTCATATTACTATCGCGCCAGCTTCTCCTGCGCCACTC

GCCTTAACTTCTCTACTTGTTCACTCACGAAATCTGTATGTATACATGATGAGGATTGTGGCGATTTCTT

GCCCTGGTTGAGGCGAAAATCCGGAACTGACATTTCGTCTTTTTTATCTATAGGAAACTCTTCTACTTAT

GGCAGGTATTTGTTTGCTTCCAAGTGCATATACACAGGGGATTCCATACTCAAGGTTCCTTGCAGCGTGC

AATTGTCGCCTGATAATCTCGTACCACAAATAAAATCATTGTTAAAAGATGAAGTCAGCGATAATGCAAA

GCTTGCTGTTGTCATTCTAGTTGAACAGAAATTAGGCCAGGACTCCGAATGGGCCCCATACATTAGCCGT

CTTCCGCGGTTGGGAGAATTGAATAGCACGATATTTTGGAGTAATGATGATCTGGAGATGATTGAGAAAA

GCTCTATATATGAAGAAACAATCAGACATAACATTCAGATTGAAGAAGATTTTTTGGCCTTAAAACCTGC

TCTTCAGCACTTCCCTTCGTTTTTTCAAGAAGTCACGCTGAAGGAGTTCCGCCATGCATATGGTTTAGTT

TCTTCTCGGGCATGGACTAGCATGAAGGGTGTATCCATGATACCATTTGCAGATTTTCTAAATCATGATG

GCATTTCTGAGGCGGATGTATTATGTGATGAAACCAAACAACTGTCCGAGGTCATTGCAGATCGTGACTA

CGCTCCAGGTGATCAGGTTATGATAAGATATGGAAAATTTTCCAACCGCACCCTCCTGTTAGATTTTGGT

TTCACACTTGCTGACAACGTCTATGAACAGGTCAGTATCTGCCTTCTGGTCAAGTCTATAAAGGTATTTG

ATGATGAATGAAGTGCATTTAAATCTTCG

>MSTRG.4037.2 gene=MSTRG.4037

CAAATAAATTACTTGTTGGTTCAGGGATAATGCAGCAGAGATACAATTCAAGAGCACCGGAAAATCAAGG

GGTGAGTTAGAGCTGATGACTGGGTGAGTAACTATGATAGGTCGATGCTCATATTACTATCGCGCCAGCT

TCTCCTGCGCCACTCGCCTTAACTTCTCTACTTGTTCACTCACGAAATCTGTATGTATACATGATGAGGA

TTGTGGCGATTTCTTGCCCTGGTTGAGGCGAAAATCCGGAACTGACATTTCGTCTTTTTTATCTATAGGA

AACTCTTCTACTTATGGCAGGTATTTGTTTGCTTCCAAGTGCATATACACAGGGGATTCCATACTCAAGG

TTCCTTGCAGCGTGCAATTGTCGCCTGATAATCTCGTACCACAAATAAAATCATTGTTAAAAGATGAAGT

CAGCGATAATGCAAAGCTTGCTGTTGTCATTCTAGTTGAACAGAAATTAGGCCAGGACTCCGAATGGGCC

CCATACATTAGCCGTCTTCCGCGGTTGGGAGAATTGAATAGCACGATATTTTGGAGTAATGATGATCTGG

AGATGATTGAGAAAAGCTCTATATATGAAGAAACAATCAGACATAACATTCAGATTGAAGAAGATTTTTT

GGCCTTAAAACCTTTAAAATTCTTGGAGCTTCTTGATTCCAGGCTCTTCAGCACTTCCCTTCGTTTTTTC

AAGAAGTCACGCTGAAGGAGTTCCGCCATGCATATGGTTTAGTTTCTTCTCGGGCATGGACTAGCATGAA

GGGTGTATCCATGATACCATTTGCAGATTTTCTAAATCATGATGGCATTTCTGAGGCGGATGTATTATGT

GATGAAACCAAACAACTGTCCGAGGTCATTGCAGATCGTGACTACGCTCCAGGTGATCAGGTTATGATAA

GATATGGAAAATTTTCCAACCGCACCCTCCTGTTAGATTTTGGTTTCACACTTGCTGACAACGTCTATGA

ACAGGTCA

>MSTRG.4038.1 gene=MSTRG.4038

GTTGGGAATAAATGGGTATTCATAAGAAAACGAAATGAAAAGAATGAAATTGTGAGATATAAAGCCCGAC

TTGTAGCACAAGGGTTTTCTCAAAGGCCTGGCATTGATTATCACGAGACATACTCGCCAGTAATGGATGG

AATTACTTTTCGTTTTATATTAGGTATGGCATCGAAAGAAAAATTGGAAACACGTCTTATGGACGTTGTT

ACTGCATACCTATATGGATCACTTGATAGTGATATTTTTATGAAAATCCCAGAAGGATTAAAAATGGATG

AGTTCAAGAAACCTCGTCATATATACTCCATTAAACTTCAACGATCATTGTATGGATTGAAACAATCTGG

TCGTATGTGGTATAACAGACTTAGTC

>MSTRG.4039.1 gene=MSTRG.4039

GGGAACCTCAGCCTCTGGGACAGTAGGGACATCATCCACATAGATCCCAGGCATCCAAGTATGTGGGTCG

TCGCCCTCCACCATATTTGGGGAATTAAATTGAAAAAGTAAAAACAAAACATTTAAAAGTGAATTTCCAA

CCCCTGGGGGTTTTCCAAAGCTGCACCCCTGAGGGTCTTCCTGATGGTTTGCCGGAAGTACGGGCCATCA

GGGCATGGGAATTCCTTCGAACCGTCTCGTGGAGACAGTTCTCCGTGCATCCCGAGGGTGTCAAGCATAT

GTCTCGATTACATATGTCGACGGGTTCCTAAGCAGCAGGGAGACTGTCCCTGGTCTTCATGTTCCAAGCA

ATACAAAATATACAGATATAATAAAAAGTTCGAAATTTATTGATAGTAGTACGCTTACATGATGTCTTTG

CCAGAGGGCACGTACAACCCCTACCCCCCGAGGCTTGGGAGTAATTTTATTGGTTCCAAGGCTTGTGCTT

GAATACTTTAAACAGGAAGCTAAAACAAACAGGAAAACATAAACATCTATCCCCCTTCTTGCGAGTCGAT

CCTTCAACATGCCTCGTTAAAACCTTCCTAGGTAAAACCCAGTGGGAAAAACCCCAGTGAAGGAAAAAGA

GTACATGAGTCGGACGATGAGGGGGTCTTCTGGTCTACTGGTAATACTTCTTGAGGTGCTCGGCATTCCA

TGCTCGGGGGATGTCTCTCCCTTCCAGGTCTGCCAAGTAATATGTTCCGGAAAACAACACCACTCTCACC

TTGTACGGTCCTTCCCAGTTAGCTCCAAAGACTCCATGGCTCGCCACCTTGGTGTTGGGCATGACTCTTC

GTAGGACTAGATCGCCCACCTGGAATGGTCTAGCCTTGACCTTCGTGTTGTAGTGCCTTGCCACTCTTTG

CTGGTACGATGCAATCCTTAGTTGAGCTCCTGATCTGGTTTCTTCGAGCATATCTAAGTGTAGGCGGTGG

TTGACCTCATTCACTTCCGGGTTGAAATTGTCTCTCCTGAATGACCCAGCTCCGAGCTCCACCGGTATCA

TAGCTTCGCATCCGTATGTAAGGTTGAAGGGTGACTCCTCGGTGGTCGTCCTAGGAGTCGTGTTGTAAGA

CCACAGCACATTAGGCAATTCTTCAGGCCAATTGCCCTTCCTGTCTTCGAGTTTCGCCTTCAGGGTGTGT

TTTATGATCTTGTTAACCGCTTCCGTCTGGCCATTCGACTGTGGGTGGCTGACAGCCGAGAAGCCTTTCT

TGATCCCGAGGTTGTCACATAACTCCCTCATTTCCTTGCAATCGAACTGCTTTCTGTTGTCGGAGATCAA

CTTATAAGGAATGCCGAATCGGCAAACAATAGCTCTGTACACAAATTCAACCAATTTCTTCGCTGTGATG

GTAGCTAGTGGCTCCGCTTCTGTCCACTTTGTGAAATAATCAACTGCCACCACGGCATACTTAACTCCTC

CTTTGCCCTTAGGTAGTTCTCCAATCAGGTCTATTCCCCACATAGCGAAGGGCCATGGGCTCATCAATGA

TGTAAGCGGGGCTGCAGGGCTGTTGACGTAGTTGGCGAACCTCTGAC

>MSTRG.4040.1 gene=MSTRG.4040

GTCAGAGGTTCGCCAACTACGTCAACAGCCCTGCAGCCCCGCTTACATCATTGATGAGCCCATGGCCCTT

CGCTATGTGGGGAATAGACCTGATTGGAGAACTACCTAAGGGCAAAGGAGGAGTTAAGTATGCCGTGGTG

GCAGTTGATTATTTCACAAAGTGGACAGAAGCGGAGCCACTAGCTACCATCACAGCGAAGAAATTGGTTG

AATTTGTGTACAGAGCTATTGTTTGCCGATTCGGCATTCCTTATAAGTTGATCTCCGACAACAGAAAGCA

GTTCGATTGCAAGGAAATGAGGGAGTTATGTGACAACCTCGGGATCAAGAAAGGCTTCTCGGCTGTCAGC

CACCCACAGTCGAATGGCCAGACGGAAGCGGTTAACAAGATCATAAAACACACCCTGAAGGCGAAACTCG

AAGACAGGAAGGGCAATTGGCCTGAAGAATTGCCTAATGTGCTGTGGTCTTACAACACGACTCCTAGGAC

GACCACCGAGGAGTCACCCTTCAACCTTACATACGGATGCGAAGCTATGATACCGGTGGAGCTCGGAGCT

GGGTCATTCAGGAGAGACAATTTCAACCCGGAAGTGAATGAGGTCAACCACCGCCTACACTTAGATATGC

TCGAAGAAACCAGATCAGGAGCTCAACTAAGGATTGCATCGTACCAGCAAAGAGTGGCAAGGCACTACAA

CACGAAGGTCAAGGCTAGACCATTCCAGGTGGGCGATCTAGTCCTACGAAGAGTCATGCCCAACACCAAG

GTGGCGAGCCATGGAGTCTTTGGAGCTAACTGGGAAGGACCGTACAAGGTGAGAGTGGTGTTGTTTTCCG

GAACATATTACTTGGCAGACCTGGAAGGGAGAGACATCCCCCGAGCATGGAATGCCGAGCACCTCAAGAA

GTATTACCAGTAGACCAGAAGACCCCCTCATCGTCCGACTCATGTACTCTTTTTCCTTCACTGGGGTTTT

TCCCACTGGGTTTTACCTAGGAAGGTTTTAACGAGGCATGTTGAAGGATCGACTCGCAAGAAGGGGGATA

GATGTTTATGTTTTCCTGTTTGTTTTAGCTTCCTGTTTAAAGTATTCAAGCACAAGCCTTGGAACCAATA

AAATTACTCCCAAGCCTCGGGGGGTAGGGGTTGTACGTGCCCTCTGGCAAAGACATCATGTAAGCGTACT

ACTATCAATAAATTTCGAACTTTTTATTATATCTGTATATTTTGTATTGCTTGGAAC

>MSTRG.4042.1 gene=MSTRG.4042

CCAAATTCAAATTGGGAGCCCTAATCTCCAAATCAACTTCAAATCTGCAAATACTTCAGTACAGATGGCA

CACGCAGAGAATCTAATCCAGTCCAAGCTTCGAAAATAGCCAAAAACACAAGCTAATCACTACTAAAAAC

GGGTTAATTTTTCTTGAAAAATCGGCAGAGTTTCGACTTAAATACAGAGATTTAATCGAAACCAGCAAGC

AAGAGCTTGAATAACCGTATATCAGCAAGTTTTCGTGGTGAAACTTGAAAACCGAGAACGATCGTGTGTG

TA

>MSTRG.4044.1 gene=MSTRG.4044

CAAATATTTTAGAGATACTATTATTTTTAAAATATTTCAAAAAAAAAAAAACTCAGCAAATCGTTTACTT

GGGACAGGCCAGCTTGACCATCTGAATATAGACCAACATTTGGGCTAATTAGGAGGACATTTGGAATAAA

TAACTAGAGCTCTGTCGTCAAGTTTTCAGAGGGGAATGACACACCATCAACCTCAAAATCTTCTTCCAAA

AAAATGAAAGTGGTTGGTTTATTCTTGATGCCTAATATTCCCTCATTCATTACACGCATGTGTTTATTTA

AAACTTAACCGATTACATCCACTCTATAAAAATGCAG

>MSTRG.4045.1 gene=MSTRG.4045

GATACCATGAAGCATGTTCTGAATGAGTAATCATGAATCTCAGCTAAGTTCCTCAATCCTGATCTCAATT

TTGTGCTACATTAGTTCACTATGCATGTTGATATCATTGAGACTCTACTACTTCACATATCTTAATTATA

AGTGAGACTGATTAATTTAAATTTTCTCTCATATACTAGACAGTGATTATAAACTCTGAT

>MSTRG.4046.1 gene=MSTRG.4046

TTTGACATCATCAATTATCTGTTAACTTGCACATAATTTATTCATGCACAAGTTGGGGGAGATTGTTAGA

TATAATTGATGATTACTGATGTTCTTAAGGTTTGTTTTAGAACAGGAATCATCAGAGTTTAAACTGGAAG

CTGATCAGAGTTTAGCAAAGTCTGACAGAGTTTAGCAAAGTCTGACAGAGTTTAATAAAGTCTGATCAGA

GTTTACATAGTCAAGACTCGACAGAGTTTACACGTGGAAAGAGCTCAGAAGCGG

>MSTRG.4048.1 gene=MSTRG.4048

AAATCTATTTTTACCATAATTGCTGAACGATGCCACGTCAGGAAATCCACCTAATTCGTCAAATGCCACG

TCATAAAGCCACATATTTTAATTATGCCACATCAGCAGCGCCAACTCATAAGTTCATTCATATATGTCAA

GTTCATTATAATCCCACAAATTACGCATGCATTCATGATTTTGTTTTTAATTGTCTTAACAAAAAATCAT

TCTCTTCCTAATTTTTAGCTTATTTTTTAAATGTAATTATTGACTTCTTATATCTCTAATTTAAATATGA

AAGATTTCTCAA

>MSTRG.4053.1 gene=MSTRG.4053

GTCCTATAAAATCTATGACCCCAAAGCTAAAGAAGTACAGAACTCATAAAGGAAAATCAAAGAACAGCCA

TTGAAGACCCTTTATCAATCTCATGGCTCTCACCTAAAAAGGTTAAGTTTTTTTCTTACAAAACTTTTGA

AAGCAGCAACATTTAAAGCCCTATTGGAGCTTCATTGTTAATTACAAACAAAAATACTGTAAAGGGTCAC

ATGCAGGCTGCTTTATTTTCCTTTTGTTGTTGAGTTTTGGTCATATTCCTTATGTTTTCTGCAAAAAGGA

CAGGCTCCAAACAGATGCGGAGGTACTAACTGGTTTCCTTTACACCTTGTATATGTTGATATGTAAGTTG

AGTGCCTGTTTTTTCTTTTTTGTAAGGTGTTGAGTGCGTGTTGTTACTAAAGAATATTTTATTGTCATGA

AGGGAAGTTGAACTCTCAGCTCTTGTGTGCTTTTTTATGATTTATCATTTATGTGATTTTCATGGTGAAC

GGGATAATAATAGTATAATTCCAGGTGTCTAAGTGGATTTATGTGCTTTATTTTCCTTTTGTTGTTGAGT

TTTGGTCAAATTCCTTATGTTTTCTGCAAAAAGGACAGGCTCCAAACAGATGCGGAGGTACTAATTGGTA

TCCTTTATACCTTGTATATGTTGATATGTAAGTTGAGTGCCTGTTTTTTCTTTTTTGTAAGGTGTTTAGT

GCGTGTTGTTACTAAAGAATATTTTATTGTCATGAAGGGAAGTTGAACTTTCAGCTCTTGTGTGCTTTTT

TATGATTTATCATTTATGTGATTTTCATGGTGAACGGGATAATAATAGTATAATTCCAGGTGTCTCATTG

GTTCATAAACGCCC

>MSTRG.4055.1 gene=MSTRG.4055

ATTTTCTTATGTGTTTACTGCAAGTCAAGGCAGATTTTTATAGTAATAATTCATGTGTTAGCAATAGTAT

TGACTGGTTGTTGGTATTGCAAACAGGTGGATAAGACTATAAACAGAGGAGATGCTGCCAAATTTTCGTT

AGAAATTATATGTATTTTAGTAGCTATAAAAAGGGTTATTTGTGCAATTATGTGCTATGTGTTTATCTGA

CTATGTGGTATGTATAAGCGAAGGGTAGCTATATATAGGTCCTAACGCTTTTCGGAATTAGTTAATCCGT

CTGATCCGACGACTGAAAAACGAGACGTTGATCGAGGCATTGATCGAGACGTGATTGAAATGGGTATTCA

GAAATAGCCAAGTGGTGTTGCAAGTTGAATTAAGGCGAAACTTAGTTATAAATAAAATCTTATCAGAATT

TGGTAGAAGGGGATGATATGGTGGTATATAGTGTCATTAGAAGTCTGAGCTAG

>MSTRG.4059.1 gene=MSTRG.4059

CATCAGGTCCGTCCAAAGATGGAGCTTTGCAAGTTGTCGAAATTATATCCACCATACAATGGTTATCTGA

TACAATATTTTTTTTGGAGCTTTGCACGTTGTCGAAATTATCATTGAGCTCTTTCTCCACAGCAATGTAC

ATTGGAGTTTTGCCTTCATCATTTTGAAGATGTGTATCAGTCGGGTCGGCCTCTACTAAAAGCTTAACAA

TAGAAGGGTGACCTTCCATAACTGCTCTATGTAAGGCAGTGTTCAACAATTTATCAGCTTTCCTGAGAAA

CGCTTGAAAAGAAGTAGGCAAATGTCTAGCTGCATCGATAAGGATCTCAGCCAATTCGGTGTGTCCATTT

ATTGCTGCCAAGTGAAGTGCAGTTTGTTTCTCTGCAGTCAGCGTGGCCAAAAGATTTTTTTTTGCAAACT

CCCTCAAAATAACTCGCACACGTTCTGCATATTTATTTTGAGATTCACCGTGAAGGATAGTTTCTTCACC

TCGGTTGAGCATATTAGCTCGCATCTCCAAATTATGAATGGCATCAGCATCTCCTTCAATGGCTGCCACA

TACAAAGCGTATCTGTCCATAATTTTGTATAATGTTCCCCCTGGGCTCTTACCTATATATGTGTATATGT

GAAAAAACTTTTAGCAAAACAGATAAAGAATTTAACTTGAAAGATGCAAGGAACTACCAAAAGAATCTTT

AAAGAGTATACCTTGGTCGAGATTCTTAATACACACAACAGTACTCCCATGAGGTCCGTCCAAAGATGGA

GTAATGCAAGTTGTAGAGATTATATCTGCTATATCGTTGAGCCCTTTTTCCACAGCTATGTACATTGGTG

TTTTACCATCATTGTTTTGGATATGTGTATCACTTGGATCCGCCTCTACTAGTAGCTTAACAATACTGAC

ATTTCCTTCCATGACTGCGGCATGTAAGGCGGTGTCCATATCTTTATCGCCCTGTCTAAGGAAAGCTTGA

AAAGAAGTAACTGATTTATCATTATCATCTGCAGGAGGAGGCAAATGTCGAGCTGCATCAATGAGGATCT

CAGCCACTTCAGTGTGTCCCTCGTATATAGCCATGTGAAGTGCTGTATGTTTATAATGACTAAGCTTGAC

CAGAAGATTTTTGTTTGCAAACTCCCTGAGAATGAAACGCACACGTTCTGTGTTTCCATTTCGAGATGCA

ATGTGAAGGATTGTTTCCCCATCTTCGTCGACTCTATCAGCGTCCATCTCCAATTCAGCATCTATATAAA

ACACTATAGTACTAGCCAGATCCATACCTACTATATGTTCTTATATATTTATGTATAGAAGAATTAATAG

AAGAAACTGCATCCCCTGTCTCTATATATAAAAGCTTGCTGTATTTTAAGATGCAGTCACTTGAACTAGC

TTTGTTACTGGAATCATACTTTTTG

>MSTRG.4060.1 gene=MSTRG.4060

ATTTTTTTTTTTGCATTACCCTGACCCTGAATTTACATCCTTGGAAGAAAAGTATTAGTTGTAATCTTTA

CATTTTCTTCTAGGAAATATAATTCTTCCTTGCCCTGAATTATCTTGGAGTCACCTTTATTTGCCGCATA

CTTTTCCCGTTAAAATAATATAATTCCGAAAAGTATCATCAAATAAATTGTTGAACTGTTGCATCTCCCT

GGACTCTGGAGTGACTTGTACGATACATATTATATGTCCTCACACAAGTTAATGAAATAACATTTGATTA

AGTAGCCTATTTGAACATTAAAATTGAGCACAGCCAGAAACATGGAGGCCCATCAATATTTATCAAATTT

TGAGCTATCATATATGAACTTATATATGTATATATATAGAAAGAGATGGTTCAATTGCTTCTTATATATA

TATTTTTTCGATGATATTGTGCTATTCCAGTTGGGCATAACTTGATAACCATCTCATTTTTTCTGTTCAG

GTTTTTGAATCATTTGGCTCTGTAGAGCTCGTGTAGTCGACTGGTAATGGCAAAAGTTATGATTTTGTTT

AGTTTGCACGTCTTGAAGATGCCAGAGCTGCAGAAAGTTTAAATGGACAACTGGAAATTGCAGGTCGAGC

AATAAAGGTTTCCATGTCAGGTTTTGATGATAGATTGGTAAATGGAGGAAGCATATTGGAATTGATTGGT

AAAGAGAGGAGGAATGGAAAGAGTCCCGAAGAAGTTGAAAAGATTTATACTTGAGGCATTTGAAGTTGCT

AGTATGCTCAAGGCATTTGAAGTTGCTAGTTTCTCATCTTTTTGCTATTACTGTTCATGTTTGGTTAAAT

TTGTTAGCTACTTTGATTTGGTAGTTAGTTTAGTTTGGTTGGATTTGTTAAGTACTTTGATTTGTATGAA

AGAATAGTTTATTGGGA

>MSTRG.4061.4 gene=MSTRG.4061

ATTTAGGTTGGTCAGCATCTTCTTAATATACCAAGCTTCTAGTTACATAACTAGCTCCTGGTTGCTAGCA

AGTTTGTTGTATTATAATTTATAATCCCAGATATTCCACCAGGCAGAGTAAAATATAGTTATGGATCTGG

ATACTCTGTTCGATGCTGCCATGGCCGGGGATGCTGATGCCACTGCCAAATTGGAGATGGAAGCTGATAG

ACTGTTCAAATATGATGAAGAAACTATCCTTCATGGTGAATCTGAAAATAAATATGCAGAACGTGTGCGA

GTTATTCTGAGGGAGTTTGCAAAAAAAAATCTTTTGGCTACGCTGACTGCAGAGAAACAAACTGCACTTC

ACTTGGCAGCAATAAATGGACACACTGACTTGGCTGAGATCCTCATCGATGCAGCTAGACATTTGCCTTC

TTCAGATGATGATAATACTCGTGATAAACCAGTTACTTCTTTTCAAGCGTTTCTTAGGCAATCTGATCTG

AATGAGGACACTGCCTTACATAAAGCAGTTTTGAAAGGGCACGCTCCTATTGTTAAGCTATTAGTAGAGG

CGGATCCAAGTGATACACATATTCAAAATTGGTCTGGTGAAACACCAATGTACATAGCAGTGGAGAAAGG

GTTGAATGATATAGCAGAGATCATCTCAACAACTTGCACTGCTCCATCTTTGCTGGGACCTGATTGTAGT

AGTGTTGTGCCTAACAAGAATTTGGACCAAGCTAAAAGCCCTGGTGGAACCCTCTATAAAATTATGGACA

AAGATGCTCTGTATTCAGCTGCCATTGCTGGAGATGATGATGCCATAGCTGCATTGGAGATGCATGCTGA

TAAACTGAACAAATACGAAGAAACTATTCTTCACACAGAATCCGAAAACGGAAATACAGAACACGTGCAA

TTTATATTAAAGGAGTTTGCTGAGAAAAAGCTTCTGGTCAAGCTTAACAAATATAAACAGACAGCACTTC

ACTTAGCATCGTGTGAAGGACATACAGAAGTGGCTGAGATCCTCATTAATACTGCTAGACAGTTGCTCCC

TCCTTCTCTAGATGATGATAATCCAGTGACTACTTTTCAAGCTTTCCTTTGGCAAGCTGATGATAAAATG

GAGACTGCCTTACATGATGCAGTTAGGAAGGGTAATGTGGGCCTTGTTAAGCTATTAGTGGAGGCGGATC

CAAGTCATCCGCACACACAGAACAGTGAAGGTGAAACACCGATCTACATAGCTGTGGAAAGAGGGTATTA

CGATATAGCAAAGATGATCTGTACAACTTCCACCGCTGCTCCTAATTTGGATGCTCCTGCTGGTAGGACG

ACTGTTTTACATATTCTTATCAACAATCTCGACAAAGGTATGTAGGATATATGTTGTATATAACAATTTA

ATATGCTTTAAGATACTTTTTTTTTGTTTTTTTAGATGTACAGGCCCTCTTAAAACCCCTAAGCTAAGAA

TTCTTACAACCTTGTCATATTAATAACTGTTATGGATCAGAACTGCA

>MSTRG.4062.1 gene=MSTRG.4062

GTGGTCAAGGGAGTTATGTCAAACAATCTGTCTACTTCAATCTTCCCAGACAAAAGAGAAAAAGCCGAAC

AATTGAAGAGAAGATACAGGAAGGTATTCAAAAGTTTATGAGTGAAGAGACTGATAGGATTATTCGAGAT

AGAGATGCATTTTGGGCTGCAGAAATGGCCAAGCTCAAAGAAGCTTTAGTTGGAAAGAAATTGTGTTTTG

ATGGGAGTCCAAATATTGGATCACAACAAGGAAGTTGTTCTAAAGGCGTACTTGATAATGTGGAGTTGGA

TTTAGCAGCGAGAAAGAAATTGGATCTCAATGAAGATCTTGGTTCTGCTGAAGGGAAAGATTCGGAAAAA

GATGAGTTTGGTTTTAAAGATGATGTTGAAGGGAAGAAAAATATTGAAGAAACAGATGGTTTTCAAGGGA

AGAGAAATGGTGAACAGAAGGGAGCTGAGGTAGTAGTTGTGGATGACCCTATGGACACTGCATTGAAAAA

TGAGTGTCCTTCTCTATGGGAATTGTCTATTGAAACGCCCAACAATGTTGTTGCACATGCAACAGTTGAC

ATGGATTGTACTGTTCTTCATGGAAAGCCAATTGGAAAAGATAATGCACGAGTCTCCATTACTCATGTAA

TTCAAGGAGCTGCTCAGATCCCGTTTCCAATTGAAGATGAGATCACCACTGTTGAACAAGCTGTTGGGAC

ATTTATTGCGTGGCCGAGAAACCACTTACGGGAAGTGAAGGCAACAAGTGATGATCTGGTGAAATCTAGA

ACTGTGAAGGGAGGCAAGAAAAATGGCGCAAGGAAAATGAAGAAGATGAATGACTTGGTCTCGGAACCAG

AGTATGCGGTTACCATGGGCCCTGACTTTCCTGTTGATTTGAAGAAATTATGGCTTTGGGCAAGCGATTC

TTTGAAAGATGGTCGCTCACACACTTTTCAGTTATCCGAAGAGGCTTTTGGATCGAAGGATAGGAAGCGT

TTGTTTCAATCAGATATGAAAGCTTTGTGCTTTGGAGGTGCAATTTCAGGAGCGGTCATTTGCATGTTTA

TCAATGTTCTCCAAGATAATTTGAGAAAACACAAAATGACAGACATGATCACTTTTGTTAATCCGGCCAA

GATTGGCGCCATTGGTTGTGGTACTCCGGCAGAGAGGTCACGTGCCCTGTCTCTTCGATTCAAAAATGCC

AAGCCAGGCCAAGTTTTTCTTTTGCCATATCACCACACGAACCACTGGGCACTTACTGTGGTTAATCCAG

ATGCACAAATGGTTTACCATATGGACCCGTTAAAGAGACGAATCGCAAATGAAGAATGGACCGAAGTGGT

TAACAATGGCATCAAAATTTACAAGGAGGACCTGAAGAGGTTTGTTAAGAAGAAAATAAATTGGGAGAAT

TTAGCGGGAGTTCCTGCCCAGACTGGGACCACTGATTGTGGTCTTTTTCTAATGCATTATATGAAGGAAA

TCTGTCTCGACAAGGAACTGAAGTTTGCAACAAAGTGGGCGCGGAGGACCAACATTGTTTGTGATACCGA

TGACCTCAATGAAATGAAGTGTACGTGGGCGAAGTATTTCATGAGGCAGCATGCTAGTTAAGCCGCTTAC

ATGACTGGTTTATTTTAGGATGCACATATATATATATTTTGGGAGTTGTCGTTGACTTAGATTGTTGTTC

TTGTTGCATGCAATCAGACTTGTCTCTCTTTTTTGGATATGTGCATCCAAAAATATATTAACATTGGCCA

GTCAGGAGTTGTTTATTTTCATGTAAAAATTTTTTGTTGGATGGAATTGTTGGTTCTTGGGTGAATATTG

GTTGGTGAATATTATTACTTGGTGGATATATAGTTTGGTAGTTGTTGGTTGGTATTTTGTTTGTACTGGT

TGGTTTGTTGGT

>MSTRG.4065.1 gene=MSTRG.4065

TTTCAATAAAATCGAGATCTAAATCCTTTTGGGTGTGATCTTGTTACCGTACCCATCTTTTTGGGTTGTT

GATTGTCCCCAATTTTTGGGTGTATGTCTTGTTGCGTTTTTGATTCGTTGTCATGTTGCGTGAAGAATGA

TTTCTACGGTGTATTGGGAGATCTGGGCAACCCGCTTCAAATCTGAAATTGTGGTACTTATCGCGAGAGC

TTACCTCACACAACAAAGGATGGTTCAATATATGGGGCATCTGTATCCCCAA

>MSTRG.4067.1 gene=MSTRG.4067

AATGGATCTATCGGAAAGTTTAGTGCTCCCAAGTAAAAGACAGAGAAAGGATGTGATGTTTAATAAAGCT

ACAAAATTATTAATAAATGAAAGGGATGCACGGATGAAAGCAGATCTAGAAAGGTATAAAAATACTCGTG

CTGTTGCCCATTGGTTTACAGATGCAATTGCTGGAGATGCTAATGCCAGAGCTGCATTGGAGATGGAAGC

CGAAAGACTGAACGAAGATGGAGAAACTATCCTTCACGTTGAATCCAAGAGAGGAGACACAGAACGCGTG

CGATTCATTGTGACAGAATTTTCAAACAAAGACCTATTGGTCAAGCAGGATAAATTAAAACAAACTGCAC

TTCACCTTGCAGCACATCATGGACACACTCAAGTGGTTGAGGTCCTCATTGATGCCGCCAGACGACATTT

GCCTTCTTCTTCAGCACGTGATGATACCCATAATCCATTTAGTGCTTTTAAAGCTTTTATTAGGCTAGCT

ACTAGTCCAATGCG

>MSTRG.4068.1 gene=MSTRG.4068

TAACACCATTTCATTTAGCGGTCGAGGGTGGATATACATCTACAGTTATACAACTTATGCAAATATGGCC

AGTTTCATCTTCTGAATACACCGCTGTTAATAAGAGGGGTCAAAACGTACTACATTTGGCGGCCACCAAA

AACAAAAAAGAGATGATACAAGGTATTTTGAAATATTGTCCGCAACAATATAAGGACAAGATATTGAACG

AGCAAGATGATAATGGTGATACACCTCTCCACATACTTGTTTCCCGTGGTTGCTTCATCCCGGAACTTAT

GAAACATAACGGGATTAACACAATGGTAAAAAATAACAAATATTGGACTCCTCGTGACATGTTATATCTT

GAAGATGAAATTATTGCTGATCAGGTAC

>MSTRG.4069.1 gene=MSTRG.4069

GGAAATTATATCAAGTACTCCATACCTTTGTGGACATTCTTAAGAGCAGCTTGTAGAGCAGCATGTAAAG

CAGTACTACCGCGAGGACCGTCCAAAGATGGGGCCGTGCAAGTTGTACAGATTATCTCTACTATATCGTT

GTACCCTTCTTCCGCGGCTAAATACAGTGGCGTTTTACCCACTAGGTTTTGAAGATGTGCATCACTTGAA

TCGGCCTCTACTAATAGCTTAACAACAGAAGCATGACCCATCTCAGCAGCTAAATGTAAGGCGGTGTTCA

ACCATGAATCAGCATGCCTAAGGAAAGCTTGAAGAGAAGTAAATGGATTATCACTAGCTGATGAAGAATA

CTGCAGATGTCTAGCTGCATCAATGAGGACCTCAGCCACTTGGGTGCGTCCATAATACGCAGCCCATGTA

AGAGCAGTCTTTTGTTGATTTTCCAGCTCCACCAAAAGGTTTTTGTGTGCAAACTGCTTCAAAATGAATC

GAACACGATCTGTGTTTCCATTTATGGATTCAGTGTGAAGGATTGTTCTGCGATCTTTGTCCAGTCTATC

AGCTTTCAACTCTAATTTAGCAATCGCATCAGCGTCTCCGGCAATGGCAGCATCGTACAAAGTATTCAGA

TCCATAACTGTGTATATAATCTGTAGTCGATTATTAGAAGAAACTGTATTCAGATGCTAGTCATTCATCA

CAAATTCCAAACCAAAATATGTTATCGAC

>MSTRG.4070.1 gene=MSTRG.4070

GTAGGATTTACCATGCCGGGTGGTCTACATCAAAGTGGAGAGGTTGATGAAGGATTGGTAATTCTCTCTA

AAAAGACTGCATTTAATGCATTTATGGTATCAGATGCACTGGCTTTGCTACTATCAACATGTTCTTTGTT

TCTCTACTTTCTTGAATCTATGTACGAAGATCCCCATCAAGTGTCCAAACTAAATGCTGCATCAACTGGG

CTCAACATTGTTTCTGTCATGGCAATGATGTTGACTTTCATTACTGGAACGTATGTGGTGTTATCTCACT

CACCGGCCATCGCCATTACTGTTTGCCTCATTGGTTCCTTCTTCTTTCTTTTCATCATTGTTTTATTGAT

CAAGATGATATACGACCGTCATGTGAAGAGGAATGCGGATTGAATATGCATTCCTGTGCTAATTATTTCC

ATCTGTCAT

>MSTRG.4070.2 gene=MSTRG.4070

GTAGGATTTACCATGCCGGGTGGTCTACATCAAAGTGGAGAGGTTGATGAAGGATTGGTAATTCTCTCTA

AAAAGACTGCATTTAATGCATTTATGGTATCAGATGTTCTTTGTTTCTCTACTTTCTTGAATCTATGTAC

GAAGATCCCCATCAAGTGTCCAAACTAAATGCTGCATCAACTGGGCTCAACATTGTTTCTGTCATGGCAA

TGATGTTGACTTTCATTACTGGAACGTATGTGGTGTTATCTCACTCACCGGCCATCGCCATTACTGTTTG

CCTCATTGGTTCCTTCTTCTTTCTTTTCATCATTGTTTTATTGATCAAGATGATATACGACCGTCATGTG

AAGAGGAATGCGGATTGAATATGCATTCCTGTGCTAATTATTTCCATCTG

>MSTRG.4071.1 gene=MSTRG.4071

TTAAGGATTTGGACGAGACGAGGATGACCAAAGACGTTCAGAAAGTCAAATGATATTGCAAAGCGAATAA

GGCACGAATCGAGGAAGCGAAGCGATGTGGCGAATAAAGTATAACTAGAGATGGTCTAGAGATTACGAGA

GGCATAGATTGTGAAAGTTGAAAGATGAGATGGAGATGTGAGACTGGAGTCAGATTTAAGGCAAGCTTAA

AGATGAGATGATTGATTGAGATTGTGAGTAAGGCTAATGCTAGGCGAGGAGACAGAGTTGAGGATCCAGT

GGTCGAGGAGTGTTCAGAAAAGTTGATGAGATTGGTGCAAGCTGGAATAGTTGAC

>MSTRG.4071.2 gene=MSTRG.4071

TTAAGGATTTGGACGAGACGAGGATGACCAAAGACGTTCAGAAAGTCAAATGATATTGCAAAGCGAATAA

GGCACGAATCGAGGAAGCGAAGCGATGTGGCGAATAAAGTATAACTAGAGATGGTCTAGAGATTACGAGA

GGCATAGATTGTGAAAGTTGAAAGATGAGATGGAGATGTGAGACTGGAGTCAGATTTAAGGCAAGCTTAA

AGATGAGATGATGGAGTGAGATTATGAATAAGGCTAATGCTAGGCGAGAAGACAGAGTTGAGGATCCAGT

GGTCAAGGAGTGTTCAGGAAAGGGATGAGGTTGGTGCAAGCTGAAATAGTTGACAGTGATTCTTCTTATC

GAAGATGATCTGATTTGGTAAGAGATGTTGTGTAATAAATAGTGGTAGATTCTAATTTTAATACTGTAAC

CTGGAAGCGATCCTGTTGTTTTTGGGGGTTTAAATGTTCTATTTCGAATCCTTTATTAAAGTTTAAATGT

TCTCTGG

>MSTRG.4072.1 gene=MSTRG.4072

TGTTAATAAGAGGGGTCAAAACGTACTACATTTGGCGGCCAGCAAAAACAAAAAAGAGATGATACAAGGT

ATTTTGAAATATTGTCCTCAACAATATAAGGACAAGATATTGAACGAGCAAGATGATAATGGTGATACAC

CTCTCCACATACTTGTTTCCCGTGGTTGCTTCATCCCGGAACTTATGAAACATAACGGGATTAACACAAT

GGTAAAAAATAACCAATATTGGACTCCTCGTGATATGTTATATCTTGAAGATGAAATTATTGCTGATCAG

GTACGTTTATTCCTTACATTTCTATTTTCTACTAAAT

>MSTRG.4073.1 gene=MSTRG.4073

TTTATTTAATAAAACTGATGAATCGGGATGCACGGTCTTGGAACTGGCAGTAACGGGTAATCACGTGAAC

GTGGTTGAACTGTTATTAAAAGAAGATCCAGCCTATCAACATGGTAGAGGAAGTAAATCAAAAAGTCTTA

TCCGTTTAATCAATAGAGCCATGGATGAGGAGTATAACGAAATTGTCGAAATACTTTCTAAACCTTACGA

AGCTGGTGGTCACAAAG

>MSTRG.4075.1 gene=MSTRG.4075

GTAATCCCGAAGTCACTTCTTTTCAAGCTTTTCTCAGGCAAGGTGATAAAGATTTGGACACTGCCTTACA

TGCAGCAGTCTTGAGAGGAAATTTTACTATCGTTAAGCTACTAGTAGAGGCCGATCGGACTGATACACAT

ATTCAAAACGACAACGGTAAAACACCAATGTACATAGCTGTGGAAAAAGGTCTCCACGATATAGCAGAGA

TTATCTCTACAACTTGCACTGCTCCATCTTTGCTCGGACCTGATTGTAGTACTGTTGTGCGTATTAAAAA

TCTGGACCAAGCTAAGAGCCCTGGTGGAAGCCTCTACAAGATTATGGATAAAGATGCTCTGTATGCGGCT

GCCATTGCTAGAGATGCTGATGCCATAGCTGCAATGGAAATGCATGCTGATAAATTGAACGGATACGAAG

AAACTATTCTTCACACAGAGTCCGAAAACGGAAATACAGAACACGTGCGATTTATATTAAGGGAGTTTGC

AGACAAAAACCTTCTGGTCAAGCTTAACAAATATAAACAGACTGCACTTCACTTGGCTTCGTCTCAAGGA

CACACAGAAGTGGCGGAGGTCCTCATTAATGCTGCTAGACATTTGCTGCCTCCTTCTCCACATGATGATA

ATATTGATGATGATAATCCAGTTACTTCATTTCAAGATTTCCTTAGGCAAGCTGATAAATATATGGACAC

TGCCTTACATGAAGCAGTTAAGAAGGGTAATATGGCCCTTGTTAAGCTTTTGGTAGAGGCTGATACAAGT

GGTAAACATATTCAAAATGATGAAGGCAAAACACCAATGTACATTGCTGTGGAAGAAGGGTTGAATGATA

TAGCAGAGGTAATCTCTACAACTTGCATTTCTCCATCTTTGCTCGGATCTGATTGTAGTACTGTTGTGCG

TAACAAGAATTTTGACCAAGCTAAAAGCCCAGGTGGAACCCTCTATAAAATTATGCAGAAAGATGCTCTG

TATGCAGCTGCCATTGCCAGAGATGCTGATGCCATAGCTGCATTGGAGATGCATGCTGATAAGCTGGACA

TGAATGACAGAACTATCCTTCACATTGAATCCAAAGAAGGAAATATAGAACATGTGCG

>MSTRG.4077.1 gene=MSTRG.4077

TGTGGACATTCTTTGGGACCTTGGCTTTTAGTTGGACTTGAGTGACGGGGTGTTAGTTTTAAACTTTTAC

GATTCTTGATGGGGATTCTGTGGGAGTAGTAGTTTTTGTCTTGTCTCAATAAAAGAGCATGCTTGCACAC

ACTGGCACTCCACACTTGTAAATAGAAGAGTAATTGACTTAGTAGCGAGAGAGATGAAATTCGAGAACAT

TAGCACAATCTGAGGTATTATAATTGGCAAGGCAATTACTTCATAGTTTACTCATTC

>MSTRG.4080.1 gene=MSTRG.4080

CAACAATGCCGACCCAGAATAAGAAGAAGATAGAAGAGATACTATTAGGCAAATCACGGAAACTGTTATA

AATAATTACCGGGATGAGCAGTTGAGGAACATGATTCAGTTGTTCTTGGAAAGAACTGATGAGAAAAGTT

GCATCATTTTAGAGTTGGCCGTGGAGAGAAATTATTCGGACACAGTTGAATTGATATTAGCACCGCTATA

TGTGGATATAAACCAGTGGAATACGTTTGGATTGAGGTCAGCCTCCTATCGATCAATGCCTTCTTGTTTG

ATGAGTATCAAGCCTTTAATGTACAAAGTCATGGATAAAGAATACACTAATATGTTCAACTTACTCACAA

GAACATATGAAACATACGAAAGATGGGCCAAACTGATGGCAGCTGATGTGGCTATATGCAGAATGCCGCA

GTTGATATCTGCTATTAGCCGCCGTCAAGAAGGTA

>MSTRG.4081.1 gene=MSTRG.4081

AGCTAAGAGCCATGGTGGAAGCCTCTACAAGATTATGGATAAAGATGCTCTGTATGCGGCTGCCATGGGT

AGAAATGCTGATGCCATAGCTGCACTGGAGATGCATGCTGATAAAGTGAGCAGATACTCCGAAGAAACCA

TCCTTCACACAGAGTCCGAAAACGGAAATACAGAACACGTGCGATTTATATTAAACAAGTTTGCGGAGAA

AAAGCTTCTGGTCAAGCTTAACAATCGTAAACAGACTGCACTTCACTTGGCTTTGTCTGAAGGATACACA

GAATTGGCTGAAATTCTCATTGGTGCTGCTAGACATTTGCTGCCTACTGATGATGATAATCCAGGCACTT

CATTCCAAACTTTCCTTAGGCAAGCTGAT

>MSTRG.4081.2 gene=MSTRG.4081

AGCTAAGAGCCATGGTGGAAGCCTCTACAAGATTATGGATAAAGATGCTCTGTATGCGGCTGCCATGGGT

AGAAATGCTGATGCCATAGCTGCACTGGAGATGCATGCTGATAAAGTGAGCAGATACTCCGAAGAAACCA

TCCTTCACACTGAGTCCAAAACCAGAAATACAGAACAAGTGCGATTTATATTGAGGGAGTTTGCAGACAA

AAACCTTCTGGTCAAGCTTAACAAATTTAAACAGACCGCACTTCACTTGGCATCGTATGAAGGACACACA

GAAGTGGCTGAGATTCTCATTGGTGCTGCTAGACATTTGCTGCCTACTGATGATGATAATCCAGGCACTT

CATTCCAAACTTTCCTTAGGCAAGCTGAT

>MSTRG.4081.3 gene=MSTRG.4081

AGCTAAGAGCCATGGTGGAAGCCTCTACAAGATTATGGATAAAGATGCTCTGTATGCGGCTGCCATGGGT

AGAAATGCTGATGCCATAGCTGCACTGGAGATGCATGCTGATAAAGTGAGCAGATACTCCGAAGAAACCA

TCCTTCACACTGAGTCCAAAACCAGAAATACAGAACAAGTGCGATTTATATTGAGGGAGTTTGCAGACAA

AAACCTTCTGGTCAAGCTTAACAAATTTAAACAGACTGCACTTCACTTGGCTTTGTCTGAAGGATACACA

GAATTGGCTGAAATTCTCATTGGTGCTGCTAGACATTTGCTGCCTACTGATGATGATAATCCAGGCACTT

CATTCCAAACTTTCCTTAGGCAAGCTGAT

>MSTRG.4083.1 gene=MSTRG.4083

ATCTGTGTATATACATATATATATCTATATATGCTTATTTGTGATTGTGCTGTGCGAGAAAACAGAGGCG

ACGGGAGGTAGGTGGATTGGAGGTTGAAGGAGGAGGCTGATGGGGCGTTCTTCGCCGGTGAAGGAGGCGG

CGTGTGGTGGCGCTGCTATGGATTCCAGTTTTAGCTATGGTTGCAAGTTGGGCAGGTTGTTGTAGCTGCT

GTTGTTGCAGCGCAGAGAGAGATGATGGAGGCGAGCAGTTGTAGAGCTATTTGGTGGTGGCTGAAGTCGA

CCAAGATGTGCATATAGAGGATGTTCAGAAGTAGTTGAATAGTATCGTGGAGCGAGGATGGAACAAACCA

ATAAAGCGAAGCTATGCTATGGATAGTCAGTTAAGTTCTGTCAGAAACTAAGCAGAAGGGAACAATTGTC

AGTAGATAGTATTGTTAGAAATCTGAGAAGCAGGAGTTAAGGCAAGTACTTCTATCCTTCTCTTA

>MSTRG.4084.1 gene=MSTRG.4084

TGATGATGTTAATATTTATCACAGGAACATATGTGGTATTATCCCAATCAGCGGCCCTCGCCATTACAGT

TTGCGTCATTGGTTCACTTTTCTTTATTTTCGTCATTGTTCACCTGATGAAGTTTGTACGTTCTCGTCGA

GCAAAGAAGAATAGAGATTGAGTATGTCTGACGCTATCTGTTATTTTCAAATATTTGTGTAGTTCAAGTT

ATGATGTATTGTAAAAGGAACGATGCTATCTAGCAG

>MSTRG.4085.1 gene=MSTRG.4085

GTAATTACATAACATAACATTATTGTTATTATTTAATGGAATTGAATTAAACAAGATTGGTGCAACTTGT

TAATCAAATTGGGCAAACGACGGTTGTTAGAGATGGGGGTGGCCTTTCTTCATTCTGGGATATGTTTATT

CTTTGTGTTCCATTCATCTAGCTATATATATCAAATATTCTTTGTATAATGCAACTATTCAATATGCTAC

ATGCATGCGCACTAATATCATGGACAAAATGGTCACTAGGCCTGCAGATTTTGGAAATTAAAGGAGTTCT

GCATGCGCTCAAGTATATATTATTTAAGGGTTCGTCCTGGTGTGTGCCCATGGACCCATGCTCA

>MSTRG.4086.1 gene=MSTRG.4086

TAATTACATAACATAACATTATTGTTATTATTTAATGGAATTGAATTAAACAAGATTGGTGCAACTTGTT

AGTCAAATTGGGCAAACGACGGTTGTTAGAGATGGTGGTGGCCTTTCTTCATTGTGGGAAATGTTTATTC

TTTGTGTTCCATTCATCTATATCAAATACTCTTTGTATAATGCAACTATTCAATATGCTACATGCATGCG

CACTAATATCATGGACAAAATGGTCACTAGGCCTGCAGATTTTGGAAATTAAAGGAGTTCTGTAATAAAT

TATAGTCGTCATCTCCCATCAAATAACTGGACTGAATGTTAGAGATGGCAGCTT

>MSTRG.4087.2 gene=MSTRG.4087

ATTTGTTTTCGGGGAAACTGATGTATTTCAGAGGTTTCATGAACTATATATCAGCCACACGTTCTTCAGT

CTTGCAACAACCTAACAGATATCTTCCTCTCCCTATTCTTGACCAAGTACCAAATGAACTAATGTTTGAT

ACATATGTGCTTGCCTCTATCATAAAAAATGATGTTAATTGTTTTGGGCAATCGCATATTTCTTTCTCAA

TAAACTTGTGAAAAATTCTTAGAGAATGTTTAGTTCCTTTGCACCTATTCCTTTCAAGTGCTTGACATCC

TTCCAAACCTGAGGATATATACTTTGCTTAAACAGATGAAAGAGCAAATATAGCTTGTTTTCTTGATATC

CAGGTTACAAGTATAAAAACTCTGATTTAATAATTAAGAATACGTTCCCTATCATTATCACAATATGTTT

ACACGCATAATCTCATCATAAACCACAAACAAATATAACCCATAATAAAACATAAACCTAACTTCTCCAT

ATTCTATGAATGCAAACCTAACTTCTCCATATTCTATGTACATCGGTGTTTTGTACTAGTAGCTTAACAA

TATCCAGGTGACCATACTTGACTGCTGTGTGTAAGGCGGTGTCCATGACTTTATCACCTTGCCTGAGAAA

AGATTGAATAGAAGTAACTTGATCATTTGAAGGAGGCAACTGGCGAGCTGCATCAACAAGGACCTCAGCC

ACTTCAGTGTGTCCGAAATATATAGCACGGTGAAGGATAGTTTCTTCATCTTGGAACAGTCTATCAGCTT

CCATCTCCAATTCAGCAATGGCATCAGCATCTCTGGCCATGGCAGCATGGAAGAGAGTATCCAGCAGATC

CATAACTATATTCTACTCTACCACTGCTCCTGCGTAATGGATATATATCTCGGATAATACCACAAACTCG

ATAGCATCCAGGAGCTAGTTATATAAATGGAAGCTTGGTAAGGAGATGTTGACCAACCACAGTGCTACAT

ACGCTGTGCTGATGTAAATTAATGGTTGTTGAGCTTGCTGGTAATGATAATGATGCATTATAATTTTGAA

AACTAAAAAGCTGTGAAATAATATATATTATTTATATCTTTCGTTGAAATAAATCAATCATAAGTTAAAT

TGGCTCCCAGCTTTAAGTCCCGAATATATTATAGAACTATTTTAGACCATCTCTGTTTTGGTTGAGGAGA

GTATAACTAATACATGTAAACAAAGAGTCAAATTACTTTGTATCAGAATACTTTTTATCATATTAAGC

>MSTRG.4087.1 gene=MSTRG.4087

ATTTGTTTTCGGGGAAACTGATGTATTTCAGAGGTTTCATGAACTATATATCAGCCACACGTTCTTCAGT

CTTGCAACAACCTAACAGATATCTTCCTCTCCCTATTCTTGACCAAGTACCAAATGAACTAATGTTTGAT

ACATATGTGCTTGCCTCTATCATAAAAAATGATGTTAATTGTTTTGGGCAATCGCATATTTCTTTCTCAA

TAAACTTGTGAAAAATTCTTAGAGAATGTTTAGTTCCTTTGCACCTATTCCTTTCAAGTGCTTGACATCC

TTCCAAACCTGAGGATATATACTTTGCTTAAACAGATGAAAGAGCAAATATAGCTTGTTTTCTTGATATC

CAGGTTACAAGTATAAAAACTCTGATTTAATAATTAAGAATACGTTCCCTATCATTATCACAATATGTTT

ACACGCATAATCTCATCATAAACCACAAACAAATATAACCCATAATAAAACATAAACCTAACTTCTCCAT

ATTCTATGAATGCAAACCTAACTTCTCCATATTCTATGTACATCGGTGTTTTGTACTAGTAGCTTAACAA

TATCCAGGTGACCATACTTGACTGCTGTGTGTAAGGCGGTGTCCATGACTTTATCACCTTGCCTGAGAAA

AGATTGAATAGAAGTAACTTGATCATTTGAAGGAGGCAACTGGCGAGCTGCATCAACAAGGACCTCAGCC

ACTTCAGTGTGTCCGAAATATATAGCACGGTGAAGTGCTGAATGTTTATATCTAGTGAAGGATAGTTTCT

TCATCTTGGAACAGTCTATCAGCTTCCATCTCCAATTCAGCAATGGCATCAGCATCTCTGGCCATGGCAG

CATGGAAGAGAGTATCCAGCAGATCCATAACTATATTCTACTCTACCACTGCTCCTGCGTAATGGATATA

TATCTCGGATAATACCACAAACTCGATAGCATCCAGGAGCTAGTTATATAAATGGAAGCTTGGTAAGGAG

ATGTTGACCAACCACAGTGCTACATACGCTGTGCTGATGTAAATTAATGGTTGTTGAGCTTGCTGGTAAT

GATAATGATGCATTATAATTTTGAAAACTAAAAAGCTGTGAAATAATATATATTATTTATATCTTTCGTT

GAAATAAATCAATCATAAGTTAAATTGGCTCCCAGCTTTAAGTCCCGAATATATTATAGAACTATTTTAG

ACCATCTCTGTTTTGGTTGAGGAGAGTATAACTAATACATGTAAACAAAGAGTCAAATTACTTTGTATCA

GAATACTTTTTATCATATTAAGCAC

>MSTRG.4087.3 gene=MSTRG.4087

CTCATCACACTACAAGAAAAACCGGATGACATCCGATCGTAGTTGGGTTGGTCATAGTCGATACAATGAG

CTGAAATATATAACGGAAGAATACAAGAATGGTGTGGATAGTTTTCTTAAATTTGCTTGTGATAATCTTA

AAGATGAAGATGAAGGCCTTATGAGATGTCCATGTCAAAAGTGTAAAAATAGGTATTTCAAAGATCCTAG

TGGTGTGAAAGTTGATTTGTATCTTCATGGGATTATGCAATGGTATACCAAATGGGATTTACATGGGGAG

AAAGATATGCCCCGAGTTGAAGTTGGAATGAGTTCCGGTCGTAATGATGATGATGATGACATGCATGATG

CTCATAGCATGCTAAGAGATTTTGCAGATGCAAATAGATATTTTGAGAATTTTGAGGAGGAACCAAATGC

CGATGCCAAAGAATTTTACAATATGGTGAATGATGCCTCTGAACCAATCTATCCAAATAACAAGGATTAC

ACAAGATTGTCATTTGTAAACAAGCTACTACATTTCAAGAACAAGCATCATTGTAGTAATAATGGCTTCG

ATGAGTTACTTTGTCTTATTGGATCGGTGTTACCTGATGGTCACAAACTTCCCAGAAGCTATTATGAGGT

GCGAAAGATGATATCGGCATTGCATATGGGATATGAAAAAATTGATGCTTGTGAAAATGATTGCATGTTA

TTTTACAAGGAAGACATCAACAAGACACATTGTGACATATGTTATACAAGCCGATACAAGGAGCAAAAGG

ATCCAAAGAAGAAGAAGATCGCACGAAAGATCTTGCGCTACTTTCCTCTCACCCAAAGGTTGCAACGCTT

GTTTATGGCTGAGAAGACAGCTGAGTGTATGAGGTGGCACCAGAACAGAGTTGTAGTTGAAGGCCAATTA

TCTCACCCCGCAGATGGTGATGAGTGAAAGCAGTTTGATAGTAGATTCCCGCACTTTTCAAAGGAGATTC

GAAATGTTAGACTTGGCCTTTCCACTGATGGATTTGACCCATTTCGTGATGCACATGCAAGGGAATATAC

AGTCTGGCCTGTGGTGGTTGTTGTGTACAACCTTCCTCCATCTATGTGCACGAAGGCTCCGTACATGTTC

ATGCCTCTTCTCATTCCTGGACCGAGTGATCCTACAAAAGATCTTCATGTTTATCTTCGACCACTGATTG

ATGAATTGAAATTATTGTGGCACACCGGGGTGGAAACATATGATAGATTCTCACGTACCAATTTTATGAT

GAAGGCGGCACTAATGTGGACAATCAGTGACTTTCCTGCACTTGCCATGCTTAGTGGATGGTCAACTAAA

GGCAAGTTGTCTTGTCCAGTTTGTATCGGAGAAGTTAAAGCCACTCAACTAAAACATGGTGGTAAACCTA

GCTTTTATGGCACTGCTCGATATTTCTTAGACCAAGATGATCCACTTAGAAGGAGTACAAAGTTCGGAAG

AACTAAGGTACTATCAGTCACATATCGACATTCTGGAATAATAGCAAAGACAATGTGCGAGCAGATACAG

TTTCCCAATCCGGGAAAGACATCTTGGAGAAAACCCAGGGATTATGGTGTGACACATAATTGGACTCACT

TTTCGCCATTTTTTGAGCTCCCGTATTGGGAGACACTTAGACTTCGTCATTGCATCGATGTCATGCACAC

CGAGAAAAATGTTTTTGACAACATATTTTGCACATTGATTAATGACAAGAAGAAGTCGAAAGACAACTTG

AAGGCAAGACGTGATTGCATGGAATTGGGAATACATCGTGAGTTATGGATAAAAGATGACGGCACAATAC

CAAGCGCGCCATACGTGC

>MSTRG.4089.1 gene=MSTRG.4089

CAGACTCTGTATTAGGTCTCTTGAAAGGGAGAAAACACCTTGTAACTTTTTCTGAAGCAGAAGGTTGGAC

TCCACTTCACTATGCTGCTTATCATGAATTTGTTTCAATACTTGGTCCACTAATTGAAGCACAAAAGGAT

GTTGGATATCAATTTGTGTACCGAGATATGGTATCAACACCGTTTCATGTAGCAGTTGAGCATGGATATA

CTTCTACAGTTATACAACTTGTGAAATTGTGGCCATCTACATCGTCTGCATACACAGCTGTTAACAAAGA

TAGTCAGAATATACTACATTTGGCAGCAGCTAAAAATAATAAAAATATGGTACAAGGTATTCTAAAACAT

TGTCCCCAAAAATACAAGGACCAGCTTTTACAACAGCAAGATGTCAATGGTGATACACCTCTCCATCTAC

TTATCTCTAATGGTTGTTTCGTACCAGAACTTATTGAACATAAGGGACTTAATACAATGGTAAAAAATAA

GAAAAGTTGGACTCCTAGGGACATGCTGTATTCTAAAGATGATATCATTGCGGAGCAGGTACTTAAAAAA

CCTTCTTTTCTTCTTCTTTTTTAGTTTTTACATAAACTAGTTCTTTGATCCTTTCGTAACAGATCTTTGC

AGAGTATTGAGTAAAGTTATTAGTCATATGTATAGGATGTCTTCTTATAATTACTAATTATTGGTTGTTC

TTTTATTTATAGGTGAAAATTAAAATTGCACTTGATGATGTGCAGTCTAATTCGTCCAGAAAATTTTGGC

GTAAAAGCATGAAGAAGGATACGGATATCTTAGAAAGTTCAGTGCTTCCAAGTAAAAGAGAGAAAAAGGA

TTTCATTTTTGATAAATATACAAAAATATTGATCGATGAAAAGAATGCACAGATGAAAAA

>MSTRG.4090.1 gene=MSTRG.4090

ATAAATGAAAGGGATGCACGGATGAAAGCAGATCTAGAAAGGTATAAAAATACTCGTGCTGTTGCCCATT

GGTTTACAGATGCAATTGCTGGAGATGCTAATGCCAGAGCTGCATTGGAGATGGAAGCCGAAAGACTGAA

CGAAGATGGAGAAACTATCCTTCACGTTGAATCCAAGAGAGGAGACACAGAACGCGTGCGATTCATTGTG

ACAGAATTTTCAAACAAAGACCTATTGGTCAAGCAGGATAAATTAAAACAAACTGCACTTCACCTTGCAG

CACATCATGGACACACTCAAGTGGTTGAGGTCCTCATTGATGCCGCCAGACGACATTTGCCTTCTTCTTC

AGCACGTGATGATACCC

>MSTRG.4091.1 gene=MSTRG.4091

AAAGGCGCGGTTGCTTTGATTCTTGCTATTAAAAGGCGTGATAAAGTGTCTGTATTGCATTTGTTGGGAG

AAAACAAGCACCTAGCAAGCTTTGTGGACCATCAAGGCTGGACACCACTTCACCACGCCGCCTATTATGA

ATTTGATTCTATACTTGATGTCATAATAACAGCACTAAAAGATGTTGGATACCAATTTGTATATGAAGAT

ATGATAACACCATTTCATTTAGCGGCCGAGGGTGGATATACATCTACAGTTATACGCCTTATGCAAATGT

GGCCGGTTTCATCTTCTGAATACACCGCTGTTAACAAGAGTGGTCAAAACATACTACATTTGGCGGC

>MSTRG.4092.1 gene=MSTRG.4092

GGCTATTATAACAATCAAATTTTGAACATATTTATCACGAAAATTCAACATATTACAGCTCGAGTTCAAA

ATTAACACTGTGATACATCATTTTCTTGTAAAAAATTAGCAAATCTGTCAGTAGATTTTGTCAACTAATA

AGGTTGTTAGAGATGATGGTGGCCTTTTTGGACCTCTGTGGTATTGTTTATTCGGACTTCCTCCTCTTAA

GTACCTGGCCACATGTAGCATGTGAATTCCAGAATAAAAATTGCTGCAGTAAGCAGGCCCTTGTTCCAGA

AAAAAATTGCTACGAGTAATTGTGATTTGATCCCAGGTATATATATACACACATACATATAGCTACGAAT

GCTTTTAAATTTAATAGATAATAATGGGTACAGATACTTTGTATGCTGCTGCTATTGCCGGAGATGCTGA

TGCCATAGCTCAACTGGCGACGAAAGCAGATCGACTGAACTTAAAGGGAGAAACTATCCTCCACACTGAA

TCCAGAAATGGAAATGCAGAACATGTGCGATTTATTTTGAGGGAGTTTGCAGACAAAAATCTTTTGAGCA

AGCTTTGTGAGAGTAAATATACTGCACTTTCGACTGCAATTGACCATGGACGAACTGAAGTGGCTGGGAT

CCTAATTGATGCAGCTCGACAATGGCCAATAAATTCCTTTCAAGATTTTCTTAGGTTAGGTAATTATTGG

ATGGATACTGCATTACATATAGCAGTTACTAGGAAAAACGTGGATGTTGTTAAGTTATTAGTAGAGGCCG

ATCCAAGTGATACACATACACACAACAAAGCAGGTGAAACACCAATGTACATAGCTGTGAAAGCAGGGTT

CAACGATATAGCTGAAATAATCTCTACAACTTGCTCAGCTCCATATTTGTACGGACGTTATGGTAGTAGT

GCTGTGCTTATTAAGAATGTCGACCACGCTGACAGCTCTGGTGGAACCCTGTATGAAATTATGAATAAAG

ATGCTCTGTATGTGGCTGCCATTGCCGGAAATGCTGGTGCAAGAGCTGCACTAGAGATGCAAGCAGATAA

GGTGGACAATGTGCAACCTATCCTTCAAGTTGAATCAAGTGAAGGAAATACAGAACATGTGAGATTCATT

TTGAGTGAGTTTGCAAACAAAAATCTTTTGACCAAGCTTAATTCAATTAATGAGACTGCACTGCACTGGG

CAGCATATAAAGGACACACTGAAGTGGCTGAGATCCTCATTGATGCAGCTTCACGTTTGCCTCCTTCAGA

TGATGATAGTCAAGTTACTTCTTCTCAAGCTTTCCTTAGGCAAGCTGATAAACATTTAAGAACTGCGTTA

CATAATGCAGTTATGAAGGATAATGTGGCCATTGTTAAGCTATTACTCGAGGCGGATCCAAGTGATACAC

ATTCAAAAAATGATTGGCATTTCACACTTAAGGGTCATTAAGCCATATATAGGAGGGAAGAAGTTATAGA

TGTGATTGGAATAGTCTTTGATGTGGTGAAACGTGCAAATGATCTTAGGGGATACGAAAGCACCCAATGG

TTGGTTGAGTTGTTTAATAAAATTGATGAGAATAAATGCATCATTATGGAACTCGCAGTAGAGGGAAATT

ATCTGAACTTGGTTCAACTCATATCAGAAGAACATCCAGCTTATTTCGTACTCAACATGTTTCGGCGTGT

GGAGCATCCCGATTTGATTGGTCTAGTGCCATTAATTTACAAAGCTATGGATAAAGAATATGGCGATATG

GTCAACTTACTCACCAAAATATATGAAAAAGGAGTTCAAGTTGTGTACAAATATTGCTCACCTACAAGCA

TGCCC

>MSTRG.4095.1 gene=MSTRG.4095

ACTGCCTTACATGAAGCAGTTAAGAAGGGTAATGTGGCCCTTTTTAAGCTATTAGTAGAGGCCGATCCAA

GTGATACACATACACGGAACAATGGAGGTGAAACGCCAATCTACATTGCTGCGGAAAGTGGGTACTGTGA

TATAGTGAAGATGATCTGTACAACTCGCACTGCTCCACTTGATTTGGATGGTCCTGGTGGTAGTACCACT

GTTTTACATATTCTTATCAACAATGCCGACCCA

>MSTRG.4096.1 gene=MSTRG.4096

GAATAGGGCTGTCAAACGGATAATTCGGATACGGATATGCCTATATCCGTATCCTTATCCGCAACTGTCG

GATTCGGATACGGATAATATCCGCTTTATTTCGGATACGGATAATATCCGCTTTTTCCCGGATACGGGTG

CGGGTATTTCGGACGGATGTCGGATATTTTTGATGATCATACCGTATTGATGATGATTTTTGAACAATTT

TTTTATGATTAAGCT

>MSTRG.4097.1 gene=MSTRG.4097

GCACTTCACTTGGCAATTTATAAAGGACGAATTGAAGTGGCTGAGGTCCTCATTGATGCAGCTCGCCAAT

TGCCTCCTCTTTCAGATGATGATGATGATCATCCAGTAACTTCTTTTCAAGATTTTCTCAGGCAAGCTGA

TCACAAAAAGAACACCGCCTTACATGCAGCACTTAAGAAAGGTCACGTTCCTATTGTTAAGCTACTAGTA

GAGGCCGACCCAACTGATACACATATTCAAAACGATGAGGGTAAAACACCAATGTACATAGCTGTGGA

>MSTRG.4097.2 gene=MSTRG.4097

GCACTTCACTTGGCAATTTATAAAGGACGAATTGAAGTGGCTGAGGTCCTCATTGATGCAGCTCGCCAAT

TGCCTCCTCTTTCAGATGATGATGATGATCATCCAGTAACTTCTTTTCAAGATTTTCTCAGGCAAGCTGA

TCACAAAAAGAACACCGCCTTACATGCAGCACTTAAGAAAGGTCACGTTCCTATTGTTAAGCTACTAGTA

GAGGCCGACCCAACTGATACACATATTCAAAACGATGAGGGTAAAACACCCATGTACATAGCTGTGGAAG

AAGGGTTCAACGATATAGCAGAGATTATCTCTACAACTTGCACTGCTCCATCTTTTCTTGGACCTGATTC

TAGTACTGTTGTGCGTATTAAAGATATGGAGCAAGCTAAGAGCCCTGGTGGAAGCCTCTACAAGATTATG

GATAAAGATGCTCTGTATGCGGCTGCCATTGCTAGAGATGTTGTTGCCATTGCTGCATTGGGGATGCATG

CTGATAAAGTGAGCGGATACTGGGAAGAAAC

>MSTRG.4099.1 gene=MSTRG.4099

ATTCATGTAGTTGCACTCCATGCTTGTGTTCTTGTGGTCATCTATATTATTATTACATGAGCTAGATGGA

TTTTGTGGGTACATTCGCTATAGGCCCTCACGAGACCTTACTCGTCCACTAGGGCTGCCTAGGGGTTTAA

AGGGCTTGTTGCATATGCCAAATGCAACCGTGATCACCTACGGAAGTGGTATATATCTATTAGGTGTTAG

TGTTATTTATTTTATTTGCCCGAGGACGTGCAAAAGACTAAGTGTGGGGATAATTGATAGAGCATATATT

TATATATGTTTTTATATGATTTTATTCCCATAATTTTTAGTATTTTGTAAATAATCGGGATGTTACTAAT

TGATTTTAAGTGTTTAATTTCAGGAAAATAAATATTCCTAAAGTTGGATTAAATCAAGCTTATTTGGGCT

TAATCAGATGAAAATTCGGACTTTATGGTTAGCCAGCGAATCAAATCTTGTTTGGGCCGTAACTTGAGTT

CTGGACGTCAGAATTGGGCGATTCAACAGCCCACGCGAAGATATTGAAATTCTCTACAAGTTTAAGGTGG

TCCAGATGTCAAAATCCAACTGGATCAACCCAGAATCAGGCCTGAACAGAAGCCTACGAATTTCAGCATC

AGAATCCAACCTGTTTTAGGATATTATTTTATTTTCTTGTAAATTAAGAAAACTCTTATATAT

>MSTRG.4100.1 gene=MSTRG.4100

ATCACATATTAGCGATGTTACATGTGTAGTGTCACTAGTGCAGAGCTTATTTATTGACACAAGCTTATTT

TGCTCTTAAATATTGAATTGAGACTTAGATATACTTGCTTCTTGAGGGGTAGCCGCTTGTTGATGATTAG

AATTTTGACTATTCCCTTTTGAGCTTAGTACGTAAATGCTTGAGTTTGAGGTGATTGTGGGGTGTAAATG

TTTTCTTCAAAAAAAA

>MSTRG.4101.1 gene=MSTRG.4101

ATAATTTTTAGTATTTTGTAAATAATCGGGATGTTACTAATTGATTTTAAGTGTTTAATTTCAGGAAAAT

AAATATTCCTAAAGTTGGATTAAATCAAGCTTATTTGGGCTTAATCAGATGAAAATTCGGACTTTATGGT

TAGCCAGCGAATCAAATCTTGTTTGGGCCGTAACTTGAGTTCTGGACGTCAGAATTGGGCGATTCAACAG

CCCACGCGAAGATATTGAAATTCTCTACAAGTTTAAGGTGGTCCAGATGTCAAAATCCAACTGGATCAAC

CCAGAATCAGGCCTGAACAGAAGCCTACGAATTTCAGCATCAGAATCCAACCTGTTTTAGGATATTATTT

TATTTTCTTGTAAATTAAGAAAACTCTTATATAT

>MSTRG.4105.1 gene=MSTRG.4105

GTGCAGTTTAATTTTAATATTAATTCAGGGTGGTCAGTTACACTTAGCCTCTACTATAAATAGAGAACTA

AGTTGCTGGGTTTTATACACCACACATTACATTCTCTTCTTCACACTTCCTCTCTGCTCTCTCTCCCAAA

CACAAGTCAGTAGTGGTTTTTCCAGCGAGTGAGGAGTCAGTCGTAGTTGAGCTTTGGAGGTGCTGCTTAA

CCAAAGCTCGGAGCTGTTTTATCCTGGAGCAGGTGATGCAAGCATCCCAACGCAGCAGGTGGGGGCAACT

ATCTCTTCAAGAGCAGTCAGGTCTTCGTATAAGGCCTGACGACTCAGCTCCTCTTTCGTTTTCTTTGTTG

TACCTGTTGTTGGCTTCTGTTTCATTATCAGTCTGCACAAGCTATGGTTAC

>MSTRG.4107.1 gene=MSTRG.4107

AGATGTCTAGAACAAGACCTCCCAGACGACTAGTGAGCAGCCGTGGTCAGGGTCCCATCCCTCCACCTAT

CTTTGGCACTTATGAGGTAATTGAGTCAGATATTATGGGTTATATATGTATGTTGTAATAATCAGCGAGT

GTATTCAGACTTAAGATATTTTGGTATTGTAATATTCAGGTGACCTAAATCATACTTAAACCTGGCAAGA

TCCTGGTCAGGGGTTGTATTGTTTATTATTATATTCCGGATGTATTTCCTTTTTATTATTATTAGTGACG

TCAACTCTTACCCCGGGGTTGAGGCCGTCACCGTAGATGGTGG

>MSTRG.4108.1 gene=MSTRG.4108

GCCACCAGTGCTTTATTCAGTGCTGCTTAGTCTACTCCAAATAGTCAGATGGTGAAAGAGAGGAAATTAT

TGATGATGGCAATTTTGGCCTCATCATGTTCATTGAGTTGACGCAGCTAGATCTGGTCGATTTGTCAAAA

TTACTAAAATGGTATAGCCAGAGTGGAGAAGCTACGACCTACAAGGATTGCATTCCACAATCACACCTAA

GATCACGCTTGGTCTTACAAAAGGCATAATGTCATGTAAGAGCGTGAGATCTAATAGCAGTGGAAAGAAT

GTCATTGATAGTGTTGAAGTCTATCCACGCTCTTAAACTCTATCCAGTTTATGTTATTAAACATGCAGGC

CCAGATTCTTAAAATGGAGCGAGGAATCTATAGATGCGCAAAGGATATTCAGGCAGTAGTTTTAACCTAC

CGTGTGGCACTCAGGACTTCACAAGCAATTGATCACCTTGACGCCAGAGACATCAGAACCAGATAGCTTT

GCTCTAGCAGTGATGTGAATTTTGCTTACCTCCATAAAACTCAGCCTAAGCACACAACCTATGGCCGAAC

AGAATCAAACTTTCACCGACAAATGCAGATAAACTGTCCTTTATTTGAAGAAGACCTTGATGCAATAATC

AAATCACTATAATTCTGCTGTGTTTCGTCTAAGAAAGCACTGAGTGTTTTTCTTCCTGTAATAGTTGTTG

GTAATAGTTAAGTGTCTAATGTTTTTTTACATAAAAATATTTGATCGTTATAAAACAAAGGTTGTTTATT

CTTTGTGAG

>MSTRG.4108.2 gene=MSTRG.4108

CCGGAAGTTTTGAAACTGCAGAACAAGTCACGCCACGTTTTTAGCTCCCATTTACGTTGTAAAACTACTT

CCCATGTCCNCACTCTTTCGATTCAAAATTCAAATTTTTCTTAAACCTAATCAGCATAACTGTAATCATA

TGTCTTCAACGGAGGCTAAAGGTGTGTTTGTGTTGCTCTATGCTTTGTGATAGATTTTGCTTTCCCTTTG

TAATCATTACTTAATGTTTTATGTTTGGGGTCTGAGTCACTGCATCATGGCAGGCCAATCTACTCTTTGA

GCCTCTTCAGTTCGAGGGGTTAGATGTGGTACCTTTATGAAGGGGATTGTTTTGAATCGGGGAAAGATGA

AGGCCTGTTTTTTGCGGAAATTTCGAGGTCGAAATTCGCCAATCGGACCTGGAGCGTTTGTTTTCCAAGT

ACGGGAGAGTAAAGCGCGTCGACATGAAATCTGAGGCATGGTGGCCCTTGGTTCCCTTAATTATGAATTT

GCTGGTTTTTGCTGGGAGATTGAATGAGAGGGAGCTGGGAGGTTGTGCTCTTACTCTGAGATGTCAGTAA

ATAGTTTGTTGAGTTTTTAAATCAAACTATTTCCAGATGCACTCTCAACACCAAATGCAATTCTCTCAGC

CAATGGGATCACAACAATTTCAGAGTAGGCAGCTACCTTCTGGATCTATGCAACATGGCCTCGGGCAAAG

CCAACTTAACGAGGAAATCAGTTGAATCAATTCTCTAGCCTTGCCACCAGTGCTTTATTCAGTGCTGCTT

AGTCTACTCCAAATAGTCAGATGGTCAGTTTAGCGTTGTTTAATAAATATTTTATTCATTTGGATTTGCA

AACTAGCATAATCTGATATTTGATAAGTATCATTTCTTCACAAATTATGTTCTGATGCCCTCATATGAAG

TACGTATCTTGCAAATATTTCTTTAGTAATTTATTAATATTTATTGGGACAGGTGAAAGAGAGGAAATTA

TTGATGATGGCAATTTTGGCCTCATCATGTTCATTGAGTTGACGCAGCTAGATCTGGTCGATTTGTCAAA

ATTACTAAAATGGTATAGCCAGAGTGGAGAAGCTACGACCTACAAGTTTATGTTATTAAACATGCAGGCC

CAGATTCTTAAAATGGAGCGAGGAATCTATAGATGCGCAAAGGATATTCAGGCAGTAGTTTTAACCTACC

GTGTGGCACTCAGGACTTCACAAGCAATTGATCACCTTGACGCCAGAGACATCAGAACCAGATAGCTTTG

CTCTAGCAGTGATGTGAATTTTGCTTACCTCCATAAAACTCAGCCTAAGCACACAACCTATGGCCGAACA

GAATCAAACTTTCACCGACAAATGCAGATAAACTGTCCTTTATTTGAAGAAGACCTTGATGCAATAATCA

AATCACTATAATTCTGCTGTGTTTCGTCTAAGAAAGCACTGAGTGTTTTTCTTCCTGTAATAGTTGTTGG

TAATAGTTAAGTGTCTAATGTTTTTTTACATAAAAATATTTGATCGTTATAAAACAAAGGTTGTTTATTC

TTTGTG

>MSTRG.4108.3 gene=MSTRG.4108

CCACGTTTTTAGCTCCCATTTACGTTGTAAAACTACTTCCCATGTCCNCACTCTTTCGATTCAAAATTCA

AATTTTTCTTAAACCTAATCAGCATAACTGTAATCATATGTCTTCAACGGAGGCTAAAGGCCAATCTACT

CTTTGAGCCTCTTCAGTTCGAGGGGTTAGATGTGGTACCTTTATGAAGGGGATTGTTTTGAATCGGGGAA

AGATGAAGGCCTGTTTTTTGCGGAAATTTCGAGGTCGAAATTCGCCAATCGGACCTGGAGCGTTTGTTTT

CCAAGTACGGGAGAGTAAAGCGCGTCGACATGAAATCTGAGGCATGGTGGCCCTTGGTTCCCTTAATTAT

GAATTTGCTGGTTTTTGCTGGGAGATTGAATGAGAGGGAGCTGGGAGGTTGTGCTCTTACTCTGAGATGT

CAGTAAATAGTTTGTTGAGTTTTTAAATCAAACTATTTCCAGATGCACTCTCAACACCAAATGCAATTCT

CTCAGCCAATGGGATCACAACAATTTCAGAGTAGGCAGCTACCTTCTGGATCTATGCAACATGGCCTCGG

GCAAAGCCAACTTAACGAGGAAATCAGTTGAATCAATTCTCTAGCCTTGCCACCAGTGCTTTATTCAGTG

CTGCTTAGTCTACTCCAAATAGTCAGATGGTGAAAGAGAGGAAATTATTGATGATGGCAATTTTGGCCTC

ATCATGTTCATTGAGTTGACGCAGCTAGATCTGGTCGATTTGTCAAAATTACTAAAATGGTATAGCCAGA

GTGGAGAAGCTACGACCTACAAGGTAGTTCAGGCAGTAGTTTTAACCTACCGTGTGGCACTCAGGACTTC

ACAAGCAATTGATCACCTTGACGCCAGAGACATCAGAACCAGATAGCTTTGCTCTAGCAGTGATGTGAAT

TTTGCTTACCTCCATAAAACTCAGCCTAAGCACACAACCTATGGCCGAACAGAATCAAACTTTCACCGAC

AAATGCAGATAAACTGTCCTTTATTTGAAGAAGACCTTGATGCAATAATCAAATCACTATAATTCTGCTG

TGTTTCGTCTAAGAAAGCACTGAGTGTTTTTCTTCCTGTAATAGTTGTTGGTAATAGTTAAGTGTCTAAT

GTTTTTTTACATAAAAATATTTGATCGTTATAAAACAAAGGTTGTTTATT

>MSTRG.4108.4 gene=MSTRG.4108

TTTTAGCTCCCATTTACGTTGTAAAACTACTTCCCATGTCCNCACTCTTTCGATTCAAAATTCAAATTTT

TCTTAAACCTAATCAGCATAACTGTAATCATATGTCTTCAACGGAGGCTAAAGGTGTGTTTGTGTTGCTC

TATGCTTTGTGATAGATTTTGCTTTCCCTTTGTAATCATTACTTAATGTTTTATGTTTGGGGTCTGAGTC

ACTGCATCATGGCAGGCCAATCTACTCTTTGAGCCTCTTCAGTTCGAGGGGTTAGATGTGGTACCTTTAT

GAAGGGGATTGTTTTGAATCGGGGAAAGATGAAGGCCTGTTTTTTGCGGAAATTTCGAGGTCGAAATTCG

CCAATCGGACCTGGAGCGTTTGTTTTCCAAGTACGGGAGAGTAAAGCGCGTCGACATGAAATCTGAGGCA

TGGTGGCCCTTGGTTCCCTTAATTATGAATTTGCTGGTTTTTGCTGGGAGATTGAATGAGAGGGAGCTGG

GAGGTTGTGCTCTTACTCTGAGATGTCAGTAAATAGTTTGTTGAGTTTTTAAATCAAACTATTTCCAGAT

GCACTCTCAACACCAAATGCAATTCTCTCAGCCAATGGGATCACAACAATTTCAGAGTAGGCAGCTACCT

TCTGGATCTATGCAACATGGCCTCGGGCAAAGCCAACTTAACGAGGAAATCAGTTGAATCAATTCTCTAG

CCTTGCCACCAGTGCTTTATTCAGTGCTGCTTAGTCTACTCCAAATAGTCAGATGGTGAAAGAGAGGAAA

TTATTGATGATGGCAATTTTGGCCTCATCATGTTCATTGAGTTGACGCAGCTAGATCTGGTCGATTTGTC

AAAATTACTAAAATGGTATAGCCAGAGTGGAGAAGCTACGACCTACAAGGCCCAGATTCTTAAAATGGAG

CGAGGAATCTATAGATGCGCAAAGGATATTCAGGCAGTAGTTTTAACCTACCGTGTGGCACTCAGGACTT

CACAAGCAATTGATCACCTTGACGCCAGAGACATCAGAACCAGATAGCTTTGCTCTAGCAGTGATGTGAA

TTTTGCTTACCTCCATAAAACTCAGCCTAAGCACACAACCTATGGCCGAACAGAATCAAACTTTCACCGA

CAAATGCAGATAAACTGTCCTTTATTTGAAGAAGACCTTGATGCAATAATCAAATCACTATAATTCTGCT

GTGTTTCGTCTAAGAAAGCACTGAGTGTTTTTCTTCCTGTAATAGTTGTTGGTAATAGTTAAGT

>MSTRG.4108.5 gene=MSTRG.4108

AGATGCACAAAGGTAGTTCAGGCAGTAGTTTTAACCTACCGTGTGGCACTCAGGACTTCACAAGCAATTG

ATCACCTTGACGCCAGAGACATCAGAACCAGATAGCTTTGCTCTAGCAGTGATGTGAATTTTGCTTACCT

CCATAAAACTCAGCCTAAGCACACAACCTATGGCCGAACAGAATCAAACTTTCACCGACAAATGCAGATA

AACTGTCCTTTATTTGAAGAAGACCTTGATGCAATAATCAAATCACTATAATTCTGCTGTGTTTCGTCTA

AGAAAGCACTGAGTGTTTTTCTTCCTGTAATAGTTGTTGGTAATAGTTAAGT

>MSTRG.4109.1 gene=MSTRG.4109

CAAAGATCAGAGACATGATCGGCACCAGAGTACGCTTTTGATATTGAGATAAGAAATGATAATGTCGTGG

CCAAGAGTAAGATATTCTATCTTAATGATGCTTTTGATACTTCTAATTCATCTGGAGCTTCTTTATCATA

GGATGCAATAAGATCGGATTTGAGTAATTCAAGCTTTGCTGAGAGCATGATCGACTTATCCAAATCAGGA

GACACTCCAACTTCTGATAAATCAGTCTACAAAAAGATTAAGGTTGCACGT

>MSTRG.4110.2 gene=MSTRG.4110

CAAACTCTAAACACAGTGTCTCTCTCAAACTCTAAACACAGTCTCTCGAACCCTCTCCCTCTCAAACCAC

GTACCTTCTCTCGCACATATCTCTTGCAATCCCTCCCGCACATCTCTCGCTCTTTCTCTCTCGCAACTTT

CTCACGCTCAGGACTCACTTTCGCTCAATTCTCAAGCGGGTTTAAATTGAAGATCTCTAGCAGCGGGTTG

TAAGCGAGGCCGTGTTCTGAAAAGGGGTTTCAAGTACAGGATTAATTGGGTTCAAGCTACTGAGTTTGGA

TTAGGGTTCTTCAGCTGGTAATTGGGGCGGTGGGTTCTTCAGCGACTGAGTTTGGATTAGGGTTCTTCAG

CGGTTAATTTGGGGCTTTTTCAGGCTGGTAAATTGACTGAGTTGCAGGGCAATTTTGGTGAAAACGTGAA

GTTGATGAGCAGAAGAGTCATCGATGATCCTGGCCTACCACATGGCAACCTGCTAAGTGGTTCGAGCTCT

TTTTTCCTACCAATTCCACCTAAAATGGTGTTTAATTTCTTCAGCAGTAACAGATCACGAACGGAGTGGG

ACTTGCAAAGAGGAATGCATATGATCCGACACACGACACTCCTCTTCAGCGAGAACCTAGAAGTGGTGTA

CCGTATTTAAAATTTTGAAGCTACTAGTGCAGGAAAGTTGGAGTGATCCAGTAGCATCACATATCGTGTT

TGGCCTTGTCGAAAATGAGGCAATGAATTGGATAATAACCAGGGGAGACCTAGGTTATATACCTGTTATT

GGTTGAGGTTTCACAGTATTTCCGTTGGACGTGTATCGGGCTCCGCTGAAACTCTTGGCATACTTAATAA

AGACAAATGCCAAAAAATAAAGTTGGCATTGGGTGCAGTTGGCACACCAACATCTTAAGGGTGCAGTATT

AAGAGGGTGTATTCAAGTAGGATTTCATAGATTATTTTTAATCTATGAATTTTAAGAAATTGTAT

>MSTRG.4110.1 gene=MSTRG.4110

CCGGTCCAAAGGCTGGTAAATTGACTGAGTTGCAGGGCAATTTTGGTGAAAACGTGAAGTTGATGAGCAG

AAGAGTCATCGATGATCCTGGCCTACCACATGGCAACCTGCTAAGTGGTTCGAGCTCTTTTTTCCTACCA

ATTCCACCTAAAATGGTGTTTAATTTCTTCAGCATAACAGATCACGAACGGAGTGGGACTTGCAAAGAGG

AATGCATATGATCCGACACACGACACTCCTCTTCAGCGAGAACCTAGAAGTGGTGTACCGTATTTAAAAT

TTTGAAGCTACTAGTGCAGGAAAGTTGGAGTGATCCAGTAGCATCACATATCGTGTTTGGCCTTGTCGAA

AATGAGGCAATGAATTGGATAATAACCAGGGGAGACCTAGGTTATATACCTGTTATTGGTTGAGGTTTCA

CAGTATTTCCGTTGGACGTGTATCGGGCTCCGCTGAAACTCTTGGCATACTTAATAAAGACAAATGCCAA

AAAATAAAGTTGGCATTGGGTGCAGTTGGCACACCAACATCTTAAGGGTGCAGTATTAAGAGGGTGTATT

CAAGTAGGATTTCATAGATTATTTTTAATCTATGAATTTTAAGAAATTGTAT

>MSTRG.4112.1 gene=MSTRG.4112

TGGAGAATGATTTCTTGTTAAATGGGGAGCTCGACGGGGAGGATGTACATTGATTCATTATCCTCTGTGT

CCATAGTCAGTTAAAGTATGTATTAGTTAGACTTTGTCGGATGGTTATTTATTAGCTTTAGTTTTATGTA

ATCTTTTGGTATGTACTGTTTGATTGAGATCAAACAGGAGATGATGTATAGGGTTGTTGGACAGGTTGTG

TAAAATGTTATATCTATTTTTATGCAGCTTTGATATATTTATTGTTTTAA

>MSTRG.4114.1 gene=MSTRG.4114

GCGCATACGCTTGTGGGTGAAATTCAGACAAGTTAAGAAGGATTTTACTTTTCACTAACCCGCCCCTTTT

TTCACTCACTTCATTCACAACACTGTCTCTTTTACTCCTGTATCTCTCCATTCACAGCTCTCTCCCTCAC

TGCCGCTCTCGTGCTCTCTCAGCTCTGCCTCGTGCTCCATCCTCAGCGCTGTCTTCTCTCTATCGTGCTT

AATAGGGATTCTTCATATCGGGGTTGGATTCAAGGATTTGTGTGCAGATTTTCTATTGATGCTCTTTATG

AGTATAGATTTCTGACATATTTAAATCATGGAGAATATTGAAGGATATGGAGATGGAATGTATTGATCGT

ATCACCGAGAATAATTTTGTAGAATTGATTATATGAATACTTGCAGCCCTGGAGATTGATAAATTTTATG

ATGGTGATGATGCTATCTTGCATCCAATTGAAAGTCTACCACCTTTCTGCTTCTCGGTTTCTTGATTCCT

TATGGATTTCCAAGTCTTC

>MSTRG.4114.2 gene=MSTRG.4114

CTCACTTCATTCACAACACTGTCTCTTTTACTCCTGTATCTCTCCATTCACAGCTCTCTCCCTCACTGCC

GCTCTCGTGCTCTCTCAGCTCTGCCTCGTGCTCCATCCTCAGCGCTGTCTTCTCTCTATCGTGCTTAATA

GGGATTCTTCATATCGGGGTTGGATTCAAGGATTTGTGTGCAGGAAGCCGGTGTTTAAACTTAAGTGAGA

AAGAGTTAACAACACTTTGGTGGATTTGGGAATATCTTTAGTGTAAACTTAGTGGAGATTTCGGTGCTTC

AAATTCAGATTTTCTATTGATGCTCTTTATGAGTATAGATTTCTGACATATTTAAATCATGGAGAATATT

GAAGGATATGGAGATGGAATGTATTGATCGTATCACCGAGAATAATTTTGTAGAATTGATTATATGAATA

CTTGCAGCCCTGGAGATTGATAAATTTTATGATGGTGATGATGCTATCTTGCATCCAATTGAAAGTCTAC

CACCTTTCTGCTTCTCGGTTTCTTGATTCCTTATGGATTTCCAAGTCTTCGGCTTCTTTTCAAGGTT

>MSTRG.4115.1 gene=MSTRG.4115

TTTTGCCTGCTTTCTCTCTCGACGCTTTCTCAGCACAGCATATCTCGACATTCTCTCTCCCGGCGACATC

TCTCTCCACCTCTTTGCTAACCATCTCTGAGTTCATAACATTAGGACCACCCTTCAAGCTTCATTCAGTT

GGAGATCCGGCTCACATACGTTCGATTGGAGACCCAATTGTAAAATACAATTTAAAGAATCAATTTCACG

ACCCAACTTGAAGAAACCCAGCTCATGAGGAAACAGAGTCTGATTTAGGGGTTTAGGGTTCAATTGGTTT

GATTTTGCGTTTCAAGGTTCGATTGGTTAGACTGGGTTTTAGGGATTGATTCTTTATAGTTTTCTTCAAG

ATTAAAAAAGATCTATTTTTGCTAGCTTATTTTGGCGGTTCAATAGAGGGAGCACCAGATGTGGTAATTT

GGTTGTAGACTTGCAAGACTTTGTTAAAGACAATATTGAAGCAGCCGTATCAGAATCTGCATGTCTAATG

CCACCATGGAGCAGAGCCTCACGTTCATGCATAAACATTCCTTGCCCTCTGAATGTAGCTTGTTAATGTA

GTTGGCTTGTGTTGATGGTTTTATGTCGTTATAGAATTAGCCCTCCTCATTTTGATGATGGACATGTATT

TTGTAGTATCAGGCAATAAATGTATGCTAATATTTTGTGATTAATATATGAGAATATTCCTCTTTG

>MSTRG.4115.2 gene=MSTRG.4115

AAATCTTTTATATTATTTCACCCTCTCATCTTTTGCCTGCTTTCTCTCTCGACGCTTTCTCAGCACAGCA

TATCTCGACATTCTCTCTCCCGGCGACATCTCTCTCCACCTCTTTGCTAACCATCTCTGAGGTACTCTCA

CCATCTCAACTCTCTCTTAATCTCACCTCTCTCACAATATTACATCTCTTACCATCCTCTCATTTTATCA

GTTCATAACATTAGGACCACCCTTCAAGCTTCATTCAGTTGGAGATCCGGCTCACATACGTTCGATTGGA

GACCCAATTGTAAAATACAATTTAAAGAATCAATTTCACGACCCAACTTGAAGAAACCCAGCTCATGAGG

AAACAGAGTCTGATTTAGGGGTTTAGGGTTCAATTGGTTTGATTTTGCGTTTCAAGGTTCGATTGGTTAG

ACTGGGTTTTAGGGATTGATTCTTTATAGTTTTCTTCAAGATTAAAAAAGATCTATTTTTGCTAGCTTAT

TTTGGCGGTTCAATAGAGGGAGCACCAGATGTGGTAATTTGGTTGTAGACTTGCAAGACTTTGTTAAAGA

CAATATTGAAGCAGCCGTATCAGAATCTGCATGTCTAATGCCACCATGGAGCAGAGCCTCACGTTCATGC

ATAAACATTCCTTGCCCTCTGAATGTAGCTTGTTAATGTAGTTGGCTTGTGTTGATGGTTTTATGTCGTT

ATAGAATTAGCCCTCCTCATTTTGATGATGGACATGTATTTTGTAGTATCAGGCAATAAATGTATGCTAA

TATTTTGTGATTAATATATGAG

>MSTRG.4122.1 gene=MSTRG.4122

TGTCAGGACATGAAATTAAAGCTGTTGGCCAAGGCAAGTTAGTTTTTGTTAAGATTTAAATGGAAAAAAA

ATTGCGGTCATGTCACAAGTTTTAGCCAATTAAGTAAATATTAGTTATGAGGGGTATGAGCAGCAAGTAG

AATTTGAGATATAATAGGAAGTTGAAAATAACTTCCATCTTGAAACTCATTTGGTTAGTGGCTTCAAAGA

TTTGAATTCCAGGATATGTAACTTTGCCATGATGCCTTAAAAGATTTGAATTCGAAGACATGTTTTTGCC

AGTCTGGACAAGGAGGCATCTTTATTTGCTTTTATTAGTTTTCCTAAAATATTATCATCTTTTTATTCAA

CAGGCATGCTGATCTTGCAGAGTAGCCAGGCAGACATCAGGGAAGTAAACCAGCGGCCATGAAATCTTGA

CAATATCCACGGTTACAGAACTCTAACATGATCGAGATCATGCTTGGACTTGGATTTGGGCTTTCTCTAC

TCAAATCTATGTGAACATGTAGAAATCTAAGCAAACCCGGATGACTTGGATGATGCAGGCATGGGAATGA

AGACATTGTTGGGGTGAACCATGGTGCTCAAGGAAAAGGATGATAGAAGATTTTGAAAATTCATTACATG

ATTATAATGTATGATTTCATTTTACTTTTATTTTGTTATGGACAAATCACAAAAGGCAAGC

>MSTRG.4123.1 gene=MSTRG.4123

AACTCTTTTCAGCTCCCCCCAAAATTTCACTAACTCCCTCGATTCACCTCACCCTCACACCCCCGATCCT

CTGCTCACTTCACAAATATCACACATCACAGCTCTTAATCACCGATCTCACTGGTCCCTTCATCCCGATT

CTCAGCTCACATATGGATATTCATCTTGATCTTAGATTAATCAAGGGTTTAATCTCATGGGTTTTCTTCA

AATTGCTTTTAATTACTAATAACTTCTTCTAATTTATCTTATTTGTGTCGATTTGGTTTGCACATCTCCA

TTTTTGTGTGTGTTTAGGGTTTTTACAATTTCTTAAGATTAGGGCTTATAATTGGGGCTTGAAATCAAGA

TAAATGGGGTTGTATTTTGGGGGTTTAGTTGATTTATTTTAGATATTTCGATTTATATATAGTATTTATT

TGCTGATATTTTTGGTTTTGTAGAAACTAAAAGATCTGAGGCTTTTGTTCTTTCCTGTTTCAGTTCTGTG

CGAGCTCTATGATCTCAAGTTAATTTGTTTGCATTAAGGTGTATCATGTTTTGTATTTAATCTCTGATCT

GTGATGTGATGCTCAATTACAGGGAGATGACAGAAGTAGATGAGGCATATGATTTTGTTAAGGTAGGAGT

CTTAGTTTGGTCCCTACTGCAGACATTATTAATTTTGCAAGCAAACTGCTAGAAGATTTCCGAAGCTGGT

AAACGTCGGGCTAGCATCAGTAGTGTAGCTTCTCTTGACATGTTAGGTGCAGCCTCATTAATTGTACATC

ATGTCAATGACAAGTTAACTGACCAGCAGAAATGTCCAGGGAACGTATTTGGCAAGTGTACTCCCAAAAC

ATGGGATGATCATGGAATTTGTAAGAAGTCAAGTAGTCCTTACGTATTGGGCAATGAAAGAGATTGTATA

GTAGCTGATTTTTGGCTGGTATATGTGATTGTATGTTGTTACTATAGTTGATAATAAAAACTTTTGTTAC

TAATTTATG

>MSTRG.4123.2 gene=MSTRG.4123

CCGATCTCACTGGTCCCTTCATCCCGATTCTCAGCTCACATATGGATATTCATCTTGATCTTAGATTAAT

CAAGGGTTTAATCTCATGGGTTTTCTTCAAATTGCTTTTAATTACTAATAACTTCTTCTAATTTATCTTA

TTTGTGTCGATTTGGTTTGCACATCTCCATTTTTGTGTGTGTTTAGGGTTTTTACAATTTCTTAAGATTA

GGGCTTATAATTGGGGCTTGAAATCAAGATAAATGGGGTTGTATTTTGGGGGTTTAGTTGATTTATTTTA

GATATTTCGATTTATATATAGTATTTATTTGCTGATATTTTTGGTTTTGTAGAAACTAAAAGATCTGAGG

CTTTTGTTCTTTCCTGTTTCAGTTCTGTGCGAGCTCTATGATCTCAAGTTAATTTGTTTGCATTAAGGTG

TATCATATATGCAATGCAGTTTTTGTTGCAAGAATTATGAATGTTTTGTTGAGGTTTGTTAATGAATTTG

TTGCTTGAAATTGGAACTCGTGTACACAGAGAAAGACTCTTCAGGAAACTTGACTCTCTAATATAATTTT

TAGTTCATAAATCTTGTATGATGCTGAGTATTGCATTACAGGAGTGGTAAATTTACAAGTAGTTCGAAAT

ATATTTGTGTAGATATATAGTTTAGTGACCTTATACTATGCACAAAAAGTCGGAATGCTTTAGTTTATAA

GTTGGCCTTCTTTGGCTTCTGCAGGTTTTGTATTTAATCTCTGATCTGTGATGTGATGCTCAATTACAGG

GAGATGACAGAAGTAGATGAGGCATATGATTTTGTTAAGG

>MSTRG.4126.1 gene=MSTRG.4126

GAGTGCGAGGCACCTTGTGACTTCAATAGTGCTATGGTAACCTATCAACTCGGAAACCGAACACGGCTTC

CCGTTCATGAAATCCTGGGTACGCAACCCCAAGAGGGTGGCAACAAAGAGGTTACCGTTGAGATCGACCT

CGACCCAAGGATGCCGGAAACCCAAGTGAAGACTGGAGCAGCTGGGGATACCCTGTCCATCTTGGTGGAT

GAGTCCGACCCCACTAAAGAACTAAAAATTGGGGAGCAATTGGGGCCGGACCTAAGGGAAACCCTGACCG

CATTCTTAAAGAATAACCTGGACGTATTCGCTTGGAACCATTCGGATATGATCGGGATTCACCCGGAAGT

CATGTGCCACCACCTCAATATTGACCCAGACAGGAAGGGAGTCAGACAGAAGAGAAGACCAATTAGTGGG

GAAAGG

>MSTRG.4129.1 gene=MSTRG.4129

CTTTTCTTTTGATTTTTACTCGAGGACGAGCAAAAGATAGGTATGGGGAAGTTTGATTAGTCCATATTTT

TGCATATTTAAATGCTTATCTTTTGTTTATTTTCGTAGTATTAATCGCTTATTTATTTTATTTCAGGAAA

ATGTAGATAAATAAAGAAAGAAGAGAAAAGGCAAGAAAATGGAAGAAAAAGAAAAGAAAAGAAGAAAATC

AAAATGAGTTGGCAAGTAAGGGGCACATGGAGGGCTATGATCTACCCAACACACAAGGCACATGGATGGT

CAAGATTTAGCCACCCTACCCCTAAACCCTAAATCTTTAGCTATAAATTCCCCCATCTCTTTACTTGTAA

T

>MSTRG.4131.1 gene=MSTRG.4131

CGATATCCCCCGGAGTTATAAATCTTTTACAAGTTTCCATTCAACATTCAGCTACTAGTCACCCATTGAA

ACTCATATTCTGAGCTTCTTTTATCTTCTCAGTCGATATTGAAATTTGGACTCATTGGTGGTCTGTTTTT

AACTTTATAATTTGCTAAAGAGATTGCGGTGAAGGAGTTGGAAAGAGTTAGTGATGGATGAAGGTCAGGG

TAATTTCAGTGCACTGCCTCCATTTATAGCCAAGACTTATGAAATGGTGGATGATTTTTCAACGGATCTC

ATTGTATCTTGGAGTATGAACAATAGAAGTTTTGTTGTGTGGAATCCGCCAGAGTTTTCTAGGATTTTGT

TGCCGAGATTCTTCAAGCACAATAACTTTTCCAGCTTTATCCGACAGCTTAACACATATGGTTTTAGAAA

AGTTGATCCTGAACAATGGGAATTCGCTAATGAGGATTTTGTTCGAGGTGAACCACACCTTTTGAAGAAC

ATTCACAGACGCAAGCCAGTTCACAGTCATTCAGCGCAAAATCTTCATGGACATGGAGCTTCTTCATCTC

CCTTGACTGATTCAGAAAGACGTGGCTACAGGGAAGACATTGAAAGATTGAGATATGAGAAGGAATCCCT

TCACTTGGAGCTTCAAAGGCGTAATGAAGAGAAACATATATTTGAATTGCAGATGCAGATTCTGACTGAG

CGTGTGCAACATCTAGAACGACGACAGACAGATGCAATATCATCTTTGGCGGAAACTTTAAAGAAACCAG

CGCTTGCTTTTAGTCTCGTGCCAAAGTCAGAAGAAGTACATGAGAGAAAGAGAAGGGTACCCAGAAAAAG

TTCCTGGCAGGAGGATGGAAGCTCCGAAGATGATCAGACGAATGCTTTCCAAGGTTTCCTTGCAGGGACT

TCAGGCTTCAACGTGGGGTTGTTGGAGAAATTGGAGTCTTCTCTAACATTTTGGGAGAATGTAGCCAGAG

ATGCTACAATGGCTACTGTGCCAGATGATCCATCAATGGATTTTGATCAATCTACAAGCCTCGCTGGGAG

CCCTGCTATATCTTACCCTGATCTTGGTATTGAAACTTTAGCGATTGACATGAACTCCGAGCCTACTAAG

GCAATTGTGCCTGAGATTACCGTTTTAAAAGAAGAGGCAAGAGATGGAAGCTTGTCTGATTCTCCAACTG

CAGCCAATGACGGATTCTGGGAACAATTTCTAACCGATAGCTTGGAATCAACTGATGGTCTTGAACTTCA

TTCCAAAAGAAAACATCTTGAGGATAGAAAGGATAGAAGTCAAGTAGTTGATCATGAAAATTTTTGGTCG

AATATGAGGAGTATAAATAACCTCACCGACAAGCTGGGACAGCTTGCTCCAGCTCAGAGCACTTGATGCT

GGTTTCTTTCAGCCTTTACTTATATGTATCAAATTTTGATAGTTTCACTACCTACTTGTAAATATAGGAT

GCTTAATTCAATAACAGCTACAGATTTTATGCG

>MSTRG.4132.1 gene=MSTRG.4132

ATCAGACAAGCTTCCATCTCTTGCCTCTTCTTTTAAAACGGTAATCTCAGGCACAATTGCCTTAGTAGGC

TCGGAGTTCATGTCAATCGCTAAAGTTTCAATACCAAGATCAGGGTAAGATATAGCAGGGCTCCCAGCGA

GGCTTGTAGATTGATCAAAATCCATTGATGGATCATCTGGCACAGTAGCCATTGTAGCATCTCTGGCTAC

ATTCTCCCAAAATGTTAGAGAAGACTCCAATTTCTCCAACAACCCCACGTTGAAGCCTGAAGTCCCTGCA

AGGAAACCTTGGAAAGCATTCGTCT

>MSTRG.4133.1 gene=MSTRG.4133

TCTCACATCTCTGTCACATCTCTCTGTCTATAACTCGGCCCAATTTCAATTAGGGGTCCAATTGAACCCC

AAATTGGATGAATTTTCAATTAATCAAGCTCCAGCTTCGAGAATGCCTAATTTCTTGACCATGCTTTATT

TATTTCGATTGGGTTTTGTTTATTTGTAATTAGGGTTCCTTCTTTGTTCAAATTTTTTATCTGGGTTTTT

CATTTTTGAACAGATCTGGAGGTTTGGAGGCTTCAAGAGGTTTCAGCGAGCTTGAAGAGGTT

>MSTRG.4137.1 gene=MSTRG.4137

ATTGCTTCACTTCAATTTCCCTAAACAGCTTTTTTTGTACTACCCTCAATCTGTATACACAATGTTGTAC

AACTCAAAGTGTTCTGCCACAGCTTTCTTCTGTTACAACATTATCTTGTTCTTCAGCGTCCAAGTTGGTG

GACAGTTTCCCCCTTTTCCATTGTTTCCACCTTCGACTCCCGACACGCCAACTATTCCAGGCTTGTCACC

ACCCAGCCCGAACACGTTACCACCAATTCCAGTATTGCCACCGCCTAGCCCAAATACGTTACCAGGCATT

CCAGGACTAACCCCGCCTATCACAAATCCGTTACCTCCAATTCCAGGCCAACTAACACCACCTTCCGTAC

CTGCAACTTGCCCGAAAAATAGAGTGACAATGAAGATTGTATGTGCAAACGTGCTAGGCATACCCGTGAT

GCCTGTGAGCAGAGCCATTAATCCACTAGGAACACCATGCTGCAAATTCATCGACGGCCTACTCGATATC

GAGGTTGCGGTTTGCATTTGCCGTTCAATCACATTCAATACCAGTAATCCTAGAACCAACATTCCAGTTT

CTTTGGTTTTGCTGCTCAATTATTGCAAGAAAGAACTTACCCATGACTTCCGGTGTGCAACTAGTTAGTT

TCATGATACCTCTCGTTTCCATTAGTAATTATGTATATGTTGATACTTCTAGTTTCGTGCAGTGTGACTG

TTGAAATAAAGTGTTTGCTAGGTGAATAAAGCATGAGTATATTTCTGTTGAGAGACAGTTAAGATTAATC

ATATTATTATATGATTTGTAAGTTTCACACAGGTTGGTAATAAAGTATGGCTCGTTTGGTCTTTCCCTTC

CTCTTCCTCTATTCTATATTTC

>MSTRG.4138.1 gene=MSTRG.4138

TTCTGGATCATAGGGGCTATTCTTTTTCCAACATCATCCAGAGCTACTGTCCATCCCCGATACATACAGT

TGCTGCAGAAGACGGATGAAATCATTGATTATGCCTGGGGTGCTGGTGTTTTGGCAAATCTGTACAGGGA

GTTGGGTAAAGCTGTGAAGAAGACCAGCACTTCCATTAACGGGTGCGTGATGCTTCTTCAGCTCTGGGCT

TATGAGAGGTTGTTGCCTGGTCGTCCTGTGATTGCACGTGGATTAGAGCGTGTATGGCCGAGGGCACGCG

CATGGGCTGAGCCAGTACAGGGCAGGAGGGTTAACCCTCACCATCATGTACGCTAGTATCATGGGGATTT

TGATAGTTTTGATCCAGAGTGGGTTACTTGGCAGCCGTATGCTCATTTTTATCAGCTAGATGATGTCGAT

CAGGAGGAGGATGTTGATTTGTATGTTGCTTGTCATATGGCTCTGGGTCGTATCCCTATGTTTGCTTTCG

AGATTGTTGAGTATCAGATGCCAGACAGGGTATTGAGACAGTTTGGCATTTTACAGCATATACCAGAGCG

GCCTGTTGACATGAGATTTTTGAGGGCGGAGAGGAACTCCGCGTCACAGCGGGATGATCACATTGCTAGG

CTGAGTGCTTATAGAGATGAGTGGGATAGCTTCGTATCTACGGGTATGCCCATTATCACGGAGGGTGCTT

ATGTACCTATATCGGAGTATATGCATTGGTTTAGACTCCACAGCAAATTGAGGATTGTTCCGTGCCTAGC

CCCTCCTCTTGATATCATACAACCCCGTGATTGGTTCTCGCAGCAGAGTATGGTCGATGCGTTAGAGAGT

GCTATGAGGATGGCGCACGCCATCCATGTGCGGGGT

>MSTRG.4140.1 gene=MSTRG.4140

GGATCTGCCAGACAAGGATCTGCATCAATATGATGTTTGAGTGATTGGCCTTTTCTGCAGAACTCCTCAG

AAAAATAGCAGAAGTGAGTTACTTAGACGAGTTGCTGCATGTGGTGGTGCATTTAAAAGGCCTGGATGGC

ACAAAAGTTCATCTTCCAGGAACAAACCTAAACGACATAGATTTGATTATGTACTTTATAGGCTTTGATT

ATGTACTTGATATTTAGATTAGATGGATGTGTAGTAGTTGATCGATATTGTATTTGATGGATGTGTAGTC

GGTGAAATTAGAATTTGATGGATGTATGTACAATTGGTTTATATATGCATTAATAAGATTTAGCTGTTGA

TTTG

>MSTRG.4141.1 gene=MSTRG.4141

CACTACCAGAAAAAAGGCCTACGACATCGGTTTTTAGCCGATGTCTATCTAAAAACCGAGCGATGTCTTC

GCATGTGATGTTAAAGGTACTCGTGCAAAGACATCGGTTAAAACCCGATGTCTAAACTGGGATTACACAG

CGCTTTTGGGTGTACCTGATGTCTAATATCCCATTTTTGTTATAAAATTAACTACTTCAAGATCAATATT

ATGTGCATTAGGTGTAGTAAAACATATCATACACAAGTAGTTTACAGGCTTTAACATCGTGAGTTACAAA

GAAAACGATGTCTTTCTGGGTTTTTTAAAATCGG

>MSTRG.4142.1 gene=MSTRG.4142

GCGTGCTTTATCGGATGATATAGAATACACACAGAGAAAACTTTTAAAGCTACCAAGCCTGAGAATTGAT

GTTGGTGACAAATAAATGCTTACTCTTCAGTGTATTAATAATTTGCTAAAACAATACGGAAAGAAAGTAG

GAGATTATCCTGGACTCCCAGCCTTGAACCCGACTACAGCTAACAGATACAGAAATGAGCTGTTGGTGGA

ATAGATGCTGTATGATCGAGAGGCACTTAGAGTCAGGGCAGAAAAAATAATTCGGTCCTTGAACCACATG

CAATCTACTGTCTTCCAAACAATTATTTAGGCTGTGGAATTGAATTTGGGTGGCCTTTATTTTGTATATG

GTCCAGGGGGTACTGGTAAAA

>MSTRG.4144.1 gene=MSTRG.4144

TCACGAATGGGCCAAATGTTATTAGGTTCGAGTTTGTTATTGGGCCTGATTACAAAGTGAGATAACAATA

AAAATCTAGGCCCAAGAATAGTCACCTCTGGCCATAGTCTTGTACTATATAATGCACATCATGTCATTTT

CAGCATTATAGAATGAATGAATGTTTTTATCTTACATTTTCTCTGCATTTTCCTGTCCACCACTTCTTAT

TTAATGTTCAGTAGCTTAAATCGCTTATTTTAATCGTCAAGATCATGACAAATGGTATAAGAGCCACCGA

TTTCTCGGATATGGCCAGCACAATTACTGTTGATTTTCCTATCCCTTCGTACTCTTCAAACCCCTTTCAT

CACCCTAATTTTACCCCTGATTTCTATCATCCGTACCAACACAGCCCACCTCTAACTCCTCTGCACCAGT

ATACACCACCTCCGACACCTCCGTATGTTCACCACCATTTTCAACCACCTCCAGCACCACCAGATTATCT

CTACACTACTTCCGAAACACCTTCTTTTACCACATCTCCAATGCTAATTACTGATACTACCAATGGTTTT

CTGCATCAATTAGATAGTAAGGTCACTCAGTTAGATGAAAAACTAGCTGAAATCAGTCAGTTGATCAAGA

AGAAGAAAAATAGTGGCATCTTTACCAAATCAGGTTCAGTTGACTGTTTTTGAAGGTTGTCCTGAAGTTT

GTGAAGGCAGTTCTCAGTATCAAAATAGGGAAGTTGAAGCAAATGAGTAGTGCGCGGATAAACATATTCA

AATCATTCACGTGCTTGAGAGAATGGAGAAGCGCCTTGCTGATTTGCCAAATCCAGTTAAGGAAATGTCC

TCATATAAGGTAAATCAGTCTTCAAATACACTTGCTCTTGACTCGATTGTTGATGGTTTAGATGATTTTT

GTAGAGTTAAGAAAATAGTTGATGATGATAGTGAGGATTTTAAACAATTTTAAGTATTAGAAAAAGATT

>MSTRG.4145.1 gene=MSTRG.4145

TAGCAAGACTGACATAAAAGGGTCATGGAGGTACATCTGATTCAGCATCAGGCATAGGCAGCTGCTAGGG

TCGTAGTTCAATTCCTGTTCTGTCGCGTCCCCCGTGATTCTTCAGCTGTTCGCGGACCGTGTATCGGGTA

GTTGTGGTCATATTCGTGATTGGGGAAGGAGTATTGGAGGAAGGAGGATTCGGGTGTAGCAAAGATAAAG

AGGGAAATTGGCACATGAAATCTACTCAGATATCCTAATGTGCTCCGGTGACACGAGATGAATGCCAAGG

TAAAAAGATGGAGACTCTGGAAACAGGCCATTCAGGATATGACAAAGCTGTTTCTTTTTCATCTCGATCC

AGAGAAGCCAACATGTTGCTCTATGAGTGTTATACAGGCTACTAACAGGGCAAGCTGAAGCTAGGCAGCT

AATACAAGTTCGTGAGTTTAAACTGATTTAAGAAATTTGGTTTTTTGATTGAGAGCTAAGATAGATAGAG

TGCATTTCTTAGTCCGACAAAAGAGGAAATTCTCCCTTTTCATTCAGAGACTAAGGTTTGTTGATTTAGT

ACATTTCTACAAAAGCTGAGATAATGTAATTCTATTACCATTTTGTTTTAGAAATGTACTATTACCGCTC

TCTACAGAGGGTAAATACATGCGG

>MSTRG.4145.2 gene=MSTRG.4145

GTCTTTGCGCCTTATTTCCGTTAGCAAGACTGACATAAAAGGGTCATGGAGGTACATCTGATTCAGCATC

AGGCATAGGCAGCTGCTAGGGTCGTAGTTCAATTCCTGTTCTGTCGCGTCCCCCGTGATTCTTCAGCTGT

TCGCGGACCGTGTATCGGGTAGTTGTGGTCATATTCGTGATTGGGGAAGGAGTATTGGAGGAAGGAGGAT

TCGGGTGTAGCAAAGGAAAAGAGTGAAATTAGCACATGAAATCTACTCAGATATCCTAGCGTGCTCCGGT

GACACGAGGTGTGTTTCCTCTGTGGCCTTAACTATTGTTTCCTCTTGATATTCCTTCACAAAATCGTGCT

ATTATAAGTATATACCACAAATTTGTGCATATCTTCAGTTAGGCTGGTCATGCGACATTAAACGTGTGAC

CTCTATAACTTATATGATATGATGTATGATTGCTTCATCATTGGCTATTTGTTGTTGACACTCGGT

>MSTRG.4145.3 gene=MSTRG.4145

GTCTTTGCGCCTTATTTCCGTTAGCAAGACTGACATAAAAGGGTCATGGAGGTACATCTGATTCAGCATC

AGGCATAGGCAGCTGCTAGGGTCGTAGTTCAATTCCTGTTCTGTCGCGTCCCCCGTGATTCTTCAGCTGT

TCGCGGACCGTGTATCGGGTAGTTGTGGTCATATTCGTGATTGGGGAAGGAGTATTGGAGGAAGGAGGAT

TCGGGTGTAGCAAAGGTATGGCCTGCCTCACGGAAAAGAGTGAAATTAGCACATGAAATCTACTCAGATA

TCCTAGCGTGCTCCGGTGACACGAGGTGTGTTTCCTCTGTGGCCTTAACTATTGTTTCCTCTTGATATTC

CTTCACAAAATCGTGCTATTATAAGTATATACCACAAATTTGTGCATAT

>MSTRG.4147.1 gene=MSTRG.4147

ATTCTCTGATGCTTATAACACTCTGAATCTTGATAAATGATCTCATTTTTATCTTCTCTGAGACAAAACA

CCTATGAAGATACCATGAAGCATGTTCTGAATGAGTAATCATGAATCTCAGCTAAGTTCCTCAATCCTGA

TCTCAATTTTGTGCTACATTAGTTCACTATGCATGTTGATATCATTGAGACTCTACTACTTCACATATCT

TAATTATAAGTGAGACTGATTAATTTAAATTTTCTCTCATATACTAGACAGTGATTATAAACTCTGATGT

TTTTATTACCTGAGACAAAACCCCCTGAAAAATAAGCATGGTACTCCTTTGAGATCTTAGATGTTAATAT

GACAGTGACTGGGAAGAACCAAACATTTGCTGGTATTAAGTAATTGCTAAGATTTTATAATCTATGGTGA

CCAGTCACAATCTCTGATTACTGGAGAAATACTAATGCAATAGTATATTTAAAG

>MSTRG.4150.1 gene=MSTRG.4150

GGGAGGCAAGAAAAATGGCGCAAGGAAAATGAAGAAGATGAATGACTTGGTCTCGGAACCAGAGTATGCG

GTTACCATGGGCCCTGACTTTCCTGCTGATTTGAAGAAATTATGGCTTTGGGCAAGCGATTCTTTGAAAG

ATGGTCGCTCACACACTTTTCAGTTATCCGAAGAAGCTTTTGGATCGAAGGATAAGAAGCGTTTGTTTCA

ATCAGATATGAAAGCTTTGTGCTTTGGAGGTGAAATTTCAGGAGCGGTCATTTGCATGTTTATCAATGTT

CTCCAAGATAATTTGAGAAAACACAAAATGACATACATGATCACTTTTGTTAATCTGGCCAAGATTGGCG

CCATTGGTTGTGGTACTCCGGCAGAGAGGTCACGTGCCCTGTCTCTTCGATTCAAAAATGCCAAGCCAGG

CCAAGTTTTTCTTTTGCCATATCACCACACGAACCACTGGGCACTTACTGTGGTTAATCCAGATGCACAA

ATGGTTTACCATATGGACCCGTTAAAGAGACGAATCGCAAATGAAGAATGGACCGAAGTGGTTAACAATG

GCATCAAAATTTACAAGGAGGACCTGAAGAGGTTTGTTAAGAAGAAAATAAATTGGGAGAATTTAGCGGG

AGTTCCTGCCCAGACTGGGACCACTGATTGTGGTCTTTTTCTAATGCATTATATGAAGGAAATCTGTCTC

GACAAGGAACTGAAGTTTGCAACAAAGTGGGCGCGGAGGACCAACATTGTTTGTGATACCGATGACCTCA

ATGAAATGAAGTGTACGTGGGCGAAGTATTTCATGAGGCAGCATGCTAGTTAAGCCGCTTACATGACTGG

TTTATTTTAGGATGCACATATATATATATTTTGGGAGTTGTCGTTGACTTAGATTGTTGTTCTTGTTGCA

TGCAATCAGACTTGTCTCTCTTTTTTGGATATGTGCATCCAAAAATATATTAACATTGGCCAGTCAGGAG

TTGTTTATTTTCATGTAAAAATTTTTTGTTGGATGGAATTGTTGGTTCTTGGGTGAATATTGGTTGGTGA

ATATTATTACTTGGT

>MSTRG.4151.1 gene=MSTRG.4151

GAAATTAATCATGGCACCTTCAAAGAAAGCAACAAAGACAAAATCAGCTCCTAATACCACTGCCGCCAAC

CTTTTAAAGGCATTAAAAAGAAACAATTCAGTGGAGACGGGAGGGGTAAACAAAAAGGCGAAGAAGAAGC

TGGTGCTGCAGCCTTCACCTCAAGATTCCTCCCCACAGTCTATAAAGAAAAAGGCAATTTCTACCAACCT

ACAACAGGAACTGAATAAGAAGAGGAAACGGGTGGAAGAACCGGTGGCTAGTAAAGAAGGAGTTAAGCTT

TTAAAACGGGGCGCAGTTACGATGCATAGGATTATTAGGAGAAAGATGCTAGGAATCAAAAAGACCGTGA

GTTTCAACGCAAAGGGCGAGCCATACGGTACTGTTGCCACAGAGATGCAGTCTTACATTGGGGTTCTTGC

GAGAACTAAGGCCCCTATTTGGTACGAAAATTGGAAACGAGTTCCAAAAGAAAGAAAAAACAAAATCTGG

GATTGTGTTCAGATGGCATTTGATGTGCCATTAACAGCAAGGAGGATGGTGTTAAAATCTGCTTGCCAGA

AGTGGAGGGAGTTCAAGATCCGTTTAACAACAGCTTATATTCTTCCTTTTCGTGAACAACCTGAGTTGCT

TGCACATCCACCGGCAGACTATACATTTATTGAGAAGGCACATTGGGACATTTTTGTTGCTGATCGTTTA

TCTAAAGATTTTCTAGAAATACACAATAAGCAGAAATCTAGAAGGAGTTTGAATGTATATGGTCACTACA

TGTCTCGGAAAGGTTATGCCAACCTGGAACAAGAAATTTCTGGTGGTGATCCAGAAATTGAGATAGACCG

GTCAAGCACATGGCTTCGTGCACGGTGCGATAAAGCTGGAAATTATAAGACAGATAAAATTAAGACCATT

GCTGAAGATATTGTGAAGATGAAGCAGAAAGTTCATGAAGGGGATGTGATAGAGTTAGGCACAGATGATG

TGTTGACCAAAGTTCTTGGGAATCCTGAGCACCGCGGACGGGTCCGTGGTCAAGGGAGTTATGTCAAACA

ATCTGTCTACTTCAATCTTCCCAGACAAAAGAGAAAAAGCCGAACAATTGAAGAGAAGATACAGGAAGGT

ATTCAAAAGTTTATGAGTGAAGAGACTGATAGGATTATTCGAGATAGAGATGCATTTTGGGCTGCAGAAA

TGGCCAAGCTCAAAGAAGCTTTAGTTGGAAAGAAATTGTGTTTTGATGGGAGTCCAAATATTGGATCACA

ACAAGGAAGTTGTTCTAAAGGCGTACTTGATAATGTGGAGTTGGATTTAGCAGCGAGAAAGAAATTGGAT

CTCAATGAAGATCTTGGTTCTGCTGAAGGGAAAGATTCGGAAAAAGATGAGTTTGGTTTTAAAGATGATG

TTGAAGGGAAGAAAAATATTGAAGAAACAGATGGTTTTCAAGGGAAGAGAAATGGTGAACAGAAGGGAGC

TGAGGTAGTAGTTGTGGATGACCCTATGGACACTGCATTGAAAAATGAGTGTCCTTCTCTATG

>MSTRG.4152.1 gene=MSTRG.4152

TTGAAGTGAGTGAAGCATTAAGGAAGGCATCCTTTTGTTCTTTGAATACCGAGGGAAAATCACCACAAAC

TTTTCAAGGCTTTATCCTTGAGGACAAGGATTTTCTGAAAAAGGATGCATTCACTATAATGACCAGCCAT

CCTGGTTATATTATTAAATCGTTGATAGCTAACCAAAGTTCTAAGAATTCATCTACCTTCAGTTGGACAT

GGATATTGTGTGATCAAATACCTGAAGGGAATGCAATTATGTGGGCAGTGAGAATTTCTAATTGGAGAAA

TGATCTTGCAGAAATCATCGTGAATGCTCA

>MSTRG.4153.1 gene=MSTRG.4153

AGTTGAGGAAACTATTCATGTTGAGAGTGATGATATAATCCAATTACATGTGTCCCCACTTAAAAAGTCA

GACCCCCAAATGTTTGATGACAATTGTGAAAGAGTGTATGTTGGTAAAGCCCCAGTGCTAACTACACCTA

CCATCATCTCCTTGGTTATTCTGGATGAATTGATTAAGAATGCTTTGAAAACATATTCTGCGTGTAAACT

TACTGGAGCTAATTTAGTGATGGCCTATAGGATTCATGAAAATGAGTATGGTAAACTCTACAATAAATCA

TATGA

>MSTRG.4154.1 gene=MSTRG.4154

TACAAATATCAAGATTTCTTTTCCTTTTTTTTTTAAAATTTTTAGTGGCACTACATGATCTGTAAAGTTG

TTTATAAAGATTGATCACTATTAAACTTGTTTTTGTTGTCTGATACATATTCGTCATGGTCCATGTCCAT

AAACCATATATATGTGTGCTTTACCATAATATATTATCCCCAGGAAATGATATCCATTTTATCAAAAAAT

ATTCTTGTTGAGCTGCATAATTATTGATCTTACCAGCAGAAGTTCCAGATTTTTGTACACGAATTCATGG

CTTCCACAAGTTATAATCAAACCCCATCATTTTCCACTCCAGCTCCACCTACTACATGGGATGTTTTCTT

GAGTTTCAGAGGTAAAGATACAAGATACACATTCACCGATCATCTTTACAAAGCCTTGCTCCGCACTGGC

ATTCGAACGTTTAAGGATGATCCTGAGCTACTCAGCGGAGAAGTAATCTCTCGCGCATTACCTCAAGCTA

TTCAGGAATCCAAGACTTACATTGTTGTCCTCTCGGAAAACTATGCCACTTCATCTTGGTGCCTAGACGA

GCTAGTAGAGATCCTCAATTGTTACAATCGAATGAAAAGATTAGTTATTCCCGTGTTTTACTAACTTGAT

CCATCTGTTGTTCGACACCAGATCAAGAGTTTTAAAGAAGTTTTTGAACAACATCAAACTCATTATGATG

AGGAGAAAATGAAAGAATGGCGCCTCACACTAAATCAAGTTGCTAACTTTTCCGGATATCATATATCTCA

AAACAGGTCTCAAGCTGATATTATAGAAGAAGTTGTTGAAAGGGTTTTACTCGAAATAAATCCCATAAAT

TTAAATGTTGCCAAATATCCAGTTGGGTTGGATTCTCGTGTTAATGGCGTAACAACATTGTTCAAAAGTG

AAACAGGTGTCATCAGAATTGGTATATATGGTATGGGTGGAGTTGGCAAAACGACGCTTGCCAAAGCTGT

GTATAATCTGAACTATCGTCGATTTCAGGGCAACTGTTTCCTTGCAAATGTTAGGGAGGTTTCAAAAACA

AAAGGCCTAGTATGTTTACAGCAGCAACTTATAGCCGTTGTTCTGAAATGTAACAACATAAACATTGGCA

ATGTTGATCAGGGAATTGAGTTGATAAAAGCAAGAGTCTGTTCCAAGAAAGTCCTCATTGTTGTTGATGA

TTTATATGACCCAAAACCACTAGAACATCTAGAGGGCTCATTTGCTTCGGGAAGCATGGTCATAATAACA

ACGAGAAATGAAGATTTGCTGGATAGAATTAATGTAAAAGCAAGATACAAGGTAGATAAATTGGATAAAA

ATGAGTCATACCAACTTTTTATCCAACATGCATTCGGTTGGAATACCATATCGGATACCTTCCCACAATT

GTCCAAAGTAATTCTTGAACGTGCTGGAGGGCTTCCATTAGCTCTTCAGGTTTTTGGCTCAAACTTGCTT

AACCAATCTGAAGAAGGATGGAGATGGTTCATTGACAAATTAAATCGAGTTCCCATTGGCGATATTGAGC

AAAACCTTATGATTAGCTTTGATGCCTTGAAGTTGGTTGATCCTTTTTTACAAGATATTTTCCTGGATAT

TGCTTGTTTTTACATCGGATGGAATAAAGACGAGGTTGCTAAAATGATGGAGACTTGCTATACATTTGTG

AATCATAATATTGACATTCTAAAGAAGATATGTCTAATAACGATTAGTGACAAAGATAGGCTGGGGATGC

ATGATCTGCTTCGTGATATGGGAAGGAAAATTGCGCGTAACAACTGTCCTGCAGAGCCGGAAAAACATAG

TAGATTGTGGATACCAGAAGCTATAGAAACTGTGTTAACAAATCACCAGGTAA

>MSTRG.4156.1 gene=MSTRG.4156

GTAAGTTCCTTGGCGACTGGAATCCTTGCTCATGTAAATGATATAAGATGATTATGTCACAACGATCCTC

TTCGCTCTTTGCGACTTCTCAGTAAATTAGGGATTTCCCTACNATTAGGGGGTTTTCGATTTTAATTTTC

TCAATCCGTCAGTCGCCGCCCACCTCCCATGCGGACTACAAAGGTGCGATTAATAGGTGGTTATGGTCCA

CTCCTACTTCCCCAAAGTGTGATGCCTGTTCCAAACTGGAGTCCTTAATTGGGACCTGTAAGCCCTACAA

TGTCTCCTGGTGTGCAACCTTCAGTCAAAGTTGCTTCTTTGTATGGCATGACACCACTATCTTCTTCCAC

TCCTGCATTTGGAGGACCTTGTGCTCCTCTACCTTCATATGCTGGTCCTTCAACTAATAGCCAGATGGAA

CAACTGTTTTTGCAGAGACCAGGCCAACCGGAGTGTCAATATTACATGAAGAACGAGGGCTGCATCCGAG

CTCTCTTGCAAGTATCACCAGCCAGCAGACTGGATGGCATCAAAGAAAAATTGTGTTTTTAGCCACTTAG

GGCTACCATTATGCCAGGTATGATCATTGTGCTCATCTGAAATGATTTATTACATACATTGTAAGTTTCC

GTCAAAACTTGGTAACTTTTAGCCAAAAATTTATTTGTTCACTGAAAAAGAAATGCATAAACT

>MSTRG.4156.2 gene=MSTRG.4156

GTAAGTTCCTTGGCGACTGGAATCCTTGCTCATGTAAATGATATAAGATGATTATGTCACAACGATCCTC

TTCGCTCTTTGCGACTTCTCAGTAAATTAGGGATTTCCCTACNATTAGGGGGTTTTCGATTTTAATTTTC

TCAATCCGTCAGTCGCCGCCCACCTCCCATGCGGACTACAAAGGTGCGATTAATAGATTTGTTCGTCCGC

TCAGTGAGCTTGTTCGTTTTCAGATGTGCATTGGAAATTTATTCGCTTTTTTGGAGTTATTGTGTTGATT

TTGGTAGACATTTTAGGGCTTTGTGGATGAAGTTGTATGTTCATTGCCGGTTTCATATGTATCAGGTGGT

TATGGTCCACTCCTACTTCCCCAAAGTGTGATGCCTGTTCCAAACTGGAGTCCTTAATTGGGACCTGTAA

GCCCTACAATGTCTCCTGGTGTGCAACCTTCAGTCAAAGTTGCTTCTTTGTATGGCATGACACCACTATC

TTCTTCCACTCCTGCATTTGGAGGACCTTGTGCTCCTCTACCTTCATATGCTGGTCCTTCAACTAATAGC

CAGATGGAACAACTGTTTTTGCAGAGACCAGGCCAACCGGAGTGTCAATATTACATGAAGAACGAGGGCT

GCATCCGAGCTCTCTTGCAAGTATCACCAGCCAGCAGACTGGATGGCATCAAAGAAAAATTGTGTTTTTA

GCCACTTAGGGCTACCATTATGCCAGGTATGATCATTGTGCTCATCTGAAATGATTTATTACATACATTG

TAAGTTTCCGTCAAAACTTGGTAACTTTTAGCCAAAAATTTATTTGTTCACTGAAAAAGAAATGCATAAA

CT

>MSTRG.4156.3 gene=MSTRG.4156

GTAAGTTCCTTGGCGACTGGAATCCTTGCTCATGTAAATGATATAAGATGATTATGTCACAACGATCCTC

TTCGCTCTTTGCGACTTCTCAGTAAATTAGGGATTTCCCTACNATTAGGGGGTTTTCGATTTTAATTTTC

TCAATCCGTCAGTCGCCGCCCACCTCCCATGCGGACTACAAAGGTGCGATTAATAGGGCTTTGTGGATGA

AGTTGTATGTTCATTGCCGGTTTCATATGTATCAGGTGGTTATGGTCCACTCCTACTTCCCCAAAGTGTG

ATGCCTGTTCCAAACTGGAGTCCTTAATTGGGACCTGTAAGCCCTACAATGTCTCCTGGTGTGCAACCTT

CAGTCAAAGTTGCTTCTTTGTATGGCATGACACCACTATCTTCTTCCACTCCTGCATTTGGAGGACCTTG

TGCTCCTCTACCTTCATATGCTGGTCCTTCAACTAATAGCCAGATGGAACAACTGTTTTTGCAGAGACCA

GGCCAACCGGAGTGTCAATATTACATGAAGAACGAGGGCTGCATCCGAGCTCTCTTGCAAGTATCACCAG

CCAGCAGACTGGATGGCATCAAAGAAAAATTGTGTTTTTAGCCACTTAGGGCTACCATTATGCCAGGTAT

GATCATTGTGCTCATCTGAAATGATTTATTACATACATTGTAAGTTTCCGTCAAAACTTGGTAACTTTTA

GCCAAAAATTTATTTGTTCACTGAAAAAGAAATGCATAAACT

>MSTRG.4157.1 gene=MSTRG.4157

CAAGTGGTGAAAGCTCTGCTTGGGGATCATGATTCTTGCCATTTTGCTTAGCTATTTTTAAGTTCTATAC

CCTATCCTGAATATGTCTTGTTTGGGATTACTTTTATCATGAAAACCAAAAGAATCTCATTAGCTTTAAG

CTAGCTATTAATATACCCGTAATATAAATCTTGGATATTTTAATTGTAACACATGAGGACATGGAATTTT

GCAATATAACTTCTGAAACTAGACACATTACTTATGACAAGTCAAGATATCACCTACTTACAGCTGCAGG

TTTGCCAATGGTAGAAGTAGTTGTAGTTCCCATTAGTTTCTGTATTATAGGCACGTGAATTCATATTTTC

TTATATAACATATGTGGACTGTGGTCATAAGACTTTTTTCTACAAACCACAACACCAATTCAGTCCCTCC

TTATTCAATTTCTCAATTTTTTTTGTGCTCCAGAGTTCGGATCCTTACTTTGCCCTTTTTTAATTCTTTT

CTCCTTTTATGTTATAGTTTGTTTCACATTTG

>MSTRG.4158.1 gene=MSTRG.4158

ATGTAGAAGAAGAAAGAAGGAACCCCTCGAACAATGACATAGAAAGTGACAAGAACTGGTTATCACTTGA

CAGGAGTGGCACCACTATCAATGTAGGCGAAAAAAGAAGCAACCCCTTGGTCAGTGATTTGGAAAGTAAC

AAGAACCGGTTATCTTCTTGGAGCAACACTGTCAATGTAGAAGAAGAAAGAAGGAAACCCTCGGACAATG

ACATGGAAAGTAACGAGGACTGTTTTTTACTTGCATTGAGAGGAAACAAGAAGGTACCCCTCGGAGAATG

ACATGGAAAGTAATAAGAATTGGTTATCTCTTGGTATGAGTGG

>MSTRG.4164.1 gene=MSTRG.4164

AGCACATCAGGTGCAAGTACTTTCGCATCTCCACATTTCTGACTGGAGTCGCTTTCTGTTCCAGCTTTAC

TACACTGCCTGGACCCTGGGGTGACTAATAGAACTTCCCTGAACTTTTCAGTTCTGGAAATCAAATCCTA

TAACTGGTCTTCATAAAGTTCTCCTGCTCAAAATAAATATCATGCCTTTCTTTGAGTTGCTCTATATTTT

CATCAAGTTTGGGCCATAAAATTGGCCCGCATGATACAGAGAGTGAATAAAGTTAAGAATGCCACTACTG

TCAATCTATTAAATCGTCATACAATTGCAAATACTAATTCTCGAGATAGCCCGCAGTCGGACATTACTGT

CCTGGTTCCTGATTTGGACAGAATAGGTGTTCGCCGTCCGGCCTGGTACCCTGATGAATCAAATACCAGC

CCGAAGAATTTGTTGGATTTTTTCAACAAAGTGGGGTAAAACAATGCTTCAGAAAGTCAGTTTAAAGCAG

GTGATTGTTATGGCACCAGCTGCGCATCGACGTCAGGTACCAAGGACAGTCCGGGGAAAACCAGCAATCC

AGATACATGTAACATGGATAATGGAGCTGATGATGAAACTAACACCGAGGCTGAGGAAGTAGAAGAGGCC

GGATGCGGACAGTACACAGCCCATGACGAGAATGATACCAAATCAGAGCTGCCTGATTTCTGGAAAGGCT

ACATGGATCTTGGTCCACCAACAAAACTCTATCCTAAATGTAAAGGCTGTCATGTGGAATGAGGAACGGA

ATAACAAAGGTGCCTGGTTGTTTACAAGGAAAAGTAGCTTTAATTTTGTTGATCCTGGAAGTCAGCTGCG

CATGATCACTCTGTGATCAAACTTGATGTGGAGCTCATAGTTGAATCCTCA

>MSTRG.4165.1 gene=MSTRG.4165

TGGGGGACAAGTTGTGCTACTTGGTGCCCAAGTAATACTACTTGGGGCCCAAGAAATCACATAATTAGGC

TTAAATTAATAAGTACTTAAGGAGATAATTACATTTAATAGTGTAGTATAAAAGTTAAAGAAATGAGTCA

AGCATCTCTTTTACGAAAGACACTGAAAGAGATTGTGAATGTGAAGGATAGCCAGGAGCAATTTGGGTGC

ACTATCAATCATATTAATAAGATTGTTTATCTGGAAGGCGAGCTTGATTAGTCAACTTAAATTGGAACGT

GAGTCAACTTAAATTGGAACGTGGTGCGCTGAAAGAATTCTGGAGAAGAATATTCAGGCAAGGGTAAAAG

TGAGATGAACTGAAGTGCTAGACTTGGTTGGATTTGCAAGTAGAAGTATTATGTAATCGACAGAAGTGTT

TTAAATGTGTACTTCGTAGTAGACTTGTGTCCTGGACGTTTATAGAAGCTATGTAAGACTTCTGATATAA

GGTAAAATATACATTTAAGTTATATTAAGTTTGAGATTATGAAGATATAGTTCGTTGTGTTCAGTAGAAT

GGCACCCCCGGGTCGGGTTGCTGAGGTCATCCCGGGCCGGGGGCGTCACA

>MSTRG.4166.1 gene=MSTRG.4166

AAACGGCAAAGATTGGGTGCCGACAGAGCAGGAGGTTGAGGACGAGGGGAGACGTTATCCTGAAGTTGAT

AAACTTTTAGAAGACAGACTTCGTCCGTTGGATAAAGTAGTTTCTGATGCTCGCTACAATCGATACATGG

CTGGAATTAAGCAAGAATGCGATGCATTGCAGGCTAAACTTGATGCACGTGATGAAGCTGATTATTGGGC

AATTTGGGGAGATGTTGACGCTGTGGATGATGAAGATGATTATGAAGAATATGGTTCCGACTATATTGTA

GACATCACCAAGGATTGAGAACTTAAGTCATGTTCTCATTTTAATCATATGTAATTTCGTTTTAGTTTAT

GTAATGCTTCTGTTTCTTTTAAGTTTATTTATTGCTTTAACACCATTACATTAAATTAATTAACGAAAAT

GGATTAACAACAATAAAAAAC

>MSTRG.4167.1 gene=MSTRG.4167

ATTTATGATGATGTTCTGTTAATTTACATTGGCTTCTTGAATTGAATTAGAGAATTGGATTATTGTTTTT

ATATAAACCTGTGGACCAGATTCGTGGTCATGTTAGGCCAATGAGTGCCTTGGATCCAGTGATAGAGCAT

GGCGGGGCCTGCTCGGGGTTAGGTATGACTGATCAGCATCCTAACCTTTGTTTTAATTAAAATGTGATAG

TCCAATTCGATAATTCTCTTGGAAAGATTGAATTACTCCAGTATTCCATTGATGCAAGGTATTGTGAATC

ATTCTATACAATACCTTAATCTTATGAATTTATGTCGAATCTTTTACTAAAGTATCTTGAGCCTTTGAAC

TTCTATCATAATGATTCAGAATTGTTATTGTTTATTATTACTTGCTGAG

>MSTRG.4168.10 gene=MSTRG.4168

GAAATCTGTTCAGATTTTTGGTAAGTCATCCCTAACCAACACTTTGTTACAGTCCCTTAGCCTTAGGCGC

ACAATAATCGAACGGCCTAATCTTGGTCCGGGAGTGCCTACAAATGTTTCGGTCACTGCAATTTTGGAAA

AGGCAAATGCGATTCGCCAGGCTGTTGGAAGCGATGATGGGGAAGATAACGATAGTTGGAGTGAGTGAGC

GAGCCAACATGATGTGTTGTAATTTAAAGTTTATCGGTGATGCACTGATGCTCTGGTATATGTGTGAATA

GTTACACCTCTATATATTCATCTGTACTTATTTTGTAGGGTATATGTTGTATCGTTTTCAGGTTTATTTT

CTAAAATAGCATGACTTTATGTTGTAATAAGTTGGATCAGACAGCTAGTTCCACTACATTGTAATTCCTA

ATTCTGCATGGTGCCTAAAATAATAGATAGCACCTAGCTTCGCACTTATTGAATTCATCCCTTCTGCCGT

TGCAAGTTATACAATGACTAACAGAATTTAAAGTTTCCGG

>MSTRG.4168.11 gene=MSTRG.4168

CTTAGCCTTAGGCGCACAATAATCGAACGGCCTAATCTTGGTCCGGGAGTGCCTACAAATGTTTCGGTCA

CTGCAATTTTGGAAAAGGCAAATGCGATTCGCCAGATCTATTTTACAGGCTGTTGGAAGCGATGATGGGG

AAGATAACGATAGTTGGAGTGAGTGAGCGAGCCAACATGATGTGTTGTAATTTAAAGTTTATCGGTGATG

CACTGATGCTCTGGTATATGTGTGAATAGTTACACCTCTATATATTCATCTGTACTTATTTTGTAGGGTA

TATGTTGTATCGTTTTCAGGTTTATTTTCTAAAATAGCATGACTTTATGTTGTAATAAGTTGGATCAGAC

AGCTAGTTCCACTACATTGTAATTCCTAATTCTGCATGGTGCCTAAAATAATAGATAGCACCTAGCTTCG

CACTTATTGAATTCATCCCTTCTGCCGTTGCAAGTTATACAATGACTAACAGAATTTAAAGTTTCCGG

>MSTRG.4171.1 gene=MSTRG.4171

CTTGGCTGTTCACTTAATGCTTTTGATGCAAATAATTAGCAAATTCGCTGGCAAATATGGGTCTCACACA

ATTTTAATTTCTTTGTTCAGATTGTGGAGATGATGGTGCACATGGATTGTCCTGGATGTGAAAAAAAGAT

TAAGAAAGCTCTTTTAAAATTAGATGGGGTTGAAAATGTTGAAATTGACATGGAAATGCAGAAGGTAACA

GTTACAGGATTTGCAGATCAAGAAGATATCCTAAAGACAGTTGCAAGAAATGGGAGGCAAGCGGAGCTGT

GGCCATATCAGTACAATGCTGATTACCGAGATTTTGATCAGTACTACAACTATGAAAAGAACCAGTTTCA

GAGCTATCCCACAGCTTATTCTTCGTCCGACTATAAGCCTGCGTGCAACTATTACAATGGGTACAATGTT

CACGAATACAGTTACTATCGTCAACCAACCTACCCAAGGACGATTCTGGATCAACCAGCTTTTGCTTTGT

TTAGTGATGAAAATACTAGTGCTTGTTTGATAATGTGATAATACTGGATCCAATGTATAGATCCTTACTT

CAATAATTATTTTGCAGTACAATTAGGCCGTCTTCAT

>MSTRG.4172.2 gene=MSTRG.4172

ATCTTTTTTGCCTCATATTAAACGAGGTTCACCTGATCATTTTGGTCGCTAAAATGGACATGCCATTGCT

CAAATAATGCCAGCTCTTTCTCTGTCTGCTCAAAATGTGGTGCAAGAAAAAGCCTTGGAAACTCCTACCG

AGAACTAGATTGATACTACATATACAAATAAAAGCACCGGCAAAGTGCAAAAGATTAAAGCTCCTGTTGC

AGTGGCAAACAATATTCCAGCAACTTCTGGAGGGGATATTAATTATGTGGTGTTATGACGGGATCACGAG

GTTAACTTTTGGAGGGAGGAGACATTAGTTTTTCAGGACAATGTCATTGCTCAAATGATTCCAACAATTT

TGTGTTTTTGAACTTGGTACAAGACAACATATATGTAATTTTTCAACCTGGCATACAAAATTACAAAGAA

TGAAATTAGAGCATGCCCGTGCCTTGCACGGGCTATAAAGCTAGTTATAAGATATAAATATAACCGGAGA

ATGTCTAAGCTTATATTAGTAGTTTTAGCTCTCAATATGTTGATCACTTAACCATAATTAAGGTAACTGA

CATTGGTGGTACCATTAGTAGATCCAGAGATCTTGATTATCTTTTTCTGTCGTTGTGTCATGATGCAAAA

ACTTGTAGTTGTATGGGAC

>MSTRG.4172.3 gene=MSTRG.4172

GTTGTGATTAATAATGCACTTGGAGTCTCTTGGTCAGTTCAATTTCCGCGAAAACATCATTTGTTCTCCG

CTAATATTTTTTGGGGGCATCATCATTAATAACTGGTTTCCATTTGATAGATTGACAGTTATTTATAATG

TTTTTCTTTGTCACATACTTGATAGATTTTAACATTGTGGCGAATGACGGCAAGCCACAAGAAAGTGCAA

TAGTATATAAACGACTTTGGAGGGTTTAAAGCTTGTATCGGTCCATAACTCATGGCTAATTCTATATTGA

CTGAATATGAACTCATATGCTCTCATCTTTTTTGCCTCATATTAAACGAGGTTCACCTGATCATTTTGGT

CGCTAAAATGGTAACTATCTCAGGTCATGCATGCTCCTAGATTTTACCAATAAATAACATATCTATATAT

TTTCTAAATAATCAATACAGGTGCTTGGAGTTTGAAATTAAAACCCTCGCCTCAAATTTAATGTTGGTAT

GGTGGATCTCACAGTAGTTTATCAATTTTTTTCAGGACATGCCATTGCTCAAATAATGCCAGCTCTTTCT

CTGTCTGCTCAAAATGTGGTGCAAGAAAAAGCCTTGGAAACTCCTACCGAGAACTAGATTGATACTACAT

ATACAAATAAAAGCACCGGCAAAGTGCAAAAGATTAAAGCTCCTGTTGCAGTGGCAAACAATATTCCAGC

AACTTCTGGAGGGGATATTAATTATGTGGTGTTATGACGGGATCACGAGGTTAACTTTTGGAGGGAGGAG

ACATTAGTTTTTCAGGACAATGTCATTGCTCAAATGATTCCAACAATTTTGTGTTTTTGAACTTGGTACA

AGACAACATATATGTAATTTTTCAACCTGGCATACAAAATTACAAAGAATGAAATTAGAGCATGCCCGTG

CCTTGCACGGGCTATAAAGCTAGTTATAAGATATAAATATAACCGGAGAATGTCTAAGCTTATATTAGTA

GTTTTAGCTCTCAATATGTTGATCACTTAACCATAATTAAGGTAACTGACATTGGTGGTACCATTAGTAG

ATCCAGAGATCTTGATTATCTTTTTCTGTCGTTGTGTCATGATGCAAAAACTTGTAGTTGT

>MSTRG.4172.4 gene=MSTRG.4172

TTTACAACCTATGAAGTCCATCGTTTGCCTCCAGTGATGCTCTGTCATATTGTTATTTTGCTCTCTCATA

TTCTATTCACTTTCTTTCTTATCGTCCCTTTTTTGATAAATATCAAGTTCTCTATTCAAGAAACAAGAAA

TTTAATTGTAGCCGAATAAATAGACACATCATTTCATGCTTGGTAATCGGGATGTTTGACGCTAGGCAGT

ATTAACAAGAGCTCAAGCTGGAGCTTACATGGTTTAATTAGAGCCTTCTTTTTATTAGAAGTACAACCAT

ATTCTTCCTTTTGTTTTATATTAGGGGCGCTTTTTCTAATTAATATTGGATTTCCTCTGTCAGTATTTAT

AGTACCGGAAACTGATGCATTTGTTTATTTTTTTCCTTAAAATACTATGTGAAGGTGGAAAATTACTAGA

TCGGATTTTGTCAAGGGGTGGAGGATATGCAGAGGACGATGCTAAAACCATTATTGTGCAGATACTAGGA

GTGGTTGCCTTCGGTCATCTACAAGGTGTTGTGCACCGTGATCTGAAATCGGAGAACTTTCTTTTCGCAG

CAAAAGACGGGGATGCCCCATTGAAGATCATAGATTTTGGGCCTATCTCATTTCAGCAGGCCAGATGAAC

ACCTCAATGATATTGTTGACAGTGCATACTATGTCGCGCCTGAAGTGCTCCACAGATCATATAGCATTGA

AGCTGATATTTGGAGTATAGGTGTCATAGCATATATGCTCTTATGTACCATCTGCCGAGGCAACTAGGCA

GGACATGCCATTGCTCAAATAATGCCAGCTCTTTCTCTGTCTGCTCAAAATGTGGTGCAAGAAAAAGCCT

TGGAAACTCCTACCGAGAACTAGATTGATACTACATATACAAATAAAAGCACCGGCAAAGTGCAAAAGAT

TAAAGCTCCTGTTGCAGTGGCAAACAATATTCCAGCAACTTCTGGAGGGGATATTAATTATGTGGTGTTA

TGACGGGATCACGAGGTTAACTTTTGGAGGGAGGAGACATTAGTTTTTCAGGACAATGTCATTGCTCAAA

TGATTCCAACAATTTTGTGTTTTTGAACTTGGTACAAGACAACATATATGTAATTTTTCAACCTGGCATA

CAAAATTACAAAGAATGAAATTAGAGCATGCCCGTGCCTTGCACGGGCTATAAAGCTAGTTATAAGATAT

AAATATAACCGGAGAATGTCTAAGCTTATATTAGTAGTTTTAGCTCTCAATATGTTGATCACTTAACCAT

AATTAAGGTAACTGACATTGGTGGTACCATTAGTAGATCCAGAG

>MSTRG.4172.6 gene=MSTRG.4172

TGTATTTCAATTTGAGGCAAACCAATATGATTCTCTTGAAGTGCAGATGAACACCTCAATGATATTGTTG

ACAGTGCATACTATGTCGCGCCTGAAGTGCTCCACAGATCATATAGCATTGAAGCTGATATTTGGAGTAT

AGGTGTCATAGCATATATGCTCTTATGTACCATCTGCCGAGGCAACTAGGCAGGATTAAGGTTTCATGTC

TGATCAGTCTGACAGAGAACTAGCTAGAGCTAGGCCGAGTTCATGGTTCCAAGAGCAGAATGATGCAACT

GATATAATTATATTAATGGATAATTTCGATTTCCCCATGTTGTTTAAGAGTAAGAGCACCAGAGGCTTCT

CTTTCTAATCGCAAGGTATGCTCTATGTTCTATATGAATTATGATAGAAAAATTTGTATAAAATATTTAT

TTTCA

>MSTRG.4179.1 gene=MSTRG.4179

GGCTCCTGCGGATGTCATCTCTGATTTCAAGAAGTCCCCGGAGTATGGGGATGCTTTGTCGAAGGCCGCA

GCCGCTGAAGTTATGCGCTGCTGGACCGTGGCCGAAAAGCATCTCAAGACGGACCCGGGTGCCAATGCTC

AGAGCTTCATCGACCTCTATATCGCCGCAAAGGACCAAATCGCTGCAGGTGGGCCGGAGCCCGAACCATA

TGTCGCCCCGGGTCAGGAAGTGGACTCGGATTCCTCTGCTGAGTCCGAACCTGTGGATG

>MSTRG.4181.1 gene=MSTRG.4181

TGTCATTTAGATGAGTTCCATGGAACTGTAGGTTACTGCGTTGAGTTACAGCGTACCAAACGAGGCCTTT

ATTTGGCATGCAAATAAATGATTATGGATGTGGTAAACTAAGATTTGCCCTTACTGATATTTCTACAGGA

CCTTGTGTTTTCTCCCACGCCTAGAGGTAAAAGGAAGGTGGTGCTATCAACAAATATTGCCGAAACATCA

TTAACTTTGGAGGGTGTTGTATATGTTGTTGATAGTGGGTTCTCAAAACAGCGCTTCTACAACCCGATTA

CAGATGTTGAAAATTTGATAGTAGCACCAATATCTAAAGCATCAGCTAGGCAAAGAGCTGGAAGGGCTGG

ACGAGTACGTCCTGGAAAATGTTATAGGCTATACACAGAAGAGTATTACCTCAAGGAAATGTCAACAGAG

GGGATTCCAGAGATCCAGAGAACAAGTCTTGTGTCCTCTGTCATTCAGTTAAAAGCGCTGGGCATAGATA

ATATCCTAGGCTTTGATTGGCCTGCATCGCCGTCGCCTGAAGCAATGATCAGAGCACTTGAAGTGCTATA

TTCTCTTGGAGTTGTTGATGATGACGCCAAACTTACTTCACCAGCTGGGTTCCAAATCGCGGAGATTCCT

TTAGACCCAATGATCTCGAAAATGATTTTAGCTTCAAATGAGCTTGGATGTTCTGAAGAGATCTTAACAA

TCGCTGCTGTTCTTTCCATACAGTCTATCTGGGTATCTTCTAGGGGATCACAGAAAGAGCTAGATGAAGC

AAAGATGAGATTTGCTGCTGCCGAGGGTGATCATGTGACCTTCCTGAATGTCTACAAAGGATTTCTTAAG

TGTAATAAATCTCCAACGTGGTGTCACAAGAACTTTGTGAACTATCAAGCCATGAAAAAGGTGATTGAAA

TCCGAGAACAACTCAGAAGAACAGCACAGAGAATTGGCATAATCTTGAAATCCTGTGATGGTGATACACA

GGTAGTGAGAAAGGCAGTAACTGCTGGATATTTTGCCAATGCTTGTCGTTTAGAAGCCTTCAGCCATTCT

GGAATGTATAAGACACTTAGAGGTTCTCAAGAAGTTTATATTCACCCATCATCTATCTTATTTAGAGTTA

ATCCTAAATGGATATTATTTCATTCACTAGTCTCAACTGATCGCCAGTATATGCGCAATGTAATTACCAT

CGATCCTGCTTGGCTGATAGAGGCTGCTCCACACTTCTACCGACACCGACAACCAGATTCCTTTCATCAA

TAAGATGTCCTCAAAAGATAAAGTCATGCCACTTATTGGACAAGAACCCAGCACAATTGTTTTGTTCCTG

AATCGAAAGAAAAAGAAAACTTATCAATATTGTGCAGATCATGAAGCCCTGAAGACCTGTACATTGAAAA

CATTTTGCTGATTGTTTATGGGGAAGCCCGTCCACATATTGAGATTGAAAACAAGCTGAGATGAATGAGA

TATTACAGCTGCAGCTAGTTGGAAAGATGTGTACATCTCAGCCAATGTATGGAAATTGAAACTATTAACA

TCTATACCAAATCCTACCAAGTGGATTGTAATGTAAAGTATTAATTATTTAGATTGTCACCACATTTTTC

CCTTCCTTAGTAATAGTTTTCTACTTTAAGATCTCCAGAAGGCAGTTTTTCGTTTTCTGGGTATGGTCGA

TTACCGAAATTAAATTTGTATTACTCTACCAATTGCTTGAAC

>MSTRG.4181.2 gene=MSTRG.4181

CAGGACCTTGTGTTTTCTCCCACGCCTAGAGGTAAAAGGAAGGTGGTGCTATCAACAAATATTGCCGAAA

CATCATTAACTTTGGAGGGTGTTGTATATGTTGTTGATAGTGGGTTCTCAAAACAGCGCTTCTACAACCC

GGTATCTTAAGTTTCCTTTATGGTGTGCATTGTGCATGCTGTCTTGTGTGTCTTGCATATCCTCATTTTG

TCAAATTCTCTTATAAATTTACCTAAGCTATGTCCTATATTCTAGTCCTATAATTATATATTCAGATATT

CTCAGACTCTCAGATTTACTTGGCAGAATATCTGTTCCAGTGTCTAACAGAAGCTACCTGGCTATTGAAA

CACAGTTGTAAATTGTATCTTGAACAATAGGTTTCTTTTTGTTACATACTTGCTTAGAGCTACCCCTGGT

TTCTGGATGTAATATTGAAATTATATCTAAATACGCAGCATTTTCATGGCCTTTTTAACTTGATTCTCGT

TACAACTTGCAGATTACAGATGTTGAAAATTTGATAGTAGCACCAATATCTAAAGCATCAGCTAGGCAAA

GAGCTGGAAGGGCTGGACGAGCTATACACAGAAGAGTATTACCTCAAGGAAATGTCAACAGAGGGGATTC

CAGAGATCCAGAGAACAAGTCTTGTGTCCTCTGTCATTCAGTTAAAAGCGCTGGGCATAGATAATATCCT

AGGCTTTGATTGGCCTGCATCGCCGTCGCCTGAAGCAATGATCAGAGCACTTGAAGTGCTATATTCTCTT

GGAGTTGTTGATGATGACGCCAAACTTACTTCACCAGCTGGGTTCCAAATCGCGGAGATTCCTTTAGACC

CAATGATCTCGAAAATGATTTTAGCTTCAAATGAGCTTGGATGTTCTGAAGAGATCTTAACAATCGCTGC

TGTTCTTTCCATACAGTCTATCTGGGTATCTTCTAGGGGATCACAGAAAGAGCTAGATGAAGCAAAGATG

AGATTTGCTGCTGCCGAGGGTGATCATGTGACCTTCCTGAATGTCTACAAAGGATTTCTTAAGTGTAATA

AATCTCCAACGTGGTGTCACAAGAACTTTGTGAACTATCAAGCCATGAAAAAGGTGATTGAAATCCGAGA

ACAACTCAGAAGAACAGCACAGAGAATTGGCATAATCTTGAAATCCTGTGATGGTGATACACAGGTAGTG

AGAAAGGCAGTAACTGCTGGATATTTTGCCAATGCTTGTCGTTTAGAAGCCTTCAGCCATTCTGGAATGT

ATAAGACACTTAGAGGTTCTCAAGAAGTTTATATTCACCCATCATCTATCTTATTTAGAGTTAATCCTAA

ATGGATATTATTTCATTCACTAGTCTCAACTGATCGCCAGTATATGCGCAATGTAATTACCATCGATCCT

GCTTGGCTGATAGAGGCTGCTCCACACTTCTACCGACACCGACAACCAGATTCCTTTCATCAATAAGATG

TCCTCAAAAGATAAAGTCATGCCACTTATTGGACAAGAACCCAGCACAATTGTTTTGTTCCTGAATCGAA

AGAAAAAGAAAACTTATCAATATTGTGCAGATCATGAAGCCCTGAAGACCTGTACATTGAAAACATTTTG

CTGATTGTTTATGGGGAAGCCCGTCCACATATTGAGATTGAAAACAAGCTGAGATGAATGAGATATTACA

GCTGCAGCTAGTTGGAAAGATGTGTACATCTCAGCCAATGTATGGAAATTGAAACTATTAACATCTATAC

CAAATCCTACCAAGTGGATTGTAATGTAAAGTATTAATTATTTAGATTGTCACCACATTTTTCCCTTCCT

TAGTAATAGTTTTCTACTTTAAGATCTCCAGAAGGCAGT

>MSTRG.4183.1 gene=MSTRG.4183

GTAAAATACCATCAATTCTGAATTGGCTGTTTATATTAAAATATGCAAAAGTGGATTTTGTCCTAATGTT

TTTACTTACACTACTTTAGTTTATGGGCTTCTCAAAAGCAACAACTATGATCTGTATATGATTTTCAACT

GCTTAAAGGTGCTCGCCAGTCCAAATGAGCATAACGATGAAACACTTTGCGGCTCAAACAAGAGATTGGC

ATGTTGGAGATCATTATGTATAGACAATTTCAT

>MSTRG.4184.1 gene=MSTRG.4184

GGATTGTTGACCCAACGCTCGCTACATGGGTACTAGTGCATAAATCTTTAGGGATCTCAACCATTGCACG

GTTAAATAAACCACTAGAATAAAGTGAATAATTGGTGTGTGAAGTCTTGTGGTGTTCGTAACGCATTTAC

ACTGCGAAGCGCTCTGACCTTAGACCGAGCAATTAATCACCAATAAAAAGAAAAAAAAAAAAAAAATTAG

AGTGAATAAAATAGAAGGGTGTGGACATTCTTTGGGACCTTGGCTTTTAGTTGGACTTGAGTGACGG

>MSTRG.4187.1 gene=MSTRG.4187

CATCTTTCCAATCTCCTTCGCTTTCTTCCTCATCTCCTCACCATTCATCAGCACGCATTGTACAGCCACC

TCAATCCTTTCTTTCTTCAAAACAGGACTCTTAATATTAAAGAATTCAGAGTTCCATACCTCATTTCCAA

CCGCGACTCCAATTCCAAGAACTTGAGTAATCAGCTTTTCGTTATAAAACTGTTCAGCAAACAATGGCCA

AGTAATCATAGGCACCCCAGCCATAACACTTTCAATAACAGAGTTCCATCCGCAATGAGTCATATAGCCT

CCTATAGCTGGATGGTTTAGAATATCAACTTGAGGTGCCCACTCCTTTATCACCAAACCTCGCTCTGATT

CCACTACCATTTTCTTGAATCCATCAGGCATCCAATCATCCTCGACTCTATTTTTTATCACCCAAATGAA

TGGAAATTTCATGTTTTCAAGGGCTAACGCAATCTCAATCAGTTGGGGCTCCGGAAACATGGTCATGCTA

CCAAAACAAATGTAAACCACACTATTGGCTTTTTGGGTATCCAGCCAATTCAAGGAGTAATGTGCTGATG

TGGATGTTGTTCTTTTCGGGTTGGAATTGGGATAGTGAGAGATGGGACCTACAGTCCAAATCTTCATGTT

CTTGATTTTCTTGAAATGATTGCTGTATGGAGCTTCAAGATCATAAAAAGTGTCAAAAATTATGCCATAG

CTCCGCGTTTCTGCTTCTTTTATCCTTTCTAGCACCTCACCATACCTCGTCTTAAATTTGTCATGGTCAG

GCAATTGAGATCTTTTCATTTCAATCTTATCAGGAAGGCCAGGAATCAAGAACAGTTCCGAATCAGACTC

CACCCTTTCATGAGGTCCATAAGCTTTCAAACTCTCAGGAACACAATGACAAAAAAAATTTGATGGATAA

AAAACAAGCCTTGGTATCTTCAGATGCTGAGCAGTGTCCACAGTCCAAGGAAAAAACATCCCAGAAACAA

TGCAACGTGGCGAGAGCTTGGTAATCAGATCTTCCATTGGGTTTTTAAGCAATGACAGAGCATGGCTAAC

TTTAAGGACCATCTCAGGTGATTTAACCGCGCTGTAGTTCTCAATTCCAGGTGGTAATCCAGCTTCTTCC

GAGGGAAACTCTAGAGTGGCGACAGATATTTGGTGGTTGGAAGCAATGCTGTCGTCGATTAACGACTGGA

AGACAGCTGCATTGGCAGGGGTGGTGATGATAGTGACTGTTCCACCATGAGCTGCCAATTGGCAGCCAAG

ATGAATCATTGGCATCATGTGGCTAGTAGTGAGGAAAGGAAGCAGAACAAAATCAGGCTTGGGAGGTGTT

TGATCAGACAAGGCCATATTTTTCATGTTGAGCACTTGTTTTATTGCTAATGGCTGATTTCTATAGCCTA

TAGTTGCACTTGTGTCTGCCTCTACTCTGTATAT

>MSTRG.4192.1 gene=MSTRG.4192

CTTAGCAGTATACACATCAACATAGGTGAAAGCTGCCGAGGGGAGGTCGGCTCGAAGCTGATCAACGCCC

TTCTTTAGTTCCGAATTGAAGTATTGAGTTAAATCATTGTAGCGTTTTGCACAACCAGCACTATCAAGCT

GATCGTCTGGGACTGGAGCCAGTGTCAAAAGAATTGGAAGACAGCCATTAGGTCCGAGGTTATGGATCCA

GAATGATCTACCTCCCAGGCTATACAAAATCTGTAATGCAATGGTGGATAAAATGTTTGCAAAGTTTGCA

CCATGAGAGAAGTTGGCACCCAAGGAATCCATGTATGGATGCAGGAATGGCAAGCCCAAGTCAGTTGCTG

AAAAAAATAAAATCATTAGCGATCCGTTTAGGCAAAAATATGGCAAGGGAGCTGGAGTTTAAACGTACCC

ATGAAATCGATCAATAGGCGTCCATCGGAGACTCTTCCTGCTGATCCATTGAAGTAGGACTGACCAAAGA

AAGGTGGATTGCCGAAAAACCAAGCAGCGAATGCACCCGTGTCGGAGTTTGCATCACCGAAGTTGAAGAT

TGCTGGAAAATCACAAGTCTGTGAAGATGCAGAGACATGTGAATGAGCTACAAGGATGGAAAATAAGAAA

ACAGAGAATGCAGAGTGTTTTCCCATGCTTAAAGACTTGTTTGGTCTATTATGATAGGTTGGTATTTGAT

ACTACTTGATATGAATCCAAGATGAGGAGTTAGTTGTGTACTTATAGGATTGGCATGTTGAGTCATGTGG

G

>MSTRG.4200.2 gene=MSTRG.4200

GTTTGGGCAGCGCCCCTGTTTGCAACGCTTGATGCTTGTTTTTATGAAACCCGAACCGTTTCATCCCCCT

GTATATTGCAGAATTAATTTGCAGTGGCATTATATGTGATTTATCATTTTTTAATCGTTAGCCAAAAACA

CTTGTTGGAGGGCTATATTAAGCAAGGTGATATATATATTTGCTGACAGATTTGTAGACTGTTCATAGAC

TGTTATCTTCACAGATCAGTACCAAACTTCTGCATTTTTATCACTATCCATGGATCTAAATCATGGTGCA

GGGGAGAATTATGCACACCCGCAGATATGTTTCTTCCATGTGATCTTCAAGGCTTCAGCTTTGGCATTTT

ACATACTGTCCGCTCTCTTTTTTGACAGTTTTGTCATAATTTTTGTGGTAACGGTTCTTCTGGCTGCACT

AGATTTTTGGGTAGTCAAGAATGTCAGTGGACGGATTTTGGTTGGTTTAAGGTGGTGGAATGAAATAAAT

GACGAGGGTGAGAGCATCTGGAAATTTGAATGCCTTGACCAGGAGTCACTGTCTCGAATGAACAAGAAGG

ATTCATGGCTGTTCTGGTGGACTATTTACCTTACTGCGGCCTTGTGGCTTATCTTTGGAATATTCTCCCT

CATAAGGTTTCAAGCTGATTATGTTCTTGTTGTAGGAGTTTGTTTGACACTAAGTGTTGCAAACATTGTT

GGCTTTACTAAGTGTCGTAAAGATGCTAAGAAGCAGATTCAGGCATTTGCCTCTCAAACGATTGCATCTC

GATTCTCATCCACTTTGCAGTCTGCATTCACTATTATCTAAGTAACACAGAGAGATTGGAACAGGATCAG

ATGTGTAGGAACTTTACATGCCCAAGAAATGGCGAAGAGATATGTTGGATGATACTTCAGTTTATTGAGA

GTACTTTTTGGAAGAAATTACCTTAAGCATTTAGTTATTGCTCAGGCAAGGGTACATTCTGTATGCATGA

ACCTAGCGAATACCCCCTCTTGATATATGACTTTGTATATGCTTAATACGAGAACCTAGGCACTTATACG

TGGTTAGTAATGCTGAGAAACATGGAGTATTATCCAGTCTTCTGTTGCACTGTATTGTGTTTTTCTTCTC

CTTGCATTTTCTTTGTGTTTT

>MSTRG.4200.1 gene=MSTRG.4200

GTTTGGGCAGCGCCCCTGTTTGCAACGCTTGATGCTTGTTTTTATGAAACCCGAACCGTTTCATCCCCCT

GTATATTGCAGAATTAATTTGCAGTGGCATTATATGTGATTTATCATTTTTTAATCGTTAGCCAAAAACA

CTTGTTGGAGGGCTATATTAAGCAAGGTGATATATATATTTGCTGACAGATTTGTAGACTGTTCATAGAC

TGTTATCTTCACAGATCAGTACCAAACTTCTGCATTTTTATCACTATCCATGGATCTAAATCATGGTGCA

GGGGAGAATTATGCACACCCGCAGATATGTTTCTTCCATGTGATCTTCAAGGCTTCAGCTTTGGCATTTT

ACATACTGTCCGCTCTCTTTTTTGACAGTTTTGTCATAATTTTTGTGGTAACGGTTCTTCTGGCTGCACT

AGATTTTTGGGTAGTCAAGAATGTCAGTGGACGGATTTTGGTTGGTTTAAGGTGGTGGAATGAAATAAAT

GACGAGGGTGAGAGCATCTGGAAATTTGAATGCCTTGACCAGGAGTCACTGTCTCGAATGAACAAGAAGG

ATTCATGGCTGTTCTGGTGGACTATTTACCTTACTGCGGCCTTGTGGCTTATCTTTGGAATATTCTCCCT

CATAAGGTTTCAAGCTGATTATGTTCTTGTTGTAGGAGTTTGTTTGACACTAAGTGTTGCAAACATTGTT

GGCTTTACTAAGTGTCGTAAAGATGCTAAGAAGCAGATTCAGGCATTTGCCTCTCAAACGATTGCATCTC

GATTCTCATCCACTTTGCAGTCTGCATTCACTATTATCTAAGTAACACAGAGAGATTGGAACAGGATCAG

ATGTGTAGGAACTTTACATGCCCAAGAAATGGCGAAGAGATATGTTGGATGATACTTCAGTTTATTGAGA

GTACTTTTTGGAAGAAATTACCTTAAGCATTTAGTTATTGCTCAGGCAAGGGTACATTCTGTATGCATGA

ACCTAGCGAATACCCCCTCTTGATATATGACTTTGTATATGCTTAATACGAGAACCTAGGCACTTATACG

TGGTTAGTAATGCTGAGAAACATGGAGTATTATCCAGTCTTCTGTTGCACTGTATTGTGTTTTTCTTCTC

CTTGCATTTTCTTTGTGTTTTTGTAGTAGTGAGTGAAACAATTTATTTTGTTTA

>MSTRG.4200.3 gene=MSTRG.4200

ATTTGCTGACAGATTTGTAGACTGTTCATAGACTGTTATCTTCACAGATCAGTACCAAACTTCTGCATTT

TTATCACTATCCATGGATCTAAATCATGGTGCAGGGGAGAATTATGCACACCCGCAGATATGTTTCTTCC

ATGTGATCTTCAAGGCTTCAGCTTTGGCATTTTACATACTGTCCGCTCTCTTTTTTGACAGTTTTGTCAT

AATTTTTGTGGTAACGGTTCTTCTGGCTGCACTAGATTTTTGGGTAGTCAAGAATGTCAGTGGACGGATT

TTGGTTGGTTTAAGGTGGTGGAATGAAATAAATGACGAGGGTGAGAGCATCTGGAAATTTGAATGCCTTG

ACCAGGAGTCACTGTCTCGAATGAACAAGAAGGATTCATGGCTGTTCTGGTGGACTATTTACCTTACTTG

CAGGCGGCCTTGTGGCTTATCTTTGGAATATTCTCCCTCATAAGGTTTCAAGCTGATTATGTTCTTGTTG

TAGGAGTTTGTTTGACACTAAGTGTTGCAAACATTGTTGGCTTTACTAAGTGTCGTAAAGATGCTAAGAA

GCAGATTCAGGCATTTGCCTCTCAAACGATTGCATCTCGATTCTCATCCACTTTGCAGTCTGCATTCACT

ATTATCTAAGTAACACAGAGAGATTGGAACAGGATCAGATGTGTAGGAACTTTACATGCCCAAGAAATGG

CGAAGAGATATGTTGGATGATACTTCAGTTTATTGAGAGTACTTTTTGGAAGAAATTACCTTAAGCATTT

AGTTATTGCTCAGGCAAGGGTACATTCTGTATGCATGAACCTAGCGAATACCCCCTCTTGATATATGACT

TTGTATATGCTTAATACGAGAACCTAGGCACTTATACGTGGTTAGTAATGC

>MSTRG.4200.4 gene=MSTRG.4200

CCTCTCTCGCATGCTCGCGCACACACACAAACACACAGAGTCTTTTAATACATTGGTACACAGGGTGCAG

GGGAGAATTATGCACACCCGCAGATATGTTTCTTCCATGTGATCTTCAAGGTACACAGTTTGATATTAAT

TTTTCTAATATTCTTGTTTTCTGGGTAGCATAATTTGCTCTCTGACTTGAATTTATATATTTGGCAGGCT

TCAGCTTTGGCATTTTACATACTGTCCGCTCTCTTTTTTGACAGTTTTGTCATAATTTTTGTGGTAACGG

TTCTTCTGGCTGCACTAGATTTTTGGGTAGTCAAGAATGTCAGTGGACGGATTTTGGTTGGTTTAAGGTG

GTGGAATGAAATAAATGACGAGGGTGAGAGCATCTGGAAATTTGAATGCCTTGACCAGGAGTCACTGTCT

CGAATGAACAAGAAGGATTCATGGCTGTTCTGGTGGACTATTTACCTTACTTGCAGGCGGCCTTGTGGCT

TATCTTTGGAATATTCTCCCTCATAAGGTTTCAAGCTGATTATGTTCTTGTTGTAGGAGTTTGTTTGACA

CTAAGTGTTGCAAACATTGTTGGCTTTACTAAGTGTCGTAAAGATGCTAAGAAGCAGATTCAGGCATTTG

CCTCTCAAACGATTGCATCTCGATTCTCATCCACTTTGCAGTCTGCATTCACTATTATCTAAGTAACACA

GAGAGATTGGAACAGGATCAGATGTGTAGGAACTTTACATGCCCAAGAAATGGCGAAGAGATATGTTGGA

TGATACTTCAGTTTATTGAGAGTACTTTTTGGAAGAAATTACCTTAAGCATTTAGTTATTGCTCAGGCAA

GGGTACATTCTGTATGCATGAACCTAGCGAATACCCCCTCTTGATATATGACTTTGTATATGCTTAATAC

GAGAACCTAGGCACTTATACGTGGTTAGTAATGCTGAGAAACATGGAGTATTATCCAGTCTTCTGTTGCA

CTGTATTGTGTTTTTCTTCTCCTT

>MSTRG.4204.1 gene=MSTRG.4204

AAATCCTAATCAATAATCAGAATCAAAGATCCAGCAATTAAGCTTGAAGAAATGGAGATCCAAGCAATCA

AGTTTGACAAATCGAGTTCCCATGTACCAATTTGCTGAATCATATTCTGCAGGTGAGTAGTGTGCGTTGC

GGAGAGAGAGGTCCAAGCTGTGAACACTGGTCTAGTAGCACTGCACTGTGACGAATGATTTTGG

>MSTRG.4206.1 gene=MSTRG.4206

GTTGATGTTTCAGGCTCTTCATGAAAGTGACAGAAGCCTTCATATGTGATAATATATCAGGATTAGAGAC

AAAAATTTTAGTTGTCTCCAAATGCTTGAATGTCGTAACTGACAGAAGTCGAAGTCATGCTCCTTCCATT

TAAAGCCCATTTAACCACTTGACATGTTCATCATTCTTTTTGATAATCATTGCTCAAAAGTCCCTCCCTC

CGGAGCATTATCAGATTTAGGATTTGGTTCAAAATATTTTAAACAGCTAGATACTACTTCATCATCAACA

TATGTAGTGGCTTCTTTGCCCAAATTTTAGTTGGTGCCTGACTATTTGAGTCACCTCTTCTTCATTGACA

ATAATATTATCTTTTTAAGGAGACACATCTAGAGTTAGTGTATGAGATGCCAGTTACTCCTCAACAATTT

TTTCATGAGTAATATCAGCCCTGTCAATCTTATGAACTATGGCTTCAGAAGGAATATATAAGAACTGGAT

TGCGAATCTTTGTCTTTGCTC

>MSTRG.4208.1 gene=MSTRG.4208

TACTCCCTCTGTCCCAGTTTAAGAGTCGTTTTTGTCATTTTTTGATGTCCCAGTTTAAGAGTCGTTTTTT

ATTTTCAAGACACTTAAACTACTCTTATTTATGATACCCTAATAACTCTTAGTATCTGTTTCCAAGTTTT

ATTAATCACCTATATTATTTTTCAACGCGTACACCTACTACCTTTCACCTTCTCTTTCACCTGTATATAT

ACTCTATATTATACTTACATTTTTCATCTCAGTTATATGTATGTTAAACACAAGATAAATATAGCGGTCA

TAAAAAATGTCTGGAGAGGGGAAAAACAAAGTTGAGAAAGATAACGAGCAAGGAGTAGAAAATGAGATTA

ACGGTGATGCAAGAGTACAAGCGGAGATTAATGGGGATGCAAGGGCACAAGCTGATTTTGATCTGAATGA

GATTCCACAAGATGAAGGAATACCTCTCCCTCGAGAATCAATGTTCCAGGTCATTTACATGATGTGGAGC

ATCTACTTTCCCATTGTTTTGAAATTATTTGAATAATTTCTATTATAATTTTCAGATTACATGTGATGTA

TGTCCAGAATATGTGATTTGGTTTTTATATGTATGAGCAATGAAAAGCTTTTGATTATTG

>MSTRG.4213.1 gene=MSTRG.4213

ACTACATTTTTATTATTATCAAACTTAGTTTTGTGTAGAAATTGGGGAGTAATTTATATTTTGTTAAATC

AAGAAGCTGCTGCATTGGGGACACGCAGCTCTATCACAGTTTAGTATAAAAAAAAGAAAACAGATGGACT

CGACACAACATATACAGCAACAGAGGCGGCGAGAGAAATTAGGGGTTAGGGTTTCGAGGAAATATCAACC

TCTTTGATCAATCCAACACTGCAGGTGAGGATTGATTATATACATGTATTAATTGGGTGTATGATTACAT

ATGTCTATATATGAATTATGCGGCTATGATTGTGATTATTTGGAAACGGAGGCTGGAAGTTGTGTGGGTG

GCGATTGATGGGTTGCCGGTGTGGCGTCTTGACGGTGGAGGCGGCGAGTCTGCTGCCGACCGGAGATGAT

ACAGGGATCGGAGGCGGCGCTGTGACCGCTCGGCGTGCAACGCAGCTGCTCGGAACGGCGAAGGCAGGTG

CGCGAAGAAGAGATGGGGG

>MSTRG.4217.1 gene=MSTRG.4217

TTATAATTATACCGCTACTTTCGTCTTAACCCCTTCTTTCTTCCAGCCAACCTAAAACTACGTGACAACA

CCTTCACCCACTAGACCAACCACGGGTTCCATTCATCTTTTACGTTCATGAGGTCAATTCTATTGATCGG

TTCATATGAGCTCCATTGGACACTAACATACTGCAGTTCGACAAATATAAAAGATGATGTATATATGTGC

CCAACTGTCCATACTCTGGGTTGCCTGTGTTCTTTGGGTATGTACTTACACAAATTCAGATAAAGGAAAA

AGAAGCACGGATGATTAAGATTTTAGAGCGGAATGATGTGAGATGTATATATGTACACATAATATCTTGT

TCATAAATTTATGTGTTAGGCATTCTCAAGAGAGATACGCTACTAAATGTTGATTCTTTGATTTGATTTT

CAAGCTCTTTCTGTTATTGCGGAGTCTTGGGTTCTATCTTAGAGCTTTATTGTTAATGGGTTTTATTGAA

ATTTTGAGTTTAAGTCATAGATGATTCTTTTTGTTGAGTTTGCAGAAAATGATCCTGTTATTTGAAACCC

TTACGGAGTTTGGTATTCTTAAAGTGTTGGTGAGGGGAAGCTCTCCAAAGTTGACGTGATTAATGATTTA

TGTTCTTTGTTGTGGTCGGTTTTTACAATATTTTATGTAATTTAATTCTTGTTTAAATTTATTTGGTTTT

GGAGTATCTTGGGGCATGTTGTGTGTCAGGTTTTTTATGTTCCAATACTGAGTTTGATACTGTTGACGCA

GCTCGAAAGATTTTTAATCTCCCTTTTGTGTTCTTGTCTCTCCTATTTTTTGTTGTGTGTGTGTGTTGAT

TGAAGTGCGGCTTAGCTTGTTTTGCAACAATTTTTTTTTGTGTATAGAGTTTTTTCACATTTATTTATTT

AAAGTGATATGTTATATGTGCATACAACATTTTGCCTAATTTTGGTACTTGAGAGAAGTTATTTTTGGGA

TTGATATTGTGCTTCAAACTCTTAATGCATTTCTTTATTTTATAATCAGAATGTGAATTTAAAATTGCGT

CTTCCATGTCATAGTCATATCCCATTTGGTAAATATTTTTGTGGGATGAATTTTTTCACCAATTGCATAT

ACTTTTGGTTTGGGTCATAATATTTGTACGCGTTTGTTTGCTATAATATTTGGAAAAAAGACCAAGTAGC

TTGCAGTATAAAATGTATTTTGTTGCCTATATCACTATACTTAATTGTCTTGTCATATTCAAACTCTCCA

CAAGGTGGATATAATGATCATACAAGCCACTAGTTTATTGGATGATCTTGATAAAAAGCTTAACTCCTAT

ATATGCTAAGCGAGTTTGGGAATGATATGGTTGGCATTTCCGGAGCTTGTAAAAATTGTACAGGATAACA

TACTATATTCAAAGTCTGTGGAGCTGATGGATGATCGTGCAAATGCTGCGAAGCTTGATTTTTCAGAGGA

TTATAGGAGAAGCTGCATTTCAGTCTTCAGTGCTATCTGCATCACCAAGGTCTTGTTACGTTGCACCAGA

AAATGGATACTGTAAAGCTACATAACAAACGGACACATACAACTTTGGTGTAATGCTGTTAGAACTGGTG

ACAAGCGGGGGAGCTTATGAACATATATAATCATGGGACGGAGAGTTTTTGGACATTGTTAGTAAAATCA

ACAAAACTAATGAAGCAATCCAAGTAGTTGATCCTAAGATCTCAAACTCATCCTTCTAGCAAACAATATT

ATGAGCTATAGATATTGTGGCCAGATGTGCATCTGTGGCATCAGAGAAGAGACCCCCAATATGTGAGGTT

GTGACGGCTTTGCAGAGTTTTTTTTTCAACTTCTCTCCTTTCCGATAGTGTTCCTGCTTCGCGTGGAGTG

TTTGATATGATCCACGTTCAACTTTTGGGCAATGAGTATGAATTGTTGCACTTTGTACATACACGGTAAA

TTGAGCTTGGAGAGAAAATATGAAGCCAGTAAAAAAAATTGGTCAGACTTGGCTTGAGGAGGAAAAGAAA

CATGTAGAATAGTAATAATGTGTTTGGAAATGTATCTCTTTATTCTCTCCCATGCATATATAATCATACA

AGTAAAACGTGCAAGATGATCCTTCTATAGCATTTAAAACTTGCAGAAAACTAATAGGTTGGTGAGAAGT

GTACCATTTGATTACGTTTGAGCACATTATGTTCTTTCTATGAGCCTAAAAATATTTTTATTTTATATTT

GTATTGTATGATTATCATGTACTATATAAGTAGTTAAATAACATGAGATGGGAAGGGGCTGCTATGGCTT

CAACATGTATGGCTTCAATATTTGTGAGGAAAAGACTGGTGATTATAATTTTGGCTACGATTATCATGTA

TACATTATGTACTAGATAAGTAGTTGAATAACATGAGATGGGAAGGGGCTGCTATGGCTTCAACATTTGT

GAGGAAAAGGCCGGTGATTATAATATTGACTACGGGCCAATATCAAATTTAAAGAAAGCTGTACTCTTCT

ATTGTAATAAAATATCATTTATCAAG

>MSTRG.4222.1 gene=MSTRG.4222

TAGGTACAATATGGTGTTTGCCCCCTTTACTGGTGTTGATAAGCACAATAAGTGCGTCACATTTGCATCC

ACTCTTCTTTCGAGAGAAGATGTGACAAATTTTACTTGGGCGTTTAATGCAATGTTGAAAGCTATGAGGA

GAAATCCCGTTGTTTTAGTCACGGATCAGTGTCCTGCTATGAAACAAGCTGTACCTGAATGTTTTAAAGC

AACCGATGAATTTCCTGCGTCCAGGCACCGTTTGTGCATGTGGCATATCACTGAAAAGTTTCCTGCTAAG

CTTGGCCACTTCTTATGCAAAGAAACAGATTTTATGGAAAAGATAAAAAAGTATATATGGTCATCAACGA

TAGAGCCTGCTGAGTTTGAGGAGGGTTGGAAATCCGTAATGGATGAGTTTAAGTTGGAAGATCATGCTTG

GTTATCATATTTGTATGCTATGAGGGAGTCATGGATCCCTGCATATTTTCGTGATAAACCAATGTATGGG

CTGATTAGGACGACGTCGAGGTCGGAGAGTGAAAACTTTTTTTTTAGCCAGTTTCATCAAAGTGGAAGTA

CATTGTC

>MSTRG.4225.1 gene=MSTRG.4225

ACAAGACACACACATCTAGTATGGAAAAAAGAACCATTTAGCATTGGATTTAAATCTCTAATTAATATAC

CAATAGCCGACGTATTATTACTAGAATTGTCAAAAGATATACTAATTAATTTATCACGAATCTTAAATTC

TCTAAGACATGTCATCATGTTATAATAAATAGCATTTCCGAAACGAGGGAGTTCAAATTGATCAAAAAAA

ATTATTATTTTATTCATTTGCCAAGTATCCATATCTATCCAGTGACATGTAATACACATAAAAGACATAA

ATACTGTCCAGCAGACCACACATCACTTGTTAAGCTAACTTTACCCTCATAATTTTCAAAAAAATTTATC

ATGGATTTTTTTACTTTTACATAACCTTTAATTGCATCATGACGAATAGTAATTCTACTAACCTTTTGAT

ATTTTGGTTGAAGGTTGTGATGTATGTAGCTGGTCATTCCATCATCATCAAAATTATTGAATGGCAATCC

CTGTGAG

>MSTRG.4226.1 gene=MSTRG.4226

ATTTTACAACTCCAAAAAGCTGAAATTAAGGAGCCATAAGGACTGAATCGCAAGAATGGAAGTTGACATG

AATTAAGTTGATCCTTTCAAACTGGAGCTGCAGAAAGCAGAGCAGAAGCACATGGATTGATTGAAGCAAG

GCAGAGGACGACGCTAAAACCATTATTGTGCAGATACTAAGAGTGGTTGCCTTCTGTCATCTACAAGGTG

TTGTGCACCGTGATCTGAAACCAGAGAACTTTCTTTTCGCAGCAAGAGACGAGGATGCCCCATTGAAGAT

CATAGATTTTGGCCTATCTGATTTCAGCAGGCCAGGACTAAACAATATGTTATTGGCATATGAGGATAAA

GCAGCCTATGCATTATGTGTTTTCTCACTTGCTCTTGGACCAAACGACGAGCCAATTAGGAGAGACACCG

GTATCCATTTTGATATACACTTTTTAGTGTGTTAAAAGACTCTTTTTCATGTTTATACATGCGGGACTGA

CTAATGCCTTAACATTTTAGGCTGAAATGCTTAAGGAAGAAAAGCACAAGATTTCTCAACGCTCTAGGCC

TCTTGTATTGGACAAAGATCATATTGCTGAAACTTATTGCTGCACTGTTATTACTGTTAGCTTAGTTGAT

CATATTTTGCAAGATCTTACTGTTAGCTTAGTTGATTATATCTGCGCGATGATCATTTGTATTGATTGTT

GTAAAGTATCGATAATTTGTTGATGCAGGTGATGTATGTTTTGGTTTTTATATTTGGTTTCTGCAACTTG

GAC

>MSTRG.4226.2 gene=MSTRG.4226

ATTTTACAACTCCAAAAAGCTGAAATTAAGGAGCCATAAGGACTGAATCGCAAGAATGGAAGTTGACATG

AATTAAGTTGATCCTTTCAAACTGGAGCTGCAGAAAGCAGAGCAGAAGCACATGGATTGATTGAAGCAAG

GCAGAGGACGACGCTAAAACCATTATTGTGCAGATACTAAGAGTGGTTGCCTTCTGTCATCTACAAGGTG

TTGTGCACCGTGATCTGAAACCAGAGAACTTTCTTTTCGCAGCAAGAGACGAGGATGCCCCATTGAAGAT

CATAGATTTTGGCCTATCTGATTTCAGCAGGCCAGTTCCTTAGACTCGCAACCTCCTGGATTGTGTATAG

TCTAGTGATATATCCACTGGCCATGACTTCTGTGTCCGGAAGTTGTTTGAAGCAAATAAAGGAAATATAC

CATGGAATAAAGGCAGGAAACATACTACAGACATTTATAGAAACACTACGGAAGATCAGAGAGAAGGCTA

GGCTTGCAATGCAGAATCCTAAGGACTAAACAATATGTTATTGGCATATGAGGATAAAGCAGCCTATGCA

TTATGTGTTTTCTCACTTGCTCTTGGACCAAACGACGAGCCAATTAGGAGAGACACCGGTATCCATTTTG

ATATACACTTTTTAGTGTGTTAAAAGACTCTTTTTCATGTTTATACATGCGGGACTGACTAATGCCTTAA

CATTTTAGGCTGAAATGCTTAAGGAAGAAAAGCACAAGATTTCTCAACGCTCTAGGCCTCTTGTATTGGA

CAAAGATCATATTGCTGAAACTTATTGCTGCACTGTTATTACTGTTAGCTTAGTTGATCATATTTTGCAA

GATCTTACTGTTAGCTTAGTTGATTATATCTGCGCGATGATCATTTGTATTGATTGTTGTAAAGTATCGA

TAATTTGTTGATGCAGGTGATGTATGTTTTGGTTTTTATATTTGGTTTCTGCAACTTGGAC

>MSTRG.4226.3 gene=MSTRG.4226

ATTTTACAACTCCAAAAAGCTGAAATTAAGGAGCCATAAGGACTGAATCGCAAGAATGGAAGTTGACATG

AATTAAGTTGATCCTTTCAAACTGGAGCTGCAGAAAGCAGAGCAGAAGCACATGGATTGATTGAAGCAAG

GCAGAGGACGACGCTAAAACCATTATTGTGCAGATACTAAGAGTGGTTGCCTTCTGTCATCTACAAGGTG

TTGTGCACCGTGATCTGAAACCAGAGAACTTTCTTTTCGCAGCAAGAGACGAGGATGCCCCATTGAAGAT

CATAGATTTTGGCCTATCTGATTTCAGCAGGCCAGTTCCTTAGACTCGCAACCTCCTGGATTGTGTATAG

TCTAGTGATATATCCACTGGCCATGACTTCTGTGTCCGGAAGTTGTTTGAAGCAAATAAAGGAAATATAC

CATGGAATAAAGGCAGGAAACATACTACAGACATTTATAGAAACACTACGGAAGATCAGAGAGAAGGCTA

GGCTTGCAATGCAGAATCCTAAGGTGATGATGTCATGACTAAACAATATGTTATTGGCATATGAGGATAA

AGCAGCCTATGCATTATGTGTTTTCTCACTTGCTCTTGGACCAAACGACGAGCCAATTAGGAGAGACACC

GGTATCCATTTTGATATACACTTTTTAGTGTGTTAAAAGACTCTTTTTCATGTTTATACATGCGGGACTG

ACTAATGCCTTAACATTTTAGGCTGAAATGCTTAAGGAAGAAAAGCACAAGATTTCTCAACGCTCTAGGC

CTCTTGTATTGGACAAAGATCATATTGCTGAAACTTATTGCTGCACTGTTATTACTGTTAGCTTAGTTGA

TCATATTTTGCAAGATCTTACTGTTAGCTTAGTTGATTATATCTGCGCGATGATCATTTGTATTGATTGT

TGTAAAGTATCGATAATTTGTTGATGCAGG

>MSTRG.4228.1 gene=MSTRG.4228

TGCTGTGGGAGTATGTCAAGGACAAGTATATTATTCCGGAAGAGGCTAAATCATGGGTTTTTACAACCAT

GAGTGATCAGTTTAGGAGCCACAAAAGCCGAATTAAGCGAAATTATTACACCAAGTATGCAACGGATGAG

GAGAGGTTGGAGAATAGGCCGAGGGACGTTTCTCTTGAAGACTGGAAGATTCTCTTAGCTTATTGGAGTG

ACGAAAAAATTAAGG

>MSTRG.4229.1 gene=MSTRG.4229

GTTTCTTTCTTTTAATTAAAACAAACCCAGTTCCTTTTCTTTTTAAGACAGCCGAAACCCACCTTTTCTT

AGTTTTGTCTAAACCAAACCCAGGGTCCCCTGTTTTTCCTGCTTTCCTTCTCGATTTGTTCGACCGAGTT

GCTATGGTTTGAAGCTTAGTTTAGTTTTGTGTATTGCTGCGTTTTGTCTGAGTTTAGCTCCGAGTTCTGT

TCTTGTTGGCTTGATTTATGTCGAAAACTGAGTCATTTTCTGAGTCCCATTTGCCGTCGAGTTCGTTGGT

CGAGCTGTTTCTGGCCTGAGTTCTGCCGAGTCGCAACGTTGTCTGGTTTCGATTTGTTTTTATATGTTCT

GTTTGCTGCGTTTTTCTTTCTGTTGCTGAGTCTGATGTACTGAGTTACCGAGTTGGCTGTGCTTTTGCTG

CGATTTTAGTTCCGTTTTTGTTTAGTTAGAGTCTGAGTTGAGTTATGTTTGCTGCGATTTATGCCGAGTT

TCGTCCTCTGCTCTGCTGAGTCTATTCGCCGAGCTCGAGTCTGGGTTTAGTTGTGTGTACTGAGTTTGTT

CTGTTTTTGTTGGAGTCGAGTTCTTGCTGCTGCTGCGTTTGTCGAGTCAGTTTCGTAGTTGCAGAGTCTT

AGTTGCTAGTGAGTCGCGAGTCACCAGAGTTTACAGCTTGAGTCGTTAGTCCCTGTTTTCGCCGAGTTCT

TTCTGGCGAGTCACTGCTGAGTCGAGTCAAGTATGTCTTCAGTTTTTGAGTTTTCTTGCTGCGTTTTTGT

TTTAATTTCTGCTGTGTTTCAATTGTAATTTGCGGAGTAGTTGTGTCGAGCCTAGGCTGAGATGTTTATC

TGAGTCGCTTGTTTTCCTTTCTGTTGCTTCTCTGTTTTGCTGATTTGTTGAGTCAAATCGCTGAGTTTGA

GTTCTTTTACTCCTGAGTCGTCGAGTTAGTCACTGAGTTTTGGGTCATAGTACAGAGTTGTTTCCCCAGA

GTTTTGATTTCATAAACATGTAGTATATCTAAGTTTTAGCCTTGAGTCGTAGCTGTAGGAAGAGTCAGAC

TGATTTTTATATCAAGTCATGTTTTGGACTTTTAGTTTGTTTTATTTTAATGAAATAGGGGTACATTATT

GTTA

>MSTRG.4236.1 gene=MSTRG.4236

CTTCCCTTCTCTTTGCTTTTTGCTTCCTTCTTATGCACAGACCTAATTCTTTTTTCCCTGTATGGGTTGA

ATCCACATCATTCTAGTCGACTAATTGGACTCTGGCATCAGAAAATAGAAAATCAACGGGAGTAATCATA

ATGTTTAGTGTGCTGGTCTGCGCCGCTTGAATATGTGTGTCTGGTGGAGTCGATTGTAGCTGAGGATTTT

TCAAGCGAGTGGAAGGACACTCTGAAATGCAATTACAAAGAGGCGTTTTGTTCAATCTGTTGTTGAATCG

AATAGTACGTAGCACTTTTGTATTTAGTGC

>MSTRG.4239.1 gene=MSTRG.4239

CTATGTACACAGTGGTTGCAACAAACCAGACACCAACAGACAAAGAAAATACACAACAAATATCACCAGA

TTATGTGAAATGGAGGAAAGAAGATCGTCTCATTTGTGGTTGGATTATCGGAACACTTTCAGAAGAAGCC

TTAAGTTTAGTTATTGGCCTGGAATCATCCCAATTAGTTTGGAATTCCTTGAAAGAAGCATATGCTCAAG

ACTCACAAGAACGAGAGTTTACTCTCCGACAACAACTCACATACCTCAAAAAAGAACCAAACCAATCTCT

AGCAGAACATTTGAGAAAATTCAAAACCATATGTGATAGTCTCGCAGCCATAGGAAACCCAGTACGTGAC

AAAGCAAAGGCTTACTCCTTACTTGCAAATTTAGGACCTAAATATGAATCATTCACAACCACTATGCTAA

AACCACCCATGCCATCATACTCTGAAGTTGTATCACTCTTGCAAGGTTTTGAGCAAAGAAACTCATGGAT

TGAAGCACCCAGTTCTGTCTCATATGCGTTCTATGGACAACGCAACCAAGGCCGGCAACAACAGAATTAT

AATTCTCGCACCAATCTCAACAACCAGTTCACATCAAAAGGATGTGGTTTTCAAGCTACAAATAACAGTT

TTCATTTCTCAAATTTTGCAGGTAATGGTTCAGGAATGCAGCAACAAAGCAACAGTCGTACTCCACGACC

ACCACCACCTGGTAAACGCCGAATGACCACTACAGATATTGTACTTCTTATATAGCTAAACATTTACAAG

GTAGTTGTTTGTT

>MSTRG.4241.1 gene=MSTRG.4241

CGGCCACAGGTGTATACTTCTCGCCAACCACCATGAAGGTACTTCAGTTTATCTGCATTGAACTTGCATG

ATACTTCTCTAAATCAGCGTCAAGATCATCTGAAGATAACTTCTCACCACGTCCTCTTCCACTTCCACGT

CCCCTCCCAGATCCACGACCCCGACCACCACCACCACGCGGTCTTCCAAATCCACCTCTCTCTTGGTTAC

T

>MSTRG.4245.1 gene=MSTRG.4245

CTTGAAAAAGACTCAATCTTTGGAGGGTTACAATCACCAGGGATTCAATCCTTTCTTCGAATCTTCTACA

GATGCAGAGATTAACAAGTTGAGATTGTCACCACCTCCAAAGCTCAAGTTTCTCAAGGATGCTGAGATGA

AACTGATGAATAAATTGATGGAAGATGCTGTGAGAAAGTGCCATGAGAAAGATGCTACATTTCTTACAGC

TTCAGAGTCAAATTCTACTGCTTCTTCACAG

>MSTRG.4251.2 gene=MSTRG.4251

ATTATTCTTGCATTTTGCTTATCACGCAGGTCTTGCTGCAGACAAAGGTAGATCCTGTGCAAACAATTGT

CGACAAAATATATGACAGACTGTGTGAGAAGCCTTCTGACATTGAATACATTCGTGAGCGCGCGATATTA

ACTCCCCTCAATGAATATGTTGAGACTGTTAACAATGAGGTCCTGCAGAGGCTGCCAGGAGAGGTTAAAA

TTTATAGAAGCTATGACAGTATTTCTAAAGGAAGCACAAACATGGTCGCAGACGAGATCTTATACCCTCC

GGAGTATTTGAACTCCTTGAAATTCAGCGGCCTCCCTAATCATGAACTGAAGATTAAGGTGGGTGCGCCT

ATAACGTTGCTGCGGAACCTTAATCCCAAAAAGGGCTTGTGTAACGGGACACGGCTTATCGTGACCAGGT

GTTATCAATTTCTGATAGAGGCGCAGATCATAACTGGCAATAAGATCGGGGGGATCACATTCATACCGAG

AATCAGTATGTCGCCATCTGACAATACATTCCCAGCACCATTGAAGCGCAAACAGTTTCCGGTATCAGTA

TGTTATGCTATGACAATAAATAAAAGTCAGGGTCAAACGGTTAAGAACGTCGGCCTTTACTTGCCAAAGC

CTGTATTTAGCCATGGTCAGCTTTATGTGGTGGTTTCAAGGGTGGCATCCCCGCATGGCTTAAGAATTCT

TTGTGTCGACGATGAGGAGGCACATGCAGGGGACACGAAGAATATAGTGTACCGCGAAATATTTGATGAG

TTGAGACTACAGTGATGCCACCATGTCAGCACGGCTAATTGGAGGGGCTCCTCACACACATCATTTTGAA

TAGTAATGTATGCAACTGTTCTATGCCCGGG

>MSTRG.4255.1 gene=MSTRG.4255

ACGAGTTGAGGTGATGAAGTGATTTTCTATCATTTTAGGTTGTTTTTGGTGCATTATTTGCAAGAATAGA

GAATTGTGGATTCATCACGACTTGGGATTTTGTGGGTACATCTTCTACAAACCCTCACGAGAACACACTC

GTCCACTAGAGCTATCTAGGGGTTTAAAGGGCTTGTTGCATGTGCTAAATGCAACCGTGATCACCTACGG

AAGTGGTACTAGAGCGAGGAAGGACATGCACGCGAGAACGAGCTGAAAAACGAAGAATAGGGCTGCCAGT

GGCAGACAGTAGCGCGCCCGCGCTAACCTCTAGCGCGCCCGCGCTACCTACTGCTGCCACTAGCGCGCCT

GCGCTAGTTTAGCGCGGCCGCGCCCTCCAGAAGTGCTCCGGGCCGATTTTTACAGCTTTTCTGATGGATT

TTCGCAGCGGACGAGGCCCGGTTGACTTAGTCAAAGCATTTTAATTTCTCAAGTGTAAAAACCCTAGCCT

CGAAGTAGTT

>MSTRG.4256.1 gene=MSTRG.4256

GATAAGTGGTATTTATACACACTTATAGGCCTTCATTTCCACTTAAATTGGTTGGTTGTACTTAAGTGTT

TAGTGTCTTTTGATGTGTTTTTAGTGTTTTTCTGTGCAGGTCACGAGTTGAGGTGATGAAGTGATTTTCT

ATCATTTTAGGTTGTTTTTGGTGCATTATTTGCAAGAATAGAGAATTGTGGATTCCTCACGACTTGGGAT

TTTGTGGGTACATCTTCTACAAACCCTCACGAGAGCACACTCGTCCACTAGAGCTATCTAGGGGTTTAAA

GGGCTTGTTGCATATGCTAAATGCAACCGTGATCACC

>MSTRG.4261.1 gene=MSTRG.4261

TAAGAATGCCTTCCATGAATTTTCAAATTTTTTTAGGCTCATACTTAGTTACAACTTTGGACATATCTGG

ATCAATTGAGAGATGGCTTTGTCAATATTTTGCAAAAGTGCAGATACTGCAAGGTTTTCTAATATTTAGA

TCAAATTGAGCTCAAGAACTTTGTCGTTTCGGAGGGTTGCCTTTAGCTCAGGGCTAATCCACCAAACTCC

ATAGACGCCAACTTCGGGAATGTTATGTGAATAAACAGAG

>MSTRG.4262.1 gene=MSTRG.4262

TCCATGTAGTTAATTGGTTTGTGGTAGCCATTTAGTACTCTGCAAACTTTGCAACAAATGGTTTAAGAGA

TAAGGGAGAAGGTGAAAAAAGAATATATGTGGGAAGGAAGAACGAGATTCTACAACCGAACTCTTCCATG

TGGATCAGCTGCTGGGGAATCTTTTGTCATTTGAACAAGGGTAATCTATAATGCTACCCGTGAATGGCGC

GACTGTAGATTATATAAACCCTGTTTTAAATCATGTACTCATGTTAATTTTATGGCCTAGCAATTTGTAC

ACGACACTTTGTACATGACACGAATACTTAGTATTTGGGTTTTTTATACAAAAAACAAGAAAGTACATCA

TGAATTTAGTGTCGGTTATAGGTCTGGTGGCGAGCCTTATTTCAGTTATAGTAGTTGTTCACATGAGAGT

AAATGCAATGATTAAAACGACAAAAAGTAATGAGTTCCACTATTCTCCTTCTAGATACTTGCTCCTAAAA

AGTTGC

>MSTRG.4270.1 gene=MSTRG.4270

ATTTCATTGAAGAGAAGCACTCCTAAAGAAGAAAGCAAGAGATATTGCGATGTCAATAACATTTCCCTGG

AAATATCATGTATCAAAATTAAATTTTGCTATGTTTCCTTGGGATGATAAAAGAAAAGGTTATTATTTGG

TGTAATGTTAAGGTTTTAGATTGACAACTAAATATTTGATTTGGCACTCTTTGCCTTGGTGGTTTCGGGT

AAGTTTGTTTAGCATTCTCTTAACTATTTCTGTTACTTTGCTATCTGGAATGGATAAGTGTAGGAACTAT

GCCTATATTATTCAGTGTAAAAGTTGAGATCTATAAACTGGATTAGGGACTTAAACTTGGCTGTAAGTTG

TAACTGTTGCAACTACTTTGTTAAGAATCTCGCATTTTGACTTCTAGAGCTTGCTCTTGGTTTCTCAATC

CCTAGCAGATTATAGTCTCCAACTACTTCTATTTGGTGCTATTACCTTGCATAAAAAACTAAATCACTCT

ATTTTG

>MSTRG.4271.1 gene=MSTRG.4271

GGGAGAAAGTGGGATTGAAGAGAGGGATATGGACTGCTAGAGAAGACGAAATCTTGACAAAATTTTATTA

CCCTTGGGGCCTTATTGCATTTCTTTACCACTTTTTACATTTTATAGAGAAGTTTGTGCATGTTTCTGCG

TAGAAGTACAGAGAGTACTGCGCGGGTGCGACTGCTGTTGCTGATTTGTCACCTCCATTAGCATGTTCAA

TACTGCTTCATAGCGTTCAACTAGTTCTTTCTGGTAAAGTTGTGTGAAGTGTTGTGTTAGTTTCAAAGAC

TTGG

>MSTRG.4273.1 gene=MSTRG.4273

TGGAGTTTCAAATCTTGAGCTACAAGTAGAGGCATATGTTACCCGTGAATTAGAACCACAAATAGAAAAG

TCAACCCCCTTGCGGACAATATGGTTCAAGGCAAACACCTCCGATGCACCTATATCCACAAATTTAGGAA

AATTCAGTTGGTTATGGCTCGGATTAATAACATCGTTGTCATTCATAATCTTCCTTCTTACCATGGGATT

GCTTACGCGCTTCTACATATATCCAATAGATCATAATAGTTTTATGGTCTTCTCATACTCATTACAGGCT

CTGCTAAATATGCTACTCCTATGCATATCCCTACTTGTGACAGCCACTGGAGCTTTTCTGTGGAACAAGA

AACAACAGATCGAGGAACTTAAAGGATTGTCACCTAAATTGCATAATTCAGTGGCCGTCAATCCTATTGA

GCAAGAACTCGAAAGTCTTCCTCAGCAGTGTCTTTCAAAAACTACCAAGCTGCATTATGGCCGAAGGCCT

GATCTGAGGGAGTTGATTATGTAATGTTGTAGGAATTCTAGTGGAACAAGAAGGATCAAGTGTTGGTGTT

CTAGTTTGTGGTCCGAAGAGTATGCGCCATGATGTCGCAAACATATGTTCATCAGGCTCTGCAGACAATT

TACATTTCGAATCCATAAGTTTTAGCTGGTGATCAAATATTCTCAGGGATGCAGTGCTGCTATCCACTCC

TTTGTAGTTACTGTGCAATTGCCTTTTAATATAAAAAAATTTCAGGCAATTAATTATTTAATAGTAATAT

ATACTGGTGGTTTTCATTGAGTATTCATAATTTTTGGGCTTAAATTATTTCTAAAACCTTGTTACTCCAA

GCTGTATCAATTTAATTTAACAGTTTAG

>MSTRG.4273.2 gene=MSTRG.4273

TGGAGTTTCAAATCTTGAGCTACAAGTAGAGGCATATGTTACCCGTGAATTAGAACCACAAATAGAAAAG

TCAACCCCCTTGCGGACAATATGGTTCAAGGCAAACACCTCCGATGCACCTATATCCACAAATTTAGGAA

AATTCAGTTGGTTATGGCTCGGATTAATAACATCGTTGTCATTCATAATCTTCCTTCTTACCATGGGATT

GCTTACGCGCTTCTACATATATCCAATAGATCATAATAGTTTTATGGTCTTCTCATACTCATTACAGGCT

CTGCTAAATATGCTACTCCTATGCATATCCCTACTTGTGACAGCCACTGGAGCTTTTCTGTGGAACAAGA

AACAACAGATCGAGGAACTTAAAGGATTGTCACCTAAATTGCATAATTCAGTGGCCGTCAATCCTATTGA

GCAAGAACTCGAAAGTCTTCCTCAGCAGTGTCTTTCAAAAACTACCAAGCTGCATTATGGCCGAAGGCCT

GATCTGAGGGGAATTCTAGTGGAACAAGAAGGATCAAGTGTTGGTGTTCTAGTTTGTGGTCCGAAGAGTA

TGCGCCATGATGTCGCAAACATATGTTCATCAGGCTCTGCAGACAATTTACATTTCGAATCCATAAGTTT

TAGCTGGTGATCAAATATTCTCAGGGATGCAGTGCTGCTATCCACTCCTTTGTAGTTACTGTGCAATTGC

CTTTTAATATAAAAAAATTTCAGGCAATTAATTATTTAATA

>MSTRG.4276.1 gene=MSTRG.4276

CACAATCTACCTTTGTAAATTTGGCAGTGCTCCAAACTTGACAGACCATATCAATCAAACTTACAGTCTT

AACTAACATTCGTTTAACCGCAAAATGTTGCCACTCATCTCATAGTACTAGAGTAATCAAGTGTTTTACG

ATAACTTCTTATTCCATTCTTTGATTTGAGCTTCACAAAGTGATATGGGACGAGACATTTAATTTTGGCC

GATTGAATATTTACCTTTGATGATAAACGATTTGTGACAGCCCAGTAGACGATATAACACAGGCCGTGAG

CTGTAAAAAGGAACATGGTTATATGTCCAAGCCAAATATGATACTTGATGCTGGCCTCTGAGGTCAAACC

AAGGAGAGGCAACACCGATGAAGTTCGTGTCACCGGAAAGAAGAGAAAGGTAAGACATATGTTGCCAACA

ATTCCTAGAGTTAGACCTATAATGCTCAGTTTAGCTTCCCACCTGCAATTAATTACATGCAAATGCATCA

TGAACAATTACTCCCATGTGATGATAGAACTGCTTAAAGAATGTTATGAGGTTTGTTTGATATGGCGTGT

CTCGATCAACTCCTCTAGAATAAGAAAACACCTACCTCTTGCCCTGCCCTATTTCTTTGAAAATGTCCAT

GTGATTAAAGAAGTAATGCAAGTAAGCTGAGAAAGTCCAAACCAGGAGAGCAATAAACATCACGAAAATG

AAGAGTTCTATCCAAGAAACAATCCCGAGGCATCTGATGATCATCGGCCTCTTCCACCAACTGAACATAG

ACTTCCTACAAAGAAAAGTACTAGTTCATAGTTCATGTAGCCATGTATGTGTCAAGTATATGTATGTGTG

CATATGTACCTTTTTATTTCACAGCTATTAGTACTTGCAAGTTTCAGTCCCAAGTGAAGATATACAGAGC

CCAGAACAGATATCAACAGCACTGGAAATGTGTAAACAAGAATTGTTGTACCTGCAAAAACAGGGAAAGA

GGCATATGCATAGAAATCCAAATTAAAAGAAAAACATATAAGCTGAAAACAAGTATAGTACCTTGTGTTC

CAAAGAAAGTGGAATTTGTCTGAGCCCGAATTTTCGGGTAGAAATATTCTCTATAAGCAGAGGTTGGCAT

CATAATCCACATGAAACTATACCCGAGAAAGATTATCACTACTATTCCCAATACCAGTGTCTGGATAAGT

CTAGTTTTGCCATTAGAACAAGTAACATGAGCTGCTGTTTTCTCCTCCATCAGAGAAACTCTATTTCCTA

ATTT

>MSTRG.4280.1 gene=MSTRG.4280

TTTTTTTTTAGTTACAGTTAGATGAAATAACTTGAAGTAGCAAGTTACCTTATCTAAAATGAGGCAAATA

TGCTTTAAGTCATGTTTTTGCTAATTTCCTGAATTACTAACGGCTCTCTTAGGTGTCCCATGACTTTTTG

CACGTATATTGAGTTGCTTAAAAATGGCACTAATTAAACATTCTTTTACTTGTTTTTCTATTAAAAAGAA

AATAAAAGAAAATTATAATAGTAAGTTTTAGAAGTATCATTTTTATGCACCTTAATGGGGATAAAGGGAG

TATTAAGTAATCTGATCCCTATGCTTACTTATATATGCAGGTACACCATGTATTACTGATTGTGTGA

>MSTRG.4281.1 gene=MSTRG.4281

AAGAGTCCAAATCGATGAACTGGTAGAAAACTCGAAAGCAAAATCCAACCAAATCACAAATCCTAACAGC

CAATTGGTTCTCAAAACACAGGGAAGAGGGAATCAGTCCAACAGTTAGGGTTTTAAGAAAAATGGGCAGA

GTAACGGCAGGAACAGGGGTTAATCAACACACACAAGCAAGCATAGCGCTTGGGATTAGTAGATCAGCAA

GCTTGAAGCTTGAAAAAGGGTTCGAAAATGGCAGATATGGAGGTCTGTACGTGTGTAGCGGTGTGTGTAT

ATGCAGCGTGTTGAGGACGATAAAGAAAAGAGGGGCATAGACAATCTCAGATTGGGTCGATGAGGTAGCT

AAACTTGCGGGGATGTATTCGATTGG

>MSTRG.4284.1 gene=MSTRG.4284

CGTCCTCAAACTCGATGTAATCATCAGCAGCATTCTCATCCTCCTCATCAAGTCCCCCAGTAATACCTTC

ATTAAGCCTGGTATTCTCCGGCAACTCACCGTAAGCCTTCAACAACCTCGCCTCATCCGGCATATACTTG

AGAATCACATCGGCCTTATCGTCCTGATAATCACGCAATCCGACGAGAATAATATCGCCGGCGGCGATCC

AGACCTTCTTGTGCATCTTGCCGCGAATATGACAGAGACGCTTGGTGCCGTCGATGCACATGGCTTCGCA

GCGGCCGTTACCGAGCATACGCATGACTTGAGCGTATTCCTGGCCGTCTTCTTTGAAGACGAGCTCGCGC

TTCTCGTCGTCGGCCTCGTTCTTGCCTCGCTTGCGGTTCTTGCCTCCCTTTCCCTTGTTCTTCGGCATGG

TGGAACCCTAGATCGGATTTCTTTTTGGTTTGTGGATTCAGATTATTTCAATTTGATTGAATTCAATTAG

GGGTGTGTG

>MSTRG.4289.1 gene=MSTRG.4289

CAAAAAGCTGAAAATCATGCAGTTGCTGTGCACTCTGTAACCAAATTACATAAATATAAGCACTTACAAA

TGTTTCATGTATAACAAAAGGAAAACAAACAAGGAAACTAAAATATAATGACAGAAACAGAGAAAAGGAG

CCTTAACTAGTCCATTTGCTAGCTATCTGAGCTATTGCTTTTGTTGATGATCCTTCATCACTAAGAGCCC

TGGAAGCTGCCTCTTTCAGGTCTCTCATTCGGGTTCGGAGACGCTTTCCTTCCTCGCCTTCCATTAGGTG

CTTAACAACCGTTGCAATTTCCACACGCTCCACCAGGCCATTTTCGCCAACTTGAGGCCTCAATGCAACT

TTAAGGCCTTGAGTGAGCATTACAGCATTCATCTTCTGTTCTGCATAGAGCGGCCACGCGATCAGGGGAA

CGCCACTCACTACGCTCTCCAGAATCGAGTTCCAGCCGCAATGAGTCATGAAGCCACCTGTCGAGCCATG

GCTCAGAATTTGGGCTTGAGGGGCCCAATTAGGCACCACAAAACCTTGGCCTTTTGTCCTCTGAACGAAT

CCTTCTGGCAAAAAATCAATAGGGTTGTTTTTGCTGCGGATGCCGAAGAAAGTTGCATTAGCACTCTCAT

CATTCGGAGTTCTGACAATCCATAAGAACCTCTGCTCACTCATTTCCAAGCCCAAGGCCAGCTCAGTGAT

CTGCTTGGATGACAGAGTCCCTCCACTCCCGAAAGAAATGAACAAGACCGAGCCGCGGGGCTGATCCTCC

AACCACCTCATACAATCCAACCTATCAGCAGCACTGCTTGAATGATCCATTTGTACAAGAGGTCCTACAG

GGTAAATTGGTGGGCTGCCCAACTCTTGTAAAGCTTTAATAGCTCCTTCCTCCAAGTCCTCAAAGCTATT

CACCATTATACCATGAGCCAATCTGTAATTTTTTGTATGATGAAGCACCCACTTATATGCATCATTCTTC

CTGTCCTGAACAGGGTCAAGTAGATCCTCTCCATCGATGGGTATACAACCAGGAATCTGGACTGGGGCAG

GCAAGTCCCTGTACTCGCAAGACGTTTCAGCATCAAGCTTAGGCAAGTAAAGAAACAATGACAAAGCCAT

AGCATTTGTGGGGTAAAAAATGTAAGGTTGTAACTTAAATTCAAGAGCTATATCAAGTGCATCAGTCGCG

AAAAGATCAACCACCAAAGTAACCAACTTCTTGGTAGCCATCATCGACTTAAGCACATCACGAAGCGAAG

GCAGAGATCTTGATATCATGATCGACATACGTGTTTCGGCCTTAACATCCTCTGACAAATCATCCAATTT

AAGTTGAGGAAGAACAACATAGTCCAAACCACTAGGAAGTGTGTCAAGAAATATCTTTTGAGCTTCGGAA

AGAGGTTCTCCAATAGGAATGATAAAAGTTGCCGTAAATTTGTGAAGACTAGTGAACCTTTTGGCAAATT

GAACTAGAGGAATGAGATGACCCATTCCAGGAGTGGGGAAAATAGCAATATGAGGTGTGGTTTCCATGGG

AGCAGAACTGAGAAGCAGGAAAAAAGAGTAGAGTTAGATAAATGAGAAACTGAGGCTGTATGTTTGTGCA

GACAAAATACAAGTATAAATAATGTGAAGGGGGTTGAATAGATGAAACTGATGGAGAGAATTCTAGATAG

GTTAAGTGAGCACACATAAGTGGTGATGTATGGATGATAGATGAGGAATCAATTTAGCACAGTAGTCTCT

CCACCTGGTGTGGTGATGTCAAAACAACCATAAGCCTTATGTCACCATTTCTAAAGGCAGTAATGAGTGA

GTAATCACTATTTCTTGCCTCCTACAAGTAACTACCAATGCTCTGCGAGCACGATTGATTAGCCCAGTGG

TGGGTTTATCCTCTGTTGTCCTGGGACAGTCGGGTTCGACTCGGGCTCACCCCGAGAAGTATTTGAACAG

ATACTCATCTGTAAGGCACATAGCCTAAATTGTCAGAAAACTACCTATGCTCTGTATTATTTCGTGCATT

TTAAGCATTAAAAACACACAAGATTCTGCAATTCCACATACATTTCCGCACATAAAGGTCTGAATATATG

CTCTGAAC

>MSTRG.4296.1 gene=MSTRG.4296

TTTAAAGAACGAGAAAACCAACTTAGACTGAAAGGTAGATAACAAACACGATGGTACTACGACCACTTAT

TCGTCCTCGGACGACATTACACTATCAGAATAATAATCCGGACTGGTGACCAGTGCAAGGTATTGTTGTT

CCTTAAACCAGTCCCCTGCTTCCCGAAACATCCTAGCATCATCGTATAGGCTCTGACGCTGCAGACGCAG

ATGCATGCGATTCTCCTCTTCTCTCGCTCTGAATAGGGCGGCCTCCATCTCCTGCGGCGTCTCTCGAGGC

ATAGTACGAGCCTTTTGTTTAGGAATTGGAATCCCAGCTAGTGCACAACGTTCCCTGCACACTTCAAGCG

AACGCACACGCCACGTCCACTCATCCAACATGGTGGTAATCGACTCAGGGGAGTAGCGTAAATTAGGATT

TAAAGAGTACCTAAACTCATCCTCACACCACCATTTCCTACCCATGGCAACAAACAGATCACCTGGTGTG

TAACTCATGA

>MSTRG.4307.1 gene=MSTRG.4307

TTTATAAGTCCTTCTGTTTGGTGCCTTCTATAATTCTGGTCGATATGTAATCCAAGTCGGTGATAGAAAT

ATTACAAGATTTACCGTTAGACTCCTAACGGTCATGTGTAAGTTTCCTTGATACCAGTGTGTAGTTCTAT

TTAACTAAATCCATAACCAAATAAAGGAAAGATAAATTCCTTTAACATGTACTAAACATGCAAATAAAAT

TGTATACATTTATAATAAAAGAACCAGACCTAGAGGTTGGCATGAATAAATATTGACACATATTGACATT

GATAAAAATATTTTTTTCCCTACTGATGCCCATCAATAACAATACAAATTTCAGCCATTTTCTACAAAAT

GACCTGTGCCTCGCACACTCCTGTTTATCCTGTTCAAGATAAAGAAGATATATGCATCAATGATGAAGCT

TTCATGATCATGAAAGAAAGACGCAGATTCCTCCACTTTTATAAATGCTTTTCGTCTAATTAGCATGGCC

TCAAGCCTGGATCTTTCAGGCTTCTTCGAGAAAGAGGATGTTTCTGAGAGGACCATCAGATTTACATCTC

ATCATTCTTCAAAAGAATTGACTGAGGGGATTGAAGACATTGTTACACAGATGGGTTTTTTATTCCGGAA

GAGAAGTGGACAGGTCGGTTATTATCAGGCTACTCGCTGATCATTTGTTTTCTTACAAATTTATTCTTCC

TGTTCCCTTCTCTGACCCGAGTCGCCTCCTAAACCGAGCTCTGCTACCATCTTAAGTTTTCAAAACACAC

CAACAAGATCCTATGCTACAACAAACATTATTCAACACAAACAACATGTACTGAAATATCTCTTGCTTAT

TGAATATTAACTCAAATACCCAGAAATTTAACTAATTGTCGCTTAATTTGGGCCAATGTGGTTAGCTGAA

AGTGCTGCATTTCATAAAGATCACAACAGTCTAGTCAGTCTTTCAGTTGCAGTAGAGGTTGGCCTTAAAC

TCAAATTCTACATTTATTTTAATCTACTCAGCAGCTCAATTGTCGCGATATGATTCTTATTCGAACTCAT

GCTCCTCTGCAACATAGAATCACGTGGATCATTAAAGAACAACCCCATTTCTCTGTCTTCCAGAATTAAA

CATAACTCTTTGAGTTCAATTTTTAATTACTTCCGCTTTCATGGACTACCCAGACGCTCTAACAAGGTTA

GACATATCTTTACTTTTACTTGTTGTCAGTATCTATGTCATTTCCCTGCTTTTAAGTCTTATTTTAAGCC

CTATGTGTATGTTTATAAATATTTCCACTTGGAAAGAGACATTAATTAAACATTAGGTAATTTATTTAAT

CTACAAATTGCTTAAAGATGAATAAAAAGGCTTCATATAAATTACAAGCTGATTCAAAACCATTTTCTGA

GAAATAAGGAACCAATATAATCTTGTATCCAGTTGGAAGGAATTCAATTGGGAGGAAAGGGGTGTAGAAC

TAAAGATAAATATTAGTACCAATGTGGAGATTTATGCATGATACTGATTCCTTCCTCAATTCTACTCTAC

ATATATGAACTAGATTTCAATCTCATTATTCTAACATGCTCAAATCTCATCGTAGAAAGAAGTTAGCCAC

ACAGAGTTTTTTCTTTTTAATGCTTACCTCGCTCTGAACTTTTTACTTGCACTTGTTATTTTTTTCTATT

CCATCTAAATCGTATTTGCATTATATTGTATTTTATTATTCAGTTGCATTATCCTGAAATTAACCTTAGC

GCGTATTTTATAATTAACATCATTCTGTTCTTTTGATACAAGCGCAAGACTTTTGACGAGAACTTTTTGA

GAAATACAAGTTGATTAATGCCGCACACTACTTGCAGTTGAGACTATTGTATAAAAAAAACCTGTTCTGC

TGTTGCTCGAAAGATTGAGGATGAATCTCCTGAGGAGATATTTAGTTTATTTAATTTTCCCGATGTCTTG

AAAGAGGTTTCATTCCGGCCAATGTGCTTAATCTCTATAATCGTATAGGTTTACTAAGTTAATTGGTACA

GGCTGAAAAGTTGGAGCCTTGGAACAACGCACCTGATGATGTTTTGTTTAGTATAGAGAAAGATTAGCTT

TAAACATTTTTGTATCAGTAGTTTACTCATACTATGTCTAGACCACGAAAATCTTACAAGACTTGATGTG

AG

>MSTRG.4310.1 gene=MSTRG.4310

GGCATAGGCCGTGAGAACCCCTCACCTTGTTTCAAGACATCAAAACACTTGAAATCCATACCATCAAAAC

ATGTACAAGATGGAGAATCTGAAGTCTTTGACAGTTCCGTGGCTTCGCCAAGATGGACGAGTCTGACGGT

TCCAACACCACCATCGCCCTCAAGAAACTCGACACTCTTGATAAGCTCGGAAACCATTGAGCTCCCTCGA

TTTGATGTCCT

>MSTRG.4317.1 gene=MSTRG.4317

TACACAAACCCCAAATCTTTCCCACGCCACCTTCACTGTCAACGTCCTCCAGTTTCGACTGCGCGCCATG

ACCACGCCGCCACCATCGACTATCAGCTCTCGCTTTGATTCTTCATCTTCCGTTTGTGCATCTGCTGATC

TCCCTATTTTTCTATCGACGCCATGGCGGCTCTTTCCCAATCAAACATGCGACCTAGTTCTATTAAAAAT

CTTGGGTCTCAAAACAAGCTCTTCGGGAAGTGCTCTTTCTCCTTTCCCTTGAAGTTCTTAATTTAATTTT

CCCTGGTTTATCTTTGCATTTCTTTTTGAATTCTGTCTATTCTTATCTTTTAATAATTTTGGATCTTTGT

AAACATTCAAATCAATTATAGTGGTGTATATATATATATATTTGTGTTGGTTCTGGTACTACTAATCGCT

TTCAATACAGGAGCTTCTTGCACAATAATGGATTGTAACATTCATGGTGGTAGTGGAGAAGTCTAAAAGG

GGCGGAGAAGAAGGATGCTAATTGGGCGATAATTGTGCCTTAATTAAGGATATTAATTATTCTTAACTAA

CGGTTTTTGTACCATTTAT

>MSTRG.4318.1 gene=MSTRG.4318

TGATATACAAGACAATTGAGTTACAAATATTGATAGAAGGGAATAAAGAAGACATCAGTTTTTAGTCTGT

TTTGTAAAAGATGTAATTTTGTAGGGCACTCTTTTTCCCTCTTGAGTTTTTTTCCCAAAAGGTTTTACTC

TTGAGGGGTTTTAACGAGGCCTTTTTTTGTGGGTTATCTCTTCTGACTTGTAAGGTTATGTTATCTAAAC

ATCCAGA

>MSTRG.4328.1 gene=MSTRG.4328

CCTATATCTATCCTGCCTTACAGTGAATGACTATTAGAAGATATGAGCACCACAACCACAACTGAGCTAC

CTTATCATCTCACAGCCAAACAACTCCCACAAAATACCAACTCATCTTCTTCTAGCTTAAACAATCTTTC

CCAGAAAAAGTACTTCTTGCACCTATTAGCCACAAAATGCACCACTTCACTTCAGAACTTAAAGCAAGTT

CATGCTCTGATACTGAAAACCCACCATGTCCAGGACCACTTTGTAGCTGGAACTTTGATTGGATCCTATG

CAATTCCACATTTTGGCAGTTTTAGCACTTCCATCGAGATTCTTGAACGAGTGCCTGAGCCAAATGTGTT

TGTTTGGAATGCTGTTATCAAGGGATGTTTAGATAACAATGAGCCTGAGTTGGCTCTCTCTTTTCACCAT

GAGATGGTGGCTACGGACTCCAGGGCCAATAAGTACACATATCCTCCATTGTTTAAGGCTTGTCTGGTCG

CGGGAGCTAGTAAAGAAGGTGCTCAGATTCACGCCCGTGTGGTGAAAAATGGACTTATGGGAGATGGATA

TATAAAGAGTTCTGGGATTCAGATGTATTCTTCTTTCAAGTGTTTGAAGGAGGCCAGGAATATTCTTTGT

ACAAATGGTGACTCGGATGTTATTTGTTATAATGCTATGATTGATGGGTACATGAAATTTGGAAATGTTG

AGGATGCTAAATTGTTATTCGAGGGATTTGAAGACAAGAATGTCAGTTCTTGGAATGCAATGGTTAGTGG

TTTTGCACGGTGTGGTATGATTGAGGAAGCTAAGGAATTCTTTGGTAGAATGCCTATCAAGGATGAGGTG

TCTTGGAGTGCTATGATTGATGGTTATAATCAAGGGGGTTATTATAAAGAGGGGATAGAGATTTTTGAAG

TAATGCAGAAAGCAAAAATCAGGCCCAAGAAATTCGTTCTGTCTAGTGTGGTTTCCGCCTGTGGTAATGT

TGGGGCACTTGATCAAGGTAGATGGATTCATGGGTACGTGAAGAAGAATAAAATTCCTATTAATGCAGTT

ATGGGAACTTCTTTCGTTGATATGTATGCTAAATGTGGGCGGCTGGATTTGGCATGGGAGGTTTTTGAAA

CCATGAAAGTGAAAGAAGTTTTTTCTTGGAATGCCATGATCAATGGGTTGGCTATCCATGGTAGGGCTAG

AGATGCAATCGATCTTTTGGAGAAGATGGAGAAAGAAAGAATAAAACCAGACGATATCACATTTTTGAAT

ATTTTGAATGCTTGTGCTCATGGAGGCTTAGTGGATGAAGGGATGTGGTATATGAACAATATGAAGGATG

TTTATGGAGTTGCACCTAAAGTGGAGCATTATGGTTGCGCGGTTGATCTTTTAGGACGGGCAGGGTTATT

CTTTGAAGCAGAGAAACTTATAGGCTCCATGCCAATGGAACCAAATGGTGCTATTCTAGGAGCATTATTA

GGTGCATGTTGGAAGCATGGAAATCTTGAATTAGGCGAAAAAGTTGGAGACCTTTTACTAGAAATGGAGC

CACAAAATGGCGGAAGGTACACTTTACTATCAAACATCTACGCCAAGGCCGGGAGATGGGAAGATGCTTC

AAGGGTGAGAAAATTGATGAAAGAAAGAGGAGTGAAAACAAATGGTGGTTGCAGTTCGATAGATTTGGAA

GGAACTTGGCATGAATTTAAAGTAGGTGATTCGACACACCCTCAATCAGAGGAAATATATTTTATGCTGG

ACAAGATTAAGGAGAGGTTGCAGAATGAAGGATACTTAGCCAACACAAGTCAAATATTACATGACATTGA

CGAGGAAGACAAAGAAAATGCAGTATGGCACCATAGCGAACGACTAGCTATAGCCTATGGCCTCATCAAC

TCGACACCCGGGACAACTATTCGATTGATGAAAAATCTCAGAGTCTGTGAGGACTGCCATTTTGTGACAA

AGCTCATTTCACAGATTTATGATAGGGAGATTGTTGTTAGGGATCGTTCTCGATTTCACCACTTTAAAAA

TGGGAAATGTTCATGTAATGATTTCTGGTAATTATAGAATGTGTATGTTTAGACTTTAGGACATAGGTTT

TCTTAAGGTTTCTTCTCACGGATTTGGAGCCCTTGGAAGAATATTTTAAGTTAATATGTAATTTATGATT

TATTAAAAAATAATAATAATATTTTC

>MSTRG.4333.1 gene=MSTRG.4333

GTATGCCCCTGCGGGGCCGATAGCAATAACACACAAACAACCACAAAATCAACTACAAAATCAACCATAC

TCAAACTTAAACACTTGCTACTCACCACCATCATCAACCCATAGTCTTTTAATCTGCTTCGATCTACTCA

AACCCAACACAATCAACCACACTCCAACACCCATCACACCCATCGGGAGAGGATGGAGGGGGAGAGGGTT

TCGATGAGGCGGGAGATGACGGCGTCGCGAGTGCGTTCAGAAGGGGGCAGATGGAGAAGGAGATGCTTGG

CTTGTCGGACTTTGAGATCTCGGTGTTCGGAGCTTGTTCTCTGAAATCTGAAATCCAATTCATCGGAGCG

ATTGAGAGAGATTGAAGGAGAGAGGCAGAGAAAAAGAGAGCGTAATTTAATTCAGATCGAGAGAGATTTG

GGGAAGGAGCGGGGAAAGAGAGACGAGAGAGGCGTAAAGAATGGAACTGTGGAAAGGAGGGGACCGTATA

TTTGCATAATAGAAGTAGGTAGTAACGTAAATCTGTAAGATTTTTAATGTGATAGTATTATTGCAATTAA

TATGCTAAAAAGAGGTATATTTAATAAATTCCCAAAAAATTTTGACCCCTTCTAAGTTTACCTTCTTCCA

GGTTGCTTGTCTCTGATTTTGACTTCAAAACATTATCCCCTCTTGAGGAAACCTTCATATTAGTTTTTGA

TCGAGACTCTTTGTTATTTCAAGTATTTAC

>MSTRG.4334.1 gene=MSTRG.4334

TATCACATTAAAAATCTTACAGATTTACGTTACTACCTACTTCTATTATGCAAATATACGGTCCCCTCCT

TTCCACAGTTCCATTCTTTACGCCTCTCTCGTCTCTCTTTCCCCGCTCCTTCCCCAAATCTCTCTCGATC

TGAATTAAATTACGCTCTCTTTTTCTCTGCCTCTCTCCTTCAATCTCTCTCAATCGCTCCGATGAATTGG

ATTTCAGATTTCAGAGAACAAGCTCCGAACACCGAGATCTCAAAGTCCGACAAGCCAAGCATCTCCTTCT

CCATCTGCCCCCTTCTGAACGCACTCGCGACGCCGTCATCTCCCGCCTCATCGAAACCCTCTCCCCCTCC

ATCCTCTCCCGATGGGTGTGATGGGTGTTGGAGTGTGGTTGATTGTGTTGGGTTTGAGTAGATCGAAGCA

GATTAAAAGACTATGGGTTGATGATGGTGGTGAGTAGCAAGTGTTTAAGTTTGAGTATGGTTGATTTTGT

AGTTGATTTTGTGGTTGTTTGTGTGTTATTGCTATCGGCCCCGCAGGGGCATACATTTTTAACCATTTGC

AACTAGTTACAACTTATCATGCAACTAAATGTGATATGAACATTGGTCTATCACCCGTTTTTTATGTGCG

CAGGTCAATTGTGATTCTACTATTTTCAGTGATGTTGGTGTCGACTGGATTGCTCGAGAAACTTTTGTAA

TCATTTTTATTATCAAGAATATTTATATTCTATCTGTTAAGCAATCTGAAAACAAAGCGGCTATTTGTTT

AGCTCGATTTTTGTTTCACACTTAGGTTGTAATTGTCAGTTTGGGGGTTTGTCCCGACTGTGTTCTTATG

TTTTATGTTCAATAAAATCTTAC

>MSTRG.4334.2 gene=MSTRG.4334

CTTCTATTATGCAAATATACGGTCCCCTCCTTTCCACAGTTCCATTCTTTACGCCTCTCTCGTCTCTCTT

TCCCCGCTCCTTCCCCAAATCTCTCTCGATCTGAATTAAATTACGCTCTCTTTTTCTCTGCCTCTCTCCT

TCAATCTCTCTCAATCGCTCCGATGAATTGGATTTCAGATTTCAGAGAACAAGCTCCGAACACCGAGATC

TCAAAGTCCGACAAGCCAAGCATCTCCTTCTCCATCTGCCCCCTTCTGAACGCACTCGCGACGCCGTCAT

CTCCCGCCTCATCGAAACCCTCTCCCCCTCCATCCTCTCCCGATGGGTGTGATGGGTGTTGGAGTGTGGT

TGATTGTGTTGGGTTTGAGTAGATCGAAGCAGATTAAAAGACTATGGGTTGATGATGGTGGTCAATTGTG

ATTCTACTATTTTCAGTGATGTTGGTGTCGACTGGATTGCTCGAGAAACTTTTGTAATCATTTTTATTAT

CAAGAATATTTATATTCTATCTGTTAAGCAATCTGAAAACAAAGCGGCTATTTGTTTAGCTCGATTTTTG

TTTCACACTTAGGTTGTAATTGTCAGTTTGGGGGTTTGTCCCGACTGTGTTCTTATGTTTTATGTTCAAT

AAAATCTTACTGTTCGTCAAAAAAAA

>MSTRG.4335.1 gene=MSTRG.4335

CTTGCAAAATAGATTGTTGAAATGGGTTATCACAGAATTTTTCCACTGGTATATCTTTTGATTGAGTTAA

TTTTAGTTTTGCCGGTTGCAACTGCTACAGTGGAACGAGCCTTTTCTGTTATGAATATAATCAAGTCCAA

TCTGCGCAACAAAATGGGTGATGAGTTTCTCACAGATTGTCTAGTATGCTACATTGAAAAAGACCTCTTC

ACAGCTATTGATAATGAAGTAATTCTGCAACACTTCCAAAATATGAAGACTCGACGAACTAATTTGCCAA

GGTTACGCATTGAATAAGGTGTCATTTCTTTTTATCATTTACTGATTTTGGTATATGTAATCTGGACCCC

TTTGAAAAAAGCTTCTGGCTCCGTCACTG

>MSTRG.4337.1 gene=MSTRG.4337

CATATTCTCCTGCTACTTGCCTCTTCCTCTGTTTCGTTGAAAAATAAGCACTTGTAGTTGGTACTTGGGT

GCAAAATTCAACTTTATTCCAAAGAATTGTATTGATTTGTATTGTGACTTTGTGCGATGCCTATATAGTT

GACTTCCCTCGTCAACTTCTTTGGAAGGTAATTAAATTTTTCTTTTGCTAAGAAACTACTTGGAAGGCAC

CTAATTTCACTTCCAAGTTCCTATCATTTTATATTAATCTGCCAATTTTGTGTTAGAAATACATCTTCTC

AATAATGTCTAAAGTATAGATTAACTAACAATGTTTTAATTAGTTGCAGGTTTCTATCCCTCAAACTAAT

TTAATTCATGACAAGCTAATTCACCATGGATCAGAATGCCTCGTTTTGTCATGTCTTCTTGAGTTTCAGA

GGGGAAACCCGCAACAAGTTCACATGCTTCTTGTACGAGGCACTGAAAGCCGAAGGATTTACAGCTTTCA

TGGACAGAACTGCCATACGTGTTGGAGATGAAGTAGATTCAACAATCAAAGAAGGGATCAGAAACTCTAT

GAGCGCCATAATTATATTCTCCCAAAATTATGCCTATTCTACTTGGTGTCTTGATGAACTTGTTCTTATC

CTTGAACGCAAAAAGAACTCGAGGTACTTTATTATACCAATTTTTTATGAGGTCGAAATTCGAGATATTA

AACACCAGCTTGGAAATTATGGCCTTGCGCTTGAAAAGCATAGGGCTAGGCACAATGACAAGGTAGACAA

GTGGAAAGAAGCTCTTGCAGAAGTTGGTAATATTCTTGGACAGCATGTAGAAGGGCTGCAGAGCACCTTT

ATCCAAAACATTGTAAAATTGTTTCGAGAGAAATTGGCTGCCAAGTTCCCAGAATGCCGCCTTCCTACAC

TAGAAGGTCCAAGTTCACAAGGCAGTTCTTCATCGTCAATGTTGAGACCTAAGCCATCCAACGACAAGGT

GATTCTTTACACAATTAGTGTTAATAATGGCCTAGGTGCAAATGGGTTGGTAAAGAAACTTCTGATGAGT

GGGAATGTTGTATTTGAGGAGAGGAATTGTTCAACAGAATCAGAGTATATGAGAGAGCTAGAAGAGCTGG

TCGGTAATGACAGGGTAAGGTTCCCTATGGTGATTGTGAATGGGAAGGACTTGTGCGGAGAAGAAGAGGT

GGAGGGTTTAGATGATTTCGAGAACAAGCAAAAGCTTATATCAGTACTGAATCATCTCTATGTTTCTGCA

AAAACAATAGAAAACCTTCATGACATGTTTAGTAGGAGTTTAACATTTCAAAGTAGTATTCAGGACCAGT

GGATCAATCGGAGATACTCTTAGTGGTTGTGCTACTTCTTTTCCTGTTTCTTTACCATGTTTCTTTTAAT

ATAAATGTAACTAGCTCTCATTCTTTGACTCTTCATCTTATTTAGTTTTGTGTATGTCGATTAGGCTTCT

GTCCTTTTAGGACTTGCTAGGCGAGCCTAACTCCGGCTCATCTGATAGTTTTTGGTTTTAGTTTGGTGCT

CATCTGATAGTTTTTTCTTCAAAATTGCTTGGTGTATTTAAGTATCATGTAATAGAAGCAACAGCTGATA

TAAGCTGTCATATCTATGAATCCACAGTAAGGGTACAATTCCTCTTTTAATTTGATCTCTATAACAATGT

ATAATGCAAAACAAATTGTGCGCAAAATTCAACTTTATTCCAAAGAATTGTATTGATTTGTATTATGACT

TTGTGCGAGGCCTATATAGTTGACTTCCCTCGTCAACTTCTTTGGAAGGTAACTAACTTTTTCTTTTGCT

AAGCAACTACTTGGAAGGTACCTAATTTCACTTCCAAGTTCCTATCATTTTATATTAATCTGCCAATTTT

GTGTTAGAAATACATCTTCTCAATAATGTCTAAAGTATAGATTAACCAACAATGTTTTGATTAGTTGCAA

GTTTCTATCCCTCATACTAATTTAATTCATAATAAGCTAATTCACCATGGCTCAGAATGCCTCGTTTTGT

CATGTCTTCTTGAGTTTCAGAGGGGAAACCCGCAACAAGTTCACATGCTTCTTGTACGAGGCACTGAAAG

CCGAAGGATTTACAGCTTTCATGGACAGAACTGCCATACGTGTTGGAGATGAAGTAGATTCAACAATCAA

AGAAGGGATCAGAAACTCTATGAGCGCCATAATTATATTCTCCCCAAATTATGCCTATTCTACTTGGTGT

CTTGATGAACTTGTTCTTATCCTTGAACGCAAAAAGAACTCGAGGTACTTTATTATACCAATTTTTTATG

AGGTCGAAATTCGAGATATTATAAACTCCAGCTCGGGAATTATGGCCTTGCGCTTGAAAAGCATAAGGCT

AGGTACAATGACAAGGTAGACAAGTGGAAAGAAGCTCTTGCAGAAGTTGGTAATATTCTTGGAGAGCATG

TAGAAGGGTAATTTTTCTACCTTTTCCTCACTGTAATTCTTGGTTTGGTTTCATCACTTTCAGGCTGCAG

AGCACCTTTATCCAAAACATTGTAAAATTGTTTCGAGAGAAATTGGCTGCCAAGTTCCCAGAATGCCACC

TTCCTACACTAGAAGGTCCTAGTTCACTAGGCAGTTCTTCATCGTCAATGTTGAGACCTAAGCCATCCAA

CGACAAGGTGATTCTTTACACAATTAGTGTTAATAATGGCCTAGGTGCAAATGGGTTGGTAAAGAAACTT

CTGATGAATGGGAATGTTGTATTTGAGGAGAGAAACTTCTTTCTTGTTTCTTTACCATGTTTTTTTTTTT

GGTGAAATTACC

>MSTRG.4337.2 gene=MSTRG.4337

CATATTCTCCTGCTACTTGCCTCTTCCTCTGTTTCGTTGAAAAATAAGCACTTGTAGTTGGTACTTGGGT

GCAAAATTCAACTTTATTCCAAAGAATTGTATTGATTTGTATTGTGACTTTGTGCGATGCCTATATAGTT

GACTTCCCTCGTCAACTTCTTTGGAAGGTAATTAAATTTTTCTTTTGCTAAGAAACTACTTGGAAGGCAC

CTAATTTCACTTCCAAGTTCCTATCATTTTATATTAATCTGCCAATTTTGTGTTAGAAATACATCTTCTC

AATAATGTCTAAAGTATAGATTAACTAACAATGTTTTAATTAGTTGCAGGTTTCTATCCCTCATACTAAT

TTAATTCACCTAGTCTAAAGTATAGAATGCCTCGTTTTGTCATGCCTTCTTGAGTTTCAGAGGGGAAACC

CGCAACAAGTTCACATGCCTCTTGTACGAGGGACTGAAAACCGTAGGATTTACAGCTTTCATGGACAGAA

CTGCCATACGTGTTGGAGATGAAGTAGATTCAACAATCAAAGAAGGGATCAGAAACTCTATGAGCGCCAT

AATTATATTCTCCCAAAATTATGCCTATTCTACTTGGTGTCTTGATGAACTTGTTCTTATCCTTGAACGC

AAAACGAACTCAAGGTATTTAATTATACCAATTTTTTATGAGGTCGAAATTCGAGATATTAAACACCAGC

TTGGAAATTATGGCCTTGCGCTTGAAAAGCATAGGGCTAGGCACAATGACAAGGTAGACAAGTGGAAAGA

AGCTCTTGCAGAAGTTGGTAATATTTTTGGACAGCATGTAGAAGGGCTGCAGAGCACCTTTATCCAAAAC

ATTGTAAAATTGTTTTGTGTGGAATTGGCTGCCAAGTTCCCAGATTAAAGTCACCTTCCTACACTAGAAG

GTCCAAGTTCACAAGGCAGTTCTTCATCGTCAATGTTGAGACCTAAGCCATCCAACAACAAGGTGATTCT

TTACATGATTAGTGTTAATAATGGCCTAGGTGCAAATGGGTTGGTAAAGAAACTTCTGATGAGTGGGAAT

GTTGTATTTGAGGAGAGGAATTGTTCAACAGAATCAGAGTATATGAGAGAGCTAGAAGAGCTGGTCGGTA

ATGACAGGGTAAGGTTCCCTATGGTGATTGTGAATGGGAAGGACTTGTGCGGAGAAGAAGAGGTGGAGGG

TTTAGATGATTTCGAGAACAAGCAAAAGCTTATATCAGTACTGAATCATCTCTATGTTTCTGCAAAAACA

ATAGAAAACCTTCATGACATGTTTAGTAGGAGTTTAACATTTCAAAGTAGTATTCAGGACCAGTGGATCA

ATCGGAGATACTCTTAGTGGTTGTGCTACTTCTTTTCCTGTTTCTTTACCATGTTTCTTTTAATATAAAT

GTAACTAGCTCTCATTCTTTGACTCTTCATCTTATTTAGTTTTGTGTATGTCGATTAGGCTTCTGTCCTT

TTAGGACTTGCTAGGCGAGCCTAACTCCGGCTCATCTGATAGTTTTTGGTTTTAGTTTGGTGCTCATCTG

ATAGTTTTTTCTTCAAAATTGCTTGGTGTATTTAAGTATCATGTAATAGAAGCAACAGCTGATATAAGCT

GTCATATCTATGAATCCACAGTAAGGGTACAATTCCTCTTTTAATTTGATCTCTATAACAATGTATAATG

CAAAACAAATTGTGCGCAAAATTCAACTTTATTCCAAAGAATTGTATTGATTTGTATTATGACTTTGTGC

GAGGCCTATATAGTTGACTTCCCTCGTCAACTTCTTTGGAAGGTAACTAACTTTTTCTTTTGCTAAGCAA

CTACTTGGAAGGTACCTAATTTCACTTCCAAGTTCCTATCATTTTATATTAATCTGCCAATTTTGTGTTA

GAAATACATCTTCTCAATAATGTCTAAAGTATAGATTAACCAACAATGTTTTGATTAGTTGCAAGTTTCT

ATCCCTCATACTAATTTAATTCATAATAAGCTAATTCACCATGGCTCAGAATGCCTCGTTTTGTCATGTC

TTCTTGAGTTTCAGAGGGGAAACCCGCAACAAGTTCACATGCTTCTTGTACGAGGCACTGAAAGCCGAAG

GATTTACAGCTTTCATGGACAGAACTGCCATACGTGTTGGAGATGAAGTAGATTCAACAATCAAAGAAGG

GATCAGAAACTCTATGAGCGCCATAATTATATTCTCCCCAAATTATGCCTATTCTACTTGGTGTCTTGAT

GAACTTGTTCTTATCCTTGAACGCAAAAAGAACTCGAGGTACTTTATTATACCAATTTTTTATGAGGTCG

AAATTCGAGATATTATAAACTCCAGCTCGGGAATTATGGCCTTGCGCTTGAAAAGCATAAGGCTAGGTAC

AATGACAAGGTAGACAAGTGGAAAGAAGCTCTTGCAGAAGTTGGTAATATTCTTGGAGAGCATGTAGAAG

GGTAATTTTTCTACCTTTTCCTCACTGTAATTCTTGGTTTGGTTTCATCACTTTCAGGCTGCAGAGCACC

TTTATCCAAAACATTGTAAAATTGTTTCGAGAGAAATTGGCTGCCAAGTTCCCAGAATGCCACCTTCCTA

CACTAGAAGGTCCTAGTTCACTAGGCAGTTCTTCATCGTCAATGTTGAGACCTAAGCCATCCAACGACAA

GGTGATTCTTTACACAATTAGTGTTAATAATGGCCTAGGTGCAAATGGGTTGGTAAAGAAACTTCTGATG

AATGGGAATGTTGTATTTGAGGAGAGAAACTTCTTTCTTGTTTCTTTACCATGTTTTTTTTTTTGGTGAA

ATTACC

>MSTRG.4337.3 gene=MSTRG.4337
