## Supplementary information_Chialva_etal_2020_R1 for "Insights into long non-coding RNA regulation of anthocyanin carrot root pigmentation"

### Supplementary Figure S1

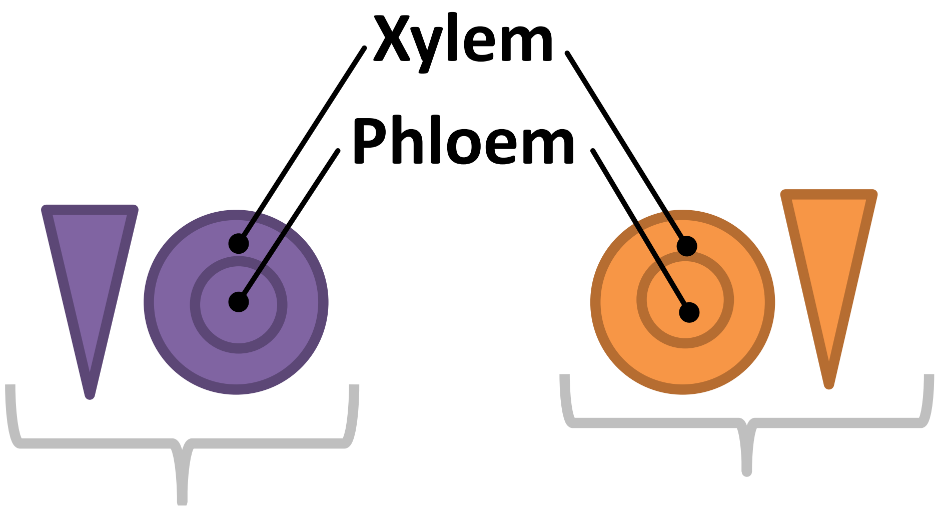


***D. carota***
**cv. Nightbird**
**(purple)**

***D. carota***
**cv. Musica**
**(orange)**

### Supplementary Figure S2

a)

| geneID | product |
|---|---|
| DCAR_010745 | DcMyb7 |
| DCAR_000385 | DcMyb6 |
| DCAR_002739 | DcbHLH3 |
| MSTRG.8365 | DcSAT1 |
| DCAR_021269 | DcUCGXT1 |
| DCAR_001218 | Myb_DNA-bind_3 domain-containing protein |
| DCAR_012866 | transcription factor bHLH145 |
| DCAR_019332 | transcription factor bHLH93-like |
| DCAR_005631 | transcription factor MYC2-like |

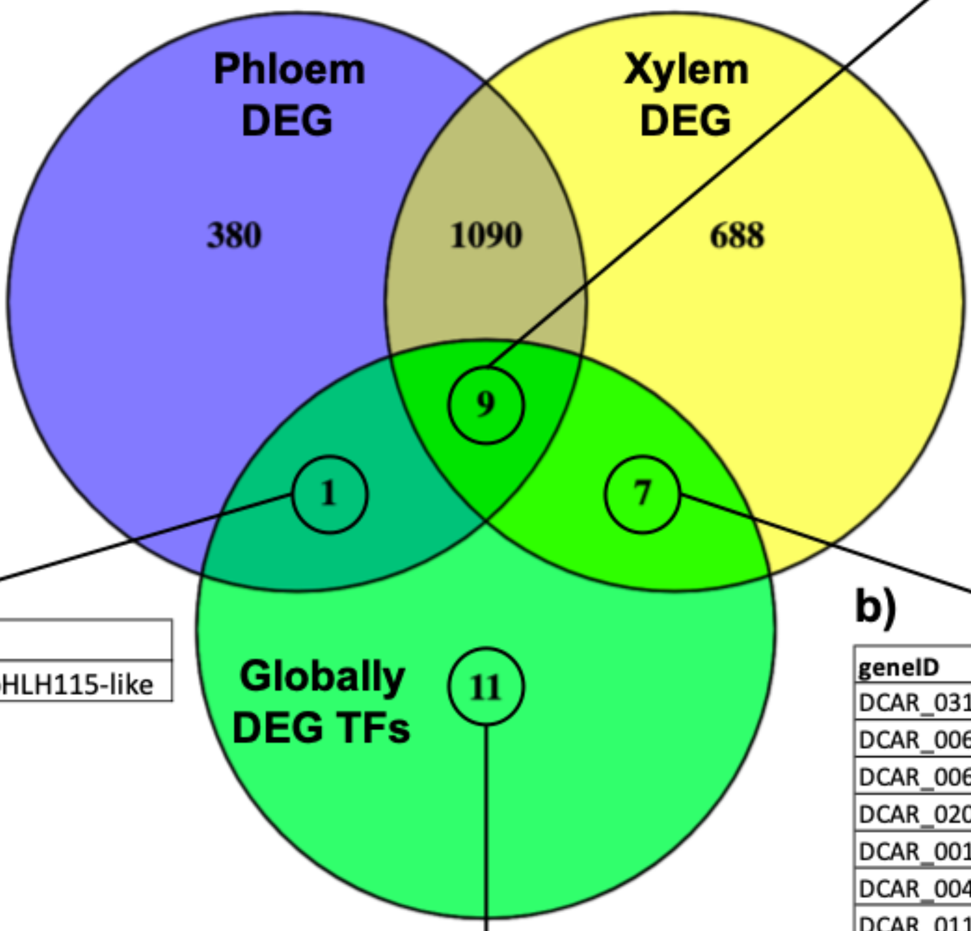


c)

| geneID | product |
|---|---|
| DCAR_02119 | transcription factor bHLH115-like |

b)

| geneID | product |
|---|---|
| DCAR_031036 | myb-related protein 3R-1-like |
| DCAR_006528 | transcription factor bHLH157-like |
| DCAR_006004 | transcription factor MYB59 |
| DCAR_020937 | transcription factor MYB44-like |
| DCAR_001572 | transcription factor bHLH63-like |
| DCAR_004664 | myb-related protein B-like |
| DCAR_011175 | transcription factor MYB35 |

d)

| geneID | product |
|---|---|
| DCAR_010746 | DcMyb8 |
| DCAR_006910 | ANTHOCYANINLESS 2 (ANL2) |
| DCAR_024963 | transcription factor MYC4 |
| DCAR_007287 | transcription factor MYB3-like |
| DCAR_019472 | myb-related protein B-like |
| DCAR_004166 | transcription factor bHLH112-like |
| DCAR_004752 | transcription factor bHLH78-like |
| DCAR_002128 | F-box/WD-40 repeat-containing protein At5g21040 |
| DCAR_022979 | transcription factor bHLH63 |
| DCAR_013994 | WD-40 repeat-containing protein MSI1-like |
| DCAR_023502 | WD40 repeat-containing subunit B1 homolog |

### Supplementary Figure S3

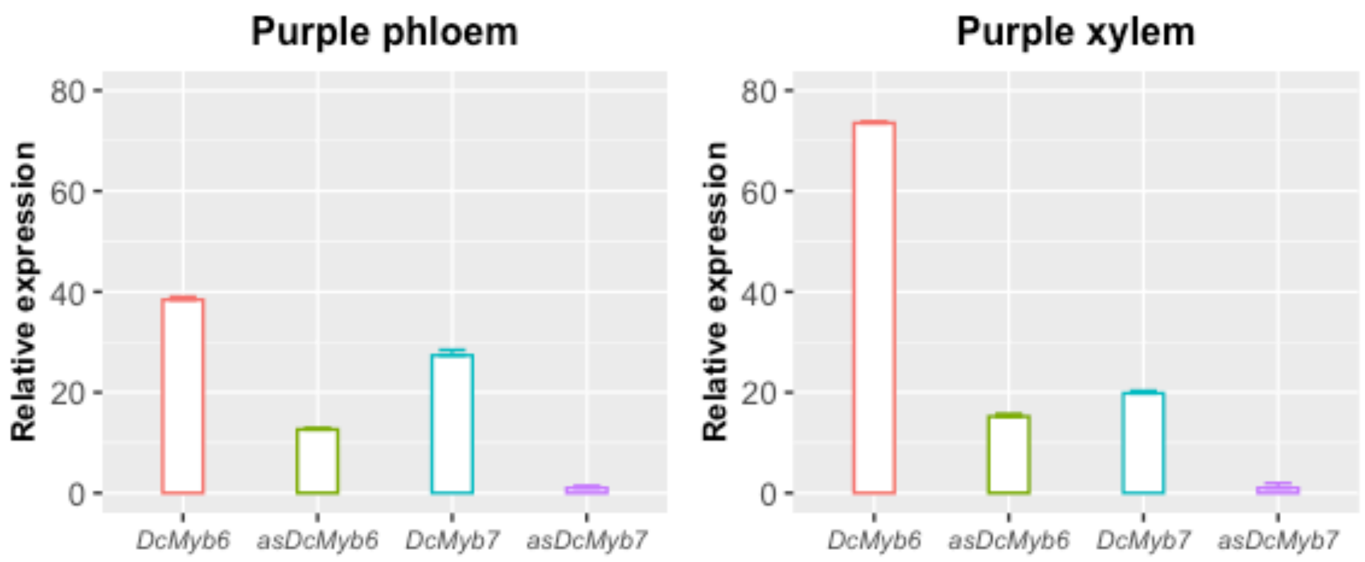

### Supplementary Table S1

| Library ID | Sample ID | Total read bases (bp) | Total read bases (Gbp) | Total reads | GC(%) | AT(%) | Q20(%) | Q30(%) | Uniquely mapped reads (%) |
|---|---|---|---|---|---|---|---|---|---|
| MORADA_M1F1 | Pur_F1 | 5699079126 | 5.7 | 56426526 | 44.83 | 55.17 | 98.1 | 94.2 | 90.9 |
| MORADA_M3F2 | Pur_F2 | 5263944462 | 5.26 | 52118262 | 44.81 | 55.19 | 98.1 | 94.1 | 91.5 |
| MORADA_M3F3 | Pur_F3 | 4935182998 | 4.94 | 48863198 | 45.26 | 54.74 | 97.9 | 93.9 | 87.8 |
| MORADA_M1X1 | Pur_X1 | 4390776232 | 4.39 | 43473032 | 44.78 | 55.22 | 98.1 | 94.2 | 92 |
| MORADA_M3X2 | Pur_X2 | 5964969302 | 5.96 | 59059102 | 44.75 | 55.25 | 98.1 | 94.1 | 91.7 |
| MORADA_M3X3 | Pur_X3 | 4845934146 | 4.85 | 47979546 | 44.91 | 55.09 | 98 | 93.9 | 86.5 |
| NARANJA_M3F1 | Org_F1 | 4704372748 | 4.7 | 46577948 | 44.67 | 55.33 | 98 | 93.9 | 91.9 |
| NARANJA_M3F2 | Org_F2 | 4703444760 | 4.7 | 46568760 | 44.79 | 55.21 | 97.9 | 93.7 | 94.1 |
| NARANJA_M3F4 | Org_F4 | 4779354946 | 4.78 | 47320346 | 44.47 | 55.53 | 98 | 94.1 | 92.8 |
| NARANJA_M3X1 | Org_X1 | 6093490994 | 6.09 | 60331594 | 45.11 | 54.89 | 98 | 94 | 92.6 |
| NARANJA_M3X2 | Org_X2 | 5819580004 | 5.82 | 57619604 | 44.67 | 55.33 | 98 | 94.2 | 86.7 |
| NARANJA_M3X4 | Org_X4 | 5069705100 | 5.07 | 50195100 | 44.28 | 55.72 | 98.1 | 94.3 | 92.8 |
| | | **Average** | 5.19 | 51377752 | 44.78 | 55.22 | 98 | 94.1 | 90.9 |
| | | **STD** | 0.569 | | 0.258 | 0.258 | 0.08 | 0.18 | 2.53 |

### Supplementary Table S3

| Transcript type | Known | New | Total |
|---|---|---|---|
| coding | 32109 | 2095 | 34204 |
| noncoding | 915 | 6373 | 7288 |
| NAT | 0 | 1521 | 1521 |
| lincRNA | 915 | 4852 | 5767 |
| structural | 1239 | 16 | 1255 |
| total | 34263 | 8484 | 42747 |

### Supplementary Table S6

| | | Correlation | | | |
|---|---|---|---|---|---|
| | | Pearson *r* | | Spearman's Rho | |
| **coding** | **lncNAT** | *r* | *p* -value | $r_s$ | *p* -value |
| ***DcMyb7*** | ***asDcMyb7*** | 0.81 | < 0.01 | 0.89 | <0.001 |
| ***DcMyb6*** | ***asDcMyb6*** | 0.86 | <0.001 | 0.79 | <0.01 |
| **DCAR_007914** | **MSTRG.6643** | -0.75 | <0.01 | -0.83 | <0.001 |
| **DCAR_010712** | **MSTRG.9085** | 0.87 | <0.001 | 0.85 | <0.001 |
| **DCAR_014717** | **MSTRG.11960** | 0.91 | <0.0001 | 0.89 | <0.001 |
| **DCAR_015415** | **MSTRG.11340** | -0.89 | <0.0001 | -0.84 | <0.001 |
| **DCAR_015459** | **MSTRG.11308** | 0.83 | <0.001 | 0.85 | <0.001 |
| **DCAR_021597** | **MSTRG.18539** | 0.71 | <0.01 | 0.74 | <0.01 |
| **DCAR_022107** | **MSTRG.18052** | 0.84 | <0.001 | 0.91 | <0.0001 |
| **DCAR_022871** | **MSTRG.17454** | 0.91 | <0.0001 | 0.90 | <0.0001 |
| **DCAR_023986** | **MSTRG.20882** | 0.71 | <0.01 | 0.81 | <0.01 |
| **DCAR_029820** | **MSTRG.26086** | 0.75 | <0.01 | <0.75 | <0.01 |
| **DCAR_031718** | **MSTRG.27809** | -0.88 | <0.001 | -0.88 | <0.001 |
| **DCAR_032195** | **MSTRG.28149** | 0.75 | <0.01 | 0.80 | <0.01 |
| **DCAR_009536** | **MSTRG.8070** | -0.82 | <0.01 | -0.80 | <0.01 |
| **DCAR_006881** | **MSTRG.5651** | 0.96 | <0 .00001 | 0.95 | <0 .00001 |
| **DCAR_007044** | **MSTRG.5819** | 0.98 | < 0.00001 | 0.90 | <0.001 |
| **DCAR_004853** | **MSTRG.3846** | 0.95 | <0.00001 | 0.92 | <0.0001 |
| **DCAR_006773** | **MSTRG.5544** | 0.81 | <0.01 | 0.85 | <0.001 |

### Supplementary Table S7

| Name | Sense | Sequence (5'->3') | Length | Tm | Product length |
|---|---|---|---|---|---|
| **DcActin-7** | **Forward** | GCTACTTGGATGTGGCTGT | 19 | 52.63 | 93 |
| | **Reverse** | CCGACTGAAGAGCTGATCC | 19 | 57.8 | |
| **asDcMyb7-1** | **Forward** | GTGTTGCTGGCAAAGGAGAGT | 21 | 61.09 | 145 |
| | **Reverse** | AAACAAACAAGCAGTGCGACTA | 22 | 59.32 | |
| **DcMyb7-1** | **Forward** | TGTTGTTAATGTCGTTGCCG | 20 | 57.33 | 128 |
| | **Reverse** | CCTTCCCCGGACCTTATCAAA | 21 | 59.44 | |
| **asDcMyb6-2** | **Forward** | TGGTCGAAGATAGTTGAGCCAC | 22 | 60.09 | 146 |
| | **Reverse** | GCAGGTACACAATTCTTCCCACAA | 24 | 61.53 | |
| **DcMyb6-1** | **Forward** | CACCACTTCGGTCTGTCCAT | 20 | 55.00 | 265 |
| | **Reverse** | CCCTCGGAGAGCAGGGTTG | 19 | 62.02 | |
